# Bioinformatic characterization of angiotensin-converting enzyme 2, the entry receptor for SARS-CoV-2

**DOI:** 10.1101/2020.04.13.038752

**Authors:** Harlan Barker, Seppo Parkkila

## Abstract

The World Health Organization declared the COVID-19 epidemic a public health emergency of international concern on March 11th, 2020, and the pandemic is rapidly spreading worldwide. COVID-19 is caused by a novel coronavirus SARS-CoV-2, which enters human target cells via angiotensin converting enzyme 2 (ACE2). We used a number of bioinformatics tools to computationally characterize ACE2 by determining its cell-specific expression in trachea, lung, and small intestine, derive its putative functions, and predict transcriptional regulation. The small intestine expressed higher levels of ACE2 than any other organ. The large intestine, kidney and testis showed moderate signals, whereas the signal was weak in the lung. Single cell RNA-Seq data from trachea indicated positive signals along the respiratory tract in key protective cell types including club, goblet, proliferating, and ciliary epithelial cells; while in lung the ratio of ACE2-expressing cells was low in all cell types (<2.6%), but was highest in vascular endothelial and goblet cells. Gene ontology analysis suggested that, besides its classical role in renin-angiotensin system, ACE2 may be functionally associated with angiogenesis/blood vessel morphogenesis. Using a novel tool for the prediction of transcription factor binding sites we identified several putative binding sites within two tissue-specific promoters of the *ACE2* gene. Our results also confirmed that age and gender play no significant role in the regulation of ACE2 mRNA expression in the lung.

**IMPORTANCE:** Vaccines and new medicines are urgently needed to prevent spread of COVID-19 pandemic, reduce the symptoms, shorten the duration of disease, prevent virus spread in the body, and most importantly to save lives. One of the key drug targets could be angiotensin-converting enzyme 2 (ACE2), which is a crucial receptor for the corona virus (SARS-CoV-2). It is known that SARS coronavirus infections lead to worse outcome in the elderly and in males. Therefore, one aim of the present study was to investigate whether age or sex could contribute to the regulation of ACE2 expression. We also decided to explore the transcriptional regulation of *ACE2* gene expression. Since data on ACE2 distribution is still conflicting, we aimed to get a more comprehensive view of the cell types expressing the receptor of SARS-CoV-2. Finally, we studied the coexpression of ACE2 with other genes and explored its putative functions using gene ontology enrichment analysis.

## INTRODUCTION

A zinc metalloenzyme, angiotensin-converting enzyme (ACE) was discovered 64 years ago and first named as a hypertension-converting enzyme [1]. Classically, ACE is well known for its roles in the regulation of arterial pressure through conversion of angiotensin I to active angiotensin II and cleavage of bradykinin and neurotensin [2]. As a zinc metalloenzyme, ACE belongs to a large cluster of zinc-binding proteins. The first zinc metalloenzyme, carbonic anhydrase was discovered in 1932 by Meldrum and Roughton [3] and thereafter thousands of such metalloenzymes have been reported in different species of all phyla [4, 5].

Angiotensin-converting enzyme 2 (ACE2) was first discovered in 2000 when a novel homologue of ACE was cloned [2, 6, 7]. Although ACE and ACE2 share significant sequence similarity in their catalytic domains, they appear to act on different peptide substrates of angiotensins [8, 9]. Previous studies identified ACE2 as a functional receptor for severe acute respiratory syndrome corona virus 1 (SARS-CoV-1) which led to an outbreak of SARS infection in 2003 [10]. ACE2 is also a crucial receptor for the novel corona virus (SARS-CoV-2), which has caused a large global outbreak of COVID-19 infection with rapidly growing numbers of patients (5,488,825 confirmed cases as of May 27^th^, 2020, https://www.who.int/emergencies/diseases/novel-coronavirus-2019). A recent report suggested that soluble ACE2 fused to the Fc portion of immunoglobulin can neutralize SARS-CoV-2 *in vitro* [11]. This result was further confirmed by showing that human recombinant soluble ACE2 reduced SARS-CoV-2 infection on cultured Vero-E6 cells in a dose dependent manner [12]. Therefore, ACE2 also holds promise for treating patients with coronavirus infection.

The structural key for target cell infection by coronavirus is the viral spike (S) protein of SARS-CoV. ACE2 acts as a locking device for the virus, whereby the binding of the surface unit S1 facilitates viral attachment to the surface of target cells [13]. The cellular serine protease (TMPRSS2) promotes SARS-CoV entry via a dual mechanism. It cleaves both the SARS-CoV S protein and the virus receptor, ACE2, promoting both the viral uptake and the viral and cellular membrane fusion events [13–15]. The critical residues contributing to the receptor-spike protein interaction were first determined for SARS-CoV-1 [16] and recently in three independent studies for SARS-CoV-2 [17–19]. It has been proposed by biolayer interferometry studies that the receptor-binding domains of SARS-CoV-1 and SARS-CoV-2 S proteins bind with comparable affinities to human ACE2 [20]. In contrast, a modelling study suggested that binding of SARS-CoV-2 is stronger [21], which was convincingly confirmed by structural and biochemical data [17, 18].

The clinical characteristics of COVID-19 infection have recently been described based on data from 1,099 patients from mainland China [22]. It was found that the clinical characteristics of COVID-19 mimic those of SARS-CoV-1 infection. The most dominant symptoms include fever, cough, fatigue, and sputum production, whereas gastrointestinal symptoms are less common. In laboratory parameters, lymphopenia was detected in 83.2% of patients on admission. According to another recent survey of 278 patients with pneumonia caused by SARS-CoV-2, fever was the most common symptom, followed by cough [23]. Bilateral pneumonia has been detected by computed tomography scans in 67.0% of patients [24]. A recent study from Wuhan, China listed the most common clinical complications determined in critically ill COVID-19 patients [25]. The complications during clinical worsening included acute respiratory distress syndrome and respiratory failure, sepsis, acute cardiac injury, and heart failure.

Data on the localization of virus receptors can provide insight into mechanisms of virus entry, tissue tropism, and pathogenesis of the disease. Therefore, it is of particular interest to correlate COVID-19 symptoms with the distribution pattern of ACE2. The first studies performed by northern blotting indicated that ACE2 is located in the human heart, kidney, and testis [2]. By immunohistochemistry, the expression of the ACE2 protein was identified in the human lung alveolar epithelial cells (type I and II pneumocytes), enterocytes of the small intestine, the brush border of the renal proximal tubules, and the endothelial cells of arteries and veins and arterial smooth muscle cells in several organs [26]. It was proposed that this distribution pattern of ACE2 could explain the tissue tropism of SARS-CoV-1 for the lung, small intestine, and kidney [27]. On the other hand, the symptoms of COVID-19, in contrast to SARS-CoV-1 infection, are not associated to the same extent with the gastrointestinal tract in spite of the high expression of ACE2 in the intestinal enterocytes [28]. In COVID-19, diarrhea has been reported in just 3.8% of patients, in contrast to 40-70% in SARS-CoV-1 infection [22, 29]. A recent preprint report indicated diarrhea in 18.1% of 254 COVID-19 patients [30].

There are conflicting reports on the expression of ACE2 in the upper respiratory tract [29]. Hamming and coworkers found that only the basal layer of nonkeratinized airway squamous epithelium shows positive signal [26], whereas Sims and colleagues demonstrated ACE2 expression on the luminal surface of ciliated cells in freshly excised human nasal and tracheobronchial tissue [31]. Ren and coworkers showed weak ACE2-positive signal in the epithelial cells of trachea and main bronchus [32]. Although lymphopenia is a typical feature of SARS [22, 29], ACE2 is not highly expressed on T or B cells or macrophages in the spleen or lymphoid organs [26].

It is known that both SARS-CoV and SARS-CoV-2 infections lead to worse outcome in the elderly [29, 33]. Therefore, one aim of the present study was to investigate whether age could contribute to the regulation of ACE2 expression. We also decided to explore the transcriptional regulation of *ACE2* gene expression using a novel computational tool recently developed by the first author of this article. Notably, data on ACE2 distribution is still conflicting, and thus we aimed to get a more comprehensive view of the cell types expressing the receptor of SARS-CoV-2. Finally, we studied the coexpression of ACE2 with other genes and explored its putative functions using a gene ontology enrichment analysis.

## RESULTS

### ACE2 is weakly expressed in the lung

The first aim of our study was to investigate different human tissues using publicly available datasets for the distribution of ACE2 mRNA and protein. In the FANTOM5 dataset, the highest values for ACE2 mRNA, ranked according to signal intensity, were seen for the small intestine, dura mater, colon, testis, thalamus, and rectum (Fig. 1).

**Figure 1.**
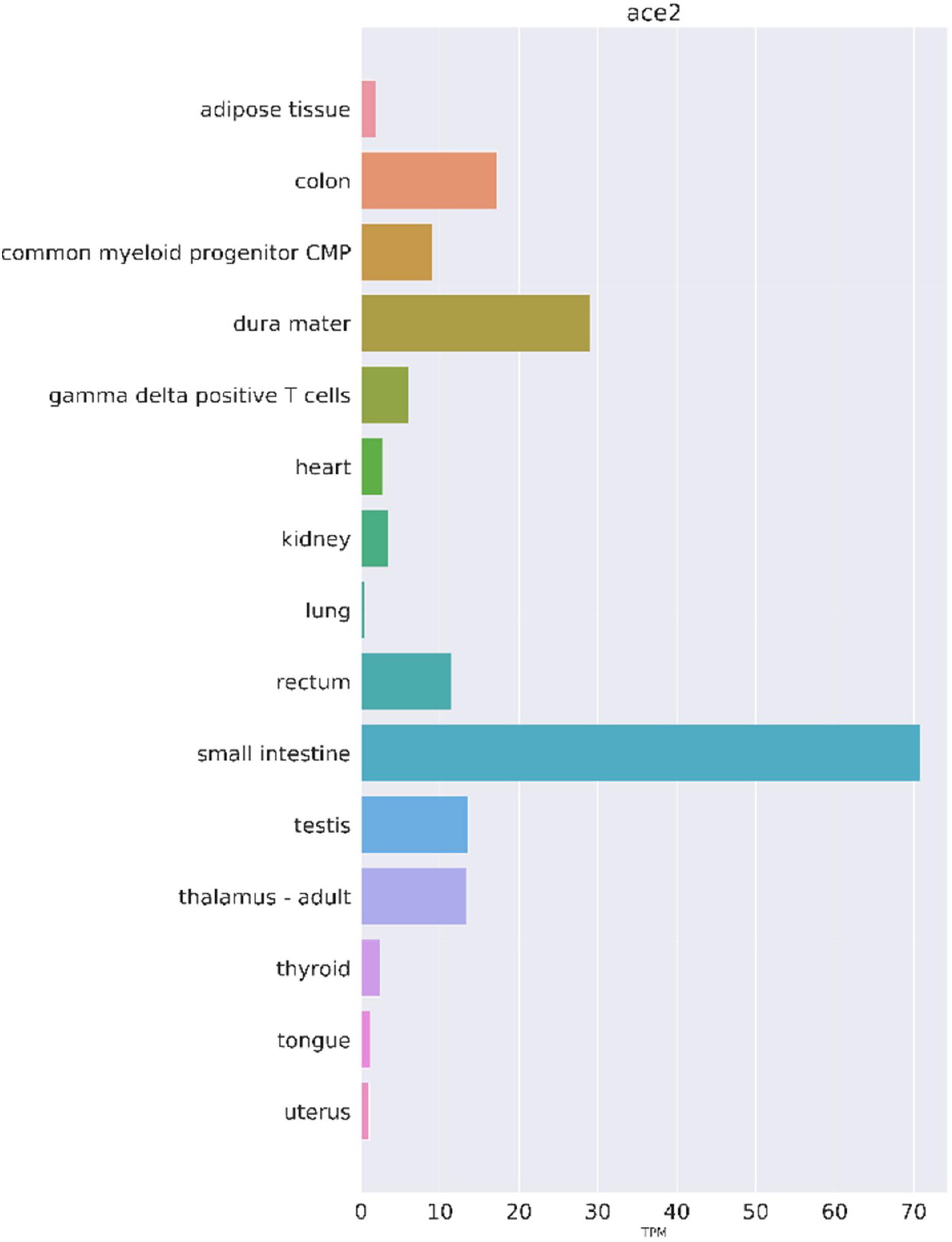
Expression of ACE2 mRNA in selected human tissues. Expression values as TPM have been extracted from the FANTOM5 dataset.

Figure 2 shows the expression of ACE2 protein in selected human tissues. Representative example images of the ACE2 immunostaining were prepared from tissue specimens of the Human Protein Atlas database (https://www.proteinatlas.org/). The results indicate a strong signal for ACE2 protein in the brush border of small intestinal enterocytes. In the kidney, prominent immunostaining reactions were present in the epithelial cells of proximal convoluted tubules and Bowman’s capsule. The seminiferous tubules and interstitial cells of testis also demonstrated strong immunostaining. No immunoreactions for ACE2 were observed in the lung specimens. Very weak signal, associated with apical membranes, was detected in sporadic ciliary cells of a nasopharyngeal mucosa sample. Although the evaluation of immunostaining reaction is generally considered semiquantitative at most, the results seem to correlate fairly well with the corresponding mRNA expression levels.

**Figure 2.**
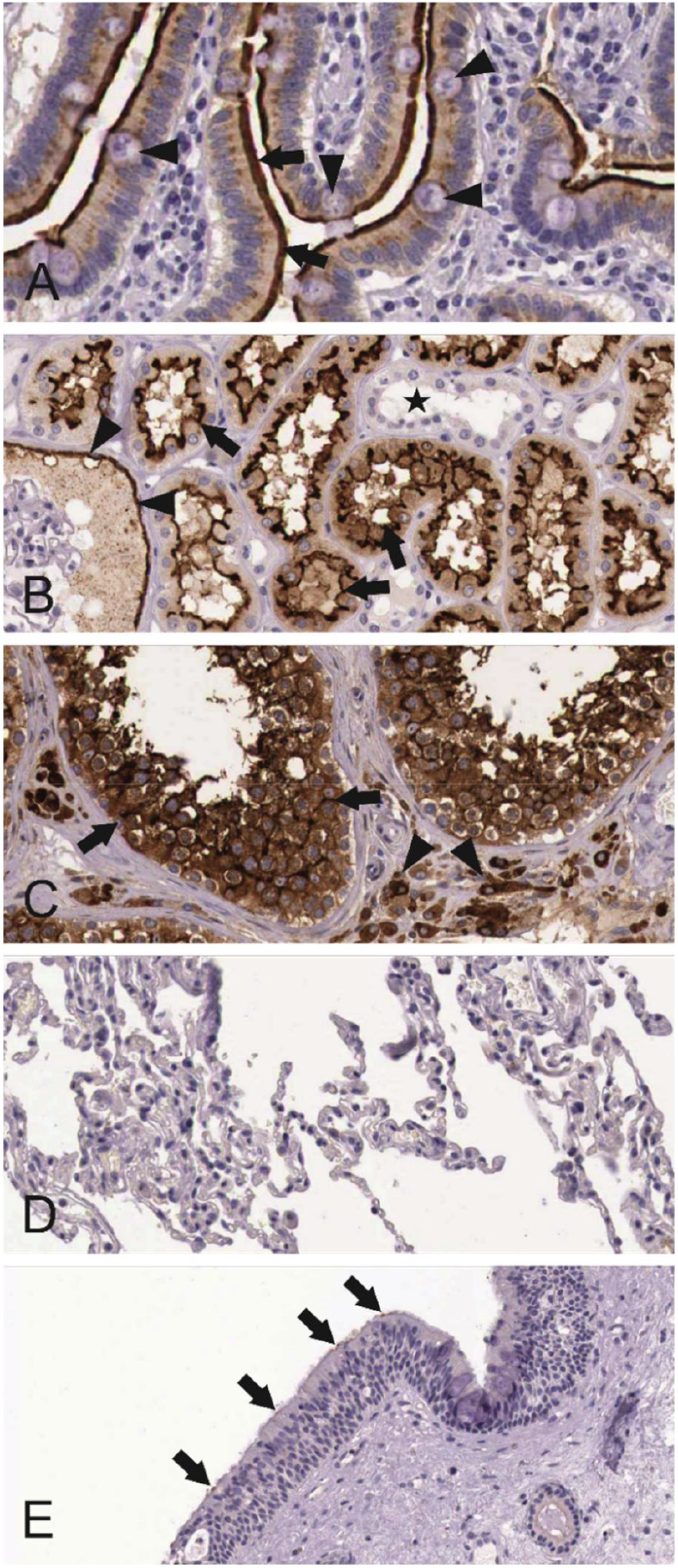
Immunohistochemical localization of ACE2 protein in selected human tissues. In the duodenum (A), the protein is most strongly localized to the apical plasma membrane of absorptive enterocytes (arrows). The goblet cells (arrowheads) show weaker apical staining. Intracellular staining is confined to the absorptive enterocytes. In the kidney (B), ACE2 shows prominent apical staining in the epithelial cells of the proximal convoluted tubules (arrows) and Bowman’s capsule epithelium (arrowheads). The distal convoluted tubules are negative (asterisk). The testis specimen (C) shows strong immunostaining in the seminiferous tubules (arrows) and interstitial cells (arrowheads). The lung sample (D) is negative. In the nasopharyngeal mucosa (E), ACE2 signal is very weak and only occasional epithelial cells show weak signals (arrows). Immunostained specimens were taken from the Protein Expression Atlas (https://www.proteinatlas.org/).

### Single cell RNA-Seq analysis indicates cell-specific expression for ACE2 mRNA

The respiratory tract is the main target region that is affected by COVID-19 infection. Bulk RNA-Seq data from lung specimens showed low expression levels for ACE2 (Fig. 1). Therefore, we performed an analysis of single cell RNA-Seq using both human lung and mouse trachea datasets, representing the breadth of the lower respiratory tract. Figures 3 and 4 show the expression of ACE2 mRNA in identified cell types of lung and trachea, respectively. In lung, ACE-2 expressing cells are generally uncommon with no cell type having a ratio of ACE2-expressing cells greater than 2.6%. The cell types with the greatest proportion of ACE2 expression are those of arterial vascular endothelial cells (2.55%), goblet cells (2.02%), and venous vascular endothelial cells (1.33%). In trachea, the highest ratio of ACE2-expressing cells included the club cells (16.62%), goblet cells (13.84%), and ciliary epithelial cells (6.63%).

**Figure 3.**
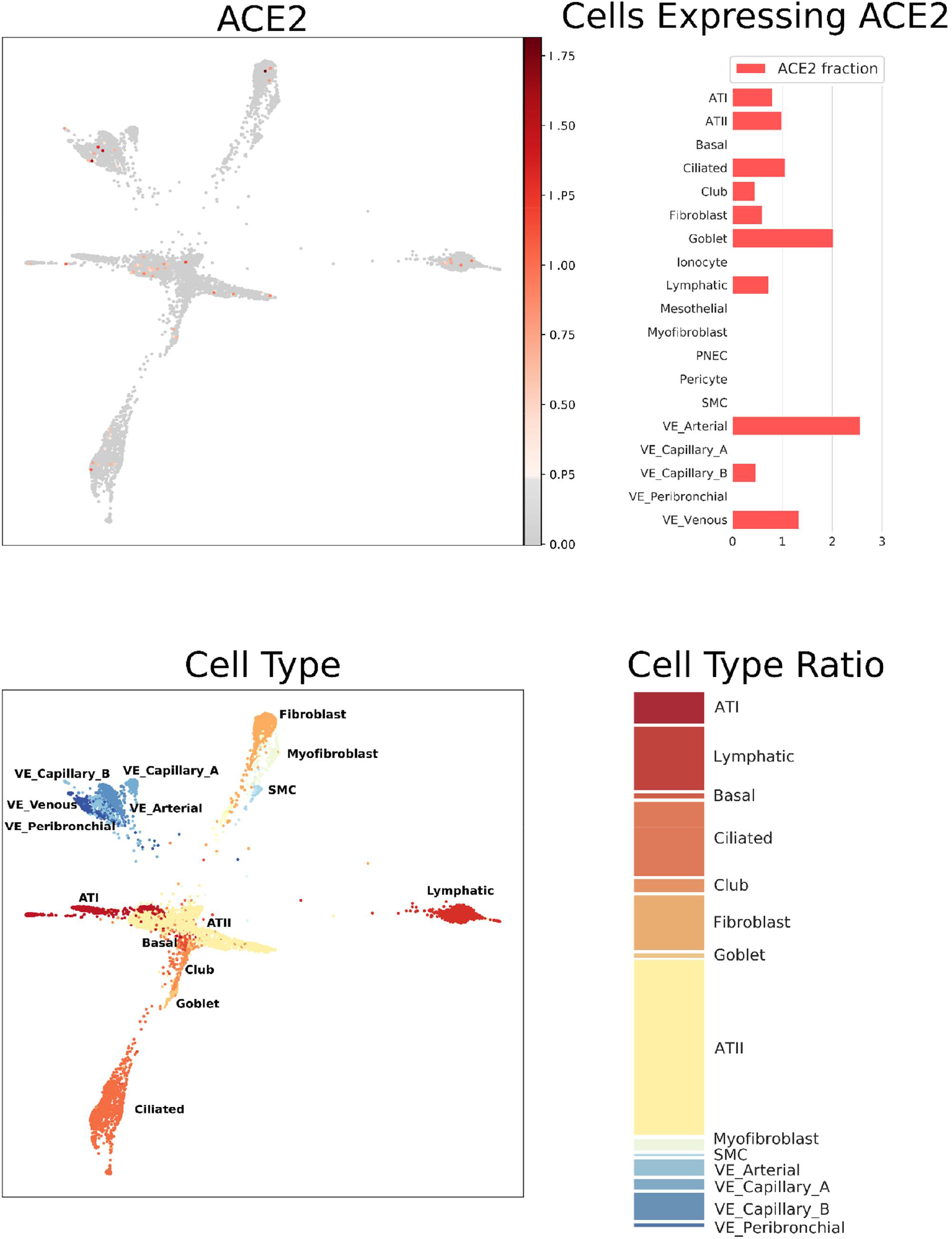
Single cell RNA-Seq analysis of different cell types from the lung (human), derived from data from GEO dataset GSE136831 [70]. ACE2 mRNA expression as normalized, batch-corrected counts is shown for comparison in upper panel. The force directed layout plot was computed and visualized in ScanPy [74]. For each cell type the ratio of cells expressing ACE2 is presented in addition to a stacked barplot of the relative cell type frequencies in the whole dataset. Alveolar type I (ATI), alveolar type II (ATII), pulmonary neuroendocrine cells (PNEC), smooth muscle cells (SMC), vascular endothelia (VE).

**Figure 4.**
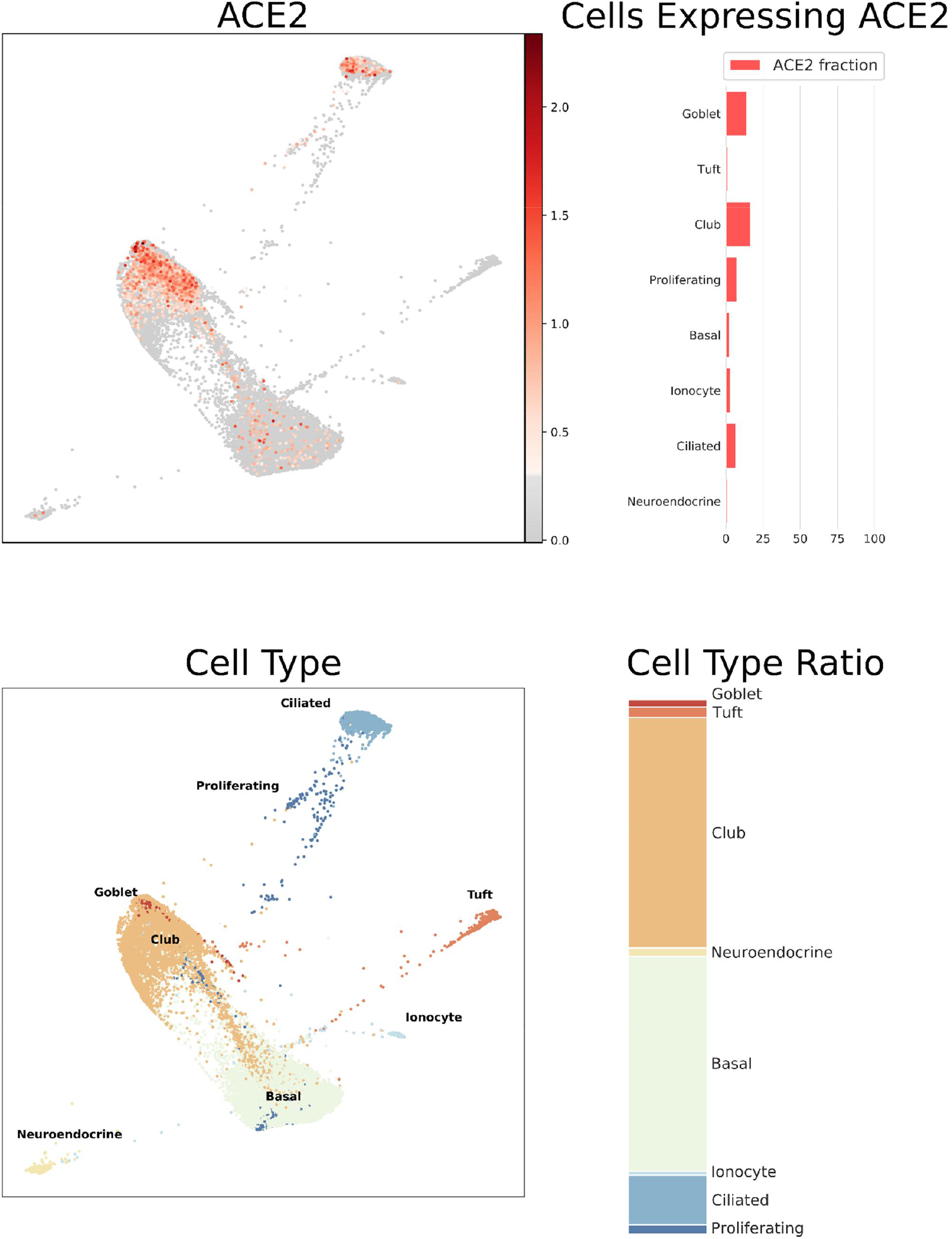
Single cell RNA-Seq analysis of different cell types from the respiratory tract (mouse tracheal epithelium), derived from data from GEO dataset GSE103354 [71]. ACE2 mRNA expression as normalized, batch-corrected counts is shown for comparison in upper panel. The force directed layout plot was computed and visualized in ScanPy [74]. For each cell type the ratio of cells expressing ACE2 is presented in addition to a stacked barplot of the relative cell type frequencies in the whole dataset.

Since both the airways and intestine contain goblet cells, SARS-CoV-1 affects gastrointestinal tract, and bulk RNA-Seq data shows high expression in small intestine and colon, we decided to analyze another single cell RNA-Seq dataset covering mouse intestinal epithelial cells. Figure 5 indicates the highest levels of ACE2 mRNA signal in the absorptive enterocytes (44.09%), whereas the intestinal goblet cells (1.35%) remain mostly negative.

**Figure 5.**
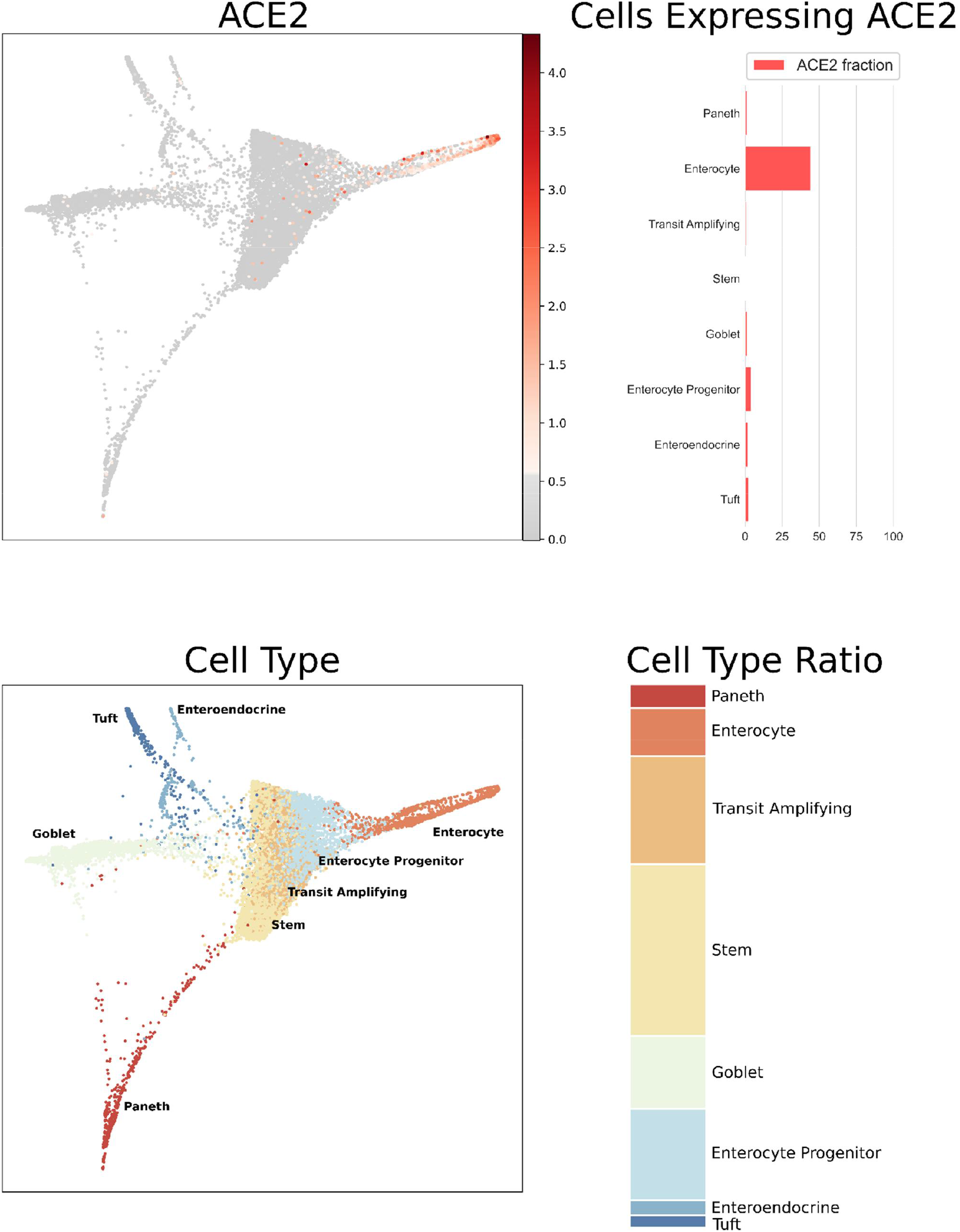
Single cell RNA-Seq analysis of mouse intestinal epithelial cells. Data is from GEO dataset GSE92332 [72]. ACE2 mRNA expression as normalized, batch-corrected counts is shown for comparison in upper panel. The force directed layout plot was computed and visualized in ScanPy [74]. For each cell type the ratio of cells expressing ACE2 is presented in addition to a stacked barplot of the relative cell type frequencies in the whole dataset.

### ACE2 mRNA expression levels are unrelated to age and gender in the lung

Since both age and gender may contribute to onset and severity of COVID-19 symptoms we aimed to investigate the effect of these variables on the expression levels of ACE2 mRNA. Figure 6 indicates that some tissues showed a slight trend to lower expression in older age categories. Among all tested tissues, statistically significant differences between the age categories were seen in the tibial nerve (p=8.58 x 10^-6^), minor salivary gland (p=0.002), aorta (p=0.003), whole blood (p=0.005), transverse colon (p=0.010), hypothalamus (p=0.039), and sun exposed skin (p=0.046). Importantly, the lung specimens showed no significant difference of ACE2 mRNA expression between different age categories (p=0.681). Complete data on ACE mRNA expression levels in different age categories are shown in Supplementary Table 1. To make a binary comparison of expression by age, samples were divided into groups of ≤45 and >45 years of age. In comparison of these younger and older age groups, significant differences in expression were found in tibial nerve (p=2.47 x 10^-7^), whole blood (p=3.21 x 10^-4^), minor salivary gland (p=4.89 x 10^-4^), sun exposed skin (p=0.003), transverse colon (p=0.022), testis (p=0.025), esophageal muscle layer (p=0.040), and subcutaneous adipose tissue (p=0.045).

**Figure 6.**
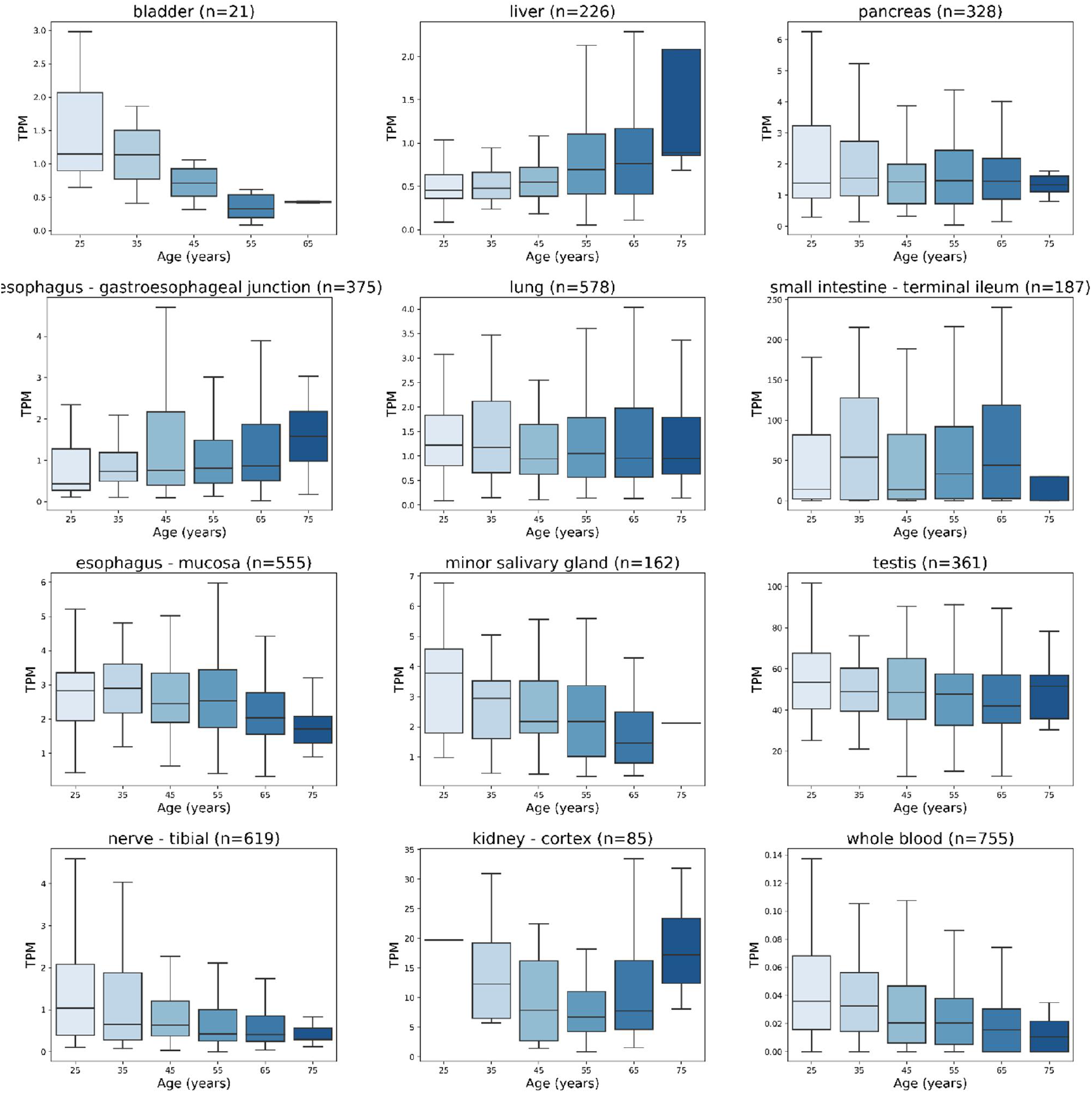
Effect of age on ACE2 mRNA expression levels. Data is extracted from the GTEx dataset as TPM. In these organs, ANOVA revealed significant differences between age categories in tibial nerve (p=8.58 x 10^-6^), minor salivary gland (p=0.002), and whole blood (p=0.005). In other tissues, the differences did not reach statistical significance. The highest TPM values are seen the small intestine, testis, and kidney.

ACE2 mRNA levels largely overlapped between male and female sexes as shown in Figure 7. In the lung, no statistically significant difference was observed in the expression levels between the male and female subjects (p=0.908). Statistically significant differences were observed in the adipose tissue (p=0.0001), whole blood (p=0.0002), amygdala (p=0.0006), transverse colon (p=0.0008), muscle layer of esophagus (p=0.002), left ventricle of heart (p=0.005), Epstein-Barr virus-transformed lymphocytes (p=0.015), and esophagus-gastroesophageal junction (p=0.024). Notably, there was no clear sex-specific trend pointing to one direction in all these cases. ACE mRNA expression levels in all studied tissues sorted according to subjects’ gender are shown in Supplementary Table 2.

**Figure 7.**
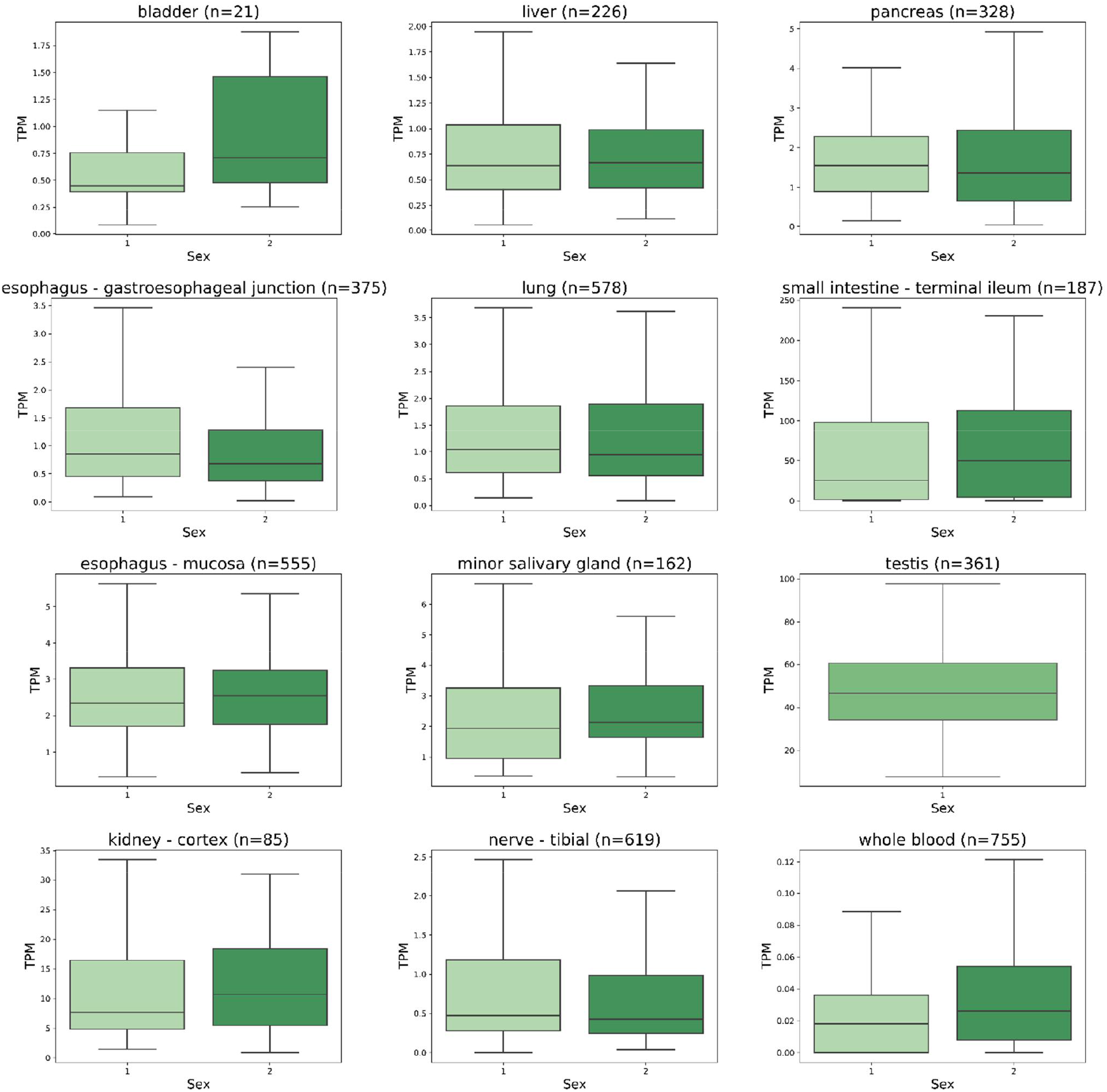
Effect of gender on ACE2 mRNA expression levels. Data is extracted from the GTEx dataset as TPM. The expression levels in males and females overlap in all tissue categories. Statistically significant differences studied by ANOVA analysis were determined in esophagus-gastroesophageal junction (p=0.024) and whole blood (p=0.0002). The ACE2 mRNA expression levels in testis specimens are shown here for comparison. 1=male, 2=female.

**Figure 8.**
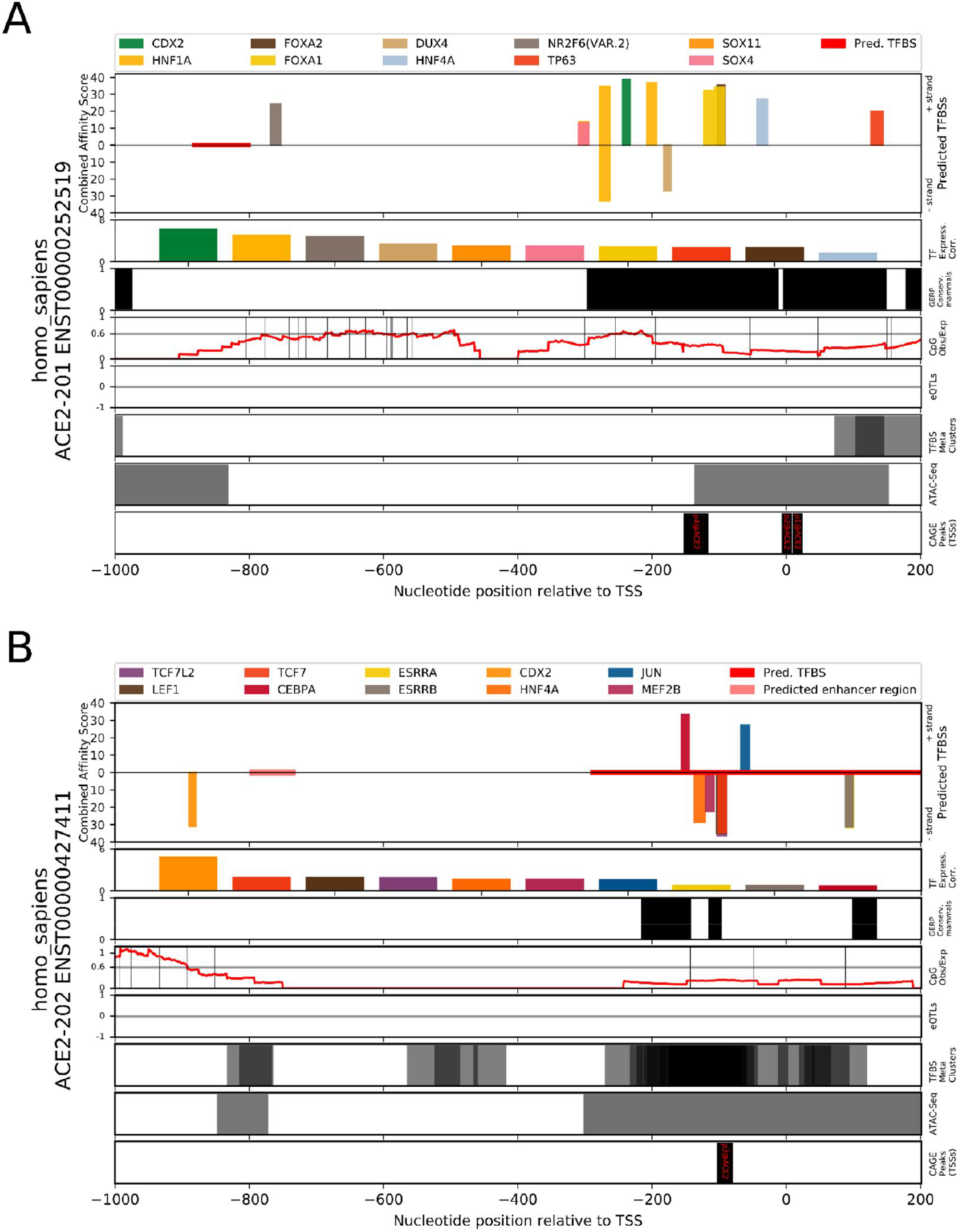
Prediction of transcription factor binding sites in the human *ACE2* gene promoter regions of Ensembl transcripts ENST00000252519 and ENST00000427411 using TFBSfootprinter. A) Promoter region of the intestine specific ACE2 transcript. The results show putative binding sites for several transcription factors which have a strong correlation of expression with ACE2 in colon, kidney, and ileum. The predicted binding sites overlap regions of conservation in mammal species (Ensembl GERP), and cluster within 400 base pairs (bp) of the transcription start site. B) Promoter region of the lung specific ACE2 transcript. The predicted binding sites cluster within 200 bp of the TSS and overlap regions of conservation in mammal species, ATAC-Seq peaks (ENCODE), and TFBS metaclusters (GTRD).

### Proximal promoter contains putative TFBSs for ileum, colon, and kidney expression

TFBS analysis of the *ACE2* intestinal transcript promoter (ENST00000252519) revealed several candidate binding sites which occur in a cluster extending from 400 bp upstream of the transcription start site; CDX2, HNF1A, FOXA1, SOX4, TP63, HNF4A, DUX4, FOXA2, NR2F6, and SOX11 (Fig. 7A). These predicted sites overlap an evolutionarily conserved region in mammals and are proximal to several ATAC-Seq peaks. In several tissues these TFs are found to be highly positively correlated (>0.7) with expression of ACE2: CDX2 (colon, terminal ileum), HNF1A (colon, kidney, terminal ileum), FOXA1 (cervix, colon, terminal ileum), HNF4A (colon, terminal ileum), FOXA2 (colon, kidney), NR2F6 (colon, kidney, terminal ileum), and SOX11 (kidney). In addition, two of the TFs are highly negatively correlated with ACE2 expression DUX4 (kidney) and FOXA1 (kidney). Full prediction results are included as Supplementary Table 3 and TF correlations by tissue are present in Supplementary Table 4.

Analysis of the *ACE2* lung transcript promoter (ENST00000427411) produced putative TFBS predictions for ESRRA, HNF4A, CDX2, CEBPA, ESRRB, MEF2B, TCF7, TCF7L2, JUN, and LEF1 (Fig. 7B). The predicted TFBSs clustered within 200 base pairs of the TSS, and overlap with evolutionarily conserved regions, TFBS metaclusters, and ATAC-Seq peaks. The TFs corresponding to predicted TFBSs, which are positively correlated (>0.7) with ACE2 expression, are ESRRA (terminal ileum, colon), HNF4A (terminal ileum, colon), CDX2 (colon, terminal ileum), CEBPA (colon, terminal ileum), ESRRB (cervix), TCF7L2 (testis). Those TFBSs with TFs which strongly (<−0.7) negatively correlate with ACE2 are ESRRA (kidney) and TCFL72 (kidney). The lung-specific transcript TSS aligns with the p3@ACE2 FANTOM5 dataset CAGE peak, which indicates that the expression of this transcript is much lower than the intestinal transcript, which corresponds with p1@ACE2 and p2@ACE2 FANTOM5 CAGE peaks. Common between the two tissue-specific transcripts, are predictions for CDX2 and HNF-family transcription factors.

### ACE2 mRNA expression correlates with metalloproteases and transporter genes

Coexpression analysis identified numerous genes in ileum, testis, colon, and kidney which are highly correlated (>0.8) with ACE2 (Table 1; Supplementary Table 5). In particular, in the ileum there are a number of genes with correlation values greater than 0.95. In contrast, analysis of the lung shows a maximum correlation of expression of 0.6275. The genes with which ACE2 mRNA expression shows the highest levels of coexpression code for metalloprotease and transporter proteins. Selected tissue-specific ACE2-correlated genes, determined with bulk RNA-Seq data, are presented with their expression levels within the scRNA-Seq trachea and intestinal datasets in Figure 9.

**Figure 9.**
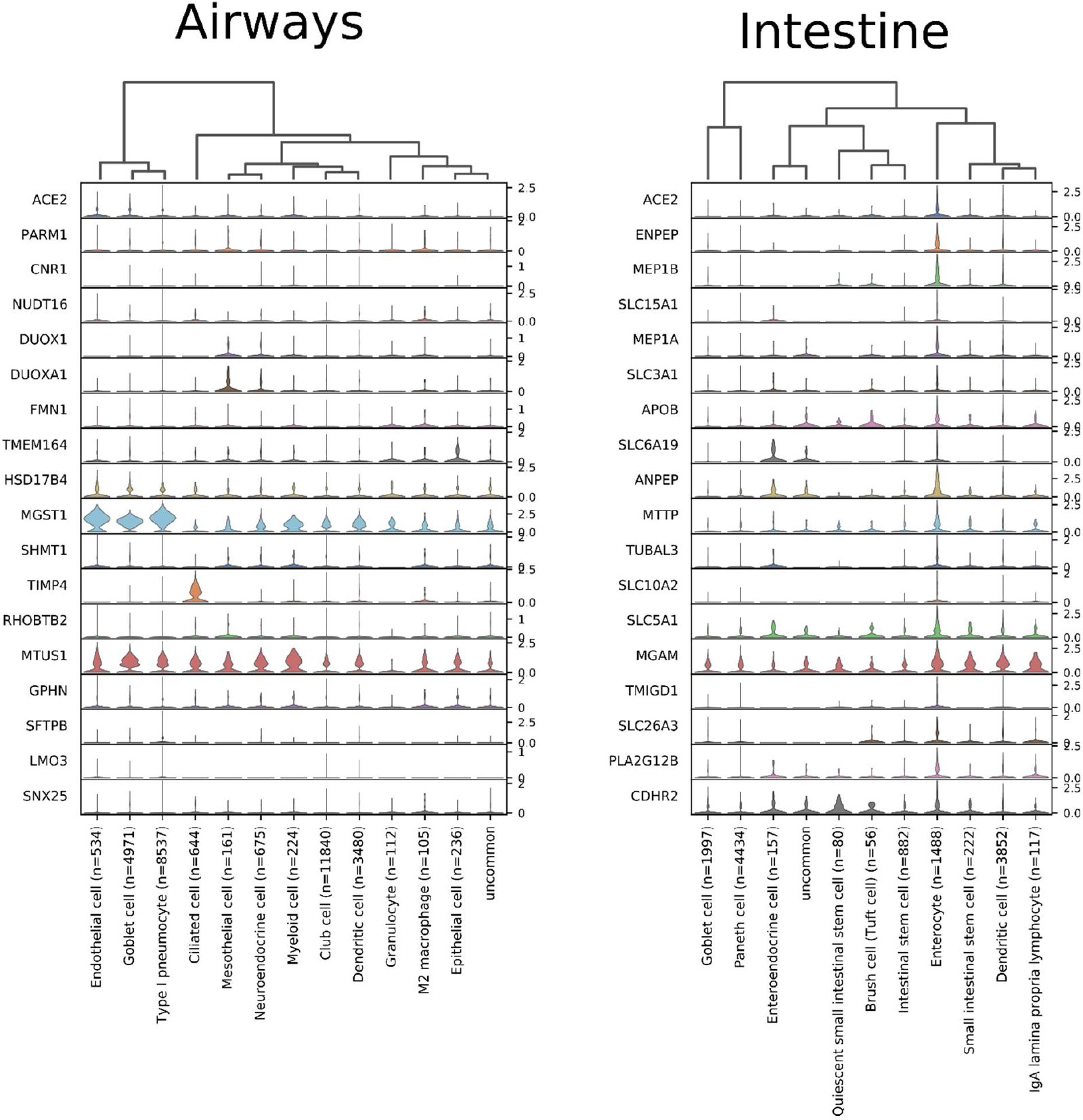
Expression of genes most highly correlated with ACE2 in single cell datasets of trachea and intestinal epithelia. Trachea expression data is taken from GSE103354 [71] and intestinal epithelia data is derived from GSE92332 [72]. Visualized in ScanPy [74].

**Table 1.** Genes associated with ACE2 mRNA expression in selected human tissues. Derived from GTEx bulk RNA-Seq data [66].

| Tissue | Correlated_<br>gene | Correlation | p-value | HGNC | UniProt | Description | Panther protein class |
| --- | --- | --- | --- | --- | --- | --- | --- |
| terminal ileum | *ENPEP* | 0.9653 | 8.29E-110 | 3355 | Q07075 | Glutamyl aminopeptidase | metalloprotease (PC00153) |
| terminal ileum | *MEP1B* | 0.9653 | 8.35E-110 | 7020 | Q16820 | Meprin A subunit beta | metalloprotease (PC00153) |
| terminal ileum | *SLC15A1* | 0.9621 | 2.19E-106 | 10920 | P46059 | Solute carrier family 15 member 1 | transporter (PC00227) |
| terminal ileum | *MEP1A* | 0.9619 | 4.4E-106 | 7015 | Q16819 | Meprin A subunit alpha | metalloprotease (PC00153) |
| terminal ileum | *SLC3A1* | 0.9608 | 5.63E-105 | 11025 | Q07837 | Neutral and basic amino acid transport protein<br>rBAT | amylase (PC00048) |
| terminal ileum | *APOB* | 0.9587 | 6.45E-103 | 603 | P04114 | Apolipoprotein B-100 |  |
| terminal ileum | *SLC6A19* | 0.9581 | 2.33E-102 | 27960 | Q695T7 | Sodium-dependent neutral amino acid<br>transporter B(0)AT1 | primary active transporter<br>(PC00068) |
| terminal ileum | *ANPEP* | 0.9562 | 1.16E-100 | 500 | P15144 | Aminopeptidase N | metalloprotease (PC00153) |
| terminal ileum | *MTTP* | 0.9559 | 2.45E-100 | 7467 | P55157 | Microsomal triglyceride transfer protein large<br>subunit | transporter (PC00227) |
| terminal ileum | *TUBAL3* | 0.9554 | 6.24E-100 | 23534 | A6NHL2 | Tubulin alpha chain-like 3 | tubulin (PC00228) |
| testis | *PLP1* | 0.9092 | 1.07E-138 | 9086 | P60201 | Myelin proteolipid protein | myelin protein (PC00161) |
| colon-transverse | *MEP1A* | 0.9056 | 1.52E-152 | 7015 | Q16819 | Meprin A subunit alpha | metalloprotease (PC00153) |
| kidney | *TINAG* | 0.9012 | 7.07E-32 | 14599 | Q9UJW2 | Tubulointerstitial nephritis antigen | cysteine protease (PC00081) |
| colon-transverse | *GDA* | 0.8964 | 8.19E-145 | 4212 | Q9Y2T3 | Guanine deaminase | deaminase (PC00088) |
| colon-transverse | *MGAM2* | 0.8963 | 9E-145 | 28101 | Q2M2H8 | Probable maltase-glucoamylase 2 | glucosidase (PC00108) |
| colon-transverse | *TMEM236* | 0.894 | 6.66E-143 | 23473 | Q5W0B7 | Transmembrane protein 236 |  |
| colon-transverse | *GDPD2* | 0.8935 | 1.58E-142 | 25974 | Q9HCC8 | Glycerophosphoinositol<br>inositolphosphodiesterase GDPD2 |  |
| colon-transverse | *PLS1* | 0.8932 | 2.81E-142 | 9090 | Q14651 | Plastin-1 | non-motor actin binding protein<br>(PC00165) |
| colon-transverse | *HHLA2* | 0.893 | 3.59E-142 | 4905 | Q9UM44 | HERV-H LTR-associating protein 2 | immunoglobulin receptor<br>superfamily (PC00124) |
| colon-transverse | *SLC3A1* | 0.8928 | 5.27E-142 | 11025 | Q07837 | Neutral and basic amino acid transport protein<br>rBAT | amylase (PC00048) |
| colon-transverse | *COL17A1* | 0.8914 | 6.12E-141 | 2194 | Q9UMD9 | Collagen alpha-1(XVII) chain | extracellular matrix structural<br>protein (PC00103) |
| testis | *NDFIP1* | 0.8911 | 2.91E-125 | 17592 | Q9BT67 | NEDD4 family-interacting protein 1 |  |
| testis | *SPARC* | 0.8908 | 4.67E-125 | 11219 | P09486 | SPARC | extracellular matrix glycoprotein<br>(PC00100) |
| **colon-transverse** | *TINAG* | 0.8905 | 2.94E-140 | 14599 | Q9UJW2 | Tubulointerstitial nephritis antigen | cysteine protease (PC00081) |
| **testis** | *SCP2* | 0.8886 | 1.34E-123 | 10606 | P22307 | Non-specific lipid-transfer protein | transfer/carrier protein (PC00219) |
| **testis** | *SMARCA1* | 0.8852 | 2.44E-121 | 11097 | P28370 | Probable global transcription activator SNF2L1 | DNA helicase (PC00011) |
| **testis** | *ATP2B1* | 0.8829 | 6.45E-120 | 814 | P20020 | Plasma membrane calcium-transporting ATPase 1 | primary active transporter (PC00068) |
| **testis** | *VSIG1* | 0.8815 | 4.6E-119 | 28675 | Q86XK7 | V-set and immunoglobulin domain-containing protein 1 |  |
| **testis** | *HTATSF1* | 0.8809 | 1.09E-118 | 5276 | O43719 | HIV Tat-specific factor 1 | RNA splicing factor (PC00148) |
| **testis** | *ACO1* | 0.8808 | 1.26E-118 | 117 | P21399 | Cytoplasmic aconitate hydratase | RNA binding protein (PC00031) |
| **testis** | *ENPP5* | 0.8802 | 3.01E-118 | 13717 | Q9UJA9 | Ectonucleotide pyrophosphatase/phosphodiesterase family member 5 | nucleotide phosphatase (PC00173) |
| **kidney** | *METTL7B* | 0.8705 | 2.78E-27 | 28276 | Q6UX53 | Methyltransferase-like protein 7B | methyltransferase (PC00155) |
| **kidney** | *ANPEP* | 0.8683 | 5.3E-27 | 500 | P15144 | Aminopeptidase N | metalloprotease (PC00153) |
| **kidney** | *LRP2* | 0.8671 | 7.59E-27 | 6694 | P98164 | Low-density lipoprotein receptor-related protein 2 |  |
| **kidney** | *SLC13A1* | 0.8667 | 8.5E-27 | 10916 | Q9BZW2 | Solute carrier family 13 member 1 | secondary carrier transporter (PC00258) |
| **kidney** | *UGT3A1* | 0.8608 | 4.52E-26 | 26625 | Q6NUS8 | UDP-glucuronosyltransferase 3A1 |  |
| **kidney** | *SLC27A2* | 0.8535 | 3.26E-25 | 10996 | O14975 | Very long-chain acyl-CoA synthetase | secondary carrier transporter (PC00258) |
| **kidney** | *TMEM27* | 0.8526 | 4.1E-25 | 29437 | Q9HBJ8 | Collectrin |  |
| **kidney** | *CLRN3* | 0.8443 | 3.36E-24 | 20795 | Q8NCR9 | Clarin-3 |  |
| **kidney** | *ACP5* | 0.8367 | 2.05E-23 | 124 | P13686 | Tartrate-resistant acid phosphatase type 5 |  |
| **lung** | *PARM1* | 0.6275 | 1.32E-64 | 24536 | Q6UWI2 | Prostate androgen-regulated mucin-like protein 1 |  |
| **lung** | *CNR1* | 0.6055 | 4.21E-59 | 2159 | P21554 | Cannabinoid receptor 1 | G-protein coupled receptor (PC00021) |
| **lung** | *NUDT16* | 0.5918 | 6.52E-56 | 26442 | Q96DE0 | U8 snoRNA-decapping enzyme |  |
| **lung** | *DUOX1* | 0.5817 | 1.23E-53 | 3062 | Q9NRD9 | Dual oxidase 1 | oxidase (PC00175) |
| **lung** | *DUOXA1* | 0.5739 | 6.2E-52 | 26507 | Q1HG43 | Dual oxidase maturation factor 1 |  |
| **lung** | *FMN1* | 0.571 | 2.53E-51 | 3768 | Q68DA7 | Formin-1 |  |
| **lung** | *TMEM164* | 0.5692 | 6.05E-51 | 26217 | Q5U3C3 | Transmembrane protein 164 |  |
| **lung** | *HSD17B4* | 0.5673 | 1.54E-50 | 5213 | P51659 | Peroxisomal multifunctional enzyme type 2 |  |
| **lung** | *MGST1* | 0.5631 | 1.16E-49 | 7061 | P10620 | Microsomal glutathione S-transferase 1 |  |
| **lung** | *SHMT1* | 0.5594 | 6.77E-49 | 10850 | P34896 | Serine hydroxymethyltransferase, cytosolic | methyltransferase (PC00155) |

### ACE2 is associated with vascular growth

GO enrichment analysis of ACE2 mRNA expression in all tissues produced 22 terms which were enriched in BP, CC, HP, KEGG, and WP ontologies (Table 2). A total of 12 of these terms were related to blood vessel growth, including the three most strongly enriched terms, ‘angiogenesis [GO:0001525]’, ‘blood vessel morphogenesis [GO:0048514]’, and ‘vasculature development [GO:0072358]’. Full GO enrichment results for relevant tissues (lung, small intestine, kidney, colon, and testis) are included as Supplementary Table 6.

**Table 2.**
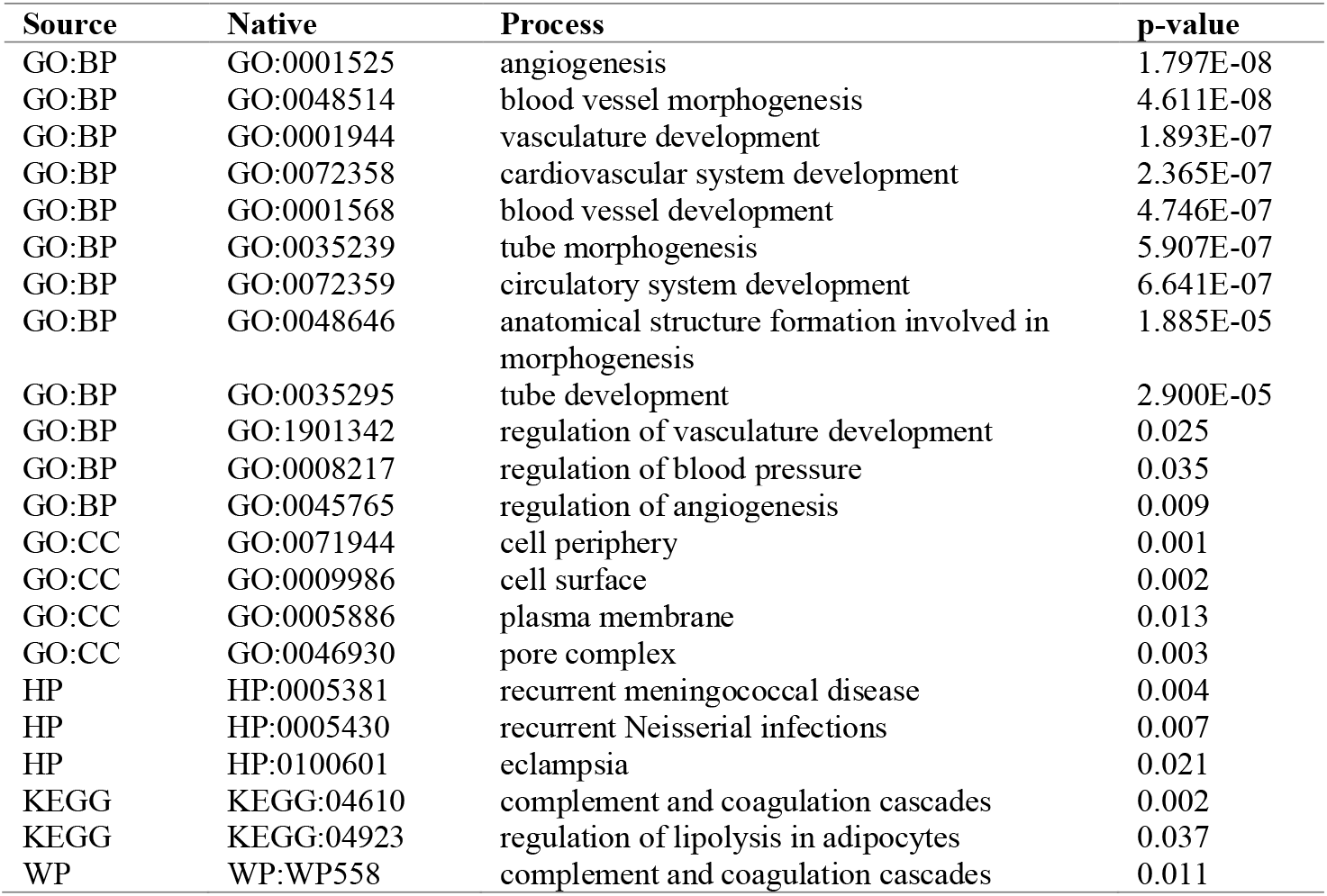
Gene ontology annotation results for the processes associated with genes strongly coexpressed (≥0.5) with ACE2 across all tissues in GTEx dataset. Calculated with GProfiler Python library [63].

| Source | Native | Process | p-value |
| --- | --- | --- | --- |
| GO:BP | GO:0001525 | angiogenesis | 1.797E-08 |
| GO:BP | GO:0048514 | blood vessel morphogenesis | 4.611E-08 |
| GO:BP | GO:0001944 | vasculature development | 1.893E-07 |
| GO:BP | GO:0072358 | cardiovascular system development | 2.365E-07 |
| GO:BP | GO:0001568 | blood vessel development | 4.746E-07 |
| GO:BP | GO:0035239 | tube morphogenesis | 5.907E-07 |
| GO:BP | GO:0072359 | circulatory system development | 6.641E-07 |
| GO:BP | GO:0048646 | anatomical structure formation involved in morphogenesis | 1.885E-05 |
| GO:BP | GO:0035295 | tube development | 2.900E-05 |
| GO:BP | GO:1901342 | regulation of vasculature development | 0.025 |
| GO:BP | GO:0008217 | regulation of blood pressure | 0.035 |
| GO:BP | GO:0045765 | regulation of angiogenesis | 0.009 |
| GO:CC | GO:0071944 | cell periphery | 0.001 |
| GO:CC | GO:0009986 | cell surface | 0.002 |
| GO:CC | GO:0005886 | plasma membrane | 0.013 |
| GO:CC | GO:0046930 | pore complex | 0.003 |
| HP | HP:0005381 | recurrent meningococcal disease | 0.004 |
| HP | HP:0005430 | recurrent Neisserial infections | 0.007 |
| HP | HP:0100601 | eclampsia | 0.021 |
| KEGG | KEGG:04610 | complement and coagulation cascades | 0.002 |
| KEGG | KEGG:04923 | regulation of lipolysis in adipocytes | 0.037 |
| WP | WP:WP558 | complement and coagulation cascades | 0.011 |

## DISCUSSION

The predominant pathological features of COVID-19 infection largely mimic those previously reported for SARS-CoV-1 infection. They include dry cough, persistent fever, progressive dyspnea, and in some cases acute exacerbation of lung function with bilateral pneumonia [31]. Major lung lesions include several pathological signs, such as diffuse alveolar damage, inflammatory exudation in the alveoli and interstitial tissue, hyperplasia of fibrous tissue, and eventually lung fibrosis [34–36]. It has been shown by fluorescence *in situ* hybridization technique that SARS-CoV-1 RNA locates to the alveolar pneumocytes and alveolar space [37, 38]. Mossel and colleagues demonstrated that SARS-CoV-1 replicates in type 2 (AT2) pneumocytes, but not in type 1 (AT1) cells [39]. Considering all of these facts, it is not surprising that most histopathological analyses have been focused on distal parts of the respiratory airways, while the regions other than the alveolus have been less systematically studied.

To understand better the pathogenesis of COVID-19 we need to know where ACE2, the receptor for SARS-CoV, is located within the human respiratory tract and elsewhere. Overall, different studies including ours have convincingly shown that several organs, such as the small intestine, colon, kidney, and testis, express higher levels of ACE2 than the lung and other parts of the respiratory tract. Our analysis of ACE2 expression in the human lung show low levels of expression in all cell types, with arterial vascular endothelial cells achieving the highest overall ratio of just ~2.5%. The present results based on mouse tracheal dataset suggested that ACE2 mRNA is predominantly expressed in the club cells, goblet cells, and ciliated epithelial cells, and at significantly higher frequency than found in the lung. The mouse dataset used in our study contained no secretory3 cells, which Lukassen and colleagues recently reported to express the highest levels of ACE2 mRNA along the human respiratory tract [40]. Another study reported positive expression in the type AT2 pneumocytes [41], which is in line with the results of Lukassen et al. [40], but only a few cells appeared positive. A third study based on single cell expression data demonstrated the strongest positive signal in the lung AT2 cells, while other cells including AT1 cells, club cells, ciliated cells, and macrophages showed weaker expression [42]. A fourth single cell expression analysis using Gene Expression Omnibus (GEO) database recently demonstrated ACE2-positive signal in 1% of AT2 cells and in 2% of respiratory tract epithelial cells [43]. This correlates with our own findings for lung. For comparison about 30% of ileal epithelial cells were ACE2-positive, and 44% of enterocytes in small intestine of mouse. Immunohistochemical analysis of mouse tissues has shown positive signal in the club cells, AT2 cells, endothelial cells, and smooth muscle cells [44]. In spite of the obvious discrepancies between different datasets, that highlights the need for large numbers of thoroughly characterized cells for single cell RNA-Seq analyses, we can now make some conclusions of the expression of ACE2 mRNA in the respiratory tract. First, ACE2 is positively though weakly expressed in the AT2 cells of the lung and less so in AT1 cells. Second, ACE2 also shows a weak positive signal, but at significantly higher proportions of cells, in several other cell types of the trachea, including goblet cells, club cells, and ciliated cells. Third, based on the findings of Lukassen et al. [40] secretory3 cells, a transient cell type of the bronchial tree, may express the highest levels of ACE2. These ACE2-positive cell types may represent the main host cells for SARS-CoV-2 along the whole respiratory tract. However, the median percentage of ACEexpressing secretory3 cells in the study was less than 6%, significantly less than that observed in club (16.62%), goblet (13.84%), and ciliated (6.63%) cells of trachea we have identified in the GSE103354 dataset.

Goblet cells, ciliated epithelial cells, and club cells are considered important cell types for the protection of airway mucosa. Lukassen and coworkers [40] described secretory3 cells as intermediate cells between goblet, ciliated, and club cells. If SARS-coronaviruses predominantly attack these cells, locating along the airway segments including the trachea, bronchi, and bronchioles until the last segment that is the respiratory bronchioles, it would be obvious that physiological protective mechanisms are severely affected. Defective mucosal protection and inefficient removal of pathogens due to viral infection may contribute to onset of severe bilateral pneumonia that is common for SARS-diseases [45]. This pathogenic mechanism is supported by previous findings, showing that early disease is manifested as a bronchiolar disease with respiratory epithelial cell necrosis, loss of cilia, squamous cell metaplasia, and intrabronchiolar fibrin deposits [31]. In fact, it has been suggested that early diffuse damage as a result of SARS-CoV-1 infection may actually initiate at the level of the respiratory bronchioles [46, 47].

Our findings confirm that the respiratory tract tissues have quite limited expression levels of ACE2 compared to several other tissues that show much more prominent signal. Because ACE2 is highly expressed in the intestine [28], as also confirmed by our bioinformatics study, it would be obvious to predict that both SARS-CoV-1 and −2 infections cause significant gastrointestinal pathology and symptoms including diarrhea. Interestingly, the patients with COVID-19 have reported less gastrointestinal symptoms than the SARS-CoV-1-infected patients [22, 29]. The pathophysiological basis for this phenomenon is not understood at this point, and thus further investigations on this topic are warranted.

When we initiated the present study, we hypothesized that understanding better the transcriptional regulation of the *ACE2* gene might help to explain the peculiar distribution pattern of ACE2 in tissues. Since upregulation of ACE2 would reflect an increased number of SARS-coronavirus receptors on cell surfaces, it could possibly help us to understand the mechanisms why certain patients (males more than females, old more than young, smokers more than non-smokers) are more susceptible for the most detrimental effects of the COVID-19 infection. In our study, the signals for ACE2 mRNA in the lung specimens did not vary much in different age groups nor did they show significant differences between males and females, which is in line with previous findings [40]. Therefore, different expression levels of lung ACE2 may not explain the variable outcome of the disease concerning age groups and genders.

To investigate the transcriptional regulation of *ACE2* gene we made predictions for the binding sites of transcription factors within the proximal promoter region of the intestine-specific and lung-specific human *ACE2* transcript promoters. Our findings introduced several putative binding sites in the *ACE2* promoter for known transcription factors, which showed high levels of coexpression with ACE2 in several tissues including the ileum, colon, and kidney. The identified transcription factors could represent potential candidate target molecules which regulate ACE2 expression. Two of our predictions, for HNF1A and HNF1B have been previously identified experimentally to drive ACE2 expression in pancreatic islet cells and insulinoma cells, respectively [48]. Later work by the same group has shown that our prediction of FOXA binding sites in the ACE2 promoter are also likely correct [49]. It is of interest that ACE2 might be regulated by oxygen status. Zhang and coworkers previously demonstrated that ACE2 mRNA and protein levels increased during the early stages of hypoxia and decreased to near-baseline levels at later stages after hypoxia inducible factor (HIF)-1α accumulation [50]. Based on these findings *ACE2* has been listed as a HIF1α-target gene [51], although it does not follow the typical HIF1α regulated expression pattern, nor is there any predicted HIF1α binding site in our analyses. However, HNF1B has been identified as upregulated in hypoxia in kidney, independent of HIF1α [52], and in hypoxic embryonic stem cells HIF1α has been shown to increase expression of transcription factors TCF7/LEF1 (predicted to bind the promoter of the lung-specific ACE2 transcript) through Wnt/β-catenin signaling [53].

The regulation of ACE2 expression remains an enigma and there may be multiple factors involved. There has been concern that the use of ACE inhibitors and angiotensin receptor blockers could increase the expression of ACE2 and increase patient susceptibility to viral host cell entry [54, 55]. Previous studies have suggested that both ACE inhibitor and angiotensin II receptor type I antagonist therapies increase ACE2 mRNA expression in the rat heart [56]. There has also been some evidence in humans showing increased expression of ACE2 in the heart, brain, and even in urine after treatment with angiotensin receptor blockers [54]. Since these drugs are widely used for treatment of hypertension and heart failure, it would be important to determine in COVID-19 patients whether these medications have any significant effects on symptoms or outcome of the disease.

Gene ontology investigations revealed interesting novel data on potential physiological roles of ACE2. The five most significant gene ontology terms included angiogenesis, blood vessel morphogenesis, vasculature development, cardiovascular system development, and blood vessel development. Our study of lung showed that the arterial and venous vascular endothelial cell types had the highest ratios of ACE2-expressing cells, first and third highest, respectively. In another study, ACE2 expression was previously detected in blood vessels [26], and a recent study showed that SARS-CoV-2 is capable of directly infecting blood vessel cells [12]. Endothelial ACE2 expression may be linked to clotting and multi-organ dysfunction reported in many patients with COVID-19. Based on the present finding angiogenesis/blood vessel morphogenesis may be considered a putative function for ACE2 in addition to its classical role as the key angiotensin-(1-7) forming enzyme [57].

Conclusions: Our bioinformatics study confirmed the low expression of ACE2 in the respiratory tract. In lung it was lowest of all, while significantly higher in the trachea. Bulk RNA-Seq analyses indicated the highest expression levels in the small intestine, colon, testis, and kidney. In the human lung scRNA-Seq dataset, the strongest positive signals for ACE2 mRNA were observed in vascular endothelial cells, goblet cells, ciliated cells, and AT2 and AT1 pneumocytes. In the mouse trachea dataset, positive signals were most common in club cells, goblet cells and ciliated epithelial cells. The results suggest that SARS-CoV infection may target the cell types that are important for the protection of airway mucosa and their damage may lead to deterioration of epithelial cell function, finally leading to a more severe lung disease with accumulation of alveolar exudate and inflammatory cells and lung edema, the signs of pneumonia recently described in the lung specimens of two patients with COVID-19 infection [58]. Gene ontology analysis based on expression in all tissues suggested that ACE2 is involved in angiogenesis/blood vessel morphogenesis processes in addition to its classical function in renin-angiotensin system.

## METHODS

### ACE2 mRNA expression

From the FANTOM5 project [59], cap analysis of gene expression (CAGE) sequencing of cDNA has been performed in 1,839 human samples from 875 different primary cells, tissues, and cell lines. Expression of transcription start sites (TSSs) was extracted and combined for all genes in all samples as tags per million (TPM). From this compiled set, *ACE2* gene expression was extracted and presented as barplot using the Matplotlib [60] and Seaborn [61] Python libraries. Similarly, human gene expression data (as TPM) was extracted from the GTEx database, along with metadata on the samples. *ACE2* gene expression values were separated by tissue and compared among 10-year interval age groups to determine if the values showed any differences throughout the lifecycle. Boxplots for tissues of relevance were generated using Matplotlib and Seaborn libraries.

### Coexpression and gene ontology enrichment analysis

In each of the tissues present in the GTEx dataset, expression values for ACE2 were compared with expression of all other genes by Spearman correlation analysis using the SciPy [62] Python library to identify those genes with concordant expression patterns. Bonferroni correction was used to derive an adjusted p-value threshold of 9.158E-07. For each tissue, those genes which both satisfied the Bonferroni-adjusted p-value threshold and had a correlation of expression of 0.50 or greater were analyzed using the Gprofiler gene ontology (GO) enrichment analysis [63] Python library to identify possible enriched terms in biological process (BP), molecular function (MF), cellular component (CC), human phenotype (HP), KEGG pathway, and WikiPathways (WP) ontologies.

### ACE2 protein expression

Immunohistochemical localization of human ACE2 was evaluated from immunostained specimens provided by Protein Expression Atlas (https://www.proteinatlas.org/). The images of the Figure 2 represent duodenum from 77-years-old female, kidney from 36-years-old male, testis from 38-years-old male, lung from 61-years-old female, and nasopharyngeal mucosa from 78-years-old female. According to Protein Expression Atlas the immunostainings were performed with the rabbit anti-human polyclonal antibody (HPA000288; Sigma Aldrich, St. Louis, MO) raised against 111 N-terminal amino acids of ACE2 and diluted 1:250 for the staining.

### Promoter Analysis

Analysis of *ACE2* promoter regions was performed using the TFBSfootprinter tool (https://github.com/thirtysix/TFBS_footprinting) which uses transcription-relevant data from several major databases to enhance prediction of putative TFBSs, including: ATAC-Seq data from ENCODE [64], transcription start sites and expression data from FANTOM5 [65], expression quantitative trail loci from GTEx [66], TFBS metacluster data from GTRD [67], TFBS binding profile data from JASPAR [68], and sequence and conservation data from Ensembl [69]. Detailed description of this novel tool is under preparation (Barker et al. manuscript under preparation). Previous studies identified two distinct tissue-specific transcription start sites (TSS) for intestine and lung expression [48], which correspond to primary protein-coding Ensembl transcripts ENST00000252519 and ENST00000427411, respectively. These two transcripts were targeted for transcription factor binding site (TFBS) analysis; input parameters of 1,000 base pairs (bp) upstream and 200 bp downstream, relative to the TSS.

### Single-Cell RNA-Seq

Single-cell expression datasets were identified for relevant tissues/cells of lung (human) [70], trachea (mouse) [71], and small intestine (mouse) [72]. Using a modified workflow described previously in [73], for each dataset the samples were filtered by Gaussian fit of read count, expressed gene count, and number of cells in which a gene is expressed. Counts were normalized by cell, log transformed, principle component analysis performed with 15 components, and k-nearest neighbors computed using SCANPY [74], and then the full data set normalized with R package ‘scran’ [75]. Batch correction by individual and sample region was performed with SCANPY using the ComBat function. The top 1,000 genes with highly differential expression were identified for cluster analysis which was performed with Uniform Manifold Approximation and Projection (UMAP) and force directed graph models. The top 100 marker genes were identified as those with higher expression unique to each cluster by Welch t-test in SCANPY. Expression of the *ACE2* gene was mapped onto cluster figures to determine overlap with previously identified cell types or cell type marker genes identified in the literature. Cell type was mapped by expression of known marker genes of cell types expressed in the lung and small intestine, as defined by *de novo* prediction in the original articles.

### Statistics

Comparisons of ACE2 expression values in different tissues and between groups delineated by age or sex, were carried out by one-way ANOVA using the stats package in the SciPy [62] Python library. Only groups with 20 or more observations and a 2-sided chi squared probability of normality of <=0.1 (due to the robustness of ANOVA to non-normal distributions) were used for comparison. Correlation of gene expression values was calculated by two-sided Spearman rank-order analysis, where a Bonferroni-corrected p-value threshold was computed using α=0.05/number of comparisons. Gene ontology enrichment analyses performed using the GProfiler tool utilize a custom algorithm for multiple testing of dependent results, which corresponds to an experiment-wide threshold of α=0.05. TFBSfootprinter analysis of the *ACE2* promoter limits results for individual TFBSs whose score satisfies a genome-wide threshold of α=0.01.

## Supporting information

Supplementary table 5 lung

Supplementary table 5 intestine

Supplementary table 5 colon

Supplementary table 5 kidney

Supplementary table 5 testis

Supplementary table 6 lung

Supplementary table 6 intestine

Supplementary table 6 colon

Supplementary table 6 kidney

Supplementary table 6 testis

