## Supplementary table 1 for "Bioinformatic characterization of angiotensin-converting enzyme 2, the entry receptor for SARS-CoV-2"

| Tissue | f-value | p-value | 25 | 35 | 45 | 55 | 65 | 75 |
|---|---|---|---|---|---|---|---|---|
| nerve - tibial | 6,384 | 8,58E-06 | 1.887±2.5 (n=50) | 1.402±1.715 (n=54) | 1.509±2.838 (n=96) | 0.935±1.402 (n=190) | 0.732±0.862 (n=206) | 0.489±0.386 (n=23) |
| minor salivary gland | 4,562 | 0,00164 | 3.405±1.868 (n=14) | 2.761±1.387 (n=16) | 2.963±2.2 (n=30) | 2.403±1.554 (n=50) | 1.736±1.139 (n=51) | 2.128±0.0 (n=1) |
| artery - aorta | 3,677 | 0,002869 | 2.611±4.705 (n=37) | 0.678±0.838 (n=38) | 0.611±0.8 (n=68) | 1.644±4.104 (n=149) | 0.774±1.907 (n=131) | 0.346±0.358 (n=9) |
| whole blood | 3,409 | 0,004722 | 0.048±0.046 (n=68) | 0.04±0.035 (n=68) | 0.039±0.097 (n=113) | 0.03±0.047 (n=234) | 0.024±0.031 (n=249) | 0.017±0.021 (n=23) |
| colon - transverse | 3,057 | 0,01012 | 9.105±10.841 (n=43) | 8.574±9.955 (n=47) | 6.938±8.003 (n=75) | 7.173±8.109 (n=136) | 4.643±8.621 (n=94) | 1.312±1.516 (n=11) |
| brain - hypothalamus | 2,571 | 0,03926 | 0.047±0.038 (n=4) | 0.075±0.044 (n=5) | 0.25±0.22 (n=16) | 0.137±0.106 (n=60) | 0.192±0.189 (n=105) | 0.13±0.063 (n=12) |
| skin - sun exposed (lower leg) | 2,271 | 0,04594 | 0.669±0.621 (n=59) | 0.775±1.212 (n=57) | 0.623±0.608 (n=107) | 0.527±0.422 (n=218) | 0.534±0.49 (n=232) | 0.539±0.313 (n=28) |
| esophagus - mucosa | 1,931 | 0,08752 | 3.05±2.241 (n=56) | 3.074±1.22 (n=51) | 2.718±1.285 (n=101) | 2.909±2.586 (n=181) | 2.464±2.011 (n=151) | 1.786±0.688 (n=15) |
| liver | 1,731 | 0,1286 | 0.511±0.279 (n=7) | 0.791±0.952 (n=17) | 0.711±0.694 (n=35) | 0.947±1.016 (n=83) | 0.986±0.966 (n=79) | 1.888±1.597 (n=5) |
| adipose - subcutaneous | 1,693 | 0,1342 | 8.275±10.025 (n=52) | 8.808±12.689 (n=58) | 6.563±8.211 (n=103) | 5.411±7.901 (n=213) | 6.648±10.922 (n=216) | 4.829±5.731 (n=21) |
| pituitary | 1,465 | 0,2015 | 0.198±0.077 (n=9) | 0.242±0.123 (n=8) | 0.448±0.709 (n=24) | 0.295±0.142 (n=87) | 0.305±0.27 (n=138) | 0.267±0.222 (n=17) |
| pancreas | 1,452 | 0,2052 | 2.494±2.913 (n=29) | 2.016±1.397 (n=31) | 1.657±1.28 (n=66) | 1.764±1.289 (n=118) | 1.836±1.45 (n=79) | 1.332±0.349 (n=5) |
| testis | 1,451 | 0,2053 | 56.634±20.505 (n=36) | 51.352±20.309 (n=31) | 50.851±19.26 (n=52) | 48.262±20.15 (n=117) | 46.784±21.317 (n=116) | 49.194±14.533 (n=9) |
| cells - ebv-transformed lymphocytes | 1,431 | 0,2261 | 0.016±0.014 (n=17) | 0.021±0.014 (n=12) | 0.03±0.026 (n=41) | 0.024±0.023 (n=54) | 0.021±0.021 (n=46) | 0.013±0.009 (n=4) |
| brain - putamen (basal ganglia) | 1,361 | 0,2407 | 0.026±0.04 (n=5) | 0.099±0.11 (n=5) | 0.12±0.103 (n=19) | 0.071±0.061 (n=67) | 0.099±0.133 (n=100) | 0.123±0.093 (n=9) |
| artery - tibial | 1,331 | 0,2493 | 0.276±0.255 (n=59) | 0.346±0.478 (n=59) | 0.425±0.687 (n=114) | 0.358±0.65 (n=216) | 0.344±0.538 (n=195) | 0.633±1.022 (n=20) |
| muscle - skeletal | 1,128 | 0,3437 | 0.697±1.123 (n=67) | 0.487±0.538 (n=65) | 0.673±1.014 (n=124) | 0.515±0.74 (n=255) | 0.652±1.172 (n=264) | 0.755±0.82 (n=28) |
| bladder | 0,959 | 0,3467 | 1.592±1.004 (n=3) | 1.137±0.728 (n=2) | 0.848±0.533 (n=7) | 0.535±0.573 (n=7) | 0.428±0.017 (n=2) | NA |
| esophagus - gastroesophageal junctio | 1,111 | 0,3538 | 0.802±0.775 (n=33) | 1.16±1.229 (n=33) | 1.617±1.917 (n=69) | 1.485±2.392 (n=137) | 1.753±2.682 (n=91) | 1.531±0.846 (n=12) |
| skin - not sun exposed (suprapubic) | 1,081 | 0,3697 | 0.627±0.416 (n=45) | 0.525±0.333 (n=52) | 0.51±0.319 (n=91) | 0.562±0.472 (n=190) | 0.504±0.303 (n=202) | 0.53±0.365 (n=24) |
| esophagus - muscularis | 1,008 | 0,4123 | 1.406±2.821 (n=60) | 1.424±1.331 (n=50) | 1.271±1.138 (n=96) | 1.804±3.517 (n=166) | 1.976±3.002 (n=133) | 1.309±0.881 (n=10) |
| uterus | 0,989 | 0,416 | 0.601±0.31 (n=21) | 1.265±1.954 (n=15) | 1.15±1.054 (n=33) | 1.14±1.361 (n=45) | 1.216±1.083 (n=27) | 4.965±0.0 (n=1) |
| brain - spinal cord (cervical c-1) | 0,946 | 0,4202 | 0.144±0.048 (n=4) | 0.123±0.101 (n=2) | 0.136±0.153 (n=16) | 0.142±0.151 (n=47) | 0.197±0.3 (n=78) | 0.105±0.058 (n=12) |
| small intestine - terminal ileum | 0,979 | 0,4204 | 50.504±60.967 (n=28) | 70.544±71.845 (n=22) | 51.135±66.283 (n=35) | 54.859±58.159 (n=56) | 75.469±86.499 (n=42) | 29.935±51.248 (n=4) |
| prostate | 0,98 | 0,4309 | 0.84±0.992 (n=30) | 1.49±3.526 (n=26) | 1.289±1.884 (n=37) | 1.003±1.51 (n=72) | 0.767±0.692 (n=73) | 1.254±1.829 (n=7) |
| brain - cerebellum | 0,949 | 0,45 | 0.068±0.04 (n=7) | 0.115±0.096 (n=9) | 0.126±0.078 (n=23) | 0.095±0.069 (n=75) | 0.1±0.08 (n=115) | 0.094±0.059 (n=12) |
| brain - anterior cingulate cortex (ba24) | 0,841 | 0,4731 | 0.022±0.022 (n=4) | 0.136±0.111 (n=4) | 0.113±0.1 (n=24) | 0.075±0.054 (n=41) | 0.115±0.194 (n=90) | 0.075±0.073 (n=13) |
| adrenal gland | 0,89 | 0,4882 | 0.609±0.331 (n=21) | 0.599±0.243 (n=21) | 0.768±0.559 (n=46) | 0.784±0.659 (n=89) | 0.887±0.94 (n=75) | 0.761±0.343 (n=6) |
| heart - atrial appendage | 0,879 | 0,4947 | 6.304±2.947 (n=15) | 5.793±3.506 (n=18) | 6.205±3.21 (n=62) | 6.272±4.394 (n=154) | 7.011±4.447 (n=163) | 5.702±2.993 (n=17) |
| heart - left ventricle | 0,877 | 0,4964 | 11.171±9.951 (n=22) | 11.633±14.459 (n=26) | 10.561±8.631 (n=66) | 10.356±8.073 (n=154) | 9.104±8.98 (n=150) | 7.185±9.831 (n=14) |
| colon - sigmoid | 0,851 | 0,5146 | 1.611±4.277 (n=36) | 1.362±2.523 (n=39) | 1.526±2.314 (n=54) | 1.295±3.862 (n=109) | 0.808±1.569 (n=123) | 0.648±0.603 (n=12) |
| adipose - visceral (omentum) | 0,789 | 0,5581 | 14.915±17.273 (n=45) | 15.282±15.495 (n=44) | 15.953±18.058 (n=86) | 12.975±13.145 (n=184) | 13.239±15.862 (n=163) | 10.498±10.153 (n=19) |
| stomach | 0,768 | 0,5735 | 0.514±0.628 (n=44) | 0.472±0.377 (n=39) | 0.713±1.295 (n=64) | 0.877±2.333 (n=128) | 0.559±0.557 (n=79) | 0.468±0.576 (n=5) |
| brain - cortex | 0,764 | 0,5762 | 0.06±0.039 (n=6) | 0.08±0.047 (n=9) | 0.104±0.078 (n=27) | 0.087±0.069 (n=77) | 0.111±0.151 (n=122) | 0.071±0.048 (n=14) |
| kidney - cortex | 0,652 | 0,5842 | 19.73±0.0 (n=1) | 14.37±8.748 (n=8) | 9.663±7.445 (n=8) | 9.73±8.237 (n=29) | 11.92±10.645 (n=35) | 18.597±8.82 (n=4) |
| brain - amygdala | 0,615 | 0,6522 | 0.062±0.018 (n=5) | 0.117±0.096 (n=4) | 0.111±0.07 (n=13) | 0.096±0.07 (n=45) | 0.122±0.178 (n=75) | 0.072±0.055 (n=10) |
| ovary | 0,606 | 0,6587 | 1.808±1.688 (n=22) | 2.071±2.153 (n=14) | 3.646±7.082 (n=37) | 5.374±16.106 (n=61) | 4.83±9.052 (n=42) | 3.921±3.324 (n=4) |
| artery - coronary | 0,652 | 0,6603 | 3.275±6.976 (n=14) | 2.816±4.178 (n=12) | 2.91±4.819 (n=37) | 4.3±6.1 (n=95) | 3.778±5.419 (n=76) | 1.28±0.941 (n=6) |
| lung | 0,624 | 0,6815 | 1.493±1.277 (n=33) | 1.474±1.121 (n=40) | 1.34±1.246 (n=93) | 1.544±1.883 (n=200) | 1.752±2.519 (n=190) | 1.498±1.395 (n=22) |
| brain - substantia nigra | 0,563 | 0,6898 | 0.058±0.045 (n=5) | 0.08±0.068 (n=4) | 0.251±0.234 (n=14) | 0.111±0.107 (n=35) | 0.35±1.122 (n=76) | 0.137±0.047 (n=5) |
| vagina | 0,61 | 0,6926 | 2.166±1.392 (n=16) | 2.693±2.382 (n=12) | 2.536±1.447 (n=36) | 2.449±2.365 (n=50) | 2.024±1.743 (n=37) | 3.416±3.899 (n=5) |
| spleen | 0,52 | 0,7213 | 0.043±0.038 (n=22) | 0.037±0.028 (n=30) | 0.046±0.102 (n=49) | 0.055±0.085 (n=86) | 0.04±0.044 (n=50) | 0.024±0.03 (n=4) |

| | | | | | | | | |
|---|---|---|---|---|---|---|---|---|
| **thyroid** | 0,529 | 0,7544 | 10.217±8.372 (n=47) | 8.326±7.022 (n=51) | 10.033±12.06 (n=110) | 9.313±10.944 (n=211) | 10.24±10.635 (n=212) | 11.767±10.92 (n=22) |
| **brain - cerebellar hemisphere** | 0,461 | 0,8052 | 0.081±0.071 (n=8) | 0.097±0.066 (n=5) | 0.13±0.077 (n=18) | 0.105±0.093 (n=65) | 0.106±0.083 (n=106) | 0.098±0.064 (n=13) |
| **brain - nucleus accumbens (basal gan** | 0,439 | 0,8213 | 0.109±0.049 (n=7) | 0.137±0.073 (n=5) | 0.133±0.09 (n=24) | 0.132±0.076 (n=75) | 0.166±0.252 (n=120) | 0.137±0.104 (n=15) |
| **cells - cultured fibroblasts** | 0,431 | 0,8271 | 0.028±0.028 (n=45) | 0.031±0.039 (n=38) | 0.043±0.053 (n=90) | 0.026±0.03 (n=169) | 0.058±0.378 (n=149) | 0.033±0.029 (n=13) |
| **brain - caudate (basal ganglia)** | 0,385 | 0,8591 | 0.122±0.082 (n=5) | 0.143±0.08 (n=6) | 0.124±0.097 (n=27) | 0.112±0.084 (n=73) | 0.153±0.28 (n=122) | 0.151±0.08 (n=13) |
| **brain - frontal cortex (ba9)** | 0,317 | 0,8661 | 0.059±0.022 (n=5) | 0.091±0.071 (n=4) | 0.14±0.102 (n=17) | 0.106±0.136 (n=63) | 0.117±0.237 (n=108) | 0.076±0.061 (n=12) |
| **brain - hippocampus** | 0,18 | 0,9699 | 0.07±0.026 (n=5) | 0.093±0.126 (n=5) | 0.131±0.162 (n=21) | 0.122±0.219 (n=52) | 0.15±0.307 (n=100) | 0.129±0.116 (n=14) |
| **breast - mammary tissue** | 0,157 | 0,9779 | 7.02±8.422 (n=39) | 6.986±11.822 (n=50) | 6.131±13.392 (n=76) | 6.657±8.906 (n=139) | 7.458±13.459 (n=138) | 6.215±7.084 (n=17) |
