## Supplementary table 2 for "Bioinformatic characterization of angiotensin-converting enzyme 2, the entry receptor for SARS-CoV-2"

| Tissue | f-value | p-value | Male | Female |
|---|---|---|---|---|
| **breast - mammary tissue** | 19,569 | 1,21E-05 | 8.627±13.377 (n=291) | 3.803±5.883 (n=168) |
| **adipose - subcutaneous** | 14,783 | 0,000132 | 5.495±8.151 (n=445) | 8.541±11.95 (n=218) |
| **whole blood** | 13,875 | 0,00021 | 0.027±0.033 (n=501) | 0.042±0.077 (n=254) |
| **brain - amygdala** | 12,233 | 0,000618 | 0.084±0.063 (n=107) | 0.165±0.217 (n=45) |
| **colon - transverse** | 11,36 | 0,000823 | 5.652±7.048 (n=259) | 8.689±11.036 (n=147) |
| **esophagus - muscularis** | 9,801 | 0,001844 | 1.932±3.308 (n=338) | 1.13±1.102 (n=177) |
| **heart - left ventricle** | 8,123 | 0,004582 | 9.112±7.813 (n=294) | 11.792±11.347 (n=138) |
| **cells - ebv-transformed lymphocytes** | 6,082 | 0,01464 | 0.02±0.019 (n=112) | 0.029±0.026 (n=62) |
| **esophagus - gastroesophageal junctio** | 5,173 | 0,02351 | 1.667±2.514 (n=251) | 1.123±1.225 (n=124) |
| **nerve - tibial** | 3,634 | 0,05708 | 1.149±1.922 (n=419) | 0.865±1.241 (n=200) |
| **muscle - skeletal** | 3,23 | 0,07269 | 0.648±1.045 (n=543) | 0.517±0.789 (n=260) |
| **heart - atrial appendage** | 2,711 | 0,1004 | 6.276±3.709 (n=293) | 6.986±4.954 (n=136) |
| **adipose - visceral (omentum)** | 2,642 | 0,1047 | 14.516±17.1 (n=371) | 12.205±10.468 (n=170) |
| **brain - hypothalamus** | 2,604 | 0,1082 | 0.16±0.169 (n=147) | 0.202±0.156 (n=55) |
| **artery - coronary** | 2,443 | 0,1194 | 3.255±4.855 (n=146) | 4.419±6.611 (n=94) |
| **artery - tibial** | 2,081 | 0,1496 | 0.388±0.645 (n=454) | 0.316±0.501 (n=209) |
| **skin - sun exposed (lower leg)** | 1,894 | 0,1692 | 0.555±0.46 (n=467) | 0.62±0.795 (n=234) |
| **brain - putamen (basal ganglia)** | 1,831 | 0,1775 | 0.085±0.104 (n=156) | 0.109±0.124 (n=49) |
| **adrenal gland** | 1,706 | 0,1927 | 0.827±0.831 (n=157) | 0.711±0.404 (n=101) |
| **pituitary** | 1,512 | 0,2199 | 0.32±0.34 (n=204) | 0.271±0.148 (n=79) |
| **kidney - cortex** | 1,444 | 0,2329 | 10.927±8.499 (n=66) | 13.926±12.266 (n=19) |
| **brain - spinal cord (cervical c-1)** | 1,303 | 0,2554 | 0.15±0.197 (n=102) | 0.194±0.286 (n=57) |
| **brain - frontal cortex (ba9)** | 0,935 | 0,3347 | 0.103±0.199 (n=153) | 0.132±0.159 (n=56) |
| **brain - caudate (basal ganglia)** | 0,846 | 0,3587 | 0.13±0.208 (n=183) | 0.158±0.204 (n=63) |
| **brain - hippocampus** | 0,845 | 0,359 | 0.125±0.247 (n=143) | 0.163±0.274 (n=54) |
| **brain - nucleus accumbens (basal gan** | 0,776 | 0,3791 | 0.142±0.198 (n=182) | 0.166±0.147 (n=64) |
| **brain - cortex** | 0,636 | 0,4259 | 0.095±0.078 (n=181) | 0.107±0.177 (n=74) |
| **artery - aorta** | 0,603 | 0,4378 | 1.273±3.182 (n=279) | 1.034±2.817 (n=153) |
| **small intestine - terminal ileum** | 0,541 | 0,463 | 56.639±67.5 (n=120) | 64.487±73.198 (n=67) |
| **brain - cerebellar hemisphere** | 0,468 | 0,4949 | 0.108±0.089 (n=157) | 0.1±0.071 (n=58) |
| **brain - anterior cingulate cortex (ba24)** | 0,467 | 0,4952 | 0.096±0.126 (n=128) | 0.114±0.199 (n=48) |
| **thyroid** | 0,451 | 0,5023 | 9.607±9.735 (n=434) | 10.2±12.238 (n=219) |
| **brain - substantia nigra** | 0,403 | 0,5265 | 0.282±0.978 (n=101) | 0.18±0.213 (n=38) |
| **cells - cultured fibroblasts** | 0,277 | 0,5988 | 0.043±0.256 (n=330) | 0.033±0.039 (n=174) |
| **bladder** | 0,272 | 0,6083 | 0.776±0.771 (n=14) | 0.96±0.626 (n=7) |
| **esophagus - mucosa** | 0,221 | 0,6382 | 2.722±1.834 (n=363) | 2.81±2.469 (n=192) |
| **colon - sigmoid** | 0,202 | 0,6537 | 1.235±2.978 (n=240) | 1.093±2.791 (n=133) |
| **stomach** | 0,187 | 0,6657 | 0.711±1.794 (n=227) | 0.637±0.997 (n=132) |
| **liver** | 0,104 | 0,7477 | 0.906±0.925 (n=161) | 0.953±1.077 (n=65) |
| **skin - not sun exposed (suprapubic)** | 0,094 | 0,7598 | 0.538±0.359 (n=411) | 0.528±0.425 (n=193) |
| **spleen** | 0,031 | 0,8613 | 0.047±0.072 (n=154) | 0.045±0.076 (n=87) |
| **pancreas** | 0,026 | 0,871 | 1.853±1.37 (n=207) | 1.823±1.833 (n=121) |
| **lung** | 0,013 | 0,9077 | 1.577±1.863 (n=395) | 1.556±2.151 (n=183) |
| **minor salivary gland** | 0,009 | 0,9231 | 2.409±1.835 (n=115) | 2.437±1.245 (n=47) |
| **brain - cerebellum** | 0,009 | 0,9242 | 0.1±0.074 (n=174) | 0.101±0.08 (n=67) |
