## Supplementary table 3 for "Bioinformatic characterization of angiotensin-converting enzyme 2, the entry receptor for SARS-CoV-2"

| binding prot. | motif | strand | start | end | TSS-relative start | TSS-relative end | PWM score | p-value | combined affinity score | combined affinity score p-value | species weights sum | cage weights sum | eqtls weights sum | atac weights sum | metacluster weights sum | cpg weight | corr. weight sum |
|---|---|---|---|---|---|---|---|---|---|---|---|---|---|---|---|---|---|
| CDX2 | TGGCCATAAAG | 1 | 756 | 767 | -244 | -233 | 12,87 | 0,0001 | 38,53 | 0,00375 | 5,6 | 5,75 | 0 | 1,31 | 0 | 6,32 | 6,67 |
| HNF1A | GATTAAAGAATAACG | 1 | 792 | 807 | -208 | -193 | 11,38 | 0,0001 | 37,01 | 0,00788 | 5,54 | 5,95 | 0 | 1,48 | 0 | 5,17 | 7,48 |
| HNF1A | AGGAAAATATTAACC | 1 | 722 | 737 | -278 | -263 | 9,8 | 0,0001 | 34,84 | 0,0136 | 5,54 | 5,63 | 0 | 1,22 | 0 | 5,17 | 7,48 |
| FOXA1 | AGATTGTTTACTGTG | 1 | 893 | 908 | -107 | -92 | 11,64 | 0,001 | 34,2 | 0,0155 | 5,54 | 6,69 | 0 | 1,89 | 0 | 2,95 | 5,49 |
| SOX4 | TAACATTTTCCGTGTA | 1 | 690 | 706 | -310 | -294 | -2,34 | 0,0001 | 12,92 | 0,0189 | 5,53 | 5,51 | 0 | 1,15 | 0 | 3,07 | 0 |
| TP63 | GACATTTTTGGACAAGTT | 1 | 1126 | 1144 | 126 | 144 | 0,02 | 0,0001 | 19,95 | 0,02 | 5,51 | 6,49 | 0 | 1,84 | 3,22 | 2,86 | 0 |
| HNF4A | AAGTTCAAAGGCTGAT | 1 | 956 | 972 | -44 | -28 | 4,53 | 0,0001 | 26,98 | 0,0206 | 5,53 | 7,68 | 0 | 1,88 | 0 | 1,7 | 5,67 |
| DUX4 | AAAGTGAATCA | -1 | 817 | 828 | -183 | -172 | 10,18 | 0,001 | 27,01 | 0,0207 | 5,6 | 6,08 | 0 | 1,6 | 0 | 3,55 | 0 |
| HNF1A | GGTTAATATTTTCCT | -1 | 722 | 737 | -278 | -263 | 7,84 | 0,001 | 32,88 | 0,0212 | 5,54 | 5,63 | 0 | 1,22 | 0 | 5,17 | 7,48 |
| FOXA2 | TGTTTACTGTGT | 1 | 897 | 909 | -103 | -91 | 13,23 | 0,0001 | 35,44 | 0,0231 | 5,58 | 6,7 | 0 | 1,89 | 0 | 2,86 | 5,18 |
| FOXA1 | TTTATATTGGCTCAG | 1 | 877 | 892 | -123 | -108 | 9,55 | 0,001 | 32,21 | 0,0232 | 5,54 | 6,55 | 0 | 1,9 | 0 | 3,19 | 5,49 |
| NR2F6(VAR.2) | CAGATCACAAGGTCA | 1 | 232 | 247 | -768 | -753 | 10,48 | 0,0001 | 24,24 | 0,0233 | 2,9 | 4,52 | 0 | 1,45 | 0 | 4,9 | 0 |
| SOX11 | AACATTTTCCGTGTA | 1 | 691 | 706 | -309 | -294 | -1,51 | 0,0001 | 13,76 | 0,0234 | 5,54 | 5,51 | 0 | 1,15 | 0 | 3,07 | 0 |
| GATA1 | CTCTTATCAGC | -1 | 966 | 977 | -34 | -23 | 12,21 | 0,0001 | 29,19 | 0,0244 | 5,6 | 7,81 | 0 | 1,87 | 0 | 1,7 | 0 |
| TP63 | AACTTGTCCAAAAATGTC | -1 | 1126 | 1144 | 126 | 144 | -1,57 | 0,0001 | 18,36 | 0,0259 | 5,51 | 6,49 | 0 | 1,84 | 3,22 | 2,86 | 0 |
| HNF1B | GTTATTCTTTAAT | -1 | 793 | 806 | -207 | -194 | 8,7 | 0,001 | 31,27 | 0,0262 | 5,57 | 5,95 | 0 | 1,48 | 0 | 5,17 | 4,41 |
| HNF1B | GGAAAATATTAAC | 1 | 723 | 736 | -277 | -264 | 8,45 | 0,001 | 30,44 | 0,0312 | 5,57 | 5,62 | 0 | 1,22 | 0 | 5,17 | 4,41 |
| FOXI1 | GTAAACA | -1 | 897 | 904 | -103 | -96 | 12,74 | 0,0001 | 34,27 | 0,0312 | 5,65 | 6,65 | 0 | 1,89 | 0 | 2,95 | 4,39 |
| HNF1A | CGTTATTCTTTAATC | -1 | 792 | 807 | -208 | -193 | 5,33 | 0,001 | 30,96 | 0,0315 | 5,54 | 5,95 | 0 | 1,48 | 0 | 5,17 | 7,48 |
| GATA4 | TCTTATCAGCC | -1 | 965 | 976 | -35 | -24 | 13,66 | 0,0001 | 30,61 | 0,0387 | 5,6 | 7,78 | 0 | 1,87 | 0 | 1,7 | 0 |
| HNF1B | ATTAAAGAATAAC | 1 | 793 | 806 | -207 | -194 | 6,75 | 0,001 | 29,32 | 0,0389 | 5,57 | 5,95 | 0 | 1,48 | 0 | 5,17 | 4,41 |
| HNF1B | GTTAATATTTTCC | -1 | 723 | 736 | -277 | -264 | 6,97 | 0,001 | 28,96 | 0,0415 | 5,57 | 5,62 | 0 | 1,22 | 0 | 5,17 | 4,41 |
| FOXG1 | GTAAACAA | -1 | 896 | 904 | -104 | -96 | 13,48 | 0,0001 | 30,61 | 0,0429 | 5,64 | 6,65 | 0 | 1,89 | 0 | 2,95 | 0 |
| FOXA2 | TATTGGCTCAGC | 1 | 881 | 893 | -119 | -107 | 9,97 | 0,001 | 32,25 | 0,045 | 5,58 | 6,56 | 0 | 1,89 | 0 | 3,07 | 5,18 |
| RARG | AAGTTCAAAGGCTGAT | 1 | 956 | 972 | -44 | -28 | -0,02 | 0,001 | 16,76 | 0,0461 | 5,53 | 7,68 | 0 | 1,88 | 0 | 1,7 | 0 |
| STAT1 | TTTCTAGGAAA | 1 | 717 | 728 | -283 | -272 | 16,32 | 1E-05 | 33,87 | 0,0474 | 5,6 | 5,59 | 0 | 1,2 | 0 | 5,17 | 0 |
| BCL6B | AGCTTTCTAGGAAAATA | 1 | 714 | 731 | -286 | -269 | -9,82 | 0,001 | 7,68 | 0,0488 | 5,52 | 5,6 | 0 | 1,21 | 0 | 5,17 | 0 |
| IRF9 | ATCAAAACCGGTACT | -1 | 740 | 755 | -260 | -245 | -1,43 | 0,0001 | 17 | 0,0503 | 5,54 | 5,7 | 0 | 1,27 | 0 | 5,92 | 0 |
| GATA1::TAL1 | CATATCTGTCCTCTCCAG | 1 | 852 | 870 | -148 | -130 | 13,4 | 0,0001 | 30,13 | 0,0524 | 5,51 | 6,37 | 0 | 1,9 | 0 | 2,95 | 0 |
| RARB | GCAGATCACAAGGTCA | 1 | 231 | 247 | -769 | -753 | 2,04 | 0,0001 | 15,81 | 0,0553 | 2,9 | 4,52 | 0 | 1,45 | 0 | 4,9 | 0 |
| NR5A2 | GATCACAAGGTCAGG | 1 | 234 | 249 | -766 | -751 | 14,25 | 0,0001 | 33,21 | 0,057 | 2,89 | 4,52 | 0 | 1,44 | 0 | 4,63 | 5,49 |
| FOXP2 | CAGTAAACAAT | -1 | 895 | 906 | -105 | -94 | 12,64 | 0,0001 | 29,74 | 0,0611 | 5,6 | 6,67 | 0 | 1,89 | 0 | 2,95 | 0 |
| IRF8 | TCAAAACCGGTACT | -1 | 740 | 754 | -260 | -246 | -3,13 | 0,0001 | 15,31 | 0,0626 | 5,56 | 5,7 | 0 | 1,27 | 0 | 5,92 | 0 |
| TP73 | GACAAGTTTAACCACGAA | 1 | 1136 | 1154 | 136 | 154 | 9,92 | 0,0001 | 30,2 | 0,0627 | 5,51 | 6,41 | 0 | 1,84 | 3,22 | 3,29 | 0 |
| STAT3 | CTTCTTGGAAT | 1 | 1185 | 1196 | 185 | 196 | 14,51 | 0,0001 | 33,13 | 0,064 | 5,6 | 6,06 | 0 | 1,55 | 1,79 | 3,63 | 0 |
| FOXH1 | CACAGTAAACA | -1 | 897 | 908 | -103 | -92 | 6,58 | 0,001 | 23,61 | 0,0652 | 5,6 | 6,69 | 0 | 1,89 | 0 | 2,86 | 0 |
| TCF7L1 | CAAGTTCAAAGG | 1 | 955 | 967 | -45 | -33 | 15,68 | 1E-05 | 32,39 | 0,0658 | 5,58 | 7,55 | 0 | 1,88 | 0 | 1,7 | 0 |
| TP63 | GACAAGTTTAACCACGAA | 1 | 1136 | 1154 | 136 | 154 | -8,11 | 0,001 | 12,17 | 0,0685 | 5,51 | 6,41 | 0 | 1,84 | 3,22 | 3,29 | 0 |
| RXRA | AAGTTCAAAGGCTG | 1 | 956 | 970 | -44 | -30 | 3,54 | 0,0001 | 20,33 | 0,0699 | 5,56 | 7,62 | 0 | 1,88 | 0 | 1,74 | 0 |
| IRF4 | TCAAAACCGGTACTT | -1 | 739 | 754 | -261 | -246 | -0,77 | 0,0001 | 17,84 | 0,0704 | 5,54 | 5,7 | 0 | 1,27 | 0 | 6,1 | 0 |
| TP53 | AAAATGTCTTGGCCTGTT | -1 | 1116 | 1134 | 116 | 134 | 8,72 | 0,0001 | 28,58 | 0,0705 | 5,51 | 6,59 | 0 | 1,85 | 3,22 | 2,7 | 0 |
| HNF1A | AATAATTTTTTAAAG | -1 | 824 | 839 | -176 | -161 | 2,16 | 0,01 | 26,57 | 0,0708 | 5,54 | 6,15 | 0 | 1,68 | 0 | 3,55 | 7,48 |
| RARG | CAGATCACAAGGTCAG | 1 | 232 | 248 | -768 | -752 | 0,22 | 0,001 | 13,83 | 0,0733 | 2,9 | 4,52 | 0 | 1,45 | 0 | 4,74 | 0 |
| HNF1A | TGTTAAAGATACACG | -1 | 700 | 715 | -300 | -285 | 1,73 | 0,01 | 26,28 | 0,0745 | 5,54 | 5,54 | 0 | 1,17 | 0 | 4,81 | 7,48 |
| BCL6 | TTTCCTAGAAAGCT | -1 | 714 | 728 | -286 | -272 | 19,06 | 1E-06 | 36,57 | 0,0747 | 5,56 | 5,59 | 0 | 1,2 | 0 | 5,17 | 0 |
| GRHL1 | AAAACCGGTACT | -1 | 740 | 752 | -260 | -248 | 12,73 | 0,0001 | 31,36 | 0,0748 | 5,58 | 5,69 | 0 | 1,26 | 0 | 6,1 | 0 |
| TCF7 | CAAGTTCAAAGG | 1 | 955 | 967 | -45 | -33 | 13,68 | 0,0001 | 30,39 | 0,0761 | 5,58 | 7,55 | 0 | 1,88 | 0 | 1,7 | 0 |
| GATA4 | CCTTCTCAGCC | 1 | 1069 | 1080 | 69 | 80 | 8,2 | 0,001 | 26,85 | 0,0777 | 5,6 | 7,19 | 0 | 1,85 | 1,79 | 2,21 | 0 |
| CDX2 | ATTCCATAAAG | 1 | 495 | 506 | -505 | -494 | 0,86 | 0,01 | 23,04 | 0,0789 | 2,94 | 4,96 | 0 | 0,98 | 0 | 6,62 | 6,67 |
| HNF1A | GAATAATTTTTTAAA | -1 | 825 | 840 | -175 | -160 | 1,6 | 0,01 | 25,9 | 0,0791 | 5,54 | 6,16 | 0 | 1,69 | 0 | 3,43 | 7,48 |
| HNF4A | TCTATCAAAGTTCACT | 1 | 1167 | 1183 | 167 | 183 | -4,4 | 0,001 | 19,37 | 0,0791 | 5,53 | 6,17 | 0 | 1,69 | 1,79 | 2,91 | 5,67 |
| STAT3 | TTTCTAGGAAA | 1 | 717 | 728 | -283 | -272 | 13,94 | 0,0001 | 31,49 | 0,0809 | 5,6 | 5,59 | 0 | 1,2 | 0 | 5,17 | 0 |
| TCF7L2 | CAAGTTCAAAGGCT | 1 | 955 | 969 | -45 | -31 | 15,18 | 1E-05 | 31,95 | 0,0816 | 5,56 | 7,6 | 0 | 1,88 | 0 | 1,74 | 0 |
| FOS | TTTGATTCACT | 1 | 815 | 826 | -185 | -174 | 7,24 | 0,001 | 23,92 | 0,0822 | 5,6 | 6,06 | 0 | 1,59 | 0 | 3,43 | 0 |
| CDX2 | AGCCAATATAA | -1 | 878 | 889 | -122 | -111 | -1,01 | 0,01 | 22,74 | 0,0826 | 5,6 | 6,52 | 0 | 1,9 | 0 | 3,07 | 6,67 |

| | | | | | | | | | | | | | | | | | |
|---|---|---|---|---|---|---|---|---|---|---|---|---|---|---|---|---|---|
| GATA1 | TCTCTATCTCC | -1 | 247 | 258 | -753 | -742 | 8,55 | 0,001 | 22,5 | 0,0831 | 2,85 | 4,53 | 0 | 1,38 | 0 | 5,19 | 0 |
| CDX1 | GCCATAAAG | 1 | 758 | 767 | -242 | -233 | 11,14 | 0,0001 | 29,93 | 0,0837 | 5,62 | 5,75 | 0 | 1,31 | 0 | 6,1 | 0 |
| CDX2 | AGACCAAAAAA | -1 | 924 | 935 | -76 | -65 | 0,43 | 0,01 | 22,64 | 0,0839 | 5,6 | 6,99 | 0 | 1,88 | 0 | 1,06 | 6,67 |
| HOXD13 | GCCATAAAGT | 1 | 758 | 768 | -242 | -232 | 8,3 | 0,001 | 27,08 | 0,0855 | 5,61 | 5,76 | 0 | 1,31 | 0 | 6,1 | 0 |
| HOXD12 | CGTGGTTAAAC | -1 | 1141 | 1152 | 141 | 152 | 4,94 | 0,001 | 25,37 | 0,0857 | 5,6 | 6,37 | 0 | 1,84 | 3,22 | 3,4 | 0 |
| RARB | CAAGTTCAAAGGCTGA | 1 | 955 | 971 | -45 | -29 | -3,73 | 0,001 | 13,06 | 0,0858 | 5,53 | 7,65 | 0 | 1,88 | 0 | 1,74 | 0 |
| TP53 | AACAGGCCAAGACATTTT | 1 | 1116 | 1134 | 116 | 134 | 7,42 | 0,0001 | 27,28 | 0,0871 | 5,51 | 6,59 | 0 | 1,85 | 3,22 | 2,7 | 0 |
| STAT1 | CTTCTTGGAAT | 1 | 1185 | 1196 | 185 | 196 | 11,38 | 0,0001 | 30 | 0,0873 | 5,6 | 6,06 | 0 | 1,55 | 1,79 | 3,63 | 0 |
| HNF1A | CGTGTATCTTTAACA | 1 | 700 | 715 | -300 | -285 | 0,61 | 0,01 | 25,16 | 0,0895 | 5,54 | 5,54 | 0 | 1,17 | 0 | 4,81 | 7,48 |
| HNF1A | AGTAAACAATCTGCT | -1 | 890 | 905 | -110 | -95 | 0,72 | 0,01 | 25,16 | 0,0895 | 5,54 | 6,66 | 0 | 1,89 | 0 | 2,86 | 7,48 |
| HNF4A | AAGGGCAAAGTCATGT | 1 | 508 | 524 | -492 | -476 | 0,95 | 0,001 | 18,47 | 0,0916 | 3 | 5 | 0 | 0,99 | 0 | 2,86 | 5,67 |
| SOX8 | AACAATCTGCTGAGCC | -1 | 885 | 901 | -115 | -99 | -14,66 | 0,001 | 2,46 | 0,092 | 5,53 | 6,63 | 0 | 1,89 | 0 | 3,07 | 0 |
| HNF1A | TGTAAAATTATAACA | 1 | 680 | 695 | -320 | -305 | 3,45 | 0,01 | 24,95 | 0,0928 | 4,35 | 5,47 | 0 | 1,13 | 0 | 3,07 | 7,48 |
| JUN | CTGCTGAGCCAAT | -1 | 882 | 895 | -118 | -105 | 3,98 | 0,001 | 21,09 | 0,093 | 5,57 | 6,57 | 0 | 1,89 | 0 | 3,07 | 0 |
| FOXQ1 | GATTGTTTACT | 1 | 894 | 905 | -106 | -95 | 11,59 | 0,0001 | 28,6 | 0,0934 | 5,6 | 6,66 | 0 | 1,89 | 0 | 2,86 | 0 |
| HNF1B | GTTAAAGATACAC | -1 | 701 | 714 | -299 | -286 | 2,75 | 0,01 | 24,24 | 0,0934 | 5,57 | 5,54 | 0 | 1,16 | 0 | 4,81 | 4,41 |
| HOXC12 | CGTGGTTAAAC | -1 | 1141 | 1152 | 141 | 152 | 3,2 | 0,001 | 23,63 | 0,0935 | 5,6 | 6,37 | 0 | 1,84 | 3,22 | 3,4 | 0 |
| HNF1A | ATCCACTGAATGACT | -1 | 1011 | 1026 | 11 | 26 | -0,38 | 0,01 | 24,88 | 0,094 | 5,54 | 8,95 | 0 | 1,86 | 0 | 1,42 | 7,48 |
| PDX1 | AAAATTAT | 1 | 830 | 838 | -170 | -162 | 3,59 | 0,01 | 26,33 | 0,0941 | 5,64 | 6,14 | 0 | 1,67 | 0 | 3,28 | 6,01 |
| SPI1 | AAAAAGAAGAAGAA | -1 | 908 | 922 | -92 | -78 | 6,99 | 0,0001 | 22,42 | 0,0944 | 5,56 | 6,83 | 0 | 1,89 | 0 | 1,15 | 0 |
| GRHL1 | AGTACCGGTTTT | 1 | 740 | 752 | -260 | -248 | 11,31 | 0,0001 | 29,94 | 0,0947 | 5,58 | 5,69 | 0 | 1,26 | 0 | 6,1 | 0 |
| FOXJ2 | GTAAACAA | -1 | 896 | 904 | -104 | -96 | 13,01 | >0.1 | 30,14 | 0,0951 | 5,64 | 6,65 | 0 | 1,89 | 0 | 2,95 | 0 |
| HOXA13 | CCAATATAAA | -1 | 877 | 887 | -123 | -113 | 10,63 | 0,001 | 27,59 | 0,0952 | 5,61 | 6,5 | 0 | 1,9 | 0 | 2,95 | 0 |
| FOXO6 | GTAAACA | -1 | 897 | 904 | -103 | -96 | 12,77 | >0.1 | 29,91 | 0,0956 | 5,65 | 6,65 | 0 | 1,89 | 0 | 2,95 | 0 |
| HNF1A | GGCCATAAAGTGACA | 1 | 757 | 772 | -243 | -228 | -1,61 | 0,01 | 24,63 | 0,098 | 5,54 | 5,78 | 0 | 1,33 | 0 | 6,1 | 7,48 |
| RXRG | AAGTTCAAAGGCTG | 1 | 956 | 970 | -44 | -30 | 1,84 | 0,0001 | 18,63 | 0,0984 | 5,56 | 7,62 | 0 | 1,88 | 0 | 1,74 | 0 |
| HOXC12 | GGCCATAAAGT | 1 | 757 | 768 | -243 | -232 | 4,44 | 0,001 | 23,21 | 0,0993 | 5,6 | 5,76 | 0 | 1,31 | 0 | 6,1 | 0 |
| HOXA10 | GGCCATAAAGT | 1 | 757 | 768 | -243 | -232 | 10,65 | 0,0001 | 29,42 | 0,0997 | 5,6 | 5,76 | 0 | 1,31 | 0 | 6,1 | 0 |
| HNF1A | AGCAGATTGTTTACT | 1 | 890 | 905 | -110 | -95 | 0,05 | 0,01 | 24,49 | 0,1 | 5,54 | 6,66 | 0 | 1,89 | 0 | 2,86 | 7,48 |
| NR1H3::RXRA | TGATCAGAAATGGCTGGGC | 1 | 165 | 184 | -835 | -816 | 2,32 | 0,01 | 15,35 | >0.1 | 3,18 | 4,44 | 0 | 1,9 | 0 | 3,51 | 0 |
| NR1H3::RXRA | TAACACGGTGAAACCCCGT | 1 | 268 | 287 | -732 | -713 | 7,4 | 0,001 | 21,03 | >0.1 | 2,77 | 4,57 | 0 | 1,3 | 0 | 4,99 | 0 |
| NR1H3::RXRA | GCACCTGTAGTCCCAGCTA | 1 | 328 | 347 | -672 | -653 | 0,09 | 0,01 | 14,4 | >0.1 | 2,59 | 4,66 | 0 | 1,15 | 0 | 5,92 | 0 |
| NR1H3::RXRA | TGAACCTGGGAGGCAGAGC | 1 | 375 | 394 | -625 | -606 | 2,92 | 0,01 | 17,96 | >0.1 | 2,57 | 4,74 | 0 | 1,07 | 0 | 6,66 | 0 |
| NR1H3::RXRA | GAGCTTGCAGTGAGCCGAG | 1 | 390 | 409 | -610 | -591 | 1,88 | 0,01 | 16,22 | >0.1 | 2,61 | 4,77 | 0 | 1,05 | 0 | 5,91 | 0 |
| NR1H3::RXRA | TGAAATAAAAATAAATGTG | 1 | 575 | 594 | -425 | -406 | 0,58 | 0,01 | 10,1 | >0.1 | 3,34 | 5,17 | 0 | 1,01 | 0 | 0 | 0 |
| NR1H3::RXRA | TAAATGTGAGATAACCTAT | 1 | 586 | 605 | -414 | -395 | 0,76 | 0,01 | 10,38 | >0.1 | 3,41 | 5,19 | 0 | 1,02 | 0 | 0 | 0 |
| NR1H3::RXRA | TATCTTCATGATCCCTAGT | 1 | 642 | 661 | -358 | -339 | 0,79 | 0,01 | 14,91 | >0.1 | 3,86 | 5,36 | 0 | 1,08 | 0 | 3,83 | 0 |
| NR1H3::RXRA | TGTAAAATTATAACATTTT | 1 | 680 | 699 | -320 | -301 | -0,15 | 0,01 | 13,81 | >0.1 | 4,39 | 5,48 | 0 | 1,14 | 0 | 2,95 | 0 |
| NR1H3::RXRA | GGCCATAAAGTGACAGGAG | 1 | 757 | 776 | -243 | -224 | 2,99 | 0,01 | 21,73 | >0.1 | 5,5 | 5,8 | 0 | 1,35 | 0 | 6,1 | 0 |
| NR1H3::RXRA | TGATCTTGGCTCACAGGGG | 1 | 1027 | 1046 | 27 | 46 | 0,68 | 0,01 | 17,74 | >0.1 | 5,5 | 8,39 | 0 | 1,86 | 0 | 1,31 | 0 |
| NR1H3::RXRA | TGGCTCACAGGGGACGATG | 1 | 1033 | 1052 | 33 | 52 | -1,07 | 0,01 | 15,73 | >0.1 | 5,5 | 8,1 | 0 | 1,86 | 0 | 1,34 | 0 |
| NR1H3::RXRA | TGGCTCCTTCTCAGCCTTG | 1 | 1064 | 1083 | 64 | 83 | 0,02 | 0,01 | 18,65 | >0.1 | 5,5 | 7,28 | 0 | 1,85 | 1,79 | 2,21 | 0 |
| NR1H3::RXRA | TGGACAAGTTTAACCACGA | 1 | 1134 | 1153 | 134 | 153 | 1,63 | 0,01 | 21,75 | >0.1 | 5,5 | 6,42 | 0 | 1,84 | 3,22 | 3,13 | 0 |
| NR1H3::RXRA | TCTATCAAAGTTCACTTGC | 1 | 1167 | 1186 | 167 | 186 | 1,5 | 0,01 | 19,68 | >0.1 | 5,5 | 6,17 | 0 | 1,69 | 1,79 | 3,02 | 0 |
| NR1H3::RXRA | TGAACTTTGATAGAACAGG | -1 | 1162 | 1181 | 162 | 181 | 6,39 | 0,001 | 24,51 | >0.1 | 5,5 | 6,21 | 0 | 1,74 | 1,79 | 2,88 | 0 |
| NR1H3::RXRA | GGCTTCGTGGTTAAACTTG | -1 | 1138 | 1157 | 138 | 157 | -1,21 | 0,01 | 19,11 | >0.1 | 5,5 | 6,39 | 0 | 1,84 | 3,22 | 3,37 | 0 |
| NR1H3::RXRA | TCCTCAATGGTGGACTGAG | -1 | 1098 | 1117 | 98 | 117 | 1,42 | 0,01 | 21,18 | >0.1 | 5,5 | 6,78 | 0 | 1,85 | 3,22 | 2,41 | 0 |
| NR1H3::RXRA | TCGTCCCCTGTGAGCCAAG | -1 | 1031 | 1050 | 31 | 50 | -0,35 | 0,01 | 16,5 | >0.1 | 5,5 | 8,19 | 0 | 1,86 | 0 | 1,31 | 0 |
| NR1H3::RXRA | TGAGCCAAGATCACATCCA | -1 | 1021 | 1040 | 21 | 40 | -1,27 | 0,01 | 16,04 | >0.1 | 5,5 | 8,7 | 0 | 1,86 | 0 | 1,25 | 0 |
| NR1H3::RXRA | TTCCCTAGACTAAAACCTC | -1 | 992 | 1011 | -8 | 11 | 2,65 | 0,01 | 20,63 | >0.1 | 5,5 | 9,12 | 0 | 1,87 | 0 | 1,5 | 0 |
| NR1H3::RXRA | TATCAGCCTTTGAACTTGG | -1 | 954 | 973 | -46 | -27 | 1,11 | 0,01 | 17,92 | >0.1 | 5,5 | 7,7 | 0 | 1,88 | 0 | 1,74 | 0 |
| NR1H3::RXRA | TTAATCCTAGAGAACCTTA | -1 | 779 | 798 | -221 | -202 | 0,5 | 0,01 | 19,26 | >0.1 | 5,5 | 5,91 | 0 | 1,44 | 0 | 5,92 | 0 |
| NR1H3::RXRA | TAATCCTAGAGAACCTTAC | -1 | 778 | 797 | -222 | -203 | 0,72 | 0,01 | 19,65 | >0.1 | 5,5 | 5,9 | 0 | 1,43 | 0 | 6,1 | 0 |
| NR1H3::RXRA | TGGCCAAATCAAAACCGGT | -1 | 743 | 762 | -257 | -238 | 1,02 | 0,01 | 19,65 | >0.1 | 5,5 | 5,73 | 0 | 1,3 | 0 | 6,1 | 0 |
| NR1H3::RXRA | TCCCTTCCAAATACATGAC | -1 | 517 | 536 | -483 | -464 | -0,7 | 0,01 | 11,43 | >0.1 | 3,05 | 5,02 | 0 | 0,99 | 0 | 3,07 | 0 |
| NR1H3::RXRA | TACATGACTTTGCCCTTGT | -1 | 506 | 525 | -494 | -475 | 0,17 | 0,01 | 12,02 | >0.1 | 3 | 5 | 0 | 0,99 | 0 | 2,86 | 0 |
| NR1H3::RXRA | GGGACTACAGGTGCCCACC | -1 | 323 | 342 | -677 | -658 | -1,15 | 0,01 | 12,83 | >0.1 | 2,6 | 4,65 | 0 | 1,16 | 0 | 5,56 | 0 |
| NR1H3::RXRA | GGACTACAGGTGCCCACCA | -1 | 322 | 341 | -678 | -659 | 0,84 | 0,01 | 14,71 | >0.1 | 2,6 | 4,65 | 0 | 1,16 | 0 | 5,46 | 0 |
| NR1H3::RXRA | TCTCCTGATCTGCCTTTGC | -1 | 87 | 106 | -913 | -894 | 1,21 | 0,01 | 11,97 | >0.1 | 3,68 | 4,35 | 0 | 1,88 | 0 | 0,85 | 0 |

| | | | | | | | | | | | | | | | | | |
|---|---|---|---|---|---|---|---|---|---|---|---|---|---|---|---|---|---|
| NR1H3::RXRA | CTTCTTTGAGTGCCTCCTT | -1 | 18 | 37 | -982 | -963 | -0,92 | 0,01 | 10,72 | >0.1 | 5,5 | 4,28 | 0 | 1,87 | 0 | 0 | 0 |
| MNT | AACACGGTGA | 1 | 269 | 279 | -731 | -721 | 2,09 | 0,01 | 15,81 | >0.1 | 2,78 | 4,56 | 0 | 1,3 | 0 | 5,09 | 0 |
| MNT | GGCACCTGTA | 1 | 327 | 337 | -673 | -663 | 0,14 | 0,01 | 14,1 | >0.1 | 2,6 | 4,65 | 0 | 1,14 | 0 | 5,56 | 0 |
| MNT | TACTCGGGAG | 1 | 345 | 355 | -655 | -645 | 0,33 | 0,01 | 15,19 | >0.1 | 2,55 | 4,67 | 0 | 1,11 | 0 | 6,52 | 0 |
| MNT | AAAATGTTGC | 1 | 535 | 545 | -465 | -455 | -2,94 | 0,01 | 7,81 | >0.1 | 3,11 | 5,04 | 0 | 0,98 | 0 | 1,61 | 0 |
| MNT | TAAATGTGAG | 1 | 586 | 596 | -414 | -404 | -0,7 | 0,01 | 8,86 | >0.1 | 3,38 | 5,17 | 0 | 1,01 | 0 | 0 | 0 |
| MNT | ATCTCATGAG | 1 | 982 | 992 | -18 | -8 | -1,11 | 0,01 | 16,33 | >0.1 | 5,61 | 8,43 | 0 | 1,87 | 0 | 1,53 | 0 |
| MNT | CTCACAGGGG | 1 | 1036 | 1046 | 36 | 46 | -1,42 | 0,01 | 15,37 | >0.1 | 5,61 | 7,99 | 0 | 1,86 | 0 | 1,34 | 0 |
| MNT | CACAGGGGAC | 1 | 1038 | 1048 | 38 | 48 | 0,87 | 0,01 | 17,59 | >0.1 | 5,61 | 7,92 | 0 | 1,86 | 0 | 1,34 | 0 |
| MNT | GCCTTGTTGC | 1 | 1077 | 1087 | 77 | 87 | -2,46 | 0,01 | 16,07 | >0.1 | 5,61 | 7,06 | 0 | 1,85 | 1,79 | 2,21 | 0 |
| MNT | GGAACAGGCC | 1 | 1114 | 1124 | 114 | 124 | -1,5 | 0,01 | 18,35 | >0.1 | 5,61 | 6,61 | 0 | 1,85 | 3,22 | 2,56 | 0 |
| MNT | TTCACTTGCT | 1 | 1177 | 1187 | 177 | 187 | -1,02 | 0,01 | 17,36 | >0.1 | 5,61 | 6,11 | 0 | 1,61 | 1,79 | 3,26 | 0 |
| MNT | AGCAAGTGAA | -1 | 1177 | 1187 | 177 | 187 | 0,41 | 0,01 | 18,79 | >0.1 | 5,61 | 6,11 | 0 | 1,61 | 1,79 | 3,26 | 0 |
| MNT | AGAACAGGTC | -1 | 1160 | 1170 | 160 | 170 | -2,77 | 0,01 | 13,86 | >0.1 | 4,4 | 6,22 | 0 | 1,76 | 1,79 | 2,46 | 0 |
| MNT | GCTTCGTGGT | -1 | 1146 | 1156 | 146 | 156 | 0,59 | 0,01 | 18,32 | >0.1 | 5,61 | 6,33 | 0 | 1,84 | 1,79 | 2,16 | 0 |
| MNT | TCCTCATGAG | -1 | 984 | 994 | -16 | -6 | 0,12 | 0,01 | 17,7 | >0.1 | 5,61 | 8,57 | 0 | 1,87 | 0 | 1,53 | 0 |
| MNT | AATACGTTAT | -1 | 801 | 811 | -199 | -189 | -0,29 | 0,01 | 16,22 | >0.1 | 5,61 | 5,98 | 0 | 1,5 | 0 | 3,43 | 0 |
| MNT | CTCACATTTA | -1 | 586 | 596 | -414 | -404 | -1,72 | 0,01 | 7,84 | >0.1 | 3,38 | 5,17 | 0 | 1,01 | 0 | 0 | 0 |
| MNT | AATACATGAC | -1 | 517 | 527 | -483 | -473 | -0,46 | 0,01 | 11,5 | >0.1 | 3,03 | 5 | 0 | 0,98 | 0 | 2,95 | 0 |
| MNT | CCCAGGTTCA | -1 | 375 | 385 | -625 | -615 | 1,77 | 0,01 | 16,51 | >0.1 | 2,56 | 4,72 | 0 | 1,07 | 0 | 6,4 | 0 |
| MNT | TACAGGTGCC | -1 | 327 | 337 | -673 | -663 | 6,77 | 0,001 | 20,73 | >0.1 | 2,6 | 4,65 | 0 | 1,14 | 0 | 5,56 | 0 |
| MNT | ACCTTGTGAT | -1 | 235 | 245 | -765 | -755 | 2,37 | 0,01 | 15,95 | >0.1 | 2,9 | 4,51 | 0 | 1,43 | 0 | 4,74 | 0 |
| MNT | TCCACAGGTC | -1 | 109 | 119 | -891 | -881 | 1,43 | 0,01 | 11,96 | >0.1 | 3,54 | 4,36 | 0 | 1,89 | 0 | 0,75 | 0 |
| MNT | AGCTCATTTC | -1 | 65 | 75 | -935 | -925 | -2,9 | 0,01 | 7,23 | >0.1 | 3,94 | 4,31 | 0 | 1,88 | 0 | 0 | 0 |
| SOX1 | CACAGAACCAGTGTG | 1 | 42 | 57 | -958 | -943 | -2,88 | 0,001 | 7,5 | >0.1 | 4,21 | 4,3 | 0 | 1,87 | 0 | 0 | 0 |
| SOX1 | TACAAAAACTTAGCT | 1 | 299 | 314 | -701 | -686 | -16,17 | 0,01 | -2,69 | >0.1 | 2,67 | 4,61 | 0 | 1,21 | 0 | 4,99 | 0 |
| SOX1 | CCCAAGTAGAGAGTT | 1 | 544 | 559 | -456 | -441 | -18,7 | 0,01 | -9,47 | >0.1 | 3,17 | 5,08 | 0 | 0,99 | 0 | 0 | 0 |
| SOX1 | ATTAATGAAATTGTC | 1 | 603 | 618 | -397 | -382 | -8,03 | 0,01 | 3,15 | >0.1 | 3,51 | 5,23 | 0 | 1,03 | 0 | 1,41 | 0 |
| SOX1 | AACACCAACATTATC | 1 | 631 | 646 | -369 | -354 | -7,27 | 0,001 | 4,44 | >0.1 | 3,73 | 5,31 | 0 | 1,05 | 0 | 1,61 | 0 |
| SOX1 | AATTATAACATTTTC | 1 | 685 | 700 | -315 | -300 | -14,26 | 0,01 | -0,24 | >0.1 | 4,45 | 5,49 | 0 | 1,14 | 0 | 2,95 | 0 |
| SOX1 | AACATTTTCCGTGTA | 1 | 691 | 706 | -309 | -294 | -10,3 | 0,01 | 4,97 | >0.1 | 5,54 | 5,51 | 0 | 1,15 | 0 | 3,07 | 0 |
| SOX1 | ACCAAAAGTACCGGT | 1 | 734 | 749 | -266 | -251 | -20,08 | 0,01 | -1,69 | >0.1 | 5,54 | 5,68 | 0 | 1,25 | 0 | 5,92 | 0 |
| SOX1 | AAGTCATTCAGTGGA | 1 | 1010 | 1025 | 10 | 25 | -20,24 | 0,01 | -2,42 | >0.1 | 5,54 | 8,99 | 0 | 1,86 | 0 | 1,42 | 0 |
| SOX1 | GGCCAAGACATTTTT | 1 | 1120 | 1135 | 120 | 135 | -13,45 | 0,01 | 6,45 | >0.1 | 5,54 | 6,55 | 0 | 1,85 | 3,22 | 2,74 | 0 |
| SOX1 | ATCACATCCACTGAA | -1 | 1016 | 1031 | 16 | 31 | -12,97 | 0,01 | 4,58 | >0.1 | 5,54 | 8,81 | 0 | 1,86 | 0 | 1,34 | 0 |
| SOX1 | TACACGGAAAATGTT | -1 | 691 | 706 | -309 | -294 | -19,23 | 0,01 | -3,96 | >0.1 | 5,54 | 5,51 | 0 | 1,15 | 0 | 3,07 | 0 |
| SOX1 | GATAATGTTGGTGTT | -1 | 631 | 646 | -369 | -354 | -16,43 | 0,01 | -4,72 | >0.1 | 3,73 | 5,31 | 0 | 1,05 | 0 | 1,61 | 0 |
| SOX1 | GACAATTTCATTAAT | -1 | 603 | 618 | -397 | -382 | 1,39 | 0,001 | 12,57 | >0.1 | 3,51 | 5,23 | 0 | 1,03 | 0 | 1,41 | 0 |
| SOX1 | CTCACATTTATTTTT | -1 | 581 | 596 | -419 | -404 | -14,76 | 0,01 | -5,21 | >0.1 | 3,37 | 5,17 | 0 | 1,01 | 0 | 0 | 0 |
| SOX1 | GACTTTGCCCTTGTC | -1 | 505 | 520 | -495 | -480 | -16,08 | 0,01 | -2,92 | >0.1 | 2,99 | 4,99 | 0 | 0,99 | 0 | 4,19 | 0 |
| SOX1 | TGCCCAGCCATTTCT | -1 | 170 | 185 | -830 | -815 | -19,94 | 0,01 | -7,11 | >0.1 | 3,17 | 4,44 | 0 | 1,88 | 0 | 3,34 | 0 |
| SOX1 | CACACTGGTTCTGTG | -1 | 42 | 57 | -958 | -943 | -16,81 | 0,01 | -6,43 | >0.1 | 4,21 | 4,3 | 0 | 1,87 | 0 | 0 | 0 |
| SOX1 | ACTTCTTTGAGTGCC | -1 | 23 | 38 | -977 | -962 | -20,56 | 0,01 | -8,87 | >0.1 | 5,54 | 4,28 | 0 | 1,87 | 0 | 0 | 0 |
| HSF2 | TCCTAGCACTTTG | 1 | 204 | 217 | -796 | -783 | -0,66 | 0,01 | 13,14 | >0.1 | 3,01 | 4,48 | 0 | 1,6 | 0 | 4,7 | 0 |
| HSF2 | TTCTGGGAATATG | 1 | 558 | 571 | -442 | -429 | -3,75 | 0,01 | 5,58 | >0.1 | 3,23 | 5,11 | 0 | 1 | 0 | 0 | 0 |
| HSF2 | TTCTAGACCTCTT | 1 | 660 | 673 | -340 | -327 | 0,08 | 0,01 | 14,76 | >0.1 | 4,03 | 5,4 | 0 | 1,09 | 0 | 4,16 | 0 |
| HSF2 | TTATAACATTTTC | 1 | 687 | 700 | -313 | -300 | -6,02 | 0,01 | 7,94 | >0.1 | 4,47 | 5,49 | 0 | 1,14 | 0 | 2,86 | 0 |
| HSF2 | TTCTAGGAAAATA | 1 | 718 | 731 | -282 | -269 | -2,16 | 0,01 | 15,2 | >0.1 | 5,57 | 5,6 | 0 | 1,2 | 0 | 4,99 | 0 |
| HSF2 | TTCTCTAGGATTA | 1 | 784 | 797 | -216 | -203 | -6,69 | 0,01 | 11,9 | >0.1 | 5,57 | 5,9 | 0 | 1,43 | 0 | 5,69 | 0 |
| HSF2 | TTCTAAAATCTGT | 1 | 837 | 850 | -163 | -150 | -1,35 | 0,01 | 15,41 | >0.1 | 5,57 | 6,23 | 0 | 1,77 | 0 | 3,19 | 0 |
| HSF2 | CTCCAGGATGAAC | 1 | 864 | 877 | -136 | -123 | -6,94 | 0,01 | 9,62 | >0.1 | 5,57 | 6,42 | 0 | 1,9 | 0 | 2,67 | 0 |
| HSF2 | TCCTGGCTCCTTC | 1 | 1061 | 1074 | 61 | 74 | -1,92 | 0,01 | 16,85 | >0.1 | 5,57 | 7,33 | 0 | 1,86 | 1,79 | 2,21 | 0 |
| HSF2 | TTCTATCAAAGTT | 1 | 1166 | 1179 | 166 | 179 | -8,64 | 0,01 | 9,47 | >0.1 | 5,57 | 6,18 | 0 | 1,7 | 1,79 | 2,86 | 0 |
| HSF2 | TGATAGAACAGGT | -1 | 1161 | 1174 | 161 | 174 | -7,11 | 0,01 | 9,7 | >0.1 | 4,45 | 6,22 | 0 | 1,75 | 1,79 | 2,61 | 0 |
| HSF2 | GTCCAAAAATGTC | -1 | 1126 | 1139 | 126 | 139 | -5,89 | 0,01 | 14,06 | >0.1 | 5,57 | 6,49 | 0 | 1,84 | 3,22 | 2,82 | 0 |
| HSF2 | GACTAAAACCTCC | -1 | 991 | 1004 | -9 | 4 | -7,39 | 0,01 | 10,46 | >0.1 | 5,57 | 8,84 | 0 | 1,87 | 0 | 1,57 | 0 |
| HSF2 | TTTTAGAATAATT | -1 | 832 | 845 | -168 | -155 | -2,99 | 0,01 | 13,93 | >0.1 | 5,57 | 6,19 | 0 | 1,73 | 0 | 3,43 | 0 |
| HSF2 | CTAGAGAACCTTA | -1 | 779 | 792 | -221 | -208 | -2,64 | 0,01 | 16,76 | >0.1 | 5,57 | 5,87 | 0 | 1,41 | 0 | 6,54 | 0 |
| HSF2 | AACCGGTACTTTT | -1 | 737 | 750 | -263 | -250 | -5,3 | 0,01 | 13,12 | >0.1 | 5,57 | 5,68 | 0 | 1,26 | 0 | 5,92 | 0 |

| | | | | | | | | | | | | | | | | | |
|---|---|---|---|---|---|---|---|---|---|---|---|---|---|---|---|---|---|
| HSF2 | TCCTAGAAAGCTG | -1 | 713 | 726 | -287 | -274 | -1,99 | 0,01 | 15,34 | >0.1 | 5,57 | 5,58 | 0 | 1,19 | 0 | 4,99 | 0 |
| HSF2 | GTCTAGAACTAGG | -1 | 655 | 668 | -345 | -332 | 0,04 | 0,01 | 14,17 | >0.1 | 3,97 | 5,38 | 0 | 1,08 | 0 | 3,7 | 0 |
| HSF2 | TTCAAGATCATAT | -1 | 566 | 579 | -434 | -421 | -5,42 | 0,01 | 3,98 | >0.1 | 3,27 | 5,13 | 0 | 1 | 0 | 0 | 0 |
| HSF2 | TCCCAGAAACTCT | -1 | 553 | 566 | -447 | -434 | -3,5 | 0,01 | 5,79 | >0.1 | 3,21 | 5,09 | 0 | 0,99 | 0 | 0 | 0 |
| HSF2 | TTATGGAATTCAT | -1 | 491 | 504 | -509 | -496 | -5,83 | 0,01 | 9,31 | >0.1 | 2,93 | 4,95 | 0 | 0,99 | 0 | 6,27 | 0 |
| HSF2 | GTCTCGGTCTGTC | -1 | 436 | 449 | -564 | -551 | -5,63 | 0,01 | 9 | >0.1 | 2,74 | 4,84 | 0 | 1,01 | 0 | 6,05 | 0 |
| LHX2 | GGAAATGAGC | 1 | 64 | 74 | -936 | -926 | 0,71 | 0,01 | 10,85 | >0.1 | 3,95 | 4,31 | 0 | 1,88 | 0 | 0 | 0 |
| LHX2 | GGAAATTAAA | 1 | 153 | 163 | -847 | -837 | 2,5 | 0,01 | 13,72 | >0.1 | 3,27 | 4,41 | 0 | 1,9 | 0 | 1,65 | 0 |
| LHX2 | CTCTATTAAA | 1 | 287 | 297 | -713 | -703 | 0,71 | 0,01 | 14,45 | >0.1 | 2,72 | 4,59 | 0 | 1,24 | 0 | 5,19 | 0 |
| LHX2 | ACCTATTAAT | 1 | 599 | 609 | -401 | -391 | 4,45 | 0,01 | 15,5 | >0.1 | 3,47 | 5,21 | 0 | 1,02 | 0 | 1,36 | 0 |
| LHX2 | ATTAATGAAA | 1 | 603 | 613 | -397 | -387 | 4,61 | 0,01 | 15,75 | >0.1 | 3,49 | 5,22 | 0 | 1,02 | 0 | 1,41 | 0 |
| LHX2 | CTTCATGATC | 1 | 645 | 655 | -355 | -345 | 2,21 | 0,01 | 16,29 | >0.1 | 3,84 | 5,34 | 0 | 1,06 | 0 | 3,83 | 0 |
| LHX2 | TCTCATGAGG | 1 | 983 | 993 | -17 | -7 | 5,08 | 0,01 | 22,63 | >0.1 | 5,61 | 8,5 | 0 | 1,87 | 0 | 1,57 | 0 |
| LHX2 | CACCATTGAG | 1 | 1105 | 1115 | 105 | 115 | 2,39 | 0,01 | 22,29 | >0.1 | 5,61 | 6,7 | 0 | 1,85 | 3,22 | 2,52 | 0 |
| LHX2 | CTCAATGGTG | -1 | 1105 | 1115 | 105 | 115 | 1,01 | 0,01 | 20,91 | >0.1 | 5,61 | 6,7 | 0 | 1,85 | 3,22 | 2,52 | 0 |
| LHX2 | CCTCATGAGA | -1 | 983 | 993 | -17 | -7 | 5,19 | 0,01 | 22,74 | >0.1 | 5,61 | 8,5 | 0 | 1,87 | 0 | 1,57 | 0 |
| LHX2 | TCTTATCAGC | -1 | 966 | 976 | -34 | -24 | 0,45 | 0,01 | 17,41 | >0.1 | 5,61 | 7,78 | 0 | 1,87 | 0 | 1,7 | 0 |
| LHX2 | GATCATGAAG | -1 | 645 | 655 | -355 | -345 | 2,54 | 0,01 | 16,62 | >0.1 | 3,84 | 5,34 | 0 | 1,06 | 0 | 3,83 | 0 |
| LHX2 | TTTCATTAAT | -1 | 603 | 613 | -397 | -387 | 4,99 | 0,01 | 16,13 | >0.1 | 3,49 | 5,22 | 0 | 1,02 | 0 | 1,41 | 0 |
| LHX2 | TTTAATTTCC | -1 | 153 | 163 | -847 | -837 | 0,95 | 0,01 | 12,17 | >0.1 | 3,27 | 4,41 | 0 | 1,9 | 0 | 1,65 | 0 |
| LHX2 | GCTCATTTCC | -1 | 64 | 74 | -936 | -926 | 3,28 | 0,01 | 13,42 | >0.1 | 3,95 | 4,31 | 0 | 1,88 | 0 | 0 | 0 |
| LHX3 | TAATGAAATTGTC | 1 | 605 | 618 | -395 | -382 | 4,89 | 0,01 | 16,11 | >0.1 | 3,52 | 5,23 | 0 | 1,02 | 0 | 1,44 | 0 |
| LHX3 | ATATTAACCAAAA | 1 | 728 | 741 | -272 | -259 | 0,47 | 0,01 | 18,44 | >0.1 | 5,57 | 5,64 | 0 | 1,23 | 0 | 5,52 | 0 |
| LHX3 | GGATTAAAGAATA | 1 | 791 | 804 | -209 | -196 | 2,53 | 0,01 | 20,82 | >0.1 | 5,57 | 5,94 | 0 | 1,47 | 0 | 5,32 | 0 |
| LHX3 | TATTTGATTCACT | 1 | 813 | 826 | -187 | -174 | 3,07 | 0,01 | 19,72 | >0.1 | 5,57 | 6,06 | 0 | 1,59 | 0 | 3,43 | 0 |
| LHX3 | ACTTTAAAAAATT | 1 | 823 | 836 | -177 | -164 | 0,56 | 0,01 | 17,62 | >0.1 | 5,57 | 6,13 | 0 | 1,66 | 0 | 3,7 | 0 |
| LHX3 | TAAAAAATTATTC | 1 | 827 | 840 | -173 | -160 | 3,82 | 0,01 | 20,66 | >0.1 | 5,57 | 6,16 | 0 | 1,69 | 0 | 3,43 | 0 |
| LHX3 | AAATTATTCTAAA | 1 | 831 | 844 | -169 | -156 | 0,2 | 0,01 | 17,22 | >0.1 | 5,57 | 6,19 | 0 | 1,72 | 0 | 3,55 | 0 |
| LHX3 | ACAGTAAACAATC | -1 | 894 | 907 | -106 | -93 | 2,54 | 0,01 | 19,63 | >0.1 | 5,57 | 6,68 | 0 | 1,89 | 0 | 2,95 | 0 |
| LHX3 | AGAATAATTTTTT | -1 | 828 | 841 | -172 | -159 | 4,86 | 0,01 | 21,57 | >0.1 | 5,57 | 6,16 | 0 | 1,7 | 0 | 3,28 | 0 |
| LHX3 | TAATTTTTTAAAG | -1 | 824 | 837 | -176 | -163 | 2,91 | 0,01 | 19,98 | >0.1 | 5,57 | 6,14 | 0 | 1,67 | 0 | 3,7 | 0 |
| LHX3 | TCTTTAATCCTAG | -1 | 788 | 801 | -212 | -199 | 0,26 | 0,01 | 18,72 | >0.1 | 5,57 | 5,92 | 0 | 1,45 | 0 | 5,52 | 0 |
| LHX3 | AATTTCATTAATA | -1 | 602 | 615 | -398 | -385 | 4,82 | 0,01 | 15,97 | >0.1 | 3,5 | 5,22 | 0 | 1,02 | 0 | 1,41 | 0 |
| LHX3 | TCATTAATAGGTT | -1 | 598 | 611 | -402 | -389 | 0,4 | 0,01 | 11,46 | >0.1 | 3,47 | 5,21 | 0 | 1,02 | 0 | 1,36 | 0 |
| LHX3 | ACATTTATTTTTA | -1 | 580 | 593 | -420 | -407 | 6,02 | 0,001 | 15,54 | >0.1 | 3,35 | 5,16 | 0 | 1,01 | 0 | 0 | 0 |
| LHX3 | GAATTCATATTTC | -1 | 486 | 499 | -514 | -501 | 1,26 | 0,01 | 16,23 | >0.1 | 2,91 | 4,94 | 0 | 0,99 | 0 | 6,13 | 0 |
| LHX3 | AGTTTTAATTTCC | -1 | 153 | 166 | -847 | -834 | 1,88 | 0,01 | 13,1 | >0.1 | 3,26 | 4,41 | 0 | 1,9 | 0 | 1,65 | 0 |
| LHX3 | TCATTTCCTATTT | -1 | 59 | 72 | -941 | -928 | 1,14 | 0,01 | 11,32 | >0.1 | 3,99 | 4,31 | 0 | 1,88 | 0 | 0 | 0 |
| LHX4 | GAAATTAA | 1 | 154 | 162 | -846 | -838 | 5,85 | 0,01 | 17,07 | >0.1 | 3,27 | 4,41 | 0 | 1,9 | 0 | 1,65 | 0 |
| LHX4 | CTGATCAG | 1 | 164 | 172 | -836 | -828 | 5,25 | 0,01 | 17,18 | >0.1 | 3,21 | 4,42 | 0 | 1,9 | 0 | 2,4 | 0 |
| LHX4 | TCTATTAA | 1 | 288 | 296 | -712 | -704 | 5,64 | 0,01 | 19,37 | >0.1 | 2,72 | 4,58 | 0 | 1,24 | 0 | 5,19 | 0 |
| LHX4 | CTGATCAG | 1 | 479 | 487 | -521 | -513 | 5,25 | 0,01 | 19,39 | >0.1 | 2,88 | 4,92 | 0 | 0,98 | 0 | 5,36 | 0 |
| LHX4 | CCTATTAA | 1 | 600 | 608 | -400 | -392 | 5,63 | 0,01 | 16,68 | >0.1 | 3,47 | 5,2 | 0 | 1,01 | 0 | 1,36 | 0 |
| LHX4 | TTAATGAA | 1 | 604 | 612 | -396 | -388 | 5,15 | 0,01 | 16,29 | >0.1 | 3,49 | 5,21 | 0 | 1,02 | 0 | 1,41 | 0 |
| LHX4 | TTATTTGA | 1 | 812 | 820 | -188 | -180 | 3,4 | 0,01 | 20,05 | >0.1 | 5,64 | 6,03 | 0 | 1,55 | 0 | 3,43 | 0 |
| LHX4 | TCTATCAA | 1 | 1167 | 1175 | 167 | 175 | 3,44 | 0,01 | 20,51 | >0.1 | 4,53 | 6,17 | 0 | 1,69 | 1,79 | 2,88 | 0 |
| LHX4 | CTTATCAG | -1 | 967 | 975 | -33 | -25 | 4,48 | 0,01 | 21,44 | >0.1 | 5,64 | 7,76 | 0 | 1,87 | 0 | 1,7 | 0 |
| LHX4 | TTCTTTAA | -1 | 794 | 802 | -206 | -198 | 3,83 | 0,01 | 22,17 | >0.1 | 5,64 | 5,93 | 0 | 1,45 | 0 | 5,32 | 0 |
| LHX4 | CAAATCAA | -1 | 750 | 758 | -250 | -242 | 4,37 | 0,01 | 22,92 | >0.1 | 5,64 | 5,71 | 0 | 1,28 | 0 | 5,92 | 0 |
| LHX4 | TTCATTAA | -1 | 604 | 612 | -396 | -388 | 8,05 | 0,001 | 19,19 | >0.1 | 3,49 | 5,21 | 0 | 1,02 | 0 | 1,41 | 0 |
| LHX4 | CTGATCAG | -1 | 479 | 487 | -521 | -513 | 5,25 | 0,01 | 19,39 | >0.1 | 2,88 | 4,92 | 0 | 0,98 | 0 | 5,36 | 0 |
| LHX4 | CTGATCAG | -1 | 164 | 172 | -836 | -828 | 5,25 | 0,01 | 17,18 | >0.1 | 3,21 | 4,42 | 0 | 1,9 | 0 | 2,4 | 0 |
| LHX4 | TTAATTTC | -1 | 154 | 162 | -846 | -838 | 4,84 | 0,01 | 16,06 | >0.1 | 3,27 | 4,41 | 0 | 1,9 | 0 | 1,65 | 0 |
| LHX6 | ACTGATCAGA | 1 | 163 | 173 | -837 | -827 | 7,24 | 0,001 | 19,18 | >0.1 | 3,21 | 4,42 | 0 | 1,9 | 0 | 2,4 | 0 |
| LHX6 | CCTGACTAAC | 1 | 262 | 272 | -738 | -728 | 4,37 | 0,01 | 17,57 | >0.1 | 2,8 | 4,55 | 0 | 1,32 | 0 | 4,53 | 0 |
| LHX6 | GCTACTCGGG | 1 | 343 | 353 | -657 | -647 | -0,41 | 0,01 | 14,46 | >0.1 | 2,56 | 4,67 | 0 | 1,11 | 0 | 6,52 | 0 |
| LHX6 | ACTGATCAGA | 1 | 478 | 488 | -522 | -512 | 7,24 | 0,001 | 21,38 | >0.1 | 2,88 | 4,92 | 0 | 0,99 | 0 | 5,36 | 0 |
| LHX6 | CCCAAGTAGA | 1 | 544 | 554 | -456 | -446 | -0,13 | 0,01 | 9,07 | >0.1 | 3,15 | 5,06 | 0 | 0,99 | 0 | 0 | 0 |
| LHX6 | ATTAATGAAA | 1 | 603 | 613 | -397 | -387 | 4,3 | 0,01 | 15,44 | >0.1 | 3,49 | 5,22 | 0 | 1,02 | 0 | 1,41 | 0 |

| | | | | | | | | | | | | | | | | | |
|---|---|---|---|---|---|---|---|---|---|---|---|---|---|---|---|---|---|
| LHX6 | ATTAACCAAA | 1 | 730 | 740 | -270 | -260 | 2,13 | 0,01 | 20,13 | >0.1 | 5,61 | 5,64 | 0 | 1,23 | 0 | 5,52 | 0 |
| LHX6 | CCTGCTCAGC | 1 | 937 | 947 | -63 | -53 | 0,77 | 0,01 | 16,33 | >0.1 | 5,61 | 7,17 | 0 | 1,88 | 0 | 0,9 | 0 |
| LHX6 | TCTCATGAGG | 1 | 983 | 993 | -17 | -7 | 0,08 | 0,01 | 17,63 | >0.1 | 5,61 | 8,5 | 0 | 1,87 | 0 | 1,57 | 0 |
| LHX6 | GCTGCTCAGT | 1 | 1094 | 1104 | 94 | 104 | 1,15 | 0,01 | 21 | >0.1 | 5,61 | 6,83 | 0 | 1,85 | 3,22 | 2,34 | 0 |
| LHX6 | TTTAACCACG | 1 | 1142 | 1152 | 142 | 152 | -0,67 | 0,01 | 19,73 | >0.1 | 5,61 | 6,36 | 0 | 1,84 | 3,22 | 3,37 | 0 |
| LHX6 | ACTGAGCAGC | -1 | 1094 | 1104 | 94 | 104 | 1,64 | 0,01 | 21,49 | >0.1 | 5,61 | 6,83 | 0 | 1,85 | 3,22 | 2,34 | 0 |
| LHX6 | CCTCATGAGA | -1 | 983 | 993 | -17 | -7 | 1,21 | 0,01 | 18,76 | >0.1 | 5,61 | 8,5 | 0 | 1,87 | 0 | 1,57 | 0 |
| LHX6 | GCTGAGCAGG | -1 | 937 | 947 | -63 | -53 | 0,69 | 0,01 | 16,25 | >0.1 | 5,61 | 7,17 | 0 | 1,88 | 0 | 0,9 | 0 |
| LHX6 | GTGAATCAAA | -1 | 815 | 825 | -185 | -175 | 0,56 | 0,01 | 17,24 | >0.1 | 5,61 | 6,06 | 0 | 1,58 | 0 | 3,43 | 0 |
| LHX6 | AGTGACCAAA | -1 | 671 | 681 | -329 | -319 | -0,92 | 0,01 | 13,31 | >0.1 | 4,16 | 5,42 | 0 | 1,1 | 0 | 3,55 | 0 |
| LHX6 | GATCATGAAG | -1 | 645 | 655 | -355 | -345 | -1,11 | 0,01 | 12,97 | >0.1 | 3,84 | 5,34 | 0 | 1,06 | 0 | 3,83 | 0 |
| LHX6 | TTTCATTAAT | -1 | 603 | 613 | -397 | -387 | 2,69 | 0,01 | 13,83 | >0.1 | 3,49 | 5,22 | 0 | 1,02 | 0 | 1,41 | 0 |
| LHX6 | TCTACTTGGG | -1 | 544 | 554 | -456 | -446 | -1,42 | 0,01 | 7,78 | >0.1 | 3,15 | 5,06 | 0 | 0,99 | 0 | 0 | 0 |
| LHX6 | TCTGATCAGT | -1 | 478 | 488 | -522 | -512 | 5,89 | 0,01 | 20,03 | >0.1 | 2,88 | 4,92 | 0 | 0,99 | 0 | 5,36 | 0 |
| LHX6 | CCCGAGTAGC | -1 | 343 | 353 | -657 | -647 | -0,73 | 0,01 | 14,14 | >0.1 | 2,56 | 4,67 | 0 | 1,11 | 0 | 6,52 | 0 |
| LHX6 | TCTGATCAGT | -1 | 163 | 173 | -837 | -827 | 5,89 | 0,01 | 17,83 | >0.1 | 3,21 | 4,42 | 0 | 1,9 | 0 | 2,4 | 0 |
| LHX8 | GCCATGGA | 1 | 148 | 156 | -852 | -844 | -3,16 | 0,01 | 8,05 | >0.1 | 3,3 | 4,4 | 0 | 1,9 | 0 | 1,61 | 0 |
| LHX8 | CTGATCAG | 1 | 164 | 172 | -836 | -828 | 9,18 | 0,001 | 21,11 | >0.1 | 3,21 | 4,42 | 0 | 1,9 | 0 | 2,4 | 0 |
| LHX8 | CTACTCGG | 1 | 344 | 352 | -656 | -648 | -4,75 | 0,01 | 10,11 | >0.1 | 2,56 | 4,67 | 0 | 1,11 | 0 | 6,52 | 0 |
| LHX8 | CTGATCAG | 1 | 479 | 487 | -521 | -513 | 9,18 | 0,001 | 23,32 | >0.1 | 2,88 | 4,92 | 0 | 0,98 | 0 | 5,36 | 0 |
| LHX8 | CCAAGTAG | 1 | 545 | 553 | -455 | -447 | -3,91 | 0,01 | 5,29 | >0.1 | 3,15 | 5,06 | 0 | 0,98 | 0 | 0 | 0 |
| LHX8 | TTAATGAA | 1 | 604 | 612 | -396 | -388 | 2,86 | 0,01 | 14 | >0.1 | 3,49 | 5,21 | 0 | 1,02 | 0 | 1,41 | 0 |
| LHX8 | TTCATGAT | 1 | 646 | 654 | -354 | -346 | -3,36 | 0,01 | 10,71 | >0.1 | 3,84 | 5,34 | 0 | 1,06 | 0 | 3,83 | 0 |
| LHX8 | CTGCTCAG | 1 | 938 | 946 | -62 | -54 | -4,96 | 0,01 | 10,61 | >0.1 | 5,64 | 7,15 | 0 | 1,88 | 0 | 0,9 | 0 |
| LHX8 | CTGATAAG | 1 | 967 | 975 | -33 | -25 | -4,66 | 0,01 | 12,3 | >0.1 | 5,64 | 7,76 | 0 | 1,87 | 0 | 1,7 | 0 |
| LHX8 | CTCATGAG | 1 | 984 | 992 | -16 | -8 | 1,24 | 0,01 | 18,76 | >0.1 | 5,64 | 8,43 | 0 | 1,87 | 0 | 1,57 | 0 |
| LHX8 | CTGCTCAG | 1 | 1095 | 1103 | 95 | 103 | -4,96 | 0,01 | 13,48 | >0.1 | 5,64 | 6,82 | 0 | 1,85 | 1,79 | 2,34 | 0 |
| LHX8 | ACCATTGA | 1 | 1106 | 1114 | 106 | 114 | 0,73 | 0,01 | 20,65 | >0.1 | 5,64 | 6,69 | 0 | 1,85 | 3,22 | 2,52 | 0 |
| LHX8 | TCAATGGT | -1 | 1106 | 1114 | 106 | 114 | -0,62 | 0,01 | 19,3 | >0.1 | 5,64 | 6,69 | 0 | 1,85 | 3,22 | 2,52 | 0 |
| LHX8 | CTGAGCAG | -1 | 1095 | 1103 | 95 | 103 | -4,96 | 0,01 | 13,48 | >0.1 | 5,64 | 6,82 | 0 | 1,85 | 1,79 | 2,34 | 0 |
| LHX8 | CTCATGAG | -1 | 984 | 992 | -16 | -8 | 1,24 | 0,01 | 18,76 | >0.1 | 5,64 | 8,43 | 0 | 1,87 | 0 | 1,57 | 0 |
| LHX8 | CTTATCAG | -1 | 967 | 975 | -33 | -25 | -4,34 | 0,01 | 12,62 | >0.1 | 5,64 | 7,76 | 0 | 1,87 | 0 | 1,7 | 0 |
| LHX8 | CTGAGCAG | -1 | 938 | 946 | -62 | -54 | -4,96 | 0,01 | 10,61 | >0.1 | 5,64 | 7,15 | 0 | 1,88 | 0 | 0,9 | 0 |
| LHX8 | ATCATGAA | -1 | 646 | 654 | -354 | -346 | -2,87 | 0,01 | 11,2 | >0.1 | 3,84 | 5,34 | 0 | 1,06 | 0 | 3,83 | 0 |
| LHX8 | TTCATTAA | -1 | 604 | 612 | -396 | -388 | 3,72 | 0,01 | 14,86 | >0.1 | 3,49 | 5,21 | 0 | 1,02 | 0 | 1,41 | 0 |
| LHX8 | CTACTTGG | -1 | 545 | 553 | -455 | -447 | -3,96 | 0,01 | 5,24 | >0.1 | 3,15 | 5,06 | 0 | 0,98 | 0 | 0 | 0 |
| LHX8 | CTGATCAG | -1 | 479 | 487 | -521 | -513 | 9,18 | 0,001 | 23,32 | >0.1 | 2,88 | 4,92 | 0 | 0,98 | 0 | 5,36 | 0 |
| LHX8 | CCGAGTAG | -1 | 344 | 352 | -656 | -648 | -5,02 | 0,01 | 9,84 | >0.1 | 2,56 | 4,67 | 0 | 1,11 | 0 | 6,52 | 0 |
| LHX8 | CTGATCAG | -1 | 164 | 172 | -836 | -828 | 9,18 | 0,001 | 21,11 | >0.1 | 3,21 | 4,42 | 0 | 1,9 | 0 | 2,4 | 0 |
| LHX8 | TCCATGGC | -1 | 148 | 156 | -852 | -844 | -3,14 | 0,01 | 8,07 | >0.1 | 3,3 | 4,4 | 0 | 1,9 | 0 | 1,61 | 0 |
| LHX9 | GAAATGAG | 1 | 65 | 73 | -935 | -927 | 4,8 | 0,01 | 14,94 | >0.1 | 3,95 | 4,31 | 0 | 1,88 | 0 | 0 | 0 |
| LHX9 | GAAATTAA | 1 | 154 | 162 | -846 | -838 | 6,21 | 0,01 | 17,43 | >0.1 | 3,27 | 4,41 | 0 | 1,9 | 0 | 1,65 | 0 |
| LHX9 | CTGATCAG | 1 | 164 | 172 | -836 | -828 | 3,55 | 0,01 | 15,48 | >0.1 | 3,21 | 4,42 | 0 | 1,9 | 0 | 2,4 | 0 |
| LHX9 | TCTATTAA | 1 | 288 | 296 | -712 | -704 | 5,67 | 0,01 | 19,4 | >0.1 | 2,72 | 4,58 | 0 | 1,24 | 0 | 5,19 | 0 |
| LHX9 | CTGATCAG | 1 | 479 | 487 | -521 | -513 | 3,55 | 0,01 | 17,69 | >0.1 | 2,88 | 4,92 | 0 | 0,98 | 0 | 5,36 | 0 |
| LHX9 | CCTATTAA | 1 | 600 | 608 | -400 | -392 | 6,87 | 0,01 | 17,92 | >0.1 | 3,47 | 5,2 | 0 | 1,01 | 0 | 1,36 | 0 |
| LHX9 | TTAATGAA | 1 | 604 | 612 | -396 | -388 | 6,64 | 0,01 | 17,78 | >0.1 | 3,49 | 5,21 | 0 | 1,02 | 0 | 1,41 | 0 |
| LHX9 | AAAATTAT | 1 | 683 | 691 | -317 | -309 | 4,09 | 0,01 | 18,07 | >0.1 | 4,34 | 5,46 | 0 | 1,12 | 0 | 3,07 | 0 |
| LHX9 | AAAATTAT | 1 | 830 | 838 | -170 | -162 | 4,09 | 0,01 | 20,82 | >0.1 | 5,64 | 6,14 | 0 | 1,67 | 0 | 3,28 | 0 |
| LHX9 | CTCATGAG | 1 | 984 | 992 | -16 | -8 | 6,38 | 0,01 | 23,9 | >0.1 | 5,64 | 8,43 | 0 | 1,87 | 0 | 1,57 | 0 |
| LHX9 | ACCATTGA | 1 | 1106 | 1114 | 106 | 114 | 3,66 | 0,01 | 23,58 | >0.1 | 5,64 | 6,69 | 0 | 1,85 | 3,22 | 2,52 | 0 |
| LHX9 | CTCATGAG | -1 | 984 | 992 | -16 | -8 | 6,38 | 0,01 | 23,9 | >0.1 | 5,64 | 8,43 | 0 | 1,87 | 0 | 1,57 | 0 |
| LHX9 | CTTATCAG | -1 | 967 | 975 | -33 | -25 | 3,66 | 0,01 | 20,62 | >0.1 | 5,64 | 7,76 | 0 | 1,87 | 0 | 1,7 | 0 |
| LHX9 | CAAATCAA | -1 | 750 | 758 | -250 | -242 | 4,12 | 0,01 | 22,67 | >0.1 | 5,64 | 5,71 | 0 | 1,28 | 0 | 5,92 | 0 |
| LHX9 | ATCATGAA | -1 | 646 | 654 | -354 | -346 | 4,46 | 0,01 | 18,53 | >0.1 | 3,84 | 5,34 | 0 | 1,06 | 0 | 3,83 | 0 |
| LHX9 | ACAATTTC | -1 | 609 | 617 | -391 | -383 | 3,55 | 0,01 | 14,87 | >0.1 | 3,53 | 5,23 | 0 | 1,02 | 0 | 1,55 | 0 |
| LHX9 | TTCATTAA | -1 | 604 | 612 | -396 | -388 | 6,59 | 0,01 | 17,73 | >0.1 | 3,49 | 5,21 | 0 | 1,02 | 0 | 1,41 | 0 |
| LHX9 | CTGATCAG | -1 | 479 | 487 | -521 | -513 | 3,55 | 0,01 | 17,69 | >0.1 | 2,88 | 4,92 | 0 | 0,98 | 0 | 5,36 | 0 |
| LHX9 | CTGATCAG | -1 | 164 | 172 | -836 | -828 | 3,55 | 0,01 | 15,48 | >0.1 | 3,21 | 4,42 | 0 | 1,9 | 0 | 2,4 | 0 |

| | | | | | | | | | | | | | | | | | |
|---|---|---|---|---|---|---|---|---|---|---|---|---|---|---|---|---|---|
| LHX9 | AAAATTAT | -1 | 132 | 140 | -868 | -860 | 4,09 | 0,01 | 15,33 | >0.1 | 3,4 | 4,38 | 0 | 1,89 | 0 | 1,57 | 0 |
| FOXQ1 | TGAACTTTATA | 1 | 872 | 883 | -128 | -117 | 3,25 | 0,01 | 20,08 | >0.1 | 5,6 | 6,47 | 0 | 1,9 | 0 | 2,86 | 0 |
| FOXQ1 | TTTAGTCTAGG | 1 | 997 | 1008 | -3 | 8 | 1,32 | 0,01 | 19,29 | >0.1 | 5,6 | 9,01 | 0 | 1,87 | 0 | 1,5 | 0 |
| FOXQ1 | CAAAGTTCACT | 1 | 1172 | 1183 | 172 | 183 | 2,62 | 0,01 | 20,77 | >0.1 | 5,6 | 6,14 | 0 | 1,65 | 1,79 | 2,97 | 0 |
| FOXQ1 | TAAAGTTCATC | -1 | 870 | 881 | -130 | -119 | 0,83 | 0,01 | 17,64 | >0.1 | 5,6 | 6,45 | 0 | 1,9 | 0 | 2,86 | 0 |
| FOXQ1 | AATTTTTTAAA | -1 | 825 | 836 | -175 | -164 | 0,83 | 0,01 | 17,91 | >0.1 | 5,6 | 6,13 | 0 | 1,66 | 0 | 3,7 | 0 |
| FOXQ1 | TATTCTTTAAT | -1 | 793 | 804 | -207 | -196 | 2,67 | 0,01 | 20,99 | >0.1 | 5,6 | 5,94 | 0 | 1,46 | 0 | 5,32 | 0 |
| FOXQ1 | TAATGTTGGTG | -1 | 633 | 644 | -367 | -356 | 3,74 | 0,01 | 15,44 | >0.1 | 3,73 | 5,3 | 0 | 1,05 | 0 | 1,61 | 0 |
| FOXQ1 | TGGTGTTTGTA | -1 | 627 | 638 | -373 | -362 | 4,45 | 0,01 | 16,01 | >0.1 | 3,68 | 5,29 | 0 | 1,04 | 0 | 1,55 | 0 |
| FOXQ1 | AATAGGTTATC | -1 | 595 | 606 | -405 | -394 | 1,16 | 0,01 | 10,81 | >0.1 | 3,44 | 5,2 | 0 | 1,01 | 0 | 0 | 0 |
| FOXQ1 | TTATTTTTATT | -1 | 578 | 589 | -422 | -411 | 3,66 | 0,01 | 13,15 | >0.1 | 3,34 | 5,15 | 0 | 1 | 0 | 0 | 0 |
| FOXQ1 | CTAAGTTTTTG | -1 | 301 | 312 | -699 | -688 | 0,68 | 0,01 | 14,15 | >0.1 | 2,67 | 4,61 | 0 | 1,2 | 0 | 4,99 | 0 |
| FOXQ1 | TATTTTTAATA | -1 | 290 | 301 | -710 | -699 | 1,15 | 0,01 | 14,77 | >0.1 | 2,71 | 4,59 | 0 | 1,23 | 0 | 5,09 | 0 |
| FOXQ1 | CATTTTTTAAA | -1 | 137 | 148 | -863 | -852 | 1,33 | 0,01 | 12,62 | >0.1 | 3,35 | 4,39 | 0 | 1,89 | 0 | 1,65 | 0 |
| GSC2 | CATAATTTTA | 1 | 131 | 141 | -869 | -859 | 3 | 0,01 | 14,25 | >0.1 | 3,4 | 4,38 | 0 | 1,89 | 0 | 1,57 | 0 |
| GSC2 | TGTAATCCTA | 1 | 199 | 209 | -801 | -791 | 7,9 | 0,001 | 21,52 | >0.1 | 3,04 | 4,46 | 0 | 1,64 | 0 | 4,48 | 0 |
| GSC2 | CATTATCTTC | 1 | 639 | 649 | -361 | -351 | 4,64 | 0,01 | 16,46 | >0.1 | 3,79 | 5,32 | 0 | 1,06 | 0 | 1,66 | 0 |
| GSC2 | CATGATCCCT | 1 | 648 | 658 | -352 | -342 | 5,95 | 0,01 | 20,07 | >0.1 | 3,87 | 5,35 | 0 | 1,07 | 0 | 3,83 | 0 |
| GSC2 | CATTTTCCGT | 1 | 693 | 703 | -307 | -297 | 3,43 | 0,01 | 17,72 | >0.1 | 4,58 | 5,5 | 0 | 1,14 | 0 | 3,07 | 0 |
| GSC2 | GCTCAGCGCC | 1 | 940 | 950 | -60 | -50 | 3,26 | 0,01 | 18,79 | >0.1 | 5,61 | 7,22 | 0 | 1,88 | 0 | 0,83 | 0 |
| GSC2 | GAAAATCTCA | 1 | 978 | 988 | -22 | -12 | 3,13 | 0,01 | 20,36 | >0.1 | 5,61 | 8,22 | 0 | 1,87 | 0 | 1,53 | 0 |
| GSC2 | TCTCAGCCTT | 1 | 1072 | 1082 | 72 | 82 | 2,75 | 0,01 | 21,42 | >0.1 | 5,61 | 7,14 | 0 | 1,85 | 1,79 | 2,27 | 0 |
| GSC2 | TATAATTCCA | -1 | 1190 | 1200 | 190 | 200 | 5,6 | 0,01 | 24,36 | >0.1 | 5,61 | 6,03 | 0 | 1,51 | 1,79 | 3,82 | 0 |
| GSC2 | CCTGTTCCTC | -1 | 1112 | 1122 | 112 | 122 | 3,8 | 0,01 | 23,65 | >0.1 | 5,61 | 6,63 | 0 | 1,85 | 3,22 | 2,54 | 0 |
| GSC2 | TCTTATCAGC | -1 | 966 | 976 | -34 | -24 | 5,08 | 0,01 | 22,04 | >0.1 | 5,61 | 7,78 | 0 | 1,87 | 0 | 1,7 | 0 |
| GSC2 | GTTCATCCTG | -1 | 867 | 877 | -133 | -123 | 4,54 | 0,01 | 21,22 | >0.1 | 5,61 | 6,42 | 0 | 1,9 | 0 | 2,75 | 0 |
| GSC2 | GTTATTCTTT | -1 | 796 | 806 | -204 | -194 | 3,1 | 0,01 | 19,56 | >0.1 | 5,61 | 5,95 | 0 | 1,47 | 0 | 3,43 | 0 |
| GSC2 | TTTAATCCTA | -1 | 789 | 799 | -211 | -201 | 9,13 | 0,001 | 27,61 | >0.1 | 5,61 | 5,91 | 0 | 1,44 | 0 | 5,52 | 0 |
| GSC2 | CTTTATGGCC | -1 | 757 | 767 | -243 | -233 | 3,5 | 0,01 | 22,28 | >0.1 | 5,61 | 5,75 | 0 | 1,31 | 0 | 6,1 | 0 |
| GSC2 | GTTAATATTT | -1 | 726 | 736 | -274 | -264 | 3,55 | 0,01 | 21,32 | >0.1 | 5,61 | 5,62 | 0 | 1,22 | 0 | 5,32 | 0 |
| GSC2 | ATTAATAGGT | -1 | 599 | 609 | -401 | -391 | 4,7 | 0,01 | 15,75 | >0.1 | 3,47 | 5,21 | 0 | 1,02 | 0 | 1,36 | 0 |
| GSC2 | GGTTATCTCA | -1 | 592 | 602 | -408 | -398 | 3,51 | 0,01 | 13,13 | >0.1 | 3,42 | 5,19 | 0 | 1,01 | 0 | 0 | 0 |
| GSC2 | CATATTCCCA | -1 | 561 | 571 | -439 | -429 | 6,64 | 0,01 | 15,98 | >0.1 | 3,24 | 5,11 | 0 | 0,99 | 0 | 0 | 0 |
| GSC2 | CATTTTCCCT | -1 | 531 | 541 | -469 | -459 | 3,36 | 0,01 | 14,08 | >0.1 | 3,09 | 5,03 | 0 | 0,98 | 0 | 1,61 | 0 |
| GSC2 | CCTCAGCCTC | -1 | 352 | 362 | -648 | -638 | 5,06 | 0,01 | 19,78 | >0.1 | 2,54 | 4,69 | 0 | 1,1 | 0 | 6,4 | 0 |
| GSC2 | TGTGATCTGC | -1 | 231 | 241 | -769 | -759 | 2,86 | 0,01 | 16,74 | >0.1 | 2,91 | 4,51 | 0 | 1,45 | 0 | 5,01 | 0 |
| GSC2 | TTTAATTTCC | -1 | 153 | 163 | -847 | -837 | 4,9 | 0,01 | 16,12 | >0.1 | 3,27 | 4,41 | 0 | 1,9 | 0 | 1,65 | 0 |
| GSC2 | CCTGATCTGC | -1 | 93 | 103 | -907 | -897 | 6,18 | 0,01 | 16,88 | >0.1 | 3,67 | 4,34 | 0 | 1,88 | 0 | 0,81 | 0 |
| GSC2 | CCTATTTCTC | -1 | 56 | 66 | -944 | -934 | 4,67 | 0,01 | 14,9 | >0.1 | 4,05 | 4,3 | 0 | 1,87 | 0 | 0 | 0 |
| SP1 | ACCCCGTCTCT | 1 | 280 | 291 | -720 | -709 | -1,98 | 0,01 | 11,69 | >0.1 | 2,74 | 4,58 | 0 | 1,26 | 0 | 5,09 | 0 |
| SP1 | GTCCCAGCTAC | 1 | 337 | 348 | -663 | -652 | -4,69 | 0,01 | 9,59 | >0.1 | 2,57 | 4,66 | 0 | 1,13 | 0 | 5,92 | 0 |
| SP1 | GCTCCAGCCTG | 1 | 422 | 433 | -578 | -567 | -4,99 | 0,01 | 9,08 | >0.1 | 2,69 | 4,81 | 0 | 1,02 | 0 | 5,56 | 0 |
| SP1 | TGTCCTCTCCA | 1 | 858 | 869 | -142 | -131 | -7,37 | 0,01 | 9,35 | >0.1 | 5,6 | 6,36 | 0 | 1,9 | 0 | 2,86 | 0 |
| SP1 | TTTCCTGCTCA | 1 | 934 | 945 | -66 | -55 | -8,02 | 0,01 | 7,49 | >0.1 | 5,6 | 7,14 | 0 | 1,88 | 0 | 0,9 | 0 |
| SP1 | TCAGCGCCCAA | 1 | 942 | 953 | -58 | -47 | -9,73 | 0,01 | 5,84 | >0.1 | 5,6 | 7,27 | 0 | 1,88 | 0 | 0,83 | 0 |
| SP1 | CGCCCAACCCA | 1 | 946 | 957 | -54 | -43 | -5,63 | 0,01 | 10,99 | >0.1 | 5,6 | 7,34 | 0 | 1,88 | 0 | 1,8 | 0 |
| SP1 | GCCCAACCCAA | 1 | 947 | 958 | -53 | -42 | -4,18 | 0,01 | 12,39 | >0.1 | 5,6 | 7,36 | 0 | 1,88 | 0 | 1,74 | 0 |
| SP1 | TTCCTGGCTCC | 1 | 1060 | 1071 | 60 | 71 | -8,44 | 0,01 | 8,53 | >0.1 | 5,6 | 7,35 | 0 | 1,86 | 0 | 2,16 | 0 |
| SP1 | GCTCCTTCTCA | 1 | 1066 | 1077 | 66 | 77 | -4,57 | 0,01 | 14,19 | >0.1 | 5,6 | 7,24 | 0 | 1,85 | 1,79 | 2,27 | 0 |
| SP1 | TCTCAGCCTTG | 1 | 1072 | 1083 | 72 | 83 | -7,13 | 0,01 | 11,53 | >0.1 | 5,6 | 7,14 | 0 | 1,85 | 1,79 | 2,27 | 0 |
| SP1 | TTACCTCTCCT | -1 | 771 | 782 | -229 | -218 | -8,74 | 0,01 | 10,15 | >0.1 | 5,6 | 5,83 | 0 | 1,37 | 0 | 6,1 | 0 |
| SP1 | TTCCCTTCCAA | -1 | 526 | 537 | -474 | -463 | -7,12 | 0,01 | 5,39 | >0.1 | 3,07 | 5,03 | 0 | 0,98 | 0 | 3,43 | 0 |
| SP1 | GCTCTGCCTCC | -1 | 383 | 394 | -617 | -606 | 8,63 | 0,001 | 23,02 | >0.1 | 2,58 | 4,74 | 0 | 1,06 | 0 | 6,02 | 0 |
| SP1 | CTGCCTCCCAG | -1 | 380 | 391 | -620 | -609 | -5,11 | 0,01 | 9,17 | >0.1 | 2,57 | 4,73 | 0 | 1,06 | 0 | 5,91 | 0 |
| SP1 | TCTCCTGCCTC | -1 | 358 | 369 | -642 | -631 | -2,94 | 0,01 | 11,62 | >0.1 | 2,52 | 4,7 | 0 | 1,09 | 0 | 6,26 | 0 |
| SP1 | CTCCTGCCTCA | -1 | 357 | 368 | -643 | -632 | 4,01 | 0,01 | 18,45 | >0.1 | 2,52 | 4,7 | 0 | 1,09 | 0 | 6,13 | 0 |
| SP1 | CCTCAGCCTCC | -1 | 351 | 362 | -649 | -638 | -1,14 | 0,01 | 13,58 | >0.1 | 2,54 | 4,69 | 0 | 1,1 | 0 | 6,4 | 0 |
| SP1 | TGCCCACCACC | -1 | 320 | 331 | -680 | -669 | -2,29 | 0,01 | 11,81 | >0.1 | 2,62 | 4,64 | 0 | 1,16 | 0 | 5,68 | 0 |
| SP1 | CCACCACCACG | -1 | 317 | 328 | -683 | -672 | -7,88 | 0,01 | 6,8 | >0.1 | 2,63 | 4,63 | 0 | 1,17 | 0 | 6,26 | 0 |

| | | | | | | | | | | | | | | | | |
|---|---|---|---|---|---|---|---|---|---|---|---|---|---|---|---|---|
| SP1 | CCACCACGCCC | -1 | 314 | 325 | -686 | -675 | -5,01 | 0,01 | 9,56 | >0.1 | 2,64 | 4,63 | 0 | 1,17 | 0 | 6,13 | 0 |
| SP1 | ACCACGCCCAG | -1 | 312 | 323 | -688 | -677 | -4,07 | 0,01 | 10,41 | >0.1 | 2,64 | 4,62 | 0 | 1,18 | 0 | 6,03 | 0 |
| SP1 | TGCCCAACTCG | -1 | 223 | 234 | -777 | -766 | -7,63 | 0,01 | 6,68 | >0.1 | 2,94 | 4,5 | 0 | 1,49 | 0 | 5,38 | 0 |
| SP1 | ACTCGGCCTCC | -1 | 217 | 228 | -783 | -772 | -2,2 | 0,01 | 12,3 | >0.1 | 2,96 | 4,49 | 0 | 1,52 | 0 | 5,52 | 0 |
| SP1 | CGGCCTCCCAA | -1 | 214 | 225 | -786 | -775 | -6,82 | 0,01 | 7,71 | >0.1 | 2,98 | 4,49 | 0 | 1,54 | 0 | 5,52 | 0 |
| SP1 | TGCCCAGCCAT | -1 | 174 | 185 | -826 | -815 | -6,83 | 0,01 | 5,84 | >0.1 | 3,15 | 4,44 | 0 | 1,84 | 0 | 3,24 | 0 |
| SP1 | TCTCCACTCCA | -1 | 115 | 126 | -885 | -874 | -5,12 | 0,01 | 5,38 | >0.1 | 3,5 | 4,37 | 0 | 1,89 | 0 | 0,75 | 0 |
| SP1 | GTGCCTCCTTC | -1 | 17 | 28 | -983 | -972 | -2,96 | 0,01 | 8,77 | >0.1 | 5,6 | 4,27 | 0 | 1,87 | 0 | 0 | 0 |
| SP1 | CCTCCTTCTCC | -1 | 14 | 25 | -986 | -975 | -2,4 | 0,01 | 9,33 | >0.1 | 5,6 | 4,27 | 0 | 1,87 | 0 | 0 | 0 |
| SP1 | CTCCTTCTCCT | -1 | 13 | 24 | -987 | -976 | -9,08 | 0,01 | 2,65 | >0.1 | 5,6 | 4,27 | 0 | 1,87 | 0 | 0 | 0 |
| SP1 | CCTTCTCCTTC | -1 | 11 | 22 | -989 | -978 | -5,16 | 0,01 | 6,56 | >0.1 | 5,6 | 4,26 | 0 | 1,86 | 0 | 0 | 0 |
| SP2 | ACCCCGTCTCTATTA | 1 | 280 | 295 | -720 | -705 | -3,17 | 0,01 | 10,5 | >0.1 | 2,73 | 4,58 | 0 | 1,26 | 0 | 5,09 | 0 |
| SP2 | GTCCCAGCTACTCGG | 1 | 337 | 352 | -663 | -648 | 0,58 | 0,01 | 15,47 | >0.1 | 2,57 | 4,67 | 0 | 1,13 | 0 | 6,52 | 0 |
| SP2 | GCTCCAGCCTGGGCG | 1 | 422 | 437 | -578 | -563 | -3,13 | 0,01 | 11,07 | >0.1 | 2,7 | 4,82 | 0 | 1,02 | 0 | 5,67 | 0 |
| SP2 | TGTCCTCTCCAGGAT | 1 | 858 | 873 | -142 | -127 | -1,09 | 0,01 | 15,61 | >0.1 | 5,54 | 6,39 | 0 | 1,9 | 0 | 2,86 | 0 |
| SP2 | GTTCTTCTTCTTTTT | 1 | 907 | 922 | -93 | -78 | -3,03 | 0,01 | 12,44 | >0.1 | 5,54 | 6,83 | 0 | 1,89 | 0 | 1,21 | 0 |
| SP2 | TTTCCTGCTCAGCGC | 1 | 934 | 949 | -66 | -51 | -4,71 | 0,01 | 10,81 | >0.1 | 5,54 | 7,2 | 0 | 1,88 | 0 | 0,9 | 0 |
| SP2 | TTCCTGCTCAGCGCC | 1 | 935 | 950 | -65 | -50 | 1,83 | 0,01 | 17,37 | >0.1 | 5,54 | 7,22 | 0 | 1,88 | 0 | 0,9 | 0 |
| SP2 | TCAGCGCCCAACCCA | 1 | 942 | 957 | -58 | -43 | -0,07 | 0,01 | 16,5 | >0.1 | 5,54 | 7,34 | 0 | 1,88 | 0 | 1,8 | 0 |
| SP2 | GCCCAACCCAAGTTC | 1 | 947 | 962 | -53 | -38 | -3,38 | 0,01 | 13,22 | >0.1 | 5,54 | 7,44 | 0 | 1,88 | 0 | 1,74 | 0 |
| SP2 | TTCCTGGCTCCTTCT | 1 | 1060 | 1075 | 60 | 75 | -2,84 | 0,01 | 15,92 | >0.1 | 5,54 | 7,35 | 0 | 1,86 | 1,79 | 2,21 | 0 |
| SP2 | GCTCCTTCTCAGCCT | 1 | 1066 | 1081 | 66 | 81 | -2,43 | 0,01 | 16,22 | >0.1 | 5,54 | 7,24 | 0 | 1,85 | 1,79 | 2,21 | 0 |
| SP2 | CTCCTTCTCAGCCTT | 1 | 1067 | 1082 | 67 | 82 | -3,61 | 0,01 | 15,02 | >0.1 | 5,54 | 7,22 | 0 | 1,85 | 1,79 | 2,21 | 0 |
| SP2 | TCTCAGCCTTGTTGC | 1 | 1072 | 1087 | 72 | 87 | -4,64 | 0,01 | 14,03 | >0.1 | 5,54 | 7,14 | 0 | 1,85 | 1,79 | 2,34 | 0 |
| SP2 | AAACCTCCTCATGAG | -1 | 984 | 999 | -16 | -1 | -0,97 | 0,01 | 16,71 | >0.1 | 5,54 | 8,69 | 0 | 1,87 | 0 | 1,57 | 0 |
| SP2 | TTACCTCTCCTGTCA | -1 | 767 | 782 | -233 | -218 | 0,4 | 0,01 | 18,83 | >0.1 | 5,54 | 5,83 | 0 | 1,37 | 0 | 5,69 | 0 |
| SP2 | GCTCTGCCTCCCAGG | -1 | 379 | 394 | -621 | -606 | 10,45 | 0,001 | 24,84 | >0.1 | 2,57 | 4,74 | 0 | 1,06 | 0 | 6,02 | 0 |
| SP2 | CTGCCTCCCAGGTTC | -1 | 376 | 391 | -624 | -609 | -2,36 | 0,01 | 12,54 | >0.1 | 2,57 | 4,74 | 0 | 1,07 | 0 | 6,53 | 0 |
| SP2 | TCTCCTGCCTCAGCC | -1 | 354 | 369 | -646 | -631 | -2,46 | 0,01 | 11,99 | >0.1 | 2,52 | 4,7 | 0 | 1,1 | 0 | 6,13 | 0 |
| SP2 | CTCCTGCCTCAGCCT | -1 | 353 | 368 | -647 | -632 | 8,3 | 0,001 | 22,66 | >0.1 | 2,53 | 4,7 | 0 | 1,1 | 0 | 6,03 | 0 |
| SP2 | CCTCAGCCTCCCGAG | -1 | 347 | 362 | -653 | -638 | 0,66 | 0,01 | 15,4 | >0.1 | 2,54 | 4,69 | 0 | 1,11 | 0 | 6,4 | 0 |
| SP2 | TGCCCACCACCACGC | -1 | 316 | 331 | -684 | -669 | 1,01 | 0,01 | 15,7 | >0.1 | 2,63 | 4,64 | 0 | 1,17 | 0 | 6,26 | 0 |
| SP2 | CCACCACCACGCCCA | -1 | 313 | 328 | -687 | -672 | -4,66 | 0,01 | 9,91 | >0.1 | 2,63 | 4,63 | 0 | 1,18 | 0 | 6,13 | 0 |
| SP2 | CCACCACGCCCAGCT | -1 | 310 | 325 | -690 | -675 | -3,71 | 0,01 | 10,78 | >0.1 | 2,64 | 4,63 | 0 | 1,18 | 0 | 6,03 | 0 |
| SP2 | ACCACGCCCAGCTAA | -1 | 308 | 323 | -692 | -677 | -4,71 | 0,01 | 9,88 | >0.1 | 2,65 | 4,62 | 0 | 1,19 | 0 | 6,13 | 0 |
| SP2 | GATCTGCCCAACTCG | -1 | 223 | 238 | -777 | -762 | 1,97 | 0,01 | 16,15 | >0.1 | 2,93 | 4,5 | 0 | 1,49 | 0 | 5,26 | 0 |
| SP2 | TGCCCAACTCGGCCT | -1 | 219 | 234 | -781 | -766 | -3,8 | 0,01 | 10,68 | >0.1 | 2,95 | 4,5 | 0 | 1,51 | 0 | 5,52 | 0 |
| SP2 | ACTCGGCCTCCCAAA | -1 | 213 | 228 | -787 | -772 | -2,87 | 0,01 | 11,78 | >0.1 | 2,97 | 4,49 | 0 | 1,55 | 0 | 5,63 | 0 |
| SP2 | CGGCCTCCCAAAGTG | -1 | 210 | 225 | -790 | -775 | -4,71 | 0,01 | 8,92 | >0.1 | 2,98 | 4,49 | 0 | 1,57 | 0 | 4,59 | 0 |
| SP2 | TGCCCAGCCATTTCT | -1 | 170 | 185 | -830 | -815 | -4,19 | 0,01 | 8,64 | >0.1 | 3,17 | 4,44 | 0 | 1,88 | 0 | 3,34 | 0 |
| SP2 | TCTCCACTCCACAGG | -1 | 111 | 126 | -889 | -874 | 1,85 | 0,01 | 12,38 | >0.1 | 3,51 | 4,37 | 0 | 1,89 | 0 | 0,77 | 0 |
| SP2 | GATCTGCCTTTGCAA | -1 | 85 | 100 | -915 | -900 | 0,07 | 0,01 | 10,01 | >0.1 | 3,72 | 4,34 | 0 | 1,88 | 0 | 0 | 0 |
| SP2 | TGAGTGCCTCCTTCT | -1 | 16 | 31 | -984 | -969 | -3,57 | 0,01 | 8,11 | >0.1 | 5,54 | 4,27 | 0 | 1,87 | 0 | 0 | 0 |
| SP2 | GTGCCTCCTTCTCCT | -1 | 13 | 28 | -987 | -972 | 4,37 | 0,01 | 16,05 | >0.1 | 5,54 | 4,27 | 0 | 1,87 | 0 | 0 | 0 |
| SP2 | CCTCCTTCTCCTTCA | -1 | 10 | 25 | -990 | -975 | -2,65 | 0,01 | 10,82 | >0.1 | 5,54 | 4,27 | 0 | 1,87 | 1,79 | 0 | 0 |
| SP2 | CTCCTTCTCCTTCAC | -1 | 9 | 24 | -991 | -976 | 2,5 | 0,01 | 15,97 | >0.1 | 5,54 | 4,27 | 0 | 1,87 | 1,79 | 0 | 0 |
| SP3 | CTCACGCCTGT | 1 | 191 | 202 | -809 | -798 | 5,49 | 0,01 | 18,85 | >0.1 | 3,08 | 4,46 | 0 | 1,7 | 0 | 4,13 | 0 |
| SP3 | ACCCCGTCTCT | 1 | 280 | 291 | -720 | -709 | 4,71 | 0,01 | 18,38 | >0.1 | 2,74 | 4,58 | 0 | 1,26 | 0 | 5,09 | 0 |
| SP3 | CAAACACCAAC | 1 | 629 | 640 | -371 | -360 | 2,85 | 0,01 | 14,55 | >0.1 | 3,7 | 5,29 | 0 | 1,05 | 0 | 1,66 | 0 |
| SP3 | TCAGCGCCCAA | 1 | 942 | 953 | -58 | -47 | 5,14 | 0,01 | 20,71 | >0.1 | 5,6 | 7,27 | 0 | 1,88 | 0 | 0,83 | 0 |
| SP3 | CGCCCAACCCA | 1 | 946 | 957 | -54 | -43 | 2,78 | 0,01 | 19,4 | >0.1 | 5,6 | 7,34 | 0 | 1,88 | 0 | 1,8 | 0 |
| SP3 | GCCCAACCCAA | 1 | 947 | 958 | -53 | -42 | 6,7 | 0,01 | 23,27 | >0.1 | 5,6 | 7,36 | 0 | 1,88 | 0 | 1,74 | 0 |
| SP3 | ATCACATCCAC | -1 | 1020 | 1031 | 20 | 31 | 4,47 | 0,01 | 21,96 | >0.1 | 5,6 | 8,72 | 0 | 1,86 | 0 | 1,31 | 0 |
| SP3 | GCTCTGCCTCC | -1 | 383 | 394 | -617 | -606 | 4,43 | 0,01 | 18,82 | >0.1 | 2,58 | 4,74 | 0 | 1,06 | 0 | 6,02 | 0 |
| SP3 | CTGCCTCCCAG | -1 | 380 | 391 | -620 | -609 | 2,94 | 0,01 | 17,22 | >0.1 | 2,57 | 4,73 | 0 | 1,06 | 0 | 5,91 | 0 |
| SP3 | CTCCTGCCTCA | -1 | 357 | 368 | -643 | -632 | 2,5 | 0,01 | 16,94 | >0.1 | 2,52 | 4,7 | 0 | 1,09 | 0 | 6,13 | 0 |
| SP3 | CCTCAGCCTCC | -1 | 351 | 362 | -649 | -638 | 3,68 | 0,01 | 18,4 | >0.1 | 2,54 | 4,69 | 0 | 1,1 | 0 | 6,4 | 0 |
| SP3 | TGCCCACCACC | -1 | 320 | 331 | -680 | -669 | 2,72 | 0,01 | 16,82 | >0.1 | 2,62 | 4,64 | 0 | 1,16 | 0 | 5,68 | 0 |
| SP3 | CCCACCACCAC | -1 | 318 | 329 | -682 | -671 | 7,76 | 0,01 | 22,44 | >0.1 | 2,63 | 4,63 | 0 | 1,16 | 0 | 6,26 | 0 |

| | | | | | | | | | | | | | | | | | |
|---|---|---|---|---|---|---|---|---|---|---|---|---|---|---|---|---|---|
| SP3 | CCACCACGCCC | -1 | 314 | 325 | -686 | -675 | 4,76 | 0,01 | 19,33 | >0.1 | 2,64 | 4,63 | 0 | 1,17 | 0 | 6,13 | 0 |
| SP3 | ACCACGCCCAG | -1 | 312 | 323 | -688 | -677 | 14,08 | 0,0001 | 28,56 | >0.1 | 2,64 | 4,62 | 0 | 1,18 | 0 | 6,03 | 0 |
| SP3 | ACTCGGCCTCC | -1 | 217 | 228 | -783 | -772 | 2,35 | 0,01 | 16,85 | >0.1 | 2,96 | 4,49 | 0 | 1,52 | 0 | 5,52 | 0 |
| SP3 | CGGCCTCCCAA | -1 | 214 | 225 | -786 | -775 | 2,68 | 0,01 | 17,21 | >0.1 | 2,98 | 4,49 | 0 | 1,54 | 0 | 5,52 | 0 |
| SP3 | TCCACTCCACA | -1 | 113 | 124 | -887 | -876 | 4,4 | 0,01 | 14,93 | >0.1 | 3,51 | 4,36 | 0 | 1,89 | 0 | 0,77 | 0 |
| SP4 | TGGCTCACGCCTGTAAT | 1 | 188 | 205 | -812 | -795 | -18,61 | 0,01 | -5,22 | >0.1 | 3,08 | 4,46 | 0 | 1,72 | 0 | 4,13 | 0 |
| SP4 | GAAACCCCGTCTCTATT | 1 | 277 | 294 | -723 | -706 | -3,3 | 0,001 | 10,38 | >0.1 | 2,74 | 4,58 | 0 | 1,27 | 0 | 5,09 | 0 |
| SP4 | GTAGTCCCAGCTACTCG | 1 | 334 | 351 | -666 | -649 | -14,15 | 0,01 | 0,14 | >0.1 | 2,57 | 4,67 | 0 | 1,13 | 0 | 5,92 | 0 |
| SP4 | GAGATCGCGCCACTGTG | 1 | 406 | 423 | -594 | -577 | -18,51 | 0,01 | -4,51 | >0.1 | 2,65 | 4,79 | 0 | 1,03 | 0 | 5,52 | 0 |
| SP4 | CCTGCTCAGCGCCCAAC | 1 | 937 | 954 | -63 | -46 | -12,54 | 0,01 | 2,98 | >0.1 | 5,52 | 7,29 | 0 | 1,88 | 0 | 0,83 | 0 |
| SP4 | AGCGCCCAACCCAAGTT | 1 | 944 | 961 | -56 | -39 | -15,09 | 0,01 | 1,47 | >0.1 | 5,52 | 7,42 | 0 | 1,88 | 0 | 1,74 | 0 |
| SP4 | CTGGCTCCTTCTCAGCC | 1 | 1063 | 1080 | 63 | 80 | -17,97 | 0,01 | 0,77 | >0.1 | 5,52 | 7,3 | 0 | 1,85 | 1,79 | 2,27 | 0 |
| SP4 | GCTGCTCAGTCCACCAT | 1 | 1094 | 1111 | 94 | 111 | -18,14 | 0,01 | 1,64 | >0.1 | 5,52 | 6,83 | 0 | 1,85 | 3,22 | 2,36 | 0 |
| SP4 | TTAACCACGAAGCCGAA | 1 | 1143 | 1160 | 143 | 160 | -15,32 | 0,01 | 3,78 | >0.1 | 5,52 | 6,35 | 0 | 1,84 | 3,22 | 2,16 | 0 |
| SP4 | TGAGCCAAGATCACATC | -1 | 1023 | 1040 | 23 | 40 | -18,3 | 0,01 | -0,98 | >0.1 | 5,52 | 8,67 | 0 | 1,86 | 0 | 1,28 | 0 |
| SP4 | AAGATCACATCCACTGA | -1 | 1017 | 1034 | 17 | 34 | -16,51 | 0,01 | 0,96 | >0.1 | 5,52 | 8,79 | 0 | 1,86 | 0 | 1,31 | 0 |
| SP4 | CTAAAACCTCCTCATGA | -1 | 985 | 1002 | -15 | 2 | -15,47 | 0,01 | 2,3 | >0.1 | 5,52 | 8,77 | 0 | 1,87 | 0 | 1,61 | 0 |
| SP4 | CAAGCTCTGCCTCCCAG | -1 | 380 | 397 | -620 | -603 | -5,37 | 0,01 | 9,03 | >0.1 | 2,58 | 4,75 | 0 | 1,06 | 0 | 6,02 | 0 |
| SP4 | CATTCTCCTGCCTCAGC | -1 | 355 | 372 | -645 | -628 | -18,42 | 0,01 | -3,84 | >0.1 | 2,52 | 4,7 | 0 | 1,1 | 0 | 6,26 | 0 |
| SP4 | CTGCCTCAGCCTCCCGA | -1 | 348 | 365 | -652 | -635 | -11,81 | 0,01 | 2,93 | >0.1 | 2,54 | 4,69 | 0 | 1,11 | 0 | 6,4 | 0 |
| SP4 | GTGCCCACCACCACGCC | -1 | 315 | 332 | -685 | -668 | -8,33 | 0,01 | 6,36 | >0.1 | 2,63 | 4,64 | 0 | 1,17 | 0 | 6,26 | 0 |
| SP4 | CCCACCACCACGCCCAG | -1 | 312 | 329 | -688 | -671 | -17,64 | 0,01 | -3,06 | >0.1 | 2,63 | 4,63 | 0 | 1,18 | 0 | 6,13 | 0 |
| SP4 | ACCACCACGCCCAGCTA | -1 | 309 | 326 | -691 | -674 | 4,91 | 0,001 | 19,4 | >0.1 | 2,64 | 4,63 | 0 | 1,19 | 0 | 6,03 | 0 |
| SP4 | TGTGATCTGCCCAACTC | -1 | 224 | 241 | -776 | -759 | -15,72 | 0,01 | -1,54 | >0.1 | 2,92 | 4,51 | 0 | 1,49 | 0 | 5,26 | 0 |
| SP4 | CCAACTCGGCCTCCCAA | -1 | 214 | 231 | -786 | -769 | -10,71 | 0,01 | 3,82 | >0.1 | 2,96 | 4,49 | 0 | 1,54 | 0 | 5,52 | 0 |
| SP4 | TGTGCCCAGCCATTTCT | -1 | 170 | 187 | -830 | -813 | -17,82 | 0,01 | -5,1 | >0.1 | 3,16 | 4,44 | 0 | 1,88 | 0 | 3,24 | 0 |
| SP4 | ATTTCCATGGCCATTTT | -1 | 142 | 159 | -858 | -841 | -16,02 | 0,01 | -4,8 | >0.1 | 3,31 | 4,41 | 0 | 1,9 | 0 | 1,61 | 0 |
| SP4 | ATGACTACTCTCCACTC | -1 | 117 | 134 | -883 | -866 | -12,66 | 0,01 | -1,28 | >0.1 | 3,46 | 4,38 | 0 | 1,89 | 0 | 1,65 | 0 |
| EHF | GAGAAGGAGGCA | 1 | 15 | 27 | -985 | -973 | -0,39 | 0,01 | 16,24 | >0.1 | 5,58 | 4,27 | 0 | 1,87 | 0 | 0 | 4,91 |
| EHF | AAGGAGGCACTC | 1 | 18 | 30 | -982 | -970 | -2,73 | 0,01 | 13,9 | >0.1 | 5,58 | 4,27 | 0 | 1,87 | 0 | 0 | 4,91 |
| EHF | CTCAAAGAAGTC | 1 | 27 | 39 | -973 | -961 | -2,81 | 0,01 | 12,78 | >0.1 | 4,53 | 4,28 | 0 | 1,87 | 0 | 0 | 4,91 |
| EHF | AAATAGGAAATG | 1 | 59 | 71 | -941 | -929 | 1,8 | 0,01 | 16,89 | >0.1 | 4 | 4,31 | 0 | 1,88 | 0 | 0 | 4,91 |
| EHF | GATCAGGAGAGT | 1 | 96 | 108 | -904 | -892 | -3,86 | 0,01 | 11,69 | >0.1 | 3,64 | 4,35 | 0 | 1,88 | 0 | 0,77 | 4,91 |
| EHF | GCCATGGAAATT | 1 | 148 | 160 | -852 | -840 | -1,92 | 0,01 | 14,23 | >0.1 | 3,29 | 4,41 | 0 | 1,9 | 0 | 1,65 | 4,91 |
| EHF | GGTCAGGAGATA | 1 | 242 | 254 | -758 | -746 | -1,67 | 0,01 | 16,36 | >0.1 | 2,87 | 4,52 | 0 | 1,4 | 0 | 4,33 | 4,91 |
| EHF | TACTCGGGAGGC | 1 | 345 | 357 | -655 | -643 | -4,15 | 0,01 | 15,63 | >0.1 | 2,55 | 4,68 | 0 | 1,11 | 0 | 6,52 | 4,91 |
| EHF | ACTCGGGAGGCT | 1 | 346 | 358 | -654 | -642 | -5,17 | 0,01 | 14,6 | >0.1 | 2,55 | 4,68 | 0 | 1,11 | 0 | 6,52 | 4,91 |
| EHF | AACCTGGGAGGC | 1 | 377 | 389 | -623 | -611 | -3,74 | 0,01 | 16,06 | >0.1 | 2,56 | 4,73 | 0 | 1,06 | 0 | 6,53 | 4,91 |
| EHF | TATTTGGAAGGG | 1 | 523 | 535 | -477 | -465 | -5,1 | 0,01 | 12,16 | >0.1 | 3,06 | 5,02 | 0 | 0,98 | 0 | 3,28 | 4,91 |
| EHF | TTCTAGGAAAAT | 1 | 718 | 730 | -282 | -270 | -4,38 | 0,01 | 17,9 | >0.1 | 5,58 | 5,6 | 0 | 1,2 | 0 | 4,99 | 4,91 |
| EHF | CTCCAGGATGAA | 1 | 864 | 876 | -136 | -124 | 0,2 | 0,01 | 21,68 | >0.1 | 5,58 | 6,41 | 0 | 1,9 | 0 | 2,67 | 4,91 |
| EHF | CATGAGGAGGTT | 1 | 986 | 998 | -14 | -2 | 2,33 | 0,01 | 24,97 | >0.1 | 5,58 | 8,66 | 0 | 1,87 | 0 | 1,61 | 4,91 |
| EHF | ATTGAGGAACAG | 1 | 1109 | 1121 | 109 | 121 | -2,9 | 0,01 | 21,89 | >0.1 | 5,58 | 6,66 | 0 | 1,85 | 3,22 | 2,56 | 4,91 |
| EHF | AACCACGAAGCC | 1 | 1145 | 1157 | 145 | 157 | 0,98 | 0,01 | 25,04 | >0.1 | 5,58 | 6,33 | 0 | 1,84 | 3,22 | 2,16 | 4,91 |
| EHF | GAAGCCGAAGAC | 1 | 1151 | 1163 | 151 | 163 | -3,74 | 0,01 | 17,87 | >0.1 | 4,51 | 6,29 | 0 | 1,84 | 1,79 | 2,26 | 4,91 |
| EHF | TTCCAAGAAGCA | -1 | 1183 | 1195 | 183 | 195 | -0,05 | 0,01 | 23,54 | >0.1 | 5,58 | 6,07 | 0 | 1,56 | 1,79 | 3,67 | 4,91 |
| EHF | AAGAAGCAAGTG | -1 | 1179 | 1191 | 179 | 191 | -2,99 | 0,01 | 20,28 | >0.1 | 5,58 | 6,1 | 0 | 1,59 | 1,79 | 3,29 | 4,91 |
| EHF | AGCCAGGAAGAG | -1 | 1057 | 1069 | 57 | 69 | 6,93 | 0,001 | 28,91 | >0.1 | 5,58 | 7,42 | 0 | 1,86 | 0 | 2,21 | 4,91 |
| EHF | GAGCAGGAAAGA | -1 | 932 | 944 | -68 | -56 | 5,07 | 0,001 | 25,49 | >0.1 | 5,58 | 7,12 | 0 | 1,88 | 0 | 0,93 | 4,91 |
| EHF | AAAGAAGAAGAA | -1 | 908 | 920 | -92 | -80 | -3,55 | 0,01 | 16,85 | >0.1 | 5,58 | 6,81 | 0 | 1,89 | 0 | 1,21 | 4,91 |
| EHF | AAACCGGTACTT | -1 | 739 | 751 | -261 | -249 | 0,98 | 0,01 | 24,33 | >0.1 | 5,58 | 5,68 | 0 | 1,26 | 0 | 5,92 | 4,91 |
| EHF | TACACGGAAAAT | -1 | 694 | 706 | -306 | -294 | 5,88 | 0,001 | 25,98 | >0.1 | 5,58 | 5,51 | 0 | 1,15 | 0 | 2,95 | 4,91 |
| EHF | TTCCCAGAAACT | -1 | 555 | 567 | -445 | -433 | -1,51 | 0,01 | 12,71 | >0.1 | 3,21 | 5,1 | 0 | 0,99 | 0 | 0 | 4,91 |
| EHF | AGTCAGGACGGT | -1 | 257 | 269 | -743 | -731 | -3,53 | 0,01 | 14,61 | >0.1 | 2,81 | 4,55 | 0 | 1,34 | 0 | 4,53 | 4,91 |
| POU5F1::SOX2 | TTAAGTTTTGCAAAG | 1 | 78 | 93 | -922 | -907 | 2,99 | 0,01 | 12,98 | >0.1 | 3,78 | 4,33 | 0 | 1,88 | 0 | 0 | 0 |
| POU5F1::SOX2 | TTTGCAAAGGCAGAT | 1 | 84 | 99 | -916 | -901 | 5,68 | 0,01 | 15,63 | >0.1 | 3,73 | 4,34 | 0 | 1,88 | 0 | 0 | 0 |
| POU5F1::SOX2 | CTGTGGAGTGGAGAG | 1 | 112 | 127 | -888 | -873 | 0,24 | 0,01 | 10,75 | >0.1 | 3,5 | 4,37 | 0 | 1,89 | 0 | 0,75 | 0 |
| POU5F1::SOX2 | AATGGCCATGGAAAT | 1 | 144 | 159 | -856 | -841 | 5,23 | 0,01 | 16,44 | >0.1 | 3,3 | 4,4 | 0 | 1,9 | 0 | 1,61 | 0 |
| POU5F1::SOX2 | CATGGAAATTAAAAC | 1 | 150 | 165 | -850 | -835 | 2,1 | 0,01 | 13,33 | >0.1 | 3,27 | 4,41 | 0 | 1,9 | 0 | 1,65 | 0 |

| | | | | | | | | | | | | | | | | | |
|---|---|---|---|---|---|---|---|---|---|---|---|---|---|---|---|---|---|
| POU5F1::SOX2 | CTTTGGGAGGCCGAG | 1 | 212 | 227 | -788 | -773 | 1,92 | 0,01 | 16,46 | >0.1 | 2,98 | 4,49 | 0 | 1,55 | 0 | 5,52 | 0 |
| POU5F1::SOX2 | CATGTATTTGGAAGG | 1 | 519 | 534 | -481 | -466 | 0,57 | 0,01 | 12,69 | >0.1 | 3,05 | 5,02 | 0 | 0,99 | 0 | 3,07 | 0 |
| POU5F1::SOX2 | ATTTGGAAGGGAAAA | 1 | 524 | 539 | -476 | -461 | 0,3 | 0,01 | 12,82 | >0.1 | 3,07 | 5,03 | 0 | 0,99 | 0 | 3,43 | 0 |
| POU5F1::SOX2 | TTTTGATTTGGCCAT | 1 | 748 | 763 | -252 | -237 | 3,19 | 0,01 | 21,87 | >0.1 | 5,54 | 5,74 | 0 | 1,3 | 0 | 6,1 | 0 |
| POU5F1::SOX2 | TATTCTTATTTGATT | 1 | 807 | 822 | -193 | -178 | 2,44 | 0,01 | 18,87 | >0.1 | 5,54 | 6,04 | 0 | 1,57 | 0 | 3,28 | 0 |
| POU5F1::SOX2 | TTTTCTTTTTTTGGT | 1 | 918 | 933 | -82 | -67 | 1,43 | 0,01 | 16,95 | >0.1 | 5,54 | 6,97 | 0 | 1,89 | 0 | 1,12 | 0 |
| POU5F1::SOX2 | AGTTCAAAGGCTGAT | 1 | 957 | 972 | -43 | -28 | 0,42 | 0,01 | 17,21 | >0.1 | 5,54 | 7,68 | 0 | 1,88 | 0 | 1,7 | 0 |
| POU5F1::SOX2 | CTTGTTGCTGTAACT | 1 | 1079 | 1094 | 79 | 94 | 0,29 | 0,01 | 18,67 | >0.1 | 5,54 | 7,03 | 0 | 1,85 | 1,79 | 2,16 | 0 |
| POU5F1::SOX2 | CTTGCTTCTTGGAAT | 1 | 1181 | 1196 | 181 | 196 | 0,41 | 0,01 | 18,93 | >0.1 | 5,54 | 6,08 | 0 | 1,58 | 1,79 | 3,52 | 0 |
| POU5F1::SOX2 | TTTTTTAAAGTGAAT | -1 | 819 | 834 | -181 | -166 | 0,42 | 0,01 | 17,42 | >0.1 | 5,54 | 6,12 | 0 | 1,65 | 0 | 3,7 | 0 |
| POU5F1::SOX2 | TATTTTTATTTCAAG | -1 | 573 | 588 | -427 | -412 | 1,5 | 0,01 | 10,97 | >0.1 | 3,32 | 5,15 | 0 | 1 | 0 | 0 | 0 |
| POU5F1::SOX2 | TTTTCCCTTCCAAAT | -1 | 524 | 539 | -476 | -461 | 3,2 | 0,01 | 15,72 | >0.1 | 3,07 | 5,03 | 0 | 0,99 | 0 | 3,43 | 0 |
| POU5F1::SOX2 | TTTTTTTTTTTTGAG | -1 | 454 | 469 | -546 | -531 | 0,4 | 0,01 | 14,6 | >0.1 | 2,8 | 4,88 | 0 | 1 | 0 | 5,52 | 0 |
| POU5F1::SOX2 | TTTTTTTTTTGAGAC | -1 | 452 | 467 | -548 | -533 | 1,46 | 0,01 | 15,65 | >0.1 | 2,79 | 4,88 | 0 | 1 | 0 | 5,52 | 0 |
| POU5F1::SOX2 | TTTTGTATTTTTAAT | -1 | 291 | 306 | -709 | -694 | 5,54 | 0,01 | 18,52 | >0.1 | 2,7 | 4,6 | 0 | 1,23 | 0 | 4,45 | 0 |
| POU5F1::SOX2 | TCTGCCTTTGCAAAA | -1 | 83 | 98 | -917 | -902 | 1,07 | 0,01 | 11,02 | >0.1 | 3,73 | 4,34 | 0 | 1,88 | 0 | 0 | 0 |
| POU5F1::SOX2 | CCTTCTCCTTCACTT | -1 | 7 | 22 | -993 | -978 | 0,44 | 0,01 | 13,91 | >0.1 | 5,54 | 4,27 | 0 | 1,86 | 1,79 | 0 | 0 |
| GATA6 | AATAGGTAAGTGA | 1 | 0 | 13 | -1000 | -987 | 3,27 | 0,01 | 16,75 | >0.1 | 5,57 | 4,26 | 0 | 1,86 | 1,79 | 0 | 0 |
| GATA6 | GAAGGAGAAGGAG | 1 | 11 | 24 | -989 | -976 | 1,69 | 0,01 | 13,39 | >0.1 | 5,57 | 4,27 | 0 | 1,87 | 0 | 0 | 0 |
| GATA6 | GAGAAATAGGAAA | 1 | 56 | 69 | -944 | -931 | 3,04 | 0,01 | 13,25 | >0.1 | 4,03 | 4,31 | 0 | 1,87 | 0 | 0 | 0 |
| GATA6 | GGCAGATCAGGAG | 1 | 92 | 105 | -908 | -895 | 7,45 | 0,001 | 18,15 | >0.1 | 3,67 | 4,34 | 0 | 1,88 | 0 | 0,81 | 0 |
| GATA6 | AACTGATCAGAAA | 1 | 162 | 175 | -838 | -825 | 5,97 | 0,01 | 17,91 | >0.1 | 3,21 | 4,42 | 0 | 1,9 | 0 | 2,4 | 0 |
| GATA6 | GGCAGATCACAAG | 1 | 230 | 243 | -770 | -757 | 5,66 | 0,01 | 19,54 | >0.1 | 2,91 | 4,51 | 0 | 1,46 | 0 | 5,01 | 0 |
| GATA6 | AGGAGATAGAGAC | 1 | 246 | 259 | -754 | -741 | 3,5 | 0,01 | 17,46 | >0.1 | 2,85 | 4,53 | 0 | 1,38 | 0 | 5,19 | 0 |
| GATA6 | AACTGATCAGAAA | 1 | 477 | 490 | -523 | -510 | 5,97 | 0,01 | 20,12 | >0.1 | 2,88 | 4,92 | 0 | 0,99 | 0 | 5,36 | 0 |
| GATA6 | CAGAAATATGAAT | 1 | 484 | 497 | -516 | -503 | 1,86 | 0,01 | 16,82 | >0.1 | 2,9 | 4,94 | 0 | 0,99 | 0 | 6,13 | 0 |
| GATA6 | TAAAGACAAGGGC | 1 | 501 | 514 | -499 | -486 | 6,07 | 0,01 | 20,68 | >0.1 | 2,97 | 4,97 | 0 | 0,99 | 0 | 5,68 | 0 |
| GATA6 | TTGAAATAAAAAT | 1 | 574 | 587 | -426 | -413 | 2,58 | 0,01 | 12,05 | >0.1 | 3,32 | 5,15 | 0 | 1 | 0 | 0 | 0 |
| GATA6 | GTGAGATAACCTA | 1 | 591 | 604 | -409 | -396 | 9,11 | 0,001 | 18,74 | >0.1 | 3,42 | 5,19 | 0 | 1,01 | 0 | 0 | 0 |
| GATA6 | GAGAGGTAAGGTT | 1 | 773 | 786 | -227 | -214 | 6,43 | 0,01 | 25,33 | >0.1 | 5,57 | 5,84 | 0 | 1,38 | 0 | 6,1 | 0 |
| GATA6 | GGCTGATAAGAGA | 1 | 965 | 978 | -35 | -22 | 12,28 | 0,0001 | 29,26 | >0.1 | 5,57 | 7,85 | 0 | 1,87 | 0 | 1,7 | 0 |
| GATA6 | AAGAGAGAAAATC | 1 | 972 | 985 | -28 | -15 | 3,93 | 0,01 | 21,11 | >0.1 | 5,57 | 8,09 | 0 | 1,87 | 0 | 1,65 | 0 |
| GATA6 | GGCTGAGAAGGAG | -1 | 1067 | 1080 | 67 | 80 | 4,99 | 0,01 | 23,64 | >0.1 | 5,57 | 7,22 | 0 | 1,85 | 1,79 | 2,21 | 0 |
| GATA6 | CCAAGATCACATC | -1 | 1023 | 1036 | 23 | 36 | 3,88 | 0,01 | 21,23 | >0.1 | 5,57 | 8,67 | 0 | 1,86 | 0 | 1,25 | 0 |
| GATA6 | AGCAGGAAAGACC | -1 | 930 | 943 | -70 | -57 | 1,8 | 0,01 | 17,31 | >0.1 | 5,57 | 7,11 | 0 | 1,88 | 0 | 0,95 | 0 |
| GATA6 | CCAAAAAAAGAAA | -1 | 919 | 932 | -81 | -68 | 1,3 | 0,01 | 16,83 | >0.1 | 5,57 | 6,95 | 0 | 1,88 | 0 | 1,12 | 0 |
| GATA6 | AAAAGAAAAAGAA | -1 | 914 | 927 | -86 | -73 | 2,89 | 0,01 | 18,39 | >0.1 | 5,57 | 6,89 | 0 | 1,89 | 0 | 1,15 | 0 |
| GATA6 | GAAAAAGAAGAAG | -1 | 910 | 923 | -90 | -77 | 1,76 | 0,01 | 17,21 | >0.1 | 5,57 | 6,84 | 0 | 1,89 | 0 | 1,15 | 0 |
| GATA6 | GACAGATATGTAA | -1 | 849 | 862 | -151 | -138 | 5,3 | 0,01 | 21,81 | >0.1 | 5,57 | 6,31 | 0 | 1,88 | 0 | 2,75 | 0 |
| GATA6 | ATCAAATAAGAAT | -1 | 808 | 821 | -192 | -179 | 7,1 | 0,001 | 23,54 | >0.1 | 5,57 | 6,03 | 0 | 1,56 | 0 | 3,28 | 0 |
| GATA6 | CCTAGAGAACCTT | -1 | 780 | 793 | -220 | -207 | 1,38 | 0,01 | 20,56 | >0.1 | 5,57 | 5,88 | 0 | 1,41 | 0 | 6,32 | 0 |
| GATA6 | TGAAGATAATGTT | -1 | 637 | 650 | -363 | -350 | 6,5 | 0,01 | 18,27 | >0.1 | 3,78 | 5,32 | 0 | 1,06 | 0 | 1,61 | 0 |
| GATA6 | TCAAGATCATATT | -1 | 565 | 578 | -435 | -422 | 1,13 | 0,01 | 10,52 | >0.1 | 3,27 | 5,12 | 0 | 1 | 0 | 0 | 0 |
| GATA6 | CCCAGCTAAGTTT | -1 | 304 | 317 | -696 | -683 | 2,5 | 0,01 | 15,96 | >0.1 | 2,66 | 4,62 | 0 | 1,2 | 0 | 4,99 | 0 |
| GATA4 | TCCTAGCACTT | 1 | 204 | 215 | -796 | -785 | -4,92 | 0,01 | 8,77 | >0.1 | 3,02 | 4,47 | 0 | 1,6 | 0 | 4,59 | 0 |
| GATA4 | ACTTAGCTGGG | 1 | 306 | 317 | -694 | -683 | -7,72 | 0,01 | 5,83 | >0.1 | 2,66 | 4,62 | 0 | 1,19 | 0 | 5,09 | 0 |
| GATA4 | CATTATCTTCA | 1 | 639 | 650 | -361 | -350 | -1,07 | 0,01 | 10,76 | >0.1 | 3,79 | 5,32 | 0 | 1,06 | 0 | 1,66 | 0 |
| GATA4 | GTGTATCTTTA | 1 | 701 | 712 | -299 | -288 | -5,85 | 0,01 | 11,42 | >0.1 | 5,6 | 5,53 | 0 | 1,16 | 0 | 4,99 | 0 |
| GATA4 | CTTTAACAGCT | 1 | 707 | 718 | -293 | -282 | -4,22 | 0,01 | 12,91 | >0.1 | 5,6 | 5,55 | 0 | 1,17 | 0 | 4,81 | 0 |
| GATA4 | TCTTATTTGAT | 1 | 810 | 821 | -190 | -179 | -0,62 | 0,01 | 15,85 | >0.1 | 5,6 | 6,03 | 0 | 1,56 | 0 | 3,28 | 0 |
| GATA4 | ACATATCTGTC | 1 | 851 | 862 | -149 | -138 | -5,57 | 0,01 | 10,89 | >0.1 | 5,6 | 6,31 | 0 | 1,88 | 0 | 2,67 | 0 |
| GATA4 | TGTCCTCTCCA | 1 | 858 | 869 | -142 | -131 | -7,38 | 0,01 | 9,34 | >0.1 | 5,6 | 6,36 | 0 | 1,9 | 0 | 2,86 | 0 |
| GATA4 | CTTTATATTGG | 1 | 876 | 887 | -124 | -113 | -6,25 | 0,01 | 10,7 | >0.1 | 5,6 | 6,5 | 0 | 1,9 | 0 | 2,95 | 0 |
| GATA4 | TCTTCTTTTTC | 1 | 912 | 923 | -88 | -77 | -5,94 | 0,01 | 9,54 | >0.1 | 5,6 | 6,84 | 0 | 1,89 | 0 | 1,15 | 0 |
| GATA4 | CTTTTTCTTTT | 1 | 916 | 927 | -84 | -73 | -3,94 | 0,01 | 11,56 | >0.1 | 5,6 | 6,89 | 0 | 1,89 | 0 | 1,12 | 0 |
| GATA4 | CCTGCTCAGCG | 1 | 937 | 948 | -63 | -52 | -5,5 | 0,01 | 10,06 | >0.1 | 5,6 | 7,18 | 0 | 1,88 | 0 | 0,9 | 0 |
| GATA4 | TGTGATCTTGG | 1 | 1025 | 1036 | 25 | 36 | -4,75 | 0,01 | 12,47 | >0.1 | 5,6 | 8,52 | 0 | 1,86 | 0 | 1,25 | 0 |
| GATA4 | GCTGCTCAGTC | 1 | 1094 | 1105 | 94 | 105 | -7,05 | 0,01 | 12,79 | >0.1 | 5,6 | 6,83 | 0 | 1,85 | 3,22 | 2,34 | 0 |
| GATA4 | TTTTCTCTCTT | -1 | 972 | 983 | -28 | -17 | 5,65 | 0,001 | 22,83 | >0.1 | 5,6 | 8,01 | 0 | 1,87 | 0 | 1,7 | 0 |

| | | | | | | | | | | | | | | | | | |
|---|---|---|---|---|---|---|---|---|---|---|---|---|---|---|---|---|---|
| GATA4 | TTCTCTCTTAT | -1 | 970 | 981 | -30 | -19 | -1,09 | 0,01 | 16,02 | >0.1 | 5,6 | 7,94 | 0 | 1,87 | 0 | 1,7 | 0 |
| GATA4 | CCTTACCTCTC | -1 | 773 | 784 | -227 | -216 | 2,44 | 0,01 | 21,35 | >0.1 | 5,6 | 5,83 | 0 | 1,37 | 0 | 6,1 | 0 |
| GATA4 | TCCTGTCACTT | -1 | 764 | 775 | -236 | -225 | -4,9 | 0,01 | 13,93 | >0.1 | 5,6 | 5,79 | 0 | 1,34 | 0 | 6,1 | 0 |
| GATA4 | TGTTATAATTT | -1 | 684 | 695 | -316 | -305 | -5,96 | 0,01 | 7,97 | >0.1 | 4,39 | 5,47 | 0 | 1,13 | 0 | 2,95 | 0 |
| GATA4 | GGTTTTCAGAC | -1 | 615 | 626 | -385 | -374 | -5,45 | 0,01 | 5,96 | >0.1 | 3,58 | 5,25 | 0 | 1,03 | 0 | 1,55 | 0 |
| GATA4 | GGTTATCTCAC | -1 | 591 | 602 | -409 | -398 | 10,68 | 0,001 | 20,29 | >0.1 | 3,42 | 5,19 | 0 | 1,01 | 0 | 0 | 0 |
| GATA4 | TTTTATTTCAA | -1 | 574 | 585 | -426 | -415 | -4,65 | 0,01 | 4,81 | >0.1 | 3,31 | 5,14 | 0 | 1 | 0 | 0 | 0 |
| GATA4 | CCTTGTCTTTA | -1 | 501 | 512 | -499 | -488 | -0,93 | 0,01 | 13,51 | >0.1 | 2,96 | 4,97 | 0 | 0,98 | 0 | 5,52 | 0 |
| GATA4 | TCTGATCAGTT | -1 | 477 | 488 | -523 | -512 | -0,92 | 0,01 | 13,06 | >0.1 | 2,87 | 4,92 | 0 | 0,99 | 0 | 5,2 | 0 |
| GATA4 | CGCGATCTCGG | -1 | 404 | 415 | -596 | -585 | -7,38 | 0,01 | 6,83 | >0.1 | 2,64 | 4,78 | 0 | 1,03 | 0 | 5,76 | 0 |
| GATA4 | CTCTATCTCCT | -1 | 246 | 257 | -754 | -743 | 7,54 | 0,001 | 21,61 | >0.1 | 2,85 | 4,53 | 0 | 1,38 | 0 | 5,3 | 0 |
| GATA4 | TGTGATCTGCC | -1 | 230 | 241 | -770 | -759 | -0,4 | 0,01 | 13,49 | >0.1 | 2,91 | 4,51 | 0 | 1,45 | 0 | 5,01 | 0 |
| GATA4 | TCTGATCAGTT | -1 | 162 | 173 | -838 | -827 | -0,92 | 0,01 | 11,08 | >0.1 | 3,22 | 4,42 | 0 | 1,9 | 0 | 2,46 | 0 |
| GATA4 | CCTGATCTGCC | -1 | 92 | 103 | -908 | -897 | 2,57 | 0,01 | 13,3 | >0.1 | 3,67 | 4,34 | 0 | 1,88 | 0 | 0,83 | 0 |
| GATA4 | TCCTATTTCTC | -1 | 56 | 67 | -944 | -933 | -3,1 | 0,01 | 7,12 | >0.1 | 4,04 | 4,31 | 0 | 1,87 | 0 | 0 | 0 |
| GATA4 | ATTTCTCACAC | -1 | 52 | 63 | -948 | -937 | -3,28 | 0,01 | 6,99 | >0.1 | 4,09 | 4,3 | 0 | 1,87 | 0 | 0 | 0 |
| GATA4 | CCTTCTCCTTC | -1 | 11 | 22 | -989 | -978 | -4,48 | 0,01 | 7,24 | >0.1 | 5,6 | 4,26 | 0 | 1,86 | 0 | 0 | 0 |
| GATA4 | CCTTCACTTAC | -1 | 5 | 16 | -995 | -984 | -6,01 | 0,01 | 7,5 | >0.1 | 5,6 | 4,26 | 0 | 1,86 | 1,79 | 0 | 0 |
| GATA5 | AGATCAGG | 1 | 95 | 103 | -905 | -897 | 7,57 | 0,001 | 18,24 | >0.1 | 3,66 | 4,34 | 0 | 1,88 | 0 | 0,79 | 0 |
| GATA5 | TGATCAGA | 1 | 165 | 173 | -835 | -827 | 7 | 0,01 | 18,93 | >0.1 | 3,21 | 4,42 | 0 | 1,9 | 0 | 2,4 | 0 |
| GATA5 | AGATCACA | 1 | 233 | 241 | -767 | -759 | 6,96 | 0,01 | 20,82 | >0.1 | 2,91 | 4,51 | 0 | 1,44 | 0 | 5,01 | 0 |
| GATA5 | AGATAGAG | 1 | 249 | 257 | -751 | -743 | 6,73 | 0,01 | 20,56 | >0.1 | 2,85 | 4,53 | 0 | 1,37 | 0 | 5,09 | 0 |
| GATA5 | AGATCGCG | 1 | 407 | 415 | -593 | -585 | 4,85 | 0,01 | 18,94 | >0.1 | 2,64 | 4,78 | 0 | 1,03 | 0 | 5,65 | 0 |
| GATA5 | TGATCAGA | 1 | 480 | 488 | -520 | -512 | 7 | 0,01 | 21,3 | >0.1 | 2,88 | 4,92 | 0 | 0,98 | 0 | 5,52 | 0 |
| GATA5 | TGATCTTG | 1 | 569 | 577 | -431 | -423 | 1,97 | 0,01 | 11,36 | >0.1 | 3,28 | 5,12 | 0 | 0,99 | 0 | 0 | 0 |
| GATA5 | AGATAACC | 1 | 594 | 602 | -406 | -398 | 9,45 | 0,001 | 19,07 | >0.1 | 3,43 | 5,19 | 0 | 1,01 | 0 | 0 | 0 |
| GATA5 | TGATCCCT | 1 | 650 | 658 | -350 | -342 | 1,26 | 0,01 | 15,39 | >0.1 | 3,88 | 5,35 | 0 | 1,07 | 0 | 3,83 | 0 |
| GATA5 | TGATTTGG | 1 | 751 | 759 | -249 | -241 | 5,44 | 0,01 | 24,18 | >0.1 | 5,64 | 5,72 | 0 | 1,28 | 0 | 6,1 | 0 |
| GATA5 | AGGTAAGG | 1 | 776 | 784 | -224 | -216 | 2,43 | 0,01 | 21,6 | >0.1 | 5,64 | 5,83 | 0 | 1,37 | 0 | 6,32 | 0 |
| GATA5 | GGATTAAA | 1 | 791 | 799 | -209 | -201 | 3,36 | 0,01 | 21,67 | >0.1 | 5,64 | 5,91 | 0 | 1,44 | 0 | 5,32 | 0 |
| GATA5 | TGATTCAC | 1 | 817 | 825 | -183 | -175 | 2,57 | 0,01 | 19,4 | >0.1 | 5,64 | 6,06 | 0 | 1,58 | 0 | 3,55 | 0 |
| GATA5 | AGATTGTT | 1 | 893 | 901 | -107 | -99 | 1,57 | 0,01 | 18,58 | >0.1 | 5,64 | 6,62 | 0 | 1,89 | 0 | 2,86 | 0 |
| GATA5 | TGATAAGA | 1 | 968 | 976 | -32 | -24 | 10,19 | 0,0001 | 27,18 | >0.1 | 5,64 | 7,78 | 0 | 1,87 | 0 | 1,7 | 0 |
| GATA5 | TGATCTTG | 1 | 1027 | 1035 | 27 | 35 | 1,97 | 0,01 | 19,14 | >0.1 | 5,64 | 8,39 | 0 | 1,86 | 0 | 1,28 | 0 |
| GATA5 | CGATGTCA | 1 | 1047 | 1055 | 47 | 55 | 0,9 | 0,01 | 17,51 | >0.1 | 5,64 | 7,65 | 0 | 1,86 | 0 | 1,46 | 0 |
| GATA5 | TGATAGAA | -1 | 1166 | 1174 | 166 | 174 | 6,16 | 0,01 | 23,14 | >0.1 | 4,51 | 6,18 | 0 | 1,7 | 1,79 | 2,79 | 0 |
| GATA5 | AGATCACA | -1 | 1025 | 1033 | 25 | 33 | 6,96 | 0,01 | 24,23 | >0.1 | 5,64 | 8,52 | 0 | 1,86 | 0 | 1,25 | 0 |
| GATA5 | AGATTTTC | -1 | 978 | 986 | -22 | -14 | 2,63 | 0,01 | 19,85 | >0.1 | 5,64 | 8,13 | 0 | 1,87 | 0 | 1,57 | 0 |
| GATA5 | AGATATGT | -1 | 851 | 859 | -149 | -141 | 5,62 | 0,01 | 22,15 | >0.1 | 5,64 | 6,29 | 0 | 1,85 | 0 | 2,75 | 0 |
| GATA5 | AGATTTTA | -1 | 840 | 848 | -160 | -152 | 3,33 | 0,01 | 20 | >0.1 | 5,64 | 6,22 | 0 | 1,75 | 0 | 3,07 | 0 |
| GATA5 | AGATACAC | -1 | 701 | 709 | -299 | -291 | 5,15 | 0,01 | 20,65 | >0.1 | 5,64 | 5,52 | 0 | 1,15 | 0 | 3,19 | 0 |
| GATA5 | GGATCATG | -1 | 648 | 656 | -352 | -344 | 1,82 | 0,01 | 15,92 | >0.1 | 3,86 | 5,34 | 0 | 1,06 | 0 | 3,83 | 0 |
| GATA5 | AGATAATG | -1 | 639 | 647 | -361 | -353 | 8,34 | 0,001 | 20,09 | >0.1 | 3,77 | 5,31 | 0 | 1,05 | 0 | 1,61 | 0 |
| GATA5 | AGATCATA | -1 | 567 | 575 | -433 | -425 | 5,02 | 0,01 | 14,4 | >0.1 | 3,27 | 5,12 | 0 | 0,99 | 0 | 0 | 0 |
| GATA5 | TGATCAGT | -1 | 478 | 486 | -522 | -514 | 4,73 | 0,01 | 18,7 | >0.1 | 2,87 | 4,91 | 0 | 0,99 | 0 | 5,2 | 0 |
| GATA5 | CGATCTCG | -1 | 405 | 413 | -595 | -587 | 3,63 | 0,01 | 17,83 | >0.1 | 2,64 | 4,77 | 0 | 1,03 | 0 | 5,76 | 0 |
| GATA5 | TGATCTGC | -1 | 231 | 239 | -769 | -761 | 3,56 | 0,01 | 17,44 | >0.1 | 2,92 | 4,5 | 0 | 1,45 | 0 | 5,01 | 0 |
| GATA5 | GGATTACA | -1 | 199 | 207 | -801 | -793 | 4,68 | 0,01 | 18,3 | >0.1 | 3,05 | 4,46 | 0 | 1,64 | 0 | 4,48 | 0 |
| GATA5 | TGATCAGT | -1 | 163 | 171 | -837 | -829 | 4,73 | 0,01 | 16,72 | >0.1 | 3,22 | 4,42 | 0 | 1,9 | 0 | 2,46 | 0 |
| GATA5 | TGATCTGC | -1 | 93 | 101 | -907 | -899 | 3,56 | 0,01 | 14,29 | >0.1 | 3,68 | 4,34 | 0 | 1,88 | 0 | 0,83 | 0 |
| GATA2 | AACTTAGCTGG | 1 | 305 | 316 | -695 | -684 | 3,24 | 0,01 | 16,7 | >0.1 | 2,66 | 4,61 | 0 | 1,19 | 0 | 4,99 | 0 |
| GATA2 | ACATTATCTTC | 1 | 638 | 649 | -362 | -351 | 8,88 | 0,001 | 20,65 | >0.1 | 3,78 | 5,32 | 0 | 1,06 | 0 | 1,61 | 0 |
| GATA2 | CGTGTATCTTT | 1 | 700 | 711 | -300 | -289 | 3,23 | 0,01 | 18,7 | >0.1 | 5,6 | 5,53 | 0 | 1,16 | 0 | 3,19 | 0 |
| GATA2 | TCTTTAACAGC | 1 | 706 | 717 | -294 | -283 | 1,62 | 0,01 | 18,92 | >0.1 | 5,6 | 5,55 | 0 | 1,17 | 0 | 4,99 | 0 |
| GATA2 | TTCTTATTTGA | 1 | 809 | 820 | -191 | -180 | 6,32 | 0,001 | 22,78 | >0.1 | 5,6 | 6,03 | 0 | 1,55 | 0 | 3,28 | 0 |
| GATA2 | TACATATCTGT | 1 | 850 | 861 | -150 | -139 | 7,16 | 0,001 | 23,68 | >0.1 | 5,6 | 6,3 | 0 | 1,87 | 0 | 2,75 | 0 |
| GATA2 | ACTTTATATTG | 1 | 875 | 886 | -125 | -114 | 1,04 | 0,01 | 17,98 | >0.1 | 5,6 | 6,5 | 0 | 1,9 | 0 | 2,95 | 0 |
| GATA2 | TCTTTTTCTTT | 1 | 915 | 926 | -85 | -74 | 1,93 | 0,01 | 17,44 | >0.1 | 5,6 | 6,88 | 0 | 1,89 | 0 | 1,15 | 0 |
| GATA2 | TCCTTCTCAGC | 1 | 1068 | 1079 | 68 | 79 | 2,12 | 0,01 | 20,78 | >0.1 | 5,6 | 7,21 | 0 | 1,85 | 1,79 | 2,21 | 0 |

| | | | | | | | | | | | | | | | | | |
|---|---|---|---|---|---|---|---|---|---|---|---|---|---|---|---|---|---|
| GATA2 | GTTCTATCAAA | 1 | 1165 | 1176 | 165 | 176 | 1,22 | 0,01 | 18,22 | >0.1 | 4,52 | 6,19 | 0 | 1,71 | 1,79 | 2,79 | 0 |
| GATA2 | AGATTTTCTCT | -1 | 975 | 986 | -25 | -14 | 0,74 | 0,01 | 17,92 | >0.1 | 5,6 | 8,13 | 0 | 1,87 | 0 | 1,57 | 0 |
| GATA2 | CTCTTATCAGC | -1 | 966 | 977 | -34 | -23 | 11,24 | 0,0001 | 28,22 | >0.1 | 5,6 | 7,81 | 0 | 1,87 | 0 | 1,7 | 0 |
| GATA2 | ACCTTACCTCT | -1 | 774 | 785 | -226 | -215 | 6,12 | 0,01 | 25,04 | >0.1 | 5,6 | 5,84 | 0 | 1,38 | 0 | 6,1 | 0 |
| GATA2 | AGGTTATCTCA | -1 | 592 | 603 | -408 | -397 | 8,97 | 0,001 | 18,59 | >0.1 | 3,42 | 5,19 | 0 | 1,01 | 0 | 0 | 0 |
| GATA2 | CATTTATTTTT | -1 | 581 | 592 | -419 | -408 | 1,57 | 0,01 | 11,09 | >0.1 | 3,35 | 5,16 | 0 | 1 | 0 | 0 | 0 |
| GATA2 | TTTTTATTTCA | -1 | 575 | 586 | -425 | -414 | 2,97 | 0,01 | 12,44 | >0.1 | 3,32 | 5,14 | 0 | 1 | 0 | 0 | 0 |
| GATA2 | CCCTTGTCTTT | -1 | 502 | 513 | -498 | -487 | 5,2 | 0,01 | 19,81 | >0.1 | 2,97 | 4,97 | 0 | 0,98 | 0 | 5,68 | 0 |
| GATA2 | TTCATATTTCT | -1 | 485 | 496 | -515 | -504 | 1,99 | 0,01 | 16,94 | >0.1 | 2,9 | 4,93 | 0 | 0,99 | 0 | 6,13 | 0 |
| GATA2 | TTCTGATCAGT | -1 | 478 | 489 | -522 | -511 | 2,39 | 0,01 | 16,54 | >0.1 | 2,88 | 4,92 | 0 | 0,99 | 0 | 5,36 | 0 |
| GATA2 | TCTCTATCTCC | -1 | 247 | 258 | -753 | -742 | 5,24 | 0,01 | 19,19 | >0.1 | 2,85 | 4,53 | 0 | 1,38 | 0 | 5,19 | 0 |
| GATA2 | TTGTGATCTGC | -1 | 231 | 242 | -769 | -758 | 1,66 | 0,01 | 15,54 | >0.1 | 2,91 | 4,51 | 0 | 1,45 | 0 | 5,01 | 0 |
| GATA2 | TTCTGATCAGT | -1 | 163 | 174 | -837 | -826 | 2,39 | 0,01 | 14,32 | >0.1 | 3,21 | 4,42 | 0 | 1,9 | 0 | 2,4 | 0 |
| GATA2 | TCCTGATCTGC | -1 | 93 | 104 | -907 | -896 | 4,94 | 0,01 | 15,64 | >0.1 | 3,67 | 4,34 | 0 | 1,88 | 0 | 0,81 | 0 |
| GATA2 | TTCCTATTTCT | -1 | 57 | 68 | -943 | -932 | 1,71 | 0,01 | 11,92 | >0.1 | 4,03 | 4,31 | 0 | 1,87 | 0 | 0 | 0 |
| GATA2 | CACTTACCTAT | -1 | 1 | 12 | -999 | -988 | 4,07 | 0,01 | 17,58 | >0.1 | 5,6 | 4,26 | 0 | 1,86 | 1,79 | 0 | 0 |
| GATA3 | AGATCAGG | 1 | 95 | 103 | -905 | -897 | 3,44 | 0,01 | 14,11 | >0.1 | 3,66 | 4,34 | 0 | 1,88 | 0 | 0,79 | 0 |
| GATA3 | TGATCAGA | 1 | 165 | 173 | -835 | -827 | 2,45 | 0,01 | 14,38 | >0.1 | 3,21 | 4,42 | 0 | 1,9 | 0 | 2,4 | 0 |
| GATA3 | AGATCACA | 1 | 233 | 241 | -767 | -759 | 2,68 | 0,01 | 16,54 | >0.1 | 2,91 | 4,51 | 0 | 1,44 | 0 | 5,01 | 0 |
| GATA3 | AGATAGAG | 1 | 249 | 257 | -751 | -743 | 6,08 | 0,01 | 19,91 | >0.1 | 2,85 | 4,53 | 0 | 1,37 | 0 | 5,09 | 0 |
| GATA3 | AGATCGCG | 1 | 407 | 415 | -593 | -585 | -0,22 | 0,01 | 13,87 | >0.1 | 2,64 | 4,78 | 0 | 1,03 | 0 | 5,65 | 0 |
| GATA3 | TGATCAGA | 1 | 480 | 488 | -520 | -512 | 2,45 | 0,01 | 16,75 | >0.1 | 2,88 | 4,92 | 0 | 0,98 | 0 | 5,52 | 0 |
| GATA3 | AGACAAGG | 1 | 504 | 512 | -496 | -488 | -0,15 | 0,01 | 14,45 | >0.1 | 2,97 | 4,97 | 0 | 0,98 | 0 | 5,68 | 0 |
| GATA3 | AGATAACC | 1 | 594 | 602 | -406 | -398 | 10,21 | 0,001 | 19,83 | >0.1 | 3,43 | 5,19 | 0 | 1,01 | 0 | 0 | 0 |
| GATA3 | TGATTTGG | 1 | 751 | 759 | -249 | -241 | 3,11 | 0,01 | 21,85 | >0.1 | 5,64 | 5,72 | 0 | 1,28 | 0 | 6,1 | 0 |
| GATA3 | AGGTAAGG | 1 | 776 | 784 | -224 | -216 | -0,15 | 0,01 | 19,02 | >0.1 | 5,64 | 5,83 | 0 | 1,37 | 0 | 6,32 | 0 |
| GATA3 | GGATTAAA | 1 | 791 | 799 | -209 | -201 | -2,68 | 0,01 | 15,63 | >0.1 | 5,64 | 5,91 | 0 | 1,44 | 0 | 5,32 | 0 |
| GATA3 | TGATTCAC | 1 | 817 | 825 | -183 | -175 | -1,36 | 0,01 | 15,47 | >0.1 | 5,64 | 6,06 | 0 | 1,58 | 0 | 3,55 | 0 |
| GATA3 | AGATTGTT | 1 | 893 | 901 | -107 | -99 | -0,17 | 0,01 | 16,84 | >0.1 | 5,64 | 6,62 | 0 | 1,89 | 0 | 2,86 | 0 |
| GATA3 | TGATAAGA | 1 | 968 | 976 | -32 | -24 | 10,66 | 0,001 | 27,65 | >0.1 | 5,64 | 7,78 | 0 | 1,87 | 0 | 1,7 | 0 |
| GATA3 | TGATAGAA | -1 | 1166 | 1174 | 166 | 174 | 5,1 | 0,01 | 22,08 | >0.1 | 4,51 | 6,18 | 0 | 1,7 | 1,79 | 2,79 | 0 |
| GATA3 | TGAGAAGG | -1 | 1069 | 1077 | 69 | 77 | -1,04 | 0,01 | 17,65 | >0.1 | 5,64 | 7,19 | 0 | 1,85 | 1,79 | 2,21 | 0 |
| GATA3 | AGATCACA | -1 | 1025 | 1033 | 25 | 33 | 2,68 | 0,01 | 19,95 | >0.1 | 5,64 | 8,52 | 0 | 1,86 | 0 | 1,25 | 0 |
| GATA3 | AGATTTTC | -1 | 978 | 986 | -22 | -14 | 0,6 | 0,01 | 17,82 | >0.1 | 5,64 | 8,13 | 0 | 1,87 | 0 | 1,57 | 0 |
| GATA3 | AGATATGT | -1 | 851 | 859 | -149 | -141 | 4,72 | 0,01 | 21,25 | >0.1 | 5,64 | 6,29 | 0 | 1,85 | 0 | 2,75 | 0 |
| GATA3 | AGATTTTA | -1 | 840 | 848 | -160 | -152 | 1,28 | 0,01 | 17,95 | >0.1 | 5,64 | 6,22 | 0 | 1,75 | 0 | 3,07 | 0 |
| GATA3 | AAATAAGA | -1 | 810 | 818 | -190 | -182 | -1,24 | 0,01 | 15,24 | >0.1 | 5,64 | 6,02 | 0 | 1,54 | 0 | 3,28 | 0 |
| GATA3 | AGAGAACC | -1 | 782 | 790 | -218 | -210 | -1,58 | 0,01 | 17,64 | >0.1 | 5,64 | 5,86 | 0 | 1,4 | 0 | 6,32 | 0 |
| GATA3 | AGATACAC | -1 | 701 | 709 | -299 | -291 | 2,99 | 0,01 | 18,49 | >0.1 | 5,64 | 5,52 | 0 | 1,15 | 0 | 3,19 | 0 |
| GATA3 | AGATAATG | -1 | 639 | 647 | -361 | -353 | 9,02 | 0,001 | 20,77 | >0.1 | 3,77 | 5,31 | 0 | 1,05 | 0 | 1,61 | 0 |
| GATA3 | AGATCATA | -1 | 567 | 575 | -433 | -425 | 0,72 | 0,01 | 10,1 | >0.1 | 3,27 | 5,12 | 0 | 0,99 | 0 | 0 | 0 |
| GATA3 | TGATCAGT | -1 | 478 | 486 | -522 | -514 | -0,18 | 0,01 | 13,79 | >0.1 | 2,87 | 4,91 | 0 | 0,99 | 0 | 5,2 | 0 |
| GATA3 | CGATCTCG | -1 | 405 | 413 | -595 | -587 | -2,06 | 0,01 | 12,14 | >0.1 | 2,64 | 4,77 | 0 | 1,03 | 0 | 5,76 | 0 |
| GATA3 | TGATCTGC | -1 | 231 | 239 | -769 | -761 | -2,41 | 0,01 | 11,47 | >0.1 | 2,92 | 4,5 | 0 | 1,45 | 0 | 5,01 | 0 |
| GATA3 | GGATTACA | -1 | 199 | 207 | -801 | -793 | -0,79 | 0,01 | 12,83 | >0.1 | 3,05 | 4,46 | 0 | 1,64 | 0 | 4,48 | 0 |
| GATA3 | TGATCAGT | -1 | 163 | 171 | -837 | -829 | -0,18 | 0,01 | 11,81 | >0.1 | 3,22 | 4,42 | 0 | 1,9 | 0 | 2,46 | 0 |
| GATA3 | TGATCTGC | -1 | 93 | 101 | -907 | -899 | -2,41 | 0,01 | 8,32 | >0.1 | 3,68 | 4,34 | 0 | 1,88 | 0 | 0,83 | 0 |
| GATA1 | ATCCTAGCACT | 1 | 203 | 214 | -797 | -786 | -9,19 | 0,01 | 4,51 | >0.1 | 3,02 | 4,47 | 0 | 1,61 | 0 | 4,59 | 0 |
| GATA1 | AACTTAGCTGG | 1 | 305 | 316 | -695 | -684 | -8,16 | 0,01 | 5,3 | >0.1 | 2,66 | 4,61 | 0 | 1,19 | 0 | 4,99 | 0 |
| GATA1 | ACATTATCTTC | 1 | 638 | 649 | -362 | -351 | -4,91 | 0,01 | 6,86 | >0.1 | 3,78 | 5,32 | 0 | 1,06 | 0 | 1,61 | 0 |
| GATA1 | CGTGTATCTTT | 1 | 700 | 711 | -300 | -289 | -6,34 | 0,01 | 9,13 | >0.1 | 5,6 | 5,53 | 0 | 1,16 | 0 | 3,19 | 0 |
| GATA1 | TCTTTAACAGC | 1 | 706 | 717 | -294 | -283 | -6,52 | 0,01 | 10,78 | >0.1 | 5,6 | 5,55 | 0 | 1,17 | 0 | 4,99 | 0 |
| GATA1 | TTCTTATTTGA | 1 | 809 | 820 | -191 | -180 | -3,22 | 0,01 | 13,24 | >0.1 | 5,6 | 6,03 | 0 | 1,55 | 0 | 3,28 | 0 |
| GATA1 | TACATATCTGT | 1 | 850 | 861 | -150 | -139 | -4,38 | 0,01 | 12,14 | >0.1 | 5,6 | 6,3 | 0 | 1,87 | 0 | 2,75 | 0 |
| GATA1 | ACTTTATATTG | 1 | 875 | 886 | -125 | -114 | -9,84 | 0,01 | 7,1 | >0.1 | 5,6 | 6,5 | 0 | 1,9 | 0 | 2,95 | 0 |
| GATA1 | TCTTTTTCTTT | 1 | 915 | 926 | -85 | -74 | -5,12 | 0,01 | 10,39 | >0.1 | 5,6 | 6,88 | 0 | 1,89 | 0 | 1,15 | 0 |
| GATA1 | TCCTTCTCAGC | 1 | 1068 | 1079 | 68 | 79 | -3,04 | 0,01 | 15,62 | >0.1 | 5,6 | 7,21 | 0 | 1,85 | 1,79 | 2,21 | 0 |
| GATA1 | GTTCTATCAAA | 1 | 1165 | 1176 | 165 | 176 | 0,48 | 0,001 | 17,48 | >0.1 | 4,52 | 6,19 | 0 | 1,71 | 1,79 | 2,79 | 0 |
| GATA1 | ATTTTCTCTCT | -1 | 973 | 984 | -27 | -16 | -5,17 | 0,01 | 12 | >0.1 | 5,6 | 8,05 | 0 | 1,87 | 0 | 1,65 | 0 |

| | | | | | | | | | | | | | | | | | |
|---|---|---|---|---|---|---|---|---|---|---|---|---|---|---|---|---|---|
| GATA1 | ACCTTACCTCT | -1 | 774 | 785 | -226 | -215 | -2,06 | 0,01 | 16,86 | >0.1 | 5,6 | 5,84 | 0 | 1,38 | 0 | 6,1 | 0 |
| GATA1 | CTCCTGTCACT | -1 | 765 | 776 | -235 | -224 | -7,6 | 0,01 | 11,05 | >0.1 | 5,6 | 5,8 | 0 | 1,34 | 0 | 5,92 | 0 |
| GATA1 | AGGTTATCTCA | -1 | 592 | 603 | -408 | -397 | 5,3 | 0,001 | 14,92 | >0.1 | 3,42 | 5,19 | 0 | 1,01 | 0 | 0 | 0 |
| GATA1 | CATTTATTTTT | -1 | 581 | 592 | -419 | -408 | -10,35 | 0,01 | -0,83 | >0.1 | 3,35 | 5,16 | 0 | 1 | 0 | 0 | 0 |
| GATA1 | TTTTTATTTCA | -1 | 575 | 586 | -425 | -414 | -6,33 | 0,01 | 3,14 | >0.1 | 3,32 | 5,14 | 0 | 1 | 0 | 0 | 0 |
| GATA1 | CCCTTGTCTTT | -1 | 502 | 513 | -498 | -487 | -2,22 | 0,01 | 12,39 | >0.1 | 2,97 | 4,97 | 0 | 0,98 | 0 | 5,68 | 0 |
| GATA1 | TTCTGATCAGT | -1 | 478 | 489 | -522 | -511 | -2,78 | 0,01 | 11,37 | >0.1 | 2,88 | 4,92 | 0 | 0,99 | 0 | 5,36 | 0 |
| GATA1 | TTGTGATCTGC | -1 | 231 | 242 | -769 | -758 | -5,4 | 0,01 | 8,48 | >0.1 | 2,91 | 4,51 | 0 | 1,45 | 0 | 5,01 | 0 |
| GATA1 | TTCTGATCAGT | -1 | 163 | 174 | -837 | -826 | -2,78 | 0,01 | 9,15 | >0.1 | 3,21 | 4,42 | 0 | 1,9 | 0 | 2,4 | 0 |
| GATA1 | TCCTGATCTGC | -1 | 93 | 104 | -907 | -896 | -0,18 | 0,001 | 10,52 | >0.1 | 3,67 | 4,34 | 0 | 1,88 | 0 | 0,81 | 0 |
| GATA1 | TTCCTATTTCT | -1 | 57 | 68 | -943 | -932 | -5,16 | 0,01 | 5,05 | >0.1 | 4,03 | 4,31 | 0 | 1,87 | 0 | 0 | 0 |
| GATA1 | CACTTACCTAT | -1 | 1 | 12 | -999 | -988 | -6,43 | 0,01 | 7,08 | >0.1 | 5,6 | 4,26 | 0 | 1,86 | 1,79 | 0 | 0 |
| MAX | CTCACGCCTG | 1 | 191 | 201 | -809 | -799 | -3,03 | 0,01 | 10,33 | >0.1 | 3,08 | 4,45 | 0 | 1,7 | 0 | 4,13 | 0 |
| MAX | CACGCCTGTA | 1 | 193 | 203 | -807 | -797 | -2,1 | 0,01 | 11,37 | >0.1 | 3,07 | 4,46 | 0 | 1,68 | 0 | 4,26 | 0 |
| MAX | ATCACAAGGT | 1 | 235 | 245 | -765 | -755 | 0,51 | 0,01 | 14,09 | >0.1 | 2,9 | 4,51 | 0 | 1,43 | 0 | 4,74 | 0 |
| MAX | GGCACCTGTA | 1 | 327 | 337 | -673 | -663 | 3,6 | 0,01 | 17,56 | >0.1 | 2,6 | 4,65 | 0 | 1,14 | 0 | 5,56 | 0 |
| MAX | ATCTCATGAG | 1 | 982 | 992 | -18 | -8 | -2,01 | 0,01 | 15,43 | >0.1 | 5,61 | 8,43 | 0 | 1,87 | 0 | 1,53 | 0 |
| MAX | CTCATGAGGA | 1 | 984 | 994 | -16 | -6 | -1,79 | 0,01 | 15,79 | >0.1 | 5,61 | 8,57 | 0 | 1,87 | 0 | 1,53 | 0 |
| MAX | CTCACAGGGG | 1 | 1036 | 1046 | 36 | 46 | -2,24 | 0,01 | 14,55 | >0.1 | 5,61 | 7,99 | 0 | 1,86 | 0 | 1,34 | 0 |
| MAX | TTCACTTGCT | 1 | 1177 | 1187 | 177 | 187 | 1,42 | 0,01 | 19,8 | >0.1 | 5,61 | 6,11 | 0 | 1,61 | 1,79 | 3,26 | 0 |
| MAX | ATCACATCCA | -1 | 1021 | 1031 | 21 | 31 | -1,44 | 0,01 | 16,04 | >0.1 | 5,61 | 8,7 | 0 | 1,86 | 0 | 1,31 | 0 |
| MAX | TCCTCATGAG | -1 | 984 | 994 | -16 | -6 | -1,57 | 0,01 | 16,01 | >0.1 | 5,61 | 8,57 | 0 | 1,87 | 0 | 1,53 | 0 |
| MAX | CTCATGAGAT | -1 | 982 | 992 | -18 | -8 | -1,39 | 0,01 | 16,05 | >0.1 | 5,61 | 8,43 | 0 | 1,87 | 0 | 1,53 | 0 |
| MAX | TACAGGTGCC | -1 | 327 | 337 | -673 | -663 | 2,39 | 0,01 | 16,35 | >0.1 | 2,6 | 4,65 | 0 | 1,14 | 0 | 5,56 | 0 |
| MAX | ACCACGCCCA | -1 | 313 | 323 | -687 | -677 | 1,14 | 0,01 | 15,71 | >0.1 | 2,64 | 4,62 | 0 | 1,17 | 0 | 6,13 | 0 |
| MAX | ACCTTGTGAT | -1 | 235 | 245 | -765 | -755 | 0,98 | 0,01 | 14,56 | >0.1 | 2,9 | 4,51 | 0 | 1,43 | 0 | 4,74 | 0 |
| MAX | TCCACAGGTC | -1 | 109 | 119 | -891 | -881 | -0,71 | 0,01 | 9,82 | >0.1 | 3,54 | 4,36 | 0 | 1,89 | 0 | 0,75 | 0 |
| TCF12 | GGCAGATCAGG | 1 | 92 | 103 | -908 | -897 | -8,47 | 0,01 | 2,26 | >0.1 | 3,67 | 4,34 | 0 | 1,88 | 0 | 0,83 | 0 |
| TCF12 | GGCAGATCACA | 1 | 230 | 241 | -770 | -759 | -10 | 0,01 | 3,89 | >0.1 | 2,91 | 4,51 | 0 | 1,45 | 0 | 5,01 | 0 |
| TCF12 | GGCACCTGTAG | 1 | 327 | 338 | -673 | -662 | -2,33 | 0,01 | 11,63 | >0.1 | 2,6 | 4,65 | 0 | 1,14 | 0 | 5,56 | 0 |
| TCF12 | CCCAGCTACTC | 1 | 339 | 350 | -661 | -650 | -8,09 | 0,01 | 6,79 | >0.1 | 2,57 | 4,67 | 0 | 1,12 | 0 | 6,52 | 0 |
| TCF12 | GGCAGGAGAAT | 1 | 360 | 371 | -640 | -629 | -6,33 | 0,01 | 8,34 | >0.1 | 2,52 | 4,7 | 0 | 1,09 | 0 | 6,36 | 0 |
| TCF12 | AAAAACTGATC | 1 | 474 | 485 | -526 | -515 | -14,14 | 0,01 | -0,43 | >0.1 | 2,86 | 4,91 | 0 | 0,99 | 0 | 4,95 | 0 |
| TCF12 | GACAAGGGCAA | 1 | 505 | 516 | -495 | -484 | -13,85 | 0,01 | 0,77 | >0.1 | 2,98 | 4,98 | 0 | 0,98 | 0 | 5,68 | 0 |
| TCF12 | AACAGCTTTCT | 1 | 711 | 722 | -289 | -278 | -4,23 | 0,01 | 13,1 | >0.1 | 5,6 | 5,57 | 0 | 1,18 | 0 | 4,99 | 0 |
| TCF12 | GACAGGAGAGG | 1 | 768 | 779 | -232 | -221 | -3,96 | 0,01 | 14,72 | >0.1 | 5,6 | 5,81 | 0 | 1,35 | 0 | 5,92 | 0 |
| TCF12 | AGCAGATTGTT | 1 | 890 | 901 | -110 | -99 | -9,5 | 0,01 | 7,56 | >0.1 | 5,6 | 6,62 | 0 | 1,89 | 0 | 2,95 | 0 |
| TCF12 | CACAGGGGACG | 1 | 1038 | 1049 | 38 | 49 | -5,43 | 0,01 | 11,28 | >0.1 | 5,6 | 7,92 | 0 | 1,86 | 0 | 1,34 | 0 |
| TCF12 | AACTGCTGCTC | 1 | 1090 | 1101 | 90 | 101 | -0,48 | 0,001 | 17,91 | >0.1 | 5,6 | 6,88 | 0 | 1,85 | 1,79 | 2,27 | 0 |
| TCF12 | AGCAAGTGAAC | -1 | 1176 | 1187 | 176 | 187 | -0,75 | 0,01 | 17,65 | >0.1 | 5,6 | 6,11 | 0 | 1,62 | 1,79 | 3,28 | 0 |
| TCF12 | AACAGGTCTTC | -1 | 1157 | 1168 | 157 | 168 | -5,21 | 0,01 | 11,35 | >0.1 | 4,39 | 6,24 | 0 | 1,79 | 1,79 | 2,34 | 0 |
| TCF12 | AGCAGCAGTTA | -1 | 1089 | 1100 | 89 | 100 | -3,92 | 0,01 | 14,43 | >0.1 | 5,6 | 6,89 | 0 | 1,85 | 1,79 | 2,21 | 0 |
| TCF12 | GACAGATATGT | -1 | 851 | 862 | -149 | -138 | -10,1 | 0,01 | 6,36 | >0.1 | 5,6 | 6,31 | 0 | 1,88 | 0 | 2,67 | 0 |
| TCF12 | AACAGATTTTA | -1 | 840 | 851 | -160 | -149 | -10,12 | 0,01 | 6,68 | >0.1 | 5,6 | 6,24 | 0 | 1,78 | 0 | 3,19 | 0 |
| TCF12 | GAAAGCTGTTA | -1 | 710 | 721 | -290 | -279 | -3,52 | 0,01 | 13,63 | >0.1 | 5,6 | 5,56 | 0 | 1,18 | 0 | 4,81 | 0 |
| TCF12 | CCCAGGTTCAC | -1 | 374 | 385 | -626 | -615 | -10,51 | 0,01 | 4,37 | >0.1 | 2,56 | 4,72 | 0 | 1,07 | 0 | 6,53 | 0 |
| TCF12 | AGTAGCTGGGA | -1 | 338 | 349 | -662 | -651 | -2,56 | 0,01 | 11,72 | >0.1 | 2,57 | 4,67 | 0 | 1,12 | 0 | 5,92 | 0 |
| TCF12 | TACAGGTGCCC | -1 | 326 | 337 | -674 | -663 | 4,4 | 0,001 | 18,25 | >0.1 | 2,6 | 4,65 | 0 | 1,15 | 0 | 5,46 | 0 |
| TCF12 | CCCAGCTAAGT | -1 | 306 | 317 | -694 | -683 | -10,92 | 0,01 | 2,63 | >0.1 | 2,66 | 4,62 | 0 | 1,19 | 0 | 5,09 | 0 |
| TCF12 | TACAGGCGTGA | -1 | 192 | 203 | -808 | -797 | -13,58 | 0,01 | -0,1 | >0.1 | 3,07 | 4,46 | 0 | 1,69 | 0 | 4,26 | 0 |
| TCF12 | AGCCACTGTGC | -1 | 182 | 193 | -818 | -807 | -13,79 | 0,01 | -0,6 | >0.1 | 3,12 | 4,45 | 0 | 1,77 | 0 | 3,86 | 0 |
| TCF12 | CACAGGTCAAC | -1 | 106 | 117 | -894 | -883 | -6,41 | 0,01 | 4,14 | >0.1 | 3,56 | 4,36 | 0 | 1,89 | 0 | 0,75 | 0 |
| FOSB::JUN | AAAGAAGTCAG | 1 | 30 | 41 | -970 | -959 | 2,39 | 0,01 | 13,01 | >0.1 | 4,47 | 4,28 | 0 | 1,87 | 0 | 0 | 0 |
| FOSB::JUN | AATGGCGTGAA | 1 | 368 | 379 | -632 | -621 | 3,28 | 0,01 | 18,33 | >0.1 | 2,54 | 4,71 | 0 | 1,08 | 0 | 6,73 | 0 |
| FOSB::JUN | GATCGCGCCAC | 1 | 408 | 419 | -592 | -581 | 2,56 | 0,01 | 16,67 | >0.1 | 2,65 | 4,78 | 0 | 1,03 | 0 | 5,65 | 0 |
| FOSB::JUN | GGCAAAGTCAT | 1 | 511 | 522 | -489 | -478 | 0,43 | 0,01 | 12,27 | >0.1 | 3,01 | 4,99 | 0 | 0,98 | 0 | 2,86 | 0 |
| FOSB::JUN | TGTGAGATAAC | 1 | 590 | 601 | -410 | -399 | 3,15 | 0,01 | 12,75 | >0.1 | 3,41 | 5,18 | 0 | 1,01 | 0 | 0 | 0 |
| FOSB::JUN | GATTTGGCCAT | 1 | 752 | 763 | -248 | -237 | 0,64 | 0,01 | 19,81 | >0.1 | 5,6 | 5,74 | 0 | 1,3 | 0 | 6,54 | 0 |
| FOSB::JUN | GGAGAGGTAAG | 1 | 772 | 783 | -228 | -217 | 0,94 | 0,01 | 19,65 | >0.1 | 5,6 | 5,83 | 0 | 1,37 | 0 | 5,92 | 0 |

| | | | | | | | | | | | | | | | | | |
|---|---|---|---|---|---|---|---|---|---|---|---|---|---|---|---|---|---|
| FOSB::JUN | AATAACGTATT | 1 | 800 | 811 | -200 | -189 | 2,74 | 0,01 | 19,36 | >0.1 | 5,6 | 5,98 | 0 | 1,5 | 0 | 3,55 | 0 |
| FOSB::JUN | TGTTACATATC | 1 | 847 | 858 | -153 | -142 | -0,82 | 0,01 | 15,76 | >0.1 | 5,6 | 6,28 | 0 | 1,84 | 0 | 2,86 | 0 |
| FOSB::JUN | GGGAAAGTCAT | 1 | 1006 | 1017 | 6 | 17 | 0,69 | 0,01 | 18,74 | >0.1 | 5,6 | 9,12 | 0 | 1,87 | 0 | 1,46 | 0 |
| FOSB::JUN | GACGATGTCAA | 1 | 1045 | 1056 | 45 | 56 | 4,8 | 0,001 | 20,61 | >0.1 | 5,6 | 7,7 | 0 | 1,86 | 0 | 0,65 | 0 |
| FOSB::JUN | GTTGCTGTAAC | 1 | 1082 | 1093 | 82 | 93 | 0,76 | 0,01 | 19,2 | >0.1 | 5,6 | 6,99 | 0 | 1,85 | 1,79 | 2,21 | 0 |
| FOSB::JUN | AGTTACAGCAA | -1 | 1083 | 1094 | 83 | 94 | -0,59 | 0,01 | 17,84 | >0.1 | 5,6 | 6,97 | 0 | 1,85 | 1,79 | 2,21 | 0 |
| FOSB::JUN | CTTGACATCGT | -1 | 1046 | 1057 | 46 | 57 | 2,02 | 0,01 | 18,62 | >0.1 | 5,6 | 7,68 | 0 | 1,86 | 0 | 1,46 | 0 |
| FOSB::JUN | GATCACATCCA | -1 | 1021 | 1032 | 21 | 32 | -0,11 | 0,01 | 17,36 | >0.1 | 5,6 | 8,7 | 0 | 1,86 | 0 | 1,31 | 0 |
| FOSB::JUN | GCTGAGCCAAT | -1 | 882 | 893 | -118 | -107 | -0,55 | 0,01 | 16,56 | >0.1 | 5,6 | 6,56 | 0 | 1,89 | 0 | 3,07 | 0 |
| FOSB::JUN | GAATACGTTAT | -1 | 801 | 812 | -199 | -188 | 3,76 | 0,01 | 20,27 | >0.1 | 5,6 | 5,98 | 0 | 1,51 | 0 | 3,43 | 0 |
| FOSB::JUN | AATGTTATAAT | -1 | 686 | 697 | -314 | -303 | -0,14 | 0,01 | 13,84 | >0.1 | 4,43 | 5,48 | 0 | 1,13 | 0 | 2,95 | 0 |
| FOSB::JUN | AGTGACCAAAG | -1 | 670 | 681 | -330 | -319 | -0,84 | 0,01 | 13,53 | >0.1 | 4,15 | 5,42 | 0 | 1,1 | 0 | 3,7 | 0 |
| FOSB::JUN | GGTTATCTCAC | -1 | 591 | 602 | -409 | -398 | 2,18 | 0,01 | 11,79 | >0.1 | 3,42 | 5,19 | 0 | 1,01 | 0 | 0 | 0 |
| FOSB::JUN | AGTGGCGCGAT | -1 | 409 | 420 | -591 | -580 | 1,39 | 0,01 | 15,38 | >0.1 | 2,65 | 4,79 | 0 | 1,03 | 0 | 5,52 | 0 |
| FOSB::JUN | GTTCACGCCAT | -1 | 369 | 380 | -631 | -620 | 4,79 | 0,01 | 19,27 | >0.1 | 2,54 | 4,72 | 0 | 1,07 | 0 | 6,15 | 0 |
| FOSB::JUN | GCTGACTTCTT | -1 | 31 | 42 | -969 | -958 | 4,08 | 0,01 | 14,68 | >0.1 | 4,45 | 4,28 | 0 | 1,87 | 0 | 0 | 0 |
| CTCF | AGTGAAGGAGAAGGAGGCA | 1 | 8 | 27 | -992 | -973 | -0,92 | 0,01 | 12,51 | >0.1 | 5,5 | 4,27 | 0 | 1,87 | 1,79 | 0 | 0 |
| CTCF | GAAGGAGAAGGAGGCACTC | 1 | 11 | 30 | -989 | -970 | -2,14 | 0,01 | 9,49 | >0.1 | 5,5 | 4,27 | 0 | 1,87 | 0 | 0 | 0 |
| CTCF | TGACCTGTGGAGTGGAGAG | 1 | 108 | 127 | -892 | -873 | -0,31 | 0,01 | 10,23 | >0.1 | 3,52 | 4,37 | 0 | 1,89 | 0 | 0,77 | 0 |
| CTCF | AGGTCAGGAGATAGAGACC | 1 | 241 | 260 | -759 | -740 | 1,79 | 0,01 | 15,89 | >0.1 | 2,86 | 4,53 | 0 | 1,41 | 0 | 5,3 | 0 |
| CTCF | TGGGCACCTGTAGTCCCAG | 1 | 325 | 344 | -675 | -656 | -3,2 | 0,01 | 10,66 | >0.1 | 2,6 | 4,66 | 0 | 1,15 | 0 | 5,46 | 0 |
| CTCF | TACTCGGGAGGCTGAGGCA | 1 | 345 | 364 | -655 | -636 | -4,4 | 0,01 | 10,35 | >0.1 | 2,54 | 4,69 | 0 | 1,12 | 0 | 6,4 | 0 |
| CTCF | GTGAACCTGGGAGGCAGAG | 1 | 374 | 393 | -626 | -607 | -4,17 | 0,01 | 10,74 | >0.1 | 2,57 | 4,74 | 0 | 1,07 | 0 | 6,53 | 0 |
| CTCF | TGGCCATAAAGTGACAGGA | 1 | 756 | 775 | -244 | -225 | 3,42 | 0,001 | 21,97 | >0.1 | 5,5 | 5,79 | 0 | 1,34 | 0 | 5,92 | 0 |
| CTCF | GTGACAGGAGAGGTAAGGT | 1 | 766 | 785 | -234 | -215 | 2,09 | 0,01 | 20,72 | >0.1 | 5,5 | 5,84 | 0 | 1,38 | 0 | 5,92 | 0 |
| CTCF | TCCTCTCCAGGATGAACTT | 1 | 860 | 879 | -140 | -121 | -1,89 | 0,01 | 14,7 | >0.1 | 5,5 | 6,44 | 0 | 1,9 | 0 | 2,75 | 0 |
| CTCF | TGGCTCACAGGGGACGATG | 1 | 1033 | 1052 | 33 | 52 | 2,66 | 0,01 | 19,46 | >0.1 | 5,5 | 8,1 | 0 | 1,86 | 0 | 1,34 | 0 |
| CTCF | TGAGCAGCAGTTACAGCAA | -1 | 1083 | 1102 | 83 | 102 | -4,09 | 0,01 | 14,24 | >0.1 | 5,5 | 6,97 | 0 | 1,85 | 1,79 | 2,21 | 0 |
| CTCF | CAGCAACAAGGCTGAGAAG | -1 | 1070 | 1089 | 70 | 89 | -3,8 | 0,01 | 14,86 | >0.1 | 5,5 | 7,17 | 0 | 1,85 | 1,79 | 2,34 | 0 |
| CTCF | CATCCACTGAATGACTTTC | -1 | 1008 | 1027 | 8 | 27 | -5,31 | 0,01 | 12,54 | >0.1 | 5,5 | 9,07 | 0 | 1,87 | 0 | 1,42 | 0 |
| CTCF | CATCCTGGAGAGGACAGAT | -1 | 855 | 874 | -145 | -126 | 6,08 | 0,001 | 22,74 | >0.1 | 5,5 | 6,4 | 0 | 1,9 | 0 | 2,86 | 0 |
| CTCF | CTGTCACTTTATGGCCAAA | -1 | 754 | 773 | -246 | -227 | -5,46 | 0,01 | 13,25 | >0.1 | 5,5 | 5,78 | 0 | 1,33 | 0 | 6,1 | 0 |
| CTCF | CTAGAACTAGGGATCATGA | -1 | 647 | 666 | -353 | -334 | -5,31 | 0,01 | 9,05 | >0.1 | 3,91 | 5,37 | 0 | 1,08 | 0 | 3,99 | 0 |
| CTCF | CGCCCAGGCTGGAGCACAG | -1 | 418 | 437 | -582 | -563 | -2,26 | 0,01 | 11,83 | >0.1 | 2,69 | 4,82 | 0 | 1,03 | 0 | 5,56 | 0 |
| CTCF | CTGCCTCCCAGGTTCACGC | -1 | 372 | 391 | -628 | -609 | -4,46 | 0,01 | 10,31 | >0.1 | 2,56 | 4,74 | 0 | 1,07 | 0 | 6,4 | 0 |
| CTCF | GGGACTACAGGTGCCCACC | -1 | 323 | 342 | -677 | -658 | -1,26 | 0,01 | 12,72 | >0.1 | 2,6 | 4,65 | 0 | 1,16 | 0 | 5,56 | 0 |
| HINFP | CAAGGTCAGGAG | 1 | 239 | 251 | -761 | -749 | -0,22 | 0,01 | 12,92 | >0.1 | 2,88 | 4,52 | 0 | 1,41 | 0 | 4,33 | 0 |
| HINFP | GACCGTCCTGAC | 1 | 256 | 268 | -744 | -732 | 3,65 | 0,001 | 16,98 | >0.1 | 2,82 | 4,54 | 0 | 1,34 | 0 | 4,63 | 0 |
| HINFP | CCCCGTCTCTAT | 1 | 281 | 293 | -719 | -707 | -2,71 | 0,01 | 10,95 | >0.1 | 2,73 | 4,58 | 0 | 1,26 | 0 | 5,09 | 0 |
| HINFP | CCAGCTACTCGG | 1 | 340 | 352 | -660 | -648 | -4,54 | 0,01 | 10,34 | >0.1 | 2,56 | 4,67 | 0 | 1,12 | 0 | 6,52 | 0 |
| HINFP | CACTGTGCTCCA | 1 | 416 | 428 | -584 | -572 | -1,91 | 0,01 | 11,89 | >0.1 | 2,67 | 4,8 | 0 | 1,02 | 0 | 5,3 | 0 |
| HINFP | CAGCCTGGGCGA | 1 | 426 | 438 | -574 | -562 | -2,76 | 0,01 | 11,58 | >0.1 | 2,7 | 4,82 | 0 | 1,01 | 0 | 5,8 | 0 |
| HINFP | CATTTTCCGTGT | 1 | 693 | 705 | -307 | -295 | -3,23 | 0,01 | 12,07 | >0.1 | 5,58 | 5,5 | 0 | 1,15 | 0 | 3,07 | 0 |
| HINFP | TAAGGTTCTCTA | 1 | 779 | 791 | -221 | -209 | -4,02 | 0,01 | 15,38 | >0.1 | 5,58 | 5,87 | 0 | 1,4 | 0 | 6,54 | 0 |
| HINFP | ATCTGTCCTCTC | 1 | 855 | 867 | -145 | -133 | -2,83 | 0,01 | 13,95 | >0.1 | 5,58 | 6,35 | 0 | 1,9 | 0 | 2,95 | 0 |
| HINFP | CCTGCTCAGCGC | 1 | 937 | 949 | -63 | -51 | -4,47 | 0,01 | 11,07 | >0.1 | 5,58 | 7,2 | 0 | 1,88 | 0 | 0,88 | 0 |
| HINFP | CAAGCTCTTCCT | 1 | 1053 | 1065 | 53 | 65 | -2,73 | 0,01 | 14,49 | >0.1 | 5,58 | 7,5 | 0 | 1,86 | 0 | 2,27 | 0 |
| HINFP | GCTCTTCCTGGC | 1 | 1056 | 1068 | 56 | 68 | -3,29 | 0,01 | 13,86 | >0.1 | 5,58 | 7,44 | 0 | 1,86 | 0 | 2,27 | 0 |
| HINFP | CTGGCTCCTTCT | 1 | 1063 | 1075 | 63 | 75 | -3,01 | 0,01 | 15,73 | >0.1 | 5,58 | 7,3 | 0 | 1,85 | 1,79 | 2,21 | 0 |
| HINFP | ACAGGTCTTCGG | -1 | 1155 | 1167 | 155 | 167 | 0,15 | 0,01 | 16,78 | >0.1 | 4,42 | 6,26 | 0 | 1,81 | 1,79 | 2,35 | 0 |
| HINFP | CCTGTTCCTCAA | -1 | 1110 | 1122 | 110 | 122 | -1,17 | 0,01 | 18,67 | >0.1 | 5,58 | 6,65 | 0 | 1,85 | 3,22 | 2,54 | 0 |
| HINFP | CATCGTCCCCTG | -1 | 1040 | 1052 | 40 | 52 | 10,25 | 0,0001 | 26,94 | >0.1 | 5,58 | 7,85 | 0 | 1,86 | 0 | 1,39 | 0 |
| HINFP | AAACCTCCTCAT | -1 | 987 | 999 | -13 | -1 | 2,53 | 0,01 | 20,28 | >0.1 | 5,58 | 8,69 | 0 | 1,87 | 0 | 1,61 | 0 |
| HINFP | TTACCTCTCCTG | -1 | 770 | 782 | -230 | -218 | -3,02 | 0,01 | 15,86 | >0.1 | 5,58 | 5,83 | 0 | 1,37 | 0 | 6,1 | 0 |
| HINFP | CTCGGTCTGTCG | -1 | 435 | 447 | -565 | -553 | -1,7 | 0,01 | 12,8 | >0.1 | 2,73 | 4,84 | 0 | 1,01 | 0 | 5,92 | 0 |
| HINFP | CAAGCTCTGCCT | -1 | 385 | 397 | -615 | -603 | 2,24 | 0,01 | 16,75 | >0.1 | 2,59 | 4,75 | 0 | 1,05 | 0 | 6,13 | 0 |
| HINFP | CTGCCTCCCAGG | -1 | 379 | 391 | -621 | -609 | -0,52 | 0,01 | 13,76 | >0.1 | 2,57 | 4,73 | 0 | 1,06 | 0 | 5,91 | 0 |
| HINFP | CTGCCTCAGCCT | -1 | 353 | 365 | -647 | -635 | -2,04 | 0,01 | 12,8 | >0.1 | 2,53 | 4,69 | 0 | 1,1 | 0 | 6,52 | 0 |
| HINFP | CAGCCTCCCGAG | -1 | 347 | 359 | -653 | -641 | 4,43 | 0,001 | 19,17 | >0.1 | 2,55 | 4,68 | 0 | 1,11 | 0 | 6,4 | 0 |

| | | | | | | | | | | | | | | | | | |
|---|---|---|---|---|---|---|---|---|---|---|---|---|---|---|---|---|---|
| HINFP | ACAGGTGCCCAC | -1 | 324 | 336 | -676 | -664 | 1,27 | 0,01 | 15,13 | >0.1 | 2,61 | 4,64 | 0 | 1,15 | 0 | 5,46 | 0 |
| HINFP | CGGCCTCCCAAA | -1 | 213 | 225 | -787 | -775 | -2,06 | 0,01 | 12,48 | >0.1 | 2,98 | 4,49 | 0 | 1,55 | 0 | 5,52 | 0 |
| HINFP | CACTGTGCCCAG | -1 | 178 | 190 | -822 | -810 | 0,77 | 0,01 | 13,24 | >0.1 | 3,13 | 4,44 | 0 | 1,81 | 0 | 3,09 | 0 |
| HINFP | CTGGTTCTGTGG | -1 | 41 | 53 | -959 | -947 | -3,73 | 0,01 | 6,68 | >0.1 | 4,25 | 4,29 | 0 | 1,87 | 0 | 0 | 0 |
| HINFP | GTGCCTCCTTCT | -1 | 16 | 28 | -984 | -972 | -2,69 | 0,01 | 9,03 | >0.1 | 5,58 | 4,27 | 0 | 1,87 | 0 | 0 | 0 |
| ATF7 | CAAAGAAGTCAGCC | 1 | 29 | 43 | -971 | -957 | -5,6 | 0,01 | 5,01 | >0.1 | 4,46 | 4,28 | 0 | 1,87 | 0 | 0 | 0 |
| ATF7 | GAATGGCGTGAACC | 1 | 367 | 381 | -633 | -619 | -4,47 | 0,01 | 10,02 | >0.1 | 2,54 | 4,72 | 0 | 1,08 | 0 | 6,15 | 0 |
| ATF7 | AGATCGCGCCACTG | 1 | 407 | 421 | -593 | -579 | -11,1 | 0,01 | 2,89 | >0.1 | 2,65 | 4,79 | 0 | 1,03 | 0 | 5,52 | 0 |
| ATF7 | ATGTGAGATAACCT | 1 | 589 | 603 | -411 | -397 | -5,94 | 0,01 | 3,68 | >0.1 | 3,41 | 5,19 | 0 | 1,01 | 0 | 0 | 0 |
| ATF7 | CTCTTTGGTCACTG | 1 | 668 | 682 | -332 | -318 | -11,05 | 0,01 | 3,32 | >0.1 | 4,14 | 5,43 | 0 | 1,1 | 0 | 3,7 | 0 |
| ATF7 | CCATAAAGTGACAG | 1 | 759 | 773 | -241 | -227 | -10,71 | 0,01 | 8,06 | >0.1 | 5,56 | 5,78 | 0 | 1,33 | 0 | 6,1 | 0 |
| ATF7 | AAGTGACAGGAGAG | 1 | 764 | 778 | -236 | -222 | -12,78 | 0,01 | 5,85 | >0.1 | 5,56 | 5,81 | 0 | 1,35 | 0 | 5,92 | 0 |
| ATF7 | AGGAGAGGTAAGGT | 1 | 771 | 785 | -229 | -215 | -13,05 | 0,01 | 5,83 | >0.1 | 5,56 | 5,84 | 0 | 1,38 | 0 | 6,1 | 0 |
| ATF7 | GGATTAAAGAATAA | 1 | 791 | 805 | -209 | -195 | -14,05 | 0,01 | 4,24 | >0.1 | 5,56 | 5,94 | 0 | 1,47 | 0 | 5,32 | 0 |
| ATF7 | GAATAACGTATTCT | 1 | 799 | 813 | -201 | -187 | -5,56 | 0,01 | 10,92 | >0.1 | 5,56 | 5,99 | 0 | 1,51 | 0 | 3,43 | 0 |
| ATF7 | CTGTTACATATCTG | 1 | 846 | 860 | -154 | -140 | -10,39 | 0,01 | 6,28 | >0.1 | 5,56 | 6,3 | 0 | 1,86 | 0 | 2,95 | 0 |
| ATF7 | AGGGAAAGTCATTC | 1 | 1005 | 1019 | 5 | 19 | -12,05 | 0,01 | 5,96 | >0.1 | 5,56 | 9,12 | 0 | 1,87 | 0 | 1,46 | 0 |
| ATF7 | GGACGATGTCAAGC | 1 | 1044 | 1058 | 44 | 58 | -9,27 | 0,01 | 7,34 | >0.1 | 5,56 | 7,73 | 0 | 1,86 | 0 | 1,46 | 0 |
| ATF7 | TGTTGCTGTAACTG | 1 | 1081 | 1095 | 81 | 95 | -13,42 | 0,01 | 5 | >0.1 | 5,56 | 7 | 0 | 1,85 | 1,79 | 2,21 | 0 |
| ATF7 | CAGTTACAGCAACA | -1 | 1081 | 1095 | 81 | 95 | -9,37 | 0,01 | 9,05 | >0.1 | 5,56 | 7 | 0 | 1,85 | 1,79 | 2,21 | 0 |
| ATF7 | GCTTGACATCGTCC | -1 | 1044 | 1058 | 44 | 58 | -6,15 | 0,01 | 10,46 | >0.1 | 5,56 | 7,73 | 0 | 1,86 | 0 | 1,46 | 0 |
| ATF7 | CAGATATGTAACAG | -1 | 846 | 860 | -154 | -140 | -9,71 | 0,01 | 6,96 | >0.1 | 5,56 | 6,3 | 0 | 1,86 | 0 | 2,95 | 0 |
| ATF7 | AGAATACGTTATTC | -1 | 799 | 813 | -201 | -187 | -6,43 | 0,01 | 10,05 | >0.1 | 5,56 | 5,99 | 0 | 1,51 | 0 | 3,43 | 0 |
| ATF7 | TTATGGCCAAATCA | -1 | 751 | 765 | -249 | -235 | -12,82 | 0,01 | 6,11 | >0.1 | 5,56 | 5,75 | 0 | 1,3 | 0 | 6,32 | 0 |
| ATF7 | CAGTGACCAAAGAG | -1 | 668 | 682 | -332 | -318 | -11,03 | 0,01 | 3,34 | >0.1 | 4,14 | 5,43 | 0 | 1,1 | 0 | 3,7 | 0 |
| ATF7 | AGGTTATCTCACAT | -1 | 589 | 603 | -411 | -397 | -8,45 | 0,01 | 1,17 | >0.1 | 3,41 | 5,19 | 0 | 1,01 | 0 | 0 | 0 |
| ATF7 | ACATGACTTTGCCC | -1 | 510 | 524 | -490 | -476 | -13,27 | 0,01 | -1,33 | >0.1 | 3,01 | 5 | 0 | 0,99 | 0 | 2,95 | 0 |
| ATF7 | CAGTGGCGCGATCT | -1 | 407 | 421 | -593 | -579 | -8,8 | 0,01 | 5,19 | >0.1 | 2,65 | 4,79 | 0 | 1,03 | 0 | 5,52 | 0 |
| ATF7 | GGTTCACGCCATTC | -1 | 367 | 381 | -633 | -619 | -10,86 | 0,01 | 3,63 | >0.1 | 2,54 | 4,72 | 0 | 1,08 | 0 | 6,15 | 0 |
| ATF7 | GGCTGACTTCTTTG | -1 | 29 | 43 | -971 | -957 | -9,25 | 0,01 | 1,36 | >0.1 | 4,46 | 4,28 | 0 | 1,87 | 0 | 0 | 0 |
| ARNTL | GATCAGGAGA | 1 | 96 | 106 | -904 | -894 | 2,68 | 0,01 | 13,32 | >0.1 | 3,64 | 4,35 | 0 | 1,88 | 0 | 0,77 | 0 |
| ARNTL | TGACCTGTGG | 1 | 108 | 118 | -892 | -882 | -0,59 | 0,01 | 9,95 | >0.1 | 3,55 | 4,36 | 0 | 1,89 | 0 | 0,75 | 0 |
| ARNTL | GGGCACAGTG | 1 | 180 | 190 | -820 | -810 | -0,3 | 0,01 | 13,01 | >0.1 | 3,13 | 4,44 | 0 | 1,79 | 0 | 3,95 | 0 |
| ARNTL | TGGCTCACGC | 1 | 188 | 198 | -812 | -802 | -0,15 | 0,01 | 13,06 | >0.1 | 3,09 | 4,45 | 0 | 1,72 | 0 | 3,95 | 0 |
| ARNTL | GCTCACGCCT | 1 | 190 | 200 | -810 | -800 | 1,78 | 0,01 | 15,15 | >0.1 | 3,08 | 4,45 | 0 | 1,7 | 0 | 4,13 | 0 |
| ARNTL | GATCACAAGG | 1 | 234 | 244 | -766 | -756 | 4,29 | 0,01 | 18,04 | >0.1 | 2,9 | 4,51 | 0 | 1,44 | 0 | 4,9 | 0 |
| ARNTL | GGTCAGGAGA | 1 | 242 | 252 | -758 | -748 | 4,35 | 0,01 | 17,47 | >0.1 | 2,87 | 4,52 | 0 | 1,4 | 0 | 4,33 | 0 |
| ARNTL | TAACACGGTG | 1 | 268 | 278 | -732 | -722 | 0,61 | 0,01 | 13,7 | >0.1 | 2,78 | 4,56 | 0 | 1,3 | 0 | 4,45 | 0 |
| ARNTL | GGGCACCTGT | 1 | 326 | 336 | -674 | -664 | 4,98 | 0,01 | 18,83 | >0.1 | 2,6 | 4,64 | 0 | 1,15 | 0 | 5,46 | 0 |
| ARNTL | AGGCAGGAGA | 1 | 359 | 369 | -641 | -631 | 1,9 | 0,01 | 16,57 | >0.1 | 2,52 | 4,7 | 0 | 1,09 | 0 | 6,36 | 0 |
| ARNTL | AGTCATGTAT | 1 | 516 | 526 | -484 | -474 | 1,55 | 0,01 | 13,51 | >0.1 | 3,02 | 5 | 0 | 0,98 | 0 | 2,95 | 0 |
| ARNTL | TGACAGGAGA | 1 | 767 | 777 | -233 | -223 | 0,25 | 0,01 | 18,92 | >0.1 | 5,61 | 5,8 | 0 | 1,35 | 0 | 5,92 | 0 |
| ARNTL | AATCTCATGA | 1 | 981 | 991 | -19 | -9 | 2,5 | 0,01 | 19,89 | >0.1 | 5,61 | 8,37 | 0 | 1,87 | 0 | 1,53 | 0 |
| ARNTL | TCTCATGAGG | 1 | 983 | 993 | -17 | -7 | 2,28 | 0,01 | 19,83 | >0.1 | 5,61 | 8,5 | 0 | 1,87 | 0 | 1,57 | 0 |
| ARNTL | GCTCACAGGG | 1 | 1035 | 1045 | 35 | 45 | 4,22 | 0,01 | 21,02 | >0.1 | 5,61 | 8,03 | 0 | 1,86 | 0 | 1,31 | 0 |
| ARNTL | GTTCACTTGC | 1 | 1176 | 1186 | 176 | 186 | 3,72 | 0,01 | 22,13 | >0.1 | 5,61 | 6,11 | 0 | 1,62 | 1,79 | 3,28 | 0 |
| ARNTL | AAGCAAGTGA | -1 | 1178 | 1188 | 178 | 188 | 1,61 | 0,01 | 20,09 | >0.1 | 5,61 | 6,1 | 0 | 1,6 | 1,79 | 3,37 | 0 |
| ARNTL | CGTCCCCTGT | -1 | 1039 | 1049 | 39 | 49 | 1,31 | 0,01 | 18,03 | >0.1 | 5,61 | 7,88 | 0 | 1,86 | 0 | 1,36 | 0 |
| ARNTL | GATCACATCC | -1 | 1022 | 1032 | 22 | 32 | 1,61 | 0,01 | 19,04 | >0.1 | 5,61 | 8,68 | 0 | 1,86 | 0 | 1,28 | 0 |
| ARNTL | CCTCATGAGA | -1 | 983 | 993 | -17 | -7 | 2,91 | 0,01 | 20,46 | >0.1 | 5,61 | 8,5 | 0 | 1,87 | 0 | 1,57 | 0 |
| ARNTL | TGTCACTTTA | -1 | 762 | 772 | -238 | -228 | 1,02 | 0,01 | 19,65 | >0.1 | 5,61 | 5,78 | 0 | 1,33 | 0 | 5,92 | 0 |
| ARNTL | TCTCACATTT | -1 | 587 | 597 | -413 | -403 | 2,85 | 0,01 | 12,42 | >0.1 | 3,39 | 5,17 | 0 | 1,01 | 0 | 0 | 0 |
| ARNTL | AGTCTCGGTC | -1 | 440 | 450 | -560 | -550 | 1,53 | 0,01 | 16,17 | >0.1 | 2,75 | 4,84 | 0 | 1 | 0 | 6,05 | 0 |
| ARNTL | GTTCACGCCA | -1 | 370 | 380 | -630 | -620 | 0,39 | 0,01 | 14,99 | >0.1 | 2,54 | 4,72 | 0 | 1,07 | 0 | 6,27 | 0 |
| ARNTL | CCTCCCGAGT | -1 | 346 | 356 | -654 | -644 | 3,41 | 0,01 | 18,27 | >0.1 | 2,55 | 4,68 | 0 | 1,11 | 0 | 6,52 | 0 |
| ARNTL | CTACAGGTGC | -1 | 328 | 338 | -672 | -662 | 1,92 | 0,01 | 15,87 | >0.1 | 2,6 | 4,65 | 0 | 1,14 | 0 | 5,56 | 0 |
| ARNTL | CACCACGCCC | -1 | 314 | 324 | -686 | -676 | -0,73 | 0,01 | 13,84 | >0.1 | 2,64 | 4,63 | 0 | 1,17 | 0 | 6,13 | 0 |
| ARNTL | TATCTCCTGA | -1 | 244 | 254 | -756 | -746 | -0,87 | 0,01 | 12,33 | >0.1 | 2,86 | 4,52 | 0 | 1,39 | 0 | 4,42 | 0 |
| ARNTL | CTCCACAGGT | -1 | 110 | 120 | -890 | -880 | -0,49 | 0,01 | 10,06 | >0.1 | 3,54 | 4,36 | 0 | 1,89 | 0 | 0,77 | 0 |

| | | | | | | | | | | | | | | | | | |
|---|---|---|---|---|---|---|---|---|---|---|---|---|---|---|---|---|---|
| ARNTL | TCTCACACTG | -1 | 50 | 60 | -950 | -940 | -0,88 | 0,01 | 9,42 | >0.1 | 4,13 | 4,3 | 0 | 1,87 | 0 | 0 | 0 |
| RXRA::VDR | TGGCCATGGAAATTA | 1 | 146 | 161 | -854 | -839 | -3,27 | 0,01 | 7,97 | >0.1 | 3,29 | 4,41 | 0 | 1,9 | 0 | 1,65 | 0 |
| RXRA::VDR | GGGAGGCCGAGTTGG | 1 | 216 | 231 | -784 | -769 | 2,96 | 0,01 | 17,58 | >0.1 | 2,96 | 4,49 | 0 | 1,53 | 0 | 5,63 | 0 |
| RXRA::VDR | AGATCACAAGGTCAG | 1 | 233 | 248 | -767 | -752 | 0,07 | 0,01 | 13,67 | >0.1 | 2,9 | 4,52 | 0 | 1,44 | 0 | 4,74 | 0 |
| RXRA::VDR | AGGTCAGGAGATAGA | 1 | 241 | 256 | -759 | -744 | 0,75 | 0,01 | 13,88 | >0.1 | 2,87 | 4,53 | 0 | 1,41 | 0 | 4,33 | 0 |
| RXRA::VDR | AGGAGATAGAGACCG | 1 | 246 | 261 | -754 | -739 | -2,43 | 0,01 | 11,42 | >0.1 | 2,85 | 4,53 | 0 | 1,38 | 0 | 5,09 | 0 |
| RXRA::VDR | GGGAGGCAGAGCTTG | 1 | 382 | 397 | -618 | -603 | -1,55 | 0,01 | 12,85 | >0.1 | 2,58 | 4,75 | 0 | 1,06 | 0 | 6,02 | 0 |
| RXRA::VDR | AGTGAGCCGAGATCG | 1 | 398 | 413 | -602 | -587 | -2,29 | 0,01 | 12,06 | >0.1 | 2,63 | 4,77 | 0 | 1,04 | 0 | 5,91 | 0 |
| RXRA::VDR | GCGCCACTGTGCTCC | 1 | 412 | 427 | -588 | -573 | -0,9 | 0,01 | 12,79 | >0.1 | 2,67 | 4,8 | 0 | 1,03 | 0 | 5,19 | 0 |
| RXRA::VDR | AAGTCATGTATTTGG | 1 | 515 | 530 | -485 | -470 | -3,75 | 0,01 | 8,23 | >0.1 | 3,03 | 5,01 | 0 | 0,99 | 0 | 2,95 | 0 |
| RXRA::VDR | GGTTCTCTAGGATTA | 1 | 782 | 797 | -218 | -203 | 1,95 | 0,01 | 20,74 | >0.1 | 5,54 | 5,9 | 0 | 1,43 | 0 | 5,92 | 0 |
| RXRA::VDR | TGTTTACTGTGTTCT | 1 | 897 | 912 | -103 | -88 | -0,5 | 0,01 | 16,41 | >0.1 | 5,54 | 6,73 | 0 | 1,89 | 0 | 2,75 | 0 |
| RXRA::VDR | AAGTCATTCAGTGGA | 1 | 1010 | 1025 | 10 | 25 | -1,4 | 0,01 | 16,42 | >0.1 | 5,54 | 8,99 | 0 | 1,86 | 0 | 1,42 | 0 |
| RXRA::VDR | GGCTCACAGGGGACG | 1 | 1034 | 1049 | 34 | 49 | 7,55 | 0,001 | 24,35 | >0.1 | 5,54 | 8,06 | 0 | 1,86 | 0 | 1,34 | 0 |
| RXRA::VDR | ATGTCAAGCTCTTCC | 1 | 1049 | 1064 | 49 | 64 | -2,81 | 0,01 | 13,65 | >0.1 | 5,54 | 7,6 | 0 | 1,86 | 0 | 1,46 | 0 |
| RXRA::VDR | AGTTCACTTGCTTCT | 1 | 1175 | 1190 | 175 | 190 | 0,61 | 0,01 | 18,96 | >0.1 | 5,54 | 6,12 | 0 | 1,62 | 1,79 | 3,26 | 0 |
| RXRA::VDR | AGGTCTTCGGCTTCG | -1 | 1150 | 1165 | 150 | 165 | 10,2 | 0,0001 | 26,89 | >0.1 | 4,5 | 6,3 | 0 | 1,84 | 1,79 | 2,26 | 0 |
| RXRA::VDR | GGTTAAACTTGTCCA | -1 | 1134 | 1149 | 134 | 149 | -1,26 | 0,01 | 18,86 | >0.1 | 5,54 | 6,42 | 0 | 1,84 | 3,22 | 3,09 | 0 |
| RXRA::VDR | ATGTAACAGATTTTA | -1 | 840 | 855 | -160 | -145 | -1,55 | 0,01 | 15,26 | >0.1 | 5,54 | 6,26 | 0 | 1,82 | 0 | 3,19 | 0 |
| RXRA::VDR | GGTACTTTTGGTTAA | -1 | 731 | 746 | -269 | -254 | -3,66 | 0,01 | 14,71 | >0.1 | 5,54 | 5,66 | 0 | 1,25 | 0 | 5,92 | 0 |
| RXRA::VDR | TGGTTAATATTTTCC | -1 | 723 | 738 | -277 | -262 | -2,11 | 0,01 | 15,61 | >0.1 | 5,54 | 5,63 | 0 | 1,22 | 0 | 5,32 | 0 |
| RXRA::VDR | AGGTCTAGAACTAGG | -1 | 655 | 670 | -345 | -330 | -2,09 | 0,01 | 12,06 | >0.1 | 3,98 | 5,39 | 0 | 1,09 | 0 | 3,7 | 0 |
| RXRA::VDR | GGATCATGAAGATAA | -1 | 641 | 656 | -359 | -344 | 2,74 | 0,01 | 16,67 | >0.1 | 3,83 | 5,34 | 0 | 1,07 | 0 | 3,7 | 0 |
| RXRA::VDR | TGGGCAACATTTTCC | -1 | 533 | 548 | -467 | -452 | -2,42 | 0,01 | 8,34 | >0.1 | 3,11 | 5,05 | 0 | 0,99 | 0 | 1,61 | 0 |
| RXRA::VDR | TCTTTATGGAATTCA | -1 | 492 | 507 | -508 | -493 | -1,53 | 0,01 | 13,79 | >0.1 | 2,94 | 4,96 | 0 | 0,99 | 0 | 6,44 | 0 |
| RXRA::VDR | TGCCTCCCAGGTTCA | -1 | 375 | 390 | -625 | -610 | -3,39 | 0,01 | 11,37 | >0.1 | 2,56 | 4,73 | 0 | 1,07 | 0 | 6,4 | 0 |
| RXRA::VDR | GGTTCACGCCATTCT | -1 | 366 | 381 | -634 | -619 | -1,93 | 0,01 | 13,13 | >0.1 | 2,54 | 4,72 | 0 | 1,08 | 0 | 6,73 | 0 |
| RXRA::VDR | GGGACTACAGGTGCC | -1 | 327 | 342 | -673 | -658 | 3,57 | 0,001 | 17,42 | >0.1 | 2,6 | 4,65 | 0 | 1,15 | 0 | 5,46 | 0 |
| RXRA::VDR | GTTTCACCGTGTTAG | -1 | 267 | 282 | -733 | -718 | 1,8 | 0,01 | 15,53 | >0.1 | 2,78 | 4,56 | 0 | 1,31 | 0 | 5,09 | 0 |
| RXRA::VDR | AGTGCTAGGATTACA | -1 | 199 | 214 | -801 | -786 | -2,06 | 0,01 | 11,68 | >0.1 | 3,03 | 4,47 | 0 | 1,64 | 0 | 4,59 | 0 |
| RXRA::VDR | GGATTACAGGCGTGA | -1 | 192 | 207 | -808 | -793 | -3,42 | 0,01 | 10,06 | >0.1 | 3,06 | 4,46 | 0 | 1,69 | 0 | 4,26 | 0 |
| RXRA::VDR | ACTTAAAAAAGCTCA | -1 | 69 | 84 | -931 | -916 | -2,02 | 0,01 | 8,05 | >0.1 | 3,87 | 4,32 | 0 | 1,88 | 0 | 0 | 0 |
| RXRA::VDR | ACTTCTTTGAGTGCC | -1 | 23 | 38 | -977 | -962 | -1,07 | 0,01 | 10,62 | >0.1 | 5,54 | 4,28 | 0 | 1,87 | 0 | 0 | 0 |
| TFEC | GGCAGATCAG | 1 | 92 | 102 | -908 | -898 | 0,23 | 0,01 | 16,09 | >0.1 | 3,68 | 4,34 | 0 | 1,88 | 0 | 0,83 | 5,13 |
| TFEC | ATCAGGAGAG | 1 | 97 | 107 | -903 | -893 | 7,76 | 0,001 | 23,52 | >0.1 | 3,64 | 4,35 | 0 | 1,88 | 0 | 0,77 | 5,13 |
| TFEC | GGCTCACGCC | 1 | 189 | 199 | -811 | -801 | 2,26 | 0,01 | 20,68 | >0.1 | 3,09 | 4,45 | 0 | 1,71 | 0 | 4,04 | 5,13 |
| TFEC | GGCAGATCAC | 1 | 230 | 240 | -770 | -760 | 0,97 | 0,01 | 19,98 | >0.1 | 2,92 | 4,51 | 0 | 1,45 | 0 | 5,01 | 5,13 |
| TFEC | ATCACAAGGT | 1 | 235 | 245 | -765 | -755 | 7,93 | 0,001 | 26,64 | >0.1 | 2,9 | 4,51 | 0 | 1,43 | 0 | 4,74 | 5,13 |
| TFEC | GTCAGGAGAT | 1 | 243 | 253 | -757 | -747 | 7,83 | 0,001 | 26,07 | >0.1 | 2,87 | 4,52 | 0 | 1,39 | 0 | 4,33 | 5,13 |
| TFEC | CTAACACGGT | 1 | 267 | 277 | -733 | -723 | 0,59 | 0,01 | 18,89 | >0.1 | 2,78 | 4,56 | 0 | 1,3 | 0 | 4,53 | 5,13 |
| TFEC | GGAATATGAT | 1 | 563 | 573 | -437 | -427 | 2,71 | 0,01 | 17,19 | >0.1 | 3,25 | 5,11 | 0 | 0,99 | 0 | 0 | 5,13 |
| TFEC | ATAAAGTGAC | 1 | 761 | 771 | -239 | -229 | 1,96 | 0,01 | 25,9 | >0.1 | 5,61 | 5,77 | 0 | 1,32 | 0 | 6,1 | 5,13 |
| TFEC | GTGACAGGAG | 1 | 766 | 776 | -234 | -224 | 0,38 | 0,01 | 24,17 | >0.1 | 5,61 | 5,8 | 0 | 1,34 | 0 | 5,92 | 5,13 |
| TFEC | GACAGGAGAG | 1 | 768 | 778 | -232 | -222 | 6,06 | 0,01 | 29,87 | >0.1 | 5,61 | 5,81 | 0 | 1,35 | 0 | 5,92 | 5,13 |
| TFEC | CTCAGCAGAT | 1 | 887 | 897 | -113 | -103 | 1,5 | 0,01 | 23,79 | >0.1 | 5,61 | 6,59 | 0 | 1,89 | 0 | 3,07 | 5,13 |
| TFEC | ATCTCATGAG | 1 | 982 | 992 | -18 | -8 | 6,74 | 0,01 | 29,31 | >0.1 | 5,61 | 8,43 | 0 | 1,87 | 0 | 1,53 | 5,13 |
| TFEC | CTCACAGGGG | 1 | 1036 | 1046 | 36 | 46 | 5,86 | 0,01 | 27,78 | >0.1 | 5,61 | 7,99 | 0 | 1,86 | 0 | 1,34 | 5,13 |
| TFEC | CACAGGGGAC | 1 | 1038 | 1048 | 38 | 48 | 5,08 | 0,01 | 26,93 | >0.1 | 5,61 | 7,92 | 0 | 1,86 | 0 | 1,34 | 5,13 |
| TFEC | GGAACAGGCC | 1 | 1114 | 1124 | 114 | 124 | 0,8 | 0,01 | 25,78 | >0.1 | 5,61 | 6,61 | 0 | 1,85 | 3,22 | 2,56 | 5,13 |
| TFEC | GCAACAAGGC | -1 | 1077 | 1087 | 77 | 87 | 0,77 | 0,01 | 24,43 | >0.1 | 5,61 | 7,06 | 0 | 1,85 | 1,79 | 2,21 | 5,13 |
| TFEC | CCCCTGTGAG | -1 | 1036 | 1046 | 36 | 46 | 0,57 | 0,01 | 22,49 | >0.1 | 5,61 | 7,99 | 0 | 1,86 | 0 | 1,34 | 5,13 |
| TFEC | CTCATGAGAT | -1 | 982 | 992 | -18 | -8 | 6,76 | 0,01 | 29,33 | >0.1 | 5,61 | 8,43 | 0 | 1,87 | 0 | 1,53 | 5,13 |
| TFEC | AATACATGAC | -1 | 517 | 527 | -483 | -473 | 2,85 | 0,01 | 19,94 | >0.1 | 3,03 | 5 | 0 | 0,98 | 0 | 2,95 | 5,13 |
| TFEC | TACAGGTGCC | -1 | 327 | 337 | -673 | -663 | 0,51 | 0,01 | 19,6 | >0.1 | 2,6 | 4,65 | 0 | 1,14 | 0 | 5,56 | 5,13 |
| TFEC | ATCTCCTGAC | -1 | 243 | 253 | -757 | -747 | 3,83 | 0,01 | 22,07 | >0.1 | 2,87 | 4,52 | 0 | 1,39 | 0 | 4,33 | 5,13 |
| TFEC | ACCTTGTGAT | -1 | 235 | 245 | -765 | -755 | 4,26 | 0,01 | 22,97 | >0.1 | 2,9 | 4,51 | 0 | 1,43 | 0 | 4,74 | 5,13 |
| TFEC | CTCTCCTGAT | -1 | 97 | 107 | -903 | -893 | 1,9 | 0,01 | 17,66 | >0.1 | 3,64 | 4,35 | 0 | 1,88 | 0 | 0,77 | 5,13 |
| GSX2 | GGAAATTAAA | 1 | 153 | 163 | -847 | -837 | 6,96 | 0,001 | 18,18 | >0.1 | 3,27 | 4,41 | 0 | 1,9 | 0 | 1,65 | 0 |
| GSX2 | CTCTATTAAA | 1 | 287 | 297 | -713 | -703 | 6,67 | 0,01 | 20,41 | >0.1 | 2,72 | 4,59 | 0 | 1,24 | 0 | 5,19 | 0 |

| | | | | | | | | | | | | | | | | | |
|---|---|---|---|---|---|---|---|---|---|---|---|---|---|---|---|---|---|
| GSX2 | ACCTATTAAT | 1 | 599 | 609 | -401 | -391 | 7,18 | 0,001 | 18,23 | >0.1 | 3,47 | 5,21 | 0 | 1,02 | 0 | 1,36 | 0 |
| GSX2 | GGCCATAAAG | 1 | 757 | 767 | -243 | -233 | 4,68 | 0,01 | 23,46 | >0.1 | 5,61 | 5,75 | 0 | 1,31 | 0 | 6,1 | 0 |
| GSX2 | AGCAGTTACA | -1 | 1087 | 1097 | 87 | 97 | 4,93 | 0,01 | 23,32 | >0.1 | 5,61 | 6,92 | 0 | 1,85 | 1,79 | 2,21 | 0 |
| GSX2 | ATCAAATAAG | -1 | 811 | 821 | -189 | -179 | 3,9 | 0,01 | 20,53 | >0.1 | 5,61 | 6,03 | 0 | 1,56 | 0 | 3,43 | 0 |
| GSX2 | CCAAATCAAA | -1 | 749 | 759 | -251 | -241 | 4,95 | 0,01 | 23,48 | >0.1 | 5,61 | 5,72 | 0 | 1,28 | 0 | 5,92 | 0 |
| GSX2 | TTTCATTAAT | -1 | 603 | 613 | -397 | -387 | 6,76 | 0,01 | 17,9 | >0.1 | 3,49 | 5,22 | 0 | 1,02 | 0 | 1,41 | 0 |
| GSX2 | TCCAAATACA | -1 | 521 | 531 | -479 | -469 | 4,86 | 0,01 | 16,97 | >0.1 | 3,05 | 5,01 | 0 | 0,98 | 0 | 3,07 | 0 |
| GSX2 | GCTCATTTCC | -1 | 64 | 74 | -936 | -926 | 4,94 | 0,01 | 15,08 | >0.1 | 3,95 | 4,31 | 0 | 1,88 | 0 | 0 | 0 |
| SOX11 | CACAGAACCAGTGTG | 1 | 42 | 57 | -958 | -943 | -22,49 | 0,01 | -12,11 | >0.1 | 4,21 | 4,3 | 0 | 1,87 | 0 | 0 | 0 |
| SOX11 | AGAGCTTGCAGTGAG | 1 | 389 | 404 | -611 | -596 | -30,11 | 0,01 | -15,68 | >0.1 | 2,6 | 4,76 | 0 | 1,05 | 0 | 6,02 | 0 |
| SOX11 | GACAAGGGCAAAGTC | 1 | 505 | 520 | -495 | -480 | -28,66 | 0,01 | -15,5 | >0.1 | 2,99 | 4,99 | 0 | 0,99 | 0 | 4,19 | 0 |
| SOX11 | CCCAAGTAGAGAGTT | 1 | 544 | 559 | -456 | -441 | -27,62 | 0,01 | -18,39 | >0.1 | 3,17 | 5,08 | 0 | 0,99 | 0 | 0 | 0 |
| SOX11 | ATTAATGAAATTGTC | 1 | 603 | 618 | -397 | -382 | -25,07 | 0,01 | -13,89 | >0.1 | 3,51 | 5,23 | 0 | 1,03 | 0 | 1,41 | 0 |
| SOX11 | AACACCAACATTATC | 1 | 631 | 646 | -369 | -354 | -24,08 | 0,01 | -12,37 | >0.1 | 3,73 | 5,31 | 0 | 1,05 | 0 | 1,61 | 0 |
| SOX11 | AATTATAACATTTTC | 1 | 685 | 700 | -315 | -300 | -25,57 | 0,01 | -11,55 | >0.1 | 4,45 | 5,49 | 0 | 1,14 | 0 | 2,95 | 0 |
| SOX11 | AAAAGTACCGGTTTT | 1 | 737 | 752 | -263 | -248 | -28,68 | 0,01 | -10,27 | >0.1 | 5,54 | 5,69 | 0 | 1,26 | 0 | 5,92 | 0 |
| SOX11 | ATTGTTTACTGTGTT | 1 | 895 | 910 | -105 | -90 | -29,11 | 0,01 | -12,1 | >0.1 | 5,54 | 6,71 | 0 | 1,89 | 0 | 2,86 | 0 |
| SOX11 | AGCAAGTGAACTTTG | -1 | 1172 | 1187 | 172 | 187 | -31,1 | 0,01 | -12,77 | >0.1 | 5,54 | 6,14 | 0 | 1,65 | 1,79 | 3,2 | 0 |
| SOX11 | AAAAATGTCTTGGCC | -1 | 1120 | 1135 | 120 | 135 | -27,33 | 0,01 | -7,43 | >0.1 | 5,54 | 6,55 | 0 | 1,85 | 3,22 | 2,74 | 0 |
| SOX11 | TATAATTTTACAGTG | -1 | 677 | 692 | -323 | -308 | -29,53 | 0,01 | -15,47 | >0.1 | 4,29 | 5,46 | 0 | 1,12 | 0 | 3,19 | 0 |
| SOX11 | GATAATGTTGGTGTT | -1 | 631 | 646 | -369 | -354 | -22,66 | 0,01 | -10,95 | >0.1 | 3,73 | 5,31 | 0 | 1,05 | 0 | 1,61 | 0 |
| SOX11 | GACAATTTCATTAAT | -1 | 603 | 618 | -397 | -382 | -6,71 | 0,001 | 4,47 | >0.1 | 3,51 | 5,23 | 0 | 1,03 | 0 | 1,41 | 0 |
| SOX11 | AACATTTTCCCTTCC | -1 | 528 | 543 | -472 | -457 | -29,17 | 0,01 | -16,36 | >0.1 | 3,09 | 5,04 | 0 | 0,99 | 0 | 3,7 | 0 |
| SOX11 | ATCAGTTTTTTTTTT | -1 | 469 | 484 | -531 | -516 | -25,77 | 0,01 | -12,07 | >0.1 | 2,85 | 4,91 | 0 | 0,99 | 0 | 4,95 | 0 |
| SOX11 | ATCAGTTTTAATTTC | -1 | 154 | 169 | -846 | -831 | -28,08 | 0,01 | -15,99 | >0.1 | 3,25 | 4,42 | 0 | 1,9 | 0 | 2,53 | 0 |
| SOX11 | AAAAAGCTCATTTCC | -1 | 64 | 79 | -936 | -921 | -28,7 | 0,01 | -18,58 | >0.1 | 3,92 | 4,32 | 0 | 1,88 | 0 | 0 | 0 |
| GSX1 | GGAAATTAAA | 1 | 153 | 163 | -847 | -837 | 5,71 | 0,01 | 16,93 | >0.1 | 3,27 | 4,41 | 0 | 1,9 | 0 | 1,65 | 0 |
| GSX1 | CCTGACTAAC | 1 | 262 | 272 | -738 | -728 | 3,92 | 0,01 | 17,12 | >0.1 | 2,8 | 4,55 | 0 | 1,32 | 0 | 4,53 | 0 |
| GSX1 | CTCTATTAAA | 1 | 287 | 297 | -713 | -703 | 6,14 | 0,01 | 19,88 | >0.1 | 2,72 | 4,59 | 0 | 1,24 | 0 | 5,19 | 0 |
| GSX1 | ACCTATTAAT | 1 | 599 | 609 | -401 | -391 | 6,99 | 0,001 | 18,04 | >0.1 | 3,47 | 5,21 | 0 | 1,02 | 0 | 1,36 | 0 |
| GSX1 | CAACATTATC | 1 | 636 | 646 | -364 | -354 | 3,93 | 0,01 | 15,71 | >0.1 | 3,76 | 5,31 | 0 | 1,05 | 0 | 1,66 | 0 |
| GSX1 | GGCCATAAAG | 1 | 757 | 767 | -243 | -233 | 3,91 | 0,01 | 22,69 | >0.1 | 5,61 | 5,75 | 0 | 1,31 | 0 | 6,1 | 0 |
| GSX1 | AGCAGTTACA | -1 | 1087 | 1097 | 87 | 97 | 5,11 | 0,01 | 23,5 | >0.1 | 5,61 | 6,92 | 0 | 1,85 | 1,79 | 2,21 | 0 |
| GSX1 | TCTTATCAGC | -1 | 966 | 976 | -34 | -24 | 4,25 | 0,01 | 21,21 | >0.1 | 5,61 | 7,78 | 0 | 1,87 | 0 | 1,7 | 0 |
| GSX1 | CCAAATCAAA | -1 | 749 | 759 | -251 | -241 | 4,69 | 0,01 | 23,22 | >0.1 | 5,61 | 5,72 | 0 | 1,28 | 0 | 5,92 | 0 |
| GSX1 | TTTCATTAAT | -1 | 603 | 613 | -397 | -387 | 6,75 | 0,01 | 17,89 | >0.1 | 3,49 | 5,22 | 0 | 1,02 | 0 | 1,41 | 0 |
| GSX1 | TCCAAATACA | -1 | 521 | 531 | -479 | -469 | 4,83 | 0,01 | 16,94 | >0.1 | 3,05 | 5,01 | 0 | 0,98 | 0 | 3,07 | 0 |
| GSX1 | AGCCATTTCT | -1 | 170 | 180 | -830 | -820 | 3,75 | 0,01 | 16,66 | >0.1 | 3,18 | 4,43 | 0 | 1,88 | 0 | 3,43 | 0 |
| GSX1 | TTTAATTTCC | -1 | 153 | 163 | -847 | -837 | 3,79 | 0,01 | 15,01 | >0.1 | 3,27 | 4,41 | 0 | 1,9 | 0 | 1,65 | 0 |
| GSX1 | GCTCATTTCC | -1 | 64 | 74 | -936 | -926 | 5,29 | 0,01 | 15,43 | >0.1 | 3,95 | 4,31 | 0 | 1,88 | 0 | 0 | 0 |
| GSX1 | TTCACTTACC | -1 | 4 | 14 | -996 | -986 | 4,12 | 0,01 | 17,64 | >0.1 | 5,61 | 4,26 | 0 | 1,86 | 1,79 | 0 | 0 |
| SOX15 | CCACTGTGCT | 1 | 415 | 425 | -585 | -575 | 3,15 | 0,01 | 16,72 | >0.1 | 2,67 | 4,8 | 0 | 1,02 | 0 | 5,09 | 0 |
| SOX15 | AAATTGTCTG | 1 | 610 | 620 | -390 | -380 | 3,11 | 0,01 | 14,46 | >0.1 | 3,54 | 5,24 | 0 | 1,02 | 0 | 1,55 | 0 |
| SOX15 | ACATTATCTT | 1 | 638 | 648 | -362 | -352 | 3,22 | 0,01 | 14,98 | >0.1 | 3,77 | 5,32 | 0 | 1,05 | 0 | 1,61 | 0 |
| SOX15 | GTTTTGATTT | 1 | 747 | 757 | -253 | -243 | 4,41 | 0,01 | 23,11 | >0.1 | 5,61 | 5,71 | 0 | 1,28 | 0 | 6,1 | 0 |
| SOX15 | GTATTCTTAT | 1 | 806 | 816 | -194 | -184 | 4,19 | 0,01 | 20,61 | >0.1 | 5,61 | 6 | 0 | 1,53 | 0 | 3,28 | 0 |
| SOX15 | AGATTGTTTA | 1 | 893 | 903 | -107 | -97 | 3,34 | 0,01 | 20,34 | >0.1 | 5,61 | 6,64 | 0 | 1,89 | 0 | 2,86 | 0 |
| SOX15 | TTTTTCTTTT | 1 | 917 | 927 | -83 | -73 | 4,71 | 0,01 | 20,22 | >0.1 | 5,61 | 6,89 | 0 | 1,89 | 0 | 1,12 | 0 |
| SOX15 | CCATTGAGGA | 1 | 1107 | 1117 | 107 | 117 | 3,69 | 0,01 | 23,52 | >0.1 | 5,61 | 6,68 | 0 | 1,85 | 3,22 | 2,47 | 0 |
| SOX15 | CCTTTGAACT | -1 | 957 | 967 | -43 | -33 | 4,82 | 0,01 | 21,59 | >0.1 | 5,61 | 7,55 | 0 | 1,88 | 0 | 1,74 | 0 |
| SOX15 | TTATTCTTTA | -1 | 795 | 805 | -205 | -195 | 4,76 | 0,01 | 23,1 | >0.1 | 5,61 | 5,94 | 0 | 1,47 | 0 | 5,32 | 0 |
| SOX15 | CTTTTGGTTA | -1 | 732 | 742 | -268 | -258 | 3,15 | 0,01 | 21,55 | >0.1 | 5,61 | 5,65 | 0 | 1,23 | 0 | 5,92 | 0 |
| SOX15 | TGTTTGTATG | -1 | 625 | 635 | -375 | -365 | 4,05 | 0,01 | 15,57 | >0.1 | 3,66 | 5,28 | 0 | 1,04 | 0 | 1,55 | 0 |
| SOX15 | CATTTATTTT | -1 | 582 | 592 | -418 | -408 | 4,73 | 0,01 | 14,25 | >0.1 | 3,36 | 5,16 | 0 | 1 | 0 | 0 | 0 |
| SOX15 | TTTTTATTTC | -1 | 576 | 586 | -424 | -414 | 5,57 | 0,01 | 15,04 | >0.1 | 3,32 | 5,14 | 0 | 1 | 0 | 0 | 0 |
| SOX15 | CCCTTGTCTT | -1 | 503 | 513 | -497 | -487 | 4,61 | 0,01 | 19,22 | >0.1 | 2,97 | 4,97 | 0 | 0,98 | 0 | 5,68 | 0 |
| SOX15 | TCTTTATGGA | -1 | 497 | 507 | -503 | -493 | 2,75 | 0,01 | 18,07 | >0.1 | 2,95 | 4,96 | 0 | 0,98 | 0 | 6,44 | 0 |
| SOX15 | CCATTCTCCT | -1 | 363 | 373 | -637 | -627 | 6,12 | 0,01 | 21,04 | >0.1 | 2,52 | 4,7 | 0 | 1,08 | 0 | 6,61 | 0 |
| SOX15 | TTTTTGTATT | -1 | 297 | 307 | -703 | -693 | 8,61 | 0,01 | 21,56 | >0.1 | 2,69 | 4,6 | 0 | 1,21 | 0 | 4,45 | 0 |

| | | | | | | | | | | | | | | | | | |
|---|---|---|---|---|---|---|---|---|---|---|---|---|---|---|---|---|---|
| SOX15 | CCATTTTTTA | -1 | 139 | 149 | -861 | -851 | 4 | 0,01 | 15,24 | >0.1 | 3,35 | 4,39 | 0 | 1,89 | 0 | 1,61 | 0 |
| SOX15 | TCTTTGAGTG | -1 | 25 | 35 | -975 | -965 | 2,67 | 0,01 | 14,42 | >0.1 | 5,61 | 4,28 | 0 | 1,87 | 0 | 0 | 0 |
| SOX17 | ACCAGTGTG | 1 | 48 | 57 | -952 | -943 | 4,36 | 0,01 | 14,69 | >0.1 | 4,16 | 4,3 | 0 | 1,87 | 0 | 0 | 0 |
| SOX17 | CACAGTGGC | 1 | 183 | 192 | -817 | -808 | 2,23 | 0,01 | 15,41 | >0.1 | 3,12 | 4,44 | 0 | 1,76 | 0 | 3,86 | 0 |
| SOX17 | CACTTTGGG | 1 | 210 | 219 | -790 | -781 | 3,56 | 0,01 | 17,3 | >0.1 | 3 | 4,48 | 0 | 1,56 | 0 | 4,7 | 0 |
| SOX17 | GCCACTGTG | 1 | 414 | 423 | -586 | -577 | 7,22 | 0,01 | 20,78 | >0.1 | 2,66 | 4,79 | 0 | 1,02 | 0 | 5,09 | 0 |
| SOX17 | GAAATTGTC | 1 | 609 | 618 | -391 | -382 | 8,63 | 0,001 | 19,96 | >0.1 | 3,53 | 5,23 | 0 | 1,02 | 0 | 1,55 | 0 |
| SOX17 | AACATTATC | 1 | 637 | 646 | -363 | -354 | 3,57 | 0,01 | 15,35 | >0.1 | 3,76 | 5,31 | 0 | 1,05 | 0 | 1,66 | 0 |
| SOX17 | CTCTTTGGT | 1 | 668 | 677 | -332 | -323 | 2,24 | 0,01 | 16,68 | >0.1 | 4,11 | 5,41 | 0 | 1,09 | 0 | 3,83 | 0 |
| SOX17 | AACATTTTC | 1 | 691 | 700 | -309 | -300 | 3,57 | 0,01 | 17,66 | >0.1 | 4,52 | 5,49 | 0 | 1,13 | 0 | 2,95 | 0 |
| SOX17 | TATATTGGC | 1 | 879 | 888 | -121 | -112 | 6,99 | 0,01 | 24,09 | >0.1 | 5,62 | 6,51 | 0 | 1,9 | 0 | 3,07 | 0 |
| SOX17 | CAGATTGTT | 1 | 892 | 901 | -108 | -99 | 5,75 | 0,01 | 22,75 | >0.1 | 5,62 | 6,62 | 0 | 1,89 | 0 | 2,86 | 0 |
| SOX17 | TTTACTGTG | 1 | 899 | 908 | -101 | -92 | 5,77 | 0,01 | 22,83 | >0.1 | 5,62 | 6,68 | 0 | 1,89 | 0 | 2,86 | 0 |
| SOX17 | ACCATTGAG | 1 | 1106 | 1115 | 106 | 115 | 3,91 | 0,01 | 23,81 | >0.1 | 5,62 | 6,69 | 0 | 1,85 | 3,22 | 2,52 | 0 |
| SOX17 | GACATCGTC | -1 | 1045 | 1054 | 45 | 54 | 4,85 | 0,01 | 20,66 | >0.1 | 5,62 | 7,7 | 0 | 1,86 | 0 | 0,62 | 0 |
| SOX17 | AATATTTTC | -1 | 724 | 733 | -276 | -267 | 2,09 | 0,01 | 19,7 | >0.1 | 5,62 | 5,61 | 0 | 1,21 | 0 | 5,17 | 0 |
| SOX17 | AACATTTTC | -1 | 534 | 543 | -466 | -457 | 3,57 | 0,01 | 14,3 | >0.1 | 3,1 | 5,04 | 0 | 0,98 | 0 | 1,61 | 0 |
| SOX17 | GCCCTTGTC | -1 | 505 | 514 | -495 | -486 | 5,39 | 0,01 | 20,18 | >0.1 | 2,98 | 4,97 | 0 | 0,98 | 0 | 5,86 | 0 |
| SOX17 | CACAGTGGC | -1 | 414 | 423 | -586 | -577 | 2,23 | 0,01 | 15,79 | >0.1 | 2,66 | 4,79 | 0 | 1,02 | 0 | 5,09 | 0 |
| SOX17 | GCCATTCTC | -1 | 365 | 374 | -635 | -626 | 4,85 | 0,01 | 19,78 | >0.1 | 2,53 | 4,71 | 0 | 1,08 | 0 | 6,61 | 0 |
| SOX17 | GACCTTGTG | -1 | 237 | 246 | -763 | -754 | 4,14 | 0,01 | 17,6 | >0.1 | 2,89 | 4,51 | 0 | 1,42 | 0 | 4,63 | 0 |
| SOX17 | GCCACTGTG | -1 | 183 | 192 | -817 | -808 | 7,22 | 0,01 | 20,4 | >0.1 | 3,12 | 4,44 | 0 | 1,76 | 0 | 3,86 | 0 |
| SOX17 | GCCATTTTT | -1 | 141 | 150 | -859 | -850 | 1,82 | 0,01 | 13,09 | >0.1 | 3,34 | 4,39 | 0 | 1,89 | 0 | 1,65 | 0 |
| SOX17 | CACACTGGT | -1 | 48 | 57 | -952 | -943 | 2,28 | 0,01 | 12,61 | >0.1 | 4,16 | 4,3 | 0 | 1,87 | 0 | 0 | 0 |
| ZIC4 | GTTGACCTGTGGAGT | 1 | 106 | 121 | -894 | -879 | -1,52 | 0,01 | 9,02 | >0.1 | 3,55 | 4,36 | 0 | 1,89 | 0 | 0,75 | 0 |
| ZIC4 | TCACGCCTGTAATCC | 1 | 192 | 207 | -808 | -793 | 0,41 | 0,01 | 13,89 | >0.1 | 3,06 | 4,46 | 0 | 1,69 | 0 | 4,26 | 0 |
| ZIC4 | GAAACCCCGTCTCTA | 1 | 277 | 292 | -723 | -708 | 2,78 | 0,01 | 16,57 | >0.1 | 2,74 | 4,58 | 0 | 1,27 | 0 | 5,19 | 0 |
| ZIC4 | GGGCACCTGTAGTCC | 1 | 326 | 341 | -674 | -659 | 6,62 | 0,001 | 20,58 | >0.1 | 2,6 | 4,65 | 0 | 1,15 | 0 | 5,56 | 0 |
| ZIC4 | CGCGCCACTGTGCTC | 1 | 411 | 426 | -589 | -574 | -0,84 | 0,01 | 12,74 | >0.1 | 2,66 | 4,8 | 0 | 1,03 | 0 | 5,09 | 0 |
| ZIC4 | AGTCCACCATTGAGG | 1 | 1101 | 1116 | 101 | 116 | -1,48 | 0,01 | 18,35 | >0.1 | 5,54 | 6,75 | 0 | 1,85 | 3,22 | 2,47 | 0 |
| ZIC4 | TCGTCCCCTGTGAGC | -1 | 1035 | 1050 | 35 | 50 | -0,65 | 0,01 | 16,12 | >0.1 | 5,54 | 8,03 | 0 | 1,86 | 0 | 1,34 | 0 |
| ZIC4 | CGTCCCCTGTGAGCC | -1 | 1034 | 1049 | 34 | 49 | 3,33 | 0,01 | 20,13 | >0.1 | 5,54 | 8,06 | 0 | 1,86 | 0 | 1,34 | 0 |
| ZIC4 | AACCTCCTCATGAGA | -1 | 983 | 998 | -17 | -2 | 0,18 | 0,01 | 17,83 | >0.1 | 5,54 | 8,66 | 0 | 1,87 | 0 | 1,57 | 0 |
| ZIC4 | AACAATCTGCTGAGC | -1 | 886 | 901 | -114 | -99 | -0,87 | 0,01 | 16,26 | >0.1 | 5,54 | 6,63 | 0 | 1,89 | 0 | 3,07 | 0 |
| ZIC4 | TTCATCCTGGAGAGG | -1 | 861 | 876 | -139 | -124 | -2,35 | 0,01 | 14,18 | >0.1 | 5,54 | 6,41 | 0 | 1,9 | 0 | 2,67 | 0 |
| ZIC4 | CCTCTCCTGTCACTT | -1 | 764 | 779 | -236 | -221 | -1,82 | 0,01 | 16,81 | >0.1 | 5,54 | 5,81 | 0 | 1,36 | 0 | 5,92 | 0 |
| ZIC4 | TGCCTCCCAGGTTCA | -1 | 375 | 390 | -625 | -610 | 0,96 | 0,01 | 15,72 | >0.1 | 2,56 | 4,73 | 0 | 1,07 | 0 | 6,4 | 0 |
| ZIC4 | AGCCTCCCGAGTAGC | -1 | 343 | 358 | -657 | -642 | 0,95 | 0,01 | 15,82 | >0.1 | 2,55 | 4,68 | 0 | 1,12 | 0 | 6,52 | 0 |
| ZIC4 | TGCCCACCACCACGC | -1 | 316 | 331 | -684 | -669 | 3,76 | 0,01 | 18,45 | >0.1 | 2,63 | 4,64 | 0 | 1,17 | 0 | 6,26 | 0 |
| ZIC4 | ACCACCACGCCCAGC | -1 | 311 | 326 | -689 | -674 | -1,13 | 0,01 | 13,45 | >0.1 | 2,64 | 4,63 | 0 | 1,18 | 0 | 6,13 | 0 |
| ZIC4 | CACGCCCAGCTAAGT | -1 | 306 | 321 | -694 | -679 | 3,28 | 0,01 | 17,43 | >0.1 | 2,65 | 4,62 | 0 | 1,19 | 0 | 5,68 | 0 |
| ZIC4 | GGCCTCCCAAAGTGC | -1 | 209 | 224 | -791 | -776 | 8,09 | 0,001 | 21,73 | >0.1 | 2,99 | 4,48 | 0 | 1,57 | 0 | 4,59 | 0 |
| ZIC4 | TGCCTCCTTCTCCTT | -1 | 12 | 27 | -988 | -973 | -1,76 | 0,01 | 9,92 | >0.1 | 5,54 | 4,27 | 0 | 1,87 | 0 | 0 | 0 |
| VSX1 | ATAATTTT | 1 | 132 | 140 | -868 | -860 | 3,25 | 0,01 | 14,49 | >0.1 | 3,4 | 4,38 | 0 | 1,89 | 0 | 1,57 | 0 |
| VSX1 | GAAATTAA | 1 | 154 | 162 | -846 | -838 | 3,23 | 0,01 | 14,45 | >0.1 | 3,27 | 4,41 | 0 | 1,9 | 0 | 1,65 | 0 |
| VSX1 | CTGATCAG | 1 | 164 | 172 | -836 | -828 | 3,22 | 0,01 | 15,15 | >0.1 | 3,21 | 4,42 | 0 | 1,9 | 0 | 2,4 | 0 |
| VSX1 | CTGATCAG | 1 | 479 | 487 | -521 | -513 | 3,22 | 0,01 | 17,36 | >0.1 | 2,88 | 4,92 | 0 | 0,98 | 0 | 5,36 | 0 |
| VSX1 | CCTATTAA | 1 | 600 | 608 | -400 | -392 | 3,8 | 0,01 | 14,85 | >0.1 | 3,47 | 5,2 | 0 | 1,01 | 0 | 1,36 | 0 |
| VSX1 | TTAATGAA | 1 | 604 | 612 | -396 | -388 | 5,52 | 0,01 | 16,66 | >0.1 | 3,49 | 5,21 | 0 | 1,02 | 0 | 1,41 | 0 |
| VSX1 | AAAATTAT | 1 | 683 | 691 | -317 | -309 | 3,67 | 0,01 | 17,65 | >0.1 | 4,34 | 5,46 | 0 | 1,12 | 0 | 3,07 | 0 |
| VSX1 | CTTATTTG | 1 | 811 | 819 | -189 | -181 | 3,08 | 0,01 | 19,57 | >0.1 | 5,64 | 6,02 | 0 | 1,55 | 0 | 3,28 | 0 |
| VSX1 | AAAATTAT | 1 | 830 | 838 | -170 | -162 | 3,67 | 0,01 | 20,4 | >0.1 | 5,64 | 6,14 | 0 | 1,67 | 0 | 3,28 | 0 |
| VSX1 | CTCATGAG | 1 | 984 | 992 | -16 | -8 | 3,77 | 0,01 | 21,29 | >0.1 | 5,64 | 8,43 | 0 | 1,87 | 0 | 1,57 | 0 |
| VSX1 | CTCATGAG | -1 | 984 | 992 | -16 | -8 | 3,77 | 0,01 | 21,29 | >0.1 | 5,64 | 8,43 | 0 | 1,87 | 0 | 1,57 | 0 |
| VSX1 | ATAATTTT | -1 | 830 | 838 | -170 | -162 | 3,25 | 0,01 | 19,98 | >0.1 | 5,64 | 6,14 | 0 | 1,67 | 0 | 3,28 | 0 |
| VSX1 | CAAATAAG | -1 | 811 | 819 | -189 | -181 | 4,2 | 0,01 | 20,69 | >0.1 | 5,64 | 6,02 | 0 | 1,55 | 0 | 3,28 | 0 |
| VSX1 | CAAATCAA | -1 | 750 | 758 | -250 | -242 | 3,48 | 0,01 | 22,03 | >0.1 | 5,64 | 5,71 | 0 | 1,28 | 0 | 5,92 | 0 |
| VSX1 | ATAATTTT | -1 | 683 | 691 | -317 | -309 | 3,25 | 0,01 | 17,23 | >0.1 | 4,34 | 5,46 | 0 | 1,12 | 0 | 3,07 | 0 |
| VSX1 | TTCATTAA | -1 | 604 | 612 | -396 | -388 | 4,87 | 0,01 | 16,01 | >0.1 | 3,49 | 5,21 | 0 | 1,02 | 0 | 1,41 | 0 |

| | | | | | | | | | | | | | | | | | |
|---|---|---|---|---|---|---|---|---|---|---|---|---|---|---|---|---|---|
| VSX1 | TTAATAGG | -1 | 600 | 608 | -400 | -392 | 4,34 | 0,01 | 15,39 | >0.1 | 3,47 | 5,2 | 0 | 1,01 | 0 | 1,36 | 0 |
| VSX1 | CTACTTGG | -1 | 545 | 553 | -455 | -447 | 3,51 | 0,01 | 12,71 | >0.1 | 3,15 | 5,06 | 0 | 0,98 | 0 | 0 | 0 |
| VSX1 | CTGATCAG | -1 | 479 | 487 | -521 | -513 | 3,22 | 0,01 | 17,36 | >0.1 | 2,88 | 4,92 | 0 | 0,98 | 0 | 5,36 | 0 |
| VSX1 | TTAATAGA | -1 | 288 | 296 | -712 | -704 | 3,37 | 0,01 | 17,1 | >0.1 | 2,72 | 4,58 | 0 | 1,24 | 0 | 5,19 | 0 |
| VSX1 | CTGATCAG | -1 | 164 | 172 | -836 | -828 | 3,22 | 0,01 | 15,15 | >0.1 | 3,21 | 4,42 | 0 | 1,9 | 0 | 2,4 | 0 |
| VSX1 | TTAATTTC | -1 | 154 | 162 | -846 | -838 | 6,44 | 0,01 | 17,66 | >0.1 | 3,27 | 4,41 | 0 | 1,9 | 0 | 1,65 | 0 |
| VSX1 | AAAATTAT | -1 | 132 | 140 | -868 | -860 | 3,67 | 0,01 | 14,91 | >0.1 | 3,4 | 4,38 | 0 | 1,89 | 0 | 1,57 | 0 |
| VSX1 | CTCATTTC | -1 | 65 | 73 | -935 | -927 | 3,72 | 0,01 | 13,86 | >0.1 | 3,95 | 4,31 | 0 | 1,88 | 0 | 0 | 0 |
| ZIC3 | GAAACCCCGTCTCTA | 1 | 277 | 292 | -723 | -708 | -0,47 | 0,01 | 13,32 | >0.1 | 2,74 | 4,58 | 0 | 1,27 | 0 | 5,19 | 0 |
| ZIC3 | GGGCACCTGTAGTCC | 1 | 326 | 341 | -674 | -659 | -3,6 | 0,01 | 10,36 | >0.1 | 2,6 | 4,65 | 0 | 1,15 | 0 | 5,56 | 0 |
| ZIC3 | ATCGCGCCACTGTGC | 1 | 409 | 424 | -591 | -576 | -7,01 | 0,01 | 7,24 | >0.1 | 2,66 | 4,79 | 0 | 1,03 | 0 | 5,76 | 0 |
| ZIC3 | TCTTTCCTGCTCAGC | 1 | 932 | 947 | -68 | -53 | -6,27 | 0,01 | 9,22 | >0.1 | 5,54 | 7,17 | 0 | 1,88 | 0 | 0,9 | 0 |
| ZIC3 | AGTCCACCATTGAGG | 1 | 1101 | 1116 | 101 | 116 | -6,66 | 0,01 | 13,17 | >0.1 | 5,54 | 6,75 | 0 | 1,85 | 3,22 | 2,47 | 0 |
| ZIC3 | TCGTCCCCTGTGAGC | -1 | 1035 | 1050 | 35 | 50 | -6,49 | 0,01 | 10,28 | >0.1 | 5,54 | 8,03 | 0 | 1,86 | 0 | 1,34 | 0 |
| ZIC3 | CGTCCCCTGTGAGCC | -1 | 1034 | 1049 | 34 | 49 | -4,41 | 0,01 | 12,39 | >0.1 | 5,54 | 8,06 | 0 | 1,86 | 0 | 1,34 | 0 |
| ZIC3 | AACCTCCTCATGAGA | -1 | 983 | 998 | -17 | -2 | 3,07 | 0,001 | 20,72 | >0.1 | 5,54 | 8,66 | 0 | 1,87 | 0 | 1,57 | 0 |
| ZIC3 | AACAATCTGCTGAGC | -1 | 886 | 901 | -114 | -99 | -0,29 | 0,01 | 16,84 | >0.1 | 5,54 | 6,63 | 0 | 1,89 | 0 | 3,07 | 0 |
| ZIC3 | CCTCTCCTGTCACTT | -1 | 764 | 779 | -236 | -221 | -5,05 | 0,01 | 13,58 | >0.1 | 5,54 | 5,81 | 0 | 1,36 | 0 | 5,92 | 0 |
| ZIC3 | TGCCTCCCAGGTTCA | -1 | 375 | 390 | -625 | -610 | -4,95 | 0,01 | 9,81 | >0.1 | 2,56 | 4,73 | 0 | 1,07 | 0 | 6,4 | 0 |
| ZIC3 | AGCCTCCCGAGTAGC | -1 | 343 | 358 | -657 | -642 | 1,99 | 0,001 | 16,86 | >0.1 | 2,55 | 4,68 | 0 | 1,12 | 0 | 6,52 | 0 |
| ZIC3 | TGCCCACCACCACGC | -1 | 316 | 331 | -684 | -669 | 0,5 | 0,01 | 15,19 | >0.1 | 2,63 | 4,64 | 0 | 1,17 | 0 | 6,26 | 0 |
| ZIC3 | CACCACGCCCAGCTA | -1 | 309 | 324 | -691 | -676 | -1,79 | 0,01 | 12,7 | >0.1 | 2,65 | 4,63 | 0 | 1,19 | 0 | 6,03 | 0 |
| ZIC3 | CACGCCCAGCTAAGT | -1 | 306 | 321 | -694 | -679 | -1,01 | 0,01 | 13,14 | >0.1 | 2,65 | 4,62 | 0 | 1,19 | 0 | 5,68 | 0 |
| ZIC3 | GGCCTCCCAAAGTGC | -1 | 209 | 224 | -791 | -776 | 7,92 | 0,001 | 21,56 | >0.1 | 2,99 | 4,48 | 0 | 1,57 | 0 | 4,59 | 0 |
| ZIC3 | CGTGAGCCACTGTGC | -1 | 182 | 197 | -818 | -803 | -6,63 | 0,01 | 6,65 | >0.1 | 3,11 | 4,45 | 0 | 1,77 | 0 | 3,95 | 0 |
| ZIC3 | TCCACTCCACAGGTC | -1 | 109 | 124 | -891 | -876 | -6,1 | 0,01 | 4,46 | >0.1 | 3,53 | 4,37 | 0 | 1,89 | 0 | 0,79 | 0 |
| ZIC3 | ACTCTCCTGATCTGC | -1 | 93 | 108 | -907 | -892 | -7,01 | 0,01 | 3,66 | >0.1 | 3,65 | 4,35 | 0 | 1,88 | 0 | 0,79 | 0 |
| ZIC3 | TGCCTCCTTCTCCTT | -1 | 12 | 27 | -988 | -973 | -3,95 | 0,01 | 7,73 | >0.1 | 5,54 | 4,27 | 0 | 1,87 | 0 | 0 | 0 |
| NFIL3 | ATAGGTAAGTG | 1 | 1 | 12 | -999 | -988 | 1,81 | 0,01 | 15,32 | >0.1 | 5,6 | 4,26 | 0 | 1,86 | 1,79 | 0 | 0 |
| NFIL3 | TTTTTTAAGTT | 1 | 74 | 85 | -926 | -915 | 1,51 | 0,01 | 11,55 | >0.1 | 3,84 | 4,32 | 0 | 1,88 | 0 | 0 | 0 |
| NFIL3 | TCATGTATTTG | 1 | 518 | 529 | -482 | -471 | -0,79 | 0,01 | 11,3 | >0.1 | 3,04 | 5,01 | 0 | 0,98 | 0 | 3,07 | 0 |
| NFIL3 | TGAGATAACCT | 1 | 592 | 603 | -408 | -397 | 2,88 | 0,01 | 12,5 | >0.1 | 3,42 | 5,19 | 0 | 1,01 | 0 | 0 | 0 |
| NFIL3 | AGAGGTAAGGT | 1 | 774 | 785 | -226 | -215 | 1,49 | 0,01 | 20,41 | >0.1 | 5,6 | 5,84 | 0 | 1,38 | 0 | 6,1 | 0 |
| NFIL3 | TAACGTATTCT | 1 | 802 | 813 | -198 | -187 | 0,3 | 0,01 | 16,82 | >0.1 | 5,6 | 5,99 | 0 | 1,51 | 0 | 3,43 | 0 |
| NFIL3 | TTATTTGATTC | 1 | 812 | 823 | -188 | -177 | -0,65 | 0,01 | 15,99 | >0.1 | 5,6 | 6,05 | 0 | 1,57 | 0 | 3,43 | 0 |
| NFIL3 | TTACATATCTG | 1 | 849 | 860 | -151 | -140 | 2,23 | 0,01 | 18,85 | >0.1 | 5,6 | 6,3 | 0 | 1,86 | 0 | 2,86 | 0 |
| NFIL3 | TTATATTGGCT | 1 | 878 | 889 | -122 | -111 | -0,59 | 0,01 | 16,49 | >0.1 | 5,6 | 6,52 | 0 | 1,9 | 0 | 3,07 | 0 |
| NFIL3 | GGATGTGATCT | 1 | 1022 | 1033 | 22 | 33 | -1,21 | 0,01 | 16,21 | >0.1 | 5,6 | 8,68 | 0 | 1,86 | 0 | 1,28 | 0 |
| NFIL3 | TGCTGTAACTG | 1 | 1084 | 1095 | 84 | 95 | 4,11 | 0,01 | 22,47 | >0.1 | 5,6 | 6,96 | 0 | 1,85 | 1,79 | 2,16 | 0 |
| NFIL3 | TGATAGAACAG | -1 | 1163 | 1174 | 163 | 174 | 1,06 | 0,01 | 17,98 | >0.1 | 4,47 | 6,2 | 0 | 1,73 | 1,79 | 2,72 | 0 |
| NFIL3 | ATATGTAACAG | -1 | 846 | 857 | -154 | -143 | 9,75 | 0,0001 | 26,32 | >0.1 | 5,6 | 6,28 | 0 | 1,83 | 0 | 2,86 | 0 |
| NFIL3 | TTTTAGAATAA | -1 | 834 | 845 | -166 | -155 | -1,12 | 0,01 | 15,68 | >0.1 | 5,6 | 6,19 | 0 | 1,73 | 0 | 3,28 | 0 |
| NFIL3 | ATACGTTATTC | -1 | 799 | 810 | -201 | -190 | 1,48 | 0,01 | 18,09 | >0.1 | 5,6 | 5,97 | 0 | 1,5 | 0 | 3,55 | 0 |
| NFIL3 | TTCTTTAATCC | -1 | 791 | 802 | -209 | -198 | 0,22 | 0,01 | 18,52 | >0.1 | 5,6 | 5,93 | 0 | 1,45 | 0 | 5,32 | 0 |
| NFIL3 | TGTTATAATTT | -1 | 684 | 695 | -316 | -305 | 2,67 | 0,01 | 16,6 | >0.1 | 4,39 | 5,47 | 0 | 1,13 | 0 | 2,95 | 0 |
| NFIL3 | TTATCTCACAT | -1 | 589 | 600 | -411 | -400 | 2,05 | 0,01 | 11,65 | >0.1 | 3,4 | 5,18 | 0 | 1,01 | 0 | 0 | 0 |
| NFIL3 | ATACATGACTT | -1 | 515 | 526 | -485 | -474 | -0,37 | 0,01 | 11,5 | >0.1 | 3,02 | 5 | 0 | 0,98 | 0 | 2,86 | 0 |
| NFIL3 | TTATGGAATTC | -1 | 493 | 504 | -507 | -496 | 6,58 | 0,001 | 21,89 | >0.1 | 2,93 | 4,95 | 0 | 0,98 | 0 | 6,44 | 0 |
| NFIL3 | TTTTGTATTTT | -1 | 295 | 306 | -705 | -694 | 2,41 | 0,01 | 15,37 | >0.1 | 2,69 | 4,6 | 0 | 1,22 | 0 | 4,45 | 0 |
| NR1H4 | GAAATGAGCTT | 1 | 65 | 76 | -935 | -924 | 1,22 | 0,01 | 11,34 | >0.1 | 3,93 | 4,31 | 0 | 1,88 | 0 | 0 | 0 |
| NR1H4 | GAGTTGACCTG | 1 | 104 | 115 | -896 | -885 | 3,94 | 0,01 | 14,52 | >0.1 | 3,58 | 4,35 | 0 | 1,89 | 0 | 0,77 | 0 |
| NR1H4 | AAAATGGCCAT | 1 | 142 | 153 | -858 | -847 | 5,2 | 0,01 | 16,47 | >0.1 | 3,33 | 4,4 | 0 | 1,9 | 0 | 1,65 | 0 |
| NR1H4 | GAAATGGCTGG | 1 | 171 | 182 | -829 | -818 | 1,22 | 0,01 | 14,04 | >0.1 | 3,17 | 4,43 | 0 | 1,87 | 0 | 3,34 | 0 |
| NR1H4 | AGAATGGCGTG | 1 | 366 | 377 | -634 | -623 | 1,52 | 0,01 | 16,46 | >0.1 | 2,53 | 4,71 | 0 | 1,08 | 0 | 6,61 | 0 |
| NR1H4 | CGACAGACCGA | 1 | 435 | 446 | -565 | -554 | 1,24 | 0,01 | 15,74 | >0.1 | 2,73 | 4,83 | 0 | 1,01 | 0 | 5,92 | 0 |
| NR1H4 | AATATGATCTT | 1 | 565 | 576 | -435 | -424 | 1,03 | 0,01 | 10,41 | >0.1 | 3,26 | 5,12 | 0 | 1 | 0 | 0 | 0 |
| NR1H4 | GAGATAACCTA | 1 | 593 | 604 | -407 | -396 | 2,92 | 0,01 | 12,55 | >0.1 | 3,43 | 5,19 | 0 | 1,01 | 0 | 0 | 0 |
| NR1H4 | TTAATGAAATT | 1 | 604 | 615 | -396 | -385 | 3,13 | 0,01 | 14,29 | >0.1 | 3,5 | 5,22 | 0 | 1,02 | 0 | 1,41 | 0 |
| NR1H4 | TTCATGATCCC | 1 | 646 | 657 | -354 | -343 | 2,96 | 0,01 | 17,06 | >0.1 | 3,86 | 5,34 | 0 | 1,07 | 0 | 3,83 | 0 |

| | | | | | | | | | | | | | | | | | |
|---|---|---|---|---|---|---|---|---|---|---|---|---|---|---|---|---|---|
| NR1H4 | ATATTAACCAA | 1 | 728 | 739 | -272 | -261 | 1,78 | 0,01 | 19,55 | >0.1 | 5,6 | 5,63 | 0 | 1,22 | 0 | 5,32 | 0 |
| NR1H4 | AAAGTGACAGG | 1 | 763 | 774 | -237 | -226 | 1,71 | 0,01 | 20,34 | >0.1 | 5,6 | 5,79 | 0 | 1,34 | 0 | 5,92 | 0 |
| NR1H4 | AGAATAACGTA | 1 | 798 | 809 | -202 | -191 | 1,14 | 0,01 | 17,74 | >0.1 | 5,6 | 5,97 | 0 | 1,49 | 0 | 3,55 | 0 |
| NR1H4 | AGGATGAACTT | 1 | 868 | 879 | -132 | -121 | 2,74 | 0,01 | 19,54 | >0.1 | 5,6 | 6,44 | 0 | 1,9 | 0 | 2,86 | 0 |
| NR1H4 | TCAGCGCCCAA | 1 | 942 | 953 | -58 | -47 | 2,25 | 0,01 | 17,82 | >0.1 | 5,6 | 7,27 | 0 | 1,88 | 0 | 0,83 | 0 |
| NR1H4 | CCGAAGACCTG | 1 | 1155 | 1166 | 155 | 166 | 2,4 | 0,01 | 19,07 | >0.1 | 4,43 | 6,26 | 0 | 1,81 | 1,79 | 2,38 | 0 |
| NR1H4 | CAAGTGAACTT | -1 | 1174 | 1185 | 174 | 185 | 4,24 | 0,01 | 22,59 | >0.1 | 5,6 | 6,13 | 0 | 1,63 | 1,79 | 3,2 | 0 |
| NR1H4 | TGAATGACTTT | -1 | 1009 | 1020 | 9 | 20 | 7,83 | 0,001 | 25,78 | >0.1 | 5,6 | 9,03 | 0 | 1,86 | 0 | 1,46 | 0 |
| NR1H4 | CCTTTGAACTT | -1 | 956 | 967 | -44 | -33 | 1,32 | 0,01 | 18,04 | >0.1 | 5,6 | 7,55 | 0 | 1,88 | 0 | 1,7 | 0 |
| NR1H4 | GGAAAGACCAA | -1 | 928 | 939 | -72 | -61 | 4,45 | 0,01 | 20 | >0.1 | 5,6 | 7,05 | 0 | 1,88 | 0 | 1,03 | 0 |
| NR1H4 | TTTATGGCCAA | -1 | 755 | 766 | -245 | -234 | 3,11 | 0,01 | 22,08 | >0.1 | 5,6 | 5,75 | 0 | 1,31 | 0 | 6,32 | 0 |
| NR1H4 | ACAGTGACCAA | -1 | 672 | 683 | -328 | -317 | 9,01 | 0,001 | 23,42 | >0.1 | 4,18 | 5,43 | 0 | 1,1 | 0 | 3,7 | 0 |
| NR1H4 | TACATGACTTT | -1 | 514 | 525 | -486 | -475 | 1,97 | 0,01 | 13,92 | >0.1 | 3,02 | 5 | 0 | 0,98 | 0 | 2,95 | 0 |
| NR1H4 | ACTTTGCCCTT | -1 | 508 | 519 | -492 | -481 | 1,49 | 0,01 | 14,82 | >0.1 | 2,99 | 4,98 | 0 | 0,98 | 0 | 4,37 | 0 |
| NR1H4 | CTCCTGACCTT | -1 | 240 | 251 | -760 | -749 | 4,23 | 0,01 | 17,36 | >0.1 | 2,88 | 4,52 | 0 | 1,41 | 0 | 4,33 | 0 |
| NR1H4 | TTAATTTCCAT | -1 | 151 | 162 | -849 | -838 | 1,24 | 0,01 | 12,47 | >0.1 | 3,27 | 4,41 | 0 | 1,9 | 0 | 1,65 | 0 |
| NR1H4 | TCCATGGCCAT | -1 | 145 | 156 | -855 | -844 | 4,1 | 0,01 | 15,31 | >0.1 | 3,31 | 4,4 | 0 | 1,9 | 0 | 1,61 | 0 |
| NR1H4 | TGAGTGCCTCC | -1 | 20 | 31 | -980 | -969 | 1,22 | 0,01 | 12,95 | >0.1 | 5,6 | 4,27 | 0 | 1,87 | 0 | 0 | 0 |
| NR1H4 | TCACTTACCTA | -1 | 2 | 13 | -998 | -987 | 6,09 | 0,01 | 19,6 | >0.1 | 5,6 | 4,26 | 0 | 1,86 | 1,79 | 0 | 0 |
| FOXJ2 | AAAAATAC | 1 | 294 | 302 | -706 | -698 | 1,47 | 0,01 | 14,43 | >0.1 | 2,7 | 4,59 | 0 | 1,22 | 0 | 4,45 | 0 |
| FOXJ2 | AAATACAA | 1 | 296 | 304 | -704 | -696 | 2,62 | 0,01 | 15,57 | >0.1 | 2,69 | 4,6 | 0 | 1,22 | 0 | 4,45 | 0 |
| FOXJ2 | ATAAAGAC | 1 | 500 | 508 | -500 | -492 | -1,68 | 0,01 | 13,84 | >0.1 | 2,96 | 4,96 | 0 | 0,98 | 0 | 6,62 | 0 |
| FOXJ2 | ATAAAAAT | 1 | 579 | 587 | -421 | -413 | -2,36 | 0,01 | 7,12 | >0.1 | 3,33 | 5,15 | 0 | 1 | 0 | 0 | 0 |
| FOXJ2 | AAAAATAA | 1 | 581 | 589 | -419 | -411 | 3,22 | 0,01 | 12,72 | >0.1 | 3,35 | 5,15 | 0 | 1 | 0 | 0 | 0 |
| FOXJ2 | ACAAACAC | 1 | 628 | 636 | -372 | -364 | 5,84 | 0,01 | 17,38 | >0.1 | 3,68 | 5,28 | 0 | 1,04 | 0 | 1,55 | 0 |
| FOXJ2 | ACCAACAT | 1 | 634 | 642 | -366 | -358 | 4,2 | 0,01 | 15,88 | >0.1 | 3,73 | 5,3 | 0 | 1,05 | 0 | 1,61 | 0 |
| FOXJ2 | GAAAATAT | 1 | 724 | 732 | -276 | -268 | 2,01 | 0,01 | 19,63 | >0.1 | 5,64 | 5,61 | 0 | 1,2 | 0 | 5,17 | 0 |
| FOXJ2 | AGCAACAA | -1 | 1080 | 1088 | 80 | 88 | -0,85 | 0,01 | 17,66 | >0.1 | 5,64 | 7,01 | 0 | 1,85 | 1,79 | 2,21 | 0 |
| FOXJ2 | GCCAATAT | -1 | 880 | 888 | -120 | -112 | 0,06 | 0,01 | 17,29 | >0.1 | 5,64 | 6,51 | 0 | 1,9 | 0 | 3,19 | 0 |
| FOXJ2 | GGCAACAT | -1 | 538 | 546 | -462 | -454 | -2,06 | 0,01 | 8,75 | >0.1 | 3,12 | 5,05 | 0 | 0,98 | 0 | 1,66 | 0 |
| FOXJ2 | AAATACAT | -1 | 520 | 528 | -480 | -472 | 0,7 | 0,01 | 12,91 | >0.1 | 3,04 | 5 | 0 | 0,98 | 0 | 3,19 | 0 |
| FOXJ2 | GACTACAG | -1 | 332 | 340 | -668 | -660 | 0,14 | 0,01 | 14,55 | >0.1 | 2,59 | 4,65 | 0 | 1,13 | 0 | 6,03 | 0 |
| FOXJ2 | GTCAACTC | -1 | 104 | 112 | -896 | -888 | -2,49 | 0,01 | 8,1 | >0.1 | 3,59 | 4,35 | 0 | 1,88 | 0 | 0,77 | 0 |
| FOXJ3 | AGGAGGCACTCAAAGAA | 1 | 19 | 36 | -981 | -964 | -0,15 | 0,01 | 11,51 | >0.1 | 5,52 | 4,28 | 0 | 1,87 | 0 | 0 | 0 |
| FOXJ3 | ATTAAAAATACAAAAAC | 1 | 291 | 308 | -709 | -692 | 1,87 | 0,01 | 14,93 | >0.1 | 2,7 | 4,6 | 0 | 1,23 | 0 | 4,53 | 0 |
| FOXJ3 | TAAAAATACAAAAACTT | 1 | 293 | 310 | -707 | -690 | 0,36 | 0,01 | 13,41 | >0.1 | 2,69 | 4,61 | 0 | 1,23 | 0 | 4,53 | 0 |
| FOXJ3 | GAAATAAAAATAAATGT | 1 | 576 | 593 | -424 | -407 | 4,09 | 0,01 | 13,61 | >0.1 | 3,34 | 5,16 | 0 | 1,01 | 0 | 0 | 0 |
| FOXJ3 | GAAAACCATACAAACAC | 1 | 619 | 636 | -381 | -364 | 2,45 | 0,01 | 13,86 | >0.1 | 3,64 | 5,28 | 0 | 1,04 | 0 | 1,44 | 0 |
| FOXJ3 | ACCATACAAACACCAAC | 1 | 623 | 640 | -377 | -360 | 2,65 | 0,01 | 14,21 | >0.1 | 3,67 | 5,29 | 0 | 1,05 | 0 | 1,55 | 0 |
| FOXJ3 | CAAACACCAACATTATC | 1 | 629 | 646 | -371 | -354 | 2,33 | 0,01 | 14,03 | >0.1 | 3,73 | 5,31 | 0 | 1,06 | 0 | 1,61 | 0 |
| FOXJ3 | TCTAGGAAAATATTAAC | 1 | 719 | 736 | -281 | -264 | 1,72 | 0,01 | 19,25 | >0.1 | 5,52 | 5,62 | 0 | 1,22 | 0 | 5,17 | 0 |
| FOXJ3 | ACACAGTAAACAATCTG | -1 | 892 | 909 | -108 | -91 | 13,13 | 0,0001 | 30,19 | >0.1 | 5,52 | 6,7 | 0 | 1,89 | 0 | 2,95 | 0 |
| FOXJ3 | CACAGGTCAACTCTCCT | -1 | 100 | 117 | -900 | -883 | 3,63 | 0,01 | 14,22 | >0.1 | 3,58 | 4,36 | 0 | 1,89 | 0 | 0,77 | 0 |
| HOXB2 | AGTCATAATT | 1 | 128 | 138 | -872 | -862 | 3,54 | 0,01 | 14,8 | >0.1 | 3,41 | 4,38 | 0 | 1,89 | 0 | 1,57 | 0 |
| HOXB2 | GGAAATTAAA | 1 | 153 | 163 | -847 | -837 | 7,3 | 0,001 | 18,52 | >0.1 | 3,27 | 4,41 | 0 | 1,9 | 0 | 1,65 | 0 |
| HOXB2 | ACTGATCAGA | 1 | 163 | 173 | -837 | -827 | 4,52 | 0,01 | 16,46 | >0.1 | 3,21 | 4,42 | 0 | 1,9 | 0 | 2,4 | 0 |
| HOXB2 | ACTGATCAGA | 1 | 478 | 488 | -522 | -512 | 4,52 | 0,01 | 18,66 | >0.1 | 2,88 | 4,92 | 0 | 0,99 | 0 | 5,36 | 0 |
| HOXB2 | ACCTATTAAT | 1 | 599 | 609 | -401 | -391 | 3,47 | 0,01 | 14,52 | >0.1 | 3,47 | 5,21 | 0 | 1,02 | 0 | 1,36 | 0 |
| HOXB2 | CAACATTATC | 1 | 636 | 646 | -364 | -354 | 5,47 | 0,01 | 17,25 | >0.1 | 3,76 | 5,31 | 0 | 1,05 | 0 | 1,66 | 0 |
| HOXB2 | TAAAATTATA | 1 | 682 | 692 | -318 | -308 | 4,48 | 0,01 | 18,46 | >0.1 | 4,34 | 5,46 | 0 | 1,12 | 0 | 3,07 | 0 |
| HOXB2 | GGCCATAAAG | 1 | 757 | 767 | -243 | -233 | 4,94 | 0,01 | 23,72 | >0.1 | 5,61 | 5,75 | 0 | 1,31 | 0 | 6,1 | 0 |
| HOXB2 | AAAAATTATT | 1 | 829 | 839 | -171 | -161 | 4,57 | 0,01 | 21,29 | >0.1 | 5,61 | 6,15 | 0 | 1,68 | 0 | 3,28 | 0 |
| HOXB2 | GCTGATAAGA | 1 | 966 | 976 | -34 | -24 | 3,75 | 0,01 | 20,71 | >0.1 | 5,61 | 7,78 | 0 | 1,87 | 0 | 1,7 | 0 |
| HOXB2 | AGTCATTCAG | 1 | 1011 | 1021 | 11 | 21 | 4,35 | 0,01 | 22,17 | >0.1 | 5,61 | 8,95 | 0 | 1,86 | 0 | 1,39 | 0 |
| HOXB2 | CGTGGTTAAA | -1 | 1142 | 1152 | 142 | 152 | 3,46 | 0,01 | 23,86 | >0.1 | 5,61 | 6,36 | 0 | 1,84 | 3,22 | 3,37 | 0 |
| HOXB2 | AGCAGTTACA | -1 | 1087 | 1097 | 87 | 97 | 4,45 | 0,01 | 22,84 | >0.1 | 5,61 | 6,92 | 0 | 1,85 | 1,79 | 2,21 | 0 |
| HOXB2 | CCAAATCAAA | -1 | 749 | 759 | -251 | -241 | 4,19 | 0,01 | 22,72 | >0.1 | 5,61 | 5,72 | 0 | 1,28 | 0 | 5,92 | 0 |
| HOXB2 | TTTCATTAAT | -1 | 603 | 613 | -397 | -387 | 6,61 | 0,01 | 17,75 | >0.1 | 3,49 | 5,22 | 0 | 1,02 | 0 | 1,41 | 0 |
| HOXB2 | TCTGATCAGT | -1 | 478 | 488 | -522 | -512 | 3,94 | 0,01 | 18,08 | >0.1 | 2,88 | 4,92 | 0 | 0,99 | 0 | 5,36 | 0 |

| | | | | | | | | | | | | | | | | | |
|---|---|---|---|---|---|---|---|---|---|---|---|---|---|---|---|---|---|
| HOXB2 | TCTGATCAGT | -1 | 163 | 173 | -837 | -827 | 3,94 | 0,01 | 15,88 | >0.1 | 3,21 | 4,42 | 0 | 1,9 | 0 | 2,4 | 0 |
| HOXB2 | TTTAATTTCC | -1 | 153 | 163 | -847 | -837 | 3,8 | 0,01 | 15,02 | >0.1 | 3,27 | 4,41 | 0 | 1,9 | 0 | 1,65 | 0 |
| HOXB2 | TAAAATTATG | -1 | 131 | 141 | -869 | -859 | 4,96 | 0,01 | 16,21 | >0.1 | 3,4 | 4,38 | 0 | 1,89 | 0 | 1,57 | 0 |
| HOXB2 | GCTCATTTCC | -1 | 64 | 74 | -936 | -926 | 5,34 | 0,01 | 15,48 | >0.1 | 3,95 | 4,31 | 0 | 1,88 | 0 | 0 | 0 |
| HOXB3 | GGAAATTAAA | 1 | 153 | 163 | -847 | -837 | 7,12 | 0,01 | 18,34 | >0.1 | 3,27 | 4,41 | 0 | 1,9 | 0 | 1,65 | 0 |
| HOXB3 | ACTGATCAGA | 1 | 163 | 173 | -837 | -827 | 3,76 | 0,01 | 15,7 | >0.1 | 3,21 | 4,42 | 0 | 1,9 | 0 | 2,4 | 0 |
| HOXB3 | CTCTATTAAA | 1 | 287 | 297 | -713 | -703 | 3,96 | 0,01 | 17,7 | >0.1 | 2,72 | 4,59 | 0 | 1,24 | 0 | 5,19 | 0 |
| HOXB3 | AAAACTTAGC | 1 | 303 | 313 | -697 | -687 | 3,29 | 0,01 | 16,76 | >0.1 | 2,67 | 4,61 | 0 | 1,2 | 0 | 4,99 | 0 |
| HOXB3 | ACTGATCAGA | 1 | 478 | 488 | -522 | -512 | 3,76 | 0,01 | 17,9 | >0.1 | 2,88 | 4,92 | 0 | 0,99 | 0 | 5,36 | 0 |
| HOXB3 | ACCTATTAAT | 1 | 599 | 609 | -401 | -391 | 5,48 | 0,01 | 16,53 | >0.1 | 3,47 | 5,21 | 0 | 1,02 | 0 | 1,36 | 0 |
| HOXB3 | CAACATTATC | 1 | 636 | 646 | -364 | -354 | 4,76 | 0,01 | 16,54 | >0.1 | 3,76 | 5,31 | 0 | 1,05 | 0 | 1,66 | 0 |
| HOXB3 | TAAAATTATA | 1 | 682 | 692 | -318 | -308 | 3,67 | 0,01 | 17,65 | >0.1 | 4,34 | 5,46 | 0 | 1,12 | 0 | 3,07 | 0 |
| HOXB3 | AAATATTAAC | 1 | 726 | 736 | -274 | -264 | 3,32 | 0,01 | 21,09 | >0.1 | 5,61 | 5,62 | 0 | 1,22 | 0 | 5,32 | 0 |
| HOXB3 | GGCCATAAAG | 1 | 757 | 767 | -243 | -233 | 4,54 | 0,01 | 23,32 | >0.1 | 5,61 | 5,75 | 0 | 1,31 | 0 | 6,1 | 0 |
| HOXB3 | AAAAATTATT | 1 | 829 | 839 | -171 | -161 | 4,16 | 0,01 | 20,88 | >0.1 | 5,61 | 6,15 | 0 | 1,68 | 0 | 3,28 | 0 |
| HOXB3 | AGTCATTCAG | 1 | 1011 | 1021 | 11 | 21 | 3,34 | 0,01 | 21,16 | >0.1 | 5,61 | 8,95 | 0 | 1,86 | 0 | 1,39 | 0 |
| HOXB3 | AGCAGTTACA | -1 | 1087 | 1097 | 87 | 97 | 4,7 | 0,01 | 23,09 | >0.1 | 5,61 | 6,92 | 0 | 1,85 | 1,79 | 2,21 | 0 |
| HOXB3 | TCTTATCAGC | -1 | 966 | 976 | -34 | -24 | 3,09 | 0,01 | 20,05 | >0.1 | 5,61 | 7,78 | 0 | 1,87 | 0 | 1,7 | 0 |
| HOXB3 | CCAAATCAAA | -1 | 749 | 759 | -251 | -241 | 3,4 | 0,01 | 21,93 | >0.1 | 5,61 | 5,72 | 0 | 1,28 | 0 | 5,92 | 0 |
| HOXB3 | TTTCATTAAT | -1 | 603 | 613 | -397 | -387 | 6,88 | 0,01 | 18,02 | >0.1 | 3,49 | 5,22 | 0 | 1,02 | 0 | 1,41 | 0 |
| HOXB3 | AGCCATTTCT | -1 | 170 | 180 | -830 | -820 | 3,71 | 0,01 | 16,62 | >0.1 | 3,18 | 4,43 | 0 | 1,88 | 0 | 3,43 | 0 |
| HOXB3 | TAAAATTATG | -1 | 131 | 141 | -869 | -859 | 4,07 | 0,01 | 15,32 | >0.1 | 3,4 | 4,38 | 0 | 1,89 | 0 | 1,57 | 0 |
| HOXB3 | GCTCATTTCC | -1 | 64 | 74 | -936 | -926 | 4,96 | 0,01 | 15,1 | >0.1 | 3,95 | 4,31 | 0 | 1,88 | 0 | 0 | 0 |
| HOXB3 | TTCACTTACC | -1 | 4 | 14 | -996 | -986 | 3,2 | 0,01 | 16,72 | >0.1 | 5,61 | 4,26 | 0 | 1,86 | 1,79 | 0 | 0 |
| HOXB5 | TAGGAAATGAGCTTTT | 1 | 62 | 78 | -938 | -922 | 3,76 | 0,01 | 13,89 | >0.1 | 3,94 | 4,32 | 0 | 1,88 | 0 | 0 | 0 |
| HOXB5 | ATGGAAATTAAAACTG | 1 | 151 | 167 | -849 | -833 | 3,73 | 0,01 | 14,95 | >0.1 | 3,26 | 4,41 | 0 | 1,9 | 0 | 1,65 | 0 |
| HOXB5 | AAGGTCAGGAGATAGA | 1 | 240 | 256 | -760 | -744 | 0,51 | 0,01 | 13,64 | >0.1 | 2,87 | 4,53 | 0 | 1,41 | 0 | 4,33 | 0 |
| HOXB5 | CTATTAATGAAATTGT | 1 | 601 | 617 | -399 | -383 | 3,76 | 0,01 | 14,93 | >0.1 | 3,5 | 5,23 | 0 | 1,03 | 0 | 1,41 | 0 |
| HOXB5 | AATGAAATTGTCTGAA | 1 | 606 | 622 | -394 | -378 | 1,07 | 0,01 | 12,42 | >0.1 | 3,54 | 5,24 | 0 | 1,03 | 0 | 1,55 | 0 |
| HOXB5 | ATCTTCATGATCCCTA | 1 | 643 | 659 | -357 | -341 | 3,84 | 0,01 | 17,95 | >0.1 | 3,85 | 5,35 | 0 | 1,07 | 0 | 3,83 | 0 |
| HOXB5 | AATCTCATGAGGAGGT | 1 | 981 | 997 | -19 | -3 | 2,13 | 0,01 | 19,7 | >0.1 | 5,53 | 8,64 | 0 | 1,87 | 0 | 1,53 | 0 |
| HOXB5 | CTCCTCATGAGATTTT | -1 | 979 | 995 | -21 | -5 | 0,94 | 0,01 | 18,46 | >0.1 | 5,53 | 8,59 | 0 | 1,87 | 0 | 1,53 | 0 |
| HOXB5 | CAGATATGTAACAGAT | -1 | 844 | 860 | -156 | -140 | 0,19 | 0,01 | 16,74 | >0.1 | 5,53 | 6,3 | 0 | 1,86 | 0 | 2,86 | 0 |
| HOXB5 | AGAATAATTTTTTAAA | -1 | 825 | 841 | -175 | -159 | 1,27 | 0,01 | 18,09 | >0.1 | 5,53 | 6,16 | 0 | 1,7 | 0 | 3,43 | 0 |
| HOXB5 | GGGATCATGAAGATAA | -1 | 641 | 657 | -359 | -343 | 2,28 | 0,01 | 16,22 | >0.1 | 3,83 | 5,34 | 0 | 1,07 | 0 | 3,7 | 0 |
| HOXB5 | CAGACAATTTCATTAA | -1 | 604 | 620 | -396 | -380 | 0,25 | 0,01 | 11,53 | >0.1 | 3,52 | 5,24 | 0 | 1,03 | 0 | 1,5 | 0 |
| HOXB5 | AATTTCATTAATAGGT | -1 | 599 | 615 | -401 | -385 | 4,45 | 0,01 | 15,6 | >0.1 | 3,49 | 5,22 | 0 | 1,02 | 0 | 1,41 | 0 |
| HOXB5 | GTTTTAATTTCCATGG | -1 | 149 | 165 | -851 | -835 | 1,48 | 0,01 | 12,71 | >0.1 | 3,27 | 4,41 | 0 | 1,9 | 0 | 1,65 | 0 |
| HOXB5 | AAGCTCATTTCCTATT | -1 | 60 | 76 | -940 | -924 | 4,87 | 0,01 | 15,02 | >0.1 | 3,96 | 4,32 | 0 | 1,88 | 0 | 0 | 0 |
| HOXC11 | AGTCATAATTT | 1 | 128 | 139 | -872 | -861 | 3,99 | 0,01 | 15,25 | >0.1 | 3,41 | 4,38 | 0 | 1,89 | 0 | 1,57 | 0 |
| HOXC11 | GGAAATTAAAA | 1 | 153 | 164 | -847 | -836 | 0,24 | 0,01 | 11,46 | >0.1 | 3,26 | 4,41 | 0 | 1,9 | 0 | 1,65 | 0 |
| HOXC11 | TGGCGTGAACC | 1 | 370 | 381 | -630 | -619 | 0,6 | 0,01 | 15,2 | >0.1 | 2,54 | 4,72 | 0 | 1,07 | 0 | 6,27 | 0 |
| HOXC11 | TCTCAAAAAAA | 1 | 453 | 464 | -547 | -536 | 1,61 | 0,01 | 15,64 | >0.1 | 2,79 | 4,87 | 0 | 1 | 0 | 5,38 | 0 |
| HOXC11 | GATCAGAAATA | 1 | 481 | 492 | -519 | -508 | -2,92 | 0,01 | 11,53 | >0.1 | 2,89 | 4,93 | 0 | 0,99 | 0 | 5,65 | 0 |
| HOXC11 | AGTCATGTATT | 1 | 516 | 527 | -484 | -473 | -0,73 | 0,01 | 11,23 | >0.1 | 3,03 | 5 | 0 | 0,98 | 0 | 2,95 | 0 |
| HOXC11 | TGAAATAAAAA | 1 | 575 | 586 | -425 | -414 | -0,3 | 0,01 | 9,17 | >0.1 | 3,32 | 5,14 | 0 | 1 | 0 | 0 | 0 |
| HOXC11 | CACTGTAAAAT | 1 | 677 | 688 | -323 | -312 | 1,87 | 0,01 | 16,11 | >0.1 | 4,26 | 5,45 | 0 | 1,11 | 0 | 3,43 | 0 |
| HOXC11 | AATTATAACAT | 1 | 685 | 696 | -315 | -304 | 0,23 | 0,01 | 14,19 | >0.1 | 4,41 | 5,47 | 0 | 1,13 | 0 | 2,95 | 0 |
| HOXC11 | GGCCATAAAGT | 1 | 757 | 768 | -243 | -232 | 5,6 | 0,001 | 24,37 | >0.1 | 5,6 | 5,76 | 0 | 1,31 | 0 | 6,1 | 0 |
| HOXC11 | GATTGTTTACT | 1 | 894 | 905 | -106 | -95 | 0,09 | 0,01 | 17,1 | >0.1 | 5,6 | 6,66 | 0 | 1,89 | 0 | 2,86 | 0 |
| HOXC11 | CGTGGTTAAAC | -1 | 1141 | 1152 | 141 | 152 | 5,36 | 0,001 | 25,79 | >0.1 | 5,6 | 6,37 | 0 | 1,84 | 3,22 | 3,4 | 0 |
| HOXC11 | CCTCATGAGAT | -1 | 982 | 993 | -18 | -7 | -3,12 | 0,01 | 14,37 | >0.1 | 5,6 | 8,5 | 0 | 1,87 | 0 | 1,53 | 0 |
| HOXC11 | GACCAAAAAAA | -1 | 923 | 934 | -77 | -66 | -1,22 | 0,01 | 14,33 | >0.1 | 5,6 | 6,98 | 0 | 1,88 | 0 | 1,09 | 0 |
| HOXC11 | CACAGTAAACA | -1 | 897 | 908 | -103 | -92 | -0,99 | 0,01 | 16,04 | >0.1 | 5,6 | 6,69 | 0 | 1,89 | 0 | 2,86 | 0 |
| HOXC11 | AGCTGTTAAAG | -1 | 707 | 718 | -293 | -282 | 2,66 | 0,01 | 19,79 | >0.1 | 5,6 | 5,55 | 0 | 1,17 | 0 | 4,81 | 0 |
| HOXC11 | TGTTATAATTT | -1 | 684 | 695 | -316 | -305 | -1,9 | 0,01 | 12,03 | >0.1 | 4,39 | 5,47 | 0 | 1,13 | 0 | 2,95 | 0 |
| HOXC11 | GATCATGAAGA | -1 | 644 | 655 | -356 | -345 | 3,54 | 0,01 | 17,48 | >0.1 | 3,84 | 5,34 | 0 | 1,06 | 0 | 3,7 | 0 |
| HOXC11 | GATCATATTCC | -1 | 563 | 574 | -437 | -426 | -2,51 | 0,01 | 6,85 | >0.1 | 3,25 | 5,11 | 0 | 0,99 | 0 | 0 | 0 |
| HOXC11 | ACTTAAAAAAG | -1 | 73 | 84 | -927 | -916 | -3,17 | 0,01 | 6,88 | >0.1 | 3,85 | 4,32 | 0 | 1,88 | 0 | 0 | 0 |

| | | | | | | | | | | | | | | | | | |
|---|---|---|---|---|---|---|---|---|---|---|---|---|---|---|---|---|---|
| CUX1 | AAACTGATCA | 1 | 161 | 171 | -839 | -829 | -1,69 | 0,01 | 10,31 | >0.1 | 3,22 | 4,42 | 0 | 1,9 | 0 | 2,46 | 0 |
| CUX1 | AAACTGATCA | 1 | 476 | 486 | -524 | -514 | -1,69 | 0,01 | 12,28 | >0.1 | 2,87 | 4,91 | 0 | 0,99 | 0 | 5,2 | 0 |
| CUX1 | AATTCCATAA | 1 | 494 | 504 | -506 | -496 | -2,05 | 0,01 | 13,26 | >0.1 | 2,94 | 4,95 | 0 | 0,98 | 0 | 6,44 | 0 |
| CUX1 | AAATAAATGT | 1 | 583 | 593 | -417 | -407 | -1,31 | 0,01 | 8,22 | >0.1 | 3,36 | 5,16 | 0 | 1 | 0 | 0 | 0 |
| CUX1 | TAACCTATTA | 1 | 597 | 607 | -403 | -393 | -1,47 | 0,01 | 9,61 | >0.1 | 3,45 | 5,2 | 0 | 1,01 | 0 | 1,41 | 0 |
| CUX1 | CTATTAATGA | 1 | 601 | 611 | -399 | -389 | 1,62 | 0,01 | 12,74 | >0.1 | 3,48 | 5,21 | 0 | 1,02 | 0 | 1,41 | 0 |
| CUX1 | TGATCCCTAG | 1 | 650 | 660 | -350 | -340 | -1,57 | 0,01 | 12,74 | >0.1 | 3,9 | 5,35 | 0 | 1,07 | 0 | 3,99 | 0 |
| CUX1 | TAACCAAAAG | 1 | 732 | 742 | -268 | -258 | -1,38 | 0,01 | 17,02 | >0.1 | 5,61 | 5,65 | 0 | 1,23 | 0 | 5,92 | 0 |
| CUX1 | TATTTGATTC | 1 | 813 | 823 | -187 | -177 | -0,86 | 0,01 | 15,8 | >0.1 | 5,61 | 6,05 | 0 | 1,57 | 0 | 3,43 | 0 |
| CUX1 | CTATCAAAGT | 1 | 1168 | 1178 | 168 | 178 | -0,34 | 0,01 | 16,72 | >0.1 | 4,58 | 6,17 | 0 | 1,68 | 1,79 | 2,84 | 0 |
| CUX1 | TCCTCAATGG | -1 | 1107 | 1117 | 107 | 117 | 0,88 | 0,01 | 20,71 | >0.1 | 5,61 | 6,68 | 0 | 1,85 | 3,22 | 2,47 | 0 |
| CUX1 | TAAACAATCT | -1 | 893 | 903 | -107 | -97 | 2,63 | 0,01 | 19,63 | >0.1 | 5,61 | 6,64 | 0 | 1,89 | 0 | 2,86 | 0 |
| CUX1 | GAATCAAATA | -1 | 813 | 823 | -187 | -177 | 0,34 | 0,01 | 17 | >0.1 | 5,61 | 6,05 | 0 | 1,57 | 0 | 3,43 | 0 |
| CUX1 | AAATCAAAAC | -1 | 747 | 757 | -253 | -243 | 2,88 | 0,01 | 21,58 | >0.1 | 5,61 | 5,71 | 0 | 1,28 | 0 | 6,1 | 0 |
| CUX1 | TGGTTAATAT | -1 | 728 | 738 | -272 | -262 | -0,65 | 0,01 | 17,13 | >0.1 | 5,61 | 5,63 | 0 | 1,22 | 0 | 5,32 | 0 |
| CUX1 | TCATTAATAG | -1 | 601 | 611 | -399 | -389 | 3,6 | 0,01 | 14,72 | >0.1 | 3,48 | 5,21 | 0 | 1,02 | 0 | 1,41 | 0 |
| CUX1 | GAATTCATAT | -1 | 489 | 499 | -511 | -501 | -1,46 | 0,01 | 13,65 | >0.1 | 2,92 | 4,94 | 0 | 0,98 | 0 | 6,27 | 0 |
| CUX1 | TGATCAGTTT | -1 | 476 | 486 | -524 | -514 | 2,25 | 0,01 | 16,22 | >0.1 | 2,87 | 4,91 | 0 | 0,99 | 0 | 5,2 | 0 |
| CUX1 | TGATCAGTTT | -1 | 161 | 171 | -839 | -829 | 2,25 | 0,01 | 14,25 | >0.1 | 3,22 | 4,42 | 0 | 1,9 | 0 | 2,46 | 0 |
| HOXC13 | AATTTTAAAAA | 1 | 134 | 145 | -866 | -855 | -2,67 | 0,01 | 8,6 | >0.1 | 3,37 | 4,39 | 0 | 1,89 | 0 | 1,61 | 0 |
| HOXC13 | GATCAGAAATG | 1 | 166 | 177 | -834 | -823 | -1,88 | 0,01 | 10,1 | >0.1 | 3,19 | 4,43 | 0 | 1,9 | 0 | 2,46 | 0 |
| HOXC13 | GCCTGTAATCC | 1 | 196 | 207 | -804 | -793 | -3,95 | 0,01 | 9,6 | >0.1 | 3,05 | 4,46 | 0 | 1,66 | 0 | 4,37 | 0 |
| HOXC13 | TCTATTAAAAA | 1 | 288 | 299 | -712 | -701 | 1,23 | 0,01 | 14,86 | >0.1 | 2,71 | 4,59 | 0 | 1,24 | 0 | 5,09 | 0 |
| HOXC13 | TCTCAAAAAAA | 1 | 453 | 464 | -547 | -536 | 0,59 | 0,01 | 14,62 | >0.1 | 2,79 | 4,87 | 0 | 1 | 0 | 5,38 | 0 |
| HOXC13 | GATCAGAAATA | 1 | 481 | 492 | -519 | -508 | -0,97 | 0,01 | 13,48 | >0.1 | 2,89 | 4,93 | 0 | 0,99 | 0 | 5,65 | 0 |
| HOXC13 | TTCCATAAAGA | 1 | 496 | 507 | -504 | -493 | -2,57 | 0,01 | 12,75 | >0.1 | 2,94 | 4,96 | 0 | 0,98 | 0 | 6,44 | 0 |
| HOXC13 | ACCTATTAATG | 1 | 599 | 610 | -401 | -390 | -3,28 | 0,01 | 7,78 | >0.1 | 3,47 | 5,21 | 0 | 1,02 | 0 | 1,36 | 0 |
| HOXC13 | TGTCTGAAAAC | 1 | 614 | 625 | -386 | -375 | -4,17 | 0,01 | 7,19 | >0.1 | 3,58 | 5,25 | 0 | 1,03 | 0 | 1,5 | 0 |
| HOXC13 | CACTGTAAAAT | 1 | 677 | 688 | -323 | -312 | 1,86 | 0,01 | 16,1 | >0.1 | 4,26 | 5,45 | 0 | 1,11 | 0 | 3,43 | 0 |
| HOXC13 | TATCTTTAACA | 1 | 704 | 715 | -296 | -285 | -3,31 | 0,01 | 13,8 | >0.1 | 5,6 | 5,54 | 0 | 1,17 | 0 | 4,81 | 0 |
| HOXC13 | TCTAGGAAAAT | 1 | 719 | 730 | -281 | -270 | 2,43 | 0,01 | 19,81 | >0.1 | 5,6 | 5,6 | 0 | 1,2 | 0 | 4,99 | 0 |
| HOXC13 | GGCCATAAAGT | 1 | 757 | 768 | -243 | -232 | 1,14 | 0,01 | 19,91 | >0.1 | 5,6 | 5,76 | 0 | 1,31 | 0 | 6,1 | 0 |
| HOXC13 | TCACTTTAAAA | 1 | 821 | 832 | -179 | -168 | -3,84 | 0,01 | 13,19 | >0.1 | 5,6 | 6,1 | 0 | 1,63 | 0 | 3,7 | 0 |
| HOXC13 | TATTCTAAAAT | 1 | 835 | 846 | -165 | -154 | -1,37 | 0,01 | 15,44 | >0.1 | 5,6 | 6,2 | 0 | 1,74 | 0 | 3,28 | 0 |
| HOXC13 | GATTGTTTACT | 1 | 894 | 905 | -106 | -95 | -4,52 | 0,01 | 12,49 | >0.1 | 5,6 | 6,66 | 0 | 1,89 | 0 | 2,86 | 0 |
| HOXC13 | CGTGGTTAAAC | -1 | 1141 | 1152 | 141 | 152 | 2,5 | 0,01 | 22,93 | >0.1 | 5,6 | 6,37 | 0 | 1,84 | 3,22 | 3,4 | 0 |
| HOXC13 | CACAGTAAACA | -1 | 897 | 908 | -103 | -92 | -2,74 | 0,01 | 14,29 | >0.1 | 5,6 | 6,69 | 0 | 1,89 | 0 | 2,86 | 0 |
| HOXC13 | GCCAATATAAA | -1 | 877 | 888 | -123 | -112 | -4,35 | 0,01 | 12,6 | >0.1 | 5,6 | 6,51 | 0 | 1,9 | 0 | 2,95 | 0 |
| HOXC13 | TTTGGTTAATA | -1 | 729 | 740 | -271 | -260 | -3,53 | 0,01 | 14,45 | >0.1 | 5,6 | 5,64 | 0 | 1,23 | 0 | 5,52 | 0 |
| HOXC13 | AGCTGTTAAAG | -1 | 707 | 718 | -293 | -282 | -2,28 | 0,01 | 14,85 | >0.1 | 5,6 | 5,55 | 0 | 1,17 | 0 | 4,81 | 0 |
| HOXC13 | GCTGTTAAAGA | -1 | 706 | 717 | -294 | -283 | -1,78 | 0,01 | 15,52 | >0.1 | 5,6 | 5,55 | 0 | 1,17 | 0 | 4,99 | 0 |
| HOXC13 | ACACGGAAAAT | -1 | 694 | 705 | -306 | -295 | -0,62 | 0,01 | 14,69 | >0.1 | 5,6 | 5,5 | 0 | 1,14 | 0 | 3,07 | 0 |
| HOXC13 | GATCATGAAGA | -1 | 644 | 655 | -356 | -345 | -2,92 | 0,01 | 11,02 | >0.1 | 3,84 | 5,34 | 0 | 1,06 | 0 | 3,7 | 0 |
| HOXC13 | TTTCATTAATA | -1 | 602 | 613 | -398 | -387 | 0,6 | 0,01 | 11,74 | >0.1 | 3,49 | 5,22 | 0 | 1,02 | 0 | 1,41 | 0 |
| HOXC13 | TCCCAGAAACT | -1 | 555 | 566 | -445 | -434 | -2,07 | 0,01 | 7,23 | >0.1 | 3,21 | 5,09 | 0 | 0,99 | 0 | 0 | 0 |
| HOXC13 | TTTTTTAAAAT | -1 | 135 | 146 | -865 | -854 | -3,61 | 0,01 | 7,61 | >0.1 | 3,37 | 4,39 | 0 | 1,89 | 0 | 1,57 | 0 |
| HOXC12 | AATAGGAAATG | 1 | 60 | 71 | -940 | -929 | -6,57 | 0,01 | 3,61 | >0.1 | 3,99 | 4,31 | 0 | 1,88 | 0 | 0 | 0 |
| HOXC12 | AGTAGTCATAA | 1 | 125 | 136 | -875 | -864 | -3,42 | 0,01 | 7,97 | >0.1 | 3,43 | 4,38 | 0 | 1,89 | 0 | 1,7 | 0 |
| HOXC12 | AGTCATAATTT | 1 | 128 | 139 | -872 | -861 | 2,08 | 0,001 | 13,34 | >0.1 | 3,41 | 4,38 | 0 | 1,89 | 0 | 1,57 | 0 |
| HOXC12 | AATTTTAAAAA | 1 | 134 | 145 | -866 | -855 | -5,92 | 0,01 | 5,35 | >0.1 | 3,37 | 4,39 | 0 | 1,89 | 0 | 1,61 | 0 |
| HOXC12 | GGAAATTAAAA | 1 | 153 | 164 | -847 | -836 | -2,95 | 0,01 | 8,27 | >0.1 | 3,26 | 4,41 | 0 | 1,9 | 0 | 1,65 | 0 |
| HOXC12 | GATCAGAAATG | 1 | 166 | 177 | -834 | -823 | -4,57 | 0,01 | 7,41 | >0.1 | 3,19 | 4,43 | 0 | 1,9 | 0 | 2,46 | 0 |
| HOXC12 | GCCTGTAATCC | 1 | 196 | 207 | -804 | -793 | -5,07 | 0,01 | 8,48 | >0.1 | 3,05 | 4,46 | 0 | 1,66 | 0 | 4,37 | 0 |
| HOXC12 | TCTCAAAAAAA | 1 | 453 | 464 | -547 | -536 | -6,35 | 0,01 | 7,68 | >0.1 | 2,79 | 4,87 | 0 | 1 | 0 | 5,38 | 0 |
| HOXC12 | GATCAGAAATA | 1 | 481 | 492 | -519 | -508 | -3,12 | 0,01 | 11,33 | >0.1 | 2,89 | 4,93 | 0 | 0,99 | 0 | 5,65 | 0 |
| HOXC12 | TGAAATAAAAA | 1 | 575 | 586 | -425 | -414 | -2,87 | 0,01 | 6,6 | >0.1 | 3,32 | 5,14 | 0 | 1 | 0 | 0 | 0 |
| HOXC12 | CACTGTAAAAT | 1 | 677 | 688 | -323 | -312 | 1,4 | 0,01 | 15,64 | >0.1 | 4,26 | 5,45 | 0 | 1,11 | 0 | 3,43 | 0 |
| HOXC12 | AATTATAACAT | 1 | 685 | 696 | -315 | -304 | -2,99 | 0,01 | 10,97 | >0.1 | 4,41 | 5,47 | 0 | 1,13 | 0 | 2,95 | 0 |
| HOXC12 | TCTAGGAAAAT | 1 | 719 | 730 | -281 | -270 | -4,29 | 0,01 | 13,09 | >0.1 | 5,6 | 5,6 | 0 | 1,2 | 0 | 4,99 | 0 |

| | | | | | | | | | | | | | | | | | |
|---|---|---|---|---|---|---|---|---|---|---|---|---|---|---|---|---|---|
| HOXC12 | TATTCTAAAAT | 1 | 835 | 846 | -165 | -154 | -3,86 | 0,01 | 12,95 | >0.1 | 5,6 | 6,2 | 0 | 1,74 | 0 | 3,28 | 0 |
| HOXC12 | GATTGTTTACT | 1 | 894 | 905 | -106 | -95 | -2,68 | 0,01 | 14,33 | >0.1 | 5,6 | 6,66 | 0 | 1,89 | 0 | 2,86 | 0 |
| HOXC12 | AGCAGTTACAG | -1 | 1086 | 1097 | 86 | 97 | -6,22 | 0,01 | 12,17 | >0.1 | 5,6 | 6,93 | 0 | 1,85 | 1,79 | 2,21 | 0 |
| HOXC12 | CACAGTAAACA | -1 | 897 | 908 | -103 | -92 | -0,47 | 0,01 | 16,56 | >0.1 | 5,6 | 6,69 | 0 | 1,89 | 0 | 2,86 | 0 |
| HOXC12 | AGCTGTTAAAG | -1 | 707 | 718 | -293 | -282 | 1,34 | 0,01 | 18,47 | >0.1 | 5,6 | 5,55 | 0 | 1,17 | 0 | 4,81 | 0 |
| HOXC12 | ACACGGAAAAT | -1 | 694 | 705 | -306 | -295 | -6,15 | 0,01 | 9,16 | >0.1 | 5,6 | 5,5 | 0 | 1,14 | 0 | 3,07 | 0 |
| HOXC12 | TGTTATAATTT | -1 | 684 | 695 | -316 | -305 | -4,94 | 0,01 | 8,99 | >0.1 | 4,39 | 5,47 | 0 | 1,13 | 0 | 2,95 | 0 |
| HOXC12 | GATCATGAAGA | -1 | 644 | 655 | -356 | -345 | -1,44 | 0,01 | 12,5 | >0.1 | 3,84 | 5,34 | 0 | 1,06 | 0 | 3,7 | 0 |
| HOXC12 | GATCATATTCC | -1 | 563 | 574 | -437 | -426 | -5,47 | 0,01 | 3,89 | >0.1 | 3,25 | 5,11 | 0 | 0,99 | 0 | 0 | 0 |
| FOSB::JUNB | AAGGAGGCAC | 1 | 18 | 28 | -982 | -972 | -1,44 | 0,01 | 10,3 | >0.1 | 5,61 | 4,27 | 0 | 1,87 | 0 | 0 | 0 |
| FOSB::JUNB | TGTGAGAAAT | 1 | 53 | 63 | -947 | -937 | 1,97 | 0,01 | 12,23 | >0.1 | 4,09 | 4,3 | 0 | 1,87 | 0 | 0 | 0 |
| FOSB::JUNB | GAGTAGTCAT | 1 | 124 | 134 | -876 | -866 | -0,49 | 0,01 | 10,91 | >0.1 | 3,44 | 4,38 | 0 | 1,89 | 0 | 1,7 | 0 |
| FOSB::JUNB | AGTGGCTCAC | 1 | 186 | 196 | -814 | -804 | 3,22 | 0,01 | 16,36 | >0.1 | 3,1 | 4,45 | 0 | 1,74 | 0 | 3,86 | 0 |
| FOSB::JUNB | CCTGACTAAC | 1 | 262 | 272 | -738 | -728 | 8,56 | 0,001 | 21,76 | >0.1 | 2,8 | 4,55 | 0 | 1,32 | 0 | 4,53 | 0 |
| FOSB::JUNB | GCTGAGGCAG | 1 | 355 | 365 | -645 | -635 | 6,27 | 0,001 | 20,62 | >0.1 | 2,53 | 4,69 | 0 | 1,09 | 0 | 6,03 | 0 |
| FOSB::JUNB | TGGGAGGCAG | 1 | 381 | 391 | -619 | -609 | -1,7 | 0,01 | 12,68 | >0.1 | 2,57 | 4,73 | 0 | 1,06 | 0 | 6,02 | 0 |
| FOSB::JUNB | CGAGACTCAG | 1 | 443 | 453 | -557 | -547 | 2,79 | 0,01 | 17,32 | >0.1 | 2,75 | 4,85 | 0 | 1 | 0 | 5,92 | 0 |
| FOSB::JUNB | TTTGATTCAC | 1 | 815 | 825 | -185 | -175 | 5,33 | 0,001 | 22,01 | >0.1 | 5,61 | 6,06 | 0 | 1,58 | 0 | 3,43 | 0 |
| FOSB::JUNB | ATTGGCTCAG | 1 | 882 | 892 | -118 | -108 | -2,31 | 0,01 | 14,81 | >0.1 | 5,61 | 6,55 | 0 | 1,89 | 0 | 3,07 | 0 |
| FOSB::JUNB | CTTGGCTCAC | 1 | 1031 | 1041 | 31 | 41 | -0,16 | 0,01 | 16,81 | >0.1 | 5,61 | 8,19 | 0 | 1,86 | 0 | 1,31 | 0 |
| FOSB::JUNB | ATTGAGGAAC | 1 | 1109 | 1119 | 109 | 119 | 0,29 | 0,01 | 20,15 | >0.1 | 5,61 | 6,66 | 0 | 1,85 | 3,22 | 2,52 | 0 |
| FOSB::JUNB | TGTGAGCCAA | -1 | 1032 | 1042 | 32 | 42 | 0,21 | 0,01 | 17,1 | >0.1 | 5,61 | 8,15 | 0 | 1,86 | 0 | 1,28 | 0 |
| FOSB::JUNB | GCTGAGCCAA | -1 | 883 | 893 | -117 | -107 | -1,02 | 0,01 | 16,11 | >0.1 | 5,61 | 6,56 | 0 | 1,89 | 0 | 3,07 | 0 |
| FOSB::JUNB | AGTGAATCAA | -1 | 816 | 826 | -184 | -174 | 4,5 | 0,01 | 21,31 | >0.1 | 5,61 | 6,06 | 0 | 1,59 | 0 | 3,55 | 0 |
| FOSB::JUNB | AGTGACCAAA | -1 | 671 | 681 | -329 | -319 | -2,6 | 0,01 | 11,63 | >0.1 | 4,16 | 5,42 | 0 | 1,1 | 0 | 3,55 | 0 |
| FOSB::JUNB | ACTGAGTCTC | -1 | 444 | 454 | -556 | -546 | -0,62 | 0,01 | 13,91 | >0.1 | 2,76 | 4,85 | 0 | 1 | 0 | 5,92 | 0 |
| FOSB::JUNB | CCTGCCTCAG | -1 | 356 | 366 | -644 | -634 | 0,9 | 0,01 | 15,34 | >0.1 | 2,52 | 4,69 | 0 | 1,09 | 0 | 6,13 | 0 |
| FOSB::JUNB | TGTTAGTCAG | -1 | 263 | 273 | -737 | -727 | 5,45 | 0,001 | 18,56 | >0.1 | 2,8 | 4,55 | 0 | 1,32 | 0 | 4,45 | 0 |
| FOSB::JUNB | CGTGAGCCAC | -1 | 187 | 197 | -813 | -803 | 3,44 | 0,01 | 16,57 | >0.1 | 3,1 | 4,45 | 0 | 1,73 | 0 | 3,86 | 0 |
| FOSB::JUNB | TATGACTACT | -1 | 125 | 135 | -875 | -865 | 0,3 | 0,01 | 11,7 | >0.1 | 3,43 | 4,38 | 0 | 1,89 | 0 | 1,7 | 0 |
| FOSB::JUNB | TATTTCTCAC | -1 | 54 | 64 | -946 | -936 | -1,87 | 0,01 | 8,38 | >0.1 | 4,07 | 4,3 | 0 | 1,87 | 0 | 0 | 0 |
| ZEB1 | TTGACCTGTGG | 1 | 107 | 118 | -893 | -882 | 4,81 | 0,01 | 15,38 | >0.1 | 3,56 | 4,36 | 0 | 1,89 | 0 | 0,77 | 0 |
| ZEB1 | CACGCCTGTAA | 1 | 193 | 204 | -807 | -796 | 4,98 | 0,01 | 18,45 | >0.1 | 3,07 | 4,46 | 0 | 1,68 | 0 | 4,26 | 0 |
| ZEB1 | CTTAGCTGGGC | 1 | 307 | 318 | -693 | -682 | 1,67 | 0,01 | 15,81 | >0.1 | 2,66 | 4,62 | 0 | 1,19 | 0 | 5,68 | 0 |
| ZEB1 | GGCACCTGTAG | 1 | 327 | 338 | -673 | -662 | 12,23 | 0,001 | 26,19 | >0.1 | 2,6 | 4,65 | 0 | 1,14 | 0 | 5,56 | 0 |
| ZEB1 | CCCAGCTACTC | 1 | 339 | 350 | -661 | -650 | 1,12 | 0,01 | 16 | >0.1 | 2,57 | 4,67 | 0 | 1,12 | 0 | 6,52 | 0 |
| ZEB1 | TGAACCTGGGA | 1 | 375 | 386 | -625 | -614 | 4,34 | 0,01 | 19,09 | >0.1 | 2,56 | 4,73 | 0 | 1,07 | 0 | 6,4 | 0 |
| ZEB1 | CCAGCCTGGGC | 1 | 425 | 436 | -575 | -564 | 1,56 | 0,01 | 15,76 | >0.1 | 2,7 | 4,82 | 0 | 1,01 | 0 | 5,67 | 0 |
| ZEB1 | AGTACCGGTTT | 1 | 740 | 751 | -260 | -249 | -1,11 | 0,01 | 17,34 | >0.1 | 5,6 | 5,68 | 0 | 1,26 | 0 | 5,92 | 0 |
| ZEB1 | CATATCTGTCC | 1 | 852 | 863 | -148 | -137 | 0,28 | 0,01 | 16,76 | >0.1 | 5,6 | 6,32 | 0 | 1,89 | 0 | 2,67 | 0 |
| ZEB1 | CTCACAGGGGA | 1 | 1036 | 1047 | 36 | 47 | 0,8 | 0,01 | 17,58 | >0.1 | 5,6 | 7,99 | 0 | 1,86 | 0 | 1,34 | 0 |
| ZEB1 | AAGACCTGTTC | 1 | 1158 | 1169 | 158 | 169 | 3,29 | 0,01 | 19,87 | >0.1 | 4,37 | 6,24 | 0 | 1,78 | 1,79 | 2,4 | 0 |
| ZEB1 | TTCACTTGCTT | 1 | 1177 | 1188 | 177 | 188 | 1,25 | 0,01 | 19,62 | >0.1 | 5,6 | 6,11 | 0 | 1,61 | 1,79 | 3,26 | 0 |
| ZEB1 | CTTACCTCTCC | -1 | 772 | 783 | -228 | -217 | 1,37 | 0,01 | 20,08 | >0.1 | 5,6 | 5,83 | 0 | 1,37 | 0 | 5,92 | 0 |
| ZEB1 | AGTAGCTGGGA | -1 | 338 | 349 | -662 | -651 | -1,01 | 0,01 | 13,27 | >0.1 | 2,57 | 4,67 | 0 | 1,12 | 0 | 5,92 | 0 |
| ZEB1 | TACAGGTGCCC | -1 | 326 | 337 | -674 | -663 | 1,28 | 0,01 | 15,13 | >0.1 | 2,6 | 4,65 | 0 | 1,15 | 0 | 5,46 | 0 |
| ZEB1 | GTGATCTGCCC | -1 | 229 | 240 | -771 | -760 | 0,09 | 0,01 | 14,1 | >0.1 | 2,92 | 4,51 | 0 | 1,46 | 0 | 5,12 | 0 |
| ZEB1 | CCCAACTCGGC | -1 | 221 | 232 | -779 | -768 | -1,1 | 0,01 | 13,37 | >0.1 | 2,95 | 4,49 | 0 | 1,5 | 0 | 5,52 | 0 |
| ZEB1 | CTGATCTGCCT | -1 | 91 | 102 | -909 | -898 | -0,64 | 0,01 | 10,12 | >0.1 | 3,68 | 4,34 | 0 | 1,88 | 0 | 0,85 | 0 |
| TFAP4 | ATCAGGAGAG | 1 | 97 | 107 | -903 | -893 | -4,08 | 0,01 | 6,55 | >0.1 | 3,64 | 4,35 | 0 | 1,88 | 0 | 0,77 | 0 |
| TFAP4 | GTCAGGAGAT | 1 | 243 | 253 | -757 | -747 | -4,44 | 0,01 | 8,67 | >0.1 | 2,87 | 4,52 | 0 | 1,39 | 0 | 4,33 | 0 |
| TFAP4 | CTTAGCTGGG | 1 | 307 | 317 | -693 | -683 | -5,26 | 0,01 | 8,88 | >0.1 | 2,66 | 4,62 | 0 | 1,19 | 0 | 5,68 | 0 |
| TFAP4 | GGCACCTGTA | 1 | 327 | 337 | -673 | -663 | 0,3 | 0,01 | 14,26 | >0.1 | 2,6 | 4,65 | 0 | 1,14 | 0 | 5,56 | 0 |
| TFAP4 | CCCAGCTACT | 1 | 339 | 349 | -661 | -651 | -5,03 | 0,01 | 9,85 | >0.1 | 2,57 | 4,66 | 0 | 1,12 | 0 | 6,52 | 0 |
| TFAP4 | AACAGCTTTC | 1 | 711 | 721 | -289 | -279 | 0,22 | 0,01 | 17,56 | >0.1 | 5,61 | 5,56 | 0 | 1,18 | 0 | 4,99 | 0 |
| TFAP4 | GACAGGAGAG | 1 | 768 | 778 | -232 | -222 | -6,15 | 0,01 | 12,53 | >0.1 | 5,61 | 5,81 | 0 | 1,35 | 0 | 5,92 | 0 |
| TFAP4 | AAAATCTGTT | 1 | 841 | 851 | -159 | -149 | -6,35 | 0,01 | 8,69 | >0.1 | 5,61 | 6,24 | 0 | 1,78 | 0 | 1,41 | 0 |
| TFAP4 | CTCAGCAGAT | 1 | 887 | 897 | -113 | -103 | 2,25 | 0,01 | 19,41 | >0.1 | 5,61 | 6,59 | 0 | 1,89 | 0 | 3,07 | 0 |
| TFAP4 | CACAGGGGAC | 1 | 1038 | 1048 | 38 | 48 | -7,02 | 0,01 | 9,7 | >0.1 | 5,61 | 7,92 | 0 | 1,86 | 0 | 1,34 | 0 |

| | | | | | | | | | | | | | | | | | |
|---|---|---|---|---|---|---|---|---|---|---|---|---|---|---|---|---|---|
| TFAP4 | AACTGCTGCT | 1 | 1090 | 1100 | 90 | 100 | -0,77 | 0,01 | 17,64 | >0.1 | 5,61 | 6,88 | 0 | 1,85 | 1,79 | 2,27 | 0 |
| TFAP4 | AAGACCTGTT | 1 | 1158 | 1168 | 158 | 168 | -2,78 | 0,01 | 13,81 | >0.1 | 4,38 | 6,24 | 0 | 1,78 | 1,79 | 2,4 | 0 |
| TFAP4 | TTCACTTGCT | 1 | 1177 | 1187 | 177 | 187 | -6,32 | 0,01 | 12,06 | >0.1 | 5,61 | 6,11 | 0 | 1,61 | 1,79 | 3,26 | 0 |
| TFAP4 | AACAGGTCTT | -1 | 1158 | 1168 | 158 | 168 | -0,98 | 0,01 | 15,61 | >0.1 | 4,38 | 6,24 | 0 | 1,78 | 1,79 | 2,4 | 0 |
| TFAP4 | AGCAGCAGTT | -1 | 1090 | 1100 | 90 | 100 | 0,51 | 0,01 | 18,92 | >0.1 | 5,61 | 6,88 | 0 | 1,85 | 1,79 | 2,27 | 0 |
| TFAP4 | GTCCCCTGTG | -1 | 1038 | 1048 | 38 | 48 | -4,93 | 0,01 | 11,79 | >0.1 | 5,61 | 7,92 | 0 | 1,86 | 0 | 1,34 | 0 |
| TFAP4 | ATCTGCTGAG | -1 | 887 | 897 | -113 | -103 | -0,54 | 0,01 | 16,62 | >0.1 | 5,61 | 6,59 | 0 | 1,89 | 0 | 3,07 | 0 |
| TFAP4 | AACAGATTTT | -1 | 841 | 851 | -159 | -149 | -5,78 | 0,01 | 9,26 | >0.1 | 5,61 | 6,24 | 0 | 1,78 | 0 | 1,41 | 0 |
| TFAP4 | CTCTCCTGTC | -1 | 768 | 778 | -232 | -222 | -7,66 | 0,01 | 11,02 | >0.1 | 5,61 | 5,81 | 0 | 1,35 | 0 | 5,92 | 0 |
| TFAP4 | GAAAGCTGTT | -1 | 711 | 721 | -289 | -279 | -0,79 | 0,01 | 16,55 | >0.1 | 5,61 | 5,56 | 0 | 1,18 | 0 | 4,99 | 0 |
| TFAP4 | AGTAGCTGGG | -1 | 339 | 349 | -661 | -651 | -6,83 | 0,01 | 8,05 | >0.1 | 2,57 | 4,66 | 0 | 1,12 | 0 | 6,52 | 0 |
| TFAP4 | TACAGGTGCC | -1 | 327 | 337 | -673 | -663 | 1,43 | 0,01 | 15,39 | >0.1 | 2,6 | 4,65 | 0 | 1,14 | 0 | 5,56 | 0 |
| TFAP4 | CCCAGCTAAG | -1 | 307 | 317 | -693 | -683 | -4,03 | 0,01 | 10,11 | >0.1 | 2,66 | 4,62 | 0 | 1,19 | 0 | 5,68 | 0 |
| TFAP4 | ATCTCCTGAC | -1 | 243 | 253 | -757 | -747 | -5,21 | 0,01 | 7,9 | >0.1 | 2,87 | 4,52 | 0 | 1,39 | 0 | 4,33 | 0 |
| TFAP4 | TACAGGCGTG | -1 | 193 | 203 | -807 | -797 | -6,88 | 0,01 | 6,59 | >0.1 | 3,07 | 4,46 | 0 | 1,68 | 0 | 4,26 | 0 |
| TFAP4 | CACAGGTCAA | -1 | 107 | 117 | -893 | -883 | -6,61 | 0,01 | 3,96 | >0.1 | 3,56 | 4,36 | 0 | 1,89 | 0 | 0,77 | 0 |
| TFAP4 | CTCTCCTGAT | -1 | 97 | 107 | -903 | -893 | -5,66 | 0,01 | 4,97 | >0.1 | 3,64 | 4,35 | 0 | 1,88 | 0 | 0,77 | 0 |
| SPDEF | AGAAGGAGGCA | 1 | 16 | 27 | -984 | -973 | -4,87 | 0,01 | 12,29 | >0.1 | 5,6 | 4,27 | 0 | 1,87 | 0 | 0 | 5,43 |
| SPDEF | AATAGGAAATG | 1 | 60 | 71 | -940 | -929 | -7,3 | 0,01 | 8,31 | >0.1 | 3,99 | 4,31 | 0 | 1,88 | 0 | 0 | 5,43 |
| SPDEF | GTCAGGAGATA | 1 | 243 | 254 | -757 | -746 | -6,19 | 0,01 | 12,36 | >0.1 | 2,87 | 4,52 | 0 | 1,4 | 0 | 4,33 | 5,43 |
| SPDEF | AACCAAAAGTA | 1 | 733 | 744 | -267 | -256 | -7,02 | 0,01 | 16,82 | >0.1 | 5,6 | 5,66 | 0 | 1,24 | 0 | 5,92 | 5,43 |
| SPDEF | TACCGGTTTTG | 1 | 742 | 753 | -258 | -247 | -2,34 | 0,01 | 21,56 | >0.1 | 5,6 | 5,69 | 0 | 1,26 | 0 | 5,92 | 5,43 |
| SPDEF | GACAGGAGAGG | 1 | 768 | 779 | -232 | -221 | -8,46 | 0,01 | 15,65 | >0.1 | 5,6 | 5,81 | 0 | 1,35 | 0 | 5,92 | 5,43 |
| SPDEF | TCCAGGATGAA | 1 | 865 | 876 | -135 | -124 | 5,27 | 0,001 | 27,28 | >0.1 | 5,6 | 6,41 | 0 | 1,9 | 0 | 2,67 | 5,43 |
| SPDEF | ATGAGGAGGTT | 1 | 987 | 998 | -13 | -2 | -6,8 | 0,01 | 16,37 | >0.1 | 5,6 | 8,66 | 0 | 1,87 | 0 | 1,61 | 5,43 |
| SPDEF | CAGTGGATGTG | 1 | 1018 | 1029 | 18 | 29 | -5,42 | 0,01 | 17,57 | >0.1 | 5,6 | 8,76 | 0 | 1,86 | 0 | 1,34 | 5,43 |
| SPDEF | ACCACGAAGCC | 1 | 1146 | 1157 | 146 | 157 | -2,77 | 0,01 | 20,38 | >0.1 | 5,6 | 6,33 | 0 | 1,84 | 1,79 | 2,16 | 5,43 |
| SPDEF | AGCCGAAGACC | 1 | 1153 | 1164 | 153 | 164 | -6,48 | 0,01 | 15,67 | >0.1 | 4,47 | 6,27 | 0 | 1,83 | 1,79 | 2,34 | 5,43 |
| SPDEF | TCCAAGAAGCA | -1 | 1183 | 1194 | 183 | 194 | -4,36 | 0,01 | 19,62 | >0.1 | 5,6 | 6,07 | 0 | 1,56 | 1,79 | 3,52 | 5,43 |
| SPDEF | AGAAGGAGCCA | -1 | 1064 | 1075 | 64 | 75 | -7,82 | 0,01 | 16,35 | >0.1 | 5,6 | 7,28 | 0 | 1,85 | 1,79 | 2,21 | 5,43 |
| SPDEF | GCCAGGAAGAG | -1 | 1057 | 1068 | 57 | 68 | 0,26 | 0,01 | 22,84 | >0.1 | 5,6 | 7,42 | 0 | 1,86 | 0 | 2,27 | 5,43 |
| SPDEF | AGCAGGAAAGA | -1 | 932 | 943 | -68 | -57 | -0,47 | 0,01 | 20,47 | >0.1 | 5,6 | 7,11 | 0 | 1,88 | 0 | 0,93 | 5,43 |
| SPDEF | CACAGTAAACA | -1 | 897 | 908 | -103 | -92 | -8,23 | 0,01 | 14,24 | >0.1 | 5,6 | 6,69 | 0 | 1,89 | 0 | 2,86 | 5,43 |
| SPDEF | GGACAGATATG | -1 | 852 | 863 | -148 | -137 | -8,44 | 0,01 | 13,47 | >0.1 | 5,6 | 6,32 | 0 | 1,89 | 0 | 2,67 | 5,43 |
| SPDEF | AACCGGTACTT | -1 | 739 | 750 | -261 | -250 | -0,66 | 0,01 | 23,22 | >0.1 | 5,6 | 5,68 | 0 | 1,26 | 0 | 5,92 | 5,43 |
| SPDEF | ACACGGAAAAT | -1 | 694 | 705 | -306 | -295 | -0,99 | 0,01 | 19,76 | >0.1 | 5,6 | 5,5 | 0 | 1,14 | 0 | 3,07 | 5,43 |
| SPDEF | TCCCAGAAACT | -1 | 555 | 566 | -445 | -434 | -2,87 | 0,01 | 11,86 | >0.1 | 3,21 | 5,09 | 0 | 0,99 | 0 | 0 | 5,43 |
| SPDEF | CCCAGAAACTC | -1 | 554 | 565 | -446 | -435 | -8,45 | 0,01 | 6,27 | >0.1 | 3,21 | 5,09 | 0 | 0,99 | 0 | 0 | 5,43 |
| ZNF423 | GCACTCAAAGAAGTC | 1 | 24 | 39 | -976 | -961 | -6,13 | 0,01 | 5,56 | >0.1 | 5,54 | 4,28 | 0 | 1,87 | 0 | 0 | 0 |
| ZNF423 | AGCACTTTGGGAGGC | 1 | 208 | 223 | -792 | -777 | 4,25 | 0,001 | 18,01 | >0.1 | 2,99 | 4,48 | 0 | 1,58 | 0 | 4,7 | 0 |
| ZNF423 | GGCACCTGTAGTCCC | 1 | 327 | 342 | -673 | -658 | -2,08 | 0,01 | 11,77 | >0.1 | 2,6 | 4,65 | 0 | 1,15 | 0 | 5,46 | 0 |
| ZNF423 | AGCTACTCGGGAGGC | 1 | 342 | 357 | -658 | -643 | -7,2 | 0,01 | 7,67 | >0.1 | 2,56 | 4,68 | 0 | 1,12 | 0 | 6,52 | 0 |
| ZNF423 | GTGAACCTGGGAGGC | 1 | 374 | 389 | -626 | -611 | -6,3 | 0,01 | 8,46 | >0.1 | 2,56 | 4,73 | 0 | 1,07 | 0 | 6,4 | 0 |
| ZNF423 | TGAACCTGGGAGGCA | 1 | 375 | 390 | -625 | -610 | -2,07 | 0,01 | 12,69 | >0.1 | 2,56 | 4,73 | 0 | 1,07 | 0 | 6,4 | 0 |
| ZNF423 | TCCAGCCTGGGCGAC | 1 | 424 | 439 | -576 | -561 | -6,17 | 0,01 | 8,04 | >0.1 | 2,7 | 4,82 | 0 | 1,02 | 0 | 5,67 | 0 |
| ZNF423 | AGACCTCTTTGGTCA | 1 | 664 | 679 | -336 | -321 | -1,51 | 0,01 | 12,93 | >0.1 | 4,09 | 5,42 | 0 | 1,1 | 0 | 3,83 | 0 |
| ZNF423 | CCCAACCCAAGTTCA | 1 | 948 | 963 | -52 | -37 | -5,67 | 0,01 | 10,95 | >0.1 | 5,54 | 7,46 | 0 | 1,88 | 0 | 1,74 | 0 |
| ZNF423 | CCAACCCAAGTTCAA | 1 | 949 | 964 | -51 | -36 | -3,74 | 0,01 | 12,9 | >0.1 | 5,54 | 7,48 | 0 | 1,88 | 0 | 1,74 | 0 |
| ZNF423 | GGCTCACAGGGGACG | 1 | 1034 | 1049 | 34 | 49 | -5,67 | 0,01 | 11,13 | >0.1 | 5,54 | 8,06 | 0 | 1,86 | 0 | 1,34 | 0 |
| ZNF423 | TTCCTCAATGGTGGA | -1 | 1103 | 1118 | 103 | 118 | -7,29 | 0,01 | 12,57 | >0.1 | 5,54 | 6,72 | 0 | 1,85 | 3,22 | 2,52 | 0 |
| ZNF423 | CGTCCCCTGTGAGCC | -1 | 1034 | 1049 | 34 | 49 | -2,85 | 0,01 | 13,95 | >0.1 | 5,54 | 8,06 | 0 | 1,86 | 0 | 1,34 | 0 |
| ZNF423 | AGAACCTTACCTCTC | -1 | 773 | 788 | -227 | -212 | -5,31 | 0,01 | 13,8 | >0.1 | 5,54 | 5,85 | 0 | 1,39 | 0 | 6,32 | 0 |
| ZNF423 | GGTACTTTTGGTTAA | -1 | 731 | 746 | -269 | -254 | -5,38 | 0,01 | 12,99 | >0.1 | 5,54 | 5,66 | 0 | 1,25 | 0 | 5,92 | 0 |
| ZNF423 | GTGACCAAAGAGGTC | -1 | 665 | 680 | -335 | -320 | -4,42 | 0,01 | 10,04 | >0.1 | 4,11 | 5,42 | 0 | 1,1 | 0 | 3,83 | 0 |
| ZNF423 | GTCGCCCAGGCTGGA | -1 | 424 | 439 | -576 | -561 | 0,94 | 0,01 | 15,15 | >0.1 | 2,7 | 4,82 | 0 | 1,02 | 0 | 5,67 | 0 |
| ZNF423 | GGAGCACAGTGGCGC | -1 | 412 | 427 | -588 | -573 | -5,36 | 0,01 | 8,33 | >0.1 | 2,67 | 4,8 | 0 | 1,03 | 0 | 5,19 | 0 |
| ZNF423 | TGCCTCCCAGGTTCA | -1 | 375 | 390 | -625 | -610 | -2,19 | 0,01 | 12,57 | >0.1 | 2,56 | 4,73 | 0 | 1,07 | 0 | 6,4 | 0 |
| ZNF423 | GCCTCCCAGGTTCAC | -1 | 374 | 389 | -626 | -611 | -4,91 | 0,01 | 9,85 | >0.1 | 2,56 | 4,73 | 0 | 1,07 | 0 | 6,4 | 0 |
| ZNF423 | GGGACTACAGGTGCC | -1 | 327 | 342 | -673 | -658 | -4,18 | 0,01 | 9,67 | >0.1 | 2,6 | 4,65 | 0 | 1,15 | 0 | 5,46 | 0 |

| | | | | | | | | | | | | | | | | | |
|---|---|---|---|---|---|---|---|---|---|---|---|---|---|---|---|---|---|
| ZNF423 | CCCAGCTAAGTTTTT | -1 | 302 | 317 | -698 | -683 | -5,56 | 0,01 | 7,82 | >0.1 | 2,67 | 4,62 | 0 | 1,2 | 0 | 4,9 | 0 |
| ZNF423 | GGCCTCCCAAAGTGC | -1 | 209 | 224 | -791 | -776 | -7,59 | 0,01 | 6,05 | >0.1 | 2,99 | 4,48 | 0 | 1,57 | 0 | 4,59 | 0 |
| ZNF423 | GCCTCCCAAAGTGCT | -1 | 208 | 223 | -792 | -777 | -0,79 | 0,01 | 12,97 | >0.1 | 2,99 | 4,48 | 0 | 1,58 | 0 | 4,7 | 0 |
| ZNF423 | CCACTCCACAGGTCA | -1 | 108 | 123 | -892 | -877 | -4,38 | 0,01 | 6,17 | >0.1 | 3,53 | 4,36 | 0 | 1,89 | 0 | 0,77 | 0 |
| HES1 | GGCACAGTGG | 1 | 181 | 191 | -819 | -809 | 3,47 | 0,01 | 16,67 | >0.1 | 3,12 | 4,44 | 0 | 1,78 | 0 | 3,86 | 0 |
| HES1 | GGCTCACGCC | 1 | 189 | 199 | -811 | -801 | 7,53 | 0,001 | 20,82 | >0.1 | 3,09 | 4,45 | 0 | 1,71 | 0 | 4,04 | 0 |
| HES1 | CTCACGCCTG | 1 | 191 | 201 | -809 | -799 | 6,42 | 0,01 | 19,78 | >0.1 | 3,08 | 4,45 | 0 | 1,7 | 0 | 4,13 | 0 |
| HES1 | ACCCCGTCTC | 1 | 280 | 290 | -720 | -710 | 3,63 | 0,01 | 17,3 | >0.1 | 2,74 | 4,57 | 0 | 1,26 | 0 | 5,09 | 0 |
| HES1 | GGCACCTGTA | 1 | 327 | 337 | -673 | -663 | 6,54 | 0,01 | 20,5 | >0.1 | 2,6 | 4,65 | 0 | 1,14 | 0 | 5,56 | 0 |
| HES1 | GGCAGGAGAA | 1 | 360 | 370 | -640 | -630 | 4,78 | 0,01 | 19,45 | >0.1 | 2,52 | 4,7 | 0 | 1,09 | 0 | 6,36 | 0 |
| HES1 | AGATCGCGCC | 1 | 407 | 417 | -593 | -583 | 3,79 | 0,01 | 17,89 | >0.1 | 2,65 | 4,78 | 0 | 1,03 | 0 | 5,65 | 0 |
| HES1 | ATCGCGCCAC | 1 | 409 | 419 | -591 | -581 | 4,01 | 0,01 | 18 | >0.1 | 2,65 | 4,78 | 0 | 1,03 | 0 | 5,52 | 0 |
| HES1 | GACAGGAGAG | 1 | 768 | 778 | -232 | -222 | 3,73 | 0,01 | 22,41 | >0.1 | 5,61 | 5,81 | 0 | 1,35 | 0 | 5,92 | 0 |
| HES1 | CACAGGGGAC | 1 | 1038 | 1048 | 38 | 48 | 3,84 | 0,01 | 20,56 | >0.1 | 5,61 | 7,92 | 0 | 1,86 | 0 | 1,34 | 0 |
| HES1 | ACCACGAAGC | 1 | 1146 | 1156 | 146 | 156 | 3,99 | 0,01 | 21,72 | >0.1 | 5,61 | 6,33 | 0 | 1,84 | 1,79 | 2,16 | 0 |
| HES1 | CCCAGGCTGG | -1 | 425 | 435 | -575 | -565 | 4,15 | 0,01 | 18,35 | >0.1 | 2,7 | 4,81 | 0 | 1,01 | 0 | 5,67 | 0 |
| HES1 | GGCGCGATCT | -1 | 407 | 417 | -593 | -583 | 5,1 | 0,01 | 19,2 | >0.1 | 2,65 | 4,78 | 0 | 1,03 | 0 | 5,65 | 0 |
| HES1 | GGCTCACTGC | -1 | 396 | 406 | -604 | -594 | 4,06 | 0,01 | 18,24 | >0.1 | 2,61 | 4,76 | 0 | 1,04 | 0 | 5,76 | 0 |
| HES1 | CCCAGGTTCA | -1 | 375 | 385 | -625 | -615 | 3,86 | 0,01 | 18,6 | >0.1 | 2,56 | 4,72 | 0 | 1,07 | 0 | 6,4 | 0 |
| HES1 | TTCACGCCAT | -1 | 369 | 379 | -631 | -621 | 4,4 | 0,01 | 18,88 | >0.1 | 2,54 | 4,71 | 0 | 1,07 | 0 | 6,15 | 0 |
| HES1 | GCCTCCCGAG | -1 | 347 | 357 | -653 | -643 | 3,75 | 0,01 | 18,61 | >0.1 | 2,55 | 4,68 | 0 | 1,11 | 0 | 6,52 | 0 |
| HES1 | TACAGGTGCC | -1 | 327 | 337 | -673 | -663 | 5,6 | 0,01 | 19,56 | >0.1 | 2,6 | 4,65 | 0 | 1,14 | 0 | 5,56 | 0 |
| HES1 | ACCACGCCCA | -1 | 313 | 323 | -687 | -677 | 7,64 | 0,001 | 22,21 | >0.1 | 2,64 | 4,62 | 0 | 1,17 | 0 | 6,13 | 0 |
| HES1 | TACAGGCGTG | -1 | 193 | 203 | -807 | -797 | 5,17 | 0,01 | 18,64 | >0.1 | 3,07 | 4,46 | 0 | 1,68 | 0 | 4,26 | 0 |
| HES1 | GGCGTGAGCC | -1 | 189 | 199 | -811 | -801 | 5,17 | 0,01 | 18,46 | >0.1 | 3,09 | 4,45 | 0 | 1,71 | 0 | 4,04 | 0 |
| HES1 | TCCACAGGTC | -1 | 109 | 119 | -891 | -881 | 5,5 | 0,01 | 16,03 | >0.1 | 3,54 | 4,36 | 0 | 1,89 | 0 | 0,75 | 0 |
| HES2 | ATAGGAAATGAGC | 1 | 61 | 74 | -939 | -926 | 3,51 | 0,01 | 13,67 | >0.1 | 3,97 | 4,31 | 0 | 1,88 | 0 | 0 | 0 |
| HES2 | GGTGGGCACCTGT | 1 | 323 | 336 | -677 | -664 | 8,6 | 0,001 | 22,46 | >0.1 | 2,61 | 4,64 | 0 | 1,15 | 0 | 5,46 | 0 |
| HES2 | CTGAGGCAGGAGA | 1 | 356 | 369 | -644 | -631 | 4,11 | 0,01 | 18,55 | >0.1 | 2,52 | 4,7 | 0 | 1,09 | 0 | 6,13 | 0 |
| HES2 | GGCGTGAACCTGG | 1 | 371 | 384 | -629 | -616 | 3,8 | 0,01 | 18,54 | >0.1 | 2,55 | 4,72 | 0 | 1,07 | 0 | 6,4 | 0 |
| HES2 | TAAAGACAAGGGC | 1 | 501 | 514 | -499 | -486 | 8,73 | 0,001 | 23,34 | >0.1 | 2,97 | 4,97 | 0 | 0,99 | 0 | 5,68 | 0 |
| HES2 | GAAGGGAAAATGT | 1 | 529 | 542 | -471 | -458 | 5,53 | 0,01 | 18,34 | >0.1 | 3,09 | 5,04 | 0 | 0,99 | 0 | 3,7 | 0 |
| HES2 | TCTGGGAATATGA | 1 | 559 | 572 | -441 | -428 | 3,35 | 0,01 | 12,69 | >0.1 | 3,24 | 5,11 | 0 | 1 | 0 | 0 | 0 |
| HES2 | TAACCAAAAGTAC | 1 | 732 | 745 | -268 | -255 | 6,34 | 0,01 | 24,73 | >0.1 | 5,57 | 5,66 | 0 | 1,24 | 0 | 5,92 | 0 |
| HES2 | AAGTGACAGGAGA | 1 | 764 | 777 | -236 | -223 | 3,62 | 0,01 | 22,25 | >0.1 | 5,57 | 5,8 | 0 | 1,35 | 0 | 5,92 | 0 |
| HES2 | CTAGGGAAAGTCA | 1 | 1003 | 1016 | 3 | 16 | 5,53 | 0,01 | 23,5 | >0.1 | 5,57 | 9,12 | 0 | 1,87 | 0 | 1,42 | 0 |
| HES2 | TTGAGGAACAGGC | 1 | 1110 | 1123 | 110 | 123 | 5,59 | 0,01 | 25,42 | >0.1 | 5,57 | 6,65 | 0 | 1,85 | 3,22 | 2,54 | 0 |
| HES2 | TTTGGACAAGTTT | 1 | 1132 | 1145 | 132 | 145 | 6,35 | 0,01 | 26,39 | >0.1 | 5,57 | 6,44 | 0 | 1,84 | 3,22 | 2,97 | 0 |
| HES2 | AAGAAGCAAGTGA | -1 | 1178 | 1191 | 178 | 191 | 6,21 | 0,01 | 24,62 | >0.1 | 5,57 | 6,1 | 0 | 1,6 | 1,79 | 3,34 | 0 |
| HES2 | TGTCCAAAAATGT | -1 | 1127 | 1140 | 127 | 140 | 3,49 | 0,01 | 23,41 | >0.1 | 5,57 | 6,48 | 0 | 1,84 | 3,22 | 2,79 | 0 |
| HES2 | ACACGGAAAATGT | -1 | 692 | 705 | -308 | -295 | 5,18 | 0,01 | 20,47 | >0.1 | 5,57 | 5,5 | 0 | 1,15 | 0 | 3,07 | 0 |
| HES2 | TTCAGACAATTTC | -1 | 609 | 622 | -391 | -378 | 3,42 | 0,01 | 14,78 | >0.1 | 3,55 | 5,24 | 0 | 1,03 | 0 | 1,55 | 0 |
| HES2 | CAGTGGCGCGATC | -1 | 408 | 421 | -592 | -579 | 2,73 | 0,01 | 16,72 | >0.1 | 2,65 | 4,79 | 0 | 1,03 | 0 | 5,52 | 0 |
| HES2 | GGACTACAGGTGC | -1 | 328 | 341 | -672 | -659 | 7,32 | 0,001 | 21,17 | >0.1 | 2,6 | 4,65 | 0 | 1,14 | 0 | 5,46 | 0 |
| HES2 | ACTCCACAGGTCA | -1 | 108 | 121 | -892 | -879 | 3,2 | 0,01 | 13,73 | >0.1 | 3,54 | 4,36 | 0 | 1,89 | 0 | 0,75 | 0 |
| HES5 | TGGCTCACGCCT | 1 | 188 | 200 | -812 | -800 | -10,7 | 0,01 | 2,6 | >0.1 | 3,09 | 4,45 | 0 | 1,72 | 0 | 4,04 | 0 |
| HES5 | AACCCCGTCTCT | 1 | 279 | 291 | -721 | -709 | -19,55 | 0,01 | -5,88 | >0.1 | 2,74 | 4,58 | 0 | 1,27 | 0 | 5,09 | 0 |
| HES5 | GGGCACCTGTAG | 1 | 326 | 338 | -674 | -662 | -11,46 | 0,01 | 2,5 | >0.1 | 2,6 | 4,65 | 0 | 1,15 | 0 | 5,56 | 0 |
| HES5 | GAGATCGCGCCA | 1 | 406 | 418 | -594 | -582 | -21,26 | 0,01 | -7,15 | >0.1 | 2,65 | 4,78 | 0 | 1,03 | 0 | 5,65 | 0 |
| HES5 | AGACAAGGGCAA | 1 | 504 | 516 | -496 | -484 | -17,98 | 0,01 | -3,36 | >0.1 | 2,98 | 4,98 | 0 | 0,98 | 0 | 5,68 | 0 |
| HES5 | AGGAACAGGCCA | 1 | 1113 | 1125 | 113 | 125 | -16,46 | 0,01 | 3,37 | >0.1 | 5,58 | 6,62 | 0 | 1,85 | 3,22 | 2,56 | 0 |
| HES5 | TTAAACTTGTCC | -1 | 1135 | 1147 | 135 | 147 | -22,3 | 0,01 | -2,15 | >0.1 | 5,58 | 6,41 | 0 | 1,84 | 3,22 | 3,09 | 0 |
| HES5 | TGGCCTGTTCCT | -1 | 1113 | 1125 | 113 | 125 | -16,37 | 0,01 | 3,46 | >0.1 | 5,58 | 6,62 | 0 | 1,85 | 3,22 | 2,56 | 0 |
| HES5 | CGGAAAATGTTA | -1 | 690 | 702 | -310 | -298 | -22,84 | 0,01 | -8,61 | >0.1 | 4,53 | 5,49 | 0 | 1,14 | 0 | 3,07 | 0 |
| HES5 | TTGCCCTTGTCT | -1 | 504 | 516 | -496 | -484 | -19,41 | 0,01 | -4,79 | >0.1 | 2,98 | 4,98 | 0 | 0,98 | 0 | 5,68 | 0 |
| HES5 | CTACAGGTGCCC | -1 | 326 | 338 | -674 | -662 | -4,3 | 0,001 | 9,66 | >0.1 | 2,6 | 4,65 | 0 | 1,15 | 0 | 5,56 | 0 |
| HES5 | CACCACGCCCAG | -1 | 312 | 324 | -688 | -676 | -21,16 | 0,01 | -6,59 | >0.1 | 2,64 | 4,63 | 0 | 1,18 | 0 | 6,13 | 0 |
| HES5 | AGAGACGGGGTT | -1 | 279 | 291 | -721 | -709 | -21,92 | 0,01 | -8,25 | >0.1 | 2,74 | 4,58 | 0 | 1,27 | 0 | 5,09 | 0 |
| HES5 | CCCAAAGTGCTA | -1 | 207 | 219 | -793 | -781 | -21,55 | 0,01 | -7,78 | >0.1 | 3 | 4,48 | 0 | 1,58 | 0 | 4,7 | 0 |

| | | | | | | | | | | | | | | | | | |
|---|---|---|---|---|---|---|---|---|---|---|---|---|---|---|---|---|---|
| HES5 | AGGCGTGAGCCA | -1 | 188 | 200 | -812 | -800 | -13,38 | 0,01 | -0,08 | >0.1 | 3,09 | 4,45 | 0 | 1,72 | 0 | 4,04 | 0 |
| HES5 | CTCCACAGGTCA | -1 | 108 | 120 | -892 | -880 | -19,19 | 0,01 | -8,65 | >0.1 | 3,54 | 4,36 | 0 | 1,89 | 0 | 0,75 | 0 |
| FOXC1 | ATTAAAAATAC | 1 | 291 | 302 | -709 | -698 | 2,96 | 0,01 | 15,94 | >0.1 | 2,71 | 4,59 | 0 | 1,23 | 0 | 4,45 | 0 |
| FOXC1 | AGGGAAAATGT | 1 | 531 | 542 | -469 | -458 | 2,9 | 0,01 | 13,62 | >0.1 | 3,09 | 5,04 | 0 | 0,98 | 0 | 1,61 | 0 |
| FOXC1 | AATAAAAATAA | 1 | 578 | 589 | -422 | -411 | 7,94 | 0,001 | 17,43 | >0.1 | 3,34 | 5,15 | 0 | 1 | 0 | 0 | 0 |
| FOXC1 | AAAATAAATGT | 1 | 582 | 593 | -418 | -407 | 6,3 | 0,01 | 15,83 | >0.1 | 3,36 | 5,16 | 0 | 1,01 | 0 | 0 | 0 |
| FOXC1 | CATACAAACAC | 1 | 625 | 636 | -375 | -364 | 4,01 | 0,01 | 15,54 | >0.1 | 3,67 | 5,28 | 0 | 1,04 | 0 | 1,55 | 0 |
| FOXC1 | AACACCAACAT | 1 | 631 | 642 | -369 | -358 | 6,31 | 0,01 | 18,03 | >0.1 | 3,72 | 5,3 | 0 | 1,05 | 0 | 1,66 | 0 |
| FOXC1 | TAGGAAAATAT | 1 | 721 | 732 | -279 | -268 | 10 | 0,001 | 27,58 | >0.1 | 5,6 | 5,61 | 0 | 1,21 | 0 | 5,17 | 0 |
| FOXC1 | TAAAAAATTAT | 1 | 827 | 838 | -173 | -162 | 1,86 | 0,01 | 18,7 | >0.1 | 5,6 | 6,14 | 0 | 1,67 | 0 | 3,43 | 0 |
| FOXC1 | TACAGCAACAA | -1 | 1080 | 1091 | 80 | 91 | 4,31 | 0,01 | 22,78 | >0.1 | 5,6 | 7,01 | 0 | 1,85 | 1,79 | 2,21 | 0 |
| FOXC1 | TGAGCCAAGAT | -1 | 1029 | 1040 | 29 | 40 | 3,65 | 0,01 | 20,73 | >0.1 | 5,6 | 8,29 | 0 | 1,86 | 0 | 1,34 | 0 |
| FOXC1 | ACAGTAAACAA | -1 | 896 | 907 | -104 | -93 | 8,94 | 0,001 | 26,05 | >0.1 | 5,6 | 6,68 | 0 | 1,89 | 0 | 2,95 | 0 |
| FOXC1 | TGAGCCAATAT | -1 | 880 | 891 | -120 | -109 | 9,04 | 0,001 | 26,14 | >0.1 | 5,6 | 6,54 | 0 | 1,9 | 0 | 3,07 | 0 |
| FOXC1 | TTGGGCAACAT | -1 | 538 | 549 | -462 | -451 | 4,01 | 0,01 | 14,91 | >0.1 | 3,13 | 5,05 | 0 | 0,99 | 0 | 1,73 | 0 |
| HES7 | TGGCTCACGCCT | 1 | 188 | 200 | -812 | -800 | -11,06 | 0,01 | 2,24 | >0.1 | 3,09 | 4,45 | 0 | 1,72 | 0 | 4,04 | 0 |
| HES7 | AACCCCGTCTCT | 1 | 279 | 291 | -721 | -709 | -19,12 | 0,01 | -5,45 | >0.1 | 2,74 | 4,58 | 0 | 1,27 | 0 | 5,09 | 0 |
| HES7 | GGGCACCTGTAG | 1 | 326 | 338 | -674 | -662 | -9,9 | 0,01 | 4,06 | >0.1 | 2,6 | 4,65 | 0 | 1,15 | 0 | 5,56 | 0 |
| HES7 | TGGCGTGAACCT | 1 | 370 | 382 | -630 | -618 | -20,5 | 0,01 | -5,76 | >0.1 | 2,55 | 4,72 | 0 | 1,07 | 0 | 6,4 | 0 |
| HES7 | GAGATCGCGCCA | 1 | 406 | 418 | -594 | -582 | -20,67 | 0,01 | -6,56 | >0.1 | 2,65 | 4,78 | 0 | 1,03 | 0 | 5,65 | 0 |
| HES7 | AGACAAGGGCAA | 1 | 504 | 516 | -496 | -484 | -20,97 | 0,01 | -6,35 | >0.1 | 2,98 | 4,98 | 0 | 0,98 | 0 | 5,68 | 0 |
| HES7 | TGACAGGAGAGG | 1 | 767 | 779 | -233 | -221 | -19,37 | 0,01 | -0,71 | >0.1 | 5,58 | 5,81 | 0 | 1,35 | 0 | 5,92 | 0 |
| HES7 | TCACAGGGGACG | 1 | 1037 | 1049 | 37 | 49 | -19,23 | 0,01 | -2,5 | >0.1 | 5,58 | 7,95 | 0 | 1,86 | 0 | 1,34 | 0 |
| HES7 | AGGAACAGGCCA | 1 | 1113 | 1125 | 113 | 125 | -9,96 | 0,01 | 9,87 | >0.1 | 5,58 | 6,62 | 0 | 1,85 | 3,22 | 2,56 | 0 |
| HES7 | TGGCCTGTTCCT | -1 | 1113 | 1125 | 113 | 125 | -15,95 | 0,01 | 3,88 | >0.1 | 5,58 | 6,62 | 0 | 1,85 | 3,22 | 2,56 | 0 |
| HES7 | GAGAAGGAGCCA | -1 | 1064 | 1076 | 64 | 76 | -21,16 | 0,01 | -2,38 | >0.1 | 5,58 | 7,28 | 0 | 1,85 | 1,79 | 2,27 | 0 |
| HES7 | CGTCCCCTGTGA | -1 | 1037 | 1049 | 37 | 49 | -18,73 | 0,01 | -2 | >0.1 | 5,58 | 7,95 | 0 | 1,86 | 0 | 1,34 | 0 |
| HES7 | AAATACATGACT | -1 | 516 | 528 | -484 | -472 | -20,35 | 0,01 | -8,38 | >0.1 | 3,03 | 5 | 0 | 0,98 | 0 | 2,95 | 0 |
| HES7 | TTGCCCTTGTCT | -1 | 504 | 516 | -496 | -484 | -14,18 | 0,01 | 0,44 | >0.1 | 2,98 | 4,98 | 0 | 0,98 | 0 | 5,68 | 0 |
| HES7 | TCGCCCAGGCTG | -1 | 426 | 438 | -574 | -562 | -19,82 | 0,01 | -5,48 | >0.1 | 2,7 | 4,82 | 0 | 1,01 | 0 | 5,8 | 0 |
| HES7 | TGGCGCGATCTC | -1 | 406 | 418 | -594 | -582 | -19,48 | 0,01 | -5,37 | >0.1 | 2,65 | 4,78 | 0 | 1,03 | 0 | 5,65 | 0 |
| HES7 | AGGTTCACGCCA | -1 | 370 | 382 | -630 | -618 | -20,31 | 0,01 | -5,57 | >0.1 | 2,55 | 4,72 | 0 | 1,07 | 0 | 6,4 | 0 |
| HES7 | CTACAGGTGCCC | -1 | 326 | 338 | -674 | -662 | -7,47 | 0,001 | 6,49 | >0.1 | 2,6 | 4,65 | 0 | 1,15 | 0 | 5,56 | 0 |
| HES7 | AGGCGTGAGCCA | -1 | 188 | 200 | -812 | -800 | -9,85 | 0,01 | 3,45 | >0.1 | 3,09 | 4,45 | 0 | 1,72 | 0 | 4,04 | 0 |
| HES7 | CTCCACAGGTCA | -1 | 108 | 120 | -892 | -880 | -16,63 | 0,01 | -6,09 | >0.1 | 3,54 | 4,36 | 0 | 1,89 | 0 | 0,75 | 0 |
| HES7 | CTTCACTTACCT | -1 | 3 | 15 | -997 | -985 | -20,75 | 0,01 | -7,25 | >0.1 | 5,58 | 4,26 | 0 | 1,86 | 1,79 | 0 | 0 |
| SMAD4 | TGGCTGGG | 1 | 175 | 183 | -825 | -817 | -0,2 | 0,01 | 12,46 | >0.1 | 3,16 | 4,43 | 0 | 1,83 | 0 | 3,24 | 0 |
| SMAD4 | GGTCAGGA | 1 | 242 | 250 | -758 | -750 | -1,56 | 0,01 | 11,56 | >0.1 | 2,88 | 4,52 | 0 | 1,4 | 0 | 4,33 | 0 |
| SMAD4 | CGTCCTGA | 1 | 259 | 267 | -741 | -733 | -10,91 | 0,01 | 2,31 | >0.1 | 2,81 | 4,54 | 0 | 1,33 | 0 | 4,53 | 0 |
| SMAD4 | CGTCTCTA | 1 | 284 | 292 | -716 | -708 | -11,41 | 0,01 | 2,24 | >0.1 | 2,73 | 4,58 | 0 | 1,25 | 0 | 5,09 | 0 |
| SMAD4 | AGCCTGGG | 1 | 427 | 435 | -573 | -565 | -2,92 | 0,01 | 11,28 | >0.1 | 2,7 | 4,81 | 0 | 1,01 | 0 | 5,67 | 0 |
| SMAD4 | TTTCTGGG | 1 | 557 | 565 | -443 | -435 | -1,19 | 0,01 | 8,1 | >0.1 | 3,21 | 5,09 | 0 | 0,99 | 0 | 0 | 0 |
| SMAD4 | TGTCTGAA | 1 | 614 | 622 | -386 | -378 | 0,29 | 0,01 | 11,62 | >0.1 | 3,57 | 5,24 | 0 | 1,03 | 0 | 1,5 | 0 |
| SMAD4 | GTTCTAGA | 1 | 659 | 667 | -341 | -333 | -1,99 | 0,01 | 12,28 | >0.1 | 3,98 | 5,38 | 0 | 1,08 | 0 | 3,83 | 0 |
| SMAD4 | TTTCTAGG | 1 | 717 | 725 | -283 | -275 | -1,13 | 0,01 | 16,44 | >0.1 | 5,64 | 5,58 | 0 | 1,19 | 0 | 5,17 | 0 |
| SMAD4 | TCTCTAGG | 1 | 785 | 793 | -215 | -207 | -0,63 | 0,01 | 18,21 | >0.1 | 5,64 | 5,88 | 0 | 1,41 | 0 | 5,92 | 0 |
| SMAD4 | AGTCTAGG | 1 | 1000 | 1008 | 0 | 8 | 8,94 | 0,001 | 26,95 | >0.1 | 5,64 | 9,01 | 0 | 1,87 | 0 | 1,5 | 0 |
| SMAD4 | TGTCAAGC | 1 | 1050 | 1058 | 50 | 58 | 0,49 | 0,001 | 17,02 | >0.1 | 5,64 | 7,57 | 0 | 1,86 | 0 | 1,46 | 0 |
| SMAD4 | TGTCCAAA | -1 | 1132 | 1140 | 132 | 140 | -11,44 | 0,1 | 8,61 | >0.1 | 5,64 | 6,44 | 0 | 1,84 | 3,22 | 2,91 | 0 |
| SMAD4 | TGTCTTGG | -1 | 1122 | 1130 | 122 | 130 | 0,12 | 0,01 | 20,03 | >0.1 | 5,64 | 6,53 | 0 | 1,85 | 3,22 | 2,67 | 0 |
| SMAD4 | TCCCTAGA | -1 | 1002 | 1010 | 2 | 10 | -10,94 | 0,01 | 7,15 | >0.1 | 5,64 | 9,12 | 0 | 1,87 | 0 | 1,46 | 0 |
| SMAD4 | TTCCTAGA | -1 | 719 | 727 | -281 | -273 | -11,44 | 0,1 | 5,96 | >0.1 | 5,64 | 5,59 | 0 | 1,19 | 0 | 4,99 | 0 |
| SMAD4 | GGTCTAGA | -1 | 661 | 669 | -339 | -331 | 10,8 | 0,001 | 25,44 | >0.1 | 4,01 | 5,38 | 0 | 1,08 | 0 | 4,16 | 0 |
| SMAD4 | TGTATGGT | -1 | 623 | 631 | -377 | -369 | -1,87 | 0,01 | 9,51 | >0.1 | 3,64 | 5,27 | 0 | 1,03 | 0 | 1,44 | 0 |
| SMAD4 | TGTCTTTA | -1 | 501 | 509 | -499 | -491 | -11,19 | 0,01 | 4,53 | >0.1 | 2,96 | 4,96 | 0 | 0,98 | 0 | 6,81 | 0 |
| SMAD4 | AGTCTCGG | -1 | 442 | 450 | -558 | -550 | -2,11 | 0,01 | 12,4 | >0.1 | 2,75 | 4,84 | 0 | 1 | 0 | 5,92 | 0 |
| SMAD4 | GGTCTGTC | -1 | 436 | 444 | -564 | -556 | -2,41 | 0,01 | 12,08 | >0.1 | 2,73 | 4,83 | 0 | 1 | 0 | 5,92 | 0 |
| SMAD4 | AGGCTGGA | -1 | 424 | 432 | -576 | -568 | -1,44 | 0,01 | 12,74 | >0.1 | 2,69 | 4,81 | 0 | 1,01 | 0 | 5,67 | 0 |
| SMAD4 | AGTCAGGA | -1 | 261 | 269 | -739 | -731 | -1,93 | 0,01 | 11,27 | >0.1 | 2,81 | 4,54 | 0 | 1,32 | 0 | 4,53 | 0 |

| | | | | | | | | | | | | | | | | | |
|---|---|---|---|---|---|---|---|---|---|---|---|---|---|---|---|---|---|
| SMAD4 | TGCCCAGC | -1 | 177 | 185 | -823 | -815 | -10,8 | 0,01 | 1,83 | >0.1 | 3,15 | 4,43 | 0 | 1,81 | 0 | 3,24 | 0 |
| RORB | AAAGAAGTCAG | 1 | 30 | 41 | -970 | -959 | 5,59 | 0,01 | 16,21 | >0.1 | 4,47 | 4,28 | 0 | 1,87 | 0 | 0 | 0 |
| RORB | AAATAGGAAAT | 1 | 59 | 70 | -941 | -930 | 3,95 | 0,01 | 14,14 | >0.1 | 4 | 4,31 | 0 | 1,87 | 0 | 0 | 0 |
| RORB | AGGCAGATCAG | 1 | 91 | 102 | -909 | -898 | 2,38 | 0,01 | 13,14 | >0.1 | 3,68 | 4,34 | 0 | 1,88 | 0 | 0,85 | 0 |
| RORB | AAAATGGCCAT | 1 | 142 | 153 | -858 | -847 | 3,48 | 0,01 | 14,75 | >0.1 | 3,33 | 4,4 | 0 | 1,9 | 0 | 1,65 | 0 |
| RORB | CAGTGGCTCAC | 1 | 185 | 196 | -815 | -804 | 2,39 | 0,01 | 15,54 | >0.1 | 3,1 | 4,45 | 0 | 1,75 | 0 | 3,86 | 0 |
| RORB | CACAAGGTCAG | 1 | 237 | 248 | -763 | -752 | 3,94 | 0,01 | 17,29 | >0.1 | 2,89 | 4,52 | 0 | 1,42 | 0 | 4,53 | 0 |
| RORB | TCTTTGGTCAC | 1 | 669 | 680 | -331 | -320 | 4,03 | 0,01 | 18,38 | >0.1 | 4,13 | 5,42 | 0 | 1,1 | 0 | 3,7 | 0 |
| RORB | GATTTGGCCAT | 1 | 752 | 763 | -248 | -237 | 3,38 | 0,01 | 22,55 | >0.1 | 5,6 | 5,74 | 0 | 1,3 | 0 | 6,54 | 0 |
| RORB | ATTTGATTCAC | 1 | 814 | 825 | -186 | -175 | 2,12 | 0,01 | 18,79 | >0.1 | 5,6 | 6,06 | 0 | 1,58 | 0 | 3,43 | 0 |
| RORB | TATTGGCTCAG | 1 | 881 | 892 | -119 | -108 | 4,83 | 0,01 | 21,94 | >0.1 | 5,6 | 6,55 | 0 | 1,89 | 0 | 3,07 | 0 |
| RORB | TCTTGGCTCAC | 1 | 1030 | 1041 | 30 | 41 | 2,17 | 0,01 | 19,2 | >0.1 | 5,6 | 8,24 | 0 | 1,86 | 0 | 1,34 | 0 |
| RORB | GACGATGTCAA | 1 | 1045 | 1056 | 45 | 56 | 3 | 0,01 | 18,81 | >0.1 | 5,6 | 7,7 | 0 | 1,86 | 0 | 0,65 | 0 |
| RORB | GAACAGGCCAA | 1 | 1115 | 1126 | 115 | 126 | 5,23 | 0,01 | 25,03 | >0.1 | 5,6 | 6,6 | 0 | 1,85 | 3,22 | 2,54 | 0 |
| RORB | GAACAGGTCTT | -1 | 1158 | 1169 | 158 | 169 | 2,7 | 0,01 | 19,28 | >0.1 | 4,37 | 6,24 | 0 | 1,78 | 1,79 | 2,4 | 0 |
| RORB | ACTTGGGTTGG | -1 | 949 | 960 | -51 | -40 | 3,17 | 0,01 | 19,78 | >0.1 | 5,6 | 7,4 | 0 | 1,88 | 0 | 1,74 | 0 |
| RORB | ATAAAGTTCAT | -1 | 871 | 882 | -129 | -118 | 4,02 | 0,01 | 20,93 | >0.1 | 5,6 | 6,46 | 0 | 1,9 | 0 | 2,95 | 0 |
| RORB | AAAGAGGTCTA | -1 | 663 | 674 | -337 | -326 | 6,8 | 0,01 | 21,34 | >0.1 | 4,05 | 5,4 | 0 | 1,09 | 0 | 3,99 | 0 |
| RORB | TAATAGGTTAT | -1 | 596 | 607 | -404 | -393 | 8,27 | 0,001 | 19,38 | >0.1 | 3,45 | 5,2 | 0 | 1,01 | 0 | 1,44 | 0 |
| RORB | TACTTGGGCAA | -1 | 541 | 552 | -459 | -448 | 2,43 | 0,01 | 11,62 | >0.1 | 3,14 | 5,06 | 0 | 0,99 | 0 | 0 | 0 |
| RORB | ACTTGGGCAAC | -1 | 540 | 551 | -460 | -449 | 4,03 | 0,01 | 13,21 | >0.1 | 3,14 | 5,06 | 0 | 0,99 | 0 | 0 | 0 |
| RORB | CCACAGGTCAA | -1 | 107 | 118 | -893 | -882 | 2,27 | 0,01 | 12,84 | >0.1 | 3,56 | 4,36 | 0 | 1,89 | 0 | 0,77 | 0 |
| RORB | AAAAAGCTCAT | -1 | 68 | 79 | -932 | -921 | 3,78 | 0,01 | 13,87 | >0.1 | 3,9 | 4,32 | 0 | 1,88 | 0 | 0 | 0 |
| RARB(VAR.2) | AGGTAAGTGAAGGAGAA | 1 | 3 | 20 | -997 | -980 | -23,72 | 0,01 | -10,28 | >0.1 | 5,52 | 4,26 | 0 | 1,86 | 1,79 | 0 | 0 |
| RARB(VAR.2) | GGCACTCAAAGAAGTCA | 1 | 23 | 40 | -977 | -960 | -21,57 | 0,01 | -9,9 | >0.1 | 5,52 | 4,28 | 0 | 1,87 | 0 | 0 | 0 |
| RARB(VAR.2) | ATTTTAAAAAATGGCCA | 1 | 135 | 152 | -865 | -848 | -19,04 | 0,01 | -7,75 | >0.1 | 3,35 | 4,4 | 0 | 1,9 | 0 | 1,65 | 0 |
| RARB(VAR.2) | AGGCCGAGTTGGGCAGA | 1 | 219 | 236 | -781 | -764 | -24,91 | 0,01 | -10,43 | >0.1 | 2,94 | 4,5 | 0 | 1,51 | 0 | 5,52 | 0 |
| RARB(VAR.2) | GGCAGATCACAAGGTCA | 1 | 230 | 247 | -770 | -753 | -11,22 | 0,001 | 2,55 | >0.1 | 2,9 | 4,52 | 0 | 1,46 | 0 | 4,9 | 0 |
| RARB(VAR.2) | AGGTCAGGAGATAGAGA | 1 | 241 | 258 | -759 | -742 | -24,75 | 0,01 | -11,53 | >0.1 | 2,86 | 4,53 | 0 | 1,41 | 0 | 4,42 | 0 |
| RARB(VAR.2) | CCATAAAGACAAGGGCA | 1 | 498 | 515 | -502 | -485 | -20,43 | 0,01 | -5,98 | >0.1 | 2,96 | 4,98 | 0 | 0,99 | 0 | 5,52 | 0 |
| RARB(VAR.2) | AGACAAGGGCAAAGTCA | 1 | 504 | 521 | -496 | -479 | -16,44 | 0,01 | -3,28 | >0.1 | 2,99 | 4,99 | 0 | 0,99 | 0 | 4,19 | 0 |
| RARB(VAR.2) | AGGGCAAAGTCATGTAT | 1 | 509 | 526 | -491 | -474 | -25,91 | 0,01 | -13,96 | >0.1 | 3,01 | 5 | 0 | 0,99 | 0 | 2,95 | 0 |
| RARB(VAR.2) | TGGTCACTGTAAAATTA | 1 | 673 | 690 | -327 | -310 | -18,13 | 0,01 | -3,89 | >0.1 | 4,24 | 5,45 | 0 | 1,12 | 0 | 3,43 | 0 |
| RARB(VAR.2) | AAGTGACAGGAGAGGTA | 1 | 764 | 781 | -236 | -219 | -24,87 | 0,01 | -6,25 | >0.1 | 5,52 | 5,82 | 0 | 1,36 | 0 | 5,92 | 0 |
| RARB(VAR.2) | AGTGACAGGAGAGGTAA | 1 | 765 | 782 | -235 | -218 | -24,06 | 0,01 | -5,43 | >0.1 | 5,52 | 5,83 | 0 | 1,37 | 0 | 5,92 | 0 |
| RARB(VAR.2) | AGGAGAGGTAAGGTTCT | 1 | 771 | 788 | -229 | -212 | -14,62 | 0,01 | 4,25 | >0.1 | 5,52 | 5,85 | 0 | 1,39 | 0 | 6,1 | 0 |
| RARB(VAR.2) | AGGTTCTCTAGGATTAA | 1 | 781 | 798 | -219 | -202 | -24,71 | 0,01 | -5,93 | >0.1 | 5,52 | 5,91 | 0 | 1,44 | 0 | 5,92 | 0 |
| RARB(VAR.2) | CGCCCAACCCAAGTTCA | 1 | 946 | 963 | -54 | -37 | -4,09 | 0,001 | 12,51 | >0.1 | 5,52 | 7,46 | 0 | 1,88 | 0 | 1,74 | 0 |
| RARB(VAR.2) | AGTTCAAAGGCTGATAA | 1 | 957 | 974 | -43 | -26 | -21,33 | 0,01 | -4,47 | >0.1 | 5,52 | 7,73 | 0 | 1,88 | 0 | 1,74 | 0 |
| RARB(VAR.2) | ATCTCATGAGGAGGTTT | 1 | 982 | 999 | -18 | -1 | -18,13 | 0,01 | -0,48 | >0.1 | 5,52 | 8,69 | 0 | 1,87 | 0 | 1,57 | 0 |
| RARB(VAR.2) | TAGTCTAGGGAAAGTCA | 1 | 999 | 1016 | -1 | 16 | -21,9 | 0,01 | -3,93 | >0.1 | 5,52 | 9,12 | 0 | 1,87 | 0 | 1,46 | 0 |
| RARB(VAR.2) | AAGTCATTCAGTGGATG | 1 | 1010 | 1027 | 10 | 27 | -23,99 | 0,01 | -6,2 | >0.1 | 5,52 | 8,99 | 0 | 1,86 | 0 | 1,42 | 0 |
| RARB(VAR.2) | GGACGATGTCAAGCTCT | 1 | 1044 | 1061 | 44 | 61 | -22,26 | 0,01 | -5,73 | >0.1 | 5,52 | 7,73 | 0 | 1,86 | 0 | 1,42 | 0 |
| RARB(VAR.2) | CATTGAGGAACAGGCCA | 1 | 1108 | 1125 | 108 | 125 | -25,86 | 0,01 | -6,06 | >0.1 | 5,52 | 6,67 | 0 | 1,85 | 3,22 | 2,54 | 0 |
| RARB(VAR.2) | TGTTCTATCAAAGTTCA | 1 | 1164 | 1181 | 164 | 181 | -8,18 | 0,001 | 9,91 | >0.1 | 5,52 | 6,19 | 0 | 1,72 | 1,79 | 2,86 | 0 |
| RARB(VAR.2) | AGGCTGAGAAGGAGCCA | -1 | 1064 | 1081 | 64 | 81 | -20,99 | 0,01 | -2,27 | >0.1 | 5,52 | 7,28 | 0 | 1,85 | 1,79 | 2,27 | 0 |
| RARB(VAR.2) | AGATCACATCCACTGAA | -1 | 1016 | 1033 | 16 | 33 | -24,03 | 0,01 | -6,53 | >0.1 | 5,52 | 8,81 | 0 | 1,86 | 0 | 1,31 | 0 |
| RARB(VAR.2) | AGCCAATATAAAGTTCA | -1 | 872 | 889 | -128 | -111 | -4,78 | 0,001 | 12,11 | >0.1 | 5,52 | 6,52 | 0 | 1,9 | 0 | 2,95 | 0 |
| RARB(VAR.2) | AGTTCATCCTGGAGAGG | -1 | 861 | 878 | -139 | -122 | -24,19 | 0,01 | -7,59 | >0.1 | 5,52 | 6,43 | 0 | 1,9 | 0 | 2,75 | 0 |
| RARB(VAR.2) | CTGTCACTTTATGGCCA | -1 | 756 | 773 | -244 | -227 | -11,59 | 0,001 | 7,15 | >0.1 | 5,52 | 5,78 | 0 | 1,33 | 0 | 6,1 | 0 |
| RARB(VAR.2) | AGTGACCAAAGAGGTCT | -1 | 664 | 681 | -336 | -319 | -22,59 | 0,01 | -8,13 | >0.1 | 4,11 | 5,42 | 0 | 1,1 | 0 | 3,83 | 0 |
| RARB(VAR.2) | ATTTCATTAATAGGTTA | -1 | 597 | 614 | -403 | -386 | -10,15 | 0,001 | 0,93 | >0.1 | 3,48 | 5,22 | 0 | 1,02 | 0 | 1,36 | 0 |
| RARB(VAR.2) | AGGTTATCTCACATTTA | -1 | 586 | 603 | -414 | -397 | -19,33 | 0,01 | -9,72 | >0.1 | 3,4 | 5,19 | 0 | 1,02 | 0 | 0 | 0 |
| RARB(VAR.2) | GGCGCGATCTCGGCTCA | -1 | 400 | 417 | -600 | -583 | -24,38 | 0,01 | -10,16 | >0.1 | 2,63 | 4,78 | 0 | 1,04 | 0 | 5,76 | 0 |
| RARB(VAR.2) | GGCTCACTGCAAGCTCT | -1 | 389 | 406 | -611 | -594 | -8,41 | 0,001 | 6,03 | >0.1 | 2,6 | 4,76 | 0 | 1,05 | 0 | 6,02 | 0 |
| RARB(VAR.2) | GGTTCACGCCATTCTCC | -1 | 364 | 381 | -636 | -619 | -25,41 | 0,01 | -10,35 | >0.1 | 2,54 | 4,72 | 0 | 1,08 | 0 | 6,73 | 0 |
| RARB(VAR.2) | ACGCCCAGCTAAGTTTT | -1 | 303 | 320 | -697 | -680 | -24,76 | 0,01 | -11,19 | >0.1 | 2,66 | 4,62 | 0 | 1,2 | 0 | 5,09 | 0 |
| RARB(VAR.2) | CGGCCTCCCAAAGTGCT | -1 | 208 | 225 | -792 | -775 | -19,61 | 0,01 | -5,96 | >0.1 | 2,99 | 4,49 | 0 | 1,58 | 0 | 4,59 | 0 |
| REL | CGGTGAAACC | 1 | 273 | 283 | -727 | -717 | 3,97 | 0,01 | 17,67 | >0.1 | 2,76 | 4,56 | 0 | 1,28 | 0 | 5,09 | 0 |

| | | | | | | | | | | | | | | | | | |
|---|---|---|---|---|---|---|---|---|---|---|---|---|---|---|---|---|---|
| REL | TGAGATAACC | 1 | 592 | 602 | -408 | -398 | 3 | 0,01 | 12,62 | >0.1 | 3,42 | 5,19 | 0 | 1,01 | 0 | 0 | 0 |
| REL | TTGGTCTTTC | 1 | 928 | 938 | -72 | -62 | 2,71 | 0,01 | 18,26 | >0.1 | 5,61 | 7,03 | 0 | 1,88 | 0 | 1,03 | 0 |
| REL | TGGTCTTTCC | 1 | 929 | 939 | -71 | -61 | 11,38 | 0,0001 | 26,95 | >0.1 | 5,61 | 7,05 | 0 | 1,88 | 0 | 1,03 | 0 |
| REL | GGAGGTTTTA | 1 | 991 | 1001 | -9 | 1 | 3,01 | 0,01 | 20,81 | >0.1 | 5,61 | 8,74 | 0 | 1,87 | 0 | 1,57 | 0 |
| REL | GGGAAAGTCA | 1 | 1006 | 1016 | 6 | 16 | 2,77 | 0,01 | 20,83 | >0.1 | 5,61 | 9,12 | 0 | 1,87 | 0 | 1,46 | 0 |
| REL | GGCCTGTTCC | -1 | 1114 | 1124 | 114 | 124 | 2,43 | 0,01 | 22,28 | >0.1 | 5,61 | 6,61 | 0 | 1,85 | 3,22 | 2,56 | 0 |
| REL | ATGACTTTCC | -1 | 1007 | 1017 | 7 | 17 | 4,07 | 0,01 | 22,13 | >0.1 | 5,61 | 9,12 | 0 | 1,87 | 0 | 1,46 | 0 |
| REL | TGAGATTTTC | -1 | 978 | 988 | -22 | -12 | 6,25 | 0,01 | 23,48 | >0.1 | 5,61 | 8,22 | 0 | 1,87 | 0 | 1,53 | 0 |
| REL | TGGGACTACA | -1 | 333 | 343 | -667 | -657 | 4,07 | 0,01 | 18,48 | >0.1 | 2,59 | 4,66 | 0 | 1,13 | 0 | 6,03 | 0 |
| REL | CGGGGTTTCA | -1 | 276 | 286 | -724 | -714 | 9,79 | 0,001 | 23,47 | >0.1 | 2,75 | 4,57 | 0 | 1,27 | 0 | 5,09 | 0 |
| REL | GGGTTTCACC | -1 | 274 | 284 | -726 | -716 | 3,01 | 0,01 | 16,81 | >0.1 | 2,76 | 4,57 | 0 | 1,28 | 0 | 5,19 | 0 |
| REL | TTTAATTTCC | -1 | 153 | 163 | -847 | -837 | 4,06 | 0,01 | 15,28 | >0.1 | 3,27 | 4,41 | 0 | 1,9 | 0 | 1,65 | 0 |
| REL | GCTCATTTCC | -1 | 64 | 74 | -936 | -926 | 4,4 | 0,01 | 14,54 | >0.1 | 3,95 | 4,31 | 0 | 1,88 | 0 | 0 | 0 |
| SPI1 | GTGAAGGAGAAGGA | 1 | 9 | 23 | -991 | -977 | -10,1 | 0,01 | 3,38 | >0.1 | 5,56 | 4,27 | 0 | 1,86 | 1,79 | 0 | 0 |
| SPI1 | AGAAGGAGGCACTC | 1 | 16 | 30 | -984 | -970 | -6,35 | 0,001 | 5,34 | >0.1 | 5,56 | 4,27 | 0 | 1,87 | 0 | 0 | 0 |
| SPI1 | AGAAATAGGAAATG | 1 | 57 | 71 | -943 | -929 | -8,55 | 0,01 | 1,64 | >0.1 | 4,01 | 4,31 | 0 | 1,88 | 0 | 0 | 0 |
| SPI1 | CAAAGGCAGATCAG | 1 | 88 | 102 | -912 | -898 | -17,56 | 0,01 | -7,64 | >0.1 | 3,69 | 4,34 | 0 | 1,88 | 0 | 0 | 0 |
| SPI1 | AAATACAAAAACTT | 1 | 296 | 310 | -704 | -690 | -16,07 | 0,01 | -3,11 | >0.1 | 2,69 | 4,6 | 0 | 1,22 | 0 | 4,45 | 0 |
| SPI1 | AAAAAAAAAACTG | 1 | 468 | 482 | -532 | -518 | -13,7 | 0,01 | -0,01 | >0.1 | 2,85 | 4,91 | 0 | 0,99 | 0 | 4,95 | 0 |
| SPI1 | AAAAAACTGATCAG | 1 | 473 | 487 | -527 | -513 | -18,37 | 0,01 | -4,51 | >0.1 | 2,86 | 4,92 | 0 | 0,99 | 0 | 5,09 | 0 |
| SPI1 | AGAAATATGAATTC | 1 | 485 | 499 | -515 | -501 | -19,04 | 0,01 | -4,07 | >0.1 | 2,91 | 4,94 | 0 | 0,99 | 0 | 6,13 | 0 |
| SPI1 | ATTAACCAAAAGTA | 1 | 730 | 744 | -270 | -256 | -16,38 | 0,01 | 1,99 | >0.1 | 5,56 | 5,66 | 0 | 1,24 | 0 | 5,92 | 0 |
| SPI1 | TGACAGGAGAGGTA | 1 | 767 | 781 | -233 | -219 | -17,91 | 0,01 | 0,52 | >0.1 | 5,56 | 5,82 | 0 | 1,36 | 0 | 5,69 | 0 |
| SPI1 | CTCATGAGGAGGTT | 1 | 984 | 998 | -16 | -2 | -11,66 | 0,01 | 6 | >0.1 | 5,56 | 8,66 | 0 | 1,87 | 0 | 1,57 | 0 |
| SPI1 | GTCTAGGGAAAGTC | 1 | 1001 | 1015 | 1 | 15 | -19,33 | 0,01 | -1,32 | >0.1 | 5,56 | 9,12 | 0 | 1,87 | 0 | 1,46 | 0 |
| SPI1 | CCATTGAGGAACAG | 1 | 1107 | 1121 | 107 | 121 | -9,42 | 0,01 | 10,4 | >0.1 | 5,56 | 6,68 | 0 | 1,85 | 3,22 | 2,52 | 0 |
| SPI1 | ACGAAGCCGAAGAC | 1 | 1149 | 1163 | 149 | 163 | -13,92 | 0,01 | 2,76 | >0.1 | 4,53 | 6,3 | 0 | 1,84 | 1,79 | 2,21 | 0 |
| SPI1 | AATTCCAAGAAGCA | -1 | 1183 | 1197 | 183 | 197 | -17,96 | 0,01 | 0,65 | >0.1 | 5,56 | 6,07 | 0 | 1,56 | 1,79 | 3,63 | 0 |
| SPI1 | CCAAGAAGCAAGTG | -1 | 1179 | 1193 | 179 | 193 | -15,02 | 0,01 | 3,45 | >0.1 | 5,56 | 6,1 | 0 | 1,59 | 1,79 | 3,43 | 0 |
| SPI1 | AGAAGGAGCCAGGA | -1 | 1061 | 1075 | 61 | 75 | -19,16 | 0,01 | -0,35 | >0.1 | 5,56 | 7,33 | 0 | 1,86 | 1,79 | 2,27 | 0 |
| SPI1 | GGAGCCAGGAAGAG | -1 | 1057 | 1071 | 57 | 71 | -19,05 | 0,01 | -2,06 | >0.1 | 5,56 | 7,42 | 0 | 1,86 | 0 | 2,16 | 0 |
| SPI1 | CAAAAAAAGAAAAA | -1 | 917 | 931 | -83 | -69 | -17,75 | 0,01 | -2,28 | >0.1 | 5,56 | 6,94 | 0 | 1,89 | 0 | 1,09 | 0 |
| SPI1 | AAAAAGAAAAAGAA | -1 | 914 | 928 | -86 | -72 | -8,21 | 0,01 | 7,26 | >0.1 | 5,56 | 6,9 | 0 | 1,89 | 0 | 1,12 | 0 |
| SPI1 | AAGAAAAAGAAGAA | -1 | 911 | 925 | -89 | -75 | -9,02 | 0,01 | 6,44 | >0.1 | 5,56 | 6,87 | 0 | 1,89 | 0 | 1,15 | 0 |
| SPI1 | AAGAAGAAGAACAC | -1 | 905 | 919 | -95 | -81 | -9,25 | 0,01 | 6,23 | >0.1 | 5,56 | 6,8 | 0 | 1,89 | 0 | 1,24 | 0 |
| SPI1 | CAAAACCGGTACTT | -1 | 739 | 753 | -261 | -247 | 0,09 | 0,001 | 18,71 | >0.1 | 5,56 | 5,69 | 0 | 1,27 | 0 | 6,1 | 0 |
| SPI1 | GGGATCATGAAGAT | -1 | 643 | 657 | -357 | -343 | -18,48 | 0,01 | -4,39 | >0.1 | 3,84 | 5,34 | 0 | 1,07 | 0 | 3,83 | 0 |
| BSX | GAAATTAA | 1 | 154 | 162 | -846 | -838 | 5,93 | 0,01 | 17,15 | >0.1 | 3,27 | 4,41 | 0 | 1,9 | 0 | 1,65 | 0 |
| BSX | TCTATTAA | 1 | 288 | 296 | -712 | -704 | 2,85 | 0,01 | 16,58 | >0.1 | 2,72 | 4,58 | 0 | 1,24 | 0 | 5,19 | 0 |
| BSX | TCCATAAA | 1 | 497 | 505 | -503 | -495 | 1,25 | 0,01 | 16,56 | >0.1 | 2,94 | 4,95 | 0 | 0,98 | 0 | 6,44 | 0 |
| BSX | CCTATTAA | 1 | 600 | 608 | -400 | -392 | 5,11 | 0,01 | 16,16 | >0.1 | 3,47 | 5,2 | 0 | 1,01 | 0 | 1,36 | 0 |
| BSX | AACATTAT | 1 | 637 | 645 | -363 | -355 | 3,45 | 0,01 | 15,22 | >0.1 | 3,76 | 5,31 | 0 | 1,05 | 0 | 1,66 | 0 |
| BSX | AAAATTAT | 1 | 683 | 691 | -317 | -309 | 3,65 | 0,01 | 17,63 | >0.1 | 4,34 | 5,46 | 0 | 1,12 | 0 | 3,07 | 0 |
| BSX | GCCATAAA | 1 | 758 | 766 | -242 | -234 | 2,72 | 0,01 | 21,52 | >0.1 | 5,64 | 5,75 | 0 | 1,31 | 0 | 6,1 | 0 |
| BSX | AGGATTAA | 1 | 790 | 798 | -210 | -202 | 2,15 | 0,01 | 20,65 | >0.1 | 5,64 | 5,91 | 0 | 1,43 | 0 | 5,52 | 0 |
| BSX | AAAATTAT | 1 | 830 | 838 | -170 | -162 | 3,65 | 0,01 | 20,38 | >0.1 | 5,64 | 6,14 | 0 | 1,67 | 0 | 3,28 | 0 |
| BSX | CTGATAAG | 1 | 967 | 975 | -33 | -25 | 2,34 | 0,01 | 19,3 | >0.1 | 5,64 | 7,76 | 0 | 1,87 | 0 | 1,7 | 0 |
| BSX | GGAATTAT | 1 | 1191 | 1199 | 191 | 199 | 2,57 | 0,01 | 21,35 | >0.1 | 5,64 | 6,02 | 0 | 1,5 | 1,79 | 3,82 | 0 |
| BSX | GTGGTTAA | -1 | 1143 | 1151 | 143 | 151 | 5,25 | 0,01 | 25,67 | >0.1 | 5,64 | 6,35 | 0 | 1,84 | 3,22 | 3,37 | 0 |
| BSX | GCAGTTAC | -1 | 1088 | 1096 | 88 | 96 | 7,18 | 0,001 | 25,58 | >0.1 | 5,64 | 6,9 | 0 | 1,85 | 1,79 | 2,21 | 0 |
| BSX | TTGGTTAA | -1 | 731 | 739 | -269 | -261 | 3,78 | 0,01 | 21,8 | >0.1 | 5,64 | 5,63 | 0 | 1,22 | 0 | 5,52 | 0 |
| BSX | GCTGTTAA | -1 | 709 | 717 | -291 | -283 | 1,79 | 0,01 | 18,96 | >0.1 | 5,64 | 5,55 | 0 | 1,17 | 0 | 4,81 | 0 |
| BSX | TTCATTAA | -1 | 604 | 612 | -396 | -388 | 6,49 | 0,01 | 17,63 | >0.1 | 3,49 | 5,21 | 0 | 1,02 | 0 | 1,41 | 0 |
| BSX | AGGATTAC | -1 | 200 | 208 | -800 | -792 | 2,42 | 0,01 | 16,03 | >0.1 | 3,04 | 4,46 | 0 | 1,63 | 0 | 4,48 | 0 |
| BSX | AAAATTAT | -1 | 132 | 140 | -868 | -860 | 3,65 | 0,01 | 14,89 | >0.1 | 3,4 | 4,38 | 0 | 1,89 | 0 | 1,57 | 0 |
| BSX | TCACTTAC | -1 | 5 | 13 | -995 | -987 | 3,23 | 0,01 | 16,78 | >0.1 | 5,64 | 4,26 | 0 | 1,86 | 1,79 | 0 | 0 |
| HIC1 | GTCCCAGCT | 1 | 337 | 346 | -663 | -654 | 7,88 | 0,01 | 22,16 | >0.1 | 2,58 | 4,66 | 0 | 1,12 | 0 | 5,92 | 0 |
| HIC1 | GCGCCACTG | 1 | 412 | 421 | -588 | -579 | 4,3 | 0,01 | 18,53 | >0.1 | 2,66 | 4,79 | 0 | 1,02 | 0 | 5,76 | 0 |
| HIC1 | GCTCCAGCC | 1 | 422 | 431 | -578 | -569 | 2,83 | 0,01 | 16,76 | >0.1 | 2,69 | 4,81 | 0 | 1,02 | 0 | 5,42 | 0 |

| | | | | | | | | | | | | | | | | | |
|---|---|---|---|---|---|---|---|---|---|---|---|---|---|---|---|---|---|
| HIC1 | TTGCCCAAG | 1 | 541 | 550 | -459 | -450 | 4,38 | 0,01 | 13,56 | >0.1 | 3,14 | 5,06 | 0 | 0,98 | 0 | 0 | 0 |
| HIC1 | ACACCAACA | 1 | 632 | 641 | -368 | -359 | 4,01 | 0,01 | 15,72 | >0.1 | 3,72 | 5,3 | 0 | 1,05 | 0 | 1,66 | 0 |
| HIC1 | GCGCCCAAC | 1 | 945 | 954 | -55 | -46 | 4,83 | 0,01 | 21,42 | >0.1 | 5,62 | 7,29 | 0 | 1,88 | 0 | 1,8 | 0 |
| HIC1 | AGGCCAAGA | 1 | 1119 | 1128 | 119 | 128 | 2,76 | 0,01 | 22,68 | >0.1 | 5,62 | 6,56 | 0 | 1,85 | 3,22 | 2,67 | 0 |
| HIC1 | ATTCCAAGA | -1 | 1187 | 1196 | 187 | 196 | 3,09 | 0,01 | 21,69 | >0.1 | 5,62 | 6,05 | 0 | 1,53 | 1,79 | 3,61 | 0 |
| HIC1 | GAGCCAAGA | -1 | 1030 | 1039 | 30 | 39 | 2,08 | 0,01 | 19,14 | >0.1 | 5,62 | 8,24 | 0 | 1,86 | 0 | 1,34 | 0 |
| HIC1 | CTGCCTCCC | -1 | 382 | 391 | -618 | -609 | 5,72 | 0,01 | 20,1 | >0.1 | 2,57 | 4,73 | 0 | 1,06 | 0 | 6,02 | 0 |
| HIC1 | CTCCCAGGT | -1 | 378 | 387 | -622 | -613 | 2,42 | 0,01 | 17,17 | >0.1 | 2,56 | 4,73 | 0 | 1,06 | 0 | 6,4 | 0 |
| HIC1 | ACGCCATTC | -1 | 367 | 376 | -633 | -624 | 3,41 | 0,01 | 18,34 | >0.1 | 2,53 | 4,71 | 0 | 1,08 | 0 | 6,61 | 0 |
| HIC1 | GTGCCCACC | -1 | 323 | 332 | -677 | -668 | 12,71 | 0,0001 | 26,68 | >0.1 | 2,62 | 4,64 | 0 | 1,15 | 0 | 5,56 | 0 |
| HIC1 | ACGCCCAGC | -1 | 311 | 320 | -689 | -680 | 5,69 | 0,01 | 20,27 | >0.1 | 2,65 | 4,62 | 0 | 1,18 | 0 | 6,13 | 0 |
| HIC1 | CTGCCCAAC | -1 | 226 | 235 | -774 | -765 | 6,34 | 0,01 | 20,5 | >0.1 | 2,93 | 4,5 | 0 | 1,47 | 0 | 5,26 | 0 |
| HIC1 | CTCCCAAAG | -1 | 212 | 221 | -788 | -779 | 2,43 | 0,01 | 16,04 | >0.1 | 2,99 | 4,48 | 0 | 1,55 | 0 | 4,59 | 0 |
| HIC1 | GTGCCCAGC | -1 | 177 | 186 | -823 | -814 | 9,95 | 0,001 | 22,58 | >0.1 | 3,15 | 4,44 | 0 | 1,81 | 0 | 3,24 | 0 |
| HIC1 | GTGCCTCCT | -1 | 19 | 28 | -981 | -972 | 7,09 | 0,01 | 18,85 | >0.1 | 5,62 | 4,27 | 0 | 1,87 | 0 | 0 | 0 |
| HIC2 | TTGCAAAGG | 1 | 85 | 94 | -915 | -906 | 3,1 | 0,01 | 13,06 | >0.1 | 3,74 | 4,33 | 0 | 1,88 | 0 | 0 | 0 |
| HIC2 | GTCCCAGCT | 1 | 337 | 346 | -663 | -654 | 6,79 | 0,01 | 21,07 | >0.1 | 2,58 | 4,66 | 0 | 1,12 | 0 | 5,92 | 0 |
| HIC2 | GCGCCACTG | 1 | 412 | 421 | -588 | -579 | 2,67 | 0,01 | 16,9 | >0.1 | 2,66 | 4,79 | 0 | 1,02 | 0 | 5,76 | 0 |
| HIC2 | GTGCTCCAG | 1 | 420 | 429 | -580 | -571 | 2,58 | 0,01 | 16,38 | >0.1 | 2,68 | 4,8 | 0 | 1,02 | 0 | 5,3 | 0 |
| HIC2 | TTGCCCAAG | 1 | 541 | 550 | -459 | -450 | 7,63 | 0,01 | 16,81 | >0.1 | 3,14 | 5,06 | 0 | 0,98 | 0 | 0 | 0 |
| HIC2 | CTGCTCAGC | 1 | 938 | 947 | -62 | -53 | 3,33 | 0,01 | 18,9 | >0.1 | 5,62 | 7,17 | 0 | 1,88 | 0 | 0,9 | 0 |
| HIC2 | GCGCCCAAC | 1 | 945 | 954 | -55 | -46 | 6,66 | 0,01 | 23,25 | >0.1 | 5,62 | 7,29 | 0 | 1,88 | 0 | 1,8 | 0 |
| HIC2 | CTGCTCAGT | 1 | 1095 | 1104 | 95 | 104 | 2,53 | 0,01 | 22,39 | >0.1 | 5,62 | 6,82 | 0 | 1,85 | 3,22 | 2,34 | 0 |
| HIC2 | ATTCCAAGA | -1 | 1187 | 1196 | 187 | 196 | 3,51 | 0,01 | 22,11 | >0.1 | 5,62 | 6,05 | 0 | 1,53 | 1,79 | 3,61 | 0 |
| HIC2 | GTCCCCTGT | -1 | 1039 | 1048 | 39 | 48 | 5,5 | 0,01 | 22,2 | >0.1 | 5,62 | 7,88 | 0 | 1,86 | 0 | 1,34 | 0 |
| HIC2 | ATACACGGA | -1 | 698 | 707 | -302 | -293 | 2,5 | 0,01 | 17,97 | >0.1 | 5,62 | 5,51 | 0 | 1,15 | 0 | 3,19 | 0 |
| HIC2 | ATTCCCAGA | -1 | 559 | 568 | -441 | -432 | 5,23 | 0,01 | 14,55 | >0.1 | 3,23 | 5,1 | 0 | 0,99 | 0 | 0 | 0 |
| HIC2 | TTGCCCTTG | -1 | 507 | 516 | -493 | -484 | 4,81 | 0,01 | 19,44 | >0.1 | 2,98 | 4,98 | 0 | 0,98 | 0 | 5,68 | 0 |
| HIC2 | TCGCCCAGG | -1 | 429 | 438 | -571 | -562 | 3,92 | 0,01 | 18,26 | >0.1 | 2,71 | 4,82 | 0 | 1,01 | 0 | 5,8 | 0 |
| HIC2 | CTGCAAGCT | -1 | 391 | 400 | -609 | -600 | 4,06 | 0,01 | 18,47 | >0.1 | 2,6 | 4,75 | 0 | 1,05 | 0 | 6,02 | 0 |
| HIC2 | CTCCCAGGT | -1 | 378 | 387 | -622 | -613 | 3,43 | 0,01 | 18,18 | >0.1 | 2,56 | 4,73 | 0 | 1,06 | 0 | 6,4 | 0 |
| HIC2 | GTTCACGCC | -1 | 371 | 380 | -629 | -620 | 3,61 | 0,01 | 18,21 | >0.1 | 2,54 | 4,72 | 0 | 1,07 | 0 | 6,27 | 0 |
| HIC2 | GTGCCCACC | -1 | 323 | 332 | -677 | -668 | 13,56 | 1E-05 | 27,53 | >0.1 | 2,62 | 4,64 | 0 | 1,15 | 0 | 5,56 | 0 |
| HIC2 | ACGCCCAGC | -1 | 311 | 320 | -689 | -680 | 7,54 | 0,01 | 22,12 | >0.1 | 2,65 | 4,62 | 0 | 1,18 | 0 | 6,13 | 0 |
| HIC2 | CTGCCCAAC | -1 | 226 | 235 | -774 | -765 | 9,02 | 0,001 | 23,18 | >0.1 | 2,93 | 4,5 | 0 | 1,47 | 0 | 5,26 | 0 |
| HIC2 | CTCCCAAAG | -1 | 212 | 221 | -788 | -779 | 2,72 | 0,01 | 16,33 | >0.1 | 2,99 | 4,48 | 0 | 1,55 | 0 | 4,59 | 0 |
| HIC2 | GTGCCCAGC | -1 | 177 | 186 | -823 | -814 | 12,73 | 0,0001 | 25,36 | >0.1 | 3,15 | 4,44 | 0 | 1,81 | 0 | 3,24 | 0 |
| HIC2 | TTGCAAAAC | -1 | 82 | 91 | -918 | -909 | 2,71 | 0,01 | 12,69 | >0.1 | 3,77 | 4,33 | 0 | 1,88 | 0 | 0 | 0 |
| NR2F6 | AGATCACAAGGTCA | 1 | 233 | 247 | -767 | -753 | -1,53 | 0,01 | 12,07 | >0.1 | 2,9 | 4,52 | 0 | 1,44 | 0 | 4,74 | 0 |
| NR2F6 | AAGGTCAGGAGATA | 1 | 240 | 254 | -760 | -746 | -1,49 | 0,01 | 11,64 | >0.1 | 2,87 | 4,52 | 0 | 1,41 | 0 | 4,33 | 0 |
| NR2F6 | GGAGGCAGAGCTTG | 1 | 383 | 397 | -617 | -603 | -2,16 | 0,01 | 12,36 | >0.1 | 2,58 | 4,75 | 0 | 1,06 | 0 | 6,13 | 0 |
| NR2F6 | CAAGGGCAAAGTCA | 1 | 507 | 521 | -493 | -479 | -4,1 | 0,01 | 7,62 | >0.1 | 3 | 4,99 | 0 | 0,99 | 0 | 2,75 | 0 |
| NR2F6 | AAGGGCAAAGTCAT | 1 | 508 | 522 | -492 | -478 | -4,88 | 0,01 | 6,96 | >0.1 | 3 | 4,99 | 0 | 0,99 | 0 | 2,86 | 0 |
| NR2F6 | AAGGGAAAATGTTG | 1 | 530 | 544 | -470 | -456 | 1,01 | 0,01 | 11,75 | >0.1 | 3,1 | 5,04 | 0 | 0,99 | 0 | 1,61 | 0 |
| NR2F6 | AAAGTGACAGGAGA | 1 | 763 | 777 | -237 | -223 | -3,8 | 0,01 | 14,82 | >0.1 | 5,56 | 5,8 | 0 | 1,35 | 0 | 5,92 | 0 |
| NR2F6 | GAGAGGTAAGGTTC | 1 | 773 | 787 | -227 | -213 | -4,42 | 0,01 | 14,69 | >0.1 | 5,56 | 5,85 | 0 | 1,39 | 0 | 6,32 | 0 |
| NR2F6 | AGAGGTAAGGTTCT | 1 | 774 | 788 | -226 | -212 | -2,8 | 0,01 | 16,32 | >0.1 | 5,56 | 5,85 | 0 | 1,39 | 0 | 6,32 | 0 |
| NR2F6 | AGGATTAAAGAATA | 1 | 790 | 804 | -210 | -196 | -3,45 | 0,01 | 14,83 | >0.1 | 5,56 | 5,94 | 0 | 1,47 | 0 | 5,32 | 0 |
| NR2F6 | AAGTTCAAAGGCTG | 1 | 956 | 970 | -44 | -30 | 7,92 | 0,001 | 24,71 | >0.1 | 5,56 | 7,62 | 0 | 1,88 | 0 | 1,74 | 0 |
| NR2F6 | GAGAGAAAATCTCA | 1 | 974 | 988 | -26 | -12 | -4,96 | 0,01 | 12,26 | >0.1 | 5,56 | 8,22 | 0 | 1,87 | 0 | 1,57 | 0 |
| NR2F6 | GAGGTTTTAGTCTA | 1 | 992 | 1006 | -8 | 6 | -2,36 | 0,01 | 15,51 | >0.1 | 5,56 | 8,92 | 0 | 1,87 | 0 | 1,53 | 0 |
| NR2F6 | TGGCTCACAGGGGA | 1 | 1033 | 1047 | 33 | 47 | 3,12 | 0,001 | 19,95 | >0.1 | 5,56 | 8,1 | 0 | 1,86 | 0 | 1,31 | 0 |
| NR2F6 | AGGGGACGATGTCA | 1 | 1041 | 1055 | 41 | 55 | -0,75 | 0,01 | 15,11 | >0.1 | 5,56 | 7,82 | 0 | 1,86 | 0 | 0,62 | 0 |
| NR2F6 | TGAGGAACAGGCCA | 1 | 1111 | 1125 | 111 | 125 | -2,29 | 0,01 | 17,49 | >0.1 | 5,56 | 6,64 | 0 | 1,85 | 3,22 | 2,52 | 0 |
| NR2F6 | TCTATCAAAGTTCA | 1 | 1167 | 1181 | 167 | 181 | 4 | 0,001 | 22,16 | >0.1 | 5,56 | 6,17 | 0 | 1,69 | 1,79 | 2,94 | 0 |
| NR2F6 | GTGAGCCAAGATCA | -1 | 1027 | 1041 | 27 | 41 | -1,99 | 0,01 | 15,16 | >0.1 | 5,56 | 8,39 | 0 | 1,86 | 0 | 1,34 | 0 |
| NR2F6 | CAATATAAAGTTCA | -1 | 872 | 886 | -128 | -114 | -1,04 | 0,01 | 15,86 | >0.1 | 5,56 | 6,5 | 0 | 1,9 | 0 | 2,95 | 0 |
| NR2F6 | CACGGAAAATGTTA | -1 | 690 | 704 | -310 | -296 | -2,24 | 0,01 | 13,03 | >0.1 | 5,56 | 5,5 | 0 | 1,14 | 0 | 3,07 | 0 |
| NR2F6 | TCATTAATAGGTTA | -1 | 597 | 611 | -403 | -389 | -4,68 | 0,01 | 6,38 | >0.1 | 3,47 | 5,21 | 0 | 1,02 | 0 | 1,36 | 0 |

| | | | | | | | | | | | | | | | | | |
|---|---|---|---|---|---|---|---|---|---|---|---|---|---|---|---|---|---|
| NR2F6 | GGGACTACAGGTGC | -1 | 328 | 342 | -672 | -658 | -3,66 | 0,01 | 10,19 | >0.1 | 2,59 | 4,65 | 0 | 1,14 | 0 | 5,46 | 0 |
| NR2F6 | AAGTGCTAGGATTA | -1 | 201 | 215 | -799 | -785 | -3,5 | 0,01 | 10,22 | >0.1 | 3,02 | 4,47 | 0 | 1,63 | 0 | 4,59 | 0 |
| NR2F6 | CACTCCACAGGTCA | -1 | 108 | 122 | -892 | -878 | 4,75 | 0,001 | 15,3 | >0.1 | 3,54 | 4,36 | 0 | 1,89 | 0 | 0,77 | 0 |
| NR2F6 | CAGGTCAACTCTCC | -1 | 101 | 115 | -899 | -885 | -4,13 | 0,01 | 6,47 | >0.1 | 3,59 | 4,36 | 0 | 1,89 | 0 | 0,77 | 0 |
| NR2F1 | AATAGGTAAGTGA | 1 | 0 | 13 | -1000 | -987 | -9,73 | 0,01 | 3,75 | >0.1 | 5,57 | 4,26 | 0 | 1,86 | 1,79 | 0 | 0 |
| NR2F1 | AAGAAGTCAGCCA | 1 | 31 | 44 | -969 | -956 | -5,01 | 0,01 | 5,57 | >0.1 | 4,43 | 4,28 | 0 | 1,87 | 0 | 0 | 0 |
| NR2F1 | GGCAGATCAGGAG | 1 | 92 | 105 | -908 | -895 | -7,2 | 0,01 | 3,5 | >0.1 | 3,67 | 4,34 | 0 | 1,88 | 0 | 0,81 | 0 |
| NR2F1 | AGTGGCTCACGCC | 1 | 186 | 199 | -814 | -801 | -10,19 | 0,01 | 2,95 | >0.1 | 3,09 | 4,45 | 0 | 1,74 | 0 | 3,86 | 0 |
| NR2F1 | GGGAGGCCGAGTT | 1 | 216 | 229 | -784 | -771 | -12,28 | 0,01 | 2,23 | >0.1 | 2,96 | 4,49 | 0 | 1,53 | 0 | 5,52 | 0 |
| NR2F1 | GGCAGATCACAAG | 1 | 230 | 243 | -770 | -757 | -7,23 | 0,01 | 6,65 | >0.1 | 2,91 | 4,51 | 0 | 1,46 | 0 | 5,01 | 0 |
| NR2F1 | ACAAGGTCAGGAG | 1 | 238 | 251 | -762 | -749 | 13,32 | 0,0001 | 26,56 | >0.1 | 2,88 | 4,52 | 0 | 1,42 | 0 | 4,42 | 0 |
| NR2F1 | CCGAGATCGCGCC | 1 | 404 | 417 | -596 | -583 | -10,4 | 0,01 | 3,82 | >0.1 | 2,64 | 4,78 | 0 | 1,03 | 0 | 5,76 | 0 |
| NR2F1 | ACAAGGGCAAAGT | 1 | 506 | 519 | -494 | -481 | -5,01 | 0,01 | 8,14 | >0.1 | 2,99 | 4,98 | 0 | 0,99 | 0 | 4,19 | 0 |
| NR2F1 | GCAAAGTCATGTA | 1 | 512 | 525 | -488 | -475 | -6,75 | 0,01 | 5,11 | >0.1 | 3,01 | 5 | 0 | 0,99 | 0 | 2,86 | 0 |
| NR2F1 | CTTTGGTCACTGT | 1 | 670 | 683 | -330 | -317 | -8,68 | 0,01 | 5,57 | >0.1 | 4,16 | 5,43 | 0 | 1,11 | 0 | 3,55 | 0 |
| NR2F1 | GAGAGGTAAGGTT | 1 | 773 | 786 | -227 | -214 | -5,73 | 0,01 | 13,17 | >0.1 | 5,57 | 5,84 | 0 | 1,38 | 0 | 6,1 | 0 |
| NR2F1 | ATTGGCTCAGCAG | 1 | 882 | 895 | -118 | -105 | -7,84 | 0,01 | 9,27 | >0.1 | 5,57 | 6,57 | 0 | 1,89 | 0 | 3,07 | 0 |
| NR2F1 | CCAAGTTCAAAGG | 1 | 954 | 967 | -46 | -33 | -0,34 | 0,001 | 16,35 | >0.1 | 5,57 | 7,55 | 0 | 1,88 | 0 | 1,7 | 0 |
| NR2F1 | GGAAAGTCATTCA | 1 | 1007 | 1020 | 7 | 20 | -8,86 | 0,01 | 9,11 | >0.1 | 5,57 | 9,12 | 0 | 1,87 | 0 | 1,42 | 0 |
| NR2F1 | CTTGGCTCACAGG | 1 | 1031 | 1044 | 31 | 44 | -4,81 | 0,01 | 12,09 | >0.1 | 5,57 | 8,19 | 0 | 1,86 | 0 | 1,28 | 0 |
| NR2F1 | CAGGGGACGATGT | 1 | 1040 | 1053 | 40 | 53 | -10,38 | 0,01 | 6,29 | >0.1 | 5,57 | 7,85 | 0 | 1,86 | 0 | 1,39 | 0 |
| NR2F1 | ACGATGTCAAGCT | 1 | 1046 | 1059 | 46 | 59 | -5,59 | 0,01 | 10,93 | >0.1 | 5,57 | 7,68 | 0 | 1,86 | 0 | 1,42 | 0 |
| NR2F1 | AACAGGCCAAGAC | 1 | 1116 | 1129 | 116 | 129 | -6,49 | 0,01 | 13,34 | >0.1 | 5,57 | 6,59 | 0 | 1,85 | 3,22 | 2,6 | 0 |
| NR2F1 | CAAAGTTCACTTG | 1 | 1172 | 1185 | 172 | 185 | -1,29 | 0,001 | 16,95 | >0.1 | 5,57 | 6,14 | 0 | 1,65 | 1,79 | 3,09 | 0 |
| NR2F1 | AACAGGTCTTCGG | -1 | 1155 | 1168 | 155 | 168 | -4,32 | 0,01 | 12,3 | >0.1 | 4,41 | 6,26 | 0 | 1,81 | 1,79 | 2,35 | 0 |
| NR2F1 | CCAAGATCACATC | -1 | 1023 | 1036 | 23 | 36 | -5,43 | 0,01 | 11,92 | >0.1 | 5,57 | 8,67 | 0 | 1,86 | 0 | 1,25 | 0 |
| NR2F1 | CTTGGGTTGGGCG | -1 | 946 | 959 | -54 | -41 | -12,49 | 0,01 | 4,08 | >0.1 | 5,57 | 7,38 | 0 | 1,88 | 0 | 1,74 | 0 |
| NR2F1 | TAAAGTTCATCCT | -1 | 868 | 881 | -132 | -119 | -6,12 | 0,01 | 10,75 | >0.1 | 5,57 | 6,45 | 0 | 1,9 | 0 | 2,95 | 0 |
| NR2F1 | GAGAGGACAGATA | -1 | 854 | 867 | -146 | -133 | -5,46 | 0,01 | 11,22 | >0.1 | 5,57 | 6,35 | 0 | 1,9 | 0 | 2,86 | 0 |
| NR2F1 | AAGAGGTCTAGAA | -1 | 660 | 673 | -340 | -327 | -2,87 | 0,01 | 11,81 | >0.1 | 4,03 | 5,4 | 0 | 1,09 | 0 | 4,16 | 0 |
| NR2F1 | TAGGGATCATGAA | -1 | 646 | 659 | -354 | -341 | -5,48 | 0,01 | 8,64 | >0.1 | 3,87 | 5,35 | 0 | 1,07 | 0 | 3,83 | 0 |
| NR2F1 | AATAGGTTATCTC | -1 | 593 | 606 | -407 | -394 | -9,88 | 0,01 | -0,23 | >0.1 | 3,44 | 5,2 | 0 | 1,02 | 0 | 0 | 0 |
| NR2F1 | TCAAGATCATATT | -1 | 565 | 578 | -435 | -422 | -8,62 | 0,01 | 0,77 | >0.1 | 3,27 | 5,12 | 0 | 1 | 0 | 0 | 0 |
| NR2F1 | CTCGGCTCACTGC | -1 | 396 | 409 | -604 | -591 | -9,39 | 0,01 | 4,95 | >0.1 | 2,62 | 4,77 | 0 | 1,04 | 0 | 5,91 | 0 |
| NR2F1 | CCAGGTTCACGCC | -1 | 371 | 384 | -629 | -616 | -4,63 | 0,01 | 10,11 | >0.1 | 2,55 | 4,72 | 0 | 1,07 | 0 | 6,4 | 0 |
| NR2F1 | CAGGCGTGAGCCA | -1 | 188 | 201 | -812 | -799 | -12,07 | 0,01 | 1,23 | >0.1 | 3,08 | 4,46 | 0 | 1,72 | 0 | 4,04 | 0 |
| NR2F1 | CACAGGTCAACTC | -1 | 104 | 117 | -896 | -883 | 9,64 | 0,001 | 20,22 | >0.1 | 3,57 | 4,36 | 0 | 1,89 | 0 | 0,77 | 0 |
| NR2F1 | AAAAGCTCATTTC | -1 | 65 | 78 | -935 | -922 | -7,13 | 0,01 | 2,99 | >0.1 | 3,92 | 4,32 | 0 | 1,88 | 0 | 0 | 0 |
| NR2F2 | AATAGGTAAGT | 1 | 0 | 11 | -1000 | -989 | 3,36 | 0,01 | 16,87 | >0.1 | 5,6 | 4,25 | 0 | 1,86 | 1,79 | 0 | 0 |
| NR2F2 | AAGAAGTCAGC | 1 | 31 | 42 | -969 | -958 | 4,25 | 0,01 | 14,85 | >0.1 | 4,45 | 4,28 | 0 | 1,87 | 0 | 0 | 0 |
| NR2F2 | GCAAAGGCAGA | 1 | 87 | 98 | -913 | -902 | 1,71 | 0,01 | 11,64 | >0.1 | 3,72 | 4,34 | 0 | 1,88 | 0 | 0 | 0 |
| NR2F2 | GGCAGATCAGG | 1 | 92 | 103 | -908 | -897 | 1,51 | 0,01 | 12,24 | >0.1 | 3,67 | 4,34 | 0 | 1,88 | 0 | 0,83 | 0 |
| NR2F2 | AAATGGCCATG | 1 | 143 | 154 | -857 | -846 | 0,88 | 0,01 | 12,1 | >0.1 | 3,32 | 4,4 | 0 | 1,9 | 0 | 1,61 | 0 |
| NR2F2 | GGCAGATCACA | 1 | 230 | 241 | -770 | -759 | 1,47 | 0,01 | 15,36 | >0.1 | 2,91 | 4,51 | 0 | 1,45 | 0 | 5,01 | 0 |
| NR2F2 | ACAAGGTCAGG | 1 | 238 | 249 | -762 | -751 | 13,34 | 0,0001 | 26,69 | >0.1 | 2,88 | 4,52 | 0 | 1,42 | 0 | 4,53 | 0 |
| NR2F2 | ACAAGGGCAAA | 1 | 506 | 517 | -494 | -483 | 7,48 | 0,001 | 22,11 | >0.1 | 2,98 | 4,98 | 0 | 0,98 | 0 | 5,68 | 0 |
| NR2F2 | GCAAAGTCATG | 1 | 512 | 523 | -488 | -477 | 6,74 | 0,01 | 18,68 | >0.1 | 3,01 | 4,99 | 0 | 0,98 | 0 | 2,95 | 0 |
| NR2F2 | GGAAGGGAAAA | 1 | 528 | 539 | -472 | -461 | 0,72 | 0,01 | 13,36 | >0.1 | 3,08 | 5,03 | 0 | 0,98 | 0 | 3,55 | 0 |
| NR2F2 | TAAATGTGAGA | 1 | 586 | 597 | -414 | -403 | 2,05 | 0,01 | 11,62 | >0.1 | 3,39 | 5,17 | 0 | 1,01 | 0 | 0 | 0 |
| NR2F2 | CTTTGGTCACT | 1 | 670 | 681 | -330 | -319 | 2,93 | 0,01 | 17,3 | >0.1 | 4,15 | 5,42 | 0 | 1,1 | 0 | 3,7 | 0 |
| NR2F2 | GAGAGGTAAGG | 1 | 773 | 784 | -227 | -216 | 4,89 | 0,01 | 23,8 | >0.1 | 5,6 | 5,83 | 0 | 1,37 | 0 | 6,1 | 0 |
| NR2F2 | GTAAGGTTCTC | 1 | 778 | 789 | -222 | -211 | 1,24 | 0,01 | 20,63 | >0.1 | 5,6 | 5,86 | 0 | 1,4 | 0 | 6,54 | 0 |
| NR2F2 | CCAAGTTCAAA | 1 | 954 | 965 | -46 | -35 | 7,34 | 0,001 | 24,01 | >0.1 | 5,6 | 7,51 | 0 | 1,88 | 0 | 1,7 | 0 |
| NR2F2 | GGAAAGTCATT | 1 | 1007 | 1018 | 7 | 18 | 5,22 | 0,01 | 23,27 | >0.1 | 5,6 | 9,12 | 0 | 1,87 | 0 | 1,46 | 0 |
| NR2F2 | ACGATGTCAAG | 1 | 1046 | 1057 | 46 | 57 | 4,08 | 0,01 | 20,68 | >0.1 | 5,6 | 7,68 | 0 | 1,86 | 0 | 1,46 | 0 |
| NR2F2 | AACAGGCCAAG | 1 | 1116 | 1127 | 116 | 127 | 2,55 | 0,01 | 22,41 | >0.1 | 5,6 | 6,59 | 0 | 1,85 | 3,22 | 2,6 | 0 |
| NR2F2 | ACAAGTTTAAC | 1 | 1137 | 1148 | 137 | 148 | 1,66 | 0,01 | 21,79 | >0.1 | 5,6 | 6,4 | 0 | 1,84 | 3,22 | 3,07 | 0 |
| NR2F2 | CAAAGTTCACT | 1 | 1172 | 1183 | 172 | 183 | 7,1 | 0,001 | 25,25 | >0.1 | 5,6 | 6,14 | 0 | 1,65 | 1,79 | 2,97 | 0 |
| NR2F2 | AACAGGTCTTC | -1 | 1157 | 1168 | 157 | 168 | 1,85 | 0,01 | 18,41 | >0.1 | 4,39 | 6,24 | 0 | 1,79 | 1,79 | 2,34 | 0 |

| | | | | | | | | | | | | | | | | | |
|---|---|---|---|---|---|---|---|---|---|---|---|---|---|---|---|---|---|
| NR2F2 | AAAATGTCTTG | -1 | 1123 | 1134 | 123 | 134 | 1,41 | 0,01 | 21,32 | >0.1 | 5,6 | 6,52 | 0 | 1,85 | 3,22 | 2,72 | 0 |
| NR2F2 | CCAAGATCACA | -1 | 1025 | 1036 | 25 | 36 | 7,8 | 0,001 | 25,02 | >0.1 | 5,6 | 8,52 | 0 | 1,86 | 0 | 1,25 | 0 |
| NR2F2 | GAAAGACCAAA | -1 | 927 | 938 | -73 | -62 | 1,49 | 0,01 | 17,03 | >0.1 | 5,6 | 7,03 | 0 | 1,88 | 0 | 1,03 | 0 |
| NR2F2 | TAAAGTTCATC | -1 | 870 | 881 | -130 | -119 | 6,41 | 0,01 | 23,22 | >0.1 | 5,6 | 6,45 | 0 | 1,9 | 0 | 2,86 | 0 |
| NR2F2 | GAGAGGACAGA | -1 | 856 | 867 | -144 | -133 | 4,53 | 0,01 | 21,32 | >0.1 | 5,6 | 6,35 | 0 | 1,9 | 0 | 2,95 | 0 |
| NR2F2 | AAAATGTTATA | -1 | 688 | 699 | -312 | -301 | 3,39 | 0,01 | 17,34 | >0.1 | 4,47 | 5,48 | 0 | 1,13 | 0 | 2,86 | 0 |
| NR2F2 | AAGAGGTCTAG | -1 | 662 | 673 | -338 | -327 | 4,92 | 0,01 | 19,44 | >0.1 | 4,04 | 5,4 | 0 | 1,09 | 0 | 3,99 | 0 |
| NR2F2 | AATAGGTTATC | -1 | 595 | 606 | -405 | -394 | 5,74 | 0,01 | 15,39 | >0.1 | 3,44 | 5,2 | 0 | 1,01 | 0 | 0 | 0 |
| NR2F2 | TCAAGATCATA | -1 | 567 | 578 | -433 | -422 | 6,67 | 0,01 | 16,07 | >0.1 | 3,27 | 5,12 | 0 | 1 | 0 | 0 | 0 |
| NR2F2 | CCAGGTTCACG | -1 | 373 | 384 | -627 | -616 | 3,27 | 0,01 | 18,15 | >0.1 | 2,55 | 4,72 | 0 | 1,07 | 0 | 6,53 | 0 |
| NR2F2 | CACAGGTCAAC | -1 | 106 | 117 | -894 | -883 | 9,07 | 0,001 | 19,62 | >0.1 | 3,56 | 4,36 | 0 | 1,89 | 0 | 0,75 | 0 |
| NR2F2 | AAAAGCTCATT | -1 | 67 | 78 | -933 | -922 | 5,01 | 0,01 | 15,11 | >0.1 | 3,91 | 4,32 | 0 | 1,88 | 0 | 0 | 0 |
| RORA(VAR.2) | AATAGGTAAGTGAA | 1 | 0 | 14 | -1000 | -986 | -3,13 | 0,01 | 10,34 | >0.1 | 5,56 | 4,26 | 0 | 1,86 | 1,79 | 0 | 0 |
| RORA(VAR.2) | CTCAAAGAAGTCAG | 1 | 27 | 41 | -973 | -959 | -5,87 | 0,01 | 4,79 | >0.1 | 4,51 | 4,28 | 0 | 1,87 | 0 | 0 | 0 |
| RORA(VAR.2) | AAGAAGTCAGCCAC | 1 | 31 | 45 | -969 | -955 | -5,73 | 0,01 | 4,84 | >0.1 | 4,42 | 4,29 | 0 | 1,87 | 0 | 0 | 0 |
| RORA(VAR.2) | GAGAAATAGGAAAT | 1 | 56 | 70 | -944 | -930 | -3,08 | 0,01 | 7,13 | >0.1 | 4,02 | 4,31 | 0 | 1,87 | 0 | 0 | 0 |
| RORA(VAR.2) | CAAAGGCAGATCAG | 1 | 88 | 102 | -912 | -898 | -2,65 | 0,01 | 7,27 | >0.1 | 3,69 | 4,34 | 0 | 1,88 | 0 | 0 | 0 |
| RORA(VAR.2) | GCACAGTGGCTCAC | 1 | 182 | 196 | -818 | -804 | 2,36 | 0,001 | 15,64 | >0.1 | 3,11 | 4,45 | 0 | 1,77 | 0 | 3,95 | 0 |
| RORA(VAR.2) | GATCACAAGGTCAG | 1 | 234 | 248 | -766 | -752 | 6,98 | 0,001 | 20,46 | >0.1 | 2,89 | 4,52 | 0 | 1,44 | 0 | 4,63 | 0 |
| RORA(VAR.2) | ACTAACACGGTGAA | 1 | 266 | 280 | -734 | -720 | -5,5 | 0,01 | 7,6 | >0.1 | 2,78 | 4,56 | 0 | 1,31 | 0 | 4,45 | 0 |
| RORA(VAR.2) | AAAGACAAGGGCAA | 1 | 502 | 516 | -498 | -484 | -3,12 | 0,01 | 11,68 | >0.1 | 2,98 | 4,98 | 0 | 0,99 | 0 | 5,86 | 0 |
| RORA(VAR.2) | AAAATGTTGCCCAA | 1 | 535 | 549 | -465 | -451 | -3,99 | 0,01 | 6,83 | >0.1 | 3,12 | 5,05 | 0 | 0,99 | 0 | 1,66 | 0 |
| RORA(VAR.2) | TTTGATTTGGCCAT | 1 | 749 | 763 | -251 | -237 | -5,42 | 0,01 | 13,49 | >0.1 | 5,56 | 5,74 | 0 | 1,3 | 0 | 6,32 | 0 |
| RORA(VAR.2) | CTTATTTGATTCAC | 1 | 811 | 825 | -189 | -175 | -1,6 | 0,01 | 15,03 | >0.1 | 5,56 | 6,06 | 0 | 1,59 | 0 | 3,43 | 0 |
| RORA(VAR.2) | TTATATTGGCTCAG | 1 | 878 | 892 | -122 | -108 | 0,65 | 0,01 | 17,72 | >0.1 | 5,56 | 6,55 | 0 | 1,9 | 0 | 3,07 | 0 |
| RORA(VAR.2) | TTAGTCTAGGGAAA | 1 | 998 | 1012 | -2 | 12 | -4,9 | 0,01 | 13,11 | >0.1 | 5,56 | 9,12 | 0 | 1,87 | 0 | 1,46 | 0 |
| RORA(VAR.2) | GGAAAGTCATTCAG | 1 | 1007 | 1021 | 7 | 21 | -4,68 | 0,01 | 13,33 | >0.1 | 5,56 | 9,12 | 0 | 1,87 | 0 | 1,46 | 0 |
| RORA(VAR.2) | GGGGACGATGTCAA | 1 | 1042 | 1056 | 42 | 56 | -5,79 | 0,01 | 10,04 | >0.1 | 5,56 | 7,79 | 0 | 1,86 | 0 | 0,62 | 0 |
| RORA(VAR.2) | AAGAAGCAAGTGAA | -1 | 1177 | 1191 | 177 | 191 | -5,79 | 0,01 | 12,62 | >0.1 | 5,56 | 6,11 | 0 | 1,61 | 1,79 | 3,34 | 0 |
| RORA(VAR.2) | TTAAACTTGTCCAA | -1 | 1133 | 1147 | 133 | 147 | -4,26 | 0,01 | 15,82 | >0.1 | 5,56 | 6,43 | 0 | 1,84 | 3,22 | 3,02 | 0 |
| RORA(VAR.2) | TGAACTTGGGTTGG | -1 | 949 | 963 | -51 | -37 | -5,75 | 0,01 | 10,88 | >0.1 | 5,56 | 7,46 | 0 | 1,88 | 0 | 1,74 | 0 |
| RORA(VAR.2) | AATATAAAGTTCAT | -1 | 871 | 885 | -129 | -115 | -2,21 | 0,01 | 14,68 | >0.1 | 5,56 | 6,49 | 0 | 1,9 | 0 | 2,95 | 0 |
| RORA(VAR.2) | TTAAAGTGAATCAA | -1 | 816 | 830 | -184 | -170 | 0,72 | 0,01 | 17,41 | >0.1 | 5,56 | 6,09 | 0 | 1,62 | 0 | 3,43 | 0 |
| RORA(VAR.2) | GTACTTTTGGTTAA | -1 | 731 | 745 | -269 | -255 | -3,3 | 0,01 | 15,07 | >0.1 | 5,56 | 5,66 | 0 | 1,24 | 0 | 5,92 | 0 |
| RORA(VAR.2) | CATTAATAGGTTAT | -1 | 596 | 610 | -404 | -390 | -1,03 | 0,01 | 10,07 | >0.1 | 3,46 | 5,21 | 0 | 1,02 | 0 | 1,41 | 0 |
| RORA(VAR.2) | TGAGTCTCGGTCTG | -1 | 438 | 452 | -562 | -548 | -4,12 | 0,01 | 10,53 | >0.1 | 2,75 | 4,85 | 0 | 1,01 | 0 | 6,05 | 0 |
| RORA(VAR.2) | AGTAGCTGGGACTA | -1 | 335 | 349 | -665 | -651 | -4,8 | 0,01 | 9,49 | >0.1 | 2,58 | 4,67 | 0 | 1,13 | 0 | 5,92 | 0 |
| RORA(VAR.2) | TAAAATTATGACTA | -1 | 127 | 141 | -873 | -859 | -5,26 | 0,01 | 6,03 | >0.1 | 3,41 | 4,38 | 0 | 1,89 | 0 | 1,61 | 0 |
| RORA(VAR.2) | ACTCCACAGGTCAA | -1 | 107 | 121 | -893 | -879 | -4,57 | 0,01 | 5,97 | >0.1 | 3,54 | 4,36 | 0 | 1,89 | 0 | 0,75 | 0 |
| RORA(VAR.2) | TTAAAAAAGCTCAT | -1 | 68 | 82 | -932 | -918 | -1,63 | 0,01 | 8,45 | >0.1 | 3,88 | 4,32 | 0 | 1,88 | 0 | 0 | 0 |
| RORA(VAR.2) | TCACACTGGTTCTG | -1 | 44 | 58 | -956 | -942 | -5,07 | 0,01 | 5,29 | >0.1 | 4,19 | 4,3 | 0 | 1,87 | 0 | 0 | 0 |
| RORA(VAR.2) | GTTCTGTGGCTGAC | -1 | 36 | 50 | -964 | -950 | -3,97 | 0,01 | 6,51 | >0.1 | 4,32 | 4,29 | 0 | 1,87 | 0 | 0 | 0 |
| ATOH1 | AAAAAATGGC | 1 | 140 | 150 | -860 | -850 | 3,53 | 0,01 | 14,81 | >0.1 | 3,34 | 4,39 | 0 | 1,89 | 0 | 1,65 | 0 |
| ATOH1 | GGCACCTGTA | 1 | 327 | 337 | -673 | -663 | 0,53 | 0,01 | 14,49 | >0.1 | 2,6 | 4,65 | 0 | 1,14 | 0 | 5,56 | 0 |
| ATOH1 | GAAATATGAA | 1 | 486 | 496 | -514 | -504 | 2,71 | 0,01 | 17,8 | >0.1 | 2,9 | 4,93 | 0 | 0,98 | 0 | 6,27 | 0 |
| ATOH1 | GGAAAATGTT | 1 | 533 | 543 | -467 | -457 | 1,04 | 0,01 | 11,77 | >0.1 | 3,1 | 5,04 | 0 | 0,98 | 0 | 1,61 | 0 |
| ATOH1 | GGAATATGAT | 1 | 563 | 573 | -437 | -427 | 3,45 | 0,01 | 12,8 | >0.1 | 3,25 | 5,11 | 0 | 0,99 | 0 | 0 | 0 |
| ATOH1 | AACAGCTTTC | 1 | 711 | 721 | -289 | -279 | 2,81 | 0,01 | 20,15 | >0.1 | 5,61 | 5,56 | 0 | 1,18 | 0 | 4,99 | 0 |
| ATOH1 | ACCAAAAGTA | 1 | 734 | 744 | -266 | -256 | 0,04 | 0,01 | 18,46 | >0.1 | 5,61 | 5,66 | 0 | 1,24 | 0 | 5,92 | 0 |
| ATOH1 | AACGTATTCT | 1 | 803 | 813 | -197 | -187 | 3,51 | 0,01 | 20,05 | >0.1 | 5,61 | 5,99 | 0 | 1,51 | 0 | 3,43 | 0 |
| ATOH1 | AAAATCTGTT | 1 | 841 | 851 | -159 | -149 | 5,28 | 0,01 | 20,32 | >0.1 | 5,61 | 6,24 | 0 | 1,78 | 0 | 1,41 | 0 |
| ATOH1 | AGCAGATTGT | 1 | 890 | 900 | -110 | -100 | 4,87 | 0,01 | 21,93 | >0.1 | 5,61 | 6,61 | 0 | 1,89 | 0 | 2,95 | 0 |
| ATOH1 | GGACGATGTC | 1 | 1044 | 1054 | 44 | 54 | -0,26 | 0,01 | 15,56 | >0.1 | 5,61 | 7,73 | 0 | 1,86 | 0 | 0,62 | 0 |
| ATOH1 | CAAAAATGTC | -1 | 1126 | 1136 | 126 | 136 | 0,96 | 0,01 | 20,88 | >0.1 | 5,61 | 6,49 | 0 | 1,84 | 3,22 | 2,75 | 0 |
| ATOH1 | AGCAGCAGTT | -1 | 1090 | 1100 | 90 | 100 | 1,12 | 0,01 | 19,53 | >0.1 | 5,61 | 6,88 | 0 | 1,85 | 1,79 | 2,27 | 0 |
| ATOH1 | ACAATCTGCT | -1 | 890 | 900 | -110 | -100 | 4,91 | 0,01 | 21,97 | >0.1 | 5,61 | 6,61 | 0 | 1,89 | 0 | 2,95 | 0 |
| ATOH1 | GACAGATATG | -1 | 852 | 862 | -148 | -138 | -0,01 | 0,01 | 16,46 | >0.1 | 5,61 | 6,31 | 0 | 1,88 | 0 | 2,67 | 0 |
| ATOH1 | CAGATATGTA | -1 | 850 | 860 | -150 | -140 | 2,55 | 0,01 | 19,07 | >0.1 | 5,61 | 6,3 | 0 | 1,86 | 0 | 2,75 | 0 |
| ATOH1 | AACAGATTTT | -1 | 841 | 851 | -159 | -149 | 5,72 | 0,01 | 20,76 | >0.1 | 5,61 | 6,24 | 0 | 1,78 | 0 | 1,41 | 0 |

| | | | | | | | | | | | | | | | | | |
|---|---|---|---|---|---|---|---|---|---|---|---|---|---|---|---|---|---|
| ATOH1 | AGAATACGTT | -1 | 803 | 813 | -197 | -187 | -0,18 | 0,01 | 16,36 | >0.1 | 5,61 | 5,99 | 0 | 1,51 | 0 | 3,43 | 0 |
| ATOH1 | GAAAGCTGTT | -1 | 711 | 721 | -289 | -279 | 2,54 | 0,01 | 19,88 | >0.1 | 5,61 | 5,56 | 0 | 1,18 | 0 | 4,99 | 0 |
| ATOH1 | GGAAAATGTT | -1 | 691 | 701 | -309 | -299 | 1,04 | 0,01 | 15,27 | >0.1 | 4,53 | 5,49 | 0 | 1,14 | 0 | 3,07 | 0 |
| ATOH1 | ATCATATTCC | -1 | 563 | 573 | -437 | -427 | 3,68 | 0,01 | 13,03 | >0.1 | 3,25 | 5,11 | 0 | 0,99 | 0 | 0 | 0 |
| ATOH1 | TTCATATTTC | -1 | 486 | 496 | -514 | -504 | 0,43 | 0,01 | 15,52 | >0.1 | 2,9 | 4,93 | 0 | 0,98 | 0 | 6,27 | 0 |
| TCFL5 | ATCAGGAGAG | 1 | 97 | 107 | -903 | -893 | 2,7 | 0,01 | 13,33 | >0.1 | 3,64 | 4,35 | 0 | 1,88 | 0 | 0,77 | 0 |
| TCFL5 | GGCTCACGCC | 1 | 189 | 199 | -811 | -801 | 7,36 | 0,001 | 20,65 | >0.1 | 3,09 | 4,45 | 0 | 1,71 | 0 | 4,04 | 0 |
| TCFL5 | ATCACAAGGT | 1 | 235 | 245 | -765 | -755 | 4,08 | 0,01 | 17,66 | >0.1 | 2,9 | 4,51 | 0 | 1,43 | 0 | 4,74 | 0 |
| TCFL5 | GTCAGGAGAT | 1 | 243 | 253 | -757 | -747 | 3,3 | 0,01 | 16,41 | >0.1 | 2,87 | 4,52 | 0 | 1,39 | 0 | 4,33 | 0 |
| TCFL5 | TGGGCGTGGT | 1 | 313 | 323 | -687 | -677 | 4,22 | 0,01 | 18,79 | >0.1 | 2,64 | 4,62 | 0 | 1,17 | 0 | 6,13 | 0 |
| TCFL5 | GGCACCTGTA | 1 | 327 | 337 | -673 | -663 | 3,2 | 0,01 | 17,16 | >0.1 | 2,6 | 4,65 | 0 | 1,14 | 0 | 5,56 | 0 |
| TCFL5 | TACTCGGGAG | 1 | 345 | 355 | -655 | -645 | 4,61 | 0,01 | 19,47 | >0.1 | 2,55 | 4,67 | 0 | 1,11 | 0 | 6,52 | 0 |
| TCFL5 | CTCGGGAGGC | 1 | 347 | 357 | -653 | -643 | 3,88 | 0,01 | 18,74 | >0.1 | 2,55 | 4,68 | 0 | 1,11 | 0 | 6,52 | 0 |
| TCFL5 | GGCAGGAGAA | 1 | 360 | 370 | -640 | -630 | 3,87 | 0,01 | 18,54 | >0.1 | 2,52 | 4,7 | 0 | 1,09 | 0 | 6,36 | 0 |
| TCFL5 | ATGGCGTGAA | 1 | 369 | 379 | -631 | -621 | 3,31 | 0,01 | 17,79 | >0.1 | 2,54 | 4,71 | 0 | 1,07 | 0 | 6,15 | 0 |
| TCFL5 | GGCGTGAACC | 1 | 371 | 381 | -629 | -619 | 2,77 | 0,01 | 17,5 | >0.1 | 2,55 | 4,72 | 0 | 1,07 | 0 | 6,4 | 0 |
| TCFL5 | AGATCGCGCC | 1 | 407 | 417 | -593 | -583 | 6,39 | 0,001 | 20,49 | >0.1 | 2,65 | 4,78 | 0 | 1,03 | 0 | 5,65 | 0 |
| TCFL5 | ATCGCGCCAC | 1 | 409 | 419 | -591 | -581 | 4,6 | 0,01 | 18,59 | >0.1 | 2,65 | 4,78 | 0 | 1,03 | 0 | 5,52 | 0 |
| TCFL5 | TGCTCCAGCC | 1 | 421 | 431 | -579 | -569 | 3,96 | 0,01 | 17,89 | >0.1 | 2,69 | 4,81 | 0 | 1,02 | 0 | 5,42 | 0 |
| TCFL5 | AGACCGAGAC | 1 | 439 | 449 | -561 | -551 | 3,21 | 0,01 | 17,84 | >0.1 | 2,74 | 4,84 | 0 | 1 | 0 | 6,05 | 0 |
| TCFL5 | ATCTCATGAG | 1 | 982 | 992 | -18 | -8 | 4,09 | 0,01 | 21,53 | >0.1 | 5,61 | 8,43 | 0 | 1,87 | 0 | 1,53 | 0 |
| TCFL5 | CTCATGAGGA | 1 | 984 | 994 | -16 | -6 | 3,99 | 0,01 | 21,57 | >0.1 | 5,61 | 8,57 | 0 | 1,87 | 0 | 1,53 | 0 |
| TCFL5 | CTCACAGGGG | 1 | 1036 | 1046 | 36 | 46 | 3 | 0,01 | 19,79 | >0.1 | 5,61 | 7,99 | 0 | 1,86 | 0 | 1,34 | 0 |
| TCFL5 | ACCACGAAGC | 1 | 1146 | 1156 | 146 | 156 | 4,49 | 0,01 | 22,22 | >0.1 | 5,61 | 6,33 | 0 | 1,84 | 1,79 | 2,16 | 0 |
| TCFL5 | GCTTCGTGGT | -1 | 1146 | 1156 | 146 | 156 | 4,49 | 0,01 | 22,22 | >0.1 | 5,61 | 6,33 | 0 | 1,84 | 1,79 | 2,16 | 0 |
| TCFL5 | CCCCTGTGAG | -1 | 1036 | 1046 | 36 | 46 | 3 | 0,01 | 19,79 | >0.1 | 5,61 | 7,99 | 0 | 1,86 | 0 | 1,34 | 0 |
| TCFL5 | TCCTCATGAG | -1 | 984 | 994 | -16 | -6 | 3,99 | 0,01 | 21,57 | >0.1 | 5,61 | 8,57 | 0 | 1,87 | 0 | 1,53 | 0 |
| TCFL5 | CTCATGAGAT | -1 | 982 | 992 | -18 | -8 | 4,09 | 0,01 | 21,53 | >0.1 | 5,61 | 8,43 | 0 | 1,87 | 0 | 1,53 | 0 |
| TCFL5 | GTCTCGGTCT | -1 | 439 | 449 | -561 | -551 | 3,21 | 0,01 | 17,84 | >0.1 | 2,74 | 4,84 | 0 | 1 | 0 | 6,05 | 0 |
| TCFL5 | GGCTGGAGCA | -1 | 421 | 431 | -579 | -569 | 3,96 | 0,01 | 17,89 | >0.1 | 2,69 | 4,81 | 0 | 1,02 | 0 | 5,42 | 0 |
| TCFL5 | GTGGCGCGAT | -1 | 409 | 419 | -591 | -581 | 4,6 | 0,01 | 18,59 | >0.1 | 2,65 | 4,78 | 0 | 1,03 | 0 | 5,52 | 0 |
| TCFL5 | GGCGCGATCT | -1 | 407 | 417 | -593 | -583 | 6,39 | 0,001 | 20,49 | >0.1 | 2,65 | 4,78 | 0 | 1,03 | 0 | 5,65 | 0 |
| TCFL5 | GGTTCACGCC | -1 | 371 | 381 | -629 | -619 | 2,77 | 0,01 | 17,5 | >0.1 | 2,55 | 4,72 | 0 | 1,07 | 0 | 6,4 | 0 |
| TCFL5 | TTCACGCCAT | -1 | 369 | 379 | -631 | -621 | 3,31 | 0,01 | 17,79 | >0.1 | 2,54 | 4,71 | 0 | 1,07 | 0 | 6,15 | 0 |
| TCFL5 | TTCTCCTGCC | -1 | 360 | 370 | -640 | -630 | 3,87 | 0,01 | 18,54 | >0.1 | 2,52 | 4,7 | 0 | 1,09 | 0 | 6,36 | 0 |
| TCFL5 | GCCTCCCGAG | -1 | 347 | 357 | -653 | -643 | 3,88 | 0,01 | 18,74 | >0.1 | 2,55 | 4,68 | 0 | 1,11 | 0 | 6,52 | 0 |
| TCFL5 | CTCCCGAGTA | -1 | 345 | 355 | -655 | -645 | 4,61 | 0,01 | 19,47 | >0.1 | 2,55 | 4,67 | 0 | 1,11 | 0 | 6,52 | 0 |
| TCFL5 | TACAGGTGCC | -1 | 327 | 337 | -673 | -663 | 3,2 | 0,01 | 17,16 | >0.1 | 2,6 | 4,65 | 0 | 1,14 | 0 | 5,56 | 0 |
| TCFL5 | ACCACGCCCA | -1 | 313 | 323 | -687 | -677 | 4,22 | 0,01 | 18,79 | >0.1 | 2,64 | 4,62 | 0 | 1,17 | 0 | 6,13 | 0 |
| TCFL5 | ATCTCCTGAC | -1 | 243 | 253 | -757 | -747 | 3,3 | 0,01 | 16,41 | >0.1 | 2,87 | 4,52 | 0 | 1,39 | 0 | 4,33 | 0 |
| TCFL5 | ACCTTGTGAT | -1 | 235 | 245 | -765 | -755 | 4,08 | 0,01 | 17,66 | >0.1 | 2,9 | 4,51 | 0 | 1,43 | 0 | 4,74 | 0 |
| TCFL5 | GGCGTGAGCC | -1 | 189 | 199 | -811 | -801 | 7,36 | 0,001 | 20,65 | >0.1 | 3,09 | 4,45 | 0 | 1,71 | 0 | 4,04 | 0 |
| TCFL5 | CTCTCCTGAT | -1 | 97 | 107 | -903 | -893 | 2,7 | 0,01 | 13,33 | >0.1 | 3,64 | 4,35 | 0 | 1,88 | 0 | 0,77 | 0 |
| RXRG | AAGGTCAGGAGATA | 1 | 240 | 254 | -760 | -746 | -15,78 | 0,01 | -2,65 | >0.1 | 2,87 | 4,52 | 0 | 1,41 | 0 | 4,33 | 0 |
| RXRG | GGAGGCAGAGCTTG | 1 | 383 | 397 | -617 | -603 | -17,91 | 0,01 | -3,39 | >0.1 | 2,58 | 4,75 | 0 | 1,06 | 0 | 6,13 | 0 |
| RXRG | AAGGGCAAAGTCAT | 1 | 508 | 522 | -492 | -478 | -18,45 | 0,01 | -6,61 | >0.1 | 3 | 4,99 | 0 | 0,99 | 0 | 2,86 | 0 |
| RXRG | AAGGGAAAATGTTG | 1 | 530 | 544 | -470 | -456 | -14,84 | 0,01 | -4,1 | >0.1 | 3,1 | 5,04 | 0 | 0,99 | 0 | 1,61 | 0 |
| RXRG | AAGTAGAGAGTTTC | 1 | 547 | 561 | -453 | -439 | -20,45 | 0,01 | -11,2 | >0.1 | 3,18 | 5,08 | 0 | 0,99 | 0 | 0 | 0 |
| RXRG | AGAGGTAAGGTTCT | 1 | 774 | 788 | -226 | -212 | -14,58 | 0,01 | 4,54 | >0.1 | 5,56 | 5,85 | 0 | 1,39 | 0 | 6,32 | 0 |
| RXRG | AGGATTAAAGAATA | 1 | 790 | 804 | -210 | -196 | -18,37 | 0,01 | -0,09 | >0.1 | 5,56 | 5,94 | 0 | 1,47 | 0 | 5,32 | 0 |
| RXRG | GAGGTTTTAGTCTA | 1 | 992 | 1006 | -8 | 6 | -16,46 | 0,01 | 1,41 | >0.1 | 5,56 | 8,92 | 0 | 1,87 | 0 | 1,53 | 0 |
| RXRG | TGGCTCACAGGGGA | 1 | 1033 | 1047 | 33 | 47 | -12,9 | 0,01 | 3,93 | >0.1 | 5,56 | 8,1 | 0 | 1,86 | 0 | 1,31 | 0 |
| RXRG | TCTATCAAAGTTCA | 1 | 1167 | 1181 | 167 | 181 | -6,25 | 0,001 | 11,91 | >0.1 | 5,56 | 6,17 | 0 | 1,69 | 1,79 | 2,94 | 0 |
| RXRG | GGCTTCGTGGTTAA | -1 | 1143 | 1157 | 143 | 157 | -20,34 | 0,01 | -1,21 | >0.1 | 5,56 | 6,35 | 0 | 1,84 | 3,22 | 2,16 | 0 |
| RXRG | CAATATAAAGTTCA | -1 | 872 | 886 | -128 | -114 | -10,03 | 0,01 | 6,87 | >0.1 | 5,56 | 6,5 | 0 | 1,9 | 0 | 2,95 | 0 |
| RXRG | TCATTAATAGGTTA | -1 | 597 | 611 | -403 | -389 | -20,23 | 0,01 | -9,17 | >0.1 | 3,47 | 5,21 | 0 | 1,02 | 0 | 1,36 | 0 |
| RXRG | GAATTCATATTTCT | -1 | 485 | 499 | -515 | -501 | -17,62 | 0,01 | -2,65 | >0.1 | 2,91 | 4,94 | 0 | 0,99 | 0 | 6,13 | 0 |
| RXRG | GGGACTACAGGTGC | -1 | 328 | 342 | -672 | -658 | -21,49 | 0,01 | -7,64 | >0.1 | 2,59 | 4,65 | 0 | 1,14 | 0 | 5,46 | 0 |
| RXRG | AAGTGCTAGGATTA | -1 | 201 | 215 | -799 | -785 | -20,5 | 0,01 | -6,78 | >0.1 | 3,02 | 4,47 | 0 | 1,63 | 0 | 4,59 | 0 |

| | | | | | | | | | | | | | | | | | |
|---|---|---|---|---|---|---|---|---|---|---|---|---|---|---|---|---|---|
| RXRG | CAGTTTTAATTTCC | -1 | 153 | 167 | -847 | -833 | -21,14 | 0,01 | -8,97 | >0.1 | 3,25 | 4,41 | 0 | 1,9 | 0 | 2,6 | 0 |
| RXRG | CACTCCACAGGTCA | -1 | 108 | 122 | -892 | -878 | -12,9 | 0,01 | -2,35 | >0.1 | 3,54 | 4,36 | 0 | 1,89 | 0 | 0,77 | 0 |
| RXRA | GGAGGCCGAGTTGG | 1 | 217 | 231 | -783 | -769 | -17,83 | 0,01 | -3,33 | >0.1 | 2,96 | 4,49 | 0 | 1,52 | 0 | 5,52 | 0 |
| RXRA | AGATCACAAGGTCA | 1 | 233 | 247 | -767 | -753 | -14,33 | 0,01 | -0,73 | >0.1 | 2,9 | 4,52 | 0 | 1,44 | 0 | 4,74 | 0 |
| RXRA | AAGGTCAGGAGATA | 1 | 240 | 254 | -760 | -746 | -11,39 | 0,01 | 1,74 | >0.1 | 2,87 | 4,52 | 0 | 1,41 | 0 | 4,33 | 0 |
| RXRA | GGAGGCAGAGCTTG | 1 | 383 | 397 | -617 | -603 | -11,45 | 0,01 | 3,07 | >0.1 | 2,58 | 4,75 | 0 | 1,06 | 0 | 6,13 | 0 |
| RXRA | AAAGACAAGGGCAA | 1 | 502 | 516 | -498 | -484 | -17,27 | 0,01 | -2,47 | >0.1 | 2,98 | 4,98 | 0 | 0,99 | 0 | 5,86 | 0 |
| RXRA | AAGGGCAAAGTCAT | 1 | 508 | 522 | -492 | -478 | -14,54 | 0,01 | -2,7 | >0.1 | 3 | 4,99 | 0 | 0,99 | 0 | 2,86 | 0 |
| RXRA | AAGGGAAAATGTTG | 1 | 530 | 544 | -470 | -456 | -10,66 | 0,01 | 0,08 | >0.1 | 3,1 | 5,04 | 0 | 0,99 | 0 | 1,61 | 0 |
| RXRA | AAGTAGAGAGTTTC | 1 | 547 | 561 | -453 | -439 | -17,76 | 0,01 | -8,51 | >0.1 | 3,18 | 5,08 | 0 | 0,99 | 0 | 0 | 0 |
| RXRA | AAAGTGACAGGAGA | 1 | 763 | 777 | -237 | -223 | -16,63 | 0,01 | 1,99 | >0.1 | 5,56 | 5,8 | 0 | 1,35 | 0 | 5,92 | 0 |
| RXRA | AGAGGTAAGGTTCT | 1 | 774 | 788 | -226 | -212 | -13,68 | 0,01 | 5,44 | >0.1 | 5,56 | 5,85 | 0 | 1,39 | 0 | 6,32 | 0 |
| RXRA | AGGATTAAAGAATA | 1 | 790 | 804 | -210 | -196 | -12,96 | 0,01 | 5,32 | >0.1 | 5,56 | 5,94 | 0 | 1,47 | 0 | 5,32 | 0 |
| RXRA | CAGGATGAACTTTA | 1 | 867 | 881 | -133 | -119 | -17,61 | 0,01 | -0,75 | >0.1 | 5,56 | 6,45 | 0 | 1,9 | 0 | 2,95 | 0 |
| RXRA | GAGAGAAAATCTCA | 1 | 974 | 988 | -26 | -12 | -17,47 | 0,01 | -0,25 | >0.1 | 5,56 | 8,22 | 0 | 1,87 | 0 | 1,57 | 0 |
| RXRA | GAGGTTTTAGTCTA | 1 | 992 | 1006 | -8 | 6 | -14,02 | 0,01 | 3,85 | >0.1 | 5,56 | 8,92 | 0 | 1,87 | 0 | 1,53 | 0 |
| RXRA | TGGCTCACAGGGGA | 1 | 1033 | 1047 | 33 | 47 | -8,9 | 0,01 | 7,93 | >0.1 | 5,56 | 8,1 | 0 | 1,86 | 0 | 1,31 | 0 |
| RXRA | AGGGGACGATGTCA | 1 | 1041 | 1055 | 41 | 55 | -15,44 | 0,01 | 0,42 | >0.1 | 5,56 | 7,82 | 0 | 1,86 | 0 | 0,62 | 0 |
| RXRA | TCTATCAAAGTTCA | 1 | 1167 | 1181 | 167 | 181 | -6,01 | 0,001 | 12,15 | >0.1 | 5,56 | 6,17 | 0 | 1,69 | 1,79 | 2,94 | 0 |
| RXRA | GTGGTTAAACTTGT | -1 | 1137 | 1151 | 137 | 151 | -17,38 | 0,01 | 2,85 | >0.1 | 5,56 | 6,4 | 0 | 1,84 | 3,22 | 3,21 | 0 |
| RXRA | GTGAGCCAAGATCA | -1 | 1027 | 1041 | 27 | 41 | -16,44 | 0,01 | 0,71 | >0.1 | 5,56 | 8,39 | 0 | 1,86 | 0 | 1,34 | 0 |
| RXRA | CAATATAAAGTTCA | -1 | 872 | 886 | -128 | -114 | -7,77 | 0,01 | 9,13 | >0.1 | 5,56 | 6,5 | 0 | 1,9 | 0 | 2,95 | 0 |
| RXRA | CACGGAAAATGTTA | -1 | 690 | 704 | -310 | -296 | -16,75 | 0,01 | -1,48 | >0.1 | 5,56 | 5,5 | 0 | 1,14 | 0 | 3,07 | 0 |
| RXRA | GGGACTACAGGTGC | -1 | 328 | 342 | -672 | -658 | -17,64 | 0,01 | -3,79 | >0.1 | 2,59 | 4,65 | 0 | 1,14 | 0 | 5,46 | 0 |
| RXRA | AAGTGCTAGGATTA | -1 | 201 | 215 | -799 | -785 | -16,47 | 0,01 | -2,75 | >0.1 | 3,02 | 4,47 | 0 | 1,63 | 0 | 4,59 | 0 |
| RXRA | AATTTCCATGGCCA | -1 | 146 | 160 | -854 | -840 | -16,59 | 0,01 | -5,34 | >0.1 | 3,29 | 4,41 | 0 | 1,9 | 0 | 1,65 | 0 |
| RXRA | CACTCCACAGGTCA | -1 | 108 | 122 | -892 | -878 | -6,02 | 0,001 | 4,53 | >0.1 | 3,54 | 4,36 | 0 | 1,89 | 0 | 0,77 | 0 |
| RXRA | CAGGTCAACTCTCC | -1 | 101 | 115 | -899 | -885 | -12,29 | 0,01 | -1,69 | >0.1 | 3,59 | 4,36 | 0 | 1,89 | 0 | 0,77 | 0 |
| RXRB | AGATCACAAGGTCA | 1 | 233 | 247 | -767 | -753 | -19,6 | 0,01 | -6 | >0.1 | 2,9 | 4,52 | 0 | 1,44 | 0 | 4,74 | 0 |
| RXRB | AAGGTCAGGAGATA | 1 | 240 | 254 | -760 | -746 | -15,44 | 0,01 | -2,31 | >0.1 | 2,87 | 4,52 | 0 | 1,41 | 0 | 4,33 | 0 |
| RXRB | GGAGGCAGAGCTTG | 1 | 383 | 397 | -617 | -603 | -18,66 | 0,01 | -4,14 | >0.1 | 2,58 | 4,75 | 0 | 1,06 | 0 | 6,13 | 0 |
| RXRB | AAGGGCAAAGTCAT | 1 | 508 | 522 | -492 | -478 | -19,15 | 0,01 | -7,31 | >0.1 | 3 | 4,99 | 0 | 0,99 | 0 | 2,86 | 0 |
| RXRB | TATTTGGAAGGGAA | 1 | 523 | 537 | -477 | -463 | -22,51 | 0,01 | -10,15 | >0.1 | 3,06 | 5,03 | 0 | 0,99 | 0 | 3,28 | 0 |
| RXRB | AAGGGAAAATGTTG | 1 | 530 | 544 | -470 | -456 | -11,67 | 0,01 | -0,93 | >0.1 | 3,1 | 5,04 | 0 | 0,99 | 0 | 1,61 | 0 |
| RXRB | AAAGTGACAGGAGA | 1 | 763 | 777 | -237 | -223 | -18,55 | 0,01 | 0,07 | >0.1 | 5,56 | 5,8 | 0 | 1,35 | 0 | 5,92 | 0 |
| RXRB | AGAGGTAAGGTTCT | 1 | 774 | 788 | -226 | -212 | -15,38 | 0,01 | 3,74 | >0.1 | 5,56 | 5,85 | 0 | 1,39 | 0 | 6,32 | 0 |
| RXRB | AGGATTAAAGAATA | 1 | 790 | 804 | -210 | -196 | -20,43 | 0,01 | -2,15 | >0.1 | 5,56 | 5,94 | 0 | 1,47 | 0 | 5,32 | 0 |
| RXRB | AAGTTCAAAGGCTG | 1 | 956 | 970 | -44 | -30 | 0,25 | 0,0001 | 17,04 | >0.1 | 5,56 | 7,62 | 0 | 1,88 | 0 | 1,74 | 0 |
| RXRB | GAGGTTTTAGTCTA | 1 | 992 | 1006 | -8 | 6 | -19,66 | 0,01 | -1,79 | >0.1 | 5,56 | 8,92 | 0 | 1,87 | 0 | 1,53 | 0 |
| RXRB | TGGCTCACAGGGGA | 1 | 1033 | 1047 | 33 | 47 | -14,62 | 0,01 | 2,21 | >0.1 | 5,56 | 8,1 | 0 | 1,86 | 0 | 1,31 | 0 |
| RXRB | AGGGGACGATGTCA | 1 | 1041 | 1055 | 41 | 55 | -19,92 | 0,01 | -4,06 | >0.1 | 5,56 | 7,82 | 0 | 1,86 | 0 | 0,62 | 0 |
| RXRB | TGAGGAACAGGCCA | 1 | 1111 | 1125 | 111 | 125 | -22,32 | 0,01 | -2,54 | >0.1 | 5,56 | 6,64 | 0 | 1,85 | 3,22 | 2,52 | 0 |
| RXRB | TCTATCAAAGTTCA | 1 | 1167 | 1181 | 167 | 181 | -11,45 | 0,01 | 6,71 | >0.1 | 5,56 | 6,17 | 0 | 1,69 | 1,79 | 2,94 | 0 |
| RXRB | GGCTTCGTGGTTAA | -1 | 1143 | 1157 | 143 | 157 | -21,62 | 0,01 | -2,49 | >0.1 | 5,56 | 6,35 | 0 | 1,84 | 3,22 | 2,16 | 0 |
| RXRB | CAATATAAAGTTCA | -1 | 872 | 886 | -128 | -114 | -13,3 | 0,01 | 3,6 | >0.1 | 5,56 | 6,5 | 0 | 1,9 | 0 | 2,95 | 0 |
| RXRB | CACGGAAAATGTTA | -1 | 690 | 704 | -310 | -296 | -21,15 | 0,01 | -5,88 | >0.1 | 5,56 | 5,5 | 0 | 1,14 | 0 | 3,07 | 0 |
| RXRB | TCATTAATAGGTTA | -1 | 597 | 611 | -403 | -389 | -19,57 | 0,01 | -8,51 | >0.1 | 3,47 | 5,21 | 0 | 1,02 | 0 | 1,36 | 0 |
| RXRB | GAATTCATATTTCT | -1 | 485 | 499 | -515 | -501 | -18,73 | 0,01 | -3,76 | >0.1 | 2,91 | 4,94 | 0 | 0,99 | 0 | 6,13 | 0 |
| RXRB | AAGTGCTAGGATTA | -1 | 201 | 215 | -799 | -785 | -21,33 | 0,01 | -7,61 | >0.1 | 3,02 | 4,47 | 0 | 1,63 | 0 | 4,59 | 0 |
| RXRB | CAGTTTTAATTTCC | -1 | 153 | 167 | -847 | -833 | -20,31 | 0,01 | -8,14 | >0.1 | 3,25 | 4,41 | 0 | 1,9 | 0 | 2,6 | 0 |
| RXRB | CACTCCACAGGTCA | -1 | 108 | 122 | -892 | -878 | -13,13 | 0,01 | -2,58 | >0.1 | 3,54 | 4,36 | 0 | 1,89 | 0 | 0,77 | 0 |
| RXRB | CAGGTCAACTCTCC | -1 | 101 | 115 | -899 | -885 | -21,69 | 0,01 | -11,09 | >0.1 | 3,59 | 4,36 | 0 | 1,89 | 0 | 0,77 | 0 |
| SPIB | AGGAGAA | 1 | 13 | 20 | -987 | -980 | 4,98 | 0,01 | 16,76 | >0.1 | 5,65 | 4,26 | 0 | 1,86 | 0 | 0 | 0 |
| SPIB | TGAGAAA | 1 | 55 | 62 | -945 | -938 | 4,58 | 0,01 | 14,83 | >0.1 | 4,08 | 4,3 | 0 | 1,87 | 0 | 0 | 0 |
| SPIB | ATAGGAA | 1 | 61 | 68 | -939 | -932 | 6,93 | 0,01 | 17,11 | >0.1 | 4 | 4,31 | 0 | 1,87 | 0 | 0 | 0 |
| SPIB | TCAGAAA | 1 | 168 | 175 | -832 | -825 | 3,54 | 0,01 | 15,52 | >0.1 | 3,19 | 4,42 | 0 | 1,9 | 0 | 2,46 | 0 |
| SPIB | TGAGGCA | 1 | 357 | 364 | -643 | -636 | 2,96 | 0,01 | 17,3 | >0.1 | 2,53 | 4,69 | 0 | 1,09 | 0 | 6,03 | 0 |
| SPIB | AGGAGAA | 1 | 363 | 370 | -637 | -630 | 4,98 | 0,01 | 19,77 | >0.1 | 2,52 | 4,7 | 0 | 1,08 | 0 | 6,49 | 0 |
| SPIB | TCAGAAA | 1 | 483 | 490 | -517 | -510 | 3,54 | 0,01 | 17,98 | >0.1 | 2,89 | 4,92 | 0 | 0,98 | 0 | 5,65 | 0 |

| | | | | | | | | | | | | | | | | | |
|---|---|---|---|---|---|---|---|---|---|---|---|---|---|---|---|---|---|
| SPIB | AAGGGAA | 1 | 530 | 537 | -470 | -463 | 6,61 | 0,01 | 19,24 | >0.1 | 3,08 | 5,02 | 0 | 0,98 | 0 | 3,55 | 0 |
| SPIB | AGGGAAA | 1 | 531 | 538 | -469 | -462 | 4,11 | 0,01 | 16,75 | >0.1 | 3,08 | 5,03 | 0 | 0,98 | 0 | 3,55 | 0 |
| SPIB | ACAGGAG | 1 | 769 | 776 | -231 | -224 | 2,73 | 0,01 | 21,44 | >0.1 | 5,65 | 5,8 | 0 | 1,34 | 0 | 5,92 | 0 |
| SPIB | AGAGGTA | 1 | 774 | 781 | -226 | -219 | 5,83 | 0,01 | 24,58 | >0.1 | 5,65 | 5,82 | 0 | 1,36 | 0 | 5,92 | 0 |
| SPIB | AGAGAAA | 1 | 975 | 982 | -25 | -18 | 5,36 | 0,01 | 22,51 | >0.1 | 5,65 | 7,98 | 0 | 1,87 | 0 | 1,65 | 0 |
| SPIB | TGAGGAG | 1 | 988 | 995 | -12 | -5 | 2,99 | 0,01 | 20,67 | >0.1 | 5,65 | 8,59 | 0 | 1,87 | 0 | 1,57 | 0 |
| SPIB | TAGGGAA | 1 | 1004 | 1011 | 4 | 11 | 5,83 | 0,01 | 23,93 | >0.1 | 5,65 | 9,12 | 0 | 1,87 | 0 | 1,46 | 0 |
| SPIB | AGGGAAA | 1 | 1005 | 1012 | 5 | 12 | 4,11 | 0,01 | 22,21 | >0.1 | 5,65 | 9,12 | 0 | 1,87 | 0 | 1,46 | 0 |
| SPIB | AGTGGAT | 1 | 1019 | 1026 | 19 | 26 | 3,27 | 0,01 | 20,86 | >0.1 | 5,65 | 8,74 | 0 | 1,86 | 0 | 1,34 | 0 |
| SPIB | TGAGGAA | 1 | 1111 | 1118 | 111 | 118 | 10,38 | 0,001 | 30,26 | >0.1 | 5,65 | 6,64 | 0 | 1,85 | 3,22 | 2,52 | 0 |
| SPIB | AGCCGAA | 1 | 1153 | 1160 | 153 | 160 | 3,19 | 0,01 | 19,82 | >0.1 | 4,52 | 6,27 | 0 | 1,83 | 1,79 | 2,21 | 0 |
| SPIB | CCAGGAA | -1 | 1060 | 1067 | 60 | 67 | 3,33 | 0,01 | 20,4 | >0.1 | 5,65 | 7,35 | 0 | 1,86 | 0 | 2,21 | 0 |
| SPIB | GCAGGAA | -1 | 935 | 942 | -65 | -58 | 3,33 | 0,01 | 18,88 | >0.1 | 5,65 | 7,09 | 0 | 1,88 | 0 | 0,93 | 0 |
| SPIB | AAAAGAA | -1 | 920 | 927 | -80 | -73 | 2,94 | 0,01 | 18,49 | >0.1 | 5,65 | 6,89 | 0 | 1,88 | 0 | 1,12 | 0 |
| SPIB | AAAAGAA | -1 | 914 | 921 | -86 | -79 | 2,94 | 0,01 | 18,45 | >0.1 | 5,65 | 6,82 | 0 | 1,89 | 0 | 1,15 | 0 |
| SPIB | AGAAGAA | -1 | 911 | 918 | -89 | -82 | 6,24 | 0,01 | 21,77 | >0.1 | 5,65 | 6,79 | 0 | 1,89 | 0 | 1,21 | 0 |
| SPIB | AGAAGAA | -1 | 908 | 915 | -92 | -85 | 6,24 | 0,01 | 21,74 | >0.1 | 5,65 | 6,75 | 0 | 1,89 | 0 | 1,21 | 0 |
| SPIB | ACAGTAA | -1 | 900 | 907 | -100 | -93 | 4,32 | 0,01 | 21,4 | >0.1 | 5,65 | 6,67 | 0 | 1,89 | 0 | 2,86 | 0 |
| SPIB | AGAGGAC | -1 | 859 | 866 | -141 | -134 | 3,77 | 0,01 | 20,52 | >0.1 | 5,65 | 6,34 | 0 | 1,9 | 0 | 2,86 | 0 |
| SPIB | ACCGGTA | -1 | 742 | 749 | -258 | -251 | 4,7 | 0,01 | 23,19 | >0.1 | 5,65 | 5,68 | 0 | 1,25 | 0 | 5,92 | 0 |
| SPIB | ACGGAAA | -1 | 696 | 703 | -304 | -297 | 3,07 | 0,01 | 17,4 | >0.1 | 4,62 | 5,5 | 0 | 1,14 | 0 | 3,07 | 0 |
| SPIB | TATGGAA | -1 | 496 | 503 | -504 | -497 | 3,62 | 0,01 | 18,92 | >0.1 | 2,94 | 4,95 | 0 | 0,98 | 0 | 6,44 | 0 |
| SPIC | GTGAAGGAGAAGGA | 1 | 9 | 23 | -991 | -977 | -0,23 | 0,01 | 13,25 | >0.1 | 5,56 | 4,27 | 0 | 1,86 | 1,79 | 0 | 0 |
| SPIC | AGAAATAGGAAATG | 1 | 57 | 71 | -943 | -929 | 6,88 | 0,001 | 17,07 | >0.1 | 4,01 | 4,31 | 0 | 1,88 | 0 | 0 | 0 |
| SPIC | CAGATCAGGAGAGT | 1 | 94 | 108 | -906 | -892 | -0,01 | 0,01 | 10,63 | >0.1 | 3,64 | 4,35 | 0 | 1,88 | 0 | 0,77 | 0 |
| SPIC | CAGATCACAAGGTC | 1 | 232 | 246 | -768 | -754 | -0,33 | 0,01 | 13,43 | >0.1 | 2,9 | 4,51 | 0 | 1,45 | 0 | 4,9 | 0 |
| SPIC | AGATAGAGACCGTC | 1 | 249 | 263 | -751 | -737 | 1,14 | 0,01 | 14,97 | >0.1 | 2,84 | 4,54 | 0 | 1,37 | 0 | 5,09 | 0 |
| SPIC | CTACTCGGGAGGCT | 1 | 344 | 358 | -656 | -642 | 2,14 | 0,01 | 17,01 | >0.1 | 2,55 | 4,68 | 0 | 1,11 | 0 | 6,52 | 0 |
| SPIC | AGGCTGAGGCAGGA | 1 | 353 | 367 | -647 | -633 | 3,19 | 0,01 | 17,55 | >0.1 | 2,53 | 4,69 | 0 | 1,1 | 0 | 6,03 | 0 |
| SPIC | ATAAAGACAAGGGC | 1 | 500 | 514 | -500 | -486 | 0,73 | 0,01 | 15,34 | >0.1 | 2,97 | 4,97 | 0 | 0,99 | 0 | 5,68 | 0 |
| SPIC | TGGAAGGGAAAATG | 1 | 527 | 541 | -473 | -459 | 1,82 | 0,01 | 14,47 | >0.1 | 3,08 | 5,03 | 0 | 0,99 | 0 | 3,55 | 0 |
| SPIC | TGACAGGAGAGGTA | 1 | 767 | 781 | -233 | -219 | -0,3 | 0,01 | 18,13 | >0.1 | 5,56 | 5,82 | 0 | 1,36 | 0 | 5,69 | 0 |
| SPIC | CTCATGAGGAGGTT | 1 | 984 | 998 | -16 | -2 | 7,8 | 0,001 | 25,46 | >0.1 | 5,56 | 8,66 | 0 | 1,87 | 0 | 1,57 | 0 |
| SPIC | GTCTAGGGAAAGTC | 1 | 1001 | 1015 | 1 | 15 | 2,94 | 0,01 | 20,95 | >0.1 | 5,56 | 9,12 | 0 | 1,87 | 0 | 1,46 | 0 |
| SPIC | TCACAGGGGACGAT | 1 | 1037 | 1051 | 37 | 51 | 5,67 | 0,01 | 22,4 | >0.1 | 5,56 | 7,95 | 0 | 1,86 | 0 | 1,36 | 0 |
| SPIC | ACGAAGCCGAAGAC | 1 | 1149 | 1163 | 149 | 163 | 6,35 | 0,001 | 23,03 | >0.1 | 4,53 | 6,3 | 0 | 1,84 | 1,79 | 2,21 | 0 |
| SPIC | AAAAAAAGAAAAAG | -1 | 916 | 930 | -84 | -70 | 2,05 | 0,01 | 17,54 | >0.1 | 5,56 | 6,93 | 0 | 1,89 | 0 | 1,12 | 0 |
| SPIC | AAAAAGAAAAAGAA | -1 | 914 | 928 | -86 | -72 | 2,32 | 0,01 | 17,79 | >0.1 | 5,56 | 6,9 | 0 | 1,89 | 0 | 1,12 | 0 |
| SPIC | AAGAAAAAGAAGAA | -1 | 911 | 925 | -89 | -75 | 0,55 | 0,01 | 16,01 | >0.1 | 5,56 | 6,87 | 0 | 1,89 | 0 | 1,15 | 0 |
| SPIC | AAAAAGAAGAAGAA | -1 | 908 | 922 | -92 | -78 | 6,96 | 0,001 | 22,39 | >0.1 | 5,56 | 6,83 | 0 | 1,89 | 0 | 1,15 | 0 |
| SPIC | GAACACAGTAAACA | -1 | 897 | 911 | -103 | -89 | 3,11 | 0,01 | 20,02 | >0.1 | 5,56 | 6,72 | 0 | 1,89 | 0 | 2,75 | 0 |
| SPIC | CAAAACCGGTACTT | -1 | 739 | 753 | -261 | -247 | 1,58 | 0,01 | 20,2 | >0.1 | 5,56 | 5,69 | 0 | 1,27 | 0 | 6,1 | 0 |
| SPIC | ATACACGGAAAATG | -1 | 693 | 707 | -307 | -293 | -0,33 | 0,01 | 14,84 | >0.1 | 5,56 | 5,51 | 0 | 1,15 | 0 | 2,95 | 0 |
| SPIC | GGGATCATGAAGAT | -1 | 643 | 657 | -357 | -343 | 1,3 | 0,01 | 15,39 | >0.1 | 3,84 | 5,34 | 0 | 1,07 | 0 | 3,83 | 0 |
| BARX1 | GAAATTAA | 1 | 154 | 162 | -846 | -838 | 6,47 | 0,01 | 17,69 | >0.1 | 3,27 | 4,41 | 0 | 1,9 | 0 | 1,65 | 0 |
| BARX1 | TCTATTAA | 1 | 288 | 296 | -712 | -704 | 2,98 | 0,01 | 16,71 | >0.1 | 2,72 | 4,58 | 0 | 1,24 | 0 | 5,19 | 0 |
| BARX1 | AAACTTAG | 1 | 304 | 312 | -696 | -688 | 2 | 0,01 | 15,46 | >0.1 | 2,67 | 4,61 | 0 | 1,19 | 0 | 4,99 | 0 |
| BARX1 | CCTATTAA | 1 | 600 | 608 | -400 | -392 | 4,53 | 0,01 | 15,58 | >0.1 | 3,47 | 5,2 | 0 | 1,01 | 0 | 1,36 | 0 |
| BARX1 | AACATTAT | 1 | 637 | 645 | -363 | -355 | 4,63 | 0,01 | 16,4 | >0.1 | 3,76 | 5,31 | 0 | 1,05 | 0 | 1,66 | 0 |
| BARX1 | AAAATTAT | 1 | 683 | 691 | -317 | -309 | 4,88 | 0,01 | 18,86 | >0.1 | 4,34 | 5,46 | 0 | 1,12 | 0 | 3,07 | 0 |
| BARX1 | AATATTAA | 1 | 727 | 735 | -273 | -265 | 0,45 | 0,01 | 18,24 | >0.1 | 5,64 | 5,62 | 0 | 1,21 | 0 | 5,32 | 0 |
| BARX1 | AGGATTAA | 1 | 790 | 798 | -210 | -202 | 4,18 | 0,01 | 22,68 | >0.1 | 5,64 | 5,91 | 0 | 1,43 | 0 | 5,52 | 0 |
| BARX1 | AAAATTAT | 1 | 830 | 838 | -170 | -162 | 4,88 | 0,01 | 21,61 | >0.1 | 5,64 | 6,14 | 0 | 1,67 | 0 | 3,28 | 0 |
| BARX1 | GGAATTAT | 1 | 1191 | 1199 | 191 | 199 | 4,87 | 0,01 | 23,65 | >0.1 | 5,64 | 6,02 | 0 | 1,5 | 1,79 | 3,82 | 0 |
| BARX1 | GTGGTTAA | -1 | 1143 | 1151 | 143 | 151 | 2,54 | 0,01 | 22,96 | >0.1 | 5,64 | 6,35 | 0 | 1,84 | 3,22 | 3,37 | 0 |
| BARX1 | GCAGTTAC | -1 | 1088 | 1096 | 88 | 96 | 5,47 | 0,01 | 23,87 | >0.1 | 5,64 | 6,9 | 0 | 1,85 | 1,79 | 2,21 | 0 |
| BARX1 | TACGTTAT | -1 | 801 | 809 | -199 | -191 | -1,11 | 0,01 | 15,53 | >0.1 | 5,64 | 5,96 | 0 | 1,49 | 0 | 3,55 | 0 |
| BARX1 | AACCTTAC | -1 | 778 | 786 | -222 | -214 | 0,96 | 0,01 | 20,37 | >0.1 | 5,64 | 5,84 | 0 | 1,38 | 0 | 6,54 | 0 |
| BARX1 | TTGGTTAA | -1 | 731 | 739 | -269 | -261 | 0,7 | 0,01 | 18,72 | >0.1 | 5,64 | 5,63 | 0 | 1,22 | 0 | 5,52 | 0 |

| | | | | | | | | | | | | | | | | | |
|---|---|---|---|---|---|---|---|---|---|---|---|---|---|---|---|---|---|
| BARX1 | GCTGTTAA | -1 | 709 | 717 | -291 | -283 | 0,18 | 0,01 | 17,35 | >0.1 | 5,64 | 5,55 | 0 | 1,17 | 0 | 4,81 | 0 |
| BARX1 | TTCATTAA | -1 | 604 | 612 | -396 | -388 | 5,87 | 0,01 | 17,01 | >0.1 | 3,49 | 5,21 | 0 | 1,02 | 0 | 1,41 | 0 |
| BARX1 | AGGATTAC | -1 | 200 | 208 | -800 | -792 | 4,21 | 0,01 | 17,82 | >0.1 | 3,04 | 4,46 | 0 | 1,63 | 0 | 4,48 | 0 |
| BARX1 | AAAATTAT | -1 | 132 | 140 | -868 | -860 | 4,88 | 0,01 | 16,12 | >0.1 | 3,4 | 4,38 | 0 | 1,89 | 0 | 1,57 | 0 |
| BARX1 | AAACTTAA | -1 | 78 | 86 | -922 | -914 | 1,17 | 0,01 | 11,19 | >0.1 | 3,81 | 4,32 | 0 | 1,88 | 0 | 0 | 0 |
| BARX1 | TCACTTAC | -1 | 5 | 13 | -995 | -987 | 3,73 | 0,01 | 17,28 | >0.1 | 5,64 | 4,26 | 0 | 1,86 | 1,79 | 0 | 0 |
| FOSL1::JUN(VAR.2) | AAGAAGTCAG | 1 | 31 | 41 | -969 | -959 | 2,56 | 0,01 | 13,17 | >0.1 | 4,46 | 4,28 | 0 | 1,87 | 0 | 0 | 0 |
| FOSL1::JUN(VAR.2) | ATGGCGTGAA | 1 | 369 | 379 | -631 | -621 | 5,3 | 0,01 | 19,78 | >0.1 | 2,54 | 4,71 | 0 | 1,07 | 0 | 6,15 | 0 |
| FOSL1::JUN(VAR.2) | ATCGCGCCAC | 1 | 409 | 419 | -591 | -581 | 6,71 | 0,001 | 20,7 | >0.1 | 2,65 | 4,78 | 0 | 1,03 | 0 | 5,52 | 0 |
| FOSL1::JUN(VAR.2) | GTGAGATAAC | 1 | 591 | 601 | -409 | -399 | 4,33 | 0,01 | 13,94 | >0.1 | 3,41 | 5,18 | 0 | 1,01 | 0 | 0 | 0 |
| FOSL1::JUN(VAR.2) | CTTTGGTCAC | 1 | 670 | 680 | -330 | -320 | 1,81 | 0,01 | 16,17 | >0.1 | 4,14 | 5,42 | 0 | 1,1 | 0 | 3,7 | 0 |
| FOSL1::JUN(VAR.2) | ATTATAACAT | 1 | 686 | 696 | -314 | -304 | 2,38 | 0,01 | 16,35 | >0.1 | 4,42 | 5,47 | 0 | 1,13 | 0 | 2,95 | 0 |
| FOSL1::JUN(VAR.2) | ATTTGGCCAT | 1 | 753 | 763 | -247 | -237 | 2,83 | 0,01 | 21,79 | >0.1 | 5,61 | 5,74 | 0 | 1,3 | 0 | 6,32 | 0 |
| FOSL1::JUN(VAR.2) | ATAACGTATT | 1 | 801 | 811 | -199 | -189 | 4,26 | 0,01 | 20,77 | >0.1 | 5,61 | 5,98 | 0 | 1,5 | 0 | 3,43 | 0 |
| FOSL1::JUN(VAR.2) | GTTACATATC | 1 | 848 | 858 | -152 | -142 | 1,89 | 0,01 | 18,58 | >0.1 | 5,61 | 6,28 | 0 | 1,84 | 0 | 2,95 | 0 |
| FOSL1::JUN(VAR.2) | ACGATGTCAA | 1 | 1046 | 1056 | 46 | 56 | 3,21 | 0,01 | 19,82 | >0.1 | 5,61 | 7,68 | 0 | 1,86 | 0 | 1,46 | 0 |
| FOSL1::JUN(VAR.2) | TTGACATCGT | -1 | 1046 | 1056 | 46 | 56 | 3,34 | 0,01 | 19,95 | >0.1 | 5,61 | 7,68 | 0 | 1,86 | 0 | 1,46 | 0 |
| FOSL1::JUN(VAR.2) | GATATGTAAC | -1 | 848 | 858 | -152 | -142 | 1,82 | 0,01 | 18,51 | >0.1 | 5,61 | 6,28 | 0 | 1,84 | 0 | 2,95 | 0 |
| FOSL1::JUN(VAR.2) | AATACGTTAT | -1 | 801 | 811 | -199 | -189 | 4,26 | 0,01 | 20,77 | >0.1 | 5,61 | 5,98 | 0 | 1,5 | 0 | 3,43 | 0 |
| FOSL1::JUN(VAR.2) | GTCACTTTAT | -1 | 761 | 771 | -239 | -229 | 1,29 | 0,01 | 20,1 | >0.1 | 5,61 | 5,77 | 0 | 1,32 | 0 | 6,1 | 0 |
| FOSL1::JUN(VAR.2) | ATGGCCAAAT | -1 | 753 | 763 | -247 | -237 | 2,69 | 0,01 | 21,65 | >0.1 | 5,61 | 5,74 | 0 | 1,3 | 0 | 6,32 | 0 |
| FOSL1::JUN(VAR.2) | ATGTTATAAT | -1 | 686 | 696 | -314 | -304 | 2,44 | 0,01 | 16,41 | >0.1 | 4,42 | 5,47 | 0 | 1,13 | 0 | 2,95 | 0 |
| FOSL1::JUN(VAR.2) | GTGACCAAAG | -1 | 670 | 680 | -330 | -320 | 1,71 | 0,01 | 16,07 | >0.1 | 4,14 | 5,42 | 0 | 1,1 | 0 | 3,7 | 0 |
| FOSL1::JUN(VAR.2) | GTTATCTCAC | -1 | 591 | 601 | -409 | -399 | 4,29 | 0,01 | 13,9 | >0.1 | 3,41 | 5,18 | 0 | 1,01 | 0 | 0 | 0 |
| FOSL1::JUN(VAR.2) | GTGGCGCGAT | -1 | 409 | 419 | -591 | -581 | 6,78 | 0,001 | 20,77 | >0.1 | 2,65 | 4,78 | 0 | 1,03 | 0 | 5,52 | 0 |
| FOSL1::JUN(VAR.2) | TTCACGCCAT | -1 | 369 | 379 | -631 | -621 | 5,35 | 0,01 | 19,83 | >0.1 | 2,54 | 4,71 | 0 | 1,07 | 0 | 6,15 | 0 |
| FOSL1::JUN(VAR.2) | CTGACTTCTT | -1 | 31 | 41 | -969 | -959 | 2,75 | 0,01 | 13,36 | >0.1 | 4,46 | 4,28 | 0 | 1,87 | 0 | 0 | 0 |
| LEF1 | GCAGATCAGGAGAGT | 1 | 93 | 108 | -907 | -892 | -2,89 | 0,01 | 7,78 | >0.1 | 3,65 | 4,35 | 0 | 1,88 | 0 | 0,79 | 0 |
| LEF1 | GCAGATCACAAGGTC | 1 | 231 | 246 | -769 | -754 | -0,74 | 0,01 | 13,03 | >0.1 | 2,9 | 4,51 | 0 | 1,45 | 0 | 4,9 | 0 |
| LEF1 | CAAGGTCAGGAGATA | 1 | 239 | 254 | -761 | -746 | -4,87 | 0,01 | 8,27 | >0.1 | 2,87 | 4,52 | 0 | 1,41 | 0 | 4,33 | 0 |
| LEF1 | ACTGATCAGAAATAT | 1 | 478 | 493 | -522 | -507 | -4,66 | 0,01 | 9,79 | >0.1 | 2,88 | 4,93 | 0 | 0,99 | 0 | 5,65 | 0 |
| LEF1 | CAAGGGCAAAGTCAT | 1 | 507 | 522 | -493 | -478 | -2,35 | 0,01 | 9,38 | >0.1 | 3 | 4,99 | 0 | 0,99 | 0 | 2,75 | 0 |
| LEF1 | TGAAATAAAAATAAA | 1 | 575 | 590 | -425 | -410 | -5,49 | 0,01 | 4 | >0.1 | 3,33 | 5,15 | 0 | 1,01 | 0 | 0 | 0 |
| LEF1 | AAAAATAAATGTGAG | 1 | 581 | 596 | -419 | -404 | -4,5 | 0,01 | 5,05 | >0.1 | 3,37 | 5,17 | 0 | 1,01 | 0 | 0 | 0 |
| LEF1 | TAGGATTAAAGAATA | 1 | 789 | 804 | -211 | -196 | 4,72 | 0,001 | 22,99 | >0.1 | 5,54 | 5,94 | 0 | 1,47 | 0 | 5,32 | 0 |
| LEF1 | CAAGTTCAAAGGCTG | 1 | 955 | 970 | -45 | -30 | 11,98 | 0,0001 | 28,76 | >0.1 | 5,54 | 7,62 | 0 | 1,88 | 0 | 1,74 | 0 |
| LEF1 | GCTGATAAGAGAGAA | 1 | 966 | 981 | -34 | -19 | -4,18 | 0,01 | 12,88 | >0.1 | 5,54 | 7,94 | 0 | 1,87 | 0 | 1,7 | 0 |
| LEF1 | TTCTATCAAAGTTCA | 1 | 1166 | 1181 | 166 | 181 | -5,85 | 0,01 | 12,21 | >0.1 | 5,54 | 6,18 | 0 | 1,7 | 1,79 | 2,84 | 0 |
| LEF1 | CAAGAAGCAAGTGAA | -1 | 1177 | 1192 | 177 | 192 | -4,42 | 0,01 | 13,98 | >0.1 | 5,54 | 6,11 | 0 | 1,61 | 1,79 | 3,34 | 0 |
| LEF1 | CAGCAACAAGGCTGA | -1 | 1074 | 1089 | 74 | 89 | -3,31 | 0,01 | 15,26 | >0.1 | 5,54 | 7,11 | 0 | 1,85 | 1,79 | 2,27 | 0 |
| LEF1 | AAAGACCAAAAAAAG | -1 | 922 | 937 | -78 | -63 | 3,77 | 0,001 | 19,28 | >0.1 | 5,54 | 7,02 | 0 | 1,88 | 0 | 1,06 | 0 |
| LEF1 | AAAAAAGAAAAAGAA | -1 | 914 | 929 | -86 | -71 | -4,51 | 0,01 | 10,96 | >0.1 | 5,54 | 6,92 | 0 | 1,89 | 0 | 1,12 | 0 |
| LEF1 | AAAGAAAAAGAAGAA | -1 | 911 | 926 | -89 | -74 | -3,89 | 0,01 | 11,57 | >0.1 | 5,54 | 6,88 | 0 | 1,89 | 0 | 1,15 | 0 |
| LEF1 | AAAGAAGAAGAACAC | -1 | 905 | 920 | -95 | -80 | -4,82 | 0,01 | 10,66 | >0.1 | 5,54 | 6,81 | 0 | 1,89 | 0 | 1,24 | 0 |
| LEF1 | GAAGAACACAGTAAA | -1 | 899 | 914 | -101 | -86 | -0,89 | 0,01 | 14,57 | >0.1 | 5,54 | 6,75 | 0 | 1,89 | 0 | 1,28 | 0 |
| LEF1 | CCAAATCAAAACCGG | -1 | 744 | 759 | -256 | -241 | -4,09 | 0,01 | 14,37 | >0.1 | 5,54 | 5,72 | 0 | 1,29 | 0 | 5,92 | 0 |
| LEF1 | AGTGACCAAAGAGGT | -1 | 666 | 681 | -334 | -319 | 4,24 | 0,001 | 18,58 | >0.1 | 4,12 | 5,42 | 0 | 1,1 | 0 | 3,7 | 0 |
| LEF1 | CTAGAACTAGGGATC | -1 | 651 | 666 | -349 | -334 | -4,42 | 0,01 | 9,96 | >0.1 | 3,93 | 5,37 | 0 | 1,08 | 0 | 3,99 | 0 |
| LEF1 | GATCATGAAGATAAT | -1 | 640 | 655 | -360 | -345 | -5,53 | 0,01 | 8,52 | >0.1 | 3,82 | 5,34 | 0 | 1,07 | 0 | 3,83 | 0 |
| LEF1 | TAGGATTACAGGCGT | -1 | 194 | 209 | -806 | -791 | -5,76 | 0,01 | 7,81 | >0.1 | 3,05 | 4,47 | 0 | 1,68 | 0 | 4,37 | 0 |
| LEF1 | AAACTTAAAAAAGCT | -1 | 71 | 86 | -929 | -914 | -1,51 | 0,01 | 8,54 | >0.1 | 3,85 | 4,33 | 0 | 1,88 | 0 | 0 | 0 |
| OLIG2 | AAAATGTTGC | 1 | 535 | 545 | -465 | -455 | -0,71 | 0,01 | 10,04 | >0.1 | 3,11 | 5,04 | 0 | 0,98 | 0 | 1,61 | 0 |
| OLIG2 | GGAATATGAT | 1 | 563 | 573 | -437 | -427 | -1,69 | 0,01 | 7,66 | >0.1 | 3,25 | 5,11 | 0 | 0,99 | 0 | 0 | 0 |
| OLIG2 | AACATTTTCC | 1 | 691 | 701 | -309 | -299 | 0,49 | 0,01 | 14,72 | >0.1 | 4,53 | 5,49 | 0 | 1,14 | 0 | 3,07 | 0 |
| OLIG2 | GCCATAAAGT | 1 | 758 | 768 | -242 | -232 | 0,09 | 0,01 | 18,87 | >0.1 | 5,61 | 5,76 | 0 | 1,31 | 0 | 6,1 | 0 |
| OLIG2 | AACGTATTCT | 1 | 803 | 813 | -197 | -187 | 5,37 | 0,001 | 21,91 | >0.1 | 5,61 | 5,99 | 0 | 1,51 | 0 | 3,43 | 0 |
| OLIG2 | AAATTATTCT | 1 | 831 | 841 | -169 | -159 | -1,32 | 0,01 | 15,58 | >0.1 | 5,61 | 6,16 | 0 | 1,7 | 0 | 3,43 | 0 |
| OLIG2 | AAAATCTGTT | 1 | 841 | 851 | -159 | -149 | 3,86 | 0,01 | 18,9 | >0.1 | 5,61 | 6,24 | 0 | 1,78 | 0 | 1,41 | 0 |
| OLIG2 | TACATATCTG | 1 | 850 | 860 | -150 | -140 | -2,7 | 0,01 | 13,82 | >0.1 | 5,61 | 6,3 | 0 | 1,86 | 0 | 2,75 | 0 |

| | | | | | | | | | | | | | | | | | |
|---|---|---|---|---|---|---|---|---|---|---|---|---|---|---|---|---|---|
| OLIG2 | GCCTTGTTGC | 1 | 1077 | 1087 | 77 | 87 | -3,02 | 0,01 | 15,51 | >0.1 | 5,61 | 7,06 | 0 | 1,85 | 1,79 | 2,21 | 0 |
| OLIG2 | ACAATCTGCT | -1 | 890 | 900 | -110 | -100 | 4,4 | 0,01 | 21,46 | >0.1 | 5,61 | 6,61 | 0 | 1,89 | 0 | 2,95 | 0 |
| OLIG2 | CAGATATGTA | -1 | 850 | 860 | -150 | -140 | -1,22 | 0,01 | 15,3 | >0.1 | 5,61 | 6,3 | 0 | 1,86 | 0 | 2,75 | 0 |
| OLIG2 | AACAGATTTT | -1 | 841 | 851 | -159 | -149 | -2,27 | 0,01 | 12,77 | >0.1 | 5,61 | 6,24 | 0 | 1,78 | 0 | 1,41 | 0 |
| OLIG2 | AGAATAATTT | -1 | 831 | 841 | -169 | -159 | -2,74 | 0,01 | 14,16 | >0.1 | 5,61 | 6,16 | 0 | 1,7 | 0 | 3,43 | 0 |
| OLIG2 | AGAATACGTT | -1 | 803 | 813 | -197 | -187 | 4,95 | 0,001 | 21,49 | >0.1 | 5,61 | 5,99 | 0 | 1,51 | 0 | 3,43 | 0 |
| OLIG2 | ATCATATTCC | -1 | 563 | 573 | -437 | -427 | 3,68 | 0,01 | 13,03 | >0.1 | 3,25 | 5,11 | 0 | 0,99 | 0 | 0 | 0 |
| OLIG2 | AACATTTTCC | -1 | 533 | 543 | -467 | -457 | 0,49 | 0,01 | 11,22 | >0.1 | 3,1 | 5,04 | 0 | 0,98 | 0 | 1,61 | 0 |
| OLIG2 | TTCATATTTC | -1 | 486 | 496 | -514 | -504 | 0,19 | 0,01 | 15,28 | >0.1 | 2,9 | 4,93 | 0 | 0,98 | 0 | 6,27 | 0 |
| OLIG2 | GCCATTTTTT | -1 | 140 | 150 | -860 | -850 | 0,56 | 0,01 | 11,84 | >0.1 | 3,34 | 4,39 | 0 | 1,89 | 0 | 1,65 | 0 |
| OLIG3 | GAAATATGAA | 1 | 486 | 496 | -514 | -504 | 0,6 | 0,01 | 15,69 | >0.1 | 2,9 | 4,93 | 0 | 0,98 | 0 | 6,27 | 0 |
| OLIG3 | AAAATGTTGC | 1 | 535 | 545 | -465 | -455 | -0,09 | 0,01 | 10,66 | >0.1 | 3,11 | 5,04 | 0 | 0,98 | 0 | 1,61 | 0 |
| OLIG3 | GGAATATGAT | 1 | 563 | 573 | -437 | -427 | 2 | 0,01 | 11,35 | >0.1 | 3,25 | 5,11 | 0 | 0,99 | 0 | 0 | 0 |
| OLIG3 | AACAGCTTTC | 1 | 711 | 721 | -289 | -279 | 2,15 | 0,01 | 19,49 | >0.1 | 5,61 | 5,56 | 0 | 1,18 | 0 | 4,99 | 0 |
| OLIG3 | AACGTATTCT | 1 | 803 | 813 | -197 | -187 | 6,7 | 0,001 | 23,24 | >0.1 | 5,61 | 5,99 | 0 | 1,51 | 0 | 3,43 | 0 |
| OLIG3 | AAATTATTCT | 1 | 831 | 841 | -169 | -159 | 0,44 | 0,01 | 17,34 | >0.1 | 5,61 | 6,16 | 0 | 1,7 | 0 | 3,43 | 0 |
| OLIG3 | AAAATCTGTT | 1 | 841 | 851 | -159 | -149 | 6,56 | 0,001 | 21,6 | >0.1 | 5,61 | 6,24 | 0 | 1,78 | 0 | 1,41 | 0 |
| OLIG3 | AGCAGATTGT | 1 | 890 | 900 | -110 | -100 | 5,25 | 0,01 | 22,31 | >0.1 | 5,61 | 6,61 | 0 | 1,89 | 0 | 2,95 | 0 |
| OLIG3 | ACAATCTGCT | -1 | 890 | 900 | -110 | -100 | 7,43 | 0,001 | 24,49 | >0.1 | 5,61 | 6,61 | 0 | 1,89 | 0 | 2,95 | 0 |
| OLIG3 | AACAGATTTT | -1 | 841 | 851 | -159 | -149 | 6,36 | 0,001 | 21,4 | >0.1 | 5,61 | 6,24 | 0 | 1,78 | 0 | 1,41 | 0 |
| OLIG3 | AGAATAATTT | -1 | 831 | 841 | -169 | -159 | 0,45 | 0,01 | 17,35 | >0.1 | 5,61 | 6,16 | 0 | 1,7 | 0 | 3,43 | 0 |
| OLIG3 | AGAATACGTT | -1 | 803 | 813 | -197 | -187 | 6,24 | 0,01 | 22,78 | >0.1 | 5,61 | 5,99 | 0 | 1,51 | 0 | 3,43 | 0 |
| OLIG3 | GAAAGCTGTT | -1 | 711 | 721 | -289 | -279 | 0,66 | 0,01 | 18 | >0.1 | 5,61 | 5,56 | 0 | 1,18 | 0 | 4,99 | 0 |
| OLIG3 | ATCATATTCC | -1 | 563 | 573 | -437 | -427 | 2,97 | 0,01 | 12,32 | >0.1 | 3,25 | 5,11 | 0 | 0,99 | 0 | 0 | 0 |
| OLIG3 | GCAACATTTT | -1 | 535 | 545 | -465 | -455 | 0,6 | 0,01 | 11,35 | >0.1 | 3,11 | 5,04 | 0 | 0,98 | 0 | 1,61 | 0 |
| OLIG1 | AAAAAATGGC | 1 | 140 | 150 | -860 | -850 | -3,41 | 0,01 | 7,87 | >0.1 | 3,34 | 4,39 | 0 | 1,89 | 0 | 1,65 | 0 |
| OLIG1 | GGAAAATGTT | 1 | 533 | 543 | -467 | -457 | -6,41 | 0,01 | 4,32 | >0.1 | 3,1 | 5,04 | 0 | 0,98 | 0 | 1,61 | 0 |
| OLIG1 | GGAATATGAT | 1 | 563 | 573 | -437 | -427 | -5,14 | 0,01 | 4,21 | >0.1 | 3,25 | 5,11 | 0 | 0,99 | 0 | 0 | 0 |
| OLIG1 | AACATTTTCC | 1 | 691 | 701 | -309 | -299 | -4,29 | 0,01 | 9,94 | >0.1 | 4,53 | 5,49 | 0 | 1,14 | 0 | 3,07 | 0 |
| OLIG1 | AACAGCTTTC | 1 | 711 | 721 | -289 | -279 | -6,29 | 0,01 | 11,05 | >0.1 | 5,61 | 5,56 | 0 | 1,18 | 0 | 4,99 | 0 |
| OLIG1 | ACCAAAAGTA | 1 | 734 | 744 | -266 | -256 | -4,5 | 0,01 | 13,92 | >0.1 | 5,61 | 5,66 | 0 | 1,24 | 0 | 5,92 | 0 |
| OLIG1 | AACGTATTCT | 1 | 803 | 813 | -197 | -187 | 1,49 | 0,01 | 18,03 | >0.1 | 5,61 | 5,99 | 0 | 1,51 | 0 | 3,43 | 0 |
| OLIG1 | AAAATCTGTT | 1 | 841 | 851 | -159 | -149 | 0,88 | 0,01 | 15,92 | >0.1 | 5,61 | 6,24 | 0 | 1,78 | 0 | 1,41 | 0 |
| OLIG1 | AGCAGATTGT | 1 | 890 | 900 | -110 | -100 | 0,66 | 0,01 | 17,72 | >0.1 | 5,61 | 6,61 | 0 | 1,89 | 0 | 2,95 | 0 |
| OLIG1 | AACTGCTGCT | 1 | 1090 | 1100 | 90 | 100 | -4,85 | 0,01 | 13,56 | >0.1 | 5,61 | 6,88 | 0 | 1,85 | 1,79 | 2,27 | 0 |
| OLIG1 | AGCAGCAGTT | -1 | 1090 | 1100 | 90 | 100 | -4,55 | 0,01 | 13,86 | >0.1 | 5,61 | 6,88 | 0 | 1,85 | 1,79 | 2,27 | 0 |
| OLIG1 | ACAATCTGCT | -1 | 890 | 900 | -110 | -100 | 0,17 | 0,01 | 17,23 | >0.1 | 5,61 | 6,61 | 0 | 1,89 | 0 | 2,95 | 0 |
| OLIG1 | AACAGATTTT | -1 | 841 | 851 | -159 | -149 | 1,88 | 0,01 | 16,92 | >0.1 | 5,61 | 6,24 | 0 | 1,78 | 0 | 1,41 | 0 |
| OLIG1 | AGAATACGTT | -1 | 803 | 813 | -197 | -187 | 0,34 | 0,01 | 16,88 | >0.1 | 5,61 | 5,99 | 0 | 1,51 | 0 | 3,43 | 0 |
| OLIG1 | TACTTTTGGT | -1 | 734 | 744 | -266 | -256 | -6,83 | 0,01 | 11,59 | >0.1 | 5,61 | 5,66 | 0 | 1,24 | 0 | 5,92 | 0 |
| OLIG1 | GGAAAATGTT | -1 | 691 | 701 | -309 | -299 | -6,41 | 0,01 | 7,82 | >0.1 | 4,53 | 5,49 | 0 | 1,14 | 0 | 3,07 | 0 |
| OLIG1 | AACATTTTCC | -1 | 533 | 543 | -467 | -457 | -4,29 | 0,01 | 6,44 | >0.1 | 3,1 | 5,04 | 0 | 0,98 | 0 | 1,61 | 0 |
| OLIG1 | TACAGGTGCC | -1 | 327 | 337 | -673 | -663 | -7,24 | 0,01 | 6,72 | >0.1 | 2,6 | 4,65 | 0 | 1,14 | 0 | 5,56 | 0 |
| OLIG1 | GCCATTTTTT | -1 | 140 | 150 | -860 | -850 | -4,51 | 0,01 | 6,77 | >0.1 | 3,34 | 4,39 | 0 | 1,89 | 0 | 1,65 | 0 |
| TFAP2B(VAR.3) | GGCCGAGTTGGGC | 1 | 220 | 233 | -780 | -767 | -6,18 | 0,01 | 8,3 | >0.1 | 2,95 | 4,5 | 0 | 1,51 | 0 | 5,52 | 0 |
| TFAP2B(VAR.3) | GCCGAGTTGGGCA | 1 | 221 | 234 | -779 | -766 | -9,25 | 0,01 | 5,22 | >0.1 | 2,94 | 4,5 | 0 | 1,5 | 0 | 5,52 | 0 |
| TFAP2B(VAR.3) | TACTCGGGAGGCT | 1 | 345 | 358 | -655 | -642 | -10,09 | 0,01 | 4,77 | >0.1 | 2,55 | 4,68 | 0 | 1,11 | 0 | 6,52 | 0 |
| TFAP2B(VAR.3) | AACCTGGGAGGCA | 1 | 377 | 390 | -623 | -610 | -0,28 | 0,01 | 14,62 | >0.1 | 2,57 | 4,73 | 0 | 1,06 | 0 | 6,53 | 0 |
| TFAP2B(VAR.3) | AGCCGAGATCGCG | 1 | 402 | 415 | -598 | -585 | -10,55 | 0,01 | 3,66 | >0.1 | 2,63 | 4,78 | 0 | 1,04 | 0 | 5,76 | 0 |
| TFAP2B(VAR.3) | TCCAGCCTGGGCG | 1 | 424 | 437 | -576 | -563 | -3,86 | 0,01 | 10,34 | >0.1 | 2,7 | 4,82 | 0 | 1,02 | 0 | 5,67 | 0 |
| TFAP2B(VAR.3) | AGCCTGGGCGACA | 1 | 427 | 440 | -573 | -560 | -2,58 | 0,01 | 11,77 | >0.1 | 2,71 | 4,82 | 0 | 1,01 | 0 | 5,8 | 0 |
| TFAP2B(VAR.3) | GGCCATAAAGTGA | 1 | 757 | 770 | -243 | -230 | -9,1 | 0,01 | 9,66 | >0.1 | 5,57 | 5,77 | 0 | 1,32 | 0 | 6,1 | 0 |
| TFAP2B(VAR.3) | TGGCTCACAGGGG | 1 | 1033 | 1046 | 33 | 46 | -3,06 | 0,01 | 13,78 | >0.1 | 5,57 | 8,1 | 0 | 1,86 | 0 | 1,31 | 0 |
| TFAP2B(VAR.3) | GGCTCACAGGGGA | 1 | 1034 | 1047 | 34 | 47 | -1,17 | 0,01 | 15,63 | >0.1 | 5,57 | 8,06 | 0 | 1,86 | 0 | 1,31 | 0 |
| TFAP2B(VAR.3) | GCTCTTCCTGGCT | 1 | 1056 | 1069 | 56 | 69 | -8,63 | 0,01 | 8,51 | >0.1 | 5,57 | 7,44 | 0 | 1,86 | 0 | 2,27 | 0 |
| TFAP2B(VAR.3) | AGCCAGGAAGAGC | -1 | 1056 | 1069 | 56 | 69 | -9,73 | 0,01 | 7,41 | >0.1 | 5,57 | 7,44 | 0 | 1,86 | 0 | 2,27 | 0 |
| TFAP2B(VAR.3) | TCCCCTGTGAGCC | -1 | 1034 | 1047 | 34 | 47 | -0,03 | 0,01 | 16,77 | >0.1 | 5,57 | 8,06 | 0 | 1,86 | 0 | 1,31 | 0 |
| TFAP2B(VAR.3) | CCCCTGTGAGCCA | -1 | 1033 | 1046 | 33 | 46 | -2,84 | 0,01 | 14 | >0.1 | 5,57 | 8,1 | 0 | 1,86 | 0 | 1,31 | 0 |
| TFAP2B(VAR.3) | TCACTTTATGGCC | -1 | 757 | 770 | -243 | -230 | -6,03 | 0,01 | 12,73 | >0.1 | 5,57 | 5,77 | 0 | 1,32 | 0 | 6,1 | 0 |

| | | | | | | | | | | | | | | | | | |
|---|---|---|---|---|---|---|---|---|---|---|---|---|---|---|---|---|---|
| TFAP2B(VAR.3) | TGTCGCCCAGGCT | -1 | 427 | 440 | -573 | -560 | -1,66 | 0,01 | 12,69 | >0.1 | 2,71 | 4,82 | 0 | 1,01 | 0 | 5,8 | 0 |
| TFAP2B(VAR.3) | CGCCCAGGCTGGA | -1 | 424 | 437 | -576 | -563 | -4,95 | 0,01 | 9,25 | >0.1 | 2,7 | 4,82 | 0 | 1,02 | 0 | 5,67 | 0 |
| TFAP2B(VAR.3) | CGCGATCTCGGCT | -1 | 402 | 415 | -598 | -585 | -10,41 | 0,01 | 3,8 | >0.1 | 2,63 | 4,78 | 0 | 1,04 | 0 | 5,76 | 0 |
| TFAP2B(VAR.3) | TGCCTCCCAGGTT | -1 | 377 | 390 | -623 | -610 | 0,51 | 0,01 | 15,41 | >0.1 | 2,57 | 4,73 | 0 | 1,06 | 0 | 6,53 | 0 |
| TFAP2B(VAR.3) | AGCCTCCCGAGTA | -1 | 345 | 358 | -655 | -642 | -10,34 | 0,01 | 4,52 | >0.1 | 2,55 | 4,68 | 0 | 1,11 | 0 | 6,52 | 0 |
| TFAP2B(VAR.3) | TGCCCAACTCGGC | -1 | 221 | 234 | -779 | -766 | -10,3 | 0,01 | 4,17 | >0.1 | 2,94 | 4,5 | 0 | 1,5 | 0 | 5,52 | 0 |
| TFAP2B(VAR.3) | GCCCAACTCGGCC | -1 | 220 | 233 | -780 | -767 | -5,44 | 0,01 | 9,04 | >0.1 | 2,95 | 4,5 | 0 | 1,51 | 0 | 5,52 | 0 |
| TFAP2B(VAR.3) | AGCCACTGTGCCC | -1 | 180 | 193 | -820 | -807 | -10,35 | 0,01 | 2,86 | >0.1 | 3,12 | 4,45 | 0 | 1,79 | 0 | 3,86 | 0 |
| ARID5A | ATAATTTTAAAAAA | 1 | 132 | 146 | -868 | -854 | 6,94 | 0,01 | 18,21 | >0.1 | 3,38 | 4,39 | 0 | 1,89 | 0 | 1,61 | 0 |
| ARID5A | GAAATATGAATTCC | 1 | 486 | 500 | -514 | -500 | 4,34 | 0,01 | 19,45 | >0.1 | 2,91 | 4,94 | 0 | 0,99 | 0 | 6,27 | 0 |
| ARID5A | AAAATGTTGCCCAA | 1 | 535 | 549 | -465 | -451 | 2,74 | 0,01 | 13,56 | >0.1 | 3,12 | 5,05 | 0 | 0,99 | 0 | 1,66 | 0 |
| ARID5A | ATGATCTTGAAATA | 1 | 568 | 582 | -432 | -418 | 3,56 | 0,01 | 12,98 | >0.1 | 3,29 | 5,13 | 0 | 1 | 0 | 0 | 0 |
| ARID5A | CTATTAATGAAATT | 1 | 601 | 615 | -399 | -385 | 2,92 | 0,01 | 14,07 | >0.1 | 3,49 | 5,22 | 0 | 1,02 | 0 | 1,41 | 0 |
| ARID5A | GTAAAATTATAACA | 1 | 681 | 695 | -319 | -305 | 6,11 | 0,01 | 20,13 | >0.1 | 4,36 | 5,47 | 0 | 1,13 | 0 | 3,07 | 0 |
| ARID5A | AAAATATTAACCAA | 1 | 725 | 739 | -275 | -261 | 6,32 | 0,01 | 24,06 | >0.1 | 5,56 | 5,63 | 0 | 1,23 | 0 | 5,32 | 0 |
| ARID5A | CTAAAATCTGTTAC | 1 | 839 | 853 | -161 | -147 | 3,74 | 0,01 | 18,76 | >0.1 | 5,56 | 6,25 | 0 | 1,8 | 0 | 1,41 | 0 |
| ARID5A | TTTATATTGGCTCA | 1 | 877 | 891 | -123 | -109 | 5,33 | 0,01 | 22,51 | >0.1 | 5,56 | 6,54 | 0 | 1,9 | 0 | 3,19 | 0 |
| ARID5A | CCAATATAAAGTTC | -1 | 873 | 887 | -127 | -113 | 6,61 | 0,01 | 23,52 | >0.1 | 5,56 | 6,5 | 0 | 1,9 | 0 | 2,95 | 0 |
| ARID5A | CAGATTTTAGAATA | -1 | 835 | 849 | -165 | -151 | 3,52 | 0,01 | 20,25 | >0.1 | 5,56 | 6,22 | 0 | 1,76 | 0 | 3,19 | 0 |
| ARID5A | ATAATTTTTTAAAG | -1 | 824 | 838 | -176 | -162 | 4,71 | 0,01 | 21,63 | >0.1 | 5,56 | 6,14 | 0 | 1,68 | 0 | 3,55 | 0 |
| ARID5A | TTAATATTTTCCTA | -1 | 721 | 735 | -279 | -265 | 6,16 | 0,01 | 23,72 | >0.1 | 5,56 | 5,62 | 0 | 1,22 | 0 | 5,17 | 0 |
| ARID5A | ATAATTTTACAGTG | -1 | 677 | 691 | -323 | -309 | 5,62 | 0,01 | 19,67 | >0.1 | 4,28 | 5,46 | 0 | 1,12 | 0 | 3,19 | 0 |
| ARID5A | ATAATGTTGGTGTT | -1 | 631 | 645 | -369 | -355 | 4,9 | 0,01 | 16,6 | >0.1 | 3,73 | 5,31 | 0 | 1,05 | 0 | 1,61 | 0 |
| ARID5A | ATCATATTCCCAGA | -1 | 559 | 573 | -441 | -427 | 4,48 | 0,01 | 13,83 | >0.1 | 3,24 | 5,11 | 0 | 1 | 0 | 0 | 0 |
| ARID5A | TTCATATTTCTGAT | -1 | 482 | 496 | -518 | -504 | 5,69 | 0,01 | 20,64 | >0.1 | 2,9 | 4,94 | 0 | 0,99 | 0 | 6,13 | 0 |
| ARID5A | GTATTTTTAATAGA | -1 | 288 | 302 | -712 | -698 | 2,77 | 0,01 | 16,4 | >0.1 | 2,71 | 4,59 | 0 | 1,24 | 0 | 5,09 | 0 |
| ARID5A | CCATTTTTTAAAAT | -1 | 135 | 149 | -865 | -851 | 2,75 | 0,01 | 14,04 | >0.1 | 3,36 | 4,39 | 0 | 1,89 | 0 | 1,65 | 0 |
| ARID5A | TTAAAATTATGACT | -1 | 128 | 142 | -872 | -858 | 5,25 | 0,01 | 16,5 | >0.1 | 3,4 | 4,39 | 0 | 1,89 | 0 | 1,57 | 0 |
| EMX2 | GGAAATGAGC | 1 | 64 | 74 | -936 | -926 | 5,18 | 0,01 | 15,32 | >0.1 | 3,95 | 4,31 | 0 | 1,88 | 0 | 0 | 0 |
| EMX2 | GGAAATTAAA | 1 | 153 | 163 | -847 | -837 | 4,82 | 0,01 | 16,04 | >0.1 | 3,27 | 4,41 | 0 | 1,9 | 0 | 1,65 | 0 |
| EMX2 | CTCTATTAAA | 1 | 287 | 297 | -713 | -703 | 2,05 | 0,01 | 15,79 | >0.1 | 2,72 | 4,59 | 0 | 1,24 | 0 | 5,19 | 0 |
| EMX2 | AAATATGAAT | 1 | 487 | 497 | -513 | -503 | -4,55 | 0,01 | 10,41 | >0.1 | 2,91 | 4,94 | 0 | 0,98 | 0 | 6,13 | 0 |
| EMX2 | GAATATGATC | 1 | 564 | 574 | -436 | -426 | -4,03 | 0,01 | 5,33 | >0.1 | 3,25 | 5,11 | 0 | 0,99 | 0 | 0 | 0 |
| EMX2 | ACCTATTAAT | 1 | 599 | 609 | -401 | -391 | 3,6 | 0,01 | 14,65 | >0.1 | 3,47 | 5,21 | 0 | 1,02 | 0 | 1,36 | 0 |
| EMX2 | ATTAATGAAA | 1 | 603 | 613 | -397 | -387 | 5,5 | 0,01 | 16,64 | >0.1 | 3,49 | 5,22 | 0 | 1,02 | 0 | 1,41 | 0 |
| EMX2 | CAACATTATC | 1 | 636 | 646 | -364 | -354 | 0,58 | 0,01 | 12,36 | >0.1 | 3,76 | 5,31 | 0 | 1,05 | 0 | 1,66 | 0 |
| EMX2 | CTTCATGATC | 1 | 645 | 655 | -355 | -345 | 3,43 | 0,01 | 17,51 | >0.1 | 3,84 | 5,34 | 0 | 1,06 | 0 | 3,83 | 0 |
| EMX2 | TAAAATTATA | 1 | 682 | 692 | -318 | -308 | 0,28 | 0,01 | 14,26 | >0.1 | 4,34 | 5,46 | 0 | 1,12 | 0 | 3,07 | 0 |
| EMX2 | AAATATTAAC | 1 | 726 | 736 | -274 | -264 | -0,64 | 0,01 | 17,13 | >0.1 | 5,61 | 5,62 | 0 | 1,22 | 0 | 5,32 | 0 |
| EMX2 | AAAAATTATT | 1 | 829 | 839 | -171 | -161 | 1,62 | 0,01 | 18,34 | >0.1 | 5,61 | 6,15 | 0 | 1,68 | 0 | 3,28 | 0 |
| EMX2 | TCTCATGAGG | 1 | 983 | 993 | -17 | -7 | 6,18 | 0,01 | 23,73 | >0.1 | 5,61 | 8,5 | 0 | 1,87 | 0 | 1,57 | 0 |
| EMX2 | CCTCATGAGA | -1 | 983 | 993 | -17 | -7 | 6,01 | 0,01 | 23,56 | >0.1 | 5,61 | 8,5 | 0 | 1,87 | 0 | 1,57 | 0 |
| EMX2 | GATCATGAAG | -1 | 645 | 655 | -355 | -345 | 4,71 | 0,01 | 18,79 | >0.1 | 3,84 | 5,34 | 0 | 1,06 | 0 | 3,83 | 0 |
| EMX2 | TTTCATTAAT | -1 | 603 | 613 | -397 | -387 | 5,31 | 0,01 | 16,45 | >0.1 | 3,49 | 5,22 | 0 | 1,02 | 0 | 1,41 | 0 |
| EMX2 | ATACATGACT | -1 | 516 | 526 | -484 | -474 | -1,5 | 0,01 | 10,46 | >0.1 | 3,02 | 5 | 0 | 0,98 | 0 | 2,95 | 0 |
| EMX2 | TAAAATTATG | -1 | 131 | 141 | -869 | -859 | 1,55 | 0,01 | 12,8 | >0.1 | 3,4 | 4,38 | 0 | 1,89 | 0 | 1,57 | 0 |
| EMX2 | AATTATGACT | -1 | 128 | 138 | -872 | -862 | 0,73 | 0,01 | 11,99 | >0.1 | 3,41 | 4,38 | 0 | 1,89 | 0 | 1,57 | 0 |
| TP73 | GGCACAGTGGCTCACGCC | 1 | 181 | 199 | -819 | -801 | -8,55 | 0,01 | 4,65 | >0.1 | 3,1 | 4,45 | 0 | 1,78 | 0 | 3,86 | 0 |
| TP73 | AACACGGTGAAACCCCGT | 1 | 269 | 287 | -731 | -713 | -10,01 | 0,01 | 3,71 | >0.1 | 2,76 | 4,57 | 0 | 1,3 | 0 | 5,09 | 0 |
| TP73 | CACCTGTAGTCCCAGCTA | 1 | 329 | 347 | -671 | -653 | -11,35 | 0,01 | 3,08 | >0.1 | 2,59 | 4,66 | 0 | 1,14 | 0 | 6,03 | 0 |
| TP73 | TGGGAGGCAGAGCTTGCA | 1 | 381 | 399 | -619 | -601 | -10,72 | 0,01 | 3,8 | >0.1 | 2,58 | 4,75 | 0 | 1,06 | 0 | 6,13 | 0 |
| TP73 | GACAGACCGAGACTCAGT | 1 | 436 | 454 | -564 | -546 | -10,96 | 0,01 | 3,69 | >0.1 | 2,75 | 4,85 | 0 | 1,01 | 0 | 6,05 | 0 |
| TP73 | TACAAACACCAACATTAT | 1 | 627 | 645 | -373 | -355 | -8,56 | 0,01 | 3,17 | >0.1 | 3,71 | 5,31 | 0 | 1,06 | 0 | 1,66 | 0 |
| TP73 | GACCTCTTTGGTCACTGT | 1 | 665 | 683 | -335 | -317 | -11,43 | 0,01 | 2,93 | >0.1 | 4,13 | 5,43 | 0 | 1,11 | 0 | 3,7 | 0 |
| TP73 | TACCGGTTTTGATTTGGC | 1 | 742 | 760 | -258 | -240 | -11,11 | 0,01 | 7,33 | >0.1 | 5,51 | 5,72 | 0 | 1,29 | 0 | 5,92 | 0 |
| TP73 | TACATATCTGTCCTCTCC | 1 | 850 | 868 | -150 | -132 | -6,32 | 0,01 | 10,2 | >0.1 | 5,51 | 6,35 | 0 | 1,9 | 0 | 2,75 | 0 |
| TP73 | GTCAAGCTCTTCCTGGCT | 1 | 1051 | 1069 | 51 | 69 | -4,41 | 0,01 | 12,78 | >0.1 | 5,51 | 7,55 | 0 | 1,86 | 0 | 2,27 | 0 |
| TP73 | ACCATTGAGGAACAGGCC | 1 | 1106 | 1124 | 106 | 124 | -9,46 | 0,01 | 10,37 | >0.1 | 5,51 | 6,69 | 0 | 1,85 | 3,22 | 2,56 | 0 |
| TP73 | AACAGGCCAAGACATTTT | 1 | 1116 | 1134 | 116 | 134 | 7,09 | 0,001 | 26,95 | >0.1 | 5,51 | 6,59 | 0 | 1,85 | 3,22 | 2,7 | 0 |

| | | | | | | | | | | | | | | | | | |
|---|---|---|---|---|---|---|---|---|---|---|---|---|---|---|---|---|---|
| TP73 | GACATTTTTGGACAAGTT | 1 | 1126 | 1144 | 126 | 144 | 2,22 | 0,001 | 22,15 | >0.1 | 5,51 | 6,49 | 0 | 1,84 | 3,22 | 2,86 | 0 |
| TP73 | TTCGTGGTTAAACTTGTC | -1 | 1136 | 1154 | 136 | 154 | 3,12 | 0,001 | 23,4 | >0.1 | 5,51 | 6,41 | 0 | 1,84 | 3,22 | 3,29 | 0 |
| TP73 | AACTTGTCCAAAAATGTC | -1 | 1126 | 1144 | 126 | 144 | 0,37 | 0,001 | 20,3 | >0.1 | 5,51 | 6,49 | 0 | 1,84 | 3,22 | 2,86 | 0 |
| TP73 | AAAATGTCTTGGCCTGTT | -1 | 1116 | 1134 | 116 | 134 | -2,68 | 0,01 | 17,18 | >0.1 | 5,51 | 6,59 | 0 | 1,85 | 3,22 | 2,7 | 0 |
| TP73 | AGCCAGGAAGAGCTTGAC | -1 | 1051 | 1069 | 51 | 69 | -2,7 | 0,01 | 14,49 | >0.1 | 5,51 | 7,55 | 0 | 1,86 | 0 | 2,27 | 0 |
| TP73 | TACCTCTCCTGTCACTTT | -1 | 763 | 781 | -237 | -219 | -6,22 | 0,01 | 12,39 | >0.1 | 5,51 | 5,82 | 0 | 1,36 | 0 | 5,92 | 0 |
| TP73 | GCCAAATCAAAACCGGTA | -1 | 742 | 760 | -258 | -240 | -6,12 | 0,01 | 12,32 | >0.1 | 5,51 | 5,72 | 0 | 1,29 | 0 | 5,92 | 0 |
| TP73 | CCCTTCCAAATACATGAC | -1 | 517 | 535 | -483 | -465 | -8,91 | 0,01 | 3,22 | >0.1 | 3,05 | 5,02 | 0 | 0,99 | 0 | 3,07 | 0 |
| TP73 | TACATGACTTTGCCCTTG | -1 | 507 | 525 | -493 | -475 | -6,89 | 0,01 | 4,96 | >0.1 | 3 | 5 | 0 | 0,99 | 0 | 2,86 | 0 |
| TP73 | CCCAGGCTGGAGCACAGT | -1 | 417 | 435 | -583 | -565 | -10,1 | 0,01 | 3,85 | >0.1 | 2,69 | 4,81 | 0 | 1,03 | 0 | 5,42 | 0 |
| TP73 | TGCAAGCTCTGCCTCCCA | -1 | 381 | 399 | -619 | -601 | 3,06 | 0,001 | 17,58 | >0.1 | 2,58 | 4,75 | 0 | 1,06 | 0 | 6,13 | 0 |
| TP73 | GCCTCCCAGGTTCACGCC | -1 | 371 | 389 | -629 | -611 | -10,16 | 0,01 | 4,6 | >0.1 | 2,56 | 4,73 | 0 | 1,08 | 0 | 6,4 | 0 |
| TP73 | CTCCTGCCTCAGCCTCCC | -1 | 350 | 368 | -650 | -632 | -6,8 | 0,01 | 8,06 | >0.1 | 2,53 | 4,7 | 0 | 1,11 | 0 | 6,52 | 0 |
| TP73 | TACAGGTGCCCACCACCA | -1 | 319 | 337 | -681 | -663 | -4,55 | 0,01 | 9,44 | >0.1 | 2,61 | 4,65 | 0 | 1,16 | 0 | 5,56 | 0 |
| TP73 | TACAGGCGTGAGCCACTG | -1 | 185 | 203 | -815 | -797 | -8,2 | 0,01 | 5,13 | >0.1 | 3,09 | 4,46 | 0 | 1,75 | 0 | 4,04 | 0 |
| TP73 | GACTACTCTCCACTCCAC | -1 | 114 | 132 | -886 | -868 | -6,89 | 0,01 | 3,58 | >0.1 | 3,48 | 4,37 | 0 | 1,89 | 0 | 0,73 | 0 |
| TP73 | CACAGGTCAACTCTCCTG | -1 | 99 | 117 | -901 | -883 | -3,98 | 0,01 | 6,62 | >0.1 | 3,59 | 4,36 | 0 | 1,89 | 0 | 0,77 | 0 |
| EMX1 | GGAAATGAGC | 1 | 64 | 74 | -936 | -926 | 6,08 | 0,01 | 16,22 | >0.1 | 3,95 | 4,31 | 0 | 1,88 | 0 | 0 | 0 |
| EMX1 | AAAAATGGCC | 1 | 141 | 151 | -859 | -849 | 4,05 | 0,01 | 15,28 | >0.1 | 3,33 | 4,39 | 0 | 1,89 | 0 | 1,61 | 0 |
| EMX1 | GGAAATTAAA | 1 | 153 | 163 | -847 | -837 | 3,55 | 0,01 | 14,77 | >0.1 | 3,27 | 4,41 | 0 | 1,9 | 0 | 1,65 | 0 |
| EMX1 | AGAAATGGCT | 1 | 170 | 180 | -830 | -820 | 3,73 | 0,01 | 16,64 | >0.1 | 3,18 | 4,43 | 0 | 1,88 | 0 | 3,43 | 0 |
| EMX1 | ATTAATGAAA | 1 | 603 | 613 | -397 | -387 | 6,89 | 0,01 | 18,03 | >0.1 | 3,49 | 5,22 | 0 | 1,02 | 0 | 1,41 | 0 |
| EMX1 | CTTCATGATC | 1 | 645 | 655 | -355 | -345 | 3,74 | 0,01 | 17,82 | >0.1 | 3,84 | 5,34 | 0 | 1,06 | 0 | 3,83 | 0 |
| EMX1 | AAAAATTATT | 1 | 829 | 839 | -171 | -161 | 3,17 | 0,01 | 19,89 | >0.1 | 5,61 | 6,15 | 0 | 1,68 | 0 | 3,28 | 0 |
| EMX1 | TCTCATGAGG | 1 | 983 | 993 | -17 | -7 | 3,85 | 0,01 | 21,4 | >0.1 | 5,61 | 8,5 | 0 | 1,87 | 0 | 1,57 | 0 |
| EMX1 | CTCAATGGTG | -1 | 1105 | 1115 | 105 | 115 | 4,03 | 0,01 | 23,93 | >0.1 | 5,61 | 6,7 | 0 | 1,85 | 3,22 | 2,52 | 0 |
| EMX1 | CTGAATGACT | -1 | 1011 | 1021 | 11 | 21 | 3,87 | 0,01 | 21,69 | >0.1 | 5,61 | 8,95 | 0 | 1,86 | 0 | 1,39 | 0 |
| EMX1 | CCTCATGAGA | -1 | 983 | 993 | -17 | -7 | 4,59 | 0,01 | 22,14 | >0.1 | 5,61 | 8,5 | 0 | 1,87 | 0 | 1,57 | 0 |
| EMX1 | CTTTATGGCC | -1 | 757 | 767 | -243 | -233 | 3,7 | 0,01 | 22,48 | >0.1 | 5,61 | 5,75 | 0 | 1,31 | 0 | 6,1 | 0 |
| EMX1 | ATTAATAGGT | -1 | 599 | 609 | -401 | -391 | 5,35 | 0,01 | 16,4 | >0.1 | 3,47 | 5,21 | 0 | 1,02 | 0 | 1,36 | 0 |
| EMX1 | TTTAATAGAG | -1 | 287 | 297 | -713 | -703 | 3,06 | 0,01 | 16,8 | >0.1 | 2,72 | 4,59 | 0 | 1,24 | 0 | 5,19 | 0 |
| EMX1 | TTTAATTTCC | -1 | 153 | 163 | -847 | -837 | 5,6 | 0,01 | 16,82 | >0.1 | 3,27 | 4,41 | 0 | 1,9 | 0 | 1,65 | 0 |
| BATF::JUN | GAGAGTAGTCA | 1 | 122 | 133 | -878 | -867 | -6,01 | 0,01 | 5,35 | >0.1 | 3,45 | 4,37 | 0 | 1,89 | 0 | 1,65 | 0 |
| BATF::JUN | GTCCTGACTAA | 1 | 260 | 271 | -740 | -729 | 5,86 | 0,001 | 19,07 | >0.1 | 2,81 | 4,55 | 0 | 1,33 | 0 | 4,53 | 0 |
| BATF::JUN | AGGCTGAGGCA | 1 | 353 | 364 | -647 | -636 | -2,84 | 0,01 | 11,88 | >0.1 | 2,53 | 4,69 | 0 | 1,1 | 0 | 6,4 | 0 |
| BATF::JUN | AGACTCAGTCT | 1 | 445 | 456 | -555 | -544 | -8,09 | 0,01 | 5,54 | >0.1 | 2,76 | 4,85 | 0 | 1 | 0 | 5,01 | 0 |
| BATF::JUN | CCTATTAATGA | 1 | 600 | 611 | -400 | -389 | -3,96 | 0,01 | 7,11 | >0.1 | 3,48 | 5,21 | 0 | 1,02 | 0 | 1,36 | 0 |
| BATF::JUN | AAATTGTCTGA | 1 | 610 | 621 | -390 | -379 | -7,69 | 0,01 | 3,67 | >0.1 | 3,55 | 5,24 | 0 | 1,03 | 0 | 1,55 | 0 |
| BATF::JUN | TTAAAGAATAA | 1 | 794 | 805 | -206 | -195 | -9,19 | 0,01 | 8,99 | >0.1 | 5,6 | 5,94 | 0 | 1,47 | 0 | 5,17 | 0 |
| BATF::JUN | TATTTGATTCA | 1 | 813 | 824 | -187 | -176 | -4,46 | 0,01 | 12,19 | >0.1 | 5,6 | 6,05 | 0 | 1,58 | 0 | 3,43 | 0 |
| BATF::JUN | ATATTGGCTCA | 1 | 880 | 891 | -120 | -109 | -4,51 | 0,01 | 12,59 | >0.1 | 5,6 | 6,54 | 0 | 1,9 | 0 | 3,07 | 0 |
| BATF::JUN | TTGTTTACTGT | 1 | 896 | 907 | -104 | -93 | -6,29 | 0,01 | 10,82 | >0.1 | 5,6 | 6,68 | 0 | 1,89 | 0 | 2,95 | 0 |
| BATF::JUN | GGTTTTAGTCT | 1 | 994 | 1005 | -6 | 5 | 0,45 | 0,01 | 18,32 | >0.1 | 5,6 | 8,88 | 0 | 1,87 | 0 | 1,53 | 0 |
| BATF::JUN | ATCTTGGCTCA | 1 | 1029 | 1040 | 29 | 40 | -8,74 | 0,01 | 8,34 | >0.1 | 5,6 | 8,29 | 0 | 1,86 | 0 | 1,34 | 0 |
| BATF::JUN | TGGTGGACTGA | -1 | 1099 | 1110 | 99 | 110 | -8,71 | 0,01 | 11,13 | >0.1 | 5,6 | 6,77 | 0 | 1,85 | 3,22 | 2,4 | 0 |
| BATF::JUN | CCACTGAATGA | -1 | 1013 | 1024 | 13 | 24 | 1,01 | 0,01 | 18,78 | >0.1 | 5,6 | 8,89 | 0 | 1,86 | 0 | 1,42 | 0 |
| BATF::JUN | TGAATGACTTT | -1 | 1009 | 1020 | 9 | 20 | -7,06 | 0,01 | 10,89 | >0.1 | 5,6 | 9,03 | 0 | 1,86 | 0 | 1,46 | 0 |
| BATF::JUN | CTGCTGAGCCA | -1 | 884 | 895 | -116 | -105 | -5,84 | 0,01 | 11,41 | >0.1 | 5,6 | 6,57 | 0 | 1,89 | 0 | 3,19 | 0 |
| BATF::JUN | AAAGTGAATCA | -1 | 817 | 828 | -183 | -172 | -3,3 | 0,01 | 13,53 | >0.1 | 5,6 | 6,08 | 0 | 1,6 | 0 | 3,55 | 0 |
| BATF::JUN | AGACTGAGTCT | -1 | 445 | 456 | -555 | -544 | 7,57 | 0,001 | 21,2 | >0.1 | 2,76 | 4,85 | 0 | 1 | 0 | 5,01 | 0 |
| BATF::JUN | CTCCTGCCTCA | -1 | 357 | 368 | -643 | -632 | -9,52 | 0,01 | 4,92 | >0.1 | 2,52 | 4,7 | 0 | 1,09 | 0 | 6,13 | 0 |
| BATF::JUN | CGTGTTAGTCA | -1 | 264 | 275 | -736 | -725 | -9,1 | 0,01 | 4,01 | >0.1 | 2,79 | 4,55 | 0 | 1,31 | 0 | 4,45 | 0 |
| BATF::JUN | ATTATGACTAC | -1 | 126 | 137 | -874 | -863 | -4,63 | 0,01 | 6,71 | >0.1 | 3,42 | 4,38 | 0 | 1,89 | 0 | 1,65 | 0 |
| SP8 | CTCACGCCTGTA | 1 | 191 | 203 | -809 | -797 | 1,71 | 0,01 | 15,2 | >0.1 | 3,07 | 4,46 | 0 | 1,7 | 0 | 4,26 | 0 |
| SP8 | GAAACCCCGTCT | 1 | 277 | 289 | -723 | -711 | 2,7 | 0,01 | 16,49 | >0.1 | 2,75 | 4,57 | 0 | 1,27 | 0 | 5,19 | 0 |
| SP8 | ACCCCGTCTCTA | 1 | 280 | 292 | -720 | -708 | 2,79 | 0,01 | 16,56 | >0.1 | 2,74 | 4,58 | 0 | 1,26 | 0 | 5,19 | 0 |
| SP8 | GTCCCAGCTACT | 1 | 337 | 349 | -663 | -651 | 1,1 | 0,01 | 15,38 | >0.1 | 2,57 | 4,67 | 0 | 1,13 | 0 | 5,92 | 0 |
| SP8 | GCTCAGCGCCCA | 1 | 940 | 952 | -60 | -48 | 2,25 | 0,01 | 17,75 | >0.1 | 5,58 | 7,25 | 0 | 1,88 | 0 | 0,79 | 0 |
| SP8 | TCAGCGCCCAAC | 1 | 942 | 954 | -58 | -46 | 2,51 | 0,01 | 19,06 | >0.1 | 5,58 | 7,29 | 0 | 1,88 | 0 | 1,8 | 0 |

| | | | | | | | | | | | | | | | | | |
|---|---|---|---|---|---|---|---|---|---|---|---|---|---|---|---|---|---|
| SP8 | GCTCAGTCCACC | 1 | 1097 | 1109 | 97 | 109 | 2,19 | 0,01 | 21,99 | >0.1 | 5,58 | 6,79 | 0 | 1,85 | 3,22 | 2,35 | 0 |
| SP8 | ACATCGTCCCCT | -1 | 1041 | 1053 | 41 | 53 | 3,09 | 0,01 | 18,97 | >0.1 | 5,58 | 7,82 | 0 | 1,86 | 0 | 0,62 | 0 |
| SP8 | ATCACATCCACT | -1 | 1019 | 1031 | 19 | 31 | 3,49 | 0,01 | 20,98 | >0.1 | 5,58 | 8,74 | 0 | 1,86 | 0 | 1,31 | 0 |
| SP8 | AAACCTCCTCAT | -1 | 987 | 999 | -13 | -1 | 1,98 | 0,01 | 19,73 | >0.1 | 5,58 | 8,69 | 0 | 1,87 | 0 | 1,61 | 0 |
| SP8 | GCTCTGCCTCCC | -1 | 382 | 394 | -618 | -606 | 4,78 | 0,01 | 19,17 | >0.1 | 2,58 | 4,74 | 0 | 1,06 | 0 | 6,02 | 0 |
| SP8 | GCCTCAGCCTCC | -1 | 351 | 363 | -649 | -637 | 0,91 | 0,01 | 15,64 | >0.1 | 2,54 | 4,69 | 0 | 1,1 | 0 | 6,4 | 0 |
| SP8 | CCTCAGCCTCCC | -1 | 350 | 362 | -650 | -638 | 3,72 | 0,01 | 18,45 | >0.1 | 2,54 | 4,69 | 0 | 1,1 | 0 | 6,4 | 0 |
| SP8 | CCCACCACCACG | -1 | 317 | 329 | -683 | -671 | 4,99 | 0,01 | 19,67 | >0.1 | 2,63 | 4,63 | 0 | 1,17 | 0 | 6,26 | 0 |
| SP8 | ACCACCACGCCC | -1 | 314 | 326 | -686 | -674 | 5,84 | 0,01 | 20,41 | >0.1 | 2,64 | 4,63 | 0 | 1,17 | 0 | 6,13 | 0 |
| SP8 | CCACCACGCCCA | -1 | 313 | 325 | -687 | -675 | 3,47 | 0,01 | 18,04 | >0.1 | 2,64 | 4,63 | 0 | 1,18 | 0 | 6,13 | 0 |
| SP8 | ACCACGCCCAGC | -1 | 311 | 323 | -689 | -677 | 15,57 | 0,0001 | 30,05 | >0.1 | 2,65 | 4,62 | 0 | 1,18 | 0 | 6,03 | 0 |
| SP8 | GATCTGCCCAAC | -1 | 226 | 238 | -774 | -762 | 2,74 | 0,01 | 16,9 | >0.1 | 2,93 | 4,5 | 0 | 1,48 | 0 | 5,26 | 0 |
| SP8 | CCAACTCGGCCT | -1 | 219 | 231 | -781 | -769 | 1,7 | 0,01 | 16,18 | >0.1 | 2,96 | 4,49 | 0 | 1,51 | 0 | 5,52 | 0 |
| SP8 | AACTCGGCCTCC | -1 | 217 | 229 | -783 | -771 | 1,5 | 0,01 | 16,11 | >0.1 | 2,96 | 4,49 | 0 | 1,52 | 0 | 5,63 | 0 |
| SP8 | ACTCGGCCTCCC | -1 | 216 | 228 | -784 | -772 | 2,08 | 0,01 | 16,59 | >0.1 | 2,97 | 4,49 | 0 | 1,53 | 0 | 5,52 | 0 |
| SP8 | TCCATGGCCATT | -1 | 144 | 156 | -856 | -844 | 3,51 | 0,01 | 14,73 | >0.1 | 3,31 | 4,4 | 0 | 1,9 | 0 | 1,61 | 0 |
| MYF6 | AACCAGTGTG | 1 | 47 | 57 | -953 | -943 | 0,94 | 0,01 | 11,28 | >0.1 | 4,17 | 4,3 | 0 | 1,87 | 0 | 0 | 0 |
| MYF6 | GAGAGTAGTC | 1 | 122 | 132 | -878 | -868 | -5,21 | 0,01 | 6,15 | >0.1 | 3,45 | 4,37 | 0 | 1,89 | 0 | 1,65 | 0 |
| MYF6 | CACAGTGGCT | 1 | 183 | 193 | -817 | -807 | -4,77 | 0,01 | 8,41 | >0.1 | 3,11 | 4,44 | 0 | 1,76 | 0 | 3,86 | 0 |
| MYF6 | GGCACCTGTA | 1 | 327 | 337 | -673 | -663 | -2,55 | 0,01 | 11,41 | >0.1 | 2,6 | 4,65 | 0 | 1,14 | 0 | 5,56 | 0 |
| MYF6 | CGCCACTGTG | 1 | 413 | 423 | -587 | -577 | -4,13 | 0,01 | 9,43 | >0.1 | 2,66 | 4,79 | 0 | 1,02 | 0 | 5,09 | 0 |
| MYF6 | AAAAACTGAT | 1 | 474 | 484 | -526 | -516 | -2,94 | 0,01 | 10,76 | >0.1 | 2,86 | 4,91 | 0 | 0,99 | 0 | 4,95 | 0 |
| MYF6 | AACAGCTTTC | 1 | 711 | 721 | -289 | -279 | 4,92 | 0,001 | 22,26 | >0.1 | 5,61 | 5,56 | 0 | 1,18 | 0 | 4,99 | 0 |
| MYF6 | AAAATCTGTT | 1 | 841 | 851 | -159 | -149 | 1,02 | 0,01 | 16,06 | >0.1 | 5,61 | 6,24 | 0 | 1,78 | 0 | 1,41 | 0 |
| MYF6 | CATATCTGTC | 1 | 852 | 862 | -148 | -138 | -4,66 | 0,01 | 11,81 | >0.1 | 5,61 | 6,31 | 0 | 1,88 | 0 | 2,67 | 0 |
| MYF6 | AACCCAAGTT | 1 | 951 | 961 | -49 | -39 | -2,85 | 0,01 | 13,79 | >0.1 | 5,61 | 7,42 | 0 | 1,88 | 0 | 1,74 | 0 |
| MYF6 | AACTGCTGCT | 1 | 1090 | 1100 | 90 | 100 | 2,57 | 0,01 | 20,98 | >0.1 | 5,61 | 6,88 | 0 | 1,85 | 1,79 | 2,27 | 0 |
| MYF6 | AAGACCTGTT | 1 | 1158 | 1168 | 158 | 168 | 3,32 | 0,01 | 19,91 | >0.1 | 4,38 | 6,24 | 0 | 1,78 | 1,79 | 2,4 | 0 |
| MYF6 | AACAGGTCTT | -1 | 1158 | 1168 | 158 | 168 | 6,53 | 0,001 | 23,12 | >0.1 | 4,38 | 6,24 | 0 | 1,78 | 1,79 | 2,4 | 0 |
| MYF6 | AGCAGCAGTT | -1 | 1090 | 1100 | 90 | 100 | 8,51 | 0,001 | 26,92 | >0.1 | 5,61 | 6,88 | 0 | 1,85 | 1,79 | 2,27 | 0 |
| MYF6 | AACAGATTTT | -1 | 841 | 851 | -159 | -149 | 0,55 | 0,01 | 15,59 | >0.1 | 5,61 | 6,24 | 0 | 1,78 | 0 | 1,41 | 0 |
| MYF6 | GAAAGCTGTT | -1 | 711 | 721 | -289 | -279 | 3,74 | 0,001 | 21,08 | >0.1 | 5,61 | 5,56 | 0 | 1,18 | 0 | 4,99 | 0 |
| MYF6 | TACAGGTGCC | -1 | 327 | 337 | -673 | -663 | 3,43 | 0,01 | 17,39 | >0.1 | 2,6 | 4,65 | 0 | 1,14 | 0 | 5,56 | 0 |
| MYF6 | AGCCACTGTG | -1 | 183 | 193 | -817 | -807 | 0,56 | 0,01 | 13,74 | >0.1 | 3,11 | 4,44 | 0 | 1,76 | 0 | 3,86 | 0 |
| MYF6 | GTCAACTCTC | -1 | 102 | 112 | -898 | -888 | -3 | 0,01 | 7,6 | >0.1 | 3,6 | 4,35 | 0 | 1,88 | 0 | 0,77 | 0 |
| MYF6 | CACACTGGTT | -1 | 47 | 57 | -953 | -943 | -4,03 | 0,01 | 6,31 | >0.1 | 4,17 | 4,3 | 0 | 1,87 | 0 | 0 | 0 |
| HOXD12 | AGCCACAGAAC | 1 | 39 | 50 | -961 | -950 | -5,21 | 0,01 | 5,24 | >0.1 | 4,29 | 4,29 | 0 | 1,87 | 0 | 0 | 0 |
| HOXD12 | AGTCATAATTT | 1 | 128 | 139 | -872 | -861 | 5,52 | 0,001 | 16,78 | >0.1 | 3,41 | 4,38 | 0 | 1,89 | 0 | 1,57 | 0 |
| HOXD12 | GATCAGAAATG | 1 | 166 | 177 | -834 | -823 | -4,84 | 0,01 | 7,14 | >0.1 | 3,19 | 4,43 | 0 | 1,9 | 0 | 2,46 | 0 |
| HOXD12 | TGGCGTGAACC | 1 | 370 | 381 | -630 | -619 | -4,37 | 0,01 | 10,23 | >0.1 | 2,54 | 4,72 | 0 | 1,07 | 0 | 6,27 | 0 |
| HOXD12 | AGTCTCAAAAA | 1 | 451 | 462 | -549 | -538 | -3,02 | 0,01 | 11 | >0.1 | 2,78 | 4,87 | 0 | 1 | 0 | 5,38 | 0 |
| HOXD12 | TCTCAAAAAAA | 1 | 453 | 464 | -547 | -536 | -1,84 | 0,01 | 12,19 | >0.1 | 2,79 | 4,87 | 0 | 1 | 0 | 5,38 | 0 |
| HOXD12 | GATCAGAAATA | 1 | 481 | 492 | -519 | -508 | -2,54 | 0,01 | 11,91 | >0.1 | 2,89 | 4,93 | 0 | 0,99 | 0 | 5,65 | 0 |
| HOXD12 | TTCCATAAAGA | 1 | 496 | 507 | -504 | -493 | -5,8 | 0,01 | 9,52 | >0.1 | 2,94 | 4,96 | 0 | 0,98 | 0 | 6,44 | 0 |
| HOXD12 | AACCATACAAA | 1 | 622 | 633 | -378 | -367 | -4,97 | 0,01 | 6,42 | >0.1 | 3,64 | 5,27 | 0 | 1,04 | 0 | 1,44 | 0 |
| HOXD12 | CACTGTAAAAT | 1 | 677 | 688 | -323 | -312 | -5,25 | 0,01 | 8,99 | >0.1 | 4,26 | 5,45 | 0 | 1,11 | 0 | 3,43 | 0 |
| HOXD12 | TATCTTTAACA | 1 | 704 | 715 | -296 | -285 | -6,88 | 0,01 | 10,23 | >0.1 | 5,6 | 5,54 | 0 | 1,17 | 0 | 4,81 | 0 |
| HOXD12 | GGCCATAAAGT | 1 | 757 | 768 | -243 | -232 | 4,18 | 0,001 | 22,95 | >0.1 | 5,6 | 5,76 | 0 | 1,31 | 0 | 6,1 | 0 |
| HOXD12 | GCTGATAAGAG | 1 | 966 | 977 | -34 | -23 | -5,65 | 0,01 | 11,33 | >0.1 | 5,6 | 7,81 | 0 | 1,87 | 0 | 1,7 | 0 |
| HOXD12 | AGTCATTCAGT | 1 | 1011 | 1022 | 11 | 22 | -6,61 | 0,01 | 11,2 | >0.1 | 5,6 | 8,95 | 0 | 1,86 | 0 | 1,39 | 0 |
| HOXD12 | TGTCCAAAAAT | -1 | 1129 | 1140 | 129 | 140 | -7,07 | 0,01 | 12,85 | >0.1 | 5,6 | 6,47 | 0 | 1,84 | 3,22 | 2,79 | 0 |
| HOXD12 | CCTCATGAGAT | -1 | 982 | 993 | -18 | -7 | -6,84 | 0,01 | 10,65 | >0.1 | 5,6 | 8,5 | 0 | 1,87 | 0 | 1,53 | 0 |
| HOXD12 | GACCAAAAAAA | -1 | 923 | 934 | -77 | -66 | -2,64 | 0,01 | 12,91 | >0.1 | 5,6 | 6,98 | 0 | 1,88 | 0 | 1,09 | 0 |
| HOXD12 | CACAGTAAACA | -1 | 897 | 908 | -103 | -92 | -2,65 | 0,01 | 14,38 | >0.1 | 5,6 | 6,69 | 0 | 1,89 | 0 | 2,86 | 0 |
| HOXD12 | AATCCTAGAGA | -1 | 785 | 796 | -215 | -204 | -7,06 | 0,01 | 11,55 | >0.1 | 5,6 | 5,9 | 0 | 1,43 | 0 | 5,69 | 0 |
| HOXD12 | TTTGGTTAATA | -1 | 729 | 740 | -271 | -260 | -6,72 | 0,01 | 11,26 | >0.1 | 5,6 | 5,64 | 0 | 1,23 | 0 | 5,52 | 0 |
| HOXD12 | AGCTGTTAAAG | -1 | 707 | 718 | -293 | -282 | -5,68 | 0,01 | 11,45 | >0.1 | 5,6 | 5,55 | 0 | 1,17 | 0 | 4,81 | 0 |
| HOXD12 | GATCATGAAGA | -1 | 644 | 655 | -356 | -345 | -1,3 | 0,01 | 12,64 | >0.1 | 3,84 | 5,34 | 0 | 1,06 | 0 | 3,7 | 0 |
| HOXD12 | TTTCATTAATA | -1 | 602 | 613 | -398 | -387 | -1,9 | 0,01 | 9,24 | >0.1 | 3,49 | 5,22 | 0 | 1,02 | 0 | 1,41 | 0 |

| | | | | | | | | | | | | | | | | | |
|---|---|---|---|---|---|---|---|---|---|---|---|---|---|---|---|---|---|
| TFAP2A(VAR.3) | GGGCACAGTGGCT | 1 | 180 | 193 | -820 | -807 | -7,3 | 0,01 | 5,91 | >0.1 | 3,12 | 4,45 | 0 | 1,79 | 0 | 3,86 | 0 |
| TFAP2A(VAR.3) | GGCCGAGTTGGGC | 1 | 220 | 233 | -780 | -767 | -0,57 | 0,01 | 13,91 | >0.1 | 2,95 | 4,5 | 0 | 1,51 | 0 | 5,52 | 0 |
| TFAP2A(VAR.3) | GCCGAGTTGGGCA | 1 | 221 | 234 | -779 | -766 | -6,76 | 0,01 | 7,71 | >0.1 | 2,94 | 4,5 | 0 | 1,5 | 0 | 5,52 | 0 |
| TFAP2A(VAR.3) | AACCTGGGAGGCA | 1 | 377 | 390 | -623 | -610 | -1,82 | 0,01 | 13,08 | >0.1 | 2,57 | 4,73 | 0 | 1,06 | 0 | 6,53 | 0 |
| TFAP2A(VAR.3) | AGCCGAGATCGCG | 1 | 402 | 415 | -598 | -585 | -7,89 | 0,01 | 6,32 | >0.1 | 2,63 | 4,78 | 0 | 1,04 | 0 | 5,76 | 0 |
| TFAP2A(VAR.3) | TCCAGCCTGGGCG | 1 | 424 | 437 | -576 | -563 | -2,18 | 0,01 | 12,02 | >0.1 | 2,7 | 4,82 | 0 | 1,02 | 0 | 5,67 | 0 |
| TFAP2A(VAR.3) | AGCCTGGGCGACA | 1 | 427 | 440 | -573 | -560 | -3,35 | 0,01 | 11 | >0.1 | 2,71 | 4,82 | 0 | 1,01 | 0 | 5,8 | 0 |
| TFAP2A(VAR.3) | GGCCATAAAGTGA | 1 | 757 | 770 | -243 | -230 | -7,17 | 0,01 | 11,59 | >0.1 | 5,57 | 5,77 | 0 | 1,32 | 0 | 6,1 | 0 |
| TFAP2A(VAR.3) | TGGCTCACAGGGG | 1 | 1033 | 1046 | 33 | 46 | -4,61 | 0,01 | 12,23 | >0.1 | 5,57 | 8,1 | 0 | 1,86 | 0 | 1,31 | 0 |
| TFAP2A(VAR.3) | GGCTCACAGGGGA | 1 | 1034 | 1047 | 34 | 47 | 0,07 | 0,01 | 16,87 | >0.1 | 5,57 | 8,06 | 0 | 1,86 | 0 | 1,31 | 0 |
| TFAP2A(VAR.3) | GCTCTTCCTGGCT | 1 | 1056 | 1069 | 56 | 69 | -7,81 | 0,01 | 9,33 | >0.1 | 5,57 | 7,44 | 0 | 1,86 | 0 | 2,27 | 0 |
| TFAP2A(VAR.3) | AGCCAGGAAGAGC | -1 | 1056 | 1069 | 56 | 69 | -8,11 | 0,01 | 9,03 | >0.1 | 5,57 | 7,44 | 0 | 1,86 | 0 | 2,27 | 0 |
| TFAP2A(VAR.3) | TCCCCTGTGAGCC | -1 | 1034 | 1047 | 34 | 47 | -0,14 | 0,01 | 16,66 | >0.1 | 5,57 | 8,06 | 0 | 1,86 | 0 | 1,31 | 0 |
| TFAP2A(VAR.3) | CCCCTGTGAGCCA | -1 | 1033 | 1046 | 33 | 46 | -5,63 | 0,01 | 11,21 | >0.1 | 5,57 | 8,1 | 0 | 1,86 | 0 | 1,31 | 0 |
| TFAP2A(VAR.3) | TCACTTTATGGCC | -1 | 757 | 770 | -243 | -230 | -7,21 | 0,01 | 11,55 | >0.1 | 5,57 | 5,77 | 0 | 1,32 | 0 | 6,1 | 0 |
| TFAP2A(VAR.3) | TGTCGCCCAGGCT | -1 | 427 | 440 | -573 | -560 | -1,66 | 0,01 | 12,69 | >0.1 | 2,71 | 4,82 | 0 | 1,01 | 0 | 5,8 | 0 |
| TFAP2A(VAR.3) | CGCCCAGGCTGGA | -1 | 424 | 437 | -576 | -563 | -4,07 | 0,01 | 10,13 | >0.1 | 2,7 | 4,82 | 0 | 1,02 | 0 | 5,67 | 0 |
| TFAP2A(VAR.3) | GCCCAGGCTGGAG | -1 | 423 | 436 | -577 | -564 | -9,84 | 0,01 | 4,36 | >0.1 | 2,7 | 4,82 | 0 | 1,02 | 0 | 5,67 | 0 |
| TFAP2A(VAR.3) | CGCGATCTCGGCT | -1 | 402 | 415 | -598 | -585 | -7,89 | 0,01 | 6,32 | >0.1 | 2,63 | 4,78 | 0 | 1,04 | 0 | 5,76 | 0 |
| TFAP2A(VAR.3) | TGCCTCCCAGGTT | -1 | 377 | 390 | -623 | -610 | -1,38 | 0,01 | 13,52 | >0.1 | 2,57 | 4,73 | 0 | 1,06 | 0 | 6,53 | 0 |
| TFAP2A(VAR.3) | TGCCCAACTCGGC | -1 | 221 | 234 | -779 | -766 | -8,06 | 0,01 | 6,41 | >0.1 | 2,94 | 4,5 | 0 | 1,5 | 0 | 5,52 | 0 |
| TFAP2A(VAR.3) | GCCCAACTCGGCC | -1 | 220 | 233 | -780 | -767 | -1,69 | 0,01 | 12,79 | >0.1 | 2,95 | 4,5 | 0 | 1,51 | 0 | 5,52 | 0 |
| TFAP2A(VAR.3) | AGCCACTGTGCCC | -1 | 180 | 193 | -820 | -807 | -7,75 | 0,01 | 5,46 | >0.1 | 3,12 | 4,45 | 0 | 1,79 | 0 | 3,86 | 0 |
| ERF | GGAGGCACTC | 1 | 20 | 30 | -980 | -970 | -3,47 | 0,01 | 8,28 | >0.1 | 5,61 | 4,27 | 0 | 1,87 | 0 | 0 | 0 |
| ERF | ATAGGAAATG | 1 | 61 | 71 | -939 | -929 | 9,2 | 0,0001 | 19,37 | >0.1 | 3,98 | 4,31 | 0 | 1,88 | 0 | 0 | 0 |
| ERF | ATCAGAAATG | 1 | 167 | 177 | -833 | -823 | 2,22 | 0,01 | 14,14 | >0.1 | 3,19 | 4,43 | 0 | 1,9 | 0 | 2,4 | 0 |
| ERF | ACCTGTAGTC | 1 | 330 | 340 | -670 | -660 | -1,13 | 0,01 | 12,71 | >0.1 | 2,59 | 4,65 | 0 | 1,14 | 0 | 5,46 | 0 |
| ERF | CTCGGGAGGC | 1 | 347 | 357 | -653 | -643 | -1,49 | 0,01 | 13,37 | >0.1 | 2,55 | 4,68 | 0 | 1,11 | 0 | 6,52 | 0 |
| ERF | ATCAGAAATA | 1 | 482 | 492 | -518 | -508 | 1,16 | 0,01 | 15,76 | >0.1 | 2,89 | 4,93 | 0 | 0,99 | 0 | 5,8 | 0 |
| ERF | ACCGGTTTTG | 1 | 743 | 753 | -257 | -247 | -2,19 | 0,01 | 16,29 | >0.1 | 5,61 | 5,69 | 0 | 1,26 | 0 | 5,92 | 0 |
| ERF | ACAGGAGAGG | 1 | 769 | 779 | -231 | -221 | -0,93 | 0,01 | 17,54 | >0.1 | 5,61 | 5,81 | 0 | 1,35 | 0 | 5,69 | 0 |
| ERF | AGAGGTAAGG | 1 | 774 | 784 | -226 | -216 | -2,61 | 0,01 | 16,31 | >0.1 | 5,61 | 5,83 | 0 | 1,37 | 0 | 6,1 | 0 |
| ERF | AGTGGATGTG | 1 | 1019 | 1029 | 19 | 29 | -0,29 | 0,01 | 17,23 | >0.1 | 5,61 | 8,74 | 0 | 1,86 | 0 | 1,31 | 0 |
| ERF | GAAGCAAGTG | -1 | 1179 | 1189 | 179 | 189 | 0,63 | 0,01 | 19,07 | >0.1 | 5,61 | 6,1 | 0 | 1,59 | 1,79 | 3,34 | 0 |
| ERF | CCAGGAAGAG | -1 | 1057 | 1067 | 57 | 67 | -1,02 | 0,01 | 16,14 | >0.1 | 5,61 | 7,42 | 0 | 1,86 | 0 | 2,27 | 0 |
| ERF | GCAGGAAAGA | -1 | 932 | 942 | -68 | -58 | 5,38 | 0,001 | 20,89 | >0.1 | 5,61 | 7,09 | 0 | 1,88 | 0 | 0,93 | 0 |
| ERF | ACAGTAAACA | -1 | 897 | 907 | -103 | -93 | -1,1 | 0,01 | 15,94 | >0.1 | 5,61 | 6,68 | 0 | 1,89 | 0 | 2,86 | 0 |
| ERF | ACCGGTACTT | -1 | 739 | 749 | -261 | -251 | 4,95 | 0,001 | 23,4 | >0.1 | 5,61 | 5,68 | 0 | 1,25 | 0 | 5,92 | 0 |
| ERF | ACGGAAAATG | -1 | 693 | 703 | -307 | -297 | 0,25 | 0,01 | 14,54 | >0.1 | 4,58 | 5,5 | 0 | 1,14 | 0 | 3,07 | 0 |
| ERF | ATAGGTTATC | -1 | 595 | 605 | -405 | -395 | -1,82 | 0,01 | 7,83 | >0.1 | 3,44 | 5,19 | 0 | 1,01 | 0 | 0 | 0 |
| ERF | ACTGCAAGCT | -1 | 391 | 401 | -609 | -599 | -3,25 | 0,01 | 11,17 | >0.1 | 2,6 | 4,75 | 0 | 1,05 | 0 | 6,02 | 0 |
| ERG | GAAGGAGGCA | 1 | 17 | 27 | -983 | -973 | -3,23 | 0,01 | 8,51 | >0.1 | 5,61 | 4,27 | 0 | 1,87 | 0 | 0 | 0 |
| ERG | ATAGGAAATG | 1 | 61 | 71 | -939 | -929 | 6,17 | 0,001 | 16,34 | >0.1 | 3,98 | 4,31 | 0 | 1,88 | 0 | 0 | 0 |
| ERG | TCAGGAGAGT | 1 | 98 | 108 | -902 | -892 | -3,65 | 0,01 | 6,97 | >0.1 | 3,63 | 4,35 | 0 | 1,88 | 0 | 0,77 | 0 |
| ERG | CATGGAAATT | 1 | 150 | 160 | -850 | -840 | -2,94 | 0,01 | 8,29 | >0.1 | 3,28 | 4,41 | 0 | 1,9 | 0 | 1,65 | 0 |
| ERG | ATCAGAAATG | 1 | 167 | 177 | -833 | -823 | -2,88 | 0,01 | 9,04 | >0.1 | 3,19 | 4,43 | 0 | 1,9 | 0 | 2,4 | 0 |
| ERG | TCAGGAGATA | 1 | 244 | 254 | -756 | -746 | -0,12 | 0,01 | 13,08 | >0.1 | 2,86 | 4,52 | 0 | 1,39 | 0 | 4,42 | 0 |
| ERG | AAGGGAAAAT | 1 | 530 | 540 | -470 | -460 | -2,07 | 0,01 | 10,73 | >0.1 | 3,09 | 5,03 | 0 | 0,98 | 0 | 3,7 | 0 |
| ERG | ACCGGTTTTG | 1 | 743 | 753 | -257 | -247 | -1,76 | 0,01 | 16,72 | >0.1 | 5,61 | 5,69 | 0 | 1,26 | 0 | 5,92 | 0 |
| ERG | ACAGGAGAGG | 1 | 769 | 779 | -231 | -221 | 0,22 | 0,01 | 18,69 | >0.1 | 5,61 | 5,81 | 0 | 1,35 | 0 | 5,69 | 0 |
| ERG | AGAGGTAAGG | 1 | 774 | 784 | -226 | -216 | -3,8 | 0,01 | 15,12 | >0.1 | 5,61 | 5,83 | 0 | 1,37 | 0 | 6,1 | 0 |
| ERG | CCAGGATGAA | 1 | 866 | 876 | -134 | -124 | 1,34 | 0,01 | 18,01 | >0.1 | 5,61 | 6,41 | 0 | 1,9 | 0 | 2,75 | 0 |
| ERG | TGAGGAGGTT | 1 | 988 | 998 | -12 | -2 | -3,11 | 0,01 | 14,64 | >0.1 | 5,61 | 8,66 | 0 | 1,87 | 0 | 1,61 | 0 |
| ERG | TAGGGAAAGT | 1 | 1004 | 1014 | 4 | 14 | -2,5 | 0,01 | 15,51 | >0.1 | 5,61 | 9,12 | 0 | 1,87 | 0 | 1,42 | 0 |
| ERG | AGTGGATGTG | 1 | 1019 | 1029 | 19 | 29 | 1,01 | 0,01 | 18,53 | >0.1 | 5,61 | 8,74 | 0 | 1,86 | 0 | 1,31 | 0 |
| ERG | TGAGGAACAG | 1 | 1111 | 1121 | 111 | 121 | -2,47 | 0,01 | 17,39 | >0.1 | 5,61 | 6,64 | 0 | 1,85 | 3,22 | 2,54 | 0 |
| ERG | CCAGGAAGAG | -1 | 1057 | 1067 | 57 | 67 | 4,44 | 0,01 | 21,6 | >0.1 | 5,61 | 7,42 | 0 | 1,86 | 0 | 2,27 | 0 |
| ERG | GCAGGAAAGA | -1 | 932 | 942 | -68 | -58 | 5,87 | 0,001 | 21,38 | >0.1 | 5,61 | 7,09 | 0 | 1,88 | 0 | 0,93 | 0 |
| ERG | ACAGTAAACA | -1 | 897 | 907 | -103 | -93 | 2,18 | 0,01 | 19,22 | >0.1 | 5,61 | 6,68 | 0 | 1,89 | 0 | 2,86 | 0 |

| | | | | | | | | | | | | | | | | | |
|---|---|---|---|---|---|---|---|---|---|---|---|---|---|---|---|---|---|
| ERG | AGAGGACAGA | -1 | 856 | 866 | -144 | -134 | -0,48 | 0,01 | 16,32 | >0.1 | 5,61 | 6,34 | 0 | 1,9 | 0 | 2,95 | 0 |
| ERG | ACCGGTACTT | -1 | 739 | 749 | -261 | -251 | 0,86 | 0,01 | 19,31 | >0.1 | 5,61 | 5,68 | 0 | 1,25 | 0 | 5,92 | 0 |
| ERG | CACGGAAAAT | -1 | 694 | 704 | -306 | -296 | 2,02 | 0,01 | 17,34 | >0.1 | 5,61 | 5,5 | 0 | 1,14 | 0 | 3,07 | 0 |
| ERG | CCCAGAAACT | -1 | 555 | 565 | -445 | -435 | -3,02 | 0,01 | 6,27 | >0.1 | 3,21 | 5,09 | 0 | 0,99 | 0 | 0 | 0 |
| ERG | TCAGGACGGT | -1 | 257 | 267 | -743 | -733 | 1,27 | 0,01 | 14,6 | >0.1 | 2,82 | 4,54 | 0 | 1,34 | 0 | 4,63 | 0 |
| MITF | GTCAGCCACAGAACCAGT | 1 | 36 | 54 | -964 | -946 | -3,32 | 0,01 | 7,13 | >0.1 | 4,28 | 4,29 | 0 | 1,87 | 0 | 0 | 0 |
| MITF | GCAGATCAGGAGAGTTGA | 1 | 93 | 111 | -907 | -889 | 2,98 | 0,01 | 13,62 | >0.1 | 3,64 | 4,35 | 0 | 1,88 | 0 | 0,77 | 0 |
| MITF | GTTGGGCAGATCACAAGG | 1 | 226 | 244 | -774 | -756 | -0,88 | 0,01 | 13,03 | >0.1 | 2,92 | 4,51 | 0 | 1,48 | 0 | 5,01 | 0 |
| MITF | GCAGATCACAAGGTCAGG | 1 | 231 | 249 | -769 | -751 | 5,12 | 0,001 | 18,73 | >0.1 | 2,9 | 4,52 | 0 | 1,45 | 0 | 4,74 | 0 |
| MITF | CAAGGTCAGGAGATAGAG | 1 | 239 | 257 | -761 | -743 | 4,03 | 0,01 | 17,17 | >0.1 | 2,87 | 4,53 | 0 | 1,42 | 0 | 4,33 | 0 |
| MITF | GGTGGGCACCTGTAGTCC | 1 | 323 | 341 | -677 | -659 | -0,67 | 0,01 | 13,31 | >0.1 | 2,6 | 4,65 | 0 | 1,16 | 0 | 5,56 | 0 |
| MITF | CAAAGTCATGTATTTGGA | 1 | 513 | 531 | -487 | -469 | -0,87 | 0,01 | 11,11 | >0.1 | 3,03 | 5,01 | 0 | 0,99 | 0 | 2,95 | 0 |
| MITF | TCTGGGAATATGATCTTG | 1 | 559 | 577 | -441 | -423 | 0,83 | 0,01 | 10,2 | >0.1 | 3,25 | 5,12 | 0 | 1 | 0 | 0 | 0 |
| MITF | TAAAGTGACAGGAGAGGT | 1 | 762 | 780 | -238 | -220 | -3,16 | 0,01 | 15,44 | >0.1 | 5,51 | 5,82 | 0 | 1,36 | 0 | 5,92 | 0 |
| MITF | GAAAATCTCATGAGGAGG | 1 | 978 | 996 | -22 | -4 | 3,05 | 0,01 | 20,57 | >0.1 | 5,51 | 8,61 | 0 | 1,87 | 0 | 1,53 | 0 |
| MITF | TCAGTGGATGTGATCTTG | 1 | 1017 | 1035 | 17 | 35 | -0,27 | 0,01 | 17,19 | >0.1 | 5,51 | 8,79 | 0 | 1,86 | 0 | 1,31 | 0 |
| MITF | TTGGCTCACAGGGGACGA | 1 | 1032 | 1050 | 32 | 50 | -0,14 | 0,01 | 16,71 | >0.1 | 5,51 | 8,15 | 0 | 1,86 | 0 | 1,34 | 0 |
| MITF | AAAGTTCACTTGCTTCTT | 1 | 1173 | 1191 | 173 | 191 | -1,48 | 0,01 | 16,86 | >0.1 | 5,51 | 6,13 | 0 | 1,64 | 1,79 | 3,26 | 0 |
| MITF | AAGAAGCAAGTGAACTTT | -1 | 1173 | 1191 | 173 | 191 | -1,48 | 0,01 | 16,86 | >0.1 | 5,51 | 6,13 | 0 | 1,64 | 1,79 | 3,26 | 0 |
| MITF | TCGTCCCCTGTGAGCCAA | -1 | 1032 | 1050 | 32 | 50 | -0,14 | 0,01 | 16,71 | >0.1 | 5,51 | 8,15 | 0 | 1,86 | 0 | 1,34 | 0 |
| MITF | CAAGATCACATCCACTGA | -1 | 1017 | 1035 | 17 | 35 | -0,27 | 0,01 | 17,19 | >0.1 | 5,51 | 8,79 | 0 | 1,86 | 0 | 1,31 | 0 |
| MITF | CCTCCTCATGAGATTTTC | -1 | 978 | 996 | -22 | -4 | 3,05 | 0,01 | 20,57 | >0.1 | 5,51 | 8,61 | 0 | 1,87 | 0 | 1,53 | 0 |
| MITF | ACCTCTCCTGTCACTTTA | -1 | 762 | 780 | -238 | -220 | -3,16 | 0,01 | 15,44 | >0.1 | 5,51 | 5,82 | 0 | 1,36 | 0 | 5,92 | 0 |
| MITF | CAAGATCATATTCCCAGA | -1 | 559 | 577 | -441 | -423 | 0,8 | 0,01 | 10,17 | >0.1 | 3,25 | 5,12 | 0 | 1 | 0 | 0 | 0 |
| MITF | TCCAAATACATGACTTTG | -1 | 513 | 531 | -487 | -469 | -0,86 | 0,01 | 11,12 | >0.1 | 3,03 | 5,01 | 0 | 0,99 | 0 | 2,95 | 0 |
| MITF | GGACTACAGGTGCCCACC | -1 | 323 | 341 | -677 | -659 | -0,67 | 0,01 | 13,31 | >0.1 | 2,6 | 4,65 | 0 | 1,16 | 0 | 5,56 | 0 |
| MITF | CTCTATCTCCTGACCTTG | -1 | 239 | 257 | -761 | -743 | 4,03 | 0,01 | 17,17 | >0.1 | 2,87 | 4,53 | 0 | 1,42 | 0 | 4,33 | 0 |
| MITF | CCTGACCTTGTGATCTGC | -1 | 231 | 249 | -769 | -751 | 5,13 | 0,001 | 18,74 | >0.1 | 2,9 | 4,52 | 0 | 1,45 | 0 | 4,74 | 0 |
| MITF | CCTTGTGATCTGCCCAAC | -1 | 226 | 244 | -774 | -756 | -0,88 | 0,01 | 13,03 | >0.1 | 2,92 | 4,51 | 0 | 1,48 | 0 | 5,01 | 0 |
| MITF | TCAACTCTCCTGATCTGC | -1 | 93 | 111 | -907 | -889 | 2,97 | 0,01 | 13,61 | >0.1 | 3,64 | 4,35 | 0 | 1,88 | 0 | 0,77 | 0 |
| MITF | ACTGGTTCTGTGGCTGAC | -1 | 36 | 54 | -964 | -946 | -3,32 | 0,01 | 7,13 | >0.1 | 4,28 | 4,29 | 0 | 1,87 | 0 | 0 | 0 |
| RARA::RXRA | AAGTGAAGGAGAAGGAG | 1 | 7 | 24 | -993 | -976 | -1,27 | 0,01 | 12,18 | >0.1 | 5,52 | 4,27 | 0 | 1,87 | 1,79 | 0 | 0 |
| RARA::RXRA | AGTGAAGGAGAAGGAGG | 1 | 8 | 25 | -992 | -975 | -0,6 | 0,01 | 12,85 | >0.1 | 5,52 | 4,27 | 0 | 1,87 | 1,79 | 0 | 0 |
| RARA::RXRA | GGCAGATCACAAGGTCA | 1 | 230 | 247 | -770 | -753 | 2,42 | 0,01 | 16,19 | >0.1 | 2,9 | 4,52 | 0 | 1,46 | 0 | 4,9 | 0 |
| RARA::RXRA | AGGTCAGGAGATAGAGA | 1 | 241 | 258 | -759 | -742 | 3,79 | 0,01 | 17,01 | >0.1 | 2,86 | 4,53 | 0 | 1,41 | 0 | 4,42 | 0 |
| RARA::RXRA | GAGGCTGAGGCAGGAGA | 1 | 352 | 369 | -648 | -631 | -1,67 | 0,01 | 12,69 | >0.1 | 2,53 | 4,7 | 0 | 1,1 | 0 | 6,03 | 0 |
| RARA::RXRA | GAGGCAGGAGAATGGCG | 1 | 358 | 375 | -642 | -625 | -0,66 | 0,01 | 14,15 | >0.1 | 2,52 | 4,71 | 0 | 1,09 | 0 | 6,49 | 0 |
| RARA::RXRA | GGGAGGCAGAGCTTGCA | 1 | 382 | 399 | -618 | -601 | 0,08 | 0,01 | 14,6 | >0.1 | 2,58 | 4,75 | 0 | 1,06 | 0 | 6,13 | 0 |
| RARA::RXRA | AGGGCAAAGTCATGTAT | 1 | 509 | 526 | -491 | -474 | 1,44 | 0,01 | 13,39 | >0.1 | 3,01 | 5 | 0 | 0,99 | 0 | 2,95 | 0 |
| RARA::RXRA | AAGTGACAGGAGAGGTA | 1 | 764 | 781 | -236 | -219 | -0,83 | 0,01 | 17,79 | >0.1 | 5,52 | 5,82 | 0 | 1,36 | 0 | 5,92 | 0 |
| RARA::RXRA | AGTGACAGGAGAGGTAA | 1 | 765 | 782 | -235 | -218 | 0,71 | 0,01 | 19,34 | >0.1 | 5,52 | 5,83 | 0 | 1,37 | 0 | 5,92 | 0 |
| RARA::RXRA | AGGAGAGGTAAGGTTCT | 1 | 771 | 788 | -229 | -212 | 2,6 | 0,01 | 21,47 | >0.1 | 5,52 | 5,85 | 0 | 1,39 | 0 | 6,1 | 0 |
| RARA::RXRA | AGGTTCTCTAGGATTAA | 1 | 781 | 798 | -219 | -202 | -1,17 | 0,01 | 17,61 | >0.1 | 5,52 | 5,91 | 0 | 1,44 | 0 | 5,92 | 0 |
| RARA::RXRA | AGTTCAAAGGCTGATAA | 1 | 957 | 974 | -43 | -26 | 1,79 | 0,01 | 18,65 | >0.1 | 5,52 | 7,73 | 0 | 1,88 | 0 | 1,74 | 0 |
| RARA::RXRA | ATCTCATGAGGAGGTTT | 1 | 982 | 999 | -18 | -1 | 0,87 | 0,01 | 18,52 | >0.1 | 5,52 | 8,69 | 0 | 1,87 | 0 | 1,57 | 0 |
| RARA::RXRA | CATTGAGGAACAGGCCA | 1 | 1108 | 1125 | 108 | 125 | 1,65 | 0,01 | 21,45 | >0.1 | 5,52 | 6,67 | 0 | 1,85 | 3,22 | 2,54 | 0 |
| RARA::RXRA | TGTTCTATCAAAGTTCA | 1 | 1164 | 1181 | 164 | 181 | 0,88 | 0,01 | 18,97 | >0.1 | 5,52 | 6,19 | 0 | 1,72 | 1,79 | 2,86 | 0 |
| RARA::RXRA | AGATCACATCCACTGAA | -1 | 1016 | 1033 | 16 | 33 | 0,61 | 0,01 | 18,11 | >0.1 | 5,52 | 8,81 | 0 | 1,86 | 0 | 1,31 | 0 |
| RARA::RXRA | GGGTTGGGCGCTGAGCA | -1 | 939 | 956 | -61 | -44 | -0,96 | 0,01 | 14,59 | >0.1 | 5,52 | 7,32 | 0 | 1,88 | 0 | 0,83 | 0 |
| RARA::RXRA | GGGCGCTGAGCAGGAAA | -1 | 934 | 951 | -66 | -49 | -1,14 | 0,01 | 14,39 | >0.1 | 5,52 | 7,24 | 0 | 1,88 | 0 | 0,9 | 0 |
| RARA::RXRA | AGTTCATCCTGGAGAGG | -1 | 861 | 878 | -139 | -122 | 3,13 | 0,01 | 19,73 | >0.1 | 5,52 | 6,43 | 0 | 1,9 | 0 | 2,75 | 0 |
| RARA::RXRA | AGTGACCAAAGAGGTCT | -1 | 664 | 681 | -336 | -319 | -0,74 | 0,01 | 13,72 | >0.1 | 4,11 | 5,42 | 0 | 1,1 | 0 | 3,83 | 0 |
| RARA::RXRA | AGGTCTAGAACTAGGGA | -1 | 653 | 670 | -347 | -330 | -1,65 | 0,01 | 12,49 | >0.1 | 3,97 | 5,39 | 0 | 1,09 | 0 | 3,7 | 0 |
| RARA::RXRA | ATTTCATTAATAGGTTA | -1 | 597 | 614 | -403 | -386 | 0,56 | 0,01 | 11,64 | >0.1 | 3,48 | 5,22 | 0 | 1,02 | 0 | 1,36 | 0 |
| RARA::RXRA | AGGTTATCTCACATTTA | -1 | 586 | 603 | -414 | -397 | 1,31 | 0,01 | 10,92 | >0.1 | 3,4 | 5,19 | 0 | 1,02 | 0 | 0 | 0 |
| RARA::RXRA | GGAGCACAGTGGCGCGA | -1 | 410 | 427 | -590 | -573 | 1,38 | 0,01 | 14,96 | >0.1 | 2,66 | 4,8 | 0 | 1,03 | 0 | 5,09 | 0 |
| RARA::RXRA | GGCTCACTGCAAGCTCT | -1 | 389 | 406 | -611 | -594 | 2,66 | 0,01 | 17,1 | >0.1 | 2,6 | 4,76 | 0 | 1,05 | 0 | 6,02 | 0 |
| MYOG | GGCAGATCAGG | 1 | 92 | 103 | -908 | -897 | -10,3 | 0,01 | 0,43 | >0.1 | 3,67 | 4,34 | 0 | 1,88 | 0 | 0,83 | 0 |
| MYOG | AATGGCTGGGC | 1 | 173 | 184 | -827 | -816 | -18,06 | 0,01 | -5,38 | >0.1 | 3,16 | 4,43 | 0 | 1,85 | 0 | 3,24 | 0 |

| | | | | | | | | | | | | | | | | | |
|---|---|---|---|---|---|---|---|---|---|---|---|---|---|---|---|---|---|
| MYOG | GGCAGATCACA | 1 | 230 | 241 | -770 | -759 | -12,14 | 0,01 | 1,75 | >0.1 | 2,91 | 4,51 | 0 | 1,45 | 0 | 5,01 | 0 |
| MYOG | GGCACCTGTAG | 1 | 327 | 338 | -673 | -662 | -2,76 | 0,01 | 11,2 | >0.1 | 2,6 | 4,65 | 0 | 1,14 | 0 | 5,56 | 0 |
| MYOG | CCCAGCTACTC | 1 | 339 | 350 | -661 | -650 | -7,56 | 0,01 | 7,32 | >0.1 | 2,57 | 4,67 | 0 | 1,12 | 0 | 6,52 | 0 |
| MYOG | GGAGGCTGAGG | 1 | 351 | 362 | -649 | -638 | -18,45 | 0,01 | -3,73 | >0.1 | 2,54 | 4,69 | 0 | 1,1 | 0 | 6,4 | 0 |
| MYOG | GGCAGGAGAAT | 1 | 360 | 371 | -640 | -629 | -6,59 | 0,01 | 8,08 | >0.1 | 2,52 | 4,7 | 0 | 1,09 | 0 | 6,36 | 0 |
| MYOG | AACAGCTTTCT | 1 | 711 | 722 | -289 | -278 | -5,58 | 0,01 | 11,75 | >0.1 | 5,6 | 5,57 | 0 | 1,18 | 0 | 4,99 | 0 |
| MYOG | GACAGGAGAGG | 1 | 768 | 779 | -232 | -221 | -4,18 | 0,01 | 14,5 | >0.1 | 5,6 | 5,81 | 0 | 1,35 | 0 | 5,92 | 0 |
| MYOG | AGCAGATTGTT | 1 | 890 | 901 | -110 | -99 | -12,23 | 0,01 | 4,83 | >0.1 | 5,6 | 6,62 | 0 | 1,89 | 0 | 2,95 | 0 |
| MYOG | CACAGGGGACG | 1 | 1038 | 1049 | 38 | 49 | -5,73 | 0,01 | 10,98 | >0.1 | 5,6 | 7,92 | 0 | 1,86 | 0 | 1,34 | 0 |
| MYOG | AACTGCTGCTC | 1 | 1090 | 1101 | 90 | 101 | -1,48 | 0,001 | 16,91 | >0.1 | 5,6 | 6,88 | 0 | 1,85 | 1,79 | 2,27 | 0 |
| MYOG | AGCAAGTGAAC | -1 | 1176 | 1187 | 176 | 187 | -6,55 | 0,01 | 11,85 | >0.1 | 5,6 | 6,11 | 0 | 1,62 | 1,79 | 3,28 | 0 |
| MYOG | AACAGGTCTTC | -1 | 1157 | 1168 | 157 | 168 | -6,41 | 0,01 | 10,15 | >0.1 | 4,39 | 6,24 | 0 | 1,79 | 1,79 | 2,34 | 0 |
| MYOG | AGCAGCAGTTA | -1 | 1089 | 1100 | 89 | 100 | -4,85 | 0,01 | 13,5 | >0.1 | 5,6 | 6,89 | 0 | 1,85 | 1,79 | 2,21 | 0 |
| MYOG | AGCAGTTACAG | -1 | 1086 | 1097 | 86 | 97 | -18,19 | 0,01 | 0,2 | >0.1 | 5,6 | 6,93 | 0 | 1,85 | 1,79 | 2,21 | 0 |
| MYOG | GGGCGCTGAGC | -1 | 940 | 951 | -60 | -49 | -18,61 | 0,01 | -3,07 | >0.1 | 5,6 | 7,23 | 0 | 1,88 | 0 | 0,83 | 0 |
| MYOG | GACAGATATGT | -1 | 851 | 862 | -149 | -138 | -12,9 | 0,01 | 3,56 | >0.1 | 5,6 | 6,31 | 0 | 1,88 | 0 | 2,67 | 0 |
| MYOG | AACAGATTTTA | -1 | 840 | 851 | -160 | -149 | -13,62 | 0,01 | 3,18 | >0.1 | 5,6 | 6,24 | 0 | 1,78 | 0 | 3,19 | 0 |
| MYOG | GAAAGCTGTTA | -1 | 710 | 721 | -290 | -279 | -4,57 | 0,01 | 12,58 | >0.1 | 5,6 | 5,56 | 0 | 1,18 | 0 | 4,81 | 0 |
| MYOG | CACAGTGGCGC | -1 | 412 | 423 | -588 | -577 | -18,22 | 0,01 | -3,83 | >0.1 | 2,66 | 4,79 | 0 | 1,03 | 0 | 5,91 | 0 |
| MYOG | CCCAGGTTCAC | -1 | 374 | 385 | -626 | -615 | -9,91 | 0,01 | 4,97 | >0.1 | 2,56 | 4,72 | 0 | 1,07 | 0 | 6,53 | 0 |
| MYOG | AGTAGCTGGGA | -1 | 338 | 349 | -662 | -651 | -2,86 | 0,01 | 11,42 | >0.1 | 2,57 | 4,67 | 0 | 1,12 | 0 | 5,92 | 0 |
| MYOG | TACAGGTGCCC | -1 | 326 | 337 | -674 | -663 | -2,54 | 0,01 | 11,31 | >0.1 | 2,6 | 4,65 | 0 | 1,15 | 0 | 5,46 | 0 |
| MYOG | CCCAGCTAAGT | -1 | 306 | 317 | -694 | -683 | -10,29 | 0,01 | 3,26 | >0.1 | 2,66 | 4,62 | 0 | 1,19 | 0 | 5,09 | 0 |
| MYOG | CACAGGTCAAC | -1 | 106 | 117 | -894 | -883 | -6,87 | 0,01 | 3,68 | >0.1 | 3,56 | 4,36 | 0 | 1,89 | 0 | 0,75 | 0 |
| RARA::RXRG | TAGGTAAGTGAAGGAGAA | 1 | 2 | 20 | -998 | -980 | 0,91 | 0,01 | 14,34 | >0.1 | 5,51 | 4,26 | 0 | 1,86 | 1,79 | 0 | 0 |
| RARA::RXRG | AAGTGAAGGAGAAGGAGG | 1 | 7 | 25 | -993 | -975 | 0,02 | 0,01 | 13,46 | >0.1 | 5,51 | 4,27 | 0 | 1,87 | 1,79 | 0 | 0 |
| RARA::RXRG | GAAGTCAGCCACAGAACC | 1 | 33 | 51 | -967 | -949 | -0,16 | 0,01 | 10,34 | >0.1 | 4,34 | 4,29 | 0 | 1,87 | 0 | 0 | 0 |
| RARA::RXRG | AGTGTGAGAAATAGGAAA | 1 | 51 | 69 | -949 | -931 | 1,02 | 0,01 | 11,26 | >0.1 | 4,06 | 4,31 | 0 | 1,87 | 0 | 0 | 0 |
| RARA::RXRG | GAGTTGACCTGTGGAGTG | 1 | 104 | 122 | -896 | -878 | 1,39 | 0,01 | 11,94 | >0.1 | 3,55 | 4,36 | 0 | 1,89 | 0 | 0,75 | 0 |
| RARA::RXRG | AATTTTAAAAAATGGCCA | 1 | 134 | 152 | -866 | -848 | 0,62 | 0,01 | 11,91 | >0.1 | 3,35 | 4,4 | 0 | 1,9 | 0 | 1,65 | 0 |
| RARA::RXRG | GAGGCCGAGTTGGCAGA | 1 | 218 | 236 | -782 | -764 | 0,12 | 0,01 | 14,61 | >0.1 | 2,95 | 4,5 | 0 | 1,52 | 0 | 5,52 | 0 |
| RARA::RXRG | GGGCAGATCACAAGGTCA | 1 | 229 | 247 | -771 | -753 | 1,41 | 0,01 | 15,19 | >0.1 | 2,9 | 4,52 | 0 | 1,46 | 0 | 4,9 | 0 |
| RARA::RXRG | AAGGTCAGGAGATAGAGA | 1 | 240 | 258 | -760 | -742 | 3,43 | 0,01 | 16,66 | >0.1 | 2,86 | 4,53 | 0 | 1,41 | 0 | 4,42 | 0 |
| RARA::RXRG | GTGGGCACCTGTAGTCCC | 1 | 324 | 342 | -676 | -658 | 2,15 | 0,01 | 16,12 | >0.1 | 2,6 | 4,65 | 0 | 1,15 | 0 | 5,56 | 0 |
| RARA::RXRG | TGAGGCAGGAGAATGGCG | 1 | 357 | 375 | -643 | -625 | 1,54 | 0,01 | 16,35 | >0.1 | 2,52 | 4,71 | 0 | 1,1 | 0 | 6,49 | 0 |
| RARA::RXRG | GCGTGAACCTGGGAGGCA | 1 | 372 | 390 | -628 | -610 | 0,09 | 0,01 | 14,85 | >0.1 | 2,56 | 4,73 | 0 | 1,07 | 0 | 6,4 | 0 |
| RARA::RXRG | AAGGGCAAAGTCATGTAT | 1 | 508 | 526 | -492 | -474 | 2,26 | 0,01 | 14,21 | >0.1 | 3,01 | 5 | 0 | 0,99 | 0 | 2,95 | 0 |
| RARA::RXRG | AAAGTGACAGGAGAGGTA | 1 | 763 | 781 | -237 | -219 | 0,18 | 0,01 | 18,79 | >0.1 | 5,51 | 5,82 | 0 | 1,36 | 0 | 5,92 | 0 |
| RARA::RXRG | CAGGAGAGGTAAGGTTCT | 1 | 770 | 788 | -230 | -212 | 1 | 0,01 | 19,86 | >0.1 | 5,51 | 5,86 | 0 | 1,39 | 0 | 6,1 | 0 |
| RARA::RXRG | GCGCCCAACCCAAGTTCA | 1 | 945 | 963 | -55 | -37 | 1,14 | 0,01 | 17,73 | >0.1 | 5,51 | 7,46 | 0 | 1,88 | 0 | 1,74 | 0 |
| RARA::RXRG | AAGTTCAAAGGCTGATAA | 1 | 956 | 974 | -44 | -26 | 1,27 | 0,01 | 18,12 | >0.1 | 5,51 | 7,73 | 0 | 1,88 | 0 | 1,74 | 0 |
| RARA::RXRG | AATCTCATGAGGAGGTTT | 1 | 981 | 999 | -19 | -1 | 0,35 | 0,01 | 17,99 | >0.1 | 5,51 | 8,69 | 0 | 1,87 | 0 | 1,57 | 0 |
| RARA::RXRG | AGGTTTTAGTCTAGGGAA | 1 | 993 | 1011 | -7 | 11 | 1,41 | 0,01 | 19,41 | >0.1 | 5,51 | 9,12 | 0 | 1,87 | 0 | 1,5 | 0 |
| RARA::RXRG | GGTTTTAGTCTAGGGAAA | 1 | 994 | 1012 | -6 | 12 | 0,65 | 0,01 | 18,61 | >0.1 | 5,51 | 9,12 | 0 | 1,87 | 0 | 1,46 | 0 |
| RARA::RXRG | AAAGTCATTCAGTGGATG | 1 | 1009 | 1027 | 9 | 27 | 1,92 | 0,01 | 19,74 | >0.1 | 5,51 | 9,03 | 0 | 1,86 | 0 | 1,42 | 0 |
| RARA::RXRG | CTGTTCTATCAAAGTTCA | 1 | 1163 | 1181 | 163 | 181 | -0,14 | 0,01 | 17,96 | >0.1 | 5,51 | 6,2 | 0 | 1,73 | 1,79 | 2,86 | 0 |
| RARA::RXRG | AAGGCTGAGAAGGAGCCA | -1 | 1064 | 1082 | 64 | 82 | 1,66 | 0,01 | 20,31 | >0.1 | 5,51 | 7,28 | 0 | 1,85 | 1,79 | 2,21 | 0 |
| RARA::RXRG | TGGGTTGGGCGCTGAGCA | -1 | 939 | 957 | -61 | -43 | 0,21 | 0,01 | 16,74 | >0.1 | 5,51 | 7,34 | 0 | 1,88 | 0 | 1,8 | 0 |
| RARA::RXRG | AAGTTCATCCTGGAGAGG | -1 | 861 | 879 | -139 | -121 | 4,65 | 0,01 | 21,17 | >0.1 | 5,51 | 6,44 | 0 | 1,9 | 0 | 2,67 | 0 |
| RARA::RXRG | CCTGTCACTTTATGGCCA | -1 | 756 | 774 | -244 | -226 | 3,09 | 0,01 | 21,64 | >0.1 | 5,51 | 5,79 | 0 | 1,34 | 0 | 5,92 | 0 |
| RARA::RXRG | GAGGTCTAGAACTAGGGA | -1 | 653 | 671 | -347 | -329 | -0,32 | 0,01 | 13,83 | >0.1 | 3,97 | 5,39 | 0 | 1,09 | 0 | 3,7 | 0 |
| RARA::RXRG | AATTTCATTAATAGGTTA | -1 | 597 | 615 | -403 | -385 | 6,41 | 0,001 | 17,55 | >0.1 | 3,48 | 5,22 | 0 | 1,03 | 0 | 1,41 | 0 |
| RARA::RXRG | TAGGTTATCTCACATTTA | -1 | 586 | 604 | -414 | -396 | 1,8 | 0,01 | 11,42 | >0.1 | 3,41 | 5,19 | 0 | 1,02 | 0 | 0 | 0 |
| RARA::RXRG | CGGCTCACTGCAAGCTCT | -1 | 389 | 407 | -611 | -593 | -0,42 | 0,01 | 14,02 | >0.1 | 2,6 | 4,76 | 0 | 1,05 | 0 | 6,02 | 0 |
| GFI1B | CACTCAAAGAA | 1 | 25 | 36 | -975 | -964 | -5,16 | 0,01 | 6,58 | >0.1 | 5,6 | 4,28 | 0 | 1,87 | 0 | 0 | 0 |
| GFI1B | CAGCCACAGAA | 1 | 38 | 49 | -962 | -951 | 1,67 | 0,01 | 12,14 | >0.1 | 4,31 | 4,29 | 0 | 1,87 | 0 | 0 | 0 |
| GFI1B | GAACCAGTGTG | 1 | 46 | 57 | -954 | -943 | 4,78 | 0,01 | 15,13 | >0.1 | 4,18 | 4,3 | 0 | 1,87 | 0 | 0 | 0 |
| GFI1B | AAGGCAGATCA | 1 | 90 | 101 | -910 | -899 | -5,53 | 0,1 | 4,38 | >0.1 | 3,69 | 4,34 | 0 | 1,88 | 0 | 0 | 0 |
| GFI1B | AAAACTGATCA | 1 | 160 | 171 | -840 | -829 | -4,56 | 0,01 | 7,44 | >0.1 | 3,23 | 4,42 | 0 | 1,9 | 0 | 2,46 | 0 |

| | | | | | | | | | | | | | | | | | |
|---|---|---|---|---|---|---|---|---|---|---|---|---|---|---|---|---|---|
| GFI1B | TAATCCTAGCA | 1 | 201 | 212 | -799 | -788 | -4,9 | 0,01 | 8,82 | >0.1 | 3,03 | 4,47 | 0 | 1,62 | 0 | 4,59 | 0 |
| GFI1B | TAGTCCCAGCT | 1 | 335 | 346 | -665 | -654 | -1,56 | 0,01 | 12,84 | >0.1 | 2,58 | 4,66 | 0 | 1,13 | 0 | 6,03 | 0 |
| GFI1B | GAACCTGGGAG | 1 | 376 | 387 | -624 | -613 | -4,93 | 0,01 | 9,82 | >0.1 | 2,56 | 4,73 | 0 | 1,06 | 0 | 6,4 | 0 |
| GFI1B | GAGGCAGAGCT | 1 | 384 | 395 | -616 | -605 | -2,71 | 0,01 | 11,69 | >0.1 | 2,58 | 4,74 | 0 | 1,05 | 0 | 6,02 | 0 |
| GFI1B | CAGTCTCAAAA | 1 | 450 | 461 | -550 | -539 | -2,7 | 0,01 | 11,2 | >0.1 | 2,78 | 4,86 | 0 | 1 | 0 | 5,26 | 0 |
| GFI1B | AAAACTGATCA | 1 | 475 | 486 | -525 | -514 | -4,56 | 0,01 | 9,3 | >0.1 | 2,87 | 4,91 | 0 | 0,99 | 0 | 5,09 | 0 |
| GFI1B | AAATGTGAGAT | 1 | 587 | 598 | -413 | -402 | -2,87 | 0,01 | 6,71 | >0.1 | 3,39 | 5,18 | 0 | 1,01 | 0 | 0 | 0 |
| GFI1B | TAACCTATTAA | 1 | 597 | 608 | -403 | -392 | -4,58 | 0,01 | 6,51 | >0.1 | 3,46 | 5,2 | 0 | 1,02 | 0 | 1,41 | 0 |
| GFI1B | TAACCAAAAGT | 1 | 732 | 743 | -268 | -257 | -3,26 | 0,01 | 15,14 | >0.1 | 5,6 | 5,65 | 0 | 1,24 | 0 | 5,92 | 0 |
| GFI1B | AAATCTGTTAC | 1 | 842 | 853 | -158 | -147 | 3,41 | 0,01 | 20,24 | >0.1 | 5,6 | 6,25 | 0 | 1,8 | 0 | 3,19 | 0 |
| GFI1B | CAACCCAAGTT | 1 | 950 | 961 | -50 | -39 | 0,69 | 0,01 | 17,32 | >0.1 | 5,6 | 7,42 | 0 | 1,88 | 0 | 1,74 | 0 |
| GFI1B | AAATCTCATGA | 1 | 980 | 991 | -20 | -9 | 5,84 | 0,001 | 23,21 | >0.1 | 5,6 | 8,37 | 0 | 1,87 | 0 | 1,53 | 0 |
| GFI1B | CATTCAGTGGA | 1 | 1014 | 1025 | 14 | 25 | -3,69 | 0,01 | 14,02 | >0.1 | 5,6 | 8,86 | 0 | 1,86 | 0 | 1,39 | 0 |
| GFI1B | CAAGCTCTTCC | 1 | 1053 | 1064 | 53 | 64 | 2,9 | 0,01 | 20,07 | >0.1 | 5,6 | 7,5 | 0 | 1,86 | 0 | 2,21 | 0 |
| GFI1B | AAGGCTGAGAA | -1 | 1071 | 1082 | 71 | 82 | -3,35 | 0,01 | 15,32 | >0.1 | 5,6 | 7,15 | 0 | 1,85 | 1,79 | 2,27 | 0 |
| GFI1B | AGATCACATCC | -1 | 1022 | 1033 | 22 | 33 | -2,38 | 0,01 | 15,04 | >0.1 | 5,6 | 8,68 | 0 | 1,86 | 0 | 1,28 | 0 |
| GFI1B | CATCCACTGAA | -1 | 1016 | 1027 | 16 | 27 | -3,08 | 0,01 | 14,55 | >0.1 | 5,6 | 8,81 | 0 | 1,86 | 0 | 1,36 | 0 |
| GFI1B | AAACCTCCTCA | -1 | 988 | 999 | -12 | -1 | -3,54 | 0,01 | 14,22 | >0.1 | 5,6 | 8,69 | 0 | 1,87 | 0 | 1,61 | 0 |
| GFI1B | GAATCAAATAA | -1 | 812 | 823 | -188 | -177 | -1,07 | 0,01 | 15,57 | >0.1 | 5,6 | 6,05 | 0 | 1,57 | 0 | 3,43 | 0 |
| GFI1B | AAATCAAAACC | -1 | 746 | 757 | -254 | -243 | 6,38 | 0,001 | 24,88 | >0.1 | 5,6 | 5,71 | 0 | 1,28 | 0 | 5,92 | 0 |
| GFI1B | AAACCGGTACT | -1 | 740 | 751 | -260 | -249 | -5,05 | 0,01 | 13,4 | >0.1 | 5,6 | 5,68 | 0 | 1,26 | 0 | 5,92 | 0 |
| GFI1B | AAAGCTGTTAA | -1 | 709 | 720 | -291 | -280 | -1,44 | 0,01 | 15,7 | >0.1 | 5,6 | 5,56 | 0 | 1,18 | 0 | 4,81 | 0 |
| GFI1B | AACTCTCTACT | -1 | 548 | 559 | -452 | -441 | -3,06 | 0,01 | 6,18 | >0.1 | 3,17 | 5,08 | 0 | 0,99 | 0 | 0 | 0 |
| GFI1B | CAAGCTCTGCC | -1 | 386 | 397 | -614 | -603 | 9,34 | 0,001 | 23,85 | >0.1 | 2,59 | 4,75 | 0 | 1,05 | 0 | 6,13 | 0 |
| GFI1B | GAGCCACTGTG | -1 | 183 | 194 | -817 | -806 | -2,61 | 0,01 | 10,57 | >0.1 | 3,11 | 4,45 | 0 | 1,76 | 0 | 3,86 | 0 |
| GFI1B | AAAGCTCATTT | -1 | 66 | 77 | -934 | -923 | 1,01 | 0,01 | 11,12 | >0.1 | 3,92 | 4,32 | 0 | 1,88 | 0 | 0 | 0 |
| MYOD1 | TTGACCTGTGGAG | 1 | 107 | 120 | -893 | -880 | -9,18 | 0,01 | 1,36 | >0.1 | 3,55 | 4,36 | 0 | 1,89 | 0 | 0,75 | 0 |
| MYOD1 | CTTAGCTGGGCGT | 1 | 307 | 320 | -693 | -680 | -9,59 | 0,01 | 4,56 | >0.1 | 2,65 | 4,62 | 0 | 1,19 | 0 | 5,68 | 0 |
| MYOD1 | GGCACCTGTAGTC | 1 | 327 | 340 | -673 | -660 | 4,77 | 0,001 | 18,73 | >0.1 | 2,6 | 4,65 | 0 | 1,15 | 0 | 5,56 | 0 |
| MYOD1 | CCCAGCTACTCGG | 1 | 339 | 352 | -661 | -648 | -3,15 | 0,01 | 11,61 | >0.1 | 2,57 | 4,67 | 0 | 1,12 | 0 | 6,4 | 0 |
| MYOD1 | TGAACCTGGGAGG | 1 | 375 | 388 | -625 | -612 | -10,08 | 0,01 | 4,67 | >0.1 | 2,56 | 4,73 | 0 | 1,07 | 0 | 6,4 | 0 |
| MYOD1 | TGCTCCAGCCTGG | 1 | 421 | 434 | -579 | -566 | -11,3 | 0,01 | 2,78 | >0.1 | 2,69 | 4,81 | 0 | 1,02 | 0 | 5,56 | 0 |
| MYOD1 | AACAGCTTTCTAG | 1 | 711 | 724 | -289 | -276 | -6,48 | 0,01 | 10,84 | >0.1 | 5,57 | 5,58 | 0 | 1,19 | 0 | 4,99 | 0 |
| MYOD1 | CATATCTGTCCTC | 1 | 852 | 865 | -148 | -135 | -13 | 0,01 | 3,56 | >0.1 | 5,57 | 6,33 | 0 | 1,9 | 0 | 2,75 | 0 |
| MYOD1 | GGCTCCTTCTCAG | 1 | 1065 | 1078 | 65 | 78 | -12,96 | 0,01 | 5,79 | >0.1 | 5,57 | 7,26 | 0 | 1,85 | 1,79 | 2,27 | 0 |
| MYOD1 | AACTGCTGCTCAG | 1 | 1090 | 1103 | 90 | 103 | 3,99 | 0,001 | 22,42 | >0.1 | 5,57 | 6,88 | 0 | 1,85 | 1,79 | 2,34 | 0 |
| MYOD1 | AAGACCTGTTCTA | 1 | 1158 | 1171 | 158 | 171 | -5,76 | 0,01 | 10,91 | >0.1 | 4,39 | 6,24 | 0 | 1,78 | 1,79 | 2,48 | 0 |
| MYOD1 | TTCACTTGCTTCT | 1 | 1177 | 1190 | 177 | 190 | -5,71 | 0,01 | 12,74 | >0.1 | 5,57 | 6,11 | 0 | 1,61 | 1,79 | 3,37 | 0 |
| MYOD1 | AGCAGCAGTTACA | -1 | 1087 | 1100 | 87 | 100 | -1,93 | 0,01 | 16,41 | >0.1 | 5,57 | 6,92 | 0 | 1,85 | 1,79 | 2,21 | 0 |
| MYOD1 | GTCCCCTGTGAGC | -1 | 1035 | 1048 | 35 | 48 | -6,73 | 0,01 | 10,06 | >0.1 | 5,57 | 8,03 | 0 | 1,86 | 0 | 1,34 | 0 |
| MYOD1 | ATCTGCTGAGCCA | -1 | 884 | 897 | -116 | -103 | -5,86 | 0,01 | 11,26 | >0.1 | 5,57 | 6,59 | 0 | 1,89 | 0 | 3,07 | 0 |
| MYOD1 | CTCTCCTGTCACT | -1 | 765 | 778 | -235 | -222 | 5,56 | 0,001 | 24,2 | >0.1 | 5,57 | 5,81 | 0 | 1,35 | 0 | 5,92 | 0 |
| MYOD1 | GAAAGCTGTTAAA | -1 | 708 | 721 | -292 | -279 | -7,4 | 0,01 | 9,72 | >0.1 | 5,57 | 5,56 | 0 | 1,18 | 0 | 4,81 | 0 |
| MYOD1 | TTCTCCTGCCTCA | -1 | 357 | 370 | -643 | -630 | 3,99 | 0,001 | 18,56 | >0.1 | 2,52 | 4,7 | 0 | 1,09 | 0 | 6,26 | 0 |
| MYOD1 | AGTAGCTGGGACT | -1 | 336 | 349 | -664 | -651 | -6,79 | 0,01 | 7,49 | >0.1 | 2,57 | 4,67 | 0 | 1,13 | 0 | 5,92 | 0 |
| MYOD1 | TACAGGTGCCCAC | -1 | 324 | 337 | -676 | -663 | -4,29 | 0,01 | 9,57 | >0.1 | 2,61 | 4,65 | 0 | 1,15 | 0 | 5,46 | 0 |
| MYOD1 | ATCTCCTGACCTT | -1 | 240 | 253 | -760 | -747 | -5,72 | 0,01 | 7,41 | >0.1 | 2,87 | 4,52 | 0 | 1,41 | 0 | 4,33 | 0 |
| MYOD1 | CTCTCCTGATCTG | -1 | 94 | 107 | -906 | -893 | -6,49 | 0,01 | 4,17 | >0.1 | 3,65 | 4,35 | 0 | 1,88 | 0 | 0,79 | 0 |
| MYOD1 | GCCTCCTTCTCCT | -1 | 13 | 26 | -987 | -974 | -10,2 | 0,01 | 1,5 | >0.1 | 5,57 | 4,27 | 0 | 1,87 | 0 | 0 | 0 |
| FOSL1::JUND(VAR.2) | CTCACGCCTG | 1 | 191 | 201 | -809 | -799 | 3,36 | 0,01 | 16,72 | >0.1 | 3,08 | 4,45 | 0 | 1,7 | 0 | 4,13 | 0 |
| FOSL1::JUND(VAR.2) | ATGGCGTGAA | 1 | 369 | 379 | -631 | -621 | 5,33 | 0,01 | 19,81 | >0.1 | 2,54 | 4,71 | 0 | 1,07 | 0 | 6,15 | 0 |
| FOSL1::JUND(VAR.2) | ATCGCGCCAC | 1 | 409 | 419 | -591 | -581 | 5,92 | 0,01 | 19,91 | >0.1 | 2,65 | 4,78 | 0 | 1,03 | 0 | 5,52 | 0 |
| FOSL1::JUND(VAR.2) | GAGACTCAGT | 1 | 444 | 454 | -556 | -546 | 3,03 | 0,01 | 17,56 | >0.1 | 2,76 | 4,85 | 0 | 1 | 0 | 5,92 | 0 |
| FOSL1::JUND(VAR.2) | GTCATGTATT | 1 | 517 | 527 | -483 | -473 | 4,01 | 0,01 | 15,97 | >0.1 | 3,03 | 5 | 0 | 0,98 | 0 | 2,95 | 0 |
| FOSL1::JUND(VAR.2) | GTGAGATAAC | 1 | 591 | 601 | -409 | -399 | 6,68 | 0,001 | 16,29 | >0.1 | 3,41 | 5,18 | 0 | 1,01 | 0 | 0 | 0 |
| FOSL1::JUND(VAR.2) | ATAACCTATT | 1 | 596 | 606 | -404 | -394 | 3,09 | 0,01 | 14,19 | >0.1 | 3,45 | 5,2 | 0 | 1,01 | 0 | 1,44 | 0 |
| FOSL1::JUND(VAR.2) | ATAACGTATT | 1 | 801 | 811 | -199 | -189 | 6,55 | 0,001 | 23,06 | >0.1 | 5,61 | 5,98 | 0 | 1,5 | 0 | 3,43 | 0 |
| FOSL1::JUND(VAR.2) | GTTACATATC | 1 | 848 | 858 | -152 | -142 | 4,87 | 0,01 | 21,56 | >0.1 | 5,61 | 6,28 | 0 | 1,84 | 0 | 2,95 | 0 |
| FOSL1::JUND(VAR.2) | ACGATGTCAA | 1 | 1046 | 1056 | 46 | 56 | 3,49 | 0,01 | 20,1 | >0.1 | 5,61 | 7,68 | 0 | 1,86 | 0 | 1,46 | 0 |

| | | | | | | | | | | | | | | | | | |
|---|---|---|---|---|---|---|---|---|---|---|---|---|---|---|---|---|---|
| FOSL1::JUND(VAR.2) | TTGACATCGT | -1 | 1046 | 1056 | 46 | 56 | 3,93 | 0,01 | 20,54 | >0.1 | 5,61 | 7,68 | 0 | 1,86 | 0 | 1,46 | 0 |
| FOSL1::JUND(VAR.2) | GTGAGCCAAG | -1 | 1031 | 1041 | 31 | 41 | 3,26 | 0,01 | 20,23 | >0.1 | 5,61 | 8,19 | 0 | 1,86 | 0 | 1,31 | 0 |
| FOSL1::JUND(VAR.2) | ATGACTTTCC | -1 | 1007 | 1017 | 7 | 17 | 4,76 | 0,01 | 22,82 | >0.1 | 5,61 | 9,12 | 0 | 1,87 | 0 | 1,46 | 0 |
| FOSL1::JUND(VAR.2) | CTGAGCCAAT | -1 | 882 | 892 | -118 | -108 | 2,92 | 0,01 | 20,04 | >0.1 | 5,61 | 6,55 | 0 | 1,89 | 0 | 3,07 | 0 |
| FOSL1::JUND(VAR.2) | GTGAATCAAA | -1 | 815 | 825 | -185 | -175 | 2,93 | 0,01 | 19,61 | >0.1 | 5,61 | 6,06 | 0 | 1,58 | 0 | 3,43 | 0 |
| FOSL1::JUND(VAR.2) | GTCACTTTAT | -1 | 761 | 771 | -239 | -229 | 3,79 | 0,01 | 22,6 | >0.1 | 5,61 | 5,77 | 0 | 1,32 | 0 | 6,1 | 0 |
| FOSL1::JUND(VAR.2) | ATGGCCAAAT | -1 | 753 | 763 | -247 | -237 | 3,46 | 0,01 | 22,42 | >0.1 | 5,61 | 5,74 | 0 | 1,3 | 0 | 6,32 | 0 |
| FOSL1::JUND(VAR.2) | GTGACCAAAG | -1 | 670 | 680 | -330 | -320 | 5,3 | 0,01 | 19,66 | >0.1 | 4,14 | 5,42 | 0 | 1,1 | 0 | 3,7 | 0 |
| FOSL1::JUND(VAR.2) | ATGACTTTGC | -1 | 512 | 522 | -488 | -478 | 5,36 | 0,01 | 17,29 | >0.1 | 3,01 | 4,99 | 0 | 0,98 | 0 | 2,95 | 0 |
| FOSL1::JUND(VAR.2) | GTGGCGCGAT | -1 | 409 | 419 | -591 | -581 | 6,06 | 0,01 | 20,05 | >0.1 | 2,65 | 4,78 | 0 | 1,03 | 0 | 5,52 | 0 |
| FOSL1::JUND(VAR.2) | TTCACGCCAT | -1 | 369 | 379 | -631 | -621 | 5,5 | 0,01 | 19,98 | >0.1 | 2,54 | 4,71 | 0 | 1,07 | 0 | 6,15 | 0 |
| FOSL1::JUND(VAR.2) | ATGACTACTC | -1 | 124 | 134 | -876 | -866 | 6,96 | 0,001 | 18,36 | >0.1 | 3,44 | 4,38 | 0 | 1,89 | 0 | 1,7 | 0 |
| FOSL1::JUND(VAR.2) | CTGACTTCTT | -1 | 31 | 41 | -969 | -959 | 6,34 | 0,01 | 16,95 | >0.1 | 4,46 | 4,28 | 0 | 1,87 | 0 | 0 | 0 |
| FOSL1::JUND(VAR.2) | GTGCCTCCTT | -1 | 18 | 28 | -982 | -972 | 4,03 | 0,01 | 15,77 | >0.1 | 5,61 | 4,27 | 0 | 1,87 | 0 | 0 | 0 |
| TWIST1 | AAGGCAGATCAGG | 1 | 90 | 103 | -910 | -897 | 2,57 | 0,01 | 13,33 | >0.1 | 3,68 | 4,34 | 0 | 1,88 | 0 | 0,85 | 0 |
| TWIST1 | TGGGCAGATCACA | 1 | 228 | 241 | -772 | -759 | 1,73 | 0,01 | 15,74 | >0.1 | 2,92 | 4,51 | 0 | 1,47 | 0 | 5,12 | 0 |
| TWIST1 | TGGGCACCTGTAG | 1 | 325 | 338 | -675 | -662 | 3,02 | 0,01 | 16,88 | >0.1 | 2,6 | 4,65 | 0 | 1,15 | 0 | 5,46 | 0 |
| TWIST1 | GTCCCAGCTACTC | 1 | 337 | 350 | -663 | -650 | 2,48 | 0,01 | 16,76 | >0.1 | 2,57 | 4,67 | 0 | 1,13 | 0 | 5,92 | 0 |
| TWIST1 | TTAACAGCTTTCT | 1 | 709 | 722 | -291 | -278 | 2,15 | 0,01 | 19,28 | >0.1 | 5,57 | 5,57 | 0 | 1,18 | 0 | 4,81 | 0 |
| TWIST1 | GTTACATATCTGT | 1 | 848 | 861 | -152 | -139 | 1,91 | 0,01 | 18,52 | >0.1 | 5,57 | 6,3 | 0 | 1,87 | 0 | 2,86 | 0 |
| TWIST1 | TCAGCAGATTGTT | 1 | 888 | 901 | -112 | -99 | 3,44 | 0,01 | 20,59 | >0.1 | 5,57 | 6,62 | 0 | 1,89 | 0 | 3,07 | 0 |
| TWIST1 | AGAACAGGTCTTC | -1 | 1157 | 1170 | 157 | 170 | 1,77 | 0,01 | 18,37 | >0.1 | 4,37 | 6,24 | 0 | 1,79 | 1,79 | 2,4 | 0 |
| TWIST1 | TGAGCAGCAGTTA | -1 | 1089 | 1102 | 89 | 102 | 1,19 | 0,01 | 19,57 | >0.1 | 5,57 | 6,89 | 0 | 1,85 | 1,79 | 2,27 | 0 |
| TWIST1 | AGGACAGATATGT | -1 | 851 | 864 | -149 | -136 | 5,56 | 0,01 | 22,03 | >0.1 | 5,57 | 6,33 | 0 | 1,9 | 0 | 2,67 | 0 |
| TWIST1 | GTAACAGATTTTA | -1 | 840 | 853 | -160 | -147 | 5,76 | 0,01 | 20,79 | >0.1 | 5,57 | 6,25 | 0 | 1,8 | 0 | 1,41 | 0 |
| TWIST1 | TAGAAAGCTGTTA | -1 | 710 | 723 | -290 | -277 | 1,03 | 0,01 | 18,34 | >0.1 | 5,57 | 5,57 | 0 | 1,18 | 0 | 4,99 | 0 |
| TWIST1 | ACTACAGGTGCCC | -1 | 326 | 339 | -674 | -661 | 6,18 | 0,01 | 20,14 | >0.1 | 2,6 | 4,65 | 0 | 1,15 | 0 | 5,56 | 0 |
| TWIST1 | CGCCCAGCTAAGT | -1 | 306 | 319 | -694 | -681 | 0,91 | 0,01 | 15,06 | >0.1 | 2,66 | 4,62 | 0 | 1,19 | 0 | 5,68 | 0 |
| TWIST1 | ATTACAGGCGTGA | -1 | 192 | 205 | -808 | -795 | 2,49 | 0,01 | 15,97 | >0.1 | 3,07 | 4,46 | 0 | 1,69 | 0 | 4,26 | 0 |
| TWIST1 | ACTCCACAGGTCA | -1 | 108 | 121 | -892 | -879 | 4,76 | 0,01 | 15,29 | >0.1 | 3,54 | 4,36 | 0 | 1,89 | 0 | 0,75 | 0 |
| TWIST1 | TCTCCTGATCTGC | -1 | 93 | 106 | -907 | -894 | 2,66 | 0,01 | 13,33 | >0.1 | 3,66 | 4,35 | 0 | 1,88 | 0 | 0,79 | 0 |
| TWIST2 | GACCTGTGGA | 1 | 109 | 119 | -891 | -881 | -2,88 | 0,01 | 7,65 | >0.1 | 3,54 | 4,36 | 0 | 1,89 | 0 | 0,75 | 0 |
| TWIST2 | ATCACAAGGT | 1 | 235 | 245 | -765 | -755 | -4,82 | 0,01 | 8,76 | >0.1 | 2,9 | 4,51 | 0 | 1,43 | 0 | 4,74 | 0 |
| TWIST2 | GGCACCTGTA | 1 | 327 | 337 | -673 | -663 | -2,25 | 0,01 | 11,71 | >0.1 | 2,6 | 4,65 | 0 | 1,14 | 0 | 5,56 | 0 |
| TWIST2 | GAAATATGAA | 1 | 486 | 496 | -514 | -504 | -4,59 | 0,01 | 10,5 | >0.1 | 2,9 | 4,93 | 0 | 0,98 | 0 | 6,27 | 0 |
| TWIST2 | GTCATGTATT | 1 | 517 | 527 | -483 | -473 | -3,12 | 0,01 | 8,84 | >0.1 | 3,03 | 5 | 0 | 0,98 | 0 | 2,95 | 0 |
| TWIST2 | TACATATCTG | 1 | 850 | 860 | -150 | -140 | -0,19 | 0,01 | 16,33 | >0.1 | 5,61 | 6,3 | 0 | 1,86 | 0 | 2,75 | 0 |
| TWIST2 | CTCACAGGGG | 1 | 1036 | 1046 | 36 | 46 | -4,35 | 0,01 | 12,44 | >0.1 | 5,61 | 7,99 | 0 | 1,86 | 0 | 1,34 | 0 |
| TWIST2 | CCCCTGTGAG | -1 | 1036 | 1046 | 36 | 46 | -2,96 | 0,01 | 13,83 | >0.1 | 5,61 | 7,99 | 0 | 1,86 | 0 | 1,34 | 0 |
| TWIST2 | TCCTCATGAG | -1 | 984 | 994 | -16 | -6 | -3,9 | 0,01 | 13,68 | >0.1 | 5,61 | 8,57 | 0 | 1,87 | 0 | 1,53 | 0 |
| TWIST2 | CAGATATGTA | -1 | 850 | 860 | -150 | -140 | -0,78 | 0,01 | 15,74 | >0.1 | 5,61 | 6,3 | 0 | 1,86 | 0 | 2,75 | 0 |
| TWIST2 | CTCACATTTA | -1 | 586 | 596 | -414 | -404 | -4,65 | 0,01 | 4,91 | >0.1 | 3,38 | 5,17 | 0 | 1,01 | 0 | 0 | 0 |
| TWIST2 | TTCATATTTC | -1 | 486 | 496 | -514 | -504 | -3,3 | 0,01 | 11,79 | >0.1 | 2,9 | 4,93 | 0 | 0,98 | 0 | 6,27 | 0 |
| TWIST2 | TACAGGTGCC | -1 | 327 | 337 | -673 | -663 | -4,73 | 0,01 | 9,23 | >0.1 | 2,6 | 4,65 | 0 | 1,14 | 0 | 5,56 | 0 |
| TWIST2 | ACCTTGTGAT | -1 | 235 | 245 | -765 | -755 | -3,43 | 0,01 | 10,15 | >0.1 | 2,9 | 4,51 | 0 | 1,43 | 0 | 4,74 | 0 |
| TWIST2 | TCCACAGGTC | -1 | 109 | 119 | -891 | -881 | -0,1 | 0,01 | 10,43 | >0.1 | 3,54 | 4,36 | 0 | 1,89 | 0 | 0,75 | 0 |
| HNF1B | TTTAAAAAATGGC | 1 | 137 | 150 | -863 | -850 | 0,26 | 0,01 | 15,96 | >0.1 | 3,35 | 4,39 | 0 | 1,89 | 0 | 1,65 | 4,41 |
| HNF1B | GGAAAATGTTGCC | 1 | 533 | 546 | -467 | -454 | -2,68 | 0,01 | 12,53 | >0.1 | 3,11 | 5,05 | 0 | 0,99 | 0 | 1,66 | 4,41 |
| HNF1B | ATAACCTATTAAT | 1 | 596 | 609 | -404 | -391 | 1,73 | 0,01 | 17,23 | >0.1 | 3,46 | 5,21 | 0 | 1,02 | 0 | 1,41 | 4,41 |
| HNF1B | GTAAAATTATAAC | 1 | 681 | 694 | -319 | -306 | 2,96 | 0,01 | 21,37 | >0.1 | 4,35 | 5,47 | 0 | 1,12 | 0 | 3,07 | 4,41 |
| HNF1B | GTGTATCTTTAAC | 1 | 701 | 714 | -299 | -286 | 1,06 | 0,01 | 22,55 | >0.1 | 5,57 | 5,54 | 0 | 1,16 | 0 | 4,81 | 4,41 |
| HNF1B | GCCATAAAGTGAC | 1 | 758 | 771 | -242 | -229 | -2,19 | 0,01 | 20,99 | >0.1 | 5,57 | 5,77 | 0 | 1,33 | 0 | 6,1 | 4,41 |
| HNF1B | TTTAAAAAATTAT | 1 | 825 | 838 | -175 | -162 | 1,65 | 0,01 | 22,99 | >0.1 | 5,57 | 6,14 | 0 | 1,68 | 0 | 3,55 | 4,41 |
| HNF1B | TTAAAAAATTATT | 1 | 826 | 839 | -174 | -161 | 0,83 | 0,01 | 22,07 | >0.1 | 5,57 | 6,15 | 0 | 1,68 | 0 | 3,43 | 4,41 |
| HNF1B | GTTACATATCTGT | 1 | 848 | 861 | -152 | -139 | -2,05 | 0,01 | 18,97 | >0.1 | 5,57 | 6,3 | 0 | 1,87 | 0 | 2,86 | 4,41 |
| HNF1B | GCAGATTGTTTAC | 1 | 891 | 904 | -109 | -96 | -1 | 0,01 | 20,38 | >0.1 | 5,57 | 6,65 | 0 | 1,89 | 0 | 2,86 | 4,41 |
| HNF1B | GTCATTCAGTGGA | 1 | 1012 | 1025 | 12 | 25 | -1,32 | 0,01 | 20,86 | >0.1 | 5,57 | 8,92 | 0 | 1,86 | 0 | 1,42 | 4,41 |
| HNF1B | TCCACTGAATGAC | -1 | 1012 | 1025 | 12 | 25 | -1,22 | 0,01 | 20,96 | >0.1 | 5,57 | 8,92 | 0 | 1,86 | 0 | 1,42 | 4,41 |
| HNF1B | GTAAACAATCTGC | -1 | 891 | 904 | -109 | -96 | 0,05 | 0,01 | 21,43 | >0.1 | 5,57 | 6,65 | 0 | 1,89 | 0 | 2,86 | 4,41 |

| | | | | | | | | | | | | | | | | | |
|---|---|---|---|---|---|---|---|---|---|---|---|---|---|---|---|---|---|
| HNF1B | ATAATTTTTTAAA | -1 | 825 | 838 | -175 | -162 | -2,75 | 0,01 | 18,59 | >0.1 | 5,57 | 6,14 | 0 | 1,68 | 0 | 3,55 | 4,41 |
| HNF1B | GTTATAATTTTAC | -1 | 681 | 694 | -319 | -306 | 1,22 | 0,01 | 19,63 | >0.1 | 4,35 | 5,47 | 0 | 1,12 | 0 | 3,07 | 4,41 |
| HNF1B | ATTAATAGGTTAT | -1 | 596 | 609 | -404 | -391 | 1,12 | 0,01 | 16,62 | >0.1 | 3,46 | 5,21 | 0 | 1,02 | 0 | 1,41 | 4,41 |
| HNF1B | GCCATTTTTTAAA | -1 | 137 | 150 | -863 | -850 | -1,25 | 0,01 | 14,45 | >0.1 | 3,35 | 4,39 | 0 | 1,89 | 0 | 1,65 | 4,41 |
| KLF5 | CTCACGCCTG | 1 | 191 | 201 | -809 | -799 | -8,94 | 0,01 | 4,42 | >0.1 | 3,08 | 4,45 | 0 | 1,7 | 0 | 4,13 | 0 |
| KLF5 | ACCCCGTCTC | 1 | 280 | 290 | -720 | -710 | -6,47 | 0,01 | 7,2 | >0.1 | 2,74 | 4,57 | 0 | 1,26 | 0 | 5,09 | 0 |
| KLF5 | GTCCCAGCTA | 1 | 337 | 347 | -663 | -653 | -7,91 | 0,01 | 6,37 | >0.1 | 2,58 | 4,66 | 0 | 1,12 | 0 | 5,92 | 0 |
| KLF5 | GCTCCAGCCT | 1 | 422 | 432 | -578 | -568 | -2,67 | 0,01 | 11,4 | >0.1 | 2,69 | 4,81 | 0 | 1,02 | 0 | 5,56 | 0 |
| KLF5 | GCCCAACCCA | 1 | 947 | 957 | -53 | -43 | 0,17 | 0,01 | 16,74 | >0.1 | 5,61 | 7,34 | 0 | 1,88 | 0 | 1,74 | 0 |
| KLF5 | GCTCCTTCTC | 1 | 1066 | 1076 | 66 | 76 | -8,08 | 0,01 | 10,69 | >0.1 | 5,61 | 7,24 | 0 | 1,85 | 1,79 | 2,27 | 0 |
| KLF5 | TTCTCAGCCT | 1 | 1071 | 1081 | 71 | 81 | -9,18 | 0,01 | 9,51 | >0.1 | 5,61 | 7,15 | 0 | 1,85 | 1,79 | 2,27 | 0 |
| KLF5 | TCTCAGCCTT | 1 | 1072 | 1082 | 72 | 82 | -9,96 | 0,01 | 8,71 | >0.1 | 5,61 | 7,14 | 0 | 1,85 | 1,79 | 2,27 | 0 |
| KLF5 | ATCACATCCA | -1 | 1021 | 1031 | 21 | 31 | -7,98 | 0,01 | 9,5 | >0.1 | 5,61 | 8,7 | 0 | 1,86 | 0 | 1,31 | 0 |
| KLF5 | TTCCCTTCCA | -1 | 527 | 537 | -473 | -463 | -5,75 | 0,01 | 6,76 | >0.1 | 3,07 | 5,03 | 0 | 0,98 | 0 | 3,43 | 0 |
| KLF5 | ACTTTGCCCT | -1 | 509 | 519 | -491 | -481 | -10,11 | 0,01 | 1,6 | >0.1 | 3 | 4,98 | 0 | 0,98 | 0 | 2,75 | 0 |
| KLF5 | CTGTCGCCCA | -1 | 431 | 441 | -569 | -559 | -8,39 | 0,01 | 5,83 | >0.1 | 2,72 | 4,83 | 0 | 1,01 | 0 | 5,67 | 0 |
| KLF5 | GCTCTGCCTC | -1 | 384 | 394 | -616 | -606 | -6,97 | 0,01 | 7,42 | >0.1 | 2,58 | 4,74 | 0 | 1,05 | 0 | 6,02 | 0 |
| KLF5 | CTGCCTCCCA | -1 | 381 | 391 | -619 | -609 | -4,93 | 0,01 | 9,45 | >0.1 | 2,57 | 4,73 | 0 | 1,06 | 0 | 6,02 | 0 |
| KLF5 | TTCACGCCAT | -1 | 369 | 379 | -631 | -621 | -6,47 | 0,01 | 8,01 | >0.1 | 2,54 | 4,71 | 0 | 1,07 | 0 | 6,15 | 0 |
| KLF5 | TCTCCTGCCT | -1 | 359 | 369 | -641 | -631 | -5,29 | 0,01 | 9,38 | >0.1 | 2,52 | 4,7 | 0 | 1,09 | 0 | 6,36 | 0 |
| KLF5 | CTCCTGCCTC | -1 | 358 | 368 | -642 | -632 | -8,76 | 0,01 | 5,8 | >0.1 | 2,52 | 4,7 | 0 | 1,09 | 0 | 6,26 | 0 |
| KLF5 | GCCTCAGCCT | -1 | 353 | 363 | -647 | -637 | -3,84 | 0,01 | 10,88 | >0.1 | 2,53 | 4,69 | 0 | 1,1 | 0 | 6,4 | 0 |
| KLF5 | CCTCAGCCTC | -1 | 352 | 362 | -648 | -638 | -8,87 | 0,01 | 5,85 | >0.1 | 2,54 | 4,69 | 0 | 1,1 | 0 | 6,4 | 0 |
| KLF5 | ACCACGCCCA | -1 | 313 | 323 | -687 | -677 | 12,4 | 0,001 | 26,97 | >0.1 | 2,64 | 4,62 | 0 | 1,17 | 0 | 6,13 | 0 |
| KLF5 | ACTCGGCCTC | -1 | 218 | 228 | -782 | -772 | -9,37 | 0,01 | 5,23 | >0.1 | 2,96 | 4,49 | 0 | 1,52 | 0 | 5,63 | 0 |
| KLF5 | ACTACTCTCC | -1 | 121 | 131 | -879 | -869 | -7,66 | 0,01 | 3,76 | >0.1 | 3,46 | 4,37 | 0 | 1,89 | 0 | 1,7 | 0 |
| KLF5 | TCTCCACTCC | -1 | 116 | 126 | -884 | -874 | -4,26 | 0,01 | 6,23 | >0.1 | 3,49 | 4,37 | 0 | 1,89 | 0 | 0,75 | 0 |
| KLF5 | TCCACTCCAC | -1 | 114 | 124 | -886 | -876 | -3,35 | 0,01 | 7,16 | >0.1 | 3,51 | 4,36 | 0 | 1,89 | 0 | 0,75 | 0 |
| KLF5 | TTCTCACACT | -1 | 51 | 61 | -949 | -939 | -9,68 | 0,01 | 0,61 | >0.1 | 4,11 | 4,3 | 0 | 1,87 | 0 | 0 | 0 |
| KLF5 | GTGCCTCCTT | -1 | 18 | 28 | -982 | -972 | -8,24 | 0,01 | 3,5 | >0.1 | 5,61 | 4,27 | 0 | 1,87 | 0 | 0 | 0 |
| KLF5 | CCTCCTTCTC | -1 | 15 | 25 | -985 | -975 | -9,99 | 0,01 | 1,75 | >0.1 | 5,61 | 4,27 | 0 | 1,87 | 0 | 0 | 0 |
| KLF5 | CCTTCTCCTT | -1 | 12 | 22 | -988 | -978 | 1,04 | 0,01 | 12,78 | >0.1 | 5,61 | 4,26 | 0 | 1,86 | 0 | 0 | 0 |
| HNF1A | TTTTAAAAAATGGCC | 1 | 136 | 151 | -864 | -849 | 0,77 | 0,01 | 19,54 | >0.1 | 3,35 | 4,4 | 0 | 1,89 | 0 | 1,65 | 7,48 |
| HNF1A | GGGAAAATGTTGCCC | 1 | 532 | 547 | -468 | -453 | -1,92 | 0,01 | 16,37 | >0.1 | 3,11 | 5,05 | 0 | 0,99 | 0 | 1,66 | 7,48 |
| HNF1A | GATAACCTATTAATG | 1 | 595 | 610 | -405 | -390 | 2,54 | 0,01 | 21,12 | >0.1 | 3,46 | 5,21 | 0 | 1,02 | 0 | 1,41 | 7,48 |
| HNF1A | CTTTAAAAAATTATT | 1 | 824 | 839 | -176 | -161 | -0,61 | 0,01 | 23,8 | >0.1 | 5,54 | 6,15 | 0 | 1,68 | 0 | 3,55 | 7,48 |
| HNF1A | TTTAAAAAATTATTC | 1 | 825 | 840 | -175 | -160 | -1,89 | 0,01 | 22,41 | >0.1 | 5,54 | 6,16 | 0 | 1,69 | 0 | 3,43 | 7,48 |
| HNF1A | AGTCATTCAGTGGAT | 1 | 1011 | 1026 | 11 | 26 | -0,92 | 0,01 | 24,34 | >0.1 | 5,54 | 8,95 | 0 | 1,86 | 0 | 1,42 | 7,48 |
| HNF1A | AAGAACACAGTAAAC | -1 | 898 | 913 | -102 | -87 | -0,96 | 0,01 | 23,45 | >0.1 | 5,54 | 6,74 | 0 | 1,89 | 0 | 2,75 | 7,48 |
| HNF1A | GACAGATATGTAACA | -1 | 847 | 862 | -153 | -138 | -0,34 | 0,01 | 23,75 | >0.1 | 5,54 | 6,31 | 0 | 1,88 | 0 | 2,86 | 7,48 |
| HNF1A | TGTTATAATTTTACA | -1 | 680 | 695 | -320 | -305 | 2,06 | 0,01 | 23,56 | >0.1 | 4,35 | 5,47 | 0 | 1,13 | 0 | 3,07 | 7,48 |
| HNF1A | CATTAATAGGTTATC | -1 | 595 | 610 | -405 | -390 | 2,29 | 0,01 | 20,87 | >0.1 | 3,46 | 5,21 | 0 | 1,02 | 0 | 1,41 | 7,48 |
| HNF1A | GGGCAACATTTTCCC | -1 | 532 | 547 | -468 | -453 | -2,3 | 0,01 | 15,99 | >0.1 | 3,11 | 5,05 | 0 | 0,99 | 0 | 1,66 | 7,48 |
| HNF1A | GGCCATTTTTTAAAA | -1 | 136 | 151 | -864 | -849 | -0,25 | 0,01 | 18,52 | >0.1 | 3,35 | 4,4 | 0 | 1,89 | 0 | 1,65 | 7,48 |
| PROP1 | TAATTTTAAAA | 1 | 133 | 144 | -867 | -856 | -0,99 | 0,01 | 10,24 | >0.1 | 3,38 | 4,39 | 0 | 1,89 | 0 | 1,57 | 0 |
| PROP1 | GAATTCCATAA | 1 | 493 | 504 | -507 | -496 | 0,74 | 0,01 | 16,05 | >0.1 | 2,93 | 4,95 | 0 | 0,98 | 0 | 6,44 | 0 |
| PROP1 | AAATAAAAATA | 1 | 577 | 588 | -423 | -412 | 1,18 | 0,01 | 10,66 | >0.1 | 3,33 | 5,15 | 0 | 1 | 0 | 0 | 0 |
| PROP1 | AAAATAAATGT | 1 | 582 | 593 | -418 | -407 | -0,53 | 0,01 | 9 | >0.1 | 3,36 | 5,16 | 0 | 1,01 | 0 | 0 | 0 |
| PROP1 | TAACCTATTAA | 1 | 597 | 608 | -403 | -392 | 0,92 | 0,01 | 12,01 | >0.1 | 3,46 | 5,2 | 0 | 1,02 | 0 | 1,41 | 0 |
| PROP1 | TAATGAAATTG | 1 | 605 | 616 | -395 | -384 | 7,67 | 0,001 | 18,84 | >0.1 | 3,51 | 5,22 | 0 | 1,02 | 0 | 1,41 | 0 |
| PROP1 | TAAAATTATAA | 1 | 682 | 693 | -318 | -307 | -0,47 | 0,01 | 13,53 | >0.1 | 4,35 | 5,46 | 0 | 1,12 | 0 | 3,07 | 0 |
| PROP1 | TTATAACATTT | 1 | 687 | 698 | -313 | -302 | -0,31 | 0,01 | 13,7 | >0.1 | 4,45 | 5,48 | 0 | 1,13 | 0 | 2,95 | 0 |
| PROP1 | TTAAAAAATTA | 1 | 826 | 837 | -174 | -163 | 2,91 | 0,01 | 19,86 | >0.1 | 5,6 | 6,14 | 0 | 1,67 | 0 | 3,55 | 0 |
| PROP1 | AAAATCTGTTA | 1 | 841 | 852 | -159 | -148 | 0,67 | 0,01 | 15,71 | >0.1 | 5,6 | 6,24 | 0 | 1,79 | 0 | 1,41 | 0 |
| PROP1 | AAATCTCATGA | 1 | 980 | 991 | -20 | -9 | 3,64 | 0,01 | 21,01 | >0.1 | 5,6 | 8,37 | 0 | 1,87 | 0 | 1,53 | 0 |
| PROP1 | TAACAGATTTT | -1 | 841 | 852 | -159 | -148 | -1,86 | 0,01 | 13,18 | >0.1 | 5,6 | 6,24 | 0 | 1,79 | 0 | 1,41 | 0 |
| PROP1 | TAATTTTTTAA | -1 | 826 | 837 | -174 | -163 | 0,5 | 0,01 | 17,45 | >0.1 | 5,6 | 6,14 | 0 | 1,67 | 0 | 3,55 | 0 |
| PROP1 | AAAGTGAATCA | -1 | 817 | 828 | -183 | -172 | 3,21 | 0,01 | 20,04 | >0.1 | 5,6 | 6,08 | 0 | 1,6 | 0 | 3,55 | 0 |
| PROP1 | GAATCAAATAA | -1 | 812 | 823 | -188 | -177 | 5,02 | 0,01 | 21,66 | >0.1 | 5,6 | 6,05 | 0 | 1,57 | 0 | 3,43 | 0 |

| | | | | | | | | | | | | | | | | | |
|---|---|---|---|---|---|---|---|---|---|---|---|---|---|---|---|---|---|
| PROP1 | AAATGTTATAA | -1 | 687 | 698 | -313 | -302 | 0,16 | 0,01 | 14,17 | >0.1 | 4,45 | 5,48 | 0 | 1,13 | 0 | 2,95 | 0 |
| PROP1 | CAATTTCATTA | -1 | 605 | 616 | -395 | -384 | 8,12 | 0,001 | 19,29 | >0.1 | 3,51 | 5,22 | 0 | 1,02 | 0 | 1,41 | 0 |
| PROP1 | TAAAATTATGA | -1 | 130 | 141 | -870 | -859 | -1,06 | 0,01 | 10,19 | >0.1 | 3,4 | 4,38 | 0 | 1,89 | 0 | 1,57 | 0 |
| PROP1 | AAAGCTCATTT | -1 | 66 | 77 | -934 | -923 | 1,59 | 0,01 | 11,7 | >0.1 | 3,92 | 4,32 | 0 | 1,88 | 0 | 0 | 0 |
| KLF1 | GCTCACGCCTG | 1 | 190 | 201 | -810 | -799 | -4,9 | 0,01 | 8,47 | >0.1 | 3,08 | 4,46 | 0 | 1,71 | 0 | 4,13 | 0 |
| KLF1 | AGTCCCAGCTA | 1 | 336 | 347 | -664 | -653 | -4,04 | 0,01 | 10,24 | >0.1 | 2,58 | 4,66 | 0 | 1,13 | 0 | 5,92 | 0 |
| KLF1 | TGCTCCAGCCT | 1 | 421 | 432 | -579 | -568 | 0,76 | 0,01 | 14,69 | >0.1 | 2,69 | 4,81 | 0 | 1,02 | 0 | 5,42 | 0 |
| KLF1 | ACAAACACCAA | 1 | 628 | 639 | -372 | -361 | -5,35 | 0,01 | 6,28 | >0.1 | 3,69 | 5,29 | 0 | 1,04 | 0 | 1,61 | 0 |
| KLF1 | CTCAGCGCCCA | 1 | 941 | 952 | -59 | -48 | -4,99 | 0,01 | 10,52 | >0.1 | 5,6 | 7,25 | 0 | 1,88 | 0 | 0,79 | 0 |
| KLF1 | GCGCCCAACCC | 1 | 945 | 956 | -55 | -44 | -3,19 | 0,01 | 13,41 | >0.1 | 5,6 | 7,32 | 0 | 1,88 | 0 | 1,8 | 0 |
| KLF1 | CGCCCAACCCA | 1 | 946 | 957 | -54 | -43 | 4,44 | 0,01 | 21,06 | >0.1 | 5,6 | 7,34 | 0 | 1,88 | 0 | 1,8 | 0 |
| KLF1 | CAGTCCACCAT | 1 | 1100 | 1111 | 100 | 111 | -4,54 | 0,01 | 15,26 | >0.1 | 5,6 | 6,76 | 0 | 1,85 | 3,22 | 2,38 | 0 |
| KLF1 | GATCACATCCA | -1 | 1021 | 1032 | 21 | 32 | 1,6 | 0,01 | 19,07 | >0.1 | 5,6 | 8,7 | 0 | 1,86 | 0 | 1,31 | 0 |
| KLF1 | AAAACCTCCTC | -1 | 989 | 1000 | -11 | 0 | -4,83 | 0,01 | 12,96 | >0.1 | 5,6 | 8,71 | 0 | 1,87 | 0 | 1,61 | 0 |
| KLF1 | GTTCACGCCAT | -1 | 369 | 380 | -631 | -620 | -3,86 | 0,01 | 10,62 | >0.1 | 2,54 | 4,72 | 0 | 1,07 | 0 | 6,15 | 0 |
| KLF1 | TGCCTCAGCCT | -1 | 353 | 364 | -647 | -636 | -0,91 | 0,01 | 13,81 | >0.1 | 2,53 | 4,69 | 0 | 1,1 | 0 | 6,4 | 0 |
| KLF1 | GTGCCCACCAC | -1 | 321 | 332 | -679 | -668 | -5,14 | 0,01 | 8,84 | >0.1 | 2,62 | 4,64 | 0 | 1,16 | 0 | 5,56 | 0 |
| KLF1 | CACCACGCCCA | -1 | 313 | 324 | -687 | -676 | 14,67 | 0,0001 | 29,24 | >0.1 | 2,64 | 4,63 | 0 | 1,17 | 0 | 6,13 | 0 |
| KLF1 | TGATCTGCCCA | -1 | 228 | 239 | -772 | -761 | -3,93 | 0,01 | 10,08 | >0.1 | 2,92 | 4,5 | 0 | 1,46 | 0 | 5,12 | 0 |
| KLF1 | TCGGCCTCCCA | -1 | 215 | 226 | -785 | -774 | -5,32 | 0,01 | 9,31 | >0.1 | 2,97 | 4,49 | 0 | 1,54 | 0 | 5,63 | 0 |
| KLF1 | GTGCCCAGCCA | -1 | 175 | 186 | -825 | -814 | -2,19 | 0,01 | 10,47 | >0.1 | 3,15 | 4,44 | 0 | 1,83 | 0 | 3,24 | 0 |
| KLF1 | GACTACTCTCC | -1 | 121 | 132 | -879 | -868 | -4,15 | 0,01 | 7,27 | >0.1 | 3,46 | 4,37 | 0 | 1,89 | 0 | 1,7 | 0 |
| KLF1 | CTCCACTCCAC | -1 | 114 | 125 | -886 | -875 | -5,42 | 0,01 | 5,08 | >0.1 | 3,5 | 4,37 | 0 | 1,89 | 0 | 0,75 | 0 |
| KLF1 | TCCTTCTCCTT | -1 | 12 | 23 | -988 | -977 | -5,19 | 0,01 | 6,54 | >0.1 | 5,6 | 4,26 | 0 | 1,86 | 0 | 0 | 0 |
| KLF9 | AGCCACAGAACCA | 1 | 39 | 52 | -961 | -948 | 1,8 | 0,01 | 12,24 | >0.1 | 4,28 | 4,29 | 0 | 1,87 | 0 | 0 | 0 |
| KLF9 | CCACAGAACCAGT | 1 | 41 | 54 | -959 | -946 | 0,87 | 0,01 | 11,28 | >0.1 | 4,24 | 4,29 | 0 | 1,87 | 0 | 0 | 0 |
| KLF9 | GCTCACGCCTGTA | 1 | 190 | 203 | -810 | -797 | 1,6 | 0,01 | 14,97 | >0.1 | 3,08 | 4,46 | 0 | 1,71 | 0 | 4,13 | 0 |
| KLF9 | AGTCCCAGCTACT | 1 | 336 | 349 | -664 | -651 | -0,22 | 0,01 | 14,06 | >0.1 | 2,57 | 4,67 | 0 | 1,13 | 0 | 5,92 | 0 |
| KLF9 | GGCGACAGACCGA | 1 | 433 | 446 | -567 | -554 | 0,62 | 0,01 | 15 | >0.1 | 2,73 | 4,83 | 0 | 1,01 | 0 | 5,8 | 0 |
| KLF9 | CGACAGACCGAGA | 1 | 435 | 448 | -565 | -552 | 2,42 | 0,01 | 16,92 | >0.1 | 2,73 | 4,84 | 0 | 1,01 | 0 | 5,92 | 0 |
| KLF9 | ACAAACACCAACA | 1 | 628 | 641 | -372 | -359 | 0,62 | 0,01 | 12,32 | >0.1 | 3,7 | 5,3 | 0 | 1,05 | 0 | 1,66 | 0 |
| KLF9 | GCGCCCAACCCAA | 1 | 945 | 958 | -55 | -42 | -0,27 | 0,01 | 16,34 | >0.1 | 5,57 | 7,36 | 0 | 1,88 | 0 | 1,8 | 0 |
| KLF9 | CGCCCAACCCAAG | 1 | 946 | 959 | -54 | -41 | 3,55 | 0,01 | 20,12 | >0.1 | 5,57 | 7,38 | 0 | 1,88 | 0 | 1,74 | 0 |
| KLF9 | GATCACATCCACT | -1 | 1019 | 1032 | 19 | 32 | 6,67 | 0,01 | 24,15 | >0.1 | 5,57 | 8,74 | 0 | 1,86 | 0 | 1,31 | 0 |
| KLF9 | GTTCACGCCATTC | -1 | 367 | 380 | -633 | -620 | -0,05 | 0,01 | 15,01 | >0.1 | 2,54 | 4,72 | 0 | 1,08 | 0 | 6,73 | 0 |
| KLF9 | TGCCTCAGCCTCC | -1 | 351 | 364 | -649 | -636 | 4,52 | 0,01 | 19,25 | >0.1 | 2,53 | 4,69 | 0 | 1,1 | 0 | 6,4 | 0 |
| KLF9 | GTGCCCACCACCA | -1 | 319 | 332 | -681 | -668 | 1,43 | 0,01 | 15,53 | >0.1 | 2,62 | 4,64 | 0 | 1,16 | 0 | 5,68 | 0 |
| KLF9 | GCCCACCACCACG | -1 | 317 | 330 | -683 | -670 | 6,52 | 0,01 | 21,2 | >0.1 | 2,63 | 4,64 | 0 | 1,17 | 0 | 6,26 | 0 |
| KLF9 | CACCACGCCCAGC | -1 | 311 | 324 | -689 | -676 | 14,84 | 0,0001 | 29,32 | >0.1 | 2,64 | 4,63 | 0 | 1,18 | 0 | 6,03 | 0 |
| KLF9 | GTGCCCAGCCATT | -1 | 173 | 186 | -827 | -814 | 2,06 | 0,01 | 14,74 | >0.1 | 3,15 | 4,44 | 0 | 1,85 | 0 | 3,24 | 0 |
| KLF9 | CTCCACTCCACAG | -1 | 112 | 125 | -888 | -875 | -0,41 | 0,01 | 10,12 | >0.1 | 3,51 | 4,37 | 0 | 1,89 | 0 | 0,77 | 0 |
| EVX1 | GGAAATGAGC | 1 | 64 | 74 | -936 | -926 | 4,26 | 0,01 | 14,4 | >0.1 | 3,95 | 4,31 | 0 | 1,88 | 0 | 0 | 0 |
| EVX1 | GCAGATCAGG | 1 | 93 | 103 | -907 | -897 | 4,32 | 0,01 | 15,02 | >0.1 | 3,67 | 4,34 | 0 | 1,88 | 0 | 0,81 | 0 |
| EVX1 | GGAAATTAAA | 1 | 153 | 163 | -847 | -837 | 6,06 | 0,01 | 17,28 | >0.1 | 3,27 | 4,41 | 0 | 1,9 | 0 | 1,65 | 0 |
| EVX1 | ACTGATCAGA | 1 | 163 | 173 | -837 | -827 | 4,94 | 0,01 | 16,88 | >0.1 | 3,21 | 4,42 | 0 | 1,9 | 0 | 2,4 | 0 |
| EVX1 | GCAGATCACA | 1 | 231 | 241 | -769 | -759 | 3 | 0,01 | 16,88 | >0.1 | 2,91 | 4,51 | 0 | 1,45 | 0 | 5,01 | 0 |
| EVX1 | ACTGATCAGA | 1 | 478 | 488 | -522 | -512 | 4,94 | 0,01 | 19,08 | >0.1 | 2,88 | 4,92 | 0 | 0,99 | 0 | 5,36 | 0 |
| EVX1 | CCCAAGTAGA | 1 | 544 | 554 | -456 | -446 | 3,82 | 0,01 | 13,02 | >0.1 | 3,15 | 5,06 | 0 | 0,99 | 0 | 0 | 0 |
| EVX1 | TGAAATTGTC | 1 | 608 | 618 | -392 | -382 | 3,14 | 0,01 | 14,47 | >0.1 | 3,53 | 5,23 | 0 | 1,02 | 0 | 1,55 | 0 |
| EVX1 | CAACATTATC | 1 | 636 | 646 | -364 | -354 | 5,86 | 0,01 | 17,64 | >0.1 | 3,76 | 5,31 | 0 | 1,05 | 0 | 1,66 | 0 |
| EVX1 | TAAAATTATA | 1 | 682 | 692 | -318 | -308 | 3,52 | 0,01 | 17,5 | >0.1 | 4,34 | 5,46 | 0 | 1,12 | 0 | 3,07 | 0 |
| EVX1 | CAAAAGTACC | 1 | 736 | 746 | -264 | -254 | 2,9 | 0,01 | 21,33 | >0.1 | 5,61 | 5,66 | 0 | 1,24 | 0 | 5,92 | 0 |
| EVX1 | AAAAATTATT | 1 | 829 | 839 | -171 | -161 | 4,16 | 0,01 | 20,88 | >0.1 | 5,61 | 6,15 | 0 | 1,68 | 0 | 3,28 | 0 |
| EVX1 | GCTGATAAGA | 1 | 966 | 976 | -34 | -24 | 4,6 | 0,01 | 21,56 | >0.1 | 5,61 | 7,78 | 0 | 1,87 | 0 | 1,7 | 0 |
| EVX1 | TCTCATGAGG | 1 | 983 | 993 | -17 | -7 | 3,43 | 0,01 | 20,98 | >0.1 | 5,61 | 8,5 | 0 | 1,87 | 0 | 1,57 | 0 |
| EVX1 | CACCATTGAG | 1 | 1105 | 1115 | 105 | 115 | 3,24 | 0,01 | 23,14 | >0.1 | 5,61 | 6,7 | 0 | 1,85 | 3,22 | 2,52 | 0 |
| EVX1 | CGTGGTTAAA | -1 | 1142 | 1152 | 142 | 152 | 2,89 | 0,01 | 23,29 | >0.1 | 5,61 | 6,36 | 0 | 1,84 | 3,22 | 3,37 | 0 |
| EVX1 | AGCAGTTACA | -1 | 1087 | 1097 | 87 | 97 | 2,96 | 0,01 | 21,35 | >0.1 | 5,61 | 6,92 | 0 | 1,85 | 1,79 | 2,21 | 0 |
| EVX1 | CCTCATGAGA | -1 | 983 | 993 | -17 | -7 | 3,78 | 0,01 | 21,33 | >0.1 | 5,61 | 8,5 | 0 | 1,87 | 0 | 1,57 | 0 |

| | | | | | | | | | | | | | | | | | |
|---|---|---|---|---|---|---|---|---|---|---|---|---|---|---|---|---|---|
| EVX1 | TCTTATCAGC | -1 | 966 | 976 | -34 | -24 | 2,84 | 0,01 | 19,8 | >0.1 | 5,61 | 7,78 | 0 | 1,87 | 0 | 1,7 | 0 |
| EVX1 | GATCATGAAG | -1 | 645 | 655 | -355 | -345 | 3,18 | 0,01 | 17,26 | >0.1 | 3,84 | 5,34 | 0 | 1,06 | 0 | 3,83 | 0 |
| EVX1 | TTTCATTAAT | -1 | 603 | 613 | -397 | -387 | 3,41 | 0,01 | 14,55 | >0.1 | 3,49 | 5,22 | 0 | 1,02 | 0 | 1,41 | 0 |
| EVX1 | TCTGATCAGT | -1 | 478 | 488 | -522 | -512 | 3,75 | 0,01 | 17,89 | >0.1 | 2,88 | 4,92 | 0 | 0,99 | 0 | 5,36 | 0 |
| EVX1 | CCCGAGTAGC | -1 | 343 | 353 | -657 | -647 | 4,98 | 0,01 | 19,85 | >0.1 | 2,56 | 4,67 | 0 | 1,11 | 0 | 6,52 | 0 |
| EVX1 | TCTGATCAGT | -1 | 163 | 173 | -837 | -827 | 3,75 | 0,01 | 15,69 | >0.1 | 3,21 | 4,42 | 0 | 1,9 | 0 | 2,4 | 0 |
| EVX1 | TAAAATTATG | -1 | 131 | 141 | -869 | -859 | 4,46 | 0,01 | 15,71 | >0.1 | 3,4 | 4,38 | 0 | 1,89 | 0 | 1,57 | 0 |
| EVX1 | GCTCATTTCC | -1 | 64 | 74 | -936 | -926 | 5,4 | 0,01 | 15,54 | >0.1 | 3,95 | 4,31 | 0 | 1,88 | 0 | 0 | 0 |
| REST | GTCAGCCACAGAACCAGTGT | 1 | 36 | 57 | -964 | -943 | -11,15 | 0,01 | -0,72 | >0.1 | 4,26 | 4,3 | 0 | 1,87 | 0 | 0 | 0 |
| REST | CACAGAACCAGTGTGAGAAAT | 1 | 42 | 63 | -958 | -937 | -1,29 | 0,001 | 9,05 | >0.1 | 4,16 | 4,3 | 0 | 1,87 | 0 | 0 | 0 |
| REST | ACCTGTGGAGTGGAGAGTAG | 1 | 110 | 131 | -890 | -869 | -9,05 | 0,01 | 1,46 | >0.1 | 3,5 | 4,37 | 0 | 1,89 | 0 | 0,75 | 0 |
| REST | CTGGGCACAGTGGCTCACG | 1 | 178 | 199 | -822 | -801 | -2,74 | 0,01 | 10,49 | >0.1 | 3,11 | 4,45 | 0 | 1,81 | 0 | 3,86 | 0 |
| REST | CCTAGCACTTTGGGAGGCCG | 1 | 205 | 226 | -795 | -774 | -6,79 | 0,01 | 7 | >0.1 | 2,99 | 4,49 | 0 | 1,6 | 0 | 4,7 | 0 |
| REST | ACAAGGTCAGGAGATAGAGA | 1 | 238 | 259 | -762 | -741 | -7,39 | 0,01 | 5,76 | >0.1 | 2,87 | 4,53 | 0 | 1,42 | 0 | 4,33 | 0 |
| REST | CTGACTAACACGGTGAAACC | 1 | 263 | 284 | -737 | -716 | -9,43 | 0,01 | 3,69 | >0.1 | 2,78 | 4,57 | 0 | 1,32 | 0 | 4,45 | 0 |
| REST | GGCAGGAGAATGGCGTGAAC | 1 | 360 | 381 | -640 | -619 | 0,76 | 0,001 | 15,72 | >0.1 | 2,53 | 4,72 | 0 | 1,09 | 0 | 6,61 | 0 |
| REST | TGCTCCAGCCTGGGCGACAC | 1 | 421 | 442 | -579 | -558 | -10,46 | 0,01 | 3,76 | >0.1 | 2,7 | 4,83 | 0 | 1,02 | 0 | 5,67 | 0 |
| REST | GCCTGGGCGACAGACCGAGA | 1 | 428 | 449 | -572 | -551 | -5,77 | 0,01 | 8,62 | >0.1 | 2,72 | 4,84 | 0 | 1,02 | 0 | 5,8 | 0 |
| REST | ATAAAGACAAGGGCAAAGTCA | 1 | 500 | 521 | -500 | -479 | -9,41 | 0,01 | 5,24 | >0.1 | 2,98 | 4,99 | 0 | 0,99 | 0 | 5,68 | 0 |
| REST | CTCAGCGCCCAACCCAAGTT | 1 | 941 | 962 | -59 | -38 | -6,16 | 0,01 | 10,44 | >0.1 | 5,47 | 7,44 | 0 | 1,88 | 0 | 1,8 | 0 |
| REST | CAGGGGACGATGTCAAGCTC | 1 | 1040 | 1061 | 40 | 61 | -10,32 | 0,01 | 5,51 | >0.1 | 5,47 | 7,85 | 0 | 1,86 | 0 | 0,65 | 0 |
| REST | TCCACCATTGAGGAACAGGC | 1 | 1103 | 1124 | 103 | 124 | -10,99 | 0,01 | 8,81 | >0.1 | 5,47 | 6,72 | 0 | 1,85 | 3,22 | 2,53 | 0 |
| REST | TTAACCACGAAGCCGAAGAC | 1 | 1143 | 1164 | 143 | 164 | -5,88 | 0,01 | 13,19 | >0.1 | 5,47 | 6,35 | 0 | 1,84 | 3,22 | 2,18 | 0 |
| REST | TCCTCAATGGTGGACTGAGC | -1 | 1096 | 1117 | 96 | 117 | -10,6 | 0,01 | 9,13 | >0.1 | 5,47 | 6,8 | 0 | 1,85 | 3,22 | 2,38 | 0 |
| REST | CTGAGCAGCAGTTACAGCAA | -1 | 1082 | 1103 | 82 | 103 | -2,67 | 0,01 | 15,65 | >0.1 | 5,47 | 6,99 | 0 | 1,85 | 1,79 | 2,21 | 0 |
| REST | GAAGGAGCCAGGAAGAGCTT | -1 | 1053 | 1074 | 53 | 74 | -8,42 | 0,01 | 10,42 | >0.1 | 5,47 | 7,5 | 0 | 1,86 | 1,79 | 2,21 | 0 |
| REST | CTGTGAGCCAAGATCACATC | -1 | 1022 | 1043 | 22 | 43 | -11,02 | 0,01 | 6,3 | >0.1 | 5,47 | 8,68 | 0 | 1,86 | 0 | 1,31 | 0 |
| REST | AAGTTCATCCTGGAGAGGACA | -1 | 858 | 879 | -142 | -121 | -8,93 | 0,01 | 7,55 | >0.1 | 5,47 | 6,44 | 0 | 1,9 | 0 | 2,67 | 0 |
| REST | TCTAGAACTAGGGATCATGAA | -1 | 646 | 667 | -354 | -333 | -6,81 | 0,01 | 7,56 | >0.1 | 3,91 | 5,38 | 0 | 1,09 | 0 | 3,99 | 0 |
| REST | CCCAGGCTGGAGCACAGTG | -1 | 414 | 435 | -586 | -565 | -8,31 | 0,01 | 5,52 | >0.1 | 2,68 | 4,81 | 0 | 1,03 | 0 | 5,3 | 0 |
| REST | TGGAGCACAGTGGCGCGATC | -1 | 407 | 428 | -593 | -572 | 1,54 | 0,001 | 15,95 | >0.1 | 2,66 | 4,8 | 0 | 1,04 | 0 | 5,91 | 0 |
| REST | CTCAGCCTCCCGAGTAGCTG | -1 | 340 | 361 | -660 | -639 | -8,38 | 0,01 | 6,51 | >0.1 | 2,55 | 4,69 | 0 | 1,12 | 0 | 6,52 | 0 |
| REST | CCGAGTAGCTGGGACTACAG | -1 | 331 | 352 | -669 | -648 | -7,17 | 0,01 | 7,13 | >0.1 | 2,58 | 4,67 | 0 | 1,14 | 0 | 5,92 | 0 |
| REST | CCCACCACCACGCCCAGCTA | -1 | 308 | 329 | -692 | -671 | 8,22 | 0,0001 | 22,82 | >0.1 | 2,64 | 4,63 | 0 | 1,19 | 0 | 6,13 | 0 |
| REST | GTCAGGACGGTCTCTATCTC | -1 | 247 | 268 | -753 | -732 | -9,3 | 0,01 | 4,44 | >0.1 | 2,83 | 4,55 | 0 | 1,38 | 0 | 4,98 | 0 |
| REST | TGAGCCACTGTGCCCAGCCA | -1 | 174 | 195 | -826 | -805 | -8,14 | 0,01 | 4,38 | >0.1 | 3,13 | 4,45 | 0 | 1,85 | 0 | 3,09 | 0 |
| REST | TTAATTTCCATGGCCATTTTT | -1 | 141 | 162 | -859 | -838 | -7,61 | 0,01 | 3,61 | >0.1 | 3,3 | 4,41 | 0 | 1,9 | 0 | 1,61 | 0 |
| REST | TCCACTCCACAGGTCAACTC | -1 | 103 | 124 | -897 | -876 | -4,95 | 0,01 | 5,6 | >0.1 | 3,55 | 4,37 | 0 | 1,89 | 0 | 0,75 | 0 |
| REST | TGCAAAACTTAAAAAAGCTCA | -1 | 69 | 90 | -931 | -910 | -8,55 | 0,01 | 1,5 | >0.1 | 3,84 | 4,33 | 0 | 1,88 | 0 | 0 | 0 |
| NKX2-5(VAR.2) | AGGCACTCAAA | 1 | 22 | 33 | -978 | -967 | 14,32 | 0,0001 | 26,06 | >0.1 | 5,6 | 4,27 | 0 | 1,87 | 0 | 0 | 0 |
| NKX2-5(VAR.2) | TAGCACTTTGG | 1 | 207 | 218 | -793 | -782 | 1,82 | 0,01 | 15,59 | >0.1 | 3,01 | 4,48 | 0 | 1,58 | 0 | 4,7 | 0 |
| NKX2-5(VAR.2) | CGCCACTGTGC | 1 | 413 | 424 | -587 | -576 | -5,11 | 0,01 | 8,46 | >0.1 | 2,66 | 4,79 | 0 | 1,02 | 0 | 5,09 | 0 |
| NKX2-5(VAR.2) | CGAGACTCAGT | 1 | 443 | 454 | -557 | -546 | -6,58 | 0,01 | 7,95 | >0.1 | 2,76 | 4,85 | 0 | 1 | 0 | 5,92 | 0 |
| NKX2-5(VAR.2) | GACCTCTTTGG | 1 | 665 | 676 | -335 | -324 | 2,15 | 0,01 | 16,56 | >0.1 | 4,08 | 5,41 | 0 | 1,09 | 0 | 3,83 | 0 |
| NKX2-5(VAR.2) | CAGCTTTCTAG | 1 | 713 | 724 | -287 | -276 | -6,27 | 0,01 | 11,25 | >0.1 | 5,6 | 5,58 | 0 | 1,19 | 0 | 5,17 | 0 |
| NKX2-5(VAR.2) | AGGTTCTCTAG | 1 | 781 | 792 | -219 | -208 | -3,62 | 0,01 | 15,58 | >0.1 | 5,6 | 5,87 | 0 | 1,41 | 0 | 6,32 | 0 |
| NKX2-5(VAR.2) | ATTCACTTTAA | 1 | 819 | 830 | -181 | -170 | -5,9 | 0,01 | 10,95 | >0.1 | 5,6 | 6,09 | 0 | 1,62 | 0 | 3,55 | 0 |
| NKX2-5(VAR.2) | GTCCTCTCCAG | 1 | 859 | 870 | -141 | -130 | 10,8 | 0,001 | 27,53 | >0.1 | 5,6 | 6,37 | 0 | 1,9 | 0 | 2,86 | 0 |
| NKX2-5(VAR.2) | AAAATCTCATG | 1 | 979 | 990 | -21 | -10 | -5,12 | 0,01 | 12,2 | >0.1 | 5,6 | 8,32 | 0 | 1,87 | 0 | 1,53 | 0 |
| NKX2-5(VAR.2) | TGGCTCACAGG | 1 | 1033 | 1044 | 33 | 44 | -1,6 | 0,01 | 15,27 | >0.1 | 5,6 | 8,1 | 0 | 1,86 | 0 | 1,31 | 0 |
| NKX2-5(VAR.2) | TCCTTCTCAGC | 1 | 1068 | 1079 | 68 | 79 | -5,48 | 0,01 | 13,18 | >0.1 | 5,6 | 7,21 | 0 | 1,85 | 1,79 | 2,21 | 0 |
| NKX2-5(VAR.2) | ATCCACTGAAT | -1 | 1015 | 1026 | 15 | 26 | -2,86 | 0,01 | 14,83 | >0.1 | 5,6 | 8,84 | 0 | 1,86 | 0 | 1,39 | 0 |
| NKX2-5(VAR.2) | TTTCTCTCTTA | -1 | 971 | 982 | -29 | -18 | -4,91 | 0,01 | 12,18 | >0.1 | 5,6 | 7,98 | 0 | 1,87 | 0 | 1,65 | 0 |
| NKX2-5(VAR.2) | TACCTCTCCTG | -1 | 770 | 781 | -230 | -219 | 8,21 | 0,001 | 26,9 | >0.1 | 5,6 | 5,82 | 0 | 1,36 | 0 | 5,92 | 0 |
| NKX2-5(VAR.2) | AAACTCTCTAC | -1 | 549 | 560 | -451 | -440 | 6,93 | 0,001 | 16,18 | >0.1 | 3,18 | 5,08 | 0 | 0,99 | 0 | 0 | 0 |
| NKX2-5(VAR.2) | TGCCTCCCAGG | -1 | 379 | 390 | -621 | -610 | -0,65 | 0,01 | 14,38 | >0.1 | 2,57 | 4,73 | 0 | 1,06 | 0 | 6,66 | 0 |
| NKX2-5(VAR.2) | GGCCTCCCAAA | -1 | 213 | 224 | -787 | -776 | -0,78 | 0,01 | 13,87 | >0.1 | 2,98 | 4,48 | 0 | 1,55 | 0 | 5,63 | 0 |
| NKX2-5(VAR.2) | AGCCACTGTGC | -1 | 182 | 193 | -818 | -807 | -3,15 | 0,01 | 10,04 | >0.1 | 3,12 | 4,45 | 0 | 1,77 | 0 | 3,86 | 0 |
| NKX2-5(VAR.2) | CTACTCTCCAC | -1 | 119 | 130 | -881 | -870 | 5,7 | 0,001 | 17,08 | >0.1 | 3,47 | 4,37 | 0 | 1,89 | 0 | 1,65 | 0 |

| | | | | | | | | | | | | | | | | | |
|---|---|---|---|---|---|---|---|---|---|---|---|---|---|---|---|---|---|
| NKX2-5(VAR.2) | CTCCACTCCAC | -1 | 114 | 125 | -886 | -875 | 9,93 | 0,001 | 20,43 | >0.1 | 3,5 | 4,37 | 0 | 1,89 | 0 | 0,75 | 0 |
| NKX2-5(VAR.2) | CAACTCTCCTG | -1 | 99 | 110 | -901 | -890 | 5,25 | 0,001 | 15,87 | >0.1 | 3,62 | 4,35 | 0 | 1,88 | 0 | 0,77 | 0 |
| PHOX2A | AAAACTGATCA | 1 | 160 | 171 | -840 | -829 | 4,33 | 0,01 | 16,33 | >0.1 | 3,23 | 4,42 | 0 | 1,9 | 0 | 2,46 | 0 |
| PHOX2A | AAAACTGATCA | 1 | 475 | 486 | -525 | -514 | 4,33 | 0,01 | 18,19 | >0.1 | 2,87 | 4,91 | 0 | 0,99 | 0 | 5,09 | 0 |
| PHOX2A | GAATTCCATAA | 1 | 493 | 504 | -507 | -496 | 4,04 | 0,01 | 19,35 | >0.1 | 2,93 | 4,95 | 0 | 0,98 | 0 | 6,44 | 0 |
| PHOX2A | ATAACCTATTA | 1 | 596 | 607 | -404 | -393 | 1,49 | 0,01 | 12,6 | >0.1 | 3,45 | 5,2 | 0 | 1,01 | 0 | 1,44 | 0 |
| PHOX2A | TAACCTATTAA | 1 | 597 | 608 | -403 | -392 | 4,63 | 0,01 | 15,72 | >0.1 | 3,46 | 5,2 | 0 | 1,02 | 0 | 1,41 | 0 |
| PHOX2A | TAATGAAATTG | 1 | 605 | 616 | -395 | -384 | 8,31 | 0,001 | 19,48 | >0.1 | 3,51 | 5,22 | 0 | 1,02 | 0 | 1,41 | 0 |
| PHOX2A | TTATTTGATTC | 1 | 812 | 823 | -188 | -177 | 4,23 | 0,01 | 20,87 | >0.1 | 5,6 | 6,05 | 0 | 1,57 | 0 | 3,43 | 0 |
| PHOX2A | TGATTCACTTT | 1 | 817 | 828 | -183 | -172 | 4,37 | 0,01 | 21,2 | >0.1 | 5,6 | 6,08 | 0 | 1,6 | 0 | 3,55 | 0 |
| PHOX2A | TTAAAAAATTA | 1 | 826 | 837 | -174 | -163 | 2,7 | 0,01 | 19,65 | >0.1 | 5,6 | 6,14 | 0 | 1,67 | 0 | 3,55 | 0 |
| PHOX2A | AAATCTCATGA | 1 | 980 | 991 | -20 | -9 | 2,73 | 0,01 | 20,1 | >0.1 | 5,6 | 8,37 | 0 | 1,87 | 0 | 1,53 | 0 |
| PHOX2A | TCATGAGATTT | -1 | 980 | 991 | -20 | -9 | 1,57 | 0,01 | 18,94 | >0.1 | 5,6 | 8,37 | 0 | 1,87 | 0 | 1,53 | 0 |
| PHOX2A | TGAGCCAATAT | -1 | 880 | 891 | -120 | -109 | 2,98 | 0,01 | 20,08 | >0.1 | 5,6 | 6,54 | 0 | 1,9 | 0 | 3,07 | 0 |
| PHOX2A | AAAGTGAATCA | -1 | 817 | 828 | -183 | -172 | 2,92 | 0,01 | 19,75 | >0.1 | 5,6 | 6,08 | 0 | 1,6 | 0 | 3,55 | 0 |
| PHOX2A | GAATCAAATAA | -1 | 812 | 823 | -188 | -177 | 5,47 | 0,01 | 22,11 | >0.1 | 5,6 | 6,05 | 0 | 1,57 | 0 | 3,43 | 0 |
| PHOX2A | AAATGTTATAA | -1 | 687 | 698 | -313 | -302 | 3,62 | 0,01 | 17,63 | >0.1 | 4,45 | 5,48 | 0 | 1,13 | 0 | 2,95 | 0 |
| PHOX2A | CAATTTCATTA | -1 | 605 | 616 | -395 | -384 | 8,78 | 0,001 | 19,95 | >0.1 | 3,51 | 5,22 | 0 | 1,02 | 0 | 1,41 | 0 |
| PHOX2A | TTATGGAATTC | -1 | 493 | 504 | -507 | -496 | 2,82 | 0,01 | 18,13 | >0.1 | 2,93 | 4,95 | 0 | 0,98 | 0 | 6,44 | 0 |
| PHOX2A | AAAGCTCATTT | -1 | 66 | 77 | -934 | -923 | 2,9 | 0,01 | 13,01 | >0.1 | 3,92 | 4,32 | 0 | 1,88 | 0 | 0 | 0 |
| HOXD3 | AATAGGTAAGTGAAGG | 1 | 0 | 16 | -1000 | -984 | 2,2 | 0,01 | 15,65 | >0.1 | 5,53 | 4,26 | 0 | 1,86 | 1,79 | 0 | 0 |
| HOXD3 | AATAGGAAATGAGCTT | 1 | 60 | 76 | -940 | -924 | 1,76 | 0,01 | 11,91 | >0.1 | 3,96 | 4,32 | 0 | 1,88 | 0 | 0 | 0 |
| HOXD3 | CCATGGAAATTAAAAC | 1 | 149 | 165 | -851 | -835 | 1,93 | 0,01 | 13,16 | >0.1 | 3,27 | 4,41 | 0 | 1,9 | 0 | 1,65 | 0 |
| HOXD3 | ACCTATTAATGAAATT | 1 | 599 | 615 | -401 | -385 | 4,03 | 0,01 | 15,18 | >0.1 | 3,49 | 5,22 | 0 | 1,02 | 0 | 1,41 | 0 |
| HOXD3 | TTAATGAAATTGTCTG | 1 | 604 | 620 | -396 | -380 | 1,74 | 0,01 | 13,02 | >0.1 | 3,52 | 5,24 | 0 | 1,03 | 0 | 1,5 | 0 |
| HOXD3 | TTATCTTCATGATCCC | 1 | 641 | 657 | -359 | -343 | 4,62 | 0,01 | 18,56 | >0.1 | 3,83 | 5,34 | 0 | 1,07 | 0 | 3,7 | 0 |
| HOXD3 | TTTAAAAAATTATTCT | 1 | 825 | 841 | -175 | -159 | 3,1 | 0,01 | 19,92 | >0.1 | 5,53 | 6,16 | 0 | 1,7 | 0 | 3,43 | 0 |
| HOXD3 | TTGCTGTAACTGCTGC | 1 | 1083 | 1099 | 83 | 99 | 2,62 | 0,01 | 20,98 | >0.1 | 5,53 | 6,97 | 0 | 1,85 | 1,79 | 2,21 | 0 |
| HOXD3 | AAGAAGCAAGTGAACT | -1 | 1175 | 1191 | 175 | 191 | 2,09 | 0,01 | 20,53 | >0.1 | 5,53 | 6,12 | 0 | 1,62 | 1,79 | 3,37 | 0 |
| HOXD3 | ATGTTATAATTTTACA | -1 | 680 | 696 | -320 | -304 | 1,26 | 0,01 | 15,29 | >0.1 | 4,36 | 5,47 | 0 | 1,13 | 0 | 3,07 | 0 |
| HOXD3 | TAGGGATCATGAAGAT | -1 | 643 | 659 | -357 | -341 | 5,6 | 0,01 | 19,71 | >0.1 | 3,85 | 5,35 | 0 | 1,07 | 0 | 3,83 | 0 |
| HOXD3 | TTCAGACAATTTCATT | -1 | 606 | 622 | -394 | -378 | 3,14 | 0,01 | 14,49 | >0.1 | 3,54 | 5,24 | 0 | 1,03 | 0 | 1,55 | 0 |
| HOXD3 | ACAATTTCATTAATAG | -1 | 601 | 617 | -399 | -383 | 3,34 | 0,01 | 14,51 | >0.1 | 3,5 | 5,23 | 0 | 1,03 | 0 | 1,41 | 0 |
| HOXD3 | CAGTTTTAATTTCCAT | -1 | 151 | 167 | -849 | -833 | 6,05 | 0,01 | 17,27 | >0.1 | 3,26 | 4,41 | 0 | 1,9 | 0 | 1,65 | 0 |
| HOXD3 | CTCCTTCACTTACCTA | -1 | 2 | 18 | -998 | -982 | 1,3 | 0,01 | 14,75 | >0.1 | 5,53 | 4,26 | 0 | 1,86 | 1,79 | 0 | 0 |
| HOXB13 | ATTTTAAAAA | 1 | 135 | 145 | -865 | -855 | 1,56 | 0,01 | 12,79 | >0.1 | 3,37 | 4,39 | 0 | 1,89 | 0 | 1,57 | 0 |
| HOXB13 | GAAATTAAAA | 1 | 154 | 164 | -846 | -836 | 4,35 | 0,01 | 15,57 | >0.1 | 3,26 | 4,41 | 0 | 1,9 | 0 | 1,65 | 0 |
| HOXB13 | AAATTAAAAC | 1 | 155 | 165 | -845 | -835 | 1,94 | 0,01 | 14,11 | >0.1 | 3,25 | 4,41 | 0 | 1,9 | 0 | 2,6 | 0 |
| HOXB13 | TCTATTAAAA | 1 | 288 | 298 | -712 | -702 | 4,83 | 0,01 | 18,46 | >0.1 | 2,72 | 4,59 | 0 | 1,24 | 0 | 5,09 | 0 |
| HOXB13 | CTATTAAAAA | 1 | 289 | 299 | -711 | -701 | 7,8 | 0,001 | 21,42 | >0.1 | 2,71 | 4,59 | 0 | 1,24 | 0 | 5,09 | 0 |
| HOXB13 | TCCATAAAGA | 1 | 497 | 507 | -503 | -493 | 5,91 | 0,01 | 21,23 | >0.1 | 2,95 | 4,96 | 0 | 0,98 | 0 | 6,44 | 0 |
| HOXB13 | GAAATAAAAA | 1 | 576 | 586 | -424 | -414 | 7,54 | 0,001 | 17,01 | >0.1 | 3,32 | 5,14 | 0 | 1 | 0 | 0 | 0 |
| HOXB13 | AAAATAAATG | 1 | 582 | 592 | -418 | -408 | 1,26 | 0,01 | 10,78 | >0.1 | 3,36 | 5,16 | 0 | 1 | 0 | 0 | 0 |
| HOXB13 | CCTATTAATG | 1 | 600 | 610 | -400 | -390 | 4,56 | 0,01 | 15,62 | >0.1 | 3,47 | 5,21 | 0 | 1,02 | 0 | 1,36 | 0 |
| HOXB13 | TTAATGAAAT | 1 | 604 | 614 | -396 | -386 | 0,95 | 0,01 | 12,1 | >0.1 | 3,5 | 5,22 | 0 | 1,02 | 0 | 1,41 | 0 |
| HOXB13 | ACTGTAAAAT | 1 | 678 | 688 | -322 | -312 | 2,36 | 0,01 | 16,47 | >0.1 | 4,27 | 5,45 | 0 | 1,11 | 0 | 3,28 | 0 |
| HOXB13 | ATTATAACAT | 1 | 686 | 696 | -314 | -304 | 0,87 | 0,01 | 14,84 | >0.1 | 4,42 | 5,47 | 0 | 1,13 | 0 | 2,95 | 0 |
| HOXB13 | GCCATAAAGT | 1 | 758 | 768 | -242 | -232 | 7,34 | 0,001 | 26,12 | >0.1 | 5,61 | 5,76 | 0 | 1,31 | 0 | 6,1 | 0 |
| HOXB13 | CACTTTAAAA | 1 | 822 | 832 | -178 | -168 | 1,05 | 0,01 | 18,09 | >0.1 | 5,61 | 6,1 | 0 | 1,63 | 0 | 3,7 | 0 |
| HOXB13 | ACTTTAAAAA | 1 | 823 | 833 | -177 | -167 | 4,06 | 0,01 | 21,11 | >0.1 | 5,61 | 6,11 | 0 | 1,64 | 0 | 3,7 | 0 |
| HOXB13 | ACAGTAAACA | -1 | 897 | 907 | -103 | -93 | 3,69 | 0,01 | 20,73 | >0.1 | 5,61 | 6,68 | 0 | 1,89 | 0 | 2,86 | 0 |
| HOXB13 | CCAATATAAA | -1 | 877 | 887 | -123 | -113 | 7,7 | 0,001 | 24,66 | >0.1 | 5,61 | 6,5 | 0 | 1,9 | 0 | 2,95 | 0 |
| HOXB13 | CAAATCAAAA | -1 | 748 | 758 | -252 | -242 | 3,49 | 0,01 | 22,01 | >0.1 | 5,61 | 5,71 | 0 | 1,28 | 0 | 5,92 | 0 |
| HOXB13 | TTCATTAATA | -1 | 602 | 612 | -398 | -388 | 0,75 | 0,01 | 11,88 | >0.1 | 3,49 | 5,21 | 0 | 1,02 | 0 | 1,41 | 0 |
| HOXB13 | TTTTTAAAAT | -1 | 135 | 145 | -865 | -855 | 0,03 | 0,01 | 11,26 | >0.1 | 3,37 | 4,39 | 0 | 1,89 | 0 | 1,57 | 0 |
| HOXB13 | AACTTAAAAA | -1 | 75 | 85 | -925 | -915 | 0,42 | 0,01 | 10,45 | >0.1 | 3,83 | 4,32 | 0 | 1,88 | 0 | 0 | 0 |
| E2F7 | GGCTCACGCCTGTA | 1 | 189 | 203 | -811 | -797 | -20,07 | 0,01 | -6,69 | >0.1 | 3,08 | 4,46 | 0 | 1,71 | 0 | 4,13 | 0 |
| E2F7 | TAGTCCCAGCTACT | 1 | 335 | 349 | -665 | -651 | -28,35 | 0,01 | -14,06 | >0.1 | 2,58 | 4,67 | 0 | 1,13 | 0 | 5,92 | 0 |
| E2F7 | AGATCGCGCCACTG | 1 | 407 | 421 | -593 | -579 | -20,18 | 0,01 | -6,19 | >0.1 | 2,65 | 4,79 | 0 | 1,03 | 0 | 5,52 | 0 |

| | | | | | | | | | | | | | | | | | |
|---|---|---|---|---|---|---|---|---|---|---|---|---|---|---|---|---|---|
| E2F7 | AGTTTCTGGGAATA | 1 | 555 | 569 | -445 | -431 | -28,22 | 0,01 | -18,91 | >0.1 | 3,22 | 5,1 | 0 | 1 | 0 | 0 | 0 |
| E2F7 | ATTTTCCGTGTATC | 1 | 694 | 708 | -306 | -292 | -25,94 | 0,01 | -10,65 | >0.1 | 5,56 | 5,52 | 0 | 1,15 | 0 | 3,07 | 0 |
| E2F7 | CTTTCTAGGAAAAT | 1 | 716 | 730 | -284 | -270 | -27,9 | 0,01 | -10,56 | >0.1 | 5,56 | 5,6 | 0 | 1,2 | 0 | 4,99 | 0 |
| E2F7 | GATTCACTTTAAAA | 1 | 818 | 832 | -182 | -168 | -28,01 | 0,01 | -11,02 | >0.1 | 5,56 | 6,1 | 0 | 1,63 | 0 | 3,7 | 0 |
| E2F7 | GCTCAGCGCCCAAC | 1 | 940 | 954 | -60 | -46 | -29,18 | 0,01 | -13,63 | >0.1 | 5,56 | 7,29 | 0 | 1,88 | 0 | 0,83 | 0 |
| E2F7 | AGTTACAGCAACAA | -1 | 1080 | 1094 | 80 | 94 | -27,67 | 0,01 | -9,24 | >0.1 | 5,56 | 7,01 | 0 | 1,85 | 1,79 | 2,21 | 0 |
| E2F7 | CTTTCCCTAGACTA | -1 | 999 | 1013 | -1 | 13 | -21,15 | 0,01 | -3,14 | >0.1 | 5,56 | 9,12 | 0 | 1,87 | 0 | 1,46 | 0 |
| E2F7 | TTTTCTCTCTTATC | -1 | 969 | 983 | -31 | -17 | -28,91 | 0,01 | -11,82 | >0.1 | 5,56 | 8,02 | 0 | 1,87 | 0 | 1,65 | 0 |
| E2F7 | CTCTCCTGTCACTT | -1 | 764 | 778 | -236 | -222 | -24,07 | 0,01 | -5,44 | >0.1 | 5,56 | 5,81 | 0 | 1,35 | 0 | 5,92 | 0 |
| E2F7 | CTTTATGGCCAAAT | -1 | 753 | 767 | -247 | -233 | -15,11 | 0,001 | 3,83 | >0.1 | 5,56 | 5,75 | 0 | 1,31 | 0 | 6,32 | 0 |
| E2F7 | ATTTTCCTAGAAAG | -1 | 716 | 730 | -284 | -270 | -26,95 | 0,01 | -9,61 | >0.1 | 5,56 | 5,6 | 0 | 1,2 | 0 | 4,99 | 0 |
| E2F7 | GTTTTCAGACAATT | -1 | 611 | 625 | -389 | -375 | -18,62 | 0,01 | -7,27 | >0.1 | 3,57 | 5,25 | 0 | 1,03 | 0 | 1,5 | 0 |
| E2F7 | TATTCCCAGAAACT | -1 | 555 | 569 | -445 | -431 | -21,91 | 0,01 | -12,6 | >0.1 | 3,22 | 5,1 | 0 | 1 | 0 | 0 | 0 |
| E2F7 | TTTTCCCTTCCAAA | -1 | 525 | 539 | -475 | -461 | -11,06 | 0,001 | 1,46 | >0.1 | 3,07 | 5,03 | 0 | 0,99 | 0 | 3,43 | 0 |
| E2F7 | TTTCCCTTCCAAAT | -1 | 524 | 538 | -476 | -462 | -9,57 | 0,001 | 2,94 | >0.1 | 3,07 | 5,03 | 0 | 0,99 | 0 | 3,43 | 0 |
| E2F7 | GGTTCACGCCATTC | -1 | 367 | 381 | -633 | -619 | -12,03 | 0,001 | 2,46 | >0.1 | 2,54 | 4,72 | 0 | 1,08 | 0 | 6,15 | 0 |
| E2F7 | TTCTCCTGCCTCAG | -1 | 356 | 370 | -644 | -630 | -18,5 | 0,01 | -3,93 | >0.1 | 2,52 | 4,7 | 0 | 1,09 | 0 | 6,26 | 0 |
| E2F7 | GGTGCCCACCACCA | -1 | 319 | 333 | -681 | -667 | -18,41 | 0,01 | -4,42 | >0.1 | 2,62 | 4,64 | 0 | 1,16 | 0 | 5,56 | 0 |
| E2F7 | CTCGGCCTCCCAAA | -1 | 213 | 227 | -787 | -773 | -29,03 | 0,01 | -14,38 | >0.1 | 2,98 | 4,49 | 0 | 1,55 | 0 | 5,63 | 0 |
| E2F7 | TGTGCCCAGCCATT | -1 | 173 | 187 | -827 | -813 | -29,32 | 0,01 | -16,64 | >0.1 | 3,15 | 4,44 | 0 | 1,85 | 0 | 3,24 | 0 |
| E2F7 | GTGCCCAGCCATTT | -1 | 172 | 186 | -828 | -814 | -21,24 | 0,01 | -8,54 | >0.1 | 3,16 | 4,44 | 0 | 1,86 | 0 | 3,24 | 0 |
| THAP1 | CCGTCCTGA | 1 | 258 | 267 | -742 | -733 | 5,52 | 0,01 | 18,84 | >0.1 | 2,82 | 4,54 | 0 | 1,33 | 0 | 4,63 | 0 |
| THAP1 | GCTCCAGCC | 1 | 422 | 431 | -578 | -569 | 4,9 | 0,01 | 18,83 | >0.1 | 2,69 | 4,81 | 0 | 1,02 | 0 | 5,42 | 0 |
| THAP1 | TTGCCCAAG | 1 | 541 | 550 | -459 | -450 | 6,48 | 0,01 | 15,66 | >0.1 | 3,14 | 5,06 | 0 | 0,98 | 0 | 0 | 0 |
| THAP1 | TGGCCATAA | 1 | 756 | 765 | -244 | -235 | 4,45 | 0,01 | 23,44 | >0.1 | 5,62 | 5,75 | 0 | 1,3 | 0 | 6,32 | 0 |
| THAP1 | CTGTCCTCT | 1 | 857 | 866 | -143 | -134 | 5,37 | 0,01 | 22,19 | >0.1 | 5,62 | 6,34 | 0 | 1,9 | 0 | 2,95 | 0 |
| THAP1 | TCTCCAGGA | 1 | 863 | 872 | -137 | -128 | 6,06 | 0,01 | 22,72 | >0.1 | 5,62 | 6,38 | 0 | 1,9 | 0 | 2,75 | 0 |
| THAP1 | GCGCCCAAC | 1 | 945 | 954 | -55 | -46 | 5,88 | 0,01 | 22,47 | >0.1 | 5,62 | 7,29 | 0 | 1,88 | 0 | 1,8 | 0 |
| THAP1 | GAGCCAGGA | -1 | 1061 | 1070 | 61 | 70 | 5,45 | 0,01 | 22,42 | >0.1 | 5,62 | 7,33 | 0 | 1,86 | 0 | 2,16 | 0 |
| THAP1 | TTTCCCTAG | -1 | 1003 | 1012 | 3 | 12 | 4,58 | 0,01 | 22,66 | >0.1 | 5,62 | 9,12 | 0 | 1,87 | 0 | 1,46 | 0 |
| THAP1 | ATTCCCAGA | -1 | 559 | 568 | -441 | -432 | 5,73 | 0,01 | 15,05 | >0.1 | 3,23 | 5,1 | 0 | 0,99 | 0 | 0 | 0 |
| THAP1 | TTTCCCTTC | -1 | 529 | 538 | -471 | -462 | 6,49 | 0,01 | 19,13 | >0.1 | 3,08 | 5,03 | 0 | 0,98 | 0 | 3,55 | 0 |
| THAP1 | TTGCCCTTG | -1 | 507 | 516 | -493 | -484 | 7,7 | 0,01 | 22,33 | >0.1 | 2,98 | 4,98 | 0 | 0,98 | 0 | 5,68 | 0 |
| THAP1 | TCGCCCAGG | -1 | 429 | 438 | -571 | -562 | 6,51 | 0,01 | 20,85 | >0.1 | 2,71 | 4,82 | 0 | 1,01 | 0 | 5,8 | 0 |
| THAP1 | CCTCCCAGG | -1 | 379 | 388 | -621 | -612 | 5,42 | 0,01 | 20,31 | >0.1 | 2,57 | 4,73 | 0 | 1,06 | 0 | 6,53 | 0 |
| THAP1 | ACGCCATTC | -1 | 367 | 376 | -633 | -624 | 4,31 | 0,01 | 19,24 | >0.1 | 2,53 | 4,71 | 0 | 1,08 | 0 | 6,61 | 0 |
| THAP1 | CCTCCCGAG | -1 | 347 | 356 | -653 | -644 | 5,68 | 0,01 | 20,53 | >0.1 | 2,55 | 4,68 | 0 | 1,11 | 0 | 6,52 | 0 |
| THAP1 | GTGCCCACC | -1 | 323 | 332 | -677 | -668 | 8,53 | 0,001 | 22,5 | >0.1 | 2,62 | 4,64 | 0 | 1,15 | 0 | 5,56 | 0 |
| THAP1 | CCACCACCA | -1 | 319 | 328 | -681 | -672 | 4,72 | 0,01 | 19,39 | >0.1 | 2,63 | 4,63 | 0 | 1,16 | 0 | 6,26 | 0 |
| THAP1 | ACGCCCAGC | -1 | 311 | 320 | -689 | -680 | 5,75 | 0,01 | 20,33 | >0.1 | 2,65 | 4,62 | 0 | 1,18 | 0 | 6,13 | 0 |
| THAP1 | CTGCCCAAC | -1 | 226 | 235 | -774 | -765 | 8,57 | 0,001 | 22,73 | >0.1 | 2,93 | 4,5 | 0 | 1,47 | 0 | 5,26 | 0 |
| THAP1 | CCTCCCAAA | -1 | 213 | 222 | -787 | -778 | 6,73 | 0,01 | 20,34 | >0.1 | 2,98 | 4,48 | 0 | 1,55 | 0 | 4,59 | 0 |
| THAP1 | GTGCCCAGC | -1 | 177 | 186 | -823 | -814 | 7,74 | 0,01 | 20,37 | >0.1 | 3,15 | 4,44 | 0 | 1,81 | 0 | 3,24 | 0 |
| THAP1 | CAGCCATTT | -1 | 172 | 181 | -828 | -819 | 4,45 | 0,01 | 17,26 | >0.1 | 3,17 | 4,43 | 0 | 1,86 | 0 | 3,34 | 0 |
| E2F4 | ACTCGGGAGGC | 1 | 346 | 357 | -654 | -643 | 0,11 | 0,001 | 14,97 | >0.1 | 2,55 | 4,68 | 0 | 1,11 | 0 | 6,52 | 0 |
| E2F4 | AGGCAGGAGAA | 1 | 359 | 370 | -641 | -630 | 0,7 | 0,001 | 15,37 | >0.1 | 2,52 | 4,7 | 0 | 1,09 | 0 | 6,36 | 0 |
| E2F4 | TGGCGTGAACC | 1 | 370 | 381 | -630 | -619 | 1,52 | 0,001 | 16,12 | >0.1 | 2,54 | 4,72 | 0 | 1,07 | 0 | 6,27 | 0 |
| E2F4 | ACCTGGGAGGC | 1 | 378 | 389 | -622 | -611 | -7,28 | 0,01 | 7,61 | >0.1 | 2,57 | 4,73 | 0 | 1,06 | 0 | 6,53 | 0 |
| E2F4 | GGGAGGCAGAG | 1 | 382 | 393 | -618 | -607 | -1,85 | 0,01 | 12,43 | >0.1 | 2,58 | 4,74 | 0 | 1,06 | 0 | 5,91 | 0 |
| E2F4 | TCGCGCCACTG | 1 | 410 | 421 | -590 | -579 | -8,41 | 0,01 | 5,71 | >0.1 | 2,65 | 4,79 | 0 | 1,03 | 0 | 5,65 | 0 |
| E2F4 | GGGCGACAGAC | 1 | 432 | 443 | -568 | -557 | 4,79 | 0,001 | 19,02 | >0.1 | 2,72 | 4,83 | 0 | 1,01 | 0 | 5,67 | 0 |
| E2F4 | GGAAGGGAAAA | 1 | 528 | 539 | -472 | -461 | -10,35 | 0,01 | 2,29 | >0.1 | 3,08 | 5,03 | 0 | 0,98 | 0 | 3,55 | 0 |
| E2F4 | CAGCGCCCAAC | 1 | 943 | 954 | -57 | -46 | -8,5 | 0,01 | 8,06 | >0.1 | 5,6 | 7,29 | 0 | 1,88 | 0 | 1,8 | 0 |
| E2F4 | AAGAGAGAAAA | 1 | 972 | 983 | -28 | -17 | -8,66 | 0,01 | 8,52 | >0.1 | 5,6 | 8,01 | 0 | 1,87 | 0 | 1,7 | 0 |
| E2F4 | AAGCCGAAGAC | 1 | 1152 | 1163 | 152 | 163 | -9,53 | 0,01 | 7,14 | >0.1 | 4,5 | 6,28 | 0 | 1,84 | 1,79 | 2,26 | 0 |
| E2F4 | AGCCGAAGACC | 1 | 1153 | 1164 | 153 | 164 | -10,77 | 0,01 | 5,94 | >0.1 | 4,47 | 6,27 | 0 | 1,83 | 1,79 | 2,34 | 0 |
| E2F4 | GAGCAGCAGTT | -1 | 1090 | 1101 | 90 | 101 | -9,43 | 0,01 | 8,96 | >0.1 | 5,6 | 6,88 | 0 | 1,85 | 1,79 | 2,27 | 0 |
| E2F4 | AGGCTGAGAAG | -1 | 1070 | 1081 | 70 | 81 | -9,5 | 0,01 | 9,19 | >0.1 | 5,6 | 7,17 | 0 | 1,85 | 1,79 | 2,27 | 0 |
| E2F4 | GGCTGAGAAGG | -1 | 1069 | 1080 | 69 | 80 | -9,15 | 0,01 | 9,5 | >0.1 | 5,6 | 7,19 | 0 | 1,85 | 1,79 | 2,21 | 0 |

| | | | | | | | | | | | | | | | | | |
|---|---|---|---|---|---|---|---|---|---|---|---|---|---|---|---|---|---|
| E2F4 | AGGAGCCAGGA | -1 | 1061 | 1072 | 61 | 72 | -7,16 | 0,01 | 11,58 | >0.1 | 5,6 | 7,33 | 0 | 1,86 | 1,79 | 2,16 | 0 |
| E2F4 | GAGCCAGGAAG | -1 | 1059 | 1070 | 59 | 70 | -9,33 | 0,01 | 7,66 | >0.1 | 5,6 | 7,37 | 0 | 1,86 | 0 | 2,16 | 0 |
| E2F4 | AGCCAGGAAGA | -1 | 1058 | 1069 | 58 | 69 | -5,67 | 0,01 | 11,39 | >0.1 | 5,6 | 7,39 | 0 | 1,86 | 0 | 2,21 | 0 |
| E2F4 | GGGCGCTGAGC | -1 | 940 | 951 | -60 | -49 | -2,96 | 0,01 | 12,58 | >0.1 | 5,6 | 7,23 | 0 | 1,88 | 0 | 0,83 | 0 |
| E2F4 | GAGCAGGAAAG | -1 | 933 | 944 | -67 | -56 | 0,66 | 0,001 | 16,18 | >0.1 | 5,6 | 7,12 | 0 | 1,88 | 0 | 0,93 | 0 |
| E2F4 | TGGAGAGGACA | -1 | 858 | 869 | -142 | -131 | -9,31 | 0,01 | 7,41 | >0.1 | 5,6 | 6,36 | 0 | 1,9 | 0 | 2,86 | 0 |
| E2F4 | ACACGGAAAAT | -1 | 694 | 705 | -306 | -295 | -7,9 | 0,01 | 7,41 | >0.1 | 5,6 | 5,5 | 0 | 1,14 | 0 | 3,07 | 0 |
| E2F4 | AGGCTGGAGCA | -1 | 421 | 432 | -579 | -568 | 0,16 | 0,001 | 14,09 | >0.1 | 2,69 | 4,81 | 0 | 1,02 | 0 | 5,42 | 0 |
| E2F4 | TGGCGCGATCT | -1 | 407 | 418 | -593 | -582 | -1,1 | 0,01 | 13,01 | >0.1 | 2,65 | 4,78 | 0 | 1,03 | 0 | 5,65 | 0 |
| E2F4 | TAGCTGGGACT | -1 | 336 | 347 | -664 | -653 | -6,83 | 0,01 | 7,45 | >0.1 | 2,58 | 4,66 | 0 | 1,13 | 0 | 5,92 | 0 |
| E2F4 | AGGCGTGAGCC | -1 | 189 | 200 | -811 | -800 | 0,51 | 0,001 | 13,8 | >0.1 | 3,08 | 4,45 | 0 | 1,71 | 0 | 4,04 | 0 |
| E2F3 | AGGAAATGAGCTTTTTTA | 1 | 63 | 81 | -937 | -919 | -12,14 | 0,001 | -2,02 | >0.1 | 3,92 | 4,32 | 0 | 1,88 | 0 | 0 | 0 |
| E2F3 | GAGAATGGCGTGAACCTG | 1 | 365 | 383 | -635 | -617 | -12,17 | 0,001 | 2,33 | >0.1 | 2,54 | 4,72 | 0 | 1,08 | 0 | 6,15 | 0 |
| E2F3 | CGAGATCGCGCCACTGTG | 1 | 405 | 423 | -595 | -577 | -5,4 | 0,001 | 8,6 | >0.1 | 2,65 | 4,79 | 0 | 1,04 | 0 | 5,52 | 0 |
| E2F3 | GAAAATGTTGCCCAAGTA | 1 | 534 | 552 | -466 | -448 | -14,38 | 0,01 | -3,47 | >0.1 | 3,12 | 5,06 | 0 | 0,99 | 0 | 1,73 | 0 |
| E2F3 | AAATGTTGCCCAAGTAGA | 1 | 536 | 554 | -464 | -446 | -16,47 | 0,01 | -7,28 | >0.1 | 3,13 | 5,07 | 0 | 0,99 | 0 | 0 | 0 |
| E2F3 | AATAAATGTGAGATAACC | 1 | 584 | 602 | -416 | -398 | -18,86 | 0,01 | -9,26 | >0.1 | 3,4 | 5,19 | 0 | 1,02 | 0 | 0 | 0 |
| E2F3 | ACATTTTCCGTGTATCTT | 1 | 692 | 710 | -308 | -290 | -23,9 | 0,01 | -8,64 | >0.1 | 5,51 | 5,52 | 0 | 1,16 | 0 | 3,07 | 0 |
| E2F3 | TGATTTGGCCATAAAGTG | 1 | 751 | 769 | -249 | -231 | -22,72 | 0,01 | -3,81 | >0.1 | 5,51 | 5,76 | 0 | 1,32 | 0 | 6,32 | 0 |
| E2F3 | GATAAGAGAGAAAATCTC | 1 | 969 | 987 | -31 | -13 | -23,54 | 0,01 | -6,33 | >0.1 | 5,51 | 8,18 | 0 | 1,87 | 0 | 1,65 | 0 |
| E2F3 | AAGATACACGGAAAATGT | -1 | 692 | 710 | -308 | -290 | -24,65 | 0,01 | -9,39 | >0.1 | 5,51 | 5,52 | 0 | 1,16 | 0 | 3,07 | 0 |
| E2F3 | GGTTATCTCACATTTATT | -1 | 584 | 602 | -416 | -398 | -23,23 | 0,01 | -13,63 | >0.1 | 3,4 | 5,19 | 0 | 1,02 | 0 | 0 | 0 |
| E2F3 | TACTTGGGCAACATTTTC | -1 | 534 | 552 | -466 | -448 | -21,92 | 0,01 | -11,01 | >0.1 | 3,12 | 5,06 | 0 | 0,99 | 0 | 1,73 | 0 |
| E2F3 | CACAGTGGCGCGATCTCG | -1 | 405 | 423 | -595 | -577 | -5,01 | 0,001 | 8,99 | >0.1 | 2,65 | 4,79 | 0 | 1,04 | 0 | 5,52 | 0 |
| E2F3 | CAGGTTCACGCCATTCTC | -1 | 365 | 383 | -635 | -617 | -20,39 | 0,01 | -5,89 | >0.1 | 2,54 | 4,72 | 0 | 1,08 | 0 | 6,15 | 0 |
| E2F3 | TAAAAAAGCTCATTTCCT | -1 | 63 | 81 | -937 | -919 | -12,2 | 0,001 | -2,08 | >0.1 | 3,92 | 4,32 | 0 | 1,88 | 0 | 0 | 0 |
| E2F3 | TCTTTGAGTGCCTCCTTC | -1 | 17 | 35 | -983 | -965 | -24,72 | 0,01 | -13,07 | >0.1 | 5,51 | 4,28 | 0 | 1,87 | 0 | 0 | 0 |
| E2F2 | GGAAATGAGCTTTTTT | 1 | 64 | 80 | -936 | -920 | -11,18 | 0,01 | -1,07 | >0.1 | 3,92 | 4,32 | 0 | 1,88 | 0 | 0 | 0 |
| E2F2 | AGAATGGCGTGAACCT | 1 | 366 | 382 | -634 | -618 | -6,68 | 0,01 | 7,81 | >0.1 | 2,54 | 4,72 | 0 | 1,08 | 0 | 6,15 | 0 |
| E2F2 | GAGATCGCGCCACTGT | 1 | 406 | 422 | -594 | -578 | -2,23 | 0,001 | 11,77 | >0.1 | 2,65 | 4,79 | 0 | 1,03 | 0 | 5,52 | 0 |
| E2F2 | AAAATGTTGCCCAAGT | 1 | 535 | 551 | -465 | -449 | -3,09 | 0,001 | 7,82 | >0.1 | 3,12 | 5,06 | 0 | 0,99 | 0 | 1,73 | 0 |
| E2F2 | ATAAATGTGAGATAAC | 1 | 585 | 601 | -415 | -399 | -13,17 | 0,01 | -3,58 | >0.1 | 3,4 | 5,18 | 0 | 1,01 | 0 | 0 | 0 |
| E2F2 | AAAATTATAACATTTT | 1 | 683 | 699 | -317 | -301 | -11,13 | 0,01 | 2,86 | >0.1 | 4,42 | 5,48 | 0 | 1,14 | 0 | 2,95 | 0 |
| E2F2 | AAAAGTACCGGTTTTG | 1 | 737 | 753 | -263 | -247 | -14,24 | 0,01 | 4,17 | >0.1 | 5,53 | 5,69 | 0 | 1,27 | 0 | 5,92 | 0 |
| E2F2 | AAATCTGTTACATATC | 1 | 842 | 858 | -158 | -142 | -13,88 | 0,01 | 2,73 | >0.1 | 5,53 | 6,28 | 0 | 1,84 | 0 | 2,95 | 0 |
| E2F2 | TTCCTGGCTCCTTCTC | 1 | 1060 | 1076 | 60 | 76 | -13,3 | 0,01 | 5,51 | >0.1 | 5,53 | 7,35 | 0 | 1,86 | 1,79 | 2,27 | 0 |
| E2F2 | TAAAGTGAATCAAATA | -1 | 813 | 829 | -187 | -171 | -13,5 | 0,01 | 3,28 | >0.1 | 5,53 | 6,08 | 0 | 1,61 | 0 | 3,55 | 0 |
| E2F2 | AAAACCGGTACTTTTG | -1 | 736 | 752 | -264 | -248 | -11,1 | 0,01 | 7,3 | >0.1 | 5,53 | 5,69 | 0 | 1,26 | 0 | 5,92 | 0 |
| E2F2 | AAAATGTTATAATTTT | -1 | 683 | 699 | -317 | -301 | -13,46 | 0,01 | 0,53 | >0.1 | 4,42 | 5,48 | 0 | 1,14 | 0 | 2,95 | 0 |
| E2F2 | ACTTGGGCAACATTTT | -1 | 535 | 551 | -465 | -449 | -4,65 | 0,001 | 6,26 | >0.1 | 3,12 | 5,06 | 0 | 0,99 | 0 | 1,73 | 0 |
| E2F2 | ACAGTGGCGCGATCTC | -1 | 406 | 422 | -594 | -578 | 3,8 | 0,0001 | 17,8 | >0.1 | 2,65 | 4,79 | 0 | 1,03 | 0 | 5,52 | 0 |
| E2F2 | AGGTTCACGCCATTCT | -1 | 366 | 382 | -634 | -618 | -7,91 | 0,01 | 6,58 | >0.1 | 2,54 | 4,72 | 0 | 1,08 | 0 | 6,15 | 0 |
| E2F2 | ATAGAGACGGGGTTTC | -1 | 277 | 293 | -723 | -707 | -10,35 | 0,01 | 3,33 | >0.1 | 2,74 | 4,58 | 0 | 1,27 | 0 | 5,09 | 0 |
| E2F2 | AAAAAAGCTCATTTCC | -1 | 64 | 80 | -936 | -920 | -12,04 | 0,01 | -1,93 | >0.1 | 3,92 | 4,32 | 0 | 1,88 | 0 | 0 | 0 |
| E2F1 | GAAGGAGGCACT | 1 | 17 | 29 | -983 | -971 | -6,33 | 0,01 | 5,39 | >0.1 | 5,58 | 4,27 | 0 | 1,87 | 0 | 0 | 0 |
| E2F1 | GGAGGCACTCAA | 1 | 20 | 32 | -980 | -968 | -4,55 | 0,01 | 7,17 | >0.1 | 5,58 | 4,27 | 0 | 1,87 | 0 | 0 | 0 |
| E2F1 | TTGGGAGGCCGA | 1 | 214 | 226 | -786 | -774 | -3,19 | 0,01 | 11,45 | >0.1 | 2,98 | 4,49 | 0 | 1,54 | 0 | 5,63 | 0 |
| E2F1 | GGTGGTGGGCAC | 1 | 320 | 332 | -680 | -668 | -6,03 | 0,01 | 7,95 | >0.1 | 2,62 | 4,64 | 0 | 1,16 | 0 | 5,56 | 0 |
| E2F1 | GTGGGCACCTGT | 1 | 324 | 336 | -676 | -664 | -3,21 | 0,01 | 10,65 | >0.1 | 2,61 | 4,64 | 0 | 1,15 | 0 | 5,46 | 0 |
| E2F1 | AATGGCGTGAAC | 1 | 368 | 380 | -632 | -620 | -1,47 | 0,01 | 13,01 | >0.1 | 2,54 | 4,72 | 0 | 1,08 | 0 | 6,15 | 0 |
| E2F1 | CTGGGAGGCAGA | 1 | 380 | 392 | -620 | -608 | -3,87 | 0,01 | 10,52 | >0.1 | 2,57 | 4,74 | 0 | 1,06 | 0 | 6,02 | 0 |
| E2F1 | TGAGCCGAGATC | 1 | 400 | 412 | -600 | -588 | -5,58 | 0,01 | 8,62 | >0.1 | 2,63 | 4,77 | 0 | 1,04 | 0 | 5,76 | 0 |
| E2F1 | GATCGCGCCACT | 1 | 408 | 420 | -592 | -580 | 8,76 | 0,0001 | 22,75 | >0.1 | 2,65 | 4,79 | 0 | 1,03 | 0 | 5,52 | 0 |
| E2F1 | CTGGGCGACAGA | 1 | 430 | 442 | -570 | -558 | 4,71 | 0,001 | 18,93 | >0.1 | 2,72 | 4,83 | 0 | 1,01 | 0 | 5,67 | 0 |
| E2F1 | AATGTTGCCCAA | 1 | 537 | 549 | -463 | -451 | -5,66 | 0,01 | 5,24 | >0.1 | 3,12 | 5,05 | 0 | 0,99 | 0 | 1,73 | 0 |
| E2F1 | TTTGGCCATAAA | 1 | 754 | 766 | -246 | -234 | -2,98 | 0,01 | 15,98 | >0.1 | 5,58 | 5,75 | 0 | 1,31 | 0 | 6,32 | 0 |
| E2F1 | TTTCCTGCTCAG | 1 | 934 | 946 | -66 | -54 | -4,57 | 0,01 | 10,97 | >0.1 | 5,58 | 7,15 | 0 | 1,88 | 0 | 0,93 | 0 |
| E2F1 | CTCAGCGCCCAA | 1 | 941 | 953 | -59 | -47 | 8,25 | 0,0001 | 23,81 | >0.1 | 5,58 | 7,27 | 0 | 1,88 | 0 | 0,83 | 0 |
| E2F1 | CCTGGCTCCTTC | 1 | 1062 | 1074 | 62 | 74 | -4,76 | 0,01 | 14,06 | >0.1 | 5,58 | 7,31 | 0 | 1,86 | 1,79 | 2,27 | 0 |

| | | | | | | | | | | | | | | | | | |
|---|---|---|---|---|---|---|---|---|---|---|---|---|---|---|---|---|---|
| E2F1 | ACTGCTGCTCAG | 1 | 1091 | 1103 | 91 | 103 | -5,38 | 0,01 | 13,05 | >0.1 | 5,58 | 6,87 | 0 | 1,85 | 1,79 | 2,34 | 0 |
| E2F1 | CTTCGTGGTTAA | -1 | 1143 | 1155 | 143 | 155 | -5,96 | 0,01 | 13,2 | >0.1 | 5,58 | 6,35 | 0 | 1,84 | 3,22 | 2,16 | 0 |
| E2F1 | GAAGGAGCCAGG | -1 | 1062 | 1074 | 62 | 74 | -2,17 | 0,01 | 16,65 | >0.1 | 5,58 | 7,31 | 0 | 1,86 | 1,79 | 2,27 | 0 |
| E2F1 | TTGGGCGCTGAG | -1 | 941 | 953 | -59 | -47 | 7,84 | 0,0001 | 23,4 | >0.1 | 5,58 | 7,27 | 0 | 1,88 | 0 | 0,83 | 0 |
| E2F1 | TTGGGCAACATT | -1 | 537 | 549 | -463 | -451 | -6,31 | 0,01 | 4,59 | >0.1 | 3,12 | 5,05 | 0 | 0,99 | 0 | 1,73 | 0 |
| E2F1 | TCTGTCGCCCAG | -1 | 430 | 442 | -570 | -558 | 0,55 | 0,01 | 14,77 | >0.1 | 2,72 | 4,83 | 0 | 1,01 | 0 | 5,67 | 0 |
| E2F1 | AGTGGCGCGATC | -1 | 408 | 420 | -592 | -580 | 8,3 | 0,0001 | 22,29 | >0.1 | 2,65 | 4,79 | 0 | 1,03 | 0 | 5,52 | 0 |
| E2F1 | TGGCGCGATCTC | -1 | 406 | 418 | -594 | -582 | -5,86 | 0,01 | 8,25 | >0.1 | 2,65 | 4,78 | 0 | 1,03 | 0 | 5,65 | 0 |
| E2F1 | TCTGCCTCCCAG | -1 | 380 | 392 | -620 | -608 | -4,42 | 0,01 | 9,97 | >0.1 | 2,57 | 4,74 | 0 | 1,06 | 0 | 6,02 | 0 |
| E2F1 | GTTCACGCCATT | -1 | 368 | 380 | -632 | -620 | -1,86 | 0,01 | 12,62 | >0.1 | 2,54 | 4,72 | 0 | 1,08 | 0 | 6,15 | 0 |
| E2F1 | TCTCCTGCCTCA | -1 | 357 | 369 | -643 | -631 | -5,4 | 0,01 | 9,17 | >0.1 | 2,52 | 4,7 | 0 | 1,09 | 0 | 6,26 | 0 |
| E2F1 | ACAGGTGCCCAC | -1 | 324 | 336 | -676 | -664 | 1,01 | 0,01 | 14,87 | >0.1 | 2,61 | 4,64 | 0 | 1,15 | 0 | 5,46 | 0 |
| E2F1 | TCGGCCTCCCAA | -1 | 214 | 226 | -786 | -774 | -4,39 | 0,01 | 10,25 | >0.1 | 2,98 | 4,49 | 0 | 1,54 | 0 | 5,63 | 0 |
| E2F1 | TTGAGTGCCTCC | -1 | 20 | 32 | -980 | -968 | -2,29 | 0,01 | 9,43 | >0.1 | 5,58 | 4,27 | 0 | 1,87 | 0 | 0 | 0 |
| HMBOX1 | AATAGGTAAG | 1 | 0 | 10 | -1000 | -990 | 2,49 | 0,01 | 16,01 | >0.1 | 5,61 | 4,25 | 0 | 1,86 | 1,79 | 0 | 0 |
| HMBOX1 | AGAAGTCAGC | 1 | 32 | 42 | -968 | -958 | 1,35 | 0,01 | 11,94 | >0.1 | 4,44 | 4,28 | 0 | 1,87 | 0 | 0 | 0 |
| HMBOX1 | AGTAGTCATA | 1 | 125 | 135 | -875 | -865 | 3,07 | 0,01 | 14,47 | >0.1 | 3,43 | 4,38 | 0 | 1,89 | 0 | 1,7 | 0 |
| HMBOX1 | CCGAGTTGGG | 1 | 222 | 232 | -778 | -768 | 1,75 | 0,01 | 16,21 | >0.1 | 2,95 | 4,49 | 0 | 1,5 | 0 | 5,52 | 0 |
| HMBOX1 | GTGAGATAAC | 1 | 591 | 601 | -409 | -399 | 2,42 | 0,01 | 12,03 | >0.1 | 3,41 | 5,18 | 0 | 1,01 | 0 | 0 | 0 |
| HMBOX1 | CCTAGTTCTA | 1 | 655 | 665 | -345 | -335 | 1,75 | 0,01 | 15,84 | >0.1 | 3,95 | 5,37 | 0 | 1,08 | 0 | 3,7 | 0 |
| HMBOX1 | TTTGGTCACT | 1 | 671 | 681 | -329 | -319 | 2,09 | 0,01 | 16,32 | >0.1 | 4,16 | 5,42 | 0 | 1,1 | 0 | 3,55 | 0 |
| HMBOX1 | AGAGGTAAGG | 1 | 774 | 784 | -226 | -216 | 1,29 | 0,01 | 20,21 | >0.1 | 5,61 | 5,83 | 0 | 1,37 | 0 | 6,1 | 0 |
| HMBOX1 | CCTGCTCAGC | 1 | 937 | 947 | -63 | -53 | 1,8 | 0,01 | 17,36 | >0.1 | 5,61 | 7,17 | 0 | 1,88 | 0 | 0,9 | 0 |
| HMBOX1 | TTTAGTCTAG | 1 | 997 | 1007 | -3 | 7 | 2,07 | 0,01 | 20,01 | >0.1 | 5,61 | 8,96 | 0 | 1,87 | 0 | 1,5 | 0 |
| HMBOX1 | ACAAGTTTAA | 1 | 1137 | 1147 | 137 | 147 | 3,87 | 0,01 | 24,01 | >0.1 | 5,61 | 6,4 | 0 | 1,84 | 3,22 | 3,07 | 0 |
| HMBOX1 | GCAAGTGAAC | -1 | 1176 | 1186 | 176 | 186 | 1,81 | 0,01 | 20,22 | >0.1 | 5,61 | 6,11 | 0 | 1,62 | 1,79 | 3,28 | 0 |
| HMBOX1 | CGTGGTTAAA | -1 | 1142 | 1152 | 142 | 152 | 8,75 | 0,001 | 29,15 | >0.1 | 5,61 | 6,36 | 0 | 1,84 | 3,22 | 3,37 | 0 |
| HMBOX1 | AGCAGTTACA | -1 | 1087 | 1097 | 87 | 97 | 4,34 | 0,01 | 22,73 | >0.1 | 5,61 | 6,92 | 0 | 1,85 | 1,79 | 2,21 | 0 |
| HMBOX1 | CACAGTAAAC | -1 | 898 | 908 | -102 | -92 | 3,32 | 0,01 | 20,37 | >0.1 | 5,61 | 6,68 | 0 | 1,89 | 0 | 2,86 | 0 |
| HMBOX1 | CCTAGAGAAC | -1 | 783 | 793 | -217 | -207 | 1,5 | 0,01 | 20,32 | >0.1 | 5,61 | 5,88 | 0 | 1,41 | 0 | 5,92 | 0 |
| HMBOX1 | TTTGGTTAAT | -1 | 730 | 740 | -270 | -260 | 6,37 | 0,001 | 24,37 | >0.1 | 5,61 | 5,64 | 0 | 1,23 | 0 | 5,52 | 0 |
| HMBOX1 | CCTAGAAAGC | -1 | 715 | 725 | -285 | -275 | 3,82 | 0,01 | 21,36 | >0.1 | 5,61 | 5,58 | 0 | 1,19 | 0 | 5,17 | 0 |
| HMBOX1 | ATAGGTTATC | -1 | 595 | 605 | -405 | -395 | 4,89 | 0,01 | 14,54 | >0.1 | 3,44 | 5,19 | 0 | 1,01 | 0 | 0 | 0 |
| HMBOX1 | GTTAGTCAGG | -1 | 262 | 272 | -738 | -728 | 5,06 | 0,01 | 18,26 | >0.1 | 2,8 | 4,55 | 0 | 1,32 | 0 | 4,53 | 0 |
| HMBOX1 | ACAGGTCAAC | -1 | 106 | 116 | -894 | -884 | 4,01 | 0,01 | 14,56 | >0.1 | 3,57 | 4,36 | 0 | 1,89 | 0 | 0,75 | 0 |
| HMBOX1 | ACTGGTTCTG | -1 | 44 | 54 | -956 | -946 | 1,49 | 0,01 | 11,87 | >0.1 | 4,22 | 4,29 | 0 | 1,87 | 0 | 0 | 0 |
| E2F8 | TTTCTGGGAATA | 1 | 557 | 569 | -443 | -431 | -11,77 | 0,01 | -2,45 | >0.1 | 3,22 | 5,1 | 0 | 0,99 | 0 | 0 | 0 |
| E2F8 | TTTTCCGTGTAT | 1 | 695 | 707 | -305 | -293 | -4,95 | 0,01 | 10,36 | >0.1 | 5,58 | 5,51 | 0 | 1,15 | 0 | 3,07 | 0 |
| E2F8 | TTTCTAGGAAAA | 1 | 717 | 729 | -283 | -271 | -6,15 | 0,01 | 11,21 | >0.1 | 5,58 | 5,59 | 0 | 1,2 | 0 | 4,99 | 0 |
| E2F8 | TTTCCTGCTCAG | 1 | 934 | 946 | -66 | -54 | -6,4 | 0,01 | 9,14 | >0.1 | 5,58 | 7,15 | 0 | 1,88 | 0 | 0,93 | 0 |
| E2F8 | TTTTTGGACAAG | 1 | 1130 | 1142 | 130 | 142 | -10,06 | 0,01 | 9,96 | >0.1 | 5,58 | 6,46 | 0 | 1,84 | 3,22 | 2,91 | 0 |
| E2F8 | TTAACCACGAAG | 1 | 1143 | 1155 | 143 | 155 | -10,14 | 0,01 | 9,02 | >0.1 | 5,58 | 6,35 | 0 | 1,84 | 3,22 | 2,16 | 0 |
| E2F8 | TTTCTCTCTTAT | -1 | 970 | 982 | -30 | -18 | -11,29 | 0,01 | 5,79 | >0.1 | 5,58 | 7,98 | 0 | 1,87 | 0 | 1,65 | 0 |
| E2F8 | TCTCCTGTCACT | -1 | 765 | 777 | -235 | -223 | -9,51 | 0,01 | 9,14 | >0.1 | 5,58 | 5,8 | 0 | 1,35 | 0 | 5,92 | 0 |
| E2F8 | TTTATGGCCAAA | -1 | 754 | 766 | -246 | -234 | 2,31 | 0,001 | 21,27 | >0.1 | 5,58 | 5,75 | 0 | 1,31 | 0 | 6,32 | 0 |
| E2F8 | TTTTCCTAGAAA | -1 | 717 | 729 | -283 | -271 | -6,95 | 0,01 | 10,41 | >0.1 | 5,58 | 5,59 | 0 | 1,2 | 0 | 4,99 | 0 |
| E2F8 | TTTCCTAGAAAG | -1 | 716 | 728 | -284 | -272 | -11,61 | 0,01 | 5,93 | >0.1 | 5,58 | 5,59 | 0 | 1,2 | 0 | 5,17 | 0 |
| E2F8 | TTTTCAGACAAT | -1 | 612 | 624 | -388 | -376 | -6,42 | 0,01 | 4,92 | >0.1 | 3,57 | 5,25 | 0 | 1,03 | 0 | 1,5 | 0 |
| E2F8 | TATTCCCAGAAA | -1 | 557 | 569 | -443 | -431 | -10,1 | 0,01 | -0,78 | >0.1 | 3,22 | 5,1 | 0 | 0,99 | 0 | 0 | 0 |
| E2F8 | ATTCCCAGAAAC | -1 | 556 | 568 | -444 | -432 | -6,08 | 0,01 | 3,23 | >0.1 | 3,22 | 5,1 | 0 | 0,99 | 0 | 0 | 0 |
| E2F8 | TTTCCCTTCCAA | -1 | 526 | 538 | -474 | -462 | 1,39 | 0,001 | 13,9 | >0.1 | 3,07 | 5,03 | 0 | 0,99 | 0 | 3,43 | 0 |
| E2F8 | TTCCCTTCCAAA | -1 | 525 | 537 | -475 | -463 | -4 | 0,01 | 8,51 | >0.1 | 3,07 | 5,03 | 0 | 0,99 | 0 | 3,43 | 0 |
| E2F8 | TCTGTCGCCCAG | -1 | 430 | 442 | -570 | -558 | -6,29 | 0,01 | 7,93 | >0.1 | 2,72 | 4,83 | 0 | 1,01 | 0 | 5,67 | 0 |
| E2F8 | TCTGCCTCCCAG | -1 | 380 | 392 | -620 | -608 | -10,6 | 0,01 | 3,79 | >0.1 | 2,57 | 4,74 | 0 | 1,06 | 0 | 6,02 | 0 |
| E2F8 | GTTCACGCCATT | -1 | 368 | 380 | -632 | -620 | -4,54 | 0,01 | 9,94 | >0.1 | 2,54 | 4,72 | 0 | 1,08 | 0 | 6,15 | 0 |
| E2F8 | TCTCCTGCCTCA | -1 | 357 | 369 | -643 | -631 | -4,52 | 0,01 | 10,05 | >0.1 | 2,52 | 4,7 | 0 | 1,09 | 0 | 6,26 | 0 |
| E2F8 | CCTCCCGAGTAG | -1 | 344 | 356 | -656 | -644 | -10,59 | 0,01 | 4,28 | >0.1 | 2,55 | 4,68 | 0 | 1,11 | 0 | 6,52 | 0 |
| E2F8 | GTGCCCACCACC | -1 | 320 | 332 | -680 | -668 | -6,16 | 0,01 | 7,82 | >0.1 | 2,62 | 4,64 | 0 | 1,16 | 0 | 5,56 | 0 |
| E2F8 | ACGCCCAGCTAA | -1 | 308 | 320 | -692 | -680 | -11,66 | 0,01 | 3,06 | >0.1 | 2,65 | 4,62 | 0 | 1,19 | 0 | 6,26 | 0 |

| | | | | | | | | | | | | | | | | | |
|---|---|---|---|---|---|---|---|---|---|---|---|---|---|---|---|---|---|
| E2F8 | GATCTGCCCAAC | -1 | 226 | 238 | -774 | -762 | -11,48 | 0,01 | 2,68 | >0.1 | 2,93 | 4,5 | 0 | 1,48 | 0 | 5,26 | 0 |
| E2F8 | TCGGCCTCCCAA | -1 | 214 | 226 | -786 | -774 | -10,82 | 0,01 | 3,82 | >0.1 | 2,98 | 4,49 | 0 | 1,54 | 0 | 5,63 | 0 |
| E2F8 | CGGCCTCCCAAA | -1 | 213 | 225 | -787 | -775 | -8,34 | 0,01 | 6,2 | >0.1 | 2,98 | 4,49 | 0 | 1,55 | 0 | 5,52 | 0 |
| E2F8 | GTGCCCAGCCAT | -1 | 174 | 186 | -826 | -814 | -12 | 0,01 | 0,67 | >0.1 | 3,15 | 4,44 | 0 | 1,84 | 0 | 3,24 | 0 |
| E2F8 | TCTCCACTCCAC | -1 | 114 | 126 | -886 | -874 | -11,36 | 0,01 | -0,86 | >0.1 | 3,5 | 4,37 | 0 | 1,89 | 0 | 0,75 | 0 |
| SCRT1 | ACAGAACCAGTGTGA | 1 | 43 | 58 | -957 | -942 | -10,99 | 0,01 | -0,63 | >0.1 | 4,19 | 4,3 | 0 | 1,87 | 0 | 0 | 0 |
| SCRT1 | GATCACAAGGTCAGG | 1 | 234 | 249 | -766 | -751 | -11,46 | 0,01 | 2,02 | >0.1 | 2,89 | 4,52 | 0 | 1,44 | 0 | 4,63 | 0 |
| SCRT1 | GACTAACACGGTGAA | 1 | 265 | 280 | -735 | -720 | -10,98 | 0,01 | 2,2 | >0.1 | 2,78 | 4,56 | 0 | 1,31 | 0 | 4,53 | 0 |
| SCRT1 | AGGGAAAATGTTGCC | 1 | 531 | 546 | -469 | -454 | -10,79 | 0,01 | -0,04 | >0.1 | 3,1 | 5,05 | 0 | 0,99 | 0 | 1,61 | 0 |
| SCRT1 | CTTTAACAGCTTTCT | 1 | 707 | 722 | -293 | -278 | -4,74 | 0,01 | 12,37 | >0.1 | 5,54 | 5,57 | 0 | 1,18 | 0 | 4,81 | 0 |
| SCRT1 | TAACCAAAAGTACCG | 1 | 732 | 747 | -268 | -253 | -11,84 | 0,01 | 6,54 | >0.1 | 5,54 | 5,67 | 0 | 1,25 | 0 | 5,92 | 0 |
| SCRT1 | GCTCAGCAGATTGTT | 1 | 886 | 901 | -114 | -99 | -7,53 | 0,01 | 9,6 | >0.1 | 5,54 | 6,63 | 0 | 1,89 | 0 | 3,07 | 0 |
| SCRT1 | GGCTCACAGGGGACG | 1 | 1034 | 1049 | 34 | 49 | -8,43 | 0,01 | 8,37 | >0.1 | 5,54 | 8,06 | 0 | 1,86 | 0 | 1,34 | 0 |
| SCRT1 | CTGTAACTGCTGCTC | 1 | 1086 | 1101 | 86 | 101 | -13,16 | 0,01 | 5,17 | >0.1 | 5,54 | 6,93 | 0 | 1,85 | 1,79 | 2,21 | 0 |
| SCRT1 | GAGGAACAGGCCAAG | 1 | 1112 | 1127 | 112 | 127 | -4,82 | 0,01 | 14,99 | >0.1 | 5,54 | 6,63 | 0 | 1,85 | 3,22 | 2,56 | 0 |
| SCRT1 | TTTGGACAAGTTTAA | 1 | 1132 | 1147 | 132 | 147 | -12,43 | 0,01 | 7,66 | >0.1 | 5,54 | 6,44 | 0 | 1,84 | 3,22 | 3,03 | 0 |
| SCRT1 | AAGAAGCAAGTGAAC | -1 | 1176 | 1191 | 176 | 191 | -5,91 | 0,01 | 12,53 | >0.1 | 5,54 | 6,11 | 0 | 1,62 | 1,79 | 3,37 | 0 |
| SCRT1 | ATAGAACAGGTCTTC | -1 | 1157 | 1172 | 157 | 172 | 0,8 | 0,001 | 17,49 | >0.1 | 4,39 | 6,24 | 0 | 1,79 | 1,79 | 2,48 | 0 |
| SCRT1 | GAGCAGCAGTTACAG | -1 | 1086 | 1101 | 86 | 101 | -7,44 | 0,01 | 10,89 | >0.1 | 5,54 | 6,93 | 0 | 1,85 | 1,79 | 2,21 | 0 |
| SCRT1 | CAGCAACAAGGCTGA | -1 | 1074 | 1089 | 74 | 89 | 1,95 | 0,001 | 20,52 | >0.1 | 5,54 | 7,11 | 0 | 1,85 | 1,79 | 2,27 | 0 |
| SCRT1 | ATGTAACAGATTTTA | -1 | 840 | 855 | -160 | -145 | -3,13 | 0,01 | 13,68 | >0.1 | 5,54 | 6,26 | 0 | 1,82 | 0 | 3,19 | 0 |
| SCRT1 | TCAAAACCGGTACTT | -1 | 739 | 754 | -261 | -246 | -12,14 | 0,01 | 6,47 | >0.1 | 5,54 | 5,7 | 0 | 1,27 | 0 | 6,1 | 0 |
| SCRT1 | CATTAATAGGTTATC | -1 | 595 | 610 | -405 | -390 | -6,92 | 0,01 | 4,18 | >0.1 | 3,46 | 5,21 | 0 | 1,02 | 0 | 1,41 | 0 |
| SCRT1 | GGGCAACATTTTCCC | -1 | 532 | 547 | -468 | -453 | -2,68 | 0,01 | 8,12 | >0.1 | 3,11 | 5,05 | 0 | 0,99 | 0 | 1,66 | 0 |
| SCRT1 | GCCTCCCAGGTTCAC | -1 | 374 | 389 | -626 | -611 | -12,09 | 0,01 | 2,67 | >0.1 | 2,56 | 4,73 | 0 | 1,07 | 0 | 6,4 | 0 |
| SCRT1 | GGACTACAGGTGCCC | -1 | 326 | 341 | -674 | -659 | 7,36 | 0,0001 | 21,32 | >0.1 | 2,6 | 4,65 | 0 | 1,15 | 0 | 5,56 | 0 |
| SCRT1 | GGATTACAGGCGTGA | -1 | 192 | 207 | -808 | -793 | -7,31 | 0,01 | 6,17 | >0.1 | 3,06 | 4,46 | 0 | 1,69 | 0 | 4,26 | 0 |
| SCRT1 | ACTCCACAGGTCAAC | -1 | 106 | 121 | -894 | -879 | 5,64 | 0,001 | 16,18 | >0.1 | 3,55 | 4,36 | 0 | 1,89 | 0 | 0,75 | 0 |
| SCRT2 | CAGCCACAGAACC | 1 | 38 | 51 | -962 | -949 | -13,13 | 0,01 | -2,68 | >0.1 | 4,29 | 4,29 | 0 | 1,87 | 0 | 0 | 0 |
| SCRT2 | ACAGAACCAGTGT | 1 | 43 | 56 | -957 | -944 | -12,1 | 0,01 | -1,72 | >0.1 | 4,21 | 4,3 | 0 | 1,87 | 0 | 0 | 0 |
| SCRT2 | GAGAAATAGGAAA | 1 | 56 | 69 | -944 | -931 | -12,84 | 0,01 | -2,63 | >0.1 | 4,03 | 4,31 | 0 | 1,87 | 0 | 0 | 0 |
| SCRT2 | GATCACAAGGTCA | 1 | 234 | 247 | -766 | -753 | -6,33 | 0,01 | 7,26 | >0.1 | 2,9 | 4,52 | 0 | 1,44 | 0 | 4,74 | 0 |
| SCRT2 | ATACAAAAACTTA | 1 | 298 | 311 | -702 | -689 | -8,61 | 0,01 | 4,26 | >0.1 | 2,68 | 4,61 | 0 | 1,21 | 0 | 4,37 | 0 |
| SCRT2 | CTTTAACAGCTTT | 1 | 707 | 720 | -293 | -280 | -6,53 | 0,01 | 10,59 | >0.1 | 5,57 | 5,56 | 0 | 1,18 | 0 | 4,81 | 0 |
| SCRT2 | TAACCAAAAGTAC | 1 | 732 | 745 | -268 | -255 | -10,33 | 0,01 | 8,06 | >0.1 | 5,57 | 5,66 | 0 | 1,24 | 0 | 5,92 | 0 |
| SCRT2 | GGCCATAAAGTGA | 1 | 757 | 770 | -243 | -230 | -12,75 | 0,01 | 6,01 | >0.1 | 5,57 | 5,77 | 0 | 1,32 | 0 | 6,1 | 0 |
| SCRT2 | AAGTGACAGGAGA | 1 | 764 | 777 | -236 | -223 | -10,66 | 0,01 | 7,97 | >0.1 | 5,57 | 5,8 | 0 | 1,35 | 0 | 5,92 | 0 |
| SCRT2 | GCTCAGCAGATTG | 1 | 886 | 899 | -114 | -101 | -8,88 | 0,01 | 8,26 | >0.1 | 5,57 | 6,61 | 0 | 1,89 | 0 | 3,07 | 0 |
| SCRT2 | GTTCAAAGGCTGA | 1 | 958 | 971 | -42 | -29 | -8,33 | 0,01 | 8,46 | >0.1 | 5,57 | 7,65 | 0 | 1,88 | 0 | 1,7 | 0 |
| SCRT2 | TCTCATGAGGAGG | 1 | 983 | 996 | -17 | -4 | -14,04 | 0,01 | 3,54 | >0.1 | 5,57 | 8,61 | 0 | 1,87 | 0 | 1,53 | 0 |
| SCRT2 | CTGTAACTGCTGC | 1 | 1086 | 1099 | 86 | 99 | -13,3 | 0,01 | 5,06 | >0.1 | 5,57 | 6,93 | 0 | 1,85 | 1,79 | 2,21 | 0 |
| SCRT2 | GAGGAACAGGCCA | 1 | 1112 | 1125 | 112 | 125 | -6,07 | 0,01 | 13,71 | >0.1 | 5,57 | 6,63 | 0 | 1,85 | 3,22 | 2,52 | 0 |
| SCRT2 | TTTGGACAAGTTT | 1 | 1132 | 1145 | 132 | 145 | -11,72 | 0,01 | 8,32 | >0.1 | 5,57 | 6,44 | 0 | 1,84 | 3,22 | 2,97 | 0 |
| SCRT2 | TTCCAAGAAGCAA | -1 | 1182 | 1195 | 182 | 195 | -13,97 | 0,01 | 4,56 | >0.1 | 5,57 | 6,08 | 0 | 1,57 | 1,79 | 3,52 | 0 |
| SCRT2 | AAGAAGCAAGTGA | -1 | 1178 | 1191 | 178 | 191 | -7,37 | 0,01 | 11,04 | >0.1 | 5,57 | 6,1 | 0 | 1,6 | 1,79 | 3,34 | 0 |
| SCRT2 | ATAGAACAGGTCT | -1 | 1159 | 1172 | 159 | 172 | -4,38 | 0,01 | 12,28 | >0.1 | 4,41 | 6,23 | 0 | 1,77 | 1,79 | 2,46 | 0 |
| SCRT2 | GAGCAGCAGTTAC | -1 | 1088 | 1101 | 88 | 101 | -8,17 | 0,01 | 10,16 | >0.1 | 5,57 | 6,9 | 0 | 1,85 | 1,79 | 2,21 | 0 |
| SCRT2 | CAGCAACAAGGCT | -1 | 1076 | 1089 | 76 | 89 | -1,99 | 0,001 | 16,51 | >0.1 | 5,57 | 7,07 | 0 | 1,85 | 1,79 | 2,21 | 0 |
| SCRT2 | CAGGAAGAGCTTG | -1 | 1053 | 1066 | 53 | 66 | -11,82 | 0,01 | 5,38 | >0.1 | 5,57 | 7,5 | 0 | 1,86 | 0 | 2,27 | 0 |
| SCRT2 | ATGTAACAGATTT | -1 | 842 | 855 | -158 | -145 | -8,84 | 0,01 | 8 | >0.1 | 5,57 | 6,26 | 0 | 1,82 | 0 | 3,19 | 0 |
| SCRT2 | TCAAAACCGGTAC | -1 | 741 | 754 | -259 | -246 | -11,92 | 0,01 | 6,53 | >0.1 | 5,57 | 5,7 | 0 | 1,27 | 0 | 5,92 | 0 |
| SCRT2 | ACCAAAGAGGTCT | -1 | 664 | 677 | -336 | -323 | -10,02 | 0,01 | 4,4 | >0.1 | 4,08 | 5,41 | 0 | 1,1 | 0 | 3,83 | 0 |
| SCRT2 | CATTAATAGGTTA | -1 | 597 | 610 | -403 | -390 | -10,53 | 0,01 | 0,57 | >0.1 | 3,46 | 5,21 | 0 | 1,02 | 0 | 1,41 | 0 |
| SCRT2 | GGGCAACATTTTC | -1 | 534 | 547 | -466 | -453 | -11,42 | 0,01 | -0,66 | >0.1 | 3,11 | 5,05 | 0 | 0,99 | 0 | 1,61 | 0 |
| SCRT2 | GGACTACAGGTGC | -1 | 328 | 341 | -672 | -659 | 0,04 | 0,001 | 13,89 | >0.1 | 2,6 | 4,65 | 0 | 1,14 | 0 | 5,46 | 0 |
| SCRT2 | CCTCCCAAAGTGC | -1 | 209 | 222 | -791 | -778 | -12,85 | 0,01 | 0,9 | >0.1 | 2,99 | 4,48 | 0 | 1,57 | 0 | 4,7 | 0 |
| SCRT2 | ACTCCACAGGTCA | -1 | 108 | 121 | -892 | -879 | 6,48 | 0,0001 | 17,01 | >0.1 | 3,54 | 4,36 | 0 | 1,89 | 0 | 0,75 | 0 |
| FOXI1 | AAAAATA | 1 | 294 | 301 | -706 | -699 | -0,11 | 0,01 | 17,25 | >0.1 | 2,7 | 4,59 | 0 | 1,22 | 0 | 4,45 | 4,39 |
| FOXI1 | ATAAAGA | 1 | 500 | 507 | -500 | -493 | 2,44 | 0,01 | 22,16 | >0.1 | 2,95 | 4,96 | 0 | 0,98 | 0 | 6,44 | 4,39 |

| | | | | | | | | | | | | | | | | | |
|---|---|---|---|---|---|---|---|---|---|---|---|---|---|---|---|---|---|
| FOXI1 | ATAAAAA | 1 | 579 | 586 | -421 | -414 | 3,65 | 0,01 | 17,52 | >0.1 | 3,33 | 5,14 | 0 | 1 | 0 | 0 | 4,39 |
| FOXI1 | AAAAATA | 1 | 581 | 588 | -419 | -412 | -0,11 | 0,01 | 13,77 | >0.1 | 3,34 | 5,15 | 0 | 1 | 0 | 0 | 4,39 |
| FOXI1 | ACAAACA | 1 | 628 | 635 | -372 | -365 | 3,25 | 0,01 | 19,18 | >0.1 | 3,67 | 5,28 | 0 | 1,04 | 0 | 1,55 | 4,39 |
| FOXI1 | ACCAACA | 1 | 634 | 641 | -366 | -359 | 0,49 | 0,01 | 16,56 | >0.1 | 3,73 | 5,3 | 0 | 1,04 | 0 | 1,61 | 4,39 |
| FOXI1 | GAAAATA | 1 | 724 | 731 | -276 | -269 | 3,45 | 0,01 | 25,47 | >0.1 | 5,65 | 5,6 | 0 | 1,2 | 0 | 5,17 | 4,39 |
| FOXI1 | GCCAATA | -1 | 881 | 888 | -119 | -112 | 0,88 | 0,01 | 22,52 | >0.1 | 5,65 | 6,51 | 0 | 1,89 | 0 | 3,19 | 4,39 |
| FOXI1 | ATCAAAA | -1 | 748 | 755 | -252 | -245 | 0,88 | 0,01 | 23,81 | >0.1 | 5,65 | 5,7 | 0 | 1,27 | 0 | 5,92 | 4,39 |
| FOXI1 | GGCAACA | -1 | 539 | 546 | -461 | -454 | 3,28 | 0,01 | 18,48 | >0.1 | 3,12 | 5,05 | 0 | 0,98 | 0 | 1,66 | 4,39 |
| GCM2 | GAAGGAGGCA | 1 | 17 | 27 | -983 | -973 | 0,81 | 0,01 | 12,55 | >0.1 | 5,61 | 4,27 | 0 | 1,87 | 0 | 0 | 0 |
| GCM2 | CGTGGTGGTG | 1 | 317 | 327 | -683 | -673 | 5,72 | 0,01 | 20,4 | >0.1 | 2,63 | 4,63 | 0 | 1,17 | 0 | 6,26 | 0 |
| GCM2 | GGTGGTGGGC | 1 | 320 | 330 | -680 | -670 | 1,22 | 0,01 | 15,32 | >0.1 | 2,62 | 4,63 | 0 | 1,16 | 0 | 5,68 | 0 |
| GCM2 | GTGGTGGGCA | 1 | 321 | 331 | -679 | -669 | 1,07 | 0,01 | 15,05 | >0.1 | 2,62 | 4,64 | 0 | 1,16 | 0 | 5,56 | 0 |
| GCM2 | GAGGCAGGAG | 1 | 358 | 368 | -642 | -632 | 1,28 | 0,01 | 15,84 | >0.1 | 2,52 | 4,7 | 0 | 1,09 | 0 | 6,26 | 0 |
| GCM2 | GACAAGGGCA | 1 | 505 | 515 | -495 | -485 | 3,28 | 0,01 | 17,9 | >0.1 | 2,98 | 4,98 | 0 | 0,98 | 0 | 5,68 | 0 |
| GCM2 | TATATTGGCT | 1 | 879 | 889 | -121 | -111 | 1,94 | 0,01 | 19,16 | >0.1 | 5,61 | 6,52 | 0 | 1,9 | 0 | 3,19 | 0 |
| GCM2 | AATGGTGGAC | -1 | 1102 | 1112 | 102 | 112 | 0,82 | 0,01 | 20,65 | >0.1 | 5,61 | 6,73 | 0 | 1,85 | 3,22 | 2,41 | 0 |
| GCM2 | CATCCTGGAG | -1 | 864 | 874 | -136 | -126 | 1,42 | 0,01 | 18,08 | >0.1 | 5,61 | 6,4 | 0 | 1,9 | 0 | 2,75 | 0 |
| GCM2 | AATGTTGGTG | -1 | 633 | 643 | -367 | -357 | 5,33 | 0,01 | 17,02 | >0.1 | 3,73 | 5,3 | 0 | 1,05 | 0 | 1,61 | 0 |
| GCM2 | CAGGCTGGAG | -1 | 423 | 433 | -577 | -567 | 1,51 | 0,01 | 15,7 | >0.1 | 2,69 | 4,81 | 0 | 1,02 | 0 | 5,67 | 0 |
| GCM2 | ACTACAGGTG | -1 | 329 | 339 | -671 | -661 | 2,4 | 0,01 | 16,24 | >0.1 | 2,6 | 4,65 | 0 | 1,14 | 0 | 5,46 | 0 |
| GCM2 | GAGACGGGGT | -1 | 280 | 290 | -720 | -710 | 3,12 | 0,01 | 16,79 | >0.1 | 2,74 | 4,57 | 0 | 1,26 | 0 | 5,09 | 0 |
| GCM2 | ATTACAGGCG | -1 | 195 | 205 | -805 | -795 | 0,32 | 0,01 | 13,88 | >0.1 | 3,06 | 4,46 | 0 | 1,67 | 0 | 4,37 | 0 |
| GCM2 | CACACTGGTT | -1 | 47 | 57 | -953 | -943 | 2,29 | 0,01 | 12,63 | >0.1 | 4,17 | 4,3 | 0 | 1,87 | 0 | 0 | 0 |
| RARA | AAGGAGAAGGAGGCACTC | 1 | 12 | 30 | -988 | -970 | -27 | 0,01 | -15,35 | >0.1 | 5,51 | 4,27 | 0 | 1,87 | 0 | 0 | 0 |
| RARA | GGGCACAGTGGCTCACGC | 1 | 180 | 198 | -820 | -802 | -25,22 | 0,01 | -11,92 | >0.1 | 3,11 | 4,45 | 0 | 1,79 | 0 | 3,95 | 0 |
| RARA | CAGATCACAAGGTCAGGA | 1 | 232 | 250 | -768 | -750 | -11,97 | 0,001 | 1,52 | >0.1 | 2,89 | 4,52 | 0 | 1,45 | 0 | 4,63 | 0 |
| RARA | AAGGTCAGGAGATAGAGA | 1 | 240 | 258 | -760 | -742 | -24,03 | 0,01 | -10,8 | >0.1 | 2,86 | 4,53 | 0 | 1,41 | 0 | 4,42 | 0 |
| RARA | GAGGCAGAGCTTGCAGTG | 1 | 384 | 402 | -616 | -598 | -30,61 | 0,01 | -16,08 | >0.1 | 2,59 | 4,75 | 0 | 1,06 | 0 | 6,13 | 0 |
| RARA | AATTCCATAAAGACAAGG | 1 | 494 | 512 | -506 | -488 | -15,71 | 0,001 | -0,36 | >0.1 | 2,95 | 4,97 | 0 | 0,99 | 0 | 6,44 | 0 |
| RARA | TATTTGGAAGGGAAAATG | 1 | 523 | 541 | -477 | -459 | -31,22 | 0,01 | -18,69 | >0.1 | 3,07 | 5,03 | 0 | 0,99 | 0 | 3,43 | 0 |
| RARA | AAGTAGAGAGTTTCTGGG | 1 | 547 | 565 | -453 | -435 | -30,25 | 0,01 | -20,97 | >0.1 | 3,19 | 5,09 | 0 | 1 | 0 | 0 | 0 |
| RARA | TTGGCCATAAAGTGACAG | 1 | 755 | 773 | -245 | -227 | -29,74 | 0,01 | -11,01 | >0.1 | 5,51 | 5,78 | 0 | 1,33 | 0 | 6,1 | 0 |
| RARA | GAGGTAAGGTTCTCTAGG | 1 | 775 | 793 | -225 | -207 | -32,14 | 0,01 | -12,79 | >0.1 | 5,51 | 5,88 | 0 | 1,42 | 0 | 6,54 | 0 |
| RARA | AAGTTCAAAGGCTGATAA | 1 | 956 | 974 | -44 | -26 | -25,13 | 0,01 | -8,28 | >0.1 | 5,51 | 7,73 | 0 | 1,88 | 0 | 1,74 | 0 |
| RARA | AATCTCATGAGGAGGTTT | 1 | 981 | 999 | -19 | -1 | -32,26 | 0,01 | -14,62 | >0.1 | 5,51 | 8,69 | 0 | 1,87 | 0 | 1,57 | 0 |
| RARA | AGGGAAAGTCATTCAGTG | 1 | 1005 | 1023 | 5 | 23 | -26,56 | 0,01 | -8,6 | >0.1 | 5,51 | 9,12 | 0 | 1,87 | 0 | 1,46 | 0 |
| RARA | AAAGTCATTCAGTGGATG | 1 | 1009 | 1027 | 9 | 27 | -26,32 | 0,01 | -8,5 | >0.1 | 5,51 | 9,03 | 0 | 1,86 | 0 | 1,42 | 0 |
| RARA | CAGGGGACGATGTCAAGC | 1 | 1040 | 1058 | 40 | 58 | -18,66 | 0,01 | -2,82 | >0.1 | 5,51 | 7,85 | 0 | 1,86 | 0 | 0,62 | 0 |
| RARA | AATGTTATAATTTTACAG | -1 | 679 | 697 | -321 | -303 | -28,77 | 0,01 | -14,74 | >0.1 | 4,36 | 5,48 | 0 | 1,13 | 0 | 3,07 | 0 |
| RARA | GGGATCATGAAGATAATG | -1 | 639 | 657 | -361 | -343 | -18,58 | 0,01 | -4,64 | >0.1 | 3,82 | 5,35 | 0 | 1,07 | 0 | 3,7 | 0 |
| RARA | ACTTAAAAAAGCTCATTT | -1 | 66 | 84 | -934 | -916 | -30,47 | 0,01 | -20,38 | >0.1 | 3,88 | 4,32 | 0 | 1,88 | 0 | 0 | 0 |
| RARB | GAAGGAGAAGGAGGCA | 1 | 11 | 27 | -989 | -973 | -15,17 | 0,01 | -3,5 | >0.1 | 5,53 | 4,27 | 0 | 1,87 | 0 | 0 | 0 |
| RARB | ATAGGAAATGAGCTTT | 1 | 61 | 77 | -939 | -923 | -27,99 | 0,01 | -17,85 | >0.1 | 3,95 | 4,32 | 0 | 1,88 | 0 | 0 | 0 |
| RARB | AGTAGTCATAATTTTA | 1 | 125 | 141 | -875 | -859 | -28,18 | 0,01 | -16,91 | >0.1 | 3,41 | 4,38 | 0 | 1,89 | 0 | 1,57 | 0 |
| RARB | TGGGCACAGTGGCTCA | 1 | 179 | 195 | -821 | -805 | -24,85 | 0,01 | -11,63 | >0.1 | 3,12 | 4,45 | 0 | 1,8 | 0 | 3,86 | 0 |
| RARB | CGAGTTGGGCAGATCA | 1 | 223 | 239 | -777 | -761 | -24,17 | 0,01 | -9,99 | >0.1 | 2,93 | 4,5 | 0 | 1,49 | 0 | 5,26 | 0 |
| RARB | CAAGGTCAGGAGATAG | 1 | 239 | 255 | -761 | -745 | -15,98 | 0,01 | -2,84 | >0.1 | 2,87 | 4,53 | 0 | 1,41 | 0 | 4,33 | 0 |
| RARB | AAAATACAAAAACTTA | 1 | 295 | 311 | -705 | -689 | -28,66 | 0,01 | -15,7 | >0.1 | 2,69 | 4,61 | 0 | 1,22 | 0 | 4,45 | 0 |
| RARB | GAATTCCATAAAGACA | 1 | 493 | 509 | -507 | -491 | -24,48 | 0,01 | -9,15 | >0.1 | 2,94 | 4,96 | 0 | 0,99 | 0 | 6,44 | 0 |
| RARB | GAAGGGAAAATGTTGC | 1 | 529 | 545 | -471 | -455 | -27,59 | 0,01 | -16,85 | >0.1 | 3,1 | 5,04 | 0 | 0,99 | 0 | 1,61 | 0 |
| RARB | TTTGGCCATAAAGTGA | 1 | 754 | 770 | -246 | -230 | -21,56 | 0,01 | -2,84 | >0.1 | 5,53 | 5,77 | 0 | 1,32 | 0 | 6,1 | 0 |
| RARB | AAGAGAGAAAATCTCA | 1 | 972 | 988 | -28 | -12 | -25,96 | 0,01 | -8,76 | >0.1 | 5,53 | 8,22 | 0 | 1,87 | 0 | 1,57 | 0 |
| RARB | AAATCTCATGAGGAGG | 1 | 980 | 996 | -20 | -4 | -27,5 | 0,01 | -9,91 | >0.1 | 5,53 | 8,61 | 0 | 1,87 | 0 | 1,57 | 0 |
| RARB | ACAGGGGACGATGTCA | 1 | 1039 | 1055 | 39 | 55 | -11,98 | 0,001 | 3,92 | >0.1 | 5,53 | 7,88 | 0 | 1,86 | 0 | 0,62 | 0 |
| RARB | TGAGCAGGAAAGACCA | -1 | 929 | 945 | -71 | -55 | -27,2 | 0,01 | -11,72 | >0.1 | 5,53 | 7,14 | 0 | 1,88 | 0 | 0,93 | 0 |
| RARB | AAATGTTATAATTTTA | -1 | 682 | 698 | -318 | -302 | -18,74 | 0,01 | -4,78 | >0.1 | 4,4 | 5,48 | 0 | 1,13 | 0 | 2,95 | 0 |
| RARB | AGGGATCATGAAGATA | -1 | 642 | 658 | -358 | -342 | -24,68 | 0,01 | -10,59 | >0.1 | 3,84 | 5,35 | 0 | 1,07 | 0 | 3,83 | 0 |
| RARB | CTGGGACTACAGGTGC | -1 | 328 | 344 | -672 | -656 | -28,9 | 0,01 | -14,47 | >0.1 | 2,59 | 4,66 | 0 | 1,15 | 0 | 6,03 | 0 |
| RARB | AACTTAAAAAAGCTCA | -1 | 69 | 85 | -931 | -915 | -26,29 | 0,01 | -16,22 | >0.1 | 3,86 | 4,32 | 0 | 1,88 | 0 | 0 | 0 |

| | | | | | | | | | | | | | | | | | |
|---|---|---|---|---|---|---|---|---|---|---|---|---|---|---|---|---|---|
| RARG | AAGGAGAAGGAGGCAC | 1 | 12 | 28 | -988 | -972 | -18,7 | 0,01 | -7,03 | >0.1 | 5,53 | 4,27 | 0 | 1,87 | 0 | 0 | 0 |
| RARG | GTAGTCATAATTTTAA | 1 | 126 | 142 | -874 | -858 | -22,12 | 0,01 | -10,82 | >0.1 | 3,41 | 4,39 | 0 | 1,89 | 0 | 1,61 | 0 |
| RARG | GGGCACAGTGGCTCAC | 1 | 180 | 196 | -820 | -804 | -20,76 | 0,01 | -7,55 | >0.1 | 3,11 | 4,45 | 0 | 1,79 | 0 | 3,86 | 0 |
| RARG | GAGTTGGGCAGATCAC | 1 | 224 | 240 | -776 | -760 | -23,62 | 0,01 | -9,44 | >0.1 | 2,93 | 4,51 | 0 | 1,49 | 0 | 5,26 | 0 |
| RARG | AAGGTCAGGAGATAGA | 1 | 240 | 256 | -760 | -744 | -20,09 | 0,01 | -6,96 | >0.1 | 2,87 | 4,53 | 0 | 1,41 | 0 | 4,33 | 0 |
| RARG | AATTCCATAAAGACAA | 1 | 494 | 510 | -506 | -490 | -29,31 | 0,01 | -13,97 | >0.1 | 2,95 | 4,97 | 0 | 0,99 | 0 | 6,44 | 0 |
| RARG | GAAGGGAAAATGTTGC | 1 | 529 | 545 | -471 | -455 | -26,36 | 0,01 | -15,62 | >0.1 | 3,1 | 5,04 | 0 | 0,99 | 0 | 1,61 | 0 |
| RARG | AAGGGAAAATGTTGCC | 1 | 530 | 546 | -470 | -454 | -28,06 | 0,01 | -17,31 | >0.1 | 3,1 | 5,05 | 0 | 0,99 | 0 | 1,61 | 0 |
| RARG | TTGGCCATAAAGTGAC | 1 | 755 | 771 | -245 | -229 | -20,86 | 0,01 | -2,13 | >0.1 | 5,53 | 5,77 | 0 | 1,33 | 0 | 6,1 | 0 |
| RARG | CAAGTTCAAAGGCTGA | 1 | 955 | 971 | -45 | -29 | -27,48 | 0,01 | -10,69 | >0.1 | 5,53 | 7,65 | 0 | 1,88 | 0 | 1,74 | 0 |
| RARG | AGAGAGAAAATCTCAT | 1 | 973 | 989 | -27 | -11 | -28,34 | 0,01 | -11,09 | >0.1 | 5,53 | 8,27 | 0 | 1,87 | 0 | 1,57 | 0 |
| RARG | CAGGGGACGATGTCAA | 1 | 1040 | 1056 | 40 | 56 | -12,77 | 0,001 | 3,09 | >0.1 | 5,53 | 7,85 | 0 | 1,86 | 0 | 0,62 | 0 |
| RARG | AGGCTGAGAAGGAGCC | -1 | 1065 | 1081 | 65 | 81 | -28,44 | 0,01 | -9,79 | >0.1 | 5,53 | 7,26 | 0 | 1,85 | 1,79 | 2,21 | 0 |
| RARG | AATGTTATAATTTTAC | -1 | 681 | 697 | -319 | -303 | -24,53 | 0,01 | -10,6 | >0.1 | 4,38 | 5,48 | 0 | 1,13 | 0 | 2,95 | 0 |
| RARG | GGGATCATGAAGATAA | -1 | 641 | 657 | -359 | -343 | -29,38 | 0,01 | -15,44 | >0.1 | 3,83 | 5,34 | 0 | 1,07 | 0 | 3,7 | 0 |
| RARG | TGGGACTACAGGTGCC | -1 | 327 | 343 | -673 | -657 | -28,29 | 0,01 | -14,44 | >0.1 | 2,59 | 4,66 | 0 | 1,15 | 0 | 5,46 | 0 |
| RARG | CCACTCCACAGGTCAA | -1 | 107 | 123 | -893 | -877 | -24,52 | 0,01 | -13,97 | >0.1 | 3,54 | 4,36 | 0 | 1,89 | 0 | 0,77 | 0 |
| NFE2L2 | GTAAGTGAAGGAGAA | 1 | 5 | 20 | -995 | -980 | -5,16 | 0,01 | 8,31 | >0.1 | 5,54 | 4,26 | 0 | 1,86 | 1,79 | 0 | 0 |
| NFE2L2 | AGAAATGGCTGGGCA | 1 | 170 | 185 | -830 | -815 | 1,37 | 0,01 | 14,2 | >0.1 | 3,17 | 4,44 | 0 | 1,88 | 0 | 3,34 | 0 |
| NFE2L2 | CACAGTGGCTCACGC | 1 | 183 | 198 | -817 | -802 | -4,99 | 0,01 | 8,19 | >0.1 | 3,1 | 4,45 | 0 | 1,76 | 0 | 3,86 | 0 |
| NFE2L2 | CGTCCTGACTAACAC | 1 | 259 | 274 | -741 | -726 | -4,68 | 0,01 | 8,46 | >0.1 | 2,8 | 4,55 | 0 | 1,33 | 0 | 4,45 | 0 |
| NFE2L2 | CACGGTGAAACCCCG | 1 | 271 | 286 | -729 | -714 | -3,2 | 0,01 | 10,51 | >0.1 | 2,76 | 4,57 | 0 | 1,29 | 0 | 5,09 | 0 |
| NFE2L2 | CCCAGCTACTCGGGA | 1 | 339 | 354 | -661 | -646 | -5,04 | 0,01 | 9,84 | >0.1 | 2,56 | 4,67 | 0 | 1,12 | 0 | 6,52 | 0 |
| NFE2L2 | GAGGCTGAGGCAGGA | 1 | 352 | 367 | -648 | -633 | 5,27 | 0,001 | 20,12 | >0.1 | 2,53 | 4,69 | 0 | 1,1 | 0 | 6,52 | 0 |
| NFE2L2 | TGGCGTGAACCTGGG | 1 | 370 | 385 | -630 | -615 | -2,36 | 0,01 | 12,39 | >0.1 | 2,55 | 4,73 | 0 | 1,07 | 0 | 6,4 | 0 |
| NFE2L2 | TGCAGTGAGCCGAGA | 1 | 395 | 410 | -605 | -590 | -1,89 | 0,01 | 12,45 | >0.1 | 2,62 | 4,77 | 0 | 1,04 | 0 | 5,91 | 0 |
| NFE2L2 | CACTGTGCTCCAGCC | 1 | 416 | 431 | -584 | -569 | -2,73 | 0,01 | 10,97 | >0.1 | 2,68 | 4,81 | 0 | 1,02 | 0 | 5,19 | 0 |
| NFE2L2 | GACCGAGACTCAGTC | 1 | 440 | 455 | -560 | -545 | -3,06 | 0,01 | 11,36 | >0.1 | 2,75 | 4,85 | 0 | 1,01 | 0 | 5,8 | 0 |
| NFE2L2 | GAATATGATCTTGAA | 1 | 564 | 579 | -436 | -421 | -4,5 | 0,01 | 4,9 | >0.1 | 3,27 | 5,13 | 0 | 1 | 0 | 0 | 0 |
| NFE2L2 | AAATGTGAGATAACC | 1 | 587 | 602 | -413 | -398 | -1,42 | 0,01 | 8,19 | >0.1 | 3,4 | 5,19 | 0 | 1,01 | 0 | 0 | 0 |
| NFE2L2 | GGTTTTGATTTGGCC | 1 | 746 | 761 | -254 | -239 | -2,82 | 0,01 | 15,66 | >0.1 | 5,54 | 5,73 | 0 | 1,29 | 0 | 5,92 | 0 |
| NFE2L2 | TAAAGTGACAGGAGA | 1 | 762 | 777 | -238 | -223 | -2,4 | 0,01 | 16,4 | >0.1 | 5,54 | 5,8 | 0 | 1,35 | 0 | 6,1 | 0 |
| NFE2L2 | TATATTGGCTCAGCA | 1 | 879 | 894 | -121 | -106 | 0,52 | 0,01 | 17,59 | >0.1 | 5,54 | 6,57 | 0 | 1,9 | 0 | 3,07 | 0 |
| NFE2L2 | AACTTTGATAGAACA | -1 | 1164 | 1179 | 164 | 179 | -3,24 | 0,01 | 14,89 | >0.1 | 5,54 | 6,19 | 0 | 1,72 | 1,79 | 2,88 | 0 |
| NFE2L2 | AAAAATGTCTTGGCC | -1 | 1120 | 1135 | 120 | 135 | -2,34 | 0,01 | 17,56 | >0.1 | 5,54 | 6,55 | 0 | 1,85 | 3,22 | 2,74 | 0 |
| NFE2L2 | ATGGTGGACTGAGCA | -1 | 1096 | 1111 | 96 | 111 | -2,56 | 0,01 | 17,21 | >0.1 | 5,54 | 6,8 | 0 | 1,85 | 3,22 | 2,35 | 0 |
| NFE2L2 | AAGGCTGAGAAGGAG | -1 | 1067 | 1082 | 67 | 82 | -5,01 | 0,01 | 13,62 | >0.1 | 5,54 | 7,22 | 0 | 1,85 | 1,79 | 2,21 | 0 |
| NFE2L2 | GAGAAGGAGCCAGGA | -1 | 1061 | 1076 | 61 | 76 | -2,21 | 0,01 | 16,59 | >0.1 | 5,54 | 7,33 | 0 | 1,86 | 1,79 | 2,27 | 0 |
| NFE2L2 | CCCTGTGAGCCAAGA | -1 | 1030 | 1045 | 30 | 45 | 0,63 | 0,01 | 17,55 | >0.1 | 5,54 | 8,24 | 0 | 1,86 | 0 | 1,28 | 0 |
| NFE2L2 | CTGAATGACTTTCCC | -1 | 1006 | 1021 | 6 | 21 | 3,39 | 0,001 | 21,34 | >0.1 | 5,54 | 9,12 | 0 | 1,87 | 0 | 1,42 | 0 |
| NFE2L2 | GGCGCTGAGCAGGAA | -1 | 935 | 950 | -65 | -50 | -3,9 | 0,01 | 11,64 | >0.1 | 5,54 | 7,22 | 0 | 1,88 | 0 | 0,9 | 0 |
| NFE2L2 | TAAAGTGAATCAAAT | -1 | 814 | 829 | -186 | -171 | -5,46 | 0,01 | 11,33 | >0.1 | 5,54 | 6,08 | 0 | 1,61 | 0 | 3,55 | 0 |
| NFE2L2 | TACAGTGACCAAAGA | -1 | 669 | 684 | -331 | -316 | -0,27 | 0,01 | 13,98 | >0.1 | 4,16 | 5,43 | 0 | 1,11 | 0 | 3,55 | 0 |
| NFE2L2 | ATACATGACTTTGCC | -1 | 511 | 526 | -489 | -474 | 6,9 | 0,001 | 18,76 | >0.1 | 3,01 | 5 | 0 | 0,99 | 0 | 2,86 | 0 |
| NFE2L2 | GAGACTGAGTCTCGG | -1 | 442 | 457 | -558 | -543 | -3,23 | 0,01 | 11,31 | >0.1 | 2,76 | 4,86 | 0 | 1,01 | 0 | 5,92 | 0 |
| NFE2L2 | TCTCCTGCCTCAGCC | -1 | 354 | 369 | -646 | -631 | -1,7 | 0,01 | 12,75 | >0.1 | 2,52 | 4,7 | 0 | 1,1 | 0 | 6,13 | 0 |
| NFE2L2 | CCGTGTTAGTCAGGA | -1 | 261 | 276 | -739 | -724 | 5,79 | 0,001 | 18,92 | >0.1 | 2,8 | 4,56 | 0 | 1,33 | 0 | 4,45 | 0 |
| NFE2L2 | AGGCGTGAGCCACTG | -1 | 185 | 200 | -815 | -800 | -4,05 | 0,01 | 9,1 | >0.1 | 3,09 | 4,45 | 0 | 1,75 | 0 | 3,86 | 0 |
| NFE2L2 | AATTATGACTACTCT | -1 | 123 | 138 | -877 | -862 | -3,17 | 0,01 | 8,23 | >0.1 | 3,43 | 4,38 | 0 | 1,89 | 0 | 1,7 | 0 |
| NFE2L2 | TGATCTGCCTTTGCA | -1 | 86 | 101 | -914 | -899 | -4,67 | 0,01 | 5,26 | >0.1 | 3,71 | 4,34 | 0 | 1,88 | 0 | 0 | 0 |
| SIX1 | GAAATAGGAAA | 1 | 58 | 69 | -942 | -931 | 2,89 | 0,01 | 13,09 | >0.1 | 4,01 | 4,31 | 0 | 1,87 | 0 | 0 | 0 |
| SIX1 | TAAAACTGATC | 1 | 159 | 170 | -841 | -830 | 7,94 | 0,001 | 19,89 | >0.1 | 3,23 | 4,42 | 0 | 1,9 | 0 | 2,4 | 0 |
| SIX1 | GTAATCCTAGC | 1 | 200 | 211 | -800 | -789 | 2,99 | 0,01 | 16,72 | >0.1 | 3,04 | 4,47 | 0 | 1,63 | 0 | 4,59 | 0 |
| SIX1 | AAAAACTTAGC | 1 | 302 | 313 | -698 | -687 | 1,68 | 0,01 | 15,15 | >0.1 | 2,67 | 4,61 | 0 | 1,2 | 0 | 4,99 | 0 |
| SIX1 | TGAACCTGGGA | 1 | 375 | 386 | -625 | -614 | 5 | 0,01 | 19,75 | >0.1 | 2,56 | 4,73 | 0 | 1,07 | 0 | 6,4 | 0 |
| SIX1 | AAAAACTGATC | 1 | 474 | 485 | -526 | -515 | 7,75 | 0,001 | 21,46 | >0.1 | 2,86 | 4,91 | 0 | 0,99 | 0 | 4,95 | 0 |
| SIX1 | GAAATATGAAT | 1 | 486 | 497 | -514 | -503 | 5,34 | 0,01 | 20,44 | >0.1 | 2,91 | 4,94 | 0 | 0,99 | 0 | 6,27 | 0 |
| SIX1 | GGAATATGATC | 1 | 563 | 574 | -437 | -426 | 7,42 | 0,001 | 16,78 | >0.1 | 3,25 | 5,11 | 0 | 0,99 | 0 | 0 | 0 |
| SIX1 | TAAATGTGAGA | 1 | 586 | 597 | -414 | -403 | 4,09 | 0,01 | 13,66 | >0.1 | 3,39 | 5,17 | 0 | 1,01 | 0 | 0 | 0 |

| | | | | | | | | | | | | | | | | | |
|---|---|---|---|---|---|---|---|---|---|---|---|---|---|---|---|---|---|
| SIX1 | ATAACCTATTA | 1 | 596 | 607 | -404 | -393 | 2,62 | 0,01 | 13,73 | >0.1 | 3,45 | 5,2 | 0 | 1,01 | 0 | 1,44 | 0 |
| SIX1 | AAAACCATACA | 1 | 620 | 631 | -380 | -369 | 1,81 | 0,01 | 13,18 | >0.1 | 3,62 | 5,27 | 0 | 1,04 | 0 | 1,44 | 0 |
| SIX1 | CCAACATTATC | 1 | 635 | 646 | -365 | -354 | 0,94 | 0,01 | 12,66 | >0.1 | 3,75 | 5,31 | 0 | 1,05 | 0 | 1,61 | 0 |
| SIX1 | GTAAAATTATA | 1 | 681 | 692 | -319 | -308 | 2,7 | 0,01 | 16,8 | >0.1 | 4,33 | 5,46 | 0 | 1,12 | 0 | 3,19 | 0 |
| SIX1 | ATAAAGTGACA | 1 | 761 | 772 | -239 | -228 | 2,52 | 0,01 | 21,32 | >0.1 | 5,6 | 5,78 | 0 | 1,33 | 0 | 6,1 | 0 |
| SIX1 | GTGACAGGAGA | 1 | 766 | 777 | -234 | -223 | 1,75 | 0,01 | 20,41 | >0.1 | 5,6 | 5,8 | 0 | 1,35 | 0 | 5,92 | 0 |
| SIX1 | AAAATCTGTTA | 1 | 841 | 852 | -159 | -148 | 3,25 | 0,01 | 18,29 | >0.1 | 5,6 | 6,24 | 0 | 1,79 | 0 | 1,41 | 0 |
| SIX1 | TGAACTTTATA | 1 | 872 | 883 | -128 | -117 | 1,87 | 0,01 | 18,7 | >0.1 | 5,6 | 6,47 | 0 | 1,9 | 0 | 2,86 | 0 |
| SIX1 | GAAGCCGAAGA | 1 | 1151 | 1162 | 151 | 162 | 0,54 | 0,01 | 17,19 | >0.1 | 4,52 | 6,29 | 0 | 1,84 | 1,79 | 2,21 | 0 |
| SIX1 | GAACTTTGATA | -1 | 1169 | 1180 | 169 | 180 | 2,39 | 0,01 | 20,56 | >0.1 | 5,6 | 6,16 | 0 | 1,68 | 1,79 | 2,94 | 0 |
| SIX1 | TAAACTTGTCC | -1 | 1135 | 1146 | 135 | 146 | 2,51 | 0,01 | 22,61 | >0.1 | 5,6 | 6,41 | 0 | 1,84 | 3,22 | 3,02 | 0 |
| SIX1 | AGAACCTTACC | -1 | 777 | 788 | -223 | -212 | 4,31 | 0,01 | 23,7 | >0.1 | 5,6 | 5,85 | 0 | 1,39 | 0 | 6,54 | 0 |
| SIX1 | AAAACCGGTAC | -1 | 741 | 752 | -259 | -248 | 0,71 | 0,01 | 19,36 | >0.1 | 5,6 | 5,69 | 0 | 1,26 | 0 | 6,1 | 0 |
| SIX1 | GAAAGCTGTTA | -1 | 710 | 721 | -290 | -279 | 4,45 | 0,01 | 21,6 | >0.1 | 5,6 | 5,56 | 0 | 1,18 | 0 | 4,81 | 0 |
| SIX1 | GGAAAATGTTA | -1 | 690 | 701 | -310 | -299 | 0,67 | 0,01 | 14,77 | >0.1 | 4,52 | 5,49 | 0 | 1,14 | 0 | 2,95 | 0 |
| SIX1 | GTGACCAAAGA | -1 | 669 | 680 | -331 | -320 | 0,56 | 0,01 | 14,91 | >0.1 | 4,13 | 5,42 | 0 | 1,1 | 0 | 3,7 | 0 |
| SIX1 | GGATCATGAAG | -1 | 645 | 656 | -355 | -344 | 0,96 | 0,01 | 15,05 | >0.1 | 3,85 | 5,34 | 0 | 1,06 | 0 | 3,83 | 0 |
| SIX1 | AATACATGACT | -1 | 516 | 527 | -484 | -473 | 1,42 | 0,01 | 13,38 | >0.1 | 3,03 | 5 | 0 | 0,98 | 0 | 2,95 | 0 |
| SIX1 | ATCTCCTGACC | -1 | 242 | 253 | -758 | -747 | 0,59 | 0,01 | 13,71 | >0.1 | 2,87 | 4,52 | 0 | 1,4 | 0 | 4,33 | 0 |
| SIX1 | CTCTCCTGATC | -1 | 96 | 107 | -904 | -893 | 0,9 | 0,01 | 11,54 | >0.1 | 3,64 | 4,35 | 0 | 1,88 | 0 | 0,77 | 0 |
| SIX1 | AAAACTTAAAA | -1 | 76 | 87 | -924 | -913 | 2,45 | 0,01 | 12,47 | >0.1 | 3,82 | 4,33 | 0 | 1,88 | 0 | 0 | 0 |
| SIX2 | GTGAGAAATAGGAAAT | 1 | 54 | 70 | -946 | -930 | 2,71 | 0,01 | 12,93 | >0.1 | 4,03 | 4,31 | 0 | 1,87 | 0 | 0 | 0 |
| SIX2 | CCATGGAAATTAAAAC | 1 | 149 | 165 | -851 | -835 | 0,73 | 0,01 | 11,96 | >0.1 | 3,27 | 4,41 | 0 | 1,9 | 0 | 1,65 | 0 |
| SIX2 | AAATTAAAACTGATCA | 1 | 155 | 171 | -845 | -829 | 6,81 | 0,001 | 18,83 | >0.1 | 3,24 | 4,42 | 0 | 1,9 | 0 | 2,46 | 0 |
| SIX2 | GCCTGTAATCCTAGCA | 1 | 196 | 212 | -804 | -788 | 5,31 | 0,01 | 18,96 | >0.1 | 3,04 | 4,47 | 0 | 1,66 | 0 | 4,48 | 0 |
| SIX2 | CGGTGAAACCCCGTCT | 1 | 273 | 289 | -727 | -711 | 4,23 | 0,01 | 17,93 | >0.1 | 2,75 | 4,57 | 0 | 1,29 | 0 | 5,09 | 0 |
| SIX2 | CTCGGGAGGCTGAGGC | 1 | 347 | 363 | -653 | -637 | 2,83 | 0,01 | 17,57 | >0.1 | 2,54 | 4,69 | 0 | 1,11 | 0 | 6,4 | 0 |
| SIX2 | GGCGTGAACCTGGGAG | 1 | 371 | 387 | -629 | -613 | 1,84 | 0,01 | 16,73 | >0.1 | 2,55 | 4,73 | 0 | 1,07 | 0 | 6,53 | 0 |
| SIX2 | TGCAGTGAGCCGAGAT | 1 | 395 | 411 | -605 | -589 | 1,55 | 0,01 | 15,89 | >0.1 | 2,62 | 4,77 | 0 | 1,05 | 0 | 5,91 | 0 |
| SIX2 | AAAAAAAAACTGATCA | 1 | 470 | 486 | -530 | -514 | 4,6 | 0,01 | 18,31 | >0.1 | 2,86 | 4,91 | 0 | 0,99 | 0 | 4,95 | 0 |
| SIX2 | ATCAGAAATATGAATT | 1 | 482 | 498 | -518 | -502 | 6,95 | 0,001 | 21,91 | >0.1 | 2,9 | 4,94 | 0 | 0,99 | 0 | 6,13 | 0 |
| SIX2 | TCTGGGAATATGATCT | 1 | 559 | 575 | -441 | -425 | 5,72 | 0,01 | 15,08 | >0.1 | 3,25 | 5,12 | 0 | 1 | 0 | 0 | 0 |
| SIX2 | AAAATAAATGTGAGAT | 1 | 582 | 598 | -418 | -402 | 2,05 | 0,01 | 11,61 | >0.1 | 3,38 | 5,18 | 0 | 1,01 | 0 | 0 | 0 |
| SIX2 | CACTGTAAAATTATAA | 1 | 677 | 693 | -323 | -307 | 4,31 | 0,01 | 18,39 | >0.1 | 4,3 | 5,46 | 0 | 1,12 | 0 | 3,19 | 0 |
| SIX2 | TTCTAAAATCTGTTAC | 1 | 837 | 853 | -163 | -147 | 6,48 | 0,001 | 23,25 | >0.1 | 5,53 | 6,25 | 0 | 1,8 | 0 | 3,19 | 0 |
| SIX2 | GGCTGATAAGAGAGAA | 1 | 965 | 981 | -35 | -19 | 0,81 | 0,01 | 17,86 | >0.1 | 5,53 | 7,94 | 0 | 1,87 | 0 | 1,7 | 0 |
| SIX2 | CCACGAAGCCGAAGAC | 1 | 1147 | 1163 | 147 | 163 | 0,56 | 0,01 | 18,26 | >0.1 | 5,53 | 6,32 | 0 | 1,84 | 1,79 | 2,21 | 0 |
| SIX2 | GCCGAAGACCTGTTCT | 1 | 1154 | 1170 | 154 | 170 | 1,02 | 0,01 | 17,64 | >0.1 | 4,4 | 6,27 | 0 | 1,82 | 1,79 | 2,34 | 0 |
| SIX2 | AAGTGAACTTTGATAG | -1 | 1168 | 1184 | 168 | 184 | 3,98 | 0,01 | 22,18 | >0.1 | 5,53 | 6,17 | 0 | 1,68 | 1,79 | 3,02 | 0 |
| SIX2 | TGGTTAAACTTGTCCA | -1 | 1134 | 1150 | 134 | 150 | 0,74 | 0,01 | 20,83 | >0.1 | 5,53 | 6,42 | 0 | 1,84 | 3,22 | 3,07 | 0 |
| SIX2 | GACTAAAACCTCCTCA | -1 | 988 | 1004 | -12 | 4 | 2,83 | 0,01 | 20,65 | >0.1 | 5,53 | 8,84 | 0 | 1,87 | 0 | 1,57 | 0 |
| SIX2 | TCCTGTCACTTTATGG | -1 | 759 | 775 | -241 | -225 | 3,35 | 0,01 | 21,93 | >0.1 | 5,53 | 5,79 | 0 | 1,34 | 0 | 5,92 | 0 |
| SIX2 | CCTAGAAAGCTGTTAA | -1 | 709 | 725 | -291 | -275 | 4,28 | 0,01 | 21,57 | >0.1 | 5,53 | 5,58 | 0 | 1,19 | 0 | 4,99 | 0 |
| SIX2 | CCCAGAAACTCTCTAC | -1 | 549 | 565 | -451 | -435 | 1,47 | 0,01 | 10,75 | >0.1 | 3,19 | 5,09 | 0 | 1 | 0 | 0 | 0 |
| SIX2 | TCCAAATACATGACTT | -1 | 515 | 531 | -485 | -469 | 0,98 | 0,01 | 13,08 | >0.1 | 3,03 | 5,01 | 0 | 0,99 | 0 | 3,07 | 0 |
| SIX2 | CACTGCAAGCTCTGCC | -1 | 386 | 402 | -614 | -598 | 1,66 | 0,01 | 16,08 | >0.1 | 2,59 | 4,75 | 0 | 1,06 | 0 | 6,02 | 0 |
| SIX2 | CTCTATCTCCTGACCT | -1 | 241 | 257 | -759 | -743 | 1,24 | 0,01 | 14,46 | >0.1 | 2,86 | 4,53 | 0 | 1,41 | 0 | 4,42 | 0 |
| SIX3 | AATAGGAAATGAGCTTT | 1 | 60 | 77 | -940 | -923 | -2,16 | 0,01 | 7,99 | >0.1 | 3,96 | 4,32 | 0 | 1,88 | 0 | 0 | 0 |
| SIX3 | AAAGGCAGATCAGGAGA | 1 | 89 | 106 | -911 | -894 | 1,88 | 0,01 | 12,61 | >0.1 | 3,67 | 4,35 | 0 | 1,88 | 0 | 0,83 | 0 |
| SIX3 | CACAGTGGCTCACGCCT | 1 | 183 | 200 | -817 | -800 | -1,83 | 0,01 | 11,34 | >0.1 | 3,1 | 4,45 | 0 | 1,76 | 0 | 3,86 | 0 |
| SIX3 | TTGGGCAGATCACAAGG | 1 | 227 | 244 | -773 | -756 | 2,91 | 0,01 | 16,82 | >0.1 | 2,91 | 4,51 | 0 | 1,47 | 0 | 5,01 | 0 |
| SIX3 | GAGCCGAGATCGCGCCA | 1 | 401 | 418 | -599 | -582 | -0,82 | 0,01 | 13,4 | >0.1 | 2,64 | 4,78 | 0 | 1,04 | 0 | 5,76 | 0 |
| SIX3 | AATGTGAGATAACCTAT | 1 | 588 | 605 | -412 | -395 | 2,6 | 0,01 | 12,23 | >0.1 | 3,42 | 5,19 | 0 | 1,02 | 0 | 0 | 0 |
| SIX3 | TTCCGTGTATCTTTAAC | 1 | 697 | 714 | -303 | -286 | 0,64 | 0,01 | 16,05 | >0.1 | 5,52 | 5,54 | 0 | 1,17 | 0 | 3,19 | 0 |
| SIX3 | AATAACGTATTCTTATT | 1 | 800 | 817 | -200 | -183 | -1 | 0,01 | 15,49 | >0.1 | 5,52 | 6,01 | 0 | 1,54 | 0 | 3,43 | 0 |
| SIX3 | GAGAAAATCTCATGAGG | 1 | 976 | 993 | -24 | -7 | -3,18 | 0,01 | 14,24 | >0.1 | 5,52 | 8,5 | 0 | 1,87 | 0 | 1,53 | 0 |
| SIX3 | CAGTGGATGTGATCTTG | 1 | 1018 | 1035 | 18 | 35 | -2,15 | 0,01 | 15,3 | >0.1 | 5,52 | 8,76 | 0 | 1,86 | 0 | 1,31 | 0 |
| SIX3 | GGGACGATGTCAAGCTC | 1 | 1043 | 1060 | 43 | 60 | 1,2 | 0,01 | 17,8 | >0.1 | 5,52 | 7,76 | 0 | 1,86 | 0 | 1,46 | 0 |
| SIX3 | CCTGTTCTATCAAAGTT | 1 | 1162 | 1179 | 162 | 179 | -1,74 | 0,01 | 16,31 | >0.1 | 5,52 | 6,21 | 0 | 1,74 | 1,79 | 2,79 | 0 |

| | | | | | | | | | | | | | | | | | |
|---|---|---|---|---|---|---|---|---|---|---|---|---|---|---|---|---|---|
| SIX3 | GAGCCAAGATCACATCC | -1 | 1022 | 1039 | 22 | 39 | 0,25 | 0,01 | 17,56 | >0.1 | 5,52 | 8,68 | 0 | 1,86 | 0 | 1,25 | 0 |
| SIX3 | TAAAGTGAATCAAATAA | -1 | 812 | 829 | -188 | -171 | 1,27 | 0,01 | 17,91 | >0.1 | 5,52 | 6,08 | 0 | 1,61 | 0 | 3,43 | 0 |
| SIX3 | AACTAGGGATCATGAAG | -1 | 645 | 662 | -355 | -338 | 5,8 | 0,001 | 19,95 | >0.1 | 3,88 | 5,36 | 0 | 1,08 | 0 | 3,83 | 0 |
| SIX3 | AATAGGTTATCTCACAT | -1 | 589 | 606 | -411 | -394 | 4,38 | 0,001 | 14,02 | >0.1 | 3,42 | 5,2 | 0 | 1,02 | 0 | 0 | 0 |
| SIX3 | TAGGTTATCTCACATTT | -1 | 587 | 604 | -413 | -396 | -1,67 | 0,01 | 7,95 | >0.1 | 3,41 | 5,19 | 0 | 1,02 | 0 | 0 | 0 |
| SIX3 | GTGGCGCGATCTCGGCT | -1 | 402 | 419 | -598 | -581 | -2,74 | 0,01 | 11,49 | >0.1 | 2,64 | 4,79 | 0 | 1,04 | 0 | 5,76 | 0 |
| SIX3 | GACGGGGTTTCACCGTG | -1 | 271 | 288 | -729 | -712 | 7,42 | 0,001 | 21,24 | >0.1 | 2,76 | 4,57 | 0 | 1,29 | 0 | 5,19 | 0 |
| FOSL2::JUN(VAR.2) | AAAGAAGTCAG | 1 | 30 | 41 | -970 | -959 | 0,98 | 0,01 | 11,6 | >0.1 | 4,47 | 4,28 | 0 | 1,87 | 0 | 0 | 0 |
| FOSL2::JUN(VAR.2) | AATGGCGTGAA | 1 | 368 | 379 | -632 | -621 | 1,02 | 0,01 | 16,07 | >0.1 | 2,54 | 4,71 | 0 | 1,08 | 0 | 6,73 | 0 |
| FOSL2::JUN(VAR.2) | GATCGCGCCAC | 1 | 408 | 419 | -592 | -581 | 1,63 | 0,01 | 15,74 | >0.1 | 2,65 | 4,78 | 0 | 1,03 | 0 | 5,65 | 0 |
| FOSL2::JUN(VAR.2) | TGTGAGATAAC | 1 | 590 | 601 | -410 | -399 | 4,68 | 0,001 | 14,28 | >0.1 | 3,41 | 5,18 | 0 | 1,01 | 0 | 0 | 0 |
| FOSL2::JUN(VAR.2) | GATAACCTATT | 1 | 595 | 606 | -405 | -394 | -1,06 | 0,01 | 8,59 | >0.1 | 3,44 | 5,2 | 0 | 1,01 | 0 | 0 | 0 |
| FOSL2::JUN(VAR.2) | GATTTGGCCAT | 1 | 752 | 763 | -248 | -237 | -0,34 | 0,01 | 18,83 | >0.1 | 5,6 | 5,74 | 0 | 1,3 | 0 | 6,54 | 0 |
| FOSL2::JUN(VAR.2) | AGTGACAGGAG | 1 | 765 | 776 | -235 | -224 | -0,45 | 0,01 | 18,2 | >0.1 | 5,6 | 5,8 | 0 | 1,34 | 0 | 5,92 | 0 |
| FOSL2::JUN(VAR.2) | GGAGAGGTAAG | 1 | 772 | 783 | -228 | -217 | 2,21 | 0,01 | 20,92 | >0.1 | 5,6 | 5,83 | 0 | 1,37 | 0 | 5,92 | 0 |
| FOSL2::JUN(VAR.2) | AATAACGTATT | 1 | 800 | 811 | -200 | -189 | 1,62 | 0,01 | 18,24 | >0.1 | 5,6 | 5,98 | 0 | 1,5 | 0 | 3,55 | 0 |
| FOSL2::JUN(VAR.2) | TGTTACATATC | 1 | 847 | 858 | -153 | -142 | -0,33 | 0,01 | 16,25 | >0.1 | 5,6 | 6,28 | 0 | 1,84 | 0 | 2,86 | 0 |
| FOSL2::JUN(VAR.2) | GACGATGTCAA | 1 | 1045 | 1056 | 45 | 56 | 3,26 | 0,01 | 19,07 | >0.1 | 5,6 | 7,7 | 0 | 1,86 | 0 | 0,65 | 0 |
| FOSL2::JUN(VAR.2) | GTTGCTGTAAC | 1 | 1082 | 1093 | 82 | 93 | -0,82 | 0,01 | 17,62 | >0.1 | 5,6 | 6,99 | 0 | 1,85 | 1,79 | 2,21 | 0 |
| FOSL2::JUN(VAR.2) | CTTGACATCGT | -1 | 1046 | 1057 | 46 | 57 | 0,91 | 0,01 | 17,51 | >0.1 | 5,6 | 7,68 | 0 | 1,86 | 0 | 1,46 | 0 |
| FOSL2::JUN(VAR.2) | TGTGAGCCAAG | -1 | 1031 | 1042 | 31 | 42 | 0,55 | 0,01 | 17,5 | >0.1 | 5,6 | 8,19 | 0 | 1,86 | 0 | 1,31 | 0 |
| FOSL2::JUN(VAR.2) | GCTGAGCCAAT | -1 | 882 | 893 | -118 | -107 | -0,03 | 0,01 | 17,08 | >0.1 | 5,6 | 6,56 | 0 | 1,89 | 0 | 3,07 | 0 |
| FOSL2::JUN(VAR.2) | GAATACGTTAT | -1 | 801 | 812 | -199 | -188 | -0,47 | 0,01 | 16,04 | >0.1 | 5,6 | 5,98 | 0 | 1,51 | 0 | 3,43 | 0 |
| FOSL2::JUN(VAR.2) | TATGGCCAAAT | -1 | 753 | 764 | -247 | -236 | -1,33 | 0,01 | 17,63 | >0.1 | 5,6 | 5,74 | 0 | 1,3 | 0 | 6,32 | 0 |
| FOSL2::JUN(VAR.2) | AATGTTATAAT | -1 | 686 | 697 | -314 | -303 | 1,64 | 0,01 | 15,62 | >0.1 | 4,43 | 5,48 | 0 | 1,13 | 0 | 2,95 | 0 |
| FOSL2::JUN(VAR.2) | AGTGACCAAAG | -1 | 670 | 681 | -330 | -319 | 3,09 | 0,01 | 17,46 | >0.1 | 4,15 | 5,42 | 0 | 1,1 | 0 | 3,7 | 0 |
| FOSL2::JUN(VAR.2) | GGTTATCTCAC | -1 | 591 | 602 | -409 | -398 | 3,59 | 0,01 | 13,2 | >0.1 | 3,42 | 5,19 | 0 | 1,01 | 0 | 0 | 0 |
| FOSL2::JUN(VAR.2) | AGTGGCGCGAT | -1 | 409 | 420 | -591 | -580 | 1,26 | 0,01 | 15,25 | >0.1 | 2,65 | 4,79 | 0 | 1,03 | 0 | 5,52 | 0 |
| FOSL2::JUN(VAR.2) | GTTCACGCCAT | -1 | 369 | 380 | -631 | -620 | 3,37 | 0,01 | 17,85 | >0.1 | 2,54 | 4,72 | 0 | 1,07 | 0 | 6,15 | 0 |
| FOSL2::JUN(VAR.2) | GCTGACTTCTT | -1 | 31 | 42 | -969 | -958 | 1,97 | 0,01 | 12,57 | >0.1 | 4,45 | 4,28 | 0 | 1,87 | 0 | 0 | 0 |
| POU5F1B | TTTGCAAAG | 1 | 84 | 93 | -916 | -907 | 4,09 | 0,01 | 14,05 | >0.1 | 3,75 | 4,33 | 0 | 1,88 | 0 | 0 | 0 |
| POU5F1B | AAGGCAGAT | 1 | 90 | 99 | -910 | -901 | 3,22 | 0,01 | 13,14 | >0.1 | 3,7 | 4,34 | 0 | 1,88 | 0 | 0 | 0 |
| POU5F1B | AATTTTAAA | 1 | 134 | 143 | -866 | -857 | 2,94 | 0,01 | 14,17 | >0.1 | 3,38 | 4,39 | 0 | 1,89 | 0 | 1,57 | 0 |
| POU5F1B | CATGGAAAT | 1 | 150 | 159 | -850 | -841 | 9,27 | 0,001 | 20,51 | >0.1 | 3,29 | 4,4 | 0 | 1,9 | 0 | 1,65 | 0 |
| POU5F1B | TATTAAAAA | 1 | 290 | 299 | -710 | -701 | 3,48 | 0,01 | 17,1 | >0.1 | 2,71 | 4,59 | 0 | 1,23 | 0 | 5,09 | 0 |
| POU5F1B | AATACAAAA | 1 | 297 | 306 | -703 | -694 | 3,24 | 0,01 | 16,27 | >0.1 | 2,69 | 4,6 | 0 | 1,21 | 0 | 4,53 | 0 |
| POU5F1B | TAATGAAAT | 1 | 605 | 614 | -395 | -386 | 2,49 | 0,01 | 13,64 | >0.1 | 3,5 | 5,22 | 0 | 1,02 | 0 | 1,41 | 0 |
| POU5F1B | CATACAAAC | 1 | 625 | 634 | -375 | -366 | 3,12 | 0,01 | 14,59 | >0.1 | 3,66 | 5,28 | 0 | 1,04 | 0 | 1,5 | 0 |
| POU5F1B | TAGGAAAAT | 1 | 721 | 730 | -279 | -270 | 3,97 | 0,01 | 21,38 | >0.1 | 5,62 | 5,6 | 0 | 1,2 | 0 | 4,99 | 0 |
| POU5F1B | TATTCTTAT | 1 | 807 | 816 | -193 | -184 | 6,79 | 0,01 | 23,23 | >0.1 | 5,62 | 6 | 0 | 1,53 | 0 | 3,28 | 0 |
| POU5F1B | TATTCTAAA | 1 | 835 | 844 | -165 | -156 | 7,59 | 0,001 | 24,4 | >0.1 | 5,62 | 6,18 | 0 | 1,72 | 0 | 3,28 | 0 |
| POU5F1B | TAGGGAAAG | 1 | 1004 | 1013 | 4 | 13 | 3 | 0,01 | 21,08 | >0.1 | 5,62 | 9,12 | 0 | 1,87 | 0 | 1,46 | 0 |
| POU5F1B | TATGTAACA | -1 | 847 | 856 | -153 | -144 | 2,96 | 0,01 | 19,54 | >0.1 | 5,62 | 6,27 | 0 | 1,82 | 0 | 2,86 | 0 |
| POU5F1B | GAATCAAAT | -1 | 814 | 823 | -186 | -177 | 2,31 | 0,01 | 18,98 | >0.1 | 5,62 | 6,05 | 0 | 1,57 | 0 | 3,43 | 0 |
| POU5F1B | CACGGAAAA | -1 | 695 | 704 | -305 | -296 | 2,45 | 0,01 | 17,78 | >0.1 | 5,62 | 5,5 | 0 | 1,14 | 0 | 3,07 | 0 |
| POU5F1B | CATGAAGAT | -1 | 643 | 652 | -357 | -348 | 4,1 | 0,01 | 18,14 | >0.1 | 3,82 | 5,33 | 0 | 1,06 | 0 | 3,83 | 0 |
| POU5F1B | TATTTTTAT | -1 | 579 | 588 | -421 | -412 | 4,98 | 0,01 | 14,47 | >0.1 | 3,34 | 5,15 | 0 | 1 | 0 | 0 | 0 |
| POU5F1B | TATGGAATT | -1 | 494 | 503 | -506 | -497 | 5,65 | 0,01 | 20,95 | >0.1 | 2,93 | 4,95 | 0 | 0,98 | 0 | 6,44 | 0 |
| POU5F1B | AATTCATAT | -1 | 489 | 498 | -511 | -502 | 4,92 | 0,01 | 20,03 | >0.1 | 2,91 | 4,94 | 0 | 0,98 | 0 | 6,27 | 0 |
| POU5F1B | TATTTTTAA | -1 | 292 | 301 | -708 | -699 | 2,37 | 0,01 | 15,34 | >0.1 | 2,71 | 4,59 | 0 | 1,23 | 0 | 4,45 | 0 |
| POU5F1B | TTTGCAAAA | -1 | 83 | 92 | -917 | -908 | 4,65 | 0,01 | 14,62 | >0.1 | 3,76 | 4,33 | 0 | 1,88 | 0 | 0 | 0 |
| FOXF2 | TAAAAATACAAAAA | 1 | 293 | 307 | -707 | -693 | -1,37 | 0,01 | 11,6 | >0.1 | 2,69 | 4,6 | 0 | 1,23 | 0 | 4,45 | 0 |
| FOXF2 | CAAAAAAAAAAAAA | 1 | 456 | 470 | -544 | -530 | -1,23 | 0,01 | 13,09 | >0.1 | 2,8 | 4,88 | 0 | 1 | 0 | 5,63 | 0 |
| FOXF2 | TGATCAGAAATATG | 1 | 480 | 494 | -520 | -506 | 1,21 | 0,01 | 15,82 | >0.1 | 2,89 | 4,93 | 0 | 0,99 | 0 | 5,8 | 0 |
| FOXF2 | ATTCCATAAAGACA | 1 | 495 | 509 | -505 | -491 | -0,23 | 0,01 | 15,1 | >0.1 | 2,95 | 4,96 | 0 | 0,99 | 0 | 6,44 | 0 |
| FOXF2 | TTGAAATAAAAATA | 1 | 574 | 588 | -426 | -412 | 0,14 | 0,01 | 9,62 | >0.1 | 3,32 | 5,15 | 0 | 1 | 0 | 0 | 0 |
| FOXF2 | TAAAAATAAATGTG | 1 | 580 | 594 | -420 | -406 | 0,73 | 0,01 | 10,26 | >0.1 | 3,36 | 5,17 | 0 | 1,01 | 0 | 0 | 0 |
| FOXF2 | CAAACACCAACATT | 1 | 629 | 643 | -371 | -357 | 1,32 | 0,01 | 13,04 | >0.1 | 3,71 | 5,3 | 0 | 1,05 | 0 | 1,66 | 0 |
| FOXF2 | TCTAGGAAAATATT | 1 | 719 | 733 | -281 | -267 | 2,25 | 0,01 | 19,79 | >0.1 | 5,56 | 5,61 | 0 | 1,21 | 0 | 5,17 | 0 |

| | | | | | | | | | | | | | | | | | |
|---|---|---|---|---|---|---|---|---|---|---|---|---|---|---|---|---|---|
| FOXF2 | CCAGGATGAACTTT | 1 | 866 | 880 | -134 | -120 | -0,48 | 0,01 | 16,28 | >0.1 | 5,56 | 6,45 | 0 | 1,9 | 0 | 2,86 | 0 |
| FOXF2 | GTTACAGCAACAAG | -1 | 1079 | 1093 | 79 | 93 | -0,77 | 0,01 | 17,62 | >0.1 | 5,56 | 7,03 | 0 | 1,85 | 1,79 | 2,16 | 0 |
| FOXF2 | ACACAGTAAACAAT | -1 | 895 | 909 | -105 | -91 | 11,39 | 0,0001 | 28,4 | >0.1 | 5,56 | 6,7 | 0 | 1,89 | 0 | 2,86 | 0 |
| FOXF2 | TTTTGGTTAATATT | -1 | 727 | 741 | -273 | -259 | -0,06 | 0,01 | 17,89 | >0.1 | 5,56 | 5,64 | 0 | 1,23 | 0 | 5,52 | 0 |
| FOXF2 | TAAAGATACACGGA | -1 | 698 | 712 | -302 | -288 | -1,02 | 0,01 | 14,42 | >0.1 | 5,56 | 5,53 | 0 | 1,16 | 0 | 3,19 | 0 |
| FOXF2 | TTACAGTGACCAAA | -1 | 671 | 685 | -329 | -315 | -0,33 | 0,01 | 13,95 | >0.1 | 4,19 | 5,44 | 0 | 1,11 | 0 | 3,55 | 0 |
| FOXF2 | CACAGGTCAACTCT | -1 | 103 | 117 | -897 | -883 | 1,01 | 0,01 | 11,59 | >0.1 | 3,57 | 4,36 | 0 | 1,89 | 0 | 0,77 | 0 |
| FOXF2 | AAAACTTAAAAAAG | -1 | 73 | 87 | -927 | -913 | -0,31 | 0,01 | 9,73 | >0.1 | 3,83 | 4,33 | 0 | 1,88 | 0 | 0 | 0 |
| ZNF263 | TAGGTAAGTGAAGGAGAAGG | 1 | 2 | 23 | -998 | -977 | 3,84 | 0,01 | 17,24 | >0.1 | 5,47 | 4,27 | 0 | 1,86 | 1,79 | 0 | 0 |
| ZNF263 | AGGTAAGTGAAGGAGAAGGA | 1 | 3 | 24 | -997 | -976 | 3,42 | 0,01 | 16,82 | >0.1 | 5,47 | 4,27 | 0 | 1,87 | 1,79 | 0 | 0 |
| ZNF263 | GGTAAGTGAAGGAGAAGGAG | 1 | 4 | 25 | -996 | -975 | 1,94 | 0,01 | 15,34 | >0.1 | 5,47 | 4,27 | 0 | 1,87 | 1,79 | 0 | 0 |
| ZNF263 | AAGTGAAGGAGAAGGAGGCA | 1 | 7 | 28 | -993 | -972 | 2,7 | 0,01 | 16,1 | >0.1 | 5,47 | 4,27 | 0 | 1,87 | 1,79 | 0 | 0 |
| ZNF263 | GAAGGAGGCACTCAAAGAAG | 1 | 17 | 38 | -983 | -962 | 8,34 | 0,01 | 19,96 | >0.1 | 5,47 | 4,28 | 0 | 1,87 | 0 | 0 | 0 |
| ZNF263 | AAAGGCAGATCAGGAGAGTT | 1 | 89 | 110 | -911 | -890 | 2,53 | 0,01 | 13,21 | >0.1 | 3,66 | 4,35 | 0 | 1,88 | 0 | 0,79 | 0 |
| ZNF263 | AGTTGACCTGTGGAGTGGAG | 1 | 105 | 126 | -895 | -874 | 1,42 | 0,01 | 11,98 | >0.1 | 3,53 | 4,37 | 0 | 1,89 | 0 | 0,77 | 0 |
| ZNF263 | GGAGGCTGAGGCAGGAGAAT | 1 | 351 | 372 | -649 | -628 | 11,13 | 0,01 | 25,59 | >0.1 | 2,52 | 4,7 | 0 | 1,11 | 0 | 6,13 | 0 |
| ZNF263 | GAGGCTGAGGCAGGAGAATG | 1 | 352 | 373 | -648 | -627 | 7,73 | 0,01 | 22,19 | >0.1 | 2,52 | 4,71 | 0 | 1,1 | 0 | 6,13 | 0 |
| ZNF263 | GAGGCAGGAGAATGGCGTGA | 1 | 358 | 379 | -642 | -621 | 1,44 | 0,01 | 16,39 | >0.1 | 2,53 | 4,72 | 0 | 1,1 | 0 | 6,61 | 0 |
| ZNF263 | GGAGGCAGAGCTTGCAGTGA | 1 | 383 | 404 | -617 | -596 | 5,68 | 0,01 | 20,22 | >0.1 | 2,59 | 4,76 | 0 | 1,06 | 0 | 6,13 | 0 |
| ZNF263 | GAGGCAGAGCTTGCAGTGAG | 1 | 384 | 405 | -616 | -595 | 2,67 | 0,01 | 17,1 | >0.1 | 2,6 | 4,76 | 0 | 1,06 | 0 | 6,02 | 0 |
| ZNF263 | TAAAGTGACAGGAGAGGTAA | 1 | 762 | 783 | -238 | -217 | 2,06 | 0,01 | 20,65 | >0.1 | 5,47 | 5,83 | 0 | 1,37 | 0 | 5,92 | 0 |
| ZNF263 | AAGTGACAGGAGAGGTAAGG | 1 | 764 | 785 | -236 | -215 | 1,45 | 0,01 | 19,84 | >0.1 | 5,47 | 5,84 | 0 | 1,38 | 0 | 5,69 | 0 |
| ZNF263 | AAAGGCTGATAAGAGAGAAA | 1 | 962 | 983 | -38 | -17 | 7,11 | 0,01 | 24,17 | >0.1 | 5,47 | 8,02 | 0 | 1,87 | 0 | 1,7 | 0 |
| ZNF263 | TGAGAAGGAGCCAGGAAGAG | -1 | 1056 | 1077 | 56 | 77 | 7,17 | 0,01 | 25,89 | >0.1 | 5,47 | 7,44 | 0 | 1,86 | 1,79 | 2,16 | 0 |
| ZNF263 | TGAGCAGGAAAGACCAAAAA | -1 | 924 | 945 | -76 | -55 | 7,96 | 0,01 | 23,48 | >0.1 | 5,47 | 7,14 | 0 | 1,88 | 0 | 1,03 | 0 |
| ZNF263 | GAAAGACCAAAAAAGAAAAA | -1 | 917 | 938 | -83 | -62 | 5,03 | 0,01 | 20,51 | >0.1 | 5,47 | 7,04 | 0 | 1,89 | 0 | 1,09 | 0 |
| ZNF263 | AAAAAAAGAAAAAGAAGAAGA | -1 | 909 | 930 | -91 | -70 | 5,53 | 0,01 | 20,97 | >0.1 | 5,47 | 6,93 | 0 | 1,89 | 0 | 1,15 | 0 |
| ZNF263 | AAAAAAGAAAAAGAAGAAGAA | -1 | 908 | 929 | -92 | -71 | 4,41 | 0,01 | 19,84 | >0.1 | 5,47 | 6,92 | 0 | 1,89 | 0 | 1,15 | 0 |
| ZNF263 | AAAAGAAAAAGAAGAAGAACA | -1 | 906 | 927 | -94 | -73 | 4,46 | 0,01 | 19,87 | >0.1 | 5,47 | 6,89 | 0 | 1,89 | 0 | 1,15 | 0 |
| ZNF263 | TAGGGATCATGAAGATAATGT | -1 | 638 | 659 | -362 | -341 | 1,21 | 0,01 | 15,16 | >0.1 | 3,83 | 5,35 | 0 | 1,07 | 0 | 3,7 | 0 |
| ZNF263 | GGATCATGAAGATAATGTTGG | -1 | 635 | 656 | -365 | -344 | 4,9 | 0,01 | 16,77 | >0.1 | 3,8 | 5,34 | 0 | 1,07 | 0 | 1,66 | 0 |
| RELB | TGAAACCCCGT | 1 | 276 | 287 | -724 | -713 | 3,83 | 0,01 | 17,51 | >0.1 | 2,75 | 4,57 | 0 | 1,27 | 0 | 5,09 | 0 |
| RELB | GAAACCCCGTC | 1 | 277 | 288 | -723 | -712 | 2,29 | 0,01 | 16,08 | >0.1 | 2,75 | 4,57 | 0 | 1,27 | 0 | 5,19 | 0 |
| RELB | GTAGTCCCAGC | 1 | 334 | 345 | -666 | -655 | 5,69 | 0,01 | 19,98 | >0.1 | 2,58 | 4,66 | 0 | 1,13 | 0 | 5,92 | 0 |
| RELB | GAATTCCATAA | 1 | 493 | 504 | -507 | -496 | 3,41 | 0,01 | 18,72 | >0.1 | 2,93 | 4,95 | 0 | 0,98 | 0 | 6,44 | 0 |
| RELB | ATGTTGCCCAA | 1 | 538 | 549 | -462 | -451 | 2,08 | 0,01 | 12,98 | >0.1 | 3,13 | 5,05 | 0 | 0,99 | 0 | 1,73 | 0 |
| RELB | ACATTTTCCGT | 1 | 692 | 703 | -308 | -297 | 2,35 | 0,01 | 16,63 | >0.1 | 4,57 | 5,5 | 0 | 1,14 | 0 | 3,07 | 0 |
| RELB | TCAGTCCACCA | 1 | 1099 | 1110 | 99 | 110 | 3,1 | 0,01 | 22,94 | >0.1 | 5,6 | 6,77 | 0 | 1,85 | 3,22 | 2,4 | 0 |
| RELB | TAATTCCAAGA | -1 | 1187 | 1198 | 187 | 198 | 2,27 | 0,01 | 20,79 | >0.1 | 5,6 | 6,05 | 0 | 1,53 | 1,79 | 3,55 | 0 |
| RELB | ATCGTCCCCTG | -1 | 1040 | 1051 | 40 | 51 | 5,44 | 0,01 | 22,11 | >0.1 | 5,6 | 7,85 | 0 | 1,86 | 0 | 1,36 | 0 |
| RELB | GACTTTCCCTA | -1 | 1004 | 1015 | 4 | 15 | 5,86 | 0,01 | 23,86 | >0.1 | 5,6 | 9,12 | 0 | 1,87 | 0 | 1,42 | 0 |
| RELB | ACTTTCCCTAG | -1 | 1003 | 1014 | 3 | 14 | 3,11 | 0,01 | 21,16 | >0.1 | 5,6 | 9,12 | 0 | 1,87 | 0 | 1,46 | 0 |
| RELB | ATATTCCCAGA | -1 | 559 | 570 | -441 | -430 | 6,98 | 0,01 | 16,31 | >0.1 | 3,23 | 5,1 | 0 | 0,99 | 0 | 0 | 0 |
| RELB | CATTTTCCCTT | -1 | 530 | 541 | -470 | -459 | 3,78 | 0,01 | 16,59 | >0.1 | 3,09 | 5,03 | 0 | 0,98 | 0 | 3,7 | 0 |
| RELB | ATTTTCCCTTC | -1 | 529 | 540 | -471 | -460 | 2,97 | 0,01 | 15,62 | >0.1 | 3,08 | 5,03 | 0 | 0,98 | 0 | 3,55 | 0 |
| RELB | CCATTCTCCTG | -1 | 362 | 373 | -638 | -627 | 6,87 | 0,01 | 21,67 | >0.1 | 2,52 | 4,7 | 0 | 1,08 | 0 | 6,49 | 0 |
| RELB | GGGTTTCACCG | -1 | 273 | 284 | -727 | -716 | 2,15 | 0,01 | 15,85 | >0.1 | 2,76 | 4,57 | 0 | 1,28 | 0 | 5,09 | 0 |
| RELB | GGTTTCACCGT | -1 | 272 | 283 | -728 | -717 | 5,19 | 0,01 | 18,8 | >0.1 | 2,77 | 4,57 | 0 | 1,29 | 0 | 4,99 | 0 |
| RELB | TAATTTCCATG | -1 | 150 | 161 | -850 | -839 | 2,93 | 0,01 | 14,16 | >0.1 | 3,28 | 4,41 | 0 | 1,9 | 0 | 1,65 | 0 |
| RELB | CTACTCTCCAC | -1 | 119 | 130 | -881 | -870 | 2,22 | 0,01 | 13,6 | >0.1 | 3,47 | 4,37 | 0 | 1,89 | 0 | 1,65 | 0 |
| RELB | CCACTCCACAG | -1 | 112 | 123 | -888 | -877 | 3,56 | 0,01 | 14,1 | >0.1 | 3,52 | 4,36 | 0 | 1,89 | 0 | 0,77 | 0 |
| RELB | CAACTCTCCTG | -1 | 99 | 110 | -901 | -890 | 3,29 | 0,01 | 13,91 | >0.1 | 3,62 | 4,35 | 0 | 1,88 | 0 | 0,77 | 0 |
| RELA | GTGAAACCCC | 1 | 275 | 285 | -725 | -715 | 1,51 | 0,01 | 15,2 | >0.1 | 2,76 | 4,57 | 0 | 1,28 | 0 | 5,09 | 0 |
| RELA | GAGAGTTTCT | 1 | 552 | 562 | -448 | -438 | 2,41 | 0,01 | 11,68 | >0.1 | 3,19 | 5,08 | 0 | 0,99 | 0 | 0 | 0 |
| RELA | TGGTCTTTCC | 1 | 929 | 939 | -71 | -61 | 7,83 | 0,001 | 23,4 | >0.1 | 5,61 | 7,05 | 0 | 1,88 | 0 | 1,03 | 0 |
| RELA | GGGAAAGTCA | 1 | 1006 | 1016 | 6 | 16 | 1,22 | 0,01 | 19,28 | >0.1 | 5,61 | 9,12 | 0 | 1,87 | 0 | 1,46 | 0 |
| RELA | ATGACTTTCC | -1 | 1007 | 1017 | 7 | 17 | 5,3 | 0,001 | 23,36 | >0.1 | 5,61 | 9,12 | 0 | 1,87 | 0 | 1,46 | 0 |
| RELA | TGAGATTTTC | -1 | 978 | 988 | -22 | -12 | 0,67 | 0,01 | 17,9 | >0.1 | 5,61 | 8,22 | 0 | 1,87 | 0 | 1,53 | 0 |
| RELA | GAGATTTTCT | -1 | 977 | 987 | -23 | -13 | 0,11 | 0,01 | 17,34 | >0.1 | 5,61 | 8,18 | 0 | 1,87 | 0 | 1,57 | 0 |

| | | | | | | | | | | | | | | | | | |
|---|---|---|---|---|---|---|---|---|---|---|---|---|---|---|---|---|---|
| RELA | ATGACTTTGC | -1 | 512 | 522 | -488 | -478 | -1,21 | 0,01 | 10,72 | >0.1 | 3,01 | 4,99 | 0 | 0,98 | 0 | 2,95 | 0 |
| RELA | CGGGGTTTCA | -1 | 276 | 286 | -724 | -714 | 7,78 | 0,001 | 21,46 | >0.1 | 2,75 | 4,57 | 0 | 1,27 | 0 | 5,09 | 0 |
| RELA | GGGGTTTCAC | -1 | 275 | 285 | -725 | -715 | 4,5 | 0,01 | 18,19 | >0.1 | 2,76 | 4,57 | 0 | 1,28 | 0 | 5,09 | 0 |
| RELA | TCGGCCTCCC | -1 | 216 | 226 | -784 | -774 | -1,16 | 0,01 | 13,35 | >0.1 | 2,97 | 4,49 | 0 | 1,53 | 0 | 5,52 | 0 |
| RELA | TTTAATTTCC | -1 | 153 | 163 | -847 | -837 | 3,47 | 0,01 | 14,69 | >0.1 | 3,27 | 4,41 | 0 | 1,9 | 0 | 1,65 | 0 |
| RAX | GGAAATGAGC | 1 | 64 | 74 | -936 | -926 | 3,43 | 0,01 | 13,57 | >0.1 | 3,95 | 4,31 | 0 | 1,88 | 0 | 0 | 0 |
| RAX | GGAAATTAAA | 1 | 153 | 163 | -847 | -837 | 7,5 | 0,001 | 18,72 | >0.1 | 3,27 | 4,41 | 0 | 1,9 | 0 | 1,65 | 0 |
| RAX | CTCTATTAAA | 1 | 287 | 297 | -713 | -703 | 3,58 | 0,01 | 17,32 | >0.1 | 2,72 | 4,59 | 0 | 1,24 | 0 | 5,19 | 0 |
| RAX | ACCTATTAAT | 1 | 599 | 609 | -401 | -391 | 5,4 | 0,01 | 16,45 | >0.1 | 3,47 | 5,21 | 0 | 1,02 | 0 | 1,36 | 0 |
| RAX | ATTAATGAAA | 1 | 603 | 613 | -397 | -387 | 4,27 | 0,01 | 15,41 | >0.1 | 3,49 | 5,22 | 0 | 1,02 | 0 | 1,41 | 0 |
| RAX | TAAAATTATA | 1 | 682 | 692 | -318 | -308 | 3,31 | 0,01 | 17,29 | >0.1 | 4,34 | 5,46 | 0 | 1,12 | 0 | 3,07 | 0 |
| RAX | AAAAATTATT | 1 | 829 | 839 | -171 | -161 | 4,74 | 0,01 | 21,46 | >0.1 | 5,61 | 6,15 | 0 | 1,68 | 0 | 3,28 | 0 |
| RAX | GCCAATATAA | -1 | 878 | 888 | -122 | -112 | 2,44 | 0,01 | 19,53 | >0.1 | 5,61 | 6,51 | 0 | 1,9 | 0 | 3,07 | 0 |
| RAX | AATAATTTTT | -1 | 829 | 839 | -171 | -161 | 2,89 | 0,01 | 19,61 | >0.1 | 5,61 | 6,15 | 0 | 1,68 | 0 | 3,28 | 0 |
| RAX | ATCAAATAAG | -1 | 811 | 821 | -189 | -179 | 5,77 | 0,01 | 22,4 | >0.1 | 5,61 | 6,03 | 0 | 1,56 | 0 | 3,43 | 0 |
| RAX | CCAAATCAAA | -1 | 749 | 759 | -251 | -241 | 2,8 | 0,01 | 21,33 | >0.1 | 5,61 | 5,72 | 0 | 1,28 | 0 | 5,92 | 0 |
| RAX | GACAATTTCA | -1 | 608 | 618 | -392 | -382 | 4,13 | 0,01 | 15,46 | >0.1 | 3,53 | 5,23 | 0 | 1,02 | 0 | 1,55 | 0 |
| RAX | TCCAAATACA | -1 | 521 | 531 | -479 | -469 | 4,19 | 0,01 | 16,3 | >0.1 | 3,05 | 5,01 | 0 | 0,98 | 0 | 3,07 | 0 |
| RAX | TTTAATTTCC | -1 | 153 | 163 | -847 | -837 | 3,71 | 0,01 | 14,93 | >0.1 | 3,27 | 4,41 | 0 | 1,9 | 0 | 1,65 | 0 |
| RAX | TAAAATTATG | -1 | 131 | 141 | -869 | -859 | 3,31 | 0,01 | 14,56 | >0.1 | 3,4 | 4,38 | 0 | 1,89 | 0 | 1,57 | 0 |
| RAX | TTCACTTACC | -1 | 4 | 14 | -996 | -986 | 3,16 | 0,01 | 16,68 | >0.1 | 5,61 | 4,26 | 0 | 1,86 | 1,79 | 0 | 0 |
| CENPB | CCAGTGTGAGAAATA | 1 | 49 | 64 | -951 | -936 | -4,27 | 0,01 | 6,01 | >0.1 | 4,11 | 4,3 | 0 | 1,87 | 0 | 0 | 0 |
| CENPB | CCCGTCTCTATTAAA | 1 | 282 | 297 | -718 | -703 | 5,29 | 0,001 | 18,85 | >0.1 | 2,73 | 4,59 | 0 | 1,26 | 0 | 4,99 | 0 |
| CENPB | TCCAGCCTGGGCGAC | 1 | 424 | 439 | -576 | -561 | -0,45 | 0,01 | 13,76 | >0.1 | 2,7 | 4,82 | 0 | 1,02 | 0 | 5,67 | 0 |
| CENPB | CTCAGTCTCAAAAAA | 1 | 448 | 463 | -552 | -537 | -4,39 | 0,01 | 9,52 | >0.1 | 2,78 | 4,87 | 0 | 1 | 0 | 5,26 | 0 |
| CENPB | ACCTCTTTGGTCACT | 1 | 666 | 681 | -334 | -319 | -3,36 | 0,01 | 10,98 | >0.1 | 4,12 | 5,42 | 0 | 1,1 | 0 | 3,7 | 0 |
| CENPB | CCTGCTCAGCGCCCA | 1 | 937 | 952 | -63 | -48 | -2,05 | 0,01 | 13,48 | >0.1 | 5,54 | 7,25 | 0 | 1,88 | 0 | 0,85 | 0 |
| CENPB | ACCTGTTCTATCAAA | 1 | 1161 | 1176 | 161 | 176 | 3,33 | 0,001 | 20,28 | >0.1 | 4,47 | 6,22 | 0 | 1,75 | 1,79 | 2,72 | 0 |
| CENPB | CCCCTGTGAGCCAAG | -1 | 1031 | 1046 | 31 | 46 | 0,99 | 0,01 | 17,89 | >0.1 | 5,54 | 8,19 | 0 | 1,86 | 0 | 1,31 | 0 |
| CENPB | ATCACATCCACTGAA | -1 | 1016 | 1031 | 16 | 31 | -1,42 | 0,01 | 16,13 | >0.1 | 5,54 | 8,81 | 0 | 1,86 | 0 | 1,34 | 0 |
| CENPB | ACCTCCTCATGAGAT | -1 | 982 | 997 | -18 | -3 | 0,28 | 0,01 | 17,86 | >0.1 | 5,54 | 8,64 | 0 | 1,87 | 0 | 1,53 | 0 |
| CENPB | CACTTTATGGCCAAA | -1 | 754 | 769 | -246 | -231 | -3,72 | 0,01 | 15,23 | >0.1 | 5,54 | 5,76 | 0 | 1,32 | 0 | 6,32 | 0 |
| CENPB | ATCATATTCCCAGAA | -1 | 558 | 573 | -442 | -427 | -3,54 | 0,01 | 5,81 | >0.1 | 3,24 | 5,11 | 0 | 1 | 0 | 0 | 0 |
| CENPB | CCAGGCTGGAGCACA | -1 | 419 | 434 | -581 | -566 | -0,67 | 0,01 | 13,27 | >0.1 | 2,69 | 4,81 | 0 | 1,02 | 0 | 5,42 | 0 |
| CENPB | TCAGCCTCCCGAGTA | -1 | 345 | 360 | -655 | -640 | -2,67 | 0,01 | 12,2 | >0.1 | 2,55 | 4,68 | 0 | 1,11 | 0 | 6,52 | 0 |
| CENPB | TCCTTCTCCTTCACT | -1 | 8 | 23 | -992 | -977 | -3,61 | 0,01 | 9,86 | >0.1 | 5,54 | 4,27 | 0 | 1,86 | 1,79 | 0 | 0 |
| CENPB | TCCTTCACTTACCTA | -1 | 2 | 17 | -998 | -983 | -3,32 | 0,01 | 10,14 | >0.1 | 5,54 | 4,26 | 0 | 1,86 | 1,79 | 0 | 0 |
| ESX1 | GGAAATTAAA | 1 | 153 | 163 | -847 | -837 | 5,53 | 0,01 | 16,75 | >0.1 | 3,27 | 4,41 | 0 | 1,9 | 0 | 1,65 | 0 |
| ESX1 | ACTGATCAGA | 1 | 163 | 173 | -837 | -827 | 4,22 | 0,01 | 16,16 | >0.1 | 3,21 | 4,42 | 0 | 1,9 | 0 | 2,4 | 0 |
| ESX1 | CCTGACTAAC | 1 | 262 | 272 | -738 | -728 | 3,32 | 0,01 | 16,52 | >0.1 | 2,8 | 4,55 | 0 | 1,32 | 0 | 4,53 | 0 |
| ESX1 | CTCTATTAAA | 1 | 287 | 297 | -713 | -703 | 4,74 | 0,01 | 18,48 | >0.1 | 2,72 | 4,59 | 0 | 1,24 | 0 | 5,19 | 0 |
| ESX1 | AAAACTTAGC | 1 | 303 | 313 | -697 | -687 | 3,3 | 0,01 | 16,77 | >0.1 | 2,67 | 4,61 | 0 | 1,2 | 0 | 4,99 | 0 |
| ESX1 | ACTGATCAGA | 1 | 478 | 488 | -522 | -512 | 4,22 | 0,01 | 18,36 | >0.1 | 2,88 | 4,92 | 0 | 0,99 | 0 | 5,36 | 0 |
| ESX1 | ACCTATTAAT | 1 | 599 | 609 | -401 | -391 | 5,88 | 0,01 | 16,93 | >0.1 | 3,47 | 5,21 | 0 | 1,02 | 0 | 1,36 | 0 |
| ESX1 | ATTAATGAAA | 1 | 603 | 613 | -397 | -387 | 3,69 | 0,01 | 14,83 | >0.1 | 3,49 | 5,22 | 0 | 1,02 | 0 | 1,41 | 0 |
| ESX1 | TAAAATTATA | 1 | 682 | 692 | -318 | -308 | 3,35 | 0,01 | 17,33 | >0.1 | 4,34 | 5,46 | 0 | 1,12 | 0 | 3,07 | 0 |
| ESX1 | AAAAATTATT | 1 | 829 | 839 | -171 | -161 | 3,82 | 0,01 | 20,54 | >0.1 | 5,61 | 6,15 | 0 | 1,68 | 0 | 3,28 | 0 |
| ESX1 | GCTGATAAGA | 1 | 966 | 976 | -34 | -24 | 3,76 | 0,01 | 20,72 | >0.1 | 5,61 | 7,78 | 0 | 1,87 | 0 | 1,7 | 0 |
| ESX1 | AGCAGTTACA | -1 | 1087 | 1097 | 87 | 97 | 3,32 | 0,01 | 21,71 | >0.1 | 5,61 | 6,92 | 0 | 1,85 | 1,79 | 2,21 | 0 |
| ESX1 | TCTTATCAGC | -1 | 966 | 976 | -34 | -24 | 4,11 | 0,01 | 21,07 | >0.1 | 5,61 | 7,78 | 0 | 1,87 | 0 | 1,7 | 0 |
| ESX1 | AACAATCTGC | -1 | 891 | 901 | -109 | -99 | 3,5 | 0,01 | 20,49 | >0.1 | 5,61 | 6,62 | 0 | 1,89 | 0 | 2,86 | 0 |
| ESX1 | CCAAATCAAA | -1 | 749 | 759 | -251 | -241 | 4,34 | 0,01 | 22,87 | >0.1 | 5,61 | 5,72 | 0 | 1,28 | 0 | 5,92 | 0 |
| ESX1 | GACAATTTCA | -1 | 608 | 618 | -392 | -382 | 4,45 | 0,01 | 15,78 | >0.1 | 3,53 | 5,23 | 0 | 1,02 | 0 | 1,55 | 0 |
| ESX1 | TTTCATTAAT | -1 | 603 | 613 | -397 | -387 | 4,32 | 0,01 | 15,46 | >0.1 | 3,49 | 5,22 | 0 | 1,02 | 0 | 1,41 | 0 |
| ESX1 | TCCAAATACA | -1 | 521 | 531 | -479 | -469 | 3,58 | 0,01 | 15,69 | >0.1 | 3,05 | 5,01 | 0 | 0,98 | 0 | 3,07 | 0 |
| ESX1 | TCTGATCAGT | -1 | 478 | 488 | -522 | -512 | 3,6 | 0,01 | 17,74 | >0.1 | 2,88 | 4,92 | 0 | 0,99 | 0 | 5,36 | 0 |
| ESX1 | TCTGATCAGT | -1 | 163 | 173 | -837 | -827 | 3,6 | 0,01 | 15,54 | >0.1 | 3,21 | 4,42 | 0 | 1,9 | 0 | 2,4 | 0 |
| ESX1 | TTTAATTTCC | -1 | 153 | 163 | -847 | -837 | 4,9 | 0,01 | 16,12 | >0.1 | 3,27 | 4,41 | 0 | 1,9 | 0 | 1,65 | 0 |
| ESX1 | TAAAATTATG | -1 | 131 | 141 | -869 | -859 | 3,56 | 0,01 | 14,81 | >0.1 | 3,4 | 4,38 | 0 | 1,89 | 0 | 1,57 | 0 |

| | | | | | | | | | | | | | | | | | |
|---|---|---|---|---|---|---|---|---|---|---|---|---|---|---|---|---|---|
| ESX1 | GCTCATTTCC | -1 | 64 | 74 | -936 | -926 | 4,03 | 0,01 | 14,17 | >0.1 | 3,95 | 4,31 | 0 | 1,88 | 0 | 0 | 0 |
| ESX1 | TTCACTTACC | -1 | 4 | 14 | -996 | -986 | 5,88 | 0,01 | 19,4 | >0.1 | 5,61 | 4,26 | 0 | 1,86 | 1,79 | 0 | 0 |
| SHOX2 | GAAATGAG | 1 | 65 | 73 | -935 | -927 | 2,62 | 0,01 | 12,76 | >0.1 | 3,95 | 4,31 | 0 | 1,88 | 0 | 0 | 0 |
| SHOX2 | ATAATTTT | 1 | 132 | 140 | -868 | -860 | 2,46 | 0,01 | 13,7 | >0.1 | 3,4 | 4,38 | 0 | 1,89 | 0 | 1,57 | 0 |
| SHOX2 | GAAATTAA | 1 | 154 | 162 | -846 | -838 | 6,06 | 0,01 | 17,28 | >0.1 | 3,27 | 4,41 | 0 | 1,9 | 0 | 1,65 | 0 |
| SHOX2 | TCTATTAA | 1 | 288 | 296 | -712 | -704 | 5,24 | 0,01 | 18,97 | >0.1 | 2,72 | 4,58 | 0 | 1,24 | 0 | 5,19 | 0 |
| SHOX2 | CCTATTAA | 1 | 600 | 608 | -400 | -392 | 6,42 | 0,01 | 17,47 | >0.1 | 3,47 | 5,2 | 0 | 1,01 | 0 | 1,36 | 0 |
| SHOX2 | TTAATGAA | 1 | 604 | 612 | -396 | -388 | 5,45 | 0,01 | 16,59 | >0.1 | 3,49 | 5,21 | 0 | 1,02 | 0 | 1,41 | 0 |
| SHOX2 | AAAATTAT | 1 | 683 | 691 | -317 | -309 | 3,58 | 0,01 | 17,56 | >0.1 | 4,34 | 5,46 | 0 | 1,12 | 0 | 3,07 | 0 |
| SHOX2 | CTTATTTG | 1 | 811 | 819 | -189 | -181 | 2,05 | 0,01 | 18,54 | >0.1 | 5,64 | 6,02 | 0 | 1,55 | 0 | 3,28 | 0 |
| SHOX2 | AAAATTAT | 1 | 830 | 838 | -170 | -162 | 3,58 | 0,01 | 20,31 | >0.1 | 5,64 | 6,14 | 0 | 1,67 | 0 | 3,28 | 0 |
| SHOX2 | CTCATGAG | 1 | 984 | 992 | -16 | -8 | 2,71 | 0,01 | 20,23 | >0.1 | 5,64 | 8,43 | 0 | 1,87 | 0 | 1,57 | 0 |
| SHOX2 | GGAATTAT | 1 | 1191 | 1199 | 191 | 199 | 2,93 | 0,01 | 21,71 | >0.1 | 5,64 | 6,02 | 0 | 1,5 | 1,79 | 3,82 | 0 |
| SHOX2 | CTCATGAG | -1 | 984 | 992 | -16 | -8 | 2,71 | 0,01 | 20,23 | >0.1 | 5,64 | 8,43 | 0 | 1,87 | 0 | 1,57 | 0 |
| SHOX2 | ATAATTTT | -1 | 830 | 838 | -170 | -162 | 2,46 | 0,01 | 19,19 | >0.1 | 5,64 | 6,14 | 0 | 1,67 | 0 | 3,28 | 0 |
| SHOX2 | CAAATAAG | -1 | 811 | 819 | -189 | -181 | 2,81 | 0,01 | 19,3 | >0.1 | 5,64 | 6,02 | 0 | 1,55 | 0 | 3,28 | 0 |
| SHOX2 | CAAATCAA | -1 | 750 | 758 | -250 | -242 | 2,72 | 0,01 | 21,27 | >0.1 | 5,64 | 5,71 | 0 | 1,28 | 0 | 5,92 | 0 |
| SHOX2 | ATAATTTT | -1 | 683 | 691 | -317 | -309 | 2,46 | 0,01 | 16,44 | >0.1 | 4,34 | 5,46 | 0 | 1,12 | 0 | 3,07 | 0 |
| SHOX2 | ACAATTTC | -1 | 609 | 617 | -391 | -383 | 3,84 | 0,01 | 15,16 | >0.1 | 3,53 | 5,23 | 0 | 1,02 | 0 | 1,55 | 0 |
| SHOX2 | TTCATTAA | -1 | 604 | 612 | -396 | -388 | 4,97 | 0,01 | 16,11 | >0.1 | 3,49 | 5,21 | 0 | 1,02 | 0 | 1,41 | 0 |
| SHOX2 | TTAATTTC | -1 | 154 | 162 | -846 | -838 | 4,37 | 0,01 | 15,59 | >0.1 | 3,27 | 4,41 | 0 | 1,9 | 0 | 1,65 | 0 |
| SHOX2 | AAAATTAT | -1 | 132 | 140 | -868 | -860 | 3,58 | 0,01 | 14,82 | >0.1 | 3,4 | 4,38 | 0 | 1,89 | 0 | 1,57 | 0 |
| SHOX2 | TCACTTAC | -1 | 5 | 13 | -995 | -987 | 2,65 | 0,01 | 16,2 | >0.1 | 5,64 | 4,26 | 0 | 1,86 | 1,79 | 0 | 0 |
| SNAI2 | AATAGGTAA | 1 | 0 | 9 | -1000 | -991 | 2,19 | 0,01 | 15,72 | >0.1 | 5,62 | 4,25 | 0 | 1,86 | 1,79 | 0 | 0 |
| SNAI2 | GGTAAGTGA | 1 | 4 | 13 | -996 | -987 | 3 | 0,01 | 16,54 | >0.1 | 5,62 | 4,26 | 0 | 1,86 | 1,79 | 0 | 0 |
| SNAI2 | AACCAGTGT | 1 | 47 | 56 | -953 | -944 | 4,06 | 0,01 | 14,41 | >0.1 | 4,18 | 4,29 | 0 | 1,87 | 0 | 0 | 0 |
| SNAI2 | GACCTGTGG | 1 | 109 | 118 | -891 | -882 | 2,3 | 0,01 | 12,84 | >0.1 | 3,55 | 4,36 | 0 | 1,89 | 0 | 0,75 | 0 |
| SNAI2 | TGGTGGTGG | 1 | 319 | 328 | -681 | -672 | 1,85 | 0,01 | 16,52 | >0.1 | 2,63 | 4,63 | 0 | 1,16 | 0 | 6,26 | 0 |
| SNAI2 | GTCATGTAT | 1 | 517 | 526 | -483 | -474 | 2,68 | 0,01 | 14,64 | >0.1 | 3,03 | 5 | 0 | 0,98 | 0 | 2,95 | 0 |
| SNAI2 | CCCAAGTAG | 1 | 544 | 553 | -456 | -447 | 2,19 | 0,01 | 11,39 | >0.1 | 3,15 | 5,06 | 0 | 0,99 | 0 | 0 | 0 |
| SNAI2 | TACCGGTTT | 1 | 742 | 751 | -258 | -249 | 3,66 | 0,01 | 22,33 | >0.1 | 5,62 | 5,68 | 0 | 1,26 | 0 | 6,1 | 0 |
| SNAI2 | GAGAGGTAA | 1 | 773 | 782 | -227 | -218 | 2,49 | 0,01 | 21,22 | >0.1 | 5,62 | 5,82 | 0 | 1,37 | 0 | 5,92 | 0 |
| SNAI2 | GTAAGGTTC | 1 | 778 | 787 | -222 | -213 | 3,99 | 0,01 | 23,39 | >0.1 | 5,62 | 5,85 | 0 | 1,39 | 0 | 6,54 | 0 |
| SNAI2 | CCCAAGTTC | 1 | 953 | 962 | -47 | -38 | 3,17 | 0,01 | 19,81 | >0.1 | 5,62 | 7,44 | 0 | 1,88 | 0 | 1,7 | 0 |
| SNAI2 | AGGAGGTTT | 1 | 990 | 999 | -10 | -1 | 4,87 | 0,01 | 22,66 | >0.1 | 5,62 | 8,69 | 0 | 1,87 | 0 | 1,61 | 0 |
| SNAI2 | GACAAGTTT | 1 | 1136 | 1145 | 136 | 145 | 6,94 | 0,01 | 27,06 | >0.1 | 5,62 | 6,41 | 0 | 1,84 | 3,22 | 3,02 | 0 |
| SNAI2 | AGCAAGTGA | -1 | 1178 | 1187 | 178 | 187 | 7,89 | 0,001 | 26,27 | >0.1 | 5,62 | 6,1 | 0 | 1,6 | 1,79 | 3,26 | 0 |
| SNAI2 | AACAGGTCT | -1 | 1159 | 1168 | 159 | 168 | 8,4 | 0,001 | 24,96 | >0.1 | 4,37 | 6,23 | 0 | 1,77 | 1,79 | 2,4 | 0 |
| SNAI2 | CAATGGTGG | -1 | 1104 | 1113 | 104 | 113 | 1,91 | 0,01 | 21,79 | >0.1 | 5,62 | 6,71 | 0 | 1,85 | 3,22 | 2,47 | 0 |
| SNAI2 | AACCGGTAC | -1 | 741 | 750 | -259 | -250 | 4,53 | 0,01 | 23 | >0.1 | 5,62 | 5,68 | 0 | 1,25 | 0 | 5,92 | 0 |
| SNAI2 | AAGAGGTCT | -1 | 664 | 673 | -336 | -327 | 3,03 | 0,01 | 17,56 | >0.1 | 4,05 | 5,4 | 0 | 1,09 | 0 | 3,99 | 0 |
| SNAI2 | TGTTGGTGT | -1 | 632 | 641 | -368 | -359 | 2,02 | 0,01 | 13,73 | >0.1 | 3,72 | 5,3 | 0 | 1,05 | 0 | 1,66 | 0 |
| SNAI2 | AATAGGTTA | -1 | 597 | 606 | -403 | -394 | 2,6 | 0,01 | 13,7 | >0.1 | 3,45 | 5,2 | 0 | 1,01 | 0 | 1,44 | 0 |
| SNAI2 | CCCAGGTTC | -1 | 376 | 385 | -624 | -615 | 6,45 | 0,01 | 21,19 | >0.1 | 2,56 | 4,72 | 0 | 1,06 | 0 | 6,4 | 0 |
| SNAI2 | TACAGGTGC | -1 | 328 | 337 | -672 | -663 | 12,37 | 0,001 | 26,32 | >0.1 | 2,6 | 4,65 | 0 | 1,14 | 0 | 5,56 | 0 |
| SNAI2 | TACAGGCGT | -1 | 194 | 203 | -806 | -797 | 4,67 | 0,01 | 18,13 | >0.1 | 3,07 | 4,46 | 0 | 1,67 | 0 | 4,26 | 0 |
| SNAI2 | CACAGGTCA | -1 | 108 | 117 | -892 | -883 | 5,68 | 0,01 | 16,24 | >0.1 | 3,56 | 4,36 | 0 | 1,89 | 0 | 0,77 | 0 |
| SNAI2 | CACTGGTTC | -1 | 46 | 55 | -954 | -945 | 4,53 | 0,01 | 14,89 | >0.1 | 4,19 | 4,29 | 0 | 1,87 | 0 | 0 | 0 |
| RFX4 | AGTAGTCATAATTTTA | 1 | 125 | 141 | -875 | -859 | -3,31 | 0,01 | 7,96 | >0.1 | 3,41 | 4,38 | 0 | 1,89 | 0 | 1,57 | 0 |
| RFX4 | AATGGCCATGGAAATT | 1 | 144 | 160 | -856 | -840 | 3,92 | 0,001 | 15,13 | >0.1 | 3,3 | 4,41 | 0 | 1,9 | 0 | 1,61 | 0 |
| RFX4 | TGTAATCCTAGCACTT | 1 | 199 | 215 | -801 | -785 | -0,5 | 0,01 | 13,35 | >0.1 | 3,03 | 4,47 | 0 | 1,64 | 0 | 4,7 | 0 |
| RFX4 | GACCGTCCTGACTAAC | 1 | 256 | 272 | -744 | -728 | -3,65 | 0,01 | 9,58 | >0.1 | 2,81 | 4,55 | 0 | 1,34 | 0 | 4,53 | 0 |
| RFX4 | TGTAGTCCCAGCTACT | 1 | 333 | 349 | -667 | -651 | 6,73 | 0,001 | 21,02 | >0.1 | 2,58 | 4,67 | 0 | 1,14 | 0 | 5,92 | 0 |
| RFX4 | AGCTACTCGGGAGGCT | 1 | 342 | 358 | -658 | -642 | -3,69 | 0,01 | 11,19 | >0.1 | 2,55 | 4,68 | 0 | 1,12 | 0 | 6,52 | 0 |
| RFX4 | TCCAGCCTGGGCGACA | 1 | 424 | 440 | -576 | -560 | -3,47 | 0,01 | 10,88 | >0.1 | 2,7 | 4,82 | 0 | 1,02 | 0 | 5,8 | 0 |
| RFX4 | GAATTCCATAAAGACA | 1 | 493 | 509 | -507 | -491 | 2,36 | 0,01 | 17,69 | >0.1 | 2,94 | 4,96 | 0 | 0,99 | 0 | 6,44 | 0 |
| RFX4 | TGTTGCCCAAGTAGAG | 1 | 539 | 555 | -461 | -445 | -1,56 | 0,01 | 7,64 | >0.1 | 3,14 | 5,07 | 0 | 0,99 | 0 | 0 | 0 |
| RFX4 | AGTTTCTGGGAATATG | 1 | 555 | 571 | -445 | -429 | 0,94 | 0,01 | 10,27 | >0.1 | 3,22 | 5,11 | 0 | 1 | 0 | 0 | 0 |
| RFX4 | GAAAACCATACAAACA | 1 | 619 | 635 | -381 | -365 | 1,87 | 0,01 | 13,27 | >0.1 | 3,64 | 5,28 | 0 | 1,04 | 0 | 1,44 | 0 |

| | | | | | | | | | | | | | | | | | |
|---|---|---|---|---|---|---|---|---|---|---|---|---|---|---|---|---|---|
| RFX4 | TATCTTCATGATCCCT | 1 | 642 | 658 | -358 | -342 | -2,98 | 0,01 | 11,11 | >0.1 | 3,84 | 5,35 | 0 | 1,07 | 0 | 3,83 | 0 |
| RFX4 | GACCTCTTTGGTCACT | 1 | 665 | 681 | -335 | -319 | -1,4 | 0,01 | 12,94 | >0.1 | 4,11 | 5,42 | 0 | 1,1 | 0 | 3,7 | 0 |
| RFX4 | TGTATCTTTAACAGCT | 1 | 702 | 718 | -298 | -282 | 3,09 | 0,01 | 20,34 | >0.1 | 5,53 | 5,55 | 0 | 1,17 | 0 | 4,99 | 0 |
| RFX4 | ATTAACCAAAAGTACC | 1 | 730 | 746 | -270 | -254 | -1,52 | 0,01 | 16,84 | >0.1 | 5,53 | 5,66 | 0 | 1,25 | 0 | 5,92 | 0 |
| RFX4 | GCTCTTCCTGGCTCCT | 1 | 1056 | 1072 | 56 | 72 | -3,88 | 0,01 | 14,9 | >0.1 | 5,53 | 7,44 | 0 | 1,86 | 1,79 | 2,16 | 0 |
| RFX4 | TGTTGCTGTAACTGCT | 1 | 1081 | 1097 | 81 | 97 | 5,41 | 0,001 | 23,75 | >0.1 | 5,53 | 7 | 0 | 1,85 | 1,79 | 2,16 | 0 |
| RFX4 | TGCTGCTCAGTCCACC | 1 | 1093 | 1109 | 93 | 109 | -4,1 | 0,01 | 15,67 | >0.1 | 5,53 | 6,84 | 0 | 1,85 | 3,22 | 2,32 | 0 |
| RFX4 | TTTAACCACGAAGCCG | 1 | 1142 | 1158 | 142 | 158 | -1,33 | 0,01 | 17,79 | >0.1 | 5,53 | 6,36 | 0 | 1,84 | 3,22 | 2,16 | 0 |
| RFX4 | CGGCTTCGTGGTTAAA | -1 | 1142 | 1158 | 142 | 158 | -2,83 | 0,01 | 16,29 | >0.1 | 5,53 | 6,36 | 0 | 1,84 | 3,22 | 2,16 | 0 |
| RFX4 | AGCAGTTACAGCAACA | -1 | 1081 | 1097 | 81 | 97 | 4,13 | 0,001 | 22,47 | >0.1 | 5,53 | 7 | 0 | 1,85 | 1,79 | 2,16 | 0 |
| RFX4 | AGGAGCCAGGAAGAGC | -1 | 1056 | 1072 | 56 | 72 | -4,29 | 0,01 | 14,49 | >0.1 | 5,53 | 7,44 | 0 | 1,86 | 1,79 | 2,16 | 0 |
| RFX4 | GTGAGCCAAGATCACA | -1 | 1025 | 1041 | 25 | 41 | -3,46 | 0,01 | 13,76 | >0.1 | 5,53 | 8,52 | 0 | 1,86 | 0 | 1,31 | 0 |
| RFX4 | GGTACTTTTGGTTAAT | -1 | 730 | 746 | -270 | -254 | -1,23 | 0,01 | 17,13 | >0.1 | 5,53 | 5,66 | 0 | 1,25 | 0 | 5,92 | 0 |
| RFX4 | TATTTTCCTAGAAAGC | -1 | 715 | 731 | -285 | -269 | -3,86 | 0,01 | 13,47 | >0.1 | 5,53 | 5,6 | 0 | 1,21 | 0 | 4,99 | 0 |
| RFX4 | AGCTGTTAAAGATACA | -1 | 702 | 718 | -298 | -282 | 1,41 | 0,01 | 18,66 | >0.1 | 5,53 | 5,55 | 0 | 1,17 | 0 | 4,99 | 0 |
| RFX4 | TAATTTTACAGTGACC | -1 | 674 | 690 | -326 | -310 | -2,35 | 0,01 | 11,9 | >0.1 | 4,25 | 5,45 | 0 | 1,12 | 0 | 3,43 | 0 |
| RFX4 | CATATTCCCAGAAACT | -1 | 555 | 571 | -445 | -429 | 0,92 | 0,01 | 10,25 | >0.1 | 3,22 | 5,11 | 0 | 1 | 0 | 0 | 0 |
| RFX4 | CTCTACTTGGGCAACA | -1 | 539 | 555 | -461 | -445 | 0,66 | 0,01 | 9,86 | >0.1 | 3,14 | 5,07 | 0 | 0,99 | 0 | 0 | 0 |
| RFX4 | TGTCTTTATGGAATTC | -1 | 493 | 509 | -507 | -491 | -2,27 | 0,01 | 13,06 | >0.1 | 2,94 | 4,96 | 0 | 0,99 | 0 | 6,44 | 0 |
| RFX4 | TGTCGCCCAGGCTGGA | -1 | 424 | 440 | -576 | -560 | -2,87 | 0,01 | 11,48 | >0.1 | 2,7 | 4,82 | 0 | 1,02 | 0 | 5,8 | 0 |
| RFX4 | AGCCTCCCGAGTAGCT | -1 | 342 | 358 | -658 | -642 | -3,64 | 0,01 | 11,24 | >0.1 | 2,55 | 4,68 | 0 | 1,12 | 0 | 6,52 | 0 |
| RFX4 | AGTAGCTGGGACTACA | -1 | 333 | 349 | -667 | -651 | 8,38 | 0,001 | 22,67 | >0.1 | 2,58 | 4,67 | 0 | 1,14 | 0 | 5,92 | 0 |
| RFX4 | AAGTGCTAGGATTACA | -1 | 199 | 215 | -801 | -785 | 2,09 | 0,01 | 15,94 | >0.1 | 3,03 | 4,47 | 0 | 1,64 | 0 | 4,7 | 0 |
| RFX4 | AATTTCCATGGCCATT | -1 | 144 | 160 | -856 | -840 | 4,79 | 0,001 | 16 | >0.1 | 3,3 | 4,41 | 0 | 1,9 | 0 | 1,61 | 0 |
| RFX4 | TAAAATTATGACTACT | -1 | 125 | 141 | -875 | -859 | 3,25 | 0,001 | 14,52 | >0.1 | 3,41 | 4,38 | 0 | 1,89 | 0 | 1,57 | 0 |
| RFX4 | TGGTTCTGTGGCTGAC | -1 | 36 | 52 | -964 | -948 | -4,37 | 0,01 | 6,09 | >0.1 | 4,3 | 4,29 | 0 | 1,87 | 0 | 0 | 0 |
| RFX5 | AATGGCCATGGAAATT | 1 | 144 | 160 | -856 | -840 | 0,13 | 0,001 | 11,34 | >0.1 | 3,3 | 4,41 | 0 | 1,9 | 0 | 1,61 | 0 |
| RFX5 | TGTAATCCTAGCACTT | 1 | 199 | 215 | -801 | -785 | -8,41 | 0,01 | 5,44 | >0.1 | 3,03 | 4,47 | 0 | 1,64 | 0 | 4,7 | 0 |
| RFX5 | CGTCTCTATTAAAAAT | 1 | 284 | 300 | -716 | -700 | -9,46 | 0,01 | 4,3 | >0.1 | 2,72 | 4,59 | 0 | 1,25 | 0 | 5,19 | 0 |
| RFX5 | CGTGGTGGTGGGCACC | 1 | 317 | 333 | -683 | -667 | -10,91 | 0,01 | 3,2 | >0.1 | 2,62 | 4,64 | 0 | 1,17 | 0 | 5,68 | 0 |
| RFX5 | TGTAGTCCCAGCTACT | 1 | 333 | 349 | -667 | -651 | -5,77 | 0,01 | 8,52 | >0.1 | 2,58 | 4,67 | 0 | 1,14 | 0 | 5,92 | 0 |
| RFX5 | GAATTCCATAAAGACA | 1 | 493 | 509 | -507 | -491 | -8,12 | 0,01 | 7,21 | >0.1 | 2,94 | 4,96 | 0 | 0,99 | 0 | 6,44 | 0 |
| RFX5 | TGTTGCCCAAGTAGAG | 1 | 539 | 555 | -461 | -445 | -0,65 | 0,001 | 8,55 | >0.1 | 3,14 | 5,07 | 0 | 0,99 | 0 | 0 | 0 |
| RFX5 | GAAAACCATACAAACA | 1 | 619 | 635 | -381 | -365 | -8,63 | 0,01 | 2,77 | >0.1 | 3,64 | 5,28 | 0 | 1,04 | 0 | 1,44 | 0 |
| RFX5 | GACCTCTTTGGTCACT | 1 | 665 | 681 | -335 | -319 | -9,08 | 0,01 | 5,26 | >0.1 | 4,11 | 5,42 | 0 | 1,1 | 0 | 3,7 | 0 |
| RFX5 | TGTATCTTTAACAGCT | 1 | 702 | 718 | -298 | -282 | -5,18 | 0,01 | 12,07 | >0.1 | 5,53 | 5,55 | 0 | 1,17 | 0 | 4,99 | 0 |
| RFX5 | TGTGATCTTGGCTCAC | 1 | 1025 | 1041 | 25 | 41 | -11,17 | 0,01 | 6,05 | >0.1 | 5,53 | 8,52 | 0 | 1,86 | 0 | 1,31 | 0 |
| RFX5 | TGTTGCTGTAACTGCT | 1 | 1081 | 1097 | 81 | 97 | -2,26 | 0,001 | 16,08 | >0.1 | 5,53 | 7 | 0 | 1,85 | 1,79 | 2,16 | 0 |
| RFX5 | GGTGGACTGAGCAGCA | -1 | 1093 | 1109 | 93 | 109 | -10,76 | 0,01 | 9,01 | >0.1 | 5,53 | 6,84 | 0 | 1,85 | 3,22 | 2,32 | 0 |
| RFX5 | AGCAGTTACAGCAACA | -1 | 1081 | 1097 | 81 | 97 | -3,17 | 0,01 | 15,17 | >0.1 | 5,53 | 7 | 0 | 1,85 | 1,79 | 2,16 | 0 |
| RFX5 | GGTACTTTTGGTTAAT | -1 | 730 | 746 | -270 | -254 | -10,45 | 0,01 | 7,91 | >0.1 | 5,53 | 5,66 | 0 | 1,25 | 0 | 5,92 | 0 |
| RFX5 | TATTTTCCTAGAAAGC | -1 | 715 | 731 | -285 | -269 | -8,15 | 0,01 | 9,18 | >0.1 | 5,53 | 5,6 | 0 | 1,21 | 0 | 4,99 | 0 |
| RFX5 | AGCTGTTAAAGATACA | -1 | 702 | 718 | -298 | -282 | -5,98 | 0,01 | 11,27 | >0.1 | 5,53 | 5,55 | 0 | 1,17 | 0 | 4,99 | 0 |
| RFX5 | AGTGACCAAAGAGGTC | -1 | 665 | 681 | -335 | -319 | -10,73 | 0,01 | 3,61 | >0.1 | 4,11 | 5,42 | 0 | 1,1 | 0 | 3,7 | 0 |
| RFX5 | TGTTTGTATGGTTTTC | -1 | 619 | 635 | -381 | -365 | -6,67 | 0,01 | 4,73 | >0.1 | 3,64 | 5,28 | 0 | 1,04 | 0 | 1,44 | 0 |
| RFX5 | CTCTACTTGGGCAACA | -1 | 539 | 555 | -461 | -445 | -2,27 | 0,001 | 6,93 | >0.1 | 3,14 | 5,07 | 0 | 0,99 | 0 | 0 | 0 |
| RFX5 | TGTCTTTATGGAATTC | -1 | 493 | 509 | -507 | -491 | -7,78 | 0,01 | 7,55 | >0.1 | 2,94 | 4,96 | 0 | 0,99 | 0 | 6,44 | 0 |
| RFX5 | TGTCGCCCAGGCTGGA | -1 | 424 | 440 | -576 | -560 | -9,19 | 0,01 | 5,16 | >0.1 | 2,7 | 4,82 | 0 | 1,02 | 0 | 5,8 | 0 |
| RFX5 | AGTAGCTGGGACTACA | -1 | 333 | 349 | -667 | -651 | -6,16 | 0,01 | 8,13 | >0.1 | 2,58 | 4,67 | 0 | 1,14 | 0 | 5,92 | 0 |
| RFX5 | GGTGCCCACCACCACG | -1 | 317 | 333 | -683 | -667 | -9,72 | 0,01 | 4,39 | >0.1 | 2,62 | 4,64 | 0 | 1,17 | 0 | 5,68 | 0 |
| RFX5 | AAGTGCTAGGATTACA | -1 | 199 | 215 | -801 | -785 | -9,15 | 0,01 | 4,7 | >0.1 | 3,03 | 4,47 | 0 | 1,64 | 0 | 4,7 | 0 |
| RFX5 | AATTTCCATGGCCATT | -1 | 144 | 160 | -856 | -840 | 0,5 | 0,001 | 11,71 | >0.1 | 3,3 | 4,41 | 0 | 1,9 | 0 | 1,61 | 0 |
| RFX5 | ATCTGCCTTTGCAAAA | -1 | 83 | 99 | -917 | -901 | -10,24 | 0,01 | -0,29 | >0.1 | 3,73 | 4,34 | 0 | 1,88 | 0 | 0 | 0 |
| RFX1 | ATGGCCATGGAAAT | 1 | 145 | 159 | -855 | -841 | 0,35 | 0,01 | 11,56 | >0.1 | 3,3 | 4,4 | 0 | 1,9 | 0 | 1,61 | 0 |
| RFX1 | CTGGGCACAGTGGC | 1 | 178 | 192 | -822 | -808 | -3,89 | 0,01 | 9,44 | >0.1 | 3,13 | 4,44 | 0 | 1,81 | 0 | 3,95 | 0 |
| RFX1 | GTAGTCCCAGCTAC | 1 | 334 | 348 | -666 | -652 | 5,99 | 0,001 | 20,28 | >0.1 | 2,58 | 4,66 | 0 | 1,13 | 0 | 5,92 | 0 |
| RFX1 | GCTACTCGGGAGGC | 1 | 343 | 357 | -657 | -643 | -0,95 | 0,01 | 13,92 | >0.1 | 2,55 | 4,68 | 0 | 1,12 | 0 | 6,52 | 0 |
| RFX1 | TGAACCTGGGAGGC | 1 | 375 | 389 | -625 | -611 | -2,25 | 0,01 | 12,51 | >0.1 | 2,56 | 4,73 | 0 | 1,07 | 0 | 6,4 | 0 |
| RFX1 | CCAGCCTGGGCGAC | 1 | 425 | 439 | -575 | -561 | 6,6 | 0,001 | 20,95 | >0.1 | 2,7 | 4,82 | 0 | 1,02 | 0 | 5,8 | 0 |

| | | | | | | | | | | | | | | | | | |
|---|---|---|---|---|---|---|---|---|---|---|---|---|---|---|---|---|---|
| RFX1 | AATTCCATAAAGAC | 1 | 494 | 508 | -506 | -492 | 2,89 | 0,01 | 18,22 | >0.1 | 2,94 | 4,96 | 0 | 0,99 | 0 | 6,44 | 0 |
| RFX1 | GTTGCCCAAGTAGA | 1 | 540 | 554 | -460 | -446 | -6,83 | 0,01 | 2,37 | >0.1 | 3,14 | 5,07 | 0 | 0,99 | 0 | 0 | 0 |
| RFX1 | AATTGTCTGAAAAC | 1 | 611 | 625 | -389 | -375 | -3,22 | 0,01 | 8,13 | >0.1 | 3,57 | 5,25 | 0 | 1,03 | 0 | 1,5 | 0 |
| RFX1 | ACCTCTTTGGTCAC | 1 | 666 | 680 | -334 | -320 | -6,79 | 0,01 | 7,54 | >0.1 | 4,11 | 5,42 | 0 | 1,1 | 0 | 3,7 | 0 |
| RFX1 | GTATCTTTAACAGC | 1 | 703 | 717 | -297 | -283 | 1,74 | 0,01 | 19 | >0.1 | 5,56 | 5,55 | 0 | 1,17 | 0 | 4,99 | 0 |
| RFX1 | GCTTTCTAGGAAAA | 1 | 715 | 729 | -285 | -271 | -6,13 | 0,01 | 11,39 | >0.1 | 5,56 | 5,6 | 0 | 1,2 | 0 | 5,17 | 0 |
| RFX1 | TTAACCAAAAGTAC | 1 | 731 | 745 | -269 | -255 | -5,97 | 0,01 | 12,4 | >0.1 | 5,56 | 5,66 | 0 | 1,24 | 0 | 5,92 | 0 |
| RFX1 | GGCCATAAAGTGAC | 1 | 757 | 771 | -243 | -229 | -2,62 | 0,01 | 16,14 | >0.1 | 5,56 | 5,77 | 0 | 1,33 | 0 | 6,1 | 0 |
| RFX1 | CTCTTCCTGGCTCC | 1 | 1057 | 1071 | 57 | 71 | -2,87 | 0,01 | 14,12 | >0.1 | 5,56 | 7,42 | 0 | 1,86 | 0 | 2,16 | 0 |
| RFX1 | GTTGCTGTAACTGC | 1 | 1082 | 1096 | 82 | 96 | -0,52 | 0,01 | 17,83 | >0.1 | 5,56 | 6,99 | 0 | 1,85 | 1,79 | 2,16 | 0 |
| RFX1 | GTGGACTGAGCAGC | -1 | 1094 | 1108 | 94 | 108 | 2,25 | 0,01 | 22,03 | >0.1 | 5,56 | 6,83 | 0 | 1,85 | 3,22 | 2,32 | 0 |
| RFX1 | GCAGTTACAGCAAC | -1 | 1082 | 1096 | 82 | 96 | 4,04 | 0,001 | 22,39 | >0.1 | 5,56 | 6,99 | 0 | 1,85 | 1,79 | 2,16 | 0 |
| RFX1 | ATCCACTGAATGAC | -1 | 1012 | 1026 | 12 | 26 | -4,57 | 0,01 | 13,16 | >0.1 | 5,56 | 8,92 | 0 | 1,86 | 0 | 1,39 | 0 |
| RFX1 | TTCATCCTGGAGAG | -1 | 862 | 876 | -138 | -124 | -5,49 | 0,01 | 11,13 | >0.1 | 5,56 | 6,41 | 0 | 1,9 | 0 | 2,75 | 0 |
| RFX1 | TAATCCTAGAGAAC | -1 | 783 | 797 | -217 | -203 | -5,96 | 0,01 | 12,62 | >0.1 | 5,56 | 5,9 | 0 | 1,43 | 0 | 5,69 | 0 |
| RFX1 | ATTTTCCTAGAAAG | -1 | 716 | 730 | -284 | -270 | -3,97 | 0,01 | 13,37 | >0.1 | 5,56 | 5,6 | 0 | 1,2 | 0 | 4,99 | 0 |
| RFX1 | TTTTCCTAGAAAGC | -1 | 715 | 729 | -285 | -271 | 0,88 | 0,01 | 18,4 | >0.1 | 5,56 | 5,6 | 0 | 1,2 | 0 | 5,17 | 0 |
| RFX1 | GCTGTTAAAGATAC | -1 | 703 | 717 | -297 | -283 | -0,58 | 0,01 | 16,68 | >0.1 | 5,56 | 5,55 | 0 | 1,17 | 0 | 4,99 | 0 |
| RFX1 | AATTTTACAGTGAC | -1 | 675 | 689 | -325 | -311 | -6,1 | 0,01 | 8,14 | >0.1 | 4,25 | 5,45 | 0 | 1,12 | 0 | 3,43 | 0 |
| RFX1 | ATATTCCCAGAAAC | -1 | 556 | 570 | -444 | -430 | 3,13 | 0,01 | 12,45 | >0.1 | 3,22 | 5,1 | 0 | 1 | 0 | 0 | 0 |
| RFX1 | TCTACTTGGGCAAC | -1 | 540 | 554 | -460 | -446 | 4,98 | 0,001 | 14,18 | >0.1 | 3,14 | 5,07 | 0 | 0,99 | 0 | 0 | 0 |
| RFX1 | GTCGCCCAGGCTGG | -1 | 425 | 439 | -575 | -561 | -4,22 | 0,01 | 10,13 | >0.1 | 2,7 | 4,82 | 0 | 1,02 | 0 | 5,8 | 0 |
| RFX1 | GCCTCCCGAGTAGC | -1 | 343 | 357 | -657 | -643 | 3,94 | 0,001 | 18,81 | >0.1 | 2,55 | 4,68 | 0 | 1,12 | 0 | 6,52 | 0 |
| RFX1 | GTAGCTGGGACTAC | -1 | 334 | 348 | -666 | -652 | -0,51 | 0,01 | 13,78 | >0.1 | 2,58 | 4,66 | 0 | 1,13 | 0 | 5,92 | 0 |
| RFX1 | TTTTTAATAGAGAC | -1 | 285 | 299 | -715 | -701 | -6,5 | 0,01 | 7,25 | >0.1 | 2,72 | 4,59 | 0 | 1,25 | 0 | 5,19 | 0 |
| RFX1 | AGTGCTAGGATTAC | -1 | 200 | 214 | -800 | -786 | -2,68 | 0,01 | 11,16 | >0.1 | 3,03 | 4,47 | 0 | 1,63 | 0 | 4,7 | 0 |
| RFX1 | AAAATTATGACTAC | -1 | 126 | 140 | -874 | -860 | -5,37 | 0,01 | 5,89 | >0.1 | 3,41 | 4,38 | 0 | 1,89 | 0 | 1,57 | 0 |
| RFX1 | CTCCACAGGTCAAC | -1 | 106 | 120 | -894 | -880 | -4,76 | 0,01 | 5,78 | >0.1 | 3,55 | 4,36 | 0 | 1,89 | 0 | 0,75 | 0 |
| RFX2 | AGTAGTCATAATTTTA | 1 | 125 | 141 | -875 | -859 | -8,3 | 0,01 | 2,97 | >0.1 | 3,41 | 4,38 | 0 | 1,89 | 0 | 1,57 | 0 |
| RFX2 | AATGGCCATGGAAATT | 1 | 144 | 160 | -856 | -840 | -1,99 | 0,001 | 9,22 | >0.1 | 3,3 | 4,41 | 0 | 1,9 | 0 | 1,61 | 0 |
| RFX2 | TGTAATCCTAGCACTT | 1 | 199 | 215 | -801 | -785 | -9,4 | 0,01 | 4,45 | >0.1 | 3,03 | 4,47 | 0 | 1,64 | 0 | 4,7 | 0 |
| RFX2 | CGTCTCTATTAAAAAT | 1 | 284 | 300 | -716 | -700 | -14,49 | 0,01 | -0,73 | >0.1 | 2,72 | 4,59 | 0 | 1,25 | 0 | 5,19 | 0 |
| RFX2 | TGTAGTCCCAGCTACT | 1 | 333 | 349 | -667 | -651 | -2,64 | 0,001 | 11,65 | >0.1 | 2,58 | 4,67 | 0 | 1,14 | 0 | 5,92 | 0 |
| RFX2 | TCCAGCCTGGGCGACA | 1 | 424 | 440 | -576 | -560 | -13,58 | 0,01 | 0,77 | >0.1 | 2,7 | 4,82 | 0 | 1,02 | 0 | 5,8 | 0 |
| RFX2 | GAATTCCATAAAGACA | 1 | 493 | 509 | -507 | -491 | -4,54 | 0,01 | 10,79 | >0.1 | 2,94 | 4,96 | 0 | 0,99 | 0 | 6,44 | 0 |
| RFX2 | TGTTGCCCAAGTAGAG | 1 | 539 | 555 | -461 | -445 | -11,15 | 0,01 | -1,95 | >0.1 | 3,14 | 5,07 | 0 | 0,99 | 0 | 0 | 0 |
| RFX2 | AGTTTCTGGGAATATG | 1 | 555 | 571 | -445 | -429 | -10,98 | 0,01 | -1,65 | >0.1 | 3,22 | 5,11 | 0 | 1 | 0 | 0 | 0 |
| RFX2 | GAAAACCATACAAACA | 1 | 619 | 635 | -381 | -365 | -6,14 | 0,01 | 5,26 | >0.1 | 3,64 | 5,28 | 0 | 1,04 | 0 | 1,44 | 0 |
| RFX2 | CATTATCTTCATGATC | 1 | 639 | 655 | -361 | -345 | -14,29 | 0,01 | -0,24 | >0.1 | 3,81 | 5,34 | 0 | 1,07 | 0 | 3,83 | 0 |
| RFX2 | TATCTTCATGATCCCT | 1 | 642 | 658 | -358 | -342 | -14,66 | 0,01 | -0,57 | >0.1 | 3,84 | 5,35 | 0 | 1,07 | 0 | 3,83 | 0 |
| RFX2 | GACCTCTTTGGTCACT | 1 | 665 | 681 | -335 | -319 | -13,65 | 0,01 | 0,69 | >0.1 | 4,11 | 5,42 | 0 | 1,1 | 0 | 3,7 | 0 |
| RFX2 | GGTCACTGTAAAATTA | 1 | 674 | 690 | -326 | -310 | -12,56 | 0,01 | 1,69 | >0.1 | 4,25 | 5,45 | 0 | 1,12 | 0 | 3,43 | 0 |
| RFX2 | TGTATCTTTAACAGCT | 1 | 702 | 718 | -298 | -282 | -4,43 | 0,01 | 12,82 | >0.1 | 5,53 | 5,55 | 0 | 1,17 | 0 | 4,99 | 0 |
| RFX2 | GCTTTCTAGGAAAATA | 1 | 715 | 731 | -285 | -269 | -11,07 | 0,01 | 6,26 | >0.1 | 5,53 | 5,6 | 0 | 1,21 | 0 | 4,99 | 0 |
| RFX2 | ATTAACCAAAAGTACC | 1 | 730 | 746 | -270 | -254 | -10,09 | 0,01 | 8,27 | >0.1 | 5,53 | 5,66 | 0 | 1,25 | 0 | 5,92 | 0 |
| RFX2 | TGTTGCTGTAACTGCT | 1 | 1081 | 1097 | 81 | 97 | -2,47 | 0,001 | 15,87 | >0.1 | 5,53 | 7 | 0 | 1,85 | 1,79 | 2,16 | 0 |
| RFX2 | TTTAACCACGAAGCCG | 1 | 1142 | 1158 | 142 | 158 | -14,42 | 0,01 | 4,7 | >0.1 | 5,53 | 6,36 | 0 | 1,84 | 3,22 | 2,16 | 0 |
| RFX2 | AGCAGTTACAGCAACA | -1 | 1081 | 1097 | 81 | 97 | -2,56 | 0,001 | 15,78 | >0.1 | 5,53 | 7 | 0 | 1,85 | 1,79 | 2,16 | 0 |
| RFX2 | GGTACTTTTGGTTAAT | -1 | 730 | 746 | -270 | -254 | -10,69 | 0,01 | 7,67 | >0.1 | 5,53 | 5,66 | 0 | 1,25 | 0 | 5,92 | 0 |
| RFX2 | TATTTTCCTAGAAAGC | -1 | 715 | 731 | -285 | -269 | -14,51 | 0,01 | 2,82 | >0.1 | 5,53 | 5,6 | 0 | 1,21 | 0 | 4,99 | 0 |
| RFX2 | AGCTGTTAAAGATACA | -1 | 702 | 718 | -298 | -282 | -5,18 | 0,01 | 12,07 | >0.1 | 5,53 | 5,55 | 0 | 1,17 | 0 | 4,99 | 0 |
| RFX2 | TAATTTTACAGTGACC | -1 | 674 | 690 | -326 | -310 | -12,67 | 0,01 | 1,58 | >0.1 | 4,25 | 5,45 | 0 | 1,12 | 0 | 3,43 | 0 |
| RFX2 | TGTTTGTATGGTTTTC | -1 | 619 | 635 | -381 | -365 | -7,25 | 0,01 | 4,15 | >0.1 | 3,64 | 5,28 | 0 | 1,04 | 0 | 1,44 | 0 |
| RFX2 | CATATTCCCAGAAACT | -1 | 555 | 571 | -445 | -429 | -10,48 | 0,01 | -1,15 | >0.1 | 3,22 | 5,11 | 0 | 1 | 0 | 0 | 0 |
| RFX2 | CTCTACTTGGGCAACA | -1 | 539 | 555 | -461 | -445 | -11,33 | 0,01 | -2,13 | >0.1 | 3,14 | 5,07 | 0 | 0,99 | 0 | 0 | 0 |
| RFX2 | TGTCTTTATGGAATTC | -1 | 493 | 509 | -507 | -491 | -5,8 | 0,01 | 9,53 | >0.1 | 2,94 | 4,96 | 0 | 0,99 | 0 | 6,44 | 0 |
| RFX2 | AGTAGCTGGGACTACA | -1 | 333 | 349 | -667 | -651 | -2,92 | 0,001 | 11,37 | >0.1 | 2,58 | 4,67 | 0 | 1,14 | 0 | 5,92 | 0 |
| RFX2 | ATTTTTAATAGAGACG | -1 | 284 | 300 | -716 | -700 | -14,44 | 0,01 | -0,68 | >0.1 | 2,72 | 4,59 | 0 | 1,25 | 0 | 5,19 | 0 |
| RFX2 | AAGTGCTAGGATTACA | -1 | 199 | 215 | -801 | -785 | -10,6 | 0,01 | 3,25 | >0.1 | 3,03 | 4,47 | 0 | 1,64 | 0 | 4,7 | 0 |

| | | | | | | | | | | | | | | | | | |
|---|---|---|---|---|---|---|---|---|---|---|---|---|---|---|---|---|---|
| RFX2 | AATTTCCATGGCCATT | -1 | 144 | 160 | -856 | -840 | -0,84 | 0,001 | 10,37 | >0.1 | 3,3 | 4,41 | 0 | 1,9 | 0 | 1,61 | 0 |
| RFX2 | TAAAATTATGACTACT | -1 | 125 | 141 | -875 | -859 | -8,14 | 0,01 | 3,13 | >0.1 | 3,41 | 4,38 | 0 | 1,89 | 0 | 1,57 | 0 |
| RFX3 | AGTAGTCATAATTTTA | 1 | 125 | 141 | -875 | -859 | -4,3 | 0,01 | 6,97 | >0.1 | 3,41 | 4,38 | 0 | 1,89 | 0 | 1,57 | 0 |
| RFX3 | AATGGCCATGGAAATT | 1 | 144 | 160 | -856 | -840 | 3,21 | 0,001 | 14,42 | >0.1 | 3,3 | 4,41 | 0 | 1,9 | 0 | 1,61 | 0 |
| RFX3 | TGTAATCCTAGCACTT | 1 | 199 | 215 | -801 | -785 | -3,92 | 0,01 | 9,93 | >0.1 | 3,03 | 4,47 | 0 | 1,64 | 0 | 4,7 | 0 |
| RFX3 | TGTAGTCCCAGCTACT | 1 | 333 | 349 | -667 | -651 | 0,66 | 0,01 | 14,95 | >0.1 | 2,58 | 4,67 | 0 | 1,14 | 0 | 5,92 | 0 |
| RFX3 | TCCAGCCTGGGCGACA | 1 | 424 | 440 | -576 | -560 | -4,33 | 0,01 | 10,02 | >0.1 | 2,7 | 4,82 | 0 | 1,02 | 0 | 5,8 | 0 |
| RFX3 | GAATTCCATAAAGACA | 1 | 493 | 509 | -507 | -491 | 2,47 | 0,001 | 17,8 | >0.1 | 2,94 | 4,96 | 0 | 0,99 | 0 | 6,44 | 0 |
| RFX3 | TGTTGCCCAAGTAGAG | 1 | 539 | 555 | -461 | -445 | -0,41 | 0,01 | 8,79 | >0.1 | 3,14 | 5,07 | 0 | 0,99 | 0 | 0 | 0 |
| RFX3 | AGTTTCTGGGAATATG | 1 | 555 | 571 | -445 | -429 | -3,25 | 0,01 | 6,08 | >0.1 | 3,22 | 5,11 | 0 | 1 | 0 | 0 | 0 |
| RFX3 | GAAAACCATACAAACA | 1 | 619 | 635 | -381 | -365 | 1,13 | 0,01 | 12,53 | >0.1 | 3,64 | 5,28 | 0 | 1,04 | 0 | 1,44 | 0 |
| RFX3 | GACCTCTTTGGTCACT | 1 | 665 | 681 | -335 | -319 | -4,67 | 0,01 | 9,67 | >0.1 | 4,11 | 5,42 | 0 | 1,1 | 0 | 3,7 | 0 |
| RFX3 | GGTCACTGTAAAATTA | 1 | 674 | 690 | -326 | -310 | -5,76 | 0,01 | 8,49 | >0.1 | 4,25 | 5,45 | 0 | 1,12 | 0 | 3,43 | 0 |
| RFX3 | TGTATCTTTAACAGCT | 1 | 702 | 718 | -298 | -282 | 1,44 | 0,01 | 18,69 | >0.1 | 5,53 | 5,55 | 0 | 1,17 | 0 | 4,99 | 0 |
| RFX3 | GCTTTCTAGGAAAATA | 1 | 715 | 731 | -285 | -269 | -5,72 | 0,01 | 11,61 | >0.1 | 5,53 | 5,6 | 0 | 1,21 | 0 | 4,99 | 0 |
| RFX3 | ATTAACCAAAAGTACC | 1 | 730 | 746 | -270 | -254 | -2,03 | 0,01 | 16,33 | >0.1 | 5,53 | 5,66 | 0 | 1,25 | 0 | 5,92 | 0 |
| RFX3 | TGTTGCTGTAACTGCT | 1 | 1081 | 1097 | 81 | 97 | 2,68 | 0,001 | 21,02 | >0.1 | 5,53 | 7 | 0 | 1,85 | 1,79 | 2,16 | 0 |
| RFX3 | TTTAACCACGAAGCCG | 1 | 1142 | 1158 | 142 | 158 | -5,33 | 0,01 | 13,79 | >0.1 | 5,53 | 6,36 | 0 | 1,84 | 3,22 | 2,16 | 0 |
| RFX3 | CGGCTTCGTGGTTAAA | -1 | 1142 | 1158 | 142 | 158 | -5,96 | 0,01 | 13,16 | >0.1 | 5,53 | 6,36 | 0 | 1,84 | 3,22 | 2,16 | 0 |
| RFX3 | AGCAGTTACAGCAACA | -1 | 1081 | 1097 | 81 | 97 | 1,82 | 0,01 | 20,16 | >0.1 | 5,53 | 7 | 0 | 1,85 | 1,79 | 2,16 | 0 |
| RFX3 | AGGAGCCAGGAAGAGC | -1 | 1056 | 1072 | 56 | 72 | -5,87 | 0,01 | 12,91 | >0.1 | 5,53 | 7,44 | 0 | 1,86 | 1,79 | 2,16 | 0 |
| RFX3 | GGTACTTTTGGTTAAT | -1 | 730 | 746 | -270 | -254 | -4,31 | 0,01 | 14,05 | >0.1 | 5,53 | 5,66 | 0 | 1,25 | 0 | 5,92 | 0 |
| RFX3 | TATTTTCCTAGAAAGC | -1 | 715 | 731 | -285 | -269 | -4,81 | 0,01 | 12,52 | >0.1 | 5,53 | 5,6 | 0 | 1,21 | 0 | 4,99 | 0 |
| RFX3 | AGCTGTTAAAGATACA | -1 | 702 | 718 | -298 | -282 | 0,97 | 0,01 | 18,22 | >0.1 | 5,53 | 5,55 | 0 | 1,17 | 0 | 4,99 | 0 |
| RFX3 | TAATTTTACAGTGACC | -1 | 674 | 690 | -326 | -310 | -5,38 | 0,01 | 8,87 | >0.1 | 4,25 | 5,45 | 0 | 1,12 | 0 | 3,43 | 0 |
| RFX3 | AGTGACCAAAGAGGTC | -1 | 665 | 681 | -335 | -319 | -4,26 | 0,01 | 10,08 | >0.1 | 4,11 | 5,42 | 0 | 1,1 | 0 | 3,7 | 0 |
| RFX3 | TGTTTGTATGGTTTTC | -1 | 619 | 635 | -381 | -365 | -1,59 | 0,01 | 9,81 | >0.1 | 3,64 | 5,28 | 0 | 1,04 | 0 | 1,44 | 0 |
| RFX3 | CATATTCCCAGAAACT | -1 | 555 | 571 | -445 | -429 | -4,39 | 0,01 | 4,94 | >0.1 | 3,22 | 5,11 | 0 | 1 | 0 | 0 | 0 |
| RFX3 | CTCTACTTGGGCAACA | -1 | 539 | 555 | -461 | -445 | -1,54 | 0,01 | 7,66 | >0.1 | 3,14 | 5,07 | 0 | 0,99 | 0 | 0 | 0 |
| RFX3 | TGTCTTTATGGAATTC | -1 | 493 | 509 | -507 | -491 | -1,13 | 0,01 | 14,2 | >0.1 | 2,94 | 4,96 | 0 | 0,99 | 0 | 6,44 | 0 |
| RFX3 | TGTCGCCCAGGCTGGA | -1 | 424 | 440 | -576 | -560 | -4,19 | 0,01 | 10,16 | >0.1 | 2,7 | 4,82 | 0 | 1,02 | 0 | 5,8 | 0 |
| RFX3 | AGTAGCTGGGACTACA | -1 | 333 | 349 | -667 | -651 | 2,76 | 0,001 | 17,05 | >0.1 | 2,58 | 4,67 | 0 | 1,14 | 0 | 5,92 | 0 |
| RFX3 | AAGTGCTAGGATTACA | -1 | 199 | 215 | -801 | -785 | -1,61 | 0,01 | 12,24 | >0.1 | 3,03 | 4,47 | 0 | 1,64 | 0 | 4,7 | 0 |
| RFX3 | AATTTCCATGGCCATT | -1 | 144 | 160 | -856 | -840 | 2,87 | 0,001 | 14,08 | >0.1 | 3,3 | 4,41 | 0 | 1,9 | 0 | 1,61 | 0 |
| RFX3 | TAAAATTATGACTACT | -1 | 125 | 141 | -875 | -859 | -2,86 | 0,01 | 8,41 | >0.1 | 3,41 | 4,38 | 0 | 1,89 | 0 | 1,57 | 0 |
| RFX3 | TGGTTCTGTGGCTGAC | -1 | 36 | 52 | -964 | -948 | -5,91 | 0,01 | 4,55 | >0.1 | 4,3 | 4,29 | 0 | 1,87 | 0 | 0 | 0 |
| RFX3 | GACTTCTTTGAGTGCC | -1 | 23 | 39 | -977 | -961 | -4,96 | 0,01 | 6,72 | >0.1 | 5,53 | 4,28 | 0 | 1,87 | 0 | 0 | 0 |
| ELK3 | TAGGTAAGTG | 1 | 2 | 12 | -998 | -988 | -1,85 | 0,01 | 11,67 | >0.1 | 5,61 | 4,26 | 0 | 1,86 | 1,79 | 0 | 0 |
| ELK3 | ATAGGAAATG | 1 | 61 | 71 | -939 | -929 | 3,9 | 0,01 | 14,07 | >0.1 | 3,98 | 4,31 | 0 | 1,88 | 0 | 0 | 0 |
| ELK3 | CATGGAAATT | 1 | 150 | 160 | -850 | -840 | -1,5 | 0,01 | 9,73 | >0.1 | 3,28 | 4,41 | 0 | 1,9 | 0 | 1,65 | 0 |
| ELK3 | ATCAGAAATG | 1 | 167 | 177 | -833 | -823 | -2,75 | 0,01 | 9,17 | >0.1 | 3,19 | 4,43 | 0 | 1,9 | 0 | 2,4 | 0 |
| ELK3 | ATCAGAAATA | 1 | 482 | 492 | -518 | -508 | -2,99 | 0,01 | 11,61 | >0.1 | 2,89 | 4,93 | 0 | 0,99 | 0 | 5,8 | 0 |
| ELK3 | TTTGGAAGGG | 1 | 525 | 535 | -475 | -465 | -2,85 | 0,01 | 9,5 | >0.1 | 3,06 | 5,02 | 0 | 0,98 | 0 | 3,28 | 0 |
| ELK3 | AAGGGAAAAT | 1 | 530 | 540 | -470 | -460 | -1,67 | 0,01 | 11,13 | >0.1 | 3,09 | 5,03 | 0 | 0,98 | 0 | 3,7 | 0 |
| ELK3 | ACCGGTTTTG | 1 | 743 | 753 | -257 | -247 | -2,66 | 0,01 | 15,82 | >0.1 | 5,61 | 5,69 | 0 | 1,26 | 0 | 5,92 | 0 |
| ELK3 | CCAGGATGAA | 1 | 866 | 876 | -134 | -124 | 2,7 | 0,01 | 19,37 | >0.1 | 5,61 | 6,41 | 0 | 1,9 | 0 | 2,75 | 0 |
| ELK3 | AGTGGATGTG | 1 | 1019 | 1029 | 19 | 29 | 2,05 | 0,01 | 19,57 | >0.1 | 5,61 | 8,74 | 0 | 1,86 | 0 | 1,31 | 0 |
| ELK3 | GCTGTAACTG | 1 | 1085 | 1095 | 85 | 95 | -1,64 | 0,01 | 16,77 | >0.1 | 5,61 | 6,94 | 0 | 1,85 | 1,79 | 2,21 | 0 |
| ELK3 | TGAGGAACAG | 1 | 1111 | 1121 | 111 | 121 | -0,78 | 0,01 | 19,08 | >0.1 | 5,61 | 6,64 | 0 | 1,85 | 3,22 | 2,54 | 0 |
| ELK3 | GAAGCAAGTG | -1 | 1179 | 1189 | 179 | 189 | -1,85 | 0,01 | 16,59 | >0.1 | 5,61 | 6,1 | 0 | 1,59 | 1,79 | 3,34 | 0 |
| ELK3 | CCAGGAAGAG | -1 | 1057 | 1067 | 57 | 67 | 5,59 | 0,001 | 22,75 | >0.1 | 5,61 | 7,42 | 0 | 1,86 | 0 | 2,27 | 0 |
| ELK3 | GCAGGAAAGA | -1 | 932 | 942 | -68 | -58 | 4,62 | 0,001 | 20,13 | >0.1 | 5,61 | 7,09 | 0 | 1,88 | 0 | 0,93 | 0 |
| ELK3 | ACAGTAAACA | -1 | 897 | 907 | -103 | -93 | 1,98 | 0,01 | 19,02 | >0.1 | 5,61 | 6,68 | 0 | 1,89 | 0 | 2,86 | 0 |
| ELK3 | AGAGGACAGA | -1 | 856 | 866 | -144 | -134 | -2,75 | 0,01 | 14,05 | >0.1 | 5,61 | 6,34 | 0 | 1,9 | 0 | 2,95 | 0 |
| ELK3 | ACCGGTACTT | -1 | 739 | 749 | -261 | -251 | 1,68 | 0,01 | 20,13 | >0.1 | 5,61 | 5,68 | 0 | 1,25 | 0 | 5,92 | 0 |
| ELK3 | CACGGAAAAT | -1 | 694 | 704 | -306 | -296 | 3,2 | 0,01 | 18,52 | >0.1 | 5,61 | 5,5 | 0 | 1,14 | 0 | 3,07 | 0 |
| ELK3 | CCCAGAAACT | -1 | 555 | 565 | -445 | -435 | -2,43 | 0,01 | 6,86 | >0.1 | 3,21 | 5,09 | 0 | 0,99 | 0 | 0 | 0 |
| ELK3 | TCAGGACGGT | -1 | 257 | 267 | -743 | -733 | 0,29 | 0,01 | 13,62 | >0.1 | 2,82 | 4,54 | 0 | 1,34 | 0 | 4,63 | 0 |
| ELK1 | TAGGTAAGTG | 1 | 2 | 12 | -998 | -988 | -6,68 | 0,01 | 6,84 | >0.1 | 5,61 | 4,26 | 0 | 1,86 | 1,79 | 0 | 0 |

| | | | | | | | | | | | | | | | | | |
|---|---|---|---|---|---|---|---|---|---|---|---|---|---|---|---|---|---|
| ELK1 | ATAGGAAATG | 1 | 61 | 71 | -939 | -929 | -1,86 | 0,01 | 8,31 | >0.1 | 3,98 | 4,31 | 0 | 1,88 | 0 | 0 | 0 |
| ELK1 | TCAGGAGATA | 1 | 244 | 254 | -756 | -746 | -6,72 | 0,01 | 6,48 | >0.1 | 2,86 | 4,52 | 0 | 1,39 | 0 | 4,42 | 0 |
| ELK1 | ACCTGTAGTC | 1 | 330 | 340 | -670 | -660 | -5,32 | 0,01 | 8,52 | >0.1 | 2,59 | 4,65 | 0 | 1,14 | 0 | 5,46 | 0 |
| ELK1 | TCGGGAGGCT | 1 | 348 | 358 | -652 | -642 | -7,39 | 0,01 | 7,34 | >0.1 | 2,55 | 4,68 | 0 | 1,11 | 0 | 6,4 | 0 |
| ELK1 | ACCGGTTTTG | 1 | 743 | 753 | -257 | -247 | -4,86 | 0,01 | 13,62 | >0.1 | 5,61 | 5,69 | 0 | 1,26 | 0 | 5,92 | 0 |
| ELK1 | ACAGGAGAGG | 1 | 769 | 779 | -231 | -221 | -7,69 | 0,01 | 10,78 | >0.1 | 5,61 | 5,81 | 0 | 1,35 | 0 | 5,69 | 0 |
| ELK1 | CCAGGATGAA | 1 | 866 | 876 | -134 | -124 | -3,83 | 0,01 | 12,84 | >0.1 | 5,61 | 6,41 | 0 | 1,9 | 0 | 2,75 | 0 |
| ELK1 | AGTGGATGTG | 1 | 1019 | 1029 | 19 | 29 | -2,78 | 0,01 | 14,74 | >0.1 | 5,61 | 8,74 | 0 | 1,86 | 0 | 1,31 | 0 |
| ELK1 | GCTGTAACTG | 1 | 1085 | 1095 | 85 | 95 | -5,53 | 0,01 | 12,88 | >0.1 | 5,61 | 6,94 | 0 | 1,85 | 1,79 | 2,21 | 0 |
| ELK1 | TGAGGAACAG | 1 | 1111 | 1121 | 111 | 121 | -7,22 | 0,01 | 12,64 | >0.1 | 5,61 | 6,64 | 0 | 1,85 | 3,22 | 2,54 | 0 |
| ELK1 | CCACGAAGCC | 1 | 1147 | 1157 | 147 | 157 | -7,72 | 0,01 | 10,05 | >0.1 | 5,61 | 6,32 | 0 | 1,84 | 1,79 | 2,21 | 0 |
| ELK1 | AGCCGAAGAC | 1 | 1153 | 1163 | 153 | 163 | -4,9 | 0,01 | 11,82 | >0.1 | 4,48 | 6,27 | 0 | 1,83 | 1,79 | 2,34 | 0 |
| ELK1 | GAAGCAAGTG | -1 | 1179 | 1189 | 179 | 189 | -6,52 | 0,01 | 11,92 | >0.1 | 5,61 | 6,1 | 0 | 1,59 | 1,79 | 3,34 | 0 |
| ELK1 | CCAGGAAGAG | -1 | 1057 | 1067 | 57 | 67 | 1,45 | 0,001 | 18,61 | >0.1 | 5,61 | 7,42 | 0 | 1,86 | 0 | 2,27 | 0 |
| ELK1 | GCAGGAAAGA | -1 | 932 | 942 | -68 | -58 | -0,82 | 0,01 | 14,69 | >0.1 | 5,61 | 7,09 | 0 | 1,88 | 0 | 0,93 | 0 |
| ELK1 | ACAGTAAACA | -1 | 897 | 907 | -103 | -93 | -3,5 | 0,01 | 13,54 | >0.1 | 5,61 | 6,68 | 0 | 1,89 | 0 | 2,86 | 0 |
| ELK1 | ACCGGTACTT | -1 | 739 | 749 | -261 | -251 | 1,54 | 0,001 | 19,99 | >0.1 | 5,61 | 5,68 | 0 | 1,25 | 0 | 5,92 | 0 |
| ELK1 | CACGGAAAAT | -1 | 694 | 704 | -306 | -296 | -2,71 | 0,01 | 12,61 | >0.1 | 5,61 | 5,5 | 0 | 1,14 | 0 | 3,07 | 0 |
| ELK1 | ACTGCAAGCT | -1 | 391 | 401 | -609 | -599 | -7,41 | 0,01 | 7,01 | >0.1 | 2,6 | 4,75 | 0 | 1,05 | 0 | 6,02 | 0 |
| ELK1 | TCAGGACGGT | -1 | 257 | 267 | -743 | -733 | -5,31 | 0,01 | 8,02 | >0.1 | 2,82 | 4,54 | 0 | 1,34 | 0 | 4,63 | 0 |
| ELK1 | GGCGTGAGCC | -1 | 189 | 199 | -811 | -801 | -7,39 | 0,01 | 5,9 | >0.1 | 3,09 | 4,45 | 0 | 1,71 | 0 | 4,04 | 0 |
| ELK1 | TCCTGATCTG | -1 | 94 | 104 | -906 | -896 | -4,8 | 0,01 | 5,87 | >0.1 | 3,66 | 4,34 | 0 | 1,88 | 0 | 0,79 | 0 |
| ELK4 | GACCGTCCTGA | 1 | 256 | 267 | -744 | -733 | -7,62 | 0,01 | 5,82 | >0.1 | 2,82 | 4,54 | 0 | 1,34 | 0 | 4,73 | 0 |
| ELK4 | GAGTTTCTGGG | 1 | 554 | 565 | -446 | -435 | -3,47 | 0,01 | 5,82 | >0.1 | 3,21 | 5,09 | 0 | 0,99 | 0 | 0 | 0 |
| ELK4 | TATCTTCATGA | 1 | 642 | 653 | -358 | -347 | -9,25 | 0,01 | 4,79 | >0.1 | 3,82 | 5,33 | 0 | 1,06 | 0 | 3,83 | 0 |
| ELK4 | CATTTTCCGTG | 1 | 693 | 704 | -307 | -296 | 3,42 | 0,001 | 18,73 | >0.1 | 5,6 | 5,5 | 0 | 1,14 | 0 | 3,07 | 0 |
| ELK4 | AAAGTACCGGT | 1 | 738 | 749 | -262 | -251 | -9,18 | 0,01 | 9,26 | >0.1 | 5,6 | 5,68 | 0 | 1,25 | 0 | 5,92 | 0 |
| ELK4 | TTGTTTACTGT | 1 | 896 | 907 | -104 | -93 | -6,26 | 0,01 | 10,85 | >0.1 | 5,6 | 6,68 | 0 | 1,89 | 0 | 2,95 | 0 |
| ELK4 | GTTCTTCTTCT | 1 | 907 | 918 | -93 | -82 | -8,5 | 0,01 | 7,01 | >0.1 | 5,6 | 6,79 | 0 | 1,89 | 0 | 1,24 | 0 |
| ELK4 | GTCTTTCCTGC | 1 | 931 | 942 | -69 | -58 | 6,68 | 0,001 | 22,2 | >0.1 | 5,6 | 7,09 | 0 | 1,88 | 0 | 0,95 | 0 |
| ELK4 | GCTCTTCCTGG | 1 | 1056 | 1067 | 56 | 67 | 10,46 | 0,0001 | 27,56 | >0.1 | 5,6 | 7,44 | 0 | 1,86 | 0 | 2,21 | 0 |
| ELK4 | TCACTTGCTTC | 1 | 1178 | 1189 | 178 | 189 | -5,69 | 0,01 | 12,77 | >0.1 | 5,6 | 6,1 | 0 | 1,6 | 1,79 | 3,37 | 0 |
| ELK4 | TTGCTTCTTGG | 1 | 1182 | 1193 | 182 | 193 | -4,23 | 0,01 | 14,38 | >0.1 | 5,6 | 6,08 | 0 | 1,57 | 1,79 | 3,57 | 0 |
| ELK4 | GGTCTTCGGCT | -1 | 1153 | 1164 | 153 | 164 | -4,48 | 0,01 | 12,23 | >0.1 | 4,47 | 6,27 | 0 | 1,83 | 1,79 | 2,34 | 0 |
| ELK4 | CGGCTTCGTGG | -1 | 1147 | 1158 | 147 | 158 | -1,83 | 0,01 | 15,93 | >0.1 | 5,6 | 6,32 | 0 | 1,84 | 1,79 | 2,21 | 0 |
| ELK4 | CCTGTTCCTCA | -1 | 1111 | 1122 | 111 | 122 | -0,4 | 0,01 | 19,45 | >0.1 | 5,6 | 6,64 | 0 | 1,85 | 3,22 | 2,54 | 0 |
| ELK4 | TCACATCCACT | -1 | 1019 | 1030 | 19 | 30 | -9,1 | 0,01 | 8,41 | >0.1 | 5,6 | 8,74 | 0 | 1,86 | 0 | 1,31 | 0 |
| ELK4 | AAACCTCCTCA | -1 | 988 | 999 | -12 | -1 | -8,88 | 0,01 | 8,88 | >0.1 | 5,6 | 8,69 | 0 | 1,87 | 0 | 1,61 | 0 |
| ELK4 | AGATTTTCTCT | -1 | 975 | 986 | -25 | -14 | -9,16 | 0,01 | 8,02 | >0.1 | 5,6 | 8,13 | 0 | 1,87 | 0 | 1,57 | 0 |
| ELK4 | GTTCATCCTGG | -1 | 866 | 877 | -134 | -123 | -0,51 | 0,01 | 16,16 | >0.1 | 5,6 | 6,42 | 0 | 1,9 | 0 | 2,75 | 0 |
| ELK4 | ACCTCTCCTGT | -1 | 769 | 780 | -231 | -220 | -7,24 | 0,01 | 11,22 | >0.1 | 5,6 | 5,82 | 0 | 1,36 | 0 | 5,69 | 0 |
| ELK4 | GCTCTGCCTCC | -1 | 383 | 394 | -617 | -606 | -4,71 | 0,01 | 9,68 | >0.1 | 2,58 | 4,74 | 0 | 1,06 | 0 | 6,02 | 0 |
| ELK4 | CATTCTCCTGC | -1 | 361 | 372 | -639 | -628 | -6,65 | 0,01 | 8,15 | >0.1 | 2,52 | 4,7 | 0 | 1,09 | 0 | 6,49 | 0 |
| ELK4 | CTCCTGCCTCA | -1 | 357 | 368 | -643 | -632 | -8,89 | 0,01 | 5,55 | >0.1 | 2,52 | 4,7 | 0 | 1,09 | 0 | 6,13 | 0 |
| ELK4 | CTATCTCCTGA | -1 | 244 | 255 | -756 | -745 | -5,46 | 0,01 | 7,74 | >0.1 | 2,86 | 4,53 | 0 | 1,39 | 0 | 4,42 | 0 |
| ELK4 | TAATTTCCATG | -1 | 150 | 161 | -850 | -839 | -9,02 | 0,01 | 2,21 | >0.1 | 3,28 | 4,41 | 0 | 1,9 | 0 | 1,65 | 0 |
| ELK4 | TCATTTCCTAT | -1 | 61 | 72 | -939 | -928 | -3,25 | 0,01 | 6,91 | >0.1 | 3,98 | 4,31 | 0 | 1,88 | 0 | 0 | 0 |
| ELK4 | TGACTTCTTTG | -1 | 29 | 40 | -971 | -960 | -8,83 | 0,01 | 1,81 | >0.1 | 4,5 | 4,28 | 0 | 1,87 | 0 | 0 | 0 |
| ELK4 | GTGCCTCCTTC | -1 | 17 | 28 | -983 | -972 | -7,25 | 0,01 | 4,48 | >0.1 | 5,6 | 4,27 | 0 | 1,87 | 0 | 0 | 0 |
| ELK4 | CCTTCTCCTTC | -1 | 11 | 22 | -989 | -978 | -9,46 | 0,01 | 2,26 | >0.1 | 5,6 | 4,26 | 0 | 1,86 | 0 | 0 | 0 |
| DBP | AGTTTTGCAAAG | 1 | 81 | 93 | -919 | -907 | -5,25 | 0,01 | 4,73 | >0.1 | 3,77 | 4,33 | 0 | 1,88 | 0 | 0 | 0 |
| DBP | AATGGCGTGAAC | 1 | 368 | 380 | -632 | -620 | -5,7 | 0,01 | 8,78 | >0.1 | 2,54 | 4,72 | 0 | 1,08 | 0 | 6,15 | 0 |
| DBP | GATCGCGCCACT | 1 | 408 | 420 | -592 | -580 | -5,41 | 0,01 | 8,58 | >0.1 | 2,65 | 4,79 | 0 | 1,03 | 0 | 5,52 | 0 |
| DBP | AATTCCATAAAG | 1 | 494 | 506 | -506 | -494 | -6,04 | 0,01 | 9,46 | >0.1 | 2,94 | 4,96 | 0 | 0,99 | 0 | 6,62 | 0 |
| DBP | TGTTGCCCAAGT | 1 | 539 | 551 | -461 | -449 | -1,68 | 0,01 | 7,5 | >0.1 | 3,13 | 5,06 | 0 | 0,99 | 0 | 0 | 0 |
| DBP | TGTGAGATAACC | 1 | 590 | 602 | -410 | -398 | 2,78 | 0,001 | 12,39 | >0.1 | 3,41 | 5,19 | 0 | 1,01 | 0 | 0 | 0 |
| DBP | AATTATAACATT | 1 | 685 | 697 | -315 | -303 | -5,29 | 0,01 | 8,68 | >0.1 | 4,42 | 5,48 | 0 | 1,13 | 0 | 2,95 | 0 |
| DBP | GGAGAGGTAAGG | 1 | 772 | 784 | -228 | -216 | -6,69 | 0,01 | 12,2 | >0.1 | 5,58 | 5,83 | 0 | 1,37 | 0 | 6,1 | 0 |
| DBP | GATTAAAGAATA | 1 | 792 | 804 | -208 | -196 | -7,75 | 0,01 | 10,55 | >0.1 | 5,58 | 5,94 | 0 | 1,47 | 0 | 5,32 | 0 |

| | | | | | | | | | | | | | | | | |
|---|---|---|---|---|---|---|---|---|---|---|---|---|---|---|---|---|
| DBP | TGTTACATATCT | 1 | 847 | 859 | -153 | -141 | 1,51 | 0,01 | 18,19 | >0.1 | 5,58 | 6,29 | 0 | 1,85 | 0 | 2,95 | 0 |
| DBP | TTTGATAGAACA | -1 | 1164 | 1176 | 164 | 176 | -5,74 | 0,01 | 11,27 | >0.1 | 4,51 | 6,19 | 0 | 1,72 | 1,79 | 2,79 | 0 |
| DBP | AGTTACAGCAAC | -1 | 1082 | 1094 | 82 | 94 | -4,71 | 0,01 | 13,72 | >0.1 | 5,58 | 6,99 | 0 | 1,85 | 1,79 | 2,21 | 0 |
| DBP | AGATATGTAACA | -1 | 847 | 859 | -153 | -141 | 3,38 | 0,001 | 20,06 | >0.1 | 5,58 | 6,29 | 0 | 1,85 | 0 | 2,95 | 0 |
| DBP | GAATACGTTATT | -1 | 800 | 812 | -200 | -188 | -6,27 | 0,01 | 10,23 | >0.1 | 5,58 | 5,98 | 0 | 1,51 | 0 | 3,43 | 0 |
| DBP | AATGTTATAATT | -1 | 685 | 697 | -315 | -303 | -3,83 | 0,01 | 10,14 | >0.1 | 4,42 | 5,48 | 0 | 1,13 | 0 | 2,95 | 0 |
| DBP | GGTTATCTCACA | -1 | 590 | 602 | -410 | -398 | 1,77 | 0,01 | 11,38 | >0.1 | 3,41 | 5,19 | 0 | 1,01 | 0 | 0 | 0 |
| DBP | ACTTGGGCAACA | -1 | 539 | 551 | -461 | -449 | -0,82 | 0,01 | 8,36 | >0.1 | 3,13 | 5,06 | 0 | 0,99 | 0 | 0 | 0 |
| DBP | AAATACATGACT | -1 | 516 | 528 | -484 | -472 | -7,38 | 0,01 | 4,59 | >0.1 | 3,03 | 5 | 0 | 0,98 | 0 | 2,95 | 0 |
| DBP | CTTTATGGAATT | -1 | 494 | 506 | -506 | -494 | -2,52 | 0,01 | 12,98 | >0.1 | 2,94 | 4,96 | 0 | 0,99 | 0 | 6,62 | 0 |
| DBP | AGTGGCGCGATC | -1 | 408 | 420 | -592 | -580 | -4,12 | 0,01 | 9,87 | >0.1 | 2,65 | 4,79 | 0 | 1,03 | 0 | 5,52 | 0 |
| DBP | CTTTGCAAAACT | -1 | 81 | 93 | -919 | -907 | -6,64 | 0,01 | 3,34 | >0.1 | 3,77 | 4,33 | 0 | 1,88 | 0 | 0 | 0 |
| FOXP2 | TTAAAAATACA | 1 | 292 | 303 | -708 | -697 | 0,18 | 0,01 | 13,15 | >0.1 | 2,7 | 4,59 | 0 | 1,23 | 0 | 4,45 | 0 |
| FOXP2 | AAAAATACAAA | 1 | 294 | 305 | -706 | -695 | -2,81 | 0,01 | 10,23 | >0.1 | 2,7 | 4,6 | 0 | 1,22 | 0 | 4,53 | 0 |
| FOXP2 | CAAAAAAAAAA | 1 | 456 | 467 | -544 | -533 | -4,34 | 0,01 | 9,85 | >0.1 | 2,8 | 4,88 | 0 | 1 | 0 | 5,52 | 0 |
| FOXP2 | AAAAAAAAAAA | 1 | 457 | 468 | -543 | -532 | -3,29 | 0,01 | 11,02 | >0.1 | 2,8 | 4,88 | 0 | 1 | 0 | 5,63 | 0 |
| FOXP2 | AAAAAAAAAAA | 1 | 458 | 469 | -542 | -531 | -3,29 | 0,01 | 11,02 | >0.1 | 2,81 | 4,88 | 0 | 0,99 | 0 | 5,63 | 0 |
| FOXP2 | AAAAAAAAAAA | 1 | 459 | 470 | -541 | -530 | -3,29 | 0,01 | 10,92 | >0.1 | 2,81 | 4,88 | 0 | 0,99 | 0 | 5,52 | 0 |
| FOXP2 | AAAAAAAAAAA | 1 | 460 | 471 | -540 | -529 | -3,29 | 0,01 | 10,92 | >0.1 | 2,81 | 4,88 | 0 | 0,99 | 0 | 5,52 | 0 |
| FOXP2 | AAAAAAAAAAA | 1 | 461 | 472 | -539 | -528 | -3,29 | 0,01 | 11,04 | >0.1 | 2,82 | 4,89 | 0 | 0,99 | 0 | 5,63 | 0 |
| FOXP2 | AAAAAAAAAAA | 1 | 462 | 473 | -538 | -527 | -3,29 | 0,01 | 11,04 | >0.1 | 2,82 | 4,89 | 0 | 0,99 | 0 | 5,63 | 0 |
| FOXP2 | AAAAAAAAAAA | 1 | 463 | 474 | -537 | -526 | -3,29 | 0,01 | 11,19 | >0.1 | 2,82 | 4,89 | 0 | 0,99 | 0 | 5,78 | 0 |
| FOXP2 | AAAAAAAAAAA | 1 | 464 | 475 | -536 | -525 | -3,29 | 0,01 | 11,2 | >0.1 | 2,83 | 4,89 | 0 | 0,99 | 0 | 5,78 | 0 |
| FOXP2 | AAAAAAAAAAA | 1 | 465 | 476 | -535 | -524 | -3,29 | 0,01 | 11,2 | >0.1 | 2,83 | 4,89 | 0 | 0,99 | 0 | 5,78 | 0 |
| FOXP2 | AAAAAAAAAAA | 1 | 466 | 477 | -534 | -523 | -3,29 | 0,01 | 11,07 | >0.1 | 2,83 | 4,9 | 0 | 0,99 | 0 | 5,63 | 0 |
| FOXP2 | AAAAAAAAAAA | 1 | 467 | 478 | -533 | -522 | -3,29 | 0,01 | 11,21 | >0.1 | 2,84 | 4,9 | 0 | 0,99 | 0 | 5,78 | 0 |
| FOXP2 | AAAAAAAAAAA | 1 | 468 | 479 | -532 | -521 | -3,29 | 0,01 | 11,36 | >0.1 | 2,84 | 4,9 | 0 | 0,99 | 0 | 5,92 | 0 |
| FOXP2 | AAAAAAAAAAC | 1 | 469 | 480 | -531 | -520 | -4,21 | 0,01 | 9,35 | >0.1 | 2,84 | 4,9 | 0 | 0,99 | 0 | 4,83 | 0 |
| FOXP2 | AAAAAAACTGA | 1 | 472 | 483 | -528 | -517 | -2,33 | 0,01 | 11,37 | >0.1 | 2,85 | 4,91 | 0 | 0,99 | 0 | 4,95 | 0 |
| FOXP2 | ATAAAGACAAG | 1 | 500 | 511 | -500 | -489 | -2,96 | 0,01 | 12,77 | >0.1 | 2,96 | 4,97 | 0 | 0,98 | 0 | 6,81 | 0 |
| FOXP2 | AAATAAAAATA | 1 | 577 | 588 | -423 | -412 | -3,45 | 0,01 | 6,03 | >0.1 | 3,33 | 5,15 | 0 | 1 | 0 | 0 | 0 |
| FOXP2 | ATAAAAATAAA | 1 | 579 | 590 | -421 | -410 | 1,94 | 0,01 | 11,44 | >0.1 | 3,34 | 5,15 | 0 | 1 | 0 | 0 | 0 |
| FOXP2 | CTGAAAACCAT | 1 | 617 | 628 | -383 | -372 | -2,64 | 0,01 | 8,75 | >0.1 | 3,6 | 5,26 | 0 | 1,03 | 0 | 1,5 | 0 |
| FOXP2 | ATACAAACACC | 1 | 626 | 637 | -374 | -363 | -0,29 | 0,01 | 11,25 | >0.1 | 3,67 | 5,28 | 0 | 1,04 | 0 | 1,55 | 0 |
| FOXP2 | AGGAAAATATT | 1 | 722 | 733 | -278 | -267 | -1,25 | 0,01 | 16,33 | >0.1 | 5,6 | 5,61 | 0 | 1,21 | 0 | 5,17 | 0 |
| FOXP2 | AAAAAAAGAAA | -1 | 919 | 930 | -81 | -70 | -2,79 | 0,01 | 12,71 | >0.1 | 5,6 | 6,93 | 0 | 1,88 | 0 | 1,09 | 0 |
| FOXP2 | AGAAAAAGAAG | -1 | 913 | 924 | -87 | -76 | -3,91 | 0,01 | 11,58 | >0.1 | 5,6 | 6,85 | 0 | 1,89 | 0 | 1,15 | 0 |
| FOXP2 | AGAAGAACACA | -1 | 904 | 915 | -96 | -85 | -3,53 | 0,01 | 11,92 | >0.1 | 5,6 | 6,76 | 0 | 1,89 | 0 | 1,21 | 0 |
| FOXP2 | AAGAACACAGT | -1 | 902 | 913 | -98 | -87 | -0,82 | 0,01 | 14,68 | >0.1 | 5,6 | 6,73 | 0 | 1,89 | 0 | 1,28 | 0 |
| NFATC1 | GTGCTCCAGC | 1 | 420 | 430 | -580 | -570 | 3,16 | 0,01 | 17,08 | >0.1 | 2,68 | 4,8 | 0 | 1,02 | 0 | 5,42 | 0 |
| NFATC1 | GAATTCCATA | 1 | 493 | 503 | -507 | -497 | 7,12 | 0,01 | 22,42 | >0.1 | 2,93 | 4,95 | 0 | 0,98 | 0 | 6,44 | 0 |
| NFATC1 | ATTTTCCGTG | 1 | 694 | 704 | -306 | -296 | 10,89 | 0,0001 | 26,21 | >0.1 | 5,61 | 5,5 | 0 | 1,14 | 0 | 3,07 | 0 |
| NFATC1 | CCTCTCCAGG | 1 | 861 | 871 | -139 | -129 | 4,03 | 0,01 | 20,67 | >0.1 | 5,61 | 6,38 | 0 | 1,9 | 0 | 2,75 | 0 |
| NFATC1 | TCTTTCCTGC | 1 | 932 | 942 | -68 | -58 | 6,61 | 0,01 | 22,12 | >0.1 | 5,61 | 7,09 | 0 | 1,88 | 0 | 0,93 | 0 |
| NFATC1 | CTCTTCCTGG | 1 | 1057 | 1067 | 57 | 67 | 3,21 | 0,01 | 20,37 | >0.1 | 5,61 | 7,42 | 0 | 1,86 | 0 | 2,27 | 0 |
| NFATC1 | TAATTCCAAG | -1 | 1188 | 1198 | 188 | 198 | 7,06 | 0,01 | 25,73 | >0.1 | 5,61 | 6,04 | 0 | 1,53 | 1,79 | 3,7 | 0 |
| NFATC1 | CTGTTCCTCA | -1 | 1111 | 1121 | 111 | 121 | 5,03 | 0,01 | 24,89 | >0.1 | 5,61 | 6,64 | 0 | 1,85 | 3,22 | 2,54 | 0 |
| NFATC1 | ACTTTCCCTA | -1 | 1004 | 1014 | 4 | 14 | 7,13 | 0,01 | 25,14 | >0.1 | 5,61 | 9,12 | 0 | 1,87 | 0 | 1,42 | 0 |
| NFATC1 | ATTTTCCTAG | -1 | 720 | 730 | -280 | -270 | 7,55 | 0,01 | 24,95 | >0.1 | 5,61 | 5,6 | 0 | 1,2 | 0 | 4,99 | 0 |
| NFATC1 | ATTTTACAGT | -1 | 678 | 688 | -322 | -312 | 3,57 | 0,01 | 17,68 | >0.1 | 4,27 | 5,45 | 0 | 1,11 | 0 | 3,28 | 0 |
| NFATC1 | ATATTCCCAG | -1 | 560 | 570 | -440 | -430 | 5,11 | 0,01 | 14,44 | >0.1 | 3,23 | 5,1 | 0 | 0,99 | 0 | 0 | 0 |
| NFATC1 | TATTCCCAGA | -1 | 559 | 569 | -441 | -431 | 3,99 | 0,01 | 13,31 | >0.1 | 3,23 | 5,1 | 0 | 0,99 | 0 | 0 | 0 |
| NFATC1 | ATTTTCCCTT | -1 | 530 | 540 | -470 | -460 | 8,26 | 0,001 | 21,06 | >0.1 | 3,09 | 5,03 | 0 | 0,98 | 0 | 3,7 | 0 |
| NFATC1 | TTTTCCCTTC | -1 | 529 | 539 | -471 | -461 | 3,73 | 0,01 | 16,37 | >0.1 | 3,08 | 5,03 | 0 | 0,98 | 0 | 3,55 | 0 |
| NFATC1 | CCCTTCCAAA | -1 | 525 | 535 | -475 | -465 | 5,44 | 0,01 | 17,79 | >0.1 | 3,06 | 5,02 | 0 | 0,98 | 0 | 3,28 | 0 |
| NFATC1 | TATTTCTGAT | -1 | 482 | 492 | -518 | -508 | 3,2 | 0,01 | 17,8 | >0.1 | 2,89 | 4,93 | 0 | 0,99 | 0 | 5,8 | 0 |
| NFATC1 | ATTCTCCTGC | -1 | 361 | 371 | -639 | -629 | 3,17 | 0,01 | 17,96 | >0.1 | 2,52 | 4,7 | 0 | 1,08 | 0 | 6,49 | 0 |
| NFATC1 | AATTTCCATG | -1 | 150 | 160 | -850 | -840 | 10,13 | 0,001 | 21,36 | >0.1 | 3,28 | 4,41 | 0 | 1,9 | 0 | 1,65 | 0 |
| NFATC1 | ACTCTCCACT | -1 | 118 | 128 | -882 | -872 | 6,46 | 0,01 | 16,92 | >0.1 | 3,48 | 4,37 | 0 | 1,89 | 0 | 0,73 | 0 |

| | | | | | | | | | | | | | | | | |
|---|---|---|---|---|---|---|---|---|---|---|---|---|---|---|---|---|
| NFATC1 | CATTTCCTAT | -1 | 61 | 71 | -939 | -929 | 6,09 | 0,01 | 16,26 | >0.1 | 3,98 | 4,31 | 0 | 1,88 | 0 | 0 | 0 |
| NFATC1 | CTTCTCCTTC | -1 | 11 | 21 | -989 | -979 | 3,26 | 0,01 | 15 | >0.1 | 5,61 | 4,26 | 0 | 1,86 | 0 | 0 | 0 |
| NFATC2 | TTTTGCA | 1 | 83 | 90 | -917 | -910 | 4,84 | 0,01 | 14,82 | >0.1 | 3,77 | 4,33 | 0 | 1,88 | 0 | 0 | 0 |
| NFATC2 | TGCTCCA | 1 | 421 | 428 | -579 | -572 | 5,21 | 0,01 | 19,01 | >0.1 | 2,68 | 4,8 | 0 | 1,02 | 0 | 5,3 | 0 |
| NFATC2 | AATTCCA | 1 | 494 | 501 | -506 | -499 | 5,02 | 0,01 | 20,14 | >0.1 | 2,93 | 4,95 | 0 | 0,98 | 0 | 6,27 | 0 |
| NFATC2 | TTTTCCG | 1 | 695 | 702 | -305 | -298 | 8,39 | 0,01 | 22,68 | >0.1 | 4,6 | 5,49 | 0 | 1,14 | 0 | 3,07 | 0 |
| NFATC2 | TGTTACA | 1 | 847 | 854 | -153 | -146 | 4,16 | 0,01 | 20,82 | >0.1 | 5,65 | 6,26 | 0 | 1,81 | 0 | 2,95 | 0 |
| NFATC2 | CTCTCCA | 1 | 862 | 869 | -138 | -131 | 4,4 | 0,01 | 21,18 | >0.1 | 5,65 | 6,36 | 0 | 1,9 | 0 | 2,86 | 0 |
| NFATC2 | CTTTCCT | 1 | 933 | 940 | -67 | -60 | 6,13 | 0,01 | 21,67 | >0.1 | 5,65 | 7,06 | 0 | 1,88 | 0 | 0,95 | 0 |
| NFATC2 | TCTTCCT | 1 | 1058 | 1065 | 58 | 65 | 6,93 | 0,01 | 24,05 | >0.1 | 5,65 | 7,39 | 0 | 1,86 | 0 | 2,21 | 0 |
| NFATC2 | AATTCCA | -1 | 1190 | 1197 | 190 | 197 | 5,02 | 0,01 | 23,7 | >0.1 | 5,65 | 6,03 | 0 | 1,51 | 1,79 | 3,7 | 0 |
| NFATC2 | TTGTCCA | -1 | 1134 | 1141 | 134 | 141 | 5,4 | 0,01 | 25,52 | >0.1 | 5,65 | 6,42 | 0 | 1,84 | 3,22 | 2,98 | 0 |
| NFATC2 | TGTTCCT | -1 | 1113 | 1120 | 113 | 120 | 6,93 | 0,01 | 26,81 | >0.1 | 5,65 | 6,62 | 0 | 1,85 | 3,22 | 2,54 | 0 |
| NFATC2 | CTTTCCC | -1 | 1006 | 1013 | 6 | 13 | 6,23 | 0,01 | 24,29 | >0.1 | 5,65 | 9,12 | 0 | 1,87 | 0 | 1,42 | 0 |
| NFATC2 | TTTCCCT | -1 | 1005 | 1012 | 5 | 12 | 3,17 | 0,01 | 21,27 | >0.1 | 5,65 | 9,12 | 0 | 1,87 | 0 | 1,46 | 0 |
| NFATC2 | TTTTCCT | -1 | 722 | 729 | -278 | -271 | 9,7 | 0,001 | 27,13 | >0.1 | 5,65 | 5,59 | 0 | 1,2 | 0 | 4,99 | 0 |
| NFATC2 | TTTTACA | -1 | 680 | 687 | -320 | -313 | 6,93 | 0,01 | 21,04 | >0.1 | 4,28 | 5,44 | 0 | 1,11 | 0 | 3,28 | 0 |
| NFATC2 | TATTCCC | -1 | 562 | 569 | -438 | -431 | 5,66 | 0,01 | 14,99 | >0.1 | 3,24 | 5,1 | 0 | 0,99 | 0 | 0 | 0 |
| NFATC2 | TTTTCCC | -1 | 532 | 539 | -468 | -461 | 9,81 | 0,001 | 22,61 | >0.1 | 3,09 | 5,03 | 0 | 0,98 | 0 | 3,7 | 0 |
| NFATC2 | TTTCCCT | -1 | 531 | 538 | -469 | -462 | 3,17 | 0,01 | 15,81 | >0.1 | 3,08 | 5,03 | 0 | 0,98 | 0 | 3,55 | 0 |
| NFATC2 | CCTTCCA | -1 | 527 | 534 | -473 | -466 | 5,47 | 0,01 | 17,82 | >0.1 | 3,07 | 5,02 | 0 | 0,98 | 0 | 3,28 | 0 |
| NFATC2 | TTTGCCC | -1 | 510 | 517 | -490 | -483 | 3,27 | 0,01 | 16,6 | >0.1 | 2,99 | 4,98 | 0 | 0,98 | 0 | 4,37 | 0 |
| NFATC2 | TTCTCCT | -1 | 363 | 370 | -637 | -630 | 5,87 | 0,01 | 20,66 | >0.1 | 2,52 | 4,7 | 0 | 1,08 | 0 | 6,49 | 0 |
| NFATC2 | ATCTCCT | -1 | 246 | 253 | -754 | -747 | 3,21 | 0,01 | 16,4 | >0.1 | 2,86 | 4,52 | 0 | 1,38 | 0 | 4,42 | 0 |
| NFATC2 | ATTTCCA | -1 | 152 | 159 | -848 | -841 | 9,16 | 0,001 | 20,39 | >0.1 | 3,28 | 4,4 | 0 | 1,9 | 0 | 1,65 | 0 |
| NFATC2 | CTCTCCA | -1 | 120 | 127 | -880 | -873 | 4,4 | 0,01 | 14,86 | >0.1 | 3,48 | 4,37 | 0 | 1,89 | 0 | 0,73 | 0 |
| NFATC2 | ATTTCCT | -1 | 63 | 70 | -937 | -930 | 7,04 | 0,01 | 17,2 | >0.1 | 3,98 | 4,31 | 0 | 1,87 | 0 | 0 | 0 |
| NFATC2 | TTCTCCT | -1 | 13 | 20 | -987 | -980 | 5,87 | 0,01 | 17,65 | >0.1 | 5,65 | 4,26 | 0 | 1,86 | 0 | 0 | 0 |
| FOXP1 | AAATAAAAATAA | 1 | 577 | 589 | -423 | -411 | 1,03 | 0,01 | 10,52 | >0.1 | 3,33 | 5,15 | 0 | 1 | 0 | 0 | 0 |
| FOXP1 | ATACAAACACCA | 1 | 626 | 638 | -374 | -362 | 4,62 | 0,01 | 16,17 | >0.1 | 3,68 | 5,29 | 0 | 1,04 | 0 | 1,55 | 0 |
| FOXP1 | AGGAAAATATTA | 1 | 722 | 734 | -278 | -266 | 2,1 | 0,01 | 19,68 | >0.1 | 5,58 | 5,61 | 0 | 1,21 | 0 | 5,17 | 0 |
| FOXP1 | CAGGAAAGACCA | -1 | 929 | 941 | -71 | -59 | 1,02 | 0,01 | 16,56 | >0.1 | 5,58 | 7,08 | 0 | 1,88 | 0 | 1 | 0 |
| FOXP1 | AAGAACACAGTA | -1 | 901 | 913 | -99 | -87 | 5,2 | 0,01 | 20,68 | >0.1 | 5,58 | 6,74 | 0 | 1,89 | 0 | 1,28 | 0 |
| FOXP1 | CAGTAAACAATC | -1 | 894 | 906 | -106 | -94 | 12,4 | 0,0001 | 29,49 | >0.1 | 5,58 | 6,67 | 0 | 1,89 | 0 | 2,95 | 0 |
| FOXP1 | TGGGCAACATTT | -1 | 536 | 548 | -464 | -452 | 1,38 | 0,01 | 12,19 | >0.1 | 3,12 | 5,05 | 0 | 0,99 | 0 | 1,66 | 0 |
| FOXP1 | AGGTCAACTCTC | -1 | 102 | 114 | -898 | -886 | 2,84 | 0,01 | 13,43 | >0.1 | 3,59 | 4,35 | 0 | 1,89 | 0 | 0,77 | 0 |
| NOBOX | TAAGTGAA | 1 | 6 | 14 | -994 | -986 | 2,45 | 0,01 | 16 | >0.1 | 5,64 | 4,26 | 0 | 1,86 | 1,79 | 0 | 0 |
| NOBOX | TAAATGTG | 1 | 586 | 594 | -414 | -406 | 3,07 | 0,01 | 12,62 | >0.1 | 3,38 | 5,16 | 0 | 1 | 0 | 0 | 0 |
| NOBOX | TAATGAAA | 1 | 605 | 613 | -395 | -387 | 2,43 | 0,01 | 13,58 | >0.1 | 3,5 | 5,22 | 0 | 1,02 | 0 | 1,41 | 0 |
| NOBOX | TAACAGCT | 1 | 710 | 718 | -290 | -282 | 2,71 | 0,01 | 19,88 | >0.1 | 5,64 | 5,55 | 0 | 1,17 | 0 | 4,81 | 0 |
| NOBOX | TAACTGCT | 1 | 1089 | 1097 | 89 | 97 | 6,58 | 0,01 | 24,97 | >0.1 | 5,64 | 6,89 | 0 | 1,85 | 1,79 | 2,21 | 0 |
| NOBOX | TAATTTTT | -1 | 829 | 837 | -171 | -163 | 4,18 | 0,01 | 21,05 | >0.1 | 5,64 | 6,14 | 0 | 1,67 | 0 | 3,43 | 0 |
| NOBOX | TTATGGCC | -1 | 757 | 765 | -243 | -235 | 4,55 | 0,01 | 23,56 | >0.1 | 5,64 | 5,75 | 0 | 1,3 | 0 | 6,32 | 0 |
| NOBOX | TAATAGGT | -1 | 599 | 607 | -401 | -393 | 7,62 | 0,001 | 18,7 | >0.1 | 3,46 | 5,2 | 0 | 1,01 | 0 | 1,41 | 0 |
| NOBOX | TACTTGGG | -1 | 544 | 552 | -456 | -448 | 3,44 | 0,01 | 12,63 | >0.1 | 3,15 | 5,06 | 0 | 0,98 | 0 | 0 | 0 |
| NOBOX | TAATAGAG | -1 | 287 | 295 | -713 | -705 | 4,04 | 0,01 | 17,67 | >0.1 | 2,72 | 4,58 | 0 | 1,24 | 0 | 5,09 | 0 |
| NOBOX | TAATTTCC | -1 | 153 | 161 | -847 | -839 | 6,33 | 0,01 | 17,55 | >0.1 | 3,27 | 4,41 | 0 | 1,9 | 0 | 1,65 | 0 |
| NOBOX | TTATGACT | -1 | 128 | 136 | -872 | -864 | 3,39 | 0,01 | 14,69 | >0.1 | 3,42 | 4,38 | 0 | 1,89 | 0 | 1,61 | 0 |
| ZBTB33 | AGATCAGGAGAGTTG | 1 | 95 | 110 | -905 | -890 | -6,32 | 0,01 | 4,31 | >0.1 | 3,63 | 4,35 | 0 | 1,88 | 0 | 0,77 | 0 |
| ZBTB33 | GGCTCACGCCTGTAA | 1 | 189 | 204 | -811 | -796 | -0,48 | 0,01 | 12,9 | >0.1 | 3,08 | 4,46 | 0 | 1,71 | 0 | 4,13 | 0 |
| ZBTB33 | CTAACACGGTGAAAC | 1 | 267 | 282 | -733 | -718 | -6,54 | 0,01 | 7,19 | >0.1 | 2,78 | 4,56 | 0 | 1,31 | 0 | 5,09 | 0 |
| ZBTB33 | TGGGCGTGGTGGTGG | 1 | 313 | 328 | -687 | -672 | -5,93 | 0,01 | 8,64 | >0.1 | 2,63 | 4,63 | 0 | 1,18 | 0 | 6,13 | 0 |
| ZBTB33 | TACTCGGGAGGCTGA | 1 | 345 | 360 | -655 | -640 | -1,61 | 0,01 | 13,26 | >0.1 | 2,55 | 4,68 | 0 | 1,11 | 0 | 6,52 | 0 |
| ZBTB33 | ATGGCGTGAACCTGG | 1 | 369 | 384 | -631 | -616 | -6,51 | 0,01 | 8,23 | >0.1 | 2,55 | 4,72 | 0 | 1,08 | 0 | 6,4 | 0 |
| ZBTB33 | GAGCCGAGATCGCGC | 1 | 401 | 416 | -599 | -584 | -7,55 | 0,01 | 6,67 | >0.1 | 2,63 | 4,78 | 0 | 1,04 | 0 | 5,76 | 0 |
| ZBTB33 | AGATCGCGCCACTGT | 1 | 407 | 422 | -593 | -578 | 2,92 | 0,001 | 16,92 | >0.1 | 2,65 | 4,79 | 0 | 1,03 | 0 | 5,52 | 0 |
| ZBTB33 | ATCGCGCCACTGTGC | 1 | 409 | 424 | -591 | -576 | -6,63 | 0,01 | 7,62 | >0.1 | 2,66 | 4,79 | 0 | 1,03 | 0 | 5,76 | 0 |
| ZBTB33 | TCTTCATGATCCCTA | 1 | 644 | 659 | -356 | -341 | -2,98 | 0,01 | 11,13 | >0.1 | 3,86 | 5,35 | 0 | 1,07 | 0 | 3,83 | 0 |
| ZBTB33 | TTTCCGTGTATCTTT | 1 | 696 | 711 | -304 | -289 | -7,15 | 0,01 | 8,27 | >0.1 | 5,54 | 5,53 | 0 | 1,16 | 0 | 3,19 | 0 |

| | | | | | | | | | | | | | | | | | |
|---|---|---|---|---|---|---|---|---|---|---|---|---|---|---|---|---|---|
| ZBTB33 | ATCTCATGAGGAGGT | 1 | 982 | 997 | -18 | -3 | 1,84 | 0,001 | 19,42 | >0.1 | 5,54 | 8,64 | 0 | 1,87 | 0 | 1,53 | 0 |
| ZBTB33 | ATGTCAAGCTCTTCC | 1 | 1049 | 1064 | 49 | 64 | -6,54 | 0,01 | 9,92 | >0.1 | 5,54 | 7,6 | 0 | 1,86 | 0 | 1,46 | 0 |
| ZBTB33 | TAACCACGAAGCCGA | 1 | 1144 | 1159 | 144 | 159 | -7,66 | 0,01 | 11,45 | >0.1 | 5,54 | 6,34 | 0 | 1,84 | 3,22 | 2,16 | 0 |
| ZBTB33 | TTCACTTGCTTCTTG | 1 | 1177 | 1192 | 177 | 192 | -4,44 | 0,01 | 13,96 | >0.1 | 5,54 | 6,11 | 0 | 1,61 | 1,79 | 3,34 | 0 |
| ZBTB33 | GTCCCCTGTGAGCCA | -1 | 1033 | 1048 | 33 | 48 | -4,17 | 0,01 | 12,64 | >0.1 | 5,54 | 8,1 | 0 | 1,86 | 0 | 1,31 | 0 |
| ZBTB33 | TCCTCATGAGATTTT | -1 | 979 | 994 | -21 | -6 | 6,85 | 0,0001 | 24,36 | >0.1 | 5,54 | 8,57 | 0 | 1,87 | 0 | 1,53 | 0 |
| ZBTB33 | TCCTGGAGAGGACAG | -1 | 857 | 872 | -143 | -128 | -6,81 | 0,01 | 9,88 | >0.1 | 5,54 | 6,38 | 0 | 1,9 | 0 | 2,86 | 0 |
| ZBTB33 | CTCTCCTGTCACTTT | -1 | 763 | 778 | -237 | -222 | -2,64 | 0,01 | 15,98 | >0.1 | 5,54 | 5,81 | 0 | 1,35 | 0 | 5,92 | 0 |
| ZBTB33 | GAGTCTCGGTCTGTC | -1 | 436 | 451 | -564 | -549 | -7,48 | 0,01 | 7,16 | >0.1 | 2,74 | 4,84 | 0 | 1,01 | 0 | 6,05 | 0 |
| ZBTB33 | CTGTCGCCCAGGCTG | -1 | 426 | 441 | -574 | -559 | -5,44 | 0,01 | 8,92 | >0.1 | 2,71 | 4,83 | 0 | 1,02 | 0 | 5,8 | 0 |
| ZBTB33 | CAGTGGCGCGATCTC | -1 | 406 | 421 | -594 | -579 | -4,5 | 0,01 | 9,62 | >0.1 | 2,65 | 4,79 | 0 | 1,03 | 0 | 5,65 | 0 |
| ZBTB33 | GTGGCGCGATCTCGG | -1 | 404 | 419 | -596 | -581 | 4,27 | 0,001 | 18,38 | >0.1 | 2,64 | 4,78 | 0 | 1,04 | 0 | 5,65 | 0 |
| ZBTB33 | CTCCCAGGTTCACGC | -1 | 372 | 387 | -628 | -613 | -6,46 | 0,01 | 8,43 | >0.1 | 2,56 | 4,73 | 0 | 1,07 | 0 | 6,53 | 0 |
| ZBTB33 | GGTTCACGCCATTCT | -1 | 366 | 381 | -634 | -619 | -2,43 | 0,01 | 12,63 | >0.1 | 2,54 | 4,72 | 0 | 1,08 | 0 | 6,73 | 0 |
| ZBTB33 | TTCACGCCATTCTCC | -1 | 364 | 379 | -636 | -621 | -2,31 | 0,01 | 12,63 | >0.1 | 2,53 | 4,71 | 0 | 1,08 | 0 | 6,61 | 0 |
| ZBTB33 | GCCTCCCGAGTAGCT | -1 | 342 | 357 | -658 | -643 | -3,21 | 0,01 | 11,66 | >0.1 | 2,56 | 4,68 | 0 | 1,12 | 0 | 6,52 | 0 |
| ZBTB33 | CTCCCGAGTAGCTGG | -1 | 340 | 355 | -660 | -645 | -1,13 | 0,01 | 13,63 | >0.1 | 2,56 | 4,68 | 0 | 1,12 | 0 | 6,4 | 0 |
| ZBTB33 | CCACCACGCCCAGCT | -1 | 310 | 325 | -690 | -675 | -6,45 | 0,01 | 8,04 | >0.1 | 2,64 | 4,63 | 0 | 1,18 | 0 | 6,03 | 0 |
| ZBTB33 | TCACCGTGTTAGTCA | -1 | 264 | 279 | -736 | -721 | -2,39 | 0,01 | 10,8 | >0.1 | 2,79 | 4,56 | 0 | 1,32 | 0 | 4,53 | 0 |
| ZBTB33 | CTATCTCCTGACCTT | -1 | 240 | 255 | -760 | -745 | -7,69 | 0,01 | 5,44 | >0.1 | 2,87 | 4,53 | 0 | 1,41 | 0 | 4,33 | 0 |
| ZBTB33 | ATCTCCTGACCTTGT | -1 | 238 | 253 | -762 | -747 | -4,75 | 0,01 | 8,39 | >0.1 | 2,88 | 4,52 | 0 | 1,42 | 0 | 4,33 | 0 |
| ZBTB33 | TGACCTTGTGATCTG | -1 | 232 | 247 | -768 | -753 | -6,02 | 0,01 | 7,74 | >0.1 | 2,9 | 4,52 | 0 | 1,45 | 0 | 4,9 | 0 |
| ZBTB33 | ACCTTGTGATCTGCC | -1 | 230 | 245 | -770 | -755 | -7,23 | 0,01 | 6,65 | >0.1 | 2,91 | 4,51 | 0 | 1,46 | 0 | 5,01 | 0 |
| ZBTB33 | CCAACTCGGCCTCCC | -1 | 216 | 231 | -784 | -769 | -6,98 | 0,01 | 7,64 | >0.1 | 2,96 | 4,49 | 0 | 1,53 | 0 | 5,63 | 0 |
| ZBTB33 | TTTCCATGGCCATTT | -1 | 143 | 158 | -857 | -842 | -7,39 | 0,01 | 3,83 | >0.1 | 3,31 | 4,4 | 0 | 1,9 | 0 | 1,61 | 0 |
| ZBTB33 | CTCTCCTGATCTGCC | -1 | 92 | 107 | -908 | -893 | -1,83 | 0,01 | 8,84 | >0.1 | 3,66 | 4,35 | 0 | 1,88 | 0 | 0,79 | 0 |
| ZBTB33 | TTCTCACACTGGTTC | -1 | 46 | 61 | -954 | -939 | -1,72 | 0,01 | 8,6 | >0.1 | 4,15 | 4,3 | 0 | 1,87 | 0 | 0 | 0 |
| ZBTB33 | CTCACACTGGTTCTG | -1 | 44 | 59 | -956 | -941 | -5,33 | 0,01 | 5,02 | >0.1 | 4,18 | 4,3 | 0 | 1,87 | 0 | 0 | 0 |
| POU3F1 | TTTTGCAAAGGC | 1 | 83 | 95 | -917 | -905 | 3,98 | 0,01 | 13,94 | >0.1 | 3,75 | 4,33 | 0 | 1,88 | 0 | 0 | 0 |
| POU3F1 | AAAGGCAGATCA | 1 | 89 | 101 | -911 | -899 | 1,86 | 0,01 | 11,78 | >0.1 | 3,69 | 4,34 | 0 | 1,88 | 0 | 0 | 0 |
| POU3F1 | TAATTTTAAAAA | 1 | 133 | 145 | -867 | -855 | 3,06 | 0,01 | 14,33 | >0.1 | 3,38 | 4,39 | 0 | 1,89 | 0 | 1,61 | 0 |
| POU3F1 | CCATGGAAATTA | 1 | 149 | 161 | -851 | -839 | 7,37 | 0,001 | 18,61 | >0.1 | 3,28 | 4,41 | 0 | 1,9 | 0 | 1,65 | 0 |
| POU3F1 | CTATTAAAAATA | 1 | 289 | 301 | -711 | -699 | 1,67 | 0,01 | 15,29 | >0.1 | 2,71 | 4,59 | 0 | 1,24 | 0 | 5,09 | 0 |
| POU3F1 | TTAATGAAATTG | 1 | 604 | 616 | -396 | -384 | 2,66 | 0,01 | 13,83 | >0.1 | 3,51 | 5,22 | 0 | 1,02 | 0 | 1,41 | 0 |
| POU3F1 | TAATGAAATTGT | 1 | 605 | 617 | -395 | -383 | 3,2 | 0,01 | 14,41 | >0.1 | 3,52 | 5,23 | 0 | 1,02 | 0 | 1,44 | 0 |
| POU3F1 | TTATCTTCATGA | 1 | 641 | 653 | -359 | -347 | 2,07 | 0,01 | 16,11 | >0.1 | 3,81 | 5,33 | 0 | 1,06 | 0 | 3,83 | 0 |
| POU3F1 | GTATTCTTATTT | 1 | 806 | 818 | -194 | -182 | 6,42 | 0,01 | 22,84 | >0.1 | 5,58 | 6,02 | 0 | 1,54 | 0 | 3,28 | 0 |
| POU3F1 | TTATTCTAAAAT | 1 | 834 | 846 | -166 | -154 | 6,33 | 0,01 | 23,13 | >0.1 | 5,58 | 6,2 | 0 | 1,74 | 0 | 3,28 | 0 |
| POU3F1 | ATATGTAACAGA | -1 | 845 | 857 | -155 | -143 | 3,98 | 0,01 | 20,53 | >0.1 | 5,58 | 6,28 | 0 | 1,83 | 0 | 2,86 | 0 |
| POU3F1 | TCATGAAGATAA | -1 | 641 | 653 | -359 | -347 | 3,25 | 0,01 | 17,29 | >0.1 | 3,81 | 5,33 | 0 | 1,06 | 0 | 3,83 | 0 |
| POU3F1 | GTATGGTTTTCA | -1 | 618 | 630 | -382 | -370 | 1,65 | 0,01 | 13,06 | >0.1 | 3,61 | 5,26 | 0 | 1,04 | 0 | 1,5 | 0 |
| POU3F1 | TTATTTTTATTT | -1 | 577 | 589 | -423 | -411 | 6,44 | 0,01 | 15,93 | >0.1 | 3,33 | 5,15 | 0 | 1 | 0 | 0 | 0 |
| POU3F1 | TTATGGAATTCA | -1 | 492 | 504 | -508 | -496 | 6,88 | 0,01 | 22,18 | >0.1 | 2,93 | 4,95 | 0 | 0,99 | 0 | 6,44 | 0 |
| POU3F1 | GAATTCATATTT | -1 | 487 | 499 | -513 | -501 | 4,57 | 0,01 | 19,68 | >0.1 | 2,91 | 4,94 | 0 | 0,99 | 0 | 6,27 | 0 |
| POU3F1 | GTATTTTTAATA | -1 | 290 | 302 | -710 | -698 | 2,18 | 0,01 | 15,16 | >0.1 | 2,71 | 4,59 | 0 | 1,23 | 0 | 4,45 | 0 |
| POU3F1 | AAAAGCTCATTT | -1 | 66 | 78 | -934 | -922 | 2,45 | 0,01 | 12,56 | >0.1 | 3,92 | 4,32 | 0 | 1,88 | 0 | 0 | 0 |
| SPZ1 | AGTGAAGGAGA | 1 | 8 | 19 | -992 | -981 | 2,39 | 0,01 | 15,91 | >0.1 | 5,6 | 4,26 | 0 | 1,86 | 1,79 | 0 | 0 |
| SPZ1 | GGAGAAGGAGG | 1 | 14 | 25 | -986 | -975 | 5,32 | 0,01 | 17,05 | >0.1 | 5,6 | 4,27 | 0 | 1,87 | 0 | 0 | 0 |
| SPZ1 | AGGAAATGAGC | 1 | 63 | 74 | -937 | -926 | 5,08 | 0,01 | 15,22 | >0.1 | 3,96 | 4,31 | 0 | 1,88 | 0 | 0 | 0 |
| SPZ1 | AGGTCAGGAGA | 1 | 241 | 252 | -759 | -748 | 2,75 | 0,01 | 15,88 | >0.1 | 2,87 | 4,52 | 0 | 1,4 | 0 | 4,33 | 0 |
| SPZ1 | GGAGATAGAGA | 1 | 247 | 258 | -753 | -742 | 4,85 | 0,01 | 18,8 | >0.1 | 2,85 | 4,53 | 0 | 1,38 | 0 | 5,19 | 0 |
| SPZ1 | ACGGTGAAACC | 1 | 272 | 283 | -728 | -717 | 3,8 | 0,01 | 17,41 | >0.1 | 2,77 | 4,57 | 0 | 1,29 | 0 | 4,99 | 0 |
| SPZ1 | GGAGGCAGAGC | 1 | 383 | 394 | -617 | -606 | 3,85 | 0,01 | 18,24 | >0.1 | 2,58 | 4,74 | 0 | 1,06 | 0 | 6,02 | 0 |
| SPZ1 | AGGGCAAAGTC | 1 | 509 | 520 | -491 | -480 | 4,21 | 0,01 | 15,93 | >0.1 | 3 | 4,99 | 0 | 0,98 | 0 | 2,75 | 0 |
| SPZ1 | AGAGTTTCTGG | 1 | 553 | 564 | -447 | -436 | 5,29 | 0,01 | 14,57 | >0.1 | 3,2 | 5,09 | 0 | 0,99 | 0 | 0 | 0 |
| SPZ1 | AGGATTAAAGA | 1 | 790 | 801 | -210 | -199 | 4 | 0,01 | 22,29 | >0.1 | 5,6 | 5,92 | 0 | 1,45 | 0 | 5,32 | 0 |
| SPZ1 | AGGTTTTAGTC | 1 | 993 | 1004 | -7 | 4 | 4,14 | 0,01 | 21,98 | >0.1 | 5,6 | 8,84 | 0 | 1,87 | 0 | 1,53 | 0 |
| SPZ1 | AGGGGACGATG | 1 | 1041 | 1052 | 41 | 52 | 3,53 | 0,01 | 20,2 | >0.1 | 5,6 | 7,82 | 0 | 1,86 | 0 | 1,39 | 0 |
| SPZ1 | GCTGTAACTGC | 1 | 1085 | 1096 | 85 | 96 | 2,17 | 0,01 | 20,57 | >0.1 | 5,6 | 6,94 | 0 | 1,85 | 1,79 | 2,21 | 0 |

| | | | | | | | | | | | | | | | | | |
|---|---|---|---|---|---|---|---|---|---|---|---|---|---|---|---|---|---|
| SPZ1 | TGAGGAACAGG | 1 | 1111 | 1122 | 111 | 122 | 4,38 | 0,01 | 24,23 | >0.1 | 5,6 | 6,64 | 0 | 1,85 | 3,22 | 2,54 | 0 |
| SPZ1 | AGGTCTTCGGC | -1 | 1154 | 1165 | 154 | 165 | 4,96 | 0,01 | 21,6 | >0.1 | 4,45 | 6,27 | 0 | 1,82 | 1,79 | 2,31 | 0 |
| SPZ1 | GTGGACTGAGC | -1 | 1097 | 1108 | 97 | 108 | 2,06 | 0,01 | 21,88 | >0.1 | 5,6 | 6,79 | 0 | 1,85 | 3,22 | 2,36 | 0 |
| SPZ1 | GCAGTTACAGC | -1 | 1085 | 1096 | 85 | 96 | 10,34 | 0,001 | 28,74 | >0.1 | 5,6 | 6,94 | 0 | 1,85 | 1,79 | 2,21 | 0 |
| SPZ1 | TGAGAAGGAGC | -1 | 1066 | 1077 | 66 | 77 | 2,72 | 0,01 | 21,48 | >0.1 | 5,6 | 7,24 | 0 | 1,85 | 1,79 | 2,27 | 0 |
| SPZ1 | CCAGGAAGAGC | -1 | 1056 | 1067 | 56 | 67 | 2,65 | 0,01 | 19,75 | >0.1 | 5,6 | 7,44 | 0 | 1,86 | 0 | 2,21 | 0 |
| SPZ1 | AGACTAAAACC | -1 | 994 | 1005 | -6 | 5 | 2,21 | 0,01 | 20,08 | >0.1 | 5,6 | 8,88 | 0 | 1,87 | 0 | 1,53 | 0 |
| SPZ1 | TGGTTTTCAGA | -1 | 616 | 627 | -384 | -373 | 3,94 | 0,01 | 15,32 | >0.1 | 3,59 | 5,26 | 0 | 1,03 | 0 | 1,5 | 0 |
| SPZ1 | AGGTTCACGCC | -1 | 371 | 382 | -629 | -618 | 3,87 | 0,01 | 18,61 | >0.1 | 2,55 | 4,72 | 0 | 1,07 | 0 | 6,4 | 0 |
| SPZ1 | GGGGTTTCACC | -1 | 274 | 285 | -726 | -715 | 11,61 | 0,0001 | 25,41 | >0.1 | 2,76 | 4,57 | 0 | 1,28 | 0 | 5,19 | 0 |
| SPZ1 | AGGATTACAGG | -1 | 197 | 208 | -803 | -792 | 7,93 | 0,001 | 21,47 | >0.1 | 3,05 | 4,46 | 0 | 1,65 | 0 | 4,37 | 0 |
| ZBTB7A | AATAGGAAATGAG | 1 | 60 | 73 | -940 | -927 | 1,74 | 0,01 | 11,91 | >0.1 | 3,98 | 4,31 | 0 | 1,88 | 0 | 0 | 0 |
| ZBTB7A | GTCAGGAGATAGA | 1 | 243 | 256 | -757 | -744 | 0,6 | 0,01 | 13,81 | >0.1 | 2,86 | 4,53 | 0 | 1,4 | 0 | 4,42 | 0 |
| ZBTB7A | AGCTGGGCGTGGT | 1 | 310 | 323 | -690 | -677 | 1,2 | 0,01 | 15,69 | >0.1 | 2,65 | 4,62 | 0 | 1,18 | 0 | 6,03 | 0 |
| ZBTB7A | CACCTGTAGTCCC | 1 | 329 | 342 | -671 | -658 | 1,82 | 0,01 | 15,67 | >0.1 | 2,59 | 4,65 | 0 | 1,14 | 0 | 5,46 | 0 |
| ZBTB7A | ACTCGGGAGGCTG | 1 | 346 | 359 | -654 | -641 | 1,41 | 0,01 | 16,27 | >0.1 | 2,55 | 4,68 | 0 | 1,11 | 0 | 6,52 | 0 |
| ZBTB7A | CTCGGGAGGCTGA | 1 | 347 | 360 | -653 | -640 | 2,47 | 0,01 | 17,21 | >0.1 | 2,54 | 4,68 | 0 | 1,11 | 0 | 6,4 | 0 |
| ZBTB7A | CTGAGGCAGGAGA | 1 | 356 | 369 | -644 | -631 | 0,35 | 0,01 | 14,79 | >0.1 | 2,52 | 4,7 | 0 | 1,09 | 0 | 6,13 | 0 |
| ZBTB7A | ACCTGGGAGGCAG | 1 | 378 | 391 | -622 | -609 | 2,47 | 0,01 | 17,5 | >0.1 | 2,57 | 4,74 | 0 | 1,06 | 0 | 6,66 | 0 |
| ZBTB7A | GGGAGGCAGAGCT | 1 | 382 | 395 | -618 | -605 | 3,28 | 0,01 | 17,68 | >0.1 | 2,58 | 4,74 | 0 | 1,06 | 0 | 6,02 | 0 |
| ZBTB7A | GCCATAAAGTGAC | 1 | 758 | 771 | -242 | -229 | 0,34 | 0,01 | 19,11 | >0.1 | 5,57 | 5,77 | 0 | 1,33 | 0 | 6,1 | 0 |
| ZBTB7A | TCCAGGATGAACT | 1 | 865 | 878 | -135 | -122 | 2,42 | 0,01 | 19,07 | >0.1 | 5,57 | 6,43 | 0 | 1,9 | 0 | 2,75 | 0 |
| ZBTB7A | TTGAGGAACAGGC | 1 | 1110 | 1123 | 110 | 123 | 3,64 | 0,01 | 23,47 | >0.1 | 5,57 | 6,65 | 0 | 1,85 | 3,22 | 2,54 | 0 |
| ZBTB7A | ACCACGAAGCCGA | 1 | 1146 | 1159 | 146 | 159 | 6,17 | 0,01 | 23,91 | >0.1 | 5,57 | 6,33 | 0 | 1,84 | 1,79 | 2,21 | 0 |
| ZBTB7A | AAGCCGAAGACCT | 1 | 1152 | 1165 | 152 | 165 | 1,51 | 0,01 | 18,24 | >0.1 | 4,47 | 6,28 | 0 | 1,84 | 1,79 | 2,34 | 0 |
| ZBTB7A | TCCAAGAAGCAAG | -1 | 1181 | 1194 | 181 | 194 | 3,43 | 0,01 | 22,02 | >0.1 | 5,57 | 6,08 | 0 | 1,58 | 1,79 | 3,57 | 0 |
| ZBTB7A | AGAAGCAAGTGAA | -1 | 1177 | 1190 | 177 | 190 | 1,37 | 0,01 | 19,82 | >0.1 | 5,57 | 6,11 | 0 | 1,61 | 1,79 | 3,37 | 0 |
| ZBTB7A | GCCAGGAAGAGCT | -1 | 1055 | 1068 | 55 | 68 | 11,79 | 0,0001 | 28,89 | >0.1 | 5,57 | 7,46 | 0 | 1,86 | 0 | 2,21 | 0 |
| ZBTB7A | AGCAGGAAAGACC | -1 | 930 | 943 | -70 | -57 | 5,84 | 0,01 | 21,35 | >0.1 | 5,57 | 7,11 | 0 | 1,88 | 0 | 0,95 | 0 |
| ZBTB7A | ATCCTGGAGAGGA | -1 | 860 | 873 | -140 | -127 | 0,71 | 0,01 | 17,32 | >0.1 | 5,57 | 6,39 | 0 | 1,9 | 0 | 2,75 | 0 |
| ZBTB7A | ACACGGAAAATGT | -1 | 692 | 705 | -308 | -295 | 0,95 | 0,01 | 16,24 | >0.1 | 5,57 | 5,5 | 0 | 1,15 | 0 | 3,07 | 0 |
| ZBTB7A | TCCCAGAAACTCT | -1 | 553 | 566 | -447 | -434 | 0,43 | 0,01 | 9,72 | >0.1 | 3,21 | 5,09 | 0 | 0,99 | 0 | 0 | 0 |
| ZBTB7A | CCCAGAAACTCTC | -1 | 552 | 565 | -448 | -435 | 0,61 | 0,01 | 9,89 | >0.1 | 3,2 | 5,09 | 0 | 0,99 | 0 | 0 | 0 |
| ZBTB7A | CCCAGGCTGGAGC | -1 | 422 | 435 | -578 | -565 | 1,08 | 0,01 | 15,27 | >0.1 | 2,69 | 4,81 | 0 | 1,02 | 0 | 5,67 | 0 |
| ZBTB7A | GTCAGGACGGTCT | -1 | 255 | 268 | -745 | -732 | 2,78 | 0,01 | 16,22 | >0.1 | 2,82 | 4,54 | 0 | 1,35 | 0 | 4,73 | 0 |
| ZBTB7A | CTCCCAAAGTGCT | -1 | 208 | 221 | -792 | -779 | 1,74 | 0,01 | 15,5 | >0.1 | 3 | 4,48 | 0 | 1,58 | 0 | 4,7 | 0 |
| MNX1 | GGCACTCAAA | 1 | 23 | 33 | -977 | -967 | 3,45 | 0,01 | 15,2 | >0.1 | 5,61 | 4,27 | 0 | 1,87 | 0 | 0 | 0 |
| MNX1 | AGTCATAATT | 1 | 128 | 138 | -872 | -862 | 4,72 | 0,01 | 15,98 | >0.1 | 3,41 | 4,38 | 0 | 1,89 | 0 | 1,57 | 0 |
| MNX1 | GGAAATTAAA | 1 | 153 | 163 | -847 | -837 | 3,38 | 0,01 | 14,6 | >0.1 | 3,27 | 4,41 | 0 | 1,9 | 0 | 1,65 | 0 |
| MNX1 | CTCTATTAAA | 1 | 287 | 297 | -713 | -703 | 3,74 | 0,01 | 17,48 | >0.1 | 2,72 | 4,59 | 0 | 1,24 | 0 | 5,19 | 0 |
| MNX1 | TTCCATAAAG | 1 | 496 | 506 | -504 | -494 | 4,6 | 0,01 | 19,92 | >0.1 | 2,94 | 4,96 | 0 | 0,98 | 0 | 6,44 | 0 |
| MNX1 | ACCTATTAAT | 1 | 599 | 609 | -401 | -391 | 6,04 | 0,01 | 17,09 | >0.1 | 3,47 | 5,21 | 0 | 1,02 | 0 | 1,36 | 0 |
| MNX1 | ATTAATGAAA | 1 | 603 | 613 | -397 | -387 | 7,4 | 0,01 | 18,54 | >0.1 | 3,49 | 5,22 | 0 | 1,02 | 0 | 1,41 | 0 |
| MNX1 | GGCCATAAAG | 1 | 757 | 767 | -243 | -233 | 6,64 | 0,01 | 25,42 | >0.1 | 5,61 | 5,75 | 0 | 1,31 | 0 | 6,1 | 0 |
| MNX1 | TCTCATGAGG | 1 | 983 | 993 | -17 | -7 | 4,26 | 0,01 | 21,81 | >0.1 | 5,61 | 8,5 | 0 | 1,87 | 0 | 1,57 | 0 |
| MNX1 | AGCAACAAGG | -1 | 1078 | 1088 | 78 | 88 | 3,64 | 0,01 | 22,15 | >0.1 | 5,61 | 7,04 | 0 | 1,85 | 1,79 | 2,21 | 0 |
| MNX1 | TCCACTGAAT | -1 | 1015 | 1025 | 15 | 25 | 2,75 | 0,01 | 20,45 | >0.1 | 5,61 | 8,84 | 0 | 1,86 | 0 | 1,39 | 0 |
| MNX1 | CCTCATGAGA | -1 | 983 | 993 | -17 | -7 | 2,88 | 0,01 | 20,43 | >0.1 | 5,61 | 8,5 | 0 | 1,87 | 0 | 1,57 | 0 |
| MNX1 | ATCAAATAAG | -1 | 811 | 821 | -189 | -179 | 2,77 | 0,01 | 19,4 | >0.1 | 5,61 | 6,03 | 0 | 1,56 | 0 | 3,43 | 0 |
| MNX1 | TGTTATAATT | -1 | 685 | 695 | -315 | -305 | 2,68 | 0,01 | 16,62 | >0.1 | 4,4 | 5,47 | 0 | 1,12 | 0 | 2,95 | 0 |
| MNX1 | GATCATGAAG | -1 | 645 | 655 | -355 | -345 | 5,31 | 0,01 | 19,39 | >0.1 | 3,84 | 5,34 | 0 | 1,06 | 0 | 3,83 | 0 |
| MNX1 | TTTCATTAAT | -1 | 603 | 613 | -397 | -387 | 6,66 | 0,01 | 17,8 | >0.1 | 3,49 | 5,22 | 0 | 1,02 | 0 | 1,41 | 0 |
| MNX1 | TTCACTTACC | -1 | 4 | 14 | -996 | -986 | 3,09 | 0,01 | 16,61 | >0.1 | 5,61 | 4,26 | 0 | 1,86 | 1,79 | 0 | 0 |
| RUNX2 | CAGCCACAG | 1 | 38 | 47 | -962 | -953 | -4,04 | 0,01 | 6,44 | >0.1 | 4,33 | 4,29 | 0 | 1,87 | 0 | 0 | 0 |
| RUNX2 | AGATCACAA | 1 | 233 | 242 | -767 | -758 | -4,26 | 0,01 | 9,6 | >0.1 | 2,91 | 4,51 | 0 | 1,44 | 0 | 5,01 | 0 |
| RUNX2 | AAACCCCGT | 1 | 278 | 287 | -722 | -713 | 1,83 | 0,01 | 15,61 | >0.1 | 2,75 | 4,57 | 0 | 1,27 | 0 | 5,19 | 0 |
| RUNX2 | AGACCGAGA | 1 | 439 | 448 | -561 | -552 | -5,82 | 0,01 | 8,81 | >0.1 | 2,74 | 4,84 | 0 | 1 | 0 | 6,05 | 0 |
| RUNX2 | AAACCATAC | 1 | 621 | 630 | -379 | -370 | -4,44 | 0,01 | 6,92 | >0.1 | 3,62 | 5,26 | 0 | 1,03 | 0 | 1,44 | 0 |
| RUNX2 | CAAACACCA | 1 | 629 | 638 | -371 | -362 | -6,94 | 0,01 | 4,69 | >0.1 | 3,69 | 5,29 | 0 | 1,04 | 0 | 1,61 | 0 |

| | | | | | | | | | | | | | | | | | |
|---|---|---|---|---|---|---|---|---|---|---|---|---|---|---|---|---|---|
| RUNX2 | AGACCTCTT | 1 | 664 | 673 | -336 | -327 | -0,64 | 0,01 | 13,89 | >0.1 | 4,05 | 5,4 | 0 | 1,09 | 0 | 3,99 | 0 |
| RUNX2 | TAACCAAAA | 1 | 732 | 741 | -268 | -259 | -2,8 | 0,01 | 15,39 | >0.1 | 5,62 | 5,64 | 0 | 1,23 | 0 | 5,69 | 0 |
| RUNX2 | AGTCCACCA | 1 | 1101 | 1110 | 101 | 110 | -7,06 | 0,01 | 12,76 | >0.1 | 5,62 | 6,75 | 0 | 1,85 | 3,22 | 2,38 | 0 |
| RUNX2 | TAACCACGA | 1 | 1144 | 1153 | 144 | 153 | 9,35 | 0,001 | 28,49 | >0.1 | 5,62 | 6,34 | 0 | 1,84 | 3,22 | 2,11 | 0 |
| RUNX2 | TTACAGCAA | -1 | 1083 | 1092 | 83 | 92 | -4,51 | 0,01 | 13,94 | >0.1 | 5,62 | 6,97 | 0 | 1,85 | 1,79 | 2,21 | 0 |
| RUNX2 | AGATCACAT | -1 | 1024 | 1033 | 24 | 33 | -7,05 | 0,01 | 10,3 | >0.1 | 5,62 | 8,59 | 0 | 1,86 | 0 | 1,28 | 0 |
| RUNX2 | AAACCTCCT | -1 | 990 | 999 | -10 | -1 | 3,12 | 0,01 | 20,91 | >0.1 | 5,62 | 8,69 | 0 | 1,87 | 0 | 1,61 | 0 |
| RUNX2 | AGACCAAAA | -1 | 926 | 935 | -74 | -65 | -4,26 | 0,01 | 11,3 | >0.1 | 5,62 | 6,99 | 0 | 1,88 | 0 | 1,06 | 0 |
| RUNX2 | AGAACACAG | -1 | 903 | 912 | -97 | -88 | -5,65 | 0,01 | 9,86 | >0.1 | 5,62 | 6,72 | 0 | 1,89 | 0 | 1,28 | 0 |
| RUNX2 | TTACCTCTC | -1 | 773 | 782 | -227 | -218 | -2,04 | 0,01 | 16,69 | >0.1 | 5,62 | 5,82 | 0 | 1,37 | 0 | 5,92 | 0 |
| RUNX2 | AAACCGGTA | -1 | 742 | 751 | -258 | -249 | -5,13 | 0,01 | 13,54 | >0.1 | 5,62 | 5,68 | 0 | 1,26 | 0 | 6,1 | 0 |
| RUNX2 | TGACCAAAG | -1 | 670 | 679 | -330 | -321 | -6,14 | 0,01 | 8,2 | >0.1 | 4,13 | 5,41 | 0 | 1,1 | 0 | 3,7 | 0 |
| RUNX2 | TCACTGCAA | -1 | 394 | 403 | -606 | -597 | -4,08 | 0,01 | 10,34 | >0.1 | 2,61 | 4,76 | 0 | 1,04 | 0 | 6,02 | 0 |
| RUNX2 | TGCCCACCA | -1 | 322 | 331 | -678 | -669 | 3,61 | 0,01 | 17,58 | >0.1 | 2,62 | 4,64 | 0 | 1,15 | 0 | 5,56 | 0 |
| RUNX2 | CCACCACCA | -1 | 319 | 328 | -681 | -672 | 5,5 | 0,001 | 20,17 | >0.1 | 2,63 | 4,63 | 0 | 1,16 | 0 | 6,26 | 0 |
| RUNX2 | CCACCACGC | -1 | 316 | 325 | -684 | -675 | 3,3 | 0,01 | 17,86 | >0.1 | 2,63 | 4,63 | 0 | 1,17 | 0 | 6,13 | 0 |
| RUNX2 | ACTCCACAG | -1 | 112 | 121 | -888 | -879 | -6,28 | 0,01 | 4,28 | >0.1 | 3,53 | 4,36 | 0 | 1,89 | 0 | 0,79 | 0 |
| ZBTB7B | TGAAACCCCGTC | 1 | 276 | 288 | -724 | -712 | -0,23 | 0,01 | 13,56 | >0.1 | 2,75 | 4,57 | 0 | 1,27 | 0 | 5,19 | 0 |
| ZBTB7B | GAAACCCCGTCT | 1 | 277 | 289 | -723 | -711 | -7,38 | 0,01 | 6,41 | >0.1 | 2,75 | 4,57 | 0 | 1,27 | 0 | 5,19 | 0 |
| ZBTB7B | ACAAACACCAAC | 1 | 628 | 640 | -372 | -360 | 4,68 | 0,001 | 16,37 | >0.1 | 3,69 | 5,29 | 0 | 1,05 | 0 | 1,66 | 0 |
| ZBTB7B | AACACCAACATT | 1 | 631 | 643 | -369 | -357 | -6,23 | 0,01 | 5,45 | >0.1 | 3,72 | 5,3 | 0 | 1,05 | 0 | 1,61 | 0 |
| ZBTB7B | GCGCCCAACCCA | 1 | 945 | 957 | -55 | -43 | 4,8 | 0,001 | 21,4 | >0.1 | 5,58 | 7,34 | 0 | 1,88 | 0 | 1,8 | 0 |
| ZBTB7B | CAGTCCACCATT | 1 | 1100 | 1112 | 100 | 112 | -1,09 | 0,01 | 18,7 | >0.1 | 5,58 | 6,76 | 0 | 1,85 | 3,22 | 2,38 | 0 |
| ZBTB7B | TTAACCACGAAG | 1 | 1143 | 1155 | 143 | 155 | -5,99 | 0,01 | 13,17 | >0.1 | 5,58 | 6,35 | 0 | 1,84 | 3,22 | 2,16 | 0 |
| ZBTB7B | TCGTCCCCTGTG | -1 | 1038 | 1050 | 38 | 50 | -3,5 | 0,01 | 13,22 | >0.1 | 5,58 | 7,92 | 0 | 1,86 | 0 | 1,36 | 0 |
| ZBTB7B | ACATCCACTGAA | -1 | 1016 | 1028 | 16 | 28 | -1,13 | 0,01 | 16,46 | >0.1 | 5,58 | 8,81 | 0 | 1,86 | 0 | 1,34 | 0 |
| ZBTB7B | AAAACCTCCTCA | -1 | 988 | 1000 | -12 | 0 | -2,06 | 0,01 | 15,72 | >0.1 | 5,58 | 8,71 | 0 | 1,87 | 0 | 1,61 | 0 |
| ZBTB7B | GGAAAGACCAAA | -1 | 927 | 939 | -73 | -61 | -5,82 | 0,01 | 9,72 | >0.1 | 5,58 | 7,05 | 0 | 1,88 | 0 | 1,03 | 0 |
| ZBTB7B | CAGCCTCCCGAG | -1 | 347 | 359 | -653 | -641 | -4,49 | 0,01 | 10,25 | >0.1 | 2,55 | 4,68 | 0 | 1,11 | 0 | 6,4 | 0 |
| ZBTB7B | GTGCCCACCACC | -1 | 320 | 332 | -680 | -668 | 7,25 | 0,001 | 21,23 | >0.1 | 2,62 | 4,64 | 0 | 1,16 | 0 | 5,56 | 0 |
| ZBTB7B | CCCACCACCACG | -1 | 317 | 329 | -683 | -671 | 6,06 | 0,001 | 20,74 | >0.1 | 2,63 | 4,63 | 0 | 1,17 | 0 | 6,26 | 0 |
| ZBTB7B | ACCACCACGCCC | -1 | 314 | 326 | -686 | -674 | -5,8 | 0,01 | 8,77 | >0.1 | 2,64 | 4,63 | 0 | 1,17 | 0 | 6,13 | 0 |
| ZBTB7B | ACCACGCCCAGC | -1 | 311 | 323 | -689 | -677 | -3,17 | 0,01 | 11,31 | >0.1 | 2,65 | 4,62 | 0 | 1,18 | 0 | 6,03 | 0 |
| ZBTB7B | ACGCCCAGCTAA | -1 | 308 | 320 | -692 | -680 | 0,54 | 0,01 | 15,26 | >0.1 | 2,65 | 4,62 | 0 | 1,19 | 0 | 6,26 | 0 |
| ZBTB7B | CTGCCCAACTCG | -1 | 223 | 235 | -777 | -765 | -5,77 | 0,01 | 8,41 | >0.1 | 2,94 | 4,5 | 0 | 1,49 | 0 | 5,26 | 0 |
| ZBTB7B | TCGGCCTCCCAA | -1 | 214 | 226 | -786 | -774 | -5,09 | 0,01 | 9,55 | >0.1 | 2,98 | 4,49 | 0 | 1,54 | 0 | 5,63 | 0 |
| ZBTB7B | CGGCCTCCCAAA | -1 | 213 | 225 | -787 | -775 | -3,53 | 0,01 | 11,01 | >0.1 | 2,98 | 4,49 | 0 | 1,55 | 0 | 5,52 | 0 |
| ZBTB7B | GTGCCCAGCCAT | -1 | 174 | 186 | -826 | -814 | -4,57 | 0,01 | 8,1 | >0.1 | 3,15 | 4,44 | 0 | 1,84 | 0 | 3,24 | 0 |
| EN1 | GAAATGAG | 1 | 65 | 73 | -935 | -927 | 3,04 | 0,01 | 13,18 | >0.1 | 3,95 | 4,31 | 0 | 1,88 | 0 | 0 | 0 |
| EN1 | ATAATTTT | 1 | 132 | 140 | -868 | -860 | 2,31 | 0,01 | 13,55 | >0.1 | 3,4 | 4,38 | 0 | 1,89 | 0 | 1,57 | 0 |
| EN1 | GAAATTAA | 1 | 154 | 162 | -846 | -838 | 5,38 | 0,01 | 16,6 | >0.1 | 3,27 | 4,41 | 0 | 1,9 | 0 | 1,65 | 0 |
| EN1 | CTGATCAG | 1 | 164 | 172 | -836 | -828 | 4,29 | 0,01 | 16,22 | >0.1 | 3,21 | 4,42 | 0 | 1,9 | 0 | 2,4 | 0 |
| EN1 | CTGATCAG | 1 | 479 | 487 | -521 | -513 | 4,29 | 0,01 | 18,43 | >0.1 | 2,88 | 4,92 | 0 | 0,98 | 0 | 5,36 | 0 |
| EN1 | TTAATGAA | 1 | 604 | 612 | -396 | -388 | 4,59 | 0,01 | 15,73 | >0.1 | 3,49 | 5,21 | 0 | 1,02 | 0 | 1,41 | 0 |
| EN1 | AAAATTAT | 1 | 683 | 691 | -317 | -309 | 2,85 | 0,01 | 16,83 | >0.1 | 4,34 | 5,46 | 0 | 1,12 | 0 | 3,07 | 0 |
| EN1 | AAAATTAT | 1 | 830 | 838 | -170 | -162 | 2,85 | 0,01 | 19,58 | >0.1 | 5,64 | 6,14 | 0 | 1,67 | 0 | 3,28 | 0 |
| EN1 | CTCATGAG | 1 | 984 | 992 | -16 | -8 | 3,19 | 0,01 | 20,71 | >0.1 | 5,64 | 8,43 | 0 | 1,87 | 0 | 1,57 | 0 |
| EN1 | CTCATGAG | -1 | 984 | 992 | -16 | -8 | 3,19 | 0,01 | 20,71 | >0.1 | 5,64 | 8,43 | 0 | 1,87 | 0 | 1,57 | 0 |
| EN1 | ACAATCTG | -1 | 892 | 900 | -108 | -100 | 2,99 | 0,01 | 19,99 | >0.1 | 5,64 | 6,61 | 0 | 1,89 | 0 | 2,86 | 0 |
| EN1 | ATAATTTT | -1 | 830 | 838 | -170 | -162 | 2,31 | 0,01 | 19,04 | >0.1 | 5,64 | 6,14 | 0 | 1,67 | 0 | 3,28 | 0 |
| EN1 | CAAATAAG | -1 | 811 | 819 | -189 | -181 | 1,65 | 0,01 | 18,14 | >0.1 | 5,64 | 6,02 | 0 | 1,55 | 0 | 3,28 | 0 |
| EN1 | CAAATCAA | -1 | 750 | 758 | -250 | -242 | 3,91 | 0,01 | 22,46 | >0.1 | 5,64 | 5,71 | 0 | 1,28 | 0 | 5,92 | 0 |
| EN1 | ATAATTTT | -1 | 683 | 691 | -317 | -309 | 2,31 | 0,01 | 16,29 | >0.1 | 4,34 | 5,46 | 0 | 1,12 | 0 | 3,07 | 0 |
| EN1 | ACAATTTC | -1 | 609 | 617 | -391 | -383 | 4,24 | 0,01 | 15,56 | >0.1 | 3,53 | 5,23 | 0 | 1,02 | 0 | 1,55 | 0 |
| EN1 | TTCATTAA | -1 | 604 | 612 | -396 | -388 | 3,88 | 0,01 | 15,02 | >0.1 | 3,49 | 5,21 | 0 | 1,02 | 0 | 1,41 | 0 |
| EN1 | CTGATCAG | -1 | 479 | 487 | -521 | -513 | 4,29 | 0,01 | 18,43 | >0.1 | 2,88 | 4,92 | 0 | 0,98 | 0 | 5,36 | 0 |
| EN1 | CTGATCAG | -1 | 164 | 172 | -836 | -828 | 4,29 | 0,01 | 16,22 | >0.1 | 3,21 | 4,42 | 0 | 1,9 | 0 | 2,4 | 0 |
| EN1 | TTAATTTC | -1 | 154 | 162 | -846 | -838 | 4,28 | 0,01 | 15,5 | >0.1 | 3,27 | 4,41 | 0 | 1,9 | 0 | 1,65 | 0 |
| EN1 | AAAATTAT | -1 | 132 | 140 | -868 | -860 | 2,85 | 0,01 | 14,09 | >0.1 | 3,4 | 4,38 | 0 | 1,89 | 0 | 1,57 | 0 |

| | | | | | | | | | | | | | | | | | |
|---|---|---|---|---|---|---|---|---|---|---|---|---|---|---|---|---|---|
| EN1 | CTCATTTC | -1 | 65 | 73 | -935 | -927 | 1,57 | 0,01 | 11,71 | >0.1 | 3,95 | 4,31 | 0 | 1,88 | 0 | 0 | 0 |
| EN1 | TCACTTAC | -1 | 5 | 13 | -995 | -987 | 2,81 | 0,01 | 16,36 | >0.1 | 5,64 | 4,26 | 0 | 1,86 | 1,79 | 0 | 0 |
| EN2 | GGCACTCAAA | 1 | 23 | 33 | -977 | -967 | 3,31 | 0,01 | 15,06 | >0.1 | 5,61 | 4,27 | 0 | 1,87 | 0 | 0 | 0 |
| EN2 | GGAAATGAGC | 1 | 64 | 74 | -936 | -926 | 2,4 | 0,01 | 12,54 | >0.1 | 3,95 | 4,31 | 0 | 1,88 | 0 | 0 | 0 |
| EN2 | ACTGATCAGA | 1 | 163 | 173 | -837 | -827 | 4,13 | 0,01 | 16,07 | >0.1 | 3,21 | 4,42 | 0 | 1,9 | 0 | 2,4 | 0 |
| EN2 | CTCTATTAAA | 1 | 287 | 297 | -713 | -703 | 2,85 | 0,01 | 16,59 | >0.1 | 2,72 | 4,59 | 0 | 1,24 | 0 | 5,19 | 0 |
| EN2 | ACTGATCAGA | 1 | 478 | 488 | -522 | -512 | 4,13 | 0,01 | 18,27 | >0.1 | 2,88 | 4,92 | 0 | 0,99 | 0 | 5,36 | 0 |
| EN2 | ACCTATTAAT | 1 | 599 | 609 | -401 | -391 | 4 | 0,01 | 15,05 | >0.1 | 3,47 | 5,21 | 0 | 1,02 | 0 | 1,36 | 0 |
| EN2 | ATTAATGAAA | 1 | 603 | 613 | -397 | -387 | 3,09 | 0,01 | 14,23 | >0.1 | 3,49 | 5,22 | 0 | 1,02 | 0 | 1,41 | 0 |
| EN2 | CTTCATGATC | 1 | 645 | 655 | -355 | -345 | 3,58 | 0,01 | 17,66 | >0.1 | 3,84 | 5,34 | 0 | 1,06 | 0 | 3,83 | 0 |
| EN2 | CCTGCTCAGC | 1 | 937 | 947 | -63 | -53 | 4,11 | 0,01 | 19,67 | >0.1 | 5,61 | 7,17 | 0 | 1,88 | 0 | 0,9 | 0 |
| EN2 | TCTCATGAGG | 1 | 983 | 993 | -17 | -7 | 4,99 | 0,01 | 22,54 | >0.1 | 5,61 | 8,5 | 0 | 1,87 | 0 | 1,57 | 0 |
| EN2 | CCTTCTCAGC | 1 | 1069 | 1079 | 69 | 79 | 1,98 | 0,01 | 20,64 | >0.1 | 5,61 | 7,19 | 0 | 1,85 | 1,79 | 2,21 | 0 |
| EN2 | GCTGCTCAGT | 1 | 1094 | 1104 | 94 | 104 | 1,98 | 0,01 | 21,83 | >0.1 | 5,61 | 6,83 | 0 | 1,85 | 3,22 | 2,34 | 0 |
| EN2 | CACCATTGAG | 1 | 1105 | 1115 | 105 | 115 | 2,33 | 0,01 | 22,23 | >0.1 | 5,61 | 6,7 | 0 | 1,85 | 3,22 | 2,52 | 0 |
| EN2 | TCCACTGAAT | -1 | 1015 | 1025 | 15 | 25 | 2,12 | 0,01 | 19,82 | >0.1 | 5,61 | 8,84 | 0 | 1,86 | 0 | 1,39 | 0 |
| EN2 | CTGAATGACT | -1 | 1011 | 1021 | 11 | 21 | 2,05 | 0,01 | 19,87 | >0.1 | 5,61 | 8,95 | 0 | 1,86 | 0 | 1,39 | 0 |
| EN2 | CCTCCTCATG | -1 | 986 | 996 | -14 | -4 | 2,06 | 0,01 | 19,73 | >0.1 | 5,61 | 8,61 | 0 | 1,87 | 0 | 1,57 | 0 |
| EN2 | CCTCATGAGA | -1 | 983 | 993 | -17 | -7 | 4,92 | 0,01 | 22,47 | >0.1 | 5,61 | 8,5 | 0 | 1,87 | 0 | 1,57 | 0 |
| EN2 | TCTTATCAGC | -1 | 966 | 976 | -34 | -24 | 3,64 | 0,01 | 20,6 | >0.1 | 5,61 | 7,78 | 0 | 1,87 | 0 | 1,7 | 0 |
| EN2 | GGCGCTGAGC | -1 | 940 | 950 | -60 | -50 | 3,24 | 0,01 | 18,77 | >0.1 | 5,61 | 7,22 | 0 | 1,88 | 0 | 0,83 | 0 |
| EN2 | GATCATGAAG | -1 | 645 | 655 | -355 | -345 | 2,99 | 0,01 | 17,07 | >0.1 | 3,84 | 5,34 | 0 | 1,06 | 0 | 3,83 | 0 |
| EN2 | TTTCATTAAT | -1 | 603 | 613 | -397 | -387 | 3,44 | 0,01 | 14,58 | >0.1 | 3,49 | 5,22 | 0 | 1,02 | 0 | 1,41 | 0 |
| EN2 | TCTGATCAGT | -1 | 478 | 488 | -522 | -512 | 3,94 | 0,01 | 18,08 | >0.1 | 2,88 | 4,92 | 0 | 0,99 | 0 | 5,36 | 0 |
| EN2 | TCTCCTGACC | -1 | 242 | 252 | -758 | -748 | 1,73 | 0,01 | 14,85 | >0.1 | 2,87 | 4,52 | 0 | 1,4 | 0 | 4,33 | 0 |
| EN2 | TCTGATCAGT | -1 | 163 | 173 | -837 | -827 | 3,94 | 0,01 | 15,88 | >0.1 | 3,21 | 4,42 | 0 | 1,9 | 0 | 2,4 | 0 |
| EN2 | TTCACTTACC | -1 | 4 | 14 | -996 | -986 | 5,96 | 0,01 | 19,48 | >0.1 | 5,61 | 4,26 | 0 | 1,86 | 1,79 | 0 | 0 |
| RUNX1 | ACCTGTGGAGT | 1 | 110 | 121 | -890 | -879 | 3,12 | 0,01 | 13,67 | >0.1 | 3,53 | 4,36 | 0 | 1,89 | 0 | 0,77 | 0 |
| RUNX1 | CACAGTGGCTC | 1 | 183 | 194 | -817 | -806 | 4,86 | 0,01 | 18,04 | >0.1 | 3,11 | 4,45 | 0 | 1,76 | 0 | 3,86 | 0 |
| RUNX1 | GGTGGTGGGCA | 1 | 320 | 331 | -680 | -669 | 2,61 | 0,01 | 16,71 | >0.1 | 2,62 | 4,64 | 0 | 1,16 | 0 | 5,68 | 0 |
| RUNX1 | GGCTGAGGCAG | 1 | 354 | 365 | -646 | -635 | 5,15 | 0,01 | 19,99 | >0.1 | 2,53 | 4,69 | 0 | 1,1 | 0 | 6,52 | 0 |
| RUNX1 | CACTGTGCTCC | 1 | 416 | 427 | -584 | -573 | 4,71 | 0,01 | 18,4 | >0.1 | 2,67 | 4,8 | 0 | 1,02 | 0 | 5,19 | 0 |
| RUNX1 | GTTTCTGGGAA | 1 | 556 | 567 | -444 | -433 | 3,54 | 0,01 | 12,84 | >0.1 | 3,22 | 5,1 | 0 | 0,99 | 0 | 0 | 0 |
| RUNX1 | CTCTTTGGTCA | 1 | 668 | 679 | -332 | -321 | 7,49 | 0,01 | 21,82 | >0.1 | 4,12 | 5,42 | 0 | 1,1 | 0 | 3,7 | 0 |
| RUNX1 | TACTGTGTTCT | 1 | 901 | 912 | -99 | -88 | 7,17 | 0,01 | 22,66 | >0.1 | 5,6 | 6,72 | 0 | 1,89 | 0 | 1,28 | 0 |
| RUNX1 | TTTTTTGGTCT | 1 | 924 | 935 | -76 | -65 | 6,94 | 0,01 | 22,47 | >0.1 | 5,6 | 6,99 | 0 | 1,88 | 0 | 1,06 | 0 |
| RUNX1 | GGATGTGATCT | 1 | 1022 | 1033 | 22 | 33 | 3,52 | 0,01 | 20,94 | >0.1 | 5,6 | 8,68 | 0 | 1,86 | 0 | 1,28 | 0 |
| RUNX1 | CCTTGTTGCTG | 1 | 1078 | 1089 | 78 | 89 | 3,42 | 0,01 | 21,92 | >0.1 | 5,6 | 7,04 | 0 | 1,85 | 1,79 | 2,21 | 0 |
| RUNX1 | GTCTTCGGCTT | -1 | 1152 | 1163 | 152 | 163 | 5,57 | 0,01 | 22,24 | >0.1 | 4,5 | 6,28 | 0 | 1,84 | 1,79 | 2,26 | 0 |
| RUNX1 | CCCTGTGAGCC | -1 | 1034 | 1045 | 34 | 45 | 2,55 | 0,01 | 19,38 | >0.1 | 5,6 | 8,06 | 0 | 1,86 | 0 | 1,31 | 0 |
| RUNX1 | ACTTTTGGTTA | -1 | 732 | 743 | -268 | -257 | 5,77 | 0,01 | 24,17 | >0.1 | 5,6 | 5,65 | 0 | 1,24 | 0 | 5,92 | 0 |
| RUNX1 | CCTTGTGATCT | -1 | 233 | 244 | -767 | -756 | 5,5 | 0,01 | 19,25 | >0.1 | 2,9 | 4,51 | 0 | 1,44 | 0 | 4,9 | 0 |
| RUNX1 | CACTGTGCCCA | -1 | 179 | 190 | -821 | -810 | 4,63 | 0,01 | 17,09 | >0.1 | 3,13 | 4,44 | 0 | 1,8 | 0 | 3,09 | 0 |
| RUNX1 | TTCTGTGGCTG | -1 | 38 | 49 | -962 | -951 | 12,18 | 0,001 | 22,65 | >0.1 | 4,31 | 4,29 | 0 | 1,87 | 0 | 0 | 0 |
| TCF4 | CACGCCTGTA | 1 | 193 | 203 | -807 | -797 | -2,48 | 0,01 | 10,99 | >0.1 | 3,07 | 4,46 | 0 | 1,68 | 0 | 4,26 | 0 |
| TCF4 | CTTAGCTGGG | 1 | 307 | 317 | -693 | -683 | -3,96 | 0,01 | 10,18 | >0.1 | 2,66 | 4,62 | 0 | 1,19 | 0 | 5,68 | 0 |
| TCF4 | GGCACCTGTA | 1 | 327 | 337 | -673 | -663 | 12,49 | 0,001 | 26,45 | >0.1 | 2,6 | 4,65 | 0 | 1,14 | 0 | 5,56 | 0 |
| TCF4 | CCCAGCTACT | 1 | 339 | 349 | -661 | -651 | 0,04 | 0,01 | 14,92 | >0.1 | 2,57 | 4,66 | 0 | 1,12 | 0 | 6,52 | 0 |
| TCF4 | TGAACCTGGG | 1 | 375 | 385 | -625 | -615 | 1,97 | 0,01 | 16,71 | >0.1 | 2,56 | 4,72 | 0 | 1,07 | 0 | 6,4 | 0 |
| TCF4 | AACACCAACA | 1 | 631 | 641 | -369 | -359 | -4,41 | 0,01 | 7,3 | >0.1 | 3,71 | 5,3 | 0 | 1,05 | 0 | 1,66 | 0 |
| TCF4 | AACAGCTTTC | 1 | 711 | 721 | -289 | -279 | -0,58 | 0,01 | 16,76 | >0.1 | 5,61 | 5,56 | 0 | 1,18 | 0 | 4,99 | 0 |
| TCF4 | AAAATCTGTT | 1 | 841 | 851 | -159 | -149 | -4,23 | 0,01 | 10,81 | >0.1 | 5,61 | 6,24 | 0 | 1,78 | 0 | 1,41 | 0 |
| TCF4 | CATATCTGTC | 1 | 852 | 862 | -148 | -138 | -1,5 | 0,01 | 14,97 | >0.1 | 5,61 | 6,31 | 0 | 1,88 | 0 | 2,67 | 0 |
| TCF4 | CTCAGCAGAT | 1 | 887 | 897 | -113 | -103 | -5,23 | 0,01 | 11,93 | >0.1 | 5,61 | 6,59 | 0 | 1,89 | 0 | 3,07 | 0 |
| TCF4 | AAGACCTGTT | 1 | 1158 | 1168 | 158 | 168 | -4,26 | 0,01 | 12,33 | >0.1 | 4,38 | 6,24 | 0 | 1,78 | 1,79 | 2,4 | 0 |
| TCF4 | AGCAGCAGTT | -1 | 1090 | 1100 | 90 | 100 | 2,6 | 0,01 | 21,01 | >0.1 | 5,61 | 6,88 | 0 | 1,85 | 1,79 | 2,27 | 0 |
| TCF4 | ACAATCTGCT | -1 | 890 | 900 | -110 | -100 | -4,68 | 0,01 | 12,38 | >0.1 | 5,61 | 6,61 | 0 | 1,89 | 0 | 2,95 | 0 |
| TCF4 | GAAAGCTGTT | -1 | 711 | 721 | -289 | -279 | -3,6 | 0,01 | 13,74 | >0.1 | 5,61 | 5,56 | 0 | 1,18 | 0 | 4,99 | 0 |
| TCF4 | AGTAGCTGGG | -1 | 339 | 349 | -661 | -651 | -0,2 | 0,01 | 14,68 | >0.1 | 2,57 | 4,66 | 0 | 1,12 | 0 | 6,52 | 0 |

| | | | | | | | | | | | | | | | | | |
|---|---|---|---|---|---|---|---|---|---|---|---|---|---|---|---|---|---|
| TCF4 | TACAGGTGCC | -1 | 327 | 337 | -673 | -663 | 4,7 | 0,01 | 18,66 | >0.1 | 2,6 | 4,65 | 0 | 1,14 | 0 | 5,56 | 0 |
| TCF4 | CCCAGCTAAG | -1 | 307 | 317 | -693 | -683 | -4,4 | 0,01 | 9,74 | >0.1 | 2,66 | 4,62 | 0 | 1,19 | 0 | 5,68 | 0 |
| TCF7 | GGAAATTAAAAC | 1 | 153 | 165 | -847 | -835 | -4,05 | 0,01 | 7,17 | >0.1 | 3,26 | 4,41 | 0 | 1,9 | 0 | 1,65 | 0 |
| TCF7 | ACTGATCAGAAA | 1 | 163 | 175 | -837 | -825 | -2,02 | 0,01 | 9,91 | >0.1 | 3,21 | 4,42 | 0 | 1,9 | 0 | 2,4 | 0 |
| TCF7 | GCAGATCACAAG | 1 | 231 | 243 | -769 | -757 | 2,8 | 0,01 | 16,68 | >0.1 | 2,91 | 4,51 | 0 | 1,45 | 0 | 5,01 | 0 |
| TCF7 | ACTGATCAGAAA | 1 | 478 | 490 | -522 | -510 | -2,02 | 0,01 | 12,29 | >0.1 | 2,88 | 4,92 | 0 | 0,99 | 0 | 5,52 | 0 |
| TCF7 | CAAGGGCAAAGT | 1 | 507 | 519 | -493 | -481 | 2,51 | 0,01 | 15,84 | >0.1 | 2,99 | 4,98 | 0 | 0,98 | 0 | 4,37 | 0 |
| TCF7 | AAAAATAAATGT | 1 | 581 | 593 | -419 | -407 | -4,12 | 0,01 | 5,41 | >0.1 | 3,36 | 5,16 | 0 | 1,01 | 0 | 0 | 0 |
| TCF7 | TGACAGGAGAGG | 1 | 767 | 779 | -233 | -221 | -1,52 | 0,01 | 17,14 | >0.1 | 5,58 | 5,81 | 0 | 1,35 | 0 | 5,92 | 0 |
| TCF7 | GGAGAGGTAAGG | 1 | 772 | 784 | -228 | -216 | -0,54 | 0,01 | 18,35 | >0.1 | 5,58 | 5,83 | 0 | 1,37 | 0 | 6,1 | 0 |
| TCF7 | TAGGATTAAAGA | 1 | 789 | 801 | -211 | -199 | 5,08 | 0,001 | 23,35 | >0.1 | 5,58 | 5,92 | 0 | 1,45 | 0 | 5,32 | 0 |
| TCF7 | GCTGATAAGAGA | 1 | 966 | 978 | -34 | -22 | -4,63 | 0,01 | 12,37 | >0.1 | 5,58 | 7,85 | 0 | 1,87 | 0 | 1,7 | 0 |
| TCF7 | TTGGCTCACAGG | 1 | 1032 | 1044 | 32 | 44 | -4,18 | 0,01 | 12,71 | >0.1 | 5,58 | 8,15 | 0 | 1,86 | 0 | 1,31 | 0 |
| TCF7 | TTCTATCAAAGT | 1 | 1166 | 1178 | 166 | 178 | -2,61 | 0,01 | 14,49 | >0.1 | 4,56 | 6,18 | 0 | 1,7 | 1,79 | 2,86 | 0 |
| TCF7 | CAAGATCACATC | -1 | 1023 | 1035 | 23 | 35 | -3,18 | 0,01 | 14,18 | >0.1 | 5,58 | 8,67 | 0 | 1,86 | 0 | 1,25 | 0 |
| TCF7 | CTAGACTAAAAC | -1 | 995 | 1007 | -5 | 7 | -4,94 | 0,01 | 12,97 | >0.1 | 5,58 | 8,96 | 0 | 1,87 | 0 | 1,5 | 0 |
| TCF7 | GAGCAGGAAAGA | -1 | 932 | 944 | -68 | -56 | 0,24 | 0,01 | 15,75 | >0.1 | 5,58 | 7,12 | 0 | 1,88 | 0 | 0,93 | 0 |
| TCF7 | AAAGACCAAAAA | -1 | 925 | 937 | -75 | -63 | 6,12 | 0,001 | 21,67 | >0.1 | 5,58 | 7,02 | 0 | 1,88 | 0 | 1,06 | 0 |
| TCF7 | AAAAAGAAAAAG | -1 | 916 | 928 | -84 | -72 | -3,41 | 0,01 | 12,08 | >0.1 | 5,58 | 6,9 | 0 | 1,89 | 0 | 1,12 | 0 |
| TCF7 | GAAGAACACAGT | -1 | 902 | 914 | -98 | -86 | -3,16 | 0,01 | 12,29 | >0.1 | 5,58 | 6,75 | 0 | 1,89 | 0 | 1,24 | 0 |
| TCF7 | GGAGAGGACAGA | -1 | 856 | 868 | -144 | -132 | -2,46 | 0,01 | 14,24 | >0.1 | 5,58 | 6,35 | 0 | 1,9 | 0 | 2,86 | 0 |
| TCF7 | CCAAATCAAAAC | -1 | 747 | 759 | -253 | -241 | -0,96 | 0,01 | 17,54 | >0.1 | 5,58 | 5,72 | 0 | 1,28 | 0 | 5,92 | 0 |
| TCF7 | AGTGACCAAAGA | -1 | 669 | 681 | -331 | -319 | 4,51 | 0,001 | 18,87 | >0.1 | 4,14 | 5,42 | 0 | 1,1 | 0 | 3,7 | 0 |
| TCF7 | CTGGAGCACAGT | -1 | 417 | 429 | -583 | -571 | -4,33 | 0,01 | 9,36 | >0.1 | 2,68 | 4,8 | 0 | 1,02 | 0 | 5,19 | 0 |
| TCF7 | TAGGATTACAGG | -1 | 197 | 209 | -803 | -791 | 0,77 | 0,01 | 14,41 | >0.1 | 3,05 | 4,47 | 0 | 1,65 | 0 | 4,48 | 0 |
| TCF7 | AAACTTAAAAAA | -1 | 74 | 86 | -926 | -914 | 1,07 | 0,01 | 11,11 | >0.1 | 3,83 | 4,32 | 0 | 1,88 | 0 | 0 | 0 |
| TCF3 | CACGCCTGTA | 1 | 193 | 203 | -807 | -797 | -1,48 | 0,01 | 11,99 | >0.1 | 3,07 | 4,46 | 0 | 1,68 | 0 | 4,26 | 0 |
| TCF3 | GGCACCTGTA | 1 | 327 | 337 | -673 | -663 | 12,43 | 0,001 | 26,39 | >0.1 | 2,6 | 4,65 | 0 | 1,14 | 0 | 5,56 | 0 |
| TCF3 | CCCAGCTACT | 1 | 339 | 349 | -661 | -651 | 1,95 | 0,01 | 16,83 | >0.1 | 2,57 | 4,66 | 0 | 1,12 | 0 | 6,52 | 0 |
| TCF3 | TGAACCTGGG | 1 | 375 | 385 | -625 | -615 | -2,31 | 0,01 | 12,43 | >0.1 | 2,56 | 4,72 | 0 | 1,07 | 0 | 6,4 | 0 |
| TCF3 | TGCTCCAGCC | 1 | 421 | 431 | -579 | -569 | -2,7 | 0,01 | 11,23 | >0.1 | 2,69 | 4,81 | 0 | 1,02 | 0 | 5,42 | 0 |
| TCF3 | AACACCAACA | 1 | 631 | 641 | -369 | -359 | -2,24 | 0,01 | 9,47 | >0.1 | 3,71 | 5,3 | 0 | 1,05 | 0 | 1,66 | 0 |
| TCF3 | AACAGCTTTC | 1 | 711 | 721 | -289 | -279 | -0,57 | 0,01 | 16,77 | >0.1 | 5,61 | 5,56 | 0 | 1,18 | 0 | 4,99 | 0 |
| TCF3 | AACTGCTGCT | 1 | 1090 | 1100 | 90 | 100 | 2,74 | 0,01 | 21,15 | >0.1 | 5,61 | 6,88 | 0 | 1,85 | 1,79 | 2,27 | 0 |
| TCF3 | AAGACCTGTT | 1 | 1158 | 1168 | 158 | 168 | -1,94 | 0,01 | 14,65 | >0.1 | 4,38 | 6,24 | 0 | 1,78 | 1,79 | 2,4 | 0 |
| TCF3 | AACAGGTCTT | -1 | 1158 | 1168 | 158 | 168 | -1,92 | 0,01 | 14,67 | >0.1 | 4,38 | 6,24 | 0 | 1,78 | 1,79 | 2,4 | 0 |
| TCF3 | AGCAGCAGTT | -1 | 1090 | 1100 | 90 | 100 | 3,4 | 0,01 | 21,81 | >0.1 | 5,61 | 6,88 | 0 | 1,85 | 1,79 | 2,27 | 0 |
| TCF3 | GTCCCCTGTG | -1 | 1038 | 1048 | 38 | 48 | -1,34 | 0,01 | 15,38 | >0.1 | 5,61 | 7,92 | 0 | 1,86 | 0 | 1,34 | 0 |
| TCF3 | CTCTCCTGTC | -1 | 768 | 778 | -232 | -222 | 0,53 | 0,01 | 19,21 | >0.1 | 5,61 | 5,81 | 0 | 1,35 | 0 | 5,92 | 0 |
| TCF3 | TTCTCCTGCC | -1 | 360 | 370 | -640 | -630 | 0,56 | 0,01 | 15,23 | >0.1 | 2,52 | 4,7 | 0 | 1,09 | 0 | 6,36 | 0 |
| TCF3 | TACAGGTGCC | -1 | 327 | 337 | -673 | -663 | 6,52 | 0,01 | 20,48 | >0.1 | 2,6 | 4,65 | 0 | 1,14 | 0 | 5,56 | 0 |
| TCF3 | CCCACCACCA | -1 | 319 | 329 | -681 | -671 | -2,89 | 0,01 | 11,09 | >0.1 | 2,63 | 4,63 | 0 | 1,16 | 0 | 5,56 | 0 |
| TCF3 | CCCAGCTAAG | -1 | 307 | 317 | -693 | -683 | -1,73 | 0,01 | 12,41 | >0.1 | 2,66 | 4,62 | 0 | 1,19 | 0 | 5,68 | 0 |
| TCF3 | ATCTCCTGAC | -1 | 243 | 253 | -757 | -747 | -2,89 | 0,01 | 10,22 | >0.1 | 2,87 | 4,52 | 0 | 1,39 | 0 | 4,33 | 0 |
| TCF3 | TACAGGCGTG | -1 | 193 | 203 | -807 | -797 | -0,92 | 0,01 | 12,55 | >0.1 | 3,07 | 4,46 | 0 | 1,68 | 0 | 4,26 | 0 |
| TCF3 | CTCTCCTGAT | -1 | 97 | 107 | -903 | -893 | -2,61 | 0,01 | 8,02 | >0.1 | 3,64 | 4,35 | 0 | 1,88 | 0 | 0,77 | 0 |
| GBX2 | GGAAATGAGC | 1 | 64 | 74 | -936 | -926 | 2,93 | 0,01 | 13,07 | >0.1 | 3,95 | 4,31 | 0 | 1,88 | 0 | 0 | 0 |
| GBX2 | CATAATTTTA | 1 | 131 | 141 | -869 | -859 | 0,97 | 0,01 | 12,22 | >0.1 | 3,4 | 4,38 | 0 | 1,89 | 0 | 1,57 | 0 |
| GBX2 | GGAAATTAAA | 1 | 153 | 163 | -847 | -837 | 5,65 | 0,01 | 16,87 | >0.1 | 3,27 | 4,41 | 0 | 1,9 | 0 | 1,65 | 0 |
| GBX2 | ACTGATCAGA | 1 | 163 | 173 | -837 | -827 | 1,56 | 0,01 | 13,5 | >0.1 | 3,21 | 4,42 | 0 | 1,9 | 0 | 2,4 | 0 |
| GBX2 | CTCTATTAAA | 1 | 287 | 297 | -713 | -703 | 1,69 | 0,01 | 15,43 | >0.1 | 2,72 | 4,59 | 0 | 1,24 | 0 | 5,19 | 0 |
| GBX2 | ACTGATCAGA | 1 | 478 | 488 | -522 | -512 | 1,56 | 0,01 | 15,7 | >0.1 | 2,88 | 4,92 | 0 | 0,99 | 0 | 5,36 | 0 |
| GBX2 | CCCAAGTAGA | 1 | 544 | 554 | -456 | -446 | 2,93 | 0,01 | 12,13 | >0.1 | 3,15 | 5,06 | 0 | 0,99 | 0 | 0 | 0 |
| GBX2 | ACCTATTAAT | 1 | 599 | 609 | -401 | -391 | 3,97 | 0,01 | 15,02 | >0.1 | 3,47 | 5,21 | 0 | 1,02 | 0 | 1,36 | 0 |
| GBX2 | ATTAATGAAA | 1 | 603 | 613 | -397 | -387 | 3,38 | 0,01 | 14,52 | >0.1 | 3,49 | 5,22 | 0 | 1,02 | 0 | 1,41 | 0 |
| GBX2 | TAAAATTATA | 1 | 682 | 692 | -318 | -308 | 1,33 | 0,01 | 15,31 | >0.1 | 4,34 | 5,46 | 0 | 1,12 | 0 | 3,07 | 0 |
| GBX2 | AAAAATTATT | 1 | 829 | 839 | -171 | -161 | 2,89 | 0,01 | 19,61 | >0.1 | 5,61 | 6,15 | 0 | 1,68 | 0 | 3,28 | 0 |
| GBX2 | AGCAGTTACA | -1 | 1087 | 1097 | 87 | 97 | 2,68 | 0,01 | 21,07 | >0.1 | 5,61 | 6,92 | 0 | 1,85 | 1,79 | 2,21 | 0 |
| GBX2 | AATAATTTTT | -1 | 829 | 839 | -171 | -161 | 1,57 | 0,01 | 18,29 | >0.1 | 5,61 | 6,15 | 0 | 1,68 | 0 | 3,28 | 0 |

| | | | | | | | | | | | | | | | | | |
|---|---|---|---|---|---|---|---|---|---|---|---|---|---|---|---|---|---|
| GBX2 | GACAATTTCA | -1 | 608 | 618 | -392 | -382 | 3,19 | 0,01 | 14,52 | >0.1 | 3,53 | 5,23 | 0 | 1,02 | 0 | 1,55 | 0 |
| GBX2 | TTTCATTAAT | -1 | 603 | 613 | -397 | -387 | 1,77 | 0,01 | 12,91 | >0.1 | 3,49 | 5,22 | 0 | 1,02 | 0 | 1,41 | 0 |
| GBX2 | TTTAATTTCC | -1 | 153 | 163 | -847 | -837 | 2,26 | 0,01 | 13,48 | >0.1 | 3,27 | 4,41 | 0 | 1,9 | 0 | 1,65 | 0 |
| GBX2 | TAAAATTATG | -1 | 131 | 141 | -869 | -859 | 2,22 | 0,01 | 13,47 | >0.1 | 3,4 | 4,38 | 0 | 1,89 | 0 | 1,57 | 0 |
| GBX2 | TTCACTTACC | -1 | 4 | 14 | -996 | -986 | 2,26 | 0,01 | 15,78 | >0.1 | 5,61 | 4,26 | 0 | 1,86 | 1,79 | 0 | 0 |
| ZNF24 | AAGTCATGTATTT | 1 | 515 | 528 | -485 | -472 | -0,2 | 0,01 | 11,77 | >0.1 | 3,03 | 5 | 0 | 0,99 | 0 | 2,95 | 0 |
| ZNF24 | ATTTGATTCACTT | 1 | 814 | 827 | -186 | -173 | 1,91 | 0,01 | 18,58 | >0.1 | 5,57 | 6,07 | 0 | 1,6 | 0 | 3,43 | 0 |
| ZNF24 | AAGTCATTCAGTG | 1 | 1010 | 1023 | 10 | 23 | 6,33 | 0,001 | 24,15 | >0.1 | 5,57 | 8,99 | 0 | 1,86 | 0 | 1,39 | 0 |
| ZNF24 | CATCCACTGAATG | -1 | 1014 | 1027 | 14 | 27 | -2,95 | 0,01 | 14,74 | >0.1 | 5,57 | 8,86 | 0 | 1,86 | 0 | 1,39 | 0 |
| ZNF24 | TATTCTTTAATCC | -1 | 791 | 804 | -209 | -196 | 3,1 | 0,01 | 21,39 | >0.1 | 5,57 | 5,94 | 0 | 1,47 | 0 | 5,32 | 0 |
| ZNF24 | ATTTCATTAATAG | -1 | 601 | 614 | -399 | -386 | 0,31 | 0,01 | 11,45 | >0.1 | 3,49 | 5,22 | 0 | 1,02 | 0 | 1,41 | 0 |
| ZNF24 | CTCACATTTATTT | -1 | 583 | 596 | -417 | -404 | 2,07 | 0,01 | 11,62 | >0.1 | 3,37 | 5,17 | 0 | 1,01 | 0 | 0 | 0 |
| ZNF24 | CATTTATTTTTAT | -1 | 579 | 592 | -421 | -408 | -1,98 | 0,01 | 7,53 | >0.1 | 3,35 | 5,16 | 0 | 1,01 | 0 | 0 | 0 |
| ZNF24 | TTCTCCTTCACTT | -1 | 7 | 20 | -993 | -980 | 1,03 | 0,01 | 14,52 | >0.1 | 5,57 | 4,26 | 0 | 1,86 | 1,79 | 0 | 0 |
| ZNF24 | CCTTCACTTACCT | -1 | 3 | 16 | -997 | -984 | 1,66 | 0,01 | 15,15 | >0.1 | 5,57 | 4,26 | 0 | 1,86 | 1,79 | 0 | 0 |
| NKX2-8 | GCACTCAAA | 1 | 24 | 33 | -976 | -967 | 9,7 | 0,001 | 21,46 | >0.1 | 5,62 | 4,27 | 0 | 1,87 | 0 | 0 | 0 |
| NKX2-8 | GCACTTTGG | 1 | 209 | 218 | -791 | -782 | 5,02 | 0,01 | 18,77 | >0.1 | 3 | 4,48 | 0 | 1,57 | 0 | 4,7 | 0 |
| NKX2-8 | GCACCTGTA | 1 | 328 | 337 | -672 | -663 | 4,38 | 0,01 | 18,33 | >0.1 | 2,6 | 4,65 | 0 | 1,14 | 0 | 5,56 | 0 |
| NKX2-8 | CTACTCGGG | 1 | 344 | 353 | -656 | -647 | 6,4 | 0,01 | 21,26 | >0.1 | 2,56 | 4,67 | 0 | 1,11 | 0 | 6,52 | 0 |
| NKX2-8 | TCACTTTAA | 1 | 821 | 830 | -179 | -170 | 3,94 | 0,01 | 20,97 | >0.1 | 5,62 | 6,09 | 0 | 1,62 | 0 | 3,7 | 0 |
| NKX2-8 | CCTCTCCAG | 1 | 861 | 870 | -139 | -130 | 7,04 | 0,01 | 23,8 | >0.1 | 5,62 | 6,37 | 0 | 1,9 | 0 | 2,86 | 0 |
| NKX2-8 | GCTCTTCCT | 1 | 1056 | 1065 | 56 | 65 | 2,74 | 0,01 | 19,93 | >0.1 | 5,62 | 7,44 | 0 | 1,86 | 0 | 2,27 | 0 |
| NKX2-8 | CTCCTTCTC | 1 | 1067 | 1076 | 67 | 76 | 3,79 | 0,01 | 22,56 | >0.1 | 5,62 | 7,22 | 0 | 1,85 | 1,79 | 2,27 | 0 |
| NKX2-8 | TCACTTGCT | 1 | 1178 | 1187 | 178 | 187 | 3,7 | 0,01 | 22,08 | >0.1 | 5,62 | 6,1 | 0 | 1,6 | 1,79 | 3,26 | 0 |
| NKX2-8 | CTTCTTGGA | 1 | 1185 | 1194 | 185 | 194 | 3,07 | 0,01 | 21,77 | >0.1 | 5,62 | 6,06 | 0 | 1,55 | 1,79 | 3,67 | 0 |
| NKX2-8 | CTCCTCATG | -1 | 986 | 995 | -14 | -5 | 2,38 | 0,01 | 20,04 | >0.1 | 5,62 | 8,59 | 0 | 1,87 | 0 | 1,57 | 0 |
| NKX2-8 | GCCTTTGAA | -1 | 959 | 968 | -41 | -32 | 2,3 | 0,01 | 19,11 | >0.1 | 5,62 | 7,57 | 0 | 1,88 | 0 | 1,74 | 0 |
| NKX2-8 | GAACTTGGG | -1 | 953 | 962 | -47 | -38 | 2,07 | 0,01 | 18,71 | >0.1 | 5,62 | 7,44 | 0 | 1,88 | 0 | 1,7 | 0 |
| NKX2-8 | CCTCTCCTG | -1 | 770 | 779 | -230 | -221 | 5,05 | 0,01 | 23,53 | >0.1 | 5,62 | 5,81 | 0 | 1,35 | 0 | 5,69 | 0 |
| NKX2-8 | TCACTTTAT | -1 | 761 | 770 | -239 | -230 | 1,96 | 0,01 | 20,59 | >0.1 | 5,62 | 5,77 | 0 | 1,32 | 0 | 5,92 | 0 |
| NKX2-8 | GTACTTTTG | -1 | 736 | 745 | -264 | -255 | 3,67 | 0,01 | 22,11 | >0.1 | 5,62 | 5,66 | 0 | 1,24 | 0 | 5,92 | 0 |
| NKX2-8 | CTACTTGGG | -1 | 544 | 553 | -456 | -447 | 7,32 | 0,01 | 16,52 | >0.1 | 3,15 | 5,06 | 0 | 0,99 | 0 | 0 | 0 |
| NKX2-8 | GCCCTTGTC | -1 | 505 | 514 | -495 | -486 | 6,88 | 0,01 | 21,67 | >0.1 | 2,98 | 4,97 | 0 | 0,98 | 0 | 5,86 | 0 |
| NKX2-8 | CCTCCCGAG | -1 | 347 | 356 | -653 | -644 | 1,9 | 0,01 | 16,75 | >0.1 | 2,55 | 4,68 | 0 | 1,11 | 0 | 6,52 | 0 |
| NKX2-8 | CCATTTCTG | -1 | 169 | 178 | -831 | -822 | 3,25 | 0,01 | 15,21 | >0.1 | 3,19 | 4,43 | 0 | 1,89 | 0 | 2,46 | 0 |
| NKX2-8 | CTACTCTCC | -1 | 121 | 130 | -879 | -870 | 1,9 | 0,01 | 13,27 | >0.1 | 3,46 | 4,37 | 0 | 1,89 | 0 | 1,65 | 0 |
| NKX2-8 | ACTCTCCAC | -1 | 119 | 128 | -881 | -872 | 5,28 | 0,01 | 15,74 | >0.1 | 3,48 | 4,37 | 0 | 1,89 | 0 | 0,73 | 0 |
| NKX2-8 | CCACTCCAC | -1 | 114 | 123 | -886 | -877 | 10,69 | 0,001 | 21,22 | >0.1 | 3,51 | 4,36 | 0 | 1,89 | 0 | 0,77 | 0 |
| NKX2-8 | ACTCTCCTG | -1 | 99 | 108 | -901 | -892 | 3,69 | 0,01 | 14,31 | >0.1 | 3,62 | 4,35 | 0 | 1,88 | 0 | 0,77 | 0 |
| NKX2-8 | AAACTTAAA | -1 | 77 | 86 | -923 | -914 | 2,15 | 0,01 | 12,17 | >0.1 | 3,82 | 4,32 | 0 | 1,88 | 0 | 0 | 0 |
| NKX2-8 | CTCCTTCTC | -1 | 15 | 24 | -985 | -976 | 3,79 | 0,01 | 15,54 | >0.1 | 5,62 | 4,27 | 0 | 1,87 | 0 | 0 | 0 |
| NKX2-8 | CTCCTTCAC | -1 | 9 | 18 | -991 | -982 | 5,78 | 0,01 | 19,32 | >0.1 | 5,62 | 4,26 | 0 | 1,86 | 1,79 | 0 | 0 |
| NKX2-8 | TCACTTACC | -1 | 4 | 13 | -996 | -987 | 3,67 | 0,01 | 17,21 | >0.1 | 5,62 | 4,26 | 0 | 1,86 | 1,79 | 0 | 0 |
| RARG(VAR.2) | TAGGTAAGTGAAGGAGA | 1 | 2 | 19 | -998 | -981 | -16,07 | 0,001 | -2,63 | >0.1 | 5,52 | 4,26 | 0 | 1,86 | 1,79 | 0 | 0 |
| RARG(VAR.2) | AAAGGCAGATCAGGAGA | 1 | 89 | 106 | -911 | -894 | -27,09 | 0,01 | -16,36 | >0.1 | 3,67 | 4,35 | 0 | 1,88 | 0 | 0,83 | 0 |
| RARG(VAR.2) | CAGATCAGGAGAGTTGA | 1 | 94 | 111 | -906 | -889 | -11,93 | 0,001 | -1,3 | >0.1 | 3,63 | 4,35 | 0 | 1,88 | 0 | 0,77 | 0 |
| RARG(VAR.2) | ATGGCCATGGAAATTAA | 1 | 145 | 162 | -855 | -838 | -26,87 | 0,01 | -15,62 | >0.1 | 3,29 | 4,41 | 0 | 1,9 | 0 | 1,65 | 0 |
| RARG(VAR.2) | CTGGGCACAGTGGCTCA | 1 | 178 | 195 | -822 | -805 | -22,68 | 0,01 | -9,45 | >0.1 | 3,12 | 4,45 | 0 | 1,81 | 0 | 3,86 | 0 |
| RARG(VAR.2) | GAGGCCGAGTTGGGCAG | 1 | 218 | 235 | -782 | -765 | -28,97 | 0,01 | -14,48 | >0.1 | 2,95 | 4,5 | 0 | 1,52 | 0 | 5,52 | 0 |
| RARG(VAR.2) | GAGTTGGGCAGATCACA | 1 | 224 | 241 | -776 | -759 | -33,55 | 0,01 | -19,37 | >0.1 | 2,92 | 4,51 | 0 | 1,49 | 0 | 5,26 | 0 |
| RARG(VAR.2) | AAGGTCAGGAGATAGAG | 1 | 240 | 257 | -760 | -743 | -33,77 | 0,01 | -20,64 | >0.1 | 2,87 | 4,53 | 0 | 1,41 | 0 | 4,33 | 0 |
| RARG(VAR.2) | GTGGGCACCTGTAGTCC | 1 | 324 | 341 | -676 | -659 | -32,87 | 0,01 | -18,9 | >0.1 | 2,6 | 4,65 | 0 | 1,15 | 0 | 5,56 | 0 |
| RARG(VAR.2) | GAGGCTGAGGCAGGAGA | 1 | 352 | 369 | -648 | -631 | -31,49 | 0,01 | -17,13 | >0.1 | 2,53 | 4,7 | 0 | 1,1 | 0 | 6,03 | 0 |
| RARG(VAR.2) | AATGTTGCCCAAGTAGA | 1 | 537 | 554 | -463 | -446 | -32,08 | 0,01 | -22,89 | >0.1 | 3,14 | 5,07 | 0 | 0,99 | 0 | 0 | 0 |
| RARG(VAR.2) | TTGGCCATAAAGTGACA | 1 | 755 | 772 | -245 | -228 | -28,15 | 0,01 | -9,42 | >0.1 | 5,52 | 5,78 | 0 | 1,33 | 0 | 6,1 | 0 |
| RARG(VAR.2) | AGTGACAGGAGAGGTAA | 1 | 765 | 782 | -235 | -218 | -12,7 | 0,001 | 5,93 | >0.1 | 5,52 | 5,83 | 0 | 1,37 | 0 | 5,92 | 0 |
| RARG(VAR.2) | CAGGAGAGGTAAGGTTC | 1 | 770 | 787 | -230 | -213 | -30,92 | 0,01 | -12,06 | >0.1 | 5,52 | 5,85 | 0 | 1,39 | 0 | 6,1 | 0 |
| RARG(VAR.2) | AAGGTTCTCTAGGATTA | 1 | 780 | 797 | -220 | -203 | -26 | 0,01 | -7,23 | >0.1 | 5,52 | 5,9 | 0 | 1,43 | 0 | 5,92 | 0 |
| RARG(VAR.2) | AGGTTCTCTAGGATTAA | 1 | 781 | 798 | -219 | -202 | -27,19 | 0,01 | -8,41 | >0.1 | 5,52 | 5,91 | 0 | 1,44 | 0 | 5,92 | 0 |

| | | | | | | | | | | | | | | | | | |
|---|---|---|---|---|---|---|---|---|---|---|---|---|---|---|---|---|---|
| RARG(VAR.2) | AACTTTATATTGGCTCA | 1 | 874 | 891 | -126 | -109 | -26,26 | 0,01 | -9,36 | >0.1 | 5,52 | 6,54 | 0 | 1,9 | 0 | 2,95 | 0 |
| RARG(VAR.2) | AAGTTCAAAGGCTGATA | 1 | 956 | 973 | -44 | -27 | -18,03 | 0,001 | -1,24 | >0.1 | 5,52 | 7,7 | 0 | 1,88 | 0 | 1,7 | 0 |
| RARG(VAR.2) | TAGTCTAGGGAAAGTCA | 1 | 999 | 1016 | -1 | 16 | -26,16 | 0,01 | -8,19 | >0.1 | 5,52 | 9,12 | 0 | 1,87 | 0 | 1,46 | 0 |
| RARG(VAR.2) | GATGTGATCTTGGCTCA | 1 | 1023 | 1040 | 23 | 40 | -20,75 | 0,01 | -3,43 | >0.1 | 5,52 | 8,67 | 0 | 1,86 | 0 | 1,28 | 0 |
| RARG(VAR.2) | AAGCTCTTCCTGGCTCC | 1 | 1054 | 1071 | 54 | 71 | -30 | 0,01 | -12,87 | >0.1 | 5,52 | 7,48 | 0 | 1,86 | 0 | 2,27 | 0 |
| RARG(VAR.2) | GCTGTAACTGCTGCTCA | 1 | 1085 | 1102 | 85 | 102 | -34,22 | 0,01 | -15,9 | >0.1 | 5,52 | 6,94 | 0 | 1,85 | 1,79 | 2,21 | 0 |
| RARG(VAR.2) | AACTGCTGCTCAGTCCA | 1 | 1090 | 1107 | 90 | 107 | -34,23 | 0,01 | -14,42 | >0.1 | 5,52 | 6,88 | 0 | 1,85 | 3,22 | 2,34 | 0 |
| RARG(VAR.2) | CAGGCCAAGACATTTTT | 1 | 1118 | 1135 | 118 | 135 | -33,06 | 0,01 | -13,23 | >0.1 | 5,52 | 6,57 | 0 | 1,85 | 3,22 | 2,67 | 0 |
| RARG(VAR.2) | TGTTCTATCAAAGTTCA | 1 | 1164 | 1181 | 164 | 181 | -28,67 | 0,01 | -10,58 | >0.1 | 5,52 | 6,19 | 0 | 1,72 | 1,79 | 2,86 | 0 |
| RARG(VAR.2) | AAGACCAAAAAAAGAAA | -1 | 919 | 936 | -81 | -64 | -32,08 | 0,01 | -16,58 | >0.1 | 5,52 | 7,01 | 0 | 1,88 | 0 | 1,09 | 0 |
| RARG(VAR.2) | AGTGACCAAAGAGGTCT | -1 | 664 | 681 | -336 | -319 | -29,12 | 0,01 | -14,66 | >0.1 | 4,11 | 5,42 | 0 | 1,1 | 0 | 3,83 | 0 |
| RARG(VAR.2) | GTTTTCAGACAATTTCA | -1 | 608 | 625 | -392 | -375 | -29,37 | 0,01 | -18,03 | >0.1 | 3,56 | 5,25 | 0 | 1,03 | 0 | 1,5 | 0 |
| RARG(VAR.2) | GGCGCGATCTCGGCTCA | -1 | 400 | 417 | -600 | -583 | -30,48 | 0,01 | -16,26 | >0.1 | 2,63 | 4,78 | 0 | 1,04 | 0 | 5,76 | 0 |
| RARG(VAR.2) | TCTGCCTCCCAGGTTCA | -1 | 375 | 392 | -625 | -608 | -29,05 | 0,01 | -14,15 | >0.1 | 2,57 | 4,74 | 0 | 1,07 | 0 | 6,53 | 0 |
| RARG(VAR.2) | AGGTTCACGCCATTCTC | -1 | 365 | 382 | -635 | -618 | -31,84 | 0,01 | -16,77 | >0.1 | 2,54 | 4,72 | 0 | 1,08 | 0 | 6,73 | 0 |
| RARG(VAR.2) | CAGGTCAACTCTCCTGA | -1 | 98 | 115 | -902 | -885 | -34,04 | 0,01 | -23,41 | >0.1 | 3,6 | 4,36 | 0 | 1,89 | 0 | 0,79 | 0 |
| RARG(VAR.2) | AAACTTAAAAAAGCTCA | -1 | 69 | 86 | -931 | -914 | -29,83 | 0,01 | -19,77 | >0.1 | 3,86 | 4,33 | 0 | 1,88 | 0 | 0 | 0 |
| IRF3 | AGGTAAGTGAAGGAGAAGGA | 1 | 3 | 24 | -997 | -976 | -8,16 | 0,01 | 5,24 | >0.1 | 5,47 | 4,27 | 0 | 1,87 | 1,79 | 0 | 0 |
| IRF3 | AAGTGAAGGAGAAGGAGGCA | 1 | 7 | 28 | -993 | -972 | -14,65 | 0,01 | -1,25 | >0.1 | 5,47 | 4,27 | 0 | 1,87 | 1,79 | 0 | 0 |
| IRF3 | CAGTGTGAGAAATAGGAAATG | 1 | 50 | 71 | -950 | -929 | -12,95 | 0,01 | -2,71 | >0.1 | 4,05 | 4,31 | 0 | 1,88 | 0 | 0 | 0 |
| IRF3 | AGTGTGAGAAATAGGAAATGA | 1 | 51 | 72 | -949 | -928 | -12,13 | 0,01 | -1,9 | >0.1 | 4,04 | 4,31 | 0 | 1,88 | 0 | 0 | 0 |
| IRF3 | AAAAAATGGCCATGGAAATTA | 1 | 140 | 161 | -860 | -839 | -6,92 | 0,01 | 4,3 | >0.1 | 3,31 | 4,41 | 0 | 1,9 | 0 | 1,61 | 0 |
| IRF3 | TGGCCATGGAAATTAAAACTG | 1 | 146 | 167 | -854 | -833 | -1,83 | 0,001 | 9,41 | >0.1 | 3,27 | 4,42 | 0 | 1,9 | 0 | 1,65 | 0 |
| IRF3 | TGGAAATTAAAACTGATCAGA | 1 | 152 | 173 | -848 | -827 | -6,14 | 0,01 | 5,96 | >0.1 | 3,24 | 4,42 | 0 | 1,9 | 0 | 2,53 | 0 |
| IRF3 | ATTAAAACTGATCAGAAATGG | 1 | 157 | 178 | -843 | -822 | -13,84 | 0,01 | -1,84 | >0.1 | 3,22 | 4,43 | 0 | 1,9 | 0 | 2,46 | 0 |
| IRF3 | AAGGTCAGGAGATAGAGACC | 1 | 240 | 261 | -760 | -739 | -6,92 | 0,01 | 7,19 | >0.1 | 2,86 | 4,54 | 0 | 1,41 | 0 | 5,3 | 0 |
| IRF3 | TCAAAAAAAAAAAAAAAAAAA | 1 | 455 | 476 | -545 | -524 | -12,41 | 0,01 | 1,82 | >0.1 | 2,81 | 4,89 | 0 | 1 | 0 | 5,52 | 0 |
| IRF3 | CAAAAAAAAAAAAAAAAAAAA | 1 | 456 | 477 | -544 | -523 | -12,82 | 0,01 | 1,53 | >0.1 | 2,82 | 4,9 | 0 | 1 | 0 | 5,63 | 0 |
| IRF3 | AAAAAAAAAAAAAAAAAAAAA | 1 | 457 | 478 | -543 | -522 | -13,73 | 0,01 | 0,63 | >0.1 | 2,82 | 4,9 | 0 | 1 | 0 | 5,63 | 0 |
| IRF3 | AAAAAAAAAAAAAAAAAAAAA | 1 | 458 | 479 | -542 | -521 | -13,73 | 0,01 | 0,77 | >0.1 | 2,82 | 4,9 | 0 | 1 | 0 | 5,78 | 0 |
| IRF3 | AAAAAAAAAAAAAAAAAAACT | 1 | 460 | 481 | -540 | -519 | -13,53 | 0,01 | 0,98 | >0.1 | 2,83 | 4,9 | 0 | 1 | 0 | 5,78 | 0 |
| IRF3 | AAAAAAAAAAAAAAAAAACTG | 1 | 461 | 482 | -539 | -518 | -5,25 | 0,01 | 9,13 | >0.1 | 2,83 | 4,91 | 0 | 1 | 0 | 5,63 | 0 |
| IRF3 | AAAAAAAAAAAAACTGATCAGA | 1 | 467 | 488 | -533 | -512 | -11,5 | 0,01 | 2,22 | >0.1 | 2,85 | 4,92 | 0 | 1 | 0 | 4,95 | 0 |
| IRF3 | ATGTATTTGGAAGGGAAAATG | 1 | 520 | 541 | -480 | -459 | -5,97 | 0,01 | 6,41 | >0.1 | 3,07 | 5,04 | 0 | 0,99 | 0 | 3,28 | 0 |
| IRF3 | TGTATTTGGAAGGGAAAATGT | 1 | 521 | 542 | -479 | -458 | -13,77 | 0,01 | -1,24 | >0.1 | 3,07 | 5,04 | 0 | 0,99 | 0 | 3,43 | 0 |
| IRF3 | TTGGAAGGGAAAATGTTGCCC | 1 | 526 | 547 | -474 | -453 | -7,16 | 0,01 | 3,59 | >0.1 | 3,09 | 5,05 | 0 | 0,99 | 0 | 1,61 | 0 |
| IRF3 | GGGAATATGATCTTGAAATAA | 1 | 562 | 583 | -438 | -417 | -12,06 | 0,01 | -2,64 | >0.1 | 3,27 | 5,14 | 0 | 1,01 | 0 | 0 | 0 |
| IRF3 | ATGATCTTGAAATAAAAATAA | 1 | 568 | 589 | -432 | -411 | -10,54 | 0,01 | -1,07 | >0.1 | 3,31 | 5,15 | 0 | 1,01 | 0 | 0 | 0 |
| IRF3 | CTTGAAATAAAAATAAATGTG | 1 | 573 | 594 | -427 | -406 | -8,58 | 0,01 | 0,94 | >0.1 | 3,34 | 5,17 | 0 | 1,01 | 0 | 0 | 0 |
| IRF3 | CTGAAAACCATACAAACACCA | 1 | 617 | 638 | -383 | -362 | -12,74 | 0,01 | -1,32 | >0.1 | 3,64 | 5,29 | 0 | 1,05 | 0 | 1,44 | 0 |
| IRF3 | GGAAAATATTAACCAAAAGTA | 1 | 723 | 744 | -277 | -256 | -9,24 | 0,01 | 8,45 | >0.1 | 5,47 | 5,66 | 0 | 1,24 | 0 | 5,32 | 0 |
| IRF3 | TTAACCACGAAGCCGAAGAC | 1 | 1143 | 1164 | 143 | 164 | -12,27 | 0,01 | 6,8 | >0.1 | 5,47 | 6,35 | 0 | 1,84 | 3,22 | 2,18 | 0 |
| IRF3 | CTGAGAAGGAGCCAGGAAGA | -1 | 1057 | 1078 | 57 | 78 | -6,71 | 0,01 | 12,04 | >0.1 | 5,47 | 7,42 | 0 | 1,86 | 1,79 | 2,21 | 0 |
| IRF3 | CTGAGCAGGAAAGACCAAAA | -1 | 925 | 946 | -75 | -54 | -14,45 | 0,01 | 1,06 | >0.1 | 5,47 | 7,15 | 0 | 1,88 | 0 | 1 | 0 |
| IRF3 | AAAGACCAAAAAAAGAAAAAG | -1 | 916 | 937 | -84 | -63 | -11,7 | 0,01 | 3,8 | >0.1 | 5,47 | 7,02 | 0 | 1,89 | 0 | 1,12 | 0 |
| IRF3 | CAAAAAAAGAAAAAGAAGAAG | -1 | 910 | 931 | -90 | -69 | -5,26 | 0,01 | 10,2 | >0.1 | 5,47 | 6,94 | 0 | 1,89 | 0 | 1,15 | 0 |
| IRF3 | AAAAAAAGAAAAAGAAGAAGA | -1 | 909 | 930 | -91 | -70 | -11,92 | 0,01 | 3,52 | >0.1 | 5,47 | 6,93 | 0 | 1,89 | 0 | 1,15 | 0 |
| IRF3 | AAGAAAAAGAAGAAGAACACA | -1 | 904 | 925 | -96 | -75 | -11,46 | 0,01 | 3,98 | >0.1 | 5,47 | 6,87 | 0 | 1,89 | 0 | 1,21 | 0 |
| IRF3 | AGAAAAAGAAGAAGAACACAG | -1 | 903 | 924 | -97 | -76 | -11,69 | 0,01 | 3,74 | >0.1 | 5,47 | 6,86 | 0 | 1,89 | 0 | 1,21 | 0 |
| IRF3 | AGAAGAAGAACACAGTAAACA | -1 | 897 | 918 | -103 | -82 | -11,5 | 0,01 | 3,93 | >0.1 | 5,47 | 6,79 | 0 | 1,89 | 0 | 1,28 | 0 |
| IRF3 | ACAGATATGTAACAGATTTTA | -1 | 840 | 861 | -160 | -139 | -12,74 | 0,01 | 3,86 | >0.1 | 5,47 | 6,3 | 0 | 1,88 | 0 | 2,95 | 0 |
| IRF3 | TTTAAAGTGAATCAAATAAGA | -1 | 810 | 831 | -190 | -169 | -10,32 | 0,01 | 6,3 | >0.1 | 5,47 | 6,1 | 0 | 1,63 | 0 | 3,43 | 0 |
| IRF3 | AGTGAATCAAATAAGAATACG | -1 | 805 | 826 | -195 | -174 | -10,81 | 0,01 | 5,6 | >0.1 | 5,47 | 6,07 | 0 | 1,59 | 0 | 3,28 | 0 |
| IRF3 | TTTATGGCCAAATCAAAACCG | -1 | 745 | 766 | -255 | -234 | -9,52 | 0,01 | 9,12 | >0.1 | 5,47 | 5,75 | 0 | 1,31 | 0 | 6,1 | 0 |
| IRF3 | GCCAAATCAAAACCGGTACTT | -1 | 739 | 760 | -261 | -240 | 0,96 | 0,001 | 19,36 | >0.1 | 5,47 | 5,72 | 0 | 1,29 | 0 | 5,92 | 0 |
| IRF3 | GTTAAAGATACACGGAAAATG | -1 | 693 | 714 | -307 | -286 | -10,08 | 0,01 | 5,29 | >0.1 | 5,47 | 5,54 | 0 | 1,17 | 0 | 3,19 | 0 |
| IRF3 | GATACACGGAAAATGTTATAA | -1 | 687 | 708 | -313 | -292 | -12,8 | 0,01 | 2,41 | >0.1 | 5,47 | 5,52 | 0 | 1,16 | 0 | 3,07 | 0 |
| IRF2 | GGTAAGTGAAGGAGAAGG | 1 | 4 | 22 | -996 | -978 | 3,47 | 0,001 | 16,9 | >0.1 | 5,51 | 4,27 | 0 | 1,86 | 1,79 | 0 | 0 |
| IRF2 | TGAAGGAGAAGGAGGCAC | 1 | 10 | 28 | -990 | -972 | -2,96 | 0,01 | 10,48 | >0.1 | 5,51 | 4,27 | 0 | 1,87 | 1,79 | 0 | 0 |

| | | | | | | | | | | | | | | | | | |
|---|---|---|---|---|---|---|---|---|---|---|---|---|---|---|---|---|---|
| IRF2 | GAAATAGGAAATGAGCTT | 1 | 58 | 76 | -942 | -924 | -3,45 | 0,01 | 6,72 | >0.1 | 3,97 | 4,32 | 0 | 1,88 | 0 | 0 | 0 |
| IRF2 | GGCCATGGAAATTAAAAC | 1 | 147 | 165 | -853 | -835 | -3,8 | 0,01 | 7,44 | >0.1 | 3,28 | 4,41 | 0 | 1,9 | 0 | 1,65 | 0 |
| IRF2 | TGGAAATTAAAACTGATC | 1 | 152 | 170 | -848 | -830 | -0,96 | 0,01 | 11,14 | >0.1 | 3,25 | 4,42 | 0 | 1,9 | 0 | 2,53 | 0 |
| IRF2 | GGAAATTAAAACTGATCA | 1 | 153 | 171 | -847 | -829 | 7,47 | 0,001 | 19,57 | >0.1 | 3,25 | 4,42 | 0 | 1,9 | 0 | 2,53 | 0 |
| IRF2 | GGAGATAGAGACCGTCCT | 1 | 247 | 265 | -753 | -735 | -2,55 | 0,01 | 11,3 | >0.1 | 2,84 | 4,54 | 0 | 1,38 | 0 | 5,09 | 0 |
| IRF2 | AAAAAAAAAAACTGATCA | 1 | 468 | 486 | -532 | -514 | -2,4 | 0,01 | 11,31 | >0.1 | 2,85 | 4,91 | 0 | 1 | 0 | 4,95 | 0 |
| IRF2 | GAAATATGAATTCCATAA | 1 | 486 | 504 | -514 | -496 | -0,8 | 0,01 | 14,33 | >0.1 | 2,92 | 4,95 | 0 | 0,99 | 0 | 6,27 | 0 |
| IRF2 | TGGAAGGGAAAATGTTGC | 1 | 527 | 545 | -473 | -455 | 1,65 | 0,01 | 12,39 | >0.1 | 3,09 | 5,04 | 0 | 0,99 | 0 | 1,61 | 0 |
| IRF2 | GGAAGGGAAAATGTTGCC | 1 | 528 | 546 | -472 | -454 | -3,21 | 0,01 | 7,53 | >0.1 | 3,1 | 5,05 | 0 | 0,99 | 0 | 1,61 | 0 |
| IRF2 | GGAAAATGTTGCCCAAGT | 1 | 533 | 551 | -467 | -449 | 0,33 | 0,01 | 11,16 | >0.1 | 3,12 | 5,06 | 0 | 0,99 | 0 | 1,66 | 0 |
| IRF2 | TAAAAATAAATGTGAGAT | 1 | 580 | 598 | -420 | -402 | -1,9 | 0,01 | 7,66 | >0.1 | 3,37 | 5,18 | 0 | 1,01 | 0 | 0 | 0 |
| IRF2 | TGAAAACCATACAAACAC | 1 | 618 | 636 | -382 | -364 | 0 | 0,01 | 11,41 | >0.1 | 3,64 | 5,28 | 0 | 1,05 | 0 | 1,44 | 0 |
| IRF2 | GGAAAATATTAACCAAAA | 1 | 723 | 741 | -277 | -259 | -3,95 | 0,01 | 13,75 | >0.1 | 5,51 | 5,64 | 0 | 1,23 | 0 | 5,32 | 0 |
| IRF2 | GAAAATATTAACCAAAAG | 1 | 724 | 742 | -276 | -258 | -3,87 | 0,01 | 13,84 | >0.1 | 5,51 | 5,65 | 0 | 1,24 | 0 | 5,32 | 0 |
| IRF2 | ATAAAGTGACAGGAGAGG | 1 | 761 | 779 | -239 | -221 | 0,72 | 0,01 | 19,31 | >0.1 | 5,51 | 5,81 | 0 | 1,36 | 0 | 5,92 | 0 |
| IRF2 | TAAGAGAGAAAATCTCAT | 1 | 971 | 989 | -29 | -11 | -3,65 | 0,01 | 13,57 | >0.1 | 5,51 | 8,27 | 0 | 1,87 | 0 | 1,57 | 0 |
| IRF2 | GTCTAGGGAAAGTCATTC | 1 | 1001 | 1019 | 1 | 19 | 2,13 | 0,01 | 20,04 | >0.1 | 5,51 | 9,12 | 0 | 1,87 | 0 | 1,42 | 0 |
| IRF2 | GACAAGTTTAACCACGAA | 1 | 1136 | 1154 | 136 | 154 | -3,59 | 0,01 | 16,69 | >0.1 | 5,51 | 6,41 | 0 | 1,84 | 3,22 | 3,29 | 0 |
| IRF2 | TAACCACGAAGCCGAAGA | 1 | 1144 | 1162 | 144 | 162 | -2,77 | 0,01 | 16,32 | >0.1 | 5,51 | 6,34 | 0 | 1,84 | 3,22 | 2,18 | 0 |
| IRF2 | CGAAGCCGAAGACCTGTT | 1 | 1150 | 1168 | 150 | 168 | -3,06 | 0,01 | 13,65 | >0.1 | 4,46 | 6,3 | 0 | 1,84 | 1,79 | 2,31 | 0 |
| IRF2 | AGCAAGTGAACTTTGATA | -1 | 1169 | 1187 | 169 | 187 | -2,96 | 0,01 | 15,27 | >0.1 | 5,51 | 6,16 | 0 | 1,68 | 1,79 | 3,09 | 0 |
| IRF2 | AAAAAGAAAAAGAAGAAG | -1 | 910 | 928 | -90 | -72 | -4,1 | 0,01 | 11,35 | >0.1 | 5,51 | 6,91 | 0 | 1,89 | 0 | 1,15 | 0 |
| IRF2 | TTAAAGTGAATCAAATAA | -1 | 812 | 830 | -188 | -170 | 1,83 | 0,01 | 18,6 | >0.1 | 5,51 | 6,09 | 0 | 1,62 | 0 | 3,55 | 0 |
| IRF2 | TTATGGCCAAATCAAAAC | -1 | 747 | 765 | -253 | -235 | -4,1 | 0,01 | 14,78 | >0.1 | 5,51 | 5,75 | 0 | 1,31 | 0 | 6,32 | 0 |
| IRF2 | CCAAATCAAAACCGGTAC | -1 | 741 | 759 | -259 | -241 | 4,38 | 0,001 | 22,81 | >0.1 | 5,51 | 5,72 | 0 | 1,29 | 0 | 5,92 | 0 |
| IRF2 | CAAAACCGGTACTTTTGG | -1 | 735 | 753 | -265 | -247 | -3,36 | 0,01 | 15,02 | >0.1 | 5,51 | 5,69 | 0 | 1,27 | 0 | 5,92 | 0 |
| IRF2 | GGATCATGAAGATAATGT | -1 | 638 | 656 | -362 | -344 | -3,4 | 0,01 | 10,66 | >0.1 | 3,81 | 5,34 | 0 | 1,07 | 0 | 3,83 | 0 |
| IRF2 | TAATAGAGACGGGGTTTC | -1 | 277 | 295 | -723 | -705 | -3,49 | 0,01 | 10,3 | >0.1 | 2,74 | 4,58 | 0 | 1,27 | 0 | 5,19 | 0 |
| IRF2 | CCAAAGTGCTAGGATTAC | -1 | 200 | 218 | -800 | -782 | 4,04 | 0,001 | 17,76 | >0.1 | 3,02 | 4,48 | 0 | 1,63 | 0 | 4,59 | 0 |
| IRF1 | TAACATTTTCCGTGTATCTTT | 1 | 690 | 711 | -310 | -289 | -1,82 | 0,01 | 13,29 | >0.1 | 5,47 | 5,53 | 0 | 1,16 | 0 | 2,95 | 0 |
| IRF1 | TACCGGTTTTGATTTGGCCAT | 1 | 742 | 763 | -258 | -237 | 2,15 | 0,01 | 20,76 | >0.1 | 5,47 | 5,74 | 0 | 1,3 | 0 | 6,1 | 0 |
| IRF1 | TATTTGATTCACTTTAAAAAA | 1 | 813 | 834 | -187 | -166 | 0,24 | 0,01 | 16,9 | >0.1 | 5,47 | 6,12 | 0 | 1,65 | 0 | 3,43 | 0 |
| IRF1 | ACTGTGTTCTTCTTCTTTTTC | 1 | 902 | 923 | -98 | -77 | 1,78 | 0,01 | 17,22 | >0.1 | 5,47 | 6,85 | 0 | 1,89 | 0 | 1,24 | 0 |
| IRF1 | GTGTTCTTCTTCTTTTTCTTT | 1 | 905 | 926 | -95 | -74 | 9,04 | 0,001 | 24,44 | >0.1 | 5,47 | 6,88 | 0 | 1,89 | 0 | 1,15 | 0 |
| IRF1 | TGTTCTTCTTCTTTTTCTTTT | 1 | 906 | 927 | -94 | -73 | -0,9 | 0,01 | 14,51 | >0.1 | 5,47 | 6,89 | 0 | 1,89 | 0 | 1,15 | 0 |
| IRF1 | GTTCTTCTTCTTTTTCTTTTT | 1 | 907 | 928 | -93 | -72 | 0,2 | 0,01 | 15,62 | >0.1 | 5,47 | 6,91 | 0 | 1,89 | 0 | 1,15 | 0 |
| IRF1 | TTCTTCTTCTTTTTCTTTTTT | 1 | 908 | 929 | -92 | -71 | 2,82 | 0,01 | 18,25 | >0.1 | 5,47 | 6,92 | 0 | 1,89 | 0 | 1,15 | 0 |
| IRF1 | TTCTTCTTTTTCTTTTTTTGG | 1 | 911 | 932 | -89 | -68 | 8,85 | 0,001 | 24,29 | >0.1 | 5,47 | 6,96 | 0 | 1,89 | 0 | 1,12 | 0 |
| IRF1 | TCTTCTTTTTCTTTTTTTGGT | 1 | 912 | 933 | -88 | -67 | -0,97 | 0,01 | 14,48 | >0.1 | 5,47 | 6,97 | 0 | 1,89 | 0 | 1,12 | 0 |
| IRF1 | CTTCTTTTTCTTTTTTTGGTC | 1 | 913 | 934 | -87 | -66 | 7,35 | 0,001 | 22,81 | >0.1 | 5,47 | 6,98 | 0 | 1,89 | 0 | 1,12 | 0 |
| IRF1 | TTCTTTTTCTTTTTTTGGTCT | 1 | 914 | 935 | -86 | -65 | 0,52 | 0,01 | 15,96 | >0.1 | 5,47 | 7 | 0 | 1,89 | 0 | 1,09 | 0 |
| IRF1 | TTCTTTTTTTGGTCTTTCCTG | 1 | 920 | 941 | -80 | -59 | -0,64 | 0,01 | 14,86 | >0.1 | 5,47 | 7,08 | 0 | 1,88 | 0 | 1,06 | 0 |
| IRF1 | CATGAGATTTTCTCTCTTATC | -1 | 969 | 990 | -31 | -10 | -1,42 | 0,01 | 15,86 | >0.1 | 5,47 | 8,32 | 0 | 1,87 | 0 | 1,61 | 0 |
| IRF1 | TGAGATTTTCTCTCTTATCAG | -1 | 967 | 988 | -33 | -12 | -0,33 | 0,01 | 16,93 | >0.1 | 5,47 | 8,22 | 0 | 1,87 | 0 | 1,7 | 0 |
| IRF1 | CTCTCCTGTCACTTTATGGC( | -1 | 757 | 778 | -243 | -222 | -1,66 | 0,01 | 16,89 | >0.1 | 5,47 | 5,81 | 0 | 1,35 | 0 | 5,92 | 0 |
| IRF1 | CGGTACTTTTGGTTAATATTT | -1 | 726 | 747 | -274 | -253 | 2,04 | 0,01 | 20,12 | >0.1 | 5,47 | 5,67 | 0 | 1,25 | 0 | 5,69 | 0 |
| IRF1 | AGGTTATCTCACATTTATTTT | -1 | 582 | 603 | -418 | -397 | 1,51 | 0,01 | 11,11 | >0.1 | 3,39 | 5,19 | 0 | 1,02 | 0 | 0 | 0 |
| IRF1 | TCTCACATTTATTTTTATTTC | -1 | 576 | 597 | -424 | -403 | 8,48 | 0,001 | 18,02 | >0.1 | 3,35 | 5,17 | 0 | 1,01 | 0 | 0 | 0 |
| IRF1 | CATTTATTTTTATTTCAAGAT | -1 | 571 | 592 | -429 | -408 | -1,38 | 0,01 | 8,12 | >0.1 | 3,33 | 5,16 | 0 | 1,01 | 0 | 0 | 0 |
| IRF1 | ATTTATTTTTATTTCAAGATC | -1 | 570 | 591 | -430 | -409 | -0,11 | 0,01 | 9,38 | >0.1 | 3,32 | 5,16 | 0 | 1,01 | 0 | 0 | 0 |
| IRF1 | CAACATTTTCCCTTCCAAATA | -1 | 523 | 544 | -477 | -456 | 3,51 | 0,01 | 16,17 | >0.1 | 3,08 | 5,04 | 0 | 0,99 | 0 | 3,55 | 0 |
| IRF1 | GATCAGTTTTTTTTTTTTTTT | -1 | 464 | 485 | -536 | -515 | 11,27 | 0,001 | 24,85 | >0.1 | 2,84 | 4,91 | 0 | 1 | 0 | 4,83 | 0 |
| IRF1 | ATCAGTTTTTTTTTTTTTTTT | -1 | 463 | 484 | -537 | -516 | 4,89 | 0,01 | 19,56 | >0.1 | 2,84 | 4,91 | 0 | 1 | 0 | 5,92 | 0 |
| IRF1 | TCAGTTTTTTTTTTTTTTTTT | -1 | 462 | 483 | -538 | -517 | 2,48 | 0,01 | 17 | >0.1 | 2,84 | 4,91 | 0 | 1 | 0 | 5,78 | 0 |
| IRF1 | CAGTTTTTTTTTTTTTTTTTT | -1 | 461 | 482 | -539 | -518 | 4,2 | 0,01 | 18,58 | >0.1 | 2,83 | 4,91 | 0 | 1 | 0 | 5,63 | 0 |
| IRF1 | AGTTTTTTTTTTTTTTTTTTT | -1 | 460 | 481 | -540 | -519 | 5,31 | 0,01 | 19,82 | >0.1 | 2,83 | 4,9 | 0 | 1 | 0 | 5,78 | 0 |
| IRF1 | GTTTTTTTTTTTTTTTTTTTT | -1 | 459 | 480 | -541 | -520 | 6,26 | 0,01 | 20,77 | >0.1 | 2,83 | 4,9 | 0 | 1 | 0 | 5,78 | 0 |
| IRF1 | TTTTTTTTTTTTTTTTTTTTT | -1 | 458 | 479 | -542 | -521 | 6,55 | 0,01 | 21,05 | >0.1 | 2,82 | 4,9 | 0 | 1 | 0 | 5,78 | 0 |
| IRF1 | TTTTTTTTTTTTTTTTTTTTT | -1 | 457 | 478 | -543 | -522 | 6,55 | 0,01 | 20,91 | >0.1 | 2,82 | 4,9 | 0 | 1 | 0 | 5,63 | 0 |

| | | | | | | | | | | | | | | | | | |
|---|---|---|---|---|---|---|---|---|---|---|---|---|---|---|---|---|---|
| IRF1 | TTTTTTTTTTTTTTTTTTTTG | -1 | 456 | 477 | -544 | -523 | 4,81 | 0,01 | 19,16 | >0.1 | 2,82 | 4,9 | 0 | 1 | 0 | 5,63 | 0 |
| IRF1 | TTTTTTTTTTTTTTTTTTTGA | -1 | 455 | 476 | -545 | -524 | 1,81 | 0,01 | 16,04 | >0.1 | 2,81 | 4,89 | 0 | 1 | 0 | 5,52 | 0 |
| IRF1 | TTTTTTTTTTTTTTTTTTGAG | -1 | 454 | 475 | -546 | -525 | 0,16 | 0,01 | 14,39 | >0.1 | 2,81 | 4,89 | 0 | 1 | 0 | 5,52 | 0 |
| IRF1 | TTTTTTTTTTTTTTTTTGAGA | -1 | 453 | 474 | -547 | -526 | -0,77 | 0,01 | 13,57 | >0.1 | 2,81 | 4,89 | 0 | 1,01 | 0 | 5,63 | 0 |
| IRF1 | TTTTTTTTTTTTTTTTGAGAC | -1 | 452 | 473 | -548 | -527 | 0,58 | 0,01 | 14,91 | >0.1 | 2,8 | 4,89 | 0 | 1,01 | 0 | 5,63 | 0 |
| IRF1 | TTTTTTTTTTTTTTTGAGACT | -1 | 451 | 472 | -549 | -528 | -0,01 | 0,01 | 14,2 | >0.1 | 2,8 | 4,89 | 0 | 1,01 | 0 | 5,52 | 0 |
| IRF1 | ACTGAGTCTCGGTCTGTCGC | -1 | 433 | 454 | -567 | -546 | -1,62 | 0,01 | 13,03 | >0.1 | 2,74 | 4,85 | 0 | 1,02 | 0 | 6,05 | 0 |
| IRF1 | GCTAAGTTTTTGTATTTTTAA | -1 | 292 | 313 | -708 | -687 | -0,75 | 0,01 | 12,23 | >0.1 | 2,69 | 4,61 | 0 | 1,23 | 0 | 4,45 | 0 |
| IRF1 | GATCAGTTTTAATTTCCATGG | -1 | 149 | 170 | -851 | -830 | 7,4 | 0,001 | 18,63 | >0.1 | 3,26 | 4,42 | 0 | 1,9 | 0 | 1,65 | 0 |
| IRF1 | GCTCATTTCCTATTTCTCACA | -1 | 53 | 74 | -947 | -926 | 0,94 | 0,01 | 11,15 | >0.1 | 4,01 | 4,31 | 0 | 1,88 | 0 | 0 | 0 |
| IRF7 | AGGTAAGTGAAGGA | 1 | 3 | 17 | -997 | -983 | -4,5 | 0,01 | 8,97 | >0.1 | 5,56 | 4,26 | 0 | 1,86 | 1,79 | 0 | 0 |
| IRF7 | GAGAAATAGGAAAT | 1 | 56 | 70 | -944 | -930 | -3,61 | 0,01 | 6,6 | >0.1 | 4,02 | 4,31 | 0 | 1,87 | 0 | 0 | 0 |
| IRF7 | ATGGAAATTAAAAC | 1 | 151 | 165 | -849 | -835 | -4,51 | 0,01 | 6,71 | >0.1 | 3,27 | 4,41 | 0 | 1,9 | 0 | 1,65 | 0 |
| IRF7 | TGGAAATTAAAACT | 1 | 152 | 166 | -848 | -834 | 4,42 | 0,001 | 15,64 | >0.1 | 3,26 | 4,41 | 0 | 1,9 | 0 | 1,65 | 0 |
| IRF7 | TTAAAAATACAAAA | 1 | 292 | 306 | -708 | -694 | -5,3 | 0,01 | 7,75 | >0.1 | 2,7 | 4,6 | 0 | 1,23 | 0 | 4,53 | 0 |
| IRF7 | CTCAAAAAAAAAAA | 1 | 454 | 468 | -546 | -532 | -2,79 | 0,01 | 11,41 | >0.1 | 2,8 | 4,88 | 0 | 1 | 0 | 5,52 | 0 |
| IRF7 | TCAAAAAAAAAAAA | 1 | 455 | 469 | -545 | -531 | -0,56 | 0,01 | 13,75 | >0.1 | 2,8 | 4,88 | 0 | 1 | 0 | 5,63 | 0 |
| IRF7 | CAAAAAAAAAAAAA | 1 | 456 | 470 | -544 | -530 | -3,1 | 0,01 | 11,22 | >0.1 | 2,8 | 4,88 | 0 | 1 | 0 | 5,63 | 0 |
| IRF7 | AAAAAAAAAAAAAA | 1 | 457 | 471 | -543 | -529 | -3,39 | 0,01 | 10,82 | >0.1 | 2,81 | 4,88 | 0 | 1 | 0 | 5,52 | 0 |
| IRF7 | AAAAAAAAAAAAAA | 1 | 458 | 472 | -542 | -528 | -3,39 | 0,01 | 10,83 | >0.1 | 2,81 | 4,89 | 0 | 1 | 0 | 5,52 | 0 |
| IRF7 | AAAAAAAAAAAAAA | 1 | 459 | 473 | -541 | -527 | -3,39 | 0,01 | 10,94 | >0.1 | 2,81 | 4,89 | 0 | 1 | 0 | 5,63 | 0 |
| IRF7 | AAAAAAAAAAAAAA | 1 | 460 | 474 | -540 | -526 | -3,39 | 0,01 | 10,95 | >0.1 | 2,82 | 4,89 | 0 | 1 | 0 | 5,63 | 0 |
| IRF7 | AAAAAAAAAAAAAA | 1 | 461 | 475 | -539 | -525 | -3,39 | 0,01 | 11,1 | >0.1 | 2,82 | 4,89 | 0 | 1 | 0 | 5,78 | 0 |
| IRF7 | AAAAAAAAAAAAAA | 1 | 462 | 476 | -538 | -524 | -3,39 | 0,01 | 11,1 | >0.1 | 2,83 | 4,89 | 0 | 1 | 0 | 5,78 | 0 |
| IRF7 | AAAAAAAAAAAAAA | 1 | 463 | 477 | -537 | -523 | -3,39 | 0,01 | 11,11 | >0.1 | 2,83 | 4,9 | 0 | 1 | 0 | 5,78 | 0 |
| IRF7 | AAAAAAAAAAAAAA | 1 | 464 | 478 | -536 | -522 | -3,39 | 0,01 | 10,97 | >0.1 | 2,83 | 4,9 | 0 | 0,99 | 0 | 5,63 | 0 |
| IRF7 | AAAAAAAAAAAAAA | 1 | 465 | 479 | -535 | -521 | -3,39 | 0,01 | 11,12 | >0.1 | 2,83 | 4,9 | 0 | 0,99 | 0 | 5,78 | 0 |
| IRF7 | AAAAAAAAAAAAAC | 1 | 466 | 480 | -534 | -520 | -2,96 | 0,01 | 11,7 | >0.1 | 2,84 | 4,9 | 0 | 0,99 | 0 | 5,92 | 0 |
| IRF7 | AAAAAAAAAAAACT | 1 | 467 | 481 | -533 | -519 | -0,87 | 0,01 | 12,69 | >0.1 | 2,84 | 4,9 | 0 | 0,99 | 0 | 4,83 | 0 |
| IRF7 | TTGGAAGGGAAAAT | 1 | 526 | 540 | -474 | -460 | -1,59 | 0,01 | 11,06 | >0.1 | 3,08 | 5,03 | 0 | 0,99 | 0 | 3,55 | 0 |
| IRF7 | CTTGAAATAAAAAT | 1 | 573 | 587 | -427 | -413 | -4,16 | 0,01 | 5,31 | >0.1 | 3,32 | 5,15 | 0 | 1 | 0 | 0 | 0 |
| IRF7 | ATAAAAATAAATGT | 1 | 579 | 593 | -421 | -407 | -0,6 | 0,01 | 8,92 | >0.1 | 3,35 | 5,16 | 0 | 1,01 | 0 | 0 | 0 |
| IRF7 | CTGAAAACCATACA | 1 | 617 | 631 | -383 | -369 | -4,5 | 0,01 | 6,91 | >0.1 | 3,61 | 5,27 | 0 | 1,04 | 0 | 1,5 | 0 |
| IRF7 | CATAAAGTGACAGG | 1 | 760 | 774 | -240 | -226 | 1,53 | 0,01 | 20,13 | >0.1 | 5,56 | 5,79 | 0 | 1,34 | 0 | 5,92 | 0 |
| IRF7 | ACGAAGCCGAAGAC | 1 | 1149 | 1163 | 149 | 163 | 2,14 | 0,01 | 18,82 | >0.1 | 4,53 | 6,3 | 0 | 1,84 | 1,79 | 2,21 | 0 |
| IRF7 | TCCAAGAAGCAAGT | -1 | 1180 | 1194 | 180 | 194 | -5,33 | 0,01 | 13,27 | >0.1 | 5,56 | 6,09 | 0 | 1,58 | 1,79 | 3,57 | 0 |
| IRF7 | AAGCAAGTGAACTT | -1 | 1174 | 1188 | 174 | 188 | 0,75 | 0,01 | 19,14 | >0.1 | 5,56 | 6,13 | 0 | 1,63 | 1,79 | 3,28 | 0 |
| IRF7 | CCTAGACTAAAACC | -1 | 994 | 1008 | -6 | 8 | -3,92 | 0,01 | 14,01 | >0.1 | 5,56 | 9,01 | 0 | 1,87 | 0 | 1,5 | 0 |
| IRF7 | AGGAAAGACCAAAA | -1 | 926 | 940 | -74 | -60 | -5,4 | 0,01 | 10,13 | >0.1 | 5,56 | 7,06 | 0 | 1,88 | 0 | 1,03 | 0 |
| IRF7 | CAAAAAAAGAAAAA | -1 | 917 | 931 | -83 | -69 | 2,38 | 0,01 | 17,85 | >0.1 | 5,56 | 6,94 | 0 | 1,89 | 0 | 1,09 | 0 |
| IRF7 | AAAAAAGAAAAAGA | -1 | 915 | 929 | -85 | -71 | -1,48 | 0,01 | 14 | >0.1 | 5,56 | 6,92 | 0 | 1,89 | 0 | 1,12 | 0 |
| IRF7 | AAGAAAAAGAAGAA | -1 | 911 | 925 | -89 | -75 | 0,76 | 0,01 | 16,22 | >0.1 | 5,56 | 6,87 | 0 | 1,89 | 0 | 1,15 | 0 |
| IRF7 | TTTAAAGTGAATCA | -1 | 817 | 831 | -183 | -169 | 3,89 | 0,001 | 20,72 | >0.1 | 5,56 | 6,1 | 0 | 1,63 | 0 | 3,55 | 0 |
| IRF7 | GCCAAATCAAAACC | -1 | 746 | 760 | -254 | -240 | 3,15 | 0,001 | 21,63 | >0.1 | 5,56 | 5,72 | 0 | 1,29 | 0 | 5,92 | 0 |
| IRF7 | TCAAAACCGGTACT | -1 | 740 | 754 | -260 | -246 | -3,23 | 0,01 | 15,21 | >0.1 | 5,56 | 5,7 | 0 | 1,27 | 0 | 5,92 | 0 |
| IRF7 | CCCAAAGTGCTAGG | -1 | 205 | 219 | -795 | -781 | -1,9 | 0,01 | 11,89 | >0.1 | 3,01 | 4,48 | 0 | 1,6 | 0 | 4,7 | 0 |
| IRF7 | GCAAAACTTAAAAA | -1 | 75 | 89 | -925 | -911 | 1,62 | 0,01 | 11,64 | >0.1 | 3,81 | 4,33 | 0 | 1,88 | 0 | 0 | 0 |
| IRF5 | TGGAAATTAAAACT | 1 | 152 | 166 | -848 | -834 | 6,44 | 0,001 | 17,66 | >0.1 | 3,26 | 4,41 | 0 | 1,9 | 0 | 1,65 | 0 |
| IRF5 | TTAAAACTGATCAG | 1 | 158 | 172 | -842 | -828 | 1,68 | 0,01 | 13,69 | >0.1 | 3,23 | 4,42 | 0 | 1,9 | 0 | 2,46 | 0 |
| IRF5 | GTGAAACCCCGTCT | 1 | 275 | 289 | -725 | -711 | 3,83 | 0,001 | 17,63 | >0.1 | 2,75 | 4,57 | 0 | 1,28 | 0 | 5,19 | 0 |
| IRF5 | CCGAGATCGCGCCA | 1 | 404 | 418 | -596 | -582 | 2,05 | 0,01 | 16,16 | >0.1 | 2,64 | 4,78 | 0 | 1,03 | 0 | 5,65 | 0 |
| IRF5 | GACAGACCGAGACT | 1 | 436 | 450 | -564 | -550 | 1,18 | 0,01 | 15,82 | >0.1 | 2,74 | 4,84 | 0 | 1,01 | 0 | 6,05 | 0 |
| IRF5 | CCGAGACTCAGTCT | 1 | 442 | 456 | -558 | -544 | 5,27 | 0,001 | 19,81 | >0.1 | 2,76 | 4,85 | 0 | 1,01 | 0 | 5,92 | 0 |
| IRF5 | AAAAAACTGATCAG | 1 | 473 | 487 | -527 | -513 | -0,49 | 0,01 | 13,37 | >0.1 | 2,86 | 4,92 | 0 | 0,99 | 0 | 5,09 | 0 |
| IRF5 | CTGAAAACCATACA | 1 | 617 | 631 | -383 | -369 | 4,09 | 0,001 | 15,5 | >0.1 | 3,61 | 5,27 | 0 | 1,04 | 0 | 1,5 | 0 |
| IRF5 | CAAACACCAACATT | 1 | 629 | 643 | -371 | -357 | -1,59 | 0,01 | 10,13 | >0.1 | 3,71 | 5,3 | 0 | 1,05 | 0 | 1,66 | 0 |
| IRF5 | TATTAACCAAAAGT | 1 | 729 | 743 | -271 | -257 | 1,61 | 0,01 | 19,75 | >0.1 | 5,56 | 5,65 | 0 | 1,24 | 0 | 5,69 | 0 |
| IRF5 | ACGATGTCAAGCTC | 1 | 1046 | 1060 | 46 | 60 | -2,31 | 0,01 | 14,2 | >0.1 | 5,56 | 7,68 | 0 | 1,86 | 0 | 1,42 | 0 |
| IRF5 | CTGTAACTGCTGCT | 1 | 1086 | 1100 | 86 | 100 | 2,97 | 0,01 | 21,32 | >0.1 | 5,56 | 6,93 | 0 | 1,85 | 1,79 | 2,21 | 0 |

| | | | | | | | | | | | | | | | | | |
|---|---|---|---|---|---|---|---|---|---|---|---|---|---|---|---|---|---|
| IRF5 | ACGAAGCCGAAGAC | 1 | 1149 | 1163 | 149 | 163 | 8,38 | 0,001 | 25,06 | >0.1 | 4,53 | 6,3 | 0 | 1,84 | 1,79 | 2,21 | 0 |
| IRF5 | CCTAGACTAAAACC | -1 | 994 | 1008 | -6 | 8 | 7,94 | 0,001 | 25,87 | >0.1 | 5,56 | 9,01 | 0 | 1,87 | 0 | 1,5 | 0 |
| IRF5 | GAAAGACCAAAAAA | -1 | 924 | 938 | -76 | -62 | -2,7 | 0,01 | 12,83 | >0.1 | 5,56 | 7,03 | 0 | 1,88 | 0 | 1,06 | 0 |
| IRF5 | GCCAAATCAAAACC | -1 | 746 | 760 | -254 | -240 | 4,89 | 0,001 | 23,37 | >0.1 | 5,56 | 5,72 | 0 | 1,29 | 0 | 5,92 | 0 |
| IRF5 | TCAAAACCGGTACT | -1 | 740 | 754 | -260 | -246 | 8,93 | 0,0001 | 27,37 | >0.1 | 5,56 | 5,7 | 0 | 1,27 | 0 | 5,92 | 0 |
| IRF5 | CAGTGACCAAAGAG | -1 | 668 | 682 | -332 | -318 | -2,81 | 0,01 | 11,56 | >0.1 | 4,14 | 5,43 | 0 | 1,1 | 0 | 3,7 | 0 |
| IRF5 | TTGAGACTGAGTCT | -1 | 445 | 459 | -555 | -541 | 3,65 | 0,001 | 17,54 | >0.1 | 2,77 | 4,86 | 0 | 1 | 0 | 5,26 | 0 |
| IRF5 | CCACCACCACGCCC | -1 | 314 | 328 | -686 | -672 | -2,89 | 0,01 | 11,68 | >0.1 | 2,63 | 4,63 | 0 | 1,17 | 0 | 6,13 | 0 |
| IRF5 | GCAAAACTTAAAAA | -1 | 75 | 89 | -925 | -911 | 0,69 | 0,01 | 10,71 | >0.1 | 3,81 | 4,33 | 0 | 1,88 | 0 | 0 | 0 |
| IRF4 | GAGAAATAGGAAATG | 1 | 56 | 71 | -944 | -929 | -22,22 | 0,01 | -12,02 | >0.1 | 4,01 | 4,31 | 0 | 1,88 | 0 | 0 | 0 |
| IRF4 | ATGGAAATTAAAACT | 1 | 151 | 166 | -849 | -834 | -21,14 | 0,01 | -9,92 | >0.1 | 3,26 | 4,41 | 0 | 1,9 | 0 | 1,65 | 0 |
| IRF4 | TGGAAATTAAAACTG | 1 | 152 | 167 | -848 | -833 | -6,17 | 0,001 | 5,05 | >0.1 | 3,26 | 4,41 | 0 | 1,9 | 0 | 1,65 | 0 |
| IRF4 | TTAAAACTGATCAGA | 1 | 158 | 173 | -842 | -827 | -18,38 | 0,01 | -6,37 | >0.1 | 3,23 | 4,42 | 0 | 1,9 | 0 | 2,46 | 0 |
| IRF4 | AAAATACAAAAACTT | 1 | 295 | 310 | -705 | -690 | -19,01 | 0,01 | -6,05 | >0.1 | 2,69 | 4,61 | 0 | 1,22 | 0 | 4,45 | 0 |
| IRF4 | GACAGACCGAGACTC | 1 | 436 | 451 | -564 | -549 | -16,07 | 0,01 | -1,43 | >0.1 | 2,74 | 4,84 | 0 | 1,01 | 0 | 6,05 | 0 |
| IRF4 | TCAAAAAAAAAAAAA | 1 | 455 | 470 | -545 | -530 | -22,13 | 0,01 | -7,81 | >0.1 | 2,8 | 4,88 | 0 | 1 | 0 | 5,63 | 0 |
| IRF4 | AAAAAAAAAAAACTG | 1 | 467 | 482 | -533 | -518 | -21,06 | 0,01 | -7,49 | >0.1 | 2,84 | 4,91 | 0 | 0,99 | 0 | 4,83 | 0 |
| IRF4 | AAAAAACTGATCAGA | 1 | 473 | 488 | -527 | -512 | -22,82 | 0,01 | -8,96 | >0.1 | 2,87 | 4,92 | 0 | 0,99 | 0 | 5,09 | 0 |
| IRF4 | TTGGAAGGGAAAATG | 1 | 526 | 541 | -474 | -459 | -13,75 | 0,01 | -1,1 | >0.1 | 3,08 | 5,03 | 0 | 0,99 | 0 | 3,55 | 0 |
| IRF4 | ATGATCTTGAAATAA | 1 | 568 | 583 | -432 | -417 | -22,18 | 0,01 | -12,75 | >0.1 | 3,29 | 5,14 | 0 | 1 | 0 | 0 | 0 |
| IRF4 | TTGAAATAAAAATAA | 1 | 574 | 589 | -426 | -411 | -20,01 | 0,01 | -10,53 | >0.1 | 3,33 | 5,15 | 0 | 1,01 | 0 | 0 | 0 |
| IRF4 | CTATTAATGAAATTG | 1 | 601 | 616 | -399 | -384 | -23,4 | 0,01 | -12,24 | >0.1 | 3,5 | 5,22 | 0 | 1,02 | 0 | 1,41 | 0 |
| IRF4 | CTGAAAACCATACAA | 1 | 617 | 632 | -383 | -368 | -7,85 | 0,001 | 3,57 | >0.1 | 3,62 | 5,27 | 0 | 1,04 | 0 | 1,5 | 0 |
| IRF4 | ACCATACAAACACCA | 1 | 623 | 638 | -377 | -362 | -23,75 | 0,01 | -12,21 | >0.1 | 3,67 | 5,29 | 0 | 1,05 | 0 | 1,55 | 0 |
| IRF4 | TATTAACCAAAAGTA | 1 | 729 | 744 | -271 | -256 | -14,93 | 0,01 | 3,2 | >0.1 | 5,54 | 5,66 | 0 | 1,24 | 0 | 5,69 | 0 |
| IRF4 | CATAAAGTGACAGGA | 1 | 760 | 775 | -240 | -225 | -23,3 | 0,01 | -4,71 | >0.1 | 5,54 | 5,79 | 0 | 1,34 | 0 | 5,92 | 0 |
| IRF4 | ATAAGAGAGAAAATC | 1 | 970 | 985 | -30 | -15 | -21,61 | 0,01 | -4,4 | >0.1 | 5,54 | 8,09 | 0 | 1,87 | 0 | 1,7 | 0 |
| IRF4 | CTGTAACTGCTGCTC | 1 | 1086 | 1101 | 86 | 101 | -19,28 | 0,01 | -0,95 | >0.1 | 5,54 | 6,93 | 0 | 1,85 | 1,79 | 2,21 | 0 |
| IRF4 | ACGAAGCCGAAGACC | 1 | 1149 | 1164 | 149 | 164 | -8,87 | 0,001 | 7,8 | >0.1 | 4,52 | 6,3 | 0 | 1,84 | 1,79 | 2,21 | 0 |
| IRF4 | AAGCAAGTGAACTTT | -1 | 1173 | 1188 | 173 | 188 | -22,53 | 0,01 | -4,12 | >0.1 | 5,54 | 6,13 | 0 | 1,64 | 1,79 | 3,29 | 0 |
| IRF4 | CAGTTACAGCAACAA | -1 | 1080 | 1095 | 80 | 95 | -21,1 | 0,01 | -2,68 | >0.1 | 5,54 | 7,01 | 0 | 1,85 | 1,79 | 2,21 | 0 |
| IRF4 | CCTAGACTAAAACCT | -1 | 993 | 1008 | -7 | 8 | -10,37 | 0,001 | 7,58 | >0.1 | 5,54 | 9,01 | 0 | 1,87 | 0 | 1,53 | 0 |
| IRF4 | GAAAGACCAAAAAAA | -1 | 923 | 938 | -77 | -62 | -22,51 | 0,01 | -6,99 | >0.1 | 5,54 | 7,03 | 0 | 1,88 | 0 | 1,06 | 0 |
| IRF4 | CAAAAAAAGAAAAAG | -1 | 916 | 931 | -84 | -69 | -20,18 | 0,01 | -4,69 | >0.1 | 5,54 | 6,94 | 0 | 1,89 | 0 | 1,12 | 0 |
| IRF4 | TTTAAAGTGAATCAA | -1 | 816 | 831 | -184 | -169 | -15,11 | 0,01 | 1,58 | >0.1 | 5,54 | 6,1 | 0 | 1,63 | 0 | 3,43 | 0 |
| IRF4 | GCCAAATCAAAACCG | -1 | 745 | 760 | -255 | -240 | -4,88 | 0,001 | 13,78 | >0.1 | 5,54 | 5,72 | 0 | 1,29 | 0 | 6,1 | 0 |
| IRF4 | CTGTTAAAGATACAC | -1 | 701 | 716 | -299 | -284 | -23,49 | 0,01 | -6,42 | >0.1 | 5,54 | 5,55 | 0 | 1,17 | 0 | 4,81 | 0 |
| IRF4 | TTGAGACTGAGTCTC | -1 | 444 | 459 | -556 | -541 | -16,22 | 0,01 | -2,47 | >0.1 | 2,77 | 4,86 | 0 | 1 | 0 | 5,12 | 0 |
| IRF4 | CCCAAAGTGCTAGGA | -1 | 204 | 219 | -796 | -781 | -17,72 | 0,01 | -3,92 | >0.1 | 3,01 | 4,48 | 0 | 1,61 | 0 | 4,7 | 0 |
| IRF4 | GCAAAACTTAAAAAA | -1 | 74 | 89 | -926 | -911 | -12,97 | 0,01 | -2,94 | >0.1 | 3,82 | 4,33 | 0 | 1,88 | 0 | 0 | 0 |
| TBP | ATTTTAAAAAATGGC | 1 | 135 | 150 | -865 | -850 | 2,15 | 0,01 | 13,44 | >0.1 | 3,35 | 4,39 | 0 | 1,89 | 0 | 1,65 | 0 |
| TBP | TTTTAAAAAATGGCC | 1 | 136 | 151 | -864 | -849 | 4,73 | 0,01 | 16,02 | >0.1 | 3,35 | 4,4 | 0 | 1,89 | 0 | 1,65 | 0 |
| TBP | TCAGAAATGGCTGGG | 1 | 168 | 183 | -832 | -817 | 2,24 | 0,01 | 15,18 | >0.1 | 3,17 | 4,43 | 0 | 1,9 | 0 | 3,43 | 0 |
| TBP | CTATTAAAAATACAA | 1 | 289 | 304 | -711 | -696 | 3,93 | 0,01 | 16,92 | >0.1 | 2,71 | 4,6 | 0 | 1,24 | 0 | 4,45 | 0 |
| TBP | ATACAAAAACTTAGC | 1 | 298 | 313 | -702 | -687 | 3,54 | 0,01 | 16,32 | >0.1 | 2,68 | 4,61 | 0 | 1,21 | 0 | 4,28 | 0 |
| TBP | CCATAAAGACAAGGG | 1 | 498 | 513 | -502 | -487 | 9,58 | 0,001 | 25,32 | >0.1 | 2,96 | 4,97 | 0 | 0,99 | 0 | 6,81 | 0 |
| TBP | AAATAAAAATAAATG | 1 | 577 | 592 | -423 | -408 | 3,16 | 0,01 | 12,67 | >0.1 | 3,34 | 5,16 | 0 | 1,01 | 0 | 0 | 0 |
| TBP | AAATAAATGTGAGAT | 1 | 583 | 598 | -417 | -402 | 3,44 | 0,01 | 13,01 | >0.1 | 3,38 | 5,18 | 0 | 1,01 | 0 | 0 | 0 |
| TBP | CTATTAATGAAATTG | 1 | 601 | 616 | -399 | -384 | 5,04 | 0,01 | 16,2 | >0.1 | 3,5 | 5,22 | 0 | 1,02 | 0 | 1,41 | 0 |
| TBP | CCATAAAGTGACAGG | 1 | 759 | 774 | -241 | -226 | 7,5 | 0,001 | 26,27 | >0.1 | 5,54 | 5,79 | 0 | 1,34 | 0 | 6,1 | 0 |
| TBP | CTTTAAAAAATTATT | 1 | 824 | 839 | -176 | -161 | 2,51 | 0,01 | 19,44 | >0.1 | 5,54 | 6,15 | 0 | 1,68 | 0 | 3,55 | 0 |
| TBP | CTTTATATTGGCTCA | 1 | 876 | 891 | -124 | -109 | 5,19 | 0,01 | 22,24 | >0.1 | 5,54 | 6,54 | 0 | 1,9 | 0 | 3,07 | 0 |
| TBP | CAATATAAAGTTCAT | -1 | 871 | 886 | -129 | -114 | 3,22 | 0,01 | 20,11 | >0.1 | 5,54 | 6,5 | 0 | 1,9 | 0 | 2,95 | 0 |
| TBP | ATATAAAGTTCATCC | -1 | 869 | 884 | -131 | -116 | 4,78 | 0,01 | 21,65 | >0.1 | 5,54 | 6,48 | 0 | 1,9 | 0 | 2,95 | 0 |
| TBP | TTTTTAATAGAGACG | -1 | 284 | 299 | -716 | -701 | 2,97 | 0,01 | 16,62 | >0.1 | 2,72 | 4,59 | 0 | 1,25 | 0 | 5,09 | 0 |
| TBP | ACTTAAAAAAGCTCA | -1 | 69 | 84 | -931 | -916 | 3,53 | 0,01 | 13,6 | >0.1 | 3,87 | 4,32 | 0 | 1,88 | 0 | 0 | 0 |
| IRF9 | CATGGAAATTAAAAC | 1 | 150 | 165 | -850 | -835 | -26,56 | 0,01 | -15,33 | >0.1 | 3,27 | 4,41 | 0 | 1,9 | 0 | 1,65 | 0 |
| IRF9 | ATGGAAATTAAAACT | 1 | 151 | 166 | -849 | -834 | -3,1 | 0,001 | 8,12 | >0.1 | 3,26 | 4,41 | 0 | 1,9 | 0 | 1,65 | 0 |
| IRF9 | AAAAATACAAAAACT | 1 | 294 | 309 | -706 | -691 | -15,41 | 0,01 | -2,36 | >0.1 | 2,69 | 4,6 | 0 | 1,22 | 0 | 4,53 | 0 |

| | | | | | | | | | | | | | | | | | |
|---|---|---|---|---|---|---|---|---|---|---|---|---|---|---|---|---|---|
| IRF9 | CGACAGACCGAGACT | 1 | 435 | 450 | -565 | -550 | -25,01 | 0,01 | -10,37 | >0.1 | 2,74 | 4,84 | 0 | 1,01 | 0 | 6,05 | 0 |
| IRF9 | AAAAAAAAAAAAAAC | 1 | 465 | 480 | -535 | -520 | -22,89 | 0,01 | -8,38 | >0.1 | 2,84 | 4,9 | 0 | 1 | 0 | 5,78 | 0 |
| IRF9 | AAAAAAAAAAAAACT | 1 | 466 | 481 | -534 | -519 | -12,15 | 0,001 | 2,51 | >0.1 | 2,84 | 4,9 | 0 | 0,99 | 0 | 5,92 | 0 |
| IRF9 | TTTGGAAGGGAAAAT | 1 | 525 | 540 | -475 | -460 | -23,42 | 0,01 | -10,9 | >0.1 | 3,08 | 5,03 | 0 | 0,99 | 0 | 3,43 | 0 |
| IRF9 | AATAAAAATAAATGT | 1 | 578 | 593 | -422 | -407 | -19,15 | 0,01 | -9,63 | >0.1 | 3,35 | 5,16 | 0 | 1,01 | 0 | 0 | 0 |
| IRF9 | TCTGAAAACCATACA | 1 | 616 | 631 | -384 | -369 | -17,45 | 0,01 | -5,99 | >0.1 | 3,61 | 5,27 | 0 | 1,04 | 0 | 1,55 | 0 |
| IRF9 | AACCATACAAACACC | 1 | 622 | 637 | -378 | -363 | -23,59 | 0,01 | -12,11 | >0.1 | 3,66 | 5,28 | 0 | 1,04 | 0 | 1,5 | 0 |
| IRF9 | ATATTAACCAAAAGT | 1 | 728 | 743 | -272 | -257 | -19,24 | 0,01 | -1,28 | >0.1 | 5,54 | 5,65 | 0 | 1,24 | 0 | 5,52 | 0 |
| IRF9 | CCATAAAGTGACAGG | 1 | 759 | 774 | -241 | -226 | -25,75 | 0,01 | -6,98 | >0.1 | 5,54 | 5,79 | 0 | 1,34 | 0 | 6,1 | 0 |
| IRF9 | GATAAGAGAGAAAAT | 1 | 969 | 984 | -31 | -16 | -26,68 | 0,01 | -9,56 | >0.1 | 5,54 | 8,05 | 0 | 1,87 | 0 | 1,65 | 0 |
| IRF9 | AGGGAAAGTCATTCA | 1 | 1005 | 1020 | 5 | 20 | -25,04 | 0,01 | -7,04 | >0.1 | 5,54 | 9,12 | 0 | 1,87 | 0 | 1,46 | 0 |
| IRF9 | CACGAAGCCGAAGAC | 1 | 1148 | 1163 | 148 | 163 | -15,98 | 0,01 | 1,72 | >0.1 | 5,54 | 6,31 | 0 | 1,84 | 1,79 | 2,21 | 0 |
| IRF9 | GAAGCAAGTGAACTT | -1 | 1174 | 1189 | 174 | 189 | -25,26 | 0,01 | -6,89 | >0.1 | 5,54 | 6,13 | 0 | 1,63 | 1,79 | 3,28 | 0 |
| IRF9 | CCCTAGACTAAAACC | -1 | 994 | 1009 | -6 | 9 | -17,53 | 0,01 | 0,44 | >0.1 | 5,54 | 9,06 | 0 | 1,87 | 0 | 1,5 | 0 |
| IRF9 | CCAAAAAAAGAAAAA | -1 | 917 | 932 | -83 | -68 | -23,1 | 0,01 | -7,63 | >0.1 | 5,54 | 6,95 | 0 | 1,89 | 0 | 1,09 | 0 |
| IRF9 | AAAAAAAGAAAAAGA | -1 | 915 | 930 | -85 | -70 | -17,88 | 0,01 | -2,4 | >0.1 | 5,54 | 6,93 | 0 | 1,89 | 0 | 1,12 | 0 |
| IRF9 | AAAGAAAAAGAAGAA | -1 | 911 | 926 | -89 | -74 | -22,54 | 0,01 | -7,08 | >0.1 | 5,54 | 6,88 | 0 | 1,89 | 0 | 1,15 | 0 |
| IRF9 | TTTTAAAGTGAATCA | -1 | 817 | 832 | -183 | -168 | -15,23 | 0,01 | 1,6 | >0.1 | 5,54 | 6,1 | 0 | 1,63 | 0 | 3,55 | 0 |
| IRF9 | GGCCAAATCAAAACC | -1 | 746 | 761 | -254 | -239 | -12,21 | 0,001 | 6,27 | >0.1 | 5,54 | 5,73 | 0 | 1,29 | 0 | 5,92 | 0 |
| IRF9 | TTTGAGACTGAGTCT | -1 | 445 | 460 | -555 | -540 | -23,82 | 0,01 | -9,93 | >0.1 | 2,77 | 4,86 | 0 | 1 | 0 | 5,26 | 0 |
| IRF9 | TCCCAAAGTGCTAGG | -1 | 205 | 220 | -795 | -780 | -25,22 | 0,01 | -11,43 | >0.1 | 3,01 | 4,48 | 0 | 1,6 | 0 | 4,7 | 0 |
| IRF9 | TGCAAAACTTAAAAA | -1 | 75 | 90 | -925 | -910 | -16,66 | 0,01 | -6,64 | >0.1 | 3,81 | 4,33 | 0 | 1,88 | 0 | 0 | 0 |
| IRF8 | AGGTAAGTGAAGGA | 1 | 3 | 17 | -997 | -983 | -25,83 | 0,01 | -12,36 | >0.1 | 5,56 | 4,26 | 0 | 1,86 | 1,79 | 0 | 0 |
| IRF8 | GTGAAGGAGAAGGA | 1 | 9 | 23 | -991 | -977 | -28,64 | 0,01 | -15,16 | >0.1 | 5,56 | 4,27 | 0 | 1,86 | 1,79 | 0 | 0 |
| IRF8 | GAGAAATAGGAAAT | 1 | 56 | 70 | -944 | -930 | -24,01 | 0,01 | -13,8 | >0.1 | 4,02 | 4,31 | 0 | 1,87 | 0 | 0 | 0 |
| IRF8 | ATGGAAATTAAAAC | 1 | 151 | 165 | -849 | -835 | -29,61 | 0,01 | -18,39 | >0.1 | 3,27 | 4,41 | 0 | 1,9 | 0 | 1,65 | 0 |
| IRF8 | TGGAAATTAAAACT | 1 | 152 | 166 | -848 | -834 | -11,75 | 0,001 | -0,53 | >0.1 | 3,26 | 4,41 | 0 | 1,9 | 0 | 1,65 | 0 |
| IRF8 | TTAAAACTGATCAG | 1 | 158 | 172 | -842 | -828 | -26,94 | 0,01 | -14,93 | >0.1 | 3,23 | 4,42 | 0 | 1,9 | 0 | 2,46 | 0 |
| IRF8 | AAAATACAAAAACT | 1 | 295 | 309 | -705 | -691 | -27,2 | 0,01 | -14,24 | >0.1 | 2,69 | 4,6 | 0 | 1,22 | 0 | 4,45 | 0 |
| IRF8 | GTGAACCTGGGAGG | 1 | 374 | 388 | -626 | -612 | -29,35 | 0,01 | -14,59 | >0.1 | 2,56 | 4,73 | 0 | 1,07 | 0 | 6,4 | 0 |
| IRF8 | GACAGACCGAGACT | 1 | 436 | 450 | -564 | -550 | -23,44 | 0,01 | -8,8 | >0.1 | 2,74 | 4,84 | 0 | 1,01 | 0 | 6,05 | 0 |
| IRF8 | AAAAAAAAAAAACT | 1 | 467 | 481 | -533 | -519 | -26,98 | 0,01 | -13,42 | >0.1 | 2,84 | 4,9 | 0 | 0,99 | 0 | 4,83 | 0 |
| IRF8 | TTGGAAGGGAAAAT | 1 | 526 | 540 | -474 | -460 | -20,31 | 0,01 | -7,66 | >0.1 | 3,08 | 5,03 | 0 | 0,99 | 0 | 3,55 | 0 |
| IRF8 | ATGATCTTGAAATA | 1 | 568 | 582 | -432 | -418 | -27,1 | 0,01 | -17,68 | >0.1 | 3,29 | 5,13 | 0 | 1 | 0 | 0 | 0 |
| IRF8 | TTGAAATAAAAATA | 1 | 574 | 588 | -426 | -412 | -27,64 | 0,01 | -18,16 | >0.1 | 3,32 | 5,15 | 0 | 1 | 0 | 0 | 0 |
| IRF8 | CTGAAAACCATACA | 1 | 617 | 631 | -383 | -369 | -10,48 | 0,001 | 0,93 | >0.1 | 3,61 | 5,27 | 0 | 1,04 | 0 | 1,5 | 0 |
| IRF8 | TATTAACCAAAAGT | 1 | 729 | 743 | -271 | -257 | -26,58 | 0,01 | -8,44 | >0.1 | 5,56 | 5,65 | 0 | 1,24 | 0 | 5,69 | 0 |
| IRF8 | CATAAAGTGACAGG | 1 | 760 | 774 | -240 | -226 | -25,34 | 0,01 | -6,74 | >0.1 | 5,56 | 5,79 | 0 | 1,34 | 0 | 5,92 | 0 |
| IRF8 | ATAAGAGAGAAAAT | 1 | 970 | 984 | -30 | -16 | -27,8 | 0,01 | -10,62 | >0.1 | 5,56 | 8,05 | 0 | 1,87 | 0 | 1,7 | 0 |
| IRF8 | ACGAAGCCGAAGAC | 1 | 1149 | 1163 | 149 | 163 | -15,37 | 0,01 | 1,31 | >0.1 | 4,53 | 6,3 | 0 | 1,84 | 1,79 | 2,21 | 0 |
| IRF8 | AAGCAAGTGAACTT | -1 | 1174 | 1188 | 174 | 188 | -18,38 | 0,01 | 0,01 | >0.1 | 5,56 | 6,13 | 0 | 1,63 | 1,79 | 3,28 | 0 |
| IRF8 | CCTAGACTAAAACC | -1 | 994 | 1008 | -6 | 8 | -20,36 | 0,01 | -2,43 | >0.1 | 5,56 | 9,01 | 0 | 1,87 | 0 | 1,5 | 0 |
| IRF8 | CAAAAAAAGAAAAA | -1 | 917 | 931 | -83 | -69 | -24,27 | 0,01 | -8,8 | >0.1 | 5,56 | 6,94 | 0 | 1,89 | 0 | 1,09 | 0 |
| IRF8 | AAAAAAGAAAAAGA | -1 | 915 | 929 | -85 | -71 | -29,73 | 0,01 | -14,25 | >0.1 | 5,56 | 6,92 | 0 | 1,89 | 0 | 1,12 | 0 |
| IRF8 | AAGAAAAAGAAGAA | -1 | 911 | 925 | -89 | -75 | -28,38 | 0,01 | -12,92 | >0.1 | 5,56 | 6,87 | 0 | 1,89 | 0 | 1,15 | 0 |
| IRF8 | TTTAAAGTGAATCA | -1 | 817 | 831 | -183 | -169 | -15,26 | 0,01 | 1,57 | >0.1 | 5,56 | 6,1 | 0 | 1,63 | 0 | 3,55 | 0 |
| IRF8 | GCCAAATCAAAACC | -1 | 746 | 760 | -254 | -240 | -18,41 | 0,01 | 0,07 | >0.1 | 5,56 | 5,72 | 0 | 1,29 | 0 | 5,92 | 0 |
| IRF8 | CTGTTAAAGATACA | -1 | 702 | 716 | -298 | -284 | -28,33 | 0,01 | -11,25 | >0.1 | 5,56 | 5,55 | 0 | 1,17 | 0 | 4,81 | 0 |
| IRF8 | CGGAAAATGTTATA | -1 | 688 | 702 | -312 | -298 | -27,35 | 0,01 | -13,26 | >0.1 | 4,51 | 5,49 | 0 | 1,14 | 0 | 2,95 | 0 |
| IRF8 | TTGAGACTGAGTCT | -1 | 445 | 459 | -555 | -541 | -21,08 | 0,01 | -7,19 | >0.1 | 2,77 | 4,86 | 0 | 1 | 0 | 5,26 | 0 |
| IRF8 | CCCAAAGTGCTAGG | -1 | 205 | 219 | -795 | -781 | -27,7 | 0,01 | -13,91 | >0.1 | 3,01 | 4,48 | 0 | 1,6 | 0 | 4,7 | 0 |
| IRF8 | GCAAAACTTAAAAA | -1 | 75 | 89 | -925 | -911 | -19,49 | 0,01 | -9,47 | >0.1 | 3,81 | 4,33 | 0 | 1,88 | 0 | 0 | 0 |
| MEF2C | GTGTGAGAAATAGGA | 1 | 52 | 67 | -948 | -933 | -0,57 | 0,01 | 9,68 | >0.1 | 4,07 | 4,31 | 0 | 1,87 | 0 | 0 | 0 |
| MEF2C | TGATCAGAAATGGCT | 1 | 165 | 180 | -835 | -820 | 5,69 | 0,01 | 17,61 | >0.1 | 3,19 | 4,43 | 0 | 1,9 | 0 | 2,4 | 0 |
| MEF2C | CTATTAAAAATACAA | 1 | 289 | 304 | -711 | -696 | 9,77 | 0,001 | 22,76 | >0.1 | 2,71 | 4,6 | 0 | 1,24 | 0 | 4,45 | 0 |
| MEF2C | CAGTCTCAAAAAAAA | 1 | 450 | 465 | -550 | -535 | -0,53 | 0,01 | 13,51 | >0.1 | 2,79 | 4,87 | 0 | 1 | 0 | 5,38 | 0 |
| MEF2C | AGTCTCAAAAAAAAA | 1 | 451 | 466 | -549 | -534 | 7,71 | 0,001 | 21,75 | >0.1 | 2,79 | 4,87 | 0 | 1 | 0 | 5,38 | 0 |
| MEF2C | GTCTCAAAAAAAAAA | 1 | 452 | 467 | -548 | -533 | 6,36 | 0,01 | 20,55 | >0.1 | 2,79 | 4,88 | 0 | 1 | 0 | 5,52 | 0 |
| MEF2C | TGATCAGAAATATGA | 1 | 480 | 495 | -520 | -505 | 3,13 | 0,01 | 17,75 | >0.1 | 2,89 | 4,93 | 0 | 0,99 | 0 | 5,8 | 0 |

| | | | | | | | | | | | | | | | | | |
|---|---|---|---|---|---|---|---|---|---|---|---|---|---|---|---|---|---|
| MEF2C | GATCTTGAAATAAAA | 1 | 570 | 585 | -430 | -415 | 10,44 | 0,001 | 19,89 | >0.1 | 3,3 | 5,14 | 0 | 1 | 0 | 0 | 0 |
| MEF2C | GAAATAAAAATAAAT | 1 | 576 | 591 | -424 | -409 | 11,49 | 0,001 | 20,99 | >0.1 | 3,34 | 5,16 | 0 | 1,01 | 0 | 0 | 0 |
| MEF2C | ACTGTAAAATTATAA | 1 | 678 | 693 | -322 | -307 | 0,33 | 0,01 | 14,42 | >0.1 | 4,31 | 5,46 | 0 | 1,12 | 0 | 3,19 | 0 |
| MEF2C | TAACCAAAAGTACCG | 1 | 732 | 747 | -268 | -253 | 1,92 | 0,01 | 20,3 | >0.1 | 5,54 | 5,67 | 0 | 1,25 | 0 | 5,92 | 0 |
| MEF2C | TGTCCAAAAATGTCT | -1 | 1125 | 1140 | 125 | 140 | 5,06 | 0,01 | 24,99 | >0.1 | 5,54 | 6,5 | 0 | 1,84 | 3,22 | 2,82 | 0 |
| MEF2C | GAGCCAGGAAGAGCT | -1 | 1055 | 1070 | 55 | 70 | -0,93 | 0,01 | 16,2 | >0.1 | 5,54 | 7,46 | 0 | 1,86 | 0 | 2,27 | 0 |
| MEF2C | GAGCCAAGATCACAT | -1 | 1024 | 1039 | 24 | 39 | 0,68 | 0,01 | 17,95 | >0.1 | 5,54 | 8,59 | 0 | 1,86 | 0 | 1,28 | 0 |
| MEF2C | AAGACCAAAAAAAGA | -1 | 921 | 936 | -79 | -64 | 3,76 | 0,01 | 19,28 | >0.1 | 5,54 | 7,01 | 0 | 1,88 | 0 | 1,09 | 0 |
| MEF2C | AGACCAAAAAAAGAA | -1 | 920 | 935 | -80 | -65 | 11,83 | 0,001 | 27,34 | >0.1 | 5,54 | 6,99 | 0 | 1,88 | 0 | 1,09 | 0 |
| MEF2C | TGAGCCAATATAAAG | -1 | 876 | 891 | -124 | -109 | 0,33 | 0,01 | 17,38 | >0.1 | 5,54 | 6,54 | 0 | 1,9 | 0 | 3,07 | 0 |
| MEF2C | GATTTTAGAATAATT | -1 | 832 | 847 | -168 | -153 | 3,45 | 0,01 | 20,23 | >0.1 | 5,54 | 6,21 | 0 | 1,75 | 0 | 3,28 | 0 |
| MEF2C | TGAATCAAATAAGAA | -1 | 809 | 824 | -191 | -176 | 1,71 | 0,01 | 18,31 | >0.1 | 5,54 | 6,05 | 0 | 1,58 | 0 | 3,43 | 0 |
| MEF2C | CAAATAAGAATACGT | -1 | 804 | 819 | -196 | -181 | 2,9 | 0,01 | 19,3 | >0.1 | 5,54 | 6,02 | 0 | 1,55 | 0 | 3,28 | 0 |
| MEF2C | ATGGCCAAATCAAAA | -1 | 748 | 763 | -252 | -237 | -0,06 | 0,01 | 18,62 | >0.1 | 5,54 | 5,74 | 0 | 1,3 | 0 | 6,1 | 0 |
| MEF2C | CTGTTAAAGATACAC | -1 | 701 | 716 | -299 | -284 | 1,1 | 0,01 | 18,17 | >0.1 | 5,54 | 5,55 | 0 | 1,17 | 0 | 4,81 | 0 |
| MEF2C | ATTTCATTAATAGGT | -1 | 599 | 614 | -401 | -386 | 2,28 | 0,01 | 13,42 | >0.1 | 3,48 | 5,22 | 0 | 1,02 | 0 | 1,41 | 0 |
| MEF2C | CCCTTCCAAATACAT | -1 | 520 | 535 | -480 | -465 | -0,78 | 0,01 | 11,47 | >0.1 | 3,05 | 5,02 | 0 | 0,99 | 0 | 3,19 | 0 |
| MEF2C | CTTCCAAATACATGA | -1 | 518 | 533 | -482 | -467 | -0,81 | 0,01 | 11,31 | >0.1 | 3,04 | 5,02 | 0 | 0,99 | 0 | 3,07 | 0 |
| MEF2C | GTATTTTTAATAGAG | -1 | 287 | 302 | -713 | -698 | -0,49 | 0,01 | 13,15 | >0.1 | 2,71 | 4,59 | 0 | 1,24 | 0 | 5,09 | 0 |
| MEF2C | GTGCTAGGATTACAG | -1 | 198 | 213 | -802 | -787 | 0,98 | 0,01 | 14,72 | >0.1 | 3,04 | 4,47 | 0 | 1,65 | 0 | 4,59 | 0 |
| MEF2C | TTTTTAAAATTATGA | -1 | 130 | 145 | -870 | -855 | 1,06 | 0,01 | 12,34 | >0.1 | 3,39 | 4,39 | 0 | 1,89 | 0 | 1,61 | 0 |
| MEF2C | AAACTTAAAAAAGCT | -1 | 71 | 86 | -929 | -914 | 8,47 | 0,001 | 18,52 | >0.1 | 3,85 | 4,33 | 0 | 1,88 | 0 | 0 | 0 |
| MEF2B | CATAATTTTAAA | 1 | 131 | 143 | -869 | -857 | -12,5 | 0,01 | -1,22 | >0.1 | 3,39 | 4,39 | 0 | 1,89 | 0 | 1,61 | 0 |
| MEF2B | TCTATTAAAAAT | 1 | 288 | 300 | -712 | -700 | -6,47 | 0,01 | 7,16 | >0.1 | 2,71 | 4,59 | 0 | 1,24 | 0 | 5,09 | 0 |
| MEF2B | ATTAAAAATACA | 1 | 291 | 303 | -709 | -697 | 1,57 | 0,001 | 14,55 | >0.1 | 2,7 | 4,59 | 0 | 1,23 | 0 | 4,45 | 0 |
| MEF2B | TCCATAAAGACA | 1 | 497 | 509 | -503 | -491 | -11,15 | 0,01 | 4,18 | >0.1 | 2,95 | 4,96 | 0 | 0,98 | 0 | 6,44 | 0 |
| MEF2B | TCTTGAAATAAA | 1 | 572 | 584 | -428 | -416 | -9,59 | 0,01 | -0,14 | >0.1 | 3,31 | 5,14 | 0 | 1 | 0 | 0 | 0 |
| MEF2B | AATAAAAATAAA | 1 | 578 | 590 | -422 | -410 | 4 | 0,001 | 13,5 | >0.1 | 3,34 | 5,15 | 0 | 1 | 0 | 0 | 0 |
| MEF2B | CCTATTAATGAA | 1 | 600 | 612 | -400 | -388 | -3,82 | 0,01 | 7,3 | >0.1 | 3,48 | 5,21 | 0 | 1,02 | 0 | 1,41 | 0 |
| MEF2B | ACCAAAAGTACC | 1 | 734 | 746 | -266 | -254 | -8,27 | 0,01 | 10,14 | >0.1 | 5,58 | 5,66 | 0 | 1,24 | 0 | 5,92 | 0 |
| MEF2B | GCCATAAAGTGA | 1 | 758 | 770 | -242 | -230 | -11,43 | 0,01 | 7,34 | >0.1 | 5,58 | 5,77 | 0 | 1,32 | 0 | 6,1 | 0 |
| MEF2B | ACTTTAAAAAAT | 1 | 823 | 835 | -177 | -165 | -10,56 | 0,01 | 6,5 | >0.1 | 5,58 | 6,12 | 0 | 1,65 | 0 | 3,7 | 0 |
| MEF2B | ACTTTATATTGG | 1 | 875 | 887 | -125 | -113 | -7,37 | 0,01 | 9,56 | >0.1 | 5,58 | 6,5 | 0 | 1,9 | 0 | 2,95 | 0 |
| MEF2B | GCTGATAAGAGA | 1 | 966 | 978 | -34 | -22 | -10,76 | 0,01 | 6,24 | >0.1 | 5,58 | 7,85 | 0 | 1,87 | 0 | 1,7 | 0 |
| MEF2B | TCCAAAAATGTC | -1 | 1126 | 1138 | 126 | 138 | -9,35 | 0,01 | 10,61 | >0.1 | 5,58 | 6,49 | 0 | 1,84 | 3,22 | 2,82 | 0 |
| MEF2B | ACCAAAAAAAGA | -1 | 921 | 933 | -79 | -67 | -2,98 | 0,01 | 12,54 | >0.1 | 5,58 | 6,97 | 0 | 1,88 | 0 | 1,09 | 0 |
| MEF2B | AATAAGAATACG | -1 | 805 | 817 | -195 | -183 | -8,23 | 0,01 | 8,18 | >0.1 | 5,58 | 6,01 | 0 | 1,54 | 0 | 3,28 | 0 |
| MEF2B | GTTAAAGATACA | -1 | 702 | 714 | -298 | -286 | -7,58 | 0,01 | 9,51 | >0.1 | 5,58 | 5,54 | 0 | 1,16 | 0 | 4,81 | 0 |
| MEF2B | TATAATTTTACA | -1 | 680 | 692 | -320 | -308 | -12,04 | 0,01 | 2,05 | >0.1 | 4,32 | 5,46 | 0 | 1,12 | 0 | 3,19 | 0 |
| MEF2B | TCTAGAACTAGG | -1 | 655 | 667 | -345 | -333 | -12,7 | 0,01 | 1,42 | >0.1 | 3,96 | 5,38 | 0 | 1,08 | 0 | 3,7 | 0 |
| MEF2B | TTCATTAATAGG | -1 | 600 | 612 | -400 | -388 | -3,86 | 0,01 | 7,26 | >0.1 | 3,48 | 5,21 | 0 | 1,02 | 0 | 1,41 | 0 |
| MEF2B | ATTTTTAATAGA | -1 | 288 | 300 | -712 | -700 | -0,77 | 0,001 | 12,86 | >0.1 | 2,71 | 4,59 | 0 | 1,24 | 0 | 5,09 | 0 |
| MEF2B | TTTAAAATTATG | -1 | 131 | 143 | -869 | -857 | -11,95 | 0,01 | -0,67 | >0.1 | 3,39 | 4,39 | 0 | 1,89 | 0 | 1,61 | 0 |
| MEF2B | ACTTAAAAAAGC | -1 | 72 | 84 | -928 | -916 | -2,63 | 0,01 | 7,42 | >0.1 | 3,85 | 4,32 | 0 | 1,88 | 0 | 0 | 0 |
| MEF2A | CATAATTTTAAA | 1 | 131 | 143 | -869 | -857 | -2,92 | 0,01 | 8,36 | >0.1 | 3,39 | 4,39 | 0 | 1,89 | 0 | 1,61 | 0 |
| MEF2A | TCTATTAAAAAT | 1 | 288 | 300 | -712 | -700 | -2,37 | 0,01 | 11,26 | >0.1 | 2,71 | 4,59 | 0 | 1,24 | 0 | 5,09 | 0 |
| MEF2A | ATTAAAAATACA | 1 | 291 | 303 | -709 | -697 | 4,56 | 0,001 | 17,54 | >0.1 | 2,7 | 4,59 | 0 | 1,23 | 0 | 4,45 | 0 |
| MEF2A | TCTCAAAAAAAA | 1 | 453 | 465 | -547 | -535 | -4,62 | 0,01 | 9,56 | >0.1 | 2,79 | 4,87 | 0 | 1 | 0 | 5,52 | 0 |
| MEF2A | TCCATAAAGACA | 1 | 497 | 509 | -503 | -491 | -7,28 | 0,01 | 8,05 | >0.1 | 2,95 | 4,96 | 0 | 0,98 | 0 | 6,44 | 0 |
| MEF2A | TCTTGAAATAAA | 1 | 572 | 584 | -428 | -416 | 0,14 | 0,01 | 9,59 | >0.1 | 3,31 | 5,14 | 0 | 1 | 0 | 0 | 0 |
| MEF2A | AATAAAAATAAA | 1 | 578 | 590 | -422 | -410 | 9,05 | 0,001 | 18,55 | >0.1 | 3,34 | 5,15 | 0 | 1 | 0 | 0 | 0 |
| MEF2A | CCTATTAATGAA | 1 | 600 | 612 | -400 | -388 | -1,24 | 0,01 | 9,88 | >0.1 | 3,48 | 5,21 | 0 | 1,02 | 0 | 1,41 | 0 |
| MEF2A | CCTAGTTCTAGA | 1 | 655 | 667 | -345 | -333 | -5,23 | 0,01 | 8,89 | >0.1 | 3,96 | 5,38 | 0 | 1,08 | 0 | 3,7 | 0 |
| MEF2A | TGTAAAATTATA | 1 | 680 | 692 | -320 | -308 | -1,26 | 0,01 | 12,83 | >0.1 | 4,32 | 5,46 | 0 | 1,12 | 0 | 3,19 | 0 |
| MEF2A | TCTAAAATCTGT | 1 | 838 | 850 | -162 | -150 | -6,82 | 0,01 | 9,83 | >0.1 | 5,58 | 6,23 | 0 | 1,77 | 0 | 3,07 | 0 |
| MEF2A | GCTGATAAGAGA | 1 | 966 | 978 | -34 | -22 | -7,29 | 0,01 | 9,71 | >0.1 | 5,58 | 7,85 | 0 | 1,87 | 0 | 1,7 | 0 |
| MEF2A | TCCAAAAATGTC | -1 | 1126 | 1138 | 126 | 138 | -6,58 | 0,01 | 13,38 | >0.1 | 5,58 | 6,49 | 0 | 1,84 | 3,22 | 2,82 | 0 |
| MEF2A | TCTCTTATCAGC | -1 | 966 | 978 | -34 | -22 | -5,29 | 0,01 | 11,71 | >0.1 | 5,58 | 7,85 | 0 | 1,87 | 0 | 1,7 | 0 |
| MEF2A | ACCAAAAAAAGA | -1 | 921 | 933 | -79 | -67 | -1,62 | 0,01 | 13,9 | >0.1 | 5,58 | 6,97 | 0 | 1,88 | 0 | 1,09 | 0 |

| | | | | | | | | | | | | | | | | | |
|---|---|---|---|---|---|---|---|---|---|---|---|---|---|---|---|---|---|
| MEF2A | GTTAAAGATACA | -1 | 702 | 714 | -298 | -286 | -5,18 | 0,01 | 11,91 | >0.1 | 5,58 | 5,54 | 0 | 1,16 | 0 | 4,81 | 0 |
| MEF2A | GTTATAATTTTA | -1 | 682 | 694 | -318 | -306 | -7,17 | 0,01 | 6,84 | >0.1 | 4,36 | 5,47 | 0 | 1,12 | 0 | 3,07 | 0 |
| MEF2A | TATAATTTTACA | -1 | 680 | 692 | -320 | -308 | -5,38 | 0,01 | 8,71 | >0.1 | 4,32 | 5,46 | 0 | 1,12 | 0 | 3,19 | 0 |
| MEF2A | TCTAGAACTAGG | -1 | 655 | 667 | -345 | -333 | 0,31 | 0,01 | 14,43 | >0.1 | 3,96 | 5,38 | 0 | 1,08 | 0 | 3,7 | 0 |
| MEF2A | TTCATTAATAGG | -1 | 600 | 612 | -400 | -388 | 1,25 | 0,01 | 12,37 | >0.1 | 3,48 | 5,21 | 0 | 1,02 | 0 | 1,41 | 0 |
| MEF2A | TTTATTTTTATT | -1 | 578 | 590 | -422 | -410 | -7,5 | 0,01 | 2 | >0.1 | 3,34 | 5,15 | 0 | 1 | 0 | 0 | 0 |
| MEF2A | TGTATTTTTAAT | -1 | 291 | 303 | -709 | -697 | -7,25 | 0,01 | 5,73 | >0.1 | 2,7 | 4,59 | 0 | 1,23 | 0 | 4,45 | 0 |
| MEF2A | ATTTTTAATAGA | -1 | 288 | 300 | -712 | -700 | 1,42 | 0,01 | 15,05 | >0.1 | 2,71 | 4,59 | 0 | 1,24 | 0 | 5,09 | 0 |
| MEF2A | TTTAAAATTATG | -1 | 131 | 143 | -869 | -857 | -1,13 | 0,01 | 10,15 | >0.1 | 3,39 | 4,39 | 0 | 1,89 | 0 | 1,61 | 0 |
| MEF2A | ACTTAAAAAAGC | -1 | 72 | 84 | -928 | -916 | -3,47 | 0,01 | 6,58 | >0.1 | 3,85 | 4,32 | 0 | 1,88 | 0 | 0 | 0 |
| CREM | AAAGAAGTCA | 1 | 30 | 40 | -970 | -960 | -11,5 | 0,01 | -0,87 | >0.1 | 4,48 | 4,28 | 0 | 1,87 | 0 | 0 | 0 |
| CREM | TTTTAAGTTT | 1 | 76 | 86 | -924 | -914 | -7,39 | 0,01 | 2,64 | >0.1 | 3,82 | 4,32 | 0 | 1,88 | 0 | 0 | 0 |
| CREM | GTTGACCTGT | 1 | 106 | 116 | -894 | -884 | -4,43 | 0,01 | 6,12 | >0.1 | 3,57 | 4,36 | 0 | 1,89 | 0 | 0,75 | 0 |
| CREM | AATGGCGTGA | 1 | 368 | 378 | -632 | -622 | -0,46 | 0,001 | 14,59 | >0.1 | 2,54 | 4,71 | 0 | 1,07 | 0 | 6,73 | 0 |
| CREM | TATGAATTCC | 1 | 490 | 500 | -510 | -500 | -12,16 | 0,01 | 2,96 | >0.1 | 2,92 | 4,94 | 0 | 0,98 | 0 | 6,27 | 0 |
| CREM | TGTGAGATAA | 1 | 590 | 600 | -410 | -400 | -11,57 | 0,01 | -1,97 | >0.1 | 3,41 | 5,18 | 0 | 1,01 | 0 | 0 | 0 |
| CREM | GATAACCTAT | 1 | 595 | 605 | -405 | -395 | -11,33 | 0,01 | -1,68 | >0.1 | 3,44 | 5,19 | 0 | 1,01 | 0 | 0 | 0 |
| CREM | TTTTCCGTGT | 1 | 695 | 705 | -305 | -295 | -5,9 | 0,01 | 9,31 | >0.1 | 5,61 | 5,5 | 0 | 1,14 | 0 | 2,95 | 0 |
| CREM | GGAGAGGTAA | 1 | 772 | 782 | -228 | -218 | -10,99 | 0,01 | 7,73 | >0.1 | 5,61 | 5,83 | 0 | 1,37 | 0 | 5,92 | 0 |
| CREM | AATAACGTAT | 1 | 800 | 810 | -200 | -190 | -0,11 | 0,001 | 16,51 | >0.1 | 5,61 | 5,97 | 0 | 1,5 | 0 | 3,55 | 0 |
| CREM | TGTTACATAT | 1 | 847 | 857 | -153 | -143 | -2,82 | 0,01 | 13,76 | >0.1 | 5,61 | 6,28 | 0 | 1,83 | 0 | 2,86 | 0 |
| CREM | GACGATGTCA | 1 | 1045 | 1055 | 45 | 55 | -9,94 | 0,01 | 5,88 | >0.1 | 5,61 | 7,7 | 0 | 1,86 | 0 | 0,65 | 0 |
| CREM | GTTGCTGTAA | 1 | 1082 | 1092 | 82 | 92 | -12,54 | 0,01 | 5,92 | >0.1 | 5,61 | 6,99 | 0 | 1,85 | 1,79 | 2,21 | 0 |
| CREM | CTTGACATCG | -1 | 1047 | 1057 | 47 | 57 | -2,75 | 0,01 | 13,79 | >0.1 | 5,61 | 7,65 | 0 | 1,86 | 0 | 1,42 | 0 |
| CREM | GATCACATCC | -1 | 1022 | 1032 | 22 | 32 | -11,12 | 0,01 | 6,31 | >0.1 | 5,61 | 8,68 | 0 | 1,86 | 0 | 1,28 | 0 |
| CREM | AATGACTTTC | -1 | 1008 | 1018 | 8 | 18 | -3,1 | 0,01 | 14,86 | >0.1 | 5,61 | 9,07 | 0 | 1,87 | 0 | 1,42 | 0 |
| CREM | GAATACGTTA | -1 | 802 | 812 | -198 | -188 | -2,29 | 0,01 | 14,24 | >0.1 | 5,61 | 5,98 | 0 | 1,51 | 0 | 3,43 | 0 |
| CREM | CCTTACCTCT | -1 | 774 | 784 | -226 | -216 | -4,15 | 0,01 | 14,77 | >0.1 | 5,61 | 5,83 | 0 | 1,37 | 0 | 6,1 | 0 |
| CREM | AGTGACCAAA | -1 | 671 | 681 | -329 | -319 | -12,06 | 0,01 | 2,17 | >0.1 | 4,16 | 5,42 | 0 | 1,1 | 0 | 3,55 | 0 |
| CREM | CATGACTTTG | -1 | 513 | 523 | -487 | -477 | -2,61 | 0,01 | 9,24 | >0.1 | 3,01 | 4,99 | 0 | 0,98 | 0 | 2,86 | 0 |
| CREM | GTTCACGCCA | -1 | 370 | 380 | -630 | -620 | -11,87 | 0,01 | 2,73 | >0.1 | 2,54 | 4,72 | 0 | 1,07 | 0 | 6,27 | 0 |
| CREM | CCTGACCTTG | -1 | 239 | 249 | -761 | -751 | -4,5 | 0,01 | 8,74 | >0.1 | 2,88 | 4,52 | 0 | 1,41 | 0 | 4,42 | 0 |
| CREM | TATGACTACT | -1 | 125 | 135 | -875 | -865 | -12,16 | 0,01 | -0,76 | >0.1 | 3,43 | 4,38 | 0 | 1,89 | 0 | 1,7 | 0 |
| CREM | GCTGACTTCT | -1 | 32 | 42 | -968 | -958 | -2,19 | 0,01 | 8,4 | >0.1 | 4,44 | 4,28 | 0 | 1,87 | 0 | 0 | 0 |
| CREM | ACTTACCTAT | -1 | 1 | 11 | -999 | -989 | -4,36 | 0,01 | 9,16 | >0.1 | 5,61 | 4,25 | 0 | 1,86 | 1,79 | 0 | 0 |
| MEF2D | CATAATTTTAAA | 1 | 131 | 143 | -869 | -857 | -12,18 | 0,01 | -0,9 | >0.1 | 3,39 | 4,39 | 0 | 1,89 | 0 | 1,61 | 0 |
| MEF2D | TTAAAAAATGGC | 1 | 138 | 150 | -862 | -850 | -9,8 | 0,01 | 1,44 | >0.1 | 3,35 | 4,39 | 0 | 1,89 | 0 | 1,61 | 0 |
| MEF2D | TCTATTAAAAAT | 1 | 288 | 300 | -712 | -700 | -6,61 | 0,01 | 7,02 | >0.1 | 2,71 | 4,59 | 0 | 1,24 | 0 | 5,09 | 0 |
| MEF2D | ATTAAAAATACA | 1 | 291 | 303 | -709 | -697 | 2,74 | 0,001 | 15,72 | >0.1 | 2,7 | 4,59 | 0 | 1,23 | 0 | 4,45 | 0 |
| MEF2D | TCTCAAAAAAAA | 1 | 453 | 465 | -547 | -535 | -10,35 | 0,01 | 3,83 | >0.1 | 2,79 | 4,87 | 0 | 1 | 0 | 5,52 | 0 |
| MEF2D | TCAAAAAAAAAA | 1 | 455 | 467 | -545 | -533 | -12,2 | 0,01 | 1,99 | >0.1 | 2,8 | 4,88 | 0 | 1 | 0 | 5,52 | 0 |
| MEF2D | TCCATAAAGACA | 1 | 497 | 509 | -503 | -491 | -12,1 | 0,01 | 3,23 | >0.1 | 2,95 | 4,96 | 0 | 0,98 | 0 | 6,44 | 0 |
| MEF2D | TCTTGAAATAAA | 1 | 572 | 584 | -428 | -416 | -6,23 | 0,01 | 3,22 | >0.1 | 3,31 | 5,14 | 0 | 1 | 0 | 0 | 0 |
| MEF2D | AATAAAAATAAA | 1 | 578 | 590 | -422 | -410 | 5,63 | 0,001 | 15,13 | >0.1 | 3,34 | 5,15 | 0 | 1 | 0 | 0 | 0 |
| MEF2D | CCTATTAATGAA | 1 | 600 | 612 | -400 | -388 | -2,69 | 0,01 | 8,43 | >0.1 | 3,48 | 5,21 | 0 | 1,02 | 0 | 1,41 | 0 |
| MEF2D | CCTAGTTCTAGA | 1 | 655 | 667 | -345 | -333 | -13,69 | 0,01 | 0,43 | >0.1 | 3,96 | 5,38 | 0 | 1,08 | 0 | 3,7 | 0 |
| MEF2D | TGTAAAATTATA | 1 | 680 | 692 | -320 | -308 | -13,8 | 0,01 | 0,29 | >0.1 | 4,32 | 5,46 | 0 | 1,12 | 0 | 3,19 | 0 |
| MEF2D | TAAAATTATAAC | 1 | 682 | 694 | -318 | -306 | -13,01 | 0,01 | 1 | >0.1 | 4,36 | 5,47 | 0 | 1,12 | 0 | 3,07 | 0 |
| MEF2D | GAAAATATTAAC | 1 | 724 | 736 | -276 | -264 | -13,93 | 0,01 | 3,81 | >0.1 | 5,58 | 5,62 | 0 | 1,22 | 0 | 5,32 | 0 |
| MEF2D | ACCAAAAGTACC | 1 | 734 | 746 | -266 | -254 | -9,08 | 0,01 | 9,33 | >0.1 | 5,58 | 5,66 | 0 | 1,24 | 0 | 5,92 | 0 |
| MEF2D | ACTTTAAAAAAT | 1 | 823 | 835 | -177 | -165 | -13,22 | 0,01 | 3,84 | >0.1 | 5,58 | 6,12 | 0 | 1,65 | 0 | 3,7 | 0 |
| MEF2D | GCTGATAAGAGA | 1 | 966 | 978 | -34 | -22 | -9,33 | 0,01 | 7,67 | >0.1 | 5,58 | 7,85 | 0 | 1,87 | 0 | 1,7 | 0 |
| MEF2D | TCCAAAAATGTC | -1 | 1126 | 1138 | 126 | 138 | -9,86 | 0,01 | 10,1 | >0.1 | 5,58 | 6,49 | 0 | 1,84 | 3,22 | 2,82 | 0 |
| MEF2D | TCTCTTATCAGC | -1 | 966 | 978 | -34 | -22 | -14,24 | 0,01 | 2,76 | >0.1 | 5,58 | 7,85 | 0 | 1,87 | 0 | 1,7 | 0 |
| MEF2D | ACCAAAAAAAGA | -1 | 921 | 933 | -79 | -67 | -3,95 | 0,01 | 11,57 | >0.1 | 5,58 | 6,97 | 0 | 1,88 | 0 | 1,09 | 0 |
| MEF2D | GTTAAAGATACA | -1 | 702 | 714 | -298 | -286 | -7,76 | 0,01 | 9,33 | >0.1 | 5,58 | 5,54 | 0 | 1,16 | 0 | 4,81 | 0 |
| MEF2D | TATAATTTTACA | -1 | 680 | 692 | -320 | -308 | -12,57 | 0,01 | 1,52 | >0.1 | 4,32 | 5,46 | 0 | 1,12 | 0 | 3,19 | 0 |
| MEF2D | TCTAGAACTAGG | -1 | 655 | 667 | -345 | -333 | -7,66 | 0,01 | 6,46 | >0.1 | 3,96 | 5,38 | 0 | 1,08 | 0 | 3,7 | 0 |
| MEF2D | TTCATTAATAGG | -1 | 600 | 612 | -400 | -388 | -4,56 | 0,01 | 6,56 | >0.1 | 3,48 | 5,21 | 0 | 1,02 | 0 | 1,41 | 0 |

| | | | | | | | | | | | | | | | | | |
|---|---|---|---|---|---|---|---|---|---|---|---|---|---|---|---|---|---|
| MEF2D | ATTTTTAATAGA | -1 | 288 | 300 | -712 | -700 | -1,51 | 0,01 | 12,12 | >0.1 | 2,71 | 4,59 | 0 | 1,24 | 0 | 5,09 | 0 |
| MEF2D | TTTAAAATTATG | -1 | 131 | 143 | -869 | -857 | -11,26 | 0,01 | 0,02 | >0.1 | 3,39 | 4,39 | 0 | 1,89 | 0 | 1,61 | 0 |
| MEF2D | ACTTAAAAAAGC | -1 | 72 | 84 | -928 | -916 | -6,1 | 0,01 | 3,95 | >0.1 | 3,85 | 4,32 | 0 | 1,88 | 0 | 0 | 0 |
| PDX1 | GAAATTAA | 1 | 154 | 162 | -846 | -838 | 6,16 | 0,01 | 23,39 | >0.1 | 3,27 | 4,41 | 0 | 1,9 | 0 | 1,65 | 6,01 |
| PDX1 | TCTATTAA | 1 | 288 | 296 | -712 | -704 | 2 | 0,01 | 21,74 | >0.1 | 2,72 | 4,58 | 0 | 1,24 | 0 | 5,19 | 6,01 |
| PDX1 | AAACTTAG | 1 | 304 | 312 | -696 | -688 | -0,96 | 0,01 | 18,51 | >0.1 | 2,67 | 4,61 | 0 | 1,19 | 0 | 4,99 | 6,01 |
| PDX1 | CCAAGTAG | 1 | 545 | 553 | -455 | -447 | -4,14 | 0,01 | 11,07 | >0.1 | 3,15 | 5,06 | 0 | 0,98 | 0 | 0 | 6,01 |
| PDX1 | CCTATTAA | 1 | 600 | 608 | -400 | -392 | 3,64 | 0,01 | 20,69 | >0.1 | 3,47 | 5,2 | 0 | 1,01 | 0 | 1,36 | 6,01 |
| PDX1 | AACATTAT | 1 | 637 | 645 | -363 | -355 | 3,01 | 0,01 | 20,79 | >0.1 | 3,76 | 5,31 | 0 | 1,05 | 0 | 1,66 | 6,01 |
| PDX1 | AAAATTAT | 1 | 683 | 691 | -317 | -309 | 3,59 | 0,01 | 23,57 | >0.1 | 4,34 | 5,46 | 0 | 1,12 | 0 | 3,07 | 6,01 |
| PDX1 | AATATTAA | 1 | 727 | 735 | -273 | -265 | -1,01 | 0,01 | 22,78 | >0.1 | 5,64 | 5,62 | 0 | 1,21 | 0 | 5,32 | 6,01 |
| PDX1 | ATTCTTAT | 1 | 808 | 816 | -192 | -184 | -5,17 | 0,01 | 17,29 | >0.1 | 5,64 | 6 | 0 | 1,53 | 0 | 3,28 | 6,01 |
| PDX1 | CTCATGAG | 1 | 984 | 992 | -16 | -8 | -4,3 | 0,01 | 19,22 | >0.1 | 5,64 | 8,43 | 0 | 1,87 | 0 | 1,57 | 6,01 |
| PDX1 | GTCATTCA | 1 | 1012 | 1020 | 12 | 20 | -5,05 | 0,01 | 18,77 | >0.1 | 5,64 | 8,92 | 0 | 1,86 | 0 | 1,39 | 6,01 |
| PDX1 | GGAATTAT | 1 | 1191 | 1199 | 191 | 199 | -5,21 | 0,01 | 19,57 | >0.1 | 5,64 | 6,02 | 0 | 1,5 | 1,79 | 3,82 | 6,01 |
| PDX1 | GCAGTTAC | -1 | 1088 | 1096 | 88 | 96 | -5,24 | 0,01 | 19,17 | >0.1 | 5,64 | 6,9 | 0 | 1,85 | 1,79 | 2,21 | 6,01 |
| PDX1 | CTCATGAG | -1 | 984 | 992 | -16 | -8 | -4,3 | 0,01 | 19,22 | >0.1 | 5,64 | 8,43 | 0 | 1,87 | 0 | 1,57 | 6,01 |
| PDX1 | AACCTTAC | -1 | 778 | 786 | -222 | -214 | -2,79 | 0,01 | 22,62 | >0.1 | 5,64 | 5,84 | 0 | 1,38 | 0 | 6,54 | 6,01 |
| PDX1 | TTCATTAA | -1 | 604 | 612 | -396 | -388 | 7,79 | 0,001 | 24,93 | >0.1 | 3,49 | 5,21 | 0 | 1,02 | 0 | 1,41 | 6,01 |
| PDX1 | AAAATTAT | -1 | 132 | 140 | -868 | -860 | 3,59 | 0,01 | 20,84 | >0.1 | 3,4 | 4,38 | 0 | 1,89 | 0 | 1,57 | 6,01 |
| PDX1 | AAACTTAA | -1 | 78 | 86 | -922 | -914 | -1,86 | 0,01 | 14,16 | >0.1 | 3,81 | 4,32 | 0 | 1,88 | 0 | 0 | 6,01 |
| PDX1 | TCACTTAC | -1 | 5 | 13 | -995 | -987 | 0,8 | 0,01 | 20,36 | >0.1 | 5,64 | 4,26 | 0 | 1,86 | 1,79 | 0 | 6,01 |
| ISX | ATAATTTT | 1 | 132 | 140 | -868 | -860 | 2,16 | 0,01 | 19,86 | >0.1 | 3,4 | 4,38 | 0 | 1,89 | 0 | 1,57 | 6,45 |
| ISX | GAAATTAA | 1 | 154 | 162 | -846 | -838 | 2,99 | 0,01 | 20,66 | >0.1 | 3,27 | 4,41 | 0 | 1,9 | 0 | 1,65 | 6,45 |
| ISX | GCACTTTG | 1 | 209 | 217 | -791 | -783 | -1,42 | 0,01 | 18,79 | >0.1 | 3 | 4,47 | 0 | 1,57 | 0 | 4,7 | 6,45 |
| ISX | TCTATTAA | 1 | 288 | 296 | -712 | -704 | 3,7 | 0,01 | 23,89 | >0.1 | 2,72 | 4,58 | 0 | 1,24 | 0 | 5,19 | 6,45 |
| ISX | CCAAGTAG | 1 | 545 | 553 | -455 | -447 | -1,12 | 0,01 | 14,53 | >0.1 | 3,15 | 5,06 | 0 | 0,98 | 0 | 0 | 6,45 |
| ISX | CCTATTAA | 1 | 600 | 608 | -400 | -392 | 4,93 | 0,01 | 22,43 | >0.1 | 3,47 | 5,2 | 0 | 1,01 | 0 | 1,36 | 6,45 |
| ISX | TTAATGAA | 1 | 604 | 612 | -396 | -388 | 3,79 | 0,01 | 21,38 | >0.1 | 3,49 | 5,21 | 0 | 1,02 | 0 | 1,41 | 6,45 |
| ISX | AAAATTAT | 1 | 683 | 691 | -317 | -309 | 0,85 | 0,01 | 21,28 | >0.1 | 4,34 | 5,46 | 0 | 1,12 | 0 | 3,07 | 6,45 |
| ISX | CTTATTTG | 1 | 811 | 819 | -189 | -181 | -0,51 | 0,01 | 22,43 | >0.1 | 5,64 | 6,02 | 0 | 1,55 | 0 | 3,28 | 6,45 |
| ISX | AAAATTAT | 1 | 830 | 838 | -170 | -162 | 0,85 | 0,01 | 24,04 | >0.1 | 5,64 | 6,14 | 0 | 1,67 | 0 | 3,28 | 6,45 |
| ISX | CTCATGAG | 1 | 984 | 992 | -16 | -8 | 0,46 | 0,01 | 24,43 | >0.1 | 5,64 | 8,43 | 0 | 1,87 | 0 | 1,57 | 6,45 |
| ISX | CCACTGAA | -1 | 1016 | 1024 | 16 | 24 | -1,07 | 0,01 | 23,09 | >0.1 | 5,64 | 8,81 | 0 | 1,86 | 0 | 1,39 | 6,45 |
| ISX | CTCATGAG | -1 | 984 | 992 | -16 | -8 | 0,46 | 0,01 | 24,43 | >0.1 | 5,64 | 8,43 | 0 | 1,87 | 0 | 1,57 | 6,45 |
| ISX | ATAATTTT | -1 | 830 | 838 | -170 | -162 | 2,16 | 0,01 | 25,35 | >0.1 | 5,64 | 6,14 | 0 | 1,67 | 0 | 3,28 | 6,45 |
| ISX | ATAATTTT | -1 | 683 | 691 | -317 | -309 | 2,16 | 0,01 | 22,59 | >0.1 | 4,34 | 5,46 | 0 | 1,12 | 0 | 3,07 | 6,45 |
| ISX | ACAATTTC | -1 | 609 | 617 | -391 | -383 | 3,19 | 0,01 | 20,97 | >0.1 | 3,53 | 5,23 | 0 | 1,02 | 0 | 1,55 | 6,45 |
| ISX | TTCATTAA | -1 | 604 | 612 | -396 | -388 | 4,78 | 0,01 | 22,37 | >0.1 | 3,49 | 5,21 | 0 | 1,02 | 0 | 1,41 | 6,45 |
| ISX | ACATTTAT | -1 | 585 | 593 | -415 | -407 | 0,31 | 0,01 | 16,3 | >0.1 | 3,37 | 5,16 | 0 | 1 | 0 | 0 | 6,45 |
| ISX | GCCATTTC | -1 | 171 | 179 | -829 | -821 | -1,04 | 0,01 | 18,32 | >0.1 | 3,18 | 4,43 | 0 | 1,87 | 0 | 3,43 | 6,45 |
| ISX | TTAATTTC | -1 | 154 | 162 | -846 | -838 | 3,48 | 0,01 | 21,15 | >0.1 | 3,27 | 4,41 | 0 | 1,9 | 0 | 1,65 | 6,45 |
| ISX | AAAATTAT | -1 | 132 | 140 | -868 | -860 | 0,85 | 0,01 | 18,55 | >0.1 | 3,4 | 4,38 | 0 | 1,89 | 0 | 1,57 | 6,45 |
| ISX | CTCATTTC | -1 | 65 | 73 | -935 | -927 | -0,21 | 0,01 | 16,38 | >0.1 | 3,95 | 4,31 | 0 | 1,88 | 0 | 0 | 6,45 |
| ISX | CCTATTTC | -1 | 58 | 66 | -942 | -934 | -1,29 | 0,01 | 15,38 | >0.1 | 4,03 | 4,3 | 0 | 1,87 | 0 | 0 | 6,45 |
| ISX | TCACTTAC | -1 | 5 | 13 | -995 | -987 | 2,79 | 0,01 | 22,79 | >0.1 | 5,64 | 4,26 | 0 | 1,86 | 1,79 | 0 | 6,45 |
| CLOCK | AACCAGTGTG | 1 | 47 | 57 | -953 | -943 | 3,07 | 0,01 | 13,41 | >0.1 | 4,17 | 4,3 | 0 | 1,87 | 0 | 0 | 0 |
| CLOCK | GACCTGTGGA | 1 | 109 | 119 | -891 | -881 | 2,31 | 0,01 | 12,84 | >0.1 | 3,54 | 4,36 | 0 | 1,89 | 0 | 0,75 | 0 |
| CLOCK | AAAAAATGGC | 1 | 140 | 150 | -860 | -850 | -0,48 | 0,01 | 10,8 | >0.1 | 3,34 | 4,39 | 0 | 1,89 | 0 | 1,65 | 0 |
| CLOCK | ATCACAAGGT | 1 | 235 | 245 | -765 | -755 | -1,65 | 0,01 | 11,93 | >0.1 | 2,9 | 4,51 | 0 | 1,43 | 0 | 4,74 | 0 |
| CLOCK | AACACGGTGA | 1 | 269 | 279 | -731 | -721 | 1,48 | 0,01 | 15,2 | >0.1 | 2,78 | 4,56 | 0 | 1,3 | 0 | 5,09 | 0 |
| CLOCK | ACCCCGTCTC | 1 | 280 | 290 | -720 | -710 | 1,1 | 0,01 | 14,77 | >0.1 | 2,74 | 4,57 | 0 | 1,26 | 0 | 5,09 | 0 |
| CLOCK | TGGGCGTGGT | 1 | 313 | 323 | -687 | -677 | -0,94 | 0,01 | 13,63 | >0.1 | 2,64 | 4,62 | 0 | 1,17 | 0 | 6,13 | 0 |
| CLOCK | GGCACCTGTA | 1 | 327 | 337 | -673 | -663 | 1,23 | 0,01 | 15,19 | >0.1 | 2,6 | 4,65 | 0 | 1,14 | 0 | 5,56 | 0 |
| CLOCK | GACAAGGGCA | 1 | 505 | 515 | -495 | -485 | 1,98 | 0,01 | 16,6 | >0.1 | 2,98 | 4,98 | 0 | 0,98 | 0 | 5,68 | 0 |
| CLOCK | GGAAAATGTT | 1 | 533 | 543 | -467 | -457 | -1,63 | 0,01 | 9,1 | >0.1 | 3,1 | 5,04 | 0 | 0,98 | 0 | 1,61 | 0 |
| CLOCK | AACCCAAGTT | 1 | 951 | 961 | -49 | -39 | -1,03 | 0,01 | 15,61 | >0.1 | 5,61 | 7,42 | 0 | 1,88 | 0 | 1,74 | 0 |
| CLOCK | CCCAAGTTCA | 1 | 953 | 963 | -47 | -37 | -1,43 | 0,01 | 15,21 | >0.1 | 5,61 | 7,46 | 0 | 1,88 | 0 | 1,7 | 0 |
| CLOCK | AAGACATTTT | 1 | 1124 | 1134 | 124 | 134 | -1,74 | 0,01 | 18,24 | >0.1 | 5,61 | 6,51 | 0 | 1,84 | 3,22 | 2,79 | 0 |

| | | | | | | | | | | | | | | | | | |
|---|---|---|---|---|---|---|---|---|---|---|---|---|---|---|---|---|---|
| CLOCK | GACAAGTTTA | 1 | 1136 | 1146 | 136 | 146 | 2,81 | 0,01 | 22,98 | >0.1 | 5,61 | 6,41 | 0 | 1,84 | 3,22 | 3,09 | 0 |
| CLOCK | AGCAAGTGAA | -1 | 1177 | 1187 | 177 | 187 | -0,62 | 0,01 | 17,76 | >0.1 | 5,61 | 6,11 | 0 | 1,61 | 1,79 | 3,26 | 0 |
| CLOCK | AGAACAGGTC | -1 | 1160 | 1170 | 160 | 170 | -1,44 | 0,01 | 15,19 | >0.1 | 4,4 | 6,22 | 0 | 1,76 | 1,79 | 2,46 | 0 |
| CLOCK | CAAAAATGTC | -1 | 1126 | 1136 | 126 | 136 | -1,21 | 0,01 | 18,71 | >0.1 | 5,61 | 6,49 | 0 | 1,84 | 3,22 | 2,75 | 0 |
| CLOCK | AAAATGTCTT | -1 | 1124 | 1134 | 124 | 134 | -1,43 | 0,01 | 18,55 | >0.1 | 5,61 | 6,51 | 0 | 1,84 | 3,22 | 2,79 | 0 |
| CLOCK | GGAAAATGTT | -1 | 691 | 701 | -309 | -299 | -1,63 | 0,01 | 12,6 | >0.1 | 4,53 | 5,49 | 0 | 1,14 | 0 | 3,07 | 0 |
| CLOCK | TACAGGTGCC | -1 | 327 | 337 | -673 | -663 | 2,22 | 0,01 | 16,18 | >0.1 | 2,6 | 4,65 | 0 | 1,14 | 0 | 5,56 | 0 |
| CLOCK | ACCACGCCCA | -1 | 313 | 323 | -687 | -677 | -0,92 | 0,01 | 13,65 | >0.1 | 2,64 | 4,62 | 0 | 1,17 | 0 | 6,13 | 0 |
| CLOCK | GAGACGGGGT | -1 | 280 | 290 | -720 | -710 | 0,09 | 0,01 | 13,76 | >0.1 | 2,74 | 4,57 | 0 | 1,26 | 0 | 5,09 | 0 |
| CLOCK | TCACCGTGTT | -1 | 269 | 279 | -731 | -721 | 0,77 | 0,01 | 14,49 | >0.1 | 2,78 | 4,56 | 0 | 1,3 | 0 | 5,09 | 0 |
| CLOCK | CCAAAGTGCT | -1 | 208 | 218 | -792 | -782 | -0,18 | 0,01 | 13,58 | >0.1 | 3 | 4,48 | 0 | 1,58 | 0 | 4,7 | 0 |
| CLOCK | TCCACAGGTC | -1 | 109 | 119 | -891 | -881 | 2,67 | 0,01 | 13,2 | >0.1 | 3,54 | 4,36 | 0 | 1,89 | 0 | 0,75 | 0 |
| ONECUT3 | TAAAAATACAAAAA | 1 | 293 | 307 | -707 | -693 | -4,97 | 0,01 | 8 | >0.1 | 2,69 | 4,6 | 0 | 1,23 | 0 | 4,45 | 0 |
| ONECUT3 | AAAAATACAAAAAC | 1 | 294 | 308 | -706 | -692 | -4,25 | 0,01 | 8,8 | >0.1 | 2,69 | 4,6 | 0 | 1,22 | 0 | 4,53 | 0 |
| ONECUT3 | CAAAAAAAAAAAAA | 1 | 456 | 470 | -544 | -530 | -4,62 | 0,01 | 9,7 | >0.1 | 2,8 | 4,88 | 0 | 1 | 0 | 5,63 | 0 |
| ONECUT3 | AAAAAAAAAAAAAA | 1 | 457 | 471 | -543 | -529 | -3,64 | 0,01 | 10,57 | >0.1 | 2,81 | 4,88 | 0 | 1 | 0 | 5,52 | 0 |
| ONECUT3 | AAAAAAAAAAAAAA | 1 | 458 | 472 | -542 | -528 | -3,64 | 0,01 | 10,58 | >0.1 | 2,81 | 4,89 | 0 | 1 | 0 | 5,52 | 0 |
| ONECUT3 | AAAAAAAAAAAAAA | 1 | 459 | 473 | -541 | -527 | -3,64 | 0,01 | 10,69 | >0.1 | 2,81 | 4,89 | 0 | 1 | 0 | 5,63 | 0 |
| ONECUT3 | AAAAAAAAAAAAAA | 1 | 460 | 474 | -540 | -526 | -3,64 | 0,01 | 10,7 | >0.1 | 2,82 | 4,89 | 0 | 1 | 0 | 5,63 | 0 |
| ONECUT3 | AAAAAAAAAAAAAA | 1 | 461 | 475 | -539 | -525 | -3,64 | 0,01 | 10,85 | >0.1 | 2,82 | 4,89 | 0 | 1 | 0 | 5,78 | 0 |
| ONECUT3 | AAAAAAAAAAAAAA | 1 | 462 | 476 | -538 | -524 | -3,64 | 0,01 | 10,85 | >0.1 | 2,83 | 4,89 | 0 | 1 | 0 | 5,78 | 0 |
| ONECUT3 | AAAAAAAAAAAAAA | 1 | 463 | 477 | -537 | -523 | -3,64 | 0,01 | 10,86 | >0.1 | 2,83 | 4,9 | 0 | 1 | 0 | 5,78 | 0 |
| ONECUT3 | AAAAAAAAAAAAAA | 1 | 464 | 478 | -536 | -522 | -3,64 | 0,01 | 10,72 | >0.1 | 2,83 | 4,9 | 0 | 0,99 | 0 | 5,63 | 0 |
| ONECUT3 | AAAAAAAAAAAAAA | 1 | 465 | 479 | -535 | -521 | -3,64 | 0,01 | 10,87 | >0.1 | 2,83 | 4,9 | 0 | 0,99 | 0 | 5,78 | 0 |
| ONECUT3 | AAAAAAAAAAAAAC | 1 | 466 | 480 | -534 | -520 | -3,7 | 0,01 | 10,96 | >0.1 | 2,84 | 4,9 | 0 | 0,99 | 0 | 5,92 | 0 |
| ONECUT3 | AAAAAAAAAAAACT | 1 | 467 | 481 | -533 | -519 | -4,9 | 0,01 | 8,66 | >0.1 | 2,84 | 4,9 | 0 | 0,99 | 0 | 4,83 | 0 |
| ONECUT3 | AAAAAACTGATCAG | 1 | 473 | 487 | -527 | -513 | -3,3 | 0,01 | 10,56 | >0.1 | 2,86 | 4,92 | 0 | 0,99 | 0 | 5,09 | 0 |
| ONECUT3 | GAAATATGAATTCC | 1 | 486 | 500 | -514 | -500 | -4,42 | 0,01 | 10,69 | >0.1 | 2,91 | 4,94 | 0 | 0,99 | 0 | 6,27 | 0 |
| ONECUT3 | ATGAATTCCATAAA | 1 | 491 | 505 | -509 | -495 | -2,17 | 0,01 | 13,14 | >0.1 | 2,93 | 4,95 | 0 | 0,99 | 0 | 6,44 | 0 |
| ONECUT3 | TTGAAATAAAAATA | 1 | 574 | 588 | -426 | -412 | -5,02 | 0,01 | 4,46 | >0.1 | 3,32 | 5,15 | 0 | 1 | 0 | 0 | 0 |
| ONECUT3 | GAAATAAAAATAAA | 1 | 576 | 590 | -424 | -410 | -3,85 | 0,01 | 5,65 | >0.1 | 3,33 | 5,15 | 0 | 1,01 | 0 | 0 | 0 |
| ONECUT3 | TAAAAATAAATGTG | 1 | 580 | 594 | -420 | -406 | 2,08 | 0,01 | 11,61 | >0.1 | 3,36 | 5,17 | 0 | 1,01 | 0 | 0 | 0 |
| ONECUT3 | AGATAACCTATTAA | 1 | 594 | 608 | -406 | -392 | -1,97 | 0,01 | 9,14 | >0.1 | 3,45 | 5,2 | 0 | 1,02 | 0 | 1,44 | 0 |
| ONECUT3 | CTGAAAACCATACA | 1 | 617 | 631 | -383 | -369 | -3,2 | 0,01 | 8,21 | >0.1 | 3,61 | 5,27 | 0 | 1,04 | 0 | 1,5 | 0 |
| ONECUT3 | CAAACACCAACATT | 1 | 629 | 643 | -371 | -357 | -4,77 | 0,01 | 6,95 | >0.1 | 3,71 | 5,3 | 0 | 1,05 | 0 | 1,66 | 0 |
| ONECUT3 | TATTAACCAAAAGT | 1 | 729 | 743 | -271 | -257 | -2,12 | 0,01 | 16,02 | >0.1 | 5,56 | 5,65 | 0 | 1,24 | 0 | 5,69 | 0 |
| ONECUT3 | GATTAAAGAATAAC | 1 | 792 | 806 | -208 | -194 | -4,57 | 0,01 | 13,58 | >0.1 | 5,56 | 5,95 | 0 | 1,48 | 0 | 5,17 | 0 |
| ONECUT3 | CTAAAATCTGTTAC | 1 | 839 | 853 | -161 | -147 | -5,36 | 0,01 | 9,66 | >0.1 | 5,56 | 6,25 | 0 | 1,8 | 0 | 1,41 | 0 |
| ONECUT3 | GGAAAGTCATTCAG | 1 | 1007 | 1021 | 7 | 21 | -3,43 | 0,01 | 14,58 | >0.1 | 5,56 | 9,12 | 0 | 1,87 | 0 | 1,46 | 0 |
| ONECUT3 | GAAAGACCAAAAAA | -1 | 924 | 938 | -76 | -62 | 1,13 | 0,01 | 16,66 | >0.1 | 5,56 | 7,03 | 0 | 1,88 | 0 | 1,06 | 0 |
| ONECUT3 | CAAAAAAAGAAAAA | -1 | 917 | 931 | -83 | -69 | -5,01 | 0,01 | 10,46 | >0.1 | 5,56 | 6,94 | 0 | 1,89 | 0 | 1,09 | 0 |
| ONECUT3 | CAGTAAACAATCTG | -1 | 892 | 906 | -108 | -94 | -2,41 | 0,01 | 14,57 | >0.1 | 5,56 | 6,67 | 0 | 1,89 | 0 | 2,86 | 0 |
| ONECUT3 | AGTGAATCAAATAA | -1 | 812 | 826 | -188 | -174 | -1 | 0,01 | 15,64 | >0.1 | 5,56 | 6,06 | 0 | 1,59 | 0 | 3,43 | 0 |
| ONECUT3 | GCCAAATCAAAACC | -1 | 746 | 760 | -254 | -240 | 3,46 | 0,001 | 21,94 | >0.1 | 5,56 | 5,72 | 0 | 1,29 | 0 | 5,92 | 0 |
| ONECUT3 | TCAAAACCGGTACT | -1 | 740 | 754 | -260 | -246 | -0,95 | 0,01 | 17,49 | >0.1 | 5,56 | 5,7 | 0 | 1,27 | 0 | 5,92 | 0 |
| ONECUT2 | TGGAAATTAAAACT | 1 | 152 | 166 | -848 | -834 | -3,51 | 0,01 | 7,71 | >0.1 | 3,26 | 4,41 | 0 | 1,9 | 0 | 1,65 | 0 |
| ONECUT2 | TAAAAATACAAAAA | 1 | 293 | 307 | -707 | -693 | -5,11 | 0,01 | 7,86 | >0.1 | 2,69 | 4,6 | 0 | 1,23 | 0 | 4,45 | 0 |
| ONECUT2 | AAAAAACTGATCAG | 1 | 473 | 487 | -527 | -513 | -3,44 | 0,01 | 10,42 | >0.1 | 2,86 | 4,92 | 0 | 0,99 | 0 | 5,09 | 0 |
| ONECUT2 | ATGAATTCCATAAA | 1 | 491 | 505 | -509 | -495 | -0,56 | 0,01 | 14,75 | >0.1 | 2,93 | 4,95 | 0 | 0,99 | 0 | 6,44 | 0 |
| ONECUT2 | TTGAAATAAAAATA | 1 | 574 | 588 | -426 | -412 | -5,03 | 0,01 | 4,45 | >0.1 | 3,32 | 5,15 | 0 | 1 | 0 | 0 | 0 |
| ONECUT2 | TAAAAATAAATGTG | 1 | 580 | 594 | -420 | -406 | 1,2 | 0,01 | 10,73 | >0.1 | 3,36 | 5,17 | 0 | 1,01 | 0 | 0 | 0 |
| ONECUT2 | CTGAAAACCATACA | 1 | 617 | 631 | -383 | -369 | -1,8 | 0,01 | 9,61 | >0.1 | 3,61 | 5,27 | 0 | 1,04 | 0 | 1,5 | 0 |
| ONECUT2 | CAAACACCAACATT | 1 | 629 | 643 | -371 | -357 | -5,1 | 0,01 | 6,62 | >0.1 | 3,71 | 5,3 | 0 | 1,05 | 0 | 1,66 | 0 |
| ONECUT2 | TATTAACCAAAAGT | 1 | 729 | 743 | -271 | -257 | -4,29 | 0,01 | 13,85 | >0.1 | 5,56 | 5,65 | 0 | 1,24 | 0 | 5,69 | 0 |
| ONECUT2 | AAAAAATTATTCTA | 1 | 828 | 842 | -172 | -158 | -3,67 | 0,01 | 13,04 | >0.1 | 5,56 | 6,17 | 0 | 1,71 | 0 | 3,28 | 0 |
| ONECUT2 | GGAAAGTCATTCAG | 1 | 1007 | 1021 | 7 | 21 | -2,27 | 0,01 | 15,74 | >0.1 | 5,56 | 9,12 | 0 | 1,87 | 0 | 1,46 | 0 |
| ONECUT2 | GAAAGACCAAAAAA | -1 | 924 | 938 | -76 | -62 | -1,25 | 0,01 | 14,28 | >0.1 | 5,56 | 7,03 | 0 | 1,88 | 0 | 1,06 | 0 |
| ONECUT2 | CAGTAAACAATCTG | -1 | 892 | 906 | -108 | -94 | -1,61 | 0,01 | 15,37 | >0.1 | 5,56 | 6,67 | 0 | 1,89 | 0 | 2,86 | 0 |
| ONECUT2 | AGTGAATCAAATAA | -1 | 812 | 826 | -188 | -174 | -0,26 | 0,01 | 16,38 | >0.1 | 5,56 | 6,06 | 0 | 1,59 | 0 | 3,43 | 0 |

| | | | | | | | | | | | | | | | | | |
|---|---|---|---|---|---|---|---|---|---|---|---|---|---|---|---|---|---|
| ONECUT2 | GCCAAATCAAAACC | -1 | 746 | 760 | -254 | -240 | 4,52 | 0,001 | 23 | >0.1 | 5,56 | 5,72 | 0 | 1,29 | 0 | 5,92 | 0 |
| ONECUT2 | TCAAAACCGGTACT | -1 | 740 | 754 | -260 | -246 | -3,46 | 0,01 | 14,98 | >0.1 | 5,56 | 5,7 | 0 | 1,27 | 0 | 5,92 | 0 |
| ONECUT1 | TGGAAATTAAAACT | 1 | 152 | 166 | -848 | -834 | -8,97 | 0,01 | 2,25 | >0.1 | 3,26 | 4,41 | 0 | 1,9 | 0 | 1,65 | 0 |
| ONECUT1 | TTAAAACTGATCAG | 1 | 158 | 172 | -842 | -828 | -12,98 | 0,01 | -0,97 | >0.1 | 3,23 | 4,42 | 0 | 1,9 | 0 | 2,46 | 0 |
| ONECUT1 | AAAAAACTGATCAG | 1 | 473 | 487 | -527 | -513 | -9,89 | 0,01 | 3,97 | >0.1 | 2,86 | 4,92 | 0 | 0,99 | 0 | 5,09 | 0 |
| ONECUT1 | GAAATATGAATTCC | 1 | 486 | 500 | -514 | -500 | -13,51 | 0,01 | 1,6 | >0.1 | 2,91 | 4,94 | 0 | 0,99 | 0 | 6,27 | 0 |
| ONECUT1 | ATGAATTCCATAAA | 1 | 491 | 505 | -509 | -495 | -12,92 | 0,01 | 2,39 | >0.1 | 2,93 | 4,95 | 0 | 0,99 | 0 | 6,44 | 0 |
| ONECUT1 | TAAAAATAAATGTG | 1 | 580 | 594 | -420 | -406 | -5,79 | 0,01 | 3,74 | >0.1 | 3,36 | 5,17 | 0 | 1,01 | 0 | 0 | 0 |
| ONECUT1 | AGATAACCTATTAA | 1 | 594 | 608 | -406 | -392 | -11,33 | 0,01 | -0,22 | >0.1 | 3,45 | 5,2 | 0 | 1,02 | 0 | 1,44 | 0 |
| ONECUT1 | TATTAACCAAAAGT | 1 | 729 | 743 | -271 | -257 | -12,74 | 0,01 | 5,4 | >0.1 | 5,56 | 5,65 | 0 | 1,24 | 0 | 5,69 | 0 |
| ONECUT1 | AAAAAATTATTCTA | 1 | 828 | 842 | -172 | -158 | -9,81 | 0,01 | 6,9 | >0.1 | 5,56 | 6,17 | 0 | 1,71 | 0 | 3,28 | 0 |
| ONECUT1 | CTAAAATCTGTTAC | 1 | 839 | 853 | -161 | -147 | -13,12 | 0,01 | 1,9 | >0.1 | 5,56 | 6,25 | 0 | 1,8 | 0 | 1,41 | 0 |
| ONECUT1 | AGCAGATTGTTTAC | 1 | 890 | 904 | -110 | -96 | -12,47 | 0,01 | 4,49 | >0.1 | 5,56 | 6,65 | 0 | 1,89 | 0 | 2,86 | 0 |
| ONECUT1 | GGAAAGTCATTCAG | 1 | 1007 | 1021 | 7 | 21 | -10,18 | 0,01 | 7,83 | >0.1 | 5,56 | 9,12 | 0 | 1,87 | 0 | 1,46 | 0 |
| ONECUT1 | GAAAGACCAAAAAA | -1 | 924 | 938 | -76 | -62 | -10,61 | 0,01 | 4,92 | >0.1 | 5,56 | 7,03 | 0 | 1,88 | 0 | 1,06 | 0 |
| ONECUT1 | CAGTAAACAATCTG | -1 | 892 | 906 | -108 | -94 | -8,69 | 0,01 | 8,29 | >0.1 | 5,56 | 6,67 | 0 | 1,89 | 0 | 2,86 | 0 |
| ONECUT1 | AGTGAATCAAATAA | -1 | 812 | 826 | -188 | -174 | -7,05 | 0,01 | 9,59 | >0.1 | 5,56 | 6,06 | 0 | 1,59 | 0 | 3,43 | 0 |
| ONECUT1 | GCCAAATCAAAACC | -1 | 746 | 760 | -254 | -240 | 0,96 | 0,001 | 19,44 | >0.1 | 5,56 | 5,72 | 0 | 1,29 | 0 | 5,92 | 0 |
| ONECUT1 | TCAAAACCGGTACT | -1 | 740 | 754 | -260 | -246 | -10,21 | 0,01 | 8,23 | >0.1 | 5,56 | 5,7 | 0 | 1,27 | 0 | 5,92 | 0 |
| ONECUT1 | TTCTGATCAGTTTT | -1 | 475 | 489 | -525 | -511 | -12,57 | 0,01 | 1,42 | >0.1 | 2,87 | 4,92 | 0 | 0,99 | 0 | 5,2 | 0 |
| ONECUT1 | TTCTGATCAGTTTT | -1 | 160 | 174 | -840 | -826 | -12,57 | 0,01 | -0,57 | >0.1 | 3,22 | 4,42 | 0 | 1,9 | 0 | 2,46 | 0 |
| ONECUT1 | AAAAGCTCATTTCC | -1 | 64 | 78 | -936 | -922 | -11,78 | 0,01 | -1,66 | >0.1 | 3,93 | 4,32 | 0 | 1,88 | 0 | 0 | 0 |
| SMAD2::SMAD3::SMAD4 | AAGTCAGCCACAG | 1 | 34 | 47 | -966 | -953 | -2,3 | 0,01 | 8,22 | >0.1 | 4,37 | 4,29 | 0 | 1,87 | 0 | 0 | 0 |
| SMAD2::SMAD3::SMAD4 | ATGGCTGGGCACA | 1 | 174 | 187 | -826 | -813 | -3,8 | 0,01 | 8,87 | >0.1 | 3,15 | 4,44 | 0 | 1,84 | 0 | 3,24 | 0 |
| SMAD2::SMAD3::SMAD4 | GTGGCTCACGCCT | 1 | 187 | 200 | -813 | -800 | 10,04 | 0,001 | 23,26 | >0.1 | 3,09 | 4,45 | 0 | 1,73 | 0 | 3,95 | 0 |
| SMAD2::SMAD3::SMAD4 | CTGACTAACACGG | 1 | 263 | 276 | -737 | -724 | 5,47 | 0,01 | 18,58 | >0.1 | 2,79 | 4,56 | 0 | 1,32 | 0 | 4,45 | 0 |
| SMAD2::SMAD3::SMAD4 | CAGCCTGGGCGAC | 1 | 426 | 439 | -574 | -561 | -3,43 | 0,01 | 10,92 | >0.1 | 2,71 | 4,82 | 0 | 1,02 | 0 | 5,8 | 0 |
| SMAD2::SMAD3::SMAD4 | GAGACTCAGTCTC | 1 | 444 | 457 | -556 | -543 | -3,18 | 0,01 | 10,45 | >0.1 | 2,76 | 4,86 | 0 | 1 | 0 | 5,01 | 0 |
| SMAD2::SMAD3::SMAD4 | TTGTCTGAAAACC | 1 | 613 | 626 | -387 | -374 | -0,94 | 0,01 | 10,42 | >0.1 | 3,58 | 5,25 | 0 | 1,03 | 0 | 1,5 | 0 |
| SMAD2::SMAD3::SMAD4 | TTGGCTCACAGGG | 1 | 1032 | 1045 | 32 | 45 | 4,57 | 0,01 | 21,45 | >0.1 | 5,57 | 8,15 | 0 | 1,86 | 0 | 1,31 | 0 |
| SMAD2::SMAD3::SMAD4 | ATGTCAAGCTCTT | 1 | 1049 | 1062 | 49 | 62 | -1,54 | 0,01 | 14,95 | >0.1 | 5,57 | 7,6 | 0 | 1,86 | 0 | 1,46 | 0 |
| SMAD2::SMAD3::SMAD4 | CTTCCTGGCTCCT | 1 | 1059 | 1072 | 59 | 72 | 1,66 | 0,01 | 20,41 | >0.1 | 5,57 | 7,37 | 0 | 1,86 | 1,79 | 2,16 | 0 |
| SMAD2::SMAD3::SMAD4 | CTGGCTCCTTCTC | 1 | 1063 | 1076 | 63 | 76 | -2,84 | 0,01 | 15,89 | >0.1 | 5,57 | 7,3 | 0 | 1,85 | 1,79 | 2,21 | 0 |
| SMAD2::SMAD3::SMAD4 | ATGACTTTCCCTA | -1 | 1004 | 1017 | 4 | 17 | -0,65 | 0,01 | 17,32 | >0.1 | 5,57 | 9,12 | 0 | 1,87 | 0 | 1,42 | 0 |
| SMAD2::SMAD3::SMAD4 | CAGCCTTTGAACT | -1 | 957 | 970 | -43 | -30 | -3,12 | 0,01 | 13,69 | >0.1 | 5,57 | 7,62 | 0 | 1,88 | 0 | 1,74 | 0 |
| SMAD2::SMAD3::SMAD4 | TCTCCTGTCACTT | -1 | 764 | 777 | -236 | -223 | -2,96 | 0,01 | 15,67 | >0.1 | 5,57 | 5,8 | 0 | 1,35 | 0 | 5,92 | 0 |
| SMAD2::SMAD3::SMAD4 | GAAACTCTCTACT | -1 | 548 | 561 | -452 | -439 | -2,9 | 0,01 | 6,35 | >0.1 | 3,18 | 5,08 | 0 | 0,99 | 0 | 0 | 0 |
| SMAD2::SMAD3::SMAD4 | GAGACTGAGTCTC | -1 | 444 | 457 | -556 | -543 | -1,95 | 0,01 | 11,68 | >0.1 | 2,76 | 4,86 | 0 | 1 | 0 | 5,01 | 0 |
| SMAD2::SMAD3::SMAD4 | GAGTCTCGGTCTG | -1 | 438 | 451 | -562 | -549 | -0,44 | 0,01 | 14,2 | >0.1 | 2,74 | 4,84 | 0 | 1,01 | 0 | 6,05 | 0 |
| SMAD2::SMAD3::SMAD4 | CGGTCTGTCGCCC | -1 | 432 | 445 | -568 | -555 | 10,89 | 0,001 | 25,26 | >0.1 | 2,72 | 4,83 | 0 | 1,01 | 0 | 5,8 | 0 |
| SMAD2::SMAD3::SMAD4 | CTGTCGCCCAGGC | -1 | 428 | 441 | -572 | -559 | -0,23 | 0,01 | 13,99 | >0.1 | 2,71 | 4,83 | 0 | 1,01 | 0 | 5,67 | 0 |
| SMAD2::SMAD3::SMAD4 | TCGGCTCACTGCA | -1 | 395 | 408 | -605 | -592 | 3,88 | 0,01 | 18,07 | >0.1 | 2,62 | 4,76 | 0 | 1,04 | 0 | 5,76 | 0 |
| SMAD2::SMAD3::SMAD4 | AGCTCTGCCTCCC | -1 | 382 | 395 | -618 | -605 | 3,51 | 0,01 | 17,91 | >0.1 | 2,58 | 4,74 | 0 | 1,06 | 0 | 6,02 | 0 |
| SMAD2::SMAD3::SMAD4 | CTGCCTCCCAGGT | -1 | 378 | 391 | -622 | -609 | 8,75 | 0,001 | 23,78 | >0.1 | 2,57 | 4,74 | 0 | 1,06 | 0 | 6,66 | 0 |
| SMAD2::SMAD3::SMAD4 | CTGCCTCAGCCTC | -1 | 352 | 365 | -648 | -635 | -1,12 | 0,01 | 13,6 | >0.1 | 2,53 | 4,69 | 0 | 1,1 | 0 | 6,4 | 0 |
| SMAD2::SMAD3::SMAD4 | CAGCCTCCCGAGT | -1 | 346 | 359 | -654 | -641 | -0,38 | 0,01 | 14,48 | >0.1 | 2,55 | 4,68 | 0 | 1,11 | 0 | 6,52 | 0 |
| SMAD2::SMAD3::SMAD4 | GTAGCTGGGACTA | -1 | 335 | 348 | -665 | -652 | -2,24 | 0,01 | 12,05 | >0.1 | 2,58 | 4,66 | 0 | 1,13 | 0 | 5,92 | 0 |
| SMAD2::SMAD3::SMAD4 | GTGCCCACCACCA | -1 | 319 | 332 | -681 | -668 | -2,06 | 0,01 | 12,04 | >0.1 | 2,62 | 4,64 | 0 | 1,16 | 0 | 5,68 | 0 |
| SMAD2::SMAD3::SMAD4 | CGGTCTCTATCTC | -1 | 248 | 261 | -752 | -739 | -3,48 | 0,01 | 10,47 | >0.1 | 2,84 | 4,53 | 0 | 1,38 | 0 | 5,19 | 0 |
| SMAD2::SMAD3::SMAD4 | GTCTCTATCTCCT | -1 | 246 | 259 | -754 | -741 | 6,32 | 0,001 | 20,28 | >0.1 | 2,85 | 4,53 | 0 | 1,38 | 0 | 5,19 | 0 |
| SMAD2::SMAD3::SMAD4 | CGGCCTCCCAAAG | -1 | 212 | 225 | -788 | -775 | -0,13 | 0,01 | 14,52 | >0.1 | 2,98 | 4,49 | 0 | 1,55 | 0 | 5,63 | 0 |
| SMAD2::SMAD3::SMAD4 | CTATTTCTCACAC | -1 | 52 | 65 | -948 | -935 | -3,7 | 0,01 | 6,56 | >0.1 | 4,08 | 4,3 | 0 | 1,87 | 0 | 0 | 0 |
| SMAD2::SMAD3::SMAD4 | ATTTCTCACACTG | -1 | 50 | 63 | -950 | -937 | -1,75 | 0,01 | 8,53 | >0.1 | 4,11 | 4,3 | 0 | 1,87 | 0 | 0 | 0 |
| SMAD2::SMAD3::SMAD4 | GTGGCTGACTTCT | -1 | 32 | 45 | -968 | -955 | 0,84 | 0,01 | 11,4 | >0.1 | 4,41 | 4,29 | 0 | 1,87 | 0 | 0 | 0 |
| SMAD2::SMAD3::SMAD4 | TTGAGTGCCTCCT | -1 | 19 | 32 | -981 | -968 | -2,18 | 0,01 | 9,53 | >0.1 | 5,57 | 4,27 | 0 | 1,87 | 0 | 0 | 0 |
| NHLH1 | GGCAGATCAG | 1 | 92 | 102 | -908 | -898 | -2,2 | 0,01 | 8,53 | >0.1 | 3,68 | 4,34 | 0 | 1,88 | 0 | 0,83 | 0 |
| NHLH1 | CACGCCTGTA | 1 | 193 | 203 | -807 | -797 | -3,58 | 0,01 | 9,89 | >0.1 | 3,07 | 4,46 | 0 | 1,68 | 0 | 4,26 | 0 |
| NHLH1 | GGCAGATCAC | 1 | 230 | 240 | -770 | -760 | -3,76 | 0,01 | 10,13 | >0.1 | 2,92 | 4,51 | 0 | 1,45 | 0 | 5,01 | 0 |
| NHLH1 | CTTAGCTGGG | 1 | 307 | 317 | -693 | -683 | -3,98 | 0,01 | 10,16 | >0.1 | 2,66 | 4,62 | 0 | 1,19 | 0 | 5,68 | 0 |

| | | | | | | | | | | | | | | | | |
|---|---|---|---|---|---|---|---|---|---|---|---|---|---|---|---|---|
| NHLH1 | GGCACCTGTA | 1 | 327 | 337 | -673 | -663 | 5,89 | 0,001 | 19,85 | >0.1 | 2,6 | 4,65 | 0 | 1,14 | 0 | 5,56 | 0 |
| NHLH1 | CCCAGCTACT | 1 | 339 | 349 | -661 | -651 | 3,67 | 0,01 | 18,55 | >0.1 | 2,57 | 4,66 | 0 | 1,12 | 0 | 6,52 | 0 |
| NHLH1 | TGAACCTGGG | 1 | 375 | 385 | -625 | -615 | -3,19 | 0,01 | 11,55 | >0.1 | 2,56 | 4,72 | 0 | 1,07 | 0 | 6,4 | 0 |
| NHLH1 | AACAGCTTTC | 1 | 711 | 721 | -289 | -279 | -1,93 | 0,01 | 15,41 | >0.1 | 5,61 | 5,56 | 0 | 1,18 | 0 | 4,99 | 0 |
| NHLH1 | AAAATCTGTT | 1 | 841 | 851 | -159 | -149 | -4,74 | 0,01 | 10,3 | >0.1 | 5,61 | 6,24 | 0 | 1,78 | 0 | 1,41 | 0 |
| NHLH1 | CTCAGCAGAT | 1 | 887 | 897 | -113 | -103 | -3,49 | 0,01 | 13,67 | >0.1 | 5,61 | 6,59 | 0 | 1,89 | 0 | 3,07 | 0 |
| NHLH1 | CACAGGGGAC | 1 | 1038 | 1048 | 38 | 48 | -1,54 | 0,01 | 15,18 | >0.1 | 5,61 | 7,92 | 0 | 1,86 | 0 | 1,34 | 0 |
| NHLH1 | AACTGCTGCT | 1 | 1090 | 1100 | 90 | 100 | -0,38 | 0,01 | 18,03 | >0.1 | 5,61 | 6,88 | 0 | 1,85 | 1,79 | 2,27 | 0 |
| NHLH1 | AACAGGTCTT | -1 | 1158 | 1168 | 158 | 168 | -3,23 | 0,01 | 13,36 | >0.1 | 4,38 | 6,24 | 0 | 1,78 | 1,79 | 2,4 | 0 |
| NHLH1 | AGCAGCAGTT | -1 | 1090 | 1100 | 90 | 100 | -0,77 | 0,01 | 17,64 | >0.1 | 5,61 | 6,88 | 0 | 1,85 | 1,79 | 2,27 | 0 |
| NHLH1 | GTCCCCTGTG | -1 | 1038 | 1048 | 38 | 48 | -4,14 | 0,01 | 12,58 | >0.1 | 5,61 | 7,92 | 0 | 1,86 | 0 | 1,34 | 0 |
| NHLH1 | ACAATCTGCT | -1 | 890 | 900 | -110 | -100 | -2,48 | 0,01 | 14,58 | >0.1 | 5,61 | 6,61 | 0 | 1,89 | 0 | 2,95 | 0 |
| NHLH1 | ATCTGCTGAG | -1 | 887 | 897 | -113 | -103 | -4,4 | 0,01 | 12,76 | >0.1 | 5,61 | 6,59 | 0 | 1,89 | 0 | 3,07 | 0 |
| NHLH1 | GAAAGCTGTT | -1 | 711 | 721 | -289 | -279 | -1,62 | 0,01 | 15,72 | >0.1 | 5,61 | 5,56 | 0 | 1,18 | 0 | 4,99 | 0 |
| NHLH1 | CCCAGGTTCA | -1 | 375 | 385 | -625 | -615 | -3,6 | 0,01 | 11,14 | >0.1 | 2,56 | 4,72 | 0 | 1,07 | 0 | 6,4 | 0 |
| NHLH1 | AGTAGCTGGG | -1 | 339 | 349 | -661 | -651 | -1,51 | 0,01 | 13,37 | >0.1 | 2,57 | 4,66 | 0 | 1,12 | 0 | 6,52 | 0 |
| NHLH1 | TACAGGTGCC | -1 | 327 | 337 | -673 | -663 | 5,56 | 0,01 | 19,52 | >0.1 | 2,6 | 4,65 | 0 | 1,14 | 0 | 5,56 | 0 |
| NHLH1 | CCCAGCTAAG | -1 | 307 | 317 | -693 | -683 | 1,34 | 0,01 | 15,48 | >0.1 | 2,66 | 4,62 | 0 | 1,19 | 0 | 5,68 | 0 |
| NHLH1 | TACAGGCGTG | -1 | 193 | 203 | -807 | -797 | -4,53 | 0,01 | 8,94 | >0.1 | 3,07 | 4,46 | 0 | 1,68 | 0 | 4,26 | 0 |
| NHLH1 | CACAGGTCAA | -1 | 107 | 117 | -893 | -883 | -3,79 | 0,01 | 6,78 | >0.1 | 3,56 | 4,36 | 0 | 1,89 | 0 | 0,77 | 0 |
| PHOX2B | AAAACTGATCA | 1 | 160 | 171 | -840 | -829 | 3,28 | 0,01 | 15,28 | >0.1 | 3,23 | 4,42 | 0 | 1,9 | 0 | 2,46 | 0 |
| PHOX2B | AAAACTGATCA | 1 | 475 | 486 | -525 | -514 | 3,28 | 0,01 | 17,14 | >0.1 | 2,87 | 4,91 | 0 | 0,99 | 0 | 5,09 | 0 |
| PHOX2B | GAATTCCATAA | 1 | 493 | 504 | -507 | -496 | 2,77 | 0,01 | 18,08 | >0.1 | 2,93 | 4,95 | 0 | 0,98 | 0 | 6,44 | 0 |
| PHOX2B | ATAACCTATTA | 1 | 596 | 607 | -404 | -393 | -0,32 | 0,01 | 10,79 | >0.1 | 3,45 | 5,2 | 0 | 1,01 | 0 | 1,44 | 0 |
| PHOX2B | TAACCTATTAA | 1 | 597 | 608 | -403 | -392 | 2,9 | 0,01 | 13,99 | >0.1 | 3,46 | 5,2 | 0 | 1,02 | 0 | 1,41 | 0 |
| PHOX2B | TAATGAAATTG | 1 | 605 | 616 | -395 | -384 | 8,54 | 0,001 | 19,71 | >0.1 | 3,51 | 5,22 | 0 | 1,02 | 0 | 1,41 | 0 |
| PHOX2B | TTATTTGATTC | 1 | 812 | 823 | -188 | -177 | 2,54 | 0,01 | 19,18 | >0.1 | 5,6 | 6,05 | 0 | 1,57 | 0 | 3,43 | 0 |
| PHOX2B | TGATTCACTTT | 1 | 817 | 828 | -183 | -172 | 3,46 | 0,01 | 20,29 | >0.1 | 5,6 | 6,08 | 0 | 1,6 | 0 | 3,55 | 0 |
| PHOX2B | TTAAAAAATTA | 1 | 826 | 837 | -174 | -163 | 1,4 | 0,01 | 18,35 | >0.1 | 5,6 | 6,14 | 0 | 1,67 | 0 | 3,55 | 0 |
| PHOX2B | AAATCTCATGA | 1 | 980 | 991 | -20 | -9 | 0,56 | 0,01 | 17,93 | >0.1 | 5,6 | 8,37 | 0 | 1,87 | 0 | 1,53 | 0 |
| PHOX2B | TCATGAGATTT | -1 | 980 | 991 | -20 | -9 | 1,54 | 0,01 | 18,91 | >0.1 | 5,6 | 8,37 | 0 | 1,87 | 0 | 1,53 | 0 |
| PHOX2B | TGAGCCAATAT | -1 | 880 | 891 | -120 | -109 | 1,91 | 0,01 | 19,01 | >0.1 | 5,6 | 6,54 | 0 | 1,9 | 0 | 3,07 | 0 |
| PHOX2B | AAAGTGAATCA | -1 | 817 | 828 | -183 | -172 | 2,34 | 0,01 | 19,17 | >0.1 | 5,6 | 6,08 | 0 | 1,6 | 0 | 3,55 | 0 |
| PHOX2B | GAATCAAATAA | -1 | 812 | 823 | -188 | -177 | 4,77 | 0,01 | 21,41 | >0.1 | 5,6 | 6,05 | 0 | 1,57 | 0 | 3,43 | 0 |
| PHOX2B | AAATGTTATAA | -1 | 687 | 698 | -313 | -302 | 2,58 | 0,01 | 16,59 | >0.1 | 4,45 | 5,48 | 0 | 1,13 | 0 | 2,95 | 0 |
| PHOX2B | CAATTTCATTA | -1 | 605 | 616 | -395 | -384 | 8,03 | 0,001 | 19,2 | >0.1 | 3,51 | 5,22 | 0 | 1,02 | 0 | 1,41 | 0 |
| PHOX2B | TTATGGAATTC | -1 | 493 | 504 | -507 | -496 | 1,7 | 0,01 | 17,01 | >0.1 | 2,93 | 4,95 | 0 | 0,98 | 0 | 6,44 | 0 |
| PHOX2B | AAAGCTCATTT | -1 | 66 | 77 | -934 | -923 | 1,73 | 0,01 | 11,84 | >0.1 | 3,92 | 4,32 | 0 | 1,88 | 0 | 0 | 0 |
| MECOM | TAGGTAAGTGAAGG | 1 | 2 | 16 | -998 | -984 | -2,99 | 0,01 | 10,48 | >0.1 | 5,56 | 4,26 | 0 | 1,86 | 1,79 | 0 | 0 |
| MECOM | ATCACAAGGTCAGG | 1 | 235 | 249 | -765 | -751 | 1,3 | 0,01 | 14,67 | >0.1 | 2,89 | 4,52 | 0 | 1,43 | 0 | 4,53 | 0 |
| MECOM | GAGGCAGGAGAATG | 1 | 358 | 372 | -642 | -628 | 0,23 | 0,01 | 14,91 | >0.1 | 2,52 | 4,7 | 0 | 1,09 | 0 | 6,36 | 0 |
| MECOM | GAGCCGAGATCGCG | 1 | 401 | 415 | -599 | -585 | -1,3 | 0,01 | 12,91 | >0.1 | 2,63 | 4,78 | 0 | 1,04 | 0 | 5,76 | 0 |
| MECOM | AAGACAAGGGCAAA | 1 | 503 | 517 | -497 | -483 | 2,57 | 0,01 | 17,2 | >0.1 | 2,98 | 4,98 | 0 | 0,99 | 0 | 5,68 | 0 |
| MECOM | AATGTGAGATAACC | 1 | 588 | 602 | -412 | -398 | 1,44 | 0,01 | 11,05 | >0.1 | 3,41 | 5,19 | 0 | 1,01 | 0 | 0 | 0 |
| MECOM | GAGATAACCTATTA | 1 | 593 | 607 | -407 | -393 | -1,42 | 0,01 | 8,24 | >0.1 | 3,44 | 5,2 | 0 | 1,02 | 0 | 0 | 0 |
| MECOM | AGGAAAATATTAAC | 1 | 722 | 736 | -278 | -264 | 1,3 | 0,01 | 18,86 | >0.1 | 5,56 | 5,62 | 0 | 1,22 | 0 | 5,17 | 0 |
| MECOM | GTGACAGGAGAGGT | 1 | 766 | 780 | -234 | -220 | 2,58 | 0,01 | 21,23 | >0.1 | 5,56 | 5,82 | 0 | 1,36 | 0 | 5,92 | 0 |
| MECOM | AACCCAAGTTCAAA | 1 | 951 | 965 | -49 | -35 | -2,48 | 0,01 | 14,16 | >0.1 | 5,56 | 7,51 | 0 | 1,88 | 0 | 1,7 | 0 |
| MECOM | CTGATAAGAGAGAA | 1 | 967 | 981 | -33 | -19 | 6,75 | 0,001 | 23,82 | >0.1 | 5,56 | 7,94 | 0 | 1,87 | 0 | 1,7 | 0 |
| MECOM | GATAAGAGAGAAAA | 1 | 969 | 983 | -31 | -17 | -2,76 | 0,01 | 14,33 | >0.1 | 5,56 | 8,02 | 0 | 1,87 | 0 | 1,65 | 0 |
| MECOM | AGGCCAAGACATTT | 1 | 1119 | 1133 | 119 | 133 | -1,18 | 0,01 | 18,68 | >0.1 | 5,56 | 6,56 | 0 | 1,85 | 3,22 | 2,67 | 0 |
| MECOM | TGGACAAGTTTAAC | 1 | 1134 | 1148 | 134 | 148 | 3,93 | 0,01 | 24,07 | >0.1 | 5,56 | 6,42 | 0 | 1,84 | 3,22 | 3,09 | 0 |
| MECOM | TTGATAGAACAGGT | -1 | 1161 | 1175 | 161 | 175 | -1,58 | 0,01 | 15,36 | >0.1 | 4,46 | 6,22 | 0 | 1,75 | 1,79 | 2,72 | 0 |
| MECOM | GAGCCAAGATCACA | -1 | 1025 | 1039 | 25 | 39 | 9,01 | 0,001 | 26,25 | >0.1 | 5,56 | 8,52 | 0 | 1,86 | 0 | 1,31 | 0 |
| MECOM | CAGGAAAGACCAAA | -1 | 927 | 941 | -73 | -59 | 0,8 | 0,01 | 16,34 | >0.1 | 5,56 | 7,08 | 0 | 1,88 | 0 | 1,03 | 0 |
| MECOM | AAAAAAAGAAAAAG | -1 | 916 | 930 | -84 | -70 | -0,3 | 0,01 | 15,19 | >0.1 | 5,56 | 6,93 | 0 | 1,89 | 0 | 1,12 | 0 |
| MECOM | GAGCCAATATAAAG | -1 | 876 | 890 | -124 | -110 | 2,8 | 0,01 | 19,85 | >0.1 | 5,56 | 6,53 | 0 | 1,9 | 0 | 3,07 | 0 |
| MECOM | TGGAGAGGACAGAT | -1 | 855 | 869 | -145 | -131 | -0,99 | 0,01 | 15,69 | >0.1 | 5,56 | 6,36 | 0 | 1,9 | 0 | 2,86 | 0 |
| MECOM | CAAATAAGAATACG | -1 | 805 | 819 | -195 | -181 | -1,49 | 0,01 | 14,92 | >0.1 | 5,56 | 6,02 | 0 | 1,55 | 0 | 3,28 | 0 |

| | | | | | | | | | | | | | | | | | |
|---|---|---|---|---|---|---|---|---|---|---|---|---|---|---|---|---|---|
| MECOM | AAGATAATGTTGGT | -1 | 634 | 648 | -366 | -352 | -1,54 | 0,01 | 10,25 | >0.1 | 3,76 | 5,32 | 0 | 1,06 | 0 | 1,66 | 0 |
| MECOM | ATTTCAAGATCATA | -1 | 567 | 581 | -433 | -419 | 1,56 | 0,01 | 10,98 | >0.1 | 3,28 | 5,13 | 0 | 1 | 0 | 0 | 0 |
| MECOM | GTGCTAGGATTACA | -1 | 199 | 213 | -801 | -787 | 3,34 | 0,01 | 17,08 | >0.1 | 3,03 | 4,47 | 0 | 1,64 | 0 | 4,59 | 0 |
| ZBTB18 | AAGGCAGATCAGG | 1 | 90 | 103 | -910 | -897 | -6,81 | 0,01 | 3,95 | >0.1 | 3,68 | 4,34 | 0 | 1,88 | 0 | 0,85 | 0 |
| ZBTB18 | GGCACAGTGGCTC | 1 | 181 | 194 | -819 | -806 | -4,21 | 0,01 | 8,99 | >0.1 | 3,12 | 4,45 | 0 | 1,78 | 0 | 3,86 | 0 |
| ZBTB18 | TGGGCACCTGTAG | 1 | 325 | 338 | -675 | -662 | -0,45 | 0,01 | 13,41 | >0.1 | 2,6 | 4,65 | 0 | 1,15 | 0 | 5,46 | 0 |
| ZBTB18 | GTCCCAGCTACTC | 1 | 337 | 350 | -663 | -650 | -5,46 | 0,01 | 8,82 | >0.1 | 2,57 | 4,67 | 0 | 1,13 | 0 | 5,92 | 0 |
| ZBTB18 | GAGGCAGAGCTTG | 1 | 384 | 397 | -616 | -603 | -5,3 | 0,01 | 9,22 | >0.1 | 2,58 | 4,75 | 0 | 1,06 | 0 | 6,13 | 0 |
| ZBTB18 | GTAACTGCTGCTC | 1 | 1088 | 1101 | 88 | 101 | -2,59 | 0,01 | 15,74 | >0.1 | 5,57 | 6,9 | 0 | 1,85 | 1,79 | 2,21 | 0 |
| ZBTB18 | AGAACAGGTCTTC | -1 | 1157 | 1170 | 157 | 170 | -6,85 | 0,01 | 9,75 | >0.1 | 4,37 | 6,24 | 0 | 1,79 | 1,79 | 2,4 | 0 |
| ZBTB18 | TGAGCAGCAGTTA | -1 | 1089 | 1102 | 89 | 102 | -3,59 | 0,01 | 14,79 | >0.1 | 5,57 | 6,89 | 0 | 1,85 | 1,79 | 2,27 | 0 |
| ZBTB18 | CCTCCTCATGAGA | -1 | 983 | 996 | -17 | -4 | -5,11 | 0,01 | 12,47 | >0.1 | 5,57 | 8,61 | 0 | 1,87 | 0 | 1,53 | 0 |
| ZBTB18 | AGGACAGATATGT | -1 | 851 | 864 | -149 | -136 | 1,27 | 0,01 | 17,74 | >0.1 | 5,57 | 6,33 | 0 | 1,9 | 0 | 2,67 | 0 |
| ZBTB18 | GTAACAGATTTTA | -1 | 840 | 853 | -160 | -147 | -4,08 | 0,01 | 10,95 | >0.1 | 5,57 | 6,25 | 0 | 1,8 | 0 | 1,41 | 0 |
| ZBTB18 | TAGAAAGCTGTTA | -1 | 710 | 723 | -290 | -277 | 0,55 | 0,01 | 17,86 | >0.1 | 5,57 | 5,57 | 0 | 1,18 | 0 | 4,99 | 0 |
| ZBTB18 | AATGTTGGTGTTT | -1 | 630 | 643 | -370 | -357 | -7,04 | 0,01 | 4,69 | >0.1 | 3,72 | 5,3 | 0 | 1,05 | 0 | 1,66 | 0 |
| ZBTB18 | AGCACAGTGGCGC | -1 | 412 | 425 | -588 | -575 | -4,26 | 0,01 | 9,31 | >0.1 | 2,66 | 4,8 | 0 | 1,03 | 0 | 5,09 | 0 |
| ZBTB18 | ACTACAGGTGCCC | -1 | 326 | 339 | -674 | -661 | -1,56 | 0,01 | 12,4 | >0.1 | 2,6 | 4,65 | 0 | 1,15 | 0 | 5,56 | 0 |
| ZBTB18 | ATTACAGGCGTGA | -1 | 192 | 205 | -808 | -795 | -1,05 | 0,01 | 12,43 | >0.1 | 3,07 | 4,46 | 0 | 1,69 | 0 | 4,26 | 0 |
| ZBTB18 | ACTCCACAGGTCA | -1 | 108 | 121 | -892 | -879 | -3,29 | 0,01 | 7,24 | >0.1 | 3,54 | 4,36 | 0 | 1,89 | 0 | 0,75 | 0 |
| ZBTB18 | TCTCCTGATCTGC | -1 | 93 | 106 | -907 | -894 | -3,7 | 0,01 | 6,97 | >0.1 | 3,66 | 4,35 | 0 | 1,88 | 0 | 0,79 | 0 |
| ZBTB18 | CTCACACTGGTTC | -1 | 46 | 59 | -954 | -941 | -4,43 | 0,01 | 5,9 | >0.1 | 4,16 | 4,3 | 0 | 1,87 | 0 | 0 | 0 |
| USF2 | GCAGATCAGGAGAGTT | 1 | 93 | 109 | -907 | -891 | 1,06 | 0,01 | 11,7 | >0.1 | 3,64 | 4,35 | 0 | 1,88 | 0 | 0,77 | 0 |
| USF2 | GCTGGGCACAGTGGCT | 1 | 177 | 193 | -823 | -807 | -4,42 | 0,01 | 8,92 | >0.1 | 3,13 | 4,45 | 0 | 1,82 | 0 | 3,95 | 0 |
| USF2 | GTGGCTCACGCCTGTA | 1 | 187 | 203 | -813 | -797 | -3,41 | 0,01 | 9,99 | >0.1 | 3,08 | 4,46 | 0 | 1,73 | 0 | 4,13 | 0 |
| USF2 | GCAGATCACAAGGTCA | 1 | 231 | 247 | -769 | -753 | 5,95 | 0,001 | 19,72 | >0.1 | 2,9 | 4,52 | 0 | 1,45 | 0 | 4,9 | 0 |
| USF2 | CAAGGTCAGGAGATAG | 1 | 239 | 255 | -761 | -745 | 3,89 | 0,001 | 17,03 | >0.1 | 2,87 | 4,53 | 0 | 1,41 | 0 | 4,33 | 0 |
| USF2 | GGTGGGCACCTGTAGT | 1 | 323 | 339 | -677 | -661 | 0,85 | 0,01 | 14,72 | >0.1 | 2,6 | 4,65 | 0 | 1,16 | 0 | 5,46 | 0 |
| USF2 | CTGAGGCAGGAGAATG | 1 | 356 | 372 | -644 | -628 | -3,9 | 0,01 | 10,78 | >0.1 | 2,52 | 4,7 | 0 | 1,1 | 0 | 6,36 | 0 |
| USF2 | GAGAATGGCGTGAACC | 1 | 365 | 381 | -635 | -619 | -2,07 | 0,01 | 12,99 | >0.1 | 2,54 | 4,72 | 0 | 1,08 | 0 | 6,73 | 0 |
| USF2 | CAAAGTCATGTATTTG | 1 | 513 | 529 | -487 | -471 | -1,26 | 0,01 | 10,71 | >0.1 | 3,02 | 5,01 | 0 | 0,99 | 0 | 2,95 | 0 |
| USF2 | TAAAGTGACAGGAGAG | 1 | 762 | 778 | -238 | -222 | -3,52 | 0,01 | 15,09 | >0.1 | 5,53 | 5,81 | 0 | 1,35 | 0 | 5,92 | 0 |
| USF2 | AAGTGACAGGAGAGGT | 1 | 764 | 780 | -236 | -220 | -4,16 | 0,01 | 14,46 | >0.1 | 5,53 | 5,82 | 0 | 1,36 | 0 | 5,92 | 0 |
| USF2 | GAAAATCTCATGAGGA | 1 | 978 | 994 | -22 | -6 | -0,66 | 0,01 | 16,84 | >0.1 | 5,53 | 8,57 | 0 | 1,87 | 0 | 1,53 | 0 |
| USF2 | AAATCTCATGAGGAGG | 1 | 980 | 996 | -20 | -4 | -4,52 | 0,01 | 13,07 | >0.1 | 5,53 | 8,61 | 0 | 1,87 | 0 | 1,57 | 0 |
| USF2 | TTGGCTCACAGGGGAC | 1 | 1032 | 1048 | 32 | 48 | 2,79 | 0,01 | 19,63 | >0.1 | 5,53 | 8,15 | 0 | 1,86 | 0 | 1,31 | 0 |
| USF2 | TTTAACCACGAAGCCG | 1 | 1142 | 1158 | 142 | 158 | -4,39 | 0,01 | 14,73 | >0.1 | 5,53 | 6,36 | 0 | 1,84 | 3,22 | 2,16 | 0 |
| USF2 | AAAGTTCACTTGCTTC | 1 | 1173 | 1189 | 173 | 189 | -2,03 | 0,01 | 16,35 | >0.1 | 5,53 | 6,13 | 0 | 1,64 | 1,79 | 3,28 | 0 |
| USF2 | AAGAAGCAAGTGAACT | -1 | 1175 | 1191 | 175 | 191 | -4,05 | 0,01 | 14,39 | >0.1 | 5,53 | 6,12 | 0 | 1,62 | 1,79 | 3,37 | 0 |
| USF2 | TTCGGCTTCGTGGTTA | -1 | 1144 | 1160 | 144 | 160 | -3,76 | 0,01 | 15,39 | >0.1 | 5,53 | 6,34 | 0 | 1,84 | 3,22 | 2,21 | 0 |
| USF2 | CATCGTCCCCTGTGAG | -1 | 1036 | 1052 | 36 | 52 | -2,93 | 0,01 | 13,81 | >0.1 | 5,53 | 7,99 | 0 | 1,86 | 0 | 1,36 | 0 |
| USF2 | TCGTCCCCTGTGAGCC | -1 | 1034 | 1050 | 34 | 50 | -3,17 | 0,01 | 13,62 | >0.1 | 5,53 | 8,06 | 0 | 1,86 | 0 | 1,34 | 0 |
| USF2 | CAAGATCACATCCACT | -1 | 1019 | 1035 | 19 | 35 | -2,35 | 0,01 | 15,06 | >0.1 | 5,53 | 8,74 | 0 | 1,86 | 0 | 1,28 | 0 |
| USF2 | CCTCCTCATGAGATTT | -1 | 980 | 996 | -20 | -4 | -1,53 | 0,01 | 16,06 | >0.1 | 5,53 | 8,61 | 0 | 1,87 | 0 | 1,57 | 0 |
| USF2 | TCCTGTCACTTTATGG | -1 | 759 | 775 | -241 | -225 | -2,65 | 0,01 | 15,93 | >0.1 | 5,53 | 5,79 | 0 | 1,34 | 0 | 5,92 | 0 |
| USF2 | CAAGATCATATTCCCA | -1 | 561 | 577 | -439 | -423 | -4,33 | 0,01 | 5,05 | >0.1 | 3,25 | 5,12 | 0 | 1 | 0 | 0 | 0 |
| USF2 | CACAGTGGCGCGATCT | -1 | 407 | 423 | -593 | -577 | -3,39 | 0,01 | 10,74 | >0.1 | 2,65 | 4,79 | 0 | 1,03 | 0 | 5,65 | 0 |
| USF2 | CAGGTTCACGCCATTC | -1 | 367 | 383 | -633 | -617 | -3,35 | 0,01 | 11,26 | >0.1 | 2,54 | 4,72 | 0 | 1,08 | 0 | 6,27 | 0 |
| USF2 | GGACTACAGGTGCCCA | -1 | 325 | 341 | -675 | -659 | -3,49 | 0,01 | 10,48 | >0.1 | 2,6 | 4,65 | 0 | 1,15 | 0 | 5,56 | 0 |
| USF2 | AATAGAGACGGGGTTT | -1 | 278 | 294 | -722 | -706 | -3,6 | 0,01 | 10,18 | >0.1 | 2,74 | 4,58 | 0 | 1,27 | 0 | 5,19 | 0 |
| USF2 | CTCTATCTCCTGACCT | -1 | 241 | 257 | -759 | -743 | -2,35 | 0,01 | 10,87 | >0.1 | 2,86 | 4,53 | 0 | 1,41 | 0 | 4,42 | 0 |
| USF2 | CCTGACCTTGTGATCT | -1 | 233 | 249 | -767 | -751 | -1,33 | 0,01 | 12,15 | >0.1 | 2,89 | 4,52 | 0 | 1,44 | 0 | 4,63 | 0 |
| USF2 | TCAACTCTCCTGATCT | -1 | 95 | 111 | -905 | -889 | -4,19 | 0,01 | 6,44 | >0.1 | 3,63 | 4,35 | 0 | 1,88 | 0 | 0,77 | 0 |
| USF2 | ACTGGTTCTGTGGCTG | -1 | 38 | 54 | -962 | -946 | -3,14 | 0,01 | 7,29 | >0.1 | 4,27 | 4,29 | 0 | 1,87 | 0 | 0 | 0 |
| FOSL2::JUNB(VAR.2) | AAAGAAGTCAGC | 1 | 30 | 42 | -970 | -958 | 1,07 | 0,01 | 11,68 | >0.1 | 4,46 | 4,28 | 0 | 1,87 | 0 | 0 | 0 |
| FOSL2::JUNB(VAR.2) | AATGGCGTGAAC | 1 | 368 | 380 | -632 | -620 | 2,39 | 0,01 | 16,87 | >0.1 | 2,54 | 4,72 | 0 | 1,08 | 0 | 6,15 | 0 |
| FOSL2::JUNB(VAR.2) | GATCGCGCCACT | 1 | 408 | 420 | -592 | -580 | 0,19 | 0,01 | 14,18 | >0.1 | 2,65 | 4,79 | 0 | 1,03 | 0 | 5,52 | 0 |
| FOSL2::JUNB(VAR.2) | GGCAAAGTCATG | 1 | 511 | 523 | -489 | -477 | -1,94 | 0,01 | 10 | >0.1 | 3,01 | 4,99 | 0 | 0,98 | 0 | 2,95 | 0 |
| FOSL2::JUNB(VAR.2) | TGTGAGATAACC | 1 | 590 | 602 | -410 | -398 | 4 | 0,01 | 13,61 | >0.1 | 3,41 | 5,19 | 0 | 1,01 | 0 | 0 | 0 |

| | | | | | | | | | | | | | | | | | |
|---|---|---|---|---|---|---|---|---|---|---|---|---|---|---|---|---|---|
| FOSL2::JUNB(VAR.2) | TCTTTGGTCACT | 1 | 669 | 681 | -331 | -319 | -1,66 | 0,01 | 12,7 | >0.1 | 4,14 | 5,42 | 0 | 1,1 | 0 | 3,7 | 0 |
| FOSL2::JUNB(VAR.2) | AATTATAACATT | 1 | 685 | 697 | -315 | -303 | -0,8 | 0,01 | 13,17 | >0.1 | 4,42 | 5,48 | 0 | 1,13 | 0 | 2,95 | 0 |
| FOSL2::JUNB(VAR.2) | GGAGAGGTAAGG | 1 | 772 | 784 | -228 | -216 | -1,08 | 0,01 | 17,81 | >0.1 | 5,58 | 5,83 | 0 | 1,37 | 0 | 6,1 | 0 |
| FOSL2::JUNB(VAR.2) | AATAACGTATTC | 1 | 800 | 812 | -200 | -188 | 1,73 | 0,01 | 18,23 | >0.1 | 5,58 | 5,98 | 0 | 1,51 | 0 | 3,43 | 0 |
| FOSL2::JUNB(VAR.2) | TGTTACATATCT | 1 | 847 | 859 | -153 | -141 | -0,88 | 0,01 | 15,8 | >0.1 | 5,58 | 6,29 | 0 | 1,85 | 0 | 2,95 | 0 |
| FOSL2::JUNB(VAR.2) | GTGGATGTGATC | 1 | 1020 | 1032 | 20 | 32 | -1,58 | 0,01 | 15,89 | >0.1 | 5,58 | 8,72 | 0 | 1,86 | 0 | 1,31 | 0 |
| FOSL2::JUNB(VAR.2) | GACGATGTCAAG | 1 | 1045 | 1057 | 45 | 57 | 3,67 | 0,01 | 20,28 | >0.1 | 5,58 | 7,7 | 0 | 1,86 | 0 | 1,46 | 0 |
| FOSL2::JUNB(VAR.2) | GTTGCTGTAACT | 1 | 1082 | 1094 | 82 | 94 | -1,24 | 0,01 | 17,19 | >0.1 | 5,58 | 6,99 | 0 | 1,85 | 1,79 | 2,21 | 0 |
| FOSL2::JUNB(VAR.2) | AGTTACAGCAAC | -1 | 1082 | 1094 | 82 | 94 | -1,04 | 0,01 | 17,39 | >0.1 | 5,58 | 6,99 | 0 | 1,85 | 1,79 | 2,21 | 0 |
| FOSL2::JUNB(VAR.2) | CTTGACATCGTC | -1 | 1045 | 1057 | 45 | 57 | 3,86 | 0,01 | 20,47 | >0.1 | 5,58 | 7,7 | 0 | 1,86 | 0 | 1,46 | 0 |
| FOSL2::JUNB(VAR.2) | GATCACATCCAC | -1 | 1020 | 1032 | 20 | 32 | -2,02 | 0,01 | 15,45 | >0.1 | 5,58 | 8,72 | 0 | 1,86 | 0 | 1,31 | 0 |
| FOSL2::JUNB(VAR.2) | AGATATGTAACA | -1 | 847 | 859 | -153 | -141 | -0,89 | 0,01 | 15,79 | >0.1 | 5,58 | 6,29 | 0 | 1,85 | 0 | 2,95 | 0 |
| FOSL2::JUNB(VAR.2) | GAATACGTTATT | -1 | 800 | 812 | -200 | -188 | 1,86 | 0,01 | 18,36 | >0.1 | 5,58 | 5,98 | 0 | 1,51 | 0 | 3,43 | 0 |
| FOSL2::JUNB(VAR.2) | CCTTACCTCTCC | -1 | 772 | 784 | -228 | -216 | -1,23 | 0,01 | 17,66 | >0.1 | 5,58 | 5,83 | 0 | 1,37 | 0 | 6,1 | 0 |
| FOSL2::JUNB(VAR.2) | AATGTTATAATT | -1 | 685 | 697 | -315 | -303 | -0,8 | 0,01 | 13,17 | >0.1 | 4,42 | 5,48 | 0 | 1,13 | 0 | 2,95 | 0 |
| FOSL2::JUNB(VAR.2) | AGTGACCAAAGA | -1 | 669 | 681 | -331 | -319 | -1,71 | 0,01 | 12,65 | >0.1 | 4,14 | 5,42 | 0 | 1,1 | 0 | 3,7 | 0 |
| FOSL2::JUNB(VAR.2) | GGTTATCTCACA | -1 | 590 | 602 | -410 | -398 | 3,92 | 0,01 | 13,53 | >0.1 | 3,41 | 5,19 | 0 | 1,01 | 0 | 0 | 0 |
| FOSL2::JUNB(VAR.2) | CATGACTTTGCC | -1 | 511 | 523 | -489 | -477 | -1,7 | 0,01 | 10,24 | >0.1 | 3,01 | 4,99 | 0 | 0,98 | 0 | 2,95 | 0 |
| FOSL2::JUNB(VAR.2) | AGTGGCGCGATC | -1 | 408 | 420 | -592 | -580 | 0,32 | 0,01 | 14,31 | >0.1 | 2,65 | 4,79 | 0 | 1,03 | 0 | 5,52 | 0 |
| FOSL2::JUNB(VAR.2) | GTTCACGCCATT | -1 | 368 | 380 | -632 | -620 | 2,21 | 0,01 | 16,69 | >0.1 | 2,54 | 4,72 | 0 | 1,08 | 0 | 6,15 | 0 |
| FOSL2::JUNB(VAR.2) | GCTGACTTCTTT | -1 | 30 | 42 | -970 | -958 | 1,04 | 0,01 | 11,65 | >0.1 | 4,46 | 4,28 | 0 | 1,87 | 0 | 0 | 0 |
| USF1 | AAAAAATGGCC | 1 | 140 | 151 | -860 | -849 | -7,03 | 0,01 | 4,24 | >0.1 | 3,34 | 4,4 | 0 | 1,89 | 0 | 1,65 | 0 |
| USF1 | TAAAACTGATC | 1 | 159 | 170 | -841 | -830 | -9,55 | 0,01 | 2,4 | >0.1 | 3,23 | 4,42 | 0 | 1,9 | 0 | 2,4 | 0 |
| USF1 | GGCAGATCACA | 1 | 230 | 241 | -770 | -759 | -8,93 | 0,01 | 4,96 | >0.1 | 2,91 | 4,51 | 0 | 1,45 | 0 | 5,01 | 0 |
| USF1 | ATCACAAGGTC | 1 | 235 | 246 | -765 | -754 | -8,62 | 0,01 | 4,96 | >0.1 | 2,9 | 4,51 | 0 | 1,43 | 0 | 4,74 | 0 |
| USF1 | GTCAGGAGATA | 1 | 243 | 254 | -757 | -746 | -7,35 | 0,01 | 5,76 | >0.1 | 2,87 | 4,52 | 0 | 1,4 | 0 | 4,33 | 0 |
| USF1 | AAAAACTGATC | 1 | 474 | 485 | -526 | -515 | -7,37 | 0,01 | 6,34 | >0.1 | 2,86 | 4,91 | 0 | 0,99 | 0 | 4,95 | 0 |
| USF1 | GTCATGTATTT | 1 | 517 | 528 | -483 | -472 | -9,05 | 0,01 | 2,92 | >0.1 | 3,03 | 5 | 0 | 0,98 | 0 | 2,95 | 0 |
| USF1 | GGAATATGATC | 1 | 563 | 574 | -437 | -426 | 0,99 | 0,01 | 10,35 | >0.1 | 3,25 | 5,11 | 0 | 0,99 | 0 | 0 | 0 |
| USF1 | ATAAAGTGACA | 1 | 761 | 772 | -239 | -228 | -1,61 | 0,01 | 17,19 | >0.1 | 5,6 | 5,78 | 0 | 1,33 | 0 | 6,1 | 0 |
| USF1 | AAAATCTGTTA | 1 | 841 | 852 | -159 | -148 | -9,85 | 0,01 | 5,19 | >0.1 | 5,6 | 6,24 | 0 | 1,79 | 0 | 1,41 | 0 |
| USF1 | TGGATGTGATC | 1 | 1021 | 1032 | 21 | 32 | -5,82 | 0,01 | 11,65 | >0.1 | 5,6 | 8,7 | 0 | 1,86 | 0 | 1,31 | 0 |
| USF1 | CACAGGGGACG | 1 | 1038 | 1049 | 38 | 49 | -7,4 | 0,01 | 9,31 | >0.1 | 5,6 | 7,92 | 0 | 1,86 | 0 | 1,34 | 0 |
| USF1 | GCCTTGTTGCT | 1 | 1077 | 1088 | 77 | 88 | -10,11 | 0,01 | 8,41 | >0.1 | 5,6 | 7,06 | 0 | 1,85 | 1,79 | 2,21 | 0 |
| USF1 | AGCAAGTGAAC | -1 | 1176 | 1187 | 176 | 187 | -6,3 | 0,01 | 12,1 | >0.1 | 5,6 | 6,11 | 0 | 1,62 | 1,79 | 3,28 | 0 |
| USF1 | CAAAAATGTCT | -1 | 1125 | 1136 | 125 | 136 | -10,17 | 0,01 | 9,76 | >0.1 | 5,6 | 6,5 | 0 | 1,84 | 3,22 | 2,77 | 0 |
| USF1 | CTCATGAGATT | -1 | 981 | 992 | -19 | -8 | -5,1 | 0,01 | 12,33 | >0.1 | 5,6 | 8,43 | 0 | 1,87 | 0 | 1,53 | 0 |
| USF1 | CTCTCCTGTCA | -1 | 767 | 778 | -233 | -222 | -1,07 | 0,01 | 17,6 | >0.1 | 5,6 | 5,81 | 0 | 1,35 | 0 | 5,92 | 0 |
| USF1 | AATACATGACT | -1 | 516 | 527 | -484 | -473 | -7,4 | 0,01 | 4,56 | >0.1 | 3,03 | 5 | 0 | 0,98 | 0 | 2,95 | 0 |
| USF1 | TACAGGTGCCC | -1 | 326 | 337 | -674 | -663 | -6,37 | 0,01 | 7,48 | >0.1 | 2,6 | 4,65 | 0 | 1,15 | 0 | 5,46 | 0 |
| USF1 | ATCTCCTGACC | -1 | 242 | 253 | -758 | -747 | 6,13 | 0,001 | 19,25 | >0.1 | 2,87 | 4,52 | 0 | 1,4 | 0 | 4,33 | 0 |
| USF1 | ACCTTGTGATC | -1 | 234 | 245 | -766 | -755 | 8,09 | 0,001 | 21,84 | >0.1 | 2,9 | 4,51 | 0 | 1,44 | 0 | 4,9 | 0 |
| USF1 | AAATTATGACT | -1 | 128 | 139 | -872 | -861 | -7,01 | 0,01 | 4,25 | >0.1 | 3,41 | 4,38 | 0 | 1,89 | 0 | 1,57 | 0 |
| USF1 | CTCTCCTGATC | -1 | 96 | 107 | -904 | -893 | 3,93 | 0,01 | 14,57 | >0.1 | 3,64 | 4,35 | 0 | 1,88 | 0 | 0,77 | 0 |
| TBX15 | AGGAGGCA | 1 | 19 | 27 | -981 | -973 | -0,92 | 0,01 | 10,85 | >0.1 | 5,64 | 4,27 | 0 | 1,87 | 0 | 0 | 0 |
| TBX15 | CAGTGTGA | 1 | 50 | 58 | -950 | -942 | 3,89 | 0,01 | 14,2 | >0.1 | 4,14 | 4,3 | 0 | 1,87 | 0 | 0 | 0 |
| TBX15 | GTGTGAGA | 1 | 52 | 60 | -948 | -940 | -1,32 | 0,01 | 8,96 | >0.1 | 4,11 | 4,3 | 0 | 1,87 | 0 | 0 | 0 |
| TBX15 | TGGAGTGG | 1 | 115 | 123 | -885 | -877 | -3,02 | 0,01 | 7,48 | >0.1 | 3,51 | 4,36 | 0 | 1,89 | 0 | 0,75 | 0 |
| TBX15 | AGGAGATA | 1 | 246 | 254 | -754 | -746 | -2,08 | 0,01 | 11,99 | >0.1 | 2,86 | 4,52 | 0 | 1,38 | 0 | 5,3 | 0 |
| TBX15 | GGGCGTGG | 1 | 314 | 322 | -686 | -678 | 2,06 | 0,01 | 16,63 | >0.1 | 2,64 | 4,62 | 0 | 1,17 | 0 | 6,13 | 0 |
| TBX15 | TGGTGGGC | 1 | 322 | 330 | -678 | -670 | -2,22 | 0,01 | 11,75 | >0.1 | 2,62 | 4,63 | 0 | 1,15 | 0 | 5,56 | 0 |
| TBX15 | TGGCGTGA | 1 | 370 | 378 | -630 | -622 | 3,6 | 0,01 | 18,07 | >0.1 | 2,54 | 4,71 | 0 | 1,07 | 0 | 6,15 | 0 |
| TBX15 | AAATGTGA | 1 | 587 | 595 | -413 | -405 | 0,95 | 0,01 | 10,5 | >0.1 | 3,38 | 5,17 | 0 | 1 | 0 | 0 | 0 |
| TBX15 | ATGTGAGA | 1 | 589 | 597 | -411 | -403 | 2,16 | 0,01 | 11,73 | >0.1 | 3,4 | 5,17 | 0 | 1,01 | 0 | 0 | 0 |
| TBX15 | AAGTGACA | 1 | 764 | 772 | -236 | -228 | -0,22 | 0,01 | 18,44 | >0.1 | 5,64 | 5,78 | 0 | 1,33 | 0 | 5,92 | 0 |
| TBX15 | AGGAGAGG | 1 | 771 | 779 | -229 | -221 | -2,7 | 0,01 | 16,02 | >0.1 | 5,64 | 5,81 | 0 | 1,35 | 0 | 5,92 | 0 |
| TBX15 | GGATGTGA | 1 | 1022 | 1030 | 22 | 30 | 2,48 | 0,01 | 19,97 | >0.1 | 5,64 | 8,68 | 0 | 1,86 | 0 | 1,31 | 0 |
| TBX15 | AGGAGCCA | -1 | 1064 | 1072 | 64 | 72 | -0,49 | 0,01 | 18,35 | >0.1 | 5,64 | 7,28 | 0 | 1,85 | 1,79 | 2,27 | 0 |
| TBX15 | AGCTGTTA | -1 | 710 | 718 | -290 | -282 | 1,06 | 0,01 | 18,23 | >0.1 | 5,64 | 5,55 | 0 | 1,17 | 0 | 4,81 | 0 |

| | | | | | | | | | | | | | | | | |
|---|---|---|---|---|---|---|---|---|---|---|---|---|---|---|---|---|
| TBX15 | TGGTGTTT | -1 | 630 | 638 | -370 | -362 | -1,75 | 0,01 | 9,93 | >0.1 | 3,69 | 5,29 | 0 | 1,04 | 0 | 1,66 | 0 |
| TBX15 | TGGCGCGA | -1 | 410 | 418 | -590 | -582 | -0,5 | 0,01 | 13,48 | >0.1 | 2,65 | 4,78 | 0 | 1,03 | 0 | 5,52 | 0 |
| TBX15 | AGCTGGGA | -1 | 338 | 346 | -662 | -654 | 0,52 | 0,01 | 14,79 | >0.1 | 2,58 | 4,66 | 0 | 1,12 | 0 | 5,92 | 0 |
| TBX15 | AGGTGCCC | -1 | 326 | 334 | -674 | -666 | 0,22 | 0,01 | 14,07 | >0.1 | 2,61 | 4,64 | 0 | 1,15 | 0 | 5,46 | 0 |
| TBX15 | CCGTGTTA | -1 | 268 | 276 | -732 | -724 | -1,77 | 0,01 | 11,4 | >0.1 | 2,78 | 4,55 | 0 | 1,3 | 0 | 4,53 | 0 |
| TBX15 | AAGTGCTA | -1 | 207 | 215 | -793 | -785 | 2,46 | 0,01 | 16,23 | >0.1 | 3,01 | 4,47 | 0 | 1,58 | 0 | 4,7 | 0 |
| TBX15 | AGGCGTGA | -1 | 192 | 200 | -808 | -800 | 10,15 | 0,001 | 23,5 | >0.1 | 3,08 | 4,45 | 0 | 1,69 | 0 | 4,13 | 0 |
| MAX::MYC | GAGAAGGAGGC | 1 | 15 | 26 | -985 | -974 | -0,24 | 0,01 | 11,49 | >0.1 | 5,6 | 4,27 | 0 | 1,87 | 0 | 0 | 0 |
| MAX::MYC | TGACCTGTGGA | 1 | 108 | 119 | -892 | -881 | 0,1 | 0,01 | 10,64 | >0.1 | 3,55 | 4,36 | 0 | 1,89 | 0 | 0,75 | 0 |
| MAX::MYC | GATCACAAGGT | 1 | 234 | 245 | -766 | -755 | 3,83 | 0,01 | 17,58 | >0.1 | 2,9 | 4,51 | 0 | 1,44 | 0 | 4,9 | 0 |
| MAX::MYC | TAACACGGTGA | 1 | 268 | 279 | -732 | -721 | -1,12 | 0,01 | 11,97 | >0.1 | 2,78 | 4,56 | 0 | 1,3 | 0 | 4,45 | 0 |
| MAX::MYC | AACCCCGTCTC | 1 | 279 | 290 | -721 | -710 | -0,58 | 0,01 | 13,2 | >0.1 | 2,74 | 4,58 | 0 | 1,26 | 0 | 5,19 | 0 |
| MAX::MYC | CTGGGCGTGGT | 1 | 312 | 323 | -688 | -677 | -1 | 0,01 | 13,48 | >0.1 | 2,64 | 4,62 | 0 | 1,18 | 0 | 6,03 | 0 |
| MAX::MYC | GGGCACCTGTA | 1 | 326 | 337 | -674 | -663 | 4,01 | 0,01 | 17,86 | >0.1 | 2,6 | 4,65 | 0 | 1,15 | 0 | 5,46 | 0 |
| MAX::MYC | GAGATCGCGCC | 1 | 406 | 417 | -594 | -583 | -1,15 | 0,01 | 12,95 | >0.1 | 2,64 | 4,78 | 0 | 1,03 | 0 | 5,65 | 0 |
| MAX::MYC | GAAAATGTTGC | 1 | 534 | 545 | -466 | -455 | -1,28 | 0,01 | 9,51 | >0.1 | 3,11 | 5,04 | 0 | 0,99 | 0 | 1,66 | 0 |
| MAX::MYC | GCCCAAGTAGA | 1 | 543 | 554 | -457 | -446 | -1,09 | 0,01 | 8,11 | >0.1 | 3,15 | 5,06 | 0 | 0,99 | 0 | 0 | 0 |
| MAX::MYC | GGGAATATGAT | 1 | 562 | 573 | -438 | -427 | -0,99 | 0,01 | 8,36 | >0.1 | 3,25 | 5,11 | 0 | 0,99 | 0 | 0 | 0 |
| MAX::MYC | GTGGATGTGAT | 1 | 1020 | 1031 | 20 | 31 | -0,81 | 0,01 | 16,68 | >0.1 | 5,6 | 8,72 | 0 | 1,86 | 0 | 1,31 | 0 |
| MAX::MYC | AACCACGAAGC | 1 | 1145 | 1156 | 145 | 156 | 2,92 | 0,01 | 22,08 | >0.1 | 5,6 | 6,33 | 0 | 1,84 | 3,22 | 2,16 | 0 |
| MAX::MYC | GTTCACTTGCT | 1 | 1176 | 1187 | 176 | 187 | 0,32 | 0,01 | 18,72 | >0.1 | 5,6 | 6,11 | 0 | 1,62 | 1,79 | 3,28 | 0 |
| MAX::MYC | AAGCAAGTGAA | -1 | 1177 | 1188 | 177 | 188 | 4,03 | 0,01 | 22,4 | >0.1 | 5,6 | 6,11 | 0 | 1,61 | 1,79 | 3,26 | 0 |
| MAX::MYC | GGCTTCGTGGT | -1 | 1146 | 1157 | 146 | 157 | 3,25 | 0,01 | 20,97 | >0.1 | 5,6 | 6,33 | 0 | 1,84 | 1,79 | 2,16 | 0 |
| MAX::MYC | AGCAACAAGGC | -1 | 1077 | 1088 | 77 | 88 | -0,36 | 0,01 | 18,16 | >0.1 | 5,6 | 7,06 | 0 | 1,85 | 1,79 | 2,21 | 0 |
| MAX::MYC | CACCACGCCCA | -1 | 313 | 324 | -687 | -676 | -0,71 | 0,01 | 13,86 | >0.1 | 2,64 | 4,63 | 0 | 1,17 | 0 | 6,13 | 0 |
| MAX::MYC | AGAGACGGGGT | -1 | 280 | 291 | -720 | -709 | 0,91 | 0,01 | 14,58 | >0.1 | 2,74 | 4,58 | 0 | 1,26 | 0 | 5,09 | 0 |
| MAX::MYC | GACCTTGTGAT | -1 | 235 | 246 | -765 | -754 | 2,78 | 0,01 | 16,36 | >0.1 | 2,9 | 4,51 | 0 | 1,43 | 0 | 4,74 | 0 |
| FOXB1 | AATAAAAATAA | 1 | 578 | 589 | -422 | -411 | 5,24 | 0,01 | 14,73 | >0.1 | 3,34 | 5,15 | 0 | 1 | 0 | 0 | 0 |
| FOXB1 | AAAATAAATGT | 1 | 582 | 593 | -418 | -407 | 5,48 | 0,01 | 15,01 | >0.1 | 3,36 | 5,16 | 0 | 1,01 | 0 | 0 | 0 |
| FOXB1 | CATACAAACAC | 1 | 625 | 636 | -375 | -364 | 1,9 | 0,01 | 13,43 | >0.1 | 3,67 | 5,28 | 0 | 1,04 | 0 | 1,55 | 0 |
| FOXB1 | AACACCAACAT | 1 | 631 | 642 | -369 | -358 | 2,04 | 0,01 | 13,76 | >0.1 | 3,72 | 5,3 | 0 | 1,05 | 0 | 1,66 | 0 |
| FOXB1 | TAGGAAAATAT | 1 | 721 | 732 | -279 | -268 | 7,35 | 0,01 | 24,93 | >0.1 | 5,6 | 5,61 | 0 | 1,21 | 0 | 5,17 | 0 |
| FOXB1 | TGAGCCAAGAT | -1 | 1029 | 1040 | 29 | 40 | 2,91 | 0,01 | 19,99 | >0.1 | 5,6 | 8,29 | 0 | 1,86 | 0 | 1,34 | 0 |
| FOXB1 | ACAGTAAACAA | -1 | 896 | 907 | -104 | -93 | 7,87 | 0,001 | 24,98 | >0.1 | 5,6 | 6,68 | 0 | 1,89 | 0 | 2,95 | 0 |
| FOXB1 | TGAGCCAATAT | -1 | 880 | 891 | -120 | -109 | 7,88 | 0,001 | 24,98 | >0.1 | 5,6 | 6,54 | 0 | 1,9 | 0 | 3,07 | 0 |
| FOXB1 | TTGGTTAATAT | -1 | 728 | 739 | -272 | -261 | 3,29 | 0,01 | 21,06 | >0.1 | 5,6 | 5,63 | 0 | 1,22 | 0 | 5,32 | 0 |
| FOXB1 | TATTTCAAGAT | -1 | 571 | 582 | -429 | -418 | 3,6 | 0,01 | 13,03 | >0.1 | 3,3 | 5,13 | 0 | 1 | 0 | 0 | 0 |
| FOXB1 | TTGGGCAACAT | -1 | 538 | 549 | -462 | -451 | 2,57 | 0,01 | 13,47 | >0.1 | 3,13 | 5,05 | 0 | 0,99 | 0 | 1,73 | 0 |
| FOS::JUN | GTGAGAAATA | 1 | 54 | 64 | -946 | -936 | 0,85 | 0,01 | 11,1 | >0.1 | 4,07 | 4,3 | 0 | 1,87 | 0 | 0 | 0 |
| FOS::JUN | AGTAGTCATA | 1 | 125 | 135 | -875 | -865 | 3,51 | 0,01 | 14,91 | >0.1 | 3,43 | 4,38 | 0 | 1,89 | 0 | 1,7 | 0 |
| FOS::JUN | GTGGCTCACG | 1 | 187 | 197 | -813 | -803 | 4,24 | 0,01 | 17,37 | >0.1 | 3,1 | 4,45 | 0 | 1,73 | 0 | 3,86 | 0 |
| FOS::JUN | CTGACTAACA | 1 | 263 | 273 | -737 | -727 | 4,91 | 0,01 | 18,02 | >0.1 | 2,8 | 4,55 | 0 | 1,32 | 0 | 4,45 | 0 |
| FOS::JUN | CTGAGGCAGG | 1 | 356 | 366 | -644 | -634 | 2,04 | 0,01 | 16,48 | >0.1 | 2,52 | 4,69 | 0 | 1,09 | 0 | 6,13 | 0 |
| FOS::JUN | GAGACTCAGT | 1 | 444 | 454 | -556 | -546 | 0,85 | 0,01 | 15,38 | >0.1 | 2,76 | 4,85 | 0 | 1 | 0 | 5,92 | 0 |
| FOS::JUN | TTTGGTCACT | 1 | 671 | 681 | -329 | -319 | -0,3 | 0,01 | 13,93 | >0.1 | 4,16 | 5,42 | 0 | 1,1 | 0 | 3,55 | 0 |
| FOS::JUN | TTGATTCACT | 1 | 816 | 826 | -184 | -174 | 5,44 | 0,01 | 22,25 | >0.1 | 5,61 | 6,06 | 0 | 1,59 | 0 | 3,55 | 0 |
| FOS::JUN | TTGGCTCAGC | 1 | 883 | 893 | -117 | -107 | 0,63 | 0,01 | 17,76 | >0.1 | 5,61 | 6,56 | 0 | 1,89 | 0 | 3,07 | 0 |
| FOS::JUN | TTGGCTCACA | 1 | 1032 | 1042 | 32 | 42 | 1,33 | 0,01 | 18,22 | >0.1 | 5,61 | 8,15 | 0 | 1,86 | 0 | 1,28 | 0 |
| FOS::JUN | GTTCCTCAAT | -1 | 1109 | 1119 | 109 | 119 | 1,71 | 0,01 | 21,57 | >0.1 | 5,61 | 6,66 | 0 | 1,85 | 3,22 | 2,52 | 0 |
| FOS::JUN | GTGAGCCAAG | -1 | 1031 | 1041 | 31 | 41 | 0,78 | 0,01 | 17,75 | >0.1 | 5,61 | 8,19 | 0 | 1,86 | 0 | 1,31 | 0 |
| FOS::JUN | CTGAATGACT | -1 | 1011 | 1021 | 11 | 21 | -0,3 | 0,01 | 17,52 | >0.1 | 5,61 | 8,95 | 0 | 1,86 | 0 | 1,39 | 0 |
| FOS::JUN | GTGAATCAAA | -1 | 815 | 825 | -185 | -175 | 5,72 | 0,01 | 22,4 | >0.1 | 5,61 | 6,06 | 0 | 1,58 | 0 | 3,43 | 0 |
| FOS::JUN | CTGAGTCTCG | -1 | 443 | 453 | -557 | -547 | 4,2 | 0,01 | 18,73 | >0.1 | 2,75 | 4,85 | 0 | 1 | 0 | 5,92 | 0 |
| FOS::JUN | CTGCCTCCCA | -1 | 381 | 391 | -619 | -609 | -0,06 | 0,01 | 14,32 | >0.1 | 2,57 | 4,73 | 0 | 1,06 | 0 | 6,02 | 0 |
| FOS::JUN | CTGCCTCAGC | -1 | 355 | 365 | -645 | -635 | 7,55 | 0,001 | 21,9 | >0.1 | 2,53 | 4,69 | 0 | 1,09 | 0 | 6,03 | 0 |
| FOS::JUN | GTTAGTCAGG | -1 | 262 | 272 | -738 | -728 | 8,78 | 0,001 | 21,98 | >0.1 | 2,8 | 4,55 | 0 | 1,32 | 0 | 4,53 | 0 |
| FOS::JUN | GTGAGCCACT | -1 | 186 | 196 | -814 | -804 | 4,46 | 0,01 | 17,6 | >0.1 | 3,1 | 4,45 | 0 | 1,74 | 0 | 3,86 | 0 |
| FOS::JUN | ATGACTACTC | -1 | 124 | 134 | -876 | -866 | -0,11 | 0,01 | 11,29 | >0.1 | 3,44 | 4,38 | 0 | 1,89 | 0 | 1,7 | 0 |
| FOS::JUN | ATTTCTCACA | -1 | 53 | 63 | -947 | -937 | 5,35 | 0,01 | 15,61 | >0.1 | 4,09 | 4,3 | 0 | 1,87 | 0 | 0 | 0 |

| | | | | | | | | | | | | | | | | | |
|---|---|---|---|---|---|---|---|---|---|---|---|---|---|---|---|---|---|
| FOS::JUN | GTGCCTCCTT | -1 | 18 | 28 | -982 | -972 | 0,74 | 0,01 | 12,48 | >0.1 | 5,61 | 4,27 | 0 | 1,87 | 0 | 0 | 0 |
| POU1F1 | AAATAGGAAATGAG | 1 | 59 | 73 | -941 | -927 | 1,04 | 0,01 | 11,21 | >0.1 | 3,98 | 4,31 | 0 | 1,88 | 0 | 0 | 0 |
| POU1F1 | GTTTTGCAAAGGCA | 1 | 82 | 96 | -918 | -904 | 1,05 | 0,01 | 11,01 | >0.1 | 3,75 | 4,34 | 0 | 1,88 | 0 | 0 | 0 |
| POU1F1 | CAAAGGCAGATCAG | 1 | 88 | 102 | -912 | -898 | 3,45 | 0,01 | 13,37 | >0.1 | 3,69 | 4,34 | 0 | 1,88 | 0 | 0 | 0 |
| POU1F1 | ATAATTTTAAAAAA | 1 | 132 | 146 | -868 | -854 | 0,54 | 0,01 | 11,81 | >0.1 | 3,38 | 4,39 | 0 | 1,89 | 0 | 1,61 | 0 |
| POU1F1 | GCCATGGAAATTAA | 1 | 148 | 162 | -852 | -838 | 5,9 | 0,001 | 17,14 | >0.1 | 3,28 | 4,41 | 0 | 1,9 | 0 | 1,65 | 0 |
| POU1F1 | TCTATTAAAAATAC | 1 | 288 | 302 | -712 | -698 | -0,13 | 0,01 | 13,5 | >0.1 | 2,71 | 4,59 | 0 | 1,24 | 0 | 5,09 | 0 |
| POU1F1 | GATCTTGAAATAAA | 1 | 570 | 584 | -430 | -416 | -0,58 | 0,01 | 8,86 | >0.1 | 3,3 | 5,14 | 0 | 1 | 0 | 0 | 0 |
| POU1F1 | GAAATAAAAATAAA | 1 | 576 | 590 | -424 | -410 | -0,67 | 0,01 | 8,83 | >0.1 | 3,33 | 5,15 | 0 | 1,01 | 0 | 0 | 0 |
| POU1F1 | ATTAATGAAATTGT | 1 | 603 | 617 | -397 | -383 | 1,22 | 0,01 | 12,39 | >0.1 | 3,51 | 5,23 | 0 | 1,02 | 0 | 1,41 | 0 |
| POU1F1 | ATTATCTTCATGAT | 1 | 640 | 654 | -360 | -346 | -0,28 | 0,01 | 13,76 | >0.1 | 3,81 | 5,34 | 0 | 1,06 | 0 | 3,83 | 0 |
| POU1F1 | CGTATTCTTATTTG | 1 | 805 | 819 | -195 | -181 | 4,01 | 0,01 | 20,42 | >0.1 | 5,56 | 6,02 | 0 | 1,55 | 0 | 3,28 | 0 |
| POU1F1 | ATTATTCTAAAATC | 1 | 833 | 847 | -167 | -153 | 2,88 | 0,01 | 19,67 | >0.1 | 5,56 | 6,21 | 0 | 1,75 | 0 | 3,28 | 0 |
| POU1F1 | CAAAGGCTGATAAG | 1 | 961 | 975 | -39 | -25 | 2,28 | 0,01 | 19,16 | >0.1 | 5,56 | 7,76 | 0 | 1,88 | 0 | 1,7 | 0 |
| POU1F1 | AAAAAGAAAAAGAA | -1 | 914 | 928 | -86 | -72 | -0,23 | 0,01 | 15,24 | >0.1 | 5,56 | 6,9 | 0 | 1,89 | 0 | 1,12 | 0 |
| POU1F1 | GATATGTAACAGAT | -1 | 844 | 858 | -156 | -142 | 1,43 | 0,01 | 17,98 | >0.1 | 5,56 | 6,28 | 0 | 1,84 | 0 | 2,86 | 0 |
| POU1F1 | GTGAATCAAATAAG | -1 | 811 | 825 | -189 | -175 | 2,35 | 0,01 | 18,98 | >0.1 | 5,56 | 6,06 | 0 | 1,59 | 0 | 3,43 | 0 |
| POU1F1 | AAGCTGTTAAAGAT | -1 | 705 | 719 | -295 | -281 | -0,8 | 0,01 | 16,3 | >0.1 | 5,56 | 5,56 | 0 | 1,18 | 0 | 4,81 | 0 |
| POU1F1 | AATGTTATAATTTT | -1 | 683 | 697 | -317 | -303 | -0,39 | 0,01 | 13,56 | >0.1 | 4,4 | 5,48 | 0 | 1,13 | 0 | 2,95 | 0 |
| POU1F1 | ATCATGAAGATAAT | -1 | 640 | 654 | -360 | -346 | 1,68 | 0,01 | 15,72 | >0.1 | 3,81 | 5,34 | 0 | 1,06 | 0 | 3,83 | 0 |
| POU1F1 | ATGAAGATAATGTT | -1 | 637 | 651 | -363 | -349 | -0,41 | 0,01 | 11,42 | >0.1 | 3,79 | 5,33 | 0 | 1,06 | 0 | 1,66 | 0 |
| POU1F1 | TTTATTTTTATTTC | -1 | 576 | 590 | -424 | -410 | 1,48 | 0,01 | 10,98 | >0.1 | 3,33 | 5,15 | 0 | 1,01 | 0 | 0 | 0 |
| POU1F1 | TTTATGGAATTCAT | -1 | 491 | 505 | -509 | -495 | 0,92 | 0,01 | 16,23 | >0.1 | 2,93 | 4,95 | 0 | 0,99 | 0 | 6,44 | 0 |
| POU1F1 | GGAATTCATATTTC | -1 | 486 | 500 | -514 | -500 | -0,38 | 0,01 | 14,73 | >0.1 | 2,91 | 4,94 | 0 | 0,99 | 0 | 6,27 | 0 |
| POU1F1 | ATATTTCTGATCAG | -1 | 479 | 493 | -521 | -507 | -0,52 | 0,01 | 13,93 | >0.1 | 2,89 | 4,93 | 0 | 0,99 | 0 | 5,65 | 0 |
| POU1F1 | TGTATTTTTAATAG | -1 | 289 | 303 | -711 | -697 | -0,13 | 0,01 | 12,86 | >0.1 | 2,71 | 4,59 | 0 | 1,24 | 0 | 4,45 | 0 |
| POU1F1 | TTTTTTAAAATTAT | -1 | 132 | 146 | -868 | -854 | 0,1 | 0,01 | 11,37 | >0.1 | 3,38 | 4,39 | 0 | 1,89 | 0 | 1,61 | 0 |
| DLX6 | ATAATTTT | 1 | 132 | 140 | -868 | -860 | 4,39 | 0,01 | 15,63 | >0.1 | 3,4 | 4,38 | 0 | 1,89 | 0 | 1,57 | 0 |
| DLX6 | GAAATTAA | 1 | 154 | 162 | -846 | -838 | 5,85 | 0,01 | 17,07 | >0.1 | 3,27 | 4,41 | 0 | 1,9 | 0 | 1,65 | 0 |
| DLX6 | TCTATTAA | 1 | 288 | 296 | -712 | -704 | 4,67 | 0,01 | 18,4 | >0.1 | 2,72 | 4,58 | 0 | 1,24 | 0 | 5,19 | 0 |
| DLX6 | CCTATTAA | 1 | 600 | 608 | -400 | -392 | 6,32 | 0,01 | 17,37 | >0.1 | 3,47 | 5,2 | 0 | 1,01 | 0 | 1,36 | 0 |
| DLX6 | TTAATGAA | 1 | 604 | 612 | -396 | -388 | 3,02 | 0,01 | 14,16 | >0.1 | 3,49 | 5,21 | 0 | 1,02 | 0 | 1,41 | 0 |
| DLX6 | AAAATTAT | 1 | 683 | 691 | -317 | -309 | 5,01 | 0,01 | 18,99 | >0.1 | 4,34 | 5,46 | 0 | 1,12 | 0 | 3,07 | 0 |
| DLX6 | AAAATTAT | 1 | 830 | 838 | -170 | -162 | 5,01 | 0,01 | 21,74 | >0.1 | 5,64 | 6,14 | 0 | 1,67 | 0 | 3,28 | 0 |
| DLX6 | GGAATTAT | 1 | 1191 | 1199 | 191 | 199 | 3,49 | 0,01 | 22,27 | >0.1 | 5,64 | 6,02 | 0 | 1,5 | 1,79 | 3,82 | 0 |
| DLX6 | ATAATTCC | -1 | 1191 | 1199 | 191 | 199 | 3,11 | 0,01 | 21,89 | >0.1 | 5,64 | 6,02 | 0 | 1,5 | 1,79 | 3,82 | 0 |
| DLX6 | ATAATTTT | -1 | 830 | 838 | -170 | -162 | 4,39 | 0,01 | 21,12 | >0.1 | 5,64 | 6,14 | 0 | 1,67 | 0 | 3,28 | 0 |
| DLX6 | ATAATTTT | -1 | 683 | 691 | -317 | -309 | 4,39 | 0,01 | 18,37 | >0.1 | 4,34 | 5,46 | 0 | 1,12 | 0 | 3,07 | 0 |
| DLX6 | ACAATTTC | -1 | 609 | 617 | -391 | -383 | 6 | 0,01 | 17,32 | >0.1 | 3,53 | 5,23 | 0 | 1,02 | 0 | 1,55 | 0 |
| DLX6 | TTCATTAA | -1 | 604 | 612 | -396 | -388 | 5,02 | 0,01 | 16,16 | >0.1 | 3,49 | 5,21 | 0 | 1,02 | 0 | 1,41 | 0 |
| DLX6 | ACATTTAT | -1 | 585 | 593 | -415 | -407 | 3,32 | 0,01 | 12,85 | >0.1 | 3,37 | 5,16 | 0 | 1 | 0 | 0 | 0 |
| DLX6 | CCAAATAC | -1 | 522 | 530 | -478 | -470 | 3,13 | 0,01 | 15,23 | >0.1 | 3,05 | 5,01 | 0 | 0,98 | 0 | 3,07 | 0 |
| DLX6 | GCCATTTC | -1 | 171 | 179 | -829 | -821 | 3,91 | 0,01 | 16,81 | >0.1 | 3,18 | 4,43 | 0 | 1,87 | 0 | 3,43 | 0 |
| DLX6 | TTAATTTC | -1 | 154 | 162 | -846 | -838 | 4,75 | 0,01 | 15,97 | >0.1 | 3,27 | 4,41 | 0 | 1,9 | 0 | 1,65 | 0 |
| DLX6 | GCCATTTT | -1 | 142 | 150 | -858 | -850 | 3,01 | 0,01 | 14,24 | >0.1 | 3,33 | 4,39 | 0 | 1,89 | 0 | 1,61 | 0 |
| DLX6 | AAAATTAT | -1 | 132 | 140 | -868 | -860 | 5,01 | 0,01 | 16,25 | >0.1 | 3,4 | 4,38 | 0 | 1,89 | 0 | 1,57 | 0 |
| DLX6 | CTCATTTC | -1 | 65 | 73 | -935 | -927 | 3,64 | 0,01 | 13,78 | >0.1 | 3,95 | 4,31 | 0 | 1,88 | 0 | 0 | 0 |
| DLX6 | CCTATTTC | -1 | 58 | 66 | -942 | -934 | 3,29 | 0,01 | 13,5 | >0.1 | 4,03 | 4,3 | 0 | 1,87 | 0 | 0 | 0 |
| DLX6 | TCACTTAC | -1 | 5 | 13 | -995 | -987 | 5,32 | 0,01 | 18,87 | >0.1 | 5,64 | 4,26 | 0 | 1,86 | 1,79 | 0 | 0 |
| ASCL2 | GGCACCTGTA | 1 | 327 | 337 | -673 | -663 | 8,01 | 0,001 | 21,97 | >0.1 | 2,6 | 4,65 | 0 | 1,14 | 0 | 5,56 | 0 |
| ASCL2 | CCCAGCTACT | 1 | 339 | 349 | -661 | -651 | 1,96 | 0,01 | 16,84 | >0.1 | 2,57 | 4,66 | 0 | 1,12 | 0 | 6,52 | 0 |
| ASCL2 | TGCTCCAGCC | 1 | 421 | 431 | -579 | -569 | -1,23 | 0,01 | 12,7 | >0.1 | 2,69 | 4,81 | 0 | 1,02 | 0 | 5,42 | 0 |
| ASCL2 | AACAGCTTTC | 1 | 711 | 721 | -289 | -279 | 6,66 | 0,001 | 24 | >0.1 | 5,61 | 5,56 | 0 | 1,18 | 0 | 4,99 | 0 |
| ASCL2 | AAAATCTGTT | 1 | 841 | 851 | -159 | -149 | 3,15 | 0,01 | 18,19 | >0.1 | 5,61 | 6,24 | 0 | 1,78 | 0 | 1,41 | 0 |
| ASCL2 | CATATCTGTC | 1 | 852 | 862 | -148 | -138 | -0,97 | 0,01 | 15,5 | >0.1 | 5,61 | 6,31 | 0 | 1,88 | 0 | 2,67 | 0 |
| ASCL2 | AGCAGATTGT | 1 | 890 | 900 | -110 | -100 | -1,12 | 0,01 | 15,94 | >0.1 | 5,61 | 6,61 | 0 | 1,89 | 0 | 2,95 | 0 |
| ASCL2 | GGCTCCTTCT | 1 | 1065 | 1075 | 65 | 75 | 0,28 | 0,01 | 19,07 | >0.1 | 5,61 | 7,26 | 0 | 1,85 | 1,79 | 2,27 | 0 |
| ASCL2 | TGTAACTGCT | 1 | 1087 | 1097 | 87 | 97 | -0,93 | 0,01 | 17,46 | >0.1 | 5,61 | 6,92 | 0 | 1,85 | 1,79 | 2,21 | 0 |
| ASCL2 | AACTGCTGCT | 1 | 1090 | 1100 | 90 | 100 | 12,06 | 0,0001 | 30,47 | >0.1 | 5,61 | 6,88 | 0 | 1,85 | 1,79 | 2,27 | 0 |

| | | | | | | | | | | | | | | | | | |
|---|---|---|---|---|---|---|---|---|---|---|---|---|---|---|---|---|---|
| ASCL2 | AAGACCTGTT | 1 | 1158 | 1168 | 158 | 168 | 0,49 | 0,01 | 17,08 | >0.1 | 4,38 | 6,24 | 0 | 1,78 | 1,79 | 2,4 | 0 |
| ASCL2 | AACAGGTCTT | -1 | 1158 | 1168 | 158 | 168 | -0,33 | 0,01 | 16,26 | >0.1 | 4,38 | 6,24 | 0 | 1,78 | 1,79 | 2,4 | 0 |
| ASCL2 | AGCAGCAGTT | -1 | 1090 | 1100 | 90 | 100 | 9,5 | 0,001 | 27,91 | >0.1 | 5,61 | 6,88 | 0 | 1,85 | 1,79 | 2,27 | 0 |
| ASCL2 | ACAATCTGCT | -1 | 890 | 900 | -110 | -100 | 3,63 | 0,01 | 20,69 | >0.1 | 5,61 | 6,61 | 0 | 1,89 | 0 | 2,95 | 0 |
| ASCL2 | AACAGATTTT | -1 | 841 | 851 | -159 | -149 | 0,1 | 0,01 | 15,14 | >0.1 | 5,61 | 6,24 | 0 | 1,78 | 0 | 1,41 | 0 |
| ASCL2 | GAAAGCTGTT | -1 | 711 | 721 | -289 | -279 | 6,27 | 0,01 | 23,61 | >0.1 | 5,61 | 5,56 | 0 | 1,18 | 0 | 4,99 | 0 |
| ASCL2 | AGTAGCTGGG | -1 | 339 | 349 | -661 | -651 | 2,73 | 0,01 | 17,61 | >0.1 | 2,57 | 4,66 | 0 | 1,12 | 0 | 6,52 | 0 |
| ASCL2 | TACAGGTGCC | -1 | 327 | 337 | -673 | -663 | 4,92 | 0,01 | 18,88 | >0.1 | 2,6 | 4,65 | 0 | 1,14 | 0 | 5,56 | 0 |
| ASCL1 | GGAGAAGGAGGCA | 1 | 14 | 27 | -986 | -973 | -0,71 | 0,01 | 10,99 | >0.1 | 5,57 | 4,27 | 0 | 1,87 | 0 | 0 | 0 |
| ASCL1 | CGTGGTGGTGGGC | 1 | 317 | 330 | -683 | -670 | -0,66 | 0,01 | 14,02 | >0.1 | 2,63 | 4,64 | 0 | 1,17 | 0 | 6,26 | 0 |
| ASCL1 | TGGGCACCTGTAG | 1 | 325 | 338 | -675 | -662 | 8,51 | 0,001 | 22,37 | >0.1 | 2,6 | 4,65 | 0 | 1,15 | 0 | 5,46 | 0 |
| ASCL1 | GTCCCAGCTACTC | 1 | 337 | 350 | -663 | -650 | -0,71 | 0,01 | 13,57 | >0.1 | 2,57 | 4,67 | 0 | 1,13 | 0 | 5,92 | 0 |
| ASCL1 | CGGGAGGCTGAGG | 1 | 349 | 362 | -651 | -638 | -1,02 | 0,01 | 13,71 | >0.1 | 2,54 | 4,69 | 0 | 1,11 | 0 | 6,4 | 0 |
| ASCL1 | GAGGCAGGAGAAT | 1 | 358 | 371 | -642 | -629 | 0,09 | 0,01 | 14,76 | >0.1 | 2,52 | 4,7 | 0 | 1,09 | 0 | 6,36 | 0 |
| ASCL1 | CGTGAACCTGGGA | 1 | 373 | 386 | -627 | -614 | 1,83 | 0,01 | 16,71 | >0.1 | 2,56 | 4,73 | 0 | 1,07 | 0 | 6,53 | 0 |
| ASCL1 | TCAGCAGATTGTT | 1 | 888 | 901 | -112 | -99 | 1,41 | 0,01 | 18,56 | >0.1 | 5,57 | 6,62 | 0 | 1,89 | 0 | 3,07 | 0 |
| ASCL1 | GTAACTGCTGCTC | 1 | 1088 | 1101 | 88 | 101 | 4,17 | 0,01 | 22,5 | >0.1 | 5,57 | 6,9 | 0 | 1,85 | 1,79 | 2,21 | 0 |
| ASCL1 | CGAAGACCTGTTC | 1 | 1156 | 1169 | 156 | 169 | -1,21 | 0,01 | 15,36 | >0.1 | 4,39 | 6,25 | 0 | 1,8 | 1,79 | 2,34 | 0 |
| ASCL1 | GAAGCAAGTGAAC | -1 | 1176 | 1189 | 176 | 189 | 0,52 | 0,01 | 18,88 | >0.1 | 5,57 | 6,11 | 0 | 1,62 | 1,79 | 3,26 | 0 |
| ASCL1 | TGAGCAGCAGTTA | -1 | 1089 | 1102 | 89 | 102 | 6,44 | 0,01 | 24,82 | >0.1 | 5,57 | 6,89 | 0 | 1,85 | 1,79 | 2,27 | 0 |
| ASCL1 | GCAGCAGTTACAG | -1 | 1086 | 1099 | 86 | 99 | 0,4 | 0,01 | 18,76 | >0.1 | 5,57 | 6,93 | 0 | 1,85 | 1,79 | 2,21 | 0 |
| ASCL1 | CGAGTAGCTGGGA | -1 | 338 | 351 | -662 | -649 | 8,88 | 0,001 | 23,76 | >0.1 | 2,57 | 4,67 | 0 | 1,12 | 0 | 6,52 | 0 |
| ASCL1 | ACTACAGGTGCCC | -1 | 326 | 339 | -674 | -661 | 5,85 | 0,01 | 19,81 | >0.1 | 2,6 | 4,65 | 0 | 1,15 | 0 | 5,56 | 0 |
| ASCL1 | TGAGCCACTGTGC | -1 | 182 | 195 | -818 | -805 | -0,48 | 0,01 | 12,71 | >0.1 | 3,11 | 4,45 | 0 | 1,77 | 0 | 3,86 | 0 |
| TBX19 | CACAGAACCAGTGTGAGAAA | 1 | 42 | 62 | -958 | -938 | -0,44 | 0,01 | 9,91 | >0.1 | 4,17 | 4,3 | 0 | 1,87 | 0 | 0 | 0 |
| TBX19 | AGTGGCTCACGCCTGTAATC | 1 | 186 | 206 | -814 | -794 | -5,46 | 0,01 | 7,95 | >0.1 | 3,08 | 4,46 | 0 | 1,74 | 0 | 4,13 | 0 |
| TBX19 | GATCACAAGGTCAGGAGATA | 1 | 234 | 254 | -766 | -746 | -1,57 | 0,01 | 11,7 | >0.1 | 2,88 | 4,53 | 0 | 1,44 | 0 | 4,42 | 0 |
| TBX19 | AGTCCCAGCTACTCGGGAGG | 1 | 336 | 356 | -664 | -644 | 0,87 | 0,01 | 15,77 | >0.1 | 2,56 | 4,68 | 0 | 1,13 | 0 | 6,52 | 0 |
| TBX19 | GGCAGGAGAATGGCGTGAAC | 1 | 360 | 380 | -640 | -620 | -4,21 | 0,01 | 10,74 | >0.1 | 2,53 | 4,72 | 0 | 1,09 | 0 | 6,61 | 0 |
| TBX19 | AATGGCGTGAACCTGGGAGG | 1 | 368 | 388 | -632 | -612 | 0,09 | 0,01 | 14,98 | >0.1 | 2,55 | 4,73 | 0 | 1,08 | 0 | 6,53 | 0 |
| TBX19 | GCTTGCAGTGAGCCGAGATC | 1 | 392 | 412 | -608 | -588 | -2,6 | 0,01 | 11,75 | >0.1 | 2,62 | 4,77 | 0 | 1,05 | 0 | 5,91 | 0 |
| TBX19 | GATCGCGCCACTGTGCTCCA | 1 | 408 | 428 | -592 | -572 | 2,02 | 0,001 | 15,6 | >0.1 | 2,66 | 4,8 | 0 | 1,03 | 0 | 5,09 | 0 |
| TBX19 | GCTCCAGCCTGGGCGACAGA | 1 | 422 | 442 | -578 | -558 | -5,4 | 0,01 | 8,96 | >0.1 | 2,7 | 4,83 | 0 | 1,02 | 0 | 5,8 | 0 |
| TBX19 | ATAAAAATAAATGTGAGATA | 1 | 579 | 599 | -421 | -401 | -4,15 | 0,01 | 5,41 | >0.1 | 3,37 | 5,18 | 0 | 1,02 | 0 | 0 | 0 |
| TBX19 | TTTAACAGCTTTCTAGGAAA | 1 | 708 | 728 | -292 | -272 | 0,55 | 0,01 | 17,99 | >0.1 | 5,48 | 5,59 | 0 | 1,2 | 0 | 5,17 | 0 |
| TBX19 | TTTGGCCATAAAGTGACAGG | 1 | 754 | 774 | -246 | -226 | -2,71 | 0,01 | 16 | >0.1 | 5,48 | 5,79 | 0 | 1,34 | 0 | 6,1 | 0 |
| TBX19 | GAGGACAGATATGTAACAGA | -1 | 845 | 865 | -155 | -135 | -5,44 | 0,01 | 11,03 | >0.1 | 5,48 | 6,33 | 0 | 1,91 | 0 | 2,75 | 0 |
| TBX19 | CCTGTCACTTTATGGCCAAA | -1 | 754 | 774 | -246 | -226 | -1,6 | 0,01 | 17,11 | >0.1 | 5,48 | 5,79 | 0 | 1,34 | 0 | 6,1 | 0 |
| TBX19 | TTTCCTAGAAAGCTGTTAAA | -1 | 708 | 728 | -292 | -272 | -0,16 | 0,01 | 17,28 | >0.1 | 5,48 | 5,59 | 0 | 1,2 | 0 | 5,17 | 0 |
| TBX19 | AATTTCATTAATAGGTTATC | -1 | 595 | 615 | -405 | -385 | -6,07 | 0,1 | 5,02 | >0.1 | 3,47 | 5,22 | 0 | 1,03 | 0 | 1,36 | 0 |
| TBX19 | TATCTCACATTTATTTTTAT | -1 | 579 | 599 | -421 | -401 | -5,21 | 0,01 | 4,35 | >0.1 | 3,37 | 5,18 | 0 | 1,02 | 0 | 0 | 0 |
| TBX19 | TCTGTCGCCCAGGCTGGAGC | -1 | 422 | 442 | -578 | -558 | -4,86 | 0,01 | 9,5 | >0.1 | 2,7 | 4,83 | 0 | 1,02 | 0 | 5,8 | 0 |
| TBX19 | TGGAGCACAGTGGCGCGATC | -1 | 408 | 428 | -592 | -572 | 3,7 | 0,001 | 17,28 | >0.1 | 2,66 | 4,8 | 0 | 1,03 | 0 | 5,09 | 0 |
| TBX19 | GATCTCGGCTCACTGCAAGC | -1 | 392 | 412 | -608 | -588 | -3,76 | 0,01 | 10,59 | >0.1 | 2,62 | 4,77 | 0 | 1,05 | 0 | 5,91 | 0 |
| TBX19 | CCTCCCAGGTTCACGCCATT | -1 | 368 | 388 | -632 | -612 | -2,17 | 0,01 | 12,72 | >0.1 | 2,55 | 4,73 | 0 | 1,08 | 0 | 6,53 | 0 |
| TBX19 | CCTCCCGAGTAGCTGGGACT | -1 | 336 | 356 | -664 | -644 | -0,44 | 0,01 | 14,46 | >0.1 | 2,56 | 4,68 | 0 | 1,13 | 0 | 6,52 | 0 |
| TBX19 | TATCTCCTGACCTTGTGATC | -1 | 234 | 254 | -766 | -746 | -4,76 | 0,01 | 8,51 | >0.1 | 2,88 | 4,53 | 0 | 1,44 | 0 | 4,42 | 0 |
| TBX19 | GATTACAGGCGTGAGCCACT | -1 | 186 | 206 | -814 | -794 | -4,67 | 0,01 | 8,74 | >0.1 | 3,08 | 4,46 | 0 | 1,74 | 0 | 4,13 | 0 |
| TBX19 | TTTCTCACACTGGTTCTGTG | -1 | 42 | 62 | -958 | -938 | -1,95 | 0,01 | 8,4 | >0.1 | 4,17 | 4,3 | 0 | 1,87 | 0 | 0 | 0 |
| FOXL1 | ACAAACA | 1 | 628 | 635 | -372 | -365 | 7,92 | 0,001 | 19,45 | >0.1 | 3,67 | 5,28 | 0 | 1,04 | 0 | 1,55 | 0 |
| FOXL1 | ACCAACA | 1 | 634 | 641 | -366 | -359 | 5,49 | 0,01 | 17,16 | >0.1 | 3,73 | 5,3 | 0 | 1,04 | 0 | 1,61 | 0 |
| FOXL1 | GAAAATA | 1 | 724 | 731 | -276 | -269 | 0,6 | 0,01 | 18,22 | >0.1 | 5,65 | 5,6 | 0 | 1,2 | 0 | 5,17 | 0 |
| FOXL1 | TTAACCA | 1 | 731 | 738 | -269 | -262 | 1,67 | 0,01 | 19,7 | >0.1 | 5,65 | 5,63 | 0 | 1,22 | 0 | 5,52 | 0 |
| FOXL1 | TTAACCA | 1 | 1143 | 1150 | 143 | 150 | 1,67 | 0,01 | 22,14 | >0.1 | 5,65 | 6,35 | 0 | 1,84 | 3,22 | 3,4 | 0 |
| FOXL1 | GTAAACA | -1 | 897 | 904 | -103 | -96 | 11,79 | >0.1 | 28,93 | >0.1 | 5,65 | 6,65 | 0 | 1,89 | 0 | 2,95 | 0 |
| FOXL1 | GCCAATA | -1 | 881 | 888 | -119 | -112 | 4,02 | 0,01 | 21,27 | >0.1 | 5,65 | 6,51 | 0 | 1,89 | 0 | 3,19 | 0 |
| FOXL1 | TCAAATA | -1 | 813 | 820 | -187 | -180 | 1,7 | 0,01 | 18,36 | >0.1 | 5,65 | 6,03 | 0 | 1,55 | 0 | 3,43 | 0 |
| FOXL1 | GTGACCA | -1 | 673 | 680 | -327 | -320 | 0,53 | 0,01 | 14,76 | >0.1 | 4,16 | 5,42 | 0 | 1,1 | 0 | 3,55 | 0 |
| FOXL1 | GTCAACT | -1 | 105 | 112 | -895 | -888 | 1,24 | 0,01 | 11,83 | >0.1 | 3,58 | 4,35 | 0 | 1,88 | 0 | 0,77 | 0 |

| | | | | | | | | | | | | | | | | | |
|---|---|---|---|---|---|---|---|---|---|---|---|---|---|---|---|---|---|
| MZF1(VAR.2) | GAAGGAGAAG | 1 | 11 | 21 | -989 | -979 | 3,47 | 0,01 | 15,21 | >0.1 | 5,61 | 4,26 | 0 | 1,86 | 0 | 0 | 0 |
| MZF1(VAR.2) | GGAGTGGAGA | 1 | 116 | 126 | -884 | -874 | 7,84 | 0,01 | 18,33 | >0.1 | 3,49 | 4,37 | 0 | 1,89 | 0 | 0,75 | 0 |
| MZF1(VAR.2) | GGCTGGGCAC | 1 | 176 | 186 | -824 | -814 | 5,7 | 0,01 | 18,35 | >0.1 | 3,15 | 4,44 | 0 | 1,82 | 0 | 3,24 | 0 |
| MZF1(VAR.2) | GGAGGCCGAG | 1 | 217 | 227 | -783 | -773 | 5,33 | 0,01 | 19,83 | >0.1 | 2,97 | 4,49 | 0 | 1,52 | 0 | 5,52 | 0 |
| MZF1(VAR.2) | GGCGTGGTGG | 1 | 315 | 325 | -685 | -675 | 4,6 | 0,01 | 19,16 | >0.1 | 2,64 | 4,63 | 0 | 1,17 | 0 | 6,13 | 0 |
| MZF1(VAR.2) | GGAGGCTGAG | 1 | 351 | 361 | -649 | -639 | 8,3 | 0,001 | 23,02 | >0.1 | 2,54 | 4,68 | 0 | 1,1 | 0 | 6,4 | 0 |
| MZF1(VAR.2) | TGAGGCAGGA | 1 | 357 | 367 | -643 | -633 | 3,25 | 0,01 | 17,69 | >0.1 | 2,52 | 4,69 | 0 | 1,09 | 0 | 6,13 | 0 |
| MZF1(VAR.2) | GGAGGCAGAG | 1 | 383 | 393 | -617 | -607 | 4,84 | 0,01 | 19,23 | >0.1 | 2,58 | 4,74 | 0 | 1,06 | 0 | 6,02 | 0 |
| MZF1(VAR.2) | CAAGGGCAAA | 1 | 507 | 517 | -493 | -483 | 3,65 | 0,01 | 16,79 | >0.1 | 2,99 | 4,98 | 0 | 0,98 | 0 | 4,19 | 0 |
| MZF1(VAR.2) | GGAAGGGAAA | 1 | 528 | 538 | -472 | -462 | 6,48 | 0,01 | 19,12 | >0.1 | 3,08 | 5,03 | 0 | 0,98 | 0 | 3,55 | 0 |
| MZF1(VAR.2) | GAAGGGAAAA | 1 | 529 | 539 | -471 | -461 | 3,93 | 0,01 | 16,57 | >0.1 | 3,08 | 5,03 | 0 | 0,98 | 0 | 3,55 | 0 |
| MZF1(VAR.2) | TTCTGGGAAT | 1 | 558 | 568 | -442 | -432 | 4,35 | 0,01 | 13,66 | >0.1 | 3,22 | 5,1 | 0 | 0,99 | 0 | 0 | 0 |
| MZF1(VAR.2) | GGAGAGGTAA | 1 | 772 | 782 | -228 | -218 | 6,44 | 0,01 | 25,16 | >0.1 | 5,61 | 5,83 | 0 | 1,37 | 0 | 5,92 | 0 |
| MZF1(VAR.2) | CTAGGGAAAG | 1 | 1003 | 1013 | 3 | 13 | 4,4 | 0,01 | 22,46 | >0.1 | 5,61 | 9,12 | 0 | 1,87 | 0 | 1,46 | 0 |
| MZF1(VAR.2) | CAATGGTGGA | -1 | 1103 | 1113 | 103 | 113 | 6,08 | 0,01 | 25,96 | >0.1 | 5,61 | 6,72 | 0 | 1,85 | 3,22 | 2,47 | 0 |
| MZF1(VAR.2) | CAAGGCTGAG | -1 | 1073 | 1083 | 73 | 83 | 5,31 | 0,01 | 23,96 | >0.1 | 5,61 | 7,12 | 0 | 1,85 | 1,79 | 2,27 | 0 |
| MZF1(VAR.2) | GTATGGTTTT | -1 | 620 | 630 | -380 | -370 | 3,78 | 0,01 | 15,14 | >0.1 | 3,62 | 5,26 | 0 | 1,03 | 0 | 1,44 | 0 |
| MZF1(VAR.2) | GACGGGGTTT | -1 | 278 | 288 | -722 | -712 | 3,69 | 0,01 | 17,47 | >0.1 | 2,75 | 4,57 | 0 | 1,27 | 0 | 5,19 | 0 |
| ARNT | AGTGTG | 1 | 51 | 57 | -949 | -943 | -1,11 | 0,1 | 9,2 | >0.1 | 4,14 | 4,29 | 0 | 1,87 | 0 | 0 | 0 |
| ARNT | GACCTG | 1 | 109 | 115 | -891 | -885 | -1 | 0,01 | 9,56 | >0.1 | 3,56 | 4,35 | 0 | 1,89 | 0 | 0,77 | 0 |
| ARNT | CACGCC | 1 | 193 | 199 | -807 | -801 | -0,82 | 0,01 | 12,52 | >0.1 | 3,08 | 4,45 | 0 | 1,68 | 0 | 4,13 | 0 |
| ARNT | CGCCTG | 1 | 195 | 201 | -805 | -799 | 1,83 | 0,01 | 15,28 | >0.1 | 3,07 | 4,45 | 0 | 1,67 | 0 | 4,26 | 0 |
| ARNT | CCCGTC | 1 | 282 | 288 | -718 | -712 | -0,75 | 0,01 | 12,9 | >0.1 | 2,74 | 4,57 | 0 | 1,25 | 0 | 5,09 | 0 |
| ARNT | GGCGTG | 1 | 315 | 321 | -685 | -679 | 2,15 | 0,01 | 16,71 | >0.1 | 2,64 | 4,62 | 0 | 1,17 | 0 | 6,13 | 0 |
| ARNT | CACCTG | 1 | 329 | 335 | -671 | -665 | 5,34 | 0,001 | 19,29 | >0.1 | 2,6 | 4,64 | 0 | 1,14 | 0 | 5,56 | 0 |
| ARNT | CTCGGG | 1 | 347 | 353 | -653 | -647 | -0,75 | 0,01 | 14,1 | >0.1 | 2,55 | 4,67 | 0 | 1,11 | 0 | 6,52 | 0 |
| ARNT | GGCGTG | 1 | 371 | 377 | -629 | -623 | 2,15 | 0,01 | 16,62 | >0.1 | 2,54 | 4,71 | 0 | 1,07 | 0 | 6,15 | 0 |
| ARNT | AACCTG | 1 | 377 | 383 | -623 | -617 | 2,89 | 0,01 | 17,62 | >0.1 | 2,56 | 4,72 | 0 | 1,06 | 0 | 6,4 | 0 |
| ARNT | AGCTTG | 1 | 391 | 397 | -609 | -603 | -1,11 | 0,1 | 13,29 | >0.1 | 2,59 | 4,74 | 0 | 1,04 | 0 | 6,02 | 0 |
| ARNT | AGCCTG | 1 | 427 | 433 | -573 | -567 | -0,61 | 0,01 | 13,58 | >0.1 | 2,7 | 4,81 | 0 | 1,01 | 0 | 5,67 | 0 |
| ARNT | AATGTG | 1 | 588 | 594 | -412 | -406 | 2,4 | 0,01 | 11,95 | >0.1 | 3,38 | 5,16 | 0 | 1 | 0 | 0 | 0 |
| ARNT | TCCGTG | 1 | 698 | 704 | -302 | -296 | -0,93 | 0,01 | 14,45 | >0.1 | 5,67 | 5,5 | 0 | 1,14 | 0 | 3,07 | 0 |
| ARNT | AAAGTG | 1 | 763 | 769 | -237 | -231 | 2,4 | 0,01 | 21,25 | >0.1 | 5,67 | 5,76 | 0 | 1,32 | 0 | 6,1 | 0 |
| ARNT | AACGTA | 1 | 803 | 809 | -197 | -191 | 2,4 | 0,01 | 18,95 | >0.1 | 5,67 | 5,96 | 0 | 1,49 | 0 | 3,43 | 0 |
| ARNT | CAGGGG | 1 | 1040 | 1046 | 40 | 46 | -0,82 | 0,01 | 15,89 | >0.1 | 5,67 | 7,85 | 0 | 1,86 | 0 | 1,34 | 0 |
| ARNT | GACCTG | 1 | 1160 | 1166 | 160 | 166 | -1 | 0,01 | 15,55 | >0.1 | 4,38 | 6,22 | 0 | 1,76 | 1,79 | 2,4 | 0 |
| ARNT | CACTTG | 1 | 1179 | 1185 | 179 | 185 | 4,84 | 0,01 | 23,25 | >0.1 | 5,67 | 6,1 | 0 | 1,59 | 1,79 | 3,26 | 0 |
| ARNT | CAAGTG | -1 | 1179 | 1185 | 179 | 185 | 4,84 | 0,01 | 23,25 | >0.1 | 5,67 | 6,1 | 0 | 1,59 | 1,79 | 3,26 | 0 |
| ARNT | AACTTG | -1 | 1138 | 1144 | 138 | 144 | 2,4 | 0,01 | 22,61 | >0.1 | 5,67 | 6,39 | 0 | 1,84 | 3,22 | 3,09 | 0 |
| ARNT | AGCTTG | -1 | 1053 | 1059 | 53 | 59 | -1,11 | 0,1 | 15,38 | >0.1 | 5,67 | 7,5 | 0 | 1,86 | 0 | 1,46 | 0 |
| ARNT | CCCCTG | -1 | 1040 | 1046 | 40 | 46 | -0,75 | 0,01 | 15,96 | >0.1 | 5,67 | 7,85 | 0 | 1,86 | 0 | 1,34 | 0 |
| ARNT | AACTTG | -1 | 955 | 961 | -45 | -39 | 2,4 | 0,01 | 19,06 | >0.1 | 5,67 | 7,42 | 0 | 1,88 | 0 | 1,7 | 0 |
| ARNT | AAAGTG | -1 | 822 | 828 | -178 | -172 | 2,4 | 0,01 | 19,44 | >0.1 | 5,67 | 6,08 | 0 | 1,6 | 0 | 3,7 | 0 |
| ARNT | CACGCC | -1 | 371 | 377 | -629 | -623 | -0,82 | 0,01 | 13,65 | >0.1 | 2,54 | 4,71 | 0 | 1,07 | 0 | 6,15 | 0 |
| ARNT | CCCGAG | -1 | 347 | 353 | -653 | -647 | -0,75 | 0,01 | 14,1 | >0.1 | 2,55 | 4,67 | 0 | 1,11 | 0 | 6,52 | 0 |
| ARNT | CAGGTG | -1 | 329 | 335 | -671 | -665 | 5,34 | 0,001 | 19,29 | >0.1 | 2,6 | 4,64 | 0 | 1,14 | 0 | 5,56 | 0 |
| ARNT | CACGCC | -1 | 315 | 321 | -685 | -679 | -0,82 | 0,01 | 13,74 | >0.1 | 2,64 | 4,62 | 0 | 1,17 | 0 | 6,13 | 0 |
| ARNT | GACGGG | -1 | 282 | 288 | -718 | -712 | -0,51 | 0,01 | 13,14 | >0.1 | 2,74 | 4,57 | 0 | 1,25 | 0 | 5,09 | 0 |
| ARNT | ACCGTG | -1 | 271 | 277 | -729 | -723 | 3,46 | 0,01 | 17,17 | >0.1 | 2,78 | 4,56 | 0 | 1,29 | 0 | 5,09 | 0 |
| ARNT | AAAGTG | -1 | 210 | 216 | -790 | -784 | 2,4 | 0,01 | 16,14 | >0.1 | 3 | 4,47 | 0 | 1,56 | 0 | 4,7 | 0 |
| ARNT | CAGGCG | -1 | 195 | 201 | -805 | -799 | -0,82 | 0,01 | 12,63 | >0.1 | 3,07 | 4,45 | 0 | 1,67 | 0 | 4,26 | 0 |
| ARNT | GGCGTG | -1 | 193 | 199 | -807 | -801 | 2,15 | 0,01 | 15,49 | >0.1 | 3,08 | 4,45 | 0 | 1,68 | 0 | 4,13 | 0 |
| VENTX | GGAAATTAA | 1 | 153 | 162 | -847 | -838 | 1,71 | 0,01 | 12,93 | >0.1 | 3,27 | 4,41 | 0 | 1,9 | 0 | 1,65 | 0 |
| VENTX | CTCTATTAA | 1 | 287 | 296 | -713 | -704 | 2,27 | 0,01 | 15,9 | >0.1 | 2,72 | 4,58 | 0 | 1,24 | 0 | 5,09 | 0 |
| VENTX | AAAACTTAG | 1 | 303 | 312 | -697 | -688 | -0,71 | 0,01 | 12,75 | >0.1 | 2,67 | 4,61 | 0 | 1,2 | 0 | 4,99 | 0 |
| VENTX | ACCTATTAA | 1 | 599 | 608 | -401 | -392 | 7,65 | 0,001 | 18,74 | >0.1 | 3,46 | 5,2 | 0 | 1,01 | 0 | 1,41 | 0 |
| VENTX | CAACATTAT | 1 | 636 | 645 | -364 | -355 | -1,22 | 0,01 | 10,5 | >0.1 | 3,75 | 5,31 | 0 | 1,05 | 0 | 1,61 | 0 |
| VENTX | TAAAATTAT | 1 | 682 | 691 | -318 | -309 | 0,15 | 0,01 | 14,24 | >0.1 | 4,33 | 5,46 | 0 | 1,12 | 0 | 3,19 | 0 |
| VENTX | TATCTTTAA | 1 | 704 | 713 | -296 | -287 | -3,32 | 0,01 | 13,81 | >0.1 | 5,62 | 5,53 | 0 | 1,16 | 0 | 4,81 | 0 |

| | | | | | | | | | | | | | | | | | |
|---|---|---|---|---|---|---|---|---|---|---|---|---|---|---|---|---|---|
| VENTX | AAATATTAA | 1 | 726 | 735 | -274 | -265 | 0,76 | 0,01 | 18,53 | >0.1 | 5,62 | 5,62 | 0 | 1,21 | 0 | 5,32 | 0 |
| VENTX | AAAAATTAT | 1 | 829 | 838 | -171 | -162 | 2,39 | 0,01 | 19,26 | >0.1 | 5,62 | 6,14 | 0 | 1,67 | 0 | 3,43 | 0 |
| VENTX | ATTGTTTAC | 1 | 895 | 904 | -105 | -96 | -3,47 | 0,01 | 13,55 | >0.1 | 5,62 | 6,65 | 0 | 1,89 | 0 | 2,86 | 0 |
| VENTX | ATTCTTTAA | -1 | 794 | 803 | -206 | -197 | -1,84 | 0,01 | 16,49 | >0.1 | 5,62 | 5,93 | 0 | 1,46 | 0 | 5,32 | 0 |
| VENTX | TTTCATTAA | -1 | 604 | 613 | -396 | -387 | 4,25 | 0,01 | 15,39 | >0.1 | 3,5 | 5,22 | 0 | 1,02 | 0 | 1,41 | 0 |
| VENTX | CACATTTAT | -1 | 585 | 594 | -415 | -406 | -3,68 | 0,01 | 5,86 | >0.1 | 3,37 | 5,16 | 0 | 1 | 0 | 0 | 0 |
| VENTX | TAAAATTAT | -1 | 132 | 141 | -868 | -859 | 0,15 | 0,01 | 11,39 | >0.1 | 3,39 | 4,38 | 0 | 1,89 | 0 | 1,57 | 0 |
| VENTX | AAAACTTAA | -1 | 78 | 87 | -922 | -913 | -1,98 | 0,01 | 8,03 | >0.1 | 3,81 | 4,33 | 0 | 1,88 | 0 | 0 | 0 |
| VENTX | TTCACTTAC | -1 | 5 | 14 | -995 | -986 | -2,22 | 0,01 | 11,32 | >0.1 | 5,62 | 4,26 | 0 | 1,86 | 1,79 | 0 | 0 |
| HOXA5 | AGGAAATG | 1 | 63 | 71 | -937 | -929 | 6,05 | 0,01 | 16,21 | >0.1 | 3,97 | 4,31 | 0 | 1,88 | 0 | 0 | 0 |
| HOXA5 | CAGAAATG | 1 | 169 | 177 | -831 | -823 | 9,11 | 0,001 | 21,07 | >0.1 | 3,19 | 4,43 | 0 | 1,89 | 0 | 2,46 | 0 |
| HOXA5 | CTGTAATC | 1 | 198 | 206 | -802 | -794 | 6,53 | 0,01 | 20,06 | >0.1 | 3,05 | 4,46 | 0 | 1,64 | 0 | 4,37 | 0 |
| HOXA5 | CTGTAGTC | 1 | 332 | 340 | -668 | -660 | 4,34 | 0,01 | 18,75 | >0.1 | 2,59 | 4,65 | 0 | 1,13 | 0 | 6,03 | 0 |
| HOXA5 | CAGAAATA | 1 | 484 | 492 | -516 | -508 | 3,66 | 0,01 | 18,39 | >0.1 | 2,89 | 4,93 | 0 | 0,98 | 0 | 5,93 | 0 |
| HOXA5 | CATGTATT | 1 | 519 | 527 | -481 | -473 | 3,81 | 0,01 | 15,89 | >0.1 | 3,03 | 5 | 0 | 0,98 | 0 | 3,07 | 0 |
| HOXA5 | AATAAATG | 1 | 584 | 592 | -416 | -408 | 5,22 | 0,01 | 14,75 | >0.1 | 3,36 | 5,16 | 0 | 1 | 0 | 0 | 0 |
| HOXA5 | ATGAAATT | 1 | 607 | 615 | -393 | -385 | 5,32 | 0,01 | 16,52 | >0.1 | 3,52 | 5,22 | 0 | 1,02 | 0 | 1,44 | 0 |
| HOXA5 | CCCTAGTT | 1 | 654 | 662 | -346 | -338 | 3,35 | 0,01 | 17,7 | >0.1 | 3,93 | 5,36 | 0 | 1,07 | 0 | 3,99 | 0 |
| HOXA5 | AGCAAGTG | -1 | 1179 | 1187 | 179 | 187 | 4,33 | 0,01 | 22,82 | >0.1 | 5,64 | 6,1 | 0 | 1,59 | 1,79 | 3,37 | 0 |
| HOXA5 | CAAAAATG | -1 | 1128 | 1136 | 128 | 136 | 3,91 | 0,01 | 23,91 | >0.1 | 5,64 | 6,48 | 0 | 1,84 | 3,22 | 2,82 | 0 |
| HOXA5 | CTCTTATC | -1 | 969 | 977 | -31 | -23 | 4,31 | 0,01 | 21,33 | >0.1 | 5,64 | 7,81 | 0 | 1,87 | 0 | 1,7 | 0 |
| HOXA5 | CAGATATG | -1 | 852 | 860 | -148 | -140 | 6,42 | 0,01 | 22,89 | >0.1 | 5,64 | 6,3 | 0 | 1,86 | 0 | 2,67 | 0 |
| HOXA5 | CTTTAATC | -1 | 792 | 800 | -208 | -200 | 5,73 | 0,01 | 24,05 | >0.1 | 5,64 | 5,92 | 0 | 1,44 | 0 | 5,32 | 0 |
| HOXA5 | CGGTACTT | -1 | 739 | 747 | -261 | -253 | 4,13 | 0,01 | 22,6 | >0.1 | 5,64 | 5,67 | 0 | 1,25 | 0 | 5,92 | 0 |
| HOXA5 | CATTAATA | -1 | 602 | 610 | -398 | -390 | 3,42 | 0,01 | 14,53 | >0.1 | 3,48 | 5,21 | 0 | 1,01 | 0 | 1,41 | 0 |
| HOXA5 | CATTTATT | -1 | 584 | 592 | -416 | -408 | 6,26 | 0,01 | 15,79 | >0.1 | 3,36 | 5,16 | 0 | 1 | 0 | 0 | 0 |
| HOXA5 | CAGAAACT | -1 | 555 | 563 | -445 | -437 | 3,34 | 0,01 | 12,62 | >0.1 | 3,2 | 5,09 | 0 | 0,99 | 0 | 0 | 0 |
| HOXA5 | CTCTACTT | -1 | 547 | 555 | -453 | -445 | 3,79 | 0,01 | 13 | >0.1 | 3,16 | 5,07 | 0 | 0,99 | 0 | 0 | 0 |
| HOXA5 | ATGGAATT | -1 | 494 | 502 | -506 | -498 | 3,45 | 0,01 | 18,75 | >0.1 | 2,93 | 4,95 | 0 | 0,98 | 0 | 6,44 | 0 |
| HOXA2 | GCAGATCAGG | 1 | 93 | 103 | -907 | -897 | 2,9 | 0,01 | 13,6 | >0.1 | 3,67 | 4,34 | 0 | 1,88 | 0 | 0,81 | 0 |
| HOXA2 | GGAAATTAAA | 1 | 153 | 163 | -847 | -837 | 5,89 | 0,01 | 17,11 | >0.1 | 3,27 | 4,41 | 0 | 1,9 | 0 | 1,65 | 0 |
| HOXA2 | ACTGATCAGA | 1 | 163 | 173 | -837 | -827 | 4,26 | 0,01 | 16,2 | >0.1 | 3,21 | 4,42 | 0 | 1,9 | 0 | 2,4 | 0 |
| HOXA2 | ACTGATCAGA | 1 | 478 | 488 | -522 | -512 | 4,26 | 0,01 | 18,4 | >0.1 | 2,88 | 4,92 | 0 | 0,99 | 0 | 5,36 | 0 |
| HOXA2 | ACCTATTAAT | 1 | 599 | 609 | -401 | -391 | 2,79 | 0,01 | 13,84 | >0.1 | 3,47 | 5,21 | 0 | 1,02 | 0 | 1,36 | 0 |
| HOXA2 | CAACATTATC | 1 | 636 | 646 | -364 | -354 | 5,09 | 0,01 | 16,87 | >0.1 | 3,76 | 5,31 | 0 | 1,05 | 0 | 1,66 | 0 |
| HOXA2 | TAAAATTATA | 1 | 682 | 692 | -318 | -308 | 2,76 | 0,01 | 16,74 | >0.1 | 4,34 | 5,46 | 0 | 1,12 | 0 | 3,07 | 0 |
| HOXA2 | AAAAATTATT | 1 | 829 | 839 | -171 | -161 | 2,94 | 0,01 | 19,66 | >0.1 | 5,61 | 6,15 | 0 | 1,68 | 0 | 3,28 | 0 |
| HOXA2 | AGTCATTCAG | 1 | 1011 | 1021 | 11 | 21 | 2,91 | 0,01 | 20,73 | >0.1 | 5,61 | 8,95 | 0 | 1,86 | 0 | 1,39 | 0 |
| HOXA2 | AGCAGTTACA | -1 | 1087 | 1097 | 87 | 97 | 4,57 | 0,01 | 22,96 | >0.1 | 5,61 | 6,92 | 0 | 1,85 | 1,79 | 2,21 | 0 |
| HOXA2 | TCTTATCAGC | -1 | 966 | 976 | -34 | -24 | 3,51 | 0,01 | 20,47 | >0.1 | 5,61 | 7,78 | 0 | 1,87 | 0 | 1,7 | 0 |
| HOXA2 | CCAAATCAAA | -1 | 749 | 759 | -251 | -241 | 4,03 | 0,01 | 22,56 | >0.1 | 5,61 | 5,72 | 0 | 1,28 | 0 | 5,92 | 0 |
| HOXA2 | GACAATTTCA | -1 | 608 | 618 | -392 | -382 | 2,88 | 0,01 | 14,21 | >0.1 | 3,53 | 5,23 | 0 | 1,02 | 0 | 1,55 | 0 |
| HOXA2 | TTTCATTAAT | -1 | 603 | 613 | -397 | -387 | 5,79 | 0,01 | 16,93 | >0.1 | 3,49 | 5,22 | 0 | 1,02 | 0 | 1,41 | 0 |
| HOXA2 | TCTGATCAGT | -1 | 478 | 488 | -522 | -512 | 3,86 | 0,01 | 18 | >0.1 | 2,88 | 4,92 | 0 | 0,99 | 0 | 5,36 | 0 |
| HOXA2 | CGCCATTCTC | -1 | 365 | 375 | -635 | -625 | 2,89 | 0,01 | 17,82 | >0.1 | 2,53 | 4,71 | 0 | 1,08 | 0 | 6,61 | 0 |
| HOXA2 | GTTAGTCAGG | -1 | 262 | 272 | -738 | -728 | 2,83 | 0,01 | 16,03 | >0.1 | 2,8 | 4,55 | 0 | 1,32 | 0 | 4,53 | 0 |
| HOXA2 | AGCCATTTCT | -1 | 170 | 180 | -830 | -820 | 3,01 | 0,01 | 15,92 | >0.1 | 3,18 | 4,43 | 0 | 1,88 | 0 | 3,43 | 0 |
| HOXA2 | TCTGATCAGT | -1 | 163 | 173 | -837 | -827 | 3,86 | 0,01 | 15,8 | >0.1 | 3,21 | 4,42 | 0 | 1,9 | 0 | 2,4 | 0 |
| HOXA2 | TTTAATTTCC | -1 | 153 | 163 | -847 | -837 | 4,15 | 0,01 | 15,37 | >0.1 | 3,27 | 4,41 | 0 | 1,9 | 0 | 1,65 | 0 |
| HOXA2 | TAAAATTATG | -1 | 131 | 141 | -869 | -859 | 3,57 | 0,01 | 14,82 | >0.1 | 3,4 | 4,38 | 0 | 1,89 | 0 | 1,57 | 0 |
| HOXA2 | GCTCATTTCC | -1 | 64 | 74 | -936 | -926 | 6,45 | 0,01 | 16,59 | >0.1 | 3,95 | 4,31 | 0 | 1,88 | 0 | 0 | 0 |
| HOXA2 | TTCACTTACC | -1 | 4 | 14 | -996 | -986 | 4 | 0,01 | 17,52 | >0.1 | 5,61 | 4,26 | 0 | 1,86 | 1,79 | 0 | 0 |
| HOXA9 | GTCATAATTTT | 1 | 129 | 140 | -871 | -860 | -0,34 | 0,01 | 10,95 | >0.1 | 3,4 | 4,38 | 0 | 1,89 | 0 | 1,61 | 0 |
| HOXA9 | AATATGAATTC | 1 | 488 | 499 | -512 | -501 | -1,57 | 0,01 | 13,54 | >0.1 | 2,91 | 4,94 | 0 | 0,99 | 0 | 6,27 | 0 |
| HOXA9 | TCCATAAAGAC | 1 | 497 | 508 | -503 | -492 | -1,78 | 0,01 | 13,55 | >0.1 | 2,95 | 4,96 | 0 | 0,98 | 0 | 6,44 | 0 |
| HOXA9 | GTCATGTATTT | 1 | 517 | 528 | -483 | -472 | 2,17 | 0,01 | 14,14 | >0.1 | 3,03 | 5 | 0 | 0,98 | 0 | 2,95 | 0 |
| HOXA9 | GAAATAAAAAT | 1 | 576 | 587 | -424 | -413 | -0,24 | 0,01 | 9,23 | >0.1 | 3,33 | 5,15 | 0 | 1 | 0 | 0 | 0 |
| HOXA9 | AAAATAAATGT | 1 | 582 | 593 | -418 | -407 | -1,66 | 0,01 | 7,87 | >0.1 | 3,36 | 5,16 | 0 | 1,01 | 0 | 0 | 0 |
| HOXA9 | CCTATTAATGA | 1 | 600 | 611 | -400 | -389 | -0,17 | 0,01 | 10,9 | >0.1 | 3,48 | 5,21 | 0 | 1,02 | 0 | 1,36 | 0 |

| | | | | | | | | | | | | | | | | | |
|---|---|---|---|---|---|---|---|---|---|---|---|---|---|---|---|---|---|
| HOXA9 | ACCATACAAAC | 1 | 623 | 634 | -377 | -366 | -1,3 | 0,01 | 10,1 | >0.1 | 3,65 | 5,28 | 0 | 1,04 | 0 | 1,44 | 0 |
| HOXA9 | TCCGTGTATCT | 1 | 698 | 709 | -302 | -291 | -0,34 | 0,01 | 15,12 | >0.1 | 5,6 | 5,52 | 0 | 1,15 | 0 | 3,19 | 0 |
| HOXA9 | GTAAACAATCT | -1 | 893 | 904 | -107 | -96 | 1,51 | 0,01 | 18,51 | >0.1 | 5,6 | 6,65 | 0 | 1,89 | 0 | 2,86 | 0 |
| HOXA9 | GATATGTAACA | -1 | 847 | 858 | -153 | -142 | -1,58 | 0,01 | 15 | >0.1 | 5,6 | 6,28 | 0 | 1,84 | 0 | 2,86 | 0 |
| HOXA9 | AAAGTGAATCA | -1 | 817 | 828 | -183 | -172 | 3,12 | 0,01 | 19,95 | >0.1 | 5,6 | 6,08 | 0 | 1,6 | 0 | 3,55 | 0 |
| HOXA9 | CACATTTATTT | -1 | 583 | 594 | -417 | -406 | -1,48 | 0,01 | 8,06 | >0.1 | 3,37 | 5,16 | 0 | 1,01 | 0 | 0 | 0 |
| HOXA9 | GGTCTCTATCT | -1 | 249 | 260 | -751 | -740 | -1,77 | 0,01 | 12,17 | >0.1 | 2,84 | 4,53 | 0 | 1,37 | 0 | 5,19 | 0 |
| HOXA9 | CCTATTTCTCA | -1 | 55 | 66 | -945 | -934 | -1,34 | 0,01 | 8,89 | >0.1 | 4,05 | 4,3 | 0 | 1,87 | 0 | 0 | 0 |
| NRF1 | ACAGTGGCTCA | 1 | 184 | 195 | -816 | -805 | -11,17 | 0,01 | 2,09 | >0.1 | 3,11 | 4,45 | 0 | 1,75 | 0 | 3,95 | 0 |
| NRF1 | GAACCTGGGAG | 1 | 376 | 387 | -624 | -613 | -8,75 | 0,01 | 6 | >0.1 | 2,56 | 4,73 | 0 | 1,06 | 0 | 6,4 | 0 |
| NRF1 | GAGATCGCGCC | 1 | 406 | 417 | -594 | -583 | -4,51 | 0,01 | 9,59 | >0.1 | 2,64 | 4,78 | 0 | 1,03 | 0 | 5,65 | 0 |
| NRF1 | GATCGCGCCAC | 1 | 408 | 419 | -592 | -581 | -12,56 | 0,01 | 1,55 | >0.1 | 2,65 | 4,78 | 0 | 1,03 | 0 | 5,65 | 0 |
| NRF1 | CAGCCTGGGCG | 1 | 426 | 437 | -574 | -563 | -0,67 | 0,01 | 13,53 | >0.1 | 2,7 | 4,82 | 0 | 1,01 | 0 | 5,67 | 0 |
| NRF1 | GCCTGGGCGAC | 1 | 428 | 439 | -572 | -561 | -0,33 | 0,001 | 14,02 | >0.1 | 2,71 | 4,82 | 0 | 1,01 | 0 | 5,8 | 0 |
| NRF1 | AATGTTGCCCA | 1 | 537 | 548 | -463 | -452 | -10,71 | 0,01 | 0,11 | >0.1 | 3,12 | 5,05 | 0 | 0,99 | 0 | 1,66 | 0 |
| NRF1 | GACGATGTCAA | 1 | 1045 | 1056 | 45 | 56 | -5,81 | 0,01 | 10 | >0.1 | 5,6 | 7,7 | 0 | 1,86 | 0 | 0,65 | 0 |
| NRF1 | GAACAGGCCAA | 1 | 1115 | 1126 | 115 | 126 | -1,05 | 0,01 | 18,75 | >0.1 | 5,6 | 6,6 | 0 | 1,85 | 3,22 | 2,54 | 0 |
| NRF1 | AAGCAAGTGAA | -1 | 1177 | 1188 | 177 | 188 | -11,88 | 0,01 | 6,49 | >0.1 | 5,6 | 6,11 | 0 | 1,61 | 1,79 | 3,26 | 0 |
| NRF1 | AAACTTGTCCA | -1 | 1134 | 1145 | 134 | 145 | -2,09 | 0,01 | 18,03 | >0.1 | 5,6 | 6,42 | 0 | 1,84 | 3,22 | 3,03 | 0 |
| NRF1 | AAGGCTGAGAA | -1 | 1071 | 1082 | 71 | 82 | -10,5 | 0,01 | 8,17 | >0.1 | 5,6 | 7,15 | 0 | 1,85 | 1,79 | 2,27 | 0 |
| NRF1 | TCCCCTGTGAG | -1 | 1036 | 1047 | 36 | 47 | -9,79 | 0,01 | 6,99 | >0.1 | 5,6 | 7,99 | 0 | 1,86 | 0 | 1,34 | 0 |
| NRF1 | CCCTGTGAGCC | -1 | 1034 | 1045 | 34 | 45 | -9,77 | 0,01 | 7,06 | >0.1 | 5,6 | 8,06 | 0 | 1,86 | 0 | 1,31 | 0 |
| NRF1 | CAGCCTTTGAA | -1 | 959 | 970 | -41 | -30 | -12,09 | 0,01 | 4,7 | >0.1 | 5,6 | 7,62 | 0 | 1,88 | 0 | 1,7 | 0 |
| NRF1 | ACAGTGGCGCG | -1 | 411 | 422 | -589 | -578 | 3,7 | 0,001 | 17,94 | >0.1 | 2,66 | 4,79 | 0 | 1,03 | 0 | 5,76 | 0 |
| NRF1 | CTACAGGTGCC | -1 | 327 | 338 | -673 | -662 | -11,29 | 0,01 | 2,67 | >0.1 | 2,6 | 4,65 | 0 | 1,14 | 0 | 5,56 | 0 |
| NRF1 | ACAGGTGCCCA | -1 | 325 | 336 | -675 | -664 | 4,48 | 0,001 | 18,34 | >0.1 | 2,61 | 4,64 | 0 | 1,15 | 0 | 5,46 | 0 |
| NRF1 | CACCACGCCCA | -1 | 313 | 324 | -687 | -676 | 2,16 | 0,001 | 16,73 | >0.1 | 2,64 | 4,63 | 0 | 1,17 | 0 | 6,13 | 0 |
| NRF1 | ACAGGCGTGAG | -1 | 191 | 202 | -809 | -798 | -5,47 | 0,01 | 7,89 | >0.1 | 3,08 | 4,46 | 0 | 1,7 | 0 | 4,13 | 0 |
| NRF1 | GCGTGAGCCAC | -1 | 187 | 198 | -813 | -802 | -11,32 | 0,01 | 1,81 | >0.1 | 3,09 | 4,45 | 0 | 1,73 | 0 | 3,86 | 0 |
| NRF1 | GCCACTGTGCC | -1 | 181 | 192 | -819 | -808 | -4,47 | 0,01 | 8,73 | >0.1 | 3,12 | 4,44 | 0 | 1,78 | 0 | 3,86 | 0 |
| NRF1 | CACTGTGCCCA | -1 | 179 | 190 | -821 | -810 | -0,68 | 0,01 | 11,78 | >0.1 | 3,13 | 4,44 | 0 | 1,8 | 0 | 3,09 | 0 |
| NRF1 | CCACAGGTCAA | -1 | 107 | 118 | -893 | -882 | -5,57 | 0,01 | 5 | >0.1 | 3,56 | 4,36 | 0 | 1,89 | 0 | 0,77 | 0 |
| NFYB | CACAGAACCAGTGTG | 1 | 42 | 57 | -958 | -943 | -2,29 | 0,01 | 8,09 | >0.1 | 4,21 | 4,3 | 0 | 1,87 | 0 | 0 | 0 |
| NFYB | GAGTTGACCTGTGGA | 1 | 104 | 119 | -896 | -881 | -12,07 | 0,01 | -1,52 | >0.1 | 3,56 | 4,36 | 0 | 1,89 | 0 | 0,75 | 0 |
| NFYB | GATCGCGCCACTGTG | 1 | 408 | 423 | -592 | -577 | -7,75 | 0,01 | 6,38 | >0.1 | 2,65 | 4,79 | 0 | 1,03 | 0 | 5,65 | 0 |
| NFYB | ATGTTGCCCAAGTAG | 1 | 538 | 553 | -462 | -447 | -8,14 | 0,01 | 1,05 | >0.1 | 3,14 | 5,06 | 0 | 0,99 | 0 | 0 | 0 |
| NFYB | GAGATAACCTATTAA | 1 | 593 | 608 | -407 | -392 | -9,63 | 0,01 | 0,03 | >0.1 | 3,44 | 5,2 | 0 | 1,02 | 0 | 0 | 0 |
| NFYB | ATATTAACCAAAAGT | 1 | 728 | 743 | -272 | -257 | -8,95 | 0,01 | 9,01 | >0.1 | 5,54 | 5,65 | 0 | 1,24 | 0 | 5,52 | 0 |
| NFYB | TCAGCGCCCAACCCA | 1 | 942 | 957 | -58 | -43 | -11,99 | 0,01 | 4,58 | >0.1 | 5,54 | 7,34 | 0 | 1,88 | 0 | 1,8 | 0 |
| NFYB | CAGTCCACCATTGAG | 1 | 1100 | 1115 | 100 | 115 | -0,04 | 0,001 | 19,75 | >0.1 | 5,54 | 6,76 | 0 | 1,85 | 3,22 | 2,41 | 0 |
| NFYB | GAACAGGCCAAGACA | 1 | 1115 | 1130 | 115 | 130 | -10,02 | 0,01 | 9,8 | >0.1 | 5,54 | 6,6 | 0 | 1,85 | 3,22 | 2,6 | 0 |
| NFYB | CAAGAAGCAAGTGAA | -1 | 1177 | 1192 | 177 | 192 | -13,26 | 0,01 | 5,14 | >0.1 | 5,54 | 6,11 | 0 | 1,61 | 1,79 | 3,34 | 0 |
| NFYB | AACTTGTCCAAAAAT | -1 | 1129 | 1144 | 129 | 144 | -9,08 | 0,01 | 10,91 | >0.1 | 5,54 | 6,47 | 0 | 1,84 | 3,22 | 2,91 | 0 |
| NFYB | CTGTTCCTCAATGGT | -1 | 1106 | 1121 | 106 | 121 | -13,12 | 0,01 | 6,72 | >0.1 | 5,54 | 6,69 | 0 | 1,85 | 3,22 | 2,53 | 0 |
| NFYB | GAAGGAGCCAGGAAG | -1 | 1059 | 1074 | 59 | 74 | -8,66 | 0,01 | 10,07 | >0.1 | 5,54 | 7,37 | 0 | 1,86 | 1,79 | 2,16 | 0 |
| NFYB | CTGTGAGCCAAGATC | -1 | 1028 | 1043 | 28 | 43 | -12,89 | 0,01 | 4,19 | >0.1 | 5,54 | 8,34 | 0 | 1,86 | 0 | 1,34 | 0 |
| NFYB | ATCACATCCACTGAA | -1 | 1016 | 1031 | 16 | 31 | -7,21 | 0,01 | 10,34 | >0.1 | 5,54 | 8,81 | 0 | 1,86 | 0 | 1,34 | 0 |
| NFYB | GGAAAGACCAAAAAA | -1 | 924 | 939 | -76 | -61 | -8,59 | 0,01 | 6,95 | >0.1 | 5,54 | 7,05 | 0 | 1,88 | 0 | 1,06 | 0 |
| NFYB | ACAGTAAACAATCTG | -1 | 892 | 907 | -108 | -93 | -7,88 | 0,01 | 9,1 | >0.1 | 5,54 | 6,68 | 0 | 1,89 | 0 | 2,86 | 0 |
| NFYB | TGCTGAGCCAATATA | -1 | 879 | 894 | -121 | -106 | -0,61 | 0,001 | 16,46 | >0.1 | 5,54 | 6,57 | 0 | 1,9 | 0 | 3,07 | 0 |
| NFYB | AGTGAATCAAATAAG | -1 | 811 | 826 | -189 | -174 | -9,14 | 0,01 | 7,49 | >0.1 | 5,54 | 6,06 | 0 | 1,59 | 0 | 3,43 | 0 |
| NFYB | TTATGGCCAAATCAA | -1 | 750 | 765 | -250 | -235 | -7,16 | 0,01 | 11,98 | >0.1 | 5,54 | 5,75 | 0 | 1,31 | 0 | 6,54 | 0 |
| NFYB | ACAGTGACCAAAGAG | -1 | 668 | 683 | -332 | -317 | 0,51 | 0,001 | 14,89 | >0.1 | 4,15 | 5,43 | 0 | 1,11 | 0 | 3,7 | 0 |
| NFYB | GTTCACGCCATTCTC | -1 | 365 | 380 | -635 | -620 | -12,75 | 0,01 | 2,31 | >0.1 | 2,54 | 4,72 | 0 | 1,08 | 0 | 6,73 | 0 |
| NFYB | GATCTGCCCAACTCG | -1 | 223 | 238 | -777 | -762 | -12,85 | 0,01 | 1,33 | >0.1 | 2,93 | 4,5 | 0 | 1,49 | 0 | 5,26 | 0 |
| NFYB | GCGTGAGCCACTGTG | -1 | 183 | 198 | -817 | -802 | -6,71 | 0,01 | 6,47 | >0.1 | 3,1 | 4,45 | 0 | 1,76 | 0 | 3,86 | 0 |
| NFYB | CTACTCTCCACTCCA | -1 | 115 | 130 | -885 | -870 | -10,79 | 0,01 | -0,3 | >0.1 | 3,48 | 4,37 | 0 | 1,89 | 0 | 0,75 | 0 |
| SOX8 | AGCTTTTTTAAGTTTT | 1 | 71 | 87 | -929 | -913 | -29,8 | 0,01 | -19,75 | >0.1 | 3,84 | 4,33 | 0 | 1,88 | 0 | 0 | 0 |
| SOX8 | AAAAATGGCCATGGAA | 1 | 141 | 157 | -859 | -843 | -28,96 | 0,01 | -17,77 | >0.1 | 3,32 | 4,4 | 0 | 1,9 | 0 | 1,57 | 0 |

| | | | | | | | | | | | | | | | | | |
|---|---|---|---|---|---|---|---|---|---|---|---|---|---|---|---|---|---|
| SOX8 | AAAACTGATCAGAAAT | 1 | 160 | 176 | -840 | -824 | -29,72 | 0,01 | -17,78 | >0.1 | 3,21 | 4,43 | 0 | 1,9 | 0 | 2,4 | 0 |
| SOX8 | CACAAGGTCAGGAGAT | 1 | 237 | 253 | -763 | -747 | -34,02 | 0,01 | -20,87 | >0.1 | 2,88 | 4,52 | 0 | 1,42 | 0 | 4,33 | 0 |
| SOX8 | CAGAGCTTGCAGTGAG | 1 | 388 | 404 | -612 | -596 | -33,35 | 0,01 | -18,92 | >0.1 | 2,6 | 4,76 | 0 | 1,05 | 0 | 6,02 | 0 |
| SOX8 | TGCAGTGAGCCGAGAT | 1 | 395 | 411 | -605 | -589 | -33,76 | 0,01 | -19,42 | >0.1 | 2,62 | 4,77 | 0 | 1,05 | 0 | 5,91 | 0 |
| SOX8 | ACCGAGACTCAGTCTC | 1 | 441 | 457 | -559 | -543 | -28,02 | 0,01 | -13,48 | >0.1 | 2,76 | 4,86 | 0 | 1,01 | 0 | 5,92 | 0 |
| SOX8 | AAAACTGATCAGAAAT | 1 | 475 | 491 | -525 | -509 | -29,72 | 0,01 | -15,56 | >0.1 | 2,88 | 4,92 | 0 | 0,99 | 0 | 5,36 | 0 |
| SOX8 | TAACATTTTCCGTGTA | 1 | 690 | 706 | -310 | -294 | -23,64 | 0,01 | -8,38 | >0.1 | 5,53 | 5,51 | 0 | 1,15 | 0 | 3,07 | 0 |
| SOX8 | AACATTTTCCGTGTAT | 1 | 691 | 707 | -309 | -293 | -32,46 | 0,01 | -17,2 | >0.1 | 5,53 | 5,51 | 0 | 1,15 | 0 | 3,07 | 0 |
| SOX8 | AAGGTTCTCTAGGATT | 1 | 780 | 796 | -220 | -204 | -33,46 | 0,01 | -14,69 | >0.1 | 5,53 | 5,9 | 0 | 1,43 | 0 | 5,92 | 0 |
| SOX8 | CAGATTGTTTACTGTG | 1 | 892 | 908 | -108 | -92 | -33,02 | 0,01 | -15,96 | >0.1 | 5,53 | 6,69 | 0 | 1,89 | 0 | 2,95 | 0 |
| SOX8 | GATTGTTTACTGTGTT | 1 | 894 | 910 | -106 | -90 | -30,37 | 0,01 | -13,38 | >0.1 | 5,53 | 6,71 | 0 | 1,89 | 0 | 2,86 | 0 |
| SOX8 | AAAGTCATTCAGTGGA | 1 | 1009 | 1025 | 9 | 25 | -31,31 | 0,01 | -13,47 | >0.1 | 5,53 | 9,03 | 0 | 1,86 | 0 | 1,42 | 0 |
| SOX8 | AAAAATGTCTTGGCCT | -1 | 1119 | 1135 | 119 | 135 | -31,55 | 0,01 | -11,65 | >0.1 | 5,53 | 6,56 | 0 | 1,85 | 3,22 | 2,74 | 0 |
| SOX8 | CACAGTAAACAATCTG | -1 | 892 | 908 | -108 | -92 | -33,75 | 0,01 | -16,69 | >0.1 | 5,53 | 6,69 | 0 | 1,89 | 0 | 2,95 | 0 |
| SOX8 | AAAATGTTATAATTTT | -1 | 683 | 699 | -317 | -301 | -32,31 | 0,01 | -18,32 | >0.1 | 4,42 | 5,48 | 0 | 1,14 | 0 | 2,95 | 0 |
| SOX8 | ATAATTTTACAGTGAC | -1 | 675 | 691 | -325 | -309 | -21,31 | 0,01 | -7,19 | >0.1 | 4,27 | 5,46 | 0 | 1,12 | 0 | 3,28 | 0 |
| SOX8 | AAGTTTTTGTATTTTT | -1 | 294 | 310 | -706 | -690 | -32,49 | 0,01 | -19,52 | >0.1 | 2,69 | 4,61 | 0 | 1,22 | 0 | 4,45 | 0 |
| SOX8 | TTCCATGGCCATTTTT | -1 | 141 | 157 | -859 | -843 | -31,55 | 0,01 | -20,36 | >0.1 | 3,32 | 4,4 | 0 | 1,9 | 0 | 1,57 | 0 |
| SOX9 | CCAGTGTGA | 1 | 49 | 58 | -951 | -942 | 4,98 | 0,01 | 15,3 | >0.1 | 4,15 | 4,3 | 0 | 1,87 | 0 | 0 | 0 |
| SOX9 | CCATGGAAA | 1 | 149 | 158 | -851 | -842 | 3,95 | 0,01 | 15,19 | >0.1 | 3,29 | 4,4 | 0 | 1,9 | 0 | 1,65 | 0 |
| SOX9 | CCACTGTGC | 1 | 415 | 424 | -585 | -576 | 5,74 | 0,01 | 19,41 | >0.1 | 2,67 | 4,79 | 0 | 1,02 | 0 | 5,19 | 0 |
| SOX9 | CTATTAATG | 1 | 601 | 610 | -399 | -390 | 2,8 | 0,01 | 13,86 | >0.1 | 3,48 | 5,21 | 0 | 1,02 | 0 | 1,36 | 0 |
| SOX9 | AAATTGTCT | 1 | 610 | 619 | -390 | -381 | 3,16 | 0,01 | 14,5 | >0.1 | 3,54 | 5,23 | 0 | 1,02 | 0 | 1,55 | 0 |
| SOX9 | CCCTAGTTC | 1 | 654 | 663 | -346 | -337 | 5,19 | 0,01 | 19,55 | >0.1 | 3,93 | 5,36 | 0 | 1,07 | 0 | 3,99 | 0 |
| SOX9 | TCTTTGGTC | 1 | 669 | 678 | -331 | -322 | 3,28 | 0,01 | 17,6 | >0.1 | 4,12 | 5,41 | 0 | 1,1 | 0 | 3,7 | 0 |
| SOX9 | GTTTTGATT | 1 | 747 | 756 | -253 | -244 | 2,91 | 0,01 | 21,43 | >0.1 | 5,62 | 5,71 | 0 | 1,27 | 0 | 5,92 | 0 |
| SOX9 | GTATTCTTA | 1 | 806 | 815 | -194 | -185 | 3,36 | 0,01 | 19,93 | >0.1 | 5,62 | 6 | 0 | 1,52 | 0 | 3,43 | 0 |
| SOX9 | AGATTGTTT | 1 | 893 | 902 | -107 | -98 | 5,67 | 0,01 | 22,68 | >0.1 | 5,62 | 6,63 | 0 | 1,89 | 0 | 2,86 | 0 |
| SOX9 | GCCTTGTTG | 1 | 1077 | 1086 | 77 | 86 | 3,67 | 0,01 | 22,27 | >0.1 | 5,62 | 7,06 | 0 | 1,85 | 1,79 | 2,27 | 0 |
| SOX9 | CCATTGAGG | 1 | 1107 | 1116 | 107 | 116 | 6,85 | 0,01 | 26,7 | >0.1 | 5,62 | 6,68 | 0 | 1,85 | 3,22 | 2,48 | 0 |
| SOX9 | CAATGGTGG | -1 | 1104 | 1113 | 104 | 113 | 2,84 | 0,01 | 22,72 | >0.1 | 5,62 | 6,71 | 0 | 1,85 | 3,22 | 2,47 | 0 |
| SOX9 | CCACTGAAT | -1 | 1015 | 1024 | 15 | 24 | 2,78 | 0,01 | 20,5 | >0.1 | 5,62 | 8,84 | 0 | 1,86 | 0 | 1,39 | 0 |
| SOX9 | CCTTTGAAC | -1 | 958 | 967 | -42 | -33 | 4,96 | 0,01 | 21,75 | >0.1 | 5,62 | 7,55 | 0 | 1,88 | 0 | 1,74 | 0 |
| SOX9 | TTATTCTTT | -1 | 796 | 805 | -204 | -195 | 3,63 | 0,01 | 21,98 | >0.1 | 5,62 | 5,94 | 0 | 1,47 | 0 | 5,32 | 0 |
| SOX9 | CTTTTGGTT | -1 | 733 | 742 | -267 | -258 | 4,34 | 0,01 | 22,76 | >0.1 | 5,62 | 5,65 | 0 | 1,23 | 0 | 5,92 | 0 |
| SOX9 | GTATGGTTT | -1 | 621 | 630 | -379 | -370 | 5,7 | 0,01 | 17,06 | >0.1 | 3,62 | 5,26 | 0 | 1,03 | 0 | 1,44 | 0 |
| SOX9 | CCCTTGTCT | -1 | 504 | 513 | -496 | -487 | 5,09 | 0,01 | 19,7 | >0.1 | 2,97 | 4,97 | 0 | 0,98 | 0 | 5,68 | 0 |
| SOX9 | CCATTCTCC | -1 | 364 | 373 | -636 | -627 | 6,31 | 0,01 | 21,23 | >0.1 | 2,53 | 4,7 | 0 | 1,08 | 0 | 6,61 | 0 |
| SOX9 | CCAAAGTGC | -1 | 209 | 218 | -791 | -782 | 2,94 | 0,01 | 16,69 | >0.1 | 3 | 4,48 | 0 | 1,57 | 0 | 4,7 | 0 |
| SOX9 | CCACTGTGC | -1 | 182 | 191 | -818 | -809 | 5,74 | 0,01 | 18,93 | >0.1 | 3,12 | 4,44 | 0 | 1,77 | 0 | 3,86 | 0 |
| SOX9 | CCATTTTTT | -1 | 140 | 149 | -860 | -851 | 3,91 | 0,01 | 15,15 | >0.1 | 3,34 | 4,39 | 0 | 1,89 | 0 | 1,61 | 0 |
| MYBL1 | GAAGTCAGCCAC | 1 | 33 | 45 | -967 | -955 | -4,92 | 0,01 | 5,63 | >0.1 | 4,4 | 4,28 | 0 | 1,87 | 0 | 0 | 0 |
| MYBL1 | AGAGTTGACCTG | 1 | 103 | 115 | -897 | -885 | -1,22 | 0,01 | 9,37 | >0.1 | 3,58 | 4,36 | 0 | 1,89 | 0 | 0,77 | 0 |
| MYBL1 | ACGGTGAAACCC | 1 | 272 | 284 | -728 | -716 | -6,76 | 0,01 | 6,94 | >0.1 | 2,76 | 4,57 | 0 | 1,29 | 0 | 5,09 | 0 |
| MYBL1 | AAACTTAGCTGG | 1 | 304 | 316 | -696 | -684 | -7,04 | 0,01 | 6,42 | >0.1 | 2,67 | 4,61 | 0 | 1,2 | 0 | 4,99 | 0 |
| MYBL1 | GGCGTGAACCTG | 1 | 371 | 383 | -629 | -617 | -5,08 | 0,01 | 9,66 | >0.1 | 2,55 | 4,72 | 0 | 1,07 | 0 | 6,4 | 0 |
| MYBL1 | GCAGTGAGCCGA | 1 | 396 | 408 | -604 | -592 | -6,15 | 0,01 | 8,18 | >0.1 | 2,62 | 4,76 | 0 | 1,04 | 0 | 5,91 | 0 |
| MYBL1 | TGAGATAACCTA | 1 | 592 | 604 | -408 | -396 | -2,55 | 0,01 | 7,08 | >0.1 | 3,43 | 5,19 | 0 | 1,01 | 0 | 0 | 0 |
| MYBL1 | AACATTATCTTC | 1 | 637 | 649 | -363 | -351 | -6,15 | 0,01 | 5,61 | >0.1 | 3,77 | 5,32 | 0 | 1,06 | 0 | 1,61 | 0 |
| MYBL1 | ATCTTTAACAGC | 1 | 705 | 717 | -295 | -283 | 1,39 | 0,001 | 18,68 | >0.1 | 5,58 | 5,55 | 0 | 1,17 | 0 | 4,99 | 0 |
| MYBL1 | AATATTAACCAA | 1 | 727 | 739 | -273 | -261 | -7,82 | 0,01 | 9,94 | >0.1 | 5,58 | 5,63 | 0 | 1,22 | 0 | 5,32 | 0 |
| MYBL1 | AGGATTAAAGAA | 1 | 790 | 802 | -210 | -198 | -5,38 | 0,01 | 12,9 | >0.1 | 5,58 | 5,93 | 0 | 1,46 | 0 | 5,32 | 0 |
| MYBL1 | AAGAATAACGTA | 1 | 797 | 809 | -203 | -191 | -3,65 | 0,01 | 12,94 | >0.1 | 5,58 | 5,97 | 0 | 1,49 | 0 | 3,55 | 0 |
| MYBL1 | ATTGTTTACTGT | 1 | 895 | 907 | -105 | -93 | -7,77 | 0,01 | 9,33 | >0.1 | 5,58 | 6,68 | 0 | 1,89 | 0 | 2,95 | 0 |
| MYBL1 | ACCATTGAGGAA | 1 | 1106 | 1118 | 106 | 118 | -8,23 | 0,01 | 11,59 | >0.1 | 5,58 | 6,69 | 0 | 1,85 | 3,22 | 2,47 | 0 |
| MYBL1 | AAGTTTAACCAC | 1 | 1139 | 1151 | 139 | 151 | 1,2 | 0,001 | 21,53 | >0.1 | 5,58 | 6,38 | 0 | 1,84 | 3,22 | 3,29 | 0 |
| MYBL1 | AAAGTTCACTTG | 1 | 1173 | 1185 | 173 | 185 | -7,51 | 0,01 | 10,84 | >0.1 | 5,58 | 6,13 | 0 | 1,64 | 1,79 | 3,2 | 0 |
| MYBL1 | GTGGTTAAACTT | -1 | 1139 | 1151 | 139 | 151 | 0,64 | 0,001 | 20,97 | >0.1 | 5,58 | 6,38 | 0 | 1,84 | 3,22 | 3,29 | 0 |
| MYBL1 | GCAGTTACAGCA | -1 | 1084 | 1096 | 84 | 96 | -3,67 | 0,01 | 14,73 | >0.1 | 5,58 | 6,96 | 0 | 1,85 | 1,79 | 2,21 | 0 |

| | | | | | | | | | | | | | | | | | |
|---|---|---|---|---|---|---|---|---|---|---|---|---|---|---|---|---|---|
| MYBL1 | GCCTTTGAACTT | -1 | 956 | 968 | -44 | -32 | -8,05 | 0,01 | 8,72 | >0.1 | 5,58 | 7,57 | 0 | 1,88 | 0 | 1,74 | 0 |
| MYBL1 | ACAGTAAACAAT | -1 | 895 | 907 | -105 | -93 | 1,7 | 0,001 | 18,8 | >0.1 | 5,58 | 6,68 | 0 | 1,89 | 0 | 2,95 | 0 |
| MYBL1 | TTCTTTAATCCT | -1 | 790 | 802 | -210 | -198 | -8,31 | 0,01 | 9,97 | >0.1 | 5,58 | 5,93 | 0 | 1,46 | 0 | 5,32 | 0 |
| MYBL1 | AACCTTACCTCT | -1 | 774 | 786 | -226 | -214 | -2,23 | 0,01 | 16,9 | >0.1 | 5,58 | 5,84 | 0 | 1,38 | 0 | 6,32 | 0 |
| MYBL1 | TTGGTTAATATT | -1 | 727 | 739 | -273 | -261 | -4,93 | 0,01 | 12,83 | >0.1 | 5,58 | 5,63 | 0 | 1,22 | 0 | 5,32 | 0 |
| MYBL1 | GCTGTTAAAGAT | -1 | 705 | 717 | -295 | -283 | -5,93 | 0,01 | 11,36 | >0.1 | 5,58 | 5,55 | 0 | 1,17 | 0 | 4,99 | 0 |
| MYBL1 | GAAGATAATGTT | -1 | 637 | 649 | -363 | -351 | -6,04 | 0,01 | 5,72 | >0.1 | 3,77 | 5,32 | 0 | 1,06 | 0 | 1,61 | 0 |
| MYBL1 | TCGGCTCACTGC | -1 | 396 | 408 | -604 | -592 | -6,63 | 0,01 | 7,7 | >0.1 | 2,62 | 4,76 | 0 | 1,04 | 0 | 5,91 | 0 |
| MYBL1 | CAGGTTCACGCC | -1 | 371 | 383 | -629 | -617 | -7,28 | 0,01 | 7,46 | >0.1 | 2,55 | 4,72 | 0 | 1,07 | 0 | 6,4 | 0 |
| MYBL1 | AGGATTACAGGC | -1 | 196 | 208 | -804 | -792 | -7,45 | 0,01 | 6,1 | >0.1 | 3,05 | 4,46 | 0 | 1,66 | 0 | 4,37 | 0 |
| MYBL1 | GGCGTGAGCCAC | -1 | 187 | 199 | -813 | -801 | -7,7 | 0,01 | 5,52 | >0.1 | 3,09 | 4,45 | 0 | 1,73 | 0 | 3,95 | 0 |
| MYBL1 | CAGGTCAACTCT | -1 | 103 | 115 | -897 | -885 | -3,65 | 0,01 | 6,94 | >0.1 | 3,58 | 4,36 | 0 | 1,89 | 0 | 0,77 | 0 |
| MYBL1 | GTGGCTGACTTC | -1 | 33 | 45 | -967 | -955 | -5,85 | 0,01 | 4,7 | >0.1 | 4,4 | 4,28 | 0 | 1,87 | 0 | 0 | 0 |
| MYBL1 | TCACTTACCTAT | -1 | 1 | 13 | -999 | -987 | -5,69 | 0,01 | 7,81 | >0.1 | 5,58 | 4,26 | 0 | 1,86 | 1,79 | 0 | 0 |
| PITX3 | TGTAATCCT | 1 | 199 | 208 | -801 | -792 | 8,12 | 0,001 | 21,74 | >0.1 | 3,04 | 4,46 | 0 | 1,64 | 0 | 4,48 | 0 |
| PITX3 | AGTGAGCCG | 1 | 398 | 407 | -602 | -593 | 2,82 | 0,01 | 17,15 | >0.1 | 2,62 | 4,76 | 0 | 1,04 | 0 | 5,91 | 0 |
| PITX3 | CATGATCCC | 1 | 648 | 657 | -352 | -343 | 7,67 | 0,001 | 21,78 | >0.1 | 3,87 | 5,34 | 0 | 1,06 | 0 | 3,83 | 0 |
| PITX3 | TAAAATCTG | 1 | 840 | 849 | -160 | -151 | 1,96 | 0,01 | 18,64 | >0.1 | 5,62 | 6,22 | 0 | 1,76 | 0 | 3,07 | 0 |
| PITX3 | GAAAATCTC | 1 | 978 | 987 | -22 | -13 | 4,67 | 0,01 | 21,92 | >0.1 | 5,62 | 8,18 | 0 | 1,87 | 0 | 1,57 | 0 |
| PITX3 | TATAATTCC | -1 | 1191 | 1200 | 191 | 200 | 5,19 | 0,01 | 23,95 | >0.1 | 5,62 | 6,02 | 0 | 1,5 | 1,79 | 3,82 | 0 |
| PITX3 | GTTAAACTT | -1 | 1139 | 1148 | 139 | 148 | 3,02 | 0,01 | 23,22 | >0.1 | 5,62 | 6,38 | 0 | 1,84 | 3,22 | 3,13 | 0 |
| PITX3 | TGTGAGCCA | -1 | 1033 | 1042 | 33 | 42 | 1,95 | 0,01 | 18,82 | >0.1 | 5,62 | 8,1 | 0 | 1,86 | 0 | 1,28 | 0 |
| PITX3 | ACTAAAACC | -1 | 994 | 1003 | -6 | 3 | 3 | 0,01 | 20,83 | >0.1 | 5,62 | 8,81 | 0 | 1,87 | 0 | 1,53 | 0 |
| PITX3 | CTAAAACCT | -1 | 993 | 1002 | -7 | 2 | 2,78 | 0,01 | 20,62 | >0.1 | 5,62 | 8,77 | 0 | 1,87 | 0 | 1,57 | 0 |
| PITX3 | GCTGAGCCA | -1 | 884 | 893 | -116 | -107 | 4,9 | 0,01 | 22,04 | >0.1 | 5,62 | 6,55 | 0 | 1,89 | 0 | 3,07 | 0 |
| PITX3 | GTTCATCCT | -1 | 868 | 877 | -132 | -123 | 3,68 | 0,01 | 20,38 | >0.1 | 5,62 | 6,42 | 0 | 1,9 | 0 | 2,75 | 0 |
| PITX3 | TTTAATCCT | -1 | 790 | 799 | -210 | -201 | 9,68 | 0,001 | 28,18 | >0.1 | 5,62 | 5,91 | 0 | 1,44 | 0 | 5,52 | 0 |
| PITX3 | CCAAATCAA | -1 | 750 | 759 | -250 | -241 | 2,3 | 0,01 | 20,84 | >0.1 | 5,62 | 5,72 | 0 | 1,28 | 0 | 5,92 | 0 |
| PITX3 | GTTAATATT | -1 | 727 | 736 | -273 | -264 | 2,81 | 0,01 | 20,59 | >0.1 | 5,62 | 5,62 | 0 | 1,22 | 0 | 5,32 | 0 |
| PITX3 | GGTTATCTC | -1 | 593 | 602 | -407 | -398 | 2,54 | 0,01 | 12,16 | >0.1 | 3,42 | 5,19 | 0 | 1,01 | 0 | 0 | 0 |
| PITX3 | CCAAATACA | -1 | 521 | 530 | -479 | -470 | 2,78 | 0,01 | 14,88 | >0.1 | 3,04 | 5,01 | 0 | 0,98 | 0 | 3,07 | 0 |
| PITX3 | CCTCAGCCT | -1 | 353 | 362 | -647 | -638 | 2,2 | 0,01 | 16,92 | >0.1 | 2,53 | 4,69 | 0 | 1,1 | 0 | 6,4 | 0 |
| PITX3 | CGTGAGCCA | -1 | 188 | 197 | -812 | -803 | 3,29 | 0,01 | 16,41 | >0.1 | 3,09 | 4,45 | 0 | 1,72 | 0 | 3,86 | 0 |
| PITX3 | TTTAATTTC | -1 | 154 | 163 | -846 | -837 | 3,07 | 0,01 | 14,29 | >0.1 | 3,26 | 4,41 | 0 | 1,9 | 0 | 1,65 | 0 |
| PITX3 | CCTGATCTG | -1 | 94 | 103 | -906 | -897 | 4,23 | 0,01 | 14,93 | >0.1 | 3,67 | 4,34 | 0 | 1,88 | 0 | 0,81 | 0 |
| PITX3 | AAAAAGCTC | -1 | 70 | 79 | -930 | -921 | 2,16 | 0,01 | 12,24 | >0.1 | 3,89 | 4,32 | 0 | 1,88 | 0 | 0 | 0 |
| MYBL2 | GAGAGTTGACCTGTG | 1 | 102 | 117 | -898 | -883 | 0,52 | 0,001 | 11,11 | >0.1 | 3,58 | 4,36 | 0 | 1,89 | 0 | 0,77 | 0 |
| MYBL2 | ACACGGTGAAACCCC | 1 | 270 | 285 | -730 | -715 | -3,39 | 0,01 | 10,23 | >0.1 | 2,77 | 4,57 | 0 | 1,3 | 0 | 4,99 | 0 |
| MYBL2 | CACGGTGAAACCCCG | 1 | 271 | 286 | -729 | -714 | -4,61 | 0,01 | 9,1 | >0.1 | 2,76 | 4,57 | 0 | 1,29 | 0 | 5,09 | 0 |
| MYBL2 | GGGCGACAGACCGAG | 1 | 432 | 447 | -568 | -553 | -7,83 | 0,01 | 6,55 | >0.1 | 2,73 | 4,84 | 0 | 1,01 | 0 | 5,8 | 0 |
| MYBL2 | GTGAGATAACCTATT | 1 | 591 | 606 | -409 | -394 | -4,37 | 0,01 | 5,27 | >0.1 | 3,43 | 5,2 | 0 | 1,02 | 0 | 0 | 0 |
| MYBL2 | GTATCTTTAACAGCT | 1 | 703 | 718 | -297 | -282 | -6,96 | 0,01 | 10,3 | >0.1 | 5,54 | 5,55 | 0 | 1,17 | 0 | 4,99 | 0 |
| MYBL2 | ATCTGTTACATATCT | 1 | 844 | 859 | -156 | -141 | -6,29 | 0,01 | 10,26 | >0.1 | 5,54 | 6,29 | 0 | 1,85 | 0 | 2,86 | 0 |
| MYBL2 | CCAAGTTCAAAGGCT | 1 | 954 | 969 | -46 | -31 | -3,51 | 0,01 | 13,21 | >0.1 | 5,54 | 7,6 | 0 | 1,88 | 0 | 1,7 | 0 |
| MYBL2 | AACTGCTGCTCAGTC | 1 | 1090 | 1105 | 90 | 105 | -5,7 | 0,01 | 14,14 | >0.1 | 5,54 | 6,88 | 0 | 1,85 | 3,22 | 2,34 | 0 |
| MYBL2 | ACAAGTTTAACCACG | 1 | 1137 | 1152 | 137 | 152 | 0,82 | 0,001 | 21,04 | >0.1 | 5,54 | 6,4 | 0 | 1,84 | 3,22 | 3,21 | 0 |
| MYBL2 | AGCCGAAGACCTGTT | 1 | 1153 | 1168 | 153 | 168 | -6,73 | 0,01 | 9,97 | >0.1 | 4,43 | 6,27 | 0 | 1,83 | 1,79 | 2,38 | 0 |
| MYBL2 | AACAGGTCTTCGGCT | -1 | 1153 | 1168 | 153 | 168 | -2,73 | 0,01 | 13,97 | >0.1 | 4,43 | 6,27 | 0 | 1,83 | 1,79 | 2,38 | 0 |
| MYBL2 | CGTGGTTAAACTTGT | -1 | 1137 | 1152 | 137 | 152 | 2,04 | 0,001 | 22,26 | >0.1 | 5,54 | 6,4 | 0 | 1,84 | 3,22 | 3,21 | 0 |
| MYBL2 | AGCAGTTACAGCAAC | -1 | 1082 | 1097 | 82 | 97 | 0,03 | 0,001 | 18,37 | >0.1 | 5,54 | 6,99 | 0 | 1,85 | 1,79 | 2,16 | 0 |
| MYBL2 | AGCCTTTGAACTTGG | -1 | 954 | 969 | -46 | -31 | -4,26 | 0,01 | 12,46 | >0.1 | 5,54 | 7,6 | 0 | 1,88 | 0 | 1,7 | 0 |
| MYBL2 | GGGCGCTGAGCAGGA | -1 | 936 | 951 | -64 | -49 | -7,52 | 0,01 | 8,01 | >0.1 | 5,54 | 7,23 | 0 | 1,88 | 0 | 0,88 | 0 |
| MYBL2 | ACACAGTAAACAATC | -1 | 894 | 909 | -106 | -91 | -5,56 | 0,01 | 11,52 | >0.1 | 5,54 | 6,7 | 0 | 1,89 | 0 | 2,95 | 0 |
| MYBL2 | ATCTGCTGAGCCAAT | -1 | 882 | 897 | -118 | -103 | -6,76 | 0,01 | 10,46 | >0.1 | 5,54 | 6,59 | 0 | 1,89 | 0 | 3,19 | 0 |
| MYBL2 | ATACGTTATTCTTTA | -1 | 795 | 810 | -205 | -190 | -4,7 | 0,01 | 11,86 | >0.1 | 5,54 | 5,97 | 0 | 1,5 | 0 | 3,55 | 0 |
| MYBL2 | TTTGGTTAATATTTT | -1 | 725 | 740 | -275 | -260 | -7,7 | 0,01 | 10,03 | >0.1 | 5,54 | 5,64 | 0 | 1,23 | 0 | 5,32 | 0 |
| MYBL2 | AGCTGTTAAAGATAC | -1 | 703 | 718 | -297 | -282 | -0,14 | 0,001 | 17,12 | >0.1 | 5,54 | 5,55 | 0 | 1,17 | 0 | 4,99 | 0 |
| MYBL2 | TAAAGATACACGGAA | -1 | 697 | 712 | -303 | -288 | -7,65 | 0,01 | 7,77 | >0.1 | 5,54 | 5,53 | 0 | 1,16 | 0 | 3,19 | 0 |
| MYBL2 | TTTCATTAATAGGTT | -1 | 598 | 613 | -402 | -387 | -8,01 | 0,01 | 3,07 | >0.1 | 3,48 | 5,22 | 0 | 1,02 | 0 | 1,36 | 0 |

| | | | | | | | | | | | | | | | | | |
|---|---|---|---|---|---|---|---|---|---|---|---|---|---|---|---|---|---|
| MYBL2 | AATAGGTTATCTCAC | -1 | 591 | 606 | -409 | -394 | -2,94 | 0,01 | 6,7 | >0.1 | 3,43 | 5,2 | 0 | 1,02 | 0 | 0 | 0 |
| MYBL2 | AACATTTTCCCTTCC | -1 | 528 | 543 | -472 | -457 | -7,5 | 0,01 | 5,31 | >0.1 | 3,09 | 5,04 | 0 | 0,99 | 0 | 3,7 | 0 |
| MYBL2 | CTCGGTCTGTCGCCC | -1 | 432 | 447 | -568 | -553 | -6,01 | 0,01 | 8,37 | >0.1 | 2,73 | 4,84 | 0 | 1,01 | 0 | 5,8 | 0 |
| MYBL2 | CCCAGGTTCACGCCA | -1 | 370 | 385 | -630 | -615 | -6,98 | 0,01 | 7,77 | >0.1 | 2,55 | 4,73 | 0 | 1,07 | 0 | 6,4 | 0 |
| MYBL2 | CGCCATTCTCCTGCC | -1 | 360 | 375 | -640 | -625 | -7,28 | 0,01 | 7,53 | >0.1 | 2,52 | 4,71 | 0 | 1,09 | 0 | 6,49 | 0 |
| MYBL2 | CGGGGTTTCACCGTG | -1 | 271 | 286 | -729 | -714 | 0,96 | 0,001 | 14,67 | >0.1 | 2,76 | 4,57 | 0 | 1,29 | 0 | 5,09 | 0 |
| MYBL2 | GACGGTCTCTATCTC | -1 | 248 | 263 | -752 | -737 | -7,85 | 0,01 | 6,1 | >0.1 | 2,84 | 4,54 | 0 | 1,38 | 0 | 5,19 | 0 |
| MYBL2 | ATCAGTTTTAATTTC | -1 | 154 | 169 | -846 | -831 | -3,99 | 0,01 | 8,1 | >0.1 | 3,25 | 4,42 | 0 | 1,9 | 0 | 2,53 | 0 |
| MYBL2 | CACAGGTCAACTCTC | -1 | 102 | 117 | -898 | -883 | 1,79 | 0,001 | 12,38 | >0.1 | 3,58 | 4,36 | 0 | 1,89 | 0 | 0,77 | 0 |
| SOX6 | CCACTGTGCT | 1 | 415 | 425 | -585 | -575 | 1,09 | 0,01 | 19,51 | >0.1 | 2,67 | 4,8 | 0 | 1,02 | 0 | 5,09 | 4,85 |
| SOX6 | AAATTGTCTG | 1 | 610 | 620 | -390 | -380 | -2,6 | 0,01 | 13,6 | >0.1 | 3,54 | 5,24 | 0 | 1,02 | 0 | 1,55 | 4,85 |
| SOX6 | ACATTATCTT | 1 | 638 | 648 | -362 | -352 | -1,23 | 0,01 | 15,37 | >0.1 | 3,77 | 5,32 | 0 | 1,05 | 0 | 1,61 | 4,85 |
| SOX6 | TCTTTGGTCA | 1 | 669 | 679 | -331 | -321 | -0,62 | 0,01 | 18,56 | >0.1 | 4,13 | 5,42 | 0 | 1,1 | 0 | 3,7 | 4,85 |
| SOX6 | ACATTTTCCG | 1 | 692 | 702 | -308 | -298 | -2,53 | 0,01 | 16,57 | >0.1 | 4,56 | 5,49 | 0 | 1,14 | 0 | 3,07 | 4,85 |
| SOX6 | GTTTTGATTT | 1 | 747 | 757 | -253 | -243 | -1,33 | 0,01 | 22,22 | >0.1 | 5,61 | 5,71 | 0 | 1,28 | 0 | 6,1 | 4,85 |
| SOX6 | ACTGTGTTCT | 1 | 902 | 912 | -98 | -88 | -1,19 | 0,01 | 19,16 | >0.1 | 5,61 | 6,72 | 0 | 1,89 | 0 | 1,28 | 4,85 |
| SOX6 | TCTTTTTCTT | 1 | 915 | 925 | -85 | -75 | 1,64 | 0,01 | 22 | >0.1 | 5,61 | 6,87 | 0 | 1,89 | 0 | 1,15 | 4,85 |
| SOX6 | TTTTTCTTTT | 1 | 917 | 927 | -83 | -73 | -1,66 | 0,01 | 18,7 | >0.1 | 5,61 | 6,89 | 0 | 1,89 | 0 | 1,12 | 4,85 |
| SOX6 | TCTTTTTTTG | 1 | 921 | 931 | -79 | -69 | -0,23 | 0,01 | 20,17 | >0.1 | 5,61 | 6,94 | 0 | 1,88 | 0 | 1,12 | 4,85 |
| SOX6 | TTTTTGGTCT | 1 | 925 | 935 | -75 | -65 | -0,92 | 0,01 | 19,47 | >0.1 | 5,61 | 6,99 | 0 | 1,88 | 0 | 1,06 | 4,85 |
| SOX6 | TTTTGGTCTT | 1 | 926 | 936 | -74 | -64 | -0,84 | 0,01 | 19,57 | >0.1 | 5,61 | 7,01 | 0 | 1,88 | 0 | 1,06 | 4,85 |
| SOX6 | ACATCGTCCC | -1 | 1043 | 1053 | 43 | 53 | -0,25 | 0,01 | 20,45 | >0.1 | 5,61 | 7,76 | 0 | 1,86 | 0 | 0,62 | 4,85 |
| SOX6 | CTTTTGGTTA | -1 | 732 | 742 | -268 | -258 | -2,54 | 0,01 | 20,71 | >0.1 | 5,61 | 5,65 | 0 | 1,23 | 0 | 5,92 | 4,85 |
| SOX6 | GTATGGTTTT | -1 | 620 | 630 | -380 | -370 | 0,04 | 0,01 | 16,25 | >0.1 | 3,62 | 5,26 | 0 | 1,03 | 0 | 1,44 | 4,85 |
| SOX6 | TTTTTATTTC | -1 | 576 | 586 | -424 | -414 | -2,41 | 0,01 | 11,9 | >0.1 | 3,32 | 5,14 | 0 | 1 | 0 | 0 | 4,85 |
| SOX6 | ACATTTTCCC | -1 | 532 | 542 | -468 | -458 | -1,24 | 0,01 | 14,33 | >0.1 | 3,1 | 5,04 | 0 | 0,98 | 0 | 1,61 | 4,85 |
| SOX6 | ACTTTGCCCT | -1 | 509 | 519 | -491 | -481 | -0,86 | 0,01 | 15,7 | >0.1 | 3 | 4,98 | 0 | 0,98 | 0 | 2,75 | 4,85 |
| SOX6 | CCCTTGTCTT | -1 | 503 | 513 | -497 | -487 | 5,94 | 0,001 | 25,39 | >0.1 | 2,97 | 4,97 | 0 | 0,98 | 0 | 5,68 | 4,85 |
| SOX6 | GTTTTTTTTT | -1 | 470 | 480 | -530 | -520 | -1,83 | 0,01 | 16,7 | >0.1 | 2,85 | 4,9 | 0 | 0,99 | 0 | 4,95 | 4,85 |
| SOX6 | TTTTTTTTTT | -1 | 469 | 479 | -531 | -521 | -2,16 | 0,01 | 16,24 | >0.1 | 2,84 | 4,9 | 0 | 0,99 | 0 | 4,83 | 4,85 |
| SOX6 | TTTTTTTTTT | -1 | 468 | 478 | -532 | -522 | -2,16 | 0,01 | 17,34 | >0.1 | 2,84 | 4,9 | 0 | 0,99 | 0 | 5,92 | 4,85 |
| SOX6 | TTTTTTTTTT | -1 | 467 | 477 | -533 | -523 | -2,16 | 0,01 | 17,18 | >0.1 | 2,83 | 4,9 | 0 | 0,99 | 0 | 5,78 | 4,85 |
| SOX6 | TTTTTTTTTT | -1 | 466 | 476 | -534 | -524 | -2,16 | 0,01 | 17,04 | >0.1 | 2,83 | 4,89 | 0 | 0,99 | 0 | 5,63 | 4,85 |
| SOX6 | TTTTTTTTTT | -1 | 465 | 475 | -535 | -525 | -2,16 | 0,01 | 17,17 | >0.1 | 2,83 | 4,89 | 0 | 0,99 | 0 | 5,78 | 4,85 |
| SOX6 | TTTTTTTTTT | -1 | 464 | 474 | -536 | -526 | -2,16 | 0,01 | 17,17 | >0.1 | 2,83 | 4,89 | 0 | 0,99 | 0 | 5,78 | 4,85 |
| SOX6 | TTTTTTTTTT | -1 | 463 | 473 | -537 | -527 | -2,16 | 0,01 | 17,16 | >0.1 | 2,82 | 4,89 | 0 | 0,99 | 0 | 5,78 | 4,85 |
| SOX6 | TTTTTTTTTT | -1 | 462 | 472 | -538 | -528 | -2,16 | 0,01 | 17,02 | >0.1 | 2,82 | 4,89 | 0 | 0,99 | 0 | 5,63 | 4,85 |
| SOX6 | TTTTTTTTTT | -1 | 461 | 471 | -539 | -529 | -2,16 | 0,01 | 17,01 | >0.1 | 2,81 | 4,88 | 0 | 0,99 | 0 | 5,63 | 4,85 |
| SOX6 | TTTTTTTTTT | -1 | 460 | 470 | -540 | -530 | -2,16 | 0,01 | 16,9 | >0.1 | 2,81 | 4,88 | 0 | 0,99 | 0 | 5,52 | 4,85 |
| SOX6 | TTTTTTTTTT | -1 | 459 | 469 | -541 | -531 | -2,16 | 0,01 | 16,89 | >0.1 | 2,81 | 4,88 | 0 | 0,99 | 0 | 5,52 | 4,85 |
| SOX6 | TTTTTTTTTT | -1 | 458 | 468 | -542 | -532 | -2,16 | 0,01 | 17 | >0.1 | 2,8 | 4,88 | 0 | 0,99 | 0 | 5,63 | 4,85 |
| SOX6 | TTTTTTTTTT | -1 | 457 | 467 | -543 | -533 | -2,16 | 0,01 | 16,99 | >0.1 | 2,8 | 4,88 | 0 | 0,99 | 0 | 5,63 | 4,85 |
| SOX6 | CCATTCTCCT | -1 | 363 | 373 | -637 | -627 | 5,06 | 0,001 | 24,83 | >0.1 | 2,52 | 4,7 | 0 | 1,08 | 0 | 6,61 | 4,85 |
| SOX6 | TTTTTGTATT | -1 | 297 | 307 | -703 | -693 | -0,72 | 0,01 | 17,08 | >0.1 | 2,69 | 4,6 | 0 | 1,21 | 0 | 4,45 | 4,85 |
| SOX6 | CCACTGTGCC | -1 | 181 | 191 | -819 | -809 | 0,83 | 0,01 | 18,88 | >0.1 | 3,12 | 4,44 | 0 | 1,78 | 0 | 3,86 | 4,85 |
| SOX6 | CCATTTTTTA | -1 | 139 | 149 | -861 | -851 | 0,81 | 0,01 | 16,9 | >0.1 | 3,35 | 4,39 | 0 | 1,89 | 0 | 1,61 | 4,85 |
| SOX4 | CCACAGAACCAGTGTG | 1 | 41 | 57 | -959 | -943 | -24,81 | 0,01 | -14,42 | >0.1 | 4,22 | 4,3 | 0 | 1,87 | 0 | 0 | 0 |
| SOX4 | AAAAAATGGCCATGGA | 1 | 140 | 156 | -860 | -844 | -30,69 | 0,01 | -19,46 | >0.1 | 3,32 | 4,4 | 0 | 1,9 | 0 | 1,61 | 0 |
| SOX4 | CAGAAATGGCTGGGCA | 1 | 169 | 185 | -831 | -815 | -29,94 | 0,01 | -17,1 | >0.1 | 3,17 | 4,44 | 0 | 1,89 | 0 | 3,34 | 0 |
| SOX4 | TCACAAGGTCAGGAGA | 1 | 236 | 252 | -764 | -748 | -33,74 | 0,01 | -20,48 | >0.1 | 2,88 | 4,52 | 0 | 1,43 | 0 | 4,42 | 0 |
| SOX4 | CAGAGCTTGCAGTGAG | 1 | 388 | 404 | -612 | -596 | -29,6 | 0,01 | -15,17 | >0.1 | 2,6 | 4,76 | 0 | 1,05 | 0 | 6,02 | 0 |
| SOX4 | AGACAAGGGCAAAGTC | 1 | 504 | 520 | -496 | -480 | -27,99 | 0,01 | -14,84 | >0.1 | 2,99 | 4,99 | 0 | 0,99 | 0 | 4,19 | 0 |
| SOX4 | GCCCAAGTAGAGAGTT | 1 | 543 | 559 | -457 | -441 | -26,94 | 0,01 | -17,71 | >0.1 | 3,16 | 5,08 | 0 | 0,99 | 0 | 0 | 0 |
| SOX4 | TATTAATGAAATTGTC | 1 | 602 | 618 | -398 | -382 | -29,72 | 0,01 | -18,54 | >0.1 | 3,51 | 5,23 | 0 | 1,03 | 0 | 1,41 | 0 |
| SOX4 | AAACACCAACATTATC | 1 | 630 | 646 | -370 | -354 | -31,18 | 0,01 | -19,47 | >0.1 | 3,73 | 5,31 | 0 | 1,06 | 0 | 1,61 | 0 |
| SOX4 | CAAAAGTACCGGTTTT | 1 | 736 | 752 | -264 | -248 | -26,3 | 0,01 | -7,9 | >0.1 | 5,53 | 5,69 | 0 | 1,26 | 0 | 5,92 | 0 |
| SOX4 | GATTGTTTACTGTGTT | 1 | 894 | 910 | -106 | -90 | -25,84 | 0,01 | -8,85 | >0.1 | 5,53 | 6,71 | 0 | 1,89 | 0 | 2,86 | 0 |
| SOX4 | AAGCAAGTGAACTTTG | -1 | 1172 | 1188 | 172 | 188 | -30,09 | 0,01 | -11,68 | >0.1 | 5,53 | 6,14 | 0 | 1,65 | 1,79 | 3,29 | 0 |
| SOX4 | CAAAAATGTCTTGGCC | -1 | 1120 | 1136 | 120 | 136 | -30,95 | 0,01 | -11,08 | >0.1 | 5,53 | 6,55 | 0 | 1,85 | 3,22 | 2,72 | 0 |

| | | | | | | | | | | | | | | | | | |
|---|---|---|---|---|---|---|---|---|---|---|---|---|---|---|---|---|---|
| SOX4 | CAGCAGTTACAGCAAC | -1 | 1082 | 1098 | 82 | 98 | -28,97 | 0,01 | -10,59 | >0.1 | 5,53 | 6,99 | 0 | 1,85 | 1,79 | 2,21 | 0 |
| SOX4 | AGATAATGTTGGTGTT | -1 | 631 | 647 | -369 | -353 | -22,06 | 0,01 | -10,34 | >0.1 | 3,74 | 5,31 | 0 | 1,06 | 0 | 1,61 | 0 |
| SOX4 | AGACAATTTCATTAAT | -1 | 603 | 619 | -397 | -381 | -10,46 | 0,001 | 0,76 | >0.1 | 3,52 | 5,23 | 0 | 1,03 | 0 | 1,44 | 0 |
| SOX4 | CAACATTTTCCCTTCC | -1 | 528 | 544 | -472 | -456 | -34,12 | 0,01 | -23,39 | >0.1 | 3,09 | 5,04 | 0 | 0,99 | 0 | 1,61 | 0 |
| SOX4 | GATCAGTTTTTTTTTT | -1 | 469 | 485 | -531 | -515 | -30,98 | 0,01 | -17,27 | >0.1 | 2,85 | 4,91 | 0 | 0,99 | 0 | 4,95 | 0 |
| SOX4 | TAATAGAGACGGGGTT | -1 | 279 | 295 | -721 | -705 | -26,32 | 0,01 | -12,65 | >0.1 | 2,73 | 4,58 | 0 | 1,27 | 0 | 5,09 | 0 |
| SOX5 | AATGTTG | 1 | 537 | 544 | -463 | -456 | 3,59 | 0,01 | 14,33 | >0.1 | 3,11 | 5,04 | 0 | 0,98 | 0 | 1,61 | 0 |
| SOX5 | ATTGTCT | 1 | 612 | 619 | -388 | -381 | 7,2 | 0,001 | 18,55 | >0.1 | 3,55 | 5,23 | 0 | 1,02 | 0 | 1,55 | 0 |
| SOX5 | ATTATCT | 1 | 640 | 647 | -360 | -353 | 2,5 | 0,01 | 14,25 | >0.1 | 3,78 | 5,31 | 0 | 1,05 | 0 | 1,61 | 0 |
| SOX5 | ATTCTTA | 1 | 808 | 815 | -192 | -185 | 4,05 | 0,01 | 20,5 | >0.1 | 5,65 | 6 | 0 | 1,52 | 0 | 3,28 | 0 |
| SOX5 | ATTATTC | 1 | 833 | 840 | -167 | -160 | 5,28 | 0,01 | 22,2 | >0.1 | 5,65 | 6,16 | 0 | 1,69 | 0 | 3,43 | 0 |
| SOX5 | ATTGTTT | 1 | 895 | 902 | -105 | -98 | 10,85 | >0.1 | 27,88 | >0.1 | 5,65 | 6,63 | 0 | 1,89 | 0 | 2,86 | 0 |
| SOX5 | CTTGTTG | 1 | 1079 | 1086 | 79 | 86 | 4,02 | 0,01 | 22,56 | >0.1 | 5,65 | 7,03 | 0 | 1,85 | 1,79 | 2,21 | 0 |
| SOX5 | ATTTTTG | 1 | 1129 | 1136 | 129 | 136 | 3,09 | 0,01 | 23,1 | >0.1 | 5,65 | 6,47 | 0 | 1,84 | 3,22 | 2,82 | 0 |
| SOX5 | ATTTTTT | -1 | 828 | 835 | -172 | -165 | 3,56 | 0,01 | 20,53 | >0.1 | 5,65 | 6,12 | 0 | 1,65 | 0 | 3,55 | 0 |
| SOX5 | ATTCTTT | -1 | 796 | 803 | -204 | -197 | 4,05 | 0,01 | 22,26 | >0.1 | 5,65 | 5,93 | 0 | 1,46 | 0 | 5,17 | 0 |
| SOX5 | AATGTTA | -1 | 690 | 697 | -310 | -303 | 4,05 | 0,01 | 17,99 | >0.1 | 4,47 | 5,48 | 0 | 1,13 | 0 | 2,86 | 0 |
| SOX5 | AATGTTG | -1 | 636 | 643 | -364 | -357 | 3,59 | 0,01 | 15,29 | >0.1 | 3,74 | 5,3 | 0 | 1,05 | 0 | 1,61 | 0 |
| SOX5 | ATGGTTT | -1 | 621 | 628 | -379 | -372 | 4,49 | 0,01 | 15,89 | >0.1 | 3,62 | 5,26 | 0 | 1,03 | 0 | 1,5 | 0 |
| SOX5 | ATTTTTA | -1 | 580 | 587 | -420 | -413 | 3,56 | 0,01 | 13,04 | >0.1 | 3,34 | 5,15 | 0 | 1 | 0 | 0 | 0 |
| SOX5 | ATTTTTA | -1 | 293 | 300 | -707 | -700 | 3,56 | 0,01 | 16,53 | >0.1 | 2,71 | 4,59 | 0 | 1,22 | 0 | 4,45 | 0 |
| SOX5 | AGTGCTA | -1 | 207 | 214 | -793 | -786 | 3,42 | 0,01 | 17,19 | >0.1 | 3,01 | 4,47 | 0 | 1,58 | 0 | 4,7 | 0 |
| SOX5 | ATTTTTT | -1 | 140 | 147 | -860 | -853 | 3,56 | 0,01 | 14,84 | >0.1 | 3,35 | 4,39 | 0 | 1,89 | 0 | 1,65 | 0 |
| ARNT::HIF1A | GTAAGTGA | 1 | 5 | 13 | -995 | -987 | 0,85 | 0,01 | 14,4 | >0.1 | 5,64 | 4,26 | 0 | 1,86 | 1,79 | 0 | 0 |
| ARNT::HIF1A | CAGTGTGA | 1 | 50 | 58 | -950 | -942 | -0,24 | 0,01 | 10,07 | >0.1 | 4,14 | 4,3 | 0 | 1,87 | 0 | 0 | 0 |
| ARNT::HIF1A | GGGCGTGG | 1 | 314 | 322 | -686 | -678 | 10,38 | 0,0001 | 24,95 | >0.1 | 2,64 | 4,62 | 0 | 1,17 | 0 | 6,13 | 0 |
| ARNT::HIF1A | GCACCTGT | 1 | 328 | 336 | -672 | -664 | 1,47 | 0,01 | 15,42 | >0.1 | 2,6 | 4,64 | 0 | 1,14 | 0 | 5,56 | 0 |
| ARNT::HIF1A | TGGCGTGA | 1 | 370 | 378 | -630 | -622 | 1,29 | 0,01 | 15,76 | >0.1 | 2,54 | 4,71 | 0 | 1,07 | 0 | 6,15 | 0 |
| ARNT::HIF1A | GAACCTGG | 1 | 376 | 384 | -624 | -616 | 0,37 | 0,01 | 15,11 | >0.1 | 2,56 | 4,72 | 0 | 1,06 | 0 | 6,4 | 0 |
| ARNT::HIF1A | GAGCTTGC | 1 | 390 | 398 | -610 | -602 | -0,04 | 0,01 | 14,36 | >0.1 | 2,59 | 4,75 | 0 | 1,05 | 0 | 6,02 | 0 |
| ARNT::HIF1A | AAATGTGA | 1 | 587 | 595 | -413 | -405 | 0,47 | 0,01 | 10,02 | >0.1 | 3,38 | 5,17 | 0 | 1 | 0 | 0 | 0 |
| ARNT::HIF1A | GGATGTGA | 1 | 1022 | 1030 | 22 | 30 | 4,07 | 0,001 | 21,56 | >0.1 | 5,64 | 8,68 | 0 | 1,86 | 0 | 1,31 | 0 |
| ARNT::HIF1A | AGACCTGT | 1 | 1159 | 1167 | 159 | 167 | 0,35 | 0,01 | 16,92 | >0.1 | 4,38 | 6,23 | 0 | 1,77 | 1,79 | 2,4 | 0 |
| ARNT::HIF1A | GCAAGTGA | -1 | 1178 | 1186 | 178 | 186 | 1,25 | 0,01 | 19,65 | >0.1 | 5,64 | 6,1 | 0 | 1,6 | 1,79 | 3,26 | 0 |
| ARNT::HIF1A | CTTCGTGG | -1 | 1147 | 1155 | 147 | 155 | 4,18 | 0,001 | 21,93 | >0.1 | 5,64 | 6,32 | 0 | 1,84 | 1,79 | 2,16 | 0 |
| ARNT::HIF1A | GAACTTGG | -1 | 954 | 962 | -46 | -38 | -0,13 | 0,01 | 16,52 | >0.1 | 5,64 | 7,44 | 0 | 1,88 | 0 | 1,7 | 0 |
| ARNT::HIF1A | ATACGTTA | -1 | 802 | 810 | -198 | -190 | -0,51 | 0,01 | 16,02 | >0.1 | 5,64 | 5,97 | 0 | 1,49 | 0 | 3,43 | 0 |
| ARNT::HIF1A | CTACTTGG | -1 | 545 | 553 | -455 | -447 | 0,68 | 0,01 | 9,88 | >0.1 | 3,15 | 5,06 | 0 | 0,98 | 0 | 0 | 0 |
| ARNT::HIF1A | ATACATGA | -1 | 518 | 526 | -482 | -474 | -0,51 | 0,01 | 11,45 | >0.1 | 3,03 | 5 | 0 | 0,98 | 0 | 2,95 | 0 |
| ARNT::HIF1A | ACAGGTGC | -1 | 328 | 336 | -672 | -664 | 2,39 | 0,01 | 16,34 | >0.1 | 2,6 | 4,64 | 0 | 1,14 | 0 | 5,56 | 0 |
| ARNT::HIF1A | AGACGGGG | -1 | 281 | 289 | -719 | -711 | 1,75 | 0,01 | 15,41 | >0.1 | 2,74 | 4,57 | 0 | 1,26 | 0 | 5,09 | 0 |
| ARNT::HIF1A | CACCGTGT | -1 | 270 | 278 | -730 | -722 | 3,03 | 0,01 | 16,74 | >0.1 | 2,78 | 4,56 | 0 | 1,29 | 0 | 5,09 | 0 |
| ARNT::HIF1A | CAAAGTGC | -1 | 209 | 217 | -791 | -783 | 0,42 | 0,01 | 14,17 | >0.1 | 3 | 4,47 | 0 | 1,57 | 0 | 4,7 | 0 |
| ARNT::HIF1A | AGGCGTGA | -1 | 192 | 200 | -808 | -800 | 8,38 | 0,001 | 21,73 | >0.1 | 3,08 | 4,45 | 0 | 1,69 | 0 | 4,13 | 0 |
| CEBPA | ATCGCGCCACT | 1 | 409 | 420 | -591 | -580 | -4,87 | 0,01 | 9,12 | >0.1 | 2,65 | 4,79 | 0 | 1,03 | 0 | 5,52 | 0 |
| CEBPA | GTTGCCCAAGT | 1 | 540 | 551 | -460 | -449 | 9,93 | 0,001 | 19,11 | >0.1 | 3,14 | 5,06 | 0 | 0,99 | 0 | 0 | 0 |
| CEBPA | CTTTGGTCACT | 1 | 670 | 681 | -330 | -319 | -11,2 | 0,01 | 3,17 | >0.1 | 4,15 | 5,42 | 0 | 1,1 | 0 | 3,7 | 0 |
| CEBPA | ATTATAACATT | 1 | 686 | 697 | -314 | -303 | -9,43 | 0,01 | 4,55 | >0.1 | 4,43 | 5,48 | 0 | 1,13 | 0 | 2,95 | 0 |
| CEBPA | ATTTGGCCATA | 1 | 753 | 764 | -247 | -236 | -7 | 0,01 | 11,96 | >0.1 | 5,6 | 5,74 | 0 | 1,3 | 0 | 6,32 | 0 |
| CEBPA | GTTACATATCT | 1 | 848 | 859 | -152 | -141 | -7,51 | 0,01 | 9,18 | >0.1 | 5,6 | 6,29 | 0 | 1,85 | 0 | 2,95 | 0 |
| CEBPA | ATTGGCTCAGC | 1 | 882 | 893 | -118 | -107 | -6,94 | 0,01 | 10,17 | >0.1 | 5,6 | 6,56 | 0 | 1,89 | 0 | 3,07 | 0 |
| CEBPA | CTTGGCTCACA | 1 | 1031 | 1042 | 31 | 42 | -8,01 | 0,01 | 8,94 | >0.1 | 5,6 | 8,19 | 0 | 1,86 | 0 | 1,31 | 0 |
| CEBPA | GTTTAACCACG | 1 | 1141 | 1152 | 141 | 152 | -7,7 | 0,01 | 12,73 | >0.1 | 5,6 | 6,37 | 0 | 1,84 | 3,22 | 3,4 | 0 |
| CEBPA | CTTGCTTCTTG | 1 | 1181 | 1192 | 181 | 192 | -11,07 | 0,01 | 7,41 | >0.1 | 5,6 | 6,08 | 0 | 1,58 | 1,79 | 3,43 | 0 |
| CEBPA | GTTCCTCAATG | -1 | 1108 | 1119 | 108 | 119 | -6,59 | 0,01 | 13,28 | >0.1 | 5,6 | 6,67 | 0 | 1,85 | 3,22 | 2,53 | 0 |
| CEBPA | CTTTCCCTAGA | -1 | 1002 | 1013 | 2 | 13 | -9,02 | 0,01 | 9,03 | >0.1 | 5,6 | 9,12 | 0 | 1,87 | 0 | 1,46 | 0 |
| CEBPA | ATGGCCAAATC | -1 | 752 | 763 | -248 | -237 | -4,25 | 0,01 | 14,92 | >0.1 | 5,6 | 5,74 | 0 | 1,3 | 0 | 6,54 | 0 |
| CEBPA | ATTTCAAGATC | -1 | 570 | 581 | -430 | -419 | -3,45 | 0,01 | 5,97 | >0.1 | 3,29 | 5,13 | 0 | 1 | 0 | 0 | 0 |
| CEBPA | CTTGGGCAACA | -1 | 539 | 550 | -461 | -450 | -5,67 | 0,01 | 3,5 | >0.1 | 3,13 | 5,06 | 0 | 0,99 | 0 | 0 | 0 |

| | | | | | | | | | | | | | | | | | |
|---|---|---|---|---|---|---|---|---|---|---|---|---|---|---|---|---|---|
| CEBPA | CCTTCCAAATA | -1 | 523 | 534 | -477 | -466 | -9,25 | 0,01 | 3 | >0.1 | 3,06 | 5,02 | 0 | 0,98 | 0 | 3,19 | 0 |
| CEBPA | CCTGCCTCAGC | -1 | 355 | 366 | -645 | -634 | -8,9 | 0,01 | 5,45 | >0.1 | 2,53 | 4,69 | 0 | 1,09 | 0 | 6,03 | 0 |
| CEBPA | GTTTCACCGTG | -1 | 271 | 282 | -729 | -718 | -6,74 | 0,01 | 6,87 | >0.1 | 2,77 | 4,56 | 0 | 1,29 | 0 | 4,99 | 0 |
| CEBPA | TTTGCAAAACT | -1 | 81 | 92 | -919 | -908 | -2,34 | 0,01 | 7,64 | >0.1 | 3,77 | 4,33 | 0 | 1,88 | 0 | 0 | 0 |
| CEBPA | CTTAAAAAAGC | -1 | 72 | 83 | -928 | -917 | -11,09 | 0,01 | -1,03 | >0.1 | 3,86 | 4,32 | 0 | 1,88 | 0 | 0 | 0 |
| CEBPA | ATTTCCTATTT | -1 | 59 | 70 | -941 | -930 | -7,03 | 0,01 | 3,16 | >0.1 | 4 | 4,31 | 0 | 1,87 | 0 | 0 | 0 |
| CEBPG | AATAGGAAAT | 1 | 60 | 70 | -940 | -930 | -6 | 0,01 | 4,18 | >0.1 | 4 | 4,31 | 0 | 1,87 | 0 | 0 | 0 |
| CEBPG | GTTTTGCAAA | 1 | 82 | 92 | -918 | -908 | -1,89 | 0,01 | 8,09 | >0.1 | 3,77 | 4,33 | 0 | 1,88 | 0 | 0 | 0 |
| CEBPG | ACGGTGAAAC | 1 | 272 | 282 | -728 | -718 | -5,94 | 0,01 | 7,67 | >0.1 | 2,77 | 4,56 | 0 | 1,29 | 0 | 4,99 | 0 |
| CEBPG | ATGGCGTGAA | 1 | 369 | 379 | -631 | -621 | -6,27 | 0,01 | 8,21 | >0.1 | 2,54 | 4,71 | 0 | 1,07 | 0 | 6,15 | 0 |
| CEBPG | GTTGCCCAAG | 1 | 540 | 550 | -460 | -450 | 4,82 | 0,001 | 13,99 | >0.1 | 3,13 | 5,06 | 0 | 0,99 | 0 | 0 | 0 |
| CEBPG | ACTGTAAAAT | 1 | 678 | 688 | -322 | -312 | -6,39 | 0,01 | 7,72 | >0.1 | 4,27 | 5,45 | 0 | 1,11 | 0 | 3,28 | 0 |
| CEBPG | ATTAACCAAA | 1 | 730 | 740 | -270 | -260 | -5,01 | 0,01 | 12,99 | >0.1 | 5,61 | 5,64 | 0 | 1,23 | 0 | 5,52 | 0 |
| CEBPG | ATTAAAGAAT | 1 | 793 | 803 | -207 | -197 | -5,25 | 0,01 | 13,07 | >0.1 | 5,61 | 5,93 | 0 | 1,46 | 0 | 5,32 | 0 |
| CEBPG | ATAACGTATT | 1 | 801 | 811 | -199 | -189 | -6,29 | 0,01 | 10,22 | >0.1 | 5,61 | 5,98 | 0 | 1,5 | 0 | 3,43 | 0 |
| CEBPG | GTTACATATC | 1 | 848 | 858 | -152 | -142 | -2,58 | 0,01 | 14,11 | >0.1 | 5,61 | 6,28 | 0 | 1,84 | 0 | 2,95 | 0 |
| CEBPG | AGCGCCCAAC | 1 | 944 | 954 | -56 | -46 | -5,48 | 0,01 | 11,1 | >0.1 | 5,61 | 7,29 | 0 | 1,88 | 0 | 1,8 | 0 |
| CEBPG | ATTGAGGAAC | 1 | 1109 | 1119 | 109 | 119 | 0,22 | 0,01 | 20,08 | >0.1 | 5,61 | 6,66 | 0 | 1,85 | 3,22 | 2,52 | 0 |
| CEBPG | GTGGTTAAAC | -1 | 1141 | 1151 | 141 | 151 | -5,78 | 0,01 | 14,67 | >0.1 | 5,61 | 6,37 | 0 | 1,84 | 3,22 | 3,4 | 0 |
| CEBPG | CTTGTCCAAA | -1 | 1132 | 1142 | 132 | 142 | -3,93 | 0,01 | 16,16 | >0.1 | 5,61 | 6,44 | 0 | 1,84 | 3,22 | 2,98 | 0 |
| CEBPG | GTTCCTCAAT | -1 | 1109 | 1119 | 109 | 119 | -2,46 | 0,01 | 17,4 | >0.1 | 5,61 | 6,66 | 0 | 1,85 | 3,22 | 2,52 | 0 |
| CEBPG | AGTAAACAAT | -1 | 895 | 905 | -105 | -95 | -2,48 | 0,01 | 14,63 | >0.1 | 5,61 | 6,66 | 0 | 1,89 | 0 | 2,95 | 0 |
| CEBPG | CTGAGCCAAT | -1 | 882 | 892 | -118 | -108 | -6,13 | 0,01 | 10,99 | >0.1 | 5,61 | 6,55 | 0 | 1,89 | 0 | 3,07 | 0 |
| CEBPG | GATATGTAAC | -1 | 848 | 858 | -152 | -142 | -2,23 | 0,01 | 14,46 | >0.1 | 5,61 | 6,28 | 0 | 1,84 | 0 | 2,95 | 0 |
| CEBPG | AATACGTTAT | -1 | 801 | 811 | -199 | -189 | -5,82 | 0,01 | 10,69 | >0.1 | 5,61 | 5,98 | 0 | 1,5 | 0 | 3,43 | 0 |
| CEBPG | ATGGCCAAAT | -1 | 753 | 763 | -247 | -237 | 0,4 | 0,01 | 19,36 | >0.1 | 5,61 | 5,74 | 0 | 1,3 | 0 | 6,32 | 0 |
| CEBPG | GTGACCAAAG | -1 | 670 | 680 | -330 | -320 | -5,98 | 0,01 | 8,38 | >0.1 | 4,14 | 5,42 | 0 | 1,1 | 0 | 3,7 | 0 |
| CEBPG | CTTGGGCAAC | -1 | 540 | 550 | -460 | -450 | 5,78 | 0,001 | 14,95 | >0.1 | 3,13 | 5,06 | 0 | 0,99 | 0 | 0 | 0 |
| CEBPG | TTTATGGAAT | -1 | 495 | 505 | -505 | -495 | -4,37 | 0,01 | 11,13 | >0.1 | 2,94 | 4,95 | 0 | 0,98 | 0 | 6,62 | 0 |
| CEBPG | GTGGCGCGAT | -1 | 409 | 419 | -591 | -581 | 3,86 | 0,001 | 17,85 | >0.1 | 2,65 | 4,78 | 0 | 1,03 | 0 | 5,52 | 0 |
| CEBPG | TCTGCCCAAC | -1 | 226 | 236 | -774 | -764 | -2,42 | 0,01 | 11,74 | >0.1 | 2,93 | 4,5 | 0 | 1,47 | 0 | 5,26 | 0 |
| CEBPG | TTTGCAAAAC | -1 | 82 | 92 | -918 | -908 | 1,44 | 0,001 | 11,42 | >0.1 | 3,77 | 4,33 | 0 | 1,88 | 0 | 0 | 0 |
| DMBX1 | GAAATGAGCTTTTTTAA | 1 | 65 | 82 | -935 | -918 | -1,07 | 0,01 | 9,03 | >0.1 | 3,9 | 4,32 | 0 | 1,88 | 0 | 0 | 0 |
| DMBX1 | AAAGGCAGATCAGGAGA | 1 | 89 | 106 | -911 | -894 | 3,22 | 0,01 | 13,95 | >0.1 | 3,67 | 4,35 | 0 | 1,88 | 0 | 0,83 | 0 |
| DMBX1 | GAGTGGAGAGTAGTCAT | 1 | 117 | 134 | -883 | -866 | 0,05 | 0,01 | 11,43 | >0.1 | 3,46 | 4,38 | 0 | 1,89 | 0 | 1,65 | 0 |
| DMBX1 | TCGGGAGGCTGAGGCAG | 1 | 348 | 365 | -652 | -635 | -0,14 | 0,01 | 14,6 | >0.1 | 2,54 | 4,69 | 0 | 1,11 | 0 | 6,4 | 0 |
| DMBX1 | TGGAAGGGAAAATGTTG | 1 | 527 | 544 | -473 | -456 | -1,85 | 0,01 | 10,97 | >0.1 | 3,09 | 5,04 | 0 | 0,99 | 0 | 3,7 | 0 |
| DMBX1 | TTTCTGGGAATATGATC | 1 | 557 | 574 | -443 | -426 | -1,79 | 0,01 | 7,56 | >0.1 | 3,24 | 5,11 | 0 | 1 | 0 | 0 | 0 |
| DMBX1 | AGGAAAATATTAACCAA | 1 | 722 | 739 | -278 | -261 | -0,08 | 0,01 | 17,62 | >0.1 | 5,52 | 5,64 | 0 | 1,23 | 0 | 5,32 | 0 |
| DMBX1 | AGTACCGGTTTTGATTT | 1 | 740 | 757 | -260 | -243 | -0,59 | 0,01 | 17,84 | >0.1 | 5,52 | 5,71 | 0 | 1,28 | 0 | 5,92 | 0 |
| DMBX1 | TCTCTAGGATTAAAGAA | 1 | 785 | 802 | -215 | -198 | 7 | 0,001 | 25,22 | >0.1 | 5,52 | 5,93 | 0 | 1,46 | 0 | 5,32 | 0 |
| DMBX1 | TTTAAAAAATTATTCTA | 1 | 825 | 842 | -175 | -158 | -2,18 | 0,01 | 14,64 | >0.1 | 5,52 | 6,17 | 0 | 1,71 | 0 | 3,43 | 0 |
| DMBX1 | TCTCCAGGATGAACTTT | 1 | 863 | 880 | -137 | -120 | -0,09 | 0,01 | 16,53 | >0.1 | 5,52 | 6,45 | 0 | 1,9 | 0 | 2,75 | 0 |
| DMBX1 | TGAGGAGGTTTTAGTCT | 1 | 988 | 1005 | -12 | 5 | 0,09 | 0,01 | 17,93 | >0.1 | 5,52 | 8,88 | 0 | 1,87 | 0 | 1,57 | 0 |
| DMBX1 | TGGACAAGTTTAACCAC | 1 | 1134 | 1151 | 134 | 151 | 0,67 | 0,01 | 20,75 | >0.1 | 5,52 | 6,42 | 0 | 1,84 | 3,22 | 3,07 | 0 |
| DMBX1 | GTCTTCGGCTTCGTGGT | -1 | 1146 | 1163 | 146 | 163 | 0,05 | 0,01 | 17,77 | >0.1 | 5,52 | 6,33 | 0 | 1,84 | 1,79 | 2,24 | 0 |
| DMBX1 | CTCATGAGATTTTCTCT | -1 | 975 | 992 | -25 | -8 | -1,2 | 0,01 | 16,16 | >0.1 | 5,52 | 8,43 | 0 | 1,87 | 0 | 1,53 | 0 |
| DMBX1 | TGTAACAGATTTTAGAA | -1 | 837 | 854 | -163 | -146 | 2,52 | 0,01 | 19,29 | >0.1 | 5,52 | 6,26 | 0 | 1,81 | 0 | 3,19 | 0 |
| DMBX1 | AACTAGGGATCATGAAG | -1 | 645 | 662 | -355 | -338 | 2,96 | 0,01 | 17,11 | >0.1 | 3,88 | 5,36 | 0 | 1,08 | 0 | 3,83 | 0 |
| DMBX1 | TCATGAAGATAATGTTG | -1 | 636 | 653 | -364 | -347 | -0,02 | 0,01 | 11,82 | >0.1 | 3,79 | 5,33 | 0 | 1,06 | 0 | 1,66 | 0 |
| DMBX1 | GATCTCGGCTCACTGCA | -1 | 395 | 412 | -605 | -588 | -1,28 | 0,01 | 13,07 | >0.1 | 2,62 | 4,77 | 0 | 1,05 | 0 | 5,91 | 0 |
| DMBX1 | TAGCTGGGACTACAGGT | -1 | 330 | 347 | -670 | -653 | -0,96 | 0,01 | 13,46 | >0.1 | 2,58 | 4,66 | 0 | 1,14 | 0 | 6,03 | 0 |
| DMBX1 | ACGCCCAGCTAAGTTTT | -1 | 303 | 320 | -697 | -680 | -1,2 | 0,01 | 12,37 | >0.1 | 2,66 | 4,62 | 0 | 1,2 | 0 | 5,09 | 0 |
| DMBX1 | GTGCTAGGATTACAGGC | -1 | 196 | 213 | -804 | -787 | 5,1 | 0,001 | 18,75 | >0.1 | 3,04 | 4,47 | 0 | 1,66 | 0 | 4,48 | 0 |
| DMBX1 | TAAAAAAGCTCATTTCC | -1 | 64 | 81 | -936 | -919 | -1,7 | 0,01 | 8,41 | >0.1 | 3,91 | 4,32 | 0 | 1,88 | 0 | 0 | 0 |
| MZF1 | TGTGGA | 1 | 113 | 119 | -887 | -881 | 3,64 | 0,01 | 14,2 | >0.1 | 3,53 | 4,36 | 0 | 1,89 | 0 | 0,79 | 0 |
| MZF1 | TGGAGA | 1 | 120 | 126 | -880 | -874 | 3,56 | 0,01 | 14,02 | >0.1 | 3,48 | 4,37 | 0 | 1,89 | 0 | 0,73 | 0 |
| MZF1 | TGGGCA | 1 | 179 | 185 | -821 | -815 | 3,98 | 0,01 | 16,51 | >0.1 | 3,14 | 4,43 | 0 | 1,8 | 0 | 3,16 | 0 |
| MZF1 | TTGGGA | 1 | 214 | 220 | -786 | -780 | 6,14 | 0,01 | 19,74 | >0.1 | 2,99 | 4,48 | 0 | 1,54 | 0 | 4,59 | 0 |

| | | | | | | | | | | | | | | | | | |
|---|---|---|---|---|---|---|---|---|---|---|---|---|---|---|---|---|---|
| MZF1 | TGGGCA | 1 | 228 | 234 | -772 | -766 | 3,98 | 0,01 | 18,13 | >0.1 | 2,93 | 4,5 | 0 | 1,46 | 0 | 5,26 | 0 |
| MZF1 | CGGTGA | 1 | 273 | 279 | -727 | -721 | 5,98 | 0,01 | 19,58 | >0.1 | 2,77 | 4,56 | 0 | 1,28 | 0 | 4,99 | 0 |
| MZF1 | TGGTGG | 1 | 319 | 325 | -681 | -675 | 3,59 | 0,01 | 18,26 | >0.1 | 2,63 | 4,63 | 0 | 1,16 | 0 | 6,26 | 0 |
| MZF1 | TGGTGG | 1 | 322 | 328 | -678 | -672 | 3,59 | 0,01 | 17,68 | >0.1 | 2,62 | 4,63 | 0 | 1,15 | 0 | 5,68 | 0 |
| MZF1 | TGGGCA | 1 | 325 | 331 | -675 | -669 | 3,98 | 0,01 | 17,94 | >0.1 | 2,61 | 4,64 | 0 | 1,15 | 0 | 5,56 | 0 |
| MZF1 | TCGGGA | 1 | 348 | 354 | -652 | -646 | 4,05 | 0,01 | 18,9 | >0.1 | 2,55 | 4,67 | 0 | 1,1 | 0 | 6,52 | 0 |
| MZF1 | CTGGGA | 1 | 380 | 386 | -620 | -614 | 5,98 | 0,01 | 20,86 | >0.1 | 2,56 | 4,73 | 0 | 1,06 | 0 | 6,53 | 0 |
| MZF1 | GGGCGA | 1 | 432 | 438 | -568 | -562 | 3,59 | 0,01 | 17,8 | >0.1 | 2,71 | 4,82 | 0 | 1,01 | 0 | 5,67 | 0 |
| MZF1 | CTGGGA | 1 | 560 | 566 | -440 | -434 | 5,98 | 0,01 | 15,28 | >0.1 | 3,22 | 5,09 | 0 | 0,99 | 0 | 0 | 0 |
| MZF1 | TGAGGA | 1 | 988 | 994 | -12 | -6 | 7,1 | 0,01 | 24,78 | >0.1 | 5,67 | 8,57 | 0 | 1,87 | 0 | 1,57 | 0 |
| MZF1 | TAGGGA | 1 | 1004 | 1010 | 4 | 10 | 3,56 | 0,01 | 21,68 | >0.1 | 5,67 | 9,12 | 0 | 1,87 | 0 | 1,46 | 0 |
| MZF1 | AGGGGA | 1 | 1041 | 1047 | 41 | 47 | 9,28 | 0,001 | 25,99 | >0.1 | 5,67 | 7,82 | 0 | 1,86 | 0 | 1,36 | 0 |
| MZF1 | TGAGGA | 1 | 1111 | 1117 | 111 | 117 | 7,1 | 0,01 | 26,99 | >0.1 | 5,67 | 6,64 | 0 | 1,85 | 3,22 | 2,52 | 0 |
| MZF1 | TGGTGG | -1 | 1104 | 1110 | 104 | 110 | 3,59 | 0,01 | 23,46 | >0.1 | 5,67 | 6,71 | 0 | 1,85 | 3,22 | 2,41 | 0 |
| MZF1 | TGGAGA | -1 | 863 | 869 | -137 | -131 | 3,56 | 0,01 | 20,24 | >0.1 | 5,67 | 6,36 | 0 | 1,9 | 0 | 2,75 | 0 |
| MZF1 | AGAGGA | -1 | 860 | 866 | -140 | -134 | 5,74 | 0,01 | 22,51 | >0.1 | 5,67 | 6,34 | 0 | 1,9 | 0 | 2,86 | 0 |
| MZF1 | TAGGGA | -1 | 653 | 659 | -347 | -341 | 3,56 | 0,01 | 17,88 | >0.1 | 3,91 | 5,35 | 0 | 1,07 | 0 | 3,99 | 0 |
| MZF1 | TGGGCA | -1 | 542 | 548 | -458 | -452 | 3,98 | 0,01 | 13,15 | >0.1 | 3,13 | 5,05 | 0 | 0,98 | 0 | 0 | 0 |
| MZF1 | CTGGGA | -1 | 338 | 344 | -662 | -656 | 5,98 | 0,01 | 20,25 | >0.1 | 2,58 | 4,66 | 0 | 1,12 | 0 | 5,92 | 0 |
| MZF1 | CGGGGT | -1 | 280 | 286 | -720 | -714 | 3,97 | 0,01 | 17,74 | >0.1 | 2,75 | 4,57 | 0 | 1,26 | 0 | 5,19 | 0 |
| CEBPD | GTTTTGCAAA | 1 | 82 | 92 | -918 | -908 | 2,49 | 0,01 | 12,47 | >0.1 | 3,77 | 4,33 | 0 | 1,88 | 0 | 0 | 0 |
| CEBPD | ATTTTAAAAA | 1 | 135 | 145 | -865 | -855 | -4,25 | 0,01 | 6,98 | >0.1 | 3,37 | 4,39 | 0 | 1,89 | 0 | 1,57 | 0 |
| CEBPD | TTTAAAAAAT | 1 | 137 | 147 | -863 | -853 | -2,6 | 0,01 | 8,69 | >0.1 | 3,36 | 4,39 | 0 | 1,89 | 0 | 1,65 | 0 |
| CEBPD | ATCGCGCCAC | 1 | 409 | 419 | -591 | -581 | -1,06 | 0,01 | 12,93 | >0.1 | 2,65 | 4,78 | 0 | 1,03 | 0 | 5,52 | 0 |
| CEBPD | GTTGCCCAAG | 1 | 540 | 550 | -460 | -450 | 5,53 | 0,001 | 14,7 | >0.1 | 3,13 | 5,06 | 0 | 0,99 | 0 | 0 | 0 |
| CEBPD | GTGAGATAAC | 1 | 591 | 601 | -409 | -399 | -2,74 | 0,01 | 6,87 | >0.1 | 3,41 | 5,18 | 0 | 1,01 | 0 | 0 | 0 |
| CEBPD | CTTTGGTCAC | 1 | 670 | 680 | -330 | -320 | -4,44 | 0,01 | 9,92 | >0.1 | 4,14 | 5,42 | 0 | 1,1 | 0 | 3,7 | 0 |
| CEBPD | ATTATAACAT | 1 | 686 | 696 | -314 | -304 | -0,92 | 0,01 | 13,05 | >0.1 | 4,42 | 5,47 | 0 | 1,13 | 0 | 2,95 | 0 |
| CEBPD | ATTAACCAAA | 1 | 730 | 740 | -270 | -260 | -3,97 | 0,01 | 14,03 | >0.1 | 5,61 | 5,64 | 0 | 1,23 | 0 | 5,52 | 0 |
| CEBPD | ATTTGGCCAT | 1 | 753 | 763 | -247 | -237 | 0,33 | 0,01 | 19,29 | >0.1 | 5,61 | 5,74 | 0 | 1,3 | 0 | 6,32 | 0 |
| CEBPD | TTTAAAAAAT | 1 | 825 | 835 | -175 | -165 | -2,6 | 0,01 | 14,48 | >0.1 | 5,61 | 6,12 | 0 | 1,65 | 0 | 3,7 | 0 |
| CEBPD | ATTCTAAAAT | 1 | 836 | 846 | -164 | -154 | -4,61 | 0,01 | 12,13 | >0.1 | 5,61 | 6,2 | 0 | 1,74 | 0 | 3,19 | 0 |
| CEBPD | ATTGAGGAAC | 1 | 1109 | 1119 | 109 | 119 | -1,8 | 0,01 | 18,06 | >0.1 | 5,61 | 6,66 | 0 | 1,85 | 3,22 | 2,52 | 0 |
| CEBPD | TTTGGACAAG | 1 | 1132 | 1142 | 132 | 142 | -2,99 | 0,01 | 17,1 | >0.1 | 5,61 | 6,44 | 0 | 1,84 | 3,22 | 2,98 | 0 |
| CEBPD | GTTTAACCAC | 1 | 1141 | 1151 | 141 | 151 | -2,8 | 0,01 | 17,65 | >0.1 | 5,61 | 6,37 | 0 | 1,84 | 3,22 | 3,4 | 0 |
| CEBPD | GTGGTTAAAC | -1 | 1141 | 1151 | 141 | 151 | -1,72 | 0,01 | 18,73 | >0.1 | 5,61 | 6,37 | 0 | 1,84 | 3,22 | 3,4 | 0 |
| CEBPD | CTTGTCCAAA | -1 | 1132 | 1142 | 132 | 142 | -1,62 | 0,01 | 18,47 | >0.1 | 5,61 | 6,44 | 0 | 1,84 | 3,22 | 2,98 | 0 |
| CEBPD | GTTCCTCAAT | -1 | 1109 | 1119 | 109 | 119 | -0,91 | 0,01 | 18,95 | >0.1 | 5,61 | 6,66 | 0 | 1,85 | 3,22 | 2,52 | 0 |
| CEBPD | CTGAGCCAAT | -1 | 882 | 892 | -118 | -108 | -2,4 | 0,01 | 14,72 | >0.1 | 5,61 | 6,55 | 0 | 1,89 | 0 | 3,07 | 0 |
| CEBPD | ATGGCCAAAT | -1 | 753 | 763 | -247 | -237 | 3,37 | 0,01 | 22,33 | >0.1 | 5,61 | 5,74 | 0 | 1,3 | 0 | 6,32 | 0 |
| CEBPD | ATGTTATAAT | -1 | 686 | 696 | -314 | -304 | -0,96 | 0,01 | 13,01 | >0.1 | 4,42 | 5,47 | 0 | 1,13 | 0 | 2,95 | 0 |
| CEBPD | GTGACCAAAG | -1 | 670 | 680 | -330 | -320 | -1,4 | 0,01 | 12,96 | >0.1 | 4,14 | 5,42 | 0 | 1,1 | 0 | 3,7 | 0 |
| CEBPD | GTTATCTCAC | -1 | 591 | 601 | -409 | -399 | -4,29 | 0,01 | 5,32 | >0.1 | 3,41 | 5,18 | 0 | 1,01 | 0 | 0 | 0 |
| CEBPD | CTTGGGCAAC | -1 | 540 | 550 | -460 | -450 | 7,39 | 0,001 | 16,56 | >0.1 | 3,13 | 5,06 | 0 | 0,99 | 0 | 0 | 0 |
| CEBPD | TTTATGGAAT | -1 | 495 | 505 | -505 | -495 | -3,71 | 0,01 | 11,79 | >0.1 | 2,94 | 4,95 | 0 | 0,98 | 0 | 6,62 | 0 |
| CEBPD | GTGGCGCGAT | -1 | 409 | 419 | -591 | -581 | -1,72 | 0,01 | 12,27 | >0.1 | 2,65 | 4,78 | 0 | 1,03 | 0 | 5,52 | 0 |
| CEBPD | TTTTTAAAAT | -1 | 135 | 145 | -865 | -855 | -2,07 | 0,01 | 9,16 | >0.1 | 3,37 | 4,39 | 0 | 1,89 | 0 | 1,57 | 0 |
| CEBPD | TTTGCAAAAC | -1 | 82 | 92 | -918 | -908 | 4,89 | 0,001 | 14,87 | >0.1 | 3,77 | 4,33 | 0 | 1,88 | 0 | 0 | 0 |
| CEBPD | CTTAAAAAAG | -1 | 73 | 83 | -927 | -917 | -3,31 | 0,01 | 6,74 | >0.1 | 3,85 | 4,32 | 0 | 1,88 | 0 | 0 | 0 |
| ALX3 | CATAATTTTA | 1 | 131 | 141 | -869 | -859 | 4,38 | 0,01 | 15,63 | >0.1 | 3,4 | 4,38 | 0 | 1,89 | 0 | 1,57 | 0 |
| ALX3 | GGAAATTAAA | 1 | 153 | 163 | -847 | -837 | 5,46 | 0,01 | 16,68 | >0.1 | 3,27 | 4,41 | 0 | 1,9 | 0 | 1,65 | 0 |
| ALX3 | ACTGATCAGA | 1 | 163 | 173 | -837 | -827 | 3,93 | 0,01 | 15,87 | >0.1 | 3,21 | 4,42 | 0 | 1,9 | 0 | 2,4 | 0 |
| ALX3 | CTCTATTAAA | 1 | 287 | 297 | -713 | -703 | 5,95 | 0,01 | 19,69 | >0.1 | 2,72 | 4,59 | 0 | 1,24 | 0 | 5,19 | 0 |
| ALX3 | ACTGATCAGA | 1 | 478 | 488 | -522 | -512 | 3,93 | 0,01 | 18,07 | >0.1 | 2,88 | 4,92 | 0 | 0,99 | 0 | 5,36 | 0 |
| ALX3 | ACCTATTAAT | 1 | 599 | 609 | -401 | -391 | 4,46 | 0,01 | 15,51 | >0.1 | 3,47 | 5,21 | 0 | 1,02 | 0 | 1,36 | 0 |
| ALX3 | ATTAATGAAA | 1 | 603 | 613 | -397 | -387 | 3,68 | 0,01 | 14,82 | >0.1 | 3,49 | 5,22 | 0 | 1,02 | 0 | 1,41 | 0 |
| ALX3 | TAAAATTATA | 1 | 682 | 692 | -318 | -308 | 5,03 | 0,01 | 19,01 | >0.1 | 4,34 | 5,46 | 0 | 1,12 | 0 | 3,07 | 0 |
| ALX3 | TTTGATTTGG | 1 | 749 | 759 | -251 | -241 | 3,88 | 0,01 | 22,41 | >0.1 | 5,61 | 5,72 | 0 | 1,28 | 0 | 5,92 | 0 |
| ALX3 | TCTTATTTGA | 1 | 810 | 820 | -190 | -180 | 4,31 | 0,01 | 20,78 | >0.1 | 5,61 | 6,03 | 0 | 1,55 | 0 | 3,28 | 0 |

| | | | | | | | | | | | | | | | | | |
|---|---|---|---|---|---|---|---|---|---|---|---|---|---|---|---|---|---|
| ALX3 | TTTGATTCAC | 1 | 815 | 825 | -185 | -175 | 3,7 | 0,01 | 20,38 | >0.1 | 5,61 | 6,06 | 0 | 1,58 | 0 | 3,43 | 0 |
| ALX3 | GCTGATAAGA | 1 | 966 | 976 | -34 | -24 | 3,63 | 0,01 | 20,59 | >0.1 | 5,61 | 7,78 | 0 | 1,87 | 0 | 1,7 | 0 |
| ALX3 | TATAATTCCA | -1 | 1190 | 1200 | 190 | 200 | 4,65 | 0,01 | 23,41 | >0.1 | 5,61 | 6,03 | 0 | 1,51 | 1,79 | 3,82 | 0 |
| ALX3 | TCTTATCAGC | -1 | 966 | 976 | -34 | -24 | 4,71 | 0,01 | 21,67 | >0.1 | 5,61 | 7,78 | 0 | 1,87 | 0 | 1,7 | 0 |
| ALX3 | TCAAATAAGA | -1 | 810 | 820 | -190 | -180 | 4,96 | 0,01 | 21,43 | >0.1 | 5,61 | 6,03 | 0 | 1,55 | 0 | 3,28 | 0 |
| ALX3 | CCAAATCAAA | -1 | 749 | 759 | -251 | -241 | 5,12 | 0,01 | 23,65 | >0.1 | 5,61 | 5,72 | 0 | 1,28 | 0 | 5,92 | 0 |
| ALX3 | TATAATTTTA | -1 | 682 | 692 | -318 | -308 | 4,53 | 0,01 | 18,51 | >0.1 | 4,34 | 5,46 | 0 | 1,12 | 0 | 3,07 | 0 |
| ALX3 | GACAATTTCA | -1 | 608 | 618 | -392 | -382 | 4,17 | 0,01 | 15,5 | >0.1 | 3,53 | 5,23 | 0 | 1,02 | 0 | 1,55 | 0 |
| ALX3 | TTTCATTAAT | -1 | 603 | 613 | -397 | -387 | 4,96 | 0,01 | 16,1 | >0.1 | 3,49 | 5,22 | 0 | 1,02 | 0 | 1,41 | 0 |
| ALX3 | TCCAAATACA | -1 | 521 | 531 | -479 | -469 | 3,95 | 0,01 | 16,06 | >0.1 | 3,05 | 5,01 | 0 | 0,98 | 0 | 3,07 | 0 |
| ALX3 | TCTGATCAGT | -1 | 478 | 488 | -522 | -512 | 4,13 | 0,01 | 18,27 | >0.1 | 2,88 | 4,92 | 0 | 0,99 | 0 | 5,36 | 0 |
| ALX3 | TCTGATCAGT | -1 | 163 | 173 | -837 | -827 | 4,13 | 0,01 | 16,07 | >0.1 | 3,21 | 4,42 | 0 | 1,9 | 0 | 2,4 | 0 |
| ALX3 | TTTAATTTCC | -1 | 153 | 163 | -847 | -837 | 6,84 | 0,01 | 18,06 | >0.1 | 3,27 | 4,41 | 0 | 1,9 | 0 | 1,65 | 0 |
| ALX3 | TAAAATTATG | -1 | 131 | 141 | -869 | -859 | 4,65 | 0,01 | 15,9 | >0.1 | 3,4 | 4,38 | 0 | 1,89 | 0 | 1,57 | 0 |
| ALX3 | TTCACTTACC | -1 | 4 | 14 | -996 | -986 | 5,79 | 0,01 | 19,31 | >0.1 | 5,61 | 4,26 | 0 | 1,86 | 1,79 | 0 | 0 |
| ALX1 | CATGGAAATTAAAACTG | 1 | 150 | 167 | -850 | -833 | -1,24 | 0,01 | 9,99 | >0.1 | 3,26 | 4,41 | 0 | 1,9 | 0 | 1,65 | 0 |
| ALX1 | CGTCCTGACTAACACGG | 1 | 259 | 276 | -741 | -724 | 0,82 | 0,01 | 14,04 | >0.1 | 2,8 | 4,56 | 0 | 1,33 | 0 | 4,53 | 0 |
| ALX1 | CGTCTCTATTAAAAATA | 1 | 284 | 301 | -716 | -699 | -1,65 | 0,01 | 12,11 | >0.1 | 2,72 | 4,59 | 0 | 1,25 | 0 | 5,19 | 0 |
| ALX1 | CCTATTAATGAAATTGT | 1 | 600 | 617 | -400 | -383 | 2,76 | 0,01 | 13,92 | >0.1 | 3,5 | 5,23 | 0 | 1,03 | 0 | 1,41 | 0 |
| ALX1 | GGTTTTGATTTGGCCAT | 1 | 746 | 763 | -254 | -237 | -0,28 | 0,01 | 18,19 | >0.1 | 5,52 | 5,74 | 0 | 1,3 | 0 | 5,92 | 0 |
| ALX1 | AGGATTAAAGAATAACG | 1 | 790 | 807 | -210 | -193 | -0,92 | 0,01 | 17,35 | >0.1 | 5,52 | 5,95 | 0 | 1,48 | 0 | 5,32 | 0 |
| ALX1 | ATTCTTATTTGATTCAC | 1 | 808 | 825 | -192 | -175 | -0,23 | 0,01 | 16,36 | >0.1 | 5,52 | 6,06 | 0 | 1,59 | 0 | 3,43 | 0 |
| ALX1 | TGAATCAAATAAGAATA | -1 | 807 | 824 | -193 | -176 | 2,43 | 0,01 | 18,86 | >0.1 | 5,52 | 6,05 | 0 | 1,58 | 0 | 3,28 | 0 |
| ALX1 | TGGCCAAATCAAAACCG | -1 | 745 | 762 | -255 | -238 | -1,45 | 0,01 | 17,01 | >0.1 | 5,52 | 5,73 | 0 | 1,3 | 0 | 5,92 | 0 |
| ALX1 | TGTTATAATTTTACAGT | -1 | 678 | 695 | -322 | -305 | -1,28 | 0,01 | 12,84 | >0.1 | 4,33 | 5,47 | 0 | 1,13 | 0 | 3,19 | 0 |
| ALX1 | CAATTTCATTAATAGGT | -1 | 599 | 616 | -401 | -384 | 2,18 | 0,01 | 13,33 | >0.1 | 3,49 | 5,22 | 0 | 1,03 | 0 | 1,41 | 0 |
| ALX1 | TTCATTAATAGGTTATC | -1 | 595 | 612 | -405 | -388 | 0,54 | 0,01 | 11,65 | >0.1 | 3,46 | 5,21 | 0 | 1,02 | 0 | 1,41 | 0 |
| ALX1 | CTCTCTACTTGGGCAAC | -1 | 540 | 557 | -460 | -443 | 0,31 | 0,01 | 9,53 | >0.1 | 3,15 | 5,07 | 0 | 0,99 | 0 | 0 | 0 |
| ALX1 | CGTGTTAGTCAGGACGG | -1 | 258 | 275 | -742 | -725 | 1,67 | 0,01 | 14,81 | >0.1 | 2,8 | 4,55 | 0 | 1,34 | 0 | 4,45 | 0 |
| ALX1 | AGTTTTAATTTCCATGG | -1 | 149 | 166 | -851 | -834 | -0,56 | 0,01 | 10,67 | >0.1 | 3,27 | 4,41 | 0 | 1,9 | 0 | 1,65 | 0 |
| ALX4 | CATGGAAATTAAAACTG | 1 | 150 | 167 | -850 | -833 | -0,04 | 0,01 | 11,19 | >0.1 | 3,26 | 4,41 | 0 | 1,9 | 0 | 1,65 | 0 |
| ALX4 | CGTCCTGACTAACACGG | 1 | 259 | 276 | -741 | -724 | 2,89 | 0,01 | 16,11 | >0.1 | 2,8 | 4,56 | 0 | 1,33 | 0 | 4,53 | 0 |
| ALX4 | CGTCTCTATTAAAAATA | 1 | 284 | 301 | -716 | -699 | 0,35 | 0,01 | 14,11 | >0.1 | 2,72 | 4,59 | 0 | 1,25 | 0 | 5,19 | 0 |
| ALX4 | CCTATTAATGAAATTGT | 1 | 600 | 617 | -400 | -383 | 7,06 | 0,001 | 18,22 | >0.1 | 3,5 | 5,23 | 0 | 1,03 | 0 | 1,41 | 0 |
| ALX4 | GGTTTTGATTTGGCCAT | 1 | 746 | 763 | -254 | -237 | -0,71 | 0,01 | 17,76 | >0.1 | 5,52 | 5,74 | 0 | 1,3 | 0 | 5,92 | 0 |
| ALX4 | AGGATTAAAGAATAACG | 1 | 790 | 807 | -210 | -193 | 2,13 | 0,01 | 20,4 | >0.1 | 5,52 | 5,95 | 0 | 1,48 | 0 | 5,32 | 0 |
| ALX4 | ATTCTTATTTGATTCAC | 1 | 808 | 825 | -192 | -175 | 1,14 | 0,01 | 17,73 | >0.1 | 5,52 | 6,06 | 0 | 1,59 | 0 | 3,43 | 0 |
| ALX4 | CCTCCTCATGAGATTTT | -1 | 979 | 996 | -21 | -4 | -1,07 | 0,01 | 16,46 | >0.1 | 5,52 | 8,61 | 0 | 1,87 | 0 | 1,53 | 0 |
| ALX4 | CACAGTAAACAATCTGC | -1 | 891 | 908 | -109 | -92 | -1,66 | 0,01 | 15,3 | >0.1 | 5,52 | 6,69 | 0 | 1,89 | 0 | 2,86 | 0 |
| ALX4 | TGAATCAAATAAGAATA | -1 | 807 | 824 | -193 | -176 | 1,39 | 0,01 | 17,82 | >0.1 | 5,52 | 6,05 | 0 | 1,58 | 0 | 3,28 | 0 |
| ALX4 | TGGCCAAATCAAAACCG | -1 | 745 | 762 | -255 | -238 | -1,02 | 0,01 | 17,44 | >0.1 | 5,52 | 5,73 | 0 | 1,3 | 0 | 5,92 | 0 |
| ALX4 | ACTTTTGGTTAATATTT | -1 | 726 | 743 | -274 | -257 | -1,84 | 0,01 | 16,09 | >0.1 | 5,52 | 5,65 | 0 | 1,24 | 0 | 5,52 | 0 |
| ALX4 | CAATTTCATTAATAGGT | -1 | 599 | 616 | -401 | -384 | 5,4 | 0,001 | 16,55 | >0.1 | 3,49 | 5,22 | 0 | 1,03 | 0 | 1,41 | 0 |
| ALX4 | TTCATTAATAGGTTATC | -1 | 595 | 612 | -405 | -388 | 0,25 | 0,01 | 11,36 | >0.1 | 3,46 | 5,21 | 0 | 1,02 | 0 | 1,41 | 0 |
| ALX4 | CTCTCTACTTGGGCAAC | -1 | 540 | 557 | -460 | -443 | 1,66 | 0,01 | 10,88 | >0.1 | 3,15 | 5,07 | 0 | 0,99 | 0 | 0 | 0 |
| ALX4 | CGTGTTAGTCAGGACGG | -1 | 258 | 275 | -742 | -725 | 3,48 | 0,01 | 16,62 | >0.1 | 2,8 | 4,55 | 0 | 1,34 | 0 | 4,45 | 0 |
| ALX4 | AGTTTTAATTTCCATGG | -1 | 149 | 166 | -851 | -834 | 0,19 | 0,01 | 11,42 | >0.1 | 3,27 | 4,41 | 0 | 1,9 | 0 | 1,65 | 0 |
| EOMES | CCAGTGTGAGAAA | 1 | 49 | 62 | -951 | -938 | 4,43 | 0,01 | 14,72 | >0.1 | 4,12 | 4,3 | 0 | 1,87 | 0 | 0 | 0 |
| EOMES | AGTGTGAGAAATA | 1 | 51 | 64 | -949 | -936 | 6,82 | 0,001 | 17,09 | >0.1 | 4,09 | 4,3 | 0 | 1,87 | 0 | 0 | 0 |
| EOMES | GACCTGTGGAGTG | 1 | 109 | 122 | -891 | -878 | -0,75 | 0,01 | 9,8 | >0.1 | 3,53 | 4,36 | 0 | 1,89 | 0 | 0,77 | 0 |
| EOMES | ACGGTGAAACCCC | 1 | 272 | 285 | -728 | -715 | -1,38 | 0,01 | 12,33 | >0.1 | 2,76 | 4,57 | 0 | 1,29 | 0 | 5,09 | 0 |
| EOMES | TGGGCGTGGTGGT | 1 | 313 | 326 | -687 | -674 | 0,55 | 0,01 | 15,12 | >0.1 | 2,64 | 4,63 | 0 | 1,18 | 0 | 6,13 | 0 |
| EOMES | ATGGCGTGAACCT | 1 | 369 | 382 | -631 | -618 | 8,44 | 0,001 | 23,05 | >0.1 | 2,54 | 4,72 | 0 | 1,07 | 0 | 6,27 | 0 |
| EOMES | GAGCCGAGATCGC | 1 | 401 | 414 | -599 | -586 | -0,36 | 0,01 | 13,85 | >0.1 | 2,63 | 4,78 | 0 | 1,04 | 0 | 5,76 | 0 |
| EOMES | TGGGCGACAGACC | 1 | 431 | 444 | -569 | -556 | -1,39 | 0,01 | 12,84 | >0.1 | 2,72 | 4,83 | 0 | 1,01 | 0 | 5,67 | 0 |
| EOMES | TAAATGTGAGATA | 1 | 586 | 599 | -414 | -401 | 0,96 | 0,01 | 10,54 | >0.1 | 3,39 | 5,18 | 0 | 1,01 | 0 | 0 | 0 |
| EOMES | AATGTGAGATAAC | 1 | 588 | 601 | -412 | -399 | 8,71 | 0,001 | 18,31 | >0.1 | 3,4 | 5,18 | 0 | 1,01 | 0 | 0 | 0 |
| EOMES | ATTGTCTGAAAAC | 1 | 612 | 625 | -388 | -375 | 0,43 | 0,01 | 11,78 | >0.1 | 3,57 | 5,25 | 0 | 1,03 | 0 | 1,5 | 0 |
| EOMES | AAAGTGACAGGAG | 1 | 763 | 776 | -237 | -224 | 2,13 | 0,01 | 20,94 | >0.1 | 5,57 | 5,8 | 0 | 1,34 | 0 | 6,1 | 0 |

| | | | | | | | | | | | | | | | | | |
|---|---|---|---|---|---|---|---|---|---|---|---|---|---|---|---|---|---|
| EOMES | AATCTGTTACATA | 1 | 843 | 856 | -157 | -144 | 1,21 | 0,01 | 17,94 | >0.1 | 5,57 | 6,27 | 0 | 1,83 | 0 | 3,07 | 0 |
| EOMES | AGGCTGATAAGAG | 1 | 964 | 977 | -36 | -23 | -1 | 0,01 | 15,95 | >0.1 | 5,57 | 7,82 | 0 | 1,87 | 0 | 1,7 | 0 |
| EOMES | TAAGAGAGAAAAT | 1 | 971 | 984 | -29 | -16 | 0,24 | 0,01 | 17,43 | >0.1 | 5,57 | 8,05 | 0 | 1,87 | 0 | 1,7 | 0 |
| EOMES | TGGATGTGATCTT | 1 | 1021 | 1034 | 21 | 34 | 2,15 | 0,01 | 19,56 | >0.1 | 5,57 | 8,7 | 0 | 1,86 | 0 | 1,28 | 0 |
| EOMES | AGGCTGAGAAGGA | -1 | 1068 | 1081 | 68 | 81 | 1,66 | 0,01 | 20,3 | >0.1 | 5,57 | 7,21 | 0 | 1,85 | 1,79 | 2,21 | 0 |
| EOMES | AAGCTGTTAAAGA | -1 | 706 | 719 | -294 | -281 | 5,86 | 0,001 | 22,97 | >0.1 | 5,57 | 5,56 | 0 | 1,18 | 0 | 4,81 | 0 |
| EOMES | AAAATGTTATAAT | -1 | 686 | 699 | -314 | -301 | 0,04 | 0,01 | 14,06 | >0.1 | 4,45 | 5,48 | 0 | 1,13 | 0 | 2,95 | 0 |
| EOMES | GTGGCGCGATCTC | -1 | 406 | 419 | -594 | -581 | 5,19 | 0,01 | 19,3 | >0.1 | 2,65 | 4,78 | 0 | 1,03 | 0 | 5,65 | 0 |
| EOMES | TAGCTGGGACTAC | -1 | 334 | 347 | -666 | -653 | 0,79 | 0,01 | 15,2 | >0.1 | 2,58 | 4,66 | 0 | 1,13 | 0 | 6,03 | 0 |
| EOMES | CAGGTGCCCACCA | -1 | 322 | 335 | -678 | -665 | -0,37 | 0,01 | 13,61 | >0.1 | 2,61 | 4,64 | 0 | 1,16 | 0 | 5,56 | 0 |
| EOMES | ACCGTGTTAGTCA | -1 | 264 | 277 | -736 | -723 | -0,31 | 0,01 | 12,88 | >0.1 | 2,79 | 4,56 | 0 | 1,31 | 0 | 4,53 | 0 |
| EOMES | ACCTTGTGATCTG | -1 | 232 | 245 | -768 | -755 | -1,04 | 0,01 | 12,72 | >0.1 | 2,9 | 4,51 | 0 | 1,45 | 0 | 4,9 | 0 |
| EOMES | AAAGTGCTAGGAT | -1 | 203 | 216 | -797 | -784 | 4,63 | 0,01 | 18,33 | >0.1 | 3,02 | 4,47 | 0 | 1,61 | 0 | 4,59 | 0 |
| EOMES | CAGGCGTGAGCCA | -1 | 188 | 201 | -812 | -799 | 7,61 | 0,001 | 20,91 | >0.1 | 3,08 | 4,46 | 0 | 1,72 | 0 | 4,04 | 0 |
| ETS1 | GAAGGAGAAG | 1 | 11 | 21 | -989 | -979 | -3,85 | 0,01 | 12,09 | >0.1 | 5,61 | 4,26 | 0 | 1,86 | 0 | 0 | 4,2 |
| ETS1 | GAAGGAGGCA | 1 | 17 | 27 | -983 | -973 | -0,27 | 0,01 | 15,68 | >0.1 | 5,61 | 4,27 | 0 | 1,87 | 0 | 0 | 4,2 |
| ETS1 | ATAGGAAATG | 1 | 61 | 71 | -939 | -929 | 5,1 | 0,001 | 19,47 | >0.1 | 3,98 | 4,31 | 0 | 1,88 | 0 | 0 | 4,2 |
| ETS1 | TCAGGAGAGT | 1 | 98 | 108 | -902 | -892 | -2,33 | 0,01 | 12,5 | >0.1 | 3,63 | 4,35 | 0 | 1,88 | 0 | 0,77 | 4,2 |
| ETS1 | TCAGGAGATA | 1 | 244 | 254 | -756 | -746 | 2,18 | 0,01 | 19,59 | >0.1 | 2,86 | 4,52 | 0 | 1,39 | 0 | 4,42 | 4,2 |
| ETS1 | TCGGGAGGCT | 1 | 348 | 358 | -652 | -642 | -4,18 | 0,01 | 14,76 | >0.1 | 2,55 | 4,68 | 0 | 1,11 | 0 | 6,4 | 4,2 |
| ETS1 | GCAGGAGAAT | 1 | 361 | 371 | -639 | -629 | -1,4 | 0,01 | 17,6 | >0.1 | 2,52 | 4,7 | 0 | 1,08 | 0 | 6,49 | 4,2 |
| ETS1 | AGCCGAGATC | 1 | 402 | 412 | -598 | -588 | -1,86 | 0,01 | 16,54 | >0.1 | 2,63 | 4,77 | 0 | 1,03 | 0 | 5,76 | 4,2 |
| ETS1 | AAGGGAAAAT | 1 | 530 | 540 | -470 | -460 | -3,68 | 0,01 | 13,32 | >0.1 | 3,09 | 5,03 | 0 | 0,98 | 0 | 3,7 | 4,2 |
| ETS1 | CTAGGAAAAT | 1 | 720 | 730 | -280 | -270 | -4,49 | 0,01 | 17,11 | >0.1 | 5,61 | 5,6 | 0 | 1,2 | 0 | 4,99 | 4,2 |
| ETS1 | ACAGGAGAGG | 1 | 769 | 779 | -231 | -221 | 3,09 | 0,01 | 25,76 | >0.1 | 5,61 | 5,81 | 0 | 1,35 | 0 | 5,69 | 4,2 |
| ETS1 | CCAGGATGAA | 1 | 866 | 876 | -134 | -124 | 3,4 | 0,01 | 24,28 | >0.1 | 5,61 | 6,41 | 0 | 1,9 | 0 | 2,75 | 4,2 |
| ETS1 | TGAGGAGGTT | 1 | 988 | 998 | -12 | -2 | -1 | 0,01 | 20,95 | >0.1 | 5,61 | 8,66 | 0 | 1,87 | 0 | 1,61 | 4,2 |
| ETS1 | AGTGGATGTG | 1 | 1019 | 1029 | 19 | 29 | -2,26 | 0,01 | 19,47 | >0.1 | 5,61 | 8,74 | 0 | 1,86 | 0 | 1,31 | 4,2 |
| ETS1 | GGACGATGTC | 1 | 1044 | 1054 | 44 | 54 | -1,2 | 0,01 | 18,83 | >0.1 | 5,61 | 7,73 | 0 | 1,86 | 0 | 0,62 | 4,2 |
| ETS1 | TGAGGAACAG | 1 | 1111 | 1121 | 111 | 121 | -3,81 | 0,01 | 20,25 | >0.1 | 5,61 | 6,64 | 0 | 1,85 | 3,22 | 2,54 | 4,2 |
| ETS1 | CCACGAAGCC | 1 | 1147 | 1157 | 147 | 157 | 0,96 | 0,01 | 22,94 | >0.1 | 5,61 | 6,32 | 0 | 1,84 | 1,79 | 2,21 | 4,2 |
| ETS1 | AGCCGAAGAC | 1 | 1153 | 1163 | 153 | 163 | 0,63 | 0,01 | 21,56 | >0.1 | 4,48 | 6,27 | 0 | 1,83 | 1,79 | 2,34 | 4,2 |
| ETS1 | CCAGGAAGAG | -1 | 1057 | 1067 | 57 | 67 | 5,88 | 0,001 | 27,24 | >0.1 | 5,61 | 7,42 | 0 | 1,86 | 0 | 2,27 | 4,2 |
| ETS1 | GCAGGAAAGA | -1 | 932 | 942 | -68 | -58 | 6,52 | 0,001 | 26,23 | >0.1 | 5,61 | 7,09 | 0 | 1,88 | 0 | 0,93 | 4,2 |
| ETS1 | ACAGTAAACA | -1 | 897 | 907 | -103 | -93 | 2,33 | 0,01 | 23,57 | >0.1 | 5,61 | 6,68 | 0 | 1,89 | 0 | 2,86 | 4,2 |
| ETS1 | AGAGGACAGA | -1 | 856 | 866 | -144 | -134 | -2,77 | 0,01 | 18,24 | >0.1 | 5,61 | 6,34 | 0 | 1,9 | 0 | 2,95 | 4,2 |
| ETS1 | ACCGGTACTT | -1 | 739 | 749 | -261 | -251 | -5,2 | 0,01 | 17,46 | >0.1 | 5,61 | 5,68 | 0 | 1,25 | 0 | 5,92 | 4,2 |
| ETS1 | CACGGAAAAT | -1 | 694 | 704 | -306 | -296 | 1,68 | 0,01 | 21,21 | >0.1 | 5,61 | 5,5 | 0 | 1,14 | 0 | 3,07 | 4,2 |
| ETS1 | CCCAGAAACT | -1 | 555 | 565 | -445 | -435 | -4,64 | 0,01 | 8,85 | >0.1 | 3,21 | 5,09 | 0 | 0,99 | 0 | 0 | 4,2 |
| ETS1 | TCAGGACGGT | -1 | 257 | 267 | -743 | -733 | -3,08 | 0,01 | 14,45 | >0.1 | 2,82 | 4,54 | 0 | 1,34 | 0 | 4,63 | 4,2 |
| AHR::ARNT | AGTGTG | 1 | 51 | 57 | -949 | -943 | 4,56 | 0,01 | 14,87 | >0.1 | 4,14 | 4,29 | 0 | 1,87 | 0 | 0 | 0 |
| AHR::ARNT | GGAGTG | 1 | 116 | 122 | -884 | -878 | 1,93 | 0,01 | 12,43 | >0.1 | 3,51 | 4,36 | 0 | 1,89 | 0 | 0,75 | 0 |
| AHR::ARNT | CGCCTG | 1 | 195 | 201 | -805 | -799 | 4,32 | 0,01 | 17,77 | >0.1 | 3,07 | 4,45 | 0 | 1,67 | 0 | 4,26 | 0 |
| AHR::ARNT | GGCGTG | 1 | 315 | 321 | -685 | -679 | 9,28 | 0,001 | 23,84 | >0.1 | 2,64 | 4,62 | 0 | 1,17 | 0 | 6,13 | 0 |
| AHR::ARNT | GGCGTG | 1 | 371 | 377 | -629 | -623 | 9,28 | 0,001 | 23,75 | >0.1 | 2,54 | 4,71 | 0 | 1,07 | 0 | 6,15 | 0 |
| AHR::ARNT | AGCTTG | 1 | 391 | 397 | -609 | -603 | 4,56 | 0,01 | 18,96 | >0.1 | 2,59 | 4,74 | 0 | 1,04 | 0 | 6,02 | 0 |
| AHR::ARNT | AGCCTG | 1 | 427 | 433 | -573 | -567 | 2,47 | 0,01 | 16,66 | >0.1 | 2,7 | 4,81 | 0 | 1,01 | 0 | 5,67 | 0 |
| AHR::ARNT | TCCGTG | 1 | 698 | 704 | -302 | -296 | 4,28 | 0,01 | 19,66 | >0.1 | 5,67 | 5,5 | 0 | 1,14 | 0 | 3,07 | 0 |
| AHR::ARNT | TTCTTG | 1 | 1186 | 1192 | 186 | 192 | 1,6 | 0,01 | 20,33 | >0.1 | 5,67 | 6,05 | 0 | 1,54 | 1,79 | 3,67 | 0 |
| AHR::ARNT | TTCGTG | -1 | 1148 | 1154 | 148 | 154 | 6,37 | 0,001 | 24,14 | >0.1 | 5,67 | 6,31 | 0 | 1,84 | 1,79 | 2,16 | 0 |
| AHR::ARNT | GGCCTG | -1 | 1118 | 1124 | 118 | 124 | 2,43 | 0,01 | 22,34 | >0.1 | 5,67 | 6,57 | 0 | 1,85 | 3,22 | 2,6 | 0 |
| AHR::ARNT | AGCTTG | -1 | 1053 | 1059 | 53 | 59 | 4,56 | 0,01 | 21,05 | >0.1 | 5,67 | 7,5 | 0 | 1,86 | 0 | 1,46 | 0 |
| AHR::ARNT | TGTTTG | -1 | 629 | 635 | -371 | -365 | 1,6 | 0,01 | 13,14 | >0.1 | 3,68 | 5,28 | 0 | 1,04 | 0 | 1,55 | 0 |
| AHR::ARNT | GGCGCG | -1 | 411 | 417 | -589 | -583 | 2,86 | 0,01 | 16,84 | >0.1 | 2,65 | 4,78 | 0 | 1,02 | 0 | 5,52 | 0 |
| AHR::ARNT | ACCGTG | -1 | 271 | 277 | -729 | -723 | 2,47 | 0,01 | 16,18 | >0.1 | 2,78 | 4,56 | 0 | 1,29 | 0 | 5,09 | 0 |
| AHR::ARNT | CTTGTG | -1 | 237 | 243 | -763 | -757 | 1,64 | 0,01 | 15,21 | >0.1 | 2,9 | 4,51 | 0 | 1,42 | 0 | 4,74 | 0 |
| AHR::ARNT | GGCGTG | -1 | 193 | 199 | -807 | -801 | 9,28 | 0,001 | 22,62 | >0.1 | 3,08 | 4,45 | 0 | 1,68 | 0 | 4,13 | 0 |
| AHR::ARNT | TGAGTG | -1 | 25 | 31 | -975 | -969 | 3,78 | 0,01 | 15,58 | >0.1 | 5,67 | 4,27 | 0 | 1,87 | 0 | 0 | 0 |
| NR4A2 | AAGGAGAA | 1 | 12 | 20 | -988 | -980 | 3,09 | 0,01 | 14,85 | >0.1 | 5,64 | 4,26 | 0 | 1,86 | 0 | 0 | 0 |

| | | | | | | | | | | | | | | | | | |
|---|---|---|---|---|---|---|---|---|---|---|---|---|---|---|---|---|---|
| NR4A2 | GAAGTCAG | 1 | 33 | 41 | -967 | -959 | 3,9 | 0,01 | 14,49 | >0.1 | 4,44 | 4,28 | 0 | 1,87 | 0 | 0 | 0 |
| NR4A2 | CAGATCAG | 1 | 94 | 102 | -906 | -898 | 6,72 | 0,01 | 17,42 | >0.1 | 3,67 | 4,34 | 0 | 1,88 | 0 | 0,81 | 0 |
| NR4A2 | GAGGCCGA | 1 | 218 | 226 | -782 | -774 | 3,11 | 0,01 | 17,6 | >0.1 | 2,97 | 4,49 | 0 | 1,52 | 0 | 5,52 | 0 |
| NR4A2 | CAGATCAC | 1 | 232 | 240 | -768 | -760 | 8,62 | 0,001 | 22,49 | >0.1 | 2,91 | 4,51 | 0 | 1,44 | 0 | 5,01 | 0 |
| NR4A2 | AAGGTCAG | 1 | 240 | 248 | -760 | -752 | 11,3 | 0,0001 | 24,53 | >0.1 | 2,88 | 4,52 | 0 | 1,41 | 0 | 4,42 | 0 |
| NR4A2 | GAGATCGC | 1 | 406 | 414 | -594 | -586 | 4,38 | 0,01 | 18,59 | >0.1 | 2,64 | 4,78 | 0 | 1,03 | 0 | 5,76 | 0 |
| NR4A2 | AAGGGCAA | 1 | 508 | 516 | -492 | -484 | 6,17 | 0,01 | 19,31 | >0.1 | 2,99 | 4,98 | 0 | 0,98 | 0 | 4,19 | 0 |
| NR4A2 | GAGATAAC | 1 | 593 | 601 | -407 | -399 | 3,49 | 0,01 | 13,1 | >0.1 | 3,42 | 5,18 | 0 | 1,01 | 0 | 0 | 0 |
| NR4A2 | TTGGTCAC | 1 | 672 | 680 | -328 | -320 | 4,38 | 0,01 | 18,6 | >0.1 | 4,16 | 5,42 | 0 | 1,1 | 0 | 3,55 | 0 |
| NR4A2 | GAGGTAAG | 1 | 775 | 783 | -225 | -217 | 3,9 | 0,01 | 22,84 | >0.1 | 5,64 | 5,83 | 0 | 1,37 | 0 | 6,1 | 0 |
| NR4A2 | AAGAATAA | 1 | 797 | 805 | -203 | -195 | 2,87 | 0,01 | 19,35 | >0.1 | 5,64 | 5,94 | 0 | 1,47 | 0 | 3,43 | 0 |
| NR4A2 | AAGTTCAA | 1 | 956 | 964 | -44 | -36 | 5,04 | 0,01 | 21,73 | >0.1 | 5,64 | 7,48 | 0 | 1,88 | 0 | 1,7 | 0 |
| NR4A2 | GATGTCAA | 1 | 1048 | 1056 | 48 | 56 | 6,53 | 0,01 | 23,07 | >0.1 | 5,64 | 7,62 | 0 | 1,86 | 0 | 1,42 | 0 |
| NR4A2 | CAGTCCAC | 1 | 1100 | 1108 | 100 | 108 | 2,9 | 0,01 | 22,77 | >0.1 | 5,64 | 6,76 | 0 | 1,85 | 3,22 | 2,4 | 0 |
| NR4A2 | AGGAACAG | 1 | 1113 | 1121 | 113 | 121 | 3,82 | 0,01 | 23,69 | >0.1 | 5,64 | 6,62 | 0 | 1,85 | 3,22 | 2,54 | 0 |
| NR4A2 | CAGGCCAA | 1 | 1118 | 1126 | 118 | 126 | 7,35 | 0,01 | 27,23 | >0.1 | 5,64 | 6,57 | 0 | 1,85 | 3,22 | 2,6 | 0 |
| NR4A2 | AAGTTCAC | 1 | 1174 | 1182 | 174 | 182 | 6,89 | 0,01 | 25,17 | >0.1 | 5,64 | 6,13 | 0 | 1,63 | 1,79 | 3,09 | 0 |
| NR4A2 | TAGAACAG | -1 | 1163 | 1171 | 163 | 171 | 4,01 | 0,01 | 20,78 | >0.1 | 4,44 | 6,2 | 0 | 1,73 | 1,79 | 2,61 | 0 |
| NR4A2 | AAGATCAC | -1 | 1026 | 1034 | 26 | 34 | 10,89 | 0,001 | 28,09 | >0.1 | 5,64 | 8,45 | 0 | 1,86 | 0 | 1,25 | 0 |
| NR4A2 | AAGACCAA | -1 | 928 | 936 | -72 | -64 | 7,32 | 0,01 | 22,87 | >0.1 | 5,64 | 7 | 0 | 1,88 | 0 | 1,03 | 0 |
| NR4A2 | AAGAACAC | -1 | 905 | 913 | -95 | -87 | 8,68 | 0,001 | 24,15 | >0.1 | 5,64 | 6,73 | 0 | 1,89 | 0 | 1,21 | 0 |
| NR4A2 | GAGGACAG | -1 | 857 | 865 | -143 | -135 | 8,22 | 0,001 | 25,04 | >0.1 | 5,64 | 6,33 | 0 | 1,9 | 0 | 2,95 | 0 |
| NR4A2 | AAGAATAC | -1 | 806 | 814 | -194 | -186 | 4,72 | 0,01 | 21,3 | >0.1 | 5,64 | 5,99 | 0 | 1,52 | 0 | 3,43 | 0 |
| NR4A2 | ATGGCCAA | -1 | 755 | 763 | -245 | -237 | 3,58 | 0,01 | 22,57 | >0.1 | 5,64 | 5,74 | 0 | 1,3 | 0 | 6,32 | 0 |
| NR4A2 | GAGGTCTA | -1 | 663 | 671 | -337 | -329 | 4,33 | 0,01 | 18,83 | >0.1 | 4,03 | 5,39 | 0 | 1,08 | 0 | 3,99 | 0 |
| NR4A2 | AAGATCAT | -1 | 568 | 576 | -432 | -424 | 5,04 | 0,01 | 14,42 | >0.1 | 3,27 | 5,12 | 0 | 0,99 | 0 | 0 | 0 |
| NR4A2 | AGGTTCAC | -1 | 374 | 382 | -626 | -618 | 3,93 | 0,01 | 18,8 | >0.1 | 2,55 | 4,72 | 0 | 1,07 | 0 | 6,53 | 0 |
| NR4A2 | AGGATTAC | -1 | 200 | 208 | -800 | -792 | 3,97 | 0,01 | 17,58 | >0.1 | 3,04 | 4,46 | 0 | 1,63 | 0 | 4,48 | 0 |
| NR4A2 | CAGGTCAA | -1 | 107 | 115 | -893 | -885 | 9,07 | 0,001 | 19,62 | >0.1 | 3,57 | 4,35 | 0 | 1,89 | 0 | 0,75 | 0 |
| PPARG | AAGGAGAAGGAGGCACTCAA | 1 | 12 | 32 | -988 | -968 | -5,38 | 0,01 | 6,25 | >0.1 | 5,48 | 4,28 | 0 | 1,87 | 0 | 0 | 0 |
| PPARG | GCTGGGCACAGTGGCTCACC | 1 | 177 | 197 | -823 | -803 | 1,04 | 0,01 | 14,29 | >0.1 | 3,12 | 4,45 | 0 | 1,82 | 0 | 3,86 | 0 |
| PPARG | TCGCGCCACTGTGCTCCAGC | 1 | 410 | 430 | -590 | -570 | -4,61 | 0,01 | 8,98 | >0.1 | 2,67 | 4,81 | 0 | 1,03 | 0 | 5,09 | 0 |
| PPARG | CAAGGGCAAAGTCATGTATT | 1 | 507 | 527 | -493 | -473 | 3,02 | 0,001 | 14,97 | >0.1 | 3,01 | 5 | 0 | 0,99 | 0 | 2,95 | 0 |
| PPARG | AAGGGAAAATGTTGCCCAAG | 1 | 530 | 550 | -470 | -450 | -5,47 | 0,01 | 5,3 | >0.1 | 3,11 | 5,06 | 0 | 0,99 | 0 | 1,61 | 0 |
| PPARG | AGGGAAAATGTTGCCCAAGT | 1 | 531 | 551 | -469 | -449 | -4,46 | 0,01 | 6,36 | >0.1 | 3,11 | 5,06 | 0 | 0,99 | 0 | 1,66 | 0 |
| PPARG | TTTGGTCACTGTAAAATTAT | 1 | 671 | 691 | -329 | -309 | -2,04 | 0,01 | 12,2 | >0.1 | 4,23 | 5,46 | 0 | 1,12 | 0 | 3,43 | 0 |
| PPARG | CTAGGAAAATATTAACCAAA | 1 | 720 | 740 | -280 | -260 | 0,7 | 0,01 | 18,37 | >0.1 | 5,48 | 5,64 | 0 | 1,23 | 0 | 5,32 | 0 |
| PPARG | TTACATATCTGTCCTCTCCA | 1 | 849 | 869 | -151 | -131 | -5,36 | 0,01 | 11,14 | >0.1 | 5,48 | 6,36 | 0 | 1,9 | 0 | 2,75 | 0 |
| PPARG | GGATGTGATCTTGGCTCACA | 1 | 1022 | 1042 | 22 | 42 | -2,75 | 0,01 | 14,58 | >0.1 | 5,48 | 8,68 | 0 | 1,86 | 0 | 1,31 | 0 |
| PPARG | CAAGCTCTTCCTGGCTCCTT | 1 | 1053 | 1073 | 53 | 73 | -3,35 | 0,01 | 15,5 | >0.1 | 5,48 | 7,5 | 0 | 1,86 | 1,79 | 2,21 | 0 |
| PPARG | CTGCTGCTCAGTCCACCATT | 1 | 1092 | 1112 | 92 | 112 | 1,92 | 0,01 | 21,69 | >0.1 | 5,48 | 6,85 | 0 | 1,85 | 3,22 | 2,36 | 0 |
| PPARG | TTTGGACAAGTTTAACCACG | 1 | 1132 | 1152 | 132 | 152 | -2,99 | 0,01 | 17,07 | >0.1 | 5,48 | 6,44 | 0 | 1,84 | 3,22 | 3,07 | 0 |
| PPARG | TTGGACAAGTTTAACCACGA | 1 | 1133 | 1153 | 133 | 153 | -0,8 | 0,01 | 19,31 | >0.1 | 5,48 | 6,43 | 0 | 1,84 | 3,22 | 3,13 | 0 |
| PPARG | GTGGTTAAACTTGTCCAAAA | -1 | 1131 | 1151 | 131 | 151 | 6,48 | 0,001 | 26,57 | >0.1 | 5,48 | 6,45 | 0 | 1,84 | 3,22 | 3,09 | 0 |
| PPARG | ATGGTGGACTGAGCAGCAGT | -1 | 1091 | 1111 | 91 | 111 | -2,24 | 0,01 | 17,51 | >0.1 | 5,48 | 6,87 | 0 | 1,85 | 3,22 | 2,32 | 0 |
| PPARG | AAGGCTGAGAAGGAGCCAGG | -1 | 1062 | 1082 | 62 | 82 | -5,78 | 0,01 | 12,94 | >0.1 | 5,48 | 7,31 | 0 | 1,86 | 1,79 | 2,27 | 0 |
| PPARG | CAGGAAGAGCTTGACATCGT | -1 | 1046 | 1066 | 46 | 66 | -3,5 | 0,01 | 12,98 | >0.1 | 5,48 | 7,68 | 0 | 1,86 | 0 | 1,46 | 0 |
| PPARG | ATCGTCCCCTGTGAGCCAAG | -1 | 1031 | 1051 | 31 | 51 | -4,12 | 0,01 | 12,75 | >0.1 | 5,48 | 8,19 | 0 | 1,86 | 0 | 1,34 | 0 |
| PPARG | GTGAGCCAAGATCACATCCA | -1 | 1021 | 1041 | 21 | 41 | 1,41 | 0,01 | 18,74 | >0.1 | 5,48 | 8,7 | 0 | 1,86 | 0 | 1,28 | 0 |
| PPARG | CAAGATCACATCCACTGAAT | -1 | 1015 | 1035 | 15 | 35 | -1,47 | 0,01 | 16,02 | >0.1 | 5,48 | 8,84 | 0 | 1,86 | 0 | 1,31 | 0 |
| PPARG | GAAGAACACAGTAAACAATC | -1 | 894 | 914 | -106 | -86 | -4,36 | 0,01 | 12,52 | >0.1 | 5,48 | 6,75 | 0 | 1,89 | 0 | 2,75 | 0 |
| PPARG | CTAGAGAACCTTACCTCTCC | -1 | 772 | 792 | -228 | -208 | -2,96 | 0,01 | 16,35 | >0.1 | 5,48 | 5,88 | 0 | 1,41 | 0 | 6,54 | 0 |
| PPARG | TTGGTTAATATTTTCCTAGA | -1 | 719 | 739 | -281 | -261 | 0,79 | 0,01 | 18,3 | >0.1 | 5,48 | 5,64 | 0 | 1,23 | 0 | 5,17 | 0 |
| PPARG | TAATTTTACAGTGACCAAAG | -1 | 670 | 690 | -330 | -310 | -3,81 | 0,01 | 10,41 | >0.1 | 4,22 | 5,45 | 0 | 1,12 | 0 | 3,43 | 0 |
| PPARG | CAAGATCATATTCCCAGAAA | -1 | 557 | 577 | -443 | -423 | -4,43 | 0,01 | 4,94 | >0.1 | 3,25 | 5,12 | 0 | 1 | 0 | 0 | 0 |
| PPARG | CTTGGGCAACATTTTCCCTT | -1 | 530 | 550 | -470 | -450 | -3,3 | 0,01 | 7,47 | >0.1 | 3,11 | 5,06 | 0 | 0,99 | 0 | 1,61 | 0 |
| PPARG | TTGGGCAACATTTTCCCTTC | -1 | 529 | 549 | -471 | -451 | -4,94 | 0,01 | 5,87 | >0.1 | 3,1 | 5,05 | 0 | 0,99 | 0 | 1,66 | 0 |
| PPARG | ATACATGACTTTGCCCTTGT | -1 | 506 | 526 | -494 | -474 | 4,95 | 0,001 | 16,81 | >0.1 | 3 | 5 | 0 | 0,99 | 0 | 2,86 | 0 |
| PPARG | CTGGAGCACAGTGGCGCGAT | -1 | 409 | 429 | -591 | -571 | -0,14 | 0,01 | 13,56 | >0.1 | 2,67 | 4,8 | 0 | 1,03 | 0 | 5,19 | 0 |

| | | | | | | | | | | | | | | | | | |
|---|---|---|---|---|---|---|---|---|---|---|---|---|---|---|---|---|---|
| PPARG | CAGGTTCACGCCATTCTCCT | -1 | 363 | 383 | -637 | -617 | -5,38 | 0,01 | 9,69 | >0.1 | 2,54 | 4,72 | 0 | 1,09 | 0 | 6,73 | 0 |
| PPARG | CTGCCCAACTCGGCCTCCCA | -1 | 215 | 235 | -785 | -765 | -5,11 | 0,01 | 9,41 | >0.1 | 2,96 | 4,5 | 0 | 1,54 | 0 | 5,52 | 0 |
| PPARG | GTGAGCCCACTGTGCCCAGCC | -1 | 176 | 196 | -824 | -804 | 1,27 | 0,01 | 14,53 | >0.1 | 3,12 | 4,45 | 0 | 1,83 | 0 | 3,86 | 0 |
| PPARG | GAGTGCCTCCTTCTCCTTCA | -1 | 10 | 30 | -990 | -970 | -3,82 | 0,01 | 9,6 | >0.1 | 5,48 | 4,27 | 0 | 1,87 | 1,79 | 0 | 0 |
| PPARG | CTCCTTCTCCTTCACTTACC | -1 | 4 | 24 | -996 | -976 | -4,97 | 0,01 | 8,44 | >0.1 | 5,48 | 4,27 | 0 | 1,87 | 1,79 | 0 | 0 |
| NR4A1 | AATAGGTAAG | 1 | 0 | 10 | -1000 | -990 | 0,3 | 0,01 | 13,82 | >0.1 | 5,61 | 4,25 | 0 | 1,86 | 1,79 | 0 | 0 |
| NR4A1 | AAGAAGTCAG | 1 | 31 | 41 | -969 | -959 | -0,53 | 0,01 | 10,08 | >0.1 | 4,46 | 4,28 | 0 | 1,87 | 0 | 0 | 0 |
| NR4A1 | AAATGGCCAT | 1 | 143 | 153 | -857 | -847 | -0,58 | 0,01 | 10,65 | >0.1 | 3,32 | 4,4 | 0 | 1,9 | 0 | 1,61 | 0 |
| NR4A1 | GAAATGGCTG | 1 | 171 | 181 | -829 | -819 | -2,6 | 0,01 | 10,22 | >0.1 | 3,17 | 4,43 | 0 | 1,87 | 0 | 3,34 | 0 |
| NR4A1 | ACAAGGTCAG | 1 | 238 | 248 | -762 | -752 | 11,6 | 0,0001 | 24,95 | >0.1 | 2,89 | 4,52 | 0 | 1,42 | 0 | 4,53 | 0 |
| NR4A1 | AGACCGTCCT | 1 | 255 | 265 | -745 | -735 | -1,38 | 0,01 | 12,18 | >0.1 | 2,83 | 4,54 | 0 | 1,35 | 0 | 4,85 | 0 |
| NR4A1 | ACAAGGGCAA | 1 | 506 | 516 | -494 | -484 | 2,99 | 0,01 | 17,61 | >0.1 | 2,98 | 4,98 | 0 | 0,98 | 0 | 5,68 | 0 |
| NR4A1 | GCAAAGTCAT | 1 | 512 | 522 | -488 | -478 | 2,15 | 0,01 | 14,08 | >0.1 | 3,01 | 4,99 | 0 | 0,98 | 0 | 2,95 | 0 |
| NR4A1 | AAAATGTTGC | 1 | 535 | 545 | -465 | -455 | -2,93 | 0,01 | 7,82 | >0.1 | 3,11 | 5,04 | 0 | 0,98 | 0 | 1,61 | 0 |
| NR4A1 | AAATTGTCTG | 1 | 610 | 620 | -390 | -380 | -1,69 | 0,01 | 9,66 | >0.1 | 3,54 | 5,24 | 0 | 1,02 | 0 | 1,55 | 0 |
| NR4A1 | CTTTGGTCAC | 1 | 670 | 680 | -330 | -320 | -1,13 | 0,01 | 13,23 | >0.1 | 4,14 | 5,42 | 0 | 1,1 | 0 | 3,7 | 0 |
| NR4A1 | GAGAGGTAAG | 1 | 773 | 783 | -227 | -217 | 0,48 | 0,01 | 19,39 | >0.1 | 5,61 | 5,83 | 0 | 1,37 | 0 | 6,1 | 0 |
| NR4A1 | GTAAGGTTCT | 1 | 778 | 788 | -222 | -212 | -1 | 0,01 | 18,4 | >0.1 | 5,61 | 5,85 | 0 | 1,39 | 0 | 6,54 | 0 |
| NR4A1 | CCAAGTTCAA | 1 | 954 | 964 | -46 | -36 | 1,61 | 0,01 | 18,28 | >0.1 | 5,61 | 7,48 | 0 | 1,88 | 0 | 1,7 | 0 |
| NR4A1 | AAAATCTCAT | 1 | 979 | 989 | -21 | -11 | -0,74 | 0,01 | 16,54 | >0.1 | 5,61 | 8,27 | 0 | 1,87 | 0 | 1,53 | 0 |
| NR4A1 | GGAAAGTCAT | 1 | 1007 | 1017 | 7 | 17 | 3,65 | 0,01 | 21,71 | >0.1 | 5,61 | 9,12 | 0 | 1,87 | 0 | 1,46 | 0 |
| NR4A1 | ACGATGTCAA | 1 | 1046 | 1056 | 46 | 56 | -1,35 | 0,01 | 15,26 | >0.1 | 5,61 | 7,68 | 0 | 1,86 | 0 | 1,46 | 0 |
| NR4A1 | TCAAGCTCTT | 1 | 1052 | 1062 | 52 | 62 | -1,03 | 0,01 | 16,24 | >0.1 | 5,61 | 7,53 | 0 | 1,86 | 0 | 2,27 | 0 |
| NR4A1 | AACAGGCCAA | 1 | 1116 | 1126 | 116 | 126 | -2,3 | 0,01 | 17,57 | >0.1 | 5,61 | 6,59 | 0 | 1,85 | 3,22 | 2,6 | 0 |
| NR4A1 | CAAAGTTCAC | 1 | 1172 | 1182 | 172 | 182 | 5,51 | 0,001 | 23,67 | >0.1 | 5,61 | 6,14 | 0 | 1,65 | 1,79 | 2,97 | 0 |
| NR4A1 | AACAGGTCTT | -1 | 1158 | 1168 | 158 | 168 | 2,23 | 0,01 | 18,82 | >0.1 | 4,38 | 6,24 | 0 | 1,78 | 1,79 | 2,4 | 0 |
| NR4A1 | AAAATGTCTT | -1 | 1124 | 1134 | 124 | 134 | 4,99 | 0,001 | 24,97 | >0.1 | 5,61 | 6,51 | 0 | 1,84 | 3,22 | 2,79 | 0 |
| NR4A1 | CCAAGATCAC | -1 | 1026 | 1036 | 26 | 36 | 3,39 | 0,01 | 20,59 | >0.1 | 5,61 | 8,45 | 0 | 1,86 | 0 | 1,28 | 0 |
| NR4A1 | GAAAGACCAA | -1 | 928 | 938 | -72 | -62 | -1,79 | 0,01 | 13,76 | >0.1 | 5,61 | 7,03 | 0 | 1,88 | 0 | 1,03 | 0 |
| NR4A1 | TAAAGTTCAT | -1 | 871 | 881 | -129 | -119 | 4,76 | 0,01 | 21,67 | >0.1 | 5,61 | 6,45 | 0 | 1,9 | 0 | 2,95 | 0 |
| NR4A1 | GAGAGGACAG | -1 | 857 | 867 | -143 | -133 | -0,42 | 0,01 | 16,3 | >0.1 | 5,61 | 6,35 | 0 | 1,9 | 0 | 2,86 | 0 |
| NR4A1 | TTATGGCCAA | -1 | 755 | 765 | -245 | -235 | -3,11 | 0,01 | 15,87 | >0.1 | 5,61 | 5,75 | 0 | 1,3 | 0 | 6,32 | 0 |
| NR4A1 | AAAATGTTAT | -1 | 689 | 699 | -311 | -301 | -1,24 | 0,01 | 12,81 | >0.1 | 4,48 | 5,48 | 0 | 1,13 | 0 | 2,95 | 0 |
| NR4A1 | AAGAGGTCTA | -1 | 663 | 673 | -337 | -327 | 4,79 | 0,01 | 19,31 | >0.1 | 4,05 | 5,4 | 0 | 1,09 | 0 | 3,99 | 0 |
| NR4A1 | AATAGGTTAT | -1 | 596 | 606 | -404 | -394 | -1,59 | 0,01 | 9,51 | >0.1 | 3,45 | 5,2 | 0 | 1,01 | 0 | 1,44 | 0 |
| NR4A1 | TCAAGATCAT | -1 | 568 | 578 | -432 | -422 | 2,64 | 0,01 | 12,04 | >0.1 | 3,28 | 5,12 | 0 | 1 | 0 | 0 | 0 |
| NR4A1 | TCTCGGTCTG | -1 | 438 | 448 | -562 | -552 | -3,24 | 0,01 | 11,39 | >0.1 | 2,74 | 4,84 | 0 | 1 | 0 | 6,05 | 0 |
| NR4A1 | GCAAGCTCTG | -1 | 388 | 398 | -612 | -602 | -0,46 | 0,01 | 14,06 | >0.1 | 2,59 | 4,75 | 0 | 1,05 | 0 | 6,13 | 0 |
| NR4A1 | GGACGGTCTC | -1 | 254 | 264 | -746 | -736 | 5,03 | 0,001 | 18,6 | >0.1 | 2,83 | 4,54 | 0 | 1,35 | 0 | 4,85 | 0 |
| NR4A1 | CACAGGTCAA | -1 | 107 | 117 | -893 | -883 | 5,41 | 0,001 | 15,98 | >0.1 | 3,56 | 4,36 | 0 | 1,89 | 0 | 0,77 | 0 |
| NR4A1 | AAAAGCTCAT | -1 | 68 | 78 | -932 | -922 | 5,39 | 0,001 | 15,49 | >0.1 | 3,91 | 4,32 | 0 | 1,88 | 0 | 0 | 0 |
| EBF1 | GATCACAAGGTCAG | 1 | 234 | 248 | -766 | -752 | -7,58 | 0,01 | 5,9 | >0.1 | 2,89 | 4,52 | 0 | 1,44 | 0 | 4,63 | 0 |
| EBF1 | CTACTCGGGAGGCT | 1 | 344 | 358 | -656 | -642 | -5,73 | 0,01 | 9,14 | >0.1 | 2,55 | 4,68 | 0 | 1,11 | 0 | 6,52 | 0 |
| EBF1 | AGTCTCAAAAAAAA | 1 | 451 | 465 | -549 | -535 | -7,7 | 0,01 | 6,34 | >0.1 | 2,79 | 4,87 | 0 | 1 | 0 | 5,38 | 0 |
| EBF1 | ATTCCATAAAGACA | 1 | 495 | 509 | -505 | -491 | -4,86 | 0,01 | 10,47 | >0.1 | 2,95 | 4,96 | 0 | 0,99 | 0 | 6,44 | 0 |
| EBF1 | GTTGCCCAAGTAGA | 1 | 540 | 554 | -460 | -446 | -6,83 | 0,01 | 2,37 | >0.1 | 3,14 | 5,07 | 0 | 0,99 | 0 | 0 | 0 |
| EBF1 | AGTGACAGGAGAGG | 1 | 765 | 779 | -235 | -221 | -8,08 | 0,01 | 10,56 | >0.1 | 5,56 | 5,81 | 0 | 1,36 | 0 | 5,92 | 0 |
| EBF1 | GTTCTCTAGGATTA | 1 | 783 | 797 | -217 | -203 | -6,01 | 0,01 | 12,57 | >0.1 | 5,56 | 5,9 | 0 | 1,43 | 0 | 5,69 | 0 |
| EBF1 | CCTCTCCAGGATGA | 1 | 861 | 875 | -139 | -125 | -7,3 | 0,01 | 9,23 | >0.1 | 5,56 | 6,41 | 0 | 1,9 | 0 | 2,67 | 0 |
| EBF1 | AATCTCATGAGGAG | 1 | 981 | 995 | -19 | -5 | 1,05 | 0,01 | 18,64 | >0.1 | 5,56 | 8,59 | 0 | 1,87 | 0 | 1,57 | 0 |
| EBF1 | TTAGTCTAGGGAAA | 1 | 998 | 1012 | -2 | 12 | -4,93 | 0,01 | 13,08 | >0.1 | 5,56 | 9,12 | 0 | 1,87 | 0 | 1,46 | 0 |
| EBF1 | GCTCACAGGGGACG | 1 | 1035 | 1049 | 35 | 49 | 4,68 | 0,001 | 21,46 | >0.1 | 5,56 | 8,03 | 0 | 1,86 | 0 | 1,34 | 0 |
| EBF1 | CGTCCCCTGTGAGC | -1 | 1035 | 1049 | 35 | 49 | 1,26 | 0,01 | 18,04 | >0.1 | 5,56 | 8,03 | 0 | 1,86 | 0 | 1,34 | 0 |
| EBF1 | TTTCCCTAGACTAA | -1 | 998 | 1012 | -2 | 12 | -6,99 | 0,01 | 11,02 | >0.1 | 5,56 | 9,12 | 0 | 1,87 | 0 | 1,46 | 0 |
| EBF1 | CTCCTCATGAGATT | -1 | 981 | 995 | -19 | -5 | 2,5 | 0,001 | 20,09 | >0.1 | 5,56 | 8,59 | 0 | 1,87 | 0 | 1,57 | 0 |
| EBF1 | TAATCCTAGAGAAC | -1 | 783 | 797 | -217 | -203 | -4,64 | 0,01 | 13,94 | >0.1 | 5,56 | 5,9 | 0 | 1,43 | 0 | 5,69 | 0 |
| EBF1 | AATCCTAGAGAACC | -1 | 782 | 796 | -218 | -204 | -5,7 | 0,01 | 13,09 | >0.1 | 5,56 | 5,9 | 0 | 1,43 | 0 | 5,92 | 0 |
| EBF1 | TTTTCCTAGAAAGC | -1 | 715 | 729 | -285 | -271 | -7,88 | 0,01 | 9,64 | >0.1 | 5,56 | 5,6 | 0 | 1,2 | 0 | 5,17 | 0 |
| EBF1 | TATTCCCAGAAACT | -1 | 555 | 569 | -445 | -431 | -6,48 | 0,01 | 2,83 | >0.1 | 3,22 | 5,1 | 0 | 1 | 0 | 0 | 0 |

| | | | | | | | | | | | | | | | | | |
|---|---|---|---|---|---|---|---|---|---|---|---|---|---|---|---|---|---|
| EBF1 | TTTCCCTTCCAAAT | -1 | 524 | 538 | -476 | -462 | -6,63 | 0,01 | 5,88 | >0.1 | 3,07 | 5,03 | 0 | 0,99 | 0 | 3,43 | 0 |
| EBF1 | AGCCTCCCGAGTAG | -1 | 344 | 358 | -656 | -642 | -4,78 | 0,01 | 10,09 | >0.1 | 2,55 | 4,68 | 0 | 1,11 | 0 | 6,52 | 0 |
| EBF1 | CCTCCCAAAGTGCT | -1 | 208 | 222 | -792 | -778 | 0,85 | 0,01 | 14,61 | >0.1 | 3 | 4,48 | 0 | 1,58 | 0 | 4,7 | 0 |
| EBF1 | ACTCCACAGGTCAA | -1 | 107 | 121 | -893 | -879 | -7,64 | 0,01 | 2,9 | >0.1 | 3,54 | 4,36 | 0 | 1,89 | 0 | 0,75 | 0 |
| MLX | GGTAAGTGAA | 1 | 4 | 14 | -996 | -986 | -5,18 | 0,01 | 8,34 | >0.1 | 5,61 | 4,26 | 0 | 1,86 | 1,79 | 0 | 0 |
| MLX | ATCAGGAGAG | 1 | 97 | 107 | -903 | -893 | -4,76 | 0,01 | 5,87 | >0.1 | 3,64 | 4,35 | 0 | 1,88 | 0 | 0,77 | 0 |
| MLX | ATCACAAGGT | 1 | 235 | 245 | -765 | -755 | -1,32 | 0,01 | 12,26 | >0.1 | 2,9 | 4,51 | 0 | 1,43 | 0 | 4,74 | 0 |
| MLX | GTCAGGAGAT | 1 | 243 | 253 | -757 | -747 | -0,2 | 0,01 | 12,91 | >0.1 | 2,87 | 4,52 | 0 | 1,39 | 0 | 4,33 | 0 |
| MLX | TGGGCGTGGT | 1 | 313 | 323 | -687 | -677 | 0,16 | 0,01 | 14,73 | >0.1 | 2,64 | 4,62 | 0 | 1,17 | 0 | 6,13 | 0 |
| MLX | GGCACCTGTA | 1 | 327 | 337 | -673 | -663 | -3,29 | 0,01 | 10,67 | >0.1 | 2,6 | 4,65 | 0 | 1,14 | 0 | 5,56 | 0 |
| MLX | GGCAGGAGAA | 1 | 360 | 370 | -640 | -630 | -5,44 | 0,01 | 9,23 | >0.1 | 2,52 | 4,7 | 0 | 1,09 | 0 | 6,36 | 0 |
| MLX | ATGGCGTGAA | 1 | 369 | 379 | -631 | -621 | 3,1 | 0,001 | 17,58 | >0.1 | 2,54 | 4,71 | 0 | 1,07 | 0 | 6,15 | 0 |
| MLX | ATCGCGCCAC | 1 | 409 | 419 | -591 | -581 | -7,23 | 0,01 | 6,76 | >0.1 | 2,65 | 4,78 | 0 | 1,03 | 0 | 5,52 | 0 |
| MLX | GTCATGTATT | 1 | 517 | 527 | -483 | -473 | -5,9 | 0,01 | 6,06 | >0.1 | 3,03 | 5 | 0 | 0,98 | 0 | 2,95 | 0 |
| MLX | ATAAAGTGAC | 1 | 761 | 771 | -239 | -229 | -1,12 | 0,01 | 17,69 | >0.1 | 5,61 | 5,77 | 0 | 1,32 | 0 | 6,1 | 0 |
| MLX | ATAACGTATT | 1 | 801 | 811 | -199 | -189 | -1,68 | 0,01 | 14,83 | >0.1 | 5,61 | 5,98 | 0 | 1,5 | 0 | 3,43 | 0 |
| MLX | TTCACTTTAA | 1 | 820 | 830 | -180 | -170 | -4,56 | 0,01 | 12,45 | >0.1 | 5,61 | 6,09 | 0 | 1,62 | 0 | 3,7 | 0 |
| MLX | ATCTCATGAG | 1 | 982 | 992 | -18 | -8 | -6,82 | 0,01 | 10,62 | >0.1 | 5,61 | 8,43 | 0 | 1,87 | 0 | 1,53 | 0 |
| MLX | TGGATGTGAT | 1 | 1021 | 1031 | 21 | 31 | -3,65 | 0,01 | 13,83 | >0.1 | 5,61 | 8,7 | 0 | 1,86 | 0 | 1,31 | 0 |
| MLX | GGGACGATGT | 1 | 1043 | 1053 | 43 | 53 | -7,58 | 0,01 | 8,27 | >0.1 | 5,61 | 7,76 | 0 | 1,86 | 0 | 0,62 | 0 |
| MLX | ACCACGAAGC | 1 | 1146 | 1156 | 146 | 156 | -5,15 | 0,01 | 12,58 | >0.1 | 5,61 | 6,33 | 0 | 1,84 | 1,79 | 2,16 | 0 |
| MLX | TTCACTTGCT | 1 | 1177 | 1187 | 177 | 187 | 1,8 | 0,001 | 20,18 | >0.1 | 5,61 | 6,11 | 0 | 1,61 | 1,79 | 3,26 | 0 |
| MLX | AGCAAGTGAA | -1 | 1177 | 1187 | 177 | 187 | 1,41 | 0,001 | 19,79 | >0.1 | 5,61 | 6,11 | 0 | 1,61 | 1,79 | 3,26 | 0 |
| MLX | GCTTCGTGGT | -1 | 1146 | 1156 | 146 | 156 | -5,31 | 0,01 | 12,42 | >0.1 | 5,61 | 6,33 | 0 | 1,84 | 1,79 | 2,16 | 0 |
| MLX | AGAGCTTGAC | -1 | 1051 | 1061 | 51 | 61 | -6,3 | 0,01 | 10,18 | >0.1 | 5,61 | 7,55 | 0 | 1,86 | 0 | 1,46 | 0 |
| MLX | CTCATGAGAT | -1 | 982 | 992 | -18 | -8 | -5,41 | 0,01 | 12,03 | >0.1 | 5,61 | 8,43 | 0 | 1,87 | 0 | 1,53 | 0 |
| MLX | TTAAAGTGAA | -1 | 820 | 830 | -180 | -170 | -5,58 | 0,01 | 11,43 | >0.1 | 5,61 | 6,09 | 0 | 1,62 | 0 | 3,7 | 0 |
| MLX | AATACGTTAT | -1 | 801 | 811 | -199 | -189 | -6,81 | 0,01 | 9,7 | >0.1 | 5,61 | 5,98 | 0 | 1,5 | 0 | 3,43 | 0 |
| MLX | GTCACTTTAT | -1 | 761 | 771 | -239 | -229 | 0,7 | 0,01 | 19,51 | >0.1 | 5,61 | 5,77 | 0 | 1,32 | 0 | 6,1 | 0 |
| MLX | GTGGCGCGAT | -1 | 409 | 419 | -591 | -581 | -1,29 | 0,01 | 12,7 | >0.1 | 2,65 | 4,78 | 0 | 1,03 | 0 | 5,52 | 0 |
| MLX | TTCACGCCAT | -1 | 369 | 379 | -631 | -621 | -3,61 | 0,01 | 10,87 | >0.1 | 2,54 | 4,71 | 0 | 1,07 | 0 | 6,15 | 0 |
| MLX | ATCTCCTGAC | -1 | 243 | 253 | -757 | -747 | -5,15 | 0,01 | 7,96 | >0.1 | 2,87 | 4,52 | 0 | 1,39 | 0 | 4,33 | 0 |
| MLX | ACCTTGTGAT | -1 | 235 | 245 | -765 | -755 | -7,27 | 0,01 | 6,31 | >0.1 | 2,9 | 4,51 | 0 | 1,43 | 0 | 4,74 | 0 |
| MLX | TTCACTTACC | -1 | 4 | 14 | -996 | -986 | -7,66 | 0,01 | 5,86 | >0.1 | 5,61 | 4,26 | 0 | 1,86 | 1,79 | 0 | 0 |
| TAL1::TCF3 | TGGGCACCTGTA | 1 | 325 | 337 | -675 | -663 | -0,6 | 0,01 | 13,26 | >0.1 | 2,6 | 4,65 | 0 | 1,15 | 0 | 5,46 | 0 |
| TAL1::TCF3 | TTAACAGCTTTC | 1 | 709 | 721 | -291 | -279 | 4,28 | 0,01 | 21,42 | >0.1 | 5,58 | 5,56 | 0 | 1,18 | 0 | 4,81 | 0 |
| TAL1::TCF3 | CTAAAATCTGTT | 1 | 839 | 851 | -161 | -149 | 6,37 | 0,001 | 23,16 | >0.1 | 5,58 | 6,24 | 0 | 1,78 | 0 | 3,19 | 0 |
| TAL1::TCF3 | TCAGCAGATTGT | 1 | 888 | 900 | -112 | -100 | 0,65 | 0,01 | 17,81 | >0.1 | 5,58 | 6,61 | 0 | 1,89 | 0 | 3,07 | 0 |
| TAL1::TCF3 | GTAACTGCTGCT | 1 | 1088 | 1100 | 88 | 100 | 2,79 | 0,01 | 21,13 | >0.1 | 5,58 | 6,9 | 0 | 1,85 | 1,79 | 2,21 | 0 |
| TAL1::TCF3 | CCAAGACATTTT | 1 | 1122 | 1134 | 122 | 134 | -0,89 | 0,01 | 19,02 | >0.1 | 5,58 | 6,53 | 0 | 1,85 | 3,22 | 2,72 | 0 |
| TAL1::TCF3 | CGAAGACCTGTT | 1 | 1156 | 1168 | 156 | 168 | 7,66 | 0,001 | 24,24 | >0.1 | 4,4 | 6,25 | 0 | 1,8 | 1,79 | 2,34 | 0 |
| TAL1::TCF3 | AGAACAGGTCTT | -1 | 1158 | 1170 | 158 | 170 | 0,36 | 0,01 | 17,02 | >0.1 | 4,38 | 6,24 | 0 | 1,78 | 1,79 | 2,48 | 0 |
| TAL1::TCF3 | TGAGCAGCAGTT | -1 | 1090 | 1102 | 90 | 102 | 0,98 | 0,01 | 19,43 | >0.1 | 5,58 | 6,88 | 0 | 1,85 | 1,79 | 2,34 | 0 |
| TAL1::TCF3 | GGAAGAGCTTGA | -1 | 1052 | 1064 | 52 | 64 | 1,82 | 0,01 | 19 | >0.1 | 5,58 | 7,53 | 0 | 1,86 | 0 | 2,21 | 0 |
| TAL1::TCF3 | AAACAATCTGCT | -1 | 890 | 902 | -110 | -98 | 5,12 | 0,01 | 22,09 | >0.1 | 5,58 | 6,63 | 0 | 1,89 | 0 | 2,86 | 0 |
| TAL1::TCF3 | AGGACAGATATG | -1 | 852 | 864 | -148 | -136 | -1,28 | 0,01 | 15,28 | >0.1 | 5,58 | 6,33 | 0 | 1,9 | 0 | 2,75 | 0 |
| TAL1::TCF3 | GTAACAGATTTT | -1 | 841 | 853 | -159 | -147 | 5,24 | 0,001 | 22,06 | >0.1 | 5,58 | 6,25 | 0 | 1,8 | 0 | 3,19 | 0 |
| TAL1::TCF3 | TAGAAAGCTGTT | -1 | 711 | 723 | -289 | -277 | 1,93 | 0,01 | 19,26 | >0.1 | 5,58 | 5,57 | 0 | 1,18 | 0 | 4,99 | 0 |
| TAL1::TCF3 | GCAAGCTCTGCC | -1 | 386 | 398 | -614 | -602 | -1,92 | 0,01 | 12,6 | >0.1 | 2,59 | 4,75 | 0 | 1,05 | 0 | 6,13 | 0 |
| TAL1::TCF3 | CGAGTAGCTGGG | -1 | 339 | 351 | -661 | -649 | -1,4 | 0,01 | 13,36 | >0.1 | 2,57 | 4,67 | 0 | 1,12 | 0 | 6,4 | 0 |
| TAL1::TCF3 | ACTACAGGTGCC | -1 | 327 | 339 | -673 | -661 | -1,78 | 0,01 | 12,18 | >0.1 | 2,6 | 4,65 | 0 | 1,15 | 0 | 5,56 | 0 |
| TAL1::TCF3 | TGGCCATTTTTT | -1 | 140 | 152 | -860 | -848 | 0,1 | 0,01 | 11,34 | >0.1 | 3,33 | 4,4 | 0 | 1,9 | 0 | 1,61 | 0 |
| PBX3 | AGGTAAGTGAAGGAGAA | 1 | 3 | 20 | -997 | -980 | 2,3 | 0,01 | 15,74 | >0.1 | 5,52 | 4,26 | 0 | 1,86 | 1,79 | 0 | 0 |
| PBX3 | AAGTGAAGGAGAAGGAG | 1 | 7 | 24 | -993 | -976 | 2,06 | 0,01 | 15,51 | >0.1 | 5,52 | 4,27 | 0 | 1,87 | 1,79 | 0 | 0 |
| PBX3 | AAGTCAGCCACAGAACC | 1 | 34 | 51 | -966 | -949 | 2,19 | 0,01 | 12,68 | >0.1 | 4,33 | 4,29 | 0 | 1,87 | 0 | 0 | 0 |
| PBX3 | GTGTGAGAAATAGGAAA | 1 | 52 | 69 | -948 | -931 | 2,44 | 0,01 | 12,68 | >0.1 | 4,05 | 4,31 | 0 | 1,87 | 0 | 0 | 0 |
| PBX3 | GGCACAGTGGCTCACGC | 1 | 181 | 198 | -819 | -802 | 1,2 | 0,01 | 14,49 | >0.1 | 3,11 | 4,45 | 0 | 1,78 | 0 | 3,95 | 0 |
| PBX3 | TCCTGACTAACACGGTG | 1 | 261 | 278 | -739 | -722 | 7,6 | 0,001 | 20,73 | >0.1 | 2,79 | 4,56 | 0 | 1,33 | 0 | 4,45 | 0 |
| PBX3 | AGGAGAATGGCGTGAAC | 1 | 363 | 380 | -637 | -620 | 1,82 | 0,01 | 16,77 | >0.1 | 2,53 | 4,72 | 0 | 1,09 | 0 | 6,61 | 0 |

| | | | | | | | | | | | | | | | | | |
|---|---|---|---|---|---|---|---|---|---|---|---|---|---|---|---|---|---|
| PBX3 | ACCTGGGAGGCAGAGCT | 1 | 378 | 395 | -622 | -605 | 5,03 | 0,01 | 19,43 | >0.1 | 2,57 | 4,74 | 0 | 1,07 | 0 | 6,02 | 0 |
| PBX3 | CTGGGAGGCAGAGCTTG | 1 | 380 | 397 | -620 | -603 | 1,53 | 0,01 | 15,93 | >0.1 | 2,58 | 4,75 | 0 | 1,06 | 0 | 6,02 | 0 |
| PBX3 | GCCTGGGCGACAGACCG | 1 | 428 | 445 | -572 | -555 | 10,82 | 0,001 | 25,06 | >0.1 | 2,72 | 4,83 | 0 | 1,02 | 0 | 5,67 | 0 |
| PBX3 | GGGCGACAGACCGAGAC | 1 | 432 | 449 | -568 | -551 | 2,07 | 0,01 | 16,58 | >0.1 | 2,73 | 4,84 | 0 | 1,01 | 0 | 5,92 | 0 |
| PBX3 | CATAAAGTGACAGGAGA | 1 | 760 | 777 | -240 | -223 | 10,08 | 0,001 | 28,67 | >0.1 | 5,52 | 5,8 | 0 | 1,35 | 0 | 5,92 | 0 |
| PBX3 | TCTTGGCTCACAGGGGA | 1 | 1030 | 1047 | 30 | 47 | 7,69 | 0,001 | 24,61 | >0.1 | 5,52 | 8,24 | 0 | 1,86 | 0 | 1,31 | 0 |
| PBX3 | CATTGAGGAACAGGCCA | 1 | 1108 | 1125 | 108 | 125 | 9,92 | 0,001 | 29,72 | >0.1 | 5,52 | 6,67 | 0 | 1,85 | 3,22 | 2,54 | 0 |
| PBX3 | CAATGGTGGACTGAGCA | -1 | 1096 | 1113 | 96 | 113 | 0,65 | 0,01 | 20,45 | >0.1 | 5,52 | 6,8 | 0 | 1,85 | 3,22 | 2,4 | 0 |
| PBX3 | GGTGGACTGAGCAGCAG | -1 | 1092 | 1109 | 92 | 109 | 1,74 | 0,01 | 21,53 | >0.1 | 5,52 | 6,85 | 0 | 1,85 | 3,22 | 2,34 | 0 |
| PBX3 | GACTGAGCAGCAGTTAC | -1 | 1088 | 1105 | 88 | 105 | 3,45 | 0,01 | 23,29 | >0.1 | 5,52 | 6,9 | 0 | 1,85 | 3,22 | 2,34 | 0 |
| PBX3 | GCAGCAGTTACAGCAAC | -1 | 1082 | 1099 | 82 | 99 | 3,08 | 0,01 | 21,44 | >0.1 | 5,52 | 6,99 | 0 | 1,85 | 1,79 | 2,21 | 0 |
| PBX3 | GGCTGAGAAGGAGCCAG | -1 | 1063 | 1080 | 63 | 80 | 2,26 | 0,01 | 21 | >0.1 | 5,52 | 7,3 | 0 | 1,85 | 1,79 | 2,27 | 0 |
| PBX3 | AAGAGCTTGACATCGTC | -1 | 1045 | 1062 | 45 | 62 | 2,44 | 0,01 | 18,94 | >0.1 | 5,52 | 7,7 | 0 | 1,86 | 0 | 1,42 | 0 |
| PBX3 | CACTGAATGACTTTCCC | -1 | 1006 | 1023 | 6 | 23 | 5,74 | 0,01 | 23,71 | >0.1 | 5,52 | 9,12 | 0 | 1,87 | 0 | 1,46 | 0 |
| PBX3 | GACTGAGTCTCGGTCTG | -1 | 438 | 455 | -562 | -545 | 4,51 | 0,01 | 19,04 | >0.1 | 2,75 | 4,85 | 0 | 1,01 | 0 | 5,92 | 0 |
| PBX3 | GGCTGGAGCACAGTGGC | -1 | 414 | 431 | -586 | -569 | 1,31 | 0,01 | 15,12 | >0.1 | 2,67 | 4,81 | 0 | 1,03 | 0 | 5,3 | 0 |
| PBX3 | AGCACAGTGGCGCGATC | -1 | 408 | 425 | -592 | -575 | 1,05 | 0,01 | 15,3 | >0.1 | 2,66 | 4,8 | 0 | 1,03 | 0 | 5,76 | 0 |
| PBX3 | CTCTGCCTCCCAGGTTC | -1 | 376 | 393 | -624 | -607 | 1,51 | 0,01 | 16,55 | >0.1 | 2,57 | 4,74 | 0 | 1,07 | 0 | 6,66 | 0 |
| PBX3 | TCCCGAGTAGCTGGGAC | -1 | 337 | 354 | -663 | -646 | 3,05 | 0,01 | 17,82 | >0.1 | 2,57 | 4,67 | 0 | 1,13 | 0 | 6,4 | 0 |
| PBX3 | TTAATAGAGACGGGGTT | -1 | 279 | 296 | -721 | -704 | 0,5 | 0,01 | 14,17 | >0.1 | 2,73 | 4,58 | 0 | 1,27 | 0 | 5,09 | 0 |
| PBX3 | CCGTGTTAGTCAGGACG | -1 | 259 | 276 | -741 | -724 | 2,97 | 0,01 | 16,19 | >0.1 | 2,8 | 4,56 | 0 | 1,33 | 0 | 4,53 | 0 |
| PBX3 | GCTAGGATTACAGGCGT | -1 | 194 | 211 | -806 | -789 | 2,44 | 0,01 | 16 | >0.1 | 3,05 | 4,47 | 0 | 1,68 | 0 | 4,37 | 0 |
| PBX3 | GCGTGAGCCACTGTGCC | -1 | 181 | 198 | -819 | -802 | 5,11 | 0,01 | 18,4 | >0.1 | 3,11 | 4,45 | 0 | 1,78 | 0 | 3,95 | 0 |
| PBX3 | GGTTCTGTGGCTGACTT | -1 | 34 | 51 | -966 | -949 | 1,41 | 0,01 | 11,9 | >0.1 | 4,33 | 4,29 | 0 | 1,87 | 0 | 0 | 0 |
| PBX3 | CTGTGGCTGACTTCTTT | -1 | 30 | 47 | -970 | -953 | 2,23 | 0,01 | 12,79 | >0.1 | 4,41 | 4,29 | 0 | 1,87 | 0 | 0 | 0 |
| PBX3 | CTTTGAGTGCCTCCTTC | -1 | 17 | 34 | -983 | -966 | 4,29 | 0,01 | 15,95 | >0.1 | 5,52 | 4,28 | 0 | 1,87 | 0 | 0 | 0 |
| PBX1 | CTATTAATGAAA | 1 | 601 | 613 | -399 | -387 | 2,19 | 0,01 | 13,33 | >0.1 | 3,49 | 5,22 | 0 | 1,02 | 0 | 1,41 | 0 |
| PBX1 | AAACCATACAAA | 1 | 621 | 633 | -379 | -367 | 1,43 | 0,01 | 12,82 | >0.1 | 3,64 | 5,27 | 0 | 1,04 | 0 | 1,44 | 0 |
| PBX1 | ATATTAACCAAA | 1 | 728 | 740 | -272 | -260 | 2,02 | 0,01 | 19,99 | >0.1 | 5,58 | 5,64 | 0 | 1,23 | 0 | 5,52 | 0 |
| PBX1 | AAATTATTCTAA | 1 | 831 | 843 | -169 | -157 | 0,35 | 0,01 | 17,37 | >0.1 | 5,58 | 6,18 | 0 | 1,71 | 0 | 3,55 | 0 |
| PBX1 | CGCCCAACCCAA | 1 | 946 | 958 | -54 | -42 | 0,49 | 0,01 | 17,05 | >0.1 | 5,58 | 7,36 | 0 | 1,88 | 0 | 1,74 | 0 |
| PBX1 | AAGTCATTCAGT | 1 | 1010 | 1022 | 10 | 22 | 4,6 | 0,01 | 22,43 | >0.1 | 5,58 | 8,99 | 0 | 1,86 | 0 | 1,39 | 0 |
| PBX1 | TGCTCAGTCCAC | 1 | 1096 | 1108 | 96 | 108 | 1,6 | 0,01 | 21,42 | >0.1 | 5,58 | 6,8 | 0 | 1,85 | 3,22 | 2,36 | 0 |
| PBX1 | CAGTCCACCATT | 1 | 1100 | 1112 | 100 | 112 | 1,53 | 0,01 | 21,32 | >0.1 | 5,58 | 6,76 | 0 | 1,85 | 3,22 | 2,38 | 0 |
| PBX1 | CCACCATTGAGG | 1 | 1104 | 1116 | 104 | 116 | -0,14 | 0,01 | 19,74 | >0.1 | 5,58 | 6,71 | 0 | 1,85 | 3,22 | 2,52 | 0 |
| PBX1 | TGTTCTATCAAA | 1 | 1164 | 1176 | 164 | 176 | 2,68 | 0,01 | 19,69 | >0.1 | 4,51 | 6,19 | 0 | 1,72 | 1,79 | 2,79 | 0 |
| PBX1 | CCAAGAAGCAAG | -1 | 1181 | 1193 | 181 | 193 | 2,71 | 0,01 | 21,32 | >0.1 | 5,58 | 6,08 | 0 | 1,58 | 1,79 | 3,57 | 0 |
| PBX1 | TAAACAATCTGC | -1 | 891 | 903 | -109 | -97 | 3,19 | 0,01 | 20,17 | >0.1 | 5,58 | 6,64 | 0 | 1,89 | 0 | 2,86 | 0 |
| PBX1 | AAGTGAATCAAA | -1 | 815 | 827 | -185 | -173 | 5,19 | 0,01 | 21,99 | >0.1 | 5,58 | 6,07 | 0 | 1,6 | 0 | 3,55 | 0 |
| PBX1 | GGCCAAATCAAA | -1 | 749 | 761 | -251 | -239 | 1,18 | 0,01 | 19,88 | >0.1 | 5,58 | 5,73 | 0 | 1,29 | 0 | 6,1 | 0 |
| PBX1 | ACGCCCAGCTAA | -1 | 308 | 320 | -692 | -680 | 0,01 | 0,01 | 14,73 | >0.1 | 2,65 | 4,62 | 0 | 1,19 | 0 | 6,26 | 0 |
| NRL | GGAGAGTTGAC | 1 | 101 | 112 | -899 | -888 | 3,33 | 0,01 | 13,95 | >0.1 | 3,6 | 4,35 | 0 | 1,88 | 0 | 0,79 | 0 |
| NRL | AAATGGCTGGG | 1 | 172 | 183 | -828 | -817 | 2,95 | 0,01 | 15,75 | >0.1 | 3,17 | 4,43 | 0 | 1,86 | 0 | 3,34 | 0 |
| NRL | GTTGGGCAGAT | 1 | 226 | 237 | -774 | -763 | 3,34 | 0,01 | 17,5 | >0.1 | 2,93 | 4,5 | 0 | 1,47 | 0 | 5,26 | 0 |
| NRL | ACCGTCCTGAC | 1 | 257 | 268 | -743 | -732 | 3,07 | 0,01 | 16,4 | >0.1 | 2,82 | 4,54 | 0 | 1,34 | 0 | 4,63 | 0 |
| NRL | GGGAGGCTGAG | 1 | 350 | 361 | -650 | -639 | 4,73 | 0,01 | 19,46 | >0.1 | 2,54 | 4,68 | 0 | 1,1 | 0 | 6,4 | 0 |
| NRL | AGTGAGCCGAG | 1 | 398 | 409 | -602 | -591 | 3,78 | 0,01 | 18,12 | >0.1 | 2,62 | 4,77 | 0 | 1,04 | 0 | 5,91 | 0 |
| NRL | AAAATGTTGCC | 1 | 535 | 546 | -465 | -454 | 3,65 | 0,01 | 14,4 | >0.1 | 3,11 | 5,05 | 0 | 0,99 | 0 | 1,61 | 0 |
| NRL | TAACAGCTTTC | 1 | 710 | 721 | -290 | -279 | 3,53 | 0,01 | 20,68 | >0.1 | 5,6 | 5,56 | 0 | 1,18 | 0 | 4,81 | 0 |
| NRL | GCTCAGCAGAT | 1 | 886 | 897 | -114 | -103 | 3,26 | 0,01 | 20,41 | >0.1 | 5,6 | 6,59 | 0 | 1,89 | 0 | 3,07 | 0 |
| NRL | AGATTGTTTAC | 1 | 893 | 904 | -107 | -96 | 4,01 | 0,01 | 21,01 | >0.1 | 5,6 | 6,65 | 0 | 1,89 | 0 | 2,86 | 0 |
| NRL | TTCCTGCTCAG | 1 | 935 | 946 | -65 | -54 | 3,96 | 0,01 | 19,51 | >0.1 | 5,6 | 7,15 | 0 | 1,88 | 0 | 0,93 | 0 |
| NRL | CAAAGGCTGAT | 1 | 961 | 972 | -39 | -28 | 6,36 | 0,01 | 23,2 | >0.1 | 5,6 | 7,67 | 0 | 1,88 | 0 | 1,7 | 0 |
| NRL | TTGTTGCTGTA | 1 | 1080 | 1091 | 80 | 91 | 3,03 | 0,01 | 21,5 | >0.1 | 5,6 | 7,01 | 0 | 1,85 | 1,79 | 2,21 | 0 |
| NRL | TAACTGCTGCT | 1 | 1089 | 1100 | 89 | 100 | 7,35 | 0,01 | 25,7 | >0.1 | 5,6 | 6,89 | 0 | 1,85 | 1,79 | 2,21 | 0 |
| NRL | CTGCTGCTCAG | 1 | 1092 | 1103 | 92 | 103 | 2,88 | 0,01 | 21,32 | >0.1 | 5,6 | 6,85 | 0 | 1,85 | 1,79 | 2,34 | 0 |
| NRL | AAGCCGAAGAC | 1 | 1152 | 1163 | 152 | 163 | 4,12 | 0,01 | 20,79 | >0.1 | 4,5 | 6,28 | 0 | 1,84 | 1,79 | 2,26 | 0 |
| NRL | CACTTGCTTCT | 1 | 1179 | 1190 | 179 | 190 | 3,44 | 0,01 | 21,86 | >0.1 | 5,6 | 6,1 | 0 | 1,59 | 1,79 | 3,34 | 0 |
| NRL | ACAAGGCTGAG | -1 | 1073 | 1084 | 73 | 84 | 5,83 | 0,01 | 24,47 | >0.1 | 5,6 | 7,12 | 0 | 1,85 | 1,79 | 2,27 | 0 |

| | | | | | | | | | | | | | | | | | |
|---|---|---|---|---|---|---|---|---|---|---|---|---|---|---|---|---|---|
| NRL | TGGGCGCTGAG | -1 | 941 | 952 | -59 | -48 | 7,32 | 0,01 | 22,83 | >0.1 | 5,6 | 7,25 | 0 | 1,88 | 0 | 0,79 | 0 |
| NRL | AATCTGCTGAG | -1 | 887 | 898 | -113 | -102 | 11,78 | 0,0001 | 28,93 | >0.1 | 5,6 | 6,6 | 0 | 1,89 | 0 | 3,07 | 0 |
| NRL | AGAAAGCTGTT | -1 | 711 | 722 | -289 | -278 | 3,34 | 0,01 | 20,67 | >0.1 | 5,6 | 5,57 | 0 | 1,18 | 0 | 4,99 | 0 |
| NRL | ATCATGAAGAT | -1 | 643 | 654 | -357 | -346 | 3,11 | 0,01 | 17,03 | >0.1 | 3,83 | 5,34 | 0 | 1,06 | 0 | 3,7 | 0 |
| NRL | TATCTCCTGAC | -1 | 243 | 254 | -757 | -746 | 5,12 | 0,01 | 18,23 | >0.1 | 2,87 | 4,52 | 0 | 1,4 | 0 | 4,33 | 0 |
| NRL | ATTATGACTAC | -1 | 126 | 137 | -874 | -863 | 2,98 | 0,01 | 14,32 | >0.1 | 3,42 | 4,38 | 0 | 1,89 | 0 | 1,65 | 0 |
| NRL | CTGTGGCTGAC | -1 | 36 | 47 | -964 | -953 | 6,77 | 0,01 | 17,27 | >0.1 | 4,35 | 4,29 | 0 | 1,87 | 0 | 0 | 0 |
| SOX10 | CACTCAAAGAA | 1 | 25 | 36 | -975 | -964 | 5,28 | 0,01 | 17,02 | >0.1 | 5,6 | 4,28 | 0 | 1,87 | 0 | 0 | 0 |
| SOX10 | TTTGCAAAGGC | 1 | 84 | 95 | -916 | -905 | 1,49 | 0,01 | 11,45 | >0.1 | 3,74 | 4,33 | 0 | 1,88 | 0 | 0 | 0 |
| SOX10 | GGCACAGTGGC | 1 | 181 | 192 | -819 | -808 | 1,75 | 0,01 | 14,95 | >0.1 | 3,12 | 4,44 | 0 | 1,78 | 0 | 3,86 | 0 |
| SOX10 | ATCACAAGGTC | 1 | 235 | 246 | -765 | -754 | 3,15 | 0,01 | 16,73 | >0.1 | 2,9 | 4,51 | 0 | 1,43 | 0 | 4,74 | 0 |
| SOX10 | GAGATAGAGAC | 1 | 248 | 259 | -752 | -741 | 0,23 | 0,01 | 14,07 | >0.1 | 2,85 | 4,53 | 0 | 1,37 | 0 | 5,09 | 0 |
| SOX10 | AATACAAAAAC | 1 | 297 | 308 | -703 | -692 | 2,99 | 0,01 | 15,94 | >0.1 | 2,69 | 4,6 | 0 | 1,21 | 0 | 4,45 | 0 |
| SOX10 | TCCATAAAGAC | 1 | 497 | 508 | -503 | -492 | 4,98 | 0,01 | 20,31 | >0.1 | 2,95 | 4,96 | 0 | 0,98 | 0 | 6,44 | 0 |
| SOX10 | AAGACAAGGGC | 1 | 503 | 514 | -497 | -486 | 5,52 | 0,01 | 20,13 | >0.1 | 2,97 | 4,97 | 0 | 0,98 | 0 | 5,68 | 0 |
| SOX10 | AGGGCAAAGTC | 1 | 509 | 520 | -491 | -480 | 1,7 | 0,01 | 13,42 | >0.1 | 3 | 4,99 | 0 | 0,98 | 0 | 2,75 | 0 |
| SOX10 | CATACAAACAC | 1 | 625 | 636 | -375 | -364 | 3,57 | 0,01 | 15,1 | >0.1 | 3,67 | 5,28 | 0 | 1,04 | 0 | 1,55 | 0 |
| SOX10 | GCCATAAAGTG | 1 | 758 | 769 | -242 | -231 | 3,96 | 0,01 | 22,74 | >0.1 | 5,6 | 5,76 | 0 | 1,32 | 0 | 6,1 | 0 |
| SOX10 | GGATTAAAGAA | 1 | 791 | 802 | -209 | -198 | 1,19 | 0,01 | 19,49 | >0.1 | 5,6 | 5,93 | 0 | 1,45 | 0 | 5,32 | 0 |
| SOX10 | AGTTCAAAGGC | 1 | 957 | 968 | -43 | -32 | 5,33 | 0,01 | 22,11 | >0.1 | 5,6 | 7,57 | 0 | 1,88 | 0 | 1,74 | 0 |
| SOX10 | GGCTCACAGGG | 1 | 1034 | 1045 | 34 | 45 | 0,77 | 0,01 | 17,6 | >0.1 | 5,6 | 8,06 | 0 | 1,86 | 0 | 1,31 | 0 |
| SOX10 | ACCACGAAGCC | 1 | 1146 | 1157 | 146 | 157 | 3,92 | 0,01 | 21,64 | >0.1 | 5,6 | 6,33 | 0 | 1,84 | 1,79 | 2,16 | 0 |
| SOX10 | CTATCAAAGTT | 1 | 1168 | 1179 | 168 | 179 | 3,33 | 0,01 | 21,41 | >0.1 | 5,6 | 6,17 | 0 | 1,68 | 1,79 | 2,84 | 0 |
| SOX10 | GCAACAAGGCT | -1 | 1076 | 1087 | 76 | 87 | 4,81 | 0,01 | 23,4 | >0.1 | 5,6 | 7,07 | 0 | 1,85 | 1,79 | 2,27 | 0 |
| SOX10 | CAAAAAAAGAA | -1 | 920 | 931 | -80 | -69 | 2,92 | 0,01 | 18,46 | >0.1 | 5,6 | 6,94 | 0 | 1,88 | 0 | 1,12 | 0 |
| SOX10 | AAGAAAAAGAA | -1 | 914 | 925 | -86 | -75 | 1,89 | 0,01 | 17,39 | >0.1 | 5,6 | 6,87 | 0 | 1,89 | 0 | 1,15 | 0 |
| SOX10 | AGAACACAGTA | -1 | 901 | 912 | -99 | -88 | 5,9 | 0,01 | 21,39 | >0.1 | 5,6 | 6,72 | 0 | 1,89 | 0 | 1,28 | 0 |
| SOX10 | AATATAAAGTT | -1 | 874 | 885 | -126 | -115 | 2,62 | 0,01 | 19,55 | >0.1 | 5,6 | 6,49 | 0 | 1,9 | 0 | 2,95 | 0 |
| SOX10 | CCTAGAAAGCT | -1 | 714 | 725 | -286 | -275 | 1,59 | 0,01 | 18,94 | >0.1 | 5,6 | 5,58 | 0 | 1,19 | 0 | 4,99 | 0 |
| SOX10 | TGACCAAAGAG | -1 | 668 | 679 | -332 | -321 | 4,28 | 0,01 | 18,61 | >0.1 | 4,12 | 5,42 | 0 | 1,1 | 0 | 3,7 | 0 |
| SOX10 | AGCACAGTGGC | -1 | 414 | 425 | -586 | -575 | 1,63 | 0,01 | 15,31 | >0.1 | 2,67 | 4,8 | 0 | 1,02 | 0 | 5,19 | 0 |
| SOX10 | TTAATAGAGAC | -1 | 285 | 296 | -715 | -704 | 0,22 | 0,01 | 13,86 | >0.1 | 2,72 | 4,58 | 0 | 1,25 | 0 | 5,09 | 0 |
| SOX10 | CTCCCAAAGTG | -1 | 210 | 221 | -790 | -779 | 1,15 | 0,01 | 14,89 | >0.1 | 2,99 | 4,48 | 0 | 1,57 | 0 | 4,7 | 0 |
| SOX10 | TTAAAAAAGCT | -1 | 71 | 82 | -929 | -918 | 1,39 | 0,01 | 11,46 | >0.1 | 3,87 | 4,32 | 0 | 1,88 | 0 | 0 | 0 |
| BACH2 | AAGAAGTCAGCCAC | 1 | 31 | 45 | -969 | -955 | -0,37 | 0,01 | 10,2 | >0.1 | 4,42 | 4,29 | 0 | 1,87 | 0 | 0 | 0 |
| BACH2 | AATGGCTGGGCACA | 1 | 173 | 187 | -827 | -813 | -0,25 | 0,01 | 12,43 | >0.1 | 3,15 | 4,44 | 0 | 1,85 | 0 | 3,24 | 0 |
| BACH2 | AGTGGCTCACGCCT | 1 | 186 | 200 | -814 | -800 | 0,05 | 0,01 | 13,28 | >0.1 | 3,09 | 4,45 | 0 | 1,74 | 0 | 3,95 | 0 |
| BACH2 | CCTGACTAACACGG | 1 | 262 | 276 | -738 | -724 | 1,28 | 0,01 | 14,4 | >0.1 | 2,79 | 4,56 | 0 | 1,32 | 0 | 4,45 | 0 |
| BACH2 | GCTGAGGCAGGAGA | 1 | 355 | 369 | -645 | -631 | 8,08 | 0,001 | 22,53 | >0.1 | 2,52 | 4,7 | 0 | 1,1 | 0 | 6,13 | 0 |
| BACH2 | CGAGACTCAGTCTC | 1 | 443 | 457 | -557 | -543 | 3,18 | 0,01 | 16,81 | >0.1 | 2,76 | 4,86 | 0 | 1 | 0 | 5,01 | 0 |
| BACH2 | ATTGGCTCAGCAGA | 1 | 882 | 896 | -118 | -104 | 9,33 | 0,001 | 26,55 | >0.1 | 5,56 | 6,58 | 0 | 1,89 | 0 | 3,19 | 0 |
| BACH2 | TCCTGCTCAGCGCC | 1 | 936 | 950 | -64 | -50 | -1,84 | 0,01 | 13,69 | >0.1 | 5,56 | 7,22 | 0 | 1,88 | 0 | 0,88 | 0 |
| BACH2 | AGTCATTCAGTGGA | 1 | 1011 | 1025 | 11 | 25 | -1,18 | 0,01 | 16,61 | >0.1 | 5,56 | 8,95 | 0 | 1,86 | 0 | 1,42 | 0 |
| BACH2 | TCCTTCTCAGCCTT | 1 | 1068 | 1082 | 68 | 82 | -1,85 | 0,01 | 16,83 | >0.1 | 5,56 | 7,21 | 0 | 1,85 | 1,79 | 2,27 | 0 |
| BACH2 | GTGGACTGAGCAGC | -1 | 1094 | 1108 | 94 | 108 | 3,15 | 0,01 | 22,93 | >0.1 | 5,56 | 6,83 | 0 | 1,85 | 3,22 | 2,32 | 0 |
| BACH2 | TGTGAGCCAAGATC | -1 | 1028 | 1042 | 28 | 42 | -0,89 | 0,01 | 16,2 | >0.1 | 5,56 | 8,34 | 0 | 1,86 | 0 | 1,34 | 0 |
| BACH2 | AATGACTTTCCCTA | -1 | 1004 | 1018 | 4 | 18 | -1,89 | 0,01 | 16,12 | >0.1 | 5,56 | 9,12 | 0 | 1,87 | 0 | 1,46 | 0 |
| BACH2 | GCTGAGCCAATATA | -1 | 879 | 893 | -121 | -107 | -0,43 | 0,01 | 16,65 | >0.1 | 5,56 | 6,56 | 0 | 1,9 | 0 | 3,07 | 0 |
| BACH2 | AGTGAATCAAATAA | -1 | 812 | 826 | -188 | -174 | -0,06 | 0,01 | 16,58 | >0.1 | 5,56 | 6,06 | 0 | 1,59 | 0 | 3,43 | 0 |
| BACH2 | CATGACTTTGCCCT | -1 | 509 | 523 | -491 | -477 | 4,8 | 0,01 | 16,65 | >0.1 | 3 | 4,99 | 0 | 0,99 | 0 | 2,86 | 0 |
| BACH2 | ACTGAGTCTCGGTC | -1 | 440 | 454 | -560 | -546 | -1,73 | 0,01 | 12,68 | >0.1 | 2,75 | 4,85 | 0 | 1,01 | 0 | 5,8 | 0 |
| BACH2 | CCTGCCTCAGCCTC | -1 | 352 | 366 | -648 | -634 | 10,32 | 0,001 | 25,17 | >0.1 | 2,53 | 4,69 | 0 | 1,1 | 0 | 6,52 | 0 |
| BACH2 | TGTTAGTCAGGACG | -1 | 259 | 273 | -741 | -727 | 5,93 | 0,001 | 19,07 | >0.1 | 2,8 | 4,55 | 0 | 1,33 | 0 | 4,45 | 0 |
| BACH2 | CCCAACTCGGCCTC | -1 | 218 | 232 | -782 | -768 | -1,89 | 0,01 | 12,6 | >0.1 | 2,96 | 4,5 | 0 | 1,52 | 0 | 5,52 | 0 |
| BACH2 | TATGACTACTCTCC | -1 | 121 | 135 | -879 | -865 | 0,15 | 0,01 | 11,56 | >0.1 | 3,45 | 4,38 | 0 | 1,89 | 0 | 1,7 | 0 |
| BACH2 | GACTACTCTCCACT | -1 | 118 | 132 | -882 | -868 | -1,58 | 0,01 | 9,8 | >0.1 | 3,47 | 4,37 | 0 | 1,89 | 0 | 1,65 | 0 |
| BACH2 | TCTGCCTTTGCAAA | -1 | 84 | 98 | -916 | -902 | -0,45 | 0,01 | 9,5 | >0.1 | 3,73 | 4,34 | 0 | 1,88 | 0 | 0 | 0 |
| POU2F2 | AAGTTTTGCAAAG | 1 | 80 | 93 | -920 | -907 | -4,43 | 0,01 | 5,55 | >0.1 | 3,77 | 4,33 | 0 | 1,88 | 0 | 0 | 0 |
| POU2F2 | GAAATATGAATTC | 1 | 486 | 499 | -514 | -501 | -3,93 | 0,01 | 11,04 | >0.1 | 2,91 | 4,94 | 0 | 0,99 | 0 | 6,13 | 0 |

| | | | | | | | | | | | | | | | | | |
|---|---|---|---|---|---|---|---|---|---|---|---|---|---|---|---|---|---|
| POU2F2 | ATTATCTTCATGA | 1 | 640 | 653 | -360 | -347 | 0,2 | 0,01 | 14,23 | >0.1 | 3,81 | 5,33 | 0 | 1,06 | 0 | 3,83 | 0 |
| POU2F2 | ATCTGTTACATAT | 1 | 844 | 857 | -156 | -143 | -3,32 | 0,01 | 13,31 | >0.1 | 5,57 | 6,28 | 0 | 1,83 | 0 | 2,95 | 0 |
| POU2F2 | GAACTTTATATTG | 1 | 873 | 886 | -127 | -114 | -6,73 | 0,01 | 10,18 | >0.1 | 5,57 | 6,5 | 0 | 1,9 | 0 | 2,95 | 0 |
| POU2F2 | TTCTTTTTCTTTT | 1 | 914 | 927 | -86 | -73 | -4,33 | 0,01 | 11,17 | >0.1 | 5,57 | 6,89 | 0 | 1,89 | 0 | 1,15 | 0 |
| POU2F2 | AATATTTTCCTAG | -1 | 720 | 733 | -280 | -267 | 1,51 | 0,01 | 19,07 | >0.1 | 5,57 | 5,61 | 0 | 1,21 | 0 | 5,17 | 0 |
| POU2F2 | GGTGTTTGTATGG | -1 | 624 | 637 | -376 | -363 | -5,51 | 0,01 | 6,03 | >0.1 | 3,67 | 5,28 | 0 | 1,04 | 0 | 1,55 | 0 |
| POU2F2 | GACAATTTCATTA | -1 | 605 | 618 | -395 | -382 | -3,7 | 0,01 | 7,52 | >0.1 | 3,52 | 5,23 | 0 | 1,02 | 0 | 1,44 | 0 |
| POU2F2 | TTTATTTTTATTT | -1 | 577 | 590 | -423 | -410 | 3,12 | 0,01 | 12,62 | >0.1 | 3,34 | 5,15 | 0 | 1 | 0 | 0 | 0 |
| POU2F2 | AACATTTTCCCTT | -1 | 530 | 543 | -470 | -457 | -4,88 | 0,01 | 5,85 | >0.1 | 3,09 | 5,04 | 0 | 0,99 | 0 | 1,61 | 0 |
| POU2F2 | TCCAAATACATGA | -1 | 518 | 531 | -482 | -469 | -5,51 | 0,01 | 6,72 | >0.1 | 3,04 | 5,01 | 0 | 0,99 | 0 | 3,19 | 0 |
| POU2F2 | TGACTTTGCCCTT | -1 | 508 | 521 | -492 | -479 | -5,96 | 0,01 | 5,76 | >0.1 | 3 | 4,99 | 0 | 0,99 | 0 | 2,75 | 0 |
| POU2F2 | TTCATATTTCTGA | -1 | 483 | 496 | -517 | -504 | -4,38 | 0,01 | 10,57 | >0.1 | 2,9 | 4,94 | 0 | 0,99 | 0 | 6,13 | 0 |
| POU2F2 | AGTTTTTGTATTT | -1 | 296 | 309 | -704 | -691 | 3,41 | 0,01 | 16,37 | >0.1 | 2,69 | 4,6 | 0 | 1,22 | 0 | 4,45 | 0 |
| POU2F2 | TTAATTTCCATGG | -1 | 149 | 162 | -851 | -838 | 0,17 | 0,01 | 11,4 | >0.1 | 3,28 | 4,41 | 0 | 1,9 | 0 | 1,65 | 0 |
| POU2F2 | CTGATCTGCCTTT | -1 | 89 | 102 | -911 | -898 | 0,58 | 0,01 | 10,49 | >0.1 | 3,69 | 4,34 | 0 | 1,88 | 0 | 0 | 0 |
| POU2F2 | TGCCTTTGCAAAA | -1 | 83 | 96 | -917 | -904 | -3,01 | 0,01 | 6,95 | >0.1 | 3,74 | 4,34 | 0 | 1,88 | 0 | 0 | 0 |
| POU2F2 | TTCACTTACCTAT | -1 | 1 | 14 | -999 | -986 | -5,03 | 0,01 | 8,45 | >0.1 | 5,57 | 4,26 | 0 | 1,86 | 1,79 | 0 | 0 |
| POU2F3 | AGGCACTCAAAGAAGT | 1 | 22 | 38 | -978 | -962 | -2,51 | 0,01 | 9,17 | >0.1 | 5,53 | 4,28 | 0 | 1,87 | 0 | 0 | 0 |
| POU2F3 | AGTTTTGCAAAGGCAG | 1 | 81 | 97 | -919 | -903 | 3,42 | 0,01 | 13,39 | >0.1 | 3,75 | 4,34 | 0 | 1,88 | 0 | 0 | 0 |
| POU2F3 | GCAAAGGCAGATCAGG | 1 | 87 | 103 | -913 | -897 | -1,33 | 0,01 | 8,59 | >0.1 | 3,69 | 4,34 | 0 | 1,88 | 0 | 0 | 0 |
| POU2F3 | GGCCATGGAAATTAAA | 1 | 147 | 163 | -853 | -837 | 5,08 | 0,001 | 16,32 | >0.1 | 3,28 | 4,41 | 0 | 1,9 | 0 | 1,65 | 0 |
| POU2F3 | CTCTATTAAAAATACA | 1 | 287 | 303 | -713 | -697 | -2,64 | 0,01 | 11 | >0.1 | 2,71 | 4,59 | 0 | 1,24 | 0 | 5,09 | 0 |
| POU2F3 | AGTCATGTATTTGGAA | 1 | 516 | 532 | -484 | -468 | -1,63 | 0,01 | 10,6 | >0.1 | 3,04 | 5,01 | 0 | 0,99 | 0 | 3,19 | 0 |
| POU2F3 | ACGTATTCTTATTTGA | 1 | 804 | 820 | -196 | -180 | 7,48 | 0,001 | 23,88 | >0.1 | 5,53 | 6,03 | 0 | 1,55 | 0 | 3,28 | 0 |
| POU2F3 | AATTATTCTAAAATCT | 1 | 832 | 848 | -168 | -152 | 0,26 | 0,01 | 17,04 | >0.1 | 5,53 | 6,22 | 0 | 1,76 | 0 | 3,28 | 0 |
| POU2F3 | TCAAAGGCTGATAAGA | 1 | 960 | 976 | -40 | -24 | -2,1 | 0,01 | 14,79 | >0.1 | 5,53 | 7,79 | 0 | 1,88 | 0 | 1,7 | 0 |
| POU2F3 | AGTGAATCAAATAAGA | -1 | 810 | 826 | -190 | -174 | -1,84 | 0,01 | 14,78 | >0.1 | 5,53 | 6,06 | 0 | 1,59 | 0 | 3,43 | 0 |
| POU2F3 | TTGTATGGTTTTCAGA | -1 | 616 | 632 | -384 | -368 | -1,08 | 0,01 | 10,34 | >0.1 | 3,61 | 5,27 | 0 | 1,04 | 0 | 1,5 | 0 |
| POU2F3 | ATTTATTTTTATTTCA | -1 | 575 | 591 | -425 | -409 | 1,7 | 0,01 | 11,2 | >0.1 | 3,33 | 5,16 | 0 | 1,01 | 0 | 0 | 0 |
| POU2F3 | CTTTATGGAATTCATA | -1 | 490 | 506 | -510 | -494 | 0,89 | 0,01 | 16,2 | >0.1 | 2,93 | 4,96 | 0 | 0,99 | 0 | 6,44 | 0 |
| POU2F3 | TGGAATTCATATTTCT | -1 | 485 | 501 | -515 | -499 | 4,39 | 0,01 | 19,51 | >0.1 | 2,91 | 4,95 | 0 | 0,99 | 0 | 6,27 | 0 |
| POU2F3 | TTGTATTTTTAATAGA | -1 | 288 | 304 | -712 | -696 | 1,1 | 0,01 | 14,09 | >0.1 | 2,71 | 4,6 | 0 | 1,24 | 0 | 4,45 | 0 |
| POU2F3 | GCCTTTGCAAAACTTA | -1 | 79 | 95 | -921 | -905 | 2,08 | 0,01 | 12,06 | >0.1 | 3,77 | 4,33 | 0 | 1,88 | 0 | 0 | 0 |
| TGIF1 | TCAAAGAAGTCA | 1 | 28 | 40 | -972 | -960 | -19,28 | 0,01 | -8,62 | >0.1 | 4,51 | 4,28 | 0 | 1,87 | 0 | 0 | 0 |
| TGIF1 | TGACCTGTGGAG | 1 | 108 | 120 | -892 | -880 | -20,88 | 0,01 | -10,34 | >0.1 | 3,54 | 4,36 | 0 | 1,89 | 0 | 0,75 | 0 |
| TGIF1 | GGAGAGTAGTCA | 1 | 121 | 133 | -879 | -867 | -24,12 | 0,01 | -12,75 | >0.1 | 3,45 | 4,37 | 0 | 1,89 | 0 | 1,65 | 0 |
| TGIF1 | TCACGCCTGTAA | 1 | 192 | 204 | -808 | -796 | -13,4 | 0,01 | 0,08 | >0.1 | 3,07 | 4,46 | 0 | 1,69 | 0 | 4,26 | 0 |
| TGIF1 | GGGCAGATCACA | 1 | 229 | 241 | -771 | -759 | -20,59 | 0,01 | -6,7 | >0.1 | 2,92 | 4,51 | 0 | 1,46 | 0 | 5,01 | 0 |
| TGIF1 | GGGCACCTGTAG | 1 | 326 | 338 | -674 | -662 | -23,41 | 0,01 | -9,45 | >0.1 | 2,6 | 4,65 | 0 | 1,15 | 0 | 5,56 | 0 |
| TGIF1 | AGACAAGGGCAA | 1 | 504 | 516 | -496 | -484 | -24,68 | 0,01 | -10,06 | >0.1 | 2,98 | 4,98 | 0 | 0,98 | 0 | 5,68 | 0 |
| TGIF1 | TAACAGCTTTCT | 1 | 710 | 722 | -290 | -278 | -6,33 | 0,001 | 10,99 | >0.1 | 5,58 | 5,57 | 0 | 1,18 | 0 | 4,99 | 0 |
| TGIF1 | TGACAGGAGAGG | 1 | 767 | 779 | -233 | -221 | -17,45 | 0,01 | 1,21 | >0.1 | 5,58 | 5,81 | 0 | 1,35 | 0 | 5,92 | 0 |
| TGIF1 | TAAAATCTGTTA | 1 | 840 | 852 | -160 | -148 | -15,51 | 0,01 | -0,48 | >0.1 | 5,58 | 6,24 | 0 | 1,79 | 0 | 1,41 | 0 |
| TGIF1 | ACATATCTGTCC | 1 | 851 | 863 | -149 | -137 | -19,42 | 0,01 | -2,96 | >0.1 | 5,58 | 6,32 | 0 | 1,89 | 0 | 2,67 | 0 |
| TGIF1 | TCACAGGGGACG | 1 | 1037 | 1049 | 37 | 49 | -14,86 | 0,01 | 1,87 | >0.1 | 5,58 | 7,95 | 0 | 1,86 | 0 | 1,34 | 0 |
| TGIF1 | GGACAAGTTTAA | 1 | 1135 | 1147 | 135 | 147 | -21,56 | 0,01 | -1,41 | >0.1 | 5,58 | 6,41 | 0 | 1,84 | 3,22 | 3,09 | 0 |
| TGIF1 | TGACATCGTCCC | -1 | 1043 | 1055 | 43 | 55 | -18,63 | 0,01 | -2,81 | >0.1 | 5,58 | 7,76 | 0 | 1,86 | 0 | 0,62 | 0 |
| TGIF1 | GGACAGATATGT | -1 | 851 | 863 | -149 | -137 | -23 | 0,01 | -6,54 | >0.1 | 5,58 | 6,32 | 0 | 1,89 | 0 | 2,67 | 0 |
| TGIF1 | AGATATGTAACA | -1 | 847 | 859 | -153 | -141 | -21,1 | 0,01 | -4,42 | >0.1 | 5,58 | 6,29 | 0 | 1,85 | 0 | 2,95 | 0 |
| TGIF1 | TAACAGATTTTA | -1 | 840 | 852 | -160 | -148 | -13,4 | 0,01 | 1,63 | >0.1 | 5,58 | 6,24 | 0 | 1,79 | 0 | 1,41 | 0 |
| TGIF1 | AGAAAGCTGTTA | -1 | 710 | 722 | -290 | -278 | -8,68 | 0,001 | 8,64 | >0.1 | 5,58 | 5,57 | 0 | 1,18 | 0 | 4,99 | 0 |
| TGIF1 | CTACAGGTGCCC | -1 | 326 | 338 | -674 | -662 | -16,46 | 0,01 | -2,5 | >0.1 | 2,6 | 4,65 | 0 | 1,15 | 0 | 5,56 | 0 |
| TGIF1 | TGTGATCTGCCC | -1 | 229 | 241 | -771 | -759 | -24,28 | 0,01 | -10,39 | >0.1 | 2,92 | 4,51 | 0 | 1,46 | 0 | 5,01 | 0 |
| TGIF1 | TTACAGGCGTGA | -1 | 192 | 204 | -808 | -796 | -7,63 | 0,001 | 5,85 | >0.1 | 3,07 | 4,46 | 0 | 1,69 | 0 | 4,26 | 0 |
| TGIF1 | TGACTACTCTCC | -1 | 121 | 133 | -879 | -867 | -16,09 | 0,01 | -4,72 | >0.1 | 3,45 | 4,37 | 0 | 1,89 | 0 | 1,65 | 0 |
| TGIF1 | TGACTTCTTTGA | -1 | 28 | 40 | -972 | -960 | -12,48 | 0,01 | -1,82 | >0.1 | 4,51 | 4,28 | 0 | 1,87 | 0 | 0 | 0 |
| POU2F1 | AATAGGTAAGTG | 1 | 0 | 12 | -1000 | -988 | 1,42 | 0,01 | 14,91 | >0.1 | 5,58 | 4,26 | 0 | 1,86 | 1,79 | 0 | 0 |
| POU2F1 | GTTTTGCAAAGG | 1 | 82 | 94 | -918 | -906 | 3,35 | 0,01 | 13,32 | >0.1 | 3,76 | 4,33 | 0 | 1,88 | 0 | 0 | 0 |
| POU2F1 | CAAAGGCAGATC | 1 | 88 | 100 | -912 | -900 | 2,52 | 0,01 | 12,44 | >0.1 | 3,7 | 4,34 | 0 | 1,88 | 0 | 0 | 0 |

| | | | | | | | | | | | | | | | | | |
|---|---|---|---|---|---|---|---|---|---|---|---|---|---|---|---|---|---|
| POU2F1 | ATAATTTTAAAA | 1 | 132 | 144 | -868 | -856 | 1,76 | 0,01 | 13 | >0.1 | 3,38 | 4,39 | 0 | 1,89 | 0 | 1,57 | 0 |
| POU2F1 | GCCATGGAAATT | 1 | 148 | 160 | -852 | -840 | 7,59 | 0,001 | 18,83 | >0.1 | 3,29 | 4,41 | 0 | 1,9 | 0 | 1,65 | 0 |
| POU2F1 | AAAATACAAAAA | 1 | 295 | 307 | -705 | -693 | 2,03 | 0,01 | 15,07 | >0.1 | 2,69 | 4,6 | 0 | 1,22 | 0 | 4,53 | 0 |
| POU2F1 | ATTAATGAAATT | 1 | 603 | 615 | -397 | -385 | 3,56 | 0,01 | 14,72 | >0.1 | 3,5 | 5,22 | 0 | 1,02 | 0 | 1,41 | 0 |
| POU2F1 | TCTAGGAAAATA | 1 | 719 | 731 | -281 | -269 | 1,14 | 0,01 | 18,52 | >0.1 | 5,58 | 5,6 | 0 | 1,2 | 0 | 4,99 | 0 |
| POU2F1 | CGTATTCTTATT | 1 | 805 | 817 | -195 | -183 | 6,28 | 0,01 | 22,69 | >0.1 | 5,58 | 6,01 | 0 | 1,54 | 0 | 3,28 | 0 |
| POU2F1 | ATTATTCTAAAA | 1 | 833 | 845 | -167 | -155 | 7,71 | 0,001 | 24,49 | >0.1 | 5,58 | 6,19 | 0 | 1,73 | 0 | 3,28 | 0 |
| POU2F1 | CAAAGGCTGATA | 1 | 961 | 973 | -39 | -27 | 1,2 | 0,01 | 18,05 | >0.1 | 5,58 | 7,7 | 0 | 1,88 | 0 | 1,7 | 0 |
| POU2F1 | TCTAGGGAAAGT | 1 | 1002 | 1014 | 2 | 14 | 1,6 | 0,01 | 19,63 | >0.1 | 5,58 | 9,12 | 0 | 1,87 | 0 | 1,46 | 0 |
| POU2F1 | GATATGTAACAG | -1 | 846 | 858 | -154 | -142 | 1,93 | 0,01 | 18,5 | >0.1 | 5,58 | 6,28 | 0 | 1,84 | 0 | 2,86 | 0 |
| POU2F1 | AATACGTTATTC | -1 | 799 | 811 | -201 | -189 | 2,41 | 0,01 | 19,02 | >0.1 | 5,58 | 5,98 | 0 | 1,5 | 0 | 3,55 | 0 |
| POU2F1 | TACACGGAAAAT | -1 | 694 | 706 | -306 | -294 | 2,63 | 0,01 | 17,82 | >0.1 | 5,58 | 5,51 | 0 | 1,15 | 0 | 2,95 | 0 |
| POU2F1 | ATCATGAAGATA | -1 | 642 | 654 | -358 | -346 | 3,82 | 0,01 | 17,74 | >0.1 | 3,82 | 5,34 | 0 | 1,06 | 0 | 3,7 | 0 |
| POU2F1 | TTTATTTTTATT | -1 | 578 | 590 | -422 | -410 | 4,53 | 0,01 | 14,03 | >0.1 | 3,34 | 5,15 | 0 | 1 | 0 | 0 | 0 |
| POU2F1 | TTTATGGAATTC | -1 | 493 | 505 | -507 | -495 | 5,1 | 0,01 | 20,41 | >0.1 | 2,94 | 4,95 | 0 | 0,99 | 0 | 6,44 | 0 |
| POU2F1 | GGAATTCATATT | -1 | 488 | 500 | -512 | -500 | 3,03 | 0,01 | 18,15 | >0.1 | 2,92 | 4,94 | 0 | 0,99 | 0 | 6,27 | 0 |
| POU2F1 | CCTTTGCAAAAC | -1 | 82 | 94 | -918 | -906 | 1,87 | 0,01 | 11,84 | >0.1 | 3,76 | 4,33 | 0 | 1,88 | 0 | 0 | 0 |
| HSF1 | TCCTAGCACTTTG | 1 | 204 | 217 | -796 | -783 | -4,1 | 0,01 | 9,7 | >0.1 | 3,01 | 4,48 | 0 | 1,6 | 0 | 4,7 | 0 |
| HSF1 | ATAGAGACCGTCC | 1 | 251 | 264 | -749 | -736 | -9,4 | 0,01 | 4,31 | >0.1 | 2,83 | 4,54 | 0 | 1,36 | 0 | 4,98 | 0 |
| HSF1 | TTCTGGGAATATG | 1 | 558 | 571 | -442 | -429 | -9,17 | 0,01 | 0,16 | >0.1 | 3,23 | 5,11 | 0 | 1 | 0 | 0 | 0 |
| HSF1 | TTCTAGACCTCTT | 1 | 660 | 673 | -340 | -327 | -4,86 | 0,01 | 9,82 | >0.1 | 4,03 | 5,4 | 0 | 1,09 | 0 | 4,16 | 0 |
| HSF1 | TTATAACATTTTC | 1 | 687 | 700 | -313 | -300 | -9,42 | 0,01 | 4,54 | >0.1 | 4,47 | 5,49 | 0 | 1,14 | 0 | 2,86 | 0 |
| HSF1 | TTCTAGGAAAATA | 1 | 718 | 731 | -282 | -269 | -6,82 | 0,01 | 10,54 | >0.1 | 5,57 | 5,6 | 0 | 1,2 | 0 | 4,99 | 0 |
| HSF1 | TTCTAAAATCTGT | 1 | 837 | 850 | -163 | -150 | -10,97 | 0,01 | 5,79 | >0.1 | 5,57 | 6,23 | 0 | 1,77 | 0 | 3,19 | 0 |
| HSF1 | CTCCAGGATGAAC | 1 | 864 | 877 | -136 | -123 | -9,07 | 0,01 | 7,49 | >0.1 | 5,57 | 6,42 | 0 | 1,9 | 0 | 2,67 | 0 |
| HSF1 | TCCTGGCTCCTTC | 1 | 1061 | 1074 | 61 | 74 | -1,88 | 0,001 | 16,89 | >0.1 | 5,57 | 7,33 | 0 | 1,86 | 1,79 | 2,21 | 0 |
| HSF1 | TCTTGGCCTGTTC | -1 | 1115 | 1128 | 115 | 128 | -12,03 | 0,01 | 7,81 | >0.1 | 5,57 | 6,6 | 0 | 1,85 | 3,22 | 2,6 | 0 |
| HSF1 | TTTTAGAATAATT | -1 | 832 | 845 | -168 | -155 | -3,31 | 0,01 | 13,61 | >0.1 | 5,57 | 6,19 | 0 | 1,73 | 0 | 3,43 | 0 |
| HSF1 | CTAGAGAACCTTA | -1 | 779 | 792 | -221 | -208 | -4,87 | 0,01 | 14,53 | >0.1 | 5,57 | 5,87 | 0 | 1,41 | 0 | 6,54 | 0 |
| HSF1 | AACCGGTACTTTT | -1 | 737 | 750 | -263 | -250 | -11,41 | 0,01 | 7,01 | >0.1 | 5,57 | 5,68 | 0 | 1,26 | 0 | 5,92 | 0 |
| HSF1 | TCCTAGAAAGCTG | -1 | 713 | 726 | -287 | -274 | -6,41 | 0,01 | 10,92 | >0.1 | 5,57 | 5,58 | 0 | 1,19 | 0 | 4,99 | 0 |
| HSF1 | GTCTAGAACTAGG | -1 | 655 | 668 | -345 | -332 | -8,33 | 0,01 | 5,8 | >0.1 | 3,97 | 5,38 | 0 | 1,08 | 0 | 3,7 | 0 |
| HSF1 | TTTCAGACAATTT | -1 | 610 | 623 | -390 | -377 | -9,79 | 0,01 | 1,54 | >0.1 | 3,56 | 5,24 | 0 | 1,03 | 0 | 1,5 | 0 |
| HSF1 | TTCAAGATCATAT | -1 | 566 | 579 | -434 | -421 | -9,92 | 0,01 | -0,52 | >0.1 | 3,27 | 5,13 | 0 | 1 | 0 | 0 | 0 |
| HSF1 | TCCCAGAAACTCT | -1 | 553 | 566 | -447 | -434 | -10,9 | 0,01 | -1,61 | >0.1 | 3,21 | 5,09 | 0 | 0,99 | 0 | 0 | 0 |
| HSF1 | TTGGGCAACATTT | -1 | 536 | 549 | -464 | -451 | -11,54 | 0,01 | -0,72 | >0.1 | 3,12 | 5,05 | 0 | 0,99 | 0 | 1,66 | 0 |
| ZFX | GTGGCTCACGCCTG | 1 | 187 | 201 | -813 | -799 | 8,76 | 0,001 | 22,07 | >0.1 | 3,09 | 4,46 | 0 | 1,73 | 0 | 4,04 | 0 |
| ZFX | TAGAGACCGTCCTG | 1 | 252 | 266 | -748 | -734 | -2,41 | 0,01 | 11,17 | >0.1 | 2,83 | 4,54 | 0 | 1,36 | 0 | 4,85 | 0 |
| ZFX | CTTAGCTGGGCGTG | 1 | 307 | 321 | -693 | -679 | 0,52 | 0,01 | 15,24 | >0.1 | 2,65 | 4,62 | 0 | 1,19 | 0 | 6,26 | 0 |
| ZFX | GGAGGCAGAGCTTG | 1 | 383 | 397 | -617 | -603 | -0,45 | 0,01 | 14,07 | >0.1 | 2,58 | 4,75 | 0 | 1,06 | 0 | 6,13 | 0 |
| ZFX | TGTGCTCCAGCCTG | 1 | 419 | 433 | -581 | -567 | 7,27 | 0,001 | 21,21 | >0.1 | 2,69 | 4,81 | 0 | 1,02 | 0 | 5,42 | 0 |
| ZFX | TAGTTCTAGACCTC | 1 | 657 | 671 | -343 | -329 | -2,92 | 0,01 | 11,55 | >0.1 | 4 | 5,39 | 0 | 1,09 | 0 | 3,99 | 0 |
| ZFX | TCCTTCTCAGCCTT | 1 | 1068 | 1082 | 68 | 82 | 2,64 | 0,01 | 21,32 | >0.1 | 5,56 | 7,21 | 0 | 1,85 | 1,79 | 2,27 | 0 |
| ZFX | CCTTCTCAGCCTTG | 1 | 1069 | 1083 | 69 | 83 | -3,89 | 0,01 | 14,78 | >0.1 | 5,56 | 7,19 | 0 | 1,85 | 1,79 | 2,27 | 0 |
| ZFX | AGGTCTTCGGCTTC | -1 | 1151 | 1165 | 151 | 165 | 3,01 | 0,01 | 19,76 | >0.1 | 4,48 | 6,29 | 0 | 1,84 | 1,79 | 2,34 | 0 |
| ZFX | AATGTCTTGGCCTG | -1 | 1118 | 1132 | 118 | 132 | 5,36 | 0,01 | 25,25 | >0.1 | 5,56 | 6,57 | 0 | 1,85 | 3,22 | 2,7 | 0 |
| ZFX | TTGGCCTGTTCCTC | -1 | 1112 | 1126 | 112 | 126 | -2,67 | 0,01 | 17,15 | >0.1 | 5,56 | 6,63 | 0 | 1,85 | 3,22 | 2,56 | 0 |
| ZFX | CTCTTATCAGCCTT | -1 | 963 | 977 | -37 | -23 | -3,55 | 0,01 | 13,39 | >0.1 | 5,56 | 7,82 | 0 | 1,87 | 0 | 1,7 | 0 |
| ZFX | ATGACTTTGCCCTT | -1 | 508 | 522 | -492 | -478 | -1,98 | 0,01 | 9,86 | >0.1 | 3 | 4,99 | 0 | 0,99 | 0 | 2,86 | 0 |
| ZFX | TGAGTCTCGGTCTG | -1 | 438 | 452 | -562 | -548 | 1,28 | 0,01 | 15,93 | >0.1 | 2,75 | 4,85 | 0 | 1,01 | 0 | 6,05 | 0 |
| ZFX | GTCGCCCAGGCTGG | -1 | 425 | 439 | -575 | -561 | -1,59 | 0,01 | 12,76 | >0.1 | 2,7 | 4,82 | 0 | 1,02 | 0 | 5,8 | 0 |
| ZFX | GCAAGCTCTGCCTC | -1 | 384 | 398 | -616 | -602 | 5,1 | 0,01 | 19,62 | >0.1 | 2,59 | 4,75 | 0 | 1,06 | 0 | 6,13 | 0 |
| ZFX | CATTCTCCTGCCTC | -1 | 358 | 372 | -642 | -628 | 0,68 | 0,01 | 15,36 | >0.1 | 2,52 | 4,7 | 0 | 1,09 | 0 | 6,36 | 0 |
| ZFX | CCTGCCTCAGCCTC | -1 | 352 | 366 | -648 | -634 | 10,15 | 0,001 | 25 | >0.1 | 2,53 | 4,69 | 0 | 1,1 | 0 | 6,52 | 0 |
| ZFX | AGTAGCTGGGACTA | -1 | 335 | 349 | -665 | -651 | -2,48 | 0,01 | 11,81 | >0.1 | 2,58 | 4,67 | 0 | 1,13 | 0 | 5,92 | 0 |
| ZFX | CCCAACTCGGCCTC | -1 | 218 | 232 | -782 | -768 | 11,11 | 0,001 | 25,6 | >0.1 | 2,96 | 4,5 | 0 | 1,52 | 0 | 5,52 | 0 |
| ZFX | TCCTGATCTGCCTT | -1 | 90 | 104 | -910 | -896 | -3,74 | 0,01 | 6,99 | >0.1 | 3,68 | 4,34 | 0 | 1,88 | 0 | 0,83 | 0 |
| HSF4 | GTGGAGAGTAGTC | 1 | 119 | 132 | -881 | -868 | -1,43 | 0,01 | 9,94 | >0.1 | 3,46 | 4,37 | 0 | 1,89 | 0 | 1,65 | 0 |
| HSF4 | TCCTAGCACTTTG | 1 | 204 | 217 | -796 | -783 | 1,77 | 0,01 | 15,57 | >0.1 | 3,01 | 4,48 | 0 | 1,6 | 0 | 4,7 | 0 |

| | | | | | | | | | | | | | | | | | |
|---|---|---|---|---|---|---|---|---|---|---|---|---|---|---|---|---|---|
| HSF4 | ATAGAGACCGTCC | 1 | 251 | 264 | -749 | -736 | -2,25 | 0,01 | 11,46 | >0.1 | 2,83 | 4,54 | 0 | 1,36 | 0 | 4,98 | 0 |
| HSF4 | CTCCAGCCTGGGC | 1 | 423 | 436 | -577 | -564 | -2,43 | 0,01 | 11,77 | >0.1 | 2,7 | 4,82 | 0 | 1,02 | 0 | 5,67 | 0 |
| HSF4 | TTCTGGGAATATG | 1 | 558 | 571 | -442 | -429 | -1,47 | 0,01 | 7,86 | >0.1 | 3,23 | 5,11 | 0 | 1 | 0 | 0 | 0 |
| HSF4 | TTCTAGACCTCTT | 1 | 660 | 673 | -340 | -327 | 2,99 | 0,01 | 17,67 | >0.1 | 4,03 | 5,4 | 0 | 1,09 | 0 | 4,16 | 0 |
| HSF4 | TTCTAGGAAAATA | 1 | 718 | 731 | -282 | -269 | -0,98 | 0,01 | 16,38 | >0.1 | 5,57 | 5,6 | 0 | 1,2 | 0 | 4,99 | 0 |
| HSF4 | CTCCAGGATGAAC | 1 | 864 | 877 | -136 | -123 | -0,38 | 0,01 | 16,18 | >0.1 | 5,57 | 6,42 | 0 | 1,9 | 0 | 2,67 | 0 |
| HSF4 | TCCTGGCTCCTTC | 1 | 1061 | 1074 | 61 | 74 | 5,94 | 0,001 | 24,71 | >0.1 | 5,57 | 7,33 | 0 | 1,86 | 1,79 | 2,21 | 0 |
| HSF4 | TTTTAGAATAATT | -1 | 832 | 845 | -168 | -155 | 0,72 | 0,01 | 17,64 | >0.1 | 5,57 | 6,19 | 0 | 1,73 | 0 | 3,43 | 0 |
| HSF4 | CTAGAGAACCTTA | -1 | 779 | 792 | -221 | -208 | 0,17 | 0,01 | 19,57 | >0.1 | 5,57 | 5,87 | 0 | 1,41 | 0 | 6,54 | 0 |
| HSF4 | TTATGGCCAAATC | -1 | 752 | 765 | -248 | -235 | -1,65 | 0,01 | 17,29 | >0.1 | 5,57 | 5,75 | 0 | 1,3 | 0 | 6,32 | 0 |
| HSF4 | AACCGGTACTTTT | -1 | 737 | 750 | -263 | -250 | 0,68 | 0,01 | 19,1 | >0.1 | 5,57 | 5,68 | 0 | 1,26 | 0 | 5,92 | 0 |
| HSF4 | TCCTAGAAAGCTG | -1 | 713 | 726 | -287 | -274 | -0,43 | 0,01 | 16,9 | >0.1 | 5,57 | 5,58 | 0 | 1,19 | 0 | 4,99 | 0 |
| HSF4 | GTCTAGAACTAGG | -1 | 655 | 668 | -345 | -332 | -0,17 | 0,01 | 13,96 | >0.1 | 3,97 | 5,38 | 0 | 1,08 | 0 | 3,7 | 0 |
| HSF4 | TTTCAGACAATTT | -1 | 610 | 623 | -390 | -377 | -2,24 | 0,01 | 9,09 | >0.1 | 3,56 | 5,24 | 0 | 1,03 | 0 | 1,5 | 0 |
| HSF4 | TTCAAGATCATAT | -1 | 566 | 579 | -434 | -421 | -2,73 | 0,01 | 6,67 | >0.1 | 3,27 | 5,13 | 0 | 1 | 0 | 0 | 0 |
| HSF4 | TCCCAGAAACTCT | -1 | 553 | 566 | -447 | -434 | -2,97 | 0,01 | 6,32 | >0.1 | 3,21 | 5,09 | 0 | 0,99 | 0 | 0 | 0 |
| HSF4 | GTCTCGGTCTGTC | -1 | 436 | 449 | -564 | -551 | 0,3 | 0,01 | 14,93 | >0.1 | 2,74 | 4,84 | 0 | 1,01 | 0 | 6,05 | 0 |
| HSF4 | GCCCAGCCATTTC | -1 | 171 | 184 | -829 | -816 | -1,03 | 0,01 | 11,79 | >0.1 | 3,17 | 4,43 | 0 | 1,87 | 0 | 3,34 | 0 |
| TBX21 | CCAGTGTGAG | 1 | 49 | 59 | -951 | -941 | 1,33 | 0,01 | 11,64 | >0.1 | 4,14 | 4,3 | 0 | 1,87 | 0 | 0 | 0 |
| TBX21 | AGTGTGAGAA | 1 | 51 | 61 | -949 | -939 | 5,66 | 0,001 | 15,95 | >0.1 | 4,11 | 4,3 | 0 | 1,87 | 0 | 0 | 0 |
| TBX21 | GACCTGTGGA | 1 | 109 | 119 | -891 | -881 | -2,06 | 0,01 | 8,47 | >0.1 | 3,54 | 4,36 | 0 | 1,89 | 0 | 0,75 | 0 |
| TBX21 | ACGGTGAAAC | 1 | 272 | 282 | -728 | -718 | -3,05 | 0,01 | 10,56 | >0.1 | 2,77 | 4,56 | 0 | 1,29 | 0 | 4,99 | 0 |
| TBX21 | TGGGCGTGGT | 1 | 313 | 323 | -687 | -677 | 0,54 | 0,01 | 15,11 | >0.1 | 2,64 | 4,62 | 0 | 1,17 | 0 | 6,13 | 0 |
| TBX21 | ATGGCGTGAA | 1 | 369 | 379 | -631 | -621 | 8,28 | 0,001 | 22,76 | >0.1 | 2,54 | 4,71 | 0 | 1,07 | 0 | 6,15 | 0 |
| TBX21 | GAGCCGAGAT | 1 | 401 | 411 | -599 | -589 | -1,96 | 0,01 | 12,24 | >0.1 | 2,63 | 4,77 | 0 | 1,04 | 0 | 5,76 | 0 |
| TBX21 | TAAATGTGAG | 1 | 586 | 596 | -414 | -404 | -2,21 | 0,01 | 7,35 | >0.1 | 3,38 | 5,17 | 0 | 1,01 | 0 | 0 | 0 |
| TBX21 | AATGTGAGAT | 1 | 588 | 598 | -412 | -402 | 7 | 0,001 | 16,58 | >0.1 | 3,4 | 5,18 | 0 | 1,01 | 0 | 0 | 0 |
| TBX21 | AAAGTGACAG | 1 | 763 | 773 | -237 | -227 | 0,98 | 0,01 | 19,62 | >0.1 | 5,61 | 5,78 | 0 | 1,33 | 0 | 5,92 | 0 |
| TBX21 | AATCTGTTAC | 1 | 843 | 853 | -157 | -147 | -1,56 | 0,01 | 15,28 | >0.1 | 5,61 | 6,25 | 0 | 1,8 | 0 | 3,19 | 0 |
| TBX21 | AGGCTGATAA | 1 | 964 | 974 | -36 | -26 | -2,47 | 0,01 | 14,44 | >0.1 | 5,61 | 7,73 | 0 | 1,87 | 0 | 1,7 | 0 |
| TBX21 | TAAGAGAGAA | 1 | 971 | 981 | -29 | -19 | -1,49 | 0,01 | 15,58 | >0.1 | 5,61 | 7,94 | 0 | 1,87 | 0 | 1,65 | 0 |
| TBX21 | GAGGTTTTAG | 1 | 992 | 1002 | -8 | 2 | -2,35 | 0,01 | 15,48 | >0.1 | 5,61 | 8,77 | 0 | 1,87 | 0 | 1,57 | 0 |
| TBX21 | TGGATGTGAT | 1 | 1021 | 1031 | 21 | 31 | -0,46 | 0,01 | 17,02 | >0.1 | 5,61 | 8,7 | 0 | 1,86 | 0 | 1,31 | 0 |
| TBX21 | AGGCTGAGAA | -1 | 1071 | 1081 | 71 | 81 | 1,62 | 0,01 | 20,31 | >0.1 | 5,61 | 7,15 | 0 | 1,85 | 1,79 | 2,27 | 0 |
| TBX21 | AAGCTGTTAA | -1 | 709 | 719 | -291 | -281 | 5,22 | 0,001 | 22,37 | >0.1 | 5,61 | 5,56 | 0 | 1,17 | 0 | 4,81 | 0 |
| TBX21 | AAAATGTTAT | -1 | 689 | 699 | -311 | -301 | -3,36 | 0,01 | 10,69 | >0.1 | 4,48 | 5,48 | 0 | 1,13 | 0 | 2,95 | 0 |
| TBX21 | GTGGCGCGAT | -1 | 409 | 419 | -591 | -581 | 2,46 | 0,01 | 16,45 | >0.1 | 2,65 | 4,78 | 0 | 1,03 | 0 | 5,52 | 0 |
| TBX21 | TAGCTGGGAC | -1 | 337 | 347 | -663 | -653 | -1,44 | 0,01 | 12,84 | >0.1 | 2,58 | 4,66 | 0 | 1,12 | 0 | 5,92 | 0 |
| TBX21 | ACCGTGTTAG | -1 | 267 | 277 | -733 | -723 | -0,97 | 0,01 | 12,2 | >0.1 | 2,78 | 4,56 | 0 | 1,3 | 0 | 4,53 | 0 |
| TBX21 | ACCTTGTGAT | -1 | 235 | 245 | -765 | -755 | -2,9 | 0,01 | 10,68 | >0.1 | 2,9 | 4,51 | 0 | 1,43 | 0 | 4,74 | 0 |
| TBX21 | AAAGTGCTAG | -1 | 206 | 216 | -794 | -784 | 3,34 | 0,01 | 17,12 | >0.1 | 3,01 | 4,47 | 0 | 1,59 | 0 | 4,7 | 0 |
| TBX21 | CAGGCGTGAG | -1 | 191 | 201 | -809 | -799 | 7,25 | 0,001 | 20,61 | >0.1 | 3,08 | 4,45 | 0 | 1,7 | 0 | 4,13 | 0 |
| TBX20 | CCAGTGTGAGA | 1 | 49 | 60 | -951 | -940 | 0,42 | 0,01 | 10,73 | >0.1 | 4,13 | 4,3 | 0 | 1,87 | 0 | 0 | 0 |
| TBX20 | AGTGTGAGAAA | 1 | 51 | 62 | -949 | -938 | 2,57 | 0,01 | 12,85 | >0.1 | 4,11 | 4,3 | 0 | 1,87 | 0 | 0 | 0 |
| TBX20 | GAGTTGGGCAG | 1 | 224 | 235 | -776 | -765 | -0,12 | 0,01 | 14,06 | >0.1 | 2,94 | 4,5 | 0 | 1,49 | 0 | 5,26 | 0 |
| TBX20 | ATGGCGTGAAC | 1 | 369 | 380 | -631 | -620 | 0,13 | 0,01 | 14,61 | >0.1 | 2,54 | 4,72 | 0 | 1,07 | 0 | 6,15 | 0 |
| TBX20 | TAAATGTGAGA | 1 | 586 | 597 | -414 | -403 | -2,03 | 0,01 | 7,54 | >0.1 | 3,39 | 5,17 | 0 | 1,01 | 0 | 0 | 0 |
| TBX20 | AATGTGAGATA | 1 | 588 | 599 | -412 | -401 | 3,72 | 0,01 | 13,31 | >0.1 | 3,4 | 5,18 | 0 | 1,01 | 0 | 0 | 0 |
| TBX20 | ATTGTCTGAAA | 1 | 612 | 623 | -388 | -377 | -1,87 | 0,01 | 9,41 | >0.1 | 3,56 | 5,24 | 0 | 1,03 | 0 | 1,44 | 0 |
| TBX20 | AAAGTGACAGG | 1 | 763 | 774 | -237 | -226 | -0,87 | 0,01 | 17,76 | >0.1 | 5,6 | 5,79 | 0 | 1,34 | 0 | 5,92 | 0 |
| TBX20 | GAGGTTTTAGT | 1 | 992 | 1003 | -8 | 3 | 1,14 | 0,01 | 18,98 | >0.1 | 5,6 | 8,81 | 0 | 1,87 | 0 | 1,57 | 0 |
| TBX20 | TGGATGTGATC | 1 | 1021 | 1032 | 21 | 32 | -0,61 | 0,01 | 16,86 | >0.1 | 5,6 | 8,7 | 0 | 1,86 | 0 | 1,31 | 0 |
| TBX20 | TCGTGGTTAAA | -1 | 1142 | 1153 | 142 | 153 | -0,18 | 0,01 | 20,21 | >0.1 | 5,6 | 6,36 | 0 | 1,84 | 3,22 | 3,37 | 0 |
| TBX20 | AAGCTGTTAAA | -1 | 708 | 719 | -292 | -281 | 1,76 | 0,01 | 18,9 | >0.1 | 5,6 | 5,56 | 0 | 1,17 | 0 | 4,81 | 0 |
| TBX20 | TTGGTGTTTGT | -1 | 628 | 639 | -372 | -361 | -1,1 | 0,01 | 10,53 | >0.1 | 3,69 | 5,29 | 0 | 1,04 | 0 | 1,61 | 0 |
| TBX20 | ACCGTGTTAGT | -1 | 266 | 277 | -734 | -723 | 0,74 | 0,01 | 13,92 | >0.1 | 2,79 | 4,56 | 0 | 1,31 | 0 | 4,53 | 0 |
| TBX20 | ACCTTGTGATC | -1 | 234 | 245 | -766 | -755 | 1,41 | 0,01 | 15,16 | >0.1 | 2,9 | 4,51 | 0 | 1,44 | 0 | 4,9 | 0 |
| TBX20 | AAAGTGCTAGG | -1 | 205 | 216 | -795 | -784 | -0,64 | 0,01 | 13,15 | >0.1 | 3,01 | 4,47 | 0 | 1,6 | 0 | 4,7 | 0 |
| STAT1::STAT2 | AAAGTACCGGTTTTG | 1 | 738 | 753 | -262 | -247 | -0,65 | 0,01 | 17,77 | >0.1 | 5,54 | 5,69 | 0 | 1,27 | 0 | 5,92 | 0 |

| | | | | | | | | | | | | | | | | | |
|---|---|---|---|---|---|---|---|---|---|---|---|---|---|---|---|---|---|
| STAT1::STAT2 | CCGGTTTTGATTTGG | 1 | 744 | 759 | -256 | -241 | 3,89 | 0,01 | 22,35 | >0.1 | 5,54 | 5,72 | 0 | 1,29 | 0 | 5,92 | 0 |
| STAT1::STAT2 | TGTGTTCTTCTTCTT | 1 | 904 | 919 | -96 | -81 | 0,08 | 0,01 | 15,52 | >0.1 | 5,54 | 6,8 | 0 | 1,89 | 0 | 1,21 | 0 |
| STAT1::STAT2 | GTTCTTCTTCTTTTT | 1 | 907 | 922 | -93 | -78 | -2,21 | 0,01 | 13,26 | >0.1 | 5,54 | 6,83 | 0 | 1,89 | 0 | 1,21 | 0 |
| STAT1::STAT2 | TCTTCTTCTTTTTCT | 1 | 909 | 924 | -91 | -76 | -0,74 | 0,01 | 14,7 | >0.1 | 5,54 | 6,86 | 0 | 1,89 | 0 | 1,15 | 0 |
| STAT1::STAT2 | CTTCTTCTTTTTCTT | 1 | 910 | 925 | -90 | -75 | -2,76 | 0,01 | 12,69 | >0.1 | 5,54 | 6,87 | 0 | 1,89 | 0 | 1,15 | 0 |
| STAT1::STAT2 | CTTCTTTTTCTTTTT | 1 | 913 | 928 | -87 | -72 | 2,51 | 0,01 | 18 | >0.1 | 5,54 | 6,9 | 0 | 1,89 | 0 | 1,15 | 0 |
| STAT1::STAT2 | TTCTTTTTCTTTTTT | 1 | 914 | 929 | -86 | -71 | -2,8 | 0,01 | 12,67 | >0.1 | 5,54 | 6,92 | 0 | 1,89 | 0 | 1,12 | 0 |
| STAT1::STAT2 | TCTTTTTCTTTTTTT | 1 | 915 | 930 | -85 | -70 | 4,21 | 0,01 | 19,69 | >0.1 | 5,54 | 6,93 | 0 | 1,89 | 0 | 1,12 | 0 |
| STAT1::STAT2 | TCACTTGCTTCTTGG | 1 | 1178 | 1193 | 178 | 193 | -2,09 | 0,01 | 16,24 | >0.1 | 5,54 | 6,1 | 0 | 1,6 | 1,79 | 3,29 | 0 |
| STAT1::STAT2 | TGACTTTCCCTAGAC | -1 | 1001 | 1016 | 1 | 16 | -0,97 | 0,01 | 17,03 | >0.1 | 5,54 | 9,12 | 0 | 1,87 | 0 | 1,46 | 0 |
| STAT1::STAT2 | GTACTTTTGGTTAAT | -1 | 730 | 745 | -270 | -255 | -1,02 | 0,01 | 17,34 | >0.1 | 5,54 | 5,66 | 0 | 1,24 | 0 | 5,92 | 0 |
| STAT1::STAT2 | TCACATTTATTTTTA | -1 | 580 | 595 | -420 | -405 | -2,86 | 0,01 | 6,68 | >0.1 | 3,36 | 5,17 | 0 | 1,01 | 0 | 0 | 0 |
| STAT1::STAT2 | TTTATTTTTATTTCA | -1 | 575 | 590 | -425 | -410 | -1,94 | 0,01 | 7,55 | >0.1 | 3,33 | 5,15 | 0 | 1,01 | 0 | 0 | 0 |
| STAT1::STAT2 | TTATTTTTATTTCAA | -1 | 574 | 589 | -426 | -411 | -2,59 | 0,01 | 6,89 | >0.1 | 3,33 | 5,15 | 0 | 1,01 | 0 | 0 | 0 |
| STAT1::STAT2 | ACATTTTCCCTTCCA | -1 | 527 | 542 | -473 | -458 | 3,9 | 0,01 | 16,56 | >0.1 | 3,08 | 5,04 | 0 | 0,99 | 0 | 3,55 | 0 |
| STAT1::STAT2 | TCAGTTTTTTTTTTT | -1 | 468 | 483 | -532 | -517 | 11,79 | 0,0001 | 25,49 | >0.1 | 2,85 | 4,91 | 0 | 0,99 | 0 | 4,95 | 0 |
| STAT1::STAT2 | CAGTTTTTTTTTTTT | -1 | 467 | 482 | -533 | -518 | -0,77 | 0,01 | 12,8 | >0.1 | 2,84 | 4,91 | 0 | 0,99 | 0 | 4,83 | 0 |
| STAT1::STAT2 | GTTTTTTTTTTTTTT | -1 | 465 | 480 | -535 | -520 | -3,15 | 0,01 | 11,36 | >0.1 | 2,84 | 4,9 | 0 | 1 | 0 | 5,78 | 0 |
| STAT1::STAT2 | TTTTTTTTTTTTTTT | -1 | 464 | 479 | -536 | -521 | -1,24 | 0,01 | 13,12 | >0.1 | 2,83 | 4,9 | 0 | 1 | 0 | 5,63 | 0 |
| STAT1::STAT2 | TTTTTTTTTTTTTTT | -1 | 463 | 478 | -537 | -522 | -1,24 | 0,01 | 13,26 | >0.1 | 2,83 | 4,9 | 0 | 1 | 0 | 5,78 | 0 |
| STAT1::STAT2 | TTTTTTTTTTTTTTT | -1 | 462 | 477 | -538 | -523 | -1,24 | 0,01 | 13,26 | >0.1 | 2,83 | 4,9 | 0 | 1 | 0 | 5,78 | 0 |
| STAT1::STAT2 | TTTTTTTTTTTTTTT | -1 | 461 | 476 | -539 | -524 | -1,24 | 0,01 | 13,25 | >0.1 | 2,82 | 4,89 | 0 | 1 | 0 | 5,78 | 0 |
| STAT1::STAT2 | TTTTTTTTTTTTTTT | -1 | 460 | 475 | -540 | -525 | -1,24 | 0,01 | 13,1 | >0.1 | 2,82 | 4,89 | 0 | 1 | 0 | 5,63 | 0 |
| STAT1::STAT2 | TTTTTTTTTTTTTTT | -1 | 459 | 474 | -541 | -526 | -1,24 | 0,01 | 13,1 | >0.1 | 2,82 | 4,89 | 0 | 1 | 0 | 5,63 | 0 |
| STAT1::STAT2 | TTTTTTTTTTTTTTT | -1 | 458 | 473 | -542 | -527 | -1,24 | 0,01 | 12,98 | >0.1 | 2,81 | 4,89 | 0 | 1 | 0 | 5,52 | 0 |
| STAT1::STAT2 | TTTTTTTTTTTTTTT | -1 | 457 | 472 | -543 | -528 | -1,24 | 0,01 | 12,98 | >0.1 | 2,81 | 4,89 | 0 | 1 | 0 | 5,52 | 0 |
| STAT1::STAT2 | TTTTTTTTTTTTTTG | -1 | 456 | 471 | -544 | -529 | -3,14 | 0,01 | 11,18 | >0.1 | 2,81 | 4,88 | 0 | 1 | 0 | 5,63 | 0 |
| STAT1::STAT2 | TGAGTCTCGGTCTGT | -1 | 437 | 452 | -563 | -548 | -2,21 | 0,01 | 12,43 | >0.1 | 2,74 | 4,85 | 0 | 1,01 | 0 | 6,05 | 0 |
| STAT1::STAT2 | TAAGTTTTTGTATTT | -1 | 296 | 311 | -704 | -689 | 4,69 | 0,01 | 17,65 | >0.1 | 2,68 | 4,61 | 0 | 1,22 | 0 | 4,45 | 0 |
| STAT1::STAT2 | ACGGTCTCTATCTCC | -1 | 247 | 262 | -753 | -738 | 0,29 | 0,01 | 14,24 | >0.1 | 2,84 | 4,54 | 0 | 1,38 | 0 | 5,19 | 0 |
| STAT1::STAT2 | TCAGTTTTAATTTCC | -1 | 153 | 168 | -847 | -832 | 16,1 | 1E-05 | 28,27 | >0.1 | 3,25 | 4,42 | 0 | 1,9 | 0 | 2,6 | 0 |
| STAT1::STAT2 | TCATTTCCTATTTCT | -1 | 57 | 72 | -943 | -928 | 4,45 | 0,01 | 14,64 | >0.1 | 4 | 4,31 | 0 | 1,88 | 0 | 0 | 0 |
| STAT1::STAT2 | TCTCCTTCACTTACC | -1 | 4 | 19 | -996 | -981 | 1,28 | 0,01 | 14,75 | >0.1 | 5,54 | 4,26 | 0 | 1,86 | 1,79 | 0 | 0 |
| ZNF384 | TTTTAAAAAATG | 1 | 136 | 148 | -864 | -852 | 8,41 | 0,01 | 19,7 | >0.1 | 3,36 | 4,39 | 0 | 1,89 | 0 | 1,65 | 0 |
| ZNF384 | ATTAAAAATACA | 1 | 291 | 303 | -709 | -697 | 4,49 | 0,01 | 17,47 | >0.1 | 2,7 | 4,59 | 0 | 1,23 | 0 | 4,45 | 0 |
| ZNF384 | TCTCAAAAAAAA | 1 | 453 | 465 | -547 | -535 | 14,08 | 0,0001 | 28,26 | >0.1 | 2,79 | 4,87 | 0 | 1 | 0 | 5,52 | 0 |
| ZNF384 | CTCAAAAAAAAA | 1 | 454 | 466 | -546 | -534 | 12,28 | 0,01 | 26,47 | >0.1 | 2,79 | 4,87 | 0 | 1 | 0 | 5,52 | 0 |
| ZNF384 | TCAAAAAAAAAA | 1 | 455 | 467 | -545 | -533 | 13,93 | 0,001 | 28,12 | >0.1 | 2,8 | 4,88 | 0 | 1 | 0 | 5,52 | 0 |
| ZNF384 | CAAAAAAAAAAA | 1 | 456 | 468 | -544 | -532 | 13,24 | 0,01 | 27,55 | >0.1 | 2,8 | 4,88 | 0 | 1 | 0 | 5,63 | 0 |
| ZNF384 | AAAAAAAAAAAA | 1 | 457 | 469 | -543 | -531 | 13,67 | 0,001 | 27,98 | >0.1 | 2,8 | 4,88 | 0 | 1 | 0 | 5,63 | 0 |
| ZNF384 | AAAAAAAAAAAA | 1 | 458 | 470 | -542 | -530 | 13,67 | 0,001 | 27,88 | >0.1 | 2,81 | 4,88 | 0 | 1 | 0 | 5,52 | 0 |
| ZNF384 | AAAAAAAAAAAA | 1 | 459 | 471 | -541 | -529 | 13,67 | 0,001 | 27,88 | >0.1 | 2,81 | 4,88 | 0 | 1 | 0 | 5,52 | 0 |
| ZNF384 | AAAAAAAAAAAA | 1 | 460 | 472 | -540 | -528 | 13,67 | 0,001 | 28 | >0.1 | 2,81 | 4,89 | 0 | 0,99 | 0 | 5,63 | 0 |
| ZNF384 | AAAAAAAAAAAA | 1 | 461 | 473 | -539 | -527 | 13,67 | 0,001 | 28 | >0.1 | 2,82 | 4,89 | 0 | 0,99 | 0 | 5,63 | 0 |
| ZNF384 | AAAAAAAAAAAA | 1 | 462 | 474 | -538 | -526 | 13,67 | 0,001 | 28,15 | >0.1 | 2,82 | 4,89 | 0 | 0,99 | 0 | 5,78 | 0 |
| ZNF384 | AAAAAAAAAAAA | 1 | 463 | 475 | -537 | -525 | 13,67 | 0,001 | 28,16 | >0.1 | 2,83 | 4,89 | 0 | 0,99 | 0 | 5,78 | 0 |
| ZNF384 | AAAAAAAAAAAA | 1 | 464 | 476 | -536 | -524 | 13,67 | 0,001 | 28,16 | >0.1 | 2,83 | 4,89 | 0 | 0,99 | 0 | 5,78 | 0 |
| ZNF384 | AAAAAAAAAAAA | 1 | 465 | 477 | -535 | -523 | 13,67 | 0,001 | 28,02 | >0.1 | 2,83 | 4,9 | 0 | 0,99 | 0 | 5,63 | 0 |
| ZNF384 | AAAAAAAAAAAA | 1 | 466 | 478 | -534 | -522 | 13,67 | 0,001 | 28,17 | >0.1 | 2,83 | 4,9 | 0 | 0,99 | 0 | 5,78 | 0 |
| ZNF384 | AAAAAAAAAAAA | 1 | 467 | 479 | -533 | -521 | 13,67 | 0,001 | 28,32 | >0.1 | 2,84 | 4,9 | 0 | 0,99 | 0 | 5,92 | 0 |
| ZNF384 | AAAAAAAAAAAC | 1 | 468 | 480 | -532 | -520 | 11,92 | 0,01 | 25,48 | >0.1 | 2,84 | 4,9 | 0 | 0,99 | 0 | 4,83 | 0 |
| ZNF384 | AAAAAAAAAACT | 1 | 469 | 481 | -531 | -519 | 9,34 | 0,01 | 23,03 | >0.1 | 2,85 | 4,9 | 0 | 0,99 | 0 | 4,95 | 0 |
| ZNF384 | CTTGAAATAAAA | 1 | 573 | 585 | -427 | -415 | 5,06 | 0,01 | 14,51 | >0.1 | 3,31 | 5,14 | 0 | 1 | 0 | 0 | 0 |
| ZNF384 | TTGAAATAAAAA | 1 | 574 | 586 | -426 | -414 | 6,71 | 0,01 | 16,17 | >0.1 | 3,32 | 5,14 | 0 | 1 | 0 | 0 | 0 |
| ZNF384 | TGAAATAAAAAT | 1 | 575 | 587 | -425 | -413 | 4,13 | 0,01 | 13,6 | >0.1 | 3,32 | 5,15 | 0 | 1 | 0 | 0 | 0 |
| ZNF384 | GAAATAAAAATA | 1 | 576 | 588 | -424 | -412 | 2,23 | 0,01 | 11,71 | >0.1 | 3,33 | 5,15 | 0 | 1 | 0 | 0 | 0 |
| ZNF384 | AATAAAAATAAA | 1 | 578 | 590 | -422 | -410 | 6,8 | 0,01 | 16,3 | >0.1 | 3,34 | 5,15 | 0 | 1 | 0 | 0 | 0 |
| ZNF384 | ATAAAAATAAAT | 1 | 579 | 591 | -421 | -409 | 4,03 | 0,01 | 13,54 | >0.1 | 3,35 | 5,16 | 0 | 1 | 0 | 0 | 0 |
| ZNF384 | TAAAAATAAATG | 1 | 580 | 592 | -420 | -408 | 2,82 | 0,01 | 12,34 | >0.1 | 3,35 | 5,16 | 0 | 1,01 | 0 | 0 | 0 |

| | | | | | | | | | | | | | | | | | |
|---|---|---|---|---|---|---|---|---|---|---|---|---|---|---|---|---|---|
| ZNF384 | ACTTTAAAAAAT | 1 | 823 | 835 | -177 | -165 | 1,77 | 0,01 | 18,83 | >0.1 | 5,58 | 6,12 | 0 | 1,65 | 0 | 3,7 | 0 |
| ZNF384 | CTTTAAAAAATT | 1 | 824 | 836 | -176 | -164 | 8,03 | 0,01 | 25,1 | >0.1 | 5,58 | 6,13 | 0 | 1,66 | 0 | 3,7 | 0 |
| ZNF384 | GACCAAAAAAAG | -1 | 922 | 934 | -78 | -66 | 9,98 | 0,01 | 25,51 | >0.1 | 5,58 | 6,98 | 0 | 1,88 | 0 | 1,09 | 0 |
| ZNF384 | ACCAAAAAAAGA | -1 | 921 | 933 | -79 | -67 | 10,45 | 0,01 | 25,97 | >0.1 | 5,58 | 6,97 | 0 | 1,88 | 0 | 1,09 | 0 |
| ZNF384 | CAAAAAAAGAAA | -1 | 919 | 931 | -81 | -69 | 2,68 | 0,01 | 18,21 | >0.1 | 5,58 | 6,94 | 0 | 1,88 | 0 | 1,12 | 0 |
| ZNF384 | AAAAAAAGAAAA | -1 | 918 | 930 | -82 | -70 | 3,21 | 0,01 | 18,69 | >0.1 | 5,58 | 6,93 | 0 | 1,89 | 0 | 1,09 | 0 |
| ZNF384 | AAAAAAGAAAAA | -1 | 917 | 929 | -83 | -71 | 4,01 | 0,01 | 19,52 | >0.1 | 5,58 | 6,91 | 0 | 1,89 | 0 | 1,12 | 0 |
| ZNF384 | AATCAAATAAGA | -1 | 810 | 822 | -190 | -178 | 3,82 | 0,01 | 20,44 | >0.1 | 5,58 | 6,04 | 0 | 1,57 | 0 | 3,43 | 0 |
| ZNF384 | ACTTAAAAAAGC | -1 | 72 | 84 | -928 | -916 | 8,66 | 0,01 | 18,71 | >0.1 | 3,85 | 4,32 | 0 | 1,88 | 0 | 0 | 0 |
| ZNF354C | AGCCAC | 1 | 39 | 45 | -961 | -955 | 4,72 | 0,01 | 15,21 | >0.1 | 4,34 | 4,28 | 0 | 1,87 | 0 | 0 | 0 |
| ZNF354C | AACCAG | 1 | 47 | 53 | -953 | -947 | 4,57 | 0,01 | 14,93 | >0.1 | 4,2 | 4,29 | 0 | 1,87 | 0 | 0 | 0 |
| ZNF354C | CGCCAC | 1 | 413 | 419 | -587 | -581 | 5,01 | 0,01 | 19,24 | >0.1 | 2,66 | 4,78 | 0 | 1,02 | 0 | 5,76 | 0 |
| ZNF354C | CTCCAG | 1 | 423 | 429 | -577 | -571 | 6,65 | 0,01 | 20,57 | >0.1 | 2,69 | 4,8 | 0 | 1,01 | 0 | 5,42 | 0 |
| ZNF354C | CTCCAG | 1 | 864 | 870 | -136 | -130 | 6,65 | 0,01 | 23,34 | >0.1 | 5,67 | 6,37 | 0 | 1,9 | 0 | 2,75 | 0 |
| ZNF354C | GTCCAC | 1 | 1102 | 1108 | 102 | 108 | 9,36 | 0,001 | 29,21 | >0.1 | 5,67 | 6,73 | 0 | 1,85 | 3,22 | 2,38 | 0 |
| ZNF354C | AACCAC | 1 | 1145 | 1151 | 145 | 151 | 8,23 | 0,01 | 27,41 | >0.1 | 5,67 | 6,33 | 0 | 1,84 | 3,22 | 2,11 | 0 |
| ZNF354C | GTCCCC | -1 | 1042 | 1048 | 42 | 48 | 3,52 | 0,01 | 20,2 | >0.1 | 5,67 | 7,79 | 0 | 1,86 | 0 | 1,36 | 0 |
| ZNF354C | ATCCAC | -1 | 1020 | 1026 | 20 | 26 | 10,04 | 0,001 | 27,63 | >0.1 | 5,67 | 8,72 | 0 | 1,86 | 0 | 1,34 | 0 |
| ZNF354C | CTCCTC | -1 | 989 | 995 | -11 | -5 | 3,98 | 0,01 | 20,69 | >0.1 | 4,64 | 8,59 | 0 | 1,87 | 0 | 1,61 | 0 |
| ZNF354C | CTCTAC | -1 | 549 | 555 | -451 | -445 | 3,48 | 0,01 | 12,7 | >0.1 | 3,17 | 5,07 | 0 | 0,98 | 0 | 0 | 0 |
| ZNF354C | GCCCAG | -1 | 430 | 436 | -570 | -564 | 3,85 | 0,01 | 18,18 | >0.1 | 2,71 | 4,82 | 0 | 1,01 | 0 | 5,8 | 0 |
| ZNF354C | GCCCAC | -1 | 324 | 330 | -676 | -670 | 7,51 | 0,01 | 21,47 | >0.1 | 2,62 | 4,63 | 0 | 1,15 | 0 | 5,56 | 0 |
| ZNF354C | CACCAC | -1 | 321 | 327 | -679 | -673 | 8,51 | 0,001 | 22,48 | >0.1 | 2,63 | 4,63 | 0 | 1,15 | 0 | 5,56 | 0 |
| ZNF354C | CACCAC | -1 | 318 | 324 | -682 | -676 | 8,51 | 0,001 | 23,06 | >0.1 | 2,63 | 4,63 | 0 | 1,16 | 0 | 6,13 | 0 |
| ZNF354C | GCCCAG | -1 | 312 | 318 | -688 | -682 | 3,85 | 0,01 | 18,42 | >0.1 | 2,65 | 4,62 | 0 | 1,17 | 0 | 6,13 | 0 |
| ZNF354C | AGCCAC | -1 | 187 | 193 | -813 | -807 | 4,72 | 0,01 | 17,85 | >0.1 | 3,1 | 4,44 | 0 | 1,73 | 0 | 3,86 | 0 |
| ZNF354C | GCCCAG | -1 | 178 | 184 | -822 | -816 | 3,85 | 0,01 | 16,47 | >0.1 | 3,15 | 4,43 | 0 | 1,8 | 0 | 3,24 | 0 |
| ZNF354C | CTCCAC | -1 | 119 | 125 | -881 | -875 | 10,32 | >0.1 | 20,81 | >0.1 | 3,49 | 4,36 | 0 | 1,89 | 0 | 0,75 | 0 |
| ZNF354C | CTCCAC | -1 | 114 | 120 | -886 | -880 | 10,32 | >0.1 | 20,85 | >0.1 | 3,52 | 4,36 | 0 | 1,89 | 0 | 0,77 | 0 |
| ZNF354C | CTTCAC | -1 | 9 | 15 | -991 | -985 | 3,48 | 0,01 | 17,06 | >0.1 | 5,67 | 4,26 | 0 | 1,86 | 1,79 | 0 | 0 |
| BHLHE40 | ATCAGGAGAG | 1 | 97 | 107 | -903 | -893 | -4,93 | 0,01 | 5,7 | >0.1 | 3,64 | 4,35 | 0 | 1,88 | 0 | 0,77 | 0 |
| BHLHE40 | GGCTCACGCC | 1 | 189 | 199 | -811 | -801 | -3,96 | 0,01 | 9,33 | >0.1 | 3,09 | 4,45 | 0 | 1,71 | 0 | 4,04 | 0 |
| BHLHE40 | CTCACGCCTG | 1 | 191 | 201 | -809 | -799 | -4,8 | 0,01 | 8,56 | >0.1 | 3,08 | 4,45 | 0 | 1,7 | 0 | 4,13 | 0 |
| BHLHE40 | ATCACAAGGT | 1 | 235 | 245 | -765 | -755 | -5,13 | 0,01 | 8,45 | >0.1 | 2,9 | 4,51 | 0 | 1,43 | 0 | 4,74 | 0 |
| BHLHE40 | GTCAGGAGAT | 1 | 243 | 253 | -757 | -747 | -4,41 | 0,01 | 8,7 | >0.1 | 2,87 | 4,52 | 0 | 1,39 | 0 | 4,33 | 0 |
| BHLHE40 | GGCACCTGTA | 1 | 327 | 337 | -673 | -663 | 1,3 | 0,01 | 15,26 | >0.1 | 2,6 | 4,65 | 0 | 1,14 | 0 | 5,56 | 0 |
| BHLHE40 | TACTCGGGAG | 1 | 345 | 355 | -655 | -645 | -3,8 | 0,01 | 11,06 | >0.1 | 2,55 | 4,67 | 0 | 1,11 | 0 | 6,52 | 0 |
| BHLHE40 | GGCAGGAGAA | 1 | 360 | 370 | -640 | -630 | -4,98 | 0,01 | 9,69 | >0.1 | 2,52 | 4,7 | 0 | 1,09 | 0 | 6,36 | 0 |
| BHLHE40 | AGATCGCGCC | 1 | 407 | 417 | -593 | -583 | -4,24 | 0,01 | 9,86 | >0.1 | 2,65 | 4,78 | 0 | 1,03 | 0 | 5,65 | 0 |
| BHLHE40 | AGACCGAGAC | 1 | 439 | 449 | -561 | -551 | -4,38 | 0,01 | 10,25 | >0.1 | 2,74 | 4,84 | 0 | 1 | 0 | 6,05 | 0 |
| BHLHE40 | GACAAGGGCA | 1 | 505 | 515 | -495 | -485 | -4,53 | 0,01 | 10,09 | >0.1 | 2,98 | 4,98 | 0 | 0,98 | 0 | 5,68 | 0 |
| BHLHE40 | ATAAAGTGAC | 1 | 761 | 771 | -239 | -229 | -1,98 | 0,01 | 16,83 | >0.1 | 5,61 | 5,77 | 0 | 1,32 | 0 | 6,1 | 0 |
| BHLHE40 | ATCTCATGAG | 1 | 982 | 992 | -18 | -8 | -1,63 | 0,01 | 15,81 | >0.1 | 5,61 | 8,43 | 0 | 1,87 | 0 | 1,53 | 0 |
| BHLHE40 | CACAGGGGAC | 1 | 1038 | 1048 | 38 | 48 | -5,33 | 0,01 | 11,39 | >0.1 | 5,61 | 7,92 | 0 | 1,86 | 0 | 1,34 | 0 |
| BHLHE40 | TTCACTTGCT | 1 | 1177 | 1187 | 177 | 187 | -3,48 | 0,01 | 14,9 | >0.1 | 5,61 | 6,11 | 0 | 1,61 | 1,79 | 3,26 | 0 |
| BHLHE40 | AGCAAGTGAA | -1 | 1177 | 1187 | 177 | 187 | 3,88 | 0,001 | 22,26 | >0.1 | 5,61 | 6,11 | 0 | 1,61 | 1,79 | 3,26 | 0 |
| BHLHE40 | GTCCCCTGTG | -1 | 1038 | 1048 | 38 | 48 | -4,47 | 0,01 | 12,25 | >0.1 | 5,61 | 7,92 | 0 | 1,86 | 0 | 1,34 | 0 |
| BHLHE40 | CCCCTGTGAG | -1 | 1036 | 1046 | 36 | 46 | -4,46 | 0,01 | 12,33 | >0.1 | 5,61 | 7,99 | 0 | 1,86 | 0 | 1,34 | 0 |
| BHLHE40 | ATCACATCCA | -1 | 1021 | 1031 | 21 | 31 | -4,3 | 0,01 | 13,18 | >0.1 | 5,61 | 8,7 | 0 | 1,86 | 0 | 1,31 | 0 |
| BHLHE40 | CTCATGAGAT | -1 | 982 | 992 | -18 | -8 | -0,99 | 0,01 | 16,45 | >0.1 | 5,61 | 8,43 | 0 | 1,87 | 0 | 1,53 | 0 |
| BHLHE40 | TTAAAGTGAA | -1 | 820 | 830 | -180 | -170 | -5,76 | 0,01 | 11,25 | >0.1 | 5,61 | 6,09 | 0 | 1,62 | 0 | 3,7 | 0 |
| BHLHE40 | CTCTCCTGTC | -1 | 768 | 778 | -232 | -222 | -3,63 | 0,01 | 15,05 | >0.1 | 5,61 | 5,81 | 0 | 1,35 | 0 | 5,92 | 0 |
| BHLHE40 | AATACATGAC | -1 | 517 | 527 | -483 | -473 | -4,69 | 0,01 | 7,27 | >0.1 | 3,03 | 5 | 0 | 0,98 | 0 | 2,95 | 0 |
| BHLHE40 | CGCCCAGGCT | -1 | 427 | 437 | -573 | -563 | -5,81 | 0,01 | 8,53 | >0.1 | 2,7 | 4,82 | 0 | 1,01 | 0 | 5,8 | 0 |
| BHLHE40 | TTCACGCCAT | -1 | 369 | 379 | -631 | -621 | -2,39 | 0,01 | 12,09 | >0.1 | 2,54 | 4,71 | 0 | 1,07 | 0 | 6,15 | 0 |
| BHLHE40 | TTCTCCTGCC | -1 | 360 | 370 | -640 | -630 | -1,29 | 0,01 | 13,38 | >0.1 | 2,52 | 4,7 | 0 | 1,09 | 0 | 6,36 | 0 |
| BHLHE40 | CTCCCGAGTA | -1 | 345 | 355 | -655 | -645 | -2,29 | 0,01 | 12,57 | >0.1 | 2,55 | 4,67 | 0 | 1,11 | 0 | 6,52 | 0 |
| BHLHE40 | TACAGGTGCC | -1 | 327 | 337 | -673 | -663 | -0,05 | 0,01 | 13,91 | >0.1 | 2,6 | 4,65 | 0 | 1,14 | 0 | 5,56 | 0 |
| BHLHE40 | ACCACGCCCA | -1 | 313 | 323 | -687 | -677 | -4,59 | 0,01 | 9,98 | >0.1 | 2,64 | 4,62 | 0 | 1,17 | 0 | 6,13 | 0 |

| | | | | | | | | | | | | | | | | | |
|---|---|---|---|---|---|---|---|---|---|---|---|---|---|---|---|---|---|
| BHLHE40 | ATCTCCTGAC | -1 | 243 | 253 | -757 | -747 | -0,22 | 0,01 | 12,89 | >0.1 | 2,87 | 4,52 | 0 | 1,39 | 0 | 4,33 | 0 |
| BHLHE40 | ACCTTGTGAT | -1 | 235 | 245 | -765 | -755 | -5,46 | 0,01 | 8,12 | >0.1 | 2,9 | 4,51 | 0 | 1,43 | 0 | 4,74 | 0 |
| BHLHE40 | GGCGTGAGCC | -1 | 189 | 199 | -811 | -801 | -5,55 | 0,01 | 7,74 | >0.1 | 3,09 | 4,45 | 0 | 1,71 | 0 | 4,04 | 0 |
| BHLHE40 | CTCTCCTGAT | -1 | 97 | 107 | -903 | -893 | -1,7 | 0,01 | 8,93 | >0.1 | 3,64 | 4,35 | 0 | 1,88 | 0 | 0,77 | 0 |
| BHLHE41 | GGTAAGTGAA | 1 | 4 | 14 | -996 | -986 | -14,61 | 0,01 | -1,09 | >0.1 | 5,61 | 4,26 | 0 | 1,86 | 1,79 | 0 | 0 |
| BHLHE41 | ATCAGGAGAG | 1 | 97 | 107 | -903 | -893 | -5,73 | 0,01 | 4,9 | >0.1 | 3,64 | 4,35 | 0 | 1,88 | 0 | 0,77 | 0 |
| BHLHE41 | CTCACGCCTG | 1 | 191 | 201 | -809 | -799 | -14,36 | 0,01 | -1 | >0.1 | 3,08 | 4,45 | 0 | 1,7 | 0 | 4,13 | 0 |
| BHLHE41 | ATCACAAGGT | 1 | 235 | 245 | -765 | -755 | -13,95 | 0,01 | -0,37 | >0.1 | 2,9 | 4,51 | 0 | 1,43 | 0 | 4,74 | 0 |
| BHLHE41 | GTCAGGAGAT | 1 | 243 | 253 | -757 | -747 | -3,79 | 0,01 | 9,32 | >0.1 | 2,87 | 4,52 | 0 | 1,39 | 0 | 4,33 | 0 |
| BHLHE41 | GGCACCTGTA | 1 | 327 | 337 | -673 | -663 | -9,35 | 0,01 | 4,61 | >0.1 | 2,6 | 4,65 | 0 | 1,14 | 0 | 5,56 | 0 |
| BHLHE41 | GGCAGGAGAA | 1 | 360 | 370 | -640 | -630 | -9,62 | 0,01 | 5,05 | >0.1 | 2,52 | 4,7 | 0 | 1,09 | 0 | 6,36 | 0 |
| BHLHE41 | ATGGCGTGAA | 1 | 369 | 379 | -631 | -621 | -7,9 | 0,01 | 6,58 | >0.1 | 2,54 | 4,71 | 0 | 1,07 | 0 | 6,15 | 0 |
| BHLHE41 | ATCGCGCCAC | 1 | 409 | 419 | -591 | -581 | -10,52 | 0,01 | 3,47 | >0.1 | 2,65 | 4,78 | 0 | 1,03 | 0 | 5,52 | 0 |
| BHLHE41 | AGACCGAGAC | 1 | 439 | 449 | -561 | -551 | -12,3 | 0,01 | 2,33 | >0.1 | 2,74 | 4,84 | 0 | 1 | 0 | 6,05 | 0 |
| BHLHE41 | GTCATGTATT | 1 | 517 | 527 | -483 | -473 | -14,57 | 0,01 | -2,61 | >0.1 | 3,03 | 5 | 0 | 0,98 | 0 | 2,95 | 0 |
| BHLHE41 | ATAAAGTGAC | 1 | 761 | 771 | -239 | -229 | -7,34 | 0,01 | 11,47 | >0.1 | 5,61 | 5,77 | 0 | 1,32 | 0 | 6,1 | 0 |
| BHLHE41 | GTGACAGGAG | 1 | 766 | 776 | -234 | -224 | -14,42 | 0,01 | 4,24 | >0.1 | 5,61 | 5,8 | 0 | 1,34 | 0 | 5,92 | 0 |
| BHLHE41 | GACAGGAGAG | 1 | 768 | 778 | -232 | -222 | -10,11 | 0,01 | 8,57 | >0.1 | 5,61 | 5,81 | 0 | 1,35 | 0 | 5,92 | 0 |
| BHLHE41 | ATCTCATGAG | 1 | 982 | 992 | -18 | -8 | -7,93 | 0,01 | 9,51 | >0.1 | 5,61 | 8,43 | 0 | 1,87 | 0 | 1,53 | 0 |
| BHLHE41 | TGGATGTGAT | 1 | 1021 | 1031 | 21 | 31 | -12,92 | 0,01 | 4,56 | >0.1 | 5,61 | 8,7 | 0 | 1,86 | 0 | 1,31 | 0 |
| BHLHE41 | CACAGGGGAC | 1 | 1038 | 1048 | 38 | 48 | -9,77 | 0,01 | 6,95 | >0.1 | 5,61 | 7,92 | 0 | 1,86 | 0 | 1,34 | 0 |
| BHLHE41 | TTCACTTGCT | 1 | 1177 | 1187 | 177 | 187 | -7,42 | 0,01 | 10,96 | >0.1 | 5,61 | 6,11 | 0 | 1,61 | 1,79 | 3,26 | 0 |
| BHLHE41 | AGCAAGTGAA | -1 | 1177 | 1187 | 177 | 187 | -3,38 | 0,01 | 15 | >0.1 | 5,61 | 6,11 | 0 | 1,61 | 1,79 | 3,26 | 0 |
| BHLHE41 | GTCCCCTGTG | -1 | 1038 | 1048 | 38 | 48 | -12,35 | 0,01 | 4,37 | >0.1 | 5,61 | 7,92 | 0 | 1,86 | 0 | 1,34 | 0 |
| BHLHE41 | CCCCTGTGAG | -1 | 1036 | 1046 | 36 | 46 | -13,08 | 0,01 | 3,71 | >0.1 | 5,61 | 7,99 | 0 | 1,86 | 0 | 1,34 | 0 |
| BHLHE41 | CTCATGAGAT | -1 | 982 | 992 | -18 | -8 | -4,58 | 0,01 | 12,86 | >0.1 | 5,61 | 8,43 | 0 | 1,87 | 0 | 1,53 | 0 |
| BHLHE41 | GTCACTTTAT | -1 | 761 | 771 | -239 | -229 | -13,95 | 0,01 | 4,86 | >0.1 | 5,61 | 5,77 | 0 | 1,32 | 0 | 6,1 | 0 |
| BHLHE41 | AATACATGAC | -1 | 517 | 527 | -483 | -473 | -13,17 | 0,01 | -1,21 | >0.1 | 3,03 | 5 | 0 | 0,98 | 0 | 2,95 | 0 |
| BHLHE41 | GTGGCGCGAT | -1 | 409 | 419 | -591 | -581 | -9,66 | 0,01 | 4,33 | >0.1 | 2,65 | 4,78 | 0 | 1,03 | 0 | 5,52 | 0 |
| BHLHE41 | TTCACGCCAT | -1 | 369 | 379 | -631 | -621 | -10,59 | 0,01 | 3,89 | >0.1 | 2,54 | 4,71 | 0 | 1,07 | 0 | 6,15 | 0 |
| BHLHE41 | TTCTCCTGCC | -1 | 360 | 370 | -640 | -630 | -13,11 | 0,01 | 1,56 | >0.1 | 2,52 | 4,7 | 0 | 1,09 | 0 | 6,36 | 0 |
| BHLHE41 | TACAGGTGCC | -1 | 327 | 337 | -673 | -663 | -8,17 | 0,01 | 5,79 | >0.1 | 2,6 | 4,65 | 0 | 1,14 | 0 | 5,56 | 0 |
| BHLHE41 | ATCTCCTGAC | -1 | 243 | 253 | -757 | -747 | -5,81 | 0,01 | 7,3 | >0.1 | 2,87 | 4,52 | 0 | 1,39 | 0 | 4,33 | 0 |
| BHLHE41 | ACCTTGTGAT | -1 | 235 | 245 | -765 | -755 | -14,04 | 0,01 | -0,46 | >0.1 | 2,9 | 4,51 | 0 | 1,43 | 0 | 4,74 | 0 |
| BHLHE41 | CAGGCGTGAG | -1 | 191 | 201 | -809 | -799 | -10,47 | 0,01 | 2,89 | >0.1 | 3,08 | 4,45 | 0 | 1,7 | 0 | 4,13 | 0 |
| BHLHE41 | GGCGTGAGCC | -1 | 189 | 199 | -811 | -801 | -10,5 | 0,01 | 2,79 | >0.1 | 3,09 | 4,45 | 0 | 1,71 | 0 | 4,04 | 0 |
| BHLHE41 | CTCTCCTGAT | -1 | 97 | 107 | -903 | -893 | -8,79 | 0,01 | 1,84 | >0.1 | 3,64 | 4,35 | 0 | 1,88 | 0 | 0,77 | 0 |
| RORC | TCAAAGAAGTCA | 1 | 28 | 40 | -972 | -960 | 2,21 | 0,01 | 18,8 | >0.1 | 4,51 | 4,28 | 0 | 1,87 | 0 | 0 | 5,94 |
| RORC | AGAAATAGGAAA | 1 | 57 | 69 | -943 | -931 | 1,91 | 0,01 | 18,05 | >0.1 | 4,02 | 4,31 | 0 | 1,87 | 0 | 0 | 5,94 |
| RORC | AAAGGCAGATCA | 1 | 89 | 101 | -911 | -899 | 3,1 | 0,01 | 18,95 | >0.1 | 3,69 | 4,34 | 0 | 1,88 | 0 | 0 | 5,94 |
| RORC | AAAAAATGGCCA | 1 | 140 | 152 | -860 | -848 | 1,68 | 0,01 | 18,85 | >0.1 | 3,33 | 4,4 | 0 | 1,9 | 0 | 1,61 | 5,94 |
| RORC | TTTGGGAGGCCG | 1 | 213 | 225 | -787 | -775 | 2,12 | 0,01 | 22,59 | >0.1 | 2,98 | 4,49 | 0 | 1,55 | 0 | 5,52 | 5,94 |
| RORC | ATCACAAGGTCA | 1 | 235 | 247 | -765 | -753 | 5,11 | 0,01 | 24,52 | >0.1 | 2,89 | 4,52 | 0 | 1,43 | 0 | 4,63 | 5,94 |
| RORC | AAGACAAGGGCA | 1 | 503 | 515 | -497 | -485 | 2,95 | 0,01 | 23,68 | >0.1 | 2,98 | 4,98 | 0 | 0,98 | 0 | 5,86 | 5,94 |
| RORC | TAAAATCTGTTA | 1 | 840 | 852 | -160 | -148 | 2,17 | 0,01 | 23,13 | >0.1 | 5,58 | 6,24 | 0 | 1,79 | 0 | 1,41 | 5,94 |
| RORC | TATATTGGCTCA | 1 | 879 | 891 | -121 | -109 | 2,14 | 0,01 | 25,16 | >0.1 | 5,58 | 6,54 | 0 | 1,9 | 0 | 3,07 | 5,94 |
| RORC | GGGACGATGTCA | 1 | 1043 | 1055 | 43 | 55 | 1,76 | 0,01 | 23,52 | >0.1 | 5,58 | 7,76 | 0 | 1,86 | 0 | 0,62 | 5,94 |
| RORC | AGGAACAGGCCA | 1 | 1113 | 1125 | 113 | 125 | 2,51 | 0,01 | 28,28 | >0.1 | 5,58 | 6,62 | 0 | 1,85 | 3,22 | 2,56 | 5,94 |
| RORC | GCAACAAGGCTG | -1 | 1075 | 1087 | 75 | 87 | 2,64 | 0,01 | 27,17 | >0.1 | 5,58 | 7,09 | 0 | 1,85 | 1,79 | 2,27 | 5,94 |
| RORC | GAACTTGGGTTG | -1 | 950 | 962 | -50 | -38 | 3,81 | 0,01 | 26,38 | >0.1 | 5,58 | 7,44 | 0 | 1,88 | 0 | 1,74 | 5,94 |
| RORC | ATATAAAGTTCA | -1 | 872 | 884 | -128 | -116 | 2,23 | 0,01 | 25,07 | >0.1 | 5,58 | 6,48 | 0 | 1,9 | 0 | 2,95 | 5,94 |
| RORC | CCAAAGAGGTCT | -1 | 664 | 676 | -336 | -324 | 5,94 | 0,01 | 26,28 | >0.1 | 4,07 | 5,41 | 0 | 1,09 | 0 | 3,83 | 5,94 |
| RORC | ATTAATAGGTTA | -1 | 597 | 609 | -403 | -391 | 8,25 | 0,001 | 25,28 | >0.1 | 3,46 | 5,21 | 0 | 1,02 | 0 | 1,41 | 5,94 |
| RORC | ATTTCAAGATCA | -1 | 569 | 581 | -431 | -419 | 2,19 | 0,01 | 17,55 | >0.1 | 3,29 | 5,13 | 0 | 1 | 0 | 0 | 5,94 |
| RORC | TCTACTTGGGCA | -1 | 542 | 554 | -458 | -446 | 2,91 | 0,01 | 18,05 | >0.1 | 3,15 | 5,07 | 0 | 0,99 | 0 | 0 | 5,94 |
| RORC | GTAGCTGGGACT | -1 | 336 | 348 | -664 | -652 | 3,31 | 0,01 | 23,53 | >0.1 | 2,58 | 4,66 | 0 | 1,13 | 0 | 5,92 | 5,94 |
| RORC | TTAGTCAGGACG | -1 | 259 | 271 | -741 | -729 | 2,47 | 0,01 | 21,62 | >0.1 | 2,81 | 4,55 | 0 | 1,33 | 0 | 4,53 | 5,94 |
| RORC | CCAACTCGGCCT | -1 | 219 | 231 | -781 | -769 | 2,53 | 0,01 | 22,95 | >0.1 | 2,96 | 4,49 | 0 | 1,51 | 0 | 5,52 | 5,94 |
| RORC | AAAATTATGACT | -1 | 128 | 140 | -872 | -860 | 1,99 | 0,01 | 19,22 | >0.1 | 3,41 | 4,38 | 0 | 1,89 | 0 | 1,61 | 5,94 |

| | | | | | | | | | | | | | | | | | |
|---|---|---|---|---|---|---|---|---|---|---|---|---|---|---|---|---|---|
| HAND1::TCF3 | CGCCTGTAAT | 1 | 195 | 205 | -805 | -795 | 5,42 | 0,01 | 23,6 | >0.1 | 3,06 | 4,46 | 0 | 1,67 | 0 | 4,37 | 4,62 |
| HAND1::TCF3 | GGTCAGGAGA | 1 | 242 | 252 | -758 | -748 | 2,92 | 0,01 | 20,66 | >0.1 | 2,87 | 4,52 | 0 | 1,4 | 0 | 4,33 | 4,62 |
| HAND1::TCF3 | AACCTGGGAG | 1 | 377 | 387 | -623 | -613 | 3,88 | 0,01 | 23,25 | >0.1 | 2,56 | 4,73 | 0 | 1,06 | 0 | 6,4 | 4,62 |
| HAND1::TCF3 | TATTTGGAAG | 1 | 523 | 533 | -477 | -467 | 5,02 | 0,01 | 21,88 | >0.1 | 3,06 | 5,02 | 0 | 0,98 | 0 | 3,19 | 4,62 |
| HAND1::TCF3 | TTTCTGGGAA | 1 | 557 | 567 | -443 | -433 | 6,16 | 0,01 | 20,08 | >0.1 | 3,22 | 5,1 | 0 | 0,99 | 0 | 0 | 4,62 |
| HAND1::TCF3 | GATCTTGAAA | 1 | 570 | 580 | -430 | -420 | 3,19 | 0,01 | 17,22 | >0.1 | 3,29 | 5,13 | 0 | 1 | 0 | 0 | 4,62 |
| HAND1::TCF3 | TGTCTGAAAA | 1 | 614 | 624 | -386 | -376 | 3,48 | 0,01 | 19,44 | >0.1 | 3,57 | 5,25 | 0 | 1,03 | 0 | 1,5 | 4,62 |
| HAND1::TCF3 | AATCTGTTAC | 1 | 843 | 853 | -157 | -147 | 3,58 | 0,01 | 25,04 | >0.1 | 5,61 | 6,25 | 0 | 1,8 | 0 | 3,19 | 4,62 |
| HAND1::TCF3 | TTCCTGGCTC | 1 | 1060 | 1070 | 60 | 70 | 6,28 | 0,01 | 27,88 | >0.1 | 5,61 | 7,35 | 0 | 1,86 | 0 | 2,16 | 4,62 |
| HAND1::TCF3 | AATCTGCTGA | -1 | 888 | 898 | -112 | -102 | 3,46 | 0,01 | 25,24 | >0.1 | 5,61 | 6,6 | 0 | 1,89 | 0 | 3,07 | 4,62 |
| HAND1::TCF3 | ATCCTGGAGA | -1 | 863 | 873 | -137 | -127 | 3,01 | 0,01 | 24,2 | >0.1 | 5,61 | 6,39 | 0 | 1,9 | 0 | 2,67 | 4,62 |
| HAND1::TCF3 | CTCCTGTCAC | -1 | 766 | 776 | -234 | -224 | 2,74 | 0,01 | 26,02 | >0.1 | 5,61 | 5,8 | 0 | 1,34 | 0 | 5,92 | 4,62 |
| HAND1::TCF3 | CTTTTGGTTA | -1 | 732 | 742 | -268 | -258 | 2,74 | 0,01 | 25,76 | >0.1 | 5,61 | 5,65 | 0 | 1,23 | 0 | 5,92 | 4,62 |
| HAND1::TCF3 | GGTCTAGAAC | -1 | 659 | 669 | -341 | -331 | 4,45 | 0,01 | 23,52 | >0.1 | 4 | 5,38 | 0 | 1,08 | 0 | 3,99 | 4,62 |
| HAND1::TCF3 | TGTATGGTTT | -1 | 621 | 631 | -379 | -369 | 3,46 | 0,01 | 19,45 | >0.1 | 3,63 | 5,27 | 0 | 1,04 | 0 | 1,44 | 4,62 |
| HAND1::TCF3 | GGTCTGTCGC | -1 | 434 | 444 | -566 | -556 | 6,53 | 0,01 | 25,52 | >0.1 | 2,73 | 4,83 | 0 | 1,01 | 0 | 5,8 | 4,62 |
| HAND1::TCF3 | AGGCTGGAGC | -1 | 422 | 432 | -578 | -568 | 4,68 | 0,01 | 23,37 | >0.1 | 2,69 | 4,81 | 0 | 1,02 | 0 | 5,56 | 4,62 |
| HAND1::TCF3 | GCTCTGCCTC | -1 | 384 | 394 | -616 | -606 | 3,37 | 0,01 | 22,38 | >0.1 | 2,58 | 4,74 | 0 | 1,05 | 0 | 6,02 | 4,62 |
| HAND1::TCF3 | CTCCTGCCTC | -1 | 358 | 368 | -642 | -632 | 3,99 | 0,01 | 23,17 | >0.1 | 2,52 | 4,7 | 0 | 1,09 | 0 | 6,26 | 4,62 |
| HAND1::TCF3 | TAGCTGGGAC | -1 | 337 | 347 | -663 | -653 | 3,79 | 0,01 | 22,69 | >0.1 | 2,58 | 4,66 | 0 | 1,12 | 0 | 5,92 | 4,62 |
| HAND1::TCF3 | GATCTGCCCA | -1 | 228 | 238 | -772 | -762 | 3,18 | 0,01 | 21,81 | >0.1 | 2,92 | 4,5 | 0 | 1,46 | 0 | 5,12 | 4,62 |
| HAND1::TCF3 | GATCTGCCTT | -1 | 90 | 100 | -910 | -900 | 9,99 | 0,001 | 24,52 | >0.1 | 3,69 | 4,34 | 0 | 1,88 | 0 | 0 | 4,62 |
| RORA | AAAGAAGTCA | 1 | 30 | 40 | -970 | -960 | 5,2 | 0,01 | 15,83 | >0.1 | 4,48 | 4,28 | 0 | 1,87 | 0 | 0 | 0 |
| RORA | AGGCAGATCA | 1 | 91 | 101 | -909 | -899 | -1,13 | 0,01 | 9,63 | >0.1 | 3,69 | 4,34 | 0 | 1,88 | 0 | 0,85 | 0 |
| RORA | AAAATGGCCA | 1 | 142 | 152 | -858 | -848 | 0,07 | 0,01 | 11,34 | >0.1 | 3,33 | 4,4 | 0 | 1,9 | 0 | 1,65 | 0 |
| RORA | CAGTGGCTCA | 1 | 185 | 195 | -815 | -805 | 0,6 | 0,01 | 13,75 | >0.1 | 3,1 | 4,45 | 0 | 1,75 | 0 | 3,86 | 0 |
| RORA | CACAAGGTCA | 1 | 237 | 247 | -763 | -753 | 11,59 | 0,0001 | 24,94 | >0.1 | 2,89 | 4,51 | 0 | 1,42 | 0 | 4,53 | 0 |
| RORA | AACACGGTGA | 1 | 269 | 279 | -731 | -721 | 1,55 | 0,01 | 15,27 | >0.1 | 2,78 | 4,56 | 0 | 1,3 | 0 | 5,09 | 0 |
| RORA | GACAAGGGCA | 1 | 505 | 515 | -495 | -485 | 5,98 | 0,001 | 20,6 | >0.1 | 2,98 | 4,98 | 0 | 0,98 | 0 | 5,68 | 0 |
| RORA | GGCAAAGTCA | 1 | 511 | 521 | -489 | -479 | -0,04 | 0,01 | 11,8 | >0.1 | 3 | 4,99 | 0 | 0,98 | 0 | 2,86 | 0 |
| RORA | ATAAATGTGA | 1 | 585 | 595 | -415 | -405 | -0,44 | 0,01 | 9,11 | >0.1 | 3,38 | 5,17 | 0 | 1,01 | 0 | 0 | 0 |
| RORA | TCTTTGGTCA | 1 | 669 | 679 | -331 | -321 | 1,44 | 0,01 | 15,78 | >0.1 | 4,13 | 5,42 | 0 | 1,1 | 0 | 3,7 | 0 |
| RORA | TTCTAGGAAA | 1 | 718 | 728 | -282 | -272 | -0,69 | 0,01 | 16,69 | >0.1 | 5,61 | 5,59 | 0 | 1,2 | 0 | 4,99 | 0 |
| RORA | TATTGGCTCA | 1 | 881 | 891 | -119 | -109 | 1,8 | 0,01 | 18,91 | >0.1 | 5,61 | 6,54 | 0 | 1,89 | 0 | 3,07 | 0 |
| RORA | TTCCTGCTCA | 1 | 935 | 945 | -65 | -55 | -1,15 | 0,01 | 14,4 | >0.1 | 5,61 | 7,14 | 0 | 1,88 | 0 | 0,93 | 0 |
| RORA | CCCAAGTTCA | 1 | 953 | 963 | -47 | -37 | 1,67 | 0,01 | 18,31 | >0.1 | 5,61 | 7,46 | 0 | 1,88 | 0 | 1,7 | 0 |
| RORA | TCTTGGCTCA | 1 | 1030 | 1040 | 30 | 40 | -0,64 | 0,01 | 16,4 | >0.1 | 5,61 | 8,24 | 0 | 1,86 | 0 | 1,34 | 0 |
| RORA | GACGATGTCA | 1 | 1045 | 1055 | 45 | 55 | 4,38 | 0,01 | 20,2 | >0.1 | 5,61 | 7,7 | 0 | 1,86 | 0 | 0,65 | 0 |
| RORA | GTCAAGCTCT | 1 | 1051 | 1061 | 51 | 61 | -0,77 | 0,01 | 15,71 | >0.1 | 5,61 | 7,55 | 0 | 1,86 | 0 | 1,46 | 0 |
| RORA | GAACAGGCCA | 1 | 1115 | 1125 | 115 | 125 | 2,66 | 0,01 | 22,48 | >0.1 | 5,61 | 6,6 | 0 | 1,85 | 3,22 | 2,54 | 0 |
| RORA | TCAAAGTTCA | 1 | 1171 | 1181 | 171 | 181 | 2,29 | 0,01 | 20,52 | >0.1 | 5,61 | 6,15 | 0 | 1,66 | 1,79 | 3,02 | 0 |
| RORA | GAACAGGTCT | -1 | 1159 | 1169 | 159 | 169 | 2,16 | 0,01 | 18,81 | >0.1 | 4,38 | 6,23 | 0 | 1,77 | 1,79 | 2,48 | 0 |
| RORA | TTCGTGGTTA | -1 | 1144 | 1154 | 144 | 154 | -0,41 | 0,01 | 18,77 | >0.1 | 5,61 | 6,34 | 0 | 1,84 | 3,22 | 2,16 | 0 |
| RORA | AAAAATGTCT | -1 | 1125 | 1135 | 125 | 135 | -1,17 | 0,01 | 18,78 | >0.1 | 5,61 | 6,5 | 0 | 1,84 | 3,22 | 2,77 | 0 |
| RORA | GCCAAGATCA | -1 | 1027 | 1037 | 27 | 37 | 2,55 | 0,01 | 19,72 | >0.1 | 5,61 | 8,39 | 0 | 1,86 | 0 | 1,31 | 0 |
| RORA | AAGAAGAACA | -1 | 906 | 916 | -94 | -84 | -1,23 | 0,01 | 14,24 | >0.1 | 5,61 | 6,77 | 0 | 1,89 | 0 | 1,21 | 0 |
| RORA | ATAAAGTTCA | -1 | 872 | 882 | -128 | -118 | 6,53 | 0,001 | 23,36 | >0.1 | 5,61 | 6,46 | 0 | 1,9 | 0 | 2,86 | 0 |
| RORA | TTTATGGCCA | -1 | 756 | 766 | -244 | -234 | -1,23 | 0,01 | 17,76 | >0.1 | 5,61 | 5,75 | 0 | 1,31 | 0 | 6,32 | 0 |
| RORA | AAAGAGGTCT | -1 | 664 | 674 | -336 | -326 | 5,7 | 0,001 | 20,08 | >0.1 | 4,06 | 5,4 | 0 | 1,09 | 0 | 3,83 | 0 |
| RORA | CTAGGGATCA | -1 | 650 | 660 | -350 | -340 | 0,92 | 0,01 | 15,23 | >0.1 | 3,9 | 5,35 | 0 | 1,07 | 0 | 3,99 | 0 |
| RORA | ATGAAGATAA | -1 | 641 | 651 | -359 | -349 | -1 | 0,01 | 13,02 | >0.1 | 3,8 | 5,33 | 0 | 1,06 | 0 | 3,83 | 0 |
| RORA | TAATAGGTTA | -1 | 597 | 607 | -403 | -393 | 4,09 | 0,01 | 15,17 | >0.1 | 3,45 | 5,2 | 0 | 1,01 | 0 | 1,41 | 0 |
| RORA | TTCAAGATCA | -1 | 569 | 579 | -431 | -421 | 6,93 | 0,001 | 16,34 | >0.1 | 3,28 | 5,13 | 0 | 1 | 0 | 0 | 0 |
| RORA | TACTTGGGCA | -1 | 542 | 552 | -458 | -448 | -0,17 | 0,01 | 9,02 | >0.1 | 3,14 | 5,06 | 0 | 0,99 | 0 | 0 | 0 |
| RORA | GTCTCGGTCT | -1 | 439 | 449 | -561 | -551 | -0,66 | 0,01 | 13,97 | >0.1 | 2,74 | 4,84 | 0 | 1 | 0 | 6,05 | 0 |
| RORA | CCCAGGTTCA | -1 | 375 | 385 | -625 | -615 | -0,17 | 0,01 | 14,57 | >0.1 | 2,56 | 4,72 | 0 | 1,07 | 0 | 6,4 | 0 |
| RORA | CCACAGGTCA | -1 | 108 | 118 | -892 | -882 | 5,82 | 0,001 | 16,36 | >0.1 | 3,55 | 4,36 | 0 | 1,89 | 0 | 0,75 | 0 |
| RORA | AAAAAGCTCA | -1 | 69 | 79 | -931 | -921 | 6,31 | 0,001 | 16,4 | >0.1 | 3,9 | 4,32 | 0 | 1,88 | 0 | 0 | 0 |
| POU6F2 | TCATAATTTT | 1 | 130 | 140 | -870 | -860 | 0,67 | 0,01 | 11,92 | >0.1 | 3,4 | 4,38 | 0 | 1,89 | 0 | 1,57 | 0 |

| | | | | | | | | | | | | | | | | | |
|---|---|---|---|---|---|---|---|---|---|---|---|---|---|---|---|---|---|
| POU6F2 | TGGAAATTAA | 1 | 152 | 162 | -848 | -838 | 2,37 | 0,01 | 13,6 | >0.1 | 3,27 | 4,41 | 0 | 1,9 | 0 | 1,65 | 0 |
| POU6F2 | AACCTATTAA | 1 | 598 | 608 | -402 | -392 | 2,37 | 0,01 | 13,46 | >0.1 | 3,46 | 5,2 | 0 | 1,01 | 0 | 1,41 | 0 |
| POU6F2 | CCAACATTAT | 1 | 635 | 645 | -365 | -355 | 2,22 | 0,01 | 13,93 | >0.1 | 3,75 | 5,31 | 0 | 1,05 | 0 | 1,61 | 0 |
| POU6F2 | TCTTCATGAT | 1 | 644 | 654 | -356 | -346 | 2,22 | 0,01 | 16,15 | >0.1 | 3,83 | 5,34 | 0 | 1,06 | 0 | 3,7 | 0 |
| POU6F2 | AAAAAATTAT | 1 | 828 | 838 | -172 | -162 | 1,71 | 0,01 | 18,56 | >0.1 | 5,61 | 6,14 | 0 | 1,67 | 0 | 3,43 | 0 |
| POU6F2 | ATCTCATGAG | 1 | 982 | 992 | -18 | -8 | 3,37 | 0,01 | 20,81 | >0.1 | 5,61 | 8,43 | 0 | 1,87 | 0 | 1,53 | 0 |
| POU6F2 | AAGTCATTCA | 1 | 1010 | 1020 | 10 | 20 | 2,41 | 0,01 | 20,29 | >0.1 | 5,61 | 8,99 | 0 | 1,86 | 0 | 1,42 | 0 |
| POU6F2 | GGCTCCTTCT | 1 | 1065 | 1075 | 65 | 75 | 2,27 | 0,01 | 21,06 | >0.1 | 5,61 | 7,26 | 0 | 1,85 | 1,79 | 2,27 | 0 |
| POU6F2 | TTGGAATTAT | 1 | 1189 | 1199 | 189 | 199 | 0 | 0,01 | 18,63 | >0.1 | 5,61 | 6,03 | 0 | 1,52 | 1,79 | 3,67 | 0 |
| POU6F2 | ACCTCCTCAT | -1 | 987 | 997 | -13 | -3 | 1,44 | 0,01 | 19,16 | >0.1 | 5,61 | 8,64 | 0 | 1,87 | 0 | 1,61 | 0 |
| POU6F2 | TCCTCATGAG | -1 | 984 | 994 | -16 | -6 | 4,26 | 0,01 | 21,84 | >0.1 | 5,61 | 8,57 | 0 | 1,87 | 0 | 1,53 | 0 |
| POU6F2 | GAATAATTTT | -1 | 830 | 840 | -170 | -160 | 0,04 | 0,01 | 16,78 | >0.1 | 5,61 | 6,16 | 0 | 1,69 | 0 | 3,28 | 0 |
| POU6F2 | GGATCATGAA | -1 | 646 | 656 | -354 | -344 | 3,54 | 0,01 | 17,63 | >0.1 | 3,85 | 5,34 | 0 | 1,06 | 0 | 3,83 | 0 |
| POU6F2 | ATTTCATTAA | -1 | 604 | 614 | -396 | -386 | 7,15 | 0,001 | 18,3 | >0.1 | 3,5 | 5,22 | 0 | 1,02 | 0 | 1,41 | 0 |
| POU6F2 | ATCTCCTGAC | -1 | 243 | 253 | -757 | -747 | -0,18 | 0,01 | 12,93 | >0.1 | 2,87 | 4,52 | 0 | 1,39 | 0 | 4,33 | 0 |
| POU6F2 | AGCTCATTTC | -1 | 65 | 75 | -935 | -925 | 8,14 | 0,001 | 18,27 | >0.1 | 3,94 | 4,31 | 0 | 1,88 | 0 | 0 | 0 |
| POU6F2 | GCCTCCTTCT | -1 | 16 | 26 | -984 | -974 | 1,43 | 0,01 | 13,17 | >0.1 | 5,61 | 4,27 | 0 | 1,87 | 0 | 0 | 0 |
| POU6F1 | GGAAATGAGC | 1 | 64 | 74 | -936 | -926 | 2,26 | 0,01 | 12,4 | >0.1 | 3,95 | 4,31 | 0 | 1,88 | 0 | 0 | 0 |
| POU6F1 | GGAAATTAAA | 1 | 153 | 163 | -847 | -837 | 3,22 | 0,01 | 14,44 | >0.1 | 3,27 | 4,41 | 0 | 1,9 | 0 | 1,65 | 0 |
| POU6F1 | ATTAATGAAA | 1 | 603 | 613 | -397 | -387 | 6,46 | 0,01 | 17,6 | >0.1 | 3,49 | 5,22 | 0 | 1,02 | 0 | 1,41 | 0 |
| POU6F1 | CTTCATGATC | 1 | 645 | 655 | -355 | -345 | 2,89 | 0,01 | 16,97 | >0.1 | 3,84 | 5,34 | 0 | 1,06 | 0 | 3,83 | 0 |
| POU6F1 | ATTAAAGAAT | 1 | 793 | 803 | -207 | -197 | 2,93 | 0,01 | 21,25 | >0.1 | 5,61 | 5,93 | 0 | 1,46 | 0 | 5,32 | 0 |
| POU6F1 | AAAAATTATT | 1 | 829 | 839 | -171 | -161 | 4,83 | 0,01 | 21,55 | >0.1 | 5,61 | 6,15 | 0 | 1,68 | 0 | 3,28 | 0 |
| POU6F1 | TCTCATGAGG | 1 | 983 | 993 | -17 | -7 | 3,35 | 0,01 | 20,9 | >0.1 | 5,61 | 8,5 | 0 | 1,87 | 0 | 1,57 | 0 |
| POU6F1 | CCTCATGAGA | -1 | 983 | 993 | -17 | -7 | 3,35 | 0,01 | 20,9 | >0.1 | 5,61 | 8,5 | 0 | 1,87 | 0 | 1,57 | 0 |
| POU6F1 | AATAATTTTT | -1 | 829 | 839 | -171 | -161 | 4,83 | 0,01 | 21,55 | >0.1 | 5,61 | 6,15 | 0 | 1,68 | 0 | 3,28 | 0 |
| POU6F1 | ATTCTTTAAT | -1 | 793 | 803 | -207 | -197 | 2,93 | 0,01 | 21,25 | >0.1 | 5,61 | 5,93 | 0 | 1,46 | 0 | 5,32 | 0 |
| POU6F1 | GATCATGAAG | -1 | 645 | 655 | -355 | -345 | 2,89 | 0,01 | 16,97 | >0.1 | 3,84 | 5,34 | 0 | 1,06 | 0 | 3,83 | 0 |
| POU6F1 | TTTCATTAAT | -1 | 603 | 613 | -397 | -387 | 6,46 | 0,01 | 17,6 | >0.1 | 3,49 | 5,22 | 0 | 1,02 | 0 | 1,41 | 0 |
| POU6F1 | TTTAATTTCC | -1 | 153 | 163 | -847 | -837 | 3,22 | 0,01 | 14,44 | >0.1 | 3,27 | 4,41 | 0 | 1,9 | 0 | 1,65 | 0 |
| POU6F1 | GCTCATTTCC | -1 | 64 | 74 | -936 | -926 | 2,26 | 0,01 | 12,4 | >0.1 | 3,95 | 4,31 | 0 | 1,88 | 0 | 0 | 0 |
| PPARA::RXRA | AGGTAAGTGAAGGAGAAG | 1 | 3 | 21 | -997 | -979 | 1,69 | 0,01 | 15,12 | >0.1 | 5,51 | 4,27 | 0 | 1,86 | 1,79 | 0 | 0 |
| PPARA::RXRA | AAGTGAAGGAGAAGGAGG | 1 | 7 | 25 | -993 | -975 | 1,72 | 0,01 | 15,16 | >0.1 | 5,51 | 4,27 | 0 | 1,87 | 1,79 | 0 | 0 |
| PPARA::RXRA | AAGTTTTGCAAAGGCAGA | 1 | 80 | 98 | -920 | -902 | 1,38 | 0,01 | 11,35 | >0.1 | 3,75 | 4,34 | 0 | 1,88 | 0 | 0 | 0 |
| PPARA::RXRA | TTGGGAGGCCGAGTTGGG | 1 | 214 | 232 | -786 | -768 | 3,63 | 0,01 | 18,27 | >0.1 | 2,96 | 4,5 | 0 | 1,54 | 0 | 5,63 | 0 |
| PPARA::RXRA | CACAAGGTCAGGAGATAG | 1 | 237 | 255 | -763 | -745 | 2,8 | 0,01 | 15,95 | >0.1 | 2,88 | 4,53 | 0 | 1,42 | 0 | 4,33 | 0 |
| PPARA::RXRA | CTGGGAGGCAGAGCTTGC | 1 | 380 | 398 | -620 | -602 | 5,32 | 0,01 | 19,73 | >0.1 | 2,58 | 4,75 | 0 | 1,06 | 0 | 6,02 | 0 |
| PPARA::RXRA | AGACAAGGGCAAAGTCAT | 1 | 504 | 522 | -496 | -478 | 2,44 | 0,01 | 15,79 | >0.1 | 2,99 | 4,99 | 0 | 0,99 | 0 | 4,37 | 0 |
| PPARA::RXRA | GACAAGGGCAAAGTCATG | 1 | 505 | 523 | -495 | -477 | 2,93 | 0,01 | 14,66 | >0.1 | 3 | 4,99 | 0 | 0,99 | 0 | 2,75 | 0 |
| PPARA::RXRA | ATGTATTTGGAAGGGAAA | 1 | 520 | 538 | -480 | -462 | 3,57 | 0,01 | 15,93 | >0.1 | 3,06 | 5,03 | 0 | 0,99 | 0 | 3,28 | 0 |
| PPARA::RXRA | TGGAAGGGAAAATGTTGC | 1 | 527 | 545 | -473 | -455 | 2,53 | 0,01 | 13,27 | >0.1 | 3,09 | 5,04 | 0 | 0,99 | 0 | 1,61 | 0 |
| PPARA::RXRA | ATTTGGCCATAAAGTGAC | 1 | 753 | 771 | -247 | -229 | 1,94 | 0,01 | 20,65 | >0.1 | 5,51 | 5,77 | 0 | 1,33 | 0 | 6,1 | 0 |
| PPARA::RXRA | AGTGACAGGAGAGGTAAG | 1 | 765 | 783 | -235 | -217 | 2,82 | 0,01 | 21,22 | >0.1 | 5,51 | 5,83 | 0 | 1,37 | 0 | 5,69 | 0 |
| PPARA::RXRA | CAGGAGAGGTAAGGTTCT | 1 | 770 | 788 | -230 | -212 | 3,58 | 0,01 | 22,44 | >0.1 | 5,51 | 5,86 | 0 | 1,39 | 0 | 6,1 | 0 |
| PPARA::RXRA | AGGAGAGGTAAGGTTCTC | 1 | 771 | 789 | -229 | -211 | 2,3 | 0,01 | 21,39 | >0.1 | 5,51 | 5,86 | 0 | 1,4 | 0 | 6,32 | 0 |
| PPARA::RXRA | CCCAAGTTCAAAGGCTGA | 1 | 953 | 971 | -47 | -29 | 3,06 | 0,01 | 19,83 | >0.1 | 5,51 | 7,65 | 0 | 1,88 | 0 | 1,74 | 0 |
| PPARA::RXRA | GAGGAGGTTTTAGTCTAG | 1 | 989 | 1007 | -11 | 7 | 2,48 | 0,01 | 20,35 | >0.1 | 5,51 | 8,96 | 0 | 1,87 | 0 | 1,53 | 0 |
| PPARA::RXRA | TAGTCTAGGGAAAGTCAT | 1 | 999 | 1017 | -1 | 17 | 2,8 | 0,01 | 20,76 | >0.1 | 5,51 | 9,12 | 0 | 1,87 | 0 | 1,46 | 0 |
| PPARA::RXRA | TCTTGGCTCACAGGGGAC | 1 | 1030 | 1048 | 30 | 48 | 3,72 | 0,01 | 20,63 | >0.1 | 5,51 | 8,24 | 0 | 1,86 | 0 | 1,31 | 0 |
| PPARA::RXRA | CATTGAGGAACAGGCCAA | 1 | 1108 | 1126 | 108 | 126 | 4,7 | 0,01 | 24,49 | >0.1 | 5,51 | 6,67 | 0 | 1,85 | 3,22 | 2,54 | 0 |
| PPARA::RXRA | TGTTCTATCAAAGTTCAC | 1 | 1164 | 1182 | 164 | 182 | 5,39 | 0,01 | 23,44 | >0.1 | 5,51 | 6,19 | 0 | 1,72 | 1,79 | 2,84 | 0 |
| PPARA::RXRA | ATACACGGAAAATGTTAT | -1 | 689 | 707 | -311 | -293 | 1,98 | 0,01 | 17,22 | >0.1 | 5,51 | 5,51 | 0 | 1,15 | 0 | 3,07 | 0 |
| PPARA::RXRA | ATTTCATTAATAGGTTAT | -1 | 596 | 614 | -404 | -386 | 3,48 | 0,01 | 14,56 | >0.1 | 3,47 | 5,22 | 0 | 1,02 | 0 | 1,36 | 0 |
| PPARA::RXRA | CTCCACTCCACAGGTCAA | -1 | 107 | 125 | -893 | -875 | 2,17 | 0,01 | 12,74 | >0.1 | 3,53 | 4,37 | 0 | 1,89 | 0 | 0,79 | 0 |
| HOXD8 | ATTTTAAAAAATGGCCA | 1 | 135 | 152 | -865 | -848 | 3,21 | 0,01 | 14,5 | >0.1 | 3,35 | 4,4 | 0 | 1,9 | 0 | 1,65 | 0 |
| HOXD8 | ATGGAAATTAAAACTGA | 1 | 151 | 168 | -849 | -832 | 2 | 0,01 | 13,22 | >0.1 | 3,26 | 4,42 | 0 | 1,9 | 0 | 1,65 | 0 |
| HOXD8 | CTGATCAGAAATGGCTG | 1 | 164 | 181 | -836 | -819 | 3,25 | 0,01 | 15,18 | >0.1 | 3,19 | 4,43 | 0 | 1,9 | 0 | 2,4 | 0 |
| HOXD8 | GTCTCTATTAAAAATAC | 1 | 285 | 302 | -715 | -698 | 2,04 | 0,01 | 15,68 | >0.1 | 2,71 | 4,59 | 0 | 1,25 | 0 | 5,09 | 0 |
| HOXD8 | ATAAAAATAAATGTGAG | 1 | 579 | 596 | -421 | -404 | 1,73 | 0,01 | 11,27 | >0.1 | 3,36 | 5,17 | 0 | 1,01 | 0 | 0 | 0 |

| | | | | | | | | | | | | | | | | | |
|---|---|---|---|---|---|---|---|---|---|---|---|---|---|---|---|---|---|
| HOXD8 | TAACCTATTAATGAAAT | 1 | 597 | 614 | -403 | -386 | 8,88 | 0,001 | 19,96 | >0.1 | 3,48 | 5,22 | 0 | 1,02 | 0 | 1,36 | 0 |
| HOXD8 | TGTATCTTTAACAGCTT | 1 | 702 | 719 | -298 | -281 | 5,52 | 0,01 | 22,76 | >0.1 | 5,52 | 5,56 | 0 | 1,18 | 0 | 4,99 | 0 |
| HOXD8 | TAGGAAAATATTAACCA | 1 | 721 | 738 | -279 | -262 | 4,52 | 0,01 | 22,06 | >0.1 | 5,52 | 5,63 | 0 | 1,22 | 0 | 5,17 | 0 |
| HOXD8 | GAAAATATTAACCAAAA | 1 | 724 | 741 | -276 | -259 | 2,35 | 0,01 | 20,06 | >0.1 | 5,52 | 5,64 | 0 | 1,23 | 0 | 5,32 | 0 |
| HOXD8 | TAAAAAATTATTCTAAA | 1 | 827 | 844 | -173 | -156 | 5,02 | 0,01 | 21,73 | >0.1 | 5,52 | 6,19 | 0 | 1,72 | 0 | 3,28 | 0 |
| HOXD8 | GAACTTTATATTGGCTC | 1 | 873 | 890 | -127 | -110 | 2,95 | 0,01 | 19,85 | >0.1 | 5,52 | 6,53 | 0 | 1,9 | 0 | 2,95 | 0 |
| HOXD8 | CAAGACATTTTTGGACA | 1 | 1123 | 1140 | 123 | 140 | 2,19 | 0,01 | 22,05 | >0.1 | 5,52 | 6,52 | 0 | 1,85 | 3,22 | 2,75 | 0 |
| HOXD8 | TTGTCCAAAAATGTCTT | -1 | 1124 | 1141 | 124 | 141 | 2,55 | 0,01 | 22,47 | >0.1 | 5,52 | 6,51 | 0 | 1,84 | 3,22 | 2,82 | 0 |
| HOXD8 | TAATTTTTTAAAGTGAA | -1 | 820 | 837 | -180 | -163 | 3,14 | 0,01 | 20,16 | >0.1 | 5,52 | 6,14 | 0 | 1,67 | 0 | 3,7 | 0 |
| HOXD8 | GAATCAAATAAGAATAC | -1 | 806 | 823 | -194 | -177 | 4,42 | 0,01 | 20,84 | >0.1 | 5,52 | 6,05 | 0 | 1,57 | 0 | 3,28 | 0 |
| HOXD8 | TTATTCTTTAATCCTAG | -1 | 788 | 805 | -212 | -195 | 1,91 | 0,01 | 20,16 | >0.1 | 5,52 | 5,94 | 0 | 1,47 | 0 | 5,32 | 0 |
| HOXD8 | CTGTCACTTTATGGCCA | -1 | 756 | 773 | -244 | -227 | 2,74 | 0,01 | 21,48 | >0.1 | 5,52 | 5,78 | 0 | 1,33 | 0 | 6,1 | 0 |
| HOXD8 | AATTTCATTAATAGGTT | -1 | 598 | 615 | -402 | -385 | 9,84 | 0,001 | 20,98 | >0.1 | 3,48 | 5,22 | 0 | 1,03 | 0 | 1,41 | 0 |
| HOXD8 | TGTATTTTTAATAGAGA | -1 | 286 | 303 | -714 | -697 | 4,71 | 0,01 | 18,35 | >0.1 | 2,71 | 4,59 | 0 | 1,25 | 0 | 5,09 | 0 |
| HOXD8 | TTTTAAAATTATGACTA | -1 | 127 | 144 | -873 | -856 | 1,77 | 0,01 | 13,02 | >0.1 | 3,4 | 4,39 | 0 | 1,89 | 0 | 1,57 | 0 |
| HOXD9 | GTGAGAAATA | 1 | 54 | 64 | -946 | -936 | 3,58 | 0,01 | 13,83 | >0.1 | 4,07 | 4,3 | 0 | 1,87 | 0 | 0 | 0 |
| HOXD9 | GTCATAATTT | 1 | 129 | 139 | -871 | -861 | 3,83 | 0,01 | 15,12 | >0.1 | 3,41 | 4,38 | 0 | 1,89 | 0 | 1,61 | 0 |
| HOXD9 | GAAATTAAAA | 1 | 154 | 164 | -846 | -836 | 8,84 | 0,001 | 20,06 | >0.1 | 3,26 | 4,41 | 0 | 1,9 | 0 | 1,65 | 0 |
| HOXD9 | CTATTAAAAA | 1 | 289 | 299 | -711 | -701 | 4,54 | 0,01 | 18,16 | >0.1 | 2,71 | 4,59 | 0 | 1,24 | 0 | 5,09 | 0 |
| HOXD9 | CTCAAAAAAA | 1 | 454 | 464 | -546 | -536 | 5,05 | 0,01 | 19,23 | >0.1 | 2,79 | 4,87 | 0 | 1 | 0 | 5,52 | 0 |
| HOXD9 | TCAAAAAAAA | 1 | 455 | 465 | -545 | -535 | 4,12 | 0,01 | 18,3 | >0.1 | 2,79 | 4,87 | 0 | 1 | 0 | 5,52 | 0 |
| HOXD9 | TCCATAAAGA | 1 | 497 | 507 | -503 | -493 | 5,39 | 0,01 | 20,71 | >0.1 | 2,95 | 4,96 | 0 | 0,98 | 0 | 6,44 | 0 |
| HOXD9 | GAAATAAAAA | 1 | 576 | 586 | -424 | -414 | 8,9 | 0,001 | 18,37 | >0.1 | 3,32 | 5,14 | 0 | 1 | 0 | 0 | 0 |
| HOXD9 | AAAATAAATG | 1 | 582 | 592 | -418 | -408 | 3,28 | 0,01 | 12,8 | >0.1 | 3,36 | 5,16 | 0 | 1 | 0 | 0 | 0 |
| HOXD9 | ACCATACAAA | 1 | 623 | 633 | -377 | -367 | 4,23 | 0,01 | 15,63 | >0.1 | 3,64 | 5,27 | 0 | 1,04 | 0 | 1,44 | 0 |
| HOXD9 | GCCATAAAGT | 1 | 758 | 768 | -242 | -232 | 9,94 | 0,001 | 28,72 | >0.1 | 5,61 | 5,76 | 0 | 1,31 | 0 | 6,1 | 0 |
| HOXD9 | GTGGTTAAAC | -1 | 1141 | 1151 | 141 | 151 | 6,71 | 0,01 | 27,16 | >0.1 | 5,61 | 6,37 | 0 | 1,84 | 3,22 | 3,4 | 0 |
| HOXD9 | GCAGTTACAG | -1 | 1086 | 1096 | 86 | 96 | 4,82 | 0,01 | 23,22 | >0.1 | 5,61 | 6,93 | 0 | 1,85 | 1,79 | 2,21 | 0 |
| HOXD9 | GCAGGAAAGA | -1 | 932 | 942 | -68 | -58 | 4,72 | 0,01 | 20,23 | >0.1 | 5,61 | 7,09 | 0 | 1,88 | 0 | 0,93 | 0 |
| HOXD9 | ACCAAAAAAA | -1 | 923 | 933 | -77 | -67 | 6,32 | 0,01 | 21,87 | >0.1 | 5,61 | 6,97 | 0 | 1,88 | 0 | 1,09 | 0 |
| HOXD9 | CCAAAAAAAG | -1 | 922 | 932 | -78 | -68 | 5,27 | 0,01 | 20,8 | >0.1 | 5,61 | 6,95 | 0 | 1,88 | 0 | 1,09 | 0 |
| HOXD9 | ACAGTAAACA | -1 | 897 | 907 | -103 | -93 | 6,5 | 0,01 | 23,54 | >0.1 | 5,61 | 6,68 | 0 | 1,89 | 0 | 2,86 | 0 |
| HOXD9 | GCCAATATAA | -1 | 878 | 888 | -122 | -112 | 3,37 | 0,01 | 20,46 | >0.1 | 5,61 | 6,51 | 0 | 1,9 | 0 | 3,07 | 0 |
| HOXD9 | CCAATATAAA | -1 | 877 | 887 | -123 | -113 | 7,27 | 0,01 | 24,23 | >0.1 | 5,61 | 6,5 | 0 | 1,9 | 0 | 2,95 | 0 |
| HOXD9 | TTCATTAATA | -1 | 602 | 612 | -398 | -388 | 3,97 | 0,01 | 15,1 | >0.1 | 3,49 | 5,21 | 0 | 1,02 | 0 | 1,41 | 0 |
| CREB5 | AAAGAAGTCAGC | 1 | 30 | 42 | -970 | -958 | -9,13 | 0,01 | 1,48 | >0.1 | 4,46 | 4,28 | 0 | 1,87 | 0 | 0 | 0 |
| CREB5 | AATGGCGTGAAC | 1 | 368 | 380 | -632 | -620 | -12,62 | 0,01 | 1,86 | >0.1 | 2,54 | 4,72 | 0 | 1,08 | 0 | 6,15 | 0 |
| CREB5 | GATCGCGCCACT | 1 | 408 | 420 | -592 | -580 | -10,24 | 0,01 | 3,75 | >0.1 | 2,65 | 4,79 | 0 | 1,03 | 0 | 5,52 | 0 |
| CREB5 | AATTCCATAAAG | 1 | 494 | 506 | -506 | -494 | -13,14 | 0,01 | 2,36 | >0.1 | 2,94 | 4,96 | 0 | 0,99 | 0 | 6,62 | 0 |
| CREB5 | TGTGAGATAACC | 1 | 590 | 602 | -410 | -398 | -6,59 | 0,01 | 3,02 | >0.1 | 3,41 | 5,19 | 0 | 1,01 | 0 | 0 | 0 |
| CREB5 | TCTTTGGTCACT | 1 | 669 | 681 | -331 | -319 | -11,03 | 0,01 | 3,33 | >0.1 | 4,14 | 5,42 | 0 | 1,1 | 0 | 3,7 | 0 |
| CREB5 | AATTATAACATT | 1 | 685 | 697 | -315 | -303 | -3,83 | 0,01 | 10,14 | >0.1 | 4,42 | 5,48 | 0 | 1,13 | 0 | 2,95 | 0 |
| CREB5 | AATAACGTATTC | 1 | 800 | 812 | -200 | -188 | -3,01 | 0,01 | 13,49 | >0.1 | 5,58 | 5,98 | 0 | 1,51 | 0 | 3,43 | 0 |
| CREB5 | TGTTACATATCT | 1 | 847 | 859 | -153 | -141 | -6,52 | 0,01 | 10,16 | >0.1 | 5,58 | 6,29 | 0 | 1,85 | 0 | 2,95 | 0 |
| CREB5 | GACGATGTCAAG | 1 | 1045 | 1057 | 45 | 57 | -2,86 | 0,001 | 13,75 | >0.1 | 5,58 | 7,7 | 0 | 1,86 | 0 | 1,46 | 0 |
| CREB5 | GTTGCTGTAACT | 1 | 1082 | 1094 | 82 | 94 | -10,43 | 0,01 | 8 | >0.1 | 5,58 | 6,99 | 0 | 1,85 | 1,79 | 2,21 | 0 |
| CREB5 | AGTTACAGCAAC | -1 | 1082 | 1094 | 82 | 94 | -12 | 0,01 | 6,43 | >0.1 | 5,58 | 6,99 | 0 | 1,85 | 1,79 | 2,21 | 0 |
| CREB5 | CTTGACATCGTC | -1 | 1045 | 1057 | 45 | 57 | -6,68 | 0,01 | 9,93 | >0.1 | 5,58 | 7,7 | 0 | 1,86 | 0 | 1,46 | 0 |
| CREB5 | AGATATGTAACA | -1 | 847 | 859 | -153 | -141 | -6,24 | 0,01 | 10,44 | >0.1 | 5,58 | 6,29 | 0 | 1,85 | 0 | 2,95 | 0 |
| CREB5 | GAATACGTTATT | -1 | 800 | 812 | -200 | -188 | -3,72 | 0,01 | 12,78 | >0.1 | 5,58 | 5,98 | 0 | 1,51 | 0 | 3,43 | 0 |
| CREB5 | CCTTACCTCTCC | -1 | 772 | 784 | -228 | -216 | -11,78 | 0,01 | 7,11 | >0.1 | 5,58 | 5,83 | 0 | 1,37 | 0 | 6,1 | 0 |
| CREB5 | AATGTTATAATT | -1 | 685 | 697 | -315 | -303 | -4,67 | 0,01 | 9,3 | >0.1 | 4,42 | 5,48 | 0 | 1,13 | 0 | 2,95 | 0 |
| CREB5 | GGTTATCTCACA | -1 | 590 | 602 | -410 | -398 | -8,36 | 0,01 | 1,25 | >0.1 | 3,41 | 5,19 | 0 | 1,01 | 0 | 0 | 0 |
| CREB5 | AAATACATGACT | -1 | 516 | 528 | -484 | -472 | -12,54 | 0,01 | -0,57 | >0.1 | 3,03 | 5 | 0 | 0,98 | 0 | 2,95 | 0 |
| CREB5 | GTTCACGCCATT | -1 | 368 | 380 | -632 | -620 | -6,18 | 0,01 | 8,3 | >0.1 | 2,54 | 4,72 | 0 | 1,08 | 0 | 6,15 | 0 |
| CREB5 | GCTGACTTCTTT | -1 | 30 | 42 | -970 | -958 | -10,63 | 0,01 | -0,02 | >0.1 | 4,46 | 4,28 | 0 | 1,87 | 0 | 0 | 0 |
| ZIC1 | GTTGACCTGTGGAG | 1 | 106 | 120 | -894 | -880 | -10,87 | 0,01 | -0,33 | >0.1 | 3,55 | 4,36 | 0 | 1,89 | 0 | 0,75 | 0 |
| ZIC1 | TCACGCCTGTAATC | 1 | 192 | 206 | -808 | -794 | -13,89 | 0,01 | -0,41 | >0.1 | 3,06 | 4,46 | 0 | 1,69 | 0 | 4,26 | 0 |
| ZIC1 | GAAACCCCGTCTCT | 1 | 277 | 291 | -723 | -709 | -10,27 | 0,01 | 3,52 | >0.1 | 2,75 | 4,58 | 0 | 1,27 | 0 | 5,19 | 0 |

| | | | | | | | | | | | | | | | | | |
|---|---|---|---|---|---|---|---|---|---|---|---|---|---|---|---|---|---|
| ZIC1 | GGGCACCTGTAGTC | 1 | 326 | 340 | -674 | -660 | -1,86 | 0,001 | 12,1 | >0.1 | 2,6 | 4,65 | 0 | 1,15 | 0 | 5,56 | 0 |
| ZIC1 | TCCCAGCTACTCGG | 1 | 338 | 352 | -662 | -648 | -13,76 | 0,01 | 1 | >0.1 | 2,57 | 4,67 | 0 | 1,12 | 0 | 6,4 | 0 |
| ZIC1 | CAGAGCTTGCAGTG | 1 | 388 | 402 | -612 | -598 | -13,92 | 0,01 | 0,5 | >0.1 | 2,6 | 4,75 | 0 | 1,05 | 0 | 6,02 | 0 |
| ZIC1 | ATCGCGCCACTGTG | 1 | 409 | 423 | -591 | -577 | -12,9 | 0,01 | 1,34 | >0.1 | 2,66 | 4,79 | 0 | 1,03 | 0 | 5,76 | 0 |
| ZIC1 | TCTTTCCTGCTCAG | 1 | 932 | 946 | -68 | -54 | -13,24 | 0,01 | 2,25 | >0.1 | 5,56 | 7,15 | 0 | 1,88 | 0 | 0,9 | 0 |
| ZIC1 | TCAGCCTTGTTGCT | 1 | 1074 | 1088 | 74 | 88 | -11,1 | 0,01 | 7,48 | >0.1 | 5,56 | 7,11 | 0 | 1,85 | 1,79 | 2,27 | 0 |
| ZIC1 | TAACTGCTGCTCAG | 1 | 1089 | 1103 | 89 | 103 | -13,48 | 0,01 | 4,95 | >0.1 | 5,56 | 6,89 | 0 | 1,85 | 1,79 | 2,34 | 0 |
| ZIC1 | CATCGTCCCCTGTG | -1 | 1038 | 1052 | 38 | 52 | -12,92 | 0,01 | 3,78 | >0.1 | 5,56 | 7,92 | 0 | 1,86 | 0 | 1,36 | 0 |
| ZIC1 | CGTCCCCTGTGAGC | -1 | 1035 | 1049 | 35 | 49 | -9,64 | 0,01 | 7,14 | >0.1 | 5,56 | 8,03 | 0 | 1,86 | 0 | 1,34 | 0 |
| ZIC1 | AACCTCCTCATGAG | -1 | 984 | 998 | -16 | -2 | -7,8 | 0,01 | 9,86 | >0.1 | 5,56 | 8,66 | 0 | 1,87 | 0 | 1,57 | 0 |
| ZIC1 | AACAATCTGCTGAG | -1 | 887 | 901 | -113 | -99 | -3,65 | 0,01 | 13,49 | >0.1 | 5,56 | 6,62 | 0 | 1,89 | 0 | 3,07 | 0 |
| ZIC1 | TTCATCCTGGAGAG | -1 | 862 | 876 | -138 | -124 | -10,02 | 0,01 | 6,6 | >0.1 | 5,56 | 6,41 | 0 | 1,9 | 0 | 2,75 | 0 |
| ZIC1 | CCTCTCCTGTCACT | -1 | 765 | 779 | -235 | -221 | -13,97 | 0,01 | 4,67 | >0.1 | 5,56 | 5,81 | 0 | 1,36 | 0 | 5,92 | 0 |
| ZIC1 | TGCCCACCACCACG | -1 | 317 | 331 | -683 | -669 | -12,21 | 0,01 | 1,78 | >0.1 | 2,63 | 4,64 | 0 | 1,17 | 0 | 5,56 | 0 |
| ZIC1 | ACCACCACGCCCAG | -1 | 312 | 326 | -688 | -674 | -10,39 | 0,01 | 4,19 | >0.1 | 2,64 | 4,63 | 0 | 1,18 | 0 | 6,13 | 0 |
| ZIC1 | CACCACGCCCAGCT | -1 | 310 | 324 | -690 | -676 | -10,29 | 0,01 | 4,2 | >0.1 | 2,65 | 4,63 | 0 | 1,18 | 0 | 6,03 | 0 |
| ZIC1 | CACGCCCAGCTAAG | -1 | 307 | 321 | -693 | -679 | -5,43 | 0,01 | 9,29 | >0.1 | 2,65 | 4,62 | 0 | 1,19 | 0 | 6,26 | 0 |
| ZIC1 | GGCCTCCCAAAGTG | -1 | 210 | 224 | -790 | -776 | 0,42 | 0,001 | 14,05 | >0.1 | 2,99 | 4,48 | 0 | 1,57 | 0 | 4,59 | 0 |
| ZIC1 | CGTGAGCCACTGTG | -1 | 183 | 197 | -817 | -803 | -13,9 | 0,01 | -0,72 | >0.1 | 3,1 | 4,45 | 0 | 1,76 | 0 | 3,86 | 0 |
| ZIC1 | TCCACTCCACAGGT | -1 | 110 | 124 | -890 | -876 | -11,53 | 0,01 | -0,99 | >0.1 | 3,52 | 4,37 | 0 | 1,89 | 0 | 0,77 | 0 |
| ZIC1 | TGCCTCCTTCTCCT | -1 | 13 | 27 | -987 | -973 | -10,21 | 0,01 | 1,48 | >0.1 | 5,56 | 4,27 | 0 | 1,87 | 0 | 0 | 0 |
| CREB3 | CAAAGAAGTCAGCC | 1 | 29 | 43 | -971 | -957 | -5,58 | 0,01 | 5,03 | >0.1 | 4,46 | 4,28 | 0 | 1,87 | 0 | 0 | 0 |
| CREB3 | CAGCCACAGAACCA | 1 | 38 | 52 | -962 | -948 | -4,8 | 0,01 | 5,65 | >0.1 | 4,28 | 4,29 | 0 | 1,87 | 0 | 0 | 0 |
| CREB3 | GAGAGTAGTCATAA | 1 | 122 | 136 | -878 | -864 | -7,77 | 0,01 | 3,64 | >0.1 | 3,44 | 4,38 | 0 | 1,89 | 0 | 1,7 | 0 |
| CREB3 | TGGGCAGATCACAA | 1 | 228 | 242 | -772 | -758 | -7,76 | 0,01 | 6,14 | >0.1 | 2,92 | 4,51 | 0 | 1,47 | 0 | 5,01 | 0 |
| CREB3 | TCACAAGGTCAGGA | 1 | 236 | 250 | -764 | -750 | -6,28 | 0,01 | 7,08 | >0.1 | 2,89 | 4,52 | 0 | 1,43 | 0 | 4,53 | 0 |
| CREB3 | AAACCCCGTCTCTA | 1 | 278 | 292 | -722 | -708 | -4,96 | 0,01 | 8,72 | >0.1 | 2,74 | 4,58 | 0 | 1,27 | 0 | 5,09 | 0 |
| CREB3 | GTGAGCCGAGATCG | 1 | 399 | 413 | -601 | -587 | -6,75 | 0,01 | 7,46 | >0.1 | 2,63 | 4,77 | 0 | 1,04 | 0 | 5,76 | 0 |
| CREB3 | GGGCAAAGTCATGT | 1 | 510 | 524 | -490 | -476 | 0,09 | 0,001 | 12,03 | >0.1 | 3,01 | 5 | 0 | 0,99 | 0 | 2,95 | 0 |
| CREB3 | TCTAGACCTCTTTG | 1 | 661 | 675 | -339 | -325 | -7,57 | 0,01 | 6,97 | >0.1 | 4,05 | 5,4 | 0 | 1,09 | 0 | 3,99 | 0 |
| CREB3 | AGGAGAGGTAAGGT | 1 | 771 | 785 | -229 | -215 | -7,45 | 0,01 | 11,43 | >0.1 | 5,56 | 5,84 | 0 | 1,38 | 0 | 6,1 | 0 |
| CREB3 | GAATAACGTATTCT | 1 | 799 | 813 | -201 | -187 | -7,6 | 0,01 | 8,88 | >0.1 | 5,56 | 5,99 | 0 | 1,51 | 0 | 3,43 | 0 |
| CREB3 | AGGGAAAGTCATTC | 1 | 1005 | 1019 | 5 | 19 | -6,44 | 0,01 | 11,57 | >0.1 | 5,56 | 9,12 | 0 | 1,87 | 0 | 1,46 | 0 |
| CREB3 | GGACGATGTCAAGC | 1 | 1044 | 1058 | 44 | 58 | -6,79 | 0,01 | 9,82 | >0.1 | 5,56 | 7,73 | 0 | 1,86 | 0 | 1,46 | 0 |
| CREB3 | AGGCCAAGACATTT | 1 | 1119 | 1133 | 119 | 133 | -3,32 | 0,01 | 16,54 | >0.1 | 5,56 | 6,56 | 0 | 1,85 | 3,22 | 2,67 | 0 |
| CREB3 | TAACCACGAAGCCG | 1 | 1144 | 1158 | 144 | 158 | -4,91 | 0,01 | 14,21 | >0.1 | 5,56 | 6,34 | 0 | 1,84 | 3,22 | 2,16 | 0 |
| CREB3 | AGAACAGGTCTTCG | -1 | 1156 | 1170 | 156 | 170 | -2,56 | 0,01 | 14,06 | >0.1 | 4,38 | 6,25 | 0 | 1,8 | 1,79 | 2,4 | 0 |
| CREB3 | CGGCTTCGTGGTTA | -1 | 1144 | 1158 | 144 | 158 | -4,91 | 0,01 | 14,21 | >0.1 | 5,56 | 6,34 | 0 | 1,84 | 3,22 | 2,16 | 0 |
| CREB3 | GTTAAACTTGTCCA | -1 | 1134 | 1148 | 134 | 148 | -6,1 | 0,01 | 14,04 | >0.1 | 5,56 | 6,42 | 0 | 1,84 | 3,22 | 3,09 | 0 |
| CREB3 | GCTTGACATCGTCC | -1 | 1044 | 1058 | 44 | 58 | -1,84 | 0,01 | 14,77 | >0.1 | 5,56 | 7,73 | 0 | 1,86 | 0 | 1,46 | 0 |
| CREB3 | TGACATCGTCCCCT | -1 | 1041 | 1055 | 41 | 55 | -7,51 | 0,01 | 8,35 | >0.1 | 5,56 | 7,82 | 0 | 1,86 | 0 | 0,62 | 0 |
| CREB3 | CAGATATGTAACAG | -1 | 846 | 860 | -154 | -140 | -6,75 | 0,01 | 9,92 | >0.1 | 5,56 | 6,3 | 0 | 1,86 | 0 | 2,95 | 0 |
| CREB3 | AGAATACGTTATTC | -1 | 799 | 813 | -201 | -187 | -1,92 | 0,01 | 14,56 | >0.1 | 5,56 | 5,99 | 0 | 1,51 | 0 | 3,43 | 0 |
| CREB3 | CTGTCACTTTATGG | -1 | 759 | 773 | -241 | -227 | 0,67 | 0,001 | 19,44 | >0.1 | 5,56 | 5,78 | 0 | 1,33 | 0 | 6,1 | 0 |
| CREB3 | TGGCCAAATCAAAA | -1 | 748 | 762 | -252 | -238 | -6,19 | 0,01 | 12,5 | >0.1 | 5,56 | 5,73 | 0 | 1,29 | 0 | 6,1 | 0 |
| CREB3 | GGGCAACATTTTCC | -1 | 533 | 547 | -467 | -453 | -5,45 | 0,01 | 5,31 | >0.1 | 3,11 | 5,05 | 0 | 0,99 | 0 | 1,61 | 0 |
| CREB3 | GGTTCACGCCATTC | -1 | 367 | 381 | -633 | -619 | 3,63 | 0,001 | 18,12 | >0.1 | 2,54 | 4,72 | 0 | 1,08 | 0 | 6,15 | 0 |
| CREB3 | ACTACAGGTGCCCA | -1 | 325 | 339 | -675 | -661 | -5,93 | 0,01 | 8,04 | >0.1 | 2,6 | 4,65 | 0 | 1,15 | 0 | 5,56 | 0 |
| CREB3 | CCACCACGCCCAGC | -1 | 311 | 325 | -689 | -675 | -6,93 | 0,01 | 7,65 | >0.1 | 2,64 | 4,63 | 0 | 1,18 | 0 | 6,13 | 0 |
| CREB3 | TCTCCACTCCACAG | -1 | 112 | 126 | -888 | -874 | -1,1 | 0,01 | 9,41 | >0.1 | 3,51 | 4,37 | 0 | 1,89 | 0 | 0,75 | 0 |
| CREB1 | AAAGAAGTCAGC | 1 | 30 | 42 | -970 | -958 | 4,62 | 0,01 | 15,23 | >0.1 | 4,46 | 4,28 | 0 | 1,87 | 0 | 0 | 0 |
| CREB1 | GCTCACGCCTGT | 1 | 190 | 202 | -810 | -798 | 0,72 | 0,01 | 14,09 | >0.1 | 3,08 | 4,46 | 0 | 1,71 | 0 | 4,13 | 0 |
| CREB1 | CACAAGGTCAGG | 1 | 237 | 249 | -763 | -751 | 0,46 | 0,01 | 13,81 | >0.1 | 2,89 | 4,52 | 0 | 1,42 | 0 | 4,53 | 0 |
| CREB1 | AATGGCGTGAAC | 1 | 368 | 380 | -632 | -620 | 3,94 | 0,01 | 18,42 | >0.1 | 2,54 | 4,72 | 0 | 1,08 | 0 | 6,15 | 0 |
| CREB1 | GATCGCGCCACT | 1 | 408 | 420 | -592 | -580 | 2,28 | 0,01 | 16,27 | >0.1 | 2,65 | 4,79 | 0 | 1,03 | 0 | 5,52 | 0 |
| CREB1 | TGTGAGATAACC | 1 | 590 | 602 | -410 | -398 | 4,22 | 0,01 | 13,83 | >0.1 | 3,41 | 5,19 | 0 | 1,01 | 0 | 0 | 0 |
| CREB1 | AGTGACAGGAGA | 1 | 765 | 777 | -235 | -223 | 0,72 | 0,01 | 19,37 | >0.1 | 5,58 | 5,8 | 0 | 1,35 | 0 | 5,92 | 0 |
| CREB1 | GGAGAGGTAAGG | 1 | 772 | 784 | -228 | -216 | 2,03 | 0,01 | 20,92 | >0.1 | 5,58 | 5,83 | 0 | 1,37 | 0 | 6,1 | 0 |
| CREB1 | TCTGTCCTCTCC | 1 | 856 | 868 | -144 | -132 | 1,57 | 0,01 | 18,27 | >0.1 | 5,58 | 6,35 | 0 | 1,9 | 0 | 2,86 | 0 |

| | | | | | | | | | | | | | | | | | |
|---|---|---|---|---|---|---|---|---|---|---|---|---|---|---|---|---|---|
| CREB1 | GTGGATGTGATC | 1 | 1020 | 1032 | 20 | 32 | 0,89 | 0,01 | 18,36 | >0.1 | 5,58 | 8,72 | 0 | 1,86 | 0 | 1,31 | 0 |
| CREB1 | GACGATGTCAAG | 1 | 1045 | 1057 | 45 | 57 | 5,83 | 0,01 | 22,44 | >0.1 | 5,58 | 7,7 | 0 | 1,86 | 0 | 1,46 | 0 |
| CREB1 | CTTGACATCGTC | -1 | 1045 | 1057 | 45 | 57 | 5,83 | 0,01 | 22,44 | >0.1 | 5,58 | 7,7 | 0 | 1,86 | 0 | 1,46 | 0 |
| CREB1 | GATCACATCCAC | -1 | 1020 | 1032 | 20 | 32 | 0,89 | 0,01 | 18,36 | >0.1 | 5,58 | 8,72 | 0 | 1,86 | 0 | 1,31 | 0 |
| CREB1 | GGAGAGGACAGA | -1 | 856 | 868 | -144 | -132 | 1,57 | 0,01 | 18,27 | >0.1 | 5,58 | 6,35 | 0 | 1,9 | 0 | 2,86 | 0 |
| CREB1 | CCTTACCTCTCC | -1 | 772 | 784 | -228 | -216 | 2,03 | 0,01 | 20,92 | >0.1 | 5,58 | 5,83 | 0 | 1,37 | 0 | 6,1 | 0 |
| CREB1 | TCTCCTGTCACT | -1 | 765 | 777 | -235 | -223 | 0,72 | 0,01 | 19,37 | >0.1 | 5,58 | 5,8 | 0 | 1,35 | 0 | 5,92 | 0 |
| CREB1 | GGTTATCTCACA | -1 | 590 | 602 | -410 | -398 | 4,22 | 0,01 | 13,83 | >0.1 | 3,41 | 5,19 | 0 | 1,01 | 0 | 0 | 0 |
| CREB1 | AGTGGCGCGATC | -1 | 408 | 420 | -592 | -580 | 2,28 | 0,01 | 16,27 | >0.1 | 2,65 | 4,79 | 0 | 1,03 | 0 | 5,52 | 0 |
| CREB1 | GTTCACGCCATT | -1 | 368 | 380 | -632 | -620 | 3,94 | 0,01 | 18,42 | >0.1 | 2,54 | 4,72 | 0 | 1,08 | 0 | 6,15 | 0 |
| CREB1 | CCTGACCTTGTG | -1 | 237 | 249 | -763 | -751 | 0,46 | 0,01 | 13,81 | >0.1 | 2,89 | 4,52 | 0 | 1,42 | 0 | 4,53 | 0 |
| CREB1 | ACAGGCGTGAGC | -1 | 190 | 202 | -810 | -798 | 0,72 | 0,01 | 14,09 | >0.1 | 3,08 | 4,46 | 0 | 1,71 | 0 | 4,13 | 0 |
| CREB1 | GCTGACTTCTTT | -1 | 30 | 42 | -970 | -958 | 4,62 | 0,01 | 15,23 | >0.1 | 4,46 | 4,28 | 0 | 1,87 | 0 | 0 | 0 |
| EVX2 | GGAAATGAGC | 1 | 64 | 74 | -936 | -926 | 4,07 | 0,01 | 14,21 | >0.1 | 3,95 | 4,31 | 0 | 1,88 | 0 | 0 | 0 |
| EVX2 | GCAGATCAGG | 1 | 93 | 103 | -907 | -897 | 3,58 | 0,01 | 14,28 | >0.1 | 3,67 | 4,34 | 0 | 1,88 | 0 | 0,81 | 0 |
| EVX2 | GGAAATTAAA | 1 | 153 | 163 | -847 | -837 | 6,14 | 0,01 | 17,36 | >0.1 | 3,27 | 4,41 | 0 | 1,9 | 0 | 1,65 | 0 |
| EVX2 | ACTGATCAGA | 1 | 163 | 173 | -837 | -827 | 4,37 | 0,01 | 16,31 | >0.1 | 3,21 | 4,42 | 0 | 1,9 | 0 | 2,4 | 0 |
| EVX2 | CCTGACTAAC | 1 | 262 | 272 | -738 | -728 | 2,8 | 0,01 | 16 | >0.1 | 2,8 | 4,55 | 0 | 1,32 | 0 | 4,53 | 0 |
| EVX2 | ACTGATCAGA | 1 | 478 | 488 | -522 | -512 | 4,37 | 0,01 | 18,51 | >0.1 | 2,88 | 4,92 | 0 | 0,99 | 0 | 5,36 | 0 |
| EVX2 | CCCAAGTAGA | 1 | 544 | 554 | -456 | -446 | 4,09 | 0,01 | 13,29 | >0.1 | 3,15 | 5,06 | 0 | 0,99 | 0 | 0 | 0 |
| EVX2 | CAACATTATC | 1 | 636 | 646 | -364 | -354 | 5,44 | 0,01 | 17,22 | >0.1 | 3,76 | 5,31 | 0 | 1,05 | 0 | 1,66 | 0 |
| EVX2 | TAAAATTATA | 1 | 682 | 692 | -318 | -308 | 3,2 | 0,01 | 17,18 | >0.1 | 4,34 | 5,46 | 0 | 1,12 | 0 | 3,07 | 0 |
| EVX2 | AAAAATTATT | 1 | 829 | 839 | -171 | -161 | 3,78 | 0,01 | 20,5 | >0.1 | 5,61 | 6,15 | 0 | 1,68 | 0 | 3,28 | 0 |
| EVX2 | GCTGATAAGA | 1 | 966 | 976 | -34 | -24 | 3,68 | 0,01 | 20,64 | >0.1 | 5,61 | 7,78 | 0 | 1,87 | 0 | 1,7 | 0 |
| EVX2 | TCTCATGAGG | 1 | 983 | 993 | -17 | -7 | 3,94 | 0,01 | 21,49 | >0.1 | 5,61 | 8,5 | 0 | 1,87 | 0 | 1,57 | 0 |
| EVX2 | CACCATTGAG | 1 | 1105 | 1115 | 105 | 115 | 3,41 | 0,01 | 23,31 | >0.1 | 5,61 | 6,7 | 0 | 1,85 | 3,22 | 2,52 | 0 |
| EVX2 | CGTGGTTAAA | -1 | 1142 | 1152 | 142 | 152 | 3,84 | 0,01 | 24,24 | >0.1 | 5,61 | 6,36 | 0 | 1,84 | 3,22 | 3,37 | 0 |
| EVX2 | AGCAGTTACA | -1 | 1087 | 1097 | 87 | 97 | 3,18 | 0,01 | 21,57 | >0.1 | 5,61 | 6,92 | 0 | 1,85 | 1,79 | 2,21 | 0 |
| EVX2 | CCTCATGAGA | -1 | 983 | 993 | -17 | -7 | 4,27 | 0,01 | 21,82 | >0.1 | 5,61 | 8,5 | 0 | 1,87 | 0 | 1,57 | 0 |
| EVX2 | GATCATGAAG | -1 | 645 | 655 | -355 | -345 | 3,56 | 0,01 | 17,64 | >0.1 | 3,84 | 5,34 | 0 | 1,06 | 0 | 3,83 | 0 |
| EVX2 | TTTCATTAAT | -1 | 603 | 613 | -397 | -387 | 4,49 | 0,01 | 15,63 | >0.1 | 3,49 | 5,22 | 0 | 1,02 | 0 | 1,41 | 0 |
| EVX2 | TCTGATCAGT | -1 | 478 | 488 | -522 | -512 | 3,91 | 0,01 | 18,05 | >0.1 | 2,88 | 4,92 | 0 | 0,99 | 0 | 5,36 | 0 |
| EVX2 | CCCGAGTAGC | -1 | 343 | 353 | -657 | -647 | 4,95 | 0,01 | 19,82 | >0.1 | 2,56 | 4,67 | 0 | 1,11 | 0 | 6,52 | 0 |
| EVX2 | TCTGATCAGT | -1 | 163 | 173 | -837 | -827 | 3,91 | 0,01 | 15,85 | >0.1 | 3,21 | 4,42 | 0 | 1,9 | 0 | 2,4 | 0 |
| EVX2 | TAAAATTATG | -1 | 131 | 141 | -869 | -859 | 4,1 | 0,01 | 15,35 | >0.1 | 3,4 | 4,38 | 0 | 1,89 | 0 | 1,57 | 0 |
| EVX2 | GCTCATTTCC | -1 | 64 | 74 | -936 | -926 | 5,23 | 0,01 | 15,37 | >0.1 | 3,95 | 4,31 | 0 | 1,88 | 0 | 0 | 0 |
| NR2C2 | AAGGAGAAGGAGGCA | 1 | 12 | 27 | -988 | -973 | -0,05 | 0,01 | 11,63 | >0.1 | 5,54 | 4,27 | 0 | 1,87 | 0 | 0 | 0 |
| NR2C2 | CAGTGTGAGAAATAG | 1 | 50 | 65 | -950 | -935 | -2,78 | 0,01 | 7,49 | >0.1 | 4,09 | 4,3 | 0 | 1,87 | 0 | 0 | 0 |
| NR2C2 | GTGGAGTGGAGAGTA | 1 | 114 | 129 | -886 | -871 | -3,81 | 0,01 | 6,69 | >0.1 | 3,49 | 4,37 | 0 | 1,89 | 0 | 0,75 | 0 |
| NR2C2 | CACTTTGGGAGGCCG | 1 | 210 | 225 | -790 | -775 | -0,42 | 0,01 | 13,21 | >0.1 | 2,98 | 4,49 | 0 | 1,57 | 0 | 4,59 | 0 |
| NR2C2 | CAAGGTCAGGAGATA | 1 | 239 | 254 | -761 | -746 | 0,23 | 0,01 | 13,37 | >0.1 | 2,87 | 4,52 | 0 | 1,41 | 0 | 4,33 | 0 |
| NR2C2 | GCGTGGTGGTGGGCA | 1 | 316 | 331 | -684 | -669 | -2,28 | 0,01 | 12,41 | >0.1 | 2,63 | 4,64 | 0 | 1,17 | 0 | 6,26 | 0 |
| NR2C2 | GGGAGGCAGAGCTTG | 1 | 382 | 397 | -618 | -603 | -4,64 | 0,01 | 9,76 | >0.1 | 2,58 | 4,75 | 0 | 1,06 | 0 | 6,02 | 0 |
| NR2C2 | CAAGGGCAAAGTCAT | 1 | 507 | 522 | -493 | -478 | 3,37 | 0,01 | 15,1 | >0.1 | 3 | 4,99 | 0 | 0,99 | 0 | 2,75 | 0 |
| NR2C2 | GTATTTGGAAGGGAA | 1 | 522 | 537 | -478 | -463 | -1,7 | 0,01 | 10,66 | >0.1 | 3,06 | 5,03 | 0 | 0,99 | 0 | 3,28 | 0 |
| NR2C2 | GAAGGGAAAATGTTG | 1 | 529 | 544 | -471 | -456 | -3,22 | 0,01 | 7,51 | >0.1 | 3,09 | 5,04 | 0 | 0,99 | 0 | 1,61 | 0 |
| NR2C2 | TGACAGGAGAGGTAA | 1 | 767 | 782 | -233 | -218 | 1,13 | 0,01 | 19,56 | >0.1 | 5,54 | 5,83 | 0 | 1,37 | 0 | 5,69 | 0 |
| NR2C2 | CAAGTTCAAAGGCTG | 1 | 955 | 970 | -45 | -30 | 9,23 | 0,001 | 26,01 | >0.1 | 5,54 | 7,62 | 0 | 1,88 | 0 | 1,74 | 0 |
| NR2C2 | TTGGCTCACAGGGGA | 1 | 1032 | 1047 | 32 | 47 | -2,61 | 0,01 | 14,25 | >0.1 | 5,54 | 8,15 | 0 | 1,86 | 0 | 1,31 | 0 |
| NR2C2 | TTCTATCAAAGTTCA | 1 | 1166 | 1181 | 166 | 181 | -0,63 | 0,01 | 17,43 | >0.1 | 5,54 | 6,18 | 0 | 1,7 | 1,79 | 2,84 | 0 |
| NR2C2 | AAGGCTGAGAAGGAG | -1 | 1067 | 1082 | 67 | 82 | -1,17 | 0,01 | 17,46 | >0.1 | 5,54 | 7,22 | 0 | 1,85 | 1,79 | 2,21 | 0 |
| NR2C2 | GAGCAGGAAAGACCA | -1 | 929 | 944 | -71 | -56 | -2,27 | 0,01 | 13,23 | >0.1 | 5,54 | 7,12 | 0 | 1,88 | 0 | 0,95 | 0 |
| NR2C2 | AGAGGTCTAGAACTA | -1 | 657 | 672 | -343 | -328 | -3,13 | 0,01 | 11,35 | >0.1 | 4 | 5,39 | 0 | 1,09 | 0 | 3,99 | 0 |
| NR2C2 | AAAGTGCTAGGATTA | -1 | 201 | 216 | -799 | -784 | -2,15 | 0,01 | 11,56 | >0.1 | 3,02 | 4,47 | 0 | 1,63 | 0 | 4,59 | 0 |
| FOXH1 | GGCAGATCACA | 1 | 230 | 241 | -770 | -759 | -11,19 | 0,01 | 2,7 | >0.1 | 2,91 | 4,51 | 0 | 1,45 | 0 | 5,01 | 0 |
| FOXH1 | TCTATTAAAAA | 1 | 288 | 299 | -712 | -701 | -10,68 | 0,01 | 2,95 | >0.1 | 2,71 | 4,59 | 0 | 1,24 | 0 | 5,09 | 0 |
| FOXH1 | AAAAATACAAA | 1 | 294 | 305 | -706 | -695 | -6,27 | 0,01 | 6,77 | >0.1 | 2,7 | 4,6 | 0 | 1,22 | 0 | 4,53 | 0 |
| FOXH1 | TGAATTCCATA | 1 | 492 | 503 | -508 | -497 | -6,02 | 0,01 | 9,11 | >0.1 | 2,93 | 4,95 | 0 | 0,98 | 0 | 6,27 | 0 |
| FOXH1 | TGAAATAAAAA | 1 | 575 | 586 | -425 | -414 | -6,45 | 0,01 | 3,02 | >0.1 | 3,32 | 5,14 | 0 | 1 | 0 | 0 | 0 |

| | | | | | | | | | | | | | | | | | |
|---|---|---|---|---|---|---|---|---|---|---|---|---|---|---|---|---|---|
| FOXH1 | AGGATTAAAGA | 1 | 790 | 801 | -210 | -199 | -9,52 | 0,01 | 8,77 | >0.1 | 5,6 | 5,92 | 0 | 1,45 | 0 | 5,32 | 0 |
| FOXH1 | CTCAGTCCACC | 1 | 1098 | 1109 | 98 | 109 | -3,05 | 0,01 | 16,75 | >0.1 | 5,6 | 6,78 | 0 | 1,85 | 3,22 | 2,35 | 0 |
| FOXH1 | CAAAGTTCACT | 1 | 1172 | 1183 | 172 | 183 | -0,84 | 0,01 | 17,31 | >0.1 | 5,6 | 6,14 | 0 | 1,65 | 1,79 | 2,97 | 0 |
| FOXH1 | GTGGTTAAACT | -1 | 1140 | 1151 | 140 | 151 | -11,07 | 0,01 | 9,26 | >0.1 | 5,6 | 6,37 | 0 | 1,84 | 3,22 | 3,29 | 0 |
| FOXH1 | TCACATCCACT | -1 | 1019 | 1030 | 19 | 30 | -3,52 | 0,01 | 13,99 | >0.1 | 5,6 | 8,74 | 0 | 1,86 | 0 | 1,31 | 0 |
| FOXH1 | AGAAGAACACA | -1 | 904 | 915 | -96 | -85 | -9,38 | 0,01 | 6,07 | >0.1 | 5,6 | 6,76 | 0 | 1,89 | 0 | 1,21 | 0 |
| FOXH1 | GTGAATCAAAT | -1 | 814 | 825 | -186 | -175 | -6,17 | 0,01 | 10,5 | >0.1 | 5,6 | 6,06 | 0 | 1,58 | 0 | 3,43 | 0 |
| FOXH1 | CCAAATCAAAA | -1 | 748 | 759 | -252 | -241 | -4,1 | 0,01 | 14,41 | >0.1 | 5,6 | 5,72 | 0 | 1,28 | 0 | 5,92 | 0 |
| FOXH1 | TGGAATTCATA | -1 | 490 | 501 | -510 | -499 | -6,7 | 0,01 | 8,42 | >0.1 | 2,92 | 4,95 | 0 | 0,98 | 0 | 6,27 | 0 |
| FOXH1 | TCCACTCCACA | -1 | 113 | 124 | -887 | -876 | 1,58 | 0,001 | 12,11 | >0.1 | 3,51 | 4,36 | 0 | 1,89 | 0 | 0,77 | 0 |
| PPARG::RXRA | GTAAGTGAAGGAGAA | 1 | 5 | 20 | -995 | -980 | 4,2 | 0,01 | 17,67 | >0.1 | 5,54 | 4,26 | 0 | 1,86 | 1,79 | 0 | 0 |
| PPARG::RXRA | GTGAAGGAGAAGGAG | 1 | 9 | 24 | -991 | -976 | 2,3 | 0,01 | 15,77 | >0.1 | 5,54 | 4,27 | 0 | 1,87 | 1,79 | 0 | 0 |
| PPARG::RXRA | GAAGGAGAAGGAGGC | 1 | 11 | 26 | -989 | -974 | 3,7 | 0,01 | 15,38 | >0.1 | 5,54 | 4,27 | 0 | 1,87 | 0 | 0 | 0 |
| PPARG::RXRA | AAGGAGAAGGAGGCA | 1 | 12 | 27 | -988 | -973 | 4,57 | 0,01 | 16,25 | >0.1 | 5,54 | 4,27 | 0 | 1,87 | 0 | 0 | 0 |
| PPARG::RXRA | GCTGGGCACAGTGGC | 1 | 177 | 192 | -823 | -808 | 6,64 | 0,01 | 19,12 | >0.1 | 3,13 | 4,44 | 0 | 1,82 | 0 | 3,09 | 0 |
| PPARG::RXRA | GGGAGGCCGAGTTGG | 1 | 216 | 231 | -784 | -769 | 3,51 | 0,01 | 18,13 | >0.1 | 2,96 | 4,49 | 0 | 1,53 | 0 | 5,63 | 0 |
| PPARG::RXRA | GCAGATCACAAGGTC | 1 | 231 | 246 | -769 | -754 | 3,05 | 0,01 | 16,82 | >0.1 | 2,9 | 4,51 | 0 | 1,45 | 0 | 4,9 | 0 |
| PPARG::RXRA | CAAGGTCAGGAGATA | 1 | 239 | 254 | -761 | -746 | 4,18 | 0,01 | 17,32 | >0.1 | 2,87 | 4,52 | 0 | 1,41 | 0 | 4,33 | 0 |
| PPARG::RXRA | GCAGGAGAATGGCGT | 1 | 361 | 376 | -639 | -624 | 3,66 | 0,01 | 18,6 | >0.1 | 2,53 | 4,71 | 0 | 1,09 | 0 | 6,61 | 0 |
| PPARG::RXRA | GGGAGGCAGAGCTTG | 1 | 382 | 397 | -618 | -603 | 4,18 | 0,01 | 18,58 | >0.1 | 2,58 | 4,75 | 0 | 1,06 | 0 | 6,02 | 0 |
| PPARG::RXRA | CTGGGCGACAGACCG | 1 | 430 | 445 | -570 | -555 | 5,8 | 0,01 | 20,04 | >0.1 | 2,72 | 4,83 | 0 | 1,01 | 0 | 5,67 | 0 |
| PPARG::RXRA | ATAAAGACAAGGGCA | 1 | 500 | 515 | -500 | -485 | 4,39 | 0,01 | 19 | >0.1 | 2,97 | 4,98 | 0 | 0,99 | 0 | 5,68 | 0 |
| PPARG::RXRA | TAAAGACAAGGGCAA | 1 | 501 | 516 | -499 | -484 | 4,52 | 0,01 | 19,14 | >0.1 | 2,97 | 4,98 | 0 | 0,99 | 0 | 5,68 | 0 |
| PPARG::RXRA | ACAAGGGCAAAGTCA | 1 | 506 | 521 | -494 | -479 | 4,66 | 0,01 | 18 | >0.1 | 2,99 | 4,99 | 0 | 0,99 | 0 | 4,37 | 0 |
| PPARG::RXRA | CAAGGGCAAAGTCAT | 1 | 507 | 522 | -493 | -478 | 7,33 | 0,01 | 19,06 | >0.1 | 3 | 4,99 | 0 | 0,99 | 0 | 2,75 | 0 |
| PPARG::RXRA | GAAGGGAAAATGTTG | 1 | 529 | 544 | -471 | -456 | 2,96 | 0,01 | 13,69 | >0.1 | 3,09 | 5,04 | 0 | 0,99 | 0 | 1,61 | 0 |
| PPARG::RXRA | GGAGAGGTAAGGTTC | 1 | 772 | 787 | -228 | -213 | 2,97 | 0,01 | 21,85 | >0.1 | 5,54 | 5,85 | 0 | 1,39 | 0 | 6,1 | 0 |
| PPARG::RXRA | GAGAGGTAAGGTTCT | 1 | 773 | 788 | -227 | -212 | 4,72 | 0,01 | 23,83 | >0.1 | 5,54 | 5,85 | 0 | 1,39 | 0 | 6,32 | 0 |
| PPARG::RXRA | CAAGTTCAAAGGCTG | 1 | 955 | 970 | -45 | -30 | 4,75 | 0,01 | 21,53 | >0.1 | 5,54 | 7,62 | 0 | 1,88 | 0 | 1,74 | 0 |
| PPARG::RXRA | TTGGCTCACAGGGGA | 1 | 1032 | 1047 | 32 | 47 | 7,57 | 0,01 | 24,43 | >0.1 | 5,54 | 8,15 | 0 | 1,86 | 0 | 1,31 | 0 |
| PPARG::RXRA | CAGGGGACGATGTCA | 1 | 1040 | 1055 | 40 | 55 | 4,31 | 0,01 | 20,19 | >0.1 | 5,54 | 7,85 | 0 | 1,86 | 0 | 0,62 | 0 |
| PPARG::RXRA | TTGAGGAACAGGCCA | 1 | 1110 | 1125 | 110 | 125 | 8,43 | 0,001 | 28,23 | >0.1 | 5,54 | 6,65 | 0 | 1,85 | 3,22 | 2,54 | 0 |
| PPARG::RXRA | TTCTATCAAAGTTCA | 1 | 1166 | 1181 | 166 | 181 | 5,07 | 0,01 | 23,13 | >0.1 | 5,54 | 6,18 | 0 | 1,7 | 1,79 | 2,84 | 0 |
| PPARG::RXRA | TGATAGAACAGGTCT | -1 | 1159 | 1174 | 159 | 174 | 2,39 | 0,01 | 19,15 | >0.1 | 4,43 | 6,23 | 0 | 1,77 | 1,79 | 2,54 | 0 |
| PPARG::RXRA | GCTGAGAAGGAGCCA | -1 | 1064 | 1079 | 64 | 79 | 3,92 | 0,01 | 22,66 | >0.1 | 5,54 | 7,28 | 0 | 1,85 | 1,79 | 2,27 | 0 |
| PPARG::RXRA | TGTGAGCCAAGATCA | -1 | 1027 | 1042 | 27 | 42 | 3,62 | 0,01 | 20,76 | >0.1 | 5,54 | 8,39 | 0 | 1,86 | 0 | 1,34 | 0 |
| PPARG::RXRA | CCAATATAAAGTTCA | -1 | 872 | 887 | -128 | -113 | 3,71 | 0,01 | 20,61 | >0.1 | 5,54 | 6,5 | 0 | 1,9 | 0 | 2,95 | 0 |
| PPARG::RXRA | CTGGAGCACAGTGGC | -1 | 414 | 429 | -586 | -571 | 7,56 | 0,01 | 21,25 | >0.1 | 2,67 | 4,8 | 0 | 1,03 | 0 | 5,19 | 0 |
| PPARG::RXRA | TGGGACTACAGGTGC | -1 | 328 | 343 | -672 | -657 | 2,6 | 0,01 | 16,45 | >0.1 | 2,59 | 4,66 | 0 | 1,14 | 0 | 5,46 | 0 |
| PPARG::RXRA | CCACTCCACAGGTCA | -1 | 108 | 123 | -892 | -877 | 3,19 | 0,01 | 13,74 | >0.1 | 3,53 | 4,36 | 0 | 1,89 | 0 | 0,77 | 0 |
| TCF7L1 | AAATAGGAAATG | 1 | 59 | 71 | -941 | -929 | -2,78 | 0,01 | 7,4 | >0.1 | 4 | 4,31 | 0 | 1,88 | 0 | 0 | 0 |
| TCF7L1 | GTTTTGCAAAGG | 1 | 82 | 94 | -918 | -906 | -3,88 | 0,01 | 6,09 | >0.1 | 3,76 | 4,33 | 0 | 1,88 | 0 | 0 | 0 |
| TCF7L1 | CAAGGGCAAAGT | 1 | 507 | 519 | -493 | -481 | 7,98 | 0,001 | 21,31 | >0.1 | 2,99 | 4,98 | 0 | 0,98 | 0 | 4,37 | 0 |
| TCF7L1 | GAAGGGAAAATG | 1 | 529 | 541 | -471 | -459 | -0,47 | 0,01 | 12,33 | >0.1 | 3,09 | 5,03 | 0 | 0,99 | 0 | 3,7 | 0 |
| TCF7L1 | TAGGATTAAAGA | 1 | 789 | 801 | -211 | -199 | -3,2 | 0,01 | 15,07 | >0.1 | 5,58 | 5,92 | 0 | 1,45 | 0 | 5,32 | 0 |
| TCF7L1 | AGAGAGAAAATC | 1 | 973 | 985 | -27 | -15 | 0,75 | 0,01 | 17,91 | >0.1 | 5,58 | 8,09 | 0 | 1,87 | 0 | 1,61 | 0 |
| TCF7L1 | CGATGTCAAGCT | 1 | 1047 | 1059 | 47 | 59 | -4,94 | 0,01 | 11,57 | >0.1 | 5,58 | 7,65 | 0 | 1,86 | 0 | 1,42 | 0 |
| TCF7L1 | TTCTATCAAAGT | 1 | 1166 | 1178 | 166 | 178 | -3,99 | 0,01 | 13,11 | >0.1 | 4,56 | 6,18 | 0 | 1,7 | 1,79 | 2,86 | 0 |
| TCF7L1 | ATAATTCCAAGA | -1 | 1187 | 1199 | 187 | 199 | -4,83 | 0,01 | 13,82 | >0.1 | 5,58 | 6,05 | 0 | 1,53 | 1,79 | 3,7 | 0 |
| TCF7L1 | TGTGAGCCAAGA | -1 | 1030 | 1042 | 30 | 42 | -0,06 | 0,01 | 16,93 | >0.1 | 5,58 | 8,24 | 0 | 1,86 | 0 | 1,31 | 0 |
| TCF7L1 | CAAGATCACATC | -1 | 1023 | 1035 | 23 | 35 | 0,62 | 0,01 | 17,98 | >0.1 | 5,58 | 8,67 | 0 | 1,86 | 0 | 1,25 | 0 |
| TCF7L1 | AAAGACCAAAAA | -1 | 925 | 937 | -75 | -63 | 3,93 | 0,001 | 19,48 | >0.1 | 5,58 | 7,02 | 0 | 1,88 | 0 | 1,06 | 0 |
| TCF7L1 | TATGGCCAAATC | -1 | 752 | 764 | -248 | -236 | -2,7 | 0,01 | 16,25 | >0.1 | 5,58 | 5,74 | 0 | 1,3 | 0 | 6,32 | 0 |
| TCF7L1 | AAAGATACACGG | -1 | 699 | 711 | -301 | -289 | 0,5 | 0,01 | 15,96 | >0.1 | 5,58 | 5,53 | 0 | 1,16 | 0 | 3,19 | 0 |
| TCF7L1 | ACACGGAAAATG | -1 | 693 | 705 | -307 | -295 | -2,72 | 0,01 | 12,58 | >0.1 | 5,58 | 5,5 | 0 | 1,15 | 0 | 3,07 | 0 |
| TCF7L1 | AGTGACCAAAGA | -1 | 669 | 681 | -331 | -319 | 7,22 | 0,001 | 21,58 | >0.1 | 4,14 | 5,42 | 0 | 1,1 | 0 | 3,7 | 0 |
| TCF7L1 | CAAGATCATATT | -1 | 565 | 577 | -435 | -423 | -0,37 | 0,01 | 9,01 | >0.1 | 3,27 | 5,12 | 0 | 1 | 0 | 0 | 0 |
| TCF7L1 | AAAGTGCTAGGA | -1 | 204 | 216 | -796 | -784 | -3,81 | 0,01 | 9,99 | >0.1 | 3,02 | 4,47 | 0 | 1,6 | 0 | 4,7 | 0 |
| TCF7L1 | TAATTTCCATGG | -1 | 149 | 161 | -851 | -839 | -4,71 | 0,01 | 6,53 | >0.1 | 3,28 | 4,41 | 0 | 1,9 | 0 | 1,65 | 0 |

| | | | | | | | | | | | | | | | | | |
|---|---|---|---|---|---|---|---|---|---|---|---|---|---|---|---|---|---|
| TCF7L1 | CCACTCCACAGG | -1 | 111 | 123 | -889 | -877 | -3,54 | 0,01 | 7 | >0.1 | 3,52 | 4,36 | 0 | 1,89 | 0 | 0,77 | 0 |
| TCF7L1 | ACAGGTCAACTC | -1 | 104 | 116 | -896 | -884 | -4,51 | 0,01 | 6,07 | >0.1 | 3,57 | 4,36 | 0 | 1,89 | 0 | 0,77 | 0 |
| TCF7L1 | AAACTTAAAAAA | -1 | 74 | 86 | -926 | -914 | -2,37 | 0,01 | 7,67 | >0.1 | 3,83 | 4,32 | 0 | 1,88 | 0 | 0 | 0 |
| TCF7L2 | GGCACTCAAAGAAG | 1 | 23 | 37 | -977 | -963 | 0,1 | 0,01 | 11,8 | >0.1 | 5,56 | 4,28 | 0 | 1,87 | 0 | 0 | 0 |
| TCF7L2 | GCAGATCAGGAGAG | 1 | 93 | 107 | -907 | -893 | -0,41 | 0,01 | 10,26 | >0.1 | 3,65 | 4,35 | 0 | 1,88 | 0 | 0,79 | 0 |
| TCF7L2 | ACTGATCAGAAATG | 1 | 163 | 177 | -837 | -823 | 0,85 | 0,01 | 12,78 | >0.1 | 3,2 | 4,43 | 0 | 1,9 | 0 | 2,4 | 0 |
| TCF7L2 | GCAGATCACAAGGT | 1 | 231 | 245 | -769 | -755 | 4,56 | 0,01 | 18,33 | >0.1 | 2,9 | 4,51 | 0 | 1,45 | 0 | 4,9 | 0 |
| TCF7L2 | AAAATACAAAAACT | 1 | 295 | 309 | -705 | -691 | 0,15 | 0,01 | 13,11 | >0.1 | 2,69 | 4,6 | 0 | 1,22 | 0 | 4,45 | 0 |
| TCF7L2 | CAGTCTCAAAAAAA | 1 | 450 | 464 | -550 | -536 | -0,93 | 0,01 | 13,1 | >0.1 | 2,78 | 4,87 | 0 | 1 | 0 | 5,38 | 0 |
| TCF7L2 | TCTCAAAAAAAAAA | 1 | 453 | 467 | -547 | -533 | -2,24 | 0,01 | 11,95 | >0.1 | 2,79 | 4,88 | 0 | 1 | 0 | 5,52 | 0 |
| TCF7L2 | ACTGATCAGAAATA | 1 | 478 | 492 | -522 | -508 | 1,59 | 0,01 | 16,04 | >0.1 | 2,88 | 4,93 | 0 | 0,99 | 0 | 5,65 | 0 |
| TCF7L2 | CAAGTAGAGAGTTT | 1 | 546 | 560 | -454 | -440 | -2,3 | 0,01 | 6,94 | >0.1 | 3,17 | 5,08 | 0 | 0,99 | 0 | 0 | 0 |
| TCF7L2 | TGAAATAAAAATAA | 1 | 575 | 589 | -425 | -411 | -0,87 | 0,01 | 8,62 | >0.1 | 3,33 | 5,15 | 0 | 1 | 0 | 0 | 0 |
| TCF7L2 | TAGGATTAAAGAAT | 1 | 789 | 803 | -211 | -197 | -2,37 | 0,01 | 15,9 | >0.1 | 5,56 | 5,93 | 0 | 1,46 | 0 | 5,32 | 0 |
| TCF7L2 | GCTGATAAGAGAGA | 1 | 966 | 980 | -34 | -20 | 2,82 | 0,01 | 19,86 | >0.1 | 5,56 | 7,91 | 0 | 1,87 | 0 | 1,7 | 0 |
| TCF7L2 | TGATAAGAGAGAAA | 1 | 968 | 982 | -32 | -18 | -1,62 | 0,01 | 15,48 | >0.1 | 5,56 | 7,98 | 0 | 1,87 | 0 | 1,7 | 0 |
| TCF7L2 | TTGGCTCACAGGGG | 1 | 1032 | 1046 | 32 | 46 | -1,79 | 0,01 | 15,08 | >0.1 | 5,56 | 8,15 | 0 | 1,86 | 0 | 1,31 | 0 |
| TCF7L2 | TTCTATCAAAGTTC | 1 | 1166 | 1180 | 166 | 180 | 0,71 | 0,01 | 18,78 | >0.1 | 5,56 | 6,18 | 0 | 1,7 | 1,79 | 2,84 | 0 |
| TCF7L2 | CAGCAACAAGGCTG | -1 | 1075 | 1089 | 75 | 89 | -0,59 | 0,01 | 17,92 | >0.1 | 5,56 | 7,09 | 0 | 1,85 | 1,79 | 2,21 | 0 |
| TCF7L2 | AAGGCTGAGAAGGA | -1 | 1068 | 1082 | 68 | 82 | -1,97 | 0,01 | 16,71 | >0.1 | 5,56 | 7,21 | 0 | 1,85 | 1,79 | 2,27 | 0 |
| TCF7L2 | GAGCAGGAAAGACC | -1 | 930 | 944 | -70 | -56 | -2,14 | 0,01 | 13,35 | >0.1 | 5,56 | 7,12 | 0 | 1,88 | 0 | 0,93 | 0 |
| TCF7L2 | AAAGACCAAAAAAA | -1 | 923 | 937 | -77 | -63 | -1,76 | 0,01 | 13,76 | >0.1 | 5,56 | 7,02 | 0 | 1,88 | 0 | 1,06 | 0 |
| TCF7L2 | GACCAAAAAAAGAA | -1 | 920 | 934 | -80 | -66 | -0,25 | 0,01 | 15,26 | >0.1 | 5,56 | 6,98 | 0 | 1,88 | 0 | 1,09 | 0 |
| TCF7L2 | AAAAAAGAAAAAGA | -1 | 915 | 929 | -85 | -71 | 0,23 | 0,01 | 15,71 | >0.1 | 5,56 | 6,92 | 0 | 1,89 | 0 | 1,12 | 0 |
| TCF7L2 | AAAGAAGAAGAACA | -1 | 906 | 920 | -94 | -80 | -2,09 | 0,01 | 13,37 | >0.1 | 5,56 | 6,81 | 0 | 1,89 | 0 | 1,21 | 0 |
| TCF7L2 | GAAGAACACAGTAA | -1 | 900 | 914 | -100 | -86 | 2,03 | 0,01 | 17,5 | >0.1 | 5,56 | 6,75 | 0 | 1,89 | 0 | 1,28 | 0 |
| TCF7L2 | CCAAATCAAAACCG | -1 | 745 | 759 | -255 | -241 | 5,52 | 0,01 | 24,18 | >0.1 | 5,56 | 5,72 | 0 | 1,28 | 0 | 6,1 | 0 |
| TCF7L2 | TTCCTAGAAAGCTG | -1 | 713 | 727 | -287 | -273 | -1,27 | 0,01 | 16,23 | >0.1 | 5,56 | 5,59 | 0 | 1,19 | 0 | 5,17 | 0 |
| TCF7L2 | AGTGACCAAAGAGG | -1 | 667 | 681 | -333 | -319 | -1,06 | 0,01 | 13,29 | >0.1 | 4,13 | 5,42 | 0 | 1,1 | 0 | 3,7 | 0 |
| TCF7L2 | GATCATGAAGATAA | -1 | 641 | 655 | -359 | -345 | -2,03 | 0,01 | 11,89 | >0.1 | 3,82 | 5,34 | 0 | 1,06 | 0 | 3,7 | 0 |
| TCF7L2 | AAACTTAAAAAAGC | -1 | 72 | 86 | -928 | -914 | 5,74 | 0,01 | 15,79 | >0.1 | 3,84 | 4,33 | 0 | 1,88 | 0 | 0 | 0 |
| FOXA1 | GGAGAGTTGACCTGT | 1 | 101 | 116 | -899 | -884 | -6 | 0,01 | 10,08 | >0.1 | 3,58 | 4,36 | 0 | 1,89 | 0 | 0,77 | 5,49 |
| FOXA1 | GTGATCTTGGCTCAC | 1 | 1026 | 1041 | 26 | 41 | -6,68 | 0,01 | 15,97 | >0.1 | 5,54 | 8,45 | 0 | 1,86 | 0 | 1,31 | 5,49 |
| FOXA1 | CAATGGTGGACTGAG | -1 | 1098 | 1113 | 98 | 113 | -4,37 | 0,01 | 20,89 | >0.1 | 5,54 | 6,78 | 0 | 1,85 | 3,22 | 2,38 | 5,49 |
| FOXA1 | TTAATATTTTCCTAG | -1 | 720 | 735 | -280 | -265 | -4,78 | 0,01 | 18,25 | >0.1 | 5,54 | 5,62 | 0 | 1,22 | 0 | 5,17 | 5,49 |
| FOXA1 | TTGGTGTTTGTATGG | -1 | 624 | 639 | -376 | -361 | 2,75 | 0,01 | 19,79 | >0.1 | 3,67 | 5,29 | 0 | 1,05 | 0 | 1,55 | 5,49 |
| FOXA1 | CTCACATTTATTTTT | -1 | 581 | 596 | -419 | -404 | -7,1 | 0,01 | 7,93 | >0.1 | 3,37 | 5,17 | 0 | 1,01 | 0 | 0 | 5,49 |
| RUNX3 | CAGCCACAGA | 1 | 38 | 48 | -962 | -952 | 7,82 | 0,001 | 18,3 | >0.1 | 4,32 | 4,29 | 0 | 1,87 | 0 | 0 | 0 |
| RUNX3 | AGACCGTCCT | 1 | 255 | 265 | -745 | -735 | -0,9 | 0,01 | 12,66 | >0.1 | 2,83 | 4,54 | 0 | 1,35 | 0 | 4,85 | 0 |
| RUNX3 | AAACCCCGTC | 1 | 278 | 288 | -722 | -712 | 0,81 | 0,01 | 14,59 | >0.1 | 2,75 | 4,57 | 0 | 1,27 | 0 | 5,19 | 0 |
| RUNX3 | AGACCGAGAC | 1 | 439 | 449 | -561 | -551 | 0,09 | 0,01 | 14,72 | >0.1 | 2,74 | 4,84 | 0 | 1 | 0 | 6,05 | 0 |
| RUNX3 | AAACCATACA | 1 | 621 | 631 | -379 | -369 | 4,52 | 0,01 | 15,89 | >0.1 | 3,63 | 5,27 | 0 | 1,04 | 0 | 1,44 | 0 |
| RUNX3 | CAAACACCAA | 1 | 629 | 639 | -371 | -361 | -1,86 | 0,01 | 9,83 | >0.1 | 3,69 | 5,29 | 0 | 1,04 | 0 | 1,66 | 0 |
| RUNX3 | ACACCAACAT | 1 | 632 | 642 | -368 | -358 | -2,21 | 0,01 | 9,47 | >0.1 | 3,72 | 5,3 | 0 | 1,05 | 0 | 1,61 | 0 |
| RUNX3 | AGACCTCTTT | 1 | 664 | 674 | -336 | -326 | -2,59 | 0,01 | 11,79 | >0.1 | 4,06 | 5,4 | 0 | 1,09 | 0 | 3,83 | 0 |
| RUNX3 | TAACCAAAAG | 1 | 732 | 742 | -268 | -258 | 3,4 | 0,01 | 21,8 | >0.1 | 5,61 | 5,65 | 0 | 1,23 | 0 | 5,92 | 0 |
| RUNX3 | TGGCCATAAA | 1 | 756 | 766 | -244 | -234 | 0,5 | 0,01 | 19,49 | >0.1 | 5,61 | 5,75 | 0 | 1,31 | 0 | 6,32 | 0 |
| RUNX3 | CAACCCAAGT | 1 | 950 | 960 | -50 | -40 | -2,66 | 0,01 | 13,96 | >0.1 | 5,61 | 7,4 | 0 | 1,88 | 0 | 1,74 | 0 |
| RUNX3 | TAACCACGAA | 1 | 1144 | 1154 | 144 | 154 | 8,91 | 0,001 | 28,09 | >0.1 | 5,61 | 6,34 | 0 | 1,84 | 3,22 | 2,16 | 0 |
| RUNX3 | AAGCCGAAGA | 1 | 1152 | 1162 | 152 | 162 | 1,36 | 0,01 | 18,05 | >0.1 | 4,51 | 6,28 | 0 | 1,84 | 1,79 | 2,26 | 0 |
| RUNX3 | AAACCTCCTC | -1 | 989 | 999 | -11 | -1 | 0,29 | 0,01 | 18,07 | >0.1 | 5,61 | 8,69 | 0 | 1,87 | 0 | 1,61 | 0 |
| RUNX3 | AGACCAAAAA | -1 | 925 | 935 | -75 | -65 | 4,99 | 0,01 | 20,54 | >0.1 | 5,61 | 6,99 | 0 | 1,88 | 0 | 1,06 | 0 |
| RUNX3 | AGAACACAGT | -1 | 902 | 912 | -98 | -88 | -1,37 | 0,01 | 14,13 | >0.1 | 5,61 | 6,72 | 0 | 1,89 | 0 | 1,28 | 0 |
| RUNX3 | AAACCGGTAC | -1 | 741 | 751 | -259 | -249 | 2,78 | 0,01 | 21,43 | >0.1 | 5,61 | 5,68 | 0 | 1,26 | 0 | 6,1 | 0 |
| RUNX3 | TGACCAAAGA | -1 | 669 | 679 | -331 | -321 | 2,4 | 0,01 | 16,74 | >0.1 | 4,13 | 5,42 | 0 | 1,1 | 0 | 3,7 | 0 |
| RUNX3 | TTCCCAGAAA | -1 | 557 | 567 | -443 | -433 | -0,82 | 0,01 | 8,49 | >0.1 | 3,22 | 5,1 | 0 | 0,99 | 0 | 0 | 0 |
| RUNX3 | TGCCCACCAC | -1 | 321 | 331 | -679 | -669 | 3,3 | 0,01 | 17,28 | >0.1 | 2,62 | 4,64 | 0 | 1,16 | 0 | 5,56 | 0 |
| RUNX3 | CCACCACCAC | -1 | 318 | 328 | -682 | -672 | 5,6 | 0,001 | 20,28 | >0.1 | 2,63 | 4,63 | 0 | 1,16 | 0 | 6,26 | 0 |
| RUNX3 | CCACCACGCC | -1 | 315 | 325 | -685 | -675 | 2,32 | 0,01 | 16,88 | >0.1 | 2,64 | 4,63 | 0 | 1,17 | 0 | 6,13 | 0 |

| | | | | | | | | | | | | | | | | | |
|---|---|---|---|---|---|---|---|---|---|---|---|---|---|---|---|---|---|
| RUNX3 | GAGCCACTGT | -1 | 184 | 194 | -816 | -806 | -2,31 | 0,01 | 10,95 | >0.1 | 3,11 | 4,45 | 0 | 1,75 | 0 | 3,95 | 0 |
| FOXA2 | TGTTGCCCAAGT | 1 | 539 | 551 | -461 | -449 | 2,58 | 0,01 | 16,94 | >0.1 | 3,13 | 5,06 | 0 | 0,99 | 0 | 0 | 5,18 |
| FOXA2 | TTTTGATTTGGC | 1 | 748 | 760 | -252 | -240 | -0,23 | 0,01 | 23,46 | >0.1 | 5,58 | 5,72 | 0 | 1,29 | 0 | 5,92 | 5,18 |
| FOXA2 | TCTTGGCTCACA | 1 | 1030 | 1042 | 30 | 42 | 0,08 | 0,01 | 22,25 | >0.1 | 5,58 | 8,24 | 0 | 1,86 | 0 | 1,31 | 5,18 |
| FOXA2 | GGTGGACTGAGC | -1 | 1097 | 1109 | 97 | 109 | -0,18 | 0,01 | 24,79 | >0.1 | 5,58 | 6,79 | 0 | 1,85 | 3,22 | 2,35 | 5,18 |
| FOXA2 | TATTTTCCTAGA | -1 | 719 | 731 | -281 | -269 | 3,72 | 0,01 | 26,28 | >0.1 | 5,58 | 5,6 | 0 | 1,2 | 0 | 4,99 | 5,18 |
| FOXA2 | TGTTGGTGTTTG | -1 | 629 | 641 | -371 | -359 | 0,07 | 0,01 | 17,03 | >0.1 | 3,7 | 5,3 | 0 | 1,05 | 0 | 1,73 | 5,18 |
| FOXA2 | TGTTTGTATGGT | -1 | 623 | 635 | -377 | -365 | 5,42 | 0,01 | 22,07 | >0.1 | 3,65 | 5,28 | 0 | 1,04 | 0 | 1,5 | 5,18 |
| FOXA2 | TATTTTTATTTC | -1 | 576 | 588 | -424 | -412 | 0,3 | 0,01 | 14,96 | >0.1 | 3,33 | 5,15 | 0 | 1 | 0 | 0 | 5,18 |
| PRRX1 | ATAATTTT | 1 | 132 | 140 | -868 | -860 | 3 | 0,01 | 14,24 | >0.1 | 3,4 | 4,38 | 0 | 1,89 | 0 | 1,57 | 0 |
| PRRX1 | GAAATTAA | 1 | 154 | 162 | -846 | -838 | 6,75 | 0,01 | 17,97 | >0.1 | 3,27 | 4,41 | 0 | 1,9 | 0 | 1,65 | 0 |
| PRRX1 | CTGATCAG | 1 | 164 | 172 | -836 | -828 | 4,03 | 0,01 | 15,96 | >0.1 | 3,21 | 4,42 | 0 | 1,9 | 0 | 2,4 | 0 |
| PRRX1 | TCTATTAA | 1 | 288 | 296 | -712 | -704 | 4,65 | 0,01 | 18,38 | >0.1 | 2,72 | 4,58 | 0 | 1,24 | 0 | 5,19 | 0 |
| PRRX1 | CTGATCAG | 1 | 479 | 487 | -521 | -513 | 4,03 | 0,01 | 18,17 | >0.1 | 2,88 | 4,92 | 0 | 0,98 | 0 | 5,36 | 0 |
| PRRX1 | CCTATTAA | 1 | 600 | 608 | -400 | -392 | 5,66 | 0,01 | 16,71 | >0.1 | 3,47 | 5,2 | 0 | 1,01 | 0 | 1,36 | 0 |
| PRRX1 | TTAATGAA | 1 | 604 | 612 | -396 | -388 | 5,16 | 0,01 | 16,3 | >0.1 | 3,49 | 5,21 | 0 | 1,02 | 0 | 1,41 | 0 |
| PRRX1 | AAAATTAT | 1 | 683 | 691 | -317 | -309 | 4,85 | 0,01 | 18,83 | >0.1 | 4,34 | 5,46 | 0 | 1,12 | 0 | 3,07 | 0 |
| PRRX1 | AAAATTAT | 1 | 830 | 838 | -170 | -162 | 4,85 | 0,01 | 21,58 | >0.1 | 5,64 | 6,14 | 0 | 1,67 | 0 | 3,28 | 0 |
| PRRX1 | GGAATTAT | 1 | 1191 | 1199 | 191 | 199 | 4,83 | 0,01 | 23,61 | >0.1 | 5,64 | 6,02 | 0 | 1,5 | 1,79 | 3,82 | 0 |
| PRRX1 | ATAATTCC | -1 | 1191 | 1199 | 191 | 199 | 3,24 | 0,01 | 22,02 | >0.1 | 5,64 | 6,02 | 0 | 1,5 | 1,79 | 3,82 | 0 |
| PRRX1 | ATAATTTT | -1 | 830 | 838 | -170 | -162 | 3 | 0,01 | 19,73 | >0.1 | 5,64 | 6,14 | 0 | 1,67 | 0 | 3,28 | 0 |
| PRRX1 | TGAATCAA | -1 | 816 | 824 | -184 | -176 | 3,34 | 0,01 | 20,03 | >0.1 | 5,64 | 6,05 | 0 | 1,58 | 0 | 3,43 | 0 |
| PRRX1 | CAAATAAG | -1 | 811 | 819 | -189 | -181 | 3,49 | 0,01 | 19,98 | >0.1 | 5,64 | 6,02 | 0 | 1,55 | 0 | 3,28 | 0 |
| PRRX1 | CAAATCAA | -1 | 750 | 758 | -250 | -242 | 4,94 | 0,01 | 23,49 | >0.1 | 5,64 | 5,71 | 0 | 1,28 | 0 | 5,92 | 0 |
| PRRX1 | ATAATTTT | -1 | 683 | 691 | -317 | -309 | 3 | 0,01 | 16,98 | >0.1 | 4,34 | 5,46 | 0 | 1,12 | 0 | 3,07 | 0 |
| PRRX1 | ACAATTTC | -1 | 609 | 617 | -391 | -383 | 3,97 | 0,01 | 15,29 | >0.1 | 3,53 | 5,23 | 0 | 1,02 | 0 | 1,55 | 0 |
| PRRX1 | TTCATTAA | -1 | 604 | 612 | -396 | -388 | 5,28 | 0,01 | 16,42 | >0.1 | 3,49 | 5,21 | 0 | 1,02 | 0 | 1,41 | 0 |
| PRRX1 | CTGATCAG | -1 | 479 | 487 | -521 | -513 | 4,03 | 0,01 | 18,17 | >0.1 | 2,88 | 4,92 | 0 | 0,98 | 0 | 5,36 | 0 |
| PRRX1 | CTGATCAG | -1 | 164 | 172 | -836 | -828 | 4,03 | 0,01 | 15,96 | >0.1 | 3,21 | 4,42 | 0 | 1,9 | 0 | 2,4 | 0 |
| PRRX1 | TTAATTTC | -1 | 154 | 162 | -846 | -838 | 4,3 | 0,01 | 15,52 | >0.1 | 3,27 | 4,41 | 0 | 1,9 | 0 | 1,65 | 0 |
| PRRX1 | AAAATTAT | -1 | 132 | 140 | -868 | -860 | 4,85 | 0,01 | 16,09 | >0.1 | 3,4 | 4,38 | 0 | 1,89 | 0 | 1,57 | 0 |
| PRRX1 | TCACTTAC | -1 | 5 | 13 | -995 | -987 | 3,8 | 0,01 | 17,35 | >0.1 | 5,64 | 4,26 | 0 | 1,86 | 1,79 | 0 | 0 |
| PRRX2 | GAAATTAA | 1 | 154 | 162 | -846 | -838 | 6,03 | 0,01 | 17,25 | >0.1 | 3,27 | 4,41 | 0 | 1,9 | 0 | 1,65 | 0 |
| PRRX2 | CTGATCAG | 1 | 164 | 172 | -836 | -828 | 4,34 | 0,01 | 16,27 | >0.1 | 3,21 | 4,42 | 0 | 1,9 | 0 | 2,4 | 0 |
| PRRX2 | TCTATTAA | 1 | 288 | 296 | -712 | -704 | 5,11 | 0,01 | 18,84 | >0.1 | 2,72 | 4,58 | 0 | 1,24 | 0 | 5,19 | 0 |
| PRRX2 | CTGATCAG | 1 | 479 | 487 | -521 | -513 | 4,34 | 0,01 | 18,48 | >0.1 | 2,88 | 4,92 | 0 | 0,98 | 0 | 5,36 | 0 |
| PRRX2 | CCTATTAA | 1 | 600 | 608 | -400 | -392 | 6,26 | 0,01 | 17,31 | >0.1 | 3,47 | 5,2 | 0 | 1,01 | 0 | 1,36 | 0 |
| PRRX2 | TTAATGAA | 1 | 604 | 612 | -396 | -388 | 4,6 | 0,01 | 15,74 | >0.1 | 3,49 | 5,21 | 0 | 1,02 | 0 | 1,41 | 0 |
| PRRX2 | AAAATTAT | 1 | 683 | 691 | -317 | -309 | 4,57 | 0,01 | 18,55 | >0.1 | 4,34 | 5,46 | 0 | 1,12 | 0 | 3,07 | 0 |
| PRRX2 | AAAATTAT | 1 | 830 | 838 | -170 | -162 | 4,57 | 0,01 | 21,3 | >0.1 | 5,64 | 6,14 | 0 | 1,67 | 0 | 3,28 | 0 |
| PRRX2 | CTGATAAG | 1 | 967 | 975 | -33 | -25 | 3,73 | 0,01 | 20,69 | >0.1 | 5,64 | 7,76 | 0 | 1,87 | 0 | 1,7 | 0 |
| PRRX2 | GGAATTAT | 1 | 1191 | 1199 | 191 | 199 | 5,01 | 0,01 | 23,79 | >0.1 | 5,64 | 6,02 | 0 | 1,5 | 1,79 | 3,82 | 0 |
| PRRX2 | ATAATTCC | -1 | 1191 | 1199 | 191 | 199 | 3,74 | 0,01 | 22,52 | >0.1 | 5,64 | 6,02 | 0 | 1,5 | 1,79 | 3,82 | 0 |
| PRRX2 | GCAGTTAC | -1 | 1088 | 1096 | 88 | 96 | 4,4 | 0,01 | 22,8 | >0.1 | 5,64 | 6,9 | 0 | 1,85 | 1,79 | 2,21 | 0 |
| PRRX2 | CCAATATA | -1 | 879 | 887 | -121 | -113 | 3,64 | 0,01 | 20,75 | >0.1 | 5,64 | 6,5 | 0 | 1,9 | 0 | 3,07 | 0 |
| PRRX2 | CAAATAAG | -1 | 811 | 819 | -189 | -181 | 4,31 | 0,01 | 20,8 | >0.1 | 5,64 | 6,02 | 0 | 1,55 | 0 | 3,28 | 0 |
| PRRX2 | CAAATCAA | -1 | 750 | 758 | -250 | -242 | 4,68 | 0,01 | 23,23 | >0.1 | 5,64 | 5,71 | 0 | 1,28 | 0 | 5,92 | 0 |
| PRRX2 | ACAATTTC | -1 | 609 | 617 | -391 | -383 | 4,95 | 0,01 | 16,27 | >0.1 | 3,53 | 5,23 | 0 | 1,02 | 0 | 1,55 | 0 |
| PRRX2 | TTCATTAA | -1 | 604 | 612 | -396 | -388 | 5,19 | 0,01 | 16,33 | >0.1 | 3,49 | 5,21 | 0 | 1,02 | 0 | 1,41 | 0 |
| PRRX2 | CTGATCAG | -1 | 479 | 487 | -521 | -513 | 4,34 | 0,01 | 18,48 | >0.1 | 2,88 | 4,92 | 0 | 0,98 | 0 | 5,36 | 0 |
| PRRX2 | CTGATCAG | -1 | 164 | 172 | -836 | -828 | 4,34 | 0,01 | 16,27 | >0.1 | 3,21 | 4,42 | 0 | 1,9 | 0 | 2,4 | 0 |
| PRRX2 | TTAATTTC | -1 | 154 | 162 | -846 | -838 | 4,85 | 0,01 | 16,07 | >0.1 | 3,27 | 4,41 | 0 | 1,9 | 0 | 1,65 | 0 |
| PRRX2 | AAAATTAT | -1 | 132 | 140 | -868 | -860 | 4,57 | 0,01 | 15,81 | >0.1 | 3,4 | 4,38 | 0 | 1,89 | 0 | 1,57 | 0 |
| PRRX2 | TCACTTAC | -1 | 5 | 13 | -995 | -987 | 5,25 | 0,01 | 18,8 | >0.1 | 5,64 | 4,26 | 0 | 1,86 | 1,79 | 0 | 0 |
| DUX4 | GGAGGCACTCA | 1 | 20 | 31 | -980 | -969 | -13,77 | 0,01 | -2,04 | >0.1 | 5,6 | 4,27 | 0 | 1,87 | 0 | 0 | 0 |
| DUX4 | AAAACTGATCA | 1 | 160 | 171 | -840 | -829 | -4,32 | 0,01 | 7,68 | >0.1 | 3,23 | 4,42 | 0 | 1,9 | 0 | 2,46 | 0 |
| DUX4 | TAATCCTAGCA | 1 | 201 | 212 | -799 | -788 | -9,98 | 0,01 | 3,74 | >0.1 | 3,03 | 4,47 | 0 | 1,62 | 0 | 4,59 | 0 |
| DUX4 | AAACCCCGTCT | 1 | 278 | 289 | -722 | -711 | -8,71 | 0,01 | 5,07 | >0.1 | 2,75 | 4,57 | 0 | 1,27 | 0 | 5,19 | 0 |
| DUX4 | AGACTCAGTCT | 1 | 445 | 456 | -555 | -544 | -10,5 | 0,01 | 3,13 | >0.1 | 2,76 | 4,85 | 0 | 1 | 0 | 5,01 | 0 |

| | | | | | | | | | | | | | | | | | |
|---|---|---|---|---|---|---|---|---|---|---|---|---|---|---|---|---|---|
| DUX4 | AAAACTGATCA | 1 | 475 | 486 | -525 | -514 | -4,32 | 0,01 | 9,54 | >0.1 | 2,87 | 4,91 | 0 | 0,99 | 0 | 5,09 | 0 |
| DUX4 | GAATTCCATAA | 1 | 493 | 504 | -507 | -496 | -9,38 | 0,01 | 5,93 | >0.1 | 2,93 | 4,95 | 0 | 0,98 | 0 | 6,44 | 0 |
| DUX4 | CAAAGTCATGT | 1 | 513 | 524 | -487 | -476 | -13,73 | 0,01 | -1,87 | >0.1 | 3,01 | 5 | 0 | 0,98 | 0 | 2,86 | 0 |
| DUX4 | TAACGTATTCT | 1 | 802 | 813 | -198 | -187 | -6,03 | 0,01 | 10,49 | >0.1 | 5,6 | 5,99 | 0 | 1,51 | 0 | 3,43 | 0 |
| DUX4 | TGATTCACTTT | 1 | 817 | 828 | -183 | -172 | -14,39 | 0,01 | 2,44 | >0.1 | 5,6 | 6,08 | 0 | 1,6 | 0 | 3,55 | 0 |
| DUX4 | AAAATTATTCT | 1 | 830 | 841 | -170 | -159 | -8,34 | 0,01 | 8,4 | >0.1 | 5,6 | 6,16 | 0 | 1,7 | 0 | 3,28 | 0 |
| DUX4 | CAACCCAAGTT | 1 | 950 | 961 | -50 | -39 | -12,68 | 0,01 | 3,95 | >0.1 | 5,6 | 7,42 | 0 | 1,88 | 0 | 1,74 | 0 |
| DUX4 | AAATCTCATGA | 1 | 980 | 991 | -20 | -9 | -6,13 | 0,01 | 11,24 | >0.1 | 5,6 | 8,37 | 0 | 1,87 | 0 | 1,53 | 0 |
| DUX4 | AAAGTCATTCA | 1 | 1009 | 1020 | 9 | 20 | -6,18 | 0,01 | 11,77 | >0.1 | 5,6 | 9,03 | 0 | 1,86 | 0 | 1,46 | 0 |
| DUX4 | GGATGTGATCT | 1 | 1022 | 1033 | 22 | 33 | -13,93 | 0,01 | 3,49 | >0.1 | 5,6 | 8,68 | 0 | 1,86 | 0 | 1,28 | 0 |
| DUX4 | AAACCTCCTCA | -1 | 988 | 999 | -12 | -1 | -5,9 | 0,01 | 11,86 | >0.1 | 5,6 | 8,69 | 0 | 1,87 | 0 | 1,61 | 0 |
| DUX4 | TGAGCCAATAT | -1 | 880 | 891 | -120 | -109 | -11,62 | 0,01 | 5,48 | >0.1 | 5,6 | 6,54 | 0 | 1,9 | 0 | 3,07 | 0 |
| DUX4 | AAAGTTCATCC | -1 | 869 | 880 | -131 | -120 | -8,73 | 0,01 | 8,16 | >0.1 | 5,6 | 6,45 | 0 | 1,9 | 0 | 2,95 | 0 |
| DUX4 | TGAATCAAATA | -1 | 813 | 824 | -187 | -176 | -13,83 | 0,01 | 2,82 | >0.1 | 5,6 | 6,05 | 0 | 1,58 | 0 | 3,43 | 0 |
| DUX4 | AAATGTTATAA | -1 | 687 | 698 | -313 | -302 | -14,7 | 0,01 | -0,69 | >0.1 | 4,45 | 5,48 | 0 | 1,13 | 0 | 2,95 | 0 |
| DUX4 | CAATTTCATTA | -1 | 605 | 616 | -395 | -384 | 3,83 | 0,001 | 15 | >0.1 | 3,51 | 5,22 | 0 | 1,02 | 0 | 1,41 | 0 |
| DUX4 | AGACTGAGTCT | -1 | 445 | 456 | -555 | -544 | -11,69 | 0,01 | 1,94 | >0.1 | 2,76 | 4,85 | 0 | 1 | 0 | 5,01 | 0 |
| DUX4 | AAAGCTCATTT | -1 | 66 | 77 | -934 | -923 | -0,35 | 0,001 | 9,76 | >0.1 | 3,92 | 4,32 | 0 | 1,88 | 0 | 0 | 0 |
| CRX | AATGAGCTTTT | 1 | 67 | 78 | -933 | -922 | 2,97 | 0,01 | 13,07 | >0.1 | 3,91 | 4,32 | 0 | 1,88 | 0 | 0 | 0 |
| CRX | AGGCAGATCAG | 1 | 91 | 102 | -909 | -898 | 6,35 | 0,001 | 17,11 | >0.1 | 3,68 | 4,34 | 0 | 1,88 | 0 | 0,85 | 0 |
| CRX | CAGGAGAGTTG | 1 | 99 | 110 | -901 | -890 | -3,48 | 0,01 | 7,14 | >0.1 | 3,62 | 4,35 | 0 | 1,88 | 0 | 0,77 | 0 |
| CRX | GTGGAGAGTAG | 1 | 119 | 130 | -881 | -870 | -1,56 | 0,01 | 9,82 | >0.1 | 3,47 | 4,37 | 0 | 1,89 | 0 | 1,65 | 0 |
| CRX | ATGGAAATTAA | 1 | 151 | 162 | -849 | -838 | -3,43 | 0,01 | 7,8 | >0.1 | 3,27 | 4,41 | 0 | 1,9 | 0 | 1,65 | 0 |
| CRX | AAACTGATCAG | 1 | 161 | 172 | -839 | -828 | -0,9 | 0,01 | 11,1 | >0.1 | 3,22 | 4,42 | 0 | 1,9 | 0 | 2,46 | 0 |
| CRX | CAGTGGCTCAC | 1 | 185 | 196 | -815 | -804 | 0,73 | 0,01 | 13,88 | >0.1 | 3,1 | 4,45 | 0 | 1,75 | 0 | 3,86 | 0 |
| CRX | GGGCAGATCAC | 1 | 229 | 240 | -771 | -760 | 0,78 | 0,01 | 14,79 | >0.1 | 2,92 | 4,51 | 0 | 1,46 | 0 | 5,12 | 0 |
| CRX | CAAAAACTTAG | 1 | 301 | 312 | -699 | -688 | -4,38 | 0,01 | 9,09 | >0.1 | 2,67 | 4,61 | 0 | 1,2 | 0 | 4,99 | 0 |
| CRX | GGGAGGCTGAG | 1 | 350 | 361 | -650 | -639 | 0,64 | 0,01 | 15,37 | >0.1 | 2,54 | 4,68 | 0 | 1,1 | 0 | 6,4 | 0 |
| CRX | AAACTGATCAG | 1 | 476 | 487 | -524 | -513 | -0,9 | 0,01 | 13,08 | >0.1 | 2,87 | 4,92 | 0 | 0,99 | 0 | 5,2 | 0 |
| CRX | TAACAGCTTTC | 1 | 710 | 721 | -290 | -279 | -2,71 | 0,01 | 14,44 | >0.1 | 5,6 | 5,56 | 0 | 1,18 | 0 | 4,81 | 0 |
| CRX | GAAAATATTAA | 1 | 724 | 735 | -276 | -265 | -4,22 | 0,01 | 13,38 | >0.1 | 5,6 | 5,62 | 0 | 1,21 | 0 | 5,17 | 0 |
| CRX | TCTAGGATTAA | 1 | 787 | 798 | -213 | -202 | 4,28 | 0,01 | 22,74 | >0.1 | 5,6 | 5,91 | 0 | 1,44 | 0 | 5,52 | 0 |
| CRX | TATTGGCTCAG | 1 | 881 | 892 | -119 | -108 | 0,56 | 0,01 | 17,67 | >0.1 | 5,6 | 6,55 | 0 | 1,89 | 0 | 3,07 | 0 |
| CRX | CAGCAGATTGT | 1 | 889 | 900 | -111 | -100 | -4,43 | 0,01 | 12,74 | >0.1 | 5,6 | 6,61 | 0 | 1,89 | 0 | 3,07 | 0 |
| CRX | CAAAGGCTGAT | 1 | 961 | 972 | -39 | -28 | -4,68 | 0,01 | 12,16 | >0.1 | 5,6 | 7,67 | 0 | 1,88 | 0 | 1,7 | 0 |
| CRX | AGGAGGTTTTA | 1 | 990 | 1001 | -10 | 1 | -3,37 | 0,01 | 14,45 | >0.1 | 5,6 | 8,74 | 0 | 1,87 | 0 | 1,61 | 0 |
| CRX | CAGTGGATGTG | 1 | 1018 | 1029 | 18 | 29 | -4,77 | 0,01 | 12,79 | >0.1 | 5,6 | 8,76 | 0 | 1,86 | 0 | 1,34 | 0 |
| CRX | CTGCTGCTCAG | 1 | 1092 | 1103 | 92 | 103 | -4,15 | 0,01 | 14,29 | >0.1 | 5,6 | 6,85 | 0 | 1,85 | 1,79 | 2,34 | 0 |
| CRX | GACAAGTTTAA | 1 | 1136 | 1147 | 136 | 147 | -5,52 | 0,01 | 14,64 | >0.1 | 5,6 | 6,41 | 0 | 1,84 | 3,22 | 3,09 | 0 |
| CRX | ACAAGGCTGAG | -1 | 1073 | 1084 | 73 | 84 | -4,31 | 0,01 | 14,33 | >0.1 | 5,6 | 7,12 | 0 | 1,85 | 1,79 | 2,27 | 0 |
| CRX | GAAGAGCTTGA | -1 | 1052 | 1063 | 52 | 63 | -4,17 | 0,01 | 13,08 | >0.1 | 5,6 | 7,53 | 0 | 1,86 | 0 | 2,27 | 0 |
| CRX | CATGAGATTTT | -1 | 979 | 990 | -21 | -10 | 2,76 | 0,01 | 20,08 | >0.1 | 5,6 | 8,32 | 0 | 1,87 | 0 | 1,53 | 0 |
| CRX | TAACAGATTTT | -1 | 841 | 852 | -159 | -148 | -0,14 | 0,01 | 14,9 | >0.1 | 5,6 | 6,24 | 0 | 1,79 | 0 | 1,41 | 0 |
| CRX | GAGAACCTTAC | -1 | 778 | 789 | -222 | -211 | -5,35 | 0,01 | 14,04 | >0.1 | 5,6 | 5,86 | 0 | 1,4 | 0 | 6,54 | 0 |
| CRX | CTAGGGATCAT | -1 | 649 | 660 | -351 | -340 | 1,14 | 0,01 | 15,29 | >0.1 | 3,89 | 5,35 | 0 | 1,07 | 0 | 3,83 | 0 |
| CRX | ATGAAGATAAT | -1 | 640 | 651 | -360 | -349 | -4,85 | 0,01 | 6,99 | >0.1 | 3,8 | 5,33 | 0 | 1,06 | 0 | 1,66 | 0 |
| CRX | TTCAAGATCAT | -1 | 568 | 579 | -432 | -421 | -5,06 | 0,01 | 4,34 | >0.1 | 3,28 | 5,13 | 0 | 1 | 0 | 0 | 0 |
| CRX | GCTAGGATTAC | -1 | 200 | 211 | -800 | -789 | 3,65 | 0,01 | 17,38 | >0.1 | 3,04 | 4,47 | 0 | 1,63 | 0 | 4,59 | 0 |
| CRX | AAAAAGCTCAT | -1 | 68 | 79 | -932 | -921 | 1,93 | 0,01 | 12,02 | >0.1 | 3,9 | 4,32 | 0 | 1,88 | 0 | 0 | 0 |
| FOSL2::JUN | GTGTGAGAAATA | 1 | 52 | 64 | -948 | -936 | -1,2 | 0,01 | 9,06 | >0.1 | 4,09 | 4,3 | 0 | 1,87 | 0 | 0 | 0 |
| FOSL2::JUN | AGAGTAGTCATA | 1 | 123 | 135 | -877 | -865 | 2,62 | 0,01 | 14,02 | >0.1 | 3,44 | 4,38 | 0 | 1,89 | 0 | 1,7 | 0 |
| FOSL2::JUN | CAGTGGCTCACG | 1 | 185 | 197 | -815 | -803 | 2,18 | 0,01 | 15,33 | >0.1 | 3,1 | 4,45 | 0 | 1,75 | 0 | 3,86 | 0 |
| FOSL2::JUN | TCCTGACTAACA | 1 | 261 | 273 | -739 | -727 | 3,77 | 0,01 | 16,98 | >0.1 | 2,8 | 4,55 | 0 | 1,32 | 0 | 4,53 | 0 |
| FOSL2::JUN | GGCTGAGGCAGG | 1 | 354 | 366 | -646 | -634 | 2,75 | 0,01 | 17,1 | >0.1 | 2,53 | 4,69 | 0 | 1,1 | 0 | 6,03 | 0 |
| FOSL2::JUN | CTGTGCTCCAGC | 1 | 418 | 430 | -582 | -570 | -2,55 | 0,01 | 11,26 | >0.1 | 2,68 | 4,8 | 0 | 1,02 | 0 | 5,3 | 0 |
| FOSL2::JUN | CCGAGACTCAGT | 1 | 442 | 454 | -558 | -546 | -0,28 | 0,01 | 14,25 | >0.1 | 2,75 | 4,85 | 0 | 1 | 0 | 5,92 | 0 |
| FOSL2::JUN | TCTTTGGTCACT | 1 | 669 | 681 | -331 | -319 | -1,7 | 0,01 | 12,66 | >0.1 | 4,14 | 5,42 | 0 | 1,1 | 0 | 3,7 | 0 |
| FOSL2::JUN | ATTTGATTCACT | 1 | 814 | 826 | -186 | -174 | 5,68 | 0,001 | 22,35 | >0.1 | 5,58 | 6,06 | 0 | 1,59 | 0 | 3,43 | 0 |
| FOSL2::JUN | TATTGGCTCAGC | 1 | 881 | 893 | -119 | -107 | -1,82 | 0,01 | 15,28 | >0.1 | 5,58 | 6,56 | 0 | 1,89 | 0 | 3,07 | 0 |

| | | | | | | | | | | | | | | | | | |
|---|---|---|---|---|---|---|---|---|---|---|---|---|---|---|---|---|---|
| FOSL2::JUN | TCTTGGCTCACA | 1 | 1030 | 1042 | 30 | 42 | -0,74 | 0,01 | 16,25 | >0.1 | 5,58 | 8,24 | 0 | 1,86 | 0 | 1,31 | 0 |
| FOSL2::JUN | CTGTTCCTCAAT | -1 | 1109 | 1121 | 109 | 121 | 1,54 | 0,01 | 21,42 | >0.1 | 5,58 | 6,66 | 0 | 1,85 | 3,22 | 2,56 | 0 |
| FOSL2::JUN | CTGTGAGCCAAG | -1 | 1031 | 1043 | 31 | 43 | 0,12 | 0,01 | 17,03 | >0.1 | 5,58 | 8,19 | 0 | 1,86 | 0 | 1,28 | 0 |
| FOSL2::JUN | CACTGAATGACT | -1 | 1011 | 1023 | 11 | 23 | -0,98 | 0,01 | 16,84 | >0.1 | 5,58 | 8,95 | 0 | 1,86 | 0 | 1,42 | 0 |
| FOSL2::JUN | TGCTGAGCCAAT | -1 | 882 | 894 | -118 | -106 | -0,94 | 0,01 | 16,17 | >0.1 | 5,58 | 6,57 | 0 | 1,89 | 0 | 3,07 | 0 |
| FOSL2::JUN | AAGTGAATCAAA | -1 | 815 | 827 | -185 | -173 | 4,93 | 0,01 | 21,73 | >0.1 | 5,58 | 6,07 | 0 | 1,6 | 0 | 3,55 | 0 |
| FOSL2::JUN | GACTGAGTCTCG | -1 | 443 | 455 | -557 | -545 | 2,78 | 0,01 | 17,32 | >0.1 | 2,76 | 4,85 | 0 | 1 | 0 | 5,92 | 0 |
| FOSL2::JUN | TCCTGCCTCAGC | -1 | 355 | 367 | -645 | -633 | 6,73 | 0,001 | 21,17 | >0.1 | 2,52 | 4,69 | 0 | 1,1 | 0 | 6,13 | 0 |
| FOSL2::JUN | GTGTTAGTCAGG | -1 | 262 | 274 | -738 | -726 | 9,44 | 0,001 | 22,56 | >0.1 | 2,8 | 4,55 | 0 | 1,32 | 0 | 4,45 | 0 |
| FOSL2::JUN | GCGTGAGCCACT | -1 | 186 | 198 | -814 | -802 | 4,45 | 0,01 | 17,59 | >0.1 | 3,1 | 4,45 | 0 | 1,74 | 0 | 3,86 | 0 |
| FOSL2::JUN | CTATTTCTCACA | -1 | 53 | 65 | -947 | -935 | 4,24 | 0,01 | 14,49 | >0.1 | 4,07 | 4,3 | 0 | 1,87 | 0 | 0 | 0 |
| FOSL2::JUN | GAGTGCCTCCTT | -1 | 18 | 30 | -982 | -970 | -2,48 | 0,01 | 9,24 | >0.1 | 5,58 | 4,27 | 0 | 1,87 | 0 | 0 | 0 |
| HIF1A | GTAAGTGAAG | 1 | 5 | 15 | -995 | -985 | -3,15 | 0,01 | 10,38 | >0.1 | 5,61 | 4,26 | 0 | 1,86 | 1,79 | 0 | 0 |
| HIF1A | TCACGCCTGT | 1 | 192 | 202 | -808 | -798 | -2,22 | 0,01 | 11,26 | >0.1 | 3,07 | 4,46 | 0 | 1,69 | 0 | 4,26 | 0 |
| HIF1A | GACCGTCCTG | 1 | 256 | 266 | -744 | -734 | -4,33 | 0,01 | 9,11 | >0.1 | 2,82 | 4,54 | 0 | 1,34 | 0 | 4,73 | 0 |
| HIF1A | GGGCGTGGTG | 1 | 314 | 324 | -686 | -676 | -0,07 | 0,01 | 14,5 | >0.1 | 2,64 | 4,63 | 0 | 1,17 | 0 | 6,13 | 0 |
| HIF1A | GCACCTGTAG | 1 | 328 | 338 | -672 | -662 | -3,83 | 0,01 | 10,12 | >0.1 | 2,6 | 4,65 | 0 | 1,14 | 0 | 5,56 | 0 |
| HIF1A | TGGCGTGAAC | 1 | 370 | 380 | -630 | -620 | -0,42 | 0,01 | 14,18 | >0.1 | 2,54 | 4,72 | 0 | 1,07 | 0 | 6,27 | 0 |
| HIF1A | GATCGCGCCA | 1 | 408 | 418 | -592 | -582 | -1,7 | 0,01 | 12,41 | >0.1 | 2,65 | 4,78 | 0 | 1,03 | 0 | 5,65 | 0 |
| HIF1A | AAATGTGAGA | 1 | 587 | 597 | -413 | -403 | -4,24 | 0,01 | 5,33 | >0.1 | 3,39 | 5,17 | 0 | 1,01 | 0 | 0 | 0 |
| HIF1A | TAACGTATTC | 1 | 802 | 812 | -198 | -188 | 1,05 | 0,01 | 17,58 | >0.1 | 5,61 | 5,98 | 0 | 1,51 | 0 | 3,43 | 0 |
| HIF1A | GGATGTGATC | 1 | 1022 | 1032 | 22 | 32 | -0,09 | 0,01 | 17,34 | >0.1 | 5,61 | 8,68 | 0 | 1,86 | 0 | 1,28 | 0 |
| HIF1A | CCACGAAGCC | 1 | 1147 | 1157 | 147 | 157 | -4,07 | 0,01 | 13,7 | >0.1 | 5,61 | 6,32 | 0 | 1,84 | 1,79 | 2,21 | 0 |
| HIF1A | TCACTTGCTT | 1 | 1178 | 1188 | 178 | 188 | -1,91 | 0,01 | 16,57 | >0.1 | 5,61 | 6,1 | 0 | 1,6 | 1,79 | 3,37 | 0 |
| HIF1A | GCAAGTGAAC | -1 | 1176 | 1186 | 176 | 186 | -2,78 | 0,01 | 15,63 | >0.1 | 5,61 | 6,11 | 0 | 1,62 | 1,79 | 3,28 | 0 |
| HIF1A | CTTCGTGGTT | -1 | 1145 | 1155 | 145 | 155 | -2,27 | 0,01 | 16,9 | >0.1 | 5,61 | 6,33 | 0 | 1,84 | 3,22 | 2,16 | 0 |
| HIF1A | CATCGTCCCC | -1 | 1042 | 1052 | 42 | 52 | -1,16 | 0,01 | 14,72 | >0.1 | 5,61 | 7,79 | 0 | 1,86 | 0 | 0,62 | 0 |
| HIF1A | TCACATCCAC | -1 | 1020 | 1030 | 20 | 30 | -2,87 | 0,01 | 14,63 | >0.1 | 5,61 | 8,72 | 0 | 1,86 | 0 | 1,31 | 0 |
| HIF1A | ATACGTTATT | -1 | 800 | 810 | -200 | -190 | 0,71 | 0,01 | 17,33 | >0.1 | 5,61 | 5,97 | 0 | 1,5 | 0 | 3,55 | 0 |
| HIF1A | ACACGGAAAA | -1 | 695 | 705 | -305 | -295 | -2,74 | 0,01 | 12,47 | >0.1 | 5,61 | 5,5 | 0 | 1,14 | 0 | 2,95 | 0 |
| HIF1A | CTACTTGGGC | -1 | 543 | 553 | -457 | -447 | -4,6 | 0,01 | 4,6 | >0.1 | 3,15 | 5,06 | 0 | 0,99 | 0 | 0 | 0 |
| HIF1A | ATACATGACT | -1 | 516 | 526 | -484 | -474 | -1,88 | 0,01 | 10,08 | >0.1 | 3,02 | 5 | 0 | 0,98 | 0 | 2,95 | 0 |
| HIF1A | TGGCGCGATC | -1 | 408 | 418 | -592 | -582 | -4,6 | 0,01 | 9,51 | >0.1 | 2,65 | 4,78 | 0 | 1,03 | 0 | 5,65 | 0 |
| HIF1A | TCACGCCATT | -1 | 368 | 378 | -632 | -622 | -1,57 | 0,01 | 13,48 | >0.1 | 2,54 | 4,71 | 0 | 1,07 | 0 | 6,73 | 0 |
| HIF1A | ACAGGTGCCC | -1 | 326 | 336 | -674 | -664 | 0,88 | 0,01 | 14,73 | >0.1 | 2,6 | 4,64 | 0 | 1,15 | 0 | 5,46 | 0 |
| HIF1A | CCACGCCCAG | -1 | 312 | 322 | -688 | -678 | 3,97 | 0,001 | 18,45 | >0.1 | 2,65 | 4,62 | 0 | 1,18 | 0 | 6,03 | 0 |
| HIF1A | AGACGGGGTT | -1 | 279 | 289 | -721 | -711 | 0,04 | 0,01 | 13,82 | >0.1 | 2,75 | 4,57 | 0 | 1,26 | 0 | 5,19 | 0 |
| HIF1A | GGACGGTCTC | -1 | 254 | 264 | -746 | -736 | 1,2 | 0,01 | 14,77 | >0.1 | 2,83 | 4,54 | 0 | 1,35 | 0 | 4,85 | 0 |
| HIF1A | CAAAGTGCTA | -1 | 207 | 217 | -793 | -783 | -3,05 | 0,01 | 10,72 | >0.1 | 3,01 | 4,48 | 0 | 1,58 | 0 | 4,7 | 0 |
| HIF1A | AGGCGTGAGC | -1 | 190 | 200 | -810 | -800 | 0,89 | 0,01 | 14,26 | >0.1 | 3,08 | 4,45 | 0 | 1,7 | 0 | 4,13 | 0 |
| GCM1 | GAAGGAGGCAC | 1 | 17 | 28 | -983 | -972 | -1,4 | 0,01 | 10,33 | >0.1 | 5,6 | 4,27 | 0 | 1,87 | 0 | 0 | 0 |
| GCM1 | CGTGGTGGTGG | 1 | 317 | 328 | -683 | -672 | 6,35 | 0,001 | 21,03 | >0.1 | 2,63 | 4,63 | 0 | 1,17 | 0 | 6,26 | 0 |
| GCM1 | GTGGTGGGCAC | 1 | 321 | 332 | -679 | -668 | -2,21 | 0,01 | 11,77 | >0.1 | 2,62 | 4,64 | 0 | 1,16 | 0 | 5,56 | 0 |
| GCM1 | GACAAGGGCAA | 1 | 505 | 516 | -495 | -484 | -2,36 | 0,01 | 12,26 | >0.1 | 2,98 | 4,98 | 0 | 0,98 | 0 | 5,68 | 0 |
| GCM1 | AGTACCGGTTT | 1 | 740 | 751 | -260 | -249 | -0,85 | 0,01 | 17,6 | >0.1 | 5,6 | 5,68 | 0 | 1,26 | 0 | 5,92 | 0 |
| GCM1 | TATATTGGCTC | 1 | 879 | 890 | -121 | -110 | 1,78 | 0,01 | 18,99 | >0.1 | 5,6 | 6,53 | 0 | 1,9 | 0 | 3,19 | 0 |
| GCM1 | GATCTTGGCTC | 1 | 1028 | 1039 | 28 | 39 | -1,23 | 0,01 | 15,87 | >0.1 | 5,6 | 8,34 | 0 | 1,86 | 0 | 1,31 | 0 |
| GCM1 | CTTCCTGGCTC | 1 | 1059 | 1070 | 59 | 70 | -1,76 | 0,01 | 15,23 | >0.1 | 5,6 | 7,37 | 0 | 1,86 | 0 | 2,16 | 0 |
| GCM1 | CATCCTGGAGA | -1 | 863 | 874 | -137 | -126 | -3,09 | 0,01 | 13,47 | >0.1 | 5,6 | 6,4 | 0 | 1,9 | 0 | 2,67 | 0 |
| GCM1 | AAAACCGGTAC | -1 | 741 | 752 | -259 | -248 | 0,4 | 0,01 | 19,05 | >0.1 | 5,6 | 5,69 | 0 | 1,26 | 0 | 6,1 | 0 |
| GCM1 | AATGTTGGTGT | -1 | 632 | 643 | -368 | -357 | 4,64 | 0,001 | 16,33 | >0.1 | 3,73 | 5,3 | 0 | 1,05 | 0 | 1,61 | 0 |
| GCM1 | CAGGCTGGAGC | -1 | 422 | 433 | -578 | -567 | 0,48 | 0,01 | 14,55 | >0.1 | 2,69 | 4,81 | 0 | 1,02 | 0 | 5,56 | 0 |
| GCM1 | CACAGTGGCGC | -1 | 412 | 423 | -588 | -577 | -1,27 | 0,01 | 13,12 | >0.1 | 2,66 | 4,79 | 0 | 1,03 | 0 | 5,91 | 0 |
| GCM1 | ACTACAGGTGC | -1 | 328 | 339 | -672 | -661 | 4,18 | 0,01 | 18,13 | >0.1 | 2,6 | 4,65 | 0 | 1,14 | 0 | 5,56 | 0 |
| GCM1 | CACACTGGTTC | -1 | 46 | 57 | -954 | -943 | 3,08 | 0,01 | 13,43 | >0.1 | 4,18 | 4,3 | 0 | 1,87 | 0 | 0 | 0 |
| TFAP2C(VAR.3) | GGGCACAGTGGCT | 1 | 180 | 193 | -820 | -807 | -12,59 | 0,01 | 0,62 | >0.1 | 3,12 | 4,45 | 0 | 1,79 | 0 | 3,86 | 0 |
| TFAP2C(VAR.3) | GGCCGAGTTGGGC | 1 | 220 | 233 | -780 | -767 | -5,27 | 0,01 | 9,21 | >0.1 | 2,95 | 4,5 | 0 | 1,51 | 0 | 5,52 | 0 |
| TFAP2C(VAR.3) | GCCGAGTTGGGCA | 1 | 221 | 234 | -779 | -766 | -10,8 | 0,01 | 3,67 | >0.1 | 2,94 | 4,5 | 0 | 1,5 | 0 | 5,52 | 0 |
| TFAP2C(VAR.3) | TACTCGGGAGGCT | 1 | 345 | 358 | -655 | -642 | -12,63 | 0,01 | 2,23 | >0.1 | 2,55 | 4,68 | 0 | 1,11 | 0 | 6,52 | 0 |

| | | | | | | | | | | | | | | | | | |
|---|---|---|---|---|---|---|---|---|---|---|---|---|---|---|---|---|---|
| TFAP2C(VAR.3) | AACCTGGGAGGCA | 1 | 377 | 390 | -623 | -610 | -1,62 | 0,01 | 13,28 | >0.1 | 2,57 | 4,73 | 0 | 1,06 | 0 | 6,53 | 0 |
| TFAP2C(VAR.3) | AGCCGAGATCGCG | 1 | 402 | 415 | -598 | -585 | -11,57 | 0,01 | 2,64 | >0.1 | 2,63 | 4,78 | 0 | 1,04 | 0 | 5,76 | 0 |
| TFAP2C(VAR.3) | TCCAGCCTGGGCG | 1 | 424 | 437 | -576 | -563 | -4,56 | 0,01 | 9,64 | >0.1 | 2,7 | 4,82 | 0 | 1,02 | 0 | 5,67 | 0 |
| TFAP2C(VAR.3) | AGCCTGGGCGACA | 1 | 427 | 440 | -573 | -560 | -3,31 | 0,01 | 11,04 | >0.1 | 2,71 | 4,82 | 0 | 1,01 | 0 | 5,8 | 0 |
| TFAP2C(VAR.3) | GGCCATAAAGTGA | 1 | 757 | 770 | -243 | -230 | -10,33 | 0,01 | 8,43 | >0.1 | 5,57 | 5,77 | 0 | 1,32 | 0 | 6,1 | 0 |
| TFAP2C(VAR.3) | TGGCTCACAGGGG | 1 | 1033 | 1046 | 33 | 46 | -5,14 | 0,01 | 11,7 | >0.1 | 5,57 | 8,1 | 0 | 1,86 | 0 | 1,31 | 0 |
| TFAP2C(VAR.3) | GGCTCACAGGGGA | 1 | 1034 | 1047 | 34 | 47 | -1,36 | 0,01 | 15,44 | >0.1 | 5,57 | 8,06 | 0 | 1,86 | 0 | 1,31 | 0 |
| TFAP2C(VAR.3) | GCTCTTCCTGGCT | 1 | 1056 | 1069 | 56 | 69 | -10,51 | 0,01 | 6,63 | >0.1 | 5,57 | 7,44 | 0 | 1,86 | 0 | 2,27 | 0 |
| TFAP2C(VAR.3) | AGCCAGGAAGAGC | -1 | 1056 | 1069 | 56 | 69 | -11,16 | 0,01 | 5,98 | >0.1 | 5,57 | 7,44 | 0 | 1,86 | 0 | 2,27 | 0 |
| TFAP2C(VAR.3) | TCCCCTGTGAGCC | -1 | 1034 | 1047 | 34 | 47 | -1,65 | 0,01 | 15,15 | >0.1 | 5,57 | 8,06 | 0 | 1,86 | 0 | 1,31 | 0 |
| TFAP2C(VAR.3) | CCCCTGTGAGCCA | -1 | 1033 | 1046 | 33 | 46 | -5,1 | 0,01 | 11,74 | >0.1 | 5,57 | 8,1 | 0 | 1,86 | 0 | 1,31 | 0 |
| TFAP2C(VAR.3) | TCACTTTATGGCC | -1 | 757 | 770 | -243 | -230 | -9,68 | 0,01 | 9,08 | >0.1 | 5,57 | 5,77 | 0 | 1,32 | 0 | 6,1 | 0 |
| TFAP2C(VAR.3) | TGTCGCCCAGGCT | -1 | 427 | 440 | -573 | -560 | -3,24 | 0,01 | 11,11 | >0.1 | 2,71 | 4,82 | 0 | 1,01 | 0 | 5,8 | 0 |
| TFAP2C(VAR.3) | CGCCCAGGCTGGA | -1 | 424 | 437 | -576 | -563 | -4,96 | 0,01 | 9,24 | >0.1 | 2,7 | 4,82 | 0 | 1,02 | 0 | 5,67 | 0 |
| TFAP2C(VAR.3) | CGCGATCTCGGCT | -1 | 402 | 415 | -598 | -585 | -11,53 | 0,01 | 2,68 | >0.1 | 2,63 | 4,78 | 0 | 1,04 | 0 | 5,76 | 0 |
| TFAP2C(VAR.3) | TGCCTCCCAGGTT | -1 | 377 | 390 | -623 | -610 | -1,71 | 0,01 | 13,19 | >0.1 | 2,57 | 4,73 | 0 | 1,06 | 0 | 6,53 | 0 |
| TFAP2C(VAR.3) | AGCCTCCCGAGTA | -1 | 345 | 358 | -655 | -642 | -12,89 | 0,01 | 1,97 | >0.1 | 2,55 | 4,68 | 0 | 1,11 | 0 | 6,52 | 0 |
| TFAP2C(VAR.3) | TGCCCAACTCGGC | -1 | 221 | 234 | -779 | -766 | -11,5 | 0,01 | 2,97 | >0.1 | 2,94 | 4,5 | 0 | 1,5 | 0 | 5,52 | 0 |
| TFAP2C(VAR.3) | GCCCAACTCGGCC | -1 | 220 | 233 | -780 | -767 | -5,13 | 0,01 | 9,35 | >0.1 | 2,95 | 4,5 | 0 | 1,51 | 0 | 5,52 | 0 |
| TFAP2C(VAR.3) | AGCCACTGTGCCC | -1 | 180 | 193 | -820 | -807 | -12,72 | 0,01 | 0,49 | >0.1 | 3,12 | 4,45 | 0 | 1,79 | 0 | 3,86 | 0 |
| ZSCAN4 | AGGAGGCACTCAAAG | 1 | 19 | 34 | -981 | -966 | -15,13 | 0,01 | -3,44 | >0.1 | 5,54 | 4,28 | 0 | 1,87 | 0 | 0 | 0 |
| ZSCAN4 | GGCACTCAAAGAAGT | 1 | 23 | 38 | -977 | -962 | -13,54 | 0,01 | -1,85 | >0.1 | 5,54 | 4,28 | 0 | 1,87 | 0 | 0 | 0 |
| ZSCAN4 | GGCTCACGCCTGTAA | 1 | 189 | 204 | -811 | -796 | -7,22 | 0,01 | 6,16 | >0.1 | 3,08 | 4,46 | 0 | 1,71 | 0 | 4,13 | 0 |
| ZSCAN4 | AGATCACAAGGTCAG | 1 | 233 | 248 | -767 | -752 | -16,11 | 0,01 | -2,51 | >0.1 | 2,9 | 4,52 | 0 | 1,44 | 0 | 4,74 | 0 |
| ZSCAN4 | CTAACACGGTGAAAC | 1 | 267 | 282 | -733 | -718 | -18,47 | 0,01 | -4,74 | >0.1 | 2,78 | 4,56 | 0 | 1,31 | 0 | 5,09 | 0 |
| ZSCAN4 | GACTCAGTCTCAAAA | 1 | 446 | 461 | -554 | -539 | -14,6 | 0,01 | -0,7 | >0.1 | 2,77 | 4,86 | 0 | 1 | 0 | 5,26 | 0 |
| ZSCAN4 | TAAAGACAAGGGCAA | 1 | 501 | 516 | -499 | -484 | -19,8 | 0,01 | -5,18 | >0.1 | 2,97 | 4,98 | 0 | 0,99 | 0 | 5,68 | 0 |
| ZSCAN4 | CATACAAACACCAAC | 1 | 625 | 640 | -375 | -360 | -19,51 | 0,01 | -7,94 | >0.1 | 3,68 | 5,29 | 0 | 1,05 | 0 | 1,55 | 0 |
| ZSCAN4 | TACAAACACCAACAT | 1 | 627 | 642 | -373 | -358 | -11,18 | 0,01 | 0,53 | >0.1 | 3,7 | 5,3 | 0 | 1,05 | 0 | 1,66 | 0 |
| ZSCAN4 | AACCCAAGTTCAAAG | 1 | 951 | 966 | -49 | -34 | -18,61 | 0,01 | -1,96 | >0.1 | 5,54 | 7,53 | 0 | 1,88 | 0 | 1,7 | 0 |
| ZSCAN4 | GGCTCACAGGGGACG | 1 | 1034 | 1049 | 34 | 49 | -18,61 | 0,01 | -1,81 | >0.1 | 5,54 | 8,06 | 0 | 1,86 | 0 | 1,34 | 0 |
| ZSCAN4 | AGCTCTTCCTGGCTC | 1 | 1055 | 1070 | 55 | 70 | -16,05 | 0,01 | 1,08 | >0.1 | 5,54 | 7,46 | 0 | 1,86 | 0 | 2,27 | 0 |
| ZSCAN4 | AGCAGTTACAGCAAC | -1 | 1082 | 1097 | 82 | 97 | -16,44 | 0,01 | 1,9 | >0.1 | 5,54 | 6,99 | 0 | 1,85 | 1,79 | 2,16 | 0 |
| ZSCAN4 | CACATCCACTGAATG | -1 | 1014 | 1029 | 14 | 29 | -15,47 | 0,01 | 2,16 | >0.1 | 5,54 | 8,86 | 0 | 1,86 | 0 | 1,36 | 0 |
| ZSCAN4 | AGTTCATCCTGGAGA | -1 | 863 | 878 | -137 | -122 | -15,57 | 0,01 | 0,97 | >0.1 | 5,54 | 6,43 | 0 | 1,9 | 0 | 2,67 | 0 |
| ZSCAN4 | AAGATACACGGAAAA | -1 | 695 | 710 | -305 | -290 | -7,84 | 0,01 | 7,57 | >0.1 | 5,54 | 5,52 | 0 | 1,16 | 0 | 3,19 | 0 |
| ZSCAN4 | TTCAGACAATTTCAT | -1 | 607 | 622 | -393 | -378 | -18,33 | 0,01 | -6,97 | >0.1 | 3,54 | 5,24 | 0 | 1,03 | 0 | 1,55 | 0 |
| ZSCAN4 | CGCCCAGGCTGGAGC | -1 | 422 | 437 | -578 | -563 | -14 | 0,01 | 0,2 | >0.1 | 2,7 | 4,82 | 0 | 1,02 | 0 | 5,67 | 0 |
| ZSCAN4 | TCGGCTCACTGCAAG | -1 | 393 | 408 | -607 | -592 | -19,64 | 0,01 | -5,45 | >0.1 | 2,61 | 4,77 | 0 | 1,05 | 0 | 5,76 | 0 |
| ZSCAN4 | TGCAAGCTCTGCCTC | -1 | 384 | 399 | -616 | -601 | -19,54 | 0,01 | -5,02 | >0.1 | 2,59 | 4,75 | 0 | 1,06 | 0 | 6,13 | 0 |
| ZSCAN4 | AGCTCTGCCTCCCAG | -1 | 380 | 395 | -620 | -605 | -19,48 | 0,01 | -5,19 | >0.1 | 2,58 | 4,74 | 0 | 1,06 | 0 | 5,91 | 0 |
| ZSCAN4 | TGCCCACCACCACGC | -1 | 316 | 331 | -684 | -669 | -19,56 | 0,01 | -4,87 | >0.1 | 2,63 | 4,64 | 0 | 1,17 | 0 | 6,26 | 0 |
| ZSCAN4 | ACCACGCCCAGCTAA | -1 | 308 | 323 | -692 | -677 | -13,87 | 0,01 | 0,72 | >0.1 | 2,65 | 4,62 | 0 | 1,19 | 0 | 6,13 | 0 |
| ZSCAN4 | TGCAAAACTTAAAAA | -1 | 75 | 90 | -925 | -910 | -17,76 | 0,01 | -7,74 | >0.1 | 3,81 | 4,33 | 0 | 1,88 | 0 | 0 | 0 |
| ZSCAN4 | TTCTCACACTGGTTC | -1 | 46 | 61 | -954 | -939 | -18,44 | 0,01 | -8,12 | >0.1 | 4,15 | 4,3 | 0 | 1,87 | 0 | 0 | 0 |
| FOSL1::JUND | AAGGAGGCAC | 1 | 18 | 28 | -982 | -972 | 4,69 | 0,01 | 16,43 | >0.1 | 5,61 | 4,27 | 0 | 1,87 | 0 | 0 | 0 |
| FOSL1::JUND | TGTGAGAAAT | 1 | 53 | 63 | -947 | -937 | 5,78 | 0,01 | 16,04 | >0.1 | 4,09 | 4,3 | 0 | 1,87 | 0 | 0 | 0 |
| FOSL1::JUND | GAGTAGTCAT | 1 | 124 | 134 | -876 | -866 | 4,29 | 0,01 | 15,69 | >0.1 | 3,44 | 4,38 | 0 | 1,89 | 0 | 1,7 | 0 |
| FOSL1::JUND | AGTGGCTCAC | 1 | 186 | 196 | -814 | -804 | 8,52 | 0,001 | 21,66 | >0.1 | 3,1 | 4,45 | 0 | 1,74 | 0 | 3,86 | 0 |
| FOSL1::JUND | CCTGACTAAC | 1 | 262 | 272 | -738 | -728 | 8,76 | 0,001 | 21,96 | >0.1 | 2,8 | 4,55 | 0 | 1,32 | 0 | 4,53 | 0 |
| FOSL1::JUND | GCTGAGGCAG | 1 | 355 | 365 | -645 | -635 | 7,33 | 0,01 | 21,68 | >0.1 | 2,53 | 4,69 | 0 | 1,09 | 0 | 6,03 | 0 |
| FOSL1::JUND | TGGGAGGCAG | 1 | 381 | 391 | -619 | -609 | 3,49 | 0,01 | 17,87 | >0.1 | 2,57 | 4,73 | 0 | 1,06 | 0 | 6,02 | 0 |
| FOSL1::JUND | CGAGACTCAG | 1 | 443 | 453 | -557 | -547 | 6,17 | 0,01 | 20,7 | >0.1 | 2,75 | 4,85 | 0 | 1 | 0 | 5,92 | 0 |
| FOSL1::JUND | TTTGATTCAC | 1 | 815 | 825 | -185 | -175 | 5,75 | 0,01 | 22,43 | >0.1 | 5,61 | 6,06 | 0 | 1,58 | 0 | 3,43 | 0 |
| FOSL1::JUND | ATTGGCTCAG | 1 | 882 | 892 | -118 | -108 | 4,65 | 0,01 | 21,77 | >0.1 | 5,61 | 6,55 | 0 | 1,89 | 0 | 3,07 | 0 |
| FOSL1::JUND | AGTCATTCAG | 1 | 1011 | 1021 | 11 | 21 | 5 | 0,01 | 22,82 | >0.1 | 5,61 | 8,95 | 0 | 1,86 | 0 | 1,39 | 0 |
| FOSL1::JUND | CTTGGCTCAC | 1 | 1031 | 1041 | 31 | 41 | 5,44 | 0,01 | 22,41 | >0.1 | 5,61 | 8,19 | 0 | 1,86 | 0 | 1,31 | 0 |
| FOSL1::JUND | CCTGGCTCCT | 1 | 1062 | 1072 | 62 | 72 | 4,24 | 0,01 | 23,03 | >0.1 | 5,61 | 7,31 | 0 | 1,86 | 1,79 | 2,21 | 0 |
| FOSL1::JUND | CGTGGTTAAA | -1 | 1142 | 1152 | 142 | 152 | 4,04 | 0,01 | 24,44 | >0.1 | 5,61 | 6,36 | 0 | 1,84 | 3,22 | 3,37 | 0 |

| | | | | | | | | | | | | | | | | | |
|---|---|---|---|---|---|---|---|---|---|---|---|---|---|---|---|---|---|
| FOSL1::JUND | TGTGAGCCAA | -1 | 1032 | 1042 | 32 | 42 | 5,46 | 0,01 | 22,35 | >0.1 | 5,61 | 8,15 | 0 | 1,86 | 0 | 1,28 | 0 |
| FOSL1::JUND | GCTGAGCCAA | -1 | 883 | 893 | -117 | -107 | 5,06 | 0,01 | 22,19 | >0.1 | 5,61 | 6,56 | 0 | 1,89 | 0 | 3,07 | 0 |
| FOSL1::JUND | AGTGAATCAA | -1 | 816 | 826 | -184 | -174 | 6,57 | 0,01 | 23,38 | >0.1 | 5,61 | 6,06 | 0 | 1,59 | 0 | 3,55 | 0 |
| FOSL1::JUND | AGTGACCAAA | -1 | 671 | 681 | -329 | -319 | 5,5 | 0,01 | 19,73 | >0.1 | 4,16 | 5,42 | 0 | 1,1 | 0 | 3,55 | 0 |
| FOSL1::JUND | ACTGAGTCTC | -1 | 444 | 454 | -556 | -546 | 3,79 | 0,01 | 18,32 | >0.1 | 2,76 | 4,85 | 0 | 1 | 0 | 5,92 | 0 |
| FOSL1::JUND | CCTGCCTCAG | -1 | 356 | 366 | -644 | -634 | 4,29 | 0,01 | 18,73 | >0.1 | 2,52 | 4,69 | 0 | 1,09 | 0 | 6,13 | 0 |
| FOSL1::JUND | TGTTAGTCAG | -1 | 263 | 273 | -737 | -727 | 6,11 | 0,01 | 19,22 | >0.1 | 2,8 | 4,55 | 0 | 1,32 | 0 | 4,45 | 0 |
| FOSL1::JUND | CGTGAGCCAC | -1 | 187 | 197 | -813 | -803 | 6,65 | 0,01 | 19,78 | >0.1 | 3,1 | 4,45 | 0 | 1,73 | 0 | 3,86 | 0 |
| FOSL1::JUND | TATGACTACT | -1 | 125 | 135 | -875 | -865 | 6,61 | 0,01 | 18,01 | >0.1 | 3,43 | 4,38 | 0 | 1,89 | 0 | 1,7 | 0 |
| FOSL1::JUND | TATTTCTCAC | -1 | 54 | 64 | -946 | -936 | 4,35 | 0,01 | 14,6 | >0.1 | 4,07 | 4,3 | 0 | 1,87 | 0 | 0 | 0 |
| FOSL1::JUND | TGTGGCTGAC | -1 | 36 | 46 | -964 | -954 | 3,67 | 0,01 | 14,18 | >0.1 | 4,36 | 4,29 | 0 | 1,87 | 0 | 0 | 0 |
| FOSL1::JUND | GCTGACTTCT | -1 | 32 | 42 | -968 | -958 | 3,66 | 0,01 | 14,25 | >0.1 | 4,44 | 4,28 | 0 | 1,87 | 0 | 0 | 0 |
| FOSL1::JUNB | GAAGGAGGCACTC | 1 | 17 | 30 | -983 | -970 | 3,03 | 0,01 | 14,74 | >0.1 | 5,57 | 4,27 | 0 | 1,87 | 0 | 0 | 0 |
| FOSL1::JUNB | GTGTGAGAAATAG | 1 | 52 | 65 | -948 | -935 | 5,88 | 0,01 | 16,14 | >0.1 | 4,08 | 4,3 | 0 | 1,87 | 0 | 0 | 0 |
| FOSL1::JUNB | AGAGTAGTCATAA | 1 | 123 | 136 | -877 | -864 | 2,74 | 0,01 | 14,14 | >0.1 | 3,44 | 4,38 | 0 | 1,89 | 0 | 1,7 | 0 |
| FOSL1::JUNB | CAGTGGCTCACGC | 1 | 185 | 198 | -815 | -802 | 7,94 | 0,001 | 21,09 | >0.1 | 3,1 | 4,45 | 0 | 1,75 | 0 | 3,86 | 0 |
| FOSL1::JUNB | TCCTGACTAACAC | 1 | 261 | 274 | -739 | -726 | 9,19 | 0,001 | 22,4 | >0.1 | 2,8 | 4,55 | 0 | 1,33 | 0 | 4,53 | 0 |
| FOSL1::JUNB | GGCTGAGGCAGGA | 1 | 354 | 367 | -646 | -633 | 7,05 | 0,01 | 21,4 | >0.1 | 2,53 | 4,69 | 0 | 1,1 | 0 | 6,03 | 0 |
| FOSL1::JUNB | CTGGGAGGCAGAG | 1 | 380 | 393 | -620 | -607 | 3,06 | 0,01 | 17,45 | >0.1 | 2,57 | 4,74 | 0 | 1,06 | 0 | 6,02 | 0 |
| FOSL1::JUNB | CCGAGACTCAGTC | 1 | 442 | 455 | -558 | -545 | 6,52 | 0,01 | 21,05 | >0.1 | 2,76 | 4,85 | 0 | 1 | 0 | 5,92 | 0 |
| FOSL1::JUNB | ATTTGATTCACTT | 1 | 814 | 827 | -186 | -173 | 5 | 0,01 | 21,67 | >0.1 | 5,57 | 6,07 | 0 | 1,6 | 0 | 3,43 | 0 |
| FOSL1::JUNB | TATTGGCTCAGCA | 1 | 881 | 894 | -119 | -106 | 3,87 | 0,01 | 20,97 | >0.1 | 5,57 | 6,57 | 0 | 1,89 | 0 | 3,07 | 0 |
| FOSL1::JUNB | AAGTCATTCAGTG | 1 | 1010 | 1023 | 10 | 23 | 2,49 | 0,01 | 20,31 | >0.1 | 5,57 | 8,99 | 0 | 1,86 | 0 | 1,39 | 0 |
| FOSL1::JUNB | TCTTGGCTCACAG | 1 | 1030 | 1043 | 30 | 43 | 4,07 | 0,01 | 21,04 | >0.1 | 5,57 | 8,24 | 0 | 1,86 | 0 | 1,31 | 0 |
| FOSL1::JUNB | CATTGAGGAACAG | 1 | 1108 | 1121 | 108 | 121 | 3,62 | 0,01 | 23,45 | >0.1 | 5,57 | 6,67 | 0 | 1,85 | 3,22 | 2,52 | 0 |
| FOSL1::JUNB | CTGTGAGCCAAGA | -1 | 1030 | 1043 | 30 | 43 | 3,53 | 0,01 | 20,5 | >0.1 | 5,57 | 8,24 | 0 | 1,86 | 0 | 1,31 | 0 |
| FOSL1::JUNB | GAATGACTTTCCC | -1 | 1006 | 1019 | 6 | 19 | 4,15 | 0,01 | 22,17 | >0.1 | 5,57 | 9,12 | 0 | 1,87 | 0 | 1,46 | 0 |
| FOSL1::JUNB | AAGTGAATCAAAT | -1 | 814 | 827 | -186 | -173 | 5 | 0,01 | 21,67 | >0.1 | 5,57 | 6,07 | 0 | 1,6 | 0 | 3,43 | 0 |
| FOSL1::JUNB | CAGTGACCAAAGA | -1 | 669 | 682 | -331 | -318 | 3,08 | 0,01 | 17,46 | >0.1 | 4,15 | 5,43 | 0 | 1,1 | 0 | 3,7 | 0 |
| FOSL1::JUNB | ACATGACTTTGCC | -1 | 511 | 524 | -489 | -476 | 3,46 | 0,01 | 15,4 | >0.1 | 3,01 | 5 | 0 | 0,99 | 0 | 2,95 | 0 |
| FOSL1::JUNB | GACTGAGTCTCGG | -1 | 442 | 455 | -558 | -545 | 4,21 | 0,01 | 18,74 | >0.1 | 2,76 | 4,85 | 0 | 1 | 0 | 5,92 | 0 |
| FOSL1::JUNB | TCCTGCCTCAGCC | -1 | 354 | 367 | -646 | -633 | 6,09 | 0,01 | 20,44 | >0.1 | 2,53 | 4,69 | 0 | 1,1 | 0 | 6,03 | 0 |
| FOSL1::JUNB | GTGTTAGTCAGGA | -1 | 261 | 274 | -739 | -726 | 5,57 | 0,01 | 18,78 | >0.1 | 2,8 | 4,55 | 0 | 1,33 | 0 | 4,53 | 0 |
| FOSL1::JUNB | GCGTGAGCCACTG | -1 | 185 | 198 | -815 | -802 | 5,11 | 0,01 | 18,26 | >0.1 | 3,1 | 4,45 | 0 | 1,75 | 0 | 3,86 | 0 |
| FOSL1::JUNB | TTATGACTACTCT | -1 | 123 | 136 | -877 | -864 | 7,03 | 0,01 | 18,43 | >0.1 | 3,44 | 4,38 | 0 | 1,89 | 0 | 1,7 | 0 |
| FOSL1::JUNB | CTATTTCTCACAC | -1 | 52 | 65 | -948 | -935 | 4,97 | 0,01 | 15,23 | >0.1 | 4,08 | 4,3 | 0 | 1,87 | 0 | 0 | 0 |
| FOSL1::JUNB | GAGTGCCTCCTTC | -1 | 17 | 30 | -983 | -970 | 2,75 | 0,01 | 14,46 | >0.1 | 5,57 | 4,27 | 0 | 1,87 | 0 | 0 | 0 |
| EGR2 | TCAGCCACAGA | 1 | 37 | 48 | -963 | -952 | -4,22 | 0,01 | 6,27 | >0.1 | 4,33 | 4,29 | 0 | 1,87 | 0 | 0 | 0 |
| EGR2 | ATGGCCATGGA | 1 | 145 | 156 | -855 | -844 | -2,78 | 0,01 | 8,43 | >0.1 | 3,31 | 4,4 | 0 | 1,9 | 0 | 1,61 | 0 |
| EGR2 | TGGCTCACGCC | 1 | 188 | 199 | -812 | -801 | 1,8 | 0,01 | 15,01 | >0.1 | 3,09 | 4,45 | 0 | 1,72 | 0 | 3,95 | 0 |
| EGR2 | CCAGCTACTCG | 1 | 340 | 351 | -660 | -649 | 1,43 | 0,01 | 16,18 | >0.1 | 2,57 | 4,67 | 0 | 1,12 | 0 | 6,4 | 0 |
| EGR2 | TTGCCCAAGTA | 1 | 541 | 552 | -459 | -448 | -4,7 | 0,01 | 4,49 | >0.1 | 3,14 | 5,06 | 0 | 0,99 | 0 | 0 | 0 |
| EGR2 | TTTCCTGCTCA | 1 | 934 | 945 | -66 | -55 | -5,36 | 0,01 | 10,15 | >0.1 | 5,6 | 7,14 | 0 | 1,88 | 0 | 0,9 | 0 |
| EGR2 | GCTCCTTCTCA | 1 | 1066 | 1077 | 66 | 77 | -4,63 | 0,01 | 14,13 | >0.1 | 5,6 | 7,24 | 0 | 1,85 | 1,79 | 2,27 | 0 |
| EGR2 | ACTGCTGCTCA | 1 | 1091 | 1102 | 91 | 102 | -3,39 | 0,01 | 15,06 | >0.1 | 5,6 | 6,87 | 0 | 1,85 | 1,79 | 2,34 | 0 |
| EGR2 | TCGGCTTCGTG | -1 | 1148 | 1159 | 148 | 159 | -6,32 | 0,01 | 11,4 | >0.1 | 5,6 | 6,31 | 0 | 1,84 | 1,79 | 2,18 | 0 |
| EGR2 | TCGTCCCCTGT | -1 | 1039 | 1050 | 39 | 50 | -5,46 | 0,01 | 11,24 | >0.1 | 5,6 | 7,88 | 0 | 1,86 | 0 | 1,36 | 0 |
| EGR2 | CGTCCCCTGTG | -1 | 1038 | 1049 | 38 | 49 | -5,5 | 0,01 | 11,21 | >0.1 | 5,6 | 7,92 | 0 | 1,86 | 0 | 1,34 | 0 |
| EGR2 | ACATCCACTGA | -1 | 1017 | 1028 | 17 | 28 | -0,79 | 0,01 | 16,79 | >0.1 | 5,6 | 8,79 | 0 | 1,86 | 0 | 1,34 | 0 |
| EGR2 | TCGCCCAGGCT | -1 | 427 | 438 | -573 | -562 | 6,43 | 0,001 | 20,77 | >0.1 | 2,71 | 4,82 | 0 | 1,01 | 0 | 5,8 | 0 |
| EGR2 | CCTCCCAGGTT | -1 | 377 | 388 | -623 | -612 | 0,34 | 0,01 | 15,09 | >0.1 | 2,56 | 4,73 | 0 | 1,06 | 0 | 6,4 | 0 |
| EGR2 | AGGTTCACGCC | -1 | 371 | 382 | -629 | -618 | -1,41 | 0,01 | 13,33 | >0.1 | 2,55 | 4,72 | 0 | 1,07 | 0 | 6,4 | 0 |
| EGR2 | ACCACCACGCC | -1 | 315 | 326 | -685 | -674 | -4,68 | 0,01 | 9,88 | >0.1 | 2,63 | 4,63 | 0 | 1,17 | 0 | 6,13 | 0 |
| EGR2 | ATGACTACTCT | -1 | 123 | 134 | -877 | -866 | -2,39 | 0,01 | 9,01 | >0.1 | 3,44 | 4,38 | 0 | 1,89 | 0 | 1,7 | 0 |
| EGR2 | CTGCCTTTGCA | -1 | 86 | 97 | -914 | -903 | -3,64 | 0,01 | 6,3 | >0.1 | 3,73 | 4,34 | 0 | 1,88 | 0 | 0 | 0 |
| EGR2 | TCTCACACTGG | -1 | 49 | 60 | -951 | -940 | -5,67 | 0,01 | 4,64 | >0.1 | 4,13 | 4,3 | 0 | 1,87 | 0 | 0 | 0 |
| EGR2 | GTGCCTCCTTC | -1 | 17 | 28 | -983 | -972 | -6,07 | 0,01 | 5,66 | >0.1 | 5,6 | 4,27 | 0 | 1,87 | 0 | 0 | 0 |
| EGR2 | CCTCCTTCTCC | -1 | 14 | 25 | -986 | -975 | -4,74 | 0,01 | 6,99 | >0.1 | 5,6 | 4,27 | 0 | 1,87 | 0 | 0 | 0 |
| EGR3 | AGTGGCTCACGCCTG | 1 | 186 | 201 | -814 | -799 | -11,07 | 0,01 | 2,16 | >0.1 | 3,09 | 4,46 | 0 | 1,74 | 0 | 3,95 | 0 |

| | | | | | | | | | | | | | | | | | |
|---|---|---|---|---|---|---|---|---|---|---|---|---|---|---|---|---|---|
| EGR3 | AAACCCCGTCTCTAT | 1 | 278 | 293 | -722 | -707 | -11,8 | 0,01 | 1,88 | >0.1 | 2,74 | 4,58 | 0 | 1,27 | 0 | 5,09 | 0 |
| EGR3 | TCCCAGCTACTCGGG | 1 | 338 | 353 | -662 | -647 | -9,74 | 0,01 | 5,02 | >0.1 | 2,57 | 4,67 | 0 | 1,13 | 0 | 6,4 | 0 |
| EGR3 | TCCAGCCTGGGCGAC | 1 | 424 | 439 | -576 | -561 | -10,1 | 0,01 | 4,11 | >0.1 | 2,7 | 4,82 | 0 | 1,02 | 0 | 5,67 | 0 |
| EGR3 | CAGCGCCCAACCCAA | 1 | 943 | 958 | -57 | -42 | -8,31 | 0,01 | 8,28 | >0.1 | 5,54 | 7,36 | 0 | 1,88 | 0 | 1,8 | 0 |
| EGR3 | TAAAACCTCCTCATG | -1 | 986 | 1001 | -14 | 1 | -7,41 | 0,01 | 10,36 | >0.1 | 5,54 | 8,74 | 0 | 1,87 | 0 | 1,61 | 0 |
| EGR3 | TGTCGCCCAGGCTGG | -1 | 425 | 440 | -575 | -560 | 0,82 | 0,001 | 15,17 | >0.1 | 2,71 | 4,82 | 0 | 1,02 | 0 | 5,8 | 0 |
| EGR3 | CTCTGCCTCCCAGGT | -1 | 378 | 393 | -622 | -607 | -10,94 | 0,01 | 3,34 | >0.1 | 2,57 | 4,74 | 0 | 1,06 | 0 | 5,91 | 0 |
| EGR3 | TGCCTCCCAGGTTCA | -1 | 375 | 390 | -625 | -610 | -4,21 | 0,01 | 10,55 | >0.1 | 2,56 | 4,73 | 0 | 1,07 | 0 | 6,4 | 0 |
| EGR3 | TCCTGCCTCAGCCTC | -1 | 352 | 367 | -648 | -633 | -9,03 | 0,01 | 5,82 | >0.1 | 2,53 | 4,69 | 0 | 1,1 | 0 | 6,52 | 0 |
| EGR3 | CTCAGCCTCCCGAGT | -1 | 346 | 361 | -654 | -639 | -11,49 | 0,01 | 3,25 | >0.1 | 2,54 | 4,68 | 0 | 1,11 | 0 | 6,4 | 0 |
| EGR3 | GCCCACCACCACGCC | -1 | 315 | 330 | -685 | -670 | -6,16 | 0,01 | 8,53 | >0.1 | 2,63 | 4,64 | 0 | 1,17 | 0 | 6,26 | 0 |
| EGR3 | CCACCACCACGCCCA | -1 | 313 | 328 | -687 | -672 | -0,26 | 0,001 | 14,31 | >0.1 | 2,63 | 4,63 | 0 | 1,18 | 0 | 6,13 | 0 |
| EGR3 | CCACGCCCAGCTAAG | -1 | 307 | 322 | -693 | -678 | 0,92 | 0,001 | 15,64 | >0.1 | 2,65 | 4,62 | 0 | 1,19 | 0 | 6,26 | 0 |
| EGR3 | ACTCGGCCTCCCAAA | -1 | 213 | 228 | -787 | -772 | -8,72 | 0,01 | 5,93 | >0.1 | 2,97 | 4,49 | 0 | 1,55 | 0 | 5,63 | 0 |
| EGR3 | CTGTGCCCAGCCATT | -1 | 173 | 188 | -827 | -812 | -4,73 | 0,01 | 7,95 | >0.1 | 3,15 | 4,44 | 0 | 1,85 | 0 | 3,24 | 0 |
| EGR3 | ATCTGCCTTTGCAAA | -1 | 84 | 99 | -916 | -901 | -10,34 | 0,01 | -0,39 | >0.1 | 3,73 | 4,34 | 0 | 1,88 | 0 | 0 | 0 |
| EGR3 | TGCCTCCTTCTCCTT | -1 | 12 | 27 | -988 | -973 | -9,08 | 0,01 | 2,6 | >0.1 | 5,54 | 4,27 | 0 | 1,87 | 0 | 0 | 0 |
| EGR1 | GTGGCTCACGCCTG | 1 | 187 | 201 | -813 | -799 | -7,03 | 0,001 | 6,28 | >0.1 | 3,09 | 4,46 | 0 | 1,73 | 0 | 4,04 | 0 |
| EGR1 | CTCACGCCTGTAAT | 1 | 191 | 205 | -809 | -795 | -21,87 | 0,01 | -8,38 | >0.1 | 3,07 | 4,46 | 0 | 1,7 | 0 | 4,26 | 0 |
| EGR1 | GAAACCCCGTCTCT | 1 | 277 | 291 | -723 | -709 | -21,62 | 0,01 | -7,83 | >0.1 | 2,75 | 4,58 | 0 | 1,27 | 0 | 5,19 | 0 |
| EGR1 | CCCAGCTACTCGGG | 1 | 339 | 353 | -661 | -647 | -19,41 | 0,01 | -4,53 | >0.1 | 2,56 | 4,67 | 0 | 1,12 | 0 | 6,52 | 0 |
| EGR1 | CGCGCCACTGTGCT | 1 | 411 | 425 | -589 | -575 | -20,26 | 0,01 | -6,69 | >0.1 | 2,66 | 4,8 | 0 | 1,03 | 0 | 5,09 | 0 |
| EGR1 | CCAGCCTGGGCGAC | 1 | 425 | 439 | -575 | -561 | -15,41 | 0,01 | -1,06 | >0.1 | 2,7 | 4,82 | 0 | 1,02 | 0 | 5,8 | 0 |
| EGR1 | GTTGCCCAAGTAGA | 1 | 540 | 554 | -460 | -446 | -15,59 | 0,01 | -6,39 | >0.1 | 3,14 | 5,07 | 0 | 0,99 | 0 | 0 | 0 |
| EGR1 | AGCGCCCAACCCAA | 1 | 944 | 958 | -56 | -42 | -18,05 | 0,01 | -1,45 | >0.1 | 5,56 | 7,36 | 0 | 1,88 | 0 | 1,8 | 0 |
| EGR1 | CCAACCCAAGTTCA | 1 | 949 | 963 | -51 | -37 | -18,31 | 0,01 | -1,68 | >0.1 | 5,56 | 7,46 | 0 | 1,88 | 0 | 1,74 | 0 |
| EGR1 | CACATCCACTGAAT | -1 | 1015 | 1029 | 15 | 29 | -19,92 | 0,01 | -2,33 | >0.1 | 5,56 | 8,84 | 0 | 1,86 | 0 | 1,34 | 0 |
| EGR1 | AAAACCTCCTCATG | -1 | 986 | 1000 | -14 | 0 | -10,22 | 0,01 | 7,53 | >0.1 | 5,56 | 8,71 | 0 | 1,87 | 0 | 1,61 | 0 |
| EGR1 | GTCGCCCAGGCTGG | -1 | 425 | 439 | -575 | -561 | -5,54 | 0,001 | 8,81 | >0.1 | 2,7 | 4,82 | 0 | 1,02 | 0 | 5,8 | 0 |
| EGR1 | GCCTCCCAGGTTCA | -1 | 375 | 389 | -625 | -611 | -18,78 | 0,01 | -4,02 | >0.1 | 2,56 | 4,73 | 0 | 1,07 | 0 | 6,4 | 0 |
| EGR1 | CAGGTTCACGCCAT | -1 | 369 | 383 | -631 | -617 | -12,2 | 0,01 | 2,54 | >0.1 | 2,55 | 4,72 | 0 | 1,08 | 0 | 6,4 | 0 |
| EGR1 | CCTGCCTCAGCCTC | -1 | 352 | 366 | -648 | -634 | -14,02 | 0,01 | 0,83 | >0.1 | 2,53 | 4,69 | 0 | 1,1 | 0 | 6,52 | 0 |
| EGR1 | TCAGCCTCCCGAGT | -1 | 346 | 360 | -654 | -640 | -20,96 | 0,01 | -6,22 | >0.1 | 2,55 | 4,68 | 0 | 1,11 | 0 | 6,4 | 0 |
| EGR1 | CCCACCACCACGCC | -1 | 315 | 329 | -685 | -671 | -16,43 | 0,01 | -1,74 | >0.1 | 2,63 | 4,63 | 0 | 1,17 | 0 | 6,26 | 0 |
| EGR1 | CACCACCACGCCCA | -1 | 313 | 327 | -687 | -673 | -9,3 | 0,001 | 5,27 | >0.1 | 2,64 | 4,63 | 0 | 1,18 | 0 | 6,13 | 0 |
| EGR1 | CACGCCCAGCTAAG | -1 | 307 | 321 | -693 | -679 | -12,23 | 0,01 | 2,49 | >0.1 | 2,65 | 4,62 | 0 | 1,19 | 0 | 6,26 | 0 |
| EGR1 | TGTGCCCAGCCATT | -1 | 173 | 187 | -827 | -813 | -19,15 | 0,01 | -6,47 | >0.1 | 3,15 | 4,44 | 0 | 1,85 | 0 | 3,24 | 0 |
| EGR1 | TATGACTACTCTCC | -1 | 121 | 135 | -879 | -865 | -21,44 | 0,01 | -10,03 | >0.1 | 3,45 | 4,38 | 0 | 1,89 | 0 | 1,7 | 0 |
| EGR1 | TCTGCCTTTGCAAA | -1 | 84 | 98 | -916 | -902 | -16,3 | 0,01 | -6,35 | >0.1 | 3,73 | 4,34 | 0 | 1,88 | 0 | 0 | 0 |
| JUN | AGAAATAGGAAAT | 1 | 57 | 70 | -943 | -930 | -13,76 | 0,01 | -3,56 | >0.1 | 4,01 | 4,31 | 0 | 1,87 | 0 | 0 | 0 |
| JUN | TAAGTTTTGCAAA | 1 | 79 | 92 | -921 | -908 | -16,34 | 0,01 | -6,35 | >0.1 | 3,78 | 4,33 | 0 | 1,88 | 0 | 0 | 0 |
| JUN | TGGCTGGGCACAG | 1 | 175 | 188 | -825 | -812 | -15,94 | 0,01 | -3,28 | >0.1 | 3,15 | 4,44 | 0 | 1,83 | 0 | 3,24 | 0 |
| JUN | AGGCTGAGGCAGG | 1 | 353 | 366 | -647 | -634 | -9,33 | 0,01 | 5,51 | >0.1 | 2,53 | 4,69 | 0 | 1,1 | 0 | 6,52 | 0 |
| JUN | GCAGTGAGCCGAG | 1 | 396 | 409 | -604 | -591 | -12,52 | 0,01 | 1,82 | >0.1 | 2,62 | 4,77 | 0 | 1,04 | 0 | 5,91 | 0 |
| JUN | AATGTGAGATAAC | 1 | 588 | 601 | -412 | -399 | -6,56 | 0,01 | 3,04 | >0.1 | 3,4 | 5,18 | 0 | 1,01 | 0 | 0 | 0 |
| JUN | TTCATGATCCCTA | 1 | 646 | 659 | -354 | -341 | -14,4 | 0,01 | -0,28 | >0.1 | 3,87 | 5,35 | 0 | 1,07 | 0 | 3,83 | 0 |
| JUN | CCTCTTTGGTCAC | 1 | 667 | 680 | -333 | -320 | -14,12 | 0,01 | 0,22 | >0.1 | 4,12 | 5,42 | 0 | 1,1 | 0 | 3,7 | 0 |
| JUN | AAAATTATAACAT | 1 | 683 | 696 | -317 | -304 | -11,64 | 0,01 | 2,3 | >0.1 | 4,39 | 5,47 | 0 | 1,13 | 0 | 2,95 | 0 |
| JUN | TTGATTTGGCCAT | 1 | 750 | 763 | -250 | -237 | -7,08 | 0,01 | 11,84 | >0.1 | 5,57 | 5,74 | 0 | 1,3 | 0 | 6,32 | 0 |
| JUN | CAGGAGAGGTAAG | 1 | 770 | 783 | -230 | -217 | -6,78 | 0,01 | 12,09 | >0.1 | 5,57 | 5,83 | 0 | 1,37 | 0 | 6,1 | 0 |
| JUN | GGGACGATGTCAA | 1 | 1043 | 1056 | 43 | 56 | -4,1 | 0,01 | 11,71 | >0.1 | 5,57 | 7,76 | 0 | 1,86 | 0 | 0,62 | 0 |
| JUN | CCTGTGAGCCAAG | -1 | 1031 | 1044 | 31 | 44 | 0,08 | 0,001 | 16,98 | >0.1 | 5,57 | 8,19 | 0 | 1,86 | 0 | 1,28 | 0 |
| JUN | GCCAAGATCACAT | -1 | 1024 | 1037 | 24 | 37 | -10,6 | 0,01 | 6,67 | >0.1 | 5,57 | 8,59 | 0 | 1,86 | 0 | 1,25 | 0 |
| JUN | CTGGAGAGGACAG | -1 | 857 | 870 | -143 | -130 | -9,66 | 0,01 | 7,04 | >0.1 | 5,57 | 6,37 | 0 | 1,9 | 0 | 2,86 | 0 |
| JUN | ACAGATATGTAAC | -1 | 848 | 861 | -152 | -139 | -12,63 | 0,01 | 3,98 | >0.1 | 5,57 | 6,3 | 0 | 1,87 | 0 | 2,86 | 0 |
| JUN | TTTATGGCCAAAT | -1 | 753 | 766 | -247 | -234 | -12,85 | 0,01 | 6,1 | >0.1 | 5,57 | 5,75 | 0 | 1,31 | 0 | 6,32 | 0 |
| JUN | AAAATGTTATAAT | -1 | 686 | 699 | -314 | -301 | -15,18 | 0,01 | -1,16 | >0.1 | 4,45 | 5,48 | 0 | 1,13 | 0 | 2,95 | 0 |
| JUN | ACAGTGACCAAAG | -1 | 670 | 683 | -330 | -317 | -0,5 | 0,001 | 13,75 | >0.1 | 4,16 | 5,43 | 0 | 1,11 | 0 | 3,55 | 0 |
| JUN | TAGGTTATCTCAC | -1 | 591 | 604 | -409 | -396 | 1,69 | 0,001 | 11,32 | >0.1 | 3,42 | 5,19 | 0 | 1,01 | 0 | 0 | 0 |

| | | | | | | | | | | | | | | | | | |
|---|---|---|---|---|---|---|---|---|---|---|---|---|---|---|---|---|---|
| JUN | GGCGTGAGCCACT | -1 | 186 | 199 | -814 | -801 | -11,15 | 0,01 | 1,99 | >0.1 | 3,09 | 4,45 | 0 | 1,74 | 0 | 3,86 | 0 |
| JUN | CCACTGTGCCCAG | -1 | 178 | 191 | -822 | -809 | -13,66 | 0,01 | -1,19 | >0.1 | 3,13 | 4,44 | 0 | 1,81 | 0 | 3,09 | 0 |
| MEIS2 | AAGTCAGC | 1 | 34 | 42 | -966 | -958 | 2,17 | 0,01 | 12,74 | >0.1 | 4,42 | 4,28 | 0 | 1,87 | 0 | 0 | 0 |
| MEIS2 | TTGACCTG | 1 | 107 | 115 | -893 | -885 | 1,78 | 0,01 | 12,33 | >0.1 | 3,57 | 4,35 | 0 | 1,89 | 0 | 0,75 | 0 |
| MEIS2 | TGGGCAGA | 1 | 228 | 236 | -772 | -764 | 1,3 | 0,01 | 15,45 | >0.1 | 2,93 | 4,5 | 0 | 1,46 | 0 | 5,26 | 0 |
| MEIS2 | AGGTCAGG | 1 | 241 | 249 | -759 | -751 | 2,92 | 0,01 | 16,04 | >0.1 | 2,88 | 4,52 | 0 | 1,4 | 0 | 4,33 | 0 |
| MEIS2 | GTGAAACC | 1 | 275 | 283 | -725 | -717 | 0,69 | 0,01 | 14,48 | >0.1 | 2,76 | 4,56 | 0 | 1,28 | 0 | 5,19 | 0 |
| MEIS2 | TGGGCACC | 1 | 325 | 333 | -675 | -667 | 0,17 | 0,01 | 14,02 | >0.1 | 2,61 | 4,64 | 0 | 1,15 | 0 | 5,46 | 0 |
| MEIS2 | GAGGCAGG | 1 | 358 | 366 | -642 | -634 | 1,37 | 0,01 | 15,8 | >0.1 | 2,52 | 4,69 | 0 | 1,09 | 0 | 6,13 | 0 |
| MEIS2 | GCGACAGA | 1 | 434 | 442 | -566 | -558 | 1,28 | 0,01 | 15,64 | >0.1 | 2,72 | 4,83 | 0 | 1,01 | 0 | 5,8 | 0 |
| MEIS2 | AAGACAAG | 1 | 503 | 511 | -497 | -489 | 1,98 | 0,01 | 16,58 | >0.1 | 2,97 | 4,97 | 0 | 0,98 | 0 | 5,68 | 0 |
| MEIS2 | TTGGAAGG | 1 | 526 | 534 | -474 | -466 | 0,72 | 0,01 | 13,06 | >0.1 | 3,06 | 5,02 | 0 | 0,98 | 0 | 3,28 | 0 |
| MEIS2 | CTGAAAAC | 1 | 617 | 625 | -383 | -375 | -0,3 | 0,01 | 11,06 | >0.1 | 3,59 | 5,25 | 0 | 1,03 | 0 | 1,5 | 0 |
| MEIS2 | TTAACAGC | 1 | 709 | 717 | -291 | -283 | 0,55 | 0,01 | 17,72 | >0.1 | 5,64 | 5,55 | 0 | 1,17 | 0 | 4,81 | 0 |
| MEIS2 | GTGACAGG | 1 | 766 | 774 | -234 | -226 | 12,05 | 0,0001 | 30,72 | >0.1 | 5,64 | 5,79 | 0 | 1,33 | 0 | 5,92 | 0 |
| MEIS2 | ATGTCAAG | 1 | 1049 | 1057 | 49 | 57 | 2,98 | 0,01 | 19,49 | >0.1 | 5,64 | 7,6 | 0 | 1,86 | 0 | 1,42 | 0 |
| MEIS2 | TGGACAAG | 1 | 1134 | 1142 | 134 | 142 | 4,57 | 0,01 | 24,66 | >0.1 | 5,64 | 6,42 | 0 | 1,84 | 3,22 | 2,97 | 0 |
| MEIS2 | TTGATAGA | -1 | 1167 | 1175 | 167 | 175 | 2,6 | 0,01 | 19,67 | >0.1 | 4,53 | 6,17 | 0 | 1,69 | 1,79 | 2,88 | 0 |
| MEIS2 | TTGACATC | -1 | 1048 | 1056 | 48 | 56 | 8,95 | 0,001 | 25,49 | >0.1 | 5,64 | 7,62 | 0 | 1,86 | 0 | 1,42 | 0 |
| MEIS2 | AGGACAGA | -1 | 856 | 864 | -144 | -136 | 4,82 | 0,01 | 21,55 | >0.1 | 5,64 | 6,32 | 0 | 1,9 | 0 | 2,86 | 0 |
| MEIS2 | CTGTCACT | -1 | 765 | 773 | -235 | -227 | 0,2 | 0,01 | 19,05 | >0.1 | 5,64 | 5,78 | 0 | 1,33 | 0 | 6,1 | 0 |
| MEIS2 | TAGAAAGC | -1 | 715 | 723 | -285 | -277 | 0,64 | 0,01 | 18,02 | >0.1 | 5,64 | 5,57 | 0 | 1,18 | 0 | 4,99 | 0 |
| MEIS2 | TAGTCAGG | -1 | 262 | 270 | -738 | -730 | 3,78 | 0,01 | 16,9 | >0.1 | 2,8 | 4,55 | 0 | 1,32 | 0 | 4,45 | 0 |
| EGR4 | AAATGGCCATGGAAAT | 1 | 143 | 159 | -857 | -841 | 0,12 | 0,01 | 11,34 | >0.1 | 3,31 | 4,41 | 0 | 1,9 | 0 | 1,61 | 0 |
| EGR4 | AGTGGCTCACGCCTGT | 1 | 186 | 202 | -814 | -798 | 1,21 | 0,01 | 14,53 | >0.1 | 3,09 | 4,46 | 0 | 1,74 | 0 | 4,04 | 0 |
| EGR4 | AAACCCCGTCTCTATT | 1 | 278 | 294 | -722 | -706 | -0,66 | 0,01 | 13,12 | >0.1 | 2,74 | 4,58 | 0 | 1,27 | 0 | 5,19 | 0 |
| EGR4 | GTAGTCCCAGCTACTC | 1 | 334 | 350 | -666 | -650 | 0,73 | 0,01 | 15,02 | >0.1 | 2,58 | 4,67 | 0 | 1,13 | 0 | 5,92 | 0 |
| EGR4 | CAGCGCCCAACCCAAG | 1 | 943 | 959 | -57 | -41 | 5,02 | 0,01 | 21,61 | >0.1 | 5,53 | 7,38 | 0 | 1,88 | 0 | 1,8 | 0 |
| EGR4 | TCACATCCACTGAATG | -1 | 1014 | 1030 | 14 | 30 | 1,46 | 0,01 | 19,05 | >0.1 | 5,53 | 8,86 | 0 | 1,86 | 0 | 1,34 | 0 |
| EGR4 | CTTTGCCCTTGTCTTT | -1 | 502 | 518 | -498 | -482 | -1,03 | 0,01 | 13,6 | >0.1 | 2,98 | 4,98 | 0 | 0,99 | 0 | 5,68 | 0 |
| EGR4 | TGTCGCCCAGGCTGGA | -1 | 424 | 440 | -576 | -560 | 4 | 0,01 | 18,35 | >0.1 | 2,7 | 4,82 | 0 | 1,02 | 0 | 5,8 | 0 |
| EGR4 | GCTCTGCCTCCCAGGT | -1 | 378 | 394 | -622 | -606 | -0,78 | 0,01 | 13,61 | >0.1 | 2,57 | 4,74 | 0 | 1,06 | 0 | 6,02 | 0 |
| EGR4 | TGCCTCCCAGGTTCAC | -1 | 374 | 390 | -626 | -610 | 2,34 | 0,01 | 17,1 | >0.1 | 2,56 | 4,73 | 0 | 1,07 | 0 | 6,4 | 0 |
| EGR4 | CCAGGTTCACGCCATT | -1 | 368 | 384 | -632 | -616 | 3,18 | 0,01 | 17,93 | >0.1 | 2,55 | 4,72 | 0 | 1,08 | 0 | 6,4 | 0 |
| EGR4 | TCCTGCCTCAGCCTCC | -1 | 351 | 367 | -649 | -633 | 0,07 | 0,01 | 14,92 | >0.1 | 2,53 | 4,69 | 0 | 1,1 | 0 | 6,52 | 0 |
| EGR4 | CCTCAGCCTCCCGAGT | -1 | 346 | 362 | -654 | -638 | -0,99 | 0,01 | 13,75 | >0.1 | 2,54 | 4,69 | 0 | 1,11 | 0 | 6,4 | 0 |
| EGR4 | CTCAGCCTCCCGAGTA | -1 | 345 | 361 | -655 | -639 | 0,17 | 0,01 | 14,91 | >0.1 | 2,55 | 4,69 | 0 | 1,11 | 0 | 6,4 | 0 |
| EGR4 | CCACCACCACGCCCAG | -1 | 312 | 328 | -688 | -672 | 5,99 | 0,001 | 20,57 | >0.1 | 2,64 | 4,63 | 0 | 1,18 | 0 | 6,13 | 0 |
| EGR4 | CCACGCCCAGCTAAGT | -1 | 306 | 322 | -694 | -678 | 8,96 | 0,001 | 23,69 | >0.1 | 2,65 | 4,62 | 0 | 1,19 | 0 | 6,26 | 0 |
| EGR4 | ACTCGGCCTCCCAAAG | -1 | 212 | 228 | -788 | -772 | 4,14 | 0,01 | 18,79 | >0.1 | 2,98 | 4,49 | 0 | 1,55 | 0 | 5,63 | 0 |
| EGR4 | CTGTGCCCAGCCATTT | -1 | 172 | 188 | -828 | -812 | 7,26 | 0,001 | 19,95 | >0.1 | 3,15 | 4,44 | 0 | 1,86 | 0 | 3,24 | 0 |
| EGR4 | ATCTGCCTTTGCAAAA | -1 | 83 | 99 | -917 | -901 | 1,49 | 0,01 | 11,44 | >0.1 | 3,73 | 4,34 | 0 | 1,88 | 0 | 0 | 0 |
| EGR4 | TGCCTCCTTCTCCTTC | -1 | 11 | 27 | -989 | -973 | 0 | 0,01 | 11,67 | >0.1 | 5,53 | 4,27 | 0 | 1,87 | 0 | 0 | 0 |
| NFE2 | AAGGAGGCACT | 1 | 18 | 29 | -982 | -971 | -15,93 | 0,01 | -4,2 | >0.1 | 5,6 | 4,27 | 0 | 1,87 | 0 | 0 | 0 |
| NFE2 | AGTCAGCCACA | 1 | 35 | 46 | -965 | -954 | -19,93 | 0,01 | -9,41 | >0.1 | 4,37 | 4,29 | 0 | 1,87 | 0 | 0 | 0 |
| NFE2 | AATGAGCTTTT | 1 | 67 | 78 | -933 | -922 | -19,78 | 0,01 | -9,68 | >0.1 | 3,91 | 4,32 | 0 | 1,88 | 0 | 0 | 0 |
| NFE2 | GAGTAGTCATA | 1 | 124 | 135 | -876 | -865 | -17,54 | 0,01 | -6,14 | >0.1 | 3,44 | 4,38 | 0 | 1,89 | 0 | 1,7 | 0 |
| NFE2 | AGTGGCTCACG | 1 | 186 | 197 | -814 | -803 | -4,51 | 0,001 | 8,63 | >0.1 | 3,1 | 4,45 | 0 | 1,74 | 0 | 3,86 | 0 |
| NFE2 | CCTGACTAACA | 1 | 262 | 273 | -738 | -727 | -13,08 | 0,01 | 0,12 | >0.1 | 2,8 | 4,55 | 0 | 1,32 | 0 | 4,53 | 0 |
| NFE2 | GCTGAGGCAGG | 1 | 355 | 366 | -645 | -634 | -16,96 | 0,01 | -2,61 | >0.1 | 2,53 | 4,69 | 0 | 1,09 | 0 | 6,03 | 0 |
| NFE2 | CGAGACTCAGT | 1 | 443 | 454 | -557 | -546 | -9,79 | 0,01 | 4,74 | >0.1 | 2,76 | 4,85 | 0 | 1 | 0 | 5,92 | 0 |
| NFE2 | CATGATCCCTA | 1 | 648 | 659 | -352 | -341 | -15,38 | 0,01 | -1,25 | >0.1 | 3,88 | 5,35 | 0 | 1,07 | 0 | 3,83 | 0 |
| NFE2 | CGTGTATCTTT | 1 | 700 | 711 | -300 | -289 | -19,41 | 0,01 | -3,94 | >0.1 | 5,6 | 5,53 | 0 | 1,16 | 0 | 3,19 | 0 |
| NFE2 | TTTGATTCACT | 1 | 815 | 826 | -185 | -174 | -11,15 | 0,01 | 5,53 | >0.1 | 5,6 | 6,06 | 0 | 1,59 | 0 | 3,43 | 0 |
| NFE2 | ATTGGCTCAGC | 1 | 882 | 893 | -118 | -107 | -20,06 | 0,01 | -2,95 | >0.1 | 5,6 | 6,56 | 0 | 1,89 | 0 | 3,07 | 0 |
| NFE2 | GGAAAGTCATT | 1 | 1007 | 1018 | 7 | 18 | -18,58 | 0,01 | -0,53 | >0.1 | 5,6 | 9,12 | 0 | 1,87 | 0 | 1,46 | 0 |
| NFE2 | CTTGGCTCACA | 1 | 1031 | 1042 | 31 | 42 | -15,99 | 0,01 | 0,96 | >0.1 | 5,6 | 8,19 | 0 | 1,86 | 0 | 1,31 | 0 |
| NFE2 | CCTGGCTCCTT | 1 | 1062 | 1073 | 62 | 73 | -19,53 | 0,01 | -0,76 | >0.1 | 5,6 | 7,31 | 0 | 1,86 | 1,79 | 2,21 | 0 |
| NFE2 | AAGGAGCCAGG | -1 | 1062 | 1073 | 62 | 73 | -20,25 | 0,01 | -1,48 | >0.1 | 5,6 | 7,31 | 0 | 1,86 | 1,79 | 2,21 | 0 |

| | | | | | | | | | | | | | | | | | |
|---|---|---|---|---|---|---|---|---|---|---|---|---|---|---|---|---|---|
| NFE2 | TGTGAGCCAAG | -1 | 1031 | 1042 | 31 | 42 | -16,68 | 0,01 | 0,27 | >0.1 | 5,6 | 8,19 | 0 | 1,86 | 0 | 1,31 | 0 |
| NFE2 | AATGACTTTCC | -1 | 1007 | 1018 | 7 | 18 | -9,01 | 0,01 | 9,04 | >0.1 | 5,6 | 9,12 | 0 | 1,87 | 0 | 1,46 | 0 |
| NFE2 | AGTGAATCAAA | -1 | 815 | 826 | -185 | -174 | -15,66 | 0,01 | 1,02 | >0.1 | 5,6 | 6,06 | 0 | 1,59 | 0 | 3,43 | 0 |
| NFE2 | CATGACTTTGC | -1 | 512 | 523 | -488 | -477 | -16,19 | 0,01 | -4,25 | >0.1 | 3,01 | 4,99 | 0 | 0,98 | 0 | 2,95 | 0 |
| NFE2 | ACTGAGTCTCG | -1 | 443 | 454 | -557 | -546 | -9,14 | 0,01 | 5,39 | >0.1 | 2,76 | 4,85 | 0 | 1 | 0 | 5,92 | 0 |
| NFE2 | CCTGCCTCAGC | -1 | 355 | 366 | -645 | -634 | -12,07 | 0,01 | 2,28 | >0.1 | 2,53 | 4,69 | 0 | 1,09 | 0 | 6,03 | 0 |
| NFE2 | TGTTAGTCAGG | -1 | 262 | 273 | -738 | -727 | -14,7 | 0,01 | -1,5 | >0.1 | 2,8 | 4,55 | 0 | 1,32 | 0 | 4,53 | 0 |
| NFE2 | CGTGAGCCACT | -1 | 186 | 197 | -814 | -803 | 0,26 | 0,001 | 13,4 | >0.1 | 3,1 | 4,45 | 0 | 1,74 | 0 | 3,86 | 0 |
| NFE2 | TATGACTACTC | -1 | 124 | 135 | -876 | -865 | -9,21 | 0,01 | 2,19 | >0.1 | 3,44 | 4,38 | 0 | 1,89 | 0 | 1,7 | 0 |
| NFE2 | GACTACTCTCC | -1 | 121 | 132 | -879 | -868 | -19,8 | 0,01 | -8,38 | >0.1 | 3,46 | 4,37 | 0 | 1,89 | 0 | 1,7 | 0 |
| NFE2 | TATTTCTCACA | -1 | 53 | 64 | -947 | -936 | -17,63 | 0,01 | -7,37 | >0.1 | 4,08 | 4,3 | 0 | 1,87 | 0 | 0 | 0 |
| NFE2 | GCTGACTTCTT | -1 | 31 | 42 | -969 | -958 | -16,97 | 0,01 | -6,37 | >0.1 | 4,45 | 4,28 | 0 | 1,87 | 0 | 0 | 0 |
| NFE2 | AGTGCCTCCTT | -1 | 18 | 29 | -982 | -971 | -7,44 | 0,01 | 4,29 | >0.1 | 5,6 | 4,27 | 0 | 1,87 | 0 | 0 | 0 |
| NKX2-5 | GTAAGTG | 1 | 5 | 12 | -995 | -988 | 6,86 | 0,01 | 20,42 | >0.1 | 5,65 | 4,25 | 0 | 1,86 | 1,79 | 0 | 0 |
| NKX2-5 | TTAAGTT | 1 | 78 | 85 | -922 | -915 | 7 | 0,01 | 17,02 | >0.1 | 3,82 | 4,32 | 0 | 1,88 | 0 | 0 | 0 |
| NKX2-5 | ATAATTT | 1 | 132 | 139 | -868 | -861 | 6,65 | 0,01 | 17,89 | >0.1 | 3,4 | 4,38 | 0 | 1,89 | 0 | 1,57 | 0 |
| NKX2-5 | ATAAGAG | 1 | 970 | 977 | -30 | -23 | 7,53 | 0,001 | 24,56 | >0.1 | 5,65 | 7,81 | 0 | 1,87 | 0 | 1,7 | 0 |
| NKX2-5 | TCAAGCT | 1 | 1052 | 1059 | 52 | 59 | 3,99 | 0,01 | 20,49 | >0.1 | 5,65 | 7,53 | 0 | 1,86 | 0 | 1,46 | 0 |
| NKX2-5 | ACAAGTT | 1 | 1137 | 1144 | 137 | 144 | 5,49 | 0,01 | 25,63 | >0.1 | 5,65 | 6,4 | 0 | 1,84 | 3,22 | 3,02 | 0 |
| NKX2-5 | ATAATTC | -1 | 1192 | 1199 | 192 | 199 | 5,73 | 0,01 | 24,51 | >0.1 | 5,65 | 6,02 | 0 | 1,5 | 1,79 | 3,82 | 0 |
| NKX2-5 | GCAAGTG | -1 | 1179 | 1186 | 179 | 186 | 5,7 | 0,01 | 24,1 | >0.1 | 5,65 | 6,1 | 0 | 1,59 | 1,79 | 3,26 | 0 |
| NKX2-5 | ACAATCT | -1 | 893 | 900 | -107 | -100 | 3,64 | 0,01 | 20,66 | >0.1 | 5,65 | 6,61 | 0 | 1,89 | 0 | 2,86 | 0 |
| NKX2-5 | ATAATTT | -1 | 831 | 838 | -169 | -162 | 6,65 | 0,01 | 23,4 | >0.1 | 5,65 | 6,14 | 0 | 1,67 | 0 | 3,28 | 0 |
| NKX2-5 | ATAAGAA | -1 | 809 | 816 | -191 | -184 | 4,69 | 0,01 | 21,16 | >0.1 | 5,65 | 6 | 0 | 1,53 | 0 | 3,28 | 0 |
| NKX2-5 | TTAATCC | -1 | 791 | 798 | -209 | -202 | 4,23 | 0,01 | 22,74 | >0.1 | 5,65 | 5,91 | 0 | 1,43 | 0 | 5,52 | 0 |
| NKX2-5 | TTAATAT | -1 | 728 | 735 | -272 | -265 | 5,59 | 0,01 | 23,39 | >0.1 | 5,65 | 5,62 | 0 | 1,21 | 0 | 5,32 | 0 |
| NKX2-5 | ATAATTT | -1 | 684 | 691 | -316 | -309 | 6,65 | 0,01 | 20,64 | >0.1 | 4,35 | 5,46 | 0 | 1,12 | 0 | 3,07 | 0 |
| NKX2-5 | ACAATTT | -1 | 610 | 617 | -390 | -383 | 5,49 | 0,01 | 16,82 | >0.1 | 3,53 | 5,23 | 0 | 1,02 | 0 | 1,55 | 0 |
| NKX2-5 | TTAATAG | -1 | 601 | 608 | -399 | -392 | 7,89 | 0,001 | 18,94 | >0.1 | 3,47 | 5,2 | 0 | 1,01 | 0 | 1,36 | 0 |
| NKX2-5 | TCAAGAT | -1 | 571 | 578 | -429 | -422 | 4,42 | 0,01 | 13,82 | >0.1 | 3,29 | 5,12 | 0 | 0,99 | 0 | 0 | 0 |
| NKX2-5 | TTAATAG | -1 | 289 | 296 | -711 | -704 | 7,89 | 0,001 | 21,62 | >0.1 | 2,72 | 4,58 | 0 | 1,23 | 0 | 5,19 | 0 |
| NKX2-5 | TTAATTT | -1 | 155 | 162 | -845 | -838 | 7 | 0,01 | 18,22 | >0.1 | 3,26 | 4,41 | 0 | 1,9 | 0 | 1,65 | 0 |
| BACH1::MAFK | CAAAGAAGTCAGCCA | 1 | 29 | 44 | -971 | -956 | -8,78 | 0,01 | 1,82 | >0.1 | 4,45 | 4,28 | 0 | 1,87 | 0 | 0 | 0 |
| BACH1::MAFK | GTGGAGAGTAGTCAT | 1 | 119 | 134 | -881 | -866 | -6,74 | 0,01 | 4,68 | >0.1 | 3,46 | 4,38 | 0 | 1,89 | 0 | 1,7 | 0 |
| BACH1::MAFK | GAAATGGCTGGGCAC | 1 | 171 | 186 | -829 | -814 | 0,67 | 0,01 | 13,38 | >0.1 | 3,16 | 4,44 | 0 | 1,87 | 0 | 3,24 | 0 |
| BACH1::MAFK | ACAGTGGCTCACGCC | 1 | 184 | 199 | -816 | -801 | -7,97 | 0,01 | 5,19 | >0.1 | 3,1 | 4,45 | 0 | 1,76 | 0 | 3,86 | 0 |
| BACH1::MAFK | GTCCTGACTAACACG | 1 | 260 | 275 | -740 | -725 | -5,01 | 0,01 | 8,2 | >0.1 | 2,8 | 4,55 | 0 | 1,33 | 0 | 4,53 | 0 |
| BACH1::MAFK | ACAAAAACTTAGCTG | 1 | 300 | 315 | -700 | -685 | -7,92 | 0,01 | 5,56 | >0.1 | 2,67 | 4,61 | 0 | 1,21 | 0 | 4,99 | 0 |
| BACH1::MAFK | CCAGCTACTCGGGAG | 1 | 340 | 355 | -660 | -645 | -5,04 | 0,01 | 9,72 | >0.1 | 2,56 | 4,68 | 0 | 1,12 | 0 | 6,4 | 0 |
| BACH1::MAFK | AGGCTGAGGCAGGAG | 1 | 353 | 368 | -647 | -632 | 6,41 | 0,001 | 20,77 | >0.1 | 2,53 | 4,7 | 0 | 1,1 | 0 | 6,03 | 0 |
| BACH1::MAFK | CGCGCCACTGTGCTC | 1 | 411 | 426 | -589 | -574 | -8,76 | 0,01 | 4,82 | >0.1 | 2,66 | 4,8 | 0 | 1,03 | 0 | 5,09 | 0 |
| BACH1::MAFK | ACCGAGACTCAGTCT | 1 | 441 | 456 | -559 | -544 | 1,59 | 0,001 | 16,13 | >0.1 | 2,76 | 4,85 | 0 | 1,01 | 0 | 5,92 | 0 |
| BACH1::MAFK | ATATTGGCTCAGCAG | 1 | 880 | 895 | -120 | -105 | 2,34 | 0,001 | 19,42 | >0.1 | 5,54 | 6,57 | 0 | 1,9 | 0 | 3,07 | 0 |
| BACH1::MAFK | TTTCCTGCTCAGCGC | 1 | 934 | 949 | -66 | -51 | -8,13 | 0,01 | 7,39 | >0.1 | 5,54 | 7,2 | 0 | 1,88 | 0 | 0,9 | 0 |
| BACH1::MAFK | GCTCCTTCTCAGCCT | 1 | 1066 | 1081 | 66 | 81 | -4,5 | 0,01 | 14,15 | >0.1 | 5,54 | 7,24 | 0 | 1,85 | 1,79 | 2,21 | 0 |
| BACH1::MAFK | AAAATGTCTTGGCCT | -1 | 1119 | 1134 | 119 | 134 | -7,04 | 0,01 | 12,8 | >0.1 | 5,54 | 6,56 | 0 | 1,85 | 3,22 | 2,67 | 0 |
| BACH1::MAFK | TGGTGGACTGAGCAG | -1 | 1095 | 1110 | 95 | 110 | -0,74 | 0,01 | 19,05 | >0.1 | 5,54 | 6,82 | 0 | 1,85 | 3,22 | 2,36 | 0 |
| BACH1::MAFK | AACAAGGCTGAGAAG | -1 | 1070 | 1085 | 70 | 85 | -8,63 | 0,01 | 10,01 | >0.1 | 5,54 | 7,17 | 0 | 1,85 | 1,79 | 2,27 | 0 |
| BACH1::MAFK | TGAATGACTTTCCCT | -1 | 1005 | 1020 | 5 | 20 | -7,55 | 0,01 | 10,45 | >0.1 | 5,54 | 9,12 | 0 | 1,87 | 0 | 1,46 | 0 |
| BACH1::MAFK | TACATGACTTTGCCC | -1 | 510 | 525 | -490 | -475 | 0,17 | 0,01 | 12,12 | >0.1 | 3,01 | 5 | 0 | 0,99 | 0 | 2,95 | 0 |
| BACH1::MAFK | AGACTGAGTCTCGGT | -1 | 441 | 456 | -559 | -544 | -8,66 | 0,01 | 5,88 | >0.1 | 2,76 | 4,85 | 0 | 1,01 | 0 | 5,92 | 0 |
| BACH1::MAFK | CTCCTGCCTCAGCCT | -1 | 353 | 368 | -647 | -632 | 4,6 | 0,001 | 18,96 | >0.1 | 2,53 | 4,7 | 0 | 1,1 | 0 | 6,03 | 0 |
| BACH1::MAFK | CGTGTTAGTCAGGAC | -1 | 260 | 275 | -740 | -725 | 2 | 0,001 | 15,21 | >0.1 | 2,8 | 4,55 | 0 | 1,33 | 0 | 4,53 | 0 |
| BACH1::MAFK | TGCCCAACTCGGCCT | -1 | 219 | 234 | -781 | -766 | -5,59 | 0,01 | 8,89 | >0.1 | 2,95 | 4,5 | 0 | 1,51 | 0 | 5,52 | 0 |
| BACH1::MAFK | GCCATTTCTGATCAG | -1 | 164 | 179 | -836 | -821 | -3,06 | 0,01 | 8,92 | >0.1 | 3,19 | 4,43 | 0 | 1,9 | 0 | 2,46 | 0 |
| BACH1::MAFK | ATTATGACTACTCTC | -1 | 122 | 137 | -878 | -863 | -6,84 | 0,01 | 4,56 | >0.1 | 3,44 | 4,38 | 0 | 1,89 | 0 | 1,7 | 0 |
| BACH1::MAFK | ATGACTACTCTCCAC | -1 | 119 | 134 | -881 | -866 | -0,78 | 0,01 | 10,64 | >0.1 | 3,46 | 4,38 | 0 | 1,89 | 0 | 1,7 | 0 |
| FOS | GAGTAGTCATA | 1 | 124 | 135 | -876 | -865 | -7,6 | 0,01 | 3,8 | >0.1 | 3,44 | 4,38 | 0 | 1,89 | 0 | 1,7 | 0 |
| FOS | AGTGGCTCACG | 1 | 186 | 197 | -814 | -803 | -2,91 | 0,01 | 10,23 | >0.1 | 3,1 | 4,45 | 0 | 1,74 | 0 | 3,86 | 0 |

| | | | | | | | | | | | | | | | | | |
|---|---|---|---|---|---|---|---|---|---|---|---|---|---|---|---|---|---|
| FOS | CCTGACTAACA | 1 | 262 | 273 | -738 | -727 | -8,44 | 0,01 | 4,76 | >0.1 | 2,8 | 4,55 | 0 | 1,32 | 0 | 4,53 | 0 |
| FOS | GCTGAGGCAGG | 1 | 355 | 366 | -645 | -634 | -4,45 | 0,01 | 9,9 | >0.1 | 2,53 | 4,69 | 0 | 1,09 | 0 | 6,03 | 0 |
| FOS | CGAGACTCAGT | 1 | 443 | 454 | -557 | -546 | -8,56 | 0,01 | 5,97 | >0.1 | 2,76 | 4,85 | 0 | 1 | 0 | 5,92 | 0 |
| FOS | CTTTGGTCACT | 1 | 670 | 681 | -330 | -319 | -14,62 | 0,01 | -0,25 | >0.1 | 4,15 | 5,42 | 0 | 1,1 | 0 | 3,7 | 0 |
| FOS | ATTGGCTCAGC | 1 | 882 | 893 | -118 | -107 | -8,85 | 0,01 | 8,26 | >0.1 | 5,6 | 6,56 | 0 | 1,89 | 0 | 3,07 | 0 |
| FOS | AGTCATTCAGT | 1 | 1011 | 1022 | 11 | 22 | -8,15 | 0,01 | 9,66 | >0.1 | 5,6 | 8,95 | 0 | 1,86 | 0 | 1,39 | 0 |
| FOS | CTTGGCTCACA | 1 | 1031 | 1042 | 31 | 42 | -11,43 | 0,01 | 5,52 | >0.1 | 5,6 | 8,19 | 0 | 1,86 | 0 | 1,31 | 0 |
| FOS | GTTTAACCACG | 1 | 1141 | 1152 | 141 | 152 | -14,57 | 0,01 | 5,86 | >0.1 | 5,6 | 6,37 | 0 | 1,84 | 3,22 | 3,4 | 0 |
| FOS | TGTTCCTCAAT | -1 | 1109 | 1120 | 109 | 120 | -11,73 | 0,01 | 8,11 | >0.1 | 5,6 | 6,66 | 0 | 1,85 | 3,22 | 2,52 | 0 |
| FOS | TGTGAGCCAAG | -1 | 1031 | 1042 | 31 | 42 | -7,73 | 0,01 | 9,22 | >0.1 | 5,6 | 8,19 | 0 | 1,86 | 0 | 1,31 | 0 |
| FOS | ACTGAATGACT | -1 | 1011 | 1022 | 11 | 22 | -8,07 | 0,01 | 9,74 | >0.1 | 5,6 | 8,95 | 0 | 1,86 | 0 | 1,39 | 0 |
| FOS | GCTGAGCCAAT | -1 | 882 | 893 | -118 | -107 | -7,79 | 0,01 | 9,32 | >0.1 | 5,6 | 6,56 | 0 | 1,89 | 0 | 3,07 | 0 |
| FOS | AGTGAATCAAA | -1 | 815 | 826 | -185 | -174 | 2,98 | 0,001 | 19,66 | >0.1 | 5,6 | 6,06 | 0 | 1,59 | 0 | 3,43 | 0 |
| FOS | CGTTATTCTTT | -1 | 796 | 807 | -204 | -193 | -11,97 | 0,01 | 4,49 | >0.1 | 5,6 | 5,95 | 0 | 1,48 | 0 | 3,43 | 0 |
| FOS | ACTGAGTCTCG | -1 | 443 | 454 | -557 | -546 | -3,82 | 0,01 | 10,71 | >0.1 | 2,76 | 4,85 | 0 | 1 | 0 | 5,92 | 0 |
| FOS | CCTGCCTCAGC | -1 | 355 | 366 | -645 | -634 | -7,83 | 0,01 | 6,52 | >0.1 | 2,53 | 4,69 | 0 | 1,09 | 0 | 6,03 | 0 |
| FOS | TGTTAGTCAGG | -1 | 262 | 273 | -738 | -727 | 7,95 | 0,001 | 21,15 | >0.1 | 2,8 | 4,55 | 0 | 1,32 | 0 | 4,53 | 0 |
| FOS | CGTGAGCCACT | -1 | 186 | 197 | -814 | -803 | -6,47 | 0,01 | 6,67 | >0.1 | 3,1 | 4,45 | 0 | 1,74 | 0 | 3,86 | 0 |
| FOS | TATTTCTCACA | -1 | 53 | 64 | -947 | -936 | -9,48 | 0,01 | 0,78 | >0.1 | 4,08 | 4,3 | 0 | 1,87 | 0 | 0 | 0 |
| LBX2 | GGAAATGAGC | 1 | 64 | 74 | -936 | -926 | 3,95 | 0,01 | 14,09 | >0.1 | 3,95 | 4,31 | 0 | 1,88 | 0 | 0 | 0 |
| LBX2 | GGAAATTAAA | 1 | 153 | 163 | -847 | -837 | 5,18 | 0,01 | 16,4 | >0.1 | 3,27 | 4,41 | 0 | 1,9 | 0 | 1,65 | 0 |
| LBX2 | ACTGATCAGA | 1 | 163 | 173 | -837 | -827 | 4,3 | 0,01 | 16,24 | >0.1 | 3,21 | 4,42 | 0 | 1,9 | 0 | 2,4 | 0 |
| LBX2 | CTCTATTAAA | 1 | 287 | 297 | -713 | -703 | 3,26 | 0,01 | 17 | >0.1 | 2,72 | 4,59 | 0 | 1,24 | 0 | 5,19 | 0 |
| LBX2 | ACTGATCAGA | 1 | 478 | 488 | -522 | -512 | 4,3 | 0,01 | 18,44 | >0.1 | 2,88 | 4,92 | 0 | 0,99 | 0 | 5,36 | 0 |
| LBX2 | CCCAAGTAGA | 1 | 544 | 554 | -456 | -446 | 7 | 0,001 | 16,2 | >0.1 | 3,15 | 5,06 | 0 | 0,99 | 0 | 0 | 0 |
| LBX2 | ACCTATTAAT | 1 | 599 | 609 | -401 | -391 | 4,03 | 0,01 | 15,08 | >0.1 | 3,47 | 5,21 | 0 | 1,02 | 0 | 1,36 | 0 |
| LBX2 | ATTAATGAAA | 1 | 603 | 613 | -397 | -387 | 4,99 | 0,01 | 16,13 | >0.1 | 3,49 | 5,22 | 0 | 1,02 | 0 | 1,41 | 0 |
| LBX2 | GCTGATAAGA | 1 | 966 | 976 | -34 | -24 | 3,56 | 0,01 | 20,52 | >0.1 | 5,61 | 7,78 | 0 | 1,87 | 0 | 1,7 | 0 |
| LBX2 | TCTCATGAGG | 1 | 983 | 993 | -17 | -7 | 4,67 | 0,01 | 22,22 | >0.1 | 5,61 | 8,5 | 0 | 1,87 | 0 | 1,57 | 0 |
| LBX2 | ACCATTGAGG | 1 | 1106 | 1116 | 106 | 116 | 3,4 | 0,01 | 23,25 | >0.1 | 5,61 | 6,69 | 0 | 1,85 | 3,22 | 2,48 | 0 |
| LBX2 | AGCAGTTACA | -1 | 1087 | 1097 | 87 | 97 | 3,8 | 0,01 | 22,19 | >0.1 | 5,61 | 6,92 | 0 | 1,85 | 1,79 | 2,21 | 0 |
| LBX2 | TCCACTGAAT | -1 | 1015 | 1025 | 15 | 25 | 3,62 | 0,01 | 21,32 | >0.1 | 5,61 | 8,84 | 0 | 1,86 | 0 | 1,39 | 0 |
| LBX2 | CCTCATGAGA | -1 | 983 | 993 | -17 | -7 | 4,9 | 0,01 | 22,45 | >0.1 | 5,61 | 8,5 | 0 | 1,87 | 0 | 1,57 | 0 |
| LBX2 | CCAAATCAAA | -1 | 749 | 759 | -251 | -241 | 3,56 | 0,01 | 22,09 | >0.1 | 5,61 | 5,72 | 0 | 1,28 | 0 | 5,92 | 0 |
| LBX2 | TTTCATTAAT | -1 | 603 | 613 | -397 | -387 | 4,44 | 0,01 | 15,58 | >0.1 | 3,49 | 5,22 | 0 | 1,02 | 0 | 1,41 | 0 |
| LBX2 | TCTGATCAGT | -1 | 478 | 488 | -522 | -512 | 3,54 | 0,01 | 17,68 | >0.1 | 2,88 | 4,92 | 0 | 0,99 | 0 | 5,36 | 0 |
| LBX2 | CCCGAGTAGC | -1 | 343 | 353 | -657 | -647 | 5,23 | 0,01 | 20,1 | >0.1 | 2,56 | 4,67 | 0 | 1,11 | 0 | 6,52 | 0 |
| LBX2 | TCTGATCAGT | -1 | 163 | 173 | -837 | -827 | 3,54 | 0,01 | 15,48 | >0.1 | 3,21 | 4,42 | 0 | 1,9 | 0 | 2,4 | 0 |
| LBX2 | TTTAATTTCC | -1 | 153 | 163 | -847 | -837 | 3,36 | 0,01 | 14,58 | >0.1 | 3,27 | 4,41 | 0 | 1,9 | 0 | 1,65 | 0 |
| LBX2 | TTCACTTACC | -1 | 4 | 14 | -996 | -986 | 5,58 | 0,01 | 19,1 | >0.1 | 5,61 | 4,26 | 0 | 1,86 | 1,79 | 0 | 0 |
| LBX1 | TTAGCTGG | 1 | 308 | 316 | -692 | -684 | -1,83 | 0,01 | 12,31 | >0.1 | 2,66 | 4,61 | 0 | 1,18 | 0 | 5,68 | 0 |
| LBX1 | GTATTTGG | 1 | 522 | 530 | -478 | -470 | -2,29 | 0,01 | 9,81 | >0.1 | 3,05 | 5,01 | 0 | 0,98 | 0 | 3,07 | 0 |
| LBX1 | TTAATGAA | 1 | 604 | 612 | -396 | -388 | 4,51 | 0,01 | 15,65 | >0.1 | 3,49 | 5,21 | 0 | 1,02 | 0 | 1,41 | 0 |
| LBX1 | TTAACAGC | 1 | 709 | 717 | -291 | -283 | 4,79 | 0,01 | 21,96 | >0.1 | 5,64 | 5,55 | 0 | 1,17 | 0 | 4,81 | 0 |
| LBX1 | GTAACTGC | 1 | 1088 | 1096 | 88 | 96 | 6,2 | 0,01 | 24,6 | >0.1 | 5,64 | 6,9 | 0 | 1,85 | 1,79 | 2,21 | 0 |
| LBX1 | TCAATGGT | -1 | 1106 | 1114 | 106 | 114 | 0,75 | 0,01 | 20,67 | >0.1 | 5,64 | 6,69 | 0 | 1,85 | 3,22 | 2,52 | 0 |
| LBX1 | GCAACAAG | -1 | 1079 | 1087 | 79 | 87 | 3,81 | 0,01 | 22,34 | >0.1 | 5,64 | 7,03 | 0 | 1,85 | 1,79 | 2,21 | 0 |
| LBX1 | GTAACAGA | -1 | 845 | 853 | -155 | -147 | 1,37 | 0,01 | 18,12 | >0.1 | 5,64 | 6,25 | 0 | 1,8 | 0 | 3,07 | 0 |
| LBX1 | AGAACTAG | -1 | 656 | 664 | -344 | -336 | -1,94 | 0,01 | 12,15 | >0.1 | 3,95 | 5,37 | 0 | 1,07 | 0 | 3,7 | 0 |
| LBX1 | TTAATAGG | -1 | 600 | 608 | -400 | -392 | 8,59 | 0,001 | 19,64 | >0.1 | 3,47 | 5,2 | 0 | 1,01 | 0 | 1,36 | 0 |
| LBX1 | CTACTTGG | -1 | 545 | 553 | -455 | -447 | -2,2 | 0,01 | 7 | >0.1 | 3,15 | 5,06 | 0 | 0,98 | 0 | 0 | 0 |
| LBX1 | GTAGCTGG | -1 | 340 | 348 | -660 | -652 | -2,79 | 0,01 | 12,08 | >0.1 | 2,57 | 4,66 | 0 | 1,12 | 0 | 6,52 | 0 |
| LBX1 | TTAATAGA | -1 | 288 | 296 | -712 | -704 | 3,32 | 0,01 | 17,05 | >0.1 | 2,72 | 4,58 | 0 | 1,24 | 0 | 5,19 | 0 |
| LBX1 | TTAATTTC | -1 | 154 | 162 | -846 | -838 | -2,73 | 0,01 | 8,49 | >0.1 | 3,27 | 4,41 | 0 | 1,9 | 0 | 1,65 | 0 |
| PAX5 | CAGTGTGAGAAA | 1 | 50 | 62 | -950 | -938 | 1,4 | 0,01 | 11,69 | >0.1 | 4,11 | 4,3 | 0 | 1,87 | 0 | 0 | 0 |
| PAX5 | CGGTGAAACCCC | 1 | 273 | 285 | -727 | -715 | 1,21 | 0,01 | 15,02 | >0.1 | 2,76 | 4,57 | 0 | 1,28 | 0 | 5,19 | 0 |
| PAX5 | GGGCGTGGTGGT | 1 | 314 | 326 | -686 | -674 | 3,43 | 0,01 | 18 | >0.1 | 2,64 | 4,63 | 0 | 1,17 | 0 | 6,13 | 0 |
| PAX5 | TGGCGTGAACCT | 1 | 370 | 382 | -630 | -618 | 8,65 | 0,001 | 23,39 | >0.1 | 2,55 | 4,72 | 0 | 1,07 | 0 | 6,4 | 0 |
| PAX5 | TGCAGTGAGCCG | 1 | 395 | 407 | -605 | -593 | 0,71 | 0,01 | 14,89 | >0.1 | 2,61 | 4,76 | 0 | 1,04 | 0 | 5,76 | 0 |

| | | | | | | | | | | | | | | | | | |
|---|---|---|---|---|---|---|---|---|---|---|---|---|---|---|---|---|---|
| PAX5 | AGCCGAGATCGC | 1 | 402 | 414 | -598 | -586 | 3,54 | 0,01 | 17,75 | >0.1 | 2,63 | 4,78 | 0 | 1,04 | 0 | 5,76 | 0 |
| PAX5 | TCGCGCCACTGT | 1 | 410 | 422 | -590 | -578 | 1 | 0,01 | 15,24 | >0.1 | 2,66 | 4,79 | 0 | 1,03 | 0 | 5,76 | 0 |
| PAX5 | CAGCCTGGGCGA | 1 | 426 | 438 | -574 | -562 | 1,71 | 0,01 | 16,05 | >0.1 | 2,7 | 4,82 | 0 | 1,01 | 0 | 5,8 | 0 |
| PAX5 | CTGGGCGACAGA | 1 | 430 | 442 | -570 | -558 | 0,93 | 0,01 | 15,15 | >0.1 | 2,72 | 4,83 | 0 | 1,01 | 0 | 5,67 | 0 |
| PAX5 | GACCGAGACTCA | 1 | 440 | 452 | -560 | -548 | 6,98 | 0,001 | 21,5 | >0.1 | 2,75 | 4,85 | 0 | 1,01 | 0 | 5,92 | 0 |
| PAX5 | GGCCAAGACATT | 1 | 1120 | 1132 | 120 | 132 | 1,33 | 0,01 | 21,2 | >0.1 | 5,58 | 6,55 | 0 | 1,85 | 3,22 | 2,67 | 0 |
| PAX5 | GAGCTTGACATC | -1 | 1048 | 1060 | 48 | 60 | 6,65 | 0,001 | 23,18 | >0.1 | 5,58 | 7,62 | 0 | 1,86 | 0 | 1,46 | 0 |
| PAX5 | CCCTGTGAGCCA | -1 | 1033 | 1045 | 33 | 45 | 1,85 | 0,01 | 18,7 | >0.1 | 5,58 | 8,1 | 0 | 1,86 | 0 | 1,31 | 0 |
| PAX5 | AGGAAAGACCAA | -1 | 928 | 940 | -72 | -60 | 1,29 | 0,01 | 16,84 | >0.1 | 5,58 | 7,06 | 0 | 1,88 | 0 | 1,03 | 0 |
| PAX5 | TACAGTGACCAA | -1 | 672 | 684 | -328 | -316 | 4 | 0,01 | 18,27 | >0.1 | 4,19 | 5,43 | 0 | 1,11 | 0 | 3,55 | 0 |
| PAX5 | ATACATGACTTT | -1 | 514 | 526 | -486 | -474 | 1,11 | 0,01 | 12,98 | >0.1 | 3,02 | 5 | 0 | 0,98 | 0 | 2,86 | 0 |
| PAX5 | AGGCTGGAGCAC | -1 | 420 | 432 | -580 | -568 | 1,11 | 0,01 | 15,04 | >0.1 | 2,69 | 4,81 | 0 | 1,02 | 0 | 5,42 | 0 |
| PAX5 | TGGCGCGATCTC | -1 | 406 | 418 | -594 | -582 | 4,27 | 0,01 | 18,38 | >0.1 | 2,65 | 4,78 | 0 | 1,03 | 0 | 5,65 | 0 |
| PAX5 | AGCTGGGACTAC | -1 | 334 | 346 | -666 | -654 | 1,01 | 0,01 | 15,41 | >0.1 | 2,58 | 4,66 | 0 | 1,13 | 0 | 6,03 | 0 |
| PAX5 | GCCCACCACCAC | -1 | 318 | 330 | -682 | -670 | 0,55 | 0,01 | 14,54 | >0.1 | 2,63 | 4,64 | 0 | 1,16 | 0 | 5,56 | 0 |
| PAX5 | CACCACGCCCAG | -1 | 312 | 324 | -688 | -676 | 0,97 | 0,01 | 15,54 | >0.1 | 2,64 | 4,63 | 0 | 1,18 | 0 | 6,13 | 0 |
| PAX5 | GGGTTTCACCGT | -1 | 272 | 284 | -728 | -716 | 1,09 | 0,01 | 14,79 | >0.1 | 2,76 | 4,57 | 0 | 1,29 | 0 | 5,09 | 0 |
| PAX5 | AGGCGTGAGCCA | -1 | 188 | 200 | -812 | -800 | 11,65 | 0,0001 | 24,95 | >0.1 | 3,09 | 4,45 | 0 | 1,72 | 0 | 4,04 | 0 |
| PAX5 | CACTGTGCCCAG | -1 | 178 | 190 | -822 | -810 | 3,8 | 0,01 | 16,27 | >0.1 | 3,13 | 4,44 | 0 | 1,81 | 0 | 3,09 | 0 |
| PAX5 | TTCCATGGCCAT | -1 | 145 | 157 | -855 | -843 | 2,43 | 0,01 | 13,64 | >0.1 | 3,31 | 4,4 | 0 | 1,9 | 0 | 1,61 | 0 |
| PAX5 | GTGGCTGACTTC | -1 | 33 | 45 | -967 | -955 | 2,45 | 0,01 | 13 | >0.1 | 4,4 | 4,28 | 0 | 1,87 | 0 | 0 | 0 |
| PAX4 | GAAATGAG | 1 | 65 | 73 | -935 | -927 | 2,46 | 0,01 | 12,6 | >0.1 | 3,95 | 4,31 | 0 | 1,88 | 0 | 0 | 0 |
| PAX4 | GAAATTAA | 1 | 154 | 162 | -846 | -838 | 5,34 | 0,01 | 16,56 | >0.1 | 3,27 | 4,41 | 0 | 1,9 | 0 | 1,65 | 0 |
| PAX4 | TGAATTCC | 1 | 492 | 500 | -508 | -500 | 5,95 | 0,01 | 20,93 | >0.1 | 2,92 | 4,94 | 0 | 0,98 | 0 | 6,13 | 0 |
| PAX4 | CCAAGTAG | 1 | 545 | 553 | -455 | -447 | 3,85 | 0,01 | 13,05 | >0.1 | 3,15 | 5,06 | 0 | 0,98 | 0 | 0 | 0 |
| PAX4 | TTAATGAA | 1 | 604 | 612 | -396 | -388 | 3,99 | 0,01 | 15,13 | >0.1 | 3,49 | 5,21 | 0 | 1,02 | 0 | 1,41 | 0 |
| PAX4 | AAAATTAT | 1 | 683 | 691 | -317 | -309 | 2,64 | 0,01 | 16,62 | >0.1 | 4,34 | 5,46 | 0 | 1,12 | 0 | 3,07 | 0 |
| PAX4 | AAAATTAT | 1 | 830 | 838 | -170 | -162 | 2,64 | 0,01 | 19,37 | >0.1 | 5,64 | 6,14 | 0 | 1,67 | 0 | 3,28 | 0 |
| PAX4 | GGAATTAT | 1 | 1191 | 1199 | 191 | 199 | 5,02 | 0,01 | 23,8 | >0.1 | 5,64 | 6,02 | 0 | 1,5 | 1,79 | 3,82 | 0 |
| PAX4 | ATAATTCC | -1 | 1191 | 1199 | 191 | 199 | 5,42 | 0,01 | 24,2 | >0.1 | 5,64 | 6,02 | 0 | 1,5 | 1,79 | 3,82 | 0 |
| PAX4 | TGAATGAC | -1 | 1012 | 1020 | 12 | 20 | 2,82 | 0,01 | 20,64 | >0.1 | 5,64 | 8,92 | 0 | 1,86 | 0 | 1,39 | 0 |
| PAX4 | TGAATCAA | -1 | 816 | 824 | -184 | -176 | 3,15 | 0,01 | 19,84 | >0.1 | 5,64 | 6,05 | 0 | 1,58 | 0 | 3,43 | 0 |
| PAX4 | CAAATAAG | -1 | 811 | 819 | -189 | -181 | 5,61 | 0,01 | 22,1 | >0.1 | 5,64 | 6,02 | 0 | 1,55 | 0 | 3,28 | 0 |
| PAX4 | AGAATACG | -1 | 805 | 813 | -195 | -187 | 2,09 | 0,01 | 18,65 | >0.1 | 5,64 | 5,99 | 0 | 1,51 | 0 | 3,43 | 0 |
| PAX4 | CAAATCAA | -1 | 750 | 758 | -250 | -242 | 3,76 | 0,01 | 22,31 | >0.1 | 5,64 | 5,71 | 0 | 1,28 | 0 | 5,92 | 0 |
| PAX4 | TTCATTAA | -1 | 604 | 612 | -396 | -388 | 1,77 | 0,01 | 12,91 | >0.1 | 3,49 | 5,21 | 0 | 1,02 | 0 | 1,41 | 0 |
| PAX4 | CAAATACA | -1 | 521 | 529 | -479 | -471 | 2,56 | 0,01 | 14,66 | >0.1 | 3,04 | 5,01 | 0 | 0,98 | 0 | 3,07 | 0 |
| PAX4 | GGAATTCA | -1 | 492 | 500 | -508 | -500 | 5,48 | 0,01 | 20,46 | >0.1 | 2,92 | 4,94 | 0 | 0,98 | 0 | 6,13 | 0 |
| PAX4 | TTAATTTC | -1 | 154 | 162 | -846 | -838 | 3,56 | 0,01 | 14,78 | >0.1 | 3,27 | 4,41 | 0 | 1,9 | 0 | 1,65 | 0 |
| PAX4 | AAAATTAT | -1 | 132 | 140 | -868 | -860 | 2,64 | 0,01 | 13,88 | >0.1 | 3,4 | 4,38 | 0 | 1,89 | 0 | 1,57 | 0 |
| PAX7 | AAACTGATCA | 1 | 161 | 171 | -839 | -829 | -1,16 | 0,01 | 10,84 | >0.1 | 3,22 | 4,42 | 0 | 1,9 | 0 | 2,46 | 0 |
| PAX7 | AAACTGATCA | 1 | 476 | 486 | -524 | -514 | -1,16 | 0,01 | 12,81 | >0.1 | 2,87 | 4,91 | 0 | 0,99 | 0 | 5,2 | 0 |
| PAX7 | ATATGAATTC | 1 | 489 | 499 | -511 | -501 | -8,09 | 0,01 | 7,02 | >0.1 | 2,92 | 4,94 | 0 | 0,98 | 0 | 6,27 | 0 |
| PAX7 | AAATAAATGT | 1 | 583 | 593 | -417 | -407 | -7,48 | 0,01 | 2,05 | >0.1 | 3,36 | 5,16 | 0 | 1 | 0 | 0 | 0 |
| PAX7 | TAACCTATTA | 1 | 597 | 607 | -403 | -393 | -3,73 | 0,01 | 7,35 | >0.1 | 3,45 | 5,2 | 0 | 1,01 | 0 | 1,41 | 0 |
| PAX7 | TAATGAAATT | 1 | 605 | 615 | -395 | -385 | -3,37 | 0,01 | 7,79 | >0.1 | 3,51 | 5,22 | 0 | 1,02 | 0 | 1,41 | 0 |
| PAX7 | TAAAGAATAA | 1 | 795 | 805 | -205 | -195 | -7,91 | 0,01 | 10,43 | >0.1 | 5,61 | 5,94 | 0 | 1,47 | 0 | 5,32 | 0 |
| PAX7 | TATTTGATTC | 1 | 813 | 823 | -187 | -177 | -4,42 | 0,01 | 12,24 | >0.1 | 5,61 | 6,05 | 0 | 1,57 | 0 | 3,43 | 0 |
| PAX7 | TAAAAAATTA | 1 | 827 | 837 | -173 | -163 | -2,2 | 0,01 | 14,64 | >0.1 | 5,61 | 6,14 | 0 | 1,67 | 0 | 3,43 | 0 |
| PAX7 | AAATTATTCT | 1 | 831 | 841 | -169 | -159 | -8,28 | 0,01 | 8,62 | >0.1 | 5,61 | 6,16 | 0 | 1,7 | 0 | 3,43 | 0 |
| PAX7 | AAATCTGTTA | 1 | 842 | 852 | -158 | -148 | -7,24 | 0,01 | 9,59 | >0.1 | 5,61 | 6,24 | 0 | 1,79 | 0 | 3,19 | 0 |
| PAX7 | AGATTGTTTA | 1 | 893 | 903 | -107 | -97 | -3,35 | 0,01 | 13,65 | >0.1 | 5,61 | 6,64 | 0 | 1,89 | 0 | 2,86 | 0 |
| PAX7 | TAAACAATCT | -1 | 893 | 903 | -107 | -97 | -4,13 | 0,01 | 12,87 | >0.1 | 5,61 | 6,64 | 0 | 1,89 | 0 | 2,86 | 0 |
| PAX7 | TAACAGATTT | -1 | 842 | 852 | -158 | -148 | -2,15 | 0,01 | 14,68 | >0.1 | 5,61 | 6,24 | 0 | 1,79 | 0 | 3,19 | 0 |
| PAX7 | TAATTTTTTA | -1 | 827 | 837 | -173 | -163 | -7,71 | 0,01 | 9,13 | >0.1 | 5,61 | 6,14 | 0 | 1,67 | 0 | 3,43 | 0 |
| PAX7 | GAATCAAATA | -1 | 813 | 823 | -187 | -177 | 2,7 | 0,001 | 19,36 | >0.1 | 5,61 | 6,05 | 0 | 1,57 | 0 | 3,43 | 0 |
| PAX7 | AAATCAAAAC | -1 | 747 | 757 | -253 | -243 | -7,64 | 0,01 | 11,06 | >0.1 | 5,61 | 5,71 | 0 | 1,28 | 0 | 6,1 | 0 |
| PAX7 | AATTTCATTA | -1 | 605 | 615 | -395 | -385 | -8,24 | 0,01 | 2,92 | >0.1 | 3,51 | 5,22 | 0 | 1,02 | 0 | 1,41 | 0 |
| PAX7 | TCATTAATAG | -1 | 601 | 611 | -399 | -389 | -7,15 | 0,01 | 3,97 | >0.1 | 3,48 | 5,21 | 0 | 1,02 | 0 | 1,41 | 0 |

| | | | | | | | | | | | | | | | | | |
|---|---|---|---|---|---|---|---|---|---|---|---|---|---|---|---|---|---|
| PAX7 | TAATAGGTTA | -1 | 597 | 607 | -403 | -393 | 0,69 | 0,01 | 11,77 | >0.1 | 3,45 | 5,2 | 0 | 1,01 | 0 | 1,41 | 0 |
| PAX7 | AAATACATGA | -1 | 518 | 528 | -482 | -472 | -8,47 | 0,01 | 3,62 | >0.1 | 3,03 | 5 | 0 | 0,98 | 0 | 3,07 | 0 |
| PAX7 | GAATTCATAT | -1 | 489 | 499 | -511 | -501 | -7,7 | 0,01 | 7,41 | >0.1 | 2,92 | 4,94 | 0 | 0,98 | 0 | 6,27 | 0 |
| PAX7 | TGATCAGTTT | -1 | 476 | 486 | -524 | -514 | -3,56 | 0,01 | 10,41 | >0.1 | 2,87 | 4,91 | 0 | 0,99 | 0 | 5,2 | 0 |
| PAX7 | TGATCAGTTT | -1 | 161 | 171 | -839 | -829 | -3,56 | 0,01 | 8,44 | >0.1 | 3,22 | 4,42 | 0 | 1,9 | 0 | 2,46 | 0 |
| PAX6 | CTCACGCCTGTAAT | 1 | 191 | 205 | -809 | -795 | 4,72 | 0,01 | 18,21 | >0.1 | 3,07 | 4,46 | 0 | 1,7 | 0 | 4,26 | 0 |
| PAX6 | CTCCAGCCTGGGCG | 1 | 423 | 437 | -577 | -563 | 3,3 | 0,01 | 17,5 | >0.1 | 2,7 | 4,82 | 0 | 1,02 | 0 | 5,67 | 0 |
| PAX6 | TTAAAGAATAACGT | 1 | 794 | 808 | -206 | -192 | 0,78 | 0,01 | 17,21 | >0.1 | 5,56 | 5,96 | 0 | 1,49 | 0 | 3,43 | 0 |
| PAX6 | TATTTGATTCACTT | 1 | 813 | 827 | -187 | -173 | -0,27 | 0,01 | 16,38 | >0.1 | 5,56 | 6,07 | 0 | 1,6 | 0 | 3,43 | 0 |
| PAX6 | CTCCAGGATGAACT | 1 | 864 | 878 | -136 | -122 | 4,3 | 0,01 | 20,94 | >0.1 | 5,56 | 6,43 | 0 | 1,9 | 0 | 2,75 | 0 |
| PAX6 | TTTCCTGCTCAGCG | 1 | 934 | 948 | -66 | -52 | 0,58 | 0,01 | 16,1 | >0.1 | 5,56 | 7,18 | 0 | 1,88 | 0 | 0,9 | 0 |
| PAX6 | GTCAAGCTCTTCCT | 1 | 1051 | 1065 | 51 | 65 | -0,2 | 0,01 | 16,98 | >0.1 | 5,56 | 7,55 | 0 | 1,86 | 0 | 2,21 | 0 |
| PAX6 | TTTTTGGACAAGTT | 1 | 1130 | 1144 | 130 | 144 | 0,81 | 0,01 | 20,87 | >0.1 | 5,56 | 6,46 | 0 | 1,84 | 3,22 | 2,98 | 0 |
| PAX6 | ATCAAAGTTCACTT | 1 | 1170 | 1184 | 170 | 184 | 2,6 | 0,01 | 20,74 | >0.1 | 5,56 | 6,15 | 0 | 1,67 | 1,79 | 2,97 | 0 |
| PAX6 | CTTCGGCTTCGTGG | -1 | 1147 | 1161 | 147 | 161 | 0,63 | 0,01 | 18,38 | >0.1 | 5,56 | 6,32 | 0 | 1,84 | 1,79 | 2,24 | 0 |
| PAX6 | TTCGTGGTTAAACT | -1 | 1140 | 1154 | 140 | 154 | 2,38 | 0,01 | 22,75 | >0.1 | 5,56 | 6,37 | 0 | 1,84 | 3,22 | 3,37 | 0 |
| PAX6 | TCAATGGTGGACTG | -1 | 1100 | 1114 | 100 | 114 | 2,33 | 0,01 | 22,13 | >0.1 | 5,56 | 6,76 | 0 | 1,85 | 3,22 | 2,41 | 0 |
| PAX6 | TTACAGCAACAAGG | -1 | 1078 | 1092 | 78 | 92 | 0,26 | 0,01 | 18,72 | >0.1 | 5,56 | 7,04 | 0 | 1,85 | 1,79 | 2,21 | 0 |
| PAX6 | TTTGAACTTGGGTT | -1 | 951 | 965 | -49 | -35 | 2,55 | 0,01 | 19,19 | >0.1 | 5,56 | 7,51 | 0 | 1,88 | 0 | 1,7 | 0 |
| PAX6 | TTCATCCTGGAGAG | -1 | 862 | 876 | -138 | -124 | -0,01 | 0,01 | 16,61 | >0.1 | 5,56 | 6,41 | 0 | 1,9 | 0 | 2,75 | 0 |
| PAX6 | TACACGGAAAATGT | -1 | 692 | 706 | -308 | -294 | 0,08 | 0,01 | 15,36 | >0.1 | 5,56 | 5,51 | 0 | 1,15 | 0 | 3,07 | 0 |
| PAX6 | TTCATTAATAGGTT | -1 | 598 | 612 | -402 | -388 | 0,13 | 0,01 | 11,2 | >0.1 | 3,47 | 5,21 | 0 | 1,02 | 0 | 1,36 | 0 |
| PAX6 | CTTGGGCAACATTT | -1 | 536 | 550 | -464 | -450 | 0,04 | 0,01 | 10,94 | >0.1 | 3,12 | 5,06 | 0 | 0,99 | 0 | 1,73 | 0 |
| PAX6 | CAAATACATGACTT | -1 | 515 | 529 | -485 | -471 | 1,63 | 0,01 | 13,6 | >0.1 | 3,03 | 5,01 | 0 | 0,99 | 0 | 2,95 | 0 |
| PAX6 | TTTGCCCTTGTCTT | -1 | 503 | 517 | -497 | -483 | 1,27 | 0,01 | 15,9 | >0.1 | 2,98 | 4,98 | 0 | 0,99 | 0 | 5,68 | 0 |
| PAX6 | ATTTCTGATCAGTT | -1 | 477 | 491 | -523 | -509 | 0,34 | 0,01 | 14,66 | >0.1 | 2,88 | 4,92 | 0 | 0,99 | 0 | 5,52 | 0 |
| PAX6 | ATCTCGGCTCACTG | -1 | 397 | 411 | -603 | -589 | 4,48 | 0,01 | 18,82 | >0.1 | 2,62 | 4,77 | 0 | 1,04 | 0 | 5,91 | 0 |
| PAX6 | CTCCCGAGTAGCTG | -1 | 341 | 355 | -659 | -645 | -0,26 | 0,01 | 14,62 | >0.1 | 2,56 | 4,68 | 0 | 1,12 | 0 | 6,52 | 0 |
| PAX6 | TACAGGCGTGAGCC | -1 | 189 | 203 | -811 | -797 | 0 | 0,01 | 13,38 | >0.1 | 3,08 | 4,46 | 0 | 1,71 | 0 | 4,13 | 0 |
| PAX6 | ATTTCTGATCAGTT | -1 | 162 | 176 | -838 | -824 | 0,34 | 0,01 | 12,28 | >0.1 | 3,21 | 4,42 | 0 | 1,9 | 0 | 2,4 | 0 |
| PAX6 | TCCACTCCACAGGT | -1 | 110 | 124 | -890 | -876 | -0,27 | 0,01 | 10,27 | >0.1 | 3,52 | 4,37 | 0 | 1,89 | 0 | 0,77 | 0 |
| PAX6 | TCCACAGGTCAACT | -1 | 105 | 119 | -895 | -881 | 1,5 | 0,01 | 12,07 | >0.1 | 3,56 | 4,36 | 0 | 1,89 | 0 | 0,77 | 0 |
| PAX6 | TTCTCACACTGGTT | -1 | 47 | 61 | -953 | -939 | 1,12 | 0,01 | 11,44 | >0.1 | 4,14 | 4,3 | 0 | 1,87 | 0 | 0 | 0 |
| PAX6 | CTCACACTGGTTCT | -1 | 45 | 59 | -955 | -941 | -0,62 | 0,01 | 9,72 | >0.1 | 4,17 | 4,3 | 0 | 1,87 | 0 | 0 | 0 |
| PAX6 | TCTGTGGCTGACTT | -1 | 34 | 48 | -966 | -952 | 1,3 | 0,01 | 11,82 | >0.1 | 4,36 | 4,29 | 0 | 1,87 | 0 | 0 | 0 |
| PAX6 | CTTCTCCTTCACTT | -1 | 7 | 21 | -993 | -979 | 0,53 | 0,01 | 14,01 | >0.1 | 5,56 | 4,26 | 0 | 1,86 | 1,79 | 0 | 0 |
| PAX1 | AGAAATGGCTGGGCACA | 1 | 170 | 187 | -830 | -813 | -8,9 | 0,01 | 3,82 | >0.1 | 3,16 | 4,44 | 0 | 1,88 | 0 | 3,24 | 0 |
| PAX1 | GCTCACGCCTGTAATCC | 1 | 190 | 207 | -810 | -793 | -16,74 | 0,01 | -3,24 | >0.1 | 3,07 | 4,46 | 0 | 1,71 | 0 | 4,26 | 0 |
| PAX1 | AGACCGTCCTGACTAAC | 1 | 255 | 272 | -745 | -728 | -18,12 | 0,01 | -4,88 | >0.1 | 2,81 | 4,55 | 0 | 1,35 | 0 | 4,53 | 0 |
| PAX1 | CTACTCGGGAGGCTGAG | 1 | 344 | 361 | -656 | -639 | -18,07 | 0,01 | -3,2 | >0.1 | 2,55 | 4,69 | 0 | 1,12 | 0 | 6,52 | 0 |
| PAX1 | GCTCCAGCCTGGGCGAC | 1 | 422 | 439 | -578 | -561 | -13,68 | 0,01 | 0,53 | >0.1 | 2,7 | 4,82 | 0 | 1,02 | 0 | 5,67 | 0 |
| PAX1 | GGTCACTGTAAAATTAT | 1 | 674 | 691 | -326 | -309 | -15,21 | 0,01 | -0,95 | >0.1 | 4,26 | 5,46 | 0 | 1,12 | 0 | 3,43 | 0 |
| PAX1 | CTTTCCTGCTCAGCGCC | 1 | 933 | 950 | -67 | -50 | -7,94 | 0,01 | 7,58 | >0.1 | 5,52 | 7,22 | 0 | 1,88 | 0 | 0,9 | 0 |
| PAX1 | GATCTTGGCTCACAGGG | 1 | 1028 | 1045 | 28 | 45 | -18,38 | 0,01 | -1,36 | >0.1 | 5,52 | 8,34 | 0 | 1,86 | 0 | 1,31 | 0 |
| PAX1 | TATCAAAGTTCACTTGC | 1 | 1169 | 1186 | 169 | 186 | -18,04 | 0,01 | 0,07 | >0.1 | 5,52 | 6,16 | 0 | 1,68 | 1,79 | 2,97 | 0 |
| PAX1 | TGTTCCTCAATGGTGGA | -1 | 1103 | 1120 | 103 | 120 | -14,04 | 0,01 | 5,75 | >0.1 | 5,52 | 6,72 | 0 | 1,85 | 3,22 | 2,48 | 0 |
| PAX1 | CTCAATGGTGGACTGAG | -1 | 1098 | 1115 | 98 | 115 | -14,34 | 0,01 | 5,41 | >0.1 | 5,52 | 6,78 | 0 | 1,85 | 3,22 | 2,38 | 0 |
| PAX1 | CAACAAGGCTGAGAAGG | -1 | 1069 | 1086 | 69 | 86 | -17,38 | 0,01 | 1,25 | >0.1 | 5,52 | 7,19 | 0 | 1,85 | 1,79 | 2,27 | 0 |
| PAX1 | GTTCATCCTGGAGAGGA | -1 | 860 | 877 | -140 | -123 | -13,18 | 0,01 | 3,33 | >0.1 | 5,52 | 6,42 | 0 | 1,9 | 0 | 2,67 | 0 |
| PAX1 | CGTTATTCTTTAATCCT | -1 | 790 | 807 | -210 | -193 | -9,37 | 0,01 | 8,9 | >0.1 | 5,52 | 5,95 | 0 | 1,48 | 0 | 5,32 | 0 |
| PAX1 | CTTTATGGCCAAATCAA | -1 | 750 | 767 | -250 | -233 | -19,2 | 0,01 | -0,29 | >0.1 | 5,52 | 5,76 | 0 | 1,31 | 0 | 6,32 | 0 |
| PAX1 | AGAAACTCTCTACTTGG | -1 | 545 | 562 | -455 | -438 | -18,31 | 0,01 | -9,06 | >0.1 | 3,17 | 5,08 | 0 | 1 | 0 | 0 | 0 |
| PAX1 | CTCTCTACTTGGGCAAC | -1 | 540 | 557 | -460 | -443 | -15,49 | 0,01 | -6,27 | >0.1 | 3,15 | 5,07 | 0 | 0,99 | 0 | 0 | 0 |
| PAX1 | AGTCTCGGTCTGTCGCC | -1 | 433 | 450 | -567 | -550 | -8,78 | 0,01 | 5,73 | >0.1 | 2,73 | 4,84 | 0 | 1,01 | 0 | 5,92 | 0 |
| PAX1 | CGCCCAGGCTGGAGCAC | -1 | 420 | 437 | -580 | -563 | -19,15 | 0,01 | -4,95 | >0.1 | 2,69 | 4,82 | 0 | 1,02 | 0 | 5,67 | 0 |
| PAX1 | GATCTCGGCTCACTGCA | -1 | 395 | 412 | -605 | -588 | -6,85 | 0,001 | 7,5 | >0.1 | 2,62 | 4,77 | 0 | 1,05 | 0 | 5,91 | 0 |
| PAX1 | AGCTCTGCCTCCCAGGT | -1 | 378 | 395 | -622 | -605 | -16,16 | 0,01 | -1,76 | >0.1 | 2,57 | 4,74 | 0 | 1,07 | 0 | 6,02 | 0 |
| PAX1 | CTCCCAGGTTCACGCCA | -1 | 370 | 387 | -630 | -613 | -18,66 | 0,01 | -3,77 | >0.1 | 2,55 | 4,73 | 0 | 1,08 | 0 | 6,53 | 0 |
| PAX1 | CACCACGCCCAGCTAAG | -1 | 307 | 324 | -693 | -676 | -16,11 | 0,01 | -1,51 | >0.1 | 2,65 | 4,63 | 0 | 1,19 | 0 | 6,13 | 0 |

| | | | | | | | | | | | | | | | | | |
|---|---|---|---|---|---|---|---|---|---|---|---|---|---|---|---|---|---|
| PAX1 | TGCCCAACTCGGCCTCC | -1 | 217 | 234 | -783 | -766 | -18,25 | 0,01 | -3,75 | >0.1 | 2,95 | 4,5 | 0 | 1,53 | 0 | 5,52 | 0 |
| PAX1 | TTACAGGCGTGAGCCAC | -1 | 187 | 204 | -813 | -796 | -19,19 | 0,01 | -5,79 | >0.1 | 3,08 | 4,46 | 0 | 1,73 | 0 | 4,13 | 0 |
| PAX1 | GACTACTCTCCACTCCA | -1 | 115 | 132 | -885 | -868 | -18,5 | 0,01 | -8,03 | >0.1 | 3,48 | 4,37 | 0 | 1,89 | 0 | 0,73 | 0 |
| PAX1 | CTCTCCACTCCACAGGT | -1 | 110 | 127 | -890 | -873 | -15,72 | 0,01 | -5,18 | >0.1 | 3,51 | 4,37 | 0 | 1,89 | 0 | 0,77 | 0 |
| PAX1 | CTCCACAGGTCAACTCT | -1 | 103 | 120 | -897 | -880 | -11,9 | 0,01 | -1,34 | >0.1 | 3,56 | 4,36 | 0 | 1,89 | 0 | 0,75 | 0 |
| PAX1 | TTCTGTGGCTGACTTCT | -1 | 32 | 49 | -968 | -951 | -15,6 | 0,01 | -5,07 | >0.1 | 4,37 | 4,29 | 0 | 1,87 | 0 | 0 | 0 |
| PAX1 | CCTTCTCCTTCACTTAC | -1 | 5 | 22 | -995 | -978 | -14,96 | 0,01 | -1,52 | >0.1 | 5,52 | 4,27 | 0 | 1,86 | 1,79 | 0 | 0 |
| SOX21 | CACAGAACCAGTGTG | 1 | 42 | 57 | -958 | -943 | -12,71 | 0,01 | -2,33 | >0.1 | 4,21 | 4,3 | 0 | 1,87 | 0 | 0 | 0 |
| SOX21 | AAAAATGGCCATGGA | 1 | 141 | 156 | -859 | -844 | -15,8 | 0,01 | -4,57 | >0.1 | 3,32 | 4,4 | 0 | 1,9 | 0 | 1,61 | 0 |
| SOX21 | AGAGCTTGCAGTGAG | 1 | 389 | 404 | -611 | -596 | -20,58 | 0,01 | -6,15 | >0.1 | 2,6 | 4,76 | 0 | 1,05 | 0 | 6,02 | 0 |
| SOX21 | GACAAGGGCAAAGTC | 1 | 505 | 520 | -495 | -480 | -7,84 | 0,001 | 5,32 | >0.1 | 2,99 | 4,99 | 0 | 0,99 | 0 | 4,19 | 0 |
| SOX21 | CCCAAGTAGAGAGTT | 1 | 544 | 559 | -456 | -441 | -13,26 | 0,01 | -4,03 | >0.1 | 3,17 | 5,08 | 0 | 0,99 | 0 | 0 | 0 |
| SOX21 | ATTAATGAAATTGTC | 1 | 603 | 618 | -397 | -382 | -4,42 | 0,001 | 6,76 | >0.1 | 3,51 | 5,23 | 0 | 1,03 | 0 | 1,41 | 0 |
| SOX21 | AACACCAACATTATC | 1 | 631 | 646 | -369 | -354 | -20,06 | 0,01 | -8,35 | >0.1 | 3,73 | 5,31 | 0 | 1,05 | 0 | 1,61 | 0 |
| SOX21 | AACATTTTCCGTGTA | 1 | 691 | 706 | -309 | -294 | -7,9 | 0,001 | 7,37 | >0.1 | 5,54 | 5,51 | 0 | 1,15 | 0 | 3,07 | 0 |
| SOX21 | AAAAGTACCGGTTTT | 1 | 737 | 752 | -263 | -248 | -19,84 | 0,01 | -1,43 | >0.1 | 5,54 | 5,69 | 0 | 1,26 | 0 | 5,92 | 0 |
| SOX21 | AGGAGAGGTAAGGTT | 1 | 771 | 786 | -229 | -214 | -20,14 | 0,01 | -1,27 | >0.1 | 5,54 | 5,84 | 0 | 1,38 | 0 | 6,1 | 0 |
| SOX21 | GACGATGTCAAGCTC | 1 | 1045 | 1060 | 45 | 60 | -21,02 | 0,01 | -4,5 | >0.1 | 5,54 | 7,7 | 0 | 1,86 | 0 | 1,42 | 0 |
| SOX21 | GGCCAAGACATTTTT | 1 | 1120 | 1135 | 120 | 135 | -19,97 | 0,01 | -0,07 | >0.1 | 5,54 | 6,55 | 0 | 1,85 | 3,22 | 2,74 | 0 |
| SOX21 | AGCAAGTGAACTTTG | -1 | 1172 | 1187 | 172 | 187 | -18,91 | 0,01 | -0,58 | >0.1 | 5,54 | 6,14 | 0 | 1,65 | 1,79 | 3,2 | 0 |
| SOX21 | CTCAATGGTGGACTG | -1 | 1100 | 1115 | 100 | 115 | -19,75 | 0,01 | 0,04 | >0.1 | 5,54 | 6,76 | 0 | 1,85 | 3,22 | 2,41 | 0 |
| SOX21 | AGCCAATATAAAGTT | -1 | 874 | 889 | -126 | -111 | -18,11 | 0,01 | -1,2 | >0.1 | 5,54 | 6,52 | 0 | 1,9 | 0 | 2,95 | 0 |
| SOX21 | TACACGGAAAATGTT | -1 | 691 | 706 | -309 | -294 | -16,45 | 0,01 | -1,18 | >0.1 | 5,54 | 5,51 | 0 | 1,15 | 0 | 3,07 | 0 |
| SOX21 | GAAAATGTTATAATT | -1 | 685 | 700 | -315 | -300 | -12,3 | 0,01 | 1,72 | >0.1 | 4,45 | 5,49 | 0 | 1,14 | 0 | 2,95 | 0 |
| SOX21 | TATAATTTTACAGTG | -1 | 677 | 692 | -323 | -308 | -16,31 | 0,01 | -2,25 | >0.1 | 4,29 | 5,46 | 0 | 1,12 | 0 | 3,19 | 0 |
| SOX21 | GATAATGTTGGTGTT | -1 | 631 | 646 | -369 | -354 | 0,47 | 0,001 | 12,18 | >0.1 | 3,73 | 5,31 | 0 | 1,05 | 0 | 1,61 | 0 |
| SOX21 | GACAATTTCATTAAT | -1 | 603 | 618 | -397 | -382 | -4,07 | 0,001 | 7,11 | >0.1 | 3,51 | 5,23 | 0 | 1,03 | 0 | 1,41 | 0 |
| SOX21 | ATCAGTTTTTTTTTT | -1 | 469 | 484 | -531 | -516 | -17,37 | 0,01 | -3,67 | >0.1 | 2,85 | 4,91 | 0 | 0,99 | 0 | 4,95 | 0 |
| SOX21 | AATAGAGACGGGGTT | -1 | 279 | 294 | -721 | -706 | -20,88 | 0,01 | -7,1 | >0.1 | 2,74 | 4,58 | 0 | 1,27 | 0 | 5,19 | 0 |
| SOX21 | CAAAGTGCTAGGATT | -1 | 202 | 217 | -798 | -783 | -20,65 | 0,01 | -6,94 | >0.1 | 3,02 | 4,48 | 0 | 1,62 | 0 | 4,59 | 0 |
| SOX21 | ATCAGTTTTAATTTC | -1 | 154 | 169 | -846 | -831 | -13,81 | 0,01 | -1,72 | >0.1 | 3,25 | 4,42 | 0 | 1,9 | 0 | 2,53 | 0 |
| SOX21 | CACACTGGTTCTGTG | -1 | 42 | 57 | -958 | -943 | -19,72 | 0,01 | -9,34 | >0.1 | 4,21 | 4,3 | 0 | 1,87 | 0 | 0 | 0 |
| PAX3 | AAACTGATCA | 1 | 161 | 171 | -839 | -829 | 1,64 | 0,01 | 13,64 | >0.1 | 3,22 | 4,42 | 0 | 1,9 | 0 | 2,46 | 0 |
| PAX3 | AAACTGATCA | 1 | 476 | 486 | -524 | -514 | 1,64 | 0,01 | 15,61 | >0.1 | 2,87 | 4,91 | 0 | 0,99 | 0 | 5,2 | 0 |
| PAX3 | ATATGAATTC | 1 | 489 | 499 | -511 | -501 | -5,69 | 0,01 | 9,42 | >0.1 | 2,92 | 4,94 | 0 | 0,98 | 0 | 6,27 | 0 |
| PAX3 | TAACCTATTA | 1 | 597 | 607 | -403 | -393 | 0,95 | 0,01 | 12,03 | >0.1 | 3,45 | 5,2 | 0 | 1,01 | 0 | 1,41 | 0 |
| PAX3 | TATTTGATTC | 1 | 813 | 823 | -187 | -177 | 1,46 | 0,01 | 18,12 | >0.1 | 5,61 | 6,05 | 0 | 1,57 | 0 | 3,43 | 0 |
| PAX3 | TAAAAAATTA | 1 | 827 | 837 | -173 | -163 | -3,71 | 0,01 | 13,13 | >0.1 | 5,61 | 6,14 | 0 | 1,67 | 0 | 3,43 | 0 |
| PAX3 | AAATTATTCT | 1 | 831 | 841 | -169 | -159 | -4,93 | 0,01 | 11,97 | >0.1 | 5,61 | 6,16 | 0 | 1,7 | 0 | 3,43 | 0 |
| PAX3 | AAATCTGTTA | 1 | 842 | 852 | -158 | -148 | -1,87 | 0,01 | 14,96 | >0.1 | 5,61 | 6,24 | 0 | 1,79 | 0 | 3,19 | 0 |
| PAX3 | AGATTGTTTA | 1 | 893 | 903 | -107 | -97 | -0,57 | 0,01 | 16,43 | >0.1 | 5,61 | 6,64 | 0 | 1,89 | 0 | 2,86 | 0 |
| PAX3 | TAAACAATCT | -1 | 893 | 903 | -107 | -97 | -3,22 | 0,01 | 13,78 | >0.1 | 5,61 | 6,64 | 0 | 1,89 | 0 | 2,86 | 0 |
| PAX3 | TAACAGATTT | -1 | 842 | 852 | -158 | -148 | -1,19 | 0,01 | 15,64 | >0.1 | 5,61 | 6,24 | 0 | 1,79 | 0 | 3,19 | 0 |
| PAX3 | TAATTTTTTA | -1 | 827 | 837 | -173 | -163 | -2,72 | 0,01 | 14,12 | >0.1 | 5,61 | 6,14 | 0 | 1,67 | 0 | 3,43 | 0 |
| PAX3 | GAATCAAATA | -1 | 813 | 823 | -187 | -177 | -3,82 | 0,01 | 12,84 | >0.1 | 5,61 | 6,05 | 0 | 1,57 | 0 | 3,43 | 0 |
| PAX3 | AATTTCATTA | -1 | 605 | 615 | -395 | -385 | -2,71 | 0,01 | 8,45 | >0.1 | 3,51 | 5,22 | 0 | 1,02 | 0 | 1,41 | 0 |
| PAX3 | TCATTAATAG | -1 | 601 | 611 | -399 | -389 | -5,56 | 0,01 | 5,56 | >0.1 | 3,48 | 5,21 | 0 | 1,02 | 0 | 1,41 | 0 |
| PAX3 | TAATAGGTTA | -1 | 597 | 607 | -403 | -393 | 0,89 | 0,01 | 11,97 | >0.1 | 3,45 | 5,2 | 0 | 1,01 | 0 | 1,41 | 0 |
| PAX3 | GAATTCATAT | -1 | 489 | 499 | -511 | -501 | -5,32 | 0,01 | 9,79 | >0.1 | 2,92 | 4,94 | 0 | 0,98 | 0 | 6,27 | 0 |
| PAX3 | TGATCAGTTT | -1 | 476 | 486 | -524 | -514 | -0,11 | 0,01 | 13,86 | >0.1 | 2,87 | 4,91 | 0 | 0,99 | 0 | 5,2 | 0 |
| PAX3 | TGATCAGTTT | -1 | 161 | 171 | -839 | -829 | -0,11 | 0,01 | 11,89 | >0.1 | 3,22 | 4,42 | 0 | 1,9 | 0 | 2,46 | 0 |
| PAX2 | AGCCACAG | 1 | 39 | 47 | -961 | -953 | 3,91 | 0,01 | 14,38 | >0.1 | 4,32 | 4,29 | 0 | 1,87 | 0 | 0 | 0 |
| PAX2 | GGCCATGG | 1 | 147 | 155 | -853 | -845 | 5,94 | 0,01 | 17,15 | >0.1 | 3,31 | 4,4 | 0 | 1,9 | 0 | 1,61 | 0 |
| PAX2 | GCTCACGC | 1 | 190 | 198 | -810 | -802 | 5,99 | 0,01 | 19,27 | >0.1 | 3,09 | 4,45 | 0 | 1,7 | 0 | 4,04 | 0 |
| PAX2 | TAACACGG | 1 | 268 | 276 | -732 | -724 | 5,38 | 0,01 | 18,55 | >0.1 | 2,78 | 4,55 | 0 | 1,3 | 0 | 4,53 | 0 |
| PAX2 | GATCGCGC | 1 | 408 | 416 | -592 | -584 | 5,54 | 0,01 | 19,64 | >0.1 | 2,65 | 4,78 | 0 | 1,03 | 0 | 5,65 | 0 |
| PAX2 | CGCCACTG | 1 | 413 | 421 | -587 | -579 | 6,12 | 0,01 | 20,5 | >0.1 | 2,66 | 4,79 | 0 | 1,02 | 0 | 5,91 | 0 |
| PAX2 | AGACAAGG | 1 | 504 | 512 | -496 | -488 | 4,12 | 0,01 | 18,72 | >0.1 | 2,97 | 4,97 | 0 | 0,98 | 0 | 5,68 | 0 |
| PAX2 | AGTCATGT | 1 | 516 | 524 | -484 | -476 | 8,55 | 0,001 | 20,41 | >0.1 | 3,02 | 5 | 0 | 0,98 | 0 | 2,86 | 0 |

| | | | | | | | | | | | | | | | | | |
|---|---|---|---|---|---|---|---|---|---|---|---|---|---|---|---|---|---|
| PAX2 | GGTCACTG | 1 | 674 | 682 | -326 | -318 | 8,73 | 0,001 | 22,99 | >0.1 | 4,19 | 5,42 | 0 | 1,1 | 0 | 3,55 | 0 |
| PAX2 | ATTCACTT | 1 | 819 | 827 | -181 | -173 | 4,49 | 0,01 | 21,22 | >0.1 | 5,64 | 6,07 | 0 | 1,6 | 0 | 3,43 | 0 |
| PAX2 | AGTCATTC | 1 | 1011 | 1019 | 11 | 19 | 8,09 | 0,001 | 25,96 | >0.1 | 5,64 | 8,95 | 0 | 1,86 | 0 | 1,42 | 0 |
| PAX2 | TGTCAAGC | 1 | 1050 | 1058 | 50 | 58 | 7 | 0,001 | 23,53 | >0.1 | 5,64 | 7,57 | 0 | 1,86 | 0 | 1,46 | 0 |
| PAX2 | AGACATTT | 1 | 1125 | 1133 | 125 | 133 | 4,11 | 0,01 | 24,11 | >0.1 | 5,64 | 6,5 | 0 | 1,84 | 3,22 | 2,79 | 0 |
| PAX2 | GTTCACTT | 1 | 1176 | 1184 | 176 | 184 | 4,02 | 0,01 | 22,48 | >0.1 | 5,64 | 6,11 | 0 | 1,62 | 1,79 | 3,29 | 0 |
| PAX2 | TGTCTTGG | -1 | 1122 | 1130 | 122 | 130 | 4,9 | 0,01 | 24,81 | >0.1 | 5,64 | 6,53 | 0 | 1,85 | 3,22 | 2,67 | 0 |
| PAX2 | CGTTATTC | -1 | 799 | 807 | -201 | -193 | 3,9 | 0,01 | 20,52 | >0.1 | 5,64 | 5,95 | 0 | 1,48 | 0 | 3,55 | 0 |
| PAX2 | TGTCACTT | -1 | 764 | 772 | -236 | -228 | 8,11 | 0,001 | 26,77 | >0.1 | 5,64 | 5,78 | 0 | 1,33 | 0 | 5,92 | 0 |
| PAX2 | ATACACGG | -1 | 699 | 707 | -301 | -293 | 3,81 | 0,01 | 19,3 | >0.1 | 5,64 | 5,51 | 0 | 1,15 | 0 | 3,19 | 0 |
| PAX2 | AGTCTCGG | -1 | 442 | 450 | -558 | -550 | 6,16 | 0,01 | 20,67 | >0.1 | 2,75 | 4,84 | 0 | 1 | 0 | 5,92 | 0 |
| PAX2 | GCTCACTG | -1 | 397 | 405 | -603 | -595 | 4,58 | 0,01 | 18,75 | >0.1 | 2,61 | 4,76 | 0 | 1,04 | 0 | 5,76 | 0 |
| PAX2 | GTTCACGC | -1 | 372 | 380 | -628 | -620 | 6,37 | 0,01 | 21,1 | >0.1 | 2,55 | 4,72 | 0 | 1,07 | 0 | 6,4 | 0 |
| PAX2 | CGCCATTC | -1 | 367 | 375 | -633 | -625 | 5 | 0,01 | 19,93 | >0.1 | 2,53 | 4,71 | 0 | 1,07 | 0 | 6,61 | 0 |
| PAX2 | CACCACGC | -1 | 316 | 324 | -684 | -676 | 5,47 | 0,01 | 20,03 | >0.1 | 2,64 | 4,63 | 0 | 1,17 | 0 | 6,13 | 0 |
| PAX2 | AGCCACTG | -1 | 185 | 193 | -815 | -807 | 6,13 | 0,01 | 19,38 | >0.1 | 3,11 | 4,44 | 0 | 1,74 | 0 | 3,95 | 0 |
| PAX2 | AGCCATTT | -1 | 172 | 180 | -828 | -820 | 4,06 | 0,01 | 16,87 | >0.1 | 3,17 | 4,43 | 0 | 1,86 | 0 | 3,34 | 0 |
| PAX2 | CTTCACTT | -1 | 7 | 15 | -993 | -985 | 4,49 | 0,01 | 18,04 | >0.1 | 5,64 | 4,26 | 0 | 1,86 | 1,79 | 0 | 0 |
| SRY | GCACTCAAA | 1 | 24 | 33 | -976 | -967 | 3,79 | 0,01 | 15,55 | >0.1 | 5,62 | 4,27 | 0 | 1,87 | 0 | 0 | 0 |
| SRY | TTAAAAAAT | 1 | 138 | 147 | -862 | -853 | 3,4 | 0,01 | 14,69 | >0.1 | 3,35 | 4,39 | 0 | 1,89 | 0 | 1,65 | 0 |
| SRY | GATCACAAG | 1 | 234 | 243 | -766 | -757 | 5,84 | 0,01 | 19,58 | >0.1 | 2,9 | 4,51 | 0 | 1,43 | 0 | 4,9 | 0 |
| SRY | AAATACAAA | 1 | 296 | 305 | -704 | -695 | 6,06 | 0,01 | 19,01 | >0.1 | 2,69 | 4,6 | 0 | 1,22 | 0 | 4,45 | 0 |
| SRY | AAAGACAAG | 1 | 502 | 511 | -498 | -489 | 4,95 | 0,01 | 19,39 | >0.1 | 2,96 | 4,97 | 0 | 0,98 | 0 | 5,52 | 0 |
| SRY | TGAAATAAA | 1 | 575 | 584 | -425 | -416 | 4,29 | 0,01 | 13,74 | >0.1 | 3,31 | 5,14 | 0 | 1 | 0 | 0 | 0 |
| SRY | AAAAATAAA | 1 | 581 | 590 | -419 | -410 | 4,1 | 0,01 | 13,61 | >0.1 | 3,35 | 5,15 | 0 | 1 | 0 | 0 | 0 |
| SRY | GAAAACCAT | 1 | 619 | 628 | -381 | -372 | 4,41 | 0,01 | 15,85 | >0.1 | 3,61 | 5,26 | 0 | 1,03 | 0 | 1,55 | 0 |
| SRY | CCATACAAA | 1 | 624 | 633 | -376 | -367 | 5,69 | 0,01 | 17,09 | >0.1 | 3,65 | 5,27 | 0 | 1,04 | 0 | 1,44 | 0 |
| SRY | TTAAAGAAT | 1 | 794 | 803 | -206 | -197 | 3,9 | 0,01 | 22,23 | >0.1 | 5,62 | 5,93 | 0 | 1,46 | 0 | 5,32 | 0 |
| SRY | TTAAAAAAT | 1 | 826 | 835 | -174 | -165 | 3,4 | 0,01 | 20,5 | >0.1 | 5,62 | 6,12 | 0 | 1,65 | 0 | 3,7 | 0 |
| SRY | TTGGACAAG | 1 | 1133 | 1142 | 133 | 142 | 4,54 | 0,01 | 24,64 | >0.1 | 5,62 | 6,43 | 0 | 1,84 | 3,22 | 2,98 | 0 |
| SRY | TCTATCAAA | 1 | 1167 | 1176 | 167 | 176 | 4,42 | 0,01 | 21,5 | >0.1 | 4,54 | 6,17 | 0 | 1,69 | 1,79 | 2,88 | 0 |
| SRY | GAACACAGT | -1 | 902 | 911 | -98 | -89 | 5,04 | 0,01 | 20,54 | >0.1 | 5,62 | 6,71 | 0 | 1,89 | 0 | 1,28 | 0 |
| SRY | GTAAACAAT | -1 | 895 | 904 | -105 | -96 | 11,74 | 1E-05 | 28,76 | >0.1 | 5,62 | 6,65 | 0 | 1,89 | 0 | 2,86 | 0 |
| SRY | TGTAACAGA | -1 | 845 | 854 | -155 | -146 | 3,78 | 0,01 | 20,53 | >0.1 | 5,62 | 6,26 | 0 | 1,81 | 0 | 3,07 | 0 |
| SRY | TAGAATAAT | -1 | 833 | 842 | -167 | -158 | 4,42 | 0,01 | 21,47 | >0.1 | 5,62 | 6,17 | 0 | 1,7 | 0 | 3,55 | 0 |
| SRY | TGAATCAAA | -1 | 815 | 824 | -185 | -176 | 5,38 | 0,01 | 22,06 | >0.1 | 5,62 | 6,05 | 0 | 1,58 | 0 | 3,43 | 0 |
| SRY | CAAATCAAA | -1 | 749 | 758 | -251 | -242 | 5,69 | 0,01 | 24,22 | >0.1 | 5,62 | 5,71 | 0 | 1,28 | 0 | 5,92 | 0 |
| SRY | TTTTACAGT | -1 | 678 | 687 | -322 | -313 | 4,23 | 0,01 | 18,47 | >0.1 | 4,26 | 5,44 | 0 | 1,11 | 0 | 3,43 | 0 |
| SRY | GAAGATAAT | -1 | 640 | 649 | -360 | -351 | 4,97 | 0,01 | 16,79 | >0.1 | 3,79 | 5,32 | 0 | 1,05 | 0 | 1,66 | 0 |
| SRY | TCAGACAAT | -1 | 612 | 621 | -388 | -379 | 7,37 | 0,01 | 18,69 | >0.1 | 3,56 | 5,24 | 0 | 1,02 | 0 | 1,5 | 0 |
| PAX9 | AGAAATGGCTGGGCACA | 1 | 170 | 187 | -830 | -813 | -17,77 | 0,01 | -5,05 | >0.1 | 3,16 | 4,44 | 0 | 1,88 | 0 | 3,24 | 0 |
| PAX9 | GCTCACGCCTGTAATCC | 1 | 190 | 207 | -810 | -793 | -19,4 | 0,01 | -5,9 | >0.1 | 3,07 | 4,46 | 0 | 1,71 | 0 | 4,26 | 0 |
| PAX9 | AGACCGTCCTGACTAAC | 1 | 255 | 272 | -745 | -728 | -25,82 | 0,01 | -12,58 | >0.1 | 2,81 | 4,55 | 0 | 1,35 | 0 | 4,53 | 0 |
| PAX9 | CTACTCGGGAGGCTGAG | 1 | 344 | 361 | -656 | -639 | -22,28 | 0,01 | -7,41 | >0.1 | 2,55 | 4,69 | 0 | 1,12 | 0 | 6,52 | 0 |
| PAX9 | GCTCCAGCCTGGGCGAC | 1 | 422 | 439 | -578 | -561 | -17,86 | 0,01 | -3,65 | >0.1 | 2,7 | 4,82 | 0 | 1,02 | 0 | 5,67 | 0 |
| PAX9 | CAGCCTGGGCGACAGAC | 1 | 426 | 443 | -574 | -557 | -28,24 | 0,01 | -14,01 | >0.1 | 2,71 | 4,83 | 0 | 1,02 | 0 | 5,67 | 0 |
| PAX9 | GGTCACTGTAAAATTAT | 1 | 674 | 691 | -326 | -309 | -20,68 | 0,01 | -6,42 | >0.1 | 4,26 | 5,46 | 0 | 1,12 | 0 | 3,43 | 0 |
| PAX9 | CTTTCCTGCTCAGCGCC | 1 | 933 | 950 | -67 | -50 | -16,66 | 0,01 | -1,14 | >0.1 | 5,52 | 7,22 | 0 | 1,88 | 0 | 0,9 | 0 |
| PAX9 | GATCTTGGCTCACAGGG | 1 | 1028 | 1045 | 28 | 45 | -22,77 | 0,01 | -5,75 | >0.1 | 5,52 | 8,34 | 0 | 1,86 | 0 | 1,31 | 0 |
| PAX9 | GGACAAGTTTAACCACG | 1 | 1135 | 1152 | 135 | 152 | -28,16 | 0,01 | -8,03 | >0.1 | 5,52 | 6,41 | 0 | 1,84 | 3,22 | 3,13 | 0 |
| PAX9 | TGTCTTGGCCTGTTCCT | -1 | 1113 | 1130 | 113 | 130 | -27,1 | 0,01 | -7,29 | >0.1 | 5,52 | 6,62 | 0 | 1,85 | 3,22 | 2,6 | 0 |
| PAX9 | TGTTCCTCAATGGTGGA | -1 | 1103 | 1120 | 103 | 120 | -26,69 | 0,01 | -6,9 | >0.1 | 5,52 | 6,72 | 0 | 1,85 | 3,22 | 2,48 | 0 |
| PAX9 | CTCAATGGTGGACTGAG | -1 | 1098 | 1115 | 98 | 115 | -25,71 | 0,01 | -5,96 | >0.1 | 5,52 | 6,78 | 0 | 1,85 | 3,22 | 2,38 | 0 |
| PAX9 | CGTCCCCTGTGAGCCAA | -1 | 1032 | 1049 | 32 | 49 | -25,81 | 0,01 | -8,98 | >0.1 | 5,52 | 8,15 | 0 | 1,86 | 0 | 1,31 | 0 |
| PAX9 | GTTCATCCTGGAGAGGA | -1 | 860 | 877 | -140 | -123 | -25,77 | 0,01 | -9,26 | >0.1 | 5,52 | 6,42 | 0 | 1,9 | 0 | 2,67 | 0 |
| PAX9 | CGTTATTCTTTAATCCT | -1 | 790 | 807 | -210 | -193 | -18,24 | 0,01 | 0,03 | >0.1 | 5,52 | 5,95 | 0 | 1,48 | 0 | 5,32 | 0 |
| PAX9 | CTTTATGGCCAAATCAA | -1 | 750 | 767 | -250 | -233 | -23,27 | 0,01 | -4,36 | >0.1 | 5,52 | 5,76 | 0 | 1,31 | 0 | 6,32 | 0 |
| PAX9 | CGGTACTTTTGGTTAAT | -1 | 730 | 747 | -270 | -253 | -26,94 | 0,01 | -8,59 | >0.1 | 5,52 | 5,67 | 0 | 1,25 | 0 | 5,92 | 0 |
| PAX9 | AGTCTCGGTCTGTCGCC | -1 | 433 | 450 | -567 | -550 | -13,63 | 0,001 | 0,88 | >0.1 | 2,73 | 4,84 | 0 | 1,01 | 0 | 5,92 | 0 |

| | | | | | | | | | | | | | | | | | |
|---|---|---|---|---|---|---|---|---|---|---|---|---|---|---|---|---|---|
| PAX9 | GATCTCGGCTCACTGCA | -1 | 395 | 412 | -605 | -588 | -5,83 | 0,001 | 8,52 | >0.1 | 2,62 | 4,77 | 0 | 1,05 | 0 | 5,91 | 0 |
| PAX9 | CCTCCCGAGTAGCTGGG | -1 | 339 | 356 | -661 | -644 | -28,01 | 0,01 | -13,25 | >0.1 | 2,56 | 4,68 | 0 | 1,12 | 0 | 6,4 | 0 |
| PAX9 | CACCACGCCCAGCTAAG | -1 | 307 | 324 | -693 | -676 | -22,85 | 0,01 | -8,25 | >0.1 | 2,65 | 4,63 | 0 | 1,19 | 0 | 6,13 | 0 |
| PAX9 | TGATCTGCCCAACTCGG | -1 | 222 | 239 | -778 | -761 | -26,71 | 0,01 | -12,52 | >0.1 | 2,93 | 4,5 | 0 | 1,5 | 0 | 5,26 | 0 |
| PAX9 | TTACAGGCGTGAGCCAC | -1 | 187 | 204 | -813 | -796 | -24,69 | 0,01 | -11,29 | >0.1 | 3,08 | 4,46 | 0 | 1,73 | 0 | 4,13 | 0 |
| PAX9 | CTCCACAGGTCAACTCT | -1 | 103 | 120 | -897 | -880 | -23,34 | 0,01 | -12,78 | >0.1 | 3,56 | 4,36 | 0 | 1,89 | 0 | 0,75 | 0 |
| PAX9 | TTCTGTGGCTGACTTCT | -1 | 32 | 49 | -968 | -951 | -22,61 | 0,01 | -12,08 | >0.1 | 4,37 | 4,29 | 0 | 1,87 | 0 | 0 | 0 |
| PAX9 | TGACTTCTTTGAGTGCC | -1 | 23 | 40 | -977 | -960 | -26,52 | 0,01 | -14,85 | >0.1 | 5,52 | 4,28 | 0 | 1,87 | 0 | 0 | 0 |
| PAX9 | CCTTCTCCTTCACTTAC | -1 | 5 | 22 | -995 | -978 | -21,85 | 0,01 | -8,41 | >0.1 | 5,52 | 4,27 | 0 | 1,86 | 1,79 | 0 | 0 |
| PKNOX1 | TCAAAGAAGTCA | 1 | 28 | 40 | -972 | -960 | -11,49 | 0,01 | -0,83 | >0.1 | 4,51 | 4,28 | 0 | 1,87 | 0 | 0 | 0 |
| PKNOX1 | TCACGCCTGTAA | 1 | 192 | 204 | -808 | -796 | -15,2 | 0,01 | -1,72 | >0.1 | 3,07 | 4,46 | 0 | 1,69 | 0 | 4,26 | 0 |
| PKNOX1 | GGGCAGATCACA | 1 | 229 | 241 | -771 | -759 | -11,89 | 0,01 | 2 | >0.1 | 2,92 | 4,51 | 0 | 1,46 | 0 | 5,01 | 0 |
| PKNOX1 | GGGCACCTGTAG | 1 | 326 | 338 | -674 | -662 | -18,94 | 0,01 | -4,98 | >0.1 | 2,6 | 4,65 | 0 | 1,15 | 0 | 5,56 | 0 |
| PKNOX1 | TAACAGCTTTCT | 1 | 710 | 722 | -290 | -278 | -5,17 | 0,001 | 12,15 | >0.1 | 5,58 | 5,57 | 0 | 1,18 | 0 | 4,99 | 0 |
| PKNOX1 | TGACAGGAGAGG | 1 | 767 | 779 | -233 | -221 | -9,5 | 0,01 | 9,16 | >0.1 | 5,58 | 5,81 | 0 | 1,35 | 0 | 5,92 | 0 |
| PKNOX1 | TAAAATCTGTTA | 1 | 840 | 852 | -160 | -148 | -14,2 | 0,01 | 0,83 | >0.1 | 5,58 | 6,24 | 0 | 1,79 | 0 | 1,41 | 0 |
| PKNOX1 | ACATATCTGTCC | 1 | 851 | 863 | -149 | -137 | -16,65 | 0,01 | -0,19 | >0.1 | 5,58 | 6,32 | 0 | 1,89 | 0 | 2,67 | 0 |
| PKNOX1 | TCACAGGGGACG | 1 | 1037 | 1049 | 37 | 49 | -7,2 | 0,001 | 9,53 | >0.1 | 5,58 | 7,95 | 0 | 1,86 | 0 | 1,34 | 0 |
| PKNOX1 | GGGACGATGTCA | 1 | 1043 | 1055 | 43 | 55 | -17,67 | 0,01 | -1,85 | >0.1 | 5,58 | 7,76 | 0 | 1,86 | 0 | 0,62 | 0 |
| PKNOX1 | GGACAAGTTTAA | 1 | 1135 | 1147 | 135 | 147 | -17,59 | 0,01 | 2,56 | >0.1 | 5,58 | 6,41 | 0 | 1,84 | 3,22 | 3,09 | 0 |
| PKNOX1 | TGACATCGTCCC | -1 | 1043 | 1055 | 43 | 55 | -18,57 | 0,01 | -2,75 | >0.1 | 5,58 | 7,76 | 0 | 1,86 | 0 | 0,62 | 0 |
| PKNOX1 | GGACAGATATGT | -1 | 851 | 863 | -149 | -137 | -8,04 | 0,01 | 8,42 | >0.1 | 5,58 | 6,32 | 0 | 1,89 | 0 | 2,67 | 0 |
| PKNOX1 | AGATATGTAACA | -1 | 847 | 859 | -153 | -141 | -14,01 | 0,01 | 2,67 | >0.1 | 5,58 | 6,29 | 0 | 1,85 | 0 | 2,95 | 0 |
| PKNOX1 | TAACAGATTTTA | -1 | 840 | 852 | -160 | -148 | -14,22 | 0,01 | 0,81 | >0.1 | 5,58 | 6,24 | 0 | 1,79 | 0 | 1,41 | 0 |
| PKNOX1 | AGAAAGCTGTTA | -1 | 710 | 722 | -290 | -278 | -6,83 | 0,001 | 10,49 | >0.1 | 5,58 | 5,57 | 0 | 1,18 | 0 | 4,99 | 0 |
| PKNOX1 | CTACAGGTGCCC | -1 | 326 | 338 | -674 | -662 | -8,87 | 0,01 | 5,09 | >0.1 | 2,6 | 4,65 | 0 | 1,15 | 0 | 5,56 | 0 |
| PKNOX1 | TTACAGGCGTGA | -1 | 192 | 204 | -808 | -796 | -4,82 | 0,001 | 8,66 | >0.1 | 3,07 | 4,46 | 0 | 1,69 | 0 | 4,26 | 0 |
| PKNOX1 | TGACTACTCTCC | -1 | 121 | 133 | -879 | -867 | -17,24 | 0,01 | -5,87 | >0.1 | 3,45 | 4,37 | 0 | 1,89 | 0 | 1,65 | 0 |
| PKNOX1 | TGACTTCTTTGA | -1 | 28 | 40 | -972 | -960 | -13,33 | 0,01 | -2,67 | >0.1 | 4,51 | 4,28 | 0 | 1,87 | 0 | 0 | 0 |
| PKNOX2 | TCAAAGAAGTCA | 1 | 28 | 40 | -972 | -960 | -14,78 | 0,01 | -4,12 | >0.1 | 4,51 | 4,28 | 0 | 1,87 | 0 | 0 | 0 |
| PKNOX2 | TGACCTGTGGAG | 1 | 108 | 120 | -892 | -880 | -12,57 | 0,01 | -2,03 | >0.1 | 3,54 | 4,36 | 0 | 1,89 | 0 | 0,75 | 0 |
| PKNOX2 | TCACGCCTGTAA | 1 | 192 | 204 | -808 | -796 | -15,08 | 0,01 | -1,6 | >0.1 | 3,07 | 4,46 | 0 | 1,69 | 0 | 4,26 | 0 |
| PKNOX2 | GGGCAGATCACA | 1 | 229 | 241 | -771 | -759 | -10,75 | 0,01 | 3,14 | >0.1 | 2,92 | 4,51 | 0 | 1,46 | 0 | 5,01 | 0 |
| PKNOX2 | GGGCACCTGTAG | 1 | 326 | 338 | -674 | -662 | -15,59 | 0,01 | -1,63 | >0.1 | 2,6 | 4,65 | 0 | 1,15 | 0 | 5,56 | 0 |
| PKNOX2 | AGACAAGGGCAA | 1 | 504 | 516 | -496 | -484 | -13,85 | 0,01 | 0,77 | >0.1 | 2,98 | 4,98 | 0 | 0,98 | 0 | 5,68 | 0 |
| PKNOX2 | TAACAGCTTTCT | 1 | 710 | 722 | -290 | -278 | -8,79 | 0,01 | 8,53 | >0.1 | 5,58 | 5,57 | 0 | 1,18 | 0 | 4,99 | 0 |
| PKNOX2 | TGACAGGAGAGG | 1 | 767 | 779 | -233 | -221 | -9,04 | 0,01 | 9,62 | >0.1 | 5,58 | 5,81 | 0 | 1,35 | 0 | 5,92 | 0 |
| PKNOX2 | TAAAATCTGTTA | 1 | 840 | 852 | -160 | -148 | -14,72 | 0,01 | 0,31 | >0.1 | 5,58 | 6,24 | 0 | 1,79 | 0 | 1,41 | 0 |
| PKNOX2 | ACATATCTGTCC | 1 | 851 | 863 | -149 | -137 | -11,66 | 0,01 | 4,8 | >0.1 | 5,58 | 6,32 | 0 | 1,89 | 0 | 2,67 | 0 |
| PKNOX2 | TCACAGGGGACG | 1 | 1037 | 1049 | 37 | 49 | -8,67 | 0,01 | 8,06 | >0.1 | 5,58 | 7,95 | 0 | 1,86 | 0 | 1,34 | 0 |
| PKNOX2 | GGACAAGTTTAA | 1 | 1135 | 1147 | 135 | 147 | -11,72 | 0,01 | 8,43 | >0.1 | 5,58 | 6,41 | 0 | 1,84 | 3,22 | 3,09 | 0 |
| PKNOX2 | GGACAGATATGT | -1 | 851 | 863 | -149 | -137 | -11,76 | 0,01 | 4,7 | >0.1 | 5,58 | 6,32 | 0 | 1,89 | 0 | 2,67 | 0 |
| PKNOX2 | ACAGATATGTAA | -1 | 849 | 861 | -151 | -139 | -18 | 0,01 | -1,49 | >0.1 | 5,58 | 6,3 | 0 | 1,87 | 0 | 2,75 | 0 |
| PKNOX2 | AGATATGTAACA | -1 | 847 | 859 | -153 | -141 | -9,27 | 0,01 | 7,41 | >0.1 | 5,58 | 6,29 | 0 | 1,85 | 0 | 2,95 | 0 |
| PKNOX2 | TAACAGATTTTA | -1 | 840 | 852 | -160 | -148 | -14,35 | 0,01 | 0,68 | >0.1 | 5,58 | 6,24 | 0 | 1,79 | 0 | 1,41 | 0 |
| PKNOX2 | AGAAAGCTGTTA | -1 | 710 | 722 | -290 | -278 | -5,09 | 0,001 | 12,23 | >0.1 | 5,58 | 5,57 | 0 | 1,18 | 0 | 4,99 | 0 |
| PKNOX2 | CTACAGGTGCCC | -1 | 326 | 338 | -674 | -662 | -16,24 | 0,01 | -2,28 | >0.1 | 2,6 | 4,65 | 0 | 1,15 | 0 | 5,56 | 0 |
| PKNOX2 | TGTGATCTGCCC | -1 | 229 | 241 | -771 | -759 | -16,21 | 0,01 | -2,32 | >0.1 | 2,92 | 4,51 | 0 | 1,46 | 0 | 5,01 | 0 |
| PKNOX2 | TTACAGGCGTGA | -1 | 192 | 204 | -808 | -796 | -5,25 | 0,001 | 8,23 | >0.1 | 3,07 | 4,46 | 0 | 1,69 | 0 | 4,26 | 0 |
| PKNOX2 | TGACTACTCTCC | -1 | 121 | 133 | -879 | -867 | -13,59 | 0,01 | -2,22 | >0.1 | 3,45 | 4,37 | 0 | 1,89 | 0 | 1,65 | 0 |
| PKNOX2 | TGACTTCTTTGA | -1 | 28 | 40 | -972 | -960 | -10,85 | 0,01 | -0,19 | >0.1 | 4,51 | 4,28 | 0 | 1,87 | 0 | 0 | 0 |
| ESR2 | AGGAGAAGGAGGCAC | 1 | 13 | 28 | -987 | -972 | 1,17 | 0,01 | 12,85 | >0.1 | 5,54 | 4,27 | 0 | 1,87 | 0 | 0 | 0 |
| ESR2 | AAGTCAGCCACAGAA | 1 | 34 | 49 | -966 | -951 | 1,44 | 0,01 | 11,95 | >0.1 | 4,35 | 4,29 | 0 | 1,87 | 0 | 0 | 0 |
| ESR2 | TGGGCACAGTGGCTC | 1 | 179 | 194 | -821 | -806 | 9,54 | 0,001 | 22,76 | >0.1 | 3,12 | 4,45 | 0 | 1,8 | 0 | 3,86 | 0 |
| ESR2 | AGATCACAAGGTCAG | 1 | 233 | 248 | -767 | -752 | 2,97 | 0,01 | 16,57 | >0.1 | 2,9 | 4,52 | 0 | 1,44 | 0 | 4,74 | 0 |
| ESR2 | AGGAGATAGAGACCG | 1 | 246 | 261 | -754 | -739 | 3,14 | 0,01 | 16,99 | >0.1 | 2,85 | 4,53 | 0 | 1,38 | 0 | 5,09 | 0 |
| ESR2 | ATGTGAGATAACCTA | 1 | 589 | 604 | -411 | -396 | 1,22 | 0,01 | 10,84 | >0.1 | 3,42 | 5,19 | 0 | 1,02 | 0 | 0 | 0 |
| ESR2 | TGGTCACTGTAAAAT | 1 | 673 | 688 | -327 | -312 | 4,29 | 0,01 | 18,5 | >0.1 | 4,22 | 5,45 | 0 | 1,11 | 0 | 3,43 | 0 |
| ESR2 | ATGTGATCTTGGCTC | 1 | 1024 | 1039 | 24 | 39 | 6,46 | 0,001 | 23,73 | >0.1 | 5,54 | 8,59 | 0 | 1,86 | 0 | 1,28 | 0 |

| | | | | | | | | | | | | | | | | | |
|---|---|---|---|---|---|---|---|---|---|---|---|---|---|---|---|---|---|
| ESR2 | AGGGGACGATGTCAA | 1 | 1041 | 1056 | 41 | 56 | 7,32 | 0,001 | 23,17 | >0.1 | 5,54 | 7,82 | 0 | 1,86 | 0 | 0,62 | 0 |
| ESR2 | ATGTCAAGCTCTTCC | 1 | 1049 | 1064 | 49 | 64 | 4,73 | 0,01 | 21,19 | >0.1 | 5,54 | 7,6 | 0 | 1,86 | 0 | 1,46 | 0 |
| ESR2 | AGGCCAAGACATTTT | 1 | 1119 | 1134 | 119 | 134 | -0,84 | 0,01 | 19 | >0.1 | 5,54 | 6,56 | 0 | 1,85 | 3,22 | 2,67 | 0 |
| ESR2 | AGTTCACTTGCTTCT | 1 | 1175 | 1190 | 175 | 190 | -0,5 | 0,01 | 17,85 | >0.1 | 5,54 | 6,12 | 0 | 1,62 | 1,79 | 3,26 | 0 |
| ESR2 | GAGCCAGGAAGAGCT | -1 | 1055 | 1070 | 55 | 70 | 3,12 | 0,01 | 20,25 | >0.1 | 5,54 | 7,46 | 0 | 1,86 | 0 | 2,27 | 0 |
| ESR2 | TTGACATCGTCCCCT | -1 | 1041 | 1056 | 41 | 56 | -1,13 | 0,01 | 14,72 | >0.1 | 5,54 | 7,82 | 0 | 1,86 | 0 | 0,62 | 0 |
| ESR2 | GAGCCAAGATCACAT | -1 | 1024 | 1039 | 24 | 39 | -0,95 | 0,01 | 16,32 | >0.1 | 5,54 | 8,59 | 0 | 1,86 | 0 | 1,28 | 0 |
| ESR2 | AGATCACATCCACTG | -1 | 1018 | 1033 | 18 | 33 | 1,04 | 0,01 | 18,52 | >0.1 | 5,54 | 8,76 | 0 | 1,86 | 0 | 1,31 | 0 |
| ESR2 | ATTTTACAGTGACCA | -1 | 673 | 688 | -327 | -312 | -0,94 | 0,01 | 13,27 | >0.1 | 4,22 | 5,45 | 0 | 1,11 | 0 | 3,43 | 0 |
| ESR2 | AGATCATATTCCCAG | -1 | 560 | 575 | -440 | -425 | -0,32 | 0,01 | 9,04 | >0.1 | 3,25 | 5,12 | 0 | 1 | 0 | 0 | 0 |
| ESR2 | GGTTCACGCCATTCT | -1 | 366 | 381 | -634 | -619 | 2,76 | 0,01 | 17,82 | >0.1 | 2,54 | 4,72 | 0 | 1,08 | 0 | 6,73 | 0 |
| ESR2 | GTTTCACCGTGTTAG | -1 | 267 | 282 | -733 | -718 | -0,67 | 0,01 | 13,06 | >0.1 | 2,78 | 4,56 | 0 | 1,31 | 0 | 5,09 | 0 |
| ESR2 | TAGTCAGGACGGTCT | -1 | 255 | 270 | -745 | -730 | 3,8 | 0,01 | 17,14 | >0.1 | 2,82 | 4,55 | 0 | 1,35 | 0 | 4,63 | 0 |
| ESR2 | CGGTCTCTATCTCCT | -1 | 246 | 261 | -754 | -739 | 1,81 | 0,01 | 15,66 | >0.1 | 2,85 | 4,53 | 0 | 1,38 | 0 | 5,09 | 0 |
| ESR2 | GAGCCACTGTGCCCA | -1 | 179 | 194 | -821 | -806 | 6,9 | 0,001 | 20,12 | >0.1 | 3,12 | 4,45 | 0 | 1,8 | 0 | 3,86 | 0 |
| ESR2 | AGGTCAACTCTCCTG | -1 | 99 | 114 | -901 | -886 | 2,95 | 0,01 | 13,58 | >0.1 | 3,6 | 4,35 | 0 | 1,89 | 0 | 0,79 | 0 |
| ESR2 | GTGCCTCCTTCTCCT | -1 | 13 | 28 | -987 | -972 | -0,66 | 0,01 | 11,02 | >0.1 | 5,54 | 4,27 | 0 | 1,87 | 0 | 0 | 0 |
| HOXC10 | GTCATAATTT | 1 | 129 | 139 | -871 | -861 | 7,67 | 0,001 | 18,96 | >0.1 | 3,41 | 4,38 | 0 | 1,89 | 0 | 1,61 | 0 |
| HOXC10 | ATTTTAAAAA | 1 | 135 | 145 | -865 | -855 | -0,05 | 0,01 | 11,18 | >0.1 | 3,37 | 4,39 | 0 | 1,89 | 0 | 1,57 | 0 |
| HOXC10 | GAAATTAAAA | 1 | 154 | 164 | -846 | -836 | 1,38 | 0,01 | 12,6 | >0.1 | 3,26 | 4,41 | 0 | 1,9 | 0 | 1,65 | 0 |
| HOXC10 | CTATTAAAAA | 1 | 289 | 299 | -711 | -701 | 0,97 | 0,01 | 14,59 | >0.1 | 2,71 | 4,59 | 0 | 1,24 | 0 | 5,09 | 0 |
| HOXC10 | CTCAAAAAAA | 1 | 454 | 464 | -546 | -536 | 4,7 | 0,01 | 18,88 | >0.1 | 2,79 | 4,87 | 0 | 1 | 0 | 5,52 | 0 |
| HOXC10 | TCCATAAAGA | 1 | 497 | 507 | -503 | -493 | -0,32 | 0,01 | 15 | >0.1 | 2,95 | 4,96 | 0 | 0,98 | 0 | 6,44 | 0 |
| HOXC10 | GAAATAAAAA | 1 | 576 | 586 | -424 | -414 | 2,43 | 0,01 | 11,9 | >0.1 | 3,32 | 5,14 | 0 | 1 | 0 | 0 | 0 |
| HOXC10 | CCTATTAATG | 1 | 600 | 610 | -400 | -390 | 0,22 | 0,01 | 11,28 | >0.1 | 3,47 | 5,21 | 0 | 1,02 | 0 | 1,36 | 0 |
| HOXC10 | ACTGTAAAAT | 1 | 678 | 688 | -322 | -312 | 4,42 | 0,01 | 18,53 | >0.1 | 4,27 | 5,45 | 0 | 1,11 | 0 | 3,28 | 0 |
| HOXC10 | ATTATAACAT | 1 | 686 | 696 | -314 | -304 | 1,96 | 0,01 | 15,93 | >0.1 | 4,42 | 5,47 | 0 | 1,13 | 0 | 2,95 | 0 |
| HOXC10 | ATCTTTAACA | 1 | 705 | 715 | -295 | -285 | 1,75 | 0,01 | 19,05 | >0.1 | 5,61 | 5,54 | 0 | 1,17 | 0 | 4,99 | 0 |
| HOXC10 | GCCATAAAGT | 1 | 758 | 768 | -242 | -232 | 8,04 | 0,001 | 26,82 | >0.1 | 5,61 | 5,76 | 0 | 1,31 | 0 | 6,1 | 0 |
| HOXC10 | ATTGTTTACT | 1 | 895 | 905 | -105 | -95 | -0,43 | 0,01 | 16,68 | >0.1 | 5,61 | 6,66 | 0 | 1,89 | 0 | 2,95 | 0 |
| HOXC10 | GTCATTCAGT | 1 | 1012 | 1022 | 12 | 22 | 0,73 | 0,01 | 18,54 | >0.1 | 5,61 | 8,92 | 0 | 1,86 | 0 | 1,42 | 0 |
| HOXC10 | GTGGTTAAAC | -1 | 1141 | 1151 | 141 | 151 | 4,86 | 0,01 | 25,31 | >0.1 | 5,61 | 6,37 | 0 | 1,84 | 3,22 | 3,4 | 0 |
| HOXC10 | GCAGTTACAG | -1 | 1086 | 1096 | 86 | 96 | 0,35 | 0,01 | 18,75 | >0.1 | 5,61 | 6,93 | 0 | 1,85 | 1,79 | 2,21 | 0 |
| HOXC10 | ACCAAAAAAA | -1 | 923 | 933 | -77 | -67 | 1,51 | 0,01 | 17,06 | >0.1 | 5,61 | 6,97 | 0 | 1,88 | 0 | 1,09 | 0 |
| HOXC10 | ACAGTAAACA | -1 | 897 | 907 | -103 | -93 | 3,28 | 0,01 | 20,32 | >0.1 | 5,61 | 6,68 | 0 | 1,89 | 0 | 2,86 | 0 |
| HOXC10 | CCAATATAAA | -1 | 877 | 887 | -123 | -113 | -0,45 | 0,01 | 16,51 | >0.1 | 5,61 | 6,5 | 0 | 1,9 | 0 | 2,95 | 0 |
| HOXC10 | GCTGTTAAAG | -1 | 707 | 717 | -293 | -283 | 4,2 | 0,01 | 21,34 | >0.1 | 5,61 | 5,55 | 0 | 1,17 | 0 | 4,81 | 0 |
| HOXC10 | GTTATAATTT | -1 | 684 | 694 | -316 | -306 | 2,77 | 0,01 | 16,68 | >0.1 | 4,38 | 5,47 | 0 | 1,12 | 0 | 2,95 | 0 |
| HOXC10 | ATCATGAAGA | -1 | 644 | 654 | -356 | -346 | 1 | 0,01 | 14,93 | >0.1 | 3,83 | 5,34 | 0 | 1,06 | 0 | 3,7 | 0 |
| HOXC10 | TTCATTAATA | -1 | 602 | 612 | -398 | -388 | 2,36 | 0,01 | 13,49 | >0.1 | 3,49 | 5,21 | 0 | 1,02 | 0 | 1,41 | 0 |
| HOXC10 | ATCATATTCC | -1 | 563 | 573 | -437 | -427 | 0,44 | 0,01 | 9,79 | >0.1 | 3,25 | 5,11 | 0 | 0,99 | 0 | 0 | 0 |
| CUX2 | AAACTGATCA | 1 | 161 | 171 | -839 | -829 | 0,16 | 0,01 | 12,16 | >0.1 | 3,22 | 4,42 | 0 | 1,9 | 0 | 2,46 | 0 |
| CUX2 | AGACCGAGAC | 1 | 439 | 449 | -561 | -551 | -1,88 | 0,01 | 12,75 | >0.1 | 2,74 | 4,84 | 0 | 1 | 0 | 6,05 | 0 |
| CUX2 | AAACTGATCA | 1 | 476 | 486 | -524 | -514 | 0,16 | 0,01 | 14,13 | >0.1 | 2,87 | 4,91 | 0 | 0,99 | 0 | 5,2 | 0 |
| CUX2 | TAACCTATTA | 1 | 597 | 607 | -403 | -393 | 1,82 | 0,01 | 12,9 | >0.1 | 3,45 | 5,2 | 0 | 1,01 | 0 | 1,41 | 0 |
| CUX2 | CTATTAATGA | 1 | 601 | 611 | -399 | -389 | -0,27 | 0,01 | 10,85 | >0.1 | 3,48 | 5,21 | 0 | 1,02 | 0 | 1,41 | 0 |
| CUX2 | TAACCAAAAG | 1 | 732 | 742 | -268 | -258 | 0,53 | 0,01 | 18,93 | >0.1 | 5,61 | 5,65 | 0 | 1,23 | 0 | 5,92 | 0 |
| CUX2 | TATTTGATTC | 1 | 813 | 823 | -187 | -177 | -1,27 | 0,01 | 15,39 | >0.1 | 5,61 | 6,05 | 0 | 1,57 | 0 | 3,43 | 0 |
| CUX2 | TGTTCTATCA | 1 | 1164 | 1174 | 164 | 174 | -1,1 | 0,01 | 15,77 | >0.1 | 4,48 | 6,19 | 0 | 1,72 | 1,79 | 2,67 | 0 |
| CUX2 | CTATCAAAGT | 1 | 1168 | 1178 | 168 | 178 | -1,5 | 0,01 | 15,56 | >0.1 | 4,58 | 6,17 | 0 | 1,68 | 1,79 | 2,84 | 0 |
| CUX2 | TCCTCAATGG | -1 | 1107 | 1117 | 107 | 117 | -0,26 | 0,01 | 19,57 | >0.1 | 5,61 | 6,68 | 0 | 1,85 | 3,22 | 2,47 | 0 |
| CUX2 | AGACCAAAAA | -1 | 925 | 935 | -75 | -65 | -0,94 | 0,01 | 14,61 | >0.1 | 5,61 | 6,99 | 0 | 1,88 | 0 | 1,06 | 0 |
| CUX2 | TAAACAATCT | -1 | 893 | 903 | -107 | -97 | 4,13 | 0,001 | 21,13 | >0.1 | 5,61 | 6,64 | 0 | 1,89 | 0 | 2,86 | 0 |
| CUX2 | GAGCCAATAT | -1 | 880 | 890 | -120 | -110 | -0,8 | 0,01 | 16,3 | >0.1 | 5,61 | 6,53 | 0 | 1,9 | 0 | 3,07 | 0 |
| CUX2 | GAATCAAATA | -1 | 813 | 823 | -187 | -177 | 1,3 | 0,01 | 17,96 | >0.1 | 5,61 | 6,05 | 0 | 1,57 | 0 | 3,43 | 0 |
| CUX2 | AAATCAAAAC | -1 | 747 | 757 | -253 | -243 | 3,09 | 0,01 | 21,79 | >0.1 | 5,61 | 5,71 | 0 | 1,28 | 0 | 6,1 | 0 |
| CUX2 | AAACCGGTAC | -1 | 741 | 751 | -259 | -249 | -0,12 | 0,01 | 18,53 | >0.1 | 5,61 | 5,68 | 0 | 1,26 | 0 | 6,1 | 0 |
| CUX2 | TGGTTAATAT | -1 | 728 | 738 | -272 | -262 | -0,65 | 0,01 | 17,13 | >0.1 | 5,61 | 5,63 | 0 | 1,22 | 0 | 5,32 | 0 |
| CUX2 | TGACCAAAGA | -1 | 669 | 679 | -331 | -321 | -0,54 | 0,01 | 13,8 | >0.1 | 4,13 | 5,42 | 0 | 1,1 | 0 | 3,7 | 0 |

| | | | | | | | | | | | | | | | | | |
|---|---|---|---|---|---|---|---|---|---|---|---|---|---|---|---|---|---|
| CUX2 | TCATTAATAG | -1 | 601 | 611 | -399 | -389 | 2,39 | 0,01 | 13,51 | >0.1 | 3,48 | 5,21 | 0 | 1,02 | 0 | 1,41 | 0 |
| CUX2 | TGATCAGTTT | -1 | 476 | 486 | -524 | -514 | 2,16 | 0,01 | 16,13 | >0.1 | 2,87 | 4,91 | 0 | 0,99 | 0 | 5,2 | 0 |
| CUX2 | TGATCAGTTT | -1 | 161 | 171 | -839 | -829 | 2,16 | 0,01 | 14,16 | >0.1 | 3,22 | 4,42 | 0 | 1,9 | 0 | 2,46 | 0 |
| INSM1 | GAGAAGGAGGCA | 1 | 15 | 27 | -985 | -973 | 0,07 | 0,01 | 11,79 | >0.1 | 5,58 | 4,27 | 0 | 1,87 | 0 | 0 | 0 |
| INSM1 | TGGCTGGGCACA | 1 | 175 | 187 | -825 | -813 | 3,39 | 0,01 | 16,05 | >0.1 | 3,15 | 4,44 | 0 | 1,83 | 0 | 3,24 | 0 |
| INSM1 | GGTCAGGAGATA | 1 | 242 | 254 | -758 | -746 | 1,59 | 0,01 | 14,71 | >0.1 | 2,87 | 4,52 | 0 | 1,4 | 0 | 4,33 | 0 |
| INSM1 | TAGCTGGGCGTG | 1 | 309 | 321 | -691 | -679 | 1,69 | 0,01 | 16,28 | >0.1 | 2,65 | 4,62 | 0 | 1,18 | 0 | 6,13 | 0 |
| INSM1 | GGCGTGGTGGTG | 1 | 315 | 327 | -685 | -673 | 1,76 | 0,01 | 16,32 | >0.1 | 2,63 | 4,63 | 0 | 1,17 | 0 | 6,13 | 0 |
| INSM1 | TGGTGGTGGGCA | 1 | 319 | 331 | -681 | -669 | 2,18 | 0,01 | 16,28 | >0.1 | 2,62 | 4,64 | 0 | 1,16 | 0 | 5,68 | 0 |
| INSM1 | AGGCTGAGGCAG | 1 | 353 | 365 | -647 | -635 | 0,83 | 0,01 | 15,67 | >0.1 | 2,53 | 4,69 | 0 | 1,1 | 0 | 6,52 | 0 |
| INSM1 | AGCCTGGGCGAC | 1 | 427 | 439 | -573 | -561 | -0,43 | 0,01 | 13,92 | >0.1 | 2,71 | 4,82 | 0 | 1,01 | 0 | 5,8 | 0 |
| INSM1 | TGGAAGGGAAAA | 1 | 527 | 539 | -473 | -461 | -0,75 | 0,01 | 11,89 | >0.1 | 3,08 | 5,03 | 0 | 0,99 | 0 | 3,55 | 0 |
| INSM1 | TTTCTGGGAATA | 1 | 557 | 569 | -443 | -431 | -0,46 | 0,01 | 8,86 | >0.1 | 3,22 | 5,1 | 0 | 0,99 | 0 | 0 | 0 |
| INSM1 | TCACAGGGGACG | 1 | 1037 | 1049 | 37 | 49 | 0,89 | 0,01 | 17,62 | >0.1 | 5,58 | 7,95 | 0 | 1,86 | 0 | 1,34 | 0 |
| INSM1 | TGTCTTGGCCTG | -1 | 1118 | 1130 | 118 | 130 | 0,1 | 0,01 | 20,04 | >0.1 | 5,58 | 6,57 | 0 | 1,85 | 3,22 | 2,72 | 0 |
| INSM1 | AGCCAGGAAGAG | -1 | 1057 | 1069 | 57 | 69 | -1,43 | 0,01 | 15,64 | >0.1 | 5,58 | 7,42 | 0 | 1,86 | 0 | 2,21 | 0 |
| INSM1 | GGGTTGGGCGCT | -1 | 944 | 956 | -56 | -44 | 3,57 | 0,01 | 20,15 | >0.1 | 5,58 | 7,32 | 0 | 1,88 | 0 | 1,8 | 0 |
| INSM1 | GTTCTGTGGCTG | -1 | 38 | 50 | -962 | -950 | 2,62 | 0,01 | 13,08 | >0.1 | 4,3 | 4,29 | 0 | 1,87 | 0 | 0 | 0 |
| E2F6 | GTGAAGGAGAA | 1 | 9 | 20 | -991 | -980 | -8,03 | 0,01 | 5,49 | >0.1 | 5,6 | 4,26 | 0 | 1,86 | 1,79 | 0 | 0 |
| E2F6 | GAGAAGGAGGC | 1 | 15 | 26 | -985 | -974 | 4,14 | 0,001 | 15,87 | >0.1 | 5,6 | 4,27 | 0 | 1,87 | 0 | 0 | 0 |
| E2F6 | AGGAGGCACTC | 1 | 19 | 30 | -981 | -970 | -7,54 | 0,01 | 4,19 | >0.1 | 5,6 | 4,27 | 0 | 1,87 | 0 | 0 | 0 |
| E2F6 | CAGTGTGAGAA | 1 | 50 | 61 | -950 | -939 | -8,59 | 0,01 | 1,7 | >0.1 | 4,12 | 4,3 | 0 | 1,87 | 0 | 0 | 0 |
| E2F6 | GATCAGGAGAG | 1 | 96 | 107 | -904 | -893 | -8,4 | 0,01 | 2,24 | >0.1 | 3,64 | 4,35 | 0 | 1,88 | 0 | 0,77 | 0 |
| E2F6 | GGGAGGCCGAG | 1 | 216 | 227 | -784 | -773 | -4,3 | 0,01 | 10,21 | >0.1 | 2,97 | 4,49 | 0 | 1,53 | 0 | 5,52 | 0 |
| E2F6 | GGTCAGGAGAT | 1 | 242 | 253 | -758 | -747 | -7,58 | 0,01 | 5,54 | >0.1 | 2,87 | 4,52 | 0 | 1,4 | 0 | 4,33 | 0 |
| E2F6 | TGGTGGGCACC | 1 | 322 | 333 | -678 | -667 | -3,06 | 0,01 | 10,91 | >0.1 | 2,62 | 4,64 | 0 | 1,15 | 0 | 5,56 | 0 |
| E2F6 | ACTCGGGAGGC | 1 | 346 | 357 | -654 | -643 | 0,09 | 0,01 | 14,95 | >0.1 | 2,55 | 4,68 | 0 | 1,11 | 0 | 6,52 | 0 |
| E2F6 | GGGAGGCTGAG | 1 | 350 | 361 | -650 | -639 | -4,8 | 0,01 | 9,93 | >0.1 | 2,54 | 4,68 | 0 | 1,1 | 0 | 6,4 | 0 |
| E2F6 | CTGAGGCAGGA | 1 | 356 | 367 | -644 | -633 | -8,93 | 0,01 | 5,51 | >0.1 | 2,52 | 4,69 | 0 | 1,09 | 0 | 6,13 | 0 |
| E2F6 | AGGCAGGAGAA | 1 | 359 | 370 | -641 | -630 | 6,43 | 0,001 | 21,1 | >0.1 | 2,52 | 4,7 | 0 | 1,09 | 0 | 6,36 | 0 |
| E2F6 | TGGCGTGAACC | 1 | 370 | 381 | -630 | -619 | 0,34 | 0,01 | 14,94 | >0.1 | 2,54 | 4,72 | 0 | 1,07 | 0 | 6,27 | 0 |
| E2F6 | ACCTGGGAGGC | 1 | 378 | 389 | -622 | -611 | -3,81 | 0,01 | 11,08 | >0.1 | 2,57 | 4,73 | 0 | 1,06 | 0 | 6,53 | 0 |
| E2F6 | GGGAGGCAGAG | 1 | 382 | 393 | -618 | -607 | 8,96 | 0,001 | 23,24 | >0.1 | 2,58 | 4,74 | 0 | 1,06 | 0 | 5,91 | 0 |
| E2F6 | GGGCGACAGAC | 1 | 432 | 443 | -568 | -557 | -3,62 | 0,01 | 10,61 | >0.1 | 2,72 | 4,83 | 0 | 1,01 | 0 | 5,67 | 0 |
| E2F6 | GGAAGGGAAAA | 1 | 528 | 539 | -472 | -461 | 0,67 | 0,01 | 13,31 | >0.1 | 3,08 | 5,03 | 0 | 0,98 | 0 | 3,55 | 0 |
| E2F6 | TGACAGGAGAG | 1 | 767 | 778 | -233 | -222 | -7,91 | 0,01 | 10,76 | >0.1 | 5,6 | 5,81 | 0 | 1,35 | 0 | 5,92 | 0 |
| E2F6 | GAGAGGTAAGG | 1 | 773 | 784 | -227 | -216 | -0,48 | 0,01 | 18,43 | >0.1 | 5,6 | 5,83 | 0 | 1,37 | 0 | 6,1 | 0 |
| E2F6 | AAGAGAGAAAA | 1 | 972 | 983 | -28 | -17 | -3,23 | 0,01 | 13,95 | >0.1 | 5,6 | 8,01 | 0 | 1,87 | 0 | 1,7 | 0 |
| E2F6 | TCTAGGGAAAG | 1 | 1002 | 1013 | 2 | 13 | -2,94 | 0,01 | 15,11 | >0.1 | 5,6 | 9,12 | 0 | 1,87 | 0 | 1,46 | 0 |
| E2F6 | AAGCCGAAGAC | 1 | 1152 | 1163 | 152 | 163 | -8,04 | 0,01 | 8,63 | >0.1 | 4,5 | 6,28 | 0 | 1,84 | 1,79 | 2,26 | 0 |
| E2F6 | AAGAAGCAAGT | -1 | 1180 | 1191 | 180 | 191 | -4,95 | 0,01 | 13,41 | >0.1 | 5,6 | 6,09 | 0 | 1,58 | 1,79 | 3,29 | 0 |
| E2F6 | GAGCAGCAGTT | -1 | 1090 | 1101 | 90 | 101 | -4,93 | 0,01 | 13,46 | >0.1 | 5,6 | 6,88 | 0 | 1,85 | 1,79 | 2,27 | 0 |
| E2F6 | GAGAAGGAGCC | -1 | 1065 | 1076 | 65 | 76 | 2,82 | 0,01 | 21,6 | >0.1 | 5,6 | 7,26 | 0 | 1,85 | 1,79 | 2,27 | 0 |
| E2F6 | AGGAGCCAGGA | -1 | 1061 | 1072 | 61 | 72 | -6,12 | 0,01 | 12,62 | >0.1 | 5,6 | 7,33 | 0 | 1,86 | 1,79 | 2,16 | 0 |
| E2F6 | AGCCAGGAAGA | -1 | 1058 | 1069 | 58 | 69 | -6,47 | 0,01 | 10,59 | >0.1 | 5,6 | 7,39 | 0 | 1,86 | 0 | 2,21 | 0 |
| E2F6 | GCCAGGAAGAG | -1 | 1057 | 1068 | 57 | 68 | -8,06 | 0,01 | 9,08 | >0.1 | 5,6 | 7,42 | 0 | 1,86 | 0 | 2,27 | 0 |
| E2F6 | GAGCAGGAAAG | -1 | 933 | 944 | -67 | -56 | 6 | 0,001 | 21,52 | >0.1 | 5,6 | 7,12 | 0 | 1,88 | 0 | 0,93 | 0 |
| E2F6 | AGCAGGAAAGA | -1 | 932 | 943 | -68 | -57 | -6,99 | 0,01 | 8,52 | >0.1 | 5,6 | 7,11 | 0 | 1,88 | 0 | 0,93 | 0 |
| E2F6 | AAGAAGAAGAA | -1 | 908 | 919 | -92 | -81 | -5,77 | 0,01 | 9,72 | >0.1 | 5,6 | 6,8 | 0 | 1,89 | 0 | 1,21 | 0 |
| E2F6 | GAGAGGACAGA | -1 | 856 | 867 | -144 | -133 | -7,81 | 0,01 | 8,98 | >0.1 | 5,6 | 6,35 | 0 | 1,9 | 0 | 2,95 | 0 |
| E2F6 | AGGCTGGAGCA | -1 | 421 | 432 | -579 | -568 | 0,64 | 0,01 | 14,57 | >0.1 | 2,69 | 4,81 | 0 | 1,02 | 0 | 5,42 | 0 |
| E2F6 | AGGCGTGAGCC | -1 | 189 | 200 | -811 | -800 | 1 | 0,01 | 14,29 | >0.1 | 3,08 | 4,45 | 0 | 1,71 | 0 | 4,04 | 0 |
| DMRT3 | TCAGTCTCAAA | 1 | 449 | 460 | -551 | -540 | -4,06 | 0,01 | 9,83 | >0.1 | 2,78 | 4,86 | 0 | 1 | 0 | 5,26 | 0 |
| DMRT3 | CATGTATTTGG | 1 | 519 | 530 | -481 | -470 | -2,96 | 0,01 | 9,26 | >0.1 | 3,04 | 5,01 | 0 | 0,98 | 0 | 3,19 | 0 |
| DMRT3 | AATGTTGCCCA | 1 | 537 | 548 | -463 | -452 | -3,33 | 0,01 | 7,49 | >0.1 | 3,12 | 5,05 | 0 | 0,99 | 0 | 1,66 | 0 |
| DMRT3 | ACTGTAAAATT | 1 | 678 | 689 | -322 | -311 | -2,91 | 0,01 | 11,21 | >0.1 | 4,28 | 5,45 | 0 | 1,11 | 0 | 3,28 | 0 |
| DMRT3 | ATTATAACATT | 1 | 686 | 697 | -314 | -303 | -2,6 | 0,01 | 11,38 | >0.1 | 4,43 | 5,48 | 0 | 1,13 | 0 | 2,95 | 0 |
| DMRT3 | CGTGTATCTTT | 1 | 700 | 711 | -300 | -289 | 8,56 | 0,001 | 24,03 | >0.1 | 5,6 | 5,53 | 0 | 1,16 | 0 | 3,19 | 0 |
| DMRT3 | AAAGTACCGGT | 1 | 738 | 749 | -262 | -251 | -3,55 | 0,01 | 14,89 | >0.1 | 5,6 | 5,68 | 0 | 1,25 | 0 | 5,92 | 0 |

| | | | | | | | | | | | | | | | | | |
|---|---|---|---|---|---|---|---|---|---|---|---|---|---|---|---|---|---|
| DMRT3 | AAAGTGACAGG | 1 | 763 | 774 | -237 | -226 | -3,66 | 0,01 | 14,97 | >0.1 | 5,6 | 5,79 | 0 | 1,34 | 0 | 5,92 | 0 |
| DMRT3 | GCTGTAACTGC | 1 | 1085 | 1096 | 85 | 96 | 5,77 | 0,001 | 24,17 | >0.1 | 5,6 | 6,94 | 0 | 1,85 | 1,79 | 2,21 | 0 |
| DMRT3 | AGTTTAACCAC | 1 | 1140 | 1151 | 140 | 151 | -2,99 | 0,01 | 17,34 | >0.1 | 5,6 | 6,37 | 0 | 1,84 | 3,22 | 3,29 | 0 |
| DMRT3 | GTTCTATCAAA | 1 | 1165 | 1176 | 165 | 176 | -0,74 | 0,01 | 16,26 | >0.1 | 4,52 | 6,19 | 0 | 1,71 | 1,79 | 2,79 | 0 |
| DMRT3 | CAAGAAGCAAG | -1 | 1181 | 1192 | 181 | 192 | -3,53 | 0,01 | 14,95 | >0.1 | 5,6 | 6,08 | 0 | 1,58 | 1,79 | 3,43 | 0 |
| DMRT3 | ACAGCAACAAG | -1 | 1079 | 1090 | 79 | 90 | -3,79 | 0,01 | 14,69 | >0.1 | 5,6 | 7,03 | 0 | 1,85 | 1,79 | 2,21 | 0 |
| DMRT3 | TATGTAACAGA | -1 | 845 | 856 | -155 | -144 | 9,08 | 0,001 | 25,72 | >0.1 | 5,6 | 6,27 | 0 | 1,82 | 0 | 2,95 | 0 |
| DMRT3 | AGTGAATCAAA | -1 | 815 | 826 | -185 | -174 | 0 | 0,01 | 16,68 | >0.1 | 5,6 | 6,06 | 0 | 1,59 | 0 | 3,43 | 0 |
| DMRT3 | TTTGTATGGTT | -1 | 622 | 633 | -378 | -367 | -4,05 | 0,01 | 7,34 | >0.1 | 3,64 | 5,27 | 0 | 1,04 | 0 | 1,44 | 0 |
| DMRT3 | GCTGGAGCACA | -1 | 419 | 430 | -581 | -570 | -4,3 | 0,01 | 9,51 | >0.1 | 2,68 | 4,8 | 0 | 1,02 | 0 | 5,3 | 0 |
| DMRT3 | CGAGTAGCTGG | -1 | 340 | 351 | -660 | -649 | 2,95 | 0,01 | 17,7 | >0.1 | 2,57 | 4,67 | 0 | 1,12 | 0 | 6,4 | 0 |
| HLTF | GGCCATGGAA | 1 | 147 | 157 | -853 | -843 | 5,63 | 0,01 | 16,84 | >0.1 | 3,3 | 4,4 | 0 | 1,9 | 0 | 1,61 | 0 |
| HLTF | GCACTTTGGG | 1 | 209 | 219 | -791 | -781 | 7,18 | 0,01 | 20,93 | >0.1 | 3 | 4,48 | 0 | 1,57 | 0 | 4,7 | 0 |
| HLTF | AAACTTAGCT | 1 | 304 | 314 | -696 | -686 | 7,82 | 0,001 | 21,19 | >0.1 | 2,67 | 4,61 | 0 | 1,2 | 0 | 4,9 | 0 |
| HLTF | TAACATTTTC | 1 | 690 | 700 | -310 | -300 | 5,81 | 0,01 | 19,89 | >0.1 | 4,51 | 5,49 | 0 | 1,13 | 0 | 2,95 | 0 |
| HLTF | TTACATATCT | 1 | 849 | 859 | -151 | -141 | 7,01 | 0,01 | 23,62 | >0.1 | 5,61 | 6,29 | 0 | 1,85 | 0 | 2,86 | 0 |
| HLTF | CTCCTTCTCA | 1 | 1067 | 1077 | 67 | 77 | 6,38 | 0,01 | 25,14 | >0.1 | 5,61 | 7,22 | 0 | 1,85 | 1,79 | 2,27 | 0 |
| HLTF | AGCCTTGTTG | 1 | 1076 | 1086 | 76 | 86 | 5,58 | 0,01 | 24,19 | >0.1 | 5,61 | 7,07 | 0 | 1,85 | 1,79 | 2,27 | 0 |
| HLTF | CACCATTGAG | 1 | 1105 | 1115 | 105 | 115 | 7,16 | 0,01 | 27,06 | >0.1 | 5,61 | 6,7 | 0 | 1,85 | 3,22 | 2,52 | 0 |
| HLTF | AGACATTTTT | 1 | 1125 | 1135 | 125 | 135 | 4,47 | 0,01 | 24,42 | >0.1 | 5,61 | 6,5 | 0 | 1,84 | 3,22 | 2,77 | 0 |
| HLTF | GAACTTTGAT | -1 | 1170 | 1180 | 170 | 180 | 7,1 | 0,01 | 25,23 | >0.1 | 5,61 | 6,15 | 0 | 1,67 | 1,79 | 2,91 | 0 |
| HLTF | AAACTTGTCC | -1 | 1135 | 1145 | 135 | 145 | 7,75 | 0,01 | 27,86 | >0.1 | 5,61 | 6,41 | 0 | 1,84 | 3,22 | 3,02 | 0 |
| HLTF | TGACATCGTC | -1 | 1045 | 1055 | 45 | 55 | 2,79 | 0,01 | 18,61 | >0.1 | 5,61 | 7,7 | 0 | 1,86 | 0 | 0,65 | 0 |
| HLTF | AGCCTTTGAA | -1 | 959 | 969 | -41 | -31 | 6,07 | 0,01 | 22,85 | >0.1 | 5,61 | 7,6 | 0 | 1,88 | 0 | 1,7 | 0 |
| HLTF | GAACTTGGGT | -1 | 952 | 962 | -48 | -38 | 7,41 | 0,01 | 24,03 | >0.1 | 5,61 | 7,44 | 0 | 1,88 | 0 | 1,7 | 0 |
| HLTF | AACCTTACCT | -1 | 776 | 786 | -224 | -214 | 1,86 | 0,01 | 21,02 | >0.1 | 5,61 | 5,84 | 0 | 1,38 | 0 | 6,32 | 0 |
| HLTF | GTACTTTTGG | -1 | 735 | 745 | -265 | -255 | 7,35 | 0,01 | 25,78 | >0.1 | 5,61 | 5,66 | 0 | 1,24 | 0 | 5,92 | 0 |
| HLTF | TCACATTTAT | -1 | 585 | 595 | -415 | -405 | 6,29 | 0,01 | 15,84 | >0.1 | 3,38 | 5,17 | 0 | 1,01 | 0 | 0 | 0 |
| HLTF | CTACTTGGGC | -1 | 543 | 553 | -457 | -447 | 7,02 | 0,01 | 16,22 | >0.1 | 3,15 | 5,06 | 0 | 0,99 | 0 | 0 | 0 |
| HLTF | CAACATTTTC | -1 | 534 | 544 | -466 | -456 | 5,94 | 0,01 | 16,73 | >0.1 | 3,1 | 5,04 | 0 | 0,98 | 0 | 1,66 | 0 |
| HLTF | TGACTTTGCC | -1 | 511 | 521 | -489 | -479 | 5,83 | 0,01 | 17,67 | >0.1 | 3 | 4,99 | 0 | 0,98 | 0 | 2,86 | 0 |
| HLTF | GCCCTTGTCT | -1 | 504 | 514 | -496 | -486 | 8,27 | 0,001 | 23,06 | >0.1 | 2,98 | 4,97 | 0 | 0,98 | 0 | 5,86 | 0 |
| HLTF | GACCTTGTGA | -1 | 236 | 246 | -764 | -754 | 8,18 | 0,001 | 21,64 | >0.1 | 2,89 | 4,51 | 0 | 1,43 | 0 | 4,63 | 0 |
| HLTF | AGCCATTTCT | -1 | 170 | 180 | -830 | -820 | 6,59 | 0,01 | 19,5 | >0.1 | 3,18 | 4,43 | 0 | 1,88 | 0 | 3,43 | 0 |
| HLTF | TTCCATGGCC | -1 | 147 | 157 | -853 | -843 | 7,29 | 0,01 | 18,5 | >0.1 | 3,3 | 4,4 | 0 | 1,9 | 0 | 1,61 | 0 |
| HLTF | GGCCATTTTT | -1 | 141 | 151 | -859 | -849 | 5,39 | 0,01 | 16,62 | >0.1 | 3,33 | 4,39 | 0 | 1,89 | 0 | 1,61 | 0 |
| HLTF | TGCCTTTGCA | -1 | 86 | 96 | -914 | -904 | 6,17 | 0,01 | 16,11 | >0.1 | 3,73 | 4,33 | 0 | 1,88 | 0 | 0 | 0 |
| HLTF | TGACTTCTTT | -1 | 30 | 40 | -970 | -960 | 3,3 | 0,01 | 13,93 | >0.1 | 4,48 | 4,28 | 0 | 1,87 | 0 | 0 | 0 |
| HLTF | CTCCTTCTCC | -1 | 14 | 24 | -986 | -976 | 6,87 | 0,01 | 18,61 | >0.1 | 5,61 | 4,27 | 0 | 1,87 | 0 | 0 | 0 |
| DUXA | AGGAGGCACTCAA | 1 | 19 | 32 | -981 | -968 | 1,87 | 0,01 | 13,58 | >0.1 | 5,57 | 4,27 | 0 | 1,87 | 0 | 0 | 0 |
| DUXA | TCAAAGAAGTCAG | 1 | 28 | 41 | -972 | -959 | -0,19 | 0,01 | 10,46 | >0.1 | 4,5 | 4,28 | 0 | 1,87 | 0 | 0 | 0 |
| DUXA | TAAAACTGATCAG | 1 | 159 | 172 | -841 | -828 | 7,75 | 0,001 | 19,75 | >0.1 | 3,23 | 4,42 | 0 | 1,9 | 0 | 2,46 | 0 |
| DUXA | GTAATCCTAGCAC | 1 | 200 | 213 | -800 | -787 | 1,53 | 0,01 | 15,26 | >0.1 | 3,03 | 4,47 | 0 | 1,63 | 0 | 4,59 | 0 |
| DUXA | GAGACTCAGTCTC | 1 | 444 | 457 | -556 | -543 | 1,55 | 0,01 | 15,18 | >0.1 | 2,76 | 4,86 | 0 | 1 | 0 | 5,01 | 0 |
| DUXA | AAAAACTGATCAG | 1 | 474 | 487 | -526 | -513 | 7,41 | 0,001 | 21,27 | >0.1 | 2,87 | 4,92 | 0 | 0,99 | 0 | 5,09 | 0 |
| DUXA | GGAATATGATCTT | 1 | 563 | 576 | -437 | -424 | -0,77 | 0,01 | 8,6 | >0.1 | 3,26 | 5,12 | 0 | 1 | 0 | 0 | 0 |
| DUXA | TTAATGAAATTGT | 1 | 604 | 617 | -396 | -383 | 0,88 | 0,01 | 12,06 | >0.1 | 3,51 | 5,23 | 0 | 1,02 | 0 | 1,41 | 0 |
| DUXA | CCAACATTATCTT | 1 | 635 | 648 | -365 | -352 | -0,47 | 0,01 | 11,32 | >0.1 | 3,76 | 5,32 | 0 | 1,06 | 0 | 1,66 | 0 |
| DUXA | ATAACGTATTCTT | 1 | 801 | 814 | -199 | -186 | 2,75 | 0,01 | 19,26 | >0.1 | 5,57 | 5,99 | 0 | 1,52 | 0 | 3,43 | 0 |
| DUXA | TTGATTCACTTTA | 1 | 816 | 829 | -184 | -171 | 0,14 | 0,01 | 16,95 | >0.1 | 5,57 | 6,08 | 0 | 1,61 | 0 | 3,55 | 0 |
| DUXA | AAAAATTATTCTA | 1 | 829 | 842 | -171 | -158 | 0,2 | 0,01 | 16,93 | >0.1 | 5,57 | 6,17 | 0 | 1,71 | 0 | 3,28 | 0 |
| DUXA | CCAACCCAAGTTC | 1 | 949 | 962 | -51 | -38 | -0,94 | 0,01 | 15,69 | >0.1 | 5,57 | 7,44 | 0 | 1,88 | 0 | 1,74 | 0 |
| DUXA | AAAATCTCATGAG | 1 | 979 | 992 | -21 | -8 | 1,54 | 0,01 | 18,95 | >0.1 | 5,57 | 8,43 | 0 | 1,87 | 0 | 1,53 | 0 |
| DUXA | GAAAGTCATTCAG | 1 | 1008 | 1021 | 8 | 21 | 3,4 | 0,01 | 21,37 | >0.1 | 5,57 | 9,07 | 0 | 1,87 | 0 | 1,46 | 0 |
| DUXA | TGGATGTGATCTT | 1 | 1021 | 1034 | 21 | 34 | 3,39 | 0,01 | 20,8 | >0.1 | 5,57 | 8,7 | 0 | 1,86 | 0 | 1,28 | 0 |
| DUXA | AAGATCACATCCA | -1 | 1021 | 1034 | 21 | 34 | -0,72 | 0,01 | 16,69 | >0.1 | 5,57 | 8,7 | 0 | 1,86 | 0 | 1,28 | 0 |
| DUXA | AAAACCTCCTCAT | -1 | 987 | 1000 | -13 | 0 | 1,97 | 0,01 | 19,73 | >0.1 | 5,57 | 8,71 | 0 | 1,87 | 0 | 1,61 | 0 |
| DUXA | TAAAGTTCATCCT | -1 | 868 | 881 | -132 | -119 | 2,27 | 0,01 | 19,14 | >0.1 | 5,57 | 6,45 | 0 | 1,9 | 0 | 2,95 | 0 |
| DUXA | TAAAGTGAATCAA | -1 | 816 | 829 | -184 | -171 | 8,88 | 0,001 | 25,69 | >0.1 | 5,57 | 6,08 | 0 | 1,61 | 0 | 3,55 | 0 |

| | | | | | | | | | | | | | | | | | |
|---|---|---|---|---|---|---|---|---|---|---|---|---|---|---|---|---|---|
| DUXA | ATGGCCAAATCAA | -1 | 750 | 763 | -250 | -237 | 3,48 | 0,01 | 22,4 | >0.1 | 5,57 | 5,74 | 0 | 1,3 | 0 | 6,32 | 0 |
| DUXA | ACAATTTCATTAA | -1 | 604 | 617 | -396 | -383 | 6,32 | 0,001 | 17,5 | >0.1 | 3,51 | 5,23 | 0 | 1,02 | 0 | 1,41 | 0 |
| DUXA | GAGACTGAGTCTC | -1 | 444 | 457 | -556 | -543 | -0,01 | 0,01 | 13,62 | >0.1 | 2,76 | 4,86 | 0 | 1 | 0 | 5,01 | 0 |
| DUXA | GTGGCGCGATCTC | -1 | 406 | 419 | -594 | -581 | -0,78 | 0,01 | 13,33 | >0.1 | 2,65 | 4,78 | 0 | 1,03 | 0 | 5,65 | 0 |
| DUXA | AAAAGCTCATTTC | -1 | 65 | 78 | -935 | -922 | 0,19 | 0,01 | 10,31 | >0.1 | 3,92 | 4,32 | 0 | 1,88 | 0 | 0 | 0 |
| TFDP1 | GTGTGAGAAAT | 1 | 52 | 63 | -948 | -937 | 0,31 | 0,01 | 10,58 | >0.1 | 4,09 | 4,3 | 0 | 1,87 | 0 | 0 | 0 |
| TFDP1 | GGCTGGGCACA | 1 | 176 | 187 | -824 | -813 | 0,49 | 0,01 | 13,13 | >0.1 | 3,15 | 4,44 | 0 | 1,82 | 0 | 3,24 | 0 |
| TFDP1 | GGGCGTGGTGG | 1 | 314 | 325 | -686 | -675 | 2,16 | 0,01 | 16,73 | >0.1 | 2,64 | 4,63 | 0 | 1,17 | 0 | 6,13 | 0 |
| TFDP1 | TGGTGGGCACC | 1 | 322 | 333 | -678 | -667 | 4,62 | 0,01 | 18,59 | >0.1 | 2,62 | 4,64 | 0 | 1,15 | 0 | 5,56 | 0 |
| TFDP1 | ACTCGGGAGGC | 1 | 346 | 357 | -654 | -643 | 5,74 | 0,001 | 20,6 | >0.1 | 2,55 | 4,68 | 0 | 1,11 | 0 | 6,52 | 0 |
| TFDP1 | AGGCAGGAGAA | 1 | 359 | 370 | -641 | -630 | 3,13 | 0,01 | 17,8 | >0.1 | 2,52 | 4,7 | 0 | 1,09 | 0 | 6,36 | 0 |
| TFDP1 | TGGCGTGAACC | 1 | 370 | 381 | -630 | -619 | 8,96 | 0,001 | 23,56 | >0.1 | 2,54 | 4,72 | 0 | 1,07 | 0 | 6,27 | 0 |
| TFDP1 | ACCTGGGAGGC | 1 | 378 | 389 | -622 | -611 | 2,45 | 0,01 | 17,34 | >0.1 | 2,57 | 4,73 | 0 | 1,06 | 0 | 6,53 | 0 |
| TFDP1 | GGGAGGCAGAG | 1 | 382 | 393 | -618 | -607 | 0,93 | 0,01 | 15,21 | >0.1 | 2,58 | 4,74 | 0 | 1,06 | 0 | 5,91 | 0 |
| TFDP1 | AGCCGAGATCG | 1 | 402 | 413 | -598 | -587 | 0,24 | 0,01 | 14,45 | >0.1 | 2,63 | 4,77 | 0 | 1,03 | 0 | 5,76 | 0 |
| TFDP1 | TCGCGCCACTG | 1 | 410 | 421 | -590 | -579 | 0,15 | 0,01 | 14,27 | >0.1 | 2,65 | 4,79 | 0 | 1,03 | 0 | 5,65 | 0 |
| TFDP1 | GGGCGACAGAC | 1 | 432 | 443 | -568 | -557 | 1,2 | 0,01 | 15,43 | >0.1 | 2,72 | 4,83 | 0 | 1,01 | 0 | 5,67 | 0 |
| TFDP1 | GGAAGGGAAAA | 1 | 528 | 539 | -472 | -461 | 2,88 | 0,01 | 15,52 | >0.1 | 3,08 | 5,03 | 0 | 0,98 | 0 | 3,55 | 0 |
| TFDP1 | GAGAGGTAAGG | 1 | 773 | 784 | -227 | -216 | 2,02 | 0,01 | 20,93 | >0.1 | 5,6 | 5,83 | 0 | 1,37 | 0 | 6,1 | 0 |
| TFDP1 | TCTAGGGAAAG | 1 | 1002 | 1013 | 2 | 13 | 1,07 | 0,01 | 19,12 | >0.1 | 5,6 | 9,12 | 0 | 1,87 | 0 | 1,46 | 0 |
| TFDP1 | GGCTGAGAAGG | -1 | 1069 | 1080 | 69 | 80 | 1,33 | 0,01 | 19,98 | >0.1 | 5,6 | 7,19 | 0 | 1,85 | 1,79 | 2,21 | 0 |
| TFDP1 | AGCCAGGAAGA | -1 | 1058 | 1069 | 58 | 69 | 2,48 | 0,01 | 19,54 | >0.1 | 5,6 | 7,39 | 0 | 1,86 | 0 | 2,21 | 0 |
| TFDP1 | GGGCGCTGAGC | -1 | 940 | 951 | -60 | -49 | 2,53 | 0,01 | 18,07 | >0.1 | 5,6 | 7,23 | 0 | 1,88 | 0 | 0,83 | 0 |
| TFDP1 | GAGCAGGAAAG | -1 | 933 | 944 | -67 | -56 | 6,56 | 0,001 | 22,08 | >0.1 | 5,6 | 7,12 | 0 | 1,88 | 0 | 0,93 | 0 |
| TFDP1 | ACACGGAAAAT | -1 | 694 | 705 | -306 | -295 | 0,91 | 0,01 | 16,22 | >0.1 | 5,6 | 5,5 | 0 | 1,14 | 0 | 3,07 | 0 |
| TFDP1 | AGGCTGGAGCA | -1 | 421 | 432 | -579 | -568 | 3,12 | 0,01 | 17,05 | >0.1 | 2,69 | 4,81 | 0 | 1,02 | 0 | 5,42 | 0 |
| TFDP1 | TGGCGCGATCT | -1 | 407 | 418 | -593 | -582 | 4,98 | 0,01 | 19,09 | >0.1 | 2,65 | 4,78 | 0 | 1,03 | 0 | 5,65 | 0 |
| TFDP1 | TAGCTGGGACT | -1 | 336 | 347 | -664 | -653 | -0,22 | 0,01 | 14,06 | >0.1 | 2,58 | 4,66 | 0 | 1,13 | 0 | 5,92 | 0 |
| TFDP1 | AGCTGGGACTA | -1 | 335 | 346 | -665 | -654 | 0,18 | 0,01 | 14,58 | >0.1 | 2,58 | 4,66 | 0 | 1,13 | 0 | 6,03 | 0 |
| TFDP1 | AGGCGTGAGCC | -1 | 189 | 200 | -811 | -800 | 6,19 | 0,001 | 19,48 | >0.1 | 3,08 | 4,45 | 0 | 1,71 | 0 | 4,04 | 0 |
| YY1 | GGAGAAGGAGGC | 1 | 14 | 26 | -986 | -974 | -5,33 | 0,01 | 6,39 | >0.1 | 5,58 | 4,27 | 0 | 1,87 | 0 | 0 | 0 |
| YY1 | AAAAATGGCCAT | 1 | 141 | 153 | -859 | -847 | 10,51 | 0,001 | 21,78 | >0.1 | 3,33 | 4,4 | 0 | 1,9 | 0 | 1,65 | 0 |
| YY1 | GGCCATGGAAAT | 1 | 147 | 159 | -853 | -841 | -5,26 | 0,01 | 5,98 | >0.1 | 3,29 | 4,4 | 0 | 1,9 | 0 | 1,65 | 0 |
| YY1 | AGAAATGGCTGG | 1 | 170 | 182 | -830 | -818 | 7,58 | 0,001 | 20,41 | >0.1 | 3,17 | 4,43 | 0 | 1,88 | 0 | 3,34 | 0 |
| YY1 | CACAGTGGCTCA | 1 | 183 | 195 | -817 | -805 | -6,49 | 0,01 | 6,78 | >0.1 | 3,11 | 4,45 | 0 | 1,76 | 0 | 3,95 | 0 |
| YY1 | GGAGATAGAGAC | 1 | 247 | 259 | -753 | -741 | -2,89 | 0,01 | 10,95 | >0.1 | 2,85 | 4,53 | 0 | 1,38 | 0 | 5,09 | 0 |
| YY1 | TAACACGGTGAA | 1 | 268 | 280 | -732 | -720 | -1,85 | 0,01 | 11,88 | >0.1 | 2,78 | 4,56 | 0 | 1,3 | 0 | 5,09 | 0 |
| YY1 | GAGAATGGCGTG | 1 | 365 | 377 | -635 | -623 | 6,13 | 0,001 | 21,07 | >0.1 | 2,53 | 4,71 | 0 | 1,08 | 0 | 6,61 | 0 |
| YY1 | CGAGATCGCGCC | 1 | 405 | 417 | -595 | -583 | -1,93 | 0,01 | 12,17 | >0.1 | 2,64 | 4,78 | 0 | 1,03 | 0 | 5,65 | 0 |
| YY1 | AAATATGAATTC | 1 | 487 | 499 | -513 | -501 | -3,35 | 0,01 | 11,76 | >0.1 | 2,91 | 4,94 | 0 | 0,99 | 0 | 6,27 | 0 |
| YY1 | GAAAATGTTGCC | 1 | 534 | 546 | -466 | -454 | -4,6 | 0,01 | 6,15 | >0.1 | 3,11 | 5,05 | 0 | 0,99 | 0 | 1,61 | 0 |
| YY1 | GAATATGATCTT | 1 | 564 | 576 | -436 | -424 | -6,11 | 0,01 | 3,27 | >0.1 | 3,26 | 5,12 | 0 | 1 | 0 | 0 | 0 |
| YY1 | CAGGATGAACTT | 1 | 867 | 879 | -133 | -121 | -3,39 | 0,01 | 13,39 | >0.1 | 5,58 | 6,44 | 0 | 1,9 | 0 | 2,86 | 0 |
| YY1 | CACCATTGAGGA | 1 | 1105 | 1117 | 105 | 117 | -7,96 | 0,01 | 11,87 | >0.1 | 5,58 | 6,7 | 0 | 1,85 | 3,22 | 2,48 | 0 |
| YY1 | AAAAATGTCTTG | -1 | 1123 | 1135 | 123 | 135 | -6,42 | 0,01 | 13,55 | >0.1 | 5,58 | 6,52 | 0 | 1,85 | 3,22 | 2,79 | 0 |
| YY1 | CTCAATGGTGGA | -1 | 1103 | 1115 | 103 | 115 | -0,76 | 0,01 | 19,15 | >0.1 | 5,58 | 6,72 | 0 | 1,85 | 3,22 | 2,53 | 0 |
| YY1 | CAACAAGGCTGA | -1 | 1074 | 1086 | 74 | 86 | 0,35 | 0,01 | 18,96 | >0.1 | 5,58 | 7,11 | 0 | 1,85 | 1,79 | 2,27 | 0 |
| YY1 | CTTTATGGCCAA | -1 | 755 | 767 | -245 | -233 | -5,18 | 0,01 | 13,79 | >0.1 | 5,58 | 5,75 | 0 | 1,31 | 0 | 6,32 | 0 |
| YY1 | CAGGCTGGAGCA | -1 | 421 | 433 | -579 | -567 | -6,39 | 0,01 | 7,68 | >0.1 | 2,69 | 4,81 | 0 | 1,02 | 0 | 5,56 | 0 |
| YY1 | CACAGTGGCGCG | -1 | 411 | 423 | -589 | -577 | -6,69 | 0,01 | 7,7 | >0.1 | 2,66 | 4,79 | 0 | 1,03 | 0 | 5,91 | 0 |
| YY1 | CAGGACGGTCTC | -1 | 254 | 266 | -746 | -734 | 0,12 | 0,01 | 13,69 | >0.1 | 2,83 | 4,54 | 0 | 1,35 | 0 | 4,85 | 0 |
| YY1 | TTCCATGGCCAT | -1 | 145 | 157 | -855 | -843 | -5,36 | 0,01 | 5,85 | >0.1 | 3,31 | 4,4 | 0 | 1,9 | 0 | 1,61 | 0 |
| YY1 | AATTATGACTAC | -1 | 126 | 138 | -874 | -862 | -6,52 | 0,01 | 4,78 | >0.1 | 3,42 | 4,38 | 0 | 1,89 | 0 | 1,61 | 0 |
| YY2 | AATGGCCATGG | 1 | 144 | 155 | -856 | -845 | 5,93 | 0,001 | 17,11 | >0.1 | 3,31 | 4,4 | 0 | 1,9 | 0 | 1,57 | 0 |
| YY2 | AAACCCCGTCT | 1 | 278 | 289 | -722 | -711 | 0,27 | 0,01 | 14,05 | >0.1 | 2,75 | 4,57 | 0 | 1,27 | 0 | 5,19 | 0 |
| YY2 | CGTCTCTATTA | 1 | 284 | 295 | -716 | -705 | 0,03 | 0,01 | 13,58 | >0.1 | 2,73 | 4,58 | 0 | 1,25 | 0 | 4,99 | 0 |
| YY2 | GAATTCCATAA | 1 | 493 | 504 | -507 | -496 | -0,52 | 0,01 | 14,79 | >0.1 | 2,93 | 4,95 | 0 | 0,98 | 0 | 6,44 | 0 |
| YY2 | AAACACCAACA | 1 | 630 | 641 | -370 | -359 | -0,95 | 0,01 | 10,83 | >0.1 | 3,71 | 5,3 | 0 | 1,05 | 0 | 1,73 | 0 |
| YY2 | CACCAACATTA | 1 | 633 | 644 | -367 | -356 | 4,25 | 0,01 | 15,95 | >0.1 | 3,73 | 5,3 | 0 | 1,05 | 0 | 1,61 | 0 |

| | | | | | | | | | | | | | | | | | |
|---|---|---|---|---|---|---|---|---|---|---|---|---|---|---|---|---|---|
| YY2 | TTTGGCCATAA | 1 | 754 | 765 | -246 | -235 | 6,01 | 0,001 | 24,98 | >0.1 | 5,6 | 5,75 | 0 | 1,3 | 0 | 6,32 | 0 |
| YY2 | CTCAGCCTTGT | 1 | 1073 | 1084 | 73 | 84 | 2,13 | 0,01 | 20,77 | >0.1 | 5,6 | 7,12 | 0 | 1,85 | 1,79 | 2,27 | 0 |
| YY2 | GTCCACCATTG | 1 | 1102 | 1113 | 102 | 113 | 10,62 | 0,0001 | 30,44 | >0.1 | 5,6 | 6,73 | 0 | 1,85 | 3,22 | 2,41 | 0 |
| YY2 | GCTTGACATCG | -1 | 1047 | 1058 | 47 | 58 | -0,53 | 0,01 | 15,99 | >0.1 | 5,6 | 7,65 | 0 | 1,86 | 0 | 1,42 | 0 |
| YY2 | GTGAGCCAAGA | -1 | 1030 | 1041 | 30 | 41 | -0,95 | 0,01 | 16,08 | >0.1 | 5,6 | 8,24 | 0 | 1,86 | 0 | 1,34 | 0 |
| YY2 | ATCAGCCTTTG | -1 | 961 | 972 | -39 | -28 | 3,59 | 0,01 | 20,43 | >0.1 | 5,6 | 7,67 | 0 | 1,88 | 0 | 1,7 | 0 |
| YY2 | CTGAGCCAATA | -1 | 881 | 892 | -119 | -108 | -0,54 | 0,01 | 16,57 | >0.1 | 5,6 | 6,55 | 0 | 1,89 | 0 | 3,07 | 0 |
| YY2 | TATGGCCAAAT | -1 | 753 | 764 | -247 | -236 | -0,55 | 0,01 | 18,41 | >0.1 | 5,6 | 5,74 | 0 | 1,3 | 0 | 6,32 | 0 |
| YY2 | ATCTCACATTT | -1 | 587 | 598 | -413 | -402 | -0,7 | 0,01 | 8,88 | >0.1 | 3,39 | 5,18 | 0 | 1,01 | 0 | 0 | 0 |
| YY2 | GGGCAACATTT | -1 | 536 | 547 | -464 | -453 | 0,64 | 0,01 | 11,45 | >0.1 | 3,12 | 5,05 | 0 | 0,99 | 0 | 1,66 | 0 |
| YY2 | CTTTGCCCTTG | -1 | 507 | 518 | -493 | -482 | 0,8 | 0,01 | 13,95 | >0.1 | 2,99 | 4,98 | 0 | 0,98 | 0 | 4,19 | 0 |
| YY2 | TCACGCCATTC | -1 | 367 | 378 | -633 | -622 | 7,34 | 0,001 | 22,39 | >0.1 | 2,54 | 4,71 | 0 | 1,08 | 0 | 6,73 | 0 |
| YY2 | TTTCACCGTGT | -1 | 270 | 281 | -730 | -719 | -0,15 | 0,01 | 13,47 | >0.1 | 2,77 | 4,56 | 0 | 1,29 | 0 | 4,99 | 0 |
| YY2 | GGTCTCTATCT | -1 | 249 | 260 | -751 | -740 | -0,69 | 0,01 | 13,25 | >0.1 | 2,84 | 4,53 | 0 | 1,37 | 0 | 5,19 | 0 |
| YY2 | CCCAGCCATTT | -1 | 172 | 183 | -828 | -817 | 7,54 | 0,001 | 20,34 | >0.1 | 3,17 | 4,43 | 0 | 1,86 | 0 | 3,34 | 0 |
| YY2 | AATTTCCATGG | -1 | 149 | 160 | -851 | -840 | -0,66 | 0,01 | 10,58 | >0.1 | 3,29 | 4,41 | 0 | 1,9 | 0 | 1,65 | 0 |
| YY2 | CATGGCCATTT | -1 | 143 | 154 | -857 | -846 | 8,1 | 0,001 | 19,32 | >0.1 | 3,32 | 4,4 | 0 | 1,9 | 0 | 1,61 | 0 |
| YY2 | ATCTGCCTTTG | -1 | 88 | 99 | -912 | -901 | 4,22 | 0,01 | 14,15 | >0.1 | 3,71 | 4,34 | 0 | 1,88 | 0 | 0 | 0 |
| YY2 | TGCCTCCTTCT | -1 | 16 | 27 | -984 | -973 | 2,23 | 0,01 | 13,96 | >0.1 | 5,6 | 4,27 | 0 | 1,87 | 0 | 0 | 0 |
| YY2 | CTTCTCCTTCA | -1 | 10 | 21 | -990 | -979 | 1,82 | 0,01 | 15,34 | >0.1 | 5,6 | 4,26 | 0 | 1,86 | 1,79 | 0 | 0 |
| POU3F4 | TTTGCAAAG | 1 | 84 | 93 | -916 | -907 | 1,7 | 0,01 | 11,66 | >0.1 | 3,75 | 4,33 | 0 | 1,88 | 0 | 0 | 0 |
| POU3F4 | CATGGAAAT | 1 | 150 | 159 | -850 | -841 | 6,48 | 0,001 | 17,72 | >0.1 | 3,29 | 4,4 | 0 | 1,9 | 0 | 1,65 | 0 |
| POU3F4 | AATACAAAA | 1 | 297 | 306 | -703 | -694 | -0,56 | 0,01 | 12,47 | >0.1 | 2,69 | 4,6 | 0 | 1,21 | 0 | 4,53 | 0 |
| POU3F4 | TAGGAAAAT | 1 | 721 | 730 | -279 | -270 | -0,01 | 0,01 | 17,4 | >0.1 | 5,62 | 5,6 | 0 | 1,2 | 0 | 4,99 | 0 |
| POU3F4 | TATTCTTAT | 1 | 807 | 816 | -193 | -184 | 5,06 | 0,01 | 21,5 | >0.1 | 5,62 | 6 | 0 | 1,53 | 0 | 3,28 | 0 |
| POU3F4 | TATTCTAAA | 1 | 835 | 844 | -165 | -156 | 4,74 | 0,01 | 21,55 | >0.1 | 5,62 | 6,18 | 0 | 1,72 | 0 | 3,28 | 0 |
| POU3F4 | TATGTAACA | -1 | 847 | 856 | -153 | -144 | 1,21 | 0,01 | 17,79 | >0.1 | 5,62 | 6,27 | 0 | 1,82 | 0 | 2,86 | 0 |
| POU3F4 | TACGTTATT | -1 | 800 | 809 | -200 | -191 | -1 | 0,01 | 15,63 | >0.1 | 5,62 | 5,97 | 0 | 1,49 | 0 | 3,55 | 0 |
| POU3F4 | CATGAAGAT | -1 | 643 | 652 | -357 | -348 | -0,81 | 0,01 | 13,23 | >0.1 | 3,82 | 5,33 | 0 | 1,06 | 0 | 3,83 | 0 |
| POU3F4 | TATGGTTTT | -1 | 620 | 629 | -380 | -371 | 0,81 | 0,01 | 12,22 | >0.1 | 3,62 | 5,26 | 0 | 1,03 | 0 | 1,5 | 0 |
| POU3F4 | TATTTTTAT | -1 | 579 | 588 | -421 | -412 | 2,45 | 0,01 | 11,94 | >0.1 | 3,34 | 5,15 | 0 | 1 | 0 | 0 | 0 |
| POU3F4 | TATGGAATT | -1 | 494 | 503 | -506 | -497 | 4,93 | 0,01 | 20,23 | >0.1 | 2,93 | 4,95 | 0 | 0,98 | 0 | 6,44 | 0 |
| POU3F4 | AATTCATAT | -1 | 489 | 498 | -511 | -502 | 1,97 | 0,01 | 17,08 | >0.1 | 2,91 | 4,94 | 0 | 0,98 | 0 | 6,27 | 0 |
| POU3F4 | TTTGCAAAA | -1 | 83 | 92 | -917 | -908 | 2,47 | 0,01 | 12,44 | >0.1 | 3,76 | 4,33 | 0 | 1,88 | 0 | 0 | 0 |
| POU3F4 | AAAGCTCAT | -1 | 68 | 77 | -932 | -923 | -0,97 | 0,01 | 9,13 | >0.1 | 3,91 | 4,32 | 0 | 1,88 | 0 | 0 | 0 |
| MEIS1 | ATCACAA | 1 | 235 | 242 | -765 | -758 | 2,98 | 0,01 | 16,72 | >0.1 | 2,9 | 4,51 | 0 | 1,43 | 0 | 4,9 | 0 |
| MEIS1 | CTGACTA | 1 | 263 | 270 | -737 | -730 | 4,77 | 0,01 | 17,88 | >0.1 | 2,8 | 4,55 | 0 | 1,32 | 0 | 4,45 | 0 |
| MEIS1 | GCGACAG | 1 | 434 | 441 | -566 | -559 | 6,14 | 0,01 | 20,36 | >0.1 | 2,72 | 4,82 | 0 | 1,01 | 0 | 5,67 | 0 |
| MEIS1 | AAGACAA | 1 | 503 | 510 | -497 | -490 | 5,78 | 0,01 | 20,21 | >0.1 | 2,96 | 4,96 | 0 | 0,98 | 0 | 5,52 | 0 |
| MEIS1 | GTGACAG | 1 | 766 | 773 | -234 | -227 | 9,43 | 0,001 | 28,3 | >0.1 | 5,65 | 5,78 | 0 | 1,33 | 0 | 6,1 | 0 |
| MEIS1 | CTGATAA | 1 | 967 | 974 | -33 | -26 | 2,53 | 0,01 | 19,48 | >0.1 | 5,65 | 7,73 | 0 | 1,87 | 0 | 1,7 | 0 |
| MEIS1 | CTCACAG | 1 | 1036 | 1043 | 36 | 43 | 4,44 | 0,01 | 21,25 | >0.1 | 5,65 | 7,99 | 0 | 1,86 | 0 | 1,31 | 0 |
| MEIS1 | AAGACAT | 1 | 1124 | 1131 | 124 | 131 | 5,59 | 0,01 | 25,56 | >0.1 | 5,65 | 6,51 | 0 | 1,84 | 3,22 | 2,74 | 0 |
| MEIS1 | TGGACAA | 1 | 1134 | 1141 | 134 | 141 | 4,89 | 0,01 | 25,01 | >0.1 | 5,65 | 6,42 | 0 | 1,84 | 3,22 | 2,98 | 0 |
| MEIS1 | TTGATAG | -1 | 1168 | 1175 | 168 | 175 | 3,74 | 0,01 | 20,81 | >0.1 | 4,54 | 6,17 | 0 | 1,68 | 1,79 | 2,88 | 0 |
| MEIS1 | TTGACAT | -1 | 1049 | 1056 | 49 | 56 | 8,81 | 0,001 | 25,34 | >0.1 | 5,65 | 7,6 | 0 | 1,86 | 0 | 1,42 | 0 |
| MEIS1 | ATCACAT | -1 | 1024 | 1031 | 24 | 31 | 2,79 | 0,01 | 20,17 | >0.1 | 5,65 | 8,59 | 0 | 1,86 | 0 | 1,28 | 0 |
| MEIS1 | ATGACTT | -1 | 1010 | 1017 | 10 | 17 | 4,42 | 0,01 | 22,34 | >0.1 | 5,65 | 8,99 | 0 | 1,86 | 0 | 1,42 | 0 |
| MEIS1 | AGGACAG | -1 | 857 | 864 | -143 | -136 | 6,12 | 0,01 | 22,86 | >0.1 | 5,65 | 6,32 | 0 | 1,9 | 0 | 2,86 | 0 |
| MEIS1 | CAGACAA | -1 | 613 | 620 | -387 | -380 | 5,94 | 0,01 | 17,25 | >0.1 | 3,56 | 5,24 | 0 | 1,02 | 0 | 1,5 | 0 |
| MEIS1 | CTCACAT | -1 | 589 | 596 | -411 | -404 | 2,95 | 0,01 | 12,52 | >0.1 | 3,39 | 5,17 | 0 | 1 | 0 | 0 | 0 |
| MEIS1 | ATGACTT | -1 | 515 | 522 | -485 | -478 | 4,42 | 0,01 | 16,27 | >0.1 | 3,01 | 4,99 | 0 | 0,98 | 0 | 2,86 | 0 |
| MEIS1 | GAGACGG | -1 | 283 | 290 | -717 | -710 | 3,72 | 0,01 | 17,48 | >0.1 | 2,74 | 4,57 | 0 | 1,25 | 0 | 5,19 | 0 |
| MEIS1 | AGGACGG | -1 | 258 | 265 | -742 | -735 | 3,56 | 0,01 | 16,98 | >0.1 | 2,82 | 4,54 | 0 | 1,33 | 0 | 4,73 | 0 |
| MEIS1 | ATGACTA | -1 | 127 | 134 | -873 | -866 | 4,61 | 0,01 | 16 | >0.1 | 3,43 | 4,38 | 0 | 1,89 | 0 | 1,7 | 0 |
| MEIS1 | CTCACAC | -1 | 52 | 59 | -948 | -941 | 3,42 | 0,01 | 13,71 | >0.1 | 4,12 | 4,3 | 0 | 1,87 | 0 | 0 | 0 |
| MEIS1 | CTGACTT | -1 | 34 | 41 | -966 | -959 | 4,58 | 0,01 | 15,16 | >0.1 | 4,43 | 4,28 | 0 | 1,87 | 0 | 0 | 0 |
| TBX2 | CCAGTGTGAGA | 1 | 49 | 60 | -951 | -940 | 4,69 | 0,01 | 15 | >0.1 | 4,13 | 4,3 | 0 | 1,87 | 0 | 0 | 0 |
| TBX2 | AGTGTGAGAAA | 1 | 51 | 62 | -949 | -938 | 7,06 | 0,001 | 17,34 | >0.1 | 4,11 | 4,3 | 0 | 1,87 | 0 | 0 | 0 |

| | | | | | | | | | | | | | | | | | |
|---|---|---|---|---|---|---|---|---|---|---|---|---|---|---|---|---|---|
| TBX2 | GACCTGTGGAG | 1 | 109 | 120 | -891 | -880 | 1,37 | 0,01 | 11,9 | >0.1 | 3,54 | 4,36 | 0 | 1,89 | 0 | 0,75 | 0 |
| TBX2 | TGGGCGTGGTG | 1 | 313 | 324 | -687 | -676 | 3,74 | 0,01 | 18,31 | >0.1 | 2,64 | 4,63 | 0 | 1,17 | 0 | 6,13 | 0 |
| TBX2 | ATGGCGTGAAC | 1 | 369 | 380 | -631 | -620 | 8,9 | 0,001 | 23,38 | >0.1 | 2,54 | 4,72 | 0 | 1,07 | 0 | 6,15 | 0 |
| TBX2 | TGGGCGACAGA | 1 | 431 | 442 | -569 | -558 | 4,24 | 0,01 | 18,46 | >0.1 | 2,72 | 4,83 | 0 | 1,01 | 0 | 5,67 | 0 |
| TBX2 | TAAATGTGAGA | 1 | 586 | 597 | -414 | -403 | 1,52 | 0,01 | 11,09 | >0.1 | 3,39 | 5,17 | 0 | 1,01 | 0 | 0 | 0 |
| TBX2 | AATGTGAGATA | 1 | 588 | 599 | -412 | -401 | 6,77 | 0,01 | 16,36 | >0.1 | 3,4 | 5,18 | 0 | 1,01 | 0 | 0 | 0 |
| TBX2 | AAAGTGACAGG | 1 | 763 | 774 | -237 | -226 | 5,59 | 0,01 | 24,22 | >0.1 | 5,6 | 5,79 | 0 | 1,34 | 0 | 5,92 | 0 |
| TBX2 | AATCTGTTACA | 1 | 843 | 854 | -157 | -146 | 2,09 | 0,01 | 18,94 | >0.1 | 5,6 | 6,26 | 0 | 1,81 | 0 | 3,19 | 0 |
| TBX2 | AGGCTGATAAG | 1 | 964 | 975 | -36 | -25 | 1,77 | 0,01 | 18,69 | >0.1 | 5,6 | 7,76 | 0 | 1,87 | 0 | 1,7 | 0 |
| TBX2 | GAGGTTTTAGT | 1 | 992 | 1003 | -8 | 3 | 2,34 | 0,01 | 20,18 | >0.1 | 5,6 | 8,81 | 0 | 1,87 | 0 | 1,57 | 0 |
| TBX2 | TGGATGTGATC | 1 | 1021 | 1032 | 21 | 32 | 1,35 | 0,01 | 18,82 | >0.1 | 5,6 | 8,7 | 0 | 1,86 | 0 | 1,31 | 0 |
| TBX2 | AGGCTGAGAAG | -1 | 1070 | 1081 | 70 | 81 | 3,44 | 0,01 | 22,13 | >0.1 | 5,6 | 7,17 | 0 | 1,85 | 1,79 | 2,27 | 0 |
| TBX2 | TGGGCGCTGAG | -1 | 941 | 952 | -59 | -48 | 1,49 | 0,01 | 17 | >0.1 | 5,6 | 7,25 | 0 | 1,88 | 0 | 0,79 | 0 |
| TBX2 | AAGCTGTTAAA | -1 | 708 | 719 | -292 | -281 | 7,63 | 0,001 | 24,77 | >0.1 | 5,6 | 5,56 | 0 | 1,17 | 0 | 4,81 | 0 |
| TBX2 | TTGGTGTTTGT | -1 | 628 | 639 | -372 | -361 | 2,95 | 0,01 | 14,58 | >0.1 | 3,69 | 5,29 | 0 | 1,04 | 0 | 1,61 | 0 |
| TBX2 | GTGGCGCGATC | -1 | 408 | 419 | -592 | -581 | 4,12 | 0,01 | 18,23 | >0.1 | 2,65 | 4,78 | 0 | 1,03 | 0 | 5,65 | 0 |
| TBX2 | CAGGTGCCCAC | -1 | 324 | 335 | -676 | -665 | 3,12 | 0,01 | 16,98 | >0.1 | 2,61 | 4,64 | 0 | 1,15 | 0 | 5,46 | 0 |
| TBX2 | GGGGTTTCACC | -1 | 274 | 285 | -726 | -715 | 0,86 | 0,01 | 14,66 | >0.1 | 2,76 | 4,57 | 0 | 1,28 | 0 | 5,19 | 0 |
| TBX2 | ACCGTGTTAGT | -1 | 266 | 277 | -734 | -723 | 3,31 | 0,01 | 16,49 | >0.1 | 2,79 | 4,56 | 0 | 1,31 | 0 | 4,53 | 0 |
| TBX2 | AAAGTGCTAGG | -1 | 205 | 216 | -795 | -784 | 6,07 | 0,01 | 19,86 | >0.1 | 3,01 | 4,47 | 0 | 1,6 | 0 | 4,7 | 0 |
| TBX2 | CAGGCGTGAGC | -1 | 190 | 201 | -810 | -799 | 8,84 | 0,001 | 22,21 | >0.1 | 3,08 | 4,46 | 0 | 1,71 | 0 | 4,13 | 0 |
| TBX1 | AGGAGGCA | 1 | 19 | 27 | -981 | -973 | -3,73 | 0,01 | 8,04 | >0.1 | 5,64 | 4,27 | 0 | 1,87 | 0 | 0 | 0 |
| TBX1 | CAGTGTGA | 1 | 50 | 58 | -950 | -942 | 5,66 | 0,001 | 15,97 | >0.1 | 4,14 | 4,3 | 0 | 1,87 | 0 | 0 | 0 |
| TBX1 | GTGTGAGA | 1 | 52 | 60 | -948 | -940 | 1,2 | 0,01 | 11,48 | >0.1 | 4,11 | 4,3 | 0 | 1,87 | 0 | 0 | 0 |
| TBX1 | AGGAGATA | 1 | 246 | 254 | -754 | -746 | -3,82 | 0,01 | 10,25 | >0.1 | 2,86 | 4,52 | 0 | 1,38 | 0 | 5,3 | 0 |
| TBX1 | CGGTGAAA | 1 | 273 | 281 | -727 | -719 | -4,19 | 0,01 | 9,41 | >0.1 | 2,77 | 4,56 | 0 | 1,28 | 0 | 4,99 | 0 |
| TBX1 | GGGCGTGG | 1 | 314 | 322 | -686 | -678 | -1,92 | 0,01 | 12,65 | >0.1 | 2,64 | 4,62 | 0 | 1,17 | 0 | 6,13 | 0 |
| TBX1 | TGGTGGGC | 1 | 322 | 330 | -678 | -670 | -3,35 | 0,01 | 10,62 | >0.1 | 2,62 | 4,63 | 0 | 1,15 | 0 | 5,56 | 0 |
| TBX1 | TGGCGTGA | 1 | 370 | 378 | -630 | -622 | 5,61 | 0,001 | 20,08 | >0.1 | 2,54 | 4,71 | 0 | 1,07 | 0 | 6,15 | 0 |
| TBX1 | GGGCGACA | 1 | 432 | 440 | -568 | -560 | -3,5 | 0,01 | 10,72 | >0.1 | 2,72 | 4,82 | 0 | 1,01 | 0 | 5,67 | 0 |
| TBX1 | AAGGGCAA | 1 | 508 | 516 | -492 | -484 | -4,7 | 0,01 | 8,44 | >0.1 | 2,99 | 4,98 | 0 | 0,98 | 0 | 4,19 | 0 |
| TBX1 | AAATGTGA | 1 | 587 | 595 | -413 | -405 | -4,47 | 0,01 | 5,08 | >0.1 | 3,38 | 5,17 | 0 | 1 | 0 | 0 | 0 |
| TBX1 | ATGTGAGA | 1 | 589 | 597 | -411 | -403 | 4,27 | 0,01 | 13,84 | >0.1 | 3,4 | 5,17 | 0 | 1,01 | 0 | 0 | 0 |
| TBX1 | AAGTGACA | 1 | 764 | 772 | -236 | -228 | 1,45 | 0,01 | 20,11 | >0.1 | 5,64 | 5,78 | 0 | 1,33 | 0 | 5,92 | 0 |
| TBX1 | AAGAGAGA | 1 | 972 | 980 | -28 | -20 | -3,23 | 0,01 | 13,84 | >0.1 | 5,64 | 7,91 | 0 | 1,87 | 0 | 1,65 | 0 |
| TBX1 | GGATGTGA | 1 | 1022 | 1030 | 22 | 30 | -4,11 | 0,01 | 13,38 | >0.1 | 5,64 | 8,68 | 0 | 1,86 | 0 | 1,31 | 0 |
| TBX1 | AGGAGCCA | -1 | 1064 | 1072 | 64 | 72 | -1,24 | 0,01 | 17,6 | >0.1 | 5,64 | 7,28 | 0 | 1,85 | 1,79 | 2,27 | 0 |
| TBX1 | AGCTGTTA | -1 | 710 | 718 | -290 | -282 | -4,56 | 0,01 | 12,61 | >0.1 | 5,64 | 5,55 | 0 | 1,17 | 0 | 4,81 | 0 |
| TBX1 | TGGCGCGA | -1 | 410 | 418 | -590 | -582 | 1,83 | 0,01 | 15,81 | >0.1 | 2,65 | 4,78 | 0 | 1,03 | 0 | 5,52 | 0 |
| TBX1 | AGGTGCCC | -1 | 326 | 334 | -674 | -666 | -0,51 | 0,01 | 13,34 | >0.1 | 2,61 | 4,64 | 0 | 1,15 | 0 | 5,46 | 0 |
| TBX1 | CCGTGTTA | -1 | 268 | 276 | -732 | -724 | 1,01 | 0,01 | 14,18 | >0.1 | 2,78 | 4,55 | 0 | 1,3 | 0 | 4,53 | 0 |
| TBX1 | AAGTGCTA | -1 | 207 | 215 | -793 | -785 | 4,08 | 0,01 | 17,85 | >0.1 | 3,01 | 4,47 | 0 | 1,58 | 0 | 4,7 | 0 |
| TBX1 | AGGCGTGA | -1 | 192 | 200 | -808 | -800 | 10,01 | 0,001 | 23,36 | >0.1 | 3,08 | 4,45 | 0 | 1,69 | 0 | 4,13 | 0 |
| TBX4 | CAGTGTGA | 1 | 50 | 58 | -950 | -942 | 5,89 | 0,01 | 16,2 | >0.1 | 4,14 | 4,3 | 0 | 1,87 | 0 | 0 | 0 |
| TBX4 | GTGTGAGA | 1 | 52 | 60 | -948 | -940 | 3,32 | 0,01 | 13,6 | >0.1 | 4,11 | 4,3 | 0 | 1,87 | 0 | 0 | 0 |
| TBX4 | ACCTGTGG | 1 | 110 | 118 | -890 | -882 | 1,34 | 0,01 | 11,87 | >0.1 | 3,54 | 4,36 | 0 | 1,89 | 0 | 0,75 | 0 |
| TBX4 | CGGTGAAA | 1 | 273 | 281 | -727 | -719 | 1,12 | 0,01 | 14,72 | >0.1 | 2,77 | 4,56 | 0 | 1,28 | 0 | 4,99 | 0 |
| TBX4 | GGGCGTGG | 1 | 314 | 322 | -686 | -678 | 5,47 | 0,01 | 20,04 | >0.1 | 2,64 | 4,62 | 0 | 1,17 | 0 | 6,13 | 0 |
| TBX4 | TGGCGTGA | 1 | 370 | 378 | -630 | -622 | 7,03 | 0,01 | 21,5 | >0.1 | 2,54 | 4,71 | 0 | 1,07 | 0 | 6,15 | 0 |
| TBX4 | GGGCGACA | 1 | 432 | 440 | -568 | -560 | 3,03 | 0,01 | 17,25 | >0.1 | 2,72 | 4,82 | 0 | 1,01 | 0 | 5,67 | 0 |
| TBX4 | AAATGTGA | 1 | 587 | 595 | -413 | -405 | 1,38 | 0,01 | 10,93 | >0.1 | 3,38 | 5,17 | 0 | 1 | 0 | 0 | 0 |
| TBX4 | ATGTGAGA | 1 | 589 | 597 | -411 | -403 | 5,81 | 0,01 | 15,38 | >0.1 | 3,4 | 5,17 | 0 | 1,01 | 0 | 0 | 0 |
| TBX4 | AAGTGACA | 1 | 764 | 772 | -236 | -228 | 5,43 | 0,01 | 24,09 | >0.1 | 5,64 | 5,78 | 0 | 1,33 | 0 | 5,92 | 0 |
| TBX4 | ATCTGTTA | 1 | 844 | 852 | -156 | -148 | 1,13 | 0,01 | 17,99 | >0.1 | 5,64 | 6,24 | 0 | 1,79 | 0 | 3,19 | 0 |
| TBX4 | GGATGTGA | 1 | 1022 | 1030 | 22 | 30 | 1,98 | 0,01 | 19,47 | >0.1 | 5,64 | 8,68 | 0 | 1,86 | 0 | 1,31 | 0 |
| TBX4 | GGCTGAGA | -1 | 1072 | 1080 | 72 | 80 | 1,2 | 0,01 | 19,9 | >0.1 | 5,64 | 7,14 | 0 | 1,85 | 1,79 | 2,27 | 0 |
| TBX4 | GGGCGCTG | -1 | 943 | 951 | -57 | -49 | 1,61 | 0,01 | 17,19 | >0.1 | 5,64 | 7,23 | 0 | 1,88 | 0 | 0,83 | 0 |
| TBX4 | AGCTGTTA | -1 | 710 | 718 | -290 | -282 | 5,63 | 0,01 | 22,8 | >0.1 | 5,64 | 5,55 | 0 | 1,17 | 0 | 4,81 | 0 |
| TBX4 | TGGTGTTT | -1 | 630 | 638 | -370 | -362 | 4,12 | 0,01 | 15,8 | >0.1 | 3,69 | 5,29 | 0 | 1,04 | 0 | 1,66 | 0 |

| | | | | | | | | | | | | | | | | | |
|---|---|---|---|---|---|---|---|---|---|---|---|---|---|---|---|---|---|
| TBX4 | TGGCGCGA | -1 | 410 | 418 | -590 | -582 | 4,82 | 0,01 | 18,8 | >0.1 | 2,65 | 4,78 | 0 | 1,03 | 0 | 5,52 | 0 |
| TBX4 | AGCTGGGA | -1 | 338 | 346 | -662 | -654 | 1,84 | 0,01 | 16,11 | >0.1 | 2,58 | 4,66 | 0 | 1,12 | 0 | 5,92 | 0 |
| TBX4 | AGGTGCCC | -1 | 326 | 334 | -674 | -666 | 5,41 | 0,01 | 19,26 | >0.1 | 2,61 | 4,64 | 0 | 1,15 | 0 | 5,46 | 0 |
| TBX4 | CCGTGTTA | -1 | 268 | 276 | -732 | -724 | 4,33 | 0,01 | 17,5 | >0.1 | 2,78 | 4,55 | 0 | 1,3 | 0 | 4,53 | 0 |
| TBX4 | AAGTGCTA | -1 | 207 | 215 | -793 | -785 | 6,95 | 0,01 | 20,72 | >0.1 | 3,01 | 4,47 | 0 | 1,58 | 0 | 4,7 | 0 |
| TBX4 | AGGCGTGA | -1 | 192 | 200 | -808 | -800 | 10,89 | 0,001 | 24,24 | >0.1 | 3,08 | 4,45 | 0 | 1,69 | 0 | 4,13 | 0 |
| TBX5 | AGGAGGCA | 1 | 19 | 27 | -981 | -973 | 3,67 | 0,01 | 15,44 | >0.1 | 5,64 | 4,27 | 0 | 1,87 | 0 | 0 | 0 |
| TBX5 | CAGTGTGA | 1 | 50 | 58 | -950 | -942 | 5,93 | 0,01 | 16,24 | >0.1 | 4,14 | 4,3 | 0 | 1,87 | 0 | 0 | 0 |
| TBX5 | GTGTGAGA | 1 | 52 | 60 | -948 | -940 | 3,83 | 0,01 | 14,11 | >0.1 | 4,11 | 4,3 | 0 | 1,87 | 0 | 0 | 0 |
| TBX5 | TGGAGTGG | 1 | 115 | 123 | -885 | -877 | 2,87 | 0,1 | 13,37 | >0.1 | 3,51 | 4,36 | 0 | 1,89 | 0 | 0,75 | 0 |
| TBX5 | AGGAGATA | 1 | 246 | 254 | -754 | -746 | 2,94 | 0,01 | 17,01 | >0.1 | 2,86 | 4,52 | 0 | 1,38 | 0 | 5,3 | 0 |
| TBX5 | GGGCGTGG | 1 | 314 | 322 | -686 | -678 | 6,38 | 0,01 | 20,95 | >0.1 | 2,64 | 4,62 | 0 | 1,17 | 0 | 6,13 | 0 |
| TBX5 | TGGTGGTG | 1 | 319 | 327 | -681 | -673 | 4,05 | 0,01 | 18,72 | >0.1 | 2,63 | 4,63 | 0 | 1,16 | 0 | 6,26 | 0 |
| TBX5 | TGGTGGGC | 1 | 322 | 330 | -678 | -670 | 3,35 | 0,01 | 17,32 | >0.1 | 2,62 | 4,63 | 0 | 1,15 | 0 | 5,56 | 0 |
| TBX5 | TGGCGTGA | 1 | 370 | 378 | -630 | -622 | 6,8 | 0,01 | 21,27 | >0.1 | 2,54 | 4,71 | 0 | 1,07 | 0 | 6,15 | 0 |
| TBX5 | GGGCGACA | 1 | 432 | 440 | -568 | -560 | 3,26 | 0,01 | 17,48 | >0.1 | 2,72 | 4,82 | 0 | 1,01 | 0 | 5,67 | 0 |
| TBX5 | ATGTGAGA | 1 | 589 | 597 | -411 | -403 | 5,48 | 0,01 | 15,05 | >0.1 | 3,4 | 5,17 | 0 | 1,01 | 0 | 0 | 0 |
| TBX5 | AAGTGACA | 1 | 764 | 772 | -236 | -228 | 4,75 | 0,01 | 23,41 | >0.1 | 5,64 | 5,78 | 0 | 1,33 | 0 | 5,92 | 0 |
| TBX5 | AGGGGACG | 1 | 1041 | 1049 | 41 | 49 | 3,34 | 0,01 | 20,02 | >0.1 | 5,64 | 7,82 | 0 | 1,86 | 0 | 1,36 | 0 |
| TBX5 | AGGAGCCA | -1 | 1064 | 1072 | 64 | 72 | 3,84 | 0,01 | 22,68 | >0.1 | 5,64 | 7,28 | 0 | 1,85 | 1,79 | 2,27 | 0 |
| TBX5 | TGGTGTTT | -1 | 630 | 638 | -370 | -362 | 5,94 | 0,01 | 17,62 | >0.1 | 3,69 | 5,29 | 0 | 1,04 | 0 | 1,66 | 0 |
| TBX5 | TGGCGCGA | -1 | 410 | 418 | -590 | -582 | 4,3 | 0,01 | 18,28 | >0.1 | 2,65 | 4,78 | 0 | 1,03 | 0 | 5,52 | 0 |
| TBX5 | AGGTGCCC | -1 | 326 | 334 | -674 | -666 | 5,09 | 0,01 | 18,94 | >0.1 | 2,61 | 4,64 | 0 | 1,15 | 0 | 5,46 | 0 |
| TBX5 | CCGTGTTA | -1 | 268 | 276 | -732 | -724 | 4,65 | 0,01 | 17,82 | >0.1 | 2,78 | 4,55 | 0 | 1,3 | 0 | 4,53 | 0 |
| TBX5 | AAGTGCTA | -1 | 207 | 215 | -793 | -785 | 5,79 | 0,01 | 19,56 | >0.1 | 3,01 | 4,47 | 0 | 1,58 | 0 | 4,7 | 0 |
| TBX5 | AGGCGTGA | -1 | 192 | 200 | -808 | -800 | 9,93 | 0,001 | 23,28 | >0.1 | 3,08 | 4,45 | 0 | 1,69 | 0 | 4,13 | 0 |
| MEOX1 | GGAAATGAGC | 1 | 64 | 74 | -936 | -926 | 3,6 | 0,01 | 13,74 | >0.1 | 3,95 | 4,31 | 0 | 1,88 | 0 | 0 | 0 |
| MEOX1 | GGCCATGGAA | 1 | 147 | 157 | -853 | -843 | 1,69 | 0,01 | 12,9 | >0.1 | 3,3 | 4,4 | 0 | 1,9 | 0 | 1,61 | 0 |
| MEOX1 | GGAAATTAAA | 1 | 153 | 163 | -847 | -837 | 5,2 | 0,01 | 16,42 | >0.1 | 3,27 | 4,41 | 0 | 1,9 | 0 | 1,65 | 0 |
| MEOX1 | ACTGATCAGA | 1 | 163 | 173 | -837 | -827 | 2,85 | 0,01 | 14,79 | >0.1 | 3,21 | 4,42 | 0 | 1,9 | 0 | 2,4 | 0 |
| MEOX1 | CCTGACTAAC | 1 | 262 | 272 | -738 | -728 | 2,22 | 0,01 | 15,42 | >0.1 | 2,8 | 4,55 | 0 | 1,32 | 0 | 4,53 | 0 |
| MEOX1 | ACTGATCAGA | 1 | 478 | 488 | -522 | -512 | 2,85 | 0,01 | 16,99 | >0.1 | 2,88 | 4,92 | 0 | 0,99 | 0 | 5,36 | 0 |
| MEOX1 | ACCTATTAAT | 1 | 599 | 609 | -401 | -391 | 1,57 | 0,01 | 12,62 | >0.1 | 3,47 | 5,21 | 0 | 1,02 | 0 | 1,36 | 0 |
| MEOX1 | ATTAATGAAA | 1 | 603 | 613 | -397 | -387 | 3,37 | 0,01 | 14,51 | >0.1 | 3,49 | 5,22 | 0 | 1,02 | 0 | 1,41 | 0 |
| MEOX1 | TGAAATTGTC | 1 | 608 | 618 | -392 | -382 | 1,98 | 0,01 | 13,31 | >0.1 | 3,53 | 5,23 | 0 | 1,02 | 0 | 1,55 | 0 |
| MEOX1 | CAACATTATC | 1 | 636 | 646 | -364 | -354 | 0,92 | 0,01 | 12,7 | >0.1 | 3,76 | 5,31 | 0 | 1,05 | 0 | 1,66 | 0 |
| MEOX1 | CTTCATGATC | 1 | 645 | 655 | -355 | -345 | 3,39 | 0,01 | 17,47 | >0.1 | 3,84 | 5,34 | 0 | 1,06 | 0 | 3,83 | 0 |
| MEOX1 | TCTCATGAGG | 1 | 983 | 993 | -17 | -7 | 5,22 | 0,01 | 22,77 | >0.1 | 5,61 | 8,5 | 0 | 1,87 | 0 | 1,57 | 0 |
| MEOX1 | CCTCATGAGA | -1 | 983 | 993 | -17 | -7 | 5,91 | 0,01 | 23,46 | >0.1 | 5,61 | 8,5 | 0 | 1,87 | 0 | 1,57 | 0 |
| MEOX1 | TCTTATCAGC | -1 | 966 | 976 | -34 | -24 | 0,94 | 0,01 | 17,9 | >0.1 | 5,61 | 7,78 | 0 | 1,87 | 0 | 1,7 | 0 |
| MEOX1 | CCAAATCAAA | -1 | 749 | 759 | -251 | -241 | 1,85 | 0,01 | 20,38 | >0.1 | 5,61 | 5,72 | 0 | 1,28 | 0 | 5,92 | 0 |
| MEOX1 | GATCATGAAG | -1 | 645 | 655 | -355 | -345 | 3,65 | 0,01 | 17,73 | >0.1 | 3,84 | 5,34 | 0 | 1,06 | 0 | 3,83 | 0 |
| MEOX1 | TTTCATTAAT | -1 | 603 | 613 | -397 | -387 | 3,41 | 0,01 | 14,55 | >0.1 | 3,49 | 5,22 | 0 | 1,02 | 0 | 1,41 | 0 |
| MEOX1 | TCTGATCAGT | -1 | 478 | 488 | -522 | -512 | 1,52 | 0,01 | 15,66 | >0.1 | 2,88 | 4,92 | 0 | 0,99 | 0 | 5,36 | 0 |
| MEOX1 | GCTCACTGCA | -1 | 395 | 405 | -605 | -595 | 1,28 | 0,01 | 15,46 | >0.1 | 2,61 | 4,76 | 0 | 1,04 | 0 | 5,76 | 0 |
| MEOX1 | TCTGATCAGT | -1 | 163 | 173 | -837 | -827 | 1,52 | 0,01 | 13,46 | >0.1 | 3,21 | 4,42 | 0 | 1,9 | 0 | 2,4 | 0 |
| MEOX1 | GCTCATTTCC | -1 | 64 | 74 | -936 | -926 | 2,89 | 0,01 | 13,03 | >0.1 | 3,95 | 4,31 | 0 | 1,88 | 0 | 0 | 0 |
| MEOX2 | GGAAATGAGC | 1 | 64 | 74 | -936 | -926 | 4,86 | 0,01 | 15 | >0.1 | 3,95 | 4,31 | 0 | 1,88 | 0 | 0 | 0 |
| MEOX2 | AGTCATAATT | 1 | 128 | 138 | -872 | -862 | 4,37 | 0,01 | 15,63 | >0.1 | 3,41 | 4,38 | 0 | 1,89 | 0 | 1,57 | 0 |
| MEOX2 | GGCCATGGAA | 1 | 147 | 157 | -853 | -843 | 4,14 | 0,01 | 15,35 | >0.1 | 3,3 | 4,4 | 0 | 1,9 | 0 | 1,61 | 0 |
| MEOX2 | GGAAATTAAA | 1 | 153 | 163 | -847 | -837 | 6,59 | 0,01 | 17,81 | >0.1 | 3,27 | 4,41 | 0 | 1,9 | 0 | 1,65 | 0 |
| MEOX2 | ACTGATCAGA | 1 | 163 | 173 | -837 | -827 | 4,9 | 0,01 | 16,84 | >0.1 | 3,21 | 4,42 | 0 | 1,9 | 0 | 2,4 | 0 |
| MEOX2 | ACTGATCAGA | 1 | 478 | 488 | -522 | -512 | 4,9 | 0,01 | 19,04 | >0.1 | 2,88 | 4,92 | 0 | 0,99 | 0 | 5,36 | 0 |
| MEOX2 | ATTAATGAAA | 1 | 603 | 613 | -397 | -387 | 5,66 | 0,01 | 16,8 | >0.1 | 3,49 | 5,22 | 0 | 1,02 | 0 | 1,41 | 0 |
| MEOX2 | TGAAATTGTC | 1 | 608 | 618 | -392 | -382 | 3,93 | 0,01 | 15,26 | >0.1 | 3,53 | 5,23 | 0 | 1,02 | 0 | 1,55 | 0 |
| MEOX2 | CTTCATGATC | 1 | 645 | 655 | -355 | -345 | 4,44 | 0,01 | 18,52 | >0.1 | 3,84 | 5,34 | 0 | 1,06 | 0 | 3,83 | 0 |
| MEOX2 | GGCCATAAAG | 1 | 757 | 767 | -243 | -233 | 4,02 | 0,01 | 22,8 | >0.1 | 5,61 | 5,75 | 0 | 1,31 | 0 | 6,1 | 0 |
| MEOX2 | TCTCATGAGG | 1 | 983 | 993 | -17 | -7 | 5,99 | 0,01 | 23,54 | >0.1 | 5,61 | 8,5 | 0 | 1,87 | 0 | 1,57 | 0 |
| MEOX2 | AGTCATTCAG | 1 | 1011 | 1021 | 11 | 21 | 4,33 | 0,01 | 22,15 | >0.1 | 5,61 | 8,95 | 0 | 1,86 | 0 | 1,39 | 0 |

| | | | | | | | | | | | | | | | | | |
|---|---|---|---|---|---|---|---|---|---|---|---|---|---|---|---|---|---|
| MEOX2 | CCTCATGAGA | -1 | 983 | 993 | -17 | -7 | 5,61 | 0,01 | 23,16 | >0.1 | 5,61 | 8,5 | 0 | 1,87 | 0 | 1,57 | 0 |
| MEOX2 | CCAAATCAAA | -1 | 749 | 759 | -251 | -241 | 3,52 | 0,01 | 22,05 | >0.1 | 5,61 | 5,72 | 0 | 1,28 | 0 | 5,92 | 0 |
| MEOX2 | GATCATGAAG | -1 | 645 | 655 | -355 | -345 | 5,68 | 0,01 | 19,76 | >0.1 | 3,84 | 5,34 | 0 | 1,06 | 0 | 3,83 | 0 |
| MEOX2 | TTTCATTAAT | -1 | 603 | 613 | -397 | -387 | 5,65 | 0,01 | 16,79 | >0.1 | 3,49 | 5,22 | 0 | 1,02 | 0 | 1,41 | 0 |
| MEOX2 | TCTGATCAGT | -1 | 478 | 488 | -522 | -512 | 4,04 | 0,01 | 18,18 | >0.1 | 2,88 | 4,92 | 0 | 0,99 | 0 | 5,36 | 0 |
| MEOX2 | TCTGATCAGT | -1 | 163 | 173 | -837 | -827 | 4,04 | 0,01 | 15,98 | >0.1 | 3,21 | 4,42 | 0 | 1,9 | 0 | 2,4 | 0 |
| MEOX2 | GCTCATTTCC | -1 | 64 | 74 | -936 | -926 | 4,19 | 0,01 | 14,33 | >0.1 | 3,95 | 4,31 | 0 | 1,88 | 0 | 0 | 0 |
| EWSR1-FLI1 | GTAAGTGAAGGAGAAGGA | 1 | 5 | 23 | -995 | -977 | -19,85 | 0,01 | -6,42 | >0.1 | 5,51 | 4,27 | 0 | 1,86 | 1,79 | 0 | 0 |
| EWSR1-FLI1 | GAAGGAGAAGGAGGCACT | 1 | 11 | 29 | -989 | -971 | -20,85 | 0,01 | -9,2 | >0.1 | 5,51 | 4,27 | 0 | 1,87 | 0 | 0 | 0 |
| EWSR1-FLI1 | GAGAAGGAGGCACTCAAA | 1 | 15 | 33 | -985 | -967 | -25,1 | 0,01 | -13,45 | >0.1 | 5,51 | 4,28 | 0 | 1,87 | 0 | 0 | 0 |
| EWSR1-FLI1 | AGAAGGAGGCACTCAAAG | 1 | 16 | 34 | -984 | -966 | -11,83 | 0,01 | -0,18 | >0.1 | 5,51 | 4,28 | 0 | 1,87 | 0 | 0 | 0 |
| EWSR1-FLI1 | GGAGGCACTCAAAGAAGT | 1 | 20 | 38 | -980 | -962 | -15,32 | 0,01 | -3,66 | >0.1 | 5,51 | 4,28 | 0 | 1,87 | 0 | 0 | 0 |
| EWSR1-FLI1 | GAGGCTGAGGCAGGAGAA | 1 | 352 | 370 | -648 | -630 | -22,07 | 0,01 | -7,61 | >0.1 | 2,52 | 4,7 | 0 | 1,1 | 0 | 6,13 | 0 |
| EWSR1-FLI1 | GAATGGCGTGAACCTGGG | 1 | 367 | 385 | -633 | -615 | -24,06 | 0,01 | -9,31 | >0.1 | 2,55 | 4,73 | 0 | 1,08 | 0 | 6,4 | 0 |
| EWSR1-FLI1 | GGAGGCAGAGCTTGCAGT | 1 | 383 | 401 | -617 | -599 | -25,93 | 0,01 | -11,4 | >0.1 | 2,59 | 4,75 | 0 | 1,06 | 0 | 6,13 | 0 |
| EWSR1-FLI1 | GCAACAAGGCTGAGAAGG | -1 | 1069 | 1087 | 69 | 87 | -25,05 | 0,01 | -6,43 | >0.1 | 5,51 | 7,19 | 0 | 1,85 | 1,79 | 2,27 | 0 |
| EWSR1-FLI1 | GGCTGAGAAGGAGCCAGG | -1 | 1062 | 1080 | 62 | 80 | -17,08 | 0,01 | 1,67 | >0.1 | 5,51 | 7,31 | 0 | 1,86 | 1,79 | 2,27 | 0 |
| EWSR1-FLI1 | GAGAAGGAGCCAGGAAGA | -1 | 1058 | 1076 | 58 | 76 | -2,91 | 0,001 | 15,85 | >0.1 | 5,51 | 7,39 | 0 | 1,86 | 1,79 | 2,21 | 0 |
| EWSR1-FLI1 | AGAAGGAGCCAGGAAGAG | -1 | 1057 | 1075 | 57 | 75 | -20,32 | 0,01 | -1,59 | >0.1 | 5,51 | 7,42 | 0 | 1,86 | 1,79 | 2,16 | 0 |
| EWSR1-FLI1 | GGGCGCTGAGCAGGAAAG | -1 | 933 | 951 | -67 | -49 | -19,18 | 0,01 | -3,66 | >0.1 | 5,51 | 7,24 | 0 | 1,88 | 0 | 0,9 | 0 |
| EWSR1-FLI1 | GGAAAGACCAAAAAAAGA | -1 | 921 | 939 | -79 | -61 | -22,33 | 0,01 | -6,83 | >0.1 | 5,51 | 7,05 | 0 | 1,88 | 0 | 1,06 | 0 |
| EWSR1-FLI1 | AAAAGAAAAAGAAGAAGA | -1 | 909 | 927 | -91 | -73 | -24,49 | 0,01 | -9,05 | >0.1 | 5,51 | 6,89 | 0 | 1,89 | 0 | 1,15 | 0 |
| EWSR1-FLI1 | GAAAAAGAAGAAGAACAC | -1 | 905 | 923 | -95 | -77 | -25,14 | 0,01 | -9,69 | >0.1 | 5,51 | 6,85 | 0 | 1,89 | 0 | 1,21 | 0 |
| STAT6 | TAATTTTAAAAAATG | 1 | 133 | 148 | -867 | -852 | -3,33 | 0,01 | 7,9 | >0.1 | 3,37 | 4,39 | 0 | 1,89 | 0 | 1,57 | 0 |
| STAT6 | GAATTCCATAAAGAC | 1 | 493 | 508 | -507 | -492 | -3,99 | 0,01 | 11,52 | >0.1 | 2,94 | 4,96 | 0 | 0,99 | 0 | 6,62 | 0 |
| STAT6 | GAGTTTCTGGGAATA | 1 | 554 | 569 | -446 | -431 | 8,97 | 0,001 | 18,28 | >0.1 | 3,22 | 5,1 | 0 | 1 | 0 | 0 | 0 |
| STAT6 | ACCTATTAATGAAAT | 1 | 599 | 614 | -401 | -386 | -2,52 | 0,01 | 8,62 | >0.1 | 3,48 | 5,22 | 0 | 1,02 | 0 | 1,41 | 0 |
| STAT6 | AGCTTTCTAGGAAAA | 1 | 714 | 729 | -286 | -271 | 6,74 | 0,001 | 24,25 | >0.1 | 5,54 | 5,6 | 0 | 1,2 | 0 | 5,17 | 0 |
| STAT6 | GCTTTCTAGGAAAAT | 1 | 715 | 730 | -285 | -270 | 7,79 | 0,001 | 25,3 | >0.1 | 5,54 | 5,6 | 0 | 1,2 | 0 | 5,17 | 0 |
| STAT6 | AAATTATTCTAAAAT | 1 | 831 | 846 | -169 | -154 | -0,86 | 0,01 | 16,05 | >0.1 | 5,54 | 6,2 | 0 | 1,74 | 0 | 3,43 | 0 |
| STAT6 | TTAGTCTAGGGAAAG | 1 | 998 | 1013 | -2 | 13 | -2,25 | 0,01 | 15,75 | >0.1 | 5,54 | 9,12 | 0 | 1,87 | 0 | 1,46 | 0 |
| STAT6 | CATTTTTGGACAAGT | 1 | 1128 | 1143 | 128 | 143 | -3,59 | 0,01 | 16,36 | >0.1 | 5,54 | 6,48 | 0 | 1,84 | 3,22 | 2,86 | 0 |
| STAT6 | CTGTTCTATCAAAGT | 1 | 1163 | 1178 | 163 | 178 | -2,8 | 0,01 | 14,24 | >0.1 | 4,52 | 6,2 | 0 | 1,73 | 1,79 | 2,79 | 0 |
| STAT6 | TAATTCCAAGAAGCA | -1 | 1183 | 1198 | 183 | 198 | -4,43 | 0,01 | 14,17 | >0.1 | 5,54 | 6,07 | 0 | 1,56 | 1,79 | 3,63 | 0 |
| STAT6 | CACATCCACTGAATG | -1 | 1014 | 1029 | 14 | 29 | -0,19 | 0,01 | 17,44 | >0.1 | 5,54 | 8,86 | 0 | 1,86 | 0 | 1,36 | 0 |
| STAT6 | GATTTTAGAATAATT | -1 | 832 | 847 | -168 | -153 | -3,24 | 0,01 | 13,54 | >0.1 | 5,54 | 6,21 | 0 | 1,75 | 0 | 3,28 | 0 |
| STAT6 | TTTTTAAAGTGAATC | -1 | 818 | 833 | -182 | -167 | 4,22 | 0,01 | 21,21 | >0.1 | 5,54 | 6,11 | 0 | 1,64 | 0 | 3,7 | 0 |
| STAT6 | TAATCCTAGAGAACC | -1 | 782 | 797 | -218 | -203 | 0,54 | 0,01 | 19,33 | >0.1 | 5,54 | 5,9 | 0 | 1,43 | 0 | 5,92 | 0 |
| STAT6 | TACTTTTGGTTAATA | -1 | 729 | 744 | -271 | -256 | -1,87 | 0,01 | 16,26 | >0.1 | 5,54 | 5,66 | 0 | 1,24 | 0 | 5,69 | 0 |
| STAT6 | TATTTTCCTAGAAAG | -1 | 716 | 731 | -284 | -269 | 1,79 | 0,01 | 19,13 | >0.1 | 5,54 | 5,6 | 0 | 1,21 | 0 | 4,99 | 0 |
| STAT6 | ATTTTCCTAGAAAGC | -1 | 715 | 730 | -285 | -270 | 6,26 | 0,001 | 23,77 | >0.1 | 5,54 | 5,6 | 0 | 1,2 | 0 | 5,17 | 0 |
| STAT6 | AAGATACACGGAAAA | -1 | 695 | 710 | -305 | -290 | -4,5 | 0,01 | 10,91 | >0.1 | 5,54 | 5,52 | 0 | 1,16 | 0 | 3,19 | 0 |
| STAT6 | ATTTTACAGTGACCA | -1 | 673 | 688 | -327 | -312 | -4,29 | 0,01 | 9,92 | >0.1 | 4,22 | 5,45 | 0 | 1,11 | 0 | 3,43 | 0 |
| STAT6 | GGTTTTCAGACAATT | -1 | 611 | 626 | -389 | -374 | -2,68 | 0,01 | 8,67 | >0.1 | 3,57 | 5,25 | 0 | 1,03 | 0 | 1,5 | 0 |
| STAT6 | ATATTCCCAGAAACT | -1 | 555 | 570 | -445 | -430 | -0,3 | 0,01 | 9,02 | >0.1 | 3,22 | 5,1 | 0 | 1 | 0 | 0 | 0 |
| STAT6 | CTCTACTTGGGCAAC | -1 | 540 | 555 | -460 | -445 | -4,35 | 0,01 | 4,85 | >0.1 | 3,15 | 5,07 | 0 | 0,99 | 0 | 0 | 0 |
| STAT6 | CCCTTCCAAATACAT | -1 | 520 | 535 | -480 | -465 | -3,44 | 0,01 | 8,81 | >0.1 | 3,05 | 5,02 | 0 | 0,99 | 0 | 3,19 | 0 |
| STAT6 | TTTTTTTTTTGAGAC | -1 | 452 | 467 | -548 | -533 | -4,88 | 0,01 | 9,31 | >0.1 | 2,79 | 4,88 | 0 | 1 | 0 | 5,52 | 0 |
| STAT6 | TTTTTTTTGAGACTG | -1 | 450 | 465 | -550 | -535 | -5,2 | 0,01 | 8,84 | >0.1 | 2,79 | 4,87 | 0 | 1 | 0 | 5,38 | 0 |
| STAT6 | CCATTTTTTAAAATT | -1 | 134 | 149 | -866 | -851 | -2,88 | 0,01 | 8,38 | >0.1 | 3,36 | 4,39 | 0 | 1,89 | 0 | 1,61 | 0 |
| STAT4 | ACTCAAAGAAGTCA | 1 | 26 | 40 | -974 | -960 | -0,78 | 0,01 | 9,9 | >0.1 | 4,53 | 4,28 | 0 | 1,87 | 0 | 0 | 0 |
| STAT4 | GAAATAGGAAATGA | 1 | 58 | 72 | -942 | -928 | -1,21 | 0,01 | 8,97 | >0.1 | 4 | 4,31 | 0 | 1,88 | 0 | 0 | 0 |
| STAT4 | CTGATCAGAAATGG | 1 | 164 | 178 | -836 | -822 | 6,17 | 0,001 | 18,16 | >0.1 | 3,2 | 4,43 | 0 | 1,9 | 0 | 2,46 | 0 |
| STAT4 | CTGATCAGAAATAT | 1 | 479 | 493 | -521 | -507 | 3,07 | 0,01 | 17,52 | >0.1 | 2,89 | 4,93 | 0 | 0,99 | 0 | 5,65 | 0 |
| STAT4 | ATTCCATAAAGACA | 1 | 495 | 509 | -505 | -491 | -2,42 | 0,01 | 12,91 | >0.1 | 2,95 | 4,96 | 0 | 0,99 | 0 | 6,44 | 0 |
| STAT4 | TGGAAGGGAAAATG | 1 | 527 | 541 | -473 | -459 | -4,16 | 0,01 | 8,49 | >0.1 | 3,08 | 5,03 | 0 | 0,99 | 0 | 3,55 | 0 |
| STAT4 | TTTCTGGGAATATG | 1 | 557 | 571 | -443 | -429 | 11,74 | 0,001 | 21,07 | >0.1 | 3,23 | 5,11 | 0 | 1 | 0 | 0 | 0 |
| STAT4 | ATTAATGAAATTGT | 1 | 603 | 617 | -397 | -383 | -3,94 | 0,01 | 7,23 | >0.1 | 3,51 | 5,23 | 0 | 1,02 | 0 | 1,41 | 0 |
| STAT4 | TTTCTAGGAAAATA | 1 | 717 | 731 | -283 | -269 | 13,73 | 0,0001 | 31,08 | >0.1 | 5,56 | 5,6 | 0 | 1,2 | 0 | 4,99 | 0 |

| | | | | | | | | | | | | | | | | | |
|---|---|---|---|---|---|---|---|---|---|---|---|---|---|---|---|---|---|
| STAT4 | GTGACAGGAGAGGT | 1 | 766 | 780 | -234 | -220 | -1,75 | 0,01 | 16,9 | >0.1 | 5,56 | 5,82 | 0 | 1,36 | 0 | 5,92 | 0 |
| STAT4 | CTGATAAGAGAGAA | 1 | 967 | 981 | -33 | -19 | -3,91 | 0,01 | 13,16 | >0.1 | 5,56 | 7,94 | 0 | 1,87 | 0 | 1,7 | 0 |
| STAT4 | CTTCTTGGAATTAT | 1 | 1185 | 1199 | 185 | 199 | 8,11 | 0,001 | 26,61 | >0.1 | 5,56 | 6,06 | 0 | 1,55 | 1,79 | 3,55 | 0 |
| STAT4 | ATTCCAAGAAGCAA | -1 | 1182 | 1196 | 182 | 196 | 11,43 | 0,001 | 30,1 | >0.1 | 5,56 | 6,08 | 0 | 1,57 | 1,79 | 3,67 | 0 |
| STAT4 | TTTGATAGAACAGG | -1 | 1162 | 1176 | 162 | 176 | -0,22 | 0,01 | 16,68 | >0.1 | 4,48 | 6,21 | 0 | 1,74 | 1,79 | 2,67 | 0 |
| STAT4 | GTTACAGCAACAAG | -1 | 1079 | 1093 | 79 | 93 | -2,88 | 0,01 | 15,51 | >0.1 | 5,56 | 7,03 | 0 | 1,85 | 1,79 | 2,16 | 0 |
| STAT4 | AGGCTGAGAAGGAG | -1 | 1067 | 1081 | 67 | 81 | -3,7 | 0,01 | 14,94 | >0.1 | 5,56 | 7,22 | 0 | 1,85 | 1,79 | 2,21 | 0 |
| STAT4 | GAGCCAGGAAGAGC | -1 | 1056 | 1070 | 56 | 70 | 5,16 | 0,001 | 22,22 | >0.1 | 5,56 | 7,44 | 0 | 1,86 | 0 | 2,21 | 0 |
| STAT4 | GAGCAGGAAAGACC | -1 | 930 | 944 | -70 | -56 | -4,34 | 0,01 | 11,15 | >0.1 | 5,56 | 7,12 | 0 | 1,88 | 0 | 0,93 | 0 |
| STAT4 | TTTCCTAGAAAGCT | -1 | 714 | 728 | -286 | -272 | 10,66 | 0,001 | 28,17 | >0.1 | 5,56 | 5,59 | 0 | 1,2 | 0 | 5,17 | 0 |
| STAT4 | TTCCTAGAAAGCTG | -1 | 713 | 727 | -287 | -273 | -4,15 | 0,01 | 13,35 | >0.1 | 5,56 | 5,59 | 0 | 1,19 | 0 | 5,17 | 0 |
| STAT4 | ATACACGGAAAATG | -1 | 693 | 707 | -307 | -293 | -1,26 | 0,01 | 13,91 | >0.1 | 5,56 | 5,51 | 0 | 1,15 | 0 | 2,95 | 0 |
| STAT4 | ATTCCCAGAAACTC | -1 | 554 | 568 | -446 | -432 | 5,8 | 0,001 | 15,11 | >0.1 | 3,21 | 5,1 | 0 | 0,99 | 0 | 0 | 0 |
| STAT4 | TTTAATAGAGACGG | -1 | 283 | 297 | -717 | -703 | -5,14 | 0,01 | 8,51 | >0.1 | 2,73 | 4,59 | 0 | 1,25 | 0 | 5,09 | 0 |
| STAT4 | GTGCTAGGATTACA | -1 | 199 | 213 | -801 | -787 | -4,49 | 0,01 | 9,25 | >0.1 | 3,03 | 4,47 | 0 | 1,64 | 0 | 4,59 | 0 |
| STAT3 | GAAATAGGAAA | 1 | 58 | 69 | -942 | -931 | -2,09 | 0,01 | 8,11 | >0.1 | 4,01 | 4,31 | 0 | 1,87 | 0 | 0 | 0 |
| STAT3 | AATTTTAAAAA | 1 | 134 | 145 | -866 | -855 | -4,34 | 0,01 | 6,93 | >0.1 | 3,37 | 4,39 | 0 | 1,89 | 0 | 1,61 | 0 |
| STAT3 | ATTTTAAAAAA | 1 | 135 | 146 | -865 | -854 | -0,27 | 0,01 | 10,95 | >0.1 | 3,37 | 4,39 | 0 | 1,89 | 0 | 1,57 | 0 |
| STAT3 | TTTTAAAAAAT | 1 | 136 | 147 | -864 | -853 | -3,27 | 0,01 | 7,99 | >0.1 | 3,36 | 4,39 | 0 | 1,89 | 0 | 1,61 | 0 |
| STAT3 | CTATTAAAAAT | 1 | 289 | 300 | -711 | -700 | -4,52 | 0,01 | 9,1 | >0.1 | 2,71 | 4,59 | 0 | 1,24 | 0 | 5,09 | 0 |
| STAT3 | GTCTCAAAAAA | 1 | 452 | 463 | -548 | -537 | -4 | 0,01 | 10,03 | >0.1 | 2,79 | 4,87 | 0 | 1 | 0 | 5,38 | 0 |
| STAT3 | GTATTTGGAAG | 1 | 522 | 533 | -478 | -467 | 3,63 | 0,01 | 15,87 | >0.1 | 3,05 | 5,02 | 0 | 0,98 | 0 | 3,19 | 0 |
| STAT3 | TTTCTGGGAAT | 1 | 557 | 568 | -443 | -432 | 15,1 | 0,0001 | 24,41 | >0.1 | 3,22 | 5,1 | 0 | 0,99 | 0 | 0 | 0 |
| STAT3 | GATCTTGAAAT | 1 | 570 | 581 | -430 | -419 | 5,07 | 0,01 | 14,49 | >0.1 | 3,29 | 5,13 | 0 | 1 | 0 | 0 | 0 |
| STAT3 | ATTAATGAAAT | 1 | 603 | 614 | -397 | -386 | -0,24 | 0,01 | 10,91 | >0.1 | 3,5 | 5,22 | 0 | 1,02 | 0 | 1,41 | 0 |
| STAT3 | TTGTCTGAAAA | 1 | 613 | 624 | -387 | -376 | 5,5 | 0,01 | 16,84 | >0.1 | 3,57 | 5,25 | 0 | 1,03 | 0 | 1,5 | 0 |
| STAT3 | GTCACTGTAAA | 1 | 675 | 686 | -325 | -314 | -1,71 | 0,01 | 12,49 | >0.1 | 4,22 | 5,44 | 0 | 1,11 | 0 | 3,43 | 0 |
| STAT3 | TTCTAGGAAAA | 1 | 718 | 729 | -282 | -271 | -0,97 | 0,01 | 16,41 | >0.1 | 5,6 | 5,59 | 0 | 1,2 | 0 | 4,99 | 0 |
| STAT3 | CTTTAAAAAAT | 1 | 824 | 835 | -176 | -165 | -2,54 | 0,01 | 14,53 | >0.1 | 5,6 | 6,12 | 0 | 1,65 | 0 | 3,7 | 0 |
| STAT3 | TATTCTAAAAT | 1 | 835 | 846 | -165 | -154 | -3,64 | 0,01 | 13,17 | >0.1 | 5,6 | 6,2 | 0 | 1,74 | 0 | 3,28 | 0 |
| STAT3 | GTCTAGGGAAA | 1 | 1001 | 1012 | 1 | 12 | 1,97 | 0,01 | 20,02 | >0.1 | 5,6 | 9,12 | 0 | 1,87 | 0 | 1,46 | 0 |
| STAT3 | ATTCCAAGAAG | -1 | 1185 | 1196 | 185 | 196 | 7,61 | 0,001 | 26,23 | >0.1 | 5,6 | 6,06 | 0 | 1,55 | 1,79 | 3,63 | 0 |
| STAT3 | GAGCCAGGAAG | -1 | 1059 | 1070 | 59 | 70 | 7,48 | 0,001 | 24,47 | >0.1 | 5,6 | 7,37 | 0 | 1,86 | 0 | 2,16 | 0 |
| STAT3 | GAGCAGGAAAG | -1 | 933 | 944 | -67 | -56 | 2,52 | 0,01 | 18,04 | >0.1 | 5,6 | 7,12 | 0 | 1,88 | 0 | 0,93 | 0 |
| STAT3 | GATTTTAGAAT | -1 | 836 | 847 | -164 | -153 | -0,46 | 0,01 | 16,28 | >0.1 | 5,6 | 6,21 | 0 | 1,75 | 0 | 3,19 | 0 |
| STAT3 | TTTCCTAGAAA | -1 | 717 | 728 | -283 | -272 | 10,34 | 0,001 | 27,89 | >0.1 | 5,6 | 5,59 | 0 | 1,2 | 0 | 5,17 | 0 |
| STAT3 | TTCCTAGAAAG | -1 | 716 | 727 | -284 | -273 | 5,6 | 0,01 | 23,14 | >0.1 | 5,6 | 5,59 | 0 | 1,19 | 0 | 5,17 | 0 |
| STAT3 | TACACGGAAAA | -1 | 695 | 706 | -305 | -294 | -2,17 | 0,01 | 13,03 | >0.1 | 5,6 | 5,51 | 0 | 1,15 | 0 | 2,95 | 0 |
| STAT3 | TTCCCAGAAAC | -1 | 556 | 567 | -444 | -433 | 2,58 | 0,01 | 11,88 | >0.1 | 3,22 | 5,1 | 0 | 0,99 | 0 | 0 | 0 |
| STAT3 | CTTTATGGAAT | -1 | 495 | 506 | -505 | -494 | 5,81 | 0,01 | 21,31 | >0.1 | 2,94 | 4,96 | 0 | 0,98 | 0 | 6,62 | 0 |
| STAT3 | TTTTTTAAAAT | -1 | 135 | 146 | -865 | -854 | 1,49 | 0,01 | 12,71 | >0.1 | 3,37 | 4,39 | 0 | 1,89 | 0 | 1,57 | 0 |
| STAT1 | ACTCAAAGAAG | 1 | 26 | 37 | -974 | -963 | -1,11 | 0,01 | 9,6 | >0.1 | 4,57 | 4,28 | 0 | 1,87 | 0 | 0 | 0 |
| STAT1 | ACGCCTGTAAT | 1 | 194 | 205 | -806 | -795 | -4,16 | 0,01 | 9,3 | >0.1 | 3,06 | 4,46 | 0 | 1,67 | 0 | 4,26 | 0 |
| STAT1 | CTTGCAGTGAG | 1 | 393 | 404 | -607 | -596 | -6,43 | 0,01 | 7,99 | >0.1 | 2,61 | 4,76 | 0 | 1,04 | 0 | 6,02 | 0 |
| STAT1 | TCTCAAAAAAA | 1 | 453 | 464 | -547 | -536 | -5,35 | 0,01 | 8,68 | >0.1 | 2,79 | 4,87 | 0 | 1 | 0 | 5,38 | 0 |
| STAT1 | TTTCTGGGAAT | 1 | 557 | 568 | -443 | -432 | 15,33 | 0,0001 | 24,64 | >0.1 | 3,22 | 5,1 | 0 | 0,99 | 0 | 0 | 0 |
| STAT1 | CTTGAAATAAA | 1 | 573 | 584 | -427 | -416 | -7,03 | 0,01 | 2,42 | >0.1 | 3,31 | 5,14 | 0 | 1 | 0 | 0 | 0 |
| STAT1 | ATTAATGAAAT | 1 | 603 | 614 | -397 | -386 | -4,95 | 0,01 | 6,2 | >0.1 | 3,5 | 5,22 | 0 | 1,02 | 0 | 1,41 | 0 |
| STAT1 | GTCACTGTAAA | 1 | 675 | 686 | -325 | -314 | -3,39 | 0,01 | 10,81 | >0.1 | 4,22 | 5,44 | 0 | 1,11 | 0 | 3,43 | 0 |
| STAT1 | GTGACAGGAGA | 1 | 766 | 777 | -234 | -223 | -5,25 | 0,01 | 13,41 | >0.1 | 5,6 | 5,8 | 0 | 1,35 | 0 | 5,92 | 0 |
| STAT1 | TCTCTAGGATT | 1 | 785 | 796 | -215 | -204 | -4,83 | 0,01 | 13,78 | >0.1 | 5,6 | 5,9 | 0 | 1,43 | 0 | 5,69 | 0 |
| STAT1 | TCTCCAGGATG | 1 | 863 | 874 | -137 | -126 | -3,4 | 0,01 | 13,16 | >0.1 | 5,6 | 6,4 | 0 | 1,9 | 0 | 2,67 | 0 |
| STAT1 | GTTGCTGTAAC | 1 | 1082 | 1093 | 82 | 93 | -2,2 | 0,01 | 16,24 | >0.1 | 5,6 | 6,99 | 0 | 1,85 | 1,79 | 2,21 | 0 |
| STAT1 | ATTCCAAGAAG | -1 | 1185 | 1196 | 185 | 196 | 9,19 | 0,001 | 27,81 | >0.1 | 5,6 | 6,06 | 0 | 1,55 | 1,79 | 3,63 | 0 |
| STAT1 | TTTGATAGAAC | -1 | 1165 | 1176 | 165 | 176 | -6,3 | 0,01 | 10,7 | >0.1 | 4,52 | 6,19 | 0 | 1,71 | 1,79 | 2,79 | 0 |
| STAT1 | GTTACAGCAAC | -1 | 1082 | 1093 | 82 | 93 | 1,9 | 0,01 | 20,34 | >0.1 | 5,6 | 6,99 | 0 | 1,85 | 1,79 | 2,21 | 0 |
| STAT1 | GAGCCAGGAAG | -1 | 1059 | 1070 | 59 | 70 | -2,94 | 0,01 | 14,05 | >0.1 | 5,6 | 7,37 | 0 | 1,86 | 0 | 2,16 | 0 |
| STAT1 | TTTAAAGTGAA | -1 | 820 | 831 | -180 | -169 | -4,12 | 0,01 | 12,89 | >0.1 | 5,6 | 6,1 | 0 | 1,62 | 0 | 3,7 | 0 |
| STAT1 | TTTCCTAGAAA | -1 | 717 | 728 | -283 | -272 | 9,54 | 0,001 | 27,09 | >0.1 | 5,6 | 5,59 | 0 | 1,2 | 0 | 5,17 | 0 |

| | | | | | | | | | | | | | | | | | |
|---|---|---|---|---|---|---|---|---|---|---|---|---|---|---|---|---|---|
| STAT1 | TTCCTAGAAAG | -1 | 716 | 727 | -284 | -273 | -0,58 | 0,01 | 16,96 | >0.1 | 5,6 | 5,59 | 0 | 1,19 | 0 | 5,17 | 0 |
| STAT1 | TTTACAGTGAC | -1 | 675 | 686 | -325 | -314 | -1,52 | 0,01 | 12,68 | >0.1 | 4,22 | 5,44 | 0 | 1,11 | 0 | 3,43 | 0 |
| STAT1 | TTTCAGACAAT | -1 | 612 | 623 | -388 | -377 | -2,26 | 0,01 | 9,02 | >0.1 | 3,56 | 5,24 | 0 | 1,03 | 0 | 1,44 | 0 |
| STAT1 | ATTCCCAGAAA | -1 | 557 | 568 | -443 | -432 | -2,53 | 0,01 | 6,78 | >0.1 | 3,22 | 5,1 | 0 | 0,99 | 0 | 0 | 0 |
| STAT1 | TTCCCAGAAAC | -1 | 556 | 567 | -444 | -433 | -0,9 | 0,01 | 8,4 | >0.1 | 3,22 | 5,1 | 0 | 0,99 | 0 | 0 | 0 |
| STAT1 | CTTTATGGAAT | -1 | 495 | 506 | -505 | -494 | -6,9 | 0,01 | 8,6 | >0.1 | 2,94 | 4,96 | 0 | 0,98 | 0 | 6,62 | 0 |
| STAT1 | GTGCTAGGATT | -1 | 202 | 213 | -798 | -787 | -4,38 | 0,01 | 9,44 | >0.1 | 3,03 | 4,47 | 0 | 1,62 | 0 | 4,7 | 0 |
| FOS::JUND | GTGTGAGAAATAG | 1 | 52 | 65 | -948 | -935 | -2,38 | 0,01 | 7,88 | >0.1 | 4,08 | 4,3 | 0 | 1,87 | 0 | 0 | 0 |
| FOS::JUND | AGAGTAGTCATAA | 1 | 123 | 136 | -877 | -864 | 0,21 | 0,01 | 11,61 | >0.1 | 3,44 | 4,38 | 0 | 1,89 | 0 | 1,7 | 0 |
| FOS::JUND | CAGTGGCTCACGC | 1 | 185 | 198 | -815 | -802 | -2,09 | 0,01 | 11,06 | >0.1 | 3,1 | 4,45 | 0 | 1,75 | 0 | 3,86 | 0 |
| FOS::JUND | TCCTGACTAACAC | 1 | 261 | 274 | -739 | -726 | 2,45 | 0,01 | 15,66 | >0.1 | 2,8 | 4,55 | 0 | 1,33 | 0 | 4,53 | 0 |
| FOS::JUND | GGCTGAGGCAGGA | 1 | 354 | 367 | -646 | -633 | 0,38 | 0,01 | 14,73 | >0.1 | 2,53 | 4,69 | 0 | 1,1 | 0 | 6,03 | 0 |
| FOS::JUND | CCGAGACTCAGTC | 1 | 442 | 455 | -558 | -545 | -4,86 | 0,01 | 9,67 | >0.1 | 2,76 | 4,85 | 0 | 1 | 0 | 5,92 | 0 |
| FOS::JUND | TCTTTGGTCACTG | 1 | 669 | 682 | -331 | -318 | -4,68 | 0,01 | 9,7 | >0.1 | 4,15 | 5,43 | 0 | 1,1 | 0 | 3,7 | 0 |
| FOS::JUND | ATTTGATTCACTT | 1 | 814 | 827 | -186 | -173 | 2,68 | 0,01 | 19,35 | >0.1 | 5,57 | 6,07 | 0 | 1,6 | 0 | 3,43 | 0 |
| FOS::JUND | TCTTGGCTCACAG | 1 | 1030 | 1043 | 30 | 43 | -5,43 | 0,01 | 11,54 | >0.1 | 5,57 | 8,24 | 0 | 1,86 | 0 | 1,31 | 0 |
| FOS::JUND | CTGTTCCTCAATG | -1 | 1108 | 1121 | 108 | 121 | -0,68 | 0,01 | 19,15 | >0.1 | 5,57 | 6,67 | 0 | 1,85 | 3,22 | 2,52 | 0 |
| FOS::JUND | CTGTGAGCCAAGA | -1 | 1030 | 1043 | 30 | 43 | -2,06 | 0,01 | 14,91 | >0.1 | 5,57 | 8,24 | 0 | 1,86 | 0 | 1,31 | 0 |
| FOS::JUND | CACTGAATGACTT | -1 | 1010 | 1023 | 10 | 23 | -5,27 | 0,01 | 12,55 | >0.1 | 5,57 | 8,99 | 0 | 1,86 | 0 | 1,39 | 0 |
| FOS::JUND | TGCTGAGCCAATA | -1 | 881 | 894 | -119 | -106 | -3,34 | 0,01 | 13,76 | >0.1 | 5,57 | 6,57 | 0 | 1,89 | 0 | 3,07 | 0 |
| FOS::JUND | AAGTGAATCAAAT | -1 | 814 | 827 | -186 | -173 | 3,58 | 0,01 | 20,25 | >0.1 | 5,57 | 6,07 | 0 | 1,6 | 0 | 3,43 | 0 |
| FOS::JUND | GACTGAGTCTCGG | -1 | 442 | 455 | -558 | -545 | 0,84 | 0,01 | 15,37 | >0.1 | 2,76 | 4,85 | 0 | 1 | 0 | 5,92 | 0 |
| FOS::JUND | CTCTGCCTCCCAG | -1 | 380 | 393 | -620 | -607 | -5,87 | 0,01 | 8,52 | >0.1 | 2,57 | 4,74 | 0 | 1,06 | 0 | 6,02 | 0 |
| FOS::JUND | TCCTGCCTCAGCC | -1 | 354 | 367 | -646 | -633 | 4,36 | 0,001 | 18,71 | >0.1 | 2,53 | 4,69 | 0 | 1,1 | 0 | 6,03 | 0 |
| FOS::JUND | GTATTTTTAATAG | -1 | 289 | 302 | -711 | -698 | -4,56 | 0,01 | 9,07 | >0.1 | 2,71 | 4,59 | 0 | 1,24 | 0 | 5,09 | 0 |
| FOS::JUND | GTGTTAGTCAGGA | -1 | 261 | 274 | -739 | -726 | 9,21 | 0,001 | 22,42 | >0.1 | 2,8 | 4,55 | 0 | 1,33 | 0 | 4,53 | 0 |
| FOS::JUND | GCGTGAGCCACTG | -1 | 185 | 198 | -815 | -802 | 2,49 | 0,01 | 15,64 | >0.1 | 3,1 | 4,45 | 0 | 1,75 | 0 | 3,86 | 0 |
| FOS::JUND | CTATTTCTCACAC | -1 | 52 | 65 | -948 | -935 | 1,46 | 0,01 | 11,72 | >0.1 | 4,08 | 4,3 | 0 | 1,87 | 0 | 0 | 0 |
| FOS::JUND | GAGTGCCTCCTTC | -1 | 17 | 30 | -983 | -970 | -4,67 | 0,01 | 7,04 | >0.1 | 5,57 | 4,27 | 0 | 1,87 | 0 | 0 | 0 |
| NFIC::TLX1 | AGCCACAGAACCAG | 1 | 39 | 53 | -961 | -947 | -2,31 | 0,01 | 8,12 | >0.1 | 4,27 | 4,29 | 0 | 1,87 | 0 | 0 | 0 |
| NFIC::TLX1 | TGGAGAGTAGTCAT | 1 | 120 | 134 | -880 | -866 | -4,78 | 0,01 | 6,59 | >0.1 | 3,45 | 4,38 | 0 | 1,89 | 0 | 1,65 | 0 |
| NFIC::TLX1 | GGGCACAGTGGCTC | 1 | 180 | 194 | -820 | -806 | -3,13 | 0,01 | 10,08 | >0.1 | 3,12 | 4,45 | 0 | 1,79 | 0 | 3,86 | 0 |
| NFIC::TLX1 | TAGCACTTTGGGAG | 1 | 207 | 221 | -793 | -779 | -2,73 | 0,01 | 11,04 | >0.1 | 3 | 4,48 | 0 | 1,59 | 0 | 4,7 | 0 |
| NFIC::TLX1 | TGGGCAGATCACAA | 1 | 228 | 242 | -772 | -758 | -2,53 | 0,01 | 11,37 | >0.1 | 2,92 | 4,51 | 0 | 1,47 | 0 | 5,01 | 0 |
| NFIC::TLX1 | GGGCAGATCACAAG | 1 | 229 | 243 | -771 | -757 | -4,06 | 0,01 | 9,83 | >0.1 | 2,91 | 4,51 | 0 | 1,46 | 0 | 5,01 | 0 |
| NFIC::TLX1 | TTGCAGTGAGCCGA | 1 | 394 | 408 | -606 | -592 | 6,93 | 0,001 | 21,12 | >0.1 | 2,61 | 4,76 | 0 | 1,04 | 0 | 5,76 | 0 |
| NFIC::TLX1 | AGCCTGGGCGACAG | 1 | 427 | 441 | -573 | -559 | -5 | 0,01 | 9,22 | >0.1 | 2,71 | 4,83 | 0 | 1,02 | 0 | 5,67 | 0 |
| NFIC::TLX1 | GGGCGACAGACCGA | 1 | 432 | 446 | -568 | -554 | -4,37 | 0,01 | 10,01 | >0.1 | 2,73 | 4,83 | 0 | 1,01 | 0 | 5,8 | 0 |
| NFIC::TLX1 | TTCCATAAAGACAA | 1 | 496 | 510 | -504 | -490 | -3,04 | 0,01 | 12,3 | >0.1 | 2,95 | 4,97 | 0 | 0,99 | 0 | 6,44 | 0 |
| NFIC::TLX1 | TGTCTGAAAACCAT | 1 | 614 | 628 | -386 | -372 | -1,22 | 0,01 | 10,16 | >0.1 | 3,59 | 5,26 | 0 | 1,03 | 0 | 1,5 | 0 |
| NFIC::TLX1 | CTGCTCAGCGCCCA | 1 | 938 | 952 | -62 | -48 | -1,71 | 0,01 | 13,81 | >0.1 | 5,56 | 7,25 | 0 | 1,88 | 0 | 0,83 | 0 |
| NFIC::TLX1 | TGGCTCACAGGGGA | 1 | 1033 | 1047 | 33 | 47 | -3,93 | 0,01 | 12,9 | >0.1 | 5,56 | 8,1 | 0 | 1,86 | 0 | 1,31 | 0 |
| NFIC::TLX1 | GGGGACGATGTCAA | 1 | 1042 | 1056 | 42 | 56 | 3,43 | 0,001 | 19,26 | >0.1 | 5,56 | 7,79 | 0 | 1,86 | 0 | 0,62 | 0 |
| NFIC::TLX1 | TGGCTCCTTCTCAG | 1 | 1064 | 1078 | 64 | 78 | 2,44 | 0,01 | 21,2 | >0.1 | 5,56 | 7,28 | 0 | 1,85 | 1,79 | 2,27 | 0 |
| NFIC::TLX1 | GAGGAACAGGCCAA | 1 | 1112 | 1126 | 112 | 126 | 0,79 | 0,01 | 20,61 | >0.1 | 5,56 | 6,63 | 0 | 1,85 | 3,22 | 2,56 | 0 |
| NFIC::TLX1 | AACCACGAAGCCGA | 1 | 1145 | 1159 | 145 | 159 | -3,69 | 0,01 | 15,48 | >0.1 | 5,56 | 6,33 | 0 | 1,84 | 3,22 | 2,21 | 0 |
| NFIC::TLX1 | TTCCAAGAAGCAAG | -1 | 1181 | 1195 | 181 | 195 | -3,8 | 0,01 | 14,73 | >0.1 | 5,56 | 6,08 | 0 | 1,58 | 1,79 | 3,52 | 0 |
| NFIC::TLX1 | TGGCCTGTTCCTCA | -1 | 1111 | 1125 | 111 | 125 | -4,08 | 0,01 | 15,7 | >0.1 | 5,56 | 6,64 | 0 | 1,85 | 3,22 | 2,52 | 0 |
| NFIC::TLX1 | TGAGAAGGAGCCAG | -1 | 1063 | 1077 | 63 | 77 | 3,06 | 0,01 | 21,84 | >0.1 | 5,56 | 7,3 | 0 | 1,85 | 1,79 | 2,27 | 0 |
| NFIC::TLX1 | TGACATCGTCCCCT | -1 | 1041 | 1055 | 41 | 55 | -2,41 | 0,01 | 13,45 | >0.1 | 5,56 | 7,82 | 0 | 1,86 | 0 | 0,62 | 0 |
| NFIC::TLX1 | CCCCTGTGAGCCAA | -1 | 1032 | 1046 | 32 | 46 | -2,48 | 0,01 | 14,39 | >0.1 | 5,56 | 8,15 | 0 | 1,86 | 0 | 1,31 | 0 |
| NFIC::TLX1 | TGGCCAAATCAAAA | -1 | 748 | 762 | -252 | -238 | -4,45 | 0,01 | 14,24 | >0.1 | 5,56 | 5,73 | 0 | 1,29 | 0 | 6,1 | 0 |
| NFIC::TLX1 | TGGTTTTCAGACAA | -1 | 613 | 627 | -387 | -373 | -2,35 | 0,01 | 9,07 | >0.1 | 3,58 | 5,26 | 0 | 1,03 | 0 | 1,55 | 0 |
| NFIC::TLX1 | CGGTCTGTCGCCCA | -1 | 431 | 445 | -569 | -555 | -4,25 | 0,01 | 10,12 | >0.1 | 2,72 | 4,83 | 0 | 1,01 | 0 | 5,8 | 0 |
| NFIC::TLX1 | CGGCTCACTGCAAG | -1 | 393 | 407 | -607 | -593 | -0,38 | 0,01 | 13,81 | >0.1 | 2,61 | 4,76 | 0 | 1,05 | 0 | 5,76 | 0 |
| NFIC::TLX1 | CACCACCACGCCCA | -1 | 313 | 327 | -687 | -673 | -2,12 | 0,01 | 12,45 | >0.1 | 2,64 | 4,63 | 0 | 1,18 | 0 | 6,13 | 0 |
| NFIC::TLX1 | TGTGATCTGCCCAA | -1 | 227 | 241 | -773 | -759 | -1,75 | 0,01 | 12,27 | >0.1 | 2,92 | 4,51 | 0 | 1,47 | 0 | 5,12 | 0 |
| NFIC::TLX1 | TCCCAAAGTGCTAG | -1 | 206 | 220 | -794 | -780 | -3,93 | 0,01 | 9,85 | >0.1 | 3 | 4,48 | 0 | 1,59 | 0 | 4,7 | 0 |
| NFIC::TLX1 | AGCCACTGTGCCCA | -1 | 179 | 193 | -821 | -807 | 2,52 | 0,01 | 15,74 | >0.1 | 3,12 | 4,45 | 0 | 1,8 | 0 | 3,86 | 0 |

| | | | | | | | | | | | | | | | | | |
|---|---|---|---|---|---|---|---|---|---|---|---|---|---|---|---|---|---|
| NFIC::TLX1 | TGACTACTCTCCAC | -1 | 119 | 133 | -881 | -867 | -3,71 | 0,01 | 7,71 | >0.1 | 3,46 | 4,38 | 0 | 1,89 | 0 | 1,7 | 0 |
| HOXA13 | ATTTTAAAAA | 1 | 135 | 145 | -865 | -855 | 5,09 | 0,01 | 16,32 | >0.1 | 3,37 | 4,39 | 0 | 1,89 | 0 | 1,57 | 0 |
| HOXA13 | GAAATTAAAA | 1 | 154 | 164 | -846 | -836 | 0,56 | 0,01 | 11,78 | >0.1 | 3,26 | 4,41 | 0 | 1,9 | 0 | 1,65 | 0 |
| HOXA13 | AAATTAAAAC | 1 | 155 | 165 | -845 | -835 | 0,87 | 0,01 | 13,04 | >0.1 | 3,25 | 4,41 | 0 | 1,9 | 0 | 2,6 | 0 |
| HOXA13 | TCTATTAAAA | 1 | 288 | 298 | -712 | -702 | 1,64 | 0,01 | 15,27 | >0.1 | 2,72 | 4,59 | 0 | 1,24 | 0 | 5,09 | 0 |
| HOXA13 | CTATTAAAAA | 1 | 289 | 299 | -711 | -701 | 10,09 | 0,001 | 23,71 | >0.1 | 2,71 | 4,59 | 0 | 1,24 | 0 | 5,09 | 0 |
| HOXA13 | CTCAAAAAAA | 1 | 454 | 464 | -546 | -536 | 1,49 | 0,01 | 15,67 | >0.1 | 2,79 | 4,87 | 0 | 1 | 0 | 5,52 | 0 |
| HOXA13 | TCCATAAAGA | 1 | 497 | 507 | -503 | -493 | 4,88 | 0,01 | 20,2 | >0.1 | 2,95 | 4,96 | 0 | 0,98 | 0 | 6,44 | 0 |
| HOXA13 | GAAATAAAAA | 1 | 576 | 586 | -424 | -414 | 4,58 | 0,01 | 14,05 | >0.1 | 3,32 | 5,14 | 0 | 1 | 0 | 0 | 0 |
| HOXA13 | CCTATTAATG | 1 | 600 | 610 | -400 | -390 | 3,49 | 0,01 | 14,55 | >0.1 | 3,47 | 5,21 | 0 | 1,02 | 0 | 1,36 | 0 |
| HOXA13 | ACCATACAAA | 1 | 623 | 633 | -377 | -367 | 0,98 | 0,01 | 12,38 | >0.1 | 3,64 | 5,27 | 0 | 1,04 | 0 | 1,44 | 0 |
| HOXA13 | ATTATAACAT | 1 | 686 | 696 | -314 | -304 | 1,19 | 0,01 | 15,16 | >0.1 | 4,42 | 5,47 | 0 | 1,13 | 0 | 2,95 | 0 |
| HOXA13 | ATCTTTAACA | 1 | 705 | 715 | -295 | -285 | 0,72 | 0,01 | 18,02 | >0.1 | 5,61 | 5,54 | 0 | 1,17 | 0 | 4,99 | 0 |
| HOXA13 | GCCATAAAGT | 1 | 758 | 768 | -242 | -232 | 8,34 | 0,001 | 27,12 | >0.1 | 5,61 | 5,76 | 0 | 1,31 | 0 | 6,1 | 0 |
| HOXA13 | GGATTAAAGA | 1 | 791 | 801 | -209 | -199 | 2,88 | 0,01 | 21,18 | >0.1 | 5,61 | 5,92 | 0 | 1,45 | 0 | 5,32 | 0 |
| HOXA13 | ACTTTAAAAA | 1 | 823 | 833 | -177 | -167 | 6,57 | 0,01 | 23,62 | >0.1 | 5,61 | 6,11 | 0 | 1,64 | 0 | 3,7 | 0 |
| HOXA13 | CCTTTGAACT | -1 | 957 | 967 | -43 | -33 | 1,33 | 0,01 | 18,1 | >0.1 | 5,61 | 7,55 | 0 | 1,88 | 0 | 1,74 | 0 |
| HOXA13 | CCAAAAAAAG | -1 | 922 | 932 | -78 | -68 | 2,62 | 0,01 | 18,15 | >0.1 | 5,61 | 6,95 | 0 | 1,88 | 0 | 1,09 | 0 |
| HOXA13 | ACAGTAAACA | -1 | 897 | 907 | -103 | -93 | 0,76 | 0,01 | 17,8 | >0.1 | 5,61 | 6,68 | 0 | 1,89 | 0 | 2,86 | 0 |
| HOXA13 | CAAATCAAAA | -1 | 748 | 758 | -252 | -242 | 2,12 | 0,01 | 20,64 | >0.1 | 5,61 | 5,71 | 0 | 1,28 | 0 | 5,92 | 0 |
| HOXA13 | CTGTTAAAGA | -1 | 706 | 716 | -294 | -284 | 3,86 | 0,01 | 21,17 | >0.1 | 5,61 | 5,55 | 0 | 1,17 | 0 | 4,99 | 0 |
| HOXA13 | ATCATGAAGA | -1 | 644 | 654 | -356 | -346 | 2,21 | 0,01 | 16,14 | >0.1 | 3,83 | 5,34 | 0 | 1,06 | 0 | 3,7 | 0 |
| HOXA13 | CCCAGAAACT | -1 | 555 | 565 | -445 | -435 | 0,81 | 0,01 | 10,1 | >0.1 | 3,21 | 5,09 | 0 | 0,99 | 0 | 0 | 0 |
| HOXA11 | AGTCATAATTTT | 1 | 128 | 140 | -872 | -860 | 7,03 | 0,001 | 18,32 | >0.1 | 3,41 | 4,38 | 0 | 1,89 | 0 | 1,61 | 0 |
| HOXA11 | AATTTTAAAAAA | 1 | 134 | 146 | -866 | -854 | -0,19 | 0,01 | 11,04 | >0.1 | 3,37 | 4,39 | 0 | 1,89 | 0 | 1,57 | 0 |
| HOXA11 | GATCAGAAATGG | 1 | 166 | 178 | -834 | -822 | 2,3 | 0,01 | 14,22 | >0.1 | 3,19 | 4,43 | 0 | 1,9 | 0 | 2,4 | 0 |
| HOXA11 | GCCTGTAATCCT | 1 | 196 | 208 | -804 | -792 | 0,23 | 0,01 | 13,78 | >0.1 | 3,05 | 4,46 | 0 | 1,66 | 0 | 4,37 | 0 |
| HOXA11 | TCTATTAAAAAT | 1 | 288 | 300 | -712 | -700 | 0,45 | 0,01 | 14,08 | >0.1 | 2,71 | 4,59 | 0 | 1,24 | 0 | 5,09 | 0 |
| HOXA11 | TGGCGTGAACCT | 1 | 370 | 382 | -630 | -618 | 3,13 | 0,01 | 17,87 | >0.1 | 2,55 | 4,72 | 0 | 1,07 | 0 | 6,4 | 0 |
| HOXA11 | TCTCAAAAAAAA | 1 | 453 | 465 | -547 | -535 | 3,68 | 0,01 | 17,86 | >0.1 | 2,79 | 4,87 | 0 | 1 | 0 | 5,52 | 0 |
| HOXA11 | GATCAGAAATAT | 1 | 481 | 493 | -519 | -507 | 3,17 | 0,01 | 17,78 | >0.1 | 2,89 | 4,93 | 0 | 0,99 | 0 | 5,8 | 0 |
| HOXA11 | AGTCATGTATTT | 1 | 516 | 528 | -484 | -472 | 1,95 | 0,01 | 13,92 | >0.1 | 3,03 | 5 | 0 | 0,98 | 0 | 2,95 | 0 |
| HOXA11 | GATCTTGAAATA | 1 | 570 | 582 | -430 | -418 | 1,4 | 0,01 | 10,83 | >0.1 | 3,29 | 5,13 | 0 | 1 | 0 | 0 | 0 |
| HOXA11 | CACTGTAAAATT | 1 | 677 | 689 | -323 | -311 | 4,09 | 0,01 | 18,2 | >0.1 | 4,27 | 5,45 | 0 | 1,12 | 0 | 3,28 | 0 |
| HOXA11 | AATTATAACATT | 1 | 685 | 697 | -315 | -303 | 2,27 | 0,01 | 16,24 | >0.1 | 4,42 | 5,48 | 0 | 1,13 | 0 | 2,95 | 0 |
| HOXA11 | TATCTTTAACAG | 1 | 704 | 716 | -296 | -284 | 1,26 | 0,01 | 18,54 | >0.1 | 5,58 | 5,55 | 0 | 1,17 | 0 | 4,99 | 0 |
| HOXA11 | GGCCATAAAGTG | 1 | 757 | 769 | -243 | -231 | 6,92 | 0,001 | 25,69 | >0.1 | 5,58 | 5,76 | 0 | 1,32 | 0 | 6,1 | 0 |
| HOXA11 | GATTGTTTACTG | 1 | 894 | 906 | -106 | -94 | 1,3 | 0,01 | 18,39 | >0.1 | 5,58 | 6,67 | 0 | 1,89 | 0 | 2,95 | 0 |
| HOXA11 | CGTGGTTAAACT | -1 | 1140 | 1152 | 140 | 152 | 1,93 | 0,01 | 22,36 | >0.1 | 5,58 | 6,37 | 0 | 1,84 | 3,22 | 3,4 | 0 |
| HOXA11 | CCTCATGAGATT | -1 | 981 | 993 | -19 | -7 | -0,69 | 0,01 | 16,79 | >0.1 | 5,58 | 8,5 | 0 | 1,87 | 0 | 1,53 | 0 |
| HOXA11 | GACCAAAAAAAG | -1 | 922 | 934 | -78 | -66 | 1,4 | 0,01 | 16,93 | >0.1 | 5,58 | 6,98 | 0 | 1,88 | 0 | 1,09 | 0 |
| HOXA11 | CACAGTAAACAA | -1 | 896 | 908 | -104 | -92 | 0,64 | 0,01 | 17,66 | >0.1 | 5,58 | 6,69 | 0 | 1,89 | 0 | 2,86 | 0 |
| HOXA11 | AATCAAATAAGA | -1 | 810 | 822 | -190 | -178 | -0,18 | 0,01 | 16,44 | >0.1 | 5,58 | 6,04 | 0 | 1,57 | 0 | 3,43 | 0 |
| HOXA11 | AGCTGTTAAAGA | -1 | 706 | 718 | -294 | -282 | 3,58 | 0,01 | 20,7 | >0.1 | 5,58 | 5,55 | 0 | 1,17 | 0 | 4,81 | 0 |
| HOXA11 | TGTTATAATTTT | -1 | 683 | 695 | -317 | -305 | 1,57 | 0,01 | 15,49 | >0.1 | 4,38 | 5,47 | 0 | 1,13 | 0 | 2,95 | 0 |
| HOXA11 | GATCATGAAGAT | -1 | 643 | 655 | -357 | -345 | 5,49 | 0,001 | 19,42 | >0.1 | 3,83 | 5,34 | 0 | 1,06 | 0 | 3,7 | 0 |
| HOXA11 | TTTCATTAATAG | -1 | 601 | 613 | -399 | -387 | -0,34 | 0,01 | 10,8 | >0.1 | 3,49 | 5,22 | 0 | 1,02 | 0 | 1,41 | 0 |
| HOXA11 | GATCATATTCCC | -1 | 562 | 574 | -438 | -426 | 1,25 | 0,01 | 10,61 | >0.1 | 3,25 | 5,11 | 0 | 1 | 0 | 0 | 0 |
| HOXA11 | ACTTAAAAAAGC | -1 | 72 | 84 | -928 | -916 | -0,02 | 0,01 | 10,03 | >0.1 | 3,85 | 4,32 | 0 | 1,88 | 0 | 0 | 0 |
| HOXA11 | GCTCATTTCCTA | -1 | 62 | 74 | -938 | -926 | -0,02 | 0,01 | 10,13 | >0.1 | 3,96 | 4,31 | 0 | 1,88 | 0 | 0 | 0 |
| HOXA10 | GGCACTCAAAG | 1 | 23 | 34 | -977 | -966 | 3,38 | 0,01 | 15,12 | >0.1 | 5,6 | 4,27 | 0 | 1,87 | 0 | 0 | 0 |
| HOXA10 | AGTCATAATTT | 1 | 128 | 139 | -872 | -861 | 4,75 | 0,01 | 16,01 | >0.1 | 3,41 | 4,38 | 0 | 1,89 | 0 | 1,57 | 0 |
| HOXA10 | GGAAATTAAAA | 1 | 153 | 164 | -847 | -836 | 7,08 | 0,01 | 18,3 | >0.1 | 3,26 | 4,41 | 0 | 1,9 | 0 | 1,65 | 0 |
| HOXA10 | CTCTATTAAAA | 1 | 287 | 298 | -713 | -702 | 4,9 | 0,01 | 18,64 | >0.1 | 2,72 | 4,59 | 0 | 1,24 | 0 | 5,19 | 0 |
| HOXA10 | TCTATTAAAAA | 1 | 288 | 299 | -712 | -701 | 4,68 | 0,01 | 18,31 | >0.1 | 2,71 | 4,59 | 0 | 1,24 | 0 | 5,09 | 0 |
| HOXA10 | TCTCAAAAAAA | 1 | 453 | 464 | -547 | -536 | 4,75 | 0,01 | 18,78 | >0.1 | 2,79 | 4,87 | 0 | 1 | 0 | 5,38 | 0 |
| HOXA10 | CTCAAAAAAAA | 1 | 454 | 465 | -546 | -535 | 4,81 | 0,01 | 18,99 | >0.1 | 2,79 | 4,87 | 0 | 1 | 0 | 5,52 | 0 |
| HOXA10 | TTCCATAAAGA | 1 | 496 | 507 | -504 | -493 | 6,72 | 0,01 | 22,04 | >0.1 | 2,94 | 4,96 | 0 | 0,98 | 0 | 6,44 | 0 |
| HOXA10 | TGAAATAAAAA | 1 | 575 | 586 | -425 | -414 | 7,03 | 0,01 | 16,5 | >0.1 | 3,32 | 5,14 | 0 | 1 | 0 | 0 | 0 |

| | | | | | | | | | | | | | | | | | |
|---|---|---|---|---|---|---|---|---|---|---|---|---|---|---|---|---|---|
| HOXA10 | ACCTATTAATG | 1 | 599 | 610 | -401 | -390 | 3,16 | 0,01 | 14,22 | >0.1 | 3,47 | 5,21 | 0 | 1,02 | 0 | 1,36 | 0 |
| HOXA10 | ATTAATGAAAT | 1 | 603 | 614 | -397 | -386 | 3,57 | 0,01 | 14,72 | >0.1 | 3,5 | 5,22 | 0 | 1,02 | 0 | 1,41 | 0 |
| HOXA10 | CACTGTAAAAT | 1 | 677 | 688 | -323 | -312 | 3,44 | 0,01 | 17,68 | >0.1 | 4,26 | 5,45 | 0 | 1,11 | 0 | 3,43 | 0 |
| HOXA10 | AATTATAACAT | 1 | 685 | 696 | -315 | -304 | 3,03 | 0,01 | 16,99 | >0.1 | 4,41 | 5,47 | 0 | 1,13 | 0 | 2,95 | 0 |
| HOXA10 | CGTGGTTAAAC | -1 | 1141 | 1152 | 141 | 152 | 5,27 | 0,01 | 25,7 | >0.1 | 5,6 | 6,37 | 0 | 1,84 | 3,22 | 3,4 | 0 |
| HOXA10 | AGCAGTTACAG | -1 | 1086 | 1097 | 86 | 97 | 4,14 | 0,01 | 22,53 | >0.1 | 5,6 | 6,93 | 0 | 1,85 | 1,79 | 2,21 | 0 |
| HOXA10 | GACCAAAAAAA | -1 | 923 | 934 | -77 | -66 | 6,34 | 0,01 | 21,89 | >0.1 | 5,6 | 6,98 | 0 | 1,88 | 0 | 1,09 | 0 |
| HOXA10 | ACCAAAAAAAG | -1 | 922 | 933 | -78 | -67 | 5,11 | 0,01 | 20,64 | >0.1 | 5,6 | 6,97 | 0 | 1,88 | 0 | 1,09 | 0 |
| HOXA10 | CACAGTAAACA | -1 | 897 | 908 | -103 | -92 | 5,62 | 0,01 | 22,65 | >0.1 | 5,6 | 6,69 | 0 | 1,89 | 0 | 2,86 | 0 |
| HOXA10 | GCCAATATAAA | -1 | 877 | 888 | -123 | -112 | 6,84 | 0,01 | 23,79 | >0.1 | 5,6 | 6,51 | 0 | 1,9 | 0 | 2,95 | 0 |
| HOXA10 | AGCTGTTAAAG | -1 | 707 | 718 | -293 | -282 | 4,39 | 0,01 | 21,52 | >0.1 | 5,6 | 5,55 | 0 | 1,17 | 0 | 4,81 | 0 |
| HOXA10 | GATCATGAAGA | -1 | 644 | 655 | -356 | -345 | 3,53 | 0,01 | 17,47 | >0.1 | 3,84 | 5,34 | 0 | 1,06 | 0 | 3,7 | 0 |
| HOXA10 | TTTCATTAATA | -1 | 602 | 613 | -398 | -387 | 5,05 | 0,01 | 16,19 | >0.1 | 3,49 | 5,22 | 0 | 1,02 | 0 | 1,41 | 0 |
| FOSL1 | TGTGAGAAATA | 1 | 53 | 64 | -947 | -936 | -6,25 | 0,01 | 4,01 | >0.1 | 4,08 | 4,3 | 0 | 1,87 | 0 | 0 | 0 |
| FOSL1 | AGTGGCTCACG | 1 | 186 | 197 | -814 | -803 | 1 | 0,001 | 14,14 | >0.1 | 3,1 | 4,45 | 0 | 1,74 | 0 | 3,86 | 0 |
| FOSL1 | CCTGACTAACA | 1 | 262 | 273 | -738 | -727 | 2,45 | 0,001 | 15,65 | >0.1 | 2,8 | 4,55 | 0 | 1,32 | 0 | 4,53 | 0 |
| FOSL1 | GCTGAGGCAGG | 1 | 355 | 366 | -645 | -634 | -0,44 | 0,001 | 13,91 | >0.1 | 2,53 | 4,69 | 0 | 1,09 | 0 | 6,03 | 0 |
| FOSL1 | TGGGAGGCAGA | 1 | 381 | 392 | -619 | -608 | -14,74 | 0,01 | -0,35 | >0.1 | 2,57 | 4,74 | 0 | 1,06 | 0 | 6,02 | 0 |
| FOSL1 | CGAGACTCAGT | 1 | 443 | 454 | -557 | -546 | -4,13 | 0,01 | 10,4 | >0.1 | 2,76 | 4,85 | 0 | 1 | 0 | 5,92 | 0 |
| FOSL1 | AAAGAATAACG | 1 | 796 | 807 | -204 | -193 | -11,46 | 0,01 | 5 | >0.1 | 5,6 | 5,95 | 0 | 1,48 | 0 | 3,43 | 0 |
| FOSL1 | TTTGATTCACT | 1 | 815 | 826 | -185 | -174 | -7,22 | 0,01 | 9,46 | >0.1 | 5,6 | 6,06 | 0 | 1,59 | 0 | 3,43 | 0 |
| FOSL1 | ATTGGCTCAGC | 1 | 882 | 893 | -118 | -107 | -2,85 | 0,01 | 14,26 | >0.1 | 5,6 | 6,56 | 0 | 1,89 | 0 | 3,07 | 0 |
| FOSL1 | AGTCATTCAGT | 1 | 1011 | 1022 | 11 | 22 | -15,14 | 0,01 | 2,67 | >0.1 | 5,6 | 8,95 | 0 | 1,86 | 0 | 1,39 | 0 |
| FOSL1 | CTTGGCTCACA | 1 | 1031 | 1042 | 31 | 42 | -8,36 | 0,01 | 8,59 | >0.1 | 5,6 | 8,19 | 0 | 1,86 | 0 | 1,31 | 0 |
| FOSL1 | ATTGAGGAACA | 1 | 1109 | 1120 | 109 | 120 | -10,91 | 0,01 | 8,93 | >0.1 | 5,6 | 6,66 | 0 | 1,85 | 3,22 | 2,52 | 0 |
| FOSL1 | AAGGAGCCAGG | -1 | 1062 | 1073 | 62 | 73 | -14,19 | 0,01 | 4,58 | >0.1 | 5,6 | 7,31 | 0 | 1,86 | 1,79 | 2,21 | 0 |
| FOSL1 | TGTGAGCCAAG | -1 | 1031 | 1042 | 31 | 42 | -5,43 | 0,01 | 11,52 | >0.1 | 5,6 | 8,19 | 0 | 1,86 | 0 | 1,31 | 0 |
| FOSL1 | ACTGAATGACT | -1 | 1011 | 1022 | 11 | 22 | -9,74 | 0,01 | 8,07 | >0.1 | 5,6 | 8,95 | 0 | 1,86 | 0 | 1,39 | 0 |
| FOSL1 | GCTGAGCCAAT | -1 | 882 | 893 | -118 | -107 | -8,8 | 0,01 | 8,31 | >0.1 | 5,6 | 6,56 | 0 | 1,89 | 0 | 3,07 | 0 |
| FOSL1 | AGTGAATCAAA | -1 | 815 | 826 | -185 | -174 | 3,95 | 0,001 | 20,63 | >0.1 | 5,6 | 6,06 | 0 | 1,59 | 0 | 3,43 | 0 |
| FOSL1 | AGTGACCAAAG | -1 | 670 | 681 | -330 | -319 | -9,28 | 0,01 | 5,09 | >0.1 | 4,15 | 5,42 | 0 | 1,1 | 0 | 3,7 | 0 |
| FOSL1 | ACTGAGTCTCG | -1 | 443 | 454 | -557 | -546 | -2,94 | 0,01 | 11,59 | >0.1 | 2,76 | 4,85 | 0 | 1 | 0 | 5,92 | 0 |
| FOSL1 | CCTGCCTCAGC | -1 | 355 | 366 | -645 | -634 | -4,3 | 0,01 | 10,05 | >0.1 | 2,53 | 4,69 | 0 | 1,09 | 0 | 6,03 | 0 |
| FOSL1 | TGTTAGTCAGG | -1 | 262 | 273 | -738 | -727 | -0,39 | 0,001 | 12,81 | >0.1 | 2,8 | 4,55 | 0 | 1,32 | 0 | 4,53 | 0 |
| FOSL1 | CGTGAGCCACT | -1 | 186 | 197 | -814 | -803 | -5,35 | 0,01 | 7,79 | >0.1 | 3,1 | 4,45 | 0 | 1,74 | 0 | 3,86 | 0 |
| FOSL1 | TATGACTACTC | -1 | 124 | 135 | -876 | -865 | -5,62 | 0,01 | 5,78 | >0.1 | 3,44 | 4,38 | 0 | 1,89 | 0 | 1,7 | 0 |
| FOSL1 | AGTGCCTCCTT | -1 | 18 | 29 | -982 | -971 | -14,27 | 0,01 | -2,54 | >0.1 | 5,6 | 4,27 | 0 | 1,87 | 0 | 0 | 0 |
| FOSL2 | GTGTGAGAAAT | 1 | 52 | 63 | -948 | -937 | -6,87 | 0,01 | 3,4 | >0.1 | 4,09 | 4,3 | 0 | 1,87 | 0 | 0 | 0 |
| FOSL2 | CAGTGGCTCAC | 1 | 185 | 196 | -815 | -804 | 10,13 | 0,001 | 23,28 | >0.1 | 3,1 | 4,45 | 0 | 1,75 | 0 | 3,86 | 0 |
| FOSL2 | TCCTGACTAAC | 1 | 261 | 272 | -739 | -728 | 2,95 | 0,001 | 16,07 | >0.1 | 2,8 | 4,55 | 0 | 1,32 | 0 | 4,45 | 0 |
| FOSL2 | GGCTGAGGCAG | 1 | 354 | 365 | -646 | -635 | -3,23 | 0,01 | 11,61 | >0.1 | 2,53 | 4,69 | 0 | 1,1 | 0 | 6,52 | 0 |
| FOSL2 | CTGTGCTCCAG | 1 | 418 | 429 | -582 | -571 | -11,72 | 0,01 | 1,97 | >0.1 | 2,68 | 4,8 | 0 | 1,02 | 0 | 5,19 | 0 |
| FOSL2 | CCGAGACTCAG | 1 | 442 | 453 | -558 | -547 | -0,75 | 0,01 | 13,66 | >0.1 | 2,75 | 4,85 | 0 | 1 | 0 | 5,8 | 0 |
| FOSL2 | TCTTTGGTCAC | 1 | 669 | 680 | -331 | -320 | -12,24 | 0,01 | 2,11 | >0.1 | 4,13 | 5,42 | 0 | 1,1 | 0 | 3,7 | 0 |
| FOSL2 | ATTTGATTCAC | 1 | 814 | 825 | -186 | -175 | 1,93 | 0,001 | 18,6 | >0.1 | 5,6 | 6,06 | 0 | 1,58 | 0 | 3,43 | 0 |
| FOSL2 | TATTGGCTCAG | 1 | 881 | 892 | -119 | -108 | 5,13 | 0,001 | 22,24 | >0.1 | 5,6 | 6,55 | 0 | 1,89 | 0 | 3,07 | 0 |
| FOSL2 | AAGTCATTCAG | 1 | 1010 | 1021 | 10 | 21 | -5,21 | 0,01 | 12,66 | >0.1 | 5,6 | 8,99 | 0 | 1,86 | 0 | 1,42 | 0 |
| FOSL2 | TCTTGGCTCAC | 1 | 1030 | 1041 | 30 | 41 | 3,79 | 0,001 | 20,82 | >0.1 | 5,6 | 8,24 | 0 | 1,86 | 0 | 1,34 | 0 |
| FOSL2 | TCCTGGCTCCT | 1 | 1061 | 1072 | 61 | 72 | -10,33 | 0,01 | 8,41 | >0.1 | 5,6 | 7,33 | 0 | 1,86 | 1,79 | 2,16 | 0 |
| FOSL2 | CATTGAGGAAC | 1 | 1108 | 1119 | 108 | 119 | -11,29 | 0,01 | 8,58 | >0.1 | 5,6 | 6,67 | 0 | 1,85 | 3,22 | 2,53 | 0 |
| FOSL2 | GAAGGAGCCAG | -1 | 1063 | 1074 | 63 | 74 | -13,55 | 0,01 | 5,27 | >0.1 | 5,6 | 7,3 | 0 | 1,85 | 1,79 | 2,27 | 0 |
| FOSL2 | TTTTGGTTAAT | -1 | 730 | 741 | -270 | -259 | -6,67 | 0,01 | 11,32 | >0.1 | 5,6 | 5,64 | 0 | 1,23 | 0 | 5,52 | 0 |
| FOSL2 | GACTGAGTCTC | -1 | 444 | 455 | -556 | -545 | -5,51 | 0,01 | 9,03 | >0.1 | 2,76 | 4,85 | 0 | 1 | 0 | 5,92 | 0 |
| FOSL2 | TCTCGGCTCAC | -1 | 399 | 410 | -601 | -590 | -10,41 | 0,01 | 3,93 | >0.1 | 2,62 | 4,77 | 0 | 1,04 | 0 | 5,91 | 0 |
| FOSL2 | CTCTGCCTCCC | -1 | 382 | 393 | -618 | -607 | -13,01 | 0,01 | 1,27 | >0.1 | 2,58 | 4,74 | 0 | 1,06 | 0 | 5,91 | 0 |
| FOSL2 | TCCTGCCTCAG | -1 | 356 | 367 | -644 | -633 | 2,42 | 0,001 | 16,86 | >0.1 | 2,52 | 4,69 | 0 | 1,09 | 0 | 6,13 | 0 |
| FOSL2 | GTGTTAGTCAG | -1 | 263 | 274 | -737 | -726 | -1,77 | 0,01 | 11,34 | >0.1 | 2,8 | 4,55 | 0 | 1,32 | 0 | 4,45 | 0 |
| FOSL2 | GCGTGAGCCAC | -1 | 187 | 198 | -813 | -802 | -0,49 | 0,01 | 12,64 | >0.1 | 3,09 | 4,45 | 0 | 1,73 | 0 | 3,86 | 0 |
| FOSL2 | TTATGACTACT | -1 | 125 | 136 | -875 | -864 | -6,8 | 0,01 | 4,59 | >0.1 | 3,43 | 4,38 | 0 | 1,89 | 0 | 1,7 | 0 |

| | | | | | | | | | | | | | | | | | |
|---|---|---|---|---|---|---|---|---|---|---|---|---|---|---|---|---|---|
| FOSL2 | CTGTGGCTGAC | -1 | 36 | 47 | -964 | -953 | -6,14 | 0,01 | 4,36 | >0.1 | 4,35 | 4,29 | 0 | 1,87 | 0 | 0 | 0 |
| FOSL2 | GAGTGCCTCCT | -1 | 19 | 30 | -981 | -970 | -4,91 | 0,01 | 6,82 | >0.1 | 5,6 | 4,27 | 0 | 1,87 | 0 | 0 | 0 |
| FOSL2::JUNB | AAGGAGGCAC | 1 | 18 | 28 | -982 | -972 | -4,28 | 0,01 | 7,46 | >0.1 | 5,61 | 4,27 | 0 | 1,87 | 0 | 0 | 0 |
| FOSL2::JUNB | TGTGAGAAAT | 1 | 53 | 63 | -947 | -937 | 0,92 | 0,01 | 11,18 | >0.1 | 4,09 | 4,3 | 0 | 1,87 | 0 | 0 | 0 |
| FOSL2::JUNB | GAGTAGTCAT | 1 | 124 | 134 | -876 | -866 | -3,48 | 0,01 | 7,92 | >0.1 | 3,44 | 4,38 | 0 | 1,89 | 0 | 1,7 | 0 |
| FOSL2::JUNB | AGTGGCTCAC | 1 | 186 | 196 | -814 | -804 | 0,91 | 0,01 | 14,05 | >0.1 | 3,1 | 4,45 | 0 | 1,74 | 0 | 3,86 | 0 |
| FOSL2::JUNB | CCTGACTAAC | 1 | 262 | 272 | -738 | -728 | 7,95 | 0,001 | 21,15 | >0.1 | 2,8 | 4,55 | 0 | 1,32 | 0 | 4,53 | 0 |
| FOSL2::JUNB | GCTGAGGCAG | 1 | 355 | 365 | -645 | -635 | 5,23 | 0,001 | 19,58 | >0.1 | 2,53 | 4,69 | 0 | 1,09 | 0 | 6,03 | 0 |
| FOSL2::JUNB | TGGGAGGCAG | 1 | 381 | 391 | -619 | -609 | -5,05 | 0,01 | 9,33 | >0.1 | 2,57 | 4,73 | 0 | 1,06 | 0 | 6,02 | 0 |
| FOSL2::JUNB | CGAGACTCAG | 1 | 443 | 453 | -557 | -547 | -0,13 | 0,01 | 14,4 | >0.1 | 2,75 | 4,85 | 0 | 1 | 0 | 5,92 | 0 |
| FOSL2::JUNB | AAAGAATAAC | 1 | 796 | 806 | -204 | -194 | -5,78 | 0,01 | 10,68 | >0.1 | 5,61 | 5,95 | 0 | 1,47 | 0 | 3,43 | 0 |
| FOSL2::JUNB | TTTGATTCAC | 1 | 815 | 825 | -185 | -175 | 5,27 | 0,001 | 21,95 | >0.1 | 5,61 | 6,06 | 0 | 1,58 | 0 | 3,43 | 0 |
| FOSL2::JUNB | ATTGGCTCAG | 1 | 882 | 892 | -118 | -108 | -5,32 | 0,01 | 11,8 | >0.1 | 5,61 | 6,55 | 0 | 1,89 | 0 | 3,07 | 0 |
| FOSL2::JUNB | AGTCATTCAG | 1 | 1011 | 1021 | 11 | 21 | -3,96 | 0,01 | 13,86 | >0.1 | 5,61 | 8,95 | 0 | 1,86 | 0 | 1,39 | 0 |
| FOSL2::JUNB | CTTGGCTCAC | 1 | 1031 | 1041 | 31 | 41 | -3,54 | 0,01 | 13,43 | >0.1 | 5,61 | 8,19 | 0 | 1,86 | 0 | 1,31 | 0 |
| FOSL2::JUNB | ATTGAGGAAC | 1 | 1109 | 1119 | 109 | 119 | -0,85 | 0,01 | 19,01 | >0.1 | 5,61 | 6,66 | 0 | 1,85 | 3,22 | 2,52 | 0 |
| FOSL2::JUNB | TGTGAGCCAA | -1 | 1032 | 1042 | 32 | 42 | -2,47 | 0,01 | 14,42 | >0.1 | 5,61 | 8,15 | 0 | 1,86 | 0 | 1,28 | 0 |
| FOSL2::JUNB | ACTGAATGAC | -1 | 1012 | 1022 | 12 | 22 | -5,77 | 0,01 | 12,04 | >0.1 | 5,61 | 8,92 | 0 | 1,86 | 0 | 1,42 | 0 |
| FOSL2::JUNB | GCTGAGCCAA | -1 | 883 | 893 | -117 | -107 | -3,98 | 0,01 | 13,15 | >0.1 | 5,61 | 6,56 | 0 | 1,89 | 0 | 3,07 | 0 |
| FOSL2::JUNB | AGTGAATCAA | -1 | 816 | 826 | -184 | -174 | 5,49 | 0,001 | 22,3 | >0.1 | 5,61 | 6,06 | 0 | 1,59 | 0 | 3,55 | 0 |
| FOSL2::JUNB | AGTGACCAAA | -1 | 671 | 681 | -329 | -319 | -5,18 | 0,01 | 9,05 | >0.1 | 4,16 | 5,42 | 0 | 1,1 | 0 | 3,55 | 0 |
| FOSL2::JUNB | ACTGAGTCTC | -1 | 444 | 454 | -556 | -546 | -3,11 | 0,01 | 11,42 | >0.1 | 2,76 | 4,85 | 0 | 1 | 0 | 5,92 | 0 |
| FOSL2::JUNB | CCTGCCTCAG | -1 | 356 | 366 | -644 | -634 | -2,2 | 0,01 | 12,24 | >0.1 | 2,52 | 4,69 | 0 | 1,09 | 0 | 6,13 | 0 |
| FOSL2::JUNB | TGTTAGTCAG | -1 | 263 | 273 | -737 | -727 | 4,84 | 0,001 | 17,95 | >0.1 | 2,8 | 4,55 | 0 | 1,32 | 0 | 4,45 | 0 |
| FOSL2::JUNB | CGTGAGCCAC | -1 | 187 | 197 | -813 | -803 | 0,61 | 0,01 | 13,74 | >0.1 | 3,1 | 4,45 | 0 | 1,73 | 0 | 3,86 | 0 |
| FOSL2::JUNB | TATGACTACT | -1 | 125 | 135 | -875 | -865 | 0,24 | 0,01 | 11,64 | >0.1 | 3,43 | 4,38 | 0 | 1,89 | 0 | 1,7 | 0 |
| FOSL2::JUNB | TATTTCTCAC | -1 | 54 | 64 | -946 | -936 | -4,83 | 0,01 | 5,42 | >0.1 | 4,07 | 4,3 | 0 | 1,87 | 0 | 0 | 0 |
| JUND(VAR.2) | GAGGCTGAGGCAGGA | 1 | 352 | 367 | -648 | -633 | -12,84 | 0,01 | 2,01 | >0.1 | 2,53 | 4,69 | 0 | 1,1 | 0 | 6,52 | 0 |
| JUND(VAR.2) | GAGAATGGCGTGAAC | 1 | 365 | 380 | -635 | -620 | -12,63 | 0,01 | 2,43 | >0.1 | 2,54 | 4,72 | 0 | 1,08 | 0 | 6,73 | 0 |
| JUND(VAR.2) | TGCAGTGAGCCGAGA | 1 | 395 | 410 | -605 | -590 | -1,5 | 0,001 | 12,84 | >0.1 | 2,62 | 4,77 | 0 | 1,04 | 0 | 5,91 | 0 |
| JUND(VAR.2) | AAATGTGAGATAACC | 1 | 587 | 602 | -413 | -398 | 7,28 | 0,001 | 16,89 | >0.1 | 3,4 | 5,19 | 0 | 1,01 | 0 | 0 | 0 |
| JUND(VAR.2) | CTTCATGATCCCTAG | 1 | 645 | 660 | -355 | -340 | -16,56 | 0,01 | -2,43 | >0.1 | 3,87 | 5,35 | 0 | 1,07 | 0 | 3,83 | 0 |
| JUND(VAR.2) | TAAAATTATAACATT | 1 | 682 | 697 | -318 | -303 | -11,91 | 0,01 | 2,03 | >0.1 | 4,39 | 5,48 | 0 | 1,13 | 0 | 2,95 | 0 |
| JUND(VAR.2) | TTTGATTTGGCCATA | 1 | 749 | 764 | -251 | -236 | -11,51 | 0,01 | 7,4 | >0.1 | 5,54 | 5,74 | 0 | 1,3 | 0 | 6,32 | 0 |
| JUND(VAR.2) | TAAAGTGACAGGAGA | 1 | 762 | 777 | -238 | -223 | -13,6 | 0,01 | 5,2 | >0.1 | 5,54 | 5,8 | 0 | 1,35 | 0 | 6,1 | 0 |
| JUND(VAR.2) | ACAGGAGAGGTAAGG | 1 | 769 | 784 | -231 | -216 | -7,09 | 0,01 | 11,77 | >0.1 | 5,54 | 5,84 | 0 | 1,38 | 0 | 6,1 | 0 |
| JUND(VAR.2) | GGGGACGATGTCAAG | 1 | 1042 | 1057 | 42 | 57 | -5,34 | 0,01 | 10,48 | >0.1 | 5,54 | 7,79 | 0 | 1,86 | 0 | 0,62 | 0 |
| JUND(VAR.2) | CCCTGTGAGCCAAGA | -1 | 1030 | 1045 | 30 | 45 | -1,08 | 0,001 | 15,84 | >0.1 | 5,54 | 8,24 | 0 | 1,86 | 0 | 1,28 | 0 |
| JUND(VAR.2) | TCTGCTGAGCCAATA | -1 | 881 | 896 | -119 | -104 | 1,02 | 0,001 | 18,11 | >0.1 | 5,54 | 6,58 | 0 | 1,89 | 0 | 3,07 | 0 |
| JUND(VAR.2) | GACAGATATGTAACA | -1 | 847 | 862 | -153 | -138 | -13,87 | 0,01 | 2,73 | >0.1 | 5,54 | 6,31 | 0 | 1,88 | 0 | 2,86 | 0 |
| JUND(VAR.2) | GAAAATGTTATAATT | -1 | 685 | 700 | -315 | -300 | -5,26 | 0,01 | 8,76 | >0.1 | 4,45 | 5,49 | 0 | 1,14 | 0 | 2,95 | 0 |
| JUND(VAR.2) | TACAGTGACCAAAGA | -1 | 669 | 684 | -331 | -316 | -10,34 | 0,01 | 3,91 | >0.1 | 4,16 | 5,43 | 0 | 1,11 | 0 | 3,55 | 0 |
| JUND(VAR.2) | ATAGGTTATCTCACA | -1 | 590 | 605 | -410 | -395 | 2,35 | 0,001 | 11,98 | >0.1 | 3,42 | 5,19 | 0 | 1,02 | 0 | 0 | 0 |
| JUND(VAR.2) | CACAGTGGCGCGATC | -1 | 408 | 423 | -592 | -577 | -11,25 | 0,01 | 2,88 | >0.1 | 2,65 | 4,79 | 0 | 1,03 | 0 | 5,65 | 0 |
| JUND(VAR.2) | AGGCGTGAGCCACTG | -1 | 185 | 200 | -815 | -800 | -11,21 | 0,01 | 1,94 | >0.1 | 3,09 | 4,45 | 0 | 1,75 | 0 | 3,86 | 0 |
| ZBED1 | TAGTCATAATTTT | 1 | 127 | 140 | -873 | -860 | -17,09 | 0,01 | -5,83 | >0.1 | 3,41 | 4,38 | 0 | 1,89 | 0 | 1,57 | 0 |
| ZBED1 | AAATGGCCATGGA | 1 | 143 | 156 | -857 | -844 | -16,57 | 0,01 | -5,38 | >0.1 | 3,31 | 4,4 | 0 | 1,9 | 0 | 1,57 | 0 |
| ZBED1 | GGCTCACGCCTGT | 1 | 189 | 202 | -811 | -798 | -17,7 | 0,01 | -4,32 | >0.1 | 3,08 | 4,46 | 0 | 1,71 | 0 | 4,13 | 0 |
| ZBED1 | CCGTCCTGACTAA | 1 | 258 | 271 | -742 | -729 | -12,61 | 0,01 | 0,61 | >0.1 | 2,81 | 4,55 | 0 | 1,34 | 0 | 4,53 | 0 |
| ZBED1 | CTAACACGGTGAA | 1 | 267 | 280 | -733 | -720 | -18,1 | 0,01 | -5,01 | >0.1 | 2,78 | 4,56 | 0 | 1,3 | 0 | 4,45 | 0 |
| ZBED1 | AGATCGCGCCACT | 1 | 407 | 420 | -593 | -580 | 0,28 | 0,0001 | 14,39 | >0.1 | 2,65 | 4,79 | 0 | 1,03 | 0 | 5,65 | 0 |
| ZBED1 | AGACCGAGACTCA | 1 | 439 | 452 | -561 | -548 | -13,58 | 0,01 | 1,06 | >0.1 | 2,75 | 4,85 | 0 | 1,01 | 0 | 6,05 | 0 |
| ZBED1 | TGATCTTGAAATA | 1 | 569 | 582 | -431 | -418 | -13,61 | 0,01 | -4,18 | >0.1 | 3,29 | 5,13 | 0 | 1 | 0 | 0 | 0 |
| ZBED1 | AAATTATAACATT | 1 | 684 | 697 | -316 | -303 | -16,68 | 0,01 | -2,72 | >0.1 | 4,41 | 5,48 | 0 | 1,13 | 0 | 2,95 | 0 |
| ZBED1 | CGGTTTTGATTTG | 1 | 745 | 758 | -255 | -242 | -18,65 | 0,01 | -0,17 | >0.1 | 5,57 | 5,71 | 0 | 1,28 | 0 | 5,92 | 0 |
| ZBED1 | ATTTGGCCATAAA | 1 | 753 | 766 | -247 | -234 | -16,25 | 0,01 | 2,7 | >0.1 | 5,57 | 5,75 | 0 | 1,31 | 0 | 6,32 | 0 |
| ZBED1 | ATAAAGTGACAGG | 1 | 761 | 774 | -239 | -226 | -12,74 | 0,01 | 5,87 | >0.1 | 5,57 | 5,79 | 0 | 1,34 | 0 | 5,92 | 0 |
| ZBED1 | ATGTCAAGCTCTT | 1 | 1049 | 1062 | 49 | 62 | -18,4 | 0,01 | -1,91 | >0.1 | 5,57 | 7,6 | 0 | 1,86 | 0 | 1,46 | 0 |
| ZBED1 | AGGCCAAGACATT | 1 | 1119 | 1132 | 119 | 132 | -13,48 | 0,01 | 6,41 | >0.1 | 5,57 | 6,56 | 0 | 1,85 | 3,22 | 2,7 | 0 |

| | | | | | | | | | | | | | | | | | |
|---|---|---|---|---|---|---|---|---|---|---|---|---|---|---|---|---|---|
| ZBED1 | ATGTCTTGGCCTG | -1 | 1118 | 1131 | 118 | 131 | -14,2 | 0,01 | 5,73 | >0.1 | 5,57 | 6,57 | 0 | 1,85 | 3,22 | 2,72 | 0 |
| ZBED1 | CTGTCACTTTATG | -1 | 760 | 773 | -240 | -227 | -16,8 | 0,01 | 1,99 | >0.1 | 5,57 | 5,78 | 0 | 1,33 | 0 | 6,1 | 0 |
| ZBED1 | TTATGGCCAAATC | -1 | 752 | 765 | -248 | -235 | -12,82 | 0,01 | 6,12 | >0.1 | 5,57 | 5,75 | 0 | 1,3 | 0 | 6,32 | 0 |
| ZBED1 | TTTTTGAGACTGA | -1 | 449 | 462 | -551 | -538 | -15,67 | 0,01 | -1,77 | >0.1 | 2,78 | 4,87 | 0 | 1 | 0 | 5,26 | 0 |
| ZBED1 | GAGTCTCGGTCTG | -1 | 438 | 451 | -562 | -549 | -14,98 | 0,01 | -0,34 | >0.1 | 2,74 | 4,84 | 0 | 1,01 | 0 | 6,05 | 0 |
| ZBED1 | CAGTGGCGCGATC | -1 | 408 | 421 | -592 | -579 | -17,09 | 0,01 | -3,1 | >0.1 | 2,65 | 4,79 | 0 | 1,03 | 0 | 5,52 | 0 |
| ZBED1 | GTGGCGCGATCTC | -1 | 406 | 419 | -594 | -581 | -8,25 | 0,001 | 5,86 | >0.1 | 2,65 | 4,78 | 0 | 1,03 | 0 | 5,65 | 0 |
| ZBED1 | CGATCTCGGCTCA | -1 | 400 | 413 | -600 | -587 | -12,37 | 0,01 | 1,84 | >0.1 | 2,63 | 4,77 | 0 | 1,04 | 0 | 5,76 | 0 |
| ZBED1 | GGTTCACGCCATT | -1 | 368 | 381 | -632 | -619 | -11,4 | 0,01 | 3,09 | >0.1 | 2,54 | 4,72 | 0 | 1,08 | 0 | 6,15 | 0 |
| ZBED1 | TAGTCAGGACGGT | -1 | 257 | 270 | -743 | -730 | -17,72 | 0,01 | -4,49 | >0.1 | 2,81 | 4,55 | 0 | 1,34 | 0 | 4,53 | 0 |
| ZBED1 | ATCTCCTGACCTT | -1 | 240 | 253 | -760 | -747 | -14,09 | 0,01 | -0,96 | >0.1 | 2,87 | 4,52 | 0 | 1,41 | 0 | 4,33 | 0 |
| ZBED1 | CCATGGCCATTTT | -1 | 142 | 155 | -858 | -845 | -15,24 | 0,01 | -4,01 | >0.1 | 3,32 | 4,4 | 0 | 1,9 | 0 | 1,61 | 0 |
| ZBED1 | AAATTATGACTAC | -1 | 126 | 139 | -874 | -861 | -16,99 | 0,01 | -5,69 | >0.1 | 3,42 | 4,38 | 0 | 1,89 | 0 | 1,61 | 0 |
| ZBED1 | CTCTCCTGATCTG | -1 | 94 | 107 | -906 | -893 | -13,11 | 0,01 | -2,45 | >0.1 | 3,65 | 4,35 | 0 | 1,88 | 0 | 0,79 | 0 |
| SRF | AAGGCAGATCAGGAGA | 1 | 90 | 106 | -910 | -894 | -14,54 | 0,01 | -3,83 | >0.1 | 3,67 | 4,35 | 0 | 1,88 | 0 | 0,81 | 0 |
| SRF | TGGCCATGGAAATTAA | 1 | 146 | 162 | -854 | -838 | -11,79 | 0,01 | -0,55 | >0.1 | 3,29 | 4,41 | 0 | 1,9 | 0 | 1,65 | 0 |
| SRF | TGATCAGAAATGGCTG | 1 | 165 | 181 | -835 | -819 | -14,98 | 0,01 | -3 | >0.1 | 3,19 | 4,43 | 0 | 1,9 | 0 | 2,46 | 0 |
| SRF | TTGCCCAAGTAGAGAG | 1 | 541 | 557 | -459 | -443 | -16,04 | 0,01 | -6,82 | >0.1 | 3,15 | 5,07 | 0 | 0,99 | 0 | 0 | 0 |
| SRF | TGCCCAAGTAGAGAGT | 1 | 542 | 558 | -458 | -442 | -13,52 | 0,01 | -4,3 | >0.1 | 3,16 | 5,07 | 0 | 0,99 | 0 | 0 | 0 |
| SRF | TCTGGGAATATGATCT | 1 | 559 | 575 | -441 | -425 | -13,78 | 0,01 | -4,42 | >0.1 | 3,25 | 5,12 | 0 | 1 | 0 | 0 | 0 |
| SRF | AAACCATACAAACACC | 1 | 621 | 637 | -379 | -363 | -12,96 | 0,01 | -1,48 | >0.1 | 3,65 | 5,28 | 0 | 1,05 | 0 | 1,5 | 0 |
| SRF | TTTCCGTGTATCTTTA | 1 | 696 | 712 | -304 | -288 | -12,81 | 0,01 | 2,6 | >0.1 | 5,53 | 5,53 | 0 | 1,16 | 0 | 3,19 | 0 |
| SRF | TGGCCATAAAGTGACA | 1 | 756 | 772 | -244 | -228 | -13,24 | 0,01 | 5,5 | >0.1 | 5,53 | 5,78 | 0 | 1,33 | 0 | 6,1 | 0 |
| SRF | GAACTTTATATTGGCT | 1 | 873 | 889 | -127 | -111 | -8,36 | 0,01 | 8,54 | >0.1 | 5,53 | 6,52 | 0 | 1,9 | 0 | 2,95 | 0 |
| SRF | TTTCTTTTTTTGGTCT | 1 | 919 | 935 | -81 | -65 | -10,07 | 0,01 | 5,43 | >0.1 | 5,53 | 6,99 | 0 | 1,88 | 0 | 1,09 | 0 |
| SRF | AAGGCTGATAAGAGAG | 1 | 963 | 979 | -37 | -21 | -15,23 | 0,01 | 1,75 | >0.1 | 5,53 | 7,88 | 0 | 1,87 | 0 | 1,7 | 0 |
| SRF | AAGACATTTTTGGACA | 1 | 1124 | 1140 | 124 | 140 | -16,35 | 0,01 | 3,58 | >0.1 | 5,53 | 6,51 | 0 | 1,84 | 3,22 | 2,82 | 0 |
| SRF | TGTCCAAAAATGTCTT | -1 | 1124 | 1140 | 124 | 140 | -8,32 | 0,01 | 11,61 | >0.1 | 5,53 | 6,51 | 0 | 1,84 | 3,22 | 2,82 | 0 |
| SRF | AGACCAAAAAAAGAAA | -1 | 919 | 935 | -81 | -65 | -5,01 | 0,001 | 10,49 | >0.1 | 5,53 | 6,99 | 0 | 1,88 | 0 | 1,09 | 0 |
| SRF | TGAGCCAATATAAAGT | -1 | 875 | 891 | -125 | -109 | -14,66 | 0,01 | 2,37 | >0.1 | 5,53 | 6,54 | 0 | 1,9 | 0 | 3,07 | 0 |
| SRF | AGCCAATATAAAGTTC | -1 | 873 | 889 | -127 | -111 | -3,12 | 0,001 | 13,78 | >0.1 | 5,53 | 6,52 | 0 | 1,9 | 0 | 2,95 | 0 |
| SRF | AGGACAGATATGTAAC | -1 | 848 | 864 | -152 | -136 | -14,72 | 0,01 | 1,71 | >0.1 | 5,53 | 6,33 | 0 | 1,9 | 0 | 2,67 | 0 |
| SRF | GAATCAAATAAGAATA | -1 | 807 | 823 | -193 | -177 | -15,78 | 0,01 | 0,65 | >0.1 | 5,53 | 6,05 | 0 | 1,57 | 0 | 3,28 | 0 |
| SRF | TGTCACTTTATGGCCA | -1 | 756 | 772 | -244 | -228 | -6,42 | 0,01 | 12,32 | >0.1 | 5,53 | 5,78 | 0 | 1,33 | 0 | 6,1 | 0 |
| SRF | TGGCCAAATCAAAACC | -1 | 746 | 762 | -254 | -238 | -15,45 | 0,01 | 3,02 | >0.1 | 5,53 | 5,73 | 0 | 1,3 | 0 | 5,92 | 0 |
| SRF | TGACCAAAGAGGTCTA | -1 | 663 | 679 | -337 | -321 | -13,16 | 0,01 | 1,27 | >0.1 | 4,09 | 5,42 | 0 | 1,1 | 0 | 3,83 | 0 |
| SRF | GGTGTTTGTATGGTTT | -1 | 621 | 637 | -379 | -363 | -12,34 | 0,01 | -0,86 | >0.1 | 3,65 | 5,28 | 0 | 1,05 | 0 | 1,5 | 0 |
| SRF | TTTCATTAATAGGTTA | -1 | 597 | 613 | -403 | -387 | -14,27 | 0,01 | -3,19 | >0.1 | 3,47 | 5,22 | 0 | 1,02 | 0 | 1,36 | 0 |
| SRF | ACTCTCTACTTGGGCA | -1 | 542 | 558 | -458 | -442 | -7,91 | 0,01 | 1,31 | >0.1 | 3,16 | 5,07 | 0 | 0,99 | 0 | 0 | 0 |
| SRF | CTTCCAAATACATGAC | -1 | 517 | 533 | -483 | -467 | -13,62 | 0,01 | -1,51 | >0.1 | 3,04 | 5,02 | 0 | 0,99 | 0 | 3,07 | 0 |
| SRF | CGCCCAGCTAAGTTTT | -1 | 303 | 319 | -697 | -681 | -11,97 | 0,01 | 1,6 | >0.1 | 2,66 | 4,62 | 0 | 1,2 | 0 | 5,09 | 0 |
| SRF | TCACCGTGTTAGTCAG | -1 | 263 | 279 | -737 | -721 | -14,17 | 0,01 | -0,97 | >0.1 | 2,79 | 4,56 | 0 | 1,32 | 0 | 4,53 | 0 |
| SRF | CTGCCCAACTCGGCCT | -1 | 219 | 235 | -781 | -765 | -15,44 | 0,01 | -0,96 | >0.1 | 2,95 | 4,5 | 0 | 1,51 | 0 | 5,52 | 0 |
| SRF | CAGCCATTTCTGATCA | -1 | 165 | 181 | -835 | -819 | -15,81 | 0,01 | -3,83 | >0.1 | 3,19 | 4,43 | 0 | 1,9 | 0 | 2,46 | 0 |
| FOXD2 | ACTAACA | 1 | 266 | 273 | -734 | -727 | 3,92 | 0,01 | 17,02 | >0.1 | 2,79 | 4,55 | 0 | 1,31 | 0 | 4,45 | 0 |
| FOXD2 | AAAAATA | 1 | 294 | 301 | -706 | -699 | 4,9 | 0,01 | 17,86 | >0.1 | 2,7 | 4,59 | 0 | 1,22 | 0 | 4,45 | 0 |
| FOXD2 | AAAAACT | 1 | 302 | 309 | -698 | -691 | 3,42 | 0,01 | 16,18 | >0.1 | 2,68 | 4,6 | 0 | 1,2 | 0 | 4,28 | 0 |
| FOXD2 | AAAAACT | 1 | 474 | 481 | -526 | -519 | 3,42 | 0,01 | 17,11 | >0.1 | 2,85 | 4,9 | 0 | 0,99 | 0 | 4,95 | 0 |
| FOXD2 | ATAAAGA | 1 | 500 | 507 | -500 | -493 | 4,07 | 0,01 | 19,4 | >0.1 | 2,95 | 4,96 | 0 | 0,98 | 0 | 6,44 | 0 |
| FOXD2 | GAAAATG | 1 | 534 | 541 | -466 | -459 | 3,89 | 0,01 | 14,61 | >0.1 | 3,1 | 5,03 | 0 | 0,98 | 0 | 1,61 | 0 |
| FOXD2 | AAAAATA | 1 | 581 | 588 | -419 | -412 | 4,9 | 0,01 | 14,39 | >0.1 | 3,34 | 5,15 | 0 | 1 | 0 | 0 | 0 |
| FOXD2 | ATAAATG | 1 | 585 | 592 | -415 | -408 | 5,17 | 0,01 | 14,7 | >0.1 | 3,37 | 5,16 | 0 | 1 | 0 | 0 | 0 |
| FOXD2 | ACAAACA | 1 | 628 | 635 | -372 | -365 | 7,88 | 0,001 | 19,41 | >0.1 | 3,67 | 5,28 | 0 | 1,04 | 0 | 1,55 | 0 |
| FOXD2 | ACCAACA | 1 | 634 | 641 | -366 | -359 | 5,64 | 0,01 | 17,31 | >0.1 | 3,73 | 5,3 | 0 | 1,04 | 0 | 1,61 | 0 |
| FOXD2 | GAAAATA | 1 | 724 | 731 | -276 | -269 | 6,45 | 0,01 | 24,07 | >0.1 | 5,65 | 5,6 | 0 | 1,2 | 0 | 5,17 | 0 |
| FOXD2 | GAACACA | -1 | 904 | 911 | -96 | -89 | 3,9 | 0,01 | 19,43 | >0.1 | 5,65 | 6,71 | 0 | 1,89 | 0 | 1,28 | 0 |
| FOXD2 | GTAAACA | -1 | 897 | 904 | -103 | -96 | 11,22 | >0.1 | 28,36 | >0.1 | 5,65 | 6,65 | 0 | 1,89 | 0 | 2,95 | 0 |
| FOXD2 | GCCAATA | -1 | 881 | 888 | -119 | -112 | 5,27 | 0,01 | 22,52 | >0.1 | 5,65 | 6,51 | 0 | 1,89 | 0 | 3,19 | 0 |
| FOXD2 | GTTAATA | -1 | 729 | 736 | -271 | -264 | 5,34 | 0,01 | 23,15 | >0.1 | 5,65 | 5,62 | 0 | 1,21 | 0 | 5,32 | 0 |

| | | | | | | | | | | | | | | | | | |
|---|---|---|---|---|---|---|---|---|---|---|---|---|---|---|---|---|---|
| FOXD2 | GAAAATG | -1 | 693 | 700 | -307 | -300 | 3,89 | 0,01 | 18,12 | >0.1 | 4,54 | 5,49 | 0 | 1,13 | 0 | 3,07 | 0 |
| FOXD2 | ATTAATA | -1 | 602 | 609 | -398 | -391 | 3,78 | 0,01 | 14,84 | >0.1 | 3,48 | 5,2 | 0 | 1,01 | 0 | 1,36 | 0 |
| FOXD2 | GTCAACT | -1 | 105 | 112 | -895 | -888 | 5,58 | 0,01 | 16,17 | >0.1 | 3,58 | 4,35 | 0 | 1,88 | 0 | 0,77 | 0 |
| FOXD3 | CAACATTATCTT | 1 | 636 | 648 | -364 | -352 | 3,12 | 0,01 | 14,87 | >0.1 | 3,77 | 5,32 | 0 | 1,06 | 0 | 1,61 | 0 |
| FOXD3 | TAACGTATTCTT | 1 | 802 | 814 | -198 | -186 | 3,14 | 0,01 | 19,66 | >0.1 | 5,58 | 5,99 | 0 | 1,52 | 0 | 3,43 | 0 |
| FOXD3 | TCTTATTTGATT | 1 | 810 | 822 | -190 | -178 | 6,43 | 0,01 | 23,05 | >0.1 | 5,58 | 6,04 | 0 | 1,57 | 0 | 3,43 | 0 |
| FOXD3 | TTATATTGGCTC | 1 | 878 | 890 | -122 | -110 | 7,36 | 0,01 | 24,56 | >0.1 | 5,58 | 6,53 | 0 | 1,9 | 0 | 3,19 | 0 |
| FOXD3 | GATTGTTTACTG | 1 | 894 | 906 | -106 | -94 | 6,73 | 0,01 | 23,82 | >0.1 | 5,58 | 6,67 | 0 | 1,89 | 0 | 2,95 | 0 |
| FOXD3 | TTTTCTTTTTTT | 1 | 918 | 930 | -82 | -70 | 4,46 | 0,01 | 19,94 | >0.1 | 5,58 | 6,93 | 0 | 1,89 | 0 | 1,09 | 0 |
| FOXD3 | TTTTTTTGGTCT | 1 | 923 | 935 | -77 | -65 | 2,98 | 0,01 | 18,5 | >0.1 | 5,58 | 6,99 | 0 | 1,88 | 0 | 1,06 | 0 |
| FOXD3 | GAATAATTTTTT | -1 | 828 | 840 | -172 | -160 | 5,48 | 0,01 | 22,19 | >0.1 | 5,58 | 6,16 | 0 | 1,69 | 0 | 3,28 | 0 |
| FOXD3 | ATACGTTATTCT | -1 | 798 | 810 | -202 | -190 | 3,48 | 0,01 | 20,08 | >0.1 | 5,58 | 5,97 | 0 | 1,5 | 0 | 3,55 | 0 |
| FOXD3 | TAATATTTTCCT | -1 | 722 | 734 | -278 | -266 | 5,56 | 0,01 | 23,14 | >0.1 | 5,58 | 5,61 | 0 | 1,21 | 0 | 5,17 | 0 |
| FOXD3 | AAATGTTATAAT | -1 | 686 | 698 | -314 | -302 | 3,47 | 0,01 | 17,47 | >0.1 | 4,44 | 5,48 | 0 | 1,13 | 0 | 2,95 | 0 |
| FOXD3 | TAATGTTGGTGT | -1 | 632 | 644 | -368 | -356 | 10,36 | 0,001 | 22,05 | >0.1 | 3,73 | 5,31 | 0 | 1,05 | 0 | 1,61 | 0 |
| FOXD3 | GTTTGTATGGTT | -1 | 622 | 634 | -378 | -366 | 3,34 | 0,01 | 14,74 | >0.1 | 3,64 | 5,28 | 0 | 1,04 | 0 | 1,44 | 0 |
| FOXD3 | TCACATTTATTT | -1 | 583 | 595 | -417 | -405 | 5,11 | 0,01 | 14,65 | >0.1 | 3,37 | 5,17 | 0 | 1,01 | 0 | 0 | 0 |
| FOXD3 | ATTTATTTTTAT | -1 | 579 | 591 | -421 | -409 | 5,46 | 0,01 | 14,97 | >0.1 | 3,35 | 5,16 | 0 | 1 | 0 | 0 | 0 |
| FOXD3 | TTATTTTTATTT | -1 | 577 | 589 | -423 | -411 | 6,89 | 0,01 | 16,38 | >0.1 | 3,33 | 5,15 | 0 | 1 | 0 | 0 | 0 |
| FOXD3 | TATTTTTATTTC | -1 | 576 | 588 | -424 | -412 | 3,87 | 0,01 | 13,35 | >0.1 | 3,33 | 5,15 | 0 | 1 | 0 | 0 | 0 |
| FOXD3 | GATCAGTTTTTT | -1 | 473 | 485 | -527 | -515 | 3,02 | 0,01 | 16,73 | >0.1 | 2,86 | 4,91 | 0 | 0,99 | 0 | 4,95 | 0 |
| FOXD3 | ATCAGTTTTTTT | -1 | 472 | 484 | -528 | -516 | 3,66 | 0,01 | 17,36 | >0.1 | 2,86 | 4,91 | 0 | 0,99 | 0 | 4,95 | 0 |
| FOXD3 | GTTTTTTTTTTT | -1 | 468 | 480 | -532 | -520 | 8,9 | 0,001 | 22,46 | >0.1 | 2,84 | 4,9 | 0 | 0,99 | 0 | 4,83 | 0 |
| FOXD3 | TTTTTTTTTTTT | -1 | 467 | 479 | -533 | -521 | 6,54 | 0,01 | 21,19 | >0.1 | 2,84 | 4,9 | 0 | 0,99 | 0 | 5,92 | 0 |
| FOXD3 | TTTTTTTTTTTT | -1 | 466 | 478 | -534 | -522 | 6,54 | 0,01 | 21,04 | >0.1 | 2,83 | 4,9 | 0 | 0,99 | 0 | 5,78 | 0 |
| FOXD3 | TTTTTTTTTTTT | -1 | 465 | 477 | -535 | -523 | 6,54 | 0,01 | 20,89 | >0.1 | 2,83 | 4,9 | 0 | 0,99 | 0 | 5,63 | 0 |
| FOXD3 | TTTTTTTTTTTT | -1 | 464 | 476 | -536 | -524 | 6,54 | 0,01 | 21,03 | >0.1 | 2,83 | 4,89 | 0 | 0,99 | 0 | 5,78 | 0 |
| FOXD3 | TTTTTTTTTTTT | -1 | 463 | 475 | -537 | -525 | 6,54 | 0,01 | 21,03 | >0.1 | 2,83 | 4,89 | 0 | 0,99 | 0 | 5,78 | 0 |
| FOXD3 | TTTTTTTTTTTT | -1 | 462 | 474 | -538 | -526 | 6,54 | 0,01 | 21,02 | >0.1 | 2,82 | 4,89 | 0 | 0,99 | 0 | 5,78 | 0 |
| FOXD3 | TTTTTTTTTTTT | -1 | 461 | 473 | -539 | -527 | 6,54 | 0,01 | 20,87 | >0.1 | 2,82 | 4,89 | 0 | 0,99 | 0 | 5,63 | 0 |
| FOXD3 | TTTTTTTTTTTT | -1 | 460 | 472 | -540 | -528 | 6,54 | 0,01 | 20,87 | >0.1 | 2,81 | 4,89 | 0 | 0,99 | 0 | 5,63 | 0 |
| FOXD3 | TTTTTTTTTTTT | -1 | 459 | 471 | -541 | -529 | 6,54 | 0,01 | 20,75 | >0.1 | 2,81 | 4,88 | 0 | 1 | 0 | 5,52 | 0 |
| FOXD3 | TTTTTTTTTTTT | -1 | 458 | 470 | -542 | -530 | 6,54 | 0,01 | 20,75 | >0.1 | 2,81 | 4,88 | 0 | 1 | 0 | 5,52 | 0 |
| FOXD3 | TTTTTTTTTTTT | -1 | 457 | 469 | -543 | -531 | 6,54 | 0,01 | 20,85 | >0.1 | 2,8 | 4,88 | 0 | 1 | 0 | 5,63 | 0 |
| FOXD3 | CTAAGTTTTTGT | -1 | 300 | 312 | -700 | -688 | 3,43 | 0,01 | 16,91 | >0.1 | 2,68 | 4,61 | 0 | 1,21 | 0 | 4,99 | 0 |
| FOXD3 | TTTTGTATTTTT | -1 | 294 | 306 | -706 | -694 | 6,61 | 0,01 | 19,57 | >0.1 | 2,69 | 4,6 | 0 | 1,22 | 0 | 4,45 | 0 |
| FOXD1 | TTAAAAAA | 1 | 138 | 146 | -862 | -854 | 4,01 | 0,01 | 15,3 | >0.1 | 3,36 | 4,39 | 0 | 1,89 | 0 | 1,65 | 0 |
| FOXD1 | TTAAAAAT | 1 | 292 | 300 | -708 | -700 | 4,2 | 0,01 | 17,17 | >0.1 | 2,71 | 4,59 | 0 | 1,23 | 0 | 4,45 | 0 |
| FOXD1 | ATAAAAAT | 1 | 579 | 587 | -421 | -413 | 4,2 | 0,01 | 13,68 | >0.1 | 3,33 | 5,15 | 0 | 1 | 0 | 0 | 0 |
| FOXD1 | GTAAAATT | 1 | 681 | 689 | -319 | -311 | 1,88 | 0,01 | 15,93 | >0.1 | 4,3 | 5,45 | 0 | 1,11 | 0 | 3,19 | 0 |
| FOXD1 | GAAAATAT | 1 | 724 | 732 | -276 | -268 | 1,88 | 0,01 | 19,5 | >0.1 | 5,64 | 5,61 | 0 | 1,2 | 0 | 5,17 | 0 |
| FOXD1 | TTAACCAA | 1 | 731 | 739 | -269 | -261 | 5 | 0,01 | 23,02 | >0.1 | 5,64 | 5,63 | 0 | 1,22 | 0 | 5,52 | 0 |
| FOXD1 | TTAAAGAA | 1 | 794 | 802 | -206 | -198 | 1,92 | 0,01 | 20,26 | >0.1 | 5,64 | 5,93 | 0 | 1,45 | 0 | 5,32 | 0 |
| FOXD1 | TTAAAAAA | 1 | 826 | 834 | -174 | -166 | 4,01 | 0,01 | 21,11 | >0.1 | 5,64 | 6,12 | 0 | 1,64 | 0 | 3,7 | 0 |
| FOXD1 | TTAACCAC | 1 | 1143 | 1151 | 143 | 151 | 3,79 | 0,01 | 24,21 | >0.1 | 5,64 | 6,35 | 0 | 1,84 | 3,22 | 3,37 | 0 |
| FOXD1 | TTAAACTT | -1 | 1139 | 1147 | 139 | 147 | 1,96 | 0,01 | 22,18 | >0.1 | 5,64 | 6,38 | 0 | 1,84 | 3,22 | 3,13 | 0 |
| FOXD1 | GTAAACAA | -1 | 896 | 904 | -104 | -96 | 12,77 | 0,0001 | 29,9 | >0.1 | 5,64 | 6,65 | 0 | 1,89 | 0 | 2,95 | 0 |
| FOXD1 | GTTAATAT | -1 | 728 | 736 | -272 | -264 | 1,95 | 0,01 | 19,74 | >0.1 | 5,64 | 5,62 | 0 | 1,22 | 0 | 5,32 | 0 |
| FOXD1 | TTAAAGAT | -1 | 705 | 713 | -295 | -287 | 2,11 | 0,01 | 19,25 | >0.1 | 5,64 | 5,53 | 0 | 1,16 | 0 | 4,81 | 0 |
| FOXD1 | GTGACCAA | -1 | 672 | 680 | -328 | -320 | 3,25 | 0,01 | 17,47 | >0.1 | 4,16 | 5,42 | 0 | 1,1 | 0 | 3,55 | 0 |
| FOXD1 | GGCAACAT | -1 | 538 | 546 | -462 | -454 | 3,29 | 0,01 | 14,1 | >0.1 | 3,12 | 5,05 | 0 | 0,98 | 0 | 1,66 | 0 |
| FOXD1 | TTAAAAAA | -1 | 74 | 82 | -926 | -918 | 4,01 | 0,01 | 14,06 | >0.1 | 3,85 | 4,32 | 0 | 1,88 | 0 | 0 | 0 |
| GMEB2 | ATAGGTAA | 1 | 1 | 9 | -999 | -991 | -0,58 | 0,01 | 12,97 | >0.1 | 5,64 | 4,25 | 0 | 1,86 | 1,79 | 0 | 0 |
| GMEB2 | GTAAGTGA | 1 | 5 | 13 | -995 | -987 | 1,69 | 0,01 | 15,24 | >0.1 | 5,64 | 4,26 | 0 | 1,86 | 1,79 | 0 | 0 |
| GMEB2 | AGAAGTCA | 1 | 32 | 40 | -968 | -960 | 0,98 | 0,01 | 11,59 | >0.1 | 4,46 | 4,28 | 0 | 1,87 | 0 | 0 | 0 |
| GMEB2 | TCACGCCT | 1 | 192 | 200 | -808 | -800 | -1,49 | 0,01 | 11,86 | >0.1 | 3,08 | 4,45 | 0 | 1,69 | 0 | 4,13 | 0 |
| GMEB2 | GGGCGTGG | 1 | 314 | 322 | -686 | -678 | -0,8 | 0,01 | 13,77 | >0.1 | 2,64 | 4,62 | 0 | 1,17 | 0 | 6,13 | 0 |
| GMEB2 | TGGCGTGA | 1 | 370 | 378 | -630 | -622 | 2,65 | 0,01 | 17,12 | >0.1 | 2,54 | 4,71 | 0 | 1,07 | 0 | 6,15 | 0 |
| GMEB2 | TCGCGCCA | 1 | 410 | 418 | -590 | -582 | -0,99 | 0,01 | 12,99 | >0.1 | 2,65 | 4,78 | 0 | 1,03 | 0 | 5,52 | 0 |

| | | | | | | | | | | | | | | | | | |
|---|---|---|---|---|---|---|---|---|---|---|---|---|---|---|---|---|---|
| GMEB2 | CAAAGTCA | 1 | 513 | 521 | -487 | -479 | 0,34 | 0,01 | 12,27 | >0.1 | 3,01 | 4,99 | 0 | 0,98 | 0 | 2,95 | 0 |
| GMEB2 | TTGCCCAA | 1 | 541 | 549 | -459 | -451 | -1,41 | 0,01 | 7,76 | >0.1 | 3,13 | 5,05 | 0 | 0,98 | 0 | 0 | 0 |
| GMEB2 | AAAAGTAC | 1 | 737 | 745 | -263 | -255 | -0,94 | 0,01 | 17,51 | >0.1 | 5,64 | 5,66 | 0 | 1,24 | 0 | 5,92 | 0 |
| GMEB2 | TAAAGTGA | 1 | 762 | 770 | -238 | -230 | -0,92 | 0,01 | 17,91 | >0.1 | 5,64 | 5,77 | 0 | 1,32 | 0 | 6,1 | 0 |
| GMEB2 | AGAGGTAA | 1 | 774 | 782 | -226 | -218 | -1,56 | 0,01 | 17,37 | >0.1 | 5,64 | 5,82 | 0 | 1,37 | 0 | 6,1 | 0 |
| GMEB2 | TAACGTAT | 1 | 802 | 810 | -198 | -190 | 3,68 | 0,01 | 20,21 | >0.1 | 5,64 | 5,97 | 0 | 1,49 | 0 | 3,43 | 0 |
| GMEB2 | CAACCCAA | 1 | 950 | 958 | -50 | -42 | -0,8 | 0,01 | 15,81 | >0.1 | 5,64 | 7,36 | 0 | 1,88 | 0 | 1,74 | 0 |
| GMEB2 | GAAAGTCA | 1 | 1008 | 1016 | 8 | 16 | 0,88 | 0,01 | 18,92 | >0.1 | 5,64 | 9,07 | 0 | 1,87 | 0 | 1,46 | 0 |
| GMEB2 | AGAAGCAA | -1 | 1182 | 1190 | 182 | 190 | -0,66 | 0,01 | 17,85 | >0.1 | 5,64 | 6,08 | 0 | 1,57 | 1,79 | 3,43 | 0 |
| GMEB2 | ATATGTAA | -1 | 849 | 857 | -151 | -143 | -1,08 | 0,01 | 15,62 | >0.1 | 5,64 | 6,28 | 0 | 1,83 | 0 | 2,95 | 0 |
| GMEB2 | TAAAGTGA | -1 | 821 | 829 | -179 | -171 | -0,92 | 0,01 | 16,11 | >0.1 | 5,64 | 6,08 | 0 | 1,61 | 0 | 3,7 | 0 |
| GMEB2 | ATACGTTA | -1 | 802 | 810 | -198 | -190 | 4,94 | 0,001 | 21,47 | >0.1 | 5,64 | 5,97 | 0 | 1,49 | 0 | 3,43 | 0 |
| GMEB2 | TTACCTCT | -1 | 774 | 782 | -226 | -218 | -0,27 | 0,01 | 18,66 | >0.1 | 5,64 | 5,82 | 0 | 1,37 | 0 | 6,1 | 0 |
| GMEB2 | TGGCGCGA | -1 | 410 | 418 | -590 | -582 | -0,35 | 0,01 | 13,63 | >0.1 | 2,65 | 4,78 | 0 | 1,03 | 0 | 5,52 | 0 |
| GMEB2 | TCACGCCA | -1 | 370 | 378 | -630 | -622 | 4,04 | 0,001 | 18,51 | >0.1 | 2,54 | 4,71 | 0 | 1,07 | 0 | 6,15 | 0 |
| GMEB2 | CTGCCTCA | -1 | 357 | 365 | -643 | -635 | -0,94 | 0,01 | 13,5 | >0.1 | 2,52 | 4,69 | 0 | 1,09 | 0 | 6,13 | 0 |
| GMEB2 | CCACGCCC | -1 | 314 | 322 | -686 | -678 | 1,85 | 0,01 | 16,42 | >0.1 | 2,64 | 4,62 | 0 | 1,17 | 0 | 6,13 | 0 |
| GMEB2 | AGGCGTGA | -1 | 192 | 200 | -808 | -800 | -0,16 | 0,01 | 13,19 | >0.1 | 3,08 | 4,45 | 0 | 1,69 | 0 | 4,13 | 0 |
| GMEB2 | GTGCCTCC | -1 | 20 | 28 | -980 | -972 | -1,41 | 0,01 | 10,36 | >0.1 | 5,64 | 4,27 | 0 | 1,87 | 0 | 0 | 0 |
| GMEB2 | TTACCTAT | -1 | 1 | 9 | -999 | -991 | 1,09 | 0,01 | 14,64 | >0.1 | 5,64 | 4,25 | 0 | 1,86 | 1,79 | 0 | 0 |
| TP53 | GTTTTGCAAAGGCAGATC | 1 | 82 | 100 | -918 | -900 | -6,46 | 0,01 | 3,49 | >0.1 | 3,73 | 4,34 | 0 | 1,88 | 0 | 0 | 0 |
| TP53 | GAAATGGCTGGGCACAGT | 1 | 171 | 189 | -829 | -811 | -8,57 | 0,01 | 4,14 | >0.1 | 3,15 | 4,44 | 0 | 1,87 | 0 | 3,24 | 0 |
| TP53 | CAGTGGCTCACGCCTGTA | 1 | 185 | 203 | -815 | -797 | -11,68 | 0,01 | 1,65 | >0.1 | 3,09 | 4,46 | 0 | 1,75 | 0 | 4,04 | 0 |
| TP53 | AACTTAGCTGGGCGTGGT | 1 | 305 | 323 | -695 | -677 | -8,05 | 0,01 | 6,68 | >0.1 | 2,65 | 4,63 | 0 | 1,2 | 0 | 6,26 | 0 |
| TP53 | GGGAGGCTGAGGCAGGAG | 1 | 350 | 368 | -650 | -632 | -11,17 | 0,01 | 3,69 | >0.1 | 2,53 | 4,7 | 0 | 1,11 | 0 | 6,52 | 0 |
| TP53 | ACTGTGCTCCAGCCTGGG | 1 | 417 | 435 | -583 | -565 | -9,15 | 0,01 | 4,8 | >0.1 | 2,69 | 4,81 | 0 | 1,03 | 0 | 5,42 | 0 |
| TP53 | CAAGGGCAAAGTCATGTA | 1 | 507 | 525 | -493 | -475 | -11,34 | 0,01 | 0,51 | >0.1 | 3 | 5 | 0 | 0,99 | 0 | 2,86 | 0 |
| TP53 | AATGTGAGATAACCTATT | 1 | 588 | 606 | -412 | -394 | -12,81 | 0,01 | -3,17 | >0.1 | 3,42 | 5,2 | 0 | 1,02 | 0 | 0 | 0 |
| TP53 | TACAAACACCAACATTAT | 1 | 627 | 645 | -373 | -355 | -11,43 | 0,01 | 0,3 | >0.1 | 3,71 | 5,31 | 0 | 1,06 | 0 | 1,66 | 0 |
| TP53 | GTCAAGCTCTTCCTGGCT | 1 | 1051 | 1069 | 51 | 69 | -8,61 | 0,01 | 8,58 | >0.1 | 5,51 | 7,55 | 0 | 1,86 | 0 | 2,27 | 0 |
| TP53 | GACATTTTTGGACAAGTT | 1 | 1126 | 1144 | 126 | 144 | 1,76 | 0,001 | 21,69 | >0.1 | 5,51 | 6,49 | 0 | 1,84 | 3,22 | 2,86 | 0 |
| TP53 | GACAAGTTTAACCACGAA | 1 | 1136 | 1154 | 136 | 154 | -7,27 | 0,01 | 13,01 | >0.1 | 5,51 | 6,41 | 0 | 1,84 | 3,22 | 3,29 | 0 |
| TP53 | GACCTGTTCTATCAAAGT | 1 | 1160 | 1178 | 160 | 178 | -8,33 | 0,01 | 8,6 | >0.1 | 4,48 | 6,22 | 0 | 1,76 | 1,79 | 2,67 | 0 |
| TP53 | ACTTTGATAGAACAGGTC | -1 | 1160 | 1178 | 160 | 178 | -9,22 | 0,01 | 7,71 | >0.1 | 4,48 | 6,22 | 0 | 1,76 | 1,79 | 2,67 | 0 |
| TP53 | TTCGTGGTTAAACTTGTC | -1 | 1136 | 1154 | 136 | 154 | -0,56 | 0,001 | 19,72 | >0.1 | 5,51 | 6,41 | 0 | 1,84 | 3,22 | 3,29 | 0 |
| TP53 | AACTTGTCCAAAAATGTC | -1 | 1126 | 1144 | 126 | 144 | 2 | 0,001 | 21,93 | >0.1 | 5,51 | 6,49 | 0 | 1,84 | 3,22 | 2,86 | 0 |
| TP53 | AGCCAGGAAGAGCTTGAC | -1 | 1051 | 1069 | 51 | 69 | -6,24 | 0,01 | 10,95 | >0.1 | 5,51 | 7,55 | 0 | 1,86 | 0 | 2,27 | 0 |
| TP53 | TACATGACTTTGCCCTTG | -1 | 507 | 525 | -493 | -475 | -11,09 | 0,01 | 0,76 | >0.1 | 3 | 5 | 0 | 0,99 | 0 | 2,86 | 0 |
| TP53 | CTCCTGCCTCAGCCTCCC | -1 | 350 | 368 | -650 | -632 | -5,87 | 0,01 | 8,99 | >0.1 | 2,53 | 4,7 | 0 | 1,11 | 0 | 6,52 | 0 |
| TP53 | ACCACGCCCAGCTAAGTT | -1 | 305 | 323 | -695 | -677 | -9,88 | 0,01 | 4,85 | >0.1 | 2,65 | 4,63 | 0 | 1,2 | 0 | 6,26 | 0 |
| TP53 | TACAGGCGTGAGCCACTG | -1 | 185 | 203 | -815 | -797 | -10,1 | 0,01 | 3,23 | >0.1 | 3,09 | 4,46 | 0 | 1,75 | 0 | 4,04 | 0 |
| TP53 | ACTGTGCCCAGCCATTTC | -1 | 171 | 189 | -829 | -811 | -3,93 | 0,01 | 8,78 | >0.1 | 3,15 | 4,44 | 0 | 1,87 | 0 | 3,24 | 0 |
| TP53 | GATCTGCCTTTGCAAAAC | -1 | 82 | 100 | -918 | -900 | -3,32 | 0,01 | 6,63 | >0.1 | 3,73 | 4,34 | 0 | 1,88 | 0 | 0 | 0 |
| GMEB1 | TGGCTCACGCCTGTAAT | 1 | 188 | 205 | -812 | -795 | 3,27 | 0,01 | 16,66 | >0.1 | 3,08 | 4,46 | 0 | 1,72 | 0 | 4,13 | 0 |
| GMEB1 | AGCTGGGCGTGGTGGTG | 1 | 310 | 327 | -690 | -673 | 2,35 | 0,01 | 16,94 | >0.1 | 2,64 | 4,63 | 0 | 1,18 | 0 | 6,13 | 0 |
| GMEB1 | GGTGGTGGGCACCTGTA | 1 | 320 | 337 | -680 | -663 | 0,36 | 0,01 | 14,35 | >0.1 | 2,61 | 4,65 | 0 | 1,16 | 0 | 5,56 | 0 |
| GMEB1 | TGGTGGGCACCTGTAGT | 1 | 322 | 339 | -678 | -661 | -0,05 | 0,01 | 13,82 | >0.1 | 2,61 | 4,65 | 0 | 1,16 | 0 | 5,46 | 0 |
| GMEB1 | AGAATGGCGTGAACCTG | 1 | 366 | 383 | -634 | -617 | 0,89 | 0,01 | 15,39 | >0.1 | 2,54 | 4,72 | 0 | 1,08 | 0 | 6,15 | 0 |
| GMEB1 | GAGATCGCGCCACTGTG | 1 | 406 | 423 | -594 | -577 | 5,03 | 0,001 | 19,03 | >0.1 | 2,65 | 4,79 | 0 | 1,03 | 0 | 5,52 | 0 |
| GMEB1 | CACTGTGCTCCAGCCTG | 1 | 416 | 433 | -584 | -567 | 0,5 | 0,01 | 14,32 | >0.1 | 2,68 | 4,81 | 0 | 1,03 | 0 | 5,3 | 0 |
| GMEB1 | GCCTGGGCGACAGACCG | 1 | 428 | 445 | -572 | -555 | 4,38 | 0,01 | 18,62 | >0.1 | 2,72 | 4,83 | 0 | 1,02 | 0 | 5,67 | 0 |
| GMEB1 | CAGGAGAGGTAAGGTTC | 1 | 770 | 787 | -230 | -213 | 1,36 | 0,01 | 20,22 | >0.1 | 5,52 | 5,85 | 0 | 1,39 | 0 | 6,1 | 0 |
| GMEB1 | TCTGTTACATATCTGTC | 1 | 845 | 862 | -155 | -138 | 1,96 | 0,01 | 18,62 | >0.1 | 5,52 | 6,31 | 0 | 1,88 | 0 | 2,95 | 0 |
| GMEB1 | TGCTCAGCGCCCAACCC | 1 | 939 | 956 | -61 | -44 | 3,24 | 0,01 | 18,79 | >0.1 | 5,52 | 7,32 | 0 | 1,88 | 0 | 0,83 | 0 |
| GMEB1 | CAGTGGATGTGATCTTG | 1 | 1018 | 1035 | 18 | 35 | 0,44 | 0,01 | 17,89 | >0.1 | 5,52 | 8,76 | 0 | 1,86 | 0 | 1,31 | 0 |
| GMEB1 | GGTTGGGCGCTGAGCAG | -1 | 938 | 955 | -62 | -45 | 3,53 | 0,01 | 19,02 | >0.1 | 5,52 | 7,31 | 0 | 1,88 | 0 | 0,79 | 0 |
| GMEB1 | ACAGATATGTAACAGAT | -1 | 844 | 861 | -156 | -139 | 0,85 | 0,01 | 17,41 | >0.1 | 5,52 | 6,3 | 0 | 1,87 | 0 | 2,86 | 0 |
| GMEB1 | AAGAATACGTTATTCTT | -1 | 797 | 814 | -203 | -186 | 2,46 | 0,01 | 19,04 | >0.1 | 5,52 | 5,99 | 0 | 1,52 | 0 | 3,55 | 0 |
| GMEB1 | GGTCTGTCGCCCAGGCT | -1 | 427 | 444 | -573 | -556 | 0,13 | 0,01 | 14,36 | >0.1 | 2,71 | 4,83 | 0 | 1,02 | 0 | 5,67 | 0 |

| | | | | | | | | | | | | | | | | | |
|---|---|---|---|---|---|---|---|---|---|---|---|---|---|---|---|---|---|
| GMEB1 | ACAGTGGCGCGATCTCG | -1 | 405 | 422 | -595 | -578 | 3 | 0,01 | 17,12 | >0.1 | 2,65 | 4,79 | 0 | 1,04 | 0 | 5,65 | 0 |
| GMEB1 | AGGTTCACGCCATTCTC | -1 | 365 | 382 | -635 | -618 | 4,42 | 0,01 | 19,49 | >0.1 | 2,54 | 4,72 | 0 | 1,08 | 0 | 6,73 | 0 |
| GMEB1 | ACAGGTGCCCACCACCA | -1 | 319 | 336 | -681 | -664 | 0 | 0,01 | 13,99 | >0.1 | 2,62 | 4,65 | 0 | 1,16 | 0 | 5,56 | 0 |
| GMEB1 | ACCACCACGCCCAGCTA | -1 | 309 | 326 | -691 | -674 | 1,58 | 0,01 | 16,07 | >0.1 | 2,64 | 4,63 | 0 | 1,19 | 0 | 6,03 | 0 |
| GMEB1 | TTACAGGCGTGAGCCAC | -1 | 187 | 204 | -813 | -796 | 2,44 | 0,01 | 15,84 | >0.1 | 3,08 | 4,46 | 0 | 1,73 | 0 | 4,13 | 0 |
| NEUROD2 | GGCTCACGCC | 1 | 189 | 199 | -811 | -801 | 2,79 | 0,01 | 16,08 | >0.1 | 3,09 | 4,45 | 0 | 1,71 | 0 | 4,04 | 0 |
| NEUROD2 | CACGCCTGTA | 1 | 193 | 203 | -807 | -797 | 0,41 | 0,01 | 13,88 | >0.1 | 3,07 | 4,46 | 0 | 1,68 | 0 | 4,26 | 0 |
| NEUROD2 | ACCGTCCTGA | 1 | 257 | 267 | -743 | -733 | 1,09 | 0,01 | 14,42 | >0.1 | 2,82 | 4,54 | 0 | 1,34 | 0 | 4,63 | 0 |
| NEUROD2 | GGCACCTGTA | 1 | 327 | 337 | -673 | -663 | 4,33 | 0,01 | 18,29 | >0.1 | 2,6 | 4,65 | 0 | 1,14 | 0 | 5,56 | 0 |
| NEUROD2 | AAGTCATGTA | 1 | 515 | 525 | -485 | -475 | -1,64 | 0,01 | 10,22 | >0.1 | 3,02 | 5 | 0 | 0,98 | 0 | 2,86 | 0 |
| NEUROD2 | AACATTTTCC | 1 | 691 | 701 | -309 | -299 | -0,74 | 0,01 | 13,49 | >0.1 | 4,53 | 5,49 | 0 | 1,14 | 0 | 3,07 | 0 |
| NEUROD2 | AACGTATTCT | 1 | 803 | 813 | -197 | -187 | 5,67 | 0,01 | 22,21 | >0.1 | 5,61 | 5,99 | 0 | 1,51 | 0 | 3,43 | 0 |
| NEUROD2 | GCCTTGTTGC | 1 | 1077 | 1087 | 77 | 87 | 1 | 0,01 | 19,53 | >0.1 | 5,61 | 7,06 | 0 | 1,85 | 1,79 | 2,21 | 0 |
| NEUROD2 | AAGACATTTT | 1 | 1124 | 1134 | 124 | 134 | 0,68 | 0,01 | 20,66 | >0.1 | 5,61 | 6,51 | 0 | 1,84 | 3,22 | 2,79 | 0 |
| NEUROD2 | AAGACCTGTT | 1 | 1158 | 1168 | 158 | 168 | 1,78 | 0,01 | 18,37 | >0.1 | 4,38 | 6,24 | 0 | 1,78 | 1,79 | 2,4 | 0 |
| NEUROD2 | CAGATATGTA | -1 | 850 | 860 | -150 | -140 | 4,29 | 0,01 | 20,81 | >0.1 | 5,61 | 6,3 | 0 | 1,86 | 0 | 2,75 | 0 |
| NEUROD2 | ACGTTATTCT | -1 | 798 | 808 | -202 | -192 | -0,22 | 0,01 | 16,38 | >0.1 | 5,61 | 5,96 | 0 | 1,49 | 0 | 3,55 | 0 |
| NEUROD2 | CCGGTACTTT | -1 | 738 | 748 | -262 | -252 | -0,73 | 0,01 | 17,72 | >0.1 | 5,61 | 5,67 | 0 | 1,25 | 0 | 5,92 | 0 |
| NEUROD2 | AACATTTTCC | -1 | 533 | 543 | -467 | -457 | -0,74 | 0,01 | 9,99 | >0.1 | 3,1 | 5,04 | 0 | 0,98 | 0 | 1,61 | 0 |
| NEUROD2 | TCGGTCTGTC | -1 | 436 | 446 | -564 | -554 | 1,6 | 0,01 | 16,09 | >0.1 | 2,73 | 4,83 | 0 | 1,01 | 0 | 5,92 | 0 |
| NEUROD2 | GGCTCACTGC | -1 | 396 | 406 | -604 | -594 | -0,61 | 0,01 | 13,57 | >0.1 | 2,61 | 4,76 | 0 | 1,04 | 0 | 5,76 | 0 |
| NEUROD2 | GCCATTCTCC | -1 | 364 | 374 | -636 | -626 | -0,45 | 0,01 | 14,48 | >0.1 | 2,53 | 4,71 | 0 | 1,08 | 0 | 6,61 | 0 |
| NEUROD2 | GCCATTTTTT | -1 | 140 | 150 | -860 | -850 | 1,1 | 0,01 | 12,38 | >0.1 | 3,34 | 4,39 | 0 | 1,89 | 0 | 1,65 | 0 |
| NEUROD2 | AGCTCATTTC | -1 | 65 | 75 | -935 | -925 | -0,54 | 0,01 | 9,59 | >0.1 | 3,94 | 4,31 | 0 | 1,88 | 0 | 0 | 0 |
| NEUROD2 | GACTTCTTTG | -1 | 29 | 39 | -971 | -961 | -1,98 | 0,01 | 8,67 | >0.1 | 4,51 | 4,28 | 0 | 1,87 | 0 | 0 | 0 |
| NEUROD2 | GCCTCCTTCT | -1 | 16 | 26 | -984 | -974 | -1,4 | 0,01 | 10,34 | >0.1 | 5,61 | 4,27 | 0 | 1,87 | 0 | 0 | 0 |
| NEUROD1 | AAGGAGAAGGAGG | 1 | 12 | 25 | -988 | -975 | 1 | 0,01 | 12,7 | >0.1 | 5,57 | 4,27 | 0 | 1,87 | 0 | 0 | 0 |
| NEUROD1 | AGGCAGATCAGGA | 1 | 91 | 104 | -909 | -896 | 0,57 | 0,01 | 11,3 | >0.1 | 3,67 | 4,34 | 0 | 1,88 | 0 | 0,83 | 0 |
| NEUROD1 | TAAAAAATGGCCA | 1 | 139 | 152 | -861 | -848 | 0,44 | 0,01 | 11,72 | >0.1 | 3,34 | 4,4 | 0 | 1,9 | 0 | 1,65 | 0 |
| NEUROD1 | GGGCAGATCACAA | 1 | 229 | 242 | -771 | -758 | 2,27 | 0,01 | 16,16 | >0.1 | 2,91 | 4,51 | 0 | 1,46 | 0 | 5,01 | 0 |
| NEUROD1 | GGGCACCTGTAGT | 1 | 326 | 339 | -674 | -661 | 0,79 | 0,01 | 14,75 | >0.1 | 2,6 | 4,65 | 0 | 1,15 | 0 | 5,56 | 0 |
| NEUROD1 | CGACAGACCGAGA | 1 | 435 | 448 | -565 | -552 | 1,55 | 0,01 | 16,05 | >0.1 | 2,73 | 4,84 | 0 | 1,01 | 0 | 5,92 | 0 |
| NEUROD1 | TAACAGCTTTCTA | 1 | 710 | 723 | -290 | -277 | 0,79 | 0,01 | 18,1 | >0.1 | 5,57 | 5,57 | 0 | 1,18 | 0 | 4,99 | 0 |
| NEUROD1 | TGACAGGAGAGGT | 1 | 767 | 780 | -233 | -220 | 0,2 | 0,01 | 18,86 | >0.1 | 5,57 | 5,82 | 0 | 1,36 | 0 | 5,92 | 0 |
| NEUROD1 | CAGCAGATTGTTT | 1 | 889 | 902 | -111 | -98 | 6,12 | 0,01 | 23,16 | >0.1 | 5,57 | 6,63 | 0 | 1,89 | 0 | 2,95 | 0 |
| NEUROD1 | GAGCAGCAGTTAC | -1 | 1088 | 1101 | 88 | 101 | 2,23 | 0,01 | 20,56 | >0.1 | 5,57 | 6,9 | 0 | 1,85 | 1,79 | 2,21 | 0 |
| NEUROD1 | GGACAGATATGTA | -1 | 850 | 863 | -150 | -137 | 3,3 | 0,01 | 19,75 | >0.1 | 5,57 | 6,32 | 0 | 1,89 | 0 | 2,67 | 0 |
| NEUROD1 | TAACAGATTTTAG | -1 | 839 | 852 | -161 | -148 | 4,24 | 0,01 | 21,03 | >0.1 | 5,57 | 6,24 | 0 | 1,79 | 0 | 3,19 | 0 |
| NEUROD1 | AGAAAGCTGTTAA | -1 | 709 | 722 | -291 | -278 | 0,14 | 0,01 | 17,27 | >0.1 | 5,57 | 5,57 | 0 | 1,18 | 0 | 4,81 | 0 |
| NEUROD1 | GAGTAGCTGGGAC | -1 | 337 | 350 | -663 | -650 | 3,57 | 0,01 | 17,85 | >0.1 | 2,57 | 4,67 | 0 | 1,13 | 0 | 5,92 | 0 |
| NEUROD1 | CTACAGGTGCCCA | -1 | 325 | 338 | -675 | -662 | 5,64 | 0,01 | 19,5 | >0.1 | 2,6 | 4,65 | 0 | 1,15 | 0 | 5,46 | 0 |
| NEUROD1 | ATAGAGACGGGGT | -1 | 280 | 293 | -720 | -707 | 0,84 | 0,01 | 14,61 | >0.1 | 2,74 | 4,58 | 0 | 1,26 | 0 | 5,19 | 0 |
| NEUROD1 | TTACAGGCGTGAG | -1 | 191 | 204 | -809 | -796 | 0,46 | 0,01 | 13,95 | >0.1 | 3,07 | 4,46 | 0 | 1,7 | 0 | 4,26 | 0 |
| JUNB(VAR.2) | AATAGGTAAG | 1 | 0 | 10 | -1000 | -990 | 2,04 | 0,01 | 15,56 | >0.1 | 5,61 | 4,25 | 0 | 1,86 | 1,79 | 0 | 0 |
| JUNB(VAR.2) | AAGAAGTCAG | 1 | 31 | 41 | -969 | -959 | 3,64 | 0,01 | 14,25 | >0.1 | 4,46 | 4,28 | 0 | 1,87 | 0 | 0 | 0 |
| JUNB(VAR.2) | ATGGCGTGAA | 1 | 369 | 379 | -631 | -621 | 6,39 | 0,001 | 20,87 | >0.1 | 2,54 | 4,71 | 0 | 1,07 | 0 | 6,15 | 0 |
| JUNB(VAR.2) | ATCGCGCCAC | 1 | 409 | 419 | -591 | -581 | 5,07 | 0,01 | 19,06 | >0.1 | 2,65 | 4,78 | 0 | 1,03 | 0 | 5,52 | 0 |
| JUNB(VAR.2) | GTGAGATAAC | 1 | 591 | 601 | -409 | -399 | 5,08 | 0,01 | 14,69 | >0.1 | 3,41 | 5,18 | 0 | 1,01 | 0 | 0 | 0 |
| JUNB(VAR.2) | CTTTGGTCAC | 1 | 670 | 680 | -330 | -320 | 3,74 | 0,01 | 18,1 | >0.1 | 4,14 | 5,42 | 0 | 1,1 | 0 | 3,7 | 0 |
| JUNB(VAR.2) | ATTATAACAT | 1 | 686 | 696 | -314 | -304 | 2,18 | 0,01 | 16,15 | >0.1 | 4,42 | 5,47 | 0 | 1,13 | 0 | 2,95 | 0 |
| JUNB(VAR.2) | ATTTGGCCAT | 1 | 753 | 763 | -247 | -237 | 2,82 | 0,01 | 21,78 | >0.1 | 5,61 | 5,74 | 0 | 1,3 | 0 | 6,32 | 0 |
| JUNB(VAR.2) | GAGAGGTAAG | 1 | 773 | 783 | -227 | -217 | 3,15 | 0,01 | 22,06 | >0.1 | 5,61 | 5,83 | 0 | 1,37 | 0 | 6,1 | 0 |
| JUNB(VAR.2) | ATAACGTATT | 1 | 801 | 811 | -199 | -189 | 4,47 | 0,01 | 20,98 | >0.1 | 5,61 | 5,98 | 0 | 1,5 | 0 | 3,43 | 0 |
| JUNB(VAR.2) | GTTACATATC | 1 | 848 | 858 | -152 | -142 | 2,97 | 0,01 | 19,66 | >0.1 | 5,61 | 6,28 | 0 | 1,84 | 0 | 2,95 | 0 |
| JUNB(VAR.2) | ACGATGTCAA | 1 | 1046 | 1056 | 46 | 56 | 4,2 | 0,01 | 20,81 | >0.1 | 5,61 | 7,68 | 0 | 1,86 | 0 | 1,46 | 0 |
| JUNB(VAR.2) | GTTACAGCAA | -1 | 1083 | 1093 | 83 | 93 | 2,12 | 0,01 | 20,56 | >0.1 | 5,61 | 6,97 | 0 | 1,85 | 1,79 | 2,21 | 0 |
| JUNB(VAR.2) | TTGACATCGT | -1 | 1046 | 1056 | 46 | 56 | 4,41 | 0,01 | 21,02 | >0.1 | 5,61 | 7,68 | 0 | 1,86 | 0 | 1,46 | 0 |
| JUNB(VAR.2) | GATATGTAAC | -1 | 848 | 858 | -152 | -142 | 2,92 | 0,01 | 19,61 | >0.1 | 5,61 | 6,28 | 0 | 1,84 | 0 | 2,95 | 0 |
| JUNB(VAR.2) | AATACGTTAT | -1 | 801 | 811 | -199 | -189 | 4,5 | 0,01 | 21,01 | >0.1 | 5,61 | 5,98 | 0 | 1,5 | 0 | 3,43 | 0 |

| | | | | | | | | | | | | | | | | |
|---|---|---|---|---|---|---|---|---|---|---|---|---|---|---|---|---|
| JUNB(VAR.2) | CTTACCTCTC | -1 | 773 | 783 | -227 | -217 | 3,08 | 0,01 | 21,99 | >0.1 | 5,61 | 5,83 | 0 | 1,37 | 0 | 6,1 | 0 |
| JUNB(VAR.2) | ATGGCCAAAT | -1 | 753 | 763 | -247 | -237 | 2,77 | 0,01 | 21,73 | >0.1 | 5,61 | 5,74 | 0 | 1,3 | 0 | 6,32 | 0 |
| JUNB(VAR.2) | ATGTTATAAT | -1 | 686 | 696 | -314 | -304 | 2,23 | 0,01 | 16,2 | >0.1 | 4,42 | 5,47 | 0 | 1,13 | 0 | 2,95 | 0 |
| JUNB(VAR.2) | GTGACCAAAG | -1 | 670 | 680 | -330 | -320 | 3,67 | 0,01 | 18,03 | >0.1 | 4,14 | 5,42 | 0 | 1,1 | 0 | 3,7 | 0 |
| JUNB(VAR.2) | GTTATCTCAC | -1 | 591 | 601 | -409 | -399 | 5,02 | 0,01 | 14,63 | >0.1 | 3,41 | 5,18 | 0 | 1,01 | 0 | 0 | 0 |
| JUNB(VAR.2) | GTGGCGCGAT | -1 | 409 | 419 | -591 | -581 | 5,16 | 0,01 | 19,15 | >0.1 | 2,65 | 4,78 | 0 | 1,03 | 0 | 5,52 | 0 |
| JUNB(VAR.2) | TTCACGCCAT | -1 | 369 | 379 | -631 | -621 | 6,27 | 0,001 | 20,75 | >0.1 | 2,54 | 4,71 | 0 | 1,07 | 0 | 6,15 | 0 |
| JUNB(VAR.2) | CTGACTTCTT | -1 | 31 | 41 | -969 | -959 | 3,76 | 0,01 | 14,37 | >0.1 | 4,46 | 4,28 | 0 | 1,87 | 0 | 0 | 0 |
| ESR1 | TCAGGAGAGTTGACCTG | 1 | 98 | 115 | -902 | -885 | -16,98 | 0,01 | -6,35 | >0.1 | 3,6 | 4,36 | 0 | 1,89 | 0 | 0,79 | 0 |
| ESR1 | CTGGGCACAGTGGCTCA | 1 | 178 | 195 | -822 | -805 | -16,81 | 0,01 | -3,58 | >0.1 | 3,12 | 4,45 | 0 | 1,81 | 0 | 3,86 | 0 |
| ESR1 | CCTGTAATCCTAGCACT | 1 | 197 | 214 | -803 | -786 | -22,79 | 0,01 | -9,04 | >0.1 | 3,04 | 4,47 | 0 | 1,65 | 0 | 4,59 | 0 |
| ESR1 | GAGGCCGAGTTGGGCAG | 1 | 218 | 235 | -782 | -765 | -20,79 | 0,01 | -6,3 | >0.1 | 2,95 | 4,5 | 0 | 1,52 | 0 | 5,52 | 0 |
| ESR1 | CAGATCACAAGGTCAGG | 1 | 232 | 249 | -768 | -751 | -14,67 | 0,01 | -1,06 | >0.1 | 2,9 | 4,52 | 0 | 1,45 | 0 | 4,74 | 0 |
| ESR1 | CAGGAGATAGAGACCGT | 1 | 245 | 262 | -755 | -738 | -19,1 | 0,01 | -5,24 | >0.1 | 2,85 | 4,54 | 0 | 1,39 | 0 | 5,09 | 0 |
| ESR1 | ACTAACACGGTGAAACC | 1 | 266 | 283 | -734 | -717 | -21,75 | 0,01 | -8,01 | >0.1 | 2,78 | 4,57 | 0 | 1,31 | 0 | 5,09 | 0 |
| ESR1 | GTGGTGGTGGGCACCTG | 1 | 318 | 335 | -682 | -665 | -21,96 | 0,01 | -7,97 | >0.1 | 2,62 | 4,64 | 0 | 1,17 | 0 | 5,56 | 0 |
| ESR1 | GAGAATGGCGTGAACCT | 1 | 365 | 382 | -635 | -618 | -20,68 | 0,01 | -5,61 | >0.1 | 2,54 | 4,72 | 0 | 1,08 | 0 | 6,73 | 0 |
| ESR1 | GAGCTTGCAGTGAGCCG | 1 | 390 | 407 | -610 | -593 | -6,38 | 0,001 | 8,06 | >0.1 | 2,61 | 4,76 | 0 | 1,05 | 0 | 6,02 | 0 |
| ESR1 | GGCGACAGACCGAGACT | 1 | 433 | 450 | -567 | -550 | -20,84 | 0,01 | -6,33 | >0.1 | 2,73 | 4,84 | 0 | 1,01 | 0 | 5,92 | 0 |
| ESR1 | AAGGGCAAAGTCATGTA | 1 | 508 | 525 | -492 | -475 | -19,94 | 0,01 | -8,08 | >0.1 | 3,01 | 5 | 0 | 0,99 | 0 | 2,86 | 0 |
| ESR1 | AGGGAAAATGTTGCCCA | 1 | 531 | 548 | -469 | -452 | -22,1 | 0,01 | -11,29 | >0.1 | 3,11 | 5,05 | 0 | 0,99 | 0 | 1,66 | 0 |
| ESR1 | TCTGGGAATATGATCTT | 1 | 559 | 576 | -441 | -424 | -19,11 | 0,01 | -9,74 | >0.1 | 3,25 | 5,12 | 0 | 1 | 0 | 0 | 0 |
| ESR1 | AAAATAAATGTGAGATA | 1 | 582 | 599 | -418 | -401 | -22,48 | 0,01 | -12,91 | >0.1 | 3,38 | 5,18 | 0 | 1,01 | 0 | 0 | 0 |
| ESR1 | TTGGTCACTGTAAAATT | 1 | 672 | 689 | -328 | -311 | -12,96 | 0,01 | 1,26 | >0.1 | 4,22 | 5,45 | 0 | 1,12 | 0 | 3,43 | 0 |
| ESR1 | AATCTCATGAGGAGGTT | 1 | 981 | 998 | -19 | -2 | -20,53 | 0,01 | -2,95 | >0.1 | 5,52 | 8,66 | 0 | 1,87 | 0 | 1,53 | 0 |
| ESR1 | TCAGTGGATGTGATCTT | 1 | 1017 | 1034 | 17 | 34 | -22,3 | 0,01 | -4,83 | >0.1 | 5,52 | 8,79 | 0 | 1,86 | 0 | 1,31 | 0 |
| ESR1 | GATGTGATCTTGGCTCA | 1 | 1023 | 1040 | 23 | 40 | -22,08 | 0,01 | -4,76 | >0.1 | 5,52 | 8,67 | 0 | 1,86 | 0 | 1,28 | 0 |
| ESR1 | CAGGGGACGATGTCAAG | 1 | 1040 | 1057 | 40 | 57 | -19,58 | 0,01 | -3,73 | >0.1 | 5,52 | 7,85 | 0 | 1,86 | 0 | 0,62 | 0 |
| ESR1 | GATGTCAAGCTCTTCCT | 1 | 1048 | 1065 | 48 | 65 | -17,06 | 0,01 | -0,59 | >0.1 | 5,52 | 7,62 | 0 | 1,86 | 0 | 1,46 | 0 |
| ESR1 | AATGGTGGACTGAGCAG | -1 | 1095 | 1112 | 95 | 112 | -19,98 | 0,01 | -0,22 | >0.1 | 5,52 | 6,82 | 0 | 1,85 | 3,22 | 2,35 | 0 |
| ESR1 | AGGCTGAGAAGGAGCCA | -1 | 1064 | 1081 | 64 | 81 | -20,38 | 0,01 | -1,66 | >0.1 | 5,52 | 7,28 | 0 | 1,85 | 1,79 | 2,27 | 0 |
| ESR1 | GGAGCCAGGAAGAGCTT | -1 | 1054 | 1071 | 54 | 71 | -18,64 | 0,01 | -1,51 | >0.1 | 5,52 | 7,48 | 0 | 1,86 | 0 | 2,27 | 0 |
| ESR1 | AGGAAGAGCTTGACATC | -1 | 1048 | 1065 | 48 | 65 | -18,38 | 0,01 | -1,91 | >0.1 | 5,52 | 7,62 | 0 | 1,86 | 0 | 1,46 | 0 |
| ESR1 | TGAGCCAAGATCACATC | -1 | 1023 | 1040 | 23 | 40 | -17,9 | 0,01 | -0,58 | >0.1 | 5,52 | 8,67 | 0 | 1,86 | 0 | 1,28 | 0 |
| ESR1 | AATTTTACAGTGACCAA | -1 | 672 | 689 | -328 | -311 | -12,73 | 0,01 | 1,49 | >0.1 | 4,22 | 5,45 | 0 | 1,12 | 0 | 3,43 | 0 |
| ESR1 | TATTTCAAGATCATATT | -1 | 565 | 582 | -435 | -418 | -21,8 | 0,01 | -12,38 | >0.1 | 3,28 | 5,13 | 0 | 1 | 0 | 0 | 0 |
| ESR1 | TACATGACTTTGCCCTT | -1 | 508 | 525 | -492 | -475 | -20,79 | 0,01 | -8,93 | >0.1 | 3,01 | 5 | 0 | 0,99 | 0 | 2,86 | 0 |
| ESR1 | TGGAGCACAGTGGCGCG | -1 | 411 | 428 | -589 | -572 | -11,48 | 0,01 | 2,21 | >0.1 | 2,67 | 4,8 | 0 | 1,03 | 0 | 5,19 | 0 |
| ESR1 | CGGCTCACTGCAAGCTC | -1 | 390 | 407 | -610 | -593 | -9,72 | 0,01 | 4,72 | >0.1 | 2,61 | 4,76 | 0 | 1,05 | 0 | 6,02 | 0 |
| ESR1 | ACGGTCTCTATCTCCTG | -1 | 245 | 262 | -755 | -738 | -14,71 | 0,01 | -0,85 | >0.1 | 2,85 | 4,54 | 0 | 1,39 | 0 | 5,09 | 0 |
| ESR1 | AGTGCTAGGATTACAGG | -1 | 197 | 214 | -803 | -786 | -22,3 | 0,01 | -8,55 | >0.1 | 3,04 | 4,47 | 0 | 1,65 | 0 | 4,59 | 0 |
| ESR1 | TGAGCCACTGTGCCCAG | -1 | 178 | 195 | -822 | -805 | -18,7 | 0,01 | -5,47 | >0.1 | 3,12 | 4,45 | 0 | 1,81 | 0 | 3,86 | 0 |
| NFYA | AGGCACTCAAA | 1 | 22 | 33 | -978 | -967 | 1,55 | 0,01 | 13,29 | >0.1 | 5,6 | 4,27 | 0 | 1,87 | 0 | 0 | 0 |
| NFYA | AGTCAGCCACA | 1 | 35 | 46 | -965 | -954 | -0,1 | 0,01 | 10,42 | >0.1 | 4,37 | 4,29 | 0 | 1,87 | 0 | 0 | 0 |
| NFYA | AACCAGTGTGA | 1 | 47 | 58 | -953 | -942 | 0,65 | 0,01 | 10,98 | >0.1 | 4,16 | 4,3 | 0 | 1,87 | 0 | 0 | 0 |
| NFYA | GGCAGATCAGG | 1 | 92 | 103 | -908 | -897 | 0,93 | 0,01 | 11,66 | >0.1 | 3,67 | 4,34 | 0 | 1,88 | 0 | 0,83 | 0 |
| NFYA | AACTGATCAGA | 1 | 162 | 173 | -838 | -827 | 1,39 | 0,01 | 13,39 | >0.1 | 3,22 | 4,42 | 0 | 1,9 | 0 | 2,46 | 0 |
| NFYA | GGCAGATCACA | 1 | 230 | 241 | -770 | -759 | 0,23 | 0,01 | 14,12 | >0.1 | 2,91 | 4,51 | 0 | 1,45 | 0 | 5,01 | 0 |
| NFYA | AGCTACTCGGG | 1 | 342 | 353 | -658 | -647 | 1,18 | 0,01 | 15,93 | >0.1 | 2,56 | 4,67 | 0 | 1,12 | 0 | 6,4 | 0 |
| NFYA | CGCCACTGTGC | 1 | 413 | 424 | -587 | -576 | -0,07 | 0,01 | 13,5 | >0.1 | 2,66 | 4,79 | 0 | 1,02 | 0 | 5,09 | 0 |
| NFYA | AACTGATCAGA | 1 | 477 | 488 | -523 | -512 | 1,39 | 0,01 | 15,37 | >0.1 | 2,87 | 4,92 | 0 | 0,99 | 0 | 5,2 | 0 |
| NFYA | GCCCAAGTAGA | 1 | 543 | 554 | -457 | -446 | 0,85 | 0,01 | 10,05 | >0.1 | 3,15 | 5,06 | 0 | 0,99 | 0 | 0 | 0 |
| NFYA | AACCTATTAAT | 1 | 598 | 609 | -402 | -391 | 0,75 | 0,01 | 11,85 | >0.1 | 3,46 | 5,21 | 0 | 1,02 | 0 | 1,41 | 0 |
| NFYA | GCCCAACCCAA | 1 | 947 | 958 | -53 | -42 | 3,57 | 0,01 | 20,14 | >0.1 | 5,6 | 7,36 | 0 | 1,88 | 0 | 1,74 | 0 |
| NFYA | AGTCATTCAGT | 1 | 1011 | 1022 | 11 | 22 | 1,5 | 0,01 | 19,31 | >0.1 | 5,6 | 8,95 | 0 | 1,86 | 0 | 1,39 | 0 |
| NFYA | CACCATTGAGG | 1 | 1105 | 1116 | 105 | 116 | 3,2 | 0,01 | 23,09 | >0.1 | 5,6 | 6,7 | 0 | 1,85 | 3,22 | 2,52 | 0 |
| NFYA | GTTCTATCAAA | 1 | 1165 | 1176 | 165 | 176 | 0,65 | 0,01 | 17,65 | >0.1 | 4,52 | 6,19 | 0 | 1,71 | 1,79 | 2,79 | 0 |
| NFYA | CCCCTGTGAGC | -1 | 1035 | 1046 | 35 | 46 | -0,62 | 0,01 | 16,17 | >0.1 | 5,6 | 8,03 | 0 | 1,86 | 0 | 1,31 | 0 |
| NFYA | ATCCACTGAAT | -1 | 1015 | 1026 | 15 | 26 | 1,5 | 0,01 | 19,19 | >0.1 | 5,6 | 8,84 | 0 | 1,86 | 0 | 1,39 | 0 |

| | | | | | | | | | | | | | | | | | |
|---|---|---|---|---|---|---|---|---|---|---|---|---|---|---|---|---|---|
| NFYA | CTCTTATCAGC | -1 | 966 | 977 | -34 | -23 | 1,01 | 0,01 | 17,99 | >0.1 | 5,6 | 7,81 | 0 | 1,87 | 0 | 1,7 | 0 |
| NFYA | GACCAAAAAAA | -1 | 923 | 934 | -77 | -66 | 1,61 | 0,01 | 17,16 | >0.1 | 5,6 | 6,98 | 0 | 1,88 | 0 | 1,09 | 0 |
| NFYA | AAACAATCTGC | -1 | 891 | 902 | -109 | -98 | 1 | 0,01 | 17,98 | >0.1 | 5,6 | 6,63 | 0 | 1,89 | 0 | 2,86 | 0 |
| NFYA | AGCCAATATAA | -1 | 878 | 889 | -122 | -111 | 1,72 | 0,01 | 18,8 | >0.1 | 5,6 | 6,52 | 0 | 1,9 | 0 | 3,07 | 0 |
| NFYA | AGTGAATCAAA | -1 | 815 | 826 | -185 | -174 | 0,48 | 0,01 | 17,16 | >0.1 | 5,6 | 6,06 | 0 | 1,59 | 0 | 3,43 | 0 |
| NFYA | ATCAAATAAGA | -1 | 810 | 821 | -190 | -179 | 1,46 | 0,01 | 17,93 | >0.1 | 5,6 | 6,03 | 0 | 1,56 | 0 | 3,28 | 0 |
| NFYA | CTCCTGTCACT | -1 | 765 | 776 | -235 | -224 | 0,57 | 0,01 | 19,22 | >0.1 | 5,6 | 5,8 | 0 | 1,34 | 0 | 5,92 | 0 |
| NFYA | GCCAAATCAAA | -1 | 749 | 760 | -251 | -240 | 4,24 | 0,01 | 22,76 | >0.1 | 5,6 | 5,72 | 0 | 1,29 | 0 | 5,92 | 0 |
| NFYA | GACCAAAGAGG | -1 | 667 | 678 | -333 | -322 | 5,12 | 0,001 | 19,57 | >0.1 | 4,11 | 5,41 | 0 | 1,1 | 0 | 3,83 | 0 |
| NFYA | CGCCATTCTCC | -1 | 364 | 375 | -636 | -625 | -0,13 | 0,01 | 14,8 | >0.1 | 2,53 | 4,71 | 0 | 1,08 | 0 | 6,61 | 0 |
| NFYA | TGCCCACCACC | -1 | 320 | 331 | -680 | -669 | 1,31 | 0,01 | 15,41 | >0.1 | 2,62 | 4,64 | 0 | 1,16 | 0 | 5,68 | 0 |
| NFYA | GCCCAACTCGG | -1 | 222 | 233 | -778 | -767 | 0,08 | 0,01 | 14,54 | >0.1 | 2,94 | 4,5 | 0 | 1,5 | 0 | 5,52 | 0 |
| NFYA | AGCCACTGTGC | -1 | 182 | 193 | -818 | -807 | 0,76 | 0,01 | 13,95 | >0.1 | 3,12 | 4,45 | 0 | 1,77 | 0 | 3,86 | 0 |
| NFYA | GCCCAGCCATT | -1 | 173 | 184 | -827 | -816 | 0,53 | 0,01 | 13,21 | >0.1 | 3,16 | 4,43 | 0 | 1,85 | 0 | 3,24 | 0 |
| NFYA | CTCCACTCCAC | -1 | 114 | 125 | -886 | -875 | 2,06 | 0,01 | 12,56 | >0.1 | 3,5 | 4,37 | 0 | 1,89 | 0 | 0,75 | 0 |
| GABPA | GAGGCACTCAA | 1 | 21 | 32 | -979 | -968 | -5,83 | 0,01 | 5,91 | >0.1 | 5,6 | 4,27 | 0 | 1,87 | 0 | 0 | 0 |
| GABPA | TAGGAAATGAG | 1 | 62 | 73 | -938 | -927 | -0,8 | 0,01 | 9,35 | >0.1 | 3,97 | 4,31 | 0 | 1,88 | 0 | 0 | 0 |
| GABPA | ATGGAAATTAA | 1 | 151 | 162 | -849 | -838 | -5,51 | 0,01 | 5,72 | >0.1 | 3,27 | 4,41 | 0 | 1,9 | 0 | 1,65 | 0 |
| GABPA | CAGGAGATAGA | 1 | 245 | 256 | -755 | -744 | -4,51 | 0,01 | 9,56 | >0.1 | 2,86 | 4,53 | 0 | 1,39 | 0 | 5,3 | 0 |
| GABPA | CGGGAGGCTGA | 1 | 349 | 360 | -651 | -640 | -5,85 | 0,01 | 8,88 | >0.1 | 2,54 | 4,68 | 0 | 1,1 | 0 | 6,4 | 0 |
| GABPA | GAGGCAGGAGA | 1 | 358 | 369 | -642 | -631 | -3,87 | 0,01 | 10,69 | >0.1 | 2,52 | 4,7 | 0 | 1,09 | 0 | 6,26 | 0 |
| GABPA | GAGGCAGAGCT | 1 | 384 | 395 | -616 | -605 | -2,37 | 0,01 | 12,03 | >0.1 | 2,58 | 4,74 | 0 | 1,05 | 0 | 6,02 | 0 |
| GABPA | AGGGAAAATGT | 1 | 531 | 542 | -469 | -458 | -5,97 | 0,01 | 4,75 | >0.1 | 3,09 | 5,04 | 0 | 0,98 | 0 | 1,61 | 0 |
| GABPA | CAGGATGAACT | 1 | 867 | 878 | -133 | -122 | -2,42 | 0,01 | 14,26 | >0.1 | 5,6 | 6,43 | 0 | 1,9 | 0 | 2,75 | 0 |
| GABPA | AGGGAAAGTCA | 1 | 1005 | 1016 | 5 | 16 | -6,32 | 0,01 | 11,68 | >0.1 | 5,6 | 9,12 | 0 | 1,87 | 0 | 1,42 | 0 |
| GABPA | GTGGATGTGAT | 1 | 1020 | 1031 | 20 | 31 | -4,84 | 0,01 | 12,65 | >0.1 | 5,6 | 8,72 | 0 | 1,86 | 0 | 1,31 | 0 |
| GABPA | GAGGAACAGGC | 1 | 1112 | 1123 | 112 | 123 | 4,25 | 0,001 | 24,08 | >0.1 | 5,6 | 6,63 | 0 | 1,85 | 3,22 | 2,54 | 0 |
| GABPA | CACGAAGCCGA | 1 | 1148 | 1159 | 148 | 159 | -1,78 | 0,01 | 15,94 | >0.1 | 5,6 | 6,31 | 0 | 1,84 | 1,79 | 2,18 | 0 |
| GABPA | GCCGAAGACCT | 1 | 1154 | 1165 | 154 | 165 | -3,39 | 0,01 | 13,25 | >0.1 | 4,45 | 6,27 | 0 | 1,82 | 1,79 | 2,31 | 0 |
| GABPA | CAAGAAGCAAG | -1 | 1181 | 1192 | 181 | 192 | -3,31 | 0,01 | 15,17 | >0.1 | 5,6 | 6,08 | 0 | 1,58 | 1,79 | 3,43 | 0 |
| GABPA | AAGCAAGTGAA | -1 | 1177 | 1188 | 177 | 188 | -5,07 | 0,01 | 13,3 | >0.1 | 5,6 | 6,11 | 0 | 1,61 | 1,79 | 3,26 | 0 |
| GABPA | CAGGAAGAGCT | -1 | 1055 | 1066 | 55 | 66 | 8,85 | 0,001 | 26,03 | >0.1 | 5,6 | 7,46 | 0 | 1,86 | 0 | 2,27 | 0 |
| GABPA | CAGGAAAGACC | -1 | 930 | 941 | -70 | -59 | 4,11 | 0,001 | 19,66 | >0.1 | 5,6 | 7,08 | 0 | 1,88 | 0 | 1 | 0 |
| GABPA | CCTGGAGAGGA | -1 | 860 | 871 | -140 | -129 | -5,26 | 0,01 | 11,48 | >0.1 | 5,6 | 6,38 | 0 | 1,9 | 0 | 2,86 | 0 |
| GABPA | ACGGAAAATGT | -1 | 692 | 703 | -308 | -297 | 1,8 | 0,01 | 16,08 | >0.1 | 4,57 | 5,5 | 0 | 1,14 | 0 | 3,07 | 0 |
| GABPA | CATGAAGATAA | -1 | 641 | 652 | -359 | -348 | -4,56 | 0,01 | 9,47 | >0.1 | 3,81 | 5,33 | 0 | 1,06 | 0 | 3,83 | 0 |
| GABPA | CCAGAAACTCT | -1 | 553 | 564 | -447 | -436 | -3,85 | 0,01 | 5,43 | >0.1 | 3,2 | 5,09 | 0 | 0,99 | 0 | 0 | 0 |
| GABPA | CAGGACGGTCT | -1 | 255 | 266 | -745 | -734 | -2,71 | 0,01 | 10,85 | >0.1 | 2,82 | 4,54 | 0 | 1,35 | 0 | 4,85 | 0 |
| GABPA | CCTGATCTGCC | -1 | 92 | 103 | -908 | -897 | -5,68 | 0,01 | 5,05 | >0.1 | 3,67 | 4,34 | 0 | 1,88 | 0 | 0,83 | 0 |
| FOXO3 | TTAAAAAA | 1 | 138 | 146 | -862 | -854 | -3,44 | 0,1 | 7,85 | >0.1 | 3,36 | 4,39 | 0 | 1,89 | 0 | 1,65 | 0 |
| FOXO3 | GTAATCCT | 1 | 200 | 208 | -800 | -792 | -2,95 | 0,01 | 10,66 | >0.1 | 3,04 | 4,46 | 0 | 1,63 | 0 | 4,48 | 0 |
| FOXO3 | GTGAACCT | 1 | 374 | 382 | -626 | -618 | 2,25 | 0,01 | 17,12 | >0.1 | 2,55 | 4,72 | 0 | 1,07 | 0 | 6,53 | 0 |
| FOXO3 | GGGAAAAT | 1 | 532 | 540 | -468 | -460 | -3,14 | 0,01 | 7,57 | >0.1 | 3,09 | 5,03 | 0 | 0,98 | 0 | 1,61 | 0 |
| FOXO3 | ATAAAAAT | 1 | 579 | 587 | -421 | -413 | 2,73 | 0,01 | 12,21 | >0.1 | 3,33 | 5,15 | 0 | 1 | 0 | 0 | 0 |
| FOXO3 | GAAAACCA | 1 | 619 | 627 | -381 | -373 | -2,39 | 0,01 | 9,04 | >0.1 | 3,6 | 5,26 | 0 | 1,03 | 0 | 1,55 | 0 |
| FOXO3 | TTAACCAA | 1 | 731 | 739 | -269 | -261 | -3,02 | 0,01 | 15 | >0.1 | 5,64 | 5,63 | 0 | 1,22 | 0 | 5,52 | 0 |
| FOXO3 | TTAAAAAA | 1 | 826 | 834 | -174 | -166 | -3,44 | 0,1 | 13,66 | >0.1 | 5,64 | 6,12 | 0 | 1,64 | 0 | 3,7 | 0 |
| FOXO3 | AGCAACAA | -1 | 1080 | 1088 | 80 | 88 | -0,46 | 0,01 | 18,05 | >0.1 | 5,64 | 7,01 | 0 | 1,85 | 1,79 | 2,21 | 0 |
| FOXO3 | GTAAACAA | -1 | 896 | 904 | -104 | -96 | 14,06 | 0,0001 | 31,19 | >0.1 | 5,64 | 6,65 | 0 | 1,89 | 0 | 2,95 | 0 |
| FOXO3 | GTTAATAT | -1 | 728 | 736 | -272 | -264 | -3,17 | 0,01 | 14,62 | >0.1 | 5,64 | 5,62 | 0 | 1,22 | 0 | 5,32 | 0 |
| FOXO3 | GTGACCAA | -1 | 672 | 680 | -328 | -320 | 2,67 | 0,01 | 16,89 | >0.1 | 4,16 | 5,42 | 0 | 1,1 | 0 | 3,55 | 0 |
| FOXO3 | GGCAACAT | -1 | 538 | 546 | -462 | -454 | 4,12 | 0,01 | 14,93 | >0.1 | 3,12 | 5,05 | 0 | 0,98 | 0 | 1,66 | 0 |
| FOXO3 | TTAAAAAA | -1 | 74 | 82 | -926 | -918 | -3,44 | 0,1 | 6,61 | >0.1 | 3,85 | 4,32 | 0 | 1,88 | 0 | 0 | 0 |
| SOX2 | CCAGTGTG | 1 | 49 | 57 | -951 | -943 | 0,38 | 0,01 | 10,7 | >0.1 | 4,16 | 4,3 | 0 | 1,87 | 0 | 0 | 0 |
| SOX2 | CCACTGTG | 1 | 415 | 423 | -585 | -577 | 0,38 | 0,01 | 14,05 | >0.1 | 2,67 | 4,79 | 0 | 1,02 | 0 | 5,19 | 0 |
| SOX2 | CCCTAGTT | 1 | 654 | 662 | -346 | -338 | -10,47 | 0,01 | 3,88 | >0.1 | 3,93 | 5,36 | 0 | 1,07 | 0 | 3,99 | 0 |
| SOX2 | TCTTTGGT | 1 | 669 | 677 | -331 | -323 | -11,84 | 0,01 | 2,47 | >0.1 | 4,11 | 5,41 | 0 | 1,09 | 0 | 3,7 | 0 |
| SOX2 | CTTTGGTC | 1 | 670 | 678 | -330 | -322 | -12,2 | 0,1 | 2,13 | >0.1 | 4,13 | 5,41 | 0 | 1,09 | 0 | 3,7 | 0 |
| SOX2 | ACTGTGTT | 1 | 902 | 910 | -98 | -90 | -11,84 | 0,01 | 3,67 | >0.1 | 5,64 | 6,7 | 0 | 1,89 | 0 | 1,28 | 0 |

| | | | | | | | | | | | | | | | | | |
|---|---|---|---|---|---|---|---|---|---|---|---|---|---|---|---|---|---|
| SOX2 | GCCTTGTT | 1 | 1077 | 1085 | 77 | 85 | -10,47 | 0,01 | 8,15 | >0.1 | 5,64 | 7,06 | 0 | 1,85 | 1,79 | 2,27 | 0 |
| SOX2 | CCATTGAG | 1 | 1107 | 1115 | 107 | 115 | -0,12 | 0,01 | 19,75 | >0.1 | 5,64 | 6,68 | 0 | 1,85 | 3,22 | 2,48 | 0 |
| SOX2 | GCTTCGTG | -1 | 1148 | 1156 | 148 | 156 | -12,2 | 0,1 | 5,59 | >0.1 | 5,64 | 6,31 | 0 | 1,84 | 1,79 | 2,21 | 0 |
| SOX2 | CCCCTGTG | -1 | 1038 | 1046 | 38 | 46 | -10,83 | 0,01 | 5,92 | >0.1 | 5,64 | 7,92 | 0 | 1,86 | 0 | 1,34 | 0 |
| SOX2 | CCTGTGAG | -1 | 1036 | 1044 | 36 | 44 | -12,2 | 0,1 | 4,59 | >0.1 | 5,64 | 7,99 | 0 | 1,86 | 0 | 1,31 | 0 |
| SOX2 | CCTTTGAA | -1 | 959 | 967 | -41 | -33 | -10,88 | 0,01 | 5,92 | >0.1 | 5,64 | 7,55 | 0 | 1,88 | 0 | 1,74 | 0 |
| SOX2 | CTTTTGGT | -1 | 734 | 742 | -266 | -258 | -11,84 | 0,01 | 6,59 | >0.1 | 5,64 | 5,65 | 0 | 1,23 | 0 | 5,92 | 0 |
| SOX2 | ACTTTGCC | -1 | 511 | 519 | -489 | -481 | -12,2 | 0,1 | -0,37 | >0.1 | 3 | 4,98 | 0 | 0,98 | 0 | 2,86 | 0 |
| SOX2 | CCCTTGTC | -1 | 505 | 513 | -495 | -487 | 2,02 | 0,001 | 16,8 | >0.1 | 2,98 | 4,97 | 0 | 0,98 | 0 | 5,86 | 0 |
| SOX2 | CCATTCTC | -1 | 365 | 373 | -635 | -627 | 0,38 | 0,01 | 15,3 | >0.1 | 2,53 | 4,7 | 0 | 1,08 | 0 | 6,61 | 0 |
| SOX2 | ACCTTGTG | -1 | 237 | 245 | -763 | -755 | -11,82 | 0,01 | 1,64 | >0.1 | 2,89 | 4,51 | 0 | 1,42 | 0 | 4,63 | 0 |
| SOX2 | CCACTGTG | -1 | 183 | 191 | -817 | -809 | 0,38 | 0,01 | 13,56 | >0.1 | 3,12 | 4,44 | 0 | 1,76 | 0 | 3,86 | 0 |
| SOX2 | CCATTTCT | -1 | 170 | 178 | -830 | -822 | -12,11 | 0,01 | 0,89 | >0.1 | 3,18 | 4,43 | 0 | 1,88 | 0 | 3,51 | 0 |
| SOX2 | CCATGGCC | -1 | 147 | 155 | -853 | -845 | -11,48 | 0,01 | -0,27 | >0.1 | 3,31 | 4,4 | 0 | 1,9 | 0 | 1,61 | 0 |
| SOX2 | CCATTTTT | -1 | 141 | 149 | -859 | -851 | 0,25 | 0,01 | 11,52 | >0.1 | 3,34 | 4,39 | 0 | 1,89 | 0 | 1,65 | 0 |
| SOX2 | CCTTTGCA | -1 | 86 | 94 | -914 | -906 | -10,38 | 0,01 | -0,43 | >0.1 | 3,74 | 4,33 | 0 | 1,88 | 0 | 0 | 0 |
| FOXO1 | ACCTGTAGTCC | 1 | 330 | 341 | -670 | -659 | -3,27 | 0,01 | 10,57 | >0.1 | 2,59 | 4,65 | 0 | 1,14 | 0 | 5,46 | 0 |
| FOXO1 | AAATGTTGCCC | 1 | 536 | 547 | -464 | -453 | 0,94 | 0,01 | 11,75 | >0.1 | 3,12 | 5,05 | 0 | 0,99 | 0 | 1,66 | 0 |
| FOXO1 | GATTGTTTACT | 1 | 894 | 905 | -106 | -95 | 10,1 | 0,001 | 27,11 | >0.1 | 5,6 | 6,66 | 0 | 1,89 | 0 | 2,86 | 0 |
| FOXO1 | TACTGTGTTCT | 1 | 901 | 912 | -99 | -88 | -2,78 | 0,01 | 12,71 | >0.1 | 5,6 | 6,72 | 0 | 1,89 | 0 | 1,28 | 0 |
| FOXO1 | TCTTCTTTTTC | 1 | 912 | 923 | -88 | -77 | -3,27 | 0,01 | 12,21 | >0.1 | 5,6 | 6,84 | 0 | 1,89 | 0 | 1,15 | 0 |
| FOXO1 | TTCTTTTTCTT | 1 | 914 | 925 | -86 | -75 | -6 | 0,01 | 9,5 | >0.1 | 5,6 | 6,87 | 0 | 1,89 | 0 | 1,15 | 0 |
| FOXO1 | TTTTCTTTTTT | 1 | 918 | 929 | -82 | -71 | -5,77 | 0,01 | 9,75 | >0.1 | 5,6 | 6,91 | 0 | 1,89 | 0 | 1,12 | 0 |
| FOXO1 | TGGTCTTTCCT | 1 | 929 | 940 | -71 | -60 | -4,02 | 0,01 | 11,55 | >0.1 | 5,6 | 7,06 | 0 | 1,88 | 0 | 1,03 | 0 |
| FOXO1 | CCTTGTTGCTG | 1 | 1078 | 1089 | 78 | 89 | 2,13 | 0,01 | 20,63 | >0.1 | 5,6 | 7,04 | 0 | 1,85 | 1,79 | 2,21 | 0 |
| FOXO1 | ACTTGCTTCTT | 1 | 1180 | 1191 | 180 | 191 | -5,33 | 0,01 | 13,03 | >0.1 | 5,6 | 6,09 | 0 | 1,58 | 1,79 | 3,29 | 0 |
| FOXO1 | GCCTGTTCCTC | -1 | 1112 | 1123 | 112 | 123 | -3,23 | 0,01 | 16,6 | >0.1 | 5,6 | 6,63 | 0 | 1,85 | 3,22 | 2,54 | 0 |
| FOXO1 | TGTTGGTGTTT | -1 | 630 | 641 | -370 | -359 | -6,05 | 0,01 | 5,73 | >0.1 | 3,71 | 5,3 | 0 | 1,05 | 0 | 1,73 | 0 |
| FOXO1 | TGGTGTTTGTA | -1 | 627 | 638 | -373 | -362 | -2,12 | 0,01 | 9,44 | >0.1 | 3,68 | 5,29 | 0 | 1,04 | 0 | 1,55 | 0 |
| FOXO1 | TATGGTTTTCA | -1 | 618 | 629 | -382 | -371 | -3,64 | 0,01 | 7,81 | >0.1 | 3,61 | 5,26 | 0 | 1,03 | 0 | 1,55 | 0 |
| FOXO1 | CCTTGTCTTTA | -1 | 501 | 512 | -499 | -488 | -4,06 | 0,01 | 10,38 | >0.1 | 2,96 | 4,97 | 0 | 0,98 | 0 | 5,52 | 0 |
| FOXO1 | ATCAGTTTTTT | -1 | 473 | 484 | -527 | -516 | -5,88 | 0,01 | 7,82 | >0.1 | 2,86 | 4,91 | 0 | 0,99 | 0 | 4,95 | 0 |
| FOXO1 | AGTTTTTTTTT | -1 | 470 | 481 | -530 | -519 | -6,03 | 0,01 | 7,66 | >0.1 | 2,85 | 4,9 | 0 | 0,99 | 0 | 4,95 | 0 |
| FOXO1 | GTCTGTCGCCC | -1 | 432 | 443 | -568 | -557 | -5,74 | 0,01 | 8,49 | >0.1 | 2,72 | 4,83 | 0 | 1,01 | 0 | 5,67 | 0 |
| FOXO1 | TTTTGTATTTT | -1 | 295 | 306 | -705 | -694 | -5,77 | 0,01 | 7,19 | >0.1 | 2,69 | 4,6 | 0 | 1,22 | 0 | 4,45 | 0 |
| FOXO1 | TCCTATTTCTC | -1 | 56 | 67 | -944 | -933 | -3,49 | 0,01 | 6,73 | >0.1 | 4,04 | 4,31 | 0 | 1,87 | 0 | 0 | 0 |
| FOXO6 | AGAAATA | 1 | 57 | 64 | -943 | -936 | -1,97 | 0,01 | 8,26 | >0.1 | 4,05 | 4,3 | 0 | 1,87 | 0 | 0 | 0 |
| FOXO6 | AAAAATA | 1 | 294 | 301 | -706 | -699 | -0,21 | 0,01 | 12,75 | >0.1 | 2,7 | 4,59 | 0 | 1,22 | 0 | 4,45 | 0 |
| FOXO6 | AAATACA | 1 | 296 | 303 | -704 | -697 | -3,59 | 0,1 | 9,44 | >0.1 | 2,7 | 4,59 | 0 | 1,21 | 0 | 4,53 | 0 |
| FOXO6 | AGAAATA | 1 | 485 | 492 | -515 | -508 | -1,97 | 0,01 | 12,76 | >0.1 | 2,9 | 4,93 | 0 | 0,98 | 0 | 5,93 | 0 |
| FOXO6 | ATAAAGA | 1 | 500 | 507 | -500 | -493 | 2,17 | 0,01 | 17,5 | >0.1 | 2,95 | 4,96 | 0 | 0,98 | 0 | 6,44 | 0 |
| FOXO6 | AAAAATA | 1 | 581 | 588 | -419 | -412 | -0,21 | 0,01 | 9,28 | >0.1 | 3,34 | 5,15 | 0 | 1 | 0 | 0 | 0 |
| FOXO6 | ATAAATG | 1 | 585 | 592 | -415 | -408 | -3,26 | 0,01 | 6,27 | >0.1 | 3,37 | 5,16 | 0 | 1 | 0 | 0 | 0 |
| FOXO6 | GAAAATA | 1 | 724 | 731 | -276 | -269 | 2,57 | 0,01 | 20,19 | >0.1 | 5,65 | 5,6 | 0 | 1,2 | 0 | 5,17 | 0 |
| FOXO6 | AGCAACA | -1 | 1081 | 1088 | 81 | 88 | 0,6 | 0,01 | 19,11 | >0.1 | 5,65 | 7 | 0 | 1,85 | 1,79 | 2,21 | 0 |
| FOXO6 | GGAAAGA | -1 | 932 | 939 | -68 | -61 | -2,32 | 0,01 | 13,26 | >0.1 | 5,65 | 7,05 | 0 | 1,88 | 0 | 1 | 0 |
| FOXO6 | AAAAAGA | -1 | 921 | 928 | -79 | -72 | -3,34 | 0,01 | 12,18 | >0.1 | 5,65 | 6,9 | 0 | 1,88 | 0 | 1,09 | 0 |
| FOXO6 | AAAAAGA | -1 | 915 | 922 | -85 | -78 | -3,34 | 0,01 | 12,18 | >0.1 | 5,65 | 6,83 | 0 | 1,89 | 0 | 1,15 | 0 |
| FOXO6 | GAACACA | -1 | 904 | 911 | -96 | -89 | 2,58 | 0,01 | 18,11 | >0.1 | 5,65 | 6,71 | 0 | 1,89 | 0 | 1,28 | 0 |
| FOXO6 | ATACACG | -1 | 700 | 707 | -300 | -293 | -3,24 | 0,01 | 12,26 | >0.1 | 5,65 | 5,51 | 0 | 1,15 | 0 | 3,19 | 0 |
| FOXO6 | GGCAACA | -1 | 539 | 546 | -461 | -454 | 3,38 | 0,01 | 14,19 | >0.1 | 3,12 | 5,05 | 0 | 0,98 | 0 | 1,66 | 0 |
| FOXO6 | AAATACA | -1 | 521 | 528 | -479 | -472 | -3,59 | 0,1 | 8,62 | >0.1 | 3,04 | 5 | 0 | 0,98 | 0 | 3,19 | 0 |
| FOXO6 | GACTACA | -1 | 333 | 340 | -667 | -660 | -2,93 | 0,01 | 11,47 | >0.1 | 2,59 | 4,65 | 0 | 1,13 | 0 | 6,03 | 0 |
| FOXO4 | AGAAATA | 1 | 57 | 64 | -943 | -936 | 2,11 | 0,01 | 12,34 | >0.1 | 4,05 | 4,3 | 0 | 1,87 | 0 | 0 | 0 |
| FOXO4 | AAAAATA | 1 | 294 | 301 | -706 | -699 | 3,1 | 0,01 | 16,06 | >0.1 | 2,7 | 4,59 | 0 | 1,22 | 0 | 4,45 | 0 |
| FOXO4 | AGAAATA | 1 | 485 | 492 | -515 | -508 | 2,11 | 0,01 | 16,84 | >0.1 | 2,9 | 4,93 | 0 | 0,98 | 0 | 5,93 | 0 |
| FOXO4 | ATAAAGA | 1 | 500 | 507 | -500 | -493 | 2,71 | 0,01 | 18,04 | >0.1 | 2,95 | 4,96 | 0 | 0,98 | 0 | 6,44 | 0 |
| FOXO4 | ATAAAAA | 1 | 579 | 586 | -421 | -414 | 3,02 | 0,01 | 12,49 | >0.1 | 3,33 | 5,14 | 0 | 1 | 0 | 0 | 0 |
| FOXO4 | AAAAATA | 1 | 581 | 588 | -419 | -412 | 3,1 | 0,01 | 12,59 | >0.1 | 3,34 | 5,15 | 0 | 1 | 0 | 0 | 0 |

| | | | | | | | | | | | | | | | | | |
|---|---|---|---|---|---|---|---|---|---|---|---|---|---|---|---|---|---|
| FOXO4 | ACAAACA | 1 | 628 | 635 | -372 | -365 | 4,26 | 0,01 | 15,79 | >0.1 | 3,67 | 5,28 | 0 | 1,04 | 0 | 1,55 | 0 |
| FOXO4 | ACCAACA | 1 | 634 | 641 | -366 | -359 | 2,66 | 0,01 | 14,33 | >0.1 | 3,73 | 5,3 | 0 | 1,04 | 0 | 1,61 | 0 |
| FOXO4 | GAAAATA | 1 | 724 | 731 | -276 | -269 | 5,28 | 0,01 | 22,9 | >0.1 | 5,65 | 5,6 | 0 | 1,2 | 0 | 5,17 | 0 |
| FOXO4 | AGCAACA | -1 | 1081 | 1088 | 81 | 88 | 3,5 | 0,01 | 22,01 | >0.1 | 5,65 | 7 | 0 | 1,85 | 1,79 | 2,21 | 0 |
| FOXO4 | GAACACA | -1 | 904 | 911 | -96 | -89 | 3,56 | 0,01 | 19,09 | >0.1 | 5,65 | 6,71 | 0 | 1,89 | 0 | 1,28 | 0 |
| FOXO4 | GTAAACA | -1 | 897 | 904 | -103 | -96 | 12,23 | 0,0001 | 29,37 | >0.1 | 5,65 | 6,65 | 0 | 1,89 | 0 | 2,95 | 0 |
| FOXO4 | GCCAATA | -1 | 881 | 888 | -119 | -112 | 1,86 | 0,01 | 19,11 | >0.1 | 5,65 | 6,51 | 0 | 1,89 | 0 | 3,19 | 0 |
| FOXO4 | ATCAAAA | -1 | 748 | 755 | -252 | -245 | 1,42 | 0,01 | 19,96 | >0.1 | 5,65 | 5,7 | 0 | 1,27 | 0 | 5,92 | 0 |
| FOXO4 | GGCAACA | -1 | 539 | 546 | -461 | -454 | 5,68 | 0,01 | 16,49 | >0.1 | 3,12 | 5,05 | 0 | 0,98 | 0 | 1,66 | 0 |
| FOXO4 | GTCAACT | -1 | 105 | 112 | -895 | -888 | 3,57 | 0,01 | 14,16 | >0.1 | 3,58 | 4,35 | 0 | 1,88 | 0 | 0,77 | 0 |
| TEAD3 | ACAGACCG | 1 | 437 | 445 | -563 | -555 | -5,75 | 0,01 | 8,74 | >0.1 | 2,73 | 4,83 | 0 | 1 | 0 | 5,92 | 0 |
| TEAD3 | GAATTCCA | 1 | 493 | 501 | -507 | -499 | 9,76 | 0,001 | 24,88 | >0.1 | 2,93 | 4,95 | 0 | 0,98 | 0 | 6,27 | 0 |
| TEAD3 | AAACACCA | 1 | 630 | 638 | -370 | -362 | -8,34 | 0,01 | 3,34 | >0.1 | 3,69 | 5,29 | 0 | 1,04 | 0 | 1,66 | 0 |
| TEAD3 | TGATCCCT | 1 | 650 | 658 | -350 | -342 | -5,42 | 0,01 | 8,71 | >0.1 | 3,88 | 5,35 | 0 | 1,07 | 0 | 3,83 | 0 |
| TEAD3 | ACATATCT | 1 | 851 | 859 | -149 | -141 | -6,58 | 0,01 | 9,95 | >0.1 | 5,64 | 6,29 | 0 | 1,85 | 0 | 2,75 | 0 |
| TEAD3 | TCTTTCCT | 1 | 932 | 940 | -68 | -60 | -7,7 | 0,01 | 7,83 | >0.1 | 5,64 | 7,06 | 0 | 1,88 | 0 | 0,95 | 0 |
| TEAD3 | TCAGTCCA | 1 | 1099 | 1107 | 99 | 107 | -6,55 | 0,01 | 13,28 | >0.1 | 5,64 | 6,77 | 0 | 1,85 | 3,22 | 2,35 | 0 |
| TEAD3 | ACAGGCCA | 1 | 1117 | 1125 | 117 | 125 | -6,68 | 0,01 | 13,21 | >0.1 | 5,64 | 6,58 | 0 | 1,85 | 3,22 | 2,6 | 0 |
| TEAD3 | TAATTCCA | -1 | 1190 | 1198 | 190 | 198 | 6,26 | 0,001 | 24,9 | >0.1 | 5,64 | 6,03 | 0 | 1,51 | 1,79 | 3,67 | 0 |
| TEAD3 | ACTTTCCC | -1 | 1006 | 1014 | 6 | 14 | -7,42 | 0,01 | 10,62 | >0.1 | 5,64 | 9,12 | 0 | 1,87 | 0 | 1,42 | 0 |
| TEAD3 | AAAGACCA | -1 | 929 | 937 | -71 | -63 | -8,34 | 0,01 | 7,23 | >0.1 | 5,64 | 7,02 | 0 | 1,88 | 0 | 1,03 | 0 |
| TEAD3 | ATATTCCC | -1 | 562 | 570 | -438 | -430 | -7,74 | 0,01 | 1,59 | >0.1 | 3,24 | 5,1 | 0 | 0,99 | 0 | 0 | 0 |
| TEAD3 | ACATGACT | -1 | 516 | 524 | -484 | -476 | -8,32 | 0,01 | 3,54 | >0.1 | 3,02 | 5 | 0 | 0,98 | 0 | 2,86 | 0 |
| TEAD3 | GAATTCAT | -1 | 491 | 499 | -509 | -501 | -8,09 | 0,01 | 7,02 | >0.1 | 2,92 | 4,94 | 0 | 0,98 | 0 | 6,27 | 0 |
| TEAD3 | AATTTCCA | -1 | 152 | 160 | -848 | -840 | -6,95 | 0,01 | 4,28 | >0.1 | 3,28 | 4,4 | 0 | 1,9 | 0 | 1,65 | 0 |
| TEAD3 | ACTTACCT | -1 | 3 | 11 | -997 | -989 | -6,09 | 0,01 | 7,46 | >0.1 | 5,64 | 4,25 | 0 | 1,86 | 1,79 | 0 | 0 |
| TEAD2 | AGACCGTCCTGAC | 1 | 255 | 268 | -745 | -732 | 1,7 | 0,01 | 15,14 | >0.1 | 2,82 | 4,54 | 0 | 1,35 | 0 | 4,73 | 0 |
| TEAD2 | ATGAATTCCATAA | 1 | 491 | 504 | -509 | -496 | 5,52 | 0,01 | 20,66 | >0.1 | 2,93 | 4,95 | 0 | 0,99 | 0 | 6,27 | 0 |
| TEAD2 | TTACATATCTGTC | 1 | 849 | 862 | -151 | -138 | 4,17 | 0,01 | 20,68 | >0.1 | 5,57 | 6,31 | 0 | 1,88 | 0 | 2,75 | 0 |
| TEAD2 | GGTCTTTCCTGCT | 1 | 930 | 943 | -70 | -57 | 2,6 | 0,01 | 18,11 | >0.1 | 5,57 | 7,11 | 0 | 1,88 | 0 | 0,95 | 0 |
| TEAD2 | GCTCAGTCCACCA | 1 | 1097 | 1110 | 97 | 110 | 4,05 | 0,01 | 23,83 | >0.1 | 5,57 | 6,79 | 0 | 1,85 | 3,22 | 2,35 | 0 |
| TEAD2 | AGACATTTTTGGA | 1 | 1125 | 1138 | 125 | 138 | 2,95 | 0,01 | 22,84 | >0.1 | 5,57 | 6,5 | 0 | 1,84 | 3,22 | 2,75 | 0 |
| TEAD2 | TATAATTCCAAGA | -1 | 1187 | 1200 | 187 | 200 | 3,11 | 0,01 | 21,75 | >0.1 | 5,57 | 6,05 | 0 | 1,53 | 1,79 | 3,7 | 0 |
| TEAD2 | TGACTTTCCCTAG | -1 | 1003 | 1016 | 3 | 16 | 4,8 | 0,01 | 22,77 | >0.1 | 5,57 | 9,12 | 0 | 1,87 | 0 | 1,42 | 0 |
| TEAD2 | TCATATTCCCAGA | -1 | 559 | 572 | -441 | -428 | 6,56 | 0,01 | 15,9 | >0.1 | 3,24 | 5,11 | 0 | 1 | 0 | 0 | 0 |
| TEAD2 | CAACATTTTCCCT | -1 | 531 | 544 | -469 | -456 | 1,34 | 0,01 | 12,08 | >0.1 | 3,1 | 5,04 | 0 | 0,99 | 0 | 1,61 | 0 |
| TEAD2 | TTCCCTTCCAAAT | -1 | 524 | 537 | -476 | -463 | 1,37 | 0,01 | 13,73 | >0.1 | 3,07 | 5,03 | 0 | 0,99 | 0 | 3,28 | 0 |
| TEAD2 | TCATATTTCTGAT | -1 | 482 | 495 | -518 | -505 | 2,92 | 0,01 | 17,67 | >0.1 | 2,9 | 4,93 | 0 | 0,99 | 0 | 5,93 | 0 |
| TEAD2 | CGCCATTCTCCTG | -1 | 362 | 375 | -638 | -625 | 2,08 | 0,01 | 17,01 | >0.1 | 2,53 | 4,71 | 0 | 1,08 | 0 | 6,61 | 0 |
| TEAD2 | TAGGATTACAGGC | -1 | 196 | 209 | -804 | -791 | 1,41 | 0,01 | 14,96 | >0.1 | 3,05 | 4,47 | 0 | 1,66 | 0 | 4,37 | 0 |
| TEAD2 | AGCCATTTCTGAT | -1 | 167 | 180 | -833 | -820 | 2,98 | 0,01 | 14,95 | >0.1 | 3,19 | 4,43 | 0 | 1,9 | 0 | 2,46 | 0 |
| TEAD2 | TTAATTTCCATGG | -1 | 149 | 162 | -851 | -838 | 3,47 | 0,01 | 14,7 | >0.1 | 3,28 | 4,41 | 0 | 1,9 | 0 | 1,65 | 0 |
| TEAD2 | CTACTCTCCACTC | -1 | 117 | 130 | -883 | -870 | 3,37 | 0,01 | 13,83 | >0.1 | 3,48 | 4,37 | 0 | 1,89 | 0 | 0,73 | 0 |
| TEAD2 | CTCCACTCCACAG | -1 | 112 | 125 | -888 | -875 | 4,07 | 0,01 | 14,6 | >0.1 | 3,51 | 4,37 | 0 | 1,89 | 0 | 0,77 | 0 |
| TEAD2 | GCTCATTTCCTAT | -1 | 61 | 74 | -939 | -926 | 1,64 | 0,01 | 11,8 | >0.1 | 3,97 | 4,31 | 0 | 1,88 | 0 | 0 | 0 |
| TEAD1 | TGAATTCCAT | 1 | 492 | 502 | -508 | -498 | 9,44 | 0,001 | 24,57 | >0.1 | 2,93 | 4,95 | 0 | 0,98 | 0 | 6,27 | 0 |
| TEAD1 | CAAACACCAA | 1 | 629 | 639 | -371 | -361 | 2,45 | 0,01 | 14,14 | >0.1 | 3,69 | 5,29 | 0 | 1,04 | 0 | 1,66 | 0 |
| TEAD1 | AGCTTTCTAG | 1 | 714 | 724 | -286 | -276 | 3,02 | 0,01 | 20,38 | >0.1 | 5,61 | 5,58 | 0 | 1,19 | 0 | 4,99 | 0 |
| TEAD1 | CGTATTCTTA | 1 | 805 | 815 | -195 | -185 | 3,79 | 0,01 | 20,35 | >0.1 | 5,61 | 6 | 0 | 1,52 | 0 | 3,43 | 0 |
| TEAD1 | TACATATCTG | 1 | 850 | 860 | -150 | -140 | 3,49 | 0,01 | 20,01 | >0.1 | 5,61 | 6,3 | 0 | 1,86 | 0 | 2,75 | 0 |
| TEAD1 | CTCAGTCCAC | 1 | 1098 | 1108 | 98 | 108 | 2,45 | 0,01 | 22,26 | >0.1 | 5,61 | 6,78 | 0 | 1,85 | 3,22 | 2,35 | 0 |
| TEAD1 | ATAATTCCAA | -1 | 1189 | 1199 | 189 | 199 | 5,06 | 0,01 | 23,69 | >0.1 | 5,61 | 6,03 | 0 | 1,52 | 1,79 | 3,67 | 0 |
| TEAD1 | GACTTTCCCT | -1 | 1005 | 1015 | 5 | 15 | 4,69 | 0,01 | 22,7 | >0.1 | 5,61 | 9,12 | 0 | 1,87 | 0 | 1,42 | 0 |
| TEAD1 | ACCTTACCTC | -1 | 775 | 785 | -225 | -215 | 2,82 | 0,01 | 21,97 | >0.1 | 5,61 | 5,84 | 0 | 1,38 | 0 | 6,32 | 0 |
| TEAD1 | CATATTCCCA | -1 | 561 | 571 | -439 | -429 | 4,9 | 0,01 | 14,24 | >0.1 | 3,24 | 5,11 | 0 | 0,99 | 0 | 0 | 0 |
| TEAD1 | TCCCTTCCAA | -1 | 526 | 536 | -474 | -464 | 6 | 0,01 | 18,5 | >0.1 | 3,07 | 5,02 | 0 | 0,98 | 0 | 3,43 | 0 |
| TEAD1 | TGCCTCCCAG | -1 | 380 | 390 | -620 | -610 | 3,26 | 0,01 | 17,53 | >0.1 | 2,57 | 4,73 | 0 | 1,06 | 0 | 5,91 | 0 |
| TEAD1 | GCCATTCTCC | -1 | 364 | 374 | -636 | -626 | 5,4 | 0,01 | 20,33 | >0.1 | 2,53 | 4,71 | 0 | 1,08 | 0 | 6,61 | 0 |
| TEAD1 | AGCCTCCCGA | -1 | 348 | 358 | -652 | -642 | 2,16 | 0,01 | 16,89 | >0.1 | 2,55 | 4,68 | 0 | 1,11 | 0 | 6,4 | 0 |

| | | | | | | | | | | | | | | | | | |
|---|---|---|---|---|---|---|---|---|---|---|---|---|---|---|---|---|---|
| TEAD1 | GGCCTCCCAA | -1 | 214 | 224 | -786 | -776 | 3,18 | 0,01 | 17,71 | >0.1 | 2,98 | 4,48 | 0 | 1,54 | 0 | 5,52 | 0 |
| TEAD1 | GCCATTTCTG | -1 | 169 | 179 | -831 | -821 | 2,22 | 0,01 | 15,23 | >0.1 | 3,18 | 4,43 | 0 | 1,89 | 0 | 3,51 | 0 |
| TEAD1 | TAATTTCCAT | -1 | 151 | 161 | -849 | -839 | 4,69 | 0,01 | 15,92 | >0.1 | 3,28 | 4,41 | 0 | 1,9 | 0 | 1,65 | 0 |
| TEAD1 | TCCACTCCAC | -1 | 114 | 124 | -886 | -876 | 4,17 | 0,01 | 14,68 | >0.1 | 3,51 | 4,36 | 0 | 1,89 | 0 | 0,75 | 0 |
| TEAD1 | TGACTTCTTT | -1 | 30 | 40 | -970 | -960 | 3,03 | 0,01 | 13,66 | >0.1 | 4,48 | 4,28 | 0 | 1,87 | 0 | 0 | 0 |
| TEAD1 | CACTTACCTA | -1 | 2 | 12 | -998 | -988 | 5,95 | 0,01 | 19,47 | >0.1 | 5,61 | 4,26 | 0 | 1,86 | 1,79 | 0 | 0 |
| TEAD4 | AAAATACAAA | 1 | 295 | 305 | -705 | -695 | 2,58 | 0,01 | 15,54 | >0.1 | 2,69 | 4,6 | 0 | 1,22 | 0 | 4,45 | 0 |
| TEAD4 | TGAATTCCAT | 1 | 492 | 502 | -508 | -498 | 9,8 | 0,001 | 24,93 | >0.1 | 2,93 | 4,95 | 0 | 0,98 | 0 | 6,27 | 0 |
| TEAD4 | AACATTTTCC | 1 | 691 | 701 | -309 | -299 | 2,25 | 0,01 | 16,48 | >0.1 | 4,53 | 5,49 | 0 | 1,14 | 0 | 3,07 | 0 |
| TEAD4 | AGCTTTCTAG | 1 | 714 | 724 | -286 | -276 | 4,13 | 0,01 | 21,49 | >0.1 | 5,61 | 5,58 | 0 | 1,19 | 0 | 4,99 | 0 |
| TEAD4 | CGTATTCTTA | 1 | 805 | 815 | -195 | -185 | 3,57 | 0,01 | 20,13 | >0.1 | 5,61 | 6 | 0 | 1,52 | 0 | 3,43 | 0 |
| TEAD4 | TACATATCTG | 1 | 850 | 860 | -150 | -140 | 3,38 | 0,01 | 19,9 | >0.1 | 5,61 | 6,3 | 0 | 1,86 | 0 | 2,75 | 0 |
| TEAD4 | GTCTTTCCTG | 1 | 931 | 941 | -69 | -59 | 2,15 | 0,01 | 17,67 | >0.1 | 5,61 | 7,08 | 0 | 1,88 | 0 | 0,95 | 0 |
| TEAD4 | GTCATTCAGT | 1 | 1012 | 1022 | 12 | 22 | 2,21 | 0,01 | 20,02 | >0.1 | 5,61 | 8,92 | 0 | 1,86 | 0 | 1,42 | 0 |
| TEAD4 | GACATTTTTG | 1 | 1126 | 1136 | 126 | 136 | 4,09 | 0,01 | 24,01 | >0.1 | 5,61 | 6,49 | 0 | 1,84 | 3,22 | 2,75 | 0 |
| TEAD4 | ATAATTCCAA | -1 | 1189 | 1199 | 189 | 199 | 5,27 | 0,01 | 23,9 | >0.1 | 5,61 | 6,03 | 0 | 1,52 | 1,79 | 3,67 | 0 |
| TEAD4 | GACTTTCCCT | -1 | 1005 | 1015 | 5 | 15 | 4,76 | 0,01 | 22,77 | >0.1 | 5,61 | 9,12 | 0 | 1,87 | 0 | 1,42 | 0 |
| TEAD4 | AACAATCTGC | -1 | 891 | 901 | -109 | -99 | 3,85 | 0,01 | 20,84 | >0.1 | 5,61 | 6,62 | 0 | 1,89 | 0 | 2,86 | 0 |
| TEAD4 | CACATTTATT | -1 | 584 | 594 | -416 | -406 | 2,12 | 0,01 | 11,66 | >0.1 | 3,37 | 5,16 | 0 | 1 | 0 | 0 | 0 |
| TEAD4 | CATATTCCCA | -1 | 561 | 571 | -439 | -429 | 4,01 | 0,01 | 13,35 | >0.1 | 3,24 | 5,11 | 0 | 0,99 | 0 | 0 | 0 |
| TEAD4 | AACATTTTCC | -1 | 533 | 543 | -467 | -457 | 2,25 | 0,01 | 12,98 | >0.1 | 3,1 | 5,04 | 0 | 0,98 | 0 | 1,61 | 0 |
| TEAD4 | CAAATACATG | -1 | 519 | 529 | -481 | -471 | 3,76 | 0,01 | 15,98 | >0.1 | 3,04 | 5,01 | 0 | 0,98 | 0 | 3,19 | 0 |
| TEAD4 | GGAATTCATA | -1 | 490 | 500 | -510 | -500 | 4,58 | 0,01 | 19,7 | >0.1 | 2,92 | 4,94 | 0 | 0,98 | 0 | 6,27 | 0 |
| TEAD4 | GCCATTCTCC | -1 | 364 | 374 | -636 | -626 | 4,05 | 0,01 | 18,98 | >0.1 | 2,53 | 4,71 | 0 | 1,08 | 0 | 6,61 | 0 |
| TEAD4 | TAATTTCCAT | -1 | 151 | 161 | -849 | -839 | 4,85 | 0,01 | 16,08 | >0.1 | 3,28 | 4,41 | 0 | 1,9 | 0 | 1,65 | 0 |
| TEAD4 | CACTTACCTA | -1 | 2 | 12 | -998 | -988 | 4,7 | 0,01 | 18,22 | >0.1 | 5,61 | 4,26 | 0 | 1,86 | 1,79 | 0 | 0 |
| KLF13 | GTGAAACCCCGTCTCTAT | 1 | 275 | 293 | -725 | -707 | -15,49 | 0,01 | -1,69 | >0.1 | 2,75 | 4,58 | 0 | 1,28 | 0 | 5,19 | 0 |
| KLF13 | AAACCCCGTCTCTATTAA | 1 | 278 | 296 | -722 | -704 | -10,82 | 0,01 | 2,86 | >0.1 | 2,73 | 4,58 | 0 | 1,27 | 0 | 5,09 | 0 |
| KLF13 | GTGGTGGGCACCTGTAGT | 1 | 321 | 339 | -679 | -661 | -12,45 | 0,01 | 1,43 | >0.1 | 2,61 | 4,65 | 0 | 1,16 | 0 | 5,46 | 0 |
| KLF13 | AGATCGCGCCACTGTGCT | 1 | 407 | 425 | -593 | -575 | -5,22 | 0,001 | 9,03 | >0.1 | 2,66 | 4,8 | 0 | 1,03 | 0 | 5,76 | 0 |
| KLF13 | GTTCTAGACCTCTTTGGT | 1 | 659 | 677 | -341 | -323 | -8,36 | 0,01 | 6,19 | >0.1 | 4,05 | 5,41 | 0 | 1,1 | 0 | 3,99 | 0 |
| KLF13 | GCTCAGCGCCCAACCCAA | 1 | 940 | 958 | -60 | -42 | -16,39 | 0,01 | 0,16 | >0.1 | 5,51 | 7,36 | 0 | 1,88 | 0 | 1,8 | 0 |
| KLF13 | ATGTCAAGCTCTTCCTGG | 1 | 1049 | 1067 | 49 | 67 | -15,48 | 0,01 | 1,7 | >0.1 | 5,51 | 7,6 | 0 | 1,86 | 0 | 2,21 | 0 |
| KLF13 | GGAACAGGCCAAGACATT | 1 | 1114 | 1132 | 114 | 132 | -14,12 | 0,01 | 5,73 | >0.1 | 5,51 | 6,61 | 0 | 1,85 | 3,22 | 2,67 | 0 |
| KLF13 | AGAACAGGTCTTCGGCTT | -1 | 1152 | 1170 | 152 | 170 | -17,17 | 0,01 | -0,49 | >0.1 | 4,42 | 6,28 | 0 | 1,84 | 1,79 | 2,35 | 0 |
| KLF13 | TGACATCGTCCCCTGTGA | -1 | 1037 | 1055 | 37 | 55 | -8,05 | 0,01 | 8,66 | >0.1 | 5,51 | 7,95 | 0 | 1,86 | 0 | 1,39 | 0 |
| KLF13 | AGATCACATCCACTGAAT | -1 | 1015 | 1033 | 15 | 33 | -3,86 | 0,001 | 13,65 | >0.1 | 5,51 | 8,84 | 0 | 1,86 | 0 | 1,31 | 0 |
| KLF13 | CTAAAACCTCCTCATGAG | -1 | 984 | 1002 | -16 | 2 | -14,18 | 0,01 | 3,58 | >0.1 | 5,51 | 8,77 | 0 | 1,87 | 0 | 1,61 | 0 |
| KLF13 | ATGGAATTCATATTTCTG | -1 | 484 | 502 | -516 | -498 | -17,02 | 0,01 | -1,9 | >0.1 | 2,91 | 4,95 | 0 | 0,99 | 0 | 6,27 | 0 |
| KLF13 | AAGCTCTGCCTCCCAGGT | -1 | 378 | 396 | -622 | -604 | -16,14 | 0,01 | -1,84 | >0.1 | 2,58 | 4,74 | 0 | 1,07 | 0 | 5,91 | 0 |
| KLF13 | GGTTCACGCCATTCTCCT | -1 | 363 | 381 | -637 | -619 | -9,04 | 0,01 | 6,03 | >0.1 | 2,54 | 4,72 | 0 | 1,09 | 0 | 6,73 | 0 |
| KLF13 | CCACCACGCCCAGCTAAG | -1 | 307 | 325 | -693 | -675 | 4,25 | 0,0001 | 18,75 | >0.1 | 2,65 | 4,63 | 0 | 1,19 | 0 | 6,03 | 0 |
| KLF13 | GTGATCTGCCCAACTCGG | -1 | 222 | 240 | -778 | -760 | -15,04 | 0,01 | -0,85 | >0.1 | 2,93 | 4,51 | 0 | 1,5 | 0 | 5,26 | 0 |
| KLF13 | TTTCCATGGCCATTTTTT | -1 | 140 | 158 | -860 | -842 | -5,05 | 0,001 | 6,14 | >0.1 | 3,32 | 4,4 | 0 | 1,9 | 0 | 1,57 | 0 |
| KLF13 | TAAAAAAGCTCATTTCCT | -1 | 63 | 81 | -937 | -919 | -15,48 | 0,01 | -5,36 | >0.1 | 3,92 | 4,32 | 0 | 1,88 | 0 | 0 | 0 |
| KLF13 | TTTGAGTGCCTCCTTCTC | -1 | 15 | 33 | -985 | -967 | -15,41 | 0,01 | -3,76 | >0.1 | 5,51 | 4,28 | 0 | 1,87 | 0 | 0 | 0 |
| KLF12 | GGAGGCACTCAAAGA | 1 | 20 | 35 | -980 | -965 | -9,05 | 0,01 | 2,64 | >0.1 | 5,54 | 4,28 | 0 | 1,87 | 0 | 0 | 0 |
| KLF12 | GCTCACGCCTGTAAT | 1 | 190 | 205 | -810 | -795 | -7,6 | 0,01 | 5,9 | >0.1 | 3,07 | 4,46 | 0 | 1,71 | 0 | 4,26 | 0 |
| KLF12 | GAGACCGTCCTGACT | 1 | 254 | 269 | -746 | -731 | -6,36 | 0,01 | 7,09 | >0.1 | 2,82 | 4,55 | 0 | 1,35 | 0 | 4,73 | 0 |
| KLF12 | GGTGAAACCCCGTCT | 1 | 274 | 289 | -726 | -711 | -3,64 | 0,01 | 10,06 | >0.1 | 2,75 | 4,57 | 0 | 1,28 | 0 | 5,09 | 0 |
| KLF12 | AACCCCGTCTCTATT | 1 | 279 | 294 | -721 | -706 | -1,33 | 0,01 | 12,45 | >0.1 | 2,74 | 4,58 | 0 | 1,27 | 0 | 5,19 | 0 |
| KLF12 | GATCGCGCCACTGTG | 1 | 408 | 423 | -592 | -577 | -8,65 | 0,01 | 5,48 | >0.1 | 2,65 | 4,79 | 0 | 1,03 | 0 | 5,65 | 0 |
| KLF12 | TGCTCAGCGCCCAAC | 1 | 939 | 954 | -61 | -46 | -8,9 | 0,01 | 6,6 | >0.1 | 5,54 | 7,29 | 0 | 1,88 | 0 | 0,79 | 0 |
| KLF12 | CGCCCAACCCAAGTT | 1 | 946 | 961 | -54 | -39 | 0,77 | 0,001 | 17,41 | >0.1 | 5,54 | 7,42 | 0 | 1,88 | 0 | 1,8 | 0 |
| KLF12 | TGTCAAGCTCTTCCT | 1 | 1050 | 1065 | 50 | 65 | -0,56 | 0,01 | 16,69 | >0.1 | 5,54 | 7,57 | 0 | 1,86 | 0 | 2,27 | 0 |
| KLF12 | TGCTCAGTCCACCAT | 1 | 1096 | 1111 | 96 | 111 | -8,93 | 0,01 | 10,84 | >0.1 | 5,54 | 6,8 | 0 | 1,85 | 3,22 | 2,35 | 0 |
| KLF12 | GGCCAAGACATTTTT | 1 | 1120 | 1135 | 120 | 135 | -8,83 | 0,01 | 11,07 | >0.1 | 5,54 | 6,55 | 0 | 1,85 | 3,22 | 2,74 | 0 |
| KLF12 | GACATCGTCCCCTGT | -1 | 1039 | 1054 | 39 | 54 | -8,72 | 0,01 | 7,96 | >0.1 | 5,54 | 7,88 | 0 | 1,86 | 0 | 1,39 | 0 |
| KLF12 | AGCCAAGATCACATC | -1 | 1023 | 1038 | 23 | 38 | -6,75 | 0,01 | 10,57 | >0.1 | 5,54 | 8,67 | 0 | 1,86 | 0 | 1,25 | 0 |

| | | | | | | | | | | | | | | | | | |
|---|---|---|---|---|---|---|---|---|---|---|---|---|---|---|---|---|---|
| KLF12 | GATCACATCCACTGA | -1 | 1017 | 1032 | 17 | 32 | 2,01 | 0,001 | 19,51 | >0.1 | 5,54 | 8,79 | 0 | 1,86 | 0 | 1,31 | 0 |
| KLF12 | GACTAAAACCTCCTC | -1 | 989 | 1004 | -11 | 4 | -8,64 | 0,01 | 9,19 | >0.1 | 5,54 | 8,84 | 0 | 1,87 | 0 | 1,57 | 0 |
| KLF12 | AGAAACTCTCTACTT | -1 | 547 | 562 | -453 | -438 | -7,66 | 0,01 | 1,6 | >0.1 | 3,18 | 5,08 | 0 | 0,99 | 0 | 0 | 0 |
| KLF12 | GACTTTGCCCTTGTC | -1 | 505 | 520 | -495 | -480 | -7,57 | 0,01 | 5,59 | >0.1 | 2,99 | 4,99 | 0 | 0,99 | 0 | 4,19 | 0 |
| KLF12 | GTTCACGCCATTCTC | -1 | 365 | 380 | -635 | -620 | -8,38 | 0,01 | 6,68 | >0.1 | 2,54 | 4,72 | 0 | 1,08 | 0 | 6,73 | 0 |
| KLF12 | CGCCATTCTCCTGCC | -1 | 360 | 375 | -640 | -625 | -7,24 | 0,01 | 7,57 | >0.1 | 2,52 | 4,71 | 0 | 1,09 | 0 | 6,49 | 0 |
| KLF12 | TGCCTCAGCCTCCCG | -1 | 349 | 364 | -651 | -636 | -9,01 | 0,01 | 5,72 | >0.1 | 2,54 | 4,69 | 0 | 1,11 | 0 | 6,4 | 0 |
| KLF12 | GCCCACCACCACGCC | -1 | 315 | 330 | -685 | -670 | -8,31 | 0,01 | 6,38 | >0.1 | 2,63 | 4,64 | 0 | 1,17 | 0 | 6,26 | 0 |
| KLF12 | CACCACGCCCAGCTA | -1 | 309 | 324 | -691 | -676 | 15,1 | 1E-05 | 29,59 | >0.1 | 2,65 | 4,63 | 0 | 1,19 | 0 | 6,03 | 0 |
| KLF12 | TGATCTGCCCAACTC | -1 | 224 | 239 | -776 | -761 | -7,51 | 0,01 | 6,67 | >0.1 | 2,93 | 4,5 | 0 | 1,49 | 0 | 5,26 | 0 |
| KLF12 | TGCCCAGCCATTTCT | -1 | 170 | 185 | -830 | -815 | -4,63 | 0,01 | 8,2 | >0.1 | 3,17 | 4,44 | 0 | 1,88 | 0 | 3,34 | 0 |
| KLF12 | GACTACTCTCCACTC | -1 | 117 | 132 | -883 | -868 | -1,56 | 0,01 | 9,82 | >0.1 | 3,47 | 4,37 | 0 | 1,89 | 0 | 1,65 | 0 |
| KLF12 | AAAAAAGCTCATTTC | -1 | 65 | 80 | -935 | -920 | -7,77 | 0,01 | 2,34 | >0.1 | 3,91 | 4,32 | 0 | 1,88 | 0 | 0 | 0 |
| PITX1 | GTCAGCCA | 1 | 36 | 44 | -964 | -956 | 3,18 | 0,01 | 13,71 | >0.1 | 4,38 | 4,28 | 0 | 1,87 | 0 | 0 | 0 |
| PITX1 | ATAATTTT | 1 | 132 | 140 | -868 | -860 | 2,49 | 0,01 | 13,73 | >0.1 | 3,4 | 4,38 | 0 | 1,89 | 0 | 1,57 | 0 |
| PITX1 | GTAATCCT | 1 | 200 | 208 | -800 | -792 | 9,23 | 0,001 | 22,84 | >0.1 | 3,04 | 4,46 | 0 | 1,63 | 0 | 4,48 | 0 |
| PITX1 | GTGAGCCG | 1 | 399 | 407 | -601 | -593 | 3,39 | 0,01 | 17,72 | >0.1 | 2,62 | 4,76 | 0 | 1,04 | 0 | 5,91 | 0 |
| PITX1 | ATGATCTT | 1 | 568 | 576 | -432 | -424 | 3,32 | 0,01 | 12,7 | >0.1 | 3,27 | 5,12 | 0 | 0,99 | 0 | 0 | 0 |
| PITX1 | ATGATCCC | 1 | 649 | 657 | -351 | -343 | 7,53 | 0,001 | 21,64 | >0.1 | 3,87 | 5,34 | 0 | 1,06 | 0 | 3,83 | 0 |
| PITX1 | TTGATTCA | 1 | 816 | 824 | -184 | -176 | 3,12 | 0,01 | 19,81 | >0.1 | 5,64 | 6,05 | 0 | 1,58 | 0 | 3,43 | 0 |
| PITX1 | AAAATCTC | 1 | 979 | 987 | -21 | -13 | 3,81 | 0,01 | 21,03 | >0.1 | 5,64 | 8,18 | 0 | 1,87 | 0 | 1,53 | 0 |
| PITX1 | GTGATCTT | 1 | 1026 | 1034 | 26 | 34 | 2,99 | 0,01 | 20,19 | >0.1 | 5,64 | 8,45 | 0 | 1,86 | 0 | 1,25 | 0 |
| PITX1 | CTCAGCCT | 1 | 1073 | 1081 | 73 | 81 | 4,47 | 0,01 | 23,15 | >0.1 | 5,64 | 7,12 | 0 | 1,85 | 1,79 | 2,27 | 0 |
| PITX1 | ATAATTCC | -1 | 1191 | 1199 | 191 | 199 | 6,7 | 0,01 | 25,48 | >0.1 | 5,64 | 6,02 | 0 | 1,5 | 1,79 | 3,82 | 0 |
| PITX1 | GTGAGCCA | -1 | 1033 | 1041 | 33 | 41 | 3,62 | 0,01 | 20,5 | >0.1 | 5,64 | 8,1 | 0 | 1,86 | 0 | 1,28 | 0 |
| PITX1 | CACATCCA | -1 | 1021 | 1029 | 21 | 29 | 2,47 | 0,01 | 19,98 | >0.1 | 5,64 | 8,7 | 0 | 1,86 | 0 | 1,31 | 0 |
| PITX1 | ATCAGCCT | -1 | 964 | 972 | -36 | -28 | 3,49 | 0,01 | 20,37 | >0.1 | 5,64 | 7,67 | 0 | 1,87 | 0 | 1,7 | 0 |
| PITX1 | CTGAGCCA | -1 | 884 | 892 | -116 | -108 | 4,93 | 0,01 | 22,08 | >0.1 | 5,64 | 6,55 | 0 | 1,89 | 0 | 3,07 | 0 |
| PITX1 | TTCATCCT | -1 | 868 | 876 | -132 | -124 | 6,88 | 0,01 | 23,58 | >0.1 | 5,64 | 6,41 | 0 | 1,9 | 0 | 2,75 | 0 |
| PITX1 | ATAATTTT | -1 | 830 | 838 | -170 | -162 | 2,49 | 0,01 | 19,22 | >0.1 | 5,64 | 6,14 | 0 | 1,67 | 0 | 3,28 | 0 |
| PITX1 | TTAATCCT | -1 | 790 | 798 | -210 | -202 | 10,71 | 0,001 | 29,21 | >0.1 | 5,64 | 5,91 | 0 | 1,43 | 0 | 5,52 | 0 |
| PITX1 | TTAATATT | -1 | 727 | 735 | -273 | -265 | 3,56 | 0,01 | 21,35 | >0.1 | 5,64 | 5,62 | 0 | 1,21 | 0 | 5,32 | 0 |
| PITX1 | ATAATTTT | -1 | 683 | 691 | -317 | -309 | 2,49 | 0,01 | 16,47 | >0.1 | 4,34 | 5,46 | 0 | 1,12 | 0 | 3,07 | 0 |
| PITX1 | ATATTCCC | -1 | 562 | 570 | -438 | -430 | 3,42 | 0,01 | 12,75 | >0.1 | 3,24 | 5,1 | 0 | 0,99 | 0 | 0 | 0 |
| PITX1 | CTCAGCCT | -1 | 353 | 361 | -647 | -639 | 4,47 | 0,01 | 19,19 | >0.1 | 2,54 | 4,68 | 0 | 1,1 | 0 | 6,4 | 0 |
| PITX1 | GTGATCTG | -1 | 232 | 240 | -768 | -760 | 2,78 | 0,01 | 16,65 | >0.1 | 2,91 | 4,51 | 0 | 1,44 | 0 | 5,01 | 0 |
| PITX1 | GTGAGCCA | -1 | 188 | 196 | -812 | -804 | 3,62 | 0,01 | 16,74 | >0.1 | 3,1 | 4,45 | 0 | 1,72 | 0 | 3,86 | 0 |
| PITX1 | TTAATTTC | -1 | 154 | 162 | -846 | -838 | 5,01 | 0,01 | 16,23 | >0.1 | 3,27 | 4,41 | 0 | 1,9 | 0 | 1,65 | 0 |
| PITX1 | CTGATCTG | -1 | 94 | 102 | -906 | -898 | 4,1 | 0,01 | 14,8 | >0.1 | 3,67 | 4,34 | 0 | 1,88 | 0 | 0,81 | 0 |
| KLF16 | CTCACGCCTGT | 1 | 191 | 202 | -809 | -798 | 1,44 | 0,01 | 14,8 | >0.1 | 3,08 | 4,46 | 0 | 1,7 | 0 | 4,13 | 0 |
| KLF16 | ATCGCGCCACT | 1 | 409 | 420 | -591 | -580 | -0,05 | 0,01 | 13,94 | >0.1 | 2,65 | 4,79 | 0 | 1,03 | 0 | 5,52 | 0 |
| KLF16 | CAAACACCAAC | 1 | 629 | 640 | -371 | -360 | 1,06 | 0,01 | 12,76 | >0.1 | 3,7 | 5,29 | 0 | 1,05 | 0 | 1,66 | 0 |
| KLF16 | GCTCAGCGCCC | 1 | 940 | 951 | -60 | -49 | 1,23 | 0,01 | 16,77 | >0.1 | 5,6 | 7,23 | 0 | 1,88 | 0 | 0,83 | 0 |
| KLF16 | TCAGCGCCCAA | 1 | 942 | 953 | -58 | -47 | -0,74 | 0,01 | 14,83 | >0.1 | 5,6 | 7,27 | 0 | 1,88 | 0 | 0,83 | 0 |
| KLF16 | GCCCAACCCAA | 1 | 947 | 958 | -53 | -42 | 4,24 | 0,01 | 20,81 | >0.1 | 5,6 | 7,36 | 0 | 1,88 | 0 | 1,74 | 0 |
| KLF16 | CTCAGTCCACC | 1 | 1098 | 1109 | 98 | 109 | -1,63 | 0,01 | 18,17 | >0.1 | 5,6 | 6,78 | 0 | 1,85 | 3,22 | 2,35 | 0 |
| KLF16 | CATCGTCCCCT | -1 | 1041 | 1052 | 41 | 52 | -0,64 | 0,01 | 16,03 | >0.1 | 5,6 | 7,82 | 0 | 1,86 | 0 | 1,39 | 0 |
| KLF16 | AAACCTCCTCA | -1 | 988 | 999 | -12 | -1 | -1,88 | 0,01 | 15,88 | >0.1 | 5,6 | 8,69 | 0 | 1,87 | 0 | 1,61 | 0 |
| KLF16 | GCTCTGCCTCC | -1 | 383 | 394 | -617 | -606 | 2,52 | 0,01 | 16,91 | >0.1 | 2,58 | 4,74 | 0 | 1,06 | 0 | 6,02 | 0 |
| KLF16 | CCTCAGCCTCC | -1 | 351 | 362 | -649 | -638 | 2,71 | 0,01 | 17,43 | >0.1 | 2,54 | 4,69 | 0 | 1,1 | 0 | 6,4 | 0 |
| KLF16 | TGCCCACCACC | -1 | 320 | 331 | -680 | -669 | 2,65 | 0,01 | 16,75 | >0.1 | 2,62 | 4,64 | 0 | 1,16 | 0 | 5,68 | 0 |
| KLF16 | CCCACCACCAC | -1 | 318 | 329 | -682 | -671 | 5,58 | 0,01 | 20,26 | >0.1 | 2,63 | 4,63 | 0 | 1,16 | 0 | 6,26 | 0 |
| KLF16 | ACCACCACGCC | -1 | 315 | 326 | -685 | -674 | -1,66 | 0,01 | 12,9 | >0.1 | 2,63 | 4,63 | 0 | 1,17 | 0 | 6,13 | 0 |
| KLF16 | CCACCACGCCC | -1 | 314 | 325 | -686 | -675 | 3,01 | 0,01 | 17,58 | >0.1 | 2,64 | 4,63 | 0 | 1,17 | 0 | 6,13 | 0 |
| KLF16 | ACCACGCCCAG | -1 | 312 | 323 | -688 | -677 | 10,61 | 0,001 | 25,09 | >0.1 | 2,64 | 4,62 | 0 | 1,18 | 0 | 6,03 | 0 |
| KLF16 | ACTCGGCCTCC | -1 | 217 | 228 | -783 | -772 | 1,08 | 0,01 | 15,58 | >0.1 | 2,96 | 4,49 | 0 | 1,52 | 0 | 5,52 | 0 |
| KLF16 | GCCCAGCCATT | -1 | 173 | 184 | -827 | -816 | -0,62 | 0,01 | 12,06 | >0.1 | 3,16 | 4,43 | 0 | 1,85 | 0 | 3,24 | 0 |
| KLF16 | TCCACTCCACA | -1 | 113 | 124 | -887 | -876 | 1,41 | 0,01 | 11,94 | >0.1 | 3,51 | 4,36 | 0 | 1,89 | 0 | 0,77 | 0 |
| KLF14 | GCTCACGCCTGTAA | 1 | 190 | 204 | -810 | -796 | -3,34 | 0,01 | 10,16 | >0.1 | 3,07 | 4,46 | 0 | 1,71 | 0 | 4,26 | 0 |

| | | | | | | | | | | | | | | | | | |
|---|---|---|---|---|---|---|---|---|---|---|---|---|---|---|---|---|---|
| KLF14 | AACCCCGTCTCTAT | 1 | 279 | 293 | -721 | -707 | -4,82 | 0,01 | 8,96 | >0.1 | 2,74 | 4,58 | 0 | 1,27 | 0 | 5,19 | 0 |
| KLF14 | AGTCCCAGCTACTC | 1 | 336 | 350 | -664 | -650 | -4,88 | 0,01 | 9,4 | >0.1 | 2,57 | 4,67 | 0 | 1,13 | 0 | 5,92 | 0 |
| KLF14 | GATCGCGCCACTGT | 1 | 408 | 422 | -592 | -578 | -2,67 | 0,01 | 11,45 | >0.1 | 2,65 | 4,79 | 0 | 1,03 | 0 | 5,65 | 0 |
| KLF14 | TGCTCAGCGCCCAA | 1 | 939 | 953 | -61 | -47 | -5,76 | 0,01 | 9,73 | >0.1 | 5,56 | 7,27 | 0 | 1,88 | 0 | 0,79 | 0 |
| KLF14 | CGCCCAACCCAAGT | 1 | 946 | 960 | -54 | -40 | -7,42 | 0,01 | 9,22 | >0.1 | 5,56 | 7,4 | 0 | 1,88 | 0 | 1,8 | 0 |
| KLF14 | TGTCAAGCTCTTCC | 1 | 1050 | 1064 | 50 | 64 | -6,27 | 0,01 | 10,99 | >0.1 | 5,56 | 7,57 | 0 | 1,86 | 0 | 2,27 | 0 |
| KLF14 | GATCACATCCACTG | -1 | 1018 | 1032 | 18 | 32 | -2,09 | 0,01 | 15,4 | >0.1 | 5,56 | 8,76 | 0 | 1,86 | 0 | 1,31 | 0 |
| KLF14 | AGCTCTGCCTCCCA | -1 | 381 | 395 | -619 | -605 | -1,02 | 0,01 | 13,38 | >0.1 | 2,58 | 4,74 | 0 | 1,06 | 0 | 6,02 | 0 |
| KLF14 | GTTCACGCCATTCT | -1 | 366 | 380 | -634 | -620 | -5,08 | 0,01 | 9,98 | >0.1 | 2,54 | 4,72 | 0 | 1,08 | 0 | 6,73 | 0 |
| KLF14 | TGCCTCAGCCTCCC | -1 | 350 | 364 | -650 | -636 | -6,86 | 0,01 | 7,87 | >0.1 | 2,54 | 4,69 | 0 | 1,1 | 0 | 6,4 | 0 |
| KLF14 | GCCTCAGCCTCCCG | -1 | 349 | 363 | -651 | -637 | -4,87 | 0,01 | 9,86 | >0.1 | 2,54 | 4,69 | 0 | 1,11 | 0 | 6,4 | 0 |
| KLF14 | GCCCACCACCACGC | -1 | 316 | 330 | -684 | -670 | -5,64 | 0,01 | 9,05 | >0.1 | 2,63 | 4,64 | 0 | 1,17 | 0 | 6,26 | 0 |
| KLF14 | CACCACGCCCAGCT | -1 | 310 | 324 | -690 | -676 | 12,37 | 0,0001 | 26,86 | >0.1 | 2,65 | 4,63 | 0 | 1,18 | 0 | 6,03 | 0 |
| KLF14 | TGATCTGCCCAACT | -1 | 225 | 239 | -775 | -761 | -2,64 | 0,01 | 11,53 | >0.1 | 2,93 | 4,5 | 0 | 1,48 | 0 | 5,26 | 0 |
| KLF14 | CAACTCGGCCTCCC | -1 | 216 | 230 | -784 | -770 | -5,43 | 0,01 | 9,19 | >0.1 | 2,96 | 4,49 | 0 | 1,53 | 0 | 5,63 | 0 |
| KLF14 | AACTCGGCCTCCCA | -1 | 215 | 229 | -785 | -771 | -5,4 | 0,01 | 9,12 | >0.1 | 2,97 | 4,49 | 0 | 1,54 | 0 | 5,52 | 0 |
| KLF14 | CACTGTGCCCAGCC | -1 | 176 | 190 | -824 | -810 | -5,87 | 0,01 | 6,69 | >0.1 | 3,14 | 4,44 | 0 | 1,83 | 0 | 3,16 | 0 |
| KLF14 | GTGCCCAGCCATTT | -1 | 172 | 186 | -828 | -814 | -6,37 | 0,01 | 6,33 | >0.1 | 3,16 | 4,44 | 0 | 1,86 | 0 | 3,24 | 0 |
| KLF14 | TTCCATGGCCATTT | -1 | 143 | 157 | -857 | -843 | -4,47 | 0,01 | 6,75 | >0.1 | 3,31 | 4,4 | 0 | 1,9 | 0 | 1,61 | 0 |
| KLF14 | GACTACTCTCCACT | -1 | 118 | 132 | -882 | -868 | -2,7 | 0,01 | 8,68 | >0.1 | 3,47 | 4,37 | 0 | 1,89 | 0 | 1,65 | 0 |
| TFAP2A | TGGCTCACGCC | 1 | 188 | 199 | -812 | -801 | 5,7 | 0,01 | 18,91 | >0.1 | 3,09 | 4,45 | 0 | 1,72 | 0 | 3,95 | 0 |
| TFAP2A | AGGCTGAGGCA | 1 | 353 | 364 | -647 | -636 | 10,56 | 0,001 | 25,28 | >0.1 | 2,53 | 4,69 | 0 | 1,1 | 0 | 6,4 | 0 |
| TFAP2A | TGCTCCAGCCT | 1 | 421 | 432 | -579 | -568 | 3,28 | 0,01 | 17,21 | >0.1 | 2,69 | 4,81 | 0 | 1,02 | 0 | 5,42 | 0 |
| TFAP2A | CAGCCTGGGCG | 1 | 426 | 437 | -574 | -563 | 0,94 | 0,01 | 15,14 | >0.1 | 2,7 | 4,82 | 0 | 1,01 | 0 | 5,67 | 0 |
| TFAP2A | AGCCTGGGCGA | 1 | 427 | 438 | -573 | -562 | 3,32 | 0,01 | 17,66 | >0.1 | 2,71 | 4,82 | 0 | 1,01 | 0 | 5,8 | 0 |
| TFAP2A | TGCTCAGCGCC | 1 | 939 | 950 | -61 | -50 | 1,09 | 0,01 | 16,63 | >0.1 | 5,6 | 7,22 | 0 | 1,88 | 0 | 0,85 | 0 |
| TFAP2A | CGCCCAACCCA | 1 | 946 | 957 | -54 | -43 | 2,18 | 0,01 | 18,8 | >0.1 | 5,6 | 7,34 | 0 | 1,88 | 0 | 1,8 | 0 |
| TFAP2A | GGCTCACAGGG | 1 | 1034 | 1045 | 34 | 45 | -0,7 | 0,01 | 16,13 | >0.1 | 5,6 | 8,06 | 0 | 1,86 | 0 | 1,31 | 0 |
| TFAP2A | CCCTGTGAGCC | -1 | 1034 | 1045 | 34 | 45 | -0,13 | 0,01 | 16,7 | >0.1 | 5,6 | 8,06 | 0 | 1,86 | 0 | 1,31 | 0 |
| TFAP2A | TCGCCCAGGCT | -1 | 427 | 438 | -573 | -562 | 8,01 | 0,001 | 22,35 | >0.1 | 2,71 | 4,82 | 0 | 1,01 | 0 | 5,8 | 0 |
| TFAP2A | AGGCTGGAGCA | -1 | 421 | 432 | -579 | -568 | 5,61 | 0,01 | 19,54 | >0.1 | 2,69 | 4,81 | 0 | 1,02 | 0 | 5,42 | 0 |
| TFAP2A | CACTGCAAGCT | -1 | 391 | 402 | -609 | -598 | 1,98 | 0,01 | 16,4 | >0.1 | 2,6 | 4,75 | 0 | 1,05 | 0 | 6,02 | 0 |
| TFAP2A | GCCTCCCAGGT | -1 | 378 | 389 | -622 | -611 | -0,58 | 0,01 | 14,31 | >0.1 | 2,57 | 4,73 | 0 | 1,06 | 0 | 6,53 | 0 |
| TFAP2A | AGGTTCACGCC | -1 | 371 | 382 | -629 | -618 | 1,27 | 0,01 | 16,01 | >0.1 | 2,55 | 4,72 | 0 | 1,07 | 0 | 6,4 | 0 |
| TFAP2A | TGCCTCAGCCT | -1 | 353 | 364 | -647 | -636 | 8,37 | 0,001 | 23,09 | >0.1 | 2,53 | 4,69 | 0 | 1,1 | 0 | 6,4 | 0 |
| TFAP2A | AGCCTCCCGAG | -1 | 347 | 358 | -653 | -642 | 2,84 | 0,01 | 17,7 | >0.1 | 2,55 | 4,68 | 0 | 1,11 | 0 | 6,52 | 0 |
| TFAP2A | AGGCGTGAGCC | -1 | 189 | 200 | -811 | -800 | 0,83 | 0,01 | 14,12 | >0.1 | 3,08 | 4,45 | 0 | 1,71 | 0 | 4,04 | 0 |
| MSC | ATGAGCTTTT | 1 | 68 | 78 | -932 | -922 | 0,48 | 0,01 | 10,58 | >0.1 | 3,91 | 4,32 | 0 | 1,88 | 0 | 0 | 0 |
| MSC | GGCACCTGTA | 1 | 327 | 337 | -673 | -663 | 1,58 | 0,01 | 15,54 | >0.1 | 2,6 | 4,65 | 0 | 1,14 | 0 | 5,56 | 0 |
| MSC | CCCAGCTACT | 1 | 339 | 349 | -661 | -651 | -3,26 | 0,01 | 11,62 | >0.1 | 2,57 | 4,66 | 0 | 1,12 | 0 | 6,52 | 0 |
| MSC | TAGACCTCTT | 1 | 663 | 673 | -337 | -327 | -0,75 | 0,01 | 13,77 | >0.1 | 4,05 | 5,4 | 0 | 1,09 | 0 | 3,99 | 0 |
| MSC | AACAGCTTTC | 1 | 711 | 721 | -289 | -279 | 10,93 | 0,0001 | 28,27 | >0.1 | 5,61 | 5,56 | 0 | 1,18 | 0 | 4,99 | 0 |
| MSC | AGTACCGGTT | 1 | 740 | 750 | -260 | -250 | -2,84 | 0,01 | 15,62 | >0.1 | 5,61 | 5,68 | 0 | 1,26 | 0 | 5,92 | 0 |
| MSC | TACCGGTTTT | 1 | 742 | 752 | -258 | -248 | -2,29 | 0,01 | 16,19 | >0.1 | 5,61 | 5,69 | 0 | 1,26 | 0 | 5,92 | 0 |
| MSC | AAAATCTGTT | 1 | 841 | 851 | -159 | -149 | 0,02 | 0,01 | 15,06 | >0.1 | 5,61 | 6,24 | 0 | 1,78 | 0 | 1,41 | 0 |
| MSC | AGCAGATTGT | 1 | 890 | 900 | -110 | -100 | -3,07 | 0,01 | 13,99 | >0.1 | 5,61 | 6,61 | 0 | 1,89 | 0 | 2,95 | 0 |
| MSC | AAAGGCTGAT | 1 | 962 | 972 | -38 | -28 | -3,42 | 0,01 | 13,43 | >0.1 | 5,61 | 7,67 | 0 | 1,87 | 0 | 1,7 | 0 |
| MSC | AGGAGGTTTT | 1 | 990 | 1000 | -10 | 0 | 0,54 | 0,01 | 18,34 | >0.1 | 5,61 | 8,71 | 0 | 1,87 | 0 | 1,61 | 0 |
| MSC | AACTGCTGCT | 1 | 1090 | 1100 | 90 | 100 | 7,19 | 0,001 | 25,6 | >0.1 | 5,61 | 6,88 | 0 | 1,85 | 1,79 | 2,27 | 0 |
| MSC | AAGACATTTT | 1 | 1124 | 1134 | 124 | 134 | -2,52 | 0,01 | 17,46 | >0.1 | 5,61 | 6,51 | 0 | 1,84 | 3,22 | 2,79 | 0 |
| MSC | AAGACCTGTT | 1 | 1158 | 1168 | 158 | 168 | 10,95 | 0,0001 | 27,54 | >0.1 | 4,38 | 6,24 | 0 | 1,78 | 1,79 | 2,4 | 0 |
| MSC | AACAGGTCTT | -1 | 1158 | 1168 | 158 | 168 | 5,02 | 0,001 | 21,61 | >0.1 | 4,38 | 6,24 | 0 | 1,78 | 1,79 | 2,4 | 0 |
| MSC | AGCAGCAGTT | -1 | 1090 | 1100 | 90 | 100 | 5,51 | 0,001 | 23,92 | >0.1 | 5,61 | 6,88 | 0 | 1,85 | 1,79 | 2,27 | 0 |
| MSC | AAGAGCTTGA | -1 | 1052 | 1062 | 52 | 62 | -3,07 | 0,01 | 14,2 | >0.1 | 5,61 | 7,53 | 0 | 1,86 | 0 | 2,27 | 0 |
| MSC | ATCAGCCTTT | -1 | 962 | 972 | -38 | -28 | -3,02 | 0,01 | 13,83 | >0.1 | 5,61 | 7,67 | 0 | 1,87 | 0 | 1,7 | 0 |
| MSC | AACAGATTTT | -1 | 841 | 851 | -159 | -149 | 4,42 | 0,01 | 19,46 | >0.1 | 5,61 | 6,24 | 0 | 1,78 | 0 | 1,41 | 0 |
| MSC | GAAAGCTGTT | -1 | 711 | 721 | -289 | -279 | 7,53 | 0,001 | 24,87 | >0.1 | 5,61 | 5,56 | 0 | 1,18 | 0 | 4,99 | 0 |
| MSC | AATAGGTTAT | -1 | 596 | 606 | -404 | -394 | -1,87 | 0,01 | 9,23 | >0.1 | 3,45 | 5,2 | 0 | 1,01 | 0 | 1,44 | 0 |
| MSC | AGTAGCTGGG | -1 | 339 | 349 | -661 | -651 | 0,51 | 0,01 | 15,39 | >0.1 | 2,57 | 4,66 | 0 | 1,12 | 0 | 6,52 | 0 |

| | | | | | | | | | | | | | | | | | |
|---|---|---|---|---|---|---|---|---|---|---|---|---|---|---|---|---|---|
| MSC | TACAGGTGCC | -1 | 327 | 337 | -673 | -663 | 5,51 | 0,001 | 19,47 | >0.1 | 2,6 | 4,65 | 0 | 1,14 | 0 | 5,56 | 0 |
| MSC | TACAGGCGTG | -1 | 193 | 203 | -807 | -797 | -1,3 | 0,01 | 12,17 | >0.1 | 3,07 | 4,46 | 0 | 1,68 | 0 | 4,26 | 0 |
| MSC | GCCTCCTTCT | -1 | 16 | 26 | -984 | -974 | -3,29 | 0,01 | 8,45 | >0.1 | 5,61 | 4,27 | 0 | 1,87 | 0 | 0 | 0 |
| TFAP2C | TGGCTCACGCCT | 1 | 188 | 200 | -812 | -800 | 5,41 | 0,01 | 18,71 | >0.1 | 3,09 | 4,45 | 0 | 1,72 | 0 | 4,04 | 0 |
| TFAP2C | TGGTGGTGGGCA | 1 | 319 | 331 | -681 | -669 | 1,93 | 0,01 | 16,03 | >0.1 | 2,62 | 4,64 | 0 | 1,16 | 0 | 5,68 | 0 |
| TFAP2C | ACTCGGGAGGCT | 1 | 346 | 358 | -654 | -642 | 1,15 | 0,01 | 16,01 | >0.1 | 2,55 | 4,68 | 0 | 1,11 | 0 | 6,52 | 0 |
| TFAP2C | AGGCTGAGGCAG | 1 | 353 | 365 | -647 | -635 | 2,5 | 0,01 | 17,34 | >0.1 | 2,53 | 4,69 | 0 | 1,1 | 0 | 6,52 | 0 |
| TFAP2C | AACCTGGGAGGC | 1 | 377 | 389 | -623 | -611 | 1,11 | 0,01 | 16 | >0.1 | 2,56 | 4,73 | 0 | 1,06 | 0 | 6,53 | 0 |
| TFAP2C | ACCTGGGAGGCA | 1 | 378 | 390 | -622 | -610 | 4,31 | 0,01 | 19,34 | >0.1 | 2,57 | 4,73 | 0 | 1,06 | 0 | 6,66 | 0 |
| TFAP2C | CAGCCTGGGCGA | 1 | 426 | 438 | -574 | -562 | 5,35 | 0,01 | 19,69 | >0.1 | 2,7 | 4,82 | 0 | 1,01 | 0 | 5,8 | 0 |
| TFAP2C | GGCTCACAGGGG | 1 | 1034 | 1046 | 34 | 46 | 2,09 | 0,01 | 18,9 | >0.1 | 5,58 | 8,06 | 0 | 1,86 | 0 | 1,31 | 0 |
| TFAP2C | CCCCTGTGAGCC | -1 | 1034 | 1046 | 34 | 46 | 2,4 | 0,01 | 19,21 | >0.1 | 5,58 | 8,06 | 0 | 1,86 | 0 | 1,31 | 0 |
| TFAP2C | CCCTGTGAGCCA | -1 | 1033 | 1045 | 33 | 45 | 1,97 | 0,01 | 18,82 | >0.1 | 5,58 | 8,1 | 0 | 1,86 | 0 | 1,31 | 0 |
| TFAP2C | TCGCCCAGGCTG | -1 | 426 | 438 | -574 | -562 | 5,24 | 0,01 | 19,58 | >0.1 | 2,7 | 4,82 | 0 | 1,01 | 0 | 5,8 | 0 |
| TFAP2C | TGCCTCCCAGGT | -1 | 378 | 390 | -622 | -610 | 3,41 | 0,01 | 18,44 | >0.1 | 2,57 | 4,73 | 0 | 1,06 | 0 | 6,66 | 0 |
| TFAP2C | CTGCCTCAGCCT | -1 | 353 | 365 | -647 | -635 | 1,81 | 0,01 | 16,65 | >0.1 | 2,53 | 4,69 | 0 | 1,1 | 0 | 6,52 | 0 |
| TFAP2C | AGCCTCCCGAGT | -1 | 346 | 358 | -654 | -642 | 1,73 | 0,01 | 16,59 | >0.1 | 2,55 | 4,68 | 0 | 1,11 | 0 | 6,52 | 0 |
| TFAP2C | TGCCCACCACCA | -1 | 319 | 331 | -681 | -669 | 3,61 | 0,01 | 17,71 | >0.1 | 2,62 | 4,64 | 0 | 1,16 | 0 | 5,68 | 0 |
| TFAP2C | AGGCGTGAGCCA | -1 | 188 | 200 | -812 | -800 | 5,97 | 0,01 | 19,27 | >0.1 | 3,09 | 4,45 | 0 | 1,72 | 0 | 4,04 | 0 |
| TFAP2B | TGGCTCACGCCT | 1 | 188 | 200 | -812 | -800 | 5,27 | 0,01 | 18,57 | >0.1 | 3,09 | 4,45 | 0 | 1,72 | 0 | 4,04 | 0 |
| TFAP2B | TGGTGGTGGGCA | 1 | 319 | 331 | -681 | -669 | 0,25 | 0,01 | 14,35 | >0.1 | 2,62 | 4,64 | 0 | 1,16 | 0 | 5,68 | 0 |
| TFAP2B | ACTCGGGAGGCT | 1 | 346 | 358 | -654 | -642 | 0,51 | 0,01 | 15,37 | >0.1 | 2,55 | 4,68 | 0 | 1,11 | 0 | 6,52 | 0 |
| TFAP2B | AGGCTGAGGCAG | 1 | 353 | 365 | -647 | -635 | 1,73 | 0,01 | 16,57 | >0.1 | 2,53 | 4,69 | 0 | 1,1 | 0 | 6,52 | 0 |
| TFAP2B | AACCTGGGAGGC | 1 | 377 | 389 | -623 | -611 | 0,78 | 0,01 | 15,67 | >0.1 | 2,56 | 4,73 | 0 | 1,06 | 0 | 6,53 | 0 |
| TFAP2B | ACCTGGGAGGCA | 1 | 378 | 390 | -622 | -610 | 3,01 | 0,01 | 18,04 | >0.1 | 2,57 | 4,73 | 0 | 1,06 | 0 | 6,66 | 0 |
| TFAP2B | CAGCCTGGGCGA | 1 | 426 | 438 | -574 | -562 | 5,15 | 0,01 | 19,49 | >0.1 | 2,7 | 4,82 | 0 | 1,01 | 0 | 5,8 | 0 |
| TFAP2B | TGGCTCACAGGG | 1 | 1033 | 1045 | 33 | 45 | 0,71 | 0,01 | 17,56 | >0.1 | 5,58 | 8,1 | 0 | 1,86 | 0 | 1,31 | 0 |
| TFAP2B | GGCTCACAGGGG | 1 | 1034 | 1046 | 34 | 46 | 0,93 | 0,01 | 17,74 | >0.1 | 5,58 | 8,06 | 0 | 1,86 | 0 | 1,31 | 0 |
| TFAP2B | CCCCTGTGAGCC | -1 | 1034 | 1046 | 34 | 46 | 2,36 | 0,01 | 19,17 | >0.1 | 5,58 | 8,06 | 0 | 1,86 | 0 | 1,31 | 0 |
| TFAP2B | CCCTGTGAGCCA | -1 | 1033 | 1045 | 33 | 45 | 0,86 | 0,01 | 17,71 | >0.1 | 5,58 | 8,1 | 0 | 1,86 | 0 | 1,31 | 0 |
| TFAP2B | TCGCCCAGGCTG | -1 | 426 | 438 | -574 | -562 | 4,94 | 0,01 | 19,28 | >0.1 | 2,7 | 4,82 | 0 | 1,01 | 0 | 5,8 | 0 |
| TFAP2B | TGCCTCCCAGGT | -1 | 378 | 390 | -622 | -610 | 3,16 | 0,01 | 18,19 | >0.1 | 2,57 | 4,73 | 0 | 1,06 | 0 | 6,66 | 0 |
| TFAP2B | CTGCCTCAGCCT | -1 | 353 | 365 | -647 | -635 | 2,18 | 0,01 | 17,02 | >0.1 | 2,53 | 4,69 | 0 | 1,1 | 0 | 6,52 | 0 |
| TFAP2B | AGCCTCCCGAGT | -1 | 346 | 358 | -654 | -642 | 0,91 | 0,01 | 15,77 | >0.1 | 2,55 | 4,68 | 0 | 1,11 | 0 | 6,52 | 0 |
| TFAP2B | TGCCCACCACCA | -1 | 319 | 331 | -681 | -669 | 3,46 | 0,01 | 17,56 | >0.1 | 2,62 | 4,64 | 0 | 1,16 | 0 | 5,68 | 0 |
| TFAP2B | AGGCGTGAGCCA | -1 | 188 | 200 | -812 | -800 | 5,36 | 0,01 | 18,66 | >0.1 | 3,09 | 4,45 | 0 | 1,72 | 0 | 4,04 | 0 |
| UNCX | GAAATTAA | 1 | 154 | 162 | -846 | -838 | 6,11 | 0,01 | 17,33 | >0.1 | 3,27 | 4,41 | 0 | 1,9 | 0 | 1,65 | 0 |
| UNCX | CTGATCAG | 1 | 164 | 172 | -836 | -828 | 4,54 | 0,01 | 16,47 | >0.1 | 3,21 | 4,42 | 0 | 1,9 | 0 | 2,4 | 0 |
| UNCX | TCTATTAA | 1 | 288 | 296 | -712 | -704 | 5,55 | 0,01 | 19,28 | >0.1 | 2,72 | 4,58 | 0 | 1,24 | 0 | 5,19 | 0 |
| UNCX | CTGATCAG | 1 | 479 | 487 | -521 | -513 | 4,54 | 0,01 | 18,68 | >0.1 | 2,88 | 4,92 | 0 | 0,98 | 0 | 5,36 | 0 |
| UNCX | CCTATTAA | 1 | 600 | 608 | -400 | -392 | 6,1 | 0,01 | 17,15 | >0.1 | 3,47 | 5,2 | 0 | 1,01 | 0 | 1,36 | 0 |
| UNCX | TTAATGAA | 1 | 604 | 612 | -396 | -388 | 5,25 | 0,01 | 16,39 | >0.1 | 3,49 | 5,21 | 0 | 1,02 | 0 | 1,41 | 0 |
| UNCX | AAAATTAT | 1 | 683 | 691 | -317 | -309 | 3,9 | 0,01 | 17,88 | >0.1 | 4,34 | 5,46 | 0 | 1,12 | 0 | 3,07 | 0 |
| UNCX | TTGATTTG | 1 | 750 | 758 | -250 | -242 | 3,46 | 0,1 | 22,01 | >0.1 | 5,64 | 5,71 | 0 | 1,28 | 0 | 5,92 | 0 |
| UNCX | AAAATTAT | 1 | 830 | 838 | -170 | -162 | 3,9 | 0,01 | 20,63 | >0.1 | 5,64 | 6,14 | 0 | 1,67 | 0 | 3,28 | 0 |
| UNCX | CTGATAAG | 1 | 967 | 975 | -33 | -25 | 3,69 | 0,01 | 20,65 | >0.1 | 5,64 | 7,76 | 0 | 1,87 | 0 | 1,7 | 0 |
| UNCX | GGAATTAT | 1 | 1191 | 1199 | 191 | 199 | 4,16 | 0,01 | 22,94 | >0.1 | 5,64 | 6,02 | 0 | 1,5 | 1,79 | 3,82 | 0 |
| UNCX | ATAATTCC | -1 | 1191 | 1199 | 191 | 199 | 3,87 | 0,01 | 22,65 | >0.1 | 5,64 | 6,02 | 0 | 1,5 | 1,79 | 3,82 | 0 |
| UNCX | TGAATCAA | -1 | 816 | 824 | -184 | -176 | 3,63 | 0,01 | 20,32 | >0.1 | 5,64 | 6,05 | 0 | 1,58 | 0 | 3,43 | 0 |
| UNCX | CAAATAAG | -1 | 811 | 819 | -189 | -181 | 3,87 | 0,01 | 20,36 | >0.1 | 5,64 | 6,02 | 0 | 1,55 | 0 | 3,28 | 0 |
| UNCX | CAAATCAA | -1 | 750 | 758 | -250 | -242 | 4,61 | 0,01 | 23,16 | >0.1 | 5,64 | 5,71 | 0 | 1,28 | 0 | 5,92 | 0 |
| UNCX | ACAATTTC | -1 | 609 | 617 | -391 | -383 | 4,42 | 0,01 | 15,74 | >0.1 | 3,53 | 5,23 | 0 | 1,02 | 0 | 1,55 | 0 |
| UNCX | TTCATTAA | -1 | 604 | 612 | -396 | -388 | 5,76 | 0,01 | 16,9 | >0.1 | 3,49 | 5,21 | 0 | 1,02 | 0 | 1,41 | 0 |
| UNCX | CTGATCAG | -1 | 479 | 487 | -521 | -513 | 4,54 | 0,01 | 18,68 | >0.1 | 2,88 | 4,92 | 0 | 0,98 | 0 | 5,36 | 0 |
| UNCX | CTGATCAG | -1 | 164 | 172 | -836 | -828 | 4,54 | 0,01 | 16,47 | >0.1 | 3,21 | 4,42 | 0 | 1,9 | 0 | 2,4 | 0 |
| UNCX | TTAATTTC | -1 | 154 | 162 | -846 | -838 | 5,56 | 0,01 | 16,78 | >0.1 | 3,27 | 4,41 | 0 | 1,9 | 0 | 1,65 | 0 |
| UNCX | AAAATTAT | -1 | 132 | 140 | -868 | -860 | 3,9 | 0,01 | 15,14 | >0.1 | 3,4 | 4,38 | 0 | 1,89 | 0 | 1,57 | 0 |
| UNCX | TCACTTAC | -1 | 5 | 13 | -995 | -987 | 4,71 | 0,01 | 18,26 | >0.1 | 5,64 | 4,26 | 0 | 1,86 | 1,79 | 0 | 0 |
| NPAS2 | GGTAAGTGAA | 1 | 4 | 14 | -996 | -986 | -4,94 | 0,01 | 8,58 | >0.1 | 5,61 | 4,26 | 0 | 1,86 | 1,79 | 0 | 0 |

| | | | | | | | | | | | | | | | | | |
|---|---|---|---|---|---|---|---|---|---|---|---|---|---|---|---|---|---|
| NPAS2 | AACCAGTGTG | 1 | 47 | 57 | -953 | -943 | -3,61 | 0,01 | 6,73 | >0.1 | 4,17 | 4,3 | 0 | 1,87 | 0 | 0 | 0 |
| NPAS2 | GGCACCTGTA | 1 | 327 | 337 | -673 | -663 | 2,61 | 0,01 | 16,57 | >0.1 | 2,6 | 4,65 | 0 | 1,14 | 0 | 5,56 | 0 |
| NPAS2 | GACAAGGGCA | 1 | 505 | 515 | -495 | -485 | -4,18 | 0,01 | 10,44 | >0.1 | 2,98 | 4,98 | 0 | 0,98 | 0 | 5,68 | 0 |
| NPAS2 | CCCAAGTAGA | 1 | 544 | 554 | -456 | -446 | 4,81 | 0,01 | 14,01 | >0.1 | 3,15 | 5,06 | 0 | 0,99 | 0 | 0 | 0 |
| NPAS2 | CCCAAGTTCA | 1 | 953 | 963 | -47 | -37 | -3,99 | 0,01 | 12,65 | >0.1 | 5,61 | 7,46 | 0 | 1,88 | 0 | 1,7 | 0 |
| NPAS2 | GACAAGTTTA | 1 | 1136 | 1146 | 136 | 146 | -3,83 | 0,01 | 16,34 | >0.1 | 5,61 | 6,41 | 0 | 1,84 | 3,22 | 3,09 | 0 |
| NPAS2 | ACCACGAAGC | 1 | 1146 | 1156 | 146 | 156 | -2,85 | 0,01 | 14,88 | >0.1 | 5,61 | 6,33 | 0 | 1,84 | 1,79 | 2,16 | 0 |
| NPAS2 | TTCACTTGCT | 1 | 1177 | 1187 | 177 | 187 | -3,97 | 0,01 | 14,41 | >0.1 | 5,61 | 6,11 | 0 | 1,61 | 1,79 | 3,26 | 0 |
| NPAS2 | AGCAAGTGAA | -1 | 1177 | 1187 | 177 | 187 | 7,25 | 0,001 | 25,63 | >0.1 | 5,61 | 6,11 | 0 | 1,61 | 1,79 | 3,26 | 0 |
| NPAS2 | TCCTCATGAG | -1 | 984 | 994 | -16 | -6 | -5,34 | 0,01 | 12,24 | >0.1 | 5,61 | 8,57 | 0 | 1,87 | 0 | 1,53 | 0 |
| NPAS2 | ATCAAATAAG | -1 | 811 | 821 | -189 | -179 | -4,43 | 0,01 | 12,2 | >0.1 | 5,61 | 6,03 | 0 | 1,56 | 0 | 3,43 | 0 |
| NPAS2 | TCCAAATACA | -1 | 521 | 531 | -479 | -469 | 0,72 | 0,01 | 12,83 | >0.1 | 3,05 | 5,01 | 0 | 0,98 | 0 | 3,07 | 0 |
| NPAS2 | TACAGGTGCC | -1 | 327 | 337 | -673 | -663 | -0,74 | 0,01 | 13,22 | >0.1 | 2,6 | 4,65 | 0 | 1,14 | 0 | 5,56 | 0 |
| NPAS2 | CCAAAGTGCT | -1 | 208 | 218 | -792 | -782 | -4,67 | 0,01 | 9,09 | >0.1 | 3 | 4,48 | 0 | 1,58 | 0 | 4,7 | 0 |
| NPAS2 | TCCACAGGTC | -1 | 109 | 119 | -891 | -881 | -1,35 | 0,01 | 9,18 | >0.1 | 3,54 | 4,36 | 0 | 1,89 | 0 | 0,75 | 0 |
| PLAG1 | GAAGGAGAAGGAGG | 1 | 11 | 25 | -989 | -975 | -1,94 | 0,01 | 9,75 | >0.1 | 5,56 | 4,27 | 0 | 1,87 | 0 | 0 | 0 |
| PLAG1 | GCTGGGCACAGTGG | 1 | 177 | 191 | -823 | -809 | -3,08 | 0,01 | 9,4 | >0.1 | 3,13 | 4,44 | 0 | 1,82 | 0 | 3,09 | 0 |
| PLAG1 | GTGGCTCACGCCTG | 1 | 187 | 201 | -813 | -799 | 1,95 | 0,01 | 15,26 | >0.1 | 3,09 | 4,46 | 0 | 1,73 | 0 | 4,04 | 0 |
| PLAG1 | GGAGGCCGAGTTGG | 1 | 217 | 231 | -783 | -769 | -1,9 | 0,01 | 12,6 | >0.1 | 2,96 | 4,49 | 0 | 1,52 | 0 | 5,52 | 0 |
| PLAG1 | GAGGCCGAGTTGGG | 1 | 218 | 232 | -782 | -768 | 8,52 | 0,001 | 23,01 | >0.1 | 2,96 | 4,5 | 0 | 1,52 | 0 | 5,52 | 0 |
| PLAG1 | TGGGCGTGGTGGTG | 1 | 313 | 327 | -687 | -673 | -0,39 | 0,01 | 14,18 | >0.1 | 2,64 | 4,63 | 0 | 1,18 | 0 | 6,13 | 0 |
| PLAG1 | GGCGTGGTGGTGGG | 1 | 315 | 329 | -685 | -671 | -3,44 | 0,01 | 11,25 | >0.1 | 2,63 | 4,63 | 0 | 1,17 | 0 | 6,26 | 0 |
| PLAG1 | GGGAGGCTGAGGCA | 1 | 350 | 364 | -650 | -636 | -1,77 | 0,01 | 12,96 | >0.1 | 2,54 | 4,69 | 0 | 1,1 | 0 | 6,4 | 0 |
| PLAG1 | GGAGGCTGAGGCAG | 1 | 351 | 365 | -649 | -635 | -3,41 | 0,01 | 11,32 | >0.1 | 2,53 | 4,69 | 0 | 1,1 | 0 | 6,4 | 0 |
| PLAG1 | GAGGCTGAGGCAGG | 1 | 352 | 366 | -648 | -634 | 1,92 | 0,01 | 16,77 | >0.1 | 2,53 | 4,69 | 0 | 1,1 | 0 | 6,52 | 0 |
| PLAG1 | GTGAACCTGGGAGG | 1 | 374 | 388 | -626 | -612 | -1,16 | 0,01 | 13,6 | >0.1 | 2,56 | 4,73 | 0 | 1,07 | 0 | 6,4 | 0 |
| PLAG1 | GTTGCCCAAGTAGA | 1 | 540 | 554 | -460 | -446 | -2,15 | 0,01 | 7,05 | >0.1 | 3,14 | 5,07 | 0 | 0,99 | 0 | 0 | 0 |
| PLAG1 | GAGAGTTTCTGGGA | 1 | 552 | 566 | -448 | -434 | 1,37 | 0,01 | 10,66 | >0.1 | 3,2 | 5,09 | 0 | 0,99 | 0 | 0 | 0 |
| PLAG1 | TTGGCCATAAAGTG | 1 | 755 | 769 | -245 | -231 | -1,09 | 0,01 | 17,65 | >0.1 | 5,56 | 5,76 | 0 | 1,32 | 0 | 6,1 | 0 |
| PLAG1 | TTGGCTCACAGGGG | 1 | 1032 | 1046 | 32 | 46 | 8,15 | 0,001 | 25,02 | >0.1 | 5,56 | 8,15 | 0 | 1,86 | 0 | 1,31 | 0 |
| PLAG1 | GTTCCTCAATGGTG | -1 | 1105 | 1119 | 105 | 119 | -1,04 | 0,01 | 18,76 | >0.1 | 5,56 | 6,7 | 0 | 1,85 | 3,22 | 2,47 | 0 |
| PLAG1 | TGGGCGCTGAGCAG | -1 | 938 | 952 | -62 | -48 | -3,73 | 0,01 | 11,79 | >0.1 | 5,56 | 7,25 | 0 | 1,88 | 0 | 0,83 | 0 |
| PLAG1 | GGGCGCTGAGCAGG | -1 | 937 | 951 | -63 | -49 | -3 | 0,01 | 12,52 | >0.1 | 5,56 | 7,23 | 0 | 1,88 | 0 | 0,85 | 0 |
| PLAG1 | GTGACCAAAGAGGT | -1 | 666 | 680 | -334 | -320 | -0,3 | 0,01 | 14,03 | >0.1 | 4,11 | 5,42 | 0 | 1,1 | 0 | 3,7 | 0 |
| PLAG1 | GTCGCCCAGGCTGG | -1 | 425 | 439 | -575 | -561 | 0,37 | 0,01 | 14,72 | >0.1 | 2,7 | 4,82 | 0 | 1,02 | 0 | 5,8 | 0 |
| PLAG1 | GGAGCACAGTGGCG | -1 | 413 | 427 | -587 | -573 | 1,17 | 0,01 | 14,75 | >0.1 | 2,67 | 4,8 | 0 | 1,03 | 0 | 5,09 | 0 |
| PLAG1 | GGCGCGATCTCGGC | -1 | 403 | 417 | -597 | -583 | -3,27 | 0,01 | 10,95 | >0.1 | 2,64 | 4,78 | 0 | 1,04 | 0 | 5,76 | 0 |
| PLAG1 | TGGGACTACAGGTG | -1 | 329 | 343 | -671 | -657 | 3,31 | 0,01 | 17,73 | >0.1 | 2,59 | 4,66 | 0 | 1,14 | 0 | 6,03 | 0 |
| PLAG1 | GTGAGCCACTGTGC | -1 | 182 | 196 | -818 | -804 | -3,58 | 0,01 | 9,7 | >0.1 | 3,11 | 4,45 | 0 | 1,77 | 0 | 3,95 | 0 |
| VSX2 | GAAATTAA | 1 | 154 | 162 | -846 | -838 | 2,68 | 0,01 | 13,9 | >0.1 | 3,27 | 4,41 | 0 | 1,9 | 0 | 1,65 | 0 |
| VSX2 | CTGATCAG | 1 | 164 | 172 | -836 | -828 | 2,9 | 0,01 | 14,83 | >0.1 | 3,21 | 4,42 | 0 | 1,9 | 0 | 2,4 | 0 |
| VSX2 | CTGATCAG | 1 | 479 | 487 | -521 | -513 | 2,9 | 0,01 | 17,04 | >0.1 | 2,88 | 4,92 | 0 | 0,98 | 0 | 5,36 | 0 |
| VSX2 | CCAAGTAG | 1 | 545 | 553 | -455 | -447 | 3,27 | 0,01 | 12,47 | >0.1 | 3,15 | 5,06 | 0 | 0,98 | 0 | 0 | 0 |
| VSX2 | CCTATTAA | 1 | 600 | 608 | -400 | -392 | 3,83 | 0,01 | 14,88 | >0.1 | 3,47 | 5,2 | 0 | 1,01 | 0 | 1,36 | 0 |
| VSX2 | TTAATGAA | 1 | 604 | 612 | -396 | -388 | 4,66 | 0,01 | 15,8 | >0.1 | 3,49 | 5,21 | 0 | 1,02 | 0 | 1,41 | 0 |
| VSX2 | AAAATTAT | 1 | 683 | 691 | -317 | -309 | 2,35 | 0,01 | 16,33 | >0.1 | 4,34 | 5,46 | 0 | 1,12 | 0 | 3,07 | 0 |
| VSX2 | CTTATTTG | 1 | 811 | 819 | -189 | -181 | 2,97 | 0,01 | 19,46 | >0.1 | 5,64 | 6,02 | 0 | 1,55 | 0 | 3,28 | 0 |
| VSX2 | AAAATTAT | 1 | 830 | 838 | -170 | -162 | 2,35 | 0,01 | 19,08 | >0.1 | 5,64 | 6,14 | 0 | 1,67 | 0 | 3,28 | 0 |
| VSX2 | CTGATAAG | 1 | 967 | 975 | -33 | -25 | 2,54 | 0,01 | 19,5 | >0.1 | 5,64 | 7,76 | 0 | 1,87 | 0 | 1,7 | 0 |
| VSX2 | CTCATGAG | 1 | 984 | 992 | -16 | -8 | 2,77 | 0,01 | 20,29 | >0.1 | 5,64 | 8,43 | 0 | 1,87 | 0 | 1,57 | 0 |
| VSX2 | CTCATGAG | -1 | 984 | 992 | -16 | -8 | 2,77 | 0,01 | 20,29 | >0.1 | 5,64 | 8,43 | 0 | 1,87 | 0 | 1,57 | 0 |
| VSX2 | CTTATCAG | -1 | 967 | 975 | -33 | -25 | 2,63 | 0,01 | 19,59 | >0.1 | 5,64 | 7,76 | 0 | 1,87 | 0 | 1,7 | 0 |
| VSX2 | CAAATAAG | -1 | 811 | 819 | -189 | -181 | 3,7 | 0,01 | 20,19 | >0.1 | 5,64 | 6,02 | 0 | 1,55 | 0 | 3,28 | 0 |
| VSX2 | CAAATCAA | -1 | 750 | 758 | -250 | -242 | 2,99 | 0,01 | 21,54 | >0.1 | 5,64 | 5,71 | 0 | 1,28 | 0 | 5,92 | 0 |
| VSX2 | TTCATTAA | -1 | 604 | 612 | -396 | -388 | 4,49 | 0,01 | 15,63 | >0.1 | 3,49 | 5,21 | 0 | 1,02 | 0 | 1,41 | 0 |
| VSX2 | TTAATAGG | -1 | 600 | 608 | -400 | -392 | 3,5 | 0,01 | 14,55 | >0.1 | 3,47 | 5,2 | 0 | 1,01 | 0 | 1,36 | 0 |
| VSX2 | CTACTTGG | -1 | 545 | 553 | -455 | -447 | 3,33 | 0,01 | 12,53 | >0.1 | 3,15 | 5,06 | 0 | 0,98 | 0 | 0 | 0 |
| VSX2 | CCAAATAC | -1 | 522 | 530 | -478 | -470 | 2,73 | 0,01 | 14,83 | >0.1 | 3,05 | 5,01 | 0 | 0,98 | 0 | 3,07 | 0 |
| VSX2 | CTGATCAG | -1 | 479 | 487 | -521 | -513 | 2,9 | 0,01 | 17,04 | >0.1 | 2,88 | 4,92 | 0 | 0,98 | 0 | 5,36 | 0 |

| | | | | | | | | | | | | | | | | | |
|---|---|---|---|---|---|---|---|---|---|---|---|---|---|---|---|---|---|
| VSX2 | TTAATAGA | -1 | 288 | 296 | -712 | -704 | 2,44 | 0,01 | 16,17 | >0.1 | 2,72 | 4,58 | 0 | 1,24 | 0 | 5,19 | 0 |
| VSX2 | CTGATCAG | -1 | 164 | 172 | -836 | -828 | 2,9 | 0,01 | 14,83 | >0.1 | 3,21 | 4,42 | 0 | 1,9 | 0 | 2,4 | 0 |
| VSX2 | TTAATTTC | -1 | 154 | 162 | -846 | -838 | 6,01 | 0,01 | 17,23 | >0.1 | 3,27 | 4,41 | 0 | 1,9 | 0 | 1,65 | 0 |
| VSX2 | AAAATTAT | -1 | 132 | 140 | -868 | -860 | 2,35 | 0,01 | 13,59 | >0.1 | 3,4 | 4,38 | 0 | 1,89 | 0 | 1,57 | 0 |
| VSX2 | CTCATTTC | -1 | 65 | 73 | -935 | -927 | 3,05 | 0,01 | 13,19 | >0.1 | 3,95 | 4,31 | 0 | 1,88 | 0 | 0 | 0 |
| PRDM1 | GCACTTTGGG | 1 | 209 | 219 | -791 | -781 | -1,86 | 0,01 | 11,89 | >0.1 | 3 | 4,48 | 0 | 1,57 | 0 | 4,7 | 0 |
| PRDM1 | TGACTAACAC | 1 | 264 | 274 | -736 | -726 | -1,99 | 0,01 | 11,12 | >0.1 | 2,79 | 4,55 | 0 | 1,31 | 0 | 4,45 | 0 |
| PRDM1 | AGAGTTTCTG | 1 | 553 | 563 | -447 | -437 | -3,45 | 0,01 | 5,82 | >0.1 | 3,2 | 5,09 | 0 | 0,99 | 0 | 0 | 0 |
| PRDM1 | TCACTGTAAA | 1 | 676 | 686 | -324 | -314 | -5,15 | 0,01 | 9,06 | >0.1 | 4,23 | 5,44 | 0 | 1,11 | 0 | 3,43 | 0 |
| PRDM1 | ACATTTTCCG | 1 | 692 | 702 | -308 | -298 | 0,71 | 0,01 | 14,97 | >0.1 | 4,56 | 5,49 | 0 | 1,14 | 0 | 3,07 | 0 |
| PRDM1 | CAGCTTTCTA | 1 | 713 | 723 | -287 | -277 | 1,38 | 0,01 | 18,91 | >0.1 | 5,61 | 5,57 | 0 | 1,18 | 0 | 5,17 | 0 |
| PRDM1 | TAAGGTTCTC | 1 | 779 | 789 | -221 | -211 | -4,95 | 0,01 | 14,46 | >0.1 | 5,61 | 5,86 | 0 | 1,39 | 0 | 6,54 | 0 |
| PRDM1 | TCACTTTAAA | 1 | 821 | 831 | -179 | -169 | 3,6 | 0,001 | 20,63 | >0.1 | 5,61 | 6,1 | 0 | 1,62 | 0 | 3,7 | 0 |
| PRDM1 | GAACTTTATA | 1 | 873 | 883 | -127 | -117 | -0,52 | 0,01 | 16,41 | >0.1 | 5,61 | 6,47 | 0 | 1,9 | 0 | 2,95 | 0 |
| PRDM1 | GGTCTTTCCT | 1 | 930 | 940 | -70 | -60 | -2,91 | 0,01 | 12,64 | >0.1 | 5,61 | 7,06 | 0 | 1,88 | 0 | 1 | 0 |
| PRDM1 | CCAAGTTCAA | 1 | 954 | 964 | -46 | -36 | -5,36 | 0,01 | 11,31 | >0.1 | 5,61 | 7,48 | 0 | 1,88 | 0 | 1,7 | 0 |
| PRDM1 | CAAAGTTCAC | 1 | 1172 | 1182 | 172 | 182 | -4,71 | 0,01 | 13,45 | >0.1 | 5,61 | 6,14 | 0 | 1,65 | 1,79 | 2,97 | 0 |
| PRDM1 | TCACTTGCTT | 1 | 1178 | 1188 | 178 | 188 | 2,8 | 0,01 | 21,28 | >0.1 | 5,61 | 6,1 | 0 | 1,6 | 1,79 | 3,37 | 0 |
| PRDM1 | GAACTTTGAT | -1 | 1170 | 1180 | 170 | 180 | -0,6 | 0,01 | 17,53 | >0.1 | 5,61 | 6,15 | 0 | 1,67 | 1,79 | 2,91 | 0 |
| PRDM1 | TCACATCCAC | -1 | 1020 | 1030 | 20 | 30 | -1,85 | 0,01 | 15,65 | >0.1 | 5,61 | 8,72 | 0 | 1,86 | 0 | 1,31 | 0 |
| PRDM1 | TGACTTTCCC | -1 | 1006 | 1016 | 6 | 16 | 12,56 | 0,0001 | 30,62 | >0.1 | 5,61 | 9,12 | 0 | 1,87 | 0 | 1,46 | 0 |
| PRDM1 | AGATTTTCTC | -1 | 976 | 986 | -24 | -14 | 1,27 | 0,01 | 18,46 | >0.1 | 5,61 | 8,13 | 0 | 1,87 | 0 | 1,57 | 0 |
| PRDM1 | TCACTTTATG | -1 | 760 | 770 | -240 | -230 | 2,72 | 0,01 | 21,33 | >0.1 | 5,61 | 5,77 | 0 | 1,32 | 0 | 5,92 | 0 |
| PRDM1 | ATATTTTCCT | -1 | 722 | 732 | -278 | -268 | -0,83 | 0,01 | 16,76 | >0.1 | 5,61 | 5,61 | 0 | 1,21 | 0 | 5,17 | 0 |
| PRDM1 | ACAATTTCAT | -1 | 607 | 617 | -393 | -383 | 1,43 | 0,01 | 12,7 | >0.1 | 3,52 | 5,23 | 0 | 1,02 | 0 | 1,5 | 0 |
| PRDM1 | TCATATTCCC | -1 | 562 | 572 | -438 | -428 | -3,69 | 0,01 | 5,66 | >0.1 | 3,25 | 5,11 | 0 | 0,99 | 0 | 0 | 0 |
| PRDM1 | AAACTCTCTA | -1 | 550 | 560 | -450 | -440 | -1,2 | 0,01 | 8,05 | >0.1 | 3,18 | 5,08 | 0 | 0,99 | 0 | 0 | 0 |
| PRDM1 | ACATTTTCCC | -1 | 532 | 542 | -468 | -458 | 3,82 | 0,001 | 14,55 | >0.1 | 3,1 | 5,04 | 0 | 0,98 | 0 | 1,61 | 0 |
| PRDM1 | TGACTTTGCC | -1 | 511 | 521 | -489 | -479 | 1,82 | 0,01 | 13,66 | >0.1 | 3 | 4,99 | 0 | 0,98 | 0 | 2,86 | 0 |
| PRDM1 | CCAGGTTCAC | -1 | 374 | 384 | -626 | -616 | -3,45 | 0,01 | 11,43 | >0.1 | 2,55 | 4,72 | 0 | 1,07 | 0 | 6,53 | 0 |
| PRDM1 | GGACGGTCTC | -1 | 254 | 264 | -746 | -736 | -4,01 | 0,01 | 9,56 | >0.1 | 2,83 | 4,54 | 0 | 1,35 | 0 | 4,85 | 0 |
| PRDM1 | TCTCTATCTC | -1 | 248 | 258 | -752 | -742 | -4,1 | 0,01 | 9,74 | >0.1 | 2,85 | 4,53 | 0 | 1,37 | 0 | 5,09 | 0 |
| PRDM1 | CCACTGTGCC | -1 | 181 | 191 | -819 | -809 | -5,44 | 0,01 | 7,76 | >0.1 | 3,12 | 4,44 | 0 | 1,78 | 0 | 3,86 | 0 |
| PRDM1 | TTAATTTCCA | -1 | 152 | 162 | -848 | -838 | -0,32 | 0,01 | 10,91 | >0.1 | 3,27 | 4,41 | 0 | 1,9 | 0 | 1,65 | 0 |
| PRDM1 | CTACTCTCCA | -1 | 120 | 130 | -880 | -870 | -1,15 | 0,01 | 10,23 | >0.1 | 3,47 | 4,37 | 0 | 1,89 | 0 | 1,65 | 0 |
| PRDM1 | CAACTCTCCT | -1 | 100 | 110 | -900 | -890 | -0,9 | 0,01 | 9,73 | >0.1 | 3,61 | 4,35 | 0 | 1,88 | 0 | 0,79 | 0 |
| PRDM1 | TCTCCTTCAC | -1 | 9 | 19 | -991 | -981 | -5,68 | 0,01 | 7,85 | >0.1 | 5,61 | 4,26 | 0 | 1,86 | 1,79 | 0 | 0 |
| PRDM1 | TCACTTACCT | -1 | 3 | 13 | -997 | -987 | 5,47 | 0,001 | 18,99 | >0.1 | 5,61 | 4,26 | 0 | 1,86 | 1,79 | 0 | 0 |
| POU3F3 | AATAGGTAAGTGA | 1 | 0 | 13 | -1000 | -987 | 3,65 | 0,01 | 17,13 | >0.1 | 5,57 | 4,26 | 0 | 1,86 | 1,79 | 0 | 0 |
| POU3F3 | GTTTTGCAAAGGC | 1 | 82 | 95 | -918 | -905 | 1,09 | 0,01 | 11,06 | >0.1 | 3,75 | 4,33 | 0 | 1,88 | 0 | 0 | 0 |
| POU3F3 | CAAAGGCAGATCA | 1 | 88 | 101 | -912 | -899 | 1,89 | 0,01 | 11,81 | >0.1 | 3,7 | 4,34 | 0 | 1,88 | 0 | 0 | 0 |
| POU3F3 | ATAATTTTAAAAA | 1 | 132 | 145 | -868 | -855 | 1,95 | 0,01 | 13,19 | >0.1 | 3,38 | 4,39 | 0 | 1,89 | 0 | 1,57 | 0 |
| POU3F3 | GCCATGGAAATTA | 1 | 148 | 161 | -852 | -839 | 5,67 | 0,01 | 16,91 | >0.1 | 3,29 | 4,41 | 0 | 1,9 | 0 | 1,65 | 0 |
| POU3F3 | ATTAATGAAATTG | 1 | 603 | 616 | -397 | -384 | 3,16 | 0,01 | 14,32 | >0.1 | 3,5 | 5,22 | 0 | 1,02 | 0 | 1,41 | 0 |
| POU3F3 | ATTATCTTCATGA | 1 | 640 | 653 | -360 | -347 | 1,16 | 0,01 | 15,19 | >0.1 | 3,81 | 5,33 | 0 | 1,06 | 0 | 3,83 | 0 |
| POU3F3 | TCTAGGAAAATAT | 1 | 719 | 732 | -281 | -268 | 1,33 | 0,01 | 18,7 | >0.1 | 5,57 | 5,61 | 0 | 1,21 | 0 | 4,99 | 0 |
| POU3F3 | CGTATTCTTATTT | 1 | 805 | 818 | -195 | -182 | 5,69 | 0,01 | 22,1 | >0.1 | 5,57 | 6,02 | 0 | 1,54 | 0 | 3,28 | 0 |
| POU3F3 | ATTATTCTAAAAT | 1 | 833 | 846 | -167 | -154 | 7,18 | 0,001 | 23,97 | >0.1 | 5,57 | 6,2 | 0 | 1,74 | 0 | 3,28 | 0 |
| POU3F3 | CAAAGGCTGATAA | 1 | 961 | 974 | -39 | -26 | 2,34 | 0,01 | 19,21 | >0.1 | 5,57 | 7,73 | 0 | 1,88 | 0 | 1,7 | 0 |
| POU3F3 | TTTAGAATAATTT | -1 | 831 | 844 | -169 | -156 | 1,03 | 0,01 | 18,05 | >0.1 | 5,57 | 6,19 | 0 | 1,72 | 0 | 3,55 | 0 |
| POU3F3 | AATACGTTATTCT | -1 | 798 | 811 | -202 | -189 | 1,94 | 0,01 | 18,54 | >0.1 | 5,57 | 5,98 | 0 | 1,5 | 0 | 3,55 | 0 |
| POU3F3 | TTTTGGTTAATAT | -1 | 728 | 741 | -272 | -259 | 3,41 | 0,01 | 21,38 | >0.1 | 5,57 | 5,64 | 0 | 1,23 | 0 | 5,52 | 0 |
| POU3F3 | AATGTTATAATTT | -1 | 684 | 697 | -316 | -303 | 1,43 | 0,01 | 15,39 | >0.1 | 4,41 | 5,48 | 0 | 1,13 | 0 | 2,95 | 0 |
| POU3F3 | ATCATGAAGATAA | -1 | 641 | 654 | -359 | -346 | 2,66 | 0,01 | 16,71 | >0.1 | 3,82 | 5,34 | 0 | 1,06 | 0 | 3,83 | 0 |
| POU3F3 | ATGAAGATAATGT | -1 | 638 | 651 | -362 | -349 | 1,39 | 0,01 | 13,22 | >0.1 | 3,79 | 5,33 | 0 | 1,06 | 0 | 1,66 | 0 |
| POU3F3 | ATAATGTTGGTGT | -1 | 632 | 645 | -368 | -355 | 1,97 | 0,01 | 13,67 | >0.1 | 3,73 | 5,31 | 0 | 1,05 | 0 | 1,61 | 0 |
| POU3F3 | TTTATTTTTATTT | -1 | 577 | 590 | -423 | -410 | 6,78 | 0,001 | 16,28 | >0.1 | 3,34 | 5,15 | 0 | 1 | 0 | 0 | 0 |
| POU3F3 | TTTATGGAATTCA | -1 | 492 | 505 | -508 | -495 | 4,77 | 0,01 | 20,08 | >0.1 | 2,93 | 4,95 | 0 | 0,99 | 0 | 6,44 | 0 |
| POU3F3 | GGAATTCATATTT | -1 | 487 | 500 | -513 | -500 | 3,32 | 0,01 | 18,43 | >0.1 | 2,91 | 4,94 | 0 | 0,99 | 0 | 6,27 | 0 |

| | | | | | | | | | | | | | | | | | |
|---|---|---|---|---|---|---|---|---|---|---|---|---|---|---|---|---|---|
| POU3F3 | ATTTTTTAAAATT | -1 | 134 | 147 | -866 | -853 | 1,86 | 0,01 | 13,09 | >0.1 | 3,37 | 4,39 | 0 | 1,89 | 0 | 1,57 | 0 |
| POU3F3 | TTTTTTAAAATTA | -1 | 133 | 146 | -867 | -854 | 1,59 | 0,01 | 12,86 | >0.1 | 3,37 | 4,39 | 0 | 1,89 | 0 | 1,61 | 0 |
| POU3F3 | AAAAAGCTCATTT | -1 | 66 | 79 | -934 | -921 | 2,72 | 0,01 | 12,83 | >0.1 | 3,91 | 4,32 | 0 | 1,88 | 0 | 0 | 0 |
| POU3F2 | AAAGGCAGATCA | 1 | 89 | 101 | -911 | -899 | 1,05 | 0,01 | 10,97 | >0.1 | 3,69 | 4,34 | 0 | 1,88 | 0 | 0 | 0 |
| POU3F2 | TAATTTTAAAAA | 1 | 133 | 145 | -867 | -855 | -0,17 | 0,01 | 11,1 | >0.1 | 3,38 | 4,39 | 0 | 1,89 | 0 | 1,61 | 0 |
| POU3F2 | CCATGGAAATTA | 1 | 149 | 161 | -851 | -839 | 7,28 | 0,001 | 18,52 | >0.1 | 3,28 | 4,41 | 0 | 1,9 | 0 | 1,65 | 0 |
| POU3F2 | CTATTAAAAATA | 1 | 289 | 301 | -711 | -699 | 0,15 | 0,01 | 13,77 | >0.1 | 2,71 | 4,59 | 0 | 1,24 | 0 | 5,09 | 0 |
| POU3F2 | ATCTTGAAATAA | 1 | 571 | 583 | -429 | -417 | 0,51 | 0,01 | 9,95 | >0.1 | 3,3 | 5,14 | 0 | 1 | 0 | 0 | 0 |
| POU3F2 | AAATAAAAATAA | 1 | 577 | 589 | -423 | -411 | 0,03 | 0,01 | 9,52 | >0.1 | 3,33 | 5,15 | 0 | 1 | 0 | 0 | 0 |
| POU3F2 | TTAATGAAATTG | 1 | 604 | 616 | -396 | -384 | 1,86 | 0,01 | 13,03 | >0.1 | 3,51 | 5,22 | 0 | 1,02 | 0 | 1,41 | 0 |
| POU3F2 | TAATGAAATTGT | 1 | 605 | 617 | -395 | -383 | 1,25 | 0,01 | 12,46 | >0.1 | 3,52 | 5,23 | 0 | 1,02 | 0 | 1,44 | 0 |
| POU3F2 | TTATCTTCATGA | 1 | 641 | 653 | -359 | -347 | 1,35 | 0,01 | 15,39 | >0.1 | 3,81 | 5,33 | 0 | 1,06 | 0 | 3,83 | 0 |
| POU3F2 | GTATTCTTATTT | 1 | 806 | 818 | -194 | -182 | 4,5 | 0,01 | 20,92 | >0.1 | 5,58 | 6,02 | 0 | 1,54 | 0 | 3,28 | 0 |
| POU3F2 | TTATTCTAAAAT | 1 | 834 | 846 | -166 | -154 | 3,91 | 0,01 | 20,71 | >0.1 | 5,58 | 6,2 | 0 | 1,74 | 0 | 3,28 | 0 |
| POU3F2 | AAAGGCTGATAA | 1 | 962 | 974 | -38 | -26 | 0,09 | 0,01 | 16,97 | >0.1 | 5,58 | 7,73 | 0 | 1,87 | 0 | 1,7 | 0 |
| POU3F2 | ATATGTAACAGA | -1 | 845 | 857 | -155 | -143 | 2,69 | 0,01 | 19,24 | >0.1 | 5,58 | 6,28 | 0 | 1,83 | 0 | 2,86 | 0 |
| POU3F2 | ATACGTTATTCT | -1 | 798 | 810 | -202 | -190 | -0,12 | 0,01 | 16,48 | >0.1 | 5,58 | 5,97 | 0 | 1,5 | 0 | 3,55 | 0 |
| POU3F2 | ATGTTATAATTT | -1 | 684 | 696 | -316 | -304 | 0,71 | 0,01 | 14,66 | >0.1 | 4,4 | 5,47 | 0 | 1,13 | 0 | 2,95 | 0 |
| POU3F2 | TCATGAAGATAA | -1 | 641 | 653 | -359 | -347 | 1,19 | 0,01 | 15,23 | >0.1 | 3,81 | 5,33 | 0 | 1,06 | 0 | 3,83 | 0 |
| POU3F2 | TGAAGATAATGT | -1 | 638 | 650 | -362 | -350 | 0,63 | 0,01 | 12,45 | >0.1 | 3,79 | 5,32 | 0 | 1,06 | 0 | 1,66 | 0 |
| POU3F2 | TTATTTTTATTT | -1 | 577 | 589 | -423 | -411 | 3,75 | 0,01 | 13,24 | >0.1 | 3,33 | 5,15 | 0 | 1 | 0 | 0 | 0 |
| POU3F2 | TTATGGAATTCA | -1 | 492 | 504 | -508 | -496 | 5,65 | 0,01 | 20,95 | >0.1 | 2,93 | 4,95 | 0 | 0,99 | 0 | 6,44 | 0 |
| POU3F2 | GAATTCATATTT | -1 | 487 | 499 | -513 | -501 | 1,71 | 0,01 | 16,82 | >0.1 | 2,91 | 4,94 | 0 | 0,99 | 0 | 6,27 | 0 |
| POU3F2 | AAAAGCTCATTT | -1 | 66 | 78 | -934 | -922 | 1,23 | 0,01 | 11,34 | >0.1 | 3,92 | 4,32 | 0 | 1,88 | 0 | 0 | 0 |
| ESRRG | TGAAGGAGAA | 1 | 10 | 20 | -990 | -980 | 4,07 | 0,01 | 17,6 | >0.1 | 5,61 | 4,26 | 0 | 1,86 | 1,79 | 0 | 0 |
| ESRRG | AGAAGGAGGC | 1 | 16 | 26 | -984 | -974 | -0,71 | 0,01 | 11,03 | >0.1 | 5,61 | 4,27 | 0 | 1,87 | 0 | 0 | 0 |
| ESRRG | GCAAAGGCAG | 1 | 87 | 97 | -913 | -903 | -1,82 | 0,01 | 8,12 | >0.1 | 3,72 | 4,34 | 0 | 1,88 | 0 | 0 | 0 |
| ESRRG | GGGAGGCCGA | 1 | 216 | 226 | -784 | -774 | -1,97 | 0,01 | 12,54 | >0.1 | 2,97 | 4,49 | 0 | 1,53 | 0 | 5,52 | 0 |
| ESRRG | ACAAGGTCAG | 1 | 238 | 248 | -762 | -752 | 12,4 | 0,0001 | 25,75 | >0.1 | 2,89 | 4,52 | 0 | 1,42 | 0 | 4,53 | 0 |
| ESRRG | ACACGGTGAA | 1 | 270 | 280 | -730 | -720 | -1,77 | 0,01 | 11,85 | >0.1 | 2,77 | 4,56 | 0 | 1,29 | 0 | 4,99 | 0 |
| ESRRG | AGAATGGCGT | 1 | 366 | 376 | -634 | -624 | 0,73 | 0,01 | 15,66 | >0.1 | 2,53 | 4,71 | 0 | 1,08 | 0 | 6,61 | 0 |
| ESRRG | ACAAGGGCAA | 1 | 506 | 516 | -494 | -484 | 10,12 | 0,001 | 24,74 | >0.1 | 2,98 | 4,98 | 0 | 0,98 | 0 | 5,68 | 0 |
| ESRRG | GCAAAGTCAT | 1 | 512 | 522 | -488 | -478 | 3,73 | 0,01 | 15,66 | >0.1 | 3,01 | 4,99 | 0 | 0,98 | 0 | 2,95 | 0 |
| ESRRG | GGAAGGGAAA | 1 | 528 | 538 | -472 | -462 | -2,24 | 0,01 | 10,4 | >0.1 | 3,08 | 5,03 | 0 | 0,98 | 0 | 3,55 | 0 |
| ESRRG | AAAATGTTGC | 1 | 535 | 545 | -465 | -455 | -3,42 | 0,01 | 7,33 | >0.1 | 3,11 | 5,04 | 0 | 0,98 | 0 | 1,61 | 0 |
| ESRRG | TAAATGTGAG | 1 | 586 | 596 | -414 | -404 | -1,75 | 0,01 | 7,81 | >0.1 | 3,38 | 5,17 | 0 | 1,01 | 0 | 0 | 0 |
| ESRRG | GTAAGGTTCT | 1 | 778 | 788 | -222 | -212 | -2,68 | 0,01 | 16,72 | >0.1 | 5,61 | 5,85 | 0 | 1,39 | 0 | 6,54 | 0 |
| ESRRG | CCAAGTTCAA | 1 | 954 | 964 | -46 | -36 | 4,51 | 0,01 | 21,18 | >0.1 | 5,61 | 7,48 | 0 | 1,88 | 0 | 1,7 | 0 |
| ESRRG | CAAAGGCTGA | 1 | 961 | 971 | -39 | -29 | -4,08 | 0,01 | 12,75 | >0.1 | 5,61 | 7,65 | 0 | 1,88 | 0 | 1,7 | 0 |
| ESRRG | AGGAGGTTTT | 1 | 990 | 1000 | -10 | 0 | -2,85 | 0,01 | 14,95 | >0.1 | 5,61 | 8,71 | 0 | 1,87 | 0 | 1,61 | 0 |
| ESRRG | GGAAAGTCAT | 1 | 1007 | 1017 | 7 | 17 | 2,7 | 0,01 | 20,76 | >0.1 | 5,61 | 9,12 | 0 | 1,87 | 0 | 1,46 | 0 |
| ESRRG | TGGATGTGAT | 1 | 1021 | 1031 | 21 | 31 | -0,69 | 0,01 | 16,79 | >0.1 | 5,61 | 8,7 | 0 | 1,86 | 0 | 1,31 | 0 |
| ESRRG | ACGATGTCAA | 1 | 1046 | 1056 | 46 | 56 | 3,5 | 0,01 | 20,11 | >0.1 | 5,61 | 7,68 | 0 | 1,86 | 0 | 1,46 | 0 |
| ESRRG | TCAAGCTCTT | 1 | 1052 | 1062 | 52 | 62 | -1,32 | 0,01 | 15,95 | >0.1 | 5,61 | 7,53 | 0 | 1,86 | 0 | 2,27 | 0 |
| ESRRG | ACAAGTTTAA | 1 | 1137 | 1147 | 137 | 147 | -0,8 | 0,01 | 19,34 | >0.1 | 5,61 | 6,4 | 0 | 1,84 | 3,22 | 3,07 | 0 |
| ESRRG | CAAAGTTCAC | 1 | 1172 | 1182 | 172 | 182 | 0,26 | 0,01 | 18,42 | >0.1 | 5,61 | 6,14 | 0 | 1,65 | 1,79 | 2,97 | 0 |
| ESRRG | TCGTGGTTAA | -1 | 1143 | 1153 | 143 | 153 | -3,79 | 0,01 | 15,35 | >0.1 | 5,61 | 6,35 | 0 | 1,84 | 3,22 | 2,11 | 0 |
| ESRRG | AAAATGTCTT | -1 | 1124 | 1134 | 124 | 134 | -4,12 | 0,01 | 15,86 | >0.1 | 5,61 | 6,51 | 0 | 1,84 | 3,22 | 2,79 | 0 |
| ESRRG | TCAATGGTGG | -1 | 1104 | 1114 | 104 | 114 | -2,09 | 0,01 | 17,84 | >0.1 | 5,61 | 6,71 | 0 | 1,85 | 3,22 | 2,53 | 0 |
| ESRRG | ACAAGGCTGA | -1 | 1074 | 1084 | 74 | 84 | -1,03 | 0,01 | 17,68 | >0.1 | 5,61 | 7,11 | 0 | 1,85 | 1,79 | 2,34 | 0 |
| ESRRG | CCAAGATCAC | -1 | 1026 | 1036 | 26 | 36 | -1,57 | 0,01 | 15,63 | >0.1 | 5,61 | 8,45 | 0 | 1,86 | 0 | 1,28 | 0 |
| ESRRG | TGAATGACTT | -1 | 1010 | 1020 | 10 | 20 | -3,71 | 0,01 | 14,17 | >0.1 | 5,61 | 8,99 | 0 | 1,86 | 0 | 1,42 | 0 |
| ESRRG | TAAAGTTCAT | -1 | 871 | 881 | -129 | -119 | 1,54 | 0,01 | 18,45 | >0.1 | 5,61 | 6,45 | 0 | 1,9 | 0 | 2,95 | 0 |
| ESRRG | GAGAGGACAG | -1 | 857 | 867 | -143 | -133 | -0,86 | 0,01 | 15,86 | >0.1 | 5,61 | 6,35 | 0 | 1,9 | 0 | 2,86 | 0 |
| ESRRG | AAAATGTTAT | -1 | 689 | 699 | -311 | -301 | -0,2 | 0,01 | 13,85 | >0.1 | 4,48 | 5,48 | 0 | 1,13 | 0 | 2,95 | 0 |
| ESRRG | AAGAGGTCTA | -1 | 663 | 673 | -337 | -327 | -2,64 | 0,01 | 11,88 | >0.1 | 4,05 | 5,4 | 0 | 1,09 | 0 | 3,99 | 0 |
| ESRRG | ACAATTTCAT | -1 | 607 | 617 | -393 | -383 | -2,37 | 0,01 | 8,9 | >0.1 | 3,52 | 5,23 | 0 | 1,02 | 0 | 1,5 | 0 |
| ESRRG | AATAGGTTAT | -1 | 596 | 606 | -404 | -394 | -0,32 | 0,01 | 10,78 | >0.1 | 3,45 | 5,2 | 0 | 1,01 | 0 | 1,44 | 0 |
| ESRRG | TCAAGATCAT | -1 | 568 | 578 | -432 | -422 | -0,29 | 0,01 | 9,11 | >0.1 | 3,28 | 5,12 | 0 | 1 | 0 | 0 | 0 |

| | | | | | | | | | | | | | | | | | |
|---|---|---|---|---|---|---|---|---|---|---|---|---|---|---|---|---|---|
| ESRRG | CCAGGTTCAC | -1 | 374 | 384 | -626 | -616 | -3,69 | 0,01 | 11,19 | >0.1 | 2,55 | 4,72 | 0 | 1,07 | 0 | 6,53 | 0 |
| ESRRG | CCATGGCCAT | -1 | 145 | 155 | -855 | -845 | -3,92 | 0,01 | 7,3 | >0.1 | 3,31 | 4,4 | 0 | 1,9 | 0 | 1,61 | 0 |
| ESRRG | AAAAGCTCAT | -1 | 68 | 78 | -932 | -922 | 0,22 | 0,01 | 10,32 | >0.1 | 3,91 | 4,32 | 0 | 1,88 | 0 | 0 | 0 |
| ESRRA | GTGAAGGAGAA | 1 | 9 | 20 | -991 | -980 | -1,05 | 0,001 | 12,47 | >0.1 | 5,6 | 4,26 | 0 | 1,86 | 1,79 | 0 | 0 |
| ESRRA | GAGAAGGAGGC | 1 | 15 | 26 | -985 | -974 | -10,34 | 0,01 | 1,39 | >0.1 | 5,6 | 4,27 | 0 | 1,87 | 0 | 0 | 0 |
| ESRRA | TGCAAAGGCAG | 1 | 86 | 97 | -914 | -903 | -9,81 | 0,01 | 0,13 | >0.1 | 3,73 | 4,34 | 0 | 1,88 | 0 | 0 | 0 |
| ESRRA | TGGGAGGCCGA | 1 | 215 | 226 | -785 | -774 | -13,08 | 0,01 | 1,55 | >0.1 | 2,97 | 4,49 | 0 | 1,54 | 0 | 5,63 | 0 |
| ESRRA | CACAAGGTCAG | 1 | 237 | 248 | -763 | -752 | 7,08 | 0,0001 | 20,43 | >0.1 | 2,89 | 4,52 | 0 | 1,42 | 0 | 4,53 | 0 |
| ESRRA | AACACGGTGAA | 1 | 269 | 280 | -731 | -720 | -11,38 | 0,01 | 2,34 | >0.1 | 2,78 | 4,56 | 0 | 1,3 | 0 | 5,09 | 0 |
| ESRRA | GAGAATGGCGT | 1 | 365 | 376 | -635 | -624 | -8,91 | 0,01 | 6,02 | >0.1 | 2,53 | 4,71 | 0 | 1,08 | 0 | 6,61 | 0 |
| ESRRA | GACAAGGGCAA | 1 | 505 | 516 | -495 | -484 | 5,4 | 0,001 | 20,02 | >0.1 | 2,98 | 4,98 | 0 | 0,98 | 0 | 5,68 | 0 |
| ESRRA | GGCAAAGTCAT | 1 | 511 | 522 | -489 | -478 | 0,56 | 0,001 | 12,4 | >0.1 | 3,01 | 4,99 | 0 | 0,98 | 0 | 2,86 | 0 |
| ESRRA | TGGAAGGGAAA | 1 | 527 | 538 | -473 | -462 | -12,71 | 0,01 | -0,2 | >0.1 | 3,08 | 5,03 | 0 | 0,98 | 0 | 3,43 | 0 |
| ESRRA | ATAAATGTGAG | 1 | 585 | 596 | -415 | -404 | -12,57 | 0,01 | -3,01 | >0.1 | 3,38 | 5,17 | 0 | 1,01 | 0 | 0 | 0 |
| ESRRA | CCCAAGTTCAA | 1 | 953 | 964 | -47 | -36 | 1,15 | 0,001 | 17,8 | >0.1 | 5,6 | 7,48 | 0 | 1,88 | 0 | 1,7 | 0 |
| ESRRA | TCAAAGGCTGA | 1 | 960 | 971 | -40 | -29 | -12,28 | 0,01 | 4,58 | >0.1 | 5,6 | 7,65 | 0 | 1,88 | 0 | 1,74 | 0 |
| ESRRA | GGGAAAGTCAT | 1 | 1006 | 1017 | 6 | 17 | -3,01 | 0,01 | 15,04 | >0.1 | 5,6 | 9,12 | 0 | 1,87 | 0 | 1,46 | 0 |
| ESRRA | GTGGATGTGAT | 1 | 1020 | 1031 | 20 | 31 | -8,33 | 0,01 | 9,16 | >0.1 | 5,6 | 8,72 | 0 | 1,86 | 0 | 1,31 | 0 |
| ESRRA | GACGATGTCAA | 1 | 1045 | 1056 | 45 | 56 | -7,75 | 0,01 | 8,06 | >0.1 | 5,6 | 7,7 | 0 | 1,86 | 0 | 0,65 | 0 |
| ESRRA | GTCAAGCTCTT | 1 | 1051 | 1062 | 51 | 62 | -3,27 | 0,01 | 13,2 | >0.1 | 5,6 | 7,55 | 0 | 1,86 | 0 | 1,46 | 0 |
| ESRRA | GACAAGTTTAA | 1 | 1136 | 1147 | 136 | 147 | -6,94 | 0,01 | 13,22 | >0.1 | 5,6 | 6,41 | 0 | 1,84 | 3,22 | 3,09 | 0 |
| ESRRA | TCAAAGTTCAC | 1 | 1171 | 1182 | 171 | 182 | -4,81 | 0,01 | 13,4 | >0.1 | 5,6 | 6,15 | 0 | 1,66 | 1,79 | 3,02 | 0 |
| ESRRA | AACAAGGCTGA | -1 | 1074 | 1085 | 74 | 85 | -11,92 | 0,01 | 6,77 | >0.1 | 5,6 | 7,11 | 0 | 1,85 | 1,79 | 2,34 | 0 |
| ESRRA | GCCAAGATCAC | -1 | 1026 | 1037 | 26 | 37 | -2,34 | 0,01 | 14,85 | >0.1 | 5,6 | 8,45 | 0 | 1,86 | 0 | 1,28 | 0 |
| ESRRA | ATAAAGTTCAT | -1 | 871 | 882 | -129 | -118 | -4,05 | 0,01 | 12,86 | >0.1 | 5,6 | 6,46 | 0 | 1,9 | 0 | 2,95 | 0 |
| ESRRA | TTTAAAGTGAA | -1 | 820 | 831 | -180 | -169 | -9,81 | 0,01 | 7,2 | >0.1 | 5,6 | 6,1 | 0 | 1,62 | 0 | 3,7 | 0 |
| ESRRA | GAAAATGTTAT | -1 | 689 | 700 | -311 | -300 | -10,79 | 0,01 | 3,28 | >0.1 | 4,5 | 5,49 | 0 | 1,13 | 0 | 2,95 | 0 |
| ESRRA | GACAATTTCAT | -1 | 607 | 618 | -393 | -382 | -6,84 | 0,01 | 4,44 | >0.1 | 3,53 | 5,23 | 0 | 1,02 | 0 | 1,5 | 0 |
| ESRRA | TTCAAGATCAT | -1 | 568 | 579 | -432 | -421 | 5,05 | 0,001 | 14,45 | >0.1 | 3,28 | 5,13 | 0 | 1 | 0 | 0 | 0 |
| ESRRA | TGCAAGCTCTG | -1 | 388 | 399 | -612 | -601 | -13,51 | 0,01 | 1,01 | >0.1 | 2,59 | 4,75 | 0 | 1,05 | 0 | 6,13 | 0 |
| ESRRA | CCCAGGTTCAC | -1 | 374 | 385 | -626 | -615 | -9,08 | 0,01 | 5,8 | >0.1 | 2,56 | 4,72 | 0 | 1,07 | 0 | 6,53 | 0 |
| ESRRA | CCCAAAGTGCT | -1 | 208 | 219 | -792 | -781 | -11,56 | 0,01 | 2,2 | >0.1 | 3 | 4,48 | 0 | 1,58 | 0 | 4,7 | 0 |
| ESRRA | TCCATGGCCAT | -1 | 145 | 156 | -855 | -844 | -6,01 | 0,01 | 5,2 | >0.1 | 3,31 | 4,4 | 0 | 1,9 | 0 | 1,61 | 0 |
| ESRRA | CCACAGGTCAA | -1 | 107 | 118 | -893 | -882 | -9,52 | 0,01 | 1,05 | >0.1 | 3,56 | 4,36 | 0 | 1,89 | 0 | 0,77 | 0 |
| ESRRA | AAAAAGCTCAT | -1 | 68 | 79 | -932 | -921 | -9,28 | 0,01 | 0,81 | >0.1 | 3,9 | 4,32 | 0 | 1,88 | 0 | 0 | 0 |
| ZBTB7C | TCAGCCACAGAA | 1 | 37 | 49 | -963 | -951 | 3,65 | 0,01 | 14,13 | >0.1 | 4,32 | 4,29 | 0 | 1,87 | 0 | 0 | 0 |
| ZBTB7C | ACTAACACGGTG | 1 | 266 | 278 | -734 | -722 | 2,5 | 0,01 | 15,68 | >0.1 | 2,78 | 4,56 | 0 | 1,31 | 0 | 4,53 | 0 |
| ZBTB7C | TGAAACCCCGTC | 1 | 276 | 288 | -724 | -712 | 4,38 | 0,01 | 18,17 | >0.1 | 2,75 | 4,57 | 0 | 1,27 | 0 | 5,19 | 0 |
| ZBTB7C | GAAACCCCGTCT | 1 | 277 | 289 | -723 | -711 | 2,94 | 0,01 | 16,73 | >0.1 | 2,75 | 4,57 | 0 | 1,27 | 0 | 5,19 | 0 |
| ZBTB7C | ACAAACACCAAC | 1 | 628 | 640 | -372 | -360 | 8,5 | 0,001 | 20,19 | >0.1 | 3,69 | 5,29 | 0 | 1,05 | 0 | 1,66 | 0 |
| ZBTB7C | GCGCCCAACCCA | 1 | 945 | 957 | -55 | -43 | 7,51 | 0,001 | 24,11 | >0.1 | 5,58 | 7,34 | 0 | 1,88 | 0 | 1,8 | 0 |
| ZBTB7C | CAGTCCACCATT | 1 | 1100 | 1112 | 100 | 112 | 4,02 | 0,01 | 23,81 | >0.1 | 5,58 | 6,76 | 0 | 1,85 | 3,22 | 2,38 | 0 |
| ZBTB7C | TCCACCATTGAG | 1 | 1103 | 1115 | 103 | 115 | 2,5 | 0,01 | 22,41 | >0.1 | 5,58 | 6,72 | 0 | 1,85 | 3,22 | 2,53 | 0 |
| ZBTB7C | TTAACCACGAAG | 1 | 1143 | 1155 | 143 | 155 | 4,34 | 0,01 | 23,5 | >0.1 | 5,58 | 6,35 | 0 | 1,84 | 3,22 | 2,16 | 0 |
| ZBTB7C | TCGTCCCCTGTG | -1 | 1038 | 1050 | 38 | 50 | 3,77 | 0,01 | 20,49 | >0.1 | 5,58 | 7,92 | 0 | 1,86 | 0 | 1,36 | 0 |
| ZBTB7C | ACATCCACTGAA | -1 | 1016 | 1028 | 16 | 28 | 5,44 | 0,01 | 23,03 | >0.1 | 5,58 | 8,81 | 0 | 1,86 | 0 | 1,34 | 0 |
| ZBTB7C | AAAACCTCCTCA | -1 | 988 | 1000 | -12 | 0 | 1,62 | 0,01 | 19,4 | >0.1 | 5,58 | 8,71 | 0 | 1,87 | 0 | 1,61 | 0 |
| ZBTB7C | AAGACCAAAAAA | -1 | 924 | 936 | -76 | -64 | 2,04 | 0,01 | 17,57 | >0.1 | 5,58 | 7,01 | 0 | 1,88 | 0 | 1,06 | 0 |
| ZBTB7C | AAGAACACAGTA | -1 | 901 | 913 | -99 | -87 | 3,42 | 0,01 | 18,9 | >0.1 | 5,58 | 6,74 | 0 | 1,89 | 0 | 1,28 | 0 |
| ZBTB7C | GTGACCAAAGAG | -1 | 668 | 680 | -332 | -320 | 3,86 | 0,01 | 18,2 | >0.1 | 4,13 | 5,42 | 0 | 1,1 | 0 | 3,7 | 0 |
| ZBTB7C | CAGCCTCCCGAG | -1 | 347 | 359 | -653 | -641 | 3,15 | 0,01 | 17,89 | >0.1 | 2,55 | 4,68 | 0 | 1,11 | 0 | 6,4 | 0 |
| ZBTB7C | GGGACTACAGGT | -1 | 330 | 342 | -670 | -658 | 2,77 | 0,01 | 17,19 | >0.1 | 2,59 | 4,65 | 0 | 1,14 | 0 | 6,03 | 0 |
| ZBTB7C | GTGCCCACCACC | -1 | 320 | 332 | -680 | -668 | 10,9 | 0,001 | 24,88 | >0.1 | 2,62 | 4,64 | 0 | 1,16 | 0 | 5,56 | 0 |
| ZBTB7C | CCCACCACCACG | -1 | 317 | 329 | -683 | -671 | 10,64 | 0,001 | 25,32 | >0.1 | 2,63 | 4,63 | 0 | 1,17 | 0 | 6,26 | 0 |
| ZBTB7C | ACCACCACGCCC | -1 | 314 | 326 | -686 | -674 | 7,42 | 0,01 | 21,99 | >0.1 | 2,64 | 4,63 | 0 | 1,17 | 0 | 6,13 | 0 |
| ZBTB7C | ACCACGCCCAGC | -1 | 311 | 323 | -689 | -677 | 2,17 | 0,01 | 16,65 | >0.1 | 2,65 | 4,62 | 0 | 1,18 | 0 | 6,03 | 0 |
| ZBTB7C | ACGCCCAGCTAA | -1 | 308 | 320 | -692 | -680 | 5,14 | 0,01 | 19,86 | >0.1 | 2,65 | 4,62 | 0 | 1,19 | 0 | 6,26 | 0 |
| ZBTB7C | CGGCCTCCCAAA | -1 | 213 | 225 | -787 | -775 | 2,59 | 0,01 | 17,13 | >0.1 | 2,98 | 4,49 | 0 | 1,55 | 0 | 5,52 | 0 |
| ZBTB7C | GTGCCCAGCCAT | -1 | 174 | 186 | -826 | -814 | 2,73 | 0,01 | 15,4 | >0.1 | 3,15 | 4,44 | 0 | 1,84 | 0 | 3,24 | 0 |

| | | | | | | | | | | | | | | | | | |
|---|---|---|---|---|---|---|---|---|---|---|---|---|---|---|---|---|---|
| CEBPB | GTTTTGCAAA | 1 | 82 | 92 | -918 | -908 | 2,4 | 0,01 | 12,38 | >0.1 | 3,77 | 4,33 | 0 | 1,88 | 0 | 0 | 0 |
| CEBPB | TTTAAAAAAT | 1 | 137 | 147 | -863 | -853 | -1,74 | 0,01 | 9,55 | >0.1 | 3,36 | 4,39 | 0 | 1,89 | 0 | 1,65 | 0 |
| CEBPB | ATCGCGCCAC | 1 | 409 | 419 | -591 | -581 | 1,19 | 0,01 | 15,18 | >0.1 | 2,65 | 4,78 | 0 | 1,03 | 0 | 5,52 | 0 |
| CEBPB | GTTGCCCAAG | 1 | 540 | 550 | -460 | -450 | 6,89 | 0,001 | 16,06 | >0.1 | 3,13 | 5,06 | 0 | 0,99 | 0 | 0 | 0 |
| CEBPB | GTGAGATAAC | 1 | 591 | 601 | -409 | -399 | -1,47 | 0,01 | 8,14 | >0.1 | 3,41 | 5,18 | 0 | 1,01 | 0 | 0 | 0 |
| CEBPB | CTTTGGTCAC | 1 | 670 | 680 | -330 | -320 | -2,73 | 0,01 | 11,63 | >0.1 | 4,14 | 5,42 | 0 | 1,1 | 0 | 3,7 | 0 |
| CEBPB | ATTATAACAT | 1 | 686 | 696 | -314 | -304 | -0,28 | 0,01 | 13,69 | >0.1 | 4,42 | 5,47 | 0 | 1,13 | 0 | 2,95 | 0 |
| CEBPB | ATTAACCAAA | 1 | 730 | 740 | -270 | -260 | -2,9 | 0,01 | 15,1 | >0.1 | 5,61 | 5,64 | 0 | 1,23 | 0 | 5,52 | 0 |
| CEBPB | ATTTGGCCAT | 1 | 753 | 763 | -247 | -237 | 1,45 | 0,01 | 20,41 | >0.1 | 5,61 | 5,74 | 0 | 1,3 | 0 | 6,32 | 0 |
| CEBPB | TTTAAAAAAT | 1 | 825 | 835 | -175 | -165 | -1,74 | 0,01 | 15,34 | >0.1 | 5,61 | 6,12 | 0 | 1,65 | 0 | 3,7 | 0 |
| CEBPB | ATTGAGGAAC | 1 | 1109 | 1119 | 109 | 119 | -3,14 | 0,01 | 16,72 | >0.1 | 5,61 | 6,66 | 0 | 1,85 | 3,22 | 2,52 | 0 |
| CEBPB | TTTGGACAAG | 1 | 1132 | 1142 | 132 | 142 | -0,99 | 0,01 | 19,1 | >0.1 | 5,61 | 6,44 | 0 | 1,84 | 3,22 | 2,98 | 0 |
| CEBPB | GTTTAACCAC | 1 | 1141 | 1151 | 141 | 151 | -1,35 | 0,01 | 19,1 | >0.1 | 5,61 | 6,37 | 0 | 1,84 | 3,22 | 3,4 | 0 |
| CEBPB | GTGGTTAAAC | -1 | 1141 | 1151 | 141 | 151 | -0,64 | 0,01 | 19,81 | >0.1 | 5,61 | 6,37 | 0 | 1,84 | 3,22 | 3,4 | 0 |
| CEBPB | CTTGTCCAAA | -1 | 1132 | 1142 | 132 | 142 | -0,39 | 0,01 | 19,7 | >0.1 | 5,61 | 6,44 | 0 | 1,84 | 3,22 | 2,98 | 0 |
| CEBPB | GTTCCTCAAT | -1 | 1109 | 1119 | 109 | 119 | 0,96 | 0,01 | 20,82 | >0.1 | 5,61 | 6,66 | 0 | 1,85 | 3,22 | 2,52 | 0 |
| CEBPB | CTGAGCCAAT | -1 | 882 | 892 | -118 | -108 | -0,36 | 0,01 | 16,76 | >0.1 | 5,61 | 6,55 | 0 | 1,89 | 0 | 3,07 | 0 |
| CEBPB | GATATGTAAC | -1 | 848 | 858 | -152 | -142 | -2,05 | 0,01 | 14,64 | >0.1 | 5,61 | 6,28 | 0 | 1,84 | 0 | 2,95 | 0 |
| CEBPB | ATGGCCAAAT | -1 | 753 | 763 | -247 | -237 | 3,85 | 0,01 | 22,81 | >0.1 | 5,61 | 5,74 | 0 | 1,3 | 0 | 6,32 | 0 |
| CEBPB | ATGTTATAAT | -1 | 686 | 696 | -314 | -304 | -0,97 | 0,01 | 13 | >0.1 | 4,42 | 5,47 | 0 | 1,13 | 0 | 2,95 | 0 |
| CEBPB | GTGACCAAAG | -1 | 670 | 680 | -330 | -320 | -0,2 | 0,01 | 14,16 | >0.1 | 4,14 | 5,42 | 0 | 1,1 | 0 | 3,7 | 0 |
| CEBPB | GTTATCTCAC | -1 | 591 | 601 | -409 | -399 | -2,64 | 0,01 | 6,97 | >0.1 | 3,41 | 5,18 | 0 | 1,01 | 0 | 0 | 0 |
| CEBPB | CTTGGGCAAC | -1 | 540 | 550 | -460 | -450 | 8,73 | 0,001 | 17,9 | >0.1 | 3,13 | 5,06 | 0 | 0,99 | 0 | 0 | 0 |
| CEBPB | GTGGCGCGAT | -1 | 409 | 419 | -591 | -581 | 1,84 | 0,01 | 15,83 | >0.1 | 2,65 | 4,78 | 0 | 1,03 | 0 | 5,52 | 0 |
| CEBPB | TCTGCCCAAC | -1 | 226 | 236 | -774 | -764 | -1,86 | 0,01 | 12,3 | >0.1 | 2,93 | 4,5 | 0 | 1,47 | 0 | 5,26 | 0 |
| CEBPB | TTTTTAAAAT | -1 | 135 | 145 | -865 | -855 | -1,53 | 0,01 | 9,7 | >0.1 | 3,37 | 4,39 | 0 | 1,89 | 0 | 1,57 | 0 |
| CEBPB | TTTGCAAAAC | -1 | 82 | 92 | -918 | -908 | 5,53 | 0,001 | 15,51 | >0.1 | 3,77 | 4,33 | 0 | 1,88 | 0 | 0 | 0 |
| CEBPB | CTTAAAAAAG | -1 | 73 | 83 | -927 | -917 | -1,89 | 0,01 | 8,16 | >0.1 | 3,85 | 4,32 | 0 | 1,88 | 0 | 0 | 0 |
| POU5F1 | TTTTGCAAAGG | 1 | 83 | 94 | -917 | -906 | 5,42 | 0,01 | 15,39 | >0.1 | 3,75 | 4,33 | 0 | 1,88 | 0 | 0 | 0 |
| POU5F1 | AAAGGCAGATC | 1 | 89 | 100 | -911 | -900 | 3,19 | 0,01 | 13,11 | >0.1 | 3,7 | 4,34 | 0 | 1,88 | 0 | 0 | 0 |
| POU5F1 | CCATGGAAATT | 1 | 149 | 160 | -851 | -840 | 7,65 | 0,01 | 18,89 | >0.1 | 3,29 | 4,41 | 0 | 1,9 | 0 | 1,65 | 0 |
| POU5F1 | AAATTAAAACT | 1 | 155 | 166 | -845 | -834 | 2,65 | 0,01 | 14,82 | >0.1 | 3,25 | 4,41 | 0 | 1,9 | 0 | 2,6 | 0 |
| POU5F1 | AAATACAAAAA | 1 | 296 | 307 | -704 | -693 | 7,31 | 0,01 | 20,35 | >0.1 | 2,69 | 4,6 | 0 | 1,22 | 0 | 4,53 | 0 |
| POU5F1 | AAATAAAAATA | 1 | 577 | 588 | -423 | -412 | 3,86 | 0,01 | 13,34 | >0.1 | 3,33 | 5,15 | 0 | 1 | 0 | 0 | 0 |
| POU5F1 | TAATGAAATTG | 1 | 605 | 616 | -395 | -384 | 3,54 | 0,01 | 14,71 | >0.1 | 3,51 | 5,22 | 0 | 1,02 | 0 | 1,41 | 0 |
| POU5F1 | CCATACAAACA | 1 | 624 | 635 | -376 | -365 | 6,2 | 0,01 | 17,67 | >0.1 | 3,66 | 5,28 | 0 | 1,04 | 0 | 1,5 | 0 |
| POU5F1 | TTATTCTAAAA | 1 | 834 | 845 | -166 | -155 | 4,11 | 0,01 | 20,91 | >0.1 | 5,6 | 6,19 | 0 | 1,73 | 0 | 3,28 | 0 |
| POU5F1 | AAGTTCAAAGG | 1 | 956 | 967 | -44 | -33 | 2,56 | 0,01 | 19,28 | >0.1 | 5,6 | 7,55 | 0 | 1,88 | 0 | 1,7 | 0 |
| POU5F1 | CAATATAAAGT | -1 | 875 | 886 | -125 | -114 | 3,04 | 0,01 | 19,98 | >0.1 | 5,6 | 6,5 | 0 | 1,9 | 0 | 2,95 | 0 |
| POU5F1 | ATATGTAACAG | -1 | 846 | 857 | -154 | -143 | 4,68 | 0,01 | 21,25 | >0.1 | 5,6 | 6,28 | 0 | 1,83 | 0 | 2,86 | 0 |
| POU5F1 | TCATGAAGATA | -1 | 642 | 653 | -358 | -347 | 5,59 | 0,01 | 19,63 | >0.1 | 3,82 | 5,33 | 0 | 1,06 | 0 | 3,83 | 0 |
| POU5F1 | TTATGGAATTC | -1 | 493 | 504 | -507 | -496 | 3,96 | 0,01 | 19,27 | >0.1 | 2,93 | 4,95 | 0 | 0,98 | 0 | 6,44 | 0 |
| POU5F1 | GAATTCATATT | -1 | 488 | 499 | -512 | -501 | 4,04 | 0,01 | 19,15 | >0.1 | 2,91 | 4,94 | 0 | 0,99 | 0 | 6,27 | 0 |
| POU5F1 | CTTTGCAAAAC | -1 | 82 | 93 | -918 | -907 | 3,28 | 0,01 | 13,25 | >0.1 | 3,76 | 4,33 | 0 | 1,88 | 0 | 0 | 0 |
| HLF | AGTTTTGCAAAG | 1 | 81 | 93 | -919 | -907 | 0,35 | 0,01 | 10,33 | >0.1 | 3,77 | 4,33 | 0 | 1,88 | 0 | 0 | 0 |
| HLF | GATCGCGCCACT | 1 | 408 | 420 | -592 | -580 | 2,38 | 0,01 | 16,37 | >0.1 | 2,65 | 4,79 | 0 | 1,03 | 0 | 5,52 | 0 |
| HLF | TGTTGCCCAAGT | 1 | 539 | 551 | -461 | -449 | 3,91 | 0,01 | 13,09 | >0.1 | 3,13 | 5,06 | 0 | 0,99 | 0 | 0 | 0 |
| HLF | TGTGAGATAACC | 1 | 590 | 602 | -410 | -398 | 6,07 | 0,001 | 15,68 | >0.1 | 3,41 | 5,19 | 0 | 1,01 | 0 | 0 | 0 |
| HLF | TCTTTGGTCACT | 1 | 669 | 681 | -331 | -319 | -1,49 | 0,01 | 12,87 | >0.1 | 4,14 | 5,42 | 0 | 1,1 | 0 | 3,7 | 0 |
| HLF | AATTATAACATT | 1 | 685 | 697 | -315 | -303 | 1,82 | 0,01 | 15,79 | >0.1 | 4,42 | 5,48 | 0 | 1,13 | 0 | 2,95 | 0 |
| HLF | TGTTACATATCT | 1 | 847 | 859 | -153 | -141 | 5 | 0,01 | 21,68 | >0.1 | 5,58 | 6,29 | 0 | 1,85 | 0 | 2,95 | 0 |
| HLF | TCTTGGCTCACA | 1 | 1030 | 1042 | 30 | 42 | -0,91 | 0,01 | 16,08 | >0.1 | 5,58 | 8,24 | 0 | 1,86 | 0 | 1,31 | 0 |
| HLF | ACTTGTCCAAAA | -1 | 1131 | 1143 | 131 | 143 | 0,1 | 0,01 | 20,18 | >0.1 | 5,58 | 6,45 | 0 | 1,84 | 3,22 | 2,98 | 0 |
| HLF | AGTTACAGCAAC | -1 | 1082 | 1094 | 82 | 94 | 0,4 | 0,01 | 18,83 | >0.1 | 5,58 | 6,99 | 0 | 1,85 | 1,79 | 2,21 | 0 |
| HLF | TGTGAGCCAAGA | -1 | 1030 | 1042 | 30 | 42 | -1,63 | 0,01 | 15,36 | >0.1 | 5,58 | 8,24 | 0 | 1,86 | 0 | 1,31 | 0 |
| HLF | GCTGAGCCAATA | -1 | 881 | 893 | -119 | -107 | -1,08 | 0,01 | 16,02 | >0.1 | 5,58 | 6,56 | 0 | 1,89 | 0 | 3,07 | 0 |
| HLF | AGATATGTAACA | -1 | 847 | 859 | -153 | -141 | 4,07 | 0,01 | 20,75 | >0.1 | 5,58 | 6,29 | 0 | 1,85 | 0 | 2,95 | 0 |
| HLF | CCTTACCTCTCC | -1 | 772 | 784 | -228 | -216 | 0,58 | 0,01 | 19,47 | >0.1 | 5,58 | 5,83 | 0 | 1,37 | 0 | 6,1 | 0 |
| HLF | TATGGCCAAATC | -1 | 752 | 764 | -248 | -236 | -0,57 | 0,01 | 18,38 | >0.1 | 5,58 | 5,74 | 0 | 1,3 | 0 | 6,32 | 0 |

| | | | | | | | | | | | | | | | | | |
|---|---|---|---|---|---|---|---|---|---|---|---|---|---|---|---|---|---|
| HLF | AATGTTATAATT | -1 | 685 | 697 | -315 | -303 | 0,3 | 0,01 | 14,27 | >0.1 | 4,42 | 5,48 | 0 | 1,13 | 0 | 2,95 | 0 |
| HLF | AGTGACCAAAGA | -1 | 669 | 681 | -331 | -319 | -0,35 | 0,01 | 14,01 | >0.1 | 4,14 | 5,42 | 0 | 1,1 | 0 | 3,7 | 0 |
| HLF | GGTTATCTCACA | -1 | 590 | 602 | -410 | -398 | 5,69 | 0,001 | 15,3 | >0.1 | 3,41 | 5,19 | 0 | 1,01 | 0 | 0 | 0 |
| HLF | ACTTGGGCAACA | -1 | 539 | 551 | -461 | -449 | 4,38 | 0,01 | 13,56 | >0.1 | 3,13 | 5,06 | 0 | 0,99 | 0 | 0 | 0 |
| HLF | CTTTATGGAATT | -1 | 494 | 506 | -506 | -494 | 0,81 | 0,01 | 16,31 | >0.1 | 2,94 | 4,96 | 0 | 0,99 | 0 | 6,62 | 0 |
| HLF | AGTGGCGCGATC | -1 | 408 | 420 | -592 | -580 | -0,6 | 0,01 | 13,39 | >0.1 | 2,65 | 4,79 | 0 | 1,03 | 0 | 5,52 | 0 |
| HLF | GTTCACGCCATT | -1 | 368 | 380 | -632 | -620 | 0,43 | 0,01 | 14,91 | >0.1 | 2,54 | 4,72 | 0 | 1,08 | 0 | 6,15 | 0 |
| HLF | CTTTGCAAAACT | -1 | 81 | 93 | -919 | -907 | 2,84 | 0,01 | 12,82 | >0.1 | 3,77 | 4,33 | 0 | 1,88 | 0 | 0 | 0 |
| HLF | ACTTAAAAAAGC | -1 | 72 | 84 | -928 | -916 | -1,45 | 0,01 | 8,6 | >0.1 | 3,85 | 4,32 | 0 | 1,88 | 0 | 0 | 0 |
| ARID3A | ATTAAA | 1 | 157 | 163 | -843 | -837 | 9,38 | >0.1 | 21,54 | >0.1 | 3,25 | 4,41 | 0 | 1,9 | 0 | 2,6 | 0 |
| ARID3A | ATTAAA | 1 | 291 | 297 | -709 | -703 | 9,38 | >0.1 | 22,99 | >0.1 | 2,71 | 4,58 | 0 | 1,23 | 0 | 5,09 | 0 |
| ARID3A | ATAAAA | 1 | 579 | 585 | -421 | -415 | 5,54 | 0,01 | 15,01 | >0.1 | 3,33 | 5,14 | 0 | 1 | 0 | 0 | 0 |
| ARID3A | ATAAAT | 1 | 585 | 591 | -415 | -409 | 3,83 | 0,01 | 13,35 | >0.1 | 3,36 | 5,16 | 0 | 1 | 0 | 0 | 0 |
| ARID3A | ATTAAT | 1 | 603 | 609 | -397 | -391 | 7,67 | 0,01 | 18,78 | >0.1 | 3,48 | 5,2 | 0 | 1,01 | 0 | 1,41 | 0 |
| ARID3A | ATGAAA | 1 | 607 | 613 | -393 | -387 | 3,45 | 0,01 | 14,6 | >0.1 | 3,51 | 5,22 | 0 | 1,02 | 0 | 1,41 | 0 |
| ARID3A | ATTAAC | 1 | 730 | 736 | -270 | -264 | 3,21 | 0,01 | 21,03 | >0.1 | 5,67 | 5,62 | 0 | 1,21 | 0 | 5,32 | 0 |
| ARID3A | ATTAAA | 1 | 793 | 799 | -207 | -201 | 9,38 | >0.1 | 27,71 | >0.1 | 5,67 | 5,91 | 0 | 1,44 | 0 | 5,32 | 0 |
| ARID3A | ATCAAA | 1 | 1170 | 1176 | 170 | 176 | 8,97 | 0,01 | 26 | >0.1 | 4,58 | 6,15 | 0 | 1,67 | 1,79 | 2,84 | 0 |
| ARID3A | GTTAAA | -1 | 1142 | 1148 | 142 | 148 | 2,79 | 0,01 | 23,18 | >0.1 | 5,67 | 6,36 | 0 | 1,84 | 3,22 | 3,29 | 0 |
| ARID3A | ACTAAA | -1 | 997 | 1003 | -3 | 3 | 2,79 | 0,01 | 20,66 | >0.1 | 5,67 | 8,8 | 0 | 1,87 | 0 | 1,53 | 0 |
| ARID3A | AGTAAA | -1 | 899 | 905 | -101 | -95 | 2,79 | 0,01 | 19,86 | >0.1 | 5,67 | 6,65 | 0 | 1,89 | 0 | 2,86 | 0 |
| ARID3A | ATCAAA | -1 | 815 | 821 | -185 | -179 | 8,97 | 0,01 | 25,66 | >0.1 | 5,67 | 6,03 | 0 | 1,56 | 0 | 3,43 | 0 |
| ARID3A | ATCAAA | -1 | 749 | 755 | -251 | -245 | 8,97 | 0,01 | 27,71 | >0.1 | 5,67 | 5,7 | 0 | 1,27 | 0 | 6,1 | 0 |
| ARID3A | GTTAAA | -1 | 708 | 714 | -292 | -286 | 2,79 | 0,01 | 20,14 | >0.1 | 5,67 | 5,54 | 0 | 1,16 | 0 | 4,99 | 0 |
| ARID3A | ATTAAT | -1 | 603 | 609 | -397 | -391 | 7,67 | 0,01 | 18,78 | >0.1 | 3,48 | 5,2 | 0 | 1,01 | 0 | 1,41 | 0 |
| ARID3A | ATTACA | -1 | 199 | 205 | -801 | -795 | 2,79 | 0,01 | 16,31 | >0.1 | 3,05 | 4,46 | 0 | 1,64 | 0 | 4,37 | 0 |
| ARID3A | CTTAAA | -1 | 77 | 83 | -923 | -917 | 2,79 | 0,01 | 12,82 | >0.1 | 3,83 | 4,32 | 0 | 1,88 | 0 | 0 | 0 |
| ARID3B | GTCATAATTTT | 1 | 129 | 140 | -871 | -860 | 4,83 | 0,01 | 16,12 | >0.1 | 3,4 | 4,38 | 0 | 1,89 | 0 | 1,61 | 0 |
| ARID3B | ATTTTAAAAAA | 1 | 135 | 146 | -865 | -854 | 7,29 | 0,01 | 18,51 | >0.1 | 3,37 | 4,39 | 0 | 1,89 | 0 | 1,57 | 0 |
| ARID3B | AAATTAAAACT | 1 | 155 | 166 | -845 | -834 | 5,87 | 0,01 | 18,04 | >0.1 | 3,25 | 4,41 | 0 | 1,9 | 0 | 2,6 | 0 |
| ARID3B | CTATTAAAAAT | 1 | 289 | 300 | -711 | -700 | 7,04 | 0,01 | 20,66 | >0.1 | 2,71 | 4,59 | 0 | 1,24 | 0 | 5,09 | 0 |
| ARID3B | GAAATAAAAAT | 1 | 576 | 587 | -424 | -413 | 2,89 | 0,01 | 12,36 | >0.1 | 3,33 | 5,15 | 0 | 1 | 0 | 0 | 0 |
| ARID3B | AAAATAAATGT | 1 | 582 | 593 | -418 | -407 | 3,69 | 0,01 | 13,22 | >0.1 | 3,36 | 5,16 | 0 | 1,01 | 0 | 0 | 0 |
| ARID3B | CTATTAATGAA | 1 | 601 | 612 | -399 | -388 | 9,75 | 0,001 | 20,88 | >0.1 | 3,48 | 5,21 | 0 | 1,02 | 0 | 1,41 | 0 |
| ARID3B | AAATTATAACA | 1 | 684 | 695 | -316 | -305 | 3,21 | 0,01 | 17,14 | >0.1 | 4,39 | 5,47 | 0 | 1,13 | 0 | 2,95 | 0 |
| ARID3B | AAAATATTAAC | 1 | 725 | 736 | -275 | -264 | 4,29 | 0,01 | 22,04 | >0.1 | 5,6 | 5,62 | 0 | 1,22 | 0 | 5,32 | 0 |
| ARID3B | GGATTAAAGAA | 1 | 791 | 802 | -209 | -198 | 4,07 | 0,01 | 22,37 | >0.1 | 5,6 | 5,93 | 0 | 1,45 | 0 | 5,32 | 0 |
| ARID3B | TTCTTATTTGA | 1 | 809 | 820 | -191 | -180 | 5,28 | 0,01 | 21,74 | >0.1 | 5,6 | 6,03 | 0 | 1,55 | 0 | 3,28 | 0 |
| ARID3B | ACTTTAAAAAA | 1 | 823 | 834 | -177 | -166 | 4,97 | 0,01 | 22,02 | >0.1 | 5,6 | 6,12 | 0 | 1,65 | 0 | 3,7 | 0 |
| ARID3B | AAATTATTCTA | 1 | 831 | 842 | -169 | -158 | 5,93 | 0,01 | 22,83 | >0.1 | 5,6 | 6,17 | 0 | 1,71 | 0 | 3,43 | 0 |
| ARID3B | AGAATAATTTT | -1 | 830 | 841 | -170 | -159 | 5,27 | 0,01 | 22,01 | >0.1 | 5,6 | 6,16 | 0 | 1,7 | 0 | 3,28 | 0 |
| ARID3B | TTTTTAAAGTG | -1 | 822 | 833 | -178 | -167 | 2,82 | 0,01 | 19,86 | >0.1 | 5,6 | 6,11 | 0 | 1,64 | 0 | 3,7 | 0 |
| ARID3B | TCTTTAATCCT | -1 | 790 | 801 | -210 | -199 | 5,05 | 0,01 | 23,34 | >0.1 | 5,6 | 5,92 | 0 | 1,45 | 0 | 5,32 | 0 |
| ARID3B | TTAATATTTTC | -1 | 724 | 735 | -276 | -265 | 3,34 | 0,01 | 20,94 | >0.1 | 5,6 | 5,62 | 0 | 1,21 | 0 | 5,17 | 0 |
| ARID3B | GTTATAATTTT | -1 | 683 | 694 | -317 | -306 | 5,34 | 0,01 | 19,36 | >0.1 | 4,37 | 5,47 | 0 | 1,12 | 0 | 3,07 | 0 |
| ARID3B | TCATTAATAGG | -1 | 600 | 611 | -400 | -389 | 7,04 | 0,01 | 18,11 | >0.1 | 3,48 | 5,21 | 0 | 1,02 | 0 | 1,36 | 0 |
| ARID3B | CATTTATTTTT | -1 | 581 | 592 | -419 | -408 | 4,04 | 0,01 | 13,56 | >0.1 | 3,35 | 5,16 | 0 | 1 | 0 | 0 | 0 |
| ARID3B | TTTTTATTTCA | -1 | 575 | 586 | -425 | -414 | 5,79 | 0,01 | 15,26 | >0.1 | 3,32 | 5,14 | 0 | 1 | 0 | 0 | 0 |
| ARID3B | TTTTTAATAGA | -1 | 288 | 299 | -712 | -701 | 8,21 | 0,001 | 21,84 | >0.1 | 2,71 | 4,59 | 0 | 1,24 | 0 | 5,09 | 0 |
| ARID3B | GTTTTAATTTC | -1 | 154 | 165 | -846 | -835 | 8,94 | 0,001 | 20,16 | >0.1 | 3,26 | 4,41 | 0 | 1,9 | 0 | 1,65 | 0 |
| ARID3B | TTTTTAAAATT | -1 | 134 | 145 | -866 | -855 | 3,66 | 0,01 | 14,93 | >0.1 | 3,37 | 4,39 | 0 | 1,89 | 0 | 1,61 | 0 |
| ARID3B | AACTTAAAAAA | -1 | 74 | 85 | -926 | -915 | 5,74 | 0,01 | 15,78 | >0.1 | 3,84 | 4,32 | 0 | 1,88 | 0 | 0 | 0 |
| CEBPE | GTTTTGCAAA | 1 | 82 | 92 | -918 | -908 | 3,84 | 0,01 | 13,82 | >0.1 | 3,77 | 4,33 | 0 | 1,88 | 0 | 0 | 0 |
| CEBPE | TTTAAAAAAT | 1 | 137 | 147 | -863 | -853 | -0,16 | 0,01 | 11,13 | >0.1 | 3,36 | 4,39 | 0 | 1,89 | 0 | 1,65 | 0 |
| CEBPE | ACGGTGAAAC | 1 | 272 | 282 | -728 | -718 | 0,95 | 0,01 | 14,56 | >0.1 | 2,77 | 4,56 | 0 | 1,29 | 0 | 4,99 | 0 |
| CEBPE | ATCGCGCCAC | 1 | 409 | 419 | -591 | -581 | 3,51 | 0,01 | 17,5 | >0.1 | 2,65 | 4,78 | 0 | 1,03 | 0 | 5,52 | 0 |
| CEBPE | GTTGCCCAAG | 1 | 540 | 550 | -460 | -450 | 7,69 | 0,001 | 16,86 | >0.1 | 3,13 | 5,06 | 0 | 0,99 | 0 | 0 | 0 |
| CEBPE | GTGAGATAAC | 1 | 591 | 601 | -409 | -399 | 0,03 | 0,01 | 9,64 | >0.1 | 3,41 | 5,18 | 0 | 1,01 | 0 | 0 | 0 |
| CEBPE | ACTGTAAAAT | 1 | 678 | 688 | -322 | -312 | 0,15 | 0,01 | 14,26 | >0.1 | 4,27 | 5,45 | 0 | 1,11 | 0 | 3,28 | 0 |

| | | | | | | | | | | | | | | | | | |
|---|---|---|---|---|---|---|---|---|---|---|---|---|---|---|---|---|---|
| CEBPE | ATTATAACAT | 1 | 686 | 696 | -314 | -304 | 1,41 | 0,01 | 15,38 | >0.1 | 4,42 | 5,47 | 0 | 1,13 | 0 | 2,95 | 0 |
| CEBPE | ATTTGGCCAT | 1 | 753 | 763 | -247 | -237 | 3,08 | 0,01 | 22,04 | >0.1 | 5,61 | 5,74 | 0 | 1,3 | 0 | 6,32 | 0 |
| CEBPE | TTTAAAAAAT | 1 | 825 | 835 | -175 | -165 | -0,16 | 0,01 | 16,92 | >0.1 | 5,61 | 6,12 | 0 | 1,65 | 0 | 3,7 | 0 |
| CEBPE | ATTGAGGAAC | 1 | 1109 | 1119 | 109 | 119 | 1,8 | 0,01 | 21,66 | >0.1 | 5,61 | 6,66 | 0 | 1,85 | 3,22 | 2,52 | 0 |
| CEBPE | TTTGGACAAG | 1 | 1132 | 1142 | 132 | 142 | 0,72 | 0,01 | 20,81 | >0.1 | 5,61 | 6,44 | 0 | 1,84 | 3,22 | 2,98 | 0 |
| CEBPE | GTTTAACCAC | 1 | 1141 | 1151 | 141 | 151 | 0,44 | 0,01 | 20,89 | >0.1 | 5,61 | 6,37 | 0 | 1,84 | 3,22 | 3,4 | 0 |
| CEBPE | GTGGTTAAAC | -1 | 1141 | 1151 | 141 | 151 | 1,22 | 0,01 | 21,67 | >0.1 | 5,61 | 6,37 | 0 | 1,84 | 3,22 | 3,4 | 0 |
| CEBPE | CTTGTCCAAA | -1 | 1132 | 1142 | 132 | 142 | 1,74 | 0,01 | 21,83 | >0.1 | 5,61 | 6,44 | 0 | 1,84 | 3,22 | 2,98 | 0 |
| CEBPE | GTTCCTCAAT | -1 | 1109 | 1119 | 109 | 119 | 1,52 | 0,01 | 21,38 | >0.1 | 5,61 | 6,66 | 0 | 1,85 | 3,22 | 2,52 | 0 |
| CEBPE | CTGAGCCAAT | -1 | 882 | 892 | -118 | -108 | 1,45 | 0,01 | 18,57 | >0.1 | 5,61 | 6,55 | 0 | 1,89 | 0 | 3,07 | 0 |
| CEBPE | GATATGTAAC | -1 | 848 | 858 | -152 | -142 | 0,58 | 0,01 | 17,27 | >0.1 | 5,61 | 6,28 | 0 | 1,84 | 0 | 2,95 | 0 |
| CEBPE | ATGGCCAAAT | -1 | 753 | 763 | -247 | -237 | 5,24 | 0,01 | 24,2 | >0.1 | 5,61 | 5,74 | 0 | 1,3 | 0 | 6,32 | 0 |
| CEBPE | ATGTTATAAT | -1 | 686 | 696 | -314 | -304 | 0,48 | 0,01 | 14,45 | >0.1 | 4,42 | 5,47 | 0 | 1,13 | 0 | 2,95 | 0 |
| CEBPE | GTGACCAAAG | -1 | 670 | 680 | -330 | -320 | 1,42 | 0,01 | 15,78 | >0.1 | 4,14 | 5,42 | 0 | 1,1 | 0 | 3,7 | 0 |
| CEBPE | GTTATCTCAC | -1 | 591 | 601 | -409 | -399 | -0,61 | 0,01 | 9 | >0.1 | 3,41 | 5,18 | 0 | 1,01 | 0 | 0 | 0 |
| CEBPE | CTTGGGCAAC | -1 | 540 | 550 | -460 | -450 | 9,38 | 0,001 | 18,55 | >0.1 | 3,13 | 5,06 | 0 | 0,99 | 0 | 0 | 0 |
| CEBPE | TTTATGGAAT | -1 | 495 | 505 | -505 | -495 | -0,09 | 0,01 | 15,41 | >0.1 | 2,94 | 4,95 | 0 | 0,98 | 0 | 6,62 | 0 |
| CEBPE | TTCACGCCAT | -1 | 369 | 379 | -631 | -621 | -0,64 | 0,01 | 13,84 | >0.1 | 2,54 | 4,71 | 0 | 1,07 | 0 | 6,15 | 0 |
| CEBPE | TCTGCCCAAC | -1 | 226 | 236 | -774 | -764 | 2,15 | 0,01 | 16,31 | >0.1 | 2,93 | 4,5 | 0 | 1,47 | 0 | 5,26 | 0 |
| CEBPE | TTTTTAAAAT | -1 | 135 | 145 | -865 | -855 | 0,18 | 0,01 | 11,41 | >0.1 | 3,37 | 4,39 | 0 | 1,89 | 0 | 1,57 | 0 |
| CEBPE | TTTGCAAAAC | -1 | 82 | 92 | -918 | -908 | 6,43 | 0,001 | 16,41 | >0.1 | 3,77 | 4,33 | 0 | 1,88 | 0 | 0 | 0 |
| CEBPE | CTTAAAAAAG | -1 | 73 | 83 | -927 | -917 | -0,39 | 0,01 | 9,66 | >0.1 | 3,85 | 4,32 | 0 | 1,88 | 0 | 0 | 0 |
| ARX | CCATGGAAATTAAAACT | 1 | 149 | 166 | -851 | -834 | -1,73 | 0,01 | 9,5 | >0.1 | 3,27 | 4,41 | 0 | 1,9 | 0 | 1,65 | 0 |
| ARX | GATAACCTATTAATGAA | 1 | 595 | 612 | -405 | -388 | 4,06 | 0,01 | 15,17 | >0.1 | 3,46 | 5,21 | 0 | 1,02 | 0 | 1,41 | 0 |
| ARX | ACCTATTAATGAAATTG | 1 | 599 | 616 | -401 | -384 | -0,11 | 0,01 | 11,04 | >0.1 | 3,49 | 5,22 | 0 | 1,03 | 0 | 1,41 | 0 |
| ARX | TATTCTTATTTGATTCA | 1 | 807 | 824 | -193 | -176 | -0,16 | 0,01 | 16,27 | >0.1 | 5,52 | 6,05 | 0 | 1,58 | 0 | 3,28 | 0 |
| ARX | GAAAGTCATTCAGTGGA | 1 | 1008 | 1025 | 8 | 25 | -2,03 | 0,01 | 15,82 | >0.1 | 5,52 | 9,07 | 0 | 1,87 | 0 | 1,39 | 0 |
| ARX | GAGCAGCAGTTACAGCA | -1 | 1084 | 1101 | 84 | 101 | -1,94 | 0,01 | 16,4 | >0.1 | 5,52 | 6,96 | 0 | 1,85 | 1,79 | 2,21 | 0 |
| ARX | CTGAGCCAATATAAAGT | -1 | 875 | 892 | -125 | -108 | -1,6 | 0,01 | 15,43 | >0.1 | 5,52 | 6,55 | 0 | 1,9 | 0 | 3,07 | 0 |
| ARX | GTGAATCAAATAAGAAT | -1 | 808 | 825 | -192 | -175 | 3,26 | 0,01 | 19,85 | >0.1 | 5,52 | 6,06 | 0 | 1,59 | 0 | 3,43 | 0 |
| ARX | ACAATTTCATTAATAGG | -1 | 600 | 617 | -400 | -383 | -0,8 | 0,01 | 10,36 | >0.1 | 3,5 | 5,23 | 0 | 1,03 | 0 | 1,41 | 0 |
| ARX | TTTCATTAATAGGTTAT | -1 | 596 | 613 | -404 | -387 | 3,86 | 0,01 | 14,93 | >0.1 | 3,47 | 5,22 | 0 | 1,02 | 0 | 1,36 | 0 |
| ARX | GGCCATTTTTTAAAATT | -1 | 134 | 151 | -866 | -849 | 1,03 | 0,01 | 12,32 | >0.1 | 3,35 | 4,4 | 0 | 1,89 | 0 | 1,65 | 0 |
| CTCFL | AAGGAGAAGGAGGC | 1 | 12 | 26 | -988 | -974 | -1,51 | 0,01 | 10,18 | >0.1 | 5,56 | 4,27 | 0 | 1,87 | 0 | 0 | 0 |
| CTCFL | GAGAAGGAGGCACT | 1 | 15 | 29 | -985 | -971 | 8,56 | 0,001 | 20,25 | >0.1 | 5,56 | 4,27 | 0 | 1,87 | 0 | 0 | 0 |
| CTCFL | CTTTGGGAGGCCGA | 1 | 212 | 226 | -788 | -774 | -0,14 | 0,01 | 14,4 | >0.1 | 2,98 | 4,49 | 0 | 1,55 | 0 | 5,52 | 0 |
| CTCFL | TGGTGGTGGGCACC | 1 | 319 | 333 | -681 | -667 | 2,17 | 0,01 | 16,16 | >0.1 | 2,62 | 4,64 | 0 | 1,16 | 0 | 5,56 | 0 |
| CTCFL | ACTCGGGAGGCTGA | 1 | 346 | 360 | -654 | -640 | -1,95 | 0,01 | 12,79 | >0.1 | 2,55 | 4,68 | 0 | 1,11 | 0 | 6,4 | 0 |
| CTCFL | CGGGAGGCTGAGGC | 1 | 349 | 363 | -651 | -637 | 0,36 | 0,01 | 15,09 | >0.1 | 2,54 | 4,69 | 0 | 1,11 | 0 | 6,4 | 0 |
| CTCFL | ACCTGGGAGGCAGA | 1 | 378 | 392 | -622 | -608 | 2,53 | 0,01 | 16,81 | >0.1 | 2,57 | 4,74 | 0 | 1,06 | 0 | 5,91 | 0 |
| CTCFL | TTGCAGTGAGCCGA | 1 | 394 | 408 | -606 | -592 | -1,22 | 0,01 | 12,97 | >0.1 | 2,61 | 4,76 | 0 | 1,04 | 0 | 5,76 | 0 |
| CTCFL | GCCGAGATCGCGCC | 1 | 403 | 417 | -597 | -583 | 1,68 | 0,01 | 15,9 | >0.1 | 2,64 | 4,78 | 0 | 1,04 | 0 | 5,76 | 0 |
| CTCFL | CGCCACTGTGCTCC | 1 | 413 | 427 | -587 | -573 | -1,46 | 0,01 | 12,12 | >0.1 | 2,67 | 4,8 | 0 | 1,03 | 0 | 5,09 | 0 |
| CTCFL | CAGGAGAGGTAAGG | 1 | 770 | 784 | -230 | -216 | 0,5 | 0,01 | 19,18 | >0.1 | 5,56 | 5,84 | 0 | 1,38 | 0 | 5,92 | 0 |
| CTCFL | CTCCAGGATGAACT | 1 | 864 | 878 | -136 | -122 | 1,32 | 0,01 | 17,96 | >0.1 | 5,56 | 6,43 | 0 | 1,9 | 0 | 2,75 | 0 |
| CTCFL | TCACAGGGGACGAT | 1 | 1037 | 1051 | 37 | 51 | 4,73 | 0,01 | 21,46 | >0.1 | 5,56 | 7,95 | 0 | 1,86 | 0 | 1,36 | 0 |
| CTCFL | GAGAAGGAGCCAGG | -1 | 1062 | 1076 | 62 | 76 | 2,64 | 0,01 | 21,37 | >0.1 | 5,56 | 7,31 | 0 | 1,86 | 1,79 | 2,21 | 0 |
| CTCFL | AGCCAGGAAGAGCT | -1 | 1055 | 1069 | 55 | 69 | -0,3 | 0,01 | 16,84 | >0.1 | 5,56 | 7,46 | 0 | 1,86 | 0 | 2,27 | 0 |
| CTCFL | GGGTTGGGCGCTGA | -1 | 942 | 956 | -58 | -44 | -1,42 | 0,01 | 15,14 | >0.1 | 5,56 | 7,32 | 0 | 1,88 | 0 | 1,8 | 0 |
| CTCFL | CTGGAGAGGACAGA | -1 | 856 | 870 | -144 | -130 | 4,56 | 0,01 | 21,25 | >0.1 | 5,56 | 6,37 | 0 | 1,9 | 0 | 2,86 | 0 |
| CTCFL | AACTAGGGATCATG | -1 | 648 | 662 | -352 | -338 | -0,27 | 0,01 | 14,05 | >0.1 | 3,9 | 5,36 | 0 | 1,07 | 0 | 3,99 | 0 |
| CTCFL | GCCCAGGCTGGAGC | -1 | 422 | 436 | -578 | -564 | -0,62 | 0,01 | 13,58 | >0.1 | 2,69 | 4,82 | 0 | 1,02 | 0 | 5,67 | 0 |
| CTCFL | CTACAGGTGCCCAC | -1 | 324 | 338 | -676 | -662 | 3,67 | 0,01 | 17,53 | >0.1 | 2,6 | 4,65 | 0 | 1,15 | 0 | 5,46 | 0 |
| CTCFL | CTCCACAGGTCAAC | -1 | 106 | 120 | -894 | -880 | 1,96 | 0,01 | 12,5 | >0.1 | 3,55 | 4,36 | 0 | 1,89 | 0 | 0,75 | 0 |
| FOXC2 | ATTAAAAATACA | 1 | 291 | 303 | -709 | -697 | 2,57 | 0,01 | 15,55 | >0.1 | 2,7 | 4,59 | 0 | 1,23 | 0 | 4,45 | 0 |
| FOXC2 | TAAAAATACAAA | 1 | 293 | 305 | -707 | -695 | 2,89 | 0,01 | 15,94 | >0.1 | 2,7 | 4,6 | 0 | 1,22 | 0 | 4,53 | 0 |
| FOXC2 | AATAAAAATAAA | 1 | 578 | 590 | -422 | -410 | 8,04 | 0,01 | 17,54 | >0.1 | 3,34 | 5,15 | 0 | 1 | 0 | 0 | 0 |
| FOXC2 | AAAATAAATGTG | 1 | 582 | 594 | -418 | -406 | 3,78 | 0,01 | 13,32 | >0.1 | 3,36 | 5,17 | 0 | 1,01 | 0 | 0 | 0 |
| FOXC2 | CATACAAACACC | 1 | 625 | 637 | -375 | -363 | 5,38 | 0,01 | 16,92 | >0.1 | 3,67 | 5,28 | 0 | 1,04 | 0 | 1,55 | 0 |

| | | | | | | | | | | | | | | | | | |
|---|---|---|---|---|---|---|---|---|---|---|---|---|---|---|---|---|---|
| FOXC2 | AACACCAACATT | 1 | 631 | 643 | -369 | -357 | 8,09 | 0,001 | 19,77 | >0.1 | 3,72 | 5,3 | 0 | 1,05 | 0 | 1,61 | 0 |
| FOXC2 | TAGGAAAATATT | 1 | 721 | 733 | -279 | -267 | 7,87 | 0,01 | 25,44 | >0.1 | 5,58 | 5,61 | 0 | 1,21 | 0 | 5,17 | 0 |
| FOXC2 | TACAGCAACAAG | -1 | 1079 | 1091 | 79 | 91 | 4,46 | 0,01 | 22,93 | >0.1 | 5,58 | 7,03 | 0 | 1,85 | 1,79 | 2,21 | 0 |
| FOXC2 | ACAGTAAACAAT | -1 | 895 | 907 | -105 | -93 | 11,73 | 0,001 | 28,83 | >0.1 | 5,58 | 6,68 | 0 | 1,89 | 0 | 2,95 | 0 |
| FOXC2 | TGAGCCAATATA | -1 | 879 | 891 | -121 | -109 | 8,24 | 0,001 | 25,32 | >0.1 | 5,58 | 6,54 | 0 | 1,9 | 0 | 3,07 | 0 |
| FOXC2 | TTGGGCAACATT | -1 | 537 | 549 | -463 | -451 | 3,08 | 0,01 | 13,98 | >0.1 | 3,12 | 5,05 | 0 | 0,99 | 0 | 1,73 | 0 |
| TFCP2 | GAACCAGTGT | 1 | 46 | 56 | -954 | -944 | 8,1 | 0,001 | 18,45 | >0.1 | 4,19 | 4,29 | 0 | 1,87 | 0 | 0 | 0 |
| TFCP2 | AAACTGATCA | 1 | 161 | 171 | -839 | -829 | -0,34 | 0,01 | 11,66 | >0.1 | 3,22 | 4,42 | 0 | 1,9 | 0 | 2,46 | 0 |
| TFCP2 | AAACTGATCA | 1 | 476 | 486 | -524 | -514 | -0,34 | 0,01 | 13,63 | >0.1 | 2,87 | 4,91 | 0 | 0,99 | 0 | 5,2 | 0 |
| TFCP2 | GTTCTAGACC | 1 | 659 | 669 | -341 | -331 | -1,21 | 0,01 | 13,25 | >0.1 | 4 | 5,38 | 0 | 1,08 | 0 | 3,99 | 0 |
| TFCP2 | AGACCTCTTT | 1 | 664 | 674 | -336 | -326 | -0,87 | 0,01 | 13,51 | >0.1 | 4,06 | 5,4 | 0 | 1,09 | 0 | 3,83 | 0 |
| TFCP2 | TAACAGCTTT | 1 | 710 | 720 | -290 | -280 | -0,12 | 0,01 | 17,04 | >0.1 | 5,61 | 5,56 | 0 | 1,18 | 0 | 4,81 | 0 |
| TFCP2 | GTACCGGTTT | 1 | 741 | 751 | -259 | -249 | 10,93 | 0,0001 | 29,58 | >0.1 | 5,61 | 5,68 | 0 | 1,26 | 0 | 6,1 | 0 |
| TFCP2 | TAACTGCTGC | 1 | 1089 | 1099 | 89 | 99 | 0,2 | 0,01 | 18,56 | >0.1 | 5,61 | 6,89 | 0 | 1,85 | 1,79 | 2,21 | 0 |
| TFCP2 | GAACAGGCCA | 1 | 1115 | 1125 | 115 | 125 | 0,12 | 0,01 | 19,94 | >0.1 | 5,61 | 6,6 | 0 | 1,85 | 3,22 | 2,54 | 0 |
| TFCP2 | GGACAAGTTT | 1 | 1135 | 1145 | 135 | 145 | 1,07 | 0,01 | 21,18 | >0.1 | 5,61 | 6,41 | 0 | 1,84 | 3,22 | 3,02 | 0 |
| TFCP2 | AGACCTGTTC | 1 | 1159 | 1169 | 159 | 169 | 5,23 | 0,001 | 21,88 | >0.1 | 4,38 | 6,23 | 0 | 1,77 | 1,79 | 2,48 | 0 |
| TFCP2 | GAACAGGTCT | -1 | 1159 | 1169 | 159 | 169 | 5,04 | 0,01 | 21,69 | >0.1 | 4,38 | 6,23 | 0 | 1,77 | 1,79 | 2,48 | 0 |
| TFCP2 | AAACTTGTCC | -1 | 1135 | 1145 | 135 | 145 | 3,77 | 0,01 | 23,88 | >0.1 | 5,61 | 6,41 | 0 | 1,84 | 3,22 | 3,02 | 0 |
| TFCP2 | AAACCTCCTC | -1 | 989 | 999 | -11 | -1 | -0,2 | 0,01 | 17,58 | >0.1 | 5,61 | 8,69 | 0 | 1,87 | 0 | 1,61 | 0 |
| TFCP2 | AATCTGCTGA | -1 | 888 | 898 | -112 | -102 | -0,97 | 0,01 | 16,2 | >0.1 | 5,61 | 6,6 | 0 | 1,89 | 0 | 3,07 | 0 |
| TFCP2 | AAACCGGTAC | -1 | 741 | 751 | -259 | -249 | 12,48 | 1E-05 | 31,13 | >0.1 | 5,61 | 5,68 | 0 | 1,26 | 0 | 6,1 | 0 |
| TFCP2 | AAAGCTGTTA | -1 | 710 | 720 | -290 | -280 | -0,25 | 0,01 | 16,91 | >0.1 | 5,61 | 5,56 | 0 | 1,18 | 0 | 4,81 | 0 |
| TFCP2 | GATCTGCCTT | -1 | 90 | 100 | -910 | -900 | 0,33 | 0,01 | 10,24 | >0.1 | 3,69 | 4,34 | 0 | 1,88 | 0 | 0 | 0 |
| TFCP2 | ACACTGGTTC | -1 | 46 | 56 | -954 | -944 | 7,31 | 0,001 | 17,66 | >0.1 | 4,19 | 4,29 | 0 | 1,87 | 0 | 0 | 0 |
| BCL6 | TCTATTAAAAATAC | 1 | 288 | 302 | -712 | -698 | -1,62 | 0,01 | 12,01 | >0.1 | 2,71 | 4,59 | 0 | 1,24 | 0 | 5,09 | 0 |
| BCL6 | GAACCTGGGAGGCA | 1 | 376 | 390 | -624 | -610 | 1,16 | 0,01 | 16,06 | >0.1 | 2,56 | 4,73 | 0 | 1,07 | 0 | 6,53 | 0 |
| BCL6 | CAGCCTGGGCGACA | 1 | 426 | 440 | -574 | -560 | -1,63 | 0,01 | 12,72 | >0.1 | 2,71 | 4,82 | 0 | 1,02 | 0 | 5,8 | 0 |
| BCL6 | GTATTTGGAAGGGA | 1 | 522 | 536 | -478 | -464 | 0,97 | 0,01 | 13,32 | >0.1 | 3,06 | 5,02 | 0 | 0,99 | 0 | 3,28 | 0 |
| BCL6 | GTTTCTGGGAATAT | 1 | 556 | 570 | -444 | -430 | 3,32 | 0,01 | 12,64 | >0.1 | 3,22 | 5,1 | 0 | 1 | 0 | 0 | 0 |
| BCL6 | GATCTTGAAATAAA | 1 | 570 | 584 | -430 | -416 | 1,98 | 0,01 | 11,42 | >0.1 | 3,3 | 5,14 | 0 | 1 | 0 | 0 | 0 |
| BCL6 | TTGTCTGAAAACCA | 1 | 613 | 627 | -387 | -373 | 4,42 | 0,01 | 15,84 | >0.1 | 3,58 | 5,26 | 0 | 1,03 | 0 | 1,55 | 0 |
| BCL6 | TATCTTTAACAGCT | 1 | 704 | 718 | -296 | -282 | 1,7 | 0,01 | 18,97 | >0.1 | 5,56 | 5,55 | 0 | 1,17 | 0 | 4,99 | 0 |
| BCL6 | CTTTCTAGGAAAAT | 1 | 716 | 730 | -284 | -270 | 2,05 | 0,01 | 19,39 | >0.1 | 5,56 | 5,6 | 0 | 1,2 | 0 | 4,99 | 0 |
| BCL6 | TTTCTAGGAAAATA | 1 | 717 | 731 | -283 | -269 | -0,51 | 0,01 | 16,84 | >0.1 | 5,56 | 5,6 | 0 | 1,2 | 0 | 4,99 | 0 |
| BCL6 | TTCTCTAGGATTAA | 1 | 784 | 798 | -216 | -202 | -1,6 | 0,01 | 16,99 | >0.1 | 5,56 | 5,91 | 0 | 1,44 | 0 | 5,69 | 0 |
| BCL6 | TATTCTAAAATCTG | 1 | 835 | 849 | -165 | -151 | -1,4 | 0,01 | 15,33 | >0.1 | 5,56 | 6,22 | 0 | 1,76 | 0 | 3,19 | 0 |
| BCL6 | ATTTTTGGACAAGT | 1 | 1129 | 1143 | 129 | 143 | 2,51 | 0,01 | 22,51 | >0.1 | 5,56 | 6,47 | 0 | 1,84 | 3,22 | 2,91 | 0 |
| BCL6 | CTTCTTGGAATTAT | 1 | 1185 | 1199 | 185 | 199 | 8,84 | 0,001 | 27,34 | >0.1 | 5,56 | 6,06 | 0 | 1,55 | 1,79 | 3,55 | 0 |
| BCL6 | ATTCCAAGAAGCAA | -1 | 1182 | 1196 | 182 | 196 | 0,94 | 0,01 | 19,61 | >0.1 | 5,56 | 6,08 | 0 | 1,57 | 1,79 | 3,67 | 0 |
| BCL6 | GTTCCTCAATGGTG | -1 | 1105 | 1119 | 105 | 119 | 6,91 | 0,001 | 26,71 | >0.1 | 5,56 | 6,7 | 0 | 1,85 | 3,22 | 2,47 | 0 |
| BCL6 | TTCCCTAGACTAAA | -1 | 997 | 1011 | -3 | 11 | 7,3 | 0,001 | 25,34 | >0.1 | 5,56 | 9,12 | 0 | 1,87 | 0 | 1,5 | 0 |
| BCL6 | CATCCTGGAGAGGA | -1 | 860 | 874 | -140 | -126 | 11,45 | 0,0001 | 28,06 | >0.1 | 5,56 | 6,4 | 0 | 1,9 | 0 | 2,75 | 0 |
| BCL6 | GATTTTAGAATAAT | -1 | 833 | 847 | -167 | -153 | -1,58 | 0,01 | 15,21 | >0.1 | 5,56 | 6,21 | 0 | 1,75 | 0 | 3,28 | 0 |
| BCL6 | TTTTTTAAAGTGAA | -1 | 820 | 834 | -180 | -166 | 2,47 | 0,01 | 19,49 | >0.1 | 5,56 | 6,12 | 0 | 1,65 | 0 | 3,7 | 0 |
| BCL6 | AATCCTAGAGAACC | -1 | 782 | 796 | -218 | -204 | 9,23 | 0,001 | 28,02 | >0.1 | 5,56 | 5,9 | 0 | 1,43 | 0 | 5,92 | 0 |
| BCL6 | ATTCCCAGAAACTC | -1 | 554 | 568 | -446 | -432 | 1,73 | 0,01 | 11,04 | >0.1 | 3,21 | 5,1 | 0 | 0,99 | 0 | 0 | 0 |
| BCL6 | TTTTTTTGAGACTG | -1 | 450 | 464 | -550 | -536 | -0,33 | 0,01 | 13,7 | >0.1 | 2,78 | 4,87 | 0 | 1 | 0 | 5,38 | 0 |
| BCL6 | TTTTTTAAAATTAT | -1 | 132 | 146 | -868 | -854 | 0,44 | 0,01 | 11,71 | >0.1 | 3,38 | 4,39 | 0 | 1,89 | 0 | 1,61 | 0 |
| BCL6 | TCTCCTGATCTGCC | -1 | 92 | 106 | -908 | -894 | -0,54 | 0,01 | 10,14 | >0.1 | 3,66 | 4,35 | 0 | 1,88 | 0 | 0,79 | 0 |
| BCL6 | CTTCTTTGAGTGCC | -1 | 23 | 37 | -977 | -963 | 7,45 | 0,001 | 19,15 | >0.1 | 5,56 | 4,28 | 0 | 1,87 | 0 | 0 | 0 |
| CDX1 | GCACTCAAA | 1 | 24 | 33 | -976 | -967 | 2,37 | 0,01 | 14,13 | >0.1 | 5,62 | 4,27 | 0 | 1,87 | 0 | 0 | 0 |
| CDX1 | GTCATAATT | 1 | 129 | 138 | -871 | -862 | 6,37 | 0,01 | 17,63 | >0.1 | 3,41 | 4,38 | 0 | 1,89 | 0 | 1,57 | 0 |
| CDX1 | GAAATTAAA | 1 | 154 | 163 | -846 | -837 | 5,11 | 0,01 | 16,33 | >0.1 | 3,26 | 4,41 | 0 | 1,9 | 0 | 1,65 | 0 |
| CDX1 | TCTATTAAA | 1 | 288 | 297 | -712 | -703 | 3,51 | 0,01 | 17,24 | >0.1 | 2,72 | 4,59 | 0 | 1,24 | 0 | 5,19 | 0 |
| CDX1 | CTATTAAAA | 1 | 289 | 298 | -711 | -702 | 3 | 0,01 | 16,62 | >0.1 | 2,71 | 4,59 | 0 | 1,23 | 0 | 5,09 | 0 |
| CDX1 | TCCATAAAG | 1 | 497 | 506 | -503 | -494 | 7,54 | 0,001 | 22,86 | >0.1 | 2,94 | 4,96 | 0 | 0,98 | 0 | 6,44 | 0 |
| CDX1 | GAAATAAAA | 1 | 576 | 585 | -424 | -415 | 7,79 | 0,001 | 17,25 | >0.1 | 3,32 | 5,14 | 0 | 1 | 0 | 0 | 0 |
| CDX1 | AAAATAAAT | 1 | 582 | 591 | -418 | -409 | 4,63 | 0,01 | 14,14 | >0.1 | 3,35 | 5,16 | 0 | 1 | 0 | 0 | 0 |

| | | | | | | | | | | | | | | | | | |
|---|---|---|---|---|---|---|---|---|---|---|---|---|---|---|---|---|---|
| CDX1 | CCTATTAAT | 1 | 600 | 609 | -400 | -391 | 3,93 | 0,01 | 14,98 | >0.1 | 3,47 | 5,2 | 0 | 1,01 | 0 | 1,36 | 0 |
| CDX1 | ACCATACAA | 1 | 623 | 632 | -377 | -368 | 4,43 | 0,01 | 15,82 | >0.1 | 3,64 | 5,27 | 0 | 1,04 | 0 | 1,44 | 0 |
| CDX1 | ACTGTAAAA | 1 | 678 | 687 | -322 | -313 | 5,04 | 0,01 | 19,28 | >0.1 | 4,26 | 5,44 | 0 | 1,11 | 0 | 3,43 | 0 |
| CDX1 | ATTATAACA | 1 | 686 | 695 | -314 | -305 | 3,6 | 0,01 | 17,55 | >0.1 | 4,41 | 5,47 | 0 | 1,12 | 0 | 2,95 | 0 |
| CDX1 | GCAGTTACA | -1 | 1087 | 1096 | 87 | 96 | 2,9 | 0,01 | 21,3 | >0.1 | 5,62 | 6,92 | 0 | 1,85 | 1,79 | 2,21 | 0 |
| CDX1 | ACCAAAAAA | -1 | 924 | 933 | -76 | -67 | 2,54 | 0,01 | 18,1 | >0.1 | 5,62 | 6,96 | 0 | 1,88 | 0 | 1,09 | 0 |
| CDX1 | CCAAAAAAA | -1 | 923 | 932 | -77 | -68 | 2,42 | 0,01 | 17,97 | >0.1 | 5,62 | 6,95 | 0 | 1,88 | 0 | 1,09 | 0 |
| CDX1 | ACAGTAAAC | -1 | 898 | 907 | -102 | -93 | 6,77 | 0,01 | 23,82 | >0.1 | 5,62 | 6,67 | 0 | 1,89 | 0 | 2,86 | 0 |
| CDX1 | CCAATATAA | -1 | 878 | 887 | -122 | -113 | 7,31 | 0,01 | 24,28 | >0.1 | 5,62 | 6,5 | 0 | 1,9 | 0 | 2,95 | 0 |
| CDX1 | GCTGTTAAA | -1 | 708 | 717 | -292 | -283 | 3,99 | 0,01 | 21,14 | >0.1 | 5,62 | 5,55 | 0 | 1,17 | 0 | 4,81 | 0 |
| CDX1 | GTTATAATT | -1 | 685 | 694 | -315 | -306 | 3,62 | 0,01 | 17,55 | >0.1 | 4,39 | 5,47 | 0 | 1,12 | 0 | 2,95 | 0 |
| CDX1 | ATCATGAAG | -1 | 645 | 654 | -355 | -346 | 2,83 | 0,01 | 16,76 | >0.1 | 3,84 | 5,34 | 0 | 1,06 | 0 | 3,7 | 0 |
| CDX1 | TTCATTAAT | -1 | 603 | 612 | -397 | -388 | 3,92 | 0,01 | 15,05 | >0.1 | 3,49 | 5,21 | 0 | 1,02 | 0 | 1,41 | 0 |
| ISL2 | GCACTCAA | 1 | 24 | 32 | -976 | -968 | 9,32 | 0,001 | 21,1 | >0.1 | 5,64 | 4,27 | 0 | 1,87 | 0 | 0 | 0 |
| ISL2 | GAAATTAA | 1 | 154 | 162 | -846 | -838 | 4,84 | 0,01 | 16,06 | >0.1 | 3,27 | 4,41 | 0 | 1,9 | 0 | 1,65 | 0 |
| ISL2 | GCACTTTG | 1 | 209 | 217 | -791 | -783 | 9,45 | 0,001 | 23,2 | >0.1 | 3 | 4,47 | 0 | 1,57 | 0 | 4,7 | 0 |
| ISL2 | AAACTTAG | 1 | 304 | 312 | -696 | -688 | 3,55 | 0,01 | 17,01 | >0.1 | 2,67 | 4,61 | 0 | 1,19 | 0 | 4,99 | 0 |
| ISL2 | CCAAGTAG | 1 | 545 | 553 | -455 | -447 | 4,51 | 0,01 | 13,71 | >0.1 | 3,15 | 5,06 | 0 | 0,98 | 0 | 0 | 0 |
| ISL2 | CCTATTAA | 1 | 600 | 608 | -400 | -392 | 4,86 | 0,01 | 15,91 | >0.1 | 3,47 | 5,2 | 0 | 1,01 | 0 | 1,36 | 0 |
| ISL2 | CCTCTTTG | 1 | 667 | 675 | -333 | -325 | 3,84 | 0,01 | 18,25 | >0.1 | 4,09 | 5,4 | 0 | 1,09 | 0 | 3,83 | 0 |
| ISL2 | TCACTTTA | 1 | 821 | 829 | -179 | -171 | 6,89 | 0,01 | 23,92 | >0.1 | 5,64 | 6,08 | 0 | 1,61 | 0 | 3,7 | 0 |
| ISL2 | GAACTTTA | 1 | 873 | 881 | -127 | -119 | 4,11 | 0,01 | 20,96 | >0.1 | 5,64 | 6,45 | 0 | 1,9 | 0 | 2,86 | 0 |
| ISL2 | GAACTTTG | -1 | 1172 | 1180 | 172 | 180 | 3,82 | 0,01 | 22,06 | >0.1 | 5,64 | 6,14 | 0 | 1,65 | 1,79 | 3,02 | 0 |
| ISL2 | GCAGTTAC | -1 | 1088 | 1096 | 88 | 96 | 5,06 | 0,01 | 23,46 | >0.1 | 5,64 | 6,9 | 0 | 1,85 | 1,79 | 2,21 | 0 |
| ISL2 | ACAATCTG | -1 | 892 | 900 | -108 | -100 | 3,32 | 0,01 | 20,32 | >0.1 | 5,64 | 6,61 | 0 | 1,89 | 0 | 2,86 | 0 |
| ISL2 | TCACTTTA | -1 | 762 | 770 | -238 | -230 | 6,89 | 0,01 | 25,72 | >0.1 | 5,64 | 5,77 | 0 | 1,32 | 0 | 6,1 | 0 |
| ISL2 | GTACTTTT | -1 | 737 | 745 | -263 | -255 | 4,88 | 0,01 | 23,33 | >0.1 | 5,64 | 5,66 | 0 | 1,24 | 0 | 5,92 | 0 |
| ISL2 | ACAATTTC | -1 | 609 | 617 | -391 | -383 | 4,39 | 0,01 | 15,71 | >0.1 | 3,53 | 5,23 | 0 | 1,02 | 0 | 1,55 | 0 |
| ISL2 | ACATTTAT | -1 | 585 | 593 | -415 | -407 | 3,25 | 0,01 | 12,78 | >0.1 | 3,37 | 5,16 | 0 | 1 | 0 | 0 | 0 |
| ISL2 | CCACTCCA | -1 | 115 | 123 | -885 | -877 | 3,94 | 0,01 | 14,44 | >0.1 | 3,51 | 4,36 | 0 | 1,89 | 0 | 0,75 | 0 |
| ISL2 | AAACTTAA | -1 | 78 | 86 | -922 | -914 | 3,84 | 0,01 | 13,86 | >0.1 | 3,81 | 4,32 | 0 | 1,88 | 0 | 0 | 0 |
| ISL2 | TCACTTAC | -1 | 5 | 13 | -995 | -987 | 7,25 | 0,01 | 20,8 | >0.1 | 5,64 | 4,26 | 0 | 1,86 | 1,79 | 0 | 0 |
| CDX2 | AGGCACTCAAA | 1 | 22 | 33 | -978 | -967 | -9,9 | 0,01 | 8,51 | >0.1 | 5,6 | 4,27 | 0 | 1,87 | 0 | 0 | 6,67 |
| CDX2 | CAGCCACAGAA | 1 | 38 | 49 | -962 | -951 | -9,68 | 0,01 | 7,46 | >0.1 | 4,31 | 4,29 | 0 | 1,87 | 0 | 0 | 6,67 |
| CDX2 | TAGTCATAATT | 1 | 127 | 138 | -873 | -862 | -3,48 | 0,01 | 14,49 | >0.1 | 3,42 | 4,38 | 0 | 1,89 | 0 | 1,61 | 6,67 |
| CDX2 | TAATTTTAAAA | 1 | 133 | 144 | -867 | -856 | -6,93 | 0,01 | 10,98 | >0.1 | 3,38 | 4,39 | 0 | 1,89 | 0 | 1,57 | 6,67 |
| CDX2 | TGATCAGAAAT | 1 | 165 | 176 | -835 | -824 | -5,3 | 0,01 | 13,3 | >0.1 | 3,2 | 4,42 | 0 | 1,9 | 0 | 2,4 | 6,67 |
| CDX2 | CTCTATTAAAA | 1 | 287 | 298 | -713 | -702 | -3,63 | 0,01 | 16,78 | >0.1 | 2,72 | 4,59 | 0 | 1,24 | 0 | 5,19 | 6,67 |
| CDX2 | CTATTAAAAAT | 1 | 289 | 300 | -711 | -700 | -9,36 | 0,01 | 10,94 | >0.1 | 2,71 | 4,59 | 0 | 1,24 | 0 | 5,09 | 6,67 |
| CDX2 | GTCTCAAAAAA | 1 | 452 | 463 | -548 | -537 | -2,29 | 0,01 | 18,41 | >0.1 | 2,79 | 4,87 | 0 | 1 | 0 | 5,38 | 6,67 |
| CDX2 | TGATCAGAAAT | 1 | 480 | 491 | -520 | -509 | -5,3 | 0,01 | 15,82 | >0.1 | 2,88 | 4,92 | 0 | 0,99 | 0 | 5,65 | 6,67 |
| CDX2 | TTGAAATAAAA | 1 | 574 | 585 | -426 | -415 | -0,29 | 0,01 | 15,84 | >0.1 | 3,31 | 5,14 | 0 | 1 | 0 | 0 | 6,67 |
| CDX2 | TAAAAATAAAT | 1 | 580 | 591 | -420 | -409 | -5,1 | 0,01 | 11,08 | >0.1 | 3,35 | 5,16 | 0 | 1 | 0 | 0 | 6,67 |
| CDX2 | AACCTATTAAT | 1 | 598 | 609 | -402 | -391 | -7,51 | 0,01 | 10,26 | >0.1 | 3,46 | 5,21 | 0 | 1,02 | 0 | 1,41 | 6,67 |
| CDX2 | AAACCATACAA | 1 | 621 | 632 | -379 | -368 | 0,83 | 0,01 | 18,88 | >0.1 | 3,63 | 5,27 | 0 | 1,04 | 0 | 1,44 | 6,67 |
| CDX2 | AAATTATAACA | 1 | 684 | 695 | -316 | -305 | -4,7 | 0,01 | 15,91 | >0.1 | 4,39 | 5,47 | 0 | 1,13 | 0 | 2,95 | 6,67 |
| CDX2 | TTATTCTAAAA | 1 | 834 | 845 | -166 | -155 | -7,09 | 0,01 | 16,38 | >0.1 | 5,6 | 6,19 | 0 | 1,73 | 0 | 3,28 | 6,67 |
| CDX2 | GAGCCAGGAAG | -1 | 1059 | 1070 | 59 | 70 | -10,29 | 0,01 | 13,37 | >0.1 | 5,6 | 7,37 | 0 | 1,86 | 0 | 2,16 | 6,67 |
| CDX2 | GACCAAAAAAA | -1 | 923 | 934 | -77 | -66 | -0,95 | 0,01 | 21,27 | >0.1 | 5,6 | 6,98 | 0 | 1,88 | 0 | 1,09 | 6,67 |
| CDX2 | ACACAGTAAAC | -1 | 898 | 909 | -102 | -91 | -8,02 | 0,01 | 15,7 | >0.1 | 5,6 | 6,7 | 0 | 1,89 | 0 | 2,86 | 6,67 |
| CDX2 | TTCCTAGAAAG | -1 | 716 | 727 | -284 | -273 | -7,54 | 0,01 | 16,68 | >0.1 | 5,6 | 5,59 | 0 | 1,19 | 0 | 5,17 | 6,67 |
| CDX2 | ATGTTATAATT | -1 | 685 | 696 | -315 | -304 | -6,52 | 0,01 | 14,11 | >0.1 | 4,41 | 5,47 | 0 | 1,13 | 0 | 2,95 | 6,67 |
| CDX2 | GGATCATGAAG | -1 | 645 | 656 | -355 | -344 | -1,56 | 0,01 | 19,2 | >0.1 | 3,85 | 5,34 | 0 | 1,06 | 0 | 3,83 | 6,67 |
| CDX2 | TTCCCAGAAAC | -1 | 556 | 567 | -444 | -433 | -4,49 | 0,01 | 11,49 | >0.1 | 3,22 | 5,1 | 0 | 0,99 | 0 | 0 | 6,67 |
| CDX2 | AACTTAAAAAA | -1 | 74 | 85 | -926 | -915 | -5,88 | 0,01 | 10,83 | >0.1 | 3,84 | 4,32 | 0 | 1,88 | 0 | 0 | 6,67 |
| GSC | CATAATTTTA | 1 | 131 | 141 | -869 | -859 | 2,65 | 0,01 | 13,9 | >0.1 | 3,4 | 4,38 | 0 | 1,89 | 0 | 1,57 | 0 |
| GSC | ACTGATCAGA | 1 | 163 | 173 | -837 | -827 | 1,79 | 0,01 | 13,73 | >0.1 | 3,21 | 4,42 | 0 | 1,9 | 0 | 2,4 | 0 |
| GSC | TGTAATCCTA | 1 | 199 | 209 | -801 | -791 | 7,16 | 0,001 | 20,78 | >0.1 | 3,04 | 4,46 | 0 | 1,64 | 0 | 4,48 | 0 |
| GSC | ACTGATCAGA | 1 | 478 | 488 | -522 | -512 | 1,79 | 0,01 | 15,93 | >0.1 | 2,88 | 4,92 | 0 | 0,99 | 0 | 5,36 | 0 |

| | | | | | | | | | | | | | | | | | |
|---|---|---|---|---|---|---|---|---|---|---|---|---|---|---|---|---|---|
| GSC | CATTATCTTC | 1 | 639 | 649 | -361 | -351 | 1,5 | 0,01 | 13,32 | >0.1 | 3,79 | 5,32 | 0 | 1,06 | 0 | 1,66 | 0 |
| GSC | CATGATCCCT | 1 | 648 | 658 | -352 | -342 | 7,03 | 0,001 | 21,15 | >0.1 | 3,87 | 5,35 | 0 | 1,07 | 0 | 3,83 | 0 |
| GSC | TTTGATTCAC | 1 | 815 | 825 | -185 | -175 | 1,29 | 0,01 | 17,97 | >0.1 | 5,61 | 6,06 | 0 | 1,58 | 0 | 3,43 | 0 |
| GSC | TATAATTCCA | -1 | 1190 | 1200 | 190 | 200 | 5,6 | 0,01 | 24,36 | >0.1 | 5,61 | 6,03 | 0 | 1,51 | 1,79 | 3,82 | 0 |
| GSC | TCTTATCAGC | -1 | 966 | 976 | -34 | -24 | 1,31 | 0,01 | 18,27 | >0.1 | 5,61 | 7,78 | 0 | 1,87 | 0 | 1,7 | 0 |
| GSC | AACAATCTGC | -1 | 891 | 901 | -109 | -99 | 1,3 | 0,01 | 18,29 | >0.1 | 5,61 | 6,62 | 0 | 1,89 | 0 | 2,86 | 0 |
| GSC | GCTGAGCCAA | -1 | 883 | 893 | -117 | -107 | 1,31 | 0,01 | 18,44 | >0.1 | 5,61 | 6,56 | 0 | 1,89 | 0 | 3,07 | 0 |
| GSC | GTTCATCCTG | -1 | 867 | 877 | -133 | -123 | 3,73 | 0,01 | 20,41 | >0.1 | 5,61 | 6,42 | 0 | 1,9 | 0 | 2,75 | 0 |
| GSC | AATAATTTTT | -1 | 829 | 839 | -171 | -161 | 2,52 | 0,01 | 19,24 | >0.1 | 5,61 | 6,15 | 0 | 1,68 | 0 | 3,28 | 0 |
| GSC | TTTAATCCTA | -1 | 789 | 799 | -211 | -201 | 8,27 | 0,001 | 26,75 | >0.1 | 5,61 | 5,91 | 0 | 1,44 | 0 | 5,52 | 0 |
| GSC | GTTAATATTT | -1 | 726 | 736 | -274 | -264 | 3,63 | 0,01 | 21,4 | >0.1 | 5,61 | 5,62 | 0 | 1,22 | 0 | 5,32 | 0 |
| GSC | TATAATTTTA | -1 | 682 | 692 | -318 | -308 | 1,04 | 0,01 | 15,02 | >0.1 | 4,34 | 5,46 | 0 | 1,12 | 0 | 3,07 | 0 |
| GSC | GATAATGTTG | -1 | 636 | 646 | -364 | -354 | 2,11 | 0,01 | 13,89 | >0.1 | 3,76 | 5,31 | 0 | 1,05 | 0 | 1,66 | 0 |
| GSC | ATTAATAGGT | -1 | 599 | 609 | -401 | -391 | 3,64 | 0,01 | 14,69 | >0.1 | 3,47 | 5,21 | 0 | 1,02 | 0 | 1,36 | 0 |
| GSC | GGTTATCTCA | -1 | 592 | 602 | -408 | -398 | 1,41 | 0,01 | 11,03 | >0.1 | 3,42 | 5,19 | 0 | 1,01 | 0 | 0 | 0 |
| GSC | TCTGATCAGT | -1 | 478 | 488 | -522 | -512 | 2,27 | 0,01 | 16,41 | >0.1 | 2,88 | 4,92 | 0 | 0,99 | 0 | 5,36 | 0 |
| GSC | CCTCAGCCTC | -1 | 352 | 362 | -648 | -638 | 1,83 | 0,01 | 16,55 | >0.1 | 2,54 | 4,69 | 0 | 1,1 | 0 | 6,4 | 0 |
| GSC | GCTAAGTTTT | -1 | 303 | 313 | -697 | -687 | 0,91 | 0,01 | 14,38 | >0.1 | 2,67 | 4,61 | 0 | 1,2 | 0 | 4,99 | 0 |
| GSC | TGTGATCTGC | -1 | 231 | 241 | -769 | -759 | 2,21 | 0,01 | 16,09 | >0.1 | 2,91 | 4,51 | 0 | 1,45 | 0 | 5,01 | 0 |
| GSC | TCTGATCAGT | -1 | 163 | 173 | -837 | -827 | 2,27 | 0,01 | 14,21 | >0.1 | 3,21 | 4,42 | 0 | 1,9 | 0 | 2,4 | 0 |
| GSC | TTTAATTTCC | -1 | 153 | 163 | -847 | -837 | 4,96 | 0,01 | 16,18 | >0.1 | 3,27 | 4,41 | 0 | 1,9 | 0 | 1,65 | 0 |
| GSC | CCTGATCTGC | -1 | 93 | 103 | -907 | -897 | 6,2 | 0,01 | 16,9 | >0.1 | 3,67 | 4,34 | 0 | 1,88 | 0 | 0,81 | 0 |
| GSC | GCTCATTTCC | -1 | 64 | 74 | -936 | -926 | 2,01 | 0,01 | 12,15 | >0.1 | 3,95 | 4,31 | 0 | 1,88 | 0 | 0 | 0 |
| MAFK | TTTTGCAAAGGCAGATCAG | 1 | 83 | 102 | -917 | -898 | 1,72 | 0,01 | 11,66 | >0.1 | 3,72 | 4,34 | 0 | 1,88 | 0 | 0 | 0 |
| MAFK | GAGTGGAGAGTAGTCATAA | 1 | 117 | 136 | -883 | -864 | 1,58 | 0,01 | 13 | >0.1 | 3,46 | 4,38 | 0 | 1,89 | 0 | 1,7 | 0 |
| MAFK | ACCTGTAGTCCCAGCTACT | 1 | 330 | 349 | -670 | -651 | 1,22 | 0,01 | 15,53 | >0.1 | 2,58 | 4,67 | 0 | 1,14 | 0 | 5,92 | 0 |
| MAFK | GGAGGCTGAGGCAGGAGAA | 1 | 351 | 370 | -649 | -630 | 8,59 | 0,001 | 22,96 | >0.1 | 2,53 | 4,7 | 0 | 1,11 | 0 | 6,03 | 0 |
| MAFK | AAGGGCAAAGTCATGTATT | 1 | 508 | 527 | -492 | -473 | 3,2 | 0,01 | 15,15 | >0.1 | 3,01 | 5 | 0 | 0,99 | 0 | 2,95 | 0 |
| MAFK | AACAGCTTTCTAGGAAAAT | 1 | 711 | 730 | -289 | -270 | 1,13 | 0,01 | 18,6 | >0.1 | 5,5 | 5,6 | 0 | 1,2 | 0 | 5,17 | 0 |
| MAFK | TTATATTGGCTCAGCAGAT | 1 | 878 | 897 | -122 | -103 | 8,23 | 0,001 | 25,28 | >0.1 | 5,5 | 6,59 | 0 | 1,9 | 0 | 3,07 | 0 |
| MAFK | GGCTGATAAGAGAGAAAAT | 1 | 965 | 984 | -35 | -16 | -0,54 | 0,01 | 16,58 | >0.1 | 5,5 | 8,05 | 0 | 1,87 | 0 | 1,7 | 0 |
| MAFK | TGTTGCTGTAACTGCTGCT | 1 | 1081 | 1100 | 81 | 100 | 3,73 | 0,01 | 22,08 | >0.1 | 5,5 | 7 | 0 | 1,85 | 1,79 | 2,21 | 0 |
| MAFK | TGCTGCTCAGTCCACCATT | 1 | 1093 | 1112 | 93 | 112 | 2,43 | 0,01 | 22,2 | >0.1 | 5,5 | 6,84 | 0 | 1,85 | 3,22 | 2,36 | 0 |
| MAFK | AATGGTGGACTGAGCAGCA | -1 | 1093 | 1112 | 93 | 112 | 2,06 | 0,01 | 21,83 | >0.1 | 5,5 | 6,84 | 0 | 1,85 | 3,22 | 2,36 | 0 |
| MAFK | AGCAGCAGTTACAGCAACA | -1 | 1081 | 1100 | 81 | 100 | 4,14 | 0,01 | 22,49 | >0.1 | 5,5 | 7 | 0 | 1,85 | 1,79 | 2,21 | 0 |
| MAFK | ATTTTCTCTCTTATCAGCC | -1 | 965 | 984 | -35 | -16 | -0,4 | 0,01 | 16,72 | >0.1 | 5,5 | 8,05 | 0 | 1,87 | 0 | 1,7 | 0 |
| MAFK | GGGCGCTGAGCAGGAAAGA | -1 | 932 | 951 | -68 | -49 | -0,84 | 0,01 | 14,67 | >0.1 | 5,5 | 7,24 | 0 | 1,88 | 0 | 0,9 | 0 |
| MAFK | ATCTGCTGAGCCAATATAA | -1 | 878 | 897 | -122 | -103 | 8,89 | 0,001 | 25,94 | >0.1 | 5,5 | 6,59 | 0 | 1,9 | 0 | 3,07 | 0 |
| MAFK | ATTTTCCTAGAAAGCTGTT | -1 | 711 | 730 | -289 | -270 | 1,34 | 0,01 | 18,81 | >0.1 | 5,5 | 5,6 | 0 | 1,2 | 0 | 5,17 | 0 |
| MAFK | CTCTACTTGGGCAACATTT | -1 | 536 | 555 | -464 | -445 | -1,29 | 0,01 | 7,91 | >0.1 | 3,14 | 5,07 | 0 | 0,99 | 0 | 0 | 0 |
| MAFK | AATACATGACTTTGCCCTT | -1 | 508 | 527 | -492 | -473 | 0,84 | 0,01 | 12,79 | >0.1 | 3,01 | 5 | 0 | 0,99 | 0 | 2,95 | 0 |
| MAFK | TTCTCCTGCCTCAGCCTCC | -1 | 351 | 370 | -649 | -630 | 7,67 | 0,001 | 22,04 | >0.1 | 2,53 | 4,7 | 0 | 1,11 | 0 | 6,03 | 0 |
| MAFK | AGTAGCTGGGACTACAGGT | -1 | 330 | 349 | -670 | -651 | 0,52 | 0,01 | 14,83 | >0.1 | 2,58 | 4,67 | 0 | 1,14 | 0 | 5,92 | 0 |
| MAFK | TTCTGATCAGTTTTAATTT | -1 | 155 | 174 | -845 | -826 | -1,49 | 0,01 | 10,47 | >0.1 | 3,23 | 4,42 | 0 | 1,9 | 0 | 2,4 | 0 |
| MAFK | TTATGACTACTCTCCACTC | -1 | 117 | 136 | -883 | -864 | 0,06 | 0,01 | 11,48 | >0.1 | 3,46 | 4,38 | 0 | 1,89 | 0 | 1,7 | 0 |
| MAFK | CTGATCTGCCTTTGCAAAA | -1 | 83 | 102 | -917 | -898 | -0,81 | 0,01 | 9,13 | >0.1 | 3,72 | 4,34 | 0 | 1,88 | 0 | 0 | 0 |
| NKX2-3 | GGCACTCAAA | 1 | 23 | 33 | -977 | -967 | 9,09 | 0,001 | 20,84 | >0.1 | 5,61 | 4,27 | 0 | 1,87 | 0 | 0 | 0 |
| NKX2-3 | AGCACTTTGG | 1 | 208 | 218 | -792 | -782 | 7,09 | 0,001 | 20,85 | >0.1 | 3 | 4,48 | 0 | 1,58 | 0 | 4,7 | 0 |
| NKX2-3 | AAAACTTAGC | 1 | 303 | 313 | -697 | -687 | 0,63 | 0,01 | 14,1 | >0.1 | 2,67 | 4,61 | 0 | 1,2 | 0 | 4,99 | 0 |
| NKX2-3 | GCTACTCGGG | 1 | 343 | 353 | -657 | -647 | 2,72 | 0,01 | 17,59 | >0.1 | 2,56 | 4,67 | 0 | 1,11 | 0 | 6,52 | 0 |
| NKX2-3 | TGATCTTGAA | 1 | 569 | 579 | -431 | -421 | -1,86 | 0,01 | 7,55 | >0.1 | 3,28 | 5,13 | 0 | 1 | 0 | 0 | 0 |
| NKX2-3 | ACCTCTTTGG | 1 | 666 | 676 | -334 | -324 | 2,92 | 0,01 | 17,34 | >0.1 | 4,09 | 5,41 | 0 | 1,09 | 0 | 3,83 | 0 |
| NKX2-3 | TTCACTTTAA | 1 | 820 | 830 | -180 | -170 | 1,9 | 0,01 | 18,91 | >0.1 | 5,61 | 6,09 | 0 | 1,62 | 0 | 3,7 | 0 |
| NKX2-3 | TCCTCTCCAG | 1 | 860 | 870 | -140 | -130 | 3,31 | 0,01 | 20,05 | >0.1 | 5,61 | 6,37 | 0 | 1,9 | 0 | 2,86 | 0 |
| NKX2-3 | AGCTCTTCCT | 1 | 1055 | 1065 | 55 | 65 | 0,66 | 0,01 | 17,86 | >0.1 | 5,61 | 7,46 | 0 | 1,86 | 0 | 2,27 | 0 |
| NKX2-3 | GCTCCTTCTC | 1 | 1066 | 1076 | 66 | 76 | -2,44 | 0,01 | 16,33 | >0.1 | 5,61 | 7,24 | 0 | 1,85 | 1,79 | 2,27 | 0 |
| NKX2-3 | TTCACTTGCT | 1 | 1177 | 1187 | 177 | 187 | 0,94 | 0,01 | 19,32 | >0.1 | 5,61 | 6,11 | 0 | 1,61 | 1,79 | 3,26 | 0 |
| NKX2-3 | GCTTCTTGGA | 1 | 1184 | 1194 | 184 | 194 | 0 | 0,01 | 18,7 | >0.1 | 5,61 | 6,06 | 0 | 1,55 | 1,79 | 3,67 | 0 |
| NKX2-3 | TAAACTTGTC | -1 | 1136 | 1146 | 136 | 146 | 0,07 | 0,01 | 20,24 | >0.1 | 5,61 | 6,41 | 0 | 1,84 | 3,22 | 3,09 | 0 |

| | | | | | | | | | | | | | | | | | |
|---|---|---|---|---|---|---|---|---|---|---|---|---|---|---|---|---|---|
| NKX2-3 | CTCTCTTATC | -1 | 969 | 979 | -31 | -21 | -0,32 | 0,01 | 16,74 | >0.1 | 5,61 | 7,88 | 0 | 1,87 | 0 | 1,7 | 0 |
| NKX2-3 | TGAACTTGGG | -1 | 953 | 963 | -47 | -37 | 1,82 | 0,01 | 18,46 | >0.1 | 5,61 | 7,46 | 0 | 1,88 | 0 | 1,7 | 0 |
| NKX2-3 | ACCTCTCCTG | -1 | 770 | 780 | -230 | -220 | 2,6 | 0,01 | 21,3 | >0.1 | 5,61 | 5,82 | 0 | 1,36 | 0 | 5,92 | 0 |
| NKX2-3 | GTCACTTTAT | -1 | 761 | 771 | -239 | -229 | 0,29 | 0,01 | 19,1 | >0.1 | 5,61 | 5,77 | 0 | 1,32 | 0 | 6,1 | 0 |
| NKX2-3 | GGTACTTTTG | -1 | 736 | 746 | -264 | -254 | 0,02 | 0,01 | 18,45 | >0.1 | 5,61 | 5,66 | 0 | 1,24 | 0 | 5,92 | 0 |
| NKX2-3 | AACTCTCTAC | -1 | 549 | 559 | -451 | -441 | -0,76 | 0,01 | 8,48 | >0.1 | 3,18 | 5,08 | 0 | 0,99 | 0 | 0 | 0 |
| NKX2-3 | TCTACTTGGG | -1 | 544 | 554 | -456 | -446 | 4,65 | 0,01 | 13,85 | >0.1 | 3,15 | 5,06 | 0 | 0,99 | 0 | 0 | 0 |
| NKX2-3 | TGCCCTTGTC | -1 | 505 | 515 | -495 | -485 | 4,07 | 0,01 | 18,69 | >0.1 | 2,98 | 4,98 | 0 | 0,98 | 0 | 5,68 | 0 |
| NKX2-3 | ACCACGCCCA | -1 | 313 | 323 | -687 | -677 | -1,91 | 0,01 | 12,66 | >0.1 | 2,64 | 4,62 | 0 | 1,17 | 0 | 6,13 | 0 |
| NKX2-3 | CCAACTCGGC | -1 | 221 | 231 | -779 | -769 | 1,04 | 0,01 | 15,51 | >0.1 | 2,95 | 4,49 | 0 | 1,5 | 0 | 5,52 | 0 |
| NKX2-3 | ACTACTCTCC | -1 | 121 | 131 | -879 | -869 | -2,32 | 0,01 | 9,1 | >0.1 | 3,46 | 4,37 | 0 | 1,89 | 0 | 1,7 | 0 |
| NKX2-3 | TACTCTCCAC | -1 | 119 | 129 | -881 | -871 | 0,6 | 0,01 | 11,98 | >0.1 | 3,47 | 4,37 | 0 | 1,89 | 0 | 1,65 | 0 |
| NKX2-3 | TCCACTCCAC | -1 | 114 | 124 | -886 | -876 | 8,51 | 0,001 | 19,02 | >0.1 | 3,51 | 4,36 | 0 | 1,89 | 0 | 0,75 | 0 |
| NKX2-3 | AACTCTCCTG | -1 | 99 | 109 | -901 | -891 | -0,09 | 0,01 | 10,53 | >0.1 | 3,62 | 4,35 | 0 | 1,88 | 0 | 0,77 | 0 |
| NKX2-3 | AAAACTTAAA | -1 | 77 | 87 | -923 | -913 | 2,18 | 0,01 | 12,2 | >0.1 | 3,81 | 4,33 | 0 | 1,88 | 0 | 0 | 0 |
| NKX2-3 | CCTCCTTCTC | -1 | 15 | 25 | -985 | -975 | -2,3 | 0,01 | 9,44 | >0.1 | 5,61 | 4,27 | 0 | 1,87 | 0 | 0 | 0 |
| NKX2-3 | TCTCCTTCAC | -1 | 9 | 19 | -991 | -981 | -1,38 | 0,01 | 12,15 | >0.1 | 5,61 | 4,26 | 0 | 1,86 | 1,79 | 0 | 0 |
| NKX2-3 | TTCACTTACC | -1 | 4 | 14 | -996 | -986 | 2,98 | 0,01 | 16,5 | >0.1 | 5,61 | 4,26 | 0 | 1,86 | 1,79 | 0 | 0 |
| MAFB | AAACTGATCAGA | 1 | 161 | 173 | -839 | -827 | 2,46 | 0,01 | 14,46 | >0.1 | 3,22 | 4,42 | 0 | 1,9 | 0 | 2,46 | 0 |
| MAFB | ACCGTCCTGACT | 1 | 257 | 269 | -743 | -731 | 3,98 | 0,01 | 17,21 | >0.1 | 2,81 | 4,55 | 0 | 1,34 | 0 | 4,53 | 0 |
| MAFB | GGGAGGCTGAGG | 1 | 350 | 362 | -650 | -638 | 2,64 | 0,01 | 17,37 | >0.1 | 2,54 | 4,69 | 0 | 1,1 | 0 | 6,4 | 0 |
| MAFB | AGTGAGCCGAGA | 1 | 398 | 410 | -602 | -590 | 1,74 | 0,01 | 16,08 | >0.1 | 2,62 | 4,77 | 0 | 1,04 | 0 | 5,91 | 0 |
| MAFB | AAACTGATCAGA | 1 | 476 | 488 | -524 | -512 | 2,46 | 0,01 | 16,44 | >0.1 | 2,87 | 4,92 | 0 | 0,99 | 0 | 5,2 | 0 |
| MAFB | AGATTGTTTACT | 1 | 893 | 905 | -107 | -95 | 0,56 | 0,01 | 17,55 | >0.1 | 5,58 | 6,66 | 0 | 1,89 | 0 | 2,86 | 0 |
| MAFB | CAAAGGCTGATA | 1 | 961 | 973 | -39 | -27 | 4,31 | 0,01 | 21,16 | >0.1 | 5,58 | 7,7 | 0 | 1,88 | 0 | 1,7 | 0 |
| MAFB | AATCTCATGAGG | 1 | 981 | 993 | -19 | -7 | 2,59 | 0,01 | 20,07 | >0.1 | 5,58 | 8,5 | 0 | 1,87 | 0 | 1,53 | 0 |
| MAFB | TAACTGCTGCTC | 1 | 1089 | 1101 | 89 | 101 | 1,95 | 0,01 | 20,34 | >0.1 | 5,58 | 6,89 | 0 | 1,85 | 1,79 | 2,27 | 0 |
| MAFB | CTGCTGCTCAGT | 1 | 1092 | 1104 | 92 | 104 | 0,94 | 0,01 | 20,79 | >0.1 | 5,58 | 6,85 | 0 | 1,85 | 3,22 | 2,34 | 0 |
| MAFB | AAGCCGAAGACC | 1 | 1152 | 1164 | 152 | 164 | 0,88 | 0,01 | 17,62 | >0.1 | 4,48 | 6,28 | 0 | 1,84 | 1,79 | 2,34 | 0 |
| MAFB | ACAAGGCTGAGA | -1 | 1072 | 1084 | 72 | 84 | 5,1 | 0,01 | 23,74 | >0.1 | 5,58 | 7,14 | 0 | 1,85 | 1,79 | 2,27 | 0 |
| MAFB | TGGGCGCTGAGC | -1 | 940 | 952 | -60 | -48 | 4,38 | 0,01 | 19,88 | >0.1 | 5,58 | 7,25 | 0 | 1,88 | 0 | 0,79 | 0 |
| MAFB | AATCTGCTGAGC | -1 | 886 | 898 | -114 | -102 | 11,01 | 0,0001 | 28,15 | >0.1 | 5,58 | 6,6 | 0 | 1,89 | 0 | 3,07 | 0 |
| MAFB | AAATACATGACT | -1 | 516 | 528 | -484 | -472 | 2,28 | 0,01 | 14,25 | >0.1 | 3,03 | 5 | 0 | 0,98 | 0 | 2,95 | 0 |
| MAFB | ATATTTCTGATC | -1 | 481 | 493 | -519 | -507 | 0,22 | 0,01 | 14,83 | >0.1 | 2,89 | 4,93 | 0 | 0,99 | 0 | 5,8 | 0 |
| MAFB | ATTCTCCTGCCT | -1 | 359 | 371 | -641 | -629 | 0,11 | 0,01 | 14,78 | >0.1 | 2,52 | 4,7 | 0 | 1,09 | 0 | 6,36 | 0 |
| MAFB | GAGTAGCTGGGA | -1 | 338 | 350 | -662 | -650 | 0,29 | 0,01 | 15,17 | >0.1 | 2,57 | 4,67 | 0 | 1,12 | 0 | 6,52 | 0 |
| MAFB | TATCTCCTGACC | -1 | 242 | 254 | -758 | -746 | 4,43 | 0,01 | 17,55 | >0.1 | 2,87 | 4,52 | 0 | 1,4 | 0 | 4,33 | 0 |
| MAFB | AAAGTGCTAGGA | -1 | 204 | 216 | -796 | -784 | 0,74 | 0,01 | 14,54 | >0.1 | 3,02 | 4,47 | 0 | 1,6 | 0 | 4,7 | 0 |
| MAFB | AAAATTATGACT | -1 | 128 | 140 | -872 | -860 | 3,66 | 0,01 | 14,95 | >0.1 | 3,41 | 4,38 | 0 | 1,89 | 0 | 1,61 | 0 |
| MAFB | ATTATGACTACT | -1 | 125 | 137 | -875 | -863 | 1,68 | 0,01 | 13,03 | >0.1 | 3,43 | 4,38 | 0 | 1,89 | 0 | 1,65 | 0 |
| MAFB | AAAAAGCTCATT | -1 | 67 | 79 | -933 | -921 | 2,35 | 0,01 | 12,45 | >0.1 | 3,91 | 4,32 | 0 | 1,88 | 0 | 0 | 0 |
| MAFB | CTGTGGCTGACT | -1 | 35 | 47 | -965 | -953 | 4,95 | 0,01 | 15,46 | >0.1 | 4,36 | 4,29 | 0 | 1,87 | 0 | 0 | 0 |
| MAFF | GTTTTGCAAAGGCAGATCAG( | 1 | 82 | 103 | -918 | -897 | 2,76 | 0,01 | 12,7 | >0.1 | 3,72 | 4,34 | 0 | 1,88 | 0 | 0 | 0 |
| MAFF | GGAGTGGAGAGTAGTCATAAT | 1 | 116 | 137 | -884 | -863 | 2,67 | 0,01 | 14,09 | >0.1 | 3,46 | 4,38 | 0 | 1,89 | 0 | 1,7 | 0 |
| MAFF | CACCTGTAGTCCCAGCTACT | 1 | 329 | 350 | -671 | -650 | 0,07 | 0,01 | 14,38 | >0.1 | 2,58 | 4,67 | 0 | 1,15 | 0 | 5,92 | 0 |
| MAFF | GGGAGGCTGAGGCAGGAGA/ | 1 | 350 | 371 | -650 | -629 | 9,2 | 0,001 | 23,57 | >0.1 | 2,53 | 4,7 | 0 | 1,11 | 0 | 6,03 | 0 |
| MAFF | CAAGGGCAAAGTCATGTATTT | 1 | 507 | 528 | -493 | -472 | 2,6 | 0,01 | 14,56 | >0.1 | 3,01 | 5,01 | 0 | 0,99 | 0 | 2,95 | 0 |
| MAFF | TAACAGCTTTCTAGGAAAATA | 1 | 710 | 731 | -290 | -269 | 0,93 | 0,01 | 18,38 | >0.1 | 5,47 | 5,6 | 0 | 1,21 | 0 | 5,17 | 0 |
| MAFF | TTTATATTGGCTCAGCAGATT | 1 | 877 | 898 | -123 | -102 | 8,91 | 0,001 | 25,95 | >0.1 | 5,47 | 6,6 | 0 | 1,9 | 0 | 3,07 | 0 |
| MAFF | AGGCTGATAAGAGAGAAAATC | 1 | 964 | 985 | -36 | -15 | 0,57 | 0,01 | 17,71 | >0.1 | 5,47 | 8,09 | 0 | 1,87 | 0 | 1,7 | 0 |
| MAFF | TTGTTGCTGTAACTGCTGCT( | 1 | 1080 | 1101 | 80 | 101 | 3,62 | 0,01 | 21,97 | >0.1 | 5,47 | 7,01 | 0 | 1,85 | 1,79 | 2,21 | 0 |
| MAFF | CTGCTGCTCAGTCCACCATT | 1 | 1092 | 1113 | 92 | 113 | 1,42 | 0,01 | 21,18 | >0.1 | 5,47 | 6,85 | 0 | 1,85 | 3,22 | 2,36 | 0 |
| MAFF | CAATGGTGGACTGAGCAGCA | -1 | 1092 | 1113 | 92 | 113 | 0,61 | 0,01 | 20,37 | >0.1 | 5,47 | 6,85 | 0 | 1,85 | 3,22 | 2,36 | 0 |
| MAFF | GAGCAGCAGTTACAGCAACA, | -1 | 1080 | 1101 | 80 | 101 | 3,99 | 0,01 | 22,34 | >0.1 | 5,47 | 7,01 | 0 | 1,85 | 1,79 | 2,21 | 0 |
| MAFF | GATTTTCTCTCTTATCAGCCT | -1 | 964 | 985 | -36 | -15 | 0,53 | 0,01 | 17,67 | >0.1 | 5,47 | 8,09 | 0 | 1,87 | 0 | 1,7 | 0 |
| MAFF | AATCTGCTGAGCCAATATAAA | -1 | 877 | 898 | -123 | -102 | 10,49 | 0,0001 | 27,53 | >0.1 | 5,47 | 6,6 | 0 | 1,9 | 0 | 3,07 | 0 |
| MAFF | TATTTTCCTAGAAAGCTGTTA | -1 | 710 | 731 | -290 | -269 | 1,11 | 0,01 | 18,56 | >0.1 | 5,47 | 5,6 | 0 | 1,21 | 0 | 5,17 | 0 |
| MAFF | TCTCTACTTGGGCAACATTTT | -1 | 535 | 556 | -465 | -444 | -0,45 | 0,01 | 8,75 | >0.1 | 3,14 | 5,07 | 0 | 1 | 0 | 0 | 0 |
| MAFF | AAATACATGACTTTGCCCTTG | -1 | 507 | 528 | -493 | -472 | 0,07 | 0,01 | 12,03 | >0.1 | 3,01 | 5,01 | 0 | 0,99 | 0 | 2,95 | 0 |

| | | | | | | | | | | | | | | | | | |
|---|---|---|---|---|---|---|---|---|---|---|---|---|---|---|---|---|---|
| MAFF | ATTCTCCTGCCTCAGCCTCC | -1 | 350 | 371 | -650 | -629 | 8,04 | 0,001 | 22,41 | >0.1 | 2,53 | 4,7 | 0 | 1,11 | 0 | 6,03 | 0 |
| MAFF | GAGTAGCTGGGACTACAGGT | -1 | 329 | 350 | -671 | -650 | 0,17 | 0,01 | 14,48 | >0.1 | 2,58 | 4,67 | 0 | 1,15 | 0 | 5,92 | 0 |
| MAFF | ATTATGACTACTCTCCACTC | -1 | 116 | 137 | -884 | -863 | 1,15 | 0,01 | 12,57 | >0.1 | 3,46 | 4,38 | 0 | 1,89 | 0 | 1,7 | 0 |
| MAFF | CCTGATCTGCCTTTGCAAAA | -1 | 82 | 103 | -918 | -897 | -0,48 | 0,01 | 9,46 | >0.1 | 3,72 | 4,34 | 0 | 1,88 | 0 | 0 | 0 |
| RREB1 | CAGCCACAGAACCAGTGTGA | 1 | 38 | 58 | -962 | -942 | -1,97 | 0,01 | 8,43 | >0.1 | 4,23 | 4,3 | 0 | 1,87 | 0 | 0 | 0 |
| RREB1 | CTGAAAACCATACAAACACC | 1 | 617 | 637 | -383 | -363 | -1,34 | 0,01 | 10,07 | >0.1 | 3,64 | 5,28 | 0 | 1,05 | 0 | 1,44 | 0 |
| RREB1 | AAACCATACAAACACCAACA | 1 | 621 | 641 | -379 | -359 | 8,95 | 0,001 | 20,51 | >0.1 | 3,67 | 5,3 | 0 | 1,05 | 0 | 1,55 | 0 |
| RREB1 | AACCATACAAACACCAACAT | 1 | 622 | 642 | -378 | -358 | 3,83 | 0,01 | 15,41 | >0.1 | 3,68 | 5,3 | 0 | 1,05 | 0 | 1,55 | 0 |
| RREB1 | CATACAAACACCAACATTAT | 1 | 625 | 645 | -375 | -355 | -0,44 | 0,01 | 11,36 | >0.1 | 3,7 | 5,31 | 0 | 1,06 | 0 | 1,73 | 0 |
| RREB1 | GCCCAACCCAAGTTCAAAGG | 1 | 947 | 967 | -53 | -33 | 1,98 | 0,01 | 18,59 | >0.1 | 5,48 | 7,55 | 0 | 1,88 | 0 | 1,7 | 0 |
| RREB1 | CCCTAGACTAAAACCTCCTC | -1 | 989 | 1009 | -11 | 9 | -0,53 | 0,01 | 17,42 | >0.1 | 5,48 | 9,06 | 0 | 1,87 | 0 | 1,53 | 0 |
| RREB1 | GGCCAAATCAAAACCGGTAC | -1 | 741 | 761 | -259 | -239 | -2,15 | 0,01 | 16,27 | >0.1 | 5,48 | 5,73 | 0 | 1,29 | 0 | 5,92 | 0 |
| RREB1 | CAGGTGCCCACCACCACGC | -1 | 315 | 335 | -685 | -665 | -2,2 | 0,01 | 11,92 | >0.1 | 2,62 | 4,64 | 0 | 1,17 | 0 | 5,68 | 0 |
| RREB1 | GTGCCCACCACCACGCCCA | -1 | 312 | 332 | -688 | -668 | 1,09 | 0,01 | 15,8 | >0.1 | 2,63 | 4,64 | 0 | 1,18 | 0 | 6,26 | 0 |
| RREB1 | TGCCCACCACCACGCCCAG | -1 | 311 | 331 | -689 | -669 | -1,25 | 0,01 | 13,33 | >0.1 | 2,63 | 4,64 | 0 | 1,18 | 0 | 6,13 | 0 |
| RREB1 | GCCCACCACCACGCCCAGC | -1 | 310 | 330 | -690 | -670 | 2,37 | 0,01 | 16,96 | >0.1 | 2,64 | 4,64 | 0 | 1,19 | 0 | 6,13 | 0 |
| RREB1 | CCCACCACCACGCCCAGCT | -1 | 309 | 329 | -691 | -671 | 8,31 | 0,001 | 22,9 | >0.1 | 2,64 | 4,63 | 0 | 1,19 | 0 | 6,13 | 0 |
| RREB1 | CCACCACCACGCCCAGCTA | -1 | 308 | 328 | -692 | -672 | -2,27 | 0,01 | 12,33 | >0.1 | 2,64 | 4,63 | 0 | 1,19 | 0 | 6,13 | 0 |
| RREB1 | CACCACCACGCCCAGCTAA | -1 | 307 | 327 | -693 | -673 | 4,57 | 0,01 | 19,07 | >0.1 | 2,65 | 4,63 | 0 | 1,19 | 0 | 6,03 | 0 |
| RREB1 | ACCACCACGCCCAGCTAAGT | -1 | 306 | 326 | -694 | -674 | -1,2 | 0,01 | 13,31 | >0.1 | 2,65 | 4,63 | 0 | 1,2 | 0 | 6,03 | 0 |
| RREB1 | CTGCCCAACTCGGCCTCCC | -1 | 215 | 235 | -785 | -765 | -2,29 | 0,01 | 12,23 | >0.1 | 2,96 | 4,5 | 0 | 1,54 | 0 | 5,52 | 0 |
| RREB1 | GCCCAACTCGGCCTCCCAA | -1 | 213 | 233 | -787 | -767 | 1,57 | 0,01 | 16,21 | >0.1 | 2,96 | 4,5 | 0 | 1,55 | 0 | 5,63 | 0 |
| RREB1 | CCACAGGTCAACTCTCCTGA | -1 | 98 | 118 | -902 | -882 | -1,07 | 0,01 | 9,53 | >0.1 | 3,59 | 4,36 | 0 | 1,89 | 0 | 0,77 | 0 |
| ZNF143 | CAGCCACAGAACCAGT | 1 | 38 | 54 | -962 | -946 | -16,97 | 0,01 | -6,54 | >0.1 | 4,27 | 4,29 | 0 | 1,87 | 0 | 0 | 0 |
| ZNF143 | CTAACACGGTGAAACC | 1 | 267 | 283 | -733 | -717 | -18,46 | 0,01 | -4,83 | >0.1 | 2,77 | 4,57 | 0 | 1,31 | 0 | 4,99 | 0 |
| ZNF143 | ATTCCATAAAGACAAG | 1 | 495 | 511 | -505 | -489 | -19,56 | 0,01 | -4,22 | >0.1 | 2,95 | 4,97 | 0 | 0,99 | 0 | 6,44 | 0 |
| ZNF143 | TCTCCAGGATGAACTT | 1 | 863 | 879 | -137 | -121 | -19,07 | 0,01 | -2,45 | >0.1 | 5,53 | 6,44 | 0 | 1,9 | 0 | 2,75 | 0 |
| ZNF143 | CTCACAGGGGACGATG | 1 | 1036 | 1052 | 36 | 52 | -19,91 | 0,01 | -3,17 | >0.1 | 5,53 | 7,99 | 0 | 1,86 | 0 | 1,36 | 0 |
| ZNF143 | CAGCCTTGTTGCTGTA | 1 | 1075 | 1091 | 75 | 91 | -20,82 | 0,01 | -2,34 | >0.1 | 5,53 | 7,09 | 0 | 1,85 | 1,79 | 2,21 | 0 |
| ZNF143 | TAACCACGAAGCCGAA | 1 | 1144 | 1160 | 144 | 160 | -4,67 | 0,001 | 14,48 | >0.1 | 5,53 | 6,34 | 0 | 1,84 | 3,22 | 2,21 | 0 |
| ZNF143 | GTCCAAAAATGTCTTG | -1 | 1123 | 1139 | 123 | 139 | -19,38 | 0,01 | 0,49 | >0.1 | 5,53 | 6,52 | 0 | 1,85 | 3,22 | 2,75 | 0 |
| ZNF143 | TTGACATCGTCCCCTG | -1 | 1040 | 1056 | 40 | 56 | -10,18 | 0,01 | 5,68 | >0.1 | 5,53 | 7,85 | 0 | 1,86 | 0 | 0,62 | 0 |
| ZNF143 | ATCCTAGAGAACCTTA | -1 | 779 | 795 | -221 | -205 | -18,23 | 0,01 | 0,72 | >0.1 | 5,53 | 5,89 | 0 | 1,42 | 0 | 6,1 | 0 |
| ZNF143 | TTCCTAGAAAGCTGTT | -1 | 711 | 727 | -289 | -273 | -15,4 | 0,01 | 1,9 | >0.1 | 5,53 | 5,59 | 0 | 1,2 | 0 | 4,99 | 0 |
| ZNF143 | TTCCCAGAAACTCTCT | -1 | 551 | 567 | -449 | -433 | -11,76 | 0,01 | -2,46 | >0.1 | 3,2 | 5,1 | 0 | 1 | 0 | 0 | 0 |
| ZNF143 | CGCCCAGGCTGGAGCA | -1 | 421 | 437 | -579 | -563 | -18,99 | 0,01 | -4,79 | >0.1 | 2,69 | 4,82 | 0 | 1,02 | 0 | 5,67 | 0 |
| ZNF143 | CTCACTGCAAGCTCTG | -1 | 388 | 404 | -612 | -596 | -14,15 | 0,01 | 0,28 | >0.1 | 2,6 | 4,76 | 0 | 1,05 | 0 | 6,02 | 0 |
| ZNF143 | TTCACGCCATTCTCCT | -1 | 363 | 379 | -637 | -621 | -12,13 | 0,01 | 2,82 | >0.1 | 2,53 | 4,71 | 0 | 1,09 | 0 | 6,61 | 0 |
| ZNF143 | CCACCACCACGCCCAG | -1 | 312 | 328 | -688 | -672 | -14,18 | 0,01 | 0,4 | >0.1 | 2,64 | 4,63 | 0 | 1,18 | 0 | 6,13 | 0 |
| ZNF143 | CTCCCAAAGTGCTAGG | -1 | 205 | 221 | -795 | -779 | -5,4 | 0,001 | 8,39 | >0.1 | 3 | 4,48 | 0 | 1,6 | 0 | 4,7 | 0 |
| ZNF143 | GAGCCACTGTGCCCAG | -1 | 178 | 194 | -822 | -806 | -9,4 | 0,01 | 3,84 | >0.1 | 3,12 | 4,45 | 0 | 1,81 | 0 | 3,86 | 0 |
| RBPJ | TGTGAGAAAT | 1 | 53 | 63 | -947 | -937 | 3,67 | 0,01 | 18,88 | >0.1 | 4,09 | 4,3 | 0 | 1,87 | 0 | 0 | 4,95 |
| RBPJ | AATAGGAAAT | 1 | 60 | 70 | -940 | -930 | 3,58 | 0,01 | 18,71 | >0.1 | 4 | 4,31 | 0 | 1,87 | 0 | 0 | 4,95 |
| RBPJ | CCTGTGGAGT | 1 | 111 | 121 | -889 | -879 | -0,99 | 0,01 | 14,52 | >0.1 | 3,53 | 4,36 | 0 | 1,89 | 0 | 0,79 | 4,95 |
| RBPJ | CATGGAAATT | 1 | 150 | 160 | -850 | -840 | 2,25 | 0,01 | 18,43 | >0.1 | 3,28 | 4,41 | 0 | 1,9 | 0 | 1,65 | 4,95 |
| RBPJ | GCTGGGCACA | 1 | 177 | 187 | -823 | -813 | 0,09 | 0,01 | 17,59 | >0.1 | 3,14 | 4,44 | 0 | 1,82 | 0 | 3,16 | 4,95 |
| RBPJ | TTTGGGAGGC | 1 | 213 | 223 | -787 | -777 | 0,3 | 0,01 | 19,89 | >0.1 | 2,98 | 4,48 | 0 | 1,55 | 0 | 5,63 | 4,95 |
| RBPJ | GGTGGGCACC | 1 | 323 | 333 | -677 | -667 | 0,48 | 0,01 | 19,4 | >0.1 | 2,61 | 4,64 | 0 | 1,15 | 0 | 5,56 | 4,95 |
| RBPJ | GGCGTGAACC | 1 | 371 | 381 | -629 | -619 | 0,64 | 0,01 | 20,32 | >0.1 | 2,55 | 4,72 | 0 | 1,07 | 0 | 6,4 | 4,95 |
| RBPJ | CCTGGGAGGC | 1 | 379 | 389 | -621 | -611 | 3,54 | 0,01 | 23,51 | >0.1 | 2,57 | 4,73 | 0 | 1,06 | 0 | 6,66 | 4,95 |
| RBPJ | ATTTGGAAGG | 1 | 524 | 534 | -476 | -466 | -1,19 | 0,01 | 16,1 | >0.1 | 3,06 | 5,02 | 0 | 0,98 | 0 | 3,28 | 4,95 |
| RBPJ | GAAGGGAAAA | 1 | 529 | 539 | -471 | -461 | 1,4 | 0,01 | 18,99 | >0.1 | 3,08 | 5,03 | 0 | 0,98 | 0 | 3,55 | 4,95 |
| RBPJ | TCTGGGAATA | 1 | 559 | 569 | -441 | -431 | 11,49 | 0,001 | 25,76 | >0.1 | 3,23 | 5,1 | 0 | 0,99 | 0 | 0 | 4,95 |
| RBPJ | TGTCTGAAAA | 1 | 614 | 624 | -386 | -376 | -0,11 | 0,01 | 16,18 | >0.1 | 3,57 | 5,25 | 0 | 1,03 | 0 | 1,5 | 4,95 |
| RBPJ | TCTAGGAAAA | 1 | 719 | 729 | -281 | -271 | 5,78 | 0,01 | 28,12 | >0.1 | 5,61 | 5,59 | 0 | 1,2 | 0 | 4,99 | 4,95 |
| RBPJ | GCTGATAAGA | 1 | 966 | 976 | -34 | -24 | -0,87 | 0,01 | 21,04 | >0.1 | 5,61 | 7,78 | 0 | 1,87 | 0 | 1,7 | 4,95 |
| RBPJ | CTAGGGAAAG | 1 | 1003 | 1013 | 3 | 13 | 1,76 | 0,01 | 24,77 | >0.1 | 5,61 | 9,12 | 0 | 1,87 | 0 | 1,46 | 4,95 |
| RBPJ | TCTTGGAATT | 1 | 1187 | 1197 | 187 | 197 | 0,01 | 0,01 | 23,49 | >0.1 | 5,61 | 6,05 | 0 | 1,53 | 1,79 | 3,55 | 4,95 |
| RBPJ | CGTGGTTAAA | -1 | 1142 | 1152 | 142 | 152 | 1,42 | 0,01 | 26,77 | >0.1 | 5,61 | 6,36 | 0 | 1,84 | 3,22 | 3,37 | 4,95 |

| | | | | | | | | | | | | | | | | | |
|---|---|---|---|---|---|---|---|---|---|---|---|---|---|---|---|---|---|
| RBPJ | GCTGAGAAGG | -1 | 1069 | 1079 | 69 | 79 | 5,34 | 0,01 | 28,95 | >0.1 | 5,61 | 7,19 | 0 | 1,85 | 1,79 | 2,21 | 4,95 |
| RBPJ | GCCAGGAAGA | -1 | 1058 | 1068 | 58 | 68 | -0,68 | 0,01 | 21,34 | >0.1 | 5,61 | 7,39 | 0 | 1,86 | 0 | 2,21 | 4,95 |
| RBPJ | AGCAGGAAAG | -1 | 933 | 943 | -67 | -57 | -1,48 | 0,01 | 18,99 | >0.1 | 5,61 | 7,11 | 0 | 1,88 | 0 | 0,93 | 4,95 |
| RBPJ | CCTAGAAAGC | -1 | 715 | 725 | -285 | -275 | -1,38 | 0,01 | 21,11 | >0.1 | 5,61 | 5,58 | 0 | 1,19 | 0 | 5,17 | 4,95 |
| RBPJ | GCTGTTAAAG | -1 | 707 | 717 | -293 | -283 | 2,5 | 0,01 | 24,59 | >0.1 | 5,61 | 5,55 | 0 | 1,17 | 0 | 4,81 | 4,95 |
| RBPJ | CTTGGGCAAC | -1 | 540 | 550 | -460 | -450 | -1,49 | 0,01 | 12,63 | >0.1 | 3,13 | 5,06 | 0 | 0,99 | 0 | 0 | 4,95 |
| RBPJ | GCTGGGACTA | -1 | 335 | 345 | -665 | -655 | 5,17 | 0,01 | 24,52 | >0.1 | 2,58 | 4,66 | 0 | 1,13 | 0 | 6,03 | 4,95 |
| MAFG::NFE2L1 | AATGAG | 1 | 67 | 73 | -933 | -927 | 4,98 | 0,01 | 15,11 | >0.1 | 3,94 | 4,31 | 0 | 1,88 | 0 | 0 | 0 |
| MAFG::NFE2L1 | CCTGAC | 1 | 262 | 268 | -738 | -732 | 6,45 | 0,01 | 19,65 | >0.1 | 2,81 | 4,54 | 0 | 1,32 | 0 | 4,53 | 0 |
| MAFG::NFE2L1 | GCTGAG | 1 | 355 | 361 | -645 | -639 | 2,74 | 0,1 | 17,45 | >0.1 | 2,53 | 4,68 | 0 | 1,09 | 0 | 6,4 | 0 |
| MAFG::NFE2L1 | TATGAT | 1 | 567 | 573 | -433 | -427 | 5,76 | 0,01 | 15,12 | >0.1 | 3,26 | 5,11 | 0 | 0,99 | 0 | 0 | 0 |
| MAFG::NFE2L1 | GATAAC | 1 | 595 | 601 | -405 | -399 | 4,31 | 0,01 | 13,93 | >0.1 | 3,43 | 5,18 | 0 | 1,01 | 0 | 0 | 0 |
| MAFG::NFE2L1 | CATGAT | 1 | 648 | 654 | -352 | -346 | 7,11 | 0,001 | 21,19 | >0.1 | 3,85 | 5,33 | 0 | 1,06 | 0 | 3,83 | 0 |
| MAFG::NFE2L1 | TATAAC | 1 | 688 | 694 | -312 | -306 | 3,09 | 0,01 | 17,04 | >0.1 | 4,42 | 5,47 | 0 | 1,12 | 0 | 2,95 | 0 |
| MAFG::NFE2L1 | AGTGAC | 1 | 765 | 771 | -235 | -229 | 5,86 | 0,01 | 24,54 | >0.1 | 5,67 | 5,77 | 0 | 1,32 | 0 | 5,92 | 0 |
| MAFG::NFE2L1 | AATAAC | 1 | 800 | 806 | -200 | -194 | 3,31 | 0,01 | 19,95 | >0.1 | 5,67 | 5,95 | 0 | 1,47 | 0 | 3,55 | 0 |
| MAFG::NFE2L1 | GCTGAT | 1 | 966 | 972 | -34 | -28 | 3,74 | 0,01 | 20,65 | >0.1 | 5,67 | 7,67 | 0 | 1,87 | 0 | 1,7 | 0 |
| MAFG::NFE2L1 | CATGAG | 1 | 986 | 992 | -14 | -8 | 6,11 | 0,01 | 23,61 | >0.1 | 5,67 | 8,43 | 0 | 1,87 | 0 | 1,53 | 0 |
| MAFG::NFE2L1 | TGTGAT | 1 | 1025 | 1031 | 25 | 31 | 3,07 | 0,01 | 20,39 | >0.1 | 5,67 | 8,52 | 0 | 1,86 | 0 | 1,28 | 0 |
| MAFG::NFE2L1 | GCTGAG | -1 | 1073 | 1079 | 73 | 79 | 2,74 | 0,1 | 21,45 | >0.1 | 5,67 | 7,12 | 0 | 1,85 | 1,79 | 2,27 | 0 |
| MAFG::NFE2L1 | AATGAC | -1 | 1012 | 1018 | 12 | 18 | 8,55 | 0,001 | 26,42 | >0.1 | 5,67 | 8,92 | 0 | 1,86 | 0 | 1,42 | 0 |
| MAFG::NFE2L1 | CATGAG | -1 | 984 | 990 | -16 | -10 | 6,11 | 0,01 | 23,5 | >0.1 | 5,67 | 8,32 | 0 | 1,87 | 0 | 1,53 | 0 |
| MAFG::NFE2L1 | GCTGAG | -1 | 941 | 947 | -59 | -53 | 2,74 | 0,1 | 18,3 | >0.1 | 5,67 | 7,17 | 0 | 1,88 | 0 | 0,85 | 0 |
| MAFG::NFE2L1 | GCTGAG | -1 | 887 | 893 | -113 | -107 | 2,74 | 0,1 | 19,92 | >0.1 | 5,67 | 6,55 | 0 | 1,89 | 0 | 3,07 | 0 |
| MAFG::NFE2L1 | AGTGAC | -1 | 675 | 681 | -325 | -319 | 5,86 | 0,01 | 20,11 | >0.1 | 4,19 | 5,42 | 0 | 1,1 | 0 | 3,55 | 0 |
| MAFG::NFE2L1 | CATGAC | -1 | 517 | 523 | -483 | -477 | 9,68 | 0,001 | 21,53 | >0.1 | 3,02 | 4,99 | 0 | 0,98 | 0 | 2,86 | 0 |
| MAFG::NFE2L1 | CAGGAC | -1 | 260 | 266 | -740 | -734 | 2,76 | 0,01 | 15,97 | >0.1 | 2,81 | 4,54 | 0 | 1,33 | 0 | 4,53 | 0 |
| MAFG::NFE2L1 | CCTGAC | -1 | 243 | 249 | -757 | -751 | 6,45 | 0,01 | 19,56 | >0.1 | 2,88 | 4,52 | 0 | 1,39 | 0 | 4,33 | 0 |
| MAFG::NFE2L1 | TGTGAT | -1 | 235 | 241 | -765 | -759 | 3,07 | 0,01 | 16,81 | >0.1 | 2,9 | 4,51 | 0 | 1,43 | 0 | 4,9 | 0 |
| MAFG::NFE2L1 | GATTAC | -1 | 200 | 206 | -800 | -794 | 4,31 | 0,01 | 17,92 | >0.1 | 3,05 | 4,46 | 0 | 1,63 | 0 | 4,48 | 0 |
| MAFG::NFE2L1 | CGTGAG | -1 | 191 | 197 | -809 | -803 | 3,42 | 0,01 | 16,69 | >0.1 | 3,09 | 4,45 | 0 | 1,7 | 0 | 4,04 | 0 |
| MAFG::NFE2L1 | CATGGC | -1 | 148 | 154 | -852 | -846 | 2,76 | 0,01 | 13,97 | >0.1 | 3,31 | 4,4 | 0 | 1,9 | 0 | 1,61 | 0 |
| MAFG::NFE2L1 | TATGAC | -1 | 129 | 135 | -871 | -865 | 8,33 | 0,001 | 19,63 | >0.1 | 3,42 | 4,38 | 0 | 1,89 | 0 | 1,61 | 0 |
| MAFG::NFE2L1 | CCTGAT | -1 | 97 | 103 | -903 | -897 | 3,87 | 0,01 | 14,53 | >0.1 | 3,65 | 4,34 | 0 | 1,88 | 0 | 0,79 | 0 |
| MAFG::NFE2L1 | GCTGAC | -1 | 36 | 42 | -964 | -958 | 6,32 | 0,01 | 16,87 | >0.1 | 4,4 | 4,28 | 0 | 1,87 | 0 | 0 | 0 |
| SREBF2(VAR.2) | ATCAGGAGAG | 1 | 97 | 107 | -903 | -893 | 1,77 | 0,001 | 12,4 | >0.1 | 3,64 | 4,35 | 0 | 1,88 | 0 | 0,77 | 0 |
| SREBF2(VAR.2) | CTCACGCCTG | 1 | 191 | 201 | -809 | -799 | -5,8 | 0,01 | 7,56 | >0.1 | 3,08 | 4,45 | 0 | 1,7 | 0 | 4,13 | 0 |
| SREBF2(VAR.2) | ATCACAAGGT | 1 | 235 | 245 | -765 | -755 | -3,2 | 0,01 | 10,38 | >0.1 | 2,9 | 4,51 | 0 | 1,43 | 0 | 4,74 | 0 |
| SREBF2(VAR.2) | GTCAGGAGAT | 1 | 243 | 253 | -757 | -747 | 4,46 | 0,001 | 17,57 | >0.1 | 2,87 | 4,52 | 0 | 1,39 | 0 | 4,33 | 0 |
| SREBF2(VAR.2) | GGCACCTGTA | 1 | 327 | 337 | -673 | -663 | -5,97 | 0,01 | 7,99 | >0.1 | 2,6 | 4,65 | 0 | 1,14 | 0 | 5,56 | 0 |
| SREBF2(VAR.2) | CTCGGGAGGC | 1 | 347 | 357 | -653 | -643 | -8,13 | 0,01 | 6,73 | >0.1 | 2,55 | 4,68 | 0 | 1,11 | 0 | 6,52 | 0 |
| SREBF2(VAR.2) | ATGGCGTGAA | 1 | 369 | 379 | -631 | -621 | -2,18 | 0,01 | 12,3 | >0.1 | 2,54 | 4,71 | 0 | 1,07 | 0 | 6,15 | 0 |
| SREBF2(VAR.2) | GTGAGCCGAG | 1 | 399 | 409 | -601 | -591 | -7,72 | 0,01 | 6,62 | >0.1 | 2,62 | 4,77 | 0 | 1,04 | 0 | 5,91 | 0 |
| SREBF2(VAR.2) | ATCGCGCCAC | 1 | 409 | 419 | -591 | -581 | 1,19 | 0,001 | 15,18 | >0.1 | 2,65 | 4,78 | 0 | 1,03 | 0 | 5,52 | 0 |
| SREBF2(VAR.2) | GTCATGTATT | 1 | 517 | 527 | -483 | -473 | -8,21 | 0,01 | 3,75 | >0.1 | 3,03 | 5 | 0 | 0,98 | 0 | 2,95 | 0 |
| SREBF2(VAR.2) | ATAAAGTGAC | 1 | 761 | 771 | -239 | -229 | -2,18 | 0,01 | 16,63 | >0.1 | 5,61 | 5,77 | 0 | 1,32 | 0 | 6,1 | 0 |
| SREBF2(VAR.2) | GACAGGAGAG | 1 | 768 | 778 | -232 | -222 | -2,55 | 0,01 | 16,13 | >0.1 | 5,61 | 5,81 | 0 | 1,35 | 0 | 5,92 | 0 |
| SREBF2(VAR.2) | ATAACGTATT | 1 | 801 | 811 | -199 | -189 | -7,89 | 0,01 | 8,62 | >0.1 | 5,61 | 5,98 | 0 | 1,5 | 0 | 3,43 | 0 |
| SREBF2(VAR.2) | CTCAGCAGAT | 1 | 887 | 897 | -113 | -103 | -0,97 | 0,01 | 16,19 | >0.1 | 5,61 | 6,59 | 0 | 1,89 | 0 | 3,07 | 0 |
| SREBF2(VAR.2) | ATCTCATGAG | 1 | 982 | 992 | -18 | -8 | -2,54 | 0,01 | 14,9 | >0.1 | 5,61 | 8,43 | 0 | 1,87 | 0 | 1,53 | 0 |
| SREBF2(VAR.2) | CACAGGGGAC | 1 | 1038 | 1048 | 38 | 48 | -7,4 | 0,01 | 9,32 | >0.1 | 5,61 | 7,92 | 0 | 1,86 | 0 | 1,34 | 0 |
| SREBF2(VAR.2) | AAGACCTGTT | 1 | 1158 | 1168 | 158 | 168 | -2,36 | 0,01 | 14,23 | >0.1 | 4,38 | 6,24 | 0 | 1,78 | 1,79 | 2,4 | 0 |
| SREBF2(VAR.2) | AGCAAGTGAA | -1 | 1177 | 1187 | 177 | 187 | -7,23 | 0,01 | 11,15 | >0.1 | 5,61 | 6,11 | 0 | 1,61 | 1,79 | 3,26 | 0 |
| SREBF2(VAR.2) | AACAGGTCTT | -1 | 1158 | 1168 | 158 | 168 | -4,7 | 0,01 | 11,89 | >0.1 | 4,38 | 6,24 | 0 | 1,78 | 1,79 | 2,4 | 0 |
| SREBF2(VAR.2) | GTCCCCTGTG | -1 | 1038 | 1048 | 38 | 48 | -5,98 | 0,01 | 10,74 | >0.1 | 5,61 | 7,92 | 0 | 1,86 | 0 | 1,34 | 0 |
| SREBF2(VAR.2) | ACCTCCTCAT | -1 | 987 | 997 | -13 | -3 | -7,04 | 0,01 | 10,68 | >0.1 | 5,61 | 8,64 | 0 | 1,87 | 0 | 1,61 | 0 |
| SREBF2(VAR.2) | CTCATGAGAT | -1 | 982 | 992 | -18 | -8 | -0,81 | 0,01 | 16,63 | >0.1 | 5,61 | 8,43 | 0 | 1,87 | 0 | 1,53 | 0 |
| SREBF2(VAR.2) | ATCTGCTGAG | -1 | 887 | 897 | -113 | -103 | -3,62 | 0,01 | 13,54 | >0.1 | 5,61 | 6,59 | 0 | 1,89 | 0 | 3,07 | 0 |
| SREBF2(VAR.2) | AATACGTTAT | -1 | 801 | 811 | -199 | -189 | -7,47 | 0,01 | 9,04 | >0.1 | 5,61 | 5,98 | 0 | 1,5 | 0 | 3,43 | 0 |

| | | | | | | | | | | | | | | | | | |
|---|---|---|---|---|---|---|---|---|---|---|---|---|---|---|---|---|---|
| SREBF2(VAR.2) | CTCTCCTGTC | -1 | 768 | 778 | -232 | -222 | -1,56 | 0,01 | 17,12 | >0.1 | 5,61 | 5,81 | 0 | 1,35 | 0 | 5,92 | 0 |
| SREBF2(VAR.2) | GTCACTTTAT | -1 | 761 | 771 | -239 | -229 | -7,65 | 0,01 | 11,16 | >0.1 | 5,61 | 5,77 | 0 | 1,32 | 0 | 6,1 | 0 |
| SREBF2(VAR.2) | AATACATGAC | -1 | 517 | 527 | -483 | -473 | -5,4 | 0,01 | 6,56 | >0.1 | 3,03 | 5 | 0 | 0,98 | 0 | 2,95 | 0 |
| SREBF2(VAR.2) | GTGGCGCGAT | -1 | 409 | 419 | -591 | -581 | -0,17 | 0,01 | 13,82 | >0.1 | 2,65 | 4,78 | 0 | 1,03 | 0 | 5,52 | 0 |
| SREBF2(VAR.2) | CTCGGCTCAC | -1 | 399 | 409 | -601 | -591 | -7,24 | 0,01 | 7,1 | >0.1 | 2,62 | 4,77 | 0 | 1,04 | 0 | 5,91 | 0 |
| SREBF2(VAR.2) | TTCACGCCAT | -1 | 369 | 379 | -631 | -621 | -2,06 | 0,01 | 12,42 | >0.1 | 2,54 | 4,71 | 0 | 1,07 | 0 | 6,15 | 0 |
| SREBF2(VAR.2) | GCCTCCCGAG | -1 | 347 | 357 | -653 | -643 | -6,97 | 0,01 | 7,89 | >0.1 | 2,55 | 4,68 | 0 | 1,11 | 0 | 6,52 | 0 |
| SREBF2(VAR.2) | TACAGGTGCC | -1 | 327 | 337 | -673 | -663 | -6,96 | 0,01 | 7 | >0.1 | 2,6 | 4,65 | 0 | 1,14 | 0 | 5,56 | 0 |
| SREBF2(VAR.2) | ATCTCCTGAC | -1 | 243 | 253 | -757 | -747 | 6,31 | 0,001 | 19,42 | >0.1 | 2,87 | 4,52 | 0 | 1,39 | 0 | 4,33 | 0 |
| SREBF2(VAR.2) | ACCTTGTGAT | -1 | 235 | 245 | -765 | -755 | -4,24 | 0,01 | 9,34 | >0.1 | 2,9 | 4,51 | 0 | 1,43 | 0 | 4,74 | 0 |
| SREBF2(VAR.2) | CAGGCGTGAG | -1 | 191 | 201 | -809 | -799 | -5,53 | 0,01 | 7,83 | >0.1 | 3,08 | 4,45 | 0 | 1,7 | 0 | 4,13 | 0 |
| SREBF2(VAR.2) | CTCTCCTGAT | -1 | 97 | 107 | -903 | -893 | 2,76 | 0,001 | 13,39 | >0.1 | 3,64 | 4,35 | 0 | 1,88 | 0 | 0,77 | 0 |
| MAF::NFE2 | GAGAGTAGTCATAAT | 1 | 122 | 137 | -878 | -863 | -8,17 | 0,01 | 3,23 | >0.1 | 3,44 | 4,38 | 0 | 1,89 | 0 | 1,7 | 0 |
| MAF::NFE2 | ATGGCTGGGCACAGT | 1 | 174 | 189 | -826 | -811 | -1,52 | 0,01 | 11,15 | >0.1 | 3,15 | 4,44 | 0 | 1,84 | 0 | 3,24 | 0 |
| MAF::NFE2 | CTGAGGCAGGAGAAT | 1 | 356 | 371 | -644 | -629 | 3,14 | 0,001 | 17,71 | >0.1 | 2,52 | 4,7 | 0 | 1,1 | 0 | 6,26 | 0 |
| MAF::NFE2 | GAGACTCAGTCTCAA | 1 | 444 | 459 | -556 | -541 | -8,67 | 0,01 | 5,08 | >0.1 | 2,77 | 4,86 | 0 | 1 | 0 | 5,12 | 0 |
| MAF::NFE2 | GTGAGATAACCTATT | 1 | 591 | 606 | -409 | -394 | -2,17 | 0,01 | 7,47 | >0.1 | 3,43 | 5,2 | 0 | 1,02 | 0 | 0 | 0 |
| MAF::NFE2 | TTGATTTGGCCATAA | 1 | 750 | 765 | -250 | -235 | -5,78 | 0,01 | 13,36 | >0.1 | 5,54 | 5,75 | 0 | 1,31 | 0 | 6,54 | 0 |
| MAF::NFE2 | GTGACAGGAGAGGTA | 1 | 766 | 781 | -234 | -219 | -5,91 | 0,01 | 12,73 | >0.1 | 5,54 | 5,82 | 0 | 1,36 | 0 | 5,92 | 0 |
| MAF::NFE2 | GTTACATATCTGTCC | 1 | 848 | 863 | -152 | -137 | -4,67 | 0,01 | 11,84 | >0.1 | 5,54 | 6,32 | 0 | 1,89 | 0 | 2,75 | 0 |
| MAF::NFE2 | TTGGCTCAGCAGATT | 1 | 883 | 898 | -117 | -102 | 2,92 | 0,001 | 20,03 | >0.1 | 5,54 | 6,6 | 0 | 1,89 | 0 | 3,07 | 0 |
| MAF::NFE2 | ATGAGGAGGTTTTAG | 1 | 987 | 1002 | -13 | 2 | -8,81 | 0,01 | 8,99 | >0.1 | 5,54 | 8,77 | 0 | 1,87 | 0 | 1,61 | 0 |
| MAF::NFE2 | GTCATTCAGTGGATG | 1 | 1012 | 1027 | 12 | 27 | -8,86 | 0,01 | 8,86 | >0.1 | 5,54 | 8,92 | 0 | 1,86 | 0 | 1,39 | 0 |
| MAF::NFE2 | TTGATAGAACAGGTC | -1 | 1160 | 1175 | 160 | 175 | -7,5 | 0,01 | 9,33 | >0.1 | 4,45 | 6,22 | 0 | 1,76 | 1,79 | 2,61 | 0 |
| MAF::NFE2 | ATGTCTTGGCCTGTT | -1 | 1116 | 1131 | 116 | 131 | -5,69 | 0,01 | 14,18 | >0.1 | 5,54 | 6,59 | 0 | 1,85 | 3,22 | 2,67 | 0 |
| MAF::NFE2 | TGGACTGAGCAGCAG | -1 | 1092 | 1107 | 92 | 107 | -6,6 | 0,01 | 13,21 | >0.1 | 5,54 | 6,85 | 0 | 1,85 | 3,22 | 2,34 | 0 |
| MAF::NFE2 | GTGAGCCAAGATCAC | -1 | 1026 | 1041 | 26 | 41 | -6,52 | 0,01 | 10,64 | >0.1 | 5,54 | 8,45 | 0 | 1,86 | 0 | 1,31 | 0 |
| MAF::NFE2 | ATGACTTTCCCTAGA | -1 | 1002 | 1017 | 2 | 17 | -3,94 | 0,01 | 14,01 | >0.1 | 5,54 | 9,12 | 0 | 1,87 | 0 | 1,42 | 0 |
| MAF::NFE2 | GTGACCAAAGAGGTC | -1 | 665 | 680 | -335 | -320 | -5,89 | 0,01 | 8,57 | >0.1 | 4,11 | 5,42 | 0 | 1,1 | 0 | 3,83 | 0 |
| MAF::NFE2 | ATGACTTTGCCCTTG | -1 | 507 | 522 | -493 | -478 | 6,59 | 0,001 | 18,32 | >0.1 | 3 | 4,99 | 0 | 0,99 | 0 | 2,75 | 0 |
| MAF::NFE2 | ATTTCTGATCAGTTT | -1 | 476 | 491 | -524 | -509 | -7,92 | 0,01 | 6,24 | >0.1 | 2,88 | 4,92 | 0 | 0,99 | 0 | 5,36 | 0 |
| MAF::NFE2 | CTGAGTCTCGGTCTG | -1 | 438 | 453 | -562 | -547 | -9,04 | 0,01 | 5,61 | >0.1 | 2,75 | 4,85 | 0 | 1,01 | 0 | 6,05 | 0 |
| MAF::NFE2 | CTGCCTCAGCCTCCC | -1 | 350 | 365 | -650 | -635 | -3,78 | 0,01 | 10,95 | >0.1 | 2,53 | 4,69 | 0 | 1,1 | 0 | 6,4 | 0 |
| MAF::NFE2 | GTTAGTCAGGACGGT | -1 | 257 | 272 | -743 | -728 | 2,44 | 0,001 | 15,67 | >0.1 | 2,81 | 4,55 | 0 | 1,34 | 0 | 4,53 | 0 |
| MAF::NFE2 | ATTTCTGATCAGTTT | -1 | 161 | 176 | -839 | -824 | -7,92 | 0,01 | 4,02 | >0.1 | 3,21 | 4,43 | 0 | 1,9 | 0 | 2,4 | 0 |
| MAF::NFE2 | ATGACTACTCTCCAC | -1 | 119 | 134 | -881 | -866 | -5,77 | 0,01 | 5,65 | >0.1 | 3,46 | 4,38 | 0 | 1,89 | 0 | 1,7 | 0 |
| MAF::NFE2 | CTGCCTTTGCAAAAC | -1 | 82 | 97 | -918 | -903 | -4,04 | 0,01 | 5,92 | >0.1 | 3,74 | 4,34 | 0 | 1,88 | 0 | 0 | 0 |
| MYC | TGACCTGTGGAG | 1 | 108 | 120 | -892 | -880 | 0,35 | 0,01 | 10,89 | >0.1 | 3,54 | 4,36 | 0 | 1,89 | 0 | 0,75 | 0 |
| MYC | GGGCACCTGTAG | 1 | 326 | 338 | -674 | -662 | 5,99 | 0,01 | 19,95 | >0.1 | 2,6 | 4,65 | 0 | 1,15 | 0 | 5,56 | 0 |
| MYC | GCCCAAGTAGAG | 1 | 543 | 555 | -457 | -445 | 0 | 0,01 | 9,21 | >0.1 | 3,15 | 5,07 | 0 | 0,99 | 0 | 0 | 0 |
| MYC | AACCACGAAGCC | 1 | 1145 | 1157 | 145 | 157 | 0,16 | 0,01 | 19,3 | >0.1 | 5,58 | 6,33 | 0 | 1,84 | 3,22 | 2,16 | 0 |
| MYC | GTTCACTTGCTT | 1 | 1176 | 1188 | 176 | 188 | 3,36 | 0,01 | 21,73 | >0.1 | 5,58 | 6,11 | 0 | 1,62 | 1,79 | 3,26 | 0 |
| MYC | AAGCAAGTGAAC | -1 | 1176 | 1188 | 176 | 188 | 1,61 | 0,01 | 19,98 | >0.1 | 5,58 | 6,11 | 0 | 1,62 | 1,79 | 3,26 | 0 |
| MYC | GGCTTCGTGGTT | -1 | 1145 | 1157 | 145 | 157 | 1,92 | 0,01 | 21,06 | >0.1 | 5,58 | 6,33 | 0 | 1,84 | 3,22 | 2,16 | 0 |
| MYC | TCCCCTGTGAGC | -1 | 1035 | 1047 | 35 | 47 | 2,49 | 0,01 | 19,29 | >0.1 | 5,58 | 8,03 | 0 | 1,86 | 0 | 1,34 | 0 |
| MYC | GATCACATCCAC | -1 | 1020 | 1032 | 20 | 32 | -0,01 | 0,01 | 17,46 | >0.1 | 5,58 | 8,72 | 0 | 1,86 | 0 | 1,31 | 0 |
| MYC | CTCCTCATGAGA | -1 | 983 | 995 | -17 | -5 | 0,4 | 0,01 | 17,97 | >0.1 | 5,58 | 8,59 | 0 | 1,87 | 0 | 1,53 | 0 |
| MYC | TCCCAGGTTCAC | -1 | 374 | 386 | -626 | -614 | 2,14 | 0,01 | 16,89 | >0.1 | 2,56 | 4,73 | 0 | 1,07 | 0 | 6,4 | 0 |
| MYC | ATTCTCCTGCCT | -1 | 359 | 371 | -641 | -629 | -0,01 | 0,01 | 14,66 | >0.1 | 2,52 | 4,7 | 0 | 1,09 | 0 | 6,36 | 0 |
| MYC | CTACAGGTGCCC | -1 | 326 | 338 | -674 | -662 | 6,93 | 0,001 | 20,89 | >0.1 | 2,6 | 4,65 | 0 | 1,15 | 0 | 5,56 | 0 |
| MYC | CACCACGCCCAG | -1 | 312 | 324 | -688 | -676 | 4,41 | 0,01 | 18,98 | >0.1 | 2,64 | 4,63 | 0 | 1,18 | 0 | 6,13 | 0 |
| MYC | GACCTTGTGATC | -1 | 234 | 246 | -766 | -754 | 3,41 | 0,01 | 17 | >0.1 | 2,9 | 4,51 | 0 | 1,44 | 0 | 4,74 | 0 |
| MYC | CCCAAAGTGCTA | -1 | 207 | 219 | -793 | -781 | 1,09 | 0,01 | 14,86 | >0.1 | 3 | 4,48 | 0 | 1,58 | 0 | 4,7 | 0 |
| MYC | GCCACTGTGCCC | -1 | 180 | 192 | -820 | -808 | 0,37 | 0,01 | 13,58 | >0.1 | 3,12 | 4,44 | 0 | 1,79 | 0 | 3,86 | 0 |
| MYC | CTCCACAGGTCA | -1 | 108 | 120 | -892 | -880 | 0,92 | 0,01 | 11,46 | >0.1 | 3,54 | 4,36 | 0 | 1,89 | 0 | 0,75 | 0 |
| MYB | TAAAACTGAT | 1 | 159 | 169 | -841 | -831 | -2,78 | 0,01 | 9,17 | >0.1 | 3,23 | 4,42 | 0 | 1,9 | 0 | 2,4 | 0 |
| MYB | CACGCCTGTA | 1 | 193 | 203 | -807 | -797 | -1,86 | 0,01 | 11,61 | >0.1 | 3,07 | 4,46 | 0 | 1,68 | 0 | 4,26 | 0 |
| MYB | GGCACCTGTA | 1 | 327 | 337 | -673 | -663 | 3,33 | 0,01 | 17,29 | >0.1 | 2,6 | 4,65 | 0 | 1,14 | 0 | 5,56 | 0 |
| MYB | CGCCACTGTG | 1 | 413 | 423 | -587 | -577 | 3,99 | 0,01 | 17,55 | >0.1 | 2,66 | 4,79 | 0 | 1,02 | 0 | 5,09 | 0 |

| | | | | | | | | | | | | | | | | | |
|---|---|---|---|---|---|---|---|---|---|---|---|---|---|---|---|---|---|
| MYB | GGCGACAGAC | 1 | 433 | 443 | -567 | -557 | -2,15 | 0,01 | 12,21 | >0.1 | 2,72 | 4,83 | 0 | 1,01 | 0 | 5,8 | 0 |
| MYB | AAAAACTGAT | 1 | 474 | 484 | -526 | -516 | -2,62 | 0,01 | 11,08 | >0.1 | 2,86 | 4,91 | 0 | 0,99 | 0 | 4,95 | 0 |
| MYB | GGTCACTGTA | 1 | 674 | 684 | -326 | -316 | -3,95 | 0,01 | 10,22 | >0.1 | 4,2 | 5,43 | 0 | 1,11 | 0 | 3,43 | 0 |
| MYB | CATATCTGTC | 1 | 852 | 862 | -148 | -138 | -3,16 | 0,01 | 13,31 | >0.1 | 5,61 | 6,31 | 0 | 1,88 | 0 | 2,67 | 0 |
| MYB | GTTTACTGTG | 1 | 898 | 908 | -102 | -92 | -4,13 | 0,01 | 12,92 | >0.1 | 5,61 | 6,68 | 0 | 1,89 | 0 | 2,86 | 0 |
| MYB | CCCAACCCAA | 1 | 948 | 958 | -52 | -42 | -0,37 | 0,01 | 16,28 | >0.1 | 5,61 | 7,36 | 0 | 1,88 | 0 | 1,8 | 0 |
| MYB | TGTAACTGCT | 1 | 1087 | 1097 | 87 | 97 | 4,44 | 0,001 | 22,83 | >0.1 | 5,61 | 6,92 | 0 | 1,85 | 1,79 | 2,21 | 0 |
| MYB | AGCAAGTGAA | -1 | 1177 | 1187 | 177 | 187 | -0,94 | 0,01 | 17,44 | >0.1 | 5,61 | 6,11 | 0 | 1,61 | 1,79 | 3,26 | 0 |
| MYB | GTGGACTGAG | -1 | 1098 | 1108 | 98 | 108 | -1,83 | 0,01 | 17,98 | >0.1 | 5,61 | 6,78 | 0 | 1,85 | 3,22 | 2,35 | 0 |
| MYB | ATCCACTGAA | -1 | 1016 | 1026 | 16 | 26 | 3,16 | 0,01 | 20,81 | >0.1 | 5,61 | 8,81 | 0 | 1,86 | 0 | 1,36 | 0 |
| MYB | CTCTCCTGTC | -1 | 768 | 778 | -232 | -222 | -3,49 | 0,01 | 15,19 | >0.1 | 5,61 | 5,81 | 0 | 1,35 | 0 | 5,92 | 0 |
| MYB | GCTCACTGCA | -1 | 395 | 405 | -605 | -595 | -3,39 | 0,01 | 10,79 | >0.1 | 2,61 | 4,76 | 0 | 1,04 | 0 | 5,76 | 0 |
| MYB | TTCTCCTGCC | -1 | 360 | 370 | -640 | -630 | -3,68 | 0,01 | 10,99 | >0.1 | 2,52 | 4,7 | 0 | 1,09 | 0 | 6,36 | 0 |
| MYB | CAGGACGGTC | -1 | 256 | 266 | -744 | -734 | -4,15 | 0,01 | 9,29 | >0.1 | 2,82 | 4,54 | 0 | 1,34 | 0 | 4,73 | 0 |
| MYB | ATCTCCTGAC | -1 | 243 | 253 | -757 | -747 | -4,28 | 0,01 | 8,83 | >0.1 | 2,87 | 4,52 | 0 | 1,39 | 0 | 4,33 | 0 |
| MYB | GTGATCTGCC | -1 | 230 | 240 | -770 | -760 | -3,03 | 0,01 | 10,86 | >0.1 | 2,92 | 4,51 | 0 | 1,45 | 0 | 5,01 | 0 |
| MYB | CCCAACTCGG | -1 | 222 | 232 | -778 | -768 | 2,31 | 0,01 | 16,77 | >0.1 | 2,95 | 4,49 | 0 | 1,5 | 0 | 5,52 | 0 |
| MYB | AGCCACTGTG | -1 | 183 | 193 | -817 | -807 | 3,51 | 0,01 | 16,69 | >0.1 | 3,11 | 4,44 | 0 | 1,76 | 0 | 3,86 | 0 |
| MYB | GTCAACTCTC | -1 | 102 | 112 | -898 | -888 | 3,75 | 0,01 | 14,35 | >0.1 | 3,6 | 4,35 | 0 | 1,88 | 0 | 0,77 | 0 |
| MYB | CTGATCTGCC | -1 | 92 | 102 | -908 | -898 | -2,54 | 0,01 | 8,19 | >0.1 | 3,68 | 4,34 | 0 | 1,88 | 0 | 0,83 | 0 |
| GATA1::TAL1 | CTGATCAGAAATATGAAT | 1 | 479 | 497 | -521 | -503 | -7,16 | 0,01 | 7,59 | >0.1 | 2,89 | 4,94 | 0 | 0,99 | 0 | 5,93 | 0 |
| GATA1::TAL1 | ATTATCTTCATGATCCCT | 1 | 640 | 658 | -360 | -342 | -3,41 | 0,01 | 10,54 | >0.1 | 3,83 | 5,35 | 0 | 1,07 | 0 | 3,7 | 0 |
| GATA1::TAL1 | TTTATATTGGCTCAGCAG | 1 | 877 | 895 | -123 | -105 | 0,54 | 0,01 | 17,59 | >0.1 | 5,51 | 6,57 | 0 | 1,9 | 0 | 3,07 | 0 |
| GATA1::TAL1 | GATGTCAAGCTCTTCCTG | 1 | 1048 | 1066 | 48 | 66 | -8,68 | 0,01 | 8,58 | >0.1 | 5,51 | 7,62 | 0 | 1,86 | 0 | 2,27 | 0 |
| GATA1::TAL1 | TTAACCACGAAGCCGAAG | 1 | 1143 | 1161 | 143 | 161 | -7,15 | 0,01 | 11,98 | >0.1 | 5,51 | 6,35 | 0 | 1,84 | 3,22 | 2,21 | 0 |
| GATA1::TAL1 | TCTATCAAAGTTCACTTG | 1 | 1167 | 1185 | 167 | 185 | -7,83 | 0,01 | 10,36 | >0.1 | 5,51 | 6,17 | 0 | 1,69 | 1,79 | 3,02 | 0 |
| GATA1::TAL1 | CTTATCAGCCTTTGAACT | -1 | 957 | 975 | -43 | -25 | -8,13 | 0,01 | 8,71 | >0.1 | 5,51 | 7,76 | 0 | 1,88 | 0 | 1,7 | 0 |
| GATA1::TAL1 | ACAATCTGCTGAGCCAAT | -1 | 882 | 900 | -118 | -100 | 1,08 | 0,01 | 18,17 | >0.1 | 5,51 | 6,62 | 0 | 1,89 | 0 | 3,07 | 0 |
| GATA1::TAL1 | TTAATCCTAGAGAACCTT | -1 | 780 | 798 | -220 | -202 | -1,22 | 0,01 | 17,55 | >0.1 | 5,51 | 5,91 | 0 | 1,44 | 0 | 5,92 | 0 |
| GATA1::TAL1 | GTTTTCAGACAATTTCAT | -1 | 607 | 625 | -393 | -375 | -5,36 | 0,01 | 5,97 | >0.1 | 3,55 | 5,25 | 0 | 1,03 | 0 | 1,5 | 0 |
| GATA1::TAL1 | GTTATCTCACATTTATTT | -1 | 583 | 601 | -417 | -399 | -8,58 | 0,01 | 1,01 | >0.1 | 3,39 | 5,18 | 0 | 1,02 | 0 | 0 | 0 |
| GATA1::TAL1 | ACTCTCTACTTGGGCAAC | -1 | 540 | 558 | -460 | -442 | -9,12 | 0,01 | 0,1 | >0.1 | 3,15 | 5,08 | 0 | 0,99 | 0 | 0 | 0 |
| GATA1::TAL1 | GCAAGCTCTGCCTCCCAG | -1 | 380 | 398 | -620 | -602 | -3,06 | 0,01 | 11,35 | >0.1 | 2,58 | 4,75 | 0 | 1,06 | 0 | 6,02 | 0 |
| GATA1::TAL1 | ATTATGACTACTCTCCAC | -1 | 119 | 137 | -881 | -863 | -1,95 | 0,01 | 9,46 | >0.1 | 3,45 | 4,38 | 0 | 1,89 | 0 | 1,7 | 0 |
| GATA1::TAL1 | CTACTCTCCACTCCACAG | -1 | 112 | 130 | -888 | -870 | -1,43 | 0,01 | 9,07 | >0.1 | 3,49 | 4,37 | 0 | 1,89 | 0 | 0,75 | 0 |
| GATA1::TAL1 | CTGATCTGCCTTTGCAAA | -1 | 84 | 102 | -916 | -898 | -8,52 | 0,01 | 1,42 | >0.1 | 3,71 | 4,34 | 0 | 1,88 | 0 | 0 | 0 |
| GATA1::TAL1 | TTTCTCACACTGGTTCTG | -1 | 44 | 62 | -956 | -938 | -4,96 | 0,01 | 5,37 | >0.1 | 4,16 | 4,3 | 0 | 1,87 | 0 | 0 | 0 |
| JUN(VAR.2) | CCAGTGTGAGAAAT | 1 | 49 | 63 | -951 | -937 | 2,81 | 0,001 | 13,1 | >0.1 | 4,11 | 4,3 | 0 | 1,87 | 0 | 0 | 0 |
| JUN(VAR.2) | AACTGATCAGAAAT | 1 | 162 | 176 | -838 | -824 | -11,3 | 0,01 | 0,64 | >0.1 | 3,21 | 4,42 | 0 | 1,9 | 0 | 2,4 | 0 |
| JUN(VAR.2) | GCACAGTGGCTCAC | 1 | 182 | 196 | -818 | -804 | -2,33 | 0,01 | 10,95 | >0.1 | 3,11 | 4,45 | 0 | 1,77 | 0 | 3,95 | 0 |
| JUN(VAR.2) | CCGTCCTGACTAAC | 1 | 258 | 272 | -742 | -728 | 4,56 | 0,001 | 17,78 | >0.1 | 2,81 | 4,55 | 0 | 1,34 | 0 | 4,53 | 0 |
| JUN(VAR.2) | ACACGGTGAAACCC | 1 | 270 | 284 | -730 | -716 | -11,68 | 0,01 | 1,94 | >0.1 | 2,77 | 4,57 | 0 | 1,29 | 0 | 4,99 | 0 |
| JUN(VAR.2) | GGAGGCTGAGGCAG | 1 | 351 | 365 | -649 | -635 | -0,95 | 0,01 | 13,78 | >0.1 | 2,53 | 4,69 | 0 | 1,1 | 0 | 6,4 | 0 |
| JUN(VAR.2) | AGACCGAGACTCAG | 1 | 439 | 453 | -561 | -547 | -3,14 | 0,01 | 11,39 | >0.1 | 2,75 | 4,85 | 0 | 1,01 | 0 | 5,92 | 0 |
| JUN(VAR.2) | AACTGATCAGAAAT | 1 | 477 | 491 | -523 | -509 | -11,3 | 0,01 | 3,02 | >0.1 | 2,88 | 4,92 | 0 | 0,99 | 0 | 5,52 | 0 |
| JUN(VAR.2) | GATTAAAGAATAAC | 1 | 792 | 806 | -208 | -194 | -10,73 | 0,01 | 7,42 | >0.1 | 5,56 | 5,95 | 0 | 1,48 | 0 | 5,17 | 0 |
| JUN(VAR.2) | CTTATTTGATTCAC | 1 | 811 | 825 | -189 | -175 | -7,3 | 0,01 | 9,33 | >0.1 | 5,56 | 6,06 | 0 | 1,59 | 0 | 3,43 | 0 |
| JUN(VAR.2) | AAAATCTGTTACAT | 1 | 841 | 855 | -159 | -145 | -7,57 | 0,01 | 9,25 | >0.1 | 5,56 | 6,26 | 0 | 1,82 | 0 | 3,19 | 0 |
| JUN(VAR.2) | GGAAAGTCATTCAG | 1 | 1007 | 1021 | 7 | 21 | -3,01 | 0,01 | 15 | >0.1 | 5,56 | 9,12 | 0 | 1,87 | 0 | 1,46 | 0 |
| JUN(VAR.2) | CCCCTGTGAGCCAA | -1 | 1032 | 1046 | 32 | 46 | -9,47 | 0,01 | 7,4 | >0.1 | 5,56 | 8,15 | 0 | 1,86 | 0 | 1,31 | 0 |
| JUN(VAR.2) | ATCCACTGAATGAC | -1 | 1012 | 1026 | 12 | 26 | -10,28 | 0,01 | 7,45 | >0.1 | 5,56 | 8,92 | 0 | 1,86 | 0 | 1,39 | 0 |
| JUN(VAR.2) | ATCTGCTGAGCCAA | -1 | 883 | 897 | -117 | -103 | -8,37 | 0,01 | 8,74 | >0.1 | 5,56 | 6,59 | 0 | 1,89 | 0 | 3,07 | 0 |
| JUN(VAR.2) | CAGATATGTAACAG | -1 | 846 | 860 | -154 | -140 | -7,81 | 0,01 | 8,86 | >0.1 | 5,56 | 6,3 | 0 | 1,86 | 0 | 2,95 | 0 |
| JUN(VAR.2) | TTAAAGTGAATCAA | -1 | 816 | 830 | -184 | -170 | 5,56 | 0,001 | 22,25 | >0.1 | 5,56 | 6,09 | 0 | 1,62 | 0 | 3,43 | 0 |
| JUN(VAR.2) | GAAAGCTGTTAAAG | -1 | 707 | 721 | -293 | -279 | -10,71 | 0,01 | 6,4 | >0.1 | 5,56 | 5,56 | 0 | 1,18 | 0 | 4,81 | 0 |
| JUN(VAR.2) | TTACAGTGACCAAA | -1 | 671 | 685 | -329 | -315 | -10,62 | 0,01 | 3,66 | >0.1 | 4,19 | 5,44 | 0 | 1,11 | 0 | 3,55 | 0 |
| JUN(VAR.2) | TGAGACTGAGTCTC | -1 | 444 | 458 | -556 | -542 | -3,04 | 0,01 | 10,71 | >0.1 | 2,76 | 4,86 | 0 | 1 | 0 | 5,12 | 0 |
| JUN(VAR.2) | TTCTCCTGCCTCAG | -1 | 356 | 370 | -644 | -630 | -8,69 | 0,01 | 5,88 | >0.1 | 2,52 | 4,7 | 0 | 1,09 | 0 | 6,26 | 0 |
| JUN(VAR.2) | ACCGTGTTAGTCAG | -1 | 263 | 277 | -737 | -723 | -4,09 | 0,01 | 9,11 | >0.1 | 2,79 | 4,56 | 0 | 1,32 | 0 | 4,53 | 0 |

| | | | | | | | | | | | | | | | | | |
|---|---|---|---|---|---|---|---|---|---|---|---|---|---|---|---|---|---|
| JUN(VAR.2) | CAGGCGTGAGCCAC | -1 | 187 | 201 | -813 | -799 | -2,86 | 0,01 | 10,45 | >0.1 | 3,09 | 4,46 | 0 | 1,73 | 0 | 4,04 | 0 |
| JUN(VAR.2) | AAATTATGACTACT | -1 | 125 | 139 | -875 | -861 | -4,91 | 0,01 | 6,39 | >0.1 | 3,42 | 4,38 | 0 | 1,89 | 0 | 1,61 | 0 |
| FLI1 | GAAGGAGGCA | 1 | 17 | 27 | -983 | -973 | -6,74 | 0,01 | 5 | >0.1 | 5,61 | 4,27 | 0 | 1,87 | 0 | 0 | 0 |
| FLI1 | ATAGGAAATG | 1 | 61 | 71 | -939 | -929 | 4,33 | 0,001 | 14,5 | >0.1 | 3,98 | 4,31 | 0 | 1,88 | 0 | 0 | 0 |
| FLI1 | TCAGGAGAGT | 1 | 98 | 108 | -902 | -892 | -7,06 | 0,01 | 3,56 | >0.1 | 3,63 | 4,35 | 0 | 1,88 | 0 | 0,77 | 0 |
| FLI1 | ATCAGAAATG | 1 | 167 | 177 | -833 | -823 | -4,75 | 0,01 | 7,17 | >0.1 | 3,19 | 4,43 | 0 | 1,9 | 0 | 2,4 | 0 |
| FLI1 | TCAGGAGATA | 1 | 244 | 254 | -756 | -746 | -2,85 | 0,01 | 10,35 | >0.1 | 2,86 | 4,52 | 0 | 1,39 | 0 | 4,42 | 0 |
| FLI1 | ATCAGAAATA | 1 | 482 | 492 | -518 | -508 | -6,04 | 0,01 | 8,56 | >0.1 | 2,89 | 4,93 | 0 | 0,99 | 0 | 5,8 | 0 |
| FLI1 | AAGGGAAAAT | 1 | 530 | 540 | -470 | -460 | -3,96 | 0,01 | 8,84 | >0.1 | 3,09 | 5,03 | 0 | 0,98 | 0 | 3,7 | 0 |
| FLI1 | CTAGGAAAAT | 1 | 720 | 730 | -280 | -270 | -7,22 | 0,01 | 10,18 | >0.1 | 5,61 | 5,6 | 0 | 1,2 | 0 | 4,99 | 0 |
| FLI1 | ACCGGTTTTG | 1 | 743 | 753 | -257 | -247 | -6,11 | 0,01 | 12,37 | >0.1 | 5,61 | 5,69 | 0 | 1,26 | 0 | 5,92 | 0 |
| FLI1 | ACAGGAGAGG | 1 | 769 | 779 | -231 | -221 | -1,8 | 0,01 | 16,67 | >0.1 | 5,61 | 5,81 | 0 | 1,35 | 0 | 5,69 | 0 |
| FLI1 | CCAGGATGAA | 1 | 866 | 876 | -134 | -124 | -0,82 | 0,01 | 15,85 | >0.1 | 5,61 | 6,41 | 0 | 1,9 | 0 | 2,75 | 0 |
| FLI1 | TGAGGAGGTT | 1 | 988 | 998 | -12 | -2 | -7,22 | 0,01 | 10,53 | >0.1 | 5,61 | 8,66 | 0 | 1,87 | 0 | 1,61 | 0 |
| FLI1 | TAGGGAAAGT | 1 | 1004 | 1014 | 4 | 14 | -5,6 | 0,01 | 12,41 | >0.1 | 5,61 | 9,12 | 0 | 1,87 | 0 | 1,42 | 0 |
| FLI1 | TGAGGAACAG | 1 | 1111 | 1121 | 111 | 121 | -5,55 | 0,01 | 14,31 | >0.1 | 5,61 | 6,64 | 0 | 1,85 | 3,22 | 2,54 | 0 |
| FLI1 | CCAAGAAGCA | -1 | 1183 | 1193 | 183 | 193 | -7,37 | 0,01 | 11,19 | >0.1 | 5,61 | 6,07 | 0 | 1,56 | 1,79 | 3,52 | 0 |
| FLI1 | CCAGGAAGAG | -1 | 1057 | 1067 | 57 | 67 | 3,32 | 0,001 | 20,48 | >0.1 | 5,61 | 7,42 | 0 | 1,86 | 0 | 2,27 | 0 |
| FLI1 | GCAGGAAAGA | -1 | 932 | 942 | -68 | -58 | 4,63 | 0,001 | 20,14 | >0.1 | 5,61 | 7,09 | 0 | 1,88 | 0 | 0,93 | 0 |
| FLI1 | ACAGTAAACA | -1 | 897 | 907 | -103 | -93 | -0,29 | 0,01 | 16,75 | >0.1 | 5,61 | 6,68 | 0 | 1,89 | 0 | 2,86 | 0 |
| FLI1 | AGAGGACAGA | -1 | 856 | 866 | -144 | -134 | -4,4 | 0,01 | 12,4 | >0.1 | 5,61 | 6,34 | 0 | 1,9 | 0 | 2,95 | 0 |
| FLI1 | GACAGATATG | -1 | 852 | 862 | -148 | -138 | -6,86 | 0,01 | 9,61 | >0.1 | 5,61 | 6,31 | 0 | 1,88 | 0 | 2,67 | 0 |
| FLI1 | ACCGGTACTT | -1 | 739 | 749 | -261 | -251 | -1,99 | 0,01 | 16,46 | >0.1 | 5,61 | 5,68 | 0 | 1,25 | 0 | 5,92 | 0 |
| FLI1 | CACGGAAAAT | -1 | 694 | 704 | -306 | -296 | 0,56 | 0,01 | 15,88 | >0.1 | 5,61 | 5,5 | 0 | 1,14 | 0 | 3,07 | 0 |
| FLI1 | CCCAGAAACT | -1 | 555 | 565 | -445 | -435 | -4,45 | 0,01 | 4,84 | >0.1 | 3,21 | 5,09 | 0 | 0,99 | 0 | 0 | 0 |
| FLI1 | TCAGGACGGT | -1 | 257 | 267 | -743 | -733 | -2,95 | 0,01 | 10,38 | >0.1 | 2,82 | 4,54 | 0 | 1,34 | 0 | 4,63 | 0 |
| KLF4 | TCACGCCTGTA | 1 | 192 | 203 | -808 | -797 | 0,76 | 0,01 | 14,24 | >0.1 | 3,07 | 4,46 | 0 | 1,69 | 0 | 4,26 | 0 |
| KLF4 | TGAAACCCCGT | 1 | 276 | 287 | -724 | -713 | 1,1 | 0,01 | 14,78 | >0.1 | 2,75 | 4,57 | 0 | 1,27 | 0 | 5,09 | 0 |
| KLF4 | CTCCAGCCTGG | 1 | 423 | 434 | -577 | -566 | 2,24 | 0,01 | 16,43 | >0.1 | 2,69 | 4,81 | 0 | 1,02 | 0 | 5,67 | 0 |
| KLF4 | AAACACCAACA | 1 | 630 | 641 | -370 | -359 | 4,31 | 0,01 | 16,09 | >0.1 | 3,71 | 5,3 | 0 | 1,05 | 0 | 1,73 | 0 |
| KLF4 | CAGCGCCCAAC | 1 | 943 | 954 | -57 | -46 | 4,45 | 0,01 | 21,01 | >0.1 | 5,6 | 7,29 | 0 | 1,88 | 0 | 1,8 | 0 |
| KLF4 | CCCAACCCAAG | 1 | 948 | 959 | -52 | -41 | 1,42 | 0,01 | 18,08 | >0.1 | 5,6 | 7,38 | 0 | 1,88 | 0 | 1,8 | 0 |
| KLF4 | TCTCAGCCTTG | 1 | 1072 | 1083 | 72 | 83 | 1,24 | 0,01 | 19,9 | >0.1 | 5,6 | 7,14 | 0 | 1,85 | 1,79 | 2,27 | 0 |
| KLF4 | TCACATCCACT | -1 | 1019 | 1030 | 19 | 30 | 6,66 | 0,01 | 24,17 | >0.1 | 5,6 | 8,74 | 0 | 1,86 | 0 | 1,31 | 0 |
| KLF4 | TGTCGCCCAGG | -1 | 429 | 440 | -571 | -560 | 1,64 | 0,01 | 15,86 | >0.1 | 2,71 | 4,82 | 0 | 1,01 | 0 | 5,67 | 0 |
| KLF4 | TCACGCCATTC | -1 | 367 | 378 | -633 | -622 | 4,88 | 0,01 | 19,93 | >0.1 | 2,54 | 4,71 | 0 | 1,08 | 0 | 6,73 | 0 |
| KLF4 | CCTCAGCCTCC | -1 | 351 | 362 | -649 | -638 | 5,37 | 0,01 | 20,09 | >0.1 | 2,54 | 4,69 | 0 | 1,1 | 0 | 6,4 | 0 |
| KLF4 | CCACCACCACG | -1 | 317 | 328 | -683 | -672 | 0,72 | 0,01 | 15,4 | >0.1 | 2,63 | 4,63 | 0 | 1,17 | 0 | 6,26 | 0 |
| KLF4 | CCACGCCCAGC | -1 | 311 | 322 | -689 | -678 | 13,7 | 0,0001 | 28,18 | >0.1 | 2,65 | 4,62 | 0 | 1,18 | 0 | 6,03 | 0 |
| KLF4 | ACTCGGCCTCC | -1 | 217 | 228 | -783 | -772 | 1,74 | 0,01 | 16,24 | >0.1 | 2,96 | 4,49 | 0 | 1,52 | 0 | 5,52 | 0 |
| KLF4 | GCCCAGCCATT | -1 | 173 | 184 | -827 | -816 | 1,23 | 0,01 | 13,91 | >0.1 | 3,16 | 4,43 | 0 | 1,85 | 0 | 3,24 | 0 |
| KLF4 | CCACTCCACAG | -1 | 112 | 123 | -888 | -877 | 0,64 | 0,01 | 11,18 | >0.1 | 3,52 | 4,36 | 0 | 1,89 | 0 | 0,77 | 0 |
| KLF4 | TCTCACACTGG | -1 | 49 | 60 | -951 | -940 | 2,27 | 0,01 | 12,58 | >0.1 | 4,13 | 4,3 | 0 | 1,87 | 0 | 0 | 0 |
| T | CAGAACCAGTGTGAGA | 1 | 44 | 60 | -956 | -940 | -7,09 | 0,01 | 3,25 | >0.1 | 4,17 | 4,3 | 0 | 1,87 | 0 | 0 | 0 |
| T | TGGCTCACGCCTGTAA | 1 | 188 | 204 | -812 | -796 | -9,5 | 0,01 | 3,89 | >0.1 | 3,08 | 4,46 | 0 | 1,72 | 0 | 4,13 | 0 |
| T | TCACAAGGTCAGGAGA | 1 | 236 | 252 | -764 | -748 | -5,17 | 0,01 | 8,09 | >0.1 | 2,88 | 4,52 | 0 | 1,43 | 0 | 4,42 | 0 |
| T | TCCCAGCTACTCGGGA | 1 | 338 | 354 | -662 | -646 | -1,47 | 0,01 | 13,42 | >0.1 | 2,56 | 4,67 | 0 | 1,13 | 0 | 6,52 | 0 |
| T | TGGCGTGAACCTGGGA | 1 | 370 | 386 | -630 | -614 | -2,73 | 0,01 | 12,16 | >0.1 | 2,55 | 4,73 | 0 | 1,08 | 0 | 6,53 | 0 |
| T | TTGCAGTGAGCCGAGA | 1 | 394 | 410 | -606 | -590 | -5,94 | 0,01 | 8,4 | >0.1 | 2,62 | 4,77 | 0 | 1,05 | 0 | 5,91 | 0 |
| T | TCGCGCCACTGTGCTC | 1 | 410 | 426 | -590 | -574 | 0,47 | 0,001 | 14,05 | >0.1 | 2,66 | 4,8 | 0 | 1,03 | 0 | 5,09 | 0 |
| T | AGTCATGTATTTGGAA | 1 | 516 | 532 | -484 | -468 | -8,98 | 0,01 | 3,25 | >0.1 | 3,04 | 5,01 | 0 | 0,99 | 0 | 3,19 | 0 |
| T | AAAAATAAATGTGAGA | 1 | 581 | 597 | -419 | -403 | -11,26 | 0,01 | -1,71 | >0.1 | 3,37 | 5,17 | 0 | 1,01 | 0 | 0 | 0 |
| T | TAACAGCTTTCTAGGA | 1 | 710 | 726 | -290 | -274 | -6,79 | 0,01 | 10,68 | >0.1 | 5,53 | 5,58 | 0 | 1,19 | 0 | 5,17 | 0 |
| T | TGGCCATAAAGTGACA | 1 | 756 | 772 | -244 | -228 | -10,41 | 0,01 | 8,33 | >0.1 | 5,53 | 5,78 | 0 | 1,33 | 0 | 6,1 | 0 |
| T | TGTTACATATCTGTCC | 1 | 847 | 863 | -153 | -137 | -10,23 | 0,01 | 6,26 | >0.1 | 5,53 | 6,32 | 0 | 1,89 | 0 | 2,75 | 0 |
| T | GGACAGATATGTAACA | -1 | 847 | 863 | -153 | -137 | -7,36 | 0,01 | 9,13 | >0.1 | 5,53 | 6,32 | 0 | 1,89 | 0 | 2,75 | 0 |
| T | TGTCACTTTATGGCCA | -1 | 756 | 772 | -244 | -228 | -5,27 | 0,01 | 13,47 | >0.1 | 5,53 | 5,78 | 0 | 1,33 | 0 | 6,1 | 0 |
| T | TCCTAGAAAGCTGTTA | -1 | 710 | 726 | -290 | -274 | -9,42 | 0,01 | 8,05 | >0.1 | 5,53 | 5,58 | 0 | 1,19 | 0 | 5,17 | 0 |
| T | TTTCATTAATAGGTTA | -1 | 597 | 613 | -403 | -387 | -9,29 | 0,01 | 1,79 | >0.1 | 3,47 | 5,22 | 0 | 1,02 | 0 | 1,36 | 0 |

| | | | | | | | | | | | | | | | | | |
|---|---|---|---|---|---|---|---|---|---|---|---|---|---|---|---|---|---|
| T | TTCCAAATACATGACT | -1 | 516 | 532 | -484 | -468 | -9,73 | 0,01 | 2,5 | >0.1 | 3,04 | 5,01 | 0 | 0,99 | 0 | 3,19 | 0 |
| T | GAGCACAGTGGCGCGA | -1 | 410 | 426 | -590 | -574 | 1,59 | 0,001 | 15,17 | >0.1 | 2,66 | 4,8 | 0 | 1,03 | 0 | 5,09 | 0 |
| T | TCTCGGCTCACTGCAA | -1 | 394 | 410 | -606 | -590 | -6,31 | 0,01 | 8,03 | >0.1 | 2,62 | 4,77 | 0 | 1,05 | 0 | 5,91 | 0 |
| T | TCCCAGGTTCACGCCA | -1 | 370 | 386 | -630 | -614 | -2,76 | 0,01 | 12,13 | >0.1 | 2,55 | 4,73 | 0 | 1,08 | 0 | 6,53 | 0 |
| T | TCCCGAGTAGCTGGGA | -1 | 338 | 354 | -662 | -646 | -1,52 | 0,01 | 13,37 | >0.1 | 2,56 | 4,67 | 0 | 1,13 | 0 | 6,52 | 0 |
| T | TCTCCTGACCTTGTGA | -1 | 236 | 252 | -764 | -748 | -11,27 | 0,01 | 1,99 | >0.1 | 2,88 | 4,52 | 0 | 1,43 | 0 | 4,42 | 0 |
| T | TTACAGGCGTGAGCCA | -1 | 188 | 204 | -812 | -796 | -10,46 | 0,01 | 2,93 | >0.1 | 3,08 | 4,46 | 0 | 1,72 | 0 | 4,13 | 0 |
| T | TCTCACACTGGTTCTG | -1 | 44 | 60 | -956 | -940 | -9,11 | 0,01 | 1,23 | >0.1 | 4,17 | 4,3 | 0 | 1,87 | 0 | 0 | 0 |
| FOSB::JUNB(VAR.2) | AAGAAGTCAG | 1 | 31 | 41 | -969 | -959 | 3,24 | 0,01 | 13,85 | >0.1 | 4,46 | 4,28 | 0 | 1,87 | 0 | 0 | 0 |
| FOSB::JUNB(VAR.2) | GAGTAGTCAT | 1 | 124 | 134 | -876 | -866 | 1,04 | 0,01 | 12,44 | >0.1 | 3,44 | 4,38 | 0 | 1,89 | 0 | 1,7 | 0 |
| FOSB::JUNB(VAR.2) | ATGGCGTGAA | 1 | 369 | 379 | -631 | -621 | 2,65 | 0,01 | 17,13 | >0.1 | 2,54 | 4,71 | 0 | 1,07 | 0 | 6,15 | 0 |
| FOSB::JUNB(VAR.2) | ATCGCGCCAC | 1 | 409 | 419 | -591 | -581 | 5,3 | 0,01 | 19,29 | >0.1 | 2,65 | 4,78 | 0 | 1,03 | 0 | 5,52 | 0 |
| FOSB::JUNB(VAR.2) | GCAAAGTCAT | 1 | 512 | 522 | -488 | -478 | 0,52 | 0,01 | 12,45 | >0.1 | 3,01 | 4,99 | 0 | 0,98 | 0 | 2,95 | 0 |
| FOSB::JUNB(VAR.2) | GTCATGTATT | 1 | 517 | 527 | -483 | -473 | 1,18 | 0,01 | 13,14 | >0.1 | 3,03 | 5 | 0 | 0,98 | 0 | 2,95 | 0 |
| FOSB::JUNB(VAR.2) | GTGAGATAAC | 1 | 591 | 601 | -409 | -399 | 3,14 | 0,01 | 12,75 | >0.1 | 3,41 | 5,18 | 0 | 1,01 | 0 | 0 | 0 |
| FOSB::JUNB(VAR.2) | CTTTGGTCAC | 1 | 670 | 680 | -330 | -320 | 0,79 | 0,01 | 15,15 | >0.1 | 4,14 | 5,42 | 0 | 1,1 | 0 | 3,7 | 0 |
| FOSB::JUNB(VAR.2) | ATTATAACAT | 1 | 686 | 696 | -314 | -304 | 2,25 | 0,01 | 16,22 | >0.1 | 4,42 | 5,47 | 0 | 1,13 | 0 | 2,95 | 0 |
| FOSB::JUNB(VAR.2) | ATTTGGCCAT | 1 | 753 | 763 | -247 | -237 | 1,51 | 0,01 | 20,47 | >0.1 | 5,61 | 5,74 | 0 | 1,3 | 0 | 6,32 | 0 |
| FOSB::JUNB(VAR.2) | GAGAGGTAAG | 1 | 773 | 783 | -227 | -217 | 0,82 | 0,01 | 19,73 | >0.1 | 5,61 | 5,83 | 0 | 1,37 | 0 | 6,1 | 0 |
| FOSB::JUNB(VAR.2) | ATAACGTATT | 1 | 801 | 811 | -199 | -189 | 2,54 | 0,01 | 19,05 | >0.1 | 5,61 | 5,98 | 0 | 1,5 | 0 | 3,43 | 0 |
| FOSB::JUNB(VAR.2) | GTTACATATC | 1 | 848 | 858 | -152 | -142 | 1,18 | 0,01 | 17,87 | >0.1 | 5,61 | 6,28 | 0 | 1,84 | 0 | 2,95 | 0 |
| FOSB::JUNB(VAR.2) | ACGATGTCAA | 1 | 1046 | 1056 | 46 | 56 | 2,71 | 0,01 | 19,32 | >0.1 | 5,61 | 7,68 | 0 | 1,86 | 0 | 1,46 | 0 |
| FOSB::JUNB(VAR.2) | TTGACATCGT | -1 | 1046 | 1056 | 46 | 56 | 1,76 | 0,01 | 18,37 | >0.1 | 5,61 | 7,68 | 0 | 1,86 | 0 | 1,46 | 0 |
| FOSB::JUNB(VAR.2) | GATATGTAAC | -1 | 848 | 858 | -152 | -142 | 2,18 | 0,01 | 18,87 | >0.1 | 5,61 | 6,28 | 0 | 1,84 | 0 | 2,95 | 0 |
| FOSB::JUNB(VAR.2) | AATACGTTAT | -1 | 801 | 811 | -199 | -189 | 3,19 | 0,01 | 19,7 | >0.1 | 5,61 | 5,98 | 0 | 1,5 | 0 | 3,43 | 0 |
| FOSB::JUNB(VAR.2) | CTTACCTCTC | -1 | 773 | 783 | -227 | -217 | 1,13 | 0,01 | 20,04 | >0.1 | 5,61 | 5,83 | 0 | 1,37 | 0 | 6,1 | 0 |
| FOSB::JUNB(VAR.2) | CTCCTGTCAC | -1 | 766 | 776 | -234 | -224 | 1,44 | 0,01 | 20,1 | >0.1 | 5,61 | 5,8 | 0 | 1,34 | 0 | 5,92 | 0 |
| FOSB::JUNB(VAR.2) | ATGGCCAAAT | -1 | 753 | 763 | -247 | -237 | 1,74 | 0,01 | 20,7 | >0.1 | 5,61 | 5,74 | 0 | 1,3 | 0 | 6,32 | 0 |
| FOSB::JUNB(VAR.2) | ATGTTATAAT | -1 | 686 | 696 | -314 | -304 | 0,56 | 0,01 | 14,53 | >0.1 | 4,42 | 5,47 | 0 | 1,13 | 0 | 2,95 | 0 |
| FOSB::JUNB(VAR.2) | GTGACCAAAG | -1 | 670 | 680 | -330 | -320 | 2,13 | 0,01 | 16,49 | >0.1 | 4,14 | 5,42 | 0 | 1,1 | 0 | 3,7 | 0 |
| FOSB::JUNB(VAR.2) | GTTATCTCAC | -1 | 591 | 601 | -409 | -399 | 5,1 | 0,01 | 14,71 | >0.1 | 3,41 | 5,18 | 0 | 1,01 | 0 | 0 | 0 |
| FOSB::JUNB(VAR.2) | GTGGCGCGAT | -1 | 409 | 419 | -591 | -581 | 5,54 | 0,01 | 19,53 | >0.1 | 2,65 | 4,78 | 0 | 1,03 | 0 | 5,52 | 0 |
| FOSB::JUNB(VAR.2) | TTCACGCCAT | -1 | 369 | 379 | -631 | -621 | 3,99 | 0,01 | 18,47 | >0.1 | 2,54 | 4,71 | 0 | 1,07 | 0 | 6,15 | 0 |
| FOSB::JUNB(VAR.2) | ATGACTACTC | -1 | 124 | 134 | -876 | -866 | 0,72 | 0,01 | 12,12 | >0.1 | 3,44 | 4,38 | 0 | 1,89 | 0 | 1,7 | 0 |
| FOSB::JUNB(VAR.2) | CTGACTTCTT | -1 | 31 | 41 | -969 | -959 | 2,72 | 0,01 | 13,33 | >0.1 | 4,46 | 4,28 | 0 | 1,87 | 0 | 0 | 0 |
| NR2F6(VAR.2) | AAGGAGAAGGAGGCA | 1 | 12 | 27 | -988 | -973 | -25,2 | 0,01 | -13,52 | >0.1 | 5,54 | 4,27 | 0 | 1,87 | 0 | 0 | 0 |
| NR2F6(VAR.2) | GTAGTCATAATTTTA | 1 | 126 | 141 | -874 | -859 | -19,41 | 0,01 | -8,15 | >0.1 | 3,41 | 4,38 | 0 | 1,89 | 0 | 1,57 | 0 |
| NR2F6(VAR.2) | GGGCACAGTGGCTCA | 1 | 180 | 195 | -820 | -805 | -21,04 | 0,01 | -7,83 | >0.1 | 3,12 | 4,45 | 0 | 1,79 | 0 | 3,86 | 0 |
| NR2F6(VAR.2) | AAGGTCAGGAGATAG | 1 | 240 | 255 | -760 | -745 | -6,6 | 0,001 | 6,53 | >0.1 | 2,87 | 4,53 | 0 | 1,41 | 0 | 4,33 | 0 |
| NR2F6(VAR.2) | GGGAGGCAGAGCTTG | 1 | 382 | 397 | -618 | -603 | -25,53 | 0,01 | -11,13 | >0.1 | 2,58 | 4,75 | 0 | 1,06 | 0 | 6,02 | 0 |
| NR2F6(VAR.2) | GAAGGGAAAATGTTG | 1 | 529 | 544 | -471 | -456 | -24,2 | 0,01 | -13,47 | >0.1 | 3,09 | 5,04 | 0 | 0,99 | 0 | 1,61 | 0 |
| NR2F6(VAR.2) | ATGATCCCTAGTTCT | 1 | 649 | 664 | -351 | -336 | -15,29 | 0,01 | -0,94 | >0.1 | 3,91 | 5,37 | 0 | 1,08 | 0 | 3,99 | 0 |
| NR2F6(VAR.2) | CAGCTTTCTAGGAAA | 1 | 713 | 728 | -287 | -272 | -25,59 | 0,01 | -8,09 | >0.1 | 5,54 | 5,59 | 0 | 1,2 | 0 | 5,17 | 0 |
| NR2F6(VAR.2) | CAAGTTCAAAGGCTG | 1 | 955 | 970 | -45 | -30 | -17,62 | 0,01 | -0,84 | >0.1 | 5,54 | 7,62 | 0 | 1,88 | 0 | 1,74 | 0 |
| NR2F6(VAR.2) | AAGTTCAAAGGCTGA | 1 | 956 | 971 | -44 | -29 | -15,04 | 0,01 | 1,77 | >0.1 | 5,54 | 7,65 | 0 | 1,88 | 0 | 1,74 | 0 |
| NR2F6(VAR.2) | AATCTCATGAGGAGG | 1 | 981 | 996 | -19 | -4 | -23,91 | 0,01 | -6,31 | >0.1 | 5,54 | 8,61 | 0 | 1,87 | 0 | 1,57 | 0 |
| NR2F6(VAR.2) | CAGGGGACGATGTCA | 1 | 1040 | 1055 | 40 | 55 | -10,16 | 0,001 | 5,72 | >0.1 | 5,54 | 7,85 | 0 | 1,86 | 0 | 0,62 | 0 |
| NR2F6(VAR.2) | AATGTTATAATTTTA | -1 | 682 | 697 | -318 | -303 | -18,51 | 0,01 | -4,57 | >0.1 | 4,39 | 5,48 | 0 | 1,13 | 0 | 2,95 | 0 |
| NR2F6(VAR.2) | GAGGTCTAGAACTAG | -1 | 656 | 671 | -344 | -329 | -19,34 | 0,01 | -5,04 | >0.1 | 3,99 | 5,39 | 0 | 1,09 | 0 | 3,83 | 0 |
| NR2F6(VAR.2) | GGGATCATGAAGATA | -1 | 642 | 657 | -358 | -343 | -14,38 | 0,01 | -0,43 | >0.1 | 3,84 | 5,34 | 0 | 1,07 | 0 | 3,7 | 0 |
| NR2F6(VAR.2) | TTCATTAATAGGTTA | -1 | 597 | 612 | -403 | -388 | -20,57 | 0,01 | -9,5 | >0.1 | 3,47 | 5,21 | 0 | 1,02 | 0 | 1,36 | 0 |
| NR2F6(VAR.2) | CCACTCCACAGGTCA | -1 | 108 | 123 | -892 | -877 | -18,65 | 0,01 | -8,1 | >0.1 | 3,53 | 4,36 | 0 | 1,89 | 0 | 0,77 | 0 |
| SMAD3 | AGAGTAGTCA | 1 | 123 | 133 | -877 | -867 | -3,3 | 0,01 | 8,11 | >0.1 | 3,45 | 4,37 | 0 | 1,89 | 0 | 1,7 | 0 |
| SMAD3 | TGGCTGGGCA | 1 | 175 | 185 | -825 | -815 | -2,98 | 0,01 | 9,68 | >0.1 | 3,15 | 4,44 | 0 | 1,83 | 0 | 3,24 | 0 |
| SMAD3 | CGTCCTGACT | 1 | 259 | 269 | -741 | -731 | -3,79 | 0,01 | 9,43 | >0.1 | 2,81 | 4,55 | 0 | 1,33 | 0 | 4,53 | 0 |
| SMAD3 | CCTGTAGTCC | 1 | 331 | 341 | -669 | -659 | -6,26 | 0,01 | 8,15 | >0.1 | 2,59 | 4,65 | 0 | 1,14 | 0 | 6,03 | 0 |
| SMAD3 | AGCCGAGATC | 1 | 402 | 412 | -598 | -588 | -7,13 | 0,01 | 7,07 | >0.1 | 2,63 | 4,77 | 0 | 1,03 | 0 | 5,76 | 0 |
| SMAD3 | GCTCCAGCCT | 1 | 422 | 432 | -578 | -568 | -5,48 | 0,01 | 8,59 | >0.1 | 2,69 | 4,81 | 0 | 1,02 | 0 | 5,56 | 0 |
| SMAD3 | GTTCTAGACC | 1 | 659 | 669 | -341 | -331 | 3,26 | 0,001 | 17,72 | >0.1 | 4 | 5,38 | 0 | 1,08 | 0 | 3,99 | 0 |

| | | | | | | | | | | | | | | | | | |
|---|---|---|---|---|---|---|---|---|---|---|---|---|---|---|---|---|---|
| SMAD3 | TTTCTAGGAA | 1 | 717 | 727 | -283 | -273 | -7,29 | 0,01 | 10,27 | >0.1 | 5,61 | 5,59 | 0 | 1,19 | 0 | 5,17 | 0 |
| SMAD3 | AGTCTAGGGA | 1 | 1000 | 1010 | 0 | 10 | -1,29 | 0,01 | 16,77 | >0.1 | 5,61 | 9,12 | 0 | 1,87 | 0 | 1,46 | 0 |
| SMAD3 | GGCCAAGACA | 1 | 1120 | 1130 | 120 | 130 | -6,64 | 0,01 | 13,29 | >0.1 | 5,61 | 6,55 | 0 | 1,85 | 3,22 | 2,7 | 0 |
| SMAD3 | TTTTTGGACA | 1 | 1130 | 1140 | 130 | 140 | -6,9 | 0,01 | 13,1 | >0.1 | 5,61 | 6,46 | 0 | 1,84 | 3,22 | 2,86 | 0 |
| SMAD3 | TCCCTAGACT | -1 | 1000 | 1010 | 0 | 10 | 1,25 | 0,001 | 19,31 | >0.1 | 5,61 | 9,12 | 0 | 1,87 | 0 | 1,46 | 0 |
| SMAD3 | ATCCTAGAGA | -1 | 785 | 795 | -215 | -205 | -6,88 | 0,01 | 11,73 | >0.1 | 5,61 | 5,89 | 0 | 1,42 | 0 | 5,69 | 0 |
| SMAD3 | TTCCTAGAAA | -1 | 717 | 727 | -283 | -273 | -4,81 | 0,01 | 12,75 | >0.1 | 5,61 | 5,59 | 0 | 1,19 | 0 | 5,17 | 0 |
| SMAD3 | GGTCTAGAAC | -1 | 659 | 669 | -341 | -331 | 3,41 | 0,001 | 17,87 | >0.1 | 4 | 5,38 | 0 | 1,08 | 0 | 3,99 | 0 |
| SMAD3 | TTTTCAGACA | -1 | 614 | 624 | -386 | -376 | -5,85 | 0,01 | 5,5 | >0.1 | 3,57 | 5,25 | 0 | 1,03 | 0 | 1,5 | 0 |
| SMAD3 | CGCCCAGGCT | -1 | 427 | 437 | -573 | -563 | -5 | 0,01 | 9,34 | >0.1 | 2,7 | 4,82 | 0 | 1,01 | 0 | 5,8 | 0 |
| SMAD3 | AGGCTGGAGC | -1 | 422 | 432 | -578 | -568 | -6,6 | 0,01 | 7,47 | >0.1 | 2,69 | 4,81 | 0 | 1,02 | 0 | 5,56 | 0 |
| SMAD3 | CGCCCAGCTA | -1 | 309 | 319 | -691 | -681 | -5,47 | 0,01 | 9,24 | >0.1 | 2,65 | 4,62 | 0 | 1,18 | 0 | 6,26 | 0 |
| SMAD3 | AGTCAGGACG | -1 | 259 | 269 | -741 | -731 | -7,84 | 0,01 | 5,38 | >0.1 | 2,81 | 4,55 | 0 | 1,33 | 0 | 4,53 | 0 |
| SMAD3 | TGCCCAGCCA | -1 | 175 | 185 | -825 | -815 | 1,83 | 0,001 | 14,49 | >0.1 | 3,15 | 4,44 | 0 | 1,83 | 0 | 3,24 | 0 |
| SMAD3 | TGACTACTCT | -1 | 123 | 133 | -877 | -867 | -4,05 | 0,01 | 7,36 | >0.1 | 3,45 | 4,37 | 0 | 1,89 | 0 | 1,7 | 0 |
| NKX6-2 | GTCATAAT | 1 | 129 | 137 | -871 | -863 | 5,01 | 0,01 | 16,27 | >0.1 | 3,41 | 4,38 | 0 | 1,89 | 0 | 1,57 | 0 |
| NKX6-2 | GAAATTAA | 1 | 154 | 162 | -846 | -838 | 6,17 | 0,01 | 17,39 | >0.1 | 3,27 | 4,41 | 0 | 1,9 | 0 | 1,65 | 0 |
| NKX6-2 | TCTATTAA | 1 | 288 | 296 | -712 | -704 | 6,34 | 0,01 | 20,07 | >0.1 | 2,72 | 4,58 | 0 | 1,24 | 0 | 5,19 | 0 |
| NKX6-2 | TCCATAAA | 1 | 497 | 505 | -503 | -495 | 5,45 | 0,01 | 20,76 | >0.1 | 2,94 | 4,95 | 0 | 0,98 | 0 | 6,44 | 0 |
| NKX6-2 | CCTATTAA | 1 | 600 | 608 | -400 | -392 | 7,48 | 0,001 | 18,53 | >0.1 | 3,47 | 5,2 | 0 | 1,01 | 0 | 1,36 | 0 |
| NKX6-2 | TTAATGAA | 1 | 604 | 612 | -396 | -388 | 3,87 | 0,01 | 15,01 | >0.1 | 3,49 | 5,21 | 0 | 1,02 | 0 | 1,41 | 0 |
| NKX6-2 | AACATTAT | 1 | 637 | 645 | -363 | -355 | 3,73 | 0,01 | 15,5 | >0.1 | 3,76 | 5,31 | 0 | 1,05 | 0 | 1,66 | 0 |
| NKX6-2 | AAAATTAT | 1 | 683 | 691 | -317 | -309 | 3,82 | 0,01 | 17,8 | >0.1 | 4,34 | 5,46 | 0 | 1,12 | 0 | 3,07 | 0 |
| NKX6-2 | GCCATAAA | 1 | 758 | 766 | -242 | -234 | 6,38 | 0,01 | 25,18 | >0.1 | 5,64 | 5,75 | 0 | 1,31 | 0 | 6,1 | 0 |
| NKX6-2 | AAAATTAT | 1 | 830 | 838 | -170 | -162 | 3,82 | 0,01 | 20,55 | >0.1 | 5,64 | 6,14 | 0 | 1,67 | 0 | 3,28 | 0 |
| NKX6-2 | GTCATTCA | 1 | 1012 | 1020 | 12 | 20 | 4,14 | 0,01 | 21,96 | >0.1 | 5,64 | 8,92 | 0 | 1,86 | 0 | 1,39 | 0 |
| NKX6-2 | ACCATTGA | 1 | 1106 | 1114 | 106 | 114 | 4,23 | 0,01 | 24,15 | >0.1 | 5,64 | 6,69 | 0 | 1,85 | 3,22 | 2,52 | 0 |
| NKX6-2 | TCTATCAA | 1 | 1167 | 1175 | 167 | 175 | 3,75 | 0,01 | 20,82 | >0.1 | 4,53 | 6,17 | 0 | 1,69 | 1,79 | 2,88 | 0 |
| NKX6-2 | GGAATTAT | 1 | 1191 | 1199 | 191 | 199 | 4,76 | 0,01 | 23,54 | >0.1 | 5,64 | 6,02 | 0 | 1,5 | 1,79 | 3,82 | 0 |
| NKX6-2 | CAAATCAA | -1 | 750 | 758 | -250 | -242 | 3,79 | 0,01 | 22,34 | >0.1 | 5,64 | 5,71 | 0 | 1,28 | 0 | 5,92 | 0 |
| NKX6-2 | ATCATGAA | -1 | 646 | 654 | -354 | -346 | 3,75 | 0,01 | 17,82 | >0.1 | 3,84 | 5,34 | 0 | 1,06 | 0 | 3,83 | 0 |
| NKX6-2 | TTCATTAA | -1 | 604 | 612 | -396 | -388 | 8,39 | 0,001 | 19,53 | >0.1 | 3,49 | 5,21 | 0 | 1,02 | 0 | 1,41 | 0 |
| NKX6-2 | CCAAATAC | -1 | 522 | 530 | -478 | -470 | 3,57 | 0,01 | 15,67 | >0.1 | 3,05 | 5,01 | 0 | 0,98 | 0 | 3,07 | 0 |
| NKX6-2 | AAAATTAT | -1 | 132 | 140 | -868 | -860 | 3,82 | 0,01 | 15,06 | >0.1 | 3,4 | 4,38 | 0 | 1,89 | 0 | 1,57 | 0 |
| NKX6-2 | CTCATTTC | -1 | 65 | 73 | -935 | -927 | 3,7 | 0,01 | 13,84 | >0.1 | 3,95 | 4,31 | 0 | 1,88 | 0 | 0 | 0 |
| NKX6-1 | GTCATAAT | 1 | 129 | 137 | -871 | -863 | 3,26 | 0,01 | 14,52 | >0.1 | 3,41 | 4,38 | 0 | 1,89 | 0 | 1,57 | 0 |
| NKX6-1 | TCTATTAA | 1 | 288 | 296 | -712 | -704 | 5,65 | 0,01 | 19,38 | >0.1 | 2,72 | 4,58 | 0 | 1,24 | 0 | 5,19 | 0 |
| NKX6-1 | TCCATAAA | 1 | 497 | 505 | -503 | -495 | 2,94 | 0,01 | 18,25 | >0.1 | 2,94 | 4,95 | 0 | 0,98 | 0 | 6,44 | 0 |
| NKX6-1 | CCTATTAA | 1 | 600 | 608 | -400 | -392 | 7,01 | 0,01 | 18,06 | >0.1 | 3,47 | 5,2 | 0 | 1,01 | 0 | 1,36 | 0 |
| NKX6-1 | ATCTTTAA | 1 | 705 | 713 | -295 | -287 | 3,51 | 0,01 | 20,65 | >0.1 | 5,64 | 5,53 | 0 | 1,16 | 0 | 4,81 | 0 |
| NKX6-1 | GCCATAAA | 1 | 758 | 766 | -242 | -234 | 4,38 | 0,01 | 23,18 | >0.1 | 5,64 | 5,75 | 0 | 1,31 | 0 | 6,1 | 0 |
| NKX6-1 | GTCATTCA | 1 | 1012 | 1020 | 12 | 20 | 4,8 | 0,01 | 22,62 | >0.1 | 5,64 | 8,92 | 0 | 1,86 | 0 | 1,39 | 0 |
| NKX6-1 | ACCATTGA | 1 | 1106 | 1114 | 106 | 114 | 4,7 | 0,01 | 24,62 | >0.1 | 5,64 | 6,69 | 0 | 1,85 | 3,22 | 2,52 | 0 |
| NKX6-1 | TTCTTTAA | -1 | 794 | 802 | -206 | -198 | 3,43 | 0,01 | 21,77 | >0.1 | 5,64 | 5,93 | 0 | 1,45 | 0 | 5,32 | 0 |
| NKX6-1 | TTCATTAA | -1 | 604 | 612 | -396 | -388 | 9,21 | 0,001 | 20,35 | >0.1 | 3,49 | 5,21 | 0 | 1,02 | 0 | 1,41 | 0 |
| NKX6-1 | GTCTTTAT | -1 | 500 | 508 | -500 | -492 | 2,72 | 0,01 | 18,24 | >0.1 | 2,96 | 4,96 | 0 | 0,98 | 0 | 6,62 | 0 |
| NKX6-1 | TCACTTAC | -1 | 5 | 13 | -995 | -987 | 3,72 | 0,01 | 17,27 | >0.1 | 5,64 | 4,26 | 0 | 1,86 | 1,79 | 0 | 0 |
| NFAT5 | CTTTTTTAAG | 1 | 73 | 83 | -927 | -917 | 3,26 | 0,01 | 13,31 | >0.1 | 3,85 | 4,32 | 0 | 1,88 | 0 | 0 | 0 |
| NFAT5 | GCTTTCTAGG | 1 | 715 | 725 | -285 | -275 | 3,71 | 0,01 | 21,25 | >0.1 | 5,61 | 5,58 | 0 | 1,19 | 0 | 5,17 | 0 |
| NFAT5 | GGTTTTGATT | 1 | 746 | 756 | -254 | -244 | 3,33 | 0,01 | 21,84 | >0.1 | 5,61 | 5,71 | 0 | 1,27 | 0 | 5,92 | 0 |
| NFAT5 | CTTTTTCTTT | 1 | 916 | 926 | -84 | -74 | 5,71 | 0,01 | 21,21 | >0.1 | 5,61 | 6,88 | 0 | 1,89 | 0 | 1,12 | 0 |
| NFAT5 | TTTTTCTTTT | 1 | 917 | 927 | -83 | -73 | 5,42 | 0,01 | 20,93 | >0.1 | 5,61 | 6,89 | 0 | 1,89 | 0 | 1,12 | 0 |
| NFAT5 | TCTTTCCTGC | 1 | 932 | 942 | -68 | -58 | 3,03 | 0,01 | 18,54 | >0.1 | 5,61 | 7,09 | 0 | 1,88 | 0 | 0,93 | 0 |
| NFAT5 | ACTTTCCCTA | -1 | 1004 | 1014 | 4 | 14 | 7,78 | 0,01 | 25,79 | >0.1 | 5,61 | 9,12 | 0 | 1,87 | 0 | 1,42 | 0 |
| NFAT5 | CTTTCCCTAG | -1 | 1003 | 1013 | 3 | 13 | 4,81 | 0,01 | 22,87 | >0.1 | 5,61 | 9,12 | 0 | 1,87 | 0 | 1,46 | 0 |
| NFAT5 | GATTTTCTCT | -1 | 975 | 985 | -25 | -15 | 3,39 | 0,01 | 20,54 | >0.1 | 5,61 | 8,09 | 0 | 1,87 | 0 | 1,57 | 0 |
| NFAT5 | ATTTTCTCTC | -1 | 974 | 984 | -26 | -16 | 6,24 | 0,01 | 23,38 | >0.1 | 5,61 | 8,05 | 0 | 1,87 | 0 | 1,61 | 0 |
| NFAT5 | ATTTTTTAAA | -1 | 825 | 835 | -175 | -165 | 3,2 | 0,01 | 20,28 | >0.1 | 5,61 | 6,12 | 0 | 1,65 | 0 | 3,7 | 0 |
| NFAT5 | CCTTACCTCT | -1 | 774 | 784 | -226 | -216 | 3,6 | 0,01 | 22,52 | >0.1 | 5,61 | 5,83 | 0 | 1,37 | 0 | 6,1 | 0 |

| | | | | | | | | | | | | | | | | | |
|---|---|---|---|---|---|---|---|---|---|---|---|---|---|---|---|---|---|
| NFAT5 | TATTTTCCTA | -1 | 721 | 731 | -279 | -269 | 3,54 | 0,01 | 21,12 | >0.1 | 5,61 | 5,6 | 0 | 1,2 | 0 | 5,17 | 0 |
| NFAT5 | ATTTTCCTAG | -1 | 720 | 730 | -280 | -270 | 7,93 | 0,01 | 25,33 | >0.1 | 5,61 | 5,6 | 0 | 1,2 | 0 | 4,99 | 0 |
| NFAT5 | GGTTTTCAGA | -1 | 616 | 626 | -384 | -374 | 3,11 | 0,01 | 14,48 | >0.1 | 3,59 | 5,25 | 0 | 1,03 | 0 | 1,5 | 0 |
| NFAT5 | TATTTTTATT | -1 | 578 | 588 | -422 | -412 | 2,91 | 0,01 | 12,4 | >0.1 | 3,33 | 5,15 | 0 | 1 | 0 | 0 | 0 |
| NFAT5 | TATTCCCAGA | -1 | 559 | 569 | -441 | -431 | 4,23 | 0,01 | 13,55 | >0.1 | 3,23 | 5,1 | 0 | 0,99 | 0 | 0 | 0 |
| NFAT5 | CATTTTCCCT | -1 | 531 | 541 | -469 | -459 | 3,87 | 0,01 | 14,59 | >0.1 | 3,09 | 5,03 | 0 | 0,98 | 0 | 1,61 | 0 |
| NFAT5 | ATTTTCCCTT | -1 | 530 | 540 | -470 | -460 | 10,43 | 0,001 | 23,23 | >0.1 | 3,09 | 5,03 | 0 | 0,98 | 0 | 3,7 | 0 |
| NFAT5 | TTTTCCCTTC | -1 | 529 | 539 | -471 | -461 | 5,85 | 0,01 | 18,49 | >0.1 | 3,08 | 5,03 | 0 | 0,98 | 0 | 3,55 | 0 |
| NFAT5 | ATTTCTGATC | -1 | 481 | 491 | -519 | -509 | 2,76 | 0,01 | 17,2 | >0.1 | 2,89 | 4,92 | 0 | 0,99 | 0 | 5,65 | 0 |
| NFAT5 | ATTTCTGATC | -1 | 166 | 176 | -834 | -824 | 2,76 | 0,01 | 14,74 | >0.1 | 3,2 | 4,42 | 0 | 1,9 | 0 | 2,46 | 0 |
| NFAT5 | AATTTCCATG | -1 | 150 | 160 | -850 | -840 | 11,48 | 0,001 | 22,71 | >0.1 | 3,28 | 4,41 | 0 | 1,9 | 0 | 1,65 | 0 |
| NFAT5 | ATTTTTTAAA | -1 | 137 | 147 | -863 | -853 | 3,2 | 0,01 | 14,49 | >0.1 | 3,36 | 4,39 | 0 | 1,89 | 0 | 1,65 | 0 |
| NFAT5 | CATTTCCTAT | -1 | 61 | 71 | -939 | -929 | 5,61 | 0,01 | 15,78 | >0.1 | 3,98 | 4,31 | 0 | 1,88 | 0 | 0 | 0 |
| NFAT5 | ATTTCCTATT | -1 | 60 | 70 | -940 | -930 | 6,73 | 0,01 | 16,91 | >0.1 | 4 | 4,31 | 0 | 1,87 | 0 | 0 | 0 |
| NFAT5 | ATTTCTCACA | -1 | 53 | 63 | -947 | -937 | 5,82 | 0,01 | 16,08 | >0.1 | 4,09 | 4,3 | 0 | 1,87 | 0 | 0 | 0 |
| BARHL2 | GGTAAGTGAA | 1 | 4 | 14 | -996 | -986 | 4,84 | 0,01 | 18,36 | >0.1 | 5,61 | 4,26 | 0 | 1,86 | 1,79 | 0 | 0 |
| BARHL2 | AAAAAATGGC | 1 | 140 | 150 | -860 | -850 | 5,01 | 0,01 | 16,29 | >0.1 | 3,34 | 4,39 | 0 | 1,89 | 0 | 1,65 | 0 |
| BARHL2 | GGAAAATGTT | 1 | 533 | 543 | -467 | -457 | 4,43 | 0,01 | 15,16 | >0.1 | 3,1 | 5,04 | 0 | 0,98 | 0 | 1,61 | 0 |
| BARHL2 | AATAAATGTG | 1 | 584 | 594 | -416 | -406 | 6,45 | 0,01 | 15,99 | >0.1 | 3,37 | 5,16 | 0 | 1 | 0 | 0 | 0 |
| BARHL2 | TATTAATGAA | 1 | 602 | 612 | -398 | -388 | 3,78 | 0,01 | 14,91 | >0.1 | 3,49 | 5,21 | 0 | 1,02 | 0 | 1,41 | 0 |
| BARHL2 | CTTTAACAGC | 1 | 707 | 717 | -293 | -283 | 5,03 | 0,01 | 22,17 | >0.1 | 5,61 | 5,55 | 0 | 1,17 | 0 | 4,81 | 0 |
| BARHL2 | GTTAAACTTG | -1 | 1138 | 1148 | 138 | 148 | 4,95 | 0,01 | 25,15 | >0.1 | 5,61 | 6,39 | 0 | 1,84 | 3,22 | 3,13 | 0 |
| BARHL2 | CAAAAATGTC | -1 | 1126 | 1136 | 126 | 136 | 3,91 | 0,01 | 23,83 | >0.1 | 5,61 | 6,49 | 0 | 1,84 | 3,22 | 2,75 | 0 |
| BARHL2 | TCTTATCAGC | -1 | 966 | 976 | -34 | -24 | 2,49 | 0,01 | 19,45 | >0.1 | 5,61 | 7,78 | 0 | 1,87 | 0 | 1,7 | 0 |
| BARHL2 | AGTAAACAAT | -1 | 895 | 905 | -105 | -95 | 6,27 | 0,01 | 23,38 | >0.1 | 5,61 | 6,66 | 0 | 1,89 | 0 | 2,95 | 0 |
| BARHL2 | AATAATTTTT | -1 | 829 | 839 | -171 | -161 | 2,84 | 0,01 | 19,56 | >0.1 | 5,61 | 6,15 | 0 | 1,68 | 0 | 3,28 | 0 |
| BARHL2 | GGTTAATATT | -1 | 727 | 737 | -273 | -263 | 2,51 | 0,01 | 20,28 | >0.1 | 5,61 | 5,63 | 0 | 1,22 | 0 | 5,32 | 0 |
| BARHL2 | GGAAAATGTT | -1 | 691 | 701 | -309 | -299 | 4,43 | 0,01 | 18,66 | >0.1 | 4,53 | 5,49 | 0 | 1,14 | 0 | 3,07 | 0 |
| BARHL2 | CATTAATAGG | -1 | 600 | 610 | -400 | -390 | 2,42 | 0,01 | 13,48 | >0.1 | 3,47 | 5,21 | 0 | 1,02 | 0 | 1,36 | 0 |
| BARHL2 | ATTAATAGGT | -1 | 599 | 609 | -401 | -391 | 4,28 | 0,01 | 15,33 | >0.1 | 3,47 | 5,21 | 0 | 1,02 | 0 | 1,36 | 0 |
| BARHL2 | GCTAAGTTTT | -1 | 303 | 313 | -697 | -687 | 3,13 | 0,01 | 16,6 | >0.1 | 2,67 | 4,61 | 0 | 1,2 | 0 | 4,99 | 0 |
| BARHL2 | CCAAAGTGCT | -1 | 208 | 218 | -792 | -782 | 2,73 | 0,01 | 16,49 | >0.1 | 3 | 4,48 | 0 | 1,58 | 0 | 4,7 | 0 |
| BARHL1 | GGTAAGTGAA | 1 | 4 | 14 | -996 | -986 | 6,28 | 0,01 | 19,8 | >0.1 | 5,61 | 4,26 | 0 | 1,86 | 1,79 | 0 | 0 |
| BARHL1 | AAAAAATGGC | 1 | 140 | 150 | -860 | -850 | 3,84 | 0,01 | 15,12 | >0.1 | 3,34 | 4,39 | 0 | 1,89 | 0 | 1,65 | 0 |
| BARHL1 | AAAAATGGCC | 1 | 141 | 151 | -859 | -849 | 3,76 | 0,01 | 14,99 | >0.1 | 3,33 | 4,39 | 0 | 1,89 | 0 | 1,61 | 0 |
| BARHL1 | AGAAATGGCT | 1 | 170 | 180 | -830 | -820 | 4,15 | 0,01 | 17,06 | >0.1 | 3,18 | 4,43 | 0 | 1,88 | 0 | 3,43 | 0 |
| BARHL1 | GGAAAATGTT | 1 | 533 | 543 | -467 | -457 | 4,47 | 0,01 | 15,2 | >0.1 | 3,1 | 5,04 | 0 | 0,98 | 0 | 1,61 | 0 |
| BARHL1 | AATAAATGTG | 1 | 584 | 594 | -416 | -406 | 5,62 | 0,01 | 15,16 | >0.1 | 3,37 | 5,16 | 0 | 1 | 0 | 0 | 0 |
| BARHL1 | TGAAATTGTC | 1 | 608 | 618 | -392 | -382 | 4,32 | 0,01 | 15,65 | >0.1 | 3,53 | 5,23 | 0 | 1,02 | 0 | 1,55 | 0 |
| BARHL1 | TTTAACAGCT | 1 | 708 | 718 | -292 | -282 | 3,95 | 0,01 | 21,1 | >0.1 | 5,61 | 5,55 | 0 | 1,17 | 0 | 4,81 | 0 |
| BARHL1 | GATAAGAGAG | 1 | 969 | 979 | -31 | -21 | 4,83 | 0,01 | 21,89 | >0.1 | 5,61 | 7,88 | 0 | 1,87 | 0 | 1,7 | 0 |
| BARHL1 | TGTAACTGCT | 1 | 1087 | 1097 | 87 | 97 | 5,45 | 0,01 | 23,84 | >0.1 | 5,61 | 6,92 | 0 | 1,85 | 1,79 | 2,21 | 0 |
| BARHL1 | CAAAAATGTC | -1 | 1126 | 1136 | 126 | 136 | 3,69 | 0,01 | 23,61 | >0.1 | 5,61 | 6,49 | 0 | 1,84 | 3,22 | 2,75 | 0 |
| BARHL1 | CTTTATGGCC | -1 | 757 | 767 | -243 | -233 | 3,69 | 0,01 | 22,47 | >0.1 | 5,61 | 5,75 | 0 | 1,31 | 0 | 6,1 | 0 |
| BARHL1 | GTTAATATTT | -1 | 726 | 736 | -274 | -264 | 3,62 | 0,01 | 21,39 | >0.1 | 5,61 | 5,62 | 0 | 1,22 | 0 | 5,32 | 0 |
| BARHL1 | GGAAAATGTT | -1 | 691 | 701 | -309 | -299 | 4,47 | 0,01 | 18,7 | >0.1 | 4,53 | 5,49 | 0 | 1,14 | 0 | 3,07 | 0 |
| BARHL1 | CCAAAGAGGT | -1 | 666 | 676 | -334 | -324 | 4,13 | 0,01 | 18,55 | >0.1 | 4,09 | 5,41 | 0 | 1,09 | 0 | 3,83 | 0 |
| BARHL1 | ATTAATAGGT | -1 | 599 | 609 | -401 | -391 | 7,09 | 0,001 | 18,14 | >0.1 | 3,47 | 5,21 | 0 | 1,02 | 0 | 1,36 | 0 |
| BARHL1 | GCTAAGTTTT | -1 | 303 | 313 | -697 | -687 | 4,88 | 0,01 | 18,35 | >0.1 | 2,67 | 4,61 | 0 | 1,2 | 0 | 4,99 | 0 |
| BARHL1 | TTTAATAGAG | -1 | 287 | 297 | -713 | -703 | 5,54 | 0,01 | 19,28 | >0.1 | 2,72 | 4,59 | 0 | 1,24 | 0 | 5,19 | 0 |
| BARHL1 | CCAAAGTGCT | -1 | 208 | 218 | -792 | -782 | 6,02 | 0,01 | 19,78 | >0.1 | 3 | 4,48 | 0 | 1,58 | 0 | 4,7 | 0 |
| BARHL1 | TTTAATTTCC | -1 | 153 | 163 | -847 | -837 | 5,48 | 0,01 | 16,7 | >0.1 | 3,27 | 4,41 | 0 | 1,9 | 0 | 1,65 | 0 |
| HOXC9 | AGTCATAATTTTA | 1 | 128 | 141 | -872 | -859 | -2,31 | 0,01 | 8,98 | >0.1 | 3,4 | 4,38 | 0 | 1,89 | 0 | 1,61 | 0 |
| HOXC9 | GAAATTAAAACTG | 1 | 154 | 167 | -846 | -833 | -1,18 | 0,01 | 10,99 | >0.1 | 3,25 | 4,41 | 0 | 1,9 | 0 | 2,6 | 0 |
| HOXC9 | AAATATGAATTCC | 1 | 487 | 500 | -513 | -500 | 1,05 | 0,01 | 16,16 | >0.1 | 2,91 | 4,94 | 0 | 0,99 | 0 | 6,27 | 0 |
| HOXC9 | AGTCATGTATTTG | 1 | 516 | 529 | -484 | -471 | -1,15 | 0,01 | 10,82 | >0.1 | 3,03 | 5,01 | 0 | 0,99 | 0 | 2,95 | 0 |
| HOXC9 | TGAAATAAAAATA | 1 | 575 | 588 | -425 | -412 | -0,16 | 0,01 | 9,32 | >0.1 | 3,33 | 5,15 | 0 | 1 | 0 | 0 | 0 |
| HOXC9 | AAAAATAAATGTG | 1 | 581 | 594 | -419 | -406 | 1,38 | 0,01 | 10,91 | >0.1 | 3,36 | 5,17 | 0 | 1,01 | 0 | 0 | 0 |
| HOXC9 | ACCTATTAATGAA | 1 | 599 | 612 | -401 | -388 | 0,96 | 0,01 | 12,03 | >0.1 | 3,48 | 5,21 | 0 | 1,02 | 0 | 1,36 | 0 |

| | | | | | | | | | | | | | | | | | |
|---|---|---|---|---|---|---|---|---|---|---|---|---|---|---|---|---|---|
| HOXC9 | CACTGTAAAATTA | 1 | 677 | 690 | -323 | -310 | 0,39 | 0,01 | 14,52 | >0.1 | 4,28 | 5,45 | 0 | 1,12 | 0 | 3,28 | 0 |
| HOXC9 | TTCCGTGTATCTT | 1 | 697 | 710 | -303 | -290 | -0,84 | 0,01 | 14,6 | >0.1 | 5,57 | 5,52 | 0 | 1,16 | 0 | 3,19 | 0 |
| HOXC9 | GGCCATAAAGTGA | 1 | 757 | 770 | -243 | -230 | -2,81 | 0,01 | 15,95 | >0.1 | 5,57 | 5,77 | 0 | 1,32 | 0 | 6,1 | 0 |
| HOXC9 | TGTTACATATCTG | 1 | 847 | 860 | -153 | -140 | -2,75 | 0,01 | 13,93 | >0.1 | 5,57 | 6,3 | 0 | 1,86 | 0 | 2,95 | 0 |
| HOXC9 | TGCTGTAACTGCT | 1 | 1084 | 1097 | 84 | 97 | -2,48 | 0,01 | 15,91 | >0.1 | 5,57 | 6,96 | 0 | 1,85 | 1,79 | 2,21 | 0 |
| HOXC9 | AGTAAACAATCTG | -1 | 892 | 905 | -108 | -95 | -2,72 | 0,01 | 14,26 | >0.1 | 5,57 | 6,66 | 0 | 1,89 | 0 | 2,86 | 0 |
| HOXC9 | TAAAGTGAATCAA | -1 | 816 | 829 | -184 | -171 | 3,19 | 0,01 | 20 | >0.1 | 5,57 | 6,08 | 0 | 1,61 | 0 | 3,55 | 0 |
| HOXC9 | GTGAATCAAATAA | -1 | 812 | 825 | -188 | -175 | -2,42 | 0,01 | 14,22 | >0.1 | 5,57 | 6,06 | 0 | 1,58 | 0 | 3,43 | 0 |
| HOXC9 | ATTCTTTAATCCT | -1 | 790 | 803 | -210 | -197 | -1,48 | 0,01 | 16,8 | >0.1 | 5,57 | 5,93 | 0 | 1,46 | 0 | 5,32 | 0 |
| HOXC9 | TCCTGTCACTTTA | -1 | 762 | 775 | -238 | -225 | -2,39 | 0,01 | 16,23 | >0.1 | 5,57 | 5,79 | 0 | 1,34 | 0 | 5,92 | 0 |
| VDR | TAAGTGAA | 1 | 6 | 14 | -994 | -986 | 3,47 | 0,01 | 17,02 | >0.1 | 5,64 | 4,26 | 0 | 1,86 | 1,79 | 0 | 0 |
| VDR | AGAGTTGA | 1 | 103 | 111 | -897 | -889 | 5,96 | 0,01 | 16,56 | >0.1 | 3,6 | 4,35 | 0 | 1,88 | 0 | 0,77 | 0 |
| VDR | TGGGCACA | 1 | 179 | 187 | -821 | -813 | 3,17 | 0,01 | 15,7 | >0.1 | 3,14 | 4,44 | 0 | 1,8 | 0 | 3,16 | 0 |
| VDR | CGAGTTGG | 1 | 223 | 231 | -777 | -769 | 3,75 | 0,01 | 18,2 | >0.1 | 2,95 | 4,49 | 0 | 1,49 | 0 | 5,52 | 0 |
| VDR | CAAGGTCA | 1 | 239 | 247 | -761 | -753 | 7,18 | 0,01 | 20,52 | >0.1 | 2,89 | 4,51 | 0 | 1,41 | 0 | 4,53 | 0 |
| VDR | CAAGGGCA | 1 | 507 | 515 | -493 | -485 | 4,35 | 0,01 | 18,97 | >0.1 | 2,98 | 4,98 | 0 | 0,98 | 0 | 5,68 | 0 |
| VDR | AGAGTTTC | 1 | 553 | 561 | -447 | -439 | 3,06 | 0,1 | 12,32 | >0.1 | 3,19 | 5,08 | 0 | 0,99 | 0 | 0 | 0 |
| VDR | AGAGGTAA | 1 | 774 | 782 | -226 | -218 | 3,27 | 0,01 | 22,2 | >0.1 | 5,64 | 5,82 | 0 | 1,37 | 0 | 6,1 | 0 |
| VDR | AAGGTTCT | 1 | 780 | 788 | -220 | -212 | 3,96 | 0,01 | 23,39 | >0.1 | 5,64 | 5,85 | 0 | 1,39 | 0 | 6,54 | 0 |
| VDR | CAAGTTCA | 1 | 955 | 963 | -45 | -37 | 10,35 | 0,001 | 27,02 | >0.1 | 5,64 | 7,46 | 0 | 1,88 | 0 | 1,7 | 0 |
| VDR | TCAGTCCA | 1 | 1099 | 1107 | 99 | 107 | 3,84 | 0,01 | 23,67 | >0.1 | 5,64 | 6,77 | 0 | 1,85 | 3,22 | 2,35 | 0 |
| VDR | CAAGTTTA | 1 | 1138 | 1146 | 138 | 146 | 7,76 | 0,001 | 27,92 | >0.1 | 5,64 | 6,39 | 0 | 1,84 | 3,22 | 3,07 | 0 |
| VDR | AAAGTTCA | 1 | 1173 | 1181 | 173 | 181 | 9,52 | 0,001 | 27,69 | >0.1 | 5,64 | 6,13 | 0 | 1,64 | 1,79 | 2,97 | 0 |
| VDR | CAAGTGAA | -1 | 1177 | 1185 | 177 | 185 | 3,27 | 0,01 | 21,7 | >0.1 | 5,64 | 6,11 | 0 | 1,61 | 1,79 | 3,28 | 0 |
| VDR | CGGCTTCG | -1 | 1150 | 1158 | 150 | 158 | 3,26 | 0,01 | 20,01 | >0.1 | 4,58 | 6,3 | 0 | 1,84 | 1,79 | 2,24 | 0 |
| VDR | CAAGATCA | -1 | 1027 | 1035 | 27 | 35 | 4,65 | 0,01 | 21,82 | >0.1 | 5,64 | 8,39 | 0 | 1,86 | 0 | 1,28 | 0 |
| VDR | TGGGTTGG | -1 | 949 | 957 | -51 | -43 | 3,93 | 0,01 | 20,59 | >0.1 | 5,64 | 7,34 | 0 | 1,88 | 0 | 1,8 | 0 |
| VDR | AAAGTTCA | -1 | 872 | 880 | -128 | -120 | 9,52 | 0,001 | 26,45 | >0.1 | 5,64 | 6,45 | 0 | 1,9 | 0 | 2,95 | 0 |
| VDR | AGAGGACA | -1 | 858 | 866 | -142 | -134 | 3,76 | 0,01 | 20,5 | >0.1 | 5,64 | 6,34 | 0 | 1,9 | 0 | 2,86 | 0 |
| VDR | AGGGATCA | -1 | 650 | 658 | -350 | -342 | 4,96 | 0,01 | 19,09 | >0.1 | 3,88 | 5,35 | 0 | 1,07 | 0 | 3,83 | 0 |
| VDR | CAATTTCA | -1 | 608 | 616 | -392 | -384 | 5,02 | 0,01 | 16,28 | >0.1 | 3,52 | 5,22 | 0 | 1,02 | 0 | 1,5 | 0 |
| VDR | CAAGATCA | -1 | 569 | 577 | -431 | -423 | 4,65 | 0,01 | 14,04 | >0.1 | 3,28 | 5,12 | 0 | 0,99 | 0 | 0 | 0 |
| VDR | GGAATTCA | -1 | 492 | 500 | -508 | -500 | 3,07 | 0,01 | 18,05 | >0.1 | 2,92 | 4,94 | 0 | 0,98 | 0 | 6,13 | 0 |
| VDR | TCGGCTCA | -1 | 400 | 408 | -600 | -592 | 3,28 | 0,01 | 17,61 | >0.1 | 2,62 | 4,76 | 0 | 1,03 | 0 | 5,91 | 0 |
| VDR | CAGGTTCA | -1 | 375 | 383 | -625 | -617 | 10,32 | 0,001 | 25,19 | >0.1 | 2,55 | 4,72 | 0 | 1,06 | 0 | 6,53 | 0 |
| VDR | GGGTTTCA | -1 | 276 | 284 | -724 | -716 | 3,46 | 0,01 | 17,14 | >0.1 | 2,76 | 4,57 | 0 | 1,27 | 0 | 5,09 | 0 |
| VDR | ACAGGTCA | -1 | 108 | 116 | -892 | -884 | 3,87 | 0,01 | 14,44 | >0.1 | 3,56 | 4,36 | 0 | 1,89 | 0 | 0,77 | 0 |
| VDR | AAAGCTCA | -1 | 69 | 77 | -931 | -923 | 4,75 | 0,01 | 14,85 | >0.1 | 3,91 | 4,31 | 0 | 1,88 | 0 | 0 | 0 |
| VDR | TGAGTGCC | -1 | 23 | 31 | -977 | -969 | 3,85 | 0,01 | 15,63 | >0.1 | 5,64 | 4,27 | 0 | 1,87 | 0 | 0 | 0 |
| FOXK2 | CCATAAAGACA | 1 | 498 | 509 | -502 | -491 | 3,24 | 0,01 | 18,58 | >0.1 | 2,95 | 4,96 | 0 | 0,98 | 0 | 6,44 | 0 |
| FOXK2 | AAATAAAAATA | 1 | 577 | 588 | -423 | -412 | 3,41 | 0,01 | 12,89 | >0.1 | 3,33 | 5,15 | 0 | 1 | 0 | 0 | 0 |
| FOXK2 | AAATAAATGTG | 1 | 583 | 594 | -417 | -406 | 1,81 | 0,01 | 11,35 | >0.1 | 3,37 | 5,16 | 0 | 1,01 | 0 | 0 | 0 |
| FOXK2 | ATACAAACACC | 1 | 626 | 637 | -374 | -363 | 4,83 | 0,01 | 16,37 | >0.1 | 3,67 | 5,28 | 0 | 1,04 | 0 | 1,55 | 0 |
| FOXK2 | AGGAAAATATT | 1 | 722 | 733 | -278 | -267 | 1,86 | 0,01 | 19,44 | >0.1 | 5,6 | 5,61 | 0 | 1,21 | 0 | 5,17 | 0 |
| FOXK2 | GTTTAACCACG | 1 | 1141 | 1152 | 141 | 152 | 3 | 0,01 | 23,43 | >0.1 | 5,6 | 6,37 | 0 | 1,84 | 3,22 | 3,4 | 0 |
| FOXK2 | GGTTAAACTTG | -1 | 1138 | 1149 | 138 | 149 | 2,17 | 0,01 | 22,35 | >0.1 | 5,6 | 6,39 | 0 | 1,84 | 3,22 | 3,13 | 0 |
| FOXK2 | AAGAACACAGT | -1 | 902 | 913 | -98 | -87 | 2,06 | 0,01 | 17,56 | >0.1 | 5,6 | 6,73 | 0 | 1,89 | 0 | 1,28 | 0 |
| FOXK2 | CAGTAAACAAT | -1 | 895 | 906 | -105 | -94 | 10,89 | 0,0001 | 27,99 | >0.1 | 5,6 | 6,67 | 0 | 1,89 | 0 | 2,95 | 0 |
| FOXK2 | AAAGATACACG | -1 | 700 | 711 | -300 | -289 | 2,11 | 0,01 | 17,58 | >0.1 | 5,6 | 5,53 | 0 | 1,16 | 0 | 3,19 | 0 |
| FOXK2 | ACTTAAAAAAG | -1 | 73 | 84 | -927 | -916 | 2,18 | 0,01 | 12,23 | >0.1 | 3,85 | 4,32 | 0 | 1,88 | 0 | 0 | 0 |
| FOXK1 | GTGAGAAATAGGAA | 1 | 54 | 68 | -946 | -932 | 2,11 | 0,01 | 12,34 | >0.1 | 4,05 | 4,31 | 0 | 1,87 | 0 | 0 | 0 |
| FOXK1 | GGCGTGAACCTGGG | 1 | 371 | 385 | -629 | -615 | 2,19 | 0,01 | 17,07 | >0.1 | 2,55 | 4,72 | 0 | 1,07 | 0 | 6,53 | 0 |
| FOXK1 | TCCATAAAGACAAG | 1 | 497 | 511 | -503 | -489 | 1,22 | 0,01 | 16,75 | >0.1 | 2,96 | 4,97 | 0 | 0,99 | 0 | 6,62 | 0 |
| FOXK1 | CATAAAGACAAGGG | 1 | 499 | 513 | -501 | -487 | 3,21 | 0,01 | 17,65 | >0.1 | 2,96 | 4,97 | 0 | 0,99 | 0 | 5,52 | 0 |
| FOXK1 | AATAAAAATAAATG | 1 | 578 | 592 | -422 | -408 | 1,79 | 0,01 | 11,3 | >0.1 | 3,35 | 5,16 | 0 | 1,01 | 0 | 0 | 0 |
| FOXK1 | TCTGAAAACCATAC | 1 | 616 | 630 | -384 | -370 | 2,24 | 0,01 | 13,69 | >0.1 | 3,6 | 5,26 | 0 | 1,04 | 0 | 1,55 | 0 |
| FOXK1 | CATACAAACACCAA | 1 | 625 | 639 | -375 | -361 | 2,28 | 0,01 | 13,84 | >0.1 | 3,68 | 5,29 | 0 | 1,05 | 0 | 1,55 | 0 |
| FOXK1 | TAGGAAAATATTAA | 1 | 721 | 735 | -279 | -265 | 1,89 | 0,01 | 19,45 | >0.1 | 5,56 | 5,62 | 0 | 1,22 | 0 | 5,17 | 0 |
| FOXK1 | AGTTTAACCACGAA | 1 | 1140 | 1154 | 140 | 154 | 2,17 | 0,01 | 22,54 | >0.1 | 5,56 | 6,37 | 0 | 1,84 | 3,22 | 3,37 | 0 |

| | | | | | | | | | | | | | | | | | |
|---|---|---|---|---|---|---|---|---|---|---|---|---|---|---|---|---|---|
| FOXK1 | TACAGCAACAAGGC | -1 | 1077 | 1091 | 77 | 91 | 4,23 | 0,01 | 22,71 | >0.1 | 5,56 | 7,06 | 0 | 1,85 | 1,79 | 2,21 | 0 |
| FOXK1 | GCAGGAAAGACCAA | -1 | 928 | 942 | -72 | -58 | 1,23 | 0,01 | 16,76 | >0.1 | 5,56 | 7,09 | 0 | 1,88 | 0 | 1 | 0 |
| FOXK1 | AAGAAAAAGAAGAA | -1 | 911 | 925 | -89 | -75 | 1,69 | 0,01 | 17,15 | >0.1 | 5,56 | 6,87 | 0 | 1,89 | 0 | 1,15 | 0 |
| FOXK1 | GAAGAACACAGTAA | -1 | 900 | 914 | -100 | -86 | 2,54 | 0,01 | 18,01 | >0.1 | 5,56 | 6,75 | 0 | 1,89 | 0 | 1,28 | 0 |
| FOXK1 | ACAGTAAACAATCT | -1 | 893 | 907 | -107 | -93 | 10,95 | 0,0001 | 28,03 | >0.1 | 5,56 | 6,68 | 0 | 1,89 | 0 | 2,95 | 0 |
| FOXK1 | CAGGTCAACTCTCC | -1 | 101 | 115 | -899 | -885 | 1,52 | 0,01 | 12,12 | >0.1 | 3,59 | 4,36 | 0 | 1,89 | 0 | 0,77 | 0 |
| RAX2 | GAAATTAA | 1 | 154 | 162 | -846 | -838 | 6,04 | 0,01 | 17,26 | >0.1 | 3,27 | 4,41 | 0 | 1,9 | 0 | 1,65 | 0 |
| RAX2 | CTGATCAG | 1 | 164 | 172 | -836 | -828 | 4,7 | 0,01 | 16,63 | >0.1 | 3,21 | 4,42 | 0 | 1,9 | 0 | 2,4 | 0 |
| RAX2 | TCTATTAA | 1 | 288 | 296 | -712 | -704 | 5,1 | 0,01 | 18,83 | >0.1 | 2,72 | 4,58 | 0 | 1,24 | 0 | 5,19 | 0 |
| RAX2 | CTGATCAG | 1 | 479 | 487 | -521 | -513 | 4,7 | 0,01 | 18,84 | >0.1 | 2,88 | 4,92 | 0 | 0,98 | 0 | 5,36 | 0 |
| RAX2 | CCTATTAA | 1 | 600 | 608 | -400 | -392 | 6,16 | 0,01 | 17,21 | >0.1 | 3,47 | 5,2 | 0 | 1,01 | 0 | 1,36 | 0 |
| RAX2 | TTAATGAA | 1 | 604 | 612 | -396 | -388 | 5,12 | 0,01 | 16,26 | >0.1 | 3,49 | 5,21 | 0 | 1,02 | 0 | 1,41 | 0 |
| RAX2 | AAAATTAT | 1 | 683 | 691 | -317 | -309 | 3,88 | 0,01 | 17,86 | >0.1 | 4,34 | 5,46 | 0 | 1,12 | 0 | 3,07 | 0 |
| RAX2 | AAAATTAT | 1 | 830 | 838 | -170 | -162 | 3,88 | 0,01 | 20,61 | >0.1 | 5,64 | 6,14 | 0 | 1,67 | 0 | 3,28 | 0 |
| RAX2 | CTCATGAG | 1 | 984 | 992 | -16 | -8 | 3,39 | 0,1 | 20,91 | >0.1 | 5,64 | 8,43 | 0 | 1,87 | 0 | 1,57 | 0 |
| RAX2 | GGAATTAT | 1 | 1191 | 1199 | 191 | 199 | 4,2 | 0,01 | 22,98 | >0.1 | 5,64 | 6,02 | 0 | 1,5 | 1,79 | 3,82 | 0 |
| RAX2 | ATAATTCC | -1 | 1191 | 1199 | 191 | 199 | 3,85 | 0,01 | 22,63 | >0.1 | 5,64 | 6,02 | 0 | 1,5 | 1,79 | 3,82 | 0 |
| RAX2 | GCAGTTAC | -1 | 1088 | 1096 | 88 | 96 | 3,47 | 0,01 | 21,87 | >0.1 | 5,64 | 6,9 | 0 | 1,85 | 1,79 | 2,21 | 0 |
| RAX2 | CTCATGAG | -1 | 984 | 992 | -16 | -8 | 3,39 | 0,1 | 20,91 | >0.1 | 5,64 | 8,43 | 0 | 1,87 | 0 | 1,57 | 0 |
| RAX2 | CAAATAAG | -1 | 811 | 819 | -189 | -181 | 3,42 | 0,01 | 19,91 | >0.1 | 5,64 | 6,02 | 0 | 1,55 | 0 | 3,28 | 0 |
| RAX2 | CAAATCAA | -1 | 750 | 758 | -250 | -242 | 4,5 | 0,01 | 23,05 | >0.1 | 5,64 | 5,71 | 0 | 1,28 | 0 | 5,92 | 0 |
| RAX2 | ACAATTTC | -1 | 609 | 617 | -391 | -383 | 4,57 | 0,01 | 15,89 | >0.1 | 3,53 | 5,23 | 0 | 1,02 | 0 | 1,55 | 0 |
| RAX2 | TTCATTAA | -1 | 604 | 612 | -396 | -388 | 5,4 | 0,01 | 16,54 | >0.1 | 3,49 | 5,21 | 0 | 1,02 | 0 | 1,41 | 0 |
| RAX2 | CTGATCAG | -1 | 479 | 487 | -521 | -513 | 4,7 | 0,01 | 18,84 | >0.1 | 2,88 | 4,92 | 0 | 0,98 | 0 | 5,36 | 0 |
| RAX2 | CTGATCAG | -1 | 164 | 172 | -836 | -828 | 4,7 | 0,01 | 16,63 | >0.1 | 3,21 | 4,42 | 0 | 1,9 | 0 | 2,4 | 0 |
| RAX2 | TTAATTTC | -1 | 154 | 162 | -846 | -838 | 5,04 | 0,01 | 16,26 | >0.1 | 3,27 | 4,41 | 0 | 1,9 | 0 | 1,65 | 0 |
| RAX2 | AAAATTAT | -1 | 132 | 140 | -868 | -860 | 3,88 | 0,01 | 15,12 | >0.1 | 3,4 | 4,38 | 0 | 1,89 | 0 | 1,57 | 0 |
| RAX2 | TCACTTAC | -1 | 5 | 13 | -995 | -987 | 4,58 | 0,01 | 18,13 | >0.1 | 5,64 | 4,26 | 0 | 1,86 | 1,79 | 0 | 0 |
| GLIS2 | GAAACCCCGTCTCT | 1 | 277 | 291 | -723 | -709 | -7,42 | 0,01 | 6,37 | >0.1 | 2,75 | 4,58 | 0 | 1,27 | 0 | 5,19 | 0 |
| GLIS2 | GGGCACCTGTAGTC | 1 | 326 | 340 | -674 | -660 | -7,36 | 0,01 | 6,6 | >0.1 | 2,6 | 4,65 | 0 | 1,15 | 0 | 5,56 | 0 |
| GLIS2 | GCGCCCAACCCAAG | 1 | 945 | 959 | -55 | -41 | -6,62 | 0,01 | 9,93 | >0.1 | 5,56 | 7,38 | 0 | 1,88 | 0 | 1,74 | 0 |
| GLIS2 | CGCCCAACCCAAGT | 1 | 946 | 960 | -54 | -40 | -7,5 | 0,01 | 9,14 | >0.1 | 5,56 | 7,4 | 0 | 1,88 | 0 | 1,8 | 0 |
| GLIS2 | GTCCACCATTGAGG | 1 | 1102 | 1116 | 102 | 116 | -7,64 | 0,01 | 12,25 | >0.1 | 5,56 | 6,73 | 0 | 1,85 | 3,22 | 2,53 | 0 |
| GLIS2 | CGTCCCCTGTGAGC | -1 | 1035 | 1049 | 35 | 49 | -0,1 | 0,01 | 16,68 | >0.1 | 5,56 | 8,03 | 0 | 1,86 | 0 | 1,34 | 0 |
| GLIS2 | GGCTCACTGCAAGC | -1 | 392 | 406 | -608 | -594 | -6,31 | 0,01 | 8,02 | >0.1 | 2,61 | 4,76 | 0 | 1,05 | 0 | 5,91 | 0 |
| GLIS2 | TGCCTCCCAGGTTC | -1 | 376 | 390 | -624 | -610 | 2,84 | 0,001 | 17,74 | >0.1 | 2,56 | 4,73 | 0 | 1,07 | 0 | 6,53 | 0 |
| GLIS2 | AGCCTCCCGAGTAG | -1 | 344 | 358 | -656 | -642 | 1,82 | 0,001 | 16,69 | >0.1 | 2,55 | 4,68 | 0 | 1,11 | 0 | 6,52 | 0 |
| GLIS2 | TGCCCACCACCACG | -1 | 317 | 331 | -683 | -669 | 5,24 | 0,001 | 19,23 | >0.1 | 2,63 | 4,64 | 0 | 1,17 | 0 | 5,56 | 0 |
| GLIS2 | GCCCACCACCACGC | -1 | 316 | 330 | -684 | -670 | -6,31 | 0,01 | 8,38 | >0.1 | 2,63 | 4,64 | 0 | 1,17 | 0 | 6,26 | 0 |
| GLIS2 | CCACCACCACGCCC | -1 | 314 | 328 | -686 | -672 | -7,15 | 0,01 | 7,42 | >0.1 | 2,63 | 4,63 | 0 | 1,17 | 0 | 6,13 | 0 |
| GLIS2 | ACCACCACGCCCAG | -1 | 312 | 326 | -688 | -674 | -4,22 | 0,01 | 10,36 | >0.1 | 2,64 | 4,63 | 0 | 1,18 | 0 | 6,13 | 0 |
| GLIS2 | CACGCCCAGCTAAG | -1 | 307 | 321 | -693 | -679 | 0,91 | 0,01 | 15,63 | >0.1 | 2,65 | 4,62 | 0 | 1,19 | 0 | 6,26 | 0 |
| GLIS2 | TGCCCAACTCGGCC | -1 | 220 | 234 | -780 | -766 | -0,93 | 0,01 | 13,55 | >0.1 | 2,95 | 4,5 | 0 | 1,51 | 0 | 5,52 | 0 |
| GLIS2 | GGCCTCCCAAAGTG | -1 | 210 | 224 | -790 | -776 | 2,32 | 0,001 | 15,95 | >0.1 | 2,99 | 4,48 | 0 | 1,57 | 0 | 4,59 | 0 |
| GLIS2 | CACTCCACAGGTCA | -1 | 108 | 122 | -892 | -878 | -7,31 | 0,01 | 3,24 | >0.1 | 3,54 | 4,36 | 0 | 1,89 | 0 | 0,77 | 0 |
| GLIS2 | TGCCTCCTTCTCCT | -1 | 13 | 27 | -987 | -973 | -7,63 | 0,01 | 4,06 | >0.1 | 5,56 | 4,27 | 0 | 1,87 | 0 | 0 | 0 |
| GLIS3 | GGCACTCAAAGAAG | 1 | 23 | 37 | -977 | -963 | -15,07 | 0,01 | -3,37 | >0.1 | 5,56 | 4,28 | 0 | 1,87 | 0 | 0 | 0 |
| GLIS3 | TCCTCTCCAGGATG | 1 | 860 | 874 | -140 | -126 | -8,28 | 0,01 | 8,33 | >0.1 | 5,56 | 6,4 | 0 | 1,9 | 0 | 2,75 | 0 |
| GLIS3 | TGGCTCCTTCTCAG | 1 | 1064 | 1078 | 64 | 78 | -14,07 | 0,01 | 4,69 | >0.1 | 5,56 | 7,28 | 0 | 1,85 | 1,79 | 2,27 | 0 |
| GLIS3 | AGTCCACCATTGAG | 1 | 1101 | 1115 | 101 | 115 | -14,73 | 0,01 | 5,12 | >0.1 | 5,56 | 6,75 | 0 | 1,85 | 3,22 | 2,47 | 0 |
| GLIS3 | CGTCCCCTGTGAGC | -1 | 1035 | 1049 | 35 | 49 | -11,32 | 0,01 | 5,46 | >0.1 | 5,56 | 8,03 | 0 | 1,86 | 0 | 1,34 | 0 |
| GLIS3 | AACCTCCTCATGAG | -1 | 984 | 998 | -16 | -2 | -13,15 | 0,01 | 4,51 | >0.1 | 5,56 | 8,66 | 0 | 1,87 | 0 | 1,57 | 0 |
| GLIS3 | GAGCCAATATAAAG | -1 | 876 | 890 | -124 | -110 | -7,71 | 0,01 | 9,34 | >0.1 | 5,56 | 6,53 | 0 | 1,9 | 0 | 3,07 | 0 |
| GLIS3 | TATCTCACATTTAT | -1 | 585 | 599 | -415 | -401 | -12,59 | 0,01 | -3,01 | >0.1 | 3,39 | 5,18 | 0 | 1,01 | 0 | 0 | 0 |
| GLIS3 | AACTCTCTACTTGG | -1 | 545 | 559 | -455 | -441 | -14,21 | 0,01 | -4,97 | >0.1 | 3,17 | 5,08 | 0 | 0,99 | 0 | 0 | 0 |
| GLIS3 | TCCCTTCCAAATAC | -1 | 522 | 536 | -478 | -464 | -13,46 | 0,01 | -1,11 | >0.1 | 3,06 | 5,02 | 0 | 0,99 | 0 | 3,28 | 0 |
| GLIS3 | TGTCGCCCAGGCTG | -1 | 426 | 440 | -574 | -560 | -12,76 | 0,01 | 1,59 | >0.1 | 2,71 | 4,82 | 0 | 1,02 | 0 | 5,8 | 0 |
| GLIS3 | TGCCTCCCAGGTTC | -1 | 376 | 390 | -624 | -610 | 0,26 | 0,001 | 15,16 | >0.1 | 2,56 | 4,73 | 0 | 1,07 | 0 | 6,53 | 0 |
| GLIS3 | AGCCTCCCGAGTAG | -1 | 344 | 358 | -656 | -642 | -3,43 | 0,001 | 11,44 | >0.1 | 2,55 | 4,68 | 0 | 1,11 | 0 | 6,52 | 0 |

| | | | | | | | | | | | | | | | | | |
|---|---|---|---|---|---|---|---|---|---|---|---|---|---|---|---|---|---|
| GLIS3 | TGCCCACCACCACG | -1 | 317 | 331 | -683 | -669 | 3,16 | 0,001 | 17,15 | >0.1 | 2,63 | 4,64 | 0 | 1,17 | 0 | 5,56 | 0 |
| GLIS3 | CACGCCCAGCTAAG | -1 | 307 | 321 | -693 | -679 | -4,57 | 0,01 | 10,15 | >0.1 | 2,65 | 4,62 | 0 | 1,19 | 0 | 6,26 | 0 |
| GLIS3 | TGCCCAACTCGGCC | -1 | 220 | 234 | -780 | -766 | -13,08 | 0,01 | 1,4 | >0.1 | 2,95 | 4,5 | 0 | 1,51 | 0 | 5,52 | 0 |
| GLIS3 | GGCCTCCCAAAGTG | -1 | 210 | 224 | -790 | -776 | 1,1 | 0,001 | 14,73 | >0.1 | 2,99 | 4,48 | 0 | 1,57 | 0 | 4,59 | 0 |
| GLIS3 | CACTCCACAGGTCA | -1 | 108 | 122 | -892 | -878 | -14,59 | 0,01 | -4,04 | >0.1 | 3,54 | 4,36 | 0 | 1,89 | 0 | 0,77 | 0 |
| GLIS3 | TGCCTCCTTCTCCT | -1 | 13 | 27 | -987 | -973 | -15,29 | 0,1 | -3,6 | >0.1 | 5,56 | 4,27 | 0 | 1,87 | 0 | 0 | 0 |
| GLIS1 | AGGCACTCAAAGAAGT | 1 | 22 | 38 | -978 | -962 | -28,82 | 0,01 | -17,14 | >0.1 | 5,53 | 4,28 | 0 | 1,87 | 0 | 0 | 0 |
| GLIS1 | GTAGTCCCAGCTACTC | 1 | 334 | 350 | -666 | -650 | -33,01 | 0,01 | -18,72 | >0.1 | 2,58 | 4,67 | 0 | 1,13 | 0 | 5,92 | 0 |
| GLIS1 | GATCGCGCCACTGTGC | 1 | 408 | 424 | -592 | -576 | -32,11 | 0,01 | -17,86 | >0.1 | 2,66 | 4,79 | 0 | 1,03 | 0 | 5,76 | 0 |
| GLIS1 | CGACAGACCGAGACTC | 1 | 435 | 451 | -565 | -549 | -30,39 | 0,01 | -15,75 | >0.1 | 2,74 | 4,84 | 0 | 1,01 | 0 | 6,05 | 0 |
| GLIS1 | ATGTTGCCCAAGTAGA | 1 | 538 | 554 | -462 | -446 | -36,76 | 0,01 | -27,56 | >0.1 | 3,14 | 5,07 | 0 | 0,99 | 0 | 0 | 0 |
| GLIS1 | GTCCTCTCCAGGATGA | 1 | 859 | 875 | -141 | -125 | -32,06 | 0,01 | -15,47 | >0.1 | 5,53 | 6,41 | 0 | 1,9 | 0 | 2,75 | 0 |
| GLIS1 | TCACAGGGGACGATGT | 1 | 1037 | 1053 | 37 | 53 | -35,13 | 0,01 | -18,42 | >0.1 | 5,53 | 7,95 | 0 | 1,86 | 0 | 1,36 | 0 |
| GLIS1 | AGTTTAACCACGAAGC | 1 | 1140 | 1156 | 140 | 156 | -31,82 | 0,01 | -12,74 | >0.1 | 5,53 | 6,37 | 0 | 1,84 | 3,22 | 2,11 | 0 |
| GLIS1 | TTAACCACGAAGCCGA | 1 | 1143 | 1159 | 143 | 159 | -39,16 | 0,01 | -20,05 | >0.1 | 5,53 | 6,35 | 0 | 1,84 | 3,22 | 2,16 | 0 |
| GLIS1 | TTACCTCTCCTGTCAC | -1 | 766 | 782 | -234 | -218 | -38,87 | 0,01 | -20,45 | >0.1 | 5,53 | 5,83 | 0 | 1,37 | 0 | 5,69 | 0 |
| GLIS1 | AAACTCTCTACTTGGG | -1 | 544 | 560 | -456 | -440 | -40,17 | 0,01 | -30,93 | >0.1 | 3,17 | 5,08 | 0 | 0,99 | 0 | 0 | 0 |
| GLIS1 | CTGTCGCCCAGGCTGG | -1 | 425 | 441 | -575 | -559 | -32,84 | 0,01 | -18,49 | >0.1 | 2,71 | 4,83 | 0 | 1,02 | 0 | 5,8 | 0 |
| GLIS1 | CTGCCTCCCAGGTTCA | -1 | 375 | 391 | -625 | -609 | -27,67 | 0,01 | -12,77 | >0.1 | 2,56 | 4,74 | 0 | 1,07 | 0 | 6,53 | 0 |
| GLIS1 | CAGCCTCCCGAGTAGC | -1 | 343 | 359 | -657 | -641 | -28,18 | 0,01 | -13,31 | >0.1 | 2,55 | 4,68 | 0 | 1,12 | 0 | 6,52 | 0 |
| GLIS1 | GTGCCCACCACCACGC | -1 | 316 | 332 | -684 | -668 | -14,9 | 0,001 | -0,9 | >0.1 | 2,63 | 4,64 | 0 | 1,17 | 0 | 5,56 | 0 |
| GLIS1 | CCACGCCCAGCTAAGT | -1 | 306 | 322 | -694 | -678 | -14,14 | 0,001 | 0,59 | >0.1 | 2,65 | 4,62 | 0 | 1,19 | 0 | 6,26 | 0 |
| GLIS1 | CTGCCCAACTCGGCCT | -1 | 219 | 235 | -781 | -765 | -40,09 | 0,01 | -25,61 | >0.1 | 2,95 | 4,5 | 0 | 1,51 | 0 | 5,52 | 0 |
| GLIS1 | CGGCCTCCCAAAGTGC | -1 | 209 | 225 | -791 | -775 | -27,84 | 0,01 | -14,2 | >0.1 | 2,99 | 4,49 | 0 | 1,57 | 0 | 4,59 | 0 |
| GLIS1 | CTACTCTCCACTCCAC | -1 | 114 | 130 | -886 | -870 | -26,08 | 0,01 | -15,59 | >0.1 | 3,49 | 4,37 | 0 | 1,89 | 0 | 0,75 | 0 |
| GLIS1 | CCACTCCACAGGTCAA | -1 | 107 | 123 | -893 | -877 | -33,8 | 0,01 | -23,25 | >0.1 | 3,54 | 4,36 | 0 | 1,89 | 0 | 0,77 | 0 |
| OTX1 | CTGATCAG | 1 | 164 | 172 | -836 | -828 | 0,75 | 0,01 | 12,68 | >0.1 | 3,21 | 4,42 | 0 | 1,9 | 0 | 2,4 | 0 |
| OTX1 | GTAATCCT | 1 | 200 | 208 | -800 | -792 | 9,5 | 0,001 | 23,11 | >0.1 | 3,04 | 4,46 | 0 | 1,63 | 0 | 4,48 | 0 |
| OTX1 | CTTAGCTG | 1 | 307 | 315 | -693 | -685 | 1,05 | 0,01 | 14,6 | >0.1 | 2,66 | 4,61 | 0 | 1,19 | 0 | 5,09 | 0 |
| OTX1 | GTGAGCCG | 1 | 399 | 407 | -601 | -593 | 2,72 | 0,01 | 17,05 | >0.1 | 2,62 | 4,76 | 0 | 1,04 | 0 | 5,91 | 0 |
| OTX1 | CTGATCAG | 1 | 479 | 487 | -521 | -513 | 0,75 | 0,01 | 14,89 | >0.1 | 2,88 | 4,92 | 0 | 0,98 | 0 | 5,36 | 0 |
| OTX1 | ATTATCTT | 1 | 640 | 648 | -360 | -352 | 0,96 | 0,01 | 12,77 | >0.1 | 3,79 | 5,32 | 0 | 1,05 | 0 | 1,66 | 0 |
| OTX1 | ATGATCCC | 1 | 649 | 657 | -351 | -343 | 4,52 | 0,01 | 18,63 | >0.1 | 3,87 | 5,34 | 0 | 1,06 | 0 | 3,83 | 0 |
| OTX1 | AAAATCTG | 1 | 841 | 849 | -159 | -151 | 3,06 | 0,01 | 19,87 | >0.1 | 5,64 | 6,22 | 0 | 1,76 | 0 | 3,19 | 0 |
| OTX1 | AAAATCTC | 1 | 979 | 987 | -21 | -13 | 2,28 | 0,01 | 19,5 | >0.1 | 5,64 | 8,18 | 0 | 1,87 | 0 | 1,53 | 0 |
| OTX1 | ATAATTCC | -1 | 1191 | 1199 | 191 | 199 | 4,1 | 0,01 | 22,88 | >0.1 | 5,64 | 6,02 | 0 | 1,5 | 1,79 | 3,82 | 0 |
| OTX1 | TTAAACTT | -1 | 1139 | 1147 | 139 | 147 | 0,56 | 0,01 | 20,78 | >0.1 | 5,64 | 6,38 | 0 | 1,84 | 3,22 | 3,13 | 0 |
| OTX1 | CTTATCAG | -1 | 967 | 975 | -33 | -25 | 1,36 | 0,01 | 18,32 | >0.1 | 5,64 | 7,76 | 0 | 1,87 | 0 | 1,7 | 0 |
| OTX1 | ACAATCTG | -1 | 892 | 900 | -108 | -100 | 0,51 | 0,01 | 17,51 | >0.1 | 5,64 | 6,61 | 0 | 1,89 | 0 | 2,86 | 0 |
| OTX1 | CTGAGCCA | -1 | 884 | 892 | -116 | -108 | 0,75 | 0,01 | 17,9 | >0.1 | 5,64 | 6,55 | 0 | 1,89 | 0 | 3,07 | 0 |
| OTX1 | TTAATCCT | -1 | 790 | 798 | -210 | -202 | 10,44 | 0,001 | 28,94 | >0.1 | 5,64 | 5,91 | 0 | 1,43 | 0 | 5,52 | 0 |
| OTX1 | CAAAACCG | -1 | 745 | 753 | -255 | -247 | 0,61 | 0,01 | 19,12 | >0.1 | 5,64 | 5,69 | 0 | 1,26 | 0 | 5,92 | 0 |
| OTX1 | TTAATATT | -1 | 727 | 735 | -273 | -265 | 1,47 | 0,01 | 19,26 | >0.1 | 5,64 | 5,62 | 0 | 1,21 | 0 | 5,32 | 0 |
| OTX1 | GAAAGCTG | -1 | 713 | 721 | -287 | -279 | 0,48 | 0,01 | 17,85 | >0.1 | 5,64 | 5,56 | 0 | 1,18 | 0 | 4,99 | 0 |
| OTX1 | ATAATGTT | -1 | 637 | 645 | -363 | -355 | 1,24 | 0,01 | 13,01 | >0.1 | 3,76 | 5,31 | 0 | 1,05 | 0 | 1,66 | 0 |
| OTX1 | TTAATAGG | -1 | 600 | 608 | -400 | -392 | 4,98 | 0,01 | 16,03 | >0.1 | 3,47 | 5,2 | 0 | 1,01 | 0 | 1,36 | 0 |
| OTX1 | GTTATCTC | -1 | 593 | 601 | -407 | -399 | 1,64 | 0,01 | 11,25 | >0.1 | 3,42 | 5,18 | 0 | 1,01 | 0 | 0 | 0 |
| OTX1 | CTGATCAG | -1 | 479 | 487 | -521 | -513 | 0,75 | 0,01 | 14,89 | >0.1 | 2,88 | 4,92 | 0 | 0,98 | 0 | 5,36 | 0 |
| OTX1 | TTAATAGA | -1 | 288 | 296 | -712 | -704 | 1,98 | 0,01 | 15,71 | >0.1 | 2,72 | 4,58 | 0 | 1,24 | 0 | 5,19 | 0 |
| OTX1 | GTGATCTG | -1 | 232 | 240 | -768 | -760 | 1,82 | 0,01 | 15,69 | >0.1 | 2,91 | 4,51 | 0 | 1,44 | 0 | 5,01 | 0 |
| OTX1 | CTGATCAG | -1 | 164 | 172 | -836 | -828 | 0,75 | 0,01 | 12,68 | >0.1 | 3,21 | 4,42 | 0 | 1,9 | 0 | 2,4 | 0 |
| OTX1 | TTAATTTC | -1 | 154 | 162 | -846 | -838 | 1,56 | 0,01 | 12,78 | >0.1 | 3,27 | 4,41 | 0 | 1,9 | 0 | 1,65 | 0 |
| OTX1 | CTGATCTG | -1 | 94 | 102 | -906 | -898 | 2,85 | 0,01 | 13,55 | >0.1 | 3,67 | 4,34 | 0 | 1,88 | 0 | 0,81 | 0 |
| FOSL1::JUN | GAAGGAGGCACTC | 1 | 17 | 30 | -983 | -970 | -1,37 | 0,01 | 10,34 | >0.1 | 5,57 | 4,27 | 0 | 1,87 | 0 | 0 | 0 |
| FOSL1::JUN | GTGTGAGAAATAG | 1 | 52 | 65 | -948 | -935 | 3,39 | 0,01 | 13,65 | >0.1 | 4,08 | 4,3 | 0 | 1,87 | 0 | 0 | 0 |
| FOSL1::JUN | CAGTGGCTCACGC | 1 | 185 | 198 | -815 | -802 | 5,08 | 0,01 | 18,23 | >0.1 | 3,1 | 4,45 | 0 | 1,75 | 0 | 3,86 | 0 |
| FOSL1::JUN | TCCTGACTAACAC | 1 | 261 | 274 | -739 | -726 | 9,26 | 0,001 | 22,47 | >0.1 | 2,8 | 4,55 | 0 | 1,33 | 0 | 4,53 | 0 |
| FOSL1::JUN | GGCTGAGGCAGGA | 1 | 354 | 367 | -646 | -633 | 5,74 | 0,001 | 20,09 | >0.1 | 2,53 | 4,69 | 0 | 1,1 | 0 | 6,03 | 0 |
| FOSL1::JUN | CCGAGACTCAGTC | 1 | 442 | 455 | -558 | -545 | 2,92 | 0,01 | 17,45 | >0.1 | 2,76 | 4,85 | 0 | 1 | 0 | 5,92 | 0 |

| | | | | | | | | | | | | | | | | | |
|---|---|---|---|---|---|---|---|---|---|---|---|---|---|---|---|---|---|
| FOSL1::JUN | ATTTGATTCACTT | 1 | 814 | 827 | -186 | -173 | 4,71 | 0,01 | 21,38 | >0.1 | 5,57 | 6,07 | 0 | 1,6 | 0 | 3,43 | 0 |
| FOSL1::JUN | TATTGGCTCAGCA | 1 | 881 | 894 | -119 | -106 | 0,14 | 0,01 | 17,24 | >0.1 | 5,57 | 6,57 | 0 | 1,89 | 0 | 3,07 | 0 |
| FOSL1::JUN | AAGTCATTCAGTG | 1 | 1010 | 1023 | 10 | 23 | -1,14 | 0,01 | 16,68 | >0.1 | 5,57 | 8,99 | 0 | 1,86 | 0 | 1,39 | 0 |
| FOSL1::JUN | TCTTGGCTCACAG | 1 | 1030 | 1043 | 30 | 43 | 1,28 | 0,01 | 18,25 | >0.1 | 5,57 | 8,24 | 0 | 1,86 | 0 | 1,31 | 0 |
| FOSL1::JUN | CATTGAGGAACAG | 1 | 1108 | 1121 | 108 | 121 | 0,5 | 0,01 | 20,33 | >0.1 | 5,57 | 6,67 | 0 | 1,85 | 3,22 | 2,52 | 0 |
| FOSL1::JUN | CTGTGAGCCAAGA | -1 | 1030 | 1043 | 30 | 43 | -1,07 | 0,01 | 15,9 | >0.1 | 5,57 | 8,24 | 0 | 1,86 | 0 | 1,31 | 0 |
| FOSL1::JUN | GAATGACTTTCCC | -1 | 1006 | 1019 | 6 | 19 | -1,14 | 0,01 | 16,88 | >0.1 | 5,57 | 9,12 | 0 | 1,87 | 0 | 1,46 | 0 |
| FOSL1::JUN | AAGTGAATCAAAT | -1 | 814 | 827 | -186 | -173 | 3,81 | 0,01 | 20,48 | >0.1 | 5,57 | 6,07 | 0 | 1,6 | 0 | 3,43 | 0 |
| FOSL1::JUN | CAGTGACCAAAGA | -1 | 669 | 682 | -331 | -318 | -0,7 | 0,01 | 13,68 | >0.1 | 4,15 | 5,43 | 0 | 1,1 | 0 | 3,7 | 0 |
| FOSL1::JUN | GACTGAGTCTCGG | -1 | 442 | 455 | -558 | -545 | -0,39 | 0,01 | 14,14 | >0.1 | 2,76 | 4,85 | 0 | 1 | 0 | 5,92 | 0 |
| FOSL1::JUN | TCCTGCCTCAGCC | -1 | 354 | 367 | -646 | -633 | 3,34 | 0,01 | 17,69 | >0.1 | 2,53 | 4,69 | 0 | 1,1 | 0 | 6,03 | 0 |
| FOSL1::JUN | GTGTTAGTCAGGA | -1 | 261 | 274 | -739 | -726 | 3,44 | 0,01 | 16,65 | >0.1 | 2,8 | 4,55 | 0 | 1,33 | 0 | 4,53 | 0 |
| FOSL1::JUN | GCGTGAGCCACTG | -1 | 185 | 198 | -815 | -802 | 1,34 | 0,01 | 14,49 | >0.1 | 3,1 | 4,45 | 0 | 1,75 | 0 | 3,86 | 0 |
| FOSL1::JUN | TTATGACTACTCT | -1 | 123 | 136 | -877 | -864 | 4,75 | 0,01 | 16,15 | >0.1 | 3,44 | 4,38 | 0 | 1,89 | 0 | 1,7 | 0 |
| FOSL1::JUN | CTATTTCTCACAC | -1 | 52 | 65 | -948 | -935 | 1,6 | 0,01 | 11,86 | >0.1 | 4,08 | 4,3 | 0 | 1,87 | 0 | 0 | 0 |
| FOSL1::JUN | GAGTGCCTCCTTC | -1 | 17 | 30 | -983 | -970 | -1,64 | 0,01 | 10,07 | >0.1 | 5,57 | 4,27 | 0 | 1,87 | 0 | 0 | 0 |
| OTX2 | TTAAGTTT | 1 | 78 | 86 | -922 | -914 | 1,28 | 0,01 | 11,3 | >0.1 | 3,81 | 4,32 | 0 | 1,88 | 0 | 0 | 0 |
| OTX2 | ATAATTTT | 1 | 132 | 140 | -868 | -860 | 3,28 | 0,01 | 14,52 | >0.1 | 3,4 | 4,38 | 0 | 1,89 | 0 | 1,57 | 0 |
| OTX2 | GTAATCCT | 1 | 200 | 208 | -800 | -792 | 9,73 | 0,001 | 23,34 | >0.1 | 3,04 | 4,46 | 0 | 1,63 | 0 | 4,48 | 0 |
| OTX2 | CTTAGCTG | 1 | 307 | 315 | -693 | -685 | 1,77 | 0,01 | 15,32 | >0.1 | 2,66 | 4,61 | 0 | 1,19 | 0 | 5,09 | 0 |
| OTX2 | ATGATCTT | 1 | 568 | 576 | -432 | -424 | 2,51 | 0,01 | 11,89 | >0.1 | 3,27 | 5,12 | 0 | 0,99 | 0 | 0 | 0 |
| OTX2 | ATTATCTT | 1 | 640 | 648 | -360 | -352 | 3,89 | 0,01 | 15,7 | >0.1 | 3,79 | 5,32 | 0 | 1,05 | 0 | 1,66 | 0 |
| OTX2 | ATGATCCC | 1 | 649 | 657 | -351 | -343 | 4,91 | 0,01 | 19,02 | >0.1 | 3,87 | 5,34 | 0 | 1,06 | 0 | 3,83 | 0 |
| OTX2 | AAAATCTG | 1 | 841 | 849 | -159 | -151 | 3,72 | 0,01 | 20,53 | >0.1 | 5,64 | 6,22 | 0 | 1,76 | 0 | 3,19 | 0 |
| OTX2 | AAAATCTC | 1 | 979 | 987 | -21 | -13 | 3,6 | 0,01 | 20,82 | >0.1 | 5,64 | 8,18 | 0 | 1,87 | 0 | 1,53 | 0 |
| OTX2 | GTGATCTT | 1 | 1026 | 1034 | 26 | 34 | 1,7 | 0,01 | 18,9 | >0.1 | 5,64 | 8,45 | 0 | 1,86 | 0 | 1,25 | 0 |
| OTX2 | ATAATTCC | -1 | 1191 | 1199 | 191 | 199 | 5,69 | 0,01 | 24,47 | >0.1 | 5,64 | 6,02 | 0 | 1,5 | 1,79 | 3,82 | 0 |
| OTX2 | TTAAACTT | -1 | 1139 | 1147 | 139 | 147 | 3,12 | 0,01 | 23,34 | >0.1 | 5,64 | 6,38 | 0 | 1,84 | 3,22 | 3,13 | 0 |
| OTX2 | CTTATCAG | -1 | 967 | 975 | -33 | -25 | 1,92 | 0,01 | 18,88 | >0.1 | 5,64 | 7,76 | 0 | 1,87 | 0 | 1,7 | 0 |
| OTX2 | ACAATCTG | -1 | 892 | 900 | -108 | -100 | 1,38 | 0,01 | 18,38 | >0.1 | 5,64 | 6,61 | 0 | 1,89 | 0 | 2,86 | 0 |
| OTX2 | TTCATCCT | -1 | 868 | 876 | -132 | -124 | 2,76 | 0,01 | 19,46 | >0.1 | 5,64 | 6,41 | 0 | 1,9 | 0 | 2,75 | 0 |
| OTX2 | ATAATTTT | -1 | 830 | 838 | -170 | -162 | 3,28 | 0,01 | 20,01 | >0.1 | 5,64 | 6,14 | 0 | 1,67 | 0 | 3,28 | 0 |
| OTX2 | TTAATCCT | -1 | 790 | 798 | -210 | -202 | 11,6 | 0,0001 | 30,1 | >0.1 | 5,64 | 5,91 | 0 | 1,43 | 0 | 5,52 | 0 |
| OTX2 | TTAATATT | -1 | 727 | 735 | -273 | -265 | 4,14 | 0,01 | 21,93 | >0.1 | 5,64 | 5,62 | 0 | 1,21 | 0 | 5,32 | 0 |
| OTX2 | ATAATTTT | -1 | 683 | 691 | -317 | -309 | 3,28 | 0,01 | 17,26 | >0.1 | 4,34 | 5,46 | 0 | 1,12 | 0 | 3,07 | 0 |
| OTX2 | ATAATGTT | -1 | 637 | 645 | -363 | -355 | 2,97 | 0,01 | 14,74 | >0.1 | 3,76 | 5,31 | 0 | 1,05 | 0 | 1,66 | 0 |
| OTX2 | TTAATAGG | -1 | 600 | 608 | -400 | -392 | 2,56 | 0,01 | 13,61 | >0.1 | 3,47 | 5,2 | 0 | 1,01 | 0 | 1,36 | 0 |
| OTX2 | GTTATCTC | -1 | 593 | 601 | -407 | -399 | 3,25 | 0,01 | 12,86 | >0.1 | 3,42 | 5,18 | 0 | 1,01 | 0 | 0 | 0 |
| OTX2 | GTGATCTG | -1 | 232 | 240 | -768 | -760 | 1,98 | 0,01 | 15,85 | >0.1 | 2,91 | 4,51 | 0 | 1,44 | 0 | 5,01 | 0 |
| OTX2 | TTAATTTC | -1 | 154 | 162 | -846 | -838 | 4,51 | 0,01 | 15,73 | >0.1 | 3,27 | 4,41 | 0 | 1,9 | 0 | 1,65 | 0 |
| OTX2 | CTGATCTG | -1 | 94 | 102 | -906 | -898 | 3,46 | 0,01 | 14,16 | >0.1 | 3,67 | 4,34 | 0 | 1,88 | 0 | 0,81 | 0 |
| JUN::JUNB | AAGGAGGCAC | 1 | 18 | 28 | -982 | -972 | 5,45 | 0,01 | 17,19 | >0.1 | 5,61 | 4,27 | 0 | 1,87 | 0 | 0 | 0 |
| JUN::JUNB | TGTGAGAAAT | 1 | 53 | 63 | -947 | -937 | 3,66 | 0,01 | 13,92 | >0.1 | 4,09 | 4,3 | 0 | 1,87 | 0 | 0 | 0 |
| JUN::JUNB | GAGTAGTCAT | 1 | 124 | 134 | -876 | -866 | 2,87 | 0,01 | 14,27 | >0.1 | 3,44 | 4,38 | 0 | 1,89 | 0 | 1,7 | 0 |
| JUN::JUNB | AGTGGCTCAC | 1 | 186 | 196 | -814 | -804 | 6,97 | 0,001 | 20,11 | >0.1 | 3,1 | 4,45 | 0 | 1,74 | 0 | 3,86 | 0 |
| JUN::JUNB | GCTCACGCCT | 1 | 190 | 200 | -810 | -800 | 2,66 | 0,01 | 16,03 | >0.1 | 3,08 | 4,45 | 0 | 1,7 | 0 | 4,13 | 0 |
| JUN::JUNB | CCTGACTAAC | 1 | 262 | 272 | -738 | -728 | 8,24 | 0,001 | 21,44 | >0.1 | 2,8 | 4,55 | 0 | 1,32 | 0 | 4,53 | 0 |
| JUN::JUNB | GCTGAGGCAG | 1 | 355 | 365 | -645 | -635 | 7,57 | 0,001 | 21,92 | >0.1 | 2,53 | 4,69 | 0 | 1,09 | 0 | 6,03 | 0 |
| JUN::JUNB | TGGGAGGCAG | 1 | 381 | 391 | -619 | -609 | 2,63 | 0,01 | 17,01 | >0.1 | 2,57 | 4,73 | 0 | 1,06 | 0 | 6,02 | 0 |
| JUN::JUNB | CGAGACTCAG | 1 | 443 | 453 | -557 | -547 | 3,01 | 0,01 | 17,54 | >0.1 | 2,75 | 4,85 | 0 | 1 | 0 | 5,92 | 0 |
| JUN::JUNB | TGTGAGATAA | 1 | 590 | 600 | -410 | -400 | 1,85 | 0,01 | 11,45 | >0.1 | 3,41 | 5,18 | 0 | 1,01 | 0 | 0 | 0 |
| JUN::JUNB | AATAACGTAT | 1 | 800 | 810 | -200 | -190 | 3,08 | 0,01 | 19,7 | >0.1 | 5,61 | 5,97 | 0 | 1,5 | 0 | 3,55 | 0 |
| JUN::JUNB | TTTGATTCAC | 1 | 815 | 825 | -185 | -175 | 4,35 | 0,01 | 21,03 | >0.1 | 5,61 | 6,06 | 0 | 1,58 | 0 | 3,43 | 0 |
| JUN::JUNB | ATTGGCTCAG | 1 | 882 | 892 | -118 | -108 | 2,6 | 0,01 | 19,72 | >0.1 | 5,61 | 6,55 | 0 | 1,89 | 0 | 3,07 | 0 |
| JUN::JUNB | CTTGGCTCAC | 1 | 1031 | 1041 | 31 | 41 | 4,39 | 0,01 | 21,36 | >0.1 | 5,61 | 8,19 | 0 | 1,86 | 0 | 1,31 | 0 |
| JUN::JUNB | CCTGGCTCCT | 1 | 1062 | 1072 | 62 | 72 | 2,05 | 0,01 | 20,84 | >0.1 | 5,61 | 7,31 | 0 | 1,86 | 1,79 | 2,21 | 0 |
| JUN::JUNB | ATTGAGGAAC | 1 | 1109 | 1119 | 109 | 119 | 3,47 | 0,01 | 23,33 | >0.1 | 5,61 | 6,66 | 0 | 1,85 | 3,22 | 2,52 | 0 |
| JUN::JUNB | TGTGAGCCAA | -1 | 1032 | 1042 | 32 | 42 | 4,11 | 0,01 | 21 | >0.1 | 5,61 | 8,15 | 0 | 1,86 | 0 | 1,28 | 0 |
| JUN::JUNB | AATGACTTTC | -1 | 1008 | 1018 | 8 | 18 | 4,64 | 0,01 | 22,6 | >0.1 | 5,61 | 9,07 | 0 | 1,87 | 0 | 1,42 | 0 |

| | | | | | | | | | | | | | | | | |
|---|---|---|---|---|---|---|---|---|---|---|---|---|---|---|---|---|
| JUN::JUNB | GCTGAGCCAA | -1 | 883 | 893 | -117 | -107 | 4,75 | 0,01 | 21,88 | >0.1 | 5,61 | 6,56 | 0 | 1,89 | 0 | 3,07 | 0 |
| JUN::JUNB | AGTGAATCAA | -1 | 816 | 826 | -184 | -174 | 5,22 | 0,01 | 22,03 | >0.1 | 5,61 | 6,06 | 0 | 1,59 | 0 | 3,55 | 0 |
| JUN::JUNB | AGTGACCAAA | -1 | 671 | 681 | -329 | -319 | 3,96 | 0,01 | 18,19 | >0.1 | 4,16 | 5,42 | 0 | 1,1 | 0 | 3,55 | 0 |
| JUN::JUNB | CATGACTTTG | -1 | 513 | 523 | -487 | -477 | 2,52 | 0,01 | 14,37 | >0.1 | 3,01 | 4,99 | 0 | 0,98 | 0 | 2,86 | 0 |
| JUN::JUNB | ACTGAGTCTC | -1 | 444 | 454 | -556 | -546 | 3,1 | 0,01 | 17,63 | >0.1 | 2,76 | 4,85 | 0 | 1 | 0 | 5,92 | 0 |
| JUN::JUNB | GCTGGAGCAC | -1 | 420 | 430 | -580 | -570 | 2,37 | 0,01 | 16,29 | >0.1 | 2,68 | 4,8 | 0 | 1,02 | 0 | 5,42 | 0 |
| JUN::JUNB | CCTGCCTCAG | -1 | 356 | 366 | -644 | -634 | 2,88 | 0,01 | 17,32 | >0.1 | 2,52 | 4,69 | 0 | 1,09 | 0 | 6,13 | 0 |
| JUN::JUNB | TGTTAGTCAG | -1 | 263 | 273 | -737 | -727 | 2,32 | 0,01 | 15,43 | >0.1 | 2,8 | 4,55 | 0 | 1,32 | 0 | 4,45 | 0 |
| JUN::JUNB | CGTGAGCCAC | -1 | 187 | 197 | -813 | -803 | 4,47 | 0,01 | 17,6 | >0.1 | 3,1 | 4,45 | 0 | 1,73 | 0 | 3,86 | 0 |
| JUN::JUNB | TATGACTACT | -1 | 125 | 135 | -875 | -865 | 6,24 | 0,01 | 17,64 | >0.1 | 3,43 | 4,38 | 0 | 1,89 | 0 | 1,7 | 0 |
| JUN::JUNB | TATTTCTCAC | -1 | 54 | 64 | -946 | -936 | 2,65 | 0,01 | 12,9 | >0.1 | 4,07 | 4,3 | 0 | 1,87 | 0 | 0 | 0 |
| JUN::JUNB | GCTGACTTCT | -1 | 32 | 42 | -968 | -958 | 5,26 | 0,01 | 15,85 | >0.1 | 4,44 | 4,28 | 0 | 1,87 | 0 | 0 | 0 |
| SHOX | GTAAGTGA | 1 | 5 | 13 | -995 | -987 | 4,1 | 0,01 | 17,65 | >0.1 | 5,64 | 4,26 | 0 | 1,86 | 1,79 | 0 | 0 |
| SHOX | ATAATTTT | 1 | 132 | 140 | -868 | -860 | 4,76 | 0,01 | 16 | >0.1 | 3,4 | 4,38 | 0 | 1,89 | 0 | 1,57 | 0 |
| SHOX | GAAATTAA | 1 | 154 | 162 | -846 | -838 | 4,62 | 0,01 | 15,84 | >0.1 | 3,27 | 4,41 | 0 | 1,9 | 0 | 1,65 | 0 |
| SHOX | TTAATGAA | 1 | 604 | 612 | -396 | -388 | 6,15 | 0,01 | 17,29 | >0.1 | 3,49 | 5,21 | 0 | 1,02 | 0 | 1,41 | 0 |
| SHOX | GAAATTGT | 1 | 609 | 617 | -391 | -383 | 3,97 | 0,01 | 15,29 | >0.1 | 3,53 | 5,23 | 0 | 1,02 | 0 | 1,55 | 0 |
| SHOX | TTGATTTG | 1 | 750 | 758 | -250 | -242 | 4,31 | 0,01 | 22,86 | >0.1 | 5,64 | 5,71 | 0 | 1,28 | 0 | 5,92 | 0 |
| SHOX | CTTATTTG | 1 | 811 | 819 | -189 | -181 | 3,78 | 0,01 | 20,27 | >0.1 | 5,64 | 6,02 | 0 | 1,55 | 0 | 3,28 | 0 |
| SHOX | CTGATAAG | 1 | 967 | 975 | -33 | -25 | 3,5 | 0,01 | 20,46 | >0.1 | 5,64 | 7,76 | 0 | 1,87 | 0 | 1,7 | 0 |
| SHOX | CTCATGAG | 1 | 984 | 992 | -16 | -8 | 3,58 | 0,01 | 21,1 | >0.1 | 5,64 | 8,43 | 0 | 1,87 | 0 | 1,57 | 0 |
| SHOX | ATAATTCC | -1 | 1191 | 1199 | 191 | 199 | 5,04 | 0,01 | 23,82 | >0.1 | 5,64 | 6,02 | 0 | 1,5 | 1,79 | 3,82 | 0 |
| SHOX | CTCATGAG | -1 | 984 | 992 | -16 | -8 | 3,58 | 0,01 | 21,1 | >0.1 | 5,64 | 8,43 | 0 | 1,87 | 0 | 1,57 | 0 |
| SHOX | ATAATTTT | -1 | 830 | 838 | -170 | -162 | 4,76 | 0,01 | 21,49 | >0.1 | 5,64 | 6,14 | 0 | 1,67 | 0 | 3,28 | 0 |
| SHOX | CAAATAAG | -1 | 811 | 819 | -189 | -181 | 3,37 | 0,01 | 19,86 | >0.1 | 5,64 | 6,02 | 0 | 1,55 | 0 | 3,28 | 0 |
| SHOX | ATAATTTT | -1 | 683 | 691 | -317 | -309 | 4,76 | 0,01 | 18,74 | >0.1 | 4,34 | 5,46 | 0 | 1,12 | 0 | 3,07 | 0 |
| SHOX | TTCATTAA | -1 | 604 | 612 | -396 | -388 | 5,19 | 0,01 | 16,33 | >0.1 | 3,49 | 5,21 | 0 | 1,02 | 0 | 1,41 | 0 |
| SHOX | TTAATAGG | -1 | 600 | 608 | -400 | -392 | 7,22 | 0,001 | 18,27 | >0.1 | 3,47 | 5,2 | 0 | 1,01 | 0 | 1,36 | 0 |
| SHOX | TTAATAGA | -1 | 288 | 296 | -712 | -704 | 6,4 | 0,01 | 20,13 | >0.1 | 2,72 | 4,58 | 0 | 1,24 | 0 | 5,19 | 0 |
| SHOX | TTAATTTC | -1 | 154 | 162 | -846 | -838 | 6,75 | 0,01 | 17,97 | >0.1 | 3,27 | 4,41 | 0 | 1,9 | 0 | 1,65 | 0 |
| NR5A2 | TAAGTGAAGGAGAAG | 1 | 6 | 21 | -994 | -979 | 0,61 | 0,01 | 19,56 | >0.1 | 5,54 | 4,26 | 0 | 1,86 | 1,79 | 0 | 5,49 |
| NR5A2 | AAGGAGAAGGAGGCA | 1 | 12 | 27 | -988 | -973 | -3,52 | 0,01 | 13,64 | >0.1 | 5,54 | 4,27 | 0 | 1,87 | 0 | 0 | 5,49 |
| NR5A2 | TTTTGCAAAGGCAGA | 1 | 83 | 98 | -917 | -902 | -1,24 | 0,01 | 14,2 | >0.1 | 3,73 | 4,34 | 0 | 1,88 | 0 | 0 | 5,49 |
| NR5A2 | AAAGACAAGGGCAAA | 1 | 502 | 517 | -498 | -483 | 7,28 | 0,001 | 27,57 | >0.1 | 2,98 | 4,98 | 0 | 0,99 | 0 | 5,86 | 5,49 |
| NR5A2 | AAGGGCAAAGTCATG | 1 | 508 | 523 | -492 | -477 | 7,67 | 0,001 | 25 | >0.1 | 3 | 4,99 | 0 | 0,99 | 0 | 2,86 | 5,49 |
| NR5A2 | CCTCTCCAGGATGAA | 1 | 861 | 876 | -139 | -124 | -4,91 | 0,01 | 17,1 | >0.1 | 5,54 | 6,41 | 0 | 1,9 | 0 | 2,67 | 5,49 |
| NR5A2 | CAACCCAAGTTCAAA | 1 | 950 | 965 | -50 | -35 | -1,09 | 0,01 | 21,02 | >0.1 | 5,54 | 7,51 | 0 | 1,88 | 0 | 1,7 | 5,49 |
| NR5A2 | AAGTTCAAAGGCTGA | 1 | 956 | 971 | -44 | -29 | 7,28 | 0,001 | 29,57 | >0.1 | 5,54 | 7,65 | 0 | 1,88 | 0 | 1,74 | 5,49 |
| NR5A2 | AGTTCAAAGGCTGAT | 1 | 957 | 972 | -43 | -28 | -2,74 | 0,01 | 19,54 | >0.1 | 5,54 | 7,68 | 0 | 1,88 | 0 | 1,7 | 5,49 |
| NR5A2 | CTAGGGAAAGTCATT | 1 | 1003 | 1018 | 3 | 18 | -3,43 | 0,01 | 20 | >0.1 | 5,54 | 9,12 | 0 | 1,87 | 0 | 1,42 | 5,49 |
| NR5A2 | GATGTCAAGCTCTTC | 1 | 1048 | 1063 | 48 | 63 | 1,1 | 0,01 | 23,08 | >0.1 | 5,54 | 7,62 | 0 | 1,86 | 0 | 1,46 | 5,49 |
| NR5A2 | GAGGAACAGGCCAAG | 1 | 1112 | 1127 | 112 | 127 | 0,19 | 0,01 | 25,48 | >0.1 | 5,54 | 6,63 | 0 | 1,85 | 3,22 | 2,56 | 5,49 |
| NR5A2 | AAGCCGAAGACCTGT | 1 | 1152 | 1167 | 152 | 167 | -3,25 | 0,01 | 18,91 | >0.1 | 4,45 | 6,28 | 0 | 1,84 | 1,79 | 2,31 | 5,49 |
| NR5A2 | TCTATCAAAGTTCAC | 1 | 1167 | 1182 | 167 | 182 | -5,01 | 0,01 | 18,62 | >0.1 | 5,54 | 6,17 | 0 | 1,69 | 1,79 | 2,94 | 5,49 |
| NR5A2 | AGCAACAAGGCTGAG | -1 | 1073 | 1088 | 73 | 88 | 1,1 | 0,01 | 25,17 | >0.1 | 5,54 | 7,12 | 0 | 1,85 | 1,79 | 2,27 | 5,49 |
| NR5A2 | TGAGCCAAGATCACA | -1 | 1025 | 1040 | 25 | 40 | -2,02 | 0,01 | 20,69 | >0.1 | 5,54 | 8,52 | 0 | 1,86 | 0 | 1,31 | 5,49 |
| NR5A2 | GTGACCAAAGAGGTC | -1 | 665 | 680 | -335 | -320 | -4,53 | 0,01 | 15,41 | >0.1 | 4,11 | 5,42 | 0 | 1,1 | 0 | 3,83 | 5,49 |
| NR5A2 | TATTTCAAGATCATA | -1 | 567 | 582 | -433 | -418 | 0,37 | 0,01 | 15,28 | >0.1 | 3,29 | 5,13 | 0 | 1 | 0 | 0 | 5,49 |
| NR5A2 | GTCGCCCAGGCTGGA | -1 | 424 | 439 | -576 | -561 | -0,53 | 0,01 | 19,17 | >0.1 | 2,7 | 4,82 | 0 | 1,02 | 0 | 5,67 | 5,49 |
| NR5A2 | CACTGCAAGCTCTGC | -1 | 387 | 402 | -613 | -598 | -3,15 | 0,01 | 16,76 | >0.1 | 2,6 | 4,75 | 0 | 1,05 | 0 | 6,02 | 5,49 |
| NR5A2 | GCCTCCCAGGTTCAC | -1 | 374 | 389 | -626 | -611 | -1,67 | 0,01 | 18,57 | >0.1 | 2,56 | 4,73 | 0 | 1,07 | 0 | 6,4 | 5,49 |
| NR5A2 | AAGTGCTAGGATTAC | -1 | 200 | 215 | -800 | -785 | -4,3 | 0,01 | 15,02 | >0.1 | 3,03 | 4,47 | 0 | 1,63 | 0 | 4,7 | 5,49 |
| NR5A2 | ATTTCCATGGCCATT | -1 | 144 | 159 | -856 | -841 | -0,8 | 0,01 | 15,9 | >0.1 | 3,3 | 4,4 | 0 | 1,9 | 0 | 1,61 | 5,49 |
| NR5A2 | ACTCCACAGGTCAAC | -1 | 106 | 121 | -894 | -879 | -1,31 | 0,01 | 14,72 | >0.1 | 3,55 | 4,36 | 0 | 1,89 | 0 | 0,75 | 5,49 |
| TFE3 | CTCACGCC | 1 | 191 | 199 | -809 | -801 | -1,44 | 0,01 | 11,92 | >0.1 | 3,08 | 4,45 | 0 | 1,7 | 0 | 4,13 | 0 |
| TFE3 | CACAAGGT | 1 | 237 | 245 | -763 | -755 | -0,44 | 0,01 | 13,02 | >0.1 | 2,89 | 4,51 | 0 | 1,42 | 0 | 4,63 | 0 |
| TFE3 | CAGGAGAT | 1 | 245 | 253 | -755 | -747 | -1,48 | 0,01 | 11,72 | >0.1 | 2,86 | 4,52 | 0 | 1,39 | 0 | 4,42 | 0 |
| TFE3 | GGCGTGGT | 1 | 315 | 323 | -685 | -677 | -1,25 | 0,01 | 13,31 | >0.1 | 2,64 | 4,62 | 0 | 1,17 | 0 | 6,13 | 0 |
| TFE3 | CTCGGGAG | 1 | 347 | 355 | -653 | -645 | 0,89 | 0,01 | 15,74 | >0.1 | 2,55 | 4,67 | 0 | 1,11 | 0 | 6,52 | 0 |

| | | | | | | | | | | | | | | | | | |
|---|---|---|---|---|---|---|---|---|---|---|---|---|---|---|---|---|---|
| TFE3 | CTCATGAG | 1 | 984 | 992 | -16 | -8 | 5,04 | 0,01 | 22,56 | >0.1 | 5,64 | 8,43 | 0 | 1,87 | 0 | 1,57 | 0 |
| TFE3 | GATGTGAT | 1 | 1023 | 1031 | 23 | 31 | 1,18 | 0,01 | 18,62 | >0.1 | 5,64 | 8,67 | 0 | 1,86 | 0 | 1,28 | 0 |
| TFE3 | CAGGGGAC | 1 | 1040 | 1048 | 40 | 48 | -0,24 | 0,01 | 16,47 | >0.1 | 5,64 | 7,85 | 0 | 1,86 | 0 | 1,36 | 0 |
| TFE3 | CACTTGCT | 1 | 1179 | 1187 | 179 | 187 | 0,83 | 0,01 | 19,32 | >0.1 | 5,64 | 6,1 | 0 | 1,59 | 1,79 | 3,37 | 0 |
| TFE3 | CCTGTGAG | -1 | 1036 | 1044 | 36 | 44 | 3,92 | 0,01 | 20,71 | >0.1 | 5,64 | 7,99 | 0 | 1,86 | 0 | 1,31 | 0 |
| TFE3 | CTCATGAG | -1 | 984 | 992 | -16 | -8 | 5,04 | 0,01 | 22,56 | >0.1 | 5,64 | 8,43 | 0 | 1,87 | 0 | 1,57 | 0 |
| TFE3 | CATGAGAT | -1 | 982 | 990 | -18 | -10 | 3 | 0,01 | 20,36 | >0.1 | 5,64 | 8,32 | 0 | 1,87 | 0 | 1,53 | 0 |
| TFE3 | CTCCTGTC | -1 | 768 | 776 | -232 | -224 | 1,12 | 0,01 | 19,81 | >0.1 | 5,64 | 5,8 | 0 | 1,34 | 0 | 5,92 | 0 |
| TFE3 | TACATGAC | -1 | 517 | 525 | -483 | -475 | 1,04 | 0,01 | 12,99 | >0.1 | 3,02 | 5 | 0 | 0,98 | 0 | 2,95 | 0 |
| TFE3 | CTCCTGCC | -1 | 360 | 368 | -640 | -632 | 2,63 | 0,01 | 17,29 | >0.1 | 2,52 | 4,7 | 0 | 1,09 | 0 | 6,36 | 0 |
| TFE3 | CAGGTGCC | -1 | 327 | 335 | -673 | -665 | 4,24 | 0,01 | 18,09 | >0.1 | 2,6 | 4,64 | 0 | 1,14 | 0 | 5,46 | 0 |
| TFE3 | CTCCTGAC | -1 | 243 | 251 | -757 | -749 | 5,54 | 0,01 | 18,65 | >0.1 | 2,87 | 4,52 | 0 | 1,39 | 0 | 4,33 | 0 |
| TFE3 | CTTGTGAT | -1 | 235 | 243 | -765 | -757 | 8,5 | 0,001 | 22,24 | >0.1 | 2,9 | 4,51 | 0 | 1,43 | 0 | 4,9 | 0 |
| TFE3 | GGCGTGAG | -1 | 191 | 199 | -809 | -801 | -1,27 | 0,01 | 12,09 | >0.1 | 3,08 | 4,45 | 0 | 1,7 | 0 | 4,13 | 0 |
| TFE3 | CGTGAGCC | -1 | 189 | 197 | -811 | -803 | -1,38 | 0,01 | 11,82 | >0.1 | 3,09 | 4,45 | 0 | 1,71 | 0 | 3,95 | 0 |
| TFE3 | CACAGGTC | -1 | 109 | 117 | -891 | -883 | -0,57 | 0,01 | 9,97 | >0.1 | 3,55 | 4,36 | 0 | 1,89 | 0 | 0,75 | 0 |
| TFE3 | CTCCTGAT | -1 | 97 | 105 | -903 | -895 | 4,79 | 0,01 | 15,43 | >0.1 | 3,64 | 4,34 | 0 | 1,88 | 0 | 0,77 | 0 |
| DLX4 | ATAATTTT | 1 | 132 | 140 | -868 | -860 | 4,39 | 0,01 | 15,63 | >0.1 | 3,4 | 4,38 | 0 | 1,89 | 0 | 1,57 | 0 |
| DLX4 | GAAATTAA | 1 | 154 | 162 | -846 | -838 | 5,97 | 0,01 | 17,19 | >0.1 | 3,27 | 4,41 | 0 | 1,9 | 0 | 1,65 | 0 |
| DLX4 | TCTATTAA | 1 | 288 | 296 | -712 | -704 | 4,42 | 0,01 | 18,15 | >0.1 | 2,72 | 4,58 | 0 | 1,24 | 0 | 5,19 | 0 |
| DLX4 | CCTATTAA | 1 | 600 | 608 | -400 | -392 | 6,03 | 0,01 | 17,08 | >0.1 | 3,47 | 5,2 | 0 | 1,01 | 0 | 1,36 | 0 |
| DLX4 | TTAATGAA | 1 | 604 | 612 | -396 | -388 | 3,32 | 0,01 | 14,46 | >0.1 | 3,49 | 5,21 | 0 | 1,02 | 0 | 1,41 | 0 |
| DLX4 | AAAATTAT | 1 | 683 | 691 | -317 | -309 | 4,96 | 0,01 | 18,94 | >0.1 | 4,34 | 5,46 | 0 | 1,12 | 0 | 3,07 | 0 |
| DLX4 | AAAATTAT | 1 | 830 | 838 | -170 | -162 | 4,96 | 0,01 | 21,69 | >0.1 | 5,64 | 6,14 | 0 | 1,67 | 0 | 3,28 | 0 |
| DLX4 | GGAATTAT | 1 | 1191 | 1199 | 191 | 199 | 4,12 | 0,01 | 22,9 | >0.1 | 5,64 | 6,02 | 0 | 1,5 | 1,79 | 3,82 | 0 |
| DLX4 | ATAATTCC | -1 | 1191 | 1199 | 191 | 199 | 3,85 | 0,01 | 22,63 | >0.1 | 5,64 | 6,02 | 0 | 1,5 | 1,79 | 3,82 | 0 |
| DLX4 | ATAATTTT | -1 | 830 | 838 | -170 | -162 | 4,39 | 0,01 | 21,12 | >0.1 | 5,64 | 6,14 | 0 | 1,67 | 0 | 3,28 | 0 |
| DLX4 | ATAATTTT | -1 | 683 | 691 | -317 | -309 | 4,39 | 0,01 | 18,37 | >0.1 | 4,34 | 5,46 | 0 | 1,12 | 0 | 3,07 | 0 |
| DLX4 | ACAATTTC | -1 | 609 | 617 | -391 | -383 | 6,08 | 0,01 | 17,4 | >0.1 | 3,53 | 5,23 | 0 | 1,02 | 0 | 1,55 | 0 |
| DLX4 | TTCATTAA | -1 | 604 | 612 | -396 | -388 | 4,79 | 0,01 | 15,93 | >0.1 | 3,49 | 5,21 | 0 | 1,02 | 0 | 1,41 | 0 |
| DLX4 | ACATTTAT | -1 | 585 | 593 | -415 | -407 | 4,1 | 0,01 | 13,63 | >0.1 | 3,37 | 5,16 | 0 | 1 | 0 | 0 | 0 |
| DLX4 | CCAAATAC | -1 | 522 | 530 | -478 | -470 | 3,48 | 0,01 | 15,58 | >0.1 | 3,05 | 5,01 | 0 | 0,98 | 0 | 3,07 | 0 |
| DLX4 | GCCATTTC | -1 | 171 | 179 | -829 | -821 | 4,55 | 0,01 | 17,45 | >0.1 | 3,18 | 4,43 | 0 | 1,87 | 0 | 3,43 | 0 |
| DLX4 | TTAATTTC | -1 | 154 | 162 | -846 | -838 | 4,65 | 0,01 | 15,87 | >0.1 | 3,27 | 4,41 | 0 | 1,9 | 0 | 1,65 | 0 |
| DLX4 | GCCATTTT | -1 | 142 | 150 | -858 | -850 | 3,75 | 0,01 | 14,98 | >0.1 | 3,33 | 4,39 | 0 | 1,89 | 0 | 1,61 | 0 |
| DLX4 | AAAATTAT | -1 | 132 | 140 | -868 | -860 | 4,96 | 0,01 | 16,2 | >0.1 | 3,4 | 4,38 | 0 | 1,89 | 0 | 1,57 | 0 |
| DLX4 | CTCATTTC | -1 | 65 | 73 | -935 | -927 | 3,75 | 0,01 | 13,89 | >0.1 | 3,95 | 4,31 | 0 | 1,88 | 0 | 0 | 0 |
| DLX4 | CCTATTTC | -1 | 58 | 66 | -942 | -934 | 3,38 | 0,01 | 13,59 | >0.1 | 4,03 | 4,3 | 0 | 1,87 | 0 | 0 | 0 |
| DLX4 | TCACTTAC | -1 | 5 | 13 | -995 | -987 | 5,44 | 0,01 | 18,99 | >0.1 | 5,64 | 4,26 | 0 | 1,86 | 1,79 | 0 | 0 |
| NEUROG2 | AAAAAATGGC | 1 | 140 | 150 | -860 | -850 | -0,3 | 0,01 | 10,98 | >0.1 | 3,34 | 4,39 | 0 | 1,89 | 0 | 1,65 | 0 |
| NEUROG2 | GGAAAATGTT | 1 | 533 | 543 | -467 | -457 | -3,3 | 0,01 | 7,43 | >0.1 | 3,1 | 5,04 | 0 | 0,98 | 0 | 1,61 | 0 |
| NEUROG2 | GGAATATGAT | 1 | 563 | 573 | -437 | -427 | -3,69 | 0,01 | 5,66 | >0.1 | 3,25 | 5,11 | 0 | 0,99 | 0 | 0 | 0 |
| NEUROG2 | AACAGCTTTC | 1 | 711 | 721 | -289 | -279 | -4,29 | 0,01 | 13,05 | >0.1 | 5,61 | 5,56 | 0 | 1,18 | 0 | 4,99 | 0 |
| NEUROG2 | GCCATAAAGT | 1 | 758 | 768 | -242 | -232 | -5,41 | 0,01 | 13,37 | >0.1 | 5,61 | 5,76 | 0 | 1,31 | 0 | 6,1 | 0 |
| NEUROG2 | AACGTATTCT | 1 | 803 | 813 | -197 | -187 | -4,44 | 0,01 | 12,1 | >0.1 | 5,61 | 5,99 | 0 | 1,51 | 0 | 3,43 | 0 |
| NEUROG2 | AAAATCTGTT | 1 | 841 | 851 | -159 | -149 | 0,33 | 0,01 | 15,37 | >0.1 | 5,61 | 6,24 | 0 | 1,78 | 0 | 1,41 | 0 |
| NEUROG2 | CATATCTGTC | 1 | 852 | 862 | -148 | -138 | -2 | 0,01 | 14,47 | >0.1 | 5,61 | 6,31 | 0 | 1,88 | 0 | 2,67 | 0 |
| NEUROG2 | AGCAGATTGT | 1 | 890 | 900 | -110 | -100 | -4,2 | 0,01 | 12,86 | >0.1 | 5,61 | 6,61 | 0 | 1,89 | 0 | 2,95 | 0 |
| NEUROG2 | AACTGCTGCT | 1 | 1090 | 1100 | 90 | 100 | -6,49 | 0,01 | 11,92 | >0.1 | 5,61 | 6,88 | 0 | 1,85 | 1,79 | 2,27 | 0 |
| NEUROG2 | AACAGGTCTT | -1 | 1158 | 1168 | 158 | 168 | -6,32 | 0,01 | 10,27 | >0.1 | 4,38 | 6,24 | 0 | 1,78 | 1,79 | 2,4 | 0 |
| NEUROG2 | CAAAAATGTC | -1 | 1126 | 1136 | 126 | 136 | -5,58 | 0,01 | 14,34 | >0.1 | 5,61 | 6,49 | 0 | 1,84 | 3,22 | 2,75 | 0 |
| NEUROG2 | AGCAGCAGTT | -1 | 1090 | 1100 | 90 | 100 | -3,17 | 0,01 | 15,24 | >0.1 | 5,61 | 6,88 | 0 | 1,85 | 1,79 | 2,27 | 0 |
| NEUROG2 | GCAACAAGGC | -1 | 1077 | 1087 | 77 | 87 | -6,82 | 0,01 | 11,71 | >0.1 | 5,61 | 7,06 | 0 | 1,85 | 1,79 | 2,21 | 0 |
| NEUROG2 | ACAATCTGCT | -1 | 890 | 900 | -110 | -100 | -1,5 | 0,01 | 15,56 | >0.1 | 5,61 | 6,61 | 0 | 1,89 | 0 | 2,95 | 0 |
| NEUROG2 | AACAGATTTT | -1 | 841 | 851 | -159 | -149 | -0,95 | 0,01 | 14,09 | >0.1 | 5,61 | 6,24 | 0 | 1,78 | 0 | 1,41 | 0 |
| NEUROG2 | AGAATACGTT | -1 | 803 | 813 | -197 | -187 | -3,24 | 0,01 | 13,3 | >0.1 | 5,61 | 5,99 | 0 | 1,51 | 0 | 3,43 | 0 |
| NEUROG2 | ACTTTATGGC | -1 | 758 | 768 | -242 | -232 | -3,59 | 0,01 | 15,19 | >0.1 | 5,61 | 5,76 | 0 | 1,31 | 0 | 6,1 | 0 |
| NEUROG2 | GAAAGCTGTT | -1 | 711 | 721 | -289 | -279 | -3,52 | 0,01 | 13,82 | >0.1 | 5,61 | 5,56 | 0 | 1,18 | 0 | 4,99 | 0 |
| NEUROG2 | GGAAAATGTT | -1 | 691 | 701 | -309 | -299 | -3,3 | 0,01 | 10,93 | >0.1 | 4,53 | 5,49 | 0 | 1,14 | 0 | 3,07 | 0 |

| | | | | | | | | | | | | | | | | | |
|---|---|---|---|---|---|---|---|---|---|---|---|---|---|---|---|---|---|
| NEUROG2 | AATACATGAC | -1 | 517 | 527 | -483 | -473 | -3,4 | 0,01 | 8,56 | >0.1 | 3,03 | 5 | 0 | 0,98 | 0 | 2,95 | 0 |
| NEUROG2 | TACAGGTGCC | -1 | 327 | 337 | -673 | -663 | -4,49 | 0,01 | 9,47 | >0.1 | 2,6 | 4,65 | 0 | 1,14 | 0 | 5,56 | 0 |
| NEUROG2 | GGCGTGAGCC | -1 | 189 | 199 | -811 | -801 | -5,41 | 0,01 | 7,88 | >0.1 | 3,09 | 4,45 | 0 | 1,71 | 0 | 4,04 | 0 |
| NEUROG1 | GGAATATGAT | 1 | 563 | 573 | -437 | -427 | 4,89 | 0,01 | 14,24 | >0.1 | 3,25 | 5,11 | 0 | 0,99 | 0 | 0 | 0 |
| NEUROG1 | AACGTATTCT | 1 | 803 | 813 | -197 | -187 | 4,31 | 0,01 | 20,85 | >0.1 | 5,61 | 5,99 | 0 | 1,51 | 0 | 3,43 | 0 |
| NEUROG1 | AAAATCTGTT | 1 | 841 | 851 | -159 | -149 | 6,06 | 0,01 | 21,1 | >0.1 | 5,61 | 6,24 | 0 | 1,78 | 0 | 1,41 | 0 |
| NEUROG1 | AGCAGATTGT | 1 | 890 | 900 | -110 | -100 | 5,66 | 0,01 | 22,72 | >0.1 | 5,61 | 6,61 | 0 | 1,89 | 0 | 2,95 | 0 |
| NEUROG1 | GCAACAAGGC | -1 | 1077 | 1087 | 77 | 87 | 4,07 | 0,01 | 22,6 | >0.1 | 5,61 | 7,06 | 0 | 1,85 | 1,79 | 2,21 | 0 |
| NEUROG1 | ACAATCTGCT | -1 | 890 | 900 | -110 | -100 | 7,48 | 0,001 | 24,54 | >0.1 | 5,61 | 6,61 | 0 | 1,89 | 0 | 2,95 | 0 |
| NEUROG1 | AACAGATTTT | -1 | 841 | 851 | -159 | -149 | 4,54 | 0,01 | 19,58 | >0.1 | 5,61 | 6,24 | 0 | 1,78 | 0 | 1,41 | 0 |
| NEUROG1 | AGAATACGTT | -1 | 803 | 813 | -197 | -187 | 6,72 | 0,01 | 23,26 | >0.1 | 5,61 | 5,99 | 0 | 1,51 | 0 | 3,43 | 0 |
| NEUROG1 | ACTTTATGGC | -1 | 758 | 768 | -242 | -232 | 4,07 | 0,01 | 22,85 | >0.1 | 5,61 | 5,76 | 0 | 1,31 | 0 | 6,1 | 0 |
| NEUROG1 | GAAAGCTGTT | -1 | 711 | 721 | -289 | -279 | 3,73 | 0,01 | 21,07 | >0.1 | 5,61 | 5,56 | 0 | 1,18 | 0 | 4,99 | 0 |
| NEUROG1 | ATCATATTCC | -1 | 563 | 573 | -437 | -427 | 4,22 | 0,01 | 13,57 | >0.1 | 3,25 | 5,11 | 0 | 0,99 | 0 | 0 | 0 |
| NEUROG1 | TTCATATTTC | -1 | 486 | 496 | -514 | -504 | 4,18 | 0,01 | 19,27 | >0.1 | 2,9 | 4,93 | 0 | 0,98 | 0 | 6,27 | 0 |
| NEUROG1 | TCAGGACGGT | -1 | 257 | 267 | -743 | -733 | 4,91 | 0,01 | 18,24 | >0.1 | 2,82 | 4,54 | 0 | 1,34 | 0 | 4,63 | 0 |
| ZNF282 | CTGTGCTCCAGCCTGGG | 1 | 418 | 435 | -582 | -565 | -10,89 | 0,01 | 3,06 | >0.1 | 2,69 | 4,81 | 0 | 1,02 | 0 | 5,42 | 0 |
| ZNF282 | AATTCCATAAAGACAAG | 1 | 494 | 511 | -506 | -489 | -7,02 | 0,01 | 8,32 | >0.1 | 2,95 | 4,97 | 0 | 0,99 | 0 | 6,44 | 0 |
| ZNF282 | TTGTCTGAAAACCATAC | 1 | 613 | 630 | -387 | -370 | -9,49 | 0,01 | 1,9 | >0.1 | 3,59 | 5,26 | 0 | 1,04 | 0 | 1,5 | 0 |
| ZNF282 | TTTGGCCATAAAGTGAC | 1 | 754 | 771 | -246 | -229 | -8,92 | 0,01 | 9,8 | >0.1 | 5,52 | 5,77 | 0 | 1,33 | 0 | 6,1 | 0 |
| ZNF282 | TCAGCGCCCAACCCAAG | 1 | 942 | 959 | -58 | -41 | -6,47 | 0,01 | 10,11 | >0.1 | 5,52 | 7,38 | 0 | 1,88 | 0 | 1,8 | 0 |
| ZNF282 | CTTTGATAGAACAGGTC | -1 | 1160 | 1177 | 160 | 177 | -10,82 | 0,01 | 6,15 | >0.1 | 4,47 | 6,22 | 0 | 1,76 | 1,79 | 2,72 | 0 |
| ZNF282 | CTTTCCCTAGACTAAAA | -1 | 996 | 1013 | -4 | 13 | -0,84 | 0,001 | 17,17 | >0.1 | 5,52 | 9,12 | 0 | 1,87 | 0 | 1,5 | 0 |
| ZNF282 | CCTCCTCATGAGATTTT | -1 | 979 | 996 | -21 | -4 | -11,44 | 0,01 | 6,09 | >0.1 | 5,52 | 8,61 | 0 | 1,87 | 0 | 1,53 | 0 |
| ZNF282 | CAATCTGCTGAGCCAAT | -1 | 882 | 899 | -118 | -101 | -10,9 | 0,01 | 6,19 | >0.1 | 5,52 | 6,61 | 0 | 1,89 | 0 | 3,07 | 0 |
| ZNF282 | TTATTCTTTAATCCTAG | -1 | 788 | 805 | -212 | -195 | -10,59 | 0,01 | 7,66 | >0.1 | 5,52 | 5,94 | 0 | 1,47 | 0 | 5,32 | 0 |
| ZNF282 | TTTTCCTAGAAAGCTGT | -1 | 712 | 729 | -288 | -271 | -5,28 | 0,01 | 12,2 | >0.1 | 5,52 | 5,6 | 0 | 1,2 | 0 | 5,17 | 0 |
| ZNF282 | ATATTCCCAGAAACTCT | -1 | 553 | 570 | -447 | -430 | -9,05 | 0,01 | 0,27 | >0.1 | 3,22 | 5,1 | 0 | 1 | 0 | 0 | 0 |
| ZNF282 | TTTTCCCTTCCAAATAC | -1 | 522 | 539 | -478 | -461 | -10,73 | 0,01 | 1,64 | >0.1 | 3,07 | 5,03 | 0 | 0,99 | 0 | 3,28 | 0 |
| ZNF282 | CCTTCCAAATACATGAC | -1 | 517 | 534 | -483 | -466 | 0,43 | 0,001 | 12,55 | >0.1 | 3,04 | 5,02 | 0 | 0,99 | 0 | 3,07 | 0 |
| ZNF282 | TTTGCCCTTGTCTTTAT | -1 | 500 | 517 | -500 | -483 | -9,18 | 0,01 | 5,44 | >0.1 | 2,97 | 4,98 | 0 | 0,99 | 0 | 5,68 | 0 |
| ZNF282 | CATTCTCCTGCCTCAGC | -1 | 355 | 372 | -645 | -628 | -7,18 | 0,01 | 7,4 | >0.1 | 2,52 | 4,7 | 0 | 1,1 | 0 | 6,26 | 0 |
| ZNF282 | GTGCCCACCACCACGCC | -1 | 315 | 332 | -685 | -668 | -8,22 | 0,01 | 6,47 | >0.1 | 2,63 | 4,64 | 0 | 1,17 | 0 | 6,26 | 0 |
| ZNF282 | CTATCTCCTGACCTTGT | -1 | 238 | 255 | -762 | -745 | -0,76 | 0,001 | 12,39 | >0.1 | 2,87 | 4,53 | 0 | 1,42 | 0 | 4,33 | 0 |
| ZNF282 | GATCTGCCCAACTCGGC | -1 | 221 | 238 | -779 | -762 | -11,03 | 0,01 | 3,17 | >0.1 | 2,94 | 4,5 | 0 | 1,5 | 0 | 5,26 | 0 |
| ZNF282 | CTGTGCCCAGCCATTTC | -1 | 171 | 188 | -829 | -812 | -10,78 | 0,01 | 1,93 | >0.1 | 3,15 | 4,44 | 0 | 1,87 | 0 | 3,24 | 0 |
| ZNF282 | ATTTCCTATTTCTCACA | -1 | 53 | 70 | -947 | -930 | -5,77 | 0,01 | 4,45 | >0.1 | 4,04 | 4,31 | 0 | 1,87 | 0 | 0 | 0 |
| DLX1 | GGAAATGAGC | 1 | 64 | 74 | -936 | -926 | 4,99 | 0,01 | 15,13 | >0.1 | 3,95 | 4,31 | 0 | 1,88 | 0 | 0 | 0 |
| DLX1 | CATAATTTTA | 1 | 131 | 141 | -869 | -859 | 6,85 | 0,01 | 18,1 | >0.1 | 3,4 | 4,38 | 0 | 1,89 | 0 | 1,57 | 0 |
| DLX1 | GGAAATTAAA | 1 | 153 | 163 | -847 | -837 | 6,34 | 0,01 | 17,56 | >0.1 | 3,27 | 4,41 | 0 | 1,9 | 0 | 1,65 | 0 |
| DLX1 | CAACATTATC | 1 | 636 | 646 | -364 | -354 | 6,37 | 0,01 | 18,15 | >0.1 | 3,76 | 5,31 | 0 | 1,05 | 0 | 1,66 | 0 |
| DLX1 | TAAAATTATA | 1 | 682 | 692 | -318 | -308 | 6,3 | 0,01 | 20,28 | >0.1 | 4,34 | 5,46 | 0 | 1,12 | 0 | 3,07 | 0 |
| DLX1 | GGAAAATATT | 1 | 723 | 733 | -277 | -267 | 3,84 | 0,01 | 21,44 | >0.1 | 5,61 | 5,61 | 0 | 1,21 | 0 | 5,17 | 0 |
| DLX1 | AAAAATTATT | 1 | 829 | 839 | -171 | -161 | 6,82 | 0,01 | 23,54 | >0.1 | 5,61 | 6,15 | 0 | 1,68 | 0 | 3,28 | 0 |
| DLX1 | AATAATTTTT | -1 | 829 | 839 | -171 | -161 | 5,97 | 0,01 | 22,69 | >0.1 | 5,61 | 6,15 | 0 | 1,68 | 0 | 3,28 | 0 |
| DLX1 | TATAATTTTA | -1 | 682 | 692 | -318 | -308 | 5,45 | 0,01 | 19,43 | >0.1 | 4,34 | 5,46 | 0 | 1,12 | 0 | 3,07 | 0 |
| DLX1 | GACAATTTCA | -1 | 608 | 618 | -392 | -382 | 5,77 | 0,01 | 17,1 | >0.1 | 3,53 | 5,23 | 0 | 1,02 | 0 | 1,55 | 0 |
| DLX1 | CACATTTATT | -1 | 584 | 594 | -416 | -406 | 4,53 | 0,01 | 14,07 | >0.1 | 3,37 | 5,16 | 0 | 1 | 0 | 0 | 0 |
| DLX1 | CAACATTTTC | -1 | 534 | 544 | -466 | -456 | 4,26 | 0,01 | 15,05 | >0.1 | 3,1 | 5,04 | 0 | 0,98 | 0 | 1,66 | 0 |
| DLX1 | TCCAAATACA | -1 | 521 | 531 | -479 | -469 | 4,7 | 0,01 | 16,81 | >0.1 | 3,05 | 5,01 | 0 | 0,98 | 0 | 3,07 | 0 |
| DLX1 | TTTAATTTCC | -1 | 153 | 163 | -847 | -837 | 6 | 0,01 | 17,22 | >0.1 | 3,27 | 4,41 | 0 | 1,9 | 0 | 1,65 | 0 |
| DLX1 | TAAAATTATG | -1 | 131 | 141 | -869 | -859 | 7,03 | 0,01 | 18,28 | >0.1 | 3,4 | 4,38 | 0 | 1,89 | 0 | 1,57 | 0 |
| DLX1 | GCTCATTTCC | -1 | 64 | 74 | -936 | -926 | 5,57 | 0,01 | 15,71 | >0.1 | 3,95 | 4,31 | 0 | 1,88 | 0 | 0 | 0 |
| DLX2 | GAAATGAG | 1 | 65 | 73 | -935 | -927 | 2,25 | 0,01 | 12,39 | >0.1 | 3,95 | 4,31 | 0 | 1,88 | 0 | 0 | 0 |
| DLX2 | ATAATTTT | 1 | 132 | 140 | -868 | -860 | 4,03 | 0,01 | 15,27 | >0.1 | 3,4 | 4,38 | 0 | 1,89 | 0 | 1,57 | 0 |
| DLX2 | GAAATTAA | 1 | 154 | 162 | -846 | -838 | 6,14 | 0,01 | 17,36 | >0.1 | 3,27 | 4,41 | 0 | 1,9 | 0 | 1,65 | 0 |
| DLX2 | TCTATTAA | 1 | 288 | 296 | -712 | -704 | 4,13 | 0,01 | 17,86 | >0.1 | 2,72 | 4,58 | 0 | 1,24 | 0 | 5,19 | 0 |
| DLX2 | CCTATTAA | 1 | 600 | 608 | -400 | -392 | 5,28 | 0,01 | 16,33 | >0.1 | 3,47 | 5,2 | 0 | 1,01 | 0 | 1,36 | 0 |
| DLX2 | TTAATGAA | 1 | 604 | 612 | -396 | -388 | 4,02 | 0,01 | 15,16 | >0.1 | 3,49 | 5,21 | 0 | 1,02 | 0 | 1,41 | 0 |

| | | | | | | | | | | | | | | | | | |
|---|---|---|---|---|---|---|---|---|---|---|---|---|---|---|---|---|---|
| DLX2 | AAAATTAT | 1 | 683 | 691 | -317 | -309 | 5,05 | 0,01 | 19,03 | >0.1 | 4,34 | 5,46 | 0 | 1,12 | 0 | 3,07 | 0 |
| DLX2 | AAAATTAT | 1 | 830 | 838 | -170 | -162 | 5,05 | 0,01 | 21,78 | >0.1 | 5,64 | 6,14 | 0 | 1,67 | 0 | 3,28 | 0 |
| DLX2 | GGAATTAT | 1 | 1191 | 1199 | 191 | 199 | 4,62 | 0,01 | 23,4 | >0.1 | 5,64 | 6,02 | 0 | 1,5 | 1,79 | 3,82 | 0 |
| DLX2 | ATAATTTT | -1 | 830 | 838 | -170 | -162 | 4,03 | 0,01 | 20,76 | >0.1 | 5,64 | 6,14 | 0 | 1,67 | 0 | 3,28 | 0 |
| DLX2 | ATAATTTT | -1 | 683 | 691 | -317 | -309 | 4,03 | 0,01 | 18,01 | >0.1 | 4,34 | 5,46 | 0 | 1,12 | 0 | 3,07 | 0 |
| DLX2 | ACAATTTC | -1 | 609 | 617 | -391 | -383 | 5,07 | 0,01 | 16,39 | >0.1 | 3,53 | 5,23 | 0 | 1,02 | 0 | 1,55 | 0 |
| DLX2 | TTCATTAA | -1 | 604 | 612 | -396 | -388 | 4,29 | 0,01 | 15,43 | >0.1 | 3,49 | 5,21 | 0 | 1,02 | 0 | 1,41 | 0 |
| DLX2 | ACATTTAT | -1 | 585 | 593 | -415 | -407 | 3,06 | 0,01 | 12,59 | >0.1 | 3,37 | 5,16 | 0 | 1 | 0 | 0 | 0 |
| DLX2 | GCCATTTC | -1 | 171 | 179 | -829 | -821 | 2,32 | 0,01 | 15,22 | >0.1 | 3,18 | 4,43 | 0 | 1,87 | 0 | 3,43 | 0 |
| DLX2 | TTAATTTC | -1 | 154 | 162 | -846 | -838 | 3,87 | 0,01 | 15,09 | >0.1 | 3,27 | 4,41 | 0 | 1,9 | 0 | 1,65 | 0 |
| DLX2 | GCCATTTT | -1 | 142 | 150 | -858 | -850 | 2 | 0,01 | 13,23 | >0.1 | 3,33 | 4,39 | 0 | 1,89 | 0 | 1,61 | 0 |
| DLX2 | AAAATTAT | -1 | 132 | 140 | -868 | -860 | 5,05 | 0,01 | 16,29 | >0.1 | 3,4 | 4,38 | 0 | 1,89 | 0 | 1,57 | 0 |
| DLX2 | TCACTTAC | -1 | 5 | 13 | -995 | -987 | 4,09 | 0,01 | 17,64 | >0.1 | 5,64 | 4,26 | 0 | 1,86 | 1,79 | 0 | 0 |
| DLX3 | ATAATTTT | 1 | 132 | 140 | -868 | -860 | 4,3 | 0,01 | 15,54 | >0.1 | 3,4 | 4,38 | 0 | 1,89 | 0 | 1,57 | 0 |
| DLX3 | GAAATTAA | 1 | 154 | 162 | -846 | -838 | 6,18 | 0,01 | 17,4 | >0.1 | 3,27 | 4,41 | 0 | 1,9 | 0 | 1,65 | 0 |
| DLX3 | TCTATTAA | 1 | 288 | 296 | -712 | -704 | 4,74 | 0,01 | 18,47 | >0.1 | 2,72 | 4,58 | 0 | 1,24 | 0 | 5,19 | 0 |
| DLX3 | CCTATTAA | 1 | 600 | 608 | -400 | -392 | 6,19 | 0,01 | 17,24 | >0.1 | 3,47 | 5,2 | 0 | 1,01 | 0 | 1,36 | 0 |
| DLX3 | TTAATGAA | 1 | 604 | 612 | -396 | -388 | 3,86 | 0,01 | 15 | >0.1 | 3,49 | 5,21 | 0 | 1,02 | 0 | 1,41 | 0 |
| DLX3 | AAAATTAT | 1 | 683 | 691 | -317 | -309 | 5,22 | 0,01 | 19,2 | >0.1 | 4,34 | 5,46 | 0 | 1,12 | 0 | 3,07 | 0 |
| DLX3 | AAAATTAT | 1 | 830 | 838 | -170 | -162 | 5,22 | 0,01 | 21,95 | >0.1 | 5,64 | 6,14 | 0 | 1,67 | 0 | 3,28 | 0 |
| DLX3 | GGAATTAT | 1 | 1191 | 1199 | 191 | 199 | 4,71 | 0,01 | 23,49 | >0.1 | 5,64 | 6,02 | 0 | 1,5 | 1,79 | 3,82 | 0 |
| DLX3 | ATAATTCC | -1 | 1191 | 1199 | 191 | 199 | 3,75 | 0,01 | 22,53 | >0.1 | 5,64 | 6,02 | 0 | 1,5 | 1,79 | 3,82 | 0 |
| DLX3 | GCAGTTAC | -1 | 1088 | 1096 | 88 | 96 | 3,71 | 0,01 | 22,11 | >0.1 | 5,64 | 6,9 | 0 | 1,85 | 1,79 | 2,21 | 0 |
| DLX3 | ATAATTTT | -1 | 830 | 838 | -170 | -162 | 4,3 | 0,01 | 21,03 | >0.1 | 5,64 | 6,14 | 0 | 1,67 | 0 | 3,28 | 0 |
| DLX3 | ATAATTTT | -1 | 683 | 691 | -317 | -309 | 4,3 | 0,01 | 18,28 | >0.1 | 4,34 | 5,46 | 0 | 1,12 | 0 | 3,07 | 0 |
| DLX3 | ACAATTTC | -1 | 609 | 617 | -391 | -383 | 5,9 | 0,01 | 17,22 | >0.1 | 3,53 | 5,23 | 0 | 1,02 | 0 | 1,55 | 0 |
| DLX3 | TTCATTAA | -1 | 604 | 612 | -396 | -388 | 4,99 | 0,01 | 16,13 | >0.1 | 3,49 | 5,21 | 0 | 1,02 | 0 | 1,41 | 0 |
| DLX3 | ACATTTAT | -1 | 585 | 593 | -415 | -407 | 3,96 | 0,01 | 13,49 | >0.1 | 3,37 | 5,16 | 0 | 1 | 0 | 0 | 0 |
| DLX3 | CCAAATAC | -1 | 522 | 530 | -478 | -470 | 4,86 | 0,01 | 16,96 | >0.1 | 3,05 | 5,01 | 0 | 0,98 | 0 | 3,07 | 0 |
| DLX3 | GCCATTTC | -1 | 171 | 179 | -829 | -821 | 4,31 | 0,01 | 17,21 | >0.1 | 3,18 | 4,43 | 0 | 1,87 | 0 | 3,43 | 0 |
| DLX3 | TTAATTTC | -1 | 154 | 162 | -846 | -838 | 4,54 | 0,01 | 15,76 | >0.1 | 3,27 | 4,41 | 0 | 1,9 | 0 | 1,65 | 0 |
| DLX3 | AAAATTAT | -1 | 132 | 140 | -868 | -860 | 5,22 | 0,01 | 16,46 | >0.1 | 3,4 | 4,38 | 0 | 1,89 | 0 | 1,57 | 0 |
| DLX3 | CTCATTTC | -1 | 65 | 73 | -935 | -927 | 3,61 | 0,01 | 13,75 | >0.1 | 3,95 | 4,31 | 0 | 1,88 | 0 | 0 | 0 |
| DLX3 | TCACTTAC | -1 | 5 | 13 | -995 | -987 | 5,54 | 0,01 | 19,09 | >0.1 | 5,64 | 4,26 | 0 | 1,86 | 1,79 | 0 | 0 |
| DDIT3::CEBPA | GTGAGAAATAGG | 1 | 54 | 66 | -946 | -934 | 0,55 | 0,01 | 10,79 | >0.1 | 4,06 | 4,3 | 0 | 1,87 | 0 | 0 | 0 |
| DDIT3::CEBPA | TTTTGCAAAGGC | 1 | 83 | 95 | -917 | -905 | 1,14 | 0,01 | 11,1 | >0.1 | 3,75 | 4,33 | 0 | 1,88 | 0 | 0 | 0 |
| DDIT3::CEBPA | GCCTGTAATCCT | 1 | 196 | 208 | -804 | -792 | 7,27 | 0,001 | 20,82 | >0.1 | 3,05 | 4,46 | 0 | 1,66 | 0 | 4,37 | 0 |
| DDIT3::CEBPA | CGGTGAAACCCC | 1 | 273 | 285 | -727 | -715 | 11,11 | 0,0001 | 24,92 | >0.1 | 2,76 | 4,57 | 0 | 1,28 | 0 | 5,19 | 0 |
| DDIT3::CEBPA | ACCTGTAGTCCC | 1 | 330 | 342 | -670 | -658 | 4,94 | 0,01 | 19,36 | >0.1 | 2,59 | 4,65 | 0 | 1,14 | 0 | 6,03 | 0 |
| DDIT3::CEBPA | GCGTGAACCTGG | 1 | 372 | 384 | -628 | -616 | 1,01 | 0,01 | 15,89 | >0.1 | 2,55 | 4,72 | 0 | 1,07 | 0 | 6,53 | 0 |
| DDIT3::CEBPA | GCTTGCAGTGAG | 1 | 392 | 404 | -608 | -596 | 2,36 | 0,01 | 16,79 | >0.1 | 2,6 | 4,76 | 0 | 1,05 | 0 | 6,02 | 0 |
| DDIT3::CEBPA | ATGTGAGATAAC | 1 | 589 | 601 | -411 | -399 | 1,22 | 0,01 | 10,82 | >0.1 | 3,41 | 5,18 | 0 | 1,01 | 0 | 0 | 0 |
| DDIT3::CEBPA | TAATGAAATTGT | 1 | 605 | 617 | -395 | -383 | 4,63 | 0,01 | 15,84 | >0.1 | 3,52 | 5,23 | 0 | 1,02 | 0 | 1,44 | 0 |
| DDIT3::CEBPA | TGCTGTAACTGC | 1 | 1084 | 1096 | 84 | 96 | 2,96 | 0,01 | 21,36 | >0.1 | 5,58 | 6,96 | 0 | 1,85 | 1,79 | 2,21 | 0 |
| DDIT3::CEBPA | AAGTTTAACCAC | 1 | 1139 | 1151 | 139 | 151 | 1,37 | 0,01 | 21,7 | >0.1 | 5,58 | 6,38 | 0 | 1,84 | 3,22 | 3,29 | 0 |
| DDIT3::CEBPA | AAGTTCACTTGC | 1 | 1174 | 1186 | 174 | 186 | 2,87 | 0,01 | 21,3 | >0.1 | 5,58 | 6,13 | 0 | 1,63 | 1,79 | 3,29 | 0 |
| DDIT3::CEBPA | AAGTGAACTTTG | -1 | 1172 | 1184 | 172 | 184 | 0,78 | 0,01 | 19,04 | >0.1 | 5,58 | 6,14 | 0 | 1,65 | 1,79 | 3,09 | 0 |
| DDIT3::CEBPA | GCTTGACATCGT | -1 | 1046 | 1058 | 46 | 58 | 1,88 | 0,01 | 18,41 | >0.1 | 5,58 | 7,68 | 0 | 1,86 | 0 | 1,42 | 0 |
| DDIT3::CEBPA | TGGCGCGATCTC | -1 | 406 | 418 | -594 | -582 | 0,87 | 0,01 | 14,98 | >0.1 | 2,65 | 4,78 | 0 | 1,03 | 0 | 5,65 | 0 |
| DDIT3::CEBPA | CACTGCAAGCTC | -1 | 390 | 402 | -610 | -598 | 3,19 | 0,01 | 17,61 | >0.1 | 2,6 | 4,75 | 0 | 1,05 | 0 | 6,02 | 0 |
| DDIT3::CEBPA | CTTTGCAAAACT | -1 | 81 | 93 | -919 | -907 | 1,78 | 0,01 | 11,76 | >0.1 | 3,77 | 4,33 | 0 | 1,88 | 0 | 0 | 0 |
| BHLHA15 | GCACCTGT | 1 | 328 | 336 | -672 | -664 | -0,43 | 0,01 | 13,52 | >0.1 | 2,6 | 4,64 | 0 | 1,14 | 0 | 5,56 | 0 |
| BHLHA15 | CCAGCTAC | 1 | 340 | 348 | -660 | -652 | -3,17 | 0,01 | 11,7 | >0.1 | 2,57 | 4,66 | 0 | 1,12 | 0 | 6,52 | 0 |
| BHLHA15 | ACAGCTTT | 1 | 712 | 720 | -288 | -280 | -4,74 | 0,01 | 12,62 | >0.1 | 5,64 | 5,56 | 0 | 1,18 | 0 | 4,99 | 0 |
| BHLHA15 | AAATCTGT | 1 | 842 | 850 | -158 | -150 | -3,24 | 0,01 | 11,81 | >0.1 | 5,64 | 6,23 | 0 | 1,77 | 0 | 1,41 | 0 |
| BHLHA15 | ACATATCT | 1 | 851 | 859 | -149 | -141 | -1,25 | 0,01 | 15,28 | >0.1 | 5,64 | 6,29 | 0 | 1,85 | 0 | 2,75 | 0 |
| BHLHA15 | ATATCTGT | 1 | 853 | 861 | -147 | -139 | -3,24 | 0,01 | 13,24 | >0.1 | 5,64 | 6,3 | 0 | 1,87 | 0 | 2,67 | 0 |
| BHLHA15 | GCAGATTG | 1 | 891 | 899 | -109 | -101 | -3,25 | 0,01 | 13,83 | >0.1 | 5,64 | 6,6 | 0 | 1,89 | 0 | 2,95 | 0 |
| BHLHA15 | ACTGCTGC | 1 | 1091 | 1099 | 91 | 99 | -5,16 | 0,01 | 13,26 | >0.1 | 5,64 | 6,87 | 0 | 1,85 | 1,79 | 2,27 | 0 |

| | | | | | | | | | | | | | | | | | |
|---|---|---|---|---|---|---|---|---|---|---|---|---|---|---|---|---|---|
| BHLHA15 | GCAGCAGT | -1 | 1091 | 1099 | 91 | 99 | -2,75 | 0,01 | 15,67 | >0.1 | 5,64 | 6,87 | 0 | 1,85 | 1,79 | 2,27 | 0 |
| BHLHA15 | CAATCTGC | -1 | 891 | 899 | -109 | -101 | -1,66 | 0,01 | 15,42 | >0.1 | 5,64 | 6,6 | 0 | 1,89 | 0 | 2,95 | 0 |
| BHLHA15 | ACAGATAT | -1 | 853 | 861 | -147 | -139 | -3,24 | 0,01 | 13,24 | >0.1 | 5,64 | 6,3 | 0 | 1,87 | 0 | 2,67 | 0 |
| BHLHA15 | AGATATGT | -1 | 851 | 859 | -149 | -141 | -1,25 | 0,01 | 15,28 | >0.1 | 5,64 | 6,29 | 0 | 1,85 | 0 | 2,75 | 0 |
| BHLHA15 | ACAGATTT | -1 | 842 | 850 | -158 | -150 | -3,24 | 0,01 | 11,81 | >0.1 | 5,64 | 6,23 | 0 | 1,77 | 0 | 1,41 | 0 |
| BHLHA15 | AAAGCTGT | -1 | 712 | 720 | -288 | -280 | -4,74 | 0,01 | 12,62 | >0.1 | 5,64 | 5,56 | 0 | 1,18 | 0 | 4,99 | 0 |
| BHLHA15 | GTAGCTGG | -1 | 340 | 348 | -660 | -652 | -4,75 | 0,01 | 10,12 | >0.1 | 2,57 | 4,66 | 0 | 1,12 | 0 | 6,52 | 0 |
| BHLHA15 | ACAGGTGC | -1 | 328 | 336 | -672 | -664 | -2,84 | 0,01 | 11,11 | >0.1 | 2,6 | 4,64 | 0 | 1,14 | 0 | 5,56 | 0 |
| TFEB | ATCAGGAGAG | 1 | 97 | 107 | -903 | -893 | 1,55 | 0,01 | 12,18 | >0.1 | 3,64 | 4,35 | 0 | 1,88 | 0 | 0,77 | 0 |
| TFEB | GGCTCACGCC | 1 | 189 | 199 | -811 | -801 | 1,1 | 0,01 | 14,39 | >0.1 | 3,09 | 4,45 | 0 | 1,71 | 0 | 4,04 | 0 |
| TFEB | ATCACAAGGT | 1 | 235 | 245 | -765 | -755 | 5,13 | 0,01 | 18,71 | >0.1 | 2,9 | 4,51 | 0 | 1,43 | 0 | 4,74 | 0 |
| TFEB | GTCAGGAGAT | 1 | 243 | 253 | -757 | -747 | 3,49 | 0,01 | 16,6 | >0.1 | 2,87 | 4,52 | 0 | 1,39 | 0 | 4,33 | 0 |
| TFEB | GGCACCTGTA | 1 | 327 | 337 | -673 | -663 | 1,61 | 0,01 | 15,57 | >0.1 | 2,6 | 4,65 | 0 | 1,14 | 0 | 5,56 | 0 |
| TFEB | GGCAGGAGAA | 1 | 360 | 370 | -640 | -630 | -1,01 | 0,01 | 13,66 | >0.1 | 2,52 | 4,7 | 0 | 1,09 | 0 | 6,36 | 0 |
| TFEB | GACAGGAGAG | 1 | 768 | 778 | -232 | -222 | -0,13 | 0,01 | 18,55 | >0.1 | 5,61 | 5,81 | 0 | 1,35 | 0 | 5,92 | 0 |
| TFEB | ATCTCATGAG | 1 | 982 | 992 | -18 | -8 | 5,38 | 0,01 | 22,82 | >0.1 | 5,61 | 8,43 | 0 | 1,87 | 0 | 1,53 | 0 |
| TFEB | CTCATGAGGA | 1 | 984 | 994 | -16 | -6 | -0,63 | 0,01 | 16,95 | >0.1 | 5,61 | 8,57 | 0 | 1,87 | 0 | 1,53 | 0 |
| TFEB | CTCACAGGGG | 1 | 1036 | 1046 | 36 | 46 | 1 | 0,01 | 17,79 | >0.1 | 5,61 | 7,99 | 0 | 1,86 | 0 | 1,34 | 0 |
| TFEB | CACAGGGGAC | 1 | 1038 | 1048 | 38 | 48 | 0,9 | 0,01 | 17,62 | >0.1 | 5,61 | 7,92 | 0 | 1,86 | 0 | 1,34 | 0 |
| TFEB | GTCCCCTGTG | -1 | 1038 | 1048 | 38 | 48 | -0,17 | 0,01 | 16,55 | >0.1 | 5,61 | 7,92 | 0 | 1,86 | 0 | 1,34 | 0 |
| TFEB | CCCCTGTGAG | -1 | 1036 | 1046 | 36 | 46 | 3,18 | 0,01 | 19,97 | >0.1 | 5,61 | 7,99 | 0 | 1,86 | 0 | 1,34 | 0 |
| TFEB | TCCTCATGAG | -1 | 984 | 994 | -16 | -6 | 1,2 | 0,01 | 18,78 | >0.1 | 5,61 | 8,57 | 0 | 1,87 | 0 | 1,53 | 0 |
| TFEB | CTCATGAGAT | -1 | 982 | 992 | -18 | -8 | 5,52 | 0,01 | 22,96 | >0.1 | 5,61 | 8,43 | 0 | 1,87 | 0 | 1,53 | 0 |
| TFEB | CTCTCCTGTC | -1 | 768 | 778 | -232 | -222 | 1,4 | 0,01 | 20,08 | >0.1 | 5,61 | 5,81 | 0 | 1,35 | 0 | 5,92 | 0 |
| TFEB | AATACATGAC | -1 | 517 | 527 | -483 | -473 | 0,09 | 0,01 | 12,05 | >0.1 | 3,03 | 5 | 0 | 0,98 | 0 | 2,95 | 0 |
| TFEB | TTCTCCTGCC | -1 | 360 | 370 | -640 | -630 | 0,76 | 0,01 | 15,43 | >0.1 | 2,52 | 4,7 | 0 | 1,09 | 0 | 6,36 | 0 |
| TFEB | TACAGGTGCC | -1 | 327 | 337 | -673 | -663 | 2,18 | 0,01 | 16,14 | >0.1 | 2,6 | 4,65 | 0 | 1,14 | 0 | 5,56 | 0 |
| TFEB | ATCTCCTGAC | -1 | 243 | 253 | -757 | -747 | 6,02 | 0,01 | 19,13 | >0.1 | 2,87 | 4,52 | 0 | 1,39 | 0 | 4,33 | 0 |
| TFEB | ACCTTGTGAT | -1 | 235 | 245 | -765 | -755 | 5,64 | 0,01 | 19,22 | >0.1 | 2,9 | 4,51 | 0 | 1,43 | 0 | 4,74 | 0 |
| TFEB | GGCGTGAGCC | -1 | 189 | 199 | -811 | -801 | -0,87 | 0,01 | 12,42 | >0.1 | 3,09 | 4,45 | 0 | 1,71 | 0 | 4,04 | 0 |
| TFEB | TCCACAGGTC | -1 | 109 | 119 | -891 | -881 | 0,8 | 0,01 | 11,33 | >0.1 | 3,54 | 4,36 | 0 | 1,89 | 0 | 0,75 | 0 |
| TFEB | CTCTCCTGAT | -1 | 97 | 107 | -903 | -893 | 3,28 | 0,01 | 13,91 | >0.1 | 3,64 | 4,35 | 0 | 1,88 | 0 | 0,77 | 0 |
| LMX1A | GAAATTAA | 1 | 154 | 162 | -846 | -838 | 6,65 | 0,01 | 17,87 | >0.1 | 3,27 | 4,41 | 0 | 1,9 | 0 | 1,65 | 0 |
| LMX1A | TCTATTAA | 1 | 288 | 296 | -712 | -704 | 5,08 | 0,01 | 18,81 | >0.1 | 2,72 | 4,58 | 0 | 1,24 | 0 | 5,19 | 0 |
| LMX1A | GAAATAAA | 1 | 576 | 584 | -424 | -416 | 3,99 | 0,01 | 13,44 | >0.1 | 3,32 | 5,14 | 0 | 1 | 0 | 0 | 0 |
| LMX1A | CCTATTAA | 1 | 600 | 608 | -400 | -392 | 4,65 | 0,01 | 15,7 | >0.1 | 3,47 | 5,2 | 0 | 1,01 | 0 | 1,36 | 0 |
| LMX1A | TTAATGAA | 1 | 604 | 612 | -396 | -388 | 7,51 | 0,001 | 18,65 | >0.1 | 3,49 | 5,21 | 0 | 1,02 | 0 | 1,41 | 0 |
| LMX1A | CTGATAAG | 1 | 967 | 975 | -33 | -25 | 3,47 | 0,01 | 20,43 | >0.1 | 5,64 | 7,76 | 0 | 1,87 | 0 | 1,7 | 0 |
| LMX1A | TCAAATAA | -1 | 812 | 820 | -188 | -180 | 3,87 | 0,01 | 20,52 | >0.1 | 5,64 | 6,03 | 0 | 1,55 | 0 | 3,43 | 0 |
| LMX1A | CAAATAAG | -1 | 811 | 819 | -189 | -181 | 3,41 | 0,01 | 19,9 | >0.1 | 5,64 | 6,02 | 0 | 1,55 | 0 | 3,28 | 0 |
| LMX1A | CAAATCAA | -1 | 750 | 758 | -250 | -242 | 3,41 | 0,01 | 21,96 | >0.1 | 5,64 | 5,71 | 0 | 1,28 | 0 | 5,92 | 0 |
| LMX1A | TTCATTAA | -1 | 604 | 612 | -396 | -388 | 7,17 | 0,001 | 18,31 | >0.1 | 3,49 | 5,21 | 0 | 1,02 | 0 | 1,41 | 0 |
| LMX1A | TTAATAGG | -1 | 600 | 608 | -400 | -392 | 4,91 | 0,01 | 15,96 | >0.1 | 3,47 | 5,2 | 0 | 1,01 | 0 | 1,36 | 0 |
| LMX1A | TTAATAGA | -1 | 288 | 296 | -712 | -704 | 5,72 | 0,01 | 19,45 | >0.1 | 2,72 | 4,58 | 0 | 1,24 | 0 | 5,19 | 0 |
| LMX1A | TTAATTTC | -1 | 154 | 162 | -846 | -838 | 5,24 | 0,01 | 16,46 | >0.1 | 3,27 | 4,41 | 0 | 1,9 | 0 | 1,65 | 0 |
| LMX1B | GAAATTAA | 1 | 154 | 162 | -846 | -838 | 7,15 | 0,001 | 18,37 | >0.1 | 3,27 | 4,41 | 0 | 1,9 | 0 | 1,65 | 0 |
| LMX1B | TCTATTAA | 1 | 288 | 296 | -712 | -704 | 5,74 | 0,01 | 19,47 | >0.1 | 2,72 | 4,58 | 0 | 1,24 | 0 | 5,19 | 0 |
| LMX1B | TCCATAAA | 1 | 497 | 505 | -503 | -495 | 4,08 | 0,01 | 19,39 | >0.1 | 2,94 | 4,95 | 0 | 0,98 | 0 | 6,44 | 0 |
| LMX1B | GAAATAAA | 1 | 576 | 584 | -424 | -416 | 5,54 | 0,01 | 14,99 | >0.1 | 3,32 | 5,14 | 0 | 1 | 0 | 0 | 0 |
| LMX1B | AAAATAAA | 1 | 582 | 590 | -418 | -410 | 4,86 | 0,01 | 14,37 | >0.1 | 3,35 | 5,15 | 0 | 1 | 0 | 0 | 0 |
| LMX1B | CCTATTAA | 1 | 600 | 608 | -400 | -392 | 5,82 | 0,01 | 16,87 | >0.1 | 3,47 | 5,2 | 0 | 1,01 | 0 | 1,36 | 0 |
| LMX1B | TTAATGAA | 1 | 604 | 612 | -396 | -388 | 6,56 | 0,01 | 17,7 | >0.1 | 3,49 | 5,21 | 0 | 1,02 | 0 | 1,41 | 0 |
| LMX1B | AAAATTAT | 1 | 683 | 691 | -317 | -309 | 4,47 | 0,01 | 18,45 | >0.1 | 4,34 | 5,46 | 0 | 1,12 | 0 | 3,07 | 0 |
| LMX1B | GCCATAAA | 1 | 758 | 766 | -242 | -234 | 4,19 | 0,01 | 22,99 | >0.1 | 5,64 | 5,75 | 0 | 1,31 | 0 | 6,1 | 0 |
| LMX1B | AAAATTAT | 1 | 830 | 838 | -170 | -162 | 4,47 | 0,01 | 21,2 | >0.1 | 5,64 | 6,14 | 0 | 1,67 | 0 | 3,28 | 0 |
| LMX1B | CCAATATA | -1 | 879 | 887 | -121 | -113 | 4,34 | 0,01 | 21,45 | >0.1 | 5,64 | 6,5 | 0 | 1,9 | 0 | 3,07 | 0 |
| LMX1B | TCAAATAA | -1 | 812 | 820 | -188 | -180 | 5,23 | 0,01 | 21,88 | >0.1 | 5,64 | 6,03 | 0 | 1,55 | 0 | 3,43 | 0 |
| LMX1B | CAAATAAG | -1 | 811 | 819 | -189 | -181 | 4,63 | 0,01 | 21,12 | >0.1 | 5,64 | 6,02 | 0 | 1,55 | 0 | 3,28 | 0 |
| LMX1B | CAAATCAA | -1 | 750 | 758 | -250 | -242 | 4,3 | 0,01 | 22,85 | >0.1 | 5,64 | 5,71 | 0 | 1,28 | 0 | 5,92 | 0 |

| | | | | | | | | | | | | | | | | | |
|---|---|---|---|---|---|---|---|---|---|---|---|---|---|---|---|---|---|
| LMX1B | ACAATTTC | -1 | 609 | 617 | -391 | -383 | 4,12 | 0,01 | 15,44 | >0.1 | 3,53 | 5,23 | 0 | 1,02 | 0 | 1,55 | 0 |
| LMX1B | TTCATTAA | -1 | 604 | 612 | -396 | -388 | 6,11 | 0,01 | 17,25 | >0.1 | 3,49 | 5,21 | 0 | 1,02 | 0 | 1,41 | 0 |
| LMX1B | CCAAATAC | -1 | 522 | 530 | -478 | -470 | 4,14 | 0,01 | 16,24 | >0.1 | 3,05 | 5,01 | 0 | 0,98 | 0 | 3,07 | 0 |
| LMX1B | TTAATTTC | -1 | 154 | 162 | -846 | -838 | 5,12 | 0,01 | 16,34 | >0.1 | 3,27 | 4,41 | 0 | 1,9 | 0 | 1,65 | 0 |
| LMX1B | AAAATTAT | -1 | 132 | 140 | -868 | -860 | 4,47 | 0,01 | 15,71 | >0.1 | 3,4 | 4,38 | 0 | 1,89 | 0 | 1,57 | 0 |
| FOS::JUN(VAR.2) | AAAGAAGTCAGCCACA | 1 | 30 | 46 | -970 | -954 | 3,26 | 0,01 | 13,83 | >0.1 | 4,42 | 4,29 | 0 | 1,87 | 0 | 0 | 0 |
| FOS::JUN(VAR.2) | AGAGTAGTCATAATTT | 1 | 123 | 139 | -877 | -861 | 5,81 | 0,01 | 17,16 | >0.1 | 3,43 | 4,38 | 0 | 1,89 | 0 | 1,65 | 0 |
| FOS::JUN(VAR.2) | GGGCAGATCACAAGGT | 1 | 229 | 245 | -771 | -755 | 3,31 | 0,01 | 17,2 | >0.1 | 2,91 | 4,51 | 0 | 1,46 | 0 | 5,01 | 0 |
| FOS::JUN(VAR.2) | AATGGCGTGAACCTGG | 1 | 368 | 384 | -632 | -616 | 6,68 | 0,001 | 21,43 | >0.1 | 2,55 | 4,72 | 0 | 1,08 | 0 | 6,4 | 0 |
| FOS::JUN(VAR.2) | GATCGCGCCACTGTGC | 1 | 408 | 424 | -592 | -576 | 6,7 | 0,001 | 20,95 | >0.1 | 2,66 | 4,79 | 0 | 1,03 | 0 | 5,76 | 0 |
| FOS::JUN(VAR.2) | GGCAAAGTCATGTATT | 1 | 511 | 527 | -489 | -473 | 4,75 | 0,01 | 16,71 | >0.1 | 3,02 | 5 | 0 | 0,99 | 0 | 2,95 | 0 |
| FOS::JUN(VAR.2) | TGTGAGATAACCTATT | 1 | 590 | 606 | -410 | -394 | 3 | 0,01 | 12,64 | >0.1 | 3,43 | 5,2 | 0 | 1,02 | 0 | 0 | 0 |
| FOS::JUN(VAR.2) | GATTTGGCCATAAAGT | 1 | 752 | 768 | -248 | -232 | 2,03 | 0,01 | 20,96 | >0.1 | 5,53 | 5,76 | 0 | 1,32 | 0 | 6,32 | 0 |
| FOS::JUN(VAR.2) | AATAACGTATTCTTAT | 1 | 800 | 816 | -200 | -184 | 1,54 | 0,01 | 18,03 | >0.1 | 5,53 | 6,01 | 0 | 1,53 | 0 | 3,43 | 0 |
| FOS::JUN(VAR.2) | TCTGTCCTCTCCAGGA | 1 | 856 | 872 | -144 | -128 | 1,62 | 0,01 | 18,3 | >0.1 | 5,53 | 6,38 | 0 | 1,9 | 0 | 2,86 | 0 |
| FOS::JUN(VAR.2) | TATTGGCTCAGCAGAT | 1 | 881 | 897 | -119 | -103 | 1,58 | 0,01 | 18,79 | >0.1 | 5,53 | 6,59 | 0 | 1,89 | 0 | 3,19 | 0 |
| FOS::JUN(VAR.2) | GGGAAAGTCATTCAGT | 1 | 1006 | 1022 | 6 | 22 | 4,92 | 0,01 | 22,9 | >0.1 | 5,53 | 9,12 | 0 | 1,87 | 0 | 1,46 | 0 |
| FOS::JUN(VAR.2) | GATGTGATCTTGGCTC | 1 | 1023 | 1039 | 23 | 39 | 1,96 | 0,01 | 19,3 | >0.1 | 5,53 | 8,67 | 0 | 1,86 | 0 | 1,28 | 0 |
| FOS::JUN(VAR.2) | GACGATGTCAAGCTCT | 1 | 1045 | 1061 | 45 | 61 | 3,62 | 0,01 | 20,13 | >0.1 | 5,53 | 7,7 | 0 | 1,86 | 0 | 1,42 | 0 |
| FOS::JUN(VAR.2) | GGCCAAGACATTTTTG | 1 | 1120 | 1136 | 120 | 136 | 3,06 | 0,01 | 22,93 | >0.1 | 5,53 | 6,55 | 0 | 1,85 | 3,22 | 2,72 | 0 |
| FOS::JUN(VAR.2) | GAAGACCTGTTCTATC | 1 | 1157 | 1173 | 157 | 173 | 1,8 | 0,01 | 18,48 | >0.1 | 4,4 | 6,24 | 0 | 1,79 | 1,79 | 2,46 | 0 |
| FOS::JUN(VAR.2) | AGTTACAGCAACAAGG | -1 | 1078 | 1094 | 78 | 94 | 1,85 | 0,01 | 20,23 | >0.1 | 5,53 | 7,04 | 0 | 1,85 | 1,79 | 2,16 | 0 |
| FOS::JUN(VAR.2) | CTTGACATCGTCCCCT | -1 | 1041 | 1057 | 41 | 57 | 2,31 | 0,01 | 18,14 | >0.1 | 5,53 | 7,82 | 0 | 1,86 | 0 | 0,62 | 0 |
| FOS::JUN(VAR.2) | GACATCGTCCCCTGTG | -1 | 1038 | 1054 | 38 | 54 | 1,77 | 0,01 | 18,47 | >0.1 | 5,53 | 7,92 | 0 | 1,86 | 0 | 1,39 | 0 |
| FOS::JUN(VAR.2) | AACCTCCTCATGAGAT | -1 | 982 | 998 | -18 | -2 | 3,6 | 0,01 | 21,24 | >0.1 | 5,53 | 8,66 | 0 | 1,87 | 0 | 1,57 | 0 |
| FOS::JUN(VAR.2) | GAATACGTTATTCTTT | -1 | 796 | 812 | -204 | -188 | 7,41 | 0,001 | 23,98 | >0.1 | 5,53 | 5,98 | 0 | 1,51 | 0 | 3,55 | 0 |
| FOS::JUN(VAR.2) | GAGAACCTTACCTCTC | -1 | 773 | 789 | -227 | -211 | 2,26 | 0,01 | 21,37 | >0.1 | 5,53 | 5,86 | 0 | 1,4 | 0 | 6,32 | 0 |
| FOS::JUN(VAR.2) | CCTTACCTCTCCTGTC | -1 | 768 | 784 | -232 | -216 | 1,97 | 0,01 | 20,81 | >0.1 | 5,53 | 5,84 | 0 | 1,38 | 0 | 6,1 | 0 |
| FOS::JUN(VAR.2) | AGTGACCAAAGAGGTC | -1 | 665 | 681 | -335 | -319 | 2,23 | 0,01 | 16,57 | >0.1 | 4,11 | 5,42 | 0 | 1,1 | 0 | 3,7 | 0 |
| FOS::JUN(VAR.2) | GGTTATCTCACATTTA | -1 | 586 | 602 | -414 | -398 | 6,05 | 0,01 | 15,65 | >0.1 | 3,4 | 5,19 | 0 | 1,01 | 0 | 0 | 0 |
| FOS::JUN(VAR.2) | CATGACTTTGCCCTTG | -1 | 507 | 523 | -493 | -477 | 2 | 0,01 | 13,84 | >0.1 | 3 | 4,99 | 0 | 0,99 | 0 | 2,86 | 0 |
| FOS::JUN(VAR.2) | AGTGGCGCGATCTCGG | -1 | 404 | 420 | -596 | -580 | 4,61 | 0,01 | 18,72 | >0.1 | 2,65 | 4,79 | 0 | 1,04 | 0 | 5,65 | 0 |
| FOS::JUN(VAR.2) | GATCTCGGCTCACTGC | -1 | 396 | 412 | -604 | -588 | 2,65 | 0,01 | 17 | >0.1 | 2,62 | 4,77 | 0 | 1,04 | 0 | 5,91 | 0 |
| FOS::JUN(VAR.2) | GTTCACGCCATTCTCC | -1 | 364 | 380 | -636 | -620 | 4,79 | 0,01 | 19,85 | >0.1 | 2,54 | 4,72 | 0 | 1,08 | 0 | 6,73 | 0 |
| FOS::JUN(VAR.2) | TATCTCCTGACCTTGT | -1 | 238 | 254 | -762 | -746 | 2,62 | 0,01 | 15,77 | >0.1 | 2,88 | 4,53 | 0 | 1,42 | 0 | 4,33 | 0 |
| FOS::JUN(VAR.2) | GCTGACTTCTTTGAGT | -1 | 26 | 42 | -974 | -958 | 2,82 | 0,01 | 13,48 | >0.1 | 4,51 | 4,28 | 0 | 1,87 | 0 | 0 | 0 |
| SOX13 | AAAAAATGGCC | 1 | 140 | 151 | -860 | -849 | 7,11 | 0,001 | 18,38 | >0.1 | 3,34 | 4,4 | 0 | 1,89 | 0 | 1,65 | 0 |
| SOX13 | GCACAGTGGCT | 1 | 182 | 193 | -818 | -807 | 7,75 | 0,001 | 20,94 | >0.1 | 3,12 | 4,45 | 0 | 1,77 | 0 | 3,86 | 0 |
| SOX13 | GCACTTTGGGA | 1 | 209 | 220 | -791 | -780 | 1,62 | 0,01 | 15,37 | >0.1 | 3 | 4,48 | 0 | 1,57 | 0 | 4,7 | 0 |
| SOX13 | TCACAAGGTCA | 1 | 236 | 247 | -764 | -753 | 2,64 | 0,01 | 16,1 | >0.1 | 2,89 | 4,51 | 0 | 1,43 | 0 | 4,63 | 0 |
| SOX13 | ATACAAAAACT | 1 | 298 | 309 | -702 | -691 | 1,9 | 0,01 | 14,85 | >0.1 | 2,68 | 4,6 | 0 | 1,21 | 0 | 4,45 | 0 |
| SOX13 | GGAGAATGGCG | 1 | 364 | 375 | -636 | -625 | 7,71 | 0,001 | 22,64 | >0.1 | 2,53 | 4,71 | 0 | 1,08 | 0 | 6,61 | 0 |
| SOX13 | AGACAAGGGCA | 1 | 504 | 515 | -496 | -485 | 6,48 | 0,01 | 21,27 | >0.1 | 2,98 | 4,98 | 0 | 0,98 | 0 | 5,86 | 0 |
| SOX13 | GGAAAATGTTG | 1 | 533 | 544 | -467 | -456 | 3,22 | 0,01 | 13,96 | >0.1 | 3,1 | 5,04 | 0 | 0,99 | 0 | 1,61 | 0 |
| SOX13 | AAACCATACAA | 1 | 621 | 632 | -379 | -368 | 1,94 | 0,01 | 13,32 | >0.1 | 3,63 | 5,27 | 0 | 1,04 | 0 | 1,44 | 0 |
| SOX13 | AAAGAATAACG | 1 | 796 | 807 | -204 | -193 | 2,15 | 0,01 | 18,61 | >0.1 | 5,6 | 5,95 | 0 | 1,48 | 0 | 3,43 | 0 |
| SOX13 | GGACGATGTCA | 1 | 1044 | 1055 | 44 | 55 | 4,63 | 0,01 | 20,44 | >0.1 | 5,6 | 7,73 | 0 | 1,86 | 0 | 0,62 | 0 |
| SOX13 | CAAAAATGTCT | -1 | 1125 | 1136 | 125 | 136 | 2,9 | 0,01 | 22,83 | >0.1 | 5,6 | 6,5 | 0 | 1,84 | 3,22 | 2,77 | 0 |
| SOX13 | CCTCAATGGTG | -1 | 1105 | 1116 | 105 | 116 | 4,99 | 0,01 | 24,88 | >0.1 | 5,6 | 6,7 | 0 | 1,85 | 3,22 | 2,52 | 0 |
| SOX13 | CAACAAGGCTG | -1 | 1075 | 1086 | 75 | 86 | 3,43 | 0,01 | 22,04 | >0.1 | 5,6 | 7,09 | 0 | 1,85 | 1,79 | 2,27 | 0 |
| SOX13 | ACACAGTAAAC | -1 | 898 | 909 | -102 | -91 | 1,84 | 0,01 | 18,88 | >0.1 | 5,6 | 6,7 | 0 | 1,89 | 0 | 2,86 | 0 |
| SOX13 | AAACAATCTGC | -1 | 891 | 902 | -109 | -98 | 4,56 | 0,01 | 21,54 | >0.1 | 5,6 | 6,63 | 0 | 1,89 | 0 | 2,86 | 0 |
| SOX13 | GGAAAATGTTA | -1 | 690 | 701 | -310 | -299 | 2,96 | 0,01 | 17,06 | >0.1 | 4,52 | 5,49 | 0 | 1,14 | 0 | 2,95 | 0 |
| SOX13 | TTACAGTGACC | -1 | 674 | 685 | -326 | -315 | 4,04 | 0,01 | 18,22 | >0.1 | 4,21 | 5,44 | 0 | 1,11 | 0 | 3,43 | 0 |
| SOX13 | AGATAATGTTG | -1 | 636 | 647 | -364 | -353 | 4,41 | 0,01 | 16,2 | >0.1 | 3,76 | 5,31 | 0 | 1,05 | 0 | 1,66 | 0 |
| SOX13 | AGACAATTTCA | -1 | 608 | 619 | -392 | -381 | 4 | 0,01 | 15,34 | >0.1 | 3,53 | 5,23 | 0 | 1,02 | 0 | 1,55 | 0 |
| SOX13 | GCACAGTGGCG | -1 | 413 | 424 | -587 | -576 | 8,1 | 0,001 | 21,67 | >0.1 | 2,66 | 4,79 | 0 | 1,02 | 0 | 5,09 | 0 |
| SOX13 | TCACACTGGTT | -1 | 47 | 58 | -953 | -942 | 5,48 | 0,01 | 15,81 | >0.1 | 4,16 | 4,3 | 0 | 1,87 | 0 | 0 | 0 |
| MAFG | GTTTTGCAAAGGCAGATCAG( | 1 | 82 | 103 | -918 | -897 | -2,54 | 0,01 | 7,4 | >0.1 | 3,72 | 4,34 | 0 | 1,88 | 0 | 0 | 0 |

| | | | | | | | | | | | | | | | | | |
|---|---|---|---|---|---|---|---|---|---|---|---|---|---|---|---|---|---|
| MAFG | GGAGTGGAGAGTAGTCATAAT | 1 | 116 | 137 | -884 | -863 | 1,36 | 0,01 | 12,78 | >0.1 | 3,46 | 4,38 | 0 | 1,89 | 0 | 1,7 | 0 |
| MAFG | CACCTGTAGTCCCAGCTACT | 1 | 329 | 350 | -671 | -650 | -0,88 | 0,01 | 13,43 | >0.1 | 2,58 | 4,67 | 0 | 1,15 | 0 | 5,92 | 0 |
| MAFG | GGGAGGCTGAGGCAGGAGA | 1 | 350 | 371 | -650 | -629 | 8,34 | 0,001 | 22,71 | >0.1 | 2,53 | 4,7 | 0 | 1,11 | 0 | 6,03 | 0 |
| MAFG | GGGAGGCAGAGCTTGCAGTG | 1 | 382 | 403 | -618 | -597 | -2,13 | 0,01 | 12,41 | >0.1 | 2,59 | 4,76 | 0 | 1,06 | 0 | 6,13 | 0 |
| MAFG | AGTGAGCCGAGATCGCGCCA | 1 | 398 | 419 | -602 | -581 | 0,68 | 0,01 | 14,91 | >0.1 | 2,63 | 4,79 | 0 | 1,05 | 0 | 5,76 | 0 |
| MAFG | GTGCTCCAGCCTGGGCGACA | 1 | 420 | 441 | -580 | -559 | -2,15 | 0,01 | 12,07 | >0.1 | 2,7 | 4,83 | 0 | 1,03 | 0 | 5,67 | 0 |
| MAFG | CAAGGGCAAAGTCATGTATTT | 1 | 507 | 528 | -493 | -472 | -1,06 | 0,01 | 10,9 | >0.1 | 3,01 | 5,01 | 0 | 0,99 | 0 | 2,95 | 0 |
| MAFG | AAAATGTTGCCCAAGTAGAGA | 1 | 535 | 556 | -465 | -444 | -1,69 | 0,01 | 7,51 | >0.1 | 3,14 | 5,07 | 0 | 1 | 0 | 0 | 0 |
| MAFG | TAACAGCTTTCTAGGAAAATA | 1 | 710 | 731 | -290 | -269 | -1,49 | 0,01 | 15,96 | >0.1 | 5,47 | 5,6 | 0 | 1,21 | 0 | 5,17 | 0 |
| MAFG | TTTATATTGGCTCAGCAGATT | 1 | 877 | 898 | -123 | -102 | 5,28 | 0,001 | 22,32 | >0.1 | 5,47 | 6,6 | 0 | 1,9 | 0 | 3,07 | 0 |
| MAFG | GTCTTTCCTGCTCAGCGCCC | 1 | 931 | 952 | -69 | -48 | -2,24 | 0,01 | 13,27 | >0.1 | 5,47 | 7,25 | 0 | 1,88 | 0 | 0,9 | 0 |
| MAFG | TTGTTGCTGTAACTGCTGCT | 1 | 1080 | 1101 | 80 | 101 | 3,17 | 0,01 | 21,52 | >0.1 | 5,47 | 7,01 | 0 | 1,85 | 1,79 | 2,21 | 0 |
| MAFG | CTGCTGCTCAGTCCACCATT | 1 | 1092 | 1113 | 92 | 113 | 0,46 | 0,01 | 20,22 | >0.1 | 5,47 | 6,85 | 0 | 1,85 | 3,22 | 2,36 | 0 |
| MAFG | CAATGGTGGACTGAGCAGCA | -1 | 1092 | 1113 | 92 | 113 | 1,35 | 0,01 | 21,11 | >0.1 | 5,47 | 6,85 | 0 | 1,85 | 3,22 | 2,36 | 0 |
| MAFG | GAGCAGCAGTTACAGCAACA | -1 | 1080 | 1101 | 80 | 101 | 3,35 | 0,01 | 21,7 | >0.1 | 5,47 | 7,01 | 0 | 1,85 | 1,79 | 2,21 | 0 |
| MAFG | TGGGCGCTGAGCAGGAAAGA | -1 | 931 | 952 | -69 | -48 | -1,28 | 0,01 | 14,23 | >0.1 | 5,47 | 7,25 | 0 | 1,88 | 0 | 0,9 | 0 |
| MAFG | AATCTGCTGAGCCAATATAAA | -1 | 877 | 898 | -123 | -102 | 6,71 | 0,001 | 23,75 | >0.1 | 5,47 | 6,6 | 0 | 1,9 | 0 | 3,07 | 0 |
| MAFG | TATTTTCCTAGAAAGCTGTTA | -1 | 710 | 731 | -290 | -269 | -2,15 | 0,01 | 15,3 | >0.1 | 5,47 | 5,6 | 0 | 1,21 | 0 | 5,17 | 0 |
| MAFG | AAATACATGACTTTGCCCTTG | -1 | 507 | 528 | -493 | -472 | -2,42 | 0,01 | 9,54 | >0.1 | 3,01 | 5,01 | 0 | 0,99 | 0 | 2,95 | 0 |
| MAFG | CTGTCGCCCAGGCTGGAGCA | -1 | 420 | 441 | -580 | -559 | -1,74 | 0,01 | 12,48 | >0.1 | 2,7 | 4,83 | 0 | 1,03 | 0 | 5,67 | 0 |
| MAFG | GTGGCGCGATCTCGGCTCAC | -1 | 398 | 419 | -602 | -581 | 1,03 | 0,01 | 15,26 | >0.1 | 2,63 | 4,79 | 0 | 1,05 | 0 | 5,76 | 0 |
| MAFG | ATTCTCCTGCCTCAGCCTCC | -1 | 350 | 371 | -650 | -629 | 5,76 | 0,001 | 20,13 | >0.1 | 2,53 | 4,7 | 0 | 1,11 | 0 | 6,03 | 0 |
| MAFG | GAGTAGCTGGGACTACAGGT | -1 | 329 | 350 | -671 | -650 | -0,74 | 0,01 | 13,57 | >0.1 | 2,58 | 4,67 | 0 | 1,15 | 0 | 5,92 | 0 |
| MAFG | ATTATGACTACTCTCCACTCC | -1 | 116 | 137 | -884 | -863 | 1,05 | 0,01 | 12,47 | >0.1 | 3,46 | 4,38 | 0 | 1,89 | 0 | 1,7 | 0 |
| TFAP2B(VAR.2) | AGGCTGAGGCA | 1 | 353 | 364 | -647 | -636 | 7,05 | 0,001 | 21,77 | >0.1 | 2,53 | 4,69 | 0 | 1,1 | 0 | 6,4 | 0 |
| TFAP2B(VAR.2) | AGCTTGCAGTG | 1 | 391 | 402 | -609 | -598 | -3,07 | 0,01 | 11,35 | >0.1 | 2,6 | 4,75 | 0 | 1,05 | 0 | 6,02 | 0 |
| TFAP2B(VAR.2) | TGCTCCAGCCT | 1 | 421 | 432 | -579 | -568 | 1,55 | 0,01 | 15,48 | >0.1 | 2,69 | 4,81 | 0 | 1,02 | 0 | 5,42 | 0 |
| TFAP2B(VAR.2) | AGCCTGGGCGA | 1 | 427 | 438 | -573 | -562 | 2,03 | 0,01 | 16,37 | >0.1 | 2,71 | 4,82 | 0 | 1,01 | 0 | 5,8 | 0 |
| TFAP2B(VAR.2) | CTCTCCAGGAT | 1 | 862 | 873 | -138 | -127 | -3,34 | 0,01 | 13,3 | >0.1 | 5,6 | 6,39 | 0 | 1,9 | 0 | 2,75 | 0 |
| TFAP2B(VAR.2) | AGCCGAAGACC | 1 | 1153 | 1164 | 153 | 164 | 1,78 | 0,01 | 18,49 | >0.1 | 4,47 | 6,27 | 0 | 1,83 | 1,79 | 2,34 | 0 |
| TFAP2B(VAR.2) | TCGCCCAGGCT | -1 | 427 | 438 | -573 | -562 | 2,92 | 0,01 | 17,26 | >0.1 | 2,71 | 4,82 | 0 | 1,01 | 0 | 5,8 | 0 |
| TFAP2B(VAR.2) | AGGCTGGAGCA | -1 | 421 | 432 | -579 | -568 | -0,19 | 0,01 | 13,74 | >0.1 | 2,69 | 4,81 | 0 | 1,02 | 0 | 5,42 | 0 |
| TFAP2B(VAR.2) | CACTGCAAGCT | -1 | 391 | 402 | -609 | -598 | -1,83 | 0,01 | 12,59 | >0.1 | 2,6 | 4,75 | 0 | 1,05 | 0 | 6,02 | 0 |
| TFAP2B(VAR.2) | TGCCTCAGCCT | -1 | 353 | 364 | -647 | -636 | 7,57 | 0,001 | 22,29 | >0.1 | 2,53 | 4,69 | 0 | 1,1 | 0 | 6,4 | 0 |
| TFAP2B(VAR.2) | AGCCTCCCGAG | -1 | 347 | 358 | -653 | -642 | -1,63 | 0,01 | 13,23 | >0.1 | 2,55 | 4,68 | 0 | 1,11 | 0 | 6,52 | 0 |
| TFAP2C(VAR.2) | TGGCTCACGCC | 1 | 188 | 199 | -812 | -801 | 2,2 | 0,01 | 15,41 | >0.1 | 3,09 | 4,45 | 0 | 1,72 | 0 | 3,95 | 0 |
| TFAP2C(VAR.2) | CTCGGGAGGCT | 1 | 347 | 358 | -653 | -642 | 0,16 | 0,01 | 15,02 | >0.1 | 2,55 | 4,68 | 0 | 1,11 | 0 | 6,52 | 0 |
| TFAP2C(VAR.2) | AGGCTGAGGCA | 1 | 353 | 364 | -647 | -636 | 11,11 | 0,001 | 25,83 | >0.1 | 2,53 | 4,69 | 0 | 1,1 | 0 | 6,4 | 0 |
| TFAP2C(VAR.2) | AGCTTGCAGTG | 1 | 391 | 402 | -609 | -598 | 0,08 | 0,01 | 14,5 | >0.1 | 2,6 | 4,75 | 0 | 1,05 | 0 | 6,02 | 0 |
| TFAP2C(VAR.2) | TGCTCCAGCCT | 1 | 421 | 432 | -579 | -568 | 4,83 | 0,01 | 18,76 | >0.1 | 2,69 | 4,81 | 0 | 1,02 | 0 | 5,42 | 0 |
| TFAP2C(VAR.2) | CAGCCTGGGCG | 1 | 426 | 437 | -574 | -563 | 0,22 | 0,01 | 14,42 | >0.1 | 2,7 | 4,82 | 0 | 1,01 | 0 | 5,67 | 0 |
| TFAP2C(VAR.2) | AGCCTGGGCGA | 1 | 427 | 438 | -573 | -562 | 5,48 | 0,01 | 19,82 | >0.1 | 2,71 | 4,82 | 0 | 1,01 | 0 | 5,8 | 0 |
| TFAP2C(VAR.2) | CGCCCAACCCA | 1 | 946 | 957 | -54 | -43 | 1,59 | 0,01 | 18,21 | >0.1 | 5,6 | 7,34 | 0 | 1,88 | 0 | 1,8 | 0 |
| TFAP2C(VAR.2) | TCGCCCAGGCT | -1 | 427 | 438 | -573 | -562 | 6,44 | 0,01 | 20,78 | >0.1 | 2,71 | 4,82 | 0 | 1,01 | 0 | 5,8 | 0 |
| TFAP2C(VAR.2) | CGCCCAGGCTG | -1 | 426 | 437 | -574 | -563 | -0,6 | 0,01 | 13,6 | >0.1 | 2,7 | 4,82 | 0 | 1,01 | 0 | 5,67 | 0 |
| TFAP2C(VAR.2) | AGGCTGGAGCA | -1 | 421 | 432 | -579 | -568 | 5,88 | 0,01 | 19,81 | >0.1 | 2,69 | 4,81 | 0 | 1,02 | 0 | 5,42 | 0 |
| TFAP2C(VAR.2) | CACTGCAAGCT | -1 | 391 | 402 | -609 | -598 | 1,6 | 0,01 | 16,02 | >0.1 | 2,6 | 4,75 | 0 | 1,05 | 0 | 6,02 | 0 |
| TFAP2C(VAR.2) | TGCCTCAGCCT | -1 | 353 | 364 | -647 | -636 | 9,97 | 0,001 | 24,69 | >0.1 | 2,53 | 4,69 | 0 | 1,1 | 0 | 6,4 | 0 |
| TFAP2C(VAR.2) | AGCCTCCCGAG | -1 | 347 | 358 | -653 | -642 | 1,57 | 0,01 | 16,43 | >0.1 | 2,55 | 4,68 | 0 | 1,11 | 0 | 6,52 | 0 |
| TFAP2C(VAR.2) | AGGCGTGAGCC | -1 | 189 | 200 | -811 | -800 | 0 | 0,01 | 13,29 | >0.1 | 3,08 | 4,45 | 0 | 1,71 | 0 | 4,04 | 0 |
| TFAP2C(VAR.2) | GGCGTGAGCCA | -1 | 188 | 199 | -812 | -801 | 1,01 | 0,01 | 14,22 | >0.1 | 3,09 | 4,45 | 0 | 1,72 | 0 | 3,95 | 0 |
| ETV1 | GAAGGAGGCA | 1 | 17 | 27 | -983 | -973 | -0,38 | 0,01 | 11,36 | >0.1 | 5,61 | 4,27 | 0 | 1,87 | 0 | 0 | 0 |
| ETV1 | ATAGGAAATG | 1 | 61 | 71 | -939 | -929 | 5,04 | 0,001 | 15,21 | >0.1 | 3,98 | 4,31 | 0 | 1,88 | 0 | 0 | 0 |
| ETV1 | TCAGGAGATA | 1 | 244 | 254 | -756 | -746 | 1,29 | 0,01 | 14,49 | >0.1 | 2,86 | 4,52 | 0 | 1,39 | 0 | 4,42 | 0 |
| ETV1 | TCGGGAGGCT | 1 | 348 | 358 | -652 | -642 | -1,26 | 0,01 | 13,47 | >0.1 | 2,55 | 4,68 | 0 | 1,11 | 0 | 6,4 | 0 |
| ETV1 | AAGGGAAAAT | 1 | 530 | 540 | -470 | -460 | -0,79 | 0,01 | 12,01 | >0.1 | 3,09 | 5,03 | 0 | 0,98 | 0 | 3,7 | 0 |
| ETV1 | ACCGGTTTTG | 1 | 743 | 753 | -257 | -247 | 1,61 | 0,01 | 20,09 | >0.1 | 5,61 | 5,69 | 0 | 1,26 | 0 | 5,92 | 0 |
| ETV1 | ACAGGAGAGG | 1 | 769 | 779 | -231 | -221 | 1,01 | 0,01 | 19,48 | >0.1 | 5,61 | 5,81 | 0 | 1,35 | 0 | 5,69 | 0 |
| ETV1 | CCAGGATGAA | 1 | 866 | 876 | -134 | -124 | 4,16 | 0,01 | 20,83 | >0.1 | 5,61 | 6,41 | 0 | 1,9 | 0 | 2,75 | 0 |

| | | | | | | | | | | | | | | | | | |
|---|---|---|---|---|---|---|---|---|---|---|---|---|---|---|---|---|---|
| ETV1 | TGAGGAGGTT | 1 | 988 | 998 | -12 | -2 | -0,51 | 0,01 | 17,24 | >0.1 | 5,61 | 8,66 | 0 | 1,87 | 0 | 1,61 | 0 |
| ETV1 | AGTGGATGTG | 1 | 1019 | 1029 | 19 | 29 | 3,69 | 0,01 | 21,21 | >0.1 | 5,61 | 8,74 | 0 | 1,86 | 0 | 1,31 | 0 |
| ETV1 | TGAGGAACAG | 1 | 1111 | 1121 | 111 | 121 | 0,62 | 0,01 | 20,48 | >0.1 | 5,61 | 6,64 | 0 | 1,85 | 3,22 | 2,54 | 0 |
| ETV1 | CCAGGAAGAG | -1 | 1057 | 1067 | 57 | 67 | 5,68 | 0,001 | 22,84 | >0.1 | 5,61 | 7,42 | 0 | 1,86 | 0 | 2,27 | 0 |
| ETV1 | GCAGGAAAGA | -1 | 932 | 942 | -68 | -58 | 5,71 | 0,001 | 21,22 | >0.1 | 5,61 | 7,09 | 0 | 1,88 | 0 | 0,93 | 0 |
| ETV1 | ACAGTAAACA | -1 | 897 | 907 | -103 | -93 | 2,59 | 0,01 | 19,63 | >0.1 | 5,61 | 6,68 | 0 | 1,89 | 0 | 2,86 | 0 |
| ETV1 | AGAGGACAGA | -1 | 856 | 866 | -144 | -134 | 0,57 | 0,01 | 17,37 | >0.1 | 5,61 | 6,34 | 0 | 1,9 | 0 | 2,95 | 0 |
| ETV1 | ACCGGTACTT | -1 | 739 | 749 | -261 | -251 | 5,16 | 0,001 | 23,61 | >0.1 | 5,61 | 5,68 | 0 | 1,25 | 0 | 5,92 | 0 |
| ETV1 | CACGGAAAAT | -1 | 694 | 704 | -306 | -296 | 3,24 | 0,01 | 18,56 | >0.1 | 5,61 | 5,5 | 0 | 1,14 | 0 | 3,07 | 0 |
| ETV1 | CCAGGCTGGA | -1 | 424 | 434 | -576 | -566 | -1,46 | 0,01 | 12,73 | >0.1 | 2,69 | 4,81 | 0 | 1,01 | 0 | 5,67 | 0 |
| ETV1 | TCAGGACGGT | -1 | 257 | 267 | -743 | -733 | 2,06 | 0,01 | 15,39 | >0.1 | 2,82 | 4,54 | 0 | 1,34 | 0 | 4,63 | 0 |
| ETV3 | ATAGGAAATG | 1 | 61 | 71 | -939 | -929 | 6,5 | 0,001 | 16,67 | >0.1 | 3,98 | 4,31 | 0 | 1,88 | 0 | 0 | 0 |
| ETV3 | CATGGAAATT | 1 | 150 | 160 | -850 | -840 | 1,3 | 0,01 | 12,53 | >0.1 | 3,28 | 4,41 | 0 | 1,9 | 0 | 1,65 | 0 |
| ETV3 | ATCAGAAATG | 1 | 167 | 177 | -833 | -823 | 0,98 | 0,01 | 12,9 | >0.1 | 3,19 | 4,43 | 0 | 1,9 | 0 | 2,4 | 0 |
| ETV3 | GCCTGTAATC | 1 | 196 | 206 | -804 | -794 | 1,48 | 0,01 | 15,03 | >0.1 | 3,06 | 4,46 | 0 | 1,66 | 0 | 4,37 | 0 |
| ETV3 | CCCGTCTCTA | 1 | 282 | 292 | -718 | -708 | -0,64 | 0,01 | 13,02 | >0.1 | 2,73 | 4,58 | 0 | 1,26 | 0 | 5,09 | 0 |
| ETV3 | ACCTGTAGTC | 1 | 330 | 340 | -670 | -660 | 5,86 | 0,001 | 19,7 | >0.1 | 2,59 | 4,65 | 0 | 1,14 | 0 | 5,46 | 0 |
| ETV3 | GCTTGCAGTG | 1 | 392 | 402 | -608 | -598 | 2,06 | 0,01 | 16,48 | >0.1 | 2,6 | 4,75 | 0 | 1,04 | 0 | 6,02 | 0 |
| ETV3 | ATCAGAAATA | 1 | 482 | 492 | -518 | -508 | -0,43 | 0,01 | 14,17 | >0.1 | 2,89 | 4,93 | 0 | 0,99 | 0 | 5,8 | 0 |
| ETV3 | TTTGGAAGGG | 1 | 525 | 535 | -475 | -465 | 1,85 | 0,01 | 14,2 | >0.1 | 3,06 | 5,02 | 0 | 0,98 | 0 | 3,28 | 0 |
| ETV3 | TCTTGAAATA | 1 | 572 | 582 | -428 | -418 | 1,33 | 0,01 | 10,76 | >0.1 | 3,3 | 5,13 | 0 | 1 | 0 | 0 | 0 |
| ETV3 | ACCGGTTTTG | 1 | 743 | 753 | -257 | -247 | -0,63 | 0,01 | 17,85 | >0.1 | 5,61 | 5,69 | 0 | 1,26 | 0 | 5,92 | 0 |
| ETV3 | AGTGGATGTG | 1 | 1019 | 1029 | 19 | 29 | 7,17 | 0,001 | 24,69 | >0.1 | 5,61 | 8,74 | 0 | 1,86 | 0 | 1,31 | 0 |
| ETV3 | GCTGTAACTG | 1 | 1085 | 1095 | 85 | 95 | 2,54 | 0,01 | 20,95 | >0.1 | 5,61 | 6,94 | 0 | 1,85 | 1,79 | 2,21 | 0 |
| ETV3 | ACCTGTTCTA | 1 | 1161 | 1171 | 161 | 171 | 0,19 | 0,01 | 16,9 | >0.1 | 4,42 | 6,22 | 0 | 1,75 | 1,79 | 2,54 | 0 |
| ETV3 | CCAGGAAGAG | -1 | 1057 | 1067 | 57 | 67 | -0,39 | 0,01 | 16,77 | >0.1 | 5,61 | 7,42 | 0 | 1,86 | 0 | 2,27 | 0 |
| ETV3 | GCAGGAAAGA | -1 | 932 | 942 | -68 | -58 | 1,98 | 0,01 | 17,49 | >0.1 | 5,61 | 7,09 | 0 | 1,88 | 0 | 0,93 | 0 |
| ETV3 | ACAGTAAACA | -1 | 897 | 907 | -103 | -93 | 1,91 | 0,01 | 18,95 | >0.1 | 5,61 | 6,68 | 0 | 1,89 | 0 | 2,86 | 0 |
| ETV3 | ACCGGTACTT | -1 | 739 | 749 | -261 | -251 | 5,15 | 0,001 | 23,6 | >0.1 | 5,61 | 5,68 | 0 | 1,25 | 0 | 5,92 | 0 |
| ETV3 | TCCTGATCTG | -1 | 94 | 104 | -906 | -896 | 4,13 | 0,01 | 14,8 | >0.1 | 3,66 | 4,34 | 0 | 1,88 | 0 | 0,79 | 0 |
| ETV3 | ACTGGTTCTG | -1 | 44 | 54 | -956 | -946 | 1,49 | 0,01 | 11,87 | >0.1 | 4,22 | 4,29 | 0 | 1,87 | 0 | 0 | 0 |
| ETV2 | AATAGGAAATG | 1 | 60 | 71 | -940 | -929 | 0,85 | 0,01 | 11,03 | >0.1 | 3,99 | 4,31 | 0 | 1,88 | 0 | 0 | 0 |
| ETV2 | GATCAGAAATG | 1 | 166 | 177 | -834 | -823 | -9,41 | 0,01 | 2,57 | >0.1 | 3,19 | 4,43 | 0 | 1,9 | 0 | 2,46 | 0 |
| ETV2 | GTCAGGAGATA | 1 | 243 | 254 | -757 | -746 | -6,81 | 0,01 | 6,3 | >0.1 | 2,87 | 4,52 | 0 | 1,4 | 0 | 4,33 | 0 |
| ETV2 | GATCAGAAATA | 1 | 481 | 492 | -519 | -508 | -9,15 | 0,01 | 5,3 | >0.1 | 2,89 | 4,93 | 0 | 0,99 | 0 | 5,65 | 0 |
| ETV2 | GAAGGGAAAAT | 1 | 529 | 540 | -471 | -460 | -4,2 | 0,01 | 8,45 | >0.1 | 3,08 | 5,03 | 0 | 0,98 | 0 | 3,55 | 0 |
| ETV2 | AAGGGAAAATG | 1 | 530 | 541 | -470 | -459 | -7,27 | 0,01 | 5,54 | >0.1 | 3,09 | 5,03 | 0 | 0,98 | 0 | 3,7 | 0 |
| ETV2 | AACCAAAAGTA | 1 | 733 | 744 | -267 | -256 | -8,91 | 0,01 | 9,5 | >0.1 | 5,6 | 5,66 | 0 | 1,24 | 0 | 5,92 | 0 |
| ETV2 | GACAGGAGAGG | 1 | 768 | 779 | -232 | -221 | -2,9 | 0,01 | 15,78 | >0.1 | 5,6 | 5,81 | 0 | 1,35 | 0 | 5,92 | 0 |
| ETV2 | GAGAGGTAAGG | 1 | 773 | 784 | -227 | -216 | -6,83 | 0,01 | 12,08 | >0.1 | 5,6 | 5,83 | 0 | 1,37 | 0 | 6,1 | 0 |
| ETV2 | TCCAGGATGAA | 1 | 865 | 876 | -135 | -124 | -3,81 | 0,01 | 12,77 | >0.1 | 5,6 | 6,41 | 0 | 1,9 | 0 | 2,67 | 0 |
| ETV2 | GAGAGAAAATC | 1 | 974 | 985 | -26 | -15 | -5,12 | 0,01 | 12,05 | >0.1 | 5,6 | 8,09 | 0 | 1,87 | 0 | 1,61 | 0 |
| ETV2 | CAGTGGATGTG | 1 | 1018 | 1029 | 18 | 29 | -4,36 | 0,01 | 13,2 | >0.1 | 5,6 | 8,76 | 0 | 1,86 | 0 | 1,34 | 0 |
| ETV2 | ACCACGAAGCC | 1 | 1146 | 1157 | 146 | 157 | -8,23 | 0,01 | 9,49 | >0.1 | 5,6 | 6,33 | 0 | 1,84 | 1,79 | 2,16 | 0 |
| ETV2 | AAGCCGAAGAC | 1 | 1152 | 1163 | 152 | 163 | -6,55 | 0,01 | 10,12 | >0.1 | 4,5 | 6,28 | 0 | 1,84 | 1,79 | 2,26 | 0 |
| ETV2 | TCCAAGAAGCA | -1 | 1183 | 1194 | 183 | 194 | -8,7 | 0,01 | 9,84 | >0.1 | 5,6 | 6,07 | 0 | 1,56 | 1,79 | 3,52 | 0 |
| ETV2 | GCCAGGAAGAG | -1 | 1057 | 1068 | 57 | 68 | -0,41 | 0,01 | 16,73 | >0.1 | 5,6 | 7,42 | 0 | 1,86 | 0 | 2,27 | 0 |
| ETV2 | AGCAGGAAAGA | -1 | 932 | 943 | -68 | -57 | 5,29 | 0,001 | 20,8 | >0.1 | 5,6 | 7,11 | 0 | 1,88 | 0 | 0,93 | 0 |
| ETV2 | AAGAAGAAGAA | -1 | 908 | 919 | -92 | -81 | -9,19 | 0,01 | 6,3 | >0.1 | 5,6 | 6,8 | 0 | 1,89 | 0 | 1,21 | 0 |
| ETV2 | CACAGTAAACA | -1 | 897 | 908 | -103 | -92 | -1,57 | 0,01 | 15,46 | >0.1 | 5,6 | 6,69 | 0 | 1,89 | 0 | 2,86 | 0 |
| ETV2 | GAGAGGACAGA | -1 | 856 | 867 | -144 | -133 | -3,5 | 0,01 | 13,29 | >0.1 | 5,6 | 6,35 | 0 | 1,9 | 0 | 2,95 | 0 |
| ETV2 | AACCGGTACTT | -1 | 739 | 750 | -261 | -250 | -4,33 | 0,01 | 14,12 | >0.1 | 5,6 | 5,68 | 0 | 1,26 | 0 | 5,92 | 0 |
| ETV2 | ACACGGAAAAT | -1 | 694 | 705 | -306 | -295 | -0,96 | 0,01 | 14,35 | >0.1 | 5,6 | 5,5 | 0 | 1,14 | 0 | 3,07 | 0 |
| ETV2 | CACGGAAAATG | -1 | 693 | 704 | -307 | -296 | -5,57 | 0,01 | 9,74 | >0.1 | 5,6 | 5,5 | 0 | 1,14 | 0 | 3,07 | 0 |
| ETV2 | TCCCAGAAACT | -1 | 555 | 566 | -445 | -434 | -5,71 | 0,01 | 3,59 | >0.1 | 3,21 | 5,09 | 0 | 0,99 | 0 | 0 | 0 |
| ETV5 | GAAGGAGGCA | 1 | 17 | 27 | -983 | -973 | 2,29 | 0,01 | 14,03 | >0.1 | 5,61 | 4,27 | 0 | 1,87 | 0 | 0 | 0 |
| ETV5 | ATAGGAAATG | 1 | 61 | 71 | -939 | -929 | 5,66 | 0,001 | 15,83 | >0.1 | 3,98 | 4,31 | 0 | 1,88 | 0 | 0 | 0 |
| ETV5 | TCAGGAGATA | 1 | 244 | 254 | -756 | -746 | 2,08 | 0,01 | 15,28 | >0.1 | 2,86 | 4,52 | 0 | 1,39 | 0 | 4,42 | 0 |
| ETV5 | CTCGGGAGGC | 1 | 347 | 357 | -653 | -643 | 0,17 | 0,01 | 15,03 | >0.1 | 2,55 | 4,68 | 0 | 1,11 | 0 | 6,52 | 0 |

| | | | | | | | | | | | | | | | | | |
|---|---|---|---|---|---|---|---|---|---|---|---|---|---|---|---|---|---|
| ETV5 | TCGGGAGGCT | 1 | 348 | 358 | -652 | -642 | 1,27 | 0,01 | 16 | >0.1 | 2,55 | 4,68 | 0 | 1,11 | 0 | 6,4 | 0 |
| ETV5 | CTAGGAAAAT | 1 | 720 | 730 | -280 | -270 | -0,11 | 0,01 | 17,29 | >0.1 | 5,61 | 5,6 | 0 | 1,2 | 0 | 4,99 | 0 |
| ETV5 | ACCGGTTTTG | 1 | 743 | 753 | -257 | -247 | 3,39 | 0,01 | 21,87 | >0.1 | 5,61 | 5,69 | 0 | 1,26 | 0 | 5,92 | 0 |
| ETV5 | ACAGGAGAGG | 1 | 769 | 779 | -231 | -221 | 2,37 | 0,01 | 20,84 | >0.1 | 5,61 | 5,81 | 0 | 1,35 | 0 | 5,69 | 0 |
| ETV5 | CCAGGATGAA | 1 | 866 | 876 | -134 | -124 | 4,81 | 0,01 | 21,48 | >0.1 | 5,61 | 6,41 | 0 | 1,9 | 0 | 2,75 | 0 |
| ETV5 | TGAGGAGGTT | 1 | 988 | 998 | -12 | -2 | 2,35 | 0,01 | 20,1 | >0.1 | 5,61 | 8,66 | 0 | 1,87 | 0 | 1,61 | 0 |
| ETV5 | AGTGGATGTG | 1 | 1019 | 1029 | 19 | 29 | 3,48 | 0,01 | 21 | >0.1 | 5,61 | 8,74 | 0 | 1,86 | 0 | 1,31 | 0 |
| ETV5 | TGAGGAACAG | 1 | 1111 | 1121 | 111 | 121 | 2,01 | 0,01 | 21,87 | >0.1 | 5,61 | 6,64 | 0 | 1,85 | 3,22 | 2,54 | 0 |
| ETV5 | CCAGGAAGAG | -1 | 1057 | 1067 | 57 | 67 | 5,89 | 0,001 | 23,05 | >0.1 | 5,61 | 7,42 | 0 | 1,86 | 0 | 2,27 | 0 |
| ETV5 | GCAGGAAAGA | -1 | 932 | 942 | -68 | -58 | 5,48 | 0,01 | 20,99 | >0.1 | 5,61 | 7,09 | 0 | 1,88 | 0 | 0,93 | 0 |
| ETV5 | ACAGTAAACA | -1 | 897 | 907 | -103 | -93 | 0,46 | 0,01 | 17,5 | >0.1 | 5,61 | 6,68 | 0 | 1,89 | 0 | 2,86 | 0 |
| ETV5 | AGAGGACAGA | -1 | 856 | 866 | -144 | -134 | 1,26 | 0,01 | 18,06 | >0.1 | 5,61 | 6,34 | 0 | 1,9 | 0 | 2,95 | 0 |
| ETV5 | ACCGGTACTT | -1 | 739 | 749 | -261 | -251 | 5,21 | 0,01 | 23,66 | >0.1 | 5,61 | 5,68 | 0 | 1,25 | 0 | 5,92 | 0 |
| ETV5 | CACGGAAAAT | -1 | 694 | 704 | -306 | -296 | 4,02 | 0,01 | 19,34 | >0.1 | 5,61 | 5,5 | 0 | 1,14 | 0 | 3,07 | 0 |
| ETV5 | TCAGGACGGT | -1 | 257 | 267 | -743 | -733 | 3,03 | 0,01 | 16,36 | >0.1 | 2,82 | 4,54 | 0 | 1,34 | 0 | 4,63 | 0 |
| ETV5 | TCCTGATCTG | -1 | 94 | 104 | -906 | -896 | 1,32 | 0,01 | 11,99 | >0.1 | 3,66 | 4,34 | 0 | 1,88 | 0 | 0,79 | 0 |
| ETV4 | GAAGGAGGCA | 1 | 17 | 27 | -983 | -973 | 0,21 | 0,01 | 11,95 | >0.1 | 5,61 | 4,27 | 0 | 1,87 | 0 | 0 | 0 |
| ETV4 | ATAGGAAATG | 1 | 61 | 71 | -939 | -929 | 4,8 | 0,01 | 14,97 | >0.1 | 3,98 | 4,31 | 0 | 1,88 | 0 | 0 | 0 |
| ETV4 | TCAGGAGAGT | 1 | 98 | 108 | -902 | -892 | -1,61 | 0,01 | 9,01 | >0.1 | 3,63 | 4,35 | 0 | 1,88 | 0 | 0,77 | 0 |
| ETV4 | CATGGAAATT | 1 | 150 | 160 | -850 | -840 | -1,97 | 0,01 | 9,26 | >0.1 | 3,28 | 4,41 | 0 | 1,9 | 0 | 1,65 | 0 |
| ETV4 | TCAGGAGATA | 1 | 244 | 254 | -756 | -746 | 1,71 | 0,01 | 14,91 | >0.1 | 2,86 | 4,52 | 0 | 1,39 | 0 | 4,42 | 0 |
| ETV4 | TCGGGAGGCT | 1 | 348 | 358 | -652 | -642 | -2,02 | 0,01 | 12,71 | >0.1 | 2,55 | 4,68 | 0 | 1,11 | 0 | 6,4 | 0 |
| ETV4 | GCAGGAGAAT | 1 | 361 | 371 | -639 | -629 | -1,9 | 0,01 | 12,89 | >0.1 | 2,52 | 4,7 | 0 | 1,08 | 0 | 6,49 | 0 |
| ETV4 | AAGGGAAAAT | 1 | 530 | 540 | -470 | -460 | -0,6 | 0,01 | 12,2 | >0.1 | 3,09 | 5,03 | 0 | 0,98 | 0 | 3,7 | 0 |
| ETV4 | CTAGGAAAAT | 1 | 720 | 730 | -280 | -270 | -1,85 | 0,01 | 15,55 | >0.1 | 5,61 | 5,6 | 0 | 1,2 | 0 | 4,99 | 0 |
| ETV4 | ACCGGTTTTG | 1 | 743 | 753 | -257 | -247 | 0,36 | 0,01 | 18,84 | >0.1 | 5,61 | 5,69 | 0 | 1,26 | 0 | 5,92 | 0 |
| ETV4 | ACAGGAGAGG | 1 | 769 | 779 | -231 | -221 | 1,16 | 0,01 | 19,63 | >0.1 | 5,61 | 5,81 | 0 | 1,35 | 0 | 5,69 | 0 |
| ETV4 | CCAGGATGAA | 1 | 866 | 876 | -134 | -124 | 4,9 | 0,01 | 21,57 | >0.1 | 5,61 | 6,41 | 0 | 1,9 | 0 | 2,75 | 0 |
| ETV4 | TGAGGAGGTT | 1 | 988 | 998 | -12 | -2 | -0,14 | 0,01 | 17,61 | >0.1 | 5,61 | 8,66 | 0 | 1,87 | 0 | 1,61 | 0 |
| ETV4 | TAGGGAAAGT | 1 | 1004 | 1014 | 4 | 14 | -1,37 | 0,01 | 16,64 | >0.1 | 5,61 | 9,12 | 0 | 1,87 | 0 | 1,42 | 0 |
| ETV4 | AGTGGATGTG | 1 | 1019 | 1029 | 19 | 29 | 2,52 | 0,01 | 20,04 | >0.1 | 5,61 | 8,74 | 0 | 1,86 | 0 | 1,31 | 0 |
| ETV4 | GGACGATGTC | 1 | 1044 | 1054 | 44 | 54 | -1,57 | 0,01 | 14,25 | >0.1 | 5,61 | 7,73 | 0 | 1,86 | 0 | 0,62 | 0 |
| ETV4 | TGAGGAACAG | 1 | 1111 | 1121 | 111 | 121 | 1,1 | 0,01 | 20,96 | >0.1 | 5,61 | 6,64 | 0 | 1,85 | 3,22 | 2,54 | 0 |
| ETV4 | CCACGAAGCC | 1 | 1147 | 1157 | 147 | 157 | -0,34 | 0,01 | 17,43 | >0.1 | 5,61 | 6,32 | 0 | 1,84 | 1,79 | 2,21 | 0 |
| ETV4 | AGCCGAAGAC | 1 | 1153 | 1163 | 153 | 163 | 1,22 | 0,01 | 17,94 | >0.1 | 4,48 | 6,27 | 0 | 1,83 | 1,79 | 2,34 | 0 |
| ETV4 | CCAGGAAGAG | -1 | 1057 | 1067 | 57 | 67 | 5,93 | 0,001 | 23,09 | >0.1 | 5,61 | 7,42 | 0 | 1,86 | 0 | 2,27 | 0 |
| ETV4 | GCAGGAAAGA | -1 | 932 | 942 | -68 | -58 | 6,75 | 0,001 | 22,26 | >0.1 | 5,61 | 7,09 | 0 | 1,88 | 0 | 0,93 | 0 |
| ETV4 | ACAGTAAACA | -1 | 897 | 907 | -103 | -93 | 1,79 | 0,01 | 18,83 | >0.1 | 5,61 | 6,68 | 0 | 1,89 | 0 | 2,86 | 0 |
| ETV4 | AGAGGACAGA | -1 | 856 | 866 | -144 | -134 | 0,94 | 0,01 | 17,74 | >0.1 | 5,61 | 6,34 | 0 | 1,9 | 0 | 2,95 | 0 |
| ETV4 | ACCGGTACTT | -1 | 739 | 749 | -261 | -251 | 2,88 | 0,01 | 21,33 | >0.1 | 5,61 | 5,68 | 0 | 1,25 | 0 | 5,92 | 0 |
| ETV4 | CACGGAAAAT | -1 | 694 | 704 | -306 | -296 | 4,08 | 0,01 | 19,4 | >0.1 | 5,61 | 5,5 | 0 | 1,14 | 0 | 3,07 | 0 |
| ETV4 | TCAGGACGGT | -1 | 257 | 267 | -743 | -733 | 2,18 | 0,01 | 15,51 | >0.1 | 2,82 | 4,54 | 0 | 1,34 | 0 | 4,63 | 0 |
| ETV6 | TAGGTAAGTG | 1 | 2 | 12 | -998 | -988 | -2,38 | 0,01 | 11,14 | >0.1 | 5,61 | 4,26 | 0 | 1,86 | 1,79 | 0 | 0 |
| ETV6 | GAAGGAGGCA | 1 | 17 | 27 | -983 | -973 | -1,38 | 0,01 | 10,36 | >0.1 | 5,61 | 4,27 | 0 | 1,87 | 0 | 0 | 0 |
| ETV6 | ATAGGAAATG | 1 | 61 | 71 | -939 | -929 | 6,73 | 0,001 | 16,9 | >0.1 | 3,98 | 4,31 | 0 | 1,88 | 0 | 0 | 0 |
| ETV6 | CATGGAAATT | 1 | 150 | 160 | -850 | -840 | 2,03 | 0,01 | 13,26 | >0.1 | 3,28 | 4,41 | 0 | 1,9 | 0 | 1,65 | 0 |
| ETV6 | ATCAGAAATG | 1 | 167 | 177 | -833 | -823 | -1,02 | 0,01 | 10,9 | >0.1 | 3,19 | 4,43 | 0 | 1,9 | 0 | 2,4 | 0 |
| ETV6 | TCAGGAGATA | 1 | 244 | 254 | -756 | -746 | -0,91 | 0,01 | 12,29 | >0.1 | 2,86 | 4,52 | 0 | 1,39 | 0 | 4,42 | 0 |
| ETV6 | TCGGGAGGCT | 1 | 348 | 358 | -652 | -642 | -2,1 | 0,01 | 12,63 | >0.1 | 2,55 | 4,68 | 0 | 1,11 | 0 | 6,4 | 0 |
| ETV6 | CTGGGAGGCA | 1 | 380 | 390 | -620 | -610 | -1,92 | 0,01 | 12,35 | >0.1 | 2,57 | 4,73 | 0 | 1,06 | 0 | 5,91 | 0 |
| ETV6 | ATCAGAAATA | 1 | 482 | 492 | -518 | -508 | -2,16 | 0,01 | 12,44 | >0.1 | 2,89 | 4,93 | 0 | 0,99 | 0 | 5,8 | 0 |
| ETV6 | TTTGGAAGGG | 1 | 525 | 535 | -475 | -465 | 1,51 | 0,01 | 13,86 | >0.1 | 3,06 | 5,02 | 0 | 0,98 | 0 | 3,28 | 0 |
| ETV6 | AAGGGAAAAT | 1 | 530 | 540 | -470 | -460 | -2,26 | 0,01 | 10,54 | >0.1 | 3,09 | 5,03 | 0 | 0,98 | 0 | 3,7 | 0 |
| ETV6 | AGGGAAAATG | 1 | 531 | 541 | -469 | -459 | 0,32 | 0,01 | 11,04 | >0.1 | 3,09 | 5,03 | 0 | 0,98 | 0 | 1,61 | 0 |
| ETV6 | CTAGGAAAAT | 1 | 720 | 730 | -280 | -270 | -0,52 | 0,01 | 16,88 | >0.1 | 5,61 | 5,6 | 0 | 1,2 | 0 | 4,99 | 0 |
| ETV6 | ACAGGAGAGG | 1 | 769 | 779 | -231 | -221 | -1,88 | 0,01 | 16,59 | >0.1 | 5,61 | 5,81 | 0 | 1,35 | 0 | 5,69 | 0 |
| ETV6 | CCAGGATGAA | 1 | 866 | 876 | -134 | -124 | -0,4 | 0,01 | 16,27 | >0.1 | 5,61 | 6,41 | 0 | 1,9 | 0 | 2,75 | 0 |
| ETV6 | AGAGAAAATC | 1 | 975 | 985 | -25 | -15 | -1,51 | 0,01 | 15,64 | >0.1 | 5,61 | 8,09 | 0 | 1,87 | 0 | 1,57 | 0 |
| ETV6 | TGAGGAGGTT | 1 | 988 | 998 | -12 | -2 | 2,64 | 0,01 | 20,39 | >0.1 | 5,61 | 8,66 | 0 | 1,87 | 0 | 1,61 | 0 |

| | | | | | | | | | | | | | | | | | |
|---|---|---|---|---|---|---|---|---|---|---|---|---|---|---|---|---|---|
| ETV6 | AGGGAAAGTC | 1 | 1005 | 1015 | 5 | 15 | 0,71 | 0,01 | 18,72 | >0.1 | 5,61 | 9,12 | 0 | 1,87 | 0 | 1,42 | 0 |
| ETV6 | AGTGGATGTG | 1 | 1019 | 1029 | 19 | 29 | 4,53 | 0,01 | 22,05 | >0.1 | 5,61 | 8,74 | 0 | 1,86 | 0 | 1,31 | 0 |
| ETV6 | TGAGGAACAG | 1 | 1111 | 1121 | 111 | 121 | -0,29 | 0,01 | 19,57 | >0.1 | 5,61 | 6,64 | 0 | 1,85 | 3,22 | 2,54 | 0 |
| ETV6 | CCAAGAAGCA | -1 | 1183 | 1193 | 183 | 193 | -0,74 | 0,01 | 17,82 | >0.1 | 5,61 | 6,07 | 0 | 1,56 | 1,79 | 3,52 | 0 |
| ETV6 | CCAGGAAGAG | -1 | 1057 | 1067 | 57 | 67 | 8,08 | 0,001 | 25,24 | >0.1 | 5,61 | 7,42 | 0 | 1,86 | 0 | 2,27 | 0 |
| ETV6 | GCAGGAAAGA | -1 | 932 | 942 | -68 | -58 | 3,67 | 0,01 | 19,18 | >0.1 | 5,61 | 7,09 | 0 | 1,88 | 0 | 0,93 | 0 |
| ETV6 | AGAAGAAGAA | -1 | 908 | 918 | -92 | -82 | -2,55 | 0,01 | 12,94 | >0.1 | 5,61 | 6,79 | 0 | 1,89 | 0 | 1,21 | 0 |
| ETV6 | ACAGTAAACA | -1 | 897 | 907 | -103 | -93 | -2,71 | 0,01 | 14,33 | >0.1 | 5,61 | 6,68 | 0 | 1,89 | 0 | 2,86 | 0 |
| ETV6 | CACGGAAAAT | -1 | 694 | 704 | -306 | -296 | 1,72 | 0,01 | 17,04 | >0.1 | 5,61 | 5,5 | 0 | 1,14 | 0 | 3,07 | 0 |
| ETV6 | ACGGAAAATG | -1 | 693 | 703 | -307 | -297 | -0,53 | 0,01 | 13,76 | >0.1 | 4,58 | 5,5 | 0 | 1,14 | 0 | 3,07 | 0 |
| ETV6 | CCCAGAAACT | -1 | 555 | 565 | -445 | -435 | -2,66 | 0,01 | 6,63 | >0.1 | 3,21 | 5,09 | 0 | 0,99 | 0 | 0 | 0 |
| HNF4G | GTTTTGCAAAGGCAG | 1 | 82 | 97 | -918 | -903 | 0,5 | 0,01 | 10,46 | >0.1 | 3,74 | 4,34 | 0 | 1,88 | 0 | 0 | 0 |
| HNF4G | TGGGCACAGTGGCTC | 1 | 179 | 194 | -821 | -806 | 0,09 | 0,01 | 13,31 | >0.1 | 3,12 | 4,45 | 0 | 1,8 | 0 | 3,86 | 0 |
| HNF4G | AGATCACAAGGTCAG | 1 | 233 | 248 | -767 | -752 | 0,49 | 0,01 | 14,09 | >0.1 | 2,9 | 4,52 | 0 | 1,44 | 0 | 4,74 | 0 |
| HNF4G | AGGAGATAGAGACCG | 1 | 246 | 261 | -754 | -739 | 1,6 | 0,01 | 15,45 | >0.1 | 2,85 | 4,53 | 0 | 1,38 | 0 | 5,09 | 0 |
| HNF4G | GGGAGGCAGAGCTTG | 1 | 382 | 397 | -618 | -603 | 2,12 | 0,01 | 16,52 | >0.1 | 2,58 | 4,75 | 0 | 1,06 | 0 | 6,02 | 0 |
| HNF4G | AGTGAGCCGAGATCG | 1 | 398 | 413 | -602 | -587 | -0,64 | 0,01 | 13,71 | >0.1 | 2,63 | 4,77 | 0 | 1,04 | 0 | 5,91 | 0 |
| HNF4G | TAAAGACAAGGGCAA | 1 | 501 | 516 | -499 | -484 | 4,97 | 0,01 | 19,59 | >0.1 | 2,97 | 4,98 | 0 | 0,99 | 0 | 5,68 | 0 |
| HNF4G | CAAGGGCAAAGTCAT | 1 | 507 | 522 | -493 | -478 | 10,05 | 0,001 | 21,78 | >0.1 | 3 | 4,99 | 0 | 0,99 | 0 | 2,75 | 0 |
| HNF4G | CAAGTAGAGAGTTTC | 1 | 546 | 561 | -454 | -439 | 2,77 | 0,01 | 12,02 | >0.1 | 3,17 | 5,08 | 0 | 0,99 | 0 | 0 | 0 |
| HNF4G | GAGAGGTAAGGTTCT | 1 | 773 | 788 | -227 | -212 | 4,99 | 0,01 | 24,1 | >0.1 | 5,54 | 5,85 | 0 | 1,39 | 0 | 6,32 | 0 |
| HNF4G | TAGGATTAAAGAATA | 1 | 789 | 804 | -211 | -196 | -0,58 | 0,01 | 17,69 | >0.1 | 5,54 | 5,94 | 0 | 1,47 | 0 | 5,32 | 0 |
| HNF4G | CCCAACCCAAGTTCA | 1 | 948 | 963 | -52 | -37 | 1,22 | 0,01 | 17,84 | >0.1 | 5,54 | 7,46 | 0 | 1,88 | 0 | 1,74 | 0 |
| HNF4G | CAAGTTCAAAGGCTG | 1 | 955 | 970 | -45 | -30 | 11 | 0,001 | 27,78 | >0.1 | 5,54 | 7,62 | 0 | 1,88 | 0 | 1,74 | 0 |
| HNF4G | TTGGCTCACAGGGGA | 1 | 1032 | 1047 | 32 | 47 | -0,66 | 0,01 | 16,2 | >0.1 | 5,54 | 8,15 | 0 | 1,86 | 0 | 1,31 | 0 |
| HNF4G | CGAAGCCGAAGACCT | 1 | 1150 | 1165 | 150 | 165 | 3,09 | 0,01 | 19,78 | >0.1 | 4,5 | 6,3 | 0 | 1,84 | 1,79 | 2,26 | 0 |
| HNF4G | TTCTATCAAAGTTCA | 1 | 1166 | 1181 | 166 | 181 | 5,15 | 0,01 | 23,21 | >0.1 | 5,54 | 6,18 | 0 | 1,7 | 1,79 | 2,84 | 0 |
| HNF4G | TGTGAGCCAAGATCA | -1 | 1027 | 1042 | 27 | 42 | 3,62 | 0,01 | 20,76 | >0.1 | 5,54 | 8,39 | 0 | 1,86 | 0 | 1,34 | 0 |
| HNF4G | GAGCAGGAAAGACCA | -1 | 929 | 944 | -71 | -56 | 3,09 | 0,01 | 18,59 | >0.1 | 5,54 | 7,12 | 0 | 1,88 | 0 | 0,95 | 0 |
| HNF4G | GAAGAACACAGTAAA | -1 | 899 | 914 | -101 | -86 | -0,68 | 0,01 | 14,78 | >0.1 | 5,54 | 6,75 | 0 | 1,89 | 0 | 1,28 | 0 |
| HNF4G | CCAATATAAAGTTCA | -1 | 872 | 887 | -128 | -113 | 7,64 | 0,001 | 24,54 | >0.1 | 5,54 | 6,5 | 0 | 1,9 | 0 | 2,95 | 0 |
| HNF4G | AGTGACCAAAGAGGT | -1 | 666 | 681 | -334 | -319 | 1,37 | 0,01 | 15,71 | >0.1 | 4,12 | 5,42 | 0 | 1,1 | 0 | 3,7 | 0 |
| HNF4G | TAGAACTAGGGATCA | -1 | 650 | 665 | -350 | -335 | 0,01 | 0,01 | 14,54 | >0.1 | 3,92 | 5,37 | 0 | 1,08 | 0 | 4,16 | 0 |
| HNF4G | GGACTACAGGTGCCC | -1 | 326 | 341 | -674 | -659 | -0,2 | 0,01 | 13,76 | >0.1 | 2,6 | 4,65 | 0 | 1,15 | 0 | 5,56 | 0 |
| HNF4G | GCCTCCCAAAGTGCT | -1 | 208 | 223 | -792 | -777 | 2,93 | 0,01 | 16,69 | >0.1 | 2,99 | 4,48 | 0 | 1,58 | 0 | 4,7 | 0 |
| HNF4G | TAATTTCCATGGCCA | -1 | 146 | 161 | -854 | -839 | -0,69 | 0,01 | 10,55 | >0.1 | 3,29 | 4,41 | 0 | 1,9 | 0 | 1,65 | 0 |
| HNF4G | CCACTCCACAGGTCA | -1 | 108 | 123 | -892 | -877 | 5,89 | 0,01 | 16,44 | >0.1 | 3,53 | 4,36 | 0 | 1,89 | 0 | 0,77 | 0 |
| HNF4A | GAGTGGAGAGTAGTCA | 1 | 117 | 133 | -883 | -867 | -18,81 | 0,01 | -1,75 | >0.1 | 3,47 | 4,38 | 0 | 1,89 | 0 | 1,65 | 5,67 |
| HNF4A | GGAGGCCGAGTTGGGC | 1 | 217 | 233 | -783 | -767 | -18,42 | 0,01 | 1,75 | >0.1 | 2,96 | 4,5 | 0 | 1,53 | 0 | 5,52 | 5,67 |
| HNF4A | GGAGATAGAGACCGTC | 1 | 247 | 263 | -753 | -737 | -18,52 | 0,01 | 1,11 | >0.1 | 2,84 | 4,54 | 0 | 1,38 | 0 | 5,19 | 5,67 |
| HNF4A | ACGGTGAAACCCCGTC | 1 | 272 | 288 | -728 | -712 | -16,98 | 0,01 | 2,4 | >0.1 | 2,76 | 4,57 | 0 | 1,29 | 0 | 5,09 | 5,67 |
| HNF4A | GGAGGCAGAGCTTGCA | 1 | 383 | 399 | -617 | -601 | -12,74 | 0,01 | 7,46 | >0.1 | 2,59 | 4,75 | 0 | 1,06 | 0 | 6,13 | 5,67 |
| HNF4A | AAATATGAATTCCATA | 1 | 487 | 503 | -513 | -497 | -18,51 | 0,01 | 2,29 | >0.1 | 2,92 | 4,95 | 0 | 0,99 | 0 | 6,27 | 5,67 |
| HNF4A | AAGTAGAGAGTTTCTG | 1 | 547 | 563 | -453 | -437 | -16,75 | 0,01 | -1,81 | >0.1 | 3,18 | 5,09 | 0 | 0,99 | 0 | 0 | 5,67 |
| HNF4A | AGAGGTAAGGTTCTCT | 1 | 774 | 790 | -226 | -210 | -7,44 | 0,001 | 17,58 | >0.1 | 5,53 | 5,86 | 0 | 1,4 | 0 | 6,54 | 5,67 |
| HNF4A | AGGATTAAAGAATAAC | 1 | 790 | 806 | -210 | -194 | -11,36 | 0,01 | 12,59 | >0.1 | 5,53 | 5,95 | 0 | 1,48 | 0 | 5,32 | 5,67 |
| HNF4A | CAGGATGAACTTTATA | 1 | 867 | 883 | -133 | -117 | -15,22 | 0,01 | 7,22 | >0.1 | 5,53 | 6,47 | 0 | 1,9 | 0 | 2,86 | 5,67 |
| HNF4A | CCAACCCAAGTTCAAA | 1 | 949 | 965 | -51 | -35 | -18,33 | 0,01 | 3,96 | >0.1 | 5,53 | 7,51 | 0 | 1,88 | 0 | 1,7 | 5,67 |
| HNF4A | GAGGTTTTAGTCTAGG | 1 | 992 | 1008 | -8 | 8 | -10,26 | 0,01 | 13,35 | >0.1 | 5,53 | 9,01 | 0 | 1,87 | 0 | 1,53 | 5,67 |
| HNF4A | GGGGACGATGTCAAGC | 1 | 1042 | 1058 | 42 | 58 | -13,21 | 0,01 | 8,29 | >0.1 | 5,53 | 7,79 | 0 | 1,86 | 0 | 0,65 | 5,67 |
| HNF4A | GATGTCAAGCTCTTCC | 1 | 1048 | 1064 | 48 | 64 | -16,46 | 0,01 | 5,69 | >0.1 | 5,53 | 7,62 | 0 | 1,86 | 0 | 1,46 | 5,67 |
| HNF4A | GAAGCCGAAGACCTGT | 1 | 1151 | 1167 | 151 | 167 | -15,05 | 0,01 | 7,32 | >0.1 | 4,46 | 6,29 | 0 | 1,84 | 1,79 | 2,31 | 5,67 |
| HNF4A | GTGGTTAAACTTGTCC | -1 | 1135 | 1151 | 135 | 151 | -19,88 | 0,01 | 5,94 | >0.1 | 5,53 | 6,41 | 0 | 1,84 | 3,22 | 3,13 | 5,67 |
| HNF4A | GTGAGCCAAGATCACA | -1 | 1025 | 1041 | 25 | 41 | -20,32 | 0,01 | 2,57 | >0.1 | 5,53 | 8,52 | 0 | 1,86 | 0 | 1,31 | 5,67 |
| HNF4A | CAATATAAAGTTCATC | -1 | 870 | 886 | -130 | -114 | -6,37 | 0,001 | 16,18 | >0.1 | 5,53 | 6,5 | 0 | 1,9 | 0 | 2,95 | 5,67 |
| HNF4A | AAGATCATATTCCCAG | -1 | 560 | 576 | -440 | -424 | -8,08 | 0,01 | 6,96 | >0.1 | 3,25 | 5,12 | 0 | 1 | 0 | 0 | 5,67 |
| HNF4A | GAATTCATATTTCTGA | -1 | 483 | 499 | -517 | -501 | -17,45 | 0,01 | 3,33 | >0.1 | 2,9 | 4,94 | 0 | 0,99 | 0 | 6,27 | 5,67 |
| HNF4A | CAGGCTGGAGCACAGT | -1 | 417 | 433 | -583 | -567 | -14,54 | 0,01 | 5,07 | >0.1 | 2,68 | 4,81 | 0 | 1,02 | 0 | 5,42 | 5,67 |
| HOXD13 | ATTTTAAAAA | 1 | 135 | 145 | -865 | -855 | 1,71 | 0,01 | 12,94 | >0.1 | 3,37 | 4,39 | 0 | 1,89 | 0 | 1,57 | 0 |

| | | | | | | | | | | | | | | | | | |
|---|---|---|---|---|---|---|---|---|---|---|---|---|---|---|---|---|---|
| HOXD13 | GAAATTAAAA | 1 | 154 | 164 | -846 | -836 | 4,99 | 0,01 | 16,21 | >0.1 | 3,26 | 4,41 | 0 | 1,9 | 0 | 1,65 | 0 |
| HOXD13 | AAATTAAAAC | 1 | 155 | 165 | -845 | -835 | 0,3 | 0,01 | 12,47 | >0.1 | 3,25 | 4,41 | 0 | 1,9 | 0 | 2,6 | 0 |
| HOXD13 | TCTATTAAAA | 1 | 288 | 298 | -712 | -702 | 4,76 | 0,01 | 18,39 | >0.1 | 2,72 | 4,59 | 0 | 1,24 | 0 | 5,09 | 0 |
| HOXD13 | CTATTAAAAA | 1 | 289 | 299 | -711 | -701 | 8,85 | 0,001 | 22,47 | >0.1 | 2,71 | 4,59 | 0 | 1,24 | 0 | 5,09 | 0 |
| HOXD13 | CTCAAAAAAA | 1 | 454 | 464 | -546 | -536 | 1,17 | 0,01 | 15,35 | >0.1 | 2,79 | 4,87 | 0 | 1 | 0 | 5,52 | 0 |
| HOXD13 | TCCATAAAGA | 1 | 497 | 507 | -503 | -493 | 5,94 | 0,01 | 21,26 | >0.1 | 2,95 | 4,96 | 0 | 0,98 | 0 | 6,44 | 0 |
| HOXD13 | GAAATAAAAA | 1 | 576 | 586 | -424 | -414 | 7,64 | 0,001 | 17,11 | >0.1 | 3,32 | 5,14 | 0 | 1 | 0 | 0 | 0 |
| HOXD13 | AAAATAAATG | 1 | 582 | 592 | -418 | -408 | -0,26 | 0,01 | 9,26 | >0.1 | 3,36 | 5,16 | 0 | 1 | 0 | 0 | 0 |
| HOXD13 | CCTATTAATG | 1 | 600 | 610 | -400 | -390 | 4,04 | 0,01 | 15,1 | >0.1 | 3,47 | 5,21 | 0 | 1,02 | 0 | 1,36 | 0 |
| HOXD13 | ACCATACAAA | 1 | 623 | 633 | -377 | -367 | 2,39 | 0,01 | 13,79 | >0.1 | 3,64 | 5,27 | 0 | 1,04 | 0 | 1,44 | 0 |
| HOXD13 | ACTGTAAAAT | 1 | 678 | 688 | -322 | -312 | 2,54 | 0,01 | 16,65 | >0.1 | 4,27 | 5,45 | 0 | 1,11 | 0 | 3,28 | 0 |
| HOXD13 | CACTTTAAAA | 1 | 822 | 832 | -178 | -168 | 1,17 | 0,01 | 18,21 | >0.1 | 5,61 | 6,1 | 0 | 1,63 | 0 | 3,7 | 0 |
| HOXD13 | ACTTTAAAAA | 1 | 823 | 833 | -177 | -167 | 2,71 | 0,01 | 19,76 | >0.1 | 5,61 | 6,11 | 0 | 1,64 | 0 | 3,7 | 0 |
| HOXD13 | CCAAAAAAAG | -1 | 922 | 932 | -78 | -68 | 1,42 | 0,01 | 16,95 | >0.1 | 5,61 | 6,95 | 0 | 1,88 | 0 | 1,09 | 0 |
| HOXD13 | ACAGTAAACA | -1 | 897 | 907 | -103 | -93 | 3,27 | 0,01 | 20,31 | >0.1 | 5,61 | 6,68 | 0 | 1,89 | 0 | 2,86 | 0 |
| HOXD13 | CCAATATAAA | -1 | 877 | 887 | -123 | -113 | 7,05 | 0,001 | 24,01 | >0.1 | 5,61 | 6,5 | 0 | 1,9 | 0 | 2,95 | 0 |
| HOXD13 | GCTGTTAAAG | -1 | 707 | 717 | -293 | -283 | 2,27 | 0,01 | 19,41 | >0.1 | 5,61 | 5,55 | 0 | 1,17 | 0 | 4,81 | 0 |
| HOXD13 | TTCATTAATA | -1 | 602 | 612 | -398 | -388 | 2,53 | 0,01 | 13,66 | >0.1 | 3,49 | 5,21 | 0 | 1,02 | 0 | 1,41 | 0 |
| HOXD13 | CCCAGAAACT | -1 | 555 | 565 | -445 | -435 | -0,59 | 0,01 | 8,7 | >0.1 | 3,21 | 5,09 | 0 | 0,99 | 0 | 0 | 0 |
| HOXD13 | TTTTTTAAAA | -1 | 136 | 146 | -864 | -854 | -0,8 | 0,01 | 10,46 | >0.1 | 3,36 | 4,39 | 0 | 1,89 | 0 | 1,61 | 0 |
| HOXD13 | TTTTTAAAAT | -1 | 135 | 145 | -865 | -855 | 1,29 | 0,01 | 12,52 | >0.1 | 3,37 | 4,39 | 0 | 1,89 | 0 | 1,57 | 0 |
| HOXD13 | AACTTAAAAA | -1 | 75 | 85 | -925 | -915 | -0,46 | 0,01 | 9,57 | >0.1 | 3,83 | 4,32 | 0 | 1,88 | 0 | 0 | 0 |
| HOXD11 | GTCATAATTT | 1 | 129 | 139 | -871 | -861 | 5,92 | 0,001 | 17,21 | >0.1 | 3,41 | 4,38 | 0 | 1,89 | 0 | 1,61 | 0 |
| HOXD11 | ATTTTAAAAA | 1 | 135 | 145 | -865 | -855 | -0,6 | 0,01 | 10,63 | >0.1 | 3,37 | 4,39 | 0 | 1,89 | 0 | 1,57 | 0 |
| HOXD11 | ATCAGAAATG | 1 | 167 | 177 | -833 | -823 | -1,18 | 0,01 | 10,74 | >0.1 | 3,19 | 4,43 | 0 | 1,9 | 0 | 2,4 | 0 |
| HOXD11 | CTATTAAAAA | 1 | 289 | 299 | -711 | -701 | -1,86 | 0,01 | 11,76 | >0.1 | 2,71 | 4,59 | 0 | 1,24 | 0 | 5,09 | 0 |
| HOXD11 | ATTAAAAATA | 1 | 291 | 301 | -709 | -699 | 1,31 | 0,01 | 14,29 | >0.1 | 2,71 | 4,59 | 0 | 1,23 | 0 | 4,45 | 0 |
| HOXD11 | CTCAAAAAAA | 1 | 454 | 464 | -546 | -536 | 6,06 | 0,001 | 20,24 | >0.1 | 2,79 | 4,87 | 0 | 1 | 0 | 5,52 | 0 |
| HOXD11 | ATCAGAAATA | 1 | 482 | 492 | -518 | -508 | 0,26 | 0,01 | 14,86 | >0.1 | 2,89 | 4,93 | 0 | 0,99 | 0 | 5,8 | 0 |
| HOXD11 | CTTGAAATAA | 1 | 573 | 583 | -427 | -417 | -0,49 | 0,01 | 8,95 | >0.1 | 3,31 | 5,14 | 0 | 1 | 0 | 0 | 0 |
| HOXD11 | GAAATAAAAA | 1 | 576 | 586 | -424 | -414 | -1,53 | 0,01 | 7,94 | >0.1 | 3,32 | 5,14 | 0 | 1 | 0 | 0 | 0 |
| HOXD11 | ACTGTAAAAT | 1 | 678 | 688 | -322 | -312 | 0,04 | 0,01 | 14,15 | >0.1 | 4,27 | 5,45 | 0 | 1,11 | 0 | 3,28 | 0 |
| HOXD11 | ATTATAACAT | 1 | 686 | 696 | -314 | -304 | 2,18 | 0,01 | 16,15 | >0.1 | 4,42 | 5,47 | 0 | 1,13 | 0 | 2,95 | 0 |
| HOXD11 | ATCTTTAACA | 1 | 705 | 715 | -295 | -285 | -0,92 | 0,01 | 16,38 | >0.1 | 5,61 | 5,54 | 0 | 1,17 | 0 | 4,99 | 0 |
| HOXD11 | CTAGGAAAAT | 1 | 720 | 730 | -280 | -270 | -1,22 | 0,01 | 16,18 | >0.1 | 5,61 | 5,6 | 0 | 1,2 | 0 | 4,99 | 0 |
| HOXD11 | GCCATAAAGT | 1 | 758 | 768 | -242 | -232 | 0,41 | 0,01 | 19,19 | >0.1 | 5,61 | 5,76 | 0 | 1,31 | 0 | 6,1 | 0 |
| HOXD11 | ATTCTAAAAT | 1 | 836 | 846 | -164 | -154 | -0,45 | 0,01 | 16,29 | >0.1 | 5,61 | 6,2 | 0 | 1,74 | 0 | 3,19 | 0 |
| HOXD11 | ATTGTTTACT | 1 | 895 | 905 | -105 | -95 | -0,58 | 0,01 | 16,53 | >0.1 | 5,61 | 6,66 | 0 | 1,89 | 0 | 2,95 | 0 |
| HOXD11 | GTCATTCAGT | 1 | 1012 | 1022 | 12 | 22 | -1,23 | 0,01 | 16,58 | >0.1 | 5,61 | 8,92 | 0 | 1,86 | 0 | 1,42 | 0 |
| HOXD11 | GTGGTTAAAC | -1 | 1141 | 1151 | 141 | 151 | 3,12 | 0,01 | 23,57 | >0.1 | 5,61 | 6,37 | 0 | 1,84 | 3,22 | 3,4 | 0 |
| HOXD11 | GTCCAAAAAT | -1 | 1129 | 1139 | 129 | 139 | -0,12 | 0,01 | 19,82 | >0.1 | 5,61 | 6,47 | 0 | 1,84 | 3,22 | 2,79 | 0 |
| HOXD11 | ATCAAATAAG | -1 | 811 | 821 | -189 | -179 | -0,79 | 0,01 | 15,84 | >0.1 | 5,61 | 6,03 | 0 | 1,56 | 0 | 3,43 | 0 |
| HOXD11 | GCTGTTAAAG | -1 | 707 | 717 | -293 | -283 | -1,34 | 0,01 | 15,8 | >0.1 | 5,61 | 5,55 | 0 | 1,17 | 0 | 4,81 | 0 |
| HOXD11 | GTTATAATTT | -1 | 684 | 694 | -316 | -306 | 2,11 | 0,01 | 16,02 | >0.1 | 4,38 | 5,47 | 0 | 1,12 | 0 | 2,95 | 0 |
| HOXD11 | ATCATGAAGA | -1 | 644 | 654 | -356 | -346 | -0,19 | 0,01 | 13,74 | >0.1 | 3,83 | 5,34 | 0 | 1,06 | 0 | 3,7 | 0 |
| HOXD11 | TTCATTAATA | -1 | 602 | 612 | -398 | -388 | -1,04 | 0,01 | 10,09 | >0.1 | 3,49 | 5,21 | 0 | 1,02 | 0 | 1,41 | 0 |
| HOXD11 | ATCATATTCC | -1 | 563 | 573 | -437 | -427 | -2,32 | 0,01 | 7,03 | >0.1 | 3,25 | 5,11 | 0 | 0,99 | 0 | 0 | 0 |
| HOXD11 | CTTAAAAAAG | -1 | 73 | 83 | -927 | -917 | 0,83 | 0,01 | 10,88 | >0.1 | 3,85 | 4,32 | 0 | 1,88 | 0 | 0 | 0 |
| POU4F3 | AGAAATAGGAAATGAG | 1 | 57 | 73 | -943 | -927 | 3,59 | 0,01 | 13,78 | >0.1 | 4 | 4,31 | 0 | 1,88 | 0 | 0 | 0 |
| POU4F3 | CAGAAATATGAATTCC | 1 | 484 | 500 | -516 | -500 | 2,38 | 0,01 | 17,35 | >0.1 | 2,91 | 4,94 | 0 | 0,99 | 0 | 6,13 | 0 |
| POU4F3 | TTGAAATAAAAATAAA | 1 | 574 | 590 | -426 | -410 | 3,31 | 0,01 | 12,8 | >0.1 | 3,33 | 5,15 | 0 | 1,01 | 0 | 0 | 0 |
| POU4F3 | AATAAAAATAAATGTG | 1 | 578 | 594 | -422 | -406 | 4,17 | 0,01 | 13,7 | >0.1 | 3,35 | 5,17 | 0 | 1,01 | 0 | 0 | 0 |
| POU4F3 | ATAACCTATTAATGAA | 1 | 596 | 612 | -404 | -388 | 2,93 | 0,01 | 14 | >0.1 | 3,47 | 5,21 | 0 | 1,02 | 0 | 1,36 | 0 |
| POU4F3 | GTGTATCTTTAACAGC | 1 | 701 | 717 | -299 | -283 | 1,34 | 0,01 | 18,4 | >0.1 | 5,53 | 5,55 | 0 | 1,17 | 0 | 4,81 | 0 |
| POU4F3 | GGAAAATATTAACCAA | 1 | 723 | 739 | -277 | -261 | 1,52 | 0,01 | 19,23 | >0.1 | 5,53 | 5,64 | 0 | 1,23 | 0 | 5,32 | 0 |
| POU4F3 | AGGATTAAAGAATAAC | 1 | 790 | 806 | -210 | -194 | 2,26 | 0,01 | 20,54 | >0.1 | 5,53 | 5,95 | 0 | 1,48 | 0 | 5,32 | 0 |
| POU4F3 | AAGAATAACGTATTCT | 1 | 797 | 813 | -203 | -187 | 2,26 | 0,01 | 18,84 | >0.1 | 5,53 | 5,99 | 0 | 1,51 | 0 | 3,55 | 0 |
| POU4F3 | ATTCTTATTTGATTCA | 1 | 808 | 824 | -192 | -176 | 6,61 | 0,001 | 23,2 | >0.1 | 5,53 | 6,05 | 0 | 1,58 | 0 | 3,43 | 0 |
| POU4F3 | TTTAAAAAATTATTCT | 1 | 825 | 841 | -175 | -159 | 2,3 | 0,01 | 19,12 | >0.1 | 5,53 | 6,16 | 0 | 1,7 | 0 | 3,43 | 0 |

| | | | | | | | | | | | | | | | | | |
|---|---|---|---|---|---|---|---|---|---|---|---|---|---|---|---|---|---|
| POU4F3 | ATGAACTTTATATTGG | 1 | 871 | 887 | -129 | -113 | 2,59 | 0,01 | 19,47 | >0.1 | 5,53 | 6,5 | 0 | 1,9 | 0 | 2,95 | 0 |
| POU4F3 | GAGAAAATCTCATGAG | 1 | 976 | 992 | -24 | -8 | 2,24 | 0,01 | 19,61 | >0.1 | 5,53 | 8,43 | 0 | 1,87 | 0 | 1,53 | 0 |
| POU4F3 | TAGAATAATTTTTTAA | -1 | 826 | 842 | -174 | -158 | 3,59 | 0,01 | 20,28 | >0.1 | 5,53 | 6,17 | 0 | 1,71 | 0 | 3,28 | 0 |
| POU4F3 | ATAATTTTTTAAAGTG | -1 | 822 | 838 | -178 | -162 | 4,37 | 0,01 | 21,42 | >0.1 | 5,53 | 6,14 | 0 | 1,68 | 0 | 3,7 | 0 |
| POU4F3 | GTGAATCAAATAAGAA | -1 | 809 | 825 | -191 | -175 | 2,74 | 0,01 | 19,34 | >0.1 | 5,53 | 6,06 | 0 | 1,59 | 0 | 3,43 | 0 |
| POU4F3 | AAGAATACGTTATTCT | -1 | 798 | 814 | -202 | -186 | 2,56 | 0,01 | 19,03 | >0.1 | 5,53 | 5,99 | 0 | 1,52 | 0 | 3,43 | 0 |
| POU4F3 | GTTATTCTTTAATCCT | -1 | 790 | 806 | -210 | -194 | 1,84 | 0,01 | 20,12 | >0.1 | 5,53 | 5,95 | 0 | 1,48 | 0 | 5,32 | 0 |
| POU4F3 | GTTAATATTTTCCTAG | -1 | 720 | 736 | -280 | -264 | 1,84 | 0,01 | 19,38 | >0.1 | 5,53 | 5,62 | 0 | 1,22 | 0 | 5,17 | 0 |
| POU4F3 | ATTAATAGGTTATCTC | -1 | 593 | 609 | -407 | -391 | 3,05 | 0,01 | 14,16 | >0.1 | 3,45 | 5,21 | 0 | 1,02 | 0 | 1,44 | 0 |
| POU4F3 | CTTCCAAATACATGAC | -1 | 517 | 533 | -483 | -467 | 0,77 | 0,01 | 12,88 | >0.1 | 3,04 | 5,02 | 0 | 0,99 | 0 | 3,07 | 0 |
| POU4F3 | TTGTATTTTTAATAGA | -1 | 288 | 304 | -712 | -696 | 6,51 | 0,001 | 19,5 | >0.1 | 2,71 | 4,6 | 0 | 1,24 | 0 | 4,45 | 0 |
| POU4F3 | AGCCATTTCTGATCAG | -1 | 164 | 180 | -836 | -820 | 2,64 | 0,01 | 14,57 | >0.1 | 3,19 | 4,43 | 0 | 1,9 | 0 | 2,4 | 0 |
| BATF3 | CAAAGAAGTCAGCC | 1 | 29 | 43 | -971 | -957 | -7,46 | 0,001 | 3,15 | >0.1 | 4,46 | 4,28 | 0 | 1,87 | 0 | 0 | 0 |
| BATF3 | GAATGGCGTGAACC | 1 | 367 | 381 | -633 | -619 | -12,03 | 0,01 | 2,46 | >0.1 | 2,54 | 4,72 | 0 | 1,08 | 0 | 6,15 | 0 |
| BATF3 | AGATCGCGCCACTG | 1 | 407 | 421 | -593 | -579 | -21,36 | 0,01 | -7,37 | >0.1 | 2,65 | 4,79 | 0 | 1,03 | 0 | 5,52 | 0 |
| BATF3 | ATGTGAGATAACCT | 1 | 589 | 603 | -411 | -397 | -19,12 | 0,01 | -9,5 | >0.1 | 3,41 | 5,19 | 0 | 1,01 | 0 | 0 | 0 |
| BATF3 | TAATGAAATTGTCT | 1 | 605 | 619 | -395 | -381 | -16,71 | 0,01 | -5,43 | >0.1 | 3,52 | 5,23 | 0 | 1,03 | 0 | 1,5 | 0 |
| BATF3 | TTATAACATTTTCC | 1 | 687 | 701 | -313 | -299 | -17,94 | 0,01 | -3,88 | >0.1 | 4,48 | 5,49 | 0 | 1,14 | 0 | 2,95 | 0 |
| BATF3 | CCATAAAGTGACAG | 1 | 759 | 773 | -241 | -227 | -21,24 | 0,01 | -2,47 | >0.1 | 5,56 | 5,78 | 0 | 1,33 | 0 | 6,1 | 0 |
| BATF3 | AAGTGACAGGAGAG | 1 | 764 | 778 | -236 | -222 | -19,99 | 0,01 | -1,36 | >0.1 | 5,56 | 5,81 | 0 | 1,35 | 0 | 5,92 | 0 |
| BATF3 | GAATAACGTATTCT | 1 | 799 | 813 | -201 | -187 | -12,89 | 0,01 | 3,59 | >0.1 | 5,56 | 5,99 | 0 | 1,51 | 0 | 3,43 | 0 |
| BATF3 | TGAGGAGGTTTTAG | 1 | 988 | 1002 | -12 | 2 | -17,87 | 0,01 | -0,06 | >0.1 | 5,56 | 8,77 | 0 | 1,87 | 0 | 1,61 | 0 |
| BATF3 | AGTGGATGTGATCT | 1 | 1019 | 1033 | 19 | 33 | -19,9 | 0,01 | -2,43 | >0.1 | 5,56 | 8,74 | 0 | 1,86 | 0 | 1,31 | 0 |
| BATF3 | GGATGTGATCTTGG | 1 | 1022 | 1036 | 22 | 36 | -21,49 | 0,01 | -4,14 | >0.1 | 5,56 | 8,68 | 0 | 1,86 | 0 | 1,25 | 0 |
| BATF3 | GGACGATGTCAAGC | 1 | 1044 | 1058 | 44 | 58 | -17,33 | 0,01 | -0,72 | >0.1 | 5,56 | 7,73 | 0 | 1,86 | 0 | 1,46 | 0 |
| BATF3 | TGTTGCTGTAACTG | 1 | 1081 | 1095 | 81 | 95 | -21,23 | 0,01 | -2,81 | >0.1 | 5,56 | 7 | 0 | 1,85 | 1,79 | 2,21 | 0 |
| BATF3 | CCAAGACATTTTTG | 1 | 1122 | 1136 | 122 | 136 | -19,96 | 0,01 | -0,01 | >0.1 | 5,56 | 6,53 | 0 | 1,85 | 3,22 | 2,79 | 0 |
| BATF3 | CGAAGACCTGTTCT | 1 | 1156 | 1170 | 156 | 170 | -15,33 | 0,01 | 1,29 | >0.1 | 4,38 | 6,25 | 0 | 1,8 | 1,79 | 2,4 | 0 |
| BATF3 | AGAACAGGTCTTCG | -1 | 1156 | 1170 | 156 | 170 | -18,51 | 0,01 | -1,89 | >0.1 | 4,38 | 6,25 | 0 | 1,8 | 1,79 | 2,4 | 0 |
| BATF3 | TGGTGGACTGAGCA | -1 | 1096 | 1110 | 96 | 110 | -21,18 | 0,01 | -1,4 | >0.1 | 5,56 | 6,8 | 0 | 1,85 | 3,22 | 2,35 | 0 |
| BATF3 | CAGTTACAGCAACA | -1 | 1081 | 1095 | 81 | 95 | -19,53 | 0,01 | -1,11 | >0.1 | 5,56 | 7 | 0 | 1,85 | 1,79 | 2,21 | 0 |
| BATF3 | GCTTGACATCGTCC | -1 | 1044 | 1058 | 44 | 58 | -13,02 | 0,01 | 3,59 | >0.1 | 5,56 | 7,73 | 0 | 1,86 | 0 | 1,46 | 0 |
| BATF3 | AGATCACATCCACT | -1 | 1019 | 1033 | 19 | 33 | -17,06 | 0,01 | 0,41 | >0.1 | 5,56 | 8,74 | 0 | 1,86 | 0 | 1,31 | 0 |
| BATF3 | GAATGACTTTCCCT | -1 | 1005 | 1019 | 5 | 19 | -20,38 | 0,01 | -2,37 | >0.1 | 5,56 | 9,12 | 0 | 1,87 | 0 | 1,46 | 0 |
| BATF3 | TCATGAGATTTTCT | -1 | 977 | 991 | -23 | -9 | -16,39 | 0,01 | 0,94 | >0.1 | 5,56 | 8,38 | 0 | 1,87 | 0 | 1,53 | 0 |
| BATF3 | AGAATACGTTATTC | -1 | 799 | 813 | -201 | -187 | -16,07 | 0,01 | 0,41 | >0.1 | 5,56 | 5,99 | 0 | 1,51 | 0 | 3,43 | 0 |
| BATF3 | TTATGGCCAAATCA | -1 | 751 | 765 | -249 | -235 | -20,79 | 0,01 | -1,86 | >0.1 | 5,56 | 5,75 | 0 | 1,3 | 0 | 6,32 | 0 |
| BATF3 | ACATGACTTTGCCC | -1 | 510 | 524 | -490 | -476 | -21,55 | 0,01 | -9,61 | >0.1 | 3,01 | 5 | 0 | 0,99 | 0 | 2,95 | 0 |
| BATF3 | CAGTGGCGCGATCT | -1 | 407 | 421 | -593 | -579 | -17,82 | 0,01 | -3,83 | >0.1 | 2,65 | 4,79 | 0 | 1,03 | 0 | 5,52 | 0 |
| BATF3 | GGTTCACGCCATTC | -1 | 367 | 381 | -633 | -619 | -18,05 | 0,01 | -3,56 | >0.1 | 2,54 | 4,72 | 0 | 1,08 | 0 | 6,15 | 0 |
| BATF3 | TCAGGACGGTCTCT | -1 | 253 | 267 | -747 | -733 | -20,21 | 0,01 | -6,64 | >0.1 | 2,83 | 4,54 | 0 | 1,36 | 0 | 4,85 | 0 |
| BATF3 | TCCTGACCTTGTGA | -1 | 236 | 250 | -764 | -750 | -20,9 | 0,01 | -7,54 | >0.1 | 2,89 | 4,52 | 0 | 1,43 | 0 | 4,53 | 0 |
| BATF3 | TACAGGCGTGAGCC | -1 | 189 | 203 | -811 | -797 | -19,18 | 0,01 | -5,8 | >0.1 | 3,08 | 4,46 | 0 | 1,71 | 0 | 4,13 | 0 |
| BATF3 | GGCTGACTTCTTTG | -1 | 29 | 43 | -971 | -957 | -8,48 | 0,001 | 2,13 | >0.1 | 4,46 | 4,28 | 0 | 1,87 | 0 | 0 | 0 |
| LIN54 | TTTTAAGTT | 1 | 76 | 85 | -924 | -915 | -1,22 | 0,01 | 8,81 | >0.1 | 3,83 | 4,32 | 0 | 1,88 | 0 | 0 | 0 |
| LIN54 | TTTTAAAAA | 1 | 136 | 145 | -864 | -855 | 7,33 | 0,01 | 18,55 | >0.1 | 3,37 | 4,39 | 0 | 1,89 | 0 | 1,57 | 0 |
| LIN54 | ATTAAAACT | 1 | 157 | 166 | -843 | -834 | -2,05 | 0,01 | 10,04 | >0.1 | 3,25 | 4,41 | 0 | 1,9 | 0 | 2,53 | 0 |
| LIN54 | ATATGAATT | 1 | 489 | 498 | -511 | -502 | -0,33 | 0,01 | 14,78 | >0.1 | 2,91 | 4,94 | 0 | 0,98 | 0 | 6,27 | 0 |
| LIN54 | CTTGAAATA | 1 | 573 | 582 | -427 | -418 | -2,55 | 0,01 | 6,88 | >0.1 | 3,3 | 5,13 | 0 | 1 | 0 | 0 | 0 |
| LIN54 | TTTTGATTT | 1 | 748 | 757 | -252 | -243 | -0,86 | 0,01 | 17,85 | >0.1 | 5,62 | 5,71 | 0 | 1,28 | 0 | 6,1 | 0 |
| LIN54 | ATTTGATTC | 1 | 814 | 823 | -186 | -177 | -0,33 | 0,01 | 16,34 | >0.1 | 5,62 | 6,05 | 0 | 1,57 | 0 | 3,43 | 0 |
| LIN54 | CTTTAAAAA | 1 | 824 | 833 | -176 | -167 | 6,18 | 0,01 | 23,25 | >0.1 | 5,62 | 6,11 | 0 | 1,64 | 0 | 3,7 | 0 |
| LIN54 | GTTCAAAGG | 1 | 958 | 967 | -42 | -33 | -2,64 | 0,01 | 14,15 | >0.1 | 5,62 | 7,55 | 0 | 1,88 | 0 | 1,74 | 0 |
| LIN54 | TTTTGGACA | 1 | 1131 | 1140 | 131 | 140 | -2,09 | 0,01 | 17,91 | >0.1 | 5,62 | 6,45 | 0 | 1,84 | 3,22 | 2,86 | 0 |
| LIN54 | CTTGGAATT | 1 | 1188 | 1197 | 188 | 197 | -1,17 | 0,01 | 17,36 | >0.1 | 5,62 | 6,04 | 0 | 1,53 | 1,79 | 3,55 | 0 |
| LIN54 | GGTTAAACT | -1 | 1140 | 1149 | 140 | 149 | -2,35 | 0,01 | 17,92 | >0.1 | 5,62 | 6,37 | 0 | 1,84 | 3,22 | 3,21 | 0 |
| LIN54 | CTTTGAACT | -1 | 957 | 966 | -43 | -34 | 9,08 | 0,001 | 25,8 | >0.1 | 5,62 | 7,53 | 0 | 1,88 | 0 | 1,7 | 0 |
| LIN54 | TTTAGAATA | -1 | 835 | 844 | -165 | -156 | -1,04 | 0,01 | 15,77 | >0.1 | 5,62 | 6,18 | 0 | 1,72 | 0 | 3,28 | 0 |
| LIN54 | TTTTAAAGT | -1 | 823 | 832 | -177 | -168 | -0,67 | 0,01 | 16,39 | >0.1 | 5,62 | 6,1 | 0 | 1,63 | 0 | 3,7 | 0 |

| | | | | | | | | | | | | | | | | | |
|---|---|---|---|---|---|---|---|---|---|---|---|---|---|---|---|---|---|
| LIN54 | GTTTGTATG | -1 | 625 | 634 | -375 | -366 | -1,43 | 0,01 | 10,04 | >0.1 | 3,66 | 5,28 | 0 | 1,04 | 0 | 1,5 | 0 |
| LIN54 | ATTCATATT | -1 | 488 | 497 | -512 | -503 | -2,58 | 0,01 | 12,38 | >0.1 | 2,91 | 4,94 | 0 | 0,98 | 0 | 6,13 | 0 |
| LIN54 | TTTTGTATT | -1 | 297 | 306 | -703 | -694 | -0,86 | 0,01 | 12,17 | >0.1 | 2,69 | 4,6 | 0 | 1,21 | 0 | 4,53 | 0 |
| LIN54 | TTTTTAATA | -1 | 290 | 299 | -710 | -701 | -1,05 | 0,01 | 12,57 | >0.1 | 2,71 | 4,59 | 0 | 1,23 | 0 | 5,09 | 0 |
| LIN54 | GTTTTAATT | -1 | 156 | 165 | -844 | -835 | -0,46 | 0,01 | 11,71 | >0.1 | 3,25 | 4,41 | 0 | 1,9 | 0 | 2,6 | 0 |
| LIN54 | TTTTAATTT | -1 | 155 | 164 | -845 | -836 | -1,71 | 0,01 | 9,5 | >0.1 | 3,26 | 4,41 | 0 | 1,9 | 0 | 1,65 | 0 |
| LIN54 | TTTTAAAAT | -1 | 135 | 144 | -865 | -856 | 7,85 | 0,01 | 19,11 | >0.1 | 3,37 | 4,39 | 0 | 1,89 | 0 | 1,61 | 0 |
| LIN54 | TTTAAAATT | -1 | 134 | 143 | -866 | -857 | -1,37 | 0,01 | 9,86 | >0.1 | 3,38 | 4,39 | 0 | 1,89 | 0 | 1,57 | 0 |
| LIN54 | CTTTGAGTG | -1 | 25 | 34 | -975 | -966 | -2,15 | 0,01 | 9,62 | >0.1 | 5,62 | 4,27 | 0 | 1,87 | 0 | 0 | 0 |
| VAX2 | GAAATTAA | 1 | 154 | 162 | -846 | -838 | 3,43 | 0,01 | 14,65 | >0.1 | 3,27 | 4,41 | 0 | 1,9 | 0 | 1,65 | 0 |
| VAX2 | CCAAGTAG | 1 | 545 | 553 | -455 | -447 | 3,84 | 0,01 | 13,04 | >0.1 | 3,15 | 5,06 | 0 | 0,98 | 0 | 0 | 0 |
| VAX2 | TTAATGAA | 1 | 604 | 612 | -396 | -388 | 7,67 | 0,001 | 18,81 | >0.1 | 3,49 | 5,21 | 0 | 1,02 | 0 | 1,41 | 0 |
| VAX2 | TTCATGAT | 1 | 646 | 654 | -354 | -346 | 5,98 | 0,01 | 20,05 | >0.1 | 3,84 | 5,34 | 0 | 1,06 | 0 | 3,83 | 0 |
| VAX2 | CTCATGAG | 1 | 984 | 992 | -16 | -8 | 7,15 | 0,001 | 24,67 | >0.1 | 5,64 | 8,43 | 0 | 1,87 | 0 | 1,57 | 0 |
| VAX2 | TCAATGGT | -1 | 1106 | 1114 | 106 | 114 | 4,59 | 0,01 | 24,51 | >0.1 | 5,64 | 6,69 | 0 | 1,85 | 3,22 | 2,52 | 0 |
| VAX2 | TGAATGAC | -1 | 1012 | 1020 | 12 | 20 | 3,71 | 0,01 | 21,53 | >0.1 | 5,64 | 8,92 | 0 | 1,86 | 0 | 1,39 | 0 |
| VAX2 | CTCATGAG | -1 | 984 | 992 | -16 | -8 | 7,15 | 0,001 | 24,67 | >0.1 | 5,64 | 8,43 | 0 | 1,87 | 0 | 1,57 | 0 |
| VAX2 | TTCATTAA | -1 | 604 | 612 | -396 | -388 | 6,64 | 0,01 | 17,78 | >0.1 | 3,49 | 5,21 | 0 | 1,02 | 0 | 1,41 | 0 |
| VAX2 | TTAATAGG | -1 | 600 | 608 | -400 | -392 | 4,26 | 0,01 | 15,31 | >0.1 | 3,47 | 5,2 | 0 | 1,01 | 0 | 1,36 | 0 |
| VAX2 | CCAAATAC | -1 | 522 | 530 | -478 | -470 | 4,35 | 0,01 | 16,45 | >0.1 | 3,05 | 5,01 | 0 | 0,98 | 0 | 3,07 | 0 |
| VAX2 | TACATGAC | -1 | 517 | 525 | -483 | -475 | 4,02 | 0,01 | 15,97 | >0.1 | 3,02 | 5 | 0 | 0,98 | 0 | 2,95 | 0 |
| VAX2 | TTAATAGA | -1 | 288 | 296 | -712 | -704 | 3,68 | 0,01 | 17,41 | >0.1 | 2,72 | 4,58 | 0 | 1,24 | 0 | 5,19 | 0 |
| VAX2 | TTAATTTC | -1 | 154 | 162 | -846 | -838 | 5,39 | 0,01 | 16,61 | >0.1 | 3,27 | 4,41 | 0 | 1,9 | 0 | 1,65 | 0 |
| VAX2 | TCCATGGC | -1 | 148 | 156 | -852 | -844 | 4,11 | 0,01 | 15,32 | >0.1 | 3,3 | 4,4 | 0 | 1,9 | 0 | 1,61 | 0 |
| VAX2 | CTCATTTC | -1 | 65 | 73 | -935 | -927 | 4,29 | 0,01 | 14,43 | >0.1 | 3,95 | 4,31 | 0 | 1,88 | 0 | 0 | 0 |
| VAX1 | TTAATGAA | 1 | 604 | 612 | -396 | -388 | 7,5 | 0,001 | 18,64 | >0.1 | 3,49 | 5,21 | 0 | 1,02 | 0 | 1,41 | 0 |
| VAX1 | TTCATGAT | 1 | 646 | 654 | -354 | -346 | 5,94 | 0,01 | 20,01 | >0.1 | 3,84 | 5,34 | 0 | 1,06 | 0 | 3,83 | 0 |
| VAX1 | CTCATGAG | 1 | 984 | 992 | -16 | -8 | 6,48 | 0,01 | 24 | >0.1 | 5,64 | 8,43 | 0 | 1,87 | 0 | 1,57 | 0 |
| VAX1 | TCAATGGT | -1 | 1106 | 1114 | 106 | 114 | 4,67 | 0,01 | 24,59 | >0.1 | 5,64 | 6,69 | 0 | 1,85 | 3,22 | 2,52 | 0 |
| VAX1 | TGAATGAC | -1 | 1012 | 1020 | 12 | 20 | 3,76 | 0,01 | 21,58 | >0.1 | 5,64 | 8,92 | 0 | 1,86 | 0 | 1,39 | 0 |
| VAX1 | CTCATGAG | -1 | 984 | 992 | -16 | -8 | 6,48 | 0,01 | 24 | >0.1 | 5,64 | 8,43 | 0 | 1,87 | 0 | 1,57 | 0 |
| VAX1 | TTAAAGAT | -1 | 705 | 713 | -295 | -287 | 3,91 | 0,01 | 21,05 | >0.1 | 5,64 | 5,53 | 0 | 1,16 | 0 | 4,81 | 0 |
| VAX1 | TTCATTAA | -1 | 604 | 612 | -396 | -388 | 6,32 | 0,01 | 17,46 | >0.1 | 3,49 | 5,21 | 0 | 1,02 | 0 | 1,41 | 0 |
| VAX1 | TTAATAGG | -1 | 600 | 608 | -400 | -392 | 4,01 | 0,01 | 15,06 | >0.1 | 3,47 | 5,2 | 0 | 1,01 | 0 | 1,36 | 0 |
| VAX1 | CCAAATAC | -1 | 522 | 530 | -478 | -470 | 4,51 | 0,01 | 16,61 | >0.1 | 3,05 | 5,01 | 0 | 0,98 | 0 | 3,07 | 0 |
| VAX1 | TACATGAC | -1 | 517 | 525 | -483 | -475 | 3,9 | 0,01 | 15,85 | >0.1 | 3,02 | 5 | 0 | 0,98 | 0 | 2,95 | 0 |
| VAX1 | TTAATTTC | -1 | 154 | 162 | -846 | -838 | 6,17 | 0,01 | 17,39 | >0.1 | 3,27 | 4,41 | 0 | 1,9 | 0 | 1,65 | 0 |
| VAX1 | CTCATTTC | -1 | 65 | 73 | -935 | -927 | 4,72 | 0,01 | 14,86 | >0.1 | 3,95 | 4,31 | 0 | 1,88 | 0 | 0 | 0 |
| FOXP3 | GTAAGTG | 1 | 5 | 12 | -995 | -988 | 3,56 | 0,01 | 17,12 | >0.1 | 5,65 | 4,25 | 0 | 1,86 | 1,79 | 0 | 0 |
| FOXP3 | GCAAAGG | 1 | 87 | 94 | -913 | -906 | 3,85 | 0,01 | 13,8 | >0.1 | 3,73 | 4,33 | 0 | 1,88 | 0 | 0 | 0 |
| FOXP3 | ACTAACA | 1 | 266 | 273 | -734 | -727 | 5,46 | 0,01 | 18,56 | >0.1 | 2,79 | 4,55 | 0 | 1,31 | 0 | 4,45 | 0 |
| FOXP3 | AAAAATA | 1 | 294 | 301 | -706 | -699 | 3,43 | 0,01 | 16,39 | >0.1 | 2,7 | 4,59 | 0 | 1,22 | 0 | 4,45 | 0 |
| FOXP3 | ATAAAGA | 1 | 500 | 507 | -500 | -493 | 7,03 | 0,01 | 22,36 | >0.1 | 2,95 | 4,96 | 0 | 0,98 | 0 | 6,44 | 0 |
| FOXP3 | GTAGAGA | 1 | 549 | 556 | -451 | -444 | 4,39 | 0,01 | 13,61 | >0.1 | 3,17 | 5,07 | 0 | 0,98 | 0 | 0 | 0 |
| FOXP3 | AAAAATA | 1 | 581 | 588 | -419 | -412 | 3,43 | 0,01 | 12,92 | >0.1 | 3,34 | 5,15 | 0 | 1 | 0 | 0 | 0 |
| FOXP3 | ATAAATG | 1 | 585 | 592 | -415 | -408 | 3,78 | 0,01 | 13,31 | >0.1 | 3,37 | 5,16 | 0 | 1 | 0 | 0 | 0 |
| FOXP3 | ACAAACA | 1 | 628 | 635 | -372 | -365 | 8,22 | 0,001 | 19,75 | >0.1 | 3,67 | 5,28 | 0 | 1,04 | 0 | 1,55 | 0 |
| FOXP3 | GAAAATA | 1 | 724 | 731 | -276 | -269 | 4,81 | 0,01 | 22,43 | >0.1 | 5,65 | 5,6 | 0 | 1,2 | 0 | 5,17 | 0 |
| FOXP3 | AAAAAGA | -1 | 921 | 928 | -79 | -72 | 3,64 | 0,01 | 19,16 | >0.1 | 5,65 | 6,9 | 0 | 1,88 | 0 | 1,09 | 0 |
| FOXP3 | AAAAAGA | -1 | 915 | 922 | -85 | -78 | 3,64 | 0,01 | 19,16 | >0.1 | 5,65 | 6,83 | 0 | 1,89 | 0 | 1,15 | 0 |
| FOXP3 | GTAAACA | -1 | 897 | 904 | -103 | -96 | 11,13 | >0.1 | 28,27 | >0.1 | 5,65 | 6,65 | 0 | 1,89 | 0 | 2,95 | 0 |
| FOXP3 | GTTAATA | -1 | 729 | 736 | -271 | -264 | 5,44 | 0,01 | 23,25 | >0.1 | 5,65 | 5,62 | 0 | 1,21 | 0 | 5,32 | 0 |
| FOXP3 | ATTAATA | -1 | 602 | 609 | -398 | -391 | 4,06 | 0,01 | 15,12 | >0.1 | 3,48 | 5,2 | 0 | 1,01 | 0 | 1,36 | 0 |
| FOSL2::JUND(VAR.2) | CAAAGAAGTCAGCCA | 1 | 29 | 44 | -971 | -956 | 2,27 | 0,01 | 12,87 | >0.1 | 4,45 | 4,28 | 0 | 1,87 | 0 | 0 | 0 |
| FOSL2::JUND(VAR.2) | TGGGCAGATCACAAG | 1 | 228 | 243 | -772 | -757 | -1,54 | 0,01 | 12,36 | >0.1 | 2,91 | 4,51 | 0 | 1,47 | 0 | 5,01 | 0 |
| FOSL2::JUND(VAR.2) | GAATGGCGTGAACCT | 1 | 367 | 382 | -633 | -618 | 2,57 | 0,01 | 17,06 | >0.1 | 2,54 | 4,72 | 0 | 1,08 | 0 | 6,15 | 0 |
| FOSL2::JUND(VAR.2) | AGATCGCGCCACTGT | 1 | 407 | 422 | -593 | -578 | 2,94 | 0,01 | 16,94 | >0.1 | 2,65 | 4,79 | 0 | 1,03 | 0 | 5,52 | 0 |
| FOSL2::JUND(VAR.2) | ATGTGAGATAACCTA | 1 | 589 | 604 | -411 | -396 | 2,68 | 0,01 | 12,3 | >0.1 | 3,42 | 5,19 | 0 | 1,02 | 0 | 0 | 0 |
| FOSL2::JUND(VAR.2) | CTCTTTGGTCACTGT | 1 | 668 | 683 | -332 | -317 | 1,22 | 0,01 | 15,6 | >0.1 | 4,15 | 5,43 | 0 | 1,11 | 0 | 3,7 | 0 |

| | | | | | | | | | | | | | | | | | |
|---|---|---|---|---|---|---|---|---|---|---|---|---|---|---|---|---|---|
| FOSL2::JUND(VAR.2) | AAATTATAACATTTT | 1 | 684 | 699 | -316 | -301 | -0,39 | 0,01 | 13,61 | >0.1 | 4,43 | 5,48 | 0 | 1,13 | 0 | 2,95 | 0 |
| FOSL2::JUND(VAR.2) | CTGTTACATATCTGT | 1 | 846 | 861 | -154 | -139 | -0,49 | 0,01 | 16,18 | >0.1 | 5,54 | 6,3 | 0 | 1,87 | 0 | 2,95 | 0 |
| FOSL2::JUND(VAR.2) | GGATGTGATCTTGGC | 1 | 1022 | 1037 | 22 | 37 | -1,36 | 0,01 | 15,98 | >0.1 | 5,54 | 8,68 | 0 | 1,86 | 0 | 1,25 | 0 |
| FOSL2::JUND(VAR.2) | GGACGATGTCAAGCT | 1 | 1044 | 1059 | 44 | 59 | 2,44 | 0,01 | 19,04 | >0.1 | 5,54 | 7,73 | 0 | 1,86 | 0 | 1,46 | 0 |
| FOSL2::JUND(VAR.2) | AGAACAGGTCTTCGG | -1 | 1155 | 1170 | 155 | 170 | -0,97 | 0,01 | 15,62 | >0.1 | 4,39 | 6,26 | 0 | 1,81 | 1,79 | 2,34 | 0 |
| FOSL2::JUND(VAR.2) | CAGTTACAGCAACAA | -1 | 1080 | 1095 | 80 | 95 | 0,25 | 0,01 | 18,67 | >0.1 | 5,54 | 7,01 | 0 | 1,85 | 1,79 | 2,21 | 0 |
| FOSL2::JUND(VAR.2) | GCTTGACATCGTCCC | -1 | 1043 | 1058 | 43 | 58 | 6,39 | 0,001 | 22,2 | >0.1 | 5,54 | 7,76 | 0 | 1,86 | 0 | 0,65 | 0 |
| FOSL2::JUND(VAR.2) | AGATCACATCCACTG | -1 | 1018 | 1033 | 18 | 33 | -0,77 | 0,01 | 16,71 | >0.1 | 5,54 | 8,76 | 0 | 1,86 | 0 | 1,31 | 0 |
| FOSL2::JUND(VAR.2) | TGGAGAGGACAGATA | -1 | 854 | 869 | -146 | -131 | -1,57 | 0,01 | 15,19 | >0.1 | 5,54 | 6,36 | 0 | 1,9 | 0 | 2,95 | 0 |
| FOSL2::JUND(VAR.2) | CAGATATGTAACAGA | -1 | 845 | 860 | -155 | -140 | -1,3 | 0,01 | 15,27 | >0.1 | 5,54 | 6,3 | 0 | 1,86 | 0 | 2,86 | 0 |
| FOSL2::JUND(VAR.2) | AGAATACGTTATTCT | -1 | 798 | 813 | -202 | -187 | 2,48 | 0,01 | 19,07 | >0.1 | 5,54 | 5,99 | 0 | 1,51 | 0 | 3,55 | 0 |
| FOSL2::JUND(VAR.2) | ACCTTACCTCTCCTG | -1 | 770 | 785 | -230 | -215 | -0,61 | 0,01 | 18,07 | >0.1 | 5,54 | 5,84 | 0 | 1,38 | 0 | 5,92 | 0 |
| FOSL2::JUND(VAR.2) | CTGTCACTTTATGGC | -1 | 758 | 773 | -242 | -227 | 0,45 | 0,01 | 19,03 | >0.1 | 5,54 | 5,78 | 0 | 1,33 | 0 | 5,92 | 0 |
| FOSL2::JUND(VAR.2) | TTAATAGGTTATCTC | -1 | 593 | 608 | -407 | -392 | -0,83 | 0,01 | 8,83 | >0.1 | 3,44 | 5,2 | 0 | 1,02 | 0 | 0 | 0 |
| FOSL2::JUND(VAR.2) | AGGTTATCTCACATT | -1 | 588 | 603 | -412 | -397 | 2,52 | 0,01 | 12,13 | >0.1 | 3,41 | 5,19 | 0 | 1,01 | 0 | 0 | 0 |
| FOSL2::JUND(VAR.2) | CAAATACATGACTTT | -1 | 514 | 529 | -486 | -471 | -1,38 | 0,01 | 10,59 | >0.1 | 3,03 | 5,01 | 0 | 0,99 | 0 | 2,95 | 0 |
| FOSL2::JUND(VAR.2) | CAGTGGCGCGATCTC | -1 | 406 | 421 | -594 | -579 | 0,72 | 0,01 | 14,84 | >0.1 | 2,65 | 4,79 | 0 | 1,03 | 0 | 5,65 | 0 |
| FOSL2::JUND(VAR.2) | GGTTCACGCCATTCT | -1 | 366 | 381 | -634 | -619 | 4,44 | 0,001 | 19,5 | >0.1 | 2,54 | 4,72 | 0 | 1,08 | 0 | 6,73 | 0 |
| FOSL2::JUND(VAR.2) | TTATGACTACTCTCC | -1 | 121 | 136 | -879 | -864 | -0,05 | 0,01 | 11,36 | >0.1 | 3,44 | 4,38 | 0 | 1,89 | 0 | 1,7 | 0 |
| FOSL2::JUND(VAR.2) | GGCTGACTTCTTTGA | -1 | 28 | 43 | -972 | -957 | 3,51 | 0,01 | 14,13 | >0.1 | 4,47 | 4,28 | 0 | 1,87 | 0 | 0 | 0 |
| NFATC3 | GAATTCCATA | 1 | 493 | 503 | -507 | -497 | 6,32 | 0,01 | 21,62 | >0.1 | 2,93 | 4,95 | 0 | 0,98 | 0 | 6,44 | 0 |
| NFATC3 | ATTTTCCGTG | 1 | 694 | 704 | -306 | -296 | 11,52 | 0,0001 | 26,84 | >0.1 | 5,61 | 5,5 | 0 | 1,14 | 0 | 3,07 | 0 |
| NFATC3 | TCTTTCCTGC | 1 | 932 | 942 | -68 | -58 | 6,57 | 0,01 | 22,08 | >0.1 | 5,61 | 7,09 | 0 | 1,88 | 0 | 0,93 | 0 |
| NFATC3 | TAATTCCAAG | -1 | 1188 | 1198 | 188 | 198 | 5,97 | 0,01 | 24,64 | >0.1 | 5,61 | 6,04 | 0 | 1,53 | 1,79 | 3,7 | 0 |
| NFATC3 | CTGTTCCTCA | -1 | 1111 | 1121 | 111 | 121 | 3,59 | 0,01 | 23,45 | >0.1 | 5,61 | 6,64 | 0 | 1,85 | 3,22 | 2,54 | 0 |
| NFATC3 | ACTTTCCCTA | -1 | 1004 | 1014 | 4 | 14 | 7,33 | 0,01 | 25,34 | >0.1 | 5,61 | 9,12 | 0 | 1,87 | 0 | 1,42 | 0 |
| NFATC3 | ATTTTCCTAG | -1 | 720 | 730 | -280 | -270 | 7,32 | 0,01 | 24,72 | >0.1 | 5,61 | 5,6 | 0 | 1,2 | 0 | 4,99 | 0 |
| NFATC3 | ATTTTACAGT | -1 | 678 | 688 | -322 | -312 | 6,01 | 0,01 | 20,12 | >0.1 | 4,27 | 5,45 | 0 | 1,11 | 0 | 3,28 | 0 |
| NFATC3 | GGTTTTCAGA | -1 | 616 | 626 | -384 | -374 | 2,76 | 0,01 | 14,13 | >0.1 | 3,59 | 5,25 | 0 | 1,03 | 0 | 1,5 | 0 |
| NFATC3 | ATATTCCCAG | -1 | 560 | 570 | -440 | -430 | 4,33 | 0,01 | 13,66 | >0.1 | 3,23 | 5,1 | 0 | 0,99 | 0 | 0 | 0 |
| NFATC3 | TATTCCCAGA | -1 | 559 | 569 | -441 | -431 | 3,74 | 0,01 | 13,06 | >0.1 | 3,23 | 5,1 | 0 | 0,99 | 0 | 0 | 0 |
| NFATC3 | ATTTTCCCTT | -1 | 530 | 540 | -470 | -460 | 9,11 | 0,001 | 21,91 | >0.1 | 3,09 | 5,03 | 0 | 0,98 | 0 | 3,7 | 0 |
| NFATC3 | TTTTCCCTTC | -1 | 529 | 539 | -471 | -461 | 3,81 | 0,01 | 16,45 | >0.1 | 3,08 | 5,03 | 0 | 0,98 | 0 | 3,55 | 0 |
| NFATC3 | CCCTTCCAAA | -1 | 525 | 535 | -475 | -465 | 3,85 | 0,01 | 16,2 | >0.1 | 3,06 | 5,02 | 0 | 0,98 | 0 | 3,28 | 0 |
| NFATC3 | AATTTCCATG | -1 | 150 | 160 | -850 | -840 | 10,32 | 0,001 | 21,55 | >0.1 | 3,28 | 4,41 | 0 | 1,9 | 0 | 1,65 | 0 |
| NFATC3 | ACTCTCCACT | -1 | 118 | 128 | -882 | -872 | 5,14 | 0,01 | 15,6 | >0.1 | 3,48 | 4,37 | 0 | 1,89 | 0 | 0,73 | 0 |
| NFATC3 | CATTTCCTAT | -1 | 61 | 71 | -939 | -929 | 4,96 | 0,01 | 15,13 | >0.1 | 3,98 | 4,31 | 0 | 1,88 | 0 | 0 | 0 |
| FOS::JUNB | AAGGAGGCACTC | 1 | 18 | 30 | -982 | -970 | -3,46 | 0,01 | 8,26 | >0.1 | 5,58 | 4,27 | 0 | 1,87 | 0 | 0 | 0 |
| FOS::JUNB | TGTGAGAAATAG | 1 | 53 | 65 | -947 | -935 | -0,72 | 0,01 | 9,53 | >0.1 | 4,07 | 4,3 | 0 | 1,87 | 0 | 0 | 0 |
| FOS::JUNB | GAGTAGTCATAA | 1 | 124 | 136 | -876 | -864 | 3,38 | 0,01 | 14,78 | >0.1 | 3,43 | 4,38 | 0 | 1,89 | 0 | 1,7 | 0 |
| FOS::JUNB | AGTGGCTCACGC | 1 | 186 | 198 | -814 | -802 | 3,16 | 0,01 | 16,3 | >0.1 | 3,1 | 4,45 | 0 | 1,74 | 0 | 3,86 | 0 |
| FOS::JUNB | CCTGACTAACAC | 1 | 262 | 274 | -738 | -726 | 5,9 | 0,001 | 19,02 | >0.1 | 2,8 | 4,55 | 0 | 1,32 | 0 | 4,45 | 0 |
| FOS::JUNB | GCTGAGGCAGGA | 1 | 355 | 367 | -645 | -633 | 2,71 | 0,01 | 17,15 | >0.1 | 2,52 | 4,69 | 0 | 1,1 | 0 | 6,13 | 0 |
| FOS::JUNB | CGAGACTCAGTC | 1 | 443 | 455 | -557 | -545 | 0,14 | 0,01 | 14,68 | >0.1 | 2,76 | 4,85 | 0 | 1 | 0 | 5,92 | 0 |
| FOS::JUNB | CTTTGGTCACTG | 1 | 670 | 682 | -330 | -318 | -2,69 | 0,01 | 11,54 | >0.1 | 4,16 | 5,43 | 0 | 1,1 | 0 | 3,55 | 0 |
| FOS::JUNB | TTTGATTCACTT | 1 | 815 | 827 | -185 | -173 | 3,64 | 0,01 | 20,44 | >0.1 | 5,58 | 6,07 | 0 | 1,6 | 0 | 3,55 | 0 |
| FOS::JUNB | ATTGGCTCAGCA | 1 | 882 | 894 | -118 | -106 | -2,26 | 0,01 | 14,85 | >0.1 | 5,58 | 6,57 | 0 | 1,89 | 0 | 3,07 | 0 |
| FOS::JUNB | CTTGGCTCACAG | 1 | 1031 | 1043 | 31 | 43 | 0,11 | 0,01 | 17,02 | >0.1 | 5,58 | 8,19 | 0 | 1,86 | 0 | 1,28 | 0 |
| FOS::JUNB | ATTGAGGAACAG | 1 | 1109 | 1121 | 109 | 121 | -3,62 | 0,01 | 16,26 | >0.1 | 5,58 | 6,66 | 0 | 1,85 | 3,22 | 2,56 | 0 |
| FOS::JUNB | TGTTCCTCAATG | -1 | 1108 | 1120 | 108 | 120 | -1,18 | 0,01 | 18,66 | >0.1 | 5,58 | 6,67 | 0 | 1,85 | 3,22 | 2,52 | 0 |
| FOS::JUNB | TGTGAGCCAAGA | -1 | 1030 | 1042 | 30 | 42 | -1,56 | 0,01 | 15,43 | >0.1 | 5,58 | 8,24 | 0 | 1,86 | 0 | 1,31 | 0 |
| FOS::JUNB | ACTGAATGACTT | -1 | 1010 | 1022 | 10 | 22 | -3,77 | 0,01 | 14,06 | >0.1 | 5,58 | 8,99 | 0 | 1,86 | 0 | 1,39 | 0 |
| FOS::JUNB | GCTGAGCCAATA | -1 | 881 | 893 | -119 | -107 | -2,48 | 0,01 | 14,62 | >0.1 | 5,58 | 6,56 | 0 | 1,89 | 0 | 3,07 | 0 |
| FOS::JUNB | AGTGAATCAAAT | -1 | 814 | 826 | -186 | -174 | 2,02 | 0,01 | 18,69 | >0.1 | 5,58 | 6,06 | 0 | 1,59 | 0 | 3,43 | 0 |
| FOS::JUNB | ACTGAGTCTCGG | -1 | 442 | 454 | -558 | -546 | 2,03 | 0,01 | 16,56 | >0.1 | 2,75 | 4,85 | 0 | 1 | 0 | 5,92 | 0 |
| FOS::JUNB | TCTGCCTCCCAG | -1 | 380 | 392 | -620 | -608 | -2,13 | 0,01 | 12,26 | >0.1 | 2,57 | 4,74 | 0 | 1,06 | 0 | 6,02 | 0 |
| FOS::JUNB | CCTGCCTCAGCC | -1 | 354 | 366 | -646 | -634 | 5,47 | 0,001 | 19,82 | >0.1 | 2,53 | 4,69 | 0 | 1,1 | 0 | 6,03 | 0 |
| FOS::JUNB | TGTTAGTCAGGA | -1 | 261 | 273 | -739 | -727 | 7,74 | 0,001 | 20,95 | >0.1 | 2,8 | 4,55 | 0 | 1,32 | 0 | 4,53 | 0 |
| FOS::JUNB | CGTGAGCCACTG | -1 | 185 | 197 | -815 | -803 | 3,87 | 0,01 | 17,02 | >0.1 | 3,1 | 4,45 | 0 | 1,75 | 0 | 3,86 | 0 |

| | | | | | | | | | | | | | | | | | |
|---|---|---|---|---|---|---|---|---|---|---|---|---|---|---|---|---|---|
| FOS::JUNB | TATGACTACTCT | -1 | 123 | 135 | -877 | -865 | -0,88 | 0,01 | 10,52 | >0.1 | 3,44 | 4,38 | 0 | 1,89 | 0 | 1,7 | 0 |
| FOS::JUNB | TATTTCTCACAC | -1 | 52 | 64 | -948 | -936 | 2,75 | 0,01 | 13,01 | >0.1 | 4,09 | 4,3 | 0 | 1,87 | 0 | 0 | 0 |
| FOS::JUNB | AGTGCCTCCTTC | -1 | 17 | 29 | -983 | -971 | -1,12 | 0,01 | 10,6 | >0.1 | 5,58 | 4,27 | 0 | 1,87 | 0 | 0 | 0 |
| MGA | CAGTGTGA | 1 | 50 | 58 | -950 | -942 | 6,86 | 0,001 | 17,17 | >0.1 | 4,14 | 4,3 | 0 | 1,87 | 0 | 0 | 0 |
| MGA | GTGTGAGA | 1 | 52 | 60 | -948 | -940 | 3,78 | 0,01 | 14,06 | >0.1 | 4,11 | 4,3 | 0 | 1,87 | 0 | 0 | 0 |
| MGA | ACCTGTGG | 1 | 110 | 118 | -890 | -882 | 0,92 | 0,01 | 11,45 | >0.1 | 3,54 | 4,36 | 0 | 1,89 | 0 | 0,75 | 0 |
| MGA | GGGCGTGG | 1 | 314 | 322 | -686 | -678 | 5,96 | 0,01 | 20,53 | >0.1 | 2,64 | 4,62 | 0 | 1,17 | 0 | 6,13 | 0 |
| MGA | TGGCGTGA | 1 | 370 | 378 | -630 | -622 | 8,92 | 0,001 | 23,39 | >0.1 | 2,54 | 4,71 | 0 | 1,07 | 0 | 6,15 | 0 |
| MGA | ACCTGGGA | 1 | 378 | 386 | -622 | -614 | -0,35 | 0,01 | 14,4 | >0.1 | 2,56 | 4,73 | 0 | 1,06 | 0 | 6,4 | 0 |
| MGA | AGCCGAGA | 1 | 402 | 410 | -598 | -590 | 1,02 | 0,01 | 15,21 | >0.1 | 2,63 | 4,77 | 0 | 1,03 | 0 | 5,76 | 0 |
| MGA | GGGCGACA | 1 | 432 | 440 | -568 | -560 | 1,52 | 0,01 | 15,74 | >0.1 | 2,72 | 4,82 | 0 | 1,01 | 0 | 5,67 | 0 |
| MGA | AAATGTGA | 1 | 587 | 595 | -413 | -405 | -0,28 | 0,01 | 9,27 | >0.1 | 3,38 | 5,17 | 0 | 1 | 0 | 0 | 0 |
| MGA | ATGTGAGA | 1 | 589 | 597 | -411 | -403 | 5,06 | 0,01 | 14,63 | >0.1 | 3,4 | 5,17 | 0 | 1,01 | 0 | 0 | 0 |
| MGA | AAGTGACA | 1 | 764 | 772 | -236 | -228 | 2,1 | 0,01 | 20,76 | >0.1 | 5,64 | 5,78 | 0 | 1,33 | 0 | 5,92 | 0 |
| MGA | ATCTGTTA | 1 | 844 | 852 | -156 | -148 | 1,29 | 0,01 | 18,15 | >0.1 | 5,64 | 6,24 | 0 | 1,79 | 0 | 3,19 | 0 |
| MGA | GGCTGATA | 1 | 965 | 973 | -35 | -27 | -0,38 | 0,01 | 16,53 | >0.1 | 5,64 | 7,7 | 0 | 1,87 | 0 | 1,7 | 0 |
| MGA | AGGTTTTA | 1 | 993 | 1001 | -7 | 1 | -0,23 | 0,01 | 17,59 | >0.1 | 5,64 | 8,74 | 0 | 1,87 | 0 | 1,57 | 0 |
| MGA | GGATGTGA | 1 | 1022 | 1030 | 22 | 30 | 1,75 | 0,01 | 19,24 | >0.1 | 5,64 | 8,68 | 0 | 1,86 | 0 | 1,31 | 0 |
| MGA | TGCTGTAA | 1 | 1084 | 1092 | 84 | 92 | -0,33 | 0,01 | 18,12 | >0.1 | 5,64 | 6,96 | 0 | 1,85 | 1,79 | 2,21 | 0 |
| MGA | GGCTGAGA | -1 | 1072 | 1080 | 72 | 80 | 2,35 | 0,01 | 21,05 | >0.1 | 5,64 | 7,14 | 0 | 1,85 | 1,79 | 2,27 | 0 |
| MGA | CCCTGTGA | -1 | 1037 | 1045 | 37 | 45 | 1,25 | 0,01 | 18,04 | >0.1 | 5,64 | 7,95 | 0 | 1,86 | 0 | 1,34 | 0 |
| MGA | GGGCGCTG | -1 | 943 | 951 | -57 | -49 | 0,42 | 0,01 | 16 | >0.1 | 5,64 | 7,23 | 0 | 1,88 | 0 | 0,83 | 0 |
| MGA | AGCTGTTA | -1 | 710 | 718 | -290 | -282 | 5,77 | 0,01 | 22,94 | >0.1 | 5,64 | 5,55 | 0 | 1,17 | 0 | 4,81 | 0 |
| MGA | TGGTGTTT | -1 | 630 | 638 | -370 | -362 | 2,01 | 0,01 | 13,69 | >0.1 | 3,69 | 5,29 | 0 | 1,04 | 0 | 1,66 | 0 |
| MGA | TGGCGCGA | -1 | 410 | 418 | -590 | -582 | 6,12 | 0,01 | 20,1 | >0.1 | 2,65 | 4,78 | 0 | 1,03 | 0 | 5,52 | 0 |
| MGA | AGCTGGGA | -1 | 338 | 346 | -662 | -654 | 2,68 | 0,01 | 16,95 | >0.1 | 2,58 | 4,66 | 0 | 1,12 | 0 | 5,92 | 0 |
| MGA | AGGTGCCC | -1 | 326 | 334 | -674 | -666 | -0,13 | 0,01 | 13,72 | >0.1 | 2,61 | 4,64 | 0 | 1,15 | 0 | 5,46 | 0 |
| MGA | CCGTGTTA | -1 | 268 | 276 | -732 | -724 | 4,42 | 0,01 | 17,59 | >0.1 | 2,78 | 4,55 | 0 | 1,3 | 0 | 4,53 | 0 |
| MGA | AAGTGCTA | -1 | 207 | 215 | -793 | -785 | 5,55 | 0,01 | 19,32 | >0.1 | 3,01 | 4,47 | 0 | 1,58 | 0 | 4,7 | 0 |
| MGA | AGGCGTGA | -1 | 192 | 200 | -808 | -800 | 11,8 | 0,0001 | 25,15 | >0.1 | 3,08 | 4,45 | 0 | 1,69 | 0 | 4,13 | 0 |
| TP63 | GTGGAGTGGAGAGTAGTC | 1 | 114 | 132 | -886 | -868 | -28,99 | 0,01 | -18,52 | >0.1 | 3,48 | 4,37 | 0 | 1,89 | 0 | 0,73 | 0 |
| TP63 | GGCACAGTGGCTCACGCC | 1 | 181 | 199 | -819 | -801 | -28,37 | 0,01 | -15,17 | >0.1 | 3,1 | 4,45 | 0 | 1,78 | 0 | 3,86 | 0 |
| TP63 | TGGGAGGCAGAGCTTGCA | 1 | 381 | 399 | -619 | -601 | -28,01 | 0,01 | -13,49 | >0.1 | 2,58 | 4,75 | 0 | 1,06 | 0 | 6,13 | 0 |
| TP63 | CAAGGGCAAAGTCATGTA | 1 | 507 | 525 | -493 | -475 | -29,89 | 0,01 | -18,04 | >0.1 | 3 | 5 | 0 | 0,99 | 0 | 2,86 | 0 |
| TP63 | TACAAACACCAACATTAT | 1 | 627 | 645 | -373 | -355 | -26,92 | 0,01 | -15,19 | >0.1 | 3,71 | 5,31 | 0 | 1,06 | 0 | 1,66 | 0 |
| TP63 | GTAAAATTATAACATTTT | 1 | 681 | 699 | -319 | -301 | -29,46 | 0,01 | -15,49 | >0.1 | 4,4 | 5,48 | 0 | 1,14 | 0 | 2,95 | 0 |
| TP63 | GCCATAAAGTGACAGGAG | 1 | 758 | 776 | -242 | -224 | -25,95 | 0,01 | -7,38 | >0.1 | 5,51 | 5,8 | 0 | 1,35 | 0 | 5,92 | 0 |
| TP63 | AAAGTGACAGGAGAGGTA | 1 | 763 | 781 | -237 | -219 | -22,34 | 0,01 | -3,73 | >0.1 | 5,51 | 5,82 | 0 | 1,36 | 0 | 5,92 | 0 |
| TP63 | GACAGGAGAGGTAAGGTT | 1 | 768 | 786 | -232 | -214 | -28,63 | 0,01 | -9,98 | >0.1 | 5,51 | 5,85 | 0 | 1,39 | 0 | 5,92 | 0 |
| TP63 | AACGTATTCTTATTTGAT | 1 | 803 | 821 | -197 | -179 | -24,51 | 0,01 | -8,13 | >0.1 | 5,51 | 6,03 | 0 | 1,56 | 0 | 3,28 | 0 |
| TP63 | GTCAAGCTCTTCCTGGCT | 1 | 1051 | 1069 | 51 | 69 | -22,72 | 0,01 | -5,53 | >0.1 | 5,51 | 7,55 | 0 | 1,86 | 0 | 2,27 | 0 |
| TP63 | ACCATTGAGGAACAGGCC | 1 | 1106 | 1124 | 106 | 124 | -25,59 | 0,01 | -5,76 | >0.1 | 5,51 | 6,69 | 0 | 1,85 | 3,22 | 2,56 | 0 |
| TP63 | AACAGGCCAAGACATTTT | 1 | 1116 | 1134 | 116 | 134 | -11,27 | 0,001 | 8,59 | >0.1 | 5,51 | 6,59 | 0 | 1,85 | 3,22 | 2,7 | 0 |
| TP63 | GACCTGTTCTATCAAAGT | 1 | 1160 | 1178 | 160 | 178 | -26,62 | 0,01 | -9,69 | >0.1 | 4,48 | 6,22 | 0 | 1,76 | 1,79 | 2,67 | 0 |
| TP63 | ACTTTGATAGAACAGGTC | -1 | 1160 | 1178 | 160 | 178 | -22,72 | 0,01 | -5,79 | >0.1 | 4,48 | 6,22 | 0 | 1,76 | 1,79 | 2,67 | 0 |
| TP63 | TTCGTGGTTAAACTTGTC | -1 | 1136 | 1154 | 136 | 154 | -15,26 | 0,001 | 5,02 | >0.1 | 5,51 | 6,41 | 0 | 1,84 | 3,22 | 3,29 | 0 |
| TP63 | AAAATGTCTTGGCCTGTT | -1 | 1116 | 1134 | 116 | 134 | -13,75 | 0,001 | 6,11 | >0.1 | 5,51 | 6,59 | 0 | 1,85 | 3,22 | 2,7 | 0 |
| TP63 | AGCCAGGAAGAGCTTGAC | -1 | 1051 | 1069 | 51 | 69 | -23,52 | 0,01 | -6,33 | >0.1 | 5,51 | 7,55 | 0 | 1,86 | 0 | 2,27 | 0 |
| TP63 | ATCAAATAAGAATACGTT | -1 | 803 | 821 | -197 | -179 | -19,36 | 0,01 | -2,98 | >0.1 | 5,51 | 6,03 | 0 | 1,56 | 0 | 3,28 | 0 |
| TP63 | TACCTCTCCTGTCACTTT | -1 | 763 | 781 | -237 | -219 | -28,99 | 0,01 | -10,38 | >0.1 | 5,51 | 5,82 | 0 | 1,36 | 0 | 5,92 | 0 |
| TP63 | GCCAAATCAAAACCGGTA | -1 | 742 | 760 | -258 | -240 | -28,32 | 0,01 | -9,88 | >0.1 | 5,51 | 5,72 | 0 | 1,29 | 0 | 5,92 | 0 |
| TP63 | AAAATGTTATAATTTTAC | -1 | 681 | 699 | -319 | -301 | -27,44 | 0,01 | -13,47 | >0.1 | 4,4 | 5,48 | 0 | 1,14 | 0 | 2,95 | 0 |
| TP63 | ATAATGTTGGTGTTTGTA | -1 | 627 | 645 | -373 | -355 | -21,4 | 0,01 | -9,67 | >0.1 | 3,71 | 5,31 | 0 | 1,06 | 0 | 1,66 | 0 |
| TP63 | TGCAAGCTCTGCCTCCCA | -1 | 381 | 399 | -619 | -601 | -24,38 | 0,01 | -9,86 | >0.1 | 2,58 | 4,75 | 0 | 1,06 | 0 | 6,13 | 0 |
| GBX1 | GGAAATGAGC | 1 | 64 | 74 | -936 | -926 | 0,29 | 0,01 | 10,43 | >0.1 | 3,95 | 4,31 | 0 | 1,88 | 0 | 0 | 0 |
| GBX1 | GGAAATTAAA | 1 | 153 | 163 | -847 | -837 | 2,77 | 0,01 | 13,99 | >0.1 | 3,27 | 4,41 | 0 | 1,9 | 0 | 1,65 | 0 |
| GBX1 | ACTGATCAGA | 1 | 163 | 173 | -837 | -827 | 1 | 0,01 | 12,94 | >0.1 | 3,21 | 4,42 | 0 | 1,9 | 0 | 2,4 | 0 |
| GBX1 | CCTGACTAAC | 1 | 262 | 272 | -738 | -728 | -0,21 | 0,01 | 12,99 | >0.1 | 2,8 | 4,55 | 0 | 1,32 | 0 | 4,53 | 0 |
| GBX1 | CTCTATTAAA | 1 | 287 | 297 | -713 | -703 | 0 | 0,01 | 13,74 | >0.1 | 2,72 | 4,59 | 0 | 1,24 | 0 | 5,19 | 0 |

| | | | | | | | | | | | | | | | | | |
|---|---|---|---|---|---|---|---|---|---|---|---|---|---|---|---|---|---|
| GBX1 | ACTGATCAGA | 1 | 478 | 488 | -522 | -512 | 1 | 0,01 | 15,14 | >0.1 | 2,88 | 4,92 | 0 | 0,99 | 0 | 5,36 | 0 |
| GBX1 | ACCTATTAAT | 1 | 599 | 609 | -401 | -391 | 3,5 | 0,01 | 14,55 | >0.1 | 3,47 | 5,21 | 0 | 1,02 | 0 | 1,36 | 0 |
| GBX1 | ATTAATGAAA | 1 | 603 | 613 | -397 | -387 | 0,75 | 0,01 | 11,89 | >0.1 | 3,49 | 5,22 | 0 | 1,02 | 0 | 1,41 | 0 |
| GBX1 | AAAAATTATT | 1 | 829 | 839 | -171 | -161 | 0,08 | 0,01 | 16,8 | >0.1 | 5,61 | 6,15 | 0 | 1,68 | 0 | 3,28 | 0 |
| GBX1 | GCTGATAAGA | 1 | 966 | 976 | -34 | -24 | 0,16 | 0,01 | 17,12 | >0.1 | 5,61 | 7,78 | 0 | 1,87 | 0 | 1,7 | 0 |
| GBX1 | AGCAGTTACA | -1 | 1087 | 1097 | 87 | 97 | 0,2 | 0,01 | 18,59 | >0.1 | 5,61 | 6,92 | 0 | 1,85 | 1,79 | 2,21 | 0 |
| GBX1 | CCTCATGAGA | -1 | 983 | 993 | -17 | -7 | -0,27 | 0,01 | 17,28 | >0.1 | 5,61 | 8,5 | 0 | 1,87 | 0 | 1,57 | 0 |
| GBX1 | AATAATTTTT | -1 | 829 | 839 | -171 | -161 | -0,51 | 0,01 | 16,21 | >0.1 | 5,61 | 6,15 | 0 | 1,68 | 0 | 3,28 | 0 |
| GBX1 | ATCAAATAAG | -1 | 811 | 821 | -189 | -179 | -0,6 | 0,01 | 16,03 | >0.1 | 5,61 | 6,03 | 0 | 1,56 | 0 | 3,43 | 0 |
| GBX1 | GACAATTTCA | -1 | 608 | 618 | -392 | -382 | 0,75 | 0,01 | 12,08 | >0.1 | 3,53 | 5,23 | 0 | 1,02 | 0 | 1,55 | 0 |
| GBX1 | TTTCATTAAT | -1 | 603 | 613 | -397 | -387 | 1,08 | 0,01 | 12,22 | >0.1 | 3,49 | 5,22 | 0 | 1,02 | 0 | 1,41 | 0 |
| GBX1 | TCCAAATACA | -1 | 521 | 531 | -479 | -469 | -0,41 | 0,01 | 11,7 | >0.1 | 3,05 | 5,01 | 0 | 0,98 | 0 | 3,07 | 0 |
| GBX1 | TCTGATCAGT | -1 | 478 | 488 | -522 | -512 | 0,07 | 0,01 | 14,21 | >0.1 | 2,88 | 4,92 | 0 | 0,99 | 0 | 5,36 | 0 |
| GBX1 | TCTGATCAGT | -1 | 163 | 173 | -837 | -827 | 0,07 | 0,01 | 12,01 | >0.1 | 3,21 | 4,42 | 0 | 1,9 | 0 | 2,4 | 0 |
| GBX1 | TTTAATTTCC | -1 | 153 | 163 | -847 | -837 | 1,67 | 0,01 | 12,89 | >0.1 | 3,27 | 4,41 | 0 | 1,9 | 0 | 1,65 | 0 |
| GBX1 | GCTCATTTCC | -1 | 64 | 74 | -936 | -926 | 1,13 | 0,01 | 11,27 | >0.1 | 3,95 | 4,31 | 0 | 1,88 | 0 | 0 | 0 |
| GBX1 | TTCACTTACC | -1 | 4 | 14 | -996 | -986 | 1,74 | 0,01 | 15,26 | >0.1 | 5,61 | 4,26 | 0 | 1,86 | 1,79 | 0 | 0 |
| MEIS3 | AAGGCAGA | 1 | 90 | 98 | -910 | -902 | 2,11 | 0,01 | 12,03 | >0.1 | 3,7 | 4,34 | 0 | 1,88 | 0 | 0 | 0 |
| MEIS3 | ATGGCTGG | 1 | 174 | 182 | -826 | -818 | 2,52 | 0,01 | 15,19 | >0.1 | 3,16 | 4,43 | 0 | 1,84 | 0 | 3,24 | 0 |
| MEIS3 | TGGGCAGA | 1 | 228 | 236 | -772 | -764 | 2,66 | 0,01 | 16,81 | >0.1 | 2,93 | 4,5 | 0 | 1,46 | 0 | 5,26 | 0 |
| MEIS3 | GTGAAACC | 1 | 275 | 283 | -725 | -717 | 2,26 | 0,01 | 16,05 | >0.1 | 2,76 | 4,56 | 0 | 1,28 | 0 | 5,19 | 0 |
| MEIS3 | TGGGCACC | 1 | 325 | 333 | -675 | -667 | 2,52 | 0,01 | 16,37 | >0.1 | 2,61 | 4,64 | 0 | 1,15 | 0 | 5,46 | 0 |
| MEIS3 | GAGGCAGG | 1 | 358 | 366 | -642 | -634 | 4,63 | 0,01 | 19,06 | >0.1 | 2,52 | 4,69 | 0 | 1,09 | 0 | 6,13 | 0 |
| MEIS3 | ATGGCGTG | 1 | 369 | 377 | -631 | -623 | 2,43 | 0,01 | 17,48 | >0.1 | 2,54 | 4,71 | 0 | 1,07 | 0 | 6,73 | 0 |
| MEIS3 | GAGGCAGA | 1 | 384 | 392 | -616 | -608 | 2,55 | 0,01 | 16,93 | >0.1 | 2,58 | 4,74 | 0 | 1,05 | 0 | 6,02 | 0 |
| MEIS3 | GCGACAGA | 1 | 434 | 442 | -566 | -558 | 6,25 | 0,01 | 20,61 | >0.1 | 2,72 | 4,83 | 0 | 1,01 | 0 | 5,8 | 0 |
| MEIS3 | AAGACAAG | 1 | 503 | 511 | -497 | -489 | 5,1 | 0,01 | 19,7 | >0.1 | 2,97 | 4,97 | 0 | 0,98 | 0 | 5,68 | 0 |
| MEIS3 | GTGACAGG | 1 | 766 | 774 | -234 | -226 | 12,06 | 0,0001 | 30,73 | >0.1 | 5,64 | 5,79 | 0 | 1,33 | 0 | 5,92 | 0 |
| MEIS3 | ATGTCAAG | 1 | 1049 | 1057 | 49 | 57 | 1,97 | 0,01 | 18,48 | >0.1 | 5,64 | 7,6 | 0 | 1,86 | 0 | 1,42 | 0 |
| MEIS3 | AAGACATT | 1 | 1124 | 1132 | 124 | 132 | 3,35 | 0,01 | 23,29 | >0.1 | 5,64 | 6,51 | 0 | 1,84 | 3,22 | 2,72 | 0 |
| MEIS3 | TGGACAAG | 1 | 1134 | 1142 | 134 | 142 | 5,64 | 0,01 | 25,73 | >0.1 | 5,64 | 6,42 | 0 | 1,84 | 3,22 | 2,97 | 0 |
| MEIS3 | TTGATAGA | -1 | 1167 | 1175 | 167 | 175 | 4,45 | 0,01 | 21,52 | >0.1 | 4,53 | 6,17 | 0 | 1,69 | 1,79 | 2,88 | 0 |
| MEIS3 | TTGACATC | -1 | 1048 | 1056 | 48 | 56 | 9,94 | 0,001 | 26,48 | >0.1 | 5,64 | 7,62 | 0 | 1,86 | 0 | 1,42 | 0 |
| MEIS3 | ATGACTTT | -1 | 1009 | 1017 | 9 | 17 | 1,67 | 0,01 | 19,62 | >0.1 | 5,64 | 9,03 | 0 | 1,86 | 0 | 1,42 | 0 |
| MEIS3 | AGGACAGA | -1 | 856 | 864 | -144 | -136 | 5,25 | 0,01 | 21,98 | >0.1 | 5,64 | 6,32 | 0 | 1,9 | 0 | 2,86 | 0 |
| MEIS3 | TGGGCAAC | -1 | 540 | 548 | -460 | -452 | 1,86 | 0,01 | 11,02 | >0.1 | 3,13 | 5,05 | 0 | 0,98 | 0 | 0 | 0 |
| MEIS3 | ATGACTTT | -1 | 514 | 522 | -486 | -478 | 1,67 | 0,01 | 13,52 | >0.1 | 3,01 | 4,99 | 0 | 0,98 | 0 | 2,86 | 0 |
| MEIS3 | GTGGCGCG | -1 | 411 | 419 | -589 | -581 | 3,22 | 0,01 | 17,33 | >0.1 | 2,65 | 4,78 | 0 | 1,02 | 0 | 5,65 | 0 |
| MEIS3 | GAGACGGG | -1 | 282 | 290 | -718 | -710 | 4,67 | 0,01 | 18,43 | >0.1 | 2,74 | 4,57 | 0 | 1,25 | 0 | 5,19 | 0 |
| MEIS3 | AGGACGGT | -1 | 257 | 265 | -743 | -735 | 1,83 | 0,01 | 15,26 | >0.1 | 2,82 | 4,54 | 0 | 1,34 | 0 | 4,73 | 0 |
| MEIS3 | ATGACTAC | -1 | 126 | 134 | -874 | -866 | 3,12 | 0,01 | 14,52 | >0.1 | 3,43 | 4,38 | 0 | 1,89 | 0 | 1,7 | 0 |
| ZNF740 | ACCCCGTCTC | 1 | 280 | 290 | -720 | -710 | 1,42 | 0,01 | 15,09 | >0.1 | 2,74 | 4,57 | 0 | 1,26 | 0 | 5,09 | 0 |
| ZNF740 | CCCAACCCAA | 1 | 948 | 958 | -52 | -42 | 1,96 | 0,01 | 18,61 | >0.1 | 5,61 | 7,36 | 0 | 1,88 | 0 | 1,8 | 0 |
| ZNF740 | ACCTCCTCAT | -1 | 987 | 997 | -13 | -3 | 2,3 | 0,01 | 20,02 | >0.1 | 5,61 | 8,64 | 0 | 1,87 | 0 | 1,61 | 0 |
| ZNF740 | GCCTCCCGAG | -1 | 347 | 357 | -653 | -643 | 1,23 | 0,01 | 16,09 | >0.1 | 2,55 | 4,68 | 0 | 1,11 | 0 | 6,52 | 0 |
| ZNF740 | TGCCCACCAC | -1 | 321 | 331 | -679 | -669 | 1,4 | 0,01 | 15,38 | >0.1 | 2,62 | 4,64 | 0 | 1,16 | 0 | 5,56 | 0 |
| ZNF740 | GCCCACCACC | -1 | 320 | 330 | -680 | -670 | 4,36 | 0,01 | 18,46 | >0.1 | 2,62 | 4,63 | 0 | 1,16 | 0 | 5,68 | 0 |
| ZNF740 | CCCACCACCA | -1 | 319 | 329 | -681 | -671 | 1,98 | 0,01 | 15,96 | >0.1 | 2,63 | 4,63 | 0 | 1,16 | 0 | 5,56 | 0 |
| ZNF740 | CCACCACCAC | -1 | 318 | 328 | -682 | -672 | 6,25 | 0,01 | 20,93 | >0.1 | 2,63 | 4,63 | 0 | 1,16 | 0 | 6,26 | 0 |
| ZNF740 | ACCACCACGC | -1 | 316 | 326 | -684 | -674 | 1,17 | 0,01 | 15,73 | >0.1 | 2,63 | 4,63 | 0 | 1,17 | 0 | 6,13 | 0 |
| ZNF740 | CACCACGCCC | -1 | 314 | 324 | -686 | -676 | 0,78 | 0,01 | 15,35 | >0.1 | 2,64 | 4,63 | 0 | 1,17 | 0 | 6,13 | 0 |
| ZNF740 | CCACGCCCAG | -1 | 312 | 322 | -688 | -678 | 3,77 | 0,01 | 18,25 | >0.1 | 2,65 | 4,62 | 0 | 1,18 | 0 | 6,03 | 0 |
| ZNF740 | GGCCTCCCAA | -1 | 214 | 224 | -786 | -776 | 2,15 | 0,01 | 16,68 | >0.1 | 2,98 | 4,48 | 0 | 1,54 | 0 | 5,52 | 0 |
| ZNF740 | GCCTCCCAAA | -1 | 213 | 223 | -787 | -777 | 3,69 | 0,01 | 18,34 | >0.1 | 2,98 | 4,48 | 0 | 1,55 | 0 | 5,63 | 0 |
| ZNF740 | GCCCAGCCAT | -1 | 174 | 184 | -826 | -816 | 1,78 | 0,01 | 14,45 | >0.1 | 3,16 | 4,43 | 0 | 1,84 | 0 | 3,24 | 0 |
| ZNF740 | TCCACTCCAC | -1 | 114 | 124 | -886 | -876 | 2,42 | 0,01 | 12,93 | >0.1 | 3,51 | 4,36 | 0 | 1,89 | 0 | 0,75 | 0 |
| PBX2 | GTGAAGGAGAAG | 1 | 9 | 21 | -991 | -979 | 0,85 | 0,01 | 14,35 | >0.1 | 5,58 | 4,26 | 0 | 1,86 | 1,79 | 0 | 0 |
| PBX2 | GAGTTGGGCAGA | 1 | 224 | 236 | -776 | -764 | 2,12 | 0,01 | 16,3 | >0.1 | 2,94 | 4,5 | 0 | 1,49 | 0 | 5,26 | 0 |
| PBX2 | GAGATAGAGACC | 1 | 248 | 260 | -752 | -740 | 2,03 | 0,01 | 15,98 | >0.1 | 2,85 | 4,53 | 0 | 1,37 | 0 | 5,19 | 0 |

| | | | | | | | | | | | | | | | | | |
|---|---|---|---|---|---|---|---|---|---|---|---|---|---|---|---|---|---|
| PBX2 | CTGACTAACACG | 1 | 263 | 275 | -737 | -725 | 7,11 | 0,001 | 20,22 | >0.1 | 2,79 | 4,55 | 0 | 1,32 | 0 | 4,45 | 0 |
| PBX2 | GTGGTGGGCACC | 1 | 321 | 333 | -679 | -667 | 3,8 | 0,01 | 17,78 | >0.1 | 2,62 | 4,64 | 0 | 1,16 | 0 | 5,56 | 0 |
| PBX2 | GAGAATGGCGTG | 1 | 365 | 377 | -635 | -623 | 2,8 | 0,01 | 17,74 | >0.1 | 2,53 | 4,71 | 0 | 1,08 | 0 | 6,61 | 0 |
| PBX2 | CTGGGAGGCAGA | 1 | 380 | 392 | -620 | -608 | 1,75 | 0,01 | 16,14 | >0.1 | 2,57 | 4,74 | 0 | 1,06 | 0 | 6,02 | 0 |
| PBX2 | CTGGGCGACAGA | 1 | 430 | 442 | -570 | -558 | 3,86 | 0,01 | 18,08 | >0.1 | 2,72 | 4,83 | 0 | 1,01 | 0 | 5,67 | 0 |
| PBX2 | GCGACAGACCGA | 1 | 434 | 446 | -566 | -554 | 0,74 | 0,01 | 15,23 | >0.1 | 2,73 | 4,83 | 0 | 1,01 | 0 | 5,92 | 0 |
| PBX2 | ATCTTTAACAGC | 1 | 705 | 717 | -295 | -283 | 0,88 | 0,01 | 18,17 | >0.1 | 5,58 | 5,55 | 0 | 1,17 | 0 | 4,99 | 0 |
| PBX2 | TAAAGTGACAGG | 1 | 762 | 774 | -238 | -226 | 10,17 | 0,001 | 28,79 | >0.1 | 5,58 | 5,79 | 0 | 1,34 | 0 | 5,92 | 0 |
| PBX2 | TTGATTCACTTT | 1 | 816 | 828 | -184 | -172 | 2,64 | 0,01 | 19,45 | >0.1 | 5,58 | 6,08 | 0 | 1,6 | 0 | 3,55 | 0 |
| PBX2 | TATATTGGCTCA | 1 | 879 | 891 | -121 | -109 | 0,88 | 0,01 | 17,96 | >0.1 | 5,58 | 6,54 | 0 | 1,9 | 0 | 3,07 | 0 |
| PBX2 | TTGGCTCACAGG | 1 | 1032 | 1044 | 32 | 44 | 4,18 | 0,01 | 21,07 | >0.1 | 5,58 | 8,15 | 0 | 1,86 | 0 | 1,31 | 0 |
| PBX2 | TTGAGGAACAGG | 1 | 1110 | 1122 | 110 | 122 | 5,33 | 0,01 | 25,17 | >0.1 | 5,58 | 6,65 | 0 | 1,85 | 3,22 | 2,54 | 0 |
| PBX2 | TTTTTGGACAAG | 1 | 1130 | 1142 | 130 | 142 | 0,83 | 0,01 | 20,85 | >0.1 | 5,58 | 6,46 | 0 | 1,84 | 3,22 | 2,91 | 0 |
| PBX2 | ATGGTGGACTGA | -1 | 1099 | 1111 | 99 | 111 | 2,14 | 0,01 | 21,94 | >0.1 | 5,58 | 6,77 | 0 | 1,85 | 3,22 | 2,38 | 0 |
| PBX2 | AGCAGTTACAGC | -1 | 1085 | 1097 | 85 | 97 | 4,63 | 0,01 | 23,01 | >0.1 | 5,58 | 6,94 | 0 | 1,85 | 1,79 | 2,21 | 0 |
| PBX2 | GAGCTTGACATC | -1 | 1048 | 1060 | 48 | 60 | 7,52 | 0,001 | 24,05 | >0.1 | 5,58 | 7,62 | 0 | 1,86 | 0 | 1,46 | 0 |
| PBX2 | CTGAATGACTTT | -1 | 1009 | 1021 | 9 | 21 | 5,43 | 0,01 | 23,32 | >0.1 | 5,58 | 9,03 | 0 | 1,86 | 0 | 1,42 | 0 |
| PBX2 | CACAGTGGCGCG | -1 | 411 | 423 | -589 | -577 | 2,03 | 0,01 | 16,42 | >0.1 | 2,66 | 4,79 | 0 | 1,03 | 0 | 5,91 | 0 |
| PBX2 | AATAGAGACGGG | -1 | 282 | 294 | -718 | -706 | 1,11 | 0,01 | 14,77 | >0.1 | 2,73 | 4,58 | 0 | 1,26 | 0 | 5,09 | 0 |
| PBX2 | TAGGATTACAGG | -1 | 197 | 209 | -803 | -791 | 0,8 | 0,01 | 14,44 | >0.1 | 3,05 | 4,47 | 0 | 1,65 | 0 | 4,48 | 0 |
| PBX2 | GTGGCTGACTTC | -1 | 33 | 45 | -967 | -955 | 3,44 | 0,01 | 13,99 | >0.1 | 4,4 | 4,28 | 0 | 1,87 | 0 | 0 | 0 |
| PBX2 | TTGAGTGCCTCC | -1 | 20 | 32 | -980 | -968 | 2,9 | 0,01 | 14,62 | >0.1 | 5,58 | 4,27 | 0 | 1,87 | 0 | 0 | 0 |
| TEF | TGTGAGATAACC | 1 | 590 | 602 | -410 | -398 | -4,59 | 0,01 | 5,02 | >0.1 | 3,41 | 5,19 | 0 | 1,01 | 0 | 0 | 0 |
| TEF | ACCAACATTATC | 1 | 634 | 646 | -366 | -354 | -5,02 | 0,01 | 6,7 | >0.1 | 3,75 | 5,31 | 0 | 1,05 | 0 | 1,61 | 0 |
| TEF | TGTAAAATTATA | 1 | 680 | 692 | -320 | -308 | -3,09 | 0,01 | 11 | >0.1 | 4,32 | 5,46 | 0 | 1,12 | 0 | 3,19 | 0 |
| TEF | TATAACATTTTC | 1 | 688 | 700 | -312 | -300 | -3,35 | 0,01 | 10,71 | >0.1 | 4,48 | 5,49 | 0 | 1,14 | 0 | 2,95 | 0 |
| TEF | CATAAAGTGACA | 1 | 760 | 772 | -240 | -228 | -4 | 0,01 | 14,79 | >0.1 | 5,58 | 5,78 | 0 | 1,33 | 0 | 6,1 | 0 |
| TEF | GATTAAAGAATA | 1 | 792 | 804 | -208 | -196 | 0,42 | 0,01 | 18,72 | >0.1 | 5,58 | 5,94 | 0 | 1,47 | 0 | 5,32 | 0 |
| TEF | AATAACGTATTC | 1 | 800 | 812 | -200 | -188 | 1,32 | 0,01 | 17,82 | >0.1 | 5,58 | 5,98 | 0 | 1,51 | 0 | 3,43 | 0 |
| TEF | TGTTACATATCT | 1 | 847 | 859 | -153 | -141 | 5,81 | 0,001 | 22,49 | >0.1 | 5,58 | 6,29 | 0 | 1,85 | 0 | 2,95 | 0 |
| TEF | CATTGAGGAACA | 1 | 1108 | 1120 | 108 | 120 | -1,55 | 0,01 | 18,29 | >0.1 | 5,58 | 6,67 | 0 | 1,85 | 3,22 | 2,52 | 0 |
| TEF | CAGTAAACAATC | -1 | 894 | 906 | -106 | -94 | -6,38 | 0,01 | 10,71 | >0.1 | 5,58 | 6,67 | 0 | 1,89 | 0 | 2,95 | 0 |
| TEF | AGATATGTAACA | -1 | 847 | 859 | -153 | -141 | 7,04 | 0,001 | 23,72 | >0.1 | 5,58 | 6,29 | 0 | 1,85 | 0 | 2,95 | 0 |
| TEF | TGTAACAGATTT | -1 | 842 | 854 | -158 | -146 | -6,15 | 0,01 | 10,68 | >0.1 | 5,58 | 6,26 | 0 | 1,81 | 0 | 3,19 | 0 |
| TEF | GAATACGTTATT | -1 | 800 | 812 | -200 | -188 | 4,58 | 0,001 | 21,08 | >0.1 | 5,58 | 5,98 | 0 | 1,51 | 0 | 3,43 | 0 |
| TEF | GAAAATGTTATA | -1 | 688 | 700 | -312 | -300 | -2,59 | 0,01 | 11,47 | >0.1 | 4,48 | 5,49 | 0 | 1,14 | 0 | 2,95 | 0 |
| TEF | GGTTATCTCACA | -1 | 590 | 602 | -410 | -398 | -5,03 | 0,01 | 4,58 | >0.1 | 3,41 | 5,19 | 0 | 1,01 | 0 | 0 | 0 |
| TEF | AAATACATGACT | -1 | 516 | 528 | -484 | -472 | -0,19 | 0,01 | 11,78 | >0.1 | 3,03 | 5 | 0 | 0,98 | 0 | 2,95 | 0 |
| TEF | CTTTATGGAATT | -1 | 494 | 506 | -506 | -494 | -2,62 | 0,01 | 12,88 | >0.1 | 2,94 | 4,96 | 0 | 0,99 | 0 | 6,62 | 0 |
| HESX1 | GGTAAGTGAA | 1 | 4 | 14 | -996 | -986 | 2,14 | 0,01 | 15,66 | >0.1 | 5,61 | 4,26 | 0 | 1,86 | 1,79 | 0 | 0 |
| HESX1 | CATAATTTTA | 1 | 131 | 141 | -869 | -859 | 2,26 | 0,01 | 13,51 | >0.1 | 3,4 | 4,38 | 0 | 1,89 | 0 | 1,57 | 0 |
| HESX1 | GGAAATTAAA | 1 | 153 | 163 | -847 | -837 | 4,57 | 0,01 | 15,79 | >0.1 | 3,27 | 4,41 | 0 | 1,9 | 0 | 1,65 | 0 |
| HESX1 | TGAAATTGTC | 1 | 608 | 618 | -392 | -382 | 3,3 | 0,01 | 14,63 | >0.1 | 3,53 | 5,23 | 0 | 1,02 | 0 | 1,55 | 0 |
| HESX1 | TAAAATTATA | 1 | 682 | 692 | -318 | -308 | 1,58 | 0,01 | 15,56 | >0.1 | 4,34 | 5,46 | 0 | 1,12 | 0 | 3,07 | 0 |
| HESX1 | TTTGATTTGG | 1 | 749 | 759 | -251 | -241 | -1,07 | 0,01 | 17,46 | >0.1 | 5,61 | 5,72 | 0 | 1,28 | 0 | 5,92 | 0 |
| HESX1 | ATAAAGTGAC | 1 | 761 | 771 | -239 | -229 | -1,31 | 0,01 | 17,5 | >0.1 | 5,61 | 5,77 | 0 | 1,32 | 0 | 6,1 | 0 |
| HESX1 | TCTTATTTGA | 1 | 810 | 820 | -190 | -180 | -1,42 | 0,01 | 15,05 | >0.1 | 5,61 | 6,03 | 0 | 1,55 | 0 | 3,28 | 0 |
| HESX1 | TTTGATTCAC | 1 | 815 | 825 | -185 | -175 | -1,06 | 0,01 | 15,62 | >0.1 | 5,61 | 6,06 | 0 | 1,58 | 0 | 3,43 | 0 |
| HESX1 | AAAAATTATT | 1 | 829 | 839 | -171 | -161 | 2,34 | 0,01 | 19,06 | >0.1 | 5,61 | 6,15 | 0 | 1,68 | 0 | 3,28 | 0 |
| HESX1 | TTATATTGGC | 1 | 878 | 888 | -122 | -112 | 0 | 0,01 | 17,09 | >0.1 | 5,61 | 6,51 | 0 | 1,9 | 0 | 3,07 | 0 |
| HESX1 | GCAGATTGTT | 1 | 891 | 901 | -109 | -99 | 0,04 | 0,01 | 17,03 | >0.1 | 5,61 | 6,62 | 0 | 1,89 | 0 | 2,86 | 0 |
| HESX1 | TATAATTCCA | -1 | 1190 | 1200 | 190 | 200 | 2,44 | 0,01 | 21,2 | >0.1 | 5,61 | 6,03 | 0 | 1,51 | 1,79 | 3,82 | 0 |
| HESX1 | AATAATTTTT | -1 | 829 | 839 | -171 | -161 | 2,5 | 0,01 | 19,22 | >0.1 | 5,61 | 6,15 | 0 | 1,68 | 0 | 3,28 | 0 |
| HESX1 | TATAATTTTA | -1 | 682 | 692 | -318 | -308 | 1,74 | 0,01 | 15,72 | >0.1 | 4,34 | 5,46 | 0 | 1,12 | 0 | 3,07 | 0 |
| HESX1 | TCTACTTGGG | -1 | 544 | 554 | -456 | -446 | 1,65 | 0,01 | 10,85 | >0.1 | 3,15 | 5,06 | 0 | 0,99 | 0 | 0 | 0 |
| HESX1 | CCAAAGTGCT | -1 | 208 | 218 | -792 | -782 | -1,43 | 0,01 | 12,33 | >0.1 | 3 | 4,48 | 0 | 1,58 | 0 | 4,7 | 0 |
| HESX1 | TTTAATTTCC | -1 | 153 | 163 | -847 | -837 | 5,36 | 0,01 | 16,58 | >0.1 | 3,27 | 4,41 | 0 | 1,9 | 0 | 1,65 | 0 |
| HESX1 | TAAAATTATG | -1 | 131 | 141 | -869 | -859 | 1,28 | 0,01 | 12,53 | >0.1 | 3,4 | 4,38 | 0 | 1,89 | 0 | 1,57 | 0 |
| NFKB1 | GGAGAGTTGACCT | 1 | 101 | 114 | -899 | -886 | -21,43 | 0,01 | -10,83 | >0.1 | 3,59 | 4,35 | 0 | 1,89 | 0 | 0,77 | 0 |

| | | | | | | | | | | | | | | | | | |
|---|---|---|---|---|---|---|---|---|---|---|---|---|---|---|---|---|---|
| NFKB1 | CGGTGAAACCCCG | 1 | 273 | 286 | -727 | -714 | 6,14 | 0,0001 | 19,95 | >0.1 | 2,76 | 4,57 | 0 | 1,28 | 0 | 5,19 | 0 |
| NFKB1 | GGTGAAACCCCGT | 1 | 274 | 287 | -726 | -713 | -13,69 | 0,01 | 0,01 | >0.1 | 2,76 | 4,57 | 0 | 1,28 | 0 | 5,09 | 0 |
| NFKB1 | GTGGTGGGCACCT | 1 | 321 | 334 | -679 | -666 | -22,31 | 0,01 | -8,33 | >0.1 | 2,62 | 4,64 | 0 | 1,16 | 0 | 5,56 | 0 |
| NFKB1 | TGGTGGGCACCTG | 1 | 322 | 335 | -678 | -665 | -20,23 | 0,01 | -6,25 | >0.1 | 2,61 | 4,64 | 0 | 1,16 | 0 | 5,56 | 0 |
| NFKB1 | TGTGAGATAACCT | 1 | 590 | 603 | -410 | -397 | -17,15 | 0,01 | -7,53 | >0.1 | 3,42 | 5,19 | 0 | 1,01 | 0 | 0 | 0 |
| NFKB1 | AGGGAAAGTCATT | 1 | 1005 | 1018 | 5 | 18 | -17,95 | 0,01 | 0,07 | >0.1 | 5,57 | 9,12 | 0 | 1,87 | 0 | 1,46 | 0 |
| NFKB1 | AATGACTTTCCCT | -1 | 1005 | 1018 | 5 | 18 | -21,4 | 0,01 | -3,38 | >0.1 | 5,57 | 9,12 | 0 | 1,87 | 0 | 1,46 | 0 |
| NFKB1 | AAGTGAATCAAAT | -1 | 814 | 827 | -186 | -173 | -20,98 | 0,01 | -4,31 | >0.1 | 5,57 | 6,07 | 0 | 1,6 | 0 | 3,43 | 0 |
| NFKB1 | AGGTTATCTCACA | -1 | 590 | 603 | -410 | -397 | -16,23 | 0,01 | -6,61 | >0.1 | 3,42 | 5,19 | 0 | 1,01 | 0 | 0 | 0 |
| NFKB1 | CAGGTGCCCACCA | -1 | 322 | 335 | -678 | -665 | -20,72 | 0,01 | -6,74 | >0.1 | 2,61 | 4,64 | 0 | 1,16 | 0 | 5,56 | 0 |
| NFKB1 | ACGGGGTTTCACC | -1 | 274 | 287 | -726 | -713 | -16,23 | 0,01 | -2,53 | >0.1 | 2,76 | 4,57 | 0 | 1,28 | 0 | 5,09 | 0 |
| NFKB1 | CGGGGTTTCACCG | -1 | 273 | 286 | -727 | -714 | 8,29 | 0,0001 | 22,1 | >0.1 | 2,76 | 4,57 | 0 | 1,28 | 0 | 5,19 | 0 |
| NFKB1 | GGGGTTTCACCGT | -1 | 272 | 285 | -728 | -715 | -21,64 | 0,01 | -7,93 | >0.1 | 2,76 | 4,57 | 0 | 1,29 | 0 | 5,09 | 0 |
| NFKB1 | GGCGTGAGCCACT | -1 | 186 | 199 | -814 | -801 | -22,15 | 0,01 | -9,01 | >0.1 | 3,09 | 4,45 | 0 | 1,74 | 0 | 3,86 | 0 |
| NFKB1 | AGGTCAACTCTCC | -1 | 101 | 114 | -899 | -886 | -19,13 | 0,01 | -8,53 | >0.1 | 3,59 | 4,35 | 0 | 1,89 | 0 | 0,77 | 0 |
| NFKB2 | ACGGTGAAACCCC | 1 | 272 | 285 | -728 | -715 | -0,16 | 0,01 | 13,55 | >0.1 | 2,76 | 4,57 | 0 | 1,29 | 0 | 5,09 | 0 |
| NFKB2 | CGGTGAAACCCCG | 1 | 273 | 286 | -727 | -714 | 11,97 | 0,0001 | 25,78 | >0.1 | 2,76 | 4,57 | 0 | 1,28 | 0 | 5,19 | 0 |
| NFKB2 | GGTGAAACCCCGT | 1 | 274 | 287 | -726 | -713 | -8,65 | 0,01 | 5,05 | >0.1 | 2,76 | 4,57 | 0 | 1,28 | 0 | 5,09 | 0 |
| NFKB2 | TGGTGGGCACCTG | 1 | 322 | 335 | -678 | -665 | -10,85 | 0,01 | 3,13 | >0.1 | 2,61 | 4,64 | 0 | 1,16 | 0 | 5,56 | 0 |
| NFKB2 | TGTGAGATAACCT | 1 | 590 | 603 | -410 | -397 | -8,02 | 0,01 | 1,6 | >0.1 | 3,42 | 5,19 | 0 | 1,01 | 0 | 0 | 0 |
| NFKB2 | TTGGTCTTTCCTG | 1 | 928 | 941 | -72 | -59 | -12 | 0,01 | 3,55 | >0.1 | 5,57 | 7,08 | 0 | 1,88 | 0 | 1,03 | 0 |
| NFKB2 | AGGGAAAGTCATT | 1 | 1005 | 1018 | 5 | 18 | -9,29 | 0,01 | 8,73 | >0.1 | 5,57 | 9,12 | 0 | 1,87 | 0 | 1,46 | 0 |
| NFKB2 | AATGACTTTCCCT | -1 | 1005 | 1018 | 5 | 18 | -6,85 | 0,01 | 11,17 | >0.1 | 5,57 | 9,12 | 0 | 1,87 | 0 | 1,46 | 0 |
| NFKB2 | CAGGAAAGACCAA | -1 | 928 | 941 | -72 | -59 | -9,27 | 0,01 | 6,28 | >0.1 | 5,57 | 7,08 | 0 | 1,88 | 0 | 1,03 | 0 |
| NFKB2 | AGGTTATCTCACA | -1 | 590 | 603 | -410 | -397 | -7,83 | 0,01 | 1,79 | >0.1 | 3,42 | 5,19 | 0 | 1,01 | 0 | 0 | 0 |
| NFKB2 | CAGGTGCCCACCA | -1 | 322 | 335 | -678 | -665 | -9,88 | 0,01 | 4,1 | >0.1 | 2,61 | 4,64 | 0 | 1,16 | 0 | 5,56 | 0 |
| NFKB2 | AGGTGCCCACCAC | -1 | 321 | 334 | -679 | -666 | -9,97 | 0,01 | 4,01 | >0.1 | 2,62 | 4,64 | 0 | 1,16 | 0 | 5,56 | 0 |
| NFKB2 | ACGGGGTTTCACC | -1 | 274 | 287 | -726 | -713 | -4,36 | 0,01 | 9,34 | >0.1 | 2,76 | 4,57 | 0 | 1,28 | 0 | 5,09 | 0 |
| NFKB2 | CGGGGTTTCACCG | -1 | 273 | 286 | -727 | -714 | 10,88 | 0,0001 | 24,69 | >0.1 | 2,76 | 4,57 | 0 | 1,28 | 0 | 5,19 | 0 |
| NFKB2 | GGGGTTTCACCGT | -1 | 272 | 285 | -728 | -715 | -5 | 0,01 | 8,71 | >0.1 | 2,76 | 4,57 | 0 | 1,29 | 0 | 5,09 | 0 |
| TCF21 | GGCACAGTGGCTCA | 1 | 181 | 195 | -819 | -805 | -10,52 | 0,01 | 2,68 | >0.1 | 3,11 | 4,45 | 0 | 1,78 | 0 | 3,86 | 0 |
| TCF21 | TGGGCACCTGTAGT | 1 | 325 | 339 | -675 | -661 | -8,27 | 0,01 | 5,7 | >0.1 | 2,6 | 4,65 | 0 | 1,15 | 0 | 5,56 | 0 |
| TCF21 | GTCCCAGCTACTCG | 1 | 337 | 351 | -663 | -649 | -6,28 | 0,01 | 8,61 | >0.1 | 2,57 | 4,67 | 0 | 1,13 | 0 | 6,52 | 0 |
| TCF21 | TTAACAGCTTTCTA | 1 | 709 | 723 | -291 | -277 | 2,53 | 0,001 | 19,83 | >0.1 | 5,56 | 5,57 | 0 | 1,19 | 0 | 4,99 | 0 |
| TCF21 | AGTACCGGTTTTGA | 1 | 740 | 754 | -260 | -246 | -10,96 | 0,01 | 7,48 | >0.1 | 5,56 | 5,7 | 0 | 1,27 | 0 | 5,92 | 0 |
| TCF21 | CTAAAATCTGTTAC | 1 | 839 | 853 | -161 | -147 | 1,99 | 0,001 | 17,01 | >0.1 | 5,56 | 6,25 | 0 | 1,8 | 0 | 1,41 | 0 |
| TCF21 | TACATATCTGTCCT | 1 | 850 | 864 | -150 | -136 | -9,85 | 0,01 | 6,6 | >0.1 | 5,56 | 6,33 | 0 | 1,9 | 0 | 2,67 | 0 |
| TCF21 | GTAACTGCTGCTCA | 1 | 1088 | 1102 | 88 | 102 | 3,86 | 0,001 | 22,24 | >0.1 | 5,56 | 6,9 | 0 | 1,85 | 1,79 | 2,27 | 0 |
| TCF21 | CGAAGACCTGTTCT | 1 | 1156 | 1170 | 156 | 170 | -1,86 | 0,01 | 14,76 | >0.1 | 4,38 | 6,25 | 0 | 1,8 | 1,79 | 2,4 | 0 |
| TCF21 | AGAACAGGTCTTCG | -1 | 1156 | 1170 | 156 | 170 | 0,74 | 0,001 | 17,36 | >0.1 | 4,38 | 6,25 | 0 | 1,8 | 1,79 | 2,4 | 0 |
| TCF21 | TGAGCAGCAGTTAC | -1 | 1088 | 1102 | 88 | 102 | -1,8 | 0,01 | 16,58 | >0.1 | 5,56 | 6,9 | 0 | 1,85 | 1,79 | 2,27 | 0 |
| TCF21 | GGAAGAGCTTGACA | -1 | 1050 | 1064 | 50 | 64 | -9,72 | 0,01 | 7,54 | >0.1 | 5,56 | 7,57 | 0 | 1,86 | 0 | 2,27 | 0 |
| TCF21 | AAACAATCTGCTGA | -1 | 888 | 902 | -112 | -98 | -6,76 | 0,01 | 10,27 | >0.1 | 5,56 | 6,63 | 0 | 1,89 | 0 | 2,95 | 0 |
| TCF21 | GTAACAGATTTTAG | -1 | 839 | 853 | -161 | -147 | -0,82 | 0,01 | 14,2 | >0.1 | 5,56 | 6,25 | 0 | 1,8 | 0 | 1,41 | 0 |
| TCF21 | TAGAAAGCTGTTAA | -1 | 709 | 723 | -291 | -277 | 3,21 | 0,001 | 20,51 | >0.1 | 5,56 | 5,57 | 0 | 1,19 | 0 | 4,99 | 0 |
| TCF21 | GCAAGCTCTGCCTC | -1 | 384 | 398 | -616 | -602 | -10,32 | 0,01 | 4,2 | >0.1 | 2,59 | 4,75 | 0 | 1,06 | 0 | 6,13 | 0 |
| TCF21 | ACTACAGGTGCCCA | -1 | 325 | 339 | -675 | -661 | -1,31 | 0,01 | 12,66 | >0.1 | 2,6 | 4,65 | 0 | 1,15 | 0 | 5,56 | 0 |
| NR4A2::RXRA | AGGAGAGTTGACCTG | 1 | 100 | 115 | -900 | -885 | 6,01 | 0,001 | 16,61 | >0.1 | 3,59 | 4,36 | 0 | 1,89 | 0 | 0,77 | 0 |
| NR4A2::RXRA | ATGGCCATGGAAATT | 1 | 145 | 160 | -855 | -840 | -0,87 | 0,01 | 10,34 | >0.1 | 3,3 | 4,41 | 0 | 1,9 | 0 | 1,61 | 0 |
| NR4A2::RXRA | GGTGGTGGGCACCTG | 1 | 320 | 335 | -680 | -665 | -1,83 | 0,01 | 12,15 | >0.1 | 2,62 | 4,64 | 0 | 1,16 | 0 | 5,56 | 0 |
| NR4A2::RXRA | AATGGCGTGAACCTG | 1 | 368 | 383 | -632 | -617 | 1,97 | 0,01 | 16,58 | >0.1 | 2,54 | 4,72 | 0 | 1,08 | 0 | 6,27 | 0 |
| NR4A2::RXRA | GCTTGCAGTGAGCCG | 1 | 392 | 407 | -608 | -593 | -0,43 | 0,01 | 13,9 | >0.1 | 2,61 | 4,76 | 0 | 1,05 | 0 | 5,91 | 0 |
| NR4A2::RXRA | CTGGGCGACAGACCG | 1 | 430 | 445 | -570 | -555 | -2,16 | 0,01 | 12,08 | >0.1 | 2,72 | 4,83 | 0 | 1,01 | 0 | 5,67 | 0 |
| NR4A2::RXRA | GAAGCCGAAGACCTG | 1 | 1151 | 1166 | 151 | 166 | 2,75 | 0,01 | 19,49 | >0.1 | 4,47 | 6,29 | 0 | 1,84 | 1,79 | 2,34 | 0 |
| NR4A2::RXRA | AAGTTCACTTGCTTC | 1 | 1174 | 1189 | 174 | 189 | -2,29 | 0,01 | 16,08 | >0.1 | 5,54 | 6,13 | 0 | 1,63 | 1,79 | 3,28 | 0 |
| NR4A2::RXRA | GAAGCAAGTGAACTT | -1 | 1174 | 1189 | 174 | 189 | 0,45 | 0,01 | 18,82 | >0.1 | 5,54 | 6,13 | 0 | 1,63 | 1,79 | 3,28 | 0 |
| NR4A2::RXRA | CAGGTCTTCGGCTTC | -1 | 1151 | 1166 | 151 | 166 | 2,53 | 0,01 | 19,27 | >0.1 | 4,47 | 6,29 | 0 | 1,84 | 1,79 | 2,34 | 0 |
| NR4A2::RXRA | GAAGAGCTTGACATC | -1 | 1048 | 1063 | 48 | 63 | -0,82 | 0,01 | 15,67 | >0.1 | 5,54 | 7,62 | 0 | 1,86 | 0 | 1,46 | 0 |
| NR4A2::RXRA | CTTGACATCGTCCCC | -1 | 1042 | 1057 | 42 | 57 | -1,94 | 0,01 | 13,88 | >0.1 | 5,54 | 7,79 | 0 | 1,86 | 0 | 0,62 | 0 |

| | | | | | | | | | | | | | | | | | |
|---|---|---|---|---|---|---|---|---|---|---|---|---|---|---|---|---|---|
| NR4A2::RXRA | TCAGCCTTTGAACTT | -1 | 956 | 971 | -44 | -29 | 2,13 | 0,01 | 18,94 | >0.1 | 5,54 | 7,65 | 0 | 1,88 | 0 | 1,74 | 0 |
| NR4A2::RXRA | CATGACTTTGCCCTT | -1 | 508 | 523 | -492 | -477 | 3,29 | 0,01 | 15,13 | >0.1 | 3 | 4,99 | 0 | 0,99 | 0 | 2,86 | 0 |
| NR4A2::RXRA | TCGGTCTGTCGCCCA | -1 | 431 | 446 | -569 | -554 | 1,57 | 0,01 | 15,95 | >0.1 | 2,72 | 4,84 | 0 | 1,01 | 0 | 5,8 | 0 |
| NR4A2::RXRA | CGGTCTGTCGCCCAG | -1 | 430 | 445 | -570 | -555 | -1,4 | 0,01 | 12,84 | >0.1 | 2,72 | 4,83 | 0 | 1,01 | 0 | 5,67 | 0 |
| NR4A2::RXRA | CAGGTTCACGCCATT | -1 | 368 | 383 | -632 | -617 | 0,36 | 0,01 | 14,97 | >0.1 | 2,54 | 4,72 | 0 | 1,08 | 0 | 6,27 | 0 |
| NR4A2::RXRA | CACGCCATTCTCCTG | -1 | 362 | 377 | -638 | -623 | 0,36 | 0,01 | 15,3 | >0.1 | 2,53 | 4,71 | 0 | 1,09 | 0 | 6,61 | 0 |
| NR4A2::RXRA | GGTGCCCACCACCAC | -1 | 318 | 333 | -682 | -667 | -2,12 | 0,01 | 11,99 | >0.1 | 2,62 | 4,64 | 0 | 1,17 | 0 | 5,68 | 0 |
| NR4A2::RXRA | ACGGGGTTTCACCGT | -1 | 272 | 287 | -728 | -713 | 0,31 | 0,01 | 14,12 | >0.1 | 2,76 | 4,57 | 0 | 1,29 | 0 | 5,19 | 0 |
| NR4A2::RXRA | CAGGTCAACTCTCCT | -1 | 100 | 115 | -900 | -885 | -0,58 | 0,01 | 10,02 | >0.1 | 3,59 | 4,36 | 0 | 1,89 | 0 | 0,77 | 0 |
| NR4A2::RXRA | TCTGTGGCTGACTTC | -1 | 33 | 48 | -967 | -952 | 0,1 | 0,01 | 10,63 | >0.1 | 4,37 | 4,29 | 0 | 1,87 | 0 | 0 | 0 |
| ESRRB | TGAAGGAGAAG | 1 | 10 | 21 | -990 | -979 | -4,86 | 0,01 | 8,66 | >0.1 | 5,6 | 4,26 | 0 | 1,86 | 1,79 | 0 | 0 |
| ESRRB | GCAAAGGCAGA | 1 | 87 | 98 | -913 | -902 | -6,5 | 0,01 | 3,43 | >0.1 | 3,72 | 4,34 | 0 | 1,88 | 0 | 0 | 0 |
| ESRRB | ACAAGGTCAGG | 1 | 238 | 249 | -762 | -751 | 8,98 | 0,0001 | 22,33 | >0.1 | 2,88 | 4,52 | 0 | 1,42 | 0 | 4,53 | 0 |
| ESRRB | ACACGGTGAAA | 1 | 270 | 281 | -730 | -719 | -6,6 | 0,01 | 7,02 | >0.1 | 2,77 | 4,56 | 0 | 1,29 | 0 | 4,99 | 0 |
| ESRRB | TCAGTCTCAAA | 1 | 449 | 460 | -551 | -540 | -10,37 | 0,01 | 3,52 | >0.1 | 2,78 | 4,86 | 0 | 1 | 0 | 5,26 | 0 |
| ESRRB | ACAAGGGCAAA | 1 | 506 | 517 | -494 | -483 | 6,68 | 0,001 | 21,31 | >0.1 | 2,98 | 4,98 | 0 | 0,98 | 0 | 5,68 | 0 |
| ESRRB | GCAAAGTCATG | 1 | 512 | 523 | -488 | -477 | 3,5 | 0,001 | 15,44 | >0.1 | 3,01 | 4,99 | 0 | 0,98 | 0 | 2,95 | 0 |
| ESRRB | GGAAGGGAAAA | 1 | 528 | 539 | -472 | -461 | -6,25 | 0,01 | 6,39 | >0.1 | 3,08 | 5,03 | 0 | 0,98 | 0 | 3,55 | 0 |
| ESRRB | TCTAGGAAAAT | 1 | 719 | 730 | -281 | -270 | -9,4 | 0,01 | 7,98 | >0.1 | 5,6 | 5,6 | 0 | 1,2 | 0 | 4,99 | 0 |
| ESRRB | CCAAGTTCAAA | 1 | 954 | 965 | -46 | -35 | 6,61 | 0,001 | 23,28 | >0.1 | 5,6 | 7,51 | 0 | 1,88 | 0 | 1,7 | 0 |
| ESRRB | TCTAGGGAAAG | 1 | 1002 | 1013 | 2 | 13 | -9,51 | 0,01 | 8,54 | >0.1 | 5,6 | 9,12 | 0 | 1,87 | 0 | 1,46 | 0 |
| ESRRB | GGAAAGTCATT | 1 | 1007 | 1018 | 7 | 18 | 1,73 | 0,001 | 19,78 | >0.1 | 5,6 | 9,12 | 0 | 1,87 | 0 | 1,46 | 0 |
| ESRRB | TCAAGCTCTTC | 1 | 1052 | 1063 | 52 | 63 | 0,13 | 0,001 | 17,38 | >0.1 | 5,6 | 7,53 | 0 | 1,86 | 0 | 2,27 | 0 |
| ESRRB | ACAAGTTTAAC | 1 | 1137 | 1148 | 137 | 148 | -3,59 | 0,01 | 16,54 | >0.1 | 5,6 | 6,4 | 0 | 1,84 | 3,22 | 3,07 | 0 |
| ESRRB | CAAAGTTCACT | 1 | 1172 | 1183 | 172 | 183 | -2,78 | 0,01 | 15,37 | >0.1 | 5,6 | 6,14 | 0 | 1,65 | 1,79 | 2,97 | 0 |
| ESRRB | ACAAGGCTGAG | -1 | 1073 | 1084 | 73 | 84 | -6,05 | 0,01 | 12,59 | >0.1 | 5,6 | 7,12 | 0 | 1,85 | 1,79 | 2,27 | 0 |
| ESRRB | CCAAGATCACA | -1 | 1025 | 1036 | 25 | 36 | 4,24 | 0,001 | 21,46 | >0.1 | 5,6 | 8,52 | 0 | 1,86 | 0 | 1,25 | 0 |
| ESRRB | AGAAGAACACA | -1 | 904 | 915 | -96 | -85 | -8,96 | 0,01 | 6,49 | >0.1 | 5,6 | 6,76 | 0 | 1,89 | 0 | 1,21 | 0 |
| ESRRB | TAAAGTTCATC | -1 | 870 | 881 | -130 | -119 | 0,27 | 0,001 | 17,08 | >0.1 | 5,6 | 6,45 | 0 | 1,9 | 0 | 2,86 | 0 |
| ESRRB | TTATGGCCAAA | -1 | 754 | 765 | -246 | -235 | -10,37 | 0,01 | 8,6 | >0.1 | 5,6 | 5,75 | 0 | 1,3 | 0 | 6,32 | 0 |
| ESRRB | TGAAGATAATG | -1 | 639 | 650 | -361 | -350 | -6,78 | 0,01 | 5,05 | >0.1 | 3,79 | 5,32 | 0 | 1,06 | 0 | 1,66 | 0 |
| ESRRB | ACAATTTCATT | -1 | 606 | 617 | -394 | -383 | -6,47 | 0,01 | 4,74 | >0.1 | 3,52 | 5,23 | 0 | 1,02 | 0 | 1,44 | 0 |
| ESRRB | AATAGGTTATC | -1 | 595 | 606 | -405 | -394 | -10,19 | 0,01 | -0,54 | >0.1 | 3,44 | 5,2 | 0 | 1,01 | 0 | 0 | 0 |
| ESRRB | TCAAGATCATA | -1 | 567 | 578 | -433 | -422 | 7,5 | 0,001 | 16,9 | >0.1 | 3,27 | 5,12 | 0 | 1 | 0 | 0 | 0 |
| ESRRB | GCAAGCTCTGC | -1 | 387 | 398 | -613 | -602 | -8,04 | 0,01 | 6,48 | >0.1 | 2,59 | 4,75 | 0 | 1,05 | 0 | 6,13 | 0 |
| ESRRB | CCAGGTTCACG | -1 | 373 | 384 | -627 | -616 | -4,32 | 0,01 | 10,56 | >0.1 | 2,55 | 4,72 | 0 | 1,07 | 0 | 6,53 | 0 |
| ESRRB | GGACGGTCTCT | -1 | 253 | 264 | -747 | -736 | -9,31 | 0,01 | 4,39 | >0.1 | 2,83 | 4,54 | 0 | 1,35 | 0 | 4,98 | 0 |
| ESRRB | CCATGGCCATT | -1 | 144 | 155 | -856 | -845 | -3,47 | 0,01 | 7,71 | >0.1 | 3,31 | 4,4 | 0 | 1,9 | 0 | 1,57 | 0 |
| ESRRB | CACAGGTCAAC | -1 | 106 | 117 | -894 | -883 | -2,46 | 0,01 | 8,09 | >0.1 | 3,56 | 4,36 | 0 | 1,89 | 0 | 0,75 | 0 |
| ESRRB | AAAAGCTCATT | -1 | 67 | 78 | -933 | -922 | -2,64 | 0,01 | 7,46 | >0.1 | 3,91 | 4,32 | 0 | 1,88 | 0 | 0 | 0 |
| TBR1 | CAGTGTGAGA | 1 | 50 | 60 | -950 | -940 | 4,38 | 0,01 | 14,68 | >0.1 | 4,13 | 4,3 | 0 | 1,87 | 0 | 0 | 0 |
| TBR1 | GTGTGAGAAA | 1 | 52 | 62 | -948 | -938 | 6,75 | 0,001 | 17,02 | >0.1 | 4,1 | 4,3 | 0 | 1,87 | 0 | 0 | 0 |
| TBR1 | ACCTGTGGAG | 1 | 110 | 120 | -890 | -880 | -0,49 | 0,01 | 10,06 | >0.1 | 3,54 | 4,36 | 0 | 1,89 | 0 | 0,77 | 0 |
| TBR1 | GGGCGTGGTG | 1 | 314 | 324 | -686 | -676 | 0,82 | 0,01 | 15,39 | >0.1 | 2,64 | 4,63 | 0 | 1,17 | 0 | 6,13 | 0 |
| TBR1 | TGGCGTGAAC | 1 | 370 | 380 | -630 | -620 | 7,31 | 0,001 | 21,91 | >0.1 | 2,54 | 4,72 | 0 | 1,07 | 0 | 6,27 | 0 |
| TBR1 | AGCCGAGATC | 1 | 402 | 412 | -598 | -588 | -1,54 | 0,01 | 12,66 | >0.1 | 2,63 | 4,77 | 0 | 1,03 | 0 | 5,76 | 0 |
| TBR1 | GGGCGACAGA | 1 | 432 | 442 | -568 | -558 | -0,99 | 0,01 | 13,24 | >0.1 | 2,72 | 4,83 | 0 | 1,01 | 0 | 5,67 | 0 |
| TBR1 | AAATGTGAGA | 1 | 587 | 597 | -413 | -403 | 1,03 | 0,01 | 10,6 | >0.1 | 3,39 | 5,17 | 0 | 1,01 | 0 | 0 | 0 |
| TBR1 | ATGTGAGATA | 1 | 589 | 599 | -411 | -401 | 7,12 | 0,001 | 16,71 | >0.1 | 3,4 | 5,18 | 0 | 1,01 | 0 | 0 | 0 |
| TBR1 | AAGTGACAGG | 1 | 764 | 774 | -236 | -226 | 1,33 | 0,01 | 20,16 | >0.1 | 5,61 | 5,79 | 0 | 1,34 | 0 | 6,1 | 0 |
| TBR1 | ATCTGTTACA | 1 | 844 | 854 | -156 | -146 | 0,76 | 0,01 | 17,5 | >0.1 | 5,61 | 6,26 | 0 | 1,81 | 0 | 3,07 | 0 |
| TBR1 | GGCTGATAAG | 1 | 965 | 975 | -35 | -25 | -1,23 | 0,01 | 15,71 | >0.1 | 5,61 | 7,76 | 0 | 1,87 | 0 | 1,7 | 0 |
| TBR1 | AAGAGAGAAA | 1 | 972 | 982 | -28 | -18 | 0,46 | 0,01 | 17,62 | >0.1 | 5,61 | 7,98 | 0 | 1,87 | 0 | 1,7 | 0 |
| TBR1 | GGATGTGATC | 1 | 1022 | 1032 | 22 | 32 | 0,88 | 0,01 | 18,31 | >0.1 | 5,61 | 8,68 | 0 | 1,86 | 0 | 1,28 | 0 |
| TBR1 | GGCTGAGAAG | -1 | 1070 | 1080 | 70 | 80 | 1,57 | 0,01 | 20,27 | >0.1 | 5,61 | 7,17 | 0 | 1,85 | 1,79 | 2,27 | 0 |
| TBR1 | AGCTGTTAAA | -1 | 708 | 718 | -292 | -282 | 7,04 | 0,001 | 24,19 | >0.1 | 5,61 | 5,55 | 0 | 1,17 | 0 | 4,81 | 0 |
| TBR1 | AAATGTTATA | -1 | 688 | 698 | -312 | -302 | -0,64 | 0,01 | 13,29 | >0.1 | 4,46 | 5,48 | 0 | 1,13 | 0 | 2,86 | 0 |
| TBR1 | TGGCGCGATC | -1 | 408 | 418 | -592 | -582 | 2,44 | 0,01 | 16,55 | >0.1 | 2,65 | 4,78 | 0 | 1,03 | 0 | 5,65 | 0 |
| TBR1 | AGGTGCCCAC | -1 | 324 | 334 | -676 | -666 | 1,49 | 0,01 | 15,35 | >0.1 | 2,61 | 4,64 | 0 | 1,15 | 0 | 5,46 | 0 |

| | | | | | | | | | | | | | | | | | |
|---|---|---|---|---|---|---|---|---|---|---|---|---|---|---|---|---|---|
| TBR1 | CCGTGTTAGT | -1 | 266 | 276 | -734 | -724 | -0,66 | 0,01 | 12,52 | >0.1 | 2,79 | 4,55 | 0 | 1,31 | 0 | 4,53 | 0 |
| TBR1 | AAGTGCTAGG | -1 | 205 | 215 | -795 | -785 | 3,78 | 0,01 | 17,57 | >0.1 | 3,02 | 4,47 | 0 | 1,6 | 0 | 4,7 | 0 |
| TBR1 | AGGCGTGAGC | -1 | 190 | 200 | -810 | -800 | 8,27 | 0,001 | 21,64 | >0.1 | 3,08 | 4,45 | 0 | 1,7 | 0 | 4,13 | 0 |
| FEV | ATAGGAAATG | 1 | 61 | 71 | -939 | -929 | 5,37 | 0,001 | 21,32 | >0.1 | 3,98 | 4,31 | 0 | 1,88 | 0 | 0 | 5,78 |
| FEV | ATCAGAAATG | 1 | 167 | 177 | -833 | -823 | -0,17 | 0,01 | 17,53 | >0.1 | 3,19 | 4,43 | 0 | 1,9 | 0 | 2,4 | 5,78 |
| FEV | TCAGGAGATA | 1 | 244 | 254 | -756 | -746 | -1,66 | 0,01 | 17,32 | >0.1 | 2,86 | 4,52 | 0 | 1,39 | 0 | 4,42 | 5,78 |
| FEV | ATCAGAAATA | 1 | 482 | 492 | -518 | -508 | -0,75 | 0,01 | 19,63 | >0.1 | 2,89 | 4,93 | 0 | 0,99 | 0 | 5,8 | 5,78 |
| FEV | AAGGGAAAAT | 1 | 530 | 540 | -470 | -460 | -0,57 | 0,01 | 18,01 | >0.1 | 3,09 | 5,03 | 0 | 0,98 | 0 | 3,7 | 5,78 |
| FEV | CTAGGAAAAT | 1 | 720 | 730 | -280 | -270 | -2,91 | 0,01 | 20,27 | >0.1 | 5,61 | 5,6 | 0 | 1,2 | 0 | 4,99 | 5,78 |
| FEV | ACCGGTTTTG | 1 | 743 | 753 | -257 | -247 | -0,71 | 0,01 | 23,55 | >0.1 | 5,61 | 5,69 | 0 | 1,26 | 0 | 5,92 | 5,78 |
| FEV | ACAGGAGAGG | 1 | 769 | 779 | -231 | -221 | -1,97 | 0,01 | 22,28 | >0.1 | 5,61 | 5,81 | 0 | 1,35 | 0 | 5,69 | 5,78 |
| FEV | CCAGGATGAA | 1 | 866 | 876 | -134 | -124 | 2,9 | 0,01 | 25,35 | >0.1 | 5,61 | 6,41 | 0 | 1,9 | 0 | 2,75 | 5,78 |
| FEV | TAGGGAAAGT | 1 | 1004 | 1014 | 4 | 14 | -1,74 | 0,01 | 22,05 | >0.1 | 5,61 | 9,12 | 0 | 1,87 | 0 | 1,42 | 5,78 |
| FEV | AGTGGATGTG | 1 | 1019 | 1029 | 19 | 29 | -0,25 | 0,01 | 23,05 | >0.1 | 5,61 | 8,74 | 0 | 1,86 | 0 | 1,31 | 5,78 |
| FEV | TGAGGAACAG | 1 | 1111 | 1121 | 111 | 121 | -1,44 | 0,01 | 24,2 | >0.1 | 5,61 | 6,64 | 0 | 1,85 | 3,22 | 2,54 | 5,78 |
| FEV | CCAAGAAGCA | -1 | 1183 | 1193 | 183 | 193 | -2,44 | 0,01 | 21,9 | >0.1 | 5,61 | 6,07 | 0 | 1,56 | 1,79 | 3,52 | 5,78 |
| FEV | CCAGGAAGAG | -1 | 1057 | 1067 | 57 | 67 | 5,2 | 0,001 | 28,13 | >0.1 | 5,61 | 7,42 | 0 | 1,86 | 0 | 2,27 | 5,78 |
| FEV | GCAGGAAAGA | -1 | 932 | 942 | -68 | -58 | 5,42 | 0,001 | 26,71 | >0.1 | 5,61 | 7,09 | 0 | 1,88 | 0 | 0,93 | 5,78 |
| FEV | ACAGTAAACA | -1 | 897 | 907 | -103 | -93 | -1,43 | 0,01 | 21,39 | >0.1 | 5,61 | 6,68 | 0 | 1,89 | 0 | 2,86 | 5,78 |
| FEV | AGAGGACAGA | -1 | 856 | 866 | -144 | -134 | -1,25 | 0,01 | 21,33 | >0.1 | 5,61 | 6,34 | 0 | 1,9 | 0 | 2,95 | 5,78 |
| FEV | GACAGATATG | -1 | 852 | 862 | -148 | -138 | -1,36 | 0,01 | 20,89 | >0.1 | 5,61 | 6,31 | 0 | 1,88 | 0 | 2,67 | 5,78 |
| FEV | ACCGGTACTT | -1 | 739 | 749 | -261 | -251 | 2,41 | 0,01 | 26,64 | >0.1 | 5,61 | 5,68 | 0 | 1,25 | 0 | 5,92 | 5,78 |
| FEV | CACGGAAAAT | -1 | 694 | 704 | -306 | -296 | 3,41 | 0,01 | 24,51 | >0.1 | 5,61 | 5,5 | 0 | 1,14 | 0 | 3,07 | 5,78 |
| FEV | CCCAGAAACT | -1 | 555 | 565 | -445 | -435 | -0,52 | 0,01 | 14,55 | >0.1 | 3,21 | 5,09 | 0 | 0,99 | 0 | 0 | 5,78 |
| FEV | TCAGGACGGT | -1 | 257 | 267 | -743 | -733 | 1,05 | 0,01 | 20,16 | >0.1 | 2,82 | 4,54 | 0 | 1,34 | 0 | 4,63 | 5,78 |
| MLXIP | TGACCTGT | 1 | 108 | 116 | -892 | -884 | -11,32 | 0,01 | -0,75 | >0.1 | 3,56 | 4,36 | 0 | 1,89 | 0 | 0,77 | 0 |
| MLXIP | TCACGCCT | 1 | 192 | 200 | -808 | -800 | -10,82 | 0,01 | 2,53 | >0.1 | 3,08 | 4,45 | 0 | 1,69 | 0 | 4,13 | 0 |
| MLXIP | GCACTTTG | 1 | 209 | 217 | -791 | -783 | -11,63 | 0,01 | 2,12 | >0.1 | 3 | 4,47 | 0 | 1,57 | 0 | 4,7 | 0 |
| MLXIP | CCCCGTCT | 1 | 281 | 289 | -719 | -711 | -10,32 | 0,01 | 3,34 | >0.1 | 2,74 | 4,57 | 0 | 1,26 | 0 | 5,09 | 0 |
| MLXIP | GGGCGTGG | 1 | 314 | 322 | -686 | -678 | -10,14 | 0,01 | 4,43 | >0.1 | 2,64 | 4,62 | 0 | 1,17 | 0 | 6,13 | 0 |
| MLXIP | GCACCTGT | 1 | 328 | 336 | -672 | -664 | 2,15 | 0,001 | 16,1 | >0.1 | 2,6 | 4,64 | 0 | 1,14 | 0 | 5,56 | 0 |
| MLXIP | GCAGGAGA | 1 | 361 | 369 | -639 | -631 | -11,55 | 0,01 | 3,11 | >0.1 | 2,52 | 4,7 | 0 | 1,08 | 0 | 6,36 | 0 |
| MLXIP | GAACCTGG | 1 | 376 | 384 | -624 | -616 | -11,14 | 0,01 | 3,6 | >0.1 | 2,56 | 4,72 | 0 | 1,06 | 0 | 6,4 | 0 |
| MLXIP | TTCCGTGT | 1 | 697 | 705 | -303 | -295 | -11,32 | 0,01 | 4,03 | >0.1 | 5,64 | 5,5 | 0 | 1,14 | 0 | 3,07 | 0 |
| MLXIP | TCACAGGG | 1 | 1037 | 1045 | 37 | 45 | -11,82 | 0,1 | 4,97 | >0.1 | 5,64 | 7,95 | 0 | 1,86 | 0 | 1,34 | 0 |
| MLXIP | CCACGAAG | 1 | 1147 | 1155 | 147 | 155 | -11,82 | 0,1 | 5,93 | >0.1 | 5,64 | 6,32 | 0 | 1,84 | 1,79 | 2,16 | 0 |
| MLXIP | TCACTTGC | 1 | 1178 | 1186 | 178 | 186 | -2,01 | 0,01 | 16,39 | >0.1 | 5,64 | 6,1 | 0 | 1,6 | 1,79 | 3,26 | 0 |
| MLXIP | GCAAGTGA | -1 | 1178 | 1186 | 178 | 186 | 0,24 | 0,01 | 18,64 | >0.1 | 5,64 | 6,1 | 0 | 1,6 | 1,79 | 3,26 | 0 |
| MLXIP | CTTCGTGG | -1 | 1147 | 1155 | 147 | 155 | -11,82 | 0,1 | 5,93 | >0.1 | 5,64 | 6,32 | 0 | 1,84 | 1,79 | 2,16 | 0 |
| MLXIP | TCCCCTGT | -1 | 1039 | 1047 | 39 | 47 | -10,82 | 0,01 | 5,9 | >0.1 | 5,64 | 7,88 | 0 | 1,86 | 0 | 1,34 | 0 |
| MLXIP | GAACTTGG | -1 | 954 | 962 | -46 | -38 | -11,63 | 0,01 | 5,02 | >0.1 | 5,64 | 7,44 | 0 | 1,88 | 0 | 1,7 | 0 |
| MLXIP | TCTCCTGT | -1 | 769 | 777 | -231 | -223 | -11,32 | 0,01 | 7,38 | >0.1 | 5,64 | 5,8 | 0 | 1,35 | 0 | 5,92 | 0 |
| MLXIP | GCCCTTGT | -1 | 506 | 514 | -494 | -486 | -10,14 | 0,01 | 4,47 | >0.1 | 2,98 | 4,97 | 0 | 0,98 | 0 | 5,68 | 0 |
| MLXIP | TCCCGAGT | -1 | 346 | 354 | -654 | -646 | -10,82 | 0,01 | 4,04 | >0.1 | 2,55 | 4,67 | 0 | 1,11 | 0 | 6,52 | 0 |
| MLXIP | ACAGGTGC | -1 | 328 | 336 | -672 | -664 | -3,82 | 0,01 | 10,13 | >0.1 | 2,6 | 4,64 | 0 | 1,14 | 0 | 5,56 | 0 |
| MLXIP | CACCGTGT | -1 | 270 | 278 | -730 | -722 | -10,82 | 0,01 | 2,89 | >0.1 | 2,78 | 4,56 | 0 | 1,29 | 0 | 5,09 | 0 |
| MLXIP | CCACAGGT | -1 | 110 | 118 | -890 | -882 | -10,82 | 0,01 | -0,29 | >0.1 | 3,54 | 4,36 | 0 | 1,89 | 0 | 0,75 | 0 |
| JUN::JUNB(VAR.2) | AAAGAAGTCAGC | 1 | 30 | 42 | -970 | -958 | 2,12 | 0,01 | 12,73 | >0.1 | 4,46 | 4,28 | 0 | 1,87 | 0 | 0 | 0 |
| JUN::JUNB(VAR.2) | AGAGTAGTCATA | 1 | 123 | 135 | -877 | -865 | 1,58 | 0,01 | 12,98 | >0.1 | 3,44 | 4,38 | 0 | 1,89 | 0 | 1,7 | 0 |
| JUN::JUNB(VAR.2) | CACAAGGTCAGG | 1 | 237 | 249 | -763 | -751 | 1,22 | 0,01 | 14,57 | >0.1 | 2,89 | 4,52 | 0 | 1,42 | 0 | 4,53 | 0 |
| JUN::JUNB(VAR.2) | AATGGCGTGAAC | 1 | 368 | 380 | -632 | -620 | 1,57 | 0,01 | 16,05 | >0.1 | 2,54 | 4,72 | 0 | 1,08 | 0 | 6,15 | 0 |
| JUN::JUNB(VAR.2) | GATCGCGCCACT | 1 | 408 | 420 | -592 | -580 | 2,2 | 0,01 | 16,19 | >0.1 | 2,65 | 4,79 | 0 | 1,03 | 0 | 5,52 | 0 |
| JUN::JUNB(VAR.2) | GGCAAAGTCATG | 1 | 511 | 523 | -489 | -477 | 3,96 | 0,01 | 15,9 | >0.1 | 3,01 | 4,99 | 0 | 0,98 | 0 | 2,95 | 0 |
| JUN::JUNB(VAR.2) | TGTGAGATAACC | 1 | 590 | 602 | -410 | -398 | 3,28 | 0,01 | 12,89 | >0.1 | 3,41 | 5,19 | 0 | 1,01 | 0 | 0 | 0 |
| JUN::JUNB(VAR.2) | TCTTTGGTCACT | 1 | 669 | 681 | -331 | -319 | 4,12 | 0,01 | 18,48 | >0.1 | 4,14 | 5,42 | 0 | 1,1 | 0 | 3,7 | 0 |
| JUN::JUNB(VAR.2) | GATTTGGCCATA | 1 | 752 | 764 | -248 | -236 | 2,63 | 0,01 | 21,58 | >0.1 | 5,58 | 5,74 | 0 | 1,3 | 0 | 6,32 | 0 |
| JUN::JUNB(VAR.2) | GGAGAGGTAAGG | 1 | 772 | 784 | -228 | -216 | 4,05 | 0,01 | 22,94 | >0.1 | 5,58 | 5,83 | 0 | 1,37 | 0 | 6,1 | 0 |
| JUN::JUNB(VAR.2) | TCTGTCCTCTCC | 1 | 856 | 868 | -144 | -132 | 1,14 | 0,01 | 17,84 | >0.1 | 5,58 | 6,35 | 0 | 1,9 | 0 | 2,86 | 0 |
| JUN::JUNB(VAR.2) | GGGAAAGTCATT | 1 | 1006 | 1018 | 6 | 18 | 2,81 | 0,01 | 20,84 | >0.1 | 5,58 | 9,12 | 0 | 1,87 | 0 | 1,46 | 0 |

| | | | | | | | | | | | | | | | | |
|---|---|---|---|---|---|---|---|---|---|---|---|---|---|---|---|---|
| JUN::JUNB(VAR.2) | GACGATGTCAAG | 1 | 1045 | 1057 | 45 | 57 | 4,5 | 0,01 | 21,11 | >0.1 | 5,58 | 7,7 | 0 | 1,86 | 0 | 1,46 | 0 |
| JUN::JUNB(VAR.2) | CTTGACATCGTC | -1 | 1045 | 1057 | 45 | 57 | 3,16 | 0,01 | 19,77 | >0.1 | 5,58 | 7,7 | 0 | 1,86 | 0 | 1,46 | 0 |
| JUN::JUNB(VAR.2) | GGAGAGGACAGA | -1 | 856 | 868 | -144 | -132 | 2,07 | 0,01 | 18,77 | >0.1 | 5,58 | 6,35 | 0 | 1,9 | 0 | 2,86 | 0 |
| JUN::JUNB(VAR.2) | GAATACGTTATT | -1 | 800 | 812 | -200 | -188 | 5,59 | 0,01 | 22,09 | >0.1 | 5,58 | 5,98 | 0 | 1,51 | 0 | 3,43 | 0 |
| JUN::JUNB(VAR.2) | TGTCACTTTATG | -1 | 760 | 772 | -240 | -228 | 3,28 | 0,01 | 22,07 | >0.1 | 5,58 | 5,78 | 0 | 1,33 | 0 | 6,1 | 0 |
| JUN::JUNB(VAR.2) | TAATAGGTTATC | -1 | 595 | 607 | -405 | -393 | 1,62 | 0,01 | 12,72 | >0.1 | 3,45 | 5,2 | 0 | 1,02 | 0 | 1,44 | 0 |
| JUN::JUNB(VAR.2) | GGTTATCTCACA | -1 | 590 | 602 | -410 | -398 | 4,1 | 0,01 | 13,71 | >0.1 | 3,41 | 5,19 | 0 | 1,01 | 0 | 0 | 0 |
| JUN::JUNB(VAR.2) | AGTGGCGCGATC | -1 | 408 | 420 | -592 | -580 | 2,3 | 0,01 | 16,29 | >0.1 | 2,65 | 4,79 | 0 | 1,03 | 0 | 5,52 | 0 |
| JUN::JUNB(VAR.2) | GTTCACGCCATT | -1 | 368 | 380 | -632 | -620 | 3,72 | 0,01 | 18,2 | >0.1 | 2,54 | 4,72 | 0 | 1,08 | 0 | 6,15 | 0 |
| JUN::JUNB(VAR.2) | TGGGACTACAGG | -1 | 331 | 343 | -669 | -657 | 1,31 | 0,01 | 15,61 | >0.1 | 2,59 | 4,66 | 0 | 1,14 | 0 | 5,92 | 0 |
| JUN::JUNB(VAR.2) | GCTGACTTCTTT | -1 | 30 | 42 | -970 | -958 | 5,26 | 0,01 | 15,87 | >0.1 | 4,46 | 4,28 | 0 | 1,87 | 0 | 0 | 0 |
| MIXL1 | GGAAATGAGC | 1 | 64 | 74 | -936 | -926 | 4,23 | 0,01 | 14,37 | >0.1 | 3,95 | 4,31 | 0 | 1,88 | 0 | 0 | 0 |
| MIXL1 | CATAATTTTA | 1 | 131 | 141 | -869 | -859 | 3,52 | 0,01 | 14,77 | >0.1 | 3,4 | 4,38 | 0 | 1,89 | 0 | 1,57 | 0 |
| MIXL1 | GGAAATTAAA | 1 | 153 | 163 | -847 | -837 | 6,58 | 0,01 | 17,8 | >0.1 | 3,27 | 4,41 | 0 | 1,9 | 0 | 1,65 | 0 |
| MIXL1 | ACTGATCAGA | 1 | 163 | 173 | -837 | -827 | 5,15 | 0,01 | 17,09 | >0.1 | 3,21 | 4,42 | 0 | 1,9 | 0 | 2,4 | 0 |
| MIXL1 | CTCTATTAAA | 1 | 287 | 297 | -713 | -703 | 4,54 | 0,01 | 18,28 | >0.1 | 2,72 | 4,59 | 0 | 1,24 | 0 | 5,19 | 0 |
| MIXL1 | ACTGATCAGA | 1 | 478 | 488 | -522 | -512 | 5,15 | 0,01 | 19,29 | >0.1 | 2,88 | 4,92 | 0 | 0,99 | 0 | 5,36 | 0 |
| MIXL1 | ACCTATTAAT | 1 | 599 | 609 | -401 | -391 | 3,9 | 0,01 | 14,95 | >0.1 | 3,47 | 5,21 | 0 | 1,02 | 0 | 1,36 | 0 |
| MIXL1 | ATTAATGAAA | 1 | 603 | 613 | -397 | -387 | 5,58 | 0,01 | 16,72 | >0.1 | 3,49 | 5,22 | 0 | 1,02 | 0 | 1,41 | 0 |
| MIXL1 | TAAAATTATA | 1 | 682 | 692 | -318 | -308 | 4,02 | 0,01 | 18 | >0.1 | 4,34 | 5,46 | 0 | 1,12 | 0 | 3,07 | 0 |
| MIXL1 | TTTGATTTGG | 1 | 749 | 759 | -251 | -241 | 3,78 | 0,01 | 22,31 | >0.1 | 5,61 | 5,72 | 0 | 1,28 | 0 | 5,92 | 0 |
| MIXL1 | TTTGATTCAC | 1 | 815 | 825 | -185 | -175 | 3,83 | 0,01 | 20,51 | >0.1 | 5,61 | 6,06 | 0 | 1,58 | 0 | 3,43 | 0 |
| MIXL1 | GCTGATAAGA | 1 | 966 | 976 | -34 | -24 | 3,49 | 0,01 | 20,45 | >0.1 | 5,61 | 7,78 | 0 | 1,87 | 0 | 1,7 | 0 |
| MIXL1 | TATAATTCCA | -1 | 1190 | 1200 | 190 | 200 | 4,05 | 0,01 | 22,81 | >0.1 | 5,61 | 6,03 | 0 | 1,51 | 1,79 | 3,82 | 0 |
| MIXL1 | CCTCATGAGA | -1 | 983 | 993 | -17 | -7 | 3,47 | 0,01 | 21,02 | >0.1 | 5,61 | 8,5 | 0 | 1,87 | 0 | 1,57 | 0 |
| MIXL1 | TCTTATCAGC | -1 | 966 | 976 | -34 | -24 | 3,56 | 0,01 | 20,52 | >0.1 | 5,61 | 7,78 | 0 | 1,87 | 0 | 1,7 | 0 |
| MIXL1 | GTGAATCAAA | -1 | 815 | 825 | -185 | -175 | 3,99 | 0,01 | 20,67 | >0.1 | 5,61 | 6,06 | 0 | 1,58 | 0 | 3,43 | 0 |
| MIXL1 | CCAAATCAAA | -1 | 749 | 759 | -251 | -241 | 5 | 0,01 | 23,53 | >0.1 | 5,61 | 5,72 | 0 | 1,28 | 0 | 5,92 | 0 |
| MIXL1 | GACAATTTCA | -1 | 608 | 618 | -392 | -382 | 4,56 | 0,01 | 15,89 | >0.1 | 3,53 | 5,23 | 0 | 1,02 | 0 | 1,55 | 0 |
| MIXL1 | TTTCATTAAT | -1 | 603 | 613 | -397 | -387 | 4,78 | 0,01 | 15,92 | >0.1 | 3,49 | 5,22 | 0 | 1,02 | 0 | 1,41 | 0 |
| MIXL1 | TCTGATCAGT | -1 | 478 | 488 | -522 | -512 | 4,44 | 0,01 | 18,58 | >0.1 | 2,88 | 4,92 | 0 | 0,99 | 0 | 5,36 | 0 |
| MIXL1 | TCTGATCAGT | -1 | 163 | 173 | -837 | -827 | 4,44 | 0,01 | 16,38 | >0.1 | 3,21 | 4,42 | 0 | 1,9 | 0 | 2,4 | 0 |
| MIXL1 | TTTAATTTCC | -1 | 153 | 163 | -847 | -837 | 5,83 | 0,01 | 17,05 | >0.1 | 3,27 | 4,41 | 0 | 1,9 | 0 | 1,65 | 0 |
| MIXL1 | TAAAATTATG | -1 | 131 | 141 | -869 | -859 | 4,13 | 0,01 | 15,38 | >0.1 | 3,4 | 4,38 | 0 | 1,89 | 0 | 1,57 | 0 |
| MIXL1 | TTCACTTACC | -1 | 4 | 14 | -996 | -986 | 4,3 | 0,01 | 17,82 | >0.1 | 5,61 | 4,26 | 0 | 1,86 | 1,79 | 0 | 0 |
| HEY2 | GGTAAGTGAA | 1 | 4 | 14 | -996 | -986 | -3,42 | 0,01 | 10,1 | >0.1 | 5,61 | 4,26 | 0 | 1,86 | 1,79 | 0 | 0 |
| HEY2 | TTGACCTGTG | 1 | 107 | 117 | -893 | -883 | -0,38 | 0,01 | 10,19 | >0.1 | 3,56 | 4,36 | 0 | 1,89 | 0 | 0,77 | 0 |
| HEY2 | CTCACGCCTG | 1 | 191 | 201 | -809 | -799 | 0,04 | 0,01 | 13,4 | >0.1 | 3,08 | 4,45 | 0 | 1,7 | 0 | 4,13 | 0 |
| HEY2 | CACGCCTGTA | 1 | 193 | 203 | -807 | -797 | 1,83 | 0,01 | 15,3 | >0.1 | 3,07 | 4,46 | 0 | 1,68 | 0 | 4,26 | 0 |
| HEY2 | TGGGCGTGGT | 1 | 313 | 323 | -687 | -677 | -0,17 | 0,01 | 14,4 | >0.1 | 2,64 | 4,62 | 0 | 1,17 | 0 | 6,13 | 0 |
| HEY2 | GGCACCTGTA | 1 | 327 | 337 | -673 | -663 | 9,76 | 0,001 | 23,72 | >0.1 | 2,6 | 4,65 | 0 | 1,14 | 0 | 5,56 | 0 |
| HEY2 | TGAACCTGGG | 1 | 375 | 385 | -625 | -615 | -0,95 | 0,01 | 13,79 | >0.1 | 2,56 | 4,72 | 0 | 1,07 | 0 | 6,4 | 0 |
| HEY2 | GACAAGTTTA | 1 | 1136 | 1146 | 136 | 146 | -0,89 | 0,01 | 19,28 | >0.1 | 5,61 | 6,41 | 0 | 1,84 | 3,22 | 3,09 | 0 |
| HEY2 | AAGACCTGTT | 1 | 1158 | 1168 | 158 | 168 | 0,22 | 0,01 | 16,81 | >0.1 | 4,38 | 6,24 | 0 | 1,78 | 1,79 | 2,4 | 0 |
| HEY2 | AGCAAGTGAA | -1 | 1177 | 1187 | 177 | 187 | -2,54 | 0,01 | 15,84 | >0.1 | 5,61 | 6,11 | 0 | 1,61 | 1,79 | 3,26 | 0 |
| HEY2 | ATCACATCCA | -1 | 1021 | 1031 | 21 | 31 | -3,43 | 0,01 | 14,05 | >0.1 | 5,61 | 8,7 | 0 | 1,86 | 0 | 1,31 | 0 |
| HEY2 | GATACACGGA | -1 | 698 | 708 | -302 | -292 | -3,49 | 0,01 | 11,97 | >0.1 | 5,61 | 5,52 | 0 | 1,15 | 0 | 3,19 | 0 |
| HEY2 | GTGGCGCGAT | -1 | 409 | 419 | -591 | -581 | -2,75 | 0,01 | 11,24 | >0.1 | 2,65 | 4,78 | 0 | 1,03 | 0 | 5,52 | 0 |
| HEY2 | GGCGCGATCT | -1 | 407 | 417 | -593 | -583 | -0,09 | 0,01 | 14,01 | >0.1 | 2,65 | 4,78 | 0 | 1,03 | 0 | 5,65 | 0 |
| HEY2 | TACAGGTGCC | -1 | 327 | 337 | -673 | -663 | -0,74 | 0,01 | 13,22 | >0.1 | 2,6 | 4,65 | 0 | 1,14 | 0 | 5,56 | 0 |
| HEY2 | ACCACGCCCA | -1 | 313 | 323 | -687 | -677 | 0,65 | 0,01 | 15,22 | >0.1 | 2,64 | 4,62 | 0 | 1,17 | 0 | 6,13 | 0 |
| HEY2 | CCAAAGTGCT | -1 | 208 | 218 | -792 | -782 | -3,32 | 0,01 | 10,44 | >0.1 | 3 | 4,48 | 0 | 1,58 | 0 | 4,7 | 0 |
| HEY2 | CAGGCGTGAG | -1 | 191 | 201 | -809 | -799 | -0,95 | 0,01 | 12,41 | >0.1 | 3,08 | 4,45 | 0 | 1,7 | 0 | 4,13 | 0 |
| PROX1 | AAAGAAGTCAGC | 1 | 30 | 42 | -970 | -958 | -7,66 | 0,01 | 2,95 | >0.1 | 4,46 | 4,28 | 0 | 1,87 | 0 | 0 | 0 |
| PROX1 | TGAAACCCCGTC | 1 | 276 | 288 | -724 | -712 | -7,3 | 0,01 | 6,49 | >0.1 | 2,75 | 4,57 | 0 | 1,27 | 0 | 5,19 | 0 |
| PROX1 | CGAGACTCAGTC | 1 | 443 | 455 | -557 | -545 | -4,13 | 0,01 | 10,41 | >0.1 | 2,76 | 4,85 | 0 | 1 | 0 | 5,92 | 0 |
| PROX1 | CATGATCCCTAG | 1 | 648 | 660 | -352 | -340 | -8,82 | 0,01 | 5,32 | >0.1 | 3,88 | 5,35 | 0 | 1,07 | 0 | 3,83 | 0 |
| PROX1 | CTAGACCTCTTT | 1 | 662 | 674 | -338 | -326 | -1,39 | 0,01 | 13,14 | >0.1 | 4,05 | 5,4 | 0 | 1,09 | 0 | 3,99 | 0 |
| PROX1 | TATAACATTTTC | 1 | 688 | 700 | -312 | -300 | -7,67 | 0,01 | 6,39 | >0.1 | 4,48 | 5,49 | 0 | 1,14 | 0 | 2,95 | 0 |

| | | | | | | | | | | | | | | | | | |
|---|---|---|---|---|---|---|---|---|---|---|---|---|---|---|---|---|---|
| PROX1 | AATAACGTATTC | 1 | 800 | 812 | -200 | -188 | -4,61 | 0,01 | 11,89 | >0.1 | 5,58 | 5,98 | 0 | 1,51 | 0 | 3,43 | 0 |
| PROX1 | AACGTATTCTTA | 1 | 803 | 815 | -197 | -185 | -8,58 | 0,01 | 7,95 | >0.1 | 5,58 | 6 | 0 | 1,52 | 0 | 3,43 | 0 |
| PROX1 | GAGGAGGTTTTA | 1 | 989 | 1001 | -11 | 1 | -8,94 | 0,01 | 8,86 | >0.1 | 5,58 | 8,74 | 0 | 1,87 | 0 | 1,61 | 0 |
| PROX1 | CCTGGCTCCTTC | 1 | 1062 | 1074 | 62 | 74 | 0,35 | 0,001 | 19,17 | >0.1 | 5,58 | 7,31 | 0 | 1,86 | 1,79 | 2,27 | 0 |
| PROX1 | TCAGCCTTGTTG | 1 | 1074 | 1086 | 74 | 86 | -8,01 | 0,01 | 10,6 | >0.1 | 5,58 | 7,11 | 0 | 1,85 | 1,79 | 2,27 | 0 |
| PROX1 | CAAGACATTTTT | 1 | 1123 | 1135 | 123 | 135 | -0,94 | 0,01 | 19,03 | >0.1 | 5,58 | 6,52 | 0 | 1,85 | 3,22 | 2,79 | 0 |
| PROX1 | CACGAAGCCGAA | 1 | 1148 | 1160 | 148 | 160 | -1,73 | 0,01 | 16,04 | >0.1 | 5,58 | 6,31 | 0 | 1,84 | 1,79 | 2,24 | 0 |
| PROX1 | GAAGACCTGTTC | 1 | 1157 | 1169 | 157 | 169 | -1,39 | 0,01 | 15,21 | >0.1 | 4,38 | 6,24 | 0 | 1,79 | 1,79 | 2,4 | 0 |
| PROX1 | TTCGGCTTCGTG | -1 | 1148 | 1160 | 148 | 160 | -4,58 | 0,01 | 13,19 | >0.1 | 5,58 | 6,31 | 0 | 1,84 | 1,79 | 2,24 | 0 |
| PROX1 | AAAAATGTCTTG | -1 | 1123 | 1135 | 123 | 135 | 2,03 | 0,001 | 22 | >0.1 | 5,58 | 6,52 | 0 | 1,85 | 3,22 | 2,79 | 0 |
| PROX1 | CTTGACATCGTC | -1 | 1045 | 1057 | 45 | 57 | -2,93 | 0,01 | 13,68 | >0.1 | 5,58 | 7,7 | 0 | 1,86 | 0 | 1,46 | 0 |
| PROX1 | TAAAACCTCCTC | -1 | 989 | 1001 | -11 | 1 | -8,47 | 0,01 | 9,33 | >0.1 | 5,58 | 8,74 | 0 | 1,87 | 0 | 1,61 | 0 |
| PROX1 | AAAACCTCCTCA | -1 | 988 | 1000 | -12 | 0 | -3,71 | 0,01 | 14,07 | >0.1 | 5,58 | 8,71 | 0 | 1,87 | 0 | 1,61 | 0 |
| PROX1 | TAAGAATACGTT | -1 | 803 | 815 | -197 | -185 | -8,19 | 0,01 | 8,34 | >0.1 | 5,58 | 6 | 0 | 1,52 | 0 | 3,43 | 0 |
| PROX1 | AAAGAGGTCTAG | -1 | 662 | 674 | -338 | -326 | -6,35 | 0,01 | 8,18 | >0.1 | 4,05 | 5,4 | 0 | 1,09 | 0 | 3,99 | 0 |
| PROX1 | CATGACTTTGCC | -1 | 511 | 523 | -489 | -477 | -4,27 | 0,01 | 7,67 | >0.1 | 3,01 | 4,99 | 0 | 0,98 | 0 | 2,95 | 0 |
| PROX1 | TATGACTACTCT | -1 | 123 | 135 | -877 | -865 | -3,53 | 0,01 | 7,87 | >0.1 | 3,44 | 4,38 | 0 | 1,89 | 0 | 1,7 | 0 |
| PROX1 | GCTGACTTCTTT | -1 | 30 | 42 | -970 | -958 | -0,2 | 0,001 | 10,41 | >0.1 | 4,46 | 4,28 | 0 | 1,87 | 0 | 0 | 0 |
| PROX1 | AGTGCCTCCTTC | -1 | 17 | 29 | -983 | -971 | -2,79 | 0,01 | 8,93 | >0.1 | 5,58 | 4,27 | 0 | 1,87 | 0 | 0 | 0 |
| HEY1 | GGCTCACGCC | 1 | 189 | 199 | -811 | -801 | -6,65 | 0,01 | 6,64 | >0.1 | 3,09 | 4,45 | 0 | 1,71 | 0 | 4,04 | 0 |
| HEY1 | CTCACGCCTG | 1 | 191 | 201 | -809 | -799 | -1,51 | 0,01 | 11,85 | >0.1 | 3,08 | 4,45 | 0 | 1,7 | 0 | 4,13 | 0 |
| HEY1 | CACGCCTGTA | 1 | 193 | 203 | -807 | -797 | -0,04 | 0,01 | 13,43 | >0.1 | 3,07 | 4,46 | 0 | 1,68 | 0 | 4,26 | 0 |
| HEY1 | AACACGGTGA | 1 | 269 | 279 | -731 | -721 | -5,82 | 0,01 | 7,9 | >0.1 | 2,78 | 4,56 | 0 | 1,3 | 0 | 5,09 | 0 |
| HEY1 | ACCCCGTCTC | 1 | 280 | 290 | -720 | -710 | -1,17 | 0,01 | 12,5 | >0.1 | 2,74 | 4,57 | 0 | 1,26 | 0 | 5,09 | 0 |
| HEY1 | TGGGCGTGGT | 1 | 313 | 323 | -687 | -677 | -5,12 | 0,01 | 9,45 | >0.1 | 2,64 | 4,62 | 0 | 1,17 | 0 | 6,13 | 0 |
| HEY1 | GGCACCTGTA | 1 | 327 | 337 | -673 | -663 | 5,76 | 0,001 | 19,72 | >0.1 | 2,6 | 4,65 | 0 | 1,14 | 0 | 5,56 | 0 |
| HEY1 | TACTCGGGAG | 1 | 345 | 355 | -655 | -645 | -6,25 | 0,01 | 8,61 | >0.1 | 2,55 | 4,67 | 0 | 1,11 | 0 | 6,52 | 0 |
| HEY1 | AGATCGCGCC | 1 | 407 | 417 | -593 | -583 | -6,59 | 0,01 | 7,51 | >0.1 | 2,65 | 4,78 | 0 | 1,03 | 0 | 5,65 | 0 |
| HEY1 | ATCGCGCCAC | 1 | 409 | 419 | -591 | -581 | -5,9 | 0,01 | 8,09 | >0.1 | 2,65 | 4,78 | 0 | 1,03 | 0 | 5,52 | 0 |
| HEY1 | TTCACTTGCT | 1 | 1177 | 1187 | 177 | 187 | -7,08 | 0,01 | 11,3 | >0.1 | 5,61 | 6,11 | 0 | 1,61 | 1,79 | 3,26 | 0 |
| HEY1 | AGCAAGTGAA | -1 | 1177 | 1187 | 177 | 187 | -5,15 | 0,01 | 13,23 | >0.1 | 5,61 | 6,11 | 0 | 1,61 | 1,79 | 3,26 | 0 |
| HEY1 | GCTTCGTGGT | -1 | 1146 | 1156 | 146 | 156 | -4,43 | 0,01 | 13,3 | >0.1 | 5,61 | 6,33 | 0 | 1,84 | 1,79 | 2,16 | 0 |
| HEY1 | GTCCCCTGTG | -1 | 1038 | 1048 | 38 | 48 | -4,33 | 0,01 | 12,39 | >0.1 | 5,61 | 7,92 | 0 | 1,86 | 0 | 1,34 | 0 |
| HEY1 | AATACGTTAT | -1 | 801 | 811 | -199 | -189 | -7,51 | 0,01 | 9 | >0.1 | 5,61 | 5,98 | 0 | 1,5 | 0 | 3,43 | 0 |
| HEY1 | CTCTCCTGTC | -1 | 768 | 778 | -232 | -222 | -4,02 | 0,01 | 14,66 | >0.1 | 5,61 | 5,81 | 0 | 1,35 | 0 | 5,92 | 0 |
| HEY1 | TACACGGAAA | -1 | 696 | 706 | -304 | -294 | -6,29 | 0,01 | 9,04 | >0.1 | 5,61 | 5,51 | 0 | 1,15 | 0 | 3,07 | 0 |
| HEY1 | TTCACGCCAT | -1 | 369 | 379 | -631 | -621 | -5,22 | 0,01 | 9,26 | >0.1 | 2,54 | 4,71 | 0 | 1,07 | 0 | 6,15 | 0 |
| HEY1 | TTCTCCTGCC | -1 | 360 | 370 | -640 | -630 | -4,11 | 0,01 | 10,56 | >0.1 | 2,52 | 4,7 | 0 | 1,09 | 0 | 6,36 | 0 |
| HEY1 | GCCTCCCGAG | -1 | 347 | 357 | -653 | -643 | -5,95 | 0,01 | 8,91 | >0.1 | 2,55 | 4,68 | 0 | 1,11 | 0 | 6,52 | 0 |
| HEY1 | TACAGGTGCC | -1 | 327 | 337 | -673 | -663 | 0,32 | 0,01 | 14,28 | >0.1 | 2,6 | 4,65 | 0 | 1,14 | 0 | 5,56 | 0 |
| HEY1 | ACCACGCCCA | -1 | 313 | 323 | -687 | -677 | 1,64 | 0,01 | 16,21 | >0.1 | 2,64 | 4,62 | 0 | 1,17 | 0 | 6,13 | 0 |
| HEY1 | CACGCCCAGC | -1 | 311 | 321 | -689 | -679 | -7,55 | 0,01 | 6,93 | >0.1 | 2,65 | 4,62 | 0 | 1,18 | 0 | 6,03 | 0 |
| HEY1 | GAGACGGGGT | -1 | 280 | 290 | -720 | -710 | -6,17 | 0,01 | 7,5 | >0.1 | 2,74 | 4,57 | 0 | 1,26 | 0 | 5,09 | 0 |
| HEY1 | ATCTCCTGAC | -1 | 243 | 253 | -757 | -747 | -6,69 | 0,01 | 6,42 | >0.1 | 2,87 | 4,52 | 0 | 1,39 | 0 | 4,33 | 0 |
| HEY1 | TACAGGCGTG | -1 | 193 | 203 | -807 | -797 | -6,06 | 0,01 | 7,41 | >0.1 | 3,07 | 4,46 | 0 | 1,68 | 0 | 4,26 | 0 |
| HEY1 | CAGGCGTGAG | -1 | 191 | 201 | -809 | -799 | -3,77 | 0,01 | 9,59 | >0.1 | 3,08 | 4,45 | 0 | 1,7 | 0 | 4,13 | 0 |
| MTF1 | GGAGAAGGAGGCAC | 1 | 14 | 28 | -986 | -972 | -9,09 | 0,01 | 2,6 | >0.1 | 5,56 | 4,27 | 0 | 1,87 | 0 | 0 | 0 |
| MTF1 | TTTGCAAAGGCAGA | 1 | 84 | 98 | -916 | -902 | -8,28 | 0,01 | 1,67 | >0.1 | 3,73 | 4,34 | 0 | 1,88 | 0 | 0 | 0 |
| MTF1 | GTGGCTCACGCCTG | 1 | 187 | 201 | -813 | -799 | -4,32 | 0,01 | 8,99 | >0.1 | 3,09 | 4,46 | 0 | 1,73 | 0 | 4,04 | 0 |
| MTF1 | GTTGGGCAGATCAC | 1 | 226 | 240 | -774 | -760 | -7,69 | 0,01 | 6,34 | >0.1 | 2,92 | 4,51 | 0 | 1,48 | 0 | 5,12 | 0 |
| MTF1 | CGAGATCGCGCCAC | 1 | 405 | 419 | -595 | -581 | -9,35 | 0,01 | 4,76 | >0.1 | 2,65 | 4,78 | 0 | 1,03 | 0 | 5,65 | 0 |
| MTF1 | TGTGCTCCAGCCTG | 1 | 419 | 433 | -581 | -567 | -7,58 | 0,01 | 6,36 | >0.1 | 2,69 | 4,81 | 0 | 1,02 | 0 | 5,42 | 0 |
| MTF1 | TCTGAAAACCATAC | 1 | 616 | 630 | -384 | -370 | -2,76 | 0,01 | 8,69 | >0.1 | 3,6 | 5,26 | 0 | 1,04 | 0 | 1,55 | 0 |
| MTF1 | TTTGGTCACTGTAA | 1 | 671 | 685 | -329 | -315 | -2,93 | 0,01 | 11,35 | >0.1 | 4,19 | 5,44 | 0 | 1,11 | 0 | 3,55 | 0 |
| MTF1 | CCTGCTCAGCGCCC | 1 | 937 | 951 | -63 | -49 | -7,78 | 0,01 | 7,74 | >0.1 | 5,56 | 7,23 | 0 | 1,88 | 0 | 0,85 | 0 |
| MTF1 | TCAGCGCCCAACCC | 1 | 942 | 956 | -58 | -44 | -1,03 | 0,01 | 15,53 | >0.1 | 5,56 | 7,32 | 0 | 1,88 | 0 | 1,8 | 0 |
| MTF1 | TGTGATCTTGGCTC | 1 | 1025 | 1039 | 25 | 39 | -5,92 | 0,01 | 11,32 | >0.1 | 5,56 | 8,52 | 0 | 1,86 | 0 | 1,31 | 0 |
| MTF1 | TTGGCTCACAGGGG | 1 | 1032 | 1046 | 32 | 46 | -9,44 | 0,01 | 7,43 | >0.1 | 5,56 | 8,15 | 0 | 1,86 | 0 | 1,31 | 0 |
| MTF1 | GGCTCACAGGGGAC | 1 | 1034 | 1048 | 34 | 48 | -8,96 | 0,01 | 7,86 | >0.1 | 5,56 | 8,06 | 0 | 1,86 | 0 | 1,34 | 0 |

| | | | | | | | | | | | | | | | | | |
|---|---|---|---|---|---|---|---|---|---|---|---|---|---|---|---|---|---|
| MTF1 | GCTGCTCAGTCCAC | 1 | 1094 | 1108 | 94 | 108 | -3,88 | 0,01 | 15,9 | >0.1 | 5,56 | 6,83 | 0 | 1,85 | 3,22 | 2,32 | 0 |
| MTF1 | TTTGGACAAGTTTA | 1 | 1132 | 1146 | 132 | 146 | -6,92 | 0,01 | 13,18 | >0.1 | 5,56 | 6,44 | 0 | 1,84 | 3,22 | 3,03 | 0 |
| MTF1 | TTTAACCACGAAGC | 1 | 1142 | 1156 | 142 | 156 | -6,89 | 0,01 | 12,25 | >0.1 | 5,56 | 6,36 | 0 | 1,84 | 3,22 | 2,16 | 0 |
| MTF1 | TGAGCAGCAGTTAC | -1 | 1088 | 1102 | 88 | 102 | -4,19 | 0,01 | 14,19 | >0.1 | 5,56 | 6,9 | 0 | 1,85 | 1,79 | 2,27 | 0 |
| MTF1 | GAAGAACACAGTAA | -1 | 900 | 914 | -100 | -86 | -7,57 | 0,01 | 7,9 | >0.1 | 5,56 | 6,75 | 0 | 1,89 | 0 | 1,28 | 0 |
| MTF1 | TAAGAATACGTTAT | -1 | 801 | 815 | -199 | -185 | -9,06 | 0,01 | 7,45 | >0.1 | 5,56 | 6 | 0 | 1,52 | 0 | 3,43 | 0 |
| MTF1 | ATCAAAACCGGTAC | -1 | 741 | 755 | -259 | -245 | -8,46 | 0,01 | 9,99 | >0.1 | 5,56 | 5,7 | 0 | 1,27 | 0 | 5,92 | 0 |
| MTF1 | TTTTCAGACAATTT | -1 | 610 | 624 | -390 | -376 | -9,37 | 0,01 | 1,91 | >0.1 | 3,56 | 5,25 | 0 | 1,03 | 0 | 1,44 | 0 |
| MTF1 | TTTGCCCTTGTCTT | -1 | 503 | 517 | -497 | -483 | -7,46 | 0,01 | 7,17 | >0.1 | 2,98 | 4,98 | 0 | 0,99 | 0 | 5,68 | 0 |
| MTF1 | CGCGATCTCGGCTC | -1 | 401 | 415 | -599 | -585 | -9,48 | 0,01 | 4,73 | >0.1 | 2,63 | 4,78 | 0 | 1,04 | 0 | 5,76 | 0 |
| MTF1 | TCGGCTCACTGCAA | -1 | 394 | 408 | -606 | -592 | -1,02 | 0,001 | 13,17 | >0.1 | 2,61 | 4,76 | 0 | 1,04 | 0 | 5,76 | 0 |
| MTF1 | ACTGCAAGCTCTGC | -1 | 387 | 401 | -613 | -599 | -8,37 | 0,01 | 6,05 | >0.1 | 2,59 | 4,75 | 0 | 1,05 | 0 | 6,02 | 0 |
| MTF1 | GGTGCCCACCACCA | -1 | 319 | 333 | -681 | -667 | -4,13 | 0,01 | 9,86 | >0.1 | 2,62 | 4,64 | 0 | 1,16 | 0 | 5,56 | 0 |
| MTF1 | GCCCAACTCGGCCT | -1 | 219 | 233 | -781 | -767 | -8,44 | 0,01 | 6,04 | >0.1 | 2,95 | 4,5 | 0 | 1,51 | 0 | 5,52 | 0 |
| MTF1 | TCTCCACTCCACAG | -1 | 112 | 126 | -888 | -874 | -4,38 | 0,01 | 6,13 | >0.1 | 3,51 | 4,37 | 0 | 1,89 | 0 | 0,75 | 0 |
| MTF1 | ACTCCACAGGTCAA | -1 | 107 | 121 | -893 | -879 | -3,74 | 0,01 | 6,8 | >0.1 | 3,54 | 4,36 | 0 | 1,89 | 0 | 0,75 | 0 |
| MTF1 | TTTGCAAAACTTAA | -1 | 78 | 92 | -922 | -908 | -7,98 | 0,01 | 2,02 | >0.1 | 3,79 | 4,33 | 0 | 1,88 | 0 | 0 | 0 |
| ID4 | CACGCCTGTA | 1 | 193 | 203 | -807 | -797 | 5,32 | 0,01 | 18,79 | >0.1 | 3,07 | 4,46 | 0 | 1,68 | 0 | 4,26 | 0 |
| ID4 | ATCACAAGGT | 1 | 235 | 245 | -765 | -755 | -2,76 | 0,01 | 10,82 | >0.1 | 2,9 | 4,51 | 0 | 1,43 | 0 | 4,74 | 0 |
| ID4 | GGCACCTGTA | 1 | 327 | 337 | -673 | -663 | 13,64 | 0,0001 | 27,6 | >0.1 | 2,6 | 4,65 | 0 | 1,14 | 0 | 5,56 | 0 |
| ID4 | CCCAGCTACT | 1 | 339 | 349 | -661 | -651 | 1,66 | 0,01 | 16,54 | >0.1 | 2,57 | 4,66 | 0 | 1,12 | 0 | 6,52 | 0 |
| ID4 | GGCAGGAGAA | 1 | 360 | 370 | -640 | -630 | -2,38 | 0,01 | 12,29 | >0.1 | 2,52 | 4,7 | 0 | 1,09 | 0 | 6,36 | 0 |
| ID4 | TGAACCTGGG | 1 | 375 | 385 | -625 | -615 | -0,17 | 0,01 | 14,57 | >0.1 | 2,56 | 4,72 | 0 | 1,07 | 0 | 6,4 | 0 |
| ID4 | TGCTCCAGCC | 1 | 421 | 431 | -579 | -569 | -1,99 | 0,01 | 11,94 | >0.1 | 2,69 | 4,81 | 0 | 1,02 | 0 | 5,42 | 0 |
| ID4 | AACACCAACA | 1 | 631 | 641 | -369 | -359 | 3,11 | 0,01 | 14,82 | >0.1 | 3,71 | 5,3 | 0 | 1,05 | 0 | 1,66 | 0 |
| ID4 | GACAGGAGAG | 1 | 768 | 778 | -232 | -222 | -2,58 | 0,01 | 16,1 | >0.1 | 5,61 | 5,81 | 0 | 1,35 | 0 | 5,92 | 0 |
| ID4 | CTCAGCAGAT | 1 | 887 | 897 | -113 | -103 | -2,45 | 0,01 | 14,71 | >0.1 | 5,61 | 6,59 | 0 | 1,89 | 0 | 3,07 | 0 |
| ID4 | AAGACCTGTT | 1 | 1158 | 1168 | 158 | 168 | -0,2 | 0,01 | 16,39 | >0.1 | 4,38 | 6,24 | 0 | 1,78 | 1,79 | 2,4 | 0 |
| ID4 | TTCACTTGCT | 1 | 1177 | 1187 | 177 | 187 | 1,02 | 0,01 | 19,4 | >0.1 | 5,61 | 6,11 | 0 | 1,61 | 1,79 | 3,26 | 0 |
| ID4 | AACAGGTCTT | -1 | 1158 | 1168 | 158 | 168 | -1,02 | 0,01 | 15,57 | >0.1 | 4,38 | 6,24 | 0 | 1,78 | 1,79 | 2,4 | 0 |
| ID4 | AGCAGCAGTT | -1 | 1090 | 1100 | 90 | 100 | 4,97 | 0,01 | 23,38 | >0.1 | 5,61 | 6,88 | 0 | 1,85 | 1,79 | 2,27 | 0 |
| ID4 | TACAGCAACA | -1 | 1081 | 1091 | 81 | 91 | 0,29 | 0,01 | 18,71 | >0.1 | 5,61 | 7 | 0 | 1,85 | 1,79 | 2,16 | 0 |
| ID4 | GACAGATATG | -1 | 852 | 862 | -148 | -138 | -1,91 | 0,01 | 14,56 | >0.1 | 5,61 | 6,31 | 0 | 1,88 | 0 | 2,67 | 0 |
| ID4 | CTCTCCTGTC | -1 | 768 | 778 | -232 | -222 | -2,69 | 0,01 | 15,99 | >0.1 | 5,61 | 5,81 | 0 | 1,35 | 0 | 5,92 | 0 |
| ID4 | TTCTCCTGCC | -1 | 360 | 370 | -640 | -630 | -2,08 | 0,01 | 12,59 | >0.1 | 2,52 | 4,7 | 0 | 1,09 | 0 | 6,36 | 0 |
| ID4 | TACAGGTGCC | -1 | 327 | 337 | -673 | -663 | 6,68 | 0,01 | 20,64 | >0.1 | 2,6 | 4,65 | 0 | 1,14 | 0 | 5,56 | 0 |
| ID4 | CCCACCACCA | -1 | 319 | 329 | -681 | -671 | 0,84 | 0,01 | 14,82 | >0.1 | 2,63 | 4,63 | 0 | 1,16 | 0 | 5,56 | 0 |
| ID4 | ACCACCACGC | -1 | 316 | 326 | -684 | -674 | 1,18 | 0,01 | 15,74 | >0.1 | 2,63 | 4,63 | 0 | 1,17 | 0 | 6,13 | 0 |
| ID4 | CCCAGCTAAG | -1 | 307 | 317 | -693 | -683 | -1,89 | 0,01 | 12,25 | >0.1 | 2,66 | 4,62 | 0 | 1,19 | 0 | 5,68 | 0 |
| ID2 | GTAAGTGA | 1 | 5 | 13 | -995 | -987 | -10,89 | 0,01 | 2,66 | >0.1 | 5,64 | 4,26 | 0 | 1,86 | 1,79 | 0 | 0 |
| ID2 | ACCTGTGG | 1 | 110 | 118 | -890 | -882 | -11,61 | 0,01 | -1,08 | >0.1 | 3,54 | 4,36 | 0 | 1,89 | 0 | 0,75 | 0 |
| ID2 | GCACTTTG | 1 | 209 | 217 | -791 | -783 | -11,26 | 0,01 | 2,49 | >0.1 | 3 | 4,47 | 0 | 1,57 | 0 | 4,7 | 0 |
| ID2 | ACACGGTG | 1 | 270 | 278 | -730 | -722 | -11,61 | 0,01 | 2,1 | >0.1 | 2,78 | 4,56 | 0 | 1,29 | 0 | 5,09 | 0 |
| ID2 | GGGCGTGG | 1 | 314 | 322 | -686 | -678 | -9,77 | 0,01 | 4,8 | >0.1 | 2,64 | 4,62 | 0 | 1,17 | 0 | 6,13 | 0 |
| ID2 | GCACCTGT | 1 | 328 | 336 | -672 | -664 | 0,94 | 0,001 | 14,89 | >0.1 | 2,6 | 4,64 | 0 | 1,14 | 0 | 5,56 | 0 |
| ID2 | ACTCGGGA | 1 | 346 | 354 | -654 | -646 | -10,74 | 0,01 | 4,12 | >0.1 | 2,55 | 4,67 | 0 | 1,11 | 0 | 6,52 | 0 |
| ID2 | GCAGGAGA | 1 | 361 | 369 | -639 | -631 | -9,89 | 0,01 | 4,77 | >0.1 | 2,52 | 4,7 | 0 | 1,08 | 0 | 6,36 | 0 |
| ID2 | GAACCTGG | 1 | 376 | 384 | -624 | -616 | -10,77 | 0,01 | 3,97 | >0.1 | 2,56 | 4,72 | 0 | 1,06 | 0 | 6,4 | 0 |
| ID2 | ACAGGAGA | 1 | 769 | 777 | -231 | -223 | -11,23 | 0,01 | 7,47 | >0.1 | 5,64 | 5,8 | 0 | 1,35 | 0 | 5,92 | 0 |
| ID2 | GGATGTGA | 1 | 1022 | 1030 | 22 | 30 | -10,39 | 0,01 | 7,1 | >0.1 | 5,64 | 8,68 | 0 | 1,86 | 0 | 1,31 | 0 |
| ID2 | ACAGGGGA | 1 | 1039 | 1047 | 39 | 47 | -10,74 | 0,01 | 5,98 | >0.1 | 5,64 | 7,88 | 0 | 1,86 | 0 | 1,34 | 0 |
| ID2 | TCACTTGC | 1 | 1178 | 1186 | 178 | 186 | -4,69 | 0,01 | 13,71 | >0.1 | 5,64 | 6,1 | 0 | 1,6 | 1,79 | 3,26 | 0 |
| ID2 | GCAAGTGA | -1 | 1178 | 1186 | 178 | 186 | 1,9 | 0,001 | 20,3 | >0.1 | 5,64 | 6,1 | 0 | 1,6 | 1,79 | 3,26 | 0 |
| ID2 | CCCTGTGA | -1 | 1037 | 1045 | 37 | 45 | -11,47 | 0,01 | 5,32 | >0.1 | 5,64 | 7,95 | 0 | 1,86 | 0 | 1,34 | 0 |
| ID2 | CCTCATGA | -1 | 985 | 993 | -15 | -7 | -11,97 | 0,01 | 5,57 | >0.1 | 5,64 | 8,5 | 0 | 1,87 | 0 | 1,53 | 0 |
| ID2 | GAACTTGG | -1 | 954 | 962 | -46 | -38 | -11,26 | 0,01 | 5,39 | >0.1 | 5,64 | 7,44 | 0 | 1,88 | 0 | 1,7 | 0 |
| ID2 | ACACGGAA | -1 | 697 | 705 | -303 | -295 | -11,23 | 0,01 | 4,12 | >0.1 | 5,64 | 5,5 | 0 | 1,14 | 0 | 3,07 | 0 |
| ID2 | GCCCTTGT | -1 | 506 | 514 | -494 | -486 | -11,35 | 0,01 | 3,26 | >0.1 | 2,98 | 4,97 | 0 | 0,98 | 0 | 5,68 | 0 |
| ID2 | ACAGGTGC | -1 | 328 | 336 | -672 | -664 | -1,89 | 0,01 | 12,06 | >0.1 | 2,6 | 4,64 | 0 | 1,14 | 0 | 5,56 | 0 |

| | | | | | | | | | | | | | | | | | |
|---|---|---|---|---|---|---|---|---|---|---|---|---|---|---|---|---|---|
| ID2 | AGACGGGG | -1 | 281 | 289 | -719 | -711 | -11,11 | 0,01 | 2,55 | >0.1 | 2,74 | 4,57 | 0 | 1,26 | 0 | 5,09 | 0 |
| ID2 | CCTTGTGA | -1 | 236 | 244 | -764 | -756 | -11,97 | 0,01 | 1,61 | >0.1 | 2,9 | 4,51 | 0 | 1,43 | 0 | 4,74 | 0 |
| ID2 | AGGCGTGA | -1 | 192 | 200 | -808 | -800 | -10,74 | 0,01 | 2,61 | >0.1 | 3,08 | 4,45 | 0 | 1,69 | 0 | 4,13 | 0 |
| ID2 | ACAGGTCA | -1 | 108 | 116 | -892 | -884 | -11,23 | 0,01 | -0,66 | >0.1 | 3,56 | 4,36 | 0 | 1,89 | 0 | 0,77 | 0 |
| SREBF2 | ATCAGGAGAG | 1 | 97 | 107 | -903 | -893 | -0,48 | 0,01 | 10,15 | >0.1 | 3,64 | 4,35 | 0 | 1,88 | 0 | 0,77 | 0 |
| SREBF2 | GAGAGTTGAC | 1 | 102 | 112 | -898 | -888 | -2,08 | 0,01 | 8,52 | >0.1 | 3,6 | 4,35 | 0 | 1,88 | 0 | 0,77 | 0 |
| SREBF2 | GTGGAGTGGA | 1 | 114 | 124 | -886 | -876 | 1,27 | 0,01 | 11,78 | >0.1 | 3,51 | 4,36 | 0 | 1,89 | 0 | 0,75 | 0 |
| SREBF2 | CTGATCAGAA | 1 | 164 | 174 | -836 | -826 | -2,29 | 0,01 | 9,64 | >0.1 | 3,21 | 4,42 | 0 | 1,9 | 0 | 2,4 | 0 |
| SREBF2 | TTGGGCAGAT | 1 | 227 | 237 | -773 | -763 | -0,09 | 0,01 | 14,06 | >0.1 | 2,93 | 4,5 | 0 | 1,47 | 0 | 5,26 | 0 |
| SREBF2 | GTCAGGAGAT | 1 | 243 | 253 | -757 | -747 | 2,38 | 0,01 | 15,49 | >0.1 | 2,87 | 4,52 | 0 | 1,39 | 0 | 4,33 | 0 |
| SREBF2 | ATGGCGTGAA | 1 | 369 | 379 | -631 | -621 | 11,64 | 0,0001 | 26,12 | >0.1 | 2,54 | 4,71 | 0 | 1,07 | 0 | 6,15 | 0 |
| SREBF2 | GTGAGCCGAG | 1 | 399 | 409 | -601 | -591 | 1,55 | 0,01 | 15,89 | >0.1 | 2,62 | 4,77 | 0 | 1,04 | 0 | 5,91 | 0 |
| SREBF2 | CTGATCAGAA | 1 | 479 | 489 | -521 | -511 | -2,29 | 0,01 | 12,02 | >0.1 | 2,88 | 4,92 | 0 | 0,99 | 0 | 5,52 | 0 |
| SREBF2 | GTGAGATAAC | 1 | 591 | 601 | -409 | -399 | 0,88 | 0,01 | 10,49 | >0.1 | 3,41 | 5,18 | 0 | 1,01 | 0 | 0 | 0 |
| SREBF2 | ATTGTCTGAA | 1 | 612 | 622 | -388 | -378 | 0,57 | 0,01 | 11,84 | >0.1 | 3,56 | 5,24 | 0 | 1,03 | 0 | 1,44 | 0 |
| SREBF2 | ATAAAGTGAC | 1 | 761 | 771 | -239 | -229 | 1,66 | 0,01 | 20,47 | >0.1 | 5,61 | 5,77 | 0 | 1,32 | 0 | 6,1 | 0 |
| SREBF2 | GTGACAGGAG | 1 | 766 | 776 | -234 | -224 | -2,42 | 0,01 | 16,24 | >0.1 | 5,61 | 5,8 | 0 | 1,34 | 0 | 5,92 | 0 |
| SREBF2 | CTCAGCAGAT | 1 | 887 | 897 | -113 | -103 | -0,22 | 0,01 | 16,94 | >0.1 | 5,61 | 6,59 | 0 | 1,89 | 0 | 3,07 | 0 |
| SREBF2 | AAAGGCTGAT | 1 | 962 | 972 | -38 | -28 | -1,46 | 0,01 | 15,39 | >0.1 | 5,61 | 7,67 | 0 | 1,87 | 0 | 1,7 | 0 |
| SREBF2 | ATGAGGAGGT | 1 | 987 | 997 | -13 | -3 | -0,95 | 0,01 | 16,77 | >0.1 | 5,61 | 8,64 | 0 | 1,87 | 0 | 1,61 | 0 |
| SREBF2 | TGGATGTGAT | 1 | 1021 | 1031 | 21 | 31 | -2,21 | 0,01 | 15,27 | >0.1 | 5,61 | 8,7 | 0 | 1,86 | 0 | 1,31 | 0 |
| SREBF2 | TTGGCCTGTT | -1 | 1116 | 1126 | 116 | 126 | -1,51 | 0,01 | 18,36 | >0.1 | 5,61 | 6,59 | 0 | 1,85 | 3,22 | 2,6 | 0 |
| SREBF2 | GTGGACTGAG | -1 | 1098 | 1108 | 98 | 108 | 7,18 | 0,001 | 26,99 | >0.1 | 5,61 | 6,78 | 0 | 1,85 | 3,22 | 2,35 | 0 |
| SREBF2 | CTCATGAGAT | -1 | 982 | 992 | -18 | -8 | -0,47 | 0,01 | 16,97 | >0.1 | 5,61 | 8,43 | 0 | 1,87 | 0 | 1,53 | 0 |
| SREBF2 | ATCTGCTGAG | -1 | 887 | 897 | -113 | -103 | -0,25 | 0,01 | 16,91 | >0.1 | 5,61 | 6,59 | 0 | 1,89 | 0 | 3,07 | 0 |
| SREBF2 | GTGGCGCGAT | -1 | 409 | 419 | -591 | -581 | 7,61 | 0,001 | 21,6 | >0.1 | 2,65 | 4,78 | 0 | 1,03 | 0 | 5,52 | 0 |
| SREBF2 | CTCGGCTCAC | -1 | 399 | 409 | -601 | -591 | 1,89 | 0,01 | 16,23 | >0.1 | 2,62 | 4,77 | 0 | 1,04 | 0 | 5,91 | 0 |
| SREBF2 | ATCTCCTGAC | -1 | 243 | 253 | -757 | -747 | 1,22 | 0,01 | 14,33 | >0.1 | 2,87 | 4,52 | 0 | 1,39 | 0 | 4,33 | 0 |
| SREBF2 | GTGATCTGCC | -1 | 230 | 240 | -770 | -760 | 2,21 | 0,01 | 16,1 | >0.1 | 2,92 | 4,51 | 0 | 1,45 | 0 | 5,01 | 0 |
| SREBF2 | CAGGCGTGAG | -1 | 191 | 201 | -809 | -799 | 3,36 | 0,01 | 16,72 | >0.1 | 3,08 | 4,45 | 0 | 1,7 | 0 | 4,13 | 0 |
| SREBF2 | CTCTCCTGAT | -1 | 97 | 107 | -903 | -893 | 0,57 | 0,01 | 11,2 | >0.1 | 3,64 | 4,35 | 0 | 1,88 | 0 | 0,77 | 0 |
| SREBF2 | CTGATCTGCC | -1 | 92 | 102 | -908 | -898 | 1,58 | 0,01 | 12,31 | >0.1 | 3,68 | 4,34 | 0 | 1,88 | 0 | 0,83 | 0 |
| SREBF1 | GGCAGATCAG | 1 | 92 | 102 | -908 | -898 | -1,1 | 0,01 | 9,63 | >0.1 | 3,68 | 4,34 | 0 | 1,88 | 0 | 0,83 | 0 |
| SREBF1 | ATCAGGAGAG | 1 | 97 | 107 | -903 | -893 | 0,32 | 0,01 | 10,95 | >0.1 | 3,64 | 4,35 | 0 | 1,88 | 0 | 0,77 | 0 |
| SREBF1 | CTCACGCCTG | 1 | 191 | 201 | -809 | -799 | 4,76 | 0,01 | 18,12 | >0.1 | 3,08 | 4,45 | 0 | 1,7 | 0 | 4,13 | 0 |
| SREBF1 | GGCAGATCAC | 1 | 230 | 240 | -770 | -760 | 0,76 | 0,01 | 14,65 | >0.1 | 2,92 | 4,51 | 0 | 1,45 | 0 | 5,01 | 0 |
| SREBF1 | GTCAGGAGAT | 1 | 243 | 253 | -757 | -747 | 1,32 | 0,01 | 14,43 | >0.1 | 2,87 | 4,52 | 0 | 1,39 | 0 | 4,33 | 0 |
| SREBF1 | GTGAGCCGAG | 1 | 399 | 409 | -601 | -591 | 1,64 | 0,01 | 15,98 | >0.1 | 2,62 | 4,77 | 0 | 1,04 | 0 | 5,91 | 0 |
| SREBF1 | ATCGCGCCAC | 1 | 409 | 419 | -591 | -581 | 7,62 | 0,001 | 21,61 | >0.1 | 2,65 | 4,78 | 0 | 1,03 | 0 | 5,52 | 0 |
| SREBF1 | CTCAGCAGAT | 1 | 887 | 897 | -113 | -103 | 2,67 | 0,01 | 19,83 | >0.1 | 5,61 | 6,59 | 0 | 1,89 | 0 | 3,07 | 0 |
| SREBF1 | CTCAGCGCCC | 1 | 941 | 951 | -59 | -49 | -1,96 | 0,01 | 13,55 | >0.1 | 5,61 | 7,23 | 0 | 1,88 | 0 | 0,79 | 0 |
| SREBF1 | ATCTCATGAG | 1 | 982 | 992 | -18 | -8 | -0,48 | 0,01 | 16,96 | >0.1 | 5,61 | 8,43 | 0 | 1,87 | 0 | 1,53 | 0 |
| SREBF1 | CTCAGTCCAC | 1 | 1098 | 1108 | 98 | 108 | 8,03 | 0,001 | 27,84 | >0.1 | 5,61 | 6,78 | 0 | 1,85 | 3,22 | 2,35 | 0 |
| SREBF1 | AACAGGCCAA | 1 | 1116 | 1126 | 116 | 126 | -3,24 | 0,01 | 16,63 | >0.1 | 5,61 | 6,59 | 0 | 1,85 | 3,22 | 2,6 | 0 |
| SREBF1 | GTGAGCCAAG | -1 | 1031 | 1041 | 31 | 41 | -1,2 | 0,01 | 15,77 | >0.1 | 5,61 | 8,19 | 0 | 1,86 | 0 | 1,31 | 0 |
| SREBF1 | ACCTCCTCAT | -1 | 987 | 997 | -13 | -3 | -1,72 | 0,01 | 16 | >0.1 | 5,61 | 8,64 | 0 | 1,87 | 0 | 1,61 | 0 |
| SREBF1 | ATCAGCCTTT | -1 | 962 | 972 | -38 | -28 | -3,05 | 0,01 | 13,8 | >0.1 | 5,61 | 7,67 | 0 | 1,87 | 0 | 1,7 | 0 |
| SREBF1 | ATCTGCTGAG | -1 | 887 | 897 | -113 | -103 | -0,19 | 0,01 | 16,97 | >0.1 | 5,61 | 6,59 | 0 | 1,89 | 0 | 3,07 | 0 |
| SREBF1 | CTGAGCCAAT | -1 | 882 | 892 | -118 | -108 | -0,08 | 0,01 | 17,04 | >0.1 | 5,61 | 6,55 | 0 | 1,89 | 0 | 3,07 | 0 |
| SREBF1 | CTTACCTCTC | -1 | 773 | 783 | -227 | -217 | -2,66 | 0,01 | 16,25 | >0.1 | 5,61 | 5,83 | 0 | 1,37 | 0 | 6,1 | 0 |
| SREBF1 | GTCACTTTAT | -1 | 761 | 771 | -239 | -229 | 0,04 | 0,01 | 18,85 | >0.1 | 5,61 | 5,77 | 0 | 1,32 | 0 | 6,1 | 0 |
| SREBF1 | TTCAGACAAT | -1 | 612 | 622 | -388 | -378 | -1,66 | 0,01 | 9,61 | >0.1 | 3,56 | 5,24 | 0 | 1,03 | 0 | 1,44 | 0 |
| SREBF1 | GTTATCTCAC | -1 | 591 | 601 | -409 | -399 | -2,78 | 0,01 | 6,83 | >0.1 | 3,41 | 5,18 | 0 | 1,01 | 0 | 0 | 0 |
| SREBF1 | CTCGGCTCAC | -1 | 399 | 409 | -601 | -591 | 3,69 | 0,01 | 18,03 | >0.1 | 2,62 | 4,77 | 0 | 1,04 | 0 | 5,91 | 0 |
| SREBF1 | TTCACGCCAT | -1 | 369 | 379 | -631 | -621 | 7,61 | 0,001 | 22,09 | >0.1 | 2,54 | 4,71 | 0 | 1,07 | 0 | 6,15 | 0 |
| SREBF1 | CTCAGCCTCC | -1 | 351 | 361 | -649 | -639 | -2,49 | 0,01 | 12,23 | >0.1 | 2,54 | 4,68 | 0 | 1,1 | 0 | 6,4 | 0 |
| SREBF1 | GCCTCCCGAG | -1 | 347 | 357 | -653 | -643 | -3,21 | 0,01 | 11,65 | >0.1 | 2,55 | 4,68 | 0 | 1,11 | 0 | 6,52 | 0 |
| SREBF1 | CCCAGCTAAG | -1 | 307 | 317 | -693 | -683 | -3,03 | 0,01 | 11,11 | >0.1 | 2,66 | 4,62 | 0 | 1,19 | 0 | 5,68 | 0 |
| SREBF1 | ATCTCCTGAC | -1 | 243 | 253 | -757 | -747 | 4,51 | 0,01 | 17,62 | >0.1 | 2,87 | 4,52 | 0 | 1,39 | 0 | 4,33 | 0 |

| | | | | | | | | | | | | | | | | |
|---|---|---|---|---|---|---|---|---|---|---|---|---|---|---|---|---|
| SREBF1 | ATCTGCCCAA | -1 | 227 | 237 | -773 | -763 | -1,71 | 0,01 | 12,44 | >0.1 | 2,93 | 4,5 | 0 | 1,47 | 0 | 5,26 | 0 |
| SREBF1 | GTCAACTCTC | -1 | 102 | 112 | -898 | -888 | -2,28 | 0,01 | 8,32 | >0.1 | 3,6 | 4,35 | 0 | 1,88 | 0 | 0,77 | 0 |
| SREBF1 | CTCTCCTGAT | -1 | 97 | 107 | -903 | -893 | 3,46 | 0,01 | 14,09 | >0.1 | 3,64 | 4,35 | 0 | 1,88 | 0 | 0,77 | 0 |
| NKX3-2 | GGCACTCAA | 1 | 23 | 32 | -977 | -968 | 8,19 | 0,001 | 19,95 | >0.1 | 5,62 | 4,27 | 0 | 1,87 | 0 | 0 | 0 |
| NKX3-2 | AGCACTTTG | 1 | 208 | 217 | -792 | -783 | 8,88 | 0,001 | 22,64 | >0.1 | 3,01 | 4,48 | 0 | 1,58 | 0 | 4,7 | 0 |
| NKX3-2 | AAAACTTAG | 1 | 303 | 312 | -697 | -688 | 2 | 0,01 | 15,46 | >0.1 | 2,67 | 4,61 | 0 | 1,2 | 0 | 4,99 | 0 |
| NKX3-2 | ACCTATTAA | 1 | 599 | 608 | -401 | -392 | 3,56 | 0,01 | 14,65 | >0.1 | 3,46 | 5,2 | 0 | 1,01 | 0 | 1,41 | 0 |
| NKX3-2 | ACCTCTTTG | 1 | 666 | 675 | -334 | -325 | 5,5 | 0,01 | 19,9 | >0.1 | 4,08 | 5,4 | 0 | 1,09 | 0 | 3,83 | 0 |
| NKX3-2 | TTCACTTTA | 1 | 820 | 829 | -180 | -171 | 5,56 | 0,01 | 22,43 | >0.1 | 5,62 | 6,08 | 0 | 1,61 | 0 | 3,55 | 0 |
| NKX3-2 | AGCTCTTCC | 1 | 1055 | 1064 | 55 | 64 | 3,02 | 0,01 | 20,23 | >0.1 | 5,62 | 7,46 | 0 | 1,86 | 0 | 2,27 | 0 |
| NKX3-2 | TTCACTTGC | 1 | 1177 | 1186 | 177 | 186 | 2,32 | 0,01 | 20,73 | >0.1 | 5,62 | 6,11 | 0 | 1,61 | 1,79 | 3,28 | 0 |
| NKX3-2 | AGCAGTTAC | -1 | 1088 | 1097 | 88 | 97 | 3,14 | 0,01 | 21,53 | >0.1 | 5,62 | 6,9 | 0 | 1,85 | 1,79 | 2,21 | 0 |
| NKX3-2 | CTCTCTTAT | -1 | 970 | 979 | -30 | -21 | 3,22 | 0,01 | 20,29 | >0.1 | 5,62 | 7,88 | 0 | 1,87 | 0 | 1,7 | 0 |
| NKX3-2 | GTCACTTTA | -1 | 762 | 771 | -238 | -229 | 6,95 | 0,01 | 25,77 | >0.1 | 5,62 | 5,77 | 0 | 1,32 | 0 | 6,1 | 0 |
| NKX3-2 | GGTACTTTT | -1 | 737 | 746 | -263 | -254 | 4 | 0,01 | 22,45 | >0.1 | 5,62 | 5,66 | 0 | 1,24 | 0 | 5,92 | 0 |
| NKX3-2 | GACAATTTC | -1 | 609 | 618 | -391 | -382 | 1,27 | 0,01 | 12,6 | >0.1 | 3,53 | 5,23 | 0 | 1,02 | 0 | 1,55 | 0 |
| NKX3-2 | CACATTTAT | -1 | 585 | 594 | -415 | -406 | 0,87 | 0,01 | 10,41 | >0.1 | 3,37 | 5,16 | 0 | 1 | 0 | 0 | 0 |
| NKX3-2 | TCTACTTGG | -1 | 545 | 554 | -455 | -446 | 2,86 | 0,01 | 12,07 | >0.1 | 3,15 | 5,06 | 0 | 0,99 | 0 | 0 | 0 |
| NKX3-2 | TCCACTCCA | -1 | 115 | 124 | -885 | -876 | 4,31 | 0,01 | 14,81 | >0.1 | 3,5 | 4,36 | 0 | 1,89 | 0 | 0,75 | 0 |
| NKX3-2 | AAAACTTAA | -1 | 78 | 87 | -922 | -913 | 2,72 | 0,01 | 12,73 | >0.1 | 3,81 | 4,33 | 0 | 1,88 | 0 | 0 | 0 |
| NKX3-2 | ACTTCTTTG | -1 | 29 | 38 | -971 | -962 | 1,46 | 0,01 | 12,12 | >0.1 | 4,52 | 4,28 | 0 | 1,87 | 0 | 0 | 0 |
| NKX3-2 | TTCACTTAC | -1 | 5 | 14 | -995 | -986 | 6,87 | 0,01 | 20,41 | >0.1 | 5,62 | 4,26 | 0 | 1,86 | 1,79 | 0 | 0 |
| NKX3-1 | GGCACTCAA | 1 | 23 | 32 | -977 | -968 | 8,45 | 0,001 | 20,21 | >0.1 | 5,62 | 4,27 | 0 | 1,87 | 0 | 0 | 0 |
| NKX3-1 | AGCACTTTG | 1 | 208 | 217 | -792 | -783 | 8,52 | 0,001 | 22,28 | >0.1 | 3,01 | 4,48 | 0 | 1,58 | 0 | 4,7 | 0 |
| NKX3-1 | GCTACTCGG | 1 | 343 | 352 | -657 | -648 | 0,28 | 0,01 | 15,02 | >0.1 | 2,56 | 4,67 | 0 | 1,11 | 0 | 6,4 | 0 |
| NKX3-1 | ACCTATTAA | 1 | 599 | 608 | -401 | -392 | 1,37 | 0,01 | 12,46 | >0.1 | 3,46 | 5,2 | 0 | 1,01 | 0 | 1,41 | 0 |
| NKX3-1 | ACCTCTTTG | 1 | 666 | 675 | -334 | -325 | 4,18 | 0,01 | 18,58 | >0.1 | 4,08 | 5,4 | 0 | 1,09 | 0 | 3,83 | 0 |
| NKX3-1 | TTCACTTTA | 1 | 820 | 829 | -180 | -171 | 3,99 | 0,01 | 20,86 | >0.1 | 5,62 | 6,08 | 0 | 1,61 | 0 | 3,55 | 0 |
| NKX3-1 | AGCTCTTCC | 1 | 1055 | 1064 | 55 | 64 | 1,56 | 0,01 | 18,77 | >0.1 | 5,62 | 7,46 | 0 | 1,86 | 0 | 2,27 | 0 |
| NKX3-1 | TTCACTTGC | 1 | 1177 | 1186 | 177 | 186 | 1,47 | 0,01 | 19,88 | >0.1 | 5,62 | 6,11 | 0 | 1,61 | 1,79 | 3,28 | 0 |
| NKX3-1 | AGCAGTTAC | -1 | 1088 | 1097 | 88 | 97 | 2,09 | 0,01 | 20,48 | >0.1 | 5,62 | 6,9 | 0 | 1,85 | 1,79 | 2,21 | 0 |
| NKX3-1 | CTCTCTTAT | -1 | 970 | 979 | -30 | -21 | 0,96 | 0,01 | 18,03 | >0.1 | 5,62 | 7,88 | 0 | 1,87 | 0 | 1,7 | 0 |
| NKX3-1 | ACCTCTCCT | -1 | 771 | 780 | -229 | -220 | -1,15 | 0,01 | 17,56 | >0.1 | 5,62 | 5,82 | 0 | 1,36 | 0 | 5,92 | 0 |
| NKX3-1 | GTCACTTTA | -1 | 762 | 771 | -238 | -229 | 5,81 | 0,01 | 24,63 | >0.1 | 5,62 | 5,77 | 0 | 1,32 | 0 | 6,1 | 0 |
| NKX3-1 | GGTACTTTT | -1 | 737 | 746 | -263 | -254 | 2,69 | 0,01 | 21,14 | >0.1 | 5,62 | 5,66 | 0 | 1,24 | 0 | 5,92 | 0 |
| NKX3-1 | GACAATTTC | -1 | 609 | 618 | -391 | -382 | -0,77 | 0,01 | 10,56 | >0.1 | 3,53 | 5,23 | 0 | 1,02 | 0 | 1,55 | 0 |
| NKX3-1 | CACATTTAT | -1 | 585 | 594 | -415 | -406 | -0,96 | 0,01 | 8,58 | >0.1 | 3,37 | 5,16 | 0 | 1 | 0 | 0 | 0 |
| NKX3-1 | TCTACTTGG | -1 | 545 | 554 | -455 | -446 | 2,36 | 0,01 | 11,57 | >0.1 | 3,15 | 5,06 | 0 | 0,99 | 0 | 0 | 0 |
| NKX3-1 | GCCATTTCT | -1 | 170 | 179 | -830 | -821 | -1,35 | 0,01 | 11,65 | >0.1 | 3,18 | 4,43 | 0 | 1,88 | 0 | 3,51 | 0 |
| NKX3-1 | GCCATTTTT | -1 | 141 | 150 | -859 | -850 | -0,91 | 0,01 | 10,36 | >0.1 | 3,34 | 4,39 | 0 | 1,89 | 0 | 1,65 | 0 |
| NKX3-1 | ACTACTCTC | -1 | 122 | 131 | -878 | -869 | 1,12 | 0,01 | 12,53 | >0.1 | 3,46 | 4,37 | 0 | 1,89 | 0 | 1,7 | 0 |
| NKX3-1 | TCCACTCCA | -1 | 115 | 124 | -885 | -876 | 4,43 | 0,01 | 14,93 | >0.1 | 3,5 | 4,36 | 0 | 1,89 | 0 | 0,75 | 0 |
| NKX3-1 | ACTTCTTTG | -1 | 29 | 38 | -971 | -962 | -0,3 | 0,01 | 10,36 | >0.1 | 4,52 | 4,28 | 0 | 1,87 | 0 | 0 | 0 |
| NKX3-1 | TTCACTTAC | -1 | 5 | 14 | -995 | -986 | 5,92 | 0,01 | 19,46 | >0.1 | 5,62 | 4,26 | 0 | 1,86 | 1,79 | 0 | 0 |
| MXI1 | GGCTCACGCCTGT | 1 | 189 | 202 | -811 | -798 | 0,04 | 0,01 | 13,42 | >0.1 | 3,08 | 4,46 | 0 | 1,71 | 0 | 4,13 | 0 |
| MXI1 | AAACCCCGTCTCT | 1 | 278 | 291 | -722 | -709 | 1,38 | 0,01 | 15,16 | >0.1 | 2,74 | 4,58 | 0 | 1,27 | 0 | 5,19 | 0 |
| MXI1 | GCTGGGCGTGGTG | 1 | 311 | 324 | -689 | -676 | 1,61 | 0,01 | 16,09 | >0.1 | 2,64 | 4,63 | 0 | 1,18 | 0 | 6,03 | 0 |
| MXI1 | TGGGCACCTGTAG | 1 | 325 | 338 | -675 | -662 | 4,48 | 0,01 | 18,34 | >0.1 | 2,6 | 4,65 | 0 | 1,15 | 0 | 5,46 | 0 |
| MXI1 | CGTGAACCTGGGA | 1 | 373 | 386 | -627 | -614 | -0,58 | 0,01 | 14,3 | >0.1 | 2,56 | 4,73 | 0 | 1,07 | 0 | 6,53 | 0 |
| MXI1 | CGCCACTGTGCTC | 1 | 413 | 426 | -587 | -574 | -0,14 | 0,01 | 13,54 | >0.1 | 2,67 | 4,8 | 0 | 1,03 | 0 | 5,19 | 0 |
| MXI1 | TAACCACGAAGCC | 1 | 1144 | 1157 | 144 | 157 | -0,28 | 0,01 | 18,86 | >0.1 | 5,57 | 6,34 | 0 | 1,84 | 3,22 | 2,16 | 0 |
| MXI1 | CGAAGACCTGTTC | 1 | 1156 | 1169 | 156 | 169 | -0,29 | 0,01 | 16,28 | >0.1 | 4,39 | 6,25 | 0 | 1,8 | 1,79 | 2,34 | 0 |
| MXI1 | GAAGCAAGTGAAC | -1 | 1176 | 1189 | 176 | 189 | 2,33 | 0,01 | 20,69 | >0.1 | 5,57 | 6,11 | 0 | 1,62 | 1,79 | 3,26 | 0 |
| MXI1 | CGGCTTCGTGGTT | -1 | 1145 | 1158 | 145 | 158 | 1,14 | 0,01 | 20,27 | >0.1 | 5,57 | 6,33 | 0 | 1,84 | 3,22 | 2,16 | 0 |
| MXI1 | GTCCCCTGTGAGC | -1 | 1035 | 1048 | 35 | 48 | 2,52 | 0,01 | 19,31 | >0.1 | 5,57 | 8,03 | 0 | 1,86 | 0 | 1,34 | 0 |
| MXI1 | AGATCACATCCAC | -1 | 1020 | 1033 | 20 | 33 | 0,02 | 0,01 | 17,48 | >0.1 | 5,57 | 8,72 | 0 | 1,86 | 0 | 1,31 | 0 |
| MXI1 | CTCCCAGGTTCAC | -1 | 374 | 387 | -626 | -613 | 2,01 | 0,01 | 16,76 | >0.1 | 2,56 | 4,73 | 0 | 1,07 | 0 | 6,4 | 0 |
| MXI1 | ACTACAGGTGCCC | -1 | 326 | 339 | -674 | -661 | 5,98 | 0,001 | 19,94 | >0.1 | 2,6 | 4,65 | 0 | 1,15 | 0 | 5,56 | 0 |
| MXI1 | CCACCACGCCCAG | -1 | 312 | 325 | -688 | -675 | 6,79 | 0,001 | 21,37 | >0.1 | 2,64 | 4,63 | 0 | 1,18 | 0 | 6,13 | 0 |

| | | | | | | | | | | | | | | | | | |
|---|---|---|---|---|---|---|---|---|---|---|---|---|---|---|---|---|---|
| MXI1 | TGACCTTGTGATC | -1 | 234 | 247 | -766 | -753 | 0,91 | 0,01 | 14,5 | >0.1 | 2,9 | 4,52 | 0 | 1,44 | 0 | 4,74 | 0 |
| MXI1 | AGCCACTGTGCCC | -1 | 180 | 193 | -820 | -807 | -0,43 | 0,01 | 12,78 | >0.1 | 3,12 | 4,45 | 0 | 1,79 | 0 | 3,86 | 0 |
| JDP2(VAR.2) | AAAGAAGTCAGC | 1 | 30 | 42 | -970 | -958 | -10,18 | 0,01 | 0,43 | >0.1 | 4,46 | 4,28 | 0 | 1,87 | 0 | 0 | 0 |
| JDP2(VAR.2) | AATGGCGTGAAC | 1 | 368 | 380 | -632 | -620 | -13,18 | 0,01 | 1,3 | >0.1 | 2,54 | 4,72 | 0 | 1,08 | 0 | 6,15 | 0 |
| JDP2(VAR.2) | AATTCCATAAAG | 1 | 494 | 506 | -506 | -494 | -16,31 | 0,01 | -0,81 | >0.1 | 2,94 | 4,96 | 0 | 0,99 | 0 | 6,62 | 0 |
| JDP2(VAR.2) | TGTGAGATAACC | 1 | 590 | 602 | -410 | -398 | -6,68 | 0,01 | 2,93 | >0.1 | 3,41 | 5,19 | 0 | 1,01 | 0 | 0 | 0 |
| JDP2(VAR.2) | TCTTTGGTCACT | 1 | 669 | 681 | -331 | -319 | -14,89 | 0,01 | -0,53 | >0.1 | 4,14 | 5,42 | 0 | 1,1 | 0 | 3,7 | 0 |
| JDP2(VAR.2) | AATTATAACATT | 1 | 685 | 697 | -315 | -303 | -10,72 | 0,01 | 3,25 | >0.1 | 4,42 | 5,48 | 0 | 1,13 | 0 | 2,95 | 0 |
| JDP2(VAR.2) | GGAGAGGTAAGG | 1 | 772 | 784 | -228 | -216 | -17,01 | 0,01 | 1,88 | >0.1 | 5,58 | 5,83 | 0 | 1,37 | 0 | 6,1 | 0 |
| JDP2(VAR.2) | AATAACGTATTC | 1 | 800 | 812 | -200 | -188 | -13,96 | 0,01 | 2,54 | >0.1 | 5,58 | 5,98 | 0 | 1,51 | 0 | 3,43 | 0 |
| JDP2(VAR.2) | GACGATGTCAAG | 1 | 1045 | 1057 | 45 | 57 | -6,46 | 0,01 | 10,15 | >0.1 | 5,58 | 7,7 | 0 | 1,86 | 0 | 1,46 | 0 |
| JDP2(VAR.2) | GTTGCTGTAACT | 1 | 1082 | 1094 | 82 | 94 | -13,68 | 0,01 | 4,75 | >0.1 | 5,58 | 6,99 | 0 | 1,85 | 1,79 | 2,21 | 0 |
| JDP2(VAR.2) | AGTTACAGCAAC | -1 | 1082 | 1094 | 82 | 94 | -14,11 | 0,01 | 4,32 | >0.1 | 5,58 | 6,99 | 0 | 1,85 | 1,79 | 2,21 | 0 |
| JDP2(VAR.2) | CTTGACATCGTC | -1 | 1045 | 1057 | 45 | 57 | -7,56 | 0,01 | 9,05 | >0.1 | 5,58 | 7,7 | 0 | 1,86 | 0 | 1,46 | 0 |
| JDP2(VAR.2) | AGATATGTAACA | -1 | 847 | 859 | -153 | -141 | -14,31 | 0,01 | 2,37 | >0.1 | 5,58 | 6,29 | 0 | 1,85 | 0 | 2,95 | 0 |
| JDP2(VAR.2) | GAATACGTTATT | -1 | 800 | 812 | -200 | -188 | -13,37 | 0,01 | 3,13 | >0.1 | 5,58 | 5,98 | 0 | 1,51 | 0 | 3,43 | 0 |
| JDP2(VAR.2) | CCTTACCTCTCC | -1 | 772 | 784 | -228 | -216 | -16,82 | 0,01 | 2,07 | >0.1 | 5,58 | 5,83 | 0 | 1,37 | 0 | 6,1 | 0 |
| JDP2(VAR.2) | AATGTTATAATT | -1 | 685 | 697 | -315 | -303 | -12,14 | 0,01 | 1,83 | >0.1 | 4,42 | 5,48 | 0 | 1,13 | 0 | 2,95 | 0 |
| JDP2(VAR.2) | AGTGACCAAAGA | -1 | 669 | 681 | -331 | -319 | -17,29 | 0,01 | -2,93 | >0.1 | 4,14 | 5,42 | 0 | 1,1 | 0 | 3,7 | 0 |
| JDP2(VAR.2) | GGTTATCTCACA | -1 | 590 | 602 | -410 | -398 | -7,46 | 0,01 | 2,15 | >0.1 | 3,41 | 5,19 | 0 | 1,01 | 0 | 0 | 0 |
| JDP2(VAR.2) | GCTGACTTCTTT | -1 | 30 | 42 | -970 | -958 | -16,06 | 0,01 | -5,45 | >0.1 | 4,46 | 4,28 | 0 | 1,87 | 0 | 0 | 0 |
| HMX1 | GGAGGCACTCAAAGAAG | 1 | 20 | 37 | -980 | -963 | 10,18 | 0,0001 | 21,85 | >0.1 | 5,52 | 4,28 | 0 | 1,87 | 0 | 0 | 0 |
| HMX1 | CATGGAAATTAAAACTG | 1 | 150 | 167 | -850 | -833 | 4,11 | 0,01 | 15,34 | >0.1 | 3,26 | 4,41 | 0 | 1,9 | 0 | 1,65 | 0 |
| HMX1 | CCTAGCACTTTGGGAGG | 1 | 205 | 222 | -795 | -778 | 5,35 | 0,001 | 19,14 | >0.1 | 3 | 4,48 | 0 | 1,6 | 0 | 4,7 | 0 |
| HMX1 | CGTCTCTATTAAAAATA | 1 | 284 | 301 | -716 | -699 | -0,21 | 0,01 | 13,55 | >0.1 | 2,72 | 4,59 | 0 | 1,25 | 0 | 5,19 | 0 |
| HMX1 | ACAAAAACTTAGCTGGG | 1 | 300 | 317 | -700 | -683 | -0,34 | 0,01 | 13,14 | >0.1 | 2,67 | 4,62 | 0 | 1,21 | 0 | 4,99 | 0 |
| HMX1 | ATAACCTATTAATGAAA | 1 | 596 | 613 | -404 | -387 | 6,81 | 0,001 | 17,88 | >0.1 | 3,47 | 5,22 | 0 | 1,02 | 0 | 1,36 | 0 |
| HMX1 | GGAAAATATTAACCAAA | 1 | 723 | 740 | -277 | -260 | -0,45 | 0,01 | 17,26 | >0.1 | 5,52 | 5,64 | 0 | 1,23 | 0 | 5,32 | 0 |
| HMX1 | CTCTAGGATTAAAGAAT | 1 | 786 | 803 | -214 | -197 | -0,14 | 0,01 | 18,3 | >0.1 | 5,52 | 5,93 | 0 | 1,46 | 0 | 5,52 | 0 |
| HMX1 | TGATTCACTTTAAAAAA | 1 | 817 | 834 | -183 | -166 | -0,9 | 0,01 | 16,08 | >0.1 | 5,52 | 6,12 | 0 | 1,65 | 0 | 3,7 | 0 |
| HMX1 | AAAGTCATTCAGTGGAT | 1 | 1009 | 1026 | 9 | 26 | -1,21 | 0,01 | 16,62 | >0.1 | 5,52 | 9,03 | 0 | 1,86 | 0 | 1,42 | 0 |
| HMX1 | AGAAGCAAGTGAACTTT | -1 | 1173 | 1190 | 173 | 190 | -0,68 | 0,01 | 17,68 | >0.1 | 5,52 | 6,13 | 0 | 1,64 | 1,79 | 3,28 | 0 |
| HMX1 | AGCAGCAGTTACAGCAA | -1 | 1083 | 1100 | 83 | 100 | 3,2 | 0,01 | 21,55 | >0.1 | 5,52 | 6,97 | 0 | 1,85 | 1,79 | 2,21 | 0 |
| HMX1 | ACATCCACTGAATGACT | -1 | 1011 | 1028 | 11 | 28 | 1,75 | 0,01 | 19,48 | >0.1 | 5,52 | 8,95 | 0 | 1,86 | 0 | 1,39 | 0 |
| HMX1 | GTAAACAATCTGCTGAG | -1 | 887 | 904 | -113 | -96 | -1,61 | 0,01 | 15,41 | >0.1 | 5,52 | 6,65 | 0 | 1,89 | 0 | 2,95 | 0 |
| HMX1 | TGAGCCAATATAAAGTT | -1 | 874 | 891 | -126 | -109 | -0,72 | 0,01 | 16,18 | >0.1 | 5,52 | 6,54 | 0 | 1,9 | 0 | 2,95 | 0 |
| HMX1 | TGAATCAAATAAGAATA | -1 | 807 | 824 | -193 | -176 | 0,28 | 0,01 | 16,71 | >0.1 | 5,52 | 6,05 | 0 | 1,58 | 0 | 3,28 | 0 |
| HMX1 | CCTGTCACTTTATGGCC | -1 | 757 | 774 | -243 | -226 | 3,71 | 0,01 | 22,27 | >0.1 | 5,52 | 5,79 | 0 | 1,34 | 0 | 5,92 | 0 |
| HMX1 | GAAAGCTGTTAAAGATA | -1 | 704 | 721 | -296 | -279 | 1,18 | 0,01 | 18,26 | >0.1 | 5,52 | 5,56 | 0 | 1,18 | 0 | 4,81 | 0 |
| HMX1 | CCCAGCCATTTCTGATC | -1 | 166 | 183 | -834 | -817 | 0,48 | 0,01 | 13,5 | >0.1 | 3,18 | 4,43 | 0 | 1,9 | 0 | 3,51 | 0 |
| HMX1 | TGCAAAACTTAAAAAAG | -1 | 73 | 90 | -927 | -910 | 1,94 | 0,01 | 11,97 | >0.1 | 3,82 | 4,33 | 0 | 1,88 | 0 | 0 | 0 |
| HMX3 | GGAGGCACTCAAAGAAG | 1 | 20 | 37 | -980 | -963 | 9,36 | 0,001 | 21,03 | >0.1 | 5,52 | 4,28 | 0 | 1,87 | 0 | 0 | 0 |
| HMX3 | CATGGAAATTAAAACTG | 1 | 150 | 167 | -850 | -833 | 2,85 | 0,01 | 14,08 | >0.1 | 3,26 | 4,41 | 0 | 1,9 | 0 | 1,65 | 0 |
| HMX3 | CCTAGCACTTTGGGAGG | 1 | 205 | 222 | -795 | -778 | 3,72 | 0,01 | 17,51 | >0.1 | 3 | 4,48 | 0 | 1,6 | 0 | 4,7 | 0 |
| HMX3 | CGTCTCTATTAAAAATA | 1 | 284 | 301 | -716 | -699 | 1,25 | 0,01 | 15,01 | >0.1 | 2,72 | 4,59 | 0 | 1,25 | 0 | 5,19 | 0 |
| HMX3 | ACAAAAACTTAGCTGGG | 1 | 300 | 317 | -700 | -683 | 0,85 | 0,01 | 14,33 | >0.1 | 2,67 | 4,62 | 0 | 1,21 | 0 | 4,99 | 0 |
| HMX3 | ATAACCTATTAATGAAA | 1 | 596 | 613 | -404 | -387 | 7,93 | 0,001 | 19 | >0.1 | 3,47 | 5,22 | 0 | 1,02 | 0 | 1,36 | 0 |
| HMX3 | CTGTAAAATTATAACAT | 1 | 679 | 696 | -321 | -304 | 0,2 | 0,01 | 14,22 | >0.1 | 4,35 | 5,47 | 0 | 1,13 | 0 | 3,07 | 0 |
| HMX3 | GGAAAATATTAACCAAA | 1 | 723 | 740 | -277 | -260 | 1,02 | 0,01 | 18,73 | >0.1 | 5,52 | 5,64 | 0 | 1,23 | 0 | 5,32 | 0 |
| HMX3 | CTCTAGGATTAAAGAAT | 1 | 786 | 803 | -214 | -197 | 0,92 | 0,01 | 19,36 | >0.1 | 5,52 | 5,93 | 0 | 1,46 | 0 | 5,52 | 0 |
| HMX3 | TGATTCACTTTAAAAAA | 1 | 817 | 834 | -183 | -166 | 2,16 | 0,01 | 19,14 | >0.1 | 5,52 | 6,12 | 0 | 1,65 | 0 | 3,7 | 0 |
| HMX3 | AAAGTCATTCAGTGGAT | 1 | 1009 | 1026 | 9 | 26 | -0,76 | 0,01 | 17,07 | >0.1 | 5,52 | 9,03 | 0 | 1,86 | 0 | 1,42 | 0 |
| HMX3 | AGAAGCAAGTGAACTTT | -1 | 1173 | 1190 | 173 | 190 | 1,68 | 0,01 | 20,04 | >0.1 | 5,52 | 6,13 | 0 | 1,64 | 1,79 | 3,28 | 0 |
| HMX3 | AGCAGCAGTTACAGCAA | -1 | 1083 | 1100 | 83 | 100 | 3,88 | 0,01 | 22,23 | >0.1 | 5,52 | 6,97 | 0 | 1,85 | 1,79 | 2,21 | 0 |
| HMX3 | ACATCCACTGAATGACT | -1 | 1011 | 1028 | 11 | 28 | 2,67 | 0,01 | 20,4 | >0.1 | 5,52 | 8,95 | 0 | 1,86 | 0 | 1,39 | 0 |
| HMX3 | AAGACCAAAAAAAGAAA | -1 | 919 | 936 | -81 | -64 | 0,61 | 0,01 | 16,11 | >0.1 | 5,52 | 7,01 | 0 | 1,88 | 0 | 1,09 | 0 |
| HMX3 | TGAGCCAATATAAAGTT | -1 | 874 | 891 | -126 | -109 | -0,04 | 0,01 | 16,86 | >0.1 | 5,52 | 6,54 | 0 | 1,9 | 0 | 2,95 | 0 |
| HMX3 | GTAACAGATTTTAGAAT | -1 | 836 | 853 | -164 | -147 | 0,24 | 0,01 | 16,88 | >0.1 | 5,52 | 6,25 | 0 | 1,8 | 0 | 3,07 | 0 |
| HMX3 | TGAATCAAATAAGAATA | -1 | 807 | 824 | -193 | -176 | 3,27 | 0,01 | 19,7 | >0.1 | 5,52 | 6,05 | 0 | 1,58 | 0 | 3,28 | 0 |

| | | | | | | | | | | | | | | | | | |
|---|---|---|---|---|---|---|---|---|---|---|---|---|---|---|---|---|---|
| HMX3 | CCTGTCACTTTATGGCC | -1 | 757 | 774 | -243 | -226 | 2,87 | 0,01 | 21,43 | >0.1 | 5,52 | 5,79 | 0 | 1,34 | 0 | 5,92 | 0 |
| HMX3 | GAAAGCTGTTAAAGATA | -1 | 704 | 721 | -296 | -279 | 0,08 | 0,01 | 17,16 | >0.1 | 5,52 | 5,56 | 0 | 1,18 | 0 | 4,81 | 0 |
| HMX3 | CCCAGCCATTTCTGATC | -1 | 166 | 183 | -834 | -817 | -0,78 | 0,01 | 12,24 | >0.1 | 3,18 | 4,43 | 0 | 1,9 | 0 | 3,51 | 0 |
| HMX3 | TGCAAAACTTAAAAAAG | -1 | 73 | 90 | -927 | -910 | 3,95 | 0,01 | 13,98 | >0.1 | 3,82 | 4,33 | 0 | 1,88 | 0 | 0 | 0 |
| HMX2 | GGAGGCACTCAAAGAAG | 1 | 20 | 37 | -980 | -963 | 9,52 | 0,001 | 21,19 | >0.1 | 5,52 | 4,28 | 0 | 1,87 | 0 | 0 | 0 |
| HMX2 | CATGGAAATTAAAACTG | 1 | 150 | 167 | -850 | -833 | 2,05 | 0,01 | 13,28 | >0.1 | 3,26 | 4,41 | 0 | 1,9 | 0 | 1,65 | 0 |
| HMX2 | CCTAGCACTTTGGGAGG | 1 | 205 | 222 | -795 | -778 | 2,63 | 0,01 | 16,42 | >0.1 | 3 | 4,48 | 0 | 1,6 | 0 | 4,7 | 0 |
| HMX2 | ACAAAAACTTAGCTGGG | 1 | 300 | 317 | -700 | -683 | -0,23 | 0,01 | 13,25 | >0.1 | 2,67 | 4,62 | 0 | 1,21 | 0 | 4,99 | 0 |
| HMX2 | ATAACCTATTAATGAAA | 1 | 596 | 613 | -404 | -387 | 5,48 | 0,001 | 16,55 | >0.1 | 3,47 | 5,22 | 0 | 1,02 | 0 | 1,36 | 0 |
| HMX2 | CTGTAAAATTATAACAT | 1 | 679 | 696 | -321 | -304 | -0,24 | 0,01 | 13,78 | >0.1 | 4,35 | 5,47 | 0 | 1,13 | 0 | 3,07 | 0 |
| HMX2 | CTCTAGGATTAAAGAAT | 1 | 786 | 803 | -214 | -197 | 0,59 | 0,01 | 19,03 | >0.1 | 5,52 | 5,93 | 0 | 1,46 | 0 | 5,52 | 0 |
| HMX2 | TGATTCACTTTAAAAAA | 1 | 817 | 834 | -183 | -166 | 0,56 | 0,01 | 17,54 | >0.1 | 5,52 | 6,12 | 0 | 1,65 | 0 | 3,7 | 0 |
| HMX2 | AAAGTCATTCAGTGGAT | 1 | 1009 | 1026 | 9 | 26 | 0,05 | 0,01 | 17,88 | >0.1 | 5,52 | 9,03 | 0 | 1,86 | 0 | 1,42 | 0 |
| HMX2 | AGAAGCAAGTGAACTTT | -1 | 1173 | 1190 | 173 | 190 | 0,86 | 0,01 | 19,22 | >0.1 | 5,52 | 6,13 | 0 | 1,64 | 1,79 | 3,28 | 0 |
| HMX2 | AGCAGCAGTTACAGCAA | -1 | 1083 | 1100 | 83 | 100 | 4,91 | 0,001 | 23,26 | >0.1 | 5,52 | 6,97 | 0 | 1,85 | 1,79 | 2,21 | 0 |
| HMX2 | ACATCCACTGAATGACT | -1 | 1011 | 1028 | 11 | 28 | 3,44 | 0,01 | 21,17 | >0.1 | 5,52 | 8,95 | 0 | 1,86 | 0 | 1,39 | 0 |
| HMX2 | AAGACCAAAAAAAGAAA | -1 | 919 | 936 | -81 | -64 | 1,77 | 0,01 | 17,27 | >0.1 | 5,52 | 7,01 | 0 | 1,88 | 0 | 1,09 | 0 |
| HMX2 | GTAAACAATCTGCTGAG | -1 | 887 | 904 | -113 | -96 | -1,12 | 0,01 | 15,9 | >0.1 | 5,52 | 6,65 | 0 | 1,89 | 0 | 2,95 | 0 |
| HMX2 | TGAGCCAATATAAAGTT | -1 | 874 | 891 | -126 | -109 | 1,29 | 0,01 | 18,19 | >0.1 | 5,52 | 6,54 | 0 | 1,9 | 0 | 2,95 | 0 |
| HMX2 | GTAACAGATTTTAGAAT | -1 | 836 | 853 | -164 | -147 | 0,25 | 0,01 | 16,89 | >0.1 | 5,52 | 6,25 | 0 | 1,8 | 0 | 3,07 | 0 |
| HMX2 | TGAATCAAATAAGAATA | -1 | 807 | 824 | -193 | -176 | 1,61 | 0,01 | 18,04 | >0.1 | 5,52 | 6,05 | 0 | 1,58 | 0 | 3,28 | 0 |
| HMX2 | CCTGTCACTTTATGGCC | -1 | 757 | 774 | -243 | -226 | 1,81 | 0,01 | 20,37 | >0.1 | 5,52 | 5,79 | 0 | 1,34 | 0 | 5,92 | 0 |
| HMX2 | GAAAGCTGTTAAAGATA | -1 | 704 | 721 | -296 | -279 | 0,26 | 0,01 | 17,34 | >0.1 | 5,52 | 5,56 | 0 | 1,18 | 0 | 4,81 | 0 |
| HMX2 | TGCAAAACTTAAAAAAG | -1 | 73 | 90 | -927 | -910 | 1,61 | 0,01 | 11,64 | >0.1 | 3,82 | 4,33 | 0 | 1,88 | 0 | 0 | 0 |
| FOXG1 | GTAGTCAT | 1 | 126 | 134 | -874 | -866 | -6 | 0,01 | 5,4 | >0.1 | 3,43 | 4,38 | 0 | 1,89 | 0 | 1,7 | 0 |
| FOXG1 | GTAATCCT | 1 | 200 | 208 | -800 | -792 | -6 | 0,01 | 7,61 | >0.1 | 3,04 | 4,46 | 0 | 1,63 | 0 | 4,48 | 0 |
| FOXG1 | GTGAACCT | 1 | 374 | 382 | -626 | -618 | -5,5 | 0,01 | 9,37 | >0.1 | 2,55 | 4,72 | 0 | 1,07 | 0 | 6,53 | 0 |
| FOXG1 | ATAAAGAC | 1 | 500 | 508 | -500 | -492 | 1,47 | 0,01 | 16,99 | >0.1 | 2,96 | 4,96 | 0 | 0,98 | 0 | 6,62 | 0 |
| FOXG1 | ATAAAAAT | 1 | 579 | 587 | -421 | -413 | 1,47 | 0,01 | 10,95 | >0.1 | 3,33 | 5,15 | 0 | 1 | 0 | 0 | 0 |
| FOXG1 | GAAAACCA | 1 | 619 | 627 | -381 | -373 | -5,42 | 0,01 | 6,01 | >0.1 | 3,6 | 5,26 | 0 | 1,03 | 0 | 1,55 | 0 |
| FOXG1 | ACAAACAC | 1 | 628 | 636 | -372 | -364 | 1,97 | 0,01 | 13,51 | >0.1 | 3,68 | 5,28 | 0 | 1,04 | 0 | 1,55 | 0 |
| FOXG1 | TTAACCAA | 1 | 731 | 739 | -269 | -261 | -5,24 | 0,01 | 12,78 | >0.1 | 5,64 | 5,63 | 0 | 1,22 | 0 | 5,52 | 0 |
| FOXG1 | TTAAAGAA | 1 | 794 | 802 | -206 | -198 | -5,73 | 0,01 | 12,61 | >0.1 | 5,64 | 5,93 | 0 | 1,45 | 0 | 5,32 | 0 |
| FOXG1 | GTGAACTT | -1 | 1174 | 1182 | 174 | 182 | -6 | 0,01 | 12,28 | >0.1 | 5,64 | 6,13 | 0 | 1,63 | 1,79 | 3,09 | 0 |
| FOXG1 | GTGACCAA | -1 | 672 | 680 | -328 | -320 | -4,92 | 0,01 | 9,3 | >0.1 | 4,16 | 5,42 | 0 | 1,1 | 0 | 3,55 | 0 |
| FOXG1 | GGCAACAT | -1 | 538 | 546 | -462 | -454 | -5,5 | 0,01 | 5,31 | >0.1 | 3,12 | 5,05 | 0 | 0,98 | 0 | 1,66 | 0 |
| JUNB | GAAGGAGGCAC | 1 | 17 | 28 | -983 | -972 | -18,08 | 0,01 | -6,35 | >0.1 | 5,6 | 4,27 | 0 | 1,87 | 0 | 0 | 0 |
| JUNB | GTGTGAGAAAT | 1 | 52 | 63 | -948 | -937 | -6,67 | 0,01 | 3,6 | >0.1 | 4,09 | 4,3 | 0 | 1,87 | 0 | 0 | 0 |
| JUNB | AGAGTAGTCAT | 1 | 123 | 134 | -877 | -866 | -18,29 | 0,01 | -6,89 | >0.1 | 3,44 | 4,38 | 0 | 1,89 | 0 | 1,7 | 0 |
| JUNB | CAGTGGCTCAC | 1 | 185 | 196 | -815 | -804 | -1,95 | 0,001 | 11,2 | >0.1 | 3,1 | 4,45 | 0 | 1,75 | 0 | 3,86 | 0 |
| JUNB | TCCTGACTAAC | 1 | 261 | 272 | -739 | -728 | 4,01 | 0,001 | 17,13 | >0.1 | 2,8 | 4,55 | 0 | 1,32 | 0 | 4,45 | 0 |
| JUNB | GGCTGAGGCAG | 1 | 354 | 365 | -646 | -635 | -3,27 | 0,01 | 11,57 | >0.1 | 2,53 | 4,69 | 0 | 1,1 | 0 | 6,52 | 0 |
| JUNB | CTGGGAGGCAG | 1 | 380 | 391 | -620 | -609 | -18,47 | 0,01 | -4,19 | >0.1 | 2,57 | 4,73 | 0 | 1,06 | 0 | 5,91 | 0 |
| JUNB | CCGAGACTCAG | 1 | 442 | 453 | -558 | -547 | -3,16 | 0,01 | 11,25 | >0.1 | 2,75 | 4,85 | 0 | 1 | 0 | 5,8 | 0 |
| JUNB | CTAGGATTAAA | 1 | 788 | 799 | -212 | -201 | -14,1 | 0,01 | 4,17 | >0.1 | 5,6 | 5,91 | 0 | 1,44 | 0 | 5,32 | 0 |
| JUNB | ATTTGATTCAC | 1 | 814 | 825 | -186 | -175 | -7,33 | 0,01 | 9,34 | >0.1 | 5,6 | 6,06 | 0 | 1,58 | 0 | 3,43 | 0 |
| JUNB | TATTGGCTCAG | 1 | 881 | 892 | -119 | -108 | -16,92 | 0,01 | 0,19 | >0.1 | 5,6 | 6,55 | 0 | 1,89 | 0 | 3,07 | 0 |
| JUNB | GGGAAAGTCAT | 1 | 1006 | 1017 | 6 | 17 | -17,12 | 0,01 | 0,93 | >0.1 | 5,6 | 9,12 | 0 | 1,87 | 0 | 1,46 | 0 |
| JUNB | AAGTCATTCAG | 1 | 1010 | 1021 | 10 | 21 | -5,82 | 0,01 | 12,05 | >0.1 | 5,6 | 8,99 | 0 | 1,86 | 0 | 1,42 | 0 |
| JUNB | TCGTGGTTAAA | -1 | 1142 | 1153 | 142 | 153 | -14,62 | 0,01 | 5,77 | >0.1 | 5,6 | 6,36 | 0 | 1,84 | 3,22 | 3,37 | 0 |
| JUNB | GAAGGAGCCAG | -1 | 1063 | 1074 | 63 | 74 | -17,03 | 0,01 | 1,79 | >0.1 | 5,6 | 7,3 | 0 | 1,85 | 1,79 | 2,27 | 0 |
| JUNB | CTGTGAGCCAA | -1 | 1032 | 1043 | 32 | 43 | -6,15 | 0,01 | 10,73 | >0.1 | 5,6 | 8,15 | 0 | 1,86 | 0 | 1,28 | 0 |
| JUNB | GAATGACTTTC | -1 | 1008 | 1019 | 8 | 19 | -18,81 | 0,01 | -0,86 | >0.1 | 5,6 | 9,07 | 0 | 1,87 | 0 | 1,42 | 0 |
| JUNB | TGCTGAGCCAA | -1 | 883 | 894 | -117 | -106 | -7,89 | 0,01 | 9,23 | >0.1 | 5,6 | 6,57 | 0 | 1,89 | 0 | 3,07 | 0 |
| JUNB | AAGTGAATCAA | -1 | 816 | 827 | -184 | -173 | -5,06 | 0,01 | 11,75 | >0.1 | 5,6 | 6,07 | 0 | 1,6 | 0 | 3,55 | 0 |
| JUNB | CAGTGACCAAA | -1 | 671 | 682 | -329 | -318 | -8,6 | 0,01 | 5,64 | >0.1 | 4,16 | 5,43 | 0 | 1,1 | 0 | 3,55 | 0 |
| JUNB | TTGAGACTGAG | -1 | 448 | 459 | -552 | -541 | -18,94 | 0,01 | -5,05 | >0.1 | 2,77 | 4,86 | 0 | 1 | 0 | 5,26 | 0 |
| JUNB | GACTGAGTCTC | -1 | 444 | 455 | -556 | -545 | -6,05 | 0,01 | 8,49 | >0.1 | 2,76 | 4,85 | 0 | 1 | 0 | 5,92 | 0 |
| JUNB | TCCTGCCTCAG | -1 | 356 | 367 | -644 | -633 | -6,68 | 0,01 | 7,76 | >0.1 | 2,52 | 4,69 | 0 | 1,09 | 0 | 6,13 | 0 |

| | | | | | | | | | | | | | | | | | |
|---|---|---|---|---|---|---|---|---|---|---|---|---|---|---|---|---|---|
| JUNB | GTGTTAGTCAG | -1 | 263 | 274 | -737 | -726 | -2,76 | 0,001 | 10,35 | >0.1 | 2,8 | 4,55 | 0 | 1,32 | 0 | 4,45 | 0 |
| JUNB | GCGTGAGCCAC | -1 | 187 | 198 | -813 | -802 | -3,53 | 0,01 | 9,6 | >0.1 | 3,09 | 4,45 | 0 | 1,73 | 0 | 3,86 | 0 |
| JUNB | TTATGACTACT | -1 | 125 | 136 | -875 | -864 | -7,02 | 0,01 | 4,37 | >0.1 | 3,43 | 4,38 | 0 | 1,89 | 0 | 1,7 | 0 |
| JUNB | GAGTGCCTCCT | -1 | 19 | 30 | -981 | -970 | -16,22 | 0,01 | -4,49 | >0.1 | 5,6 | 4,27 | 0 | 1,87 | 0 | 0 | 0 |
| FOSL2::JUND | AAGGAGGCAC | 1 | 18 | 28 | -982 | -972 | -3,05 | 0,01 | 8,69 | >0.1 | 5,61 | 4,27 | 0 | 1,87 | 0 | 0 | 0 |
| FOSL2::JUND | TGTGAGAAAT | 1 | 53 | 63 | -947 | -937 | 2,86 | 0,01 | 13,12 | >0.1 | 4,09 | 4,3 | 0 | 1,87 | 0 | 0 | 0 |
| FOSL2::JUND | GAGTAGTCAT | 1 | 124 | 134 | -876 | -866 | -3,86 | 0,01 | 7,54 | >0.1 | 3,44 | 4,38 | 0 | 1,89 | 0 | 1,7 | 0 |
| FOSL2::JUND | AGTGGCTCAC | 1 | 186 | 196 | -814 | -804 | 0,83 | 0,01 | 13,97 | >0.1 | 3,1 | 4,45 | 0 | 1,74 | 0 | 3,86 | 0 |
| FOSL2::JUND | CCTGACTAAC | 1 | 262 | 272 | -738 | -728 | 7,79 | 0,001 | 20,99 | >0.1 | 2,8 | 4,55 | 0 | 1,32 | 0 | 4,53 | 0 |
| FOSL2::JUND | GCTGAGGCAG | 1 | 355 | 365 | -645 | -635 | 6,8 | 0,001 | 21,15 | >0.1 | 2,53 | 4,69 | 0 | 1,09 | 0 | 6,03 | 0 |
| FOSL2::JUND | TGGGAGGCAG | 1 | 381 | 391 | -619 | -609 | -3,91 | 0,01 | 10,47 | >0.1 | 2,57 | 4,73 | 0 | 1,06 | 0 | 6,02 | 0 |
| FOSL2::JUND | CGAGACTCAG | 1 | 443 | 453 | -557 | -547 | -0,34 | 0,01 | 14,19 | >0.1 | 2,75 | 4,85 | 0 | 1 | 0 | 5,92 | 0 |
| FOSL2::JUND | TTTGATTCAC | 1 | 815 | 825 | -185 | -175 | 4,9 | 0,001 | 21,58 | >0.1 | 5,61 | 6,06 | 0 | 1,58 | 0 | 3,43 | 0 |
| FOSL2::JUND | ATTGGCTCAG | 1 | 882 | 892 | -118 | -108 | -5,26 | 0,01 | 11,86 | >0.1 | 5,61 | 6,55 | 0 | 1,89 | 0 | 3,07 | 0 |
| FOSL2::JUND | AGTCATTCAG | 1 | 1011 | 1021 | 11 | 21 | -4,76 | 0,01 | 13,06 | >0.1 | 5,61 | 8,95 | 0 | 1,86 | 0 | 1,39 | 0 |
| FOSL2::JUND | CTTGGCTCAC | 1 | 1031 | 1041 | 31 | 41 | -3,47 | 0,01 | 13,5 | >0.1 | 5,61 | 8,19 | 0 | 1,86 | 0 | 1,31 | 0 |
| FOSL2::JUND | ATTGAGGAAC | 1 | 1109 | 1119 | 109 | 119 | 0,14 | 0,01 | 20 | >0.1 | 5,61 | 6,66 | 0 | 1,85 | 3,22 | 2,52 | 0 |
| FOSL2::JUND | TGTGAGCCAA | -1 | 1032 | 1042 | 32 | 42 | -2,42 | 0,01 | 14,47 | >0.1 | 5,61 | 8,15 | 0 | 1,86 | 0 | 1,28 | 0 |
| FOSL2::JUND | GCTGAGCCAA | -1 | 883 | 893 | -117 | -107 | -3,5 | 0,01 | 13,63 | >0.1 | 5,61 | 6,56 | 0 | 1,89 | 0 | 3,07 | 0 |
| FOSL2::JUND | AGTGAATCAA | -1 | 816 | 826 | -184 | -174 | 5,12 | 0,001 | 21,93 | >0.1 | 5,61 | 6,06 | 0 | 1,59 | 0 | 3,55 | 0 |
| FOSL2::JUND | AGTGACCAAA | -1 | 671 | 681 | -329 | -319 | -5,49 | 0,01 | 8,74 | >0.1 | 4,16 | 5,42 | 0 | 1,1 | 0 | 3,55 | 0 |
| FOSL2::JUND | ACTGAGTCTC | -1 | 444 | 454 | -556 | -546 | -2,99 | 0,01 | 11,54 | >0.1 | 2,76 | 4,85 | 0 | 1 | 0 | 5,92 | 0 |
| FOSL2::JUND | CCTGCCTCAG | -1 | 356 | 366 | -644 | -634 | 0,46 | 0,01 | 14,9 | >0.1 | 2,52 | 4,69 | 0 | 1,09 | 0 | 6,13 | 0 |
| FOSL2::JUND | TGTTAGTCAG | -1 | 263 | 273 | -737 | -727 | 4,24 | 0,001 | 17,35 | >0.1 | 2,8 | 4,55 | 0 | 1,32 | 0 | 4,45 | 0 |
| FOSL2::JUND | CGTGAGCCAC | -1 | 187 | 197 | -813 | -803 | 0,43 | 0,01 | 13,56 | >0.1 | 3,1 | 4,45 | 0 | 1,73 | 0 | 3,86 | 0 |
| FOSL2::JUND | TATGACTACT | -1 | 125 | 135 | -875 | -865 | -0,02 | 0,01 | 11,38 | >0.1 | 3,43 | 4,38 | 0 | 1,89 | 0 | 1,7 | 0 |
| FOSL2::JUND | TATTTCTCAC | -1 | 54 | 64 | -946 | -936 | -5,13 | 0,01 | 5,12 | >0.1 | 4,07 | 4,3 | 0 | 1,87 | 0 | 0 | 0 |
| JUND | AAGGAGGCACT | 1 | 18 | 29 | -982 | -971 | -22,84 | 0,01 | -11,11 | >0.1 | 5,6 | 4,27 | 0 | 1,87 | 0 | 0 | 0 |
| JUND | AGTCAGCCACA | 1 | 35 | 46 | -965 | -954 | -22,48 | 0,01 | -11,96 | >0.1 | 4,37 | 4,29 | 0 | 1,87 | 0 | 0 | 0 |
| JUND | TGTGAGAAATA | 1 | 53 | 64 | -947 | -936 | -8,44 | 0,01 | 1,82 | >0.1 | 4,08 | 4,3 | 0 | 1,87 | 0 | 0 | 0 |
| JUND | GAGTAGTCATA | 1 | 124 | 135 | -876 | -865 | -21,03 | 0,01 | -9,63 | >0.1 | 3,44 | 4,38 | 0 | 1,89 | 0 | 1,7 | 0 |
| JUND | AGTGGCTCACG | 1 | 186 | 197 | -814 | -803 | -2,62 | 0,001 | 10,52 | >0.1 | 3,1 | 4,45 | 0 | 1,74 | 0 | 3,86 | 0 |
| JUND | CCTGACTAACA | 1 | 262 | 273 | -738 | -727 | 2,28 | 0,001 | 15,48 | >0.1 | 2,8 | 4,55 | 0 | 1,32 | 0 | 4,53 | 0 |
| JUND | GCTGAGGCAGG | 1 | 355 | 366 | -645 | -634 | -5,79 | 0,01 | 8,56 | >0.1 | 2,53 | 4,69 | 0 | 1,09 | 0 | 6,03 | 0 |
| JUND | CGAGACTCAGT | 1 | 443 | 454 | -557 | -546 | -9,47 | 0,01 | 5,06 | >0.1 | 2,76 | 4,85 | 0 | 1 | 0 | 5,92 | 0 |
| JUND | GCAAAGTCATG | 1 | 512 | 523 | -488 | -477 | -21,08 | 0,01 | -9,14 | >0.1 | 3,01 | 4,99 | 0 | 0,98 | 0 | 2,95 | 0 |
| JUND | TTTGATTCACT | 1 | 815 | 826 | -185 | -174 | -8,46 | 0,01 | 8,22 | >0.1 | 5,6 | 6,06 | 0 | 1,59 | 0 | 3,43 | 0 |
| JUND | ATTGGCTCAGC | 1 | 882 | 893 | -118 | -107 | -20,52 | 0,01 | -3,41 | >0.1 | 5,6 | 6,56 | 0 | 1,89 | 0 | 3,07 | 0 |
| JUND | GGAAAGTCATT | 1 | 1007 | 1018 | 7 | 18 | -20,79 | 0,01 | -2,74 | >0.1 | 5,6 | 9,12 | 0 | 1,87 | 0 | 1,46 | 0 |
| JUND | AGTCATTCAGT | 1 | 1011 | 1022 | 11 | 22 | -10,12 | 0,01 | 7,69 | >0.1 | 5,6 | 8,95 | 0 | 1,86 | 0 | 1,39 | 0 |
| JUND | AAGGAGCCAGG | -1 | 1062 | 1073 | 62 | 73 | -22,34 | 0,01 | -3,57 | >0.1 | 5,6 | 7,31 | 0 | 1,86 | 1,79 | 2,21 | 0 |
| JUND | TGTGAGCCAAG | -1 | 1031 | 1042 | 31 | 42 | -18,52 | 0,01 | -1,57 | >0.1 | 5,6 | 8,19 | 0 | 1,86 | 0 | 1,31 | 0 |
| JUND | AATGACTTTCC | -1 | 1007 | 1018 | 7 | 18 | -21,65 | 0,01 | -3,6 | >0.1 | 5,6 | 9,12 | 0 | 1,87 | 0 | 1,46 | 0 |
| JUND | GCTGAGCCAAT | -1 | 882 | 893 | -118 | -107 | -21,44 | 0,01 | -4,33 | >0.1 | 5,6 | 6,56 | 0 | 1,89 | 0 | 3,07 | 0 |
| JUND | AGTGAATCAAA | -1 | 815 | 826 | -185 | -174 | -19,59 | 0,01 | -2,91 | >0.1 | 5,6 | 6,06 | 0 | 1,59 | 0 | 3,43 | 0 |
| JUND | AGTGACCAAAG | -1 | 670 | 681 | -330 | -319 | -22,84 | 0,01 | -8,47 | >0.1 | 4,15 | 5,42 | 0 | 1,1 | 0 | 3,7 | 0 |
| JUND | ACTGAGTCTCG | -1 | 443 | 454 | -557 | -546 | -5,77 | 0,01 | 8,76 | >0.1 | 2,76 | 4,85 | 0 | 1 | 0 | 5,92 | 0 |
| JUND | CCTGCCTCAGC | -1 | 355 | 366 | -645 | -634 | -9 | 0,01 | 5,35 | >0.1 | 2,53 | 4,69 | 0 | 1,09 | 0 | 6,03 | 0 |
| JUND | TGTTAGTCAGG | -1 | 262 | 273 | -738 | -727 | -5,52 | 0,01 | 7,68 | >0.1 | 2,8 | 4,55 | 0 | 1,32 | 0 | 4,53 | 0 |
| JUND | CGTGAGCCACT | -1 | 186 | 197 | -814 | -803 | -8,51 | 0,01 | 4,63 | >0.1 | 3,1 | 4,45 | 0 | 1,74 | 0 | 3,86 | 0 |
| JUND | TATGACTACTC | -1 | 124 | 135 | -876 | -865 | -6,18 | 0,01 | 5,22 | >0.1 | 3,44 | 4,38 | 0 | 1,89 | 0 | 1,7 | 0 |
| JUND | TGTGGCTGACT | -1 | 35 | 46 | -965 | -954 | -21,84 | 0,01 | -11,32 | >0.1 | 4,37 | 4,29 | 0 | 1,87 | 0 | 0 | 0 |
| JUND | GCTGACTTCTT | -1 | 31 | 42 | -969 | -958 | -21,49 | 0,01 | -10,89 | >0.1 | 4,45 | 4,28 | 0 | 1,87 | 0 | 0 | 0 |
| JUND | AGTGCCTCCTT | -1 | 18 | 29 | -982 | -971 | -19,23 | 0,01 | -7,5 | >0.1 | 5,6 | 4,27 | 0 | 1,87 | 0 | 0 | 0 |
| RHOXF1 | ATGAGCTT | 1 | 68 | 76 | -932 | -924 | 5,84 | 0,01 | 15,95 | >0.1 | 3,92 | 4,31 | 0 | 1,88 | 0 | 0 | 0 |
| RHOXF1 | GTAATCCT | 1 | 200 | 208 | -800 | -792 | 8,41 | 0,01 | 22,02 | >0.1 | 3,04 | 4,46 | 0 | 1,63 | 0 | 4,48 | 0 |
| RHOXF1 | CTTAGCTG | 1 | 307 | 315 | -693 | -685 | 4,07 | 0,01 | 17,62 | >0.1 | 2,66 | 4,61 | 0 | 1,19 | 0 | 5,09 | 0 |
| RHOXF1 | CAGAGCTT | 1 | 388 | 396 | -612 | -604 | 4,11 | 0,01 | 18,62 | >0.1 | 2,59 | 4,74 | 0 | 1,05 | 0 | 6,13 | 0 |
| RHOXF1 | GTGAGCCG | 1 | 399 | 407 | -601 | -593 | 8,79 | 0,001 | 23,12 | >0.1 | 2,62 | 4,76 | 0 | 1,04 | 0 | 5,91 | 0 |

| | | | | | | | | | | | | | | | | | |
|---|---|---|---|---|---|---|---|---|---|---|---|---|---|---|---|---|---|
| RHOXF1 | ATGATCTT | 1 | 568 | 576 | -432 | -424 | 6,4 | 0,01 | 15,78 | >0.1 | 3,27 | 5,12 | 0 | 0,99 | 0 | 0 | 0 |
| RHOXF1 | ATTATCTT | 1 | 640 | 648 | -360 | -352 | 3,93 | 0,01 | 15,74 | >0.1 | 3,79 | 5,32 | 0 | 1,05 | 0 | 1,66 | 0 |
| RHOXF1 | ATGATCCC | 1 | 649 | 657 | -351 | -343 | 10,36 | 0,0001 | 24,47 | >0.1 | 3,87 | 5,34 | 0 | 1,06 | 0 | 3,83 | 0 |
| RHOXF1 | TGTATCTT | 1 | 702 | 710 | -298 | -290 | -0,1 | 0,01 | 17,2 | >0.1 | 5,64 | 5,52 | 0 | 1,15 | 0 | 4,99 | 0 |
| RHOXF1 | AAAATCTG | 1 | 841 | 849 | -159 | -151 | 4,5 | 0,01 | 21,31 | >0.1 | 5,64 | 6,22 | 0 | 1,76 | 0 | 3,19 | 0 |
| RHOXF1 | CATATCTG | 1 | 852 | 860 | -148 | -140 | 2,69 | 0,01 | 19,16 | >0.1 | 5,64 | 6,3 | 0 | 1,86 | 0 | 2,67 | 0 |
| RHOXF1 | AAAATCTC | 1 | 979 | 987 | -21 | -13 | 5,2 | 0,01 | 22,42 | >0.1 | 5,64 | 8,18 | 0 | 1,87 | 0 | 1,53 | 0 |
| RHOXF1 | GTGATCTT | 1 | 1026 | 1034 | 26 | 34 | 6,1 | 0,01 | 23,3 | >0.1 | 5,64 | 8,45 | 0 | 1,86 | 0 | 1,25 | 0 |
| RHOXF1 | CGAAGCCG | 1 | 1150 | 1158 | 150 | 158 | 4,83 | 0,01 | 21,58 | >0.1 | 4,58 | 6,3 | 0 | 1,84 | 1,79 | 2,24 | 0 |
| RHOXF1 | AGGAGCCA | -1 | 1064 | 1072 | 64 | 72 | 5,18 | 0,01 | 24,02 | >0.1 | 5,64 | 7,28 | 0 | 1,85 | 1,79 | 2,27 | 0 |
| RHOXF1 | AAGAGCTT | -1 | 1054 | 1062 | 54 | 62 | 3,9 | 0,01 | 21,09 | >0.1 | 5,64 | 7,48 | 0 | 1,86 | 0 | 2,21 | 0 |
| RHOXF1 | GTGAGCCA | -1 | 1033 | 1041 | 33 | 41 | 8,9 | 0,001 | 25,78 | >0.1 | 5,64 | 8,1 | 0 | 1,86 | 0 | 1,28 | 0 |
| RHOXF1 | CTGAGCCA | -1 | 884 | 892 | -116 | -108 | 9,41 | 0,001 | 26,56 | >0.1 | 5,64 | 6,55 | 0 | 1,89 | 0 | 3,07 | 0 |
| RHOXF1 | TTAATCCT | -1 | 790 | 798 | -210 | -202 | 8,7 | 0,01 | 27,2 | >0.1 | 5,64 | 5,91 | 0 | 1,43 | 0 | 5,52 | 0 |
| RHOXF1 | GAAAGCTG | -1 | 713 | 721 | -287 | -279 | 3,63 | 0,01 | 21 | >0.1 | 5,64 | 5,56 | 0 | 1,18 | 0 | 4,99 | 0 |
| RHOXF1 | GTTATCTC | -1 | 593 | 601 | -407 | -399 | 4,83 | 0,01 | 14,44 | >0.1 | 3,42 | 5,18 | 0 | 1,01 | 0 | 0 | 0 |
| RHOXF1 | AGTAGCTG | -1 | 341 | 349 | -659 | -651 | -0,16 | 0,01 | 14,59 | >0.1 | 2,57 | 4,66 | 0 | 1,12 | 0 | 6,4 | 0 |
| RHOXF1 | GTGATCTG | -1 | 232 | 240 | -768 | -760 | 6,59 | 0,01 | 20,46 | >0.1 | 2,91 | 4,51 | 0 | 1,44 | 0 | 5,01 | 0 |
| RHOXF1 | GTGAGCCA | -1 | 188 | 196 | -812 | -804 | 8,9 | 0,001 | 22,02 | >0.1 | 3,1 | 4,45 | 0 | 1,72 | 0 | 3,86 | 0 |
| RHOXF1 | CTGATCTG | -1 | 94 | 102 | -906 | -898 | 7,1 | 0,01 | 17,8 | >0.1 | 3,67 | 4,34 | 0 | 1,88 | 0 | 0,81 | 0 |
| RHOXF1 | AAAAGCTC | -1 | 70 | 78 | -930 | -922 | 4,64 | 0,01 | 14,73 | >0.1 | 3,9 | 4,32 | 0 | 1,88 | 0 | 0 | 0 |
| STAT5A::STAT5B | CACTCAAAGAA | 1 | 25 | 36 | -975 | -964 | -8,81 | 0,01 | 2,93 | >0.1 | 5,6 | 4,28 | 0 | 1,87 | 0 | 0 | 0 |
| STAT5A::STAT5B | AGTCCCAGCTA | 1 | 336 | 347 | -664 | -653 | -8,3 | 0,01 | 5,98 | >0.1 | 2,58 | 4,66 | 0 | 1,13 | 0 | 5,92 | 0 |
| STAT5A::STAT5B | AATTCCATAAA | 1 | 494 | 505 | -506 | -495 | -6,6 | 0,01 | 8,71 | >0.1 | 2,94 | 4,95 | 0 | 0,98 | 0 | 6,44 | 0 |
| STAT5A::STAT5B | TTGCCCAAGTA | 1 | 541 | 552 | -459 | -448 | -6,65 | 0,01 | 2,54 | >0.1 | 3,14 | 5,06 | 0 | 0,99 | 0 | 0 | 0 |
| STAT5A::STAT5B | GTTTCTGGGAA | 1 | 556 | 567 | -444 | -433 | -1,99 | 0,01 | 7,31 | >0.1 | 3,22 | 5,1 | 0 | 0,99 | 0 | 0 | 0 |
| STAT5A::STAT5B | AATTGTCTGAA | 1 | 611 | 622 | -389 | -378 | -8,18 | 0,01 | 3,14 | >0.1 | 3,56 | 5,24 | 0 | 1,03 | 0 | 1,5 | 0 |
| STAT5A::STAT5B | TTTTCCGTGTA | 1 | 695 | 706 | -305 | -294 | -6,51 | 0,01 | 8,69 | >0.1 | 5,6 | 5,51 | 0 | 1,15 | 0 | 2,95 | 0 |
| STAT5A::STAT5B | GCTTTCTAGGA | 1 | 715 | 726 | -285 | -274 | -8,54 | 0,01 | 9 | >0.1 | 5,6 | 5,58 | 0 | 1,19 | 0 | 5,17 | 0 |
| STAT5A::STAT5B | CTTTCTAGGAA | 1 | 716 | 727 | -284 | -273 | 11,69 | 0,001 | 29,23 | >0.1 | 5,6 | 5,59 | 0 | 1,19 | 0 | 5,17 | 0 |
| STAT5A::STAT5B | TGTTTACTGTG | 1 | 897 | 908 | -103 | -92 | -8,48 | 0,01 | 8,55 | >0.1 | 5,6 | 6,69 | 0 | 1,89 | 0 | 2,86 | 0 |
| STAT5A::STAT5B | AATCTCATGAG | 1 | 981 | 992 | -19 | -8 | -2,31 | 0,01 | 15,12 | >0.1 | 5,6 | 8,43 | 0 | 1,87 | 0 | 1,53 | 0 |
| STAT5A::STAT5B | TGTTCTATCAA | 1 | 1164 | 1175 | 164 | 175 | -8,29 | 0,01 | 8,59 | >0.1 | 4,5 | 6,19 | 0 | 1,72 | 1,79 | 2,67 | 0 |
| STAT5A::STAT5B | GCTTCTTGGAA | 1 | 1184 | 1195 | 184 | 195 | 10,89 | 0,001 | 29,57 | >0.1 | 5,6 | 6,06 | 0 | 1,55 | 1,79 | 3,67 | 0 |
| STAT5A::STAT5B | AATTCCAAGAA | -1 | 1186 | 1197 | 186 | 197 | 14,04 | 0,0001 | 32,63 | >0.1 | 5,6 | 6,05 | 0 | 1,54 | 1,79 | 3,61 | 0 |
| STAT5A::STAT5B | CTTTGATAGAA | -1 | 1166 | 1177 | 166 | 177 | -8,44 | 0,01 | 8,66 | >0.1 | 4,54 | 6,18 | 0 | 1,7 | 1,79 | 2,88 | 0 |
| STAT5A::STAT5B | CGTCCCCTGTG | -1 | 1038 | 1049 | 38 | 49 | 1,89 | 0,01 | 18,6 | >0.1 | 5,6 | 7,92 | 0 | 1,86 | 0 | 1,34 | 0 |
| STAT5A::STAT5B | CATCCACTGAA | -1 | 1016 | 1027 | 16 | 27 | 0,79 | 0,01 | 18,42 | >0.1 | 5,6 | 8,81 | 0 | 1,86 | 0 | 1,36 | 0 |
| STAT5A::STAT5B | TTTCCCTAGAC | -1 | 1001 | 1012 | 1 | 12 | -4,3 | 0,01 | 13,75 | >0.1 | 5,6 | 9,12 | 0 | 1,87 | 0 | 1,46 | 0 |
| STAT5A::STAT5B | TCATCCTGGAG | -1 | 864 | 875 | -136 | -125 | -6,55 | 0,01 | 10,1 | >0.1 | 5,6 | 6,41 | 0 | 1,9 | 0 | 2,75 | 0 |
| STAT5A::STAT5B | TTTTTAAAGTG | -1 | 822 | 833 | -178 | -167 | -4,01 | 0,01 | 13,03 | >0.1 | 5,6 | 6,11 | 0 | 1,64 | 0 | 3,7 | 0 |
| STAT5A::STAT5B | TAATCCTAGAG | -1 | 786 | 797 | -214 | -203 | -6,86 | 0,01 | 11,76 | >0.1 | 5,6 | 5,9 | 0 | 1,43 | 0 | 5,69 | 0 |
| STAT5A::STAT5B | TTTTCCTAGAA | -1 | 718 | 729 | -282 | -271 | 14,11 | 0,0001 | 31,49 | >0.1 | 5,6 | 5,59 | 0 | 1,2 | 0 | 4,99 | 0 |
| STAT5A::STAT5B | TTTCCTAGAAA | -1 | 717 | 728 | -283 | -272 | -7,73 | 0,01 | 9,82 | >0.1 | 5,6 | 5,59 | 0 | 1,2 | 0 | 5,17 | 0 |
| STAT5A::STAT5B | ATTTTACAGTG | -1 | 677 | 688 | -323 | -312 | -2,93 | 0,01 | 11,31 | >0.1 | 4,26 | 5,45 | 0 | 1,11 | 0 | 3,43 | 0 |
| STAT5A::STAT5B | TATTCCCAGAA | -1 | 558 | 569 | -442 | -431 | 13,55 | 0,0001 | 22,87 | >0.1 | 3,23 | 5,1 | 0 | 0,99 | 0 | 0 | 0 |
| STAT5A::STAT5B | ATTCCCAGAAA | -1 | 557 | 568 | -443 | -432 | -4,56 | 0,01 | 4,75 | >0.1 | 3,22 | 5,1 | 0 | 0,99 | 0 | 0 | 0 |
| STAT5A::STAT5B | CCTTCCAAATA | -1 | 523 | 534 | -477 | -466 | -5,7 | 0,01 | 6,55 | >0.1 | 3,06 | 5,02 | 0 | 0,98 | 0 | 3,19 | 0 |
| STAT5A::STAT5B | TCTTTATGGAA | -1 | 496 | 507 | -504 | -493 | -0,73 | 0,01 | 14,59 | >0.1 | 2,94 | 4,96 | 0 | 0,98 | 0 | 6,44 | 0 |
| STAT5A::STAT5B | TTTTTTTTGAG | -1 | 454 | 465 | -546 | -535 | -1,9 | 0,01 | 12,28 | >0.1 | 2,79 | 4,87 | 0 | 1 | 0 | 5,52 | 0 |
| STAT5A::STAT5B | CCTCCCAGGTT | -1 | 377 | 388 | -623 | -612 | -8,76 | 0,01 | 5,99 | >0.1 | 2,56 | 4,73 | 0 | 1,06 | 0 | 6,4 | 0 |
| STAT5A::STAT5B | CCTCCCGAGTA | -1 | 345 | 356 | -655 | -644 | -6,2 | 0,01 | 8,66 | >0.1 | 2,55 | 4,68 | 0 | 1,11 | 0 | 6,52 | 0 |
| STAT5A::STAT5B | ACTTCTTTGAG | -1 | 27 | 38 | -973 | -962 | 5,35 | 0,01 | 16,04 | >0.1 | 4,54 | 4,28 | 0 | 1,87 | 0 | 0 | 0 |
| NR1A4::RXRA | GAAGTCAGCCACAGA | 1 | 33 | 48 | -967 | -952 | 0,59 | 0,01 | 11,12 | >0.1 | 4,37 | 4,29 | 0 | 1,87 | 0 | 0 | 0 |
| NR1A4::RXRA | AGGAGAGTTGACCTG | 1 | 100 | 115 | -900 | -885 | 3,69 | 0,01 | 14,29 | >0.1 | 3,59 | 4,36 | 0 | 1,89 | 0 | 0,77 | 0 |
| NR1A4::RXRA | ATGGCCATGGAAATT | 1 | 145 | 160 | -855 | -840 | 1,07 | 0,01 | 12,28 | >0.1 | 3,3 | 4,41 | 0 | 1,9 | 0 | 1,61 | 0 |
| NR1A4::RXRA | CAGGAGAATGGCGTG | 1 | 362 | 377 | -638 | -623 | -0,26 | 0,01 | 14,68 | >0.1 | 2,53 | 4,71 | 0 | 1,09 | 0 | 6,61 | 0 |
| NR1A4::RXRA | AATGGCGTGAACCTG | 1 | 368 | 383 | -632 | -617 | 3,54 | 0,01 | 18,15 | >0.1 | 2,54 | 4,72 | 0 | 1,08 | 0 | 6,27 | 0 |
| NR1A4::RXRA | GAGATCGCGCCACTG | 1 | 406 | 421 | -594 | -579 | 1,5 | 0,01 | 15,62 | >0.1 | 2,65 | 4,79 | 0 | 1,03 | 0 | 5,65 | 0 |

| | | | | | | | | | | | | | | | | | |
|---|---|---|---|---|---|---|---|---|---|---|---|---|---|---|---|---|---|
| NR1A4::RXRA | AAGGGCAAAGTCATG | 1 | 508 | 523 | -492 | -477 | 2,93 | 0,01 | 14,77 | >0.1 | 3 | 4,99 | 0 | 0,99 | 0 | 2,86 | 0 |
| NR1A4::RXRA | AAGTTCAAAGGCTGA | 1 | 956 | 971 | -44 | -29 | -0,01 | 0,01 | 16,8 | >0.1 | 5,54 | 7,65 | 0 | 1,88 | 0 | 1,74 | 0 |
| NR1A4::RXRA | GGGGACGATGTCAAG | 1 | 1042 | 1057 | 42 | 57 | 0,92 | 0,01 | 16,74 | >0.1 | 5,54 | 7,79 | 0 | 1,86 | 0 | 0,62 | 0 |
| NR1A4::RXRA | GAAGCCGAAGACCTG | 1 | 1151 | 1166 | 151 | 166 | 4,16 | 0,01 | 20,9 | >0.1 | 4,47 | 6,29 | 0 | 1,84 | 1,79 | 2,34 | 0 |
| NR1A4::RXRA | AAGTTCACTTGCTTC | 1 | 1174 | 1189 | 174 | 189 | 5,17 | 0,01 | 23,54 | >0.1 | 5,54 | 6,13 | 0 | 1,63 | 1,79 | 3,28 | 0 |
| NR1A4::RXRA | GAAGCAAGTGAACTT | -1 | 1174 | 1189 | 174 | 189 | 0,47 | 0,01 | 18,84 | >0.1 | 5,54 | 6,13 | 0 | 1,63 | 1,79 | 3,28 | 0 |
| NR1A4::RXRA | CAGGTCTTCGGCTTC | -1 | 1151 | 1166 | 151 | 166 | 5,49 | 0,01 | 22,23 | >0.1 | 4,47 | 6,29 | 0 | 1,84 | 1,79 | 2,34 | 0 |
| NR1A4::RXRA | AATTTCATTAATAGG | -1 | 600 | 615 | -400 | -385 | -0,17 | 0,01 | 10,98 | >0.1 | 3,49 | 5,22 | 0 | 1,02 | 0 | 1,41 | 0 |
| NR1A4::RXRA | CATGACTTTGCCCTT | -1 | 508 | 523 | -492 | -477 | 1,74 | 0,01 | 13,58 | >0.1 | 3 | 4,99 | 0 | 0,99 | 0 | 2,86 | 0 |
| NR1A4::RXRA | TCGGTCTGTCGCCCA | -1 | 431 | 446 | -569 | -554 | 1,23 | 0,01 | 15,61 | >0.1 | 2,72 | 4,84 | 0 | 1,01 | 0 | 5,8 | 0 |
| NR1A4::RXRA | CGGTCTGTCGCCCAG | -1 | 430 | 445 | -570 | -555 | 0,48 | 0,01 | 14,72 | >0.1 | 2,72 | 4,83 | 0 | 1,01 | 0 | 5,67 | 0 |
| NR1A4::RXRA | GAGCACAGTGGCGCG | -1 | 411 | 426 | -589 | -574 | 0,65 | 0,01 | 14,23 | >0.1 | 2,66 | 4,8 | 0 | 1,03 | 0 | 5,09 | 0 |
| NR1A4::RXRA | CGGCTCACTGCAAGC | -1 | 392 | 407 | -608 | -593 | 1,02 | 0,01 | 15,35 | >0.1 | 2,61 | 4,76 | 0 | 1,05 | 0 | 5,91 | 0 |
| NR1A4::RXRA | CAGGTTCACGCCATT | -1 | 368 | 383 | -632 | -617 | 1,23 | 0,01 | 15,84 | >0.1 | 2,54 | 4,72 | 0 | 1,08 | 0 | 6,27 | 0 |
| NR1A4::RXRA | AGGTTCACGCCATTC | -1 | 367 | 382 | -633 | -618 | 0,27 | 0,01 | 14,76 | >0.1 | 2,54 | 4,72 | 0 | 1,08 | 0 | 6,15 | 0 |
| NR1A4::RXRA | CACGCCATTCTCCTG | -1 | 362 | 377 | -638 | -623 | 3,87 | 0,01 | 18,81 | >0.1 | 2,53 | 4,71 | 0 | 1,09 | 0 | 6,61 | 0 |
| NR1A4::RXRA | CAGGTCAACTCTCCT | -1 | 100 | 115 | -900 | -885 | 4,95 | 0,01 | 15,55 | >0.1 | 3,59 | 4,36 | 0 | 1,89 | 0 | 0,77 | 0 |
| NR1A4::RXRA | TCCTTCACTTACCTA | -1 | 2 | 17 | -998 | -983 | 0,65 | 0,01 | 14,11 | >0.1 | 5,54 | 4,26 | 0 | 1,86 | 1,79 | 0 | 0 |
| ELF1 | GAGAAGGAGGCA | 1 | 15 | 27 | -985 | -973 | -12,22 | 0,01 | -0,5 | >0.1 | 5,58 | 4,27 | 0 | 1,87 | 0 | 0 | 0 |
| ELF1 | CTCAAAGAAGTC | 1 | 27 | 39 | -973 | -961 | -10,21 | 0,01 | 0,47 | >0.1 | 4,53 | 4,28 | 0 | 1,87 | 0 | 0 | 0 |
| ELF1 | AAATAGGAAATG | 1 | 59 | 71 | -941 | -929 | -4,82 | 0,01 | 5,36 | >0.1 | 4 | 4,31 | 0 | 1,88 | 0 | 0 | 0 |
| ELF1 | GCCATGGAAATT | 1 | 148 | 160 | -852 | -840 | -8,69 | 0,01 | 2,55 | >0.1 | 3,29 | 4,41 | 0 | 1,9 | 0 | 1,65 | 0 |
| ELF1 | GGTCAGGAGATA | 1 | 242 | 254 | -758 | -746 | -11,1 | 0,01 | 2,02 | >0.1 | 2,87 | 4,52 | 0 | 1,4 | 0 | 4,33 | 0 |
| ELF1 | TAACCAAAAGTA | 1 | 732 | 744 | -268 | -256 | -13,43 | 0,01 | 4,96 | >0.1 | 5,58 | 5,66 | 0 | 1,24 | 0 | 5,92 | 0 |
| ELF1 | CTCCAGGATGAA | 1 | 864 | 876 | -136 | -124 | -2,36 | 0,01 | 14,21 | >0.1 | 5,58 | 6,41 | 0 | 1,9 | 0 | 2,67 | 0 |
| ELF1 | AGAGAGAAAATC | 1 | 973 | 985 | -27 | -15 | -13,54 | 0,01 | 3,62 | >0.1 | 5,58 | 8,09 | 0 | 1,87 | 0 | 1,61 | 0 |
| ELF1 | CATGAGGAGGTT | 1 | 986 | 998 | -14 | -2 | -9,91 | 0,01 | 7,81 | >0.1 | 5,58 | 8,66 | 0 | 1,87 | 0 | 1,61 | 0 |
| ELF1 | TCAGTGGATGTG | 1 | 1017 | 1029 | 17 | 29 | -10,43 | 0,01 | 7,14 | >0.1 | 5,58 | 8,79 | 0 | 1,86 | 0 | 1,34 | 0 |
| ELF1 | AACCACGAAGCC | 1 | 1145 | 1157 | 145 | 157 | -2,26 | 0,01 | 16,88 | >0.1 | 5,58 | 6,33 | 0 | 1,84 | 3,22 | 2,16 | 0 |
| ELF1 | GAAGCCGAAGAC | 1 | 1151 | 1163 | 151 | 163 | -8,68 | 0,01 | 8,01 | >0.1 | 4,51 | 6,29 | 0 | 1,84 | 1,79 | 2,26 | 0 |
| ELF1 | TTCCAAGAAGCA | -1 | 1183 | 1195 | 183 | 195 | -6,92 | 0,01 | 11,76 | >0.1 | 5,58 | 6,07 | 0 | 1,56 | 1,79 | 3,67 | 0 |
| ELF1 | AAGAAGCAAGTG | -1 | 1179 | 1191 | 179 | 191 | -7,33 | 0,01 | 11,03 | >0.1 | 5,58 | 6,1 | 0 | 1,59 | 1,79 | 3,29 | 0 |
| ELF1 | AGCCAGGAAGAG | -1 | 1057 | 1069 | 57 | 69 | 5,3 | 0,001 | 22,37 | >0.1 | 5,58 | 7,42 | 0 | 1,86 | 0 | 2,21 | 0 |
| ELF1 | GAGCAGGAAAGA | -1 | 932 | 944 | -68 | -56 | -6,73 | 0,01 | 8,78 | >0.1 | 5,58 | 7,12 | 0 | 1,88 | 0 | 0,93 | 0 |
| ELF1 | AAAGAAGAAGAA | -1 | 908 | 920 | -92 | -80 | -12,92 | 0,01 | 2,57 | >0.1 | 5,58 | 6,81 | 0 | 1,89 | 0 | 1,21 | 0 |
| ELF1 | ACACAGTAAACA | -1 | 897 | 909 | -103 | -91 | -11,56 | 0,01 | 5,47 | >0.1 | 5,58 | 6,7 | 0 | 1,89 | 0 | 2,86 | 0 |
| ELF1 | AAACCGGTACTT | -1 | 739 | 751 | -261 | -249 | -12,11 | 0,01 | 6,33 | >0.1 | 5,58 | 5,68 | 0 | 1,26 | 0 | 5,92 | 0 |
| ELF1 | TACACGGAAAAT | -1 | 694 | 706 | -306 | -294 | 1,81 | 0,001 | 17 | >0.1 | 5,58 | 5,51 | 0 | 1,15 | 0 | 2,95 | 0 |
| ELF1 | GATCATGAAGAT | -1 | 643 | 655 | -357 | -345 | -13,04 | 0,01 | 0,89 | >0.1 | 3,83 | 5,34 | 0 | 1,06 | 0 | 3,7 | 0 |
| ELF1 | TTCCCAGAAACT | -1 | 555 | 567 | -445 | -433 | -6,11 | 0,01 | 3,19 | >0.1 | 3,21 | 5,1 | 0 | 0,99 | 0 | 0 | 0 |
| ELF3 | AATAGGTAAGTGA | 1 | 0 | 13 | -1000 | -987 | -5,7 | 0,01 | 12,26 | >0.1 | 5,57 | 4,26 | 0 | 1,86 | 1,79 | 0 | 4,47 |
| ELF3 | GAGAAGGAGGCAC | 1 | 15 | 28 | -985 | -972 | -8,09 | 0,01 | 8,09 | >0.1 | 5,57 | 4,27 | 0 | 1,87 | 0 | 0 | 4,47 |
| ELF3 | CTCAAAGAAGTCA | 1 | 27 | 40 | -973 | -960 | -4,07 | 0,01 | 11,07 | >0.1 | 4,52 | 4,28 | 0 | 1,87 | 0 | 0 | 4,47 |
| ELF3 | AAATAGGAAATGA | 1 | 59 | 72 | -941 | -928 | -5 | 0,01 | 9,65 | >0.1 | 3,99 | 4,31 | 0 | 1,88 | 0 | 0 | 4,47 |
| ELF3 | GCCATGGAAATTA | 1 | 148 | 161 | -852 | -839 | -9,54 | 0,01 | 6,17 | >0.1 | 3,29 | 4,41 | 0 | 1,9 | 0 | 1,65 | 4,47 |
| ELF3 | TAACCAAAAGTAC | 1 | 732 | 745 | -268 | -255 | -8,7 | 0,01 | 14,16 | >0.1 | 5,57 | 5,66 | 0 | 1,24 | 0 | 5,92 | 4,47 |
| ELF3 | CTCCAGGATGAAC | 1 | 864 | 877 | -136 | -123 | -1,21 | 0,01 | 19,82 | >0.1 | 5,57 | 6,42 | 0 | 1,9 | 0 | 2,67 | 4,47 |
| ELF3 | CATGAGGAGGTTT | 1 | 986 | 999 | -14 | -1 | -8,62 | 0,01 | 13,59 | >0.1 | 5,57 | 8,69 | 0 | 1,87 | 0 | 1,61 | 4,47 |
| ELF3 | AACCACGAAGCCG | 1 | 1145 | 1158 | 145 | 158 | -3,96 | 0,01 | 19,64 | >0.1 | 5,57 | 6,33 | 0 | 1,84 | 3,22 | 2,16 | 4,47 |
| ELF3 | TTCCAAGAAGCAA | -1 | 1182 | 1195 | 182 | 195 | -1 | 0,01 | 22 | >0.1 | 5,57 | 6,08 | 0 | 1,57 | 1,79 | 3,52 | 4,47 |
| ELF3 | AAGAAGCAAGTGA | -1 | 1178 | 1191 | 178 | 191 | 3,08 | 0,001 | 25,96 | >0.1 | 5,57 | 6,1 | 0 | 1,6 | 1,79 | 3,34 | 4,47 |
| ELF3 | AACAAGGCTGAGA | -1 | 1072 | 1085 | 72 | 85 | -8,09 | 0,01 | 15,01 | >0.1 | 5,57 | 7,14 | 0 | 1,85 | 1,79 | 2,27 | 4,47 |
| ELF3 | AGCCAGGAAGAGC | -1 | 1056 | 1069 | 56 | 69 | 2,22 | 0,001 | 23,83 | >0.1 | 5,57 | 7,44 | 0 | 1,86 | 0 | 2,27 | 4,47 |
| ELF3 | GAGCAGGAAAGAC | -1 | 931 | 944 | -69 | -56 | 1,42 | 0,001 | 21,39 | >0.1 | 5,57 | 7,12 | 0 | 1,88 | 0 | 0,93 | 4,47 |
| ELF3 | GAAAAAGAAGAAG | -1 | 910 | 923 | -90 | -77 | -8,29 | 0,01 | 11,64 | >0.1 | 5,57 | 6,84 | 0 | 1,89 | 0 | 1,15 | 4,47 |
| ELF3 | AAAGAAGAAGAAC | -1 | 907 | 920 | -93 | -80 | -7,35 | 0,01 | 12,6 | >0.1 | 5,57 | 6,81 | 0 | 1,89 | 0 | 1,21 | 4,47 |
| ELF3 | TACACGGAAAATG | -1 | 693 | 706 | -307 | -294 | 4,52 | 0,001 | 24,29 | >0.1 | 5,57 | 5,51 | 0 | 1,15 | 0 | 3,07 | 4,47 |
| ELF3 | TTCCCAGAAACTC | -1 | 554 | 567 | -446 | -433 | -5,52 | 0,01 | 8,25 | >0.1 | 3,21 | 5,1 | 0 | 0,99 | 0 | 0 | 4,47 |
| ELF5 | ATAGGTAAGTG | 1 | 1 | 12 | -999 | -988 | 1,91 | 0,01 | 21,37 | >0.1 | 5,6 | 4,26 | 0 | 1,86 | 1,79 | 0 | 5,95 |

| | | | | | | | | | | | | | | | | | |
|---|---|---|---|---|---|---|---|---|---|---|---|---|---|---|---|---|---|
| ELF5 | AGGAGGCACTC | 1 | 19 | 30 | -981 | -970 | 0,02 | 0,01 | 17,71 | >0.1 | 5,6 | 4,27 | 0 | 1,87 | 0 | 0 | 5,95 |
| ELF5 | TCAAAGAAGTC | 1 | 28 | 39 | -972 | -961 | -0,07 | 0,01 | 16,55 | >0.1 | 4,52 | 4,28 | 0 | 1,87 | 0 | 0 | 5,95 |
| ELF5 | AATAGGAAATG | 1 | 60 | 71 | -940 | -929 | 6,16 | 0,01 | 22,29 | >0.1 | 3,99 | 4,31 | 0 | 1,88 | 0 | 0 | 5,95 |
| ELF5 | CCATGGAAATT | 1 | 149 | 160 | -851 | -840 | 2,75 | 0,01 | 19,94 | >0.1 | 3,29 | 4,41 | 0 | 1,9 | 0 | 1,65 | 5,95 |
| ELF5 | ACGGTGAAACC | 1 | 272 | 283 | -728 | -717 | -0,15 | 0,01 | 19,41 | >0.1 | 2,77 | 4,57 | 0 | 1,29 | 0 | 4,99 | 5,95 |
| ELF5 | GGGAGGCAGAG | 1 | 382 | 393 | -618 | -607 | 0,38 | 0,01 | 20,62 | >0.1 | 2,58 | 4,74 | 0 | 1,06 | 0 | 5,91 | 5,95 |
| ELF5 | ATTTGGAAGGG | 1 | 524 | 535 | -476 | -465 | 0,66 | 0,01 | 18,96 | >0.1 | 3,06 | 5,02 | 0 | 0,98 | 0 | 3,28 | 5,95 |
| ELF5 | GAAGGGAAAAT | 1 | 529 | 540 | -471 | -460 | 2,18 | 0,01 | 20,78 | >0.1 | 3,08 | 5,03 | 0 | 0,98 | 0 | 3,55 | 5,95 |
| ELF5 | TCTAGGAAAAT | 1 | 719 | 730 | -281 | -270 | 2,89 | 0,01 | 26,23 | >0.1 | 5,6 | 5,6 | 0 | 1,2 | 0 | 4,99 | 5,95 |
| ELF5 | TCCAGGATGAA | 1 | 865 | 876 | -135 | -124 | 7,32 | 0,001 | 29,85 | >0.1 | 5,6 | 6,41 | 0 | 1,9 | 0 | 2,67 | 5,95 |
| ELF5 | AGGATGAACTT | 1 | 868 | 879 | -132 | -121 | -0,07 | 0,01 | 22,68 | >0.1 | 5,6 | 6,44 | 0 | 1,9 | 0 | 2,86 | 5,95 |
| ELF5 | CTAGGGAAAGT | 1 | 1003 | 1014 | 3 | 14 | 1,32 | 0,01 | 25,32 | >0.1 | 5,6 | 9,12 | 0 | 1,87 | 0 | 1,46 | 5,95 |
| ELF5 | CAGTGGATGTG | 1 | 1018 | 1029 | 18 | 29 | 1,64 | 0,01 | 25,15 | >0.1 | 5,6 | 8,76 | 0 | 1,86 | 0 | 1,34 | 5,95 |
| ELF5 | CAGGGGACGAT | 1 | 1040 | 1051 | 40 | 51 | 0,02 | 0,01 | 22,65 | >0.1 | 5,6 | 7,85 | 0 | 1,86 | 0 | 1,36 | 5,95 |
| ELF5 | TTGAGGAACAG | 1 | 1110 | 1121 | 110 | 121 | 3,86 | 0,01 | 29,69 | >0.1 | 5,6 | 6,65 | 0 | 1,85 | 3,22 | 2,56 | 5,95 |
| ELF5 | TCCAAGAAGCA | -1 | 1183 | 1194 | 183 | 194 | 1,33 | 0,01 | 25,83 | >0.1 | 5,6 | 6,07 | 0 | 1,56 | 1,79 | 3,52 | 5,95 |
| ELF5 | ACAAGGCTGAG | -1 | 1073 | 1084 | 73 | 84 | -0,13 | 0,01 | 24,46 | >0.1 | 5,6 | 7,12 | 0 | 1,85 | 1,79 | 2,27 | 5,95 |
| ELF5 | GCCAGGAAGAG | -1 | 1057 | 1068 | 57 | 68 | 10,02 | 0,001 | 33,12 | >0.1 | 5,6 | 7,42 | 0 | 1,86 | 0 | 2,27 | 5,95 |
| ELF5 | AGCAGGAAAGA | -1 | 932 | 943 | -68 | -57 | 8,8 | 0,001 | 30,26 | >0.1 | 5,6 | 7,11 | 0 | 1,88 | 0 | 0,93 | 5,95 |
| ELF5 | GAGAGGACAGA | -1 | 856 | 867 | -144 | -133 | 0,26 | 0,01 | 23,01 | >0.1 | 5,6 | 6,35 | 0 | 1,9 | 0 | 2,95 | 5,95 |
| ELF5 | AACCGGTACTT | -1 | 739 | 750 | -261 | -250 | 0,55 | 0,01 | 24,95 | >0.1 | 5,6 | 5,68 | 0 | 1,26 | 0 | 5,92 | 5,95 |
| ELF5 | ACACGGAAAAT | -1 | 694 | 705 | -306 | -295 | 9,13 | 0,001 | 30,4 | >0.1 | 5,6 | 5,5 | 0 | 1,14 | 0 | 3,07 | 5,95 |
| ELF5 | ATCATGAAGAT | -1 | 643 | 654 | -357 | -346 | 2,07 | 0,01 | 21,95 | >0.1 | 3,83 | 5,34 | 0 | 1,06 | 0 | 3,7 | 5,95 |
| ELF5 | TCCCAGAAACT | -1 | 555 | 566 | -445 | -434 | -0,09 | 0,01 | 15,16 | >0.1 | 3,21 | 5,09 | 0 | 0,99 | 0 | 0 | 5,95 |
| ELF5 | GTCAGGACGGT | -1 | 257 | 268 | -743 | -732 | 2,36 | 0,01 | 21,64 | >0.1 | 2,82 | 4,54 | 0 | 1,34 | 0 | 4,63 | 5,95 |
| ELF4 | AATAGGTAAGTG | 1 | 0 | 12 | -1000 | -988 | -5,56 | 0,01 | 7,93 | >0.1 | 5,58 | 4,26 | 0 | 1,86 | 1,79 | 0 | 0 |
| ELF4 | GAGAAGGAGGCA | 1 | 15 | 27 | -985 | -973 | -6,93 | 0,01 | 4,79 | >0.1 | 5,58 | 4,27 | 0 | 1,87 | 0 | 0 | 0 |
| ELF4 | CTCAAAGAAGTC | 1 | 27 | 39 | -973 | -961 | -6,85 | 0,01 | 3,83 | >0.1 | 4,53 | 4,28 | 0 | 1,87 | 0 | 0 | 0 |
| ELF4 | AAATAGGAAATG | 1 | 59 | 71 | -941 | -929 | -1,56 | 0,01 | 8,62 | >0.1 | 4 | 4,31 | 0 | 1,88 | 0 | 0 | 0 |
| ELF4 | GATCAGGAGAGT | 1 | 96 | 108 | -904 | -892 | -8,72 | 0,01 | 1,91 | >0.1 | 3,64 | 4,35 | 0 | 1,88 | 0 | 0,77 | 0 |
| ELF4 | GGTCAGGAGATA | 1 | 242 | 254 | -758 | -746 | -7,3 | 0,01 | 5,82 | >0.1 | 2,87 | 4,52 | 0 | 1,4 | 0 | 4,33 | 0 |
| ELF4 | TACTCGGGAGGC | 1 | 345 | 357 | -655 | -643 | -5,46 | 0,01 | 9,4 | >0.1 | 2,55 | 4,68 | 0 | 1,11 | 0 | 6,52 | 0 |
| ELF4 | ACTCGGGAGGCT | 1 | 346 | 358 | -654 | -642 | -6,42 | 0,01 | 8,44 | >0.1 | 2,55 | 4,68 | 0 | 1,11 | 0 | 6,52 | 0 |
| ELF4 | AACCTGGGAGGC | 1 | 377 | 389 | -623 | -611 | -6,42 | 0,01 | 8,47 | >0.1 | 2,56 | 4,73 | 0 | 1,06 | 0 | 6,53 | 0 |
| ELF4 | TAACCAAAAGTA | 1 | 732 | 744 | -268 | -256 | -7,28 | 0,01 | 11,11 | >0.1 | 5,58 | 5,66 | 0 | 1,24 | 0 | 5,92 | 0 |
| ELF4 | GTACCGGTTTTG | 1 | 741 | 753 | -259 | -247 | -8,97 | 0,01 | 9,49 | >0.1 | 5,58 | 5,69 | 0 | 1,27 | 0 | 5,92 | 0 |
| ELF4 | CTCCAGGATGAA | 1 | 864 | 876 | -136 | -124 | -3,56 | 0,01 | 13,01 | >0.1 | 5,58 | 6,41 | 0 | 1,9 | 0 | 2,67 | 0 |
| ELF4 | CATGAGGAGGTT | 1 | 986 | 998 | -14 | -2 | -2,58 | 0,01 | 15,14 | >0.1 | 5,58 | 8,66 | 0 | 1,87 | 0 | 1,61 | 0 |
| ELF4 | ATTGAGGAACAG | 1 | 1109 | 1121 | 109 | 121 | -6,36 | 0,01 | 13,52 | >0.1 | 5,58 | 6,66 | 0 | 1,85 | 3,22 | 2,56 | 0 |
| ELF4 | AACCACGAAGCC | 1 | 1145 | 1157 | 145 | 157 | -2,03 | 0,01 | 17,11 | >0.1 | 5,58 | 6,33 | 0 | 1,84 | 3,22 | 2,16 | 0 |
| ELF4 | GAAGCCGAAGAC | 1 | 1151 | 1163 | 151 | 163 | -8,93 | 0,01 | 7,76 | >0.1 | 4,51 | 6,29 | 0 | 1,84 | 1,79 | 2,26 | 0 |
| ELF4 | TTCCAAGAAGCA | -1 | 1183 | 1195 | 183 | 195 | -2,04 | 0,01 | 16,64 | >0.1 | 5,58 | 6,07 | 0 | 1,56 | 1,79 | 3,67 | 0 |
| ELF4 | AAGAAGCAAGTG | -1 | 1179 | 1191 | 179 | 191 | -5,9 | 0,01 | 12,46 | >0.1 | 5,58 | 6,1 | 0 | 1,59 | 1,79 | 3,29 | 0 |
| ELF4 | AGCCAGGAAGAG | -1 | 1057 | 1069 | 57 | 69 | 3,75 | 0,001 | 20,82 | >0.1 | 5,58 | 7,42 | 0 | 1,86 | 0 | 2,21 | 0 |
| ELF4 | GAGCAGGAAAGA | -1 | 932 | 944 | -68 | -56 | 1,01 | 0,001 | 16,52 | >0.1 | 5,58 | 7,12 | 0 | 1,88 | 0 | 0,93 | 0 |
| ELF4 | AAAGAAGAAGAA | -1 | 908 | 920 | -92 | -80 | -8,36 | 0,01 | 7,13 | >0.1 | 5,58 | 6,81 | 0 | 1,89 | 0 | 1,21 | 0 |
| ELF4 | ACACAGTAAACA | -1 | 897 | 909 | -103 | -91 | -8,08 | 0,01 | 8,95 | >0.1 | 5,58 | 6,7 | 0 | 1,89 | 0 | 2,86 | 0 |
| ELF4 | AAACCGGTACTT | -1 | 739 | 751 | -261 | -249 | 1,66 | 0,001 | 20,1 | >0.1 | 5,58 | 5,68 | 0 | 1,26 | 0 | 5,92 | 0 |
| ELF4 | TACACGGAAAAT | -1 | 694 | 706 | -306 | -294 | -2,32 | 0,01 | 12,87 | >0.1 | 5,58 | 5,51 | 0 | 1,15 | 0 | 2,95 | 0 |
| ELF4 | TTCCCAGAAACT | -1 | 555 | 567 | -445 | -433 | -4,47 | 0,01 | 4,83 | >0.1 | 3,21 | 5,1 | 0 | 0,99 | 0 | 0 | 0 |
| ELF4 | AGTCAGGACGGT | -1 | 257 | 269 | -743 | -731 | -5,81 | 0,01 | 7,42 | >0.1 | 2,81 | 4,55 | 0 | 1,34 | 0 | 4,53 | 0 |
| XBP1 | TCAAAGAAGTCAGC | 1 | 28 | 42 | -972 | -958 | -2,38 | 0,01 | 8,25 | >0.1 | 4,48 | 4,28 | 0 | 1,87 | 0 | 0 | 0 |
| XBP1 | TCAGCCACAGAACC | 1 | 37 | 51 | -963 | -949 | -6,07 | 0,01 | 4,39 | >0.1 | 4,3 | 4,29 | 0 | 1,87 | 0 | 0 | 0 |
| XBP1 | GGAGAGTAGTCATA | 1 | 121 | 135 | -879 | -865 | -7,23 | 0,01 | 4,18 | >0.1 | 3,45 | 4,38 | 0 | 1,89 | 0 | 1,7 | 0 |
| XBP1 | TGGCTCACGCCTGT | 1 | 188 | 202 | -812 | -798 | -6,94 | 0,01 | 6,45 | >0.1 | 3,08 | 4,46 | 0 | 1,72 | 0 | 4,13 | 0 |
| XBP1 | ATCACAAGGTCAGG | 1 | 235 | 249 | -765 | -751 | -4,94 | 0,01 | 8,43 | >0.1 | 2,89 | 4,52 | 0 | 1,43 | 0 | 4,53 | 0 |
| XBP1 | GAAACCCCGTCTCT | 1 | 277 | 291 | -723 | -709 | -4,71 | 0,01 | 9,08 | >0.1 | 2,75 | 4,58 | 0 | 1,27 | 0 | 5,19 | 0 |
| XBP1 | AGCTACTCGGGAGG | 1 | 342 | 356 | -658 | -644 | -8,61 | 0,01 | 6,26 | >0.1 | 2,56 | 4,68 | 0 | 1,12 | 0 | 6,52 | 0 |
| XBP1 | GAGATCGCGCCACT | 1 | 406 | 420 | -594 | -580 | -5,58 | 0,01 | 8,53 | >0.1 | 2,65 | 4,79 | 0 | 1,03 | 0 | 5,65 | 0 |

| | | | | | | | | | | | | | | | | | |
|---|---|---|---|---|---|---|---|---|---|---|---|---|---|---|---|---|---|
| XBP1 | AGGGCAAAGTCATG | 1 | 509 | 523 | -491 | -477 | 2,97 | 0,001 | 14,82 | >0.1 | 3 | 4,99 | 0 | 0,99 | 0 | 2,86 | 0 |
| XBP1 | ACACCAACATTATC | 1 | 632 | 646 | -368 | -354 | -8,62 | 0,01 | 3,09 | >0.1 | 3,74 | 5,31 | 0 | 1,05 | 0 | 1,61 | 0 |
| XBP1 | CAGGAGAGGTAAGG | 1 | 770 | 784 | -230 | -216 | -6,04 | 0,01 | 12,64 | >0.1 | 5,56 | 5,84 | 0 | 1,38 | 0 | 5,92 | 0 |
| XBP1 | AGAATAACGTATTC | 1 | 798 | 812 | -202 | -188 | -5,65 | 0,01 | 10,94 | >0.1 | 5,56 | 5,98 | 0 | 1,51 | 0 | 3,55 | 0 |
| XBP1 | TAGGGAAAGTCATT | 1 | 1004 | 1018 | 4 | 18 | -4,38 | 0,01 | 13,63 | >0.1 | 5,56 | 9,12 | 0 | 1,87 | 0 | 1,46 | 0 |
| XBP1 | GGGACGATGTCAAG | 1 | 1043 | 1057 | 43 | 57 | -6,46 | 0,01 | 9,36 | >0.1 | 5,56 | 7,76 | 0 | 1,86 | 0 | 0,65 | 0 |
| XBP1 | TAGAACAGGTCTTC | -1 | 1157 | 1171 | 157 | 171 | -1,9 | 0,01 | 14,78 | >0.1 | 4,38 | 6,24 | 0 | 1,79 | 1,79 | 2,48 | 0 |
| XBP1 | AGGTCTTCGGCTTC | -1 | 1151 | 1165 | 151 | 165 | -7,7 | 0,01 | 9,05 | >0.1 | 4,48 | 6,29 | 0 | 1,84 | 1,79 | 2,34 | 0 |
| XBP1 | TCGGCTTCGTGGTT | -1 | 1145 | 1159 | 145 | 159 | -6,37 | 0,01 | 12,8 | >0.1 | 5,56 | 6,33 | 0 | 1,84 | 3,22 | 2,21 | 0 |
| XBP1 | AGCTTGACATCGTC | -1 | 1045 | 1059 | 45 | 59 | -4,98 | 0,01 | 11,56 | >0.1 | 5,56 | 7,7 | 0 | 1,86 | 0 | 1,42 | 0 |
| XBP1 | TTGACATCGTCCCC | -1 | 1042 | 1056 | 42 | 56 | -3,94 | 0,01 | 11,89 | >0.1 | 5,56 | 7,79 | 0 | 1,86 | 0 | 0,62 | 0 |
| XBP1 | AAGATCACATCCAC | -1 | 1020 | 1034 | 20 | 34 | -6,76 | 0,01 | 10,66 | >0.1 | 5,56 | 8,72 | 0 | 1,86 | 0 | 1,28 | 0 |
| XBP1 | AAAACCTCCTCATG | -1 | 986 | 1000 | -14 | 0 | -8,37 | 0,01 | 9,38 | >0.1 | 5,56 | 8,71 | 0 | 1,87 | 0 | 1,61 | 0 |
| XBP1 | ACAGATATGTAACA | -1 | 847 | 861 | -153 | -139 | -5,57 | 0,01 | 11,03 | >0.1 | 5,56 | 6,3 | 0 | 1,87 | 0 | 2,86 | 0 |
| XBP1 | AAGAATACGTTATT | -1 | 800 | 814 | -200 | -186 | 1,55 | 0,001 | 18,05 | >0.1 | 5,56 | 5,99 | 0 | 1,52 | 0 | 3,43 | 0 |
| XBP1 | CCTGTCACTTTATG | -1 | 760 | 774 | -240 | -226 | -7,85 | 0,01 | 10,75 | >0.1 | 5,56 | 5,79 | 0 | 1,34 | 0 | 5,92 | 0 |
| XBP1 | ATGGCCAAATCAAA | -1 | 749 | 763 | -251 | -237 | -5,89 | 0,01 | 13,02 | >0.1 | 5,56 | 5,74 | 0 | 1,3 | 0 | 6,32 | 0 |
| XBP1 | AGATACACGGAAAA | -1 | 695 | 709 | -305 | -291 | -5,72 | 0,01 | 9,7 | >0.1 | 5,56 | 5,52 | 0 | 1,15 | 0 | 3,19 | 0 |
| XBP1 | ATTAATAGGTTATC | -1 | 595 | 609 | -405 | -391 | -8,12 | 0,01 | 2,97 | >0.1 | 3,45 | 5,21 | 0 | 1,02 | 0 | 1,41 | 0 |
| XBP1 | TGGGCAACATTTTC | -1 | 534 | 548 | -466 | -452 | -8,23 | 0,01 | 2,58 | >0.1 | 3,11 | 5,05 | 0 | 0,99 | 0 | 1,66 | 0 |
| XBP1 | AGGTTCACGCCATT | -1 | 368 | 382 | -632 | -618 | 3,02 | 0,001 | 17,63 | >0.1 | 2,54 | 4,72 | 0 | 1,08 | 0 | 6,27 | 0 |
| XBP1 | ACCACCACGCCCAG | -1 | 312 | 326 | -688 | -674 | -7,42 | 0,01 | 7,16 | >0.1 | 2,64 | 4,63 | 0 | 1,18 | 0 | 6,13 | 0 |
| XBP1 | TTACAGGCGTGAGC | -1 | 190 | 204 | -810 | -796 | -4,44 | 0,01 | 9,06 | >0.1 | 3,07 | 4,46 | 0 | 1,71 | 0 | 4,26 | 0 |
| XBP1 | CTCCACAGGTCAAC | -1 | 106 | 120 | -894 | -880 | -4,86 | 0,01 | 5,68 | >0.1 | 3,55 | 4,36 | 0 | 1,89 | 0 | 0,75 | 0 |
| GRHL1 | AGAACCAGTGTG | 1 | 45 | 57 | -955 | -943 | 6,15 | 0,001 | 16,5 | >0.1 | 4,19 | 4,3 | 0 | 1,87 | 0 | 0 | 0 |
| GRHL1 | TTGACCTGTGGA | 1 | 107 | 119 | -893 | -881 | -3,96 | 0,01 | 6,58 | >0.1 | 3,55 | 4,36 | 0 | 1,89 | 0 | 0,75 | 0 |
| GRHL1 | AAAACTGATCAG | 1 | 160 | 172 | -840 | -828 | -4,21 | 0,01 | 7,79 | >0.1 | 3,22 | 4,42 | 0 | 1,9 | 0 | 2,46 | 0 |
| GRHL1 | AAACCCCGTCTC | 1 | 278 | 290 | -722 | -710 | -1,69 | 0,01 | 12,09 | >0.1 | 2,75 | 4,58 | 0 | 1,27 | 0 | 5,19 | 0 |
| GRHL1 | GGCACCTGTAGT | 1 | 327 | 339 | -673 | -661 | -5,71 | 0,01 | 8,25 | >0.1 | 2,6 | 4,65 | 0 | 1,15 | 0 | 5,56 | 0 |
| GRHL1 | AAAACTGATCAG | 1 | 475 | 487 | -525 | -513 | -4,21 | 0,01 | 9,77 | >0.1 | 2,87 | 4,92 | 0 | 0,99 | 0 | 5,2 | 0 |
| GRHL1 | AAGTCATGTATT | 1 | 515 | 527 | -485 | -473 | -3,37 | 0,01 | 8,59 | >0.1 | 3,02 | 5 | 0 | 0,98 | 0 | 2,95 | 0 |
| GRHL1 | ATAACCTATTAA | 1 | 596 | 608 | -404 | -392 | -4,11 | 0,01 | 6,97 | >0.1 | 3,45 | 5,2 | 0 | 1,02 | 0 | 1,41 | 0 |
| GRHL1 | TAGACCTCTTTG | 1 | 663 | 675 | -337 | -325 | -1,95 | 0,01 | 12,44 | >0.1 | 4,06 | 5,4 | 0 | 1,09 | 0 | 3,83 | 0 |
| GRHL1 | TTAACAGCTTTC | 1 | 709 | 721 | -291 | -279 | -0,83 | 0,01 | 16,31 | >0.1 | 5,58 | 5,56 | 0 | 1,18 | 0 | 4,81 | 0 |
| GRHL1 | AAAATCTGTTAC | 1 | 841 | 853 | -159 | -147 | -2,29 | 0,01 | 14,53 | >0.1 | 5,58 | 6,25 | 0 | 1,8 | 0 | 3,19 | 0 |
| GRHL1 | GGAACAGGCCAA | 1 | 1114 | 1126 | 114 | 126 | -2,56 | 0,01 | 17,24 | >0.1 | 5,58 | 6,61 | 0 | 1,85 | 3,22 | 2,54 | 0 |
| GRHL1 | TGGACAAGTTTA | 1 | 1134 | 1146 | 134 | 146 | 4,02 | 0,001 | 24,11 | >0.1 | 5,58 | 6,42 | 0 | 1,84 | 3,22 | 3,02 | 0 |
| GRHL1 | AAGACCTGTTCT | 1 | 1158 | 1170 | 158 | 170 | 8,91 | 0,001 | 25,57 | >0.1 | 4,38 | 6,24 | 0 | 1,78 | 1,79 | 2,48 | 0 |
| GRHL1 | AGAACAGGTCTT | -1 | 1158 | 1170 | 158 | 170 | 10,14 | 0,0001 | 26,8 | >0.1 | 4,38 | 6,24 | 0 | 1,78 | 1,79 | 2,48 | 0 |
| GRHL1 | TAAACTTGTCCA | -1 | 1134 | 1146 | 134 | 146 | 3,67 | 0,001 | 23,76 | >0.1 | 5,58 | 6,42 | 0 | 1,84 | 3,22 | 3,02 | 0 |
| GRHL1 | TTGGCCTGTTCC | -1 | 1114 | 1126 | 114 | 126 | -4,34 | 0,01 | 15,46 | >0.1 | 5,58 | 6,61 | 0 | 1,85 | 3,22 | 2,54 | 0 |
| GRHL1 | GTAACAGATTTT | -1 | 841 | 853 | -159 | -147 | -3,13 | 0,01 | 13,69 | >0.1 | 5,58 | 6,25 | 0 | 1,8 | 0 | 3,19 | 0 |
| GRHL1 | GAAAGCTGTTAA | -1 | 709 | 721 | -291 | -279 | -2,99 | 0,01 | 14,15 | >0.1 | 5,58 | 5,56 | 0 | 1,18 | 0 | 4,81 | 0 |
| GRHL1 | CAAAGAGGTCTA | -1 | 663 | 675 | -337 | -325 | -2,7 | 0,01 | 11,69 | >0.1 | 4,06 | 5,4 | 0 | 1,09 | 0 | 3,83 | 0 |
| GRHL1 | TTAATAGGTTAT | -1 | 596 | 608 | -404 | -392 | -4,54 | 0,01 | 6,54 | >0.1 | 3,45 | 5,2 | 0 | 1,02 | 0 | 1,41 | 0 |
| GRHL1 | AATACATGACTT | -1 | 515 | 527 | -485 | -473 | -2,24 | 0,01 | 9,72 | >0.1 | 3,02 | 5 | 0 | 0,98 | 0 | 2,95 | 0 |
| GRHL1 | ACTACAGGTGCC | -1 | 327 | 339 | -673 | -661 | -3,82 | 0,01 | 10,14 | >0.1 | 2,6 | 4,65 | 0 | 1,15 | 0 | 5,56 | 0 |
| GRHL1 | GAGACGGGGTTT | -1 | 278 | 290 | -722 | -710 | -2,44 | 0,01 | 11,34 | >0.1 | 2,75 | 4,58 | 0 | 1,27 | 0 | 5,19 | 0 |
| GRHL1 | TCCACAGGTCAA | -1 | 107 | 119 | -893 | -881 | -4,57 | 0,01 | 5,97 | >0.1 | 3,55 | 4,36 | 0 | 1,89 | 0 | 0,75 | 0 |
| GRHL1 | CACACTGGTTCT | -1 | 45 | 57 | -955 | -943 | 2,91 | 0,01 | 13,26 | >0.1 | 4,19 | 4,3 | 0 | 1,87 | 0 | 0 | 0 |
| GRHL2 | CTCAAAGAAGTCAGC | 1 | 27 | 42 | -973 | -958 | 1,62 | 0,01 | 12,27 | >0.1 | 4,5 | 4,28 | 0 | 1,87 | 0 | 0 | 0 |
| GRHL2 | ACAGAACCAGTGTGA | 1 | 43 | 58 | -957 | -942 | 8,97 | 0,001 | 19,33 | >0.1 | 4,19 | 4,3 | 0 | 1,87 | 0 | 0 | 0 |
| GRHL2 | GCAAAGGCAGATCAG | 1 | 87 | 102 | -913 | -898 | 2,26 | 0,01 | 12,18 | >0.1 | 3,7 | 4,34 | 0 | 1,88 | 0 | 0 | 0 |
| GRHL2 | GTGAAACCCCGTCTC | 1 | 275 | 290 | -725 | -710 | 0,89 | 0,01 | 14,69 | >0.1 | 2,75 | 4,58 | 0 | 1,28 | 0 | 5,19 | 0 |
| GRHL2 | TGAAACCCCGTCTCT | 1 | 276 | 291 | -724 | -709 | 1,78 | 0,01 | 15,57 | >0.1 | 2,75 | 4,58 | 0 | 1,28 | 0 | 5,19 | 0 |
| GRHL2 | AAAAAAACTGATCAG | 1 | 472 | 487 | -528 | -513 | 1,32 | 0,01 | 15,04 | >0.1 | 2,86 | 4,92 | 0 | 0,99 | 0 | 4,95 | 0 |
| GRHL2 | AGATAACCTATTAAT | 1 | 594 | 609 | -406 | -391 | 1,07 | 0,01 | 12,19 | >0.1 | 3,45 | 5,21 | 0 | 1,02 | 0 | 1,44 | 0 |
| GRHL2 | TCTAGACCTCTTTGG | 1 | 661 | 676 | -339 | -324 | 3,51 | 0,01 | 18,06 | >0.1 | 4,05 | 5,41 | 0 | 1,1 | 0 | 3,99 | 0 |
| GRHL2 | AAAGTACCGGTTTTG | 1 | 738 | 753 | -262 | -247 | 7,03 | 0,001 | 25,45 | >0.1 | 5,54 | 5,69 | 0 | 1,27 | 0 | 5,92 | 0 |

| | | | | | | | | | | | | | | | | | |
|---|---|---|---|---|---|---|---|---|---|---|---|---|---|---|---|---|---|
| GRHL2 | CTAAAATCTGTTACA | 1 | 839 | 854 | -161 | -146 | 4,36 | 0,01 | 19,38 | >0.1 | 5,54 | 6,26 | 0 | 1,81 | 0 | 1,41 | 0 |
| GRHL2 | GCCAAGACATTTTTG | 1 | 1121 | 1136 | 121 | 136 | 0,36 | 0,01 | 20,24 | >0.1 | 5,54 | 6,54 | 0 | 1,85 | 3,22 | 2,72 | 0 |
| GRHL2 | TTTGGACAAGTTTAA | 1 | 1132 | 1147 | 132 | 147 | 3,23 | 0,01 | 23,32 | >0.1 | 5,54 | 6,44 | 0 | 1,84 | 3,22 | 3,03 | 0 |
| GRHL2 | CGAAGACCTGTTCTA | 1 | 1156 | 1171 | 156 | 171 | 9,38 | 0,001 | 25,99 | >0.1 | 4,37 | 6,25 | 0 | 1,8 | 1,79 | 2,4 | 0 |
| GRHL2 | ATAGAACAGGTCTTC | -1 | 1157 | 1172 | 157 | 172 | 3,45 | 0,01 | 20,14 | >0.1 | 4,39 | 6,24 | 0 | 1,79 | 1,79 | 2,48 | 0 |
| GRHL2 | GTTAAACTTGTCCAA | -1 | 1133 | 1148 | 133 | 148 | 5,26 | 0,01 | 25,32 | >0.1 | 5,54 | 6,43 | 0 | 1,84 | 3,22 | 3,02 | 0 |
| GRHL2 | TCTTGGCCTGTTCCT | -1 | 1113 | 1128 | 113 | 128 | 1,87 | 0,01 | 21,64 | >0.1 | 5,54 | 6,62 | 0 | 1,85 | 3,22 | 2,54 | 0 |
| GRHL2 | CAGCAACAAGGCTGA | -1 | 1074 | 1089 | 74 | 89 | 0,5 | 0,01 | 19,07 | >0.1 | 5,54 | 7,11 | 0 | 1,85 | 1,79 | 2,27 | 0 |
| GRHL2 | CTAAAACCTCCTCAT | -1 | 987 | 1002 | -13 | 2 | 2,91 | 0,01 | 20,71 | >0.1 | 5,54 | 8,77 | 0 | 1,87 | 0 | 1,61 | 0 |
| GRHL2 | TCAAAACCGGTACTT | -1 | 739 | 754 | -261 | -246 | 9,02 | 0,001 | 27,63 | >0.1 | 5,54 | 5,7 | 0 | 1,27 | 0 | 6,1 | 0 |
| GRHL2 | TAGAAAGCTGTTAAA | -1 | 708 | 723 | -292 | -277 | 4,27 | 0,01 | 21,38 | >0.1 | 5,54 | 5,57 | 0 | 1,19 | 0 | 4,81 | 0 |
| GRHL2 | ACCAAAGAGGTCTAG | -1 | 662 | 677 | -338 | -323 | 0,85 | 0,01 | 15,25 | >0.1 | 4,07 | 5,41 | 0 | 1,1 | 0 | 3,83 | 0 |
| GRHL2 | TTCAGACAATTTCAT | -1 | 607 | 622 | -393 | -378 | 0,37 | 0,01 | 11,73 | >0.1 | 3,54 | 5,24 | 0 | 1,03 | 0 | 1,55 | 0 |
| GRHL2 | TTCTGATCAGTTTTT | -1 | 474 | 489 | -526 | -511 | 2,83 | 0,01 | 16,82 | >0.1 | 2,87 | 4,92 | 0 | 0,99 | 0 | 5,2 | 0 |
| GRHL2 | GCCCAGCCATTTCTG | -1 | 169 | 184 | -831 | -816 | 5,4 | 0,01 | 18,24 | >0.1 | 3,17 | 4,43 | 0 | 1,89 | 0 | 3,34 | 0 |
| GRHL2 | TTCTGATCAGTTTTA | -1 | 159 | 174 | -841 | -826 | 2,62 | 0,01 | 14,62 | >0.1 | 3,22 | 4,42 | 0 | 1,9 | 0 | 2,46 | 0 |
| GRHL2 | CTCACACTGGTTCTG | -1 | 44 | 59 | -956 | -941 | 4,2 | 0,01 | 14,55 | >0.1 | 4,18 | 4,3 | 0 | 1,87 | 0 | 0 | 0 |
| NR1H2::RXRA | CAGATCACAAGGTCAGG | 1 | 232 | 249 | -768 | -751 | -8,92 | 0,01 | 4,69 | >0.1 | 2,9 | 4,52 | 0 | 1,45 | 0 | 4,74 | 0 |
| NR1H2::RXRA | CAAGGTCAGGAGATAGA | 1 | 239 | 256 | -761 | -744 | -15 | 0,01 | -1,86 | >0.1 | 2,87 | 4,53 | 0 | 1,41 | 0 | 4,33 | 0 |
| NR1H2::RXRA | CACGGTGAAACCCCGTC | 1 | 271 | 288 | -729 | -712 | -13,98 | 0,01 | -0,16 | >0.1 | 2,76 | 4,57 | 0 | 1,29 | 0 | 5,19 | 0 |
| NR1H2::RXRA | TAAAGACAAGGGCAAAG | 1 | 501 | 518 | -499 | -482 | -8,1 | 0,01 | 6,71 | >0.1 | 2,98 | 4,98 | 0 | 0,99 | 0 | 5,86 | 0 |
| NR1H2::RXRA | ACAAGGGCAAAGTCATG | 1 | 506 | 523 | -494 | -477 | -12,22 | 0,01 | -0,49 | >0.1 | 3 | 4,99 | 0 | 0,99 | 0 | 2,75 | 0 |
| NR1H2::RXRA | CAAGGGCAAAGTCATGT | 1 | 507 | 524 | -493 | -476 | -10,67 | 0,01 | 1,18 | >0.1 | 3 | 5 | 0 | 0,99 | 0 | 2,86 | 0 |
| NR1H2::RXRA | GAAGGGAAAATGTTGCC | 1 | 529 | 546 | -471 | -454 | -14,39 | 0,01 | -3,64 | >0.1 | 3,1 | 5,05 | 0 | 0,99 | 0 | 1,61 | 0 |
| NR1H2::RXRA | AGACCTCTTTGGTCACT | 1 | 664 | 681 | -336 | -319 | -8,84 | 0,01 | 5,62 | >0.1 | 4,11 | 5,42 | 0 | 1,1 | 0 | 3,83 | 0 |
| NR1H2::RXRA | TAAAGTGACAGGAGAGG | 1 | 762 | 779 | -238 | -221 | -10,75 | 0,01 | 7,85 | >0.1 | 5,52 | 5,81 | 0 | 1,36 | 0 | 5,92 | 0 |
| NR1H2::RXRA | TAGGATTAAAGAATAAC | 1 | 789 | 806 | -211 | -194 | -10,2 | 0,01 | 8,06 | >0.1 | 5,52 | 5,95 | 0 | 1,48 | 0 | 5,32 | 0 |
| NR1H2::RXRA | CCTGCTCAGCGCCCAAC | 1 | 937 | 954 | -63 | -46 | -14,01 | 0,01 | 1,51 | >0.1 | 5,52 | 7,29 | 0 | 1,88 | 0 | 0,83 | 0 |
| NR1H2::RXRA | CAAGTTCAAAGGCTGAT | 1 | 955 | 972 | -45 | -28 | 1,66 | 0,001 | 18,47 | >0.1 | 5,52 | 7,68 | 0 | 1,88 | 0 | 1,74 | 0 |
| NR1H2::RXRA | AGAGAGAAAATCTCATG | 1 | 973 | 990 | -27 | -10 | -11,04 | 0,01 | 6,25 | >0.1 | 5,52 | 8,32 | 0 | 1,87 | 0 | 1,57 | 0 |
| NR1H2::RXRA | TTGGCTCACAGGGGACG | 1 | 1032 | 1049 | 32 | 49 | -8,02 | 0,01 | 8,81 | >0.1 | 5,52 | 8,15 | 0 | 1,86 | 0 | 1,31 | 0 |
| NR1H2::RXRA | CAGGGGACGATGTCAAG | 1 | 1040 | 1057 | 40 | 57 | -7,81 | 0,01 | 8,04 | >0.1 | 5,52 | 7,85 | 0 | 1,86 | 0 | 0,62 | 0 |
| NR1H2::RXRA | TTGAGGAACAGGCCAAG | 1 | 1110 | 1127 | 110 | 127 | -11,98 | 0,01 | 7,77 | >0.1 | 5,52 | 6,65 | 0 | 1,85 | 3,22 | 2,52 | 0 |
| NR1H2::RXRA | TTCTATCAAAGTTCACT | 1 | 1166 | 1183 | 166 | 183 | -9,34 | 0,01 | 8,8 | >0.1 | 5,52 | 6,18 | 0 | 1,7 | 1,79 | 2,94 | 0 |
| NR1H2::RXRA | AAAAGAAAAAGAAGAAG | -1 | 910 | 927 | -90 | -73 | -14,14 | 0,01 | 1,31 | >0.1 | 5,52 | 6,89 | 0 | 1,89 | 0 | 1,15 | 0 |
| NR1H2::RXRA | CCAATATAAAGTTCATC | -1 | 870 | 887 | -130 | -113 | -11,5 | 0,01 | 5,37 | >0.1 | 5,52 | 6,5 | 0 | 1,9 | 0 | 2,95 | 0 |
| NR1H2::RXRA | TATGGCCAAATCAAAAC | -1 | 747 | 764 | -253 | -236 | -13,96 | 0,01 | 4,71 | >0.1 | 5,52 | 5,74 | 0 | 1,3 | 0 | 6,1 | 0 |
| NR1H2::RXRA | AGCTGTTAAAGATACAC | -1 | 701 | 718 | -299 | -282 | -13,26 | 0,01 | 3,8 | >0.1 | 5,52 | 5,55 | 0 | 1,18 | 0 | 4,81 | 0 |
| NR1H2::RXRA | AAAGATACACGGAAAAT | -1 | 694 | 711 | -306 | -289 | -14,15 | 0,01 | 1,24 | >0.1 | 5,52 | 5,53 | 0 | 1,16 | 0 | 3,19 | 0 |
| NR1H2::RXRA | CAAGATCATATTCCCAG | -1 | 560 | 577 | -440 | -423 | -11,68 | 0,01 | -2,3 | >0.1 | 3,25 | 5,12 | 0 | 1 | 0 | 0 | 0 |
| NR1H2::RXRA | TAATTTCCATGGCCATT | -1 | 144 | 161 | -856 | -839 | -9,72 | 0,01 | 1,49 | >0.1 | 3,3 | 4,41 | 0 | 1,9 | 0 | 1,61 | 0 |
| NR1H2::RXRA | CCACTCCACAGGTCAAC | -1 | 106 | 123 | -894 | -877 | 1,22 | 0,001 | 11,76 | >0.1 | 3,54 | 4,36 | 0 | 1,89 | 0 | 0,75 | 0 |
| NR1H2::RXRA | ACAGGTCAACTCTCCTG | -1 | 99 | 116 | -901 | -884 | -4,75 | 0,001 | 5,85 | >0.1 | 3,59 | 4,36 | 0 | 1,89 | 0 | 0,77 | 0 |
| BHLHE22 | GGCTCACGCC | 1 | 189 | 199 | -811 | -801 | -3,54 | 0,01 | 9,75 | >0.1 | 3,09 | 4,45 | 0 | 1,71 | 0 | 4,04 | 0 |
| BHLHE22 | GAAATATGAA | 1 | 486 | 496 | -514 | -504 | -3,01 | 0,01 | 12,08 | >0.1 | 2,9 | 4,93 | 0 | 0,98 | 0 | 6,27 | 0 |
| BHLHE22 | GGAATATGAT | 1 | 563 | 573 | -437 | -427 | 0,4 | 0,01 | 9,75 | >0.1 | 3,25 | 5,11 | 0 | 0,99 | 0 | 0 | 0 |
| BHLHE22 | AACATTTTCC | 1 | 691 | 701 | -309 | -299 | -1,03 | 0,01 | 13,2 | >0.1 | 4,53 | 5,49 | 0 | 1,14 | 0 | 3,07 | 0 |
| BHLHE22 | GCCATAAAGT | 1 | 758 | 768 | -242 | -232 | -3,51 | 0,01 | 15,27 | >0.1 | 5,61 | 5,76 | 0 | 1,31 | 0 | 6,1 | 0 |
| BHLHE22 | AACGTATTCT | 1 | 803 | 813 | -197 | -187 | 3,46 | 0,01 | 20 | >0.1 | 5,61 | 5,99 | 0 | 1,51 | 0 | 3,43 | 0 |
| BHLHE22 | AAAATCTGTT | 1 | 841 | 851 | -159 | -149 | 1,06 | 0,01 | 16,1 | >0.1 | 5,61 | 6,24 | 0 | 1,78 | 0 | 1,41 | 0 |
| BHLHE22 | TACATATCTG | 1 | 850 | 860 | -150 | -140 | -2,22 | 0,01 | 14,3 | >0.1 | 5,61 | 6,3 | 0 | 1,86 | 0 | 2,75 | 0 |
| BHLHE22 | ACAATCTGCT | -1 | 890 | 900 | -110 | -100 | 2,27 | 0,01 | 19,33 | >0.1 | 5,61 | 6,61 | 0 | 1,89 | 0 | 2,95 | 0 |
| BHLHE22 | AGAATACGTT | -1 | 803 | 813 | -197 | -187 | 4,14 | 0,001 | 20,68 | >0.1 | 5,61 | 5,99 | 0 | 1,51 | 0 | 3,43 | 0 |
| BHLHE22 | ACCTTACCTC | -1 | 775 | 785 | -225 | -215 | -2,17 | 0,01 | 16,98 | >0.1 | 5,61 | 5,84 | 0 | 1,38 | 0 | 6,32 | 0 |
| BHLHE22 | ACTTTATGGC | -1 | 758 | 768 | -242 | -232 | -3,4 | 0,01 | 15,38 | >0.1 | 5,61 | 5,76 | 0 | 1,31 | 0 | 6,1 | 0 |
| BHLHE22 | ATCATATTCC | -1 | 563 | 573 | -437 | -427 | 1,36 | 0,01 | 10,71 | >0.1 | 3,25 | 5,11 | 0 | 0,99 | 0 | 0 | 0 |
| BHLHE22 | AACATTTTCC | -1 | 533 | 543 | -467 | -457 | -1,03 | 0,01 | 9,7 | >0.1 | 3,1 | 5,04 | 0 | 0,98 | 0 | 1,61 | 0 |
| BHLHE22 | TTCATATTTC | -1 | 486 | 496 | -514 | -504 | -3,69 | 0,01 | 11,4 | >0.1 | 2,9 | 4,93 | 0 | 0,98 | 0 | 6,27 | 0 |
| BHLHE22 | ACCTTGTGAT | -1 | 235 | 245 | -765 | -755 | -3,78 | 0,01 | 9,8 | >0.1 | 2,9 | 4,51 | 0 | 1,43 | 0 | 4,74 | 0 |

| | | | | | | | | | | | | | | | | | |
|---|---|---|---|---|---|---|---|---|---|---|---|---|---|---|---|---|---|
| BHLHE22 | GCCATTTTTT | -1 | 140 | 150 | -860 | -850 | -1,26 | 0,01 | 10,02 | >0.1 | 3,34 | 4,39 | 0 | 1,89 | 0 | 1,65 | 0 |
| BHLHE23 | TAAAAAATGGCC | 1 | 139 | 151 | -861 | -849 | -9,02 | 0,01 | 2,26 | >0.1 | 3,34 | 4,4 | 0 | 1,89 | 0 | 1,65 | 0 |
| BHLHE23 | GATCACAAGGTC | 1 | 234 | 246 | -766 | -754 | -9,2 | 0,01 | 4,39 | >0.1 | 2,9 | 4,51 | 0 | 1,44 | 0 | 4,74 | 0 |
| BHLHE23 | AGAAATATGAAT | 1 | 485 | 497 | -515 | -503 | -5,25 | 0,01 | 9,85 | >0.1 | 2,9 | 4,94 | 0 | 0,99 | 0 | 6,27 | 0 |
| BHLHE23 | GGGAATATGATC | 1 | 562 | 574 | -438 | -426 | -3,36 | 0,01 | 6 | >0.1 | 3,25 | 5,11 | 0 | 1 | 0 | 0 | 0 |
| BHLHE23 | GGCCATAAAGTG | 1 | 757 | 769 | -243 | -231 | -8,71 | 0,01 | 10,06 | >0.1 | 5,58 | 5,76 | 0 | 1,32 | 0 | 6,1 | 0 |
| BHLHE23 | AAGAATAACGTA | 1 | 797 | 809 | -203 | -191 | -9,43 | 0,01 | 7,16 | >0.1 | 5,58 | 5,97 | 0 | 1,49 | 0 | 3,55 | 0 |
| BHLHE23 | TAACGTATTCTT | 1 | 802 | 814 | -198 | -186 | -0,65 | 0,01 | 15,87 | >0.1 | 5,58 | 5,99 | 0 | 1,52 | 0 | 3,43 | 0 |
| BHLHE23 | TAAAATCTGTTA | 1 | 840 | 852 | -160 | -148 | -2,61 | 0,01 | 12,42 | >0.1 | 5,58 | 6,24 | 0 | 1,79 | 0 | 1,41 | 0 |
| BHLHE23 | TTACATATCTGT | 1 | 849 | 861 | -151 | -139 | -7,05 | 0,01 | 9,46 | >0.1 | 5,58 | 6,3 | 0 | 1,87 | 0 | 2,75 | 0 |
| BHLHE23 | CAGCAGATTGTT | 1 | 889 | 901 | -111 | -99 | -6,89 | 0,01 | 10,16 | >0.1 | 5,58 | 6,62 | 0 | 1,89 | 0 | 2,95 | 0 |
| BHLHE23 | AACAATCTGCTG | -1 | 889 | 901 | -111 | -99 | -2,16 | 0,01 | 14,89 | >0.1 | 5,58 | 6,62 | 0 | 1,89 | 0 | 2,95 | 0 |
| BHLHE23 | ACAGATATGTAA | -1 | 849 | 861 | -151 | -139 | -8,31 | 0,01 | 8,2 | >0.1 | 5,58 | 6,3 | 0 | 1,87 | 0 | 2,75 | 0 |
| BHLHE23 | TAACAGATTTTA | -1 | 840 | 852 | -160 | -148 | -6,28 | 0,01 | 8,75 | >0.1 | 5,58 | 6,24 | 0 | 1,79 | 0 | 1,41 | 0 |
| BHLHE23 | AAGAATACGTTA | -1 | 802 | 814 | -198 | -186 | 2,26 | 0,001 | 18,78 | >0.1 | 5,58 | 5,99 | 0 | 1,52 | 0 | 3,43 | 0 |
| BHLHE23 | AACCTTACCTCT | -1 | 774 | 786 | -226 | -214 | -8,49 | 0,01 | 10,64 | >0.1 | 5,58 | 5,84 | 0 | 1,38 | 0 | 6,32 | 0 |
| BHLHE23 | CACTTTATGGCC | -1 | 757 | 769 | -243 | -231 | -9,85 | 0,01 | 8,92 | >0.1 | 5,58 | 5,76 | 0 | 1,32 | 0 | 6,1 | 0 |
| BHLHE23 | GATCATATTCCC | -1 | 562 | 574 | -438 | -426 | -4,52 | 0,01 | 4,84 | >0.1 | 3,25 | 5,11 | 0 | 1 | 0 | 0 | 0 |
| BHLHE23 | ATTCATATTTCT | -1 | 485 | 497 | -515 | -503 | -9,7 | 0,01 | 5,4 | >0.1 | 2,9 | 4,94 | 0 | 0,99 | 0 | 6,27 | 0 |
| BHLHE23 | AAAATTATGACT | -1 | 128 | 140 | -872 | -860 | -8,35 | 0,01 | 2,94 | >0.1 | 3,41 | 4,38 | 0 | 1,89 | 0 | 1,61 | 0 |
| FIGLA | ACCCCGTCTC | 1 | 280 | 290 | -720 | -710 | 0,74 | 0,01 | 14,41 | >0.1 | 2,74 | 4,57 | 0 | 1,26 | 0 | 5,09 | 0 |
| FIGLA | CTTAGCTGGG | 1 | 307 | 317 | -693 | -683 | 2,31 | 0,01 | 16,45 | >0.1 | 2,66 | 4,62 | 0 | 1,19 | 0 | 5,68 | 0 |
| FIGLA | GGCACCTGTA | 1 | 327 | 337 | -673 | -663 | 10 | 0,001 | 23,96 | >0.1 | 2,6 | 4,65 | 0 | 1,14 | 0 | 5,56 | 0 |
| FIGLA | CCCAGCTACT | 1 | 339 | 349 | -661 | -651 | 4,21 | 0,01 | 19,09 | >0.1 | 2,57 | 4,66 | 0 | 1,12 | 0 | 6,52 | 0 |
| FIGLA | AACACCAACA | 1 | 631 | 641 | -369 | -359 | 0,97 | 0,01 | 12,68 | >0.1 | 3,71 | 5,3 | 0 | 1,05 | 0 | 1,66 | 0 |
| FIGLA | AACAGCTTTC | 1 | 711 | 721 | -289 | -279 | 6,24 | 0,01 | 23,58 | >0.1 | 5,61 | 5,56 | 0 | 1,18 | 0 | 4,99 | 0 |
| FIGLA | CTCAGCAGAT | 1 | 887 | 897 | -113 | -103 | 2,25 | 0,01 | 19,41 | >0.1 | 5,61 | 6,59 | 0 | 1,89 | 0 | 3,07 | 0 |
| FIGLA | AACTGCTGCT | 1 | 1090 | 1100 | 90 | 100 | 4,07 | 0,01 | 22,48 | >0.1 | 5,61 | 6,88 | 0 | 1,85 | 1,79 | 2,27 | 0 |
| FIGLA | TCCACCATTG | 1 | 1103 | 1113 | 103 | 113 | 2,57 | 0,01 | 22,45 | >0.1 | 5,61 | 6,72 | 0 | 1,85 | 3,22 | 2,47 | 0 |
| FIGLA | AAGACCTGTT | 1 | 1158 | 1168 | 158 | 168 | 1,06 | 0,01 | 17,65 | >0.1 | 4,38 | 6,24 | 0 | 1,78 | 1,79 | 2,4 | 0 |
| FIGLA | AACAGGTCTT | -1 | 1158 | 1168 | 158 | 168 | 4,17 | 0,01 | 20,76 | >0.1 | 4,38 | 6,24 | 0 | 1,78 | 1,79 | 2,4 | 0 |
| FIGLA | AGCAGCAGTT | -1 | 1090 | 1100 | 90 | 100 | 5,83 | 0,01 | 24,24 | >0.1 | 5,61 | 6,88 | 0 | 1,85 | 1,79 | 2,27 | 0 |
| FIGLA | GTCCCCTGTG | -1 | 1038 | 1048 | 38 | 48 | 5,14 | 0,01 | 21,86 | >0.1 | 5,61 | 7,92 | 0 | 1,86 | 0 | 1,34 | 0 |
| FIGLA | ATCTGCTGAG | -1 | 887 | 897 | -113 | -103 | 0,6 | 0,01 | 17,76 | >0.1 | 5,61 | 6,59 | 0 | 1,89 | 0 | 3,07 | 0 |
| FIGLA | CTCTCCTGTC | -1 | 768 | 778 | -232 | -222 | 3,54 | 0,01 | 22,22 | >0.1 | 5,61 | 5,81 | 0 | 1,35 | 0 | 5,92 | 0 |
| FIGLA | CCCAGGTTCA | -1 | 375 | 385 | -625 | -615 | 1,81 | 0,01 | 16,55 | >0.1 | 2,56 | 4,72 | 0 | 1,07 | 0 | 6,4 | 0 |
| FIGLA | TTCTCCTGCC | -1 | 360 | 370 | -640 | -630 | 2,39 | 0,01 | 17,06 | >0.1 | 2,52 | 4,7 | 0 | 1,09 | 0 | 6,36 | 0 |
| FIGLA | AGTAGCTGGG | -1 | 339 | 349 | -661 | -651 | 3,45 | 0,01 | 18,33 | >0.1 | 2,57 | 4,66 | 0 | 1,12 | 0 | 6,52 | 0 |
| FIGLA | TACAGGTGCC | -1 | 327 | 337 | -673 | -663 | 6,63 | 0,01 | 20,59 | >0.1 | 2,6 | 4,65 | 0 | 1,14 | 0 | 5,56 | 0 |
| FIGLA | CCCACCACCA | -1 | 319 | 329 | -681 | -671 | 0,62 | 0,01 | 14,6 | >0.1 | 2,63 | 4,63 | 0 | 1,16 | 0 | 5,56 | 0 |
| FIGLA | ACCACCACGC | -1 | 316 | 326 | -684 | -674 | 2,99 | 0,01 | 17,55 | >0.1 | 2,63 | 4,63 | 0 | 1,17 | 0 | 6,13 | 0 |
| FIGLA | CCCAGCTAAG | -1 | 307 | 317 | -693 | -683 | 2,28 | 0,01 | 16,42 | >0.1 | 2,66 | 4,62 | 0 | 1,19 | 0 | 5,68 | 0 |
| FIGLA | ATCTCCTGAC | -1 | 243 | 253 | -757 | -747 | 2,32 | 0,01 | 15,43 | >0.1 | 2,87 | 4,52 | 0 | 1,39 | 0 | 4,33 | 0 |
| FIGLA | CTCTCCTGAT | -1 | 97 | 107 | -903 | -893 | 2,02 | 0,01 | 12,65 | >0.1 | 3,64 | 4,35 | 0 | 1,88 | 0 | 0,77 | 0 |
| GLI2 | TCAGCCACAGAA | 1 | 37 | 49 | -963 | -951 | -0,9 | 0,01 | 9,58 | >0.1 | 4,32 | 4,29 | 0 | 1,87 | 0 | 0 | 0 |
| GLI2 | CCTAGCACTTTG | 1 | 205 | 217 | -795 | -783 | -2,37 | 0,01 | 11,42 | >0.1 | 3,01 | 4,48 | 0 | 1,6 | 0 | 4,7 | 0 |
| GLI2 | CAGATCACAAGG | 1 | 232 | 244 | -768 | -756 | 1,01 | 0,01 | 14,77 | >0.1 | 2,9 | 4,51 | 0 | 1,45 | 0 | 4,9 | 0 |
| GLI2 | GAGACCGTCCTG | 1 | 254 | 266 | -746 | -734 | 0,51 | 0,01 | 14,08 | >0.1 | 2,83 | 4,54 | 0 | 1,35 | 0 | 4,85 | 0 |
| GLI2 | ACTAACACGGTG | 1 | 266 | 278 | -734 | -722 | -0,93 | 0,01 | 12,25 | >0.1 | 2,78 | 4,56 | 0 | 1,31 | 0 | 4,53 | 0 |
| GLI2 | TAGACCTCTTTG | 1 | 663 | 675 | -337 | -325 | 0,29 | 0,01 | 14,68 | >0.1 | 4,06 | 5,4 | 0 | 1,09 | 0 | 3,83 | 0 |
| GLI2 | GCGCCCAACCCA | 1 | 945 | 957 | -55 | -43 | 4,68 | 0,001 | 21,28 | >0.1 | 5,58 | 7,34 | 0 | 1,88 | 0 | 1,8 | 0 |
| GLI2 | TGGCTCACAGGG | 1 | 1033 | 1045 | 33 | 45 | -3,04 | 0,01 | 13,81 | >0.1 | 5,58 | 8,1 | 0 | 1,86 | 0 | 1,31 | 0 |
| GLI2 | CAGTCCACCATT | 1 | 1100 | 1112 | 100 | 112 | 1,54 | 0,01 | 21,33 | >0.1 | 5,58 | 6,76 | 0 | 1,85 | 3,22 | 2,38 | 0 |
| GLI2 | TTAACCACGAAG | 1 | 1143 | 1155 | 143 | 155 | 1,05 | 0,01 | 20,21 | >0.1 | 5,58 | 6,35 | 0 | 1,84 | 3,22 | 2,16 | 0 |
| GLI2 | TCGTCCCCTGTG | -1 | 1038 | 1050 | 38 | 50 | -1,29 | 0,01 | 15,43 | >0.1 | 5,58 | 7,92 | 0 | 1,86 | 0 | 1,36 | 0 |
| GLI2 | AAGACCAAAAAA | -1 | 924 | 936 | -76 | -64 | 0,65 | 0,01 | 16,18 | >0.1 | 5,58 | 7,01 | 0 | 1,88 | 0 | 1,06 | 0 |
| GLI2 | AAGAACACAGTA | -1 | 901 | 913 | -99 | -87 | 0,73 | 0,01 | 16,21 | >0.1 | 5,58 | 6,74 | 0 | 1,89 | 0 | 1,28 | 0 |
| GLI2 | GTGACCAAAGAG | -1 | 668 | 680 | -332 | -320 | 2,56 | 0,01 | 16,9 | >0.1 | 4,13 | 5,42 | 0 | 1,1 | 0 | 3,7 | 0 |
| GLI2 | TCGCCCAGGCTG | -1 | 426 | 438 | -574 | -562 | 6,34 | 0,001 | 20,68 | >0.1 | 2,7 | 4,82 | 0 | 1,01 | 0 | 5,8 | 0 |

| | | | | | | | | | | | | | | | | | |
|---|---|---|---|---|---|---|---|---|---|---|---|---|---|---|---|---|---|
| GLI2 | TGGAGCACAGTG | -1 | 416 | 428 | -584 | -572 | 2,07 | 0,01 | 15,87 | >0.1 | 2,67 | 4,8 | 0 | 1,02 | 0 | 5,3 | 0 |
| GLI2 | TCTGCCTCCCAG | -1 | 380 | 392 | -620 | -608 | -1,55 | 0,01 | 12,84 | >0.1 | 2,57 | 4,74 | 0 | 1,06 | 0 | 6,02 | 0 |
| GLI2 | GGGACTACAGGT | -1 | 330 | 342 | -670 | -658 | -0,63 | 0,01 | 13,79 | >0.1 | 2,59 | 4,65 | 0 | 1,14 | 0 | 6,03 | 0 |
| GLI2 | GTGCCCACCACC | -1 | 320 | 332 | -680 | -668 | 4,56 | 0,001 | 18,54 | >0.1 | 2,62 | 4,64 | 0 | 1,16 | 0 | 5,56 | 0 |
| GLI2 | CCCACCACCACG | -1 | 317 | 329 | -683 | -671 | 4,55 | 0,001 | 19,23 | >0.1 | 2,63 | 4,63 | 0 | 1,17 | 0 | 6,26 | 0 |
| GLI2 | ACCACCACGCCC | -1 | 314 | 326 | -686 | -674 | 4,23 | 0,01 | 18,8 | >0.1 | 2,64 | 4,63 | 0 | 1,17 | 0 | 6,13 | 0 |
| GLI2 | ACGCCCAGCTAA | -1 | 308 | 320 | -692 | -680 | 1,6 | 0,01 | 16,32 | >0.1 | 2,65 | 4,62 | 0 | 1,19 | 0 | 6,26 | 0 |
| GLI2 | TCGGCCTCCCAA | -1 | 214 | 226 | -786 | -774 | 2,99 | 0,01 | 17,63 | >0.1 | 2,98 | 4,49 | 0 | 1,54 | 0 | 5,63 | 0 |
| GLI2 | CCTCCCAAAGTG | -1 | 210 | 222 | -790 | -778 | -3,28 | 0,01 | 10,35 | >0.1 | 2,99 | 4,48 | 0 | 1,57 | 0 | 4,59 | 0 |
| GLI2 | GTGCCCAGCCAT | -1 | 174 | 186 | -826 | -814 | 1,97 | 0,01 | 14,64 | >0.1 | 3,15 | 4,44 | 0 | 1,84 | 0 | 3,24 | 0 |
| GLI2 | ATGACTACTCTC | -1 | 122 | 134 | -878 | -866 | -0,62 | 0,01 | 10,79 | >0.1 | 3,45 | 4,38 | 0 | 1,89 | 0 | 1,7 | 0 |
| MSX1 | GCACTCAA | 1 | 24 | 32 | -976 | -968 | 2,4 | 0,01 | 14,18 | >0.1 | 5,64 | 4,27 | 0 | 1,87 | 0 | 0 | 0 |
| MSX1 | GAAATGAG | 1 | 65 | 73 | -935 | -927 | 2,5 | 0,01 | 12,64 | >0.1 | 3,95 | 4,31 | 0 | 1,88 | 0 | 0 | 0 |
| MSX1 | GAAATTAA | 1 | 154 | 162 | -846 | -838 | 4,97 | 0,01 | 16,19 | >0.1 | 3,27 | 4,41 | 0 | 1,9 | 0 | 1,65 | 0 |
| MSX1 | CTGATCAG | 1 | 164 | 172 | -836 | -828 | 2,53 | 0,01 | 14,46 | >0.1 | 3,21 | 4,42 | 0 | 1,9 | 0 | 2,4 | 0 |
| MSX1 | CTGATCAG | 1 | 479 | 487 | -521 | -513 | 2,53 | 0,01 | 16,67 | >0.1 | 2,88 | 4,92 | 0 | 0,98 | 0 | 5,36 | 0 |
| MSX1 | CCAAGTAG | 1 | 545 | 553 | -455 | -447 | 5,42 | 0,01 | 14,62 | >0.1 | 3,15 | 5,06 | 0 | 0,98 | 0 | 0 | 0 |
| MSX1 | CCTATTAA | 1 | 600 | 608 | -400 | -392 | 3,34 | 0,01 | 14,39 | >0.1 | 3,47 | 5,2 | 0 | 1,01 | 0 | 1,36 | 0 |
| MSX1 | TTAATGAA | 1 | 604 | 612 | -396 | -388 | 4,25 | 0,01 | 15,39 | >0.1 | 3,49 | 5,21 | 0 | 1,02 | 0 | 1,41 | 0 |
| MSX1 | AAAATTAT | 1 | 683 | 691 | -317 | -309 | 2,81 | 0,01 | 16,79 | >0.1 | 4,34 | 5,46 | 0 | 1,12 | 0 | 3,07 | 0 |
| MSX1 | AAAATTAT | 1 | 830 | 838 | -170 | -162 | 2,81 | 0,01 | 19,54 | >0.1 | 5,64 | 6,14 | 0 | 1,67 | 0 | 3,28 | 0 |
| MSX1 | CTCATGAG | 1 | 984 | 992 | -16 | -8 | 4,07 | 0,01 | 21,59 | >0.1 | 5,64 | 8,43 | 0 | 1,87 | 0 | 1,57 | 0 |
| MSX1 | GGAATTAT | 1 | 1191 | 1199 | 191 | 199 | 3,27 | 0,01 | 22,05 | >0.1 | 5,64 | 6,02 | 0 | 1,5 | 1,79 | 3,82 | 0 |
| MSX1 | ATAATTCC | -1 | 1191 | 1199 | 191 | 199 | 3,47 | 0,01 | 22,25 | >0.1 | 5,64 | 6,02 | 0 | 1,5 | 1,79 | 3,82 | 0 |
| MSX1 | CCACTGAA | -1 | 1016 | 1024 | 16 | 24 | 3,44 | 0,01 | 21,14 | >0.1 | 5,64 | 8,81 | 0 | 1,86 | 0 | 1,39 | 0 |
| MSX1 | CTCATGAG | -1 | 984 | 992 | -16 | -8 | 4,07 | 0,01 | 21,59 | >0.1 | 5,64 | 8,43 | 0 | 1,87 | 0 | 1,57 | 0 |
| MSX1 | ACAATTTC | -1 | 609 | 617 | -391 | -383 | 4,24 | 0,01 | 15,56 | >0.1 | 3,53 | 5,23 | 0 | 1,02 | 0 | 1,55 | 0 |
| MSX1 | TTCATTAA | -1 | 604 | 612 | -396 | -388 | 4,65 | 0,01 | 15,79 | >0.1 | 3,49 | 5,21 | 0 | 1,02 | 0 | 1,41 | 0 |
| MSX1 | CTGATCAG | -1 | 479 | 487 | -521 | -513 | 2,53 | 0,01 | 16,67 | >0.1 | 2,88 | 4,92 | 0 | 0,98 | 0 | 5,36 | 0 |
| MSX1 | GCCATTTC | -1 | 171 | 179 | -829 | -821 | 2,79 | 0,01 | 15,69 | >0.1 | 3,18 | 4,43 | 0 | 1,87 | 0 | 3,43 | 0 |
| MSX1 | CTGATCAG | -1 | 164 | 172 | -836 | -828 | 2,53 | 0,01 | 14,46 | >0.1 | 3,21 | 4,42 | 0 | 1,9 | 0 | 2,4 | 0 |
| MSX1 | TTAATTTC | -1 | 154 | 162 | -846 | -838 | 2,72 | 0,01 | 13,94 | >0.1 | 3,27 | 4,41 | 0 | 1,9 | 0 | 1,65 | 0 |
| MSX1 | AAAATTAT | -1 | 132 | 140 | -868 | -860 | 2,81 | 0,01 | 14,05 | >0.1 | 3,4 | 4,38 | 0 | 1,89 | 0 | 1,57 | 0 |
| MSX1 | TCACTTAC | -1 | 5 | 13 | -995 | -987 | 5,09 | 0,01 | 18,64 | >0.1 | 5,64 | 4,26 | 0 | 1,86 | 1,79 | 0 | 0 |
| MSX2 | GAAATGAG | 1 | 65 | 73 | -935 | -927 | 0,81 | 0,01 | 10,95 | >0.1 | 3,95 | 4,31 | 0 | 1,88 | 0 | 0 | 0 |
| MSX2 | GAAATTAA | 1 | 154 | 162 | -846 | -838 | 5,8 | 0,01 | 17,02 | >0.1 | 3,27 | 4,41 | 0 | 1,9 | 0 | 1,65 | 0 |
| MSX2 | CTGATCAG | 1 | 164 | 172 | -836 | -828 | 0,83 | 0,01 | 12,76 | >0.1 | 3,21 | 4,42 | 0 | 1,9 | 0 | 2,4 | 0 |
| MSX2 | TCTATTAA | 1 | 288 | 296 | -712 | -704 | 3,88 | 0,01 | 17,61 | >0.1 | 2,72 | 4,58 | 0 | 1,24 | 0 | 5,19 | 0 |
| MSX2 | CTGATCAG | 1 | 479 | 487 | -521 | -513 | 0,83 | 0,01 | 14,97 | >0.1 | 2,88 | 4,92 | 0 | 0,98 | 0 | 5,36 | 0 |
| MSX2 | CCTATTAA | 1 | 600 | 608 | -400 | -392 | 6,6 | 0,01 | 17,65 | >0.1 | 3,47 | 5,2 | 0 | 1,01 | 0 | 1,36 | 0 |
| MSX2 | TTAATGAA | 1 | 604 | 612 | -396 | -388 | 1,97 | 0,01 | 13,11 | >0.1 | 3,49 | 5,21 | 0 | 1,02 | 0 | 1,41 | 0 |
| MSX2 | AAAATTAT | 1 | 683 | 691 | -317 | -309 | 3,41 | 0,01 | 17,39 | >0.1 | 4,34 | 5,46 | 0 | 1,12 | 0 | 3,07 | 0 |
| MSX2 | GCCATAAA | 1 | 758 | 766 | -242 | -234 | 1 | 0,01 | 19,8 | >0.1 | 5,64 | 5,75 | 0 | 1,31 | 0 | 6,1 | 0 |
| MSX2 | AAAATTAT | 1 | 830 | 838 | -170 | -162 | 3,41 | 0,01 | 20,14 | >0.1 | 5,64 | 6,14 | 0 | 1,67 | 0 | 3,28 | 0 |
| MSX2 | CTCATGAG | 1 | 984 | 992 | -16 | -8 | 0,87 | 0,01 | 18,39 | >0.1 | 5,64 | 8,43 | 0 | 1,87 | 0 | 1,57 | 0 |
| MSX2 | GGAATTAT | 1 | 1191 | 1199 | 191 | 199 | 3,84 | 0,01 | 22,62 | >0.1 | 5,64 | 6,02 | 0 | 1,5 | 1,79 | 3,82 | 0 |
| MSX2 | ATAATTCC | -1 | 1191 | 1199 | 191 | 199 | 2,15 | 0,01 | 20,93 | >0.1 | 5,64 | 6,02 | 0 | 1,5 | 1,79 | 3,82 | 0 |
| MSX2 | CTCATGAG | -1 | 984 | 992 | -16 | -8 | 0,87 | 0,01 | 18,39 | >0.1 | 5,64 | 8,43 | 0 | 1,87 | 0 | 1,57 | 0 |
| MSX2 | CAAATAAG | -1 | 811 | 819 | -189 | -181 | 0,87 | 0,01 | 17,36 | >0.1 | 5,64 | 6,02 | 0 | 1,55 | 0 | 3,28 | 0 |
| MSX2 | CAAATCAA | -1 | 750 | 758 | -250 | -242 | 2,99 | 0,01 | 21,54 | >0.1 | 5,64 | 5,71 | 0 | 1,28 | 0 | 5,92 | 0 |
| MSX2 | ACAATTTC | -1 | 609 | 617 | -391 | -383 | 2,31 | 0,01 | 13,63 | >0.1 | 3,53 | 5,23 | 0 | 1,02 | 0 | 1,55 | 0 |
| MSX2 | TTCATTAA | -1 | 604 | 612 | -396 | -388 | 3,13 | 0,01 | 14,27 | >0.1 | 3,49 | 5,21 | 0 | 1,02 | 0 | 1,41 | 0 |
| MSX2 | CTGATCAG | -1 | 479 | 487 | -521 | -513 | 0,83 | 0,01 | 14,97 | >0.1 | 2,88 | 4,92 | 0 | 0,98 | 0 | 5,36 | 0 |
| MSX2 | CTGATCAG | -1 | 164 | 172 | -836 | -828 | 0,83 | 0,01 | 12,76 | >0.1 | 3,21 | 4,42 | 0 | 1,9 | 0 | 2,4 | 0 |
| MSX2 | AAAATTAT | -1 | 132 | 140 | -868 | -860 | 3,41 | 0,01 | 14,65 | >0.1 | 3,4 | 4,38 | 0 | 1,89 | 0 | 1,57 | 0 |
| MSX2 | TCACTTAC | -1 | 5 | 13 | -995 | -987 | 2,69 | 0,01 | 16,24 | >0.1 | 5,64 | 4,26 | 0 | 1,86 | 1,79 | 0 | 0 |
| ZNF410 | TGAATTCCATAAAGACA | 1 | 492 | 509 | -508 | -491 | -16,58 | 0,01 | -1,06 | >0.1 | 2,94 | 4,96 | 0 | 0,99 | 0 | 6,62 | 0 |
| ZNF410 | GAGATAACCTATTAATG | 1 | 593 | 610 | -407 | -390 | -28,7 | 0,01 | -17,58 | >0.1 | 3,45 | 5,21 | 0 | 1,02 | 0 | 1,44 | 0 |
| ZNF410 | TGAAAACCATACAAACA | 1 | 618 | 635 | -382 | -365 | -23,11 | 0,01 | -11,71 | >0.1 | 3,63 | 5,28 | 0 | 1,04 | 0 | 1,44 | 0 |

| | | | | | | | | | | | | | | | | | |
|---|---|---|---|---|---|---|---|---|---|---|---|---|---|---|---|---|---|
| ZNF410 | TACAAACACCAACATTA | 1 | 627 | 644 | -373 | -356 | -29,28 | 0,01 | -17,48 | >0.1 | 3,71 | 5,31 | 0 | 1,05 | 0 | 1,73 | 0 |
| ZNF410 | CAAACACCAACATTATC | 1 | 629 | 646 | -371 | -354 | -29,01 | 0,01 | -17,31 | >0.1 | 3,73 | 5,31 | 0 | 1,06 | 0 | 1,61 | 0 |
| ZNF410 | CCTGTTCCTCAATGGTG | -1 | 1105 | 1122 | 105 | 122 | -29,16 | 0,01 | -9,34 | >0.1 | 5,52 | 6,7 | 0 | 1,85 | 3,22 | 2,53 | 0 |
| ZNF410 | GAAAGACCAAAAAAAGA | -1 | 921 | 938 | -79 | -62 | -24,03 | 0,01 | -8,53 | >0.1 | 5,52 | 7,03 | 0 | 1,88 | 0 | 1,06 | 0 |
| ZNF410 | AGAAGAACACAGTAAAC | -1 | 898 | 915 | -102 | -85 | -29,86 | 0,01 | -14,41 | >0.1 | 5,52 | 6,76 | 0 | 1,89 | 0 | 1,28 | 0 |
| ZNF410 | CACAGTAAACAATCTGC | -1 | 891 | 908 | -109 | -92 | -29,37 | 0,01 | -12,41 | >0.1 | 5,52 | 6,69 | 0 | 1,89 | 0 | 2,86 | 0 |
| ZNF410 | TGAATCAAATAAGAATA | -1 | 807 | 824 | -193 | -176 | -26,74 | 0,01 | -10,31 | >0.1 | 5,52 | 6,05 | 0 | 1,58 | 0 | 3,28 | 0 |
| ZNF410 | CCTGTCACTTTATGGCC | -1 | 757 | 774 | -243 | -226 | -29,82 | 0,01 | -11,26 | >0.1 | 5,52 | 5,79 | 0 | 1,34 | 0 | 5,92 | 0 |
| ZNF410 | TGCCCACCACCACGCCC | -1 | 314 | 331 | -686 | -669 | -29,81 | 0,01 | -15,11 | >0.1 | 2,63 | 4,64 | 0 | 1,18 | 0 | 6,26 | 0 |
| ZNF410 | GGCCTCCCAAAGTGCTA | -1 | 207 | 224 | -793 | -776 | -17,24 | 0,01 | -3,47 | >0.1 | 2,99 | 4,49 | 0 | 1,59 | 0 | 4,7 | 0 |
| ZNF410 | TGCAAAACTTAAAAAAG | -1 | 73 | 90 | -927 | -910 | -25,74 | 0,01 | -15,71 | >0.1 | 3,82 | 4,33 | 0 | 1,88 | 0 | 0 | 0 |
| ZNF410 | TATTTCTCACACTGGTT | -1 | 47 | 64 | -953 | -936 | -29,63 | 0,01 | -19,33 | >0.1 | 4,12 | 4,3 | 0 | 1,87 | 0 | 0 | 0 |
| NR3C2 | CTGGGCACAGTGGCTCA | 1 | 178 | 195 | -822 | -805 | -19,74 | 0,01 | -6,51 | >0.1 | 3,12 | 4,45 | 0 | 1,81 | 0 | 3,86 | 0 |
| NR3C2 | GGGCACAGTGGCTCACG | 1 | 180 | 197 | -820 | -803 | -18,76 | 0,01 | -5,55 | >0.1 | 3,11 | 4,45 | 0 | 1,79 | 0 | 3,86 | 0 |
| NR3C2 | CAGATCACAAGGTCAGG | 1 | 232 | 249 | -768 | -751 | -19,55 | 0,01 | -5,94 | >0.1 | 2,9 | 4,52 | 0 | 1,45 | 0 | 4,74 | 0 |
| NR3C2 | AGATAGAGACCGTCCTG | 1 | 249 | 266 | -751 | -734 | -14,66 | 0,01 | -0,94 | >0.1 | 2,83 | 4,54 | 0 | 1,37 | 0 | 4,98 | 0 |
| NR3C2 | ACTAACACGGTGAAACC | 1 | 266 | 283 | -734 | -717 | -20,65 | 0,01 | -6,91 | >0.1 | 2,78 | 4,57 | 0 | 1,31 | 0 | 5,09 | 0 |
| NR3C2 | GGGCACCTGTAGTCCCA | 1 | 326 | 343 | -674 | -657 | -0,85 | 0,001 | 13,01 | >0.1 | 2,6 | 4,66 | 0 | 1,15 | 0 | 5,46 | 0 |
| NR3C2 | GAGAATGGCGTGAACCT | 1 | 365 | 382 | -635 | -618 | -18,88 | 0,01 | -3,81 | >0.1 | 2,54 | 4,72 | 0 | 1,08 | 0 | 6,73 | 0 |
| NR3C2 | CGCGCCACTGTGCTCCA | 1 | 411 | 428 | -589 | -572 | -13,63 | 0,01 | 0,06 | >0.1 | 2,67 | 4,8 | 0 | 1,03 | 0 | 5,19 | 0 |
| NR3C2 | GGGAAAATGTTGCCCAA | 1 | 532 | 549 | -468 | -451 | -16,05 | 0,01 | -5,28 | >0.1 | 3,11 | 5,05 | 0 | 0,99 | 0 | 1,61 | 0 |
| NR3C2 | ACCAACATTATCTTCAT | 1 | 634 | 651 | -366 | -349 | -18,06 | 0,01 | -6,29 | >0.1 | 3,77 | 5,33 | 0 | 1,06 | 0 | 1,61 | 0 |
| NR3C2 | TAGACCTCTTTGGTCAC | 1 | 663 | 680 | -337 | -320 | -17,88 | 0,01 | -3,43 | >0.1 | 4,09 | 5,42 | 0 | 1,1 | 0 | 3,83 | 0 |
| NR3C2 | ATTAACCAAAAGTACCG | 1 | 730 | 747 | -270 | -253 | -13,33 | 0,01 | 5,02 | >0.1 | 5,52 | 5,67 | 0 | 1,25 | 0 | 5,92 | 0 |
| NR3C2 | TTGGCCATAAAGTGACA | 1 | 755 | 772 | -245 | -228 | -17,84 | 0,01 | 0,89 | >0.1 | 5,52 | 5,78 | 0 | 1,33 | 0 | 6,1 | 0 |
| NR3C2 | GGAGAGGTAAGGTTCTC | 1 | 772 | 789 | -228 | -211 | -16,46 | 0,01 | 2,64 | >0.1 | 5,52 | 5,86 | 0 | 1,4 | 0 | 6,32 | 0 |
| NR3C2 | CAGCAGATTGTTTACTG | 1 | 889 | 906 | -111 | -94 | -17,47 | 0,01 | -0,52 | >0.1 | 5,52 | 6,67 | 0 | 1,89 | 0 | 2,86 | 0 |
| NR3C2 | TTGTTTACTGTGTTCTT | 1 | 896 | 913 | -104 | -87 | -16,49 | 0,01 | 0,41 | >0.1 | 5,52 | 6,74 | 0 | 1,89 | 0 | 2,75 | 0 |
| NR3C2 | GATGTCAAGCTCTTCCT | 1 | 1048 | 1065 | 48 | 65 | -19,04 | 0,01 | -2,57 | >0.1 | 5,52 | 7,62 | 0 | 1,86 | 0 | 1,46 | 0 |
| NR3C2 | AAGTTCACTTGCTTCTT | 1 | 1174 | 1191 | 174 | 191 | -19,23 | 0,01 | -0,89 | >0.1 | 5,52 | 6,13 | 0 | 1,63 | 1,79 | 3,26 | 0 |
| NR3C2 | AAGAAGCAAGTGAACTT | -1 | 1174 | 1191 | 174 | 191 | -19,67 | 0,01 | -1,33 | >0.1 | 5,52 | 6,13 | 0 | 1,63 | 1,79 | 3,26 | 0 |
| NR3C2 | TGGTTAAACTTGTCCAA | -1 | 1133 | 1150 | 133 | 150 | -20,81 | 0,01 | -0,7 | >0.1 | 5,52 | 6,43 | 0 | 1,84 | 3,22 | 3,09 | 0 |
| NR3C2 | AAGAACACAGTAAACAA | -1 | 896 | 913 | -104 | -87 | -18,41 | 0,01 | -1,51 | >0.1 | 5,52 | 6,74 | 0 | 1,89 | 0 | 2,75 | 0 |
| NR3C2 | CAGTAAACAATCTGCTG | -1 | 889 | 906 | -111 | -94 | -15,14 | 0,01 | 1,81 | >0.1 | 5,52 | 6,67 | 0 | 1,89 | 0 | 2,86 | 0 |
| NR3C2 | GAGAACCTTACCTCTCC | -1 | 772 | 789 | -228 | -211 | -15,52 | 0,01 | 3,58 | >0.1 | 5,52 | 5,86 | 0 | 1,4 | 0 | 6,32 | 0 |
| NR3C2 | TGTCACTTTATGGCCAA | -1 | 755 | 772 | -245 | -228 | -20,02 | 0,01 | -1,29 | >0.1 | 5,52 | 5,78 | 0 | 1,33 | 0 | 6,1 | 0 |
| NR3C2 | CGGTACTTTTGGTTAAT | -1 | 730 | 747 | -270 | -253 | -18,04 | 0,01 | 0,31 | >0.1 | 5,52 | 5,67 | 0 | 1,25 | 0 | 5,92 | 0 |
| NR3C2 | GTGACCAAAGAGGTCTA | -1 | 663 | 680 | -337 | -320 | -16,41 | 0,01 | -1,96 | >0.1 | 4,09 | 5,42 | 0 | 1,1 | 0 | 3,83 | 0 |
| NR3C2 | ATGAAGATAATGTTGGT | -1 | 634 | 651 | -366 | -349 | -16,1 | 0,01 | -4,33 | >0.1 | 3,77 | 5,33 | 0 | 1,06 | 0 | 1,61 | 0 |
| NR3C2 | TTGGGCAACATTTTCCC | -1 | 532 | 549 | -468 | -451 | -15,93 | 0,01 | -5,16 | >0.1 | 3,11 | 5,05 | 0 | 0,99 | 0 | 1,61 | 0 |
| NR3C2 | TGGAGCACAGTGGCGCG | -1 | 411 | 428 | -589 | -572 | -14,66 | 0,01 | -0,97 | >0.1 | 2,67 | 4,8 | 0 | 1,03 | 0 | 5,19 | 0 |
| NR3C2 | AGGTTCACGCCATTCTC | -1 | 365 | 382 | -635 | -618 | -17,38 | 0,01 | -2,31 | >0.1 | 2,54 | 4,72 | 0 | 1,08 | 0 | 6,73 | 0 |
| NR3C2 | TGGGACTACAGGTGCCC | -1 | 326 | 343 | -674 | -657 | -1,43 | 0,001 | 12,43 | >0.1 | 2,6 | 4,66 | 0 | 1,15 | 0 | 5,46 | 0 |
| NR3C2 | GGTTTCACCGTGTTAGT | -1 | 266 | 283 | -734 | -717 | -17,99 | 0,01 | -4,25 | >0.1 | 2,78 | 4,57 | 0 | 1,31 | 0 | 5,09 | 0 |
| NR3C2 | CAGGACGGTCTCTATCT | -1 | 249 | 266 | -751 | -734 | -13,88 | 0,01 | -0,16 | >0.1 | 2,83 | 4,54 | 0 | 1,37 | 0 | 4,98 | 0 |
| NR3C2 | ACGGTCTCTATCTCCTG | -1 | 245 | 262 | -755 | -738 | -20,14 | 0,01 | -6,28 | >0.1 | 2,85 | 4,54 | 0 | 1,39 | 0 | 5,09 | 0 |
| NR3C2 | CGTGAGCCACTGTGCCC | -1 | 180 | 197 | -820 | -803 | -17,34 | 0,01 | -4,13 | >0.1 | 3,11 | 4,45 | 0 | 1,79 | 0 | 3,86 | 0 |
| NR3C2 | TGAGCCACTGTGCCCAG | -1 | 178 | 195 | -822 | -805 | -17,46 | 0,01 | -4,23 | >0.1 | 3,12 | 4,45 | 0 | 1,81 | 0 | 3,86 | 0 |
| NR3C1 | CTGGGCACAGTGGCTCA | 1 | 178 | 195 | -822 | -805 | -15,94 | 0,01 | -2,71 | >0.1 | 3,12 | 4,45 | 0 | 1,81 | 0 | 3,86 | 0 |
| NR3C1 | GGGCACAGTGGCTCACG | 1 | 180 | 197 | -820 | -803 | -14,89 | 0,01 | -1,68 | >0.1 | 3,11 | 4,45 | 0 | 1,79 | 0 | 3,86 | 0 |
| NR3C1 | GAGGCCGAGTTGGGCAG | 1 | 218 | 235 | -782 | -765 | -18,09 | 0,01 | -3,6 | >0.1 | 2,95 | 4,5 | 0 | 1,52 | 0 | 5,52 | 0 |
| NR3C1 | CAGATCACAAGGTCAGG | 1 | 232 | 249 | -768 | -751 | -17,35 | 0,01 | -3,74 | >0.1 | 2,9 | 4,52 | 0 | 1,45 | 0 | 4,74 | 0 |
| NR3C1 | AGATAGAGACCGTCCTG | 1 | 249 | 266 | -751 | -734 | -11,88 | 0,01 | 1,84 | >0.1 | 2,83 | 4,54 | 0 | 1,37 | 0 | 4,98 | 0 |
| NR3C1 | ACTAACACGGTGAAACC | 1 | 266 | 283 | -734 | -717 | -15,75 | 0,01 | -2,01 | >0.1 | 2,78 | 4,57 | 0 | 1,31 | 0 | 5,09 | 0 |
| NR3C1 | GGGCACCTGTAGTCCCA | 1 | 326 | 343 | -674 | -657 | -2,36 | 0,001 | 11,5 | >0.1 | 2,6 | 4,66 | 0 | 1,15 | 0 | 5,46 | 0 |
| NR3C1 | GAGAATGGCGTGAACCT | 1 | 365 | 382 | -635 | -618 | -15,03 | 0,01 | 0,04 | >0.1 | 2,54 | 4,72 | 0 | 1,08 | 0 | 6,73 | 0 |
| NR3C1 | CGCGCCACTGTGCTCCA | 1 | 411 | 428 | -589 | -572 | -14,06 | 0,01 | -0,37 | >0.1 | 2,67 | 4,8 | 0 | 1,03 | 0 | 5,19 | 0 |
| NR3C1 | GGGAAAATGTTGCCCAA | 1 | 532 | 549 | -468 | -451 | -16,02 | 0,01 | -5,25 | >0.1 | 3,11 | 5,05 | 0 | 0,99 | 0 | 1,61 | 0 |
| NR3C1 | ACCAACATTATCTTCAT | 1 | 634 | 651 | -366 | -349 | -15,58 | 0,01 | -3,81 | >0.1 | 3,77 | 5,33 | 0 | 1,06 | 0 | 1,61 | 0 |

| | | | | | | | | | | | | | | | | | |
|---|---|---|---|---|---|---|---|---|---|---|---|---|---|---|---|---|---|
| NR3C1 | TAGACCTCTTTGGTCAC | 1 | 663 | 680 | -337 | -320 | -15,06 | 0,01 | -0,61 | >0.1 | 4,09 | 5,42 | 0 | 1,1 | 0 | 3,83 | 0 |
| NR3C1 | ATTAACCAAAAGTACCG | 1 | 730 | 747 | -270 | -253 | -15,26 | 0,01 | 3,09 | >0.1 | 5,52 | 5,67 | 0 | 1,25 | 0 | 5,92 | 0 |
| NR3C1 | GGAGAGGTAAGGTTCTC | 1 | 772 | 789 | -228 | -211 | -11,91 | 0,01 | 7,19 | >0.1 | 5,52 | 5,86 | 0 | 1,4 | 0 | 6,32 | 0 |
| NR3C1 | AAGAATAACGTATTCTT | 1 | 797 | 814 | -203 | -186 | -18,49 | 0,01 | -1,91 | >0.1 | 5,52 | 5,99 | 0 | 1,52 | 0 | 3,55 | 0 |
| NR3C1 | TGTTACATATCTGTCCT | 1 | 847 | 864 | -153 | -136 | -17,71 | 0,01 | -1,21 | >0.1 | 5,52 | 6,33 | 0 | 1,9 | 0 | 2,75 | 0 |
| NR3C1 | CAGCAGATTGTTTACTG | 1 | 889 | 906 | -111 | -94 | -9,69 | 0,01 | 7,26 | >0.1 | 5,52 | 6,67 | 0 | 1,89 | 0 | 2,86 | 0 |
| NR3C1 | TTGTTTACTGTGTTCTT | 1 | 896 | 913 | -104 | -87 | -14,59 | 0,01 | 2,31 | >0.1 | 5,52 | 6,74 | 0 | 1,89 | 0 | 2,75 | 0 |
| NR3C1 | TGGCTCACAGGGGACGA | 1 | 1033 | 1050 | 33 | 50 | -16,09 | 0,01 | 0,73 | >0.1 | 5,52 | 8,1 | 0 | 1,86 | 0 | 1,34 | 0 |
| NR3C1 | GATGTCAAGCTCTTCCT | 1 | 1048 | 1065 | 48 | 65 | -17,08 | 0,01 | -0,61 | >0.1 | 5,52 | 7,62 | 0 | 1,86 | 0 | 1,46 | 0 |
| NR3C1 | TTGGACAAGTTTAACCA | 1 | 1133 | 1150 | 133 | 150 | -15,67 | 0,01 | 4,44 | >0.1 | 5,52 | 6,43 | 0 | 1,84 | 3,22 | 3,09 | 0 |
| NR3C1 | AAGTTCACTTGCTTCTT | 1 | 1174 | 1191 | 174 | 191 | -13,7 | 0,01 | 4,64 | >0.1 | 5,52 | 6,13 | 0 | 1,63 | 1,79 | 3,26 | 0 |
| NR3C1 | AAGAAGCAAGTGAACTT | -1 | 1174 | 1191 | 174 | 191 | -14,98 | 0,01 | 3,36 | >0.1 | 5,52 | 6,13 | 0 | 1,63 | 1,79 | 3,26 | 0 |
| NR3C1 | AGGAAGAGCTTGACATC | -1 | 1048 | 1065 | 48 | 65 | -17,51 | 0,01 | -1,04 | >0.1 | 5,52 | 7,62 | 0 | 1,86 | 0 | 1,46 | 0 |
| NR3C1 | AAGAACACAGTAAACAA | -1 | 896 | 913 | -104 | -87 | -14,93 | 0,01 | 1,97 | >0.1 | 5,52 | 6,74 | 0 | 1,89 | 0 | 2,75 | 0 |
| NR3C1 | CAGTAAACAATCTGCTG | -1 | 889 | 906 | -111 | -94 | -13,63 | 0,01 | 3,32 | >0.1 | 5,52 | 6,67 | 0 | 1,89 | 0 | 2,86 | 0 |
| NR3C1 | AAGAATACGTTATTCTT | -1 | 797 | 814 | -203 | -186 | -18,04 | 0,01 | -1,46 | >0.1 | 5,52 | 5,99 | 0 | 1,52 | 0 | 3,55 | 0 |
| NR3C1 | GAGAACCTTACCTCTCC | -1 | 772 | 789 | -228 | -211 | -13,87 | 0,01 | 5,23 | >0.1 | 5,52 | 5,86 | 0 | 1,4 | 0 | 6,32 | 0 |
| NR3C1 | TGTCACTTTATGGCCAA | -1 | 755 | 772 | -245 | -228 | -16,73 | 0,01 | 2 | >0.1 | 5,52 | 5,78 | 0 | 1,33 | 0 | 6,1 | 0 |
| NR3C1 | CGGTACTTTTGGTTAAT | -1 | 730 | 747 | -270 | -253 | -15,42 | 0,01 | 2,93 | >0.1 | 5,52 | 5,67 | 0 | 1,25 | 0 | 5,92 | 0 |
| NR3C1 | GTGACCAAAGAGGTCTA | -1 | 663 | 680 | -337 | -320 | -14,23 | 0,01 | 0,22 | >0.1 | 4,09 | 5,42 | 0 | 1,1 | 0 | 3,83 | 0 |
| NR3C1 | ATGAAGATAATGTTGGT | -1 | 634 | 651 | -366 | -349 | -14,26 | 0,01 | -2,49 | >0.1 | 3,77 | 5,33 | 0 | 1,06 | 0 | 1,61 | 0 |
| NR3C1 | TTGGGCAACATTTTCCC | -1 | 532 | 549 | -468 | -451 | -11,3 | 0,01 | -0,53 | >0.1 | 3,11 | 5,05 | 0 | 0,99 | 0 | 1,61 | 0 |
| NR3C1 | AAATACATGACTTTGCC | -1 | 511 | 528 | -489 | -472 | -18,01 | 0,01 | -6,05 | >0.1 | 3,02 | 5,01 | 0 | 0,99 | 0 | 2,95 | 0 |
| NR3C1 | TGGAGCACAGTGGCGCG | -1 | 411 | 428 | -589 | -572 | -11,36 | 0,01 | 2,33 | >0.1 | 2,67 | 4,8 | 0 | 1,03 | 0 | 5,19 | 0 |
| NR3C1 | AGGTTCACGCCATTCTC | -1 | 365 | 382 | -635 | -618 | -14,21 | 0,01 | 0,86 | >0.1 | 2,54 | 4,72 | 0 | 1,08 | 0 | 6,73 | 0 |
| NR3C1 | TGGGACTACAGGTGCCC | -1 | 326 | 343 | -674 | -657 | -1,59 | 0,001 | 12,27 | >0.1 | 2,6 | 4,66 | 0 | 1,15 | 0 | 5,46 | 0 |
| NR3C1 | GGTTTCACCGTGTTAGT | -1 | 266 | 283 | -734 | -717 | -14,87 | 0,01 | -1,13 | >0.1 | 2,78 | 4,57 | 0 | 1,31 | 0 | 5,09 | 0 |
| NR3C1 | CAGGACGGTCTCTATCT | -1 | 249 | 266 | -751 | -734 | -12,52 | 0,01 | 1,2 | >0.1 | 2,83 | 4,54 | 0 | 1,37 | 0 | 4,98 | 0 |
| NR3C1 | CGTGAGCCACTGTGCCC | -1 | 180 | 197 | -820 | -803 | -16,03 | 0,01 | -2,82 | >0.1 | 3,11 | 4,45 | 0 | 1,79 | 0 | 3,86 | 0 |
| RARA(VAR.2) | AGGTAAGTGAAGGAGAA | 1 | 3 | 20 | -997 | -980 | -13,6 | 0,01 | -0,16 | >0.1 | 5,52 | 4,26 | 0 | 1,86 | 1,79 | 0 | 0 |
| RARA(VAR.2) | AGTGAAGGAGAAGGAGG | 1 | 8 | 25 | -992 | -975 | -15,65 | 0,01 | -2,2 | >0.1 | 5,52 | 4,27 | 0 | 1,87 | 1,79 | 0 | 0 |
| RARA(VAR.2) | GGCACTCAAAGAAGTCA | 1 | 23 | 40 | -977 | -960 | -14,78 | 0,01 | -3,11 | >0.1 | 5,52 | 4,28 | 0 | 1,87 | 0 | 0 | 0 |
| RARA(VAR.2) | ATTTTAAAAAATGGCCA | 1 | 135 | 152 | -865 | -848 | -15,09 | 0,01 | -3,8 | >0.1 | 3,35 | 4,4 | 0 | 1,9 | 0 | 1,65 | 0 |
| RARA(VAR.2) | AGCACTTTGGGAGGCCG | 1 | 208 | 225 | -792 | -775 | -16,25 | 0,01 | -2,6 | >0.1 | 2,99 | 4,49 | 0 | 1,58 | 0 | 4,59 | 0 |
| RARA(VAR.2) | AGGCCGAGTTGGGCAGA | 1 | 219 | 236 | -781 | -764 | -11,84 | 0,01 | 2,64 | >0.1 | 2,94 | 4,5 | 0 | 1,51 | 0 | 5,52 | 0 |
| RARA(VAR.2) | GGCAGATCACAAGGTCA | 1 | 230 | 247 | -770 | -753 | -3,29 | 0,001 | 10,48 | >0.1 | 2,9 | 4,52 | 0 | 1,46 | 0 | 4,9 | 0 |
| RARA(VAR.2) | AGGTCAGGAGATAGAGA | 1 | 241 | 258 | -759 | -742 | -11,5 | 0,01 | 1,72 | >0.1 | 2,86 | 4,53 | 0 | 1,41 | 0 | 4,42 | 0 |
| RARA(VAR.2) | CGTGAACCTGGGAGGCA | 1 | 373 | 390 | -627 | -610 | -16,78 | 0,01 | -2,02 | >0.1 | 2,56 | 4,73 | 0 | 1,07 | 0 | 6,4 | 0 |
| RARA(VAR.2) | AGACAAGGGCAAAGTCA | 1 | 504 | 521 | -496 | -479 | -16,07 | 0,01 | -2,91 | >0.1 | 2,99 | 4,99 | 0 | 0,99 | 0 | 4,19 | 0 |
| RARA(VAR.2) | AGGGCAAAGTCATGTAT | 1 | 509 | 526 | -491 | -474 | -15,16 | 0,01 | -3,21 | >0.1 | 3,01 | 5 | 0 | 0,99 | 0 | 2,95 | 0 |
| RARA(VAR.2) | AGTGACAGGAGAGGTAA | 1 | 765 | 782 | -235 | -218 | -9,53 | 0,01 | 9,1 | >0.1 | 5,52 | 5,83 | 0 | 1,37 | 0 | 5,92 | 0 |
| RARA(VAR.2) | AGGAGAGGTAAGGTTCT | 1 | 771 | 788 | -229 | -212 | -4,77 | 0,001 | 14,1 | >0.1 | 5,52 | 5,85 | 0 | 1,39 | 0 | 6,1 | 0 |
| RARA(VAR.2) | AGGTTCTCTAGGATTAA | 1 | 781 | 798 | -219 | -202 | -13,67 | 0,01 | 5,11 | >0.1 | 5,52 | 5,91 | 0 | 1,44 | 0 | 5,92 | 0 |
| RARA(VAR.2) | CGCCCAACCCAAGTTCA | 1 | 946 | 963 | -54 | -37 | -1,98 | 0,001 | 14,62 | >0.1 | 5,52 | 7,46 | 0 | 1,88 | 0 | 1,74 | 0 |
| RARA(VAR.2) | ATCTCATGAGGAGGTTT | 1 | 982 | 999 | -18 | -1 | -10,59 | 0,01 | 7,06 | >0.1 | 5,52 | 8,69 | 0 | 1,87 | 0 | 1,57 | 0 |
| RARA(VAR.2) | AAGTCATTCAGTGGATG | 1 | 1010 | 1027 | 10 | 27 | -17,26 | 0,01 | 0,53 | >0.1 | 5,52 | 8,99 | 0 | 1,86 | 0 | 1,42 | 0 |
| RARA(VAR.2) | TGTTCTATCAAAGTTCA | 1 | 1164 | 1181 | 164 | 181 | -6,91 | 0,01 | 11,18 | >0.1 | 5,52 | 6,19 | 0 | 1,72 | 1,79 | 2,86 | 0 |
| RARA(VAR.2) | AGGCTGAGAAGGAGCCA | -1 | 1064 | 1081 | 64 | 81 | -13,64 | 0,01 | 5,08 | >0.1 | 5,52 | 7,28 | 0 | 1,85 | 1,79 | 2,27 | 0 |
| RARA(VAR.2) | AGCCAATATAAAGTTCA | -1 | 872 | 889 | -128 | -111 | -4,04 | 0,001 | 12,85 | >0.1 | 5,52 | 6,52 | 0 | 1,9 | 0 | 2,95 | 0 |
| RARA(VAR.2) | AGTTCATCCTGGAGAGG | -1 | 861 | 878 | -139 | -122 | -10,21 | 0,01 | 6,39 | >0.1 | 5,52 | 6,43 | 0 | 1,9 | 0 | 2,75 | 0 |
| RARA(VAR.2) | CTGTCACTTTATGGCCA | -1 | 756 | 773 | -244 | -227 | -7,55 | 0,01 | 11,19 | >0.1 | 5,52 | 5,78 | 0 | 1,33 | 0 | 6,1 | 0 |
| RARA(VAR.2) | AGTGACCAAAGAGGTCT | -1 | 664 | 681 | -336 | -319 | -9,03 | 0,01 | 5,43 | >0.1 | 4,11 | 5,42 | 0 | 1,1 | 0 | 3,83 | 0 |
| RARA(VAR.2) | ATTTCATTAATAGGTTA | -1 | 597 | 614 | -403 | -386 | -5,07 | 0,001 | 6,01 | >0.1 | 3,48 | 5,22 | 0 | 1,02 | 0 | 1,36 | 0 |
| RARA(VAR.2) | AGGTTATCTCACATTTA | -1 | 586 | 603 | -414 | -397 | -10,89 | 0,01 | -1,28 | >0.1 | 3,4 | 5,19 | 0 | 1,02 | 0 | 0 | 0 |
| RARA(VAR.2) | GGCGCGATCTCGGCTCA | -1 | 400 | 417 | -600 | -583 | -13,88 | 0,01 | 0,34 | >0.1 | 2,63 | 4,78 | 0 | 1,04 | 0 | 5,76 | 0 |
| RARA(VAR.2) | GGCTCACTGCAAGCTCT | -1 | 389 | 406 | -611 | -594 | -3,75 | 0,001 | 10,69 | >0.1 | 2,6 | 4,76 | 0 | 1,05 | 0 | 6,02 | 0 |
| RARA(VAR.2) | GGTTCACGCCATTCTCC | -1 | 364 | 381 | -636 | -619 | -15,22 | 0,01 | -0,16 | >0.1 | 2,54 | 4,72 | 0 | 1,08 | 0 | 6,73 | 0 |
| RARA(VAR.2) | CGGCCTCCCAAAGTGCT | -1 | 208 | 225 | -792 | -775 | -11,59 | 0,01 | 2,06 | >0.1 | 2,99 | 4,49 | 0 | 1,58 | 0 | 4,59 | 0 |
| SREBF1(VAR.2) | ATCAGGAGAG | 1 | 97 | 107 | -903 | -893 | 2,27 | 0,001 | 12,9 | >0.1 | 3,64 | 4,35 | 0 | 1,88 | 0 | 0,77 | 0 |

| | | | | | | | | | | | | | | | | | |
|---|---|---|---|---|---|---|---|---|---|---|---|---|---|---|---|---|---|
| SREBF1(VAR.2) | CTCACGCCTG | 1 | 191 | 201 | -809 | -799 | -4,89 | 0,01 | 8,47 | >0.1 | 3,08 | 4,45 | 0 | 1,7 | 0 | 4,13 | 0 |
| SREBF1(VAR.2) | CACGCCTGTA | 1 | 193 | 203 | -807 | -797 | -6,66 | 0,01 | 6,81 | >0.1 | 3,07 | 4,46 | 0 | 1,68 | 0 | 4,26 | 0 |
| SREBF1(VAR.2) | ATCACAAGGT | 1 | 235 | 245 | -765 | -755 | -4,54 | 0,01 | 9,04 | >0.1 | 2,9 | 4,51 | 0 | 1,43 | 0 | 4,74 | 0 |
| SREBF1(VAR.2) | GTCAGGAGAT | 1 | 243 | 253 | -757 | -747 | 4,94 | 0,001 | 18,05 | >0.1 | 2,87 | 4,52 | 0 | 1,39 | 0 | 4,33 | 0 |
| SREBF1(VAR.2) | GGCACCTGTA | 1 | 327 | 337 | -673 | -663 | -2,51 | 0,01 | 11,45 | >0.1 | 2,6 | 4,65 | 0 | 1,14 | 0 | 5,56 | 0 |
| SREBF1(VAR.2) | GGCAGGAGAA | 1 | 360 | 370 | -640 | -630 | -6,95 | 0,01 | 7,72 | >0.1 | 2,52 | 4,7 | 0 | 1,09 | 0 | 6,36 | 0 |
| SREBF1(VAR.2) | ATGGCGTGAA | 1 | 369 | 379 | -631 | -621 | -6,71 | 0,01 | 7,77 | >0.1 | 2,54 | 4,71 | 0 | 1,07 | 0 | 6,15 | 0 |
| SREBF1(VAR.2) | ATCGCGCCAC | 1 | 409 | 419 | -591 | -581 | 0,53 | 0,01 | 14,52 | >0.1 | 2,65 | 4,78 | 0 | 1,03 | 0 | 5,52 | 0 |
| SREBF1(VAR.2) | GTCATGTATT | 1 | 517 | 527 | -483 | -473 | -5,7 | 0,01 | 6,26 | >0.1 | 3,03 | 5 | 0 | 0,98 | 0 | 2,95 | 0 |
| SREBF1(VAR.2) | ATAAAGTGAC | 1 | 761 | 771 | -239 | -229 | -4,31 | 0,01 | 14,5 | >0.1 | 5,61 | 5,77 | 0 | 1,32 | 0 | 6,1 | 0 |
| SREBF1(VAR.2) | GACAGGAGAG | 1 | 768 | 778 | -232 | -222 | -1,47 | 0,01 | 17,21 | >0.1 | 5,61 | 5,81 | 0 | 1,35 | 0 | 5,92 | 0 |
| SREBF1(VAR.2) | ATAACGTATT | 1 | 801 | 811 | -199 | -189 | -6,68 | 0,01 | 9,83 | >0.1 | 5,61 | 5,98 | 0 | 1,5 | 0 | 3,43 | 0 |
| SREBF1(VAR.2) | CTCAGCAGAT | 1 | 887 | 897 | -113 | -103 | 0,04 | 0,01 | 17,2 | >0.1 | 5,61 | 6,59 | 0 | 1,89 | 0 | 3,07 | 0 |
| SREBF1(VAR.2) | ATCTCATGAG | 1 | 982 | 992 | -18 | -8 | -1,3 | 0,01 | 16,14 | >0.1 | 5,61 | 8,43 | 0 | 1,87 | 0 | 1,53 | 0 |
| SREBF1(VAR.2) | CACAGGGGAC | 1 | 1038 | 1048 | 38 | 48 | -0,74 | 0,01 | 15,98 | >0.1 | 5,61 | 7,92 | 0 | 1,86 | 0 | 1,34 | 0 |
| SREBF1(VAR.2) | AACTGCTGCT | 1 | 1090 | 1100 | 90 | 100 | -6,85 | 0,01 | 11,56 | >0.1 | 5,61 | 6,88 | 0 | 1,85 | 1,79 | 2,27 | 0 |
| SREBF1(VAR.2) | AAGACCTGTT | 1 | 1158 | 1168 | 158 | 168 | -6,14 | 0,01 | 10,45 | >0.1 | 4,38 | 6,24 | 0 | 1,78 | 1,79 | 2,4 | 0 |
| SREBF1(VAR.2) | AACAGGTCTT | -1 | 1158 | 1168 | 158 | 168 | -3,07 | 0,01 | 13,52 | >0.1 | 4,38 | 6,24 | 0 | 1,78 | 1,79 | 2,4 | 0 |
| SREBF1(VAR.2) | GTCCCCTGTG | -1 | 1038 | 1048 | 38 | 48 | -6,51 | 0,01 | 10,21 | >0.1 | 5,61 | 7,92 | 0 | 1,86 | 0 | 1,34 | 0 |
| SREBF1(VAR.2) | ACCTCCTCAT | -1 | 987 | 997 | -13 | -3 | -5,44 | 0,01 | 12,28 | >0.1 | 5,61 | 8,64 | 0 | 1,87 | 0 | 1,61 | 0 |
| SREBF1(VAR.2) | CTCATGAGAT | -1 | 982 | 992 | -18 | -8 | 0,94 | 0,01 | 18,38 | >0.1 | 5,61 | 8,43 | 0 | 1,87 | 0 | 1,53 | 0 |
| SREBF1(VAR.2) | ATCTGCTGAG | -1 | 887 | 897 | -113 | -103 | -1,29 | 0,01 | 15,87 | >0.1 | 5,61 | 6,59 | 0 | 1,89 | 0 | 3,07 | 0 |
| SREBF1(VAR.2) | CTCTCCTGTC | -1 | 768 | 778 | -232 | -222 | 0,54 | 0,01 | 19,22 | >0.1 | 5,61 | 5,81 | 0 | 1,35 | 0 | 5,92 | 0 |
| SREBF1(VAR.2) | AATACATGAC | -1 | 517 | 527 | -483 | -473 | -6,69 | 0,01 | 5,27 | >0.1 | 3,03 | 5 | 0 | 0,98 | 0 | 2,95 | 0 |
| SREBF1(VAR.2) | GTGGCGCGAT | -1 | 409 | 419 | -591 | -581 | -5,85 | 0,01 | 8,14 | >0.1 | 2,65 | 4,78 | 0 | 1,03 | 0 | 5,52 | 0 |
| SREBF1(VAR.2) | CTCGGCTCAC | -1 | 399 | 409 | -601 | -591 | -6,06 | 0,01 | 8,28 | >0.1 | 2,62 | 4,77 | 0 | 1,04 | 0 | 5,91 | 0 |
| SREBF1(VAR.2) | TTCACGCCAT | -1 | 369 | 379 | -631 | -621 | -0,74 | 0,01 | 13,74 | >0.1 | 2,54 | 4,71 | 0 | 1,07 | 0 | 6,15 | 0 |
| SREBF1(VAR.2) | TTCTCCTGCC | -1 | 360 | 370 | -640 | -630 | -4,4 | 0,01 | 10,27 | >0.1 | 2,52 | 4,7 | 0 | 1,09 | 0 | 6,36 | 0 |
| SREBF1(VAR.2) | GCCTCCCGAG | -1 | 347 | 357 | -653 | -643 | -5,44 | 0,01 | 9,42 | >0.1 | 2,55 | 4,68 | 0 | 1,11 | 0 | 6,52 | 0 |
| SREBF1(VAR.2) | TACAGGTGCC | -1 | 327 | 337 | -673 | -663 | -2,19 | 0,01 | 11,77 | >0.1 | 2,6 | 4,65 | 0 | 1,14 | 0 | 5,56 | 0 |
| SREBF1(VAR.2) | ATCTCCTGAC | -1 | 243 | 253 | -757 | -747 | 7,58 | 0,001 | 20,69 | >0.1 | 2,87 | 4,52 | 0 | 1,39 | 0 | 4,33 | 0 |
| SREBF1(VAR.2) | ACCTTGTGAT | -1 | 235 | 245 | -765 | -755 | -1,63 | 0,01 | 11,95 | >0.1 | 2,9 | 4,51 | 0 | 1,43 | 0 | 4,74 | 0 |
| SREBF1(VAR.2) | CACAGGTCAA | -1 | 107 | 117 | -893 | -883 | -6,78 | 0,01 | 3,79 | >0.1 | 3,56 | 4,36 | 0 | 1,89 | 0 | 0,77 | 0 |
| SREBF1(VAR.2) | CTCTCCTGAT | -1 | 97 | 107 | -903 | -893 | 4,37 | 0,001 | 15 | >0.1 | 3,64 | 4,35 | 0 | 1,88 | 0 | 0,77 | 0 |
| MYCN | TGACCTGTGGAG | 1 | 108 | 120 | -892 | -880 | 0,31 | 0,01 | 10,85 | >0.1 | 3,54 | 4,36 | 0 | 1,89 | 0 | 0,75 | 0 |
| MYCN | GATCACAAGGTC | 1 | 234 | 246 | -766 | -754 | 3,45 | 0,01 | 17,04 | >0.1 | 2,9 | 4,51 | 0 | 1,44 | 0 | 4,74 | 0 |
| MYCN | CTGGGCGTGGTG | 1 | 312 | 324 | -688 | -676 | -0,18 | 0,01 | 14,39 | >0.1 | 2,64 | 4,63 | 0 | 1,18 | 0 | 6,13 | 0 |
| MYCN | GGGCACCTGTAG | 1 | 326 | 338 | -674 | -662 | 4,48 | 0,01 | 18,44 | >0.1 | 2,6 | 4,65 | 0 | 1,15 | 0 | 5,56 | 0 |
| MYCN | TCTCATGAGGAG | 1 | 983 | 995 | -17 | -5 | 1,16 | 0,01 | 18,73 | >0.1 | 5,58 | 8,59 | 0 | 1,87 | 0 | 1,53 | 0 |
| MYCN | GCTCACAGGGGA | 1 | 1035 | 1047 | 35 | 47 | 0,74 | 0,01 | 17,54 | >0.1 | 5,58 | 8,03 | 0 | 1,86 | 0 | 1,34 | 0 |
| MYCN | AGCCTTGTTGCT | 1 | 1076 | 1088 | 76 | 88 | -0,82 | 0,01 | 17,7 | >0.1 | 5,58 | 7,07 | 0 | 1,85 | 1,79 | 2,21 | 0 |
| MYCN | AACCACGAAGCC | 1 | 1145 | 1157 | 145 | 157 | 1,9 | 0,01 | 21,04 | >0.1 | 5,58 | 6,33 | 0 | 1,84 | 3,22 | 2,16 | 0 |
| MYCN | GTTCACTTGCTT | 1 | 1176 | 1188 | 176 | 188 | -0,6 | 0,01 | 17,77 | >0.1 | 5,58 | 6,11 | 0 | 1,62 | 1,79 | 3,26 | 0 |
| MYCN | AAGCAAGTGAAC | -1 | 1176 | 1188 | 176 | 188 | -0,6 | 0,01 | 17,77 | >0.1 | 5,58 | 6,11 | 0 | 1,62 | 1,79 | 3,26 | 0 |
| MYCN | GGCTTCGTGGTT | -1 | 1145 | 1157 | 145 | 157 | 1,9 | 0,01 | 21,04 | >0.1 | 5,58 | 6,33 | 0 | 1,84 | 3,22 | 2,16 | 0 |
| MYCN | AGCAACAAGGCT | -1 | 1076 | 1088 | 76 | 88 | -0,82 | 0,01 | 17,7 | >0.1 | 5,58 | 7,07 | 0 | 1,85 | 1,79 | 2,21 | 0 |
| MYCN | TCCCCTGTGAGC | -1 | 1035 | 1047 | 35 | 47 | 0,74 | 0,01 | 17,54 | >0.1 | 5,58 | 8,03 | 0 | 1,86 | 0 | 1,34 | 0 |
| MYCN | CTCCTCATGAGA | -1 | 983 | 995 | -17 | -5 | 1,16 | 0,01 | 18,73 | >0.1 | 5,58 | 8,59 | 0 | 1,87 | 0 | 1,53 | 0 |
| MYCN | CTACAGGTGCCC | -1 | 326 | 338 | -674 | -662 | 4,48 | 0,01 | 18,44 | >0.1 | 2,6 | 4,65 | 0 | 1,15 | 0 | 5,56 | 0 |
| MYCN | CACCACGCCCAG | -1 | 312 | 324 | -688 | -676 | -0,18 | 0,01 | 14,39 | >0.1 | 2,64 | 4,63 | 0 | 1,18 | 0 | 6,13 | 0 |
| MYCN | GACCTTGTGATC | -1 | 234 | 246 | -766 | -754 | 3,45 | 0,01 | 17,04 | >0.1 | 2,9 | 4,51 | 0 | 1,44 | 0 | 4,74 | 0 |
| MYCN | CTCCACAGGTCA | -1 | 108 | 120 | -892 | -880 | 0,31 | 0,01 | 10,85 | >0.1 | 3,54 | 4,36 | 0 | 1,89 | 0 | 0,75 | 0 |
| MLXIPL | ATCAGGAGAG | 1 | 97 | 107 | -903 | -893 | -3,16 | 0,01 | 11,75 | >0.1 | 3,64 | 4,35 | 0 | 1,88 | 0 | 0,77 | 4,28 |
| MLXIPL | TTGACCTGTG | 1 | 107 | 117 | -893 | -883 | -3,46 | 0,01 | 11,39 | >0.1 | 3,56 | 4,36 | 0 | 1,89 | 0 | 0,77 | 4,28 |
| MLXIPL | ATCACAAGGT | 1 | 235 | 245 | -765 | -755 | -2,61 | 0,01 | 15,26 | >0.1 | 2,9 | 4,51 | 0 | 1,43 | 0 | 4,74 | 4,28 |
| MLXIPL | GTCAGGAGAT | 1 | 243 | 253 | -757 | -747 | -0,76 | 0,01 | 16,63 | >0.1 | 2,87 | 4,52 | 0 | 1,39 | 0 | 4,33 | 4,28 |
| MLXIPL | TGGGCGTGGT | 1 | 313 | 323 | -687 | -677 | -0,77 | 0,01 | 18,08 | >0.1 | 2,64 | 4,62 | 0 | 1,17 | 0 | 6,13 | 4,28 |
| MLXIPL | GGCACCTGTA | 1 | 327 | 337 | -673 | -663 | 2,88 | 0,01 | 21,12 | >0.1 | 2,6 | 4,65 | 0 | 1,14 | 0 | 5,56 | 4,28 |
| MLXIPL | TACTCGGGAG | 1 | 345 | 355 | -655 | -645 | -3,26 | 0,01 | 15,88 | >0.1 | 2,55 | 4,67 | 0 | 1,11 | 0 | 6,52 | 4,28 |

| | | | | | | | | | | | | | | | | | |
|---|---|---|---|---|---|---|---|---|---|---|---|---|---|---|---|---|---|
| MLXIPL | GGCAGGAGAA | 1 | 360 | 370 | -640 | -630 | -3,75 | 0,01 | 15,2 | >0.1 | 2,52 | 4,7 | 0 | 1,09 | 0 | 6,36 | 4,28 |
| MLXIPL | ATGGCGTGAA | 1 | 369 | 379 | -631 | -621 | 2,01 | 0,01 | 20,77 | >0.1 | 2,54 | 4,71 | 0 | 1,07 | 0 | 6,15 | 4,28 |
| MLXIPL | GACAAGGGCA | 1 | 505 | 515 | -495 | -485 | -3,99 | 0,01 | 14,91 | >0.1 | 2,98 | 4,98 | 0 | 0,98 | 0 | 5,68 | 4,28 |
| MLXIPL | ATAAAGTGAC | 1 | 761 | 771 | -239 | -229 | -4,53 | 0,01 | 18,56 | >0.1 | 5,61 | 5,77 | 0 | 1,32 | 0 | 6,1 | 4,28 |
| MLXIPL | GTGACAGGAG | 1 | 766 | 776 | -234 | -224 | -4,61 | 0,01 | 18,34 | >0.1 | 5,61 | 5,8 | 0 | 1,34 | 0 | 5,92 | 4,28 |
| MLXIPL | ATCTCATGAG | 1 | 982 | 992 | -18 | -8 | 1,52 | 0,01 | 23,25 | >0.1 | 5,61 | 8,43 | 0 | 1,87 | 0 | 1,53 | 4,28 |
| MLXIPL | TGGATGTGAT | 1 | 1021 | 1031 | 21 | 31 | -3,82 | 0,01 | 17,94 | >0.1 | 5,61 | 8,7 | 0 | 1,86 | 0 | 1,31 | 4,28 |
| MLXIPL | CTCACAGGGG | 1 | 1036 | 1046 | 36 | 46 | -3,64 | 0,01 | 17,44 | >0.1 | 5,61 | 7,99 | 0 | 1,86 | 0 | 1,34 | 4,28 |
| MLXIPL | CACAGGGGAC | 1 | 1038 | 1048 | 38 | 48 | -1,76 | 0,01 | 19,25 | >0.1 | 5,61 | 7,92 | 0 | 1,86 | 0 | 1,34 | 4,28 |
| MLXIPL | AAGACCTGTT | 1 | 1158 | 1168 | 158 | 168 | -1,19 | 0,01 | 19,68 | >0.1 | 4,38 | 6,24 | 0 | 1,78 | 1,79 | 2,4 | 4,28 |
| MLXIPL | TTCACTTGCT | 1 | 1177 | 1187 | 177 | 187 | 0,92 | 0,01 | 23,58 | >0.1 | 5,61 | 6,11 | 0 | 1,61 | 1,79 | 3,26 | 4,28 |
| MLXIPL | AGCAAGTGAA | -1 | 1177 | 1187 | 177 | 187 | 4,89 | 0,001 | 27,55 | >0.1 | 5,61 | 6,11 | 0 | 1,61 | 1,79 | 3,26 | 4,28 |
| MLXIPL | AACAGGTCTT | -1 | 1158 | 1168 | 158 | 168 | -4,52 | 0,01 | 16,35 | >0.1 | 4,38 | 6,24 | 0 | 1,78 | 1,79 | 2,4 | 4,28 |
| MLXIPL | ATCACATCCA | -1 | 1021 | 1031 | 21 | 31 | -4,32 | 0,01 | 17,44 | >0.1 | 5,61 | 8,7 | 0 | 1,86 | 0 | 1,31 | 4,28 |
| MLXIPL | TCCTCATGAG | -1 | 984 | 994 | -16 | -6 | -3,65 | 0,01 | 18,21 | >0.1 | 5,61 | 8,57 | 0 | 1,87 | 0 | 1,53 | 4,28 |
| MLXIPL | CTCTCCTGTC | -1 | 768 | 778 | -232 | -222 | -4,04 | 0,01 | 18,92 | >0.1 | 5,61 | 5,81 | 0 | 1,35 | 0 | 5,92 | 4,28 |
| MLXIPL | AATACATGAC | -1 | 517 | 527 | -483 | -473 | -3,93 | 0,01 | 12,32 | >0.1 | 3,03 | 5 | 0 | 0,98 | 0 | 2,95 | 4,28 |
| MLXIPL | TTCTCCTGCC | -1 | 360 | 370 | -640 | -630 | -2,22 | 0,01 | 16,73 | >0.1 | 2,52 | 4,7 | 0 | 1,09 | 0 | 6,36 | 4,28 |
| MLXIPL | TACAGGTGCC | -1 | 327 | 337 | -673 | -663 | 5,37 | 0,001 | 23,61 | >0.1 | 2,6 | 4,65 | 0 | 1,14 | 0 | 5,56 | 4,28 |
| MLXIPL | GAGACGGGGT | -1 | 280 | 290 | -720 | -710 | 0,2 | 0,01 | 18,15 | >0.1 | 2,74 | 4,57 | 0 | 1,26 | 0 | 5,09 | 4,28 |
| MLXIPL | ATCTCCTGAC | -1 | 243 | 253 | -757 | -747 | 0,58 | 0,01 | 17,97 | >0.1 | 2,87 | 4,52 | 0 | 1,39 | 0 | 4,33 | 4,28 |
| MLXIPL | CAGGCGTGAG | -1 | 191 | 201 | -809 | -799 | -3,73 | 0,01 | 13,91 | >0.1 | 3,08 | 4,45 | 0 | 1,7 | 0 | 4,13 | 4,28 |
| MLXIPL | TCCACAGGTC | -1 | 109 | 119 | -891 | -881 | -4,19 | 0,01 | 10,63 | >0.1 | 3,54 | 4,36 | 0 | 1,89 | 0 | 0,75 | 4,28 |
| MLXIPL | CTCTCCTGAT | -1 | 97 | 107 | -903 | -893 | -1,6 | 0,01 | 13,31 | >0.1 | 3,64 | 4,35 | 0 | 1,88 | 0 | 0,77 | 4,28 |
| SOX3 | CCACTGTGCT | 1 | 415 | 425 | -585 | -575 | -1,94 | 0,01 | 11,63 | >0.1 | 2,67 | 4,8 | 0 | 1,02 | 0 | 5,09 | 0 |
| SOX3 | GTTTTGATTT | 1 | 747 | 757 | -253 | -243 | -2,31 | 0,01 | 16,39 | >0.1 | 5,61 | 5,71 | 0 | 1,28 | 0 | 6,1 | 0 |
| SOX3 | GTATTCTTAT | 1 | 806 | 816 | -194 | -184 | -1,43 | 0,01 | 14,99 | >0.1 | 5,61 | 6 | 0 | 1,53 | 0 | 3,28 | 0 |
| SOX3 | TCTTTTTCTT | 1 | 915 | 925 | -85 | -75 | -1,69 | 0,01 | 13,82 | >0.1 | 5,61 | 6,87 | 0 | 1,89 | 0 | 1,15 | 0 |
| SOX3 | TTTTTCTTTT | 1 | 917 | 927 | -83 | -73 | 6,39 | 0,01 | 21,9 | >0.1 | 5,61 | 6,89 | 0 | 1,89 | 0 | 1,12 | 0 |
| SOX3 | TCTTTTTTTG | 1 | 921 | 931 | -79 | -69 | -2,39 | 0,01 | 13,17 | >0.1 | 5,61 | 6,94 | 0 | 1,88 | 0 | 1,12 | 0 |
| SOX3 | TTTTTGGTCT | 1 | 925 | 935 | -75 | -65 | -2,67 | 0,01 | 12,88 | >0.1 | 5,61 | 6,99 | 0 | 1,88 | 0 | 1,06 | 0 |
| SOX3 | TTTTGGTCTT | 1 | 926 | 936 | -74 | -64 | -2,89 | 0,01 | 12,67 | >0.1 | 5,61 | 7,01 | 0 | 1,88 | 0 | 1,06 | 0 |
| SOX3 | ATTTTCTCTC | -1 | 974 | 984 | -26 | -16 | -4,18 | 0,01 | 12,96 | >0.1 | 5,61 | 8,05 | 0 | 1,87 | 0 | 1,61 | 0 |
| SOX3 | TTATTCTTTA | -1 | 795 | 805 | -205 | -195 | -0,94 | 0,01 | 17,4 | >0.1 | 5,61 | 5,94 | 0 | 1,47 | 0 | 5,32 | 0 |
| SOX3 | CTTTTGGTTA | -1 | 732 | 742 | -268 | -258 | -4,19 | 0,01 | 14,21 | >0.1 | 5,61 | 5,65 | 0 | 1,23 | 0 | 5,92 | 0 |
| SOX3 | TTTTTATTTC | -1 | 576 | 586 | -424 | -414 | -3,7 | 0,01 | 5,77 | >0.1 | 3,32 | 5,14 | 0 | 1 | 0 | 0 | 0 |
| SOX3 | CCCTTGTCTT | -1 | 503 | 513 | -497 | -487 | 2,46 | 0,01 | 17,07 | >0.1 | 2,97 | 4,97 | 0 | 0,98 | 0 | 5,68 | 0 |
| SOX3 | GTTTTTTTTT | -1 | 470 | 480 | -530 | -520 | -2,7 | 0,01 | 10,98 | >0.1 | 2,85 | 4,9 | 0 | 0,99 | 0 | 4,95 | 0 |
| SOX3 | TTTTTTTTTT | -1 | 469 | 479 | -531 | -521 | -3,6 | 0,01 | 9,96 | >0.1 | 2,84 | 4,9 | 0 | 0,99 | 0 | 4,83 | 0 |
| SOX3 | TTTTTTTTTT | -1 | 468 | 478 | -532 | -522 | -3,6 | 0,01 | 11,05 | >0.1 | 2,84 | 4,9 | 0 | 0,99 | 0 | 5,92 | 0 |
| SOX3 | TTTTTTTTTT | -1 | 467 | 477 | -533 | -523 | -3,6 | 0,01 | 10,9 | >0.1 | 2,83 | 4,9 | 0 | 0,99 | 0 | 5,78 | 0 |
| SOX3 | TTTTTTTTTT | -1 | 466 | 476 | -534 | -524 | -3,6 | 0,01 | 10,75 | >0.1 | 2,83 | 4,89 | 0 | 0,99 | 0 | 5,63 | 0 |
| SOX3 | TTTTTTTTTT | -1 | 465 | 475 | -535 | -525 | -3,6 | 0,01 | 10,89 | >0.1 | 2,83 | 4,89 | 0 | 0,99 | 0 | 5,78 | 0 |
| SOX3 | TTTTTTTTTT | -1 | 464 | 474 | -536 | -526 | -3,6 | 0,01 | 10,88 | >0.1 | 2,83 | 4,89 | 0 | 0,99 | 0 | 5,78 | 0 |
| SOX3 | TTTTTTTTTT | -1 | 463 | 473 | -537 | -527 | -3,6 | 0,01 | 10,88 | >0.1 | 2,82 | 4,89 | 0 | 0,99 | 0 | 5,78 | 0 |
| SOX3 | TTTTTTTTTT | -1 | 462 | 472 | -538 | -528 | -3,6 | 0,01 | 10,73 | >0.1 | 2,82 | 4,89 | 0 | 0,99 | 0 | 5,63 | 0 |
| SOX3 | TTTTTTTTTT | -1 | 461 | 471 | -539 | -529 | -3,6 | 0,01 | 10,72 | >0.1 | 2,81 | 4,88 | 0 | 0,99 | 0 | 5,63 | 0 |
| SOX3 | TTTTTTTTTT | -1 | 460 | 470 | -540 | -530 | -3,6 | 0,01 | 10,61 | >0.1 | 2,81 | 4,88 | 0 | 0,99 | 0 | 5,52 | 0 |
| SOX3 | TTTTTTTTTT | -1 | 459 | 469 | -541 | -531 | -3,6 | 0,01 | 10,6 | >0.1 | 2,81 | 4,88 | 0 | 0,99 | 0 | 5,52 | 0 |
| SOX3 | TTTTTTTTTT | -1 | 458 | 468 | -542 | -532 | -3,6 | 0,01 | 10,71 | >0.1 | 2,8 | 4,88 | 0 | 0,99 | 0 | 5,63 | 0 |
| SOX3 | TTTTTTTTTT | -1 | 457 | 467 | -543 | -533 | -3,6 | 0,01 | 10,71 | >0.1 | 2,8 | 4,88 | 0 | 0,99 | 0 | 5,63 | 0 |
| SOX3 | CCATTCTCCT | -1 | 363 | 373 | -637 | -627 | 8,62 | 0,001 | 23,54 | >0.1 | 2,52 | 4,7 | 0 | 1,08 | 0 | 6,61 | 0 |
| SOX3 | TTTTTGTATT | -1 | 297 | 307 | -703 | -693 | -2,31 | 0,01 | 10,64 | >0.1 | 2,69 | 4,6 | 0 | 1,21 | 0 | 4,45 | 0 |
| SOX3 | CCACTGTGCC | -1 | 181 | 191 | -819 | -809 | -2,04 | 0,01 | 11,16 | >0.1 | 3,12 | 4,44 | 0 | 1,78 | 0 | 3,86 | 0 |
| SOX3 | TCTTTGAGTG | -1 | 25 | 35 | -975 | -965 | -3,63 | 0,01 | 8,12 | >0.1 | 5,61 | 4,28 | 0 | 1,87 | 0 | 0 | 0 |
| SOX3 | TCCTTCTCCT | -1 | 13 | 23 | -987 | -977 | -4,31 | 0,01 | 7,43 | >0.1 | 5,61 | 4,26 | 0 | 1,86 | 0 | 0 | 0 |
| POU4F1 | GAGCTTTTTTAAGT | 1 | 70 | 84 | -930 | -916 | 1,4 | 0,01 | 11,46 | >0.1 | 3,86 | 4,32 | 0 | 1,88 | 0 | 0 | 0 |
| POU4F1 | CAGAAATATGAATT | 1 | 484 | 498 | -516 | -502 | 2,42 | 0,01 | 17,52 | >0.1 | 2,9 | 4,94 | 0 | 0,99 | 0 | 6,27 | 0 |
| POU4F1 | TTGAAATAAAAATA | 1 | 574 | 588 | -426 | -412 | 3,04 | 0,01 | 12,52 | >0.1 | 3,32 | 5,15 | 0 | 1 | 0 | 0 | 0 |

| | | | | | | | | | | | | | | | | | |
|---|---|---|---|---|---|---|---|---|---|---|---|---|---|---|---|---|---|
| POU4F1 | AATAAAAATAAATG | 1 | 578 | 592 | -422 | -408 | 3,09 | 0,01 | 12,6 | >0.1 | 3,35 | 5,16 | 0 | 1,01 | 0 | 0 | 0 |
| POU4F1 | ATAACCTATTAATG | 1 | 596 | 610 | -404 | -390 | 2,47 | 0,01 | 13,57 | >0.1 | 3,46 | 5,21 | 0 | 1,02 | 0 | 1,41 | 0 |
| POU4F1 | GTGTATCTTTAACA | 1 | 701 | 715 | -299 | -285 | 2,33 | 0,01 | 19,41 | >0.1 | 5,56 | 5,54 | 0 | 1,17 | 0 | 4,81 | 0 |
| POU4F1 | AGGATTAAAGAATA | 1 | 790 | 804 | -210 | -196 | 1,94 | 0,01 | 20,22 | >0.1 | 5,56 | 5,94 | 0 | 1,47 | 0 | 5,32 | 0 |
| POU4F1 | ATTAAAGAATAACG | 1 | 793 | 807 | -207 | -193 | 1,49 | 0,01 | 19,8 | >0.1 | 5,56 | 5,95 | 0 | 1,48 | 0 | 5,32 | 0 |
| POU4F1 | AAGAATAACGTATT | 1 | 797 | 811 | -203 | -189 | 3,75 | 0,01 | 20,33 | >0.1 | 5,56 | 5,98 | 0 | 1,5 | 0 | 3,55 | 0 |
| POU4F1 | ATTCTTATTTGATT | 1 | 808 | 822 | -192 | -178 | 6,63 | 0,01 | 23,07 | >0.1 | 5,56 | 6,04 | 0 | 1,57 | 0 | 3,28 | 0 |
| POU4F1 | TTTAAAAAATTATT | 1 | 825 | 839 | -175 | -161 | 3,57 | 0,01 | 20,39 | >0.1 | 5,56 | 6,15 | 0 | 1,68 | 0 | 3,43 | 0 |
| POU4F1 | ATGAACTTTATATT | 1 | 871 | 885 | -129 | -115 | 2,14 | 0,01 | 19,03 | >0.1 | 5,56 | 6,49 | 0 | 1,9 | 0 | 2,95 | 0 |
| POU4F1 | TAGAATAATTTTTT | -1 | 828 | 842 | -172 | -158 | 2,23 | 0,01 | 18,94 | >0.1 | 5,56 | 6,17 | 0 | 1,71 | 0 | 3,28 | 0 |
| POU4F1 | ATAATTTTTTAAAG | -1 | 824 | 838 | -176 | -162 | 5,21 | 0,01 | 22,13 | >0.1 | 5,56 | 6,14 | 0 | 1,68 | 0 | 3,55 | 0 |
| POU4F1 | GTGAATCAAATAAG | -1 | 811 | 825 | -189 | -175 | 3,19 | 0,01 | 19,82 | >0.1 | 5,56 | 6,06 | 0 | 1,59 | 0 | 3,43 | 0 |
| POU4F1 | AAGAATACGTTATT | -1 | 800 | 814 | -200 | -186 | 3,1 | 0,01 | 19,6 | >0.1 | 5,56 | 5,99 | 0 | 1,52 | 0 | 3,43 | 0 |
| POU4F1 | GTTATTCTTTAATC | -1 | 792 | 806 | -208 | -194 | 1,98 | 0,01 | 20,13 | >0.1 | 5,56 | 5,95 | 0 | 1,48 | 0 | 5,17 | 0 |
| POU4F1 | GTTAATATTTTCCT | -1 | 722 | 736 | -278 | -264 | 1,54 | 0,01 | 19,1 | >0.1 | 5,56 | 5,62 | 0 | 1,22 | 0 | 5,17 | 0 |
| POU4F1 | ATTAATAGGTTATC | -1 | 595 | 609 | -405 | -391 | 4,21 | 0,01 | 15,3 | >0.1 | 3,45 | 5,21 | 0 | 1,02 | 0 | 1,41 | 0 |
| POU4F1 | TTGTATTTTTAATA | -1 | 290 | 304 | -710 | -696 | 7,45 | 0,001 | 20,43 | >0.1 | 2,7 | 4,6 | 0 | 1,23 | 0 | 4,45 | 0 |
| POU4F2 | AGAAATAGGAAATGAG | 1 | 57 | 73 | -943 | -927 | -0,04 | 0,01 | 10,15 | >0.1 | 4 | 4,31 | 0 | 1,88 | 0 | 0 | 0 |
| POU4F2 | GAGCTTTTTTAAGTTT | 1 | 70 | 86 | -930 | -914 | -6,68 | 0,01 | 3,38 | >0.1 | 3,85 | 4,33 | 0 | 1,88 | 0 | 0 | 0 |
| POU4F2 | AAAAAAAACTGATCAG | 1 | 471 | 487 | -529 | -513 | -8,42 | 0,01 | 5,3 | >0.1 | 2,86 | 4,92 | 0 | 0,99 | 0 | 4,95 | 0 |
| POU4F2 | TTGAAATAAAAATAAA | 1 | 574 | 590 | -426 | -410 | -7,43 | 0,01 | 2,06 | >0.1 | 3,33 | 5,15 | 0 | 1,01 | 0 | 0 | 0 |
| POU4F2 | AATAAAAATAAATGTG | 1 | 578 | 594 | -422 | -406 | -4,42 | 0,01 | 5,11 | >0.1 | 3,35 | 5,17 | 0 | 1,01 | 0 | 0 | 0 |
| POU4F2 | ATAAATGTGAGATAAC | 1 | 585 | 601 | -415 | -399 | -8,64 | 0,01 | 0,95 | >0.1 | 3,4 | 5,18 | 0 | 1,01 | 0 | 0 | 0 |
| POU4F2 | ATAACCTATTAATGAA | 1 | 596 | 612 | -404 | -388 | -8,21 | 0,01 | 2,86 | >0.1 | 3,47 | 5,21 | 0 | 1,02 | 0 | 1,36 | 0 |
| POU4F2 | GTGTATCTTTAACAGC | 1 | 701 | 717 | -299 | -283 | -9,45 | 0,01 | 7,61 | >0.1 | 5,53 | 5,55 | 0 | 1,17 | 0 | 4,81 | 0 |
| POU4F2 | AGGATTAAAGAATAAC | 1 | 790 | 806 | -210 | -194 | -8,79 | 0,01 | 9,49 | >0.1 | 5,53 | 5,95 | 0 | 1,48 | 0 | 5,32 | 0 |
| POU4F2 | AAGAATAACGTATTCT | 1 | 797 | 813 | -203 | -187 | -7,82 | 0,01 | 8,76 | >0.1 | 5,53 | 5,99 | 0 | 1,51 | 0 | 3,55 | 0 |
| POU4F2 | ATTCTTATTTGATTCA | 1 | 808 | 824 | -192 | -176 | -0,51 | 0,01 | 16,08 | >0.1 | 5,53 | 6,05 | 0 | 1,58 | 0 | 3,43 | 0 |
| POU4F2 | ATGAACTTTATATTGG | 1 | 871 | 887 | -129 | -113 | -7,92 | 0,01 | 8,96 | >0.1 | 5,53 | 6,5 | 0 | 1,9 | 0 | 2,95 | 0 |
| POU4F2 | GAGAAAATCTCATGAG | 1 | 976 | 992 | -24 | -8 | -5,51 | 0,01 | 11,86 | >0.1 | 5,53 | 8,43 | 0 | 1,87 | 0 | 1,53 | 0 |
| POU4F2 | TAGAATAATTTTTTAA | -1 | 826 | 842 | -174 | -158 | -5,72 | 0,01 | 10,97 | >0.1 | 5,53 | 6,17 | 0 | 1,71 | 0 | 3,28 | 0 |
| POU4F2 | ATAATTTTTTAAAGTG | -1 | 822 | 838 | -178 | -162 | -4,82 | 0,01 | 12,23 | >0.1 | 5,53 | 6,14 | 0 | 1,68 | 0 | 3,7 | 0 |
| POU4F2 | GTGAATCAAATAAGAA | -1 | 809 | 825 | -191 | -175 | -7,8 | 0,01 | 8,8 | >0.1 | 5,53 | 6,06 | 0 | 1,59 | 0 | 3,43 | 0 |
| POU4F2 | AAGAATACGTTATTCT | -1 | 798 | 814 | -202 | -186 | -5,41 | 0,01 | 11,06 | >0.1 | 5,53 | 5,99 | 0 | 1,52 | 0 | 3,43 | 0 |
| POU4F2 | GTTAATATTTTCCTAG | -1 | 720 | 736 | -280 | -264 | -6,39 | 0,01 | 11,15 | >0.1 | 5,53 | 5,62 | 0 | 1,22 | 0 | 5,17 | 0 |
| POU4F2 | ATTAATAGGTTATCTC | -1 | 593 | 609 | -407 | -391 | -8,2 | 0,01 | 2,91 | >0.1 | 3,45 | 5,21 | 0 | 1,02 | 0 | 1,44 | 0 |
| POU4F2 | TTGTATTTTTAATAGA | -1 | 288 | 304 | -712 | -696 | -0,58 | 0,01 | 12,41 | >0.1 | 2,71 | 4,6 | 0 | 1,24 | 0 | 4,45 | 0 |
| POU4F2 | AGCCATTTCTGATCAG | -1 | 164 | 180 | -836 | -820 | -3,55 | 0,01 | 8,38 | >0.1 | 3,19 | 4,43 | 0 | 1,9 | 0 | 2,4 | 0 |
| AR | GGGCACAGTGGCTCACG | 1 | 180 | 197 | -820 | -803 | -9,56 | 0,01 | 3,65 | >0.1 | 3,11 | 4,45 | 0 | 1,79 | 0 | 3,86 | 0 |
| AR | AGATAGAGACCGTCCTG | 1 | 249 | 266 | -751 | -734 | -7,76 | 0,01 | 5,96 | >0.1 | 2,83 | 4,54 | 0 | 1,37 | 0 | 4,98 | 0 |
| AR | ACTAACACGGTGAAACC | 1 | 266 | 283 | -734 | -717 | -7,3 | 0,01 | 6,44 | >0.1 | 2,78 | 4,57 | 0 | 1,31 | 0 | 5,09 | 0 |
| AR | GGTGAAACCCCGTCTCT | 1 | 274 | 291 | -726 | -709 | -13,66 | 0,01 | 0,14 | >0.1 | 2,75 | 4,58 | 0 | 1,28 | 0 | 5,19 | 0 |
| AR | GGGCACCTGTAGTCCCA | 1 | 326 | 343 | -674 | -657 | 2,71 | 0,001 | 16,57 | >0.1 | 2,6 | 4,66 | 0 | 1,15 | 0 | 5,46 | 0 |
| AR | GAGAATGGCGTGAACCT | 1 | 365 | 382 | -635 | -618 | 1,3 | 0,001 | 16,37 | >0.1 | 2,54 | 4,72 | 0 | 1,08 | 0 | 6,73 | 0 |
| AR | GAGCTTGCAGTGAGCCG | 1 | 390 | 407 | -610 | -593 | -11,16 | 0,01 | 3,28 | >0.1 | 2,61 | 4,76 | 0 | 1,05 | 0 | 6,02 | 0 |
| AR | GGGAAAATGTTGCCCAA | 1 | 532 | 549 | -468 | -451 | -7,84 | 0,01 | 2,93 | >0.1 | 3,11 | 5,05 | 0 | 0,99 | 0 | 1,61 | 0 |
| AR | ATTAACCAAAAGTACCG | 1 | 730 | 747 | -270 | -253 | -12,08 | 0,01 | 6,27 | >0.1 | 5,52 | 5,67 | 0 | 1,25 | 0 | 5,92 | 0 |
| AR | AAGTACCGGTTTTGATT | 1 | 739 | 756 | -261 | -244 | -13,14 | 0,01 | 5,28 | >0.1 | 5,52 | 5,71 | 0 | 1,28 | 0 | 5,92 | 0 |
| AR | GGAGAGGTAAGGTTCTC | 1 | 772 | 789 | -228 | -211 | -10,02 | 0,01 | 9,08 | >0.1 | 5,52 | 5,86 | 0 | 1,4 | 0 | 6,32 | 0 |
| AR | AAGAATAACGTATTCTT | 1 | 797 | 814 | -203 | -186 | -7,75 | 0,01 | 8,83 | >0.1 | 5,52 | 5,99 | 0 | 1,52 | 0 | 3,55 | 0 |
| AR | CAGCAGATTGTTTACTG | 1 | 889 | 906 | -111 | -94 | -6,89 | 0,01 | 10,06 | >0.1 | 5,52 | 6,67 | 0 | 1,89 | 0 | 2,86 | 0 |
| AR | TTGTTTACTGTGTTCTT | 1 | 896 | 913 | -104 | -87 | -7,92 | 0,01 | 8,98 | >0.1 | 5,52 | 6,74 | 0 | 1,89 | 0 | 2,75 | 0 |
| AR | GGGGACGATGTCAAGCT | 1 | 1042 | 1059 | 42 | 59 | -13,84 | 0,01 | 1,98 | >0.1 | 5,52 | 7,79 | 0 | 1,86 | 0 | 0,65 | 0 |
| AR | GATGTCAAGCTCTTCCT | 1 | 1048 | 1065 | 48 | 65 | -13,43 | 0,01 | 3,04 | >0.1 | 5,52 | 7,62 | 0 | 1,86 | 0 | 1,46 | 0 |
| AR | TTGGACAAGTTTAACCA | 1 | 1133 | 1150 | 133 | 150 | -13,8 | 0,01 | 6,31 | >0.1 | 5,52 | 6,43 | 0 | 1,84 | 3,22 | 3,09 | 0 |
| AR | AAGTTCACTTGCTTCTT | 1 | 1174 | 1191 | 174 | 191 | -10,95 | 0,01 | 7,39 | >0.1 | 5,52 | 6,13 | 0 | 1,63 | 1,79 | 3,26 | 0 |
| AR | AAGAAGCAAGTGAACTT | -1 | 1174 | 1191 | 174 | 191 | -4,96 | 0,01 | 13,38 | >0.1 | 5,52 | 6,13 | 0 | 1,63 | 1,79 | 3,26 | 0 |
| AR | AGGAAGAGCTTGACATC | -1 | 1048 | 1065 | 48 | 65 | -9,65 | 0,01 | 6,82 | >0.1 | 5,52 | 7,62 | 0 | 1,86 | 0 | 1,46 | 0 |
| AR | TCGTCCCCTGTGAGCCA | -1 | 1033 | 1050 | 33 | 50 | -10,47 | 0,01 | 6,35 | >0.1 | 5,52 | 8,1 | 0 | 1,86 | 0 | 1,34 | 0 |

| | | | | | | | | | | | | | | | | |
|---|---|---|---|---|---|---|---|---|---|---|---|---|---|---|---|---|
| AR | AAGAACACAGTAAACAA | -1 | 896 | 913 | -104 | -87 | -7,4 | 0,01 | 9,5 | >0.1 | 5,52 | 6,74 | 0 | 1,89 | 0 | 2,75 | 0 |
| AR | CAGTAAACAATCTGCTG | -1 | 889 | 906 | -111 | -94 | -6,17 | 0,01 | 10,78 | >0.1 | 5,52 | 6,67 | 0 | 1,89 | 0 | 2,86 | 0 |
| AR | AGGACAGATATGTAACA | -1 | 847 | 864 | -153 | -136 | -11,72 | 0,01 | 4,78 | >0.1 | 5,52 | 6,33 | 0 | 1,9 | 0 | 2,75 | 0 |
| AR | AAGAATACGTTATTCTT | -1 | 797 | 814 | -203 | -186 | -7,44 | 0,01 | 9,14 | >0.1 | 5,52 | 5,99 | 0 | 1,52 | 0 | 3,55 | 0 |
| AR | GAGAACCTTACCTCTCC | -1 | 772 | 789 | -228 | -211 | -9,97 | 0,01 | 9,13 | >0.1 | 5,52 | 5,86 | 0 | 1,4 | 0 | 6,32 | 0 |
| AR | AATCAAAACCGGTACTT | -1 | 739 | 756 | -261 | -244 | -13,87 | 0,01 | 4,55 | >0.1 | 5,52 | 5,71 | 0 | 1,28 | 0 | 5,92 | 0 |
| AR | TTGGGCAACATTTTCCC | -1 | 532 | 549 | -468 | -451 | -12,86 | 0,01 | -2,09 | >0.1 | 3,11 | 5,05 | 0 | 0,99 | 0 | 1,61 | 0 |
| AR | AGGTTCACGCCATTCTC | -1 | 365 | 382 | -635 | -618 | -6,04 | 0,01 | 9,03 | >0.1 | 2,54 | 4,72 | 0 | 1,08 | 0 | 6,73 | 0 |
| AR | TGGGACTACAGGTGCCC | -1 | 326 | 343 | -674 | -657 | -1,55 | 0,001 | 12,31 | >0.1 | 2,6 | 4,66 | 0 | 1,15 | 0 | 5,46 | 0 |
| AR | GGTTTCACCGTGTTAGT | -1 | 266 | 283 | -734 | -717 | -12,98 | 0,01 | 0,76 | >0.1 | 2,78 | 4,57 | 0 | 1,31 | 0 | 5,09 | 0 |
| AR | CAGGACGGTCTCTATCT | -1 | 249 | 266 | -751 | -734 | -8,5 | 0,01 | 5,22 | >0.1 | 2,83 | 4,54 | 0 | 1,37 | 0 | 4,98 | 0 |
| AR | CGTGAGCCACTGTGCCC | -1 | 180 | 197 | -820 | -803 | -9,8 | 0,01 | 3,41 | >0.1 | 3,11 | 4,45 | 0 | 1,79 | 0 | 3,86 | 0 |
| NR2E1 | GTAAGTGAA | 1 | 5 | 14 | -995 | -986 | 4,07 | 0,01 | 17,61 | >0.1 | 5,62 | 4,26 | 0 | 1,86 | 1,79 | 0 | 0 |
| NR2E1 | AGAAGTCAG | 1 | 32 | 41 | -968 | -959 | 10,66 | 0,001 | 21,26 | >0.1 | 4,45 | 4,28 | 0 | 1,87 | 0 | 0 | 0 |
| NR2E1 | AGTAGTCAT | 1 | 125 | 134 | -875 | -866 | 5,37 | 0,01 | 16,77 | >0.1 | 3,44 | 4,38 | 0 | 1,89 | 0 | 1,7 | 0 |
| NR2E1 | GGAAATTAA | 1 | 153 | 162 | -847 | -838 | 2,25 | 0,01 | 13,47 | >0.1 | 3,27 | 4,41 | 0 | 1,9 | 0 | 1,65 | 0 |
| NR2E1 | CAAGGTCAG | 1 | 239 | 248 | -761 | -752 | 6,36 | 0,01 | 19,7 | >0.1 | 2,88 | 4,52 | 0 | 1,41 | 0 | 4,53 | 0 |
| NR2E1 | TAAAGACAA | 1 | 501 | 510 | -499 | -490 | 0,31 | 0,01 | 16,03 | >0.1 | 2,96 | 4,96 | 0 | 0,98 | 0 | 6,81 | 0 |
| NR2E1 | CAAAGTCAT | 1 | 513 | 522 | -487 | -478 | 10,36 | 0,001 | 22,29 | >0.1 | 3,01 | 4,99 | 0 | 0,98 | 0 | 2,95 | 0 |
| NR2E1 | AAAAATAAA | 1 | 581 | 590 | -419 | -410 | 1,8 | 0,01 | 11,31 | >0.1 | 3,35 | 5,15 | 0 | 1 | 0 | 0 | 0 |
| NR2E1 | GAAAACCAT | 1 | 619 | 628 | -381 | -372 | 0,94 | 0,01 | 12,38 | >0.1 | 3,61 | 5,26 | 0 | 1,03 | 0 | 1,55 | 0 |
| NR2E1 | TAAAATTAT | 1 | 682 | 691 | -318 | -309 | 1,47 | 0,01 | 15,56 | >0.1 | 4,33 | 5,46 | 0 | 1,12 | 0 | 3,19 | 0 |
| NR2E1 | TAAAGTGAC | 1 | 762 | 771 | -238 | -229 | 3,64 | 0,01 | 22,46 | >0.1 | 5,62 | 5,77 | 0 | 1,32 | 0 | 6,1 | 0 |
| NR2E1 | AAAAATTAT | 1 | 829 | 838 | -171 | -162 | 2,95 | 0,01 | 19,82 | >0.1 | 5,62 | 6,14 | 0 | 1,67 | 0 | 3,43 | 0 |
| NR2E1 | CAAGTTCAA | 1 | 955 | 964 | -45 | -36 | 1,53 | 0,01 | 18,21 | >0.1 | 5,62 | 7,48 | 0 | 1,88 | 0 | 1,7 | 0 |
| NR2E1 | GAAAGTCAT | 1 | 1008 | 1017 | 8 | 17 | 10,01 | 0,001 | 28,04 | >0.1 | 5,62 | 9,07 | 0 | 1,87 | 0 | 1,46 | 0 |
| NR2E1 | CGATGTCAA | 1 | 1047 | 1056 | 47 | 56 | 3,97 | 0,01 | 20,57 | >0.1 | 5,62 | 7,65 | 0 | 1,86 | 0 | 1,46 | 0 |
| NR2E1 | ACAGGCCAA | 1 | 1117 | 1126 | 117 | 126 | 3,9 | 0,01 | 23,78 | >0.1 | 5,62 | 6,58 | 0 | 1,85 | 3,22 | 2,6 | 0 |
| NR2E1 | AAAGTTCAC | 1 | 1173 | 1182 | 173 | 182 | 0,64 | 0,01 | 18,8 | >0.1 | 5,62 | 6,13 | 0 | 1,64 | 1,79 | 2,97 | 0 |
| NR2E1 | GCAAGTGAA | -1 | 1177 | 1186 | 177 | 186 | 2,51 | 0,01 | 20,92 | >0.1 | 5,62 | 6,11 | 0 | 1,61 | 1,79 | 3,28 | 0 |
| NR2E1 | AGGAGCCAG | -1 | 1063 | 1072 | 63 | 72 | 3,92 | 0,01 | 22,7 | >0.1 | 5,62 | 7,3 | 0 | 1,85 | 1,79 | 2,21 | 0 |
| NR2E1 | GTGAGCCAA | -1 | 1032 | 1041 | 32 | 41 | 3,85 | 0,01 | 20,79 | >0.1 | 5,62 | 8,15 | 0 | 1,86 | 0 | 1,31 | 0 |
| NR2E1 | CAAGATCAC | -1 | 1026 | 1035 | 26 | 35 | 1,23 | 0,01 | 18,42 | >0.1 | 5,62 | 8,45 | 0 | 1,86 | 0 | 1,25 | 0 |
| NR2E1 | AAAGACCAA | -1 | 928 | 937 | -72 | -63 | 1,76 | 0,01 | 17,31 | >0.1 | 5,62 | 7,02 | 0 | 1,88 | 0 | 1,03 | 0 |
| NR2E1 | CTGAGCCAA | -1 | 883 | 892 | -117 | -108 | 4,2 | 0,01 | 21,33 | >0.1 | 5,62 | 6,55 | 0 | 1,89 | 0 | 3,07 | 0 |
| NR2E1 | AAAGTTCAT | -1 | 871 | 880 | -129 | -120 | 1,05 | 0,01 | 17,88 | >0.1 | 5,62 | 6,45 | 0 | 1,9 | 0 | 2,86 | 0 |
| NR2E1 | TAAAGTGAA | -1 | 820 | 829 | -180 | -171 | 6,24 | 0,01 | 23,11 | >0.1 | 5,62 | 6,08 | 0 | 1,61 | 0 | 3,55 | 0 |
| NR2E1 | GTGAATCAA | -1 | 816 | 825 | -184 | -175 | 2,32 | 0,01 | 19,01 | >0.1 | 5,62 | 6,06 | 0 | 1,58 | 0 | 3,43 | 0 |
| NR2E1 | CCAAATCAA | -1 | 750 | 759 | -250 | -241 | 4,09 | 0,01 | 22,63 | >0.1 | 5,62 | 5,72 | 0 | 1,28 | 0 | 5,92 | 0 |
| NR2E1 | AAATGTTAT | -1 | 689 | 698 | -311 | -302 | 0,5 | 0,01 | 14,44 | >0.1 | 4,47 | 5,48 | 0 | 1,13 | 0 | 2,86 | 0 |
| NR2E1 | ATAGGTTAT | -1 | 596 | 605 | -404 | -395 | 3,23 | 0,01 | 12,88 | >0.1 | 3,44 | 5,19 | 0 | 1,01 | 0 | 0 | 0 |
| NR2E1 | CAAGATCAT | -1 | 568 | 577 | -432 | -423 | 1,65 | 0,01 | 11,04 | >0.1 | 3,27 | 5,12 | 0 | 0,99 | 0 | 0 | 0 |
| NR2E1 | GGAATTCAT | -1 | 491 | 500 | -509 | -500 | 1,6 | 0,01 | 16,72 | >0.1 | 2,92 | 4,94 | 0 | 0,98 | 0 | 6,27 | 0 |
| NR2E1 | GTTAGTCAG | -1 | 263 | 272 | -737 | -728 | 1,96 | 0,01 | 15,15 | >0.1 | 2,8 | 4,55 | 0 | 1,32 | 0 | 4,53 | 0 |
| NR2E1 | GTGAGCCAC | -1 | 187 | 196 | -813 | -804 | 1,25 | 0,01 | 14,38 | >0.1 | 3,1 | 4,45 | 0 | 1,73 | 0 | 3,86 | 0 |
| NR2E1 | TAAAATTAT | -1 | 132 | 141 | -868 | -859 | 1,47 | 0,01 | 12,71 | >0.1 | 3,39 | 4,38 | 0 | 1,89 | 0 | 1,57 | 0 |
| NR2E1 | ACAGGTCAA | -1 | 107 | 116 | -893 | -884 | 7,67 | 0,001 | 18,22 | >0.1 | 3,56 | 4,36 | 0 | 1,89 | 0 | 0,75 | 0 |
| NR2E3 | TGAGCTT | 1 | 69 | 76 | -931 | -924 | 4,58 | 0,01 | 14,68 | >0.1 | 3,91 | 4,31 | 0 | 1,88 | 0 | 0 | 0 |
| NR2E3 | TAAGTTT | 1 | 79 | 86 | -921 | -914 | 3,59 | 0,01 | 13,6 | >0.1 | 3,81 | 4,32 | 0 | 1,88 | 0 | 0 | 0 |
| NR2E3 | CACGCCT | 1 | 193 | 200 | -807 | -800 | -0,05 | 0,01 | 13,29 | >0.1 | 3,08 | 4,45 | 0 | 1,68 | 0 | 4,13 | 0 |
| NR2E3 | CAAGGTC | 1 | 239 | 246 | -761 | -754 | -0,55 | 0,1 | 12,79 | >0.1 | 2,89 | 4,51 | 0 | 1,41 | 0 | 4,53 | 0 |
| NR2E3 | AAAACTT | 1 | 303 | 310 | -697 | -690 | 3,35 | 0,01 | 16,81 | >0.1 | 2,67 | 4,6 | 0 | 1,2 | 0 | 4,99 | 0 |
| NR2E3 | CCAGCTA | 1 | 340 | 347 | -660 | -653 | -0,55 | 0,1 | 13,72 | >0.1 | 2,57 | 4,66 | 0 | 1,12 | 0 | 5,92 | 0 |
| NR2E3 | GAACCTG | 1 | 376 | 383 | -624 | -617 | -0,37 | 0,01 | 14,5 | >0.1 | 2,56 | 4,72 | 0 | 1,06 | 0 | 6,53 | 0 |
| NR2E3 | AGAGCTT | 1 | 389 | 396 | -611 | -604 | 4,28 | 0,01 | 18,79 | >0.1 | 2,59 | 4,74 | 0 | 1,05 | 0 | 6,13 | 0 |
| NR2E3 | CCAGCCT | 1 | 425 | 432 | -575 | -568 | -0,05 | 0,01 | 14,13 | >0.1 | 2,69 | 4,81 | 0 | 1,01 | 0 | 5,67 | 0 |
| NR2E3 | TAACCTA | 1 | 597 | 604 | -403 | -396 | -0,12 | 0,01 | 9,52 | >0.1 | 3,44 | 5,19 | 0 | 1,01 | 0 | 0 | 0 |
| NR2E3 | AAACCAT | 1 | 621 | 628 | -379 | -372 | -0,43 | 0,01 | 10,97 | >0.1 | 3,62 | 5,26 | 0 | 1,03 | 0 | 1,5 | 0 |
| NR2E3 | CAACATT | 1 | 636 | 643 | -364 | -357 | 1,11 | 0,01 | 12,81 | >0.1 | 3,74 | 5,3 | 0 | 1,05 | 0 | 1,61 | 0 |

| | | | | | | | | | | | | | | | | | |
|---|---|---|---|---|---|---|---|---|---|---|---|---|---|---|---|---|---|
| NR2E3 | ACAGCTT | 1 | 712 | 719 | -288 | -281 | 4,28 | 0,01 | 21,47 | >0.1 | 5,65 | 5,56 | 0 | 1,17 | 0 | 4,81 | 0 |
| NR2E3 | TAAGGTT | 1 | 779 | 786 | -221 | -214 | 4,09 | 0,01 | 23,51 | >0.1 | 5,65 | 5,84 | 0 | 1,38 | 0 | 6,54 | 0 |
| NR2E3 | CAAGCTC | 1 | 1053 | 1060 | 53 | 60 | 6,31 | 0,001 | 22,79 | >0.1 | 5,65 | 7,5 | 0 | 1,86 | 0 | 1,46 | 0 |
| NR2E3 | CAGCCTT | 1 | 1075 | 1082 | 75 | 82 | 2,1 | 0,01 | 20,76 | >0.1 | 5,65 | 7,09 | 0 | 1,85 | 1,79 | 2,27 | 0 |
| NR2E3 | CAAGTTT | 1 | 1138 | 1145 | 138 | 145 | 5,32 | 0,01 | 25,52 | >0.1 | 5,65 | 6,39 | 0 | 1,84 | 3,22 | 3,09 | 0 |
| NR2E3 | TAAACTT | -1 | 1139 | 1146 | 139 | 146 | 3,65 | 0,01 | 23,82 | >0.1 | 5,65 | 6,38 | 0 | 1,84 | 3,22 | 3,07 | 0 |
| NR2E3 | CAAGGCT | -1 | 1076 | 1083 | 76 | 83 | -0,55 | 0,1 | 18,17 | >0.1 | 5,65 | 7,07 | 0 | 1,85 | 1,79 | 2,34 | 0 |
| NR2E3 | AGAGCTT | -1 | 1054 | 1061 | 54 | 61 | 4,28 | 0,01 | 21,54 | >0.1 | 5,65 | 7,48 | 0 | 1,86 | 0 | 2,27 | 0 |
| NR2E3 | AAACCTC | -1 | 992 | 999 | -8 | -1 | 0,07 | 0,01 | 17,89 | >0.1 | 5,65 | 8,69 | 0 | 1,87 | 0 | 1,61 | 0 |
| NR2E3 | CAGCCTT | -1 | 963 | 970 | -37 | -30 | 2,1 | 0,01 | 18,94 | >0.1 | 5,65 | 7,62 | 0 | 1,87 | 0 | 1,7 | 0 |
| NR2E3 | GAACCTT | -1 | 780 | 787 | -220 | -213 | 5,99 | 0,001 | 25,42 | >0.1 | 5,65 | 5,85 | 0 | 1,39 | 0 | 6,54 | 0 |
| NR2E3 | AAAGCTG | -1 | 713 | 720 | -287 | -280 | 4,28 | 0,01 | 21,65 | >0.1 | 5,65 | 5,56 | 0 | 1,18 | 0 | 4,99 | 0 |
| NR2E3 | CAACATT | -1 | 537 | 544 | -463 | -456 | 1,11 | 0,01 | 11,85 | >0.1 | 3,11 | 5,04 | 0 | 0,98 | 0 | 1,61 | 0 |
| NR2E3 | CAGGCTG | -1 | 426 | 433 | -574 | -567 | -0,05 | 0,01 | 14,14 | >0.1 | 2,7 | 4,81 | 0 | 1,01 | 0 | 5,67 | 0 |
| NR2E3 | CAAGCTC | -1 | 390 | 397 | -610 | -603 | 6,31 | 0,001 | 20,82 | >0.1 | 2,59 | 4,74 | 0 | 1,05 | 0 | 6,13 | 0 |
| NR2E3 | CCAGGTT | -1 | 377 | 384 | -623 | -616 | -0,55 | 0,1 | 14,19 | >0.1 | 2,56 | 4,72 | 0 | 1,06 | 0 | 6,4 | 0 |
| NR2E3 | CCAGCTA | -1 | 309 | 316 | -691 | -684 | -0,55 | 0,1 | 13,58 | >0.1 | 2,66 | 4,61 | 0 | 1,18 | 0 | 5,68 | 0 |
| NR2E3 | TAAGTTT | -1 | 304 | 311 | -696 | -689 | 3,59 | 0,01 | 17,05 | >0.1 | 2,67 | 4,61 | 0 | 1,19 | 0 | 4,99 | 0 |
| NR2E3 | TGACCTT | -1 | 240 | 247 | -760 | -753 | 0,37 | 0,01 | 13,7 | >0.1 | 2,88 | 4,51 | 0 | 1,41 | 0 | 4,53 | 0 |
| NR2E3 | CAGGCGT | -1 | 194 | 201 | -806 | -799 | -0,05 | 0,01 | 13,41 | >0.1 | 3,07 | 4,45 | 0 | 1,67 | 0 | 4,26 | 0 |
| NR2E3 | AAAACTT | -1 | 80 | 87 | -920 | -913 | 3,35 | 0,01 | 13,35 | >0.1 | 3,8 | 4,32 | 0 | 1,88 | 0 | 0 | 0 |
| NR2E3 | AAAGCTC | -1 | 70 | 77 | -930 | -923 | 4,28 | 0,01 | 14,37 | >0.1 | 3,9 | 4,31 | 0 | 1,88 | 0 | 0 | 0 |
| TFAP2A(VAR.2) | TGGCTCACGCCT | 1 | 188 | 200 | -812 | -800 | 5,49 | 0,01 | 18,79 | >0.1 | 3,09 | 4,45 | 0 | 1,72 | 0 | 4,04 | 0 |
| TFAP2A(VAR.2) | AGGCTGAGGCAG | 1 | 353 | 365 | -647 | -635 | 0,4 | 0,01 | 15,24 | >0.1 | 2,53 | 4,69 | 0 | 1,1 | 0 | 6,52 | 0 |
| TFAP2A(VAR.2) | AACCTGGGAGGC | 1 | 377 | 389 | -623 | -611 | 0,93 | 0,01 | 15,82 | >0.1 | 2,56 | 4,73 | 0 | 1,06 | 0 | 6,53 | 0 |
| TFAP2A(VAR.2) | ACCTGGGAGGCA | 1 | 378 | 390 | -622 | -610 | 2,57 | 0,01 | 17,6 | >0.1 | 2,57 | 4,73 | 0 | 1,06 | 0 | 6,66 | 0 |
| TFAP2A(VAR.2) | CAGCCTGGGCGA | 1 | 426 | 438 | -574 | -562 | 4,51 | 0,01 | 18,85 | >0.1 | 2,7 | 4,82 | 0 | 1,01 | 0 | 5,8 | 0 |
| TFAP2A(VAR.2) | TGCTCAGCGCCC | 1 | 939 | 951 | -61 | -49 | 0,72 | 0,01 | 16,24 | >0.1 | 5,58 | 7,23 | 0 | 1,88 | 0 | 0,83 | 0 |
| TFAP2A(VAR.2) | TGGCTCACAGGG | 1 | 1033 | 1045 | 33 | 45 | 0,89 | 0,01 | 17,74 | >0.1 | 5,58 | 8,1 | 0 | 1,86 | 0 | 1,31 | 0 |
| TFAP2A(VAR.2) | GGCTCACAGGGG | 1 | 1034 | 1046 | 34 | 46 | 0,14 | 0,01 | 16,95 | >0.1 | 5,58 | 8,06 | 0 | 1,86 | 0 | 1,31 | 0 |
| TFAP2A(VAR.2) | AGCCGAAGACCT | 1 | 1153 | 1165 | 153 | 165 | 0,2 | 0,01 | 16,87 | >0.1 | 4,46 | 6,27 | 0 | 1,83 | 1,79 | 2,31 | 0 |
| TFAP2A(VAR.2) | CCCCTGTGAGCC | -1 | 1034 | 1046 | 34 | 46 | 4,04 | 0,01 | 20,85 | >0.1 | 5,58 | 8,06 | 0 | 1,86 | 0 | 1,31 | 0 |
| TFAP2A(VAR.2) | CCCTGTGAGCCA | -1 | 1033 | 1045 | 33 | 45 | 1,7 | 0,01 | 18,55 | >0.1 | 5,58 | 8,1 | 0 | 1,86 | 0 | 1,31 | 0 |
| TFAP2A(VAR.2) | TCGCCCAGGCTG | -1 | 426 | 438 | -574 | -562 | 5,06 | 0,01 | 19,4 | >0.1 | 2,7 | 4,82 | 0 | 1,01 | 0 | 5,8 | 0 |
| TFAP2A(VAR.2) | TGCCTCCCAGGT | -1 | 378 | 390 | -622 | -610 | 4,6 | 0,01 | 19,63 | >0.1 | 2,57 | 4,73 | 0 | 1,06 | 0 | 6,66 | 0 |
| TFAP2A(VAR.2) | CTGCCTCAGCCT | -1 | 353 | 365 | -647 | -635 | 3,39 | 0,01 | 18,23 | >0.1 | 2,53 | 4,69 | 0 | 1,1 | 0 | 6,52 | 0 |
| TFAP2A(VAR.2) | TGCCTCAGCCTC | -1 | 352 | 364 | -648 | -636 | 0,17 | 0,01 | 14,89 | >0.1 | 2,53 | 4,69 | 0 | 1,1 | 0 | 6,4 | 0 |
| TFAP2A(VAR.2) | TGCCCACCACCA | -1 | 319 | 331 | -681 | -669 | 5,35 | 0,01 | 19,45 | >0.1 | 2,62 | 4,64 | 0 | 1,16 | 0 | 5,68 | 0 |
| TFAP2A(VAR.2) | AGGCGTGAGCCA | -1 | 188 | 200 | -812 | -800 | 4,76 | 0,01 | 18,06 | >0.1 | 3,09 | 4,45 | 0 | 1,72 | 0 | 4,04 | 0 |
| NOTO | GGAAATGAGC | 1 | 64 | 74 | -936 | -926 | 1,67 | 0,01 | 11,81 | >0.1 | 3,95 | 4,31 | 0 | 1,88 | 0 | 0 | 0 |
| NOTO | GGAAATTAAA | 1 | 153 | 163 | -847 | -837 | 4,8 | 0,01 | 16,02 | >0.1 | 3,27 | 4,41 | 0 | 1,9 | 0 | 1,65 | 0 |
| NOTO | ACCTATTAAT | 1 | 599 | 609 | -401 | -391 | 1,52 | 0,01 | 12,57 | >0.1 | 3,47 | 5,21 | 0 | 1,02 | 0 | 1,36 | 0 |
| NOTO | ATTAATGAAA | 1 | 603 | 613 | -397 | -387 | 3,61 | 0,01 | 14,75 | >0.1 | 3,49 | 5,22 | 0 | 1,02 | 0 | 1,41 | 0 |
| NOTO | CAACATTATC | 1 | 636 | 646 | -364 | -354 | 3,77 | 0,01 | 15,55 | >0.1 | 3,76 | 5,31 | 0 | 1,05 | 0 | 1,66 | 0 |
| NOTO | CTTCATGATC | 1 | 645 | 655 | -355 | -345 | 3,46 | 0,01 | 17,54 | >0.1 | 3,84 | 5,34 | 0 | 1,06 | 0 | 3,83 | 0 |
| NOTO | TAAAATTATA | 1 | 682 | 692 | -318 | -308 | 1,95 | 0,01 | 15,93 | >0.1 | 4,34 | 5,46 | 0 | 1,12 | 0 | 3,07 | 0 |
| NOTO | AAAAATTATT | 1 | 829 | 839 | -171 | -161 | 2,62 | 0,01 | 19,34 | >0.1 | 5,61 | 6,15 | 0 | 1,68 | 0 | 3,28 | 0 |
| NOTO | TCTCATGAGG | 1 | 983 | 993 | -17 | -7 | 5,14 | 0,01 | 22,69 | >0.1 | 5,61 | 8,5 | 0 | 1,87 | 0 | 1,57 | 0 |
| NOTO | CCTCATGAGA | -1 | 983 | 993 | -17 | -7 | 5,39 | 0,01 | 22,94 | >0.1 | 5,61 | 8,5 | 0 | 1,87 | 0 | 1,57 | 0 |
| NOTO | TCAAATAAGA | -1 | 810 | 820 | -190 | -180 | 2,72 | 0,01 | 19,19 | >0.1 | 5,61 | 6,03 | 0 | 1,55 | 0 | 3,28 | 0 |
| NOTO | TTTCATTAAT | -1 | 603 | 613 | -397 | -387 | 7,37 | 0,001 | 18,51 | >0.1 | 3,49 | 5,22 | 0 | 1,02 | 0 | 1,41 | 0 |
| NOTO | TCACATTTAT | -1 | 585 | 595 | -415 | -405 | 3,7 | 0,01 | 13,25 | >0.1 | 3,38 | 5,17 | 0 | 1,01 | 0 | 0 | 0 |
| NOTO | TTTAATTTCC | -1 | 153 | 163 | -847 | -837 | 5,67 | 0,01 | 16,89 | >0.1 | 3,27 | 4,41 | 0 | 1,9 | 0 | 1,65 | 0 |
| NOTO | TAAAATTATG | -1 | 131 | 141 | -869 | -859 | 2,87 | 0,01 | 14,12 | >0.1 | 3,4 | 4,38 | 0 | 1,89 | 0 | 1,57 | 0 |
| NOTO | GCTCATTTCC | -1 | 64 | 74 | -936 | -926 | 7,69 | 0,001 | 17,83 | >0.1 | 3,95 | 4,31 | 0 | 1,88 | 0 | 0 | 0 |
| NFIC | GGCTGGGCACA | 1 | 176 | 187 | -824 | -813 | 8,3 | 0,001 | 20,94 | >0.1 | 3,15 | 4,44 | 0 | 1,82 | 0 | 3,24 | 0 |
| NFIC | ATCCTAGCACT | 1 | 203 | 214 | -797 | -786 | 2,98 | 0,01 | 16,68 | >0.1 | 3,02 | 4,47 | 0 | 1,61 | 0 | 4,59 | 0 |
| NFIC | AGTTGGGCAGA | 1 | 225 | 236 | -775 | -764 | 2,47 | 0,01 | 16,64 | >0.1 | 2,93 | 4,5 | 0 | 1,48 | 0 | 5,26 | 0 |
| NFIC | AACTTAGCTGG | 1 | 305 | 316 | -695 | -684 | 4,24 | 0,01 | 17,7 | >0.1 | 2,66 | 4,61 | 0 | 1,19 | 0 | 4,99 | 0 |

| | | | | | | | | | | | | | | | | | |
|---|---|---|---|---|---|---|---|---|---|---|---|---|---|---|---|---|---|
| NFIC | AGCTGGGCGTG | 1 | 310 | 321 | -690 | -679 | 3,64 | 0,01 | 18,22 | >0.1 | 2,65 | 4,62 | 0 | 1,18 | 0 | 6,13 | 0 |
| NFIC | TGGTGGGCACC | 1 | 322 | 333 | -678 | -667 | 3,64 | 0,01 | 17,61 | >0.1 | 2,62 | 4,64 | 0 | 1,15 | 0 | 5,56 | 0 |
| NFIC | TACTCGGGAGG | 1 | 345 | 356 | -655 | -644 | 3,16 | 0,01 | 18,02 | >0.1 | 2,55 | 4,68 | 0 | 1,11 | 0 | 6,52 | 0 |
| NFIC | AACCTGGGAGG | 1 | 377 | 388 | -623 | -612 | 2,46 | 0,01 | 17,21 | >0.1 | 2,56 | 4,73 | 0 | 1,06 | 0 | 6,4 | 0 |
| NFIC | GCCTGGGCGAC | 1 | 428 | 439 | -572 | -561 | 3,95 | 0,01 | 18,3 | >0.1 | 2,71 | 4,82 | 0 | 1,01 | 0 | 5,8 | 0 |
| NFIC | TATTTGGAAGG | 1 | 523 | 534 | -477 | -466 | 3,25 | 0,01 | 15,5 | >0.1 | 3,06 | 5,02 | 0 | 0,98 | 0 | 3,19 | 0 |
| NFIC | TTCTGGGAATA | 1 | 558 | 569 | -442 | -431 | 2,3 | 0,01 | 11,62 | >0.1 | 3,23 | 5,1 | 0 | 0,99 | 0 | 0 | 0 |
| NFIC | TTCTAGGAAAA | 1 | 718 | 729 | -282 | -271 | 2,68 | 0,01 | 20,06 | >0.1 | 5,6 | 5,59 | 0 | 1,2 | 0 | 4,99 | 0 |
| NFIC | GATTTGGCCAT | 1 | 752 | 763 | -248 | -237 | 3,5 | 0,01 | 22,67 | >0.1 | 5,6 | 5,74 | 0 | 1,3 | 0 | 6,54 | 0 |
| NFIC | ATATTGGCTCA | 1 | 880 | 891 | -120 | -109 | 4,23 | 0,01 | 21,33 | >0.1 | 5,6 | 6,54 | 0 | 1,9 | 0 | 3,07 | 0 |
| NFIC | GGCTCAGCAGA | 1 | 885 | 896 | -115 | -104 | 3,66 | 0,01 | 20,8 | >0.1 | 5,6 | 6,58 | 0 | 1,89 | 0 | 3,07 | 0 |
| NFIC | ATCTTGGCTCA | 1 | 1029 | 1040 | 29 | 40 | 9,41 | 0,001 | 26,49 | >0.1 | 5,6 | 8,29 | 0 | 1,86 | 0 | 1,34 | 0 |
| NFIC | TTCCTGGCTCC | 1 | 1060 | 1071 | 60 | 71 | 5,51 | 0,01 | 22,48 | >0.1 | 5,6 | 7,35 | 0 | 1,86 | 0 | 2,16 | 0 |
| NFIC | TCCTTCTCAGC | 1 | 1068 | 1079 | 68 | 79 | 2,05 | 0,01 | 20,71 | >0.1 | 5,6 | 7,21 | 0 | 1,85 | 1,79 | 2,21 | 0 |
| NFIC | TTCTTGGAATT | 1 | 1186 | 1197 | 186 | 197 | 7,47 | 0,01 | 26,06 | >0.1 | 5,6 | 6,05 | 0 | 1,54 | 1,79 | 3,61 | 0 |
| NFIC | AACTTGTCCAA | -1 | 1133 | 1144 | 133 | 144 | 3,13 | 0,01 | 23,19 | >0.1 | 5,6 | 6,43 | 0 | 1,84 | 3,22 | 2,97 | 0 |
| NFIC | GTCTTGGCCTG | -1 | 1118 | 1129 | 118 | 129 | 9,46 | 0,001 | 29,36 | >0.1 | 5,6 | 6,57 | 0 | 1,85 | 3,22 | 2,67 | 0 |
| NFIC | GACTGAGCAGC | -1 | 1094 | 1105 | 94 | 105 | 3,35 | 0,01 | 23,19 | >0.1 | 5,6 | 6,83 | 0 | 1,85 | 3,22 | 2,34 | 0 |
| NFIC | AGCTTGACATC | -1 | 1048 | 1059 | 48 | 59 | 6,83 | 0,01 | 23,33 | >0.1 | 5,6 | 7,62 | 0 | 1,86 | 0 | 1,42 | 0 |
| NFIC | CTCTTATCAGC | -1 | 966 | 977 | -34 | -23 | 2,98 | 0,01 | 19,96 | >0.1 | 5,6 | 7,81 | 0 | 1,87 | 0 | 1,7 | 0 |
| NFIC | AACTTGGGTTG | -1 | 950 | 961 | -50 | -39 | 2,31 | 0,01 | 18,94 | >0.1 | 5,6 | 7,42 | 0 | 1,88 | 0 | 1,74 | 0 |
| NFIC | CGCTGAGCAGG | -1 | 937 | 948 | -63 | -52 | 3,07 | 0,01 | 18,63 | >0.1 | 5,6 | 7,18 | 0 | 1,88 | 0 | 0,9 | 0 |
| NFIC | CTCCTGTCACT | -1 | 765 | 776 | -235 | -224 | 3,41 | 0,01 | 22,06 | >0.1 | 5,6 | 5,8 | 0 | 1,34 | 0 | 5,92 | 0 |
| NFIC | TACTTGGGCAA | -1 | 541 | 552 | -459 | -448 | 2,5 | 0,01 | 11,69 | >0.1 | 3,14 | 5,06 | 0 | 0,99 | 0 | 0 | 0 |
| NFIC | ACTTGGGCAAC | -1 | 540 | 551 | -460 | -449 | 1,92 | 0,01 | 11,1 | >0.1 | 3,14 | 5,06 | 0 | 0,99 | 0 | 0 | 0 |
| NFIC | GACTTTGCCCT | -1 | 509 | 520 | -491 | -480 | 2,1 | 0,01 | 13,82 | >0.1 | 3 | 4,99 | 0 | 0,98 | 0 | 2,75 | 0 |
| NFIC | CCCTTGTCTTT | -1 | 502 | 513 | -498 | -487 | 3,16 | 0,01 | 17,77 | >0.1 | 2,97 | 4,97 | 0 | 0,98 | 0 | 5,68 | 0 |
| NFIC | ATCTCGGCTCA | -1 | 400 | 411 | -600 | -589 | 4,89 | 0,01 | 19,23 | >0.1 | 2,63 | 4,77 | 0 | 1,04 | 0 | 5,91 | 0 |
| NFIC | AACTCGGCCTC | -1 | 218 | 229 | -782 | -771 | 4,66 | 0,01 | 19,26 | >0.1 | 2,96 | 4,49 | 0 | 1,52 | 0 | 5,63 | 0 |
| NFIC | TCCATGGCCAT | -1 | 145 | 156 | -855 | -844 | 2,67 | 0,01 | 13,88 | >0.1 | 3,31 | 4,4 | 0 | 1,9 | 0 | 1,61 | 0 |
| NFIC | GCCTTTGCAAA | -1 | 84 | 95 | -916 | -905 | 5,5 | 0,01 | 15,46 | >0.1 | 3,74 | 4,33 | 0 | 1,88 | 0 | 0 | 0 |
| NFIA | TTTGCAAAGG | 1 | 84 | 94 | -916 | -906 | -7,26 | 0,01 | 2,7 | >0.1 | 3,75 | 4,33 | 0 | 1,88 | 0 | 0 | 0 |
| NFIA | GAGGCCGAGT | 1 | 218 | 228 | -782 | -772 | -23,63 | 0,01 | -9,03 | >0.1 | 2,96 | 4,49 | 0 | 1,52 | 0 | 5,63 | 0 |
| NFIA | AGTCCCAGCT | 1 | 336 | 346 | -664 | -654 | -23,93 | 0,01 | -9,65 | >0.1 | 2,58 | 4,66 | 0 | 1,13 | 0 | 5,92 | 0 |
| NFIA | TGAGCCGAGA | 1 | 400 | 410 | -600 | -590 | -23,71 | 0,01 | -9,37 | >0.1 | 2,63 | 4,77 | 0 | 1,04 | 0 | 5,91 | 0 |
| NFIA | CGCGCCACTG | 1 | 411 | 421 | -589 | -579 | -24,22 | 0,01 | -9,99 | >0.1 | 2,66 | 4,79 | 0 | 1,03 | 0 | 5,76 | 0 |
| NFIA | AGGGCAAAGT | 1 | 509 | 519 | -491 | -481 | -24,02 | 0,01 | -12,31 | >0.1 | 3 | 4,98 | 0 | 0,98 | 0 | 2,75 | 0 |
| NFIA | GTTGCCCAAG | 1 | 540 | 550 | -460 | -450 | -6,38 | 0,01 | 2,79 | >0.1 | 3,13 | 5,06 | 0 | 0,99 | 0 | 0 | 0 |
| NFIA | TTGCCCAAGT | 1 | 541 | 551 | -459 | -449 | -24,74 | 0,01 | -15,56 | >0.1 | 3,14 | 5,06 | 0 | 0,99 | 0 | 0 | 0 |
| NFIA | GATCCCTAGT | 1 | 651 | 661 | -349 | -339 | -24,67 | 0,01 | -10,34 | >0.1 | 3,91 | 5,36 | 0 | 1,07 | 0 | 3,99 | 0 |
| NFIA | AGTGACAGGA | 1 | 765 | 775 | -235 | -225 | -24,14 | 0,01 | -5,48 | >0.1 | 5,61 | 5,79 | 0 | 1,34 | 0 | 5,92 | 0 |
| NFIA | AGCGCCCAAC | 1 | 944 | 954 | -56 | -46 | -24,42 | 0,01 | -7,84 | >0.1 | 5,61 | 7,29 | 0 | 1,88 | 0 | 1,8 | 0 |
| NFIA | GATGTCAAGC | 1 | 1048 | 1058 | 48 | 58 | -6,98 | 0,01 | 9,53 | >0.1 | 5,61 | 7,62 | 0 | 1,86 | 0 | 1,42 | 0 |
| NFIA | GCTGCTCAGT | 1 | 1094 | 1104 | 94 | 104 | -24,64 | 0,01 | -4,79 | >0.1 | 5,61 | 6,83 | 0 | 1,85 | 3,22 | 2,34 | 0 |
| NFIA | CAGGCCAAGA | 1 | 1118 | 1128 | 118 | 128 | -5,7 | 0,001 | 14,22 | >0.1 | 5,61 | 6,57 | 0 | 1,85 | 3,22 | 2,67 | 0 |
| NFIA | AATTCCAAGA | -1 | 1187 | 1197 | 187 | 197 | -7,19 | 0,01 | 11,34 | >0.1 | 5,61 | 6,05 | 0 | 1,53 | 1,79 | 3,55 | 0 |
| NFIA | GGAGCCAGGA | -1 | 1061 | 1071 | 61 | 71 | -23,09 | 0,01 | -6,13 | >0.1 | 5,61 | 7,33 | 0 | 1,86 | 0 | 2,16 | 0 |
| NFIA | TGAGCCAAGA | -1 | 1030 | 1040 | 30 | 40 | -5,44 | 0,001 | 11,6 | >0.1 | 5,61 | 8,24 | 0 | 1,86 | 0 | 1,34 | 0 |
| NFIA | TGAGCCAATA | -1 | 881 | 891 | -119 | -109 | -6,69 | 0,01 | 10,42 | >0.1 | 5,61 | 6,54 | 0 | 1,89 | 0 | 3,07 | 0 |
| NFIA | ATGGCCAAAT | -1 | 753 | 763 | -247 | -237 | -6,6 | 0,01 | 12,36 | >0.1 | 5,61 | 5,74 | 0 | 1,3 | 0 | 6,32 | 0 |
| NFIA | CCTTCCAAAT | -1 | 524 | 534 | -476 | -466 | -7,37 | 0,01 | 4,97 | >0.1 | 3,06 | 5,02 | 0 | 0,98 | 0 | 3,28 | 0 |
| NFIA | GTCGCCCAGG | -1 | 429 | 439 | -571 | -561 | -23,94 | 0,01 | -9,73 | >0.1 | 2,71 | 4,82 | 0 | 1,01 | 0 | 5,67 | 0 |
| NFIA | CCTCCCAGGT | -1 | 378 | 388 | -622 | -612 | -24,34 | 0,01 | -9,45 | >0.1 | 2,56 | 4,73 | 0 | 1,06 | 0 | 6,53 | 0 |
| NFIA | CCTCCCGAGT | -1 | 346 | 356 | -654 | -644 | -24,33 | 0,01 | -9,47 | >0.1 | 2,55 | 4,68 | 0 | 1,11 | 0 | 6,52 | 0 |
| NFIA | GGTGCCCACC | -1 | 323 | 333 | -677 | -667 | -5,09 | 0,001 | 8,88 | >0.1 | 2,61 | 4,64 | 0 | 1,15 | 0 | 5,56 | 0 |
| NFIA | CACGCCCAGC | -1 | 311 | 321 | -689 | -679 | -24,41 | 0,01 | -9,93 | >0.1 | 2,65 | 4,62 | 0 | 1,18 | 0 | 6,03 | 0 |
| NFIA | TCTGCCCAAC | -1 | 226 | 236 | -774 | -764 | -7,39 | 0,01 | 6,77 | >0.1 | 2,93 | 4,5 | 0 | 1,47 | 0 | 5,26 | 0 |
| NFIA | CTGCCCAACT | -1 | 225 | 235 | -775 | -765 | -24,67 | 0,01 | -10,5 | >0.1 | 2,94 | 4,5 | 0 | 1,48 | 0 | 5,26 | 0 |
| NFIA | CCTCCCAAAG | -1 | 212 | 222 | -788 | -778 | -7,33 | 0,01 | 6,28 | >0.1 | 2,99 | 4,48 | 0 | 1,55 | 0 | 4,59 | 0 |

| | | | | | | | | | | | | | | | | | |
|---|---|---|---|---|---|---|---|---|---|---|---|---|---|---|---|---|---|
| NFIA | AGTGCTAGGA | -1 | 204 | 214 | -796 | -786 | -24,13 | 0,01 | -10,44 | >0.1 | 3,02 | 4,47 | 0 | 1,6 | 0 | 4,59 | 0 |
| NFIA | TGTGCCCAGC | -1 | 177 | 187 | -823 | -813 | -5,29 | 0,001 | 7,27 | >0.1 | 3,14 | 4,44 | 0 | 1,82 | 0 | 3,16 | 0 |
| NFIA | TTTGCAAAAC | -1 | 82 | 92 | -918 | -908 | -8,26 | 0,01 | 1,72 | >0.1 | 3,77 | 4,33 | 0 | 1,88 | 0 | 0 | 0 |
| NFIA | AGTGCCTCCT | -1 | 19 | 29 | -981 | -971 | -23,79 | 0,01 | -12,04 | >0.1 | 5,61 | 4,27 | 0 | 1,87 | 0 | 0 | 0 |
| GFI1 | GTAATCCTAG | 1 | 200 | 210 | -800 | -790 | 4 | 0,01 | 17,73 | >0.1 | 3,04 | 4,47 | 0 | 1,63 | 0 | 4,59 | 0 |
| GFI1 | CAGATCACAA | 1 | 232 | 242 | -768 | -758 | 3,09 | 0,01 | 16,96 | >0.1 | 2,91 | 4,51 | 0 | 1,44 | 0 | 5,01 | 0 |
| GFI1 | TGAAACCCCG | 1 | 276 | 286 | -724 | -714 | 3,42 | 0,01 | 17,1 | >0.1 | 2,75 | 4,57 | 0 | 1,27 | 0 | 5,09 | 0 |
| GFI1 | AAAAACTTAG | 1 | 302 | 312 | -698 | -688 | 2,15 | 0,01 | 15,62 | >0.1 | 2,67 | 4,61 | 0 | 1,2 | 0 | 4,99 | 0 |
| GFI1 | AAAATAAATG | 1 | 582 | 592 | -418 | -408 | 3,41 | 0,01 | 12,93 | >0.1 | 3,36 | 5,16 | 0 | 1 | 0 | 0 | 0 |
| GFI1 | ACAAACACCA | 1 | 628 | 638 | -372 | -362 | 3,02 | 0,01 | 14,65 | >0.1 | 3,69 | 5,29 | 0 | 1,04 | 0 | 1,61 | 0 |
| GFI1 | AAAATCTGTT | 1 | 841 | 851 | -159 | -149 | 6,12 | 0,01 | 21,16 | >0.1 | 5,61 | 6,24 | 0 | 1,78 | 0 | 1,41 | 0 |
| GFI1 | AAAATCTCAT | 1 | 979 | 989 | -21 | -11 | 7,31 | 0,01 | 24,59 | >0.1 | 5,61 | 8,27 | 0 | 1,87 | 0 | 1,53 | 0 |
| GFI1 | TCATTCAGTG | 1 | 1013 | 1023 | 13 | 23 | 2,66 | 0,01 | 20,44 | >0.1 | 5,61 | 8,89 | 0 | 1,86 | 0 | 1,42 | 0 |
| GFI1 | TCAAGCTCTT | 1 | 1052 | 1062 | 52 | 62 | 2,95 | 0,01 | 20,22 | >0.1 | 5,61 | 7,53 | 0 | 1,86 | 0 | 2,27 | 0 |
| GFI1 | ACAATCTGCT | -1 | 890 | 900 | -110 | -100 | 5,39 | 0,01 | 22,45 | >0.1 | 5,61 | 6,61 | 0 | 1,89 | 0 | 2,95 | 0 |
| GFI1 | TGAATCAAAT | -1 | 814 | 824 | -186 | -176 | 5,52 | 0,01 | 22,19 | >0.1 | 5,61 | 6,05 | 0 | 1,58 | 0 | 3,43 | 0 |
| GFI1 | CAAATAAGAA | -1 | 809 | 819 | -191 | -181 | 3,38 | 0,01 | 19,84 | >0.1 | 5,61 | 6,02 | 0 | 1,55 | 0 | 3,28 | 0 |
| GFI1 | TTAATCCTAG | -1 | 788 | 798 | -212 | -202 | 4,23 | 0,01 | 22,5 | >0.1 | 5,61 | 5,91 | 0 | 1,44 | 0 | 5,32 | 0 |
| GFI1 | CAAATCAAAA | -1 | 748 | 758 | -252 | -242 | 8,37 | 0,001 | 26,89 | >0.1 | 5,61 | 5,71 | 0 | 1,28 | 0 | 5,92 | 0 |
| GFI1 | AGAAACTCTC | -1 | 552 | 562 | -448 | -438 | 2,4 | 0,01 | 11,67 | >0.1 | 3,19 | 5,08 | 0 | 0,99 | 0 | 0 | 0 |
| GFI1 | CAAATACATG | -1 | 519 | 529 | -481 | -471 | 3,42 | 0,01 | 15,64 | >0.1 | 3,04 | 5,01 | 0 | 0,98 | 0 | 3,19 | 0 |
| GFI1 | GCAAGCTCTG | -1 | 388 | 398 | -612 | -602 | 5,66 | 0,01 | 20,18 | >0.1 | 2,59 | 4,75 | 0 | 1,05 | 0 | 6,13 | 0 |
| GFI1 | AAAAGCTCAT | -1 | 68 | 78 | -932 | -922 | 2,39 | 0,01 | 12,49 | >0.1 | 3,91 | 4,32 | 0 | 1,88 | 0 | 0 | 0 |
| JDP2 | AGGAGGCAC | 1 | 19 | 28 | -981 | -972 | -4,03 | 0,01 | 7,73 | >0.1 | 5,62 | 4,27 | 0 | 1,87 | 0 | 0 | 0 |
| JDP2 | GTGAGAAAT | 1 | 54 | 63 | -946 | -937 | -2,84 | 0,01 | 7,41 | >0.1 | 4,08 | 4,3 | 0 | 1,87 | 0 | 0 | 0 |
| JDP2 | AGTAGTCAT | 1 | 125 | 134 | -875 | -866 | -1,66 | 0,01 | 9,74 | >0.1 | 3,44 | 4,38 | 0 | 1,89 | 0 | 1,7 | 0 |
| JDP2 | GTGGCTCAC | 1 | 187 | 196 | -813 | -804 | 2,81 | 0,001 | 15,94 | >0.1 | 3,1 | 4,45 | 0 | 1,73 | 0 | 3,86 | 0 |
| JDP2 | CTGACTAAC | 1 | 263 | 272 | -737 | -728 | 2,29 | 0,001 | 15,48 | >0.1 | 2,8 | 4,55 | 0 | 1,32 | 0 | 4,53 | 0 |
| JDP2 | CTGAGGCAG | 1 | 356 | 365 | -644 | -635 | -6,31 | 0,01 | 8,03 | >0.1 | 2,53 | 4,69 | 0 | 1,09 | 0 | 6,03 | 0 |
| JDP2 | GAGACTCAG | 1 | 444 | 453 | -556 | -547 | -2,72 | 0,01 | 11,81 | >0.1 | 2,76 | 4,85 | 0 | 1 | 0 | 5,92 | 0 |
| JDP2 | TTGATTCAC | 1 | 816 | 825 | -184 | -175 | 0,26 | 0,01 | 16,95 | >0.1 | 5,62 | 6,06 | 0 | 1,58 | 0 | 3,43 | 0 |
| JDP2 | TTGGCTCAC | 1 | 1032 | 1041 | 32 | 41 | -1,78 | 0,01 | 15,16 | >0.1 | 5,62 | 8,15 | 0 | 1,86 | 0 | 1,31 | 0 |
| JDP2 | GTGAGCCAA | -1 | 1032 | 1041 | 32 | 41 | -6,8 | 0,01 | 10,14 | >0.1 | 5,62 | 8,15 | 0 | 1,86 | 0 | 1,31 | 0 |
| JDP2 | ATGACTTTC | -1 | 1008 | 1017 | 8 | 17 | -0,9 | 0,01 | 17,13 | >0.1 | 5,62 | 9,07 | 0 | 1,87 | 0 | 1,46 | 0 |
| JDP2 | GTGAATCAA | -1 | 816 | 825 | -184 | -175 | 0,27 | 0,01 | 16,96 | >0.1 | 5,62 | 6,06 | 0 | 1,58 | 0 | 3,43 | 0 |
| JDP2 | ATCTCACAT | -1 | 589 | 598 | -411 | -402 | -6,64 | 0,01 | 2,94 | >0.1 | 3,4 | 5,18 | 0 | 1,01 | 0 | 0 | 0 |
| JDP2 | ATTTCTGAT | -1 | 482 | 491 | -518 | -509 | -5,3 | 0,01 | 9,15 | >0.1 | 2,89 | 4,92 | 0 | 0,98 | 0 | 5,65 | 0 |
| JDP2 | CTGAGTCTC | -1 | 444 | 453 | -556 | -547 | -0,31 | 0,01 | 14,22 | >0.1 | 2,76 | 4,85 | 0 | 1 | 0 | 5,92 | 0 |
| JDP2 | GTTAGTCAG | -1 | 263 | 272 | -737 | -728 | -2,59 | 0,01 | 10,6 | >0.1 | 2,8 | 4,55 | 0 | 1,32 | 0 | 4,53 | 0 |
| JDP2 | GTGAGCCAC | -1 | 187 | 196 | -813 | -804 | -2,11 | 0,01 | 11,02 | >0.1 | 3,1 | 4,45 | 0 | 1,73 | 0 | 3,86 | 0 |
| JDP2 | ATTTCTGAT | -1 | 167 | 176 | -833 | -824 | -5,3 | 0,01 | 6,68 | >0.1 | 3,19 | 4,42 | 0 | 1,9 | 0 | 2,46 | 0 |
| JDP2 | ATGACTACT | -1 | 125 | 134 | -875 | -866 | -2,78 | 0,01 | 8,62 | >0.1 | 3,44 | 4,38 | 0 | 1,89 | 0 | 1,7 | 0 |
| JDP2 | ATTTCTCAC | -1 | 54 | 63 | -946 | -937 | 2,18 | 0,001 | 12,43 | >0.1 | 4,08 | 4,3 | 0 | 1,87 | 0 | 0 | 0 |
| JDP2 | GTGGCTGAC | -1 | 36 | 45 | -964 | -955 | -6,18 | 0,01 | 4,34 | >0.1 | 4,37 | 4,28 | 0 | 1,87 | 0 | 0 | 0 |
| TGIF2 | TCAAAGAAGTCA | 1 | 28 | 40 | -972 | -960 | -2,77 | 0,01 | 7,89 | >0.1 | 4,51 | 4,28 | 0 | 1,87 | 0 | 0 | 0 |
| TGIF2 | GGAGAGTAGTCA | 1 | 121 | 133 | -879 | -867 | -5,46 | 0,01 | 5,91 | >0.1 | 3,45 | 4,37 | 0 | 1,89 | 0 | 1,65 | 0 |
| TGIF2 | TCACGCCTGTAA | 1 | 192 | 204 | -808 | -796 | -3,89 | 0,01 | 9,59 | >0.1 | 3,07 | 4,46 | 0 | 1,69 | 0 | 4,26 | 0 |
| TGIF2 | GGGCAGATCACA | 1 | 229 | 241 | -771 | -759 | -2,15 | 0,01 | 11,74 | >0.1 | 2,92 | 4,51 | 0 | 1,46 | 0 | 5,01 | 0 |
| TGIF2 | GGGCACCTGTAG | 1 | 326 | 338 | -674 | -662 | -5,58 | 0,01 | 8,38 | >0.1 | 2,6 | 4,65 | 0 | 1,15 | 0 | 5,56 | 0 |
| TGIF2 | TAACAGCTTTCT | 1 | 710 | 722 | -290 | -278 | 2,51 | 0,001 | 19,83 | >0.1 | 5,58 | 5,57 | 0 | 1,18 | 0 | 4,99 | 0 |
| TGIF2 | TGACAGGAGAGG | 1 | 767 | 779 | -233 | -221 | -0,91 | 0,01 | 17,75 | >0.1 | 5,58 | 5,81 | 0 | 1,35 | 0 | 5,92 | 0 |
| TGIF2 | TAAAATCTGTTA | 1 | 840 | 852 | -160 | -148 | -1,9 | 0,01 | 13,13 | >0.1 | 5,58 | 6,24 | 0 | 1,79 | 0 | 1,41 | 0 |
| TGIF2 | ACATATCTGTCC | 1 | 851 | 863 | -149 | -137 | -5,15 | 0,01 | 11,31 | >0.1 | 5,58 | 6,32 | 0 | 1,89 | 0 | 2,67 | 0 |
| TGIF2 | TCACAGGGGACG | 1 | 1037 | 1049 | 37 | 49 | -2,23 | 0,01 | 14,5 | >0.1 | 5,58 | 7,95 | 0 | 1,86 | 0 | 1,34 | 0 |
| TGIF2 | GGGACGATGTCA | 1 | 1043 | 1055 | 43 | 55 | -6,01 | 0,01 | 9,81 | >0.1 | 5,58 | 7,76 | 0 | 1,86 | 0 | 0,62 | 0 |
| TGIF2 | GGACAAGTTTAA | 1 | 1135 | 1147 | 135 | 147 | -5,39 | 0,01 | 14,76 | >0.1 | 5,58 | 6,41 | 0 | 1,84 | 3,22 | 3,09 | 0 |
| TGIF2 | TGACATCGTCCC | -1 | 1043 | 1055 | 43 | 55 | -4,44 | 0,01 | 11,38 | >0.1 | 5,58 | 7,76 | 0 | 1,86 | 0 | 0,62 | 0 |
| TGIF2 | GGACAGATATGT | -1 | 851 | 863 | -149 | -137 | -4,24 | 0,01 | 12,22 | >0.1 | 5,58 | 6,32 | 0 | 1,89 | 0 | 2,67 | 0 |
| TGIF2 | AGATATGTAACA | -1 | 847 | 859 | -153 | -141 | -3,48 | 0,01 | 13,2 | >0.1 | 5,58 | 6,29 | 0 | 1,85 | 0 | 2,95 | 0 |

| | | | | | | | | | | | | | | | | | |
|---|---|---|---|---|---|---|---|---|---|---|---|---|---|---|---|---|---|
| TGIF2 | TAACAGATTTTA | -1 | 840 | 852 | -160 | -148 | -2,69 | 0,01 | 12,34 | >0.1 | 5,58 | 6,24 | 0 | 1,79 | 0 | 1,41 | 0 |
| TGIF2 | AGAAAGCTGTTA | -1 | 710 | 722 | -290 | -278 | 3,52 | 0,001 | 20,84 | >0.1 | 5,58 | 5,57 | 0 | 1,18 | 0 | 4,99 | 0 |
| TGIF2 | CTACAGGTGCCC | -1 | 326 | 338 | -674 | -662 | -0,38 | 0,01 | 13,58 | >0.1 | 2,6 | 4,65 | 0 | 1,15 | 0 | 5,56 | 0 |
| TGIF2 | TGTGATCTGCCC | -1 | 229 | 241 | -771 | -759 | -3,7 | 0,01 | 10,19 | >0.1 | 2,92 | 4,51 | 0 | 1,46 | 0 | 5,01 | 0 |
| TGIF2 | TTACAGGCGTGA | -1 | 192 | 204 | -808 | -796 | 2,5 | 0,001 | 15,98 | >0.1 | 3,07 | 4,46 | 0 | 1,69 | 0 | 4,26 | 0 |
| TGIF2 | AGGCGTGAGCCA | -1 | 188 | 200 | -812 | -800 | -6,13 | 0,01 | 7,17 | >0.1 | 3,09 | 4,45 | 0 | 1,72 | 0 | 4,04 | 0 |
| TGIF2 | TGACTACTCTCC | -1 | 121 | 133 | -879 | -867 | -4,32 | 0,01 | 7,05 | >0.1 | 3,45 | 4,37 | 0 | 1,89 | 0 | 1,65 | 0 |
| TGIF2 | TGACTTCTTTGA | -1 | 28 | 40 | -972 | -960 | -2,49 | 0,01 | 8,17 | >0.1 | 4,51 | 4,28 | 0 | 1,87 | 0 | 0 | 0 |
| CREB3L2 | TTGACCTGT | 1 | 107 | 116 | -893 | -884 | -0,96 | 0,01 | 9,59 | >0.1 | 3,56 | 4,36 | 0 | 1,89 | 0 | 0,75 | 0 |
| CREB3L2 | GACCTGTGG | 1 | 109 | 118 | -891 | -882 | -5,3 | 0,01 | 5,24 | >0.1 | 3,55 | 4,36 | 0 | 1,89 | 0 | 0,75 | 0 |
| CREB3L2 | CTCACGCCT | 1 | 191 | 200 | -809 | -800 | -6,58 | 0,01 | 6,78 | >0.1 | 3,08 | 4,45 | 0 | 1,7 | 0 | 4,13 | 0 |
| CREB3L2 | GTCAGGAGA | 1 | 243 | 252 | -757 | -748 | -6,79 | 0,01 | 6,32 | >0.1 | 2,87 | 4,52 | 0 | 1,39 | 0 | 4,33 | 0 |
| CREB3L2 | ACCCCGTCT | 1 | 280 | 289 | -720 | -711 | -4,28 | 0,01 | 9,49 | >0.1 | 2,74 | 4,57 | 0 | 1,26 | 0 | 5,19 | 0 |
| CREB3L2 | TGGGCGTGG | 1 | 313 | 322 | -687 | -678 | -7,18 | 0,01 | 7,29 | >0.1 | 2,64 | 4,62 | 0 | 1,17 | 0 | 6,03 | 0 |
| CREB3L2 | GCGTGGTGG | 1 | 316 | 325 | -684 | -675 | -6,28 | 0,01 | 8,28 | >0.1 | 2,63 | 4,63 | 0 | 1,17 | 0 | 6,13 | 0 |
| CREB3L2 | GGCACCTGT | 1 | 327 | 336 | -673 | -664 | -2,32 | 0,01 | 11,53 | >0.1 | 2,6 | 4,64 | 0 | 1,14 | 0 | 5,46 | 0 |
| CREB3L2 | TACTCGGGA | 1 | 345 | 354 | -655 | -646 | -7,37 | 0,01 | 7,49 | >0.1 | 2,56 | 4,67 | 0 | 1,11 | 0 | 6,52 | 0 |
| CREB3L2 | GCAAAGTCA | 1 | 512 | 521 | -488 | -479 | -7,36 | 0,01 | 4,48 | >0.1 | 3,01 | 4,99 | 0 | 0,98 | 0 | 2,86 | 0 |
| CREB3L2 | GTCATGTAT | 1 | 517 | 526 | -483 | -474 | -6,56 | 0,01 | 5,4 | >0.1 | 3,03 | 5 | 0 | 0,98 | 0 | 2,95 | 0 |
| CREB3L2 | TAGACCTCT | 1 | 663 | 672 | -337 | -328 | -2,54 | 0,01 | 11,97 | >0.1 | 4,04 | 5,39 | 0 | 1,09 | 0 | 3,99 | 0 |
| CREB3L2 | GACAGGAGA | 1 | 768 | 777 | -232 | -223 | -5,8 | 0,01 | 12,89 | >0.1 | 5,62 | 5,8 | 0 | 1,35 | 0 | 5,92 | 0 |
| CREB3L2 | GCCAAGACA | 1 | 1121 | 1130 | 121 | 130 | -7,06 | 0,01 | 12,87 | >0.1 | 5,62 | 6,54 | 0 | 1,85 | 3,22 | 2,7 | 0 |
| CREB3L2 | GACAAGTTT | 1 | 1136 | 1145 | 136 | 145 | -5,58 | 0,01 | 14,54 | >0.1 | 5,62 | 6,41 | 0 | 1,84 | 3,22 | 3,02 | 0 |
| CREB3L2 | AAGACCTGT | 1 | 1158 | 1167 | 158 | 167 | -6,8 | 0,01 | 9,74 | >0.1 | 4,39 | 6,24 | 0 | 1,78 | 1,79 | 2,34 | 0 |
| CREB3L2 | TTCACTTGC | 1 | 1177 | 1186 | 177 | 186 | -5,99 | 0,01 | 12,42 | >0.1 | 5,62 | 6,11 | 0 | 1,61 | 1,79 | 3,28 | 0 |
| CREB3L2 | AACAGGTCT | -1 | 1159 | 1168 | 159 | 168 | -6,59 | 0,01 | 9,97 | >0.1 | 4,37 | 6,23 | 0 | 1,77 | 1,79 | 2,4 | 0 |
| CREB3L2 | GCTTCGTGG | -1 | 1147 | 1156 | 147 | 156 | -3,79 | 0,01 | 13,95 | >0.1 | 5,62 | 6,32 | 0 | 1,84 | 1,79 | 2,16 | 0 |
| CREB3L2 | TTGACATCG | -1 | 1047 | 1056 | 47 | 56 | -4,53 | 0,01 | 12,07 | >0.1 | 5,62 | 7,65 | 0 | 1,86 | 0 | 1,46 | 0 |
| CREB3L2 | GTCCCCTGT | -1 | 1039 | 1048 | 39 | 48 | -5,3 | 0,01 | 11,4 | >0.1 | 5,62 | 7,88 | 0 | 1,86 | 0 | 1,34 | 0 |
| CREB3L2 | TCCTCATGA | -1 | 985 | 994 | -15 | -6 | -6,55 | 0,01 | 11,04 | >0.1 | 5,62 | 8,57 | 0 | 1,87 | 0 | 1,53 | 0 |
| CREB3L2 | GTCACTTTA | -1 | 762 | 771 | -238 | -229 | -7,56 | 0,01 | 11,26 | >0.1 | 5,62 | 5,77 | 0 | 1,32 | 0 | 6,1 | 0 |
| CREB3L2 | GCCAAATCA | -1 | 751 | 760 | -249 | -240 | -7,56 | 0,01 | 11,18 | >0.1 | 5,62 | 5,72 | 0 | 1,29 | 0 | 6,1 | 0 |
| CREB3L2 | TCTACTTGG | -1 | 545 | 554 | -455 | -446 | -6,36 | 0,01 | 2,85 | >0.1 | 3,15 | 5,06 | 0 | 0,99 | 0 | 0 | 0 |
| CREB3L2 | TTCACGCCA | -1 | 370 | 379 | -630 | -621 | -1,24 | 0,01 | 13,24 | >0.1 | 2,54 | 4,71 | 0 | 1,07 | 0 | 6,15 | 0 |
| CREB3L2 | TACAGGTGC | -1 | 328 | 337 | -672 | -663 | -4,51 | 0,01 | 9,44 | >0.1 | 2,6 | 4,65 | 0 | 1,14 | 0 | 5,56 | 0 |
| CREB3L2 | GAGACGGGG | -1 | 281 | 290 | -719 | -710 | 2,1 | 0,001 | 15,76 | >0.1 | 2,74 | 4,57 | 0 | 1,26 | 0 | 5,09 | 0 |
| CREB3L2 | TCACCGTGT | -1 | 270 | 279 | -730 | -721 | -4,87 | 0,01 | 8,84 | >0.1 | 2,78 | 4,56 | 0 | 1,29 | 0 | 5,09 | 0 |
| CREB3L2 | TACAGGCGT | -1 | 194 | 203 | -806 | -797 | -6,38 | 0,01 | 7,08 | >0.1 | 3,07 | 4,46 | 0 | 1,67 | 0 | 4,26 | 0 |
| CREB3L2 | CAGGCGTGA | -1 | 192 | 201 | -808 | -799 | -6,8 | 0,01 | 6,55 | >0.1 | 3,08 | 4,45 | 0 | 1,69 | 0 | 4,13 | 0 |
| CREB3L2 | GCCATTTCT | -1 | 170 | 179 | -830 | -821 | -6,56 | 0,01 | 6,44 | >0.1 | 3,18 | 4,43 | 0 | 1,88 | 0 | 3,51 | 0 |
| CREB3L2 | TCCACAGGT | -1 | 110 | 119 | -890 | -881 | -5,06 | 0,01 | 5,47 | >0.1 | 3,54 | 4,36 | 0 | 1,89 | 0 | 0,75 | 0 |
| CREB3L2 | CACAGGTCA | -1 | 108 | 117 | -892 | -883 | -7,08 | 0,01 | 3,48 | >0.1 | 3,56 | 4,36 | 0 | 1,89 | 0 | 0,77 | 0 |
| CREB3L1 | CAGCCACAGAACCA | 1 | 38 | 52 | -962 | -948 | 3,07 | 0,001 | 19,02 | >0.1 | 4,28 | 4,29 | 0 | 1,87 | 0 | 0 | 5,5 |
| CREB3L1 | TCACAAGGTCAGGA | 1 | 236 | 250 | -764 | -750 | -5,16 | 0,01 | 13,7 | >0.1 | 2,89 | 4,52 | 0 | 1,43 | 0 | 4,53 | 5,5 |
| CREB3L1 | CTAACACGGTGAAA | 1 | 267 | 281 | -733 | -719 | -8,87 | 0,01 | 10,36 | >0.1 | 2,78 | 4,56 | 0 | 1,3 | 0 | 5,09 | 5,5 |
| CREB3L1 | AAACCCCGTCTCTA | 1 | 278 | 292 | -722 | -708 | -5,42 | 0,01 | 13,76 | >0.1 | 2,74 | 4,58 | 0 | 1,27 | 0 | 5,09 | 5,5 |
| CREB3L1 | GTCCCAGCTACTCG | 1 | 337 | 351 | -663 | -649 | -8,05 | 0,01 | 12,34 | >0.1 | 2,57 | 4,67 | 0 | 1,13 | 0 | 6,52 | 5,5 |
| CREB3L1 | GGGCAAAGTCATGT | 1 | 510 | 524 | -490 | -476 | -6,83 | 0,01 | 10,61 | >0.1 | 3,01 | 5 | 0 | 0,99 | 0 | 2,95 | 5,5 |
| CREB3L1 | GTGACAGGAGAGGT | 1 | 766 | 780 | -234 | -220 | -5,42 | 0,01 | 18,73 | >0.1 | 5,56 | 5,82 | 0 | 1,36 | 0 | 5,92 | 5,5 |
| CREB3L1 | AGGCCAAGACATTT | 1 | 1119 | 1133 | 119 | 133 | -5,41 | 0,01 | 19,95 | >0.1 | 5,56 | 6,56 | 0 | 1,85 | 3,22 | 2,67 | 5,5 |
| CREB3L1 | TAACCACGAAGCCG | 1 | 1144 | 1158 | 144 | 158 | -1,57 | 0,01 | 23,06 | >0.1 | 5,56 | 6,34 | 0 | 1,84 | 3,22 | 2,16 | 5,5 |
| CREB3L1 | AGAACAGGTCTTCG | -1 | 1156 | 1170 | 156 | 170 | -3,5 | 0,01 | 18,62 | >0.1 | 4,38 | 6,25 | 0 | 1,8 | 1,79 | 2,4 | 5,5 |
| CREB3L1 | GAGCCAGGAAGAGC | -1 | 1056 | 1070 | 56 | 70 | -9,25 | 0,01 | 13,31 | >0.1 | 5,56 | 7,44 | 0 | 1,86 | 0 | 2,21 | 5,5 |
| CREB3L1 | AGATCACATCCACT | -1 | 1019 | 1033 | 19 | 33 | -9,02 | 0,01 | 13,95 | >0.1 | 5,56 | 8,74 | 0 | 1,86 | 0 | 1,31 | 5,5 |
| CREB3L1 | CTAAAACCTCCTCA | -1 | 988 | 1002 | -12 | 2 | -6,21 | 0,01 | 17,1 | >0.1 | 5,56 | 8,77 | 0 | 1,87 | 0 | 1,61 | 5,5 |
| CREB3L1 | AAACCTCCTCATGA | -1 | 985 | 999 | -15 | -1 | -9,15 | 0,01 | 14,08 | >0.1 | 5,56 | 8,69 | 0 | 1,87 | 0 | 1,61 | 5,5 |
| CREB3L1 | AGGACAGATATGTA | -1 | 850 | 864 | -150 | -136 | -9,23 | 0,01 | 12,73 | >0.1 | 5,56 | 6,33 | 0 | 1,9 | 0 | 2,67 | 5,5 |
| CREB3L1 | CTGTCACTTTATGG | -1 | 759 | 773 | -241 | -227 | -9,33 | 0,01 | 14,95 | >0.1 | 5,56 | 5,78 | 0 | 1,33 | 0 | 6,1 | 5,5 |
| CREB3L1 | TGGCCAAATCAAAA | -1 | 748 | 762 | -252 | -238 | -5,62 | 0,01 | 18,57 | >0.1 | 5,56 | 5,73 | 0 | 1,29 | 0 | 6,1 | 5,5 |

| | | | | | | | | | | | | | | | | | |
|---|---|---|---|---|---|---|---|---|---|---|---|---|---|---|---|---|---|
| CREB3L1 | GATACACGGAAAAT | -1 | 694 | 708 | -306 | -292 | -8,92 | 0,01 | 11,87 | >0.1 | 5,56 | 5,52 | 0 | 1,15 | 0 | 3,07 | 5,5 |
| CREB3L1 | GGGCAACATTTTCC | -1 | 533 | 547 | -467 | -453 | -7,82 | 0,01 | 8,44 | >0.1 | 3,11 | 5,05 | 0 | 0,99 | 0 | 1,61 | 5,5 |
| CREB3L1 | CTGCAAGCTCTGCC | -1 | 386 | 400 | -614 | -600 | -8,31 | 0,01 | 11,72 | >0.1 | 2,59 | 4,75 | 0 | 1,05 | 0 | 6,13 | 5,5 |
| CREB3L1 | CTCCCAGGTTCACG | -1 | 373 | 387 | -627 | -613 | -6,57 | 0,01 | 13,68 | >0.1 | 2,56 | 4,73 | 0 | 1,07 | 0 | 6,4 | 5,5 |
| CREB3L1 | CTGCCTCAGCCTCC | -1 | 351 | 365 | -649 | -635 | -8,06 | 0,01 | 12,17 | >0.1 | 2,53 | 4,69 | 0 | 1,1 | 0 | 6,4 | 5,5 |
| CREB3L1 | ACTACAGGTGCCCA | -1 | 325 | 339 | -675 | -661 | -3,69 | 0,01 | 15,78 | >0.1 | 2,6 | 4,65 | 0 | 1,15 | 0 | 5,56 | 5,5 |
| CREB3L1 | CCACCACCACGCCC | -1 | 314 | 328 | -686 | -672 | -4,11 | 0,01 | 15,96 | >0.1 | 2,63 | 4,63 | 0 | 1,17 | 0 | 6,13 | 5,5 |
| CREB3L1 | CCACCACGCCCAGC | -1 | 311 | 325 | -689 | -675 | -7,7 | 0,01 | 12,38 | >0.1 | 2,64 | 4,63 | 0 | 1,18 | 0 | 6,13 | 5,5 |
| CREB3L1 | ACTCCACAGGTCAA | -1 | 107 | 121 | -893 | -879 | -8,61 | 0,01 | 7,43 | >0.1 | 3,54 | 4,36 | 0 | 1,89 | 0 | 0,75 | 5,5 |
| CREB3L1 | TCCACAGGTCAACT | -1 | 105 | 119 | -895 | -881 | -3,63 | 0,01 | 12,44 | >0.1 | 3,56 | 4,36 | 0 | 1,89 | 0 | 0,77 | 5,5 |
| CREB3L1 | GTGCCTCCTTCTCC | -1 | 14 | 28 | -986 | -972 | -9,27 | 0,01 | 7,92 | >0.1 | 5,56 | 4,27 | 0 | 1,87 | 0 | 0 | 5,5 |
| ATF4 | GAAGGAGGCACTC | 1 | 17 | 30 | -983 | -970 | -1,36 | 0,01 | 10,35 | >0.1 | 5,57 | 4,27 | 0 | 1,87 | 0 | 0 | 0 |
| ATF4 | GGCTGAGGCAGGA | 1 | 354 | 367 | -646 | -633 | 1,51 | 0,001 | 15,86 | >0.1 | 2,53 | 4,69 | 0 | 1,1 | 0 | 6,03 | 0 |
| ATF4 | TGATCTTGAAATA | 1 | 569 | 582 | -431 | -418 | -4,39 | 0,01 | 5,04 | >0.1 | 3,29 | 5,13 | 0 | 1 | 0 | 0 | 0 |
| ATF4 | ATGTGAGATAACC | 1 | 589 | 602 | -411 | -398 | -4,87 | 0,01 | 4,74 | >0.1 | 3,41 | 5,19 | 0 | 1,01 | 0 | 0 | 0 |
| ATF4 | TATTAATGAAATT | 1 | 602 | 615 | -398 | -385 | -4,98 | 0,01 | 6,17 | >0.1 | 3,5 | 5,22 | 0 | 1,02 | 0 | 1,41 | 0 |
| ATF4 | TGATTTGGCCATA | 1 | 751 | 764 | -249 | -236 | -4,29 | 0,01 | 14,86 | >0.1 | 5,57 | 5,74 | 0 | 1,3 | 0 | 6,54 | 0 |
| ATF4 | CTAGGATTAAAGA | 1 | 788 | 801 | -212 | -199 | -7,19 | 0,01 | 11,27 | >0.1 | 5,57 | 5,92 | 0 | 1,45 | 0 | 5,52 | 0 |
| ATF4 | GAATAACGTATTC | 1 | 799 | 812 | -201 | -188 | 0,76 | 0,01 | 17,37 | >0.1 | 5,57 | 5,98 | 0 | 1,51 | 0 | 3,55 | 0 |
| ATF4 | CCAGGATGAACTT | 1 | 866 | 879 | -134 | -121 | -6,03 | 0,01 | 10,63 | >0.1 | 5,57 | 6,44 | 0 | 1,9 | 0 | 2,75 | 0 |
| ATF4 | TGTTGCTGTAACT | 1 | 1081 | 1094 | 81 | 94 | -7,94 | 0,01 | 10,49 | >0.1 | 5,57 | 7 | 0 | 1,85 | 1,79 | 2,21 | 0 |
| ATF4 | CTGTGAGCCAAGA | -1 | 1030 | 1043 | 30 | 43 | -6,63 | 0,01 | 10,34 | >0.1 | 5,57 | 8,24 | 0 | 1,86 | 0 | 1,31 | 0 |
| ATF4 | TGCTGAGCCAATA | -1 | 881 | 894 | -119 | -106 | -0,74 | 0,01 | 16,36 | >0.1 | 5,57 | 6,57 | 0 | 1,89 | 0 | 3,07 | 0 |
| ATF4 | AAATGTTATAATT | -1 | 685 | 698 | -315 | -302 | -4,7 | 0,01 | 9,29 | >0.1 | 4,43 | 5,48 | 0 | 1,13 | 0 | 2,95 | 0 |
| ATF4 | CAGTGACCAAAGA | -1 | 669 | 682 | -331 | -318 | -6,5 | 0,01 | 7,88 | >0.1 | 4,15 | 5,43 | 0 | 1,1 | 0 | 3,7 | 0 |
| ATF4 | TCTTTATGGAATT | -1 | 494 | 507 | -506 | -493 | -6,61 | 0,01 | 8,9 | >0.1 | 2,94 | 4,96 | 0 | 0,99 | 0 | 6,62 | 0 |
| ATF4 | GGTTCACGCCATT | -1 | 368 | 381 | -632 | -619 | -7,3 | 0,01 | 7,19 | >0.1 | 2,54 | 4,72 | 0 | 1,08 | 0 | 6,15 | 0 |
| BCL6B | GGCACTCAAAGAAGTCA | 1 | 23 | 40 | -977 | -960 | -27,97 | 0,01 | -16,3 | >0.1 | 5,52 | 4,28 | 0 | 1,87 | 0 | 0 | 0 |
| BCL6B | TGAATTCCATAAAGACA | 1 | 492 | 509 | -508 | -491 | -34,08 | 0,01 | -18,56 | >0.1 | 2,94 | 4,96 | 0 | 0,99 | 0 | 6,62 | 0 |
| BCL6B | GAGTTTCTGGGAATATG | 1 | 554 | 571 | -446 | -429 | -30,58 | 0,01 | -21,25 | >0.1 | 3,22 | 5,11 | 0 | 1 | 0 | 0 | 0 |
| BCL6B | TATGATCTTGAAATAAA | 1 | 567 | 584 | -433 | -416 | -35,13 | 0,01 | -25,7 | >0.1 | 3,29 | 5,14 | 0 | 1 | 0 | 0 | 0 |
| BCL6B | GGTTCTCTAGGATTAAA | 1 | 782 | 799 | -218 | -201 | -18,05 | 0,001 | 0,52 | >0.1 | 5,52 | 5,91 | 0 | 1,44 | 0 | 5,69 | 0 |
| BCL6B | TCCTCTCCAGGATGAAC | 1 | 860 | 877 | -140 | -123 | -23,55 | 0,01 | -7,04 | >0.1 | 5,52 | 6,42 | 0 | 1,9 | 0 | 2,67 | 0 |
| BCL6B | CGATGTCAAGCTCTTCC | 1 | 1047 | 1064 | 47 | 64 | -32,92 | 0,01 | -16,43 | >0.1 | 5,52 | 7,65 | 0 | 1,86 | 0 | 1,46 | 0 |
| BCL6B | TTGCTTCTTGGAATTAT | 1 | 1182 | 1199 | 182 | 199 | -19 | 0,001 | -0,41 | >0.1 | 5,52 | 6,08 | 0 | 1,57 | 1,79 | 3,63 | 0 |
| BCL6B | GACTTTCCCTAGACTAA | -1 | 998 | 1015 | -2 | 15 | -33,16 | 0,01 | -15,19 | >0.1 | 5,52 | 9,12 | 0 | 1,87 | 0 | 1,46 | 0 |
| BCL6B | CGCTGAGCAGGAAAGAC | -1 | 931 | 948 | -69 | -52 | -27,08 | 0,01 | -11,6 | >0.1 | 5,52 | 7,19 | 0 | 1,88 | 0 | 0,9 | 0 |
| BCL6B | TTTTTTAAAGTGAATCA | -1 | 817 | 834 | -183 | -166 | -32,91 | 0,01 | -15,93 | >0.1 | 5,52 | 6,12 | 0 | 1,65 | 0 | 3,7 | 0 |
| BCL6B | TGTCTTTATGGAATTCA | -1 | 492 | 509 | -508 | -491 | -18,3 | 0,001 | -2,78 | >0.1 | 2,94 | 4,96 | 0 | 0,99 | 0 | 6,62 | 0 |
| BCL6B | TTTTTTTTTGAGACTGA | -1 | 449 | 466 | -551 | -534 | -34,07 | 0,01 | -20,03 | >0.1 | 2,79 | 4,87 | 0 | 1 | 0 | 5,38 | 0 |
| BCL6B | TGCCTCCCAGGTTCACG | -1 | 373 | 390 | -627 | -610 | -33,02 | 0,01 | -18,26 | >0.1 | 2,56 | 4,73 | 0 | 1,07 | 0 | 6,4 | 0 |
| BCL6B | AAAGTGCTAGGATTACA | -1 | 199 | 216 | -801 | -784 | -32,85 | 0,01 | -19 | >0.1 | 3,03 | 4,47 | 0 | 1,64 | 0 | 4,7 | 0 |
| BCL6B | GCCTTTGCAAAACTTAA | -1 | 78 | 95 | -922 | -905 | -34,97 | 0,01 | -24,98 | >0.1 | 3,77 | 4,34 | 0 | 1,88 | 0 | 0 | 0 |
| BCL6B | TGACTTCTTTGAGTGCC | -1 | 23 | 40 | -977 | -960 | -29,93 | 0,01 | -18,26 | >0.1 | 5,52 | 4,28 | 0 | 1,87 | 0 | 0 | 0 |
| NFIX | TCAGCCACA | 1 | 37 | 46 | -963 | -954 | 3,94 | 0,01 | 14,44 | >0.1 | 4,35 | 4,29 | 0 | 1,87 | 0 | 0 | 0 |
| NFIX | AAAGGCAGA | 1 | 89 | 98 | -911 | -902 | 3,8 | 0,01 | 13,72 | >0.1 | 3,71 | 4,34 | 0 | 1,88 | 0 | 0 | 0 |
| NFIX | ATGGCCATG | 1 | 145 | 154 | -855 | -846 | 4,48 | 0,01 | 15,66 | >0.1 | 3,31 | 4,4 | 0 | 1,9 | 0 | 1,57 | 0 |
| NFIX | TGAGGCAGG | 1 | 357 | 366 | -643 | -634 | 5,3 | 0,01 | 19,74 | >0.1 | 2,52 | 4,69 | 0 | 1,09 | 0 | 6,13 | 0 |
| NFIX | GGAGGCAGA | 1 | 383 | 392 | -617 | -608 | 5,39 | 0,01 | 19,67 | >0.1 | 2,58 | 4,74 | 0 | 1,05 | 0 | 5,91 | 0 |
| NFIX | CGCGCCACT | 1 | 411 | 420 | -589 | -580 | 5,44 | 0,01 | 19,55 | >0.1 | 2,65 | 4,79 | 0 | 1,02 | 0 | 5,65 | 0 |
| NFIX | TGCTCCAGC | 1 | 421 | 430 | -579 | -570 | 4,71 | 0,01 | 18,63 | >0.1 | 2,68 | 4,8 | 0 | 1,02 | 0 | 5,42 | 0 |
| NFIX | AAGGGCAAA | 1 | 508 | 517 | -492 | -483 | 4,24 | 0,01 | 17,38 | >0.1 | 2,99 | 4,98 | 0 | 0,98 | 0 | 4,19 | 0 |
| NFIX | GTTGCCCAA | 1 | 540 | 549 | -460 | -451 | 6,83 | 0,01 | 16 | >0.1 | 3,13 | 5,05 | 0 | 0,98 | 0 | 0 | 0 |
| NFIX | TTGGCCATA | 1 | 755 | 764 | -245 | -236 | 3,73 | 0,01 | 22,72 | >0.1 | 5,62 | 5,74 | 0 | 1,3 | 0 | 6,32 | 0 |
| NFIX | CTCTCCAGG | 1 | 862 | 871 | -138 | -129 | 4,54 | 0,01 | 21,19 | >0.1 | 5,62 | 6,38 | 0 | 1,9 | 0 | 2,75 | 0 |
| NFIX | AGCGCCCAA | 1 | 944 | 953 | -56 | -47 | 5,87 | 0,01 | 22,44 | >0.1 | 5,62 | 7,27 | 0 | 1,88 | 0 | 1,8 | 0 |
| NFIX | CAGGCCAAG | 1 | 1118 | 1127 | 118 | 127 | 8,86 | 0,001 | 28,73 | >0.1 | 5,62 | 6,57 | 0 | 1,85 | 3,22 | 2,6 | 0 |
| NFIX | AATTCCAAG | -1 | 1188 | 1197 | 188 | 197 | 6 | 0,01 | 24,53 | >0.1 | 5,62 | 6,04 | 0 | 1,53 | 1,79 | 3,55 | 0 |
| NFIX | GGAGCCAGG | -1 | 1062 | 1071 | 62 | 71 | 9,37 | 0,001 | 26,33 | >0.1 | 5,62 | 7,31 | 0 | 1,86 | 0 | 2,16 | 0 |

| | | | | | | | | | | | | | | | | | |
|---|---|---|---|---|---|---|---|---|---|---|---|---|---|---|---|---|---|
| NFIX | TGAGCCAAG | -1 | 1031 | 1040 | 31 | 40 | 9,51 | 0,001 | 26,52 | >0.1 | 5,62 | 8,19 | 0 | 1,86 | 0 | 1,34 | 0 |
| NFIX | TGAGCCAAT | -1 | 882 | 891 | -118 | -109 | 8,51 | 0,01 | 25,63 | >0.1 | 5,62 | 6,54 | 0 | 1,89 | 0 | 3,07 | 0 |
| NFIX | ATGGCCAAA | -1 | 754 | 763 | -246 | -237 | 7,61 | 0,01 | 26,59 | >0.1 | 5,62 | 5,74 | 0 | 1,3 | 0 | 6,32 | 0 |
| NFIX | TTGGGCAAC | -1 | 540 | 549 | -460 | -451 | 4,8 | 0,01 | 13,97 | >0.1 | 3,13 | 5,05 | 0 | 0,98 | 0 | 0 | 0 |
| NFIX | CCTTCCAAA | -1 | 525 | 534 | -475 | -466 | 6,15 | 0,01 | 18,49 | >0.1 | 3,06 | 5,02 | 0 | 0,98 | 0 | 3,28 | 0 |
| NFIX | GTCGCCCAG | -1 | 430 | 439 | -570 | -561 | 6,21 | 0,01 | 20,42 | >0.1 | 2,71 | 4,82 | 0 | 1,01 | 0 | 5,67 | 0 |
| NFIX | CACGCCATT | -1 | 368 | 377 | -632 | -623 | 4,8 | 0,01 | 19,85 | >0.1 | 2,54 | 4,71 | 0 | 1,07 | 0 | 6,73 | 0 |
| NFIX | GGTGCCCAC | -1 | 324 | 333 | -676 | -667 | 8,39 | 0,01 | 22,36 | >0.1 | 2,61 | 4,64 | 0 | 1,15 | 0 | 5,56 | 0 |
| NFIX | CACGCCCAG | -1 | 312 | 321 | -688 | -679 | 6,19 | 0,01 | 20,67 | >0.1 | 2,65 | 4,62 | 0 | 1,18 | 0 | 6,03 | 0 |
| NFIX | TCTGCCCAA | -1 | 227 | 236 | -773 | -764 | 6,2 | 0,01 | 20,35 | >0.1 | 2,93 | 4,5 | 0 | 1,47 | 0 | 5,26 | 0 |
| NFIX | TGAGCCACT | -1 | 186 | 195 | -814 | -805 | 4,41 | 0,01 | 17,55 | >0.1 | 3,1 | 4,45 | 0 | 1,74 | 0 | 3,86 | 0 |
| NFIX | TGTGCCCAG | -1 | 178 | 187 | -822 | -813 | 7,67 | 0,01 | 20,21 | >0.1 | 3,14 | 4,44 | 0 | 1,81 | 0 | 3,16 | 0 |
| NFIX | CCAGCCATT | -1 | 173 | 182 | -827 | -818 | 4,66 | 0,01 | 17,45 | >0.1 | 3,17 | 4,43 | 0 | 1,85 | 0 | 3,34 | 0 |
| ATF1 | TTGACCTG | 1 | 107 | 115 | -893 | -885 | -3,52 | 0,01 | 7,03 | >0.1 | 3,57 | 4,35 | 0 | 1,89 | 0 | 0,75 | 0 |
| ATF1 | CTCACGCC | 1 | 191 | 199 | -809 | -801 | 2,89 | 0,001 | 16,25 | >0.1 | 3,08 | 4,45 | 0 | 1,7 | 0 | 4,13 | 0 |
| ATF1 | ATCACAAG | 1 | 235 | 243 | -765 | -757 | -4,18 | 0,01 | 9,56 | >0.1 | 2,9 | 4,51 | 0 | 1,43 | 0 | 4,9 | 0 |
| ATF1 | CTGACTAA | 1 | 263 | 271 | -737 | -729 | -4,72 | 0,01 | 8,47 | >0.1 | 2,8 | 4,55 | 0 | 1,32 | 0 | 4,53 | 0 |
| ATF1 | CTGAGGCA | 1 | 356 | 364 | -644 | -636 | -2,6 | 0,01 | 11,74 | >0.1 | 2,53 | 4,69 | 0 | 1,09 | 0 | 6,03 | 0 |
| ATF1 | ATGGCGTG | 1 | 369 | 377 | -631 | -623 | -0,21 | 0,01 | 14,84 | >0.1 | 2,54 | 4,71 | 0 | 1,07 | 0 | 6,73 | 0 |
| ATF1 | AAGACAAG | 1 | 503 | 511 | -497 | -489 | -4,32 | 0,01 | 10,28 | >0.1 | 2,97 | 4,97 | 0 | 0,98 | 0 | 5,68 | 0 |
| ATF1 | GTCATGTA | 1 | 517 | 525 | -483 | -475 | -4,76 | 0,01 | 7,19 | >0.1 | 3,02 | 5 | 0 | 0,98 | 0 | 2,95 | 0 |
| ATF1 | GAGAGGTA | 1 | 773 | 781 | -227 | -219 | -4,39 | 0,01 | 14,34 | >0.1 | 5,64 | 5,82 | 0 | 1,36 | 0 | 5,92 | 0 |
| ATF1 | ATAACGTA | 1 | 801 | 809 | -199 | -191 | 0,49 | 0,01 | 17,13 | >0.1 | 5,64 | 5,96 | 0 | 1,49 | 0 | 3,55 | 0 |
| ATF1 | GTTACATA | 1 | 848 | 856 | -152 | -144 | 2,91 | 0,001 | 19,5 | >0.1 | 5,64 | 6,27 | 0 | 1,82 | 0 | 2,86 | 0 |
| ATF1 | ATGAGGAG | 1 | 987 | 995 | -13 | -5 | -3,69 | 0,01 | 13,98 | >0.1 | 5,64 | 8,59 | 0 | 1,87 | 0 | 1,57 | 0 |
| ATF1 | GGGACGAT | 1 | 1043 | 1051 | 43 | 51 | -4,09 | 0,01 | 11,79 | >0.1 | 5,64 | 7,76 | 0 | 1,86 | 0 | 0,62 | 0 |
| ATF1 | AAGACATT | 1 | 1124 | 1132 | 124 | 132 | -2,53 | 0,01 | 17,41 | >0.1 | 5,64 | 6,51 | 0 | 1,84 | 3,22 | 2,72 | 0 |
| ATF1 | ACCACGAA | 1 | 1146 | 1154 | 146 | 154 | -0,59 | 0,01 | 17,17 | >0.1 | 5,64 | 6,33 | 0 | 1,84 | 1,79 | 2,16 | 0 |
| ATF1 | TTGACATC | -1 | 1048 | 1056 | 48 | 56 | 0,91 | 0,01 | 17,45 | >0.1 | 5,64 | 7,62 | 0 | 1,86 | 0 | 1,42 | 0 |
| ATF1 | ATCACATC | -1 | 1023 | 1031 | 23 | 31 | -1,44 | 0,01 | 16 | >0.1 | 5,64 | 8,67 | 0 | 1,86 | 0 | 1,28 | 0 |
| ATF1 | ATGACTTT | -1 | 1009 | 1017 | 9 | 17 | -2,39 | 0,01 | 15,56 | >0.1 | 5,64 | 9,03 | 0 | 1,86 | 0 | 1,42 | 0 |
| ATF1 | AATACGTT | -1 | 803 | 811 | -197 | -189 | 2,38 | 0,001 | 18,92 | >0.1 | 5,64 | 5,98 | 0 | 1,5 | 0 | 3,43 | 0 |
| ATF1 | GTGACCAA | -1 | 672 | 680 | -328 | -320 | -2,06 | 0,01 | 12,16 | >0.1 | 4,16 | 5,42 | 0 | 1,1 | 0 | 3,55 | 0 |
| ATF1 | CTCACATT | -1 | 588 | 596 | -412 | -404 | -4,33 | 0,01 | 5,23 | >0.1 | 3,39 | 5,17 | 0 | 1 | 0 | 0 | 0 |
| ATF1 | AATACATG | -1 | 519 | 527 | -481 | -473 | -4,03 | 0,01 | 8,05 | >0.1 | 3,03 | 5 | 0 | 0,98 | 0 | 3,07 | 0 |
| ATF1 | ATGACTTT | -1 | 514 | 522 | -486 | -478 | -2,39 | 0,01 | 9,46 | >0.1 | 3,01 | 4,99 | 0 | 0,98 | 0 | 2,86 | 0 |
| ATF1 | CTGTCGCC | -1 | 433 | 441 | -567 | -559 | -4,22 | 0,01 | 10 | >0.1 | 2,72 | 4,82 | 0 | 1,01 | 0 | 5,67 | 0 |
| ATF1 | GTGGCGCG | -1 | 411 | 419 | -589 | -581 | -1,32 | 0,01 | 12,79 | >0.1 | 2,65 | 4,78 | 0 | 1,02 | 0 | 5,65 | 0 |
| ATF1 | TTCACGCC | -1 | 371 | 379 | -629 | -621 | 1,99 | 0,01 | 16,58 | >0.1 | 2,54 | 4,71 | 0 | 1,07 | 0 | 6,27 | 0 |
| ATF1 | ACCACGCC | -1 | 315 | 323 | -685 | -677 | -0,58 | 0,01 | 13,98 | >0.1 | 2,64 | 4,62 | 0 | 1,17 | 0 | 6,13 | 0 |
| ATF1 | GAGACGGG | -1 | 282 | 290 | -718 | -710 | -4,79 | 0,01 | 8,97 | >0.1 | 2,74 | 4,57 | 0 | 1,25 | 0 | 5,19 | 0 |
| ATF1 | CTGACCTT | -1 | 240 | 248 | -760 | -752 | -3,82 | 0,01 | 9,41 | >0.1 | 2,88 | 4,52 | 0 | 1,41 | 0 | 4,42 | 0 |
| ATF1 | ATGACTAC | -1 | 126 | 134 | -874 | -866 | -4,42 | 0,01 | 6,98 | >0.1 | 3,43 | 4,38 | 0 | 1,89 | 0 | 1,7 | 0 |
| ATF1 | CTGACTTC | -1 | 33 | 41 | -967 | -959 | -3,36 | 0,01 | 7,23 | >0.1 | 4,44 | 4,28 | 0 | 1,87 | 0 | 0 | 0 |
| ATF1 | CTTACCTA | -1 | 2 | 10 | -998 | -990 | -3,93 | 0,01 | 9,62 | >0.1 | 5,64 | 4,25 | 0 | 1,86 | 1,79 | 0 | 0 |
| ATF3 | GATCACAA | 1 | 234 | 242 | -766 | -758 | -3,97 | 0,01 | 9,77 | >0.1 | 2,9 | 4,51 | 0 | 1,43 | 0 | 4,9 | 0 |
| ATF3 | GGTGAAAC | 1 | 274 | 282 | -726 | -718 | -4,81 | 0,01 | 8,88 | >0.1 | 2,76 | 4,56 | 0 | 1,28 | 0 | 5,09 | 0 |
| ATF3 | AATGGCGT | 1 | 368 | 376 | -632 | -624 | -0,83 | 0,01 | 14,21 | >0.1 | 2,54 | 4,71 | 0 | 1,07 | 0 | 6,73 | 0 |
| ATF3 | AATGAAAT | 1 | 606 | 614 | -394 | -386 | -5,22 | 0,01 | 5,94 | >0.1 | 3,51 | 5,22 | 0 | 1,02 | 0 | 1,41 | 0 |
| ATF3 | TATAACAT | 1 | 688 | 696 | -312 | -304 | -5,15 | 0,01 | 8,84 | >0.1 | 4,44 | 5,47 | 0 | 1,13 | 0 | 2,95 | 0 |
| ATF3 | AGTGACAG | 1 | 765 | 773 | -235 | -227 | 1,94 | 0,01 | 20,79 | >0.1 | 5,64 | 5,78 | 0 | 1,33 | 0 | 6,1 | 0 |
| ATF3 | AATAACGT | 1 | 800 | 808 | -200 | -192 | -1,75 | 0,01 | 14,88 | >0.1 | 5,64 | 5,96 | 0 | 1,48 | 0 | 3,55 | 0 |
| ATF3 | TGTTACAT | 1 | 847 | 855 | -153 | -145 | 0,99 | 0,01 | 17,57 | >0.1 | 5,64 | 6,26 | 0 | 1,81 | 0 | 2,86 | 0 |
| ATF3 | CATGAGGA | 1 | 986 | 994 | -14 | -6 | -3,04 | 0,01 | 14,61 | >0.1 | 5,64 | 8,57 | 0 | 1,87 | 0 | 1,57 | 0 |
| ATF3 | GGGGACGA | 1 | 1042 | 1050 | 42 | 50 | -1,91 | 0,01 | 14,77 | >0.1 | 5,64 | 7,79 | 0 | 1,86 | 0 | 1,39 | 0 |
| ATF3 | GATGTCAA | 1 | 1048 | 1056 | 48 | 56 | -4,04 | 0,01 | 12,5 | >0.1 | 5,64 | 7,62 | 0 | 1,86 | 0 | 1,42 | 0 |
| ATF3 | CAAGACAT | 1 | 1123 | 1131 | 123 | 131 | -3,64 | 0,01 | 16,33 | >0.1 | 5,64 | 6,52 | 0 | 1,85 | 3,22 | 2,74 | 0 |
| ATF3 | AATGTCTT | -1 | 1124 | 1132 | 124 | 132 | -5,14 | 0,01 | 14,8 | >0.1 | 5,64 | 6,51 | 0 | 1,84 | 3,22 | 2,72 | 0 |
| ATF3 | AGTTACAG | -1 | 1086 | 1094 | 86 | 94 | -2,86 | 0,01 | 15,57 | >0.1 | 5,64 | 6,93 | 0 | 1,85 | 1,79 | 2,21 | 0 |

| | | | | | | | | | | | | | | | | | |
|---|---|---|---|---|---|---|---|---|---|---|---|---|---|---|---|---|---|
| ATF3 | CTTGACAT | -1 | 1049 | 1057 | 49 | 57 | -3,37 | 0,01 | 13,14 | >0.1 | 5,64 | 7,6 | 0 | 1,86 | 0 | 1,42 | 0 |
| ATF3 | GATCACAT | -1 | 1024 | 1032 | 24 | 32 | -1,17 | 0,01 | 16,2 | >0.1 | 5,64 | 8,59 | 0 | 1,86 | 0 | 1,28 | 0 |
| ATF3 | AATGACTT | -1 | 1010 | 1018 | 10 | 18 | 7,15 | 0,001 | 25,11 | >0.1 | 5,64 | 8,99 | 0 | 1,86 | 0 | 1,46 | 0 |
| ATF3 | CATGAGAT | -1 | 982 | 990 | -18 | -10 | -3,64 | 0,01 | 13,72 | >0.1 | 5,64 | 8,32 | 0 | 1,87 | 0 | 1,53 | 0 |
| ATF3 | GAGGACAG | -1 | 857 | 865 | -143 | -135 | -4,59 | 0,01 | 12,23 | >0.1 | 5,64 | 6,33 | 0 | 1,9 | 0 | 2,95 | 0 |
| ATF3 | GAATACGT | -1 | 804 | 812 | -196 | -188 | -2,64 | 0,01 | 13,91 | >0.1 | 5,64 | 5,98 | 0 | 1,51 | 0 | 3,43 | 0 |
| ATF3 | CATGAAGA | -1 | 644 | 652 | -356 | -348 | -3,54 | 0,01 | 10,37 | >0.1 | 3,82 | 5,33 | 0 | 1,06 | 0 | 3,7 | 0 |
| ATF3 | CATGACTT | -1 | 515 | 523 | -485 | -477 | 8,23 | 0,001 | 20,17 | >0.1 | 3,02 | 4,99 | 0 | 0,98 | 0 | 2,95 | 0 |
| ATF3 | AGTGGCGC | -1 | 412 | 420 | -588 | -580 | -3,91 | 0,01 | 10,32 | >0.1 | 2,66 | 4,79 | 0 | 1,02 | 0 | 5,76 | 0 |
| ATF3 | CAGGACGG | -1 | 258 | 266 | -742 | -734 | -3,6 | 0,01 | 9,72 | >0.1 | 2,82 | 4,54 | 0 | 1,33 | 0 | 4,63 | 0 |
| ATF3 | GATTACAG | -1 | 198 | 206 | -802 | -794 | 2,4 | 0,01 | 15,93 | >0.1 | 3,05 | 4,46 | 0 | 1,64 | 0 | 4,37 | 0 |
| ATF3 | TATGACTA | -1 | 127 | 135 | -873 | -865 | 4,34 | 0,001 | 15,68 | >0.1 | 3,43 | 4,38 | 0 | 1,89 | 0 | 1,65 | 0 |
| ATF3 | GCTGACTT | -1 | 34 | 42 | -966 | -958 | -0,88 | 0,01 | 9,69 | >0.1 | 4,42 | 4,28 | 0 | 1,87 | 0 | 0 | 0 |
