## Supplementary table 4 for "Bioinformatic characterization of angiotensin-converting enzyme 2, the entry receptor for SARS-CoV-2"

| tissue | correlated_gene | correlation | pval |
|---|---|---|---|
| kidney - medulla | NKX6-2 | 1 | 0 |
| kidney - medulla | NFE2 | 1 | 0 |
| kidney - medulla | MIXL1 | 1 | 0 |
| kidney - medulla | HES5 | 1 | 0 |
| kidney - medulla | MYBL2 | 1 | 0 |
| kidney - medulla | E2F2 | 1 | 0 |
| kidney - medulla | SPIB | 1 | 0 |
| kidney - medulla | MYOG | 0,9487 | 0,05132 |
| small intestine - terminal ileum | HNF4G | 0,9451 | 7,983E-92 |
| small intestine - terminal ileum | GATA6 | 0,9032 | 7,405E-70 |
| fallopian tube | JUND | 0,9 | 0,0009431 |
| small intestine - terminal ileum | HNF1A | 0,9 | 1,238E-68 |
| small intestine - terminal ileum | HNF4A | 0,8997 | 1,634E-68 |
| small intestine - terminal ileum | ISX | 0,8989 | 3,156E-68 |
| small intestine - terminal ileum | NR1H4 | 0,8968 | 1,928E-67 |
| cervix - endocervix | MZF1 | 0,8909 | 0,0005421 |
| small intestine - terminal ileum | GATA5 | 0,8858 | 1,329E-63 |
| cervix - ectocervix | SREBF2 | 0,8833 | 0,001591 |
| cervix - ectocervix | PAX9 | 0,8833 | 0,001591 |
| fallopian tube | TWIST2 | 0,8833 | 0,001591 |
| small intestine - terminal ileum | ZBTB7B | 0,8818 | 2,834E-62 |
| small intestine - terminal ileum | PPARA | 0,8793 | 1,664E-61 |
| small intestine - terminal ileum | HNF1B | 0,8792 | 1,827E-61 |
| small intestine - terminal ileum | KLF5 | 0,8787 | 2,566E-61 |
| small intestine - terminal ileum | VDR | 0,8741 | 6,289E-60 |
| colon - transverse | HNF4G | 0,8691 | 1,327E-125 |
| small intestine - terminal ileum | ESRRA | 0,8674 | 5,806E-58 |
| cervix - endocervix | EMX2 | 0,8667 | 0,001174 |
| colon - transverse | VDR | 0,8624 | 1,574E-121 |
| small intestine - terminal ileum | NR5A2 | 0,8607 | 3,826E-56 |
| colon - transverse | HNF4A | 0,8588 | 1,961E-119 |
| cervix - ectocervix | ETV4 | 0,85 | 0,003705 |
| small intestine - terminal ileum | CDX2 | 0,8499 | 2,28E-53 |
| small intestine - terminal ileum | ATOH1 | 0,8492 | 3,491E-53 |
| cervix - endocervix | SCRT1 | 0,8424 | 0,00222 |
| cervix - endocervix | STAT6 | 0,8424 | 0,00222 |
| cervix - ectocervix | POU4F1 | 0,8416 | 0,004442 |
| colon - transverse | ZBTB7B | 0,8408 | 9,814E-110 |
| colon - transverse | GATA6 | 0,84 | 2,599E-109 |
| small intestine - terminal ileum | CEBPA | 0,838 | 1,536E-50 |
| colon - transverse | PPARG | 0,8374 | 4,648E-108 |
| testis | HLTF | 0,837 | 5,21E-96 |
| small intestine - terminal ileum | MLXIPL | 0,8367 | 3,007E-50 |
| bladder | NR1H4 | 0,8366 | 0,000002286 |
| cervix - ectocervix | RORC | 0,8333 | 0,005266 |
| cervix - ectocervix | HIF1A | 0,8333 | 0,005266 |
| cervix - ectocervix | MYC | 0,8333 | 0,005266 |
| fallopian tube | HIC1 | 0,8333 | 0,005266 |
| fallopian tube | REL | 0,8333 | 0,005266 |
| small intestine - terminal ileum | CDX1 | 0,8328 | 2,114E-49 |
| small intestine - terminal ileum | SPDEF | 0,8298 | 9,745E-49 |
| small intestine - terminal ileum | SOX6 | 0,8297 | 1,007E-48 |
| colon - transverse | NR1H4 | 0,8285 | 8,844E-104 |

| | | | |
|---|---|---|---|
| colon - transverse | ISX | 0,8282 | 1,246E-103 |
| small intestine - terminal ileum | TLX1 | 0,8282 | 2,073E-48 |
| testis | SMAD4 | 0,8252 | 4,429E-91 |
| colon - transverse | ESRRA | 0,8234 | 1,81E-101 |
| colon - transverse | NR5A2 | 0,8228 | 3,615E-101 |
| fallopian tube | HMX2 | 0,8218 | 0,006573 |
| bladder | ZBTB7C | 0,8182 | 0,000005853 |
| small intestine - terminal ileum | ELF3 | 0,818 | 2,656E-46 |
| colon - transverse | ZBTB7C | 0,8174 | 8,105E-99 |
| colon - transverse | KLF5 | 0,8169 | 1,402E-98 |
| cervix - ectocervix | FOS | 0,8167 | 0,007225 |
| cervix - ectocervix | CEBPG | 0,8167 | 0,007225 |
| cervix - ectocervix | E2F8 | 0,8167 | 0,007225 |
| testis | SOX9 | 0,813 | 2,454E-86 |
| colon - transverse | HNF1A | 0,8101 | 1,025E-95 |
| bladder | LIN54 | 0,8078 | 0,000009503 |
| cervix - endocervix | USF2 | 0,8061 | 0,004862 |
| cervix - endocervix | TGIF1 | 0,8061 | 0,004862 |
| bladder | SP1 | 0,8026 | 0,00001198 |
| colon - transverse | CDX1 | 0,8007 | 6,444E-92 |
| cervix - ectocervix | GRHL1 | 0,8 | 0,009628 |
| cervix - ectocervix | IRF5 | 0,8 | 0,009628 |
| kidney - medulla | HOXA13 | 0,8 | 0,2 |
| kidney - medulla | EVX1 | 0,8 | 0,2 |
| kidney - medulla | GCM1 | 0,8 | 0,2 |
| kidney - medulla | MNX1 | 0,8 | 0,2 |
| kidney - medulla | NR4A2 | 0,8 | 0,2 |
| kidney - medulla | NR4A1 | 0,8 | 0,2 |
| kidney - medulla | ETV4 | 0,8 | 0,2 |
| kidney - medulla | HMX2 | 0,8 | 0,2 |
| kidney - medulla | GFI1 | 0,8 | 0,2 |
| kidney - medulla | BATF | 0,8 | 0,2 |
| kidney - medulla | SOX11 | 0,8 | 0,2 |
| kidney - medulla | SOX9 | 0,8 | 0,2 |
| kidney - medulla | ELF3 | 0,8 | 0,2 |
| kidney - medulla | SIX1 | 0,8 | 0,2 |
| kidney - medulla | SIX3 | 0,8 | 0,2 |
| kidney - medulla | TFEC | 0,8 | 0,2 |
| kidney - medulla | MYB | 0,8 | 0,2 |
| kidney - medulla | GATA5 | 0,8 | 0,2 |
| kidney - medulla | ARX | 0,8 | 0,2 |
| colon - transverse | EHF | 0,7999 | 1,267E-91 |
| colon - transverse | CDX2 | 0,794 | 2,346E-89 |
| cervix - endocervix | ZNF282 | 0,7939 | 0,0061 |
| small intestine - terminal ileum | GRHL2 | 0,7931 | 1,093E-41 |
| bladder | NFAT5 | 0,7922 | 0,00001867 |
| colon - transverse | CEBPA | 0,792 | 1,359E-88 |
| testis | HIF1A | 0,7917 | 7,559E-79 |
| bladder | SP3 | 0,7883 | 0,00002191 |
| colon - transverse | HNF1B | 0,7879 | 4,311E-87 |
| cervix - endocervix | HIC1 | 0,7818 | 0,007547 |
| colon - transverse | ATOH1 | 0,7803 | 2,314E-84 |
| cervix - ectocervix | ISL2 | 0,7798 | 0,01321 |
| colon - transverse | GRHL2 | 0,7792 | 5,278E-84 |

| | | | |
|---|---|---|---|
| small intestine - terminal ileum | HES2 | 0,7788 | 2,549E-39 |
| testis | ZBTB33 | 0,7787 | 1,087E-74 |
| colon - transverse | EMX1 | 0,7767 | 3,88E-83 |
| kidney - medulla | GBX1 | 0,7746 | 0,2254 |
| kidney - medulla | DMRT3 | 0,7746 | 0,2254 |
| kidney - medulla | ISL2 | 0,7746 | 0,2254 |
| kidney - medulla | NEUROG1 | 0,7746 | 0,2254 |
| kidney - medulla | FIGLA | 0,7746 | 0,2254 |
| kidney - medulla | OTX1 | 0,7746 | 0,2254 |
| kidney - medulla | OTX2 | 0,7746 | 0,2254 |
| kidney - medulla | PROP1 | 0,7746 | 0,2254 |
| kidney - medulla | NKX3-2 | 0,7746 | 0,2254 |
| kidney - medulla | NKX2-3 | 0,7746 | 0,2254 |
| kidney - medulla | SPIC | 0,7746 | 0,2254 |
| colon - transverse | ELF4 | 0,7744 | 2,562E-82 |
| bladder | RREB1 | 0,7727 | 0,00004028 |
| small intestine - terminal ileum | CREB3L1 | 0,7714 | 3,593E-38 |
| testis | FOXO1 | 0,771 | 2,485E-72 |
| cervix - endocervix | TCF4 | 0,7697 | 0,009222 |
| small intestine - terminal ileum | ONECUT3 | 0,7693 | 7,608E-38 |
| bladder | TBP | 0,7688 | 0,00004656 |
| testis | FOXO3 | 0,7681 | 1,695E-71 |
| small intestine - terminal ileum | FOXA1 | 0,7678 | 1,3E-37 |
| colon - transverse | FOXA1 | 0,7677 | 4,31E-80 |
| cervix - endocervix | OTX1 | 0,7669 | 0,009638 |
| cervix - ectocervix | FOXA1 | 0,7667 | 0,01594 |
| cervix - ectocervix | PPARA | 0,7667 | 0,01594 |
| cervix - ectocervix | NFE2L2 | 0,7667 | 0,01594 |
| colon - transverse | HES2 | 0,7667 | 9,059E-80 |
| fallopian tube | NFYA | 0,7667 | 0,01594 |
| colon - transverse | FOXH1 | 0,7635 | 9,913E-79 |
| heart - left ventricle | MITF | 0,7628 | 1,934E-83 |
| colon - transverse | SOX9 | 0,7616 | 3,905E-78 |
| colon - transverse | MLX | 0,7611 | 5,95E-78 |
| small intestine - terminal ileum | MNX1 | 0,7608 | 1,417E-36 |
| colon - transverse | KLF4 | 0,7601 | 1,234E-77 |
| testis | ELF1 | 0,7596 | 4,694E-69 |
| cervix - endocervix | E2F8 | 0,7576 | 0,01114 |
| cervix - endocervix | IRF3 | 0,7576 | 0,01114 |
| small intestine - terminal ileum | MLX | 0,7558 | 7,399E-36 |
| testis | AR | 0,7551 | 8,44E-68 |
| colon - transverse | NR2F6 | 0,7548 | 5,347E-76 |
| colon - transverse | ELF3 | 0,7527 | 2,471E-75 |
| small intestine - terminal ileum | POU5F1 | 0,7526 | 2,097E-35 |
| testis | MEF2A | 0,7524 | 4,671E-67 |
| small intestine - terminal ileum | FOXH1 | 0,7517 | 2,75E-35 |
| heart - left ventricle | MEF2A | 0,7511 | 1,499E-79 |
| testis | GATA4 | 0,751 | 1,075E-66 |
| colon - transverse | SPIB | 0,7506 | 1,029E-74 |
| testis | FOXO4 | 0,7504 | 1,547E-66 |
| cervix - ectocervix | ZBTB7B | 0,75 | 0,01994 |
| cervix - ectocervix | ESRRB | 0,75 | 0,01994 |
| testis | ELK1 | 0,75 | 1,973E-66 |
| testis | NFE2L1 | 0,7499 | 2,096E-66 |

| | | | |
|---|---|---|---|
| heart - left ventricle | RARB | 0,7498 | 3,72E-79 |
| kidney - cortex | HNF1A | 0,7462 | 2,5E-16 |
| cervix - endocervix | RHOXF1 | 0,7455 | 0,01333 |
| colon - transverse | FOXD2 | 0,745 | 4,778E-73 |
| small intestine - terminal ileum | EHF | 0,7445 | 2,667E-34 |
| bladder | SP2 | 0,7442 | 0,0001098 |
| small intestine - terminal ileum | EMX1 | 0,7441 | 3,077E-34 |
| testis | MXI1 | 0,7435 | 1,066E-64 |
| colon - transverse | EVX1 | 0,7427 | 2,293E-72 |
| colon - transverse | E2F2 | 0,7417 | 4,481E-72 |
| bladder | REL | 0,7416 | 0,0001195 |
| testis | NKX3-1 | 0,7413 | 3,975E-64 |
| testis | ARNT | 0,7412 | 4,007E-64 |
| testis | ID4 | 0,7403 | 7,067E-64 |
| minor salivary gland | SP1 | 0,7392 | 2,912E-29 |
| testis | ZFX | 0,7392 | 1,347E-63 |
| testis | SP3 | 0,7365 | 6,609E-63 |
| testis | FOXJ2 | 0,7365 | 6,289E-63 |
| colon - transverse | RORC | 0,7348 | 4,296E-70 |
| heart - left ventricle | PPARA | 0,7348 | 1,686E-74 |
| small intestine - terminal ileum | NR3C2 | 0,7348 | 5,152E-33 |
| testis | TEAD1 | 0,7345 | 2,038E-62 |
| small intestine - terminal ileum | PAX6 | 0,7337 | 7,176E-33 |
| cervix - ectocervix | DLX3 | 0,7333 | 0,02455 |
| cervix - ectocervix | STAT1 | 0,7333 | 0,02455 |
| cervix - endocervix | ZFX | 0,7333 | 0,0158 |
| cervix - endocervix | E2F2 | 0,7333 | 0,0158 |
| fallopian tube | EN2 | 0,7333 | 0,02455 |
| bladder | FOXC1 | 0,7325 | 0,0001596 |
| bladder | ELF1 | 0,7325 | 0,0001596 |
| heart - left ventricle | KLF13 | 0,7318 | 1,263E-73 |
| small intestine - terminal ileum | ARX | 0,7309 | 1,627E-32 |
| colon - transverse | SPDEF | 0,7305 | 6,515E-69 |
| cervix - ectocervix | SP8 | 0,7303 | 0,02546 |
| cervix - endocervix | OLIG1 | 0,7301 | 0,01651 |
| colon - transverse | NR3C2 | 0,7298 | 1,024E-68 |
| small intestine - terminal ileum | RORC | 0,7288 | 2,956E-32 |
| small intestine - terminal ileum | FOXD1 | 0,7276 | 4,261E-32 |
| small intestine - terminal ileum | NEUROD1 | 0,7273 | 4,56E-32 |
| colon - transverse | SREBF2 | 0,7261 | 1,063E-67 |
| small intestine - terminal ileum | NR2F6 | 0,7257 | 7,224E-32 |
| colon - transverse | CREB3L1 | 0,7255 | 1,516E-67 |
| kidney - cortex | GLIS1 | 0,7242 | 4,756E-15 |
| colon - transverse | MNX1 | 0,7235 | 5,172E-67 |
| bladder | SOX13 | 0,7223 | 0,0002176 |
| cervix - endocervix | TFAP2C | 0,7212 | 0,01857 |
| cervix - endocervix | BACH1 | 0,7212 | 0,01857 |
| cervix - endocervix | ASCL2 | 0,7212 | 0,01857 |
| cervix - endocervix | POU2F3 | 0,7212 | 0,01857 |
| heart - left ventricle | MLX | 0,7209 | 1,759E-70 |
| small intestine - terminal ileum | ESRRG | 0,7209 | 2,824E-31 |
| bladder | HOXA5 | 0,7208 | 0,0002277 |
| bladder | SMAD3 | 0,7208 | 0,0002277 |
| heart - left ventricle | PROX1 | 0,7197 | 3,955E-70 |

| | | | |
|---|---|---|---|
| heart - left ventricle | HSF2 | 0,7132 | 2,447E-68 |
| heart - left ventricle | CUX1 | 0,7122 | 4,441E-68 |
| testis | TCF7L2 | 0,7096 | 1,548E-56 |
| colon - transverse | FOXA2 | 0,7092 | 2,688E-63 |
| cervix - endocervix | ETV2 | 0,7091 | 0,02167 |
| cervix - endocervix | GLI2 | 0,7091 | 0,02167 |
| minor salivary gland | BCL6 | 0,7085 | 5,363E-26 |
| bladder | RFX1 | 0,708 | 0,0003293 |
| bladder | SP4 | 0,7078 | 0,0003314 |
| small intestine - terminal ileum | FEV | 0,7073 | 1,123E-29 |
| colon - transverse | RREB1 | 0,7066 | 1,159E-62 |
| testis | NFIC | 0,7064 | 7,853E-56 |
| colon - transverse | POU5F1 | 0,7055 | 2,226E-62 |
| testis | TP53 | 0,7054 | 1,282E-55 |
| small intestine - terminal ileum | NPAS2 | 0,7047 | 2,228E-29 |
| bladder | ZNF384 | 0,7039 | 0,0003695 |
| small intestine - terminal ileum | MAF | 0,7039 | 2,741E-29 |
| heart - left ventricle | TEAD1 | 0,7029 | 1,339E-65 |
| testis | E2F6 | 0,7012 | 1,056E-54 |
| heart - left ventricle | NFYB | 0,701 | 4,06E-65 |
| colon - transverse | MYB | 0,7007 | 3,236E-61 |
| cervix - ectocervix | MTF1 | 0,7 | 0,03577 |
| fallopian tube | XBP1 | 0,7 | 0,03577 |
| fallopian tube | SMAD3 | 0,7 | 0,03577 |
| heart - left ventricle | NFE2L1 | 0,6995 | 9,935E-65 |
| heart - left ventricle | PBX1 | 0,6979 | 2,602E-64 |
| small intestine - terminal ileum | CEBPG | 0,6976 | 1,35E-28 |
| cervix - endocervix | E2F7 | 0,697 | 0,0251 |
| testis | NFE2L2 | 0,6964 | 1,113E-53 |
| heart - left ventricle | E2F6 | 0,696 | 7,597E-64 |
| testis | REL | 0,6941 | 3,366E-53 |
| heart - left ventricle | CLOCK | 0,6904 | 1,877E-62 |
| heart - left ventricle | SMAD4 | 0,6883 | 6,302E-62 |
| testis | NFKB1 | 0,688 | 6,273E-52 |
| kidney - cortex | TFEC | 0,6875 | 3,684E-13 |
| kidney - cortex | MLXIPL | 0,6867 | 4,026E-13 |
| heart - left ventricle | GATA6 | 0,6863 | 1,92E-61 |
| minor salivary gland | POU2F3 | 0,6856 | 7,952E-24 |
| colon - transverse | PPARA | 0,6853 | 1,35E-57 |
| cervix - endocervix | FOXO4 | 0,6848 | 0,02888 |
| cervix - endocervix | LBX2 | 0,6848 | 0,02888 |
| cervix - endocervix | HINFP | 0,6848 | 0,02888 |
| cervix - endocervix | MITF | 0,6848 | 0,02888 |
| cervix - endocervix | RUNX2 | 0,6848 | 0,02888 |
| cervix - endocervix | ESR1 | 0,6848 | 0,02888 |
| heart - left ventricle | FOXJ2 | 0,6845 | 5,247E-61 |
| testis | NFIA | 0,6842 | 3,787E-51 |
| cervix - ectocervix | PRDM1 | 0,6833 | 0,04244 |
| cervix - ectocervix | MLX | 0,6833 | 0,04244 |
| cervix - endocervix | LHX3 | 0,6833 | 0,02938 |
| heart - left ventricle | ETV1 | 0,6833 | 1,027E-60 |
| testis | RORA | 0,683 | 6,331E-51 |
| testis | PBX2 | 0,6828 | 7,094E-51 |
| heart - left ventricle | FOXO3 | 0,6825 | 1,577E-60 |

| | | | |
|---|---|---|---|
| colon - transverse | CEBPG | 0,6812 | 1,148E-56 |
| bladder | HOXA9 | 0,6792 | 0,000709 |
| heart - left ventricle | ESRRG | 0,678 | 1,915E-59 |
| testis | NFAT5 | 0,678 | 6,256E-50 |
| cervix - ectocervix | BHLHA15 | 0,6778 | 0,04481 |
| testis | TFCP2 | 0,6777 | 7,189E-50 |
| bladder | IRF2 | 0,674 | 0,0008071 |
| kidney - cortex | HNF4A | 0,674 | 1,556E-12 |
| colon - transverse | TLX1 | 0,6739 | 4,637E-55 |
| colon - transverse | PAX6 | 0,673 | 7,166E-55 |
| cervix - endocervix | EGR2 | 0,6727 | 0,03304 |
| cervix - endocervix | SMAD3 | 0,6727 | 0,03304 |
| cervix - endocervix | TP73 | 0,6727 | 0,03304 |
| bladder | REST | 0,6714 | 0,0008603 |
| colon - transverse | GFI1B | 0,6713 | 1,627E-54 |
| heart - left ventricle | NFATC3 | 0,6706 | 9,746E-58 |
| bladder | SPDEF | 0,6701 | 0,000888 |
| colon - transverse | FEV | 0,669 | 5,28E-54 |
| minor salivary gland | STAT6 | 0,669 | 2,248E-22 |
| minor salivary gland | TFAP2C | 0,6677 | 2,888E-22 |
| cervix - ectocervix | AHR | 0,6667 | 0,04987 |
| cervix - ectocervix | JUNB | 0,6667 | 0,04987 |
| cervix - ectocervix | SOX2 | 0,6667 | 0,04987 |
| cervix - ectocervix | ELF3 | 0,6667 | 0,04987 |
| cervix - ectocervix | E2F7 | 0,6667 | 0,04987 |
| fallopian tube | CREB3L2 | 0,6667 | 0,04987 |
| fallopian tube | MNX1 | 0,6667 | 0,04987 |
| fallopian tube | YY2 | 0,6667 | 0,04987 |
| fallopian tube | RUNX2 | 0,6667 | 0,04987 |
| heart - left ventricle | REST | 0,6653 | 1,552E-56 |
| artery - coronary | MLXIPL | 0,6646 | 5,785E-32 |
| heart - left ventricle | FOXO4 | 0,6639 | 3,196E-56 |
| testis | PBX3 | 0,6636 | 3,538E-47 |
| heart - left ventricle | MGA | 0,6625 | 6,338E-56 |
| cervix - endocervix | NEUROG1 | 0,6623 | 0,03693 |
| heart - left ventricle | ARNT | 0,6613 | 1,16E-55 |
| fallopian tube | GFI1B | 0,6611 | 0,05252 |
| cervix - endocervix | NR1H4 | 0,6606 | 0,03759 |
| testis | STAT3 | 0,6605 | 1,31E-46 |
| fallopian tube | HMX3 | 0,6573 | 0,0544 |
| heart - left ventricle | NFIC | 0,6573 | 8,655E-55 |
| bladder | CEBPA | 0,6571 | 0,001209 |
| bladder | STAT2 | 0,6571 | 0,001209 |
| testis | IRF2 | 0,6566 | 6,8E-46 |
| testis | MLX | 0,6557 | 9,758E-46 |
| heart - left ventricle | MEF2D | 0,6548 | 3,05E-54 |
| colon - transverse | NEUROD1 | 0,6515 | 2,06E-50 |
| testis | YY1 | 0,6512 | 6,279E-45 |
| heart - left ventricle | KLF9 | 0,6508 | 2,13E-53 |
| bladder | POU5F1B | 0,6506 | 0,001404 |
| heart - left ventricle | PBX3 | 0,6504 | 2,634E-53 |
| colon - transverse | PDX1 | 0,6502 | 3,725E-50 |
| cervix - ectocervix | SP1 | 0,65 | 0,05807 |
| colon - transverse | MECOM | 0,65 | 4,155E-50 |

| | | | |
|---|---|---|---|
| fallopian tube | MAFF | 0,65 | 0,05807 |
| heart - left ventricle | LIN54 | 0,6498 | 3,512E-53 |
| testis | MSX2 | 0,6495 | 1,251E-44 |
| cervix - endocervix | NRF1 | 0,6485 | 0,04254 |
| small intestine - terminal ileum | HOXC11 | 0,6484 | 1,103E-23 |
| testis | MEF2D | 0,6482 | 2,104E-44 |
| heart - left ventricle | SP2 | 0,6481 | 7,827E-53 |
| testis | MYC | 0,648 | 2,311E-44 |
| heart - left ventricle | FOXJ3 | 0,6475 | 1,066E-52 |
| colon - transverse | PAX4 | 0,6469 | 1,657E-49 |
| testis | MAF | 0,6462 | 4,718E-44 |
| testis | AHR | 0,6459 | 5,357E-44 |
| testis | TFE3 | 0,6456 | 5,982E-44 |
| bladder | TCF12 | 0,6442 | 0,001624 |
| testis | ATF7 | 0,6439 | 1,172E-43 |
| minor salivary gland | RUNX1 | 0,6431 | 2,799E-20 |
| heart - left ventricle | MXI1 | 0,6421 | 1,379E-51 |
| bladder | ZSCAN4 | 0,6416 | 0,00172 |
| bladder | STAT1 | 0,6416 | 0,00172 |
| heart - left ventricle | TFDP1 | 0,6414 | 1,933E-51 |
| muscle - skeletal | FOXO1 | 0,6414 | 2,792E-94 |
| heart - left ventricle | YY1 | 0,6406 | 2,814E-51 |
| heart - left ventricle | PPARG | 0,638 | 9,427E-51 |
| bladder | FOXH1 | 0,6366 | 0,001918 |
| bladder | HOXB3 | 0,6364 | 0,001926 |
| cervix - endocervix | NR1H3 | 0,6364 | 0,04791 |
| bladder | ZNF282 | 0,6351 | 0,001981 |
| bladder | TP53 | 0,6351 | 0,001981 |
| minor salivary gland | GLIS3 | 0,6351 | 1,127E-19 |
| heart - left ventricle | HIF1A | 0,6349 | 3,966E-50 |
| minor salivary gland | TP53 | 0,6341 | 1,327E-19 |
| heart - left ventricle | CTCF | 0,6336 | 7,069E-50 |
| heart - left ventricle | ELK1 | 0,6336 | 7,237E-50 |
| cervix - ectocervix | NPAS2 | 0,6333 | 0,06709 |
| cervix - ectocervix | GRHL2 | 0,6333 | 0,06709 |
| cervix - ectocervix | BHLHE40 | 0,6333 | 0,06709 |
| cervix - ectocervix | REL | 0,6333 | 0,06709 |
| cervix - ectocervix | E2F2 | 0,6333 | 0,06709 |
| fallopian tube | SREBF1 | 0,6333 | 0,06709 |
| fallopian tube | TFE3 | 0,6333 | 0,06709 |
| minor salivary gland | SOX4 | 0,6329 | 1,647E-19 |
| colon - transverse | ARX | 0,6323 | 1,019E-46 |
| heart - left ventricle | ESR2 | 0,6323 | 1,323E-49 |
| small intestine - terminal ileum | SOX9 | 0,6318 | 3,165E-22 |
| testis | ELK3 | 0,6313 | 1,537E-41 |
| bladder | MSX2 | 0,6312 | 0,002153 |
| heart - left ventricle | RREB1 | 0,6309 | 2,43E-49 |
| heart - left ventricle | RXRA | 0,6304 | 3,108E-49 |
| heart - left ventricle | ZEB1 | 0,6302 | 3,402E-49 |
| bladder | POU4F1 | 0,63 | 0,002205 |
| small intestine - terminal ileum | FOXA2 | 0,6298 | 4,653E-22 |
| heart - left ventricle | ZBTB7B | 0,6286 | 6,98E-49 |
| muscle - skeletal | NFIC | 0,6275 | 4,07E-89 |
| heart - left ventricle | TFEC | 0,6274 | 1,163E-48 |

| | | | |
|---|---|---|---|
| heart - left ventricle | ZFX | 0,6273 | 1,263E-48 |
| testis | GATA6 | 0,6269 | 8,098E-41 |
| kidney - cortex | HNF4G | 0,6268 | 1,387E-10 |
| minor salivary gland | STAT3 | 0,6259 | 5,353E-19 |
| heart - left ventricle | ELF1 | 0,6247 | 3,993E-48 |
| testis | GABPA | 0,6244 | 2,068E-40 |
| cervix - endocervix | NR2E3 | 0,6242 | 0,05372 |
| testis | STAT1 | 0,6242 | 2,16E-40 |
| testis | SREBF2 | 0,6237 | 2,684E-40 |
| minor salivary gland | GRHL2 | 0,6226 | 9,087E-19 |
| artery - aorta | PPARG | 0,6222 | 1,209E-47 |
| heart - left ventricle | ESRRB | 0,6222 | 1,176E-47 |
| bladder | FOSL2 | 0,6208 | 0,002674 |
| heart - left ventricle | NFIA | 0,6208 | 2,155E-47 |
| minor salivary gland | REST | 0,6197 | 1,477E-18 |
| minor salivary gland | BATF | 0,6197 | 1,482E-18 |
| bladder | SNAI2 | 0,6195 | 0,002746 |
| heart - left ventricle | NFIX | 0,6193 | 4,12E-47 |
| colon - transverse | ONECUT2 | 0,6192 | 2,447E-44 |
| heart - left ventricle | TBX20 | 0,6189 | 4,954E-47 |
| heart - left ventricle | SREBF2 | 0,6184 | 6,119E-47 |
| colon - transverse | E2F8 | 0,6183 | 3,537E-44 |
| bladder | NPAS2 | 0,6182 | 0,002819 |
| heart - left ventricle | TFCP2 | 0,6177 | 8,332E-47 |
| heart - left ventricle | NPAS2 | 0,6175 | 9,187E-47 |
| heart - left ventricle | HLTF | 0,6174 | 9,684E-47 |
| small intestine - terminal ileum | ZBTB7C | 0,617 | 5,301E-21 |
| cervix - ectocervix | ZBTB33 | 0,6167 | 0,07693 |
| cervix - ectocervix | TFAP2C | 0,6167 | 0,07693 |
| cervix - ectocervix | MYBL1 | 0,6167 | 0,07693 |
| cervix - ectocervix | MYBL2 | 0,6167 | 0,07693 |
| cervix - ectocervix | HOXB13 | 0,6167 | 0,07693 |
| heart - left ventricle | SP4 | 0,6166 | 1,339E-46 |
| heart - left ventricle | ZBTB18 | 0,6159 | 1,821E-46 |
| testis | YY2 | 0,615 | 6,25E-39 |
| minor salivary gland | NFKB1 | 0,6144 | 3,462E-18 |
| heart - left ventricle | NR2C2 | 0,6143 | 3,523E-46 |
| heart - left ventricle | NR3C2 | 0,6142 | 3,753E-46 |
| testis | ARNTL | 0,614 | 8,835E-39 |
| heart - left ventricle | MTF1 | 0,6134 | 5,297E-46 |
| bladder | MGA | 0,613 | 0,003131 |
| bladder | ELK4 | 0,6125 | 0,003159 |
| cervix - endocervix | TCF3 | 0,6121 | 0,05997 |
| cervix - endocervix | CEBPA | 0,6121 | 0,05997 |
| testis | REST | 0,6117 | 1,971E-38 |
| testis | SOX4 | 0,6116 | 2,032E-38 |
| heart - left ventricle | KLF12 | 0,6111 | 1,367E-45 |
| heart - left ventricle | ZNF24 | 0,61 | 2,178E-45 |
| bladder | HNF4A | 0,6099 | 0,00333 |
| testis | FOXC1 | 0,6094 | 4,435E-38 |
| bladder | POU3F1 | 0,6091 | 0,003382 |
| heart - left ventricle | STAT1 | 0,6091 | 3,174E-45 |
| breast - mammary tissue | PPARG | 0,6082 | 9,083E-48 |
| bladder | EVX1 | 0,6078 | 0,00347 |

| | | | |
|---|---|---|---|
| bladder | STAT6 | 0,6078 | 0,00347 |
| heart - left ventricle | HAND1 | 0,6077 | 5,756E-45 |
| minor salivary gland | TEAD3 | 0,6069 | 1,12E-17 |
| testis | KLF13 | 0,6062 | 1,373E-37 |
| heart - left ventricle | ZNF354C | 0,606 | 1,158E-44 |
| testis | FOXJ3 | 0,606 | 1,436E-37 |
| brain - hypothalamus | NFATC1 | 0,6058 | 1,263E-21 |
| heart - left ventricle | ZNF143 | 0,6056 | 1,399E-44 |
| heart - left ventricle | NFAT5 | 0,6045 | 2,133E-44 |
| artery - coronary | PPARG | 0,6041 | 2,89E-25 |
| minor salivary gland | TGIF1 | 0,604 | 1,749E-17 |
| minor salivary gland | GMEB1 | 0,6036 | 1,878E-17 |
| minor salivary gland | IRF5 | 0,6036 | 1,859E-17 |
| ALL_samples | NR5A2 | 0,6034 | 0 |
| heart - left ventricle | ESRRA | 0,6019 | 6,247E-44 |
| minor salivary gland | FOXO4 | 0,6019 | 2,442E-17 |
| testis | GATA5 | 0,6018 | 6,049E-37 |
| bladder | KLF5 | 0,6013 | 0,003937 |
| bladder | TP73 | 0,6013 | 0,003937 |
| small intestine - terminal ileum | PPARG | 0,6013 | 9,086E-20 |
| testis | ZBTB7A | 0,6005 | 9,458E-37 |
| cervix - ectocervix | HES1 | 0,6 | 0,08762 |
| cervix - ectocervix | FOSL1 | 0,6 | 0,08762 |
| cervix - endocervix | RORB | 0,6 | 0,06669 |
| cervix - endocervix | FOXP1 | 0,6 | 0,06669 |
| cervix - endocervix | BACH2 | 0,6 | 0,06669 |
| cervix - endocervix | DBP | 0,6 | 0,06669 |
| cervix - endocervix | TCF7L2 | 0,6 | 0,06669 |
| cervix - endocervix | ARID3B | 0,6 | 0,06669 |
| fallopian tube | NFKB2 | 0,6 | 0,08762 |
| fallopian tube | ZNF143 | 0,6 | 0,08762 |
| fallopian tube | ALX3 | 0,6 | 0,08762 |
| fallopian tube | IRF4 | 0,6 | 0,08762 |
| fallopian tube | CREB3L1 | 0,6 | 0,08762 |
| fallopian tube | CEBPD | 0,6 | 0,08762 |
| kidney - medulla | HNF1A | 0,6 | 0,4 |
| kidney - medulla | LMX1B | 0,6 | 0,4 |
| kidney - medulla | LHX6 | 0,6 | 0,4 |
| kidney - medulla | ISX | 0,6 | 0,4 |
| kidney - medulla | MAFB | 0,6 | 0,4 |
| kidney - medulla | CEBPA | 0,6 | 0,4 |
| kidney - medulla | NR1H4 | 0,6 | 0,4 |
| kidney - medulla | HNF4G | 0,6 | 0,4 |
| kidney - medulla | HNF4A | 0,6 | 0,4 |
| kidney - medulla | MAF | 0,6 | 0,4 |
| kidney - medulla | VDR | 0,6 | 0,4 |
| small intestine - terminal ileum | LBX2 | 0,5993 | 1,297E-19 |
| testis | PPARA | 0,5993 | 1,412E-36 |
| bladder | PPARG | 0,5987 | 0,004137 |
| heart - left ventricle | TCFL5 | 0,5969 | 4,807E-43 |
| muscle - skeletal | MAX | 0,5961 | 2,061E-78 |
| heart - left ventricle | SMAD2 | 0,5959 | 7,112E-43 |
| brain - hypothalamus | TBX19 | 0,5956 | 8,899E-21 |
| heart - left ventricle | ZNF384 | 0,5956 | 7,92E-43 |

| | | | |
|---|---|---|---|
| muscle - skeletal | ARNTL | 0,5955 | 3,19E-78 |
| muscle - skeletal | FOSL2 | 0,5954 | 3,432E-78 |
| heart - left ventricle | E2F3 | 0,5949 | 1,049E-42 |
| testis | CTCF | 0,5944 | 7,282E-36 |
| artery - aorta | MLXIPL | 0,5928 | 2,407E-42 |
| testis | ATF1 | 0,5924 | 1,411E-35 |
| heart - left ventricle | EHF | 0,5923 | 2,92E-42 |
| bladder | RFX3 | 0,5922 | 0,004677 |
| bladder | ETV3 | 0,5922 | 0,004677 |
| bladder | FOXJ3 | 0,5922 | 0,004677 |
| bladder | FOXA1 | 0,5922 | 0,004677 |
| testis | NFYB | 0,5917 | 1,802E-35 |
| bladder | TFAP2C | 0,5909 | 0,004791 |
| bladder | RORC | 0,5909 | 0,004791 |
| colon - transverse | IRF8 | 0,5905 | 1,622E-39 |
| small intestine - terminal ileum | SP8 | 0,5902 | 6,117E-19 |
| heart - left ventricle | AR | 0,5884 | 1,38E-41 |
| muscle - skeletal | RXRA | 0,5884 | 5,864E-76 |
| cervix - endocervix | HOXB3 | 0,5879 | 0,07388 |
| cervix - endocervix | TBX19 | 0,5879 | 0,07388 |
| testis | RORC | 0,5871 | 7,996E-35 |
| bladder | MECOM | 0,587 | 0,005149 |
| bladder | ARID3A | 0,5857 | 0,005273 |
| kidney - cortex | MAF | 0,5846 | 4,253E-09 |
| muscle - skeletal | TBP | 0,5845 | 9,339E-75 |
| bladder | SOX6 | 0,5844 | 0,0054 |
| cervix - ectocervix | ZBTB7A | 0,5833 | 0,09919 |
| cervix - ectocervix | KLF5 | 0,5833 | 0,09919 |
| cervix - ectocervix | EHF | 0,5833 | 0,09919 |
| cervix - ectocervix | FOXQ1 | 0,5833 | 0,09919 |
| fallopian tube | CEBPB | 0,5833 | 0,09919 |
| fallopian tube | BACH1 | 0,5833 | 0,09919 |
| fallopian tube | PRDM1 | 0,5833 | 0,09919 |
| fallopian tube | NFE2L2 | 0,5833 | 0,09919 |
| fallopian tube | PROX1 | 0,5833 | 0,09919 |
| bladder | GABPA | 0,5831 | 0,005529 |
| brain - hypothalamus | RFX3 | 0,5825 | 9,549E-20 |
| heart - left ventricle | TBP | 0,5821 | 1,508E-40 |
| heart - left ventricle | NR1H2 | 0,5821 | 1,509E-40 |
| muscle - skeletal | ZNF143 | 0,5808 | 1,323E-73 |
| bladder | LHX4 | 0,5805 | 0,005795 |
| heart - left ventricle | THAP1 | 0,5794 | 4,27E-40 |
| brain - hypothalamus | RREB1 | 0,5783 | 2,002E-19 |
| bladder | NFYA | 0,5779 | 0,006071 |
| minor salivary gland | ELF1 | 0,5774 | 8,734E-16 |
| bladder | HESX1 | 0,5766 | 0,006213 |
| heart - left ventricle | POU2F1 | 0,5766 | 1,225E-39 |
| heart - left ventricle | MEF2C | 0,5764 | 1,309E-39 |
| heart - left ventricle | SP1 | 0,576 | 1,501E-39 |
| testis | ZNF740 | 0,5759 | 2,773E-33 |
| cervix - endocervix | TCF12 | 0,5758 | 0,08155 |
| cervix - endocervix | TEF | 0,5758 | 0,08155 |
| cervix - endocervix | HOXA11 | 0,5758 | 0,08155 |
| bladder | DLX6 | 0,5753 | 0,006358 |

| | | | |
|---|---|---|---|
| bladder | FOS | 0,5753 | 0,006358 |
| testis | RELA | 0,575 | 3,723E-33 |
| heart - left ventricle | SRF | 0,5744 | 2,698E-39 |
| vagina | MAFB | 0,574 | 4,742E-15 |
| heart - left ventricle | PLAG1 | 0,5738 | 3,374E-39 |
| heart - left ventricle | ZBTB33 | 0,5724 | 5,826E-39 |
| heart - left ventricle | ZNF740 | 0,5724 | 5,778E-39 |
| brain - hippocampus | GLIS3 | 0,5717 | 1,73E-18 |
| small intestine - terminal ileum | TCF7L2 | 0,5714 | 1,339E-17 |
| muscle - skeletal | FOXK2 | 0,5711 | 1,043E-70 |
| minor salivary gland | ZNF282 | 0,5708 | 2,179E-15 |
| heart - left ventricle | CENPB | 0,5707 | 1,079E-38 |
| small intestine - terminal ileum | PDX1 | 0,5693 | 1,865E-17 |
| muscle - skeletal | SREBF2 | 0,5689 | 4,748E-70 |
| minor salivary gland | MXI1 | 0,5686 | 2,937E-15 |
| minor salivary gland | IRF2 | 0,5683 | 3,097E-15 |
| muscle - skeletal | BACH1 | 0,5678 | 1,018E-69 |
| colon - transverse | HOXB13 | 0,5676 | 5,305E-36 |
| bladder | GRHL1 | 0,5675 | 0,007288 |
| heart - left ventricle | CREB3L1 | 0,5675 | 3,473E-38 |
| heart - left ventricle | CREB1 | 0,567 | 4,099E-38 |
| testis | USF1 | 0,567 | 4,244E-32 |
| cervix - ectocervix | TFAP2A | 0,5667 | 0,1116 |
| cervix - ectocervix | NKX6-2 | 0,5667 | 0,1116 |
| fallopian tube | ID2 | 0,5667 | 0,1116 |
| fallopian tube | MYBL1 | 0,5667 | 0,1116 |
| bladder | EHF | 0,5662 | 0,007453 |
| minor salivary gland | ZNF143 | 0,5652 | 4,706E-15 |
| heart - left ventricle | FOXK1 | 0,5641 | 1,185E-37 |
| brain - hypothalamus | ATF7 | 0,5637 | 2,44E-18 |
| bladder | ZBTB33 | 0,5636 | 0,007793 |
| cervix - endocervix | EOMES | 0,5636 | 0,08972 |
| cervix - endocervix | MLXIPL | 0,5636 | 0,08972 |
| cervix - endocervix | RARG | 0,5636 | 0,08972 |
| cervix - endocervix | CUX1 | 0,5636 | 0,08972 |
| cervix - endocervix | MYCN | 0,5636 | 0,08972 |
| cervix - endocervix | HES2 | 0,5636 | 0,08972 |
| testis | SIX1 | 0,5627 | 1,564E-31 |
| bladder | TBX2 | 0,5623 | 0,007967 |
| minor salivary gland | TFE3 | 0,562 | 7,176E-15 |
| minor salivary gland | RELA | 0,5619 | 7,298E-15 |
| heart - left ventricle | FOXO1 | 0,5616 | 2,847E-37 |
| bladder | JUNB | 0,561 | 0,008145 |
| muscle - skeletal | MLX | 0,5609 | 9,956E-68 |
| cervix - ectocervix | HNF4A | 0,5607 | 0,1163 |
| testis | MEF2C | 0,5607 | 2,826E-31 |
| heart - left ventricle | TEAD2 | 0,5602 | 4,739E-37 |
| testis | MAFB | 0,5599 | 3,577E-31 |
| bladder | HNF1B | 0,5597 | 0,008326 |
| heart - left ventricle | MEIS2 | 0,5596 | 5,694E-37 |
| vagina | EHF | 0,5593 | 3,239E-14 |
| muscle - skeletal | E2F4 | 0,559 | 3,337E-67 |
| minor salivary gland | CTCF | 0,5584 | 1,156E-14 |
| cervix - endocervix | MIXL1 | 0,5583 | 0,09346 |

| | | | |
|---|---|---|---|
| minor salivary gland | HINFP | 0,5581 | 1,209E-14 |
| minor salivary gland | CEBPD | 0,5581 | 1,201E-14 |
| heart - left ventricle | SP3 | 0,5577 | 1,131E-36 |
| minor salivary gland | ATF1 | 0,5575 | 1,301E-14 |
| testis | RXRA | 0,5574 | 7,438E-31 |
| heart - left ventricle | RORB | 0,5571 | 1,367E-36 |
| muscle - skeletal | GMEB1 | 0,5568 | 1,476E-66 |
| muscle - skeletal | ARID3A | 0,5565 | 1,695E-66 |
| bladder | SOX5 | 0,5545 | 0,009083 |
| brain - hypothalamus | GLIS3 | 0,5537 | 1,273E-17 |
| heart - left ventricle | GATA4 | 0,5523 | 7,198E-36 |
| bladder | BHLHE41 | 0,5519 | 0,009482 |
| minor salivary gland | ZNF410 | 0,5518 | 2,719E-14 |
| testis | TGIF2 | 0,5517 | 3,815E-30 |
| cervix - endocervix | HIC2 | 0,5515 | 0,0984 |
| cervix - endocervix | FOXD2 | 0,5515 | 0,0984 |
| cervix - endocervix | ESR2 | 0,5515 | 0,0984 |
| heart - left ventricle | TBX5 | 0,5512 | 1,047E-35 |
| lung | RORB | 0,551 | 3,289E-47 |
| minor salivary gland | RELB | 0,5506 | 3,2E-14 |
| cervix - ectocervix | TP63 | 0,55 | 0,125 |
| fallopian tube | EOMES | 0,55 | 0,125 |
| fallopian tube | FOXF2 | 0,55 | 0,125 |
| fallopian tube | ATF1 | 0,55 | 0,125 |
| vagina | FOXA1 | 0,5499 | 1,051E-13 |
| heart - left ventricle | ZBTB7A | 0,5498 | 1,724E-35 |
| muscle - skeletal | ZBTB7B | 0,5496 | 1,428E-64 |
| ALL_samples | MECOM | 0,549 | 0 |
| muscle - skeletal | NPAS2 | 0,5486 | 2,668E-64 |
| bladder | FOXP3 | 0,5481 | 0,01011 |
| brain - hippocampus | RFX2 | 0,5481 | 7,617E-17 |
| heart - left ventricle | RFX5 | 0,548 | 3,105E-35 |
| cervix - ectocervix | SPIC | 0,5477 | 0,1269 |
| brain - hypothalamus | MTF1 | 0,5474 | 3,454E-17 |
| brain - hypothalamus | ARNT | 0,5469 | 3,729E-17 |
| heart - left ventricle | RBPJ | 0,5468 | 4,667E-35 |
| minor salivary gland | ZNF384 | 0,5468 | 5,174E-14 |
| cervix - endocervix | NOTO | 0,5462 | 0,1023 |
| brain - hippocampus | RREB1 | 0,5461 | 1,039E-16 |
| bladder | SOX15 | 0,5455 | 0,01054 |
| bladder | ELF3 | 0,5455 | 0,01054 |
| testis | ERF | 0,5453 | 2,342E-29 |
| brain - hypothalamus | TEAD1 | 0,5452 | 4,86E-17 |
| brain - hypothalamus | IRF2 | 0,5452 | 4,863E-17 |
| brain - hypothalamus | CREB3L2 | 0,5451 | 4,957E-17 |
| heart - left ventricle | ELK4 | 0,545 | 8,572E-35 |
| cervix - endocervix | NKX2-8 | 0,5449 | 0,1033 |
| heart - left ventricle | BHLHE41 | 0,5446 | 9,923E-35 |
| minor salivary gland | TFAP2A | 0,5443 | 7,11E-14 |
| bladder | PLAG1 | 0,5442 | 0,01077 |
| muscle - skeletal | STAT5A | 0,5438 | 5,545E-63 |
| heart - left ventricle | TCF12 | 0,5434 | 1,5E-34 |
| heart - left ventricle | RFX1 | 0,5434 | 1,496E-34 |
| testis | MITF | 0,5434 | 3,995E-29 |

| | | | |
|---|---|---|---|
| vagina | ZBTB7C | 0,5433 | 2,342E-13 |
| vagina | SOX9 | 0,5427 | 2,5E-13 |
| colon - transverse | NPAS2 | 0,5426 | 1,863E-32 |
| cervix - ectocervix | NEUROG1 | 0,5425 | 0,1313 |
| brain - hypothalamus | LHX4 | 0,5424 | 7,593E-17 |
| bladder | ELF5 | 0,5416 | 0,01123 |
| testis | KLF12 | 0,5416 | 6,65E-29 |
| brain - hypothalamus | ELF4 | 0,5405 | 1,015E-16 |
| muscle - skeletal | RELA | 0,5402 | 4,993E-62 |
| cervix - endocervix | LHX4 | 0,5394 | 0,1076 |
| cervix - endocervix | HOXD3 | 0,5394 | 0,1076 |
| cervix - endocervix | RFX2 | 0,5394 | 0,1076 |
| cervix - endocervix | HSF1 | 0,5394 | 0,1076 |
| cervix - endocervix | FOXP2 | 0,5394 | 0,1076 |
| cervix - endocervix | SOX4 | 0,5394 | 0,1076 |
| cervix - endocervix | BCL6B | 0,5394 | 0,1076 |
| cervix - endocervix | MYBL2 | 0,5394 | 0,1076 |
| cervix - endocervix | ZNF24 | 0,5394 | 0,1076 |
| heart - left ventricle | SOX6 | 0,5393 | 5,683E-34 |
| bladder | DLX4 | 0,539 | 0,0117 |
| bladder | POU2F1 | 0,539 | 0,0117 |
| bladder | RXRA | 0,539 | 0,0117 |
| minor salivary gland | SMAD2 | 0,5389 | 1,393E-13 |
| brain - hippocampus | EHF | 0,5387 | 3,191E-16 |
| esophagus - mucosa | SOX2 | 0,5381 | 5,587E-43 |
| heart - left ventricle | CREB3 | 0,5381 | 8,618E-34 |
| bladder | KLF14 | 0,5373 | 0,01201 |
| heart - left ventricle | MAFG | 0,5367 | 1,36E-33 |
| artery - coronary | TAL1 | 0,5361 | 2,963E-19 |
| breast - mammary tissue | MLXIPL | 0,5359 | 1,737E-35 |
| testis | TWIST1 | 0,5354 | 3,608E-28 |
| muscle - skeletal | GMEB2 | 0,5353 | 9,728E-61 |
| bladder | GRHL2 | 0,5351 | 0,01244 |
| brain - caudate (basal ganglia) | RFX3 | 0,5345 | 1,4E-19 |
| ALL_samples | GATA6 | 0,5343 | 0 |
| muscle - skeletal | ARID3B | 0,5343 | 1,744E-60 |
| colon - transverse | GFI1 | 0,5342 | 2,444E-31 |
| muscle - skeletal | VDR | 0,5342 | 1,878E-60 |
| artery - coronary | CEBPA | 0,5338 | 4,414E-19 |
| cervix - ectocervix | POU3F1 | 0,5333 | 0,1392 |
| cervix - ectocervix | PAX5 | 0,5333 | 0,1392 |
| fallopian tube | MAFG | 0,5333 | 0,1392 |
| fallopian tube | ETV4 | 0,5333 | 0,1392 |
| fallopian tube | ARID5A | 0,5333 | 0,1392 |
| heart - left ventricle | HLF | 0,5331 | 4,33E-33 |
| artery - coronary | SOX2 | 0,5329 | 5,244E-19 |
| heart - left ventricle | NFATC1 | 0,5329 | 4,582E-33 |
| bladder | NR1H3 | 0,5325 | 0,01296 |
| brain - hypothalamus | RFX2 | 0,5324 | 3,497E-16 |
| heart - left ventricle | PITX3 | 0,5322 | 5,844E-33 |
| testis | NR2F2 | 0,5322 | 8,677E-28 |
| brain - hypothalamus | GATA1 | 0,532 | 3,745E-16 |
| heart - left ventricle | RORA | 0,5311 | 8,292E-33 |
| minor salivary gland | GRHL1 | 0,5311 | 3,594E-13 |

| | | | |
|---|---|---|---|
| heart - left ventricle | SOX9 | 0,5307 | 9,421E-33 |
| vagina | GRHL2 | 0,5306 | 1,045E-12 |
| nerve - tibial | TWIST2 | 0,5304 | 3,367E-46 |
| bladder | POU4F3 | 0,5302 | 0,01341 |
| bladder | GATA3 | 0,5299 | 0,01349 |
| cervix - ectocervix | CDX2 | 0,5295 | 0,1427 |
| brain - hippocampus | NFATC1 | 0,5294 | 1,254E-15 |
| brain - hypothalamus | STAT3 | 0,5294 | 5,487E-16 |
| testis | FOXC2 | 0,5294 | 1,798E-27 |
| heart - left ventricle | AHR | 0,5293 | 1,473E-32 |
| heart - left ventricle | IRF5 | 0,5291 | 1,579E-32 |
| brain - hypothalamus | RFX1 | 0,5289 | 5,931E-16 |
| brain - hypothalamus | SP1 | 0,5286 | 6,212E-16 |
| heart - left ventricle | XBP1 | 0,5286 | 1,807E-32 |
| muscle - skeletal | ZBTB7A | 0,5284 | 5,913E-59 |
| bladder | NKX6-1 | 0,5281 | 0,01386 |
| bladder | HNF4G | 0,5281 | 0,01386 |
| brain - hypothalamus | MGA | 0,5278 | 6,935E-16 |
| brain - hypothalamus | NFATC2 | 0,5276 | 7,179E-16 |
| cervix - endocervix | GSX2 | 0,5276 | 0,117 |
| minor salivary gland | HMBOX1 | 0,5276 | 5,463E-13 |
| bladder | GATA2 | 0,5273 | 0,01404 |
| cervix - endocervix | MXI1 | 0,5273 | 0,1173 |
| cervix - endocervix | ZBTB18 | 0,5273 | 0,1173 |
| brain - hypothalamus | ZFX | 0,527 | 7,857E-16 |
| brain - hippocampus | RARG | 0,5264 | 1,961E-15 |
| heart - left ventricle | GMEB1 | 0,526 | 4,1E-32 |
| nerve - tibial | BCL6 | 0,525 | 3,779E-45 |
| kidney - cortex | E2F3 | 0,5249 | 2,508E-07 |
| testis | SNAI2 | 0,5246 | 6,491E-27 |
| brain - hypothalamus | PLAG1 | 0,5242 | 1,174E-15 |
| brain - hypothalamus | STAT5A | 0,5242 | 1,177E-15 |
| heart - left ventricle | NKX3-1 | 0,5242 | 7,247E-32 |
| kidney - cortex | NR1H4 | 0,5242 | 2,628E-07 |
| minor salivary gland | YY1 | 0,5242 | 8,112E-13 |
| heart - left ventricle | CEBPG | 0,5241 | 7,493E-32 |
| testis | PAX9 | 0,5241 | 7,316E-27 |
| brain - hypothalamus | SOX9 | 0,5236 | 1,283E-15 |
| minor salivary gland | HOXC13 | 0,5235 | 8,874E-13 |
| heart - left ventricle | HSF1 | 0,5231 | 1,039E-31 |
| testis | SMAD3 | 0,5227 | 1,054E-26 |
| muscle - skeletal | YY1 | 0,5225 | 1,878E-57 |
| muscle - skeletal | FOXQ1 | 0,5224 | 1,944E-57 |
| cervix - endocervix | SP8 | 0,5222 | 0,1215 |
| cervix - endocervix | RAX | 0,5222 | 0,1215 |
| colon - transverse | POU2F3 | 0,5219 | 9,739E-30 |
| minor salivary gland | SP2 | 0,5219 | 1,069E-12 |
| brain - hippocampus | IRF2 | 0,5216 | 3,851E-15 |
| minor salivary gland | TEAD2 | 0,5209 | 1,196E-12 |
| brain - hypothalamus | STAT2 | 0,5206 | 1,998E-15 |
| minor salivary gland | FOXK2 | 0,5194 | 1,42E-12 |
| muscle - skeletal | KLF9 | 0,5194 | 1,088E-56 |
| brain - hypothalamus | ZNF410 | 0,5189 | 2,566E-15 |
| pancreas | KLF9 | 0,5186 | 5,594E-24 |

| | | | |
|---|---|---|---|
| vagina | SOX2 | 0,5182 | 4,252E-12 |
| brain - hypothalamus | RARG | 0,5174 | 3,144E-15 |
| brain - hippocampus | NFATC2 | 0,517 | 7,404E-15 |
| bladder | JUN | 0,5169 | 0,01643 |
| bladder | ESRRG | 0,5169 | 0,01643 |
| bladder | NFE2L2 | 0,5169 | 0,01643 |
| cervix - ectocervix | FOXP3 | 0,5167 | 0,1544 |
| cervix - ectocervix | SREBF1 | 0,5167 | 0,1544 |
| cervix - ectocervix | ESRRA | 0,5167 | 0,1544 |
| cervix - ectocervix | HES2 | 0,5167 | 0,1544 |
| cervix - ectocervix | SOX9 | 0,5167 | 0,1544 |
| fallopian tube | RARA | 0,5167 | 0,1544 |
| fallopian tube | ONECUT3 | 0,5167 | 0,1544 |
| fallopian tube | CREM | 0,5167 | 0,1544 |
| fallopian tube | NR4A1 | 0,5167 | 0,1544 |
| brain - hypothalamus | ATF1 | 0,5165 | 3,617E-15 |
| minor salivary gland | STAT2 | 0,5163 | 2,03E-12 |
| brain - hypothalamus | BCL6 | 0,5161 | 3,799E-15 |
| kidney - cortex | LBX2 | 0,5159 | 4,335E-07 |
| testis | TCF12 | 0,5159 | 6,115E-26 |
| brain - hypothalamus | FOXO1 | 0,5158 | 3,947E-15 |
| testis | NR2C2 | 0,5156 | 6,566E-26 |
| testis | TCF21 | 0,5153 | 7,117E-26 |
| cervix - endocervix | TEAD1 | 0,5152 | 0,1276 |
| cervix - endocervix | ZNF740 | 0,5152 | 0,1276 |
| cervix - endocervix | IRF2 | 0,5152 | 0,1276 |
| cervix - endocervix | HOXC11 | 0,5152 | 0,1276 |
| heart - left ventricle | NRL | 0,5151 | 1,209E-30 |
| heart - left ventricle | PRRX1 | 0,5148 | 1,319E-30 |
| heart - left ventricle | FOXK2 | 0,5148 | 1,309E-30 |
| testis | ZBTB7C | 0,5146 | 8,56E-26 |
| stomach | STAT6 | 0,5144 | 1,226E-25 |
| ALL_samples | PPARG | 0,5141 | 0 |
| brain - hippocampus | TGIF1 | 0,5141 | 1,11E-14 |
| muscle - skeletal | BCL6 | 0,514 | 2,322E-55 |
| minor salivary gland | ZNF263 | 0,5138 | 2,71E-12 |
| muscle - skeletal | CENPB | 0,5134 | 3,331E-55 |
| minor salivary gland | PKNOX1 | 0,5132 | 2,896E-12 |
| brain - nucleus accumbens (basal ga | SP3 | 0,5128 | 6,705E-18 |
| minor salivary gland | HNF4G | 0,5128 | 3,013E-12 |
| minor salivary gland | FOXO3 | 0,5124 | 3,172E-12 |
| testis | RBPJ | 0,5122 | 1,553E-25 |
| brain - hypothalamus | SMAD4 | 0,512 | 6,783E-15 |
| heart - left ventricle | GABPA | 0,512 | 3,069E-30 |
| kidney - cortex | PPARA | 0,5116 | 0,000000561 |
| colon - sigmoid | MLXIPL | 0,5114 | 3,036E-26 |
| heart - left ventricle | MNT | 0,5112 | 3,874E-30 |
| brain - hypothalamus | TCF7 | 0,5111 | 7,744E-15 |
| minor salivary gland | GMEB2 | 0,5111 | 3,667E-12 |
| testis | USF2 | 0,5109 | 2,167E-25 |
| brain - nucleus accumbens (basal ga | ARNT | 0,5106 | 9,79E-18 |
| pancreas | TEAD1 | 0,5106 | 3,568E-23 |
| muscle - skeletal | USF1 | 0,5104 | 1,779E-54 |
| thyroid | TBX5 | 0,5104 | 1,356E-44 |

| | | | |
|---|---|---|---|
| cervix - endocervix | SPZ1 | 0,5103 | 0,1318 |
| cervix - endocervix | HMX1 | 0,5103 | 0,1318 |
| muscle - skeletal | CLOCK | 0,5103 | 1,922E-54 |
| minor salivary gland | TCF12 | 0,5102 | 4,052E-12 |
| testis | KLF9 | 0,5095 | 3,073E-25 |
| vagina | TFAP2A | 0,509 | 1,158E-11 |
| brain - hippocampus | TP73 | 0,5088 | 2,286E-14 |
| testis | BHLHE41 | 0,5088 | 3,612E-25 |
| heart - left ventricle | TFEB | 0,5076 | 1,146E-29 |
| heart - left ventricle | ATF7 | 0,5074 | 1,218E-29 |
| small intestine - terminal ileum | GFI1B | 0,5074 | 1,255E-13 |
| muscle - skeletal | ZBTB33 | 0,5073 | 9,82E-54 |
| testis | LHX9 | 0,5073 | 5,211E-25 |
| heart - left ventricle | ATF1 | 0,5071 | 1,306E-29 |
| brain - hypothalamus | RUNX2 | 0,5069 | 1,378E-14 |
| heart - left ventricle | RORC | 0,5068 | 1,452E-29 |
| brain - hippocampus | SOX9 | 0,5063 | 3,21E-14 |
| brain - hypothalamus | NFAT5 | 0,506 | 1,58E-14 |
| minor salivary gland | HIF1A | 0,506 | 6,482E-12 |
| minor salivary gland | MLXIP | 0,5057 | 6,704E-12 |
| heart - left ventricle | MEIS1 | 0,5055 | 2,092E-29 |
| brain - hypothalamus | VSX1 | 0,5053 | 1,724E-14 |
| colon - transverse | BHLHA15 | 0,5053 | 1,066E-27 |
| bladder | TP63 | 0,5052 | 0,01949 |
| bladder | FOXQ1 | 0,5052 | 0,01949 |
| bladder | FOXI1 | 0,5051 | 0,01952 |
| heart - left ventricle | STAT3 | 0,5051 | 2,381E-29 |
| testis | STAT5A | 0,505 | 9,177E-25 |
| brain - hypothalamus | IRF9 | 0,5048 | 1,862E-14 |
| small intestine - terminal ileum | KLF4 | 0,5047 | 1,774E-13 |
| brain - hippocampus | TCF7 | 0,5045 | 4,084E-14 |
| testis | CTCFL | 0,5044 | 1,072E-24 |
| heart - left ventricle | ETV3 | 0,5043 | 2,969E-29 |
| bladder | ZBTB7A | 0,5039 | 0,01986 |
| bladder | TGIF2 | 0,5039 | 0,01986 |
| small intestine - terminal ileum | POU2F3 | 0,5038 | 1,988E-13 |
| heart - left ventricle | ARID3A | 0,5037 | 3,638E-29 |
| minor salivary gland | DLX4 | 0,5037 | 8,36E-12 |
| muscle - skeletal | ZNF282 | 0,5037 | 6,878E-53 |
| minor salivary gland | SMAD4 | 0,5036 | 8,485E-12 |
| brain - hypothalamus | ZNF384 | 0,5031 | 2,336E-14 |
| brain - hypothalamus | TGIF2 | 0,5031 | 2,342E-14 |
| cervix - endocervix | HOXD9 | 0,503 | 0,1383 |
| brain - hypothalamus | MLXIP | 0,5028 | 2,434E-14 |
| bladder | TFCP2 | 0,5026 | 0,02023 |
| brain - hypothalamus | REL | 0,5022 | 2,638E-14 |
| minor salivary gland | THAP1 | 0,5021 | 9,952E-12 |
| brain - hippocampus | TEAD1 | 0,5019 | 5,744E-14 |
| brain - hypothalamus | ID4 | 0,5018 | 2,773E-14 |
| bladder | MLX | 0,5013 | 0,02061 |
| testis | NR3C1 | 0,5008 | 2,587E-24 |
| brain - hippocampus | CREB3L2 | 0,5005 | 6,968E-14 |
| brain - hypothalamus | SP3 | 0,5005 | 3,328E-14 |
| heart - left ventricle | NFYA | 0,5004 | 9,349E-29 |

| | | | |
|---|---|---|---|
| bladder | ARX | 0,5003 | 0,02091 |
| cervix - ectocervix | KLF16 | 0,5 | 0,1705 |
| cervix - ectocervix | FOSL2 | 0,5 | 0,1705 |
| colon - transverse | MYBL2 | 0,5 | 4,555E-27 |
| fallopian tube | RELB | 0,5 | 0,1705 |
| fallopian tube | SOX15 | 0,5 | 0,1705 |
| fallopian tube | NKX3-1 | 0,5 | 0,1705 |
| cervix - ectocervix | TEAD1 | -0,5 | 0,1705 |
| cervix - ectocervix | FOXF2 | -0,5 | 0,1705 |
| cervix - ectocervix | SP3 | -0,5 | 0,1705 |
| cervix - ectocervix | FOXJ3 | -0,5 | 0,1705 |
| cervix - ectocervix | ZNF24 | -0,5 | 0,1705 |
| cervix - ectocervix | ZNF354C | -0,5 | 0,1705 |
| cervix - ectocervix | GATA5 | -0,5 | 0,1705 |
| vagina | TCF21 | -0,5 | 3,006E-11 |
| vagina | TCFL5 | -0,5003 | 2,924E-11 |
| small intestine - terminal ileum | TEAD4 | -0,5006 | 2,954E-13 |
| vagina | DLX1 | -0,5007 | 2,805E-11 |
| nerve - tibial | CREB5 | -0,5009 | 1,254E-40 |
| adipose - subcutaneous | HIC1 | -0,5012 | 1,842E-43 |
| esophagus - mucosa | DDIT3 | -0,5014 | 1,153E-36 |
| small intestine - terminal ileum | HOXB3 | -0,5014 | 2,7E-13 |
| breast - mammary tissue | SPDEF | -0,5017 | 1,244E-30 |
| small intestine - terminal ileum | MEF2B | -0,5017 | 2,583E-13 |
| breast - mammary tissue | HOXD9 | -0,502 | 1,134E-30 |
| lung | EOMES | -0,502 | 3,123E-38 |
| breast - mammary tissue | POU2F3 | -0,5026 | 9,49E-31 |
| vagina | PKNOX2 | -0,5027 | 2,267E-11 |
| colon - transverse | REST | -0,5029 | 2,051E-27 |
| cervix - endocervix | MEOX1 | -0,503 | 0,1383 |
| cervix - endocervix | TBX1 | -0,503 | 0,1383 |
| cervix - endocervix | FOXC2 | -0,503 | 0,1383 |
| small intestine - terminal ileum | HIC2 | -0,503 | 2,207E-13 |
| breast - mammary tissue | ESR1 | -0,5031 | 8,138E-31 |
| vagina | NFATC2 | -0,5034 | 2,103E-11 |
| breast - mammary tissue | CDX1 | -0,5041 | 5,949E-31 |
| vagina | NR5A2 | -0,5047 | 1,841E-11 |
| vagina | GATA6 | -0,5052 | 1,739E-11 |
| small intestine - terminal ileum | VSX1 | -0,5053 | 1,632E-13 |
| adipose - subcutaneous | DBP | -0,5054 | 2,849E-44 |
| testis | BSX | -0,5056 | 8,087E-25 |
| vagina | ETS1 | -0,5058 | 1,631E-11 |
| breast - mammary tissue | TFAP2C | -0,5062 | 3,11E-31 |
| testis | PAX4 | -0,5067 | 6,097E-25 |
| artery - coronary | MSC | -0,507 | 4,433E-17 |
| bladder | MYF6 | -0,5072 | 0,01894 |
| colon - transverse | POU2F2 | -0,5073 | 6,147E-28 |
| vagina | CREM | -0,5073 | 1,397E-11 |
| colon - transverse | HOXC9 | -0,5074 | 6,027E-28 |
| breast - mammary tissue | GRHL1 | -0,5075 | 2,054E-31 |
| small intestine - terminal ileum | ZNF740 | -0,5081 | 1,139E-13 |
| colon - transverse | NFATC2 | -0,5083 | 4,688E-28 |
| nerve - tibial | SPIB | -0,5085 | 5,121E-42 |
| small intestine - terminal ileum | TGIF1 | -0,5087 | 1,065E-13 |

| | | | |
|---|---|---|---|
| vagina | SMAD4 | -0,5087 | 1,197E-11 |
| colon - transverse | MEF2B | -0,5092 | 3,597E-28 |
| vagina | PKNOX1 | -0,5093 | 1,126E-11 |
| nerve - tibial | DBP | -0,5094 | 3,494E-42 |
| lung | IRF1 | -0,5095 | 1,657E-39 |
| nerve - tibial | RXRG | -0,5104 | 2,257E-42 |
| colon - transverse | DLX3 | -0,5107 | 2,375E-28 |
| testis | GATA1 | -0,5107 | 2,258E-25 |
| colon - transverse | DMRT3 | -0,5108 | 2,292E-28 |
| colon - transverse | PLAG1 | -0,5108 | 2,282E-28 |
| small intestine - terminal ileum | STAT1 | -0,5108 | 8,092E-14 |
| vagina | CREB5 | -0,5108 | 9,545E-12 |
| vagina | HOXD3 | -0,511 | 9,342E-12 |
| adipose - subcutaneous | RXRG | -0,5113 | 1,934E-45 |
| colon - transverse | BACH1 | -0,5116 | 1,823E-28 |
| testis | IRF1 | -0,5118 | 1,713E-25 |
| lung | NFKB2 | -0,512 | 6,22E-40 |
| small intestine - terminal ileum | ZFX | -0,5122 | 6,738E-14 |
| breast - mammary tissue | TBX1 | -0,5134 | 3,103E-32 |
| nerve - tibial | SOX15 | -0,5137 | 5,6E-43 |
| breast - mammary tissue | DLX3 | -0,5138 | 2,794E-32 |
| vagina | RFX3 | -0,5141 | 6,714E-12 |
| vagina | NFIA | -0,5144 | 6,478E-12 |
| vagina | LBX2 | -0,5145 | 6,406E-12 |
| vagina | USF1 | -0,5146 | 6,297E-12 |
| artery - coronary | TFEB | -0,5147 | 1,233E-17 |
| esophagus - mucosa | HSF1 | -0,5147 | 7,138E-39 |
| vagina | KLF12 | -0,5149 | 6,144E-12 |
| breast - mammary tissue | TEAD3 | -0,5151 | 1,835E-32 |
| small intestine - terminal ileum | IRF9 | -0,5151 | 4,594E-14 |
| cervix - endocervix | NR4A1 | -0,5152 | 0,1276 |
| small intestine - terminal ileum | TEAD1 | -0,5152 | 4,522E-14 |
| bladder | SRF | -0,5156 | 0,01675 |
| bladder | ALX4 | -0,5164 | 0,01654 |
| breast - mammary tissue | RXRB | -0,5165 | 1,173E-32 |
| cervix - ectocervix | CUX2 | -0,5167 | 0,1544 |
| cervix - ectocervix | CENPB | -0,5167 | 0,1544 |
| cervix - ectocervix | FOSB | -0,5167 | 0,1544 |
| cervix - ectocervix | NFIC | -0,5167 | 0,1544 |
| fallopian tube | ZBTB33 | -0,5167 | 0,1544 |
| fallopian tube | ZIC1 | -0,5167 | 0,1544 |
| breast - mammary tissue | TBX19 | -0,5168 | 1,045E-32 |
| bladder | RXRG | -0,5169 | 0,01643 |
| small intestine - terminal ileum | ELK4 | -0,5169 | 3,639E-14 |
| vagina | HES7 | -0,5169 | 4,922E-12 |
| breast - mammary tissue | GATA3 | -0,517 | 9,779E-33 |
| esophagus - mucosa | PRRX2 | -0,517 | 2,98E-39 |
| small intestine - terminal ileum | MAFF | -0,5175 | 3,376E-14 |
| testis | LHX2 | -0,5175 | 3,999E-26 |
| breast - mammary tissue | PRRX2 | -0,5181 | 6,99E-33 |
| vagina | TGIF2 | -0,5184 | 4,174E-12 |
| small intestine - terminal ileum | BHLHE41 | -0,5185 | 2,929E-14 |
| breast - mammary tissue | RFX1 | -0,5186 | 5,766E-33 |
| cervix - ectocervix | PAX6 | -0,5188 | 0,1524 |

| | | | |
|---|---|---|---|
| lung | STAT4 | -0,519 | 3,538E-41 |
| esophagus - mucosa | ARID5A | -0,5191 | 1,278E-39 |
| breast - mammary tissue | FOXP3 | -0,5203 | 3,343E-33 |
| colon - transverse | FOXO3 | -0,5205 | 1,46E-29 |
| colon - transverse | ZFX | -0,5205 | 1,44E-29 |
| small intestine - terminal ileum | FOXO3 | -0,5205 | 2,247E-14 |
| small intestine - terminal ileum | HMBOX1 | -0,5205 | 2,254E-14 |
| small intestine - terminal ileum | ETV1 | -0,5206 | 2,205E-14 |
| adipose - visceral (omentum) | PKNOX2 | -0,5209 | 5,932E-39 |
| testis | KLF5 | -0,521 | 1,648E-26 |
| breast - mammary tissue | RARB | -0,5211 | 2,621E-33 |
| testis | STAT4 | -0,5214 | 1,474E-26 |
| colon - transverse | GATA5 | -0,5216 | 1,059E-29 |
| colon - transverse | GABPA | -0,5218 | 9,869E-30 |
| colon - transverse | HOXC10 | -0,5219 | 9,65E-30 |
| vagina | GATA1 | -0,5221 | 2,745E-12 |
| breast - mammary tissue | ZNF282 | -0,5223 | 1,75E-33 |
| breast - mammary tissue | SOX15 | -0,5224 | 1,665E-33 |
| vagina | HEY2 | -0,5224 | 2,671E-12 |
| colon - transverse | MAFG | -0,5229 | 7,155E-30 |
| small intestine - terminal ileum | SP3 | -0,523 | 1,612E-14 |
| vagina | MEOX2 | -0,5231 | 2,459E-12 |
| breast - mammary tissue | MYB | -0,5232 | 1,292E-33 |
| vagina | TBX19 | -0,5232 | 2,422E-12 |
| testis | FOXB1 | -0,5234 | 8,791E-27 |
| breast - mammary tissue | MYCN | -0,5235 | 1,18E-33 |
| vagina | HMBOX1 | -0,5236 | 2,319E-12 |
| small intestine - terminal ileum | NFE2L2 | -0,5239 | 1,426E-14 |
| esophagus - mucosa | HIC1 | -0,5241 | 1,787E-40 |
| small intestine - terminal ileum | GLIS3 | -0,5242 | 1,358E-14 |
| small intestine - terminal ileum | POU3F3 | -0,5247 | 1,274E-14 |
| small intestine - terminal ileum | TBP | -0,5248 | 1,252E-14 |
| cervix - ectocervix | PHOX2A | -0,5249 | 0,1468 |
| cervix - ectocervix | UNCX | -0,5249 | 0,1468 |
| nerve - tibial | BHLHE41 | -0,525 | 3,814E-45 |
| vagina | GMEB2 | -0,526 | 1,767E-12 |
| testis | NKX3-2 | -0,5265 | 3,899E-27 |
| artery - coronary | PRRX2 | -0,5272 | 1,439E-18 |
| cervix - endocervix | MYC | -0,5273 | 0,1173 |
| cervix - endocervix | STAT1 | -0,5273 | 0,1173 |
| colon - transverse | MAX | -0,5273 | 1,948E-30 |
| artery - coronary | FOXL1 | -0,5275 | 1,351E-18 |
| colon - transverse | NR4A1 | -0,5276 | 1,778E-30 |
| vagina | CEBPB | -0,5277 | 1,453E-12 |
| esophagus - mucosa | CEBPB | -0,5278 | 3,984E-41 |
| small intestine - terminal ileum | MSC | -0,5285 | 7,629E-15 |
| bladder | CENPB | -0,5286 | 0,01376 |
| adipose - subcutaneous | HOXA13 | -0,529 | 4,509E-49 |
| nerve - tibial | BHLHE40 | -0,529 | 6,302E-46 |
| artery - coronary | FOXC2 | -0,5298 | 9,09E-19 |
| vagina | HIC1 | -0,5304 | 1,067E-12 |
| fallopian tube | ONECUT1 | -0,5309 | 0,1414 |
| nerve - tibial | PKNOX2 | -0,531 | 2,465E-46 |
| vagina | TEAD2 | -0,531 | 9,92E-13 |

| | | | |
|---|---|---|---|
| breast - mammary tissue | FOXP1 | -0,5314 | 8,326E-35 |
| esophagus - mucosa | RARA | -0,5315 | 8,602E-42 |
| breast - mammary tissue | PBX2 | -0,5317 | 7,537E-35 |
| breast - mammary tissue | ELF3 | -0,5326 | 5,394E-35 |
| vagina | FOXD2 | -0,5327 | 8,218E-13 |
| small intestine - terminal ileum | CREB1 | -0,533 | 4,099E-15 |
| small intestine - terminal ileum | VENTX | -0,533 | 4,095E-15 |
| breast - mammary tissue | FOXF2 | -0,5332 | 4,513E-35 |
| cervix - ectocervix | MGA | -0,5333 | 0,1392 |
| cervix - ectocervix | MEIS1 | -0,5333 | 0,1392 |
| cervix - ectocervix | HOXA5 | -0,5333 | 0,1392 |
| cervix - ectocervix | DLX1 | -0,5333 | 0,1392 |
| cervix - ectocervix | TEAD2 | -0,5333 | 0,1392 |
| cervix - ectocervix | RFX1 | -0,5333 | 0,1392 |
| cervix - ectocervix | ATF4 | -0,5333 | 0,1392 |
| cervix - ectocervix | MEF2B | -0,5333 | 0,1392 |
| cervix - ectocervix | SIX2 | -0,5333 | 0,1392 |
| cervix - ectocervix | PKNOX2 | -0,5333 | 0,1392 |
| cervix - ectocervix | AR | -0,5333 | 0,1392 |
| fallopian tube | NPAS2 | -0,5333 | 0,1392 |
| vagina | FOXP1 | -0,5339 | 7,095E-13 |
| vagina | TCF4 | -0,5345 | 6,654E-13 |
| breast - mammary tissue | BHLHE41 | -0,5349 | 2,499E-35 |
| breast - mammary tissue | NR2F1 | -0,535 | 2,39E-35 |
| small intestine - terminal ileum | MYC | -0,535 | 3,07E-15 |
| small intestine - terminal ileum | TFDP1 | -0,5354 | 2,926E-15 |
| breast - mammary tissue | GRHL2 | -0,5355 | 1,999E-35 |
| breast - mammary tissue | HLTF | -0,5359 | 1,773E-35 |
| small intestine - terminal ileum | NFKB1 | -0,5362 | 2,605E-15 |
| adipose - subcutaneous | GLIS2 | -0,5366 | 1,051E-50 |
| vagina | BHLHE22 | -0,5367 | 5,124E-13 |
| vagina | RARA | -0,5368 | 5,065E-13 |
| vagina | RXRG | -0,5369 | 4,986E-13 |
| testis | PITX3 | -0,5374 | 2,089E-28 |
| small intestine - terminal ileum | ID4 | -0,5375 | 2,152E-15 |
| bladder | CREM | -0,5377 | 0,01194 |
| vagina | NR1H3 | -0,538 | 4,403E-13 |
| nerve - tibial | SOX4 | -0,5388 | 6,604E-48 |
| cervix - endocervix | POU6F2 | -0,5394 | 0,1076 |
| small intestine - terminal ileum | SOX2 | -0,5399 | 1,54E-15 |
| small intestine - terminal ileum | DLX4 | -0,5403 | 1,448E-15 |
| muscle - skeletal | E2F8 | -0,5404 | 4,339E-62 |
| small intestine - terminal ileum | CUX2 | -0,5408 | 1,351E-15 |
| colon - transverse | ZNF740 | -0,5411 | 2,947E-32 |
| breast - mammary tissue | TP63 | -0,5431 | 1,424E-36 |
| vagina | EMX2 | -0,5432 | 2,37E-13 |
| breast - mammary tissue | HSF4 | -0,5434 | 1,29E-36 |
| colon - transverse | FLI1 | -0,5439 | 1,245E-32 |
| vagina | ERG | -0,5442 | 2,099E-13 |
| testis | SPZ1 | -0,5444 | 3,06E-29 |
| testis | TBX1 | -0,5446 | 2,896E-29 |
| colon - transverse | HMBOX1 | -0,5452 | 8,094E-33 |
| vagina | HOXC9 | -0,5456 | 1,775E-13 |
| breast - mammary tissue | CTCF | -0,546 | 5,071E-37 |

| | | | |
|---|---|---|---|
| vagina | FOXJ2 | -0,5461 | 1,67E-13 |
| adipose - visceral (omentum) | PRRX1 | -0,5463 | 2,011E-43 |
| breast - mammary tissue | LMX1B | -0,5465 | 4,274E-37 |
| bladder | CREB3 | -0,5468 | 0,01032 |
| testis | HNF4A | -0,547 | 1,46E-29 |
| cervix - ectocervix | FOXI1 | -0,5477 | 0,1269 |
| fallopian tube | VAX1 | -0,5477 | 0,1269 |
| fallopian tube | INSM1 | -0,5477 | 0,1269 |
| small intestine - terminal ileum | CEBPD | -0,5482 | 4,62E-16 |
| colon - transverse | SOX13 | -0,5486 | 2,795E-33 |
| colon - transverse | CEBPD | -0,5487 | 2,663E-33 |
| breast - mammary tissue | ZNF263 | -0,5488 | 1,884E-37 |
| bladder | OLIG1 | -0,5491 | 0,00994 |
| colon - transverse | PRRX2 | -0,5494 | 2,178E-33 |
| cervix - ectocervix | DDIT3 | -0,55 | 0,125 |
| cervix - ectocervix | POU3F2 | -0,55 | 0,125 |
| cervix - ectocervix | NFYB | -0,55 | 0,125 |
| cervix - ectocervix | EBF1 | -0,55 | 0,125 |
| fallopian tube | SMAD2 | -0,55 | 0,125 |
| artery - coronary | ETV6 | -0,5503 | 2,102E-20 |
| colon - transverse | ZNF24 | -0,5505 | 1,508E-33 |
| vagina | NFKB2 | -0,5505 | 9,732E-14 |
| colon - transverse | EMX2 | -0,5516 | 1,059E-33 |
| breast - mammary tissue | RUNX2 | -0,5523 | 5,191E-38 |
| testis | ESRRB | -0,5523 | 3,233E-30 |
| colon - transverse | CEBPB | -0,5531 | 6,61E-34 |
| vagina | PPARG | -0,5534 | 6,787E-14 |
| breast - mammary tissue | TFAP4 | -0,5538 | 3,017E-38 |
| colon - transverse | FOXO1 | -0,5541 | 4,728E-34 |
| small intestine - terminal ileum | MNT | -0,5544 | 1,843E-16 |
| vagina | MSX1 | -0,5548 | 5,695E-14 |
| vagina | MEF2B | -0,5549 | 5,629E-14 |
| colon - transverse | YY1 | -0,5551 | 3,413E-34 |
| breast - mammary tissue | TFAP2A | -0,5557 | 1,517E-38 |
| small intestine - terminal ileum | FLI1 | -0,5567 | 1,296E-16 |
| colon - transverse | HESX1 | -0,5571 | 1,794E-34 |
| vagina | ARNT | -0,5571 | 4,267E-14 |
| breast - mammary tissue | TP73 | -0,5572 | 8,535E-39 |
| colon - transverse | DBP | -0,5579 | 1,388E-34 |
| breast - mammary tissue | PROX1 | -0,5582 | 5,802E-39 |
| bladder | E2F1 | -0,5584 | 0,00851 |
| vagina | NR2F2 | -0,5587 | 3,483E-14 |
| small intestine - terminal ileum | NFATC2 | -0,5595 | 8,471E-17 |
| vagina | RXRB | -0,5595 | 3,152E-14 |
| vagina | MEF2D | -0,5598 | 3,012E-14 |
| adipose - subcutaneous | GLI2 | -0,5606 | 3,94E-56 |
| testis | SHOX2 | -0,5618 | 2,017E-31 |
| vagina | SOX5 | -0,562 | 2,283E-14 |
| colon - transverse | BATF3 | -0,5621 | 3,428E-35 |
| small intestine - terminal ileum | POU2F1 | -0,5623 | 5,551E-17 |
| cervix - endocervix | EBF1 | -0,5636 | 0,08972 |
| colon - transverse | TBP | -0,5636 | 2,098E-35 |
| vagina | PROX1 | -0,5642 | 1,718E-14 |
| colon - transverse | STAT2 | -0,5643 | 1,61E-35 |

| | | | |
|---|---|---|---|
| colon - transverse | ELK4 | -0,5646 | 1,497E-35 |
| adipose - subcutaneous | CDX1 | -0,5653 | 2,926E-57 |
| vagina | MEIS3 | -0,5654 | 1,483E-14 |
| cervix - ectocervix | FOXP2 | -0,5667 | 0,1116 |
| cervix - ectocervix | BHLHE41 | -0,5667 | 0,1116 |
| cervix - ectocervix | HIC2 | -0,5667 | 0,1116 |
| cervix - ectocervix | GMEB2 | -0,5667 | 0,1116 |
| fallopian tube | ZNF282 | -0,5667 | 0,1116 |
| fallopian tube | HESX1 | -0,5667 | 0,1116 |
| vagina | RELA | -0,5669 | 1,21E-14 |
| vagina | MITF | -0,5672 | 1,169E-14 |
| vagina | FOXJ3 | -0,5673 | 1,157E-14 |
| bladder | SOX8 | -0,5675 | 0,007288 |
| testis | HSF1 | -0,5678 | 3,383E-32 |
| colon - transverse | CUX1 | -0,568 | 4,653E-36 |
| vagina | GLIS2 | -0,5681 | 1,038E-14 |
| breast - mammary tissue | MEIS2 | -0,5683 | 1,264E-40 |
| small intestine - terminal ileum | SOX1 | -0,5691 | 1,916E-17 |
| vagina | TFE3 | -0,5694 | 8,74E-15 |
| colon - transverse | CLOCK | -0,5698 | 2,52E-36 |
| small intestine - terminal ileum | NFATC1 | -0,5699 | 1,713E-17 |
| colon - transverse | IRF9 | -0,57 | 2,381E-36 |
| vagina | TFAP2B | -0,5701 | 8,025E-15 |
| small intestine - terminal ileum | TCF12 | -0,5703 | 1,59E-17 |
| vagina | GLIS1 | -0,5704 | 7,673E-15 |
| testis | EVX2 | -0,5705 | 1,496E-32 |
| breast - mammary tissue | HINFP | -0,5706 | 5,262E-41 |
| vagina | MLXIP | -0,5706 | 7,472E-15 |
| vagina | HSF4 | -0,5706 | 7,469E-15 |
| breast - mammary tissue | DBP | -0,5709 | 4,687E-41 |
| colon - transverse | CREB1 | -0,5719 | 1,224E-36 |
| small intestine - terminal ileum | JUNB | -0,5728 | 1,079E-17 |
| small intestine - terminal ileum | NFE2 | -0,573 | 1,035E-17 |
| vagina | BATF3 | -0,5732 | 5,257E-15 |
| colon - transverse | BCL6B | -0,5738 | 6,271E-37 |
| vagina | SHOX2 | -0,5741 | 4,718E-15 |
| colon - transverse | KLF9 | -0,5743 | 5,373E-37 |
| cervix - ectocervix | HAND1 | -0,5745 | 0,1057 |
| colon - transverse | GLIS3 | -0,5756 | 3,449E-37 |
| cervix - endocervix | STAT5A | -0,5758 | 0,08155 |
| small intestine - terminal ileum | CEBPB | -0,5782 | 4,527E-18 |
| small intestine - terminal ileum | TCF21 | -0,5786 | 4,253E-18 |
| breast - mammary tissue | ETV4 | -0,58 | 1,311E-42 |
| small intestine - terminal ileum | HES7 | -0,5804 | 3,123E-18 |
| vagina | PBX3 | -0,5808 | 1,905E-15 |
| breast - mammary tissue | TCF3 | -0,5817 | 6,503E-43 |
| vagina | FOXD3 | -0,5818 | 1,662E-15 |
| vagina | MEF2C | -0,5823 | 1,552E-15 |
| testis | CREM | -0,5826 | 3,382E-34 |
| vagina | CREB3 | -0,5826 | 1,481E-15 |
| colon - transverse | AHR | -0,5829 | 2,584E-38 |
| vagina | HOXC11 | -0,583 | 1,404E-15 |
| breast - mammary tissue | MEF2B | -0,5832 | 3,61E-43 |
| cervix - ectocervix | HOXA2 | -0,5833 | 0,09919 |

| | | | |
|---|---|---|---|
| cervix - ectocervix | TEAD3 | -0,5833 | 0,09919 |
| cervix - ectocervix | HLTF | -0,5833 | 0,09919 |
| cervix - ectocervix | HMBOX1 | -0,5833 | 0,09919 |
| cervix - ectocervix | MAFK | -0,5833 | 0,09919 |
| cervix - ectocervix | PBX2 | -0,5833 | 0,09919 |
| cervix - ectocervix | ZNF740 | -0,5833 | 0,09919 |
| cervix - ectocervix | HOXA11 | -0,5833 | 0,09919 |
| cervix - ectocervix | FOXC2 | -0,5833 | 0,09919 |
| fallopian tube | KLF12 | -0,5833 | 0,09919 |
| fallopian tube | IRF9 | -0,5833 | 0,09919 |
| fallopian tube | HEY2 | -0,5833 | 0,09919 |
| breast - mammary tissue | ALX4 | -0,5834 | 3,317E-43 |
| colon - transverse | NFE2L2 | -0,5834 | 2,158E-38 |
| colon - transverse | TBX1 | -0,5837 | 1,924E-38 |
| breast - mammary tissue | SOX4 | -0,5839 | 2,649E-43 |
| vagina | NFATC1 | -0,5839 | 1,233E-15 |
| colon - transverse | GATA2 | -0,5842 | 1,632E-38 |
| testis | RUNX2 | -0,5842 | 2,043E-34 |
| testis | NKX6-1 | -0,5848 | 1,702E-34 |
| colon - transverse | MEF2D | -0,5849 | 1,272E-38 |
| breast - mammary tissue | IRF3 | -0,5852 | 1,579E-43 |
| colon - transverse | HOXA10 | -0,5852 | 1,138E-38 |
| vagina | NRF1 | -0,5854 | 1,006E-15 |
| small intestine - terminal ileum | NR4A1 | -0,586 | 1,245E-18 |
| testis | NRL | -0,5868 | 8,852E-35 |
| artery - coronary | GLIS2 | -0,5872 | 1,225E-23 |
| colon - transverse | MGA | -0,5877 | 4,582E-39 |
| vagina | GLIS3 | -0,5877 | 7,339E-16 |
| colon - transverse | BACH2 | -0,5885 | 3,33E-39 |
| breast - mammary tissue | GLI2 | -0,5901 | 2,097E-44 |
| adipose - subcutaneous | CREB3L1 | -0,5923 | 5,038E-64 |
| vagina | STAT5B | -0,5927 | 3,606E-16 |
| vagina | POU2F1 | -0,5927 | 3,602E-16 |
| adipose - subcutaneous | PRRX2 | -0,5933 | 2,84E-64 |
| cervix - ectocervix | PITX3 | -0,5934 | 0,09212 |
| vagina | TCF7L2 | -0,5936 | 3,16E-16 |
| colon - transverse | GRHL1 | -0,594 | 4,507E-40 |
| small intestine - terminal ileum | EGR2 | -0,594 | 3,247E-19 |
| cervix - ectocervix | DLX2 | -0,5941 | 0,09158 |
| cervix - ectocervix | SOX3 | -0,5941 | 0,09158 |
| testis | ARID3A | -0,5947 | 6,722E-36 |
| colon - transverse | SCRT1 | -0,5958 | 2,3E-40 |
| muscle - skeletal | MEF2C | -0,5966 | 1,461E-78 |
| vagina | POU6F1 | -0,5966 | 2,075E-16 |
| small intestine - terminal ileum | FOXO6 | -0,5979 | 1,643E-19 |
| small intestine - terminal ileum | HLF | -0,5988 | 1,417E-19 |
| colon - transverse | HOXD3 | -0,5989 | 7,096E-41 |
| small intestine - terminal ileum | YY2 | -0,599 | 1,354E-19 |
| cervix - ectocervix | CTCF | -0,6 | 0,08762 |
| cervix - ectocervix | POU6F1 | -0,6 | 0,08762 |
| cervix - ectocervix | SMAD3 | -0,6 | 0,08762 |
| cervix - ectocervix | TCF12 | -0,6 | 0,08762 |
| cervix - ectocervix | MEF2D | -0,6 | 0,08762 |
| cervix - ectocervix | PKNOX1 | -0,6 | 0,08762 |

| | | | |
|---|---|---|---|
| cervix - ectocervix | HOXC10 | -0,6 | 0,08762 |
| cervix - ectocervix | ALX4 | -0,6 | 0,08762 |
| cervix - ectocervix | TGIF2 | -0,6 | 0,08762 |
| kidney - medulla | ZBTB7C | -0,6 | 0,4 |
| kidney - medulla | MEIS3 | -0,6 | 0,4 |
| kidney - medulla | BCL6 | -0,6 | 0,4 |
| kidney - medulla | ZNF410 | -0,6 | 0,4 |
| kidney - medulla | POU4F1 | -0,6 | 0,4 |
| kidney - medulla | NFYA | -0,6 | 0,4 |
| kidney - medulla | ERF | -0,6 | 0,4 |
| kidney - medulla | TEAD3 | -0,6 | 0,4 |
| kidney - medulla | RELA | -0,6 | 0,4 |
| kidney - medulla | NFKB1 | -0,6 | 0,4 |
| kidney - medulla | ZNF143 | -0,6 | 0,4 |
| kidney - medulla | STAT5B | -0,6 | 0,4 |
| kidney - medulla | KLF9 | -0,6 | 0,4 |
| kidney - medulla | MLXIP | -0,6 | 0,4 |
| kidney - medulla | AHR | -0,6 | 0,4 |
| kidney - medulla | GSC | -0,6 | 0,4 |
| kidney - medulla | POU1F1 | -0,6 | 0,4 |
| kidney - medulla | CEBPB | -0,6 | 0,4 |
| kidney - medulla | ID4 | -0,6 | 0,4 |
| kidney - medulla | SMAD3 | -0,6 | 0,4 |
| kidney - medulla | RFX2 | -0,6 | 0,4 |
| kidney - medulla | CUX2 | -0,6 | 0,4 |
| kidney - medulla | TFAP2C | -0,6 | 0,4 |
| kidney - medulla | FOXO1 | -0,6 | 0,4 |
| kidney - medulla | CREM | -0,6 | 0,4 |
| kidney - medulla | NR2E3 | -0,6 | 0,4 |
| kidney - medulla | FOXP1 | -0,6 | 0,4 |
| kidney - medulla | BACH1 | -0,6 | 0,4 |
| kidney - medulla | FOXO3 | -0,6 | 0,4 |
| kidney - medulla | POU5F1 | -0,6 | 0,4 |
| kidney - medulla | FOXG1 | -0,6 | 0,4 |
| kidney - medulla | MAFK | -0,6 | 0,4 |
| kidney - medulla | MAFG | -0,6 | 0,4 |
| kidney - medulla | TP53 | -0,6 | 0,4 |
| kidney - medulla | HIC1 | -0,6 | 0,4 |
| kidney - medulla | ZNF263 | -0,6 | 0,4 |
| kidney - medulla | ETV3 | -0,6 | 0,4 |
| kidney - medulla | ETV6 | -0,6 | 0,4 |
| kidney - medulla | SP1 | -0,6 | 0,4 |
| kidney - medulla | SP2 | -0,6 | 0,4 |
| kidney - medulla | NR5A2 | -0,6 | 0,4 |
| kidney - medulla | GLIS2 | -0,6 | 0,4 |
| kidney - medulla | ELK3 | -0,6 | 0,4 |
| kidney - medulla | ELK4 | -0,6 | 0,4 |
| kidney - medulla | PAX9 | -0,6 | 0,4 |
| kidney - medulla | HES7 | -0,6 | 0,4 |
| kidney - medulla | MNT | -0,6 | 0,4 |
| kidney - medulla | EN2 | -0,6 | 0,4 |
| kidney - medulla | SIX2 | -0,6 | 0,4 |
| kidney - medulla | SRF | -0,6 | 0,4 |
| kidney - medulla | PKNOX1 | -0,6 | 0,4 |

| | | | |
|---|---|---|---|
| kidney - medulla | PKNOX2 | -0,6 | 0,4 |
| kidney - medulla | GMEB2 | -0,6 | 0,4 |
| kidney - medulla | GMEB1 | -0,6 | 0,4 |
| kidney - medulla | CDX1 | -0,6 | 0,4 |
| kidney - medulla | IRF3 | -0,6 | 0,4 |
| kidney - medulla | IRF2 | -0,6 | 0,4 |
| kidney - medulla | ONECUT2 | -0,6 | 0,4 |
| kidney - medulla | MSX1 | -0,6 | 0,4 |
| kidney - medulla | TFE3 | -0,6 | 0,4 |
| kidney - medulla | LEF1 | -0,6 | 0,4 |
| kidney - medulla | ZNF24 | -0,6 | 0,4 |
| kidney - medulla | RUNX2 | -0,6 | 0,4 |
| kidney - medulla | ESR2 | -0,6 | 0,4 |
| kidney - medulla | FOXK1 | -0,6 | 0,4 |
| kidney - medulla | GATA6 | -0,6 | 0,4 |
| kidney - medulla | ETS1 | -0,6 | 0,4 |
| kidney - medulla | STAT4 | -0,6 | 0,4 |
| kidney - medulla | MTF1 | -0,6 | 0,4 |
| kidney - medulla | TGIF1 | -0,6 | 0,4 |
| vagina | STAT5A | -0,6005 | 1,175E-16 |
| colon - transverse | TFCP2 | -0,6011 | 3,098E-41 |
| vagina | NR1H2 | -0,6012 | 1,059E-16 |
| colon - transverse | VSX1 | -0,6013 | 2,873E-41 |
| small intestine - terminal ileum | DLX2 | -0,6013 | 9,034E-20 |
| vagina | HOXC10 | -0,6016 | 9,964E-17 |
| vagina | SOX8 | -0,6024 | 8,83E-17 |
| small intestine - terminal ileum | RORA | -0,6028 | 7,009E-20 |
| adipose - subcutaneous | PRRX1 | -0,6031 | 6,372E-67 |
| vagina | ZEB1 | -0,6033 | 7,813E-17 |
| small intestine - terminal ileum | PAX9 | -0,6034 | 6,215E-20 |
| vagina | EBF1 | -0,604 | 6,965E-17 |
| breast - mammary tissue | RARG | -0,6058 | 2,6E-47 |
| colon - transverse | POU2F1 | -0,6059 | 4,731E-42 |
| small intestine - terminal ileum | SMAD4 | -0,6065 | 3,617E-20 |
| testis | SCRT1 | -0,6067 | 1,153E-37 |
| small intestine - terminal ileum | GABPA | -0,6073 | 3,118E-20 |
| colon - transverse | FOXK2 | -0,6083 | 1,917E-42 |
| small intestine - terminal ileum | ZNF24 | -0,6086 | 2,465E-20 |
| small intestine - terminal ileum | RFX5 | -0,6089 | 2,333E-20 |
| colon - transverse | HOXD9 | -0,6091 | 1,362E-42 |
| colon - transverse | BHLHE40 | -0,6093 | 1,252E-42 |
| small intestine - terminal ileum | LHX6 | -0,6097 | 2,042E-20 |
| colon - transverse | NR1H2 | -0,6098 | 1,041E-42 |
| colon - transverse | ESRRG | -0,6101 | 9,354E-43 |
| colon - transverse | LHX6 | -0,611 | 6,591E-43 |
| small intestine - terminal ileum | FOXK1 | -0,6114 | 1,487E-20 |
| bladder | JDP2 | -0,6117 | 0,003213 |
| cervix - endocervix | NFYB | -0,6121 | 0,05997 |
| cervix - endocervix | HOXD13 | -0,6121 | 0,05997 |
| testis | ETV2 | -0,6125 | 1,485E-38 |
| testis | NR1H3 | -0,6128 | 1,337E-38 |
| testis | RFX1 | -0,6129 | 1,298E-38 |
| vagina | RBPJ | -0,6129 | 1,833E-17 |
| colon - transverse | NFATC3 | -0,6132 | 2,747E-43 |

| | | | |
|---|---|---|---|
| colon - transverse | ZNF410 | -0,6139 | 2,028E-43 |
| small intestine - terminal ileum | HESX1 | -0,6144 | 8,596E-21 |
| testis | LMX1A | -0,6157 | 4,798E-39 |
| small intestine - terminal ileum | CREB3L2 | -0,6159 | 6,523E-21 |
| small intestine - terminal ileum | ID2 | -0,616 | 6,396E-21 |
| vagina | HSF2 | -0,6162 | 1,105E-17 |
| small intestine - terminal ileum | SOX11 | -0,6164 | 5,967E-21 |
| small intestine - terminal ileum | EGR1 | -0,6166 | 5,709E-21 |
| small intestine - terminal ileum | SOX8 | -0,6166 | 5,688E-21 |
| cervix - ectocervix | EMX2 | -0,6167 | 0,07693 |
| cervix - ectocervix | POU1F1 | -0,6167 | 0,07693 |
| cervix - ectocervix | PBX3 | -0,6167 | 0,07693 |
| cervix - ectocervix | GLIS2 | -0,6167 | 0,07693 |
| cervix - ectocervix | GMEB1 | -0,6167 | 0,07693 |
| cervix - ectocervix | E2F4 | -0,6167 | 0,07693 |
| cervix - ectocervix | IRF3 | -0,6167 | 0,07693 |
| cervix - ectocervix | STAT5A | -0,6167 | 0,07693 |
| cervix - ectocervix | ARNT | -0,6167 | 0,07693 |
| fallopian tube | TFAP4 | -0,6167 | 0,07693 |
| fallopian tube | TCF7L2 | -0,6167 | 0,07693 |
| small intestine - terminal ileum | TWIST1 | -0,6168 | 5,548E-21 |
| testis | MEIS3 | -0,6178 | 2,246E-39 |
| small intestine - terminal ileum | BHLHE40 | -0,619 | 3,66E-21 |
| colon - transverse | TFEB | -0,6191 | 2,563E-44 |
| small intestine - terminal ileum | SP4 | -0,6191 | 3,582E-21 |
| vagina | TCF7L1 | -0,6198 | 6,357E-18 |
| colon - transverse | SP4 | -0,6208 | 1,281E-44 |
| small intestine - terminal ileum | GRHL1 | -0,6209 | 2,559E-21 |
| small intestine - terminal ileum | PBX1 | -0,6212 | 2,409E-21 |
| colon - transverse | FOXC1 | -0,6226 | 5,956E-45 |
| small intestine - terminal ileum | ZNF143 | -0,6228 | 1,792E-21 |
| small intestine - terminal ileum | ELK1 | -0,6238 | 1,485E-21 |
| cervix - endocervix | EVX2 | -0,6242 | 0,05372 |
| colon - transverse | HES7 | -0,6242 | 3,156E-45 |
| small intestine - terminal ileum | MGA | -0,6245 | 1,294E-21 |
| colon - transverse | MNT | -0,6246 | 2,619E-45 |
| colon - transverse | SIX2 | -0,6253 | 1,958E-45 |
| small intestine - terminal ileum | YY1 | -0,6253 | 1,11E-21 |
| breast - mammary tissue | FOXO6 | -0,6262 | 2,4E-51 |
| small intestine - terminal ileum | ETV5 | -0,6263 | 9,236E-22 |
| bladder | GLIS2 | -0,6273 | 0,002337 |
| colon - transverse | POU3F4 | -0,6273 | 8,355E-46 |
| colon - transverse | ESR1 | -0,6278 | 7,051E-46 |
| vagina | PRRX2 | -0,6282 | 1,657E-18 |
| testis | EVX1 | -0,6285 | 4,525E-41 |
| colon - transverse | PAX9 | -0,6292 | 3,902E-46 |
| small intestine - terminal ileum | CENPB | -0,6292 | 5,243E-22 |
| colon - transverse | VENTX | -0,6306 | 2,148E-46 |
| colon - transverse | ARNT | -0,6308 | 1,936E-46 |
| vagina | ATF4 | -0,6309 | 1,075E-18 |
| colon - transverse | HSF4 | -0,6317 | 1,351E-46 |
| colon - transverse | ID4 | -0,6319 | 1,199E-46 |
| colon - transverse | FOXC2 | -0,6322 | 1,05E-46 |
| vagina | MAFK | -0,6322 | 8,637E-19 |

| | | | |
|---|---|---|---|
| kidney - medulla | GCM2 | -0,6325 | 0,3675 |
| kidney - medulla | GSX2 | -0,6325 | 0,3675 |
| kidney - medulla | SOX21 | -0,6325 | 0,3675 |
| colon - transverse | ETV5 | -0,6329 | 7,862E-47 |
| colon - transverse | FOXK1 | -0,633 | 7,731E-47 |
| cervix - ectocervix | HOXA10 | -0,6333 | 0,06709 |
| cervix - ectocervix | TFAP4 | -0,6333 | 0,06709 |
| cervix - ectocervix | NFKB2 | -0,6333 | 0,06709 |
| cervix - ectocervix | RFX2 | -0,6333 | 0,06709 |
| cervix - ectocervix | GLIS1 | -0,6333 | 0,06709 |
| cervix - ectocervix | NKX3-2 | -0,6333 | 0,06709 |
| cervix - ectocervix | GLI2 | -0,6333 | 0,06709 |
| fallopian tube | CEBPA | -0,6333 | 0,06709 |
| small intestine - terminal ileum | DLX1 | -0,6334 | 2,313E-22 |
| small intestine - terminal ileum | NR1H2 | -0,6343 | 1,911E-22 |
| colon - transverse | BCL6 | -0,6352 | 2,938E-47 |
| small intestine - terminal ileum | BATF3 | -0,6361 | 1,339E-22 |
| colon - transverse | TFAP2C | -0,6373 | 1,164E-47 |
| testis | HNF1B | -0,6373 | 1,568E-42 |
| small intestine - terminal ileum | STAT5A | -0,6374 | 1,036E-22 |
| small intestine - terminal ileum | PRRX2 | -0,6387 | 7,928E-23 |
| fallopian tube | MYOD1 | -0,639 | 0,06392 |
| fallopian tube | HOXC12 | -0,639 | 0,06392 |
| small intestine - terminal ileum | RFX3 | -0,639 | 7,562E-23 |
| small intestine - terminal ileum | RFX2 | -0,6397 | 6,531E-23 |
| colon - transverse | SOX11 | -0,6398 | 4,049E-48 |
| colon - transverse | ZNF143 | -0,6415 | 1,919E-48 |
| small intestine - terminal ileum | EGR3 | -0,6428 | 3,49E-23 |
| small intestine - terminal ileum | RARB | -0,6433 | 3,118E-23 |
| colon - transverse | TCF12 | -0,6438 | 6,763E-49 |
| colon - transverse | SMAD4 | -0,6441 | 6,051E-49 |
| small intestine - terminal ileum | TFAP2A | -0,6444 | 2,522E-23 |
| colon - transverse | ERF | -0,6453 | 3,48E-49 |
| small intestine - terminal ileum | NRF1 | -0,6458 | 1,889E-23 |
| colon - transverse | HEY1 | -0,6468 | 1,782E-49 |
| small intestine - terminal ileum | HSF4 | -0,6479 | 1,23E-23 |
| vagina | ARID5A | -0,6493 | 4,817E-20 |
| cervix - ectocervix | TWIST1 | -0,65 | 0,05807 |
| cervix - ectocervix | MEIS3 | -0,65 | 0,05807 |
| cervix - ectocervix | DLX6 | -0,65 | 0,05807 |
| cervix - ectocervix | GFI1B | -0,65 | 0,05807 |
| cervix - ectocervix | TCF7L2 | -0,65 | 0,05807 |
| cervix - ectocervix | MITF | -0,65 | 0,05807 |
| cervix - ectocervix | FOXA2 | -0,65 | 0,05807 |
| colon - transverse | CENPB | -0,6502 | 3,77E-50 |
| colon - transverse | TAL1 | -0,651 | 2,663E-50 |
| colon - transverse | YY2 | -0,6513 | 2,302E-50 |
| vagina | TWIST1 | -0,6519 | 3,064E-20 |
| small intestine - terminal ileum | FOXC2 | -0,6525 | 4,63E-24 |
| testis | GFI1B | -0,653 | 2,95E-45 |
| small intestine - terminal ileum | TFCP2 | -0,6536 | 3,69E-24 |
| small intestine - terminal ileum | REST | -0,6555 | 2,447E-24 |
| small intestine - terminal ileum | FOXC1 | -0,6557 | 2,351E-24 |
| small intestine - terminal ileum | FOXF2 | -0,6558 | 2,287E-24 |

| | | | |
|---|---|---|---|
| small intestine - terminal ileum | TBX19 | -0,6562 | 2,103E-24 |
| small intestine - terminal ileum | RXRG | -0,6563 | 2,072E-24 |
| small intestine - terminal ileum | MEF2D | -0,6563 | 2,076E-24 |
| cervix - ectocervix | OLIG2 | -0,6573 | 0,0544 |
| colon - transverse | MSX1 | -0,6576 | 1,222E-51 |
| colon - transverse | TEAD1 | -0,6582 | 9,431E-52 |
| colon - transverse | SMAD2 | -0,659 | 6,399E-52 |
| colon - transverse | HMX1 | -0,6593 | 5,574E-52 |
| colon - transverse | ARID5A | -0,6601 | 3,86E-52 |
| cervix - ectocervix | GCM2 | -0,6611 | 0,05252 |
| cervix - ectocervix | USF2 | -0,6611 | 0,05252 |
| colon - transverse | MEOX1 | -0,6618 | 1,719E-52 |
| colon - transverse | ERG | -0,6629 | 1E-52 |
| colon - transverse | E2F6 | -0,6631 | 8,97E-53 |
| colon - transverse | USF2 | -0,6634 | 7,7E-53 |
| colon - transverse | TP63 | -0,6638 | 6,377E-53 |
| small intestine - terminal ileum | SIX2 | -0,6638 | 3,985E-25 |
| colon - transverse | CREB3 | -0,6642 | 5,397E-53 |
| colon - transverse | RFX3 | -0,6654 | 3,043E-53 |
| small intestine - terminal ileum | MLXIP | -0,6655 | 2,747E-25 |
| cervix - ectocervix | MEIS2 | -0,6667 | 0,04987 |
| cervix - ectocervix | HOXA13 | -0,6667 | 0,04987 |
| cervix - ectocervix | TEF | -0,6667 | 0,04987 |
| cervix - ectocervix | ZEB1 | -0,6667 | 0,04987 |
| cervix - ectocervix | NR1H2 | -0,6667 | 0,04987 |
| cervix - ectocervix | IRF4 | -0,6667 | 0,04987 |
| colon - transverse | JDP2 | -0,6689 | 5,479E-54 |
| small intestine - terminal ileum | DDIT3 | -0,6704 | 9,18E-26 |
| small intestine - terminal ileum | TAL1 | -0,6707 | 8,52E-26 |
| colon - transverse | STAT5A | -0,6714 | 1,591E-54 |
| cervix - endocervix | FOXL1 | -0,6727 | 0,03304 |
| small intestine - terminal ileum | FOXO1 | -0,6742 | 3,899E-26 |
| testis | CREB3 | -0,6744 | 3,124E-49 |
| small intestine - terminal ileum | FOXK2 | -0,6747 | 3,433E-26 |
| cervix - endocervix | FOXG1 | -0,6748 | 0,03231 |
| bladder | GLI2 | -0,6753 | 0,0007816 |
| small intestine - terminal ileum | TP63 | -0,6753 | 2,988E-26 |
| colon - transverse | TCF21 | -0,6756 | 1,909E-55 |
| small intestine - terminal ileum | RXRB | -0,6756 | 2,807E-26 |
| small intestine - terminal ileum | ZNF263 | -0,6763 | 2,352E-26 |
| colon - transverse | PKNOX1 | -0,6765 | 1,208E-55 |
| colon - transverse | RELA | -0,6768 | 1,046E-55 |
| testis | PRRX2 | -0,6775 | 7,824E-50 |
| colon - transverse | DDIT3 | -0,6777 | 6,883E-56 |
| colon - transverse | MAFK | -0,6777 | 6,683E-56 |
| vagina | HSF1 | -0,6779 | 2,505E-22 |
| small intestine - terminal ileum | SOX15 | -0,6786 | 1,391E-26 |
| small intestine - terminal ileum | VAX2 | -0,6795 | 1,116E-26 |
| colon - transverse | CREM | -0,6802 | 1,877E-56 |
| colon - transverse | FOXF2 | -0,6804 | 1,723E-56 |
| testis | TEAD2 | -0,6811 | 1,555E-50 |
| small intestine - terminal ileum | TGIF2 | -0,682 | 6,2E-27 |
| cervix - ectocervix | ID4 | -0,6833 | 0,04244 |
| cervix - ectocervix | TFAP2B | -0,6833 | 0,04244 |

| | | | |
|---|---|---|---|
| cervix - ectocervix | SOX4 | -0,6833 | 0,04244 |
| cervix - ectocervix | ZNF263 | -0,6833 | 0,04244 |
| cervix - ectocervix | YY1 | -0,6833 | 0,04244 |
| cervix - ectocervix | ATF1 | -0,6833 | 0,04244 |
| cervix - ectocervix | PAX3 | -0,6833 | 0,04244 |
| cervix - ectocervix | NFE2L1 | -0,6833 | 0,04244 |
| cervix - ectocervix | FOXD2 | -0,6833 | 0,04244 |
| cervix - ectocervix | CDX1 | -0,6833 | 0,04244 |
| cervix - ectocervix | MSX1 | -0,6833 | 0,04244 |
| cervix - ectocervix | PROX1 | -0,6833 | 0,04244 |
| cervix - ectocervix | STAT6 | -0,6833 | 0,04244 |
| cervix - ectocervix | RHOXF1 | -0,6833 | 0,04244 |
| cervix - ectocervix | HOXC12 | -0,6833 | 0,04244 |
| cervix - ectocervix | STAT2 | -0,6833 | 0,04244 |
| fallopian tube | LMX1A | -0,6833 | 0,04244 |
| small intestine - terminal ileum | TBX1 | -0,6836 | 4,283E-27 |
| colon - transverse | ZNF263 | -0,6839 | 2,743E-57 |
| colon - transverse | AR | -0,6842 | 2,317E-57 |
| small intestine - terminal ileum | CREB5 | -0,6849 | 3,157E-27 |
| colon - transverse | DLX6 | -0,685 | 1,565E-57 |
| small intestine - terminal ileum | ARNT | -0,686 | 2,372E-27 |
| colon - transverse | HAND1 | -0,6862 | 8,109E-58 |
| colon - transverse | EBF1 | -0,6863 | 7,895E-58 |
| colon - transverse | ATF7 | -0,6867 | 6,237E-58 |
| colon - transverse | HOXA11 | -0,687 | 5,587E-58 |
| small intestine - terminal ileum | CREB3 | -0,689 | 1,157E-27 |
| small intestine - terminal ileum | KLF12 | -0,6907 | 7,606E-28 |
| colon - transverse | PBX1 | -0,6933 | 1,922E-59 |
| bladder | ATF4 | -0,6935 | 0,0004899 |
| cervix - ectocervix | CRX | -0,6938 | 0,03817 |
| colon - transverse | SOX1 | -0,6938 | 1,409E-59 |
| colon - transverse | MEF2A | -0,6945 | 1,001E-59 |
| colon - transverse | NRF1 | -0,6949 | 8,138E-60 |
| colon - transverse | RFX5 | -0,6955 | 5,672E-60 |
| colon - transverse | DLX2 | -0,6963 | 3,675E-60 |
| colon - transverse | BHLHE22 | -0,6967 | 2,908E-60 |
| colon - transverse | VAX2 | -0,6978 | 1,606E-60 |
| small intestine - terminal ileum | FOXD3 | -0,6982 | 1,186E-28 |
| colon - transverse | RXRB | -0,6983 | 1,202E-60 |
| colon - transverse | PHOX2A | -0,6991 | 7,748E-61 |
| cervix - ectocervix | TCF3 | -0,7 | 0,03577 |
| cervix - ectocervix | USF1 | -0,7 | 0,03577 |
| cervix - ectocervix | NR1H3 | -0,7 | 0,03577 |
| cervix - ectocervix | GATA2 | -0,7 | 0,03577 |
| fallopian tube | REST | -0,7 | 0,03577 |
| fallopian tube | HOXC11 | -0,7 | 0,03577 |
| colon - transverse | NR2F2 | -0,7002 | 4,413E-61 |
| small intestine - terminal ileum | AR | -0,7004 | 6,768E-29 |
| small intestine - terminal ileum | ISL2 | -0,7006 | 6,434E-29 |
| colon - transverse | SOX17 | -0,7025 | 1,184E-61 |
| colon - transverse | NFIC | -0,7029 | 9,236E-62 |
| colon - transverse | PRRX1 | -0,7033 | 7,482E-62 |
| colon - transverse | SOX5 | -0,7033 | 7,559E-62 |
| colon - transverse | NKX3-1 | -0,7039 | 5,355E-62 |

| | | | |
|---|---|---|---|
| small intestine - terminal ileum | BCL6 | -0,7039 | 2,696E-29 |
| colon - transverse | NR2F1 | -0,7059 | 1,764E-62 |
| colon - transverse | ZNF423 | -0,706 | 1,595E-62 |
| small intestine - terminal ileum | NFIX | -0,7064 | 1,412E-29 |
| small intestine - terminal ileum | NKX3-2 | -0,7083 | 8,552E-30 |
| colon - transverse | FOXP2 | -0,7087 | 3,39E-63 |
| small intestine - terminal ileum | SMAD2 | -0,7088 | 7,576E-30 |
| colon - transverse | TEAD3 | -0,7094 | 2,361E-63 |
| small intestine - terminal ileum | ESR1 | -0,7095 | 6,291E-30 |
| colon - transverse | HOXB3 | -0,7102 | 1,485E-63 |
| bladder | MSX1 | -0,7104 | 0,000308 |
| colon - transverse | FOXJ2 | -0,7112 | 8,272E-64 |
| small intestine - terminal ileum | BHLHE22 | -0,7112 | 3,947E-30 |
| colon - transverse | PHOX2B | -0,712 | 5,15E-64 |
| fallopian tube | HOXD11 | -0,712 | 0,0314 |
| colon - transverse | RBPJ | -0,7121 | 4,956E-64 |
| colon - transverse | TFE3 | -0,7124 | 3,994E-64 |
| colon - transverse | NFYB | -0,7128 | 3,196E-64 |
| colon - transverse | RXRG | -0,7151 | 8,375E-65 |
| colon - transverse | SRF | -0,7152 | 7,632E-65 |
| colon - transverse | HOXD8 | -0,716 | 4,952E-65 |
| cervix - ectocervix | TBX2 | -0,7167 | 0,02982 |
| cervix - ectocervix | EGR2 | -0,7167 | 0,02982 |
| cervix - ectocervix | HSF2 | -0,7167 | 0,02982 |
| cervix - ectocervix | SOX8 | -0,7167 | 0,02982 |
| cervix - ectocervix | NFIX | -0,7167 | 0,02982 |
| colon - transverse | HLF | -0,7167 | 3,187E-65 |
| colon - transverse | ZNF354C | -0,7172 | 2,419E-65 |
| colon - transverse | CUX2 | -0,7174 | 2,143E-65 |
| small intestine - terminal ileum | GATA2 | -0,7175 | 7,106E-31 |
| colon - transverse | NR3C1 | -0,7176 | 1,85E-65 |
| colon - transverse | RORA | -0,7179 | 1,601E-65 |
| small intestine - terminal ileum | AHR | -0,7184 | 5,689E-31 |
| colon - transverse | CREB3L2 | -0,7186 | 1,05E-65 |
| colon - transverse | CREB5 | -0,7189 | 8,304E-66 |
| colon - transverse | KLF12 | -0,7192 | 7,019E-66 |
| colon - transverse | NKX3-2 | -0,7193 | 6,86E-66 |
| colon - transverse | HIC1 | -0,7194 | 6,152E-66 |
| colon - transverse | GLI2 | -0,7197 | 5,419E-66 |
| colon - transverse | HEY2 | -0,7197 | 5,339E-66 |
| colon - transverse | FOXP1 | -0,7199 | 4,733E-66 |
| small intestine - terminal ileum | NFIC | -0,7202 | 3,364E-31 |
| colon - transverse | TBX19 | -0,7207 | 2,833E-66 |
| small intestine - terminal ileum | NKX2-3 | -0,7223 | 1,877E-31 |
| colon - transverse | RFX2 | -0,7225 | 9,746E-67 |
| small intestine - terminal ileum | BCL6B | -0,7225 | 1,781E-31 |
| colon - transverse | MITF | -0,7227 | 8,681E-67 |
| small intestine - terminal ileum | E2F6 | -0,7236 | 1,326E-31 |
| colon - transverse | SOX2 | -0,7243 | 3,216E-67 |
| colon - transverse | RORB | -0,7246 | 2,628E-67 |
| small intestine - terminal ileum | MEF2A | -0,7248 | 9,411E-32 |
| small intestine - terminal ileum | NR3C1 | -0,7252 | 8,384E-32 |
| colon - transverse | POU3F3 | -0,726 | 1,117E-67 |
| colon - transverse | HSF1 | -0,7261 | 1,059E-67 |

| | | | |
|---|---|---|---|
| colon - transverse | HLTF | -0,7278 | 3,595E-68 |
| colon - transverse | FOXO6 | -0,7281 | 3,011E-68 |
| small intestine - terminal ileum | SOX5 | -0,7281 | 3,638E-32 |
| small intestine - terminal ileum | ARID5A | -0,7281 | 3,661E-32 |
| small intestine - terminal ileum | NFKB2 | -0,7291 | 2,736E-32 |
| small intestine - terminal ileum | ZNF354C | -0,7306 | 1,791E-32 |
| colon - transverse | NFIX | -0,7309 | 5,143E-69 |
| colon - transverse | ISL2 | -0,7311 | 4,575E-69 |
| colon - transverse | NKX2-3 | -0,7315 | 3,437E-69 |
| colon - transverse | TWIST1 | -0,7316 | 3,239E-69 |
| small intestine - terminal ileum | FOXJ2 | -0,732 | 1,187E-32 |
| colon - transverse | NFKB2 | -0,7322 | 2,288E-69 |
| colon - transverse | MEF2C | -0,7322 | 2,279E-69 |
| colon - transverse | LMX1B | -0,7326 | 1,708E-69 |
| small intestine - terminal ileum | MEF2C | -0,7332 | 8,368E-33 |
| cervix - ectocervix | ZNF282 | -0,7333 | 0,02455 |
| cervix - ectocervix | CREB3 | -0,7333 | 0,02455 |
| cervix - ectocervix | HSF1 | -0,7333 | 0,02455 |
| cervix - ectocervix | PRRX1 | -0,7333 | 0,02455 |
| cervix - ectocervix | PRRX2 | -0,7333 | 0,02455 |
| cervix - ectocervix | FOXL1 | -0,7333 | 0,02455 |
| fallopian tube | TFCP2 | -0,7333 | 0,02455 |
| small intestine - terminal ileum | SOX17 | -0,7361 | 3,489E-33 |
| small intestine - terminal ileum | HOXC10 | -0,7363 | 3,257E-33 |
| colon - transverse | SOX8 | -0,7369 | 1,082E-70 |
| colon - transverse | MEOX2 | -0,737 | 1,017E-70 |
| colon - transverse | TFAP2A | -0,7377 | 6,458E-71 |
| kidney - medulla | TBX4 | -0,7379 | 0,2621 |
| kidney - medulla | BHLHE41 | -0,7379 | 0,2621 |
| kidney - medulla | GATA4 | -0,7379 | 0,2621 |
| colon - transverse | TWIST2 | -0,7386 | 3,487E-71 |
| small intestine - terminal ileum | CREM | -0,7398 | 1,159E-33 |
| small intestine - terminal ileum | TWIST2 | -0,7398 | 1,139E-33 |
| colon - transverse | NFATC1 | -0,7418 | 4,351E-72 |
| small intestine - terminal ileum | HEY1 | -0,7427 | 4,762E-34 |
| small intestine - terminal ileum | MITF | -0,743 | 4,317E-34 |
| small intestine - terminal ileum | POU6F1 | -0,7431 | 4,167E-34 |
| small intestine - terminal ileum | HOXD3 | -0,7445 | 2,71E-34 |
| colon - transverse | ELK3 | -0,7462 | 2,215E-73 |
| colon - transverse | MEIS2 | -0,7463 | 2,036E-73 |
| colon - transverse | RARB | -0,7463 | 1,957E-73 |
| colon - transverse | ATF4 | -0,7477 | 7,982E-74 |
| colon - transverse | SOX15 | -0,7479 | 6,626E-74 |
| colon - transverse | STAT5B | -0,749 | 3,269E-74 |
| colon - transverse | TCFL5 | -0,7493 | 2,542E-74 |
| small intestine - terminal ileum | RBPJ | -0,7495 | 5,691E-35 |
| cervix - ectocervix | TCF4 | -0,75 | 0,01994 |
| cervix - ectocervix | BHLHE22 | -0,75 | 0,01994 |
| cervix - ectocervix | ARID5A | -0,75 | 0,01994 |
| colon - transverse | ETV1 | -0,75 | 1,573E-74 |
| colon - transverse | RARG | -0,752 | 3,946E-75 |
| colon - transverse | MEIS1 | -0,7544 | 7,422E-76 |
| colon - transverse | TEAD2 | -0,7544 | 7,318E-76 |
| colon - transverse | ZEB1 | -0,7545 | 6,591E-76 |

| | | | |
|---|---|---|---|
| colon - transverse | HSF2 | -0,7555 | 3,213E-76 |
| small intestine - terminal ileum | PBX2 | -0,7559 | 7,221E-36 |
| small intestine - terminal ileum | HIC1 | -0,7567 | 5,472E-36 |
| colon - transverse | PBX2 | -0,757 | 1,106E-76 |
| cervix - endocervix | TBX5 | -0,7576 | 0,01114 |
| colon - transverse | DLX1 | -0,758 | 5,426E-77 |
| small intestine - terminal ileum | FOXP1 | -0,758 | 3,542E-36 |
| small intestine - terminal ileum | EBF1 | -0,7598 | 1,938E-36 |
| small intestine - terminal ileum | RELA | -0,7599 | 1,878E-36 |
| small intestine - terminal ileum | NR2F2 | -0,7608 | 1,416E-36 |
| small intestine - terminal ileum | MSX1 | -0,7612 | 1,231E-36 |
| small intestine - terminal ileum | ATF4 | -0,7617 | 1,036E-36 |
| small intestine - terminal ileum | GLI2 | -0,7618 | 1,01E-36 |
| colon - transverse | MEIS3 | -0,7627 | 1,761E-78 |
| small intestine - terminal ileum | SRF | -0,7639 | 4,89E-37 |
| small intestine - terminal ileum | PRRX1 | -0,7662 | 2,256E-37 |
| cervix - ectocervix | SOX5 | -0,7667 | 0,01594 |
| cervix - ectocervix | FOXD3 | -0,7667 | 0,01594 |
| small intestine - terminal ileum | NR2F1 | -0,7674 | 1,46E-37 |
| small intestine - terminal ileum | TCF4 | -0,7684 | 1,059E-37 |
| small intestine - terminal ileum | HOXC9 | -0,7689 | 8,832E-38 |
| small intestine - terminal ileum | MEIS3 | -0,7698 | 6,37E-38 |
| colon - transverse | NFIA | -0,7703 | 6,087E-81 |
| colon - transverse | TCF4 | -0,7733 | 5,931E-82 |
| small intestine - terminal ileum | HSF2 | -0,7737 | 1,64E-38 |
| small intestine - terminal ileum | PKNOX1 | -0,7737 | 1,585E-38 |
| kidney - medulla | SP8 | -0,7746 | 0,2254 |
| kidney - medulla | TLX1 | -0,7746 | 0,2254 |
| kidney - medulla | ESX1 | -0,7746 | 0,2254 |
| kidney - medulla | PAX7 | -0,7746 | 0,2254 |
| kidney - medulla | DUXA | -0,7746 | 0,2254 |
| small intestine - terminal ileum | MEOX2 | -0,775 | 1,003E-38 |
| small intestine - terminal ileum | ERG | -0,7753 | 9,202E-39 |
| small intestine - terminal ileum | MEOX1 | -0,7757 | 7,804E-39 |
| colon - transverse | PBX3 | -0,7759 | 7,64E-83 |
| colon - transverse | TGIF2 | -0,7763 | 5,547E-83 |
| small intestine - terminal ileum | HSF1 | -0,7791 | 2,233E-39 |
| colon - transverse | FOXD3 | -0,7813 | 1,043E-84 |
| colon - transverse | TCF7L1 | -0,7823 | 4,637E-85 |
| small intestine - terminal ileum | ATF7 | -0,7828 | 5,764E-40 |
| cervix - ectocervix | LMX1A | -0,7833 | 0,01252 |
| cervix - ectocervix | NRL | -0,7833 | 0,01252 |
| cervix - ectocervix | ONECUT3 | -0,7833 | 0,01252 |
| cervix - ectocervix | TCF21 | -0,7833 | 0,01252 |
| cervix - ectocervix | MSC | -0,7833 | 0,01252 |
| cervix - ectocervix | KLF14 | -0,7833 | 0,01252 |
| colon - transverse | HOXB2 | -0,7844 | 7,733E-86 |
| small intestine - terminal ileum | HEY2 | -0,7849 | 2,561E-40 |
| fallopian tube | RFX4 | -0,7851 | 0,01219 |
| colon - transverse | GLIS2 | -0,7861 | 1,909E-86 |
| small intestine - terminal ileum | RORB | -0,7866 | 1,385E-40 |
| small intestine - terminal ileum | TFE3 | -0,7892 | 5,082E-41 |
| small intestine - terminal ileum | NKX3-1 | -0,7906 | 2,837E-41 |
| small intestine - terminal ileum | MEIS2 | -0,7914 | 2,113E-41 |

| | | | |
|---|---|---|---|
| small intestine - terminal ileum | STAT5B | -0,7932 | 1,035E-41 |
| small intestine - terminal ileum | HOXD8 | -0,7955 | 4,079E-42 |
| small intestine - terminal ileum | NFIA | -0,7956 | 3,999E-42 |
| small intestine - terminal ileum | HLTF | -0,7959 | 3,518E-42 |
| small intestine - terminal ileum | NFYB | -0,7969 | 2,381E-42 |
| small intestine - terminal ileum | HOXD9 | -0,7975 | 1,821E-42 |
| cervix - ectocervix | NR2F2 | -0,8 | 0,009628 |
| cervix - ectocervix | HIC1 | -0,8 | 0,009628 |
| cervix - ectocervix | RXRG | -0,8 | 0,009628 |
| cervix - ectocervix | HINFP | -0,8 | 0,009628 |
| kidney - medulla | ZBTB7A | -0,8 | 0,2 |
| kidney - medulla | ZBTB7B | -0,8 | 0,2 |
| kidney - medulla | POU3F2 | -0,8 | 0,2 |
| kidney - medulla | MGA | -0,8 | 0,2 |
| kidney - medulla | ZNF282 | -0,8 | 0,2 |
| kidney - medulla | TFAP4 | -0,8 | 0,2 |
| kidney - medulla | ZNF384 | -0,8 | 0,2 |
| kidney - medulla | NKX6-1 | -0,8 | 0,2 |
| kidney - medulla | NFYB | -0,8 | 0,2 |
| kidney - medulla | CREB3 | -0,8 | 0,2 |
| kidney - medulla | ERG | -0,8 | 0,2 |
| kidney - medulla | TEAD2 | -0,8 | 0,2 |
| kidney - medulla | TEAD1 | -0,8 | 0,2 |
| kidney - medulla | NFATC1 | -0,8 | 0,2 |
| kidney - medulla | NFATC2 | -0,8 | 0,2 |
| kidney - medulla | CTCF | -0,8 | 0,2 |
| kidney - medulla | POU6F1 | -0,8 | 0,2 |
| kidney - medulla | REST | -0,8 | 0,2 |
| kidney - medulla | RARB | -0,8 | 0,2 |
| kidney - medulla | KLF5 | -0,8 | 0,2 |
| kidney - medulla | SMAD2 | -0,8 | 0,2 |
| kidney - medulla | RFX5 | -0,8 | 0,2 |
| kidney - medulla | ZBTB33 | -0,8 | 0,2 |
| kidney - medulla | HSF1 | -0,8 | 0,2 |
| kidney - medulla | RORA | -0,8 | 0,2 |
| kidney - medulla | TCF21 | -0,8 | 0,2 |
| kidney - medulla | MXI1 | -0,8 | 0,2 |
| kidney - medulla | PRRX1 | -0,8 | 0,2 |
| kidney - medulla | NPAS2 | -0,8 | 0,2 |
| kidney - medulla | NFAT5 | -0,8 | 0,2 |
| kidney - medulla | FOXO6 | -0,8 | 0,2 |
| kidney - medulla | LIN54 | -0,8 | 0,2 |
| kidney - medulla | EHF | -0,8 | 0,2 |
| kidney - medulla | HMBOX1 | -0,8 | 0,2 |
| kidney - medulla | TCF12 | -0,8 | 0,2 |
| kidney - medulla | HOXD11 | -0,8 | 0,2 |
| kidney - medulla | NEUROD2 | -0,8 | 0,2 |
| kidney - medulla | TCF3 | -0,8 | 0,2 |
| kidney - medulla | SPDEF | -0,8 | 0,2 |
| kidney - medulla | NR2F6 | -0,8 | 0,2 |
| kidney - medulla | NR2F1 | -0,8 | 0,2 |
| kidney - medulla | TWIST2 | -0,8 | 0,2 |
| kidney - medulla | SNAI2 | -0,8 | 0,2 |
| kidney - medulla | HES1 | -0,8 | 0,2 |

| | | | |
|---|---|---|---|
| kidney - medulla | HIC2 | -0,8 | 0,2 |
| kidney - medulla | SP3 | -0,8 | 0,2 |
| kidney - medulla | SP4 | -0,8 | 0,2 |
| kidney - medulla | PBX3 | -0,8 | 0,2 |
| kidney - medulla | SMAD4 | -0,8 | 0,2 |
| kidney - medulla | FOXJ2 | -0,8 | 0,2 |
| kidney - medulla | FOXJ3 | -0,8 | 0,2 |
| kidney - medulla | TCFL5 | -0,8 | 0,2 |
| kidney - medulla | YY1 | -0,8 | 0,2 |
| kidney - medulla | YY2 | -0,8 | 0,2 |
| kidney - medulla | DLX4 | -0,8 | 0,2 |
| kidney - medulla | ZSCAN4 | -0,8 | 0,2 |
| kidney - medulla | GABPA | -0,8 | 0,2 |
| kidney - medulla | GLIS3 | -0,8 | 0,2 |
| kidney - medulla | SREBF2 | -0,8 | 0,2 |
| kidney - medulla | ELK1 | -0,8 | 0,2 |
| kidney - medulla | JDP2 | -0,8 | 0,2 |
| kidney - medulla | ATF4 | -0,8 | 0,2 |
| kidney - medulla | ATF7 | -0,8 | 0,2 |
| kidney - medulla | ATF1 | -0,8 | 0,2 |
| kidney - medulla | HINFP | -0,8 | 0,2 |
| kidney - medulla | REL | -0,8 | 0,2 |
| kidney - medulla | NR2C2 | -0,8 | 0,2 |
| kidney - medulla | FLI1 | -0,8 | 0,2 |
| kidney - medulla | ESRRA | -0,8 | 0,2 |
| kidney - medulla | TAL1 | -0,8 | 0,2 |
| kidney - medulla | MEF2C | -0,8 | 0,2 |
| kidney - medulla | MEF2A | -0,8 | 0,2 |
| kidney - medulla | MEF2D | -0,8 | 0,2 |
| kidney - medulla | TCF4 | -0,8 | 0,2 |
| kidney - medulla | USF1 | -0,8 | 0,2 |
| kidney - medulla | TCF7L1 | -0,8 | 0,2 |
| kidney - medulla | TCF7L2 | -0,8 | 0,2 |
| kidney - medulla | SOX15 | -0,8 | 0,2 |
| kidney - medulla | SOX17 | -0,8 | 0,2 |
| kidney - medulla | FOXA1 | -0,8 | 0,2 |
| kidney - medulla | FOXA2 | -0,8 | 0,2 |
| kidney - medulla | ZEB1 | -0,8 | 0,2 |
| kidney - medulla | THAP1 | -0,8 | 0,2 |
| kidney - medulla | TBP | -0,8 | 0,2 |
| kidney - medulla | NR1H2 | -0,8 | 0,2 |
| kidney - medulla | ARID3B | -0,8 | 0,2 |
| kidney - medulla | NR3C1 | -0,8 | 0,2 |
| kidney - medulla | SOX8 | -0,8 | 0,2 |
| kidney - medulla | SOX5 | -0,8 | 0,2 |
| kidney - medulla | ELF1 | -0,8 | 0,2 |
| kidney - medulla | KLF13 | -0,8 | 0,2 |
| kidney - medulla | KLF12 | -0,8 | 0,2 |
| kidney - medulla | KLF16 | -0,8 | 0,2 |
| kidney - medulla | E2F6 | -0,8 | 0,2 |
| kidney - medulla | E2F4 | -0,8 | 0,2 |
| kidney - medulla | FOXD1 | -0,8 | 0,2 |
| kidney - medulla | MAX | -0,8 | 0,2 |
| kidney - medulla | IRF7 | -0,8 | 0,2 |

| | | | |
|---|---|---|---|
| kidney - medulla | MZF1 | -0,8 | 0,2 |
| kidney - medulla | HOXC9 | -0,8 | 0,2 |
| kidney - medulla | ARID5A | -0,8 | 0,2 |
| kidney - medulla | GLI2 | -0,8 | 0,2 |
| kidney - medulla | FOXK2 | -0,8 | 0,2 |
| kidney - medulla | POU2F1 | -0,8 | 0,2 |
| kidney - medulla | ZNF354C | -0,8 | 0,2 |
| kidney - medulla | RXRA | -0,8 | 0,2 |
| kidney - medulla | CLOCK | -0,8 | 0,2 |
| kidney - medulla | FOXL1 | -0,8 | 0,2 |
| kidney - medulla | ZNF423 | -0,8 | 0,2 |
| kidney - medulla | MLX | -0,8 | 0,2 |
| kidney - medulla | STAT6 | -0,8 | 0,2 |
| kidney - medulla | ALX1 | -0,8 | 0,2 |
| kidney - medulla | NFIC | -0,8 | 0,2 |
| kidney - medulla | NFIA | -0,8 | 0,2 |
| kidney - medulla | NFIX | -0,8 | 0,2 |
| small intestine - terminal ileum | MEIS1 | -0,802 | 2,946E-43 |
| colon - transverse | PKNOX2 | -0,8054 | 8,687E-94 |
| colon - transverse | POU6F1 | -0,8107 | 5,949E-96 |
| small intestine - terminal ileum | ZEB1 | -0,8158 | 7,277E-46 |
| small intestine - terminal ileum | ZNF423 | -0,816 | 6,509E-46 |
| cervix - ectocervix | NFE2 | -0,8167 | 0,007225 |
| cervix - ectocervix | TWIST2 | -0,8167 | 0,007225 |
| cervix - ectocervix | NRF1 | -0,8167 | 0,007225 |
| cervix - ectocervix | HEY2 | -0,8167 | 0,007225 |
| small intestine - terminal ileum | TCFL5 | -0,8185 | 2,125E-46 |
| small intestine - terminal ileum | HOXB2 | -0,8195 | 1,348E-46 |
| small intestine - terminal ileum | TCF7L1 | -0,8274 | 3,078E-48 |
| small intestine - terminal ileum | PKNOX2 | -0,8282 | 2,113E-48 |
| small intestine - terminal ileum | GLIS2 | -0,8305 | 6,828E-49 |
| small intestine - terminal ileum | PBX3 | -0,8315 | 4,241E-49 |
| small intestine - terminal ileum | ELK3 | -0,8323 | 2,786E-49 |
| cervix - ectocervix | HSF4 | -0,8333 | 0,005266 |
| cervix - ectocervix | THAP1 | -0,8333 | 0,005266 |
| small intestine - terminal ileum | TEAD2 | -0,8344 | 9,662E-50 |
| cervix - ectocervix | YY2 | -0,85 | 0,003705 |
| cervix - ectocervix | JDP2 | -0,85 | 0,003705 |
| cervix - ectocervix | TCF7L1 | -0,85 | 0,003705 |
| small intestine - terminal ileum | RARG | -0,8587 | 1,342E-55 |
| cervix - ectocervix | NR2F1 | -0,8667 | 0,002495 |
| cervix - ectocervix | TFEB | -0,8667 | 0,002495 |
| cervix - ectocervix | RXRB | -0,8833 | 0,001591 |
| cervix - ectocervix | HOXC9 | -0,8833 | 0,001591 |
| cervix - ectocervix | MEOX2 | -0,9 | 0,0009431 |
| cervix - ectocervix | CREB3L1 | -0,9167 | 0,0005066 |
| cervix - ectocervix | MAF | -0,9167 | 0,0005066 |
| kidney - medulla | PHOX2A | -0,9487 | 0,05132 |
| kidney - medulla | EVX2 | -0,9487 | 0,05132 |
| kidney - medulla | NEUROD1 | -0,9487 | 0,05132 |
| kidney - medulla | OLIG3 | -0,9487 | 0,05132 |
| kidney - medulla | INSM1 | -0,9487 | 0,05132 |
| kidney - medulla | SRY | -0,9487 | 0,05132 |
| kidney - medulla | TBX5 | -0,9487 | 0,05132 |

| | | | |
|---|---|---|---|
| kidney - medulla | LHX4 | -1 | 0 |
| kidney - medulla | DLX6 | -1 | 0 |
| kidney - medulla | TBX2 | -1 | 0 |
| kidney - medulla | NRL | -1 | 0 |
| kidney - medulla | CUX1 | -1 | 0 |
| kidney - medulla | ONECUT1 | -1 | 0 |
| kidney - medulla | RORC | -1 | 0 |
| kidney - medulla | PITX3 | -1 | 0 |
| kidney - medulla | PITX1 | -1 | 0 |
| kidney - medulla | TFDP1 | -1 | 0 |
| kidney - medulla | BCL6B | -1 | 0 |
| kidney - medulla | CENPB | -1 | 0 |
| kidney - medulla | NFATC3 | -1 | 0 |
| kidney - medulla | FOXC2 | -1 | 0 |
| kidney - medulla | HOXC10 | -1 | 0 |
