## Supplementary table 5 lung for "Bioinformatic characterization of angiotensin-converting enzyme 2, the entry receptor for SARS-CoV-2"

| target_gene | correlated_gene | correlation | pval |
|---|---|---|---|
| ACE2 | ACE2 | 1 | 0 |
| ACE2 | PARM1 | 0,6275 | 1,31969E-64 |
| ACE2 | CNR1 | 0,6055 | 4,21393E-59 |
| ACE2 | NUDT16 | 0,5918 | 6,52256E-56 |
| ACE2 | DUOX1 | 0,5817 | 1,23385E-53 |
| ACE2 | DUOXA1 | 0,5739 | 6,20472E-52 |
| ACE2 | FMN1 | 0,571 | 2,5348E-51 |
| ACE2 | TMEM164 | 0,5692 | 6,05358E-51 |
| ACE2 | HSD17B4 | 0,5673 | 1,53876E-50 |
| ACE2 | MGST1 | 0,5631 | 1,15733E-49 |
| ACE2 | SHMT1 | 0,5594 | 6,76984E-49 |
| ACE2 | C9orf152 | 0,5582 | 1,19896E-48 |
| ACE2 | PGC | 0,5573 | 1,79634E-48 |
| ACE2 | TIMP4 | 0,5567 | 2,37306E-48 |
| ACE2 | RHOBTB2 | 0,5567 | 2,35674E-48 |
| ACE2 | MTUS1 | 0,5566 | 2,53496E-48 |
| ACE2 | GPHN | 0,5546 | 6,2203E-48 |
| ACE2 | SFTPB | 0,5537 | 9,42419E-48 |
| ACE2 | LMO3 | 0,5536 | 1,0025E-47 |
| ACE2 | SNX25 | 0,5533 | 1,15778E-47 |
| ACE2 | CD163 | 0,5516 | 2,49839E-47 |
| ACE2 | RORB | 0,551 | 3,28916E-47 |
| ACE2 | KCNJ11 | 0,5492 | 7,55408E-47 |
| ACE2 | ACACA | 0,5484 | 1,10053E-46 |
| ACE2 | TACC2 | 0,5464 | 2,6588E-46 |
| ACE2 | TMEM97 | 0,5441 | 7,7012E-46 |
| ACE2 | SDR16C5 | 0,543 | 1,24108E-45 |
| ACE2 | TFCP2L1 | 0,5428 | 1,33266E-45 |
| ACE2 | ALOX15B | 0,5419 | 2,03033E-45 |
| ACE2 | NUDT16P1 | 0,5398 | 5,01874E-45 |
| ACE2 | MCCC2 | 0,537 | 1,7071E-44 |
| ACE2 | HRASLS5 | 0,5347 | 4,70183E-44 |
| ACE2 | GUCY2D | 0,5344 | 5,40303E-44 |
| ACE2 | AKAP1 | 0,5341 | 6,04683E-44 |
| ACE2 | PLA2G4F | 0,534 | 6,27183E-44 |
| ACE2 | RP1-193H18.3 | 0,5336 | 7,49379E-44 |
| ACE2 | RPH3AL | 0,5334 | 8,30493E-44 |
| ACE2 | CPM | 0,5323 | 1,34329E-43 |
| ACE2 | SYBU | 0,5321 | 1,46439E-43 |
| ACE2 | SCAMP2 | 0,5317 | 1,69879E-43 |
| ACE2 | SERPINF2 | 0,531 | 2,34445E-43 |
| ACE2 | TMEM87A | 0,5308 | 2,54791E-43 |
| ACE2 | FMO5 | 0,5298 | 3,85358E-43 |
| ACE2 | PRR15L | 0,5295 | 4,33297E-43 |
| ACE2 | MAP7 | 0,5294 | 4,55736E-43 |
| ACE2 | MAOA | 0,5291 | 5,19594E-43 |
| ACE2 | ALPL | 0,5287 | 6,24149E-43 |
| ACE2 | CCDC141 | 0,5287 | 6,05019E-43 |
| ACE2 | MCCC1 | 0,5279 | 8,68858E-43 |
| ACE2 | PCDHAC2 | 0,5279 | 8,82258E-43 |
| ACE2 | RPS6KA2 | 0,527 | 1,27212E-42 |
| ACE2 | TM7SF2 | 0,526 | 1,97386E-42 |
| ACE2 | FHDC1 | 0,5255 | 2,41723E-42 |

| | | | |
|---|---|---|---|
| ACE2 | FASN | 0,5245 | 3,59517E-42 |
| ACE2 | TTC39A | 0,5238 | 4,90775E-42 |
| ACE2 | SMPDL3B | 0,5236 | 5,31204E-42 |
| ACE2 | RP11-352D13.6 | 0,5236 | 5,34564E-42 |
| ACE2 | RP3-340B19.3 | 0,5224 | 8,71886E-42 |
| ACE2 | C1orf210 | 0,5215 | 1,26572E-41 |
| ACE2 | PIP5K1B | 0,5214 | 1,32017E-41 |
| ACE2 | CTD-2651B20.1 | 0,5214 | 1,34437E-41 |
| ACE2 | CTSH | 0,5214 | 1,30749E-41 |
| ACE2 | LRRK2 | 0,5212 | 1,40947E-41 |
| ACE2 | PTPN13 | 0,5206 | 1,84261E-41 |
| ACE2 | ASL | 0,5204 | 1,98887E-41 |
| ACE2 | ECHDC3 | 0,5203 | 2,09897E-41 |
| ACE2 | SIAE | 0,5202 | 2,18008E-41 |
| ACE2 | ZNF385B | 0,5195 | 2,88973E-41 |
| ACE2 | SPTSSA | 0,5192 | 3,31615E-41 |
| ACE2 | DCXR | 0,5191 | 3,47308E-41 |
| ACE2 | TMEM53 | 0,5172 | 7,46548E-41 |
| ACE2 | AC090616.2 | 0,5167 | 9,27866E-41 |
| ACE2 | ABCD3 | 0,5166 | 9,68953E-41 |
| ACE2 | PCDHAC1 | 0,5156 | 1,42514E-40 |
| ACE2 | SCNN1A | 0,5151 | 1,7314E-40 |
| ACE2 | TLCD1 | 0,5149 | 1,86758E-40 |
| ACE2 | DHCR24 | 0,5145 | 2,20196E-40 |
| ACE2 | ERLEC1 | 0,5145 | 2,23072E-40 |
| ACE2 | C14orf1 | 0,5127 | 4,68483E-40 |
| ACE2 | SLC34A2 | 0,5121 | 5,82373E-40 |
| ACE2 | CPB2 | 0,512 | 6,13715E-40 |
| ACE2 | LANCL1-AS1 | 0,5115 | 7,45762E-40 |
| ACE2 | NAGS | 0,5113 | 7,95013E-40 |
| ACE2 | SLCO4C1 | 0,5111 | 8,7685E-40 |
| ACE2 | PRODH | 0,5108 | 9,90008E-40 |
| ACE2 | PCSK2 | 0,5103 | 1,18676E-39 |
| ACE2 | GSTA4 | 0,5101 | 1,3212E-39 |
| ACE2 | PON3 | 0,5101 | 1,2977E-39 |
| ACE2 | FUT3 | 0,5101 | 1,30129E-39 |
| ACE2 | NAPSA | 0,5086 | 2,38914E-39 |
| ACE2 | SNX30 | 0,5084 | 2,57513E-39 |
| ACE2 | SLC7A8 | 0,5072 | 4,05778E-39 |
| ACE2 | CPT2 | 0,5071 | 4,23175E-39 |
| ACE2 | CCT8P1 | 0,507 | 4,40311E-39 |
| ACE2 | ABCA3 | 0,507 | 4,48686E-39 |
| ACE2 | ACADL | 0,5069 | 4,72196E-39 |
| ACE2 | TMEM192 | 0,5067 | 5,0864E-39 |
| ACE2 | PCYT2 | 0,5066 | 5,16513E-39 |
| ACE2 | PCCB | 0,506 | 6,55894E-39 |
| ACE2 | EPDR1 | 0,5059 | 6,94704E-39 |
| ACE2 | LDHB | 0,5058 | 7,1198E-39 |
| ACE2 | SEZ6L2 | 0,5043 | 1,27478E-38 |
| ACE2 | PEBP4 | 0,5037 | 1,66025E-38 |
| ACE2 | POLR3H | 0,5033 | 1,92098E-38 |
| ACE2 | RMST | 0,5032 | 1,98743E-38 |
| ACE2 | GDE1 | 0,5025 | 2,65853E-38 |
| ACE2 | MUT | 0,5009 | 4,82406E-38 |

| | | | |
|---|---|---|---|
| ACE2 | USP13 | 0,5008 | 5,04794E-38 |
| ACE2 | ASAH1 | 0,5005 | 5,71592E-38 |
| ACE2 | TMEM105 | 0,5005 | 5,70865E-38 |
| ACE2 | MACROD2 | 0,5003 | 6,18929E-38 |
| ACE2 | MUC1 | 0,4996 | 8,01137E-38 |
| ACE2 | MLPH | 0,4995 | 8,2415E-38 |
| ACE2 | WFDC12 | 0,4994 | 8,82972E-38 |
| ACE2 | C2 | 0,4988 | 1,0849E-37 |
| ACE2 | SFTPD | 0,4982 | 1,36906E-37 |
| ACE2 | TMEM92 | 0,498 | 1,50515E-37 |
| ACE2 | TMEM41A | 0,4978 | 1,61191E-37 |
| ACE2 | ARMC9 | 0,4973 | 1,94705E-37 |
| ACE2 | RP11-1099M24.9 | 0,4973 | 1,91745E-37 |
| ACE2 | CDKL2 | 0,4971 | 2,1182E-37 |
| ACE2 | KCNS1 | 0,4971 | 2,0767E-37 |
| ACE2 | BEX2 | 0,4969 | 2,23361E-37 |
| ACE2 | PFKFB2 | 0,4964 | 2,766E-37 |
| ACE2 | ZBTB42 | 0,4962 | 2,97616E-37 |
| ACE2 | ZMYND8 | 0,4961 | 3,09124E-37 |
| ACE2 | SAMHD1 | 0,496 | 3,16068E-37 |
| ACE2 | NIM1K | 0,4958 | 3,43357E-37 |
| ACE2 | LSS | 0,4958 | 3,50747E-37 |
| ACE2 | RPS29P11 | 0,4955 | 3,83139E-37 |
| ACE2 | EBP | 0,4952 | 4,40814E-37 |
| ACE2 | OLAH | 0,495 | 4,68945E-37 |
| ACE2 | CHP1 | 0,4945 | 5,56338E-37 |
| ACE2 | POLR2C | 0,4945 | 5,74655E-37 |
| ACE2 | PAK6 | 0,4944 | 5,76263E-37 |
| ACE2 | FITM2 | 0,4941 | 6,65499E-37 |
| ACE2 | PTPRU | 0,4934 | 8,48891E-37 |
| ACE2 | RETSAT | 0,4933 | 8,74722E-37 |
| ACE2 | C6orf223 | 0,4933 | 8,87203E-37 |
| ACE2 | KIAA1324L | 0,493 | 9,80202E-37 |
| ACE2 | MBIP | 0,4929 | 1,02521E-36 |
| ACE2 | AP1M2 | 0,4928 | 1,07078E-36 |
| ACE2 | EGFR | 0,4924 | 1,24497E-36 |
| ACE2 | LRRC8E | 0,4924 | 1,25126E-36 |
| ACE2 | RP11-184M15.1 | 0,4921 | 1,37774E-36 |
| ACE2 | KDF1 | 0,4916 | 1,68965E-36 |
| ACE2 | SHC3 | 0,4913 | 1,87671E-36 |
| ACE2 | ACSM3 | 0,4912 | 1,92669E-36 |
| ACE2 | TCN2 | 0,4909 | 2,16519E-36 |
| ACE2 | CACNA2D2 | 0,4908 | 2,24475E-36 |
| ACE2 | SLPI | 0,4907 | 2,33564E-36 |
| ACE2 | SLC22A3 | 0,4901 | 2,9806E-36 |
| ACE2 | RP5-1132H15.3 | 0,4901 | 2,95663E-36 |
| ACE2 | B3GNT8 | 0,4901 | 2,99058E-36 |
| ACE2 | LGI3 | 0,49 | 3,0687E-36 |
| ACE2 | KRTCAP3 | 0,4898 | 3,31308E-36 |
| ACE2 | FREM2 | 0,4892 | 4,11613E-36 |
| ACE2 | RP11-630A13.4 | 0,4888 | 4,83708E-36 |
| ACE2 | ILDR1 | 0,4881 | 6,1092E-36 |
| ACE2 | AACS | 0,4877 | 7,305E-36 |
| ACE2 | RP11-352D13.5 | 0,4875 | 7,87675E-36 |

| | | | |
|---|---|---|---|
| ACE2 | NCMAP | 0,4874 | 7,9844E-36 |
| ACE2 | ANK3 | 0,4872 | 8,52845E-36 |
| ACE2 | HOPX | 0,4871 | 8,95046E-36 |
| ACE2 | MYO5C | 0,487 | 9,3091E-36 |
| ACE2 | MRPS23 | 0,4868 | 1,00597E-35 |
| ACE2 | KLF13 | 0,4856 | 1,56141E-35 |
| ACE2 | C19orf54 | 0,485 | 1,93426E-35 |
| ACE2 | AHCYL2 | 0,4848 | 2,08574E-35 |
| ACE2 | GKN2 | 0,4847 | 2,15817E-35 |
| ACE2 | PNMT | 0,4844 | 2,40199E-35 |
| ACE2 | CNIH4 | 0,4839 | 2,90104E-35 |
| ACE2 | IVD | 0,4839 | 2,97673E-35 |
| ACE2 | FZD5 | 0,4838 | 3,06194E-35 |
| ACE2 | NSDHL | 0,4838 | 3,05317E-35 |
| ACE2 | TMEM163 | 0,4834 | 3,47449E-35 |
| ACE2 | AZGP1 | 0,4834 | 3,57312E-35 |
| ACE2 | LACTB2 | 0,4831 | 3,96969E-35 |
| ACE2 | HPN | 0,483 | 4,04294E-35 |
| ACE2 | SFTA2 | 0,4829 | 4,2144E-35 |
| ACE2 | ANKS6 | 0,4826 | 4,74954E-35 |
| ACE2 | DDX28 | 0,4825 | 4,80481E-35 |
| ACE2 | TST | 0,4822 | 5,40369E-35 |
| ACE2 | FAM83H | 0,4816 | 6,7479E-35 |
| ACE2 | LAMP3 | 0,4814 | 7,36974E-35 |
| ACE2 | ROS1 | 0,4814 | 7,20659E-35 |
| ACE2 | PCSK9 | 0,4813 | 7,68288E-35 |
| ACE2 | BCAT2 | 0,4813 | 7,56328E-35 |
| ACE2 | KCTD14 | 0,4812 | 7,94561E-35 |
| ACE2 | CTB-55O6.4 | 0,481 | 8,3127E-35 |
| ACE2 | ZRANB2-AS1 | 0,4809 | 8,65521E-35 |
| ACE2 | BMP8B | 0,4808 | 9,0926E-35 |
| ACE2 | SYNE4 | 0,4805 | 1,01579E-34 |
| ACE2 | MALL | 0,4804 | 1,04029E-34 |
| ACE2 | ENTPD3 | 0,48 | 1,21424E-34 |
| ACE2 | RP11-76C10.6 | 0,4796 | 1,40252E-34 |
| ACE2 | MMP24 | 0,4796 | 1,40762E-34 |
| ACE2 | ARSE | 0,4789 | 1,78648E-34 |
| ACE2 | EMC3 | 0,4785 | 2,07438E-34 |
| ACE2 | DPCR1 | 0,4783 | 2,23433E-34 |
| ACE2 | RASEF | 0,4783 | 2,22923E-34 |
| ACE2 | SFXN2 | 0,478 | 2,46268E-34 |
| ACE2 | PRTG | 0,4778 | 2,66254E-34 |
| ACE2 | GPR39 | 0,4776 | 2,85881E-34 |
| ACE2 | GOLT1A | 0,4772 | 3,33439E-34 |
| ACE2 | PRKAB1 | 0,4772 | 3,28397E-34 |
| ACE2 | SLC29A2 | 0,4771 | 3,39317E-34 |
| ACE2 | RP4-568C11.4 | 0,4771 | 3,37026E-34 |
| ACE2 | AC013463.2 | 0,4765 | 4,27689E-34 |
| ACE2 | PRRG2 | 0,4764 | 4,39658E-34 |
| ACE2 | P3H2 | 0,4763 | 4,60755E-34 |
| ACE2 | FAM46B | 0,4762 | 4,73466E-34 |
| ACE2 | WNT3 | 0,4759 | 5,20616E-34 |
| ACE2 | BCKDHA | 0,4759 | 5,35344E-34 |
| ACE2 | SPSB2 | 0,4758 | 5,54028E-34 |

| | | | |
|---|---|---|---|
| ACE2 | AIFM1 | 0,4748 | 7,74643E-34 |
| ACE2 | ST3GAL5 | 0,4745 | 8,75454E-34 |
| ACE2 | SOWAHB | 0,474 | 1,03003E-33 |
| ACE2 | GCAT | 0,4739 | 1,07059E-33 |
| ACE2 | HHIP-AS1 | 0,4738 | 1,09594E-33 |
| ACE2 | EHF | 0,4725 | 1,74851E-33 |
| ACE2 | TFB2M | 0,4724 | 1,84046E-33 |
| ACE2 | DHTKD1 | 0,4724 | 1,78811E-33 |
| ACE2 | IRX6 | 0,4723 | 1,85322E-33 |
| ACE2 | RAP1GAP | 0,4719 | 2,17836E-33 |
| ACE2 | CLPTM1L | 0,4719 | 2,13775E-33 |
| ACE2 | CAMSAP3 | 0,4718 | 2,25435E-33 |
| ACE2 | CTD-2008P7.9 | 0,4717 | 2,36567E-33 |
| ACE2 | BDH1 | 0,4716 | 2,3984E-33 |
| ACE2 | RP11-424M22.3 | 0,4716 | 2,37299E-33 |
| ACE2 | C5 | 0,4715 | 2,50202E-33 |
| ACE2 | PLA2G3 | 0,4712 | 2,75994E-33 |
| ACE2 | LINC01671 | 0,4711 | 2,90407E-33 |
| ACE2 | RP11-476D10.1 | 0,471 | 2,95254E-33 |
| ACE2 | MERTK | 0,4709 | 3,11251E-33 |
| ACE2 | KLF15 | 0,4709 | 3,08863E-33 |
| ACE2 | RP11-297A16.4 | 0,4708 | 3,23919E-33 |
| ACE2 | RP11-95I16.2 | 0,4708 | 3,19E-33 |
| ACE2 | GGT5 | 0,4702 | 3,99963E-33 |
| ACE2 | CDH1 | 0,47 | 4,19267E-33 |
| ACE2 | MET | 0,4699 | 4,36918E-33 |
| ACE2 | PLS1 | 0,4696 | 4,81308E-33 |
| ACE2 | RP11-87E22.2 | 0,4696 | 4,8559E-33 |
| ACE2 | TBC1D14 | 0,4693 | 5,34727E-33 |
| ACE2 | FCHO2 | 0,4691 | 5,78117E-33 |
| ACE2 | SLC25A4 | 0,469 | 6,00587E-33 |
| ACE2 | PCLO | 0,469 | 6,03514E-33 |
| ACE2 | SDCCAG8 | 0,4687 | 6,72152E-33 |
| ACE2 | LGALSL | 0,4685 | 7,25014E-33 |
| ACE2 | SNTB1 | 0,4683 | 7,71697E-33 |
| ACE2 | ACSS2 | 0,468 | 8,42939E-33 |
| ACE2 | HKDC1 | 0,4678 | 9,00318E-33 |
| ACE2 | SLC46A2 | 0,4669 | 1,22801E-32 |
| ACE2 | PDP2 | 0,4669 | 1,23922E-32 |
| ACE2 | SEC23B | 0,4664 | 1,4693E-32 |
| ACE2 | SC5D | 0,4661 | 1,62184E-32 |
| ACE2 | LPCAT1 | 0,466 | 1,70505E-32 |
| ACE2 | PARP4 | 0,4659 | 1,76279E-32 |
| ACE2 | HOOK1 | 0,4658 | 1,80823E-32 |
| ACE2 | ICA1 | 0,4658 | 1,84602E-32 |
| ACE2 | GLB1 | 0,4656 | 1,91646E-32 |
| ACE2 | CARMIL1 | 0,4656 | 1,93718E-32 |
| ACE2 | GALNT10 | 0,4653 | 2,17383E-32 |
| ACE2 | ALDH6A1 | 0,4642 | 3,13746E-32 |
| ACE2 | C18orf63 | 0,4639 | 3,4821E-32 |
| ACE2 | MLXIPL | 0,4636 | 3,90718E-32 |
| ACE2 | IRS2 | 0,4634 | 4,09943E-32 |
| ACE2 | LL22NC03-N95F10.1 | 0,4633 | 4,30509E-32 |
| ACE2 | FDPS | 0,4631 | 4,51127E-32 |

| | | | |
|---|---|---|---|
| ACE2 | UBLCP1 | 0,463 | 4,75449E-32 |
| ACE2 | GSTO2 | 0,4628 | 5,06852E-32 |
| ACE2 | ATP6V0E1 | 0,4624 | 5,78773E-32 |
| ACE2 | DSG2 | 0,4623 | 6,07899E-32 |
| ACE2 | SORT1 | 0,462 | 6,65694E-32 |
| ACE2 | MID1IP1 | 0,4618 | 7,22142E-32 |
| ACE2 | AC079630.2 | 0,4617 | 7,44417E-32 |
| ACE2 | MFSD2A | 0,4616 | 7,53583E-32 |
| ACE2 | FAM84B | 0,4616 | 7,726E-32 |
| ACE2 | RP11-22C11.2 | 0,4612 | 8,63239E-32 |
| ACE2 | FA2H | 0,461 | 9,37394E-32 |
| ACE2 | ATP1B1 | 0,4609 | 9,69226E-32 |
| ACE2 | RP11-27G14.4 | 0,4605 | 1,11287E-31 |
| ACE2 | NUDT5 | 0,4602 | 1,24023E-31 |
| ACE2 | ARMT1 | 0,4598 | 1,39768E-31 |
| ACE2 | AC093627.10 | 0,4591 | 1,79742E-31 |
| ACE2 | SCD | 0,4586 | 2,12109E-31 |
| ACE2 | TPD52 | 0,4585 | 2,15443E-31 |
| ACE2 | DERA | 0,4585 | 2,16795E-31 |
| ACE2 | USP44 | 0,4585 | 2,21619E-31 |
| ACE2 | TMEM56 | 0,4584 | 2,28448E-31 |
| ACE2 | SFTA3 | 0,4584 | 2,28438E-31 |
| ACE2 | FGFR2 | 0,4582 | 2,42581E-31 |
| ACE2 | CA12 | 0,458 | 2,55445E-31 |
| ACE2 | FAM105A | 0,4579 | 2,65874E-31 |
| ACE2 | RAB40C | 0,4576 | 2,97951E-31 |
| ACE2 | PANK1 | 0,4573 | 3,26907E-31 |
| ACE2 | MMAB | 0,457 | 3,58846E-31 |
| ACE2 | GRHL2 | 0,4565 | 4,29611E-31 |
| ACE2 | ESRP1 | 0,4564 | 4,47004E-31 |
| ACE2 | CYP51A1 | 0,4563 | 4,49724E-31 |
| ACE2 | MTRR | 0,4562 | 4,79049E-31 |
| ACE2 | IDH1 | 0,456 | 5,00375E-31 |
| ACE2 | PNKD | 0,456 | 5,04892E-31 |
| ACE2 | FKBP5 | 0,4558 | 5,38059E-31 |
| ACE2 | MARC1 | 0,4557 | 5,63862E-31 |
| ACE2 | HMGCR | 0,4556 | 5,74906E-31 |
| ACE2 | PLA2G10 | 0,4556 | 5,76734E-31 |
| ACE2 | FASTKD5 | 0,4556 | 5,81818E-31 |
| ACE2 | MARVELD3 | 0,4553 | 6,34239E-31 |
| ACE2 | PRSS16 | 0,4552 | 6,66867E-31 |
| ACE2 | FOXA2 | 0,455 | 7,0255E-31 |
| ACE2 | SULT1B1 | 0,4546 | 8,13151E-31 |
| ACE2 | LRP2 | 0,4545 | 8,19993E-31 |
| ACE2 | SFTPA2 | 0,4545 | 8,3499E-31 |
| ACE2 | ECHDC1 | 0,4542 | 9,1208E-31 |
| ACE2 | PMM1 | 0,4541 | 9,42532E-31 |
| ACE2 | MIA2 | 0,4539 | 1,00594E-30 |
| ACE2 | PITPNM3 | 0,4535 | 1,16852E-30 |
| ACE2 | PXMP4 | 0,4531 | 1,30005E-30 |
| ACE2 | AMN | 0,453 | 1,37047E-30 |
| ACE2 | CTD-2547L16.1 | 0,4529 | 1,39305E-30 |
| ACE2 | ERP29 | 0,4528 | 1,45187E-30 |
| ACE2 | ATAD3C | 0,4525 | 1,60069E-30 |

| | | | |
|---|---|---|---|
| ACE2 | SLC25A16 | 0,4524 | 1,65567E-30 |
| ACE2 | SGPP2 | 0,4522 | 1,77117E-30 |
| ACE2 | KANK1 | 0,4522 | 1,75664E-30 |
| ACE2 | NFX1 | 0,4521 | 1,84055E-30 |
| ACE2 | GSR | 0,452 | 1,87077E-30 |
| ACE2 | HECTD1 | 0,452 | 1,91846E-30 |
| ACE2 | ACAT1 | 0,4517 | 2,06331E-30 |
| ACE2 | CTD-2589M5.4 | 0,4514 | 2,32791E-30 |
| ACE2 | CNDP2 | 0,4514 | 2,31574E-30 |
| ACE2 | ANKRD2 | 0,4512 | 2,49798E-30 |
| ACE2 | NCOA4 | 0,451 | 2,60358E-30 |
| ACE2 | SLC25A18 | 0,451 | 2,64884E-30 |
| ACE2 | KRT19 | 0,4509 | 2,72931E-30 |
| ACE2 | KCNK1 | 0,4508 | 2,79035E-30 |
| ACE2 | SCN1A | 0,4506 | 3,04297E-30 |
| ACE2 | KTI12 | 0,4505 | 3,15558E-30 |
| ACE2 | NPC2 | 0,4502 | 3,41535E-30 |
| ACE2 | SLC25A38 | 0,4501 | 3,58809E-30 |
| ACE2 | FAH | 0,45 | 3,69146E-30 |
| ACE2 | RP4-569M23.2 | 0,4499 | 3,82545E-30 |
| ACE2 | TSPAN3 | 0,4495 | 4,34532E-30 |
| ACE2 | GPAM | 0,4492 | 4,69425E-30 |
| ACE2 | KBTBD4 | 0,449 | 5,05799E-30 |
| ACE2 | ACPP | 0,4485 | 6,06698E-30 |
| ACE2 | FAM118B | 0,4484 | 6,19391E-30 |
| ACE2 | CTD-2245E15.3 | 0,4481 | 6,89434E-30 |
| ACE2 | NDUFS1 | 0,448 | 7,0802E-30 |
| ACE2 | CHMP4C | 0,448 | 6,93254E-30 |
| ACE2 | SFTPA1 | 0,4477 | 7,80695E-30 |
| ACE2 | SCIN | 0,4476 | 7,91323E-30 |
| ACE2 | CRB3 | 0,4476 | 7,88795E-30 |
| ACE2 | DNAJC14 | 0,4475 | 8,37082E-30 |
| ACE2 | RD3 | 0,4471 | 9,43715E-30 |
| ACE2 | MEGF9 | 0,4469 | 1,00939E-29 |
| ACE2 | RNF43 | 0,4467 | 1,08354E-29 |
| ACE2 | RP11-76C10.5 | 0,4466 | 1,09616E-29 |
| ACE2 | RNF26 | 0,4465 | 1,14636E-29 |
| ACE2 | NRTN | 0,4465 | 1,13248E-29 |
| ACE2 | VEPH1 | 0,4462 | 1,26275E-29 |
| ACE2 | GCDH | 0,4461 | 1,30209E-29 |
| ACE2 | BLVRB | 0,446 | 1,36326E-29 |
| ACE2 | PIGR | 0,4458 | 1,41632E-29 |
| ACE2 | SLC35F6 | 0,4457 | 1,4823E-29 |
| ACE2 | RASL10B | 0,4457 | 1,49372E-29 |
| ACE2 | KLHL26 | 0,4456 | 1,5087E-29 |
| ACE2 | ESRP2 | 0,4455 | 1,5787E-29 |
| ACE2 | ATP1A1 | 0,4454 | 1,64588E-29 |
| ACE2 | BMP3 | 0,4451 | 1,81932E-29 |
| ACE2 | GSPT2 | 0,445 | 1,87386E-29 |
| ACE2 | S100A14 | 0,4449 | 1,88671E-29 |
| ACE2 | CYB5A | 0,4449 | 1,92712E-29 |
| ACE2 | KIAA0391 | 0,4447 | 2,03541E-29 |
| ACE2 | H2AFY | 0,4445 | 2,20167E-29 |
| ACE2 | DSTNP2 | 0,4445 | 2,17518E-29 |

| | | | |
|---|---|---|---|
| ACE2 | CRTAC1 | 0,4444 | 2,24419E-29 |
| ACE2 | RP11-2B6.2 | 0,4441 | 2,44986E-29 |
| ACE2 | DHCR7 | 0,4439 | 2,62E-29 |
| ACE2 | UNC50 | 0,4438 | 2,73057E-29 |
| ACE2 | THUMPD3 | 0,4437 | 2,82013E-29 |
| ACE2 | TOM1L1 | 0,4437 | 2,7876E-29 |
| ACE2 | ALDH1L1 | 0,4436 | 2,91275E-29 |
| ACE2 | PLA2G1B | 0,4436 | 2,90092E-29 |
| ACE2 | MVK | 0,4435 | 2,99133E-29 |
| ACE2 | STARD7 | 0,4434 | 3,14913E-29 |
| ACE2 | SLC39A11 | 0,4433 | 3,21067E-29 |
| ACE2 | RASGRF1 | 0,4432 | 3,32666E-29 |
| ACE2 | ERGIC1 | 0,4431 | 3,4393E-29 |
| ACE2 | RIOX1 | 0,443 | 3,55184E-29 |
| ACE2 | PDSS2 | 0,4429 | 3,69099E-29 |
| ACE2 | CTD-2376I4.1 | 0,4426 | 4,06107E-29 |
| ACE2 | HPS6 | 0,4426 | 3,96682E-29 |
| ACE2 | ST7 | 0,4425 | 4,16969E-29 |
| ACE2 | ARHGAP35 | 0,4423 | 4,45479E-29 |
| ACE2 | ADI1 | 0,4422 | 4,62238E-29 |
| ACE2 | INTS5 | 0,4422 | 4,51535E-29 |
| ACE2 | CACFD1 | 0,4421 | 4,65717E-29 |
| ACE2 | GLOD5 | 0,4421 | 4,74202E-29 |
| ACE2 | LSR | 0,442 | 4,78296E-29 |
| ACE2 | RP11-26H16.4 | 0,4418 | 5,24186E-29 |
| ACE2 | LINC01736 | 0,4417 | 5,30263E-29 |
| ACE2 | CTC-425O23.5 | 0,4416 | 5,44766E-29 |
| ACE2 | AP001469.5 | 0,4416 | 5,5476E-29 |
| ACE2 | SNX8 | 0,4414 | 5,93074E-29 |
| ACE2 | BFAR | 0,4414 | 5,88687E-29 |
| ACE2 | SAP30 | 0,4413 | 5,98504E-29 |
| ACE2 | GANC | 0,4411 | 6,45395E-29 |
| ACE2 | LIPH | 0,4408 | 7,11805E-29 |
| ACE2 | HADH | 0,4407 | 7,37099E-29 |
| ACE2 | ROR1 | 0,4401 | 8,98592E-29 |
| ACE2 | RP11-259K15.2 | 0,44 | 9,2409E-29 |
| ACE2 | ZNF750 | 0,44 | 9,01979E-29 |
| ACE2 | C3orf36 | 0,4399 | 9,31598E-29 |
| ACE2 | SRPX | 0,4397 | 1,01192E-28 |
| ACE2 | TMEM106C | 0,4394 | 1,09154E-28 |
| ACE2 | RP11-635O16.2 | 0,4394 | 1,09648E-28 |
| ACE2 | EPCAM | 0,4393 | 1,14781E-28 |
| ACE2 | WDR81 | 0,4393 | 1,12496E-28 |
| ACE2 | SQLE | 0,4392 | 1,17779E-28 |
| ACE2 | CLP1 | 0,4392 | 1,1859E-28 |
| ACE2 | LINC01612 | 0,4391 | 1,21846E-28 |
| ACE2 | MGC27382 | 0,4387 | 1,39129E-28 |
| ACE2 | CXCL17 | 0,4384 | 1,50538E-28 |
| ACE2 | MLX | 0,4383 | 1,57744E-28 |
| ACE2 | RP11-933H2.4 | 0,438 | 1,73949E-28 |
| ACE2 | TADA2B | 0,4379 | 1,79604E-28 |
| ACE2 | PTPN3 | 0,4379 | 1,78832E-28 |
| ACE2 | VSIG4 | 0,4379 | 1,76972E-28 |
| ACE2 | CAPN8 | 0,4378 | 1,83062E-28 |

| | | | |
|---|---|---|---|
| ACE2 | DARS2 | 0,4377 | 1,8872E-28 |
| ACE2 | RAB20 | 0,4371 | 2,28725E-28 |
| ACE2 | AFG3L2 | 0,437 | 2,36402E-28 |
| ACE2 | ZNF622 | 0,4367 | 2,62293E-28 |
| ACE2 | TMEM245 | 0,4367 | 2,58553E-28 |
| ACE2 | IRX3 | 0,4367 | 2,58275E-28 |
| ACE2 | AAR2 | 0,4367 | 2,54925E-28 |
| ACE2 | MRPL46 | 0,4364 | 2,85399E-28 |
| ACE2 | CDS1 | 0,4362 | 3,02665E-28 |
| ACE2 | RMDN2 | 0,4361 | 3,15609E-28 |
| ACE2 | MFSD1 | 0,4361 | 3,09508E-28 |
| ACE2 | FLT3 | 0,4355 | 3,75041E-28 |
| ACE2 | FLRT3 | 0,4354 | 3,86509E-28 |
| ACE2 | NR0B2 | 0,4353 | 4,00901E-28 |
| ACE2 | RIDA | 0,4352 | 4,16481E-28 |
| ACE2 | DCLRE1A | 0,4351 | 4,27934E-28 |
| ACE2 | GOT1 | 0,4349 | 4,4689E-28 |
| ACE2 | RPL13P5 | 0,4349 | 4,49047E-28 |
| ACE2 | DESI2 | 0,4343 | 5,54917E-28 |
| ACE2 | PCDHA12 | 0,4339 | 6,14946E-28 |
| ACE2 | ABC13-47488600E1 | 0,4338 | 6,39966E-28 |
| ACE2 | RP11-76E17.4 | 0,4338 | 6,47271E-28 |
| ACE2 | SLC22A31 | 0,4336 | 6,82988E-28 |
| ACE2 | DPP3 | 0,4335 | 6,94203E-28 |
| ACE2 | VPS18 | 0,4335 | 6,96055E-28 |
| ACE2 | ZHX3 | 0,4333 | 7,40189E-28 |
| ACE2 | RBM47 | 0,4332 | 7,63895E-28 |
| ACE2 | BCAT1 | 0,4332 | 7,66397E-28 |
| ACE2 | AC010127.3 | 0,4328 | 8,6868E-28 |
| ACE2 | RDH11 | 0,4328 | 8,7396E-28 |
| ACE2 | PLA2G12B | 0,4327 | 9,0424E-28 |
| ACE2 | PARN | 0,4325 | 9,42305E-28 |
| ACE2 | EPRS | 0,4324 | 9,72763E-28 |
| ACE2 | NIF3L1 | 0,432 | 1,1155E-27 |
| ACE2 | DHRS7B | 0,432 | 1,12933E-27 |
| ACE2 | TBXAS1 | 0,4318 | 1,17127E-27 |
| ACE2 | LINC01932 | 0,4314 | 1,35101E-27 |
| ACE2 | ALCAM | 0,4313 | 1,36827E-27 |
| ACE2 | HMGCS1 | 0,4313 | 1,37868E-27 |
| ACE2 | ABC7-42404400C24 | 0,4313 | 1,3675E-27 |
| ACE2 | HHIP | 0,4309 | 1,55061E-27 |
| ACE2 | FBXO38 | 0,4308 | 1,61415E-27 |
| ACE2 | KCNQ1 | 0,4308 | 1,63319E-27 |
| ACE2 | SLC25A24 | 0,4307 | 1,67847E-27 |
| ACE2 | SLC27A3 | 0,4305 | 1,74115E-27 |
| ACE2 | ATP6V1H | 0,4304 | 1,84245E-27 |
| ACE2 | SLC25A5 | 0,4303 | 1,89095E-27 |
| ACE2 | TM9SF2 | 0,4302 | 1,96469E-27 |
| ACE2 | SYT3 | 0,4301 | 1,99699E-27 |
| ACE2 | ZDHHC16 | 0,4299 | 2,14554E-27 |
| ACE2 | AHCY | 0,4298 | 2,18974E-27 |
| ACE2 | LRRC31 | 0,4297 | 2,22781E-27 |
| ACE2 | GHITM | 0,4295 | 2,36625E-27 |
| ACE2 | USP27X | 0,4293 | 2,5663E-27 |

| | | | |
|---|---|---|---|
| ACE2 | RP11-1260E13.3 | 0,4292 | 2,6046E-27 |
| ACE2 | AC079630.4 | 0,4291 | 2,74777E-27 |
| ACE2 | FBXO27 | 0,4289 | 2,90052E-27 |
| ACE2 | MFAP3L | 0,4286 | 3,12819E-27 |
| ACE2 | ATP6V1E1 | 0,4286 | 3,12993E-27 |
| ACE2 | TMEM27 | 0,4284 | 3,3792E-27 |
| ACE2 | DEPTOR | 0,4283 | 3,45553E-27 |
| ACE2 | TBC1D16 | 0,4281 | 3,7291E-27 |
| ACE2 | ADH5 | 0,4279 | 3,87239E-27 |
| ACE2 | ANKEF1 | 0,4279 | 3,94809E-27 |
| ACE2 | DPP4 | 0,4278 | 3,97345E-27 |
| ACE2 | RP11-89H19.2 | 0,4278 | 3,98027E-27 |
| ACE2 | SMG8 | 0,4277 | 4,2103E-27 |
| ACE2 | ZDHHC4 | 0,4276 | 4,24097E-27 |
| ACE2 | C8orf34-AS1 | 0,4275 | 4,42654E-27 |
| ACE2 | HACD2 | 0,4274 | 4,59371E-27 |
| ACE2 | FAF2 | 0,4273 | 4,68512E-27 |
| ACE2 | ACTR3C | 0,4271 | 5,05301E-27 |
| ACE2 | KCNQ3 | 0,4269 | 5,23586E-27 |
| ACE2 | SECISBP2L | 0,4269 | 5,29318E-27 |
| ACE2 | SLC6A14 | 0,4268 | 5,53109E-27 |
| ACE2 | C1orf53 | 0,4267 | 5,62713E-27 |
| ACE2 | RP11-259K5.2 | 0,4267 | 5,6551E-27 |
| ACE2 | GPR160 | 0,4267 | 5,63548E-27 |
| ACE2 | ACLY | 0,4267 | 5,65932E-27 |
| ACE2 | VWA2 | 0,426 | 7,04908E-27 |
| ACE2 | AGPAT2 | 0,4259 | 7,1321E-27 |
| ACE2 | ACAT2 | 0,4258 | 7,30837E-27 |
| ACE2 | ATP13A4 | 0,4257 | 7,666E-27 |
| ACE2 | C2-AS1 | 0,4252 | 8,76972E-27 |
| ACE2 | C11orf1 | 0,4251 | 9,15462E-27 |
| ACE2 | RP11-867G23.8 | 0,425 | 9,53423E-27 |
| ACE2 | RP11-14I2.2 | 0,4249 | 9,76386E-27 |
| ACE2 | SLC31A2 | 0,4242 | 1,19141E-26 |
| ACE2 | MANEAL | 0,424 | 1,26209E-26 |
| ACE2 | FAM103A1 | 0,424 | 1,25935E-26 |
| ACE2 | TMEM268 | 0,4238 | 1,33478E-26 |
| ACE2 | DNASE1L1 | 0,4237 | 1,40522E-26 |
| ACE2 | COA7 | 0,4235 | 1,47104E-26 |
| ACE2 | MIRLET7D | 0,4233 | 1,56816E-26 |
| ACE2 | KLHDC7A | 0,4232 | 1,61708E-26 |
| ACE2 | RP11-532F12.5 | 0,4232 | 1,6221E-26 |
| ACE2 | BSPRY | 0,4229 | 1,78062E-26 |
| ACE2 | PHYHD1 | 0,4226 | 1,91628E-26 |
| ACE2 | RP11-172H24.4 | 0,4226 | 1,94148E-26 |
| ACE2 | PCK2 | 0,4224 | 2,05277E-26 |
| ACE2 | TTPAL | 0,4223 | 2,10825E-26 |
| ACE2 | EPS8 | 0,4222 | 2,15516E-26 |
| ACE2 | ZDHHC12 | 0,4221 | 2,26341E-26 |
| ACE2 | PLEKHA7 | 0,422 | 2,29285E-26 |
| ACE2 | CTAGE5 | 0,422 | 2,30865E-26 |
| ACE2 | PCDH20 | 0,4219 | 2,39923E-26 |
| ACE2 | MOCS3 | 0,4218 | 2,47621E-26 |
| ACE2 | MTHFD1 | 0,4213 | 2,83156E-26 |

| | | | |
|---|---|---|---|
| ACE2 | GALE | 0,4211 | 3,00026E-26 |
| ACE2 | LNX2 | 0,421 | 3,13505E-26 |
| ACE2 | HRASLS2 | 0,4209 | 3,15446E-26 |
| ACE2 | PTCSC3 | 0,4209 | 3,17739E-26 |
| ACE2 | RPL3P13 | 0,4208 | 3,32794E-26 |
| ACE2 | SLC26A9 | 0,4205 | 3,59784E-26 |
| ACE2 | CSTF1 | 0,4205 | 3,58898E-26 |
| ACE2 | RAB11FIP1 | 0,4198 | 4,37398E-26 |
| ACE2 | IL1R2 | 0,4197 | 4,63548E-26 |
| ACE2 | AGTR2 | 0,4196 | 4,72571E-26 |
| ACE2 | AC005324.6 | 0,4193 | 5,13651E-26 |
| ACE2 | FGFBP1 | 0,4191 | 5,42729E-26 |
| ACE2 | CADPS2 | 0,4191 | 5,49535E-26 |
| ACE2 | IDI1 | 0,4191 | 5,50853E-26 |
| ACE2 | LIPT2 | 0,4189 | 5,78323E-26 |
| ACE2 | DCAF11 | 0,4189 | 5,82611E-26 |
| ACE2 | EIF2AK4 | 0,4188 | 5,96038E-26 |
| ACE2 | AC007405.6 | 0,4187 | 6,17372E-26 |
| ACE2 | SLC6A20 | 0,4187 | 6,06017E-26 |
| ACE2 | LRPPRC | 0,4186 | 6,37374E-26 |
| ACE2 | F11 | 0,4182 | 7,20827E-26 |
| ACE2 | PXMP2 | 0,418 | 7,52199E-26 |
| ACE2 | LAD1 | 0,4178 | 7,99475E-26 |
| ACE2 | MFSD13A | 0,4176 | 8,56791E-26 |
| ACE2 | PSKH1 | 0,4176 | 8,40334E-26 |
| ACE2 | ARFGEF3 | 0,4171 | 9,84524E-26 |
| ACE2 | COL11A1 | 0,4169 | 1,03545E-25 |
| ACE2 | SFTPC | 0,4169 | 1,04019E-25 |
| ACE2 | RP1-172H20.4 | 0,4168 | 1,07293E-25 |
| ACE2 | AIMP2 | 0,4167 | 1,11507E-25 |
| ACE2 | C16orf89 | 0,4167 | 1,09874E-25 |
| ACE2 | MRPS35 | 0,4166 | 1,12773E-25 |
| ACE2 | PON2 | 0,4163 | 1,24331E-25 |
| ACE2 | RP3-342P20.2 | 0,416 | 1,37075E-25 |
| ACE2 | TMEM144 | 0,4158 | 1,42549E-25 |
| ACE2 | ZNF57 | 0,4153 | 1,65467E-25 |
| ACE2 | MRPS18B | 0,4149 | 1,87627E-25 |
| ACE2 | NDUFB9 | 0,4149 | 1,89092E-25 |
| ACE2 | ATP6V0A4 | 0,4147 | 1,96842E-25 |
| ACE2 | ATP5A1 | 0,4146 | 2,02229E-25 |
| ACE2 | DSCR3 | 0,4146 | 2,06584E-25 |
| ACE2 | PID1 | 0,4145 | 2,07243E-25 |
| ACE2 | MGC32805 | 0,4144 | 2,16514E-25 |
| ACE2 | CXADR | 0,4143 | 2,20708E-25 |
| ACE2 | ZNRF3 | 0,4142 | 2,31032E-25 |
| ACE2 | TTC39A-AS1 | 0,4141 | 2,37853E-25 |
| ACE2 | SFTPA3P | 0,4141 | 2,32803E-25 |
| ACE2 | SEC11C | 0,4141 | 2,32904E-25 |
| ACE2 | ST14 | 0,4138 | 2,55802E-25 |
| ACE2 | PKP3 | 0,4133 | 2,94008E-25 |
| ACE2 | IRX5 | 0,4129 | 3,30523E-25 |
| ACE2 | RP11-59N23.3 | 0,4127 | 3,56117E-25 |
| ACE2 | PREP | 0,4126 | 3,58801E-25 |
| ACE2 | COG2 | 0,4125 | 3,7288E-25 |

| | | | |
|---|---|---|---|
| ACE2 | RP11-599B13.3 | 0,4125 | 3,72198E-25 |
| ACE2 | MPP7 | 0,4124 | 3,87221E-25 |
| ACE2 | TRIM16L | 0,4122 | 4,04503E-25 |
| ACE2 | TTN | 0,4121 | 4,21677E-25 |
| ACE2 | AP5B1 | 0,412 | 4,31175E-25 |
| ACE2 | MAPK8IP2 | 0,412 | 4,37598E-25 |
| ACE2 | NEK8 | 0,4119 | 4,48197E-25 |
| ACE2 | LINC01107 | 0,4118 | 4,55178E-25 |
| ACE2 | RPUSD2 | 0,4116 | 4,79505E-25 |
| ACE2 | F11-AS1 | 0,4115 | 5,01363E-25 |
| ACE2 | CLDN23 | 0,4111 | 5,66423E-25 |
| ACE2 | COPG1 | 0,4109 | 5,88894E-25 |
| ACE2 | SENP8 | 0,4109 | 5,97645E-25 |
| ACE2 | DLD | 0,4104 | 6,77444E-25 |
| ACE2 | CYP2C18 | 0,4104 | 6,76261E-25 |
| ACE2 | GPRC5C | 0,4102 | 7,15934E-25 |
| ACE2 | MAPKAPK3 | 0,4101 | 7,47198E-25 |
| ACE2 | PRSS8 | 0,4101 | 7,46773E-25 |
| ACE2 | IL20RA | 0,41 | 7,68883E-25 |
| ACE2 | OPLAH | 0,41 | 7,60855E-25 |
| ACE2 | GTF3C4 | 0,4099 | 7,8182E-25 |
| ACE2 | RP3-325F22.5 | 0,4098 | 8,17144E-25 |
| ACE2 | TOX | 0,4095 | 8,93395E-25 |
| ACE2 | MAPK13 | 0,4094 | 9,08942E-25 |
| ACE2 | LAMA2 | 0,4093 | 9,34243E-25 |
| ACE2 | PLCH1 | 0,4091 | 9,92496E-25 |
| ACE2 | RP11-384L8.1 | 0,409 | 1,03482E-24 |
| ACE2 | ATP2C2 | 0,409 | 1,02732E-24 |
| ACE2 | MRC1 | 0,4089 | 1,04726E-24 |
| ACE2 | ADGRV1 | 0,4086 | 1,14733E-24 |
| ACE2 | TMEM238 | 0,4086 | 1,14155E-24 |
| ACE2 | ABCC13 | 0,4082 | 1,29948E-24 |
| ACE2 | LINC01124 | 0,4081 | 1,3337E-24 |
| ACE2 | GOLGA5 | 0,408 | 1,36186E-24 |
| ACE2 | CHAMP1 | 0,4079 | 1,41423E-24 |
| ACE2 | TDRKH | 0,4078 | 1,4323E-24 |
| ACE2 | PIGS | 0,4077 | 1,49256E-24 |
| ACE2 | DDX1 | 0,4076 | 1,54262E-24 |
| ACE2 | PLA2G16 | 0,4076 | 1,53025E-24 |
| ACE2 | TM7SF3 | 0,4076 | 1,53449E-24 |
| ACE2 | DANCR | 0,4075 | 1,56518E-24 |
| ACE2 | CLEC4E | 0,4075 | 1,55894E-24 |
| ACE2 | SDR42E1 | 0,4073 | 1,63528E-24 |
| ACE2 | CREB3L1 | 0,4072 | 1,69476E-24 |
| ACE2 | RLN3 | 0,407 | 1,79027E-24 |
| ACE2 | MAD2L1BP | 0,4067 | 1,98665E-24 |
| ACE2 | C1orf116 | 0,4066 | 1,99888E-24 |
| ACE2 | ARV1 | 0,4065 | 2,09632E-24 |
| ACE2 | SPINT1 | 0,4064 | 2,12519E-24 |
| ACE2 | ACOXL | 0,4062 | 2,23818E-24 |
| ACE2 | RP13-923O23.7 | 0,4062 | 2,25514E-24 |
| ACE2 | IPP | 0,4061 | 2,34217E-24 |
| ACE2 | RP11-582J16.4 | 0,406 | 2,41442E-24 |
| ACE2 | CTD-2530N21.5 | 0,4059 | 2,44722E-24 |

| | | | |
|---|---|---|---|
| ACE2 | TMEM125 | 0,4058 | 2,52331E-24 |
| ACE2 | PIGO | 0,4054 | 2,8159E-24 |
| ACE2 | FDFT1 | 0,4052 | 3,00236E-24 |
| ACE2 | CNKSR1 | 0,4051 | 3,097E-24 |
| ACE2 | ITPR3 | 0,405 | 3,13851E-24 |
| ACE2 | TMEM243 | 0,405 | 3,18828E-24 |
| ACE2 | RAB25 | 0,4046 | 3,59177E-24 |
| ACE2 | ECH1 | 0,4046 | 3,53127E-24 |
| ACE2 | CPSF3 | 0,4045 | 3,61003E-24 |
| ACE2 | MYBPHL | 0,4044 | 3,74278E-24 |
| ACE2 | RIMS4 | 0,4043 | 3,92126E-24 |
| ACE2 | NEU1 | 0,4042 | 3,97198E-24 |
| ACE2 | SH3BGRL2 | 0,4042 | 3,97482E-24 |
| ACE2 | ACOT2 | 0,404 | 4,23536E-24 |
| ACE2 | ATMIN | 0,4039 | 4,37513E-24 |
| ACE2 | SND1 | 0,4037 | 4,5825E-24 |
| ACE2 | NT5DC1 | 0,4036 | 4,66531E-24 |
| ACE2 | C15orf56 | 0,4036 | 4,65212E-24 |
| ACE2 | CTSE | 0,4034 | 4,95896E-24 |
| ACE2 | LRAT | 0,4033 | 5,12108E-24 |
| ACE2 | PTPMT1 | 0,4032 | 5,31626E-24 |
| ACE2 | RNF145 | 0,403 | 5,5099E-24 |
| ACE2 | RASD1 | 0,403 | 5,60401E-24 |
| ACE2 | AARD | 0,4029 | 5,79114E-24 |
| ACE2 | MPST | 0,4029 | 5,79464E-24 |
| ACE2 | RP11-566K19.5 | 0,4028 | 5,92604E-24 |
| ACE2 | GLS2 | 0,4027 | 6,00649E-24 |
| ACE2 | PCDHA11 | 0,4025 | 6,31713E-24 |
| ACE2 | MUL1 | 0,4023 | 6,75158E-24 |
| ACE2 | C2orf15 | 0,4023 | 6,84506E-24 |
| ACE2 | ZNF321P | 0,4022 | 7,01325E-24 |
| ACE2 | RP4-594I10.3 | 0,4021 | 7,19048E-24 |
| ACE2 | NAXE | 0,4018 | 7,6762E-24 |
| ACE2 | DBI | 0,4016 | 8,16389E-24 |
| ACE2 | NKX2-1 | 0,4016 | 8,18724E-24 |
| ACE2 | PHYH | 0,4015 | 8,39757E-24 |
| ACE2 | SORCS2 | 0,4014 | 8,77005E-24 |
| ACE2 | HACD3 | 0,4014 | 8,66212E-24 |
| ACE2 | MT-ND6 | 0,4013 | 9,05925E-24 |
| ACE2 | EML5 | 0,4011 | 9,51486E-24 |
| ACE2 | COX5A | 0,4011 | 9,3996E-24 |
| ACE2 | TFAP2C | 0,401 | 9,75003E-24 |
| ACE2 | APEH | 0,4009 | 9,96737E-24 |
| ACE2 | TMEM209 | 0,4006 | 1,08698E-23 |
| ACE2 | ASCC1 | 0,4006 | 1,08954E-23 |
| ACE2 | ALG5 | 0,4006 | 1,10089E-23 |
| ACE2 | EID2 | 0,4005 | 1,12377E-23 |
| ACE2 | PNPO | 0,4004 | 1,13979E-23 |
| ACE2 | RP11-545A16.3 | 0,4003 | 1,18469E-23 |
| ACE2 | SERINC2 | 0,4002 | 1,22186E-23 |
| ACE2 | MFSD5 | 0,4002 | 1,22455E-23 |
| ACE2 | GALNT15 | 0,4 | 1,26513E-23 |
| ACE2 | CEBPA | 0,3998 | 1,36811E-23 |
| ACE2 | MR1 | 0,3997 | 1,37985E-23 |

| | | | |
|---|---|---|---|
| ACE2 | SHISA2 | 0,3996 | 1,43049E-23 |
| ACE2 | PDPK1 | 0,3995 | 1,47754E-23 |
| ACE2 | JAGN1 | 0,3992 | 1,58274E-23 |
| ACE2 | ESYT3 | 0,3992 | 1,57837E-23 |
| ACE2 | PLIN5 | 0,3992 | 1,58967E-23 |
| ACE2 | CTD-2376I4.2 | 0,3987 | 1,84321E-23 |
| ACE2 | WIF1 | 0,3986 | 1,86998E-23 |
| ACE2 | RP11-293P20.2 | 0,3985 | 1,93071E-23 |
| ACE2 | UMPS | 0,3985 | 1,92504E-23 |
| ACE2 | SLC16A13 | 0,3985 | 1,96706E-23 |
| ACE2 | ALDH5A1 | 0,3983 | 2,02365E-23 |
| ACE2 | RP11-304L19.13 | 0,398 | 2,21883E-23 |
| ACE2 | STRBP | 0,3979 | 2,28283E-23 |
| ACE2 | FAM160A1 | 0,3978 | 2,34377E-23 |
| ACE2 | HNF1B | 0,3978 | 2,36154E-23 |
| ACE2 | ORMDL2 | 0,3976 | 2,45185E-23 |
| ACE2 | PAQR5 | 0,3976 | 2,45225E-23 |
| ACE2 | MCRIP2 | 0,3976 | 2,47704E-23 |
| ACE2 | H1F0 | 0,3976 | 2,50796E-23 |
| ACE2 | EHHADH | 0,3973 | 2,71211E-23 |
| ACE2 | AADAC | 0,3971 | 2,87629E-23 |
| ACE2 | RRAGD | 0,3971 | 2,86148E-23 |
| ACE2 | MMP28 | 0,397 | 2,96138E-23 |
| ACE2 | NAGLU | 0,397 | 2,89022E-23 |
| ACE2 | ALG1 | 0,3966 | 3,23366E-23 |
| ACE2 | LINC01806 | 0,3965 | 3,39337E-23 |
| ACE2 | MRM2 | 0,3963 | 3,55073E-23 |
| ACE2 | FUT6 | 0,3963 | 3,58954E-23 |
| ACE2 | HLCS | 0,3963 | 3,54711E-23 |
| ACE2 | SDHB | 0,3962 | 3,64706E-23 |
| ACE2 | TLN2 | 0,3962 | 3,65352E-23 |
| ACE2 | ZXDC | 0,396 | 3,88899E-23 |
| ACE2 | HACD1 | 0,3959 | 3,95201E-23 |
| ACE2 | SOX9 | 0,3959 | 3,91682E-23 |
| ACE2 | MED18 | 0,3958 | 4,06028E-23 |
| ACE2 | GPD1L | 0,3957 | 4,19051E-23 |
| ACE2 | SLC9A2 | 0,3956 | 4,31562E-23 |
| ACE2 | ADRA1B | 0,3956 | 4,34368E-23 |
| ACE2 | LDOC1L | 0,3956 | 4,32795E-23 |
| ACE2 | ALPP | 0,3955 | 4,46427E-23 |
| ACE2 | NHLRC1 | 0,3955 | 4,43467E-23 |
| ACE2 | TMEM30B | 0,3951 | 4,93189E-23 |
| ACE2 | DOLK | 0,3949 | 5,23183E-23 |
| ACE2 | SCAMP4 | 0,3946 | 5,70728E-23 |
| ACE2 | CTH | 0,3944 | 5,89661E-23 |
| ACE2 | ARHGEF16 | 0,3943 | 6,16145E-23 |
| ACE2 | ELF5 | 0,3941 | 6,37486E-23 |
| ACE2 | ETV5 | 0,3938 | 7,00474E-23 |
| ACE2 | SLC35E1 | 0,3937 | 7,18371E-23 |
| ACE2 | VKORC1L1 | 0,3936 | 7,33796E-23 |
| ACE2 | PLPPR1 | 0,3936 | 7,35927E-23 |
| ACE2 | METTL7B | 0,3936 | 7,45478E-23 |
| ACE2 | PIGBOS1 | 0,3935 | 7,68516E-23 |
| ACE2 | PHB | 0,3935 | 7,67736E-23 |

| | | | |
|---|---|---|---|
| ACE2 | TMEM141 | 0,3934 | 7,77307E-23 |
| ACE2 | PTPRG | 0,3933 | 7,94088E-23 |
| ACE2 | MT-TE | 0,3933 | 8,10161E-23 |
| ACE2 | C9orf64 | 0,3932 | 8,17675E-23 |
| ACE2 | ATP6V1A | 0,3931 | 8,39222E-23 |
| ACE2 | TMEM236 | 0,3931 | 8,44252E-23 |
| ACE2 | SAR1B | 0,393 | 8,66667E-23 |
| ACE2 | CLTC | 0,393 | 8,74988E-23 |
| ACE2 | WASHC5 | 0,3928 | 9,08425E-23 |
| ACE2 | PSAT1 | 0,3928 | 9,0959E-23 |
| ACE2 | CIT | 0,3926 | 9,68807E-23 |
| ACE2 | CEP70 | 0,3925 | 9,97875E-23 |
| ACE2 | POLR3B | 0,3925 | 9,8362E-23 |
| ACE2 | MBTPS2 | 0,3925 | 9,91779E-23 |
| ACE2 | PTER | 0,3924 | 1,01033E-22 |
| ACE2 | PRKCZ | 0,3922 | 1,09034E-22 |
| ACE2 | ASB13 | 0,3922 | 1,07312E-22 |
| ACE2 | DRAM1 | 0,3922 | 1,06534E-22 |
| ACE2 | MST1L | 0,3921 | 1,10509E-22 |
| ACE2 | SGMS2 | 0,3921 | 1,11321E-22 |
| ACE2 | JKAMP | 0,3921 | 1,09448E-22 |
| ACE2 | PRDX6 | 0,3919 | 1,16733E-22 |
| ACE2 | SLC46A1 | 0,3919 | 1,16205E-22 |
| ACE2 | HAGLR | 0,3916 | 1,26359E-22 |
| ACE2 | TRMT10C | 0,3916 | 1,26669E-22 |
| ACE2 | TGS1 | 0,3916 | 1,25448E-22 |
| ACE2 | ZDHHC3 | 0,3915 | 1,30666E-22 |
| ACE2 | EPB41L4B | 0,3915 | 1,31675E-22 |
| ACE2 | RP11-723J4.3 | 0,3913 | 1,38456E-22 |
| ACE2 | MRPS33 | 0,3911 | 1,44388E-22 |
| ACE2 | AC025335.1 | 0,3909 | 1,5416E-22 |
| ACE2 | TMEM9B | 0,3906 | 1,64748E-22 |
| ACE2 | GPN3 | 0,3906 | 1,68104E-22 |
| ACE2 | MOCOS | 0,3906 | 1,67387E-22 |
| ACE2 | TKT | 0,3904 | 1,73192E-22 |
| ACE2 | POLR3C | 0,3902 | 1,82677E-22 |
| ACE2 | SDHAP3 | 0,3902 | 1,83182E-22 |
| ACE2 | IRX2 | 0,3902 | 1,85932E-22 |
| ACE2 | P3H2-AS1 | 0,39 | 1,93587E-22 |
| ACE2 | OCLN | 0,3898 | 2,05902E-22 |
| ACE2 | RNF128 | 0,3898 | 2,06665E-22 |
| ACE2 | BEND3 | 0,3897 | 2,09413E-22 |
| ACE2 | TRIM24 | 0,3897 | 2,13039E-22 |
| ACE2 | HARBI1 | 0,3894 | 2,31648E-22 |
| ACE2 | PYM1 | 0,3894 | 2,29916E-22 |
| ACE2 | ADORA2B | 0,3894 | 2,25761E-22 |
| ACE2 | TTLL12 | 0,3894 | 2,31251E-22 |
| ACE2 | DHRS7 | 0,3892 | 2,41422E-22 |
| ACE2 | SACM1L | 0,389 | 2,57714E-22 |
| ACE2 | CACNB1 | 0,389 | 2,56123E-22 |
| ACE2 | FBXO40 | 0,3889 | 2,63153E-22 |
| ACE2 | ELAC1 | 0,3887 | 2,7797E-22 |
| ACE2 | MSMO1 | 0,3886 | 2,81439E-22 |
| ACE2 | APOA5 | 0,3886 | 2,86633E-22 |

| | | | |
|---|---|---|---|
| ACE2 | RP11-44F14.10 | 0,3882 | 3,11347E-22 |
| ACE2 | CTD-2015H6.3 | 0,3881 | 3,21954E-22 |
| ACE2 | RP11-1260E13.2 | 0,3881 | 3,22585E-22 |
| ACE2 | RN7SL268P | 0,3881 | 3,26473E-22 |
| ACE2 | MRPL45 | 0,388 | 3,29037E-22 |
| ACE2 | PRKAG1 | 0,3878 | 3,53793E-22 |
| ACE2 | INTS14 | 0,3878 | 3,48872E-22 |
| ACE2 | UQCRFS1 | 0,3877 | 3,609E-22 |
| ACE2 | BPNT1 | 0,3876 | 3,71574E-22 |
| ACE2 | NUP37 | 0,3876 | 3,65433E-22 |
| ACE2 | MT-ND5 | 0,3876 | 3,65025E-22 |
| ACE2 | PSMC1 | 0,3875 | 3,8166E-22 |
| ACE2 | FH | 0,3874 | 3,92209E-22 |
| ACE2 | MRPS36 | 0,3873 | 4,00914E-22 |
| ACE2 | CTB-11I22.1 | 0,3873 | 4,05061E-22 |
| ACE2 | TMBIM6 | 0,3872 | 4,13022E-22 |
| ACE2 | FAM153C | 0,3871 | 4,16647E-22 |
| ACE2 | KCNJ15 | 0,387 | 4,34884E-22 |
| ACE2 | ECE2 | 0,3868 | 4,51277E-22 |
| ACE2 | MRPS18C | 0,3868 | 4,59652E-22 |
| ACE2 | RP4-635A23.6 | 0,3867 | 4,65206E-22 |
| ACE2 | BTBD9 | 0,3864 | 5,07059E-22 |
| ACE2 | MS4A4A | 0,3862 | 5,29772E-22 |
| ACE2 | LINC01314 | 0,3862 | 5,30918E-22 |
| ACE2 | TMEM198 | 0,3861 | 5,5483E-22 |
| ACE2 | MRPS16 | 0,386 | 5,65219E-22 |
| ACE2 | CLN3 | 0,386 | 5,56879E-22 |
| ACE2 | CYP2B7P | 0,386 | 5,65963E-22 |
| ACE2 | SYS1 | 0,386 | 5,56868E-22 |
| ACE2 | TMEM213 | 0,3859 | 5,81466E-22 |
| ACE2 | MIPEP | 0,3859 | 5,72961E-22 |
| ACE2 | KIF16B | 0,3859 | 5,75601E-22 |
| ACE2 | FASTKD2 | 0,3858 | 6,01436E-22 |
| ACE2 | C5orf63 | 0,3855 | 6,47483E-22 |
| ACE2 | SQRDL | 0,3855 | 6,46423E-22 |
| ACE2 | ALDH4A1 | 0,3854 | 6,62909E-22 |
| ACE2 | FAM19A4 | 0,3854 | 6,55674E-22 |
| ACE2 | KIAA1549 | 0,3853 | 6,7249E-22 |
| ACE2 | CCNYL1 | 0,3851 | 7,2237E-22 |
| ACE2 | LONP1 | 0,385 | 7,39698E-22 |
| ACE2 | EGF | 0,3848 | 7,64718E-22 |
| ACE2 | LTV1 | 0,3848 | 7,82243E-22 |
| ACE2 | SLC12A8 | 0,3846 | 8,09954E-22 |
| ACE2 | CNOT11 | 0,3845 | 8,45333E-22 |
| ACE2 | SRD5A3 | 0,3845 | 8,47411E-22 |
| ACE2 | SLC7A6 | 0,3845 | 8,29803E-22 |
| ACE2 | CEBPA-AS1 | 0,3844 | 8,65991E-22 |
| ACE2 | RBMS1P1 | 0,3843 | 8,72434E-22 |
| ACE2 | PIP4K2C | 0,3842 | 9,0324E-22 |
| ACE2 | MTMR12 | 0,3841 | 9,19863E-22 |
| ACE2 | PPP1R8 | 0,384 | 9,53603E-22 |
| ACE2 | ODC1 | 0,384 | 9,44433E-22 |
| ACE2 | AOX1 | 0,3839 | 9,73754E-22 |
| ACE2 | LARGE2 | 0,3839 | 9,76288E-22 |

| | | | |
|---|---|---|---|
| ACE2 | LPCAT3 | 0,3839 | 9,66833E-22 |
| ACE2 | UTP15 | 0,3837 | 1,03028E-21 |
| ACE2 | SLC7A7 | 0,3837 | 1,02137E-21 |
| ACE2 | ITGB6 | 0,3835 | 1,08811E-21 |
| ACE2 | NECAB3 | 0,3835 | 1,09156E-21 |
| ACE2 | CDK7 | 0,3834 | 1,11091E-21 |
| ACE2 | ST6GAL1 | 0,3833 | 1,13235E-21 |
| ACE2 | ZNF552 | 0,3832 | 1,18112E-21 |
| ACE2 | GCNT2 | 0,3831 | 1,19586E-21 |
| ACE2 | DLAT | 0,383 | 1,23557E-21 |
| ACE2 | FICD | 0,3829 | 1,2855E-21 |
| ACE2 | ACSS3 | 0,3827 | 1,34936E-21 |
| ACE2 | XBP1 | 0,3826 | 1,36314E-21 |
| ACE2 | RHOXF1P1 | 0,3826 | 1,36E-21 |
| ACE2 | LINC00964 | 0,3824 | 1,46633E-21 |
| ACE2 | ADK | 0,3824 | 1,43132E-21 |
| ACE2 | RPN2 | 0,3824 | 1,45045E-21 |
| ACE2 | LINC01003 | 0,3823 | 1,49503E-21 |
| ACE2 | ENDOD1 | 0,3823 | 1,49249E-21 |
| ACE2 | EP300-AS1 | 0,3822 | 1,53198E-21 |
| ACE2 | UBIAD1 | 0,3818 | 1,68157E-21 |
| ACE2 | ARHGEF38 | 0,3818 | 1,67736E-21 |
| ACE2 | FAM153A | 0,3817 | 1,74094E-21 |
| ACE2 | LYRM2 | 0,3816 | 1,77696E-21 |
| ACE2 | CDCA7L | 0,3816 | 1,80604E-21 |
| ACE2 | RP11-370I10.2 | 0,3816 | 1,76615E-21 |
| ACE2 | CTNNBIP1 | 0,3815 | 1,80974E-21 |
| ACE2 | KIAA0895 | 0,3815 | 1,82435E-21 |
| ACE2 | WNT5A | 0,3814 | 1,90149E-21 |
| ACE2 | IRAK1BP1 | 0,3813 | 1,92956E-21 |
| ACE2 | BRI3 | 0,3813 | 1,91782E-21 |
| ACE2 | ADORA3 | 0,3812 | 1,96972E-21 |
| ACE2 | UBE3C | 0,3812 | 1,99764E-21 |
| ACE2 | ASNA1 | 0,3811 | 2,04339E-21 |
| ACE2 | GGT3P | 0,3809 | 2,1162E-21 |
| ACE2 | RP11-710E1.2 | 0,3808 | 2,17533E-21 |
| ACE2 | WNT7B | 0,3807 | 2,27416E-21 |
| ACE2 | MAP3K15 | 0,3806 | 2,31244E-21 |
| ACE2 | SEPHS2 | 0,3804 | 2,4235E-21 |
| ACE2 | CCL23 | 0,3804 | 2,46413E-21 |
| ACE2 | MANSC1 | 0,3803 | 2,49888E-21 |
| ACE2 | ABCC6 | 0,3802 | 2,56639E-21 |
| ACE2 | STK16 | 0,3801 | 2,6398E-21 |
| ACE2 | AC104667.3 | 0,3801 | 2,62067E-21 |
| ACE2 | TMEM45B | 0,3801 | 2,63039E-21 |
| ACE2 | AASS | 0,38 | 2,69475E-21 |
| ACE2 | TMEM265 | 0,3798 | 2,83944E-21 |
| ACE2 | ACOT1 | 0,3797 | 2,95392E-21 |
| ACE2 | DNASE2 | 0,3796 | 2,97166E-21 |
| ACE2 | DNAJC30 | 0,3795 | 3,10743E-21 |
| ACE2 | RP11-422N16.3 | 0,3792 | 3,35133E-21 |
| ACE2 | PCDHA4 | 0,3791 | 3,43507E-21 |
| ACE2 | PIGW | 0,3791 | 3,43125E-21 |
| ACE2 | RAB3D | 0,3791 | 3,43335E-21 |

| | | | |
|---|---|---|---|
| ACE2 | POGK | 0,379 | 3,45451E-21 |
| ACE2 | RP11-875O11.1 | 0,3789 | 3,59468E-21 |
| ACE2 | CFTR | 0,3788 | 3,64137E-21 |
| ACE2 | CRNDE | 0,3788 | 3,66354E-21 |
| ACE2 | PSME3 | 0,3788 | 3,703E-21 |
| ACE2 | TMIGD3 | 0,3787 | 3,74192E-21 |
| ACE2 | STK32A | 0,3787 | 3,75591E-21 |
| ACE2 | ITPK1 | 0,3787 | 3,77452E-21 |
| ACE2 | MROH1 | 0,3785 | 3,95144E-21 |
| ACE2 | MOAP1 | 0,3785 | 3,92618E-21 |
| ACE2 | PRKAR2A | 0,3784 | 4,05969E-21 |
| ACE2 | AC017060.1 | 0,3784 | 4,10699E-21 |
| ACE2 | STAM | 0,378 | 4,50378E-21 |
| ACE2 | SCAMP3 | 0,3776 | 5,02273E-21 |
| ACE2 | RP11-82L18.2 | 0,3775 | 5,15872E-21 |
| ACE2 | LCMT2 | 0,3775 | 5,08886E-21 |
| ACE2 | RP11-473M20.5 | 0,3775 | 5,14393E-21 |
| ACE2 | FAM50B | 0,3774 | 5,29586E-21 |
| ACE2 | PPP5C | 0,3773 | 5,34307E-21 |
| ACE2 | YIPF5 | 0,3772 | 5,54992E-21 |
| ACE2 | ACY1 | 0,377 | 5,81751E-21 |
| ACE2 | AC007182.6 | 0,377 | 5,82784E-21 |
| ACE2 | MRPL15 | 0,3769 | 5,97739E-21 |
| ACE2 | LAMP1 | 0,3766 | 6,43201E-21 |
| ACE2 | EIF2AK1 | 0,3765 | 6,65916E-21 |
| ACE2 | XXyac-YR38GF2.1 | 0,3763 | 6,9082E-21 |
| ACE2 | CYP4B1 | 0,376 | 7,58378E-21 |
| ACE2 | TRUB2 | 0,376 | 7,5446E-21 |
| ACE2 | POLR1B | 0,3758 | 7,92654E-21 |
| ACE2 | IPO9 | 0,3757 | 8,07847E-21 |
| ACE2 | SMIM4 | 0,3757 | 8,1014E-21 |
| ACE2 | MDH2 | 0,3757 | 8,05448E-21 |
| ACE2 | RNASE6 | 0,3757 | 8,07054E-21 |
| ACE2 | RTN4IP1 | 0,3756 | 8,2696E-21 |
| ACE2 | RPL26P27 | 0,3756 | 8,43149E-21 |
| ACE2 | ATP2C1 | 0,3755 | 8,59532E-21 |
| ACE2 | ACADSB | 0,3753 | 8,90435E-21 |
| ACE2 | WDR12 | 0,3752 | 9,2632E-21 |
| ACE2 | MTCH2 | 0,3752 | 9,34285E-21 |
| ACE2 | C19orf35 | 0,3751 | 9,37356E-21 |
| ACE2 | C4BPA | 0,375 | 9,76297E-21 |
| ACE2 | NDUFAF1 | 0,375 | 9,69907E-21 |
| ACE2 | SLC35C1 | 0,3748 | 1,03437E-20 |
| ACE2 | CRYM | 0,3747 | 1,04639E-20 |
| ACE2 | ZBTB8OS | 0,3745 | 1,10703E-20 |
| ACE2 | ATIC | 0,3744 | 1,12029E-20 |
| ACE2 | ACOT13 | 0,3744 | 1,13336E-20 |
| ACE2 | TALDO1 | 0,3743 | 1,16638E-20 |
| ACE2 | RPP25 | 0,3743 | 1,1642E-20 |
| ACE2 | C22orf29 | 0,3743 | 1,15514E-20 |
| ACE2 | PDE4DIP | 0,3742 | 1,19913E-20 |
| ACE2 | RTCB | 0,3742 | 1,18884E-20 |
| ACE2 | NUDT12 | 0,374 | 1,23925E-20 |
| ACE2 | CLCN4 | 0,374 | 1,2504E-20 |

| | | | |
|---|---|---|---|
| ACE2 | CREG1 | 0,3736 | 1,37686E-20 |
| ACE2 | CLCN3 | 0,3736 | 1,39292E-20 |
| ACE2 | PPL | 0,3736 | 1,39431E-20 |
| ACE2 | NDUFV3 | 0,3736 | 1,37472E-20 |
| ACE2 | SERPINA1 | 0,3735 | 1,42722E-20 |
| ACE2 | SPRYD4 | 0,3733 | 1,50407E-20 |
| ACE2 | AC008982.2 | 0,3732 | 1,5355E-20 |
| ACE2 | PDF | 0,3731 | 1,56708E-20 |
| ACE2 | TANC2 | 0,3731 | 1,56708E-20 |
| ACE2 | RPARP-AS1 | 0,373 | 1,62057E-20 |
| ACE2 | RPP25L | 0,3728 | 1,70846E-20 |
| ACE2 | KSR2 | 0,3727 | 1,72368E-20 |
| ACE2 | KRT15 | 0,3727 | 1,75335E-20 |
| ACE2 | SLC10A4 | 0,3724 | 1,85816E-20 |
| ACE2 | RP11-25K19.1 | 0,3722 | 1,99558E-20 |
| ACE2 | VPS33A | 0,3721 | 2,04523E-20 |
| ACE2 | TGFBRAP1 | 0,372 | 2,08453E-20 |
| ACE2 | MT1JP | 0,372 | 2,09777E-20 |
| ACE2 | MESP1 | 0,3719 | 2,11362E-20 |
| ACE2 | DHDDS | 0,3718 | 2,15448E-20 |
| ACE2 | PTGR2 | 0,3716 | 2,29816E-20 |
| ACE2 | MTHFD2L | 0,3715 | 2,35167E-20 |
| ACE2 | RP11-7F17.3 | 0,3715 | 2,34408E-20 |
| ACE2 | BECN1 | 0,3715 | 2,32736E-20 |
| ACE2 | GGTLC4P | 0,3715 | 2,37626E-20 |
| ACE2 | CBR1 | 0,3714 | 2,39334E-20 |
| ACE2 | SNAP29 | 0,3714 | 2,39084E-20 |
| ACE2 | CNOT8 | 0,371 | 2,69416E-20 |
| ACE2 | TMEM101 | 0,371 | 2,69564E-20 |
| ACE2 | APOH | 0,3708 | 2,7767E-20 |
| ACE2 | ACO2 | 0,3708 | 2,77293E-20 |
| ACE2 | ACAA2 | 0,3707 | 2,89532E-20 |
| ACE2 | CLPP | 0,3707 | 2,84762E-20 |
| ACE2 | GOLGA2P7 | 0,3705 | 3,05391E-20 |
| ACE2 | METTL2A | 0,3702 | 3,26595E-20 |
| ACE2 | RPTOR | 0,37 | 3,40641E-20 |
| ACE2 | GUCD1 | 0,3699 | 3,51935E-20 |
| ACE2 | RP11-475O23.2 | 0,3698 | 3,55566E-20 |
| ACE2 | ABHD12 | 0,3696 | 3,82165E-20 |
| ACE2 | FAM153B | 0,3695 | 3,83229E-20 |
| ACE2 | PEX11B | 0,3694 | 4,00489E-20 |
| ACE2 | TMBIM4 | 0,3694 | 3,9298E-20 |
| ACE2 | CLPX | 0,3694 | 3,9529E-20 |
| ACE2 | MAIP1 | 0,3693 | 4,03145E-20 |
| ACE2 | DYNC1LI1 | 0,3692 | 4,18282E-20 |
| ACE2 | MPZL2 | 0,3691 | 4,23671E-20 |
| ACE2 | EMC7 | 0,3689 | 4,49755E-20 |
| ACE2 | CERS2 | 0,3688 | 4,57042E-20 |
| ACE2 | PC | 0,3688 | 4,57665E-20 |
| ACE2 | MRPL58 | 0,3687 | 4,68467E-20 |
| ACE2 | C12orf49 | 0,3686 | 4,8569E-20 |
| ACE2 | AC092071.1 | 0,3684 | 5,08496E-20 |
| ACE2 | P4HB | 0,3682 | 5,29548E-20 |
| ACE2 | CHPF2 | 0,368 | 5,68309E-20 |

| | | | |
|---|---|---|---|
| ACE2 | UTP3 | 0,3679 | 5,72552E-20 |
| ACE2 | RP11-16K12.1 | 0,3678 | 5,89399E-20 |
| ACE2 | DDOST | 0,3676 | 6,22565E-20 |
| ACE2 | HUNK | 0,3675 | 6,40413E-20 |
| ACE2 | ANKRD28 | 0,3674 | 6,581E-20 |
| ACE2 | C10orf35 | 0,3674 | 6,52967E-20 |
| ACE2 | NCR3LG1 | 0,3674 | 6,56966E-20 |
| ACE2 | NSF | 0,367 | 7,24544E-20 |
| ACE2 | TUFM | 0,3669 | 7,40644E-20 |
| ACE2 | SLC27A4 | 0,3668 | 7,56137E-20 |
| ACE2 | SIGLEC10 | 0,3668 | 7,50789E-20 |
| ACE2 | SLC35B1 | 0,3667 | 7,76937E-20 |
| ACE2 | ATPAF1 | 0,3666 | 7,98236E-20 |
| ACE2 | IRX1 | 0,3666 | 7,97963E-20 |
| ACE2 | TRMO | 0,3666 | 7,89896E-20 |
| ACE2 | REPS2 | 0,3666 | 7,94436E-20 |
| ACE2 | EBAG9 | 0,3665 | 8,09876E-20 |
| ACE2 | CD38 | 0,3663 | 8,5685E-20 |
| ACE2 | C11orf52 | 0,3663 | 8,5609E-20 |
| ACE2 | AC013275.2 | 0,3662 | 8,68629E-20 |
| ACE2 | PCAT18 | 0,3662 | 8,76008E-20 |
| ACE2 | PEX11A | 0,3661 | 9,02277E-20 |
| ACE2 | RIMKLA | 0,366 | 9,16386E-20 |
| ACE2 | SSR3 | 0,366 | 9,11959E-20 |
| ACE2 | HOMEZ | 0,366 | 9,19096E-20 |
| ACE2 | RP11-44F14.8 | 0,3659 | 9,46135E-20 |
| ACE2 | TRIM71 | 0,3658 | 9,61202E-20 |
| ACE2 | KAT5 | 0,3658 | 9,6859E-20 |
| ACE2 | NDUFA7 | 0,3658 | 9,67138E-20 |
| ACE2 | CTR9 | 0,3657 | 1,00162E-19 |
| ACE2 | NUDT8 | 0,3657 | 9,96303E-20 |
| ACE2 | ATOH8 | 0,3656 | 1,01145E-19 |
| ACE2 | AP3B1 | 0,3656 | 1,01582E-19 |
| ACE2 | TBCEL | 0,3656 | 1,01393E-19 |
| ACE2 | TJP3 | 0,3655 | 1,03139E-19 |
| ACE2 | DPM2 | 0,3654 | 1,07904E-19 |
| ACE2 | SMAGP | 0,3654 | 1,06734E-19 |
| ACE2 | VPS25 | 0,3654 | 1,0635E-19 |
| ACE2 | ENO1-AS1 | 0,3653 | 1,08297E-19 |
| ACE2 | SLC4A1AP | 0,3651 | 1,15893E-19 |
| ACE2 | AGL | 0,365 | 1,19071E-19 |
| ACE2 | ANGPTL1 | 0,365 | 1,17631E-19 |
| ACE2 | ARL4A | 0,365 | 1,18653E-19 |
| ACE2 | ST20-AS1 | 0,365 | 1,17648E-19 |
| ACE2 | FLVCR2 | 0,3649 | 1,21031E-19 |
| ACE2 | SCTR | 0,3647 | 1,27175E-19 |
| ACE2 | ST7-AS2 | 0,3646 | 1,30148E-19 |
| ACE2 | MRPL49 | 0,3645 | 1,32606E-19 |
| ACE2 | FRAT1 | 0,3644 | 1,36841E-19 |
| ACE2 | NARS | 0,3644 | 1,37476E-19 |
| ACE2 | FAIM | 0,3643 | 1,40373E-19 |
| ACE2 | ZBED3 | 0,3643 | 1,41209E-19 |
| ACE2 | RP11-573D15.9 | 0,3641 | 1,45842E-19 |
| ACE2 | AC092171.4 | 0,3641 | 1,46586E-19 |

| | | | |
|---|---|---|---|
| ACE2 | C5orf22 | 0,364 | 1,49113E-19 |
| ACE2 | RP11-1099M24.8 | 0,364 | 1,52313E-19 |
| ACE2 | MRPS22 | 0,3639 | 1,53381E-19 |
| ACE2 | SRSF8 | 0,3639 | 1,55811E-19 |
| ACE2 | KIAA1429 | 0,3638 | 1,58314E-19 |
| ACE2 | LIN52 | 0,3638 | 1,57617E-19 |
| ACE2 | PNPLA4 | 0,3638 | 1,56349E-19 |
| ACE2 | PEG10 | 0,3636 | 1,65347E-19 |
| ACE2 | ATRN | 0,3636 | 1,66337E-19 |
| ACE2 | C1QC | 0,3634 | 1,73547E-19 |
| ACE2 | GALNT3 | 0,3634 | 1,73408E-19 |
| ACE2 | SOCS7 | 0,3634 | 1,76286E-19 |
| ACE2 | RAD50 | 0,3633 | 1,77187E-19 |
| ACE2 | FGL1 | 0,3633 | 1,77061E-19 |
| ACE2 | RP11-320N7.2 | 0,3633 | 1,78905E-19 |
| ACE2 | RP11-16P6.1 | 0,3632 | 1,83531E-19 |
| ACE2 | UGGT1 | 0,3631 | 1,87651E-19 |
| ACE2 | MID1IP1-AS1 | 0,363 | 1,92875E-19 |
| ACE2 | MRPL44 | 0,3629 | 1,98497E-19 |
| ACE2 | GRSF1 | 0,3629 | 1,99075E-19 |
| ACE2 | RP11-713M15.2 | 0,3629 | 1,94733E-19 |
| ACE2 | PXYLP1 | 0,3627 | 2,05541E-19 |
| ACE2 | CPNE3 | 0,3627 | 2,07544E-19 |
| ACE2 | SPINK5 | 0,3624 | 2,24161E-19 |
| ACE2 | CBS | 0,3624 | 2,25026E-19 |
| ACE2 | CXorf57 | 0,3622 | 2,32754E-19 |
| ACE2 | MIR22HG | 0,362 | 2,44316E-19 |
| ACE2 | TATDN3 | 0,3619 | 2,54013E-19 |
| ACE2 | FAM175B | 0,3619 | 2,50436E-19 |
| ACE2 | ZBED9 | 0,3618 | 2,55529E-19 |
| ACE2 | RP11-293M10.6 | 0,3618 | 2,57942E-19 |
| ACE2 | RSG1 | 0,3616 | 2,69524E-19 |
| ACE2 | IARS2 | 0,3616 | 2,71122E-19 |
| ACE2 | EVPL | 0,3616 | 2,69045E-19 |
| ACE2 | WFDC5 | 0,3614 | 2,82616E-19 |
| ACE2 | ZNF275 | 0,3613 | 2,91779E-19 |
| ACE2 | ERBB4 | 0,3612 | 2,94036E-19 |
| ACE2 | BCL2L13 | 0,3612 | 3,00439E-19 |
| ACE2 | HYAL1 | 0,3611 | 3,03897E-19 |
| ACE2 | CIAPIN1 | 0,3611 | 3,08261E-19 |
| ACE2 | FAM174B | 0,361 | 3,13065E-19 |
| ACE2 | ENO1 | 0,3608 | 3,26129E-19 |
| ACE2 | OXCT1 | 0,3605 | 3,51074E-19 |
| ACE2 | CTD-3074O7.5 | 0,3605 | 3,54888E-19 |
| ACE2 | PKP2 | 0,3605 | 3,50805E-19 |
| ACE2 | SUPT16H | 0,3605 | 3,56488E-19 |
| ACE2 | SYT15 | 0,3604 | 3,57817E-19 |
| ACE2 | CHRM2 | 0,3603 | 3,67661E-19 |
| ACE2 | PDK4 | 0,3602 | 3,80366E-19 |
| ACE2 | MRPL3 | 0,3601 | 3,92239E-19 |
| ACE2 | COPZ1 | 0,3601 | 3,92065E-19 |
| ACE2 | CLDN4 | 0,36 | 3,94774E-19 |
| ACE2 | DAAM2 | 0,3599 | 4,0524E-19 |
| ACE2 | HIP1R | 0,3599 | 4,08324E-19 |

| | | | |
|---|---|---|---|
| ACE2 | RILP | 0,3597 | 4,31253E-19 |
| ACE2 | ATP8B5P | 0,3596 | 4,35293E-19 |
| ACE2 | EAPP | 0,3596 | 4,33989E-19 |
| ACE2 | RP11-697K23.3 | 0,3594 | 4,62827E-19 |
| ACE2 | CHAC2 | 0,3593 | 4,7278E-19 |
| ACE2 | SLC25A10 | 0,3592 | 4,85916E-19 |
| ACE2 | UPRT | 0,3592 | 4,82436E-19 |
| ACE2 | F3 | 0,3591 | 4,93155E-19 |
| ACE2 | SUMF2 | 0,3591 | 4,89119E-19 |
| ACE2 | PPP1R14C | 0,359 | 5,04984E-19 |
| ACE2 | PRMT5 | 0,3589 | 5,18488E-19 |
| ACE2 | BRCC3 | 0,3589 | 5,15387E-19 |
| ACE2 | MAP3K21 | 0,3588 | 5,31268E-19 |
| ACE2 | CLUH | 0,3588 | 5,30589E-19 |
| ACE2 | GTF3C3 | 0,3587 | 5,48141E-19 |
| ACE2 | RAB17 | 0,3586 | 5,54343E-19 |
| ACE2 | RP11-946L16.1 | 0,3585 | 5,71134E-19 |
| ACE2 | PPA2 | 0,3584 | 5,86649E-19 |
| ACE2 | C1orf43 | 0,3582 | 6,1536E-19 |
| ACE2 | RN7SKP18 | 0,3582 | 6,10026E-19 |
| ACE2 | PDHX | 0,3581 | 6,32465E-19 |
| ACE2 | SPRY4-IT1 | 0,3579 | 6,63626E-19 |
| ACE2 | PEPD | 0,3579 | 6,60844E-19 |
| ACE2 | ALDH1L1-AS2 | 0,3578 | 6,78444E-19 |
| ACE2 | ABCC4 | 0,3577 | 6,85784E-19 |
| ACE2 | EVPLL | 0,3577 | 6,8702E-19 |
| ACE2 | EIF3A | 0,3575 | 7,18759E-19 |
| ACE2 | EPGN | 0,3574 | 7,46341E-19 |
| ACE2 | ZNF114 | 0,3574 | 7,36638E-19 |
| ACE2 | RP11-96D1.6 | 0,3572 | 7,85262E-19 |
| ACE2 | MTND6P4 | 0,3571 | 7,93485E-19 |
| ACE2 | ATP11A | 0,3571 | 7,948E-19 |
| ACE2 | ELOVL1 | 0,3569 | 8,41527E-19 |
| ACE2 | RP11-7F17.5 | 0,3569 | 8,4548E-19 |
| ACE2 | CAT | 0,3568 | 8,5455E-19 |
| ACE2 | RP11-132A1.4 | 0,3567 | 8,6757E-19 |
| ACE2 | C16orf62 | 0,3567 | 8,67908E-19 |
| ACE2 | RP11-44F14.9 | 0,3567 | 8,85014E-19 |
| ACE2 | MRPL37 | 0,3566 | 8,99472E-19 |
| ACE2 | RP11-206P5.2 | 0,3566 | 9,04156E-19 |
| ACE2 | FOXA1 | 0,3566 | 9,07503E-19 |
| ACE2 | LINC00264 | 0,3565 | 9,29475E-19 |
| ACE2 | ABCC8 | 0,3565 | 9,12565E-19 |
| ACE2 | PGAM5 | 0,3564 | 9,35727E-19 |
| ACE2 | ZNF319 | 0,3564 | 9,49374E-19 |
| ACE2 | LRPAP1 | 0,3563 | 9,65955E-19 |
| ACE2 | TRAM1 | 0,3563 | 9,53661E-19 |
| ACE2 | CCDC94 | 0,3563 | 9,67839E-19 |
| ACE2 | KRT7 | 0,3562 | 9,87426E-19 |
| ACE2 | ALG11 | 0,3562 | 9,9878E-19 |
| ACE2 | TMC4 | 0,3561 | 1,00225E-18 |
| ACE2 | PARS2 | 0,356 | 1,0447E-18 |
| ACE2 | GTF2I | 0,356 | 1,02297E-18 |
| ACE2 | YIF1A | 0,356 | 1,03186E-18 |

| | | | |
|---|---|---|---|
| ACE2 | CORO2A | 0,3559 | 1,0696E-18 |
| ACE2 | DNAH14 | 0,3558 | 1,09683E-18 |
| ACE2 | TPI1P2 | 0,3557 | 1,11571E-18 |
| ACE2 | MPDU1 | 0,3557 | 1,12155E-18 |
| ACE2 | CEACAM4 | 0,3557 | 1,11355E-18 |
| ACE2 | UQCRC2 | 0,3556 | 1,13161E-18 |
| ACE2 | ALG14 | 0,3555 | 1,17187E-18 |
| ACE2 | CDH16 | 0,3555 | 1,15545E-18 |
| ACE2 | GRM3 | 0,3554 | 1,19469E-18 |
| ACE2 | RP13-20L14.6 | 0,3554 | 1,20404E-18 |
| ACE2 | ELMO3 | 0,3553 | 1,22488E-18 |
| ACE2 | MON1B | 0,3553 | 1,22846E-18 |
| ACE2 | ERBB3 | 0,3552 | 1,25281E-18 |
| ACE2 | GATB | 0,3551 | 1,28342E-18 |
| ACE2 | CLDN3 | 0,3551 | 1,28542E-18 |
| ACE2 | BRI3BP | 0,3551 | 1,27502E-18 |
| ACE2 | PALB2 | 0,3551 | 1,28156E-18 |
| ACE2 | CD68 | 0,3551 | 1,27199E-18 |
| ACE2 | NHLH2 | 0,3549 | 1,3488E-18 |
| ACE2 | GFM1 | 0,3549 | 1,33299E-18 |
| ACE2 | C1QBP | 0,3549 | 1,33607E-18 |
| ACE2 | PRKAG2 | 0,3547 | 1,41317E-18 |
| ACE2 | RBPMS-AS1 | 0,3546 | 1,44831E-18 |
| ACE2 | SNX14 | 0,3544 | 1,49754E-18 |
| ACE2 | TRIM35 | 0,3544 | 1,50695E-18 |
| ACE2 | PDCD6IPP1 | 0,3544 | 1,50335E-18 |
| ACE2 | CDKL5 | 0,3544 | 1,50893E-18 |
| ACE2 | C1QB | 0,3543 | 1,5508E-18 |
| ACE2 | METAP2 | 0,3543 | 1,55332E-18 |
| ACE2 | GTF2H3 | 0,3543 | 1,55711E-18 |
| ACE2 | ZNF778 | 0,3543 | 1,54793E-18 |
| ACE2 | MAPKAP1 | 0,3541 | 1,6153E-18 |
| ACE2 | TMEM86A | 0,354 | 1,68141E-18 |
| ACE2 | CNPPD1 | 0,3539 | 1,69122E-18 |
| ACE2 | MRPS17 | 0,3539 | 1,72019E-18 |
| ACE2 | GOT2 | 0,3539 | 1,69011E-18 |
| ACE2 | LINC00336 | 0,3536 | 1,81205E-18 |
| ACE2 | FADS1 | 0,3536 | 1,82315E-18 |
| ACE2 | SYNDIG1L | 0,3535 | 1,86377E-18 |
| ACE2 | EXOSC7 | 0,3534 | 1,93452E-18 |
| ACE2 | RNASE1 | 0,3534 | 1,9243E-18 |
| ACE2 | GNPDA1 | 0,3533 | 1,9432E-18 |
| ACE2 | SDHAF2 | 0,3532 | 1,985E-18 |
| ACE2 | OGDH | 0,3531 | 2,04315E-18 |
| ACE2 | TP53I3 | 0,353 | 2,1231E-18 |
| ACE2 | PDGFC | 0,3529 | 2,14186E-18 |
| ACE2 | WASF3 | 0,3527 | 2,25113E-18 |
| ACE2 | ATP6V0D1 | 0,3526 | 2,31474E-18 |
| ACE2 | HIGD1A | 0,3525 | 2,35728E-18 |
| ACE2 | ASPH | 0,3525 | 2,3621E-18 |
| ACE2 | NKX2-1-AS1 | 0,3524 | 2,40137E-18 |
| ACE2 | MGAT1 | 0,3522 | 2,53156E-18 |
| ACE2 | LMTK3 | 0,3522 | 2,53414E-18 |
| ACE2 | PSMG1 | 0,3522 | 2,55375E-18 |

| | | | |
|---|---|---|---|
| ACE2 | PLD3 | 0,3519 | 2,73196E-18 |
| ACE2 | FAF1 | 0,3518 | 2,7778E-18 |
| ACE2 | C6orf47 | 0,3518 | 2,79077E-18 |
| ACE2 | RP11-295D4.3 | 0,3518 | 2,76654E-18 |
| ACE2 | ALG3 | 0,3517 | 2,8507E-18 |
| ACE2 | RNF141 | 0,3517 | 2,86984E-18 |
| ACE2 | C4orf50 | 0,3516 | 2,92471E-18 |
| ACE2 | L2HGDH | 0,3516 | 2,90634E-18 |
| ACE2 | AC009133.12 | 0,3516 | 2,94626E-18 |
| ACE2 | HOXD1 | 0,3515 | 2,9612E-18 |
| ACE2 | DCBLD2 | 0,3514 | 3,06833E-18 |
| ACE2 | RP11-357D18.1 | 0,3511 | 3,3121E-18 |
| ACE2 | TMEM186 | 0,3509 | 3,47893E-18 |
| ACE2 | DHX8 | 0,3509 | 3,40376E-18 |
| ACE2 | ENOX2 | 0,3506 | 3,72523E-18 |
| ACE2 | RAD17 | 0,3505 | 3,80601E-18 |
| ACE2 | CTD-3064M3.4 | 0,3504 | 3,86522E-18 |
| ACE2 | LINC02207 | 0,3504 | 3,88465E-18 |
| ACE2 | AP1B1 | 0,3503 | 3,9502E-18 |
| ACE2 | MCF2L-AS1 | 0,3502 | 4,06697E-18 |
| ACE2 | GBA | 0,3501 | 4,19147E-18 |
| ACE2 | C1orf226 | 0,3501 | 4,12554E-18 |
| ACE2 | ABO | 0,35 | 4,27434E-18 |
| ACE2 | AC137932.5 | 0,35 | 4,19798E-18 |
| ACE2 | STARD3NL | 0,3498 | 4,42579E-18 |
| ACE2 | CSPG5 | 0,3497 | 4,59281E-18 |
| ACE2 | RPN1 | 0,3497 | 4,54287E-18 |
| ACE2 | MMADHC | 0,3496 | 4,67105E-18 |
| ACE2 | APCDD1 | 0,3496 | 4,65619E-18 |
| ACE2 | EI24 | 0,3495 | 4,73447E-18 |
| ACE2 | FAM83E | 0,3492 | 5,09958E-18 |
| ACE2 | MYO5B | 0,3491 | 5,23891E-18 |
| ACE2 | FGGY | 0,349 | 5,35886E-18 |
| ACE2 | AQP3 | 0,349 | 5,40083E-18 |
| ACE2 | GLE1 | 0,3489 | 5,51497E-18 |
| ACE2 | CDC123 | 0,3489 | 5,50069E-18 |
| ACE2 | RP11-946L16.2 | 0,3489 | 5,49457E-18 |
| ACE2 | RP13-650J16.1 | 0,3488 | 5,6429E-18 |
| ACE2 | MTM1 | 0,3488 | 5,59078E-18 |
| ACE2 | GLMP | 0,3483 | 6,25621E-18 |
| ACE2 | GTF2H2C | 0,3483 | 6,35641E-18 |
| ACE2 | KISS1 | 0,3482 | 6,4613E-18 |
| ACE2 | KIF1BP | 0,3482 | 6,39148E-18 |
| ACE2 | DMKN | 0,3482 | 6,42357E-18 |
| ACE2 | SPHK2 | 0,3481 | 6,57806E-18 |
| ACE2 | ACTL10 | 0,3481 | 6,55328E-18 |
| ACE2 | DNAJC11 | 0,348 | 6,81675E-18 |
| ACE2 | RP11-27I1.4 | 0,348 | 6,67717E-18 |
| ACE2 | ZBTB16 | 0,348 | 6,73994E-18 |
| ACE2 | DSC2 | 0,348 | 6,77064E-18 |
| ACE2 | CERS6 | 0,3479 | 6,86191E-18 |
| ACE2 | RP11-789C1.2 | 0,3479 | 6,87311E-18 |
| ACE2 | TBC1D2 | 0,3479 | 6,95456E-18 |
| ACE2 | CEBPZ | 0,3476 | 7,44886E-18 |

| | | | |
|---|---|---|---|
| ACE2 | MAGIX | 0,3476 | 7,45948E-18 |
| ACE2 | AC009403.2 | 0,3475 | 7,47897E-18 |
| ACE2 | RS1 | 0,3475 | 7,57189E-18 |
| ACE2 | SDC1 | 0,3473 | 7,94839E-18 |
| ACE2 | PIGN | 0,3473 | 7,99943E-18 |
| ACE2 | TOB1 | 0,3472 | 8,04961E-18 |
| ACE2 | TCEAL4 | 0,3472 | 8,17429E-18 |
| ACE2 | FMC1 | 0,3471 | 8,36034E-18 |
| ACE2 | GSTZ1 | 0,3471 | 8,30316E-18 |
| ACE2 | MLEC | 0,347 | 8,43055E-18 |
| ACE2 | SLAIN1 | 0,347 | 8,52509E-18 |
| ACE2 | CHMP1B2P | 0,347 | 8,48057E-18 |
| ACE2 | DCDC2 | 0,3469 | 8,65488E-18 |
| ACE2 | LINC02012 | 0,3468 | 8,79974E-18 |
| ACE2 | MED1 | 0,3468 | 8,93524E-18 |
| ACE2 | METTL13 | 0,3467 | 9,01968E-18 |
| ACE2 | HACL1 | 0,3467 | 9,1776E-18 |
| ACE2 | RASSF10 | 0,3467 | 9,01941E-18 |
| ACE2 | PPP1R9A | 0,3464 | 9,68991E-18 |
| ACE2 | KNSTRN | 0,3464 | 9,74107E-18 |
| ACE2 | EIF4E2 | 0,3462 | 1,02702E-17 |
| ACE2 | CHCHD4 | 0,3462 | 1,02995E-17 |
| ACE2 | ALDH7A1 | 0,3462 | 1,02097E-17 |
| ACE2 | TCTA | 0,3461 | 1,04185E-17 |
| ACE2 | KEAP1 | 0,346 | 1,05953E-17 |
| ACE2 | SLC30A6 | 0,3459 | 1,09776E-17 |
| ACE2 | CMTM8 | 0,3458 | 1,11918E-17 |
| ACE2 | SLC38A11 | 0,3457 | 1,1446E-17 |
| ACE2 | SELENBP1 | 0,3456 | 1,16162E-17 |
| ACE2 | TXNDC9 | 0,3456 | 1,17423E-17 |
| ACE2 | PLLP | 0,3455 | 1,1987E-17 |
| ACE2 | CENPBD1 | 0,3455 | 1,20118E-17 |
| ACE2 | CCDC71 | 0,3454 | 1,22644E-17 |
| ACE2 | DDX52 | 0,3454 | 1,21556E-17 |
| ACE2 | NECAB1 | 0,3452 | 1,27692E-17 |
| ACE2 | C6orf203 | 0,3451 | 1,30773E-17 |
| ACE2 | RP11-597D13.7 | 0,345 | 1,35115E-17 |
| ACE2 | LRP2BP | 0,3448 | 1,41016E-17 |
| ACE2 | TMEM199 | 0,3448 | 1,39745E-17 |
| ACE2 | EAF1 | 0,3447 | 1,44026E-17 |
| ACE2 | TMEM33 | 0,3447 | 1,42517E-17 |
| ACE2 | MDM2 | 0,3447 | 1,43836E-17 |
| ACE2 | ARHGEF19 | 0,3446 | 1,46611E-17 |
| ACE2 | SHH | 0,3446 | 1,47503E-17 |
| ACE2 | IDE | 0,3446 | 1,4641E-17 |
| ACE2 | SEC24C | 0,3444 | 1,54486E-17 |
| ACE2 | PYGB | 0,3444 | 1,54744E-17 |
| ACE2 | RP11-26L20.4 | 0,3443 | 1,5697E-17 |
| ACE2 | SLC30A3 | 0,3441 | 1,63391E-17 |
| ACE2 | REN | 0,3439 | 1,74037E-17 |
| ACE2 | ANXA4 | 0,3439 | 1,71841E-17 |
| ACE2 | SERP1 | 0,3439 | 1,72744E-17 |
| ACE2 | CLPTM1 | 0,3437 | 1,78508E-17 |
| ACE2 | CLCN5 | 0,3437 | 1,78575E-17 |

| | | | |
|---|---|---|---|
| ACE2 | WDR3 | 0,3436 | 1,86575E-17 |
| ACE2 | RAD1 | 0,3436 | 1,83626E-17 |
| ACE2 | MTX1 | 0,3435 | 1,89024E-17 |
| ACE2 | FAM3B | 0,3435 | 1,88341E-17 |
| ACE2 | SREBF2 | 0,3435 | 1,89851E-17 |
| ACE2 | JADE1 | 0,3434 | 1,912E-17 |
| ACE2 | METTL7A | 0,3434 | 1,91496E-17 |
| ACE2 | GNS | 0,3432 | 2,00462E-17 |
| ACE2 | RNFT1 | 0,3432 | 2,00438E-17 |
| ACE2 | MAB21L3 | 0,3431 | 2,07264E-17 |
| ACE2 | RRAGA | 0,3431 | 2,04992E-17 |
| ACE2 | ZNF587B | 0,3431 | 2,07079E-17 |
| ACE2 | MAFB | 0,3431 | 2,05945E-17 |
| ACE2 | FAM134B | 0,343 | 2,12E-17 |
| ACE2 | RP11-793H13.11 | 0,3429 | 2,18128E-17 |
| ACE2 | AP001065.15 | 0,3429 | 2,16533E-17 |
| ACE2 | RXRA | 0,3428 | 2,22363E-17 |
| ACE2 | SLC25A1 | 0,3428 | 2,222E-17 |
| ACE2 | RP11-10C24.3 | 0,3426 | 2,30961E-17 |
| ACE2 | TMEM62 | 0,3426 | 2,3127E-17 |
| ACE2 | TEX264 | 0,3425 | 2,37209E-17 |
| ACE2 | APEX1 | 0,3425 | 2,35651E-17 |
| ACE2 | RP1-63M2.7 | 0,3425 | 2,38989E-17 |
| ACE2 | SULT1C2 | 0,3424 | 2,43936E-17 |
| ACE2 | GPR34 | 0,3424 | 2,41379E-17 |
| ACE2 | GABBR2 | 0,3423 | 2,49569E-17 |
| ACE2 | IP6K3 | 0,3422 | 2,56483E-17 |
| ACE2 | AC073254.1 | 0,3421 | 2,57003E-17 |
| ACE2 | RN7SKP51 | 0,3421 | 2,60247E-17 |
| ACE2 | ELP3 | 0,342 | 2,64772E-17 |
| ACE2 | RP11-566K19.6 | 0,342 | 2,67386E-17 |
| ACE2 | MRPS9 | 0,3419 | 2,73511E-17 |
| ACE2 | LNX1 | 0,3419 | 2,71191E-17 |
| ACE2 | FBXL18 | 0,3419 | 2,70378E-17 |
| ACE2 | PICK1 | 0,3419 | 2,7147E-17 |
| ACE2 | MARVELD2 | 0,3418 | 2,77901E-17 |
| ACE2 | MYH14 | 0,3418 | 2,7768E-17 |
| ACE2 | MPHOSPH10 | 0,3417 | 2,81512E-17 |
| ACE2 | ADCY8 | 0,3417 | 2,81353E-17 |
| ACE2 | NMU | 0,3416 | 2,87219E-17 |
| ACE2 | GPRIN2 | 0,3416 | 2,93375E-17 |
| ACE2 | CTD-3116E22.8 | 0,3416 | 2,88691E-17 |
| ACE2 | ATP6AP1 | 0,3414 | 3,04835E-17 |
| ACE2 | PACSIN2 | 0,3413 | 3,08016E-17 |
| ACE2 | KCNG3 | 0,3412 | 3,1621E-17 |
| ACE2 | C7orf25 | 0,3412 | 3,21066E-17 |
| ACE2 | RGP1 | 0,3412 | 3,18916E-17 |
| ACE2 | GRAMD4 | 0,3411 | 3,22931E-17 |
| ACE2 | SLC39A9 | 0,3409 | 3,39822E-17 |
| ACE2 | NFE2L1 | 0,3409 | 3,39115E-17 |
| ACE2 | TMEM179B | 0,3408 | 3,48068E-17 |
| ACE2 | UBE2Q2P2 | 0,3407 | 3,57278E-17 |
| ACE2 | NFATC3 | 0,3407 | 3,55077E-17 |
| ACE2 | FAM86C1 | 0,3406 | 3,66918E-17 |

| | | | |
|---|---|---|---|
| ACE2 | ST13P2 | 0,3405 | 3,69694E-17 |
| ACE2 | PMM2 | 0,3405 | 3,72364E-17 |
| ACE2 | HMGCL | 0,3404 | 3,78939E-17 |
| ACE2 | RAB27B | 0,3404 | 3,78841E-17 |
| ACE2 | POLR1A | 0,3403 | 3,87674E-17 |
| ACE2 | RP11-266L9.8 | 0,3403 | 3,89643E-17 |
| ACE2 | LINC00482 | 0,3403 | 3,87622E-17 |
| ACE2 | CTSA | 0,3403 | 3,93516E-17 |
| ACE2 | NUDT18 | 0,3402 | 3,97839E-17 |
| ACE2 | YRDC | 0,34 | 4,20824E-17 |
| ACE2 | COX11 | 0,34 | 4,14962E-17 |
| ACE2 | PEX3 | 0,3398 | 4,35805E-17 |
| ACE2 | TMEM139 | 0,3398 | 4,36464E-17 |
| ACE2 | RRP9 | 0,3397 | 4,40625E-17 |
| ACE2 | C3orf33 | 0,3397 | 4,48095E-17 |
| ACE2 | FAM201A | 0,3397 | 4,45839E-17 |
| ACE2 | RP5-1120P11.1 | 0,3396 | 4,59044E-17 |
| ACE2 | RPL23AP64 | 0,3396 | 4,59998E-17 |
| ACE2 | AGR2 | 0,3395 | 4,65401E-17 |
| ACE2 | XPO4 | 0,3395 | 4,70395E-17 |
| ACE2 | AC007009.1 | 0,3394 | 4,71752E-17 |
| ACE2 | SMCR8 | 0,3394 | 4,74117E-17 |
| ACE2 | GPATCH3 | 0,3393 | 4,84752E-17 |
| ACE2 | PNLIPRP3 | 0,3393 | 4,9147E-17 |
| ACE2 | MRPS11 | 0,3393 | 4,81511E-17 |
| ACE2 | MDH1 | 0,3391 | 5,14564E-17 |
| ACE2 | RNF13 | 0,3391 | 5,04371E-17 |
| ACE2 | KB-318B8.7 | 0,339 | 5,2291E-17 |
| ACE2 | HMBS | 0,3389 | 5,28896E-17 |
| ACE2 | ARSD | 0,3388 | 5,4029E-17 |
| ACE2 | NIPAL3 | 0,3387 | 5,55525E-17 |
| ACE2 | TMEM241 | 0,3387 | 5,51963E-17 |
| ACE2 | ZC2HC1C | 0,3386 | 5,66301E-17 |
| ACE2 | URB2 | 0,3385 | 5,82861E-17 |
| ACE2 | PPP1R1B | 0,3385 | 5,81978E-17 |
| ACE2 | SELENOS | 0,3384 | 6,01482E-17 |
| ACE2 | RTN4RL1 | 0,3384 | 5,90389E-17 |
| ACE2 | CHD8 | 0,3383 | 6,06501E-17 |
| ACE2 | RIOK2 | 0,3381 | 6,35186E-17 |
| ACE2 | ISCA2 | 0,3381 | 6,36808E-17 |
| ACE2 | ADSL | 0,3381 | 6,33343E-17 |
| ACE2 | HIF1AN | 0,338 | 6,48814E-17 |
| ACE2 | HTATIP2 | 0,338 | 6,55341E-17 |
| ACE2 | LANCL3 | 0,338 | 6,56069E-17 |
| ACE2 | MTMR4 | 0,3379 | 6,71837E-17 |
| ACE2 | RNU1-38P | 0,3376 | 7,18266E-17 |
| ACE2 | ASPSCR1 | 0,3376 | 7,13824E-17 |
| ACE2 | SDC4 | 0,3376 | 7,13551E-17 |
| ACE2 | HYAL2 | 0,3375 | 7,2127E-17 |
| ACE2 | BRAP | 0,3375 | 7,28619E-17 |
| ACE2 | PTK6 | 0,3375 | 7,31888E-17 |
| ACE2 | POR | 0,3373 | 7,67159E-17 |
| ACE2 | KIAA1522 | 0,3372 | 7,82004E-17 |
| ACE2 | TRMT10A | 0,3371 | 7,92678E-17 |

| | | | |
|---|---|---|---|
| ACE2 | WWC1 | 0,3369 | 8,36241E-17 |
| ACE2 | SHTN1 | 0,3369 | 8,23539E-17 |
| ACE2 | DGCR12 | 0,3368 | 8,4391E-17 |
| ACE2 | RP11-440D17.3 | 0,3367 | 8,73849E-17 |
| ACE2 | GOLPH3L | 0,3365 | 8,98058E-17 |
| ACE2 | MICALCL | 0,3365 | 9,07141E-17 |
| ACE2 | GDPD1 | 0,3364 | 9,34855E-17 |
| ACE2 | FAM71E1 | 0,3364 | 9,33336E-17 |
| ACE2 | TXNRD2 | 0,3364 | 9,20208E-17 |
| ACE2 | LRRC47 | 0,3363 | 9,52981E-17 |
| ACE2 | UGT8 | 0,3362 | 9,70531E-17 |
| ACE2 | PTAFR | 0,336 | 1,01634E-16 |
| ACE2 | ABCC6P2 | 0,336 | 1,02141E-16 |
| ACE2 | PES1 | 0,3359 | 1,04398E-16 |
| ACE2 | EIF2A | 0,3357 | 1,08064E-16 |
| ACE2 | ZC3H13 | 0,3357 | 1,09571E-16 |
| ACE2 | DDT | 0,3356 | 1,10947E-16 |
| ACE2 | SLC31A1 | 0,3355 | 1,13451E-16 |
| ACE2 | GGT1 | 0,3355 | 1,12438E-16 |
| ACE2 | SDHC | 0,3354 | 1,15992E-16 |
| ACE2 | RBM15B | 0,3354 | 1,15286E-16 |
| ACE2 | KCTD18 | 0,3353 | 1,18338E-16 |
| ACE2 | FDXACB1 | 0,3352 | 1,19695E-16 |
| ACE2 | EML2 | 0,3352 | 1,20709E-16 |
| ACE2 | MBL1P | 0,3351 | 1,23249E-16 |
| ACE2 | RP1-154K9.2 | 0,3349 | 1,28418E-16 |
| ACE2 | RP11-80P20.3 | 0,3348 | 1,33188E-16 |
| ACE2 | ASB8 | 0,3348 | 1,30918E-16 |
| ACE2 | HOMER2 | 0,3346 | 1,37546E-16 |
| ACE2 | TRIB3 | 0,3346 | 1,38835E-16 |
| ACE2 | SLC1A4 | 0,3345 | 1,41842E-16 |
| ACE2 | OBSL1 | 0,3345 | 1,40947E-16 |
| ACE2 | MMGT1 | 0,3345 | 1,39821E-16 |
| ACE2 | KB-1995A5.4 | 0,3344 | 1,45817E-16 |
| ACE2 | CNOT10 | 0,3343 | 1,47542E-16 |
| ACE2 | LETM1 | 0,3343 | 1,46884E-16 |
| ACE2 | ARRB1 | 0,3343 | 1,46563E-16 |
| ACE2 | MUM1L1 | 0,3343 | 1,46567E-16 |
| ACE2 | LLGL2 | 0,3341 | 1,5527E-16 |
| ACE2 | BCL2L1 | 0,3341 | 1,52655E-16 |
| ACE2 | XKRX | 0,3341 | 1,55392E-16 |
| ACE2 | WNT4 | 0,334 | 1,5766E-16 |
| ACE2 | NPR1 | 0,334 | 1,56335E-16 |
| ACE2 | ZBTB24 | 0,3339 | 1,62756E-16 |
| ACE2 | PCCA | 0,3339 | 1,60659E-16 |
| ACE2 | RP4-569M23.4 | 0,3339 | 1,60611E-16 |
| ACE2 | RP11-311D14.1 | 0,3338 | 1,63234E-16 |
| ACE2 | USP54 | 0,3338 | 1,63325E-16 |
| ACE2 | DIO3 | 0,3338 | 1,63385E-16 |
| ACE2 | SSBP1 | 0,3337 | 1,66977E-16 |
| ACE2 | PTK2B | 0,3337 | 1,6697E-16 |
| ACE2 | CHCHD5 | 0,3336 | 1,72521E-16 |
| ACE2 | PPP1CA | 0,3336 | 1,72926E-16 |
| ACE2 | CES3 | 0,3335 | 1,76597E-16 |

| | | | |
|---|---|---|---|
| ACE2 | GCC1 | 0,3334 | 1,78299E-16 |
| ACE2 | TMPRSS2 | 0,3334 | 1,80677E-16 |
| ACE2 | OXSM | 0,3333 | 1,8507E-16 |
| ACE2 | FAXC | 0,3333 | 1,84375E-16 |
| ACE2 | ZBTB45 | 0,3333 | 1,82751E-16 |
| ACE2 | AC009237.8 | 0,3331 | 1,9046E-16 |
| ACE2 | IFT57 | 0,3331 | 1,90921E-16 |
| ACE2 | UTP20 | 0,3331 | 1,93898E-16 |
| ACE2 | TLE6 | 0,3331 | 1,92567E-16 |
| ACE2 | PPT1 | 0,333 | 1,95437E-16 |
| ACE2 | RP11-259N19.1 | 0,3329 | 2,00093E-16 |
| ACE2 | ULBP3 | 0,3329 | 2,01255E-16 |
| ACE2 | CCDC87 | 0,3329 | 2,01404E-16 |
| ACE2 | C11orf54 | 0,3329 | 2,00911E-16 |
| ACE2 | SLC5A8 | 0,3328 | 2,03654E-16 |
| ACE2 | RP11-447D11.3 | 0,3327 | 2,10106E-16 |
| ACE2 | HEXB | 0,3326 | 2,12823E-16 |
| ACE2 | ARFGAP2 | 0,3326 | 2,11964E-16 |
| ACE2 | GJB1 | 0,3326 | 2,12342E-16 |
| ACE2 | OCIAD2 | 0,3325 | 2,20823E-16 |
| ACE2 | GNMT | 0,3324 | 2,23351E-16 |
| ACE2 | PRIM1 | 0,3324 | 2,21582E-16 |
| ACE2 | CHKA | 0,3323 | 2,26951E-16 |
| ACE2 | HMGB3P26 | 0,3323 | 2,27323E-16 |
| ACE2 | ACOX1 | 0,3322 | 2,33382E-16 |
| ACE2 | MRPS28 | 0,3321 | 2,39993E-16 |
| ACE2 | SLC25A15 | 0,3321 | 2,4111E-16 |
| ACE2 | PIGV | 0,332 | 2,41808E-16 |
| ACE2 | SFRP5 | 0,332 | 2,42272E-16 |
| ACE2 | RP11-452H21.4 | 0,332 | 2,46096E-16 |
| ACE2 | HOOK2 | 0,3319 | 2,51377E-16 |
| ACE2 | GPC4 | 0,3319 | 2,49956E-16 |
| ACE2 | OSBP | 0,3318 | 2,54021E-16 |
| ACE2 | CTC-429P9.5 | 0,3318 | 2,53103E-16 |
| ACE2 | TM9SF1 | 0,3317 | 2,60127E-16 |
| ACE2 | SUV39H1 | 0,3317 | 2,59145E-16 |
| ACE2 | PARD6B | 0,3314 | 2,7764E-16 |
| ACE2 | TSC22D3 | 0,3314 | 2,75816E-16 |
| ACE2 | LINC01144 | 0,3313 | 2,8685E-16 |
| ACE2 | MFSD9 | 0,3312 | 2,89157E-16 |
| ACE2 | SH3PXD2B | 0,3312 | 2,91267E-16 |
| ACE2 | LINC00261 | 0,331 | 3,00992E-16 |
| ACE2 | C5orf38 | 0,3309 | 3,13041E-16 |
| ACE2 | DHX32 | 0,3309 | 3,07862E-16 |
| ACE2 | MECOM | 0,3308 | 3,18656E-16 |
| ACE2 | NUBP1 | 0,3308 | 3,1339E-16 |
| ACE2 | COQ9 | 0,3308 | 3,14184E-16 |
| ACE2 | CTD-2186M15.3 | 0,3306 | 3,30875E-16 |
| ACE2 | RP1-140C12.2 | 0,3306 | 3,28547E-16 |
| ACE2 | KLF5 | 0,3306 | 3,30958E-16 |
| ACE2 | CFAP221 | 0,3305 | 3,38842E-16 |
| ACE2 | RP1-313I6.12 | 0,3305 | 3,39451E-16 |
| ACE2 | ALDH3B1 | 0,3305 | 3,41591E-16 |
| ACE2 | JMJD7 | 0,3304 | 3,45719E-16 |

| | | | |
|---|---|---|---|
| ACE2 | ICMT | 0,3303 | 3,55192E-16 |
| ACE2 | CYB5R1 | 0,3303 | 3,56123E-16 |
| ACE2 | CYB561D2 | 0,3302 | 3,64124E-16 |
| ACE2 | KIAA0319L | 0,33 | 3,8044E-16 |
| ACE2 | ZAR1 | 0,33 | 3,77746E-16 |
| ACE2 | EXOSC5 | 0,33 | 3,7799E-16 |
| ACE2 | GLO1 | 0,3299 | 3,84659E-16 |
| ACE2 | NAPA | 0,3299 | 3,82911E-16 |
| ACE2 | PECR | 0,3297 | 4,05662E-16 |
| ACE2 | CTA-407F11.8 | 0,3297 | 4,01511E-16 |
| ACE2 | B3GALT6 | 0,3295 | 4,22454E-16 |
| ACE2 | RP11-196G11.4 | 0,3295 | 4,15473E-16 |
| ACE2 | CCNB3 | 0,3295 | 4,21682E-16 |
| ACE2 | CTBP1-AS2 | 0,3294 | 4,26246E-16 |
| ACE2 | GHRHR | 0,3294 | 4,30683E-16 |
| ACE2 | SLC39A3 | 0,3294 | 4,29167E-16 |
| ACE2 | GRAMD2 | 0,3293 | 4,35255E-16 |
| ACE2 | CLDN7 | 0,3293 | 4,38002E-16 |
| ACE2 | MASP2 | 0,329 | 4,71939E-16 |
| ACE2 | TANGO2 | 0,329 | 4,71265E-16 |
| ACE2 | NPM1P25 | 0,3289 | 4,73296E-16 |
| ACE2 | LAMTOR1 | 0,3289 | 4,79021E-16 |
| ACE2 | RNF6 | 0,3289 | 4,73588E-16 |
| ACE2 | PHKB | 0,3288 | 4,88518E-16 |
| ACE2 | SMIM1 | 0,3287 | 5,02661E-16 |
| ACE2 | ADAMTS15 | 0,3285 | 5,15254E-16 |
| ACE2 | MALSU1 | 0,3284 | 5,31975E-16 |
| ACE2 | GOLGA7B | 0,3284 | 5,26952E-16 |
| ACE2 | FPR2 | 0,3284 | 5,2654E-16 |
| ACE2 | WFDC2 | 0,3284 | 5,29011E-16 |
| ACE2 | SLC25A33 | 0,3283 | 5,45745E-16 |
| ACE2 | SLC38A1 | 0,3283 | 5,41685E-16 |
| ACE2 | MT-TN | 0,328 | 5,8164E-16 |
| ACE2 | MLST8 | 0,3279 | 5,90852E-16 |
| ACE2 | TMEM116 | 0,3278 | 6,0907E-16 |
| ACE2 | HDDC3 | 0,3278 | 5,9998E-16 |
| ACE2 | BICDL2 | 0,3278 | 6,01066E-16 |
| ACE2 | AREG | 0,3277 | 6,22546E-16 |
| ACE2 | SLC35A2 | 0,3277 | 6,1957E-16 |
| ACE2 | SPRYD7 | 0,3275 | 6,4653E-16 |
| ACE2 | SCNN1B | 0,3275 | 6,50843E-16 |
| ACE2 | MINCR | 0,3273 | 6,69655E-16 |
| ACE2 | DDB1 | 0,3273 | 6,80497E-16 |
| ACE2 | PQLC2 | 0,3272 | 6,90839E-16 |
| ACE2 | NT5DC2 | 0,3272 | 6,95269E-16 |
| ACE2 | MRPS18A | 0,3272 | 6,90867E-16 |
| ACE2 | UFSP1 | 0,3272 | 6,8569E-16 |
| ACE2 | SEL1L3 | 0,3271 | 6,9604E-16 |
| ACE2 | TRAPPC13 | 0,3271 | 7,08341E-16 |
| ACE2 | ZKSCAN2 | 0,3271 | 7,05165E-16 |
| ACE2 | TACO1 | 0,3269 | 7,2697E-16 |
| ACE2 | AGA | 0,3268 | 7,54469E-16 |
| ACE2 | CTD-2306A12.1 | 0,3268 | 7,44511E-16 |
| ACE2 | PLA2G12A | 0,3266 | 7,7447E-16 |

| | | | |
|---|---|---|---|
| ACE2 | PDXK | 0,3266 | 7,75354E-16 |
| ACE2 | HGD | 0,3265 | 7,95022E-16 |
| ACE2 | QDPR | 0,3265 | 8,01898E-16 |
| ACE2 | SLC38A7 | 0,3265 | 7,93618E-16 |
| ACE2 | AC100802.3 | 0,3264 | 8,14077E-16 |
| ACE2 | TMEM251 | 0,3263 | 8,4133E-16 |
| ACE2 | AGR3 | 0,3262 | 8,49187E-16 |
| ACE2 | RP11-250B2.6 | 0,3261 | 8,63077E-16 |
| ACE2 | MIEF1 | 0,3261 | 8,6919E-16 |
| ACE2 | PIR | 0,3261 | 8,6584E-16 |
| ACE2 | FECH | 0,326 | 8,93271E-16 |
| ACE2 | VAPA | 0,3258 | 9,36592E-16 |
| ACE2 | AC016745.3 | 0,3257 | 9,56782E-16 |
| ACE2 | YTHDF2 | 0,3256 | 9,74555E-16 |
| ACE2 | PCDHA13 | 0,3256 | 9,74562E-16 |
| ACE2 | RFESD | 0,3255 | 9,88003E-16 |
| ACE2 | RP11-863K10.7 | 0,3255 | 9,80354E-16 |
| ACE2 | DNTTIP1 | 0,3255 | 9,82649E-16 |
| ACE2 | RP11-51F16.9 | 0,3254 | 1,00558E-15 |
| ACE2 | AXIN2 | 0,3254 | 1,00256E-15 |
| ACE2 | RAP1GDS1 | 0,3253 | 1,02323E-15 |
| ACE2 | TRAPPC11 | 0,3252 | 1,04515E-15 |
| ACE2 | MMP7 | 0,3252 | 1,06165E-15 |
| ACE2 | TXNDC11 | 0,3252 | 1,05181E-15 |
| ACE2 | ZNF287 | 0,3252 | 1,04385E-15 |
| ACE2 | PRPF19 | 0,3251 | 1,08717E-15 |
| ACE2 | TMEM208 | 0,325 | 1,11093E-15 |
| ACE2 | TTI1 | 0,325 | 1,08799E-15 |
| ACE2 | AC007277.3 | 0,3249 | 1,1163E-15 |
| ACE2 | CDKN2AIPNL | 0,3249 | 1,11648E-15 |
| ACE2 | TSFM | 0,3249 | 1,11256E-15 |
| ACE2 | ACTL6A | 0,3248 | 1,13881E-15 |
| ACE2 | MRPS2 | 0,3248 | 1,14298E-15 |
| ACE2 | AP3B2 | 0,3248 | 1,15749E-15 |
| ACE2 | CLINT1 | 0,3247 | 1,16275E-15 |
| ACE2 | RP11-238K6.1 | 0,3247 | 1,18033E-15 |
| ACE2 | CCPG1 | 0,3247 | 1,16452E-15 |
| ACE2 | AP5S1 | 0,3247 | 1,16273E-15 |
| ACE2 | RPL18AP7 | 0,3246 | 1,19979E-15 |
| ACE2 | SOAT1 | 0,3245 | 1,2327E-15 |
| ACE2 | RHOT1 | 0,3245 | 1,23451E-15 |
| ACE2 | PAK5 | 0,3243 | 1,28049E-15 |
| ACE2 | SNX2 | 0,3242 | 1,31569E-15 |
| ACE2 | TBL2 | 0,3242 | 1,3049E-15 |
| ACE2 | NIPSNAP1 | 0,3242 | 1,29464E-15 |
| ACE2 | TMEM147 | 0,3241 | 1,3332E-15 |
| ACE2 | MLYCD | 0,3239 | 1,38715E-15 |
| ACE2 | DBT | 0,3238 | 1,40504E-15 |
| ACE2 | TMPPE | 0,3238 | 1,43221E-15 |
| ACE2 | IGF2R | 0,3238 | 1,43131E-15 |
| ACE2 | CHIA | 0,3237 | 1,44624E-15 |
| ACE2 | RP11-452G18.2 | 0,3237 | 1,46109E-15 |
| ACE2 | ZDHHC5 | 0,3237 | 1,43856E-15 |
| ACE2 | FARS2 | 0,3236 | 1,48362E-15 |

| | | | |
|---|---|---|---|
| ACE2 | LENG9 | 0,3236 | 1,46356E-15 |
| ACE2 | DIEXF | 0,3235 | 1,50044E-15 |
| ACE2 | MMAA | 0,3235 | 1,51409E-15 |
| ACE2 | ZSCAN25 | 0,3235 | 1,49946E-15 |
| ACE2 | AKR1C3 | 0,3234 | 1,53605E-15 |
| ACE2 | FNDC10 | 0,3233 | 1,57882E-15 |
| ACE2 | HSDL2 | 0,3233 | 1,59037E-15 |
| ACE2 | RP11-110I1.11 | 0,3233 | 1,58126E-15 |
| ACE2 | DAG1 | 0,3232 | 1,59419E-15 |
| ACE2 | RP11-767I20.1 | 0,3231 | 1,63485E-15 |
| ACE2 | GORASP2 | 0,323 | 1,6937E-15 |
| ACE2 | MARS2 | 0,323 | 1,67722E-15 |
| ACE2 | SLC35D2 | 0,323 | 1,69181E-15 |
| ACE2 | MANEA | 0,3229 | 1,72081E-15 |
| ACE2 | SMC3 | 0,3229 | 1,71591E-15 |
| ACE2 | CLN6 | 0,3228 | 1,73859E-15 |
| ACE2 | LA16c-358B7.4 | 0,3228 | 1,76763E-15 |
| ACE2 | MORF4L2-AS1 | 0,3222 | 2,00472E-15 |
| ACE2 | MT-TL1 | 0,3221 | 2,00728E-15 |
| ACE2 | PDIA6 | 0,322 | 2,06458E-15 |
| ACE2 | CTC-277H1.6 | 0,322 | 2,06214E-15 |
| ACE2 | PIGM | 0,3219 | 2,12085E-15 |
| ACE2 | TSPYL5 | 0,3219 | 2,1286E-15 |
| ACE2 | LRRC52 | 0,3217 | 2,20446E-15 |
| ACE2 | C3orf80 | 0,3217 | 2,19235E-15 |
| ACE2 | NLN | 0,3217 | 2,18886E-15 |
| ACE2 | PPP4R4 | 0,3217 | 2,20162E-15 |
| ACE2 | RP11-309L24.10 | 0,3216 | 2,2451E-15 |
| ACE2 | CTD-2562J17.6 | 0,3216 | 2,26619E-15 |
| ACE2 | RP11-65E22.2 | 0,3215 | 2,29114E-15 |
| ACE2 | RENBP | 0,3215 | 2,3063E-15 |
| ACE2 | MTMR14 | 0,3214 | 2,34098E-15 |
| ACE2 | PEX7 | 0,3214 | 2,36239E-15 |
| ACE2 | ANKRD1 | 0,3212 | 2,44321E-15 |
| ACE2 | LLPH | 0,321 | 2,54258E-15 |
| ACE2 | C14orf119 | 0,321 | 2,53437E-15 |
| ACE2 | TTC30A | 0,3208 | 2,63606E-15 |
| ACE2 | SDAD1 | 0,3208 | 2,67301E-15 |
| ACE2 | SLC39A8 | 0,3208 | 2,66717E-15 |
| ACE2 | TSTD1 | 0,3207 | 2,69607E-15 |
| ACE2 | SIRT5 | 0,3207 | 2,70449E-15 |
| ACE2 | NRGN | 0,3207 | 2,72697E-15 |
| ACE2 | C14orf132 | 0,3206 | 2,80082E-15 |
| ACE2 | PIGP | 0,3206 | 2,75295E-15 |
| ACE2 | KDM4D | 0,3204 | 2,88192E-15 |
| ACE2 | ELOA | 0,3203 | 2,94272E-15 |
| ACE2 | PAIP2B | 0,3203 | 2,93687E-15 |
| ACE2 | MRPS30 | 0,3201 | 3,07161E-15 |
| ACE2 | UBE4A | 0,3201 | 3,07389E-15 |
| ACE2 | COMMD9 | 0,32 | 3,1643E-15 |
| ACE2 | HAS3 | 0,32 | 3,16168E-15 |
| ACE2 | TAF6L | 0,3198 | 3,27413E-15 |
| ACE2 | RPLP0 | 0,3198 | 3,29497E-15 |
| ACE2 | SGK3 | 0,3197 | 3,37653E-15 |

| | | | |
|---|---|---|---|
| ACE2 | CKMT1A | 0,3197 | 3,32903E-15 |
| ACE2 | VPS4B | 0,3197 | 3,34979E-15 |
| ACE2 | RP11-128B16.5 | 0,3196 | 3,43051E-15 |
| ACE2 | ADCK2 | 0,3195 | 3,48765E-15 |
| ACE2 | SPTLC1 | 0,3195 | 3,47445E-15 |
| ACE2 | PRKD1 | 0,3195 | 3,46496E-15 |
| ACE2 | SPCS1 | 0,3194 | 3,58132E-15 |
| ACE2 | WIPI1 | 0,3194 | 3,53616E-15 |
| ACE2 | MGAM2 | 0,3192 | 3,68944E-15 |
| ACE2 | HMGB1P44 | 0,3191 | 3,76271E-15 |
| ACE2 | GFM2 | 0,3191 | 3,83422E-15 |
| ACE2 | RP11-10C24.1 | 0,3188 | 3,99947E-15 |
| ACE2 | SLC25A3 | 0,3187 | 4,13458E-15 |
| ACE2 | ZMPSTE24 | 0,3186 | 4,18939E-15 |
| ACE2 | POPDC3 | 0,3186 | 4,22094E-15 |
| ACE2 | RP11-528M18.2 | 0,3185 | 4,32302E-15 |
| ACE2 | BTF3L4 | 0,3184 | 4,34889E-15 |
| ACE2 | CYB561D1 | 0,3184 | 4,40985E-15 |
| ACE2 | NOA1 | 0,3182 | 4,5618E-15 |
| ACE2 | ENOSF1 | 0,3182 | 4,57782E-15 |
| ACE2 | CBLC | 0,3182 | 4,54493E-15 |
| ACE2 | PRDX5 | 0,3181 | 4,67829E-15 |
| ACE2 | RP11-403A3.3 | 0,3181 | 4,66553E-15 |
| ACE2 | FBXL8 | 0,3179 | 4,90296E-15 |
| ACE2 | LRG1 | 0,3178 | 4,96564E-15 |
| ACE2 | MIOS | 0,3177 | 5,05265E-15 |
| ACE2 | STEAP3 | 0,3176 | 5,13167E-15 |
| ACE2 | CLDN12 | 0,3176 | 5,15925E-15 |
| ACE2 | DMBT1 | 0,3175 | 5,30635E-15 |
| ACE2 | MTOR | 0,3174 | 5,42274E-15 |
| ACE2 | DTNB | 0,3174 | 5,41667E-15 |
| ACE2 | SUCLG2 | 0,3173 | 5,54644E-15 |
| ACE2 | RPL7L1 | 0,3173 | 5,45835E-15 |
| ACE2 | SUOX | 0,3173 | 5,552E-15 |
| ACE2 | ZNF165 | 0,3172 | 5,65952E-15 |
| ACE2 | CXorf56 | 0,3171 | 5,72999E-15 |
| ACE2 | NDC1 | 0,317 | 5,91577E-15 |
| ACE2 | SDHAF4 | 0,3169 | 5,92556E-15 |
| ACE2 | THTPA | 0,3168 | 6,04839E-15 |
| ACE2 | C1GALT1C1 | 0,3168 | 6,0619E-15 |
| ACE2 | KLHL29 | 0,3167 | 6,18253E-15 |
| ACE2 | GALNT7 | 0,3167 | 6,19042E-15 |
| ACE2 | C6orf89 | 0,3167 | 6,27809E-15 |
| ACE2 | FZD6 | 0,3167 | 6,2142E-15 |
| ACE2 | PIK3R4 | 0,3166 | 6,3956E-15 |
| ACE2 | NUDT16L1 | 0,3164 | 6,58136E-15 |
| ACE2 | MRPS12 | 0,3164 | 6,65812E-15 |
| ACE2 | CUL3 | 0,3163 | 6,72326E-15 |
| ACE2 | CTD-2410N18.3 | 0,3163 | 6,77962E-15 |
| ACE2 | RMDN3 | 0,3162 | 6,95163E-15 |
| ACE2 | PRELID3B | 0,3162 | 6,88598E-15 |
| ACE2 | PSMC2 | 0,3161 | 7,11775E-15 |
| ACE2 | BRF2 | 0,3161 | 7,09016E-15 |
| ACE2 | GID8 | 0,3161 | 7,11527E-15 |

| | | | |
|---|---|---|---|
| ACE2 | UQCRC1 | 0,316 | 7,2057E-15 |
| ACE2 | SPOCK1 | 0,316 | 7,19909E-15 |
| ACE2 | MIR27B | 0,316 | 7,15652E-15 |
| ACE2 | SLC52A3 | 0,316 | 7,16533E-15 |
| ACE2 | BANK1 | 0,3159 | 7,39807E-15 |
| ACE2 | FBXW2 | 0,3159 | 7,40878E-15 |
| ACE2 | ATL1 | 0,3159 | 7,3668E-15 |
| ACE2 | DRG1 | 0,3159 | 7,41504E-15 |
| ACE2 | PRDX4 | 0,3159 | 7,3858E-15 |
| ACE2 | MT-TT | 0,3159 | 7,30767E-15 |
| ACE2 | CACNB4 | 0,3158 | 7,53602E-15 |
| ACE2 | POU2F3 | 0,3158 | 7,455E-15 |
| ACE2 | TMED2 | 0,3157 | 7,70784E-15 |
| ACE2 | VWDE | 0,3156 | 7,87478E-15 |
| ACE2 | POP7 | 0,3156 | 7,81306E-15 |
| ACE2 | JPH1 | 0,3156 | 7,85218E-15 |
| ACE2 | FZD8 | 0,3156 | 7,83923E-15 |
| ACE2 | TNFSF13 | 0,3156 | 7,84491E-15 |
| ACE2 | LLNLR-304A6.2 | 0,3156 | 7,73479E-15 |
| ACE2 | LARP1 | 0,3155 | 8,01317E-15 |
| ACE2 | TMEM102 | 0,3155 | 8,00838E-15 |
| ACE2 | SRPRB | 0,3154 | 8,20141E-15 |
| ACE2 | CTSB | 0,3154 | 8,08504E-15 |
| ACE2 | ERICH2 | 0,3152 | 8,4604E-15 |
| ACE2 | MRPL14 | 0,3152 | 8,45982E-15 |
| ACE2 | ZBED6CL | 0,3152 | 8,50649E-15 |
| ACE2 | PCDH9 | 0,3152 | 8,54679E-15 |
| ACE2 | ITFG1 | 0,3152 | 8,40482E-15 |
| ACE2 | DNM2 | 0,3152 | 8,55726E-15 |
| ACE2 | SH2D6 | 0,3151 | 8,63385E-15 |
| ACE2 | KDM4A | 0,315 | 8,86204E-15 |
| ACE2 | ODAM | 0,315 | 8,85148E-15 |
| ACE2 | CHUK | 0,315 | 8,80747E-15 |
| ACE2 | PLEKHD1 | 0,315 | 8,76625E-15 |
| ACE2 | ZNF324 | 0,315 | 8,91552E-15 |
| ACE2 | RP11-142E9.1 | 0,3149 | 9,05032E-15 |
| ACE2 | CNPY2 | 0,3148 | 9,12378E-15 |
| ACE2 | TTC22 | 0,3147 | 9,46545E-15 |
| ACE2 | ATP6V1E2 | 0,3146 | 9,67481E-15 |
| ACE2 | PTPRF | 0,3145 | 9,70411E-15 |
| ACE2 | ALG1L5P | 0,3145 | 9,86486E-15 |
| ACE2 | NAGPA-AS1 | 0,3145 | 9,79415E-15 |
| ACE2 | WFDC13 | 0,3145 | 9,8062E-15 |
| ACE2 | POLDIP2 | 0,3144 | 1,00847E-14 |
| ACE2 | HS1BP3 | 0,3142 | 1,04742E-14 |
| ACE2 | GANAB | 0,3142 | 1,04736E-14 |
| ACE2 | COX10 | 0,3141 | 1,05256E-14 |
| ACE2 | ZC3H4 | 0,3141 | 1,05692E-14 |
| ACE2 | SMARCA5 | 0,314 | 1,07321E-14 |
| ACE2 | TBCC | 0,314 | 1,08239E-14 |
| ACE2 | SLC30A9 | 0,3139 | 1,10212E-14 |
| ACE2 | CLGN | 0,3139 | 1,11448E-14 |
| ACE2 | EEF1E1 | 0,3139 | 1,10179E-14 |
| ACE2 | RP11-11N9.4 | 0,3139 | 1,09647E-14 |

| | | | |
|---|---|---|---|
| ACE2 | PCNA | 0,3139 | 1,10356E-14 |
| ACE2 | CGN | 0,3137 | 1,1543E-14 |
| ACE2 | RP11-63E9.1 | 0,3137 | 1,15897E-14 |
| ACE2 | KNDC1 | 0,3137 | 1,16022E-14 |
| ACE2 | WDR72 | 0,3137 | 1,15285E-14 |
| ACE2 | RP11-458J1.1 | 0,3137 | 1,14354E-14 |
| ACE2 | MIS18BP1 | 0,3136 | 1,16691E-14 |
| ACE2 | NPAS2 | 0,3135 | 1,19061E-14 |
| ACE2 | FABP5 | 0,3135 | 1,19855E-14 |
| ACE2 | RCN2 | 0,3135 | 1,20934E-14 |
| ACE2 | ZNF784 | 0,3135 | 1,20993E-14 |
| ACE2 | SNX25P1 | 0,3134 | 1,23307E-14 |
| ACE2 | NNT | 0,3134 | 1,23269E-14 |
| ACE2 | NQO1 | 0,3134 | 1,21814E-14 |
| ACE2 | MAGI3 | 0,3133 | 1,23854E-14 |
| ACE2 | CTD-2547L16.3 | 0,3133 | 1,23739E-14 |
| ACE2 | SLC39A4 | 0,3133 | 1,24318E-14 |
| ACE2 | TOR2A | 0,3133 | 1,25066E-14 |
| ACE2 | CCDC43 | 0,3133 | 1,25354E-14 |
| ACE2 | FEN1 | 0,3131 | 1,31056E-14 |
| ACE2 | MRPL40 | 0,3131 | 1,30786E-14 |
| ACE2 | SHROOM3 | 0,313 | 1,3155E-14 |
| ACE2 | RP13-476E20.1 | 0,3129 | 1,36049E-14 |
| ACE2 | MIEN1 | 0,3129 | 1,34272E-14 |
| ACE2 | AC083900.1 | 0,3128 | 1,38919E-14 |
| ACE2 | NAIF1 | 0,3128 | 1,37365E-14 |
| ACE2 | CAPN3 | 0,3128 | 1,39014E-14 |
| ACE2 | HNRNPA1P59 | 0,3127 | 1,42396E-14 |
| ACE2 | AQR | 0,3127 | 1,41736E-14 |
| ACE2 | PLPP6 | 0,3125 | 1,47442E-14 |
| ACE2 | ATP5F1 | 0,3124 | 1,48839E-14 |
| ACE2 | RP11-121C2.2 | 0,3124 | 1,51032E-14 |
| ACE2 | RP11-276H7.3 | 0,3123 | 1,52056E-14 |
| ACE2 | SPRY2 | 0,3123 | 1,52396E-14 |
| ACE2 | RCBTB1 | 0,3122 | 1,57003E-14 |
| ACE2 | CTA-221G9.12 | 0,3122 | 1,54949E-14 |
| ACE2 | LINC01814 | 0,3121 | 1,58199E-14 |
| ACE2 | METAP1 | 0,3121 | 1,601E-14 |
| ACE2 | PLRG1 | 0,3121 | 1,60585E-14 |
| ACE2 | PGD | 0,3119 | 1,66397E-14 |
| ACE2 | IMPA1 | 0,3117 | 1,73304E-14 |
| ACE2 | RP1-37N7.1 | 0,3117 | 1,73635E-14 |
| ACE2 | KB-1995A5.5 | 0,3117 | 1,7341E-14 |
| ACE2 | PTGFR | 0,3113 | 1,88764E-14 |
| ACE2 | PSMD14 | 0,3112 | 1,89636E-14 |
| ACE2 | PIK3R1 | 0,3111 | 1,96899E-14 |
| ACE2 | RP11-539I5.1 | 0,3111 | 1,96076E-14 |
| ACE2 | ESRRA | 0,3111 | 1,96086E-14 |
| ACE2 | GPLD1 | 0,311 | 2,00509E-14 |
| ACE2 | AC004019.13 | 0,311 | 1,99708E-14 |
| ACE2 | GALNT12 | 0,3109 | 2,01059E-14 |
| ACE2 | AP000695.6 | 0,3109 | 2,04761E-14 |
| ACE2 | RPL26L1 | 0,3108 | 2,05971E-14 |
| ACE2 | HINT2 | 0,3107 | 2,10754E-14 |

| | | | |
|---|---|---|---|
| ACE2 | ACADM | 0,3106 | 2,15279E-14 |
| ACE2 | ACOXL-AS1 | 0,3104 | 2,22698E-14 |
| ACE2 | SLC15A2 | 0,3104 | 2,26128E-14 |
| ACE2 | RP11-395N21.2 | 0,3104 | 2,23079E-14 |
| ACE2 | LSM14B | 0,3103 | 2,31305E-14 |
| ACE2 | DGCR6 | 0,3103 | 2,31122E-14 |
| ACE2 | LDHD | 0,3102 | 2,35987E-14 |
| ACE2 | CITF22-49E9.3 | 0,3102 | 2,35901E-14 |
| ACE2 | BMP1 | 0,3101 | 2,36531E-14 |
| ACE2 | RP11-166P13.4 | 0,3101 | 2,39415E-14 |
| ACE2 | IL10 | 0,31 | 2,43215E-14 |
| ACE2 | ADRB2 | 0,31 | 2,41977E-14 |
| ACE2 | AARS | 0,31 | 2,42437E-14 |
| ACE2 | RP11-353N4.5 | 0,3099 | 2,48062E-14 |
| ACE2 | OS9 | 0,3099 | 2,49875E-14 |
| ACE2 | GOSR1 | 0,3099 | 2,48636E-14 |
| ACE2 | TMEM17 | 0,3097 | 2,58329E-14 |
| ACE2 | UQCR10 | 0,3097 | 2,57222E-14 |
| ACE2 | DHFR2 | 0,3096 | 2,6437E-14 |
| ACE2 | RP11-714G18.1 | 0,3096 | 2,6416E-14 |
| ACE2 | STOML2 | 0,3096 | 2,65476E-14 |
| ACE2 | DPP9-AS1 | 0,3096 | 2,64807E-14 |
| ACE2 | CEACAM6 | 0,3096 | 2,63846E-14 |
| ACE2 | ABHD11 | 0,3095 | 2,71226E-14 |
| ACE2 | DBR1 | 0,3094 | 2,73305E-14 |
| ACE2 | XXbac-B135H6.18 | 0,3094 | 2,7234E-14 |
| ACE2 | ARMC6 | 0,3093 | 2,79112E-14 |
| ACE2 | PTPN20 | 0,3092 | 2,84046E-14 |
| ACE2 | IYD | 0,3091 | 2,8965E-14 |
| ACE2 | RP11-12G12.7 | 0,3091 | 2,92679E-14 |
| ACE2 | MFSD3 | 0,309 | 2,96068E-14 |
| ACE2 | MFSD6 | 0,3089 | 3,03236E-14 |
| ACE2 | C19orf73 | 0,3088 | 3,11372E-14 |
| ACE2 | MRPL39 | 0,3088 | 3,08144E-14 |
| ACE2 | HNRNPUL2 | 0,3087 | 3,16978E-14 |
| ACE2 | IRAK3 | 0,3087 | 3,12596E-14 |
| ACE2 | ZDHHC7 | 0,3087 | 3,13669E-14 |
| ACE2 | PRR22 | 0,3087 | 3,13273E-14 |
| ACE2 | HDHD5 | 0,3087 | 3,15591E-14 |
| ACE2 | CA2 | 0,3086 | 3,22692E-14 |
| ACE2 | IFT80 | 0,3085 | 3,26057E-14 |
| ACE2 | MT-TI | 0,3085 | 3,27433E-14 |
| ACE2 | RPL31P40 | 0,3084 | 3,32612E-14 |
| ACE2 | SPTY2D1 | 0,3084 | 3,31784E-14 |
| ACE2 | KCNB1 | 0,3084 | 3,37351E-14 |
| ACE2 | GRID2IP | 0,3082 | 3,49668E-14 |
| ACE2 | DHODH | 0,3082 | 3,4915E-14 |
| ACE2 | RP11-254F19.4 | 0,3082 | 3,4914E-14 |
| ACE2 | CKAP5 | 0,3081 | 3,53891E-14 |
| ACE2 | TSN | 0,308 | 3,63316E-14 |
| ACE2 | TIGD2 | 0,308 | 3,62862E-14 |
| ACE2 | CTNS | 0,308 | 3,64372E-14 |
| ACE2 | LINC00891 | 0,308 | 3,59405E-14 |
| ACE2 | CORO1B | 0,3079 | 3,69183E-14 |

| | | | |
|---|---|---|---|
| ACE2 | PABPC4 | 0,3078 | 3,80133E-14 |
| ACE2 | USP51 | 0,3078 | 3,74102E-14 |
| ACE2 | PSMD2 | 0,3076 | 3,91568E-14 |
| ACE2 | LANCL2 | 0,3076 | 3,96088E-14 |
| ACE2 | PHOSPHO2 | 0,3075 | 4,00216E-14 |
| ACE2 | RP11-826N14.2 | 0,3075 | 3,99083E-14 |
| ACE2 | PINK1-AS | 0,3074 | 4,07795E-14 |
| ACE2 | MTMR6 | 0,3074 | 4,05422E-14 |
| ACE2 | OAZ3 | 0,3072 | 4,29204E-14 |
| ACE2 | C19orf81 | 0,3072 | 4,29528E-14 |
| ACE2 | FAAH | 0,3071 | 4,29843E-14 |
| ACE2 | THEM6 | 0,3071 | 4,36603E-14 |
| ACE2 | ZNF185 | 0,3071 | 4,37113E-14 |
| ACE2 | CDK5 | 0,307 | 4,42613E-14 |
| ACE2 | NCOA6 | 0,307 | 4,43563E-14 |
| ACE2 | ZNF425 | 0,3069 | 4,51121E-14 |
| ACE2 | VAPB | 0,3069 | 4,54533E-14 |
| ACE2 | FUCA1 | 0,3068 | 4,63486E-14 |
| ACE2 | C1orf132 | 0,3068 | 4,61785E-14 |
| ACE2 | LACE1 | 0,3068 | 4,5838E-14 |
| ACE2 | BLVRA | 0,3068 | 4,64684E-14 |
| ACE2 | SERPINA3 | 0,3068 | 4,59519E-14 |
| ACE2 | RNF20 | 0,3067 | 4,72263E-14 |
| ACE2 | RP11-964E11.3 | 0,3067 | 4,70495E-14 |
| ACE2 | ASPHD1 | 0,3067 | 4,73076E-14 |
| ACE2 | HERPUD1 | 0,3067 | 4,70111E-14 |
| ACE2 | RNF225 | 0,3067 | 4,6652E-14 |
| ACE2 | RRP15 | 0,3066 | 4,78571E-14 |
| ACE2 | GDI2 | 0,3066 | 4,77162E-14 |
| ACE2 | PDCD10 | 0,3065 | 4,8741E-14 |
| ACE2 | ENDOG | 0,3065 | 4,84372E-14 |
| ACE2 | PTPRM | 0,3064 | 5,01037E-14 |
| ACE2 | PSMD1 | 0,3063 | 5,06459E-14 |
| ACE2 | RP11-405M12.4 | 0,3063 | 5,04512E-14 |
| ACE2 | RFC1 | 0,3061 | 5,27266E-14 |
| ACE2 | SLC16A6 | 0,3061 | 5,33155E-14 |
| ACE2 | ARFGEF2 | 0,3061 | 5,31477E-14 |
| ACE2 | RHOXF1-AS1 | 0,3061 | 5,27657E-14 |
| ACE2 | AL450992.2 | 0,306 | 5,4248E-14 |
| ACE2 | AC105053.3 | 0,306 | 5,36322E-14 |
| ACE2 | KLHL12 | 0,3059 | 5,46652E-14 |
| ACE2 | TOR1A | 0,3059 | 5,49086E-14 |
| ACE2 | UTP14C | 0,3059 | 5,5083E-14 |
| ACE2 | C5orf51 | 0,3058 | 5,64091E-14 |
| ACE2 | LYPLA1 | 0,3058 | 5,66504E-14 |
| ACE2 | FAM213A | 0,3058 | 5,6174E-14 |
| ACE2 | KLLN | 0,3058 | 5,66386E-14 |
| ACE2 | CYC1 | 0,3057 | 5,73787E-14 |
| ACE2 | RP5-965G21.6 | 0,3057 | 5,7061E-14 |
| ACE2 | RAB19 | 0,3056 | 5,80727E-14 |
| ACE2 | INSR | 0,3056 | 5,82814E-14 |
| ACE2 | UTP14A | 0,3056 | 5,86128E-14 |
| ACE2 | PIP5KL1 | 0,3055 | 5,98406E-14 |
| ACE2 | MT-TY | 0,3055 | 5,9657E-14 |

| | | | |
|---|---|---|---|
| ACE2 | GIPC1 | 0,3054 | 6,08732E-14 |
| ACE2 | FGG | 0,3053 | 6,23383E-14 |
| ACE2 | SLC1A1 | 0,3053 | 6,25062E-14 |
| ACE2 | RP4-681L3.2 | 0,3052 | 6,35673E-14 |
| ACE2 | B4GALNT3 | 0,3052 | 6,29549E-14 |
| ACE2 | RP11-178C3.2 | 0,3051 | 6,45817E-14 |
| ACE2 | TMEM43 | 0,305 | 6,59682E-14 |
| ACE2 | ERLIN2 | 0,305 | 6,55371E-14 |
| ACE2 | MT-TC | 0,305 | 6,52716E-14 |
| ACE2 | KIAA1211L | 0,3049 | 6,69753E-14 |
| ACE2 | ZNHIT2 | 0,3049 | 6,69738E-14 |
| ACE2 | HSD17B6 | 0,3048 | 6,82308E-14 |
| ACE2 | HAGH | 0,3048 | 6,87062E-14 |
| ACE2 | ETFB | 0,3048 | 6,83974E-14 |
| ACE2 | HMGB3 | 0,3048 | 6,78086E-14 |
| ACE2 | NUP133 | 0,3047 | 6,94421E-14 |
| ACE2 | TRIM61 | 0,3047 | 6,96082E-14 |
| ACE2 | CEP104 | 0,3046 | 7,13147E-14 |
| ACE2 | TSPYL1 | 0,3046 | 7,0755E-14 |
| ACE2 | NUDT19 | 0,3046 | 7,08921E-14 |
| ACE2 | PSMD8 | 0,3046 | 7,11838E-14 |
| ACE2 | SLC35A5 | 0,3045 | 7,19152E-14 |
| ACE2 | SCPEP1 | 0,3045 | 7,17998E-14 |
| ACE2 | RP11-343L5.2 | 0,3044 | 7,43859E-14 |
| ACE2 | INTS7 | 0,3043 | 7,52114E-14 |
| ACE2 | APOLD1 | 0,3043 | 7,55242E-14 |
| ACE2 | ACADS | 0,3043 | 7,50146E-14 |
| ACE2 | TMEM63B | 0,3042 | 7,69983E-14 |
| ACE2 | NACC1 | 0,3042 | 7,64552E-14 |
| ACE2 | HSD17B12 | 0,3041 | 7,77306E-14 |
| ACE2 | TARS2 | 0,304 | 7,93919E-14 |
| ACE2 | DGUOK-AS1 | 0,304 | 7,97612E-14 |
| ACE2 | TXNDC17 | 0,304 | 7,95343E-14 |
| ACE2 | AC007405.4 | 0,3039 | 8,15307E-14 |
| ACE2 | TES | 0,3039 | 8,20752E-14 |
| ACE2 | THNSL1 | 0,3039 | 8,11209E-14 |
| ACE2 | NDUFA9 | 0,3039 | 8,15624E-14 |
| ACE2 | LINC00704 | 0,3038 | 8,37594E-14 |
| ACE2 | CCDC155 | 0,3038 | 8,37477E-14 |
| ACE2 | MTND5P11 | 0,3037 | 8,49768E-14 |
| ACE2 | RP11-54H7.4 | 0,3037 | 8,53873E-14 |
| ACE2 | DMRTA1 | 0,3036 | 8,67937E-14 |
| ACE2 | PLEKHB1 | 0,3036 | 8,6555E-14 |
| ACE2 | ABT1 | 0,3035 | 8,77961E-14 |
| ACE2 | UNC13B | 0,3035 | 8,75202E-14 |
| ACE2 | EARS2 | 0,3035 | 8,88562E-14 |
| ACE2 | IMP3 | 0,3034 | 9,007E-14 |
| ACE2 | VAC14 | 0,3034 | 9,02704E-14 |
| ACE2 | CCDC69 | 0,3033 | 9,23963E-14 |
| ACE2 | AKAP13 | 0,3033 | 9,26314E-14 |
| ACE2 | CDH15 | 0,3033 | 9,20039E-14 |
| ACE2 | CTC-281F24.5 | 0,3033 | 9,21454E-14 |
| ACE2 | MRPS7 | 0,3033 | 9,21357E-14 |
| ACE2 | NUP214 | 0,3032 | 9,43706E-14 |

| | | | |
|---|---|---|---|
| ACE2 | MIEF2 | 0,3032 | 9,35272E-14 |
| ACE2 | EEF2KMT | 0,3031 | 9,63119E-14 |
| ACE2 | NF2 | 0,3031 | 9,46006E-14 |
| ACE2 | SIMC1 | 0,3027 | 1,02654E-13 |
| ACE2 | DACT2 | 0,3027 | 1,02846E-13 |
| ACE2 | RP11-517B11.7 | 0,3026 | 1,05597E-13 |
| ACE2 | ZDHHC24 | 0,3026 | 1,05926E-13 |
| ACE2 | ADIPOR2 | 0,3025 | 1,07036E-13 |
| ACE2 | CTC-497E21.3 | 0,3023 | 1,11634E-13 |
| ACE2 | RN7SL650P | 0,3023 | 1,11093E-13 |
| ACE2 | RRN3 | 0,3023 | 1,12453E-13 |
| ACE2 | LAPTM4B | 0,3021 | 1,16668E-13 |
| ACE2 | PDGFRA | 0,302 | 1,19322E-13 |
| ACE2 | USP5 | 0,3019 | 1,19986E-13 |
| ACE2 | PLEKHG4 | 0,3019 | 1,20742E-13 |
| ACE2 | SIPA1L3 | 0,3019 | 1,20276E-13 |
| ACE2 | PRLH | 0,3018 | 1,22421E-13 |
| ACE2 | TRAPPC8 | 0,3018 | 1,22355E-13 |
| ACE2 | ZSWIM1 | 0,3018 | 1,22571E-13 |
| ACE2 | NEXN | 0,3017 | 1,26101E-13 |
| ACE2 | RP11-462G2.1 | 0,3017 | 1,2466E-13 |
| ACE2 | CRTC1 | 0,3017 | 1,25292E-13 |
| ACE2 | CTD-2291D10.4 | 0,3017 | 1,25058E-13 |
| ACE2 | C9orf106 | 0,3015 | 1,29779E-13 |
| ACE2 | LINC00460 | 0,3015 | 1,30523E-13 |
| ACE2 | CENPS | 0,3014 | 1,32181E-13 |
| ACE2 | CLRN1-AS1 | 0,3014 | 1,33207E-13 |
| ACE2 | PROSER2 | 0,3014 | 1,32364E-13 |
| ACE2 | AC124861.1 | 0,3013 | 1,3427E-13 |
| ACE2 | RP11-120K24.3 | 0,3013 | 1,35352E-13 |
| ACE2 | PSEN1 | 0,3013 | 1,36349E-13 |
| ACE2 | ATP5S | 0,3012 | 1,37005E-13 |
| ACE2 | GLDN | 0,3012 | 1,39286E-13 |
| ACE2 | ID1 | 0,3012 | 1,39137E-13 |
| ACE2 | RRS1 | 0,3011 | 1,41658E-13 |
| ACE2 | TMEM128 | 0,301 | 1,43466E-13 |
| ACE2 | KRT19P1 | 0,301 | 1,43151E-13 |
| ACE2 | C3AR1 | 0,3009 | 1,45863E-13 |
| ACE2 | EFL1 | 0,3009 | 1,4704E-13 |
| ACE2 | BCRP3 | 0,3009 | 1,46039E-13 |
| ACE2 | RP11-690D19.3 | 0,3008 | 1,49541E-13 |
| ACE2 | CUTC | 0,3007 | 1,53059E-13 |
| ACE2 | RP13-580F15.2 | 0,3007 | 1,5157E-13 |
| ACE2 | ZNF888 | 0,3007 | 1,51152E-13 |
| ACE2 | BAIAP2 | 0,3006 | 1,56436E-13 |
| ACE2 | CASK | 0,3006 | 1,55896E-13 |
| ACE2 | FAM86EP | 0,3005 | 1,57713E-13 |
| ACE2 | GRIN3A | 0,3005 | 1,58637E-13 |
| ACE2 | SETD6 | 0,3005 | 1,5685E-13 |
| ACE2 | CCDC122 | 0,3004 | 1,61574E-13 |
| ACE2 | SLC16A8 | 0,3004 | 1,61239E-13 |
| ACE2 | LINC02006 | 0,3003 | 1,65416E-13 |
| ACE2 | RP11-203P23.2 | 0,3003 | 1,64434E-13 |
| ACE2 | CTSD | 0,3003 | 1,65898E-13 |

| | | | |
|---|---|---|---|
| ACE2 | KB-1592A4.16 | 0,3003 | 1,65946E-13 |
| ACE2 | PCP4L1 | 0,3002 | 1,68833E-13 |
| ACE2 | NHSL1 | 0,3002 | 1,66015E-13 |
| ACE2 | RP11-78O7.2 | 0,3002 | 1,69203E-13 |
| ACE2 | EP300 | 0,3001 | 1,72091E-13 |
| ACE2 | BCAR3 | 0,3 | 1,73725E-13 |
| ACE2 | RP3-483K16.4 | 0,3 | 1,72959E-13 |
| ACE2 | FAM181A | 0,2997 | 1,84755E-13 |
| ACE2 | LINC02038 | 0,2996 | 1,87864E-13 |
| ACE2 | RN7SL403P | 0,2996 | 1,88606E-13 |
| ACE2 | MRPL57 | 0,2996 | 1,86396E-13 |
| ACE2 | TOX3 | 0,2996 | 1,86541E-13 |
| ACE2 | RP5-831C21.1 | 0,2995 | 1,91816E-13 |
| ACE2 | HADHB | 0,2994 | 1,93742E-13 |
| ACE2 | FAM57A | 0,2994 | 1,97031E-13 |
| ACE2 | IFNGR1 | 0,2992 | 2,04147E-13 |
| ACE2 | FUCA2 | 0,2992 | 2,05084E-13 |
| ACE2 | RP11-325I22.3 | 0,2991 | 2,08416E-13 |
| ACE2 | PHAX | 0,2991 | 2,07438E-13 |
| ACE2 | INPP5J | 0,2991 | 2,08939E-13 |
| ACE2 | TADA1 | 0,299 | 2,12071E-13 |
| ACE2 | ALKBH2 | 0,299 | 2,13085E-13 |
| ACE2 | CDH26 | 0,299 | 2,0947E-13 |
| ACE2 | LMAN2 | 0,2989 | 2,13721E-13 |
| ACE2 | PRKAG2-AS1 | 0,2989 | 2,15254E-13 |
| ACE2 | TUBA1C | 0,2989 | 2,1655E-13 |
| ACE2 | AFF2 | 0,2989 | 2,17143E-13 |
| ACE2 | LINC01649 | 0,2988 | 2,19783E-13 |
| ACE2 | UFL1 | 0,2988 | 2,2083E-13 |
| ACE2 | MRPL13 | 0,2988 | 2,19583E-13 |
| ACE2 | POLR2B | 0,2987 | 2,25175E-13 |
| ACE2 | PSMD6 | 0,2986 | 2,26885E-13 |
| ACE2 | CASD1 | 0,2986 | 2,27878E-13 |
| ACE2 | TXNRD3 | 0,2985 | 2,30717E-13 |
| ACE2 | SUCLG1 | 0,2984 | 2,36022E-13 |
| ACE2 | LRRC20 | 0,2984 | 2,37134E-13 |
| ACE2 | MRPL19 | 0,298 | 2,56797E-13 |
| ACE2 | PIK3CB | 0,298 | 2,54098E-13 |
| ACE2 | RMDN1 | 0,298 | 2,57251E-13 |
| ACE2 | DOLPP1 | 0,298 | 2,5778E-13 |
| ACE2 | RN7SL832P | 0,2979 | 2,62724E-13 |
| ACE2 | FAM198B | 0,2979 | 2,59693E-13 |
| ACE2 | LINC01481 | 0,2979 | 2,59877E-13 |
| ACE2 | KIAA1143 | 0,2978 | 2,67662E-13 |
| ACE2 | RP11-93B14.10 | 0,2978 | 2,65332E-13 |
| ACE2 | POP1 | 0,2977 | 2,69503E-13 |
| ACE2 | TXNRD1 | 0,2977 | 2,72506E-13 |
| ACE2 | ABHD6 | 0,2976 | 2,73929E-13 |
| ACE2 | NOS1AP | 0,2975 | 2,80141E-13 |
| ACE2 | REEP6 | 0,2975 | 2,83458E-13 |
| ACE2 | PPCS | 0,2974 | 2,88102E-13 |
| ACE2 | SLC10A5 | 0,2974 | 2,88126E-13 |
| ACE2 | ASH2L | 0,2972 | 3,00116E-13 |
| ACE2 | AGPAT3 | 0,2971 | 3,01218E-13 |

| | | | |
|---|---|---|---|
| ACE2 | ASCC3 | 0,297 | 3,08481E-13 |
| ACE2 | TAOK3 | 0,297 | 3,10033E-13 |
| ACE2 | GABRB3 | 0,297 | 3,12638E-13 |
| ACE2 | PDE12 | 0,2969 | 3,14311E-13 |
| ACE2 | FASTKD3 | 0,2969 | 3,15293E-13 |
| ACE2 | SYK | 0,2969 | 3,14285E-13 |
| ACE2 | TMEM254 | 0,2969 | 3,1794E-13 |
| ACE2 | SURF1 | 0,2968 | 3,22599E-13 |
| ACE2 | RP11-571M6.17 | 0,2968 | 3,21677E-13 |
| ACE2 | TMCO1 | 0,2967 | 3,30708E-13 |
| ACE2 | CTC-295J13.3 | 0,2967 | 3,25986E-13 |
| ACE2 | ZNF239 | 0,2967 | 3,29261E-13 |
| ACE2 | GRB14 | 0,2965 | 3,37849E-13 |
| ACE2 | RNF123 | 0,2965 | 3,41789E-13 |
| ACE2 | CTC-265F19.3 | 0,2965 | 3,43187E-13 |
| ACE2 | IMP4 | 0,2964 | 3,45899E-13 |
| ACE2 | RNPEPL1 | 0,2964 | 3,44579E-13 |
| ACE2 | SMPD1 | 0,2964 | 3,44216E-13 |
| ACE2 | RAB5C | 0,2964 | 3,45017E-13 |
| ACE2 | NTN4 | 0,2963 | 3,51371E-13 |
| ACE2 | WASF2 | 0,2962 | 3,63294E-13 |
| ACE2 | TRMT61B | 0,2962 | 3,6128E-13 |
| ACE2 | ATP6V1D | 0,2961 | 3,66817E-13 |
| ACE2 | SPATC1 | 0,296 | 3,73315E-13 |
| ACE2 | KRT8 | 0,296 | 3,71491E-13 |
| ACE2 | SRP54 | 0,296 | 3,75302E-13 |
| ACE2 | USP8 | 0,2958 | 3,86347E-13 |
| ACE2 | TRAM1L1 | 0,2956 | 4,05098E-13 |
| ACE2 | NOMO2 | 0,2956 | 4,03582E-13 |
| ACE2 | SGF29 | 0,2956 | 4,03104E-13 |
| ACE2 | HPN-AS1 | 0,2956 | 4,06114E-13 |
| ACE2 | SRBD1 | 0,2955 | 4,15711E-13 |
| ACE2 | TGFA | 0,2955 | 4,10871E-13 |
| ACE2 | PSMA6 | 0,2955 | 4,09826E-13 |
| ACE2 | FAM217B | 0,2955 | 4,09293E-13 |
| ACE2 | RP11-156K23.3 | 0,2954 | 4,18571E-13 |
| ACE2 | HMOX2 | 0,2954 | 4,1707E-13 |
| ACE2 | POP4 | 0,2954 | 4,19504E-13 |
| ACE2 | PRPF6 | 0,2953 | 4,31054E-13 |
| ACE2 | NCBP2-AS2 | 0,2952 | 4,37016E-13 |
| ACE2 | CA3 | 0,2952 | 4,32698E-13 |
| ACE2 | GEMIN5 | 0,295 | 4,54086E-13 |
| ACE2 | LEO1 | 0,295 | 4,5486E-13 |
| ACE2 | ULK3 | 0,295 | 4,51366E-13 |
| ACE2 | YIPF2 | 0,295 | 4,49085E-13 |
| ACE2 | DPY19L3 | 0,295 | 4,52067E-13 |
| ACE2 | NTSR1 | 0,295 | 4,53108E-13 |
| ACE2 | BLNK | 0,2949 | 4,60081E-13 |
| ACE2 | NUFIP1 | 0,2949 | 4,58169E-13 |
| ACE2 | RP11-94C24.13 | 0,2949 | 4,64041E-13 |
| ACE2 | AC094019.4 | 0,2948 | 4,75143E-13 |
| ACE2 | HSD17B10 | 0,2948 | 4,68443E-13 |
| ACE2 | TBC1D8B | 0,2948 | 4,66828E-13 |
| ACE2 | HK2 | 0,2947 | 4,77743E-13 |

| | | | |
|---|---|---|---|
| ACE2 | RP11-180P8.1 | 0,2947 | 4,81306E-13 |
| ACE2 | AC020951.1 | 0,2947 | 4,83204E-13 |
| ACE2 | NSD1 | 0,2946 | 4,87642E-13 |
| ACE2 | RRS1-AS1 | 0,2945 | 4,95821E-13 |
| ACE2 | SLC35E3 | 0,2945 | 5,01565E-13 |
| ACE2 | RP11-347C12.10 | 0,2945 | 4,95405E-13 |
| ACE2 | TPBGL | 0,2944 | 5,03031E-13 |
| ACE2 | SF3A1 | 0,2943 | 5,20373E-13 |
| ACE2 | LINC01940 | 0,2942 | 5,28788E-13 |
| ACE2 | LRRC1 | 0,2941 | 5,40434E-13 |
| ACE2 | RP11-100L22.1 | 0,2939 | 5,53983E-13 |
| ACE2 | MTHFS | 0,2939 | 5,56429E-13 |
| ACE2 | SENP2 | 0,2938 | 5,69914E-13 |
| ACE2 | RLN2 | 0,2938 | 5,68503E-13 |
| ACE2 | PDIK1L | 0,2937 | 5,83018E-13 |
| ACE2 | PLXNA4 | 0,2937 | 5,80807E-13 |
| ACE2 | TMEM64 | 0,2937 | 5,81133E-13 |
| ACE2 | M6PR | 0,2937 | 5,80639E-13 |
| ACE2 | FAM58A | 0,2937 | 5,78233E-13 |
| ACE2 | DIS3L | 0,2936 | 5,94764E-13 |
| ACE2 | RP11-388K2.1 | 0,2936 | 5,87726E-13 |
| ACE2 | UBE4B | 0,2934 | 6,15918E-13 |
| ACE2 | USO1 | 0,2934 | 6,16156E-13 |
| ACE2 | LA16c-380H5.3 | 0,2934 | 6,10654E-13 |
| ACE2 | TMTC1 | 0,2933 | 6,22925E-13 |
| ACE2 | PDCD2L | 0,2933 | 6,29381E-13 |
| ACE2 | NLRX1 | 0,2932 | 6,39608E-13 |
| ACE2 | ABAT | 0,2932 | 6,32861E-13 |
| ACE2 | RPA1 | 0,2932 | 6,31318E-13 |
| ACE2 | BX842568.1 | 0,2932 | 6,37122E-13 |
| ACE2 | COPB2 | 0,2931 | 6,54161E-13 |
| ACE2 | TMED3 | 0,2931 | 6,4552E-13 |
| ACE2 | TRAPPC12 | 0,2929 | 6,68673E-13 |
| ACE2 | AADAT | 0,2928 | 6,91455E-13 |
| ACE2 | RTN3 | 0,2928 | 6,817E-13 |
| ACE2 | GCNT3 | 0,2927 | 6,9381E-13 |
| ACE2 | FGFBP3 | 0,2926 | 7,17661E-13 |
| ACE2 | YEATS4 | 0,2926 | 7,12973E-13 |
| ACE2 | CCDC97 | 0,2925 | 7,20762E-13 |
| ACE2 | SGO2 | 0,2924 | 7,36098E-13 |
| ACE2 | PEBP1 | 0,2924 | 7,34369E-13 |
| ACE2 | NOB1 | 0,2924 | 7,3657E-13 |
| ACE2 | SMAP2 | 0,2923 | 7,49992E-13 |
| ACE2 | AC004022.8 | 0,2923 | 7,50483E-13 |
| ACE2 | RP11-390F4.3 | 0,2923 | 7,52735E-13 |
| ACE2 | GRK1 | 0,2922 | 7,65793E-13 |
| ACE2 | C1orf56 | 0,2921 | 7,86471E-13 |
| ACE2 | NOP14 | 0,2921 | 7,76348E-13 |
| ACE2 | DLX3 | 0,2918 | 8,30427E-13 |
| ACE2 | SUSD2 | 0,2918 | 8,24941E-13 |
| ACE2 | BCDIN3D | 0,2917 | 8,40823E-13 |
| ACE2 | KCTD1 | 0,2917 | 8,47743E-13 |
| ACE2 | GYS1 | 0,2917 | 8,4719E-13 |
| ACE2 | EZR | 0,2916 | 8,5692E-13 |

| | | | |
|---|---|---|---|
| ACE2 | LHFPL3-AS2 | 0,2916 | 8,52877E-13 |
| ACE2 | RP11-110I1.5 | 0,2916 | 8,51524E-13 |
| ACE2 | ZNF441 | 0,2916 | 8,59753E-13 |
| ACE2 | CBR3-AS1 | 0,2916 | 8,65691E-13 |
| ACE2 | RP11-108M9.3 | 0,2915 | 8,80207E-13 |
| ACE2 | MAD1L1 | 0,2915 | 8,69941E-13 |
| ACE2 | EMC1 | 0,2914 | 8,8383E-13 |
| ACE2 | RP11-517P14.2 | 0,2914 | 8,97614E-13 |
| ACE2 | PIGH | 0,2914 | 8,98041E-13 |
| ACE2 | TMEM61 | 0,2912 | 9,20862E-13 |
| ACE2 | CTD-3032H12.1 | 0,2912 | 9,23699E-13 |
| ACE2 | C18orf21 | 0,2912 | 9,24622E-13 |
| ACE2 | LRRC8D | 0,291 | 9,63736E-13 |
| ACE2 | PIGC | 0,291 | 9,58014E-13 |
| ACE2 | TIMM23 | 0,291 | 9,64957E-13 |
| ACE2 | TMCO3 | 0,291 | 9,68269E-13 |
| ACE2 | PDK2 | 0,2908 | 9,93288E-13 |
| ACE2 | RP11-977G19.11 | 0,2907 | 1,02286E-12 |
| ACE2 | TRMT5 | 0,2906 | 1,04044E-12 |
| ACE2 | LINC00239 | 0,2906 | 1,03845E-12 |
| ACE2 | RPUSD3 | 0,2905 | 1,05711E-12 |
| ACE2 | ATP8A1 | 0,2905 | 1,04681E-12 |
| ACE2 | FGA | 0,2905 | 1,06061E-12 |
| ACE2 | POLR3G | 0,2905 | 1,06269E-12 |
| ACE2 | PAXIP1 | 0,2904 | 1,07194E-12 |
| ACE2 | SRPRA | 0,2904 | 1,07451E-12 |
| ACE2 | POLR3K | 0,2904 | 1,0689E-12 |
| ACE2 | IMMT | 0,2903 | 1,08508E-12 |
| ACE2 | RP11-219E7.1 | 0,2903 | 1,09423E-12 |
| ACE2 | EAF2 | 0,2902 | 1,11329E-12 |
| ACE2 | COL4A4 | 0,29 | 1,16162E-12 |
| ACE2 | SLC1A3 | 0,2899 | 1,17152E-12 |
| ACE2 | MCAT | 0,2899 | 1,17521E-12 |
| ACE2 | KPNA3 | 0,2897 | 1,22277E-12 |
| ACE2 | DSG2-AS1 | 0,2895 | 1,26548E-12 |
| ACE2 | RIC3 | 0,2894 | 1,29365E-12 |
| ACE2 | AMDHD2 | 0,2894 | 1,29096E-12 |
| ACE2 | RP11-166P13.3 | 0,2889 | 1,41972E-12 |
| ACE2 | SLC16A5 | 0,2889 | 1,41568E-12 |
| ACE2 | SELENOT | 0,2888 | 1,44969E-12 |
| ACE2 | PPP6C | 0,2888 | 1,43582E-12 |
| ACE2 | USP16 | 0,2888 | 1,44882E-12 |
| ACE2 | TMEM115 | 0,2887 | 1,47771E-12 |
| ACE2 | EIF3B | 0,2887 | 1,46485E-12 |
| ACE2 | LBX2-AS1 | 0,2886 | 1,49913E-12 |
| ACE2 | RP11-7I15.4 | 0,2886 | 1,51316E-12 |
| ACE2 | CTD-2020K17.4 | 0,2885 | 1,54173E-12 |
| ACE2 | TTC30B | 0,2884 | 1,55124E-12 |
| ACE2 | FOXA3 | 0,2884 | 1,54877E-12 |
| ACE2 | GSTP1 | 0,2883 | 1,59176E-12 |
| ACE2 | HYOU1 | 0,2883 | 1,59411E-12 |
| ACE2 | TVP23B | 0,2883 | 1,57538E-12 |
| ACE2 | TPRG1-AS1 | 0,2882 | 1,62766E-12 |
| ACE2 | RTCA | 0,2881 | 1,65703E-12 |

| | | | |
|---|---|---|---|
| ACE2 | TUBGCP5 | 0,2881 | 1,63708E-12 |
| ACE2 | MICU1 | 0,288 | 1,67634E-12 |
| ACE2 | ACSL5 | 0,288 | 1,6674E-12 |
| ACE2 | AMIGO1 | 0,2879 | 1,69673E-12 |
| ACE2 | CORIN | 0,2879 | 1,72212E-12 |
| ACE2 | ADH5P4 | 0,2879 | 1,7126E-12 |
| ACE2 | SF3B5 | 0,2879 | 1,69911E-12 |
| ACE2 | PRDM11 | 0,2879 | 1,70717E-12 |
| ACE2 | COIL | 0,2879 | 1,69301E-12 |
| ACE2 | RP11-498C9.15 | 0,2879 | 1,71226E-12 |
| ACE2 | DDX18 | 0,2878 | 1,72672E-12 |
| ACE2 | MPEG1 | 0,2878 | 1,74311E-12 |
| ACE2 | SLC25A11 | 0,2878 | 1,72474E-12 |
| ACE2 | EDEM1 | 0,2877 | 1,76654E-12 |
| ACE2 | TULP1 | 0,2877 | 1,78653E-12 |
| ACE2 | PSMB5 | 0,2876 | 1,81752E-12 |
| ACE2 | ST6GALNAC2 | 0,2876 | 1,81383E-12 |
| ACE2 | SLC30A5 | 0,2875 | 1,84826E-12 |
| ACE2 | RP11-87E22.1 | 0,2875 | 1,82967E-12 |
| ACE2 | SLC16A6P1 | 0,2875 | 1,82973E-12 |
| ACE2 | LINGO3 | 0,2875 | 1,84442E-12 |
| ACE2 | ITGA10 | 0,2874 | 1,86704E-12 |
| ACE2 | MPV17 | 0,2874 | 1,87627E-12 |
| ACE2 | MPPE1 | 0,2874 | 1,8798E-12 |
| ACE2 | SLC1A5 | 0,2874 | 1,87008E-12 |
| ACE2 | TKFC | 0,2873 | 1,90518E-12 |
| ACE2 | NKD1 | 0,2873 | 1,9018E-12 |
| ACE2 | NCEH1 | 0,2872 | 1,94277E-12 |
| ACE2 | ZNF106 | 0,2872 | 1,94064E-12 |
| ACE2 | MRS2 | 0,287 | 2,01234E-12 |
| ACE2 | PRDX3 | 0,287 | 2,02617E-12 |
| ACE2 | NRAS | 0,2869 | 2,06022E-12 |
| ACE2 | SMG7-AS1 | 0,2868 | 2,10223E-12 |
| ACE2 | IL1RL1 | 0,2868 | 2,107E-12 |
| ACE2 | RP11-1020A11.2 | 0,2867 | 2,12133E-12 |
| ACE2 | STX19 | 0,2867 | 2,12672E-12 |
| ACE2 | CTSC | 0,2867 | 2,1104E-12 |
| ACE2 | ADAM1B | 0,2867 | 2,11658E-12 |
| ACE2 | UBL7 | 0,2867 | 2,1404E-12 |
| ACE2 | RP11-473M20.16 | 0,2867 | 2,13741E-12 |
| ACE2 | ACTRT3 | 0,2866 | 2,16824E-12 |
| ACE2 | ITPR2 | 0,2866 | 2,15047E-12 |
| ACE2 | MED8 | 0,2864 | 2,23149E-12 |
| ACE2 | C3orf52 | 0,2864 | 2,23714E-12 |
| ACE2 | MUC15 | 0,2864 | 2,26002E-12 |
| ACE2 | BHLHA15 | 0,2862 | 2,34415E-12 |
| ACE2 | MTDH | 0,2862 | 2,32584E-12 |
| ACE2 | CCNDBP1 | 0,2862 | 2,35314E-12 |
| ACE2 | HDAC8 | 0,2862 | 2,32047E-12 |
| ACE2 | AGPS | 0,2861 | 2,36596E-12 |
| ACE2 | MRPL2 | 0,2861 | 2,36958E-12 |
| ACE2 | RP11-61K9.3 | 0,2861 | 2,37865E-12 |
| ACE2 | MRPL10 | 0,2861 | 2,37411E-12 |
| ACE2 | TSSK4 | 0,286 | 2,40406E-12 |

| | | | |
|---|---|---|---|
| ACE2 | LTN1 | 0,286 | 2,43661E-12 |
| ACE2 | TIRAP | 0,2859 | 2,47665E-12 |
| ACE2 | RP3-510O8.4 | 0,2857 | 2,55378E-12 |
| ACE2 | RP11-646I6.5 | 0,2855 | 2,63069E-12 |
| ACE2 | RP4-570O12.3 | 0,2855 | 2,65129E-12 |
| ACE2 | RAB3IP | 0,2855 | 2,67018E-12 |
| ACE2 | CHST7 | 0,2855 | 2,6577E-12 |
| ACE2 | YKT6 | 0,2854 | 2,68652E-12 |
| ACE2 | WRN | 0,2853 | 2,74175E-12 |
| ACE2 | EBPL | 0,2853 | 2,74119E-12 |
| ACE2 | GLYR1 | 0,2852 | 2,81073E-12 |
| ACE2 | IGFBP1 | 0,2851 | 2,82671E-12 |
| ACE2 | ZNF777 | 0,2851 | 2,83077E-12 |
| ACE2 | RP11-760H22.2 | 0,285 | 2,8886E-12 |
| ACE2 | DOK4 | 0,285 | 2,88231E-12 |
| ACE2 | NIP7 | 0,285 | 2,90164E-12 |
| ACE2 | CANX | 0,2849 | 2,9319E-12 |
| ACE2 | MTX2 | 0,2848 | 2,99398E-12 |
| ACE2 | TRMT6 | 0,2848 | 3,02652E-12 |
| ACE2 | AF186192.6 | 0,2847 | 3,05495E-12 |
| ACE2 | ZNF252P | 0,2847 | 3,04871E-12 |
| ACE2 | ETFA | 0,2847 | 3,05921E-12 |
| ACE2 | RP11-434D9.1 | 0,2845 | 3,16468E-12 |
| ACE2 | APRT | 0,2844 | 3,23222E-12 |
| ACE2 | SLC2A11 | 0,2844 | 3,25902E-12 |
| ACE2 | LRBA | 0,2843 | 3,32466E-12 |
| ACE2 | PROS1 | 0,2841 | 3,44755E-12 |
| ACE2 | SRP72 | 0,2841 | 3,42978E-12 |
| ACE2 | RARS | 0,2841 | 3,41718E-12 |
| ACE2 | TRIP11 | 0,2841 | 3,40329E-12 |
| ACE2 | SMIM14 | 0,284 | 3,45835E-12 |
| ACE2 | KIF13B | 0,284 | 3,48294E-12 |
| ACE2 | RP11-16E12.2 | 0,2839 | 3,56183E-12 |
| ACE2 | SLC24A5 | 0,2839 | 3,57382E-12 |
| ACE2 | GPATCH4 | 0,2837 | 3,66798E-12 |
| ACE2 | LINC01963 | 0,2837 | 3,67239E-12 |
| ACE2 | RP3-337H4.9 | 0,2837 | 3,68611E-12 |
| ACE2 | OSTM1 | 0,2837 | 3,69834E-12 |
| ACE2 | ARHGAP8 | 0,2837 | 3,68955E-12 |
| ACE2 | WWC3 | 0,2837 | 3,69982E-12 |
| ACE2 | RBM34 | 0,2836 | 3,73146E-12 |
| ACE2 | PPP1R1C | 0,2836 | 3,71418E-12 |
| ACE2 | ADAT1 | 0,2836 | 3,72258E-12 |
| ACE2 | GAS8-AS1 | 0,2836 | 3,70909E-12 |
| ACE2 | ZC3HC1 | 0,2834 | 3,86065E-12 |
| ACE2 | ARPC1B | 0,2833 | 3,96186E-12 |
| ACE2 | TOR4A | 0,2833 | 3,9433E-12 |
| ACE2 | RP11-79N23.1 | 0,2832 | 3,99593E-12 |
| ACE2 | FAM184A | 0,2832 | 4,01676E-12 |
| ACE2 | ACTG1P24 | 0,2832 | 4,04364E-12 |
| ACE2 | RP11-1016B18.1 | 0,2827 | 4,3804E-12 |
| ACE2 | SMIM22 | 0,2826 | 4,50703E-12 |
| ACE2 | MRPL34 | 0,2826 | 4,48089E-12 |
| ACE2 | USP18 | 0,2826 | 4,45526E-12 |

| | | | |
|---|---|---|---|
| ACE2 | GREB1 | 0,2825 | 4,53189E-12 |
| ACE2 | F13A1 | 0,2825 | 4,59542E-12 |
| ACE2 | RP11-479O17.12 | 0,2825 | 4,53628E-12 |
| ACE2 | LAMP2 | 0,2825 | 4,55917E-12 |
| ACE2 | POLR2D | 0,2824 | 4,68552E-12 |
| ACE2 | AMOTL2 | 0,2824 | 4,63842E-12 |
| ACE2 | DCAF4 | 0,2824 | 4,64279E-12 |
| ACE2 | AC005546.2 | 0,2824 | 4,63297E-12 |
| ACE2 | CTD-2012J19.3 | 0,2823 | 4,71956E-12 |
| ACE2 | ST6GALNAC4 | 0,2823 | 4,73981E-12 |
| ACE2 | RP5-901A4.1 | 0,2822 | 4,80871E-12 |
| ACE2 | TRIAP1 | 0,2822 | 4,81505E-12 |
| ACE2 | NAE1 | 0,2822 | 4,81209E-12 |
| ACE2 | RP11-162A12.2 | 0,2822 | 4,86015E-12 |
| ACE2 | RHOXF1P2 | 0,2822 | 4,82507E-12 |
| ACE2 | UBTD1 | 0,2821 | 4,93976E-12 |
| ACE2 | DPH5 | 0,282 | 5,0046E-12 |
| ACE2 | SLC35B2 | 0,282 | 4,95263E-12 |
| ACE2 | LSM1 | 0,282 | 5,00866E-12 |
| ACE2 | LDHAL6A | 0,282 | 4,98342E-12 |
| ACE2 | IFNLR1 | 0,2819 | 5,06323E-12 |
| ACE2 | LHX4 | 0,2819 | 5,05113E-12 |
| ACE2 | INSM2 | 0,2819 | 5,12383E-12 |
| ACE2 | RBMY1KP | 0,2819 | 5,04283E-12 |
| ACE2 | FAM134A | 0,2818 | 5,15028E-12 |
| ACE2 | CTD-2353F22.2 | 0,2818 | 5,19863E-12 |
| ACE2 | NKAPL | 0,2818 | 5,21985E-12 |
| ACE2 | RRM1 | 0,2818 | 5,15753E-12 |
| ACE2 | VSIG10 | 0,2818 | 5,14964E-12 |
| ACE2 | DPYD | 0,2816 | 5,40376E-12 |
| ACE2 | MTND6P22 | 0,2816 | 5,33621E-12 |
| ACE2 | TMEM60 | 0,2816 | 5,37088E-12 |
| ACE2 | CHAF1A | 0,2816 | 5,35118E-12 |
| ACE2 | FAM120A | 0,2815 | 5,44021E-12 |
| ACE2 | FAM169B | 0,2815 | 5,50969E-12 |
| ACE2 | RP5-994D16.9 | 0,2813 | 5,65981E-12 |
| ACE2 | RPRD2 | 0,2813 | 5,66278E-12 |
| ACE2 | IARS | 0,2813 | 5,65183E-12 |
| ACE2 | WDR5 | 0,2813 | 5,63667E-12 |
| ACE2 | ZSWIM5 | 0,2812 | 5,80388E-12 |
| ACE2 | NME6 | 0,2811 | 5,8356E-12 |
| ACE2 | MAP3K9 | 0,2811 | 5,90621E-12 |
| ACE2 | TPD52L1 | 0,281 | 5,9781E-12 |
| ACE2 | AP3D1 | 0,281 | 5,95849E-12 |
| ACE2 | SNRPGP15 | 0,281 | 6,02143E-12 |
| ACE2 | ATRAID | 0,2808 | 6,20669E-12 |
| ACE2 | ZNF69 | 0,2807 | 6,2916E-12 |
| ACE2 | DGCR2 | 0,2807 | 6,25219E-12 |
| ACE2 | RP11-448G15.1 | 0,2806 | 6,43023E-12 |
| ACE2 | VTA1 | 0,2806 | 6,43922E-12 |
| ACE2 | G6PC3 | 0,2806 | 6,43491E-12 |
| ACE2 | ZNF311 | 0,2805 | 6,48098E-12 |
| ACE2 | MNX1-AS1 | 0,2805 | 6,59004E-12 |
| ACE2 | CHRNA2 | 0,2805 | 6,50756E-12 |

| | | | |
|---|---|---|---|
| ACE2 | KRT87P | 0,2805 | 6,55159E-12 |
| ACE2 | KBTBD7 | 0,2804 | 6,67321E-12 |
| ACE2 | AATF | 0,2803 | 6,77918E-12 |
| ACE2 | BRWD1-AS2 | 0,2803 | 6,8208E-12 |
| ACE2 | SPR | 0,2802 | 6,88118E-12 |
| ACE2 | OARD1 | 0,2802 | 6,95077E-12 |
| ACE2 | FOXN1 | 0,2802 | 6,90726E-12 |
| ACE2 | KIAA0100 | 0,2802 | 6,92384E-12 |
| ACE2 | ATXN7L3B | 0,2801 | 7,0304E-12 |
| ACE2 | MYCL | 0,2799 | 7,26402E-12 |
| ACE2 | RP11-85G21.2 | 0,2798 | 7,40163E-12 |
| ACE2 | PARP1 | 0,2798 | 7,41419E-12 |
| ACE2 | HAGLROS | 0,2798 | 7,44842E-12 |
| ACE2 | NUDCD2 | 0,2798 | 7,41044E-12 |
| ACE2 | RP11-419I17.1 | 0,2798 | 7,39213E-12 |
| ACE2 | C10orf95 | 0,2798 | 7,41622E-12 |
| ACE2 | C1QA | 0,2797 | 7,60464E-12 |
| ACE2 | SYDE2 | 0,2797 | 7,50241E-12 |
| ACE2 | RP4-631H13.6 | 0,2796 | 7,66351E-12 |
| ACE2 | LZTFL1 | 0,2796 | 7,72702E-12 |
| ACE2 | TMEM68 | 0,2796 | 7,65526E-12 |
| ACE2 | FAHD1 | 0,2796 | 7,71905E-12 |
| ACE2 | GRN | 0,2796 | 7,62007E-12 |
| ACE2 | ESF1 | 0,2796 | 7,73336E-12 |
| ACE2 | LINC01811 | 0,2795 | 7,86218E-12 |
| ACE2 | GAR1 | 0,2795 | 7,76516E-12 |
| ACE2 | SMPDL3A | 0,2795 | 7,82924E-12 |
| ACE2 | KIAA2022 | 0,2795 | 7,77806E-12 |
| ACE2 | BMS1P21 | 0,2794 | 7,94607E-12 |
| ACE2 | ELAC2 | 0,2794 | 7,99503E-12 |
| ACE2 | RP11-427P5.3 | 0,2793 | 8,04691E-12 |
| ACE2 | IBTK | 0,2793 | 8,13082E-12 |
| ACE2 | DTD2 | 0,2793 | 8,04664E-12 |
| ACE2 | ZCCHC12 | 0,2793 | 8,13469E-12 |
| ACE2 | TPRG1L | 0,2792 | 8,21949E-12 |
| ACE2 | AC015971.2 | 0,2792 | 8,26692E-12 |
| ACE2 | CICP16 | 0,2792 | 8,28345E-12 |
| ACE2 | UNQ6494 | 0,2792 | 8,20992E-12 |
| ACE2 | FADD | 0,2791 | 8,40277E-12 |
| ACE2 | PWP1 | 0,2791 | 8,37819E-12 |
| ACE2 | LINC00885 | 0,279 | 8,59421E-12 |
| ACE2 | ERAL1 | 0,279 | 8,50576E-12 |
| ACE2 | ARSK | 0,2789 | 8,69206E-12 |
| ACE2 | TBL3 | 0,2789 | 8,74585E-12 |
| ACE2 | IMPA2 | 0,2789 | 8,73669E-12 |
| ACE2 | ZFAT | 0,2788 | 8,91849E-12 |
| ACE2 | ENTPD5 | 0,2787 | 8,98139E-12 |
| ACE2 | C15orf52 | 0,2787 | 9,04424E-12 |
| ACE2 | SP5 | 0,2785 | 9,29482E-12 |
| ACE2 | HORMAD2-AS1 | 0,2785 | 9,385E-12 |
| ACE2 | POLH | 0,2784 | 9,5257E-12 |
| ACE2 | GBAS | 0,2784 | 9,58361E-12 |
| ACE2 | PAFAH2 | 0,2783 | 9,70109E-12 |
| ACE2 | RP11-164O23.8 | 0,2782 | 9,82613E-12 |

| | | | |
|---|---|---|---|
| ACE2 | VPS11 | 0,2782 | 9,89902E-12 |
| ACE2 | URB1-AS1 | 0,2782 | 9,80128E-12 |
| ACE2 | MRPS15 | 0,2781 | 9,94502E-12 |
| ACE2 | COBLL1 | 0,278 | 1,02867E-11 |
| ACE2 | CCT5 | 0,278 | 1,01111E-11 |
| ACE2 | SMU1 | 0,278 | 1,02271E-11 |
| ACE2 | AC024592.9 | 0,2779 | 1,04542E-11 |
| ACE2 | MKRN2OS | 0,2778 | 1,04999E-11 |
| ACE2 | MELTF | 0,2778 | 1,05438E-11 |
| ACE2 | DUOXA2 | 0,2777 | 1,07919E-11 |
| ACE2 | ARL2BP | 0,2777 | 1,07581E-11 |
| ACE2 | IMPACT | 0,2777 | 1,0688E-11 |
| ACE2 | DHX57 | 0,2775 | 1,12262E-11 |
| ACE2 | MRPS25 | 0,2775 | 1,10984E-11 |
| ACE2 | FBXO4 | 0,2775 | 1,11418E-11 |
| ACE2 | APOOL | 0,2775 | 1,1051E-11 |
| ACE2 | B3GNT7 | 0,2774 | 1,12614E-11 |
| ACE2 | GMPPB | 0,2774 | 1,1334E-11 |
| ACE2 | PYROXD1 | 0,2774 | 1,13646E-11 |
| ACE2 | JMJD4 | 0,2773 | 1,14983E-11 |
| ACE2 | MTAPP2 | 0,2773 | 1,15456E-11 |
| ACE2 | IMPA1P1 | 0,2773 | 1,14897E-11 |
| ACE2 | G6PD | 0,2773 | 1,15274E-11 |
| ACE2 | LINC02015 | 0,2772 | 1,18306E-11 |
| ACE2 | C4orf19 | 0,2772 | 1,17369E-11 |
| ACE2 | UTP23 | 0,2772 | 1,17332E-11 |
| ACE2 | NARS2 | 0,2772 | 1,17335E-11 |
| ACE2 | SMDT1 | 0,2772 | 1,17391E-11 |
| ACE2 | TMEM187 | 0,2772 | 1,16871E-11 |
| ACE2 | CCDC6 | 0,2771 | 1,19558E-11 |
| ACE2 | ZDHHC13 | 0,2771 | 1,197E-11 |
| ACE2 | RP11-490O6.2 | 0,2771 | 1,19146E-11 |
| ACE2 | TIMM50 | 0,2771 | 1,18633E-11 |
| ACE2 | RP11-342M1.3 | 0,277 | 1,21632E-11 |
| ACE2 | PNMA2 | 0,277 | 1,20692E-11 |
| ACE2 | RP11-338I21.1 | 0,277 | 1,21907E-11 |
| ACE2 | GNPTAB | 0,277 | 1,21237E-11 |
| ACE2 | ANKDD1B | 0,2769 | 1,24947E-11 |
| ACE2 | MFSD6L | 0,2769 | 1,24458E-11 |
| ACE2 | HADHA | 0,2768 | 1,26909E-11 |
| ACE2 | RP5-1171I10.5 | 0,2767 | 1,28483E-11 |
| ACE2 | RCL1 | 0,2766 | 1,30615E-11 |
| ACE2 | RP11-16E12.1 | 0,2766 | 1,29949E-11 |
| ACE2 | RPL39P40 | 0,2766 | 1,3055E-11 |
| ACE2 | RP5-1065J22.8 | 0,2765 | 1,32041E-11 |
| ACE2 | LCMT1 | 0,2765 | 1,33714E-11 |
| ACE2 | RPS6P16 | 0,2764 | 1,35304E-11 |
| ACE2 | GALK2 | 0,2764 | 1,35482E-11 |
| ACE2 | GGT2 | 0,2764 | 1,35825E-11 |
| ACE2 | SUCO | 0,2763 | 1,36464E-11 |
| ACE2 | ADAMDEC1 | 0,2763 | 1,38336E-11 |
| ACE2 | PTGES2 | 0,2763 | 1,37916E-11 |
| ACE2 | PSMD9 | 0,2763 | 1,38602E-11 |
| ACE2 | ENAM | 0,2762 | 1,39232E-11 |

| | | | |
|---|---|---|---|
| ACE2 | EPHA4 | 0,2761 | 1,43162E-11 |
| ACE2 | ATP6V1G1 | 0,2761 | 1,43582E-11 |
| ACE2 | WDR74 | 0,2761 | 1,42158E-11 |
| ACE2 | TCEAL8 | 0,2761 | 1,43703E-11 |
| ACE2 | RP11-521O16.1 | 0,276 | 1,45136E-11 |
| ACE2 | PCBD1 | 0,276 | 1,45278E-11 |
| ACE2 | RP11-715J22.2 | 0,276 | 1,45195E-11 |
| ACE2 | NCSTN | 0,2759 | 1,4706E-11 |
| ACE2 | DTD1 | 0,2759 | 1,46575E-11 |
| ACE2 | SLU7 | 0,2758 | 1,5026E-11 |
| ACE2 | OIT3 | 0,2758 | 1,51163E-11 |
| ACE2 | SLC7A1 | 0,2758 | 1,50976E-11 |
| ACE2 | FIZ1 | 0,2758 | 1,51128E-11 |
| ACE2 | LINC01186 | 0,2758 | 1,49681E-11 |
| ACE2 | SEC23IP | 0,2757 | 1,5296E-11 |
| ACE2 | SYNC | 0,2756 | 1,55982E-11 |
| ACE2 | RP11-460M2.1 | 0,2756 | 1,56866E-11 |
| ACE2 | CCDC183-AS1 | 0,2756 | 1,55652E-11 |
| ACE2 | WDR35 | 0,2755 | 1,57996E-11 |
| ACE2 | SLC44A3 | 0,2754 | 1,6063E-11 |
| ACE2 | KCNJ8 | 0,2754 | 1,60802E-11 |
| ACE2 | STARD10 | 0,2753 | 1,64541E-11 |
| ACE2 | RP11-627G18.1 | 0,2753 | 1,62629E-11 |
| ACE2 | MIR3677 | 0,2752 | 1,67371E-11 |
| ACE2 | GS1-594A7.3 | 0,2752 | 1,66802E-11 |
| ACE2 | SH2D5 | 0,2751 | 1,69309E-11 |
| ACE2 | PTCD1 | 0,2751 | 1,69085E-11 |
| ACE2 | MED11 | 0,2751 | 1,7086E-11 |
| ACE2 | CTD-2666L21.2 | 0,2751 | 1,71284E-11 |
| ACE2 | MNX1 | 0,275 | 1,7383E-11 |
| ACE2 | CLCN7 | 0,275 | 1,72419E-11 |
| ACE2 | XRN2 | 0,275 | 1,71389E-11 |
| ACE2 | MIGA1 | 0,2748 | 1,80511E-11 |
| ACE2 | GUF1 | 0,2748 | 1,78537E-11 |
| ACE2 | C1RL | 0,2747 | 1,824E-11 |
| ACE2 | PTPRK | 0,2746 | 1,84283E-11 |
| ACE2 | EDRF1-AS1 | 0,2746 | 1,84332E-11 |
| ACE2 | NDUFS8 | 0,2746 | 1,8551E-11 |
| ACE2 | FNIP2 | 0,2745 | 1,87329E-11 |
| ACE2 | GPNMB | 0,2745 | 1,8863E-11 |
| ACE2 | LINC00940 | 0,2745 | 1,8777E-11 |
| ACE2 | KRT18 | 0,2745 | 1,89314E-11 |
| ACE2 | GOLGA6L9 | 0,2745 | 1,88304E-11 |
| ACE2 | RCC1 | 0,2744 | 1,91797E-11 |
| ACE2 | PRMT6 | 0,2744 | 1,9331E-11 |
| ACE2 | KRR1 | 0,2744 | 1,9191E-11 |
| ACE2 | RP11-499E18.1 | 0,2743 | 1,94686E-11 |
| ACE2 | KIF12 | 0,2743 | 1,96202E-11 |
| ACE2 | ZNF33A | 0,2743 | 1,96732E-11 |
| ACE2 | RP11-38G5.2 | 0,2743 | 1,96134E-11 |
| ACE2 | CTB-161C1.1 | 0,2742 | 1,98697E-11 |
| ACE2 | BRMS1 | 0,2742 | 2,00111E-11 |
| ACE2 | RPGR | 0,2742 | 1,97245E-11 |
| ACE2 | ADIPOR1 | 0,2741 | 2,02185E-11 |

| | | | |
|---|---|---|---|
| ACE2 | PDIA4 | 0,2741 | 2,0092E-11 |
| ACE2 | PEX12 | 0,2741 | 2,03755E-11 |
| ACE2 | CISD3 | 0,2741 | 2,00653E-11 |
| ACE2 | NRSN2-AS1 | 0,2741 | 2,01189E-11 |
| ACE2 | IL17RE | 0,274 | 2,05359E-11 |
| ACE2 | MGAT2 | 0,274 | 2,04704E-11 |
| ACE2 | RNF126P1 | 0,2739 | 2,10952E-11 |
| ACE2 | TMA16 | 0,2737 | 2,16152E-11 |
| ACE2 | STT3A | 0,2737 | 2,18285E-11 |
| ACE2 | RN7SL600P | 0,2736 | 2,20797E-11 |
| ACE2 | CISD2 | 0,2736 | 2,19713E-11 |
| ACE2 | SPATA32 | 0,2736 | 2,19431E-11 |
| ACE2 | ELFN2 | 0,2735 | 2,24211E-11 |
| ACE2 | TACSTD2 | 0,2733 | 2,31588E-11 |
| ACE2 | PIGF | 0,2733 | 2,32663E-11 |
| ACE2 | SLCO2A1 | 0,2733 | 2,31111E-11 |
| ACE2 | SIGMAR1 | 0,2733 | 2,33084E-11 |
| ACE2 | RP11-93B14.9 | 0,2733 | 2,31773E-11 |
| ACE2 | AMACR | 0,2732 | 2,36834E-11 |
| ACE2 | SQSTM1 | 0,2732 | 2,35143E-11 |
| ACE2 | SUPT16HP1 | 0,2732 | 2,34847E-11 |
| ACE2 | NUDT7 | 0,2732 | 2,38264E-11 |
| ACE2 | RP5-862K6.4 | 0,2732 | 2,37547E-11 |
| ACE2 | NOL9 | 0,2731 | 2,42003E-11 |
| ACE2 | ST3GAL6 | 0,2731 | 2,40384E-11 |
| ACE2 | SLC17A5 | 0,2731 | 2,40406E-11 |
| ACE2 | C9orf62 | 0,273 | 2,45383E-11 |
| ACE2 | PRMT3 | 0,273 | 2,44572E-11 |
| ACE2 | GAA | 0,273 | 2,45128E-11 |
| ACE2 | RP11-58A11.2 | 0,2729 | 2,50814E-11 |
| ACE2 | TM6SF1 | 0,2729 | 2,50077E-11 |
| ACE2 | CTD-2308L22.1 | 0,2728 | 2,52119E-11 |
| ACE2 | RP11-629B11.5 | 0,2728 | 2,52927E-11 |
| ACE2 | CTD-2506P8.6 | 0,2727 | 2,59958E-11 |
| ACE2 | PARD6A | 0,2727 | 2,56136E-11 |
| ACE2 | CTB-131K11.1 | 0,2727 | 2,56781E-11 |
| ACE2 | USP19 | 0,2726 | 2,64165E-11 |
| ACE2 | TAX1BP1 | 0,2726 | 2,61638E-11 |
| ACE2 | RP11-797A18.5 | 0,2726 | 2,62664E-11 |
| ACE2 | ETV5-AS1 | 0,2725 | 2,66704E-11 |
| ACE2 | ZMAT2 | 0,2725 | 2,68551E-11 |
| ACE2 | TMEM185B | 0,2724 | 2,72123E-11 |
| ACE2 | MT-TA | 0,2724 | 2,73205E-11 |
| ACE2 | HAPLN2 | 0,2723 | 2,76516E-11 |
| ACE2 | PSMA5 | 0,2722 | 2,80794E-11 |
| ACE2 | COG8 | 0,2722 | 2,81711E-11 |
| ACE2 | RP11-365F18.6 | 0,2721 | 2,84582E-11 |
| ACE2 | SNRPB2 | 0,2721 | 2,8385E-11 |
| ACE2 | RASSF6 | 0,272 | 2,92654E-11 |
| ACE2 | MAP9 | 0,272 | 2,91886E-11 |
| ACE2 | TWISTNB | 0,272 | 2,92621E-11 |
| ACE2 | SPINT3 | 0,272 | 2,91338E-11 |
| ACE2 | RFT1 | 0,2719 | 2,98792E-11 |
| ACE2 | SEPT7-AS1 | 0,2719 | 2,98275E-11 |

| | | | |
|---|---|---|---|
| ACE2 | AC005076.5 | 0,2719 | 2,95105E-11 |
| ACE2 | HAX1 | 0,2718 | 3,00166E-11 |
| ACE2 | RP13-870H17.3 | 0,2718 | 3,02991E-11 |
| ACE2 | DOPEY2 | 0,2718 | 3,00497E-11 |
| ACE2 | FARSA | 0,2717 | 3,07101E-11 |
| ACE2 | TRUB1 | 0,2716 | 3,14681E-11 |
| ACE2 | RITA1 | 0,2716 | 3,12355E-11 |
| ACE2 | MAFG-AS1 | 0,2716 | 3,11656E-11 |
| ACE2 | NUDT9 | 0,2715 | 3,15648E-11 |
| ACE2 | IBA57 | 0,2714 | 3,25625E-11 |
| ACE2 | TMEM177 | 0,2714 | 3,2391E-11 |
| ACE2 | ACVR1 | 0,2714 | 3,21452E-11 |
| ACE2 | C11orf91 | 0,2713 | 3,30698E-11 |
| ACE2 | RP1-92O14.3 | 0,2712 | 3,31545E-11 |
| ACE2 | TRAPPC1 | 0,2712 | 3,32076E-11 |
| ACE2 | RBBP4P2 | 0,2711 | 3,41295E-11 |
| ACE2 | RP11-153M7.3 | 0,2711 | 3,41307E-11 |
| ACE2 | TMEM223 | 0,2711 | 3,42199E-11 |
| ACE2 | RP11-680F8.1 | 0,2711 | 3,40582E-11 |
| ACE2 | RP11-84C10.2 | 0,271 | 3,4328E-11 |
| ACE2 | DANT2 | 0,271 | 3,46365E-11 |
| ACE2 | RP11-360A10.1 | 0,2709 | 3,52028E-11 |
| ACE2 | LINC01719 | 0,2708 | 3,5905E-11 |
| ACE2 | VAMP8 | 0,2708 | 3,56525E-11 |
| ACE2 | TMEM203 | 0,2708 | 3,57856E-11 |
| ACE2 | TGM4 | 0,2707 | 3,62721E-11 |
| ACE2 | CD180 | 0,2707 | 3,66824E-11 |
| ACE2 | C11orf98 | 0,2707 | 3,67227E-11 |
| ACE2 | MSLN | 0,2707 | 3,61221E-11 |
| ACE2 | PSMC5 | 0,2707 | 3,65725E-11 |
| ACE2 | EMC4 | 0,2706 | 3,68214E-11 |
| ACE2 | DPP9 | 0,2706 | 3,71865E-11 |
| ACE2 | SDPR | 0,2705 | 3,78753E-11 |
| ACE2 | RP11-215C7.3 | 0,2705 | 3,75221E-11 |
| ACE2 | EDEM2 | 0,2704 | 3,86088E-11 |
| ACE2 | ZMAT5 | 0,2703 | 3,90498E-11 |
| ACE2 | HPDL | 0,2702 | 3,95393E-11 |
| ACE2 | ARL8B | 0,2702 | 3,93578E-11 |
| ACE2 | PQLC2L | 0,2702 | 3,96462E-11 |
| ACE2 | USP46-AS1 | 0,2702 | 3,94115E-11 |
| ACE2 | ZNF747 | 0,2702 | 3,99166E-11 |
| ACE2 | AQP4 | 0,2702 | 3,98515E-11 |
| ACE2 | PSMD5 | 0,2701 | 4,05839E-11 |
| ACE2 | BICDL1 | 0,2701 | 4,01098E-11 |
| ACE2 | SCRN3 | 0,27 | 4,10816E-11 |
| ACE2 | DAP3 | 0,2699 | 4,20172E-11 |
| ACE2 | WDR17 | 0,2699 | 4,20228E-11 |
| ACE2 | AKTIP | 0,2699 | 4,15722E-11 |
| ACE2 | CDC23 | 0,2698 | 4,28396E-11 |
| ACE2 | METTL16 | 0,2697 | 4,35352E-11 |
| ACE2 | GTF3C6 | 0,2696 | 4,39186E-11 |
| ACE2 | LINC01139 | 0,2695 | 4,43914E-11 |
| ACE2 | SEC14L1P1 | 0,2695 | 4,44627E-11 |
| ACE2 | SLC2A3P2 | 0,2694 | 4,55011E-11 |

| | | | |
|---|---|---|---|
| ACE2 | AASDH | 0,2694 | 4,5532E-11 |
| ACE2 | KLB | 0,2693 | 4,6315E-11 |
| ACE2 | SPINK13 | 0,2693 | 4,60593E-11 |
| ACE2 | FCF1 | 0,2693 | 4,60481E-11 |
| ACE2 | CTSS | 0,2692 | 4,67908E-11 |
| ACE2 | TEX261 | 0,2692 | 4,71443E-11 |
| ACE2 | WDR31 | 0,2692 | 4,72676E-11 |
| ACE2 | ZFP3 | 0,2692 | 4,68353E-11 |
| ACE2 | RP11-863K10.4 | 0,2691 | 4,81843E-11 |
| ACE2 | WDR34 | 0,2691 | 4,76415E-11 |
| ACE2 | RP11-36D19.9 | 0,2691 | 4,82038E-11 |
| ACE2 | VANGL1 | 0,269 | 4,8679E-11 |
| ACE2 | FAM136A | 0,269 | 4,84425E-11 |
| ACE2 | LARS | 0,269 | 4,91332E-11 |
| ACE2 | RP13-213K19.1 | 0,269 | 4,8363E-11 |
| ACE2 | SSR4 | 0,269 | 4,91314E-11 |
| ACE2 | RPAP1 | 0,2689 | 4,92991E-11 |
| ACE2 | RP11-305A4.5 | 0,2689 | 4,91877E-11 |
| ACE2 | NEDD4L | 0,2689 | 4,93794E-11 |
| ACE2 | DAPK1 | 0,2688 | 5,02637E-11 |
| ACE2 | TIMELESS | 0,2688 | 5,03867E-11 |
| ACE2 | XPC | 0,2687 | 5,14268E-11 |
| ACE2 | RP1-179N16.3 | 0,2687 | 5,136E-11 |
| ACE2 | TMEM184C | 0,2686 | 5,20116E-11 |
| ACE2 | NAGPA | 0,2686 | 5,21268E-11 |
| ACE2 | DDX19B | 0,2686 | 5,23117E-11 |
| ACE2 | CTC-429P9.2 | 0,2686 | 5,19601E-11 |
| ACE2 | TOMM22 | 0,2686 | 5,18243E-11 |
| ACE2 | RWDD4 | 0,2685 | 5,31879E-11 |
| ACE2 | POLD2 | 0,2685 | 5,2783E-11 |
| ACE2 | RP11-88E10.5 | 0,2685 | 5,29889E-11 |
| ACE2 | CTD-2621I17.3 | 0,2685 | 5,29873E-11 |
| ACE2 | DUSP23 | 0,2684 | 5,35214E-11 |
| ACE2 | ADGRF1 | 0,2684 | 5,4188E-11 |
| ACE2 | ALG10 | 0,2684 | 5,40305E-11 |
| ACE2 | RP11-561C5.4 | 0,2684 | 5,35588E-11 |
| ACE2 | SON | 0,2684 | 5,44139E-11 |
| ACE2 | APOBEC3AP1 | 0,2683 | 5,50573E-11 |
| ACE2 | FADS2 | 0,2683 | 5,49169E-11 |
| ACE2 | RNF40 | 0,2683 | 5,51823E-11 |
| ACE2 | RP11-49K24.8 | 0,2683 | 5,48343E-11 |
| ACE2 | TRMT12 | 0,2681 | 5,68418E-11 |
| ACE2 | DPY19L1 | 0,2679 | 5,88609E-11 |
| ACE2 | TUBGCP3 | 0,2679 | 5,84204E-11 |
| ACE2 | NAA20 | 0,2679 | 5,89547E-11 |
| ACE2 | IGSF3 | 0,2678 | 5,94014E-11 |
| ACE2 | PEX16 | 0,2678 | 5,98771E-11 |
| ACE2 | MPV17L | 0,2678 | 5,99367E-11 |
| ACE2 | TMEM126A | 0,2677 | 6,10911E-11 |
| ACE2 | DOCK9 | 0,2677 | 6,06878E-11 |
| ACE2 | ME1 | 0,2676 | 6,22665E-11 |
| ACE2 | ZNF79 | 0,2676 | 6,16615E-11 |
| ACE2 | RP11-286N22.16 | 0,2676 | 6,13542E-11 |
| ACE2 | SLC25A23 | 0,2676 | 6,20182E-11 |

| | | | |
|---|---|---|---|
| ACE2 | ANAPC16 | 0,2675 | 6,26053E-11 |
| ACE2 | ALKBH4 | 0,2674 | 6,44216E-11 |
| ACE2 | RP11-148K1.10 | 0,2674 | 6,44607E-11 |
| ACE2 | TWNK | 0,2671 | 6,75714E-11 |
| ACE2 | DOCK9-AS2 | 0,267 | 6,83481E-11 |
| ACE2 | CCDC51 | 0,2669 | 6,93632E-11 |
| ACE2 | EXOC3-AS1 | 0,2669 | 6,93888E-11 |
| ACE2 | RP11-599B13.9 | 0,2669 | 6,90638E-11 |
| ACE2 | PMS1 | 0,2668 | 7,08721E-11 |
| ACE2 | CTD-2336O2.3 | 0,2668 | 7,12202E-11 |
| ACE2 | FOXN3-AS1 | 0,2667 | 7,2003E-11 |
| ACE2 | TLCD2 | 0,2667 | 7,15449E-11 |
| ACE2 | EIF3D | 0,2667 | 7,24086E-11 |
| ACE2 | FBXO9 | 0,2666 | 7,26774E-11 |
| ACE2 | C1orf127 | 0,2665 | 7,48265E-11 |
| ACE2 | RP11-152D8.1 | 0,2665 | 7,46515E-11 |
| ACE2 | CCND3 | 0,2665 | 7,46463E-11 |
| ACE2 | RP1-120G22.11 | 0,2664 | 7,60146E-11 |
| ACE2 | TSSC1 | 0,2664 | 7,60577E-11 |
| ACE2 | TMEM99 | 0,2664 | 7,5517E-11 |
| ACE2 | RBBP4 | 0,2663 | 7,65642E-11 |
| ACE2 | SNX17 | 0,2663 | 7,69141E-11 |
| ACE2 | PEX1 | 0,2663 | 7,69937E-11 |
| ACE2 | KRT27 | 0,2663 | 7,70319E-11 |
| ACE2 | CTD-2231E14.8 | 0,2663 | 7,70732E-11 |
| ACE2 | SEC14L2 | 0,2663 | 7,64482E-11 |
| ACE2 | PAK1IP1 | 0,2662 | 7,89979E-11 |
| ACE2 | AC046143.3 | 0,2661 | 8,00308E-11 |
| ACE2 | EIF3C | 0,2661 | 7,95515E-11 |
| ACE2 | LINC01836 | 0,2661 | 7,95122E-11 |
| ACE2 | NOP9 | 0,266 | 8,08224E-11 |
| ACE2 | LINC01637 | 0,266 | 8,1545E-11 |
| ACE2 | INHBB | 0,2659 | 8,2467E-11 |
| ACE2 | ZNF16 | 0,2659 | 8,30976E-11 |
| ACE2 | AP006621.9 | 0,2659 | 8,27493E-11 |
| ACE2 | LDAH | 0,2658 | 8,39161E-11 |
| ACE2 | ATP5G3 | 0,2658 | 8,32784E-11 |
| ACE2 | CTD-2609K8.3 | 0,2658 | 8,44283E-11 |
| ACE2 | OGFOD1 | 0,2658 | 8,39959E-11 |
| ACE2 | DMXL1 | 0,2657 | 8,46665E-11 |
| ACE2 | ENPP5 | 0,2657 | 8,5841E-11 |
| ACE2 | SIL1 | 0,2656 | 8,67403E-11 |
| ACE2 | RP11-473O4.4 | 0,2655 | 8,87685E-11 |
| ACE2 | LGMNP1 | 0,2655 | 8,85008E-11 |
| ACE2 | AAGAB | 0,2655 | 8,7594E-11 |
| ACE2 | GSTA9P | 0,2654 | 9,02916E-11 |
| ACE2 | VPS35 | 0,2654 | 9,03701E-11 |
| ACE2 | RP11-889L3.1 | 0,2653 | 9,13503E-11 |
| ACE2 | PLEKHM1 | 0,2653 | 9,05298E-11 |
| ACE2 | SLC2A13 | 0,2652 | 9,23902E-11 |
| ACE2 | SNX29 | 0,2651 | 9,49291E-11 |
| ACE2 | ARCN1 | 0,265 | 9,52032E-11 |
| ACE2 | AC002117.1 | 0,265 | 9,54116E-11 |
| ACE2 | RP11-38G5.4 | 0,2649 | 9,75407E-11 |

| | | | |
|---|---|---|---|
| ACE2 | RP11-22P6.2 | 0,2649 | 9,72994E-11 |
| ACE2 | TGIF2P1 | 0,2648 | 9,98089E-11 |
| ACE2 | VPS41 | 0,2648 | 9,88283E-11 |
| ACE2 | RP11-231P20.2 | 0,2647 | 1,00618E-10 |
| ACE2 | KCNS2 | 0,2647 | 1,00778E-10 |
| ACE2 | AMDHD1 | 0,2646 | 1,02292E-10 |
| ACE2 | XXbac-B444P24.10 | 0,2646 | 1,03019E-10 |
| ACE2 | TECR | 0,2645 | 1,03398E-10 |
| ACE2 | PROK1 | 0,2644 | 1,05894E-10 |
| ACE2 | SMARCD2 | 0,2644 | 1,05164E-10 |
| ACE2 | RPIA | 0,2643 | 1,08259E-10 |
| ACE2 | YAP1P1 | 0,2643 | 1,07259E-10 |
| ACE2 | SLAMF8 | 0,2641 | 1,1147E-10 |
| ACE2 | ICE1 | 0,2641 | 1,12154E-10 |
| ACE2 | OCIAD1 | 0,264 | 1,14065E-10 |
| ACE2 | ZNF658 | 0,264 | 1,13177E-10 |
| ACE2 | COASY | 0,264 | 1,12877E-10 |
| ACE2 | PPP4R1 | 0,264 | 1,12866E-10 |
| ACE2 | ZNF468 | 0,264 | 1,13074E-10 |
| ACE2 | LINC00998 | 0,2639 | 1,151E-10 |
| ACE2 | MFAP1 | 0,2639 | 1,15283E-10 |
| ACE2 | PDHB | 0,2638 | 1,16314E-10 |
| ACE2 | SMCO3 | 0,2638 | 1,16387E-10 |
| ACE2 | LINC00467 | 0,2637 | 1,18898E-10 |
| ACE2 | GPX1 | 0,2637 | 1,18681E-10 |
| ACE2 | COA3 | 0,2637 | 1,19885E-10 |
| ACE2 | ZBTB8A | 0,2636 | 1,20679E-10 |
| ACE2 | RP5-1042I8.7 | 0,2636 | 1,20942E-10 |
| ACE2 | ZNF669 | 0,2636 | 1,22086E-10 |
| ACE2 | EEF1B2 | 0,2636 | 1,20321E-10 |
| ACE2 | XAGE2 | 0,2636 | 1,20595E-10 |
| ACE2 | TATDN1 | 0,2635 | 1,23684E-10 |
| ACE2 | TSR1 | 0,2635 | 1,23496E-10 |
| ACE2 | PRPSAP2 | 0,2635 | 1,23939E-10 |
| ACE2 | SPIRE1 | 0,2635 | 1,22313E-10 |
| ACE2 | WNT5A-AS1 | 0,2634 | 1,26103E-10 |
| ACE2 | TTI2 | 0,2634 | 1,25722E-10 |
| ACE2 | PSAP | 0,2634 | 1,25842E-10 |
| ACE2 | GOLGA2P10 | 0,2634 | 1,24375E-10 |
| ACE2 | TP53RK | 0,2634 | 1,25584E-10 |
| ACE2 | MAP3K5 | 0,2633 | 1,26866E-10 |
| ACE2 | RP11-44F14.2 | 0,2633 | 1,27524E-10 |
| ACE2 | NUDCD3 | 0,2631 | 1,31297E-10 |
| ACE2 | MTND4P20 | 0,2631 | 1,32167E-10 |
| ACE2 | ZPR1 | 0,2631 | 1,32272E-10 |
| ACE2 | NRL | 0,2631 | 1,32668E-10 |
| ACE2 | GFOD1 | 0,263 | 1,34062E-10 |
| ACE2 | BTD | 0,2629 | 1,3514E-10 |
| ACE2 | CAPN12 | 0,2629 | 1,35258E-10 |
| ACE2 | SLC50A1 | 0,2628 | 1,38403E-10 |
| ACE2 | C16orf91 | 0,2626 | 1,42642E-10 |
| ACE2 | LA16c-313F4.1 | 0,2626 | 1,42968E-10 |
| ACE2 | RALY-AS1 | 0,2626 | 1,42831E-10 |
| ACE2 | EFCAB14 | 0,2625 | 1,45711E-10 |

| | | | |
|---|---|---|---|
| ACE2 | RP11-434E6.5 | 0,2625 | 1,46416E-10 |
| ACE2 | NLRC4 | 0,2624 | 1,47372E-10 |
| ACE2 | HSPA4 | 0,2623 | 1,49207E-10 |
| ACE2 | ZDHHC9 | 0,2623 | 1,50377E-10 |
| ACE2 | TMEM44-AS1 | 0,2622 | 1,52047E-10 |
| ACE2 | NBN | 0,2622 | 1,53455E-10 |
| ACE2 | TFAM | 0,2621 | 1,56576E-10 |
| ACE2 | TRIM26 | 0,262 | 1,58269E-10 |
| ACE2 | EXPH5 | 0,262 | 1,58302E-10 |
| ACE2 | MIA3 | 0,2619 | 1,61931E-10 |
| ACE2 | TIGD6 | 0,2619 | 1,60935E-10 |
| ACE2 | DERL2 | 0,2618 | 1,6397E-10 |
| ACE2 | MYDGF | 0,2618 | 1,64402E-10 |
| ACE2 | TPRN | 0,2617 | 1,66907E-10 |
| ACE2 | C16orf95 | 0,2617 | 1,65583E-10 |
| ACE2 | AGGF1 | 0,2616 | 1,68313E-10 |
| ACE2 | RP11-305L7.1 | 0,2616 | 1,67958E-10 |
| ACE2 | TMEM19 | 0,2616 | 1,68166E-10 |
| ACE2 | ACOT4 | 0,2615 | 1,7294E-10 |
| ACE2 | LGMN | 0,2614 | 1,75012E-10 |
| ACE2 | RP5-1148A21.3 | 0,2613 | 1,78562E-10 |
| ACE2 | ZNF268 | 0,2613 | 1,78851E-10 |
| ACE2 | CTC-429P9.1 | 0,2613 | 1,76847E-10 |
| ACE2 | CXXC4 | 0,2612 | 1,81108E-10 |
| ACE2 | AP1S1 | 0,2612 | 1,78993E-10 |
| ACE2 | TLR8 | 0,2612 | 1,81471E-10 |
| ACE2 | AC011899.9 | 0,2611 | 1,84333E-10 |
| ACE2 | RP11-173P15.9 | 0,2611 | 1,8494E-10 |
| ACE2 | RFC3 | 0,2611 | 1,82064E-10 |
| ACE2 | IREB2 | 0,261 | 1,85702E-10 |
| ACE2 | CSTF2 | 0,261 | 1,8742E-10 |
| ACE2 | MTIF2 | 0,2609 | 1,88096E-10 |
| ACE2 | CCL28 | 0,2609 | 1,90627E-10 |
| ACE2 | C6orf120 | 0,2609 | 1,89444E-10 |
| ACE2 | RIPK1 | 0,2608 | 1,91409E-10 |
| ACE2 | SSNA1 | 0,2608 | 1,93496E-10 |
| ACE2 | LL22NC03-22A12.12 | 0,2608 | 1,93259E-10 |
| ACE2 | LINC01134 | 0,2607 | 1,97269E-10 |
| ACE2 | PLAA | 0,2607 | 1,95776E-10 |
| ACE2 | MPI | 0,2607 | 1,94817E-10 |
| ACE2 | KLRG2 | 0,2606 | 1,9829E-10 |
| ACE2 | RNASE2 | 0,2606 | 1,993E-10 |
| ACE2 | KIAA0513 | 0,2606 | 2,00216E-10 |
| ACE2 | SHMT2 | 0,2605 | 2,0365E-10 |
| ACE2 | KRT8P3 | 0,2604 | 2,04916E-10 |
| ACE2 | RP11-641J8.1 | 0,2604 | 2,04364E-10 |
| ACE2 | CPNE4 | 0,2603 | 2,08352E-10 |
| ACE2 | WASF5P | 0,26 | 2,21324E-10 |
| ACE2 | CELSR1 | 0,26 | 2,1812E-10 |
| ACE2 | SEC22C | 0,2599 | 2,23245E-10 |
| ACE2 | RASSF7 | 0,2599 | 2,24319E-10 |
| ACE2 | SPARCL1 | 0,2598 | 2,25651E-10 |
| ACE2 | RP11-1026M7.2 | 0,2598 | 2,26248E-10 |
| ACE2 | BAIAP2-AS1 | 0,2598 | 2,26385E-10 |

| | | | |
|---|---|---|---|
| ACE2 | PCBD2 | 0,2597 | 2,29347E-10 |
| ACE2 | RP3-355L5.5 | 0,2597 | 2,31111E-10 |
| ACE2 | NUDT13 | 0,2597 | 2,32006E-10 |
| ACE2 | LA16c-325D7.2 | 0,2597 | 2,32387E-10 |
| ACE2 | ATXN1L | 0,2596 | 2,34534E-10 |
| ACE2 | ZNHIT3 | 0,2595 | 2,38308E-10 |
| ACE2 | NDST1 | 0,2594 | 2,44473E-10 |
| ACE2 | DNAJB12 | 0,2594 | 2,42572E-10 |
| ACE2 | CPPED1 | 0,2594 | 2,44487E-10 |
| ACE2 | NDUFA8 | 0,2593 | 2,48296E-10 |
| ACE2 | ACBD5 | 0,2593 | 2,45011E-10 |
| ACE2 | FAAH2 | 0,2593 | 2,46397E-10 |
| ACE2 | IRF6 | 0,2592 | 2,51647E-10 |
| ACE2 | RP11-184J23.2 | 0,2591 | 2,5666E-10 |
| ACE2 | LY6G6E | 0,2591 | 2,53528E-10 |
| ACE2 | CPB2-AS1 | 0,2591 | 2,53147E-10 |
| ACE2 | RP11-321F6.2 | 0,2591 | 2,54845E-10 |
| ACE2 | EPB41 | 0,259 | 2,58039E-10 |
| ACE2 | RAB11A | 0,259 | 2,59704E-10 |
| ACE2 | SNX24 | 0,2589 | 2,64263E-10 |
| ACE2 | B3GALT4 | 0,2589 | 2,63113E-10 |
| ACE2 | ERCC4 | 0,2589 | 2,64643E-10 |
| ACE2 | NAT8B | 0,2588 | 2,68147E-10 |
| ACE2 | CTD-2501M5.1 | 0,2588 | 2,6923E-10 |
| ACE2 | LINC01579 | 0,2587 | 2,70908E-10 |
| ACE2 | ARHGAP44 | 0,2587 | 2,71625E-10 |
| ACE2 | TAF5L | 0,2586 | 2,76071E-10 |
| ACE2 | TSNAX | 0,2586 | 2,76619E-10 |
| ACE2 | SCAP | 0,2586 | 2,76411E-10 |
| ACE2 | BCKDHB | 0,2586 | 2,7741E-10 |
| ACE2 | MAPK1 | 0,2586 | 2,78703E-10 |
| ACE2 | EFNA5 | 0,2585 | 2,80989E-10 |
| ACE2 | DDX54 | 0,2585 | 2,78932E-10 |
| ACE2 | SLC7A10 | 0,2585 | 2,81649E-10 |
| ACE2 | RNF139 | 0,2583 | 2,91535E-10 |
| ACE2 | RPS3AP2 | 0,2582 | 2,97629E-10 |
| ACE2 | SCGB2B2 | 0,2582 | 2,9312E-10 |
| ACE2 | AC002059.10 | 0,2582 | 2,96345E-10 |
| ACE2 | EBNA1BP2 | 0,2581 | 3,0164E-10 |
| ACE2 | AC123023.1 | 0,2581 | 3,0165E-10 |
| ACE2 | FAM221A | 0,2581 | 3,02098E-10 |
| ACE2 | FAM53B | 0,2581 | 2,98621E-10 |
| ACE2 | ZNF554 | 0,258 | 3,03485E-10 |
| ACE2 | MAPKAPK5 | 0,2579 | 3,08785E-10 |
| ACE2 | SLC35A4 | 0,2578 | 3,15021E-10 |
| ACE2 | SPPL3 | 0,2578 | 3,14559E-10 |
| ACE2 | PFAS | 0,2578 | 3,16781E-10 |
| ACE2 | ARHGEF18 | 0,2578 | 3,17358E-10 |
| ACE2 | TTC37 | 0,2576 | 3,26691E-10 |
| ACE2 | TMPRSS13 | 0,2576 | 3,27042E-10 |
| ACE2 | RP11-1055B8.2 | 0,2576 | 3,24523E-10 |
| ACE2 | F2RL3 | 0,2576 | 3,2803E-10 |
| ACE2 | SSU72 | 0,2575 | 3,33128E-10 |
| ACE2 | MIR573 | 0,2575 | 3,28358E-10 |

| | | | |
|---|---|---|---|
| ACE2 | GPAA1 | 0,2575 | 3,30531E-10 |
| ACE2 | CDK4 | 0,2575 | 3,29828E-10 |
| ACE2 | CRADD | 0,2575 | 3,32529E-10 |
| ACE2 | CTB-50E14.5 | 0,2575 | 3,29599E-10 |
| ACE2 | SUMO3 | 0,2575 | 3,28475E-10 |
| ACE2 | CSF2 | 0,2574 | 3,35555E-10 |
| ACE2 | NABP2 | 0,2574 | 3,35024E-10 |
| ACE2 | PIGT | 0,2574 | 3,34387E-10 |
| ACE2 | TOR1AIP2 | 0,2573 | 3,40616E-10 |
| ACE2 | COL4A3BP | 0,2573 | 3,44322E-10 |
| ACE2 | RPP40 | 0,2573 | 3,39774E-10 |
| ACE2 | API5 | 0,2573 | 3,41662E-10 |
| ACE2 | NDUFS3 | 0,2573 | 3,41972E-10 |
| ACE2 | WSB2 | 0,2573 | 3,40639E-10 |
| ACE2 | RP11-715J22.4 | 0,2573 | 3,43765E-10 |
| ACE2 | PMVK | 0,2572 | 3,47415E-10 |
| ACE2 | ESD | 0,2572 | 3,49335E-10 |
| ACE2 | EPPK1 | 0,2571 | 3,53622E-10 |
| ACE2 | SLC9A7 | 0,2571 | 3,52918E-10 |
| ACE2 | RP1-186E20.1 | 0,257 | 3,59539E-10 |
| ACE2 | ABCC2 | 0,257 | 3,56218E-10 |
| ACE2 | RHOD | 0,257 | 3,60086E-10 |
| ACE2 | RP1-117B12.4 | 0,257 | 3,5797E-10 |
| ACE2 | TFIP11 | 0,257 | 3,58261E-10 |
| ACE2 | RP11-350J20.12 | 0,2569 | 3,66031E-10 |
| ACE2 | AASDHPPT | 0,2569 | 3,63535E-10 |
| ACE2 | GPC5 | 0,2569 | 3,66135E-10 |
| ACE2 | RABGGTA | 0,2569 | 3,62337E-10 |
| ACE2 | RP11-8L8.2 | 0,2569 | 3,65484E-10 |
| ACE2 | TMEM267 | 0,2568 | 3,72779E-10 |
| ACE2 | KIF20B | 0,2568 | 3,71518E-10 |
| ACE2 | WNK1 | 0,2568 | 3,68065E-10 |
| ACE2 | STK4-AS1 | 0,2568 | 3,69861E-10 |
| ACE2 | ZBTB8B | 0,2567 | 3,75419E-10 |
| ACE2 | NSUN4 | 0,2566 | 3,81139E-10 |
| ACE2 | TOMM70 | 0,2566 | 3,83729E-10 |
| ACE2 | SMARCAD1 | 0,2566 | 3,80477E-10 |
| ACE2 | LINC01431 | 0,2566 | 3,8E-10 |
| ACE2 | SLC25A20 | 0,2565 | 3,86479E-10 |
| ACE2 | ORM2 | 0,2565 | 3,87722E-10 |
| ACE2 | ILVBL | 0,2565 | 3,9072E-10 |
| ACE2 | OSBPL11 | 0,2564 | 3,96404E-10 |
| ACE2 | PPIP5K2 | 0,2564 | 3,9833E-10 |
| ACE2 | CYP4Z2P | 0,2563 | 4,02104E-10 |
| ACE2 | AC003991.3 | 0,2563 | 4,01102E-10 |
| ACE2 | TMED10 | 0,2563 | 4,04928E-10 |
| ACE2 | ZSCAN5A | 0,2563 | 4,00282E-10 |
| ACE2 | LSM5 | 0,2562 | 4,1139E-10 |
| ACE2 | GFPT1 | 0,2561 | 4,11911E-10 |
| ACE2 | LCN10 | 0,2561 | 4,1648E-10 |
| ACE2 | RP11-510M2.2 | 0,2561 | 4,146E-10 |
| ACE2 | NCL | 0,256 | 4,25184E-10 |
| ACE2 | BLOC1S4 | 0,256 | 4,2098E-10 |
| ACE2 | HSDL1 | 0,2559 | 4,30037E-10 |

| | | | |
|---|---|---|---|
| ACE2 | ST3GAL1P1 | 0,2558 | 4,38327E-10 |
| ACE2 | ZDHHC2 | 0,2558 | 4,39048E-10 |
| ACE2 | PPM1G | 0,2557 | 4,43863E-10 |
| ACE2 | LINC01964 | 0,2557 | 4,39969E-10 |
| ACE2 | CHST2 | 0,2557 | 4,46316E-10 |
| ACE2 | FAM173B | 0,2557 | 4,40819E-10 |
| ACE2 | RP11-348F1.3 | 0,2557 | 4,44515E-10 |
| ACE2 | RP11-400N9.1 | 0,2556 | 4,49468E-10 |
| ACE2 | KLHDC10 | 0,2556 | 4,48483E-10 |
| ACE2 | RP11-631N16.2 | 0,2556 | 4,51776E-10 |
| ACE2 | RP3-414A15.12 | 0,2556 | 4,50987E-10 |
| ACE2 | WI2-1896O14.1 | 0,2554 | 4,65734E-10 |
| ACE2 | SEMA4F | 0,2554 | 4,68166E-10 |
| ACE2 | KLHL35 | 0,2554 | 4,65774E-10 |
| ACE2 | ALKBH8 | 0,2554 | 4,68178E-10 |
| ACE2 | MIR663AHG | 0,2554 | 4,65572E-10 |
| ACE2 | CTD-2626G11.2 | 0,2553 | 4,70597E-10 |
| ACE2 | MST1R | 0,2552 | 4,78307E-10 |
| ACE2 | EIF3M | 0,2552 | 4,78912E-10 |
| ACE2 | DNAAF2 | 0,2552 | 4,79769E-10 |
| ACE2 | EIF2AK3 | 0,2551 | 4,87937E-10 |
| ACE2 | RP11-71E19.1 | 0,2551 | 4,85729E-10 |
| ACE2 | ATP5B | 0,2551 | 4,86373E-10 |
| ACE2 | AJUBA | 0,2551 | 4,85698E-10 |
| ACE2 | RP11-505K9.4 | 0,2551 | 4,85252E-10 |
| ACE2 | AC006539.3 | 0,2551 | 4,87312E-10 |
| ACE2 | STAM2 | 0,255 | 4,97253E-10 |
| ACE2 | GLUD1P3 | 0,255 | 4,9911E-10 |
| ACE2 | RNY3P8 | 0,255 | 4,94919E-10 |
| ACE2 | UBE2Q1 | 0,2549 | 5,03392E-10 |
| ACE2 | CCDC112 | 0,2549 | 5,0171E-10 |
| ACE2 | CCDC25 | 0,2549 | 5,06412E-10 |
| ACE2 | FOXRED2 | 0,2549 | 5,03125E-10 |
| ACE2 | PCSK6 | 0,2548 | 5,1274E-10 |
| ACE2 | NDUFC2 | 0,2547 | 5,18845E-10 |
| ACE2 | LINC01714 | 0,2545 | 5,38283E-10 |
| ACE2 | LYPLAL1 | 0,2545 | 5,33883E-10 |
| ACE2 | SPTBN2 | 0,2545 | 5,37644E-10 |
| ACE2 | MBTPS1 | 0,2545 | 5,38648E-10 |
| ACE2 | NOC2L | 0,2543 | 5,53067E-10 |
| ACE2 | DNAJC19 | 0,2542 | 5,60915E-10 |
| ACE2 | TTC1 | 0,2542 | 5,6558E-10 |
| ACE2 | SNRPC | 0,2542 | 5,67935E-10 |
| ACE2 | MIRLET7BHG | 0,2542 | 5,67645E-10 |
| ACE2 | RP11-806O11.1 | 0,2541 | 5,7157E-10 |
| ACE2 | ACAD9 | 0,254 | 5,80693E-10 |
| ACE2 | PHACTR1 | 0,254 | 5,80924E-10 |
| ACE2 | RP3-460G2.2 | 0,254 | 5,8446E-10 |
| ACE2 | AC009237.16 | 0,2539 | 5,92005E-10 |
| ACE2 | RP11-528I4.2 | 0,2539 | 5,89575E-10 |
| ACE2 | RPL7P41 | 0,2539 | 5,92585E-10 |
| ACE2 | CA15P1 | 0,2539 | 5,95248E-10 |
| ACE2 | TMEM80 | 0,2538 | 6,02282E-10 |
| ACE2 | F8A1 | 0,2538 | 5,97562E-10 |

| | | | |
|---|---|---|---|
| ACE2 | MBLAC2 | 0,2537 | 6,12311E-10 |
| ACE2 | DNAJC24 | 0,2537 | 6,12037E-10 |
| ACE2 | RP11-832A4.7 | 0,2537 | 6,12153E-10 |
| ACE2 | ALOX5AP | 0,2537 | 6,10697E-10 |
| ACE2 | ASF1A | 0,2536 | 6,25208E-10 |
| ACE2 | AKAP3 | 0,2536 | 6,19904E-10 |
| ACE2 | C4BPB | 0,2535 | 6,30385E-10 |
| ACE2 | VPS50 | 0,2535 | 6,26016E-10 |
| ACE2 | RAB11B-AS1 | 0,2535 | 6,34442E-10 |
| ACE2 | RP11-152H18.4 | 0,2534 | 6,43272E-10 |
| ACE2 | RRP1B | 0,2534 | 6,41598E-10 |
| ACE2 | BCAP31 | 0,2534 | 6,45095E-10 |
| ACE2 | COQ2 | 0,2533 | 6,51885E-10 |
| ACE2 | HTR3A | 0,2533 | 6,50902E-10 |
| ACE2 | RP11-544M22.8 | 0,2532 | 6,65458E-10 |
| ACE2 | CCNJL | 0,2532 | 6,58871E-10 |
| ACE2 | ADGRF5 | 0,2532 | 6,60449E-10 |
| ACE2 | BCL2L2 | 0,2532 | 6,6162E-10 |
| ACE2 | GDPGP1 | 0,2532 | 6,65892E-10 |
| ACE2 | GPATCH8 | 0,2532 | 6,60862E-10 |
| ACE2 | SAMM50 | 0,2532 | 6,60179E-10 |
| ACE2 | MRPS27 | 0,2531 | 6,71175E-10 |
| ACE2 | GPR150 | 0,2531 | 6,74014E-10 |
| ACE2 | RP11-498P14.5 | 0,253 | 6,83762E-10 |
| ACE2 | RP11-505K9.1 | 0,253 | 6,80215E-10 |
| ACE2 | EFTUD2 | 0,253 | 6,79919E-10 |
| ACE2 | CAD | 0,2529 | 6,92288E-10 |
| ACE2 | MT1E | 0,2529 | 6,91506E-10 |
| ACE2 | ALG12 | 0,2529 | 6,96599E-10 |
| ACE2 | PNMA6A | 0,2529 | 6,99146E-10 |
| ACE2 | PDZK1IP1 | 0,2528 | 7,08557E-10 |
| ACE2 | ABHD11-AS1 | 0,2528 | 7,05894E-10 |
| ACE2 | ALG8 | 0,2528 | 7,05268E-10 |
| ACE2 | CDK5RAP2 | 0,2527 | 7,10876E-10 |
| ACE2 | SUN2 | 0,2527 | 7,1357E-10 |
| ACE2 | HIRIP3 | 0,2526 | 7,26888E-10 |
| ACE2 | RIMBP3 | 0,2526 | 7,32219E-10 |
| ACE2 | BOLA1 | 0,2525 | 7,34754E-10 |
| ACE2 | PPP1R13B | 0,2525 | 7,42018E-10 |
| ACE2 | MPP3 | 0,2525 | 7,42995E-10 |
| ACE2 | TDRD5 | 0,2524 | 7,47298E-10 |
| ACE2 | TOMM20 | 0,2524 | 7,52645E-10 |
| ACE2 | MRPL47 | 0,2524 | 7,47996E-10 |
| ACE2 | RP11-83B20.8 | 0,2524 | 7,51536E-10 |
| ACE2 | GSPT1 | 0,2524 | 7,55808E-10 |
| ACE2 | POFUT1 | 0,2524 | 7,55335E-10 |
| ACE2 | RGS7 | 0,2523 | 7,65674E-10 |
| ACE2 | COMMD5 | 0,2523 | 7,60848E-10 |
| ACE2 | RP11-597M12.2 | 0,2523 | 7,6422E-10 |
| ACE2 | TMPRSS7 | 0,2522 | 7,77886E-10 |
| ACE2 | IPO4 | 0,2522 | 7,72337E-10 |
| ACE2 | DDC | 0,2521 | 7,93747E-10 |
| ACE2 | SH2D4A | 0,2521 | 7,91783E-10 |
| ACE2 | RP11-187C18.4 | 0,2521 | 7,84263E-10 |

| | | | |
|---|---|---|---|
| ACE2 | JMJD8 | 0,2521 | 7,85133E-10 |
| ACE2 | RNF135 | 0,2521 | 7,87537E-10 |
| ACE2 | PEX11G | 0,2521 | 7,81932E-10 |
| ACE2 | APLP2 | 0,252 | 7,97413E-10 |
| ACE2 | SLC3A2 | 0,2519 | 8,08553E-10 |
| ACE2 | RP4-800M22.1 | 0,2518 | 8,21644E-10 |
| ACE2 | RNLS | 0,2518 | 8,30141E-10 |
| ACE2 | AC016747.3 | 0,2517 | 8,33692E-10 |
| ACE2 | ZNF664 | 0,2517 | 8,34463E-10 |
| ACE2 | RP11-83B20.9 | 0,2517 | 8,40829E-10 |
| ACE2 | DLEU1 | 0,2517 | 8,3639E-10 |
| ACE2 | TIMM13 | 0,2517 | 8,41456E-10 |
| ACE2 | C22orf46 | 0,2517 | 8,41797E-10 |
| ACE2 | RP11-42O15.3 | 0,2516 | 8,59537E-10 |
| ACE2 | C4A | 0,2516 | 8,51328E-10 |
| ACE2 | PITPNA | 0,2516 | 8,53247E-10 |
| ACE2 | ME2 | 0,2516 | 8,53152E-10 |
| ACE2 | PWAR1 | 0,2515 | 8,66212E-10 |
| ACE2 | EXO5 | 0,2514 | 8,77924E-10 |
| ACE2 | RAB10 | 0,2514 | 8,80988E-10 |
| ACE2 | LTA4H | 0,2514 | 8,86958E-10 |
| ACE2 | CLCN2 | 0,2513 | 8,89281E-10 |
| ACE2 | LINC02036 | 0,2513 | 8,91188E-10 |
| ACE2 | RP11-589B3.6 | 0,2513 | 8,95843E-10 |
| ACE2 | ADCK1 | 0,2513 | 8,96833E-10 |
| ACE2 | ZNF682 | 0,2513 | 9,00783E-10 |
| ACE2 | CCDC126 | 0,2512 | 9,08981E-10 |
| ACE2 | RP11-379H18.1 | 0,2512 | 9,12792E-10 |
| ACE2 | RP11-1096G20.5 | 0,2512 | 9,10268E-10 |
| ACE2 | PGAP3 | 0,2512 | 9,10819E-10 |
| ACE2 | SNF8 | 0,2512 | 9,1299E-10 |
| ACE2 | PRR34 | 0,2512 | 9,12402E-10 |
| ACE2 | ANKRD65 | 0,251 | 9,3554E-10 |
| ACE2 | MYRIP | 0,251 | 9,39647E-10 |
| ACE2 | TPP1 | 0,251 | 9,312E-10 |
| ACE2 | APPL2 | 0,251 | 9,3481E-10 |
| ACE2 | ZSCAN32 | 0,251 | 9,40737E-10 |
| ACE2 | GCSH | 0,251 | 9,36158E-10 |
| ACE2 | NANP | 0,251 | 9,41564E-10 |
| ACE2 | PGGT1B | 0,2508 | 9,64453E-10 |
| ACE2 | LPP-AS2 | 0,2507 | 9,78944E-10 |
| ACE2 | RN7SL473P | 0,2506 | 1,00415E-09 |
| ACE2 | POLR1C | 0,2506 | 1,00153E-09 |
| ACE2 | RP11-91K8.4 | 0,2505 | 1,01082E-09 |
| ACE2 | IGFBP3 | 0,2505 | 1,0123E-09 |
| ACE2 | LPIN2 | 0,2505 | 1,01381E-09 |
| ACE2 | FGF22 | 0,2505 | 1,00979E-09 |
| ACE2 | STRIP2 | 0,2504 | 1,03909E-09 |
| ACE2 | RTN4R | 0,2504 | 1,03256E-09 |
| ACE2 | PGM1 | 0,2503 | 1,05172E-09 |
| ACE2 | BRINP2 | 0,2503 | 1,05407E-09 |
| ACE2 | CTD-2165H16.3 | 0,2503 | 1,04215E-09 |
| ACE2 | MUC21 | 0,2503 | 1,05413E-09 |
| ACE2 | MTFMT | 0,2503 | 1,04637E-09 |

| | | | |
|---|---|---|---|
| ACE2 | AC000036.4 | 0,2503 | 1,0506E-09 |
| ACE2 | SORBS2 | 0,2501 | 1,08675E-09 |
| ACE2 | VARS | 0,2501 | 1,0896E-09 |
| ACE2 | COPB1 | 0,2501 | 1,08581E-09 |
| ACE2 | RPUSD1 | 0,2501 | 1,07317E-09 |
| ACE2 | ARHGAP29 | -0,25 | 1,09413E-09 |
| ACE2 | CXCL6 | -0,2501 | 1,07328E-09 |
| ACE2 | RP11-486G15.2 | -0,2502 | 1,06271E-09 |
| ACE2 | RP11-24C3.2 | -0,2502 | 1,06576E-09 |
| ACE2 | FGD6 | -0,2502 | 1,0714E-09 |
| ACE2 | AC012462.2 | -0,2503 | 1,05013E-09 |
| ACE2 | LCORL | -0,2503 | 1,04742E-09 |
| ACE2 | TNRC18P1 | -0,2504 | 1,03641E-09 |
| ACE2 | RP11-642D21.2 | -0,2504 | 1,03241E-09 |
| ACE2 | RP11-543H23.2 | -0,2504 | 1,03804E-09 |
| ACE2 | TAZ | -0,2504 | 1,02723E-09 |
| ACE2 | RP11-46F15.2 | -0,2505 | 1,01911E-09 |
| ACE2 | HSPB2 | -0,2505 | 1,0163E-09 |
| ACE2 | RP11-843B15.4 | -0,2505 | 1,01954E-09 |
| ACE2 | SNRPA1 | -0,2505 | 1,01988E-09 |
| ACE2 | RP11-159D12.8 | -0,2505 | 1,01871E-09 |
| ACE2 | RP11-318A15.8 | -0,2505 | 1,00885E-09 |
| ACE2 | TSPAN2 | -0,2506 | 9,94346E-10 |
| ACE2 | LINC02082 | -0,2506 | 9,99436E-10 |
| ACE2 | FNDC1-IT1 | -0,2506 | 1,00114E-09 |
| ACE2 | AC012314.8 | -0,2506 | 1,00378E-09 |
| ACE2 | GBP7 | -0,2507 | 9,90304E-10 |
| ACE2 | NEB | -0,2507 | 9,79488E-10 |
| ACE2 | SESN3 | -0,2507 | 9,88052E-10 |
| ACE2 | AC156455.1 | -0,2507 | 9,83551E-10 |
| ACE2 | COL4A2-AS1 | -0,2507 | 9,85617E-10 |
| ACE2 | ZNF121 | -0,2507 | 9,80398E-10 |
| ACE2 | RP1-142L7.8 | -0,2508 | 9,6639E-10 |
| ACE2 | STRADA | -0,2509 | 9,57894E-10 |
| ACE2 | RP11-543H23.1 | -0,2509 | 9,55796E-10 |
| ACE2 | RP3-354N19.3 | -0,251 | 9,3237E-10 |
| ACE2 | MROH6 | -0,251 | 9,3683E-10 |
| ACE2 | SNAP25 | -0,251 | 9,35587E-10 |
| ACE2 | RP11-334A14.8 | -0,2511 | 9,2778E-10 |
| ACE2 | C10orf10 | -0,2511 | 9,26298E-10 |
| ACE2 | RN7SL263P | -0,2511 | 9,28795E-10 |
| ACE2 | HIST1H2APS3 | -0,2512 | 9,12925E-10 |
| ACE2 | TNFRSF21 | -0,2512 | 9,09624E-10 |
| ACE2 | AP000487.4 | -0,2512 | 9,11154E-10 |
| ACE2 | AP001442.2 | -0,2512 | 9,07248E-10 |
| ACE2 | IL17RD | -0,2513 | 9,0118E-10 |
| ACE2 | RHBDF2 | -0,2513 | 8,88809E-10 |
| ACE2 | PRKD3 | -0,2514 | 8,77475E-10 |
| ACE2 | CFL1P1 | -0,2514 | 8,75472E-10 |
| ACE2 | RP11-324O2.3 | -0,2514 | 8,80914E-10 |
| ACE2 | RP11-626H12.3 | -0,2514 | 8,83361E-10 |
| ACE2 | GNB3 | -0,2514 | 8,76051E-10 |
| ACE2 | SCML1 | -0,2514 | 8,76487E-10 |
| ACE2 | PRRG1 | -0,2514 | 8,75105E-10 |

| | | | |
|---|---|---|---|
| ACE2 | MKNK1-AS1 | -0,2515 | 8,61228E-10 |
| ACE2 | RP13-516M14.1 | -0,2515 | 8,68178E-10 |
| ACE2 | RNA5SP39 | -0,2517 | 8,35141E-10 |
| ACE2 | RP11-395I6.3 | -0,2518 | 8,25029E-10 |
| ACE2 | MICU3 | -0,2518 | 8,26663E-10 |
| ACE2 | NELL2 | -0,2518 | 8,31705E-10 |
| ACE2 | KPNA1 | -0,2519 | 8,19149E-10 |
| ACE2 | LINGO1 | -0,2519 | 8,19771E-10 |
| ACE2 | GDF5 | -0,2519 | 8,18143E-10 |
| ACE2 | ZBTB37 | -0,252 | 8,03973E-10 |
| ACE2 | CNIH3 | -0,252 | 8,01123E-10 |
| ACE2 | CCDC146 | -0,252 | 7,9904E-10 |
| ACE2 | RP11-661A12.8 | -0,252 | 7,98204E-10 |
| ACE2 | KIAA1210 | -0,252 | 8,00384E-10 |
| ACE2 | RP11-521D12.5 | -0,2521 | 7,81967E-10 |
| ACE2 | REL | -0,2521 | 7,92429E-10 |
| ACE2 | RP13-514E23.2 | -0,2521 | 7,82143E-10 |
| ACE2 | RP11-544L8__B.4 | -0,2522 | 7,69416E-10 |
| ACE2 | RP11-16P20.4 | -0,2522 | 7,77758E-10 |
| ACE2 | RP11-95O2.5 | -0,2522 | 7,75958E-10 |
| ACE2 | RAB6C-AS1 | -0,2523 | 7,66992E-10 |
| ACE2 | RP11-1081L13.4 | -0,2523 | 7,59225E-10 |
| ACE2 | MTMR3 | -0,2523 | 7,57816E-10 |
| ACE2 | RAB33A | -0,2523 | 7,58545E-10 |
| ACE2 | HNRNPA2B1 | -0,2524 | 7,50312E-10 |
| ACE2 | ABL1 | -0,2524 | 7,49201E-10 |
| ACE2 | APBB1 | -0,2524 | 7,56709E-10 |
| ACE2 | MIR324 | -0,2524 | 7,47931E-10 |
| ACE2 | BRWD1 | -0,2524 | 7,48586E-10 |
| ACE2 | FAM159A | -0,2525 | 7,39695E-10 |
| ACE2 | MXRA7 | -0,2525 | 7,40557E-10 |
| ACE2 | KIF25-AS1 | -0,2526 | 7,23453E-10 |
| ACE2 | RP11-813P10.1 | -0,2526 | 7,31734E-10 |
| ACE2 | RNU6V | -0,2527 | 7,20831E-10 |
| ACE2 | CLCF1 | -0,2527 | 7,15297E-10 |
| ACE2 | THY1 | -0,2527 | 7,15906E-10 |
| ACE2 | HS3ST3B1 | -0,2527 | 7,13481E-10 |
| ACE2 | RBM15 | -0,2528 | 7,05474E-10 |
| ACE2 | ZNF692 | -0,2528 | 7,04463E-10 |
| ACE2 | AC007256.5 | -0,2528 | 7,02109E-10 |
| ACE2 | WNT1 | -0,2528 | 7,05773E-10 |
| ACE2 | CTD-2017C7.1 | -0,2528 | 7,03384E-10 |
| ACE2 | AC005954.3 | -0,2528 | 7,0207E-10 |
| ACE2 | HSPB6 | -0,2528 | 7,04383E-10 |
| ACE2 | CEACAM1 | -0,2528 | 7,04714E-10 |
| ACE2 | HAPLN1 | -0,2529 | 6,88759E-10 |
| ACE2 | SNORA13 | -0,2529 | 6,93758E-10 |
| ACE2 | TFAP2A-AS1 | -0,2529 | 6,9509E-10 |
| ACE2 | LINC00840 | -0,2529 | 6,95762E-10 |
| ACE2 | DOCK3 | -0,253 | 6,79079E-10 |
| ACE2 | PTGER3 | -0,2531 | 6,69637E-10 |
| ACE2 | RP4-671G15.2 | -0,2531 | 6,67214E-10 |
| ACE2 | CDCA7 | -0,2531 | 6,68769E-10 |
| ACE2 | HIGD1B | -0,2531 | 6,76878E-10 |

| | | | |
|---|---|---|---|
| ACE2 | TMEM35A | -0,2531 | 6,68052E-10 |
| ACE2 | NXT1 | -0,2532 | 6,59527E-10 |
| ACE2 | PHF21B | -0,2532 | 6,63044E-10 |
| ACE2 | SHISA5 | -0,2534 | 6,37044E-10 |
| ACE2 | RN7SL244P | -0,2534 | 6,43296E-10 |
| ACE2 | MRVI1 | -0,2534 | 6,42015E-10 |
| ACE2 | PPP1R3B | -0,2535 | 6,33409E-10 |
| ACE2 | RAB37 | -0,2535 | 6,26949E-10 |
| ACE2 | NASP | -0,2536 | 6,22266E-10 |
| ACE2 | GPSM3 | -0,2536 | 6,18022E-10 |
| ACE2 | RP1-67K17.4 | -0,2536 | 6,19922E-10 |
| ACE2 | OVOS2 | -0,2536 | 6,19875E-10 |
| ACE2 | LIM2 | -0,2536 | 6,24006E-10 |
| ACE2 | SELP | -0,2537 | 6,0946E-10 |
| ACE2 | SCN8A | -0,2537 | 6,11153E-10 |
| ACE2 | LA16c-349E10.1 | -0,2537 | 6,0736E-10 |
| ACE2 | RASIP1 | -0,2537 | 6,10586E-10 |
| ACE2 | RP3-470B24.5 | -0,2538 | 6,04585E-10 |
| ACE2 | RP11-748H22.1 | -0,2538 | 5,96879E-10 |
| ACE2 | RP11-296O14.3 | -0,2539 | 5,93197E-10 |
| ACE2 | RP11-265E18.1 | -0,2539 | 5,94183E-10 |
| ACE2 | NAA16 | -0,2539 | 5,88576E-10 |
| ACE2 | RP11-106M3.3 | -0,2539 | 5,89821E-10 |
| ACE2 | LINC00892 | -0,2539 | 5,92684E-10 |
| ACE2 | HIST1H3I | -0,254 | 5,77666E-10 |
| ACE2 | CDK14 | -0,254 | 5,80018E-10 |
| ACE2 | DENND1C | -0,254 | 5,83518E-10 |
| ACE2 | FBLIM1 | -0,2541 | 5,70937E-10 |
| ACE2 | RP11-553K8.5 | -0,2541 | 5,73505E-10 |
| ACE2 | MRNIP | -0,2541 | 5,76978E-10 |
| ACE2 | PXDNL | -0,2541 | 5,75266E-10 |
| ACE2 | PTPN5 | -0,2541 | 5,73225E-10 |
| ACE2 | ARAP1-AS2 | -0,2541 | 5,72346E-10 |
| ACE2 | PRRC2C | -0,2542 | 5,66537E-10 |
| ACE2 | DST | -0,2542 | 5,62406E-10 |
| ACE2 | LINC01229 | -0,2542 | 5,59433E-10 |
| ACE2 | ZNF700 | -0,2542 | 5,59725E-10 |
| ACE2 | RP4-665N4.8 | -0,2543 | 5,57631E-10 |
| ACE2 | AC103563.9 | -0,2543 | 5,58922E-10 |
| ACE2 | ZNF451 | -0,2543 | 5,57964E-10 |
| ACE2 | SCARB1 | -0,2545 | 5,40438E-10 |
| ACE2 | RP11-504I13.3 | -0,2545 | 5,41361E-10 |
| ACE2 | FRMD8 | -0,2546 | 5,30742E-10 |
| ACE2 | CTA-292E10.6 | -0,2546 | 5,2922E-10 |
| ACE2 | RP11-296O14.2 | -0,2547 | 5,24205E-10 |
| ACE2 | WBP1 | -0,2547 | 5,17849E-10 |
| ACE2 | RP11-330H6.6 | -0,2547 | 5,16259E-10 |
| ACE2 | CCNI2 | -0,2547 | 5,2263E-10 |
| ACE2 | KLC1 | -0,2547 | 5,17234E-10 |
| ACE2 | CC2D1A | -0,2547 | 5,1751E-10 |
| ACE2 | ATP2B4 | -0,2548 | 5,13158E-10 |
| ACE2 | AC093162.5 | -0,255 | 4,92293E-10 |
| ACE2 | LINC01945 | -0,255 | 4,94293E-10 |
| ACE2 | AC140912.1 | -0,255 | 4,97719E-10 |

| | | | |
|---|---|---|---|
| ACE2 | P2RX1 | -0,255 | 4,92074E-10 |
| ACE2 | KIZ-AS1 | -0,255 | 4,98959E-10 |
| ACE2 | ARNT | -0,2552 | 4,77538E-10 |
| ACE2 | KPNA5 | -0,2552 | 4,77525E-10 |
| ACE2 | ADAM11 | -0,2552 | 4,80226E-10 |
| ACE2 | RP11-73K9.3 | -0,2553 | 4,73638E-10 |
| ACE2 | VPS28 | -0,2553 | 4,68547E-10 |
| ACE2 | CTC-563A5.2 | -0,2554 | 4,61359E-10 |
| ACE2 | TRBV14 | -0,2554 | 4,67804E-10 |
| ACE2 | RP11-444D3.1 | -0,2554 | 4,66899E-10 |
| ACE2 | RP11-231I16.1 | -0,2554 | 4,62739E-10 |
| ACE2 | RNY1P4 | -0,2554 | 4,62555E-10 |
| ACE2 | ZNF254 | -0,2554 | 4,61226E-10 |
| ACE2 | SCNN1D | -0,2555 | 4,56437E-10 |
| ACE2 | CTC-558O2.2 | -0,2555 | 4,53886E-10 |
| ACE2 | C9orf172 | -0,2555 | 4,58126E-10 |
| ACE2 | TMEM235 | -0,2555 | 4,60403E-10 |
| ACE2 | CTC-453G23.8 | -0,2555 | 4,58849E-10 |
| ACE2 | TRBV25-1 | -0,2556 | 4,47218E-10 |
| ACE2 | RP11-47I22.2 | -0,2556 | 4,49488E-10 |
| ACE2 | CTD-3247H4.2 | -0,2556 | 4,48074E-10 |
| ACE2 | SMOX | -0,2556 | 4,50578E-10 |
| ACE2 | MAP3K20 | -0,2557 | 4,42089E-10 |
| ACE2 | COL21A1 | -0,2557 | 4,41593E-10 |
| ACE2 | RP11-1086F11.1 | -0,2558 | 4,33861E-10 |
| ACE2 | KCND2 | -0,2558 | 4,36729E-10 |
| ACE2 | STRC | -0,2558 | 4,32828E-10 |
| ACE2 | FSIP2-AS1 | -0,2559 | 4,28806E-10 |
| ACE2 | CPSF6 | -0,2559 | 4,2741E-10 |
| ACE2 | CSK | -0,2559 | 4,29332E-10 |
| ACE2 | CD19 | -0,2559 | 4,31584E-10 |
| ACE2 | NAA40 | -0,256 | 4,2405E-10 |
| ACE2 | PPFIBP1 | -0,256 | 4,19878E-10 |
| ACE2 | IFNG-AS1 | -0,256 | 4,24829E-10 |
| ACE2 | CTD-3099C6.9 | -0,256 | 4,1884E-10 |
| ACE2 | ZNF630 | -0,256 | 4,20824E-10 |
| ACE2 | CLCNKA | -0,2561 | 4,12456E-10 |
| ACE2 | MBOAT2 | -0,2561 | 4,16638E-10 |
| ACE2 | AMZ2P2 | -0,2561 | 4,13868E-10 |
| ACE2 | NACA | -0,2561 | 4,13652E-10 |
| ACE2 | RP11-454H19.2 | -0,2562 | 4,10496E-10 |
| ACE2 | RN7SKP30 | -0,2563 | 4,0474E-10 |
| ACE2 | TATDN2P3 | -0,2563 | 4,01402E-10 |
| ACE2 | KNOP1P2 | -0,2564 | 3,93522E-10 |
| ACE2 | PLEKHG5 | -0,2565 | 3,87969E-10 |
| ACE2 | DNAH1 | -0,2565 | 3,86722E-10 |
| ACE2 | TRBV12-3 | -0,2565 | 3,92263E-10 |
| ACE2 | RP11-1112C15.2 | -0,2565 | 3,9098E-10 |
| ACE2 | RP11-465I4.2 | -0,2565 | 3,90977E-10 |
| ACE2 | FNDC4 | -0,2566 | 3,80744E-10 |
| ACE2 | ASIC4 | -0,2566 | 3,81803E-10 |
| ACE2 | CTD-2049O4.1 | -0,2566 | 3,80574E-10 |
| ACE2 | TSPOAP1 | -0,2566 | 3,82206E-10 |
| ACE2 | MBD1 | -0,2566 | 3,82554E-10 |

| | | | |
|---|---|---|---|
| ACE2 | PIP5K1A | -0,2567 | 3,76214E-10 |
| ACE2 | FAM138B | -0,2567 | 3,77187E-10 |
| ACE2 | INPPL1 | -0,2567 | 3,73821E-10 |
| ACE2 | RP11-358B23.6 | -0,2567 | 3,78299E-10 |
| ACE2 | CTD-2192J16.21 | -0,2567 | 3,75286E-10 |
| ACE2 | RASA4DP | -0,2568 | 3,72906E-10 |
| ACE2 | TMEM140 | -0,2568 | 3,68777E-10 |
| ACE2 | CTC-378H22.2 | -0,2568 | 3,73275E-10 |
| ACE2 | NFIL3 | -0,2569 | 3,62253E-10 |
| ACE2 | LDB2 | -0,257 | 3,6107E-10 |
| ACE2 | HTR1DP1 | -0,257 | 3,56557E-10 |
| ACE2 | AC097359.2 | -0,2571 | 3,5152E-10 |
| ACE2 | ECE1 | -0,2572 | 3,47654E-10 |
| ACE2 | LINC00032 | -0,2572 | 3,49336E-10 |
| ACE2 | RP11-3K24.1 | -0,2572 | 3,45826E-10 |
| ACE2 | RP3-324O17.7 | -0,2572 | 3,47475E-10 |
| ACE2 | CRYBB2P1 | -0,2572 | 3,48573E-10 |
| ACE2 | CXorf21 | -0,2572 | 3,45799E-10 |
| ACE2 | RP11-37L2.1 | -0,2573 | 3,42503E-10 |
| ACE2 | AF124730.4 | -0,2574 | 3,38962E-10 |
| ACE2 | TYMP | -0,2574 | 3,36274E-10 |
| ACE2 | MBD5 | -0,2575 | 3,30762E-10 |
| ACE2 | MAPK6PS3 | -0,2575 | 3,33179E-10 |
| ACE2 | RPRD1A | -0,2575 | 3,31797E-10 |
| ACE2 | SDCBP2 | -0,2575 | 3,28395E-10 |
| ACE2 | CSNK1G3 | -0,2576 | 3,24514E-10 |
| ACE2 | NCAM1 | -0,2576 | 3,24131E-10 |
| ACE2 | RASSF2 | -0,2576 | 3,24249E-10 |
| ACE2 | AP000347.4 | -0,2576 | 3,25795E-10 |
| ACE2 | RP11-402L5.1 | -0,2577 | 3,2142E-10 |
| ACE2 | HIST3H2A | -0,2578 | 3,1277E-10 |
| ACE2 | AC009542.2 | -0,2578 | 3,15453E-10 |
| ACE2 | TLE4 | -0,2578 | 3,14531E-10 |
| ACE2 | RP11-122K13.12 | -0,2578 | 3,16399E-10 |
| ACE2 | BTN2A3P | -0,2579 | 3,11458E-10 |
| ACE2 | RP11-677N16.1 | -0,2579 | 3,09375E-10 |
| ACE2 | YPEL2 | -0,2579 | 3,11691E-10 |
| ACE2 | CTB-174D11.1 | -0,258 | 3,05849E-10 |
| ACE2 | CASP7 | -0,258 | 3,02883E-10 |
| ACE2 | CTNND1 | -0,258 | 3,07449E-10 |
| ACE2 | UBAP1L | -0,258 | 3,03681E-10 |
| ACE2 | ZNF286A | -0,258 | 3,03727E-10 |
| ACE2 | AC004019.10 | -0,258 | 3,02972E-10 |
| ACE2 | MTATP6P26 | -0,2581 | 3,00425E-10 |
| ACE2 | NELFE | -0,2581 | 3,006E-10 |
| ACE2 | COL10A1 | -0,2581 | 2,99263E-10 |
| ACE2 | PDCD4-AS1 | -0,2581 | 2,98088E-10 |
| ACE2 | MSRB3 | -0,2581 | 3,01046E-10 |
| ACE2 | RP13-991F5.2 | -0,2581 | 3,00407E-10 |
| ACE2 | LINC01578 | -0,2582 | 2,92908E-10 |
| ACE2 | GPR173 | -0,2582 | 2,96459E-10 |
| ACE2 | C1orf35 | -0,2583 | 2,92722E-10 |
| ACE2 | RP11-348J24.2 | -0,2583 | 2,91655E-10 |
| ACE2 | NRXN2 | -0,2583 | 2,89603E-10 |

| | | | |
|---|---|---|---|
| ACE2 | ANKRD33 | -0,2583 | 2,92043E-10 |
| ACE2 | TMEM72-AS1 | -0,2584 | 2,85802E-10 |
| ACE2 | C16orf74 | -0,2584 | 2,86657E-10 |
| ACE2 | RNF168 | -0,2585 | 2,83036E-10 |
| ACE2 | FGFBP2 | -0,2585 | 2,83143E-10 |
| ACE2 | BICD1 | -0,2585 | 2,80065E-10 |
| ACE2 | CCDC65 | -0,2585 | 2,83191E-10 |
| ACE2 | CERS5 | -0,2585 | 2,8297E-10 |
| ACE2 | TNFSF14 | -0,2586 | 2,77512E-10 |
| ACE2 | RPL5P13 | -0,2587 | 2,7104E-10 |
| ACE2 | PHTF2 | -0,2587 | 2,73585E-10 |
| ACE2 | ZBTB34 | -0,2587 | 2,70297E-10 |
| ACE2 | PRG2 | -0,2587 | 2,71126E-10 |
| ACE2 | RHOV | -0,2587 | 2,73405E-10 |
| ACE2 | SLAMF1 | -0,2588 | 2,66766E-10 |
| ACE2 | LINC02068 | -0,2588 | 2,68022E-10 |
| ACE2 | RP11-514D23.3 | -0,2588 | 2,67291E-10 |
| ACE2 | TPRG1 | -0,2589 | 2,63998E-10 |
| ACE2 | HID1 | -0,2589 | 2,62812E-10 |
| ACE2 | VCPKMT | -0,259 | 2,60414E-10 |
| ACE2 | NPAP1 | -0,259 | 2,609E-10 |
| ACE2 | HOXB6 | -0,259 | 2,59036E-10 |
| ACE2 | CTC-806A22.1 | -0,2591 | 2,56947E-10 |
| ACE2 | RIC1 | -0,2591 | 2,52833E-10 |
| ACE2 | AGAP1-IT1 | -0,2592 | 2,52566E-10 |
| ACE2 | RP11-145M9.5 | -0,2592 | 2,52229E-10 |
| ACE2 | E2F3P1 | -0,2592 | 2,52256E-10 |
| ACE2 | RP11-27M5.1 | -0,2593 | 2,46042E-10 |
| ACE2 | OGN | -0,2593 | 2,47668E-10 |
| ACE2 | RP11-455O6.2 | -0,2593 | 2,45182E-10 |
| ACE2 | RP11-395L14.18 | -0,2594 | 2,42509E-10 |
| ACE2 | ZNF385D | -0,2594 | 2,41486E-10 |
| ACE2 | EIF4E3 | -0,2594 | 2,4348E-10 |
| ACE2 | CCNL2P1 | -0,2594 | 2,41789E-10 |
| ACE2 | RP11-73M11.3 | -0,2594 | 2,42121E-10 |
| ACE2 | AE000661.37 | -0,2594 | 2,4192E-10 |
| ACE2 | RP11-446H18.6 | -0,2595 | 2,38749E-10 |
| ACE2 | SCX | -0,2595 | 2,37136E-10 |
| ACE2 | RAB23 | -0,2596 | 2,35196E-10 |
| ACE2 | COLEC10 | -0,2596 | 2,36422E-10 |
| ACE2 | RP11-469A15.2 | -0,2597 | 2,32703E-10 |
| ACE2 | LRRC37A | -0,2597 | 2,32574E-10 |
| ACE2 | CHIC1 | -0,2597 | 2,29572E-10 |
| ACE2 | RP11-1399P15.1 | -0,2598 | 2,27281E-10 |
| ACE2 | CTD-2666L21.1 | -0,2599 | 2,24089E-10 |
| ACE2 | TP53INP1 | -0,26 | 2,18457E-10 |
| ACE2 | RPS26P45 | -0,26 | 2,18126E-10 |
| ACE2 | RP4-671O14.6 | -0,26 | 2,19144E-10 |
| ACE2 | RP11-537A6.9 | -0,2601 | 2,17806E-10 |
| ACE2 | bP-2189O9.2 | -0,2601 | 2,15366E-10 |
| ACE2 | TSSK3 | -0,2602 | 2,11683E-10 |
| ACE2 | KB-1552D7.2 | -0,2602 | 2,12326E-10 |
| ACE2 | AC091814.2 | -0,2602 | 2,14174E-10 |
| ACE2 | RP11-31F15.2 | -0,2603 | 2,1038E-10 |

| | | | |
|---|---|---|---|
| ACE2 | RP11-172C16.4 | -0,2603 | 2,10958E-10 |
| ACE2 | RP11-561I11.4 | -0,2604 | 2,04872E-10 |
| ACE2 | ACSL6 | -0,2604 | 2,05654E-10 |
| ACE2 | RP11-266K4.9 | -0,2604 | 2,05382E-10 |
| ACE2 | RP11-396F22.1 | -0,2604 | 2,04514E-10 |
| ACE2 | RP11-715F3.2 | -0,2604 | 2,04842E-10 |
| ACE2 | RHOH | -0,2605 | 2,01199E-10 |
| ACE2 | RP11-506M13.3 | -0,2605 | 2,03068E-10 |
| ACE2 | BLID | -0,2605 | 2,04098E-10 |
| ACE2 | GREM2 | -0,2606 | 2,0025E-10 |
| ACE2 | RP3-336K20__B.2 | -0,2606 | 1,98255E-10 |
| ACE2 | LCNL1 | -0,2606 | 2,00325E-10 |
| ACE2 | CD4 | -0,2606 | 1,97924E-10 |
| ACE2 | RP11-136H19.1 | -0,2606 | 2,00067E-10 |
| ACE2 | RP11-568G11.4 | -0,2607 | 1,97006E-10 |
| ACE2 | ANK2 | -0,2607 | 1,94597E-10 |
| ACE2 | TAC1 | -0,2607 | 1,96187E-10 |
| ACE2 | ACSBG1 | -0,2607 | 1,96625E-10 |
| ACE2 | KAT6A | -0,2608 | 1,94233E-10 |
| ACE2 | RP11-796E2.4 | -0,2608 | 1,93554E-10 |
| ACE2 | ZNF207 | -0,2608 | 1,92463E-10 |
| ACE2 | ACSBG2 | -0,2608 | 1,92474E-10 |
| ACE2 | COX6B1P5 | -0,2609 | 1,89088E-10 |
| ACE2 | RP1-197B17.3 | -0,2609 | 1,8818E-10 |
| ACE2 | RP5-837M10.4 | -0,261 | 1,8533E-10 |
| ACE2 | KCNA3 | -0,261 | 1,87907E-10 |
| ACE2 | RP11-810P8.1 | -0,261 | 1,86956E-10 |
| ACE2 | RP11-1058N17.1 | -0,261 | 1,86511E-10 |
| ACE2 | RP3-510H16.3 | -0,261 | 1,87258E-10 |
| ACE2 | GPR137C | -0,2611 | 1,82392E-10 |
| ACE2 | DRP2 | -0,2611 | 1,84725E-10 |
| ACE2 | KCNK17 | -0,2612 | 1,81133E-10 |
| ACE2 | RP11-97N19.2 | -0,2612 | 1,8186E-10 |
| ACE2 | NPRL2 | -0,2613 | 1,78878E-10 |
| ACE2 | DCAF16 | -0,2613 | 1,76376E-10 |
| ACE2 | HNRNPA1P16 | -0,2613 | 1,78765E-10 |
| ACE2 | FAM114A1 | -0,2614 | 1,74473E-10 |
| ACE2 | RP3-465N24.5 | -0,2615 | 1,72374E-10 |
| ACE2 | SSBP2 | -0,2615 | 1,71744E-10 |
| ACE2 | PCDHB16 | -0,2615 | 1,71128E-10 |
| ACE2 | RP11-182J1.14 | -0,2615 | 1,7071E-10 |
| ACE2 | ENGASE | -0,2615 | 1,71284E-10 |
| ACE2 | ZNF567 | -0,2615 | 1,72922E-10 |
| ACE2 | TRH | -0,2616 | 1,67872E-10 |
| ACE2 | DEK | -0,2616 | 1,70123E-10 |
| ACE2 | RING1 | -0,2616 | 1,69688E-10 |
| ACE2 | NCALD | -0,2616 | 1,67721E-10 |
| ACE2 | ADGRB3 | -0,2617 | 1,67496E-10 |
| ACE2 | EGLN3 | -0,2617 | 1,65362E-10 |
| ACE2 | CTD-2358C21.5 | -0,2617 | 1,66248E-10 |
| ACE2 | JUN | -0,2618 | 1,6314E-10 |
| ACE2 | SLC25A2 | -0,2618 | 1,62633E-10 |
| ACE2 | HAS2 | -0,2618 | 1,62158E-10 |
| ACE2 | NXPH4 | -0,2618 | 1,62913E-10 |

| | | | |
|---|---|---|---|
| ACE2 | VASH2 | -0,2619 | 1,5973E-10 |
| ACE2 | RNF126 | -0,2619 | 1,61877E-10 |
| ACE2 | ABCB4 | -0,262 | 1,57412E-10 |
| ACE2 | VLDLR-AS1 | -0,262 | 1,58844E-10 |
| ACE2 | RP11-578B16.1 | -0,262 | 1,57594E-10 |
| ACE2 | CTD-2252P21.1 | -0,2621 | 1,55473E-10 |
| ACE2 | LUCAT1 | -0,2621 | 1,56223E-10 |
| ACE2 | PHF10 | -0,2621 | 1,54958E-10 |
| ACE2 | ZNF354B | -0,2623 | 1,49606E-10 |
| ACE2 | MED24 | -0,2623 | 1,50362E-10 |
| ACE2 | LSMEM2 | -0,2624 | 1,47414E-10 |
| ACE2 | RP11-354E11.2 | -0,2624 | 1,47811E-10 |
| ACE2 | ADIRF | -0,2624 | 1,48582E-10 |
| ACE2 | GRK5 | -0,2624 | 1,48928E-10 |
| ACE2 | CRYBB3 | -0,2624 | 1,47205E-10 |
| ACE2 | AC108463.1 | -0,2625 | 1,44805E-10 |
| ACE2 | SIAH1 | -0,2625 | 1,45053E-10 |
| ACE2 | KANSL3 | -0,2626 | 1,43556E-10 |
| ACE2 | TIFAB | -0,2626 | 1,42259E-10 |
| ACE2 | PPP1R2P6 | -0,2626 | 1,4384E-10 |
| ACE2 | VLDLR | -0,2626 | 1,42164E-10 |
| ACE2 | CDKN2A | -0,2627 | 1,41892E-10 |
| ACE2 | CCDC85C | -0,2627 | 1,40963E-10 |
| ACE2 | TYK2 | -0,2627 | 1,41754E-10 |
| ACE2 | PI4KAP2 | -0,2627 | 1,41842E-10 |
| ACE2 | RP11-230G5.2 | -0,2628 | 1,38455E-10 |
| ACE2 | SGK2 | -0,2628 | 1,38922E-10 |
| ACE2 | PDK1 | -0,2629 | 1,36733E-10 |
| ACE2 | C1QTNF4 | -0,2629 | 1,36399E-10 |
| ACE2 | ANKS1B | -0,2629 | 1,35069E-10 |
| ACE2 | RP11-356O24.1 | -0,2629 | 1,37204E-10 |
| ACE2 | HSD52 | -0,263 | 1,34664E-10 |
| ACE2 | RP11-446H18.1 | -0,263 | 1,34198E-10 |
| ACE2 | EIF5A2 | -0,263 | 1,33543E-10 |
| ACE2 | HCG27 | -0,263 | 1,33075E-10 |
| ACE2 | RP5-940J5.6 | -0,263 | 1,33482E-10 |
| ACE2 | NECAP2 | -0,2631 | 1,30642E-10 |
| ACE2 | RP11-70P17.1 | -0,2631 | 1,30626E-10 |
| ACE2 | DNMT3A | -0,2631 | 1,32299E-10 |
| ACE2 | RP11-631M6.2 | -0,2631 | 1,32188E-10 |
| ACE2 | ANGPTL2 | -0,2631 | 1,3254E-10 |
| ACE2 | AIRE | -0,2631 | 1,32135E-10 |
| ACE2 | MYOM2 | -0,2632 | 1,30254E-10 |
| ACE2 | PDLIM1 | -0,2632 | 1,3004E-10 |
| ACE2 | KCNN2 | -0,2633 | 1,26919E-10 |
| ACE2 | DCBLD1 | -0,2633 | 1,27257E-10 |
| ACE2 | PWRN1 | -0,2634 | 1,2571E-10 |
| ACE2 | PFDN2 | -0,2635 | 1,23119E-10 |
| ACE2 | ANO6 | -0,2635 | 1,24079E-10 |
| ACE2 | RP11-17E3.1 | -0,2635 | 1,22283E-10 |
| ACE2 | DLGAP4-AS1 | -0,2635 | 1,22243E-10 |
| ACE2 | RP11-735G4.1 | -0,2636 | 1,21532E-10 |
| ACE2 | AC093620.5 | -0,2636 | 1,21373E-10 |
| ACE2 | RP11-598F7.6 | -0,2636 | 1,20327E-10 |

| | | | |
|---|---|---|---|
| ACE2 | RP11-269F20.1 | -0,2637 | 1,19777E-10 |
| ACE2 | AC004540.5 | -0,2637 | 1,19299E-10 |
| ACE2 | IFITM3 | -0,2637 | 1,18868E-10 |
| ACE2 | INO80B | -0,2638 | 1,17819E-10 |
| ACE2 | TXK | -0,2638 | 1,17397E-10 |
| ACE2 | ULBP1 | -0,2638 | 1,1727E-10 |
| ACE2 | ILF3 | -0,2638 | 1,16594E-10 |
| ACE2 | CH17-140K24.6 | -0,2639 | 1,15122E-10 |
| ACE2 | RP1-225E12.2 | -0,2639 | 1,14544E-10 |
| ACE2 | C12orf42 | -0,2639 | 1,14653E-10 |
| ACE2 | RP11-133K1.12 | -0,2639 | 1,1464E-10 |
| ACE2 | SNORA70F | -0,264 | 1,13727E-10 |
| ACE2 | UBE2CP3 | -0,264 | 1,12597E-10 |
| ACE2 | RP11-737O24.5 | -0,264 | 1,13525E-10 |
| ACE2 | NFASC | -0,2641 | 1,10683E-10 |
| ACE2 | KCNH2 | -0,2641 | 1,11371E-10 |
| ACE2 | LINC00605 | -0,2641 | 1,12045E-10 |
| ACE2 | GNB4 | -0,2643 | 1,07254E-10 |
| ACE2 | HCG22 | -0,2643 | 1,08357E-10 |
| ACE2 | TMEM217 | -0,2643 | 1,08321E-10 |
| ACE2 | RP11-61L23.2 | -0,2643 | 1,08099E-10 |
| ACE2 | RP11-278C7.1 | -0,2643 | 1,07884E-10 |
| ACE2 | RARG | -0,2643 | 1,08351E-10 |
| ACE2 | RP11-802O23.3 | -0,2644 | 1,06517E-10 |
| ACE2 | SNORD15B | -0,2644 | 1,0514E-10 |
| ACE2 | SULF2 | -0,2644 | 1,05693E-10 |
| ACE2 | AC093787.1 | -0,2645 | 1,04418E-10 |
| ACE2 | GOLGA5P1 | -0,2645 | 1,04562E-10 |
| ACE2 | HIST1H2AH | -0,2645 | 1,04796E-10 |
| ACE2 | LINC00996 | -0,2645 | 1,0501E-10 |
| ACE2 | RP11-326C3.16 | -0,2645 | 1,03929E-10 |
| ACE2 | C20orf202 | -0,2645 | 1,04835E-10 |
| ACE2 | MRPS6 | -0,2645 | 1,04131E-10 |
| ACE2 | EVA1B | -0,2646 | 1,02971E-10 |
| ACE2 | USP49 | -0,2646 | 1,03024E-10 |
| ACE2 | AC007292.6 | -0,2646 | 1,02353E-10 |
| ACE2 | ZNF586 | -0,2646 | 1,01753E-10 |
| ACE2 | ALDOB | -0,2647 | 1,00815E-10 |
| ACE2 | RP11-454E5.4 | -0,2647 | 1,01092E-10 |
| ACE2 | CTDSPL2 | -0,2648 | 9,93456E-11 |
| ACE2 | CTB-50L17.9 | -0,2648 | 9,94238E-11 |
| ACE2 | SUN1 | -0,2649 | 9,74773E-11 |
| ACE2 | PLCB2 | -0,2649 | 9,69511E-11 |
| ACE2 | ILKAP | -0,265 | 9,60617E-11 |
| ACE2 | RP11-192H23.8 | -0,265 | 9,57347E-11 |
| ACE2 | CTD-3014M21.1 | -0,265 | 9,56624E-11 |
| ACE2 | PRDM8 | -0,2651 | 9,48286E-11 |
| ACE2 | MFRP | -0,2651 | 9,39029E-11 |
| ACE2 | RP11-64B16.2 | -0,2651 | 9,49749E-11 |
| ACE2 | STAT1 | -0,2652 | 9,26735E-11 |
| ACE2 | UBE2B | -0,2652 | 9,24904E-11 |
| ACE2 | TRAF3IP3 | -0,2653 | 9,11769E-11 |
| ACE2 | EHD1 | -0,2653 | 9,11162E-11 |
| ACE2 | PDE6G | -0,2653 | 9,15621E-11 |

| | | | |
|---|---|---|---|
| ACE2 | NDUFA9P1 | -0,2654 | 9,00918E-11 |
| ACE2 | HMP19 | -0,2656 | 8,65894E-11 |
| ACE2 | CRISP3 | -0,2656 | 8,61505E-11 |
| ACE2 | S100A4 | -0,2657 | 8,47434E-11 |
| ACE2 | SNCG | -0,2657 | 8,55054E-11 |
| ACE2 | LINC00294 | -0,2657 | 8,5699E-11 |
| ACE2 | NPIPB3 | -0,2658 | 8,34566E-11 |
| ACE2 | CTD-2095E4.4 | -0,2658 | 8,39933E-11 |
| ACE2 | RP11-244H3.1 | -0,2659 | 8,20261E-11 |
| ACE2 | TANK | -0,2659 | 8,23644E-11 |
| ACE2 | RP11-188P20.3 | -0,2659 | 8,2909E-11 |
| ACE2 | TET1 | -0,2659 | 8,29627E-11 |
| ACE2 | ZNF615 | -0,2659 | 8,2894E-11 |
| ACE2 | NTNG2 | -0,266 | 8,0583E-11 |
| ACE2 | CT45A11P | -0,266 | 8,1407E-11 |
| ACE2 | RPL5P23 | -0,2661 | 7,94697E-11 |
| ACE2 | RP5-940J5.8 | -0,2661 | 7,9773E-11 |
| ACE2 | SAV1 | -0,2661 | 8,03037E-11 |
| ACE2 | RP11-855A2.2 | -0,2661 | 7,90427E-11 |
| ACE2 | CSDC2 | -0,2662 | 7,87832E-11 |
| ACE2 | RP3-368A4.6 | -0,2662 | 7,85659E-11 |
| ACE2 | PTBP2 | -0,2663 | 7,71833E-11 |
| ACE2 | RP11-672A2.6 | -0,2663 | 7,64515E-11 |
| ACE2 | CCDC42 | -0,2663 | 7,70356E-11 |
| ACE2 | LINC00298 | -0,2664 | 7,54106E-11 |
| ACE2 | RAMP3 | -0,2664 | 7,60365E-11 |
| ACE2 | RP11-736N17.10 | -0,2664 | 7,52823E-11 |
| ACE2 | RP11-426J5.3 | -0,2664 | 7,53484E-11 |
| ACE2 | AC108142.1 | -0,2665 | 7,49993E-11 |
| ACE2 | RP11-830F9.5 | -0,2665 | 7,40869E-11 |
| ACE2 | TNRC6B | -0,2665 | 7,49071E-11 |
| ACE2 | FAM53C | -0,2666 | 7,30372E-11 |
| ACE2 | MAF | -0,2666 | 7,26649E-11 |
| ACE2 | TBCAP3 | -0,2667 | 7,22107E-11 |
| ACE2 | RP11-407N17.4 | -0,2667 | 7,18924E-11 |
| ACE2 | RNF24 | -0,2667 | 7,14654E-11 |
| ACE2 | ARVCF | -0,2667 | 7,20183E-11 |
| ACE2 | RP11-277B15.3 | -0,2668 | 7,13357E-11 |
| ACE2 | TRBJ2-3 | -0,2668 | 7,10631E-11 |
| ACE2 | NUB1 | -0,2668 | 7,12369E-11 |
| ACE2 | MAP4K5 | -0,2668 | 7,04346E-11 |
| ACE2 | RARA | -0,2668 | 7,12873E-11 |
| ACE2 | LINC00854 | -0,2668 | 7,06638E-11 |
| ACE2 | TRBV24-1 | -0,2669 | 6,97869E-11 |
| ACE2 | CBX3P4 | -0,2669 | 6,90335E-11 |
| ACE2 | CPLX3 | -0,2669 | 6,97883E-11 |
| ACE2 | RP11-54O7.3 | -0,267 | 6,85187E-11 |
| ACE2 | DEPDC7 | -0,267 | 6,80626E-11 |
| ACE2 | PAMR1 | -0,267 | 6,79855E-11 |
| ACE2 | OR5AK2 | -0,267 | 6,90201E-11 |
| ACE2 | SAMD10 | -0,267 | 6,89347E-11 |
| ACE2 | RP3-393E18.2 | -0,2671 | 6,67618E-11 |
| ACE2 | PRPH | -0,2671 | 6,69536E-11 |
| ACE2 | PLXNC1 | -0,2671 | 6,75617E-11 |

| | | | |
|---|---|---|---|
| ACE2 | PLPPR4 | -0,2672 | 6,59381E-11 |
| ACE2 | CILP2 | -0,2673 | 6,50646E-11 |
| ACE2 | SPACA6P-AS | -0,2673 | 6,53533E-11 |
| ACE2 | RP11-643C9.2 | -0,2674 | 6,43544E-11 |
| ACE2 | RNF113B | -0,2674 | 6,38801E-11 |
| ACE2 | FAM181B | -0,2676 | 6,14262E-11 |
| ACE2 | RP11-197N18.8 | -0,2676 | 6,17759E-11 |
| ACE2 | MKL1 | -0,2676 | 6,19966E-11 |
| ACE2 | VN1R108P | -0,2677 | 6,07475E-11 |
| ACE2 | SLC10A6 | -0,2678 | 5,96555E-11 |
| ACE2 | RAD9A | -0,2678 | 5,95254E-11 |
| ACE2 | ABCB9 | -0,2678 | 5,96204E-11 |
| ACE2 | DNAJB5 | -0,2679 | 5,82949E-11 |
| ACE2 | RP4-718P11.1 | -0,2679 | 5,91317E-11 |
| ACE2 | SNORA73B | -0,268 | 5,76346E-11 |
| ACE2 | NPIPB11 | -0,268 | 5,75196E-11 |
| ACE2 | RP11-49C9.2 | -0,268 | 5,79839E-11 |
| ACE2 | METTL21A | -0,2681 | 5,70441E-11 |
| ACE2 | TRGVA | -0,2681 | 5,66558E-11 |
| ACE2 | TMEM158 | -0,2682 | 5,61787E-11 |
| ACE2 | DPYSL3 | -0,2682 | 5,58681E-11 |
| ACE2 | MTA2 | -0,2682 | 5,58463E-11 |
| ACE2 | RP11-727A23.4 | -0,2682 | 5,58757E-11 |
| ACE2 | SOAT2 | -0,2682 | 5,57169E-11 |
| ACE2 | RP6-91H8.2 | -0,2682 | 5,60667E-11 |
| ACE2 | RP11-973H7.1 | -0,2682 | 5,57355E-11 |
| ACE2 | ZNF791 | -0,2682 | 5,53768E-11 |
| ACE2 | KDM5C | -0,2682 | 5,58552E-11 |
| ACE2 | MAP4 | -0,2683 | 5,4802E-11 |
| ACE2 | RP11-474P2.2 | -0,2684 | 5,40112E-11 |
| ACE2 | RP11-128N14.5 | -0,2684 | 5,40145E-11 |
| ACE2 | RPL7AP65 | -0,2684 | 5,36655E-11 |
| ACE2 | BRD7P4 | -0,2685 | 5,31076E-11 |
| ACE2 | EPB41L2 | -0,2685 | 5,2928E-11 |
| ACE2 | LRRN4CL | -0,2685 | 5,2923E-11 |
| ACE2 | RP11-288L9.1 | -0,2686 | 5,20348E-11 |
| ACE2 | EGFLAM | -0,2686 | 5,20061E-11 |
| ACE2 | RNU6-1098P | -0,2687 | 5,10702E-11 |
| ACE2 | APBB3 | -0,2687 | 5,17041E-11 |
| ACE2 | NR3C1 | -0,2687 | 5,16223E-11 |
| ACE2 | RAB44 | -0,2687 | 5,12268E-11 |
| ACE2 | RP11-661A12.9 | -0,2687 | 5,10511E-11 |
| ACE2 | MMP23B | -0,2688 | 5,0784E-11 |
| ACE2 | RPS15AP6 | -0,2689 | 4,94318E-11 |
| ACE2 | NDUFAF4P2 | -0,2689 | 4,95851E-11 |
| ACE2 | RP11-430H12.2 | -0,269 | 4,8688E-11 |
| ACE2 | RNF150 | -0,269 | 4,84068E-11 |
| ACE2 | CDO1 | -0,269 | 4,88985E-11 |
| ACE2 | HLA-L | -0,269 | 4,84498E-11 |
| ACE2 | RP11-367H1.1 | -0,2691 | 4,77092E-11 |
| ACE2 | ABI3BP | -0,2691 | 4,76267E-11 |
| ACE2 | DAPP1 | -0,2691 | 4,75256E-11 |
| ACE2 | MCTP2 | -0,2691 | 4,82265E-11 |
| ACE2 | RP11-346C20.4 | -0,2691 | 4,7654E-11 |

| | | | |
|---|---|---|---|
| ACE2 | HS3ST3A1 | -0,2691 | 4,7944E-11 |
| ACE2 | AC002398.11 | -0,2691 | 4,8129E-11 |
| ACE2 | CTA-384D8.36 | -0,2691 | 4,8071E-11 |
| ACE2 | EGOT | -0,2692 | 4,67922E-11 |
| ACE2 | LINC00839 | -0,2692 | 4,71728E-11 |
| ACE2 | RP1-68D18.2 | -0,2692 | 4,6771E-11 |
| ACE2 | GAPDHP15 | -0,2693 | 4,59819E-11 |
| ACE2 | RP11-508N22.12 | -0,2693 | 4,66259E-11 |
| ACE2 | THEM7P | -0,2693 | 4,62932E-11 |
| ACE2 | LLNLR-246C6.1 | -0,2693 | 4,61088E-11 |
| ACE2 | SP110 | -0,2694 | 4,57885E-11 |
| ACE2 | SNHG8 | -0,2694 | 4,55283E-11 |
| ACE2 | ENPP3 | -0,2694 | 4,52235E-11 |
| ACE2 | CEP152 | -0,2694 | 4,52582E-11 |
| ACE2 | RP1-41C23.4 | -0,2694 | 4,53206E-11 |
| ACE2 | OR7E100P | -0,2695 | 4,4603E-11 |
| ACE2 | KLHL30 | -0,2696 | 4,40672E-11 |
| ACE2 | MAP3K7 | -0,2696 | 4,37517E-11 |
| ACE2 | RP11-196G11.3 | -0,2696 | 4,39784E-11 |
| ACE2 | ITM2A | -0,2696 | 4,36526E-11 |
| ACE2 | ZNF678 | -0,2697 | 4,3186E-11 |
| ACE2 | AIRN | -0,2697 | 4,31349E-11 |
| ACE2 | RP5-1050D4.2 | -0,2697 | 4,31651E-11 |
| ACE2 | HSPB7 | -0,2698 | 4,24998E-11 |
| ACE2 | RP11-25K21.1 | -0,2698 | 4,2401E-11 |
| ACE2 | NENF | -0,2698 | 4,27183E-11 |
| ACE2 | POU5F1 | -0,2698 | 4,24434E-11 |
| ACE2 | C9orf50 | -0,2698 | 4,26902E-11 |
| ACE2 | CEMIP | -0,2698 | 4,26037E-11 |
| ACE2 | CNN3 | -0,2699 | 4,19766E-11 |
| ACE2 | C1orf204 | -0,2699 | 4,20008E-11 |
| ACE2 | ZSCAN18 | -0,2699 | 4,15265E-11 |
| ACE2 | RP11-536C5.7 | -0,27 | 4,13047E-11 |
| ACE2 | RP11-242D8.1 | -0,2701 | 4,01544E-11 |
| ACE2 | OLFML3 | -0,2702 | 3,99314E-11 |
| ACE2 | RP11-166O4.6 | -0,2702 | 3,94463E-11 |
| ACE2 | RP13-39P12.3 | -0,2702 | 3,96281E-11 |
| ACE2 | RP11-127I20.8 | -0,2702 | 3,97768E-11 |
| ACE2 | VWA3A | -0,2702 | 3,95524E-11 |
| ACE2 | HNRNPU | -0,2703 | 3,92499E-11 |
| ACE2 | LINC01278 | -0,2703 | 3,87879E-11 |
| ACE2 | LL0XNC01-240C2.1 | -0,2703 | 3,87937E-11 |
| ACE2 | SELENOW | -0,2704 | 3,85014E-11 |
| ACE2 | RP5-1024N4.4 | -0,2705 | 3,7813E-11 |
| ACE2 | CTB-120L21.1 | -0,2705 | 3,79425E-11 |
| ACE2 | TPCN2 | -0,2705 | 3,79479E-11 |
| ACE2 | METTL3 | -0,2705 | 3,7682E-11 |
| ACE2 | MPRIP | -0,2705 | 3,77923E-11 |
| ACE2 | FBXO24 | -0,2706 | 3,67422E-11 |
| ACE2 | RP4-563E14.1 | -0,2706 | 3,69838E-11 |
| ACE2 | CRABP2 | -0,2708 | 3,59547E-11 |
| ACE2 | TICAM1 | -0,2708 | 3,57063E-11 |
| ACE2 | GRAMD1A | -0,2708 | 3,60566E-11 |
| ACE2 | SYNPO | -0,2709 | 3,51371E-11 |

| | | | |
|---|---|---|---|
| ACE2 | LRCH4 | -0,2709 | 3,4942E-11 |
| ACE2 | SRRT | -0,2709 | 3,54247E-11 |
| ACE2 | LINC01915 | -0,2709 | 3,52518E-11 |
| ACE2 | PPP1R8P1 | -0,271 | 3,46907E-11 |
| ACE2 | CXCL9 | -0,271 | 3,47515E-11 |
| ACE2 | TFPI2 | -0,271 | 3,4581E-11 |
| ACE2 | TMEM121 | -0,271 | 3,47714E-11 |
| ACE2 | SNN | -0,271 | 3,42982E-11 |
| ACE2 | RP11-221J22.2 | -0,2711 | 3,38007E-11 |
| ACE2 | CPZ | -0,2711 | 3,37608E-11 |
| ACE2 | UBQLNL | -0,2711 | 3,38758E-11 |
| ACE2 | ST8SIA6-AS1 | -0,2712 | 3,35104E-11 |
| ACE2 | MUS81 | -0,2712 | 3,31679E-11 |
| ACE2 | RP11-589P10.5 | -0,2712 | 3,31996E-11 |
| ACE2 | LINC01358 | -0,2713 | 3,26881E-11 |
| ACE2 | NBPF14 | -0,2713 | 3,30516E-11 |
| ACE2 | EGFEM1P | -0,2713 | 3,30515E-11 |
| ACE2 | CTC-484P3.3 | -0,2713 | 3,29857E-11 |
| ACE2 | KATNBL1 | -0,2713 | 3,31103E-11 |
| ACE2 | KCNH6 | -0,2713 | 3,30792E-11 |
| ACE2 | FNDC11 | -0,2713 | 3,25997E-11 |
| ACE2 | DCST2 | -0,2714 | 3,24986E-11 |
| ACE2 | PLA2G4A | -0,2714 | 3,20924E-11 |
| ACE2 | ATG2A | -0,2714 | 3,2158E-11 |
| ACE2 | RBM7 | -0,2714 | 3,23826E-11 |
| ACE2 | PRKCB | -0,2714 | 3,25068E-11 |
| ACE2 | LINC00882 | -0,2715 | 3,16353E-11 |
| ACE2 | RP1-102E24.10 | -0,2715 | 3,18606E-11 |
| ACE2 | BARD1 | -0,2716 | 3,12819E-11 |
| ACE2 | CYP4F35P | -0,2716 | 3,10311E-11 |
| ACE2 | SH3BGRL | -0,2716 | 3,12897E-11 |
| ACE2 | ZBTB25 | -0,2717 | 3,05583E-11 |
| ACE2 | CTC-287O8.1 | -0,2718 | 2,99499E-11 |
| ACE2 | TUBB2B | -0,2718 | 3,01575E-11 |
| ACE2 | ZNF76 | -0,2718 | 3,03063E-11 |
| ACE2 | CCL19 | -0,2718 | 2,99392E-11 |
| ACE2 | COL27A1 | -0,2718 | 2,99715E-11 |
| ACE2 | DNM1 | -0,2718 | 3,03042E-11 |
| ACE2 | CALCB | -0,2719 | 2,93791E-11 |
| ACE2 | AC009120.6 | -0,2719 | 2,95372E-11 |
| ACE2 | CTD-2104P17.1 | -0,2719 | 2,9672E-11 |
| ACE2 | RP11-536C12.1 | -0,272 | 2,91391E-11 |
| ACE2 | RP11-227G15.10 | -0,272 | 2,89718E-11 |
| ACE2 | IFNGR2 | -0,2721 | 2,87838E-11 |
| ACE2 | HMCN1 | -0,2722 | 2,82345E-11 |
| ACE2 | BACH2 | -0,2722 | 2,79472E-11 |
| ACE2 | MYH11 | -0,2722 | 2,83515E-11 |
| ACE2 | ANKRD36 | -0,2723 | 2,75982E-11 |
| ACE2 | RP11-553L6.3 | -0,2723 | 2,76939E-11 |
| ACE2 | NDEL1 | -0,2723 | 2,74375E-11 |
| ACE2 | RP11-481C4.2 | -0,2723 | 2,74812E-11 |
| ACE2 | EPG5 | -0,2723 | 2,76187E-11 |
| ACE2 | SSC5D | -0,2723 | 2,7549E-11 |
| ACE2 | NFATC1 | -0,2724 | 2,70249E-11 |

| | | | |
|---|---|---|---|
| ACE2 | FGD1 | -0,2724 | 2,73715E-11 |
| ACE2 | AC092316.1 | -0,2725 | 2,6532E-11 |
| ACE2 | RP11-288I21.1 | -0,2726 | 2,64741E-11 |
| ACE2 | FBLN7 | -0,2726 | 2,62545E-11 |
| ACE2 | HINFP | -0,2726 | 2,60587E-11 |
| ACE2 | CTD-3184A7.4 | -0,2726 | 2,63492E-11 |
| ACE2 | TUB | -0,2727 | 2,57944E-11 |
| ACE2 | RP11-173A16.2 | -0,2727 | 2,5955E-11 |
| ACE2 | EMP3 | -0,2727 | 2,5763E-11 |
| ACE2 | RP11-477H21.2 | -0,2728 | 2,5238E-11 |
| ACE2 | CTXN1 | -0,2728 | 2,55004E-11 |
| ACE2 | U82695.9 | -0,2728 | 2,55171E-11 |
| ACE2 | KLHL5 | -0,2729 | 2,49075E-11 |
| ACE2 | RP11-297C4.2 | -0,273 | 2,45135E-11 |
| ACE2 | LINC01081 | -0,273 | 2,43303E-11 |
| ACE2 | CCL8 | -0,273 | 2,46277E-11 |
| ACE2 | TNRC6C-AS1 | -0,273 | 2,44387E-11 |
| ACE2 | CTB-102L5.7 | -0,273 | 2,44135E-11 |
| ACE2 | PPP1R12C | -0,273 | 2,44131E-11 |
| ACE2 | RP11-545P7.9 | -0,2731 | 2,4238E-11 |
| ACE2 | VMO1 | -0,2731 | 2,39088E-11 |
| ACE2 | RBMX | -0,2731 | 2,41026E-11 |
| ACE2 | RP11-662B19.2 | -0,2732 | 2,37979E-11 |
| ACE2 | FGFR1OP2 | -0,2732 | 2,37633E-11 |
| ACE2 | RP3-508I15.14 | -0,2733 | 2,3237E-11 |
| ACE2 | HRG | -0,2734 | 2,29899E-11 |
| ACE2 | RP11-108L7.14 | -0,2734 | 2,27497E-11 |
| ACE2 | CACNA1C | -0,2734 | 2,28333E-11 |
| ACE2 | EXOSC3P1 | -0,2735 | 2,22772E-11 |
| ACE2 | P2RY6 | -0,2736 | 2,21649E-11 |
| ACE2 | TCP10L | -0,2736 | 2,22133E-11 |
| ACE2 | RP11-567F11.2 | -0,2737 | 2,1728E-11 |
| ACE2 | TNXB | -0,2739 | 2,08802E-11 |
| ACE2 | AF131216.6 | -0,274 | 2,05468E-11 |
| ACE2 | SNORD22 | -0,274 | 2,04524E-11 |
| ACE2 | CEP128 | -0,274 | 2,0682E-11 |
| ACE2 | CDKN1B | -0,2742 | 1,99082E-11 |
| ACE2 | ARHGEF10L | -0,2743 | 1,95919E-11 |
| ACE2 | RP11-127B20.3 | -0,2743 | 1,95276E-11 |
| ACE2 | RP11-820K3.3 | -0,2743 | 1,94774E-11 |
| ACE2 | PSMB10 | -0,2743 | 1,95328E-11 |
| ACE2 | GATAD2A | -0,2743 | 1,95643E-11 |
| ACE2 | BICC1 | -0,2744 | 1,93637E-11 |
| ACE2 | PTHLH | -0,2744 | 1,92065E-11 |
| ACE2 | NCS1 | -0,2745 | 1,87821E-11 |
| ACE2 | CTD-2007L18.5 | -0,2745 | 1,87569E-11 |
| ACE2 | HIST1H4I | -0,2746 | 1,85956E-11 |
| ACE2 | SVIL-AS1 | -0,2747 | 1,80674E-11 |
| ACE2 | ENTPD1 | -0,2747 | 1,83624E-11 |
| ACE2 | CMIP | -0,2747 | 1,81215E-11 |
| ACE2 | CD300A | -0,2747 | 1,81345E-11 |
| ACE2 | CTB-50L17.5 | -0,2747 | 1,82132E-11 |
| ACE2 | NTAN1P2 | -0,2749 | 1,74893E-11 |
| ACE2 | RP11-131P10.2 | -0,2749 | 1,7711E-11 |

| | | | |
|---|---|---|---|
| ACE2 | AC005682.6 | -0,275 | 1,74163E-11 |
| ACE2 | HNRNPCP7 | -0,275 | 1,71584E-11 |
| ACE2 | AC073072.5 | -0,2751 | 1,70682E-11 |
| ACE2 | TMEM81 | -0,2752 | 1,66916E-11 |
| ACE2 | AL133243.1 | -0,2752 | 1,68085E-11 |
| ACE2 | MAB21L2 | -0,2752 | 1,66765E-11 |
| ACE2 | METTL24 | -0,2752 | 1,66259E-11 |
| ACE2 | RP11-686D22.10 | -0,2752 | 1,6589E-11 |
| ACE2 | WDR45B | -0,2752 | 1,67724E-11 |
| ACE2 | ZNF496 | -0,2753 | 1,63413E-11 |
| ACE2 | MAGI1-IT1 | -0,2753 | 1,62929E-11 |
| ACE2 | KDELC1P1 | -0,2753 | 1,65255E-11 |
| ACE2 | PDLIM4 | -0,2753 | 1,63543E-11 |
| ACE2 | RP11-530C5.1 | -0,2753 | 1,64042E-11 |
| ACE2 | SETBP1 | -0,2753 | 1,62667E-11 |
| ACE2 | CTC-360J11.4 | -0,2753 | 1,62628E-11 |
| ACE2 | DDX17 | -0,2753 | 1,64511E-11 |
| ACE2 | RRP8 | -0,2754 | 1,59903E-11 |
| ACE2 | TMEM132A | -0,2754 | 1,60735E-11 |
| ACE2 | RP11-592H6.3 | -0,2754 | 1,62039E-11 |
| ACE2 | RP11-930O11.3 | -0,2755 | 1,57534E-11 |
| ACE2 | MAMSTR | -0,2755 | 1,59585E-11 |
| ACE2 | AF127936.3 | -0,2755 | 1,57014E-11 |
| ACE2 | SLC6A9 | -0,2756 | 1,55616E-11 |
| ACE2 | RP1-283K11.3 | -0,2756 | 1,54458E-11 |
| ACE2 | RPS20 | -0,2757 | 1,52555E-11 |
| ACE2 | RP11-788M5.3 | -0,2757 | 1,52591E-11 |
| ACE2 | MOXD1 | -0,2758 | 1,5108E-11 |
| ACE2 | C1QTNF12 | -0,2759 | 1,4837E-11 |
| ACE2 | RP5-1024N4.5 | -0,2759 | 1,47224E-11 |
| ACE2 | IMMP2L | -0,2759 | 1,48785E-11 |
| ACE2 | CRTC3 | -0,2759 | 1,48821E-11 |
| ACE2 | CPED1 | -0,276 | 1,4392E-11 |
| ACE2 | CTA-963H5.5 | -0,276 | 1,45738E-11 |
| ACE2 | PLK2 | -0,2761 | 1,42721E-11 |
| ACE2 | CCDC136 | -0,2761 | 1,43274E-11 |
| ACE2 | RFPL3S | -0,2761 | 1,41429E-11 |
| ACE2 | RP11-133F8.2 | -0,2762 | 1,40147E-11 |
| ACE2 | PCDHGC4 | -0,2762 | 1,41128E-11 |
| ACE2 | ST3GAL2 | -0,2762 | 1,396E-11 |
| ACE2 | HLA-C | -0,2763 | 1,38747E-11 |
| ACE2 | FAM27E3 | -0,2763 | 1,37707E-11 |
| ACE2 | RP11-454F8.3 | -0,2763 | 1,37064E-11 |
| ACE2 | MAP3K1 | -0,2764 | 1,34613E-11 |
| ACE2 | POC1B | -0,2764 | 1,35549E-11 |
| ACE2 | RP11-999E24.3 | -0,2764 | 1,36109E-11 |
| ACE2 | RP11-326A19.4 | -0,2764 | 1,34365E-11 |
| ACE2 | UACA | -0,2765 | 1,31843E-11 |
| ACE2 | ATP5G1P4 | -0,2766 | 1,29555E-11 |
| ACE2 | RP11-252I14.2 | -0,2766 | 1,30668E-11 |
| ACE2 | SNHG6 | -0,2766 | 1,29881E-11 |
| ACE2 | SPRED1 | -0,2766 | 1,29944E-11 |
| ACE2 | PDE4A | -0,2767 | 1,28563E-11 |
| ACE2 | RALGPS2 | -0,2768 | 1,26834E-11 |

| | | | |
|---|---|---|---|
| ACE2 | KLHL3 | -0,2768 | 1,2525E-11 |
| ACE2 | TRBV5-5 | -0,2769 | 1,2296E-11 |
| ACE2 | NRIP3 | -0,2769 | 1,2326E-11 |
| ACE2 | KERA | -0,2769 | 1,23427E-11 |
| ACE2 | PLAC9P1 | -0,277 | 1,20629E-11 |
| ACE2 | TRBV5-6 | -0,277 | 1,20849E-11 |
| ACE2 | RBFOX2 | -0,277 | 1,22422E-11 |
| ACE2 | ZNF436-AS1 | -0,2771 | 1,18685E-11 |
| ACE2 | BTAF1 | -0,2771 | 1,19928E-11 |
| ACE2 | RP11-2E11.5 | -0,2772 | 1,17147E-11 |
| ACE2 | KLF3-AS1 | -0,2773 | 1,14958E-11 |
| ACE2 | PCDHB6 | -0,2773 | 1,15705E-11 |
| ACE2 | ITPKB-IT1 | -0,2774 | 1,13496E-11 |
| ACE2 | ECSCR | -0,2774 | 1,13014E-11 |
| ACE2 | DENND5B | -0,2774 | 1,14186E-11 |
| ACE2 | KCNQ4 | -0,2775 | 1,10728E-11 |
| ACE2 | RN7SL558P | -0,2775 | 1,1072E-11 |
| ACE2 | SCARNA9 | -0,2775 | 1,10528E-11 |
| ACE2 | RP3-449O17.1 | -0,2775 | 1,11071E-11 |
| ACE2 | TNN | -0,2776 | 1,08795E-11 |
| ACE2 | ACD | -0,2776 | 1,09787E-11 |
| ACE2 | POPDC2 | -0,2777 | 1,08084E-11 |
| ACE2 | RP11-291B21.2 | -0,2777 | 1,07238E-11 |
| ACE2 | DZIP1L | -0,2778 | 1,05523E-11 |
| ACE2 | TNFRSF1A | -0,2778 | 1,06013E-11 |
| ACE2 | LA16c-360H6.1 | -0,2778 | 1,05646E-11 |
| ACE2 | TNFRSF9 | -0,2779 | 1,02919E-11 |
| ACE2 | PM20D2 | -0,2779 | 1,04209E-11 |
| ACE2 | LINC01415 | -0,2779 | 1,03098E-11 |
| ACE2 | NFATC2 | -0,2779 | 1,03052E-11 |
| ACE2 | RP3-467L1.4 | -0,278 | 1,02516E-11 |
| ACE2 | RP11-397G17.1 | -0,278 | 1,01407E-11 |
| ACE2 | ROR2 | -0,278 | 1,02004E-11 |
| ACE2 | SNORD3C | -0,278 | 1,0254E-11 |
| ACE2 | IGFBP4 | -0,278 | 1,01455E-11 |
| ACE2 | LINC00570 | -0,2781 | 1,00713E-11 |
| ACE2 | GAK | -0,2781 | 1,00547E-11 |
| ACE2 | SLED1 | -0,2781 | 1,00884E-11 |
| ACE2 | KDM5B | -0,2782 | 9,7642E-12 |
| ACE2 | ZNF853 | -0,2782 | 9,82397E-12 |
| ACE2 | DDX50P1 | -0,2783 | 9,70606E-12 |
| ACE2 | RP11-79M19.2 | -0,2783 | 9,59175E-12 |
| ACE2 | LINC00869 | -0,2784 | 9,54732E-12 |
| ACE2 | AC011997.1 | -0,2784 | 9,58445E-12 |
| ACE2 | GGTA1P | -0,2784 | 9,58186E-12 |
| ACE2 | FCRL6 | -0,2785 | 9,31764E-12 |
| ACE2 | HDAC9 | -0,2785 | 9,26964E-12 |
| ACE2 | RP11-383C5.3 | -0,2785 | 9,40747E-12 |
| ACE2 | HTR1F | -0,2787 | 9,01158E-12 |
| ACE2 | RP11-484D2.5 | -0,2787 | 9,03805E-12 |
| ACE2 | USP36 | -0,2787 | 9,01504E-12 |
| ACE2 | DDIT4L | -0,2788 | 8,88199E-12 |
| ACE2 | SYNM | -0,2788 | 8,79275E-12 |
| ACE2 | FAM65A | -0,2788 | 8,83456E-12 |

| | | | |
|---|---|---|---|
| ACE2 | PPP1R15B | -0,2789 | 8,76434E-12 |
| ACE2 | TTC13 | -0,279 | 8,50153E-12 |
| ACE2 | APOC3 | -0,279 | 8,48014E-12 |
| ACE2 | RP11-313P22.1 | -0,279 | 8,56635E-12 |
| ACE2 | SCARF2 | -0,279 | 8,60328E-12 |
| ACE2 | RP11-277I20.3 | -0,2791 | 8,44893E-12 |
| ACE2 | RP13-188A5.1 | -0,2791 | 8,40319E-12 |
| ACE2 | BEND7 | -0,2792 | 8,28069E-12 |
| ACE2 | BEGAIN | -0,2792 | 8,18144E-12 |
| ACE2 | CDX1 | -0,2793 | 8,07152E-12 |
| ACE2 | PXT1 | -0,2793 | 8,13104E-12 |
| ACE2 | FNDC5 | -0,2794 | 8,02142E-12 |
| ACE2 | EOGT | -0,2794 | 7,97264E-12 |
| ACE2 | NHSL2 | -0,2794 | 7,99692E-12 |
| ACE2 | EED | -0,2796 | 7,74412E-12 |
| ACE2 | RP11-802E16.3 | -0,2797 | 7,51125E-12 |
| ACE2 | KCTD10 | -0,2797 | 7,50414E-12 |
| ACE2 | METRNL | -0,2797 | 7,50697E-12 |
| ACE2 | ZNF578 | -0,2797 | 7,57361E-12 |
| ACE2 | SNORD83B | -0,2797 | 7,53149E-12 |
| ACE2 | NR1H4 | -0,2798 | 7,42763E-12 |
| ACE2 | NFIA-AS2 | -0,2799 | 7,24943E-12 |
| ACE2 | CR1L | -0,2799 | 7,29204E-12 |
| ACE2 | RP11-759A24.1 | -0,2799 | 7,33724E-12 |
| ACE2 | STIM2 | -0,28 | 7,20019E-12 |
| ACE2 | MAP4K1 | -0,28 | 7,15751E-12 |
| ACE2 | HIST1H2AB | -0,2801 | 7,00282E-12 |
| ACE2 | TRPV2 | -0,2801 | 7,08323E-12 |
| ACE2 | RNU4ATAC | -0,2802 | 6,8375E-12 |
| ACE2 | RP11-379F4.1 | -0,2802 | 6,87992E-12 |
| ACE2 | BCL2A1 | -0,2802 | 6,95797E-12 |
| ACE2 | RAC2 | -0,2802 | 6,91813E-12 |
| ACE2 | FN1 | -0,2803 | 6,71938E-12 |
| ACE2 | ACTG1P14 | -0,2804 | 6,64364E-12 |
| ACE2 | CSF2RB | -0,2804 | 6,62368E-12 |
| ACE2 | HDAC10 | -0,2804 | 6,62566E-12 |
| ACE2 | PSD2 | -0,2805 | 6,57641E-12 |
| ACE2 | RP11-180C16.1 | -0,2805 | 6,50216E-12 |
| ACE2 | LINC01474 | -0,2805 | 6,5957E-12 |
| ACE2 | SIK3 | -0,2805 | 6,51251E-12 |
| ACE2 | PAGR1 | -0,2805 | 6,56929E-12 |
| ACE2 | BCAS3 | -0,2805 | 6,48937E-12 |
| ACE2 | TEX10 | -0,2806 | 6,3862E-12 |
| ACE2 | RALY | -0,2806 | 6,44468E-12 |
| ACE2 | UNC5A | -0,2807 | 6,33195E-12 |
| ACE2 | GFRA1 | -0,2807 | 6,28115E-12 |
| ACE2 | KLF16 | -0,2807 | 6,28512E-12 |
| ACE2 | AC011290.5 | -0,2808 | 6,16753E-12 |
| ACE2 | MAP4K2 | -0,2808 | 6,19063E-12 |
| ACE2 | BMP4 | -0,2808 | 6,18797E-12 |
| ACE2 | SH2B3 | -0,2809 | 6,12524E-12 |
| ACE2 | HNRNPA1P52 | -0,2809 | 6,12777E-12 |
| ACE2 | FUNDC2P4 | -0,2809 | 6,06964E-12 |
| ACE2 | IL1RN | -0,281 | 5,98246E-12 |

| | | | |
|---|---|---|---|
| ACE2 | CAMKK2 | -0,281 | 5,94343E-12 |
| ACE2 | ZRSR2 | -0,281 | 5,99813E-12 |
| ACE2 | VWA5B1 | -0,2811 | 5,85774E-12 |
| ACE2 | RP5-1116H23.3 | -0,2811 | 5,89485E-12 |
| ACE2 | OSBPL10 | -0,2812 | 5,7866E-12 |
| ACE2 | RPL23AP36 | -0,2813 | 5,63041E-12 |
| ACE2 | CD72 | -0,2813 | 5,663E-12 |
| ACE2 | YBX3 | -0,2813 | 5,70216E-12 |
| ACE2 | KCNC4 | -0,2814 | 5,54515E-12 |
| ACE2 | NR4A3 | -0,2814 | 5,59336E-12 |
| ACE2 | TRPC4 | -0,2814 | 5,57201E-12 |
| ACE2 | CTA-113A6.2 | -0,2814 | 5,59883E-12 |
| ACE2 | CYP4F22 | -0,2814 | 5,52687E-12 |
| ACE2 | RPS6KA3 | -0,2814 | 5,5782E-12 |
| ACE2 | USP31 | -0,2815 | 5,43427E-12 |
| ACE2 | PRDM15 | -0,2816 | 5,3417E-12 |
| ACE2 | LRCH3 | -0,2817 | 5,25015E-12 |
| ACE2 | PDSS1P1 | -0,2818 | 5,17065E-12 |
| ACE2 | RP11-430K21.2 | -0,2818 | 5,15194E-12 |
| ACE2 | PFKFB3 | -0,2818 | 5,18918E-12 |
| ACE2 | AGO1 | -0,2819 | 5,11244E-12 |
| ACE2 | BANCR | -0,2819 | 5,08227E-12 |
| ACE2 | CTD-2574D22.4 | -0,2819 | 5,11523E-12 |
| ACE2 | RP11-635N19.2 | -0,2819 | 5,04302E-12 |
| ACE2 | DEFB124 | -0,2819 | 5,07695E-12 |
| ACE2 | UPK3A | -0,282 | 4,96044E-12 |
| ACE2 | LGR6 | -0,2821 | 4,89754E-12 |
| ACE2 | OXER1 | -0,2821 | 4,86794E-12 |
| ACE2 | PCDHGA5 | -0,2821 | 4,88992E-12 |
| ACE2 | TRBV7-6 | -0,2821 | 4,87177E-12 |
| ACE2 | RP11-707M3.3 | -0,2821 | 4,87333E-12 |
| ACE2 | RP11-304L19.1 | -0,2822 | 4,86249E-12 |
| ACE2 | KIAA1586 | -0,2823 | 4,71298E-12 |
| ACE2 | EGR3 | -0,2823 | 4,72134E-12 |
| ACE2 | TBC1D20 | -0,2823 | 4,77265E-12 |
| ACE2 | CTA-268H5.14 | -0,2823 | 4,70012E-12 |
| ACE2 | MEST | -0,2824 | 4,62487E-12 |
| ACE2 | MBNL2 | -0,2824 | 4,6183E-12 |
| ACE2 | TICRR | -0,2824 | 4,64697E-12 |
| ACE2 | NOCT | -0,2826 | 4,46766E-12 |
| ACE2 | CTNNA3 | -0,2826 | 4,4633E-12 |
| ACE2 | RP5-821D11.7 | -0,2826 | 4,45923E-12 |
| ACE2 | NBDY | -0,2826 | 4,47226E-12 |
| ACE2 | RP4-621F18.2 | -0,2827 | 4,3675E-12 |
| ACE2 | RP11-118K6.3 | -0,2827 | 4,41433E-12 |
| ACE2 | RP11-727A23.7 | -0,2827 | 4,39521E-12 |
| ACE2 | BANP | -0,2827 | 4,40659E-12 |
| ACE2 | SAFB2 | -0,2827 | 4,41925E-12 |
| ACE2 | HCG4P3 | -0,2828 | 4,34004E-12 |
| ACE2 | COX7A2P1 | -0,2828 | 4,31239E-12 |
| ACE2 | AC006128.2 | -0,2828 | 4,30476E-12 |
| ACE2 | COL6A2 | -0,2828 | 4,30287E-12 |
| ACE2 | SCMH1 | -0,2829 | 4,26648E-12 |
| ACE2 | AC092614.2 | -0,2829 | 4,23161E-12 |

| | | | |
|---|---|---|---|
| ACE2 | MFSD14C | -0,2829 | 4,25679E-12 |
| ACE2 | VIM | -0,2829 | 4,28366E-12 |
| ACE2 | LINC00622 | -0,283 | 4,17647E-12 |
| ACE2 | RP11-144H23.2 | -0,283 | 4,16771E-12 |
| ACE2 | TEX14 | -0,283 | 4,208E-12 |
| ACE2 | CCDC130 | -0,283 | 4,17513E-12 |
| ACE2 | ZC3H12B | -0,283 | 4,14449E-12 |
| ACE2 | NKRF | -0,283 | 4,18755E-12 |
| ACE2 | NPIPB12 | -0,2831 | 4,07171E-12 |
| ACE2 | TTR | -0,2831 | 4,06373E-12 |
| ACE2 | HIST1H1PS1 | -0,2832 | 4,01934E-12 |
| ACE2 | RP11-52J3.2 | -0,2832 | 4,04048E-12 |
| ACE2 | RP1-28O10.1 | -0,2833 | 3,95853E-12 |
| ACE2 | ZNF738 | -0,2833 | 3,93061E-12 |
| ACE2 | CNBP | -0,2834 | 3,84927E-12 |
| ACE2 | AC005154.6 | -0,2834 | 3,84688E-12 |
| ACE2 | PCED1A | -0,2834 | 3,85473E-12 |
| ACE2 | RP3-402G11.25 | -0,2834 | 3,89085E-12 |
| ACE2 | PCNP | -0,2835 | 3,78663E-12 |
| ACE2 | MESTP1 | -0,2835 | 3,84082E-12 |
| ACE2 | ZEB1 | -0,2835 | 3,83494E-12 |
| ACE2 | CLDND2 | -0,2835 | 3,8138E-12 |
| ACE2 | CSNK1G2P1 | -0,2836 | 3,75509E-12 |
| ACE2 | RP11-380B4.3 | -0,2836 | 3,71845E-12 |
| ACE2 | RAB12 | -0,2836 | 3,77178E-12 |
| ACE2 | YPEL5 | -0,2837 | 3,65998E-12 |
| ACE2 | DOCK10 | -0,2837 | 3,66946E-12 |
| ACE2 | INHBA-AS1 | -0,2837 | 3,6641E-12 |
| ACE2 | TESK1 | -0,2837 | 3,64854E-12 |
| ACE2 | SOST | -0,2837 | 3,64317E-12 |
| ACE2 | ARHGAP31-AS1 | -0,2838 | 3,62702E-12 |
| ACE2 | LINC02100 | -0,2838 | 3,59842E-12 |
| ACE2 | CTC-575I10.1 | -0,2838 | 3,61849E-12 |
| ACE2 | RIMKLB | -0,2838 | 3,63608E-12 |
| ACE2 | LOXL1 | -0,2838 | 3,59279E-12 |
| ACE2 | RP1-152L7.5 | -0,2839 | 3,516E-12 |
| ACE2 | HADHAP1 | -0,284 | 3,45162E-12 |
| ACE2 | TCAF2 | -0,284 | 3,4904E-12 |
| ACE2 | PDE10A | -0,2841 | 3,38841E-12 |
| ACE2 | APOBEC3H | -0,2841 | 3,42291E-12 |
| ACE2 | PKD1L1 | -0,2842 | 3,35155E-12 |
| ACE2 | PGGHG | -0,2842 | 3,33051E-12 |
| ACE2 | PDGFA | -0,2843 | 3,27634E-12 |
| ACE2 | BBS9 | -0,2843 | 3,3035E-12 |
| ACE2 | ZKSCAN1 | -0,2843 | 3,2999E-12 |
| ACE2 | AC007879.3 | -0,2844 | 3,22883E-12 |
| ACE2 | KHDRBS2 | -0,2844 | 3,25397E-12 |
| ACE2 | AGAP6 | -0,2844 | 3,24877E-12 |
| ACE2 | C17orf82 | -0,2844 | 3,2226E-12 |
| ACE2 | ABL2 | -0,2845 | 3,20389E-12 |
| ACE2 | EPB42 | -0,2846 | 3,10835E-12 |
| ACE2 | C1orf159 | -0,2847 | 3,05655E-12 |
| ACE2 | RP11-91K8.5 | -0,2847 | 3,07211E-12 |
| ACE2 | AF001548.3 | -0,2847 | 3,06437E-12 |

| | | | |
|---|---|---|---|
| ACE2 | RP11-382D12.2 | -0,2848 | 3,03011E-12 |
| ACE2 | UBE2D3 | -0,2848 | 3,02289E-12 |
| ACE2 | RP11-454F8.2 | -0,2848 | 3,01571E-12 |
| ACE2 | UQCRBP1 | -0,2848 | 2,98524E-12 |
| ACE2 | TRAC | -0,2849 | 2,96306E-12 |
| ACE2 | ZNFX1 | -0,2849 | 2,9735E-12 |
| ACE2 | COL14A1 | -0,285 | 2,90804E-12 |
| ACE2 | POGZ | -0,2851 | 2,8592E-12 |
| ACE2 | GPR65 | -0,2851 | 2,84799E-12 |
| ACE2 | SLC26A6 | -0,2852 | 2,81114E-12 |
| ACE2 | SDC2 | -0,2852 | 2,81158E-12 |
| ACE2 | COX7A1 | -0,2852 | 2,78874E-12 |
| ACE2 | RP11-382A20.3 | -0,2853 | 2,75235E-12 |
| ACE2 | RBM17 | -0,2854 | 2,69327E-12 |
| ACE2 | PSMB8 | -0,2856 | 2,58504E-12 |
| ACE2 | RP11-981P6.1 | -0,2856 | 2,60652E-12 |
| ACE2 | LENG8 | -0,2856 | 2,62254E-12 |
| ACE2 | MAOB | -0,2856 | 2,59975E-12 |
| ACE2 | MIR222HG | -0,2856 | 2,60179E-12 |
| ACE2 | SLC20A1 | -0,2857 | 2,53698E-12 |
| ACE2 | FAP | -0,2857 | 2,57009E-12 |
| ACE2 | WNT10A | -0,2857 | 2,56636E-12 |
| ACE2 | MBOAT4 | -0,2857 | 2,5582E-12 |
| ACE2 | RP11-383M4.6 | -0,2857 | 2,54104E-12 |
| ACE2 | PRKG1 | -0,2857 | 2,57609E-12 |
| ACE2 | RP11-148B18.1 | -0,2858 | 2,50542E-12 |
| ACE2 | RP11-488L18.4 | -0,2858 | 2,49734E-12 |
| ACE2 | NOD1 | -0,2859 | 2,45069E-12 |
| ACE2 | RP11-85K15.3 | -0,2859 | 2,48038E-12 |
| ACE2 | NTN5 | -0,2859 | 2,46389E-12 |
| ACE2 | AC010733.5 | -0,286 | 2,43133E-12 |
| ACE2 | ARL5B | -0,286 | 2,44009E-12 |
| ACE2 | PHACTR2P1 | -0,286 | 2,41197E-12 |
| ACE2 | LINC00640 | -0,286 | 2,40272E-12 |
| ACE2 | RP11-2C24.7 | -0,286 | 2,41854E-12 |
| ACE2 | GTF3C2-AS1 | -0,2861 | 2,39378E-12 |
| ACE2 | IDO2 | -0,2861 | 2,36481E-12 |
| ACE2 | PPP1R12A | -0,2861 | 2,37922E-12 |
| ACE2 | RAMP2 | -0,2861 | 2,35403E-12 |
| ACE2 | CLEC11A | -0,2861 | 2,38282E-12 |
| ACE2 | MIAT | -0,2861 | 2,37979E-12 |
| ACE2 | L1CAM | -0,2861 | 2,36163E-12 |
| ACE2 | APOL6 | -0,2862 | 2,33155E-12 |
| ACE2 | ZNF250 | -0,2863 | 2,27112E-12 |
| ACE2 | LRFN5 | -0,2863 | 2,28578E-12 |
| ACE2 | CTD-2262B20.1 | -0,2863 | 2,28431E-12 |
| ACE2 | TMEM74B | -0,2863 | 2,2877E-12 |
| ACE2 | ZNF732 | -0,2864 | 2,26234E-12 |
| ACE2 | AGK | -0,2864 | 2,24928E-12 |
| ACE2 | CTD-2576N18.1 | -0,2865 | 2,22117E-12 |
| ACE2 | CTB-50L17.7 | -0,2865 | 2,19508E-12 |
| ACE2 | CHD1 | -0,2866 | 2,15532E-12 |
| ACE2 | PURG | -0,2866 | 2,16599E-12 |
| ACE2 | RDH5 | -0,2866 | 2,16526E-12 |

| | | | |
|---|---|---|---|
| ACE2 | RP4-800J21.3 | -0,2866 | 2,1713E-12 |
| ACE2 | CYS1 | -0,2867 | 2,14017E-12 |
| ACE2 | CTB-47B11.3 | -0,2867 | 2,13719E-12 |
| ACE2 | TSC22D4 | -0,2867 | 2,11206E-12 |
| ACE2 | AURKC | -0,2867 | 2,12559E-12 |
| ACE2 | FAM182B | -0,2867 | 2,14705E-12 |
| ACE2 | ALG13 | -0,2868 | 2,08342E-12 |
| ACE2 | MAML1 | -0,2869 | 2,05257E-12 |
| ACE2 | RP11-196B3.1 | -0,2869 | 2,04925E-12 |
| ACE2 | LINC01128 | -0,287 | 1,99713E-12 |
| ACE2 | RP11-84D1.1 | -0,287 | 2,00618E-12 |
| ACE2 | OSTF1 | -0,287 | 2,00405E-12 |
| ACE2 | RP11-351C21.2 | -0,287 | 2,02909E-12 |
| ACE2 | RP11-417L19.5 | -0,287 | 2,00867E-12 |
| ACE2 | AC004076.5 | -0,287 | 1,99929E-12 |
| ACE2 | MID1 | -0,287 | 2,01506E-12 |
| ACE2 | SLC4A9 | -0,2871 | 1,97747E-12 |
| ACE2 | RP3-325F22.3 | -0,2871 | 1,96659E-12 |
| ACE2 | LINC01451 | -0,2871 | 1,96562E-12 |
| ACE2 | DUSP5 | -0,2871 | 1,97737E-12 |
| ACE2 | SGO1-AS1 | -0,2872 | 1,92735E-12 |
| ACE2 | ATP13A3 | -0,2872 | 1,92643E-12 |
| ACE2 | RP1-244F24.1 | -0,2872 | 1,94745E-12 |
| ACE2 | RP11-319G9.5 | -0,2872 | 1,94442E-12 |
| ACE2 | TAF7 | -0,2873 | 1,91363E-12 |
| ACE2 | AC004471.9 | -0,2873 | 1,90918E-12 |
| ACE2 | PLCH2 | -0,2874 | 1,86412E-12 |
| ACE2 | AP000892.6 | -0,2875 | 1,83818E-12 |
| ACE2 | CTD-2026K11.6 | -0,2875 | 1,85292E-12 |
| ACE2 | GAMT | -0,2875 | 1,83242E-12 |
| ACE2 | RPSAP16 | -0,2876 | 1,8101E-12 |
| ACE2 | MAFK | -0,2876 | 1,80662E-12 |
| ACE2 | RP11-426C22.1 | -0,2876 | 1,78806E-12 |
| ACE2 | SLC5A4 | -0,2876 | 1,79554E-12 |
| ACE2 | AC002550.6 | -0,2877 | 1,76433E-12 |
| ACE2 | RP11-861L17.2 | -0,2877 | 1,78567E-12 |
| ACE2 | VEGFA | -0,2878 | 1,7338E-12 |
| ACE2 | RP13-93L13.1 | -0,2878 | 1,73685E-12 |
| ACE2 | TBRG1 | -0,2878 | 1,72545E-12 |
| ACE2 | CTD-2576D5.4 | -0,2878 | 1,72339E-12 |
| ACE2 | RP11-298D21.1 | -0,2878 | 1,73511E-12 |
| ACE2 | RP11-21M24.2 | -0,2879 | 1,71329E-12 |
| ACE2 | RP11-231C14.6 | -0,2879 | 1,71287E-12 |
| ACE2 | NDUFB4 | -0,288 | 1,66223E-12 |
| ACE2 | UNC5D | -0,288 | 1,67547E-12 |
| ACE2 | ARID4B | -0,2881 | 1,64486E-12 |
| ACE2 | RP3-337H4.6 | -0,2881 | 1,64023E-12 |
| ACE2 | FCER2 | -0,2881 | 1,6539E-12 |
| ACE2 | RP11-304L19.3 | -0,2882 | 1,61804E-12 |
| ACE2 | SMG1P2 | -0,2882 | 1,60377E-12 |
| ACE2 | AC090044.2 | -0,2883 | 1,57615E-12 |
| ACE2 | TRAFD1 | -0,2883 | 1,58636E-12 |
| ACE2 | NLRP1 | -0,2883 | 1,5845E-12 |
| ACE2 | RAPGEF5 | -0,2884 | 1,55437E-12 |

| | | | |
|---|---|---|---|
| ACE2 | UBA7 | -0,2885 | 1,5255E-12 |
| ACE2 | C10orf11 | -0,2885 | 1,52293E-12 |
| ACE2 | SLX1A | -0,2886 | 1,50284E-12 |
| ACE2 | TGM2 | -0,2886 | 1,49788E-12 |
| ACE2 | CAMK2N1 | -0,2887 | 1,48113E-12 |
| ACE2 | RGS13 | -0,2887 | 1,47479E-12 |
| ACE2 | IRAK2 | -0,2887 | 1,48434E-12 |
| ACE2 | HGFAC | -0,2887 | 1,46242E-12 |
| ACE2 | SBF1 | -0,2887 | 1,46433E-12 |
| ACE2 | FOXC2 | -0,2888 | 1,44152E-12 |
| ACE2 | MYL9 | -0,2888 | 1,43829E-12 |
| ACE2 | RP11-426L16.3 | -0,2889 | 1,40723E-12 |
| ACE2 | CARD17 | -0,2889 | 1,42036E-12 |
| ACE2 | STX16 | -0,2889 | 1,40851E-12 |
| ACE2 | AP4B1-AS1 | -0,289 | 1,39429E-12 |
| ACE2 | RP11-494K3.2 | -0,289 | 1,38119E-12 |
| ACE2 | ZNF460 | -0,289 | 1,3869E-12 |
| ACE2 | H3F3A | -0,2891 | 1,36205E-12 |
| ACE2 | PPM1M | -0,2891 | 1,37811E-12 |
| ACE2 | PARP12 | -0,2891 | 1,36644E-12 |
| ACE2 | C11orf95 | -0,2891 | 1,36979E-12 |
| ACE2 | TRIM9 | -0,2891 | 1,3652E-12 |
| ACE2 | RP11-703M24.5 | -0,2891 | 1,35776E-12 |
| ACE2 | RP11-461L13.3 | -0,2893 | 1,31697E-12 |
| ACE2 | LA16c-321D4.2 | -0,2893 | 1,31112E-12 |
| ACE2 | CTD-3105H18.4 | -0,2893 | 1,30918E-12 |
| ACE2 | FAM160B2 | -0,2894 | 1,30081E-12 |
| ACE2 | SAMD4B | -0,2894 | 1,30581E-12 |
| ACE2 | ZBTB46 | -0,2894 | 1,28262E-12 |
| ACE2 | RP3-370M22.8 | -0,2894 | 1,30214E-12 |
| ACE2 | RP4-583P15.16 | -0,2895 | 1,26121E-12 |
| ACE2 | ECM1 | -0,2896 | 1,24055E-12 |
| ACE2 | RP11-266K4.14 | -0,2896 | 1,24054E-12 |
| ACE2 | UPK3BP1 | -0,2897 | 1,23417E-12 |
| ACE2 | RP4-718D20.3 | -0,2897 | 1,21851E-12 |
| ACE2 | RP3-333H23.9 | -0,2897 | 1,23019E-12 |
| ACE2 | AC159540.1 | -0,2898 | 1,19224E-12 |
| ACE2 | RP11-274B21.4 | -0,2898 | 1,20692E-12 |
| ACE2 | HSPA6 | -0,2899 | 1,18536E-12 |
| ACE2 | LIMS4 | -0,2899 | 1,17085E-12 |
| ACE2 | GLI2 | -0,2899 | 1,1724E-12 |
| ACE2 | MICALL2 | -0,29 | 1,14809E-12 |
| ACE2 | RP11-872J21.5 | -0,29 | 1,15458E-12 |
| ACE2 | SLFN13 | -0,29 | 1,16033E-12 |
| ACE2 | MCTP1 | -0,2901 | 1,13771E-12 |
| ACE2 | IFIT1P1 | -0,2901 | 1,13086E-12 |
| ACE2 | THOC1 | -0,2901 | 1,14672E-12 |
| ACE2 | TSHZ2 | -0,2901 | 1,1425E-12 |
| ACE2 | ST3GAL4-AS1 | -0,2903 | 1,09279E-12 |
| ACE2 | RP11-932O9.10 | -0,2903 | 1,09228E-12 |
| ACE2 | GJC1 | -0,2903 | 1,08878E-12 |
| ACE2 | SUGP2 | -0,2903 | 1,09114E-12 |
| ACE2 | LINC01436 | -0,2904 | 1,08119E-12 |
| ACE2 | RPS12P17 | -0,2905 | 1,06145E-12 |

| | | | |
|---|---|---|---|
| ACE2 | C16orf52 | -0,2905 | 1,05132E-12 |
| ACE2 | LLNLR-271C9.1 | -0,2905 | 1,04634E-12 |
| ACE2 | ARMCX3-AS1 | -0,2906 | 1,03407E-12 |
| ACE2 | HIST2H2AB | -0,2907 | 1,01287E-12 |
| ACE2 | AC007899.3 | -0,2907 | 1,02467E-12 |
| ACE2 | TTC14 | -0,2907 | 1,0176E-12 |
| ACE2 | SNHG15 | -0,2907 | 1,02231E-12 |
| ACE2 | RP11-505K9.5 | -0,2907 | 1,00754E-12 |
| ACE2 | COL16A1 | -0,2908 | 9,91245E-13 |
| ACE2 | FBXO39 | -0,2909 | 9,7087E-13 |
| ACE2 | RP11-15K2.2 | -0,2909 | 9,83489E-13 |
| ACE2 | AC005954.4 | -0,2909 | 9,80545E-13 |
| ACE2 | B4GALT3 | -0,291 | 9,66211E-13 |
| ACE2 | AC098614.2 | -0,2911 | 9,35614E-13 |
| ACE2 | RNU4-78P | -0,2911 | 9,36139E-13 |
| ACE2 | RP1-134E15.3 | -0,2911 | 9,49415E-13 |
| ACE2 | HDX | -0,2911 | 9,3781E-13 |
| ACE2 | PPP2R2A | -0,2912 | 9,23696E-13 |
| ACE2 | TNFAIP6 | -0,2913 | 9,0456E-13 |
| ACE2 | RAMP1 | -0,2913 | 9,041E-13 |
| ACE2 | DACT3-AS1 | -0,2913 | 9,0504E-13 |
| ACE2 | AF129075.5 | -0,2913 | 9,06748E-13 |
| ACE2 | IL24 | -0,2914 | 8,95135E-13 |
| ACE2 | PKP4 | -0,2914 | 8,91473E-13 |
| ACE2 | OSM | -0,2916 | 8,61443E-13 |
| ACE2 | LY9 | -0,2918 | 8,24497E-13 |
| ACE2 | RP11-564A8.4 | -0,2918 | 8,27516E-13 |
| ACE2 | SYTL3 | -0,2918 | 8,34834E-13 |
| ACE2 | MEFV | -0,2918 | 8,26722E-13 |
| ACE2 | STX5 | -0,2919 | 8,18465E-13 |
| ACE2 | SKIL | -0,292 | 7,90155E-13 |
| ACE2 | MIR497HG | -0,2921 | 7,87146E-13 |
| ACE2 | FLI1 | -0,2922 | 7,64905E-13 |
| ACE2 | RP11-141M1.3 | -0,2922 | 7,70394E-13 |
| ACE2 | DENND4A | -0,2922 | 7,74736E-13 |
| ACE2 | RP11-47L3.1 | -0,2922 | 7,69487E-13 |
| ACE2 | ELMO2 | -0,2923 | 7,53734E-13 |
| ACE2 | RP1-257A7.4 | -0,2924 | 7,42546E-13 |
| ACE2 | RP11-445H22.3 | -0,2924 | 7,33997E-13 |
| ACE2 | CTA-384D8.34 | -0,2924 | 7,33279E-13 |
| ACE2 | KLRC1 | -0,2925 | 7,22837E-13 |
| ACE2 | TAS2R14 | -0,2925 | 7,20194E-13 |
| ACE2 | ZNF143 | -0,2926 | 7,07556E-13 |
| ACE2 | MYL6 | -0,2926 | 7,19069E-13 |
| ACE2 | RP11-116D17.3 | -0,2926 | 7,09956E-13 |
| ACE2 | TMEM86B | -0,2926 | 7,14882E-13 |
| ACE2 | BCL6B | -0,2927 | 6,9407E-13 |
| ACE2 | CIC | -0,2927 | 7,01816E-13 |
| ACE2 | ARHGDIG | -0,2928 | 6,90756E-13 |
| ACE2 | PDCD5 | -0,2928 | 6,8136E-13 |
| ACE2 | HELZ2 | -0,2928 | 6,88844E-13 |
| ACE2 | TNFRSF14 | -0,2929 | 6,69149E-13 |
| ACE2 | RP11-1B20.1 | -0,2929 | 6,76341E-13 |
| ACE2 | GRIA1 | -0,2929 | 6,73334E-13 |

| | | | |
|---|---|---|---|
| ACE2 | SBF2 | -0,2929 | 6,76202E-13 |
| ACE2 | VGLL4 | -0,293 | 6,60343E-13 |
| ACE2 | KRT18P31 | -0,2931 | 6,49197E-13 |
| ACE2 | PCDHGB9P | -0,2931 | 6,49874E-13 |
| ACE2 | NSMF | -0,2931 | 6,49678E-13 |
| ACE2 | TBXA2R | -0,2931 | 6,52018E-13 |
| ACE2 | ARHGAP42 | -0,2932 | 6,32004E-13 |
| ACE2 | RP11-464D20.6 | -0,2932 | 6,35293E-13 |
| ACE2 | CTD-3220F14.1 | -0,2932 | 6,37983E-13 |
| ACE2 | RP4-534N18.2 | -0,2933 | 6,29181E-13 |
| ACE2 | RAB21 | -0,2933 | 6,26618E-13 |
| ACE2 | RP11-603B24.1 | -0,2933 | 6,25046E-13 |
| ACE2 | MIR133A1HG | -0,2933 | 6,28901E-13 |
| ACE2 | C22orf24 | -0,2934 | 6,09822E-13 |
| ACE2 | TMEM108 | -0,2935 | 6,04537E-13 |
| ACE2 | BIRC3 | -0,2935 | 6,06536E-13 |
| ACE2 | RP11-159L20.2 | -0,2935 | 6,05891E-13 |
| ACE2 | KIR2DL1 | -0,2935 | 5,98691E-13 |
| ACE2 | RP11-338N10.3 | -0,2936 | 5,87157E-13 |
| ACE2 | NFIB | -0,2936 | 5,88199E-13 |
| ACE2 | CASP10 | -0,2937 | 5,77545E-13 |
| ACE2 | MGP | -0,2937 | 5,78804E-13 |
| ACE2 | ZNF419 | -0,2937 | 5,84063E-13 |
| ACE2 | DUSP22 | -0,2938 | 5,68765E-13 |
| ACE2 | ERG | -0,2938 | 5,64191E-13 |
| ACE2 | CBX3P3 | -0,294 | 5,51784E-13 |
| ACE2 | CRY1 | -0,294 | 5,47293E-13 |
| ACE2 | RP11-562A8.4 | -0,294 | 5,48582E-13 |
| ACE2 | AC008277.1 | -0,2941 | 5,36254E-13 |
| ACE2 | TARID | -0,2941 | 5,34748E-13 |
| ACE2 | DSCAML1 | -0,2942 | 5,24245E-13 |
| ACE2 | CD69 | -0,2942 | 5,23039E-13 |
| ACE2 | RP11-353N4.6 | -0,2943 | 5,20751E-13 |
| ACE2 | SNORA52 | -0,2943 | 5,15249E-13 |
| ACE2 | C18orf25 | -0,2944 | 5,1026E-13 |
| ACE2 | LINC01160 | -0,2945 | 5,02396E-13 |
| ACE2 | TAOK2 | -0,2945 | 4,97309E-13 |
| ACE2 | XXbac-BPG170G13.: | -0,2946 | 4,8756E-13 |
| ACE2 | HOXB8 | -0,2946 | 4,92507E-13 |
| ACE2 | ZCCHC11 | -0,2947 | 4,78216E-13 |
| ACE2 | RP11-561I11.3 | -0,2947 | 4,76564E-13 |
| ACE2 | MAP3K10 | -0,2947 | 4,83714E-13 |
| ACE2 | TMEM234 | -0,2948 | 4,71644E-13 |
| ACE2 | LINC02195 | -0,2948 | 4,74542E-13 |
| ACE2 | RP11-295P9.3 | -0,2949 | 4,62466E-13 |
| ACE2 | SMAD9-IT1 | -0,2949 | 4,64085E-13 |
| ACE2 | UCKL1 | -0,2949 | 4,62535E-13 |
| ACE2 | RP11-598F7.5 | -0,295 | 4,52187E-13 |
| ACE2 | CFD | -0,295 | 4,5416E-13 |
| ACE2 | LRRC8C | -0,2951 | 4,46147E-13 |
| ACE2 | POU5F2 | -0,2951 | 4,41199E-13 |
| ACE2 | SPTA1 | -0,2952 | 4,3266E-13 |
| ACE2 | NAB1 | -0,2953 | 4,29386E-13 |
| ACE2 | AC091133.1 | -0,2953 | 4,3092E-13 |

| | | | |
|---|---|---|---|
| ACE2 | TTC39C | -0,2954 | 4,22781E-13 |
| ACE2 | C1orf74 | -0,2955 | 4,14019E-13 |
| ACE2 | RBM12B | -0,2955 | 4,10433E-13 |
| ACE2 | RP11-426C22.8 | -0,2955 | 4,13467E-13 |
| ACE2 | CTD-3162L10.1 | -0,2955 | 4,08852E-13 |
| ACE2 | NPTN-IT1 | -0,2956 | 4,01527E-13 |
| ACE2 | SNRPA | -0,2956 | 4,0483E-13 |
| ACE2 | RBCK1 | -0,2956 | 4,0211E-13 |
| ACE2 | RP4-734G22.3 | -0,2957 | 3,98388E-13 |
| ACE2 | CTD-2002J20.1 | -0,2958 | 3,86181E-13 |
| ACE2 | RP11-20I20.4 | -0,2958 | 3,89503E-13 |
| ACE2 | RP11-290L1.4 | -0,2959 | 3,83027E-13 |
| ACE2 | SYNGR3 | -0,2959 | 3,79772E-13 |
| ACE2 | C16orf87 | -0,2959 | 3,82957E-13 |
| ACE2 | RP4-760C5.3 | -0,2959 | 3,81926E-13 |
| ACE2 | RAB8B | -0,296 | 3,77876E-13 |
| ACE2 | CTD-3128G10.6 | -0,296 | 3,75169E-13 |
| ACE2 | CTD-2001J20.1 | -0,2961 | 3,65483E-13 |
| ACE2 | RPS12P31 | -0,2961 | 3,69127E-13 |
| ACE2 | BID | -0,2961 | 3,68258E-13 |
| ACE2 | CTD-2036P10.5 | -0,2962 | 3,58421E-13 |
| ACE2 | SRRM1 | -0,2963 | 3,52083E-13 |
| ACE2 | RP11-66N24.3 | -0,2963 | 3,50984E-13 |
| ACE2 | RGAG4 | -0,2963 | 3,53088E-13 |
| ACE2 | AC092652.1 | -0,2964 | 3,46179E-13 |
| ACE2 | FAM69A | -0,2965 | 3,39365E-13 |
| ACE2 | RP11-383B4.4 | -0,2965 | 3,4147E-13 |
| ACE2 | CD81 | -0,2965 | 3,43863E-13 |
| ACE2 | RP11-176H8.3 | -0,2965 | 3,39836E-13 |
| ACE2 | POLR2F | -0,2965 | 3,42932E-13 |
| ACE2 | ARMCX2 | -0,2965 | 3,37742E-13 |
| ACE2 | SLC6A16 | -0,2966 | 3,31309E-13 |
| ACE2 | BMP2K | -0,2967 | 3,28489E-13 |
| ACE2 | DPY19L1P1 | -0,2967 | 3,29484E-13 |
| ACE2 | RP11-959F10.6 | -0,2967 | 3,29838E-13 |
| ACE2 | DAO | -0,2967 | 3,26095E-13 |
| ACE2 | AKT2 | -0,2967 | 3,29954E-13 |
| ACE2 | ADCY10P1 | -0,2968 | 3,19274E-13 |
| ACE2 | TRBV6-6 | -0,2968 | 3,21632E-13 |
| ACE2 | NADSYN1 | -0,2968 | 3,2039E-13 |
| ACE2 | LSAMP | -0,2969 | 3,15401E-13 |
| ACE2 | SCAF4 | -0,2969 | 3,15968E-13 |
| ACE2 | RP11-810P12.7 | -0,297 | 3,11587E-13 |
| ACE2 | RGS12 | -0,2971 | 3,06574E-13 |
| ACE2 | DBN1 | -0,2972 | 2,99994E-13 |
| ACE2 | RP11-73M18.7 | -0,2972 | 2,99846E-13 |
| ACE2 | PRICKLE3 | -0,2972 | 2,99918E-13 |
| ACE2 | DPP6 | -0,2974 | 2,84746E-13 |
| ACE2 | METTL9 | -0,2975 | 2,7909E-13 |
| ACE2 | PSMB9 | -0,2976 | 2,75552E-13 |
| ACE2 | ST3GAL4 | -0,2976 | 2,74947E-13 |
| ACE2 | NTF3 | -0,2976 | 2,74413E-13 |
| ACE2 | RPL21P135 | -0,2977 | 2,71909E-13 |
| ACE2 | CALB2 | -0,2977 | 2,71586E-13 |

| | | | |
|---|---|---|---|
| ACE2 | RP11-465L10.10 | -0,2977 | 2,68678E-13 |
| ACE2 | CALCRL | -0,2978 | 2,68245E-13 |
| ACE2 | LAMTOR3P2 | -0,2979 | 2,60581E-13 |
| ACE2 | C6orf3 | -0,298 | 2,56521E-13 |
| ACE2 | RP5-855D21.1 | -0,298 | 2,5707E-13 |
| ACE2 | ARID5B | -0,298 | 2,58237E-13 |
| ACE2 | ADRA2A | -0,2981 | 2,49101E-13 |
| ACE2 | DPYD-IT1 | -0,2982 | 2,46924E-13 |
| ACE2 | PROK2 | -0,2982 | 2,4727E-13 |
| ACE2 | ZNF259P1 | -0,2982 | 2,48627E-13 |
| ACE2 | RP11-119F19.5 | -0,2982 | 2,46637E-13 |
| ACE2 | PRND | -0,2982 | 2,4462E-13 |
| ACE2 | SH3RF3 | -0,2983 | 2,43767E-13 |
| ACE2 | BACH1 | -0,2983 | 2,4062E-13 |
| ACE2 | CTA-212A2.4 | -0,2983 | 2,43189E-13 |
| ACE2 | AAK1 | -0,2984 | 2,35385E-13 |
| ACE2 | LOXL1-AS1 | -0,2984 | 2,37181E-13 |
| ACE2 | RP11-459F6.3 | -0,2984 | 2,3576E-13 |
| ACE2 | BIRC2 | -0,2985 | 2,33545E-13 |
| ACE2 | CKS1BP1 | -0,2985 | 2,30779E-13 |
| ACE2 | RP11-342A23.2 | -0,2985 | 2,30449E-13 |
| ACE2 | RP11-120M18.2 | -0,2985 | 2,32575E-13 |
| ACE2 | PRSS57 | -0,2986 | 2,2872E-13 |
| ACE2 | RPS3AP38 | -0,2987 | 2,24259E-13 |
| ACE2 | CFL1P5 | -0,2988 | 2,18099E-13 |
| ACE2 | CUBN | -0,2988 | 2,19285E-13 |
| ACE2 | AC009303.1 | -0,2989 | 2,15031E-13 |
| ACE2 | MXD3 | -0,299 | 2,13261E-13 |
| ACE2 | RP11-25G10.2 | -0,299 | 2,12471E-13 |
| ACE2 | GADD45B | -0,299 | 2,12087E-13 |
| ACE2 | RP11-359K18.4 | -0,2991 | 2,06979E-13 |
| ACE2 | INPP4B | -0,2992 | 2,02144E-13 |
| ACE2 | INPP4A | -0,2993 | 2,00071E-13 |
| ACE2 | IL6 | -0,2993 | 2,00416E-13 |
| ACE2 | AC000068.9 | -0,2993 | 2,00922E-13 |
| ACE2 | PCDHB13 | -0,2994 | 1,94099E-13 |
| ACE2 | FNDC1 | -0,2994 | 1,95916E-13 |
| ACE2 | SERPINH1 | -0,2994 | 1,94745E-13 |
| ACE2 | JAK3 | -0,2994 | 1,97052E-13 |
| ACE2 | FLJ27354 | -0,2995 | 1,90886E-13 |
| ACE2 | HIVEP3 | -0,2996 | 1,88665E-13 |
| ACE2 | RPL29P11 | -0,2996 | 1,88775E-13 |
| ACE2 | CTD-3224K15.3 | -0,2996 | 1,87377E-13 |
| ACE2 | KLRC2 | -0,2996 | 1,87039E-13 |
| ACE2 | BAZ2A | -0,2996 | 1,87953E-13 |
| ACE2 | AKAP6 | -0,2996 | 1,8702E-13 |
| ACE2 | APOBEC3D | -0,2998 | 1,81025E-13 |
| ACE2 | ROBO1 | -0,2999 | 1,77207E-13 |
| ACE2 | AC005306.3 | -0,2999 | 1,75934E-13 |
| ACE2 | RP5-1116H23.4 | -0,2999 | 1,78222E-13 |
| ACE2 | NUAK2 | -0,3 | 1,75716E-13 |
| ACE2 | TRGJP1 | -0,3 | 1,74784E-13 |
| ACE2 | RP11-1149M10.1 | -0,3 | 1,73403E-13 |
| ACE2 | RP11-732A19.1 | -0,3 | 1,73842E-13 |

| | | | |
|---|---|---|---|
| ACE2 | PRPF3 | -0,3002 | 1,66379E-13 |
| ACE2 | BCHE | -0,3002 | 1,6784E-13 |
| ACE2 | RRM1-AS1 | -0,3002 | 1,68333E-13 |
| ACE2 | CYTH1 | -0,3002 | 1,69075E-13 |
| ACE2 | UNC80 | -0,3003 | 1,63648E-13 |
| ACE2 | AOAH-IT1 | -0,3003 | 1,65182E-13 |
| ACE2 | PGM2L1 | -0,3003 | 1,63581E-13 |
| ACE2 | OLR1 | -0,3003 | 1,63247E-13 |
| ACE2 | CTC-465D4.1 | -0,3003 | 1,63887E-13 |
| ACE2 | ABCA7 | -0,3003 | 1,63716E-13 |
| ACE2 | PXN | -0,3004 | 1,60757E-13 |
| ACE2 | MYL5 | -0,3005 | 1,59528E-13 |
| ACE2 | CLEC4A | -0,3005 | 1,59317E-13 |
| ACE2 | HBQ1 | -0,3005 | 1,59204E-13 |
| ACE2 | SPEG | -0,3006 | 1,55989E-13 |
| ACE2 | ACAP3 | -0,3007 | 1,52031E-13 |
| ACE2 | CTD-2014D20.1 | -0,3007 | 1,52975E-13 |
| ACE2 | AP000487.5 | -0,3007 | 1,53151E-13 |
| ACE2 | P2RX2 | -0,3007 | 1,52709E-13 |
| ACE2 | RP3-508I15.22 | -0,3007 | 1,5305E-13 |
| ACE2 | RP6-99M1.3 | -0,3007 | 1,50731E-13 |
| ACE2 | ELMO1 | -0,3008 | 1,49602E-13 |
| ACE2 | TNFAIP1 | -0,3008 | 1,49029E-13 |
| ACE2 | RP5-981O7.2 | -0,3009 | 1,47266E-13 |
| ACE2 | ADD3 | -0,301 | 1,42305E-13 |
| ACE2 | ACER3 | -0,301 | 1,42369E-13 |
| ACE2 | JAG2 | -0,301 | 1,43893E-13 |
| ACE2 | RP11-759A24.2 | -0,3011 | 1,40338E-13 |
| ACE2 | KIF9-AS1 | -0,3012 | 1,36809E-13 |
| ACE2 | IFITM1 | -0,3012 | 1,37795E-13 |
| ACE2 | DPT | -0,3013 | 1,35111E-13 |
| ACE2 | ADAMTS9-AS2 | -0,3013 | 1,34382E-13 |
| ACE2 | TRGV5P | -0,3014 | 1,33269E-13 |
| ACE2 | MEX3B | -0,3014 | 1,33706E-13 |
| ACE2 | TBL1X | -0,3014 | 1,32049E-13 |
| ACE2 | RP11-1H15.2 | -0,3015 | 1,30465E-13 |
| ACE2 | LAT | -0,3015 | 1,29498E-13 |
| ACE2 | C2CD4C | -0,3015 | 1,30572E-13 |
| ACE2 | TRBV13 | -0,3016 | 1,28665E-13 |
| ACE2 | XXbac-B562F10.11 | -0,3016 | 1,27744E-13 |
| ACE2 | TBL1XR1 | -0,3017 | 1,24538E-13 |
| ACE2 | CBLN3 | -0,3017 | 1,26279E-13 |
| ACE2 | MPRIP-AS1 | -0,3017 | 1,24396E-13 |
| ACE2 | ARHGEF2 | -0,3018 | 1,23195E-13 |
| ACE2 | COL6A4P2 | -0,3018 | 1,23977E-13 |
| ACE2 | RPLP1P13 | -0,3018 | 1,23841E-13 |
| ACE2 | RP3-395M20.7 | -0,3019 | 1,21624E-13 |
| ACE2 | WDFY1 | -0,3019 | 1,19803E-13 |
| ACE2 | DOC2GP | -0,3019 | 1,20521E-13 |
| ACE2 | RP11-554D20.2 | -0,3019 | 1,21501E-13 |
| ACE2 | TGFB1I1 | -0,3019 | 1,19941E-13 |
| ACE2 | SERPINC1 | -0,302 | 1,17278E-13 |
| ACE2 | RP11-410E4.1 | -0,302 | 1,19197E-13 |
| ACE2 | RPL7P18 | -0,302 | 1,17075E-13 |

| | | | |
|---|---|---|---|
| ACE2 | HLA-J | -0,302 | 1,18794E-13 |
| ACE2 | SLC17A9 | -0,302 | 1,18166E-13 |
| ACE2 | RP4-635E18.7 | -0,3021 | 1,15704E-13 |
| ACE2 | RBMXL2 | -0,3021 | 1,15735E-13 |
| ACE2 | ARHGEF12 | -0,3022 | 1,13028E-13 |
| ACE2 | PHF11 | -0,3023 | 1,12305E-13 |
| ACE2 | RP11-693M3.1 | -0,3023 | 1,10943E-13 |
| ACE2 | NRIR | -0,3024 | 1,1016E-13 |
| ACE2 | BASP1 | -0,3024 | 1,10325E-13 |
| ACE2 | RP4-751H13.5 | -0,3024 | 1,10122E-13 |
| ACE2 | HDAC1P1 | -0,3024 | 1,10309E-13 |
| ACE2 | DMPK | -0,3024 | 1,09364E-13 |
| ACE2 | RAVER2 | -0,3025 | 1,07024E-13 |
| ACE2 | TAS2R18 | -0,3025 | 1,06287E-13 |
| ACE2 | FCHO1 | -0,3025 | 1,07872E-13 |
| ACE2 | GPC3 | -0,3025 | 1,06917E-13 |
| ACE2 | RP11-50C13.1 | -0,3026 | 1,05901E-13 |
| ACE2 | FGFR1 | -0,3027 | 1,03793E-13 |
| ACE2 | GPR52 | -0,3028 | 1,00557E-13 |
| ACE2 | PCDHGA7 | -0,3028 | 1,01085E-13 |
| ACE2 | SRSF12 | -0,3028 | 1,00419E-13 |
| ACE2 | AC084219.4 | -0,3028 | 1,01841E-13 |
| ACE2 | SLAMF7 | -0,3031 | 9,49374E-14 |
| ACE2 | AC055764.1 | -0,3031 | 9,61844E-14 |
| ACE2 | TRAPPC2 | -0,3031 | 9,48185E-14 |
| ACE2 | S100A16 | -0,3032 | 9,39639E-14 |
| ACE2 | RC3H1 | -0,3032 | 9,43137E-14 |
| ACE2 | TRGV3 | -0,3033 | 9,19804E-14 |
| ACE2 | MAFA | -0,3033 | 9,09934E-14 |
| ACE2 | RP11-390B4.3 | -0,3033 | 9,11022E-14 |
| ACE2 | ARHGAP23 | -0,3033 | 9,14367E-14 |
| ACE2 | PARD6G-AS1 | -0,3033 | 9,20538E-14 |
| ACE2 | ATP1B3-AS1 | -0,3034 | 8,95921E-14 |
| ACE2 | ZNF394 | -0,3034 | 9,04995E-14 |
| ACE2 | AC093063.2 | -0,3034 | 8,92821E-14 |
| ACE2 | PIK3IP1-AS1 | -0,3035 | 8,76284E-14 |
| ACE2 | AC006272.2 | -0,3036 | 8,61978E-14 |
| ACE2 | C1orf140 | -0,3037 | 8,50994E-14 |
| ACE2 | TRGC2 | -0,3037 | 8,48331E-14 |
| ACE2 | APOA1-AS | -0,3037 | 8,52913E-14 |
| ACE2 | NSUN6 | -0,3038 | 8,31677E-14 |
| ACE2 | RP11-488L18.10 | -0,304 | 8,069E-14 |
| ACE2 | PHLDB2 | -0,304 | 7,92671E-14 |
| ACE2 | AC025165.8 | -0,304 | 7,94913E-14 |
| ACE2 | SAMD4A | -0,304 | 8,05317E-14 |
| ACE2 | RP11-222K16.1 | -0,3041 | 7,7915E-14 |
| ACE2 | EPB41L4A-AS1 | -0,3041 | 7,80922E-14 |
| ACE2 | CTC-444N24.7 | -0,3041 | 7,81936E-14 |
| ACE2 | RP11-783K16.14 | -0,3042 | 7,69859E-14 |
| ACE2 | SH3GL1 | -0,3042 | 7,67249E-14 |
| ACE2 | KCTD16 | -0,3043 | 7,58756E-14 |
| ACE2 | BAMBI | -0,3043 | 7,53799E-14 |
| ACE2 | RBM6 | -0,3044 | 7,34021E-14 |
| ACE2 | PIWIL2 | -0,3044 | 7,40627E-14 |

| | | | |
|---|---|---|---|
| ACE2 | KLHL25 | -0,3044 | 7,33956E-14 |
| ACE2 | RNF11 | -0,3045 | 7,19254E-14 |
| ACE2 | IGF2BP2 | -0,3045 | 7,31228E-14 |
| ACE2 | TRANK1 | -0,3046 | 7,12895E-14 |
| ACE2 | GLP1R | -0,3046 | 7,16337E-14 |
| ACE2 | ZUFSP | -0,3046 | 7,1013E-14 |
| ACE2 | TRBV10-3 | -0,3047 | 7,02222E-14 |
| ACE2 | KCTD6 | -0,3048 | 6,87242E-14 |
| ACE2 | SAFB | -0,3048 | 6,85075E-14 |
| ACE2 | RP5-999L4.2 | -0,3048 | 6,8806E-14 |
| ACE2 | RAPGEF4 | -0,3049 | 6,66471E-14 |
| ACE2 | AC004383.3 | -0,3049 | 6,76632E-14 |
| ACE2 | RP11-429G19.3 | -0,305 | 6,621E-14 |
| ACE2 | AMOTL1 | -0,305 | 6,54108E-14 |
| ACE2 | PHF6 | -0,305 | 6,60552E-14 |
| ACE2 | MIR6891 | -0,3051 | 6,38984E-14 |
| ACE2 | EIF4HP2 | -0,3051 | 6,39965E-14 |
| ACE2 | HNRNPD | -0,3052 | 6,29985E-14 |
| ACE2 | TRBV21-1 | -0,3052 | 6,34298E-14 |
| ACE2 | RP11-799M12.2 | -0,3053 | 6,20425E-14 |
| ACE2 | CARNS1 | -0,3053 | 6,18642E-14 |
| ACE2 | STRN3 | -0,3053 | 6,24813E-14 |
| ACE2 | RP11-56B16.5 | -0,3053 | 6,13954E-14 |
| ACE2 | SEPT4 | -0,3053 | 6,21183E-14 |
| ACE2 | TTC24 | -0,3054 | 6,10103E-14 |
| ACE2 | PRKRIP1 | -0,3054 | 6,0447E-14 |
| ACE2 | YPEL3 | -0,3054 | 6,0429E-14 |
| ACE2 | HOXB3 | -0,3054 | 6,10842E-14 |
| ACE2 | CD8B | -0,3055 | 5,97293E-14 |
| ACE2 | EPHA1-AS1 | -0,3055 | 5,93154E-14 |
| ACE2 | RP11-383F6.1 | -0,3056 | 5,79639E-14 |
| ACE2 | MIATNB | -0,3056 | 5,78831E-14 |
| ACE2 | ATF4 | -0,3056 | 5,79724E-14 |
| ACE2 | MAP1LC3A | -0,3057 | 5,68719E-14 |
| ACE2 | AC010524.4 | -0,3058 | 5,64721E-14 |
| ACE2 | HIST2H2BD | -0,3059 | 5,46545E-14 |
| ACE2 | TAP2 | -0,3059 | 5,53567E-14 |
| ACE2 | RUNDC3A | -0,3059 | 5,55697E-14 |
| ACE2 | RP11-277A4.4 | -0,306 | 5,41766E-14 |
| ACE2 | PCDHGB8P | -0,306 | 5,42542E-14 |
| ACE2 | EGFR-AS1 | -0,306 | 5,42515E-14 |
| ACE2 | CASP1P2 | -0,306 | 5,42936E-14 |
| ACE2 | BRD4 | -0,306 | 5,3597E-14 |
| ACE2 | RANGAP1 | -0,306 | 5,43479E-14 |
| ACE2 | VAMP4 | -0,3061 | 5,24894E-14 |
| ACE2 | SMG1P3 | -0,3061 | 5,26508E-14 |
| ACE2 | GADD45A | -0,3062 | 5,20793E-14 |
| ACE2 | RP11-834C11.4 | -0,3062 | 5,13736E-14 |
| ACE2 | LGALS2 | -0,3062 | 5,15055E-14 |
| ACE2 | FOXD2-AS1 | -0,3063 | 5,06E-14 |
| ACE2 | TCTEX1D1 | -0,3063 | 5,10453E-14 |
| ACE2 | ZFAND2B | -0,3063 | 5,04265E-14 |
| ACE2 | RP11-626H12.1 | -0,3063 | 5,05932E-14 |
| ACE2 | SDK2 | -0,3063 | 5,0887E-14 |

| | | | |
|---|---|---|---|
| ACE2 | TRAV4 | -0,3064 | 5,01122E-14 |
| ACE2 | LINC00982 | -0,3065 | 4,91E-14 |
| ACE2 | HIST1H2BC | -0,3065 | 4,91592E-14 |
| ACE2 | RP11-89K11.1 | -0,3065 | 4,92568E-14 |
| ACE2 | RP11-485M7.2 | -0,3066 | 4,81941E-14 |
| ACE2 | AC130469.2 | -0,3066 | 4,8131E-14 |
| ACE2 | MTMR8 | -0,3066 | 4,80176E-14 |
| ACE2 | EMD | -0,3066 | 4,76637E-14 |
| ACE2 | DDX18P1 | -0,3067 | 4,7115E-14 |
| ACE2 | RPL10P19 | -0,3068 | 4,56046E-14 |
| ACE2 | RP11-416N2.4 | -0,3068 | 4,63245E-14 |
| ACE2 | TTLL4 | -0,3069 | 4,50917E-14 |
| ACE2 | IDO1 | -0,3069 | 4,55919E-14 |
| ACE2 | SP140L | -0,307 | 4,44571E-14 |
| ACE2 | HIVEP2 | -0,3071 | 4,37152E-14 |
| ACE2 | DNAJA1 | -0,3071 | 4,30276E-14 |
| ACE2 | SLC8B1 | -0,3071 | 4,36173E-14 |
| ACE2 | RPS27AP12 | -0,3072 | 4,26885E-14 |
| ACE2 | RP11-168K11.5 | -0,3072 | 4,26238E-14 |
| ACE2 | TRIM3 | -0,3072 | 4,27858E-14 |
| ACE2 | DCHS1 | -0,3072 | 4,28456E-14 |
| ACE2 | UTS2 | -0,3073 | 4,15101E-14 |
| ACE2 | DRAXIN | -0,3073 | 4,20244E-14 |
| ACE2 | DCAF15 | -0,3074 | 4,12089E-14 |
| ACE2 | PPP4R2 | -0,3075 | 4,04378E-14 |
| ACE2 | RP13-204A15.5 | -0,3075 | 4,02481E-14 |
| ACE2 | TTYH2 | -0,3076 | 3,92792E-14 |
| ACE2 | ZNF836 | -0,3076 | 3,89107E-14 |
| ACE2 | CLEC9A | -0,3077 | 3,83975E-14 |
| ACE2 | KRT73 | -0,3077 | 3,84911E-14 |
| ACE2 | ATP11C | -0,3077 | 3,87351E-14 |
| ACE2 | ITK | -0,3078 | 3,78824E-14 |
| ACE2 | PPP1R17 | -0,3078 | 3,75644E-14 |
| ACE2 | PURB | -0,3078 | 3,80725E-14 |
| ACE2 | C1orf52 | -0,3079 | 3,70582E-14 |
| ACE2 | KBTBD2 | -0,3079 | 3,70811E-14 |
| ACE2 | PRSS12 | -0,308 | 3,63969E-14 |
| ACE2 | HDAC7 | -0,3081 | 3,52498E-14 |
| ACE2 | APBA2 | -0,3082 | 3,48244E-14 |
| ACE2 | PGPEP1L | -0,3082 | 3,45835E-14 |
| ACE2 | POFUT2 | -0,3082 | 3,51214E-14 |
| ACE2 | TM4SF1 | -0,3083 | 3,42155E-14 |
| ACE2 | MATR3 | -0,3083 | 3,4189E-14 |
| ACE2 | RP11-466A19.1 | -0,3083 | 3,41252E-14 |
| ACE2 | PABPC1P3 | -0,3083 | 3,42138E-14 |
| ACE2 | ILF2P1 | -0,3085 | 3,31162E-14 |
| ACE2 | RPS6KB1 | -0,3085 | 3,27471E-14 |
| ACE2 | CNTD2 | -0,3085 | 3,30185E-14 |
| ACE2 | RP11-379B18.5 | -0,3086 | 3,24627E-14 |
| ACE2 | ENTPD1-AS1 | -0,3086 | 3,20401E-14 |
| ACE2 | SLC22A10 | -0,3086 | 3,21998E-14 |
| ACE2 | MCRIP1 | -0,3086 | 3,24239E-14 |
| ACE2 | BATF3 | -0,3087 | 3,16693E-14 |
| ACE2 | AC092580.1 | -0,3087 | 3,15296E-14 |

| | | | |
|---|---|---|---|
| ACE2 | SEMA3G | -0,3087 | 3,15648E-14 |
| ACE2 | FAM229A | -0,3088 | 3,06671E-14 |
| ACE2 | RP11-18B16.2 | -0,3088 | 3,09147E-14 |
| ACE2 | RP11-677I18.3 | -0,3088 | 3,06319E-14 |
| ACE2 | GUCY1B3 | -0,3089 | 3,02163E-14 |
| ACE2 | RELL2 | -0,3089 | 3,03828E-14 |
| ACE2 | NEFM | -0,3089 | 3,02266E-14 |
| ACE2 | CTA-29F11.1 | -0,3089 | 3,05251E-14 |
| ACE2 | ATG12P2 | -0,3091 | 2,91947E-14 |
| ACE2 | WFDC1 | -0,3091 | 2,91055E-14 |
| ACE2 | FOLH1 | -0,3092 | 2,85765E-14 |
| ACE2 | RP11-849N15.3 | -0,3092 | 2,87181E-14 |
| ACE2 | JUNB | -0,3092 | 2,87593E-14 |
| ACE2 | LGALS9C | -0,3093 | 2,80434E-14 |
| ACE2 | RNF217 | -0,3094 | 2,73327E-14 |
| ACE2 | HCCAT5 | -0,3094 | 2,76543E-14 |
| ACE2 | SUDS3 | -0,3095 | 2,68913E-14 |
| ACE2 | ZDHHC20-IT1 | -0,3095 | 2,66656E-14 |
| ACE2 | FGF14-IT1 | -0,3095 | 2,67001E-14 |
| ACE2 | CD8A | -0,3096 | 2,61183E-14 |
| ACE2 | IGIP | -0,3096 | 2,61116E-14 |
| ACE2 | GAS6 | -0,3096 | 2,64928E-14 |
| ACE2 | METTL22 | -0,3096 | 2,612E-14 |
| ACE2 | NPC1L1 | -0,3097 | 2,6075E-14 |
| ACE2 | SGCA | -0,3097 | 2,59396E-14 |
| ACE2 | TREX2 | -0,3097 | 2,60028E-14 |
| ACE2 | RP11-981G7.6 | -0,3098 | 2,55345E-14 |
| ACE2 | MSC-AS1 | -0,3098 | 2,53224E-14 |
| ACE2 | TGFB2 | -0,31 | 2,43816E-14 |
| ACE2 | SEMA6A-AS1 | -0,31 | 2,41119E-14 |
| ACE2 | SMO | -0,31 | 2,43096E-14 |
| ACE2 | RNF19A | -0,31 | 2,41291E-14 |
| ACE2 | TRAV8-6 | -0,31 | 2,41522E-14 |
| ACE2 | CDKN2D | -0,31 | 2,45331E-14 |
| ACE2 | TUG1 | -0,31 | 2,43226E-14 |
| ACE2 | RP11-329B9.5 | -0,3101 | 2,4073E-14 |
| ACE2 | PLA2R1 | -0,3102 | 2,35851E-14 |
| ACE2 | RP5-1198O20.4 | -0,3103 | 2,27125E-14 |
| ACE2 | BICD2 | -0,3103 | 2,28297E-14 |
| ACE2 | RP11-158M2.2 | -0,3103 | 2,29714E-14 |
| ACE2 | SF3A2 | -0,3103 | 2,29629E-14 |
| ACE2 | VAMP1 | -0,3104 | 2,25725E-14 |
| ACE2 | AC108463.2 | -0,3105 | 2,19809E-14 |
| ACE2 | PCDHB11 | -0,3105 | 2,20429E-14 |
| ACE2 | CTB-109A12.1 | -0,3105 | 2,18558E-14 |
| ACE2 | PMS2P9 | -0,3105 | 2,20646E-14 |
| ACE2 | NPAT | -0,3105 | 2,196E-14 |
| ACE2 | SLC25A37 | -0,3106 | 2,15085E-14 |
| ACE2 | RBM4 | -0,3106 | 2,17138E-14 |
| ACE2 | RP11-212I21.4 | -0,3106 | 2,16391E-14 |
| ACE2 | ITGAV | -0,3107 | 2,11087E-14 |
| ACE2 | RP11-211G3.3 | -0,3107 | 2,12197E-14 |
| ACE2 | RP11-313E19.2 | -0,3107 | 2,10742E-14 |
| ACE2 | PICALM | -0,3107 | 2,1208E-14 |

| | | | |
|---|---|---|---|
| ACE2 | IRF9 | -0,3107 | 2,11406E-14 |
| ACE2 | SNHG23 | -0,3107 | 2,11992E-14 |
| ACE2 | FAM171A2 | -0,3107 | 2,11926E-14 |
| ACE2 | FLNC | -0,3108 | 2,06697E-14 |
| ACE2 | TCF7L2 | -0,3108 | 2,08002E-14 |
| ACE2 | LINC01355 | -0,3109 | 2,01146E-14 |
| ACE2 | RP11-524C21.2 | -0,3109 | 2,02548E-14 |
| ACE2 | LST1 | -0,311 | 1,9763E-14 |
| ACE2 | NDUFAB1P1 | -0,311 | 1,99733E-14 |
| ACE2 | EDIL3 | -0,3111 | 1,96783E-14 |
| ACE2 | RIPK2 | -0,3111 | 1,95899E-14 |
| ACE2 | RP11-15B17.1 | -0,3112 | 1,9092E-14 |
| ACE2 | GPRIN3 | -0,3113 | 1,8549E-14 |
| ACE2 | FHL1 | -0,3113 | 1,86794E-14 |
| ACE2 | TAGLN2 | -0,3114 | 1,83803E-14 |
| ACE2 | MIR3662 | -0,3114 | 1,81919E-14 |
| ACE2 | LAP3 | -0,3115 | 1,79523E-14 |
| ACE2 | STC2 | -0,3115 | 1,80159E-14 |
| ACE2 | LBX1 | -0,3116 | 1,76933E-14 |
| ACE2 | CD207 | -0,3117 | 1,73542E-14 |
| ACE2 | UNC119 | -0,3117 | 1,72204E-14 |
| ACE2 | CERS1 | -0,3117 | 1,73145E-14 |
| ACE2 | FAM127A | -0,3117 | 1,73324E-14 |
| ACE2 | STX6 | -0,3118 | 1,67654E-14 |
| ACE2 | LYST | -0,3118 | 1,69771E-14 |
| ACE2 | RP11-203L2.3 | -0,3118 | 1,67818E-14 |
| ACE2 | PTRF | -0,3118 | 1,68709E-14 |
| ACE2 | RP11-392O1.4 | -0,3118 | 1,69907E-14 |
| ACE2 | MAP4K4 | -0,3119 | 1,65323E-14 |
| ACE2 | KIR3DX1 | -0,3119 | 1,66702E-14 |
| ACE2 | OTUD4 | -0,312 | 1,60966E-14 |
| ACE2 | NRARP | -0,312 | 1,64007E-14 |
| ACE2 | GNAS | -0,312 | 1,62711E-14 |
| ACE2 | RAB39B | -0,312 | 1,62644E-14 |
| ACE2 | SPDYA | -0,3121 | 1,58717E-14 |
| ACE2 | RP11-521I2.3 | -0,3121 | 1,6016E-14 |
| ACE2 | RP11-322D14.1 | -0,3121 | 1,58268E-14 |
| ACE2 | RP11-2C24.5 | -0,3122 | 1,5716E-14 |
| ACE2 | FOXO6 | -0,3123 | 1,54353E-14 |
| ACE2 | PTBP3 | -0,3123 | 1,53796E-14 |
| ACE2 | FES | -0,3123 | 1,54348E-14 |
| ACE2 | ELF4 | -0,3123 | 1,52566E-14 |
| ACE2 | RP11-309L24.2 | -0,3124 | 1,49995E-14 |
| ACE2 | NPDC1 | -0,3124 | 1,49251E-14 |
| ACE2 | RP11-563J2.2 | -0,3124 | 1,50677E-14 |
| ACE2 | C1QTNF3 | -0,3125 | 1,45979E-14 |
| ACE2 | AC005387.3 | -0,3125 | 1,47171E-14 |
| ACE2 | ARHGEF17 | -0,3126 | 1,42728E-14 |
| ACE2 | LAYN | -0,3126 | 1,44534E-14 |
| ACE2 | ISLR | -0,3126 | 1,43456E-14 |
| ACE2 | ETHE1 | -0,3126 | 1,45069E-14 |
| ACE2 | LNCSRLR | -0,3127 | 1,39918E-14 |
| ACE2 | LINCR-0001 | -0,3127 | 1,4082E-14 |
| ACE2 | SFXN3 | -0,3127 | 1,41133E-14 |

| | | | |
|---|---|---|---|
| ACE2 | TRIM52 | -0,3128 | 1,38946E-14 |
| ACE2 | TRBV4-2 | -0,3128 | 1,39649E-14 |
| ACE2 | CTD-2102P23.1 | -0,3129 | 1,35476E-14 |
| ACE2 | GSTM5 | -0,313 | 1,32572E-14 |
| ACE2 | HMGB1P39 | -0,313 | 1,32334E-14 |
| ACE2 | RP11-841C19.1 | -0,313 | 1,32494E-14 |
| ACE2 | LINC00894 | -0,313 | 1,32584E-14 |
| ACE2 | CDC7 | -0,3131 | 1,30004E-14 |
| ACE2 | OLFML1 | -0,3131 | 1,29238E-14 |
| ACE2 | THAP2 | -0,3131 | 1,31171E-14 |
| ACE2 | CCDC144NL-AS1 | -0,3132 | 1,26841E-14 |
| ACE2 | HLX | -0,3133 | 1,26039E-14 |
| ACE2 | AC104134.2 | -0,3133 | 1,25647E-14 |
| ACE2 | RP11-347P5.1 | -0,3133 | 1,25255E-14 |
| ACE2 | PCBP2-OT1 | -0,3133 | 1,25382E-14 |
| ACE2 | RHBDF1 | -0,3133 | 1,24578E-14 |
| ACE2 | PRR5L | -0,3134 | 1,21535E-14 |
| ACE2 | RN7SKP161 | -0,3134 | 1,23109E-14 |
| ACE2 | MATN1 | -0,3136 | 1,16702E-14 |
| ACE2 | FAM126B | -0,3136 | 1,17228E-14 |
| ACE2 | PAG1 | -0,3136 | 1,18576E-14 |
| ACE2 | HOXB-AS1 | -0,3136 | 1,18661E-14 |
| ACE2 | B4GALNT4 | -0,3137 | 1,1616E-14 |
| ACE2 | AC005253.2 | -0,3137 | 1,15995E-14 |
| ACE2 | RASA2-IT1 | -0,3138 | 1,13702E-14 |
| ACE2 | CTB-152G17.6 | -0,3138 | 1,13213E-14 |
| ACE2 | PHF12 | -0,3138 | 1,13416E-14 |
| ACE2 | SDHCP1 | -0,3138 | 1,12802E-14 |
| ACE2 | OGT | -0,3138 | 1,12187E-14 |
| ACE2 | ODF2L | -0,3139 | 1,11615E-14 |
| ACE2 | PEAK1 | -0,3139 | 1,11604E-14 |
| ACE2 | CHD3 | -0,3139 | 1,10226E-14 |
| ACE2 | HAAO | -0,314 | 1,09094E-14 |
| ACE2 | ARID3B | -0,314 | 1,07673E-14 |
| ACE2 | ADCY10 | -0,3141 | 1,05512E-14 |
| ACE2 | PTMA | -0,3141 | 1,05294E-14 |
| ACE2 | RP11-214O1.2 | -0,3141 | 1,05141E-14 |
| ACE2 | PMS2P11 | -0,3142 | 1,04563E-14 |
| ACE2 | ACTN1-AS1 | -0,3142 | 1,04373E-14 |
| ACE2 | MARK2P8 | -0,3143 | 1,01289E-14 |
| ACE2 | SGCG | -0,3143 | 1,01289E-14 |
| ACE2 | FOXL1 | -0,3143 | 1,0247E-14 |
| ACE2 | HTRA2 | -0,3144 | 1,00729E-14 |
| ACE2 | FAM89B | -0,3144 | 9,90116E-15 |
| ACE2 | DPYD-AS1 | -0,3145 | 9,87667E-15 |
| ACE2 | MARK1 | -0,3145 | 9,81009E-15 |
| ACE2 | RP11-139E19.3 | -0,3145 | 9,88101E-15 |
| ACE2 | GCNT1P3 | -0,3146 | 9,5532E-15 |
| ACE2 | RP11-351M8.1 | -0,3146 | 9,68354E-15 |
| ACE2 | BX470102.3 | -0,3147 | 9,4626E-15 |
| ACE2 | AC004985.12 | -0,3147 | 9,3091E-15 |
| ACE2 | TATDN2 | -0,3148 | 9,13436E-15 |
| ACE2 | FBXL13 | -0,3148 | 9,26749E-15 |
| ACE2 | IPMK | -0,3148 | 9,15342E-15 |

| | | | |
|---|---|---|---|
| ACE2 | SIPA1 | -0,3148 | 9,16212E-15 |
| ACE2 | CSAD | -0,3148 | 9,23687E-15 |
| ACE2 | MORC3 | -0,3148 | 9,111E-15 |
| ACE2 | SNORD94 | -0,3149 | 9,02565E-15 |
| ACE2 | C17orf99 | -0,3149 | 9,03575E-15 |
| ACE2 | ATP2A3 | -0,315 | 8,90746E-15 |
| ACE2 | PTGER1 | -0,315 | 8,88161E-15 |
| ACE2 | PDGFB | -0,315 | 8,82156E-15 |
| ACE2 | RP11-1148L6.5 | -0,315 | 8,8721E-15 |
| ACE2 | SLC4A10 | -0,3151 | 8,6888E-15 |
| ACE2 | APLP1 | -0,3152 | 8,42575E-15 |
| ACE2 | MICB | -0,3153 | 8,23975E-15 |
| ACE2 | AMD1P1 | -0,3153 | 8,27358E-15 |
| ACE2 | AL109761.5 | -0,3153 | 8,29759E-15 |
| ACE2 | CRIM1 | -0,3155 | 7,89667E-15 |
| ACE2 | FAM66C | -0,3155 | 8,00912E-15 |
| ACE2 | LINC01146 | -0,3155 | 7,95551E-15 |
| ACE2 | NKAIN4 | -0,3155 | 7,92394E-15 |
| ACE2 | PKLR | -0,3157 | 7,67678E-15 |
| ACE2 | GPAA1P1 | -0,3157 | 7,67053E-15 |
| ACE2 | DCAF13P3 | -0,3157 | 7,64439E-15 |
| ACE2 | MAPK3 | -0,3157 | 7,62363E-15 |
| ACE2 | IL17B | -0,3158 | 7,48912E-15 |
| ACE2 | GIMAP4 | -0,3159 | 7,28907E-15 |
| ACE2 | PXK | -0,316 | 7,19784E-15 |
| ACE2 | HIST1H2BO | -0,316 | 7,12827E-15 |
| ACE2 | RP11-317B3.2 | -0,316 | 7,23908E-15 |
| ACE2 | RP11-295I5.4 | -0,316 | 7,18333E-15 |
| ACE2 | NAB2 | -0,3161 | 7,04748E-15 |
| ACE2 | ANGPTL6 | -0,3161 | 7,10267E-15 |
| ACE2 | RP11-468E2.5 | -0,3162 | 6,88676E-15 |
| ACE2 | CTD-3065B20.3 | -0,3162 | 6,87863E-15 |
| ACE2 | RP11-154H23.3 | -0,3163 | 6,83036E-15 |
| ACE2 | RP11-197M22.2 | -0,3164 | 6,58614E-15 |
| ACE2 | LGALS9B | -0,3164 | 6,6772E-15 |
| ACE2 | CTD-2583A14.8 | -0,3164 | 6,69401E-15 |
| ACE2 | TBC1D10A | -0,3165 | 6,52024E-15 |
| ACE2 | SIT1 | -0,3167 | 6,25815E-15 |
| ACE2 | ZBTB5 | -0,3168 | 6,0817E-15 |
| ACE2 | SPX | -0,3168 | 6,04684E-15 |
| ACE2 | TMEM132E | -0,3168 | 6,04891E-15 |
| ACE2 | PLD4 | -0,3169 | 5,96093E-15 |
| ACE2 | RP11-61L19.1 | -0,3169 | 6,02346E-15 |
| ACE2 | PIP5K1C | -0,3169 | 5,95314E-15 |
| ACE2 | DIP2A | -0,317 | 5,90336E-15 |
| ACE2 | KAZN | -0,3171 | 5,71995E-15 |
| ACE2 | LINC02044 | -0,3171 | 5,72854E-15 |
| ACE2 | RP11-282A11.3 | -0,3171 | 5,70979E-15 |
| ACE2 | MRGPRF-AS1 | -0,3172 | 5,60911E-15 |
| ACE2 | NHS | -0,3172 | 5,58904E-15 |
| ACE2 | HES1 | -0,3173 | 5,52681E-15 |
| ACE2 | TRBV5-1 | -0,3173 | 5,53184E-15 |
| ACE2 | CCDC159 | -0,3173 | 5,46693E-15 |
| ACE2 | CCDC117 | -0,3173 | 5,5524E-15 |

| | | | |
|---|---|---|---|
| ACE2 | IL12A-AS1 | -0,3174 | 5,41462E-15 |
| ACE2 | NDUFB1P2 | -0,3175 | 5,32948E-15 |
| ACE2 | AC011247.3 | -0,3175 | 5,32054E-15 |
| ACE2 | ATG4B | -0,3175 | 5,27233E-15 |
| ACE2 | RP11-379F4.8 | -0,3175 | 5,30938E-15 |
| ACE2 | FLJ31104 | -0,3175 | 5,29009E-15 |
| ACE2 | NAALAD2 | -0,3175 | 5,31236E-15 |
| ACE2 | RP11-728F11.3 | -0,3175 | 5,30588E-15 |
| ACE2 | TSPOAP1-AS1 | -0,3175 | 5,29539E-15 |
| ACE2 | PAXBP1 | -0,3175 | 5,29788E-15 |
| ACE2 | TMSB10 | -0,3176 | 5,16779E-15 |
| ACE2 | TDO2 | -0,3176 | 5,22364E-15 |
| ACE2 | DPYS | -0,3176 | 5,14494E-15 |
| ACE2 | RP11-739N20.2 | -0,3177 | 5,11329E-15 |
| ACE2 | LINC01215 | -0,3178 | 4,93441E-15 |
| ACE2 | TIMP2 | -0,3178 | 4,92238E-15 |
| ACE2 | RP11-386I23.1 | -0,3179 | 4,8368E-15 |
| ACE2 | RASGRP3 | -0,3179 | 4,85529E-15 |
| ACE2 | DVL3 | -0,318 | 4,8015E-15 |
| ACE2 | RN7SKP137 | -0,318 | 4,77368E-15 |
| ACE2 | CRIP1 | -0,318 | 4,7321E-15 |
| ACE2 | CTA-363E19.2 | -0,3181 | 4,64806E-15 |
| ACE2 | RP11-386I14.4 | -0,3182 | 4,55079E-15 |
| ACE2 | ZBTB12BP | -0,3182 | 4,5266E-15 |
| ACE2 | RP3-406P24.5 | -0,3183 | 4,48459E-15 |
| ACE2 | FBXO3 | -0,3183 | 4,43999E-15 |
| ACE2 | INTS6 | -0,3183 | 4,48572E-15 |
| ACE2 | ADD2 | -0,3184 | 4,36629E-15 |
| ACE2 | EPHB1 | -0,3184 | 4,38917E-15 |
| ACE2 | PSD3 | -0,3184 | 4,36301E-15 |
| ACE2 | CTB-1144G6.6 | -0,3185 | 4,31682E-15 |
| ACE2 | ULK1 | -0,3185 | 4,30456E-15 |
| ACE2 | HKR1 | -0,3185 | 4,28545E-15 |
| ACE2 | GPR15 | -0,3186 | 4,19177E-15 |
| ACE2 | TRIM38 | -0,3186 | 4,22288E-15 |
| ACE2 | TSSC2 | -0,3186 | 4,23805E-15 |
| ACE2 | C1QTNF5 | -0,3186 | 4,17726E-15 |
| ACE2 | RUSC1 | -0,3187 | 4,09519E-15 |
| ACE2 | ERO1B | -0,3187 | 4,13816E-15 |
| ACE2 | USP42 | -0,3187 | 4,11792E-15 |
| ACE2 | RP11-360N9.2 | -0,3187 | 4,11629E-15 |
| ACE2 | AL590762.11 | -0,3187 | 4,16038E-15 |
| ACE2 | ZDHHC18 | -0,3188 | 4,03286E-15 |
| ACE2 | SKINT1L | -0,3188 | 4,08054E-15 |
| ACE2 | ICOS | -0,3188 | 3,99963E-15 |
| ACE2 | ANXA2R | -0,3188 | 4,02788E-15 |
| ACE2 | RP11-626H12.2 | -0,3189 | 3,98668E-15 |
| ACE2 | TRBJ1-5 | -0,319 | 3,85641E-15 |
| ACE2 | LSP1 | -0,319 | 3,86474E-15 |
| ACE2 | HCST | -0,3191 | 3,75911E-15 |
| ACE2 | AC068196.1 | -0,3192 | 3,68383E-15 |
| ACE2 | KLHL30-AS1 | -0,3192 | 3,71537E-15 |
| ACE2 | RP11-473C18.3 | -0,3192 | 3,72438E-15 |
| ACE2 | SLC12A5 | -0,3192 | 3,71381E-15 |

| | | | |
|---|---|---|---|
| ACE2 | BAG2 | -0,3193 | 3,6233E-15 |
| ACE2 | USP11 | -0,3193 | 3,61799E-15 |
| ACE2 | RPS2P36 | -0,3194 | 3,59484E-15 |
| ACE2 | PRR4 | -0,3195 | 3,48195E-15 |
| ACE2 | FREM1 | -0,3196 | 3,44625E-15 |
| ACE2 | RP11-993B23.3 | -0,3196 | 3,40883E-15 |
| ACE2 | FAT3 | -0,3197 | 3,38461E-15 |
| ACE2 | CXorf40B | -0,3197 | 3,36941E-15 |
| ACE2 | FMOD | -0,3198 | 3,29938E-15 |
| ACE2 | LINC00994 | -0,3198 | 3,27658E-15 |
| ACE2 | PRPSAP1 | -0,3198 | 3,27445E-15 |
| ACE2 | RP11-314N13.9 | -0,3198 | 3,29105E-15 |
| ACE2 | LINC01099 | -0,3199 | 3,24196E-15 |
| ACE2 | TMEM138 | -0,3199 | 3,19185E-15 |
| ACE2 | CNOT9 | -0,32 | 3,1621E-15 |
| ACE2 | SMAD5 | -0,32 | 3,13392E-15 |
| ACE2 | ATP6V1G2 | -0,32 | 3,1525E-15 |
| ACE2 | SLFN11 | -0,3202 | 2,99824E-15 |
| ACE2 | LINC01353 | -0,3204 | 2,88638E-15 |
| ACE2 | CNOT2 | -0,3204 | 2,91126E-15 |
| ACE2 | RFLNA | -0,3204 | 2,91889E-15 |
| ACE2 | FBRSL1 | -0,3204 | 2,89024E-15 |
| ACE2 | TMEM94 | -0,3204 | 2,87288E-15 |
| ACE2 | ARHGAP4 | -0,3204 | 2,86522E-15 |
| ACE2 | ELF2 | -0,3205 | 2,85841E-15 |
| ACE2 | ASPN | -0,3205 | 2,814E-15 |
| ACE2 | PLEKHO2 | -0,3205 | 2,84923E-15 |
| ACE2 | MACF1 | -0,3206 | 2,75806E-15 |
| ACE2 | CAPN5 | -0,3206 | 2,77489E-15 |
| ACE2 | LINC00216 | -0,3206 | 2,79823E-15 |
| ACE2 | RCN1P2 | -0,3207 | 2,71987E-15 |
| ACE2 | PAK3 | -0,3207 | 2,71285E-15 |
| ACE2 | IKZF1 | -0,3208 | 2,68978E-15 |
| ACE2 | TRBV28 | -0,3208 | 2,68074E-15 |
| ACE2 | RP11-753H16.5 | -0,3208 | 2,68026E-15 |
| ACE2 | LINC01686 | -0,3209 | 2,62199E-15 |
| ACE2 | MARCH7 | -0,321 | 2,55869E-15 |
| ACE2 | HIST1H2AC | -0,321 | 2,55384E-15 |
| ACE2 | TBC1D22B | -0,321 | 2,54359E-15 |
| ACE2 | MS4A1 | -0,321 | 2,57206E-15 |
| ACE2 | KRT72 | -0,321 | 2,56265E-15 |
| ACE2 | CTD-2600O9.2 | -0,321 | 2,53139E-15 |
| ACE2 | RP11-423F24.3 | -0,3212 | 2,4389E-15 |
| ACE2 | AC009505.2 | -0,3212 | 2,46158E-15 |
| ACE2 | LRRC70 | -0,3212 | 2,43619E-15 |
| ACE2 | RP11-87H9.4 | -0,3212 | 2,42565E-15 |
| ACE2 | RET | -0,3212 | 2,42788E-15 |
| ACE2 | TSPAN9 | -0,3212 | 2,42552E-15 |
| ACE2 | TMEM63A | -0,3213 | 2,40584E-15 |
| ACE2 | LINC01828 | -0,3213 | 2,41594E-15 |
| ACE2 | HOXA4 | -0,3213 | 2,38433E-15 |
| ACE2 | RP5-1160K1.6 | -0,3214 | 2,34588E-15 |
| ACE2 | CCDC85B | -0,3214 | 2,34728E-15 |
| ACE2 | PCED1B-AS1 | -0,3214 | 2,33565E-15 |

| | | | |
|---|---|---|---|
| ACE2 | RP11-346C20.3 | -0,3214 | 2,33239E-15 |
| ACE2 | CTC-559E9.5 | -0,3214 | 2,36478E-15 |
| ACE2 | RP11-767N6.2 | -0,3215 | 2,30255E-15 |
| ACE2 | AC144449.1 | -0,3215 | 2,32352E-15 |
| ACE2 | RP4-756H11.4 | -0,3215 | 2,31339E-15 |
| ACE2 | STAT5A | -0,3215 | 2,31585E-15 |
| ACE2 | CTC-471J1.2 | -0,3215 | 2,30196E-15 |
| ACE2 | BMP5 | -0,3216 | 2,27143E-15 |
| ACE2 | GNA13 | -0,3216 | 2,23748E-15 |
| ACE2 | SLC45A3 | -0,3217 | 2,22155E-15 |
| ACE2 | SSBP4 | -0,3217 | 2,22444E-15 |
| ACE2 | STUM | -0,3219 | 2,1134E-15 |
| ACE2 | GSTM2 | -0,322 | 2,09059E-15 |
| ACE2 | RP11-573G6.6 | -0,322 | 2,09057E-15 |
| ACE2 | GPR35 | -0,3221 | 2,01745E-15 |
| ACE2 | RP11-478C6.2 | -0,3221 | 2,0192E-15 |
| ACE2 | NRN1 | -0,3222 | 1,97415E-15 |
| ACE2 | RBM12B-AS1 | -0,3222 | 2,00265E-15 |
| ACE2 | GRIP2 | -0,3223 | 1,94459E-15 |
| ACE2 | TFP1 | -0,3223 | 1,94284E-15 |
| ACE2 | HOXA9 | -0,3223 | 1,96192E-15 |
| ACE2 | WHAMM | -0,3223 | 1,93965E-15 |
| ACE2 | ZNF814 | -0,3223 | 1,94333E-15 |
| ACE2 | PSMB8-AS1 | -0,3224 | 1,91784E-15 |
| ACE2 | RN7SKP110 | -0,3225 | 1,85876E-15 |
| ACE2 | SLC16A14 | -0,3226 | 1,8328E-15 |
| ACE2 | ARHGEF3 | -0,3226 | 1,81575E-15 |
| ACE2 | CXCL12 | -0,3226 | 1,83263E-15 |
| ACE2 | C2CD4A | -0,3226 | 1,82494E-15 |
| ACE2 | MCOLN3 | -0,3227 | 1,77603E-15 |
| ACE2 | ANO7 | -0,3228 | 1,73888E-15 |
| ACE2 | HECA | -0,3228 | 1,74173E-15 |
| ACE2 | RP1-101K10.6 | -0,3228 | 1,75804E-15 |
| ACE2 | TRBV6-5 | -0,3228 | 1,74013E-15 |
| ACE2 | RP11-674P19.2 | -0,3228 | 1,74937E-15 |
| ACE2 | RP11-158K1.3 | -0,3229 | 1,72821E-15 |
| ACE2 | PRF1 | -0,3229 | 1,70945E-15 |
| ACE2 | CORO7 | -0,323 | 1,67017E-15 |
| ACE2 | LINC02126 | -0,323 | 1,69396E-15 |
| ACE2 | CCDC3 | -0,3231 | 1,63007E-15 |
| ACE2 | IL4I1 | -0,3232 | 1,61481E-15 |
| ACE2 | CNOT6L | -0,3233 | 1,57198E-15 |
| ACE2 | SLC6A7 | -0,3233 | 1,58208E-15 |
| ACE2 | FLOT1 | -0,3233 | 1,56856E-15 |
| ACE2 | FZR1 | -0,3233 | 1,57439E-15 |
| ACE2 | RP11-449G16.1 | -0,3234 | 1,5514E-15 |
| ACE2 | AC005754.8 | -0,3234 | 1,55028E-15 |
| ACE2 | LY6E | -0,3234 | 1,54866E-15 |
| ACE2 | ZNF654 | -0,3235 | 1,5259E-15 |
| ACE2 | RASSF3 | -0,3235 | 1,52429E-15 |
| ACE2 | SNURF | -0,3235 | 1,51726E-15 |
| ACE2 | NAXD | -0,3236 | 1,46569E-15 |
| ACE2 | LTBP3 | -0,3237 | 1,44764E-15 |
| ACE2 | SYCP2L | -0,3238 | 1,42235E-15 |

| | | | |
|---|---|---|---|
| ACE2 | PDGFRB | -0,3239 | 1,38387E-15 |
| ACE2 | KIF7 | -0,3239 | 1,37389E-15 |
| ACE2 | RP11-583F2.5 | -0,3239 | 1,37962E-15 |
| ACE2 | SMCHD1 | -0,3239 | 1,39155E-15 |
| ACE2 | XXyac-YX65C7_A.2 | -0,324 | 1,35612E-15 |
| ACE2 | CDH5 | -0,324 | 1,35503E-15 |
| ACE2 | JUND | -0,3241 | 1,33419E-15 |
| ACE2 | TRBV7-2 | -0,3242 | 1,31396E-15 |
| ACE2 | FAM122C | -0,3242 | 1,30313E-15 |
| ACE2 | AC009404.2 | -0,3243 | 1,27292E-15 |
| ACE2 | RP11-339B21.13 | -0,3243 | 1,27276E-15 |
| ACE2 | SACS | -0,3243 | 1,27604E-15 |
| ACE2 | PPP1R15A | -0,3243 | 1,27051E-15 |
| ACE2 | PCDHB7 | -0,3244 | 1,23936E-15 |
| ACE2 | GCK | -0,3246 | 1,19776E-15 |
| ACE2 | FBXO44 | -0,3247 | 1,16086E-15 |
| ACE2 | PLSCR4 | -0,3247 | 1,17329E-15 |
| ACE2 | RGS4 | -0,3248 | 1,13642E-15 |
| ACE2 | LRP4 | -0,3249 | 1,11147E-15 |
| ACE2 | KCNN4 | -0,3249 | 1,11379E-15 |
| ACE2 | RP11-373D23.3 | -0,325 | 1,09769E-15 |
| ACE2 | THSD7B | -0,3251 | 1,07019E-15 |
| ACE2 | ITGA1 | -0,3251 | 1,071E-15 |
| ACE2 | MAST4 | -0,3251 | 1,06863E-15 |
| ACE2 | AKNA | -0,3251 | 1,07299E-15 |
| ACE2 | RP11-527N22.1 | -0,3252 | 1,04533E-15 |
| ACE2 | NRP1 | -0,3252 | 1,0613E-15 |
| ACE2 | TBX2 | -0,3253 | 1,03585E-15 |
| ACE2 | ZNF90P1 | -0,3254 | 1,02053E-15 |
| ACE2 | ZFYVE27 | -0,3254 | 1,01119E-15 |
| ACE2 | RNU6-343P | -0,3254 | 1,01343E-15 |
| ACE2 | ZEB1-AS1 | -0,3255 | 9,9012E-16 |
| ACE2 | AC005363.9 | -0,3255 | 9,92169E-16 |
| ACE2 | RUBCN | -0,3256 | 9,61066E-16 |
| ACE2 | PLEKHH3 | -0,3256 | 9,64317E-16 |
| ACE2 | SNX9 | -0,3257 | 9,43771E-16 |
| ACE2 | HAS2-AS1 | -0,3257 | 9,38303E-16 |
| ACE2 | GPR182 | -0,3257 | 9,52338E-16 |
| ACE2 | RP11-319G9.3 | -0,3257 | 9,46221E-16 |
| ACE2 | AC004540.4 | -0,3258 | 9,3148E-16 |
| ACE2 | RP3-468O1.6 | -0,3258 | 9,22416E-16 |
| ACE2 | RP11-365O16.6 | -0,3259 | 9,11309E-16 |
| ACE2 | RP11-68I3.11 | -0,3259 | 9,04837E-16 |
| ACE2 | MAPK11 | -0,3259 | 9,04761E-16 |
| ACE2 | CXorf65 | -0,3259 | 9,10291E-16 |
| ACE2 | NAV1 | -0,326 | 8,92327E-16 |
| ACE2 | TRBJ1-1 | -0,326 | 8,88521E-16 |
| ACE2 | RP11-45A17.2 | -0,326 | 8,87683E-16 |
| ACE2 | RP11-977G19.5 | -0,326 | 8,8304E-16 |
| ACE2 | DNAJA4 | -0,326 | 8,8441E-16 |
| ACE2 | RP11-818O24.2 | -0,326 | 8,88285E-16 |
| ACE2 | TRBV2 | -0,3261 | 8,71255E-16 |
| ACE2 | DRD4 | -0,3261 | 8,70514E-16 |
| ACE2 | SV2B | -0,3261 | 8,69572E-16 |

| | | | |
|---|---|---|---|
| ACE2 | CDH11 | -0,3262 | 8,50064E-16 |
| ACE2 | RN7SL861P | -0,3263 | 8,36596E-16 |
| ACE2 | PCGF3 | -0,3263 | 8,38762E-16 |
| ACE2 | HLA-E | -0,3263 | 8,41101E-16 |
| ACE2 | PSTPIP1 | -0,3263 | 8,30712E-16 |
| ACE2 | DDHD1 | -0,3264 | 8,21737E-16 |
| ACE2 | CD300LB | -0,3264 | 8,09252E-16 |
| ACE2 | CSNK1A1 | -0,3265 | 7,95537E-16 |
| ACE2 | RP5-1007F24.1 | -0,3265 | 8,06566E-16 |
| ACE2 | STON2 | -0,3265 | 7,92522E-16 |
| ACE2 | PRICKLE2 | -0,3266 | 7,89614E-16 |
| ACE2 | GLIS2 | -0,3267 | 7,63514E-16 |
| ACE2 | CPNE1 | -0,3267 | 7,63723E-16 |
| ACE2 | RP11-167N5.5 | -0,3268 | 7,53956E-16 |
| ACE2 | IRGQ | -0,3268 | 7,47978E-16 |
| ACE2 | TRBV29-1 | -0,3269 | 7,29175E-16 |
| ACE2 | FHOD1 | -0,3269 | 7,26515E-16 |
| ACE2 | SERPINI1 | -0,327 | 7,22062E-16 |
| ACE2 | CELF1 | -0,327 | 7,24847E-16 |
| ACE2 | BORCS5 | -0,327 | 7,11318E-16 |
| ACE2 | RANP4 | -0,327 | 7,10608E-16 |
| ACE2 | RP11-106D4.2 | -0,3271 | 6,97565E-16 |
| ACE2 | PRDX3P4 | -0,3273 | 6,73757E-16 |
| ACE2 | TAGAP | -0,3273 | 6,73386E-16 |
| ACE2 | PLXNA3 | -0,3273 | 6,6638E-16 |
| ACE2 | C2orf40 | -0,3274 | 6,606E-16 |
| ACE2 | ATG16L2 | -0,3274 | 6,62782E-16 |
| ACE2 | AD001527.7 | -0,3274 | 6,61462E-16 |
| ACE2 | LTBP4 | -0,3274 | 6,60847E-16 |
| ACE2 | EQTN | -0,3275 | 6,46049E-16 |
| ACE2 | CTA-984G1.5 | -0,3276 | 6,35501E-16 |
| ACE2 | HOXA-AS2 | -0,3277 | 6,15911E-16 |
| ACE2 | TRAV2 | -0,3277 | 6,11911E-16 |
| ACE2 | CTC-524C5.2 | -0,3277 | 6,16897E-16 |
| ACE2 | RNASEK | -0,3277 | 6,20895E-16 |
| ACE2 | CHRNA4 | -0,3278 | 6,08313E-16 |
| ACE2 | PTTG2 | -0,328 | 5,7369E-16 |
| ACE2 | PITPNC1 | -0,328 | 5,76162E-16 |
| ACE2 | CYTH4 | -0,328 | 5,83415E-16 |
| ACE2 | CYP2D7 | -0,328 | 5,79188E-16 |
| ACE2 | RP13-228J13.8 | -0,328 | 5,78003E-16 |
| ACE2 | C1orf228 | -0,3282 | 5,54078E-16 |
| ACE2 | TMPRSS6 | -0,3282 | 5,54096E-16 |
| ACE2 | KLF12 | -0,3283 | 5,48786E-16 |
| ACE2 | NPM1P37 | -0,3284 | 5,28743E-16 |
| ACE2 | RP11-138A9.2 | -0,3285 | 5,19887E-16 |
| ACE2 | GABARAPL1 | -0,3285 | 5,26042E-16 |
| ACE2 | CTD-2341M24.1 | -0,3285 | 5,15554E-16 |
| ACE2 | PSMD6-AS1 | -0,3286 | 5,12208E-16 |
| ACE2 | CHSY1 | -0,3286 | 5,08246E-16 |
| ACE2 | STAB2 | -0,3287 | 4,93659E-16 |
| ACE2 | CADPS | -0,3288 | 4,83079E-16 |
| ACE2 | BTN3A3 | -0,3288 | 4,8789E-16 |
| ACE2 | RAB3A | -0,3288 | 4,88535E-16 |

| | | | |
|---|---|---|---|
| ACE2 | NRBF2P5 | -0,3289 | 4,74637E-16 |
| ACE2 | C15orf53 | -0,3289 | 4,79923E-16 |
| ACE2 | PLAUR | -0,3289 | 4,77646E-16 |
| ACE2 | HNRNPCP1 | -0,329 | 4,69907E-16 |
| ACE2 | RP11-114H24.6 | -0,329 | 4,64303E-16 |
| ACE2 | FRAS1 | -0,3291 | 4,54447E-16 |
| ACE2 | WNT10B | -0,3291 | 4,59566E-16 |
| ACE2 | ALPK3 | -0,3291 | 4,61895E-16 |
| ACE2 | PLEKHG2 | -0,3291 | 4,56777E-16 |
| ACE2 | AP001471.1 | -0,3291 | 4,61194E-16 |
| ACE2 | ODF3B | -0,3291 | 4,53581E-16 |
| ACE2 | SRGAP2C | -0,3292 | 4,48097E-16 |
| ACE2 | STARD4-AS1 | -0,3292 | 4,4655E-16 |
| ACE2 | ARAP3 | -0,3292 | 4,48611E-16 |
| ACE2 | MEX3C | -0,3292 | 4,4339E-16 |
| ACE2 | C16orf72 | -0,3293 | 4,42685E-16 |
| ACE2 | C1orf147 | -0,3294 | 4,31151E-16 |
| ACE2 | RBM5 | -0,3294 | 4,26781E-16 |
| ACE2 | RP11-286H14.8 | -0,3294 | 4,32038E-16 |
| ACE2 | TMEM156 | -0,3295 | 4,16852E-16 |
| ACE2 | NSD3 | -0,3295 | 4,20064E-16 |
| ACE2 | CYP2E1 | -0,3295 | 4,1642E-16 |
| ACE2 | PFDN5 | -0,3295 | 4,20117E-16 |
| ACE2 | PRR5 | -0,3295 | 4,22549E-16 |
| ACE2 | BTN3A1 | -0,3296 | 4,11812E-16 |
| ACE2 | CENPP | -0,3296 | 4,14419E-16 |
| ACE2 | SHANK1 | -0,3296 | 4,12528E-16 |
| ACE2 | SV2A | -0,3297 | 4,05504E-16 |
| ACE2 | SPRY1 | -0,3297 | 4,05798E-16 |
| ACE2 | RNY4P10 | -0,3297 | 4,0552E-16 |
| ACE2 | MCAM | -0,3297 | 4,04563E-16 |
| ACE2 | INHBA | -0,3298 | 3,96743E-16 |
| ACE2 | CBL | -0,3298 | 3,91173E-16 |
| ACE2 | RPL23AP32 | -0,3299 | 3,8582E-16 |
| ACE2 | ZHX2 | -0,3299 | 3,84077E-16 |
| ACE2 | PTPRD-AS1 | -0,3299 | 3,88267E-16 |
| ACE2 | KMT2A | -0,3299 | 3,81366E-16 |
| ACE2 | CTD-2537I9.12 | -0,3299 | 3,82229E-16 |
| ACE2 | MX2 | -0,3299 | 3,85608E-16 |
| ACE2 | ARID2 | -0,3301 | 3,68946E-16 |
| ACE2 | LINC01473 | -0,3302 | 3,59374E-16 |
| ACE2 | RP4-730K3.3 | -0,3303 | 3,51532E-16 |
| ACE2 | RP11-333I13.1 | -0,3303 | 3,51644E-16 |
| ACE2 | CTD-2235C13.1 | -0,3303 | 3,56237E-16 |
| ACE2 | KIAA1211 | -0,3304 | 3,44567E-16 |
| ACE2 | BEST1 | -0,3305 | 3,35117E-16 |
| ACE2 | MARK4 | -0,3305 | 3,36664E-16 |
| ACE2 | FCRL3 | -0,3306 | 3,28405E-16 |
| ACE2 | GZMA | -0,3306 | 3,28325E-16 |
| ACE2 | RP11-155G14.5 | -0,3307 | 3,25634E-16 |
| ACE2 | PAWR | -0,3307 | 3,23025E-16 |
| ACE2 | CTD-3032J10.3 | -0,3307 | 3,23558E-16 |
| ACE2 | RP11-503E24.2 | -0,3308 | 3,15379E-16 |
| ACE2 | ME3 | -0,3308 | 3,1541E-16 |

| | | | |
|---|---|---|---|
| ACE2 | PBX4 | -0,3308 | 3,17092E-16 |
| ACE2 | ITPKB | -0,3311 | 2,95273E-16 |
| ACE2 | AC106801.1 | -0,3311 | 2,95778E-16 |
| ACE2 | TEAD3 | -0,3311 | 2,96839E-16 |
| ACE2 | KLRD1 | -0,3311 | 2,94863E-16 |
| ACE2 | FGF14 | -0,3311 | 2,99639E-16 |
| ACE2 | RP11-79H23.3 | -0,3312 | 2,91012E-16 |
| ACE2 | SNORA23 | -0,3312 | 2,89554E-16 |
| ACE2 | KANK4 | -0,3313 | 2,85912E-16 |
| ACE2 | FAM171B | -0,3313 | 2,83645E-16 |
| ACE2 | THEMIS | -0,3313 | 2,85718E-16 |
| ACE2 | CST7 | -0,3313 | 2,82121E-16 |
| ACE2 | SOS1-IT1 | -0,3314 | 2,77844E-16 |
| ACE2 | RP11-329B9.4 | -0,3314 | 2,79643E-16 |
| ACE2 | PRSS42 | -0,3315 | 2,74956E-16 |
| ACE2 | PDE1C | -0,3315 | 2,73806E-16 |
| ACE2 | LINC00535 | -0,3316 | 2,66175E-16 |
| ACE2 | HNRNPH3 | -0,3316 | 2,68827E-16 |
| ACE2 | HIF1A-AS2 | -0,3316 | 2,66587E-16 |
| ACE2 | AOC4P | -0,3318 | 2,54994E-16 |
| ACE2 | TBC1D10B | -0,3319 | 2,5069E-16 |
| ACE2 | ASPA | -0,3319 | 2,50874E-16 |
| ACE2 | MMRN2 | -0,3321 | 2,36456E-16 |
| ACE2 | MYO15B | -0,3321 | 2,38531E-16 |
| ACE2 | PNISR | -0,3322 | 2,35686E-16 |
| ACE2 | TESPA1 | -0,3322 | 2,34561E-16 |
| ACE2 | LDHAL6B | -0,3322 | 2,34967E-16 |
| ACE2 | GRIPAP1 | -0,3322 | 2,31511E-16 |
| ACE2 | RP1-292B18.4 | -0,3323 | 2,28737E-16 |
| ACE2 | SSUH2 | -0,3325 | 2,18453E-16 |
| ACE2 | TRAT1 | -0,3327 | 2,08477E-16 |
| ACE2 | ABCA2 | -0,3327 | 2,11105E-16 |
| ACE2 | MYOZ3 | -0,3328 | 2,02694E-16 |
| ACE2 | RN7SL748P | -0,3328 | 2,06036E-16 |
| ACE2 | LILRB1 | -0,3329 | 1,98639E-16 |
| ACE2 | GDF10 | -0,333 | 1,97126E-16 |
| ACE2 | HSP90AB4P | -0,333 | 1,98079E-16 |
| ACE2 | SDC3 | -0,3331 | 1,92266E-16 |
| ACE2 | DCUN1D2-AS | -0,3331 | 1,9034E-16 |
| ACE2 | RP11-159D12.6 | -0,3331 | 1,93006E-16 |
| ACE2 | PPEF2 | -0,3332 | 1,88349E-16 |
| ACE2 | RGS17P1 | -0,3332 | 1,88885E-16 |
| ACE2 | TRBJ2-7 | -0,3333 | 1,8215E-16 |
| ACE2 | PDZD7 | -0,3333 | 1,83767E-16 |
| ACE2 | CX3CL1 | -0,3333 | 1,82615E-16 |
| ACE2 | ZC3H11A | -0,3334 | 1,80757E-16 |
| ACE2 | OSTN | -0,3334 | 1,79437E-16 |
| ACE2 | RP1-69D17.3 | -0,3335 | 1,761E-16 |
| ACE2 | NFE2L3 | -0,3336 | 1,71955E-16 |
| ACE2 | LA16c-390H2.4 | -0,3336 | 1,71739E-16 |
| ACE2 | ARMCX3 | -0,3336 | 1,70466E-16 |
| ACE2 | RLF | -0,3338 | 1,64045E-16 |
| ACE2 | NFYA | -0,3338 | 1,62923E-16 |
| ACE2 | TMEM255B | -0,3338 | 1,63422E-16 |

| | | | |
|---|---|---|---|
| ACE2 | RP11-270M14.5 | -0,3338 | 1,63993E-16 |
| ACE2 | RP11-424I19.2 | -0,3338 | 1,6298E-16 |
| ACE2 | LDHAP2 | -0,3339 | 1,6192E-16 |
| ACE2 | LINC01943 | -0,3339 | 1,61841E-16 |
| ACE2 | AC053503.6 | -0,3339 | 1,60398E-16 |
| ACE2 | IFNAR2 | -0,3339 | 1,60635E-16 |
| ACE2 | APOBEC3G | -0,3339 | 1,61929E-16 |
| ACE2 | RP11-111J6.2 | -0,334 | 1,58409E-16 |
| ACE2 | HMBOX1 | -0,334 | 1,59172E-16 |
| ACE2 | RP11-551L14.8 | -0,3341 | 1,54154E-16 |
| ACE2 | PCDHGC5 | -0,3342 | 1,49452E-16 |
| ACE2 | BCL2 | -0,3342 | 1,49731E-16 |
| ACE2 | CLEC17A | -0,3342 | 1,51829E-16 |
| ACE2 | INPP5E | -0,3343 | 1,48906E-16 |
| ACE2 | CTA-390C10.10 | -0,3343 | 1,46815E-16 |
| ACE2 | FGF18 | -0,3344 | 1,43456E-16 |
| ACE2 | C12orf75 | -0,3344 | 1,43776E-16 |
| ACE2 | RP1-102K2.8 | -0,3344 | 1,42969E-16 |
| ACE2 | LRRC4C | -0,3345 | 1,39753E-16 |
| ACE2 | STEAP2-AS1 | -0,3346 | 1,39577E-16 |
| ACE2 | CTB-31N19.3 | -0,3346 | 1,36972E-16 |
| ACE2 | ZNF516 | -0,3346 | 1,39189E-16 |
| ACE2 | IKZF2 | -0,3347 | 1,34736E-16 |
| ACE2 | LYSMD2 | -0,3347 | 1,36187E-16 |
| ACE2 | SH3BP1 | -0,3347 | 1,35823E-16 |
| ACE2 | MXRA8 | -0,3349 | 1,30295E-16 |
| ACE2 | RP3-426I6.5 | -0,3349 | 1,29459E-16 |
| ACE2 | HIC1 | -0,3349 | 1,28907E-16 |
| ACE2 | LYPD3 | -0,3349 | 1,28202E-16 |
| ACE2 | LINC00632 | -0,335 | 1,25767E-16 |
| ACE2 | HABP4 | -0,3352 | 1,22318E-16 |
| ACE2 | CH25H | -0,3352 | 1,20748E-16 |
| ACE2 | NCR1 | -0,3352 | 1,214E-16 |
| ACE2 | PTGES3P2 | -0,3353 | 1,189E-16 |
| ACE2 | RPL23AP1 | -0,3353 | 1,18243E-16 |
| ACE2 | MPP2 | -0,3353 | 1,17113E-16 |
| ACE2 | DYNLT3 | -0,3353 | 1,1874E-16 |
| ACE2 | TARDBP | -0,3354 | 1,15598E-16 |
| ACE2 | RP11-274B21.12 | -0,3354 | 1,15525E-16 |
| ACE2 | PCDH17 | -0,3354 | 1,15352E-16 |
| ACE2 | RP11-151N17.1 | -0,3354 | 1,14689E-16 |
| ACE2 | RASD2 | -0,3354 | 1,14663E-16 |
| ACE2 | RASA4B | -0,3355 | 1,14377E-16 |
| ACE2 | HOMER3 | -0,3355 | 1,13927E-16 |
| ACE2 | KCNMB2 | -0,3356 | 1,10193E-16 |
| ACE2 | CMA1 | -0,3356 | 1,1002E-16 |
| ACE2 | EBLN2 | -0,3358 | 1,05746E-16 |
| ACE2 | SUB1 | -0,3358 | 1,04944E-16 |
| ACE2 | PELO | -0,3358 | 1,04999E-16 |
| ACE2 | CALY | -0,3358 | 1,06603E-16 |
| ACE2 | IL4 | -0,3359 | 1,04182E-16 |
| ACE2 | TRBV27 | -0,3359 | 1,0417E-16 |
| ACE2 | PALM2 | -0,3359 | 1,03442E-16 |
| ACE2 | MRGPRF | -0,3359 | 1,03474E-16 |

| | | | |
|---|---|---|---|
| ACE2 | TMC6 | -0,3359 | 1,02912E-16 |
| ACE2 | RBMS1 | -0,336 | 1,01069E-16 |
| ACE2 | PRPH2 | -0,336 | 1,01389E-16 |
| ACE2 | RP11-73O6.3 | -0,3361 | 9,91111E-17 |
| ACE2 | CCL21 | -0,3361 | 9,92565E-17 |
| ACE2 | STK38L | -0,3361 | 9,95403E-17 |
| ACE2 | MIB2 | -0,3362 | 9,69896E-17 |
| ACE2 | USP20 | -0,3363 | 9,40505E-17 |
| ACE2 | POU5F1P5 | -0,3363 | 9,48603E-17 |
| ACE2 | TPM4 | -0,3363 | 9,50381E-17 |
| ACE2 | CORO2B | -0,3364 | 9,18318E-17 |
| ACE2 | TRBV6-2 | -0,3365 | 9,02109E-17 |
| ACE2 | WARS | -0,3365 | 9,03301E-17 |
| ACE2 | RP3-426I6.6 | -0,3366 | 8,87436E-17 |
| ACE2 | EFCC1 | -0,3366 | 8,82316E-17 |
| ACE2 | AC010226.4 | -0,3367 | 8,59658E-17 |
| ACE2 | FGF11 | -0,3369 | 8,31799E-17 |
| ACE2 | NBPF15 | -0,3371 | 7,90033E-17 |
| ACE2 | TAS2R19 | -0,3371 | 8,01041E-17 |
| ACE2 | RP11-63M22.1 | -0,3372 | 7,85621E-17 |
| ACE2 | MYCBP2-AS1 | -0,3373 | 7,53058E-17 |
| ACE2 | SPIB | -0,3373 | 7,64362E-17 |
| ACE2 | RP11-196H14.4 | -0,3374 | 7,48836E-17 |
| ACE2 | NUMB | -0,3374 | 7,45885E-17 |
| ACE2 | SELENOO | -0,3374 | 7,40096E-17 |
| ACE2 | PRRG3 | -0,3374 | 7,45231E-17 |
| ACE2 | GYPB | -0,3375 | 7,21189E-17 |
| ACE2 | RCAN1 | -0,3375 | 7,27354E-17 |
| ACE2 | L3MBTL3 | -0,3376 | 7,12148E-17 |
| ACE2 | AKAP8L | -0,3376 | 7,06805E-17 |
| ACE2 | FAM219A | -0,3377 | 6,93273E-17 |
| ACE2 | BMS1P3 | -0,3377 | 7,00585E-17 |
| ACE2 | RHOT2 | -0,3377 | 6,97454E-17 |
| ACE2 | PCDHB9 | -0,3378 | 6,83481E-17 |
| ACE2 | RDH16 | -0,3378 | 6,87995E-17 |
| ACE2 | RSRC2 | -0,338 | 6,57631E-17 |
| ACE2 | RP11-342K2.1 | -0,338 | 6,54392E-17 |
| ACE2 | RP11-225N10.3 | -0,3381 | 6,30497E-17 |
| ACE2 | TMSB4XP4 | -0,3381 | 6,41855E-17 |
| ACE2 | SLC5A10 | -0,3381 | 6,3114E-17 |
| ACE2 | SBNO2 | -0,3381 | 6,31149E-17 |
| ACE2 | LINC01700 | -0,3382 | 6,23331E-17 |
| ACE2 | RP11-317J9.1 | -0,3383 | 6,15029E-17 |
| ACE2 | LINC01266 | -0,3384 | 5,95292E-17 |
| ACE2 | AC024560.3 | -0,3384 | 5,99636E-17 |
| ACE2 | RP11-179A10.1 | -0,3384 | 5,88805E-17 |
| ACE2 | ZMYM2 | -0,3385 | 5,82472E-17 |
| ACE2 | MPZ | -0,3386 | 5,64369E-17 |
| ACE2 | IQSEC1 | -0,3386 | 5,75651E-17 |
| ACE2 | STARD9 | -0,3386 | 5,66736E-17 |
| ACE2 | SLC4A7 | -0,3387 | 5,5424E-17 |
| ACE2 | RP11-194N12.2 | -0,3387 | 5,57311E-17 |
| ACE2 | MFSD4A | -0,3388 | 5,39621E-17 |
| ACE2 | GADL1 | -0,3388 | 5,39683E-17 |

| | | | |
|---|---|---|---|
| ACE2 | DGAT1 | -0,3388 | 5,39441E-17 |
| ACE2 | KCNT1 | -0,3388 | 5,45914E-17 |
| ACE2 | PTPRG-AS1 | -0,3389 | 5,27283E-17 |
| ACE2 | TMEM71 | -0,3389 | 5,30564E-17 |
| ACE2 | SLCO1C1 | -0,3389 | 5,35162E-17 |
| ACE2 | ARFGAP1 | -0,3389 | 5,31059E-17 |
| ACE2 | CTC-428H11.2 | -0,339 | 5,24234E-17 |
| ACE2 | PPP1R2P1 | -0,339 | 5,19345E-17 |
| ACE2 | RP11-51J9.4 | -0,339 | 5,25655E-17 |
| ACE2 | RP11-635L1.2 | -0,3392 | 5,01707E-17 |
| ACE2 | TRPM4 | -0,3392 | 5,01492E-17 |
| ACE2 | MLC1 | -0,3392 | 5,02786E-17 |
| ACE2 | RP11-247A12.7 | -0,3393 | 4,86181E-17 |
| ACE2 | RP11-1H15.1 | -0,3393 | 4,87608E-17 |
| ACE2 | HSPE1P3 | -0,3393 | 4,89433E-17 |
| ACE2 | PITPNB | -0,3393 | 4,89946E-17 |
| ACE2 | TRBV3-1 | -0,3395 | 4,66376E-17 |
| ACE2 | HIST2H2AA3 | -0,3396 | 4,5352E-17 |
| ACE2 | RP11-478C6.6 | -0,3396 | 4,53929E-17 |
| ACE2 | R3HDM4 | -0,3396 | 4,50453E-17 |
| ACE2 | CDH24 | -0,3398 | 4,37019E-17 |
| ACE2 | TERF2IP | -0,3398 | 4,32179E-17 |
| ACE2 | RP11-423P10.2 | -0,3399 | 4,23342E-17 |
| ACE2 | COL23A1 | -0,3399 | 4,2131E-17 |
| ACE2 | CNOT4 | -0,3399 | 4,29091E-17 |
| ACE2 | RP11-278C7.4 | -0,3399 | 4,22507E-17 |
| ACE2 | OR5AK4P | -0,34 | 4,19264E-17 |
| ACE2 | ING1 | -0,34 | 4,12257E-17 |
| ACE2 | AC137932.4 | -0,34 | 4,20371E-17 |
| ACE2 | CA5B | -0,34 | 4,17912E-17 |
| ACE2 | RXRB | -0,3401 | 4,11541E-17 |
| ACE2 | MEX3D | -0,3401 | 4,07305E-17 |
| ACE2 | TMEM240 | -0,3402 | 3,9869E-17 |
| ACE2 | ZEB2 | -0,3402 | 4,01227E-17 |
| ACE2 | NPIPB4 | -0,3402 | 3,9561E-17 |
| ACE2 | VNN3 | -0,3403 | 3,93445E-17 |
| ACE2 | APLNR | -0,3403 | 3,87665E-17 |
| ACE2 | RP11-471B22.2 | -0,3403 | 3,86758E-17 |
| ACE2 | SEPT9 | -0,3403 | 3,8497E-17 |
| ACE2 | OLFM2 | -0,3403 | 3,87236E-17 |
| ACE2 | UBE2R2 | -0,3405 | 3,73286E-17 |
| ACE2 | CTSG | -0,3405 | 3,67896E-17 |
| ACE2 | RP11-384M20.1 | -0,3405 | 3,75786E-17 |
| ACE2 | PDE9A | -0,3405 | 3,70929E-17 |
| ACE2 | RP11-1084A12.1 | -0,3406 | 3,67343E-17 |
| ACE2 | PTOV1 | -0,3406 | 3,61838E-17 |
| ACE2 | C4orf3 | -0,3407 | 3,52819E-17 |
| ACE2 | RP11-135N5.3 | -0,3407 | 3,58245E-17 |
| ACE2 | ADAMTS10 | -0,3407 | 3,54896E-17 |
| ACE2 | C11orf96 | -0,3408 | 3,48817E-17 |
| ACE2 | GPR153 | -0,3409 | 3,39687E-17 |
| ACE2 | AHDC1 | -0,3409 | 3,40628E-17 |
| ACE2 | IL15 | -0,3409 | 3,40235E-17 |
| ACE2 | PEA15 | -0,341 | 3,29271E-17 |

| | | | |
|---|---|---|---|
| ACE2 | RP11-1029F8.1 | -0,341 | 3,32916E-17 |
| ACE2 | ARRDC4 | -0,341 | 3,31134E-17 |
| ACE2 | HNRNPM | -0,341 | 3,33435E-17 |
| ACE2 | BACH1-IT1 | -0,341 | 3,30923E-17 |
| ACE2 | PLCG2 | -0,3411 | 3,21626E-17 |
| ACE2 | GFI1 | -0,3413 | 3,0939E-17 |
| ACE2 | TRBV11-2 | -0,3413 | 3,09172E-17 |
| ACE2 | RP11-277P12.20 | -0,3413 | 3,11561E-17 |
| ACE2 | RP11-840I19.3 | -0,3413 | 3,12956E-17 |
| ACE2 | RP11-290C10.1 | -0,3414 | 3,02751E-17 |
| ACE2 | RP11-248J18.2 | -0,3414 | 3,00526E-17 |
| ACE2 | RP3-508I15.20 | -0,3414 | 3,05547E-17 |
| ACE2 | RASGRF2-AS1 | -0,3415 | 2,98106E-17 |
| ACE2 | AC063976.7 | -0,3415 | 2,95356E-17 |
| ACE2 | DGKB | -0,3415 | 2,9722E-17 |
| ACE2 | SNORA3B | -0,3416 | 2,88529E-17 |
| ACE2 | RP11-345J18.2 | -0,3416 | 2,89496E-17 |
| ACE2 | PATL1 | -0,3417 | 2,82422E-17 |
| ACE2 | AE000658.31 | -0,3417 | 2,81926E-17 |
| ACE2 | PHF8 | -0,3417 | 2,85645E-17 |
| ACE2 | AC067945.3 | -0,3418 | 2,78954E-17 |
| ACE2 | RP11-88H12.2 | -0,3418 | 2,76017E-17 |
| ACE2 | FBXL7 | -0,3418 | 2,7982E-17 |
| ACE2 | AK5 | -0,3419 | 2,71179E-17 |
| ACE2 | CTD-3128G10.7 | -0,3419 | 2,71407E-17 |
| ACE2 | ANKLE1 | -0,3419 | 2,7223E-17 |
| ACE2 | LMF2 | -0,3419 | 2,71077E-17 |
| ACE2 | MXRA5 | -0,3419 | 2,73089E-17 |
| ACE2 | ZFP36 | -0,342 | 2,67314E-17 |
| ACE2 | SPOPL | -0,3421 | 2,5732E-17 |
| ACE2 | PKMP3 | -0,3421 | 2,61521E-17 |
| ACE2 | NACAD | -0,3421 | 2,58371E-17 |
| ACE2 | TMEM14EP | -0,3422 | 2,53646E-17 |
| ACE2 | RPS18P9 | -0,3422 | 2,55241E-17 |
| ACE2 | ARHGEF39 | -0,3422 | 2,51213E-17 |
| ACE2 | AC090804.1 | -0,3423 | 2,50661E-17 |
| ACE2 | RP11-713P17.3 | -0,3423 | 2,49698E-17 |
| ACE2 | RP11-322D14.2 | -0,3423 | 2,45552E-17 |
| ACE2 | AP004290.1 | -0,3424 | 2,40367E-17 |
| ACE2 | PDE5A | -0,3425 | 2,37564E-17 |
| ACE2 | CTC-301O7.4 | -0,3425 | 2,391E-17 |
| ACE2 | HIST1H2AI | -0,3426 | 2,32941E-17 |
| ACE2 | RP5-1112D6.8 | -0,3426 | 2,29436E-17 |
| ACE2 | RPLP0P2 | -0,3426 | 2,30609E-17 |
| ACE2 | RP13-487P22.1 | -0,3426 | 2,30679E-17 |
| ACE2 | LINC01068 | -0,3427 | 2,28008E-17 |
| ACE2 | RASA4 | -0,3428 | 2,22498E-17 |
| ACE2 | ATXN2 | -0,3428 | 2,22932E-17 |
| ACE2 | ATXN2L | -0,3428 | 2,23671E-17 |
| ACE2 | SNTB2 | -0,3428 | 2,22365E-17 |
| ACE2 | GRAP2 | -0,3428 | 2,20247E-17 |
| ACE2 | RP11-308D16.2 | -0,3428 | 2,21483E-17 |
| ACE2 | BHLHE22 | -0,343 | 2,09504E-17 |
| ACE2 | FKBP8 | -0,343 | 2,10696E-17 |

| | | | |
|---|---|---|---|
| ACE2 | LRRC15 | -0,3431 | 2,048E-17 |
| ACE2 | RP1-292B18.1 | -0,3431 | 2,05318E-17 |
| ACE2 | KLRC3 | -0,3431 | 2,07209E-17 |
| ACE2 | ASB2 | -0,3431 | 2,05282E-17 |
| ACE2 | ADAMTS7P3 | -0,3431 | 2,04571E-17 |
| ACE2 | APLN | -0,3431 | 2,06977E-17 |
| ACE2 | ILK | -0,3432 | 1,99967E-17 |
| ACE2 | PRDM10 | -0,3433 | 1,97589E-17 |
| ACE2 | AKAP5 | -0,3433 | 1,95831E-17 |
| ACE2 | KDM2A | -0,3435 | 1,88885E-17 |
| ACE2 | RP1-228P16.1 | -0,3435 | 1,88332E-17 |
| ACE2 | RP11-676J12.6 | -0,3435 | 1,89684E-17 |
| ACE2 | RP11-495P10.10 | -0,3437 | 1,82096E-17 |
| ACE2 | AC012066.1 | -0,3437 | 1,79438E-17 |
| ACE2 | CD83 | -0,3437 | 1,79922E-17 |
| ACE2 | MTCH1 | -0,3437 | 1,80574E-17 |
| ACE2 | RP11-1102P22.2 | -0,3437 | 1,80297E-17 |
| ACE2 | TRBV20-1 | -0,3439 | 1,70599E-17 |
| ACE2 | AC068522.4 | -0,3439 | 1,72886E-17 |
| ACE2 | GNAL | -0,3439 | 1,7177E-17 |
| ACE2 | ADGRE3 | -0,3439 | 1,73654E-17 |
| ACE2 | SLC16A2 | -0,3439 | 1,7165E-17 |
| ACE2 | SEMA3A | -0,3441 | 1,66339E-17 |
| ACE2 | DDX39B | -0,3442 | 1,59828E-17 |
| ACE2 | CACUL1 | -0,3442 | 1,60753E-17 |
| ACE2 | ALAS2 | -0,3443 | 1,58751E-17 |
| ACE2 | NABP1 | -0,3444 | 1,5353E-17 |
| ACE2 | AFAP1-AS1 | -0,3444 | 1,55208E-17 |
| ACE2 | LIME1 | -0,3444 | 1,53106E-17 |
| ACE2 | FXYD1 | -0,3445 | 1,49777E-17 |
| ACE2 | MTMR9LP | -0,3446 | 1,47236E-17 |
| ACE2 | CFAP58-AS1 | -0,3446 | 1,48446E-17 |
| ACE2 | MRGBP | -0,3446 | 1,47344E-17 |
| ACE2 | B4GALT4 | -0,3447 | 1,44269E-17 |
| ACE2 | PRRX1 | -0,3448 | 1,40743E-17 |
| ACE2 | PPP2R2D | -0,3449 | 1,35811E-17 |
| ACE2 | CTC-558O2.1 | -0,345 | 1,34296E-17 |
| ACE2 | C22orf31 | -0,345 | 1,35176E-17 |
| ACE2 | UIMC1 | -0,3451 | 1,30314E-17 |
| ACE2 | MYOCD | -0,3452 | 1,26999E-17 |
| ACE2 | CCL4 | -0,3452 | 1,27837E-17 |
| ACE2 | RP11-563N6.6 | -0,3453 | 1,25813E-17 |
| ACE2 | RP11-552F3.10 | -0,3453 | 1,25729E-17 |
| ACE2 | CTD-2047H16.4 | -0,3453 | 1,26168E-17 |
| ACE2 | JAM3 | -0,3454 | 1,21615E-17 |
| ACE2 | RP11-147L13.8 | -0,3454 | 1,23715E-17 |
| ACE2 | NCAM2 | -0,3454 | 1,21794E-17 |
| ACE2 | RP11-92C4.3 | -0,3456 | 1,15691E-17 |
| ACE2 | PHF19 | -0,3458 | 1,12579E-17 |
| ACE2 | GTDC1 | -0,3459 | 1,08879E-17 |
| ACE2 | ATP2B1 | -0,3459 | 1,10258E-17 |
| ACE2 | ACTG1P3 | -0,346 | 1,05784E-17 |
| ACE2 | NXF3 | -0,346 | 1,06047E-17 |
| ACE2 | SNORD35A | -0,3461 | 1,03327E-17 |

| | | | |
|---|---|---|---|
| ACE2 | TMEM51 | -0,3462 | 1,00863E-17 |
| ACE2 | PANK4 | -0,3463 | 1,00122E-17 |
| ACE2 | TTN-AS1 | -0,3464 | 9,79643E-18 |
| ACE2 | RP11-867G23.3 | -0,3464 | 9,72323E-18 |
| ACE2 | PCDH18 | -0,3465 | 9,41556E-18 |
| ACE2 | FNBP4 | -0,3465 | 9,512E-18 |
| ACE2 | BTG1P1 | -0,3465 | 9,50338E-18 |
| ACE2 | LCAT | -0,3465 | 9,48181E-18 |
| ACE2 | GSDMD | -0,3466 | 9,27998E-18 |
| ACE2 | ADAM12 | -0,3466 | 9,28214E-18 |
| ACE2 | TAS2R15P | -0,3466 | 9,3494E-18 |
| ACE2 | RP13-20L14.10 | -0,3466 | 9,34771E-18 |
| ACE2 | HIST1H4B | -0,3467 | 9,1021E-18 |
| ACE2 | SNRPGP14 | -0,3467 | 9,02213E-18 |
| ACE2 | CTB-58E17.9 | -0,3467 | 8,99946E-18 |
| ACE2 | RP11-66H6.4 | -0,3469 | 8,71859E-18 |
| ACE2 | ACTN1 | -0,347 | 8,50057E-18 |
| ACE2 | SNX29P2 | -0,347 | 8,52589E-18 |
| ACE2 | NREP | -0,3471 | 8,26848E-18 |
| ACE2 | RP11-358B23.5 | -0,3471 | 8,24133E-18 |
| ACE2 | CYLD | -0,3472 | 8,12241E-18 |
| ACE2 | RP11-793A3.2 | -0,3472 | 8,10485E-18 |
| ACE2 | TRIM69 | -0,3473 | 7,96763E-18 |
| ACE2 | SNORD35B | -0,3473 | 7,87255E-18 |
| ACE2 | RP1-151F17.1 | -0,3474 | 7,72554E-18 |
| ACE2 | ZMYM5 | -0,3474 | 7,67236E-18 |
| ACE2 | TCF3 | -0,3474 | 7,69027E-18 |
| ACE2 | RP11-70L8.4 | -0,3475 | 7,60071E-18 |
| ACE2 | CTD-2349P21.10 | -0,3475 | 7,59297E-18 |
| ACE2 | SELENOM | -0,3475 | 7,6417E-18 |
| ACE2 | LINC02084 | -0,3476 | 7,35396E-18 |
| ACE2 | RP3-509L4.3 | -0,3476 | 7,34606E-18 |
| ACE2 | ZNF767P | -0,3476 | 7,45952E-18 |
| ACE2 | NUTM2B-AS1 | -0,3476 | 7,33726E-18 |
| ACE2 | RP11-893F2.13 | -0,3476 | 7,3903E-18 |
| ACE2 | TFE3 | -0,3476 | 7,4426E-18 |
| ACE2 | CHST1 | -0,3477 | 7,224E-18 |
| ACE2 | RRN3P3 | -0,3477 | 7,18277E-18 |
| ACE2 | CTD-2653D5.1 | -0,3477 | 7,20631E-18 |
| ACE2 | PACS1 | -0,3478 | 7,02855E-18 |
| ACE2 | KDM6B | -0,3478 | 7,06511E-18 |
| ACE2 | KIR2DP1 | -0,3478 | 7,04577E-18 |
| ACE2 | SFMBT2 | -0,3479 | 6,96096E-18 |
| ACE2 | TNFAIP2 | -0,348 | 6,72512E-18 |
| ACE2 | ITGA9-AS1 | -0,3481 | 6,53292E-18 |
| ACE2 | B3GNTL1P1 | -0,3481 | 6,63046E-18 |
| ACE2 | ZSWIM6 | -0,3482 | 6,43665E-18 |
| ACE2 | AOAH | -0,3482 | 6,45723E-18 |
| ACE2 | RPS19P3 | -0,3482 | 6,41148E-18 |
| ACE2 | EXD3 | -0,3483 | 6,26228E-18 |
| ACE2 | EPSTI1 | -0,3483 | 6,31491E-18 |
| ACE2 | DTNBP1 | -0,3484 | 6,17102E-18 |
| ACE2 | HIST1H2BJ | -0,3484 | 6,18269E-18 |
| ACE2 | PLA2G4C | -0,3485 | 6,00318E-18 |

| | | | |
|---|---|---|---|
| ACE2 | NPIPB13 | -0,3486 | 5,81125E-18 |
| ACE2 | LINC01588 | -0,3487 | 5,72423E-18 |
| ACE2 | NDE1 | -0,349 | 5,33451E-18 |
| ACE2 | TCTE3 | -0,3491 | 5,21573E-18 |
| ACE2 | DDX56 | -0,3491 | 5,25502E-18 |
| ACE2 | EMC9 | -0,3491 | 5,22699E-18 |
| ACE2 | THCAT158 | -0,3491 | 5,1672E-18 |
| ACE2 | RP11-474P2.5 | -0,3493 | 4,96732E-18 |
| ACE2 | SEPT5 | -0,3493 | 5,02273E-18 |
| ACE2 | FAM118A | -0,3494 | 4,92949E-18 |
| ACE2 | DNAJC18 | -0,3497 | 4,57038E-18 |
| ACE2 | ATAT1 | -0,3499 | 4,29031E-18 |
| ACE2 | HEY2 | -0,3499 | 4,3403E-18 |
| ACE2 | STAT5B | -0,3499 | 4,29394E-18 |
| ACE2 | CAMK2A | -0,35 | 4,20568E-18 |
| ACE2 | RP11-1007G5.2 | -0,3501 | 4,14497E-18 |
| ACE2 | MED12 | -0,3501 | 4,12556E-18 |
| ACE2 | SDHAP1 | -0,3502 | 4,03619E-18 |
| ACE2 | FAM102B | -0,3503 | 3,92663E-18 |
| ACE2 | ARHGEF7 | -0,3503 | 3,98803E-18 |
| ACE2 | CYCSP40 | -0,3503 | 3,94717E-18 |
| ACE2 | SATB1-AS1 | -0,3504 | 3,85788E-18 |
| ACE2 | RP11-3K24.3 | -0,3505 | 3,77665E-18 |
| ACE2 | BBC3 | -0,3505 | 3,74746E-18 |
| ACE2 | CDH6 | -0,3506 | 3,693E-18 |
| ACE2 | YPEL4 | -0,3506 | 3,68464E-18 |
| ACE2 | ACAD11 | -0,3507 | 3,60244E-18 |
| ACE2 | LINC00189 | -0,3507 | 3,5796E-18 |
| ACE2 | PCDHGA6 | -0,3508 | 3,52356E-18 |
| ACE2 | GXYLT2 | -0,3509 | 3,40032E-18 |
| ACE2 | ATN1 | -0,3509 | 3,45716E-18 |
| ACE2 | HIST2H3D | -0,351 | 3,33982E-18 |
| ACE2 | RP11-575A19.2 | -0,351 | 3,36806E-18 |
| ACE2 | RP11-680G24.6 | -0,351 | 3,39395E-18 |
| ACE2 | PTPRC | -0,3511 | 3,2585E-18 |
| ACE2 | ALB | -0,3511 | 3,31497E-18 |
| ACE2 | RP11-764D10.2 | -0,3511 | 3,2626E-18 |
| ACE2 | TBKBP1 | -0,3511 | 3,28606E-18 |
| ACE2 | KLHL41 | -0,3512 | 3,20359E-18 |
| ACE2 | SFI1 | -0,3512 | 3,22346E-18 |
| ACE2 | GTPBP2 | -0,3513 | 3,12657E-18 |
| ACE2 | RAB30 | -0,3513 | 3,12024E-18 |
| ACE2 | MRC2 | -0,3513 | 3,11449E-18 |
| ACE2 | RLIM | -0,3513 | 3,16648E-18 |
| ACE2 | RP11-452J21.2 | -0,3514 | 3,0712E-18 |
| ACE2 | SNORD21 | -0,3515 | 3,0165E-18 |
| ACE2 | AC019181.3 | -0,3515 | 3,01468E-18 |
| ACE2 | FLJ22447 | -0,3515 | 2,97595E-18 |
| ACE2 | MYL4 | -0,3515 | 2,97913E-18 |
| ACE2 | P3H1 | -0,3516 | 2,93107E-18 |
| ACE2 | GJA1P1 | -0,3516 | 2,89128E-18 |
| ACE2 | CLSTN3 | -0,3516 | 2,907E-18 |
| ACE2 | ITGAX | -0,3516 | 2,91901E-18 |
| ACE2 | CFLAR-AS1 | -0,3517 | 2,86213E-18 |

| | | | |
|---|---|---|---|
| ACE2 | RP11-44F14.6 | -0,3518 | 2,76997E-18 |
| ACE2 | HBD | -0,3519 | 2,74129E-18 |
| ACE2 | IL11RA | -0,352 | 2,67338E-18 |
| ACE2 | LRRC32 | -0,352 | 2,63254E-18 |
| ACE2 | MEG3 | -0,352 | 2,64642E-18 |
| ACE2 | AC093838.4 | -0,3523 | 2,45361E-18 |
| ACE2 | LINC00861 | -0,3523 | 2,46365E-18 |
| ACE2 | RP11-452L6.1 | -0,3523 | 2,48011E-18 |
| ACE2 | RP11-147L13.13 | -0,3524 | 2,44805E-18 |
| ACE2 | RP11-302L19.3 | -0,3525 | 2,34077E-18 |
| ACE2 | TRGV2 | -0,3525 | 2,37351E-18 |
| ACE2 | PTGES | -0,3525 | 2,3842E-18 |
| ACE2 | HCFC1R1 | -0,3525 | 2,37614E-18 |
| ACE2 | TBX4 | -0,3526 | 2,31255E-18 |
| ACE2 | UBE2W | -0,3527 | 2,24624E-18 |
| ACE2 | SRA1 | -0,3528 | 2,18238E-18 |
| ACE2 | RP11-218E20.5 | -0,3528 | 2,19045E-18 |
| ACE2 | HMGN2P46 | -0,3528 | 2,20154E-18 |
| ACE2 | CTB-131B5.2 | -0,3529 | 2,1625E-18 |
| ACE2 | GRIK1 | -0,3529 | 2,16975E-18 |
| ACE2 | LINC01138 | -0,353 | 2,10695E-18 |
| ACE2 | CHN1 | -0,353 | 2,08444E-18 |
| ACE2 | NR2F1 | -0,3531 | 2,04152E-18 |
| ACE2 | MYOZ1 | -0,3531 | 2,03901E-18 |
| ACE2 | PIEZO2 | -0,3531 | 2,03861E-18 |
| ACE2 | RGS6 | -0,3532 | 1,99195E-18 |
| ACE2 | ADGRL3 | -0,3533 | 1,9818E-18 |
| ACE2 | GRHL1 | -0,3534 | 1,91043E-18 |
| ACE2 | TRGV7 | -0,3534 | 1,91036E-18 |
| ACE2 | EXT1 | -0,3534 | 1,93595E-18 |
| ACE2 | RP11-59C5.3 | -0,3534 | 1,89681E-18 |
| ACE2 | SKAP1 | -0,3534 | 1,93645E-18 |
| ACE2 | RP11-36B15.1 | -0,3535 | 1,85173E-18 |
| ACE2 | PPP2R5B | -0,3535 | 1,85863E-18 |
| ACE2 | RP11-192M23.1 | -0,3535 | 1,88863E-18 |
| ACE2 | DLGAP1-AS2 | -0,3535 | 1,87954E-18 |
| ACE2 | RHOQ | -0,3536 | 1,84091E-18 |
| ACE2 | TRBV7-9 | -0,3536 | 1,83933E-18 |
| ACE2 | RP11-817I4.1 | -0,3536 | 1,84249E-18 |
| ACE2 | SSH1 | -0,3537 | 1,77654E-18 |
| ACE2 | TENM1 | -0,3537 | 1,77991E-18 |
| ACE2 | CTD-2357A8.2 | -0,3538 | 1,75761E-18 |
| ACE2 | ZNF548 | -0,3538 | 1,748E-18 |
| ACE2 | BTN2A1 | -0,3539 | 1,68992E-18 |
| ACE2 | RP5-1107A17.3 | -0,3539 | 1,70202E-18 |
| ACE2 | HIST2H2BC | -0,354 | 1,67368E-18 |
| ACE2 | RP11-894J14.2 | -0,354 | 1,67702E-18 |
| ACE2 | LRRC77P | -0,354 | 1,65747E-18 |
| ACE2 | PRSS53 | -0,354 | 1,65874E-18 |
| ACE2 | AC147651.4 | -0,3541 | 1,62996E-18 |
| ACE2 | LZTS1 | -0,3541 | 1,61534E-18 |
| ACE2 | RNF114 | -0,3541 | 1,63858E-18 |
| ACE2 | ACTN2 | -0,3542 | 1,58666E-18 |
| ACE2 | PLSCR1 | -0,3542 | 1,57829E-18 |

| | | | |
|---|---|---|---|
| ACE2 | ROBO2 | -0,3543 | 1,56251E-18 |
| ACE2 | GLT8D2 | -0,3544 | 1,51908E-18 |
| ACE2 | A4GALT | -0,3544 | 1,49664E-18 |
| ACE2 | RP4-620E11.8 | -0,3545 | 1,45964E-18 |
| ACE2 | RFX6 | -0,3546 | 1,42717E-18 |
| ACE2 | TRBV9 | -0,3546 | 1,44634E-18 |
| ACE2 | NTRK2 | -0,3546 | 1,44703E-18 |
| ACE2 | HPX | -0,3546 | 1,45269E-18 |
| ACE2 | C1QTNF2 | -0,3547 | 1,42283E-18 |
| ACE2 | DOK6 | -0,3547 | 1,425E-18 |
| ACE2 | RN7SL798P | -0,3548 | 1,36191E-18 |
| ACE2 | IFIT1B | -0,3548 | 1,38941E-18 |
| ACE2 | RGS9 | -0,3548 | 1,38991E-18 |
| ACE2 | CEP131 | -0,3548 | 1,37601E-18 |
| ACE2 | RN7SL364P | -0,3548 | 1,3643E-18 |
| ACE2 | PDIA2 | -0,3549 | 1,34533E-18 |
| ACE2 | AC058791.1 | -0,3551 | 1,27142E-18 |
| ACE2 | SCARF1 | -0,3551 | 1,2713E-18 |
| ACE2 | RBBP8P1 | -0,3551 | 1,27518E-18 |
| ACE2 | PEBP1P3 | -0,3552 | 1,25306E-18 |
| ACE2 | CD200R1 | -0,3552 | 1,26201E-18 |
| ACE2 | PSME2 | -0,3553 | 1,23525E-18 |
| ACE2 | CDK3 | -0,3553 | 1,22912E-18 |
| ACE2 | SLC23A2 | -0,3554 | 1,19383E-18 |
| ACE2 | TMEM178A | -0,3555 | 1,15809E-18 |
| ACE2 | TBC1D10C | -0,3555 | 1,16225E-18 |
| ACE2 | RP3-467L1.6 | -0,3556 | 1,14872E-18 |
| ACE2 | DCHS2 | -0,3556 | 1,13032E-18 |
| ACE2 | CTA-384D8.35 | -0,3556 | 1,13879E-18 |
| ACE2 | MEIS1 | -0,3557 | 1,10922E-18 |
| ACE2 | DLL1 | -0,3557 | 1,11315E-18 |
| ACE2 | RP11-861L17.3 | -0,3557 | 1,10999E-18 |
| ACE2 | RP11-73M7.6 | -0,3558 | 1,09334E-18 |
| ACE2 | CH17-262A2.1 | -0,3558 | 1,08073E-18 |
| ACE2 | RP11-180M15.7 | -0,3558 | 1,08166E-18 |
| ACE2 | TRABD | -0,3558 | 1,09416E-18 |
| ACE2 | TNF | -0,3559 | 1,06488E-18 |
| ACE2 | COX4I2 | -0,3559 | 1,05804E-18 |
| ACE2 | ZFP36L2 | -0,356 | 1,03077E-18 |
| ACE2 | GPAA1P2 | -0,356 | 1,02747E-18 |
| ACE2 | BAALC | -0,356 | 1,04707E-18 |
| ACE2 | TAF1D | -0,356 | 1,02881E-18 |
| ACE2 | ARNTL2-AS1 | -0,356 | 1,04198E-18 |
| ACE2 | RARA-AS1 | -0,356 | 1,04018E-18 |
| ACE2 | RYR2 | -0,3561 | 1,02066E-18 |
| ACE2 | PHF21A | -0,3561 | 1,01096E-18 |
| ACE2 | CTD-2536I1.3 | -0,3561 | 1,00682E-18 |
| ACE2 | RP11-157K17.5 | -0,3562 | 9,92453E-19 |
| ACE2 | RP11-85G20.2 | -0,3562 | 9,9543E-19 |
| ACE2 | LINC00926 | -0,3562 | 9,82943E-19 |
| ACE2 | RP11-87N3.6 | -0,3562 | 9,94625E-19 |
| ACE2 | PDCL3P5 | -0,3563 | 9,72848E-19 |
| ACE2 | JMJD6 | -0,3563 | 9,53127E-19 |
| ACE2 | RASGRF2 | -0,3564 | 9,50258E-19 |

| | | | |
|---|---|---|---|
| ACE2 | RP5-1021I20.5 | -0,3564 | 9,50559E-19 |
| ACE2 | RP5-1042K10.13 | -0,3564 | 9,45196E-19 |
| ACE2 | URAHP | -0,3566 | 9,02075E-19 |
| ACE2 | CIITA | -0,3567 | 8,86441E-19 |
| ACE2 | RP5-867C24.1 | -0,3567 | 8,67573E-19 |
| ACE2 | CEACAM19 | -0,3567 | 8,85774E-19 |
| ACE2 | CD3D | -0,3568 | 8,58651E-19 |
| ACE2 | TMEM233 | -0,3568 | 8,47305E-19 |
| ACE2 | SERTAD3 | -0,3568 | 8,58481E-19 |
| ACE2 | FARP2 | -0,3569 | 8,36296E-19 |
| ACE2 | RP11-705C15.2 | -0,3569 | 8,31261E-19 |
| ACE2 | CD96 | -0,357 | 8,11398E-19 |
| ACE2 | RP11-148K1.12 | -0,357 | 8,16304E-19 |
| ACE2 | LIF | -0,357 | 8,24825E-19 |
| ACE2 | CNKSR3 | -0,3572 | 7,73352E-19 |
| ACE2 | IRF2 | -0,3573 | 7,63278E-19 |
| ACE2 | CNN1 | -0,3573 | 7,63858E-19 |
| ACE2 | PDLIM7 | -0,3574 | 7,46962E-19 |
| ACE2 | ANKRD36C | -0,3575 | 7,31643E-19 |
| ACE2 | PPP2R2B | -0,3575 | 7,21066E-19 |
| ACE2 | AP3M2 | -0,3575 | 7,2376E-19 |
| ACE2 | GPSM1 | -0,3575 | 7,23234E-19 |
| ACE2 | RP11-93O14.3 | -0,3575 | 7,24566E-19 |
| ACE2 | IRF3 | -0,3576 | 7,02553E-19 |
| ACE2 | HMGN2P19 | -0,3578 | 6,66604E-19 |
| ACE2 | CATSPER2 | -0,3578 | 6,69377E-19 |
| ACE2 | LGALS1 | -0,3578 | 6,69404E-19 |
| ACE2 | DHX34 | -0,3581 | 6,26928E-19 |
| ACE2 | LINC01934 | -0,3582 | 6,18535E-19 |
| ACE2 | ANKRD13D | -0,3582 | 6,18741E-19 |
| ACE2 | RP11-212I21.3 | -0,3583 | 5,97723E-19 |
| ACE2 | HMGB1P24 | -0,3584 | 5,78057E-19 |
| ACE2 | DAPK3 | -0,3584 | 5,87693E-19 |
| ACE2 | BGN | -0,3584 | 5,83787E-19 |
| ACE2 | CDK6 | -0,3585 | 5,7285E-19 |
| ACE2 | PLEK | -0,3586 | 5,53861E-19 |
| ACE2 | RP11-12A2.1 | -0,3586 | 5,51883E-19 |
| ACE2 | HIST1H3H | -0,3587 | 5,49785E-19 |
| ACE2 | JAZF1 | -0,3587 | 5,38869E-19 |
| ACE2 | RP11-236L14.2 | -0,3587 | 5,44041E-19 |
| ACE2 | LINC00222 | -0,3588 | 5,29247E-19 |
| ACE2 | RP11-10J21.4 | -0,3588 | 5,3556E-19 |
| ACE2 | N4BP2L2-IT2 | -0,3588 | 5,33527E-19 |
| ACE2 | TTBK2 | -0,3588 | 5,28461E-19 |
| ACE2 | RSRP1 | -0,3589 | 5,19147E-19 |
| ACE2 | CAP2P1 | -0,3589 | 5,22099E-19 |
| ACE2 | RP11-358B23.7 | -0,3589 | 5,21327E-19 |
| ACE2 | RP11-513M16.7 | -0,359 | 5,06486E-19 |
| ACE2 | MMP17 | -0,359 | 5,06362E-19 |
| ACE2 | GK-IT1 | -0,359 | 5,02745E-19 |
| ACE2 | ANKRD16 | -0,3591 | 4,95267E-19 |
| ACE2 | LINC01197 | -0,3591 | 4,92038E-19 |
| ACE2 | USP12 | -0,3594 | 4,568E-19 |
| ACE2 | FBXO17 | -0,3594 | 4,56008E-19 |

| | | | |
|---|---|---|---|
| ACE2 | SH3BP2 | -0,3595 | 4,4843E-19 |
| ACE2 | RP3-477J10.1 | -0,3595 | 4,53016E-19 |
| ACE2 | EXOC3L4 | -0,3596 | 4,42498E-19 |
| ACE2 | ZNF571 | -0,3596 | 4,38953E-19 |
| ACE2 | B2M | -0,3597 | 4,28398E-19 |
| ACE2 | TRG-AS1 | -0,3598 | 4,14964E-19 |
| ACE2 | RBPMS2 | -0,3598 | 4,16314E-19 |
| ACE2 | RP3-330M21.5 | -0,3599 | 4,07723E-19 |
| ACE2 | ZNF227 | -0,3599 | 4,03292E-19 |
| ACE2 | AGAP2 | -0,36 | 3,9643E-19 |
| ACE2 | TMEM200B | -0,3601 | 3,87388E-19 |
| ACE2 | RP11-385F5.4 | -0,3601 | 3,91713E-19 |
| ACE2 | SNORA53 | -0,3601 | 3,86592E-19 |
| ACE2 | RP11-473M20.9 | -0,3601 | 3,85933E-19 |
| ACE2 | PDE3A | -0,3602 | 3,81917E-19 |
| ACE2 | PRR14 | -0,3602 | 3,75332E-19 |
| ACE2 | TRAV8-3 | -0,3603 | 3,66274E-19 |
| ACE2 | GPBP1 | -0,3604 | 3,59125E-19 |
| ACE2 | RP11-395I14.2 | -0,3604 | 3,61942E-19 |
| ACE2 | LUC7L | -0,3604 | 3,5742E-19 |
| ACE2 | CHTF18 | -0,3604 | 3,60238E-19 |
| ACE2 | FAM126A | -0,3605 | 3,50819E-19 |
| ACE2 | CASP12 | -0,3606 | 3,45927E-19 |
| ACE2 | LAMP5 | -0,3607 | 3,32038E-19 |
| ACE2 | PRKAA2 | -0,3609 | 3,20499E-19 |
| ACE2 | TAS2R20 | -0,3609 | 3,21476E-19 |
| ACE2 | PHLPP2 | -0,3609 | 3,18334E-19 |
| ACE2 | CDK11A | -0,3611 | 3,05822E-19 |
| ACE2 | MIR765 | -0,3611 | 3,01671E-19 |
| ACE2 | TRPC6 | -0,3611 | 3,06127E-19 |
| ACE2 | RP11-64B16.4 | -0,3611 | 3,03918E-19 |
| ACE2 | KCNQ1OT1 | -0,3613 | 2,93683E-19 |
| ACE2 | RASAL1 | -0,3613 | 2,92944E-19 |
| ACE2 | SLFN12L | -0,3614 | 2,84754E-19 |
| ACE2 | MAGED2 | -0,3614 | 2,83358E-19 |
| ACE2 | CBFA2T2 | -0,3615 | 2,78985E-19 |
| ACE2 | SNHG12 | -0,3616 | 2,70632E-19 |
| ACE2 | RP11-329B9.3 | -0,3616 | 2,6743E-19 |
| ACE2 | NINJ1 | -0,3616 | 2,66981E-19 |
| ACE2 | GRIK4 | -0,3616 | 2,70303E-19 |
| ACE2 | SNORD91A | -0,3619 | 2,538E-19 |
| ACE2 | MLLT6 | -0,362 | 2,44208E-19 |
| ACE2 | WNT11 | -0,3621 | 2,40349E-19 |
| ACE2 | RP11-521C22.2 | -0,3623 | 2,28943E-19 |
| ACE2 | ZNF609 | -0,3624 | 2,23075E-19 |
| ACE2 | CTD-2536I1.2 | -0,3624 | 2,21689E-19 |
| ACE2 | CXorf38 | -0,3624 | 2,22184E-19 |
| ACE2 | HIST2H2AA4 | -0,3625 | 2,19326E-19 |
| ACE2 | CTD-2006K23.1 | -0,3625 | 2,16896E-19 |
| ACE2 | FUBP1 | -0,3626 | 2,10553E-19 |
| ACE2 | RNU1-100P | -0,3626 | 2,12746E-19 |
| ACE2 | RP11-473O4.5 | -0,3626 | 2,13187E-19 |
| ACE2 | RP11-666F17.1 | -0,3626 | 2,1436E-19 |
| ACE2 | PTMS | -0,3629 | 1,96589E-19 |

| | | | |
|---|---|---|---|
| ACE2 | RP11-946P6.6 | -0,3629 | 1,97002E-19 |
| ACE2 | S100A13 | -0,363 | 1,914E-19 |
| ACE2 | SEMA7A | -0,363 | 1,93208E-19 |
| ACE2 | DEDD | -0,3631 | 1,88813E-19 |
| ACE2 | RP11-588H23.3 | -0,3631 | 1,87492E-19 |
| ACE2 | PARP14 | -0,3633 | 1,78726E-19 |
| ACE2 | ESR1 | -0,3633 | 1,78707E-19 |
| ACE2 | CPQ | -0,3633 | 1,77005E-19 |
| ACE2 | AC021188.4 | -0,3634 | 1,73007E-19 |
| ACE2 | TET2 | -0,3634 | 1,73789E-19 |
| ACE2 | KIAA0141 | -0,3634 | 1,74771E-19 |
| ACE2 | HAP1 | -0,3634 | 1,73721E-19 |
| ACE2 | RP4-738P11.4 | -0,3635 | 1,69866E-19 |
| ACE2 | LEF1 | -0,3636 | 1,64914E-19 |
| ACE2 | TCEAL7 | -0,3636 | 1,66127E-19 |
| ACE2 | MFAP2 | -0,3637 | 1,63697E-19 |
| ACE2 | HEATR9 | -0,3637 | 1,62236E-19 |
| ACE2 | ADAMTS8 | -0,3638 | 1,5674E-19 |
| ACE2 | HIST2H2BE | -0,364 | 1,49478E-19 |
| ACE2 | RP11-71H17.8 | -0,364 | 1,50477E-19 |
| ACE2 | RP11-96C23.11 | -0,364 | 1,51485E-19 |
| ACE2 | NPM1P26 | -0,3641 | 1,48374E-19 |
| ACE2 | RP11-423G4.10 | -0,3642 | 1,4221E-19 |
| ACE2 | PRRX2 | -0,3643 | 1,39374E-19 |
| ACE2 | ZNF529 | -0,3643 | 1,40771E-19 |
| ACE2 | RP11-460I13.6 | -0,3645 | 1,33989E-19 |
| ACE2 | RP11-231L11.3 | -0,3646 | 1,31063E-19 |
| ACE2 | WHRN | -0,3646 | 1,29263E-19 |
| ACE2 | MAST2 | -0,3647 | 1,28292E-19 |
| ACE2 | KIF25 | -0,3647 | 1,26723E-19 |
| ACE2 | PITPNM1 | -0,3647 | 1,26206E-19 |
| ACE2 | RP4-761J14.8 | -0,3647 | 1,2687E-19 |
| ACE2 | RP5-837J1.4 | -0,3648 | 1,22446E-19 |
| ACE2 | LINC01176 | -0,365 | 1,18243E-19 |
| ACE2 | FGF17 | -0,365 | 1,17047E-19 |
| ACE2 | ADGRB2 | -0,3651 | 1,14908E-19 |
| ACE2 | KIAA0907 | -0,3651 | 1,15689E-19 |
| ACE2 | ALMS1-IT1 | -0,3651 | 1,15944E-19 |
| ACE2 | ARHGAP20 | -0,3651 | 1,14808E-19 |
| ACE2 | SLC39A10 | -0,3652 | 1,13372E-19 |
| ACE2 | ARHGAP9 | -0,3652 | 1,13226E-19 |
| ACE2 | AEBP2 | -0,3654 | 1,06097E-19 |
| ACE2 | CTD-2036P10.6 | -0,3654 | 1,07282E-19 |
| ACE2 | APOA2 | -0,3655 | 1,05352E-19 |
| ACE2 | OLFML2A | -0,3655 | 1,03208E-19 |
| ACE2 | BIRC6-AS2 | -0,3656 | 1,01315E-19 |
| ACE2 | IER5L | -0,3656 | 1,02575E-19 |
| ACE2 | KRTAP5-1 | -0,3656 | 1,01204E-19 |
| ACE2 | RGS11 | -0,3656 | 1,01658E-19 |
| ACE2 | HSPA7 | -0,3657 | 9,88873E-20 |
| ACE2 | EVL | -0,3657 | 9,81241E-20 |
| ACE2 | DZIP1 | -0,3658 | 9,64964E-20 |
| ACE2 | YES1P1 | -0,3658 | 9,71168E-20 |
| ACE2 | RN7SL368P | -0,3659 | 9,4295E-20 |

| | | | |
|---|---|---|---|
| ACE2 | LINC01748 | -0,366 | 9,26996E-20 |
| ACE2 | HNRNPH1 | -0,366 | 9,20425E-20 |
| ACE2 | RP11-403P17.6 | -0,366 | 9,18496E-20 |
| ACE2 | RHOA-IT1 | -0,3661 | 8,89867E-20 |
| ACE2 | GNG11 | -0,3661 | 9,10409E-20 |
| ACE2 | RBM38 | -0,3661 | 9,01411E-20 |
| ACE2 | PLEKHA2 | -0,3662 | 8,70015E-20 |
| ACE2 | DGKE | -0,3662 | 8,78391E-20 |
| ACE2 | RP11-436K8.1 | -0,3664 | 8,4496E-20 |
| ACE2 | RP11-504P24.9 | -0,3664 | 8,37539E-20 |
| ACE2 | HOXB5 | -0,3664 | 8,40723E-20 |
| ACE2 | LMCD1-AS1 | -0,3665 | 8,18717E-20 |
| ACE2 | SERHL2 | -0,3665 | 8,19898E-20 |
| ACE2 | NTNG1 | -0,3666 | 7,91995E-20 |
| ACE2 | CCL2 | -0,3667 | 7,82221E-20 |
| ACE2 | RP11-264B14.1 | -0,3667 | 7,69576E-20 |
| ACE2 | LINC01268 | -0,3668 | 7,58229E-20 |
| ACE2 | PPP1R35 | -0,3668 | 7,60603E-20 |
| ACE2 | HSF1 | -0,3668 | 7,5644E-20 |
| ACE2 | SLC4A1 | -0,3668 | 7,56714E-20 |
| ACE2 | CTB-108O6.2 | -0,3669 | 7,31836E-20 |
| ACE2 | RP11-30L15.4 | -0,3669 | 7,37301E-20 |
| ACE2 | RN7SKP20 | -0,3669 | 7,35128E-20 |
| ACE2 | SNORA59A | -0,3671 | 7,01614E-20 |
| ACE2 | UBE2V1 | -0,3671 | 7,09155E-20 |
| ACE2 | PCDHB10 | -0,3673 | 6,64912E-20 |
| ACE2 | AP000577.2 | -0,3673 | 6,70848E-20 |
| ACE2 | GPCPD1 | -0,3673 | 6,6279E-20 |
| ACE2 | AP001610.5 | -0,3673 | 6,61852E-20 |
| ACE2 | RP3-508I15.21 | -0,3673 | 6,64388E-20 |
| ACE2 | FGF13 | -0,3673 | 6,73896E-20 |
| ACE2 | COL6A5 | -0,3674 | 6,60637E-20 |
| ACE2 | RP11-302B13.1 | -0,3674 | 6,55399E-20 |
| ACE2 | VAV2 | -0,3675 | 6,38239E-20 |
| ACE2 | ADAMTS14 | -0,3675 | 6,43874E-20 |
| ACE2 | VSIG8 | -0,3676 | 6,26858E-20 |
| ACE2 | ST6GALNAC6 | -0,3676 | 6,25626E-20 |
| ACE2 | ADAM8 | -0,3676 | 6,2573E-20 |
| ACE2 | RPL7P14 | -0,3677 | 6,06325E-20 |
| ACE2 | RP3-512E2.2 | -0,3677 | 6,09983E-20 |
| ACE2 | NUMBL | -0,3678 | 5,96926E-20 |
| ACE2 | RP11-45M22.3 | -0,3679 | 5,78165E-20 |
| ACE2 | RHOC | -0,368 | 5,67606E-20 |
| ACE2 | RP11-196G18.22 | -0,368 | 5,58792E-20 |
| ACE2 | CTD-2240E14.4 | -0,368 | 5,60596E-20 |
| ACE2 | CYTOR | -0,3681 | 5,54817E-20 |
| ACE2 | RP11-49G2.3 | -0,3682 | 5,38872E-20 |
| ACE2 | FAS | -0,3683 | 5,1965E-20 |
| ACE2 | RP11-1148L6.8 | -0,3683 | 5,21409E-20 |
| ACE2 | RP11-397P13.7 | -0,3684 | 5,04495E-20 |
| ACE2 | UBE2E2 | -0,3684 | 5,11692E-20 |
| ACE2 | CTC-490E21.11 | -0,3684 | 5,09889E-20 |
| ACE2 | ADGRG5 | -0,3686 | 4,85812E-20 |
| ACE2 | ZNF841 | -0,3687 | 4,75575E-20 |

| | | | |
|---|---|---|---|
| ACE2 | HOXB2 | -0,3688 | 4,66525E-20 |
| ACE2 | RP11-664H17.1 | -0,3689 | 4,48876E-20 |
| ACE2 | CDC34 | -0,3693 | 4,04835E-20 |
| ACE2 | VSNL1 | -0,3694 | 3,98967E-20 |
| ACE2 | MEF2C-AS1 | -0,3695 | 3,85264E-20 |
| ACE2 | HIST3H2BB | -0,3696 | 3,79995E-20 |
| ACE2 | CCNL1 | -0,3698 | 3,6228E-20 |
| ACE2 | ST8SIA4 | -0,3698 | 3,59824E-20 |
| ACE2 | IL7 | -0,3699 | 3,54017E-20 |
| ACE2 | TBX2-AS1 | -0,3699 | 3,53069E-20 |
| ACE2 | RP11-713H12.1 | -0,37 | 3,38773E-20 |
| ACE2 | DCP2 | -0,3701 | 3,32604E-20 |
| ACE2 | CRIP2 | -0,3701 | 3,37266E-20 |
| ACE2 | ITM2C | -0,3702 | 3,26731E-20 |
| ACE2 | MED28 | -0,3702 | 3,22719E-20 |
| ACE2 | TNR | -0,3703 | 3,21339E-20 |
| ACE2 | DISC1 | -0,3703 | 3,16657E-20 |
| ACE2 | TF | -0,3703 | 3,18761E-20 |
| ACE2 | HNRNPA3P9 | -0,3703 | 3,2004E-20 |
| ACE2 | CMTM1 | -0,3703 | 3,1648E-20 |
| ACE2 | CTD-2515A14.1 | -0,3704 | 3,08926E-20 |
| ACE2 | STEAP2 | -0,3705 | 2,98726E-20 |
| ACE2 | CTA-268H5.12 | -0,3705 | 3,03082E-20 |
| ACE2 | LIX1L | -0,3706 | 2,91471E-20 |
| ACE2 | LUM | -0,3706 | 2,95307E-20 |
| ACE2 | PLPPR2 | -0,3706 | 2,94367E-20 |
| ACE2 | CTD-2583A14.11 | -0,3706 | 2,93883E-20 |
| ACE2 | GABRA2 | -0,3707 | 2,85072E-20 |
| ACE2 | PIK3CD | -0,3708 | 2,82583E-20 |
| ACE2 | GATAD2B | -0,3708 | 2,81601E-20 |
| ACE2 | HIST1H1B | -0,3708 | 2,78402E-20 |
| ACE2 | GABBR1 | -0,3708 | 2,78365E-20 |
| ACE2 | TYRP1 | -0,3709 | 2,75875E-20 |
| ACE2 | OGFR-AS1 | -0,3711 | 2,6314E-20 |
| ACE2 | CEACAM22P | -0,3712 | 2,56662E-20 |
| ACE2 | TSEN54 | -0,3713 | 2,47847E-20 |
| ACE2 | RP11-1334A24.5 | -0,3714 | 2,42086E-20 |
| ACE2 | RP11-726G1.1 | -0,3714 | 2,38998E-20 |
| ACE2 | MAST3 | -0,3714 | 2,42909E-20 |
| ACE2 | RP11-752L20.3 | -0,3716 | 2,30924E-20 |
| ACE2 | RAP1B | -0,3716 | 2,28564E-20 |
| ACE2 | LAMA4 | -0,3717 | 2,23399E-20 |
| ACE2 | SNORA28 | -0,3718 | 2,18217E-20 |
| ACE2 | GABARAPL2 | -0,3718 | 2,17285E-20 |
| ACE2 | FLT4 | -0,3719 | 2,13484E-20 |
| ACE2 | OTUD5 | -0,3719 | 2,14152E-20 |
| ACE2 | ABI2 | -0,372 | 2,08739E-20 |
| ACE2 | RAB28P5 | -0,372 | 2,07284E-20 |
| ACE2 | RP11-107E5.3 | -0,3721 | 2,04234E-20 |
| ACE2 | DENND5A | -0,3721 | 2,04111E-20 |
| ACE2 | HBA1 | -0,3722 | 1,98339E-20 |
| ACE2 | TRBV19 | -0,3723 | 1,93451E-20 |
| ACE2 | RP11-58K22.5 | -0,3723 | 1,91254E-20 |
| ACE2 | CHORDC1 | -0,3723 | 1,90831E-20 |

| | | | |
|---|---|---|---|
| ACE2 | STK10 | -0,3724 | 1,85515E-20 |
| ACE2 | ECM2 | -0,3724 | 1,8663E-20 |
| ACE2 | PAPD7 | -0,3725 | 1,80899E-20 |
| ACE2 | WNT16 | -0,3726 | 1,77161E-20 |
| ACE2 | CTD-2175A23.1 | -0,3727 | 1,75682E-20 |
| ACE2 | OR52B3P | -0,3727 | 1,73034E-20 |
| ACE2 | SETD5 | -0,3729 | 1,65587E-20 |
| ACE2 | ASXL1 | -0,3729 | 1,66017E-20 |
| ACE2 | SMARCA1 | -0,3729 | 1,65779E-20 |
| ACE2 | RP11-6L6.4 | -0,373 | 1,62596E-20 |
| ACE2 | ADGRA2 | -0,373 | 1,63091E-20 |
| ACE2 | LTA | -0,3731 | 1,57234E-20 |
| ACE2 | RP11-325L12.6 | -0,3732 | 1,54507E-20 |
| ACE2 | RP11-208N14.5 | -0,3734 | 1,47362E-20 |
| ACE2 | RP5-1163L11.2 | -0,3735 | 1,41712E-20 |
| ACE2 | PCDHB18P | -0,3736 | 1,39189E-20 |
| ACE2 | GATA3 | -0,3736 | 1,39737E-20 |
| ACE2 | CDC42SE1 | -0,3737 | 1,36266E-20 |
| ACE2 | MIR4435-2HG | -0,3737 | 1,36046E-20 |
| ACE2 | AC067945.4 | -0,374 | 1,25365E-20 |
| ACE2 | RPS2P44 | -0,3741 | 1,23546E-20 |
| ACE2 | RP13-228J13.6 | -0,3741 | 1,23587E-20 |
| ACE2 | B3GAT2 | -0,3742 | 1,184E-20 |
| ACE2 | CD80 | -0,3743 | 1,16875E-20 |
| ACE2 | TVP23C | -0,3746 | 1,06257E-20 |
| ACE2 | CD70 | -0,3746 | 1,07784E-20 |
| ACE2 | ARHGEF40 | -0,3747 | 1,06091E-20 |
| ACE2 | ATP8B4 | -0,3747 | 1,06007E-20 |
| ACE2 | SNORA21 | -0,3747 | 1,04471E-20 |
| ACE2 | LEMD2 | -0,3749 | 1,00544E-20 |
| ACE2 | ENTPD4 | -0,3749 | 9,88806E-21 |
| ACE2 | GPRASP1 | -0,375 | 9,75709E-21 |
| ACE2 | LIMS1 | -0,3753 | 8,9461E-21 |
| ACE2 | CXXC5 | -0,3754 | 8,7704E-21 |
| ACE2 | CTD-2545G14.4 | -0,3755 | 8,6461E-21 |
| ACE2 | RP11-455F5.6 | -0,3756 | 8,32313E-21 |
| ACE2 | LRRC55 | -0,3757 | 8,04941E-21 |
| ACE2 | RP11-405A12.2 | -0,3757 | 8,15572E-21 |
| ACE2 | MIR6807 | -0,3757 | 8,21083E-21 |
| ACE2 | ZNF217 | -0,3757 | 8,18556E-21 |
| ACE2 | COL13A1 | -0,3758 | 8,00999E-21 |
| ACE2 | LL21NC02-21A1.1 | -0,376 | 7,57212E-21 |
| ACE2 | CD48 | -0,3761 | 7,29419E-21 |
| ACE2 | RP11-103C16.2 | -0,3761 | 7,4024E-21 |
| ACE2 | SLC2A3 | -0,3762 | 7,21154E-21 |
| ACE2 | GPR158 | -0,3763 | 7,03076E-21 |
| ACE2 | HIST1H2BM | -0,3765 | 6,6927E-21 |
| ACE2 | SHF | -0,3765 | 6,66626E-21 |
| ACE2 | SOX8 | -0,3765 | 6,63109E-21 |
| ACE2 | PGR-AS1 | -0,3768 | 6,10112E-21 |
| ACE2 | PTN | -0,3769 | 5,94827E-21 |
| ACE2 | HNRNPA3 | -0,3771 | 5,72109E-21 |
| ACE2 | OR2I1P | -0,3771 | 5,71634E-21 |
| ACE2 | AC019186.1 | -0,3772 | 5,57419E-21 |

| | | | |
|---|---|---|---|
| ACE2 | C11orf84 | -0,3772 | 5,48914E-21 |
| ACE2 | AC009299.2 | -0,3773 | 5,42856E-21 |
| ACE2 | PCDHB12 | -0,3773 | 5,46292E-21 |
| ACE2 | CREM | -0,3773 | 5,43028E-21 |
| ACE2 | CCR10 | -0,3774 | 5,32891E-21 |
| ACE2 | BCL11B | -0,3776 | 4,99645E-21 |
| ACE2 | GATM | -0,3776 | 4,95701E-21 |
| ACE2 | MAD2L2 | -0,3777 | 4,884E-21 |
| ACE2 | RPS15AP10 | -0,3777 | 4,82052E-21 |
| ACE2 | TCF4 | -0,3777 | 4,9178E-21 |
| ACE2 | RUNX3 | -0,3778 | 4,75649E-21 |
| ACE2 | CSNK2B | -0,3778 | 4,70717E-21 |
| ACE2 | TMEM133 | -0,3778 | 4,79409E-21 |
| ACE2 | RP11-352M15.2 | -0,3778 | 4,72839E-21 |
| ACE2 | SEPT1 | -0,3778 | 4,72479E-21 |
| ACE2 | LSM11 | -0,3779 | 4,65953E-21 |
| ACE2 | KRT18P4 | -0,3779 | 4,57788E-21 |
| ACE2 | TTC3-AS1 | -0,3779 | 4,67876E-21 |
| ACE2 | MXD1 | -0,378 | 4,56533E-21 |
| ACE2 | BRSK1 | -0,378 | 4,51252E-21 |
| ACE2 | RP11-274H2.2 | -0,3781 | 4,43087E-21 |
| ACE2 | AFAP1 | -0,3782 | 4,27877E-21 |
| ACE2 | RP11-288H12.3 | -0,3783 | 4,16528E-21 |
| ACE2 | AC000123.4 | -0,3783 | 4,19776E-21 |
| ACE2 | WDR86 | -0,3783 | 4,13482E-21 |
| ACE2 | RP11-638I2.8 | -0,3784 | 4,11627E-21 |
| ACE2 | MALT1 | -0,3784 | 4,027E-21 |
| ACE2 | FAM193B | -0,3785 | 3,98682E-21 |
| ACE2 | KLRC4 | -0,3785 | 3,98363E-21 |
| ACE2 | MIR1304 | -0,3786 | 3,82996E-21 |
| ACE2 | USPL1 | -0,3788 | 3,72642E-21 |
| ACE2 | CTD-2027I19.2 | -0,3788 | 3,69246E-21 |
| ACE2 | RP1-142L7.5 | -0,3789 | 3,54175E-21 |
| ACE2 | ZBTB10 | -0,3789 | 3,58758E-21 |
| ACE2 | FRZB | -0,379 | 3,50183E-21 |
| ACE2 | KCND1 | -0,379 | 3,53479E-21 |
| ACE2 | PNRC2 | -0,3791 | 3,41973E-21 |
| ACE2 | GPR155 | -0,3792 | 3,28882E-21 |
| ACE2 | RP11-564A8.8 | -0,3793 | 3,20333E-21 |
| ACE2 | CTC-529I10.1 | -0,3793 | 3,26526E-21 |
| ACE2 | ZEB2-AS1 | -0,3794 | 3,17045E-21 |
| ACE2 | RP11-199F11.2 | -0,3797 | 2,88751E-21 |
| ACE2 | FOXP1 | -0,3798 | 2,8395E-21 |
| ACE2 | SLC35G5 | -0,3798 | 2,80695E-21 |
| ACE2 | KMT5C | -0,3799 | 2,76316E-21 |
| ACE2 | RP1-20B11.2 | -0,38 | 2,70698E-21 |
| ACE2 | FAM131B | -0,38 | 2,67718E-21 |
| ACE2 | MAPK8IP3 | -0,38 | 2,68129E-21 |
| ACE2 | PARD6G | -0,38 | 2,71135E-21 |
| ACE2 | HBB | -0,3801 | 2,59861E-21 |
| ACE2 | C11orf65 | -0,3803 | 2,50364E-21 |
| ACE2 | TMEM136 | -0,3803 | 2,50365E-21 |
| ACE2 | RP11-709A23.2 | -0,3803 | 2,51827E-21 |
| ACE2 | BTBD11 | -0,3803 | 2,47243E-21 |

| | | | |
|---|---|---|---|
| ACE2 | GALNT16 | -0,3803 | 2,48706E-21 |
| ACE2 | MEF2B | -0,3803 | 2,51841E-21 |
| ACE2 | PROX1 | -0,3804 | 2,46503E-21 |
| ACE2 | ATP6V1C2 | -0,3804 | 2,4495E-21 |
| ACE2 | SLC12A7 | -0,3804 | 2,41709E-21 |
| ACE2 | LIMA1 | -0,3805 | 2,37153E-21 |
| ACE2 | PRTFDC1 | -0,3807 | 2,28112E-21 |
| ACE2 | RP1-69D17.4 | -0,3808 | 2,20878E-21 |
| ACE2 | RPL17P50 | -0,3808 | 2,21452E-21 |
| ACE2 | TMEM173 | -0,3809 | 2,15733E-21 |
| ACE2 | RP11-81H14.2 | -0,381 | 2,06749E-21 |
| ACE2 | TNFSF13B | -0,3811 | 2,0069E-21 |
| ACE2 | RP11-274B21.3 | -0,3812 | 1,984E-21 |
| ACE2 | MARCH5 | -0,3812 | 1,99758E-21 |
| ACE2 | SLC26A10 | -0,3812 | 1,98611E-21 |
| ACE2 | RP11-563J2.3 | -0,3813 | 1,91407E-21 |
| ACE2 | SNORA66 | -0,3815 | 1,83431E-21 |
| ACE2 | PLXDC1 | -0,3816 | 1,76541E-21 |
| ACE2 | LUC7L3 | -0,3818 | 1,70204E-21 |
| ACE2 | SELE | -0,382 | 1,59381E-21 |
| ACE2 | NBL1 | -0,3821 | 1,55169E-21 |
| ACE2 | DLEU2 | -0,3821 | 1,55962E-21 |
| ACE2 | LSMEM1 | -0,3823 | 1,49109E-21 |
| ACE2 | FHL3 | -0,3824 | 1,44015E-21 |
| ACE2 | C1orf198 | -0,3824 | 1,45134E-21 |
| ACE2 | BCORL1 | -0,3824 | 1,44784E-21 |
| ACE2 | CDK11B | -0,3825 | 1,42288E-21 |
| ACE2 | VEGFD | -0,3825 | 1,41728E-21 |
| ACE2 | RP11-603J24.5 | -0,3826 | 1,38907E-21 |
| ACE2 | PDLIM3 | -0,3828 | 1,32151E-21 |
| ACE2 | RP11-327J17.9 | -0,3829 | 1,25696E-21 |
| ACE2 | HSPB1 | -0,383 | 1,25061E-21 |
| ACE2 | EFHD2 | -0,3831 | 1,20304E-21 |
| ACE2 | RP4-530I15.9 | -0,3832 | 1,17338E-21 |
| ACE2 | CCND2 | -0,3833 | 1,15747E-21 |
| ACE2 | SPATS2 | -0,3834 | 1,11885E-21 |
| ACE2 | RP13-554M15.7 | -0,3834 | 1,111E-21 |
| ACE2 | RP1-240B8.3 | -0,3836 | 1,05444E-21 |
| ACE2 | DRAP1 | -0,3836 | 1,04737E-21 |
| ACE2 | CDK17 | -0,3837 | 1,04311E-21 |
| ACE2 | ARHGEF25 | -0,3838 | 1,00887E-21 |
| ACE2 | KRT8P50 | -0,3838 | 1,00817E-21 |
| ACE2 | TCIRG1 | -0,384 | 9,52697E-22 |
| ACE2 | RP5-1073O3.2 | -0,3841 | 9,17004E-22 |
| ACE2 | TRAF6 | -0,3841 | 9,21625E-22 |
| ACE2 | RP11-327J17.7 | -0,3841 | 9,36817E-22 |
| ACE2 | MSC | -0,3842 | 9,082E-22 |
| ACE2 | STMN1P1 | -0,3842 | 9,13051E-22 |
| ACE2 | PIGHP1 | -0,3843 | 8,83138E-22 |
| ACE2 | ZNF83 | -0,3843 | 8,86194E-22 |
| ACE2 | CCNH | -0,3844 | 8,49249E-22 |
| ACE2 | RP1-30M3.6 | -0,3844 | 8,50971E-22 |
| ACE2 | THAP9-AS1 | -0,3845 | 8,33008E-22 |
| ACE2 | IL18BP | -0,3845 | 8,41467E-22 |

| | | | |
|---|---|---|---|
| ACE2 | MAN1C1 | -0,3846 | 8,19758E-22 |
| ACE2 | PNRC1 | -0,3847 | 7,99505E-22 |
| ACE2 | RP11-664I21.6 | -0,3847 | 7,89913E-22 |
| ACE2 | MASP1 | -0,3849 | 7,60816E-22 |
| ACE2 | GAPDHP61 | -0,3849 | 7,51911E-22 |
| ACE2 | HTR5BP | -0,385 | 7,35547E-22 |
| ACE2 | OR52K3P | -0,385 | 7,32324E-22 |
| ACE2 | MMP25-AS1 | -0,385 | 7,42255E-22 |
| ACE2 | CMC2 | -0,385 | 7,30985E-22 |
| ACE2 | LINC00607 | -0,3851 | 7,11156E-22 |
| ACE2 | MALAT1 | -0,3853 | 6,83528E-22 |
| ACE2 | CHRNE | -0,3854 | 6,65223E-22 |
| ACE2 | EMID1 | -0,3854 | 6,66531E-22 |
| ACE2 | ARHGEF6 | -0,3855 | 6,36576E-22 |
| ACE2 | RP11-223P11.3 | -0,3856 | 6,33734E-22 |
| ACE2 | KLRG1 | -0,3856 | 6,33428E-22 |
| ACE2 | NCK2 | -0,3857 | 6,0634E-22 |
| ACE2 | ZDHHC14 | -0,3857 | 6,11268E-22 |
| ACE2 | SSPN | -0,3857 | 6,05886E-22 |
| ACE2 | NUFIP2 | -0,3858 | 5,93235E-22 |
| ACE2 | SRSF4 | -0,3861 | 5,55305E-22 |
| ACE2 | FAM131A | -0,3861 | 5,45112E-22 |
| ACE2 | LINC01140 | -0,3862 | 5,27806E-22 |
| ACE2 | DCUN1D4 | -0,3862 | 5,31136E-22 |
| ACE2 | CACNA2D3-AS1 | -0,3863 | 5,26259E-22 |
| ACE2 | RP11-23P13.6 | -0,3863 | 5,21382E-22 |
| ACE2 | CCDC9 | -0,3863 | 5,24933E-22 |
| ACE2 | ATF3 | -0,3864 | 5,1016E-22 |
| ACE2 | HOXB-AS3 | -0,3864 | 5,09376E-22 |
| ACE2 | ZSWIM4 | -0,3864 | 5,06414E-22 |
| ACE2 | IVNS1ABP | -0,3865 | 4,98595E-22 |
| ACE2 | PTGER4 | -0,3867 | 4,66837E-22 |
| ACE2 | SMAD3 | -0,3867 | 4,73454E-22 |
| ACE2 | RN7SL809P | -0,3868 | 4,55135E-22 |
| ACE2 | RP11-230F18.5 | -0,3869 | 4,47406E-22 |
| ACE2 | RP11-576I22.2 | -0,387 | 4,33529E-22 |
| ACE2 | HIST1H2AL | -0,387 | 4,36582E-22 |
| ACE2 | PPT2 | -0,3871 | 4,16324E-22 |
| ACE2 | RP11-157H4.1 | -0,3871 | 4,24782E-22 |
| ACE2 | RPA4 | -0,3871 | 4,17252E-22 |
| ACE2 | GUK1 | -0,3872 | 4,0518E-22 |
| ACE2 | TFEB | -0,3874 | 3,92754E-22 |
| ACE2 | RP11-325E14.5 | -0,3874 | 3,85212E-22 |
| ACE2 | NTRK1 | -0,3875 | 3,83565E-22 |
| ACE2 | SH3GL1P1 | -0,3875 | 3,82445E-22 |
| ACE2 | SHC2 | -0,3877 | 3,60683E-22 |
| ACE2 | BCL3 | -0,3877 | 3,59422E-22 |
| ACE2 | CXCL8 | -0,3878 | 3,53638E-22 |
| ACE2 | GNAI2 | -0,3879 | 3,40157E-22 |
| ACE2 | RP11-115N4.1 | -0,388 | 3,2972E-22 |
| ACE2 | HPSE2 | -0,388 | 3,28361E-22 |
| ACE2 | RP11-213G2.3 | -0,3881 | 3,26691E-22 |
| ACE2 | RP11-21G15.1 | -0,3881 | 3,23429E-22 |
| ACE2 | TCF15 | -0,3881 | 3,27047E-22 |

| | | | |
|---|---|---|---|
| ACE2 | ETS1 | -0,3884 | 3,01819E-22 |
| ACE2 | RP11-490B18.6 | -0,3884 | 2,95883E-22 |
| ACE2 | CTD-3035K23.3 | -0,3884 | 2,99432E-22 |
| ACE2 | RP11-434H14.1 | -0,3885 | 2,87221E-22 |
| ACE2 | IMMP1LP1 | -0,3885 | 2,89166E-22 |
| ACE2 | CLC | -0,3885 | 2,92686E-22 |
| ACE2 | AC116366.6 | -0,3886 | 2,79439E-22 |
| ACE2 | TRGV4 | -0,3886 | 2,8284E-22 |
| ACE2 | PITPNM2 | -0,3886 | 2,81831E-22 |
| ACE2 | PLCB1 | -0,3887 | 2,74372E-22 |
| ACE2 | AC069363.1 | -0,3888 | 2,69617E-22 |
| ACE2 | ZNF683 | -0,3889 | 2,63517E-22 |
| ACE2 | RP11-771K4.3 | -0,3889 | 2,6134E-22 |
| ACE2 | GLA | -0,3889 | 2,61266E-22 |
| ACE2 | XAF1 | -0,389 | 2,54775E-22 |
| ACE2 | AC064875.2 | -0,3892 | 2,43338E-22 |
| ACE2 | FNDC9 | -0,3892 | 2,38997E-22 |
| ACE2 | MYO3A | -0,3892 | 2,41654E-22 |
| ACE2 | GDI1 | -0,3893 | 2,33656E-22 |
| ACE2 | ATP2B1-AS1 | -0,3895 | 2,21965E-22 |
| ACE2 | ABCD4 | -0,3896 | 2,14766E-22 |
| ACE2 | ANO1 | -0,3897 | 2,13905E-22 |
| ACE2 | MAP7D1 | -0,3898 | 2,03978E-22 |
| ACE2 | IL15RA | -0,3898 | 2,07798E-22 |
| ACE2 | RP11-417N10.5 | -0,39 | 1,9383E-22 |
| ACE2 | EML4 | -0,3901 | 1,91986E-22 |
| ACE2 | SNHG1 | -0,3901 | 1,88143E-22 |
| ACE2 | PTPN22 | -0,3902 | 1,84858E-22 |
| ACE2 | RP11-168K11.3 | -0,3902 | 1,83232E-22 |
| ACE2 | FDX1P1 | -0,3903 | 1,77505E-22 |
| ACE2 | RTEL1P1 | -0,3904 | 1,75947E-22 |
| ACE2 | FAM196B | -0,3904 | 1,77195E-22 |
| ACE2 | LGALS17A | -0,3904 | 1,76424E-22 |
| ACE2 | NIT1 | -0,3906 | 1,68099E-22 |
| ACE2 | LMO2 | -0,3906 | 1,64788E-22 |
| ACE2 | SAP25 | -0,3907 | 1,611E-22 |
| ACE2 | RP5-894A10.2 | -0,3908 | 1,55448E-22 |
| ACE2 | RP11-817I4.2 | -0,3908 | 1,55627E-22 |
| ACE2 | GPR63 | -0,3909 | 1,539E-22 |
| ACE2 | RP11-536K7.3 | -0,391 | 1,47325E-22 |
| ACE2 | FXYD2 | -0,391 | 1,48483E-22 |
| ACE2 | RP11-139H15.5 | -0,391 | 1,47704E-22 |
| ACE2 | RP5-1000K24.2 | -0,3911 | 1,4427E-22 |
| ACE2 | SRGAP1 | -0,3912 | 1,41361E-22 |
| ACE2 | RP11-553L6.2 | -0,3913 | 1,38395E-22 |
| ACE2 | RP1-151F17.2 | -0,3915 | 1,29544E-22 |
| ACE2 | RP11-395G23.3 | -0,3915 | 1,29693E-22 |
| ACE2 | HMG20B | -0,3916 | 1,26453E-22 |
| ACE2 | TSC22D2 | -0,3919 | 1,17687E-22 |
| ACE2 | HIST1H2AD | -0,392 | 1,12955E-22 |
| ACE2 | ADGRB1 | -0,3922 | 1,08645E-22 |
| ACE2 | RP11-631N16.4 | -0,3922 | 1,0888E-22 |
| ACE2 | HRH1 | -0,3927 | 9,3718E-23 |
| ACE2 | CD244 | -0,3928 | 9,07743E-23 |

| | | | |
|---|---|---|---|
| ACE2 | NEURL1 | -0,3929 | 8,91188E-23 |
| ACE2 | ZNF821 | -0,3929 | 9,02247E-23 |
| ACE2 | HOXB4 | -0,3929 | 9,00164E-23 |
| ACE2 | CTC-444N24.8 | -0,3929 | 8,84937E-23 |
| ACE2 | CEACAM3 | -0,393 | 8,64743E-23 |
| ACE2 | VAC14-AS1 | -0,3931 | 8,44602E-23 |
| ACE2 | SOD1P3 | -0,3933 | 8,06581E-23 |
| ACE2 | LRFN1 | -0,3933 | 8,00513E-23 |
| ACE2 | RP11-64K12.8 | -0,3934 | 7,77717E-23 |
| ACE2 | RP11-16E23.4 | -0,3935 | 7,544E-23 |
| ACE2 | APOL3 | -0,3936 | 7,43413E-23 |
| ACE2 | OVGP1 | -0,3937 | 7,27921E-23 |
| ACE2 | RP11-147L13.14 | -0,3937 | 7,2718E-23 |
| ACE2 | SLC30A7 | -0,3938 | 6,99574E-23 |
| ACE2 | FXYD6 | -0,3938 | 6,97404E-23 |
| ACE2 | LRP1-AS | -0,3939 | 6,8293E-23 |
| ACE2 | ZFC3H1 | -0,3939 | 6,74165E-23 |
| ACE2 | CSRP1 | -0,394 | 6,54724E-23 |
| ACE2 | GPC2 | -0,394 | 6,6244E-23 |
| ACE2 | PLEKHA8P1 | -0,394 | 6,64833E-23 |
| ACE2 | SOCS2-AS1 | -0,394 | 6,72135E-23 |
| ACE2 | RP11-549J18.1 | -0,3941 | 6,43382E-23 |
| ACE2 | CTD-2033D15.2 | -0,3942 | 6,23774E-23 |
| ACE2 | LRMP | -0,3943 | 6,1846E-23 |
| ACE2 | RP11-203M5.8 | -0,3943 | 6,19656E-23 |
| ACE2 | RP11-638I2.10 | -0,3943 | 6,13567E-23 |
| ACE2 | RP3-394A18.1 | -0,3943 | 6,17839E-23 |
| ACE2 | RPS12P28 | -0,3945 | 5,71603E-23 |
| ACE2 | C20orf203 | -0,3945 | 5,72237E-23 |
| ACE2 | CAMTA1 | -0,3946 | 5,63383E-23 |
| ACE2 | TGFB1 | -0,3947 | 5,49012E-23 |
| ACE2 | IL33 | -0,3949 | 5,17854E-23 |
| ACE2 | RP11-876N24.4 | -0,3949 | 5,20921E-23 |
| ACE2 | SAPCD2P3 | -0,395 | 5,11502E-23 |
| ACE2 | TTC7B | -0,3951 | 4,88975E-23 |
| ACE2 | PPFIA3 | -0,3951 | 4,90641E-23 |
| ACE2 | AC006129.2 | -0,3952 | 4,74573E-23 |
| ACE2 | MEF2C | -0,3953 | 4,63217E-23 |
| ACE2 | IST1 | -0,3953 | 4,6807E-23 |
| ACE2 | AMPD2 | -0,3954 | 4,49071E-23 |
| ACE2 | RP11-272L14.2 | -0,3954 | 4,54878E-23 |
| ACE2 | GSDMB | -0,3954 | 4,49804E-23 |
| ACE2 | HBM | -0,3955 | 4,37673E-23 |
| ACE2 | LA16c-358B7.3 | -0,3955 | 4,42168E-23 |
| ACE2 | CFAP69 | -0,3957 | 4,17906E-23 |
| ACE2 | RP13-270P17.1 | -0,3958 | 4,07402E-23 |
| ACE2 | CCR3 | -0,396 | 3,87401E-23 |
| ACE2 | DLG4 | -0,3961 | 3,71981E-23 |
| ACE2 | MAL | -0,3962 | 3,64677E-23 |
| ACE2 | PCDHB14 | -0,3962 | 3,6329E-23 |
| ACE2 | LINC00342 | -0,3963 | 3,57829E-23 |
| ACE2 | ZNF141 | -0,3963 | 3,51178E-23 |
| ACE2 | FAM129C | -0,3963 | 3,57542E-23 |
| ACE2 | PCED1B | -0,3964 | 3,41942E-23 |

| | | | |
|---|---|---|---|
| ACE2 | RP11-155O18.6 | -0,3967 | 3,1712E-23 |
| ACE2 | GNL1 | -0,3967 | 3,19499E-23 |
| ACE2 | RP11-325F22.2 | -0,3967 | 3,183E-23 |
| ACE2 | GDPD3 | -0,397 | 2,92678E-23 |
| ACE2 | IFI16 | -0,3972 | 2,78472E-23 |
| ACE2 | GYPA | -0,3972 | 2,73822E-23 |
| ACE2 | RP11-299L17.3 | -0,3972 | 2,80633E-23 |
| ACE2 | FGF7 | -0,3972 | 2,7972E-23 |
| ACE2 | RN7SL566P | -0,3972 | 2,751E-23 |
| ACE2 | PLAC9 | -0,3973 | 2,73533E-23 |
| ACE2 | ZNF506 | -0,3973 | 2,69358E-23 |
| ACE2 | RP11-876N24.3 | -0,3974 | 2,62059E-23 |
| ACE2 | IFI44L | -0,3975 | 2,54343E-23 |
| ACE2 | MYH10 | -0,3975 | 2,58819E-23 |
| ACE2 | LINC00877 | -0,3976 | 2,51094E-23 |
| ACE2 | ADA | -0,3976 | 2,50685E-23 |
| ACE2 | RP11-260M2.1 | -0,3977 | 2,41773E-23 |
| ACE2 | TUBG2 | -0,3977 | 2,43454E-23 |
| ACE2 | RRP7BP | -0,3977 | 2,44112E-23 |
| ACE2 | TCEA2 | -0,3979 | 2,30618E-23 |
| ACE2 | RP11-73M18.6 | -0,398 | 2,1961E-23 |
| ACE2 | MYO1G | -0,3981 | 2,1543E-23 |
| ACE2 | AC000123.3 | -0,3982 | 2,12051E-23 |
| ACE2 | TNFSF10 | -0,3983 | 2,05645E-23 |
| ACE2 | AP000593.7 | -0,3984 | 1,98101E-23 |
| ACE2 | HIST1H2BF | -0,3985 | 1,96321E-23 |
| ACE2 | APOL1 | -0,3985 | 1,91424E-23 |
| ACE2 | PPFIA4 | -0,3986 | 1,9067E-23 |
| ACE2 | HSPA12A | -0,3986 | 1,89771E-23 |
| ACE2 | TRAF2 | -0,3987 | 1,81638E-23 |
| ACE2 | MARCKS | -0,3988 | 1,81011E-23 |
| ACE2 | NAMA | -0,3989 | 1,71899E-23 |
| ACE2 | NAP1L4P1 | -0,399 | 1,71327E-23 |
| ACE2 | C8orf4 | -0,399 | 1,69422E-23 |
| ACE2 | RP13-554M15.2 | -0,3991 | 1,63589E-23 |
| ACE2 | E2F5 | -0,3994 | 1,51802E-23 |
| ACE2 | PTGS2 | -0,3996 | 1,44132E-23 |
| ACE2 | TRGVB | -0,3997 | 1,39796E-23 |
| ACE2 | RP11-713N11.6 | -0,3997 | 1,40991E-23 |
| ACE2 | MROH7 | -0,3998 | 1,35614E-23 |
| ACE2 | CLDND1 | -0,3998 | 1,34067E-23 |
| ACE2 | HLA-DQB1-AS1 | -0,3998 | 1,34884E-23 |
| ACE2 | RP11-391L3.5 | -0,3998 | 1,37064E-23 |
| ACE2 | SNORA65 | -0,4 | 1,26681E-23 |
| ACE2 | STK11 | -0,4 | 1,26854E-23 |
| ACE2 | ITPRIPL1 | -0,4001 | 1,24786E-23 |
| ACE2 | STMN3 | -0,4001 | 1,24872E-23 |
| ACE2 | CTD-2366F13.2 | -0,4002 | 1,21329E-23 |
| ACE2 | WIPF1 | -0,4003 | 1,1906E-23 |
| ACE2 | CCDC88B | -0,4003 | 1,18345E-23 |
| ACE2 | INTS6L | -0,4003 | 1,18254E-23 |
| ACE2 | LIX1L-AS1 | -0,4004 | 1,16272E-23 |
| ACE2 | PKIA-AS1 | -0,4004 | 1,15556E-23 |
| ACE2 | CTD-2013N17.4 | -0,4007 | 1,04244E-23 |

| | | | |
|---|---|---|---|
| ACE2 | TRAF5 | -0,4008 | 1,03333E-23 |
| ACE2 | RP11-161M6.2 | -0,4008 | 1,0312E-23 |
| ACE2 | KLRF1 | -0,4009 | 9,9848E-24 |
| ACE2 | AEBP1 | -0,4012 | 9,19883E-24 |
| ACE2 | SLC22A17 | -0,4012 | 9,28405E-24 |
| ACE2 | CYTIP | -0,4013 | 8,9023E-24 |
| ACE2 | ASAP1-IT2 | -0,4013 | 9,03838E-24 |
| ACE2 | PBX2 | -0,4014 | 8,68974E-24 |
| ACE2 | RP11-471L13.3 | -0,4014 | 8,78432E-24 |
| ACE2 | PAIP1P1 | -0,4015 | 8,51505E-24 |
| ACE2 | RP4-742C19.12 | -0,4015 | 8,42682E-24 |
| ACE2 | RELA | -0,4016 | 8,1345E-24 |
| ACE2 | AC109333.10 | -0,4016 | 8,24952E-24 |
| ACE2 | MSX1 | -0,4017 | 8,06434E-24 |
| ACE2 | RP11-640N20.4 | -0,4019 | 7,6387E-24 |
| ACE2 | RP11-456P18.2 | -0,402 | 7,30521E-24 |
| ACE2 | ARL4AP5 | -0,402 | 7,29329E-24 |
| ACE2 | RP11-90B9.3 | -0,4021 | 7,16587E-24 |
| ACE2 | EIF4EP1 | -0,4021 | 7,13352E-24 |
| ACE2 | RP11-181E10.3 | -0,4022 | 6,93287E-24 |
| ACE2 | MOB3C | -0,4023 | 6,73617E-24 |
| ACE2 | ZYX | -0,4023 | 6,71902E-24 |
| ACE2 | COL6A6 | -0,4025 | 6,41867E-24 |
| ACE2 | RP11-1046B16.3 | -0,4025 | 6,35605E-24 |
| ACE2 | TLE3 | -0,4025 | 6,40601E-24 |
| ACE2 | MICAL3 | -0,4025 | 6,4653E-24 |
| ACE2 | RP11-701H16.4 | -0,4026 | 6,27622E-24 |
| ACE2 | RP11-97C16.1 | -0,4027 | 6,01366E-24 |
| ACE2 | CHRD | -0,4027 | 6,03171E-24 |
| ACE2 | HIST1H4E | -0,4028 | 5,85711E-24 |
| ACE2 | MB21D1 | -0,4028 | 5,85853E-24 |
| ACE2 | NR2F2-AS1 | -0,4028 | 5,89511E-24 |
| ACE2 | RP13-672B3.5 | -0,4029 | 5,79687E-24 |
| ACE2 | PLA2G4C-AS1 | -0,4029 | 5,72476E-24 |
| ACE2 | RFTN1 | -0,403 | 5,52194E-24 |
| ACE2 | HNRNPDL | -0,403 | 5,52768E-24 |
| ACE2 | IFRD1 | -0,403 | 5,49216E-24 |
| ACE2 | POU5F1P3 | -0,4031 | 5,4532E-24 |
| ACE2 | ATXN7 | -0,4032 | 5,2827E-24 |
| ACE2 | RP11-285F7.2 | -0,4033 | 5,16571E-24 |
| ACE2 | TIGIT | -0,4034 | 4,98651E-24 |
| ACE2 | TRAF1 | -0,4034 | 5,02843E-24 |
| ACE2 | TNIP1 | -0,4035 | 4,79864E-24 |
| ACE2 | POU6F1 | -0,4035 | 4,78163E-24 |
| ACE2 | ENAH | -0,4036 | 4,65182E-24 |
| ACE2 | CENPT | -0,4036 | 4,73818E-24 |
| ACE2 | MDFI | -0,4037 | 4,57088E-24 |
| ACE2 | FEZ1 | -0,4037 | 4,53368E-24 |
| ACE2 | GPATCH2L | -0,4037 | 4,56367E-24 |
| ACE2 | IFI27 | -0,4037 | 4,60571E-24 |
| ACE2 | C2CD4B | -0,4037 | 4,62842E-24 |
| ACE2 | RP3-395M20.9 | -0,4038 | 4,494E-24 |
| ACE2 | MUSTN1 | -0,4038 | 4,43522E-24 |
| ACE2 | GRK2 | -0,4038 | 4,3953E-24 |

| | | | |
|---|---|---|---|
| ACE2 | CTA-276F8.1 | -0,404 | 4,24861E-24 |
| ACE2 | LINC02019 | -0,4041 | 4,05353E-24 |
| ACE2 | CREB5 | -0,4041 | 4,11058E-24 |
| ACE2 | RP11-707G18.1 | -0,4042 | 3,92644E-24 |
| ACE2 | GDF11 | -0,4042 | 3,96106E-24 |
| ACE2 | TIE1 | -0,4043 | 3,8172E-24 |
| ACE2 | LMBR1L | -0,4043 | 3,90021E-24 |
| ACE2 | STK4 | -0,4043 | 3,90865E-24 |
| ACE2 | AC007283.4 | -0,4045 | 3,70742E-24 |
| ACE2 | PLEKHA8 | -0,4045 | 3,6519E-24 |
| ACE2 | TMEM176A | -0,4046 | 3,60039E-24 |
| ACE2 | HSPB8 | -0,4047 | 3,42402E-24 |
| ACE2 | HBA2 | -0,4048 | 3,3707E-24 |
| ACE2 | DCAF5 | -0,4049 | 3,30034E-24 |
| ACE2 | DLGAP4 | -0,4049 | 3,29389E-24 |
| ACE2 | RP11-599J14.2 | -0,405 | 3,22056E-24 |
| ACE2 | RP5-1057I20.5 | -0,405 | 3,18457E-24 |
| ACE2 | RP11-170L3.4 | -0,405 | 3,19781E-24 |
| ACE2 | RP11-166N17.1 | -0,4051 | 3,09669E-24 |
| ACE2 | SCARNA21 | -0,4053 | 2,93769E-24 |
| ACE2 | AC010976.2 | -0,4053 | 2,92384E-24 |
| ACE2 | RBP1 | -0,4053 | 2,92964E-24 |
| ACE2 | RP11-894P9.1 | -0,4055 | 2,7239E-24 |
| ACE2 | NBAT1 | -0,4056 | 2,65242E-24 |
| ACE2 | PRRT1 | -0,4056 | 2,68823E-24 |
| ACE2 | TRPS1 | -0,4056 | 2,70234E-24 |
| ACE2 | CD5 | -0,4057 | 2,57605E-24 |
| ACE2 | CTD-2521M24.13 | -0,4058 | 2,56938E-24 |
| ACE2 | NAPB | -0,4058 | 2,52612E-24 |
| ACE2 | UPB1 | -0,4058 | 2,50536E-24 |
| ACE2 | CDK9 | -0,4059 | 2,48793E-24 |
| ACE2 | RP1-68D18.4 | -0,4059 | 2,4882E-24 |
| ACE2 | MED15 | -0,4061 | 2,32385E-24 |
| ACE2 | RP5-1074L1.4 | -0,4062 | 2,28816E-24 |
| ACE2 | TEX41 | -0,4062 | 2,28483E-24 |
| ACE2 | DGKH | -0,4062 | 2,25448E-24 |
| ACE2 | CTD-2006C1.2 | -0,4063 | 2,17153E-24 |
| ACE2 | TNFAIP8 | -0,4064 | 2,14385E-24 |
| ACE2 | HIST1H1E | -0,4065 | 2,06899E-24 |
| ACE2 | CNTNAP1 | -0,4066 | 2,02031E-24 |
| ACE2 | SLC31A1P1 | -0,4067 | 1,95325E-24 |
| ACE2 | TEAD2 | -0,4067 | 1,9573E-24 |
| ACE2 | CIB2 | -0,4068 | 1,90172E-24 |
| ACE2 | RN7SL15P | -0,4068 | 1,9078E-24 |
| ACE2 | LINC00299 | -0,4069 | 1,83377E-24 |
| ACE2 | SNORD89 | -0,4069 | 1,85335E-24 |
| ACE2 | PGR | -0,407 | 1,82858E-24 |
| ACE2 | AHSP | -0,407 | 1,78429E-24 |
| ACE2 | RP4-738P11.3 | -0,4071 | 1,74207E-24 |
| ACE2 | PLCB4 | -0,4071 | 1,74382E-24 |
| ACE2 | GTPBP1 | -0,4072 | 1,71276E-24 |
| ACE2 | FGF2 | -0,4073 | 1,66118E-24 |
| ACE2 | VASN | -0,4073 | 1,65326E-24 |
| ACE2 | GBP4 | -0,4074 | 1,63209E-24 |

| | | | |
|---|---|---|---|
| ACE2 | SLCO5A1 | -0,4074 | 1,59559E-24 |
| ACE2 | LTBP2 | -0,4074 | 1,61084E-24 |
| ACE2 | ZFP36L1 | -0,4076 | 1,50983E-24 |
| ACE2 | AC022819.3 | -0,4076 | 1,5151E-24 |
| ACE2 | RP11-125O18.1 | -0,4078 | 1,41955E-24 |
| ACE2 | TMEM130 | -0,4078 | 1,43271E-24 |
| ACE2 | RASL12 | -0,4078 | 1,45782E-24 |
| ACE2 | MTND5P25 | -0,408 | 1,36595E-24 |
| ACE2 | RP11-382J24.2 | -0,408 | 1,3613E-24 |
| ACE2 | ADORA2A-AS1 | -0,408 | 1,36705E-24 |
| ACE2 | DPRX | -0,4081 | 1,31499E-24 |
| ACE2 | LEFTY2 | -0,4082 | 1,28166E-24 |
| ACE2 | LOXL2 | -0,4082 | 1,27053E-24 |
| ACE2 | RP11-44K6.4 | -0,4082 | 1,30003E-24 |
| ACE2 | CNRIP1 | -0,4083 | 1,25765E-24 |
| ACE2 | PCBP4 | -0,4084 | 1,20732E-24 |
| ACE2 | CTC-507E2.2 | -0,4085 | 1,18399E-24 |
| ACE2 | AC005740.6 | -0,4086 | 1,14305E-24 |
| ACE2 | SPACA6 | -0,4086 | 1,14556E-24 |
| ACE2 | SCN4B | -0,4087 | 1,10096E-24 |
| ACE2 | RNF44 | -0,4089 | 1,0434E-24 |
| ACE2 | CSNK1G2 | -0,4089 | 1,03948E-24 |
| ACE2 | ATAD2B | -0,409 | 1,01348E-24 |
| ACE2 | ITGA4 | -0,409 | 1,01779E-24 |
| ACE2 | ISG15 | -0,4091 | 9,9113E-25 |
| ACE2 | SLC8A1-AS1 | -0,4092 | 9,58108E-25 |
| ACE2 | DOCK4-AS1 | -0,4092 | 9,69085E-25 |
| ACE2 | RP5-1071N3.1 | -0,4093 | 9,36325E-25 |
| ACE2 | SRSF2 | -0,4093 | 9,48432E-25 |
| ACE2 | NOL4L | -0,4093 | 9,43386E-25 |
| ACE2 | KB-1460A1.5 | -0,4094 | 9,11078E-25 |
| ACE2 | AP000240.9 | -0,4094 | 8,99349E-25 |
| ACE2 | ARL10 | -0,4095 | 8,93314E-25 |
| ACE2 | PABPN1 | -0,4095 | 8,91942E-25 |
| ACE2 | ERF | -0,4095 | 8,84811E-25 |
| ACE2 | UBE2V1P1 | -0,4096 | 8,52503E-25 |
| ACE2 | SNORA61 | -0,4097 | 8,48371E-25 |
| ACE2 | CRMP1 | -0,4098 | 8,16048E-25 |
| ACE2 | ETV7 | -0,4098 | 8,0607E-25 |
| ACE2 | RP4-598P13.1 | -0,41 | 7,63194E-25 |
| ACE2 | OTULIN | -0,41 | 7,66253E-25 |
| ACE2 | TBL1XR1-AS1 | -0,4101 | 7,37269E-25 |
| ACE2 | TNFRSF25 | -0,4102 | 7,3503E-25 |
| ACE2 | CD28 | -0,4103 | 7,07877E-25 |
| ACE2 | AC007879.2 | -0,4103 | 7,11528E-25 |
| ACE2 | RP11-1114A5.4 | -0,4106 | 6,41971E-25 |
| ACE2 | TMSB4X | -0,4107 | 6,30935E-25 |
| ACE2 | RP11-384C4.2 | -0,4108 | 6,12908E-25 |
| ACE2 | UPF3B | -0,4108 | 6,03265E-25 |
| ACE2 | ELK4 | -0,411 | 5,83487E-25 |
| ACE2 | XXYLT1-AS2 | -0,411 | 5,73195E-25 |
| ACE2 | RP1-27K12.4 | -0,4111 | 5,59885E-25 |
| ACE2 | RP11-4B16.1 | -0,4111 | 5,60635E-25 |
| ACE2 | SENP5 | -0,4112 | 5,41451E-25 |

| | | | |
|---|---|---|---|
| ACE2 | ZNF697 | -0,4113 | 5,27723E-25 |
| ACE2 | BTBD10P2 | -0,4113 | 5,28627E-25 |
| ACE2 | RP11-1023L17.1 | -0,4114 | 5,12129E-25 |
| ACE2 | STK38 | -0,4114 | 5,1972E-25 |
| ACE2 | ZCCHC18 | -0,4114 | 5,09355E-25 |
| ACE2 | SLIT3 | -0,4115 | 5,03937E-25 |
| ACE2 | HLA-B | -0,4115 | 5,04094E-25 |
| ACE2 | AC005104.3 | -0,4116 | 4,80759E-25 |
| ACE2 | TMEM176B | -0,4116 | 4,83135E-25 |
| ACE2 | GNG2 | -0,4116 | 4,84999E-25 |
| ACE2 | LINC00670 | -0,4117 | 4,67144E-25 |
| ACE2 | RP11-728F11.4 | -0,4118 | 4,62751E-25 |
| ACE2 | TMEM259 | -0,4119 | 4,42013E-25 |
| ACE2 | SNORD10 | -0,412 | 4,34116E-25 |
| ACE2 | SIPA1L2 | -0,4121 | 4,26003E-25 |
| ACE2 | UBD | -0,4123 | 3,93441E-25 |
| ACE2 | RP11-225B17.2 | -0,4124 | 3,90736E-25 |
| ACE2 | RP11-120K18.3 | -0,4127 | 3,54416E-25 |
| ACE2 | SHISA3 | -0,413 | 3,23773E-25 |
| ACE2 | IFITM4P | -0,413 | 3,23483E-25 |
| ACE2 | LINC01163 | -0,413 | 3,2067E-25 |
| ACE2 | TMEM200A | -0,4132 | 3,05819E-25 |
| ACE2 | LINC00997 | -0,4134 | 2,85915E-25 |
| ACE2 | UGCG | -0,4134 | 2,84926E-25 |
| ACE2 | RP11-152L20.3 | -0,4134 | 2,90425E-25 |
| ACE2 | MIR155HG | -0,4134 | 2,88296E-25 |
| ACE2 | CSRP2 | -0,4135 | 2,7845E-25 |
| ACE2 | RP11-216B9.9 | -0,4136 | 2,70323E-25 |
| ACE2 | REC8 | -0,4138 | 2,58504E-25 |
| ACE2 | KSR1 | -0,4138 | 2,57046E-25 |
| ACE2 | ELL | -0,4138 | 2,60086E-25 |
| ACE2 | IL12A | -0,4139 | 2,5196E-25 |
| ACE2 | RP11-214J9.1 | -0,414 | 2,45457E-25 |
| ACE2 | CTD-2013N17.6 | -0,4141 | 2,34728E-25 |
| ACE2 | PSPC1-AS2 | -0,4143 | 2,21899E-25 |
| ACE2 | NLGN2 | -0,4143 | 2,21377E-25 |
| ACE2 | HLA-A | -0,4145 | 2,09204E-25 |
| ACE2 | CCDC102B | -0,4146 | 2,06199E-25 |
| ACE2 | ARL4C | -0,4147 | 1,9932E-25 |
| ACE2 | BACH1-IT2 | -0,4148 | 1,90323E-25 |
| ACE2 | CBARP | -0,4149 | 1,86145E-25 |
| ACE2 | RP4-737E23.6 | -0,4149 | 1,84388E-25 |
| ACE2 | INPP5F | -0,4151 | 1,7663E-25 |
| ACE2 | DTX1 | -0,4152 | 1,71211E-25 |
| ACE2 | YY1AP1 | -0,4153 | 1,65383E-25 |
| ACE2 | ANKRD10-IT1 | -0,4156 | 1,53952E-25 |
| ACE2 | SLC2A6 | -0,4158 | 1,4363E-25 |
| ACE2 | TKTL1 | -0,4158 | 1,42064E-25 |
| ACE2 | EGFL6 | -0,4159 | 1,39807E-25 |
| ACE2 | NOD2 | -0,4162 | 1,29151E-25 |
| ACE2 | HCAR2 | -0,4163 | 1,25059E-25 |
| ACE2 | TRIM47 | -0,4163 | 1,24276E-25 |
| ACE2 | HES4 | -0,4164 | 1,22167E-25 |
| ACE2 | RP11-417F21.1 | -0,4164 | 1,21137E-25 |

| | | | |
|---|---|---|---|
| ACE2 | ANTXR1 | -0,4167 | 1,09953E-25 |
| ACE2 | AC007919.18 | -0,4167 | 1,11221E-25 |
| ACE2 | MEX3A | -0,4168 | 1,0745E-25 |
| ACE2 | SNORD14E | -0,4168 | 1,07166E-25 |
| ACE2 | AC004623.3 | -0,4168 | 1,06088E-25 |
| ACE2 | RP3-405J10.2 | -0,4169 | 1,02722E-25 |
| ACE2 | LIMK2 | -0,4169 | 1,03726E-25 |
| ACE2 | C9orf3 | -0,417 | 1,00596E-25 |
| ACE2 | RP5-855D21.3 | -0,4172 | 9,66923E-26 |
| ACE2 | RP11-713H12.2 | -0,4172 | 9,54558E-26 |
| ACE2 | RP11-168J18.6 | -0,4175 | 8,62009E-26 |
| ACE2 | NLGN3 | -0,4175 | 8,63089E-26 |
| ACE2 | GPR176 | -0,4176 | 8,53668E-26 |
| ACE2 | RP11-327F22.4 | -0,4177 | 8,14598E-26 |
| ACE2 | VMP1 | -0,4181 | 7,31389E-26 |
| ACE2 | SUZ12P1 | -0,4183 | 6,85812E-26 |
| ACE2 | HES7 | -0,4184 | 6,64071E-26 |
| ACE2 | MIR6753 | -0,4185 | 6,4546E-26 |
| ACE2 | RP11-727F15.13 | -0,4187 | 6,10366E-26 |
| ACE2 | CCDC150P1 | -0,4193 | 5,17832E-26 |
| ACE2 | RP11-525G13.2 | -0,4194 | 4,95418E-26 |
| ACE2 | FOXS1 | -0,4194 | 4,98309E-26 |
| ACE2 | RP9P | -0,4196 | 4,63784E-26 |
| ACE2 | EZH2 | -0,4196 | 4,71484E-26 |
| ACE2 | GAN | -0,4196 | 4,72497E-26 |
| ACE2 | CTD-2081C10.7 | -0,4197 | 4,6345E-26 |
| ACE2 | NIPSNAP3B | -0,4199 | 4,26081E-26 |
| ACE2 | IGFBP7-AS1 | -0,4204 | 3,66076E-26 |
| ACE2 | COL11A2 | -0,4204 | 3,76287E-26 |
| ACE2 | CEP85L | -0,4206 | 3,52199E-26 |
| ACE2 | USP3-AS1 | -0,4206 | 3,50398E-26 |
| ACE2 | N4BP2L1 | -0,4207 | 3,38578E-26 |
| ACE2 | HIST1H2BG | -0,4209 | 3,21596E-26 |
| ACE2 | NLRC5 | -0,4209 | 3,17791E-26 |
| ACE2 | BMP7 | -0,4209 | 3,1562E-26 |
| ACE2 | SPNS3 | -0,4211 | 3,05807E-26 |
| ACE2 | TRGC1 | -0,4212 | 2,94407E-26 |
| ACE2 | RP11-803D5.1 | -0,4213 | 2,80425E-26 |
| ACE2 | RP11-347L15.1 | -0,4213 | 2,8702E-26 |
| ACE2 | CTD-3035K23.7 | -0,4214 | 2,74523E-26 |
| ACE2 | PPP1R12B | -0,4216 | 2,63712E-26 |
| ACE2 | IFI44 | -0,4218 | 2,47796E-26 |
| ACE2 | LARP6 | -0,4218 | 2,47988E-26 |
| ACE2 | MYO9B | -0,4218 | 2,45353E-26 |
| ACE2 | SH2D1B | -0,422 | 2,33329E-26 |
| ACE2 | RP4-681N20.5 | -0,4221 | 2,26304E-26 |
| ACE2 | RP11-168O16.1 | -0,4222 | 2,16628E-26 |
| ACE2 | AC093673.5 | -0,4222 | 2,1733E-26 |
| ACE2 | TPT1-AS1 | -0,4222 | 2,15465E-26 |
| ACE2 | ITGAL | -0,4222 | 2,17538E-26 |
| ACE2 | GNG8 | -0,4222 | 2,16441E-26 |
| ACE2 | MIR100HG | -0,4224 | 2,05876E-26 |
| ACE2 | FSCN1 | -0,4225 | 1,98023E-26 |
| ACE2 | CTD-2135J3.3 | -0,4226 | 1,93837E-26 |

| | | | |
|---|---|---|---|
| ACE2 | CASK-AS1 | -0,4226 | 1,92524E-26 |
| ACE2 | AC005550.3 | -0,4228 | 1,82578E-26 |
| ACE2 | PTGDS | -0,4228 | 1,81084E-26 |
| ACE2 | RP11-291I6.2 | -0,4229 | 1,77638E-26 |
| ACE2 | DLL4 | -0,423 | 1,7289E-26 |
| ACE2 | CCDC28B | -0,4231 | 1,67808E-26 |
| ACE2 | PCAT19 | -0,4231 | 1,67138E-26 |
| ACE2 | HAUS7 | -0,4231 | 1,65475E-26 |
| ACE2 | PHF13 | -0,4232 | 1,63655E-26 |
| ACE2 | CTB-193M12.5 | -0,4232 | 1,63117E-26 |
| ACE2 | SLC25A28 | -0,4234 | 1,51326E-26 |
| ACE2 | RP11-115L11.1 | -0,4236 | 1,41278E-26 |
| ACE2 | LRP5L | -0,4236 | 1,44133E-26 |
| ACE2 | RP11-360F5.1 | -0,4237 | 1,40625E-26 |
| ACE2 | SRSF1 | -0,4237 | 1,37233E-26 |
| ACE2 | ZNF876P | -0,4238 | 1,35508E-26 |
| ACE2 | RP5-1085F17.3 | -0,4239 | 1,31252E-26 |
| ACE2 | SPHK1 | -0,424 | 1,27972E-26 |
| ACE2 | RASA2 | -0,4241 | 1,24831E-26 |
| ACE2 | KIF21B | -0,4243 | 1,1587E-26 |
| ACE2 | GUCY1A2 | -0,4243 | 1,16515E-26 |
| ACE2 | GATA6-AS1 | -0,4243 | 1,1649E-26 |
| ACE2 | SRSF5 | -0,4244 | 1,13326E-26 |
| ACE2 | RP5-1028K7.2 | -0,4244 | 1,10993E-26 |
| ACE2 | SP100 | -0,4247 | 1,01322E-26 |
| ACE2 | PDE7A | -0,4248 | 9,91497E-27 |
| ACE2 | NAALADL1 | -0,4248 | 9,918E-27 |
| ACE2 | ATP1B1P1 | -0,4249 | 9,82351E-27 |
| ACE2 | FBXW7 | -0,4249 | 9,67962E-27 |
| ACE2 | POLB | -0,4249 | 9,60239E-27 |
| ACE2 | TRAV8-2 | -0,4251 | 9,21514E-27 |
| ACE2 | PPP4R1L | -0,4251 | 8,99797E-27 |
| ACE2 | RP11-23J18.1 | -0,4252 | 8,92283E-27 |
| ACE2 | CLASRP | -0,4252 | 8,7312E-27 |
| ACE2 | IL23R | -0,4253 | 8,60903E-27 |
| ACE2 | C1orf54 | -0,4253 | 8,4864E-27 |
| ACE2 | CXCL11 | -0,4253 | 8,50982E-27 |
| ACE2 | ZBTB20-AS1 | -0,4254 | 8,27074E-27 |
| ACE2 | TAPBP | -0,4255 | 8,07661E-27 |
| ACE2 | P2RY10 | -0,4258 | 7,43841E-27 |
| ACE2 | PPARD | -0,426 | 6,8685E-27 |
| ACE2 | CLEC4F | -0,4262 | 6,59108E-27 |
| ACE2 | HYPK | -0,4262 | 6,52635E-27 |
| ACE2 | CMTM3 | -0,4263 | 6,36418E-27 |
| ACE2 | AC006160.5 | -0,4264 | 6,1197E-27 |
| ACE2 | CLK3 | -0,4264 | 6,17639E-27 |
| ACE2 | PLEKHO1 | -0,4265 | 5,90753E-27 |
| ACE2 | RP11-326I11.5 | -0,4266 | 5,73911E-27 |
| ACE2 | HCP5 | -0,4266 | 5,73638E-27 |
| ACE2 | FBXW4P1 | -0,4268 | 5,5046E-27 |
| ACE2 | HYMAI | -0,4269 | 5,34695E-27 |
| ACE2 | RP11-73M18.8 | -0,4269 | 5,22707E-27 |
| ACE2 | SGIP1 | -0,4271 | 4,97282E-27 |
| ACE2 | MICA | -0,4271 | 5,00688E-27 |

| | | | |
|---|---|---|---|
| ACE2 | RP11-324E6.6 | -0,4271 | 4,95892E-27 |
| ACE2 | CTD-2382E5.6 | -0,4272 | 4,8058E-27 |
| ACE2 | RP3-405J10.3 | -0,4274 | 4,56838E-27 |
| ACE2 | LRRTM4 | -0,4275 | 4,40123E-27 |
| ACE2 | CYGB | -0,4275 | 4,45615E-27 |
| ACE2 | ACAP1 | -0,4276 | 4,30106E-27 |
| ACE2 | RP11-165D6.1 | -0,4278 | 4,04997E-27 |
| ACE2 | AC005387.2 | -0,4278 | 4,00309E-27 |
| ACE2 | PRDX3P1 | -0,4278 | 3,99663E-27 |
| ACE2 | ABHD17A | -0,4279 | 3,93213E-27 |
| ACE2 | SEMA4C | -0,428 | 3,77344E-27 |
| ACE2 | TRBV23-1 | -0,428 | 3,77428E-27 |
| ACE2 | IFT43 | -0,428 | 3,76131E-27 |
| ACE2 | EDNRA | -0,4281 | 3,72097E-27 |
| ACE2 | USF1 | -0,4286 | 3,16936E-27 |
| ACE2 | VEGFC | -0,4286 | 3,18283E-27 |
| ACE2 | CD40 | -0,4286 | 3,15791E-27 |
| ACE2 | NPIPB5 | -0,4287 | 3,03213E-27 |
| ACE2 | AL133243.2 | -0,4288 | 2,93247E-27 |
| ACE2 | ANKDD1A | -0,4289 | 2,91838E-27 |
| ACE2 | DONSON | -0,4289 | 2,83838E-27 |
| ACE2 | GCH1 | -0,4295 | 2,38711E-27 |
| ACE2 | CSF1 | -0,4297 | 2,2629E-27 |
| ACE2 | PDZD4 | -0,4297 | 2,2812E-27 |
| ACE2 | NOTCH3 | -0,4299 | 2,1143E-27 |
| ACE2 | HCG4P5 | -0,4302 | 1,94757E-27 |
| ACE2 | PALM | -0,4302 | 1,91421E-27 |
| ACE2 | ITIH5 | -0,4303 | 1,88064E-27 |
| ACE2 | CTA-212A2.1 | -0,4306 | 1,70232E-27 |
| ACE2 | AC104809.4 | -0,4307 | 1,65955E-27 |
| ACE2 | STX1A | -0,4309 | 1,56537E-27 |
| ACE2 | ETV6 | -0,4309 | 1,56147E-27 |
| ACE2 | XCL1 | -0,4311 | 1,47895E-27 |
| ACE2 | EFEMP2 | -0,4311 | 1,46297E-27 |
| ACE2 | RP11-356B19.11 | -0,4312 | 1,42262E-27 |
| ACE2 | ITGB7 | -0,4312 | 1,4404E-27 |
| ACE2 | HIST1H3D | -0,4313 | 1,38679E-27 |
| ACE2 | CTD-2371O3.2 | -0,4314 | 1,32565E-27 |
| ACE2 | RP11-196H14.2 | -0,4316 | 1,263E-27 |
| ACE2 | GEM | -0,4319 | 1,15952E-27 |
| ACE2 | RN7SL608P | -0,4321 | 1,09263E-27 |
| ACE2 | HIST1H2BE | -0,4322 | 1,03406E-27 |
| ACE2 | SLA2 | -0,4322 | 1,03123E-27 |
| ACE2 | GDPD5 | -0,4325 | 9,49284E-28 |
| ACE2 | ARSI | -0,4327 | 8,89949E-28 |
| ACE2 | RP11-968A15.2 | -0,4328 | 8,67424E-28 |
| ACE2 | LIMS1-AS1 | -0,4329 | 8,33084E-28 |
| ACE2 | SAMD14 | -0,4329 | 8,31032E-28 |
| ACE2 | XXbac-BPG252P9.9 | -0,433 | 8,17608E-28 |
| ACE2 | RP1-79C4.4 | -0,4331 | 7,98223E-28 |
| ACE2 | CTC-506B8.1 | -0,4331 | 7,90665E-28 |
| ACE2 | PCDH15 | -0,4332 | 7,59748E-28 |
| ACE2 | RASGRP1 | -0,4332 | 7,80396E-28 |
| ACE2 | RNF122 | -0,4334 | 7,16164E-28 |

| | | | |
|---|---|---|---|
| ACE2 | SGSM2 | -0,4335 | 7,01373E-28 |
| ACE2 | SLC6A1 | -0,4337 | 6,58842E-28 |
| ACE2 | EMSY | -0,4337 | 6,59296E-28 |
| ACE2 | ZBTB32 | -0,4337 | 6,58947E-28 |
| ACE2 | MAP3K14 | -0,4338 | 6,35167E-28 |
| ACE2 | C17orf67 | -0,4338 | 6,30315E-28 |
| ACE2 | HIST2H2AC | -0,4339 | 6,21515E-28 |
| ACE2 | XRCC3 | -0,434 | 5,93833E-28 |
| ACE2 | MARCKSL1 | -0,4342 | 5,70857E-28 |
| ACE2 | NR2F1-AS1 | -0,4342 | 5,5757E-28 |
| ACE2 | RP11-13A1.1 | -0,4342 | 5,58866E-28 |
| ACE2 | CTD-2525P14.5 | -0,4342 | 5,56072E-28 |
| ACE2 | RALGDS | -0,4343 | 5,52377E-28 |
| ACE2 | JAM2 | -0,4343 | 5,43444E-28 |
| ACE2 | AC003080.4 | -0,4345 | 5,09966E-28 |
| ACE2 | TRO | -0,4346 | 4,93486E-28 |
| ACE2 | PKP4-AS1 | -0,4347 | 4,87597E-28 |
| ACE2 | RP11-229O3.1 | -0,4348 | 4,67448E-28 |
| ACE2 | ANGPTL3 | -0,4352 | 4,14661E-28 |
| ACE2 | SNORA29 | -0,4352 | 4,15982E-28 |
| ACE2 | PCDHB15 | -0,4353 | 3,96235E-28 |
| ACE2 | PSME1 | -0,4354 | 3,83705E-28 |
| ACE2 | AC023271.1 | -0,4356 | 3,59571E-28 |
| ACE2 | RP11-762I7.4 | -0,4357 | 3,53062E-28 |
| ACE2 | ZNRF1 | -0,4357 | 3,54695E-28 |
| ACE2 | RP11-687M24.8 | -0,4358 | 3,40679E-28 |
| ACE2 | SMTN | -0,4358 | 3,45089E-28 |
| ACE2 | S100A3 | -0,4359 | 3,36079E-28 |
| ACE2 | CTD-2013N24.2 | -0,436 | 3,26195E-28 |
| ACE2 | PRPS1P2 | -0,436 | 3,21887E-28 |
| ACE2 | AC084117.3 | -0,436 | 3,26217E-28 |
| ACE2 | RP5-1029F21.2 | -0,436 | 3,22249E-28 |
| ACE2 | CTC-444N24.11 | -0,436 | 3,2615E-28 |
| ACE2 | HIST1H2BD | -0,4362 | 3,06052E-28 |
| ACE2 | MTND4P23 | -0,4364 | 2,8233E-28 |
| ACE2 | AC073052.1 | -0,4365 | 2,76345E-28 |
| ACE2 | ANGPTL8 | -0,4365 | 2,74399E-28 |
| ACE2 | CSNK1E | -0,4365 | 2,73262E-28 |
| ACE2 | C17orf107 | -0,4366 | 2,66703E-28 |
| ACE2 | RP11-399O19.9 | -0,4367 | 2,60239E-28 |
| ACE2 | MAX | -0,4367 | 2,60445E-28 |
| ACE2 | TNFRSF1B | -0,4368 | 2,47894E-28 |
| ACE2 | RP11-4B16.4 | -0,4368 | 2,54001E-28 |
| ACE2 | CDR2 | -0,437 | 2,39106E-28 |
| ACE2 | TMCC2 | -0,4371 | 2,27707E-28 |
| ACE2 | CTB-12A17.2 | -0,4371 | 2,31522E-28 |
| ACE2 | MAP3K20-AS1 | -0,4373 | 2,15741E-28 |
| ACE2 | MTMR11 | -0,4375 | 2,0103E-28 |
| ACE2 | RP11-177H13.2 | -0,4375 | 2,03596E-28 |
| ACE2 | CRTC2 | -0,4376 | 1,96346E-28 |
| ACE2 | DENND2A | -0,4376 | 1,92974E-28 |
| ACE2 | DMRT2 | -0,4376 | 1,92772E-28 |
| ACE2 | RN7SL494P | -0,4376 | 1,93505E-28 |
| ACE2 | RP1-59D14.5 | -0,4376 | 1,93678E-28 |

| | | | |
|---|---|---|---|
| ACE2 | TCEA1P4 | -0,4377 | 1,88686E-28 |
| ACE2 | DOCK6 | -0,4378 | 1,85107E-28 |
| ACE2 | RELB | -0,4378 | 1,84944E-28 |
| ACE2 | CDK2 | -0,4379 | 1,75372E-28 |
| ACE2 | PTPN7 | -0,438 | 1,71471E-28 |
| ACE2 | HLA-F-AS1 | -0,4381 | 1,64915E-28 |
| ACE2 | NARF | -0,4381 | 1,65342E-28 |
| ACE2 | C7orf61 | -0,4382 | 1,62817E-28 |
| ACE2 | RP11-311P8.2 | -0,4382 | 1,59791E-28 |
| ACE2 | MDK | -0,4383 | 1,54929E-28 |
| ACE2 | RNF165 | -0,4386 | 1,41004E-28 |
| ACE2 | PKD2L2 | -0,4387 | 1,38735E-28 |
| ACE2 | HIST1H2BI | -0,4387 | 1,37259E-28 |
| ACE2 | MIR4668 | -0,4387 | 1,36703E-28 |
| ACE2 | MTF2 | -0,4388 | 1,35268E-28 |
| ACE2 | CTB-167G5.6 | -0,4389 | 1,28229E-28 |
| ACE2 | ATP1B2 | -0,4392 | 1,18689E-28 |
| ACE2 | CLK1 | -0,4394 | 1,12104E-28 |
| ACE2 | RP11-422P24.11 | -0,4398 | 9,80648E-29 |
| ACE2 | SEPT8 | -0,4399 | 9,48241E-29 |
| ACE2 | AC130469.1 | -0,4399 | 9,40213E-29 |
| ACE2 | RP11-504A18.1 | -0,44 | 9,18437E-29 |
| ACE2 | CLIP2 | -0,4403 | 8,30672E-29 |
| ACE2 | HGF | -0,4403 | 8,31266E-29 |
| ACE2 | LTB | -0,4405 | 7,7977E-29 |
| ACE2 | RP11-542B15.1 | -0,4405 | 7,87874E-29 |
| ACE2 | BST2 | -0,4406 | 7,56138E-29 |
| ACE2 | TSHZ3 | -0,4409 | 6,93598E-29 |
| ACE2 | LITAF | -0,4412 | 6,18148E-29 |
| ACE2 | CTB-37A13.1 | -0,4413 | 5,99721E-29 |
| ACE2 | CPLX1 | -0,4417 | 5,30445E-29 |
| ACE2 | CD6 | -0,4417 | 5,38942E-29 |
| ACE2 | ERFE | -0,4418 | 5,18326E-29 |
| ACE2 | ACHE | -0,4418 | 5,17226E-29 |
| ACE2 | DACT1 | -0,4418 | 5,12129E-29 |
| ACE2 | RP11-324I22.3 | -0,4419 | 4,93036E-29 |
| ACE2 | AC068831.6 | -0,442 | 4,83339E-29 |
| ACE2 | OR2H2 | -0,4421 | 4,77232E-29 |
| ACE2 | GNA14 | -0,4422 | 4,49078E-29 |
| ACE2 | COL4A2-AS2 | -0,4426 | 4,0656E-29 |
| ACE2 | RASSF5 | -0,4431 | 3,44683E-29 |
| ACE2 | PPP1R18 | -0,4431 | 3,42557E-29 |
| ACE2 | MAP3K11 | -0,4432 | 3,26828E-29 |
| ACE2 | AC003104.1 | -0,4433 | 3,23631E-29 |
| ACE2 | AC004837.5 | -0,4435 | 2,95895E-29 |
| ACE2 | EBF4 | -0,4436 | 2,9485E-29 |
| ACE2 | CXCL10 | -0,4438 | 2,76726E-29 |
| ACE2 | C9orf47 | -0,4439 | 2,67874E-29 |
| ACE2 | RP11-513N24.1 | -0,444 | 2,54881E-29 |
| ACE2 | PRR33 | -0,4445 | 2,18492E-29 |
| ACE2 | RP11-416A14.1 | -0,4451 | 1,79888E-29 |
| ACE2 | BHLHE40-AS1 | -0,4451 | 1,79773E-29 |
| ACE2 | RP11-203B9.4 | -0,4451 | 1,8119E-29 |
| ACE2 | LINC00115 | -0,4452 | 1,74422E-29 |

| | | | |
|---|---|---|---|
| ACE2 | KLRB1 | -0,4452 | 1,76701E-29 |
| ACE2 | CTD-2527I21.14 | -0,4453 | 1,68542E-29 |
| ACE2 | HSPA1A | -0,4455 | 1,59131E-29 |
| ACE2 | SUMO4 | -0,4458 | 1,45402E-29 |
| ACE2 | CLEC2D | -0,4458 | 1,43669E-29 |
| ACE2 | HELB | -0,4458 | 1,44831E-29 |
| ACE2 | RP11-837J7.4 | -0,446 | 1,34345E-29 |
| ACE2 | ZNF80 | -0,4462 | 1,27343E-29 |
| ACE2 | RASAL3 | -0,4462 | 1,24379E-29 |
| ACE2 | RAB24 | -0,4464 | 1,17601E-29 |
| ACE2 | UBA6 | -0,4466 | 1,09397E-29 |
| ACE2 | LINC00922 | -0,4466 | 1,10036E-29 |
| ACE2 | PHF1 | -0,4473 | 8,78608E-30 |
| ACE2 | GNRH1 | -0,4473 | 8,86985E-30 |
| ACE2 | INSC | -0,4478 | 7,57572E-30 |
| ACE2 | MLKL | -0,4479 | 7,16953E-30 |
| ACE2 | AC007750.5 | -0,448 | 7,11256E-30 |
| ACE2 | SNORD33 | -0,4484 | 6,1021E-30 |
| ACE2 | AC002480.3 | -0,4487 | 5,5661E-30 |
| ACE2 | DNAJC19P5 | -0,449 | 5,15933E-30 |
| ACE2 | RP11-661O13.1 | -0,4491 | 4,86289E-30 |
| ACE2 | NCR3 | -0,4492 | 4,79985E-30 |
| ACE2 | MIR637 | -0,4494 | 4,52067E-30 |
| ACE2 | HSPA1B | -0,4495 | 4,29948E-30 |
| ACE2 | RP4-758J24.6 | -0,4496 | 4,16829E-30 |
| ACE2 | IER3 | -0,4497 | 4,04108E-30 |
| ACE2 | CHMP4BP1 | -0,4497 | 4,03335E-30 |
| ACE2 | CTD-2184D3.5 | -0,45 | 3,63828E-30 |
| ACE2 | PRKX | -0,4502 | 3,47677E-30 |
| ACE2 | AC005519.4 | -0,4503 | 3,28664E-30 |
| ACE2 | FRMD6 | -0,4505 | 3,14899E-30 |
| ACE2 | SOX7 | -0,4509 | 2,75691E-30 |
| ACE2 | RP11-274H2.5 | -0,4515 | 2,25601E-30 |
| ACE2 | RPL29P19 | -0,452 | 1,88946E-30 |
| ACE2 | CCL3 | -0,452 | 1,87438E-30 |
| ACE2 | TTLL3 | -0,4522 | 1,75523E-30 |
| ACE2 | RPS15AP30 | -0,4522 | 1,78509E-30 |
| ACE2 | RP11-284N8.3 | -0,4523 | 1,71187E-30 |
| ACE2 | IRF7 | -0,4523 | 1,6963E-30 |
| ACE2 | ATXN1 | -0,4526 | 1,5693E-30 |
| ACE2 | UBASH3A | -0,4527 | 1,51959E-30 |
| ACE2 | CTD-2020K17.3 | -0,4528 | 1,45754E-30 |
| ACE2 | CERKL | -0,4529 | 1,42831E-30 |
| ACE2 | ARGLU1 | -0,453 | 1,38577E-30 |
| ACE2 | GZMH | -0,453 | 1,35748E-30 |
| ACE2 | PKIG | -0,453 | 1,35066E-30 |
| ACE2 | USP35 | -0,4531 | 1,33161E-30 |
| ACE2 | PRAL | -0,4531 | 1,30913E-30 |
| ACE2 | RP11-575L7.8 | -0,4534 | 1,18577E-30 |
| ACE2 | CDC42EP5 | -0,4534 | 1,18001E-30 |
| ACE2 | TIAF1 | -0,4535 | 1,16851E-30 |
| ACE2 | SLC16A1-AS1 | -0,4537 | 1,09691E-30 |
| ACE2 | TGFB3 | -0,454 | 9,82932E-31 |
| ACE2 | SYNGR1 | -0,454 | 9,73715E-31 |

| | | | |
|---|---|---|---|
| ACE2 | LINC01871 | -0,4542 | 9,03775E-31 |
| ACE2 | RP11-2E11.9 | -0,4543 | 8,83312E-31 |
| ACE2 | OAZ2 | -0,4543 | 8,91508E-31 |
| ACE2 | RP5-1056L3.3 | -0,4545 | 8,27998E-31 |
| ACE2 | EHBP1L1 | -0,4545 | 8,41364E-31 |
| ACE2 | TMC8 | -0,4548 | 7,49121E-31 |
| ACE2 | RP11-492I21.1 | -0,4549 | 7,36924E-31 |
| ACE2 | INAFM1 | -0,4549 | 7,32808E-31 |
| ACE2 | RP1-111C20.4 | -0,455 | 7,06159E-31 |
| ACE2 | APOL2 | -0,4553 | 6,41349E-31 |
| ACE2 | TBX19 | -0,4554 | 6,15216E-31 |
| ACE2 | AP000692.9 | -0,4554 | 6,24341E-31 |
| ACE2 | PHKG1 | -0,4556 | 5,79404E-31 |
| ACE2 | A2M-AS1 | -0,4558 | 5,41949E-31 |
| ACE2 | RP5-1126H10.2 | -0,4559 | 5,18102E-31 |
| ACE2 | TSLP | -0,4561 | 4,80064E-31 |
| ACE2 | TRGV10 | -0,4562 | 4,77196E-31 |
| ACE2 | RP11-861L17.4 | -0,4563 | 4,52028E-31 |
| ACE2 | MYBL1 | -0,4567 | 4,00714E-31 |
| ACE2 | ADAM19 | -0,457 | 3,63285E-31 |
| ACE2 | FIGNL2 | -0,457 | 3,57986E-31 |
| ACE2 | PKNOX2 | -0,4571 | 3,50801E-31 |
| ACE2 | DGKA | -0,4571 | 3,48121E-31 |
| ACE2 | PIK3C2B | -0,4574 | 3,10535E-31 |
| ACE2 | SH2B2 | -0,4575 | 3,04805E-31 |
| ACE2 | HILPDA | -0,4577 | 2,84714E-31 |
| ACE2 | HIST2H2BF | -0,4579 | 2,66368E-31 |
| ACE2 | RHOJ | -0,4581 | 2,53635E-31 |
| ACE2 | STAMBPL1 | -0,4584 | 2,25399E-31 |
| ACE2 | IL2RG | -0,4584 | 2,27024E-31 |
| ACE2 | GBP5 | -0,4586 | 2,08474E-31 |
| ACE2 | ST8SIA1 | -0,4587 | 2,03667E-31 |
| ACE2 | PTGDR | -0,4587 | 2,03388E-31 |
| ACE2 | TBX21 | -0,4589 | 1,87571E-31 |
| ACE2 | GBP3 | -0,4591 | 1,78842E-31 |
| ACE2 | RP11-1090M7.1 | -0,4592 | 1,71425E-31 |
| ACE2 | RP11-638I2.9 | -0,4593 | 1,68785E-31 |
| ACE2 | FXYD7 | -0,4593 | 1,69E-31 |
| ACE2 | RP11-672A2.5 | -0,4594 | 1,61069E-31 |
| ACE2 | GZMB | -0,4595 | 1,56488E-31 |
| ACE2 | DEF6 | -0,4597 | 1,46476E-31 |
| ACE2 | CCL5 | -0,4597 | 1,43206E-31 |
| ACE2 | RASSF1 | -0,4598 | 1,41732E-31 |
| ACE2 | NFKBIE | -0,46 | 1,30447E-31 |
| ACE2 | RP5-1184F4.5 | -0,46 | 1,33521E-31 |
| ACE2 | RP11-53I6.4 | -0,4605 | 1,09546E-31 |
| ACE2 | RP11-327F22.1 | -0,4607 | 1,05527E-31 |
| ACE2 | TRDC | -0,4608 | 1,01062E-31 |
| ACE2 | CTD-3185P2.1 | -0,4608 | 9,98322E-32 |
| ACE2 | CARD19 | -0,4616 | 7,71847E-32 |
| ACE2 | HIST1H2BH | -0,4619 | 6,87168E-32 |
| ACE2 | RRN3P2 | -0,4624 | 5,87612E-32 |
| ACE2 | LA16c-329F2.2 | -0,4625 | 5,63406E-32 |
| ACE2 | ZBTB17 | -0,4626 | 5,42772E-32 |

| | | | |
|---|---|---|---|
| ACE2 | PPP1R9B | -0,4626 | 5,43349E-32 |
| ACE2 | FLT3LG | -0,4628 | 5,04593E-32 |
| ACE2 | FMNL1 | -0,4634 | 4,08193E-32 |
| ACE2 | RP11-799B12.2 | -0,4634 | 4,07643E-32 |
| ACE2 | PLCXD2 | -0,4636 | 3,86087E-32 |
| ACE2 | DUSP2 | -0,4637 | 3,72895E-32 |
| ACE2 | RP11-687E1.2 | -0,4637 | 3,79304E-32 |
| ACE2 | PPP1R14BP3 | -0,4638 | 3,62491E-32 |
| ACE2 | G0S2 | -0,4639 | 3,54065E-32 |
| ACE2 | RPSAP36 | -0,4639 | 3,52059E-32 |
| ACE2 | RP11-347C12.11 | -0,4641 | 3,20779E-32 |
| ACE2 | TRBC2 | -0,4642 | 3,15281E-32 |
| ACE2 | CTC-529I10.2 | -0,4642 | 3,15813E-32 |
| ACE2 | RP13-735L24.1 | -0,4643 | 3,05905E-32 |
| ACE2 | TVP23A | -0,4643 | 3,05391E-32 |
| ACE2 | CETP | -0,4647 | 2,63164E-32 |
| ACE2 | AC079584.3 | -0,4648 | 2,54361E-32 |
| ACE2 | RP4-724E16.2 | -0,4649 | 2,50398E-32 |
| ACE2 | TNFRSF4 | -0,4651 | 2,28835E-32 |
| ACE2 | SNORD87 | -0,4651 | 2,29261E-32 |
| ACE2 | MINK1 | -0,4654 | 2,11797E-32 |
| ACE2 | CD200 | -0,4655 | 2,00186E-32 |
| ACE2 | FGF1 | -0,4656 | 1,91545E-32 |
| ACE2 | SP140 | -0,4662 | 1,5822E-32 |
| ACE2 | ZBP1 | -0,4662 | 1,55837E-32 |
| ACE2 | RP11-134L10.1 | -0,4663 | 1,5412E-32 |
| ACE2 | ISYNA1 | -0,4663 | 1,54981E-32 |
| ACE2 | CTSW | -0,4668 | 1,27542E-32 |
| ACE2 | DRD1 | -0,468 | 8,58357E-33 |
| ACE2 | FAM110B | -0,4682 | 7,80821E-33 |
| ACE2 | CTC-510F12.3 | -0,4682 | 7,81395E-33 |
| ACE2 | PRKD2 | -0,4682 | 7,9488E-33 |
| ACE2 | ZMIZ2 | -0,4683 | 7,72468E-33 |
| ACE2 | ZNF267 | -0,4687 | 6,68038E-33 |
| ACE2 | CTC-137K3.1 | -0,4688 | 6,4199E-33 |
| ACE2 | NKG7 | -0,469 | 6,08209E-33 |
| ACE2 | DYRK2 | -0,4692 | 5,52035E-33 |
| ACE2 | HIST1H1D | -0,4695 | 4,97449E-33 |
| ACE2 | RP5-1057I20.4 | -0,47 | 4,27916E-33 |
| ACE2 | CASC15 | -0,4705 | 3,48759E-33 |
| ACE2 | RP11-138H8.8 | -0,4709 | 3,05291E-33 |
| ACE2 | ARID5A | -0,4717 | 2,35882E-33 |
| ACE2 | SYNJ2 | -0,4717 | 2,33533E-33 |
| ACE2 | RP11-680A11.5 | -0,4721 | 2,00361E-33 |
| ACE2 | RPL29P14 | -0,4722 | 1,98229E-33 |
| ACE2 | RP11-290D2.6 | -0,4734 | 1,28514E-33 |
| ACE2 | RP11-84A19.4 | -0,4735 | 1,21679E-33 |
| ACE2 | LINC-PINT | -0,4738 | 1,11294E-33 |
| ACE2 | FCHSD1 | -0,4742 | 9,65596E-34 |
| ACE2 | RP5-1014D13.2 | -0,4743 | 9,31192E-34 |
| ACE2 | RP11-552F3.9 | -0,4744 | 9,05672E-34 |
| ACE2 | MFAP4 | -0,4748 | 7,84247E-34 |
| ACE2 | CAMK4 | -0,4749 | 7,50146E-34 |
| ACE2 | RELT | -0,4752 | 6,71617E-34 |

| | | | |
|---|---|---|---|
| ACE2 | LIMD2 | -0,4752 | 6,67536E-34 |
| ACE2 | ITGB2-AS1 | -0,4752 | 6,72069E-34 |
| ACE2 | PTGIR | -0,4753 | 6,587E-34 |
| ACE2 | SOX4 | -0,4754 | 6,28432E-34 |
| ACE2 | RP11-642A1.2 | -0,4754 | 6,30907E-34 |
| ACE2 | TCF7 | -0,4758 | 5,41999E-34 |
| ACE2 | TAP1 | -0,4758 | 5,50203E-34 |
| ACE2 | LCK | -0,4761 | 4,92529E-34 |
| ACE2 | NCKAP5L | -0,4761 | 4,97541E-34 |
| ACE2 | RNF166 | -0,4767 | 3,97907E-34 |
| ACE2 | TRAF3IP2-AS1 | -0,4768 | 3,791E-34 |
| ACE2 | CARD11 | -0,4768 | 3,78381E-34 |
| ACE2 | OPTN | -0,4772 | 3,29886E-34 |
| ACE2 | TRGV9 | -0,4775 | 2,97388E-34 |
| ACE2 | RP11-423E7.2 | -0,4777 | 2,71761E-34 |
| ACE2 | EGFL8 | -0,4777 | 2,75092E-34 |
| ACE2 | ANKRD10 | -0,4779 | 2,56127E-34 |
| ACE2 | INPP5D | -0,478 | 2,52103E-34 |
| ACE2 | HIST1H2AE | -0,478 | 2,47195E-34 |
| ACE2 | DACT3 | -0,478 | 2,46762E-34 |
| ACE2 | NDUFA4L2 | -0,4789 | 1,80209E-34 |
| ACE2 | ACE | -0,4791 | 1,65462E-34 |
| ACE2 | ST6GALNAC4P1 | -0,4792 | 1,5905E-34 |
| ACE2 | CTRL | -0,4793 | 1,57225E-34 |
| ACE2 | RP1-224A6.9 | -0,4794 | 1,51476E-34 |
| ACE2 | OGFR | -0,4794 | 1,49687E-34 |
| ACE2 | TGIF2 | -0,4795 | 1,45411E-34 |
| ACE2 | RASA3 | -0,4796 | 1,42046E-34 |
| ACE2 | GTF2F2P1 | -0,4797 | 1,35125E-34 |
| ACE2 | ARTN | -0,4802 | 1,13172E-34 |
| ACE2 | VCAM1 | -0,4804 | 1,05955E-34 |
| ACE2 | CARHSP1 | -0,4807 | 9,52882E-35 |
| ACE2 | HUS1B | -0,4808 | 9,10527E-35 |
| ACE2 | FMNL3 | -0,481 | 8,50773E-35 |
| ACE2 | HMGN1P3 | -0,4814 | 7,16541E-35 |
| ACE2 | HSH2D | -0,482 | 5,84573E-35 |
| ACE2 | RP11-37B2.1 | -0,4822 | 5,44634E-35 |
| ACE2 | IFNG | -0,4822 | 5,36661E-35 |
| ACE2 | SYTL2 | -0,4825 | 4,85621E-35 |
| ACE2 | STX4 | -0,4832 | 3,76111E-35 |
| ACE2 | AD000864.6 | -0,4832 | 3,74845E-35 |
| ACE2 | GBP1P1 | -0,4834 | 3,56571E-35 |
| ACE2 | CDC37 | -0,4835 | 3,42725E-35 |
| ACE2 | S1PR5 | -0,4837 | 3,09768E-35 |
| ACE2 | RP11-373D23.2 | -0,4838 | 3,07733E-35 |
| ACE2 | RP11-342D11.2 | -0,4839 | 2,88348E-35 |
| ACE2 | KB-431C1.5 | -0,4842 | 2,63953E-35 |
| ACE2 | HDC | -0,4844 | 2,47837E-35 |
| ACE2 | PTGES3P1 | -0,4848 | 2,10656E-35 |
| ACE2 | RP11-570L15.1 | -0,4849 | 2,04707E-35 |
| ACE2 | BTN2A2 | -0,4849 | 2,00976E-35 |
| ACE2 | KB-1208A12.3 | -0,485 | 1,95144E-35 |
| ACE2 | AC107081.5 | -0,4855 | 1,61328E-35 |
| ACE2 | RUBCNL | -0,4856 | 1,56895E-35 |

| | | | |
|---|---|---|---|
| ACE2 | HOTAIRM1 | -0,4857 | 1,50516E-35 |
| ACE2 | LAG3 | -0,4858 | 1,47265E-35 |
| ACE2 | EID3 | -0,4858 | 1,45551E-35 |
| ACE2 | CTC-510F12.2 | -0,4865 | 1,13695E-35 |
| ACE2 | TRIP10 | -0,4868 | 9,92343E-36 |
| ACE2 | TNK2 | -0,4869 | 9,57713E-36 |
| ACE2 | HAPLN3 | -0,4869 | 9,60886E-36 |
| ACE2 | RP11-506B6.6 | -0,4877 | 7,20243E-36 |
| ACE2 | ADAMTS7 | -0,4878 | 7,00317E-36 |
| ACE2 | RP1-187B23.1 | -0,4879 | 6,66824E-36 |
| ACE2 | DDAH2 | -0,4881 | 6,25874E-36 |
| ACE2 | RP11-1094M14.5 | -0,4884 | 5,63451E-36 |
| ACE2 | C5orf56 | -0,4886 | 5,19961E-36 |
| ACE2 | RP11-147L13.15 | -0,4897 | 3,45706E-36 |
| ACE2 | MOV10 | -0,49 | 3,05072E-36 |
| ACE2 | CLEC2B | -0,4904 | 2,61082E-36 |
| ACE2 | RP11-297C4.6 | -0,4904 | 2,6119E-36 |
| ACE2 | PPM1J | -0,4905 | 2,56155E-36 |
| ACE2 | LRRC4B | -0,4905 | 2,54976E-36 |
| ACE2 | COL26A1 | -0,4906 | 2,41244E-36 |
| ACE2 | GPR174 | -0,4919 | 1,47996E-36 |
| ACE2 | PACERR | -0,4921 | 1,37217E-36 |
| ACE2 | LZTS2 | -0,4924 | 1,23257E-36 |
| ACE2 | CLEC1A | -0,4925 | 1,20748E-36 |
| ACE2 | ROBO3 | -0,4926 | 1,14288E-36 |
| ACE2 | CACNB3 | -0,4926 | 1,15343E-36 |
| ACE2 | CTNNAL1 | -0,4929 | 1,05258E-36 |
| ACE2 | HCAR3 | -0,4932 | 9,12923E-37 |
| ACE2 | BCAR1 | -0,4952 | 4,3032E-37 |
| ACE2 | PYHIN1 | -0,4954 | 3,98491E-37 |
| ACE2 | FAM162B | -0,4962 | 2,91569E-37 |
| ACE2 | ARHGAP15 | -0,4964 | 2,7562E-37 |
| ACE2 | PML | -0,4965 | 2,61612E-37 |
| ACE2 | RP13-516M14.10 | -0,4975 | 1,82566E-37 |
| ACE2 | IL2RB | -0,4976 | 1,75431E-37 |
| ACE2 | TESC | -0,4978 | 1,57921E-37 |
| ACE2 | ARHGAP17 | -0,4983 | 1,3124E-37 |
| ACE2 | AZIN2 | -0,4985 | 1,23424E-37 |
| ACE2 | SH2D2A | -0,5016 | 3,64225E-38 |
| ACE2 | EOMES | -0,502 | 3,12312E-38 |
| ACE2 | NFKBID | -0,5025 | 2,64517E-38 |
| ACE2 | RGS3 | -0,5031 | 2,08161E-38 |
| ACE2 | XCL2 | -0,5033 | 1,93196E-38 |
| ACE2 | PLD2 | -0,5034 | 1,82461E-38 |
| ACE2 | RP11-888D10.4 | -0,5035 | 1,76803E-38 |
| ACE2 | CARD16 | -0,5039 | 1,52613E-38 |
| ACE2 | CTD-2020K17.1 | -0,5041 | 1,39492E-38 |
| ACE2 | DCLK2 | -0,5058 | 7,27426E-39 |
| ACE2 | LPAR6 | -0,5058 | 7,11093E-39 |
| ACE2 | NEURL3 | -0,5067 | 4,9529E-39 |
| ACE2 | CASP1 | -0,5079 | 3,18777E-39 |
| ACE2 | GNLY | -0,5083 | 2,70679E-39 |
| ACE2 | MAP3K7CL | -0,5085 | 2,50431E-39 |
| ACE2 | IRF1 | -0,5095 | 1,65715E-39 |

| | | | |
|---|---|---|---|
| ACE2 | GPR18 | -0,5096 | 1,60589E-39 |
| ACE2 | RARRES2 | -0,5097 | 1,56401E-39 |
| ACE2 | VANGL2 | -0,51 | 1,35426E-39 |
| ACE2 | PHLDB3 | -0,5105 | 1,10731E-39 |
| ACE2 | NFKB2 | -0,512 | 6,21968E-40 |
| ACE2 | CCL11 | -0,5124 | 5,18745E-40 |
| ACE2 | FNBP1 | -0,513 | 4,08753E-40 |
| ACE2 | RP11-693N9.2 | -0,5131 | 3,88621E-40 |
| ACE2 | SH2D1A | -0,5137 | 3,06472E-40 |
| ACE2 | RP1-142L7.9 | -0,5146 | 2,16856E-40 |
| ACE2 | GZMM | -0,5148 | 1,98258E-40 |
| ACE2 | TRBC1 | -0,515 | 1,83982E-40 |
| ACE2 | IL32 | -0,5154 | 1,52537E-40 |
| ACE2 | TOMM20P2 | -0,5159 | 1,2796E-40 |
| ACE2 | AFAP1L2 | -0,516 | 1,21019E-40 |
| ACE2 | C19orf66 | -0,5163 | 1,06878E-40 |
| ACE2 | ZAP70 | -0,5166 | 9,35572E-41 |
| ACE2 | C5orf58 | -0,5172 | 7,40723E-41 |
| ACE2 | GBP2 | -0,5184 | 4,4662E-41 |
| ACE2 | STAT4 | -0,519 | 3,53828E-41 |
| ACE2 | GBP1 | -0,5195 | 2,93784E-41 |
| ACE2 | MCOLN2 | -0,5206 | 1,81543E-41 |
| ACE2 | KLRK1 | -0,521 | 1,57147E-41 |
| ACE2 | SERPINB9 | -0,5224 | 8,80664E-42 |
| ACE2 | ADORA2A | -0,5255 | 2,36612E-42 |
| ACE2 | IL12RB2 | -0,5263 | 1,68954E-42 |
| ACE2 | SEPT6 | -0,5264 | 1,63676E-42 |
| ACE2 | TNFRSF18 | -0,5273 | 1,13038E-42 |
| ACE2 | AP000442.1 | -0,5274 | 1,09082E-42 |
| ACE2 | AC073410.1 | -0,5282 | 7,58458E-43 |
| ACE2 | HLA-F | -0,5305 | 2,84519E-43 |
| ACE2 | CD247 | -0,5367 | 1,99362E-44 |
| ACE2 | CD7 | -0,5391 | 6,82817E-45 |
| ACE2 | SAMD3 | -0,5408 | 3,21747E-45 |
| ACE2 | RP11-247I13.3 | -0,5411 | 2,91061E-45 |
| ACE2 | FASLG | -0,5459 | 3,42157E-46 |
| ACE2 | RP11-1094M14.8 | -0,5533 | 1,18144E-47 |
