## Supplementary table 5 intestine for "Bioinformatic characterization of angiotensin-converting enzyme 2, the entry receptor for SARS-CoV-2"

| target_gene | correlated_gene | correlation | pval |
|---|---|---|---|
| ACE2 | ACE2 | 1 | 0 |
| ACE2 | ENPEP | 0,9653 | 8,287E-110 |
| ACE2 | MEP1B | 0,9653 | 8,3455E-110 |
| ACE2 | SLC15A1 | 0,9621 | 2,1858E-106 |
| ACE2 | MEP1A | 0,9619 | 4,4024E-106 |
| ACE2 | SLC3A1 | 0,9608 | 5,6337E-105 |
| ACE2 | APOB | 0,9587 | 6,4451E-103 |
| ACE2 | SLC6A19 | 0,9581 | 2,3281E-102 |
| ACE2 | ANPEP | 0,9562 | 1,1592E-100 |
| ACE2 | MTTP | 0,9559 | 2,4502E-100 |
| ACE2 | TUBAL3 | 0,9554 | 6,2351E-100 |
| ACE2 | SLC10A2 | 0,9554 | 5,9469E-100 |
| ACE2 | SLC5A1 | 0,9554 | 6,4235E-100 |
| ACE2 | CES2 | 0,9537 | 1,86936E-98 |
| ACE2 | HHLA2 | 0,9525 | 1,90613E-97 |
| ACE2 | MGAM | 0,9525 | 1,87318E-97 |
| ACE2 | TMIGD1 | 0,9525 | 1,94126E-97 |
| ACE2 | SLC26A3 | 0,9511 | 2,55794E-96 |
| ACE2 | PLA2G12B | 0,9509 | 3,39352E-96 |
| ACE2 | CDHR2 | 0,9503 | 1,14138E-95 |
| ACE2 | CYP2C18 | 0,9499 | 2,17212E-95 |
| ACE2 | SLC1A1 | 0,9497 | 3,47074E-95 |
| ACE2 | GDPD2 | 0,949 | 1,18026E-94 |
| ACE2 | CDHR5 | 0,9484 | 2,96067E-94 |
| ACE2 | LRRC19 | 0,9476 | 1,25291E-93 |
| ACE2 | PRAP1 | 0,9471 | 2,78854E-93 |
| ACE2 | PLS1 | 0,9469 | 3,99865E-93 |
| ACE2 | MME | 0,9468 | 5,40525E-93 |
| ACE2 | SI | 0,9462 | 1,41905E-92 |
| ACE2 | TMEM236 | 0,9461 | 1,54145E-92 |
| ACE2 | FADS6 | 0,9461 | 1,53611E-92 |
| ACE2 | TMEM253 | 0,946 | 1,9319E-92 |
| ACE2 | HNF4G | 0,9451 | 7,98344E-92 |
| ACE2 | CREB3L3 | 0,9449 | 1,09515E-91 |
| ACE2 | FLJ22763 | 0,9439 | 5,50806E-91 |
| ACE2 | SLC13A1 | 0,9439 | 5,49745E-91 |
| ACE2 | SLC2A2 | 0,9427 | 4,14441E-90 |
| ACE2 | TREH | 0,9427 | 3,89806E-90 |
| ACE2 | APOBEC1 | 0,9426 | 4,73636E-90 |
| ACE2 | SLC7A9 | 0,942 | 1,27917E-89 |
| ACE2 | TMEM92 | 0,9418 | 1,53808E-89 |
| ACE2 | MIA2 | 0,9416 | 2,23442E-89 |
| ACE2 | AOC1 | 0,9408 | 7,42337E-89 |
| ACE2 | DPP4 | 0,9406 | 1,03015E-88 |
| ACE2 | OIT3 | 0,9403 | 1,59236E-88 |
| ACE2 | SLC17A8 | 0,9403 | 1,50176E-88 |
| ACE2 | PTPRH | 0,9398 | 3,11669E-88 |
| ACE2 | TMEM45B | 0,9396 | 4,26146E-88 |
| ACE2 | CLRN3 | 0,9395 | 5,08969E-88 |
| ACE2 | SLC6A4 | 0,9394 | 5,72512E-88 |
| ACE2 | CDH17 | 0,9391 | 9,82149E-88 |
| ACE2 | GDA | 0,9391 | 9,18081E-88 |
| ACE2 | GUCA2A | 0,9387 | 1,79268E-87 |

| | | | |
|---|---|---|---|
| ACE2 | SMIM24 | 0,9386 | 1,94232E-87 |
| ACE2 | MUC3A | 0,9385 | 2,39038E-87 |
| ACE2 | UGT2A3 | 0,9384 | 2,50974E-87 |
| ACE2 | MUC17 | 0,9384 | 2,77924E-87 |
| ACE2 | ALPI | 0,9383 | 2,98491E-87 |
| ACE2 | MARVELD3 | 0,9381 | 4,17131E-87 |
| ACE2 | EPB41L4B | 0,9376 | 8,31415E-87 |
| ACE2 | TM6SF2 | 0,9376 | 8,81089E-87 |
| ACE2 | SCIN | 0,9361 | 7,06634E-86 |
| ACE2 | MYO1A | 0,936 | 8,09617E-86 |
| ACE2 | TMEM82 | 0,9358 | 1,13031E-85 |
| ACE2 | SLC6A20 | 0,9352 | 2,48138E-85 |
| ACE2 | DSC2 | 0,9351 | 3,0038E-85 |
| ACE2 | CYP2J2 | 0,9349 | 3,9127E-85 |
| ACE2 | AC007182.6 | 0,9349 | 3,86167E-85 |
| ACE2 | ALDOB | 0,9347 | 4,56533E-85 |
| ACE2 | PDZK1 | 0,9342 | 9,07595E-85 |
| ACE2 | ZG16 | 0,9342 | 1,00282E-84 |
| ACE2 | MS4A8 | 0,9331 | 4,12536E-84 |
| ACE2 | ANKS4B | 0,933 | 4,91552E-84 |
| ACE2 | GUCY2C | 0,9326 | 8,58183E-84 |
| ACE2 | CGN | 0,9325 | 9,75105E-84 |
| ACE2 | NR1I2 | 0,9325 | 9,43578E-84 |
| ACE2 | CEACAM20 | 0,9317 | 2,64701E-83 |
| ACE2 | KHK | 0,9316 | 3,25302E-83 |
| ACE2 | SDCBP2 | 0,9315 | 3,29614E-83 |
| ACE2 | GBA3 | 0,9314 | 3,84349E-83 |
| ACE2 | CXADR | 0,9313 | 4,35961E-83 |
| ACE2 | DPEP1 | 0,9312 | 5,11463E-83 |
| ACE2 | RETSAT | 0,9308 | 9,16644E-83 |
| ACE2 | BTNL8 | 0,9307 | 9,40135E-83 |
| ACE2 | TINAG | 0,9307 | 1,04614E-82 |
| ACE2 | AGMO | 0,9304 | 1,48832E-82 |
| ACE2 | MYH14 | 0,9304 | 1,54001E-82 |
| ACE2 | MS4A10 | 0,93 | 2,35136E-82 |
| ACE2 | A1CF | 0,9298 | 3,19943E-82 |
| ACE2 | PRSS8 | 0,9293 | 5,54544E-82 |
| ACE2 | RAB17 | 0,9292 | 6,92945E-82 |
| ACE2 | GPA33 | 0,9291 | 7,33683E-82 |
| ACE2 | CDKN2B | 0,9288 | 1,11252E-81 |
| ACE2 | B3GALT5 | 0,9287 | 1,18852E-81 |
| ACE2 | EPCAM | 0,9283 | 2,16071E-81 |
| ACE2 | RP11-116D2.1 | 0,9278 | 3,92286E-81 |
| ACE2 | ACY1 | 0,9275 | 5,84221E-81 |
| ACE2 | ADGRG7 | 0,927 | 1,08349E-80 |
| ACE2 | SLC28A1 | 0,9267 | 1,48687E-80 |
| ACE2 | TMEM150B | 0,9265 | 1,96937E-80 |
| ACE2 | PKIB | 0,926 | 3,23525E-80 |
| ACE2 | ENTPD8 | 0,9256 | 5,38043E-80 |
| ACE2 | CPO | 0,9252 | 9,36694E-80 |
| ACE2 | REEP6 | 0,9252 | 9,18551E-80 |
| ACE2 | MOGAT2 | 0,9248 | 1,35842E-79 |
| ACE2 | MISP | 0,924 | 3,49226E-79 |
| ACE2 | CYP4F2 | 0,924 | 3,77484E-79 |

| | | | |
|---|---|---|---|
| ACE2 | PRR15 | 0,9238 | 4,82733E-79 |
| ACE2 | FER1L6 | 0,9238 | 4,56169E-79 |
| ACE2 | MALL | 0,9235 | 6,84911E-79 |
| ACE2 | MYRFL | 0,9235 | 6,45954E-79 |
| ACE2 | GUCA2B | 0,9227 | 1,69377E-78 |
| ACE2 | DSG2 | 0,9225 | 2,17621E-78 |
| ACE2 | C1orf106 | 0,9224 | 2,36079E-78 |
| ACE2 | TM4SF20 | 0,9223 | 2,72919E-78 |
| ACE2 | GOLT1A | 0,9219 | 4,20362E-78 |
| ACE2 | MOGAT3 | 0,9213 | 8,50621E-78 |
| ACE2 | SLC13A2 | 0,9213 | 7,83617E-78 |
| ACE2 | HAPLN4 | 0,9211 | 9,96615E-78 |
| ACE2 | LINC00483 | 0,9209 | 1,33305E-77 |
| ACE2 | ABCG2 | 0,9207 | 1,66949E-77 |
| ACE2 | C1orf210 | 0,9205 | 2,05889E-77 |
| ACE2 | AMN | 0,9205 | 2,09934E-77 |
| ACE2 | SOWAHA | 0,92 | 3,62053E-77 |
| ACE2 | CDH1 | 0,92 | 3,68166E-77 |
| ACE2 | TM4SF5 | 0,9198 | 4,43128E-77 |
| ACE2 | DHRS11 | 0,9196 | 5,33872E-77 |
| ACE2 | UGT1A1 | 0,9195 | 5,8423E-77 |
| ACE2 | MALRD1 | 0,9195 | 6,09114E-77 |
| ACE2 | SULT1B1 | 0,9194 | 6,49569E-77 |
| ACE2 | SLC51B | 0,9191 | 9,98804E-77 |
| ACE2 | ATP1A1 | 0,919 | 1,05027E-76 |
| ACE2 | XPNPEP2 | 0,919 | 1,01046E-76 |
| ACE2 | SERINC2 | 0,9189 | 1,12511E-76 |
| ACE2 | TMEM37 | 0,9188 | 1,35309E-76 |
| ACE2 | SPINT1 | 0,9186 | 1,67688E-76 |
| ACE2 | EDN2 | 0,9185 | 1,82155E-76 |
| ACE2 | SLC44A4 | 0,9185 | 1,86983E-76 |
| ACE2 | IYD | 0,9184 | 2,03003E-76 |
| ACE2 | CYP2C19 | 0,9184 | 1,99828E-76 |
| ACE2 | SLC5A9 | 0,9181 | 2,93042E-76 |
| ACE2 | AADAC | 0,9181 | 2,68953E-76 |
| ACE2 | CYP2C9 | 0,918 | 3,13597E-76 |
| ACE2 | NAT2 | 0,9177 | 4,16972E-76 |
| ACE2 | INPP5J | 0,9177 | 4,184E-76 |
| ACE2 | MYO5B | 0,9176 | 4,69619E-76 |
| ACE2 | SLC23A1 | 0,9173 | 6,79745E-76 |
| ACE2 | SLC17A4 | 0,9171 | 8,50219E-76 |
| ACE2 | CYP3A5 | 0,9171 | 8,61578E-76 |
| ACE2 | IGSF23 | 0,9171 | 8,02446E-76 |
| ACE2 | HKDC1 | 0,917 | 9,15943E-76 |
| ACE2 | USH1C | 0,917 | 8,88622E-76 |
| ACE2 | SLC5A12 | 0,9167 | 1,29661E-75 |
| ACE2 | KRT20 | 0,9164 | 1,70951E-75 |
| ACE2 | PTPRF | 0,9163 | 1,94338E-75 |
| ACE2 | BTNL3 | 0,9162 | 2,06881E-75 |
| ACE2 | VIL1 | 0,9161 | 2,29639E-75 |
| ACE2 | PEPD | 0,916 | 2,72391E-75 |
| ACE2 | PBLD | 0,9159 | 2,97871E-75 |
| ACE2 | SLC5A11 | 0,9159 | 2,94984E-75 |
| ACE2 | F2RL1 | 0,9156 | 3,96774E-75 |

| | | | |
|---|---|---|---|
| ACE2 | ESPN | 0,9153 | 5,48259E-75 |
| ACE2 | MTND4P20 | 0,9153 | 5,56126E-75 |
| ACE2 | PLEKHG6 | 0,9153 | 5,63422E-75 |
| ACE2 | TMC4 | 0,9153 | 5,54027E-75 |
| ACE2 | IL22RA1 | 0,9149 | 8,47899E-75 |
| ACE2 | SLC51A | 0,9148 | 9,31493E-75 |
| ACE2 | PLB1 | 0,9146 | 1,12301E-74 |
| ACE2 | SLC9A3 | 0,9145 | 1,24836E-74 |
| ACE2 | FLVCR2 | 0,9145 | 1,30408E-74 |
| ACE2 | CLDN7 | 0,9145 | 1,32152E-74 |
| ACE2 | MYO7B | 0,9142 | 1,81261E-74 |
| ACE2 | TRIM10 | 0,914 | 2,08834E-74 |
| ACE2 | ABCC13 | 0,914 | 2,08564E-74 |
| ACE2 | C11orf86 | 0,9136 | 3,25178E-74 |
| ACE2 | CCL15 | 0,9136 | 3,07964E-74 |
| ACE2 | METTL7B | 0,9132 | 5,05425E-74 |
| ACE2 | PCK2 | 0,9129 | 6,57772E-74 |
| ACE2 | FRK | 0,9127 | 8,10975E-74 |
| ACE2 | CEACAM18 | 0,9126 | 9,19474E-74 |
| ACE2 | SYT8 | 0,9122 | 1,36063E-73 |
| ACE2 | TBX10 | 0,9121 | 1,50948E-73 |
| ACE2 | NAGS | 0,9118 | 1,92973E-73 |
| ACE2 | PDZD3 | 0,9114 | 2,81975E-73 |
| ACE2 | RNU1-70P | 0,9112 | 3,44622E-73 |
| ACE2 | NGEF | 0,9109 | 5,08661E-73 |
| ACE2 | MUC13 | 0,9108 | 5,15386E-73 |
| ACE2 | FBP1 | 0,9106 | 6,64748E-73 |
| ACE2 | LAMB3 | 0,9102 | 9,42286E-73 |
| ACE2 | UGT2B7 | 0,91 | 1,20106E-72 |
| ACE2 | PRSS3 | 0,9099 | 1,24633E-72 |
| ACE2 | NDFIP2 | 0,9097 | 1,5219E-72 |
| ACE2 | PTGR1 | 0,9096 | 1,68412E-72 |
| ACE2 | ACSL5 | 0,9096 | 1,70704E-72 |
| ACE2 | RP11-71L14.3 | 0,9094 | 2,07076E-72 |
| ACE2 | AGR3 | 0,9092 | 2,51108E-72 |
| ACE2 | CASP16P | 0,9091 | 2,72716E-72 |
| ACE2 | LIPH | 0,909 | 3,06145E-72 |
| ACE2 | APOA4 | 0,9088 | 3,94322E-72 |
| ACE2 | EPS8L3 | 0,9083 | 6,38797E-72 |
| ACE2 | TMEM139 | 0,908 | 8,29577E-72 |
| ACE2 | SMIM6 | 0,908 | 8,55147E-72 |
| ACE2 | RP11-680F8.1 | 0,9079 | 9,29625E-72 |
| ACE2 | PKLR | 0,9078 | 1,02601E-71 |
| ACE2 | COL17A1 | 0,9077 | 1,06439E-71 |
| ACE2 | SMIM2-AS1 | 0,9076 | 1,25049E-71 |
| ACE2 | SMLR1 | 0,9074 | 1,42118E-71 |
| ACE2 | RP11-452I5.2 | 0,9072 | 1,69641E-71 |
| ACE2 | PTK6 | 0,9072 | 1,73169E-71 |
| ACE2 | C4orf19 | 0,907 | 2,22299E-71 |
| ACE2 | RNF128 | 0,907 | 2,04141E-71 |
| ACE2 | KRT19 | 0,9069 | 2,38891E-71 |
| ACE2 | PHGR1 | 0,9067 | 2,83007E-71 |
| ACE2 | UGT1A5 | 0,9066 | 3,22393E-71 |
| ACE2 | PRODH | 0,9064 | 3,82535E-71 |

| | | | |
|---|---|---|---|
| ACE2 | FAM3B | 0,9063 | 4,28492E-71 |
| ACE2 | UGT1A10 | 0,9062 | 4,48422E-71 |
| ACE2 | CTB-127C13.1 | 0,906 | 5,61467E-71 |
| ACE2 | MUC2 | 0,9059 | 5,84266E-71 |
| ACE2 | FUT6 | 0,9058 | 6,72538E-71 |
| ACE2 | CHMP4C | 0,9057 | 7,46344E-71 |
| ACE2 | SLC52A1 | 0,9055 | 8,29494E-71 |
| ACE2 | CAPN9 | 0,9054 | 9,09698E-71 |
| ACE2 | SLC41A2 | 0,9054 | 9,10475E-71 |
| ACE2 | MIR194-2HG | 0,9051 | 1,26895E-70 |
| ACE2 | HRASLS2 | 0,905 | 1,44275E-70 |
| ACE2 | PRRG2 | 0,9045 | 2,1193E-70 |
| ACE2 | C1orf115 | 0,9044 | 2,48196E-70 |
| ACE2 | ZNF488 | 0,9044 | 2,4911E-70 |
| ACE2 | ACY3 | 0,9043 | 2,59963E-70 |
| ACE2 | CLDN4 | 0,9041 | 3,19825E-70 |
| ACE2 | DGAT1 | 0,9041 | 3,30465E-70 |
| ACE2 | RBM47 | 0,9037 | 4,65067E-70 |
| ACE2 | RP11-63P12.7 | 0,9036 | 4,92892E-70 |
| ACE2 | LMO7 | 0,9036 | 5,20661E-70 |
| ACE2 | TGFA | 0,9035 | 5,42732E-70 |
| ACE2 | CYP3A4 | 0,9033 | 6,69728E-70 |
| ACE2 | MTND4P19 | 0,9032 | 7,30526E-70 |
| ACE2 | NAALADL1 | 0,9032 | 7,42145E-70 |
| ACE2 | STYK1 | 0,9032 | 7,07344E-70 |
| ACE2 | GATA6 | 0,9032 | 7,40511E-70 |
| ACE2 | SLC39A5 | 0,9031 | 8,17996E-70 |
| ACE2 | NUDT16P1 | 0,9028 | 1,02086E-69 |
| ACE2 | GGT8P | 0,9025 | 1,32427E-69 |
| ACE2 | SPINT2 | 0,9024 | 1,44054E-69 |
| ACE2 | RN7SKP22 | 0,9023 | 1,57046E-69 |
| ACE2 | IHH | 0,9021 | 1,93914E-69 |
| ACE2 | FABP2 | 0,9021 | 1,90362E-69 |
| ACE2 | DNAJC22 | 0,9021 | 1,88979E-69 |
| ACE2 | EPS8L2 | 0,9014 | 3,50643E-69 |
| ACE2 | ABCC6 | 0,9014 | 3,48357E-69 |
| ACE2 | ARHGEF16 | 0,9012 | 4,44504E-69 |
| ACE2 | BCL2L15 | 0,9008 | 6,39128E-69 |
| ACE2 | CRB3 | 0,9007 | 7,00283E-69 |
| ACE2 | SULT1A2 | 0,9003 | 9,88377E-69 |
| ACE2 | CHST5 | 0,9003 | 9,55069E-69 |
| ACE2 | HSD17B2 | 0,9003 | 9,44749E-69 |
| ACE2 | RP11-486A14.1 | 0,9002 | 1,02427E-68 |
| ACE2 | C19orf33 | 0,9001 | 1,17718E-68 |
| ACE2 | HNF1A | 0,9 | 1,23771E-68 |
| ACE2 | APOC3 | 0,8997 | 1,68852E-68 |
| ACE2 | HNF4A | 0,8997 | 1,63388E-68 |
| ACE2 | LRRC66 | 0,8996 | 1,77317E-68 |
| ACE2 | KIAA1522 | 0,8994 | 2,16808E-68 |
| ACE2 | OTOP2 | 0,8994 | 2,03369E-68 |
| ACE2 | TTC38 | 0,8994 | 2,03764E-68 |
| ACE2 | SOWAHB | 0,8991 | 2,82871E-68 |
| ACE2 | ISX | 0,8989 | 3,15581E-68 |
| ACE2 | TMEM184A | 0,8984 | 4,83789E-68 |

| | | | |
|---|---|---|---|
| ACE2 | BAIAP2L2 | 0,8984 | 5,02971E-68 |
| ACE2 | FMO5 | 0,8983 | 5,2614E-68 |
| ACE2 | GLOD5 | 0,898 | 6,89839E-68 |
| ACE2 | CYP2B6 | 0,8979 | 7,84768E-68 |
| ACE2 | CLDN3 | 0,8972 | 1,39934E-67 |
| ACE2 | B3GNT8 | 0,8972 | 1,4349E-67 |
| ACE2 | DAB1 | 0,8971 | 1,54656E-67 |
| ACE2 | NR1H4 | 0,8968 | 1,92772E-67 |
| ACE2 | TJP3 | 0,8964 | 2,76581E-67 |
| ACE2 | IL17RE | 0,8961 | 3,53076E-67 |
| ACE2 | CDS1 | 0,896 | 3,97598E-67 |
| ACE2 | CYP2S1 | 0,8959 | 4,11352E-67 |
| ACE2 | TMPRSS2 | 0,8959 | 4,30679E-67 |
| ACE2 | LINC02000 | 0,8956 | 5,24391E-67 |
| ACE2 | ABHD11-AS1 | 0,8954 | 6,22424E-67 |
| ACE2 | ACE | 0,8952 | 7,43673E-67 |
| ACE2 | FAM83F | 0,8951 | 8,36755E-67 |
| ACE2 | IGSF9 | 0,8949 | 9,58094E-67 |
| ACE2 | CAPN13 | 0,8947 | 1,18168E-66 |
| ACE2 | SLC22A18 | 0,8947 | 1,15252E-66 |
| ACE2 | MARVELD2 | 0,8946 | 1,21505E-66 |
| ACE2 | CTD-3010D24.3 | 0,8946 | 1,28095E-66 |
| ACE2 | PLA2G4F | 0,8945 | 1,32579E-66 |
| ACE2 | HTR1D | 0,8942 | 1,69837E-66 |
| ACE2 | PLEKHA7 | 0,8942 | 1,81002E-66 |
| ACE2 | TRIM36 | 0,894 | 2,02938E-66 |
| ACE2 | TMC5 | 0,8935 | 3,03972E-66 |
| ACE2 | ACOX1 | 0,8935 | 3,14159E-66 |
| ACE2 | NAT8B | 0,8933 | 3,58326E-66 |
| ACE2 | TMEM30B | 0,8933 | 3,69408E-66 |
| ACE2 | C11orf52 | 0,8931 | 4,43522E-66 |
| ACE2 | B3GNT7 | 0,893 | 4,62347E-66 |
| ACE2 | SLC30A10 | 0,8929 | 5,11477E-66 |
| ACE2 | BCAS1 | 0,8929 | 5,08364E-66 |
| ACE2 | AIM1L | 0,8928 | 5,47821E-66 |
| ACE2 | LINC02177 | 0,8925 | 6,79295E-66 |
| ACE2 | TKFC | 0,8922 | 9,07279E-66 |
| ACE2 | RP11-697K23.3 | 0,8921 | 9,38262E-66 |
| ACE2 | RP11-766F14.1 | 0,8919 | 1,16455E-65 |
| ACE2 | PTPN3 | 0,8916 | 1,43183E-65 |
| ACE2 | XDH | 0,8914 | 1,69623E-65 |
| ACE2 | AKR7A3 | 0,8913 | 1,87521E-65 |
| ACE2 | ABCB1 | 0,8913 | 1,91768E-65 |
| ACE2 | LRRC75A | 0,8913 | 1,80789E-65 |
| ACE2 | HGD | 0,891 | 2,44095E-65 |
| ACE2 | CAMSAP3 | 0,8909 | 2,54529E-65 |
| ACE2 | SLC39A4 | 0,8908 | 2,77843E-65 |
| ACE2 | RP11-27G14.4 | 0,8904 | 3,73323E-65 |
| ACE2 | NEU4 | 0,8904 | 3,93685E-65 |
| ACE2 | GPT | 0,8901 | 4,85819E-65 |
| ACE2 | MGAM2 | 0,89 | 5,31369E-65 |
| ACE2 | TTC22 | 0,8899 | 5,60301E-65 |
| ACE2 | FTLP14 | 0,8899 | 5,71755E-65 |
| ACE2 | B3GNT3 | 0,8898 | 6,20502E-65 |

| | | | |
|---|---|---|---|
| ACE2 | XXyac-YM21GA2.7 | 0,8893 | 9,31688E-65 |
| ACE2 | DSP | 0,8892 | 9,49738E-65 |
| ACE2 | ST14 | 0,889 | 1,1346E-64 |
| ACE2 | MROH3P | 0,8887 | 1,42658E-64 |
| ACE2 | TPRN | 0,8887 | 1,44032E-64 |
| ACE2 | FXYD3 | 0,8881 | 2,24883E-64 |
| ACE2 | ERICH4 | 0,8881 | 2,42481E-64 |
| ACE2 | SLC9A3R1 | 0,888 | 2,49864E-64 |
| ACE2 | CMBL | 0,8879 | 2,74075E-64 |
| ACE2 | APOA1 | 0,8879 | 2,63896E-64 |
| ACE2 | CRYL1 | 0,8878 | 2,913E-64 |
| ACE2 | TSPAN1 | 0,8876 | 3,48258E-64 |
| ACE2 | SMPD3 | 0,8876 | 3,42257E-64 |
| ACE2 | MAP7 | 0,8875 | 3,81876E-64 |
| ACE2 | PRR15L | 0,8869 | 5,70858E-64 |
| ACE2 | HSD3B1 | 0,8868 | 6,31385E-64 |
| ACE2 | CTD-2085J24.5 | 0,8866 | 7,60479E-64 |
| ACE2 | RIPK4 | 0,8866 | 7,20854E-64 |
| ACE2 | PP7080 | 0,8861 | 1,09897E-63 |
| ACE2 | SLC46A3 | 0,8861 | 1,05344E-63 |
| ACE2 | FCGBP | 0,8861 | 1,09435E-63 |
| ACE2 | C1orf116 | 0,886 | 1,15145E-63 |
| ACE2 | MOCOS | 0,886 | 1,1697E-63 |
| ACE2 | PLCB3 | 0,8858 | 1,41066E-63 |
| ACE2 | GATA5 | 0,8858 | 1,32905E-63 |
| ACE2 | C6orf222 | 0,8855 | 1,71769E-63 |
| ACE2 | EGFR-AS1 | 0,8853 | 1,93751E-63 |
| ACE2 | ITGB6 | 0,8852 | 2,20842E-63 |
| ACE2 | PRLR | 0,8851 | 2,38014E-63 |
| ACE2 | ALDH18A1 | 0,885 | 2,50563E-63 |
| ACE2 | NBPF7 | 0,8848 | 2,88841E-63 |
| ACE2 | DHRS7 | 0,8848 | 2,95134E-63 |
| ACE2 | CHP2 | 0,8848 | 2,9105E-63 |
| ACE2 | PAQR8 | 0,8847 | 3,04476E-63 |
| ACE2 | LINC01133 | 0,8846 | 3,49391E-63 |
| ACE2 | RP11-255M6.1 | 0,8846 | 3,39176E-63 |
| ACE2 | AKR7L | 0,8844 | 4,04884E-63 |
| ACE2 | SCNN1A | 0,8843 | 4,4198E-63 |
| ACE2 | HNF4A-AS1 | 0,8843 | 4,34138E-63 |
| ACE2 | ANKRD9 | 0,884 | 5,26969E-63 |
| ACE2 | SEMA4G | 0,8839 | 5,63309E-63 |
| ACE2 | PCK1 | 0,8839 | 5,88E-63 |
| ACE2 | DOLPP1 | 0,8836 | 7,13017E-63 |
| ACE2 | NLRP6 | 0,8834 | 8,19956E-63 |
| ACE2 | DDC | 0,8833 | 9,07806E-63 |
| ACE2 | FAM110C | 0,8831 | 1,05107E-62 |
| ACE2 | GPRIN2 | 0,8831 | 1,0344E-62 |
| ACE2 | LINC01207 | 0,8827 | 1,4118E-62 |
| ACE2 | CLIC5 | 0,8824 | 1,79766E-62 |
| ACE2 | OAT | 0,8824 | 1,71876E-62 |
| ACE2 | TSPAN3 | 0,8823 | 1,93784E-62 |
| ACE2 | C6orf132 | 0,8822 | 2,0643E-62 |
| ACE2 | ATP1B1 | 0,8819 | 2,46515E-62 |
| ACE2 | ZBTB7B | 0,8818 | 2,83417E-62 |

| | | | |
|---|---|---|---|
| ACE2 | CTAGE5 | 0,8815 | 3,41922E-62 |
| ACE2 | CIDEC | 0,8814 | 3,63997E-62 |
| ACE2 | TMEM41A | 0,8814 | 3,78849E-62 |
| ACE2 | SYTL5 | 0,8814 | 3,59594E-62 |
| ACE2 | SLC36A1 | 0,8812 | 4,31753E-62 |
| ACE2 | MYOM3 | 0,8811 | 4,72122E-62 |
| ACE2 | TSPAN8 | 0,8809 | 5,36308E-62 |
| ACE2 | FAM151A | 0,8807 | 6,20124E-62 |
| ACE2 | UGT1A4 | 0,8807 | 6,10314E-62 |
| ACE2 | CLDN23 | 0,8805 | 7,29004E-62 |
| ACE2 | DHDH | 0,8805 | 7,13423E-62 |
| ACE2 | DQX1 | 0,8804 | 7,38934E-62 |
| ACE2 | WWC1 | 0,8801 | 9,28548E-62 |
| ACE2 | GALNT7 | 0,8798 | 1,2077E-61 |
| ACE2 | SLC25A15 | 0,8795 | 1,43406E-61 |
| ACE2 | PPP1R14D | 0,8795 | 1,42244E-61 |
| ACE2 | CDCP1 | 0,8793 | 1,6742E-61 |
| ACE2 | PPARA | 0,8793 | 1,6643E-61 |
| ACE2 | HNF1B | 0,8792 | 1,82711E-61 |
| ACE2 | FAM160A1 | 0,8791 | 1,97807E-61 |
| ACE2 | SLC2A5 | 0,879 | 2,09133E-61 |
| ACE2 | CORO2A | 0,8788 | 2,36444E-61 |
| ACE2 | HSD11B2 | 0,8788 | 2,47556E-61 |
| ACE2 | AC106876.2 | 0,8787 | 2,60278E-61 |
| ACE2 | KLF5 | 0,8787 | 2,56605E-61 |
| ACE2 | LYPD8 | 0,8786 | 2,69721E-61 |
| ACE2 | TST | 0,8786 | 2,82631E-61 |
| ACE2 | REG4 | 0,8785 | 2,91238E-61 |
| ACE2 | B3GNT6 | 0,8785 | 2,88727E-61 |
| ACE2 | LINC00479 | 0,8785 | 2,88823E-61 |
| ACE2 | FAM3D | 0,8784 | 3,24525E-61 |
| ACE2 | SLAMF6P1 | 0,8783 | 3,40025E-61 |
| ACE2 | NHSL1 | 0,878 | 4,12856E-61 |
| ACE2 | GIPC2 | 0,8779 | 4,47647E-61 |
| ACE2 | AQP11 | 0,8776 | 5,62292E-61 |
| ACE2 | SLC28A2 | 0,8776 | 5,457E-61 |
| ACE2 | ADTRP | 0,8775 | 6,11763E-61 |
| ACE2 | CA13 | 0,8773 | 6,76587E-61 |
| ACE2 | LRRC31 | 0,8772 | 7,56826E-61 |
| ACE2 | SEPHS2 | 0,8766 | 1,16135E-60 |
| ACE2 | PKP3 | 0,8763 | 1,41403E-60 |
| ACE2 | CYP4F12 | 0,8761 | 1,6148E-60 |
| ACE2 | ENTPD5 | 0,876 | 1,72335E-60 |
| ACE2 | RBP2 | 0,8758 | 2,02465E-60 |
| ACE2 | MYO1D | 0,8758 | 2,05594E-60 |
| ACE2 | NEDD4L | 0,8758 | 2,06738E-60 |
| ACE2 | CBLC | 0,8758 | 1,98168E-60 |
| ACE2 | ARL14 | 0,8757 | 2,15201E-60 |
| ACE2 | MKRN2OS | 0,8756 | 2,25908E-60 |
| ACE2 | TMEM125 | 0,8755 | 2,49691E-60 |
| ACE2 | MGAT4B | 0,8754 | 2,66637E-60 |
| ACE2 | G6PC | 0,875 | 3,58358E-60 |
| ACE2 | RAPGEFL1 | 0,8748 | 3,94215E-60 |
| ACE2 | ITPKA | 0,8746 | 4,52284E-60 |

| | | | |
|---|---|---|---|
| ACE2 | AC027612.4 | 0,8744 | 5,12142E-60 |
| ACE2 | VIPR1 | 0,8743 | 5,67484E-60 |
| ACE2 | NTS | 0,8743 | 5,58712E-60 |
| ACE2 | VDR | 0,8741 | 6,2886E-60 |
| ACE2 | AKR1B10 | 0,874 | 6,88366E-60 |
| ACE2 | SLC22A4 | 0,8737 | 8,77903E-60 |
| ACE2 | PPP1R16A | 0,8737 | 8,5735E-60 |
| ACE2 | SLC31A1 | 0,8736 | 9,31628E-60 |
| ACE2 | C1QTNF12 | 0,8733 | 1,12786E-59 |
| ACE2 | SMPDL3A | 0,8732 | 1,17515E-59 |
| ACE2 | MAOA | 0,8729 | 1,49299E-59 |
| ACE2 | BARX2 | 0,8728 | 1,52212E-59 |
| ACE2 | CTD-2385L22.2 | 0,8728 | 1,57931E-59 |
| ACE2 | EPHA10 | 0,8727 | 1,71591E-59 |
| ACE2 | RP11-276H7.3 | 0,8726 | 1,75558E-59 |
| ACE2 | CYP4F3 | 0,8724 | 2,11046E-59 |
| ACE2 | CTD-2396E7.11 | 0,8721 | 2,51836E-59 |
| ACE2 | B3GNT5 | 0,8719 | 2,96807E-59 |
| ACE2 | CHN2 | 0,8718 | 3,06713E-59 |
| ACE2 | KB-1247B1.1 | 0,8718 | 3,12054E-59 |
| ACE2 | TP53I3 | 0,8716 | 3,65566E-59 |
| ACE2 | RP11-800A3.4 | 0,8715 | 3,73839E-59 |
| ACE2 | GPD1 | 0,8708 | 5,91988E-59 |
| ACE2 | VIPR1-AS1 | 0,8707 | 6,3291E-59 |
| ACE2 | SLC37A4 | 0,8707 | 6,66655E-59 |
| ACE2 | CLCN2 | 0,8704 | 8,0255E-59 |
| ACE2 | SLC52A3 | 0,8703 | 8,63517E-59 |
| ACE2 | ACADS | 0,8702 | 9,17871E-59 |
| ACE2 | IRF6 | 0,8701 | 9,79691E-59 |
| ACE2 | TFF1 | 0,8701 | 9,60883E-59 |
| ACE2 | AGPAT2 | 0,87 | 1,00376E-58 |
| ACE2 | PNPLA1 | 0,8698 | 1,17916E-58 |
| ACE2 | GPR39 | 0,8693 | 1,6008E-58 |
| ACE2 | GRTP1 | 0,8693 | 1,65811E-58 |
| ACE2 | PRR13 | 0,8691 | 1,88608E-58 |
| ACE2 | RP11-611E13.3 | 0,869 | 1,97308E-58 |
| ACE2 | FABP1 | 0,8689 | 2,14574E-58 |
| ACE2 | SGK2 | 0,8689 | 2,14841E-58 |
| ACE2 | HEPACAM2 | 0,8687 | 2,38806E-58 |
| ACE2 | ACOT11 | 0,8685 | 2,76088E-58 |
| ACE2 | ZP2 | 0,8684 | 2,92023E-58 |
| ACE2 | PRELID2 | 0,8683 | 3,14847E-58 |
| ACE2 | TMEM164 | 0,8683 | 3,10512E-58 |
| ACE2 | RPL12P14 | 0,8681 | 3,63682E-58 |
| ACE2 | GNA11 | 0,8679 | 4,06074E-58 |
| ACE2 | ESRRA | 0,8674 | 5,80633E-58 |
| ACE2 | CNDP2 | 0,8674 | 5,67595E-58 |
| ACE2 | GPAT3 | 0,8668 | 8,33703E-58 |
| ACE2 | KDF1 | 0,8667 | 8,64395E-58 |
| ACE2 | GK | 0,8667 | 8,69016E-58 |
| ACE2 | CLDN15 | 0,8664 | 1,0881E-57 |
| ACE2 | PAPSS2 | 0,8664 | 1,11338E-57 |
| ACE2 | MRO | 0,8664 | 1,09835E-57 |
| ACE2 | POF1B | 0,8663 | 1,13893E-57 |

| | | | |
|---|---|---|---|
| ACE2 | SLC35G1 | 0,8662 | 1,19601E-57 |
| ACE2 | ELOVL7 | 0,8657 | 1,73644E-57 |
| ACE2 | TIAM2 | 0,8657 | 1,6927E-57 |
| ACE2 | TDP2 | 0,8654 | 2,03948E-57 |
| ACE2 | RASEF | 0,8654 | 2,11106E-57 |
| ACE2 | CBR1 | 0,8653 | 2,1354E-57 |
| ACE2 | GSTA1 | 0,8651 | 2,54436E-57 |
| ACE2 | SLC22A5 | 0,8649 | 2,80347E-57 |
| ACE2 | CHRNA7 | 0,8646 | 3,41793E-57 |
| ACE2 | AC007126.1 | 0,8645 | 3,57067E-57 |
| ACE2 | TOM1L1 | 0,8644 | 4,00591E-57 |
| ACE2 | SLC22A18AS | 0,8642 | 4,41217E-57 |
| ACE2 | AP000439.3 | 0,8642 | 4,38041E-57 |
| ACE2 | OVOL2 | 0,864 | 5,10143E-57 |
| ACE2 | HECTD3 | 0,8638 | 5,61477E-57 |
| ACE2 | S100A14 | 0,8638 | 5,66894E-57 |
| ACE2 | OVOL1 | 0,8638 | 5,81703E-57 |
| ACE2 | MMP15 | 0,8637 | 5,95123E-57 |
| ACE2 | RP11-77K12.5 | 0,8637 | 6,00307E-57 |
| ACE2 | STAP2 | 0,8637 | 6,235E-57 |
| ACE2 | HDHD3 | 0,8633 | 7,91183E-57 |
| ACE2 | CDC42BPG | 0,8633 | 7,90209E-57 |
| ACE2 | FAM109A | 0,8632 | 8,12102E-57 |
| ACE2 | FGFR3 | 0,8627 | 1,12755E-56 |
| ACE2 | CTD-2555C10.3 | 0,8627 | 1,09816E-56 |
| ACE2 | PLEKHS1 | 0,8624 | 1,38447E-56 |
| ACE2 | AP006621.9 | 0,8622 | 1,52493E-56 |
| ACE2 | BICDL2 | 0,8618 | 1,95952E-56 |
| ACE2 | RP11-503C24.6 | 0,8615 | 2,34444E-56 |
| ACE2 | SUCLG1 | 0,8613 | 2,73773E-56 |
| ACE2 | KCNK5 | 0,8611 | 3,06048E-56 |
| ACE2 | TMEM229A | 0,861 | 3,2393E-56 |
| ACE2 | FAM83E | 0,861 | 3,23604E-56 |
| ACE2 | CNKSR1 | 0,8607 | 3,90504E-56 |
| ACE2 | NR5A2 | 0,8607 | 3,82614E-56 |
| ACE2 | CALML4 | 0,8605 | 4,30767E-56 |
| ACE2 | SDC1 | 0,8604 | 4,76999E-56 |
| ACE2 | LINC00974 | 0,8604 | 4,73799E-56 |
| ACE2 | TRPM6 | 0,8603 | 5,07194E-56 |
| ACE2 | ABCC6P1 | 0,8603 | 4,92561E-56 |
| ACE2 | P2RY2 | 0,8601 | 5,57288E-56 |
| ACE2 | SLC25A5 | 0,8601 | 5,69867E-56 |
| ACE2 | RBKS | 0,8598 | 6,65181E-56 |
| ACE2 | HOOK1 | 0,8595 | 8,18588E-56 |
| ACE2 | RP11-99J16__A.2 | 0,8591 | 1,01448E-55 |
| ACE2 | RP11-462G2.1 | 0,8589 | 1,14697E-55 |
| ACE2 | GRAMD1B | 0,8589 | 1,17626E-55 |
| ACE2 | GPR35 | 0,8583 | 1,66338E-55 |
| ACE2 | RP5-881L22.6 | 0,8579 | 2,16534E-55 |
| ACE2 | MME-AS1 | 0,8575 | 2,66217E-55 |
| ACE2 | ERBB3 | 0,8575 | 2,78589E-55 |
| ACE2 | RP11-297L17.2 | 0,8568 | 4,06007E-55 |
| ACE2 | VNN1 | 0,8567 | 4,31824E-55 |
| ACE2 | ABCG8 | 0,8564 | 5,16432E-55 |

| | | | |
|---|---|---|---|
| ACE2 | PLLP | 0,8561 | 6,49827E-55 |
| ACE2 | MTMR11 | 0,8559 | 7,28952E-55 |
| ACE2 | LINC00704 | 0,8559 | 7,23078E-55 |
| ACE2 | CLCA1 | 0,8556 | 8,30042E-55 |
| ACE2 | VILL | 0,8556 | 8,56973E-55 |
| ACE2 | MFSD2A | 0,8552 | 1,09272E-54 |
| ACE2 | MYRF | 0,8551 | 1,15304E-54 |
| ACE2 | MYH15 | 0,855 | 1,24194E-54 |
| ACE2 | CCT8P1 | 0,8549 | 1,26815E-54 |
| ACE2 | SLC27A4 | 0,8549 | 1,29996E-54 |
| ACE2 | GLRA4 | 0,8549 | 1,30088E-54 |
| ACE2 | EFNA2 | 0,8548 | 1,36765E-54 |
| ACE2 | SGK1 | 0,8547 | 1,4694E-54 |
| ACE2 | USP43 | 0,8547 | 1,45947E-54 |
| ACE2 | FAM83H | 0,8546 | 1,56206E-54 |
| ACE2 | BPHL | 0,8545 | 1,60354E-54 |
| ACE2 | TRIM15 | 0,8539 | 2,36972E-54 |
| ACE2 | RAB11FIP1 | 0,8539 | 2,36972E-54 |
| ACE2 | P2RY4 | 0,8539 | 2,25342E-54 |
| ACE2 | LINC01843 | 0,8537 | 2,56784E-54 |
| ACE2 | PHLPP2 | 0,8535 | 2,94983E-54 |
| ACE2 | LGALS4 | 0,8533 | 3,20244E-54 |
| ACE2 | EPHX2 | 0,853 | 3,98227E-54 |
| ACE2 | RP11-227H15.5 | 0,853 | 3,91816E-54 |
| ACE2 | TMPRSS15 | 0,8529 | 4,13215E-54 |
| ACE2 | RP11-48O20.5 | 0,8528 | 4,34649E-54 |
| ACE2 | TTC36 | 0,8527 | 4,57444E-54 |
| ACE2 | AGXT2 | 0,8526 | 4,82982E-54 |
| ACE2 | RP11-532F12.5 | 0,8526 | 4,92245E-54 |
| ACE2 | PCYT2 | 0,8524 | 5,64803E-54 |
| ACE2 | FMO1 | 0,8522 | 6,22775E-54 |
| ACE2 | RP11-354P11.4 | 0,8521 | 6,39874E-54 |
| ACE2 | NIPAL1 | 0,8517 | 8,30336E-54 |
| ACE2 | GLRX | 0,8517 | 8,38135E-54 |
| ACE2 | RAP1GAP | 0,8516 | 8,81386E-54 |
| ACE2 | MLPH | 0,8516 | 8,83048E-54 |
| ACE2 | CEACAM1 | 0,8515 | 9,35952E-54 |
| ACE2 | PRKCZ | 0,8514 | 9,88267E-54 |
| ACE2 | KRT8 | 0,8514 | 1,00671E-53 |
| ACE2 | GLTPD2 | 0,8513 | 1,05172E-53 |
| ACE2 | PRKG2 | 0,8512 | 1,12138E-53 |
| ACE2 | MAPK13 | 0,851 | 1,22445E-53 |
| ACE2 | SLC25A48 | 0,8509 | 1,34587E-53 |
| ACE2 | OR7E14P | 0,8506 | 1,52565E-53 |
| ACE2 | CTB-186H2.3 | 0,8503 | 1,79047E-53 |
| ACE2 | MACC1 | 0,8502 | 1,94417E-53 |
| ACE2 | ABO | 0,8499 | 2,34122E-53 |
| ACE2 | CDX2 | 0,8499 | 2,28038E-53 |
| ACE2 | LSR | 0,8499 | 2,25223E-53 |
| ACE2 | MYO6 | 0,8495 | 2,97272E-53 |
| ACE2 | SMIM22 | 0,8495 | 2,95752E-53 |
| ACE2 | AQP3 | 0,8494 | 3,13958E-53 |
| ACE2 | ATOH1 | 0,8492 | 3,4915E-53 |
| ACE2 | IFNLR1 | 0,8491 | 3,72543E-53 |

| | | | |
|---|---|---|---|
| ACE2 | HNF1A-AS1 | 0,8491 | 3,69157E-53 |
| ACE2 | TAT | 0,8491 | 3,55417E-53 |
| ACE2 | SULT2A1 | 0,8491 | 3,71038E-53 |
| ACE2 | BEST2 | 0,8489 | 4,00645E-53 |
| ACE2 | SMPDL3B | 0,8488 | 4,31459E-53 |
| ACE2 | KBTBD11-OT1 | 0,8486 | 4,88456E-53 |
| ACE2 | KCNH6 | 0,8485 | 4,99921E-53 |
| ACE2 | PC | 0,8484 | 5,32084E-53 |
| ACE2 | ARHGEF5 | 0,8483 | 5,78409E-53 |
| ACE2 | LINC00511 | 0,8483 | 5,55319E-53 |
| ACE2 | TMEM171 | 0,8482 | 6,16452E-53 |
| ACE2 | NR6A1 | 0,8482 | 6,04571E-53 |
| ACE2 | LAD1 | 0,8478 | 7,56055E-53 |
| ACE2 | DOK4 | 0,8478 | 7,44073E-53 |
| ACE2 | TMEM53 | 0,8477 | 7,81626E-53 |
| ACE2 | SATB2 | 0,8475 | 9,09671E-53 |
| ACE2 | OCIAD2 | 0,8475 | 8,70184E-53 |
| ACE2 | C8G | 0,8475 | 9,12325E-53 |
| ACE2 | RP4-811M8.1 | 0,8475 | 8,99337E-53 |
| ACE2 | AKR1C3 | 0,8474 | 9,17274E-53 |
| ACE2 | PLPP2 | 0,8474 | 9,59494E-53 |
| ACE2 | MST1R | 0,8473 | 1,01535E-52 |
| ACE2 | KIF16B | 0,8472 | 1,05284E-52 |
| ACE2 | RP11-867O8.5 | 0,8469 | 1,2176E-52 |
| ACE2 | SEMA6D | 0,8469 | 1,22283E-52 |
| ACE2 | SLC27A2 | 0,8465 | 1,58428E-52 |
| ACE2 | B4GALNT2 | 0,8464 | 1,66512E-52 |
| ACE2 | OSGIN1 | 0,8463 | 1,69029E-52 |
| ACE2 | NXPE1 | 0,8462 | 1,82114E-52 |
| ACE2 | PLEKHH1 | 0,8462 | 1,78587E-52 |
| ACE2 | PIK3C2G | 0,8461 | 1,95036E-52 |
| ACE2 | SDHA | 0,846 | 2,048E-52 |
| ACE2 | PDZD7 | 0,846 | 2,01156E-52 |
| ACE2 | AQP7 | 0,8458 | 2,34239E-52 |
| ACE2 | SERPINA1 | 0,8458 | 2,28673E-52 |
| ACE2 | CKMT1B | 0,8457 | 2,41345E-52 |
| ACE2 | PCSK5 | 0,8456 | 2,59416E-52 |
| ACE2 | SCGB2A1 | 0,8455 | 2,67681E-52 |
| ACE2 | APPL2 | 0,8454 | 2,88193E-52 |
| ACE2 | PARD6B | 0,8454 | 2,85939E-52 |
| ACE2 | HSD17B11 | 0,8452 | 3,20736E-52 |
| ACE2 | HTATIP2 | 0,8452 | 3,15586E-52 |
| ACE2 | RPS29P12 | 0,845 | 3,61441E-52 |
| ACE2 | RP11-35P15.1 | 0,8448 | 3,96244E-52 |
| ACE2 | CMTM4 | 0,8448 | 3,99199E-52 |
| ACE2 | LINC00675 | 0,8448 | 3,95525E-52 |
| ACE2 | JUP | 0,8448 | 4,01441E-52 |
| ACE2 | CNGA1 | 0,8445 | 4,73625E-52 |
| ACE2 | NEURL1 | 0,8444 | 4,9136E-52 |
| ACE2 | CYB5A | 0,844 | 6,03E-52 |
| ACE2 | ABCG5 | 0,8438 | 6,81556E-52 |
| ACE2 | CARD10 | 0,8437 | 7,29155E-52 |
| ACE2 | BRINP3 | 0,8436 | 7,48123E-52 |
| ACE2 | STRADB | 0,8431 | 9,9682E-52 |

| | | | |
|---|---|---|---|
| ACE2 | CD2AP | 0,8431 | 1,00348E-51 |
| ACE2 | RP11-478K15.6 | 0,8431 | 1,00333E-51 |
| ACE2 | BPNT1 | 0,8426 | 1,29632E-51 |
| ACE2 | ERBB2 | 0,8426 | 1,27956E-51 |
| ACE2 | GGT1 | 0,8423 | 1,52993E-51 |
| ACE2 | FABP6 | 0,8422 | 1,57907E-51 |
| ACE2 | SH3RF2 | 0,8418 | 2,0578E-51 |
| ACE2 | LGALS2 | 0,8418 | 1,98877E-51 |
| ACE2 | RP11-276H7.2 | 0,8417 | 2,09035E-51 |
| ACE2 | C17orf78 | 0,8417 | 2,13939E-51 |
| ACE2 | AP1M2 | 0,8415 | 2,321E-51 |
| ACE2 | RP11-164P12.5 | 0,8413 | 2,66525E-51 |
| ACE2 | SELENOP | 0,8411 | 2,96919E-51 |
| ACE2 | SEC16B | 0,841 | 3,1396E-51 |
| ACE2 | SH3D21 | 0,8409 | 3,2384E-51 |
| ACE2 | SLC9A2 | 0,8409 | 3,22577E-51 |
| ACE2 | KRT89P | 0,8409 | 3,22411E-51 |
| ACE2 | KIAA1211 | 0,8406 | 3,71964E-51 |
| ACE2 | TPMT | 0,8406 | 3,8542E-51 |
| ACE2 | HEPH | 0,8406 | 3,78813E-51 |
| ACE2 | MICALCL | 0,8405 | 4,01382E-51 |
| ACE2 | FHDC1 | 0,8404 | 4,3337E-51 |
| ACE2 | GJB1 | 0,8403 | 4,42567E-51 |
| ACE2 | RP11-94C24.13 | 0,8402 | 4,82242E-51 |
| ACE2 | AC092569.2 | 0,8401 | 4,96742E-51 |
| ACE2 | CDKN2B-AS1 | 0,84 | 5,36381E-51 |
| ACE2 | SLC25A34 | 0,8396 | 6,48167E-51 |
| ACE2 | KISS1 | 0,8396 | 6,36208E-51 |
| ACE2 | TCN2 | 0,8391 | 8,38191E-51 |
| ACE2 | MPST | 0,8391 | 8,45567E-51 |
| ACE2 | DEGS2 | 0,8389 | 9,58925E-51 |
| ACE2 | TMEM54 | 0,8385 | 1,13451E-50 |
| ACE2 | KCNJ3 | 0,8384 | 1,23444E-50 |
| ACE2 | CEBPA | 0,838 | 1,53614E-50 |
| ACE2 | GALNT5 | 0,8376 | 1,87257E-50 |
| ACE2 | AL163953.2 | 0,8376 | 1,89046E-50 |
| ACE2 | CHP1 | 0,8376 | 1,82991E-50 |
| ACE2 | SLC7A7 | 0,8375 | 1,93726E-50 |
| ACE2 | UGT1A3 | 0,8374 | 1,99742E-50 |
| ACE2 | POR | 0,8374 | 2,04359E-50 |
| ACE2 | RPS4XP7 | 0,837 | 2,48603E-50 |
| ACE2 | PXMP2 | 0,8369 | 2,61146E-50 |
| ACE2 | DSG2-AS1 | 0,8369 | 2,63516E-50 |
| ACE2 | MLXIPL | 0,8367 | 3,00683E-50 |
| ACE2 | CES3 | 0,8367 | 2,94272E-50 |
| ACE2 | FAM83B | 0,8364 | 3,4861E-50 |
| ACE2 | CMAS | 0,8364 | 3,43636E-50 |
| ACE2 | VAMP8 | 0,8361 | 3,9696E-50 |
| ACE2 | FUT3 | 0,836 | 4,20936E-50 |
| ACE2 | PIP5K1B | 0,8359 | 4,56384E-50 |
| ACE2 | ARHGEF38 | 0,8355 | 5,46851E-50 |
| ACE2 | AC007255.8 | 0,8353 | 6,18023E-50 |
| ACE2 | OTOP3 | 0,8352 | 6,51422E-50 |
| ACE2 | BAIAP2L1 | 0,8351 | 6,65072E-50 |

| | | | |
|---|---|---|---|
| ACE2 | RP11-798K3.2 | 0,8351 | 6,81776E-50 |
| ACE2 | RP11-77K12.10 | 0,8351 | 6,59478E-50 |
| ACE2 | GAL3ST1 | 0,835 | 7,05594E-50 |
| ACE2 | KLC4 | 0,8349 | 7,31661E-50 |
| ACE2 | DEPDC7 | 0,8348 | 7,81111E-50 |
| ACE2 | C15orf48 | 0,8346 | 8,50576E-50 |
| ACE2 | SULT1C3 | 0,8343 | 1,02077E-49 |
| ACE2 | PCLO | 0,8343 | 1,01532E-49 |
| ACE2 | RP13-726E6.2 | 0,8343 | 1,03381E-49 |
| ACE2 | ANO7 | 0,8342 | 1,06604E-49 |
| ACE2 | CRAT8 | 0,8342 | 1,03971E-49 |
| ACE2 | LGALS3 | 0,834 | 1,1537E-49 |
| ACE2 | CAPN5 | 0,8337 | 1,38984E-49 |
| ACE2 | RP11-284F21.7 | 0,8334 | 1,55576E-49 |
| ACE2 | FUOM | 0,8329 | 2,04395E-49 |
| ACE2 | CDX1 | 0,8328 | 2,11408E-49 |
| ACE2 | RP11-867G23.8 | 0,8327 | 2,26153E-49 |
| ACE2 | ELMO3 | 0,8327 | 2,3208E-49 |
| ACE2 | PAFAH2 | 0,8325 | 2,53798E-49 |
| ACE2 | CPS1 | 0,8325 | 2,44908E-49 |
| ACE2 | TRIM31 | 0,8323 | 2,70695E-49 |
| ACE2 | MEP1AP1 | 0,8322 | 2,98875E-49 |
| ACE2 | SLC35D2 | 0,8322 | 2,96847E-49 |
| ACE2 | GGN | 0,8321 | 3,10093E-49 |
| ACE2 | RP4-724E13.2 | 0,832 | 3,20314E-49 |
| ACE2 | GCNT3 | 0,832 | 3,26339E-49 |
| ACE2 | TFF3 | 0,8318 | 3,61061E-49 |
| ACE2 | URAD | 0,8315 | 4,21468E-49 |
| ACE2 | VSIG10 | 0,8313 | 4,59286E-49 |
| ACE2 | RP11-284F21.10 | 0,831 | 5,18874E-49 |
| ACE2 | HEBP1 | 0,831 | 5,36967E-49 |
| ACE2 | CKMT1A | 0,8309 | 5,6571E-49 |
| ACE2 | CHAD | 0,8309 | 5,46516E-49 |
| ACE2 | TRPM4 | 0,8308 | 5,87665E-49 |
| ACE2 | CCRL2 | 0,8303 | 7,39294E-49 |
| ACE2 | TRIM40 | 0,8301 | 8,22227E-49 |
| ACE2 | SPDEF | 0,8298 | 9,74501E-49 |
| ACE2 | PNLIPRP2 | 0,8298 | 9,6072E-49 |
| ACE2 | SOX6 | 0,8297 | 1,00701E-48 |
| ACE2 | UGT2B15 | 0,8294 | 1,17338E-48 |
| ACE2 | LINC00570 | 0,8293 | 1,23431E-48 |
| ACE2 | SIAE | 0,8293 | 1,26341E-48 |
| ACE2 | SLC47A2 | 0,8293 | 1,2074E-48 |
| ACE2 | UGT1A8 | 0,8291 | 1,35538E-48 |
| ACE2 | PYY | 0,8289 | 1,5345E-48 |
| ACE2 | PANK1 | 0,8288 | 1,54908E-48 |
| ACE2 | BCL2L14 | 0,8286 | 1,76101E-48 |
| ACE2 | FUT2 | 0,8286 | 1,78139E-48 |
| ACE2 | RP11-96D1.6 | 0,8283 | 2,04453E-48 |
| ACE2 | TLX1 | 0,8282 | 2,07256E-48 |
| ACE2 | PLIN3 | 0,828 | 2,32578E-48 |
| ACE2 | RP11-380D23.1 | 0,8279 | 2,4409E-48 |
| ACE2 | ITM2C | 0,8278 | 2,58756E-48 |
| ACE2 | CTNND1 | 0,8278 | 2,61618E-48 |

| | | | |
|---|---|---|---|
| ACE2 | B4GALT4 | 0,8274 | 3,0839E-48 |
| ACE2 | LEXM | 0,8269 | 3,97172E-48 |
| ACE2 | ADH6 | 0,8269 | 3,92001E-48 |
| ACE2 | SSUH2 | 0,8267 | 4,36123E-48 |
| ACE2 | CHDH | 0,8266 | 4,54605E-48 |
| ACE2 | DERA | 0,8264 | 5,13658E-48 |
| ACE2 | ZZEF1 | 0,8264 | 5,04455E-48 |
| ACE2 | RP11-395B7.2 | 0,8262 | 5,50004E-48 |
| ACE2 | LINC00543 | 0,826 | 6,14455E-48 |
| ACE2 | SLC39A11 | 0,8253 | 8,42078E-48 |
| ACE2 | TMEM120A | 0,8252 | 9,13179E-48 |
| ACE2 | KRT12 | 0,8251 | 9,65179E-48 |
| ACE2 | C6orf136 | 0,8248 | 1,07009E-47 |
| ACE2 | NR0B2 | 0,8246 | 1,20381E-47 |
| ACE2 | KRT8P3 | 0,8246 | 1,19394E-47 |
| ACE2 | CA7 | 0,8245 | 1,288E-47 |
| ACE2 | RP11-311D14.1 | 0,8243 | 1,40156E-47 |
| ACE2 | EDN3 | 0,8243 | 1,35868E-47 |
| ACE2 | CLCN5 | 0,8243 | 1,4108E-47 |
| ACE2 | SAR1B | 0,8236 | 1,9058E-47 |
| ACE2 | SLC25A1 | 0,8236 | 1,95018E-47 |
| ACE2 | SGPP2 | 0,8235 | 2,07427E-47 |
| ACE2 | MGAT3 | 0,8234 | 2,1351E-47 |
| ACE2 | AC009014.3 | 0,8231 | 2,42035E-47 |
| ACE2 | TLCD2 | 0,8231 | 2,41886E-47 |
| ACE2 | TMEM106C | 0,823 | 2,54544E-47 |
| ACE2 | RPL7P16 | 0,8229 | 2,64329E-47 |
| ACE2 | ENPP7 | 0,8229 | 2,76425E-47 |
| ACE2 | SLC35A3 | 0,8228 | 2,84729E-47 |
| ACE2 | DNMBP-AS1 | 0,8228 | 2,8822E-47 |
| ACE2 | C19orf57 | 0,8228 | 2,81516E-47 |
| ACE2 | TOR1AIP2 | 0,8225 | 3,27621E-47 |
| ACE2 | SPIRE2 | 0,8225 | 3,27216E-47 |
| ACE2 | ZNF57 | 0,8225 | 3,32822E-47 |
| ACE2 | USH1G | 0,8223 | 3,64338E-47 |
| ACE2 | SSTR5-AS1 | 0,8218 | 4,6187E-47 |
| ACE2 | SPACA4 | 0,8217 | 4,79873E-47 |
| ACE2 | CTD-2021H9.3 | 0,8214 | 5,37255E-47 |
| ACE2 | ACVR1B | 0,8211 | 6,16265E-47 |
| ACE2 | ATP7B | 0,8211 | 6,35275E-47 |
| ACE2 | FAM83G | 0,8211 | 6,2543E-47 |
| ACE2 | DDR1 | 0,821 | 6,7013E-47 |
| ACE2 | BEST4 | 0,8207 | 7,52536E-47 |
| ACE2 | HTR4 | 0,8204 | 8,64119E-47 |
| ACE2 | ANXA4 | 0,8203 | 9,30803E-47 |
| ACE2 | PDXP | 0,8203 | 8,95363E-47 |
| ACE2 | KLHL34 | 0,8202 | 9,47964E-47 |
| ACE2 | C11orf24 | 0,8198 | 1,15008E-46 |
| ACE2 | PLA2G10 | 0,8198 | 1,1768E-46 |
| ACE2 | STBD1 | 0,8196 | 1,26887E-46 |
| ACE2 | ABCC2 | 0,8195 | 1,33934E-46 |
| ACE2 | ADGRF1 | 0,8192 | 1,49737E-46 |
| ACE2 | PFKFB4 | 0,819 | 1,64278E-46 |
| ACE2 | CARMIL1 | 0,8189 | 1,73391E-46 |

| | | | |
|---|---|---|---|
| ACE2 | LLGL2 | 0,8187 | 1,89546E-46 |
| ACE2 | TBC1D2 | 0,8186 | 2,0378E-46 |
| ACE2 | KRT15 | 0,8186 | 2,013E-46 |
| ACE2 | ARHGAP32 | 0,8185 | 2,10352E-46 |
| ACE2 | SP6 | 0,8185 | 2,13625E-46 |
| ACE2 | ACSS2 | 0,8185 | 2,11063E-46 |
| ACE2 | GSR | 0,8181 | 2,58127E-46 |
| ACE2 | ELF3 | 0,818 | 2,6559E-46 |
| ACE2 | RPSAP9 | 0,818 | 2,63752E-46 |
| ACE2 | HOOK2 | 0,8179 | 2,79302E-46 |
| ACE2 | STX19 | 0,8178 | 2,90708E-46 |
| ACE2 | REP15 | 0,8178 | 2,92547E-46 |
| ACE2 | RP11-480I12.5 | 0,8177 | 2,99031E-46 |
| ACE2 | RP11-473M20.5 | 0,8172 | 3,79245E-46 |
| ACE2 | GGT6 | 0,8172 | 3,84119E-46 |
| ACE2 | LINC02200 | 0,8164 | 5,43188E-46 |
| ACE2 | RP11-884K10.6 | 0,816 | 6,68328E-46 |
| ACE2 | OCLN | 0,8157 | 7,70851E-46 |
| ACE2 | LGR4 | 0,8154 | 8,76375E-46 |
| ACE2 | KIAA2013 | 0,8152 | 9,44789E-46 |
| ACE2 | SLC6A18 | 0,8152 | 9,54991E-46 |
| ACE2 | XKR9 | 0,8149 | 1,09653E-45 |
| ACE2 | CTC-558O2.1 | 0,8148 | 1,141E-45 |
| ACE2 | RAD17P1 | 0,8146 | 1,22393E-45 |
| ACE2 | RP11-336A10.5 | 0,8144 | 1,33926E-45 |
| ACE2 | SYT7 | 0,8144 | 1,36762E-45 |
| ACE2 | NAT1 | 0,8143 | 1,38609E-45 |
| ACE2 | LINC00488 | 0,8141 | 1,53185E-45 |
| ACE2 | SLC22A23 | 0,814 | 1,59618E-45 |
| ACE2 | TOX3 | 0,8136 | 1,89999E-45 |
| ACE2 | CPT2 | 0,8135 | 2,02499E-45 |
| ACE2 | SHBG | 0,8133 | 2,24623E-45 |
| ACE2 | CYP27A1 | 0,8129 | 2,66735E-45 |
| ACE2 | GSTA2 | 0,8129 | 2,63393E-45 |
| ACE2 | AATK | 0,8125 | 3,21398E-45 |
| ACE2 | SULT1A1 | 0,8122 | 3,66135E-45 |
| ACE2 | GSKIP | 0,8118 | 4,40741E-45 |
| ACE2 | RP11-62H20.1 | 0,8115 | 4,99673E-45 |
| ACE2 | CTD-2184D3.3 | 0,8115 | 4,93541E-45 |
| ACE2 | IGF2BP2 | 0,8114 | 5,06501E-45 |
| ACE2 | PDE9A | 0,8114 | 5,07321E-45 |
| ACE2 | FAAH | 0,8113 | 5,35496E-45 |
| ACE2 | HIST1H3E | 0,8113 | 5,43452E-45 |
| ACE2 | TCP10 | 0,8113 | 5,4577E-45 |
| ACE2 | KLB | 0,8108 | 6,65986E-45 |
| ACE2 | SCAMP2 | 0,8107 | 6,9458E-45 |
| ACE2 | KIF13B | 0,8105 | 7,79528E-45 |
| ACE2 | CTSD | 0,8105 | 7,55286E-45 |
| ACE2 | CACNA1D | 0,8103 | 8,39154E-45 |
| ACE2 | FA2H | 0,8103 | 8,41179E-45 |
| ACE2 | AC090616.2 | 0,8102 | 8,70527E-45 |
| ACE2 | RHPN2 | 0,8098 | 1,02205E-44 |
| ACE2 | GOT1 | 0,8097 | 1,09666E-44 |
| ACE2 | SLC16A5 | 0,8096 | 1,12249E-44 |

| | | | |
|---|---|---|---|
| ACE2 | LINC00955 | 0,8094 | 1,22319E-44 |
| ACE2 | ITGA3 | 0,8091 | 1,43419E-44 |
| ACE2 | TNNI1 | 0,809 | 1,44383E-44 |
| ACE2 | C5orf30 | 0,8088 | 1,6182E-44 |
| ACE2 | TDRD12 | 0,8085 | 1,79067E-44 |
| ACE2 | RP11-284F21.9 | 0,8082 | 2,06209E-44 |
| ACE2 | RP5-1148A21.3 | 0,8082 | 2,10306E-44 |
| ACE2 | RPS6KA1 | 0,8081 | 2,19057E-44 |
| ACE2 | TTC39A | 0,808 | 2,30548E-44 |
| ACE2 | KRT9 | 0,808 | 2,28958E-44 |
| ACE2 | CISD3 | 0,8076 | 2,68339E-44 |
| ACE2 | KBTBD11 | 0,8074 | 2,9498E-44 |
| ACE2 | NEURL3 | 0,8073 | 3,09191E-44 |
| ACE2 | UCN3 | 0,807 | 3,49891E-44 |
| ACE2 | ABHD17C | 0,807 | 3,52398E-44 |
| ACE2 | LA16c-380H5.5 | 0,8069 | 3,64967E-44 |
| ACE2 | RPL7AP52 | 0,8067 | 3,91619E-44 |
| ACE2 | RAB25 | 0,8066 | 4,15484E-44 |
| ACE2 | FMO4 | 0,8066 | 4,20875E-44 |
| ACE2 | WTH3DI | 0,8066 | 4,17284E-44 |
| ACE2 | BDH1 | 0,8066 | 4,08037E-44 |
| ACE2 | C3orf52 | 0,8062 | 4,90588E-44 |
| ACE2 | KCNK1 | 0,806 | 5,37045E-44 |
| ACE2 | CRYBA2 | 0,8057 | 6,19368E-44 |
| ACE2 | NAT8 | 0,8056 | 6,2205E-44 |
| ACE2 | GPRC5C | 0,805 | 8,11019E-44 |
| ACE2 | CTD-2547H18.1 | 0,8046 | 9,81491E-44 |
| ACE2 | LRRC26 | 0,8042 | 1,15E-43 |
| ACE2 | RP11-386E5.1 | 0,8041 | 1,21115E-43 |
| ACE2 | SLC37A1 | 0,8041 | 1,21374E-43 |
| ACE2 | GUCD1 | 0,8039 | 1,28412E-43 |
| ACE2 | RP11-187E13.1 | 0,8038 | 1,33421E-43 |
| ACE2 | CTB-78O21.1 | 0,8037 | 1,39059E-43 |
| ACE2 | CYP2D6 | 0,8032 | 1,77047E-43 |
| ACE2 | ACAA2 | 0,803 | 1,91737E-43 |
| ACE2 | LINC00337 | 0,8029 | 1,94963E-43 |
| ACE2 | GSTM4 | 0,8029 | 1,99212E-43 |
| ACE2 | XYLB | 0,8025 | 2,3761E-43 |
| ACE2 | GPR160 | 0,8021 | 2,76157E-43 |
| ACE2 | SPINK4 | 0,8021 | 2,79091E-43 |
| ACE2 | ANXA13 | 0,8016 | 3,35705E-43 |
| ACE2 | HNRNPA1P17 | 0,8014 | 3,74698E-43 |
| ACE2 | PADI2 | 0,8011 | 4,13574E-43 |
| ACE2 | PDZK1IP1 | 0,8011 | 4,21427E-43 |
| ACE2 | GALE | 0,8008 | 4,82169E-43 |
| ACE2 | HSD3B2 | 0,8008 | 4,68921E-43 |
| ACE2 | AC073283.4 | 0,8006 | 5,20934E-43 |
| ACE2 | RP3-523K23.2 | 0,8005 | 5,43594E-43 |
| ACE2 | UGT1A6 | 0,8004 | 5,57562E-43 |
| ACE2 | RP11-380P13.1 | 0,8003 | 5,90727E-43 |
| ACE2 | SCP2 | 0,8 | 6,5499E-43 |
| ACE2 | CALR4P | 0,7997 | 7,4089E-43 |
| ACE2 | RP11-757F18.5 | 0,7997 | 7,37614E-43 |
| ACE2 | SHROOM1 | 0,7996 | 7,79838E-43 |

| | | | |
|---|---|---|---|
| ACE2 | RP11-433O3.1 | 0,7991 | 9,51585E-43 |
| ACE2 | RP13-122B23.8 | 0,7988 | 1,07186E-42 |
| ACE2 | NPM1P32 | 0,7987 | 1,12549E-42 |
| ACE2 | RP11-844P9.6 | 0,7986 | 1,19421E-42 |
| ACE2 | ANO10 | 0,7979 | 1,59499E-42 |
| ACE2 | NAPRT | 0,7979 | 1,54504E-42 |
| ACE2 | UBE2V2P1 | 0,7979 | 1,58436E-42 |
| ACE2 | SUSD2 | 0,7979 | 1,58393E-42 |
| ACE2 | TMEM238 | 0,7978 | 1,62216E-42 |
| ACE2 | ACO2 | 0,7978 | 1,62787E-42 |
| ACE2 | SLC23A3 | 0,7976 | 1,77839E-42 |
| ACE2 | RP3-342P20.2 | 0,7973 | 1,96344E-42 |
| ACE2 | ABCC6P2 | 0,7971 | 2,20406E-42 |
| ACE2 | RFLNA | 0,7969 | 2,39679E-42 |
| ACE2 | RALGPS1 | 0,7967 | 2,56904E-42 |
| ACE2 | NXPE2 | 0,7966 | 2,70267E-42 |
| ACE2 | ZSWIM5 | 0,7965 | 2,76413E-42 |
| ACE2 | LASP1 | 0,7964 | 2,9083E-42 |
| ACE2 | C10orf99 | 0,7962 | 3,08993E-42 |
| ACE2 | RP11-379B8.1 | 0,7959 | 3,52545E-42 |
| ACE2 | AFDN-AS1 | 0,7958 | 3,66292E-42 |
| ACE2 | SLC4A4 | 0,7954 | 4,35226E-42 |
| ACE2 | GGT3P | 0,7954 | 4,30341E-42 |
| ACE2 | CTD-2085J24.3 | 0,7948 | 5,4443E-42 |
| ACE2 | OMP | 0,7947 | 5,80908E-42 |
| ACE2 | GEMIN8P4 | 0,7943 | 6,77471E-42 |
| ACE2 | SLC34A2 | 0,7941 | 7,19928E-42 |
| ACE2 | EPN1 | 0,7938 | 8,2806E-42 |
| ACE2 | RBP3 | 0,7937 | 8,37726E-42 |
| ACE2 | P2RX4 | 0,7937 | 8,46192E-42 |
| ACE2 | ASS1 | 0,7936 | 8,83417E-42 |
| ACE2 | RP11-1223D19.2 | 0,7935 | 9,19386E-42 |
| ACE2 | SSTR1 | 0,7935 | 9,33865E-42 |
| ACE2 | RP11-319G6.3 | 0,7934 | 9,47404E-42 |
| ACE2 | RP11-844P9.2 | 0,7932 | 1,03331E-41 |
| ACE2 | CDKN2B-AS | 0,7932 | 1,03012E-41 |
| ACE2 | CTA-363E6.6 | 0,7932 | 1,02646E-41 |
| ACE2 | CLCA4 | 0,7931 | 1,07806E-41 |
| ACE2 | GRHL2 | 0,7931 | 1,09274E-41 |
| ACE2 | RP3-434P1.6 | 0,7929 | 1,14604E-41 |
| ACE2 | LPCAT3 | 0,7925 | 1,37098E-41 |
| ACE2 | RP11-757F18.3 | 0,7924 | 1,43838E-41 |
| ACE2 | RP11-588L15.2 | 0,7921 | 1,59605E-41 |
| ACE2 | AC005281.2 | 0,7921 | 1,61593E-41 |
| ACE2 | MAP3K21 | 0,7918 | 1,82869E-41 |
| ACE2 | GLYCTK | 0,7914 | 2,11339E-41 |
| ACE2 | DEFB131 | 0,7911 | 2,3783E-41 |
| ACE2 | RP11-132E11.2 | 0,7911 | 2,37032E-41 |
| ACE2 | SLC6A7 | 0,7908 | 2,62215E-41 |
| ACE2 | GPRC5A | 0,7907 | 2,82262E-41 |
| ACE2 | ACOT4 | 0,7907 | 2,81918E-41 |
| ACE2 | RP3-417L20.4 | 0,7905 | 2,95926E-41 |
| ACE2 | MAPK6 | 0,7905 | 2,99811E-41 |
| ACE2 | TMPRSS4 | 0,7904 | 3,09295E-41 |

| | | | |
|---|---|---|---|
| ACE2 | MGC32805 | 0,7899 | 3,83768E-41 |
| ACE2 | FSIP2 | 0,7898 | 3,91151E-41 |
| ACE2 | RP11-191G24.1 | 0,7897 | 4,17518E-41 |
| ACE2 | RP11-297D21.2 | 0,7895 | 4,48886E-41 |
| ACE2 | RP11-627G18.1 | 0,7895 | 4,42436E-41 |
| ACE2 | VWA7 | 0,7894 | 4,66784E-41 |
| ACE2 | KCNQ1 | 0,7893 | 4,73388E-41 |
| ACE2 | CCDC162P | 0,789 | 5,32265E-41 |
| ACE2 | C1orf168 | 0,7883 | 7,07906E-41 |
| ACE2 | CTD-2385L22.1 | 0,7883 | 7,14808E-41 |
| ACE2 | RP11-432I5.2 | 0,7883 | 7,17333E-41 |
| ACE2 | SULT1E1 | 0,7878 | 8,46491E-41 |
| ACE2 | PRSS16 | 0,7878 | 8,73987E-41 |
| ACE2 | CELP | 0,7877 | 9,03572E-41 |
| ACE2 | RP11-1260E13.3 | 0,7877 | 8,76449E-41 |
| ACE2 | DHRS9 | 0,7875 | 9,49588E-41 |
| ACE2 | PDXK | 0,7874 | 1,00812E-40 |
| ACE2 | IDH1 | 0,787 | 1,17482E-40 |
| ACE2 | SLC7A4 | 0,7867 | 1,30698E-40 |
| ACE2 | RP11-459D22.1 | 0,7862 | 1,58383E-40 |
| ACE2 | LPIN3 | 0,7858 | 1,85261E-40 |
| ACE2 | TMEM105 | 0,7857 | 1,88638E-40 |
| ACE2 | TMEM144 | 0,7856 | 1,97546E-40 |
| ACE2 | SFN | 0,7854 | 2,12361E-40 |
| ACE2 | ST6GALNAC1 | 0,7853 | 2,20982E-40 |
| ACE2 | RTCB | 0,7852 | 2,28623E-40 |
| ACE2 | RBBP8NL | 0,785 | 2,53698E-40 |
| ACE2 | CASP5 | 0,7848 | 2,7165E-40 |
| ACE2 | AC021218.2 | 0,7847 | 2,80137E-40 |
| ACE2 | CASP7 | 0,7845 | 3,07506E-40 |
| ACE2 | SMIM5 | 0,7845 | 2,99244E-40 |
| ACE2 | CTD-2196E14.8 | 0,7842 | 3,43674E-40 |
| ACE2 | KIAA1161 | 0,7838 | 3,98832E-40 |
| ACE2 | GPRIN1 | 0,7836 | 4,26221E-40 |
| ACE2 | SEC14L2 | 0,7832 | 4,8706E-40 |
| ACE2 | DECR1 | 0,7829 | 5,52418E-40 |
| ACE2 | COMTD1 | 0,7826 | 6,28299E-40 |
| ACE2 | RP11-168F9.2 | 0,7824 | 6,7834E-40 |
| ACE2 | UQCRC1 | 0,782 | 7,74547E-40 |
| ACE2 | BCO1 | 0,782 | 7,7752E-40 |
| ACE2 | TMPRSS7 | 0,7819 | 8,10746E-40 |
| ACE2 | LCT | 0,7818 | 8,38637E-40 |
| ACE2 | OVOL1-AS1 | 0,7816 | 8,9966E-40 |
| ACE2 | WSCD1 | 0,7816 | 9,14862E-40 |
| ACE2 | SLC25A24 | 0,7812 | 1,02894E-39 |
| ACE2 | RSPH1 | 0,7812 | 1,02964E-39 |
| ACE2 | LINC01123 | 0,781 | 1,1144E-39 |
| ACE2 | CDH16 | 0,7809 | 1,16119E-39 |
| ACE2 | EHHADH | 0,7804 | 1,3929E-39 |
| ACE2 | RP11-286N22.16 | 0,78 | 1,60422E-39 |
| ACE2 | TMEM98 | 0,78 | 1,60804E-39 |
| ACE2 | ACE3P | 0,7799 | 1,67613E-39 |
| ACE2 | PLCH1 | 0,7796 | 1,86174E-39 |
| ACE2 | RP11-319C21.1 | 0,7795 | 1,95855E-39 |

| | | | |
|---|---|---|---|
| ACE2 | RP11-231P20.2 | 0,7794 | 2,00788E-39 |
| ACE2 | SLC25A20 | 0,7793 | 2,0859E-39 |
| ACE2 | NUDT16 | 0,7792 | 2,15106E-39 |
| ACE2 | C1orf226 | 0,7791 | 2,27803E-39 |
| ACE2 | FAM84A | 0,779 | 2,37166E-39 |
| ACE2 | KCNJ13 | 0,779 | 2,34769E-39 |
| ACE2 | MISP3 | 0,779 | 2,32748E-39 |
| ACE2 | HES2 | 0,7788 | 2,54875E-39 |
| ACE2 | RP11-150O12.3 | 0,7788 | 2,56187E-39 |
| ACE2 | LINC00705 | 0,7787 | 2,58895E-39 |
| ACE2 | CEP170B | 0,7787 | 2,59382E-39 |
| ACE2 | TMEM61 | 0,7786 | 2,75707E-39 |
| ACE2 | GS1-594A7.3 | 0,7786 | 2,71968E-39 |
| ACE2 | RP11-102C16.3 | 0,7784 | 2,98749E-39 |
| ACE2 | LGALS9B | 0,7784 | 2,98569E-39 |
| ACE2 | RP5-903G2.2 | 0,7782 | 3,14744E-39 |
| ACE2 | PHBP8 | 0,7781 | 3,32055E-39 |
| ACE2 | GSTA5 | 0,7779 | 3,51344E-39 |
| ACE2 | CDA | 0,7776 | 3,89031E-39 |
| ACE2 | CYP4A11 | 0,7773 | 4,46209E-39 |
| ACE2 | RP11-439E19.7 | 0,7773 | 4,38877E-39 |
| ACE2 | MELTF | 0,7773 | 4,33466E-39 |
| ACE2 | RP11-234B24.2 | 0,7772 | 4,5029E-39 |
| ACE2 | RILP | 0,7772 | 4,54846E-39 |
| ACE2 | RP13-497K6.1 | 0,7771 | 4,72766E-39 |
| ACE2 | NSDHL | 0,7768 | 5,2582E-39 |
| ACE2 | ATP8B1 | 0,7767 | 5,37914E-39 |
| ACE2 | GRAMD4 | 0,7767 | 5,4281E-39 |
| ACE2 | LINC00330 | 0,7764 | 5,98378E-39 |
| ACE2 | RP11-30P6.6 | 0,7758 | 7,54996E-39 |
| ACE2 | FOXA3 | 0,7756 | 8,05613E-39 |
| ACE2 | OTC | 0,7755 | 8,42319E-39 |
| ACE2 | NCMAP | 0,7754 | 8,74407E-39 |
| ACE2 | PLD1 | 0,7753 | 9,0805E-39 |
| ACE2 | PTMAP3 | 0,7752 | 9,55209E-39 |
| ACE2 | RP11-844P9.5 | 0,775 | 9,95347E-39 |
| ACE2 | MLIP | 0,7749 | 1,05731E-38 |
| ACE2 | LINC00365 | 0,7747 | 1,14359E-38 |
| ACE2 | RP1-117B12.4 | 0,7743 | 1,30253E-38 |
| ACE2 | GRAMD2 | 0,7738 | 1,56388E-38 |
| ACE2 | SLC34A3 | 0,7737 | 1,63411E-38 |
| ACE2 | HSD3BP2 | 0,7736 | 1,66951E-38 |
| ACE2 | SPAM1 | 0,7736 | 1,66803E-38 |
| ACE2 | RP3-406A7.7 | 0,7733 | 1,88607E-38 |
| ACE2 | WNK4 | 0,7733 | 1,8427E-38 |
| ACE2 | ABCC3 | 0,7732 | 1,90432E-38 |
| ACE2 | SLC26A6 | 0,7731 | 1,96282E-38 |
| ACE2 | PAK4 | 0,7731 | 1,96851E-38 |
| ACE2 | DENND1B | 0,7728 | 2,20989E-38 |
| ACE2 | CEACAM6 | 0,7728 | 2,26223E-38 |
| ACE2 | RP11-278L15.6 | 0,7725 | 2,47077E-38 |
| ACE2 | RP11-592N21.2 | 0,7725 | 2,45684E-38 |
| ACE2 | GMDS | 0,7723 | 2,68363E-38 |
| ACE2 | FAHD1 | 0,7723 | 2,64873E-38 |

| | | | |
|---|---|---|---|
| ACE2 | PLAGL2 | 0,7723 | 2,66157E-38 |
| ACE2 | CASP6 | 0,7722 | 2,7134E-38 |
| ACE2 | PPP1R14C | 0,772 | 2,92487E-38 |
| ACE2 | RP11-357H14.17 | 0,7717 | 3,31509E-38 |
| ACE2 | MTCH2 | 0,7715 | 3,50017E-38 |
| ACE2 | FAM160A2 | 0,7714 | 3,65151E-38 |
| ACE2 | CREB3L1 | 0,7714 | 3,59317E-38 |
| ACE2 | TBX3 | 0,7711 | 4,03247E-38 |
| ACE2 | ERN2 | 0,7705 | 5,02189E-38 |
| ACE2 | LINC01637 | 0,7705 | 4,9408E-38 |
| ACE2 | ASAH2 | 0,7703 | 5,45813E-38 |
| ACE2 | RP11-209E8.1 | 0,7703 | 5,39325E-38 |
| ACE2 | ENTPD7 | 0,7701 | 5,76733E-38 |
| ACE2 | F11R | 0,77 | 6,00251E-38 |
| ACE2 | CAPN8 | 0,7698 | 6,31012E-38 |
| ACE2 | SLC25A10 | 0,7698 | 6,31465E-38 |
| ACE2 | FAT1 | 0,7697 | 6,69047E-38 |
| ACE2 | RP11-391H12.8 | 0,7697 | 6,60709E-38 |
| ACE2 | RP13-452N2.1 | 0,7695 | 7,20046E-38 |
| ACE2 | RP11-480I12.9 | 0,7694 | 7,38833E-38 |
| ACE2 | FFAR4 | 0,7694 | 7,29776E-38 |
| ACE2 | KCNJ4 | 0,7694 | 7,51309E-38 |
| ACE2 | ONECUT3 | 0,7693 | 7,60822E-38 |
| ACE2 | CHST6 | 0,7691 | 8,21702E-38 |
| ACE2 | RP11-1223D19.1 | 0,7688 | 9,24431E-38 |
| ACE2 | FRMD1 | 0,7688 | 9,11155E-38 |
| ACE2 | SLC5A4 | 0,7687 | 9,31467E-38 |
| ACE2 | AGMAT | 0,7686 | 9,74278E-38 |
| ACE2 | GALNT3 | 0,7685 | 1,00069E-37 |
| ACE2 | NET1 | 0,7683 | 1,09824E-37 |
| ACE2 | AADACP1 | 0,768 | 1,20311E-37 |
| ACE2 | SLC35C1 | 0,768 | 1,21304E-37 |
| ACE2 | FOXA1 | 0,7678 | 1,2998E-37 |
| ACE2 | COX5A | 0,7678 | 1,27813E-37 |
| ACE2 | RP11-855O10.2 | 0,7675 | 1,42496E-37 |
| ACE2 | TM4SF4 | 0,7673 | 1,5089E-37 |
| ACE2 | MS4A18 | 0,7668 | 1,82999E-37 |
| ACE2 | CEACAM5 | 0,7668 | 1,8045E-37 |
| ACE2 | AP1S1 | 0,7667 | 1,91061E-37 |
| ACE2 | ARR3 | 0,7665 | 2,04052E-37 |
| ACE2 | NBPF4 | 0,7663 | 2,16694E-37 |
| ACE2 | C4BPB | 0,7663 | 2,19781E-37 |
| ACE2 | RAB20 | 0,7661 | 2,34911E-37 |
| ACE2 | ATRN | 0,766 | 2,43933E-37 |
| ACE2 | NMRAL2P | 0,7659 | 2,45531E-37 |
| ACE2 | BSPRY | 0,7657 | 2,66324E-37 |
| ACE2 | MIR621 | 0,7657 | 2,65678E-37 |
| ACE2 | CROT | 0,7655 | 2,84279E-37 |
| ACE2 | SELENOS | 0,7651 | 3,27869E-37 |
| ACE2 | ARSD | 0,765 | 3,39819E-37 |
| ACE2 | SLC17A5 | 0,7649 | 3,56672E-37 |
| ACE2 | DOCK5 | 0,7648 | 3,59656E-37 |
| ACE2 | RP11-1060J15.5 | 0,7648 | 3,5845E-37 |
| ACE2 | MPP5 | 0,7648 | 3,69086E-37 |

| | | | |
|---|---|---|---|
| ACE2 | CD164 | 0,7645 | 4,035E-37 |
| ACE2 | MS4A15 | 0,7642 | 4,38033E-37 |
| ACE2 | GRB7 | 0,7642 | 4,3994E-37 |
| ACE2 | CTD-2325A15.5 | 0,764 | 4,83001E-37 |
| ACE2 | PEX11G | 0,764 | 4,7872E-37 |
| ACE2 | AC024592.9 | 0,7637 | 5,19523E-37 |
| ACE2 | GDPD1 | 0,7636 | 5,51806E-37 |
| ACE2 | SRC | 0,7635 | 5,61749E-37 |
| ACE2 | GSTA7P | 0,7628 | 7,16632E-37 |
| ACE2 | RP11-296E3.2 | 0,7627 | 7,33009E-37 |
| ACE2 | MNX1-AS2 | 0,7624 | 8,26626E-37 |
| ACE2 | COBL | 0,7622 | 8,74177E-37 |
| ACE2 | TMBIM6 | 0,7622 | 8,64744E-37 |
| ACE2 | RIOK3 | 0,7622 | 8,67532E-37 |
| ACE2 | SLC31A2 | 0,7619 | 9,68296E-37 |
| ACE2 | ESRP2 | 0,7619 | 9,75098E-37 |
| ACE2 | MXD1 | 0,7618 | 9,92025E-37 |
| ACE2 | MPI | 0,7615 | 1,11449E-36 |
| ACE2 | KRTCAP3 | 0,761 | 1,33399E-36 |
| ACE2 | PRSS36 | 0,7609 | 1,34538E-36 |
| ACE2 | MNX1 | 0,7608 | 1,41708E-36 |
| ACE2 | ESRP1 | 0,7607 | 1,44622E-36 |
| ACE2 | TNFRSF11A | 0,7606 | 1,49271E-36 |
| ACE2 | PECR | 0,7604 | 1,59022E-36 |
| ACE2 | RP11-297C4.3 | 0,7603 | 1,65219E-36 |
| ACE2 | PRSS12 | 0,7601 | 1,78269E-36 |
| ACE2 | ABCD3 | 0,76 | 1,85967E-36 |
| ACE2 | AC097468.4 | 0,7598 | 1,97575E-36 |
| ACE2 | LINC01080 | 0,7597 | 2,05889E-36 |
| ACE2 | RAB19 | 0,7595 | 2,14126E-36 |
| ACE2 | DECR2 | 0,7593 | 2,28588E-36 |
| ACE2 | TMEM25 | 0,7592 | 2,42394E-36 |
| ACE2 | CAMK2N1 | 0,7591 | 2,51088E-36 |
| ACE2 | RNPEP | 0,759 | 2,5689E-36 |
| ACE2 | TPSG1 | 0,7587 | 2,79745E-36 |
| ACE2 | XKRX | 0,7585 | 3,00937E-36 |
| ACE2 | U47924.27 | 0,7579 | 3,72704E-36 |
| ACE2 | TRBV29OR9-2 | 0,7572 | 4,72765E-36 |
| ACE2 | CC2D1A | 0,7571 | 4,86311E-36 |
| ACE2 | LMTK2 | 0,757 | 5,05526E-36 |
| ACE2 | SHH | 0,757 | 4,98891E-36 |
| ACE2 | CYP2C58P | 0,7569 | 5,20273E-36 |
| ACE2 | AKR1B15 | 0,7568 | 5,30314E-36 |
| ACE2 | RP11-22C11.2 | 0,7566 | 5,6302E-36 |
| ACE2 | GCNT1 | 0,7566 | 5,61923E-36 |
| ACE2 | DFNA5 | 0,7564 | 6,03177E-36 |
| ACE2 | PRRG4 | 0,7563 | 6,33483E-36 |
| ACE2 | RP11-89H19.2 | 0,7563 | 6,35595E-36 |
| ACE2 | SPHK2 | 0,7562 | 6,44201E-36 |
| ACE2 | PGPEP1 | 0,7561 | 6,65358E-36 |
| ACE2 | SLC44A3 | 0,7559 | 7,22475E-36 |
| ACE2 | ECHS1 | 0,7558 | 7,32222E-36 |
| ACE2 | MLX | 0,7558 | 7,39913E-36 |
| ACE2 | LGALS3BP | 0,7556 | 7,93318E-36 |

| | | | |
|---|---|---|---|
| ACE2 | STXBP5-AS1 | 0,7554 | 8,37521E-36 |
| ACE2 | GRAMD1C | 0,7552 | 9,1209E-36 |
| ACE2 | MEP1AP4 | 0,7552 | 8,92023E-36 |
| ACE2 | LINC01451 | 0,755 | 9,65182E-36 |
| ACE2 | CACFD1 | 0,7548 | 1,01371E-35 |
| ACE2 | DNASE1 | 0,7548 | 1,0251E-35 |
| ACE2 | GATA6-AS1 | 0,7548 | 1,01182E-35 |
| ACE2 | NPY6R | 0,7547 | 1,04858E-35 |
| ACE2 | RP11-465B22.8 | 0,7541 | 1,27914E-35 |
| ACE2 | KRT19P1 | 0,7538 | 1,42023E-35 |
| ACE2 | ATG4D | 0,7536 | 1,49615E-35 |
| ACE2 | HN1 | 0,7535 | 1,55082E-35 |
| ACE2 | CTA-243E7.1 | 0,7535 | 1,58512E-35 |
| ACE2 | XK | 0,7535 | 1,57673E-35 |
| ACE2 | DISP2 | 0,7533 | 1,65394E-35 |
| ACE2 | DAPK2 | 0,7527 | 2,02104E-35 |
| ACE2 | RP11-418I22.2 | 0,7527 | 2,01372E-35 |
| ACE2 | POU5F1 | 0,7526 | 2,09681E-35 |
| ACE2 | FAHD2P1 | 0,7523 | 2,29396E-35 |
| ACE2 | EBP | 0,7523 | 2,27642E-35 |
| ACE2 | IL32 | 0,7522 | 2,38376E-35 |
| ACE2 | RBM22P5 | 0,7521 | 2,45904E-35 |
| ACE2 | MIR559 | 0,7519 | 2,6097E-35 |
| ACE2 | RP11-725G5.2 | 0,7518 | 2,68337E-35 |
| ACE2 | FOXH1 | 0,7517 | 2,74984E-35 |
| ACE2 | RP5-1033H22.2 | 0,7515 | 3,01785E-35 |
| ACE2 | OPLAH | 0,7511 | 3,41203E-35 |
| ACE2 | CANT1 | 0,7511 | 3,37689E-35 |
| ACE2 | UGT1A7 | 0,751 | 3,53343E-35 |
| ACE2 | RP11-119D9.1 | 0,7508 | 3,75805E-35 |
| ACE2 | SMCO4 | 0,7507 | 3,85444E-35 |
| ACE2 | CFAP65 | 0,7504 | 4,18397E-35 |
| ACE2 | MUC12 | 0,7504 | 4,26481E-35 |
| ACE2 | LINC01443 | 0,7504 | 4,16791E-35 |
| ACE2 | SDR42E1 | 0,7501 | 4,6698E-35 |
| ACE2 | UGT1A9 | 0,7499 | 4,99697E-35 |
| ACE2 | ZDHHC12 | 0,7499 | 4,98711E-35 |
| ACE2 | LINC01137 | 0,7496 | 5,50691E-35 |
| ACE2 | DCST1-AS1 | 0,7496 | 5,41099E-35 |
| ACE2 | CTSE | 0,7496 | 5,47589E-35 |
| ACE2 | RP11-64D22.5 | 0,7494 | 5,75869E-35 |
| ACE2 | GOLIM4 | 0,7494 | 5,8686E-35 |
| ACE2 | LINC00526 | 0,7493 | 6,03392E-35 |
| ACE2 | BRI3BP | 0,7492 | 6,18083E-35 |
| ACE2 | NUDT8 | 0,7491 | 6,40516E-35 |
| ACE2 | HMGCR | 0,749 | 6,57817E-35 |
| ACE2 | SLC35D1 | 0,7486 | 7,45038E-35 |
| ACE2 | MGST2 | 0,7486 | 7,44822E-35 |
| ACE2 | IFNE | 0,7484 | 7,99357E-35 |
| ACE2 | YBX2 | 0,7483 | 8,18579E-35 |
| ACE2 | KRT18 | 0,7482 | 8,57178E-35 |
| ACE2 | SOAT2 | 0,7482 | 8,40394E-35 |
| ACE2 | HAAO | 0,748 | 9,08713E-35 |
| ACE2 | PFKFB2 | 0,7478 | 9,49468E-35 |

| | | | |
|---|---|---|---|
| ACE2 | ABHD11 | 0,7478 | 9,78668E-35 |
| ACE2 | GUCY2GP | 0,7477 | 1,00959E-34 |
| ACE2 | BTD | 0,7476 | 1,04189E-34 |
| ACE2 | UGT8 | 0,7476 | 1,03048E-34 |
| ACE2 | PTPRD | 0,7475 | 1,04834E-34 |
| ACE2 | SHTN1 | 0,7471 | 1,21952E-34 |
| ACE2 | RP11-378I13.1 | 0,747 | 1,25626E-34 |
| ACE2 | SH2D4A | 0,747 | 1,23192E-34 |
| ACE2 | RPL39P40 | 0,7468 | 1,3296E-34 |
| ACE2 | MFSD9 | 0,7466 | 1,39958E-34 |
| ACE2 | LINC01106 | 0,7466 | 1,41241E-34 |
| ACE2 | ATP2C2 | 0,7463 | 1,53868E-34 |
| ACE2 | AC016738.4 | 0,7459 | 1,77643E-34 |
| ACE2 | AC005550.3 | 0,7458 | 1,80161E-34 |
| ACE2 | MGST3 | 0,7455 | 1,97101E-34 |
| ACE2 | RMDN2 | 0,7454 | 2,06131E-34 |
| ACE2 | GGT4P | 0,7454 | 2,02187E-34 |
| ACE2 | FZD5 | 0,7453 | 2,12791E-34 |
| ACE2 | S100P | 0,7453 | 2,0877E-34 |
| ACE2 | RP3-368B9.2 | 0,7451 | 2,27679E-34 |
| ACE2 | EPS8L1 | 0,745 | 2,34396E-34 |
| ACE2 | TRPC7-AS1 | 0,7448 | 2,50015E-34 |
| ACE2 | EHF | 0,7445 | 2,66652E-34 |
| ACE2 | EMX1 | 0,7441 | 3,07743E-34 |
| ACE2 | SHD | 0,7441 | 3,1051E-34 |
| ACE2 | PRELID3B | 0,744 | 3,18381E-34 |
| ACE2 | C1orf105 | 0,7439 | 3,25389E-34 |
| ACE2 | KB-1991G8.1 | 0,7431 | 4,15046E-34 |
| ACE2 | CYB561 | 0,7429 | 4,45731E-34 |
| ACE2 | AFG3L2 | 0,7428 | 4,56662E-34 |
| ACE2 | LACTB2 | 0,7426 | 4,81753E-34 |
| ACE2 | TTC39A-AS1 | 0,7425 | 5,09398E-34 |
| ACE2 | RP11-517C16.2 | 0,7422 | 5,54128E-34 |
| ACE2 | RN7SKP127 | 0,7421 | 5,67711E-34 |
| ACE2 | RP4-594I10.3 | 0,7417 | 6,36078E-34 |
| ACE2 | FAM3C | 0,7416 | 6,60951E-34 |
| ACE2 | CRAT | 0,7415 | 6,8558E-34 |
| ACE2 | EFNA1 | 0,7412 | 7,47702E-34 |
| ACE2 | GCNT4 | 0,7412 | 7,56851E-34 |
| ACE2 | CLDN8 | 0,7411 | 7,67929E-34 |
| ACE2 | XXyac-YM21GA2.4 | 0,7408 | 8,40044E-34 |
| ACE2 | LINC00501 | 0,7407 | 8,71233E-34 |
| ACE2 | CTD-2566J3.1 | 0,7407 | 8,60412E-34 |
| ACE2 | RP11-1260E13.2 | 0,7407 | 8,73752E-34 |
| ACE2 | TLDC1 | 0,7403 | 9,94651E-34 |
| ACE2 | PDXDC1 | 0,7396 | 1,20485E-33 |
| ACE2 | PLEKHJ1 | 0,7396 | 1,21644E-33 |
| ACE2 | RASSF7 | 0,7394 | 1,30143E-33 |
| ACE2 | RP4-659I19.1 | 0,739 | 1,47005E-33 |
| ACE2 | RP11-83M16.4 | 0,739 | 1,4645E-33 |
| ACE2 | LNX2 | 0,7388 | 1,54811E-33 |
| ACE2 | EZR | 0,7387 | 1,599E-33 |
| ACE2 | RP11-354M1.2 | 0,7386 | 1,63886E-33 |
| ACE2 | RP1-232L22__B.1 | 0,7384 | 1,73856E-33 |

| | | | |
|---|---|---|---|
| ACE2 | PRKCD | 0,738 | 1,95327E-33 |
| ACE2 | ATP6V0D1 | 0,738 | 1,96717E-33 |
| ACE2 | RP11-211N11.5 | 0,7379 | 2,02865E-33 |
| ACE2 | AC010745.1 | 0,7375 | 2,2756E-33 |
| ACE2 | SATB2-AS1 | 0,7374 | 2,37369E-33 |
| ACE2 | RP11-439C15.5 | 0,7374 | 2,39168E-33 |
| ACE2 | NEO1 | 0,7374 | 2,3526E-33 |
| ACE2 | PTP4A2P1 | 0,7374 | 2,35543E-33 |
| ACE2 | APOA5 | 0,737 | 2,64167E-33 |
| ACE2 | C2orf88 | 0,7369 | 2,73219E-33 |
| ACE2 | CTNNA1P1 | 0,7369 | 2,76671E-33 |
| ACE2 | KRT8P45 | 0,7366 | 2,97617E-33 |
| ACE2 | RP11-386I14.2 | 0,7365 | 3,05522E-33 |
| ACE2 | SULT2B1 | 0,7362 | 3,4307E-33 |
| ACE2 | MGAT4A | 0,736 | 3,58921E-33 |
| ACE2 | CASC16 | 0,7358 | 3,77375E-33 |
| ACE2 | PLEKHB2 | 0,7354 | 4,30118E-33 |
| ACE2 | RP11-720D4.2 | 0,7353 | 4,43582E-33 |
| ACE2 | C2orf72 | 0,7351 | 4,68937E-33 |
| ACE2 | LINC02038 | 0,7349 | 5,0523E-33 |
| ACE2 | NR3C2 | 0,7348 | 5,15241E-33 |
| ACE2 | TFCP2L1 | 0,7347 | 5,32146E-33 |
| ACE2 | GALM | 0,7344 | 5,70921E-33 |
| ACE2 | SPINK1 | 0,7344 | 5,85279E-33 |
| ACE2 | GALNT4 | 0,7344 | 5,84209E-33 |
| ACE2 | DEFB1 | 0,7343 | 5,92889E-33 |
| ACE2 | MARC2 | 0,7339 | 6,65196E-33 |
| ACE2 | MMEL1 | 0,7338 | 6,97953E-33 |
| ACE2 | IMPA1P1 | 0,7338 | 6,9486E-33 |
| ACE2 | PAX6 | 0,7337 | 7,17593E-33 |
| ACE2 | MYO15B | 0,7337 | 7,20509E-33 |
| ACE2 | ADAMTSL5 | 0,7337 | 7,16881E-33 |
| ACE2 | CTA-228A9.3 | 0,7337 | 7,11829E-33 |
| ACE2 | RP11-108L7.15 | 0,7336 | 7,35172E-33 |
| ACE2 | MVP | 0,7335 | 7,5067E-33 |
| ACE2 | C19orf54 | 0,7332 | 8,24832E-33 |
| ACE2 | LINC01595 | 0,7331 | 8,55828E-33 |
| ACE2 | TMEM89 | 0,733 | 8,87524E-33 |
| ACE2 | DNAH3 | 0,733 | 8,71381E-33 |
| ACE2 | ENPP3 | 0,7328 | 9,30792E-33 |
| ACE2 | UGT1A2P | 0,7326 | 9,92385E-33 |
| ACE2 | RP11-627G18.2 | 0,7325 | 1,02457E-32 |
| ACE2 | PERP | 0,7321 | 1,1285E-32 |
| ACE2 | PRR5L | 0,7321 | 1,14829E-32 |
| ACE2 | PSEN1 | 0,732 | 1,16649E-32 |
| ACE2 | SERPINB1 | 0,7318 | 1,23554E-32 |
| ACE2 | SDSL | 0,7318 | 1,25179E-32 |
| ACE2 | DCAF11 | 0,7316 | 1,31068E-32 |
| ACE2 | AKR1B10P1 | 0,7315 | 1,35013E-32 |
| ACE2 | MYO1E | 0,7314 | 1,39328E-32 |
| ACE2 | MBOAT7 | 0,7314 | 1,41449E-32 |
| ACE2 | TGFBR3L | 0,731 | 1,57762E-32 |
| ACE2 | ARX | 0,7309 | 1,62673E-32 |
| ACE2 | TMCO1 | 0,7307 | 1,71166E-32 |

| | | | |
|---|---|---|---|
| ACE2 | ARSE | 0,7307 | 1,69716E-32 |
| ACE2 | CTD-3051D23.4 | 0,7306 | 1,75252E-32 |
| ACE2 | RP5-908M14.5 | 0,7303 | 1,94308E-32 |
| ACE2 | SLC25A47 | 0,7302 | 1,95871E-32 |
| ACE2 | SIPA1L3 | 0,73 | 2,10552E-32 |
| ACE2 | GCHFR | 0,7297 | 2,30603E-32 |
| ACE2 | EFNA4 | 0,7295 | 2,43294E-32 |
| ACE2 | RXFP4 | 0,729 | 2,78967E-32 |
| ACE2 | AP006216.5 | 0,729 | 2,79589E-32 |
| ACE2 | RORC | 0,7288 | 2,95583E-32 |
| ACE2 | EXPH5 | 0,7287 | 3,02399E-32 |
| ACE2 | IL1RN | 0,7285 | 3,20655E-32 |
| ACE2 | CERS6 | 0,7285 | 3,25179E-32 |
| ACE2 | UGT2B17 | 0,7285 | 3,26406E-32 |
| ACE2 | AC098828.3 | 0,7283 | 3,40867E-32 |
| ACE2 | RNF152 | 0,7283 | 3,47526E-32 |
| ACE2 | FAM3C2 | 0,7283 | 3,45203E-32 |
| ACE2 | RP11-388N2.1 | 0,728 | 3,79453E-32 |
| ACE2 | KRT8P10 | 0,7278 | 3,96462E-32 |
| ACE2 | FOXD1 | 0,7276 | 4,26137E-32 |
| ACE2 | SH3RF1 | 0,7275 | 4,305E-32 |
| ACE2 | NEUROD1 | 0,7273 | 4,55973E-32 |
| ACE2 | MST1 | 0,7273 | 4,52926E-32 |
| ACE2 | HADHA | 0,7272 | 4,73332E-32 |
| ACE2 | KIF3B | 0,7267 | 5,47889E-32 |
| ACE2 | ADAP1 | 0,7264 | 5,86179E-32 |
| ACE2 | SLC25A15P3 | 0,7263 | 6,12991E-32 |
| ACE2 | RP11-49I11.1 | 0,7262 | 6,34342E-32 |
| ACE2 | MSMO1 | 0,7259 | 6,76506E-32 |
| ACE2 | RP11-160H22.5 | 0,7257 | 7,24103E-32 |
| ACE2 | NR2F6 | 0,7257 | 7,22354E-32 |
| ACE2 | UNC5CL | 0,7255 | 7,72411E-32 |
| ACE2 | SNORA44 | 0,7253 | 8,09293E-32 |
| ACE2 | CTD-3080P12.3 | 0,7249 | 9,17864E-32 |
| ACE2 | HTR3E | 0,7248 | 9,48915E-32 |
| ACE2 | RP11-597D13.7 | 0,7247 | 9,52941E-32 |
| ACE2 | SSTR5 | 0,7245 | 1,01142E-31 |
| ACE2 | HADHB | 0,7243 | 1,08912E-31 |
| ACE2 | GAPDHP32 | 0,7242 | 1,11691E-31 |
| ACE2 | DNM2 | 0,7242 | 1,10724E-31 |
| ACE2 | RP11-324O2.3 | 0,7241 | 1,14484E-31 |
| ACE2 | HOXD1 | 0,7237 | 1,27948E-31 |
| ACE2 | SRI | 0,7236 | 1,31763E-31 |
| ACE2 | RP11-794G24.1 | 0,7236 | 1,31129E-31 |
| ACE2 | TNIK | 0,7234 | 1,39849E-31 |
| ACE2 | CLCN1 | 0,7234 | 1,40285E-31 |
| ACE2 | MIR31HG | 0,7234 | 1,37682E-31 |
| ACE2 | ARL4A | 0,7231 | 1,51261E-31 |
| ACE2 | XXyac-YM21GA2.3 | 0,7231 | 1,49489E-31 |
| ACE2 | LRP5 | 0,723 | 1,57539E-31 |
| ACE2 | DNMBP | 0,7225 | 1,7684E-31 |
| ACE2 | GBA2 | 0,7221 | 2,02453E-31 |
| ACE2 | NBPF6 | 0,7219 | 2,09804E-31 |
| ACE2 | SH2D3A | 0,7218 | 2,16504E-31 |

| | | | |
|---|---|---|---|
| ACE2 | PROM1 | 0,7214 | 2,43235E-31 |
| ACE2 | XXbac-B476C20.9 | 0,7213 | 2,52023E-31 |
| ACE2 | MGAT3-AS1 | 0,7211 | 2,64463E-31 |
| ACE2 | ESRRG | 0,7209 | 2,82416E-31 |
| ACE2 | RP11-103J17.2 | 0,7207 | 2,92862E-31 |
| ACE2 | ZDHHC3 | 0,7203 | 3,28449E-31 |
| ACE2 | ETHE1 | 0,7203 | 3,33498E-31 |
| ACE2 | CDC42EP5 | 0,7201 | 3,53538E-31 |
| ACE2 | RP1-122P22.4 | 0,7199 | 3,66263E-31 |
| ACE2 | ZNF503-AS1 | 0,7198 | 3,76217E-31 |
| ACE2 | RP11-336K24.5 | 0,7195 | 4,10966E-31 |
| ACE2 | LBX2-AS1 | 0,7189 | 4,83276E-31 |
| ACE2 | AC112229.1 | 0,7189 | 4,95419E-31 |
| ACE2 | TMEM63B | 0,7189 | 4,82251E-31 |
| ACE2 | ABTB2 | 0,7189 | 4,90941E-31 |
| ACE2 | CTSLP8 | 0,7188 | 5,02018E-31 |
| ACE2 | RP11-166P13.4 | 0,7188 | 5,02155E-31 |
| ACE2 | TUBA1C | 0,7187 | 5,14439E-31 |
| ACE2 | CD68 | 0,7183 | 5,80641E-31 |
| ACE2 | TLDC2 | 0,7182 | 5,89111E-31 |
| ACE2 | NDRG1 | 0,718 | 6,26299E-31 |
| ACE2 | TACO1 | 0,7175 | 7,2918E-31 |
| ACE2 | FCGRT | 0,7174 | 7,33171E-31 |
| ACE2 | ESPL1 | 0,7172 | 7,71754E-31 |
| ACE2 | KDM8 | 0,7172 | 7,74356E-31 |
| ACE2 | ZG16B | 0,7168 | 8,63547E-31 |
| ACE2 | ARHGEF35 | 0,7166 | 9,23918E-31 |
| ACE2 | APOM | 0,7164 | 9,63712E-31 |
| ACE2 | TMEM177 | 0,7162 | 1,02568E-30 |
| ACE2 | HAO2-IT1 | 0,716 | 1,08038E-30 |
| ACE2 | GGT2 | 0,716 | 1,07827E-30 |
| ACE2 | BAK1 | 0,7154 | 1,28351E-30 |
| ACE2 | GALNT14 | 0,7151 | 1,37306E-30 |
| ACE2 | LUZP1 | 0,715 | 1,40612E-30 |
| ACE2 | CCDC152 | 0,7148 | 1,5037E-30 |
| ACE2 | CLDN2 | 0,7146 | 1,5703E-30 |
| ACE2 | RN7SL541P | 0,7145 | 1,6335E-30 |
| ACE2 | MCUR1 | 0,7141 | 1,83697E-30 |
| ACE2 | CLCA2 | 0,714 | 1,87982E-30 |
| ACE2 | RP11-486M3.3 | 0,7137 | 2,03204E-30 |
| ACE2 | SPPL2A | 0,7136 | 2,09206E-30 |
| ACE2 | RP11-369K16.1 | 0,713 | 2,42966E-30 |
| ACE2 | RPL15P21 | 0,7129 | 2,51758E-30 |
| ACE2 | FGD4 | 0,7128 | 2,61116E-30 |
| ACE2 | SLC35B1 | 0,7127 | 2,64471E-30 |
| ACE2 | RNF43 | 0,7123 | 2,98095E-30 |
| ACE2 | GPR153 | 0,7122 | 3,0078E-30 |
| ACE2 | EFHD2 | 0,7122 | 3,07287E-30 |
| ACE2 | LINC01814 | 0,7121 | 3,13087E-30 |
| ACE2 | ARHGEF34P | 0,7121 | 3,09562E-30 |
| ACE2 | PCDH1 | 0,712 | 3,18686E-30 |
| ACE2 | SGPL1 | 0,7118 | 3,36704E-30 |
| ACE2 | RP4-695O20.1 | 0,7118 | 3,40298E-30 |
| ACE2 | TMEM141 | 0,7115 | 3,64056E-30 |

| | | | |
|---|---|---|---|
| ACE2 | RP11-579D7.4 | 0,7115 | 3,65101E-30 |
| ACE2 | UGT1A13P | 0,7113 | 3,84232E-30 |
| ACE2 | RP11-43P8.2 | 0,7112 | 3,98848E-30 |
| ACE2 | RP11-783K16.5 | 0,7112 | 3,99417E-30 |
| ACE2 | FBLIM1 | 0,7111 | 4,07619E-30 |
| ACE2 | CCDC129 | 0,711 | 4,21905E-30 |
| ACE2 | AC013268.3 | 0,7108 | 4,38604E-30 |
| ACE2 | LINC01348 | 0,7107 | 4,5343E-30 |
| ACE2 | SPATA31E1 | 0,7106 | 4,61691E-30 |
| ACE2 | DHCR24 | 0,7105 | 4,79044E-30 |
| ACE2 | SDHB | 0,7104 | 4,85903E-30 |
| ACE2 | ACP2 | 0,7104 | 4,85564E-30 |
| ACE2 | P4HB | 0,7102 | 5,17506E-30 |
| ACE2 | AP001627.1 | 0,7101 | 5,27129E-30 |
| ACE2 | ABHD6 | 0,71 | 5,51227E-30 |
| ACE2 | RP3-325F22.5 | 0,71 | 5,44736E-30 |
| ACE2 | HID1 | 0,7098 | 5,81414E-30 |
| ACE2 | EPB41L3 | 0,7098 | 5,77163E-30 |
| ACE2 | RP11-47A17.2 | 0,7092 | 6,79474E-30 |
| ACE2 | PRR13P5 | 0,7092 | 6,85335E-30 |
| ACE2 | LCN15 | 0,7091 | 6,8978E-30 |
| ACE2 | TRIM31-AS1 | 0,7089 | 7,32625E-30 |
| ACE2 | LRRC28 | 0,7089 | 7,25084E-30 |
| ACE2 | KIFC3 | 0,7089 | 7,28974E-30 |
| ACE2 | ZFYVE28 | 0,7085 | 8,06654E-30 |
| ACE2 | SLC30A4 | 0,7085 | 8,15969E-30 |
| ACE2 | STPG3 | 0,7083 | 8,6786E-30 |
| ACE2 | LGALS9C | 0,7082 | 8,92962E-30 |
| ACE2 | ACTBP13 | 0,708 | 9,25272E-30 |
| ACE2 | NCKAP5 | 0,7078 | 9,79293E-30 |
| ACE2 | DLG3 | 0,7077 | 9,96377E-30 |
| ACE2 | AC106869.2 | 0,7076 | 1,02953E-29 |
| ACE2 | LINC00482 | 0,7074 | 1,07659E-29 |
| ACE2 | FEV | 0,7073 | 1,12344E-29 |
| ACE2 | KIF12 | 0,7071 | 1,1895E-29 |
| ACE2 | SECTM1 | 0,7069 | 1,2384E-29 |
| ACE2 | CYP3A7 | 0,7065 | 1,37924E-29 |
| ACE2 | EIF6 | 0,7064 | 1,41691E-29 |
| ACE2 | FLVCR1 | 0,7063 | 1,44744E-29 |
| ACE2 | LINC01559 | 0,7061 | 1,55119E-29 |
| ACE2 | FAM189A1 | 0,7059 | 1,63626E-29 |
| ACE2 | RP11-104J23.1 | 0,7059 | 1,60708E-29 |
| ACE2 | SCTR | 0,7053 | 1,88964E-29 |
| ACE2 | STPG3-AS1 | 0,7048 | 2,1318E-29 |
| ACE2 | NPAS2 | 0,7047 | 2,2277E-29 |
| ACE2 | HCN3 | 0,7041 | 2,59823E-29 |
| ACE2 | RP1-172H20.4 | 0,704 | 2,66326E-29 |
| ACE2 | RP11-350E12.4 | 0,7039 | 2,73916E-29 |
| ACE2 | MAF | 0,7039 | 2,74062E-29 |
| ACE2 | PLGRKT | 0,7036 | 2,9232E-29 |
| ACE2 | USP2 | 0,7035 | 3,04856E-29 |
| ACE2 | UNC93A | 0,7034 | 3,08593E-29 |
| ACE2 | RP1-80N2.4 | 0,7033 | 3,2099E-29 |
| ACE2 | GJB2 | 0,7032 | 3,30233E-29 |

| | | | |
|---|---|---|---|
| ACE2 | SLC25A13 | 0,7031 | 3,34441E-29 |
| ACE2 | PPP2R5D | 0,7026 | 3,78052E-29 |
| ACE2 | XXyac-YR21CG7.1 | 0,7024 | 3,9758E-29 |
| ACE2 | HMGCS1 | 0,7022 | 4,24728E-29 |
| ACE2 | RP11-627G23.1 | 0,7022 | 4,1677E-29 |
| ACE2 | LRRC1 | 0,702 | 4,46437E-29 |
| ACE2 | KALRN | 0,7016 | 4,90599E-29 |
| ACE2 | AKR1A1 | 0,7013 | 5,28462E-29 |
| ACE2 | TAAR3P | 0,7011 | 5,57893E-29 |
| ACE2 | GFER | 0,7011 | 5,55513E-29 |
| ACE2 | SESN2 | 0,7009 | 5,84267E-29 |
| ACE2 | AC104667.3 | 0,7009 | 5,89815E-29 |
| ACE2 | CPTP | 0,7004 | 6,68685E-29 |
| ACE2 | NHEJ1 | 0,7004 | 6,75046E-29 |
| ACE2 | C2CD2L | 0,7004 | 6,71098E-29 |
| ACE2 | RP11-482M8.1 | 0,7003 | 6,91116E-29 |
| ACE2 | OR2A7 | 0,7002 | 7,06646E-29 |
| ACE2 | SLC10A5 | 0,7002 | 7,04991E-29 |
| ACE2 | AC098614.2 | 0,7 | 7,43391E-29 |
| ACE2 | RP11-226B15.1 | 0,7 | 7,33144E-29 |
| ACE2 | PDE11A | 0,6991 | 9,39589E-29 |
| ACE2 | ADGRG4 | 0,6991 | 9,44569E-29 |
| ACE2 | RP11-384L8.1 | 0,699 | 9,61906E-29 |
| ACE2 | HK2 | 0,6989 | 9,71841E-29 |
| ACE2 | AC098828.2 | 0,6988 | 1,01336E-28 |
| ACE2 | RP11-595B24.2 | 0,6988 | 1,01898E-28 |
| ACE2 | STXBP5 | 0,6987 | 1,02783E-28 |
| ACE2 | SUCLG2 | 0,6983 | 1,14227E-28 |
| ACE2 | C9orf152 | 0,6983 | 1,13996E-28 |
| ACE2 | LINC00589 | 0,6981 | 1,18879E-28 |
| ACE2 | AC011523.2 | 0,6981 | 1,20255E-28 |
| ACE2 | FGF19 | 0,6978 | 1,2961E-28 |
| ACE2 | ATP10B | 0,6977 | 1,32871E-28 |
| ACE2 | GRTP1-AS1 | 0,6976 | 1,37626E-28 |
| ACE2 | BCAR1 | 0,6976 | 1,3642E-28 |
| ACE2 | CEBPG | 0,6976 | 1,34994E-28 |
| ACE2 | TUFT1 | 0,6974 | 1,44013E-28 |
| ACE2 | HAGLR | 0,6974 | 1,42885E-28 |
| ACE2 | MIR429 | 0,6973 | 1,47098E-28 |
| ACE2 | STARD10 | 0,6972 | 1,50103E-28 |
| ACE2 | CUBN | 0,6971 | 1,53511E-28 |
| ACE2 | NR1I3 | 0,6968 | 1,65807E-28 |
| ACE2 | RP11-309L24.4 | 0,6968 | 1,65782E-28 |
| ACE2 | AC114812.10 | 0,6967 | 1,69244E-28 |
| ACE2 | GBP3 | 0,6963 | 1,90674E-28 |
| ACE2 | ASL | 0,6963 | 1,87546E-28 |
| ACE2 | RP11-132F7.2 | 0,6963 | 1,90821E-28 |
| ACE2 | CTC-304I17.6 | 0,696 | 2,06376E-28 |
| ACE2 | ACBD5 | 0,6958 | 2,14718E-28 |
| ACE2 | NXPE4 | 0,6958 | 2,17574E-28 |
| ACE2 | RP5-984P4.6 | 0,6958 | 2,12422E-28 |
| ACE2 | EPN3 | 0,6957 | 2,20904E-28 |
| ACE2 | RP11-610P16.1 | 0,6955 | 2,29098E-28 |
| ACE2 | PSMD10P2 | 0,6952 | 2,49686E-28 |

| | | | |
|---|---|---|---|
| ACE2 | CHRM4 | 0,6952 | 2,46966E-28 |
| ACE2 | TRAF4 | 0,6949 | 2,68182E-28 |
| ACE2 | PDSS1 | 0,6948 | 2,74603E-28 |
| ACE2 | SLC25A47P1 | 0,6945 | 3,00889E-28 |
| ACE2 | SHPK | 0,6942 | 3,20683E-28 |
| ACE2 | KAZALD1 | 0,6941 | 3,30349E-28 |
| ACE2 | RP5-1185I7.1 | 0,6939 | 3,4632E-28 |
| ACE2 | LINC01224 | 0,6937 | 3,60204E-28 |
| ACE2 | MDK | 0,6931 | 4,26049E-28 |
| ACE2 | NOXA1 | 0,6929 | 4,41913E-28 |
| ACE2 | LINC01186 | 0,6928 | 4,58441E-28 |
| ACE2 | IMPA1 | 0,6926 | 4,74353E-28 |
| ACE2 | UCA1 | 0,6923 | 5,10736E-28 |
| ACE2 | SHROOM3 | 0,6919 | 5,62275E-28 |
| ACE2 | MAL2 | 0,6919 | 5,71522E-28 |
| ACE2 | TRIM36-IT1 | 0,6916 | 6,12652E-28 |
| ACE2 | ARG2 | 0,6908 | 7,37407E-28 |
| ACE2 | PLEK2 | 0,6905 | 7,93651E-28 |
| ACE2 | SYT13 | 0,69 | 8,95372E-28 |
| ACE2 | F11 | 0,6898 | 9,47978E-28 |
| ACE2 | CEBPA-AS1 | 0,6896 | 9,98252E-28 |
| ACE2 | LAMC2 | 0,6894 | 1,04243E-27 |
| ACE2 | RP11-783K16.13 | 0,6893 | 1,08076E-27 |
| ACE2 | MROH2A | 0,6889 | 1,18779E-27 |
| ACE2 | RP11-510N19.5 | 0,6883 | 1,38453E-27 |
| ACE2 | AIG1 | 0,6883 | 1,38412E-27 |
| ACE2 | RP11-314N13.9 | 0,6882 | 1,39394E-27 |
| ACE2 | RP11-297C4.1 | 0,6881 | 1,42875E-27 |
| ACE2 | ACOT12 | 0,6879 | 1,50281E-27 |
| ACE2 | CA3 | 0,6879 | 1,49568E-27 |
| ACE2 | RP11-422N16.3 | 0,687 | 1,86716E-27 |
| ACE2 | CLUHP6 | 0,6869 | 1,92406E-27 |
| ACE2 | PPARGC1A | 0,6868 | 1,96534E-27 |
| ACE2 | CFTR | 0,6864 | 2,18176E-27 |
| ACE2 | RNF157 | 0,686 | 2,41514E-27 |
| ACE2 | ZDHHC9 | 0,6858 | 2,52123E-27 |
| ACE2 | CASR | 0,6857 | 2,56772E-27 |
| ACE2 | FGFBP1 | 0,6853 | 2,84017E-27 |
| ACE2 | AGPAT3 | 0,6847 | 3,23502E-27 |
| ACE2 | ENPP7P11 | 0,6846 | 3,38482E-27 |
| ACE2 | RP11-407N17.5 | 0,6844 | 3,49095E-27 |
| ACE2 | AQP12A | 0,6842 | 3,69811E-27 |
| ACE2 | Z83844.1 | 0,6842 | 3,72746E-27 |
| ACE2 | DAGLA | 0,6841 | 3,7969E-27 |
| ACE2 | PKP2 | 0,6839 | 3,92829E-27 |
| ACE2 | PGAP3 | 0,6833 | 4,57063E-27 |
| ACE2 | RP11-10C24.3 | 0,6831 | 4,79112E-27 |
| ACE2 | LDHAL6B | 0,6828 | 5,21538E-27 |
| ACE2 | GSDMB | 0,6827 | 5,32566E-27 |
| ACE2 | SLC6A13 | 0,6823 | 5,82654E-27 |
| ACE2 | DGKQ | 0,6818 | 6,59397E-27 |
| ACE2 | ADH1C | 0,6818 | 6,49504E-27 |
| ACE2 | CENPV | 0,6817 | 6,7082E-27 |
| ACE2 | ANO7L1 | 0,6814 | 7,21E-27 |

| | | | |
|---|---|---|---|
| ACE2 | AP001625.6 | 0,6812 | 7,55674E-27 |
| ACE2 | FAM83A | 0,6807 | 8,51556E-27 |
| ACE2 | CCDC68 | 0,6806 | 8,66898E-27 |
| ACE2 | LINC01268 | 0,6804 | 9,18741E-27 |
| ACE2 | TMEM63C | 0,6804 | 9,16493E-27 |
| ACE2 | SCAMP5 | 0,6804 | 9,14774E-27 |
| ACE2 | HPGD | 0,6802 | 9,63441E-27 |
| ACE2 | PEX26 | 0,6802 | 9,61493E-27 |
| ACE2 | RNF103 | 0,6793 | 1,18384E-26 |
| ACE2 | RP13-379O24.3 | 0,6793 | 1,18243E-26 |
| ACE2 | UOX | 0,679 | 1,25531E-26 |
| ACE2 | MIR3131 | 0,679 | 1,27426E-26 |
| ACE2 | AC012485.2 | 0,679 | 1,26405E-26 |
| ACE2 | DGAT2L7P | 0,6787 | 1,368E-26 |
| ACE2 | PARP3 | 0,6785 | 1,41919E-26 |
| ACE2 | NAAA | 0,6782 | 1,52777E-26 |
| ACE2 | KCP | 0,6781 | 1,56945E-26 |
| ACE2 | C11orf54 | 0,6779 | 1,6478E-26 |
| ACE2 | PI3 | 0,6775 | 1,80076E-26 |
| ACE2 | RP11-1100L3.4 | 0,6774 | 1,85071E-26 |
| ACE2 | NMNAT1 | 0,6772 | 1,94771E-26 |
| ACE2 | LINC00628 | 0,6772 | 1,938E-26 |
| ACE2 | MIR135A1 | 0,6772 | 1,92248E-26 |
| ACE2 | LRRIQ4 | 0,6769 | 2,05193E-26 |
| ACE2 | LIPA | 0,6769 | 2,08713E-26 |
| ACE2 | UGDH | 0,6767 | 2,15338E-26 |
| ACE2 | IQGAP2 | 0,6767 | 2,14681E-26 |
| ACE2 | GFPT1 | 0,6766 | 2,19352E-26 |
| ACE2 | PROC | 0,6764 | 2,29687E-26 |
| ACE2 | FAM83H-AS1 | 0,6762 | 2,42193E-26 |
| ACE2 | ENKUR | 0,6761 | 2,47556E-26 |
| ACE2 | C11orf53 | 0,6758 | 2,65245E-26 |
| ACE2 | AC005262.3 | 0,6757 | 2,75213E-26 |
| ACE2 | CTD-2288F12.1 | 0,6756 | 2,80292E-26 |
| ACE2 | TRPM5 | 0,6754 | 2,92281E-26 |
| ACE2 | NUDT12 | 0,6753 | 3,00696E-26 |
| ACE2 | CRISP1 | 0,6753 | 2,97606E-26 |
| ACE2 | GALNT6 | 0,6747 | 3,41892E-26 |
| ACE2 | PIGR | 0,6746 | 3,50707E-26 |
| ACE2 | AC061961.2 | 0,6745 | 3,56412E-26 |
| ACE2 | AC084809.2 | 0,6744 | 3,67287E-26 |
| ACE2 | NABP1 | 0,6743 | 3,74404E-26 |
| ACE2 | IPMK | 0,6742 | 3,89431E-26 |
| ACE2 | TSPAN13 | 0,674 | 4,0642E-26 |
| ACE2 | TIGD2 | 0,6736 | 4,44293E-26 |
| ACE2 | SELENOI | 0,6734 | 4,64574E-26 |
| ACE2 | RP1-80N2.3 | 0,6732 | 4,9011E-26 |
| ACE2 | SCT | 0,6731 | 4,91314E-26 |
| ACE2 | SYTL2 | 0,6729 | 5,17675E-26 |
| ACE2 | CTSA | 0,6722 | 6,06146E-26 |
| ACE2 | ATP5G3 | 0,6718 | 6,74158E-26 |
| ACE2 | UGT2B29P | 0,6714 | 7,3725E-26 |
| ACE2 | ABCA5 | 0,6712 | 7,67788E-26 |
| ACE2 | PITX2 | 0,6711 | 7,81118E-26 |

| | | | |
|---|---|---|---|
| ACE2 | COX6A1 | 0,6711 | 7,83184E-26 |
| ACE2 | STXBP6 | 0,6711 | 7,90517E-26 |
| ACE2 | PLA2G4D | 0,6709 | 8,1295E-26 |
| ACE2 | ANO9 | 0,6707 | 8,59964E-26 |
| ACE2 | FAHD2A | 0,6706 | 8,82945E-26 |
| ACE2 | CPD | 0,6705 | 9,04554E-26 |
| ACE2 | RP11-731D1.4 | 0,6704 | 9,25104E-26 |
| ACE2 | ETFDH | 0,6701 | 9,73254E-26 |
| ACE2 | ANKFN1 | 0,6701 | 9,78832E-26 |
| ACE2 | ITGB4 | 0,67 | 1,00695E-25 |
| ACE2 | KIAA1211L | 0,6695 | 1,11597E-25 |
| ACE2 | RP5-1018K9.1 | 0,6694 | 1,15395E-25 |
| ACE2 | RP11-112J1.3 | 0,6689 | 1,28739E-25 |
| ACE2 | RMDN3 | 0,6685 | 1,40956E-25 |
| ACE2 | LINC01564 | 0,6684 | 1,43235E-25 |
| ACE2 | RP11-259K5.2 | 0,6682 | 1,52581E-25 |
| ACE2 | RP11-21L23.4 | 0,668 | 1,58018E-25 |
| ACE2 | MAB21L3 | 0,6679 | 1,60727E-25 |
| ACE2 | ULK3 | 0,6676 | 1,73695E-25 |
| ACE2 | RP11-150O12.5 | 0,6675 | 1,7546E-25 |
| ACE2 | DNPH1 | 0,6669 | 2,00631E-25 |
| ACE2 | SLC35F5 | 0,6668 | 2,0703E-25 |
| ACE2 | CTD-2005D20.1 | 0,6667 | 2,1023E-25 |
| ACE2 | RPL7P28 | 0,6667 | 2,1005E-25 |
| ACE2 | SERPINB5 | 0,6662 | 2,38128E-25 |
| ACE2 | LRRC75B | 0,6662 | 2,36331E-25 |
| ACE2 | BUB1P1 | 0,6661 | 2,40176E-25 |
| ACE2 | FRMD3 | 0,6659 | 2,51684E-25 |
| ACE2 | ACSF2 | 0,6658 | 2,60157E-25 |
| ACE2 | EPHA1 | 0,6657 | 2,66214E-25 |
| ACE2 | SHMT1 | 0,6657 | 2,66396E-25 |
| ACE2 | SCNN1G | 0,6656 | 2,6902E-25 |
| ACE2 | UCK2 | 0,6655 | 2,75881E-25 |
| ACE2 | SLC26A2 | 0,6653 | 2,90042E-25 |
| ACE2 | RFX6 | 0,665 | 3,05868E-25 |
| ACE2 | RN7SL272P | 0,6649 | 3,1314E-25 |
| ACE2 | TTC6 | 0,6649 | 3,15062E-25 |
| ACE2 | RP5-881L22.5 | 0,6648 | 3,23281E-25 |
| ACE2 | SEC24A | 0,6647 | 3,282E-25 |
| ACE2 | AC073842.19 | 0,6646 | 3,36294E-25 |
| ACE2 | CTC-251D13.1 | 0,6643 | 3,57053E-25 |
| ACE2 | PTPRR | 0,6642 | 3,66435E-25 |
| ACE2 | RP11-182L7.1 | 0,664 | 3,85915E-25 |
| ACE2 | AC011298.2 | 0,6638 | 4,03874E-25 |
| ACE2 | RP1-45I4.3 | 0,6635 | 4,28151E-25 |
| ACE2 | AZGP1 | 0,6635 | 4,27631E-25 |
| ACE2 | HOXB7 | 0,6635 | 4,25873E-25 |
| ACE2 | RNU6-339P | 0,6634 | 4,38377E-25 |
| ACE2 | DHCR7 | 0,6629 | 4,9529E-25 |
| ACE2 | RP11-153K11.3 | 0,6627 | 5,08412E-25 |
| ACE2 | RP11-219B4.5 | 0,6622 | 5,78783E-25 |
| ACE2 | AC005083.1 | 0,662 | 5,98474E-25 |
| ACE2 | RP11-83M16.6 | 0,6619 | 6,12275E-25 |
| ACE2 | LENG9 | 0,6619 | 6,08672E-25 |

| | | | |
|---|---|---|---|
| ACE2 | SH2D7 | 0,6614 | 6,79784E-25 |
| ACE2 | RNFT1P2 | 0,6612 | 7,09215E-25 |
| ACE2 | ESCO2 | 0,6612 | 7,18293E-25 |
| ACE2 | AC053503.4 | 0,6607 | 7,99854E-25 |
| ACE2 | CIB2 | 0,6607 | 8,04368E-25 |
| ACE2 | TMEM220-AS1 | 0,6607 | 7,92475E-25 |
| ACE2 | RPL3P13 | 0,6604 | 8,56381E-25 |
| ACE2 | MMAA | 0,6603 | 8,73711E-25 |
| ACE2 | CPA2 | 0,6598 | 9,75101E-25 |
| ACE2 | MAPK3 | 0,6598 | 9,7868E-25 |
| ACE2 | STK39 | 0,6597 | 9,87314E-25 |
| ACE2 | SELENBP1 | 0,6593 | 1,08198E-24 |
| ACE2 | DKFZP434K028 | 0,6591 | 1,12203E-24 |
| ACE2 | GAPDHP74 | 0,6587 | 1,22139E-24 |
| ACE2 | TM9SF2 | 0,6587 | 1,21912E-24 |
| ACE2 | RP11-524H19.2 | 0,6585 | 1,29079E-24 |
| ACE2 | KRTAP13-2 | 0,6577 | 1,5322E-24 |
| ACE2 | LINC02023 | 0,6567 | 1,90067E-24 |
| ACE2 | RP11-149I9.2 | 0,6567 | 1,88773E-24 |
| ACE2 | AIFM3 | 0,6567 | 1,88744E-24 |
| ACE2 | SEMA3B | 0,6566 | 1,91621E-24 |
| ACE2 | CTA-315H11.2 | 0,6563 | 2,04306E-24 |
| ACE2 | RAB3IP | 0,6557 | 2,32303E-24 |
| ACE2 | RAVER2 | 0,6556 | 2,41588E-24 |
| ACE2 | RP11-523J2.1 | 0,6556 | 2,41737E-24 |
| ACE2 | MT-CO1 | 0,6547 | 2,88361E-24 |
| ACE2 | KLHDC7A | 0,6545 | 3,06171E-24 |
| ACE2 | LINC00992 | 0,6541 | 3,26985E-24 |
| ACE2 | DLG1-AS1 | 0,654 | 3,38647E-24 |
| ACE2 | RP11-242C19.2 | 0,6539 | 3,46271E-24 |
| ACE2 | RP11-109E12.5 | 0,6538 | 3,49808E-24 |
| ACE2 | SEC13 | 0,6538 | 3,55193E-24 |
| ACE2 | KIAA1324 | 0,6537 | 3,60008E-24 |
| ACE2 | SLC39A14 | 0,6537 | 3,61767E-24 |
| ACE2 | B4GALT1-AS1 | 0,6537 | 3,60243E-24 |
| ACE2 | GK-AS1 | 0,6537 | 3,63277E-24 |
| ACE2 | HADH | 0,6535 | 3,7297E-24 |
| ACE2 | RPS3AP46 | 0,6532 | 4,00072E-24 |
| ACE2 | LINC02086 | 0,6531 | 4,12765E-24 |
| ACE2 | KPNA7 | 0,6529 | 4,26629E-24 |
| ACE2 | TRIM80P | 0,6525 | 4,61943E-24 |
| ACE2 | EPPK1 | 0,6524 | 4,76261E-24 |
| ACE2 | GSTO2 | 0,6521 | 5,05139E-24 |
| ACE2 | TTLL6 | 0,6521 | 5,02596E-24 |
| ACE2 | SORL1 | 0,652 | 5,15163E-24 |
| ACE2 | ITLN1 | 0,6519 | 5,22473E-24 |
| ACE2 | RN7SL208P | 0,6517 | 5,46983E-24 |
| ACE2 | STX3 | 0,6517 | 5,52658E-24 |
| ACE2 | RPS4XP8 | 0,6516 | 5,58456E-24 |
| ACE2 | F12 | 0,6513 | 5,95227E-24 |
| ACE2 | BEND7 | 0,6512 | 6,07083E-24 |
| ACE2 | TBC1D8 | 0,651 | 6,41701E-24 |
| ACE2 | CBR3-AS1 | 0,651 | 6,31904E-24 |
| ACE2 | RP13-128O4.3 | 0,651 | 6,37584E-24 |

| | | | |
|---|---|---|---|
| ACE2 | JHDM1D-AS1 | 0,6509 | 6,43939E-24 |
| ACE2 | RAB26 | 0,6509 | 6,56576E-24 |
| ACE2 | GABRA2 | 0,6505 | 7,07427E-24 |
| ACE2 | ENTPD6 | 0,6504 | 7,24449E-24 |
| ACE2 | BST1 | 0,6501 | 7,70087E-24 |
| ACE2 | PFKL | 0,6501 | 7,68395E-24 |
| ACE2 | VDAC1 | 0,65 | 7,79235E-24 |
| ACE2 | CHGA | 0,6498 | 8,27932E-24 |
| ACE2 | AC079779.4 | 0,6497 | 8,36073E-24 |
| ACE2 | MIR210HG | 0,6496 | 8,5308E-24 |
| ACE2 | MYBPC1 | 0,6495 | 8,72526E-24 |
| ACE2 | RP11-418I22.3 | 0,6494 | 8,99934E-24 |
| ACE2 | SH2D6 | 0,6489 | 9,85329E-24 |
| ACE2 | KRT8P32 | 0,6488 | 1,00844E-23 |
| ACE2 | HOXC11 | 0,6484 | 1,10264E-23 |
| ACE2 | NAA60 | 0,6484 | 1,10875E-23 |
| ACE2 | RP11-519G16.3 | 0,6483 | 1,12619E-23 |
| ACE2 | BCL2L10 | 0,6482 | 1,1532E-23 |
| ACE2 | AGR2 | 0,6478 | 1,25295E-23 |
| ACE2 | MAGI3 | 0,6476 | 1,29276E-23 |
| ACE2 | MPDU1 | 0,6475 | 1,33315E-23 |
| ACE2 | STXBP2 | 0,6475 | 1,3184E-23 |
| ACE2 | YAP1P1 | 0,6474 | 1,3652E-23 |
| ACE2 | NECTIN4 | 0,6469 | 1,50373E-23 |
| ACE2 | GCG | 0,6468 | 1,54528E-23 |
| ACE2 | CYP4F11 | 0,6468 | 1,53932E-23 |
| ACE2 | RP11-756A22.7 | 0,6466 | 1,59804E-23 |
| ACE2 | AIFM1 | 0,6465 | 1,64415E-23 |
| ACE2 | AC005105.2 | 0,6464 | 1,66955E-23 |
| ACE2 | RP13-225O21.2 | 0,6464 | 1,65237E-23 |
| ACE2 | ITPK1 | 0,6463 | 1,71003E-23 |
| ACE2 | GALNT12 | 0,6462 | 1,72785E-23 |
| ACE2 | SOCS6 | 0,6461 | 1,76687E-23 |
| ACE2 | RP11-644F5.15 | 0,646 | 1,80491E-23 |
| ACE2 | NUDT14 | 0,646 | 1,82548E-23 |
| ACE2 | ST18 | 0,6457 | 1,93928E-23 |
| ACE2 | AC009133.12 | 0,6457 | 1,90433E-23 |
| ACE2 | RP11-61A14.1 | 0,6456 | 1,96384E-23 |
| ACE2 | LINC01226 | 0,6455 | 2,00308E-23 |
| ACE2 | RP11-542M13.2 | 0,6455 | 1,99273E-23 |
| ACE2 | TRIM3 | 0,6441 | 2,65563E-23 |
| ACE2 | CTD-3193O13.1 | 0,644 | 2,72854E-23 |
| ACE2 | RP13-870H17.3 | 0,6439 | 2,80893E-23 |
| ACE2 | RP11-341D18.5 | 0,6439 | 2,79524E-23 |
| ACE2 | AP001610.9 | 0,6439 | 2,77781E-23 |
| ACE2 | PAX8 | 0,6433 | 3,15121E-23 |
| ACE2 | KIR2DL4 | 0,6432 | 3,22359E-23 |
| ACE2 | OPCML-IT1 | 0,643 | 3,3414E-23 |
| ACE2 | CTD-2589M5.4 | 0,6429 | 3,44126E-23 |
| ACE2 | CRADD | 0,6429 | 3,44241E-23 |
| ACE2 | CORO1B | 0,6428 | 3,51936E-23 |
| ACE2 | RP5-875H18.10 | 0,6428 | 3,48285E-23 |
| ACE2 | RP11-468N14.5 | 0,6423 | 3,83288E-23 |
| ACE2 | CDCA3 | 0,6422 | 3,97475E-23 |

| | | | |
|---|---|---|---|
| ACE2 | KIAA1671 | 0,642 | 4,10738E-23 |
| ACE2 | OR7E85P | 0,6417 | 4,39807E-23 |
| ACE2 | PPP1R1B | 0,6416 | 4,49258E-23 |
| ACE2 | RP11-223I10.1 | 0,6413 | 4,70681E-23 |
| ACE2 | GNG13 | 0,6412 | 4,87094E-23 |
| ACE2 | LMTK3 | 0,6411 | 4,90585E-23 |
| ACE2 | AC073342.12 | 0,6407 | 5,3E-23 |
| ACE2 | SULT1C2 | 0,6406 | 5,40983E-23 |
| ACE2 | QPRT | 0,6406 | 5,42209E-23 |
| ACE2 | ARHGEF38-IT1 | 0,6404 | 5,68825E-23 |
| ACE2 | RP11-49I11.4 | 0,6404 | 5,69996E-23 |
| ACE2 | NOL4L | 0,6401 | 5,98681E-23 |
| ACE2 | RP5-1180D12.1 | 0,64 | 6,15255E-23 |
| ACE2 | HOXB-AS4 | 0,6399 | 6,31198E-23 |
| ACE2 | LINC01351 | 0,6397 | 6,57264E-23 |
| ACE2 | AGAP1-IT1 | 0,6393 | 7,06421E-23 |
| ACE2 | ERVH-1 | 0,6389 | 7,71629E-23 |
| ACE2 | LINC01909 | 0,6389 | 7,70968E-23 |
| ACE2 | SLC43A2 | 0,6388 | 7,8839E-23 |
| ACE2 | MSLN | 0,6387 | 8,05828E-23 |
| ACE2 | MUC4 | 0,6385 | 8,34826E-23 |
| ACE2 | FUCA1 | 0,6377 | 9,70654E-23 |
| ACE2 | XPNPEP1 | 0,6376 | 9,96558E-23 |
| ACE2 | CAPN1 | 0,6376 | 1,00116E-22 |
| ACE2 | HTR3C | 0,6373 | 1,05571E-22 |
| ACE2 | DCST1 | 0,6367 | 1,20215E-22 |
| ACE2 | KB-1460A1.2 | 0,6363 | 1,29381E-22 |
| ACE2 | CTB-167B5.1 | 0,6361 | 1,33577E-22 |
| ACE2 | NEDD4 | 0,6357 | 1,462E-22 |
| ACE2 | TOM1 | 0,6354 | 1,55533E-22 |
| ACE2 | RP4-568C11.4 | 0,6353 | 1,5859E-22 |
| ACE2 | KRTAP5-AS1 | 0,6351 | 1,64739E-22 |
| ACE2 | SLC19A3 | 0,635 | 1,67714E-22 |
| ACE2 | LINC01444 | 0,6347 | 1,77428E-22 |
| ACE2 | RP11-459I19.1 | 0,6346 | 1,80904E-22 |
| ACE2 | C2orf54 | 0,6345 | 1,84123E-22 |
| ACE2 | SPATA31C1 | 0,6345 | 1,85653E-22 |
| ACE2 | PITPNA | 0,6344 | 1,89614E-22 |
| ACE2 | RP5-875H18.4 | 0,6343 | 1,92039E-22 |
| ACE2 | TECTB | 0,6341 | 1,98738E-22 |
| ACE2 | ING2 | 0,634 | 2,06055E-22 |
| ACE2 | OGDH | 0,634 | 2,03182E-22 |
| ACE2 | CHMP1A | 0,634 | 2,03404E-22 |
| ACE2 | AF064860.7 | 0,634 | 2,05236E-22 |
| ACE2 | C9orf64 | 0,6338 | 2,14012E-22 |
| ACE2 | CISD1 | 0,6338 | 2,14034E-22 |
| ACE2 | RNF223 | 0,6335 | 2,24855E-22 |
| ACE2 | TMEM74B | 0,6335 | 2,25885E-22 |
| ACE2 | CYP51A1 | 0,6334 | 2,29956E-22 |
| ACE2 | GCLM | 0,6332 | 2,39782E-22 |
| ACE2 | DNAJB3 | 0,6327 | 2,61638E-22 |
| ACE2 | NBEAL1 | 0,6321 | 2,97318E-22 |
| ACE2 | APOC2 | 0,632 | 3,02371E-22 |
| ACE2 | ENTPD2 | 0,6319 | 3,09133E-22 |

| | | | |
|---|---|---|---|
| ACE2 | SOX9 | 0,6318 | 3,16478E-22 |
| ACE2 | TAX1BP3 | 0,6316 | 3,28208E-22 |
| ACE2 | AC019330.1 | 0,6315 | 3,36699E-22 |
| ACE2 | RP11-443B7.1 | 0,6314 | 3,40075E-22 |
| ACE2 | ATP1B3 | 0,6309 | 3,75162E-22 |
| ACE2 | C2orf82 | 0,6308 | 3,86413E-22 |
| ACE2 | CMTM8 | 0,6308 | 3,84053E-22 |
| ACE2 | HMGN2P46 | 0,6308 | 3,79786E-22 |
| ACE2 | TIGD3 | 0,6303 | 4,19741E-22 |
| ACE2 | AFDN | 0,6302 | 4,2851E-22 |
| ACE2 | RP13-580F15.2 | 0,6302 | 4,3177E-22 |
| ACE2 | LINC00390 | 0,6299 | 4,54385E-22 |
| ACE2 | FOXA2 | 0,6298 | 4,65257E-22 |
| ACE2 | AP001626.2 | 0,6296 | 4,86935E-22 |
| ACE2 | AF001550.7 | 0,6295 | 4,92825E-22 |
| ACE2 | CTD-2186M15.3 | 0,6293 | 5,08837E-22 |
| ACE2 | KLK1 | 0,6291 | 5,35416E-22 |
| ACE2 | SERPINA7 | 0,6291 | 5,28175E-22 |
| ACE2 | MFSD3 | 0,629 | 5,45231E-22 |
| ACE2 | PNPO | 0,629 | 5,46648E-22 |
| ACE2 | RP4-799G3.2 | 0,6289 | 5,56531E-22 |
| ACE2 | ALAS1 | 0,6289 | 5,50862E-22 |
| ACE2 | LINC00336 | 0,6285 | 5,99379E-22 |
| ACE2 | AC084809.3 | 0,6284 | 6,07778E-22 |
| ACE2 | DUSP21 | 0,6282 | 6,32324E-22 |
| ACE2 | ME2 | 0,6275 | 7,24063E-22 |
| ACE2 | STPG1 | 0,6271 | 7,83335E-22 |
| ACE2 | RP11-387A1.6 | 0,6271 | 7,89341E-22 |
| ACE2 | SPATA2 | 0,6271 | 7,81181E-22 |
| ACE2 | ROS1 | 0,6263 | 9,181E-22 |
| ACE2 | RP11-545A16.1 | 0,626 | 9,7051E-22 |
| ACE2 | R3HDML | 0,626 | 9,68225E-22 |
| ACE2 | KB-1460A1.3 | 0,6259 | 9,86619E-22 |
| ACE2 | RP11-944C7.1 | 0,6256 | 1,05278E-21 |
| ACE2 | PHYHIPL | 0,6255 | 1,07202E-21 |
| ACE2 | LAMA1 | 0,6253 | 1,11181E-21 |
| ACE2 | AP000679.2 | 0,6252 | 1,13164E-21 |
| ACE2 | RP11-7K24.3 | 0,6249 | 1,18985E-21 |
| ACE2 | B4GALNT3 | 0,6247 | 1,24749E-21 |
| ACE2 | ABCA4 | 0,624 | 1,42689E-21 |
| ACE2 | ANKRD20A21P | 0,624 | 1,42221E-21 |
| ACE2 | CTD-2600O9.1 | 0,6238 | 1,48079E-21 |
| ACE2 | BCL2L2-PABPN1 | 0,6235 | 1,556E-21 |
| ACE2 | TMEM9B | 0,6234 | 1,60241E-21 |
| ACE2 | RP11-817J15.3 | 0,6233 | 1,62292E-21 |
| ACE2 | LRRC9 | 0,6227 | 1,83282E-21 |
| ACE2 | PPP1R36 | 0,6225 | 1,887E-21 |
| ACE2 | LA16c-380H5.2 | 0,6225 | 1,89741E-21 |
| ACE2 | LPGAT1 | 0,6222 | 2,01721E-21 |
| ACE2 | ETFA | 0,6222 | 2,01226E-21 |
| ACE2 | MIR200A | 0,622 | 2,08598E-21 |
| ACE2 | GALNT1 | 0,622 | 2,09237E-21 |
| ACE2 | NUP62CL | 0,622 | 2,07309E-21 |
| ACE2 | KCNK10 | 0,6219 | 2,1232E-21 |

| | | | |
|---|---|---|---|
| ACE2 | FLNB | 0,6215 | 2,28841E-21 |
| ACE2 | PATJ | 0,6208 | 2,58568E-21 |
| ACE2 | LRP4-AS1 | 0,6208 | 2,62561E-21 |
| ACE2 | GALNT8 | 0,6207 | 2,66384E-21 |
| ACE2 | RP11-807H22.5 | 0,6204 | 2,81465E-21 |
| ACE2 | CTD-3094K11.1 | 0,6203 | 2,86092E-21 |
| ACE2 | UBE2FP1 | 0,6201 | 2,9621E-21 |
| ACE2 | RP11-84H6.1 | 0,62 | 3,05066E-21 |
| ACE2 | EFNB1 | 0,6198 | 3,14984E-21 |
| ACE2 | RP5-875O13.7 | 0,6196 | 3,27393E-21 |
| ACE2 | CDH20 | 0,6196 | 3,29E-21 |
| ACE2 | GRIN1 | 0,6195 | 3,35807E-21 |
| ACE2 | CTD-2184C24.2 | 0,6195 | 3,35847E-21 |
| ACE2 | CCNG1P1 | 0,6194 | 3,41121E-21 |
| ACE2 | MED18 | 0,6193 | 3,46064E-21 |
| ACE2 | TLX1NB | 0,6189 | 3,75147E-21 |
| ACE2 | SLC9A1 | 0,6184 | 4,05958E-21 |
| ACE2 | RN7SL786P | 0,6183 | 4,16892E-21 |
| ACE2 | MCRIP2 | 0,6183 | 4,15615E-21 |
| ACE2 | GCNT2 | 0,6182 | 4,2118E-21 |
| ACE2 | RP11-418J17.3 | 0,6179 | 4,48978E-21 |
| ACE2 | ITLN2 | 0,6173 | 5,02631E-21 |
| ACE2 | RP11-115J16.2 | 0,6173 | 5,06469E-21 |
| ACE2 | PRKAB1 | 0,6173 | 4,99482E-21 |
| ACE2 | CLYBL | 0,6173 | 5,06395E-21 |
| ACE2 | RP11-575G13.3 | 0,6172 | 5,07396E-21 |
| ACE2 | TJP2 | 0,617 | 5,3412E-21 |
| ACE2 | ZBTB7C | 0,617 | 5,30064E-21 |
| ACE2 | CLDN12 | 0,6165 | 5,80376E-21 |
| ACE2 | FAM213B | 0,6162 | 6,1777E-21 |
| ACE2 | RP11-568K15.1 | 0,6158 | 6,58737E-21 |
| ACE2 | AKR1C4 | 0,6158 | 6,59905E-21 |
| ACE2 | LINC00261 | 0,6158 | 6,57799E-21 |
| ACE2 | ADGRF4 | 0,6157 | 6,74298E-21 |
| ACE2 | LINC01668 | 0,6157 | 6,76429E-21 |
| ACE2 | AC005336.4 | 0,6155 | 7,06789E-21 |
| ACE2 | RP5-921G16.1 | 0,6152 | 7,40699E-21 |
| ACE2 | TPH1 | 0,615 | 7,645E-21 |
| ACE2 | FREM1 | 0,6148 | 7,92373E-21 |
| ACE2 | PLS1-AS1 | 0,6146 | 8,30639E-21 |
| ACE2 | CTB-160O22.1 | 0,6146 | 8,33464E-21 |
| ACE2 | CCDC120 | 0,6145 | 8,40359E-21 |
| ACE2 | LHFPL3 | 0,6142 | 8,88959E-21 |
| ACE2 | RP4-657E11.10 | 0,6141 | 9,12666E-21 |
| ACE2 | CNNM4 | 0,6138 | 9,51371E-21 |
| ACE2 | C5orf52 | 0,6137 | 9,84844E-21 |
| ACE2 | GLTP | 0,6137 | 9,79556E-21 |
| ACE2 | RPSAP46 | 0,6134 | 1,03149E-20 |
| ACE2 | PFN1P9 | 0,6129 | 1,14205E-20 |
| ACE2 | SST | 0,6128 | 1,16319E-20 |
| ACE2 | RP11-933H2.4 | 0,6125 | 1,21192E-20 |
| ACE2 | RNF186 | 0,6121 | 1,32241E-20 |
| ACE2 | RP11-123J14.4 | 0,6121 | 1,30899E-20 |
| ACE2 | RP11-290F5.2 | 0,612 | 1,33408E-20 |

| | | | |
|---|---|---|---|
| ACE2 | LPIN2 | 0,6119 | 1,35648E-20 |
| ACE2 | PAK1 | 0,6118 | 1,38827E-20 |
| ACE2 | OR5BA1P | 0,6115 | 1,46508E-20 |
| ACE2 | MTCYBP41 | 0,6114 | 1,48642E-20 |
| ACE2 | NKX2-2 | 0,6113 | 1,50982E-20 |
| ACE2 | RP11-568A7.2 | 0,6107 | 1,69773E-20 |
| ACE2 | PIGS | 0,6104 | 1,77337E-20 |
| ACE2 | TXNDC17 | 0,6102 | 1,84116E-20 |
| ACE2 | RP11-45P22.2 | 0,6099 | 1,95139E-20 |
| ACE2 | EVPL | 0,6096 | 2,06153E-20 |
| ACE2 | WI2-2118C23.2 | 0,6095 | 2,09777E-20 |
| ACE2 | PIGZ | 0,6086 | 2,49584E-20 |
| ACE2 | SLC6A8 | 0,6084 | 2,54683E-20 |
| ACE2 | CRYM | 0,6082 | 2,67783E-20 |
| ACE2 | ABCC11 | 0,6075 | 3,00257E-20 |
| ACE2 | FUT4 | 0,6072 | 3,17803E-20 |
| ACE2 | TRIM2 | 0,6063 | 3,7301E-20 |
| ACE2 | CGREF1 | 0,6061 | 3,86551E-20 |
| ACE2 | KLHL32 | 0,6057 | 4,15646E-20 |
| ACE2 | AC102953.4 | 0,6056 | 4,23694E-20 |
| ACE2 | SNX18P3 | 0,6054 | 4,36929E-20 |
| ACE2 | RP11-293P20.2 | 0,6053 | 4,49836E-20 |
| ACE2 | RP11-44N21.1 | 0,605 | 4,70679E-20 |
| ACE2 | TMEM59 | 0,6044 | 5,23422E-20 |
| ACE2 | ARHGAP18 | 0,6044 | 5,23405E-20 |
| ACE2 | RNA5SP507 | 0,6044 | 5,24182E-20 |
| ACE2 | MUC20 | 0,6037 | 5,90467E-20 |
| ACE2 | SLC12A2 | 0,6036 | 6,04426E-20 |
| ACE2 | LL0XNC01-131B10.2 | 0,6035 | 6,17733E-20 |
| ACE2 | ITGA2 | 0,6034 | 6,25077E-20 |
| ACE2 | LINC00857 | 0,603 | 6,72994E-20 |
| ACE2 | RP11-40A7.2 | 0,6026 | 7,23145E-20 |
| ACE2 | HMGN1P7 | 0,602 | 8,09936E-20 |
| ACE2 | MSRA | 0,602 | 8,06243E-20 |
| ACE2 | C4orf48 | 0,6018 | 8,38021E-20 |
| ACE2 | IFNWP19 | 0,6014 | 8,91364E-20 |
| ACE2 | PPARG | 0,6013 | 9,08596E-20 |
| ACE2 | DDT | 0,6012 | 9,31255E-20 |
| ACE2 | LINC02068 | 0,6004 | 1,06091E-19 |
| ACE2 | CDHR1 | 0,6004 | 1,06399E-19 |
| ACE2 | PAG1 | 0,6002 | 1,10647E-19 |
| ACE2 | CTB-171A8.1 | 0,6001 | 1,11399E-19 |
| ACE2 | TFF2 | 0,5998 | 1,19128E-19 |
| ACE2 | ID2-AS1 | 0,5997 | 1,21041E-19 |
| ACE2 | CFL1 | 0,5995 | 1,24989E-19 |
| ACE2 | LBX2 | 0,5993 | 1,29659E-19 |
| ACE2 | RPL7P60 | 0,5993 | 1,28115E-19 |
| ACE2 | MYCL | 0,5986 | 1,46809E-19 |
| ACE2 | RP11-83M16.3 | 0,5984 | 1,51895E-19 |
| ACE2 | RP11-616M22.3 | 0,5983 | 1,53537E-19 |
| ACE2 | CTB-25B13.9 | 0,5983 | 1,5325E-19 |
| ACE2 | RP1-40E16.11 | 0,5982 | 1,55849E-19 |
| ACE2 | ASB13 | 0,5981 | 1,58943E-19 |
| ACE2 | UGT2B25P | 0,5979 | 1,6564E-19 |

| | | | |
|---|---|---|---|
| ACE2 | UQCRQ | 0,5979 | 1,65409E-19 |
| ACE2 | CCDC175 | 0,5979 | 1,65725E-19 |
| ACE2 | RP1-240K6.3 | 0,5978 | 1,67667E-19 |
| ACE2 | HMGB3 | 0,5978 | 1,67676E-19 |
| ACE2 | UGT2B10 | 0,5977 | 1,70988E-19 |
| ACE2 | CTD-2134A5.4 | 0,5977 | 1,71039E-19 |
| ACE2 | ABHD3 | 0,5977 | 1,69253E-19 |
| ACE2 | KNG1 | 0,5976 | 1,74846E-19 |
| ACE2 | TMEM229B | 0,5973 | 1,82197E-19 |
| ACE2 | DNAJC5 | 0,5971 | 1,87887E-19 |
| ACE2 | IL10RB | 0,5971 | 1,90087E-19 |
| ACE2 | RPL17P11 | 0,5968 | 1,97941E-19 |
| ACE2 | RP11-800A3.2 | 0,5968 | 1,9882E-19 |
| ACE2 | EXOC3L4 | 0,5968 | 1,99356E-19 |
| ACE2 | TUSC8 | 0,5967 | 2,02491E-19 |
| ACE2 | RP13-463N16.6 | 0,5964 | 2,12522E-19 |
| ACE2 | TSPAN6 | 0,5964 | 2,12283E-19 |
| ACE2 | KRTAP4-1 | 0,596 | 2,29158E-19 |
| ACE2 | MNX1-AS1 | 0,5959 | 2,3363E-19 |
| ACE2 | CTD-2196E14.5 | 0,5957 | 2,40757E-19 |
| ACE2 | RP3-395M20.8 | 0,5956 | 2,46405E-19 |
| ACE2 | RP11-27K13.3 | 0,5956 | 2,47055E-19 |
| ACE2 | FAM47E | 0,5956 | 2,43599E-19 |
| ACE2 | CTC-297N7.9 | 0,5956 | 2,44221E-19 |
| ACE2 | DDO | 0,5955 | 2,514E-19 |
| ACE2 | UBALD2 | 0,5951 | 2,68852E-19 |
| ACE2 | PRSS22 | 0,5948 | 2,8206E-19 |
| ACE2 | ATP5B | 0,5947 | 2,86214E-19 |
| ACE2 | RP11-497G19.1 | 0,5945 | 2,9441E-19 |
| ACE2 | TMEM256 | 0,5945 | 2,97919E-19 |
| ACE2 | KCNV1 | 0,5944 | 3,00954E-19 |
| ACE2 | ECEL1P1 | 0,5942 | 3,114E-19 |
| ACE2 | GIMD1 | 0,5934 | 3,57044E-19 |
| ACE2 | TOR4A | 0,5934 | 3,58189E-19 |
| ACE2 | RP11-390F4.3 | 0,5929 | 3,90268E-19 |
| ACE2 | ATP1A1-AS1 | 0,5928 | 3,99459E-19 |
| ACE2 | RPS29P33 | 0,5926 | 4,09811E-19 |
| ACE2 | KRT40 | 0,592 | 4,55664E-19 |
| ACE2 | XPR1 | 0,5917 | 4,76227E-19 |
| ACE2 | RP11-468N14.09 | 0,5915 | 4,97294E-19 |
| ACE2 | AC138517.6 | 0,5914 | 5,003E-19 |
| ACE2 | LINC00637 | 0,5913 | 5,07237E-19 |
| ACE2 | MAGEA5 | 0,5913 | 5,07196E-19 |
| ACE2 | RP11-240D10.4 | 0,5908 | 5,56895E-19 |
| ACE2 | SPECC1 | 0,5906 | 5,71231E-19 |
| ACE2 | KB-1184D12.1 | 0,5905 | 5,8781E-19 |
| ACE2 | SP8 | 0,5902 | 6,11663E-19 |
| ACE2 | LRRC8D | 0,59 | 6,3343E-19 |
| ACE2 | CRACR2B | 0,5898 | 6,60229E-19 |
| ACE2 | HMGCS2 | 0,5897 | 6,7325E-19 |
| ACE2 | MGST1 | 0,5892 | 7,29906E-19 |
| ACE2 | SURF4 | 0,5891 | 7,40142E-19 |
| ACE2 | RP11-396O20.1 | 0,5888 | 7,75512E-19 |
| ACE2 | HMGB1P11 | 0,5882 | 8,68268E-19 |

| | | | |
|---|---|---|---|
| ACE2 | LINC01978 | 0,5882 | 8,58039E-19 |
| ACE2 | RP11-16L21.7 | 0,5881 | 8,75249E-19 |
| ACE2 | ALPPL2 | 0,5879 | 9,02081E-19 |
| ACE2 | RP11-164P12.4 | 0,5878 | 9,21019E-19 |
| ACE2 | NIT1 | 0,5876 | 9,58315E-19 |
| ACE2 | CASC9 | 0,5871 | 1,04127E-18 |
| ACE2 | BAATP1 | 0,587 | 1,05801E-18 |
| ACE2 | REG3A | 0,5869 | 1,07199E-18 |
| ACE2 | ECEL1P2 | 0,5869 | 1,07869E-18 |
| ACE2 | DNPEP | 0,5867 | 1,10163E-18 |
| ACE2 | RP11-448G15.1 | 0,5867 | 1,10394E-18 |
| ACE2 | NCR3LG1 | 0,5867 | 1,10441E-18 |
| ACE2 | VWA2 | 0,5863 | 1,1878E-18 |
| ACE2 | VPS25 | 0,5861 | 1,22304E-18 |
| ACE2 | RAG1 | 0,586 | 1,25638E-18 |
| ACE2 | CRACR2A | 0,5858 | 1,28681E-18 |
| ACE2 | TRPC7 | 0,5857 | 1,3071E-18 |
| ACE2 | LINC01134 | 0,5855 | 1,35515E-18 |
| ACE2 | FAM105A | 0,5854 | 1,38126E-18 |
| ACE2 | PARP4 | 0,5851 | 1,4493E-18 |
| ACE2 | AC114812.8 | 0,5849 | 1,50807E-18 |
| ACE2 | LRP4 | 0,5849 | 1,50508E-18 |
| ACE2 | ASPDH | 0,5849 | 1,49411E-18 |
| ACE2 | HNRNPA1P9 | 0,5844 | 1,63014E-18 |
| ACE2 | SFXN3 | 0,5844 | 1,62235E-18 |
| ACE2 | REG3G | 0,5843 | 1,64343E-18 |
| ACE2 | RHOU | 0,5842 | 1,66978E-18 |
| ACE2 | SUGCT | 0,5842 | 1,68135E-18 |
| ACE2 | RP11-442J21.2 | 0,5837 | 1,84121E-18 |
| ACE2 | MYL12B | 0,5837 | 1,83297E-18 |
| ACE2 | PANCR | 0,5836 | 1,84916E-18 |
| ACE2 | NECTIN3 | 0,5835 | 1,89464E-18 |
| ACE2 | PI4K2B | 0,5834 | 1,92711E-18 |
| ACE2 | PSORS1C3 | 0,5833 | 1,95112E-18 |
| ACE2 | PREP | 0,5827 | 2,15219E-18 |
| ACE2 | MIR200B | 0,5826 | 2,18424E-18 |
| ACE2 | SC5D | 0,5819 | 2,4746E-18 |
| ACE2 | RP5-908M14.10 | 0,5819 | 2,44633E-18 |
| ACE2 | KRT8P8 | 0,5817 | 2,52833E-18 |
| ACE2 | RHOBTB2 | 0,5816 | 2,57549E-18 |
| ACE2 | CYP2B7P | 0,5816 | 2,58249E-18 |
| ACE2 | LCN12 | 0,5812 | 2,77956E-18 |
| ACE2 | RP1-232L24.3 | 0,5809 | 2,91203E-18 |
| ACE2 | PANK3 | 0,5803 | 3,22154E-18 |
| ACE2 | SH3BGRL2 | 0,5803 | 3,18935E-18 |
| ACE2 | FDX1 | 0,5803 | 3,19042E-18 |
| ACE2 | SLC26A1 | 0,5801 | 3,28959E-18 |
| ACE2 | TAS2R38 | 0,5801 | 3,32911E-18 |
| ACE2 | RP11-269G24.6 | 0,5801 | 3,29523E-18 |
| ACE2 | KRT18P28 | 0,5793 | 3,75499E-18 |
| ACE2 | RP11-15F12.6 | 0,5793 | 3,79605E-18 |
| ACE2 | TMEM220 | 0,5791 | 3,90372E-18 |
| ACE2 | RP11-356M20.1 | 0,5789 | 3,9892E-18 |
| ACE2 | AC010872.1 | 0,5788 | 4,07325E-18 |

| | | | |
|---|---|---|---|
| ACE2 | HSD3BP1 | 0,5787 | 4,14609E-18 |
| ACE2 | RPP25L | 0,5784 | 4,33903E-18 |
| ACE2 | DEFA6 | 0,5783 | 4,4594E-18 |
| ACE2 | MCU | 0,5783 | 4,41236E-18 |
| ACE2 | TRPV3 | 0,578 | 4,6418E-18 |
| ACE2 | RP11-63D14.1 | 0,5778 | 4,8258E-18 |
| ACE2 | NUTM1 | 0,5775 | 5,01179E-18 |
| ACE2 | ITGA6 | 0,5772 | 5,25921E-18 |
| ACE2 | KCNJ16 | 0,5772 | 5,28325E-18 |
| ACE2 | TMCC3 | 0,5771 | 5,36101E-18 |
| ACE2 | RP3-467K16.4 | 0,577 | 5,50206E-18 |
| ACE2 | KRT8P49 | 0,577 | 5,44945E-18 |
| ACE2 | AKR7A2 | 0,5767 | 5,74261E-18 |
| ACE2 | DEFA5 | 0,5765 | 5,9737E-18 |
| ACE2 | TPI1 | 0,5765 | 5,94734E-18 |
| ACE2 | RP11-549B18.1 | 0,5763 | 6,07899E-18 |
| ACE2 | MPND | 0,5763 | 6,1172E-18 |
| ACE2 | TP53AIP1 | 0,5762 | 6,2524E-18 |
| ACE2 | SLC30A2 | 0,5758 | 6,66628E-18 |
| ACE2 | LINC02012 | 0,5758 | 6,59323E-18 |
| ACE2 | RUNDC3B | 0,5757 | 6,79432E-18 |
| ACE2 | GSTZ1 | 0,5757 | 6,72365E-18 |
| ACE2 | LHFPL3-AS2 | 0,5753 | 7,15104E-18 |
| ACE2 | KRT18P34 | 0,5751 | 7,43524E-18 |
| ACE2 | AP002856.7 | 0,575 | 7,533E-18 |
| ACE2 | OSR2 | 0,5749 | 7,71475E-18 |
| ACE2 | TPRG1L | 0,5748 | 7,84223E-18 |
| ACE2 | AP1B1P1 | 0,5746 | 8,05374E-18 |
| ACE2 | CDKN1A | 0,5745 | 8,1167E-18 |
| ACE2 | AKR1C2 | 0,5745 | 8,14302E-18 |
| ACE2 | RP3-400B16.1 | 0,5744 | 8,33781E-18 |
| ACE2 | LGALSL | 0,5728 | 1,06557E-17 |
| ACE2 | LINC00668 | 0,5727 | 1,08783E-17 |
| ACE2 | RP11-8L2.1 | 0,5725 | 1,13027E-17 |
| ACE2 | ENPP1 | 0,5724 | 1,13822E-17 |
| ACE2 | RNU6-704P | 0,5723 | 1,15586E-17 |
| ACE2 | NOXO1 | 0,5716 | 1,3087E-17 |
| ACE2 | TCF7L2 | 0,5714 | 1,33905E-17 |
| ACE2 | PON3 | 0,5713 | 1,369E-17 |
| ACE2 | ILDR1 | 0,5711 | 1,41323E-17 |
| ACE2 | CHPT1 | 0,5709 | 1,45232E-17 |
| ACE2 | MFSD6L | 0,5709 | 1,44414E-17 |
| ACE2 | RP11-1223D19.6 | 0,5707 | 1,5041E-17 |
| ACE2 | PEX11A | 0,5703 | 1,6044E-17 |
| ACE2 | KRT42P | 0,5701 | 1,65528E-17 |
| ACE2 | MAIP1 | 0,5699 | 1,71044E-17 |
| ACE2 | CYBRD1 | 0,5697 | 1,75633E-17 |
| ACE2 | HNRNPA1P27 | 0,5695 | 1,81957E-17 |
| ACE2 | PDX1 | 0,5693 | 1,86488E-17 |
| ACE2 | IL17RB | 0,569 | 1,95357E-17 |
| ACE2 | CERS2 | 0,5689 | 1,98554E-17 |
| ACE2 | C4orf33 | 0,5688 | 2,02738E-17 |
| ACE2 | TUBB4B | 0,5688 | 2,02567E-17 |
| ACE2 | CTA-714B7.7 | 0,5688 | 2,00893E-17 |

| | | | |
|---|---|---|---|
| ACE2 | SLC35A2 | 0,5687 | 2,0647E-17 |
| ACE2 | NPIPB14P | 0,5685 | 2,11056E-17 |
| ACE2 | ALDOC | 0,5685 | 2,11365E-17 |
| ACE2 | RP11-44F14.2 | 0,5684 | 2,154E-17 |
| ACE2 | CTD-2012B7.1 | 0,5682 | 2,22544E-17 |
| ACE2 | RP11-569G9.7 | 0,5679 | 2,33931E-17 |
| ACE2 | KNL1 | 0,5677 | 2,39193E-17 |
| ACE2 | ANKEF1 | 0,5677 | 2,42184E-17 |
| ACE2 | REG1B | 0,5676 | 2,4505E-17 |
| ACE2 | ELMOD2 | 0,5676 | 2,43744E-17 |
| ACE2 | BMS1P1 | 0,5676 | 2,42378E-17 |
| ACE2 | RP11-336A10.4 | 0,5675 | 2,46467E-17 |
| ACE2 | ILVBL | 0,5675 | 2,47085E-17 |
| ACE2 | FAM135A | 0,5672 | 2,59784E-17 |
| ACE2 | RP5-1056H1.2 | 0,567 | 2,67812E-17 |
| ACE2 | RP11-242G20.1 | 0,5666 | 2,85755E-17 |
| ACE2 | RNF5 | 0,5665 | 2,89667E-17 |
| ACE2 | RP1-239B22.5 | 0,5663 | 2,99001E-17 |
| ACE2 | SAMD13 | 0,5662 | 3,03068E-17 |
| ACE2 | HPCAL1 | 0,5657 | 3,27892E-17 |
| ACE2 | DMBT1 | 0,5657 | 3,29864E-17 |
| ACE2 | RP11-1223D19.5 | 0,5655 | 3,36895E-17 |
| ACE2 | DOPEY2 | 0,5648 | 3,77966E-17 |
| ACE2 | ATOH7 | 0,5647 | 3,80703E-17 |
| ACE2 | RBBP8 | 0,5647 | 3,86541E-17 |
| ACE2 | H2AFY | 0,5646 | 3,88377E-17 |
| ACE2 | RP11-375H19.2 | 0,5645 | 3,92924E-17 |
| ACE2 | LINC02003 | 0,5642 | 4,1637E-17 |
| ACE2 | LILRP2 | 0,5642 | 4,1305E-17 |
| ACE2 | GGTLC3 | 0,5642 | 4,17263E-17 |
| ACE2 | TSKU | 0,5641 | 4,23639E-17 |
| ACE2 | TMEM8A | 0,564 | 4,30796E-17 |
| ACE2 | RP11-552M14.1 | 0,5639 | 4,32236E-17 |
| ACE2 | LPO | 0,5638 | 4,41752E-17 |
| ACE2 | SEZ6L2 | 0,5637 | 4,45756E-17 |
| ACE2 | RP11-138I1.3 | 0,5636 | 4,53389E-17 |
| ACE2 | PDXDC2P | 0,5635 | 4,58672E-17 |
| ACE2 | C11orf42 | 0,5633 | 4,79162E-17 |
| ACE2 | AQP12B | 0,5632 | 4,86343E-17 |
| ACE2 | LINC00864 | 0,5632 | 4,81044E-17 |
| ACE2 | LINC01659 | 0,563 | 5,02612E-17 |
| ACE2 | SMAGP | 0,5627 | 5,22207E-17 |
| ACE2 | LINC01924 | 0,5626 | 5,33604E-17 |
| ACE2 | MRAP2 | 0,5624 | 5,4451E-17 |
| ACE2 | AC114812.5 | 0,5618 | 6,00277E-17 |
| ACE2 | UGT3A1 | 0,5617 | 6,13076E-17 |
| ACE2 | PRAG1 | 0,5614 | 6,38638E-17 |
| ACE2 | ZNF552 | 0,5614 | 6,35413E-17 |
| ACE2 | FBXL6 | 0,5606 | 7,18827E-17 |
| ACE2 | MYO10 | 0,5605 | 7,32782E-17 |
| ACE2 | POLRMTP1 | 0,5603 | 7,50361E-17 |
| ACE2 | RP11-649A18.5 | 0,5603 | 7,53581E-17 |
| ACE2 | PRELID1 | 0,56 | 7,95823E-17 |
| ACE2 | MPC2 | 0,5598 | 8,16085E-17 |

| | | | |
|---|---|---|---|
| ACE2 | LRIT2 | 0,5598 | 8,15442E-17 |
| ACE2 | RP11-203P23.2 | 0,5597 | 8,2592E-17 |
| ACE2 | RP11-734K21.5 | 0,5596 | 8,36387E-17 |
| ACE2 | TMEM41B | 0,5593 | 8,79199E-17 |
| ACE2 | GABRE | 0,5592 | 8,96275E-17 |
| ACE2 | NEURL1B | 0,5591 | 9,06599E-17 |
| ACE2 | BBIP1P1 | 0,559 | 9,14682E-17 |
| ACE2 | RP11-519G16.5 | 0,5588 | 9,42948E-17 |
| ACE2 | RP11-95G17.2 | 0,5587 | 9,70017E-17 |
| ACE2 | SLC2A9 | 0,5584 | 1,0147E-16 |
| ACE2 | HYKK | 0,5584 | 1,00701E-16 |
| ACE2 | DPP3 | 0,5583 | 1,0181E-16 |
| ACE2 | ABCD1 | 0,5583 | 1,0241E-16 |
| ACE2 | SLC50A1 | 0,5582 | 1,04221E-16 |
| ACE2 | KIAA1217 | 0,5576 | 1,1413E-16 |
| ACE2 | RPL32P33 | 0,5574 | 1,16573E-16 |
| ACE2 | TMC7 | 0,5571 | 1,23057E-16 |
| ACE2 | GCLC | 0,557 | 1,24581E-16 |
| ACE2 | SLC9C1 | 0,5569 | 1,25953E-16 |
| ACE2 | RP11-84A14.4 | 0,5568 | 1,29154E-16 |
| ACE2 | RP11-44F14.8 | 0,5566 | 1,32989E-16 |
| ACE2 | FERMT1 | 0,5564 | 1,36965E-16 |
| ACE2 | RP11-845C23.2 | 0,5563 | 1,39542E-16 |
| ACE2 | GATM-AS1 | 0,5562 | 1,41288E-16 |
| ACE2 | SLC5A8 | 0,5561 | 1,41931E-16 |
| ACE2 | KLK3 | 0,5561 | 1,43289E-16 |
| ACE2 | ZNF664 | 0,5558 | 1,50017E-16 |
| ACE2 | OLFM4 | 0,5555 | 1,55243E-16 |
| ACE2 | LINC01979 | 0,5555 | 1,56916E-16 |
| ACE2 | CTAGE4 | 0,5553 | 1,61513E-16 |
| ACE2 | RPLP0P2 | 0,5553 | 1,60098E-16 |
| ACE2 | CTD-2503O16.4 | 0,5552 | 1,62431E-16 |
| ACE2 | ACTN4 | 0,5552 | 1,64595E-16 |
| ACE2 | TMEM117 | 0,5551 | 1,66101E-16 |
| ACE2 | TSPAN12 | 0,555 | 1,67843E-16 |
| ACE2 | SLC5A2 | 0,555 | 1,68458E-16 |
| ACE2 | RP11-515O17.2 | 0,5547 | 1,77635E-16 |
| ACE2 | RP11-412P11.1 | 0,5546 | 1,78411E-16 |
| ACE2 | LCORL | 0,5544 | 1,85686E-16 |
| ACE2 | HRCT1 | 0,5542 | 1,90574E-16 |
| ACE2 | ACKR4 | 0,5539 | 1,9846E-16 |
| ACE2 | RP11-1036E20.9 | 0,5537 | 2,03389E-16 |
| ACE2 | SLC18A1 | 0,5536 | 2,08515E-16 |
| ACE2 | SLC12A8 | 0,553 | 2,28075E-16 |
| ACE2 | THSD4 | 0,5529 | 2,31136E-16 |
| ACE2 | LETM1 | 0,5525 | 2,45849E-16 |
| ACE2 | RP11-235E17.5 | 0,5524 | 2,47417E-16 |
| ACE2 | LINC02057 | 0,5523 | 2,51609E-16 |
| ACE2 | AC005077.14 | 0,5523 | 2,50665E-16 |
| ACE2 | RP11-671J11.4 | 0,5523 | 2,5285E-16 |
| ACE2 | RP11-10C24.1 | 0,5522 | 2,5638E-16 |
| ACE2 | MOXD2P | 0,5519 | 2,6714E-16 |
| ACE2 | CTD-2256P15.5 | 0,5517 | 2,74511E-16 |
| ACE2 | ZNF711 | 0,5517 | 2,76825E-16 |

| | | | |
|---|---|---|---|
| ACE2 | OR2A1-AS1 | 0,5516 | 2,79314E-16 |
| ACE2 | FER1L6-AS1 | 0,5515 | 2,85183E-16 |
| ACE2 | CXCL16 | 0,5513 | 2,94295E-16 |
| ACE2 | RFK | 0,5512 | 2,96696E-16 |
| ACE2 | GS1-393G12.14 | 0,5511 | 2,99787E-16 |
| ACE2 | BMP8B | 0,551 | 3,04018E-16 |
| ACE2 | RP11-439C15.4 | 0,551 | 3,03894E-16 |
| ACE2 | SAT2 | 0,5506 | 3,26407E-16 |
| ACE2 | RP11-544M22.1 | 0,5504 | 3,36805E-16 |
| ACE2 | REG1A | 0,5504 | 3,33894E-16 |
| ACE2 | ELF3-AS1 | 0,5503 | 3,39925E-16 |
| ACE2 | TMBIM1 | 0,5503 | 3,40541E-16 |
| ACE2 | S100A16 | 0,55 | 3,54816E-16 |
| ACE2 | FAT2 | 0,55 | 3,53826E-16 |
| ACE2 | RIPK3 | 0,5498 | 3,6679E-16 |
| ACE2 | FAM129B | 0,5497 | 3,68986E-16 |
| ACE2 | RP11-1C8.4 | 0,5495 | 3,84668E-16 |
| ACE2 | GK-IT1 | 0,5492 | 3,99361E-16 |
| ACE2 | RP11-486F17.1 | 0,549 | 4,14047E-16 |
| ACE2 | RP11-467J12.4 | 0,549 | 4,08968E-16 |
| ACE2 | ADIPOR2 | 0,5489 | 4,17283E-16 |
| ACE2 | CHST13 | 0,5487 | 4,3109E-16 |
| ACE2 | PRSS2 | 0,5478 | 4,87175E-16 |
| ACE2 | PHEX | 0,5476 | 5,03846E-16 |
| ACE2 | RP11-49I11.3 | 0,5474 | 5,2264E-16 |
| ACE2 | MT-TL1 | 0,5472 | 5,34283E-16 |
| ACE2 | KB-1592A4.16 | 0,547 | 5,48258E-16 |
| ACE2 | TMEM87B | 0,5467 | 5,74094E-16 |
| ACE2 | RRBP1 | 0,5464 | 5,97733E-16 |
| ACE2 | RP11-797A18.5 | 0,546 | 6,36206E-16 |
| ACE2 | GPR119 | 0,5458 | 6,57932E-16 |
| ACE2 | SERPINA4 | 0,5455 | 6,84084E-16 |
| ACE2 | ETV7 | 0,5452 | 7,12427E-16 |
| ACE2 | S100G | 0,5451 | 7,30717E-16 |
| ACE2 | TCP10L2 | 0,545 | 7,37049E-16 |
| ACE2 | RP11-95I16.6 | 0,5444 | 8,02565E-16 |
| ACE2 | RN7SKP25 | 0,5443 | 8,22402E-16 |
| ACE2 | SLC17A1 | 0,544 | 8,51637E-16 |
| ACE2 | CER1 | 0,5439 | 8,65989E-16 |
| ACE2 | FGF9 | 0,5437 | 8,84871E-16 |
| ACE2 | MT-ND6 | 0,5437 | 8,858E-16 |
| ACE2 | RPH3AL | 0,5436 | 8,97314E-16 |
| ACE2 | CIR1P2 | 0,5435 | 9,23132E-16 |
| ACE2 | KRT8P37 | 0,5435 | 9,20971E-16 |
| ACE2 | ESPNP | 0,5432 | 9,53478E-16 |
| ACE2 | RP11-429J17.7 | 0,5431 | 9,69209E-16 |
| ACE2 | SYNGR4 | 0,5426 | 1,04285E-15 |
| ACE2 | EXT1 | 0,5423 | 1,09666E-15 |
| ACE2 | RP11-21L23.3 | 0,5422 | 1,10577E-15 |
| ACE2 | DOCK9-AS1 | 0,542 | 1,13579E-15 |
| ACE2 | RP11-329A14.1 | 0,5418 | 1,17582E-15 |
| ACE2 | RP11-290F24.6 | 0,5417 | 1,19516E-15 |
| ACE2 | VWA5B2 | 0,5413 | 1,25993E-15 |
| ACE2 | NAGA | 0,5412 | 1,27371E-15 |

| | | | |
|---|---|---|---|
| ACE2 | RP11-320N7.2 | 0,5411 | 1,28612E-15 |
| ACE2 | KRT8P24 | 0,5411 | 1,29628E-15 |
| ACE2 | KRT8P7 | 0,541 | 1,3053E-15 |
| ACE2 | RP11-963H4.3 | 0,541 | 1,30601E-15 |
| ACE2 | RPSAP75 | 0,5409 | 1,32479E-15 |
| ACE2 | SLC6A6 | 0,5408 | 1,35593E-15 |
| ACE2 | LINC01648 | 0,54 | 1,51063E-15 |
| ACE2 | PAQR5 | 0,5399 | 1,53585E-15 |
| ACE2 | RAB27B | 0,5398 | 1,55448E-15 |
| ACE2 | NDUFA9 | 0,5397 | 1,58868E-15 |
| ACE2 | RP6-149D17.1 | 0,5397 | 1,59163E-15 |
| ACE2 | GOLGA4 | 0,5392 | 1,68747E-15 |
| ACE2 | CYP2W1 | 0,5391 | 1,72613E-15 |
| ACE2 | FUT5 | 0,5389 | 1,77013E-15 |
| ACE2 | IL18 | 0,5387 | 1,82873E-15 |
| ACE2 | LINC02041 | 0,5385 | 1,87381E-15 |
| ACE2 | STARD5 | 0,5385 | 1,86564E-15 |
| ACE2 | LINC01709 | 0,5384 | 1,89073E-15 |
| ACE2 | RP11-1149M10.2 | 0,5384 | 1,89353E-15 |
| ACE2 | SLC3A2 | 0,5379 | 2,0382E-15 |
| ACE2 | DSG4 | 0,5377 | 2,10537E-15 |
| ACE2 | UMODL1 | 0,5377 | 2,08911E-15 |
| ACE2 | IGSF3 | 0,5373 | 2,22397E-15 |
| ACE2 | RP11-295G20.2 | 0,5371 | 2,27841E-15 |
| ACE2 | RP11-227H15.4 | 0,5367 | 2,43421E-15 |
| ACE2 | AC011747.3 | 0,5365 | 2,50765E-15 |
| ACE2 | SNRPGP4 | 0,5365 | 2,4778E-15 |
| ACE2 | COPG1 | 0,5364 | 2,53041E-15 |
| ACE2 | TPD52 | 0,5363 | 2,57097E-15 |
| ACE2 | STARD4 | 0,5361 | 2,62992E-15 |
| ACE2 | KRT18P63 | 0,5357 | 2,79265E-15 |
| ACE2 | AC107072.2 | 0,5357 | 2,77983E-15 |
| ACE2 | RPL7L1P3 | 0,5357 | 2,80222E-15 |
| ACE2 | PLEKHA6 | 0,5354 | 2,92001E-15 |
| ACE2 | ACPP | 0,5353 | 2,96605E-15 |
| ACE2 | LINC00964 | 0,5351 | 3,01486E-15 |
| ACE2 | KRT8P35 | 0,535 | 3,09261E-15 |
| ACE2 | ARFGEF3 | 0,5349 | 3,11707E-15 |
| ACE2 | KRT18P11 | 0,5349 | 3,12435E-15 |
| ACE2 | RP5-1142J19.1 | 0,5347 | 3,23038E-15 |
| ACE2 | HOXB9 | 0,5347 | 3,19227E-15 |
| ACE2 | RN7SKP204 | 0,5346 | 3,24191E-15 |
| ACE2 | RP11-104J23.2 | 0,5345 | 3,2898E-15 |
| ACE2 | NAT16 | 0,5344 | 3,3642E-15 |
| ACE2 | GPX2 | 0,5344 | 3,32907E-15 |
| ACE2 | MRPL22P1 | 0,5343 | 3,39532E-15 |
| ACE2 | RP1-317E23.3 | 0,5342 | 3,44023E-15 |
| ACE2 | CTD-2256P15.4 | 0,5342 | 3,44903E-15 |
| ACE2 | TMEM56 | 0,5338 | 3,64312E-15 |
| ACE2 | GULOP | 0,5337 | 3,70107E-15 |
| ACE2 | KDELR2 | 0,5334 | 3,83065E-15 |
| ACE2 | URGCP | 0,5333 | 3,92469E-15 |
| ACE2 | AC114812.9 | 0,5332 | 3,93346E-15 |
| ACE2 | IGF2BP2-AS1 | 0,5331 | 4,0409E-15 |

| | | | |
|---|---|---|---|
| ACE2 | RSPH3 | 0,533 | 4,08712E-15 |
| ACE2 | GOLM1 | 0,5328 | 4,16809E-15 |
| ACE2 | LTK | 0,5328 | 4,16049E-15 |
| ACE2 | UGT2B11 | 0,5326 | 4,27889E-15 |
| ACE2 | DOT1L | 0,5325 | 4,38994E-15 |
| ACE2 | PLXNA2 | 0,5324 | 4,45439E-15 |
| ACE2 | PLAC8 | 0,5317 | 4,9098E-15 |
| ACE2 | CDK18 | 0,5313 | 5,12747E-15 |
| ACE2 | CRYBB3 | 0,531 | 5,41069E-15 |
| ACE2 | AC011288.2 | 0,5309 | 5,48566E-15 |
| ACE2 | MBOAT1 | 0,5308 | 5,53998E-15 |
| ACE2 | TNNC2 | 0,5306 | 5,70974E-15 |
| ACE2 | CHKA | 0,5304 | 5,82526E-15 |
| ACE2 | FAM46A | 0,5298 | 6,36122E-15 |
| ACE2 | AC112229.6 | 0,5296 | 6,50224E-15 |
| ACE2 | ZNF213 | 0,5295 | 6,59449E-15 |
| ACE2 | POFUT1 | 0,5294 | 6,75308E-15 |
| ACE2 | RP11-432I5.8 | 0,5292 | 6,94163E-15 |
| ACE2 | PCSK9 | 0,5291 | 6,96373E-15 |
| ACE2 | MINPP1 | 0,5284 | 7,73736E-15 |
| ACE2 | CTAGE3P | 0,5283 | 7,8373E-15 |
| ACE2 | RP11-10N16.2 | 0,5278 | 8,39242E-15 |
| ACE2 | RP11-392P7.6 | 0,5277 | 8,46293E-15 |
| ACE2 | AC012507.3 | 0,5274 | 8,85165E-15 |
| ACE2 | CPPED1 | 0,5273 | 8,91583E-15 |
| ACE2 | PCYT1A | 0,527 | 9,29766E-15 |
| ACE2 | SERPINB6 | 0,5266 | 9,90046E-15 |
| ACE2 | SMAP1 | 0,5264 | 1,01065E-14 |
| ACE2 | GSTK1 | 0,5264 | 1,01428E-14 |
| ACE2 | TTC39B | 0,5262 | 1,0426E-14 |
| ACE2 | RP11-132E11.4 | 0,5262 | 1,03899E-14 |
| ACE2 | IMPG1 | 0,5261 | 1,05419E-14 |
| ACE2 | HNRNPA1P62 | 0,5261 | 1,06057E-14 |
| ACE2 | NQO1 | 0,5258 | 1,09334E-14 |
| ACE2 | IDNK | 0,5255 | 1,1483E-14 |
| ACE2 | RP11-44F21.5 | 0,5253 | 1,18456E-14 |
| ACE2 | RP11-57H12.2 | 0,5251 | 1,2089E-14 |
| ACE2 | RNF19B | 0,5249 | 1,23669E-14 |
| ACE2 | MOB3B | 0,5249 | 1,23808E-14 |
| ACE2 | TNIP2 | 0,5246 | 1,29996E-14 |
| ACE2 | GSG1 | 0,5246 | 1,28694E-14 |
| ACE2 | CASK | 0,5246 | 1,29979E-14 |
| ACE2 | STEAP3 | 0,5244 | 1,33468E-14 |
| ACE2 | SLC1A7 | 0,5243 | 1,33909E-14 |
| ACE2 | LZTS3 | 0,524 | 1,40036E-14 |
| ACE2 | AP006748.1 | 0,5238 | 1,43327E-14 |
| ACE2 | GRAMD3 | 0,5237 | 1,47002E-14 |
| ACE2 | KRT8P13 | 0,5235 | 1,49419E-14 |
| ACE2 | RPS3AP40 | 0,5233 | 1,54102E-14 |
| ACE2 | PSCA | 0,5231 | 1,5823E-14 |
| ACE2 | CFAP99 | 0,523 | 1,60261E-14 |
| ACE2 | ICA1 | 0,5228 | 1,65087E-14 |
| ACE2 | KLK12 | 0,5228 | 1,64978E-14 |
| ACE2 | RP5-1074L1.1 | 0,5224 | 1,73738E-14 |

| | | | |
|---|---|---|---|
| ACE2 | ABRACL | 0,5223 | 1,7609E-14 |
| ACE2 | GLCE | 0,522 | 1,83882E-14 |
| ACE2 | GPD2 | 0,5218 | 1,89436E-14 |
| ACE2 | EHMT2-AS1 | 0,5217 | 1,90196E-14 |
| ACE2 | RP11-568A7.3 | 0,5212 | 2,04505E-14 |
| ACE2 | PPP1R13B | 0,521 | 2,1077E-14 |
| ACE2 | RP11-486M3.2 | 0,5209 | 2,14463E-14 |
| ACE2 | LINC01975 | 0,5209 | 2,13017E-14 |
| ACE2 | RPS12P2 | 0,5205 | 2,23893E-14 |
| ACE2 | DBI | 0,5203 | 2,31832E-14 |
| ACE2 | TMEM135 | 0,5199 | 2,44132E-14 |
| ACE2 | BCL10 | 0,5195 | 2,57817E-14 |
| ACE2 | WNK2 | 0,5195 | 2,55656E-14 |
| ACE2 | AIMP2 | 0,5194 | 2,60164E-14 |
| ACE2 | COX8A | 0,5194 | 2,60109E-14 |
| ACE2 | BCL2L1 | 0,5194 | 2,59687E-14 |
| ACE2 | RP11-438P9.2 | 0,5193 | 2,64549E-14 |
| ACE2 | CYSTM1 | 0,5188 | 2,81111E-14 |
| ACE2 | ANKRD18A | 0,5187 | 2,85064E-14 |
| ACE2 | RP11-5O24.1 | 0,518 | 3,1377E-14 |
| ACE2 | RP11-734K21.3 | 0,5178 | 3,2129E-14 |
| ACE2 | CA12 | 0,5177 | 3,28675E-14 |
| ACE2 | DDAH1 | 0,5176 | 3,29055E-14 |
| ACE2 | ENDOG | 0,5176 | 3,32606E-14 |
| ACE2 | RP1-228P16.5 | 0,5176 | 3,31692E-14 |
| ACE2 | RP11-297C4.2 | 0,5171 | 3,54304E-14 |
| ACE2 | SLFNL1-AS1 | 0,517 | 3,59799E-14 |
| ACE2 | SULT1D1P | 0,5169 | 3,62742E-14 |
| ACE2 | TMED6 | 0,5168 | 3,6888E-14 |
| ACE2 | CCL25 | 0,5166 | 3,76753E-14 |
| ACE2 | DUSP27 | 0,5165 | 3,83751E-14 |
| ACE2 | NEK3 | 0,5165 | 3,84323E-14 |
| ACE2 | NCSTN | 0,5163 | 3,94471E-14 |
| ACE2 | ADAT3 | 0,5161 | 4,05103E-14 |
| ACE2 | RP11-28H5.2 | 0,5159 | 4,11928E-14 |
| ACE2 | RP11-361F15.5 | 0,5156 | 4,29689E-14 |
| ACE2 | RP11-24F11.2 | 0,5155 | 4,35734E-14 |
| ACE2 | AC074011.2 | 0,5154 | 4,4511E-14 |
| ACE2 | AC142119.1 | 0,5151 | 4,61988E-14 |
| ACE2 | RETNLB | 0,5151 | 4,58209E-14 |
| ACE2 | MIR210 | 0,5151 | 4,61632E-14 |
| ACE2 | AGAP13P | 0,5149 | 4,71243E-14 |
| ACE2 | FRMD8 | 0,5149 | 4,74752E-14 |
| ACE2 | RP3-446N13.5 | 0,5144 | 5,05355E-14 |
| ACE2 | MIR22HG | 0,5144 | 5,07524E-14 |
| ACE2 | LINC01214 | 0,5143 | 5,11618E-14 |
| ACE2 | SEMA6A | 0,5143 | 5,11387E-14 |
| ACE2 | KB-1410C5.3 | 0,5143 | 5,09461E-14 |
| ACE2 | AC009495.2 | 0,5142 | 5,18432E-14 |
| ACE2 | TRPV1 | 0,5137 | 5,51155E-14 |
| ACE2 | RNA5SP434 | 0,5137 | 5,54249E-14 |
| ACE2 | RP11-67L3.2 | 0,5136 | 5,61544E-14 |
| ACE2 | KRTAP5-5 | 0,5135 | 5,65459E-14 |
| ACE2 | LCN2 | 0,5134 | 5,75914E-14 |

| | | | |
|---|---|---|---|
| ACE2 | CTD-2600O9.2 | 0,5134 | 5,75136E-14 |
| ACE2 | LCOR | 0,5132 | 5,87698E-14 |
| ACE2 | OXER1 | 0,5124 | 6,55251E-14 |
| ACE2 | MB21D2 | 0,5121 | 6,86319E-14 |
| ACE2 | AC087491.2 | 0,5121 | 6,84484E-14 |
| ACE2 | ZNF326 | 0,5118 | 7,05335E-14 |
| ACE2 | CCNJL | 0,5118 | 7,12214E-14 |
| ACE2 | WFDC12 | 0,5113 | 7,53525E-14 |
| ACE2 | MYL5 | 0,5111 | 7,79776E-14 |
| ACE2 | FAM95C | 0,511 | 7,8896E-14 |
| ACE2 | ADCY6 | 0,5109 | 7,98244E-14 |
| ACE2 | AC022431.3 | 0,5108 | 8,03532E-14 |
| ACE2 | AXDND1 | 0,5106 | 8,31949E-14 |
| ACE2 | ANO3 | 0,5104 | 8,53293E-14 |
| ACE2 | OAF | 0,5104 | 8,47633E-14 |
| ACE2 | LDHA | 0,5103 | 8,67429E-14 |
| ACE2 | RHOC | 0,5096 | 9,49462E-14 |
| ACE2 | TLR3 | 0,5095 | 9,61611E-14 |
| ACE2 | LINC00867 | 0,5095 | 9,52249E-14 |
| ACE2 | ABCA13 | 0,5092 | 9,89844E-14 |
| ACE2 | MT-CO2 | 0,5092 | 9,97373E-14 |
| ACE2 | PQLC2 | 0,5088 | 1,04615E-13 |
| ACE2 | HMGCL | 0,5088 | 1,04054E-13 |
| ACE2 | FABP12 | 0,5088 | 1,04961E-13 |
| ACE2 | U47924.29 | 0,5087 | 1,05776E-13 |
| ACE2 | RP11-835E18.5 | 0,5087 | 1,05674E-13 |
| ACE2 | RP11-720N19.1 | 0,5084 | 1,10056E-13 |
| ACE2 | EFL1 | 0,5078 | 1,19572E-13 |
| ACE2 | GFI1B | 0,5074 | 1,25486E-13 |
| ACE2 | DRAIC | 0,5072 | 1,2783E-13 |
| ACE2 | ABHD12 | 0,5071 | 1,29575E-13 |
| ACE2 | FNBP1L | 0,507 | 1,32366E-13 |
| ACE2 | SDF4 | 0,5069 | 1,34348E-13 |
| ACE2 | NCOA4 | 0,5069 | 1,32928E-13 |
| ACE2 | PKP1 | 0,5067 | 1,36405E-13 |
| ACE2 | RN7SL8P | 0,5066 | 1,3961E-13 |
| ACE2 | RP11-982M15.6 | 0,5064 | 1,42261E-13 |
| ACE2 | KRT8P36 | 0,5062 | 1,45816E-13 |
| ACE2 | PSORS1C1 | 0,5055 | 1,59396E-13 |
| ACE2 | MICALL2 | 0,5055 | 1,59466E-13 |
| ACE2 | RP11-507E23.1 | 0,5054 | 1,61507E-13 |
| ACE2 | RP11-736N17.8 | 0,5051 | 1,67484E-13 |
| ACE2 | LNX1 | 0,5048 | 1,75458E-13 |
| ACE2 | KLF4 | 0,5047 | 1,77399E-13 |
| ACE2 | GAPDHP23 | 0,5046 | 1,78006E-13 |
| ACE2 | GLIS3-AS1 | 0,5046 | 1,78772E-13 |
| ACE2 | PLOD2 | 0,5045 | 1,82058E-13 |
| ACE2 | MT-TY | 0,5042 | 1,89057E-13 |
| ACE2 | RP11-440I14.3 | 0,5039 | 1,9654E-13 |
| ACE2 | RP11-67L14.1 | 0,5038 | 1,97198E-13 |
| ACE2 | POU2F3 | 0,5038 | 1,98765E-13 |
| ACE2 | CEACAM22P | 0,5033 | 2,10339E-13 |
| ACE2 | RP11-661A12.5 | 0,5031 | 2,16985E-13 |
| ACE2 | ATP6V1B2 | 0,5025 | 2,34521E-13 |

| | | | |
|---|---|---|---|
| ACE2 | CTB-39G8.3 | 0,5023 | 2,38818E-13 |
| ACE2 | PDK2 | 0,5022 | 2,42939E-13 |
| ACE2 | KRT8P11 | 0,5019 | 2,52422E-13 |
| ACE2 | AHCYL2 | 0,5014 | 2,69505E-13 |
| ACE2 | VPS13A | 0,5014 | 2,68077E-13 |
| ACE2 | DSG3 | 0,5014 | 2,66842E-13 |
| ACE2 | RBMY2TP | 0,5011 | 2,77319E-13 |
| ACE2 | KLK15 | 0,5006 | 2,97114E-13 |
| ACE2 | CYC1 | 0,5005 | 3,02154E-13 |
| ACE2 | FRRS1 | 0,5003 | 3,08318E-13 |
| ACE2 | DHRS1 | 0,5001 | 3,1784E-13 |
| ACE2 | RP11-349A8.3 | 0,5001 | 3,14407E-13 |
| ACE2 | NOCT | 0,5 | 3,18388E-13 |
| ACE2 | RP11-369C8.1 | 0,5 | 3,21498E-13 |
| ACE2 | TSTA3 | 0,4999 | 3,22124E-13 |
| ACE2 | COA3 | 0,4998 | 3,28902E-13 |
| ACE2 | METRNL | 0,4998 | 3,28807E-13 |
| ACE2 | KRT8P5 | 0,4997 | 3,33923E-13 |
| ACE2 | GHRH | 0,4997 | 3,32142E-13 |
| ACE2 | RP11-150O12.6 | 0,4996 | 3,37386E-13 |
| ACE2 | ECI1 | 0,4996 | 3,3718E-13 |
| ACE2 | GNG12 | 0,4994 | 3,45611E-13 |
| ACE2 | IL6R | 0,4992 | 3,52606E-13 |
| ACE2 | CCDC186 | 0,499 | 3,61765E-13 |
| ACE2 | PGM1 | 0,4988 | 3,72517E-13 |
| ACE2 | HAO2 | 0,4984 | 3,89608E-13 |
| ACE2 | RTKN | 0,4984 | 3,89304E-13 |
| ACE2 | ALDH3B1 | 0,4983 | 3,94964E-13 |
| ACE2 | ETFB | 0,4982 | 3,99766E-13 |
| ACE2 | CA1 | 0,4973 | 4,47277E-13 |
| ACE2 | AACS | 0,4969 | 4,72548E-13 |
| ACE2 | RP11-465N4.5 | 0,4963 | 5,04293E-13 |
| ACE2 | SLC12A7 | 0,4963 | 5,05799E-13 |
| ACE2 | PRADC1 | 0,4958 | 5,36909E-13 |
| ACE2 | CTD-2235C13.3 | 0,4958 | 5,40972E-13 |
| ACE2 | TMEM102 | 0,4956 | 5,50544E-13 |
| ACE2 | GGTLC5P | 0,4956 | 5,51046E-13 |
| ACE2 | SULT1C2P1 | 0,4955 | 5,62536E-13 |
| ACE2 | RP11-887P2.6 | 0,4953 | 5,70587E-13 |
| ACE2 | RP11-503L19.1 | 0,4952 | 5,78315E-13 |
| ACE2 | RP11-416N2.3 | 0,4952 | 5,80761E-13 |
| ACE2 | FAM134B | 0,495 | 5,95452E-13 |
| ACE2 | FN3K | 0,4948 | 6,07779E-13 |
| ACE2 | KCNE3 | 0,4943 | 6,53304E-13 |
| ACE2 | SOX9-AS1 | 0,4942 | 6,543E-13 |
| ACE2 | C5orf67 | 0,494 | 6,70345E-13 |
| ACE2 | KRT8P31 | 0,494 | 6,77805E-13 |
| ACE2 | SOCS5P4 | 0,494 | 6,70962E-13 |
| ACE2 | GAPDHP40 | 0,4938 | 6,88913E-13 |
| ACE2 | UPK3A | 0,4938 | 6,93472E-13 |
| ACE2 | RP11-46I1.2 | 0,4937 | 7,00788E-13 |
| ACE2 | RP11-407G23.2 | 0,4935 | 7,1916E-13 |
| ACE2 | RP1-95L4.4 | 0,4934 | 7,2446E-13 |
| ACE2 | RP11-231E6.1 | 0,4933 | 7,32E-13 |

| | | | |
|---|---|---|---|
| ACE2 | PYROXD1 | 0,4932 | 7,41418E-13 |
| ACE2 | ALG14 | 0,493 | 7,65314E-13 |
| ACE2 | MIR3944 | 0,4929 | 7,71652E-13 |
| ACE2 | RP4-680D5.8 | 0,4928 | 7,82917E-13 |
| ACE2 | SLC30A1 | 0,4927 | 7,87541E-13 |
| ACE2 | RNF157-AS1 | 0,4926 | 7,97031E-13 |
| ACE2 | MT-ND5 | 0,4926 | 8,02595E-13 |
| ACE2 | RP11-285E9.6 | 0,4921 | 8,52124E-13 |
| ACE2 | RP11-341G23.3 | 0,4918 | 8,81786E-13 |
| ACE2 | SLC28A3 | 0,4906 | 1,01741E-12 |
| ACE2 | NAPEPLD | 0,4902 | 1,07157E-12 |
| ACE2 | LINC00605 | 0,4899 | 1,11208E-12 |
| ACE2 | AC015971.2 | 0,4898 | 1,12448E-12 |
| ACE2 | RP11-362L22.1 | 0,4898 | 1,11974E-12 |
| ACE2 | JAGN1 | 0,4897 | 1,13918E-12 |
| ACE2 | VEGFA | 0,4897 | 1,13683E-12 |
| ACE2 | RP11-416N2.4 | 0,4894 | 1,18279E-12 |
| ACE2 | LIMA1 | 0,4893 | 1,18941E-12 |
| ACE2 | PEBP1 | 0,4893 | 1,19312E-12 |
| ACE2 | RNF121 | 0,4886 | 1,29429E-12 |
| ACE2 | NECTIN1 | 0,4886 | 1,30515E-12 |
| ACE2 | UGT2B27P | 0,4885 | 1,31959E-12 |
| ACE2 | DEPDC1B | 0,4884 | 1,33263E-12 |
| ACE2 | RP11-650P15.1 | 0,4883 | 1,34857E-12 |
| ACE2 | LINC00114 | 0,4883 | 1,34094E-12 |
| ACE2 | CTD-2280E9.1 | 0,4879 | 1,42266E-12 |
| ACE2 | RP11-386M24.3 | 0,4878 | 1,43701E-12 |
| ACE2 | ACAA1 | 0,4874 | 1,49617E-12 |
| ACE2 | PRDX5 | 0,4874 | 1,49366E-12 |
| ACE2 | GK5 | 0,4873 | 1,51775E-12 |
| ACE2 | CTA-243E7.2 | 0,4869 | 1,58836E-12 |
| ACE2 | ELOVL6 | 0,4867 | 1,62729E-12 |
| ACE2 | AP000344.4 | 0,4867 | 1,63764E-12 |
| ACE2 | PLGLA | 0,4865 | 1,66405E-12 |
| ACE2 | RRAS2 | 0,4865 | 1,67347E-12 |
| ACE2 | LINC00574 | 0,4863 | 1,71392E-12 |
| ACE2 | CTA-221G9.7 | 0,4863 | 1,71194E-12 |
| ACE2 | LY75 | 0,4859 | 1,79245E-12 |
| ACE2 | NEU1 | 0,4859 | 1,79235E-12 |
| ACE2 | RP11-479A21.1 | 0,4859 | 1,7922E-12 |
| ACE2 | CTB-161C1.1 | 0,4856 | 1,86204E-12 |
| ACE2 | CTC-756D1.3 | 0,4856 | 1,86745E-12 |
| ACE2 | USP30 | 0,4856 | 1,86677E-12 |
| ACE2 | ZNF440 | 0,4855 | 1,88637E-12 |
| ACE2 | ACVR2A | 0,4854 | 1,90868E-12 |
| ACE2 | PLIN2 | 0,4854 | 1,91271E-12 |
| ACE2 | ANAPC1P1 | 0,4853 | 1,92206E-12 |
| ACE2 | ABHD2 | 0,4853 | 1,92692E-12 |
| ACE2 | KRT8P33 | 0,4852 | 1,95948E-12 |
| ACE2 | RP11-575A19.2 | 0,4851 | 1,97189E-12 |
| ACE2 | FIBCD1 | 0,4848 | 2,04804E-12 |
| ACE2 | RP11-661A12.9 | 0,4843 | 2,16563E-12 |
| ACE2 | SPATA2L | 0,4841 | 2,24176E-12 |
| ACE2 | TDRD5 | 0,484 | 2,26345E-12 |

| | | | |
|---|---|---|---|
| ACE2 | ASS1P11 | 0,4839 | 2,28642E-12 |
| ACE2 | MT-TS1 | 0,4839 | 2,2741E-12 |
| ACE2 | MTCO1P12 | 0,4833 | 2,44229E-12 |
| ACE2 | RP11-2I17.4 | 0,4832 | 2,47796E-12 |
| ACE2 | CNTFR-AS1 | 0,4829 | 2,57073E-12 |
| ACE2 | TMED4 | 0,4828 | 2,61169E-12 |
| ACE2 | MT-TC | 0,4826 | 2,66882E-12 |
| ACE2 | ADRA1B | 0,4824 | 2,72959E-12 |
| ACE2 | AP002856.5 | 0,4824 | 2,72743E-12 |
| ACE2 | RP5-1057J7.7 | 0,4816 | 2,98351E-12 |
| ACE2 | CASZ1 | 0,4813 | 3,11817E-12 |
| ACE2 | RP11-525G13.2 | 0,4813 | 3,10816E-12 |
| ACE2 | GMFBP1 | 0,4813 | 3,11038E-12 |
| ACE2 | RREB1 | 0,4811 | 3,17295E-12 |
| ACE2 | PAX4 | 0,481 | 3,22626E-12 |
| ACE2 | CFB | 0,4809 | 3,26623E-12 |
| ACE2 | RP11-429J17.5 | 0,4809 | 3,2369E-12 |
| ACE2 | ENPP7P7 | 0,4808 | 3,28039E-12 |
| ACE2 | RFPL4A | 0,4808 | 3,31016E-12 |
| ACE2 | TET3 | 0,4807 | 3,33817E-12 |
| ACE2 | PMM1 | 0,4807 | 3,33906E-12 |
| ACE2 | EFNA3 | 0,4805 | 3,40373E-12 |
| ACE2 | TBC1D30 | 0,4805 | 3,4229E-12 |
| ACE2 | TMEM52 | 0,4804 | 3,44038E-12 |
| ACE2 | RP3-365O12.2 | 0,4803 | 3,48873E-12 |
| ACE2 | COL2A1 | 0,4801 | 3,56276E-12 |
| ACE2 | MYZAP | 0,4797 | 3,74832E-12 |
| ACE2 | PUDP | 0,4796 | 3,79436E-12 |
| ACE2 | SPSB1 | 0,4793 | 3,93097E-12 |
| ACE2 | LINC02029 | 0,4793 | 3,91213E-12 |
| ACE2 | RP5-892K4.1 | 0,4792 | 3,95958E-12 |
| ACE2 | RP11-386M24.6 | 0,4791 | 4,0001E-12 |
| ACE2 | EDF1 | 0,479 | 4,07817E-12 |
| ACE2 | RNU6-1065P | 0,4789 | 4,09631E-12 |
| ACE2 | RP11-100N20.1 | 0,4788 | 4,17636E-12 |
| ACE2 | MT-TE | 0,4788 | 4,15081E-12 |
| ACE2 | LINC01014 | 0,4787 | 4,2343E-12 |
| ACE2 | SNHG18 | 0,4785 | 4,32054E-12 |
| ACE2 | BRPF3 | 0,4783 | 4,43843E-12 |
| ACE2 | F11-AS1 | 0,4779 | 4,6011E-12 |
| ACE2 | RPL10L | 0,4779 | 4,65136E-12 |
| ACE2 | KB-1460A1.1 | 0,4778 | 4,68745E-12 |
| ACE2 | RP11-116O18.1 | 0,4776 | 4,81128E-12 |
| ACE2 | NT5DC1 | 0,4773 | 4,96294E-12 |
| ACE2 | RP1-170O19.24 | 0,477 | 5,15421E-12 |
| ACE2 | ARHGAP8 | 0,4765 | 5,47551E-12 |
| ACE2 | AC017074.2 | 0,4764 | 5,48843E-12 |
| ACE2 | TMEM181 | 0,4762 | 5,63471E-12 |
| ACE2 | RP11-115D19.1 | 0,4758 | 5,87506E-12 |
| ACE2 | CD58 | 0,4757 | 6,00082E-12 |
| ACE2 | RP11-122K13.7 | 0,4757 | 5,94545E-12 |
| ACE2 | FRYL | 0,4756 | 6,07768E-12 |
| ACE2 | C9orf24 | 0,4755 | 6,1213E-12 |
| ACE2 | LINC00244 | 0,4754 | 6,19551E-12 |

| | | | |
|---|---|---|---|
| ACE2 | SLC39A7 | 0,4753 | 6,28502E-12 |
| ACE2 | HMGA2 | 0,4753 | 6,28867E-12 |
| ACE2 | NXF3 | 0,4753 | 6,22826E-12 |
| ACE2 | GUSBP8 | 0,4748 | 6,66131E-12 |
| ACE2 | DHDDS | 0,4739 | 7,38194E-12 |
| ACE2 | MLN | 0,4733 | 7,85823E-12 |
| ACE2 | RP11-546O6.4 | 0,4732 | 8,01381E-12 |
| ACE2 | RP11-720D4.3 | 0,4732 | 8,0031E-12 |
| ACE2 | AC034220.3 | 0,4727 | 8,40992E-12 |
| ACE2 | CTB-60B18.10 | 0,4727 | 8,41523E-12 |
| ACE2 | RNU6-796P | 0,4725 | 8,62463E-12 |
| ACE2 | SPATA33 | 0,4724 | 8,72975E-12 |
| ACE2 | RP11-769O8.3 | 0,4724 | 8,7799E-12 |
| ACE2 | RP11-616M22.7 | 0,4722 | 8,95341E-12 |
| ACE2 | RP11-359I18.1 | 0,472 | 9,1865E-12 |
| ACE2 | RPL7L1P11 | 0,4712 | 1,00623E-11 |
| ACE2 | PKDREJ | 0,4711 | 1,01538E-11 |
| ACE2 | RP11-172C16.5 | 0,4708 | 1,04899E-11 |
| ACE2 | RP5-965G21.4 | 0,4707 | 1,05604E-11 |
| ACE2 | SIGLEC12 | 0,4704 | 1,09937E-11 |
| ACE2 | AC067959.1 | 0,4702 | 1,11818E-11 |
| ACE2 | RP11-497G19.2 | 0,47 | 1,15356E-11 |
| ACE2 | ADCY10 | 0,4699 | 1,15899E-11 |
| ACE2 | RP11-704M14.1 | 0,4699 | 1,15885E-11 |
| ACE2 | GNAT3 | 0,4696 | 1,20766E-11 |
| ACE2 | KRT14 | 0,4695 | 1,21405E-11 |
| ACE2 | AC022201.4 | 0,4689 | 1,29402E-11 |
| ACE2 | DUSP18 | 0,4689 | 1,30244E-11 |
| ACE2 | PSORS1C2 | 0,4685 | 1,35678E-11 |
| ACE2 | NRARP | 0,4684 | 1,3828E-11 |
| ACE2 | AK7 | 0,4684 | 1,38446E-11 |
| ACE2 | RP11-649A18.4 | 0,468 | 1,43522E-11 |
| ACE2 | SCML2P1 | 0,4679 | 1,46084E-11 |
| ACE2 | CCL28 | 0,4676 | 1,50209E-11 |
| ACE2 | ARRDC1 | 0,4676 | 1,51477E-11 |
| ACE2 | ENC1 | 0,4675 | 1,52212E-11 |
| ACE2 | MIR3646 | 0,4675 | 1,51734E-11 |
| ACE2 | RP11-739N20.2 | 0,4672 | 1,57401E-11 |
| ACE2 | SFXN5 | 0,4669 | 1,62522E-11 |
| ACE2 | TRIM29 | 0,4668 | 1,65024E-11 |
| ACE2 | ANXA2 | 0,4667 | 1,66964E-11 |
| ACE2 | MARK2 | 0,4666 | 1,67823E-11 |
| ACE2 | CIB1 | 0,4665 | 1,711E-11 |
| ACE2 | MAGIX | 0,4663 | 1,74102E-11 |
| ACE2 | MLIP-IT1 | 0,4662 | 1,76912E-11 |
| ACE2 | AP000437.3 | 0,4662 | 1,76241E-11 |
| ACE2 | AC004593.3 | 0,4661 | 1,78115E-11 |
| ACE2 | ACOX2 | 0,4659 | 1,81604E-11 |
| ACE2 | APOBR | 0,4658 | 1,84969E-11 |
| ACE2 | NPY | 0,4654 | 1,92672E-11 |
| ACE2 | ZNF3 | 0,4652 | 1,97394E-11 |
| ACE2 | CABP7 | 0,4652 | 1,9743E-11 |
| ACE2 | RP5-1024N4.4 | 0,4648 | 2,06327E-11 |
| ACE2 | MIER3 | 0,4648 | 2,06219E-11 |

| | | | |
|---|---|---|---|
| ACE2 | CPNE2 | 0,4648 | 2,05471E-11 |
| ACE2 | KRT80 | 0,4647 | 2,07916E-11 |
| ACE2 | AC090505.1 | 0,4646 | 2,10327E-11 |
| ACE2 | RP11-499O7.7 | 0,4644 | 2,16104E-11 |
| ACE2 | CLUH | 0,4644 | 2,15266E-11 |
| ACE2 | SPAG5 | 0,4643 | 2,17683E-11 |
| ACE2 | OR7E117P | 0,4639 | 2,28361E-11 |
| ACE2 | MAP2K6 | 0,4637 | 2,32649E-11 |
| ACE2 | MAP7D2 | 0,4637 | 2,33507E-11 |
| ACE2 | OR7E83P | 0,4635 | 2,38513E-11 |
| ACE2 | VWA3B | 0,4631 | 2,50525E-11 |
| ACE2 | RP11-555M1.3 | 0,4629 | 2,54066E-11 |
| ACE2 | CCNI2 | 0,4627 | 2,61589E-11 |
| ACE2 | C6orf223 | 0,4624 | 2,70249E-11 |
| ACE2 | YWHAZP5 | 0,4624 | 2,70706E-11 |
| ACE2 | FAM9C | 0,4624 | 2,68225E-11 |
| ACE2 | RP11-545D22.1 | 0,462 | 2,82104E-11 |
| ACE2 | CTA-363E6.3 | 0,4618 | 2,88142E-11 |
| ACE2 | RP11-260N14.1 | 0,4616 | 2,95735E-11 |
| ACE2 | FAR2 | 0,4616 | 2,95171E-11 |
| ACE2 | RP11-345K9.3 | 0,4615 | 2,98243E-11 |
| ACE2 | CTD-2651B20.4 | 0,4615 | 2,97362E-11 |
| ACE2 | RP11-1038A11.3 | 0,4614 | 3,01348E-11 |
| ACE2 | RP11-392P7.7 | 0,4614 | 2,99797E-11 |
| ACE2 | ARHGAP27 | 0,4613 | 3,05843E-11 |
| ACE2 | MVK | 0,4611 | 3,09656E-11 |
| ACE2 | SLFNL1 | 0,4609 | 3,18893E-11 |
| ACE2 | LNX1-AS2 | 0,4609 | 3,1745E-11 |
| ACE2 | PTGER4 | 0,4608 | 3,20484E-11 |
| ACE2 | MMP24-AS1 | 0,4604 | 3,36113E-11 |
| ACE2 | PTPRK | 0,4602 | 3,42488E-11 |
| ACE2 | ALDH3A2 | 0,4602 | 3,45055E-11 |
| ACE2 | ZNF165 | 0,4598 | 3,59582E-11 |
| ACE2 | F7 | 0,4598 | 3,57473E-11 |
| ACE2 | RP11-351J23.1 | 0,4597 | 3,6197E-11 |
| ACE2 | RP11-265B7.1 | 0,4596 | 3,66389E-11 |
| ACE2 | RP11-331F4.5 | 0,4594 | 3,73785E-11 |
| ACE2 | RP1-92C8.3 | 0,4593 | 3,81277E-11 |
| ACE2 | IGHVII-78-1 | 0,4593 | 3,79656E-11 |
| ACE2 | AC226118.1 | 0,4592 | 3,83341E-11 |
| ACE2 | LINC02113 | 0,4591 | 3,87823E-11 |
| ACE2 | UNC5D | 0,4591 | 3,89922E-11 |
| ACE2 | TRMT10A | 0,459 | 3,91317E-11 |
| ACE2 | OR2S1P | 0,459 | 3,93212E-11 |
| ACE2 | SRD5A1 | 0,4585 | 4,16327E-11 |
| ACE2 | NOS1AP | 0,4583 | 4,23004E-11 |
| ACE2 | RP11-665E10.2 | 0,4583 | 4,22129E-11 |
| ACE2 | APOBEC4 | 0,4582 | 4,27107E-11 |
| ACE2 | SH2D1B | 0,4581 | 4,33972E-11 |
| ACE2 | RPSAP69 | 0,4581 | 4,32305E-11 |
| ACE2 | CHRFAM7A | 0,4579 | 4,44157E-11 |
| ACE2 | NDFIP1P1 | 0,4578 | 4,46279E-11 |
| ACE2 | AC104534.2 | 0,4575 | 4,63313E-11 |
| ACE2 | ASCL2 | 0,4574 | 4,68637E-11 |

| | | | |
|---|---|---|---|
| ACE2 | MS4A12 | 0,4571 | 4,83671E-11 |
| ACE2 | C2orf48 | 0,4568 | 4,96797E-11 |
| ACE2 | AC128709.4 | 0,4568 | 4,96495E-11 |
| ACE2 | PLCXD2 | 0,4565 | 5,15834E-11 |
| ACE2 | VWA5A | 0,4562 | 5,35216E-11 |
| ACE2 | NOS2 | 0,4562 | 5,32096E-11 |
| ACE2 | ALKAL1 | 0,4559 | 5,49976E-11 |
| ACE2 | RP11-1267H10.3 | 0,4558 | 5,53669E-11 |
| ACE2 | RP11-503C24.4 | 0,4557 | 5,60042E-11 |
| ACE2 | RP11-384J4.2 | 0,4557 | 5,64103E-11 |
| ACE2 | PLA2G2A | 0,4555 | 5,73423E-11 |
| ACE2 | CLTB | 0,4555 | 5,74638E-11 |
| ACE2 | SNX24 | 0,4554 | 5,82525E-11 |
| ACE2 | AMBP | 0,4552 | 5,92403E-11 |
| ACE2 | LINC01555 | 0,4548 | 6,16349E-11 |
| ACE2 | UNC93B7 | 0,4543 | 6,51279E-11 |
| ACE2 | TULP4 | 0,4536 | 7,04309E-11 |
| ACE2 | HMOX1 | 0,4534 | 7,20925E-11 |
| ACE2 | CHCHD2P1 | 0,4527 | 7,76635E-11 |
| ACE2 | ZBTB45P2 | 0,4525 | 7,91072E-11 |
| ACE2 | TRHDE | 0,4521 | 8,30176E-11 |
| ACE2 | SPATA31C2 | 0,452 | 8,38583E-11 |
| ACE2 | RN7SL125P | 0,4519 | 8,51326E-11 |
| ACE2 | LRRFIP2 | 0,4513 | 8,99476E-11 |
| ACE2 | MIGA2 | 0,4512 | 9,09785E-11 |
| ACE2 | CERS6-AS1 | 0,4511 | 9,27189E-11 |
| ACE2 | LINC01267 | 0,4511 | 9,18455E-11 |
| ACE2 | NSUN7 | 0,4511 | 9,26234E-11 |
| ACE2 | PCSK1 | 0,4511 | 9,26941E-11 |
| ACE2 | LINC02159 | 0,451 | 9,28556E-11 |
| ACE2 | RAP1GAP2 | 0,4509 | 9,43595E-11 |
| ACE2 | TMEM132D | 0,4508 | 9,50093E-11 |
| ACE2 | MTCO2P12 | 0,4506 | 9,76667E-11 |
| ACE2 | TNIP1 | 0,4506 | 9,73431E-11 |
| ACE2 | CATSPERB | 0,4505 | 9,85235E-11 |
| ACE2 | SLC14A2 | 0,4505 | 9,827E-11 |
| ACE2 | MTND5P11 | 0,4504 | 9,97654E-11 |
| ACE2 | RP11-96L14.9 | 0,4503 | 1,00715E-10 |
| ACE2 | TMEM252 | 0,4503 | 1,00235E-10 |
| ACE2 | LAPTM4B | 0,4502 | 1,01185E-10 |
| ACE2 | CMPK1 | 0,45 | 1,03966E-10 |
| ACE2 | RAB8A | 0,45 | 1,03833E-10 |
| ACE2 | RP11-466A19.1 | 0,4499 | 1,04544E-10 |
| ACE2 | PSD3 | 0,4497 | 1,06678E-10 |
| ACE2 | RP11-797A18.6 | 0,4496 | 1,08025E-10 |
| ACE2 | HSD17B4 | 0,4494 | 1,1045E-10 |
| ACE2 | NPY4R | 0,4491 | 1,14126E-10 |
| ACE2 | PGAP2 | 0,4488 | 1,18054E-10 |
| ACE2 | HIST1H3G | 0,4487 | 1,1904E-10 |
| ACE2 | ERICH5 | 0,4487 | 1,19498E-10 |
| ACE2 | LINC01687 | 0,4487 | 1,1902E-10 |
| ACE2 | LINC01910 | 0,4485 | 1,21443E-10 |
| ACE2 | KRTAP3-2 | 0,4484 | 1,22489E-10 |
| ACE2 | RP11-10C24.2 | 0,4483 | 1,23789E-10 |

| | | | |
|---|---|---|---|
| ACE2 | FEM1C | 0,4477 | 1,33022E-10 |
| ACE2 | ID1 | 0,4477 | 1,31981E-10 |
| ACE2 | SLC39A9 | 0,4474 | 1,37396E-10 |
| ACE2 | RP11-49I11.2 | 0,4473 | 1,37786E-10 |
| ACE2 | AC002128.5 | 0,447 | 1,42376E-10 |
| ACE2 | JPH1 | 0,4469 | 1,44327E-10 |
| ACE2 | SERPINA2 | 0,4469 | 1,43874E-10 |
| ACE2 | NCKAP5-IT1 | 0,4468 | 1,44953E-10 |
| ACE2 | ANXA11 | 0,4468 | 1,45772E-10 |
| ACE2 | KRT8P2 | 0,4461 | 1,5594E-10 |
| ACE2 | MTND6P4 | 0,4458 | 1,62317E-10 |
| ACE2 | SEPSECS | 0,4454 | 1,6935E-10 |
| ACE2 | DAP | 0,4454 | 1,69571E-10 |
| ACE2 | ATP8A1 | 0,4452 | 1,71404E-10 |
| ACE2 | AC005281.1 | 0,4452 | 1,72266E-10 |
| ACE2 | TMEM210 | 0,4449 | 1,77325E-10 |
| ACE2 | CTC-228N24.2 | 0,4446 | 1,83062E-10 |
| ACE2 | CH17-360D5.1 | 0,4443 | 1,8854E-10 |
| ACE2 | CYP2A7 | 0,4441 | 1,9298E-10 |
| ACE2 | TLCD1 | 0,4439 | 1,98025E-10 |
| ACE2 | TFG | 0,4437 | 2,02037E-10 |
| ACE2 | MT-ATP8 | 0,4436 | 2,03825E-10 |
| ACE2 | LINC00870 | 0,4434 | 2,07822E-10 |
| ACE2 | MLIP-AS1 | 0,4431 | 2,14896E-10 |
| ACE2 | TCEA3 | 0,443 | 2,15839E-10 |
| ACE2 | DHRS3 | 0,4422 | 2,34837E-10 |
| ACE2 | ADGRB2 | 0,4419 | 2,43205E-10 |
| ACE2 | FAM166A | 0,4416 | 2,50981E-10 |
| ACE2 | AC004231.2 | 0,4414 | 2,55841E-10 |
| ACE2 | GRHL3 | 0,4412 | 2,61232E-10 |
| ACE2 | MT-ATP6 | 0,4407 | 2,74695E-10 |
| ACE2 | HAO1 | 0,4404 | 2,83879E-10 |
| ACE2 | PNPLA2 | 0,4402 | 2,88895E-10 |
| ACE2 | LIPG | 0,4399 | 2,99361E-10 |
| ACE2 | MT-CO3 | 0,4398 | 3,01468E-10 |
| ACE2 | RP11-188C12.2 | 0,4397 | 3,04414E-10 |
| ACE2 | PLXNB2 | 0,4396 | 3,07359E-10 |
| ACE2 | C1orf53 | 0,4394 | 3,13687E-10 |
| ACE2 | EPHB2 | 0,4393 | 3,16847E-10 |
| ACE2 | SLC22A20 | 0,4392 | 3,21189E-10 |
| ACE2 | PNPLA3 | 0,4392 | 3,21838E-10 |
| ACE2 | RNA5SP151 | 0,439 | 3,29013E-10 |
| ACE2 | SLC16A13 | 0,439 | 3,27028E-10 |
| ACE2 | CEACAM16 | 0,4386 | 3,42368E-10 |
| ACE2 | PAOX | 0,4385 | 3,45489E-10 |
| ACE2 | TNK1 | 0,4384 | 3,49582E-10 |
| ACE2 | PDCD4 | 0,4382 | 3,55412E-10 |
| ACE2 | ASS1P10 | 0,438 | 3,64408E-10 |
| ACE2 | RP11-719K4.3 | 0,4379 | 3,66129E-10 |
| ACE2 | DSCAS | 0,4378 | 3,72337E-10 |
| ACE2 | MAP3K11 | 0,4377 | 3,765E-10 |
| ACE2 | WSB2 | 0,4373 | 3,88486E-10 |
| ACE2 | SLC25A5P2 | 0,4372 | 3,92417E-10 |
| ACE2 | FARP2 | 0,437 | 4,03745E-10 |

| | | | |
|---|---|---|---|
| ACE2 | TSPO2 | 0,4368 | 4,12182E-10 |
| ACE2 | RP11-345K9.2 | 0,4368 | 4,12024E-10 |
| ACE2 | LAMA3 | 0,4368 | 4,1278E-10 |
| ACE2 | SLC20A1 | 0,4367 | 4,15972E-10 |
| ACE2 | RP11-222K16.1 | 0,4367 | 4,14091E-10 |
| ACE2 | B4GALT4-AS1 | 0,4367 | 4,15201E-10 |
| ACE2 | SSH3 | 0,4366 | 4,1749E-10 |
| ACE2 | LCLAT1 | 0,4361 | 4,42747E-10 |
| ACE2 | SLC25A25-AS1 | 0,4359 | 4,52246E-10 |
| ACE2 | BIRC3 | 0,4353 | 4,77558E-10 |
| ACE2 | LINC00896 | 0,4353 | 4,78246E-10 |
| ACE2 | YIF1A | 0,435 | 4,94218E-10 |
| ACE2 | PDLIM1 | 0,4349 | 5,00378E-10 |
| ACE2 | CASP3 | 0,4348 | 5,01093E-10 |
| ACE2 | DDC-AS1 | 0,4348 | 5,01378E-10 |
| ACE2 | GDPGP1 | 0,4348 | 5,04303E-10 |
| ACE2 | OCEL1 | 0,4348 | 5,05154E-10 |
| ACE2 | EGLN3-AS1 | 0,4347 | 5,10288E-10 |
| ACE2 | CCL24 | 0,4346 | 5,14938E-10 |
| ACE2 | KCNJ6 | 0,4337 | 5,65313E-10 |
| ACE2 | LYZ | 0,4327 | 6,19648E-10 |
| ACE2 | RP11-70D24.2 | 0,4327 | 6,24744E-10 |
| ACE2 | RP1-76B20.11 | 0,4327 | 6,19375E-10 |
| ACE2 | CTD-2529O21.2 | 0,4325 | 6,37286E-10 |
| ACE2 | GHITM | 0,4323 | 6,49946E-10 |
| ACE2 | MESP2 | 0,4323 | 6,47487E-10 |
| ACE2 | RP11-440I14.2 | 0,4322 | 6,51684E-10 |
| ACE2 | RP11-385F7.1 | 0,4322 | 6,51697E-10 |
| ACE2 | RP11-15F12.1 | 0,4321 | 6,61282E-10 |
| ACE2 | PTCHD3P3 | 0,4318 | 6,78111E-10 |
| ACE2 | GAREM1 | 0,4318 | 6,79567E-10 |
| ACE2 | MAST2 | 0,4316 | 6,97628E-10 |
| ACE2 | LINC00853 | 0,4314 | 7,09523E-10 |
| ACE2 | AP004550.1 | 0,4314 | 7,07436E-10 |
| ACE2 | PTPRJ | 0,4313 | 7,14583E-10 |
| ACE2 | AC000111.4 | 0,4312 | 7,21149E-10 |
| ACE2 | NOVA1-AS1 | 0,4311 | 7,31718E-10 |
| ACE2 | KRT19P4 | 0,431 | 7,38799E-10 |
| ACE2 | RNU6-140P | 0,431 | 7,37236E-10 |
| ACE2 | CNTN3 | 0,4303 | 7,89239E-10 |
| ACE2 | LSMEM1 | 0,4302 | 7,99913E-10 |
| ACE2 | BAAT | 0,4302 | 8,00058E-10 |
| ACE2 | EAPP | 0,4302 | 7,99683E-10 |
| ACE2 | LDHD | 0,43 | 8,13383E-10 |
| ACE2 | AVPI1 | 0,4298 | 8,35751E-10 |
| ACE2 | ABAT | 0,4298 | 8,35582E-10 |
| ACE2 | RP11-166P13.3 | 0,4298 | 8,31757E-10 |
| ACE2 | COQ4 | 0,4297 | 8,42584E-10 |
| ACE2 | RP11-232L2.2 | 0,4296 | 8,46772E-10 |
| ACE2 | ABHD17B | 0,4296 | 8,487E-10 |
| ACE2 | LINC01336 | 0,4289 | 9,10393E-10 |
| ACE2 | RN7SL720P | 0,4281 | 9,83324E-10 |
| ACE2 | MTND2P16 | 0,428 | 9,93952E-10 |
| ACE2 | SLC44A3-AS1 | 0,4278 | 1,01412E-09 |

| | | | |
|---|---|---|---|
| ACE2 | MGLL | 0,4272 | 1,07449E-09 |
| ACE2 | PPM1A | 0,4272 | 1,07273E-09 |
| ACE2 | RNU6-652P | 0,4271 | 1,08978E-09 |
| ACE2 | RP11-1267H10.4 | 0,427 | 1,10067E-09 |
| ACE2 | RGMB-AS1 | 0,427 | 1,09514E-09 |
| ACE2 | HNRNPA1P59 | 0,4266 | 1,14617E-09 |
| ACE2 | RP11-70D24.3 | 0,4265 | 1,15286E-09 |
| ACE2 | LINC01842 | 0,4265 | 1,15249E-09 |
| ACE2 | MB | 0,4261 | 1,20039E-09 |
| ACE2 | LINC01079 | 0,4259 | 1,22195E-09 |
| ACE2 | ATP5J2P5 | 0,4258 | 1,23403E-09 |
| ACE2 | MYO5C | 0,4256 | 1,25578E-09 |
| ACE2 | LL22NC03-N95F10.1 | 0,4254 | 1,286E-09 |
| ACE2 | TAT-AS1 | 0,4253 | 1,29946E-09 |
| ACE2 | ATP2A1 | 0,425 | 1,34413E-09 |
| ACE2 | RP11-626P14.2 | 0,4249 | 1,34835E-09 |
| ACE2 | LPCAT4 | 0,4246 | 1,3884E-09 |
| ACE2 | RP13-582O9.7 | 0,4245 | 1,39889E-09 |
| ACE2 | E2F3P2 | 0,4244 | 1,41328E-09 |
| ACE2 | FAM8A4P | 0,4242 | 1,44364E-09 |
| ACE2 | SH3BP5L | 0,4239 | 1,48577E-09 |
| ACE2 | LINC00654 | 0,4236 | 1,54033E-09 |
| ACE2 | CTD-2540B15.9 | 0,4232 | 1,59478E-09 |
| ACE2 | RP11-390N6.1 | 0,4231 | 1,60708E-09 |
| ACE2 | CCDC125 | 0,423 | 1,62668E-09 |
| ACE2 | ZBED9 | 0,4229 | 1,64798E-09 |
| ACE2 | PON2 | 0,4228 | 1,66425E-09 |
| ACE2 | SLC6A14 | 0,4227 | 1,67136E-09 |
| ACE2 | PRR22 | 0,4225 | 1,70107E-09 |
| ACE2 | RP11-156K23.3 | 0,4224 | 1,72631E-09 |
| ACE2 | ARSEP1 | 0,4223 | 1,73451E-09 |
| ACE2 | EPS8 | 0,4221 | 1,76882E-09 |
| ACE2 | PRR26 | 0,4218 | 1,83744E-09 |
| ACE2 | DHRS4 | 0,4215 | 1,89132E-09 |
| ACE2 | RP11-274B18.4 | 0,4213 | 1,92035E-09 |
| ACE2 | GOSR2 | 0,4213 | 1,92902E-09 |
| ACE2 | FAM25C | 0,4212 | 1,94512E-09 |
| ACE2 | HCCS | 0,4211 | 1,96549E-09 |
| ACE2 | MROH6 | 0,421 | 1,9821E-09 |
| ACE2 | KLK4 | 0,421 | 1,986E-09 |
| ACE2 | ABR | 0,4209 | 1,99548E-09 |
| ACE2 | BIK | 0,4208 | 2,01564E-09 |
| ACE2 | RP11-796E10.1 | 0,4207 | 2,04307E-09 |
| ACE2 | NCAPGP2 | 0,4204 | 2,09578E-09 |
| ACE2 | RP4-706G24.1 | 0,4203 | 2,11887E-09 |
| ACE2 | LGR5 | 0,4203 | 2,11089E-09 |
| ACE2 | SYT12 | 0,4201 | 2,14898E-09 |
| ACE2 | RP11-575G13.2 | 0,4201 | 2,16206E-09 |
| ACE2 | AREL1 | 0,4201 | 2,159E-09 |
| ACE2 | CRYZ | 0,4197 | 2,23635E-09 |
| ACE2 | C4BPA | 0,4196 | 2,25422E-09 |
| ACE2 | CHAT | 0,4196 | 2,27248E-09 |
| ACE2 | LINC01220 | 0,4196 | 2,27367E-09 |
| ACE2 | SQRDL | 0,4196 | 2,26161E-09 |

| | | | |
|---|---|---|---|
| ACE2 | RP11-38G5.4 | 0,4194 | 2,30855E-09 |
| ACE2 | MTHFS | 0,4193 | 2,32264E-09 |
| ACE2 | IQGAP3 | 0,419 | 2,40207E-09 |
| ACE2 | RTCA | 0,4188 | 2,45059E-09 |
| ACE2 | FAM95B1 | 0,4188 | 2,44573E-09 |
| ACE2 | TMEM127 | 0,4187 | 2,46085E-09 |
| ACE2 | MTATP6P1 | 0,4186 | 2,49463E-09 |
| ACE2 | DCST2 | 0,4185 | 2,52377E-09 |
| ACE2 | FCF1P5 | 0,4184 | 2,54811E-09 |
| ACE2 | PGD | 0,4179 | 2,6746E-09 |
| ACE2 | PKN2 | 0,4178 | 2,68936E-09 |
| ACE2 | HOXA1 | 0,4177 | 2,70941E-09 |
| ACE2 | RNF123 | 0,4175 | 2,7658E-09 |
| ACE2 | RP11-354K1.2 | 0,4172 | 2,84435E-09 |
| ACE2 | TRPA1 | 0,417 | 2,90373E-09 |
| ACE2 | GGACT | 0,4168 | 2,96822E-09 |
| ACE2 | CTA-363E6.2 | 0,4166 | 3,0119E-09 |
| ACE2 | PRSS58 | 0,4165 | 3,04839E-09 |
| ACE2 | POLR2H | 0,4162 | 3,13667E-09 |
| ACE2 | RP11-685B14.3 | 0,4162 | 3,13157E-09 |
| ACE2 | CYP3A51P | 0,4161 | 3,16035E-09 |
| ACE2 | MIR4653 | 0,4161 | 3,17639E-09 |
| ACE2 | CH17-302M23.1 | 0,4161 | 3,16668E-09 |
| ACE2 | TYMSOS | 0,4158 | 3,24467E-09 |
| ACE2 | FAM98C | 0,4158 | 3,24957E-09 |
| ACE2 | NEUROG3 | 0,4155 | 3,33747E-09 |
| ACE2 | RP11-379C10.4 | 0,4154 | 3,37922E-09 |
| ACE2 | MAP3K13 | 0,4153 | 3,43214E-09 |
| ACE2 | RNA5SP311 | 0,415 | 3,5283E-09 |
| ACE2 | RP11-710F7.2 | 0,4149 | 3,53588E-09 |
| ACE2 | PDLIM2 | 0,4148 | 3,58955E-09 |
| ACE2 | PRKCZ-AS1 | 0,4145 | 3,69399E-09 |
| ACE2 | RP13-16H11.8 | 0,4145 | 3,68534E-09 |
| ACE2 | ALOX15B | 0,4143 | 3,75166E-09 |
| ACE2 | RP11-489M13.3 | 0,4141 | 3,84088E-09 |
| ACE2 | RP3-431P23.5 | 0,414 | 3,87512E-09 |
| ACE2 | PLBD1 | 0,4139 | 3,90453E-09 |
| ACE2 | CTD-2651B20.8 | 0,4139 | 3,91028E-09 |
| ACE2 | UQCRC2 | 0,4137 | 3,97306E-09 |
| ACE2 | CTC-756D1.1 | 0,4136 | 4,02856E-09 |
| ACE2 | TPI1P1 | 0,4132 | 4,16422E-09 |
| ACE2 | ACTR3C | 0,4132 | 4,18633E-09 |
| ACE2 | RP11-36B15.1 | 0,4131 | 4,22864E-09 |
| ACE2 | NECTIN2 | 0,4131 | 4,22229E-09 |
| ACE2 | RP11-150C16.1 | 0,413 | 4,26356E-09 |
| ACE2 | TMEM213 | 0,4129 | 4,30621E-09 |
| ACE2 | UBE2L2 | 0,4126 | 4,41667E-09 |
| ACE2 | CTAGE11P | 0,4126 | 4,41814E-09 |
| ACE2 | MTCO1P2 | 0,4125 | 4,43447E-09 |
| ACE2 | C19orf73 | 0,4125 | 4,44791E-09 |
| ACE2 | PROM2 | 0,4124 | 4,51472E-09 |
| ACE2 | AC144836.1 | 0,4124 | 4,49382E-09 |
| ACE2 | RP11-542B15.1 | 0,4123 | 4,55621E-09 |
| ACE2 | RP11-1149M3.2 | 0,4122 | 4,59131E-09 |

| | | | |
|---|---|---|---|
| ACE2 | COX10 | 0,4122 | 4,58666E-09 |
| ACE2 | CCNO | 0,4121 | 4,61777E-09 |
| ACE2 | PPARGC1B | 0,412 | 4,68638E-09 |
| ACE2 | RBMY2EP | 0,4117 | 4,80644E-09 |
| ACE2 | KLK11 | 0,4116 | 4,83206E-09 |
| ACE2 | RP11-486M23.3 | 0,4115 | 4,90382E-09 |
| ACE2 | LINC01780 | 0,4113 | 4,98516E-09 |
| ACE2 | TMEM198 | 0,4111 | 5,08159E-09 |
| ACE2 | ARHGAP35 | 0,411 | 5,13353E-09 |
| ACE2 | ACTN4P1 | 0,4105 | 5,36017E-09 |
| ACE2 | DIRC2 | 0,4104 | 5,41637E-09 |
| ACE2 | TOB2P1 | 0,4104 | 5,43493E-09 |
| ACE2 | RP4-651E10.4 | 0,4103 | 5,4958E-09 |
| ACE2 | AC005019.3 | 0,41 | 5,61878E-09 |
| ACE2 | CCDC183 | 0,41 | 5,6593E-09 |
| ACE2 | C1QTNF1-AS1 | 0,41 | 5,62412E-09 |
| ACE2 | RP11-342C24.8 | 0,4098 | 5,73011E-09 |
| ACE2 | RDH13 | 0,4098 | 5,71724E-09 |
| ACE2 | RP11-376N17.4 | 0,4095 | 5,89671E-09 |
| ACE2 | ATP5C1 | 0,4089 | 6,25868E-09 |
| ACE2 | AC068134.6 | 0,4086 | 6,43947E-09 |
| ACE2 | MLEC | 0,4086 | 6,44147E-09 |
| ACE2 | RP11-963H4.5 | 0,4083 | 6,61516E-09 |
| ACE2 | RP11-670E13.6 | 0,4083 | 6,57745E-09 |
| ACE2 | HDGF | 0,4081 | 6,73256E-09 |
| ACE2 | CTD-2033D24.2 | 0,4081 | 6,70834E-09 |
| ACE2 | KRT18P10 | 0,4078 | 6,8833E-09 |
| ACE2 | FLJ31356 | 0,4077 | 6,95172E-09 |
| ACE2 | RP11-646E18.4 | 0,4077 | 6,98065E-09 |
| ACE2 | CTAGE15 | 0,4075 | 7,13323E-09 |
| ACE2 | ALDH3A1 | 0,4075 | 7,08541E-09 |
| ACE2 | RP3-326I13.1 | 0,4074 | 7,16966E-09 |
| ACE2 | AC019109.1 | 0,407 | 7,44037E-09 |
| ACE2 | RP11-470C13.4 | 0,407 | 7,4655E-09 |
| ACE2 | RP11-257O5.4 | 0,4068 | 7,58434E-09 |
| ACE2 | CTC-287O8.1 | 0,4067 | 7,67523E-09 |
| ACE2 | ARFIP2 | 0,4067 | 7,65755E-09 |
| ACE2 | COX5B | 0,4066 | 7,74621E-09 |
| ACE2 | ARPC1A | 0,4066 | 7,72783E-09 |
| ACE2 | RP13-20L14.6 | 0,4065 | 7,77694E-09 |
| ACE2 | AF196972.4 | 0,4065 | 7,78385E-09 |
| ACE2 | AF064860.5 | 0,4063 | 7,94909E-09 |
| ACE2 | CAPN11 | 0,4062 | 8,02796E-09 |
| ACE2 | RP11-523H20.3 | 0,4061 | 8,08307E-09 |
| ACE2 | RP11-426L16.3 | 0,4059 | 8,22861E-09 |
| ACE2 | FUNDC2P2 | 0,4059 | 8,25751E-09 |
| ACE2 | RAB15 | 0,4059 | 8,26644E-09 |
| ACE2 | C11orf80 | 0,4057 | 8,43339E-09 |
| ACE2 | LINC01954 | 0,4056 | 8,50687E-09 |
| ACE2 | RP11-51F16.9 | 0,4053 | 8,70111E-09 |
| ACE2 | NPHS2 | 0,4052 | 8,78031E-09 |
| ACE2 | SPACA3 | 0,4048 | 9,16084E-09 |
| ACE2 | CCDC47 | 0,4044 | 9,44347E-09 |
| ACE2 | ERAP1 | 0,404 | 9,8453E-09 |

| | | | |
|---|---|---|---|
| ACE2 | LEAP2 | 0,404 | 9,84944E-09 |
| ACE2 | LHFPL3-AS1 | 0,4036 | 1,01652E-08 |
| ACE2 | TM9SF3 | 0,4036 | 1,01836E-08 |
| ACE2 | DPP10-AS1 | 0,4035 | 1,0282E-08 |
| ACE2 | HSD17B8 | 0,4035 | 1,03117E-08 |
| ACE2 | LINC00393 | 0,4035 | 1,02945E-08 |
| ACE2 | REN | 0,4034 | 1,04132E-08 |
| ACE2 | RP11-697E22.3 | 0,4034 | 1,03411E-08 |
| ACE2 | UNC5B | 0,4033 | 1,05078E-08 |
| ACE2 | RP11-481L19.2 | 0,4033 | 1,04485E-08 |
| ACE2 | AP000640.10 | 0,4033 | 1,04967E-08 |
| ACE2 | LINC01645 | 0,4032 | 1,05219E-08 |
| ACE2 | AP1AR | 0,4027 | 1,10475E-08 |
| ACE2 | NR2E3 | 0,4025 | 1,12117E-08 |
| ACE2 | SQLE | 0,4023 | 1,1449E-08 |
| ACE2 | CYP4A22 | 0,4021 | 1,17107E-08 |
| ACE2 | RNU1-75P | 0,4019 | 1,19099E-08 |
| ACE2 | RP11-395L14.13 | 0,4015 | 1,22788E-08 |
| ACE2 | FAM201A | 0,4014 | 1,24565E-08 |
| ACE2 | NDUFS1 | 0,4011 | 1,27754E-08 |
| ACE2 | HINT1 | 0,4011 | 1,27733E-08 |
| ACE2 | MATN2 | 0,4011 | 1,28172E-08 |
| ACE2 | KRT18P1 | 0,4007 | 1,33018E-08 |
| ACE2 | CTD-2330K9.2 | 0,4006 | 1,33722E-08 |
| ACE2 | AICDA | 0,4006 | 1,34235E-08 |
| ACE2 | PRORY | 0,4006 | 1,33533E-08 |
| ACE2 | MTCO1P22 | 0,4004 | 1,35693E-08 |
| ACE2 | RP11-80H5.2 | 0,4001 | 1,40106E-08 |
| ACE2 | PMM2 | 0,3999 | 1,42311E-08 |
| ACE2 | APOL6 | 0,3998 | 1,44019E-08 |
| ACE2 | AC012360.6 | 0,3997 | 1,45053E-08 |
| ACE2 | KRT8P39 | 0,3996 | 1,46097E-08 |
| ACE2 | PTGDR2 | 0,399 | 1,54271E-08 |
| ACE2 | EXOC6B | 0,3986 | 1,59779E-08 |
| ACE2 | RP11-44N12.5 | 0,3986 | 1,59916E-08 |
| ACE2 | RP11-368L12.1 | 0,3986 | 1,60697E-08 |
| ACE2 | GP2 | 0,3984 | 1,63024E-08 |
| ACE2 | ACVR1C | 0,3982 | 1,66813E-08 |
| ACE2 | ACHE | 0,3982 | 1,66656E-08 |
| ACE2 | RP11-626G11.1 | 0,3982 | 1,65887E-08 |
| ACE2 | CTC-756D1.2 | 0,3981 | 1,67279E-08 |
| ACE2 | ENPP7P8 | 0,3979 | 1,70276E-08 |
| ACE2 | LINC00112 | 0,3978 | 1,72366E-08 |
| ACE2 | AMACR | 0,3977 | 1,73886E-08 |
| ACE2 | DYNAP | 0,3975 | 1,7764E-08 |
| ACE2 | HNRNPA1P33 | 0,3973 | 1,80036E-08 |
| ACE2 | RP11-111E14.1 | 0,3973 | 1,80459E-08 |
| ACE2 | RN7SL305P | 0,3973 | 1,79514E-08 |
| ACE2 | HOXA-AS3 | 0,3967 | 1,89526E-08 |
| ACE2 | RP11-44F14.10 | 0,3966 | 1,91552E-08 |
| ACE2 | LINC01558 | 0,3964 | 1,95071E-08 |
| ACE2 | TSC22D1-AS1 | 0,3964 | 1,9469E-08 |
| ACE2 | SIRT6 | 0,3961 | 1,99721E-08 |
| ACE2 | THEM6 | 0,3959 | 2,03752E-08 |

| | | | |
|---|---|---|---|
| ACE2 | RP11-159D23.2 | 0,3959 | 2,03369E-08 |
| ACE2 | CASP1 | 0,3958 | 2,06173E-08 |
| ACE2 | REG1CP | 0,3957 | 2,07559E-08 |
| ACE2 | RP11-235E17.3 | 0,3957 | 2,07747E-08 |
| ACE2 | MT-TI | 0,3957 | 2,07837E-08 |
| ACE2 | CYP51A1P1 | 0,3955 | 2,11934E-08 |
| ACE2 | RP11-708H21.1 | 0,3955 | 2,11164E-08 |
| ACE2 | MIR3180-1 | 0,3953 | 2,1519E-08 |
| ACE2 | DEFA11P | 0,3952 | 2,1796E-08 |
| ACE2 | HMGA1P5 | 0,3951 | 2,18871E-08 |
| ACE2 | RITA1 | 0,3946 | 2,29711E-08 |
| ACE2 | ASS1P7 | 0,3943 | 2,35267E-08 |
| ACE2 | ITPR3 | 0,394 | 2,41482E-08 |
| ACE2 | RP11-49L2.1 | 0,3939 | 2,44131E-08 |
| ACE2 | CLCN3 | 0,3938 | 2,45128E-08 |
| ACE2 | MAGI1-IT1 | 0,3936 | 2,50484E-08 |
| ACE2 | RASSF6 | 0,3936 | 2,49685E-08 |
| ACE2 | KCTD14 | 0,3935 | 2,53388E-08 |
| ACE2 | RP11-552J9.9 | 0,3933 | 2,56451E-08 |
| ACE2 | RN7SKP281 | 0,3929 | 2,65586E-08 |
| ACE2 | AC114730.3 | 0,3929 | 2,66116E-08 |
| ACE2 | MUC5B | 0,3929 | 2,66271E-08 |
| ACE2 | RP11-491F9.1 | 0,3929 | 2,6692E-08 |
| ACE2 | RP11-167J8.3 | 0,3925 | 2,75289E-08 |
| ACE2 | CCDC25 | 0,3924 | 2,7761E-08 |
| ACE2 | UGT2B24P | 0,3922 | 2,82427E-08 |
| ACE2 | IDH3A | 0,3921 | 2,85767E-08 |
| ACE2 | ZC3H3 | 0,3919 | 2,91054E-08 |
| ACE2 | KRT8P22 | 0,3918 | 2,94574E-08 |
| ACE2 | RP11-963H4.2 | 0,3912 | 3,0964E-08 |
| ACE2 | SLCO2B1 | 0,391 | 3,15872E-08 |
| ACE2 | IQSEC2 | 0,391 | 3,16076E-08 |
| ACE2 | AC011899.10 | 0,3909 | 3,16924E-08 |
| ACE2 | MIR608 | 0,3909 | 3,1875E-08 |
| ACE2 | MAN1A1 | 0,3907 | 3,22731E-08 |
| ACE2 | CA9 | 0,3905 | 3,28446E-08 |
| ACE2 | LINC01977 | 0,3904 | 3,3076E-08 |
| ACE2 | RP11-96H17.1 | 0,3901 | 3,41608E-08 |
| ACE2 | CYP4F23P | 0,3901 | 3,39916E-08 |
| ACE2 | DNASE2B | 0,39 | 3,4302E-08 |
| ACE2 | RP11-88L20.1 | 0,3898 | 3,50015E-08 |
| ACE2 | SIRT7 | 0,3898 | 3,49724E-08 |
| ACE2 | PNKD | 0,3896 | 3,5623E-08 |
| ACE2 | RP11-627G18.4 | 0,3894 | 3,63528E-08 |
| ACE2 | TCP11L1 | 0,3893 | 3,66374E-08 |
| ACE2 | BTBD9 | 0,3892 | 3,69166E-08 |
| ACE2 | RP13-644M16.4 | 0,3889 | 3,76987E-08 |
| ACE2 | LINC01658 | 0,3888 | 3,81393E-08 |
| ACE2 | MGC27382 | 0,3887 | 3,8401E-08 |
| ACE2 | VAV2 | 0,3885 | 3,90625E-08 |
| ACE2 | RP11-485O10.2 | 0,3881 | 4,06085E-08 |
| ACE2 | SPATA6L | 0,3876 | 4,23468E-08 |
| ACE2 | CYP4F36P | 0,3876 | 4,22986E-08 |
| ACE2 | CCDC17 | 0,3875 | 4,28932E-08 |

| | | | |
|---|---|---|---|
| ACE2 | OR52W1 | 0,3874 | 4,29516E-08 |
| ACE2 | TXN | 0,3872 | 4,40332E-08 |
| ACE2 | MGAT2 | 0,3872 | 4,39316E-08 |
| ACE2 | XIAPP3 | 0,3871 | 4,43255E-08 |
| ACE2 | RN7SL706P | 0,387 | 4,46584E-08 |
| ACE2 | KIAA1958 | 0,3869 | 4,52094E-08 |
| ACE2 | RP11-873E20.1 | 0,3868 | 4,56023E-08 |
| ACE2 | AK2 | 0,3867 | 4,58573E-08 |
| ACE2 | OSBPL6 | 0,3865 | 4,64669E-08 |
| ACE2 | RP11-321G12.1 | 0,3864 | 4,70812E-08 |
| ACE2 | MIR141 | 0,3863 | 4,72776E-08 |
| ACE2 | AC079779.5 | 0,3861 | 4,84813E-08 |
| ACE2 | CTAGE8 | 0,3861 | 4,84729E-08 |
| ACE2 | AC093818.1 | 0,386 | 4,87807E-08 |
| ACE2 | RP11-550A5.2 | 0,3856 | 5,03853E-08 |
| ACE2 | PLCH1-AS1 | 0,385 | 5,30554E-08 |
| ACE2 | AFP | 0,385 | 5,32942E-08 |
| ACE2 | SGMS2 | 0,385 | 5,30196E-08 |
| ACE2 | RP11-844P9.4 | 0,385 | 5,29154E-08 |
| ACE2 | CH507-42P11.6 | 0,385 | 5,30905E-08 |
| ACE2 | CCNYL1 | 0,3847 | 5,425E-08 |
| ACE2 | NOX1 | 0,3847 | 5,45992E-08 |
| ACE2 | RP11-203M5.6 | 0,3846 | 5,47384E-08 |
| ACE2 | OR8R1P | 0,3845 | 5,53922E-08 |
| ACE2 | RP11-386I14.3 | 0,3844 | 5,60254E-08 |
| ACE2 | MIR4635 | 0,3844 | 5,60187E-08 |
| ACE2 | SAPCD2 | 0,3844 | 5,59536E-08 |
| ACE2 | PDZD8 | 0,3843 | 5,6194E-08 |
| ACE2 | SLC32A1 | 0,3843 | 5,63731E-08 |
| ACE2 | SLC16A10 | 0,3842 | 5,67893E-08 |
| ACE2 | AC004490.1 | 0,3842 | 5,68947E-08 |
| ACE2 | RP1-69D17.3 | 0,3841 | 5,74684E-08 |
| ACE2 | ADCK5 | 0,3841 | 5,74641E-08 |
| ACE2 | MRPS36 | 0,384 | 5,77E-08 |
| ACE2 | CAPN15 | 0,3839 | 5,83178E-08 |
| ACE2 | ELDR | 0,3838 | 5,86708E-08 |
| ACE2 | GPS2P2 | 0,3837 | 5,94661E-08 |
| ACE2 | ACTG1P18 | 0,3836 | 6,00482E-08 |
| ACE2 | RP11-561B11.1 | 0,3835 | 6,04097E-08 |
| ACE2 | MSX2 | 0,3833 | 6,13108E-08 |
| ACE2 | STOML3 | 0,3831 | 6,24279E-08 |
| ACE2 | RP4-773N10.4 | 0,383 | 6,32131E-08 |
| ACE2 | NOSTRIN | 0,3828 | 6,40574E-08 |
| ACE2 | LMNTD2 | 0,3827 | 6,4761E-08 |
| ACE2 | RDH5 | 0,3826 | 6,53726E-08 |
| ACE2 | ALKBH7 | 0,3825 | 6,59163E-08 |
| ACE2 | TTC7A | 0,3824 | 6,61201E-08 |
| ACE2 | TBC1D8B | 0,3824 | 6,62049E-08 |
| ACE2 | TSPAN15 | 0,3823 | 6,67669E-08 |
| ACE2 | FTOP1 | 0,382 | 6,83833E-08 |
| ACE2 | OR5M6P | 0,3817 | 7,02973E-08 |
| ACE2 | AC004012.1 | 0,381 | 7,45517E-08 |
| ACE2 | RP11-285E23.2 | 0,381 | 7,47035E-08 |
| ACE2 | ELOCP19 | 0,3809 | 7,51434E-08 |

| | | | |
|---|---|---|---|
| ACE2 | RP11-407G23.5 | 0,3809 | 7,54002E-08 |
| ACE2 | RN7SKP249 | 0,3808 | 7,59392E-08 |
| ACE2 | OR5G5P | 0,3807 | 7,68475E-08 |
| ACE2 | MT1F | 0,3807 | 7,67808E-08 |
| ACE2 | AKAP1 | 0,3807 | 7,67302E-08 |
| ACE2 | GALC | 0,3806 | 7,75715E-08 |
| ACE2 | AC069257.8 | 0,3802 | 7,97291E-08 |
| ACE2 | SLC25A16 | 0,3801 | 8,06336E-08 |
| ACE2 | BNIP3P39 | 0,38 | 8,10042E-08 |
| ACE2 | BLACAT1 | 0,3798 | 8,27791E-08 |
| ACE2 | PLA2G2E | 0,3793 | 8,62025E-08 |
| ACE2 | KRT19P3 | 0,3791 | 8,77321E-08 |
| ACE2 | CTD-2651B20.5 | 0,3791 | 8,7922E-08 |
| ACE2 | VPS13A-AS1 | 0,3786 | 9,17592E-08 |
| ACE2 | CTD-2651B20.2 | 0,3783 | 9,39441E-08 |
| ACE2 | C8B | 0,3782 | 9,43092E-08 |
| ACE2 | UQCR10 | 0,378 | 9,62601E-08 |
| ACE2 | SNX30 | 0,3779 | 9,73406E-08 |
| ACE2 | LINC01022 | 0,3777 | 9,84516E-08 |
| ACE2 | SLC52A2 | 0,3777 | 9,85576E-08 |
| ACE2 | GRM7-AS2 | 0,3775 | 1,00418E-07 |
| ACE2 | VPS4B | 0,3774 | 1,01649E-07 |
| ACE2 | CCAT1 | 0,3772 | 1,02538E-07 |
| ACE2 | RP11-626G3.1 | 0,3771 | 1,03638E-07 |
| ACE2 | ZC2HC1C | 0,3769 | 1,05723E-07 |
| ACE2 | SMIM2-IT1 | 0,3766 | 1,0847E-07 |
| ACE2 | SYAP1 | 0,3766 | 1,08491E-07 |
| ACE2 | LYPD6B | 0,3765 | 1,089E-07 |
| ACE2 | PHACTR3 | 0,3763 | 1,11442E-07 |
| ACE2 | CTD-2008A1.3 | 0,3762 | 1,12332E-07 |
| ACE2 | ARPIN | 0,3762 | 1,11519E-07 |
| ACE2 | PRR34-AS1 | 0,3762 | 1,11483E-07 |
| ACE2 | ORMDL2 | 0,3759 | 1,15152E-07 |
| ACE2 | CTD-2270L9.4 | 0,3756 | 1,18096E-07 |
| ACE2 | KB-1269D1.8 | 0,3756 | 1,18012E-07 |
| ACE2 | RP11-483I13.6 | 0,3753 | 1,20171E-07 |
| ACE2 | WIPF2 | 0,3753 | 1,20329E-07 |
| ACE2 | FRAT2 | 0,3752 | 1,21424E-07 |
| ACE2 | CTD-2231E14.8 | 0,3748 | 1,26181E-07 |
| ACE2 | NKX1-2 | 0,3746 | 1,27501E-07 |
| ACE2 | BCAR3 | 0,3745 | 1,29284E-07 |
| ACE2 | SLC7A11 | 0,3744 | 1,30231E-07 |
| ACE2 | DCDC2 | 0,3743 | 1,31203E-07 |
| ACE2 | KDELR1 | 0,3739 | 1,35407E-07 |
| ACE2 | MTCO1P53 | 0,3739 | 1,35811E-07 |
| ACE2 | RP11-268J15.5 | 0,3738 | 1,36496E-07 |
| ACE2 | AC016734.2 | 0,3737 | 1,37919E-07 |
| ACE2 | RP11-639F1.2 | 0,3736 | 1,3953E-07 |
| ACE2 | MYO5BP3 | 0,3736 | 1,38941E-07 |
| ACE2 | FAM83A-AS1 | 0,3734 | 1,40912E-07 |
| ACE2 | CLYBL-AS2 | 0,3734 | 1,41589E-07 |
| ACE2 | RP11-274E7.2 | 0,3731 | 1,44363E-07 |
| ACE2 | CPT1A | 0,373 | 1,46299E-07 |
| ACE2 | PHBP3 | 0,3728 | 1,4826E-07 |

| | | | |
|---|---|---|---|
| ACE2 | MUC6 | 0,3727 | 1,49213E-07 |
| ACE2 | ATP5G1P6 | 0,3726 | 1,50345E-07 |
| ACE2 | TECR | 0,3725 | 1,52713E-07 |
| ACE2 | AP001625.4 | 0,3724 | 1,53387E-07 |
| ACE2 | WARSP1 | 0,3723 | 1,54225E-07 |
| ACE2 | RP11-93B14.10 | 0,3721 | 1,57271E-07 |
| ACE2 | RP11-44F14.9 | 0,3718 | 1,60646E-07 |
| ACE2 | RP11-148M9.1 | 0,3718 | 1,61509E-07 |
| ACE2 | SULT1A3 | 0,3717 | 1,62079E-07 |
| ACE2 | KRT18P29 | 0,3713 | 1,68152E-07 |
| ACE2 | CNPPD1 | 0,3712 | 1,69567E-07 |
| ACE2 | ANKRD22 | 0,3708 | 1,75766E-07 |
| ACE2 | RP11-700H6.1 | 0,3706 | 1,77251E-07 |
| ACE2 | MTCO1P40 | 0,3706 | 1,78626E-07 |
| ACE2 | HSPA8P3 | 0,3705 | 1,796E-07 |
| ACE2 | AP006621.8 | 0,3701 | 1,84975E-07 |
| ACE2 | RP11-649A18.3 | 0,3701 | 1,84688E-07 |
| ACE2 | RP11-288D15.1 | 0,37 | 1,86941E-07 |
| ACE2 | RP11-44F14.1 | 0,3699 | 1,87914E-07 |
| ACE2 | C20orf24 | 0,3698 | 1,89623E-07 |
| ACE2 | RP11-478K15.7 | 0,3697 | 1,9205E-07 |
| ACE2 | TICAM1 | 0,3697 | 1,90795E-07 |
| ACE2 | ARHGEF10L | 0,3695 | 1,94456E-07 |
| ACE2 | RP11-337L12.1 | 0,3694 | 1,9616E-07 |
| ACE2 | RP4-669P10.16 | 0,3694 | 1,96421E-07 |
| ACE2 | STIL | 0,369 | 2,02642E-07 |
| ACE2 | TMEM33 | 0,369 | 2,02395E-07 |
| ACE2 | EGF | 0,3689 | 2,04778E-07 |
| ACE2 | SSR1P1 | 0,3689 | 2,04654E-07 |
| ACE2 | ETV2 | 0,3689 | 2,03805E-07 |
| ACE2 | SERPINA6 | 0,3688 | 2,05455E-07 |
| ACE2 | ZNF823 | 0,3687 | 2,07289E-07 |
| ACE2 | MIR1229 | 0,3686 | 2,10269E-07 |
| ACE2 | CDC42BPB | 0,3685 | 2,11846E-07 |
| ACE2 | AC073046.25 | 0,3684 | 2,12279E-07 |
| ACE2 | POLD4 | 0,3683 | 2,15011E-07 |
| ACE2 | AC068831.3 | 0,3683 | 2,1498E-07 |
| ACE2 | NUDT22 | 0,3682 | 2,16011E-07 |
| ACE2 | ZMYND15 | 0,3682 | 2,16721E-07 |
| ACE2 | SF3A3P2 | 0,368 | 2,19709E-07 |
| ACE2 | SNX4 | 0,3677 | 2,25734E-07 |
| ACE2 | RP11-69G7.1 | 0,3677 | 2,25374E-07 |
| ACE2 | TMEM176A | 0,3674 | 2,30977E-07 |
| ACE2 | RP4-694A7.2 | 0,3673 | 2,32674E-07 |
| ACE2 | RP11-93B14.9 | 0,3673 | 2,33015E-07 |
| ACE2 | HAGLROS | 0,3671 | 2,35779E-07 |
| ACE2 | RP11-104N10.2 | 0,367 | 2,39537E-07 |
| ACE2 | RP11-375N15.2 | 0,3668 | 2,41791E-07 |
| ACE2 | TNFRSF11B | 0,3668 | 2,41978E-07 |
| ACE2 | LINC00456 | 0,3668 | 2,43147E-07 |
| ACE2 | TMEM19 | 0,3667 | 2,44269E-07 |
| ACE2 | WNT3 | 0,3664 | 2,49555E-07 |
| ACE2 | BLVRB | 0,3664 | 2,51079E-07 |
| ACE2 | AURKA | 0,3664 | 2,50008E-07 |

| | | | |
|---|---|---|---|
| ACE2 | TMEM92-AS1 | 0,3663 | 2,5327E-07 |
| ACE2 | RP11-326K13.4 | 0,3663 | 2,52944E-07 |
| ACE2 | C20orf202 | 0,3663 | 2,52126E-07 |
| ACE2 | RP11-350N15.5 | 0,3662 | 2,54158E-07 |
| ACE2 | MTUS1 | 0,3661 | 2,57106E-07 |
| ACE2 | RP11-245A18.1 | 0,3661 | 2,55656E-07 |
| ACE2 | ETS2 | 0,3661 | 2,57635E-07 |
| ACE2 | RP5-1125A11.7 | 0,3659 | 2,61583E-07 |
| ACE2 | RP11-1060J15.9 | 0,3655 | 2,69456E-07 |
| ACE2 | RP11-497D6.3 | 0,3654 | 2,70558E-07 |
| ACE2 | PRR5 | 0,3652 | 2,75603E-07 |
| ACE2 | SRXN1 | 0,365 | 2,8009E-07 |
| ACE2 | ITPK1-AS1 | 0,3648 | 2,85544E-07 |
| ACE2 | RP11-861E23.2 | 0,3646 | 2,90533E-07 |
| ACE2 | ARMC7 | 0,3645 | 2,92157E-07 |
| ACE2 | CAMKV | 0,3644 | 2,93729E-07 |
| ACE2 | ATG4A | 0,3644 | 2,93255E-07 |
| ACE2 | EGLN3P1 | 0,3642 | 2,99017E-07 |
| ACE2 | RP11-250H24.2 | 0,3639 | 3,06012E-07 |
| ACE2 | RP11-55L4.2 | 0,3635 | 3,17332E-07 |
| ACE2 | AC009303.2 | 0,3634 | 3,1946E-07 |
| ACE2 | VSTM5 | 0,363 | 3,28434E-07 |
| ACE2 | RP11-407G23.1 | 0,3628 | 3,3356E-07 |
| ACE2 | JOSD1 | 0,3628 | 3,35519E-07 |
| ACE2 | RP11-81A22.4 | 0,3626 | 3,40822E-07 |
| ACE2 | CDC14A | 0,3625 | 3,42347E-07 |
| ACE2 | RP11-310P5.2 | 0,3625 | 3,43265E-07 |
| ACE2 | XBP1 | 0,3625 | 3,43309E-07 |
| ACE2 | CDCA4P2 | 0,3622 | 3,50346E-07 |
| ACE2 | CHMP2A | 0,3622 | 3,50114E-07 |
| ACE2 | RP1-140A9.1 | 0,3621 | 3,52707E-07 |
| ACE2 | GS1-204I12.4 | 0,362 | 3,56591E-07 |
| ACE2 | RP11-670E13.3 | 0,362 | 3,56669E-07 |
| ACE2 | KRT18P38 | 0,3618 | 3,61299E-07 |
| ACE2 | ETF1P3 | 0,3617 | 3,64926E-07 |
| ACE2 | LHX9 | 0,3614 | 3,74413E-07 |
| ACE2 | SNX12 | 0,3614 | 3,75284E-07 |
| ACE2 | GLOD4 | 0,3613 | 3,7795E-07 |
| ACE2 | AC138472.4 | 0,3613 | 3,76621E-07 |
| ACE2 | VAV3 | 0,3611 | 3,82408E-07 |
| ACE2 | RN7SL386P | 0,361 | 3,87303E-07 |
| ACE2 | LINC01594 | 0,361 | 3,87173E-07 |
| ACE2 | KRT18P16 | 0,361 | 3,8655E-07 |
| ACE2 | LINC01225 | 0,3609 | 3,89032E-07 |
| ACE2 | HLA-J | 0,3609 | 3,88481E-07 |
| ACE2 | RP11-141C7.2 | 0,3607 | 3,94564E-07 |
| ACE2 | RHBG | 0,3606 | 3,98931E-07 |
| ACE2 | SERPINB8 | 0,3606 | 3,99262E-07 |
| ACE2 | RP11-304L19.1 | 0,3605 | 4,03008E-07 |
| ACE2 | TOB1 | 0,3604 | 4,03805E-07 |
| ACE2 | CCDC85C | 0,36 | 4,17993E-07 |
| ACE2 | AC128709.3 | 0,3599 | 4,22491E-07 |
| ACE2 | RP11-269G24.4 | 0,3595 | 4,34608E-07 |
| ACE2 | ISL1 | 0,3594 | 4,38842E-07 |

| | | | |
|---|---|---|---|
| ACE2 | RP11-347C12.3 | 0,3594 | 4,38816E-07 |
| ACE2 | FFAR2 | 0,3592 | 4,44213E-07 |
| ACE2 | UQCRFS1 | 0,359 | 4,51334E-07 |
| ACE2 | GBF1 | 0,3588 | 4,60442E-07 |
| ACE2 | RP11-555G19.1 | 0,3586 | 4,67642E-07 |
| ACE2 | CTB-73N10.1 | 0,3585 | 4,70759E-07 |
| ACE2 | CRISP2 | 0,3583 | 4,76072E-07 |
| ACE2 | SPRY4-IT1 | 0,3582 | 4,80801E-07 |
| ACE2 | NBPF5P | 0,3581 | 4,83601E-07 |
| ACE2 | RP11-20B24.2 | 0,358 | 4,87807E-07 |
| ACE2 | ATP9A | 0,3579 | 4,91276E-07 |
| ACE2 | STK4-AS1 | 0,3577 | 5,02666E-07 |
| ACE2 | RP11-57A1.1 | 0,3571 | 5,24723E-07 |
| ACE2 | TERT | 0,3569 | 5,31688E-07 |
| ACE2 | CA4 | 0,3569 | 5,33202E-07 |
| ACE2 | RP11-813F20.4 | 0,3568 | 5,38779E-07 |
| ACE2 | AC004066.3 | 0,3566 | 5,45915E-07 |
| ACE2 | PRKAG2 | 0,3566 | 5,47791E-07 |
| ACE2 | RN7SL745P | 0,3566 | 5,44652E-07 |
| ACE2 | HEXA-AS1 | 0,3564 | 5,52515E-07 |
| ACE2 | RP11-460I13.2 | 0,3563 | 5,59777E-07 |
| ACE2 | FASTKD1 | 0,3563 | 5,60009E-07 |
| ACE2 | KRT8P9 | 0,3559 | 5,76405E-07 |
| ACE2 | HMGN2P47 | 0,3558 | 5,79388E-07 |
| ACE2 | RPL7L1P2 | 0,3556 | 5,8876E-07 |
| ACE2 | RP13-890H12.2 | 0,3555 | 5,94879E-07 |
| ACE2 | ERLIN1 | 0,3554 | 5,97486E-07 |
| ACE2 | TM7SF2 | 0,3551 | 6,14238E-07 |
| ACE2 | CDK20 | 0,3544 | 6,47237E-07 |
| ACE2 | CTC-250P20.1 | 0,3541 | 6,65025E-07 |
| ACE2 | SLC29A4 | 0,3541 | 6,63875E-07 |
| ACE2 | CTAGE1 | 0,3541 | 6,65161E-07 |
| ACE2 | MMP20 | 0,3539 | 6,719E-07 |
| ACE2 | PMEL | 0,3539 | 6,75422E-07 |
| ACE2 | PKD1L3 | 0,3539 | 6,72669E-07 |
| ACE2 | RP1-232L24.4 | 0,3538 | 6,76283E-07 |
| ACE2 | SRSF9P1 | 0,3538 | 6,78892E-07 |
| ACE2 | AC004066.2 | 0,3536 | 6,89971E-07 |
| ACE2 | KCNG1 | 0,3536 | 6,89392E-07 |
| ACE2 | PRSS3P1 | 0,3534 | 7,00557E-07 |
| ACE2 | MPP1 | 0,3533 | 7,07181E-07 |
| ACE2 | MET | 0,353 | 7,21838E-07 |
| ACE2 | DLG3-AS1 | 0,353 | 7,23618E-07 |
| ACE2 | KLK13 | 0,3527 | 7,39129E-07 |
| ACE2 | PPP2R1B | 0,3526 | 7,46666E-07 |
| ACE2 | KRT8P20 | 0,3525 | 7,53331E-07 |
| ACE2 | RP11-432I5.4 | 0,3525 | 7,51988E-07 |
| ACE2 | RP11-240E2.2 | 0,3524 | 7,54725E-07 |
| ACE2 | CCL20 | 0,3523 | 7,59837E-07 |
| ACE2 | LL22NC03-102D1.18 | 0,3522 | 7,69648E-07 |
| ACE2 | PODXL2 | 0,3521 | 7,73735E-07 |
| ACE2 | LINC01942 | 0,352 | 7,81247E-07 |
| ACE2 | RP11-336N8.2 | 0,3519 | 7,87554E-07 |
| ACE2 | LINC01389 | 0,3517 | 7,9558E-07 |

| | | | |
|---|---|---|---|
| ACE2 | MAN2A1 | 0,3517 | 7,97825E-07 |
| ACE2 | RP11-502I4.3 | 0,3515 | 8,09923E-07 |
| ACE2 | CTD-2160D9.1 | 0,3514 | 8,18455E-07 |
| ACE2 | RP11-167N4.2 | 0,3514 | 8,14703E-07 |
| ACE2 | CHRNA2 | 0,3512 | 8,3188E-07 |
| ACE2 | GSTA3 | 0,3509 | 8,46977E-07 |
| ACE2 | MIRLET7D | 0,3508 | 8,56379E-07 |
| ACE2 | RNU2-33P | 0,3507 | 8,64947E-07 |
| ACE2 | CD164L2 | 0,3504 | 8,8188E-07 |
| ACE2 | UNC13B | 0,35 | 9,1042E-07 |
| ACE2 | CALM1 | 0,35 | 9,11548E-07 |
| ACE2 | C2orf91 | -0,3499 | 9,14092E-07 |
| ACE2 | PGBD4P3 | -0,35 | 9,08538E-07 |
| ACE2 | SSXP10 | -0,35 | 9,10672E-07 |
| ACE2 | AC003075.4 | -0,35 | 9,1154E-07 |
| ACE2 | OR2A1 | -0,35 | 9,08911E-07 |
| ACE2 | AMD1P4 | -0,3501 | 8,99682E-07 |
| ACE2 | SMARCE1P6 | -0,3502 | 8,95459E-07 |
| ACE2 | IL5RA | -0,3502 | 8,96069E-07 |
| ACE2 | AC009473.1 | -0,3502 | 8,98986E-07 |
| ACE2 | C17orf102 | -0,3502 | 8,95188E-07 |
| ACE2 | LIMD1-AS1 | -0,3503 | 8,91388E-07 |
| ACE2 | LSM12P1 | -0,3503 | 8,89394E-07 |
| ACE2 | CTAGE7P | -0,3503 | 8,88789E-07 |
| ACE2 | CTD-3203P2.1 | -0,3503 | 8,88832E-07 |
| ACE2 | RP11-426C22.8 | -0,3503 | 8,88854E-07 |
| ACE2 | C17orf64 | -0,3503 | 8,87992E-07 |
| ACE2 | FCGR3B | -0,3504 | 8,80075E-07 |
| ACE2 | RP11-355B11.2 | -0,3504 | 8,82015E-07 |
| ACE2 | GCA | -0,3504 | 8,85268E-07 |
| ACE2 | MEIG1 | -0,3504 | 8,80544E-07 |
| ACE2 | RP11-180M15.6 | -0,3504 | 8,80159E-07 |
| ACE2 | AC000036.4 | -0,3504 | 8,83095E-07 |
| ACE2 | FAM65B | -0,3505 | 8,78266E-07 |
| ACE2 | HLA-DOA | -0,3505 | 8,78328E-07 |
| ACE2 | COMMD9 | -0,3505 | 8,76425E-07 |
| ACE2 | RP11-693N9.2 | -0,3505 | 8,75925E-07 |
| ACE2 | CDK12 | -0,3505 | 8,78486E-07 |
| ACE2 | CORIN | -0,3506 | 8,7131E-07 |
| ACE2 | RP11-488P3.1 | -0,3507 | 8,61021E-07 |
| ACE2 | LINC01460 | -0,3507 | 8,62451E-07 |
| ACE2 | LAMTOR3 | -0,3507 | 8,58853E-07 |
| ACE2 | RP11-80H5.9 | -0,3507 | 8,61355E-07 |
| ACE2 | CTD-2555K7.2 | -0,3507 | 8,63214E-07 |
| ACE2 | RP11-932O9.8 | -0,3507 | 8,64554E-07 |
| ACE2 | LIPC-AS1 | -0,3507 | 8,59522E-07 |
| ACE2 | RP11-81K2.2 | -0,3507 | 8,65035E-07 |
| ACE2 | SRSF6 | -0,3507 | 8,62786E-07 |
| ACE2 | F2R | -0,3508 | 8,54784E-07 |
| ACE2 | CTD-2213F21.2 | -0,3508 | 8,5553E-07 |
| ACE2 | HIF1A | -0,3508 | 8,52685E-07 |
| ACE2 | RP11-806K15.1 | -0,3509 | 8,49077E-07 |
| ACE2 | RP11-567J20.1 | -0,3509 | 8,46494E-07 |
| ACE2 | CTD-3074O7.7 | -0,3509 | 8,47905E-07 |

| | | | |
|---|---|---|---|
| ACE2 | CBFA2T3 | -0,3509 | 8,50779E-07 |
| ACE2 | RP5-1074L1.4 | -0,351 | 8,44328E-07 |
| ACE2 | B3GALT1 | -0,351 | 8,40014E-07 |
| ACE2 | RP13-585F24.1 | -0,351 | 8,44797E-07 |
| ACE2 | RP11-367F23.1 | -0,351 | 8,43745E-07 |
| ACE2 | TMEM38B | -0,3511 | 8,39087E-07 |
| ACE2 | AC090587.4 | -0,3511 | 8,33892E-07 |
| ACE2 | RP11-893F2.5 | -0,3511 | 8,36701E-07 |
| ACE2 | MMADHC | -0,3512 | 8,28459E-07 |
| ACE2 | RP11-110I1.14 | -0,3512 | 8,32032E-07 |
| ACE2 | RP11-540B6.6 | -0,3512 | 8,27807E-07 |
| ACE2 | FBRS | -0,3512 | 8,31763E-07 |
| ACE2 | XPO6 | -0,3513 | 8,25703E-07 |
| ACE2 | GCAT | -0,3513 | 8,23923E-07 |
| ACE2 | RP1-164F3.8 | -0,3513 | 8,2499E-07 |
| ACE2 | HSPD1P10 | -0,3514 | 8,17846E-07 |
| ACE2 | THNSL1 | -0,3514 | 8,14328E-07 |
| ACE2 | GRK6 | -0,3515 | 8,11998E-07 |
| ACE2 | RP11-240G22.4 | -0,3515 | 8,09932E-07 |
| ACE2 | MTUS2-AS1 | -0,3515 | 8,08201E-07 |
| ACE2 | ALLC | -0,3516 | 8,07536E-07 |
| ACE2 | RP11-391M7.3 | -0,3516 | 8,03441E-07 |
| ACE2 | LINC01719 | -0,3517 | 7,97884E-07 |
| ACE2 | LINC02060 | -0,3517 | 8,01149E-07 |
| ACE2 | TRIM35 | -0,3517 | 7,97947E-07 |
| ACE2 | RP11-627K11.1 | -0,3517 | 8,00282E-07 |
| ACE2 | AC005306.3 | -0,3517 | 8,00242E-07 |
| ACE2 | TNNC1 | -0,3518 | 7,91893E-07 |
| ACE2 | SMCO2 | -0,3518 | 7,93677E-07 |
| ACE2 | CLCN7 | -0,3518 | 7,94112E-07 |
| ACE2 | RP11-34P13.13 | -0,3519 | 7,87578E-07 |
| ACE2 | RP3-476K8.3 | -0,3519 | 7,88435E-07 |
| ACE2 | USP42 | -0,3519 | 7,85804E-07 |
| ACE2 | RP11-727A23.7 | -0,3519 | 7,8507E-07 |
| ACE2 | RP11-677M14.3 | -0,3519 | 7,88193E-07 |
| ACE2 | GOLGA6L9 | -0,3519 | 7,87183E-07 |
| ACE2 | RPP14 | -0,352 | 7,82307E-07 |
| ACE2 | EIF4G1 | -0,352 | 7,777E-07 |
| ACE2 | CTD-2373H9.5 | -0,352 | 7,82454E-07 |
| ACE2 | AC091878.1 | -0,3521 | 7,76011E-07 |
| ACE2 | EXOC6 | -0,3521 | 7,75141E-07 |
| ACE2 | RP11-168K11.3 | -0,3522 | 7,67413E-07 |
| ACE2 | RP1-40G4P.1 | -0,3522 | 7,68895E-07 |
| ACE2 | LINC01930 | -0,3523 | 7,61219E-07 |
| ACE2 | RP11-563N6.6 | -0,3523 | 7,62055E-07 |
| ACE2 | RP11-1017G21.6 | -0,3523 | 7,63219E-07 |
| ACE2 | RP11-524C21.1 | -0,3524 | 7,59214E-07 |
| ACE2 | TMEM60 | -0,3524 | 7,57402E-07 |
| ACE2 | CTB-50E14.5 | -0,3524 | 7,55737E-07 |
| ACE2 | BET1 | -0,3525 | 7,4973E-07 |
| ACE2 | RN7SL809P | -0,3525 | 7,48148E-07 |
| ACE2 | NKPD1 | -0,3525 | 7,49449E-07 |
| ACE2 | AC010976.2 | -0,3527 | 7,39779E-07 |
| ACE2 | APOA1-AS | -0,3527 | 7,39446E-07 |

| | | | |
|---|---|---|---|
| ACE2 | SNORD35A | -0,3527 | 7,38452E-07 |
| ACE2 | HHIPL2 | -0,3528 | 7,3467E-07 |
| ACE2 | TSSC1 | -0,3528 | 7,33936E-07 |
| ACE2 | RAD50 | -0,3528 | 7,36168E-07 |
| ACE2 | TMEM143 | -0,3528 | 7,31494E-07 |
| ACE2 | HPRT1 | -0,3528 | 7,33621E-07 |
| ACE2 | MTF2 | -0,3529 | 7,30218E-07 |
| ACE2 | SNRNP48 | -0,3529 | 7,27079E-07 |
| ACE2 | RP11-57K17.1 | -0,3529 | 7,27898E-07 |
| ACE2 | AC073130.1 | -0,3529 | 7,28041E-07 |
| ACE2 | RN7SL75P | -0,3529 | 7,26697E-07 |
| ACE2 | MSX2P1 | -0,3529 | 7,29316E-07 |
| ACE2 | NFYA | -0,353 | 7,20317E-07 |
| ACE2 | AF067845.1 | -0,353 | 7,2184E-07 |
| ACE2 | RP11-64P14.7 | -0,3531 | 7,15806E-07 |
| ACE2 | BFAR | -0,3531 | 7,14233E-07 |
| ACE2 | RP3-467K16.2 | -0,3532 | 7,09596E-07 |
| ACE2 | AC007405.4 | -0,3532 | 7,10945E-07 |
| ACE2 | RP5-1159O4.1 | -0,3532 | 7,12324E-07 |
| ACE2 | MTHFSD | -0,3532 | 7,09492E-07 |
| ACE2 | ZNF799 | -0,3532 | 7,09433E-07 |
| ACE2 | CEP57 | -0,3533 | 7,07622E-07 |
| ACE2 | ST8SIA5 | -0,3533 | 7,08318E-07 |
| ACE2 | CTD-2587H19.2 | -0,3533 | 7,03305E-07 |
| ACE2 | RP11-930P14.2 | -0,3534 | 6,99619E-07 |
| ACE2 | RP13-554M15.8 | -0,3534 | 6,9781E-07 |
| ACE2 | RP11-13K12.2 | -0,3534 | 6,97625E-07 |
| ACE2 | RP11-770G2.2 | -0,3535 | 6,92898E-07 |
| ACE2 | POLG | -0,3535 | 6,94737E-07 |
| ACE2 | SMG1 | -0,3535 | 6,96423E-07 |
| ACE2 | RSG1 | -0,3536 | 6,89766E-07 |
| ACE2 | RP11-212P7.2 | -0,3536 | 6,87135E-07 |
| ACE2 | ENPP2 | -0,3536 | 6,88747E-07 |
| ACE2 | CERNA1 | -0,3536 | 6,88853E-07 |
| ACE2 | HPN | -0,3536 | 6,89736E-07 |
| ACE2 | RP5-1042K10.13 | -0,3536 | 6,91E-07 |
| ACE2 | TIGD6 | -0,3537 | 6,82425E-07 |
| ACE2 | RN7SKP150 | -0,3537 | 6,84252E-07 |
| ACE2 | RPSAP70 | -0,3538 | 6,81505E-07 |
| ACE2 | CYP51A1-AS1 | -0,3538 | 6,78676E-07 |
| ACE2 | RP11-574F11.3 | -0,3538 | 6,81144E-07 |
| ACE2 | CTC-525D6.1 | -0,3538 | 6,80038E-07 |
| ACE2 | AC009506.1 | -0,3539 | 6,75117E-07 |
| ACE2 | RP11-281O15.8 | -0,3539 | 6,73389E-07 |
| ACE2 | RP11-115C21.2 | -0,3539 | 6,73727E-07 |
| ACE2 | IL13 | -0,354 | 6,66901E-07 |
| ACE2 | C6orf120 | -0,354 | 6,7016E-07 |
| ACE2 | RP11-615I2.1 | -0,354 | 6,68046E-07 |
| ACE2 | USP9Y | -0,354 | 6,70495E-07 |
| ACE2 | RIMKLA | -0,3541 | 6,62848E-07 |
| ACE2 | RP13-977J11.2 | -0,3541 | 6,63789E-07 |
| ACE2 | TRAPPC6A | -0,3541 | 6,62988E-07 |
| ACE2 | SETP17 | -0,3543 | 6,54174E-07 |
| ACE2 | POLR3B | -0,3543 | 6,50555E-07 |

| | | | |
|---|---|---|---|
| ACE2 | RP11-329N15.3 | -0,3544 | 6,47606E-07 |
| ACE2 | AKAP7 | -0,3544 | 6,47891E-07 |
| ACE2 | RP11-762H8.1 | -0,3544 | 6,48817E-07 |
| ACE2 | RP11-85I17.2 | -0,3545 | 6,45396E-07 |
| ACE2 | LPAR3 | -0,3546 | 6,38382E-07 |
| ACE2 | TCP11 | -0,3546 | 6,36146E-07 |
| ACE2 | GTF2IP13 | -0,3546 | 6,36717E-07 |
| ACE2 | PSTPIP1 | -0,3546 | 6,39038E-07 |
| ACE2 | TMEM265 | -0,3546 | 6,35941E-07 |
| ACE2 | SYNPO | -0,3547 | 6,33271E-07 |
| ACE2 | SLC2A3P4 | -0,3547 | 6,30819E-07 |
| ACE2 | RSF1-IT2 | -0,3547 | 6,3516E-07 |
| ACE2 | RP11-843A23.1 | -0,3547 | 6,32461E-07 |
| ACE2 | LINC01973 | -0,3547 | 6,34635E-07 |
| ACE2 | RP11-815J4.6 | -0,3547 | 6,35492E-07 |
| ACE2 | JAK3 | -0,3547 | 6,34306E-07 |
| ACE2 | ARL4C | -0,3548 | 6,30377E-07 |
| ACE2 | CHCHD1 | -0,3548 | 6,27841E-07 |
| ACE2 | TADA2B | -0,3549 | 6,22896E-07 |
| ACE2 | RRM1 | -0,3549 | 6,24828E-07 |
| ACE2 | RP11-196E1.3 | -0,3549 | 6,22345E-07 |
| ACE2 | WRAP53 | -0,3549 | 6,2287E-07 |
| ACE2 | CTB-75G16.3 | -0,3549 | 6,23626E-07 |
| ACE2 | CFAP57 | -0,355 | 6,16982E-07 |
| ACE2 | AC009229.6 | -0,355 | 6,17843E-07 |
| ACE2 | SNORA29 | -0,355 | 6,19322E-07 |
| ACE2 | ANKRD12 | -0,355 | 6,17316E-07 |
| ACE2 | LL21NC02-1C16.2 | -0,355 | 6,16144E-07 |
| ACE2 | XX-FW83128A1.2 | -0,355 | 6,18291E-07 |
| ACE2 | SRGAP2-AS1 | -0,3551 | 6,15287E-07 |
| ACE2 | RASAL2-AS1 | -0,3551 | 6,13599E-07 |
| ACE2 | RP11-344A5.1 | -0,3551 | 6,14514E-07 |
| ACE2 | PSMD7 | -0,3551 | 6,11406E-07 |
| ACE2 | SUMO2P17 | -0,3551 | 6,13419E-07 |
| ACE2 | AP000282.3 | -0,3551 | 6,12247E-07 |
| ACE2 | RP11-326I11.5 | -0,3552 | 6,10605E-07 |
| ACE2 | C8orf37-AS1 | -0,3552 | 6,09723E-07 |
| ACE2 | AP006547.3 | -0,3552 | 6,06903E-07 |
| ACE2 | CHCHD2 | -0,3553 | 6,04717E-07 |
| ACE2 | AC104634.3 | -0,3554 | 6,01618E-07 |
| ACE2 | LPAL2 | -0,3554 | 6,01711E-07 |
| ACE2 | CABP4 | -0,3554 | 6,01281E-07 |
| ACE2 | IGFL2-AS1 | -0,3554 | 6,01588E-07 |
| ACE2 | EP300 | -0,3554 | 6,0134E-07 |
| ACE2 | RP11-234N17.1 | -0,3555 | 5,94722E-07 |
| ACE2 | CR1 | -0,3556 | 5,88759E-07 |
| ACE2 | GPC2 | -0,3556 | 5,92572E-07 |
| ACE2 | RP11-121E16.1 | -0,3556 | 5,92577E-07 |
| ACE2 | LINC00158 | -0,3556 | 5,91847E-07 |
| ACE2 | RP5-1000K24.2 | -0,3556 | 5,89621E-07 |
| ACE2 | KIAA0907 | -0,3558 | 5,83239E-07 |
| ACE2 | NYAP2 | -0,3558 | 5,83039E-07 |
| ACE2 | CEP192 | -0,3558 | 5,82038E-07 |
| ACE2 | ZDHHC11 | -0,3559 | 5,7533E-07 |

| | | | |
|---|---|---|---|
| ACE2 | AQP7P2 | -0,3559 | 5,78251E-07 |
| ACE2 | ULK4P2 | -0,3559 | 5,78467E-07 |
| ACE2 | ZNF341 | -0,3559 | 5,77143E-07 |
| ACE2 | RP5-1049N15.2 | -0,356 | 5,73959E-07 |
| ACE2 | RPS15AP1 | -0,356 | 5,72044E-07 |
| ACE2 | RP11-438B23.2 | -0,3561 | 5,68274E-07 |
| ACE2 | JAG2 | -0,3561 | 5,65716E-07 |
| ACE2 | RN7SKP23 | -0,3561 | 5,66088E-07 |
| ACE2 | LINC01695 | -0,3561 | 5,66029E-07 |
| ACE2 | RALGPS2 | -0,3562 | 5,61531E-07 |
| ACE2 | CLDN16 | -0,3562 | 5,64394E-07 |
| ACE2 | RP11-1198D22.1 | -0,3562 | 5,629E-07 |
| ACE2 | GDF9 | -0,3562 | 5,61476E-07 |
| ACE2 | WEE2-AS1 | -0,3562 | 5,63586E-07 |
| ACE2 | OGFR | -0,3562 | 5,62366E-07 |
| ACE2 | RP11-342M1.3 | -0,3563 | 5,58422E-07 |
| ACE2 | RP11-568J23.8 | -0,3563 | 5,57608E-07 |
| ACE2 | CSNK1G2 | -0,3563 | 5,60854E-07 |
| ACE2 | APOOL | -0,3563 | 5,59971E-07 |
| ACE2 | SPATA17 | -0,3564 | 5,55135E-07 |
| ACE2 | VAPB | -0,3564 | 5,55645E-07 |
| ACE2 | RP11-499E14.1 | -0,3565 | 5,51115E-07 |
| ACE2 | CTD-2023N9.1 | -0,3565 | 5,52224E-07 |
| ACE2 | SLC26A5 | -0,3565 | 5,51511E-07 |
| ACE2 | RP11-400F19.6 | -0,3566 | 5,43903E-07 |
| ACE2 | AC005523.3 | -0,3566 | 5,45256E-07 |
| ACE2 | RP11-108M9.6 | -0,3567 | 5,4127E-07 |
| ACE2 | CTC-444N24.13 | -0,3567 | 5,42197E-07 |
| ACE2 | ZNF334 | -0,3567 | 5,42874E-07 |
| ACE2 | RPS4XP6 | -0,3568 | 5,38122E-07 |
| ACE2 | MAPK8IP3 | -0,3568 | 5,35732E-07 |
| ACE2 | RP11-130F10.1 | -0,3569 | 5,34038E-07 |
| ACE2 | RP11-266L9.6 | -0,3569 | 5,33756E-07 |
| ACE2 | C1D | -0,357 | 5,27351E-07 |
| ACE2 | CTD-2201I18.1 | -0,357 | 5,2733E-07 |
| ACE2 | SMARCD2 | -0,357 | 5,27803E-07 |
| ACE2 | PREB | -0,3571 | 5,25084E-07 |
| ACE2 | CTD-2035E11.4 | -0,3572 | 5,19068E-07 |
| ACE2 | RP11-748H22.1 | -0,3572 | 5,22097E-07 |
| ACE2 | NFS1 | -0,3572 | 5,19526E-07 |
| ACE2 | AATK-AS1 | -0,3573 | 5,15536E-07 |
| ACE2 | ASPSCR1 | -0,3573 | 5,17452E-07 |
| ACE2 | RP11-196B3.2 | -0,3573 | 5,18539E-07 |
| ACE2 | TNR | -0,3574 | 5,13014E-07 |
| ACE2 | CNN2P9 | -0,3574 | 5,12196E-07 |
| ACE2 | ECHDC1 | -0,3574 | 5,12239E-07 |
| ACE2 | IFITM9P | -0,3574 | 5,13673E-07 |
| ACE2 | B3GNT4 | -0,3574 | 5,13345E-07 |
| ACE2 | WI2-87327B8.1 | -0,3574 | 5,11765E-07 |
| ACE2 | AF064858.11 | -0,3574 | 5,12792E-07 |
| ACE2 | SGPP1 | -0,3575 | 5,07827E-07 |
| ACE2 | CTD-2313J17.6 | -0,3575 | 5,101E-07 |
| ACE2 | RP4-569M23.2 | -0,3575 | 5,09119E-07 |
| ACE2 | SECISBP2 | -0,3576 | 5,05529E-07 |

| | | | |
|---|---|---|---|
| ACE2 | RP3-322G13.7 | -0,3576 | 5,05922E-07 |
| ACE2 | RP11-330M19.1 | -0,3577 | 5,00354E-07 |
| ACE2 | LINC01024 | -0,3577 | 5,01663E-07 |
| ACE2 | LCTL | -0,3577 | 5,01056E-07 |
| ACE2 | COBLL1 | -0,3579 | 4,93682E-07 |
| ACE2 | RP11-157L3.9 | -0,3579 | 4,93286E-07 |
| ACE2 | RP5-1077I2.3 | -0,3579 | 4,94633E-07 |
| ACE2 | KCNMA1-AS1 | -0,358 | 4,87502E-07 |
| ACE2 | RP11-90J7.2 | -0,358 | 4,90386E-07 |
| ACE2 | RPL12P27 | -0,358 | 4,90683E-07 |
| ACE2 | FOXP1-AS1 | -0,3581 | 4,85434E-07 |
| ACE2 | NPIPA3 | -0,3581 | 4,87081E-07 |
| ACE2 | CTD-3032J10.3 | -0,3581 | 4,86565E-07 |
| ACE2 | UBE2A | -0,3581 | 4,84668E-07 |
| ACE2 | ZNF800 | -0,3582 | 4,80381E-07 |
| ACE2 | FAM35DP | -0,3582 | 4,80813E-07 |
| ACE2 | KRT17P8 | -0,3582 | 4,82163E-07 |
| ACE2 | CTB-17P3.4 | -0,3583 | 4,77756E-07 |
| ACE2 | RP11-482M8.3 | -0,3583 | 4,78901E-07 |
| ACE2 | RPS7P11 | -0,3583 | 4,76658E-07 |
| ACE2 | RP5-827C21.4 | -0,3584 | 4,75558E-07 |
| ACE2 | RP11-317J9.1 | -0,3584 | 4,75499E-07 |
| ACE2 | PAPD4 | -0,3584 | 4,75419E-07 |
| ACE2 | C12orf56 | -0,3584 | 4,74998E-07 |
| ACE2 | FAM192A | -0,3584 | 4,73348E-07 |
| ACE2 | LINC01788 | -0,3585 | 4,68654E-07 |
| ACE2 | RP11-894J14.2 | -0,3585 | 4,70741E-07 |
| ACE2 | SETP11 | -0,3585 | 4,71494E-07 |
| ACE2 | RP11-1046B16.3 | -0,3585 | 4,69699E-07 |
| ACE2 | STK4 | -0,3585 | 4,70125E-07 |
| ACE2 | WFDC13 | -0,3585 | 4,70037E-07 |
| ACE2 | ATG16L2 | -0,3586 | 4,67929E-07 |
| ACE2 | TMEM154 | -0,3587 | 4,61262E-07 |
| ACE2 | TIMM8B | -0,3587 | 4,64494E-07 |
| ACE2 | HEXDC | -0,3587 | 4,63589E-07 |
| ACE2 | CTD-3099C6.5 | -0,3587 | 4,62919E-07 |
| ACE2 | ZSCAN5B | -0,3587 | 4,63235E-07 |
| ACE2 | AC083884.8 | -0,3588 | 4,61011E-07 |
| ACE2 | RP11-362J17.1 | -0,3589 | 4,56611E-07 |
| ACE2 | NALCN-AS1 | -0,3589 | 4,54445E-07 |
| ACE2 | FOXQ1 | -0,359 | 4,52339E-07 |
| ACE2 | CALB1 | -0,359 | 4,53659E-07 |
| ACE2 | NUP37 | -0,359 | 4,52905E-07 |
| ACE2 | OR2T8 | -0,3591 | 4,47623E-07 |
| ACE2 | PCGF2 | -0,3591 | 4,47834E-07 |
| ACE2 | CCDC130 | -0,3591 | 4,4975E-07 |
| ACE2 | DBNDD2 | -0,3591 | 4,48937E-07 |
| ACE2 | AC007365.3 | -0,3592 | 4,45822E-07 |
| ACE2 | SERPINB9P1 | -0,3592 | 4,44192E-07 |
| ACE2 | LRWD1 | -0,3592 | 4,43582E-07 |
| ACE2 | IGFL4 | -0,3592 | 4,44287E-07 |
| ACE2 | CYP24A1 | -0,3592 | 4,44689E-07 |
| ACE2 | RGR | -0,3593 | 4,43034E-07 |
| ACE2 | TNN | -0,3594 | 4,38034E-07 |

| | | | |
|---|---|---|---|
| ACE2 | SNORD94 | -0,3594 | 4,39675E-07 |
| ACE2 | PPIAP29 | -0,3594 | 4,39097E-07 |
| ACE2 | RP11-382A20.1 | -0,3594 | 4,36765E-07 |
| ACE2 | LINC01424 | -0,3594 | 4,38987E-07 |
| ACE2 | TIGD5 | -0,3595 | 4,33759E-07 |
| ACE2 | RPL21P120 | -0,3595 | 4,33963E-07 |
| ACE2 | RP11-778J15.1 | -0,3596 | 4,32617E-07 |
| ACE2 | CTB-50L17.8 | -0,3596 | 4,30227E-07 |
| ACE2 | RAB33A | -0,3596 | 4,30605E-07 |
| ACE2 | CDKN2C | -0,3597 | 4,26686E-07 |
| ACE2 | FST | -0,3597 | 4,28541E-07 |
| ACE2 | TRBV26OR9-2 | -0,3597 | 4,26865E-07 |
| ACE2 | FAM19A3 | -0,3598 | 4,24824E-07 |
| ACE2 | HIST1H3I | -0,3598 | 4,25655E-07 |
| ACE2 | LINC01479 | -0,3598 | 4,23841E-07 |
| ACE2 | RP11-423G4.10 | -0,3598 | 4,25416E-07 |
| ACE2 | PRORSD1P | -0,3599 | 4,22609E-07 |
| ACE2 | RP11-325L12.7 | -0,3599 | 4,20553E-07 |
| ACE2 | RP13-228J13.6 | -0,3599 | 4,22767E-07 |
| ACE2 | MPP4 | -0,36 | 4,1782E-07 |
| ACE2 | CUL7 | -0,3601 | 4,12974E-07 |
| ACE2 | RP11-203M5.7 | -0,3601 | 4,15302E-07 |
| ACE2 | RP11-371F15.3 | -0,3602 | 4,10948E-07 |
| ACE2 | GFM2 | -0,3602 | 4,10632E-07 |
| ACE2 | TAS2R5 | -0,3602 | 4,10377E-07 |
| ACE2 | PITRM1-AS1 | -0,3602 | 4,10819E-07 |
| ACE2 | HCG25 | -0,3603 | 4,09468E-07 |
| ACE2 | CCDC15 | -0,3603 | 4,06519E-07 |
| ACE2 | RP11-243M5.2 | -0,3603 | 4,0677E-07 |
| ACE2 | WASH7P | -0,3604 | 4,04917E-07 |
| ACE2 | AC005752.10 | -0,3604 | 4,03817E-07 |
| ACE2 | RP11-281O15.2 | -0,3604 | 4,0551E-07 |
| ACE2 | ABRA | -0,3604 | 4,03974E-07 |
| ACE2 | RELT | -0,3604 | 4,03434E-07 |
| ACE2 | MAGEB17 | -0,3604 | 4,04797E-07 |
| ACE2 | STAG3L2 | -0,3605 | 4,014E-07 |
| ACE2 | SEL1L2 | -0,3605 | 4,01906E-07 |
| ACE2 | RPL4P6 | -0,3605 | 4,02721E-07 |
| ACE2 | RP11-159D12.8 | -0,3606 | 3,97947E-07 |
| ACE2 | RP11-314C9.1 | -0,3607 | 3,96399E-07 |
| ACE2 | NKTR | -0,3607 | 3,96507E-07 |
| ACE2 | RP11-206L10.9 | -0,3608 | 3,93193E-07 |
| ACE2 | ITPRIPL1 | -0,3608 | 3,91856E-07 |
| ACE2 | RP11-472M19.2 | -0,3608 | 3,9145E-07 |
| ACE2 | SCNN1D | -0,3609 | 3,88183E-07 |
| ACE2 | SYT17 | -0,3609 | 3,89762E-07 |
| ACE2 | RP1-59D14.10 | -0,3609 | 3,88767E-07 |
| ACE2 | HS3ST3B1 | -0,3609 | 3,87978E-07 |
| ACE2 | SNCA-AS1 | -0,361 | 3,86919E-07 |
| ACE2 | RP1-67K17.4 | -0,361 | 3,85183E-07 |
| ACE2 | JOSD2 | -0,361 | 3,8612E-07 |
| ACE2 | CTA-217C2.2 | -0,361 | 3,86594E-07 |
| ACE2 | ESRRAP2 | -0,3611 | 3,82606E-07 |
| ACE2 | SNORA2 | -0,3612 | 3,80565E-07 |

| | | | |
|---|---|---|---|
| ACE2 | UTP14C | -0,3612 | 3,80024E-07 |
| ACE2 | RP1-56K13.5 | -0,3612 | 3,78919E-07 |
| ACE2 | RP3-417G15.1 | -0,3612 | 3,79464E-07 |
| ACE2 | RP11-421M1.8 | -0,3613 | 3,77149E-07 |
| ACE2 | AC003985.1 | -0,3613 | 3,78139E-07 |
| ACE2 | FBXO32 | -0,3613 | 3,77935E-07 |
| ACE2 | STARD13-IT1 | -0,3613 | 3,78168E-07 |
| ACE2 | AC139100.4 | -0,3613 | 3,78213E-07 |
| ACE2 | ALDH5A1 | -0,3614 | 3,74834E-07 |
| ACE2 | RP11-95G6.1 | -0,3614 | 3,74907E-07 |
| ACE2 | RP11-799D4.4 | -0,3614 | 3,73205E-07 |
| ACE2 | PKP4-AS1 | -0,3615 | 3,71965E-07 |
| ACE2 | ZDHHC16 | -0,3615 | 3,70833E-07 |
| ACE2 | TEX29 | -0,3615 | 3,7007E-07 |
| ACE2 | SLC22A14 | -0,3616 | 3,67071E-07 |
| ACE2 | CTB-193M12.3 | -0,3616 | 3,6844E-07 |
| ACE2 | ANKRD18CP | -0,3617 | 3,65385E-07 |
| ACE2 | RP11-885N19.6 | -0,3617 | 3,63743E-07 |
| ACE2 | ANP32A | -0,3617 | 3,64038E-07 |
| ACE2 | ARHGEF2 | -0,3618 | 3,62591E-07 |
| ACE2 | NSUN5P1 | -0,3618 | 3,62336E-07 |
| ACE2 | ALG1L9P | -0,3618 | 3,62474E-07 |
| ACE2 | ST13P11 | -0,3618 | 3,62694E-07 |
| ACE2 | CTC-523E23.11 | -0,3618 | 3,63601E-07 |
| ACE2 | ZNF75D | -0,3618 | 3,6253E-07 |
| ACE2 | RP11-11N5.1 | -0,3619 | 3,60659E-07 |
| ACE2 | LINC01394 | -0,3619 | 3,5997E-07 |
| ACE2 | CAP1 | -0,362 | 3,56873E-07 |
| ACE2 | MFSD8 | -0,362 | 3,55398E-07 |
| ACE2 | RP11-79P5.9 | -0,362 | 3,55862E-07 |
| ACE2 | YWHAZP4 | -0,362 | 3,56144E-07 |
| ACE2 | ZWILCH | -0,362 | 3,57342E-07 |
| ACE2 | MCM7 | -0,3621 | 3,53024E-07 |
| ACE2 | CCDC137 | -0,3621 | 3,53886E-07 |
| ACE2 | SNAP25-AS1 | -0,3621 | 3,5389E-07 |
| ACE2 | MLF2 | -0,3622 | 3,51214E-07 |
| ACE2 | CTD-2017C7.2 | -0,3622 | 3,51964E-07 |
| ACE2 | LL0XNC01-7P3.1 | -0,3622 | 3,5196E-07 |
| ACE2 | KNTC1 | -0,3623 | 3,48537E-07 |
| ACE2 | CTC-412M14.5 | -0,3623 | 3,49178E-07 |
| ACE2 | LCA10 | -0,3623 | 3,49292E-07 |
| ACE2 | NPM1P39 | -0,3624 | 3,46145E-07 |
| ACE2 | AC092155.4 | -0,3624 | 3,44691E-07 |
| ACE2 | PRPF40A | -0,3624 | 3,44419E-07 |
| ACE2 | CTD-3080F16.3 | -0,3624 | 3,45664E-07 |
| ACE2 | SPTY2D1 | -0,3624 | 3,46206E-07 |
| ACE2 | HBZ | -0,3624 | 3,44923E-07 |
| ACE2 | NPIPB7 | -0,3624 | 3,44678E-07 |
| ACE2 | TNFRSF4 | -0,3625 | 3,43607E-07 |
| ACE2 | USP40 | -0,3625 | 3,41264E-07 |
| ACE2 | PAK2 | -0,3625 | 3,42652E-07 |
| ACE2 | TRAF3IP2 | -0,3625 | 3,41866E-07 |
| ACE2 | SACM1L | -0,3626 | 3,40164E-07 |
| ACE2 | RHOA-IT1 | -0,3626 | 3,39917E-07 |

| | | | |
|---|---|---|---|
| ACE2 | LINC02188 | -0,3626 | 3,40261E-07 |
| ACE2 | PTGES3L | -0,3626 | 3,40363E-07 |
| ACE2 | TTC16 | -0,3627 | 3,3673E-07 |
| ACE2 | LINC02156 | -0,3627 | 3,38388E-07 |
| ACE2 | RP11-923I11.4 | -0,3627 | 3,36505E-07 |
| ACE2 | RP11-61K12.2 | -0,3628 | 3,35462E-07 |
| ACE2 | GALNT10 | -0,3628 | 3,34854E-07 |
| ACE2 | MRPL28 | -0,3628 | 3,35084E-07 |
| ACE2 | FBXL12 | -0,3628 | 3,34825E-07 |
| ACE2 | CTA-212A2.4 | -0,3628 | 3,35273E-07 |
| ACE2 | RP11-269F19.2 | -0,3629 | 3,30779E-07 |
| ACE2 | KIRREL3-AS1 | -0,3629 | 3,32458E-07 |
| ACE2 | RP13-43E11.1 | -0,3629 | 3,3166E-07 |
| ACE2 | FUT10 | -0,363 | 3,28268E-07 |
| ACE2 | CCL7 | -0,363 | 3,30387E-07 |
| ACE2 | ASB16 | -0,363 | 3,29221E-07 |
| ACE2 | RP11-207C16.4 | -0,3631 | 3,27891E-07 |
| ACE2 | TSSK1A | -0,3631 | 3,25887E-07 |
| ACE2 | RP11-247I13.11 | -0,3631 | 3,27099E-07 |
| ACE2 | CEP152 | -0,3632 | 3,2396E-07 |
| ACE2 | MPRIP-AS1 | -0,3632 | 3,23035E-07 |
| ACE2 | TEX261 | -0,3633 | 3,20989E-07 |
| ACE2 | AC000123.4 | -0,3633 | 3,20582E-07 |
| ACE2 | RP11-1112C15.2 | -0,3633 | 3,22004E-07 |
| ACE2 | PPP2R1A | -0,3633 | 3,22444E-07 |
| ACE2 | RP11-278C7.1 | -0,3634 | 3,188E-07 |
| ACE2 | ZDHHC22 | -0,3635 | 3,15718E-07 |
| ACE2 | GJA1P1 | -0,3636 | 3,14752E-07 |
| ACE2 | CTC-563A5.2 | -0,3636 | 3,13431E-07 |
| ACE2 | CTD-3065B20.2 | -0,3636 | 3,12638E-07 |
| ACE2 | SCML2P2 | -0,3636 | 3,14475E-07 |
| ACE2 | LDLRAD4 | -0,3636 | 3,14775E-07 |
| ACE2 | GTF2IP9 | -0,3637 | 3,11296E-07 |
| ACE2 | CTD-2184D3.7 | -0,3637 | 3,12254E-07 |
| ACE2 | GOLGA6L4 | -0,3637 | 3,10957E-07 |
| ACE2 | TDRKH-AS1 | -0,3638 | 3,09E-07 |
| ACE2 | RNVU1-15 | -0,3639 | 3,05374E-07 |
| ACE2 | RP11-579E24.2 | -0,3639 | 3,06992E-07 |
| ACE2 | ANKLE2 | -0,3639 | 3,06968E-07 |
| ACE2 | CTD-2313J17.5 | -0,3639 | 3,05365E-07 |
| ACE2 | GRIN2A | -0,3639 | 3,05763E-07 |
| ACE2 | SKAP2 | -0,364 | 3,04478E-07 |
| ACE2 | RP11-253I19.3 | -0,364 | 3,04415E-07 |
| ACE2 | USP15 | -0,364 | 3,02681E-07 |
| ACE2 | CTC-260E6.3 | -0,364 | 3,04052E-07 |
| ACE2 | NCAPGP1 | -0,3641 | 3,0069E-07 |
| ACE2 | FAM170B | -0,3641 | 3,01358E-07 |
| ACE2 | HS2ST1 | -0,3642 | 2,98491E-07 |
| ACE2 | TSACC | -0,3642 | 2,98825E-07 |
| ACE2 | RP11-154H23.4 | -0,3642 | 2,98822E-07 |
| ACE2 | RPA3 | -0,3643 | 2,96027E-07 |
| ACE2 | FAM71F1 | -0,3643 | 2,96084E-07 |
| ACE2 | RP11-342M3.5 | -0,3643 | 2,95608E-07 |
| ACE2 | PDCD6IPP1 | -0,3643 | 2,97362E-07 |

| | | | |
|---|---|---|---|
| ACE2 | AC004381.6 | -0,3643 | 2,96154E-07 |
| ACE2 | RP11-68I3.11 | -0,3643 | 2,95825E-07 |
| ACE2 | RP11-175P19.2 | -0,3644 | 2,94889E-07 |
| ACE2 | ZC2HC1B | -0,3644 | 2,94251E-07 |
| ACE2 | FSD2 | -0,3644 | 2,94408E-07 |
| ACE2 | TBX2 | -0,3644 | 2,94805E-07 |
| ACE2 | ARHGEF1 | -0,3644 | 2,95251E-07 |
| ACE2 | RP11-49O14.2 | -0,3645 | 2,91872E-07 |
| ACE2 | LMBR1L | -0,3645 | 2,91061E-07 |
| ACE2 | CES5A | -0,3645 | 2,92999E-07 |
| ACE2 | RPL14P3 | -0,3646 | 2,90661E-07 |
| ACE2 | RN7SL49P | -0,3646 | 2,88611E-07 |
| ACE2 | CCDC94 | -0,3646 | 2,89763E-07 |
| ACE2 | RAP2C | -0,3646 | 2,90266E-07 |
| ACE2 | C1orf234 | -0,3647 | 2,87487E-07 |
| ACE2 | RN7SKP16 | -0,3647 | 2,87686E-07 |
| ACE2 | DDX50P1 | -0,3647 | 2,86238E-07 |
| ACE2 | RP11-271M24.2 | -0,3648 | 2,84623E-07 |
| ACE2 | ALOX5AP | -0,3648 | 2,84189E-07 |
| ACE2 | AP001437.1 | -0,3648 | 2,84471E-07 |
| ACE2 | AC053503.11 | -0,3649 | 2,83602E-07 |
| ACE2 | STAB2 | -0,3649 | 2,82541E-07 |
| ACE2 | XYLT1 | -0,3649 | 2,83305E-07 |
| ACE2 | RP11-392O17.1 | -0,365 | 2,79764E-07 |
| ACE2 | RP11-562I5.2 | -0,365 | 2,7938E-07 |
| ACE2 | PAICS | -0,365 | 2,79355E-07 |
| ACE2 | SEPT7P3 | -0,365 | 2,79476E-07 |
| ACE2 | SAPCD2P3 | -0,365 | 2,80009E-07 |
| ACE2 | RP11-215D10.1 | -0,3651 | 2,77494E-07 |
| ACE2 | SNX5 | -0,3651 | 2,79027E-07 |
| ACE2 | IL12RB2 | -0,3652 | 2,76927E-07 |
| ACE2 | ANKRD17 | -0,3652 | 2,76512E-07 |
| ACE2 | AFG3L1P | -0,3652 | 2,7653E-07 |
| ACE2 | CTD-3193K9.4 | -0,3652 | 2,75265E-07 |
| ACE2 | RP11-856M7.2 | -0,3652 | 2,76866E-07 |
| ACE2 | RP5-1126H10.2 | -0,3653 | 2,738E-07 |
| ACE2 | LINC01894 | -0,3653 | 2,73308E-07 |
| ACE2 | SLC35C2 | -0,3653 | 2,73012E-07 |
| ACE2 | POU3F1 | -0,3655 | 2,68481E-07 |
| ACE2 | RPL7AP26 | -0,3655 | 2,69563E-07 |
| ACE2 | CPNE5 | -0,3655 | 2,70352E-07 |
| ACE2 | SLC26A7 | -0,3655 | 2,70105E-07 |
| ACE2 | STMN1P1 | -0,3655 | 2,69286E-07 |
| ACE2 | RP11-50D16.4 | -0,3655 | 2,6945E-07 |
| ACE2 | TMEM178A | -0,3657 | 2,65929E-07 |
| ACE2 | SLC22A3 | -0,3657 | 2,6463E-07 |
| ACE2 | CCL19 | -0,3657 | 2,65831E-07 |
| ACE2 | OAS3 | -0,3657 | 2,64623E-07 |
| ACE2 | SMG1P7 | -0,3657 | 2,65192E-07 |
| ACE2 | TMEM27 | -0,3657 | 2,6611E-07 |
| ACE2 | ARL4AP4 | -0,3658 | 2,63943E-07 |
| ACE2 | CPLX2 | -0,3658 | 2,6368E-07 |
| ACE2 | RP11-481A20.10 | -0,3658 | 2,62309E-07 |
| ACE2 | ARL17A | -0,3658 | 2,63111E-07 |

| | | | |
|---|---|---|---|
| ACE2 | POLR2J3 | -0,3659 | 2,60082E-07 |
| ACE2 | ADARB2 | -0,3659 | 2,61599E-07 |
| ACE2 | CTD-3184A7.4 | -0,3659 | 2,60478E-07 |
| ACE2 | RP11-298C3.2 | -0,3659 | 2,6078E-07 |
| ACE2 | RP11-504P24.3 | -0,366 | 2,58668E-07 |
| ACE2 | SETP14 | -0,366 | 2,59601E-07 |
| ACE2 | JRKL | -0,366 | 2,57822E-07 |
| ACE2 | PHBP9 | -0,3661 | 2,5767E-07 |
| ACE2 | RP11-76C10.3 | -0,3661 | 2,56315E-07 |
| ACE2 | ACTC1 | -0,3661 | 2,57056E-07 |
| ACE2 | SNORD3C | -0,3661 | 2,55823E-07 |
| ACE2 | CSF2RB | -0,3661 | 2,56074E-07 |
| ACE2 | CTB-113P19.3 | -0,3662 | 2,55442E-07 |
| ACE2 | GOLGA7B | -0,3662 | 2,53677E-07 |
| ACE2 | RP11-697E2.9 | -0,3663 | 2,53157E-07 |
| ACE2 | CETN4P | -0,3664 | 2,50876E-07 |
| ACE2 | TATDN2P3 | -0,3664 | 2,5098E-07 |
| ACE2 | DYNLL2 | -0,3664 | 2,5093E-07 |
| ACE2 | LINC00863 | -0,3665 | 2,47641E-07 |
| ACE2 | RPL17P36 | -0,3665 | 2,4749E-07 |
| ACE2 | RP11-685G9.2 | -0,3665 | 2,47559E-07 |
| ACE2 | RP11-360N9.2 | -0,3665 | 2,47873E-07 |
| ACE2 | RP11-561O4.1 | -0,3666 | 2,47073E-07 |
| ACE2 | RP11-632C17__A.1 | -0,3666 | 2,46536E-07 |
| ACE2 | COL20A1 | -0,3666 | 2,46631E-07 |
| ACE2 | RP11-115H13.1 | -0,3666 | 2,4675E-07 |
| ACE2 | RPS15AP12 | -0,3667 | 2,44008E-07 |
| ACE2 | CHEK2 | -0,3667 | 2,44343E-07 |
| ACE2 | RP11-431J24.2 | -0,3667 | 2,45055E-07 |
| ACE2 | SMC2 | -0,3668 | 2,42077E-07 |
| ACE2 | ANKRD33 | -0,3668 | 2,43315E-07 |
| ACE2 | YIPF6 | -0,3668 | 2,4284E-07 |
| ACE2 | GCNA | -0,3668 | 2,4226E-07 |
| ACE2 | RP11-297J22.1 | -0,3669 | 2,41399E-07 |
| ACE2 | RP11-91P24.7 | -0,3669 | 2,40488E-07 |
| ACE2 | RN7SL812P | -0,3669 | 2,40966E-07 |
| ACE2 | TMEM50A | -0,367 | 2,38163E-07 |
| ACE2 | GHRLOS | -0,367 | 2,38682E-07 |
| ACE2 | GMNC | -0,367 | 2,38209E-07 |
| ACE2 | RPS11P5 | -0,367 | 2,37983E-07 |
| ACE2 | CBX3P2 | -0,367 | 2,39242E-07 |
| ACE2 | RP11-53I6.4 | -0,367 | 2,38132E-07 |
| ACE2 | OSBPL11 | -0,3671 | 2,3595E-07 |
| ACE2 | GRM3 | -0,3671 | 2,37396E-07 |
| ACE2 | RP11-264E20.1 | -0,3671 | 2,36191E-07 |
| ACE2 | FBXW10 | -0,3671 | 2,36428E-07 |
| ACE2 | RP11-616M22.2 | -0,3672 | 2,3479E-07 |
| ACE2 | CTD-3222D19.9 | -0,3672 | 2,34E-07 |
| ACE2 | XXYLT1-AS2 | -0,3673 | 2,33638E-07 |
| ACE2 | LKAAEAR1 | -0,3673 | 2,32771E-07 |
| ACE2 | RP1-124C6.1 | -0,3674 | 2,30205E-07 |
| ACE2 | RP1-283K11.2 | -0,3674 | 2,31873E-07 |
| ACE2 | RP11-1E4.1 | -0,3674 | 2,31019E-07 |
| ACE2 | FICD | -0,3674 | 2,315E-07 |

| | | | |
|---|---|---|---|
| ACE2 | TMEM31 | -0,3674 | 2,30236E-07 |
| ACE2 | LINC00487 | -0,3675 | 2,29195E-07 |
| ACE2 | LINC01506 | -0,3675 | 2,29023E-07 |
| ACE2 | AC010729.2 | -0,3676 | 2,27342E-07 |
| ACE2 | RP11-271K11.1 | -0,3676 | 2,28141E-07 |
| ACE2 | GLIS1 | -0,3677 | 2,24846E-07 |
| ACE2 | RP11-288A5.2 | -0,3677 | 2,24794E-07 |
| ACE2 | PPP1R3E | -0,3677 | 2,25335E-07 |
| ACE2 | LINC00618 | -0,3677 | 2,25216E-07 |
| ACE2 | RP11-10A14.3 | -0,3678 | 2,24383E-07 |
| ACE2 | RAP1BP1 | -0,3678 | 2,2332E-07 |
| ACE2 | CALHM1 | -0,3678 | 2,23386E-07 |
| ACE2 | RP11-1017G21.5 | -0,3678 | 2,22755E-07 |
| ACE2 | RP11-383C5.3 | -0,368 | 2,20158E-07 |
| ACE2 | BNIP3P15 | -0,368 | 2,20026E-07 |
| ACE2 | LSMEM2 | -0,3681 | 2,17677E-07 |
| ACE2 | RP11-71E19.1 | -0,3681 | 2,18267E-07 |
| ACE2 | NSUN6 | -0,3681 | 2,19107E-07 |
| ACE2 | RP11-1007G5.2 | -0,3681 | 2,18947E-07 |
| ACE2 | BAHCC1 | -0,3681 | 2,17373E-07 |
| ACE2 | UBA1 | -0,3681 | 2,187E-07 |
| ACE2 | RP11-494H4.3 | -0,3682 | 2,16776E-07 |
| ACE2 | RPL7P26 | -0,3682 | 2,16923E-07 |
| ACE2 | RP11-415J8.7 | -0,3683 | 2,15272E-07 |
| ACE2 | RP11-122G18.11 | -0,3683 | 2,15245E-07 |
| ACE2 | C20orf197 | -0,3683 | 2,14665E-07 |
| ACE2 | AC009312.1 | -0,3684 | 2,12253E-07 |
| ACE2 | KATNB1 | -0,3684 | 2,13583E-07 |
| ACE2 | RP11-615I2.2 | -0,3684 | 2,13831E-07 |
| ACE2 | CAMTA2 | -0,3684 | 2,13564E-07 |
| ACE2 | C2orf16 | -0,3685 | 2,11302E-07 |
| ACE2 | DHX15 | -0,3686 | 2,09209E-07 |
| ACE2 | CHML | -0,3687 | 2,07408E-07 |
| ACE2 | LINC01305 | -0,3687 | 2,08573E-07 |
| ACE2 | AC010900.2 | -0,3687 | 2,08271E-07 |
| ACE2 | CARD6 | -0,3687 | 2,07651E-07 |
| ACE2 | CTB-31N19.3 | -0,3687 | 2,07338E-07 |
| ACE2 | LINC01393 | -0,3688 | 2,06745E-07 |
| ACE2 | GLIDR | -0,3688 | 2,06076E-07 |
| ACE2 | ALDH1A2 | -0,3688 | 2,05445E-07 |
| ACE2 | VDAC1P8 | -0,3689 | 2,03769E-07 |
| ACE2 | DDX54 | -0,3689 | 2,04863E-07 |
| ACE2 | PLEK | -0,369 | 2,02737E-07 |
| ACE2 | RP11-219G10.1 | -0,369 | 2,02292E-07 |
| ACE2 | ZNF92 | -0,369 | 2,02122E-07 |
| ACE2 | PSMB3 | -0,369 | 2,02197E-07 |
| ACE2 | SLC15A2 | -0,3691 | 2,00888E-07 |
| ACE2 | RP11-34F20.7 | -0,3691 | 2,01437E-07 |
| ACE2 | WDHD1 | -0,3691 | 2,00387E-07 |
| ACE2 | RP11-147L13.7 | -0,3691 | 2,0118E-07 |
| ACE2 | MYLK2 | -0,3691 | 2,00968E-07 |
| ACE2 | FITM2 | -0,3691 | 2,01822E-07 |
| ACE2 | MTND4P26 | -0,3692 | 1,99919E-07 |
| ACE2 | COL4A4 | -0,3692 | 1,99331E-07 |

| | | | |
|---|---|---|---|
| ACE2 | ZNRF3 | -0,3692 | 2,00084E-07 |
| ACE2 | THAP4 | -0,3693 | 1,9765E-07 |
| ACE2 | RP11-12J10.4 | -0,3693 | 1,9773E-07 |
| ACE2 | CASC18 | -0,3693 | 1,97824E-07 |
| ACE2 | TRIM73 | -0,3694 | 1,9667E-07 |
| ACE2 | CTD-2017C7.1 | -0,3694 | 1,96552E-07 |
| ACE2 | ZNF628 | -0,3694 | 1,95919E-07 |
| ACE2 | CATIP | -0,3695 | 1,94112E-07 |
| ACE2 | FAM90A20P | -0,3695 | 1,95073E-07 |
| ACE2 | GNAS-AS1 | -0,3695 | 1,94083E-07 |
| ACE2 | AC010883.5 | -0,3696 | 1,9277E-07 |
| ACE2 | SPATA1 | -0,3697 | 1,91496E-07 |
| ACE2 | TRIM67 | -0,3697 | 1,91283E-07 |
| ACE2 | ABT1 | -0,3697 | 1,92238E-07 |
| ACE2 | RP3-388M5.9 | -0,3697 | 1,92103E-07 |
| ACE2 | RP11-386I14.4 | -0,3698 | 1,90676E-07 |
| ACE2 | RP11-404O13.4 | -0,3698 | 1,90495E-07 |
| ACE2 | IL1RL2 | -0,3698 | 1,89862E-07 |
| ACE2 | CLIP2 | -0,3698 | 1,89699E-07 |
| ACE2 | RP11-536K7.5 | -0,3698 | 1,90083E-07 |
| ACE2 | RP11-453E17.4 | -0,3699 | 1,88136E-07 |
| ACE2 | HRH3 | -0,3699 | 1,88348E-07 |
| ACE2 | MAPKAPK5-AS1 | -0,37 | 1,87355E-07 |
| ACE2 | RPL9P7 | -0,37 | 1,8625E-07 |
| ACE2 | THEM7P | -0,3701 | 1,84781E-07 |
| ACE2 | RPS5P2 | -0,3701 | 1,85053E-07 |
| ACE2 | RP11-372K14.2 | -0,3702 | 1,83789E-07 |
| ACE2 | DFFB | -0,3703 | 1,82296E-07 |
| ACE2 | NT5C1A | -0,3703 | 1,82842E-07 |
| ACE2 | AC093787.1 | -0,3703 | 1,82035E-07 |
| ACE2 | RP11-445F6.2 | -0,3703 | 1,81756E-07 |
| ACE2 | RP11-482E14.1 | -0,3703 | 1,82238E-07 |
| ACE2 | FAM103A1 | -0,3703 | 1,82825E-07 |
| ACE2 | SLC25A15P5 | -0,3703 | 1,82492E-07 |
| ACE2 | RP11-333B11.1 | -0,3704 | 1,8154E-07 |
| ACE2 | MEIS1-AS3 | -0,3705 | 1,78962E-07 |
| ACE2 | AC114877.3 | -0,3705 | 1,79255E-07 |
| ACE2 | DUX4L50 | -0,3705 | 1,7988E-07 |
| ACE2 | IFNG | -0,3706 | 1,78366E-07 |
| ACE2 | PLPPR3 | -0,3706 | 1,7795E-07 |
| ACE2 | RP3-407E4.4 | -0,3707 | 1,76566E-07 |
| ACE2 | OGT | -0,3707 | 1,76736E-07 |
| ACE2 | CH17-172B3.1 | -0,3708 | 1,75081E-07 |
| ACE2 | EGOT | -0,3708 | 1,75543E-07 |
| ACE2 | CALCA | -0,3708 | 1,75709E-07 |
| ACE2 | RP11-6N17.4 | -0,3708 | 1,74688E-07 |
| ACE2 | RP5-1065J22.8 | -0,3709 | 1,73415E-07 |
| ACE2 | FAM205A | -0,3709 | 1,72965E-07 |
| ACE2 | SSNA1 | -0,3709 | 1,7402E-07 |
| ACE2 | MTDHP2 | -0,3709 | 1,74326E-07 |
| ACE2 | FCHSD1 | -0,371 | 1,72108E-07 |
| ACE2 | LAMB1 | -0,371 | 1,72539E-07 |
| ACE2 | RP11-995C19.2 | -0,371 | 1,72811E-07 |
| ACE2 | CTD-2525I3.5 | -0,371 | 1,71767E-07 |

| | | | |
|---|---|---|---|
| ACE2 | RP11-368J21.3 | -0,3711 | 1,71221E-07 |
| ACE2 | MAGEC3 | -0,3711 | 1,7117E-07 |
| ACE2 | FBXO16 | -0,3712 | 1,69748E-07 |
| ACE2 | RP11-736N17.11 | -0,3712 | 1,69838E-07 |
| ACE2 | RP11-16E12.1 | -0,3712 | 1,69432E-07 |
| ACE2 | CLIC3 | -0,3713 | 1,67414E-07 |
| ACE2 | NTRK3-AS1 | -0,3713 | 1,67646E-07 |
| ACE2 | STK35 | -0,3713 | 1,68527E-07 |
| ACE2 | TMEM52B | -0,3715 | 1,65062E-07 |
| ACE2 | PPP1R3D | -0,3716 | 1,63575E-07 |
| ACE2 | SLC25A2 | -0,3717 | 1,62943E-07 |
| ACE2 | RP11-77P6.2 | -0,3717 | 1,62562E-07 |
| ACE2 | RP11-1263C18.2 | -0,3718 | 1,60778E-07 |
| ACE2 | RP11-322D14.2 | -0,3718 | 1,61352E-07 |
| ACE2 | RP11-401O9.3 | -0,3718 | 1,60676E-07 |
| ACE2 | UBE2K | -0,3719 | 1,60079E-07 |
| ACE2 | RP5-1065J22.2 | -0,372 | 1,58457E-07 |
| ACE2 | CRK | -0,372 | 1,58774E-07 |
| ACE2 | MATN1-AS1 | -0,3722 | 1,55638E-07 |
| ACE2 | SMIM3 | -0,3722 | 1,56183E-07 |
| ACE2 | TCF19 | -0,3722 | 1,55438E-07 |
| ACE2 | TRPV5 | -0,3722 | 1,56197E-07 |
| ACE2 | GNA13 | -0,3722 | 1,56207E-07 |
| ACE2 | SPTA1 | -0,3723 | 1,54296E-07 |
| ACE2 | RP11-861A13.3 | -0,3723 | 1,55067E-07 |
| ACE2 | AGAP5 | -0,3723 | 1,55369E-07 |
| ACE2 | POU5F1P3 | -0,3723 | 1,55269E-07 |
| ACE2 | RP11-457P14.5 | -0,3724 | 1,5401E-07 |
| ACE2 | CCDC89 | -0,3724 | 1,53812E-07 |
| ACE2 | AK4P3 | -0,3724 | 1,52859E-07 |
| ACE2 | GPR183 | -0,3724 | 1,53491E-07 |
| ACE2 | AC022154.7 | -0,3724 | 1,53486E-07 |
| ACE2 | TBX18-AS1 | -0,3725 | 1,52074E-07 |
| ACE2 | CLPB | -0,3725 | 1,51632E-07 |
| ACE2 | ARHGDIB | -0,3725 | 1,5225E-07 |
| ACE2 | ADPGK | -0,3725 | 1,52592E-07 |
| ACE2 | RP11-130L8.3 | -0,3726 | 1,505E-07 |
| ACE2 | ZNF462 | -0,3726 | 1,50766E-07 |
| ACE2 | RP11-432J22.2 | -0,3726 | 1,50443E-07 |
| ACE2 | RP11-543P15.3 | -0,3726 | 1,50751E-07 |
| ACE2 | SCN9A | -0,3727 | 1,49835E-07 |
| ACE2 | RP11-514P8.10 | -0,3727 | 1,50293E-07 |
| ACE2 | OR13J1 | -0,3727 | 1,49281E-07 |
| ACE2 | TMBIM4 | -0,3727 | 1,49758E-07 |
| ACE2 | FBXO27 | -0,3727 | 1,50152E-07 |
| ACE2 | PP14571 | -0,3728 | 1,48859E-07 |
| ACE2 | TMEM251 | -0,3728 | 1,48972E-07 |
| ACE2 | SIRPB2 | -0,3728 | 1,4819E-07 |
| ACE2 | HTR1E | -0,3729 | 1,4722E-07 |
| ACE2 | TSPAN33 | -0,373 | 1,46434E-07 |
| ACE2 | MAFA-AS1 | -0,373 | 1,46123E-07 |
| ACE2 | RAB39A | -0,373 | 1,46157E-07 |
| ACE2 | AC003682.16 | -0,373 | 1,4627E-07 |
| ACE2 | RP3-467L1.6 | -0,3731 | 1,44819E-07 |

| | | | |
|---|---|---|---|
| ACE2 | MED19 | -0,3731 | 1,45251E-07 |
| ACE2 | AC009229.5 | -0,3732 | 1,43878E-07 |
| ACE2 | ABCC10 | -0,3732 | 1,43558E-07 |
| ACE2 | RP11-402L5.1 | -0,3732 | 1,43859E-07 |
| ACE2 | RND1 | -0,3733 | 1,42225E-07 |
| ACE2 | GTPBP3 | -0,3733 | 1,42014E-07 |
| ACE2 | CH17-385C13.2 | -0,3734 | 1,41345E-07 |
| ACE2 | AC016907.3 | -0,3734 | 1,41828E-07 |
| ACE2 | TREX1 | -0,3734 | 1,41698E-07 |
| ACE2 | RP11-635L1.3 | -0,3734 | 1,40738E-07 |
| ACE2 | RP11-299H22.3 | -0,3735 | 1,40489E-07 |
| ACE2 | GTF2IP4 | -0,3736 | 1,39099E-07 |
| ACE2 | DAPK1 | -0,3736 | 1,39209E-07 |
| ACE2 | RP11-17A1.3 | -0,3736 | 1,38465E-07 |
| ACE2 | FAM186A | -0,3736 | 1,38461E-07 |
| ACE2 | RP11-1348G14.6 | -0,3736 | 1,39112E-07 |
| ACE2 | TPRG1LP1 | -0,3736 | 1,39067E-07 |
| ACE2 | TAL2 | -0,3737 | 1,3759E-07 |
| ACE2 | RP11-578F21.2 | -0,3737 | 1,37554E-07 |
| ACE2 | RP11-1102P22.1 | -0,3737 | 1,38064E-07 |
| ACE2 | LINC01700 | -0,3737 | 1,37862E-07 |
| ACE2 | RP11-486G15.1 | -0,3738 | 1,36764E-07 |
| ACE2 | CTB-111H14.1 | -0,3739 | 1,35042E-07 |
| ACE2 | RP11-423C15.3 | -0,3739 | 1,35287E-07 |
| ACE2 | CXCL8 | -0,374 | 1,34594E-07 |
| ACE2 | CCDC78 | -0,374 | 1,34748E-07 |
| ACE2 | LINC01929 | -0,374 | 1,34839E-07 |
| ACE2 | DPY19L3 | -0,374 | 1,34851E-07 |
| ACE2 | U2AF1L4 | -0,374 | 1,34032E-07 |
| ACE2 | KANSL1 | -0,3741 | 1,33701E-07 |
| ACE2 | FAM90A25P | -0,3742 | 1,31906E-07 |
| ACE2 | SH2B1 | -0,3742 | 1,32143E-07 |
| ACE2 | RP11-81A22.5 | -0,3742 | 1,31735E-07 |
| ACE2 | RP11-848P1.3 | -0,3742 | 1,32676E-07 |
| ACE2 | ROR1-AS1 | -0,3743 | 1,31496E-07 |
| ACE2 | SDHC | -0,3743 | 1,3115E-07 |
| ACE2 | DNAJC5G | -0,3743 | 1,31539E-07 |
| ACE2 | RNU6-26P | -0,3743 | 1,31637E-07 |
| ACE2 | RAB44 | -0,3743 | 1,3125E-07 |
| ACE2 | RP5-1071N3.1 | -0,3744 | 1,3033E-07 |
| ACE2 | INHBB | -0,3744 | 1,30475E-07 |
| ACE2 | RPL21P5 | -0,3744 | 1,29703E-07 |
| ACE2 | CHCHD4 | -0,3745 | 1,28829E-07 |
| ACE2 | RP11-863P13.6 | -0,3745 | 1,29093E-07 |
| ACE2 | GRM1 | -0,3746 | 1,27531E-07 |
| ACE2 | STRIP2 | -0,3746 | 1,27894E-07 |
| ACE2 | HBE1 | -0,3746 | 1,28356E-07 |
| ACE2 | CTD-2027I19.3 | -0,3746 | 1,2755E-07 |
| ACE2 | DIAPH2-AS1 | -0,3746 | 1,27927E-07 |
| ACE2 | FCRLB | -0,3747 | 1,27372E-07 |
| ACE2 | RP11-362A9.3 | -0,3747 | 1,27133E-07 |
| ACE2 | RP11-160C18.2 | -0,3747 | 1,26872E-07 |
| ACE2 | LRRC38 | -0,3748 | 1,25786E-07 |
| ACE2 | RP11-92K2.2 | -0,3748 | 1,2543E-07 |

| | | | |
|---|---|---|---|
| ACE2 | AOX1 | -0,3748 | 1,25384E-07 |
| ACE2 | RP11-87F15.2 | -0,3748 | 1,26129E-07 |
| ACE2 | SNORD14A | -0,3748 | 1,26075E-07 |
| ACE2 | TMEM151A | -0,3749 | 1,24271E-07 |
| ACE2 | NME2P1 | -0,3749 | 1,24585E-07 |
| ACE2 | RPS2P4 | -0,3749 | 1,25204E-07 |
| ACE2 | HYPK | -0,3749 | 1,24581E-07 |
| ACE2 | RPL7AP11 | -0,375 | 1,24045E-07 |
| ACE2 | RP11-96L14.7 | -0,3751 | 1,22692E-07 |
| ACE2 | PCDHA9 | -0,3751 | 1,22687E-07 |
| ACE2 | RP11-398C13.8 | -0,3751 | 1,22464E-07 |
| ACE2 | BNIP3P9 | -0,3751 | 1,23073E-07 |
| ACE2 | WDR5B | -0,3752 | 1,21478E-07 |
| ACE2 | RP11-598F7.5 | -0,3753 | 1,2111E-07 |
| ACE2 | SMYD2 | -0,3754 | 1,19633E-07 |
| ACE2 | RAD51AP2 | -0,3754 | 1,19423E-07 |
| ACE2 | CPNE4 | -0,3754 | 1,20071E-07 |
| ACE2 | PROSER2 | -0,3754 | 1,19251E-07 |
| ACE2 | AC091153.4 | -0,3754 | 1,19524E-07 |
| ACE2 | MMP21 | -0,3755 | 1,18929E-07 |
| ACE2 | BNIP3P16 | -0,3755 | 1,18672E-07 |
| ACE2 | SPAG6 | -0,3756 | 1,1789E-07 |
| ACE2 | MPZL1 | -0,3757 | 1,17004E-07 |
| ACE2 | GCOM2 | -0,3757 | 1,16829E-07 |
| ACE2 | LLNLR-249E10.1 | -0,3757 | 1,16782E-07 |
| ACE2 | RP11-256I23.2 | -0,3758 | 1,15467E-07 |
| ACE2 | MELTF-AS1 | -0,3758 | 1,15839E-07 |
| ACE2 | HCN1 | -0,3759 | 1,15081E-07 |
| ACE2 | NPM1P6 | -0,3759 | 1,14346E-07 |
| ACE2 | RDH11 | -0,3759 | 1,14977E-07 |
| ACE2 | RP11-94C24.6 | -0,3759 | 1,15034E-07 |
| ACE2 | CTA-384D8.36 | -0,3759 | 1,14972E-07 |
| ACE2 | AC062017.1 | -0,376 | 1,1344E-07 |
| ACE2 | HIST1H1D | -0,376 | 1,13823E-07 |
| ACE2 | ACER2 | -0,376 | 1,13375E-07 |
| ACE2 | SLC14A1 | -0,376 | 1,13615E-07 |
| ACE2 | MRGBP | -0,376 | 1,13725E-07 |
| ACE2 | RNU5F-1 | -0,3761 | 1,13178E-07 |
| ACE2 | HNRNPA3P15 | -0,3761 | 1,12517E-07 |
| ACE2 | GTF3C5 | -0,3761 | 1,12715E-07 |
| ACE2 | ZNF267 | -0,3761 | 1,12488E-07 |
| ACE2 | MAPT-IT1 | -0,3761 | 1,13248E-07 |
| ACE2 | MZF1-AS1 | -0,3761 | 1,12777E-07 |
| ACE2 | CH17-195P21.2 | -0,3762 | 1,11612E-07 |
| ACE2 | RBBP5 | -0,3762 | 1,12154E-07 |
| ACE2 | RP11-325L7.1 | -0,3762 | 1,12278E-07 |
| ACE2 | RPS29 | -0,3762 | 1,12147E-07 |
| ACE2 | SERP1 | -0,3763 | 1,11259E-07 |
| ACE2 | PRKCQ | -0,3763 | 1,11064E-07 |
| ACE2 | INO80E | -0,3763 | 1,11434E-07 |
| ACE2 | FOXO4 | -0,3763 | 1,11332E-07 |
| ACE2 | CTD-2583P5.3 | -0,3764 | 1,09658E-07 |
| ACE2 | HRASLS | -0,3765 | 1,08997E-07 |
| ACE2 | HOTAIRM1 | -0,3765 | 1,09482E-07 |

| | | | |
|---|---|---|---|
| ACE2 | RP11-587D21.1 | -0,3765 | 1,09043E-07 |
| ACE2 | SMTNL1 | -0,3765 | 1,09534E-07 |
| ACE2 | RN7SKP161 | -0,3765 | 1,09241E-07 |
| ACE2 | CTD-2232E5.2 | -0,3766 | 1,08674E-07 |
| ACE2 | HN1L | -0,3766 | 1,07962E-07 |
| ACE2 | RP11-445P19.3 | -0,3767 | 1,07138E-07 |
| ACE2 | PLA2G15 | -0,3769 | 1,05433E-07 |
| ACE2 | GAN | -0,3769 | 1,05577E-07 |
| ACE2 | CYP4F24P | -0,3769 | 1,05847E-07 |
| ACE2 | DDX1 | -0,377 | 1,04826E-07 |
| ACE2 | RP11-120K18.3 | -0,3771 | 1,03756E-07 |
| ACE2 | RP11-362F19.3 | -0,3772 | 1,02819E-07 |
| ACE2 | ECHDC3 | -0,3773 | 1,02237E-07 |
| ACE2 | PGAM5 | -0,3773 | 1,02274E-07 |
| ACE2 | KLHL28 | -0,3773 | 1,023E-07 |
| ACE2 | RP11-547D13.1 | -0,3773 | 1,01877E-07 |
| ACE2 | RP11-632K20.7 | -0,3774 | 1,01095E-07 |
| ACE2 | TVP23CP2 | -0,3774 | 1,01109E-07 |
| ACE2 | NMUR1 | -0,3775 | 1,00149E-07 |
| ACE2 | RP11-367J11.3 | -0,3775 | 1,00387E-07 |
| ACE2 | FBXO8 | -0,3775 | 1,00018E-07 |
| ACE2 | LINC02148 | -0,3775 | 1,00146E-07 |
| ACE2 | RP11-727F15.9 | -0,3775 | 1,00737E-07 |
| ACE2 | BTBD6 | -0,3775 | 1,00775E-07 |
| ACE2 | RP11-361L15.5 | -0,3775 | 1,00596E-07 |
| ACE2 | ZBTB8B | -0,3777 | 9,8981E-08 |
| ACE2 | LUCAT1 | -0,3777 | 9,83802E-08 |
| ACE2 | XXbac-BPGBPG55C | -0,3777 | 9,8839E-08 |
| ACE2 | RP11-261P13.5 | -0,3777 | 9,88183E-08 |
| ACE2 | LMO7-AS1 | -0,3777 | 9,86528E-08 |
| ACE2 | TMLHE-AS1 | -0,3777 | 9,90717E-08 |
| ACE2 | FAM174A | -0,3779 | 9,6757E-08 |
| ACE2 | SDF2 | -0,3779 | 9,73096E-08 |
| ACE2 | LA16c-329F2.2 | -0,378 | 9,59796E-08 |
| ACE2 | LINC01132 | -0,3781 | 9,56549E-08 |
| ACE2 | VNN3 | -0,3781 | 9,53336E-08 |
| ACE2 | RASGRF1 | -0,3781 | 9,53432E-08 |
| ACE2 | ADAMTS7P4 | -0,3781 | 9,56407E-08 |
| ACE2 | COQ8B | -0,3781 | 9,55111E-08 |
| ACE2 | FAM109B | -0,3781 | 9,58312E-08 |
| ACE2 | RP11-296O14.2 | -0,3782 | 9,50283E-08 |
| ACE2 | LINC01918 | -0,3782 | 9,43626E-08 |
| ACE2 | PDCL3P4 | -0,3782 | 9,4869E-08 |
| ACE2 | FABP5P3 | -0,3782 | 9,4282E-08 |
| ACE2 | GPATCH8 | -0,3782 | 9,50351E-08 |
| ACE2 | AC009950.2 | -0,3783 | 9,41481E-08 |
| ACE2 | IFT88 | -0,3783 | 9,37134E-08 |
| ACE2 | EXOC1 | -0,3784 | 9,29826E-08 |
| ACE2 | TARID | -0,3784 | 9,31273E-08 |
| ACE2 | AHCY | -0,3784 | 9,28646E-08 |
| ACE2 | AGAP4 | -0,3785 | 9,24794E-08 |
| ACE2 | TIMM17A | -0,3786 | 9,14437E-08 |
| ACE2 | HAGHL | -0,3786 | 9,12386E-08 |
| ACE2 | SPAG9 | -0,3786 | 9,18683E-08 |

| | | | |
|---|---|---|---|
| ACE2 | ZFP36 | -0,3786 | 9,1487E-08 |
| ACE2 | AC011247.3 | -0,3787 | 9,05047E-08 |
| ACE2 | TUT1 | -0,3787 | 9,11204E-08 |
| ACE2 | RAB40B | -0,3787 | 9,06423E-08 |
| ACE2 | ZNF680P1 | -0,3788 | 9,00819E-08 |
| ACE2 | OXA1L | -0,3789 | 8,90638E-08 |
| ACE2 | CTC-246B18.8 | -0,3789 | 8,89902E-08 |
| ACE2 | CXorf57 | -0,3789 | 8,95376E-08 |
| ACE2 | RP11-423F24.3 | -0,379 | 8,836E-08 |
| ACE2 | RP11-756P10.2 | -0,379 | 8,88422E-08 |
| ACE2 | MOG | -0,379 | 8,81407E-08 |
| ACE2 | RP11-16F15.2 | -0,3791 | 8,7916E-08 |
| ACE2 | LMAN1 | -0,3791 | 8,78687E-08 |
| ACE2 | LINC01634 | -0,3791 | 8,76529E-08 |
| ACE2 | RP11-120B7.1 | -0,3793 | 8,59948E-08 |
| ACE2 | RNF14 | -0,3793 | 8,60742E-08 |
| ACE2 | TREML1 | -0,3793 | 8,63034E-08 |
| ACE2 | ZNF503-AS2 | -0,3793 | 8,64079E-08 |
| ACE2 | CCDC87 | -0,3793 | 8,61874E-08 |
| ACE2 | RP11-6N17.3 | -0,3793 | 8,60346E-08 |
| ACE2 | PEAK1 | -0,3794 | 8,57398E-08 |
| ACE2 | HSPE1P3 | -0,3794 | 8,56897E-08 |
| ACE2 | C1orf140 | -0,3795 | 8,44877E-08 |
| ACE2 | RNPS1P1 | -0,3795 | 8,46712E-08 |
| ACE2 | RP11-734J24.1 | -0,3795 | 8,46662E-08 |
| ACE2 | RP11-333E1.2 | -0,3795 | 8,45978E-08 |
| ACE2 | CTD-2373H9.3 | -0,3795 | 8,48712E-08 |
| ACE2 | HSPA9 | -0,3796 | 8,42732E-08 |
| ACE2 | RP11-4L24.4 | -0,3796 | 8,38402E-08 |
| ACE2 | ZMYND19 | -0,3796 | 8,38768E-08 |
| ACE2 | FGF11 | -0,3796 | 8,44352E-08 |
| ACE2 | TNK2-AS1 | -0,3797 | 8,34821E-08 |
| ACE2 | CTA-212D2.2 | -0,3797 | 8,33012E-08 |
| ACE2 | TMEM196 | -0,3797 | 8,35332E-08 |
| ACE2 | RP11-493L12.6 | -0,3797 | 8,35058E-08 |
| ACE2 | GPR84 | -0,3797 | 8,31701E-08 |
| ACE2 | AC002310.12 | -0,3798 | 8,28565E-08 |
| ACE2 | ANKRD26P1 | -0,3798 | 8,24702E-08 |
| ACE2 | RP4-555L14.4 | -0,3799 | 8,1806E-08 |
| ACE2 | RP11-727A23.4 | -0,3799 | 8,22719E-08 |
| ACE2 | LINC00266-1 | -0,38 | 8,163E-08 |
| ACE2 | RP6-109B7.5 | -0,38 | 8,1629E-08 |
| ACE2 | AC016910.1 | -0,3801 | 8,05045E-08 |
| ACE2 | RNFT2 | -0,3801 | 8,04736E-08 |
| ACE2 | RP11-405M12.3 | -0,3801 | 8,08077E-08 |
| ACE2 | L1TD1 | -0,3802 | 7,99487E-08 |
| ACE2 | MARC1 | -0,3802 | 7,98177E-08 |
| ACE2 | NUDT5 | -0,3802 | 8,00763E-08 |
| ACE2 | RP11-263K4.5 | -0,3802 | 7,98136E-08 |
| ACE2 | RNF219-AS1 | -0,3802 | 7,98718E-08 |
| ACE2 | MAPT-AS1 | -0,3802 | 8,0136E-08 |
| ACE2 | TRIM62 | -0,3803 | 7,94855E-08 |
| ACE2 | PPP1R42 | -0,3803 | 7,95164E-08 |
| ACE2 | S100A12 | -0,3804 | 7,85673E-08 |

| | | | |
|---|---|---|---|
| ACE2 | LARP4 | -0,3804 | 7,8451E-08 |
| ACE2 | RP11-326I19.2 | -0,3805 | 7,81908E-08 |
| ACE2 | RP11-727F15.11 | -0,3805 | 7,76983E-08 |
| ACE2 | CMB9-94B1.2 | -0,3805 | 7,82812E-08 |
| ACE2 | RP11-662I13.3 | -0,3805 | 7,77259E-08 |
| ACE2 | SMPX | -0,3805 | 7,80758E-08 |
| ACE2 | SEPHS1P6 | -0,3806 | 7,70561E-08 |
| ACE2 | GALNT2 | -0,3807 | 7,65013E-08 |
| ACE2 | TIMM23 | -0,3807 | 7,68189E-08 |
| ACE2 | GNPTAB | -0,3807 | 7,64027E-08 |
| ACE2 | RPL21P7 | -0,3807 | 7,68424E-08 |
| ACE2 | SCN1A | -0,3808 | 7,61028E-08 |
| ACE2 | RP11-480D4.6 | -0,3808 | 7,59746E-08 |
| ACE2 | CTD-2196P11.2 | -0,3808 | 7,56776E-08 |
| ACE2 | TMEM260 | -0,3808 | 7,59117E-08 |
| ACE2 | RP1-30M3.6 | -0,3809 | 7,53796E-08 |
| ACE2 | MAN2C1 | -0,3809 | 7,51072E-08 |
| ACE2 | RP11-218M11.7 | -0,3809 | 7,50691E-08 |
| ACE2 | CDH3 | -0,381 | 7,47813E-08 |
| ACE2 | AC055764.1 | -0,3811 | 7,40531E-08 |
| ACE2 | CTD-2236F14.1 | -0,3811 | 7,3892E-08 |
| ACE2 | RP4-725G10.3 | -0,3811 | 7,41982E-08 |
| ACE2 | TSNARE1 | -0,3811 | 7,42902E-08 |
| ACE2 | RP11-21A7A.2 | -0,3811 | 7,42127E-08 |
| ACE2 | CATSPER1 | -0,3811 | 7,37703E-08 |
| ACE2 | RP5-867C24.1 | -0,3811 | 7,37853E-08 |
| ACE2 | RP11-196B3.1 | -0,3811 | 7,43862E-08 |
| ACE2 | RP11-404E16.1 | -0,3812 | 7,36598E-08 |
| ACE2 | PSMD3 | -0,3812 | 7,36743E-08 |
| ACE2 | KMT2B | -0,3812 | 7,33648E-08 |
| ACE2 | BHMT | -0,3813 | 7,25631E-08 |
| ACE2 | SUOX | -0,3813 | 7,28009E-08 |
| ACE2 | RP11-255G12.3 | -0,3813 | 7,31091E-08 |
| ACE2 | ZNF441 | -0,3813 | 7,25962E-08 |
| ACE2 | RP11-106E7.1 | -0,3814 | 7,2184E-08 |
| ACE2 | FHOD1 | -0,3814 | 7,22441E-08 |
| ACE2 | WASHC2C | -0,3815 | 7,13251E-08 |
| ACE2 | LINC00398 | -0,3815 | 7,14274E-08 |
| ACE2 | HIF1A-AS2 | -0,3815 | 7,13892E-08 |
| ACE2 | BX322557.13 | -0,3815 | 7,17487E-08 |
| ACE2 | TFB2M | -0,3816 | 7,11313E-08 |
| ACE2 | DOK1 | -0,3816 | 7,071E-08 |
| ACE2 | RNU6-312P | -0,3816 | 7,08171E-08 |
| ACE2 | ZBTB38 | -0,3816 | 7,12987E-08 |
| ACE2 | SF3B5 | -0,3816 | 7,10983E-08 |
| ACE2 | RAD54B | -0,3816 | 7,07439E-08 |
| ACE2 | DBH | -0,3816 | 7,12149E-08 |
| ACE2 | B3GAT1 | -0,3816 | 7,11361E-08 |
| ACE2 | RP11-114N19.3 | -0,3816 | 7,09478E-08 |
| ACE2 | TRAPPC5 | -0,3816 | 7,07111E-08 |
| ACE2 | PROX1 | -0,3817 | 7,05698E-08 |
| ACE2 | LMX1B | -0,3817 | 7,03353E-08 |
| ACE2 | CXorf23 | -0,3817 | 7,01477E-08 |
| ACE2 | LINC01765 | -0,3818 | 6,97775E-08 |

| | | | |
|---|---|---|---|
| ACE2 | RP4-640E24.1 | -0,3818 | 6,95444E-08 |
| ACE2 | TARBP1 | -0,3818 | 6,98176E-08 |
| ACE2 | CTC-338M12.4 | -0,3818 | 6,97904E-08 |
| ACE2 | ALDH1B1 | -0,3818 | 7,00789E-08 |
| ACE2 | ATG101 | -0,3818 | 6,98407E-08 |
| ACE2 | ADNP2 | -0,3818 | 6,9699E-08 |
| ACE2 | THOC6 | -0,3819 | 6,91054E-08 |
| ACE2 | RP11-723D22.2 | -0,382 | 6,8799E-08 |
| ACE2 | RP11-286N22.14 | -0,382 | 6,86169E-08 |
| ACE2 | NXPH4 | -0,382 | 6,83899E-08 |
| ACE2 | DCDC2C | -0,3821 | 6,79373E-08 |
| ACE2 | ASB8 | -0,3821 | 6,81752E-08 |
| ACE2 | CHTF18 | -0,3821 | 6,81431E-08 |
| ACE2 | IRX5 | -0,3821 | 6,7925E-08 |
| ACE2 | KRT223P | -0,3821 | 6,78748E-08 |
| ACE2 | KLHDC9 | -0,3822 | 6,75297E-08 |
| ACE2 | AC097724.3 | -0,3822 | 6,76641E-08 |
| ACE2 | RP11-778D9.4 | -0,3822 | 6,74138E-08 |
| ACE2 | RP11-22B23.1 | -0,3822 | 6,75571E-08 |
| ACE2 | RN7SL574P | -0,3823 | 6,70346E-08 |
| ACE2 | RP3-428L16.1 | -0,3823 | 6,66341E-08 |
| ACE2 | RP13-968A2.1 | -0,3823 | 6,67205E-08 |
| ACE2 | AF131215.8 | -0,3823 | 6,66961E-08 |
| ACE2 | RP11-509J21.4 | -0,3823 | 6,6945E-08 |
| ACE2 | LINC01711 | -0,3823 | 6,66047E-08 |
| ACE2 | PVALB | -0,3823 | 6,68387E-08 |
| ACE2 | HSPA8 | -0,3824 | 6,62993E-08 |
| ACE2 | RPL21P99 | -0,3824 | 6,63165E-08 |
| ACE2 | RP11-989F5.1 | -0,3824 | 6,61579E-08 |
| ACE2 | BNIP3P17 | -0,3824 | 6,63001E-08 |
| ACE2 | RP11-14N7.2 | -0,3825 | 6,58296E-08 |
| ACE2 | KIFAP3 | -0,3825 | 6,56315E-08 |
| ACE2 | ATP1B1P1 | -0,3825 | 6,5573E-08 |
| ACE2 | HIGD2A | -0,3826 | 6,53524E-08 |
| ACE2 | RP11-726G1.1 | -0,3826 | 6,54697E-08 |
| ACE2 | HSP90AB4P | -0,3826 | 6,51336E-08 |
| ACE2 | EXOSC3P1 | -0,3826 | 6,51019E-08 |
| ACE2 | RCAN1 | -0,3826 | 6,54712E-08 |
| ACE2 | KCNC2 | -0,3827 | 6,46537E-08 |
| ACE2 | CDK2AP2P2 | -0,3828 | 6,41912E-08 |
| ACE2 | AKR1C1 | -0,3828 | 6,40563E-08 |
| ACE2 | RP11-493E12.2 | -0,3829 | 6,34216E-08 |
| ACE2 | AC073850.6 | -0,3829 | 6,3739E-08 |
| ACE2 | FAM107B | -0,3829 | 6,33228E-08 |
| ACE2 | RP11-24M17.4 | -0,3829 | 6,35325E-08 |
| ACE2 | CTD-2369P2.8 | -0,3829 | 6,3343E-08 |
| ACE2 | KLF17 | -0,383 | 6,28814E-08 |
| ACE2 | CDH18 | -0,383 | 6,29263E-08 |
| ACE2 | RP11-567J20.2 | -0,383 | 6,28626E-08 |
| ACE2 | SLC30A3 | -0,3831 | 6,24062E-08 |
| ACE2 | THAP9-AS1 | -0,3831 | 6,24678E-08 |
| ACE2 | DNHD1 | -0,3831 | 6,22935E-08 |
| ACE2 | LINC00384 | -0,3831 | 6,25703E-08 |
| ACE2 | AC007193.6 | -0,3831 | 6,26549E-08 |

| | | | |
|---|---|---|---|
| ACE2 | FRG1DP | -0,3831 | 6,25678E-08 |
| ACE2 | RP11-665C14.2 | -0,3832 | 6,1823E-08 |
| ACE2 | SCGB1D2 | -0,3832 | 6,21259E-08 |
| ACE2 | RP11-490B18.6 | -0,3832 | 6,20389E-08 |
| ACE2 | ARAF | -0,3832 | 6,17465E-08 |
| ACE2 | RPL5P1 | -0,3833 | 6,15065E-08 |
| ACE2 | RP11-474G23.3 | -0,3834 | 6,113E-08 |
| ACE2 | BTC | -0,3834 | 6,10791E-08 |
| ACE2 | RP1-223B1.1 | -0,3834 | 6,06826E-08 |
| ACE2 | SSR1 | -0,3834 | 6,09223E-08 |
| ACE2 | CCNC | -0,3834 | 6,07092E-08 |
| ACE2 | RPS27AP12 | -0,3834 | 6,07384E-08 |
| ACE2 | DLX3 | -0,3834 | 6,08246E-08 |
| ACE2 | SCARNA21 | -0,3835 | 6,04309E-08 |
| ACE2 | NUB1 | -0,3835 | 6,03173E-08 |
| ACE2 | RP11-712B9.2 | -0,3835 | 6,03447E-08 |
| ACE2 | RP11-326A13.1 | -0,3835 | 6,01551E-08 |
| ACE2 | RP11-626H12.1 | -0,3836 | 5,98494E-08 |
| ACE2 | AC141930.3 | -0,3837 | 5,95726E-08 |
| ACE2 | C7orf34 | -0,3837 | 5,92362E-08 |
| ACE2 | CKS1BP1 | -0,3837 | 5,91438E-08 |
| ACE2 | LINC01730 | -0,3837 | 5,94737E-08 |
| ACE2 | BRWD1 | -0,3837 | 5,91308E-08 |
| ACE2 | KIF25 | -0,3838 | 5,86373E-08 |
| ACE2 | PPP1R35 | -0,3838 | 5,86268E-08 |
| ACE2 | ADAT1 | -0,3838 | 5,90581E-08 |
| ACE2 | RP11-462H3.2 | -0,3838 | 5,90598E-08 |
| ACE2 | ZNF90P1 | -0,3839 | 5,83531E-08 |
| ACE2 | SNTG1 | -0,3839 | 5,84149E-08 |
| ACE2 | NDUFS8 | -0,3839 | 5,84621E-08 |
| ACE2 | RP11-845M18.6 | -0,3839 | 5,84779E-08 |
| ACE2 | POM121L9P | -0,3839 | 5,83158E-08 |
| ACE2 | HERC6 | -0,384 | 5,75972E-08 |
| ACE2 | FAM135B | -0,384 | 5,76225E-08 |
| ACE2 | RP11-363J20.1 | -0,384 | 5,76314E-08 |
| ACE2 | ATP8B4 | -0,384 | 5,76916E-08 |
| ACE2 | RP11-554D20.2 | -0,384 | 5,77905E-08 |
| ACE2 | CTD-2162K18.4 | -0,384 | 5,77828E-08 |
| ACE2 | RP1-161N10.1 | -0,384 | 5,78048E-08 |
| ACE2 | LINC00471 | -0,3841 | 5,75043E-08 |
| ACE2 | RP11-394J1.2 | -0,3841 | 5,71867E-08 |
| ACE2 | SUPT5H | -0,3841 | 5,73673E-08 |
| ACE2 | TARS2 | -0,3842 | 5,68406E-08 |
| ACE2 | PSMD14 | -0,3842 | 5,70026E-08 |
| ACE2 | LINC01011 | -0,3842 | 5,69659E-08 |
| ACE2 | RP11-46H11.12 | -0,3842 | 5,69347E-08 |
| ACE2 | RP11-822E23.7 | -0,3842 | 5,69704E-08 |
| ACE2 | NXF5 | -0,3842 | 5,70558E-08 |
| ACE2 | DACT2 | -0,3843 | 5,61877E-08 |
| ACE2 | MCM6 | -0,3844 | 5,58101E-08 |
| ACE2 | DMRT3 | -0,3844 | 5,57399E-08 |
| ACE2 | OPRK1 | -0,3845 | 5,53512E-08 |
| ACE2 | HBA2 | -0,3845 | 5,54373E-08 |
| ACE2 | RP5-1029F21.2 | -0,3845 | 5,53798E-08 |

| | | | |
|---|---|---|---|
| ACE2 | KNSTRN | -0,3846 | 5,48249E-08 |
| ACE2 | KCTD9 | -0,3847 | 5,45981E-08 |
| ACE2 | ZNF33BP1 | -0,3847 | 5,44523E-08 |
| ACE2 | TRIM34 | -0,3847 | 5,45287E-08 |
| ACE2 | ATF7IP2 | -0,3847 | 5,44154E-08 |
| ACE2 | KCNJ14 | -0,3847 | 5,45744E-08 |
| ACE2 | RP11-553D4.2 | -0,3848 | 5,39147E-08 |
| ACE2 | NXNL1 | -0,3848 | 5,40292E-08 |
| ACE2 | VSIG10L | -0,3848 | 5,4178E-08 |
| ACE2 | GLB1L | -0,3849 | 5,34948E-08 |
| ACE2 | RAB43 | -0,3849 | 5,33421E-08 |
| ACE2 | RP11-411K7.1 | -0,3849 | 5,3339E-08 |
| ACE2 | AC079613.1 | -0,385 | 5,32993E-08 |
| ACE2 | SLC37A3 | -0,385 | 5,29046E-08 |
| ACE2 | PTGER4P2-CDK2AF | -0,385 | 5,29526E-08 |
| ACE2 | PLEKHM1 | -0,385 | 5,31531E-08 |
| ACE2 | AC098820.3 | -0,3851 | 5,24068E-08 |
| ACE2 | CFAP77 | -0,3851 | 5,25333E-08 |
| ACE2 | AF064858.10 | -0,3851 | 5,26587E-08 |
| ACE2 | RP11-440L14.1 | -0,3852 | 5,23721E-08 |
| ACE2 | BTG1P1 | -0,3852 | 5,22602E-08 |
| ACE2 | RP11-255G12.2 | -0,3852 | 5,23478E-08 |
| ACE2 | RP11-327F22.4 | -0,3852 | 5,22538E-08 |
| ACE2 | EXOC3L1 | -0,3852 | 5,21945E-08 |
| ACE2 | DSC3 | -0,3852 | 5,21812E-08 |
| ACE2 | SNHG4 | -0,3853 | 5,16708E-08 |
| ACE2 | RP11-88G17.6 | -0,3853 | 5,18227E-08 |
| ACE2 | BET1L | -0,3854 | 5,14796E-08 |
| ACE2 | RP11-653J6.1 | -0,3854 | 5,10992E-08 |
| ACE2 | RP11-307C19.1 | -0,3854 | 5,14684E-08 |
| ACE2 | DNAI2 | -0,3854 | 5,10972E-08 |
| ACE2 | PRDM15 | -0,3854 | 5,11918E-08 |
| ACE2 | UBE2J2 | -0,3855 | 5,06589E-08 |
| ACE2 | LRPAP1 | -0,3855 | 5,08329E-08 |
| ACE2 | ANKRA2 | -0,3855 | 5,09674E-08 |
| ACE2 | RP11-540O11.1 | -0,3855 | 5,09213E-08 |
| ACE2 | TMC2 | -0,3855 | 5,07195E-08 |
| ACE2 | CELSR1 | -0,3855 | 5,06586E-08 |
| ACE2 | EIF4BP7 | -0,3855 | 5,07421E-08 |
| ACE2 | SF3B1 | -0,3856 | 5,05227E-08 |
| ACE2 | MSRB2 | -0,3856 | 5,03264E-08 |
| ACE2 | SPHKAP | -0,3857 | 4,99964E-08 |
| ACE2 | CTC-338M12.13 | -0,3857 | 5,00796E-08 |
| ACE2 | GABRG3 | -0,3857 | 5,01045E-08 |
| ACE2 | RP11-481F24.3 | -0,3857 | 4,98067E-08 |
| ACE2 | AP000692.9 | -0,3857 | 5,00911E-08 |
| ACE2 | ICK | -0,3858 | 4,9424E-08 |
| ACE2 | RP11-193M21.1 | -0,3858 | 4,95031E-08 |
| ACE2 | CMA1 | -0,3858 | 4,9345E-08 |
| ACE2 | KDM3A | -0,3859 | 4,92249E-08 |
| ACE2 | RP11-394I13.3 | -0,3859 | 4,91956E-08 |
| ACE2 | RP11-426A6.5 | -0,3859 | 4,92192E-08 |
| ACE2 | MAPK7 | -0,3859 | 4,9089E-08 |
| ACE2 | AKT1S1 | -0,3859 | 4,91398E-08 |

| | | | |
|---|---|---|---|
| ACE2 | IL27RA | -0,386 | 4,86447E-08 |
| ACE2 | RP11-155O18.6 | -0,3861 | 4,82043E-08 |
| ACE2 | ANKRD28 | -0,3861 | 4,8269E-08 |
| ACE2 | SEC61B | -0,3861 | 4,81048E-08 |
| ACE2 | COG8 | -0,3861 | 4,83724E-08 |
| ACE2 | ZNF576 | -0,3861 | 4,83397E-08 |
| ACE2 | RP4-620E11.8 | -0,3861 | 4,81374E-08 |
| ACE2 | RP11-458I7.1 | -0,3862 | 4,77396E-08 |
| ACE2 | RP11-212P7.1 | -0,3862 | 4,78267E-08 |
| ACE2 | RP11-1099M24.6 | -0,3862 | 4,79474E-08 |
| ACE2 | AC004623.3 | -0,3862 | 4,78548E-08 |
| ACE2 | APOBEC3D | -0,3862 | 4,79018E-08 |
| ACE2 | NLRP3 | -0,3863 | 4,72605E-08 |
| ACE2 | RP11-92A5.2 | -0,3863 | 4,72502E-08 |
| ACE2 | RP11-138A9.2 | -0,3863 | 4,73179E-08 |
| ACE2 | RP11-45A16.4 | -0,3863 | 4,74796E-08 |
| ACE2 | DDX23 | -0,3863 | 4,74731E-08 |
| ACE2 | PRIM1 | -0,3863 | 4,76058E-08 |
| ACE2 | ASB14 | -0,3864 | 4,71691E-08 |
| ACE2 | RP1-283K11.3 | -0,3864 | 4,68836E-08 |
| ACE2 | MYH3 | -0,3864 | 4,68454E-08 |
| ACE2 | SMARCA4 | -0,3864 | 4,71789E-08 |
| ACE2 | AC004893.11 | -0,3865 | 4,66749E-08 |
| ACE2 | LINC00535 | -0,3865 | 4,64899E-08 |
| ACE2 | PPM1H | -0,3867 | 4,5963E-08 |
| ACE2 | UQCRBP1 | -0,3867 | 4,57071E-08 |
| ACE2 | RP11-796E2.4 | -0,3868 | 4,53458E-08 |
| ACE2 | ZNF815P | -0,3869 | 4,49227E-08 |
| ACE2 | TDH | -0,3869 | 4,50786E-08 |
| ACE2 | HMX1 | -0,387 | 4,47394E-08 |
| ACE2 | CILP | -0,387 | 4,48421E-08 |
| ACE2 | ARMCX3-AS1 | -0,387 | 4,45782E-08 |
| ACE2 | MAGEA8-AS1 | -0,387 | 4,48182E-08 |
| ACE2 | CTD-2194A8.2 | -0,3871 | 4,42927E-08 |
| ACE2 | GALR2 | -0,3871 | 4,41283E-08 |
| ACE2 | AKAP3 | -0,3872 | 4,38366E-08 |
| ACE2 | PDE6H | -0,3872 | 4,37266E-08 |
| ACE2 | ZNF284 | -0,3872 | 4,37023E-08 |
| ACE2 | CNPY3 | -0,3873 | 4,34914E-08 |
| ACE2 | LINC01235 | -0,3873 | 4,35169E-08 |
| ACE2 | ATPAF2 | -0,3873 | 4,35795E-08 |
| ACE2 | C1orf159 | -0,3874 | 4,29664E-08 |
| ACE2 | HMGB3P10 | -0,3874 | 4,30117E-08 |
| ACE2 | CARTPT | -0,3874 | 4,31412E-08 |
| ACE2 | CTC-529P8.1 | -0,3874 | 4,32711E-08 |
| ACE2 | RPS3AP33 | -0,3874 | 4,30229E-08 |
| ACE2 | SNORA23 | -0,3874 | 4,32382E-08 |
| ACE2 | DTX1 | -0,3874 | 4,32498E-08 |
| ACE2 | LA16c-360A4.1 | -0,3874 | 4,30293E-08 |
| ACE2 | HCG22 | -0,3875 | 4,26197E-08 |
| ACE2 | PTPN20 | -0,3875 | 4,28175E-08 |
| ACE2 | SNRPGP14 | -0,3875 | 4,29256E-08 |
| ACE2 | FARP1-AS1 | -0,3875 | 4,27919E-08 |
| ACE2 | AC006548.28 | -0,3875 | 4,2727E-08 |

| | | | |
|---|---|---|---|
| ACE2 | MTND6P21 | -0,3876 | 4,2568E-08 |
| ACE2 | SEMA7A | -0,3877 | 4,20824E-08 |
| ACE2 | FMNL1 | -0,3877 | 4,20102E-08 |
| ACE2 | RN7SL262P | -0,3877 | 4,2131E-08 |
| ACE2 | DALRD3 | -0,3878 | 4,15647E-08 |
| ACE2 | GJA3 | -0,3878 | 4,17977E-08 |
| ACE2 | HHATL | -0,3879 | 4,13195E-08 |
| ACE2 | AC007566.10 | -0,3879 | 4,13788E-08 |
| ACE2 | RP11-545G3.2 | -0,3879 | 4,13351E-08 |
| ACE2 | RPSAP3 | -0,3879 | 4,11342E-08 |
| ACE2 | PROCA1 | -0,3879 | 4,11518E-08 |
| ACE2 | S100A2 | -0,388 | 4,09109E-08 |
| ACE2 | RP11-423E7.2 | -0,388 | 4,08493E-08 |
| ACE2 | ST8SIA4 | -0,388 | 4,09462E-08 |
| ACE2 | RP11-493L12.4 | -0,388 | 4,09295E-08 |
| ACE2 | PSMC3IP | -0,388 | 4,08972E-08 |
| ACE2 | AQP4 | -0,388 | 4,10817E-08 |
| ACE2 | RP11-389C8.1 | -0,3882 | 4,01793E-08 |
| ACE2 | FOLR2 | -0,3882 | 4,04043E-08 |
| ACE2 | RP11-753A21.1 | -0,3882 | 4,02878E-08 |
| ACE2 | CNN2P8 | -0,3883 | 3,98925E-08 |
| ACE2 | RP11-211G3.3 | -0,3883 | 4,0034E-08 |
| ACE2 | RP11-1277A3.2 | -0,3883 | 3,99898E-08 |
| ACE2 | RP11-673E11.2 | -0,3883 | 3,9977E-08 |
| ACE2 | CTD-2114J12.1 | -0,3885 | 3,93006E-08 |
| ACE2 | CALHM3 | -0,3885 | 3,9328E-08 |
| ACE2 | KIRREL2 | -0,3885 | 3,91187E-08 |
| ACE2 | ABCF1 | -0,3886 | 3,90107E-08 |
| ACE2 | RP11-163E9.1 | -0,3887 | 3,86906E-08 |
| ACE2 | DNAJC1 | -0,3887 | 3,86719E-08 |
| ACE2 | RP11-2N1.3 | -0,3887 | 3,84012E-08 |
| ACE2 | CTC-412M14.6 | -0,3888 | 3,83286E-08 |
| ACE2 | MIS18A | -0,3888 | 3,81134E-08 |
| ACE2 | MCOLN3 | -0,3889 | 3,79082E-08 |
| ACE2 | CX3CR1 | -0,3889 | 3,7876E-08 |
| ACE2 | NFXL1 | -0,3889 | 3,7705E-08 |
| ACE2 | RP11-467K18.2 | -0,3889 | 3,77449E-08 |
| ACE2 | RP11-752G15.10 | -0,3889 | 3,78303E-08 |
| ACE2 | ASS1P2 | -0,389 | 3,76824E-08 |
| ACE2 | LPP-AS2 | -0,389 | 3,76629E-08 |
| ACE2 | PRR34 | -0,389 | 3,76832E-08 |
| ACE2 | AC007277.3 | -0,3891 | 3,71642E-08 |
| ACE2 | RPL7P18 | -0,3891 | 3,70965E-08 |
| ACE2 | OLFM5P | -0,3891 | 3,73034E-08 |
| ACE2 | PPP1R1C | -0,3892 | 3,69036E-08 |
| ACE2 | NELFA | -0,3892 | 3,68121E-08 |
| ACE2 | ADAMTS2 | -0,3892 | 3,70214E-08 |
| ACE2 | RP11-490G8.1 | -0,3892 | 3,69092E-08 |
| ACE2 | RP1-37C10.3 | -0,3893 | 3,66279E-08 |
| ACE2 | SPRY2 | -0,3893 | 3,66786E-08 |
| ACE2 | ZNF845 | -0,3893 | 3,64647E-08 |
| ACE2 | KRT8P26 | -0,3894 | 3,627E-08 |
| ACE2 | CYP2G1P | -0,3894 | 3,62728E-08 |
| ACE2 | HNRNPH1 | -0,3895 | 3,60557E-08 |

| | | | |
|---|---|---|---|
| ACE2 | UPF3AP1 | -0,3895 | 3,5863E-08 |
| ACE2 | HGH1 | -0,3896 | 3,54774E-08 |
| ACE2 | LINC00664 | -0,3896 | 3,56366E-08 |
| ACE2 | LINC00634 | -0,3896 | 3,56463E-08 |
| ACE2 | CNOT6L | -0,3897 | 3,53969E-08 |
| ACE2 | GARS | -0,3897 | 3,52755E-08 |
| ACE2 | TUBB8P7 | -0,3897 | 3,52938E-08 |
| ACE2 | TMEM199 | -0,3897 | 3,527E-08 |
| ACE2 | LRRC8B | -0,3898 | 3,5046E-08 |
| ACE2 | RFWD2 | -0,3898 | 3,50505E-08 |
| ACE2 | CA2 | -0,3898 | 3,50269E-08 |
| ACE2 | PABPN1L | -0,3898 | 3,50283E-08 |
| ACE2 | DVL1 | -0,3899 | 3,46523E-08 |
| ACE2 | JRK | -0,3899 | 3,4726E-08 |
| ACE2 | OR1H1P | -0,3899 | 3,46139E-08 |
| ACE2 | KRCC1 | -0,39 | 3,45345E-08 |
| ACE2 | LINC00602 | -0,39 | 3,44479E-08 |
| ACE2 | AC006378.2 | -0,39 | 3,45323E-08 |
| ACE2 | MAG | -0,39 | 3,45402E-08 |
| ACE2 | RN7SKP239 | -0,3901 | 3,42391E-08 |
| ACE2 | PAN3 | -0,3901 | 3,39768E-08 |
| ACE2 | ADNP-AS1 | -0,3901 | 3,42259E-08 |
| ACE2 | RBBP8P1 | -0,3901 | 3,40311E-08 |
| ACE2 | RABGGTB | -0,3902 | 3,3932E-08 |
| ACE2 | RP11-44N11.2 | -0,3902 | 3,38535E-08 |
| ACE2 | MEGF11 | -0,3902 | 3,3759E-08 |
| ACE2 | DHFR2 | -0,3903 | 3,33819E-08 |
| ACE2 | MRPL3 | -0,3903 | 3,36156E-08 |
| ACE2 | RP11-430K21.2 | -0,3903 | 3,33582E-08 |
| ACE2 | AC079776.1 | -0,3904 | 3,33385E-08 |
| ACE2 | RPL29P14 | -0,3904 | 3,33541E-08 |
| ACE2 | RPSAP47 | -0,3904 | 3,31183E-08 |
| ACE2 | MYBPC3 | -0,3904 | 3,31364E-08 |
| ACE2 | RHOXF1P3 | -0,3904 | 3,33384E-08 |
| ACE2 | HS1BP3 | -0,3905 | 3,28182E-08 |
| ACE2 | NXPH2 | -0,3905 | 3,29738E-08 |
| ACE2 | LINC02172 | -0,3905 | 3,28438E-08 |
| ACE2 | CRNKL1 | -0,3906 | 3,27557E-08 |
| ACE2 | TTTY15 | -0,3906 | 3,25186E-08 |
| ACE2 | RPL7L1P9 | -0,3907 | 3,23171E-08 |
| ACE2 | EFTUD2 | -0,3907 | 3,22838E-08 |
| ACE2 | XKR8 | -0,3908 | 3,20569E-08 |
| ACE2 | FCER1A | -0,3908 | 3,21368E-08 |
| ACE2 | DMXL1 | -0,3908 | 3,20489E-08 |
| ACE2 | IRX6 | -0,3908 | 3,21512E-08 |
| ACE2 | RP11-65L3.2 | -0,3909 | 3,17362E-08 |
| ACE2 | KIAA2012 | -0,3909 | 3,17103E-08 |
| ACE2 | CTD-2012K14.7 | -0,3909 | 3,16764E-08 |
| ACE2 | RP13-228J13.1 | -0,3909 | 3,17482E-08 |
| ACE2 | PLGLB1 | -0,391 | 3,16117E-08 |
| ACE2 | RBMS3-AS2 | -0,391 | 3,13873E-08 |
| ACE2 | RP11-631N16.2 | -0,391 | 3,15745E-08 |
| ACE2 | TRHDE-AS1 | -0,391 | 3,15567E-08 |
| ACE2 | CORO6 | -0,3911 | 3,1152E-08 |

| | | | |
|---|---|---|---|
| ACE2 | RP11-85B7.2 | -0,3912 | 3,10228E-08 |
| ACE2 | PNKP | -0,3912 | 3,09044E-08 |
| ACE2 | C2orf80 | -0,3913 | 3,05574E-08 |
| ACE2 | PARG | -0,3913 | 3,06944E-08 |
| ACE2 | RP11-709A23.2 | -0,3913 | 3,0755E-08 |
| ACE2 | LOXHD1 | -0,3913 | 3,06513E-08 |
| ACE2 | SEC63P1 | -0,3915 | 3,01813E-08 |
| ACE2 | TDP1 | -0,3915 | 3,01764E-08 |
| ACE2 | BCYRN1 | -0,3916 | 2,99095E-08 |
| ACE2 | DOCK9-AS2 | -0,3916 | 2,99034E-08 |
| ACE2 | RP11-345J18.2 | -0,3916 | 2,98036E-08 |
| ACE2 | AC008753.6 | -0,3916 | 2,9943E-08 |
| ACE2 | ARHGAP5-AS1 | -0,3917 | 2,97523E-08 |
| ACE2 | PIK3CD | -0,3918 | 2,94601E-08 |
| ACE2 | LAMTOR5 | -0,3919 | 2,92082E-08 |
| ACE2 | SP110 | -0,3919 | 2,91199E-08 |
| ACE2 | TELO2 | -0,3919 | 2,92316E-08 |
| ACE2 | TMEM183AP1 | -0,392 | 2,87745E-08 |
| ACE2 | DDX3Y | -0,3921 | 2,85259E-08 |
| ACE2 | NAXE | -0,3922 | 2,83371E-08 |
| ACE2 | RP11-122G18.7 | -0,3922 | 2,84577E-08 |
| ACE2 | UFSP1 | -0,3922 | 2,84641E-08 |
| ACE2 | LINC01252 | -0,3922 | 2,84067E-08 |
| ACE2 | RP11-16E23.4 | -0,3922 | 2,84609E-08 |
| ACE2 | CTD-3193O13.8 | -0,3922 | 2,83289E-08 |
| ACE2 | CLVS1 | -0,3923 | 2,8046E-08 |
| ACE2 | TMEM186 | -0,3923 | 2,79929E-08 |
| ACE2 | TMEM214 | -0,3924 | 2,79347E-08 |
| ACE2 | RPSAP36 | -0,3924 | 2,77583E-08 |
| ACE2 | TMEM80 | -0,3924 | 2,79241E-08 |
| ACE2 | TPM1 | -0,3924 | 2,77524E-08 |
| ACE2 | ENPP6 | -0,3925 | 2,75213E-08 |
| ACE2 | DOCK4-AS1 | -0,3925 | 2,76172E-08 |
| ACE2 | RP11-166B2.3 | -0,3925 | 2,75148E-08 |
| ACE2 | RS1 | -0,3925 | 2,77259E-08 |
| ACE2 | SPEN | -0,3926 | 2,73045E-08 |
| ACE2 | ASH2LP3 | -0,3926 | 2,73937E-08 |
| ACE2 | NFKBIZ | -0,3928 | 2,68495E-08 |
| ACE2 | CTB-51J22.1 | -0,3928 | 2,68787E-08 |
| ACE2 | LY6H | -0,3928 | 2,67717E-08 |
| ACE2 | RP11-736K20.6 | -0,3928 | 2,69685E-08 |
| ACE2 | CLEC4M | -0,3928 | 2,70024E-08 |
| ACE2 | RP11-15A1.9 | -0,3928 | 2,67748E-08 |
| ACE2 | UBE2L3 | -0,3928 | 2,68717E-08 |
| ACE2 | IFRD2 | -0,3929 | 2,65797E-08 |
| ACE2 | SND1 | -0,3929 | 2,66577E-08 |
| ACE2 | ATE1 | -0,3929 | 2,66349E-08 |
| ACE2 | RP11-66B24.9 | -0,3929 | 2,67349E-08 |
| ACE2 | TMEM128 | -0,393 | 2,63972E-08 |
| ACE2 | SGMS1 | -0,393 | 2,64786E-08 |
| ACE2 | MRI1 | -0,3931 | 2,61419E-08 |
| ACE2 | BCRP3 | -0,3931 | 2,6152E-08 |
| ACE2 | RP11-791G16.2 | -0,3932 | 2,59898E-08 |
| ACE2 | EWSAT1 | -0,3932 | 2,58762E-08 |

| | | | |
|---|---|---|---|
| ACE2 | MIATNB | -0,3932 | 2,60415E-08 |
| ACE2 | FLG | -0,3933 | 2,56343E-08 |
| ACE2 | RP11-761N21.2 | -0,3933 | 2,5826E-08 |
| ACE2 | AC144530.1 | -0,3933 | 2,56651E-08 |
| ACE2 | RP11-572O17.1 | -0,3933 | 2,57908E-08 |
| ACE2 | BARX1 | -0,3933 | 2,57174E-08 |
| ACE2 | CCDC40 | -0,3933 | 2,56626E-08 |
| ACE2 | KRBOX1 | -0,3934 | 2,54468E-08 |
| ACE2 | AC002066.1 | -0,3934 | 2,55871E-08 |
| ACE2 | TMED3 | -0,3934 | 2,55678E-08 |
| ACE2 | IDH3B | -0,3934 | 2,55451E-08 |
| ACE2 | APOBEC3H | -0,3935 | 2,5191E-08 |
| ACE2 | CACNA1S | -0,3936 | 2,51427E-08 |
| ACE2 | AC092162.1 | -0,3936 | 2,5008E-08 |
| ACE2 | PRICKLE2-AS1 | -0,3936 | 2,49856E-08 |
| ACE2 | FAM19A4 | -0,3936 | 2,49869E-08 |
| ACE2 | RP11-210H10__A.1 | -0,3937 | 2,47977E-08 |
| ACE2 | DNAJC3-AS1 | -0,3937 | 2,48041E-08 |
| ACE2 | MTRF1L | -0,3938 | 2,46896E-08 |
| ACE2 | RP11-1180F24.3 | -0,3938 | 2,46539E-08 |
| ACE2 | RP11-97N19.2 | -0,3938 | 2,46797E-08 |
| ACE2 | CTB-33O18.2 | -0,3939 | 2,43884E-08 |
| ACE2 | RP11-527N22.1 | -0,3939 | 2,42913E-08 |
| ACE2 | AC091133.1 | -0,3939 | 2,4428E-08 |
| ACE2 | RP1-309F20.4 | -0,3939 | 2,4487E-08 |
| ACE2 | RP11-98J23.2 | -0,394 | 2,40882E-08 |
| ACE2 | OR7E7P | -0,394 | 2,41962E-08 |
| ACE2 | CFHR1 | -0,3941 | 2,40395E-08 |
| ACE2 | RP11-248J18.3 | -0,3941 | 2,39974E-08 |
| ACE2 | LINC01547 | -0,3941 | 2,40212E-08 |
| ACE2 | RP3-510D11.4 | -0,3942 | 2,38439E-08 |
| ACE2 | RP13-131K19.2 | -0,3942 | 2,37128E-08 |
| ACE2 | FAM175B | -0,3942 | 2,3795E-08 |
| ACE2 | RP5-1063M23.3 | -0,3942 | 2,38263E-08 |
| ACE2 | BCL2A1 | -0,3942 | 2,36557E-08 |
| ACE2 | CD70 | -0,3942 | 2,37541E-08 |
| ACE2 | VCP | -0,3943 | 2,36134E-08 |
| ACE2 | LINC00475 | -0,3943 | 2,34818E-08 |
| ACE2 | MKX-AS1 | -0,3943 | 2,35847E-08 |
| ACE2 | LINC00621 | -0,3943 | 2,36233E-08 |
| ACE2 | LIN9 | -0,3944 | 2,34168E-08 |
| ACE2 | RP11-218F10.3 | -0,3944 | 2,33019E-08 |
| ACE2 | RP11-438D14.3 | -0,3944 | 2,33485E-08 |
| ACE2 | AC006116.15 | -0,3944 | 2,33285E-08 |
| ACE2 | ZNF324 | -0,3944 | 2,32511E-08 |
| ACE2 | TBC1D31 | -0,3945 | 2,30391E-08 |
| ACE2 | ARAP1-AS2 | -0,3945 | 2,32268E-08 |
| ACE2 | RP11-31H5.3 | -0,3945 | 2,3129E-08 |
| ACE2 | ASH2L | -0,3946 | 2,28808E-08 |
| ACE2 | RP11-35O15.1 | -0,3946 | 2,30207E-08 |
| ACE2 | SMIM4 | -0,3947 | 2,28159E-08 |
| ACE2 | OR52K3P | -0,3947 | 2,28189E-08 |
| ACE2 | RP11-88E10.5 | -0,3947 | 2,27309E-08 |
| ACE2 | AC079610.2 | -0,3948 | 2,25848E-08 |

| | | | |
|---|---|---|---|
| ACE2 | RNU6-1039P | -0,3948 | 2,24229E-08 |
| ACE2 | MYRIP | -0,3949 | 2,22487E-08 |
| ACE2 | PI4KAP1 | -0,3949 | 2,23729E-08 |
| ACE2 | RP4-650F12.2 | -0,395 | 2,21987E-08 |
| ACE2 | THUMPD3-AS1 | -0,395 | 2,20754E-08 |
| ACE2 | RP11-104O19.4 | -0,395 | 2,21302E-08 |
| ACE2 | PPEF2 | -0,3951 | 2,19023E-08 |
| ACE2 | RP11-261P13.6 | -0,3951 | 2,19586E-08 |
| ACE2 | EEF1A1P12 | -0,3952 | 2,17785E-08 |
| ACE2 | ZFX-AS1 | -0,3952 | 2,17791E-08 |
| ACE2 | RP11-124N19.4 | -0,3953 | 2,15263E-08 |
| ACE2 | WHAMM | -0,3953 | 2,15024E-08 |
| ACE2 | MAN2A2 | -0,3953 | 2,15404E-08 |
| ACE2 | CTC-281F24.5 | -0,3953 | 2,1482E-08 |
| ACE2 | SIGLEC17P | -0,3953 | 2,1613E-08 |
| ACE2 | TIMM17B | -0,3953 | 2,1625E-08 |
| ACE2 | NDUFB2-AS1 | -0,3954 | 2,14445E-08 |
| ACE2 | OR1L8 | -0,3954 | 2,12963E-08 |
| ACE2 | FLJ21408 | -0,3954 | 2,13284E-08 |
| ACE2 | RP11-70P17.1 | -0,3955 | 2,11154E-08 |
| ACE2 | PRUNE1 | -0,3955 | 2,12104E-08 |
| ACE2 | C10orf82 | -0,3955 | 2,12525E-08 |
| ACE2 | SNORD89 | -0,3956 | 2,09958E-08 |
| ACE2 | EFCAB10 | -0,3956 | 2,09488E-08 |
| ACE2 | RNF34 | -0,3956 | 2,09373E-08 |
| ACE2 | DNAJB14 | -0,3957 | 2,07357E-08 |
| ACE2 | NPY2R | -0,3957 | 2,07336E-08 |
| ACE2 | MIR4458HG | -0,3957 | 2,07663E-08 |
| ACE2 | KB-1448A5.1 | -0,3957 | 2,07938E-08 |
| ACE2 | RP5-1119A7.17 | -0,3957 | 2,083E-08 |
| ACE2 | GMEB1 | -0,3958 | 2,05203E-08 |
| ACE2 | NMU | -0,3958 | 2,06355E-08 |
| ACE2 | RP11-110I1.13 | -0,3958 | 2,06554E-08 |
| ACE2 | AGPS | -0,3959 | 2,0373E-08 |
| ACE2 | PTMAP5 | -0,3959 | 2,0383E-08 |
| ACE2 | RP11-118F19.1 | -0,3959 | 2,04859E-08 |
| ACE2 | GGT5 | -0,3959 | 2,03323E-08 |
| ACE2 | AC012358.4 | -0,396 | 2,03107E-08 |
| ACE2 | AC007403.3 | -0,396 | 2,03263E-08 |
| ACE2 | KIAA1109 | -0,396 | 2,02928E-08 |
| ACE2 | RP11-93I21.3 | -0,396 | 2,03026E-08 |
| ACE2 | RP11-626G11.6 | -0,396 | 2,02588E-08 |
| ACE2 | CLIC6 | -0,396 | 2,0228E-08 |
| ACE2 | LY6K | -0,3961 | 2,00721E-08 |
| ACE2 | RNGTTP1 | -0,3961 | 2,01136E-08 |
| ACE2 | KCNC4-AS1 | -0,3963 | 1,97745E-08 |
| ACE2 | B3GALNT2 | -0,3963 | 1,96832E-08 |
| ACE2 | TMEM182 | -0,3963 | 1,96712E-08 |
| ACE2 | RPL5P22 | -0,3963 | 1,97657E-08 |
| ACE2 | MAEL | -0,3964 | 1,96152E-08 |
| ACE2 | RP11-309L24.2 | -0,3965 | 1,93424E-08 |
| ACE2 | RP11-253A20.1 | -0,3966 | 1,91512E-08 |
| ACE2 | C2orf69 | -0,3966 | 1,92114E-08 |
| ACE2 | TMUB2 | -0,3966 | 1,92177E-08 |

| | | | |
|---|---|---|---|
| ACE2 | SYNJ1 | -0,3966 | 1,92494E-08 |
| ACE2 | PIK3IP1-AS1 | -0,3966 | 1,92461E-08 |
| ACE2 | MPRIPP1 | -0,3967 | 1,90122E-08 |
| ACE2 | RN7SL771P | -0,3967 | 1,89383E-08 |
| ACE2 | PPP2R2D | -0,3967 | 1,907E-08 |
| ACE2 | RP11-212D19.5 | -0,3967 | 1,90378E-08 |
| ACE2 | CTC-338M12.5 | -0,3968 | 1,87667E-08 |
| ACE2 | SNHG10 | -0,3968 | 1,88626E-08 |
| ACE2 | VRK1 | -0,3968 | 1,88462E-08 |
| ACE2 | RP11-657O9.1 | -0,3969 | 1,86804E-08 |
| ACE2 | UMAD1 | -0,3969 | 1,87468E-08 |
| ACE2 | AC005009.2 | -0,3969 | 1,86686E-08 |
| ACE2 | HVCN1 | -0,3969 | 1,87554E-08 |
| ACE2 | RP11-346C4.3 | -0,3969 | 1,87531E-08 |
| ACE2 | MFSD12 | -0,3969 | 1,87046E-08 |
| ACE2 | AL590762.11 | -0,3969 | 1,87289E-08 |
| ACE2 | STXBP3 | -0,397 | 1,85295E-08 |
| ACE2 | FOLH1 | -0,397 | 1,844E-08 |
| ACE2 | EPB42 | -0,397 | 1,84904E-08 |
| ACE2 | POMC | -0,3971 | 1,84252E-08 |
| ACE2 | RP11-278A23.1 | -0,3971 | 1,8268E-08 |
| ACE2 | CTC-204F22.1 | -0,3971 | 1,84037E-08 |
| ACE2 | SPATA21 | -0,3973 | 1,80018E-08 |
| ACE2 | RPL24P8 | -0,3973 | 1,80872E-08 |
| ACE2 | ZNF816 | -0,3973 | 1,79727E-08 |
| ACE2 | RP11-114M1.3 | -0,3974 | 1,78264E-08 |
| ACE2 | HOMEZ | -0,3974 | 1,79314E-08 |
| ACE2 | TRAPPC1 | -0,3974 | 1,78187E-08 |
| ACE2 | RP11-11N5.3 | -0,3975 | 1,77132E-08 |
| ACE2 | RP11-73O6.3 | -0,3975 | 1,76743E-08 |
| ACE2 | RNASE4 | -0,3975 | 1,76317E-08 |
| ACE2 | RP11-519C12.1 | -0,3975 | 1,77091E-08 |
| ACE2 | SLC7A5P1 | -0,3975 | 1,7675E-08 |
| ACE2 | RP11-180O5.2 | -0,3976 | 1,75319E-08 |
| ACE2 | TMEM217 | -0,3976 | 1,75853E-08 |
| ACE2 | RP11-116D17.3 | -0,3976 | 1,74737E-08 |
| ACE2 | SPATA22 | -0,3976 | 1,7487E-08 |
| ACE2 | SBNO2 | -0,3976 | 1,75697E-08 |
| ACE2 | FCAR | -0,3976 | 1,74702E-08 |
| ACE2 | BMP6 | -0,3978 | 1,72886E-08 |
| ACE2 | NDUFB4 | -0,3979 | 1,71377E-08 |
| ACE2 | RP11-15L13.4 | -0,3979 | 1,70855E-08 |
| ACE2 | SRSF8 | -0,3979 | 1,71182E-08 |
| ACE2 | S1PR5 | -0,3979 | 1,70041E-08 |
| ACE2 | SDR42E2 | -0,398 | 1,69258E-08 |
| ACE2 | SLC13A5 | -0,398 | 1,6916E-08 |
| ACE2 | CTB-58E17.3 | -0,398 | 1,69951E-08 |
| ACE2 | GGA1 | -0,398 | 1,68768E-08 |
| ACE2 | HIST1H2BI | -0,3981 | 1,67323E-08 |
| ACE2 | RP11-504G3.2 | -0,3981 | 1,67204E-08 |
| ACE2 | DDX3X | -0,3981 | 1,67632E-08 |
| ACE2 | CTB-129O4.1 | -0,3982 | 1,66198E-08 |
| ACE2 | RP11-80H18.4 | -0,3983 | 1,65048E-08 |
| ACE2 | NFAT5 | -0,3983 | 1,64555E-08 |

| | | | |
|---|---|---|---|
| ACE2 | RPL23AP3 | -0,3983 | 1,64723E-08 |
| ACE2 | RP11-554E23.2 | -0,3984 | 1,63708E-08 |
| ACE2 | RP11-162G10.5 | -0,3985 | 1,62283E-08 |
| ACE2 | RP11-537A6.9 | -0,3985 | 1,62269E-08 |
| ACE2 | KLRF1 | -0,3985 | 1,61804E-08 |
| ACE2 | RP11-739G5.1 | -0,3986 | 1,60277E-08 |
| ACE2 | CNOT6 | -0,3987 | 1,58756E-08 |
| ACE2 | AAMDC | -0,3987 | 1,58774E-08 |
| ACE2 | HINFP | -0,3988 | 1,57903E-08 |
| ACE2 | RP11-65J3.2 | -0,3989 | 1,56011E-08 |
| ACE2 | NKD2 | -0,399 | 1,54606E-08 |
| ACE2 | LINC02021 | -0,3991 | 1,53922E-08 |
| ACE2 | GGT7 | -0,3991 | 1,52954E-08 |
| ACE2 | PRICKLE2-AS3 | -0,3992 | 1,51563E-08 |
| ACE2 | DPY19L2P3 | -0,3992 | 1,52242E-08 |
| ACE2 | NACC2 | -0,3992 | 1,51746E-08 |
| ACE2 | RP11-461A8.5 | -0,3992 | 1,52114E-08 |
| ACE2 | RHBDL2 | -0,3993 | 1,50707E-08 |
| ACE2 | TMEM258 | -0,3993 | 1,5007E-08 |
| ACE2 | C16orf91 | -0,3993 | 1,50315E-08 |
| ACE2 | RP11-481C4.2 | -0,3993 | 1,50328E-08 |
| ACE2 | ZNF780A | -0,3993 | 1,50742E-08 |
| ACE2 | CAMK2N2 | -0,3995 | 1,4815E-08 |
| ACE2 | HSPB9 | -0,3995 | 1,47851E-08 |
| ACE2 | KB-1742H10.3 | -0,3996 | 1,46972E-08 |
| ACE2 | LSP1 | -0,3996 | 1,46405E-08 |
| ACE2 | LINC01422 | -0,3996 | 1,46961E-08 |
| ACE2 | RPL12P30 | -0,3997 | 1,44723E-08 |
| ACE2 | RNF166 | -0,3997 | 1,44718E-08 |
| ACE2 | VN1R51P | -0,3998 | 1,44126E-08 |
| ACE2 | AC004383.3 | -0,3998 | 1,43233E-08 |
| ACE2 | PYGO2 | -0,3999 | 1,42451E-08 |
| ACE2 | AQP4-AS1 | -0,3999 | 1,42449E-08 |
| ACE2 | VMAC | -0,3999 | 1,42557E-08 |
| ACE2 | SLC39A1 | -0,4 | 1,41185E-08 |
| ACE2 | PTGES3P3 | -0,4 | 1,40784E-08 |
| ACE2 | RP11-685M7.3 | -0,4 | 1,41461E-08 |
| ACE2 | RP11-379B18.1 | -0,4001 | 1,3946E-08 |
| ACE2 | EEF1A1P9 | -0,4001 | 1,40264E-08 |
| ACE2 | ACTG1P14 | -0,4001 | 1,39725E-08 |
| ACE2 | RP11-700H6.4 | -0,4001 | 1,40409E-08 |
| ACE2 | ASAH1 | -0,4002 | 1,39255E-08 |
| ACE2 | TSSK5P | -0,4003 | 1,37702E-08 |
| ACE2 | MMP27 | -0,4003 | 1,37102E-08 |
| ACE2 | GRB2 | -0,4003 | 1,37272E-08 |
| ACE2 | PARGP1 | -0,4004 | 1,36148E-08 |
| ACE2 | HIF1AN | -0,4004 | 1,35822E-08 |
| ACE2 | ANKRD62P1 | -0,4004 | 1,36198E-08 |
| ACE2 | DIO1 | -0,4005 | 1,34862E-08 |
| ACE2 | AC008073.9 | -0,4005 | 1,34724E-08 |
| ACE2 | OR7E38P | -0,4005 | 1,353E-08 |
| ACE2 | POLR2J | -0,4005 | 1,35228E-08 |
| ACE2 | RP11-1134I14.6 | -0,4005 | 1,35008E-08 |
| ACE2 | RP11-17E13.2 | -0,4006 | 1,3329E-08 |

| | | | |
|---|---|---|---|
| ACE2 | GTF2IP20 | -0,4006 | 1,34356E-08 |
| ACE2 | ANXA2R | -0,4007 | 1,33035E-08 |
| ACE2 | AC091729.9 | -0,4007 | 1,32496E-08 |
| ACE2 | ZNRF2P2 | -0,4007 | 1,33055E-08 |
| ACE2 | RP11-365F18.3 | -0,4007 | 1,33145E-08 |
| ACE2 | CTD-2033D15.1 | -0,4007 | 1,33036E-08 |
| ACE2 | RP11-89B16.1 | -0,4008 | 1,31197E-08 |
| ACE2 | CTD-2199O4.6 | -0,4008 | 1,31648E-08 |
| ACE2 | RP5-1121E10.2 | -0,4008 | 1,3174E-08 |
| ACE2 | DHX38 | -0,4008 | 1,31635E-08 |
| ACE2 | SEPT9 | -0,4008 | 1,31801E-08 |
| ACE2 | C22orf24 | -0,4008 | 1,31029E-08 |
| ACE2 | RP11-407N8.6 | -0,4009 | 1,30466E-08 |
| ACE2 | TBL3 | -0,4009 | 1,30718E-08 |
| ACE2 | VASP | -0,401 | 1,29602E-08 |
| ACE2 | IL1RAPL2 | -0,401 | 1,28584E-08 |
| ACE2 | CDC7 | -0,4011 | 1,27455E-08 |
| ACE2 | RP11-536C12.1 | -0,4011 | 1,28405E-08 |
| ACE2 | HRH2 | -0,4011 | 1,27334E-08 |
| ACE2 | RP11-680F20.6 | -0,4012 | 1,26883E-08 |
| ACE2 | RP5-908M14.9 | -0,4012 | 1,27081E-08 |
| ACE2 | RP11-807C20.2 | -0,4013 | 1,25865E-08 |
| ACE2 | MADCAM1 | -0,4013 | 1,25484E-08 |
| ACE2 | NPEPPS | -0,4014 | 1,24644E-08 |
| ACE2 | TOR3A | -0,4015 | 1,23664E-08 |
| ACE2 | SUGP1 | -0,4015 | 1,23322E-08 |
| ACE2 | CYP26B1 | -0,4016 | 1,22617E-08 |
| ACE2 | TPTE2 | -0,4017 | 1,21265E-08 |
| ACE2 | RP11-145P16.3 | -0,4018 | 1,19738E-08 |
| ACE2 | RPL7AP10 | -0,4018 | 1,20298E-08 |
| ACE2 | DUX4L27 | -0,4019 | 1,18718E-08 |
| ACE2 | CTD-2026D20.2 | -0,4019 | 1,18475E-08 |
| ACE2 | SERHL | -0,4019 | 1,18642E-08 |
| ACE2 | RP11-245J9.5 | -0,402 | 1,18029E-08 |
| ACE2 | EXOC3 | -0,402 | 1,18278E-08 |
| ACE2 | RP11-806H10.4 | -0,402 | 1,18318E-08 |
| ACE2 | CRKL | -0,402 | 1,17458E-08 |
| ACE2 | SDE2 | -0,4021 | 1,16461E-08 |
| ACE2 | STX1A | -0,4021 | 1,16643E-08 |
| ACE2 | AAMP | -0,4022 | 1,15691E-08 |
| ACE2 | RP11-379B18.5 | -0,4022 | 1,15842E-08 |
| ACE2 | RP11-395I14.2 | -0,4022 | 1,15906E-08 |
| ACE2 | LINC00702 | -0,4022 | 1,16182E-08 |
| ACE2 | RP11-212E4.1 | -0,4022 | 1,1556E-08 |
| ACE2 | CLEC4GP1 | -0,4022 | 1,15272E-08 |
| ACE2 | RBM3 | -0,4022 | 1,15218E-08 |
| ACE2 | MYOM2 | -0,4023 | 1,14586E-08 |
| ACE2 | ZNF33B | -0,4023 | 1,14387E-08 |
| ACE2 | LMF2 | -0,4023 | 1,14325E-08 |
| ACE2 | RP11-501J20.5 | -0,4024 | 1,13538E-08 |
| ACE2 | CH17-262A2.1 | -0,4025 | 1,12519E-08 |
| ACE2 | FBXO4 | -0,4025 | 1,12543E-08 |
| ACE2 | ZNF282 | -0,4025 | 1,12739E-08 |
| ACE2 | SLC47A1 | -0,4025 | 1,1277E-08 |

| | | | |
|---|---|---|---|
| ACE2 | OR2L13 | -0,4026 | 1,11158E-08 |
| ACE2 | CTD-2366F13.2 | -0,4026 | 1,11296E-08 |
| ACE2 | CTC-351M12.1 | -0,4026 | 1,11669E-08 |
| ACE2 | SLC6A1-AS1 | -0,4027 | 1,1021E-08 |
| ACE2 | REC8 | -0,4027 | 1,10531E-08 |
| ACE2 | RP1-101D8.1 | -0,4027 | 1,10481E-08 |
| ACE2 | RP4-671O14.6 | -0,4027 | 1,10926E-08 |
| ACE2 | RP13-578N3.3 | -0,4028 | 1,09475E-08 |
| ACE2 | RP4-806M20.4 | -0,4028 | 1,09387E-08 |
| ACE2 | TCEA1P2 | -0,4029 | 1,08855E-08 |
| ACE2 | RP11-259P15.4 | -0,4029 | 1,08828E-08 |
| ACE2 | SNORD22 | -0,4029 | 1,08477E-08 |
| ACE2 | GRIN2B | -0,4029 | 1,08702E-08 |
| ACE2 | RN7SL494P | -0,4029 | 1,08829E-08 |
| ACE2 | MED8 | -0,403 | 1,07677E-08 |
| ACE2 | CD302 | -0,403 | 1,07818E-08 |
| ACE2 | AADACL2 | -0,403 | 1,07119E-08 |
| ACE2 | POU5F1B | -0,403 | 1,07672E-08 |
| ACE2 | GPR65 | -0,4031 | 1,06267E-08 |
| ACE2 | CD83 | -0,4032 | 1,05217E-08 |
| ACE2 | VIM-AS1 | -0,4032 | 1,05292E-08 |
| ACE2 | ATF1 | -0,4032 | 1,05213E-08 |
| ACE2 | AC062029.1 | -0,4033 | 1,04834E-08 |
| ACE2 | CTB-37A13.1 | -0,4033 | 1,04447E-08 |
| ACE2 | RP11-151N17.1 | -0,4033 | 1,05045E-08 |
| ACE2 | CDH15 | -0,4033 | 1,04545E-08 |
| ACE2 | IQCJ-SCHIP1-AS1 | -0,4034 | 1,03619E-08 |
| ACE2 | HIGD1AP18 | -0,4034 | 1,03476E-08 |
| ACE2 | ZNF214 | -0,4034 | 1,04018E-08 |
| ACE2 | ENDOV | -0,4034 | 1,03571E-08 |
| ACE2 | RP11-419C23.1 | -0,4035 | 1,02699E-08 |
| ACE2 | LINC01561 | -0,4035 | 1,03204E-08 |
| ACE2 | FABP5P7 | -0,4035 | 1,02505E-08 |
| ACE2 | RP11-314B1.2 | -0,4036 | 1,0175E-08 |
| ACE2 | OR52D1 | -0,4036 | 1,01665E-08 |
| ACE2 | ATP5S | -0,4036 | 1,02268E-08 |
| ACE2 | ELAVL2 | -0,4037 | 1,00559E-08 |
| ACE2 | AC005387.3 | -0,4037 | 1,00943E-08 |
| ACE2 | PLPPR5 | -0,4038 | 9,99278E-09 |
| ACE2 | DNAJC19P5 | -0,4038 | 9,96672E-09 |
| ACE2 | LINC01238 | -0,4038 | 9,98685E-09 |
| ACE2 | ZNF285B | -0,4038 | 1,00296E-08 |
| ACE2 | RP11-460I13.6 | -0,4039 | 9,94074E-09 |
| ACE2 | TMEM163 | -0,4039 | 9,86469E-09 |
| ACE2 | ANKRD20A10P | -0,4039 | 9,95076E-09 |
| ACE2 | PDPR | -0,4039 | 9,93176E-09 |
| ACE2 | MIS18A-AS1 | -0,4039 | 9,91158E-09 |
| ACE2 | AC006547.13 | -0,4039 | 9,89144E-09 |
| ACE2 | RP11-10E18.7 | -0,4041 | 9,71014E-09 |
| ACE2 | RP11-775C24.4 | -0,4041 | 9,73847E-09 |
| ACE2 | SCARF1 | -0,4041 | 9,74069E-09 |
| ACE2 | RPL36AP33 | -0,4042 | 9,64856E-09 |
| ACE2 | CTD-2561B21.4 | -0,4042 | 9,67408E-09 |
| ACE2 | CTD-2013N17.4 | -0,4042 | 9,60067E-09 |

| | | | |
|---|---|---|---|
| ACE2 | SLC38A9 | -0,4043 | 9,54595E-09 |
| ACE2 | CDC42P5 | -0,4044 | 9,42584E-09 |
| ACE2 | SAFB2 | -0,4044 | 9,42822E-09 |
| ACE2 | VN1R83P | -0,4044 | 9,4688E-09 |
| ACE2 | SATB1 | -0,4045 | 9,36789E-09 |
| ACE2 | LRRC37A15P | -0,4045 | 9,39509E-09 |
| ACE2 | UHRF1BP1L | -0,4045 | 9,41505E-09 |
| ACE2 | DUOXA1 | -0,4045 | 9,33435E-09 |
| ACE2 | RP11-127B20.3 | -0,4046 | 9,25797E-09 |
| ACE2 | RP11-1081M5.2 | -0,4046 | 9,26269E-09 |
| ACE2 | BRINP1 | -0,4046 | 9,32393E-09 |
| ACE2 | CTB-50L17.9 | -0,4046 | 9,25711E-09 |
| ACE2 | MICU1 | -0,4047 | 9,24323E-09 |
| ACE2 | POP4 | -0,4047 | 9,23077E-09 |
| ACE2 | LINC00909 | -0,4048 | 9,10235E-09 |
| ACE2 | RASD2 | -0,4049 | 9,0743E-09 |
| ACE2 | RP4-740C4.5 | -0,405 | 8,92174E-09 |
| ACE2 | CTC-428H11.2 | -0,405 | 8,94374E-09 |
| ACE2 | MCRS1 | -0,405 | 8,96353E-09 |
| ACE2 | PRPF39 | -0,405 | 8,95884E-09 |
| ACE2 | TBL1Y | -0,405 | 8,95912E-09 |
| ACE2 | OLFM3 | -0,4051 | 8,88891E-09 |
| ACE2 | LINC00174 | -0,4051 | 8,84974E-09 |
| ACE2 | ZNF689 | -0,4051 | 8,83586E-09 |
| ACE2 | SNORD33 | -0,4051 | 8,83738E-09 |
| ACE2 | AMPD2 | -0,4052 | 8,82601E-09 |
| ACE2 | RP11-504P24.9 | -0,4052 | 8,80771E-09 |
| ACE2 | GUSBP9 | -0,4052 | 8,79151E-09 |
| ACE2 | RP11-420L9.2 | -0,4052 | 8,77516E-09 |
| ACE2 | RP11-415D17.3 | -0,4053 | 8,69094E-09 |
| ACE2 | ARNTL2 | -0,4053 | 8,72273E-09 |
| ACE2 | RFX7 | -0,4053 | 8,72775E-09 |
| ACE2 | RP11-586D19.1 | -0,4054 | 8,65952E-09 |
| ACE2 | LSM6 | -0,4054 | 8,59532E-09 |
| ACE2 | CST6 | -0,4054 | 8,59268E-09 |
| ACE2 | RP11-890B15.2 | -0,4054 | 8,61963E-09 |
| ACE2 | STK17A | -0,4055 | 8,54665E-09 |
| ACE2 | GRK5-IT1 | -0,4055 | 8,54582E-09 |
| ACE2 | RP11-15A1.2 | -0,4055 | 8,58641E-09 |
| ACE2 | RPL23AP65 | -0,4056 | 8,5004E-09 |
| ACE2 | RP11-111K18.2 | -0,4057 | 8,39825E-09 |
| ACE2 | OGFOD1P1 | -0,4057 | 8,37656E-09 |
| ACE2 | C8orf89 | -0,4058 | 8,29722E-09 |
| ACE2 | CENPJ | -0,4058 | 8,28606E-09 |
| ACE2 | RP11-596C23.2 | -0,4058 | 8,28941E-09 |
| ACE2 | IGFL1P1 | -0,4058 | 8,33002E-09 |
| ACE2 | RNU1-122P | -0,4059 | 8,26179E-09 |
| ACE2 | TGDS | -0,4059 | 8,26042E-09 |
| ACE2 | GTF2F1 | -0,4059 | 8,25862E-09 |
| ACE2 | C19orf38 | -0,4059 | 8,25471E-09 |
| ACE2 | RP5-1198O20.4 | -0,406 | 8,14532E-09 |
| ACE2 | NMBR | -0,406 | 8,13375E-09 |
| ACE2 | RP11-467H10.1 | -0,406 | 8,1746E-09 |
| ACE2 | RAPGEF1 | -0,406 | 8,17973E-09 |

| | | | |
|---|---|---|---|
| ACE2 | KB-1125A3.12 | -0,406 | 8,16203E-09 |
| ACE2 | MSH2 | -0,4061 | 8,10954E-09 |
| ACE2 | MTX2 | -0,4061 | 8,06064E-09 |
| ACE2 | RP11-560I19.1 | -0,4061 | 8,1055E-09 |
| ACE2 | CTD-2371O3.2 | -0,4061 | 8,07334E-09 |
| ACE2 | SCAND3P1 | -0,4061 | 8,11094E-09 |
| ACE2 | FDX1P2 | -0,4061 | 8,05607E-09 |
| ACE2 | APOL2 | -0,4061 | 8,06119E-09 |
| ACE2 | SNX10 | -0,4062 | 8,01084E-09 |
| ACE2 | PCAT1 | -0,4062 | 7,98938E-09 |
| ACE2 | CTD-2003C8.2 | -0,4062 | 8,02066E-09 |
| ACE2 | RP11-165D6.1 | -0,4062 | 8,00517E-09 |
| ACE2 | BCL7B | -0,4063 | 7,95981E-09 |
| ACE2 | CTD-2005H7.1 | -0,4063 | 7,94965E-09 |
| ACE2 | RP11-884K10.7 | -0,4064 | 7,83838E-09 |
| ACE2 | TEAD3 | -0,4064 | 7,8905E-09 |
| ACE2 | RP11-244E17.1 | -0,4064 | 7,8478E-09 |
| ACE2 | GOLGA8Q | -0,4064 | 7,88142E-09 |
| ACE2 | RP13-638C3.2 | -0,4064 | 7,89975E-09 |
| ACE2 | LINC01697 | -0,4064 | 7,89371E-09 |
| ACE2 | G6PD | -0,4064 | 7,88904E-09 |
| ACE2 | RNU5E-1 | -0,4065 | 7,82447E-09 |
| ACE2 | POMGNT1 | -0,4065 | 7,76303E-09 |
| ACE2 | RP11-326C3.14 | -0,4065 | 7,82201E-09 |
| ACE2 | RP11-452J21.2 | -0,4066 | 7,71675E-09 |
| ACE2 | GNRH2 | -0,4066 | 7,71015E-09 |
| ACE2 | RP11-30L15.4 | -0,4068 | 7,56006E-09 |
| ACE2 | SURF1 | -0,4068 | 7,60304E-09 |
| ACE2 | RP11-422P24.9 | -0,407 | 7,46713E-09 |
| ACE2 | MAP3K5 | -0,407 | 7,41446E-09 |
| ACE2 | C1orf186 | -0,4071 | 7,36725E-09 |
| ACE2 | FEZF1-AS1 | -0,4073 | 7,25211E-09 |
| ACE2 | RP11-708L7.6 | -0,4073 | 7,24832E-09 |
| ACE2 | TP53TG3D | -0,4073 | 7,2281E-09 |
| ACE2 | KRT18P4 | -0,4073 | 7,25035E-09 |
| ACE2 | ADCY10P1 | -0,4074 | 7,17638E-09 |
| ACE2 | RP11-90B9.3 | -0,4074 | 7,1968E-09 |
| ACE2 | KLHL38 | -0,4075 | 7,12045E-09 |
| ACE2 | RP5-1107A17.4 | -0,4075 | 7,12881E-09 |
| ACE2 | SMCHD1 | -0,4075 | 7,08744E-09 |
| ACE2 | PLXNA3 | -0,4075 | 7,09421E-09 |
| ACE2 | RP5-1159O4.2 | -0,4076 | 7,04644E-09 |
| ACE2 | CYCSP40 | -0,4076 | 7,05954E-09 |
| ACE2 | PCBP1 | -0,4077 | 6,98005E-09 |
| ACE2 | CHRNA4 | -0,4077 | 6,97225E-09 |
| ACE2 | RP11-461L13.3 | -0,4078 | 6,91258E-09 |
| ACE2 | M6PR | -0,4078 | 6,92679E-09 |
| ACE2 | AC001226.7 | -0,4078 | 6,91523E-09 |
| ACE2 | HNMT | -0,4079 | 6,83338E-09 |
| ACE2 | CMTM7 | -0,4079 | 6,86525E-09 |
| ACE2 | SSPO | -0,4079 | 6,84854E-09 |
| ACE2 | TMEM75 | -0,4079 | 6,8657E-09 |
| ACE2 | OTUD7A | -0,4079 | 6,85507E-09 |
| ACE2 | CACNA1C-IT2 | -0,408 | 6,79304E-09 |

| | | | |
|---|---|---|---|
| ACE2 | RP11-12M9.3 | -0,408 | 6,78905E-09 |
| ACE2 | GAS5-AS1 | -0,4081 | 6,70622E-09 |
| ACE2 | RP11-464F9.22 | -0,4081 | 6,72868E-09 |
| ACE2 | KDM4E | -0,4081 | 6,70802E-09 |
| ACE2 | DRC1 | -0,4082 | 6,66918E-09 |
| ACE2 | PBX2P1 | -0,4082 | 6,65928E-09 |
| ACE2 | RN7SKP268 | -0,4082 | 6,65208E-09 |
| ACE2 | TNKS2-AS1 | -0,4082 | 6,65763E-09 |
| ACE2 | PER1 | -0,4082 | 6,64693E-09 |
| ACE2 | AP2A1 | -0,4082 | 6,63311E-09 |
| ACE2 | LL0XNC01-116E7.1 | -0,4082 | 6,66756E-09 |
| ACE2 | RNF11 | -0,4083 | 6,6156E-09 |
| ACE2 | STEAP3-AS1 | -0,4084 | 6,51434E-09 |
| ACE2 | FGFR2 | -0,4084 | 6,52023E-09 |
| ACE2 | FAM83D | -0,4084 | 6,52842E-09 |
| ACE2 | GLRA2 | -0,4084 | 6,5544E-09 |
| ACE2 | S100A9 | -0,4085 | 6,46362E-09 |
| ACE2 | KCNH7 | -0,4085 | 6,49109E-09 |
| ACE2 | LINC01300 | -0,4085 | 6,4704E-09 |
| ACE2 | ZPR1 | -0,4085 | 6,46712E-09 |
| ACE2 | RP11-15A1.7 | -0,4085 | 6,50434E-09 |
| ACE2 | RP11-46A10.6 | -0,4086 | 6,40846E-09 |
| ACE2 | VNN2 | -0,4087 | 6,33131E-09 |
| ACE2 | ANKRD50 | -0,4088 | 6,30265E-09 |
| ACE2 | TRIO | -0,4088 | 6,2842E-09 |
| ACE2 | NPIPB4 | -0,4088 | 6,28432E-09 |
| ACE2 | OAZ3 | -0,4089 | 6,26951E-09 |
| ACE2 | ARV1 | -0,4089 | 6,26966E-09 |
| ACE2 | UEVLD | -0,4089 | 6,2663E-09 |
| ACE2 | WASH4P | -0,4089 | 6,23495E-09 |
| ACE2 | RP13-270P17.1 | -0,4089 | 6,26716E-09 |
| ACE2 | PPP1R14BP3 | -0,409 | 6,19547E-09 |
| ACE2 | LINC01987 | -0,409 | 6,18555E-09 |
| ACE2 | ATXN7 | -0,4091 | 6,10699E-09 |
| ACE2 | VPS4A | -0,4091 | 6,15483E-09 |
| ACE2 | RP11-730A19.9 | -0,4092 | 6,06837E-09 |
| ACE2 | TSPAN19 | -0,4092 | 6,06648E-09 |
| ACE2 | RP11-848P1.7 | -0,4092 | 6,07871E-09 |
| ACE2 | RP11-335K5.2 | -0,4093 | 5,98784E-09 |
| ACE2 | RP11-152L20.3 | -0,4093 | 6,0038E-09 |
| ACE2 | TET2-AS1 | -0,4094 | 5,98503E-09 |
| ACE2 | DPY19L2P1 | -0,4094 | 5,98334E-09 |
| ACE2 | CUZD1 | -0,4094 | 5,95611E-09 |
| ACE2 | C1QTNF5 | -0,4094 | 5,98223E-09 |
| ACE2 | F10 | -0,4094 | 5,9445E-09 |
| ACE2 | SLC24A5 | -0,4094 | 5,95861E-09 |
| ACE2 | LRRIQ3 | -0,4095 | 5,89692E-09 |
| ACE2 | SLC30A6 | -0,4095 | 5,92627E-09 |
| ACE2 | SERBP1P3 | -0,4095 | 5,90129E-09 |
| ACE2 | RP11-723O4.7 | -0,4095 | 5,9035E-09 |
| ACE2 | ZCCHC3 | -0,4095 | 5,90536E-09 |
| ACE2 | CIAO1 | -0,4096 | 5,85738E-09 |
| ACE2 | MAP1LC3A | -0,4096 | 5,83575E-09 |
| ACE2 | NUTM2B-AS1 | -0,4097 | 5,77778E-09 |

| | | | |
|---|---|---|---|
| ACE2 | RP11-167N24.3 | -0,4097 | 5,79678E-09 |
| ACE2 | LINC01229 | -0,4097 | 5,81602E-09 |
| ACE2 | CTB-96E2.10 | -0,4097 | 5,81813E-09 |
| ACE2 | ARIH2OS | -0,4098 | 5,76309E-09 |
| ACE2 | SLC30A5 | -0,4098 | 5,7369E-09 |
| ACE2 | RP11-637O19.2 | -0,4098 | 5,7179E-09 |
| ACE2 | RAD23B | -0,4098 | 5,71489E-09 |
| ACE2 | RP11-73M18.7 | -0,4098 | 5,72263E-09 |
| ACE2 | RP11-462G12.1 | -0,4098 | 5,73943E-09 |
| ACE2 | AC005253.2 | -0,4098 | 5,76339E-09 |
| ACE2 | PRMT5-AS1 | -0,4099 | 5,68292E-09 |
| ACE2 | RCCD1 | -0,4099 | 5,66978E-09 |
| ACE2 | AC002550.6 | -0,4099 | 5,67825E-09 |
| ACE2 | VIT | -0,41 | 5,65833E-09 |
| ACE2 | LAMTOR1 | -0,41 | 5,63406E-09 |
| ACE2 | CDKL1 | -0,41 | 5,6151E-09 |
| ACE2 | RP11-429B14.4 | -0,41 | 5,6087E-09 |
| ACE2 | RP11-340I6.8 | -0,4101 | 5,55667E-09 |
| ACE2 | EIF3A | -0,4101 | 5,59097E-09 |
| ACE2 | C1orf174 | -0,4102 | 5,53663E-09 |
| ACE2 | RAB24 | -0,4102 | 5,52097E-09 |
| ACE2 | NDUFAF6 | -0,4102 | 5,52098E-09 |
| ACE2 | OR52N4 | -0,4102 | 5,54074E-09 |
| ACE2 | RPL41P1 | -0,4102 | 5,52397E-09 |
| ACE2 | CTTNBP2NL | -0,4103 | 5,47491E-09 |
| ACE2 | EVL | -0,4103 | 5,46861E-09 |
| ACE2 | OVGP1 | -0,4104 | 5,42233E-09 |
| ACE2 | CTD-2292P10.4 | -0,4105 | 5,38753E-09 |
| ACE2 | INTS5 | -0,4105 | 5,37474E-09 |
| ACE2 | RP4-646N3.1 | -0,4105 | 5,37399E-09 |
| ACE2 | LINC01135 | -0,4106 | 5,31303E-09 |
| ACE2 | DAPL1 | -0,4106 | 5,32309E-09 |
| ACE2 | DEFB109P3 | -0,4106 | 5,30146E-09 |
| ACE2 | LINC00865 | -0,4106 | 5,32456E-09 |
| ACE2 | PRPS1P2 | -0,4107 | 5,25298E-09 |
| ACE2 | VTI1B | -0,4107 | 5,26655E-09 |
| ACE2 | PGF | -0,4107 | 5,2837E-09 |
| ACE2 | RP11-242D8.3 | -0,4107 | 5,29106E-09 |
| ACE2 | PQBP1 | -0,4107 | 5,28216E-09 |
| ACE2 | TOPBP1 | -0,4108 | 5,2472E-09 |
| ACE2 | OR7M1P | -0,4108 | 5,20774E-09 |
| ACE2 | DRD2 | -0,4108 | 5,24757E-09 |
| ACE2 | AP000345.4 | -0,4108 | 5,22099E-09 |
| ACE2 | FANCB | -0,4108 | 5,20336E-09 |
| ACE2 | IGFN1 | -0,4109 | 5,15595E-09 |
| ACE2 | DUSP6 | -0,411 | 5,13965E-09 |
| ACE2 | AC009120.3 | -0,411 | 5,12666E-09 |
| ACE2 | ADCK1 | -0,4111 | 5,06786E-09 |
| ACE2 | CTB-32O4.2 | -0,4111 | 5,06799E-09 |
| ACE2 | DDX4 | -0,4112 | 5,0168E-09 |
| ACE2 | EPHA1-AS1 | -0,4112 | 5,01137E-09 |
| ACE2 | CRYM-AS1 | -0,4112 | 5,05679E-09 |
| ACE2 | CTD-3001H11.2 | -0,4112 | 5,04758E-09 |
| ACE2 | RP4-598P13.1 | -0,4113 | 5,01013E-09 |

| | | | |
|---|---|---|---|
| ACE2 | RP11-363D14.1 | -0,4113 | 4,98158E-09 |
| ACE2 | NF1 | -0,4113 | 4,98142E-09 |
| ACE2 | LRRC37A2 | -0,4113 | 5,00947E-09 |
| ACE2 | CTD-3224K15.3 | -0,4114 | 4,9479E-09 |
| ACE2 | CCND3 | -0,4114 | 4,95625E-09 |
| ACE2 | C6orf118 | -0,4114 | 4,95698E-09 |
| ACE2 | RP1 | -0,4114 | 4,95061E-09 |
| ACE2 | CASP9 | -0,4115 | 4,87768E-09 |
| ACE2 | RP11-359I18.5 | -0,4115 | 4,88299E-09 |
| ACE2 | CLEC4A | -0,4115 | 4,90619E-09 |
| ACE2 | RP11-567C2.1 | -0,4115 | 4,90436E-09 |
| ACE2 | RPL39P3 | -0,4116 | 4,83231E-09 |
| ACE2 | HS3ST4 | -0,4116 | 4,87007E-09 |
| ACE2 | GRIK2 | -0,4117 | 4,8214E-09 |
| ACE2 | RP11-204M4.3 | -0,4117 | 4,78674E-09 |
| ACE2 | RP11-84A19.4 | -0,4118 | 4,7681E-09 |
| ACE2 | AC024704.2 | -0,4118 | 4,75782E-09 |
| ACE2 | WWTR1-IT1 | -0,4118 | 4,75554E-09 |
| ACE2 | AP000442.4 | -0,4118 | 4,75069E-09 |
| ACE2 | DHX58 | -0,4118 | 4,75662E-09 |
| ACE2 | RP5-890E16.5 | -0,4118 | 4,7386E-09 |
| ACE2 | PDE4B | -0,4119 | 4,69878E-09 |
| ACE2 | RP11-513M16.7 | -0,4119 | 4,72252E-09 |
| ACE2 | DRC7 | -0,4119 | 4,71811E-09 |
| ACE2 | RP11-471L13.3 | -0,4119 | 4,73064E-09 |
| ACE2 | HIST1H2BF | -0,412 | 4,66019E-09 |
| ACE2 | SAMD9L | -0,412 | 4,66869E-09 |
| ACE2 | XRRA1 | -0,412 | 4,67675E-09 |
| ACE2 | CTD-2270N23.1 | -0,412 | 4,66225E-09 |
| ACE2 | RP4-728D4.2 | -0,4121 | 4,61657E-09 |
| ACE2 | RP11-569G13.2 | -0,4121 | 4,62498E-09 |
| ACE2 | PCNX1 | -0,4121 | 4,6311E-09 |
| ACE2 | FIRRE | -0,4121 | 4,64286E-09 |
| ACE2 | RP11-546K22.3 | -0,4122 | 4,58184E-09 |
| ACE2 | SFMBT2 | -0,4122 | 4,57972E-09 |
| ACE2 | RMST | -0,4122 | 4,57808E-09 |
| ACE2 | RP1-278D1.2 | -0,4122 | 4,60205E-09 |
| ACE2 | ARPP21 | -0,4123 | 4,54871E-09 |
| ACE2 | ANKRD20A17P | -0,4123 | 4,56001E-09 |
| ACE2 | RP11-526D8.11 | -0,4123 | 4,52504E-09 |
| ACE2 | NPTN-IT1 | -0,4123 | 4,5466E-09 |
| ACE2 | CTB-167G5.7 | -0,4123 | 4,55208E-09 |
| ACE2 | CCDC188 | -0,4123 | 4,54068E-09 |
| ACE2 | RP4-742C19.13 | -0,4123 | 4,54573E-09 |
| ACE2 | LINC01310 | -0,4123 | 4,5348E-09 |
| ACE2 | GNAQ | -0,4125 | 4,46093E-09 |
| ACE2 | RP11-343H19.1 | -0,4125 | 4,46569E-09 |
| ACE2 | NPTX2 | -0,4126 | 4,42444E-09 |
| ACE2 | RP11-266K4.1 | -0,4126 | 4,42641E-09 |
| ACE2 | TCAM1P | -0,4126 | 4,40179E-09 |
| ACE2 | FAUP1 | -0,4126 | 4,41801E-09 |
| ACE2 | RP11-290M5.4 | -0,4127 | 4,3843E-09 |
| ACE2 | LRRC71 | -0,4127 | 4,37783E-09 |
| ACE2 | TLR1 | -0,4127 | 4,36716E-09 |

| | | | |
|---|---|---|---|
| ACE2 | ANKRD13A | -0,4127 | 4,35734E-09 |
| ACE2 | ANAPC2 | -0,4128 | 4,34094E-09 |
| ACE2 | RPS2P55 | -0,4128 | 4,33147E-09 |
| ACE2 | RP11-22A3.2 | -0,4129 | 4,28318E-09 |
| ACE2 | RP11-700A24.1 | -0,413 | 4,26441E-09 |
| ACE2 | PTPN23 | -0,4132 | 4,15197E-09 |
| ACE2 | PKHD1L1 | -0,4132 | 4,1679E-09 |
| ACE2 | RP11-643C9.2 | -0,4132 | 4,15181E-09 |
| ACE2 | RP11-348F1.3 | -0,4132 | 4,18507E-09 |
| ACE2 | UBE2Q2P2 | -0,4133 | 4,14599E-09 |
| ACE2 | SH3KBP1 | -0,4133 | 4,13824E-09 |
| ACE2 | MIR34AHG | -0,4134 | 4,0822E-09 |
| ACE2 | EEF1A1P11 | -0,4134 | 4,08801E-09 |
| ACE2 | KLHL12 | -0,4134 | 4,10832E-09 |
| ACE2 | ACSL3 | -0,4134 | 4,07515E-09 |
| ACE2 | AC093495.4 | -0,4134 | 4,09203E-09 |
| ACE2 | RP3-337H4.10 | -0,4134 | 4,09295E-09 |
| ACE2 | SLC9A7 | -0,4134 | 4,10236E-09 |
| ACE2 | OR8S1 | -0,4135 | 4,03565E-09 |
| ACE2 | TOM1L2 | -0,4135 | 4,07101E-09 |
| ACE2 | RP11-16N11.2 | -0,4136 | 4,00852E-09 |
| ACE2 | GLCCI1 | -0,4136 | 4,01777E-09 |
| ACE2 | RPL5P23 | -0,4136 | 4,02361E-09 |
| ACE2 | SNORD68 | -0,4136 | 4,01846E-09 |
| ACE2 | CTC-484P3.3 | -0,4137 | 3,98263E-09 |
| ACE2 | ACTL6B | -0,4137 | 3,97154E-09 |
| ACE2 | RP11-3B12.4 | -0,4137 | 3,96785E-09 |
| ACE2 | IMPACT | -0,4137 | 3,98454E-09 |
| ACE2 | GRINA | -0,4138 | 3,94071E-09 |
| ACE2 | GRHPR | -0,4138 | 3,92249E-09 |
| ACE2 | RP11-592N21.1 | -0,4138 | 3,95835E-09 |
| ACE2 | KCTD13 | -0,4138 | 3,93883E-09 |
| ACE2 | CSNK2A1 | -0,4138 | 3,94139E-09 |
| ACE2 | RPL32P3 | -0,4139 | 3,91976E-09 |
| ACE2 | TMEM232 | -0,4139 | 3,89559E-09 |
| ACE2 | POLR3A | -0,4139 | 3,90047E-09 |
| ACE2 | NBPF3 | -0,414 | 3,86584E-09 |
| ACE2 | EIF4HP2 | -0,414 | 3,8719E-09 |
| ACE2 | CCDC18-AS1 | -0,4141 | 3,83139E-09 |
| ACE2 | WDR5 | -0,4141 | 3,82312E-09 |
| ACE2 | CCDC73 | -0,4141 | 3,82575E-09 |
| ACE2 | NLGN4Y | -0,4141 | 3,8133E-09 |
| ACE2 | ZFP36L2 | -0,4142 | 3,78985E-09 |
| ACE2 | SH3GL1P1 | -0,4142 | 3,77859E-09 |
| ACE2 | CNTD2 | -0,4142 | 3,79763E-09 |
| ACE2 | LL22NC03-N14H11.1 | -0,4142 | 3,78339E-09 |
| ACE2 | LGI3 | -0,4143 | 3,76462E-09 |
| ACE2 | ATP23 | -0,4143 | 3,7671E-09 |
| ACE2 | SSU72 | -0,4144 | 3,71703E-09 |
| ACE2 | PCDHA1 | -0,4144 | 3,7264E-09 |
| ACE2 | RPS3AP38 | -0,4145 | 3,70115E-09 |
| ACE2 | RP11-140A10.3 | -0,4145 | 3,68501E-09 |
| ACE2 | RP11-73E17.2 | -0,4145 | 3,67321E-09 |
| ACE2 | RP11-403B2.7 | -0,4145 | 3,69838E-09 |

| | | | |
|---|---|---|---|
| ACE2 | CTD-2529P6.3 | -0,4145 | 3,69534E-09 |
| ACE2 | AC090804.1 | -0,4146 | 3,65864E-09 |
| ACE2 | EDARADD | -0,4147 | 3,62688E-09 |
| ACE2 | RP11-410L14.1 | -0,4147 | 3,61926E-09 |
| ACE2 | RP11-295P9.3 | -0,4147 | 3,63472E-09 |
| ACE2 | CHD2 | -0,4147 | 3,6241E-09 |
| ACE2 | ADGRG3 | -0,4147 | 3,61742E-09 |
| ACE2 | BEAN1-AS1 | -0,4147 | 3,63335E-09 |
| ACE2 | RP11-181E10.3 | -0,4148 | 3,59673E-09 |
| ACE2 | SP100 | -0,4148 | 3,59723E-09 |
| ACE2 | RP4-555D20.2 | -0,4148 | 3,58747E-09 |
| ACE2 | RP11-80H5.7 | -0,4148 | 3,58135E-09 |
| ACE2 | RP11-327J17.8 | -0,4148 | 3,589E-09 |
| ACE2 | IZUMO1 | -0,4148 | 3,58502E-09 |
| ACE2 | RP13-52K8.1 | -0,4148 | 3,59459E-09 |
| ACE2 | RP11-338N10.2 | -0,4149 | 3,5636E-09 |
| ACE2 | ERMARD | -0,4149 | 3,56409E-09 |
| ACE2 | DIO3OS | -0,4149 | 3,55852E-09 |
| ACE2 | CNTN6 | -0,415 | 3,51743E-09 |
| ACE2 | SLC24A2 | -0,415 | 3,50427E-09 |
| ACE2 | GABARAPL1 | -0,415 | 3,51986E-09 |
| ACE2 | RP11-797D24.3 | -0,4151 | 3,49418E-09 |
| ACE2 | SLC12A9 | -0,4151 | 3,49133E-09 |
| ACE2 | MESTP1 | -0,4152 | 3,45155E-09 |
| ACE2 | RPL21P134 | -0,4152 | 3,45895E-09 |
| ACE2 | LINC00517 | -0,4153 | 3,42942E-09 |
| ACE2 | RP11-81K2.1 | -0,4153 | 3,4224E-09 |
| ACE2 | MFAP5 | -0,4154 | 3,39426E-09 |
| ACE2 | AGAP2 | -0,4154 | 3,40108E-09 |
| ACE2 | SNRPCP3 | -0,4155 | 3,35159E-09 |
| ACE2 | RP11-706J10.2 | -0,4155 | 3,35716E-09 |
| ACE2 | SNRPB | -0,4155 | 3,34634E-09 |
| ACE2 | SKP2 | -0,4156 | 3,31287E-09 |
| ACE2 | RP11-480N24.4 | -0,4156 | 3,33312E-09 |
| ACE2 | RP11-347C18.3 | -0,4156 | 3,31255E-09 |
| ACE2 | ASB16-AS1 | -0,4156 | 3,33007E-09 |
| ACE2 | LINC00163 | -0,4156 | 3,31204E-09 |
| ACE2 | RP11-332H14.1 | -0,4157 | 3,30126E-09 |
| ACE2 | CBX3P4 | -0,4157 | 3,28692E-09 |
| ACE2 | STOML1 | -0,4157 | 3,29707E-09 |
| ACE2 | CAPNS2 | -0,4157 | 3,27983E-09 |
| ACE2 | RP11-803P9.1 | -0,4158 | 3,26176E-09 |
| ACE2 | FAM20C | -0,4158 | 3,25094E-09 |
| ACE2 | ADAM32 | -0,4158 | 3,2543E-09 |
| ACE2 | PARD3-AS1 | -0,4158 | 3,25665E-09 |
| ACE2 | HERC2P3 | -0,4158 | 3,25731E-09 |
| ACE2 | RP11-185E8.2 | -0,4159 | 3,22172E-09 |
| ACE2 | RP11-141M1.4 | -0,4159 | 3,21482E-09 |
| ACE2 | SCAMP4 | -0,4159 | 3,23274E-09 |
| ACE2 | RBM44 | -0,416 | 3,19902E-09 |
| ACE2 | KEL | -0,416 | 3,20325E-09 |
| ACE2 | RP11-750H9.5 | -0,416 | 3,20268E-09 |
| ACE2 | ARAP1 | -0,416 | 3,20402E-09 |
| ACE2 | MRPL45P2 | -0,416 | 3,2107E-09 |

| | | | |
|---|---|---|---|
| ACE2 | LINC00997 | -0,4161 | 3,17024E-09 |
| ACE2 | LINC00921 | -0,4161 | 3,16869E-09 |
| ACE2 | PTOV1-AS2 | -0,4161 | 3,16343E-09 |
| ACE2 | MANBAL | -0,4161 | 3,18074E-09 |
| ACE2 | RN7SKP80 | -0,4161 | 3,17223E-09 |
| ACE2 | RP3-393E18.2 | -0,4162 | 3,14464E-09 |
| ACE2 | LINC01410 | -0,4162 | 3,12632E-09 |
| ACE2 | LMNTD1 | -0,4162 | 3,14433E-09 |
| ACE2 | LEMD3 | -0,4162 | 3,12554E-09 |
| ACE2 | CBX4 | -0,4162 | 3,12582E-09 |
| ACE2 | SLC12A5 | -0,4162 | 3,13202E-09 |
| ACE2 | ADAMTS5 | -0,4162 | 3,14036E-09 |
| ACE2 | LRIF1 | -0,4163 | 3,11392E-09 |
| ACE2 | CCNT2 | -0,4163 | 3,11971E-09 |
| ACE2 | ABC13-47488600E1 | -0,4163 | 3,09489E-09 |
| ACE2 | RP11-1000B6.8 | -0,4163 | 3,10956E-09 |
| ACE2 | PRPSAP2 | -0,4163 | 3,09515E-09 |
| ACE2 | RP3-402G11.28 | -0,4163 | 3,10221E-09 |
| ACE2 | MMD2 | -0,4164 | 3,0702E-09 |
| ACE2 | ZNF26 | -0,4164 | 3,08077E-09 |
| ACE2 | LA16c-360H6.1 | -0,4164 | 3,06951E-09 |
| ACE2 | KB-1995A5.5 | -0,4164 | 3,08422E-09 |
| ACE2 | TMEM14EP | -0,4165 | 3,06213E-09 |
| ACE2 | PIGG | -0,4165 | 3,05092E-09 |
| ACE2 | CIPC | -0,4165 | 3,05586E-09 |
| ACE2 | CPA3 | -0,4166 | 3,02137E-09 |
| ACE2 | UBE2R2 | -0,4167 | 3,00327E-09 |
| ACE2 | BTBD8 | -0,4168 | 2,95162E-09 |
| ACE2 | ZNF852 | -0,4168 | 2,95425E-09 |
| ACE2 | LINC02145 | -0,4168 | 2,96974E-09 |
| ACE2 | RP11-768F21.1 | -0,4168 | 2,9546E-09 |
| ACE2 | DLEU1 | -0,4168 | 2,95484E-09 |
| ACE2 | UBE2V1P1 | -0,4168 | 2,95519E-09 |
| ACE2 | WRB | -0,4168 | 2,95467E-09 |
| ACE2 | PHKA1P1 | -0,4169 | 2,94346E-09 |
| ACE2 | GS1-304P7.1 | -0,4169 | 2,92287E-09 |
| ACE2 | AC000123.3 | -0,4169 | 2,92532E-09 |
| ACE2 | NUP214 | -0,4169 | 2,92734E-09 |
| ACE2 | OR7E2P | -0,4169 | 2,93486E-09 |
| ACE2 | RP11-127B20.2 | -0,417 | 2,89208E-09 |
| ACE2 | ATF6B | -0,417 | 2,91762E-09 |
| ACE2 | ZMIZ1-AS1 | -0,417 | 2,89371E-09 |
| ACE2 | CTD-2026K11.4 | -0,417 | 2,91804E-09 |
| ACE2 | FAM96B | -0,417 | 2,89556E-09 |
| ACE2 | CTB-50L17.7 | -0,417 | 2,90353E-09 |
| ACE2 | CTC-559E9.5 | -0,417 | 2,90108E-09 |
| ACE2 | COQ10B | -0,4171 | 2,88232E-09 |
| ACE2 | RP11-577H5.1 | -0,4171 | 2,87023E-09 |
| ACE2 | FAM214A | -0,4171 | 2,86831E-09 |
| ACE2 | AC002472.16 | -0,4171 | 2,86537E-09 |
| ACE2 | SBF1 | -0,4171 | 2,88275E-09 |
| ACE2 | EHMT1 | -0,4172 | 2,84089E-09 |
| ACE2 | SLC35F3 | -0,4173 | 2,81688E-09 |
| ACE2 | PCDHB1 | -0,4173 | 2,82243E-09 |

| | | | |
|---|---|---|---|
| ACE2 | APCDD1L | -0,4173 | 2,82194E-09 |
| ACE2 | AC073052.1 | -0,4174 | 2,79981E-09 |
| ACE2 | C8orf22 | -0,4175 | 2,76703E-09 |
| ACE2 | RP11-433J8.1 | -0,4175 | 2,77653E-09 |
| ACE2 | TREX2 | -0,4175 | 2,77257E-09 |
| ACE2 | PDE7A | -0,4176 | 2,73934E-09 |
| ACE2 | FAAP100 | -0,4176 | 2,74204E-09 |
| ACE2 | AC005785.2 | -0,4176 | 2,75605E-09 |
| ACE2 | ACTG1P10 | -0,4176 | 2,74433E-09 |
| ACE2 | ALDH7A1 | -0,4177 | 2,7162E-09 |
| ACE2 | JMJD1C | -0,4177 | 2,72173E-09 |
| ACE2 | FAM187B | -0,4177 | 2,71059E-09 |
| ACE2 | MTAPP2 | -0,4178 | 2,69011E-09 |
| ACE2 | SERHL2 | -0,4178 | 2,68432E-09 |
| ACE2 | MILR1 | -0,418 | 2,62899E-09 |
| ACE2 | GCC2-AS1 | -0,4181 | 2,61806E-09 |
| ACE2 | CCDC54 | -0,4181 | 2,6183E-09 |
| ACE2 | SNRPGP2 | -0,4181 | 2,61988E-09 |
| ACE2 | IL10RB-AS1 | -0,4181 | 2,60804E-09 |
| ACE2 | AC007362.3 | -0,4182 | 2,5813E-09 |
| ACE2 | RP11-269M20.3 | -0,4182 | 2,60208E-09 |
| ACE2 | PCBP2 | -0,4182 | 2,58885E-09 |
| ACE2 | FOXJ3 | -0,4183 | 2,55791E-09 |
| ACE2 | EXOSC4 | -0,4183 | 2,5612E-09 |
| ACE2 | RP11-111M22.3 | -0,4183 | 2,55715E-09 |
| ACE2 | CTB-32O4.3 | -0,4183 | 2,55835E-09 |
| ACE2 | RP11-112L6.3 | -0,4183 | 2,56942E-09 |
| ACE2 | LINC00316 | -0,4183 | 2,57596E-09 |
| ACE2 | RPLP0P6 | -0,4184 | 2,5472E-09 |
| ACE2 | FANCL | -0,4184 | 2,53287E-09 |
| ACE2 | TTC21A | -0,4184 | 2,53597E-09 |
| ACE2 | L3MBTL4 | -0,4184 | 2,54646E-09 |
| ACE2 | AC005954.4 | -0,4184 | 2,53037E-09 |
| ACE2 | AC092535.3 | -0,4185 | 2,50709E-09 |
| ACE2 | NHLRC2 | -0,4185 | 2,51661E-09 |
| ACE2 | RP11-253E3.3 | -0,4185 | 2,52151E-09 |
| ACE2 | RP11-338C15.3 | -0,4186 | 2,49012E-09 |
| ACE2 | RP11-11N7.4 | -0,4186 | 2,48407E-09 |
| ACE2 | RP11-521B24.4 | -0,4186 | 2,50383E-09 |
| ACE2 | GABPB1-AS1 | -0,4186 | 2,48732E-09 |
| ACE2 | MTND1P8 | -0,4186 | 2,50206E-09 |
| ACE2 | KRT10 | -0,4186 | 2,49676E-09 |
| ACE2 | EPHX3 | -0,4186 | 2,48105E-09 |
| ACE2 | ZNF860 | -0,4187 | 2,47833E-09 |
| ACE2 | VCPKMT | -0,4187 | 2,46197E-09 |
| ACE2 | PARP9 | -0,4188 | 2,4392E-09 |
| ACE2 | LINC02067 | -0,4188 | 2,45407E-09 |
| ACE2 | FNIP1 | -0,4188 | 2,44983E-09 |
| ACE2 | DBNL | -0,4188 | 2,44742E-09 |
| ACE2 | INSC | -0,4188 | 2,45015E-09 |
| ACE2 | CTC-510F12.2 | -0,4188 | 2,45604E-09 |
| ACE2 | CTD-2175A23.1 | -0,4189 | 2,42325E-09 |
| ACE2 | RP11-618M23.5 | -0,4189 | 2,432E-09 |
| ACE2 | CTD-2319I12.10 | -0,4189 | 2,42786E-09 |

| | | | |
|---|---|---|---|
| ACE2 | RP1-32I10.10 | -0,4189 | 2,42683E-09 |
| ACE2 | ARHGAP31-AS1 | -0,419 | 2,3939E-09 |
| ACE2 | FAM193B | -0,419 | 2,39263E-09 |
| ACE2 | NPDC1 | -0,419 | 2,39601E-09 |
| ACE2 | SLC25A36P1 | -0,419 | 2,39076E-09 |
| ACE2 | PCGF1 | -0,4191 | 2,36556E-09 |
| ACE2 | GADL1 | -0,4191 | 2,3855E-09 |
| ACE2 | SNORA3B | -0,4191 | 2,38545E-09 |
| ACE2 | RP11-613M5.2 | -0,4191 | 2,36913E-09 |
| ACE2 | LINC00893 | -0,4191 | 2,37307E-09 |
| ACE2 | SEC24B-AS1 | -0,4192 | 2,35379E-09 |
| ACE2 | PRR33 | -0,4192 | 2,35749E-09 |
| ACE2 | CTD-2012K14.5 | -0,4192 | 2,35446E-09 |
| ACE2 | XXbac-BPG170G13.3 | -0,4193 | 2,32533E-09 |
| ACE2 | RP11-909N17.2 | -0,4193 | 2,32545E-09 |
| ACE2 | RP11-30K9.1 | -0,4193 | 2,32785E-09 |
| ACE2 | QSOX1 | -0,4194 | 2,31101E-09 |
| ACE2 | FAM96AP2 | -0,4194 | 2,31778E-09 |
| ACE2 | FGL2 | -0,4194 | 2,30967E-09 |
| ACE2 | NANOS3 | -0,4194 | 2,30844E-09 |
| ACE2 | HSP90AB2P | -0,4195 | 2,29328E-09 |
| ACE2 | SVIP | -0,4195 | 2,28967E-09 |
| ACE2 | CFAP161 | -0,4195 | 2,29298E-09 |
| ACE2 | MAZ | -0,4195 | 2,27831E-09 |
| ACE2 | TOMM40 | -0,4195 | 2,27941E-09 |
| ACE2 | PPP6R2 | -0,4195 | 2,28087E-09 |
| ACE2 | PPP1R2P1 | -0,4196 | 2,27242E-09 |
| ACE2 | AC006159.5 | -0,4196 | 2,27428E-09 |
| ACE2 | MBTD1 | -0,4196 | 2,26301E-09 |
| ACE2 | KAT14 | -0,4196 | 2,2697E-09 |
| ACE2 | RP11-344N10.5 | -0,4197 | 2,24891E-09 |
| ACE2 | DCX | -0,4197 | 2,24618E-09 |
| ACE2 | VWC2L | -0,4198 | 2,21295E-09 |
| ACE2 | CHD1 | -0,4198 | 2,22932E-09 |
| ACE2 | EPO | -0,4198 | 2,22326E-09 |
| ACE2 | GAL3ST3 | -0,4198 | 2,21067E-09 |
| ACE2 | ACSM5P1 | -0,4198 | 2,22164E-09 |
| ACE2 | CTC-559E9.8 | -0,4198 | 2,22716E-09 |
| ACE2 | BCL11A | -0,4199 | 2,19737E-09 |
| ACE2 | KLHL24 | -0,4199 | 2,19636E-09 |
| ACE2 | KANK1 | -0,4199 | 2,20457E-09 |
| ACE2 | CTSG | -0,4199 | 2,19682E-09 |
| ACE2 | TNRC6B | -0,4199 | 2,19013E-09 |
| ACE2 | JADE3 | -0,4199 | 2,19056E-09 |
| ACE2 | PRDX1P1 | -0,42 | 2,17617E-09 |
| ACE2 | RP11-417O11.5 | -0,42 | 2,17763E-09 |
| ACE2 | CTC-459F4.3 | -0,42 | 2,16812E-09 |
| ACE2 | CDH22 | -0,42 | 2,17211E-09 |
| ACE2 | LINC00476 | -0,4201 | 2,16462E-09 |
| ACE2 | DHX37 | -0,4201 | 2,16113E-09 |
| ACE2 | RP11-294N21.3 | -0,4202 | 2,13768E-09 |
| ACE2 | HCG2040054 | -0,4203 | 2,12002E-09 |
| ACE2 | MBD4 | -0,4203 | 2,12159E-09 |
| ACE2 | RP11-968A15.2 | -0,4203 | 2,11157E-09 |

| | | | |
|---|---|---|---|
| ACE2 | RP11-439L18.1 | -0,4204 | 2,08945E-09 |
| ACE2 | MRPS7 | -0,4204 | 2,10363E-09 |
| ACE2 | EPRS | -0,4205 | 2,08127E-09 |
| ACE2 | CSNK1A1L | -0,4205 | 2,08064E-09 |
| ACE2 | ZCCHC10 | -0,4206 | 2,04926E-09 |
| ACE2 | RP11-79M19.2 | -0,4206 | 2,0531E-09 |
| ACE2 | RP11-243M5.4 | -0,4206 | 2,05125E-09 |
| ACE2 | CTD-2325P2.3 | -0,4206 | 2,0638E-09 |
| ACE2 | NME6 | -0,4207 | 2,0442E-09 |
| ACE2 | RP11-43F13.3 | -0,4207 | 2,02819E-09 |
| ACE2 | GABRG2 | -0,4207 | 2,03685E-09 |
| ACE2 | RP11-855A2.3 | -0,4207 | 2,0438E-09 |
| ACE2 | RPS4Y1 | -0,4207 | 2,03942E-09 |
| ACE2 | AC007391.2 | -0,4208 | 2,02441E-09 |
| ACE2 | RP11-234G16.5 | -0,4208 | 2,00599E-09 |
| ACE2 | NAV2 | -0,4208 | 2,01794E-09 |
| ACE2 | RFPL3S | -0,4208 | 2,01603E-09 |
| ACE2 | RP4-777O23.1 | -0,4209 | 1,99582E-09 |
| ACE2 | NOSIP | -0,4209 | 1,98959E-09 |
| ACE2 | ZNF417 | -0,4209 | 2,0015E-09 |
| ACE2 | KLHL7-AS1 | -0,421 | 1,97048E-09 |
| ACE2 | LINC02193 | -0,421 | 1,9732E-09 |
| ACE2 | LINC01372 | -0,4211 | 1,95661E-09 |
| ACE2 | RP11-227G15.12 | -0,4211 | 1,95019E-09 |
| ACE2 | AGTR2 | -0,4211 | 1,96541E-09 |
| ACE2 | RP11-92G12.3 | -0,4212 | 1,93007E-09 |
| ACE2 | RP11-932O9.7 | -0,4212 | 1,94594E-09 |
| ACE2 | CTD-2012K14.2 | -0,4212 | 1,93133E-09 |
| ACE2 | KIF13A | -0,4213 | 1,9264E-09 |
| ACE2 | RP11-589P10.5 | -0,4213 | 1,92148E-09 |
| ACE2 | MEIS3P2 | -0,4213 | 1,9204E-09 |
| ACE2 | CFAP45 | -0,4214 | 1,90102E-09 |
| ACE2 | ZFYVE1 | -0,4214 | 1,90456E-09 |
| ACE2 | TIGD1 | -0,4215 | 1,88115E-09 |
| ACE2 | SKP1P1 | -0,4215 | 1,88464E-09 |
| ACE2 | C1RL-AS1 | -0,4215 | 1,87807E-09 |
| ACE2 | FNIP2 | -0,4216 | 1,85758E-09 |
| ACE2 | SMKR1 | -0,4218 | 1,82729E-09 |
| ACE2 | BTBD10P2 | -0,4219 | 1,8031E-09 |
| ACE2 | ZPLD1 | -0,4221 | 1,77618E-09 |
| ACE2 | ASPH | -0,4221 | 1,78199E-09 |
| ACE2 | ITIH2 | -0,4221 | 1,77124E-09 |
| ACE2 | FAM160B1 | -0,4221 | 1,77471E-09 |
| ACE2 | ADAMTS15 | -0,4221 | 1,78063E-09 |
| ACE2 | LA16c-359F1.1 | -0,4221 | 1,7767E-09 |
| ACE2 | AC007193.10 | -0,4221 | 1,77986E-09 |
| ACE2 | TRIM33 | -0,4222 | 1,75703E-09 |
| ACE2 | MTCO3P43 | -0,4222 | 1,75353E-09 |
| ACE2 | RBM45 | -0,4222 | 1,75983E-09 |
| ACE2 | CTD-2293H3.1 | -0,4222 | 1,7656E-09 |
| ACE2 | TCTE3 | -0,4223 | 1,74425E-09 |
| ACE2 | RP11-712C19.1 | -0,4223 | 1,73512E-09 |
| ACE2 | RP11-446E24.3 | -0,4224 | 1,73386E-09 |
| ACE2 | NPC2 | -0,4224 | 1,72279E-09 |

| | | | |
|---|---|---|---|
| ACE2 | SRSF11 | -0,4225 | 1,71339E-09 |
| ACE2 | ADAR | -0,4225 | 1,70773E-09 |
| ACE2 | MEX3A | -0,4225 | 1,7071E-09 |
| ACE2 | GORASP2 | -0,4225 | 1,71701E-09 |
| ACE2 | PTPRG | -0,4226 | 1,69986E-09 |
| ACE2 | ZNRD1 | -0,4226 | 1,6917E-09 |
| ACE2 | SDHAF2 | -0,4226 | 1,68943E-09 |
| ACE2 | MDM2 | -0,4226 | 1,698E-09 |
| ACE2 | CADPS | -0,4227 | 1,66889E-09 |
| ACE2 | RP11-111M22.5 | -0,4227 | 1,6735E-09 |
| ACE2 | ZNF266 | -0,4227 | 1,68402E-09 |
| ACE2 | AF064858.8 | -0,4227 | 1,67239E-09 |
| ACE2 | IGSF21 | -0,4228 | 1,66561E-09 |
| ACE2 | LINC00216 | -0,4228 | 1,65955E-09 |
| ACE2 | RP11-759A24.2 | -0,4228 | 1,65274E-09 |
| ACE2 | CGB7 | -0,4228 | 1,66278E-09 |
| ACE2 | EWSR1 | -0,4228 | 1,65483E-09 |
| ACE2 | RP11-216N14.5 | -0,4229 | 1,64704E-09 |
| ACE2 | PPP1R1A | -0,4229 | 1,64057E-09 |
| ACE2 | AC010504.2 | -0,4229 | 1,64881E-09 |
| ACE2 | RP4-738P15.1 | -0,4229 | 1,6428E-09 |
| ACE2 | CTB-174D11.1 | -0,423 | 1,63496E-09 |
| ACE2 | CKAP4 | -0,423 | 1,63417E-09 |
| ACE2 | CTD-2020K17.1 | -0,423 | 1,6272E-09 |
| ACE2 | IGLV5-52 | -0,423 | 1,63562E-09 |
| ACE2 | KCNIP3 | -0,4231 | 1,61079E-09 |
| ACE2 | LL09NC01-139C3.1 | -0,4231 | 1,60961E-09 |
| ACE2 | MIR6807 | -0,4231 | 1,618E-09 |
| ACE2 | RP3-340N1.2 | -0,4232 | 1,59456E-09 |
| ACE2 | LIMD1 | -0,4232 | 1,6009E-09 |
| ACE2 | RP11-166O4.6 | -0,4232 | 1,59337E-09 |
| ACE2 | SDHDP6 | -0,4233 | 1,58607E-09 |
| ACE2 | KLHL25 | -0,4233 | 1,57846E-09 |
| ACE2 | ANGPTL8 | -0,4233 | 1,58137E-09 |
| ACE2 | RP11-337N6.2 | -0,4234 | 1,56063E-09 |
| ACE2 | RP11-51J9.5 | -0,4234 | 1,56685E-09 |
| ACE2 | RNU4-2 | -0,4234 | 1,5641E-09 |
| ACE2 | RP11-490N5.3 | -0,4234 | 1,56733E-09 |
| ACE2 | RP11-536C5.7 | -0,4235 | 1,55654E-09 |
| ACE2 | WASHC4 | -0,4235 | 1,55163E-09 |
| ACE2 | SRP54-AS1 | -0,4235 | 1,54523E-09 |
| ACE2 | RP11-47L3.1 | -0,4235 | 1,54778E-09 |
| ACE2 | AC018816.3 | -0,4238 | 1,50612E-09 |
| ACE2 | PKD2L2 | -0,4238 | 1,50403E-09 |
| ACE2 | RP11-350J20.9 | -0,4238 | 1,49957E-09 |
| ACE2 | RPL23AP25 | -0,4239 | 1,48605E-09 |
| ACE2 | BAZ2B | -0,4239 | 1,48786E-09 |
| ACE2 | SESTD1 | -0,4239 | 1,49027E-09 |
| ACE2 | RHAG | -0,4239 | 1,49743E-09 |
| ACE2 | BNIP1 | -0,424 | 1,48339E-09 |
| ACE2 | LY6G6C | -0,424 | 1,47294E-09 |
| ACE2 | KCNV2 | -0,424 | 1,47144E-09 |
| ACE2 | LINC00612 | -0,424 | 1,47428E-09 |
| ACE2 | CCDC144B | -0,424 | 1,48019E-09 |

| | | | |
|---|---|---|---|
| ACE2 | AF038458.5 | -0,424 | 1,47393E-09 |
| ACE2 | ZNF480 | -0,424 | 1,47703E-09 |
| ACE2 | RIT2 | -0,4241 | 1,46491E-09 |
| ACE2 | HAR1B | -0,4241 | 1,46331E-09 |
| ACE2 | GATA1 | -0,4241 | 1,45714E-09 |
| ACE2 | RNF111 | -0,4242 | 1,4499E-09 |
| ACE2 | PITPNM3 | -0,4242 | 1,44138E-09 |
| ACE2 | RP11-462D18.4 | -0,4242 | 1,44912E-09 |
| ACE2 | CSTA | -0,4243 | 1,43614E-09 |
| ACE2 | SMN2 | -0,4243 | 1,43607E-09 |
| ACE2 | C9orf153 | -0,4243 | 1,4395E-09 |
| ACE2 | RP11-475A13.1 | -0,4243 | 1,43578E-09 |
| ACE2 | LAMA2 | -0,4244 | 1,42391E-09 |
| ACE2 | RP1-292L20.3 | -0,4244 | 1,41709E-09 |
| ACE2 | AL133243.2 | -0,4245 | 1,4062E-09 |
| ACE2 | COPS5 | -0,4245 | 1,40523E-09 |
| ACE2 | RP11-638I2.8 | -0,4246 | 1,38674E-09 |
| ACE2 | CLASRP | -0,4246 | 1,38911E-09 |
| ACE2 | LMNA | -0,4247 | 1,3809E-09 |
| ACE2 | RPL23AP49 | -0,4247 | 1,37505E-09 |
| ACE2 | INTS1 | -0,4247 | 1,37723E-09 |
| ACE2 | SEC14L1P1 | -0,4247 | 1,37301E-09 |
| ACE2 | HSPE1 | -0,4248 | 1,36877E-09 |
| ACE2 | RGS20 | -0,4248 | 1,37087E-09 |
| ACE2 | RP11-715J22.3 | -0,4248 | 1,3591E-09 |
| ACE2 | HLA-F-AS1 | -0,4249 | 1,35727E-09 |
| ACE2 | RP11-544I20.2 | -0,4249 | 1,35759E-09 |
| ACE2 | GJA9 | -0,425 | 1,3322E-09 |
| ACE2 | ZC3H15 | -0,425 | 1,34207E-09 |
| ACE2 | AC079354.3 | -0,425 | 1,33199E-09 |
| ACE2 | CCDC14 | -0,425 | 1,3354E-09 |
| ACE2 | SH3TC1 | -0,425 | 1,33389E-09 |
| ACE2 | CTC-338M12.9 | -0,425 | 1,33185E-09 |
| ACE2 | PITPNM2 | -0,425 | 1,33618E-09 |
| ACE2 | ACYP1 | -0,425 | 1,33905E-09 |
| ACE2 | E2F4 | -0,425 | 1,33783E-09 |
| ACE2 | ARRDC2 | -0,425 | 1,33335E-09 |
| ACE2 | RP11-225B17.2 | -0,4251 | 1,32762E-09 |
| ACE2 | RP11-166N17.1 | -0,4251 | 1,32828E-09 |
| ACE2 | RP11-138H8.6 | -0,4251 | 1,33045E-09 |
| ACE2 | RP5-971N18.3 | -0,4251 | 1,3298E-09 |
| ACE2 | ZIM2 | -0,4252 | 1,309E-09 |
| ACE2 | RP4-673M15.1 | -0,4253 | 1,29362E-09 |
| ACE2 | HNRNPUL2 | -0,4253 | 1,30545E-09 |
| ACE2 | RP11-89C3.3 | -0,4253 | 1,30393E-09 |
| ACE2 | RP11-847H18.3 | -0,4253 | 1,2978E-09 |
| ACE2 | RP11-620J15.4 | -0,4253 | 1,30264E-09 |
| ACE2 | RP3-466P17.1 | -0,4254 | 1,28741E-09 |
| ACE2 | RGS5 | -0,4255 | 1,27185E-09 |
| ACE2 | SERPINE1 | -0,4255 | 1,27305E-09 |
| ACE2 | QRFP | -0,4255 | 1,27082E-09 |
| ACE2 | ARMC3 | -0,4255 | 1,27761E-09 |
| ACE2 | HOXA-AS2 | -0,4256 | 1,26107E-09 |
| ACE2 | VCPIP1 | -0,4256 | 1,26583E-09 |

| | | | |
|---|---|---|---|
| ACE2 | TEX2 | -0,4256 | 1,26516E-09 |
| ACE2 | TPR | -0,4257 | 1,2461E-09 |
| ACE2 | RP11-349F21.5 | -0,4257 | 1,25085E-09 |
| ACE2 | C10orf76 | -0,4257 | 1,25409E-09 |
| ACE2 | CTC-591M7.1 | -0,4257 | 1,24623E-09 |
| ACE2 | SUPT6H | -0,4257 | 1,25312E-09 |
| ACE2 | CALB2 | -0,4258 | 1,23805E-09 |
| ACE2 | AP000593.7 | -0,4259 | 1,22327E-09 |
| ACE2 | GOLGA8B | -0,4259 | 1,2286E-09 |
| ACE2 | LINC01560 | -0,4259 | 1,22054E-09 |
| ACE2 | UBL5 | -0,426 | 1,20771E-09 |
| ACE2 | RBM34 | -0,4261 | 1,19547E-09 |
| ACE2 | SLC39A10 | -0,4261 | 1,19815E-09 |
| ACE2 | RP11-138E2.1 | -0,4261 | 1,20093E-09 |
| ACE2 | RP4-607I7.1 | -0,4261 | 1,20143E-09 |
| ACE2 | AAK1 | -0,4262 | 1,1932E-09 |
| ACE2 | UNC5B-AS1 | -0,4262 | 1,18354E-09 |
| ACE2 | RP13-941N14.1 | -0,4262 | 1,19504E-09 |
| ACE2 | LINC01482 | -0,4262 | 1,18976E-09 |
| ACE2 | FLJ38576 | -0,4263 | 1,18225E-09 |
| ACE2 | RPS4XP16 | -0,4263 | 1,17289E-09 |
| ACE2 | LINC01311 | -0,4263 | 1,1737E-09 |
| ACE2 | RP11-226L15.5 | -0,4264 | 1,16608E-09 |
| ACE2 | PAPPA2 | -0,4264 | 1,1618E-09 |
| ACE2 | RP11-120M18.5 | -0,4264 | 1,16891E-09 |
| ACE2 | CTD-2653M23.3 | -0,4265 | 1,15643E-09 |
| ACE2 | GRIN2C | -0,4265 | 1,15777E-09 |
| ACE2 | TEKT4P2 | -0,4265 | 1,1517E-09 |
| ACE2 | TTLL7-IT1 | -0,4266 | 1,14656E-09 |
| ACE2 | AC009264.1 | -0,4267 | 1,12713E-09 |
| ACE2 | ODAM | -0,4268 | 1,1231E-09 |
| ACE2 | FSBP | -0,4268 | 1,11733E-09 |
| ACE2 | RP11-563J2.3 | -0,4269 | 1,11144E-09 |
| ACE2 | OLR1 | -0,4269 | 1,1145E-09 |
| ACE2 | RP11-298D21.3 | -0,4269 | 1,11282E-09 |
| ACE2 | CTD-2189E23.1 | -0,4269 | 1,11191E-09 |
| ACE2 | RP4-635E18.7 | -0,427 | 1,09902E-09 |
| ACE2 | SMYD1 | -0,427 | 1,10258E-09 |
| ACE2 | CTD-2630F21.1 | -0,427 | 1,0988E-09 |
| ACE2 | RP11-319G6.1 | -0,4271 | 1,08963E-09 |
| ACE2 | RP11-1084A12.1 | -0,4271 | 1,09119E-09 |
| ACE2 | KYNU | -0,4272 | 1,08138E-09 |
| ACE2 | MKNK2 | -0,4272 | 1,07244E-09 |
| ACE2 | RP11-418J17.1 | -0,4273 | 1,07016E-09 |
| ACE2 | CISH | -0,4273 | 1,07021E-09 |
| ACE2 | PRR7 | -0,4273 | 1,07144E-09 |
| ACE2 | LL22NC03-2H8.5 | -0,4273 | 1,06556E-09 |
| ACE2 | OTUD3 | -0,4274 | 1,05429E-09 |
| ACE2 | LNPK | -0,4274 | 1,05593E-09 |
| ACE2 | C4orf36 | -0,4274 | 1,05475E-09 |
| ACE2 | RP1-278E11.3 | -0,4274 | 1,0538E-09 |
| ACE2 | MTCO1P28 | -0,4274 | 1,0529E-09 |
| ACE2 | LYSMD1 | -0,4275 | 1,04275E-09 |
| ACE2 | RP11-12A2.1 | -0,4275 | 1,04634E-09 |

| | | | |
|---|---|---|---|
| ACE2 | ABCB4 | -0,4275 | 1,05013E-09 |
| ACE2 | RP13-895J2.3 | -0,4275 | 1,0459E-09 |
| ACE2 | CTC-523E23.3 | -0,4275 | 1,0434E-09 |
| ACE2 | USP19 | -0,4276 | 1,03952E-09 |
| ACE2 | PILRB | -0,4276 | 1,03808E-09 |
| ACE2 | TRAPPC9 | -0,4276 | 1,03072E-09 |
| ACE2 | VEPH1 | -0,4277 | 1,03014E-09 |
| ACE2 | GTF2A2 | -0,4277 | 1,03015E-09 |
| ACE2 | ZNF175 | -0,4277 | 1,0274E-09 |
| ACE2 | PARP4P3 | -0,4277 | 1,02722E-09 |
| ACE2 | RP11-694I15.7 | -0,4278 | 1,01279E-09 |
| ACE2 | TPTE2P6 | -0,4278 | 1,01939E-09 |
| ACE2 | CTB-133G6.1 | -0,4278 | 1,01302E-09 |
| ACE2 | ERVFRD-1 | -0,4279 | 1,00465E-09 |
| ACE2 | RP11-705C15.2 | -0,4279 | 1,00541E-09 |
| ACE2 | DPYS | -0,4281 | 9,86126E-10 |
| ACE2 | KIF17 | -0,4282 | 9,80497E-10 |
| ACE2 | LINC01374 | -0,4282 | 9,78641E-10 |
| ACE2 | RP11-483C6.1 | -0,4282 | 9,77438E-10 |
| ACE2 | ADRA1D | -0,4283 | 9,69541E-10 |
| ACE2 | SERTAD1 | -0,4284 | 9,53521E-10 |
| ACE2 | DKFZP434A062 | -0,4285 | 9,46214E-10 |
| ACE2 | HAS3 | -0,4285 | 9,48372E-10 |
| ACE2 | MYT1 | -0,4285 | 9,47113E-10 |
| ACE2 | CYP4Z1 | -0,4286 | 9,41274E-10 |
| ACE2 | ATP6V1B1 | -0,4286 | 9,35153E-10 |
| ACE2 | RP11-188P20.3 | -0,4286 | 9,40472E-10 |
| ACE2 | RAET1E | -0,4286 | 9,4108E-10 |
| ACE2 | ZNF394 | -0,4286 | 9,34003E-10 |
| ACE2 | TMEM183A | -0,4287 | 9,27106E-10 |
| ACE2 | SLC35D3 | -0,4287 | 9,30554E-10 |
| ACE2 | LINC01285 | -0,4287 | 9,262E-10 |
| ACE2 | SNORA73B | -0,4288 | 9,22636E-10 |
| ACE2 | RP11-923I11.5 | -0,4288 | 9,15593E-10 |
| ACE2 | ALOX12-AS1 | -0,4288 | 9,21536E-10 |
| ACE2 | QPCTL | -0,4288 | 9,18767E-10 |
| ACE2 | SELENOT | -0,4289 | 9,08378E-10 |
| ACE2 | OSTN | -0,4289 | 9,09676E-10 |
| ACE2 | RP11-295I5.4 | -0,4289 | 9,13283E-10 |
| ACE2 | SUCLG2-AS1 | -0,429 | 8,97213E-10 |
| ACE2 | SARAF | -0,429 | 8,97453E-10 |
| ACE2 | NAMA | -0,429 | 9,03417E-10 |
| ACE2 | CEP164P1 | -0,429 | 9,02764E-10 |
| ACE2 | BDNF-AS | -0,429 | 8,96961E-10 |
| ACE2 | KIF27 | -0,4291 | 8,88965E-10 |
| ACE2 | HIST1H2BM | -0,4292 | 8,80488E-10 |
| ACE2 | MEPCE | -0,4292 | 8,83134E-10 |
| ACE2 | RP11-64B16.3 | -0,4292 | 8,85366E-10 |
| ACE2 | SETP2 | -0,4292 | 8,8442E-10 |
| ACE2 | HERC2P2 | -0,4292 | 8,82972E-10 |
| ACE2 | ELFN2 | -0,4292 | 8,80912E-10 |
| ACE2 | ZNF252P | -0,4293 | 8,79161E-10 |
| ACE2 | RP11-406H21.2 | -0,4293 | 8,73858E-10 |
| ACE2 | CCDC84 | -0,4293 | 8,71529E-10 |

| | | | |
|---|---|---|---|
| ACE2 | RPS23P8 | -0,4293 | 8,78421E-10 |
| ACE2 | RP11-229P13.25 | -0,4294 | 8,70215E-10 |
| ACE2 | MT3 | -0,4294 | 8,68848E-10 |
| ACE2 | RP11-75A9.3 | -0,4294 | 8,67313E-10 |
| ACE2 | RNF220 | -0,4295 | 8,57974E-10 |
| ACE2 | PCNX2 | -0,4295 | 8,59533E-10 |
| ACE2 | GPBAR1 | -0,4295 | 8,56569E-10 |
| ACE2 | MYOT | -0,4295 | 8,56055E-10 |
| ACE2 | PXN-AS1 | -0,4295 | 8,5854E-10 |
| ACE2 | SMAD6 | -0,4295 | 8,57168E-10 |
| ACE2 | RP3-467L1.4 | -0,4296 | 8,52233E-10 |
| ACE2 | MTX1P1 | -0,4296 | 8,51777E-10 |
| ACE2 | LINC01412 | -0,4296 | 8,5042E-10 |
| ACE2 | RP11-51J9.6 | -0,4296 | 8,47077E-10 |
| ACE2 | RP11-68L18.1 | -0,4296 | 8,49549E-10 |
| ACE2 | RP11-672L10.2 | -0,4296 | 8,51764E-10 |
| ACE2 | TBC1D9B | -0,4298 | 8,31175E-10 |
| ACE2 | CCDC171 | -0,4298 | 8,29335E-10 |
| ACE2 | ANTXRL | -0,4298 | 8,30571E-10 |
| ACE2 | OTOA | -0,4298 | 8,33313E-10 |
| ACE2 | RP11-108M9.5 | -0,4299 | 8,24809E-10 |
| ACE2 | SDHAF1 | -0,4299 | 8,21064E-10 |
| ACE2 | ZNF45 | -0,4299 | 8,20892E-10 |
| ACE2 | MCM8 | -0,4299 | 8,2049E-10 |
| ACE2 | RP11-855A2.2 | -0,43 | 8,18748E-10 |
| ACE2 | CTC-510F12.3 | -0,43 | 8,13277E-10 |
| ACE2 | C4orf22 | -0,4301 | 8,07947E-10 |
| ACE2 | TPBG | -0,4302 | 7,96919E-10 |
| ACE2 | MEI4 | -0,4303 | 7,91356E-10 |
| ACE2 | QRICH1 | -0,4304 | 7,8562E-10 |
| ACE2 | RP11-45P15.4 | -0,4304 | 7,81391E-10 |
| ACE2 | RP11-323P17.2 | -0,4305 | 7,74767E-10 |
| ACE2 | SSTR2 | -0,4305 | 7,78107E-10 |
| ACE2 | IL12A-AS1 | -0,4306 | 7,70948E-10 |
| ACE2 | DCAF13 | -0,4306 | 7,68434E-10 |
| ACE2 | RP11-147I3.1 | -0,4306 | 7,68037E-10 |
| ACE2 | VAC14 | -0,4306 | 7,65373E-10 |
| ACE2 | RP11-2N1.2 | -0,4307 | 7,62883E-10 |
| ACE2 | HM13 | -0,4307 | 7,62494E-10 |
| ACE2 | ZBTB5 | -0,4308 | 7,50918E-10 |
| ACE2 | RP11-785D18.3 | -0,4308 | 7,52519E-10 |
| ACE2 | RP11-756G20.1 | -0,4308 | 7,53388E-10 |
| ACE2 | FOXP2 | -0,4309 | 7,44088E-10 |
| ACE2 | EVI5 | -0,431 | 7,34796E-10 |
| ACE2 | RP11-1379J22.2 | -0,431 | 7,38979E-10 |
| ACE2 | RP11-523O18.5 | -0,431 | 7,40533E-10 |
| ACE2 | RPL13AP25 | -0,431 | 7,41676E-10 |
| ACE2 | USP6 | -0,431 | 7,34743E-10 |
| ACE2 | CTC1 | -0,431 | 7,41063E-10 |
| ACE2 | VSTM2B | -0,431 | 7,41303E-10 |
| ACE2 | RP11-177C12.1 | -0,4311 | 7,30175E-10 |
| ACE2 | POLR2B | -0,4311 | 7,3362E-10 |
| ACE2 | RNU6-485P | -0,4311 | 7,31682E-10 |
| ACE2 | NUDT4 | -0,4311 | 7,33923E-10 |

| | | | |
|---|---|---|---|
| ACE2 | PKD1L2 | -0,4311 | 7,32331E-10 |
| ACE2 | AC112497.1 | -0,4311 | 7,3275E-10 |
| ACE2 | RP11-468E2.11 | -0,4312 | 7,21364E-10 |
| ACE2 | AC012066.1 | -0,4313 | 7,15089E-10 |
| ACE2 | RP11-402G3.5 | -0,4313 | 7,15793E-10 |
| ACE2 | MIR1244-2 | -0,4314 | 7,07431E-10 |
| ACE2 | CCND3P1 | -0,4314 | 7,07474E-10 |
| ACE2 | PRRC2C | -0,4315 | 7,03162E-10 |
| ACE2 | COL26A1 | -0,4316 | 6,94412E-10 |
| ACE2 | COX8C | -0,4316 | 6,92275E-10 |
| ACE2 | MIR762HG | -0,4316 | 6,9666E-10 |
| ACE2 | ZNF765 | -0,4316 | 6,9379E-10 |
| ACE2 | LINC00846 | -0,4316 | 6,98327E-10 |
| ACE2 | RP11-104L21.2 | -0,4317 | 6,87266E-10 |
| ACE2 | RNF185 | -0,4318 | 6,78852E-10 |
| ACE2 | RP11-231L11.3 | -0,4319 | 6,76919E-10 |
| ACE2 | CD63 | -0,4319 | 6,77393E-10 |
| ACE2 | RP11-888D10.4 | -0,4319 | 6,72113E-10 |
| ACE2 | STAMBPL1 | -0,432 | 6,67215E-10 |
| ACE2 | KDM5A | -0,432 | 6,65801E-10 |
| ACE2 | SPTBN5 | -0,432 | 6,65421E-10 |
| ACE2 | C1orf228 | -0,4321 | 6,61994E-10 |
| ACE2 | LINC02018 | -0,4321 | 6,61838E-10 |
| ACE2 | ALKBH4 | -0,4321 | 6,62962E-10 |
| ACE2 | THOP1 | -0,4321 | 6,63286E-10 |
| ACE2 | TARS | -0,4322 | 6,5517E-10 |
| ACE2 | C7orf73 | -0,4322 | 6,55131E-10 |
| ACE2 | RP11-398C13.6 | -0,4322 | 6,53296E-10 |
| ACE2 | TAF6L | -0,4322 | 6,55882E-10 |
| ACE2 | DUS2 | -0,4322 | 6,57298E-10 |
| ACE2 | UNC119 | -0,4322 | 6,5137E-10 |
| ACE2 | CTC-308K20.1 | -0,4323 | 6,50435E-10 |
| ACE2 | BTN3A3 | -0,4323 | 6,48313E-10 |
| ACE2 | RP11-7F17.8 | -0,4323 | 6,46606E-10 |
| ACE2 | RP11-676J12.8 | -0,4323 | 6,47369E-10 |
| ACE2 | XXbac-B444P24.8 | -0,4323 | 6,4601E-10 |
| ACE2 | PROSER3 | -0,4324 | 6,40586E-10 |
| ACE2 | GREB1 | -0,4325 | 6,32966E-10 |
| ACE2 | DDX43 | -0,4325 | 6,32615E-10 |
| ACE2 | CCDC65 | -0,4325 | 6,38175E-10 |
| ACE2 | YPEL3 | -0,4325 | 6,37998E-10 |
| ACE2 | RP5-1029K10.2 | -0,4326 | 6,2806E-10 |
| ACE2 | CCDC57 | -0,4326 | 6,30129E-10 |
| ACE2 | ZNF429 | -0,4327 | 6,19824E-10 |
| ACE2 | C1QTNF6 | -0,4327 | 6,20947E-10 |
| ACE2 | HSD17B7P2 | -0,4328 | 6,13369E-10 |
| ACE2 | RP11-757G1.5 | -0,4328 | 6,18371E-10 |
| ACE2 | RP11-358B23.7 | -0,4328 | 6,18496E-10 |
| ACE2 | PHF6 | -0,4328 | 6,13685E-10 |
| ACE2 | RP11-204L24.2 | -0,4329 | 6,07447E-10 |
| ACE2 | SLC35E1 | -0,4329 | 6,12029E-10 |
| ACE2 | RP11-89N17.4 | -0,433 | 6,05524E-10 |
| ACE2 | RP11-133K1.12 | -0,433 | 6,00957E-10 |
| ACE2 | RP11-616M22.1 | -0,433 | 6,06113E-10 |

| | | | |
|---|---|---|---|
| ACE2 | RP1-41C23.4 | -0,433 | 6,06595E-10 |
| ACE2 | DAND5 | -0,433 | 6,02769E-10 |
| ACE2 | NAPA-AS1 | -0,433 | 6,06671E-10 |
| ACE2 | MIR3164 | -0,4331 | 5,98936E-10 |
| ACE2 | DCLRE1C | -0,4332 | 5,93666E-10 |
| ACE2 | AC145124.2 | -0,4334 | 5,78076E-10 |
| ACE2 | ODF3L1 | -0,4334 | 5,82254E-10 |
| ACE2 | GUCA1B | -0,4335 | 5,73254E-10 |
| ACE2 | TTC9 | -0,4335 | 5,73927E-10 |
| ACE2 | OSER1 | -0,4335 | 5,74158E-10 |
| ACE2 | RHOXF1-AS1 | -0,4335 | 5,72186E-10 |
| ACE2 | PPP2R5A | -0,4336 | 5,69258E-10 |
| ACE2 | DKK1 | -0,4336 | 5,67664E-10 |
| ACE2 | RGPD8 | -0,4337 | 5,64005E-10 |
| ACE2 | UBE2F | -0,4337 | 5,60824E-10 |
| ACE2 | CARD19 | -0,4337 | 5,6003E-10 |
| ACE2 | RP11-837J7.4 | -0,4337 | 5,62546E-10 |
| ACE2 | ADAP2 | -0,4337 | 5,64191E-10 |
| ACE2 | AC004471.10 | -0,4337 | 5,6081E-10 |
| ACE2 | TBK1 | -0,4338 | 5,54638E-10 |
| ACE2 | C1QBP | -0,4338 | 5,56836E-10 |
| ACE2 | RNF167 | -0,4339 | 5,51381E-10 |
| ACE2 | RP11-760H22.2 | -0,434 | 5,47201E-10 |
| ACE2 | RALGAPA1P1 | -0,434 | 5,44654E-10 |
| ACE2 | USP32P2 | -0,434 | 5,43961E-10 |
| ACE2 | CRLF3 | -0,4341 | 5,38319E-10 |
| ACE2 | RP11-21G15.1 | -0,4341 | 5,41602E-10 |
| ACE2 | OAZ1 | -0,4341 | 5,41092E-10 |
| ACE2 | MCM3 | -0,4342 | 5,33858E-10 |
| ACE2 | ELOA-AS1 | -0,4343 | 5,30656E-10 |
| ACE2 | AC073063.10 | -0,4343 | 5,31248E-10 |
| ACE2 | COQ7 | -0,4343 | 5,30072E-10 |
| ACE2 | EPAS1 | -0,4344 | 5,22246E-10 |
| ACE2 | GUCA1A | -0,4344 | 5,24659E-10 |
| ACE2 | SNORD124 | -0,4344 | 5,23965E-10 |
| ACE2 | LENG8 | -0,4344 | 5,25458E-10 |
| ACE2 | NECAP2 | -0,4345 | 5,21355E-10 |
| ACE2 | AC103563.8 | -0,4345 | 5,198E-10 |
| ACE2 | DYNLL1 | -0,4345 | 5,16561E-10 |
| ACE2 | OPRL1 | -0,4345 | 5,19825E-10 |
| ACE2 | RP11-104L21.3 | -0,4346 | 5,1612E-10 |
| ACE2 | AC092614.2 | -0,4346 | 5,11952E-10 |
| ACE2 | SIL1 | -0,4346 | 5,12252E-10 |
| ACE2 | C7orf49 | -0,4346 | 5,14172E-10 |
| ACE2 | PFN1 | -0,4346 | 5,11856E-10 |
| ACE2 | MIB2 | -0,4347 | 5,09597E-10 |
| ACE2 | AP002954.3 | -0,4347 | 5,10299E-10 |
| ACE2 | EI24 | -0,4347 | 5,06941E-10 |
| ACE2 | SH2D5 | -0,4348 | 5,04683E-10 |
| ACE2 | RP11-4K3__A.3 | -0,4348 | 5,01506E-10 |
| ACE2 | ZNF736 | -0,4348 | 5,02154E-10 |
| ACE2 | PHEX-AS1 | -0,4348 | 5,04818E-10 |
| ACE2 | RP1-290F12.3 | -0,4348 | 5,05509E-10 |
| ACE2 | RPL4P3 | -0,4349 | 4,98702E-10 |

| | | | |
|---|---|---|---|
| ACE2 | HAND1 | -0,4349 | 4,98813E-10 |
| ACE2 | RP11-362K14.6 | -0,435 | 4,95774E-10 |
| ACE2 | KERA | -0,435 | 4,93687E-10 |
| ACE2 | SPAAR | -0,4351 | 4,88344E-10 |
| ACE2 | LSM12 | -0,4351 | 4,90698E-10 |
| ACE2 | MEX3D | -0,4351 | 4,89629E-10 |
| ACE2 | BRAFP1 | -0,4351 | 4,85942E-10 |
| ACE2 | RP11-108M9.4 | -0,4352 | 4,82982E-10 |
| ACE2 | ING5 | -0,4352 | 4,84051E-10 |
| ACE2 | MAMDC2 | -0,4352 | 4,82295E-10 |
| ACE2 | AP006621.5 | -0,4352 | 4,84188E-10 |
| ACE2 | TPTE2P2 | -0,4352 | 4,82938E-10 |
| ACE2 | RP6-24A23.3 | -0,4352 | 4,8332E-10 |
| ACE2 | MAPKAPK2 | -0,4353 | 4,80274E-10 |
| ACE2 | DLG5-AS1 | -0,4353 | 4,77237E-10 |
| ACE2 | PPP2CA | -0,4355 | 4,69307E-10 |
| ACE2 | PROX2 | -0,4355 | 4,66631E-10 |
| ACE2 | MCM3AP | -0,4355 | 4,68598E-10 |
| ACE2 | SLC9C2 | -0,4356 | 4,62343E-10 |
| ACE2 | RP11-190P13.2 | -0,4356 | 4,61887E-10 |
| ACE2 | RP5-1098D14.1 | -0,4357 | 4,61595E-10 |
| ACE2 | RP11-76E17.2 | -0,4357 | 4,60101E-10 |
| ACE2 | SMCR5 | -0,4357 | 4,58714E-10 |
| ACE2 | TPM3P9 | -0,4357 | 4,57208E-10 |
| ACE2 | CTD-2263F21.1 | -0,4358 | 4,54129E-10 |
| ACE2 | MFAP1 | -0,4358 | 4,52523E-10 |
| ACE2 | U91319.1 | -0,4358 | 4,55465E-10 |
| ACE2 | LINC00211 | -0,4359 | 4,48909E-10 |
| ACE2 | RP11-424I19.2 | -0,4359 | 4,50395E-10 |
| ACE2 | DLEU2L | -0,436 | 4,43823E-10 |
| ACE2 | ITGAX | -0,436 | 4,461E-10 |
| ACE2 | RP11-1079K10.2 | -0,436 | 4,47678E-10 |
| ACE2 | RP3-426I6.6 | -0,4361 | 4,39556E-10 |
| ACE2 | SNRNP200 | -0,4361 | 4,41587E-10 |
| ACE2 | NEFL | -0,4361 | 4,39172E-10 |
| ACE2 | IRS1 | -0,4362 | 4,36854E-10 |
| ACE2 | RP11-706C16.7 | -0,4362 | 4,35919E-10 |
| ACE2 | RP11-231I16.1 | -0,4362 | 4,35189E-10 |
| ACE2 | C15orf52 | -0,4362 | 4,34747E-10 |
| ACE2 | TLE1P1 | -0,4362 | 4,37349E-10 |
| ACE2 | MINOS1 | -0,4363 | 4,30636E-10 |
| ACE2 | RNVU1-3 | -0,4363 | 4,31022E-10 |
| ACE2 | CTD-2270F17.1 | -0,4363 | 4,32186E-10 |
| ACE2 | CTGF | -0,4363 | 4,3438E-10 |
| ACE2 | ENO4 | -0,4363 | 4,33753E-10 |
| ACE2 | RP11-179A10.1 | -0,4363 | 4,30062E-10 |
| ACE2 | CFAP69 | -0,4364 | 4,27274E-10 |
| ACE2 | RP11-333O1.1 | -0,4365 | 4,23795E-10 |
| ACE2 | HMGXB4 | -0,4365 | 4,22039E-10 |
| ACE2 | RP3-508I15.20 | -0,4365 | 4,25616E-10 |
| ACE2 | PIKFYVE | -0,4366 | 4,18694E-10 |
| ACE2 | LSM8 | -0,4366 | 4,17984E-10 |
| ACE2 | BTAF1 | -0,4366 | 4,19159E-10 |
| ACE2 | ARMC6 | -0,4366 | 4,18794E-10 |

| | | | |
|---|---|---|---|
| ACE2 | MAP1LC3B2 | -0,4367 | 4,14745E-10 |
| ACE2 | ABHD13 | -0,4367 | 4,13991E-10 |
| ACE2 | RP11-413H22.3 | -0,4367 | 4,15247E-10 |
| ACE2 | B4GALT5 | -0,4367 | 4,15066E-10 |
| ACE2 | SPDYA | -0,4368 | 4,12809E-10 |
| ACE2 | SRGAP1 | -0,4368 | 4,11254E-10 |
| ACE2 | PTGES3P2 | -0,4369 | 4,0452E-10 |
| ACE2 | RPSAP12 | -0,4369 | 4,06311E-10 |
| ACE2 | RP11-1079K10.3 | -0,4369 | 4,05784E-10 |
| ACE2 | RP11-423P10.2 | -0,4371 | 3,98654E-10 |
| ACE2 | GTF3C3 | -0,4371 | 3,99258E-10 |
| ACE2 | PRUNE2 | -0,4371 | 3,97471E-10 |
| ACE2 | RP11-276H19.2 | -0,4371 | 3,968E-10 |
| ACE2 | PROSER2-AS1 | -0,4371 | 3,99725E-10 |
| ACE2 | RPL26P6 | -0,4371 | 3,96546E-10 |
| ACE2 | UBE2Q2P6 | -0,4371 | 3,99671E-10 |
| ACE2 | RP11-709D24.8 | -0,4371 | 3,99397E-10 |
| ACE2 | CTD-2358C21.5 | -0,4371 | 3,98494E-10 |
| ACE2 | CTC-453G23.8 | -0,4371 | 3,9891E-10 |
| ACE2 | HIST1H1PS1 | -0,4372 | 3,93547E-10 |
| ACE2 | C14orf119 | -0,4372 | 3,95532E-10 |
| ACE2 | RP11-258C19.4 | -0,4373 | 3,92065E-10 |
| ACE2 | RP1-137D17.1 | -0,4374 | 3,86435E-10 |
| ACE2 | GDAP1 | -0,4374 | 3,87483E-10 |
| ACE2 | RP11-384C4.2 | -0,4375 | 3,82853E-10 |
| ACE2 | RP11-262I2.2 | -0,4376 | 3,78753E-10 |
| ACE2 | ARNTL | -0,4376 | 3,7749E-10 |
| ACE2 | CTC-336P14.1 | -0,4377 | 3,76491E-10 |
| ACE2 | LHCGR | -0,4377 | 3,72789E-10 |
| ACE2 | LGI1 | -0,4377 | 3,75373E-10 |
| ACE2 | AP001062.7 | -0,4377 | 3,76401E-10 |
| ACE2 | RP11-298I3.4 | -0,4378 | 3,699E-10 |
| ACE2 | TNKS | -0,4379 | 3,6713E-10 |
| ACE2 | RASGRP2 | -0,4379 | 3,66211E-10 |
| ACE2 | SPRED3 | -0,4379 | 3,65607E-10 |
| ACE2 | AP4B1 | -0,438 | 3,61975E-10 |
| ACE2 | RP11-385F5.4 | -0,438 | 3,64741E-10 |
| ACE2 | RPRM | -0,438 | 3,63102E-10 |
| ACE2 | OSGEPL1-AS1 | -0,438 | 3,64744E-10 |
| ACE2 | RP11-646I6.3 | -0,438 | 3,61975E-10 |
| ACE2 | ZMIZ1 | -0,438 | 3,64604E-10 |
| ACE2 | RP11-439H8.4 | -0,4381 | 3,61298E-10 |
| ACE2 | TFEB | -0,4381 | 3,60699E-10 |
| ACE2 | RP1-187B23.1 | -0,4382 | 3,56649E-10 |
| ACE2 | RP11-485M7.2 | -0,4382 | 3,56253E-10 |
| ACE2 | LSM1 | -0,4382 | 3,55797E-10 |
| ACE2 | RP11-297D21.4 | -0,4382 | 3,54946E-10 |
| ACE2 | EXOC3L2 | -0,4382 | 3,54504E-10 |
| ACE2 | HLX-AS1 | -0,4383 | 3,53008E-10 |
| ACE2 | CUX1 | -0,4383 | 3,53731E-10 |
| ACE2 | AQP7P1 | -0,4383 | 3,52317E-10 |
| ACE2 | RP11-177G23.2 | -0,4384 | 3,49291E-10 |
| ACE2 | TTI2 | -0,4384 | 3,48263E-10 |
| ACE2 | ALG10B | -0,4384 | 3,49255E-10 |

| | | | |
|---|---|---|---|
| ACE2 | NOL3 | -0,4384 | 3,49433E-10 |
| ACE2 | RP11-489D6.2 | -0,4385 | 3,44612E-10 |
| ACE2 | CARM1 | -0,4385 | 3,45879E-10 |
| ACE2 | MS4A2 | -0,4386 | 3,42702E-10 |
| ACE2 | RINL | -0,4386 | 3,4146E-10 |
| ACE2 | CAPN10-AS1 | -0,4387 | 3,37617E-10 |
| ACE2 | RP11-133N21.7 | -0,4387 | 3,37008E-10 |
| ACE2 | HSP90B1 | -0,4387 | 3,39706E-10 |
| ACE2 | RP11-165D7.5 | -0,4387 | 3,38238E-10 |
| ACE2 | RP11-60A24.3 | -0,4387 | 3,39502E-10 |
| ACE2 | AC016739.2 | -0,4388 | 3,3356E-10 |
| ACE2 | IL20RA | -0,4388 | 3,3419E-10 |
| ACE2 | CFAP43 | -0,4388 | 3,35108E-10 |
| ACE2 | ZBTB14 | -0,4388 | 3,33051E-10 |
| ACE2 | MPHOSPH9 | -0,4389 | 3,32086E-10 |
| ACE2 | ARHGEF7-AS2 | -0,4389 | 3,30533E-10 |
| ACE2 | CTD-3051D23.1 | -0,4389 | 3,29871E-10 |
| ACE2 | SNORD116-24 | -0,4389 | 3,31721E-10 |
| ACE2 | AP000704.5 | -0,4389 | 3,29959E-10 |
| ACE2 | PCF11 | -0,439 | 3,26765E-10 |
| ACE2 | PDK3 | -0,439 | 3,27708E-10 |
| ACE2 | QTRT2 | -0,4391 | 3,26036E-10 |
| ACE2 | ZNF655 | -0,4391 | 3,26088E-10 |
| ACE2 | ITGB1 | -0,4391 | 3,25951E-10 |
| ACE2 | ITGBL1 | -0,4391 | 3,23412E-10 |
| ACE2 | TRDN | -0,4392 | 3,20048E-10 |
| ACE2 | TAS2R31 | -0,4392 | 3,22792E-10 |
| ACE2 | LEPROTL1 | -0,4393 | 3,16994E-10 |
| ACE2 | SNAI2 | -0,4393 | 3,17589E-10 |
| ACE2 | SRSF2 | -0,4393 | 3,18076E-10 |
| ACE2 | RASA2 | -0,4394 | 3,15693E-10 |
| ACE2 | WNT2 | -0,4394 | 3,15614E-10 |
| ACE2 | XX-C00717C00720L.[illegible] | -0,4394 | 3,14753E-10 |
| ACE2 | DCUN1D1 | -0,4395 | 3,11797E-10 |
| ACE2 | RPL23AP88 | -0,4396 | 3,0879E-10 |
| ACE2 | CTB-158E9.1 | -0,4396 | 3,0766E-10 |
| ACE2 | LINC01091 | -0,4397 | 3,06689E-10 |
| ACE2 | ZNF318 | -0,4397 | 3,03582E-10 |
| ACE2 | AC006160.5 | -0,4398 | 3,00821E-10 |
| ACE2 | RP11-643M14.1 | -0,4398 | 3,0088E-10 |
| ACE2 | HNRNPA1P48 | -0,4398 | 3,03431E-10 |
| ACE2 | RP11-342D14.1 | -0,4398 | 3,00639E-10 |
| ACE2 | CBY3 | -0,4399 | 2,98227E-10 |
| ACE2 | TRRAP | -0,4399 | 2,98044E-10 |
| ACE2 | RP11-563J2.2 | -0,4399 | 2,98571E-10 |
| ACE2 | AP000442.1 | -0,4399 | 2,97431E-10 |
| ACE2 | URB2 | -0,44 | 2,95424E-10 |
| ACE2 | ADGRV1 | -0,44 | 2,96551E-10 |
| ACE2 | QSOX2 | -0,44 | 2,97272E-10 |
| ACE2 | NABP2 | -0,44 | 2,96684E-10 |
| ACE2 | RP1-159A19.4 | -0,4401 | 2,92633E-10 |
| ACE2 | RP5-933K21.3 | -0,4401 | 2,93255E-10 |
| ACE2 | RGS22 | -0,4402 | 2,89675E-10 |
| ACE2 | RDH16 | -0,4402 | 2,88413E-10 |

| | | | |
|---|---|---|---|
| ACE2 | POLR3K | -0,4402 | 2,89079E-10 |
| ACE2 | C16orf96 | -0,4402 | 2,88921E-10 |
| ACE2 | MRNIP | -0,4403 | 2,87496E-10 |
| ACE2 | ETS1 | -0,4403 | 2,87577E-10 |
| ACE2 | ZNF200 | -0,4403 | 2,85758E-10 |
| ACE2 | RPL7AP66 | -0,4403 | 2,8732E-10 |
| ACE2 | C9orf84 | -0,4404 | 2,84711E-10 |
| ACE2 | RP11-788M5.3 | -0,4404 | 2,84623E-10 |
| ACE2 | OSBP2 | -0,4404 | 2,83697E-10 |
| ACE2 | RPS6KA3 | -0,4404 | 2,82984E-10 |
| ACE2 | KLF9 | -0,4405 | 2,81994E-10 |
| ACE2 | TMEM254-AS1 | -0,4405 | 2,82332E-10 |
| ACE2 | RP11-598F7.6 | -0,4405 | 2,81912E-10 |
| ACE2 | PUS7L | -0,4405 | 2,8178E-10 |
| ACE2 | CALML6 | -0,4406 | 2,77566E-10 |
| ACE2 | FAR1 | -0,4406 | 2,78383E-10 |
| ACE2 | GVQW2 | -0,4407 | 2,74947E-10 |
| ACE2 | DBH-AS1 | -0,4407 | 2,74229E-10 |
| ACE2 | OR8T1P | -0,4407 | 2,75108E-10 |
| ACE2 | RP11-120M18.2 | -0,4407 | 2,76291E-10 |
| ACE2 | RP13-270P17.2 | -0,4407 | 2,74212E-10 |
| ACE2 | FBXO36 | -0,4408 | 2,73249E-10 |
| ACE2 | SLC2A12 | -0,4408 | 2,71249E-10 |
| ACE2 | RP11-392A14.9 | -0,4408 | 2,71423E-10 |
| ACE2 | TSHZ1 | -0,4408 | 2,71548E-10 |
| ACE2 | TMEM59L | -0,4408 | 2,7301E-10 |
| ACE2 | LINC02139 | -0,4409 | 2,68569E-10 |
| ACE2 | C18orf65 | -0,4409 | 2,70845E-10 |
| ACE2 | METTL24 | -0,441 | 2,67735E-10 |
| ACE2 | CD274 | -0,441 | 2,67151E-10 |
| ACE2 | DHTKD1 | -0,441 | 2,65645E-10 |
| ACE2 | KCNA4 | -0,441 | 2,67376E-10 |
| ACE2 | MRPL49 | -0,441 | 2,65816E-10 |
| ACE2 | RP11-329B9.5 | -0,4411 | 2,62881E-10 |
| ACE2 | RP11-90C4.1 | -0,4412 | 2,61575E-10 |
| ACE2 | RPL3P2 | -0,4412 | 2,60172E-10 |
| ACE2 | CCAR2 | -0,4413 | 2,59476E-10 |
| ACE2 | RP11-849H4.4 | -0,4413 | 2,59777E-10 |
| ACE2 | DPH6 | -0,4414 | 2,55793E-10 |
| ACE2 | DYNLRB1 | -0,4414 | 2,55081E-10 |
| ACE2 | PACSIN1 | -0,4415 | 2,54438E-10 |
| ACE2 | MRPS31P5 | -0,4415 | 2,53245E-10 |
| ACE2 | CDC42EP1 | -0,4415 | 2,53304E-10 |
| ACE2 | RP11-50D9.3 | -0,4416 | 2,50232E-10 |
| ACE2 | NACAP1 | -0,4416 | 2,5156E-10 |
| ACE2 | RP11-157H4.1 | -0,4416 | 2,51927E-10 |
| ACE2 | RP11-46J23.1 | -0,4417 | 2,48269E-10 |
| ACE2 | RPS3AP29 | -0,4417 | 2,49071E-10 |
| ACE2 | BAALC-AS2 | -0,4417 | 2,4687E-10 |
| ACE2 | RBM26 | -0,4417 | 2,48251E-10 |
| ACE2 | COG1 | -0,4417 | 2,48823E-10 |
| ACE2 | LYSMD3 | -0,4418 | 2,46645E-10 |
| ACE2 | SLC35G5 | -0,4418 | 2,46252E-10 |
| ACE2 | RP11-209D14.2 | -0,4418 | 2,46634E-10 |

| | | | |
|---|---|---|---|
| ACE2 | DHRS13 | -0,4418 | 2,44924E-10 |
| ACE2 | CTC-459F4.1 | -0,4418 | 2,46647E-10 |
| ACE2 | ARL6IP6 | -0,4419 | 2,44221E-10 |
| ACE2 | SHROOM2 | -0,4419 | 2,44195E-10 |
| ACE2 | RP4-555D20.4 | -0,442 | 2,39866E-10 |
| ACE2 | GIMAP2 | -0,442 | 2,40158E-10 |
| ACE2 | CRISPLD2 | -0,442 | 2,39967E-10 |
| ACE2 | EEF1B2 | -0,4421 | 2,37293E-10 |
| ACE2 | IDE | -0,4421 | 2,37844E-10 |
| ACE2 | KLHL41 | -0,4422 | 2,35745E-10 |
| ACE2 | CHAMP1 | -0,4422 | 2,34588E-10 |
| ACE2 | BOD1L1 | -0,4423 | 2,34145E-10 |
| ACE2 | MIR137HG | -0,4424 | 2,31362E-10 |
| ACE2 | TXNIP | -0,4424 | 2,29639E-10 |
| ACE2 | A1BG-AS1 | -0,4424 | 2,30268E-10 |
| ACE2 | RP1-43E13.2 | -0,4425 | 2,28889E-10 |
| ACE2 | ODF2L | -0,4425 | 2,27448E-10 |
| ACE2 | TBL1XR1-AS1 | -0,4426 | 2,25296E-10 |
| ACE2 | TSPOAP1-AS1 | -0,4426 | 2,25627E-10 |
| ACE2 | IL11 | -0,4426 | 2,26255E-10 |
| ACE2 | GPCPD1 | -0,4426 | 2,25575E-10 |
| ACE2 | PRDX4 | -0,4426 | 2,26802E-10 |
| ACE2 | RP5-1102E8.3 | -0,4427 | 2,22561E-10 |
| ACE2 | RPP38 | -0,4427 | 2,23242E-10 |
| ACE2 | LINC01270 | -0,4427 | 2,23544E-10 |
| ACE2 | AC011380.8 | -0,4429 | 2,1974E-10 |
| ACE2 | NDUFAF1 | -0,4429 | 2,1918E-10 |
| ACE2 | RP11-798G7.6 | -0,4429 | 2,19884E-10 |
| ACE2 | CTD-2195M15.1 | -0,443 | 2,17265E-10 |
| ACE2 | RP11-76C10.4 | -0,443 | 2,17549E-10 |
| ACE2 | AC105339.1 | -0,443 | 2,17456E-10 |
| ACE2 | PTX4 | -0,443 | 2,17185E-10 |
| ACE2 | RP11-332H18.5 | -0,443 | 2,15996E-10 |
| ACE2 | RP5-965F6.2 | -0,4431 | 2,14929E-10 |
| ACE2 | RP11-234A1.1 | -0,4431 | 2,14418E-10 |
| ACE2 | ZNF100 | -0,4431 | 2,14129E-10 |
| ACE2 | AC096558.1 | -0,4432 | 2,11595E-10 |
| ACE2 | CBR4 | -0,4432 | 2,12491E-10 |
| ACE2 | PIK3R1 | -0,4432 | 2,12672E-10 |
| ACE2 | PTPN5 | -0,4433 | 2,09508E-10 |
| ACE2 | RP11-582E3.4 | -0,4433 | 2,10457E-10 |
| ACE2 | RP11-243M5.3 | -0,4433 | 2,10116E-10 |
| ACE2 | MLKL | -0,4433 | 2,10818E-10 |
| ACE2 | ACSL6 | -0,4434 | 2,08066E-10 |
| ACE2 | EFR3A | -0,4435 | 2,06559E-10 |
| ACE2 | AP000577.2 | -0,4435 | 2,0611E-10 |
| ACE2 | C3orf38 | -0,4436 | 2,04659E-10 |
| ACE2 | LINC01023 | -0,4436 | 2,04369E-10 |
| ACE2 | LINC02073 | -0,4436 | 2,0402E-10 |
| ACE2 | RP11-232D9.3 | -0,4436 | 2,02762E-10 |
| ACE2 | RASA4DP | -0,4437 | 2,00504E-10 |
| ACE2 | CHMP4BP1 | -0,4437 | 2,01982E-10 |
| ACE2 | WDR59 | -0,4437 | 2,00619E-10 |
| ACE2 | PAK5 | -0,4437 | 2,021E-10 |

| | | | |
|---|---|---|---|
| ACE2 | AC104653.1 | -0,4438 | 1,98445E-10 |
| ACE2 | YTHDC2 | -0,4438 | 2,00089E-10 |
| ACE2 | HIBADH | -0,4438 | 1,99317E-10 |
| ACE2 | PPP4R3B | -0,4439 | 1,974E-10 |
| ACE2 | HBP1 | -0,4439 | 1,97364E-10 |
| ACE2 | RP11-620J15.2 | -0,4439 | 1,97611E-10 |
| ACE2 | ALPL | -0,444 | 1,95786E-10 |
| ACE2 | HIST2H2AB | -0,444 | 1,95138E-10 |
| ACE2 | SLC41A3 | -0,444 | 1,96108E-10 |
| ACE2 | SLC25A36 | -0,444 | 1,96304E-10 |
| ACE2 | CTC-471C19.1 | -0,444 | 1,95283E-10 |
| ACE2 | AP000487.5 | -0,444 | 1,94753E-10 |
| ACE2 | PLAC9P1 | -0,4441 | 1,93393E-10 |
| ACE2 | LPL | -0,4442 | 1,90637E-10 |
| ACE2 | APOE | -0,4442 | 1,91358E-10 |
| ACE2 | RPS23P6 | -0,4443 | 1,89083E-10 |
| ACE2 | PRKCH | -0,4444 | 1,87831E-10 |
| ACE2 | LINC01139 | -0,4445 | 1,86035E-10 |
| ACE2 | DPP6 | -0,4445 | 1,84774E-10 |
| ACE2 | FREM2 | -0,4445 | 1,84499E-10 |
| ACE2 | MAGEA8 | -0,4445 | 1,85773E-10 |
| ACE2 | CD200R1 | -0,4446 | 1,82588E-10 |
| ACE2 | SAMD3 | -0,4446 | 1,8285E-10 |
| ACE2 | RP11-840I19.5 | -0,4446 | 1,82707E-10 |
| ACE2 | COIL | -0,4446 | 1,83115E-10 |
| ACE2 | RP11-15H20.6 | -0,4446 | 1,83932E-10 |
| ACE2 | LRIT3 | -0,4447 | 1,81021E-10 |
| ACE2 | RP11-161D15.1 | -0,4447 | 1,81205E-10 |
| ACE2 | RNF139-AS1 | -0,4447 | 1,81497E-10 |
| ACE2 | TMEM106A | -0,4447 | 1,81541E-10 |
| ACE2 | FBN3 | -0,4447 | 1,81134E-10 |
| ACE2 | TAF7L | -0,4447 | 1,82242E-10 |
| ACE2 | RP11-500G10.5 | -0,4448 | 1,79665E-10 |
| ACE2 | EDC3 | -0,4448 | 1,79854E-10 |
| ACE2 | PVT1 | -0,4449 | 1,78541E-10 |
| ACE2 | RP11-390K5.1 | -0,4449 | 1,77597E-10 |
| ACE2 | CDK15 | -0,445 | 1,76196E-10 |
| ACE2 | CCDC6 | -0,445 | 1,75701E-10 |
| ACE2 | RP11-666A8.8 | -0,445 | 1,76484E-10 |
| ACE2 | CTC-260E6.6 | -0,445 | 1,75242E-10 |
| ACE2 | CH17-296N19.1 | -0,4451 | 1,73261E-10 |
| ACE2 | LINC00941 | -0,4451 | 1,74248E-10 |
| ACE2 | MLYCD | -0,4451 | 1,74565E-10 |
| ACE2 | ZNF550 | -0,4451 | 1,74167E-10 |
| ACE2 | LRRC39 | -0,4453 | 1,7136E-10 |
| ACE2 | RP13-143G15.4 | -0,4453 | 1,7044E-10 |
| ACE2 | CTDNEP1 | -0,4453 | 1,69856E-10 |
| ACE2 | RP13-228J13.8 | -0,4453 | 1,70299E-10 |
| ACE2 | WDFY1 | -0,4454 | 1,68015E-10 |
| ACE2 | MRGPRD | -0,4454 | 1,6908E-10 |
| ACE2 | PPM1G | -0,4455 | 1,67322E-10 |
| ACE2 | AASS | -0,4455 | 1,6656E-10 |
| ACE2 | ETV6 | -0,4455 | 1,67413E-10 |
| ACE2 | ZADH2 | -0,4455 | 1,67029E-10 |

| | | | |
|---|---|---|---|
| ACE2 | RP11-350J20.13 | -0,4456 | 1,65644E-10 |
| ACE2 | NRN1L | -0,4456 | 1,65131E-10 |
| ACE2 | SNRPD3 | -0,4456 | 1,66031E-10 |
| ACE2 | ADGRD1 | -0,4457 | 1,62969E-10 |
| ACE2 | ALYREF | -0,4457 | 1,62787E-10 |
| ACE2 | SYCP2 | -0,4457 | 1,63993E-10 |
| ACE2 | ZNF37CP | -0,4458 | 1,62012E-10 |
| ACE2 | TMEM9B-AS1 | -0,4458 | 1,62326E-10 |
| ACE2 | AC005330.2 | -0,4458 | 1,62234E-10 |
| ACE2 | FCGR2C | -0,4459 | 1,60725E-10 |
| ACE2 | MRPS25 | -0,4459 | 1,60275E-10 |
| ACE2 | AC012442.5 | -0,446 | 1,57837E-10 |
| ACE2 | RP11-472N13.2 | -0,446 | 1,58162E-10 |
| ACE2 | BCAN | -0,4461 | 1,57219E-10 |
| ACE2 | RP1-228P16.8 | -0,4461 | 1,56893E-10 |
| ACE2 | TEKT1 | -0,4461 | 1,55994E-10 |
| ACE2 | RP11-552F3.4 | -0,4461 | 1,56827E-10 |
| ACE2 | MRPL54 | -0,4461 | 1,56164E-10 |
| ACE2 | GID8 | -0,4461 | 1,57477E-10 |
| ACE2 | CTA-38K21.2 | -0,4461 | 1,56353E-10 |
| ACE2 | SUMO1 | -0,4462 | 1,556E-10 |
| ACE2 | VN1R20P | -0,4462 | 1,55553E-10 |
| ACE2 | RP11-352G18.2 | -0,4462 | 1,55636E-10 |
| ACE2 | ZNF750 | -0,4462 | 1,55381E-10 |
| ACE2 | RP11-475C16.1 | -0,4463 | 1,53688E-10 |
| ACE2 | ADAM20 | -0,4463 | 1,53777E-10 |
| ACE2 | RP11-147L13.8 | -0,4463 | 1,54096E-10 |
| ACE2 | PIGC | -0,4464 | 1,52492E-10 |
| ACE2 | AC079807.2 | -0,4464 | 1,51563E-10 |
| ACE2 | AC012613.2 | -0,4464 | 1,51282E-10 |
| ACE2 | TUBGCP6 | -0,4464 | 1,52058E-10 |
| ACE2 | CTD-2207L17.1 | -0,4465 | 1,49931E-10 |
| ACE2 | ZNF225 | -0,4465 | 1,49994E-10 |
| ACE2 | AC079922.3 | -0,4466 | 1,49209E-10 |
| ACE2 | RPLP1P6 | -0,4466 | 1,4841E-10 |
| ACE2 | LARP1 | -0,4466 | 1,48281E-10 |
| ACE2 | RP4-633O19__A.1 | -0,4466 | 1,48253E-10 |
| ACE2 | RCAN3 | -0,4467 | 1,46433E-10 |
| ACE2 | TADA1 | -0,4467 | 1,46512E-10 |
| ACE2 | WNT16 | -0,4467 | 1,47494E-10 |
| ACE2 | SNX33 | -0,4467 | 1,46575E-10 |
| ACE2 | IMMP1LP1 | -0,4468 | 1,45841E-10 |
| ACE2 | RP11-9E17.1 | -0,4468 | 1,45874E-10 |
| ACE2 | RP11-356N1.2 | -0,4469 | 1,44465E-10 |
| ACE2 | KCNH8 | -0,4469 | 1,43401E-10 |
| ACE2 | TRPV4 | -0,4469 | 1,43332E-10 |
| ACE2 | RP11-243M5.5 | -0,4469 | 1,44035E-10 |
| ACE2 | WASHC5 | -0,447 | 1,4318E-10 |
| ACE2 | ARHGAP23P1 | -0,447 | 1,42186E-10 |
| ACE2 | RP13-20L14.1 | -0,447 | 1,42532E-10 |
| ACE2 | RP11-64C12.8 | -0,447 | 1,4189E-10 |
| ACE2 | RP11-100G15.7 | -0,4471 | 1,40795E-10 |
| ACE2 | RP4-740C4.7 | -0,4472 | 1,39388E-10 |
| ACE2 | NPIPB5 | -0,4472 | 1,39665E-10 |

| | | | |
|---|---|---|---|
| ACE2 | PPIH | -0,4473 | 1,38242E-10 |
| ACE2 | TAGLN2 | -0,4473 | 1,38035E-10 |
| ACE2 | HSPA8P4 | -0,4473 | 1,38455E-10 |
| ACE2 | OTULIN | -0,4474 | 1,36657E-10 |
| ACE2 | RP3-495K2.4 | -0,4474 | 1,36785E-10 |
| ACE2 | MS4A14 | -0,4474 | 1,36694E-10 |
| ACE2 | ULK4P3 | -0,4474 | 1,36787E-10 |
| ACE2 | SIGLEC6 | -0,4474 | 1,36058E-10 |
| ACE2 | BACH2 | -0,4475 | 1,34625E-10 |
| ACE2 | GIMAP6 | -0,4475 | 1,3477E-10 |
| ACE2 | RP11-316N24.2 | -0,4476 | 1,34159E-10 |
| ACE2 | RASA1 | -0,4476 | 1,34088E-10 |
| ACE2 | RP11-320L11.2 | -0,4476 | 1,34424E-10 |
| ACE2 | RTL1 | -0,4476 | 1,34046E-10 |
| ACE2 | FAM134C | -0,4476 | 1,34467E-10 |
| ACE2 | RHOBTB1 | -0,4477 | 1,31806E-10 |
| ACE2 | SLC47A1P1 | -0,4477 | 1,3234E-10 |
| ACE2 | RP11-1094H24.4 | -0,4477 | 1,32488E-10 |
| ACE2 | ATP8B3 | -0,4477 | 1,3225E-10 |
| ACE2 | RP4-735C1.4 | -0,4478 | 1,31659E-10 |
| ACE2 | CRIPT | -0,4478 | 1,30515E-10 |
| ACE2 | HOOK3 | -0,4478 | 1,30623E-10 |
| ACE2 | RP11-574O7.1 | -0,4478 | 1,30643E-10 |
| ACE2 | SNAPC4 | -0,4478 | 1,30867E-10 |
| ACE2 | RP11-452L6.1 | -0,4478 | 1,30363E-10 |
| ACE2 | RP11-234K24.3 | -0,4479 | 1,30223E-10 |
| ACE2 | RP4-669L17.10 | -0,448 | 1,28931E-10 |
| ACE2 | YAP1 | -0,448 | 1,28209E-10 |
| ACE2 | SPSB3 | -0,448 | 1,28108E-10 |
| ACE2 | RP11-21B21.4 | -0,4481 | 1,26676E-10 |
| ACE2 | RP11-384F7.2 | -0,4482 | 1,25022E-10 |
| ACE2 | RP11-762B21.5 | -0,4482 | 1,26223E-10 |
| ACE2 | RNY3P8 | -0,4482 | 1,25257E-10 |
| ACE2 | SLC24A1 | -0,4482 | 1,25132E-10 |
| ACE2 | LRRC37A | -0,4482 | 1,25391E-10 |
| ACE2 | RP11-135D11.2 | -0,4483 | 1,24677E-10 |
| ACE2 | GOLGA8R | -0,4483 | 1,23891E-10 |
| ACE2 | HYDIN | -0,4483 | 1,24145E-10 |
| ACE2 | RP11-1152H14.1 | -0,4484 | 1,22907E-10 |
| ACE2 | FAM86B3P | -0,4484 | 1,22434E-10 |
| ACE2 | SGCZ | -0,4484 | 1,23406E-10 |
| ACE2 | ADD3 | -0,4485 | 1,22083E-10 |
| ACE2 | RP11-305O6.3 | -0,4485 | 1,2106E-10 |
| ACE2 | RP11-326A19.4 | -0,4485 | 1,21083E-10 |
| ACE2 | LRRC37A6P | -0,4486 | 1,19906E-10 |
| ACE2 | P2RX1 | -0,4486 | 1,20547E-10 |
| ACE2 | RP11-179B2.2 | -0,4488 | 1,17967E-10 |
| ACE2 | PISD | -0,4488 | 1,18256E-10 |
| ACE2 | USP6NL | -0,4489 | 1,16716E-10 |
| ACE2 | DYRK1B | -0,4489 | 1,16408E-10 |
| ACE2 | AC074391.1 | -0,449 | 1,14814E-10 |
| ACE2 | SEPT10 | -0,449 | 1,15094E-10 |
| ACE2 | CTD-2298J14.2 | -0,449 | 1,1538E-10 |
| ACE2 | SCGB3A1 | -0,4491 | 1,13817E-10 |

| | | | |
|---|---|---|---|
| ACE2 | RP3-495K2.2 | -0,4491 | 1,14181E-10 |
| ACE2 | POC1B-AS1 | -0,4491 | 1,14171E-10 |
| ACE2 | GS1-304P7.2 | -0,4492 | 1,13367E-10 |
| ACE2 | RP11-367E12.4 | -0,4492 | 1,12724E-10 |
| ACE2 | C16orf58 | -0,4492 | 1,13045E-10 |
| ACE2 | RP11-158H5.2 | -0,4492 | 1,12493E-10 |
| ACE2 | PLTP | -0,4492 | 1,1261E-10 |
| ACE2 | DSTNP1 | -0,4492 | 1,12757E-10 |
| ACE2 | RP11-134G8.5 | -0,4493 | 1,11629E-10 |
| ACE2 | DKFZP434I0714 | -0,4493 | 1,11992E-10 |
| ACE2 | U47924.32 | -0,4493 | 1,12234E-10 |
| ACE2 | CWC25 | -0,4493 | 1,12228E-10 |
| ACE2 | RP11-973H7.1 | -0,4493 | 1,11532E-10 |
| ACE2 | GPX4 | -0,4493 | 1,11903E-10 |
| ACE2 | ZNF557 | -0,4493 | 1,1149E-10 |
| ACE2 | FCGR2B | -0,4494 | 1,10997E-10 |
| ACE2 | AC105398.3 | -0,4494 | 1,09991E-10 |
| ACE2 | RP11-122C5.3 | -0,4494 | 1,10374E-10 |
| ACE2 | POFUT2 | -0,4494 | 1,10133E-10 |
| ACE2 | ISG20L2 | -0,4495 | 1,09732E-10 |
| ACE2 | CLCN6 | -0,4496 | 1,08593E-10 |
| ACE2 | UNC13D | -0,4496 | 1,07889E-10 |
| ACE2 | FEZ2 | -0,4497 | 1,07058E-10 |
| ACE2 | AC093388.3 | -0,4497 | 1,07484E-10 |
| ACE2 | CCAR1 | -0,4497 | 1,07293E-10 |
| ACE2 | NHLRC4 | -0,4497 | 1,07009E-10 |
| ACE2 | TAZ | -0,4497 | 1,06629E-10 |
| ACE2 | GUSBP6 | -0,4499 | 1,05259E-10 |
| ACE2 | CTD-3074O7.5 | -0,4499 | 1,05095E-10 |
| ACE2 | PSMD12 | -0,4499 | 1,04625E-10 |
| ACE2 | FAM217B | -0,4499 | 1,05399E-10 |
| ACE2 | AC018890.6 | -0,45 | 1,03365E-10 |
| ACE2 | C5orf22 | -0,45 | 1,03753E-10 |
| ACE2 | CTD-2530H12.8 | -0,45 | 1,03596E-10 |
| ACE2 | UBAC1 | -0,4501 | 1,02368E-10 |
| ACE2 | PTK7 | -0,4502 | 1,01378E-10 |
| ACE2 | BRICD5 | -0,4502 | 1,01391E-10 |
| ACE2 | MTDHP3 | -0,4502 | 1,01931E-10 |
| ACE2 | RP1-257A7.4 | -0,4503 | 1,00913E-10 |
| ACE2 | TOR1A | -0,4503 | 1,00442E-10 |
| ACE2 | RP11-229O3.1 | -0,4503 | 1,00487E-10 |
| ACE2 | RP13-1032I1.11 | -0,4503 | 9,99662E-11 |
| ACE2 | BRD4 | -0,4503 | 1,00604E-10 |
| ACE2 | OSCAR | -0,4503 | 1,00813E-10 |
| ACE2 | CMTM6 | -0,4504 | 9,99119E-11 |
| ACE2 | RP11-358D17.1 | -0,4504 | 9,95953E-11 |
| ACE2 | PRPF4B | -0,4504 | 9,97632E-11 |
| ACE2 | ARL15 | -0,4506 | 9,72936E-11 |
| ACE2 | RP11-213G2.5 | -0,4506 | 9,74733E-11 |
| ACE2 | HELZ | -0,4506 | 9,72036E-11 |
| ACE2 | R3HDM4 | -0,4506 | 9,72079E-11 |
| ACE2 | ADCY3 | -0,4507 | 9,66544E-11 |
| ACE2 | CTD-2081C10.7 | -0,4507 | 9,61809E-11 |
| ACE2 | RPL26P19 | -0,4507 | 9,65648E-11 |

| | | | |
|---|---|---|---|
| ACE2 | RP11-1277A3.1 | -0,4507 | 9,59571E-11 |
| ACE2 | TXNL4B | -0,4507 | 9,66109E-11 |
| ACE2 | CNTN5 | -0,4508 | 9,54296E-11 |
| ACE2 | RP11-161H23.9 | -0,4508 | 9,47669E-11 |
| ACE2 | RP11-164O23.8 | -0,4509 | 9,43739E-11 |
| ACE2 | RP11-3D4.2 | -0,4509 | 9,44162E-11 |
| ACE2 | HIF3A | -0,4509 | 9,42191E-11 |
| ACE2 | NAP1L6 | -0,451 | 9,36585E-11 |
| ACE2 | PSMB2 | -0,4511 | 9,25072E-11 |
| ACE2 | AC114730.2 | -0,4511 | 9,19065E-11 |
| ACE2 | RP11-1012E15.2 | -0,4511 | 9,21248E-11 |
| ACE2 | NUP62 | -0,4511 | 9,22465E-11 |
| ACE2 | RP5-894A10.2 | -0,4512 | 9,15466E-11 |
| ACE2 | HMGN2P3 | -0,4512 | 9,1195E-11 |
| ACE2 | GOT2 | -0,4512 | 9,08726E-11 |
| ACE2 | C19orf81 | -0,4512 | 9,08527E-11 |
| ACE2 | ACAN | -0,4513 | 9,02446E-11 |
| ACE2 | ZNF98 | -0,4513 | 8,9952E-11 |
| ACE2 | RP11-956E11.1 | -0,4514 | 8,91541E-11 |
| ACE2 | RFC5 | -0,4514 | 8,9407E-11 |
| ACE2 | PRMT6 | -0,4515 | 8,80987E-11 |
| ACE2 | CBLB | -0,4516 | 8,76166E-11 |
| ACE2 | ADAMTS19 | -0,4516 | 8,74853E-11 |
| ACE2 | WDR11 | -0,4516 | 8,7148E-11 |
| ACE2 | NGB | -0,4516 | 8,78409E-11 |
| ACE2 | ZNF235 | -0,4516 | 8,75153E-11 |
| ACE2 | CDC40 | -0,4517 | 8,63159E-11 |
| ACE2 | ZBTB45 | -0,4517 | 8,6265E-11 |
| ACE2 | FOXN4 | -0,4518 | 8,54734E-11 |
| ACE2 | AC079250.1 | -0,4519 | 8,47714E-11 |
| ACE2 | LINC01799 | -0,4519 | 8,43347E-11 |
| ACE2 | HSPD1 | -0,4519 | 8,45799E-11 |
| ACE2 | PLA1A | -0,4519 | 8,4741E-11 |
| ACE2 | MARCH5 | -0,4519 | 8,46231E-11 |
| ACE2 | AC006116.12 | -0,4519 | 8,44514E-11 |
| ACE2 | MCM9 | -0,452 | 8,41661E-11 |
| ACE2 | RP11-54O7.17 | -0,4521 | 8,25533E-11 |
| ACE2 | FSCN3 | -0,4521 | 8,3323E-11 |
| ACE2 | RP11-51B23.3 | -0,4521 | 8,27748E-11 |
| ACE2 | AC138969.4 | -0,4521 | 8,28826E-11 |
| ACE2 | SCRT1 | -0,4522 | 8,16965E-11 |
| ACE2 | CCDC38 | -0,4522 | 8,15851E-11 |
| ACE2 | STAU1 | -0,4522 | 8,17141E-11 |
| ACE2 | RP11-100N21.1 | -0,4523 | 8,11425E-11 |
| ACE2 | TUBG1 | -0,4523 | 8,0835E-11 |
| ACE2 | ATP6V0E1 | -0,4524 | 7,98503E-11 |
| ACE2 | AC004381.8 | -0,4524 | 8,05001E-11 |
| ACE2 | USB1 | -0,4524 | 8,01548E-11 |
| ACE2 | RUNDC3A-AS1 | -0,4524 | 8,03739E-11 |
| ACE2 | KRT16P6 | -0,4525 | 7,95162E-11 |
| ACE2 | CTD-2349P21.10 | -0,4525 | 7,94198E-11 |
| ACE2 | AJAP1 | -0,4526 | 7,88517E-11 |
| ACE2 | AC108463.2 | -0,4526 | 7,87807E-11 |
| ACE2 | EMC3-AS1 | -0,4526 | 7,84416E-11 |

| | | | |
|---|---|---|---|
| ACE2 | DIDO1 | -0,4526 | 7,83279E-11 |
| ACE2 | NINJ2 | -0,4527 | 7,77464E-11 |
| ACE2 | PPP5C | -0,4527 | 7,73436E-11 |
| ACE2 | CTB-61M7.2 | -0,4527 | 7,80783E-11 |
| ACE2 | ARFGAP1 | -0,4527 | 7,78953E-11 |
| ACE2 | RP11-656D10.3 | -0,4528 | 7,7221E-11 |
| ACE2 | FGD5P1 | -0,4528 | 7,71297E-11 |
| ACE2 | MACROD1 | -0,4528 | 7,68196E-11 |
| ACE2 | HMGB1P31 | -0,4529 | 7,56586E-11 |
| ACE2 | LAGE3P1 | -0,4529 | 7,60799E-11 |
| ACE2 | TWNK | -0,4529 | 7,59861E-11 |
| ACE2 | RPL7P38 | -0,4529 | 7,58908E-11 |
| ACE2 | C16orf74 | -0,4529 | 7,64012E-11 |
| ACE2 | PRKCB | -0,453 | 7,51008E-11 |
| ACE2 | MDM4 | -0,4531 | 7,48019E-11 |
| ACE2 | CCDC174 | -0,4531 | 7,44283E-11 |
| ACE2 | FHIT | -0,4531 | 7,4545E-11 |
| ACE2 | LAP3 | -0,4531 | 7,42222E-11 |
| ACE2 | BTBD7 | -0,4531 | 7,46364E-11 |
| ACE2 | RP11-707O23.1 | -0,4531 | 7,47261E-11 |
| ACE2 | RP11-338N10.3 | -0,4532 | 7,39781E-11 |
| ACE2 | MYOC | -0,4532 | 7,33206E-11 |
| ACE2 | LINC01366 | -0,4532 | 7,37309E-11 |
| ACE2 | LINC01154 | -0,4532 | 7,35035E-11 |
| ACE2 | SBNO1 | -0,4532 | 7,37825E-11 |
| ACE2 | ZNF207 | -0,4532 | 7,33575E-11 |
| ACE2 | FCGR2A | -0,4533 | 7,31332E-11 |
| ACE2 | RP11-478C6.5 | -0,4533 | 7,29194E-11 |
| ACE2 | RP11-53M11.5 | -0,4533 | 7,26875E-11 |
| ACE2 | AP000487.6 | -0,4533 | 7,30568E-11 |
| ACE2 | CTD-2012K14.6 | -0,4533 | 7,27546E-11 |
| ACE2 | FAM19A1 | -0,4536 | 7,08126E-11 |
| ACE2 | RP11-131M11.2 | -0,4537 | 6,96122E-11 |
| ACE2 | UBA6 | -0,4537 | 6,94465E-11 |
| ACE2 | FAM149A | -0,4537 | 7,00952E-11 |
| ACE2 | C11orf71 | -0,4537 | 6,9879E-11 |
| ACE2 | CTD-2553L13.9 | -0,4537 | 6,95537E-11 |
| ACE2 | FTCDNL1 | -0,4538 | 6,91244E-11 |
| ACE2 | TPRXL | -0,4538 | 6,90172E-11 |
| ACE2 | RP11-1319K7.1 | -0,4538 | 6,86964E-11 |
| ACE2 | RP1-101K10.6 | -0,4538 | 6,94018E-11 |
| ACE2 | ACSM2B | -0,4538 | 6,91439E-11 |
| ACE2 | RP11-22P6.2 | -0,4538 | 6,89989E-11 |
| ACE2 | LINC01841 | -0,4538 | 6,92152E-11 |
| ACE2 | CTB-33G10.1 | -0,4538 | 6,9002E-11 |
| ACE2 | LINGO2 | -0,4539 | 6,82218E-11 |
| ACE2 | HCCAT5 | -0,4539 | 6,84263E-11 |
| ACE2 | PRR29-AS1 | -0,4539 | 6,85331E-11 |
| ACE2 | LRRC37A16P | -0,4539 | 6,80548E-11 |
| ACE2 | METTL4 | -0,4539 | 6,83641E-11 |
| ACE2 | RP11-12M9.4 | -0,4539 | 6,8591E-11 |
| ACE2 | CH17-469D17.1 | -0,454 | 6,77282E-11 |
| ACE2 | RP1-102K2.9 | -0,454 | 6,79278E-11 |
| ACE2 | MARCH9 | -0,4541 | 6,70724E-11 |

| | | | |
|---|---|---|---|
| ACE2 | RP11-15A1.3 | -0,4542 | 6,57793E-11 |
| ACE2 | AMY2B | -0,4543 | 6,56251E-11 |
| ACE2 | KMT2C | -0,4543 | 6,52229E-11 |
| ACE2 | MYOZ1 | -0,4543 | 6,52616E-11 |
| ACE2 | NPIPA7 | -0,4543 | 6,56496E-11 |
| ACE2 | RP4-781B1.5 | -0,4543 | 6,51806E-11 |
| ACE2 | LINC00115 | -0,4544 | 6,44659E-11 |
| ACE2 | LCN10 | -0,4544 | 6,48166E-11 |
| ACE2 | LINC00994 | -0,4545 | 6,39223E-11 |
| ACE2 | HIST1H2AD | -0,4545 | 6,3768E-11 |
| ACE2 | FNDC1-IT1 | -0,4545 | 6,43559E-11 |
| ACE2 | TLK2P2 | -0,4545 | 6,37921E-11 |
| ACE2 | LINC01684 | -0,4546 | 6,33707E-11 |
| ACE2 | RBMXP4 | -0,4547 | 6,28086E-11 |
| ACE2 | RP11-554F20.1 | -0,4547 | 6,28466E-11 |
| ACE2 | KCNMA1 | -0,4547 | 6,29686E-11 |
| ACE2 | ZCCHC12 | -0,4547 | 6,27435E-11 |
| ACE2 | CELF2-AS1 | -0,4548 | 6,22569E-11 |
| ACE2 | MNDA | -0,4549 | 6,12151E-11 |
| ACE2 | C3orf70 | -0,4549 | 6,10027E-11 |
| ACE2 | RBM26-AS1 | -0,4549 | 6,12306E-11 |
| ACE2 | GOSR1 | -0,455 | 6,06581E-11 |
| ACE2 | UBE3C | -0,4551 | 5,99712E-11 |
| ACE2 | AC022153.1 | -0,4551 | 6,0128E-11 |
| ACE2 | PLA2G2C | -0,4552 | 5,92319E-11 |
| ACE2 | ECEL1 | -0,4552 | 5,90892E-11 |
| ACE2 | HSPBAP1 | -0,4552 | 5,94663E-11 |
| ACE2 | LINC01554 | -0,4552 | 5,91408E-11 |
| ACE2 | DUSP1 | -0,4552 | 5,92218E-11 |
| ACE2 | FAM153B | -0,4552 | 5,93954E-11 |
| ACE2 | RP11-359E19.2 | -0,4552 | 5,92869E-11 |
| ACE2 | RP11-278C7.4 | -0,4552 | 5,90716E-11 |
| ACE2 | GRIPAP1 | -0,4552 | 5,9451E-11 |
| ACE2 | TSPAN32 | -0,4553 | 5,8843E-11 |
| ACE2 | PRDX3P1 | -0,4553 | 5,90004E-11 |
| ACE2 | MAMDC2-AS1 | -0,4554 | 5,83716E-11 |
| ACE2 | RPH3A | -0,4554 | 5,81296E-11 |
| ACE2 | HAUS4 | -0,4554 | 5,83756E-11 |
| ACE2 | GMEB2 | -0,4554 | 5,80742E-11 |
| ACE2 | LINC02048 | -0,4555 | 5,72847E-11 |
| ACE2 | PARM1 | -0,4555 | 5,76455E-11 |
| ACE2 | AC007792.1 | -0,4555 | 5,74775E-11 |
| ACE2 | ZBTB8OS | -0,4556 | 5,6857E-11 |
| ACE2 | RHOQP2 | -0,4556 | 5,70027E-11 |
| ACE2 | GRIA2 | -0,4557 | 5,6486E-11 |
| ACE2 | KLF10 | -0,4558 | 5,57931E-11 |
| ACE2 | ACAD8 | -0,4558 | 5,53524E-11 |
| ACE2 | SLC11A2 | -0,4558 | 5,5516E-11 |
| ACE2 | RP11-61J19.5 | -0,4559 | 5,48845E-11 |
| ACE2 | STARD7-AS1 | -0,4559 | 5,48318E-11 |
| ACE2 | GVQW1 | -0,4559 | 5,52682E-11 |
| ACE2 | SYNC | -0,456 | 5,423E-11 |
| ACE2 | C1orf61 | -0,456 | 5,46501E-11 |
| ACE2 | MIR199A2 | -0,456 | 5,41464E-11 |

| | | | |
|---|---|---|---|
| ACE2 | USP34 | -0,456 | 5,43199E-11 |
| ACE2 | PPIA | -0,456 | 5,41624E-11 |
| ACE2 | AC005009.1 | -0,456 | 5,41712E-11 |
| ACE2 | ADIRF-AS1 | -0,456 | 5,43706E-11 |
| ACE2 | ZNF814 | -0,456 | 5,45552E-11 |
| ACE2 | RP11-676M6.1 | -0,4561 | 5,36024E-11 |
| ACE2 | RP11-147L13.2 | -0,4561 | 5,38046E-11 |
| ACE2 | MBD2 | -0,4561 | 5,37033E-11 |
| ACE2 | RP4-530I15.9 | -0,4561 | 5,36921E-11 |
| ACE2 | SSBP3-AS1 | -0,4562 | 5,35286E-11 |
| ACE2 | MTERF4 | -0,4562 | 5,30843E-11 |
| ACE2 | RP11-259K5.1 | -0,4562 | 5,31311E-11 |
| ACE2 | CTU1 | -0,4562 | 5,34145E-11 |
| ACE2 | MIR181A1HG | -0,4563 | 5,24573E-11 |
| ACE2 | C22orf34 | -0,4563 | 5,26752E-11 |
| ACE2 | RP4-545K15.5 | -0,4563 | 5,27371E-11 |
| ACE2 | POLR2J4 | -0,4564 | 5,19842E-11 |
| ACE2 | SERTM1 | -0,4564 | 5,19086E-11 |
| ACE2 | COL6A6 | -0,4565 | 5,15949E-11 |
| ACE2 | AMZ1 | -0,4565 | 5,13479E-11 |
| ACE2 | AQP1 | -0,4565 | 5,1439E-11 |
| ACE2 | PHOX2A | -0,4565 | 5,14721E-11 |
| ACE2 | CTD-3092A11.2 | -0,4565 | 5,15149E-11 |
| ACE2 | ABLIM1 | -0,4566 | 5,07138E-11 |
| ACE2 | CTD-3065B20.3 | -0,4567 | 5,0245E-11 |
| ACE2 | TRIM22 | -0,4569 | 4,90752E-11 |
| ACE2 | SCAF11 | -0,4569 | 4,95723E-11 |
| ACE2 | RP11-493L12.7 | -0,4569 | 4,93054E-11 |
| ACE2 | CLPX | -0,4569 | 4,91583E-11 |
| ACE2 | NME1 | -0,4569 | 4,93894E-11 |
| ACE2 | ZNF525 | -0,4569 | 4,92154E-11 |
| ACE2 | USO1 | -0,4571 | 4,81788E-11 |
| ACE2 | GLB1L3 | -0,4571 | 4,80319E-11 |
| ACE2 | CNOT2 | -0,4571 | 4,83985E-11 |
| ACE2 | RP11-121A14.2 | -0,4572 | 4,79197E-11 |
| ACE2 | NAPG | -0,4572 | 4,75378E-11 |
| ACE2 | FBXL4 | -0,4573 | 4,73329E-11 |
| ACE2 | KCNA1 | -0,4573 | 4,70049E-11 |
| ACE2 | SLC6A2 | -0,4573 | 4,72289E-11 |
| ACE2 | LRRN2 | -0,4574 | 4,69555E-11 |
| ACE2 | KCTD8 | -0,4574 | 4,68781E-11 |
| ACE2 | RP11-677I18.3 | -0,4574 | 4,68731E-11 |
| ACE2 | ATP5L | -0,4574 | 4,66212E-11 |
| ACE2 | RP11-445H22.3 | -0,4574 | 4,67111E-11 |
| ACE2 | ESYT3 | -0,4575 | 4,59957E-11 |
| ACE2 | RP11-323F24.4 | -0,4576 | 4,58552E-11 |
| ACE2 | CFAP44 | -0,4577 | 4,53974E-11 |
| ACE2 | NTN4 | -0,4577 | 4,54601E-11 |
| ACE2 | HEXIM2 | -0,4577 | 4,50836E-11 |
| ACE2 | MED25 | -0,4577 | 4,50544E-11 |
| ACE2 | PCYOX1 | -0,4578 | 4,45055E-11 |
| ACE2 | S100A8 | -0,4579 | 4,43985E-11 |
| ACE2 | LINC01063 | -0,4579 | 4,40298E-11 |
| ACE2 | FAM86B2 | -0,4579 | 4,40939E-11 |

| | | | |
|---|---|---|---|
| ACE2 | EMX2OS | -0,4579 | 4,41874E-11 |
| ACE2 | RP11-266K4.9 | -0,4579 | 4,43965E-11 |
| ACE2 | RP11-1023L17.1 | -0,458 | 4,39073E-11 |
| ACE2 | DST | -0,458 | 4,36379E-11 |
| ACE2 | SNORD83B | -0,4581 | 4,31451E-11 |
| ACE2 | RP11-728K20.3 | -0,4582 | 4,27331E-11 |
| ACE2 | ACBD3 | -0,4583 | 4,22338E-11 |
| ACE2 | RP11-169D4.2 | -0,4583 | 4,22084E-11 |
| ACE2 | PTGFRN | -0,4584 | 4,20129E-11 |
| ACE2 | RP11-722E23.2 | -0,4584 | 4,19388E-11 |
| ACE2 | RP11-142M10.2 | -0,4584 | 4,19744E-11 |
| ACE2 | SNORD23 | -0,4584 | 4,18759E-11 |
| ACE2 | CCDC115 | -0,4585 | 4,13013E-11 |
| ACE2 | SULT4A1 | -0,4585 | 4,16095E-11 |
| ACE2 | ZMYM6 | -0,4586 | 4,10347E-11 |
| ACE2 | CTD-2562J17.6 | -0,4586 | 4,09751E-11 |
| ACE2 | HAP1 | -0,4586 | 4,09317E-11 |
| ACE2 | RP11-524C21.2 | -0,4587 | 4,06749E-11 |
| ACE2 | NPM2 | -0,4587 | 4,0498E-11 |
| ACE2 | RP11-541M12.6 | -0,4587 | 4,05508E-11 |
| ACE2 | ZNF430 | -0,4587 | 4,04004E-11 |
| ACE2 | EN2 | -0,4588 | 4,00761E-11 |
| ACE2 | ANK1 | -0,4588 | 4,0232E-11 |
| ACE2 | CTD-3025N20.3 | -0,4588 | 4,02199E-11 |
| ACE2 | RP11-309L24.6 | -0,4589 | 3,94879E-11 |
| ACE2 | CTD-2530H12.4 | -0,4589 | 3,94607E-11 |
| ACE2 | L3HYPDH | -0,4589 | 3,94687E-11 |
| ACE2 | AC004076.5 | -0,4589 | 3,95491E-11 |
| ACE2 | RP11-136C24.2 | -0,459 | 3,904E-11 |
| ACE2 | RP11-1415C14.3 | -0,459 | 3,91199E-11 |
| ACE2 | SNORA13 | -0,459 | 3,92449E-11 |
| ACE2 | MIOS | -0,459 | 3,93215E-11 |
| ACE2 | NLRP12 | -0,459 | 3,93596E-11 |
| ACE2 | AMIGO1 | -0,4591 | 3,8615E-11 |
| ACE2 | ECT2L | -0,4591 | 3,89622E-11 |
| ACE2 | SLC9A5 | -0,4591 | 3,85833E-11 |
| ACE2 | ZNF708 | -0,4591 | 3,88314E-11 |
| ACE2 | RP4-758J18.2 | -0,4592 | 3,83703E-11 |
| ACE2 | RAF1 | -0,4592 | 3,8216E-11 |
| ACE2 | RAB18 | -0,4592 | 3,85564E-11 |
| ACE2 | CTC-444N24.7 | -0,4592 | 3,82507E-11 |
| ACE2 | SNX9 | -0,4593 | 3,77597E-11 |
| ACE2 | RP11-642A1.2 | -0,4593 | 3,81073E-11 |
| ACE2 | LINC00683 | -0,4593 | 3,79731E-11 |
| ACE2 | PSMG1 | -0,4593 | 3,79735E-11 |
| ACE2 | CNOT8 | -0,4594 | 3,7716E-11 |
| ACE2 | RP1-234P15.4 | -0,4594 | 3,74547E-11 |
| ACE2 | AP000936.1 | -0,4594 | 3,74385E-11 |
| ACE2 | GTSE1-AS1 | -0,4594 | 3,74208E-11 |
| ACE2 | ZNF320 | -0,4595 | 3,70393E-11 |
| ACE2 | SEMA4A | -0,4596 | 3,68842E-11 |
| ACE2 | RP1-67A8.3 | -0,4597 | 3,62787E-11 |
| ACE2 | C9orf116 | -0,4597 | 3,62324E-11 |
| ACE2 | MTMR1 | -0,4597 | 3,61526E-11 |

| | | | |
|---|---|---|---|
| ACE2 | TMEM69 | -0,4598 | 3,59886E-11 |
| ACE2 | PRR18 | -0,4598 | 3,60527E-11 |
| ACE2 | SBF2-AS1 | -0,4598 | 3,58913E-11 |
| ACE2 | RP5-1024C24.1 | -0,4598 | 3,58089E-11 |
| ACE2 | NUP88 | -0,4598 | 3,57297E-11 |
| ACE2 | HHAT | -0,4599 | 3,55172E-11 |
| ACE2 | MTND5P28 | -0,4599 | 3,5416E-11 |
| ACE2 | LINC01322 | -0,4599 | 3,56102E-11 |
| ACE2 | RP11-889L3.1 | -0,4599 | 3,567E-11 |
| ACE2 | RP11-530A18.1 | -0,4599 | 3,54336E-11 |
| ACE2 | ATG3 | -0,46 | 3,52466E-11 |
| ACE2 | RP11-66B24.4 | -0,46 | 3,53195E-11 |
| ACE2 | NAGPA-AS1 | -0,46 | 3,51762E-11 |
| ACE2 | ADPRM | -0,46 | 3,52729E-11 |
| ACE2 | PIGA | -0,46 | 3,49555E-11 |
| ACE2 | UTS2B | -0,4601 | 3,48441E-11 |
| ACE2 | CLNS1A | -0,4601 | 3,45742E-11 |
| ACE2 | TMEM147-AS1 | -0,4601 | 3,47274E-11 |
| ACE2 | RP11-54O7.16 | -0,4602 | 3,43559E-11 |
| ACE2 | RP11-248N22.1 | -0,4602 | 3,45408E-11 |
| ACE2 | PKN2-AS1 | -0,4603 | 3,41513E-11 |
| ACE2 | C8orf4 | -0,4603 | 3,4063E-11 |
| ACE2 | NTMT1 | -0,4603 | 3,41763E-11 |
| ACE2 | RP11-326C3.7 | -0,4603 | 3,39779E-11 |
| ACE2 | CTD-2286N8.2 | -0,4603 | 3,38324E-11 |
| ACE2 | GPATCH1 | -0,4603 | 3,40375E-11 |
| ACE2 | RP1-29C18.8 | -0,4603 | 3,3929E-11 |
| ACE2 | CNGA4 | -0,4604 | 3,37914E-11 |
| ACE2 | TRPM7 | -0,4604 | 3,34768E-11 |
| ACE2 | CPSF3 | -0,4605 | 3,32118E-11 |
| ACE2 | IQCF1 | -0,4605 | 3,33481E-11 |
| ACE2 | STARD3 | -0,4605 | 3,33065E-11 |
| ACE2 | ZNF507 | -0,4605 | 3,32368E-11 |
| ACE2 | SYT14 | -0,4606 | 3,28439E-11 |
| ACE2 | RPL23AP32 | -0,4606 | 3,28224E-11 |
| ACE2 | RP11-809C18.3 | -0,4606 | 3,29555E-11 |
| ACE2 | APOL1 | -0,4606 | 3,29796E-11 |
| ACE2 | TCEA1P4 | -0,4607 | 3,25224E-11 |
| ACE2 | APOL4 | -0,4607 | 3,26916E-11 |
| ACE2 | STK26 | -0,4607 | 3,2608E-11 |
| ACE2 | RP11-739P1.3 | -0,4608 | 3,22099E-11 |
| ACE2 | FAM182A | -0,4608 | 3,21237E-11 |
| ACE2 | TCTEX1D1 | -0,4609 | 3,18427E-11 |
| ACE2 | RP11-274B21.10 | -0,4609 | 3,18908E-11 |
| ACE2 | RPS6P25 | -0,4609 | 3,18313E-11 |
| ACE2 | UBE2FP3 | -0,461 | 3,16104E-11 |
| ACE2 | RP11-251G23.5 | -0,461 | 3,15091E-11 |
| ACE2 | AC009245.3 | -0,461 | 3,15632E-11 |
| ACE2 | NPFFR1 | -0,461 | 3,16442E-11 |
| ACE2 | RP11-32B5.1 | -0,461 | 3,15789E-11 |
| ACE2 | ANKRD44 | -0,4611 | 3,111E-11 |
| ACE2 | SPATA4 | -0,4611 | 3,112E-11 |
| ACE2 | SLC25A21-AS1 | -0,4611 | 3,09859E-11 |
| ACE2 | SIRPA | -0,4611 | 3,11982E-11 |

| | | | |
|---|---|---|---|
| ACE2 | LINC00629 | -0,4611 | 3,10696E-11 |
| ACE2 | CTD-2587M2.1 | -0,4612 | 3,087E-11 |
| ACE2 | PVRIG | -0,4614 | 2,9953E-11 |
| ACE2 | INHBA-AS1 | -0,4615 | 2,96542E-11 |
| ACE2 | ZNF16 | -0,4615 | 2,96852E-11 |
| ACE2 | RP11-118B22.4 | -0,4615 | 2,98396E-11 |
| ACE2 | RP6-74O6.6 | -0,4616 | 2,94523E-11 |
| ACE2 | KLHL23 | -0,4616 | 2,932E-11 |
| ACE2 | TDO2 | -0,4616 | 2,93326E-11 |
| ACE2 | RP11-250B2.5 | -0,4616 | 2,95929E-11 |
| ACE2 | ULBP2 | -0,4616 | 2,93742E-11 |
| ACE2 | TPT1P4 | -0,4618 | 2,88706E-11 |
| ACE2 | AC093390.1 | -0,462 | 2,82778E-11 |
| ACE2 | HMCES | -0,462 | 2,81397E-11 |
| ACE2 | COL5A1-AS1 | -0,462 | 2,81747E-11 |
| ACE2 | LINC02080 | -0,4621 | 2,80097E-11 |
| ACE2 | RASAL2 | -0,4622 | 2,74427E-11 |
| ACE2 | RP11-158K1.3 | -0,4622 | 2,7643E-11 |
| ACE2 | CRYGS | -0,4623 | 2,71552E-11 |
| ACE2 | ZFAND6 | -0,4623 | 2,72631E-11 |
| ACE2 | AC092641.2 | -0,4624 | 2,70388E-11 |
| ACE2 | CPZ | -0,4624 | 2,70476E-11 |
| ACE2 | RP4-761J14.10 | -0,4624 | 2,69883E-11 |
| ACE2 | DESI1 | -0,4624 | 2,69957E-11 |
| ACE2 | LINC00640 | -0,4625 | 2,66224E-11 |
| ACE2 | AFF1 | -0,4626 | 2,63584E-11 |
| ACE2 | RP11-335L23.5 | -0,4626 | 2,64457E-11 |
| ACE2 | COL4A2-AS2 | -0,4626 | 2,64225E-11 |
| ACE2 | NNAT | -0,4626 | 2,63419E-11 |
| ACE2 | LOC100421166 | -0,4627 | 2,61249E-11 |
| ACE2 | RP11-399B17.1 | -0,4628 | 2,56556E-11 |
| ACE2 | PPIL3 | -0,4628 | 2,56557E-11 |
| ACE2 | FZD1 | -0,4628 | 2,5672E-11 |
| ACE2 | APLN | -0,4628 | 2,56662E-11 |
| ACE2 | CDKN2AIP | -0,4629 | 2,54556E-11 |
| ACE2 | RPL21P119 | -0,4629 | 2,55652E-11 |
| ACE2 | CHRNB2 | -0,463 | 2,51405E-11 |
| ACE2 | PEX2 | -0,463 | 2,5131E-11 |
| ACE2 | FNBP4 | -0,4632 | 2,47286E-11 |
| ACE2 | WDR90 | -0,4632 | 2,46908E-11 |
| ACE2 | MRPL58 | -0,4632 | 2,45779E-11 |
| ACE2 | CTBP1-AS2 | -0,4633 | 2,43818E-11 |
| ACE2 | LEMD2 | -0,4633 | 2,45128E-11 |
| ACE2 | UAP1L1 | -0,4633 | 2,43054E-11 |
| ACE2 | ZNF850 | -0,4633 | 2,43782E-11 |
| ACE2 | RP11-107N15.1 | -0,4634 | 2,42296E-11 |
| ACE2 | VPS53 | -0,4634 | 2,42541E-11 |
| ACE2 | LDOC1L | -0,4634 | 2,39978E-11 |
| ACE2 | RP11-284M14.1 | -0,4635 | 2,37929E-11 |
| ACE2 | MICA | -0,4635 | 2,37276E-11 |
| ACE2 | ZNF717 | -0,4636 | 2,35444E-11 |
| ACE2 | ZC3H8 | -0,4638 | 2,30748E-11 |
| ACE2 | RPL5P12 | -0,4638 | 2,31538E-11 |
| ACE2 | ATXN2L | -0,4638 | 2,30596E-11 |

| | | | |
|---|---|---|---|
| ACE2 | NME8 | -0,4639 | 2,26948E-11 |
| ACE2 | SLC2A6 | -0,464 | 2,26292E-11 |
| ACE2 | SLTM | -0,464 | 2,25606E-11 |
| ACE2 | SSR2 | -0,4641 | 2,2221E-11 |
| ACE2 | HIST1H4E | -0,4641 | 2,22642E-11 |
| ACE2 | TIMM10 | -0,4641 | 2,23607E-11 |
| ACE2 | RN7SL15P | -0,4641 | 2,22526E-11 |
| ACE2 | SASS6 | -0,4642 | 2,19914E-11 |
| ACE2 | TANC1 | -0,4642 | 2,20146E-11 |
| ACE2 | PMS2CL | -0,4642 | 2,20686E-11 |
| ACE2 | ADAM12 | -0,4642 | 2,19803E-11 |
| ACE2 | RP11-701H24.5 | -0,4642 | 2,20726E-11 |
| ACE2 | GTF2H2C | -0,4644 | 2,14678E-11 |
| ACE2 | ANKRD10-IT1 | -0,4644 | 2,16751E-11 |
| ACE2 | ARL17B | -0,4644 | 2,14878E-11 |
| ACE2 | HNRNPCP1 | -0,4645 | 2,13607E-11 |
| ACE2 | C2CD5 | -0,4646 | 2,11469E-11 |
| ACE2 | LINC00519 | -0,4646 | 2,10076E-11 |
| ACE2 | RP11-112N19.2 | -0,4646 | 2,12199E-11 |
| ACE2 | ZCCHC9 | -0,4647 | 2,07886E-11 |
| ACE2 | RP11-797H7.1 | -0,4647 | 2,09445E-11 |
| ACE2 | PLA2G4A | -0,4648 | 2,0633E-11 |
| ACE2 | HSPA1B | -0,4648 | 2,06224E-11 |
| ACE2 | OIP5-AS1 | -0,4648 | 2,07487E-11 |
| ACE2 | CD300E | -0,4648 | 2,06112E-11 |
| ACE2 | BPIFB9P | -0,4648 | 2,06983E-11 |
| ACE2 | RP4-730K3.3 | -0,4649 | 2,04335E-11 |
| ACE2 | PTCHD1-AS | -0,4649 | 2,04097E-11 |
| ACE2 | LINC02015 | -0,465 | 2,02434E-11 |
| ACE2 | ZBED3 | -0,465 | 2,01394E-11 |
| ACE2 | RP5-981O7.2 | -0,465 | 2,01327E-11 |
| ACE2 | PPP1R15A | -0,465 | 2,00915E-11 |
| ACE2 | FBXL3 | -0,4651 | 1,98962E-11 |
| ACE2 | RPL9P9 | -0,4651 | 2,00122E-11 |
| ACE2 | RP11-212I21.3 | -0,4651 | 1,99349E-11 |
| ACE2 | RP5-1014D13.2 | -0,4651 | 2,00642E-11 |
| ACE2 | RAD1P2 | -0,4653 | 1,94111E-11 |
| ACE2 | MCCC1 | -0,4653 | 1,94978E-11 |
| ACE2 | RP11-394O4.3 | -0,4653 | 1,95806E-11 |
| ACE2 | RP11-154D6.2 | -0,4654 | 1,92271E-11 |
| ACE2 | QRSL1 | -0,4654 | 1,91972E-11 |
| ACE2 | MBLAC1 | -0,4654 | 1,94004E-11 |
| ACE2 | SNORA52 | -0,4654 | 1,93159E-11 |
| ACE2 | OR11H7 | -0,4654 | 1,93046E-11 |
| ACE2 | STX12 | -0,4655 | 1,90897E-11 |
| ACE2 | CHMP7 | -0,4655 | 1,90156E-11 |
| ACE2 | MTA2 | -0,4655 | 1,91494E-11 |
| ACE2 | RP11-605F22.2 | -0,4655 | 1,90479E-11 |
| ACE2 | LINC01138 | -0,4656 | 1,88651E-11 |
| ACE2 | TRIM27 | -0,4656 | 1,88318E-11 |
| ACE2 | MOSPD2 | -0,4656 | 1,89673E-11 |
| ACE2 | ZC3HAV1 | -0,4657 | 1,86662E-11 |
| ACE2 | RP11-29A19.2 | -0,4657 | 1,86499E-11 |
| ACE2 | CCDC116 | -0,4657 | 1,86147E-11 |

| | | | |
|---|---|---|---|
| ACE2 | PLEKHG4B | -0,4658 | 1,83637E-11 |
| ACE2 | RP11-448A19.1 | -0,4658 | 1,84058E-11 |
| ACE2 | ANAPC15 | -0,4658 | 1,84617E-11 |
| ACE2 | RP11-187O7.3 | -0,4658 | 1,83637E-11 |
| ACE2 | SIKE1 | -0,466 | 1,79834E-11 |
| ACE2 | THEGL | -0,466 | 1,79806E-11 |
| ACE2 | NUDT6 | -0,466 | 1,80937E-11 |
| ACE2 | LILRA6 | -0,466 | 1,81443E-11 |
| ACE2 | ATF4P4 | -0,4661 | 1,79009E-11 |
| ACE2 | RP11-314A20.1 | -0,4661 | 1,79019E-11 |
| ACE2 | MAT2A | -0,4662 | 1,76271E-11 |
| ACE2 | OAS2 | -0,4662 | 1,76511E-11 |
| ACE2 | THOC2 | -0,4662 | 1,76216E-11 |
| ACE2 | TRIM45 | -0,4663 | 1,74761E-11 |
| ACE2 | HSP90AB3P | -0,4663 | 1,74225E-11 |
| ACE2 | SEM1 | -0,4664 | 1,71591E-11 |
| ACE2 | ZNF699 | -0,4664 | 1,7316E-11 |
| ACE2 | SLC35B4 | -0,4665 | 1,71368E-11 |
| ACE2 | PPP1R26P1 | -0,4665 | 1,69824E-11 |
| ACE2 | NBPF12 | -0,4666 | 1,68781E-11 |
| ACE2 | AC022182.3 | -0,4666 | 1,69079E-11 |
| ACE2 | RBP5 | -0,4669 | 1,62438E-11 |
| ACE2 | RPL10P15 | -0,4669 | 1,62572E-11 |
| ACE2 | H6PD | -0,467 | 1,62082E-11 |
| ACE2 | JADE2 | -0,467 | 1,60717E-11 |
| ACE2 | AC005519.4 | -0,467 | 1,61226E-11 |
| ACE2 | SLC16A6P1 | -0,467 | 1,61435E-11 |
| ACE2 | DR1 | -0,4671 | 1,58662E-11 |
| ACE2 | GIMAP5 | -0,4671 | 1,59573E-11 |
| ACE2 | RMDN1 | -0,4671 | 1,60193E-11 |
| ACE2 | SLC18A3 | -0,4671 | 1,60094E-11 |
| ACE2 | SRGAP2D | -0,4672 | 1,58202E-11 |
| ACE2 | WFDC3 | -0,4672 | 1,58117E-11 |
| ACE2 | NSD1 | -0,4673 | 1,55418E-11 |
| ACE2 | LINC00271 | -0,4673 | 1,56628E-11 |
| ACE2 | RP11-399K21.14 | -0,4673 | 1,55115E-11 |
| ACE2 | HOXC-AS3 | -0,4673 | 1,5663E-11 |
| ACE2 | Z69720.2 | -0,4673 | 1,55119E-11 |
| ACE2 | PTPN2 | -0,4673 | 1,5615E-11 |
| ACE2 | TMEM189 | -0,4673 | 1,56517E-11 |
| ACE2 | RBMS2P1 | -0,4674 | 1,54059E-11 |
| ACE2 | SPNS1 | -0,4674 | 1,53731E-11 |
| ACE2 | RP5-906A24.1 | -0,4674 | 1,54016E-11 |
| ACE2 | RCHY1 | -0,4675 | 1,52861E-11 |
| ACE2 | PUS3 | -0,4675 | 1,52945E-11 |
| ACE2 | RP11-83B20.1 | -0,4675 | 1,51971E-11 |
| ACE2 | KCNH5 | -0,4675 | 1,53024E-11 |
| ACE2 | CTCF | -0,4675 | 1,52485E-11 |
| ACE2 | RP11-183G22.1 | -0,4676 | 1,51271E-11 |
| ACE2 | ANKRD36B | -0,4676 | 1,50926E-11 |
| ACE2 | CPLX3 | -0,4676 | 1,51211E-11 |
| ACE2 | CCDC43 | -0,4676 | 1,51024E-11 |
| ACE2 | CTB-66B24.1 | -0,4676 | 1,50828E-11 |
| ACE2 | REEP1 | -0,4677 | 1,49032E-11 |

| | | | |
|---|---|---|---|
| ACE2 | ENOX1-AS1 | -0,4677 | 1,4953E-11 |
| ACE2 | AC007228.5 | -0,4677 | 1,48383E-11 |
| ACE2 | LETM1P2 | -0,4677 | 1,49103E-11 |
| ACE2 | SH3PXD2B | -0,4678 | 1,47692E-11 |
| ACE2 | NELFB | -0,4678 | 1,47231E-11 |
| ACE2 | CREBZF | -0,4678 | 1,48182E-11 |
| ACE2 | GPR19 | -0,4678 | 1,46736E-11 |
| ACE2 | EXOC3-AS1 | -0,4679 | 1,45304E-11 |
| ACE2 | RP11-578O24.2 | -0,4679 | 1,45225E-11 |
| ACE2 | PSTK | -0,4679 | 1,45667E-11 |
| ACE2 | RP6-99M1.3 | -0,4679 | 1,45814E-11 |
| ACE2 | RP11-305F18.1 | -0,468 | 1,43533E-11 |
| ACE2 | GPR3 | -0,4681 | 1,42667E-11 |
| ACE2 | CAMK2A | -0,4681 | 1,41727E-11 |
| ACE2 | RP11-645C24.5 | -0,4681 | 1,42224E-11 |
| ACE2 | PPP1R9B | -0,4681 | 1,4168E-11 |
| ACE2 | RP4-686C3.7 | -0,4681 | 1,42088E-11 |
| ACE2 | RP11-1060G2.2 | -0,4682 | 1,41317E-11 |
| ACE2 | CTNNA2 | -0,4683 | 1,38883E-11 |
| ACE2 | RWDD2A | -0,4683 | 1,39353E-11 |
| ACE2 | NPL | -0,4684 | 1,37807E-11 |
| ACE2 | RP11-161D15.3 | -0,4684 | 1,38481E-11 |
| ACE2 | C9orf139 | -0,4684 | 1,3774E-11 |
| ACE2 | SYCE1 | -0,4684 | 1,37564E-11 |
| ACE2 | AMDHD1 | -0,4684 | 1,37048E-11 |
| ACE2 | FAM184B | -0,4685 | 1,35424E-11 |
| ACE2 | DDHD2 | -0,4685 | 1,36484E-11 |
| ACE2 | TPCN2 | -0,4685 | 1,35871E-11 |
| ACE2 | ZNF785 | -0,4685 | 1,3605E-11 |
| ACE2 | CHERP | -0,4685 | 1,36658E-11 |
| ACE2 | CTC-559E9.6 | -0,4685 | 1,35662E-11 |
| ACE2 | ZNF234 | -0,4685 | 1,35563E-11 |
| ACE2 | UNKL | -0,4686 | 1,34E-11 |
| ACE2 | SLED1 | -0,4687 | 1,32881E-11 |
| ACE2 | TBC1D29 | -0,4687 | 1,33577E-11 |
| ACE2 | LINC01271 | -0,4687 | 1,33185E-11 |
| ACE2 | RN7SL861P | -0,4688 | 1,32216E-11 |
| ACE2 | FAM210A | -0,4688 | 1,31632E-11 |
| ACE2 | RP11-396C23.4 | -0,4689 | 1,30322E-11 |
| ACE2 | POLR2L | -0,469 | 1,28684E-11 |
| ACE2 | H19 | -0,469 | 1,28864E-11 |
| ACE2 | RP11-820I16.1 | -0,469 | 1,28427E-11 |
| ACE2 | PCED1A | -0,469 | 1,29129E-11 |
| ACE2 | YIPF7 | -0,4691 | 1,27907E-11 |
| ACE2 | RP11-363E6.4 | -0,4691 | 1,27692E-11 |
| ACE2 | RP11-599J14.2 | -0,4691 | 1,26953E-11 |
| ACE2 | FSTL5 | -0,4692 | 1,25829E-11 |
| ACE2 | RPS13P2 | -0,4693 | 1,2376E-11 |
| ACE2 | RP11-1415C14.4 | -0,4693 | 1,24031E-11 |
| ACE2 | HSPA4 | -0,4694 | 1,23261E-11 |
| ACE2 | MESDC1 | -0,4694 | 1,22684E-11 |
| ACE2 | C19orf70 | -0,4694 | 1,23643E-11 |
| ACE2 | RP11-138B9.1 | -0,4695 | 1,22149E-11 |
| ACE2 | WBP2 | -0,4695 | 1,21249E-11 |

| | | | |
|---|---|---|---|
| ACE2 | RAP1GDS1 | -0,4696 | 1,20513E-11 |
| ACE2 | RP11-498P14.3 | -0,4696 | 1,20775E-11 |
| ACE2 | ABCA2 | -0,4696 | 1,20286E-11 |
| ACE2 | C12orf54 | -0,4696 | 1,19576E-11 |
| ACE2 | RP11-530C5.4 | -0,4696 | 1,19746E-11 |
| ACE2 | ISG15 | -0,4697 | 1,18331E-11 |
| ACE2 | CEMIP | -0,4697 | 1,18563E-11 |
| ACE2 | APCDD1L-AS1 | -0,4697 | 1,18857E-11 |
| ACE2 | NOA1 | -0,4698 | 1,18035E-11 |
| ACE2 | TEX15 | -0,4698 | 1,17017E-11 |
| ACE2 | RP11-10L7.1 | -0,4699 | 1,16143E-11 |
| ACE2 | SYNPO2L | -0,4699 | 1,16114E-11 |
| ACE2 | RP11-838N2.5 | -0,47 | 1,1502E-11 |
| ACE2 | HYAL2 | -0,4702 | 1,12339E-11 |
| ACE2 | CTC-232P5.1 | -0,4702 | 1,11658E-11 |
| ACE2 | RP11-196G18.22 | -0,4703 | 1,11186E-11 |
| ACE2 | LYPLAL1 | -0,4703 | 1,10846E-11 |
| ACE2 | ZNF669 | -0,4703 | 1,11169E-11 |
| ACE2 | RP11-145M9.5 | -0,4703 | 1,11077E-11 |
| ACE2 | PIP5K1A | -0,4704 | 1,10131E-11 |
| ACE2 | DAZL | -0,4704 | 1,09995E-11 |
| ACE2 | ABHD16A | -0,4704 | 1,09665E-11 |
| ACE2 | LILRB3 | -0,4704 | 1,09554E-11 |
| ACE2 | HCRTR1 | -0,4706 | 1,06745E-11 |
| ACE2 | CTC-512J14.7 | -0,4706 | 1,07822E-11 |
| ACE2 | MKLN1 | -0,4706 | 1,06828E-11 |
| ACE2 | CTC-429P9.3 | -0,4706 | 1,07035E-11 |
| ACE2 | KCNMB2-AS1 | -0,4707 | 1,06049E-11 |
| ACE2 | RP3-331H24.6 | -0,4707 | 1,06619E-11 |
| ACE2 | CTD-2292M16.8 | -0,4707 | 1,05897E-11 |
| ACE2 | RP1-253P7.4 | -0,4707 | 1,06616E-11 |
| ACE2 | MIR1-1HG | -0,4707 | 1,06245E-11 |
| ACE2 | PROKR1 | -0,4708 | 1,04641E-11 |
| ACE2 | SQSTM1 | -0,4708 | 1,04339E-11 |
| ACE2 | WDR24 | -0,4708 | 1,04994E-11 |
| ACE2 | GYPE | -0,4709 | 1,03541E-11 |
| ACE2 | ZFPM2-AS1 | -0,4709 | 1,03579E-11 |
| ACE2 | RP11-932O9.9 | -0,4709 | 1,03139E-11 |
| ACE2 | SPDYE12P | -0,471 | 1,03087E-11 |
| ACE2 | RNASEH2B-AS1 | -0,471 | 1,02452E-11 |
| ACE2 | AB019441.29 | -0,4711 | 1,00926E-11 |
| ACE2 | RP11-158H5.8 | -0,4711 | 1,01859E-11 |
| ACE2 | RP11-20I20.1 | -0,4712 | 1,0007E-11 |
| ACE2 | RBM22 | -0,4712 | 1,00357E-11 |
| ACE2 | SLU7 | -0,4712 | 1,0037E-11 |
| ACE2 | RPS3AP26 | -0,4712 | 1,00177E-11 |
| ACE2 | ATXN7L3B | -0,4712 | 9,96895E-12 |
| ACE2 | ACTG1P17 | -0,4712 | 1,00349E-11 |
| ACE2 | FLII | -0,4712 | 1,00087E-11 |
| ACE2 | DNAH17 | -0,4712 | 1,00757E-11 |
| ACE2 | CTD-2342J14.6 | -0,4712 | 1,00717E-11 |
| ACE2 | LL22NC03-2H8.4 | -0,4712 | 1,0043E-11 |
| ACE2 | KHDRBS2 | -0,4713 | 9,91403E-12 |
| ACE2 | RP11-532N4.2 | -0,4713 | 9,85331E-12 |

| | | | |
|---|---|---|---|
| ACE2 | SNRPG | -0,4714 | 9,78256E-12 |
| ACE2 | PCDHA11 | -0,4714 | 9,81297E-12 |
| ACE2 | SERF2 | -0,4714 | 9,76762E-12 |
| ACE2 | STRA6 | -0,4714 | 9,79192E-12 |
| ACE2 | RP11-247I13.3 | -0,4714 | 9,80942E-12 |
| ACE2 | CTNS | -0,4715 | 9,6884E-12 |
| ACE2 | NLK | -0,4715 | 9,70876E-12 |
| ACE2 | CTD-3193K9.11 | -0,4715 | 9,65249E-12 |
| ACE2 | TYW1B | -0,4717 | 9,46631E-12 |
| ACE2 | TRAV18 | -0,4717 | 9,43966E-12 |
| ACE2 | SEZ6L | -0,4717 | 9,50111E-12 |
| ACE2 | ZMIZ2 | -0,4718 | 9,36023E-12 |
| ACE2 | GTSF1 | -0,4718 | 9,38677E-12 |
| ACE2 | GOLGA8T | -0,4718 | 9,31538E-12 |
| ACE2 | RP11-192H23.6 | -0,4718 | 9,34402E-12 |
| ACE2 | MFSD10 | -0,4719 | 9,24708E-12 |
| ACE2 | CTD-2248H3.1 | -0,4719 | 9,23254E-12 |
| ACE2 | RP11-131L12.4 | -0,4719 | 9,25067E-12 |
| ACE2 | RP11-368J22.2 | -0,4719 | 9,28205E-12 |
| ACE2 | SLC35E4 | -0,4719 | 9,29932E-12 |
| ACE2 | CCZ1B | -0,472 | 9,12078E-12 |
| ACE2 | CBLN2 | -0,472 | 9,13419E-12 |
| ACE2 | CITF22-92A6.2 | -0,472 | 9,11702E-12 |
| ACE2 | AC092597.3 | -0,4721 | 9,03038E-12 |
| ACE2 | BISPR | -0,4721 | 9,02576E-12 |
| ACE2 | TATDN2 | -0,4722 | 8,93341E-12 |
| ACE2 | AC073130.3 | -0,4722 | 8,96609E-12 |
| ACE2 | CREBBP | -0,4722 | 8,90157E-12 |
| ACE2 | KCNMB2 | -0,4723 | 8,84122E-12 |
| ACE2 | ASB5 | -0,4723 | 8,80627E-12 |
| ACE2 | EIF3LP2 | -0,4723 | 8,80489E-12 |
| ACE2 | HCAR1 | -0,4723 | 8,87006E-12 |
| ACE2 | CTD-3193K9.3 | -0,4723 | 8,86115E-12 |
| ACE2 | SLC7A14 | -0,4724 | 8,73294E-12 |
| ACE2 | RP11-98D18.3 | -0,4725 | 8,61E-12 |
| ACE2 | RP11-258F22.1 | -0,4725 | 8,62695E-12 |
| ACE2 | USP33 | -0,4726 | 8,58623E-12 |
| ACE2 | RP11-110G21.1 | -0,4726 | 8,54729E-12 |
| ACE2 | IFIT1 | -0,4726 | 8,56256E-12 |
| ACE2 | GLYR1 | -0,4726 | 8,5718E-12 |
| ACE2 | TACSTD2 | -0,4727 | 8,45584E-12 |
| ACE2 | RP11-384K6.6 | -0,4727 | 8,43786E-12 |
| ACE2 | CTD-2201E18.3 | -0,4727 | 8,40997E-12 |
| ACE2 | RP11-1081M5.3 | -0,4727 | 8,47833E-12 |
| ACE2 | ARHGEF39 | -0,4727 | 8,47765E-12 |
| ACE2 | LOC440461 | -0,4727 | 8,40174E-12 |
| ACE2 | SPOCK3 | -0,4728 | 8,35524E-12 |
| ACE2 | TNFAIP8 | -0,4728 | 8,34863E-12 |
| ACE2 | LINC01301 | -0,4728 | 8,36497E-12 |
| ACE2 | IPPK | -0,4729 | 8,26625E-12 |
| ACE2 | ORAOV1 | -0,4729 | 8,25239E-12 |
| ACE2 | RP1-292B18.1 | -0,473 | 8,1847E-12 |
| ACE2 | VN1R42P | -0,473 | 8,1478E-12 |
| ACE2 | C16orf47 | -0,473 | 8,11487E-12 |

| | | | |
|---|---|---|---|
| ACE2 | ZNF606 | -0,473 | 8,11963E-12 |
| ACE2 | AFF2 | -0,473 | 8,13115E-12 |
| ACE2 | FRZB | -0,4732 | 7,96274E-12 |
| ACE2 | SERINC1 | -0,4732 | 7,958E-12 |
| ACE2 | RGS9BP | -0,4732 | 7,97818E-12 |
| ACE2 | WASH2P | -0,4733 | 7,90192E-12 |
| ACE2 | UNC5C | -0,4734 | 7,78255E-12 |
| ACE2 | VIP | -0,4734 | 7,79294E-12 |
| ACE2 | PHACTR1 | -0,4735 | 7,74298E-12 |
| ACE2 | MTDH | -0,4735 | 7,70192E-12 |
| ACE2 | TAS2R20 | -0,4735 | 7,66926E-12 |
| ACE2 | KDM2B | -0,4735 | 7,66688E-12 |
| ACE2 | TEFM | -0,4735 | 7,69639E-12 |
| ACE2 | GPC3 | -0,4735 | 7,65852E-12 |
| ACE2 | RP11-165A20.3 | -0,4736 | 7,60763E-12 |
| ACE2 | RP11-429A20.3 | -0,4736 | 7,64438E-12 |
| ACE2 | PLK5 | -0,4736 | 7,59638E-12 |
| ACE2 | CTD-3131K8.2 | -0,4736 | 7,61492E-12 |
| ACE2 | LYPD1 | -0,4737 | 7,4901E-12 |
| ACE2 | NMRK2 | -0,4737 | 7,49903E-12 |
| ACE2 | DUXAP8 | -0,4737 | 7,5351E-12 |
| ACE2 | TRMU | -0,4737 | 7,48563E-12 |
| ACE2 | GUSBP2 | -0,4738 | 7,43961E-12 |
| ACE2 | TAF5 | -0,4738 | 7,4254E-12 |
| ACE2 | VARS | -0,4739 | 7,34934E-12 |
| ACE2 | RP11-571M6.18 | -0,4739 | 7,36217E-12 |
| ACE2 | MYBBP1A | -0,4739 | 7,38379E-12 |
| ACE2 | U2AF2 | -0,4739 | 7,34258E-12 |
| ACE2 | ALG1L | -0,474 | 7,23262E-12 |
| ACE2 | RP11-375I20.6 | -0,474 | 7,24227E-12 |
| ACE2 | IER3 | -0,474 | 7,30038E-12 |
| ACE2 | RP11-339B21.10 | -0,474 | 7,27991E-12 |
| ACE2 | KDM4D | -0,474 | 7,2977E-12 |
| ACE2 | RP11-1007O24.2 | -0,474 | 7,26273E-12 |
| ACE2 | ABCB7 | -0,474 | 7,27012E-12 |
| ACE2 | RP11-495P10.1 | -0,4741 | 7,20582E-12 |
| ACE2 | FBLN7 | -0,4741 | 7,18185E-12 |
| ACE2 | ARPC5L | -0,4741 | 7,17866E-12 |
| ACE2 | SMG1P5 | -0,4741 | 7,16147E-12 |
| ACE2 | RP11-1069G10.2 | -0,4742 | 7,12652E-12 |
| ACE2 | BRCA1 | -0,4742 | 7,13815E-12 |
| ACE2 | PTDSS1 | -0,4743 | 7,0172E-12 |
| ACE2 | RP11-110I1.12 | -0,4743 | 6,98621E-12 |
| ACE2 | CTD-2378E12.1 | -0,4743 | 6,98457E-12 |
| ACE2 | CNEP1R1 | -0,4743 | 6,99472E-12 |
| ACE2 | CASKIN2 | -0,4743 | 7,00627E-12 |
| ACE2 | DLGAP1-AS2 | -0,4743 | 7,02883E-12 |
| ACE2 | FBXL17 | -0,4744 | 6,90617E-12 |
| ACE2 | RBAK | -0,4744 | 6,97053E-12 |
| ACE2 | SMUG1 | -0,4744 | 6,91731E-12 |
| ACE2 | CA14 | -0,4746 | 6,80388E-12 |
| ACE2 | MGAT1 | -0,4746 | 6,79794E-12 |
| ACE2 | MRPL42 | -0,4746 | 6,81917E-12 |
| ACE2 | P2RX2 | -0,4746 | 6,82065E-12 |

| | | | |
|---|---|---|---|
| ACE2 | RP11-467L19.16 | -0,4746 | 6,76213E-12 |
| ACE2 | BCAS4 | -0,4746 | 6,79434E-12 |
| ACE2 | C22orf15 | -0,4746 | 6,81615E-12 |
| ACE2 | DISP1 | -0,4747 | 6,71461E-12 |
| ACE2 | FOXD4L1 | -0,4747 | 6,68864E-12 |
| ACE2 | GS1-72M22.1 | -0,4747 | 6,68752E-12 |
| ACE2 | KCTD21-AS1 | -0,4747 | 6,7285E-12 |
| ACE2 | CARD8 | -0,4747 | 6,70024E-12 |
| ACE2 | OTOF | -0,4748 | 6,59974E-12 |
| ACE2 | AC013474.4 | -0,4748 | 6,61933E-12 |
| ACE2 | DDX41 | -0,4748 | 6,6617E-12 |
| ACE2 | ACKR3 | -0,4749 | 6,53745E-12 |
| ACE2 | SETD3 | -0,4749 | 6,57277E-12 |
| ACE2 | MATN1 | -0,475 | 6,45745E-12 |
| ACE2 | RP11-1191J2.5 | -0,475 | 6,51256E-12 |
| ACE2 | TMC1 | -0,475 | 6,49094E-12 |
| ACE2 | TMEFF2 | -0,4751 | 6,41752E-12 |
| ACE2 | RP11-339B21.8 | -0,4751 | 6,40716E-12 |
| ACE2 | RP11-731J8.2 | -0,4753 | 6,25743E-12 |
| ACE2 | GGA2 | -0,4753 | 6,22115E-12 |
| ACE2 | RP3-453C12.15 | -0,4753 | 6,28718E-12 |
| ACE2 | DLEU2 | -0,4754 | 6,16302E-12 |
| ACE2 | RP11-358B23.5 | -0,4754 | 6,19986E-12 |
| ACE2 | RP11-45A17.4 | -0,4755 | 6,13137E-12 |
| ACE2 | BEAN1 | -0,4755 | 6,0897E-12 |
| ACE2 | AC002398.11 | -0,4755 | 6,1007E-12 |
| ACE2 | GIGYF1 | -0,4756 | 6,04911E-12 |
| ACE2 | C14orf39 | -0,4756 | 6,01531E-12 |
| ACE2 | NARF-IT1 | -0,4757 | 5,96095E-12 |
| ACE2 | RP4-616B8.5 | -0,4757 | 5,94592E-12 |
| ACE2 | LCMT2 | -0,4758 | 5,88857E-12 |
| ACE2 | RP11-395N3.1 | -0,4759 | 5,8558E-12 |
| ACE2 | RP1-197B17.3 | -0,4759 | 5,82821E-12 |
| ACE2 | USP7 | -0,4759 | 5,80995E-12 |
| ACE2 | RPL23AP2 | -0,4759 | 5,86577E-12 |
| ACE2 | ZNF253 | -0,4759 | 5,80854E-12 |
| ACE2 | PCP4 | -0,4759 | 5,84153E-12 |
| ACE2 | DENND4B | -0,476 | 5,74391E-12 |
| ACE2 | LDAH | -0,476 | 5,80363E-12 |
| ACE2 | MCPH1-AS1 | -0,476 | 5,75596E-12 |
| ACE2 | ODF2-AS1 | -0,476 | 5,76954E-12 |
| ACE2 | TNFSF13B | -0,476 | 5,75644E-12 |
| ACE2 | CTD-3088G3.8 | -0,476 | 5,73903E-12 |
| ACE2 | RP11-666A8.12 | -0,476 | 5,75445E-12 |
| ACE2 | CTC-471F3.5 | -0,476 | 5,75445E-12 |
| ACE2 | C15orf57 | -0,4761 | 5,70344E-12 |
| ACE2 | RP11-35G9.3 | -0,4761 | 5,71635E-12 |
| ACE2 | FAM168B | -0,4762 | 5,62974E-12 |
| ACE2 | PDZRN3-AS1 | -0,4763 | 5,56848E-12 |
| ACE2 | IL20RB | -0,4763 | 5,54315E-12 |
| ACE2 | RP11-864N7.2 | -0,4763 | 5,54635E-12 |
| ACE2 | FOXRED1 | -0,4763 | 5,5714E-12 |
| ACE2 | ADSS | -0,4764 | 5,48036E-12 |
| ACE2 | PAPD7 | -0,4764 | 5,51621E-12 |

| | | | |
|---|---|---|---|
| ACE2 | INSRR | -0,4765 | 5,46256E-12 |
| ACE2 | AF131216.6 | -0,4765 | 5,42906E-12 |
| ACE2 | SCX | -0,4765 | 5,41579E-12 |
| ACE2 | RP11-371A19.2 | -0,4765 | 5,43331E-12 |
| ACE2 | RP11-672A2.4 | -0,4765 | 5,42532E-12 |
| ACE2 | PLCD3 | -0,4765 | 5,46633E-12 |
| ACE2 | SNAP25 | -0,4765 | 5,45644E-12 |
| ACE2 | RNU6-8 | -0,4766 | 5,37096E-12 |
| ACE2 | TMEM151B | -0,4767 | 5,3295E-12 |
| ACE2 | RP11-154D17.1 | -0,4767 | 5,32794E-12 |
| ACE2 | AC069257.6 | -0,4768 | 5,28543E-12 |
| ACE2 | STMN4 | -0,4768 | 5,23338E-12 |
| ACE2 | RP11-399E6.1 | -0,4769 | 5,22328E-12 |
| ACE2 | AC107081.5 | -0,4769 | 5,19225E-12 |
| ACE2 | H1FX-AS1 | -0,477 | 5,12131E-12 |
| ACE2 | ATG14 | -0,477 | 5,13623E-12 |
| ACE2 | COMT | -0,477 | 5,13023E-12 |
| ACE2 | MSL3 | -0,477 | 5,13858E-12 |
| ACE2 | RWDD4P2 | -0,4771 | 5,06835E-12 |
| ACE2 | RBM20 | -0,4771 | 5,08566E-12 |
| ACE2 | NLRP1 | -0,4771 | 5,08269E-12 |
| ACE2 | RN7SL566P | -0,4771 | 5,09165E-12 |
| ACE2 | DRAXIN | -0,4772 | 4,99491E-12 |
| ACE2 | AHSA2 | -0,4772 | 5,02202E-12 |
| ACE2 | MAP1S | -0,4772 | 5,01325E-12 |
| ACE2 | RP11-686O6.2 | -0,4773 | 4,96246E-12 |
| ACE2 | TMEM175 | -0,4773 | 4,96723E-12 |
| ACE2 | CTC-340A15.2 | -0,4773 | 4,95512E-12 |
| ACE2 | SPDL1 | -0,4773 | 4,95309E-12 |
| ACE2 | SYT16 | -0,4774 | 4,91493E-12 |
| ACE2 | RP11-488C13.5 | -0,4774 | 4,91448E-12 |
| ACE2 | P2RY12 | -0,4775 | 4,85903E-12 |
| ACE2 | PHOX2B | -0,4776 | 4,78048E-12 |
| ACE2 | ACSBG1 | -0,4776 | 4,81111E-12 |
| ACE2 | SH3GLB1 | -0,4777 | 4,74968E-12 |
| ACE2 | PPM1B | -0,4777 | 4,75559E-12 |
| ACE2 | RP11-44N22.3 | -0,4777 | 4,72844E-12 |
| ACE2 | GP5 | -0,4777 | 4,75921E-12 |
| ACE2 | RP11-484O2.1 | -0,4777 | 4,71137E-12 |
| ACE2 | RP5-994D16.3 | -0,4778 | 4,67399E-12 |
| ACE2 | PSMD2 | -0,4778 | 4,67837E-12 |
| ACE2 | PSMG4 | -0,4778 | 4,66166E-12 |
| ACE2 | AC135178.7 | -0,4778 | 4,70277E-12 |
| ACE2 | PER3 | -0,4779 | 4,61857E-12 |
| ACE2 | EPHX1 | -0,4779 | 4,6436E-12 |
| ACE2 | CNIH2 | -0,4779 | 4,62514E-12 |
| ACE2 | TGFB2-AS1 | -0,478 | 4,59146E-12 |
| ACE2 | MTATP6P26 | -0,478 | 4,58472E-12 |
| ACE2 | RP11-791G15.2 | -0,4781 | 4,54626E-12 |
| ACE2 | CTSO | -0,4781 | 4,50869E-12 |
| ACE2 | ZFP62 | -0,4781 | 4,53778E-12 |
| ACE2 | GIMAP7 | -0,4781 | 4,50327E-12 |
| ACE2 | GOLGA1 | -0,4781 | 4,51225E-12 |
| ACE2 | GAS2 | -0,4781 | 4,49631E-12 |

| | | | |
|---|---|---|---|
| ACE2 | RP11-677M14.7 | -0,4781 | 4,50568E-12 |
| ACE2 | RPL24P2 | -0,4781 | 4,50605E-12 |
| ACE2 | GTF2B | -0,4782 | 4,47197E-12 |
| ACE2 | LINC01759 | -0,4783 | 4,39346E-12 |
| ACE2 | HSPD1P4 | -0,4783 | 4,42943E-12 |
| ACE2 | FBXO5 | -0,4784 | 4,36602E-12 |
| ACE2 | ZNF778 | -0,4784 | 4,35796E-12 |
| ACE2 | PFAS | -0,4784 | 4,38184E-12 |
| ACE2 | RP1-228H13.5 | -0,4785 | 4,33096E-12 |
| ACE2 | CTC-260E6.4 | -0,4785 | 4,28984E-12 |
| ACE2 | AP006222.2 | -0,4786 | 4,24557E-12 |
| ACE2 | LINC01783 | -0,4786 | 4,27539E-12 |
| ACE2 | UBXN10-AS1 | -0,4786 | 4,28479E-12 |
| ACE2 | KIF9 | -0,4786 | 4,25781E-12 |
| ACE2 | RP11-43F13.4 | -0,4786 | 4,28512E-12 |
| ACE2 | C5orf58 | -0,4786 | 4,28144E-12 |
| ACE2 | RP11-168P13.1 | -0,4786 | 4,25484E-12 |
| ACE2 | RP11-490K7.4 | -0,4787 | 4,22095E-12 |
| ACE2 | ZCCHC4 | -0,4787 | 4,23252E-12 |
| ACE2 | PCDHA6 | -0,4787 | 4,22129E-12 |
| ACE2 | PPHLN1 | -0,4787 | 4,20122E-12 |
| ACE2 | ARF4-AS1 | -0,4789 | 4,11635E-12 |
| ACE2 | FAM86JP | -0,4789 | 4,133E-12 |
| ACE2 | MTRNR2L4 | -0,4789 | 4,13764E-12 |
| ACE2 | KCNQ2 | -0,4789 | 4,10813E-12 |
| ACE2 | SLC16A1 | -0,479 | 4,07181E-12 |
| ACE2 | LRRTM3 | -0,479 | 4,09282E-12 |
| ACE2 | RP11-114F10.3 | -0,479 | 4,08954E-12 |
| ACE2 | NCK1-AS1 | -0,4791 | 4,03222E-12 |
| ACE2 | GPX3 | -0,4791 | 4,01223E-12 |
| ACE2 | ALKBH3-AS1 | -0,4791 | 4,01041E-12 |
| ACE2 | LONP2 | -0,4791 | 4,03125E-12 |
| ACE2 | LINC01534 | -0,4791 | 4,03937E-12 |
| ACE2 | TWF2 | -0,4792 | 3,95746E-12 |
| ACE2 | FAM153C | -0,4792 | 3,9909E-12 |
| ACE2 | SNORA65 | -0,4792 | 3,96661E-12 |
| ACE2 | RP11-1081L13.4 | -0,4792 | 3,95204E-12 |
| ACE2 | PUS1 | -0,4792 | 3,97398E-12 |
| ACE2 | FUCA1P1 | -0,4793 | 3,93169E-12 |
| ACE2 | ALCAM | -0,4793 | 3,95162E-12 |
| ACE2 | CTB-1144G6.6 | -0,4793 | 3,93138E-12 |
| ACE2 | RPS2P5 | -0,4793 | 3,92211E-12 |
| ACE2 | RP11-266L9.3 | -0,4793 | 3,92015E-12 |
| ACE2 | C5orf56 | -0,4794 | 3,87861E-12 |
| ACE2 | RIC1 | -0,4794 | 3,88123E-12 |
| ACE2 | PSAP | -0,4794 | 3,88958E-12 |
| ACE2 | CHD9 | -0,4794 | 3,87049E-12 |
| ACE2 | FKBP9 | -0,4795 | 3,83797E-12 |
| ACE2 | SEPHS1P1 | -0,4795 | 3,84706E-12 |
| ACE2 | RP11-180C16.1 | -0,4795 | 3,83187E-12 |
| ACE2 | TNKS2 | -0,4796 | 3,77512E-12 |
| ACE2 | HIPK3 | -0,4796 | 3,78447E-12 |
| ACE2 | ATAD3B | -0,4797 | 3,74428E-12 |
| ACE2 | C12orf66 | -0,4797 | 3,72961E-12 |

| | | | |
|---|---|---|---|
| ACE2 | COQ6 | -0,4798 | 3,68473E-12 |
| ACE2 | CSNK1G1 | -0,4798 | 3,71099E-12 |
| ACE2 | RP5-826L7.1 | -0,4799 | 3,65138E-12 |
| ACE2 | ZFC3H1 | -0,4799 | 3,66593E-12 |
| ACE2 | AEN | -0,4799 | 3,64524E-12 |
| ACE2 | TEX13D | -0,4799 | 3,66854E-12 |
| ACE2 | CTDSPL | -0,48 | 3,63519E-12 |
| ACE2 | HOGA1 | -0,48 | 3,62186E-12 |
| ACE2 | DCAF4 | -0,48 | 3,61651E-12 |
| ACE2 | ZNF75A | -0,48 | 3,60564E-12 |
| ACE2 | HNRNPL | -0,48 | 3,60679E-12 |
| ACE2 | TMEM257 | -0,48 | 3,6394E-12 |
| ACE2 | DPH6-AS1 | -0,4801 | 3,58855E-12 |
| ACE2 | BACE2 | -0,4801 | 3,5672E-12 |
| ACE2 | SOAT1 | -0,4802 | 3,5342E-12 |
| ACE2 | PFKFB3 | -0,4802 | 3,54534E-12 |
| ACE2 | MAX | -0,4802 | 3,55097E-12 |
| ACE2 | RPL12P4 | -0,4802 | 3,53189E-12 |
| ACE2 | RP11-263K19.6 | -0,4803 | 3,50345E-12 |
| ACE2 | CCDC13 | -0,4803 | 3,50307E-12 |
| ACE2 | OCIAD1 | -0,4803 | 3,48838E-12 |
| ACE2 | RP11-9G1.3 | -0,4803 | 3,48878E-12 |
| ACE2 | RP11-727F15.13 | -0,4803 | 3,50572E-12 |
| ACE2 | RFC3 | -0,4803 | 3,48959E-12 |
| ACE2 | FKBP4 | -0,4804 | 3,44204E-12 |
| ACE2 | GABRD | -0,4805 | 3,43101E-12 |
| ACE2 | ABCB5 | -0,4805 | 3,4056E-12 |
| ACE2 | RP11-371A22.1 | -0,4805 | 3,42493E-12 |
| ACE2 | DICER1-AS1 | -0,4805 | 3,39594E-12 |
| ACE2 | CNOT3 | -0,4805 | 3,43001E-12 |
| ACE2 | PCYT1B | -0,4805 | 3,40952E-12 |
| ACE2 | PID1 | -0,4806 | 3,38808E-12 |
| ACE2 | GATB | -0,4806 | 3,37123E-12 |
| ACE2 | C10orf107 | -0,4806 | 3,3744E-12 |
| ACE2 | TCEANC | -0,4806 | 3,37505E-12 |
| ACE2 | SAMD15 | -0,4807 | 3,32854E-12 |
| ACE2 | RP11-894P9.1 | -0,4807 | 3,3398E-12 |
| ACE2 | RP11-666O2.2 | -0,4807 | 3,3218E-12 |
| ACE2 | UTP18 | -0,4807 | 3,3238E-12 |
| ACE2 | ZNF805 | -0,4807 | 3,32137E-12 |
| ACE2 | CTD-2541M15.3 | -0,4808 | 3,29737E-12 |
| ACE2 | TMEM5-AS1 | -0,4808 | 3,28068E-12 |
| ACE2 | RSBN1 | -0,4809 | 3,23894E-12 |
| ACE2 | TSPYL1 | -0,4809 | 3,24426E-12 |
| ACE2 | KRT222 | -0,4809 | 3,24838E-12 |
| ACE2 | NPPC | -0,481 | 3,23379E-12 |
| ACE2 | CTA-223H9.9 | -0,481 | 3,23202E-12 |
| ACE2 | AC009404.2 | -0,4811 | 3,18634E-12 |
| ACE2 | CTD-2308L22.1 | -0,4811 | 3,16731E-12 |
| ACE2 | RP3-337H4.6 | -0,4811 | 3,17841E-12 |
| ACE2 | BUB3 | -0,4811 | 3,16291E-12 |
| ACE2 | MTFMT | -0,4811 | 3,16539E-12 |
| ACE2 | ADGRL1 | -0,4811 | 3,17697E-12 |
| ACE2 | B4GALT3 | -0,4812 | 3,15174E-12 |

| | | | |
|---|---|---|---|
| ACE2 | KLF14 | -0,4812 | 3,14443E-12 |
| ACE2 | JMJD7 | -0,4812 | 3,1526E-12 |
| ACE2 | HNRNPA2B1 | -0,4813 | 3,09658E-12 |
| ACE2 | KRT86 | -0,4813 | 3,09766E-12 |
| ACE2 | CTD-2561B21.5 | -0,4813 | 3,09595E-12 |
| ACE2 | LIMK1 | -0,4814 | 3,07946E-12 |
| ACE2 | FAS | -0,4814 | 3,07929E-12 |
| ACE2 | GAD1 | -0,4815 | 3,03614E-12 |
| ACE2 | SIAH1 | -0,4815 | 3,03604E-12 |
| ACE2 | LINC00670 | -0,4815 | 3,0346E-12 |
| ACE2 | CTD-3113P16.11 | -0,4815 | 3,04544E-12 |
| ACE2 | ITCH | -0,4815 | 3,03072E-12 |
| ACE2 | LRRC7 | -0,4816 | 2,98342E-12 |
| ACE2 | RP11-692D12.1 | -0,4816 | 3,00086E-12 |
| ACE2 | RP11-332H14.2 | -0,4817 | 2,94937E-12 |
| ACE2 | RP11-208N14.4 | -0,4817 | 2,96978E-12 |
| ACE2 | CUL9 | -0,4817 | 2,97625E-12 |
| ACE2 | THAP1 | -0,4817 | 2,96698E-12 |
| ACE2 | TTLL9 | -0,4817 | 2,97511E-12 |
| ACE2 | RP11-474O21.5 | -0,4819 | 2,88435E-12 |
| ACE2 | RP11-673E1.1 | -0,4819 | 2,8771E-12 |
| ACE2 | PPP4R2 | -0,482 | 2,85954E-12 |
| ACE2 | FAM193A | -0,482 | 2,84901E-12 |
| ACE2 | MANEA | -0,482 | 2,84653E-12 |
| ACE2 | FOXN3 | -0,482 | 2,84677E-12 |
| ACE2 | HMGB1P10 | -0,4821 | 2,81667E-12 |
| ACE2 | TPRKB | -0,4822 | 2,78028E-12 |
| ACE2 | OR7E13P | -0,4822 | 2,78724E-12 |
| ACE2 | HIST2H3D | -0,4823 | 2,76796E-12 |
| ACE2 | CNOT10 | -0,4823 | 2,74528E-12 |
| ACE2 | OR5K2 | -0,4823 | 2,76844E-12 |
| ACE2 | ARL1 | -0,4823 | 2,77288E-12 |
| ACE2 | DCAF6 | -0,4825 | 2,70254E-12 |
| ACE2 | RP11-15A1.4 | -0,4825 | 2,69146E-12 |
| ACE2 | TNFRSF6B | -0,4825 | 2,68892E-12 |
| ACE2 | BLZF1 | -0,4826 | 2,67465E-12 |
| ACE2 | PRSS57 | -0,4826 | 2,65456E-12 |
| ACE2 | ZNF350 | -0,4826 | 2,66566E-12 |
| ACE2 | ADCYAP1 | -0,4827 | 2,61879E-12 |
| ACE2 | RP11-343J3.2 | -0,4828 | 2,59135E-12 |
| ACE2 | DPY19L1P1 | -0,4829 | 2,56882E-12 |
| ACE2 | BAG4 | -0,4829 | 2,57693E-12 |
| ACE2 | KIN | -0,4829 | 2,55594E-12 |
| ACE2 | DCAF4L1 | -0,483 | 2,52897E-12 |
| ACE2 | CACNG4 | -0,483 | 2,55413E-12 |
| ACE2 | CTC-459F4.9 | -0,483 | 2,55336E-12 |
| ACE2 | C5AR1 | -0,483 | 2,55309E-12 |
| ACE2 | ATG13 | -0,4831 | 2,50735E-12 |
| ACE2 | SVIL-AS1 | -0,4832 | 2,47905E-12 |
| ACE2 | CACNG7 | -0,4832 | 2,48568E-12 |
| ACE2 | UBQLN1 | -0,4833 | 2,44255E-12 |
| ACE2 | ZNF414 | -0,4833 | 2,44404E-12 |
| ACE2 | ZNF530 | -0,4833 | 2,46359E-12 |
| ACE2 | FANCG | -0,4834 | 2,42257E-12 |

| | | | |
|---|---|---|---|
| ACE2 | ELAVL3 | -0,4834 | 2,42672E-12 |
| ACE2 | ADORA1 | -0,4835 | 2,39182E-12 |
| ACE2 | LMOD3 | -0,4835 | 2,39215E-12 |
| ACE2 | RP11-809O17.1 | -0,4835 | 2,38955E-12 |
| ACE2 | PHB2 | -0,4835 | 2,39288E-12 |
| ACE2 | RP11-753H16.3 | -0,4835 | 2,38844E-12 |
| ACE2 | RNF144A-AS1 | -0,4836 | 2,36319E-12 |
| ACE2 | RP11-1072C15.6 | -0,4836 | 2,37831E-12 |
| ACE2 | CTD-2561B21.8 | -0,4836 | 2,37055E-12 |
| ACE2 | RPS28 | -0,4836 | 2,35715E-12 |
| ACE2 | LRRC3-AS1 | -0,4836 | 2,37964E-12 |
| ACE2 | RNPC3 | -0,4837 | 2,33752E-12 |
| ACE2 | ROPN1B | -0,4837 | 2,32444E-12 |
| ACE2 | HSP90AA5P | -0,4837 | 2,33428E-12 |
| ACE2 | RP11-638I2.9 | -0,4837 | 2,33885E-12 |
| ACE2 | RP11-803D5.1 | -0,4839 | 2,27708E-12 |
| ACE2 | RP11-570L15.1 | -0,4839 | 2,29391E-12 |
| ACE2 | MKRN3 | -0,4839 | 2,2919E-12 |
| ACE2 | PARD6G-AS1 | -0,4839 | 2,27174E-12 |
| ACE2 | LINC01985 | -0,484 | 2,25548E-12 |
| ACE2 | PLPP4 | -0,484 | 2,24646E-12 |
| ACE2 | NOC4L | -0,4841 | 2,23622E-12 |
| ACE2 | ZNF575 | -0,4841 | 2,24159E-12 |
| ACE2 | POLR1B | -0,4842 | 2,19448E-12 |
| ACE2 | PRRT2 | -0,4842 | 2,21598E-12 |
| ACE2 | MTND5P1 | -0,4842 | 2,19203E-12 |
| ACE2 | PERM1 | -0,4844 | 2,16137E-12 |
| ACE2 | LINC00452 | -0,4844 | 2,15026E-12 |
| ACE2 | GON4L | -0,4845 | 2,12694E-12 |
| ACE2 | ARHGAP15 | -0,4845 | 2,11875E-12 |
| ACE2 | CSRNP1 | -0,4845 | 2,12612E-12 |
| ACE2 | LGR6 | -0,4846 | 2,10261E-12 |
| ACE2 | CTD-2035E11.5 | -0,4846 | 2,10156E-12 |
| ACE2 | PLXDC2 | -0,4846 | 2,10255E-12 |
| ACE2 | COX18 | -0,4847 | 2,08441E-12 |
| ACE2 | COPG2 | -0,4847 | 2,08321E-12 |
| ACE2 | BLOC1S2 | -0,4847 | 2,06829E-12 |
| ACE2 | PABPN1 | -0,4847 | 2,06587E-12 |
| ACE2 | RP3-414A15.10 | -0,4847 | 2,06904E-12 |
| ACE2 | CLK3 | -0,4847 | 2,07747E-12 |
| ACE2 | KLHDC4 | -0,4847 | 2,07191E-12 |
| ACE2 | IQCC | -0,4848 | 2,0497E-12 |
| ACE2 | MCEE | -0,4848 | 2,0555E-12 |
| ACE2 | RP11-121C2.2 | -0,4848 | 2,05537E-12 |
| ACE2 | MAN1B1 | -0,4848 | 2,04377E-12 |
| ACE2 | RP1-261D10.2 | -0,4848 | 2,0592E-12 |
| ACE2 | RP11-173M1.4 | -0,4849 | 2,03237E-12 |
| ACE2 | CTD-3138B18.6 | -0,4849 | 2,02612E-12 |
| ACE2 | RP11-181C21.4 | -0,485 | 2,00712E-12 |
| ACE2 | NT5C2 | -0,485 | 2,00129E-12 |
| ACE2 | C11orf65 | -0,485 | 2,00435E-12 |
| ACE2 | RP11-438D14.2 | -0,485 | 2,00281E-12 |
| ACE2 | CACNB3 | -0,485 | 2,00441E-12 |
| ACE2 | LRBA | -0,4851 | 1,97322E-12 |

| | | | |
|---|---|---|---|
| ACE2 | RP5-855D21.3 | -0,4851 | 1,98779E-12 |
| ACE2 | AC005082.12 | -0,4852 | 1,94728E-12 |
| ACE2 | CCDC144NL | -0,4852 | 1,9619E-12 |
| ACE2 | URB1-AS1 | -0,4852 | 1,9523E-12 |
| ACE2 | GPSM2 | -0,4854 | 1,91669E-12 |
| ACE2 | NPIPB9 | -0,4854 | 1,9111E-12 |
| ACE2 | TVP23C | -0,4854 | 1,91759E-12 |
| ACE2 | RP11-427M20.1 | -0,4855 | 1,8868E-12 |
| ACE2 | MIR222HG | -0,4855 | 1,88213E-12 |
| ACE2 | RP11-455J20.3 | -0,4856 | 1,86028E-12 |
| ACE2 | RP11-539G18.3 | -0,4856 | 1,87246E-12 |
| ACE2 | PRAL | -0,4856 | 1,87526E-12 |
| ACE2 | ADGRE5 | -0,4856 | 1,86835E-12 |
| ACE2 | PLVAP | -0,4856 | 1,86774E-12 |
| ACE2 | AP000254.8 | -0,4856 | 1,86294E-12 |
| ACE2 | SEMA5B | -0,4857 | 1,85208E-12 |
| ACE2 | RP11-147L13.13 | -0,4857 | 1,8358E-12 |
| ACE2 | CBFA2T2 | -0,4857 | 1,85093E-12 |
| ACE2 | AL022344.7 | -0,4859 | 1,78859E-12 |
| ACE2 | NTF3 | -0,4859 | 1,80117E-12 |
| ACE2 | TUBA1B | -0,4859 | 1,80509E-12 |
| ACE2 | SUMO4 | -0,486 | 1,76959E-12 |
| ACE2 | GJC3 | -0,486 | 1,76665E-12 |
| ACE2 | RPUSD2 | -0,486 | 1,77268E-12 |
| ACE2 | GLTSCR2-AS1 | -0,486 | 1,77248E-12 |
| ACE2 | CDC26 | -0,4861 | 1,7547E-12 |
| ACE2 | AC083843.4 | -0,4862 | 1,73792E-12 |
| ACE2 | ZNF830 | -0,4862 | 1,74457E-12 |
| ACE2 | RP11-4B16.1 | -0,4862 | 1,7327E-12 |
| ACE2 | INTS6-AS1 | -0,4863 | 1,72204E-12 |
| ACE2 | CTDSPL2 | -0,4863 | 1,71873E-12 |
| ACE2 | KCNB1 | -0,4863 | 1,72114E-12 |
| ACE2 | OSBPL9 | -0,4864 | 1,6998E-12 |
| ACE2 | LOX | -0,4864 | 1,69434E-12 |
| ACE2 | CTD-2527I21.5 | -0,4864 | 1,69588E-12 |
| ACE2 | WARS2 | -0,4865 | 1,67447E-12 |
| ACE2 | ATP6V1C2 | -0,4865 | 1,66533E-12 |
| ACE2 | KIAA1257 | -0,4865 | 1,66419E-12 |
| ACE2 | KY | -0,4865 | 1,6837E-12 |
| ACE2 | ERLEC1 | -0,4866 | 1,66218E-12 |
| ACE2 | RP11-637A17.2 | -0,4866 | 1,65616E-12 |
| ACE2 | SETDB1 | -0,4867 | 1,63731E-12 |
| ACE2 | XXbac-BPG154L12.5 | -0,4867 | 1,64369E-12 |
| ACE2 | LINC00565 | -0,4867 | 1,64322E-12 |
| ACE2 | RP4-565E6.1 | -0,4868 | 1,62443E-12 |
| ACE2 | SH2B2 | -0,4869 | 1,60491E-12 |
| ACE2 | RP11-848P1.5 | -0,4869 | 1,58677E-12 |
| ACE2 | ZNF41 | -0,4869 | 1,59865E-12 |
| ACE2 | STEAP2 | -0,487 | 1,57137E-12 |
| ACE2 | KIAA0087 | -0,4871 | 1,55364E-12 |
| ACE2 | GSC | -0,4871 | 1,55039E-12 |
| ACE2 | MRPL47 | -0,4872 | 1,53572E-12 |
| ACE2 | MTO1 | -0,4872 | 1,5303E-12 |
| ACE2 | ZGPAT | -0,4872 | 1,53187E-12 |

| | | | |
|---|---|---|---|
| ACE2 | DCP2 | -0,4873 | 1,51607E-12 |
| ACE2 | SNHG20 | -0,4873 | 1,5295E-12 |
| ACE2 | CTC-425O23.2 | -0,4873 | 1,52845E-12 |
| ACE2 | CACNG8 | -0,4873 | 1,52378E-12 |
| ACE2 | MOB1A | -0,4874 | 1,51052E-12 |
| ACE2 | CRHBP | -0,4874 | 1,49704E-12 |
| ACE2 | LINC01600 | -0,4874 | 1,50946E-12 |
| ACE2 | LINC00092 | -0,4874 | 1,50416E-12 |
| ACE2 | AC005104.3 | -0,4875 | 1,49061E-12 |
| ACE2 | CLVS2 | -0,4875 | 1,47845E-12 |
| ACE2 | HCAR2 | -0,4875 | 1,49156E-12 |
| ACE2 | KBTBD7 | -0,4875 | 1,47902E-12 |
| ACE2 | FAM212B-AS1 | -0,4876 | 1,46053E-12 |
| ACE2 | HIST1H3D | -0,4876 | 1,47332E-12 |
| ACE2 | RP11-771K4.3 | -0,4876 | 1,47535E-12 |
| ACE2 | MYO1H | -0,4876 | 1,46785E-12 |
| ACE2 | BMP1 | -0,4877 | 1,4484E-12 |
| ACE2 | ZNF589 | -0,4878 | 1,4315E-12 |
| ACE2 | SNRPC | -0,4878 | 1,42614E-12 |
| ACE2 | BRF2 | -0,4878 | 1,43285E-12 |
| ACE2 | RHPN1 | -0,4878 | 1,43055E-12 |
| ACE2 | MCTS2P | -0,4878 | 1,43237E-12 |
| ACE2 | ZNF621 | -0,4879 | 1,40809E-12 |
| ACE2 | RACGAP1 | -0,4879 | 1,41437E-12 |
| ACE2 | ZNF410 | -0,4879 | 1,41837E-12 |
| ACE2 | ZNF738 | -0,4879 | 1,41718E-12 |
| ACE2 | ZNFX1 | -0,4879 | 1,41727E-12 |
| ACE2 | RP11-159L20.2 | -0,488 | 1,39848E-12 |
| ACE2 | RPL7AP6 | -0,488 | 1,39031E-12 |
| ACE2 | RP11-1055B8.3 | -0,4881 | 1,3763E-12 |
| ACE2 | KIF1A | -0,4882 | 1,35689E-12 |
| ACE2 | AC004967.7 | -0,4882 | 1,36835E-12 |
| ACE2 | RP11-644N4.1 | -0,4882 | 1,36379E-12 |
| ACE2 | FAM199X | -0,4882 | 1,36122E-12 |
| ACE2 | IFI6 | -0,4883 | 1,34567E-12 |
| ACE2 | LGALS8 | -0,4883 | 1,34152E-12 |
| ACE2 | RP11-2H3.6 | -0,4883 | 1,34092E-12 |
| ACE2 | RP11-887P2.5 | -0,4883 | 1,34814E-12 |
| ACE2 | ACTN2 | -0,4884 | 1,32763E-12 |
| ACE2 | PTPRN | -0,4884 | 1,33219E-12 |
| ACE2 | RP11-830F9.6 | -0,4884 | 1,32899E-12 |
| ACE2 | ST8SIA3 | -0,4884 | 1,3264E-12 |
| ACE2 | FGF22 | -0,4884 | 1,3371E-12 |
| ACE2 | FGFBP3 | -0,4885 | 1,31757E-12 |
| ACE2 | RP11-154J22.1 | -0,4885 | 1,30949E-12 |
| ACE2 | RP11-758M4.4 | -0,4886 | 1,29939E-12 |
| ACE2 | BAMBI | -0,4886 | 1,30132E-12 |
| ACE2 | RP11-770J1.4 | -0,4886 | 1,30308E-12 |
| ACE2 | RP11-167B3.3 | -0,4886 | 1,30257E-12 |
| ACE2 | ZNF526 | -0,4886 | 1,30025E-12 |
| ACE2 | YARS | -0,4887 | 1,28699E-12 |
| ACE2 | RNGTT | -0,4887 | 1,28831E-12 |
| ACE2 | CCND1 | -0,4887 | 1,28863E-12 |
| ACE2 | CTD-3234P18.6 | -0,4887 | 1,284E-12 |

| | | | |
|---|---|---|---|
| ACE2 | RP11-48B3.3 | -0,4888 | 1,27447E-12 |
| ACE2 | AC007038.7 | -0,4889 | 1,24649E-12 |
| ACE2 | LRRC69 | -0,4889 | 1,25009E-12 |
| ACE2 | RBM15B | -0,489 | 1,24032E-12 |
| ACE2 | RP11-1070N10.3 | -0,489 | 1,24445E-12 |
| ACE2 | RP11-227G15.10 | -0,489 | 1,23562E-12 |
| ACE2 | AC005387.2 | -0,489 | 1,2363E-12 |
| ACE2 | NR0B1 | -0,489 | 1,24345E-12 |
| ACE2 | AC074289.1 | -0,4891 | 1,22838E-12 |
| ACE2 | FIP1L1 | -0,4891 | 1,22165E-12 |
| ACE2 | CFAP206 | -0,4891 | 1,22754E-12 |
| ACE2 | CA8 | -0,4891 | 1,23114E-12 |
| ACE2 | SH3GLB2 | -0,4891 | 1,21975E-12 |
| ACE2 | RP11-111M22.2 | -0,4892 | 1,2067E-12 |
| ACE2 | C14orf93 | -0,4892 | 1,2159E-12 |
| ACE2 | KRBA2 | -0,4892 | 1,20579E-12 |
| ACE2 | LRRC75A-AS1 | -0,4892 | 1,21214E-12 |
| ACE2 | MRPL39 | -0,4892 | 1,20466E-12 |
| ACE2 | STON1-GTF2A1L | -0,4893 | 1,19217E-12 |
| ACE2 | SRPRA | -0,4893 | 1,18809E-12 |
| ACE2 | NBEAP1 | -0,4893 | 1,1894E-12 |
| ACE2 | RP11-464D20.2 | -0,4893 | 1,19282E-12 |
| ACE2 | SNIP1 | -0,4894 | 1,17929E-12 |
| ACE2 | WDFY3 | -0,4894 | 1,17987E-12 |
| ACE2 | ZNF346-IT1 | -0,4894 | 1,17857E-12 |
| ACE2 | NFIL3 | -0,4894 | 1,18086E-12 |
| ACE2 | TINAGL1 | -0,4895 | 1,16772E-12 |
| ACE2 | KRT18P5 | -0,4895 | 1,1637E-12 |
| ACE2 | SOHLH2 | -0,4896 | 1,15523E-12 |
| ACE2 | KHSRP | -0,4896 | 1,15469E-12 |
| ACE2 | CTD-2623N2.11 | -0,4897 | 1,13943E-12 |
| ACE2 | MPV17L2 | -0,4897 | 1,13756E-12 |
| ACE2 | LINC00222 | -0,4898 | 1,12774E-12 |
| ACE2 | TADA2A | -0,4898 | 1,12587E-12 |
| ACE2 | WDR18 | -0,4898 | 1,11887E-12 |
| ACE2 | C1orf50 | -0,4899 | 1,11008E-12 |
| ACE2 | MAP3K14 | -0,4899 | 1,11583E-12 |
| ACE2 | LIAS | -0,49 | 1,09557E-12 |
| ACE2 | RP11-2C24.5 | -0,4901 | 1,08414E-12 |
| ACE2 | RP11-456P18.2 | -0,4902 | 1,06648E-12 |
| ACE2 | RP11-6I2.3 | -0,4902 | 1,07634E-12 |
| ACE2 | NCBP1 | -0,4903 | 1,05717E-12 |
| ACE2 | PRPF19 | -0,4903 | 1,06213E-12 |
| ACE2 | C19orf66 | -0,4903 | 1,06239E-12 |
| ACE2 | IFNAR2 | -0,4903 | 1,06092E-12 |
| ACE2 | UBL4A | -0,4903 | 1,06343E-12 |
| ACE2 | RP11-353N4.6 | -0,4904 | 1,04995E-12 |
| ACE2 | RP11-230C9.4 | -0,4904 | 1,0449E-12 |
| ACE2 | ADHFE1 | -0,4904 | 1,04724E-12 |
| ACE2 | FMR1 | -0,4905 | 1,03584E-12 |
| ACE2 | RP5-864K19.4 | -0,4906 | 1,01569E-12 |
| ACE2 | HIST2H2BF | -0,4906 | 1,01767E-12 |
| ACE2 | PRSS44 | -0,4906 | 1,01649E-12 |
| ACE2 | RP11-442O1.3 | -0,4906 | 1,01813E-12 |

| | | | |
|---|---|---|---|
| ACE2 | MEIS3P1 | -0,4906 | 1,02496E-12 |
| ACE2 | WDR4 | -0,4906 | 1,02627E-12 |
| ACE2 | RP11-665G4.1 | -0,4907 | 1,00849E-12 |
| ACE2 | PKIA-AS1 | -0,4907 | 1,0044E-12 |
| ACE2 | KCTD19 | -0,4907 | 1,00968E-12 |
| ACE2 | ZNF350-AS1 | -0,4907 | 1,01148E-12 |
| ACE2 | ELMOD3 | -0,4908 | 9,97426E-13 |
| ACE2 | MED20 | -0,4908 | 9,93038E-13 |
| ACE2 | CTD-2571L23.8 | -0,4908 | 9,95467E-13 |
| ACE2 | RHOXF1 | -0,4908 | 9,9812E-13 |
| ACE2 | RP11-470M17.2 | -0,4909 | 9,88413E-13 |
| ACE2 | VSIG8 | -0,491 | 9,70089E-13 |
| ACE2 | CTD-2334D19.1 | -0,491 | 9,66901E-13 |
| ACE2 | CTD-2017D11.2 | -0,491 | 9,67143E-13 |
| ACE2 | RP5-1039K5.16 | -0,491 | 9,74629E-13 |
| ACE2 | RAD17 | -0,4911 | 9,55929E-13 |
| ACE2 | TMEM248 | -0,4911 | 9,59537E-13 |
| ACE2 | RP11-64C12.1 | -0,4911 | 9,55084E-13 |
| ACE2 | SP5 | -0,4912 | 9,54006E-13 |
| ACE2 | TAS1R1 | -0,4913 | 9,41908E-13 |
| ACE2 | IBA57 | -0,4913 | 9,37631E-13 |
| ACE2 | KIAA0368 | -0,4913 | 9,3981E-13 |
| ACE2 | RNA5SP449 | -0,4913 | 9,37299E-13 |
| ACE2 | ZSCAN4 | -0,4913 | 9,37264E-13 |
| ACE2 | AC005076.5 | -0,4914 | 9,23929E-13 |
| ACE2 | RAB11FIP3 | -0,4914 | 9,26335E-13 |
| ACE2 | TFAP2C | -0,4914 | 9,31235E-13 |
| ACE2 | DUSP11 | -0,4915 | 9,092E-13 |
| ACE2 | NSUN2 | -0,4915 | 9,14143E-13 |
| ACE2 | VPS37A | -0,4915 | 9,19909E-13 |
| ACE2 | SUDS3 | -0,4915 | 9,18116E-13 |
| ACE2 | ZSCAN25 | -0,4916 | 9,07828E-13 |
| ACE2 | ADAMTS7P3 | -0,4917 | 8,95399E-13 |
| ACE2 | RP11-68I3.10 | -0,4917 | 8,88229E-13 |
| ACE2 | KIF5B | -0,4918 | 8,86106E-13 |
| ACE2 | PMS2P5 | -0,4919 | 8,71576E-13 |
| ACE2 | UNC45B | -0,4919 | 8,72558E-13 |
| ACE2 | UBE2E3 | -0,492 | 8,6504E-13 |
| ACE2 | AC007285.6 | -0,492 | 8,64518E-13 |
| ACE2 | TNPO3 | -0,492 | 8,59417E-13 |
| ACE2 | LPAR6 | -0,492 | 8,6175E-13 |
| ACE2 | CTD-3064H18.4 | -0,492 | 8,58794E-13 |
| ACE2 | RNU4ATAC | -0,4921 | 8,45964E-13 |
| ACE2 | PSMD11 | -0,4921 | 8,52584E-13 |
| ACE2 | C2orf50 | -0,4922 | 8,38083E-13 |
| ACE2 | RP11-536K7.3 | -0,4922 | 8,3664E-13 |
| ACE2 | PDHX | -0,4922 | 8,39393E-13 |
| ACE2 | TRIM72 | -0,4922 | 8,40422E-13 |
| ACE2 | RP11-159D12.2 | -0,4923 | 8,34324E-13 |
| ACE2 | SERPINB9 | -0,4924 | 8,21146E-13 |
| ACE2 | CDKN1C | -0,4924 | 8,1784E-13 |
| ACE2 | BCKDHA | -0,4924 | 8,19425E-13 |
| ACE2 | AP001065.15 | -0,4924 | 8,20717E-13 |
| ACE2 | SEPT3 | -0,4924 | 8,23197E-13 |

| | | | |
|---|---|---|---|
| ACE2 | WI2-89031B12.1 | -0,4924 | 8,22433E-13 |
| ACE2 | PDIA5 | -0,4925 | 8,09583E-13 |
| ACE2 | HLCS | -0,4925 | 8,09263E-13 |
| ACE2 | XXbac-BPG299F13.1 | -0,4926 | 7,96226E-13 |
| ACE2 | RRAGC | -0,4927 | 7,86215E-13 |
| ACE2 | ADH1A | -0,4927 | 7,8754E-13 |
| ACE2 | RP11-265E18.1 | -0,4927 | 7,9333E-13 |
| ACE2 | PELI2 | -0,4927 | 7,90862E-13 |
| ACE2 | SYNDIG1L | -0,4927 | 7,87694E-13 |
| ACE2 | KLHL11 | -0,4927 | 7,8754E-13 |
| ACE2 | NDRG3 | -0,4927 | 7,87253E-13 |
| ACE2 | NCF2 | -0,4928 | 7,85002E-13 |
| ACE2 | RP11-258F1.1 | -0,4928 | 7,81818E-13 |
| ACE2 | APC2 | -0,4928 | 7,84915E-13 |
| ACE2 | APMAP | -0,4928 | 7,79777E-13 |
| ACE2 | KIAA0922 | -0,4929 | 7,71546E-13 |
| ACE2 | GLIPR1L1 | -0,4929 | 7,74476E-13 |
| ACE2 | RP1-267D11.6 | -0,4929 | 7,75193E-13 |
| ACE2 | SEMA6B | -0,4929 | 7,69015E-13 |
| ACE2 | LINC01963 | -0,493 | 7,63313E-13 |
| ACE2 | RP11-498E2.9 | -0,493 | 7,60661E-13 |
| ACE2 | GLTSCR1 | -0,493 | 7,64411E-13 |
| ACE2 | SETD2 | -0,4931 | 7,52226E-13 |
| ACE2 | AC093616.4 | -0,4932 | 7,39772E-13 |
| ACE2 | CEP135 | -0,4933 | 7,36805E-13 |
| ACE2 | UTP3 | -0,4933 | 7,35977E-13 |
| ACE2 | SLC7A6OS | -0,4933 | 7,34117E-13 |
| ACE2 | RNF138P1 | -0,4934 | 7,2142E-13 |
| ACE2 | PALM2 | -0,4934 | 7,22242E-13 |
| ACE2 | CTC-490E21.10 | -0,4934 | 7,21185E-13 |
| ACE2 | MZT2A | -0,4935 | 7,20477E-13 |
| ACE2 | RP11-438L19.1 | -0,4935 | 7,20638E-13 |
| ACE2 | ADGRA1 | -0,4935 | 7,14843E-13 |
| ACE2 | ABCA6 | -0,4935 | 7,16959E-13 |
| ACE2 | RP11-674N23.4 | -0,4935 | 7,17639E-13 |
| ACE2 | RP11-573D15.1 | -0,4936 | 7,07963E-13 |
| ACE2 | GUSBP5 | -0,4936 | 7,05432E-13 |
| ACE2 | TMEM203 | -0,4936 | 7,05949E-13 |
| ACE2 | RP11-441F2.5 | -0,4936 | 7,0601E-13 |
| ACE2 | REV1 | -0,4937 | 6,99345E-13 |
| ACE2 | AC098617.1 | -0,4937 | 6,95872E-13 |
| ACE2 | ZRSR1 | -0,4937 | 6,99855E-13 |
| ACE2 | STC2 | -0,4937 | 6,98734E-13 |
| ACE2 | GLS2 | -0,4937 | 6,96435E-13 |
| ACE2 | BRD1 | -0,4937 | 6,97948E-13 |
| ACE2 | NFYC-AS1 | -0,4939 | 6,79158E-13 |
| ACE2 | GNAI3 | -0,4939 | 6,77974E-13 |
| ACE2 | RP11-1191J2.2 | -0,4939 | 6,78031E-13 |
| ACE2 | CHD8 | -0,4939 | 6,79141E-13 |
| ACE2 | CTD-2026K11.6 | -0,494 | 6,72248E-13 |
| ACE2 | TRAPPC12-AS1 | -0,4941 | 6,67246E-13 |
| ACE2 | ACTB | -0,4941 | 6,62487E-13 |
| ACE2 | GDF5 | -0,4941 | 6,66517E-13 |
| ACE2 | DENND6B | -0,4941 | 6,62188E-13 |

| | | | |
|---|---|---|---|
| ACE2 | RP11-342K6.2 | -0,4942 | 6,58781E-13 |
| ACE2 | RP3-422G23.4 | -0,4942 | 6,56038E-13 |
| ACE2 | RP11-680C21.1 | -0,4942 | 6,53618E-13 |
| ACE2 | CFH | -0,4943 | 6,50253E-13 |
| ACE2 | LINC00680 | -0,4943 | 6,52918E-13 |
| ACE2 | ESRRB | -0,4943 | 6,48526E-13 |
| ACE2 | RSL24D1P11 | -0,4943 | 6,51599E-13 |
| ACE2 | ALG13 | -0,4943 | 6,46396E-13 |
| ACE2 | PPIE | -0,4944 | 6,42595E-13 |
| ACE2 | RP11-641D5.1 | -0,4944 | 6,4412E-13 |
| ACE2 | TPPP3 | -0,4944 | 6,40188E-13 |
| ACE2 | SMURF1 | -0,4946 | 6,29547E-13 |
| ACE2 | RPS2P7 | -0,4946 | 6,2466E-13 |
| ACE2 | RP11-384O8.1 | -0,4948 | 6,11581E-13 |
| ACE2 | PRICKLE4 | -0,4948 | 6,10912E-13 |
| ACE2 | CRHR2 | -0,4948 | 6,12311E-13 |
| ACE2 | CCDC159 | -0,4948 | 6,07882E-13 |
| ACE2 | AP000255.6 | -0,4948 | 6,09514E-13 |
| ACE2 | DYNC1I2 | -0,4949 | 6,05157E-13 |
| ACE2 | AC145676.2 | -0,4949 | 5,99913E-13 |
| ACE2 | DLX6 | -0,4949 | 6,04606E-13 |
| ACE2 | PUF60 | -0,4949 | 6,05277E-13 |
| ACE2 | TBC1D2B | -0,4949 | 6,06475E-13 |
| ACE2 | HAS1 | -0,4949 | 6,04763E-13 |
| ACE2 | RBM17 | -0,495 | 5,94577E-13 |
| ACE2 | WAPL | -0,495 | 5,97782E-13 |
| ACE2 | XRCC3 | -0,495 | 5,9707E-13 |
| ACE2 | CTD-2033D15.3 | -0,495 | 5,98506E-13 |
| ACE2 | RP11-647K16.1 | -0,4951 | 5,91242E-13 |
| ACE2 | GAS2L2 | -0,4951 | 5,87626E-13 |
| ACE2 | SYN2 | -0,4952 | 5,78619E-13 |
| ACE2 | USF2 | -0,4952 | 5,78866E-13 |
| ACE2 | ACR | -0,4953 | 5,7159E-13 |
| ACE2 | FOXD4 | -0,4954 | 5,65667E-13 |
| ACE2 | RP5-1116H23.5 | -0,4954 | 5,6764E-13 |
| ACE2 | SPOCK1 | -0,4955 | 5,58838E-13 |
| ACE2 | NRAV | -0,4955 | 5,58238E-13 |
| ACE2 | NR4A2 | -0,4956 | 5,56579E-13 |
| ACE2 | CH17-472G23.4 | -0,4957 | 5,44524E-13 |
| ACE2 | LIMS1-AS1 | -0,4957 | 5,44308E-13 |
| ACE2 | PWWP2A | -0,4957 | 5,44385E-13 |
| ACE2 | TGS1 | -0,4957 | 5,49624E-13 |
| ACE2 | SCG3 | -0,4957 | 5,4434E-13 |
| ACE2 | RPS27AP16 | -0,4957 | 5,45961E-13 |
| ACE2 | LINC02035 | -0,4958 | 5,42523E-13 |
| ACE2 | ZNF559-ZNF177 | -0,4958 | 5,38002E-13 |
| ACE2 | RGS7 | -0,4959 | 5,34584E-13 |
| ACE2 | PLSCR1 | -0,4959 | 5,346E-13 |
| ACE2 | CNTRL | -0,4959 | 5,32391E-13 |
| ACE2 | SEC14L5 | -0,496 | 5,24167E-13 |
| ACE2 | SMG1P3 | -0,496 | 5,24539E-13 |
| ACE2 | PAQR6 | -0,4961 | 5,16931E-13 |
| ACE2 | LINC01829 | -0,4961 | 5,17835E-13 |
| ACE2 | RP11-562I5.1 | -0,4961 | 5,1918E-13 |

| | | | |
|---|---|---|---|
| ACE2 | RP4-605O3.4 | -0,4961 | 5,19387E-13 |
| ACE2 | AC007796.1 | -0,4961 | 5,19687E-13 |
| ACE2 | ZNF436 | -0,4962 | 5,1114E-13 |
| ACE2 | RP3-509L4.3 | -0,4962 | 5,13577E-13 |
| ACE2 | RP11-624M8.1 | -0,4962 | 5,13844E-13 |
| ACE2 | RP11-298I3.6 | -0,4962 | 5,13571E-13 |
| ACE2 | CTD-2576N18.1 | -0,4962 | 5,11195E-13 |
| ACE2 | YTHDF1 | -0,4962 | 5,13248E-13 |
| ACE2 | LINC01990 | -0,4963 | 5,07197E-13 |
| ACE2 | CYGB | -0,4963 | 5,09923E-13 |
| ACE2 | TP53TG5 | -0,4963 | 5,06029E-13 |
| ACE2 | LINC00599 | -0,4964 | 5,01096E-13 |
| ACE2 | FIGNL1 | -0,4965 | 4,96534E-13 |
| ACE2 | LINC01563 | -0,4965 | 4,9462E-13 |
| ACE2 | DKC1 | -0,4965 | 4,94353E-13 |
| ACE2 | GNG3 | -0,4966 | 4,9045E-13 |
| ACE2 | CTD-2588E21.1 | -0,4966 | 4,88003E-13 |
| ACE2 | RNF214 | -0,4967 | 4,85459E-13 |
| ACE2 | EIF4A3 | -0,4967 | 4,83035E-13 |
| ACE2 | SRRM5 | -0,4967 | 4,82857E-13 |
| ACE2 | RP11-498P14.5 | -0,4968 | 4,78498E-13 |
| ACE2 | DPYD | -0,4969 | 4,6836E-13 |
| ACE2 | RP11-227G15.9 | -0,4969 | 4,69399E-13 |
| ACE2 | RP11-230C9.2 | -0,497 | 4,64233E-13 |
| ACE2 | SORBS3 | -0,497 | 4,67016E-13 |
| ACE2 | BOP1 | -0,497 | 4,67654E-13 |
| ACE2 | RAB39B | -0,497 | 4,66033E-13 |
| ACE2 | ARID4A | -0,4971 | 4,61797E-13 |
| ACE2 | RP11-147L13.14 | -0,4971 | 4,61323E-13 |
| ACE2 | CTB-31O20.2 | -0,4971 | 4,60956E-13 |
| ACE2 | MMGT1 | -0,4971 | 4,58653E-13 |
| ACE2 | RP11-499F19.3 | -0,4972 | 4,53597E-13 |
| ACE2 | G2E3 | -0,4972 | 4,51398E-13 |
| ACE2 | FFAR3 | -0,4972 | 4,53666E-13 |
| ACE2 | GSN-AS1 | -0,4973 | 4,47902E-13 |
| ACE2 | RP11-313P22.1 | -0,4973 | 4,45615E-13 |
| ACE2 | MTMR3 | -0,4973 | 4,49077E-13 |
| ACE2 | OPHN1 | -0,4973 | 4,49385E-13 |
| ACE2 | XXbac-BPG154L12.4 | -0,4974 | 4,39948E-13 |
| ACE2 | MFSD14B | -0,4974 | 4,42765E-13 |
| ACE2 | UBOX5 | -0,4974 | 4,4221E-13 |
| ACE2 | HMP19 | -0,4975 | 4,38758E-13 |
| ACE2 | SLC25A17 | -0,4975 | 4,37198E-13 |
| ACE2 | AC022210.2 | -0,4976 | 4,29257E-13 |
| ACE2 | CREBRF | -0,4976 | 4,34118E-13 |
| ACE2 | ZNF395 | -0,4977 | 4,23941E-13 |
| ACE2 | RP11-632L2.2 | -0,4977 | 4,24811E-13 |
| ACE2 | COMP | -0,4977 | 4,28829E-13 |
| ACE2 | TRMT6 | -0,4977 | 4,24813E-13 |
| ACE2 | KCNK15 | -0,4977 | 4,27559E-13 |
| ACE2 | KCNK3 | -0,4978 | 4,19226E-13 |
| ACE2 | RP11-397E7.4 | -0,4978 | 4,19849E-13 |
| ACE2 | AK3P3 | -0,4978 | 4,21914E-13 |
| ACE2 | UBR5-AS1 | -0,4978 | 4,23204E-13 |

| | | | |
|---|---|---|---|
| ACE2 | NPAT | -0,4978 | 4,22999E-13 |
| ACE2 | AP000695.4 | -0,4978 | 4,20259E-13 |
| ACE2 | USP24 | -0,4979 | 4,18057E-13 |
| ACE2 | SLITRK3 | -0,4979 | 4,18557E-13 |
| ACE2 | SLC8A2 | -0,4979 | 4,17901E-13 |
| ACE2 | PLCB4 | -0,4979 | 4,16817E-13 |
| ACE2 | PLRG1 | -0,498 | 4,10712E-13 |
| ACE2 | CTD-2373J6.1 | -0,4981 | 4,06423E-13 |
| ACE2 | SPDYE3 | -0,4981 | 4,04116E-13 |
| ACE2 | IFNAR1 | -0,4981 | 4,05654E-13 |
| ACE2 | HDC | -0,4982 | 4,002E-13 |
| ACE2 | HSPA7 | -0,4983 | 3,94591E-13 |
| ACE2 | BSN-AS2 | -0,4983 | 3,97175E-13 |
| ACE2 | MINCR | -0,4983 | 3,96127E-13 |
| ACE2 | OMD | -0,4983 | 3,94613E-13 |
| ACE2 | RP11-64D24.4 | -0,4984 | 3,89724E-13 |
| ACE2 | RP4-694B14.8 | -0,4984 | 3,90094E-13 |
| ACE2 | EPHX4 | -0,4985 | 3,85615E-13 |
| ACE2 | MTDHP1 | -0,4985 | 3,85153E-13 |
| ACE2 | FAM166B | -0,4985 | 3,86078E-13 |
| ACE2 | KLHL35 | -0,4985 | 3,8725E-13 |
| ACE2 | PPCDC | -0,4985 | 3,8766E-13 |
| ACE2 | CTD-2339F6.1 | -0,4986 | 3,80141E-13 |
| ACE2 | ADGRG2 | -0,4986 | 3,7939E-13 |
| ACE2 | CTD-2517O10.6 | -0,4987 | 3,77334E-13 |
| ACE2 | NLRX1 | -0,4987 | 3,75181E-13 |
| ACE2 | SRRM4 | -0,4987 | 3,76775E-13 |
| ACE2 | WDR25 | -0,4987 | 3,76138E-13 |
| ACE2 | LLNLF-187D8.1 | -0,4987 | 3,75971E-13 |
| ACE2 | DNER | -0,4988 | 3,71303E-13 |
| ACE2 | MRPS28 | -0,4988 | 3,74034E-13 |
| ACE2 | DRD1 | -0,4989 | 3,67507E-13 |
| ACE2 | DUXAP10 | -0,4989 | 3,66846E-13 |
| ACE2 | NUFIP2 | -0,4989 | 3,65221E-13 |
| ACE2 | WASF2 | -0,499 | 3,62325E-13 |
| ACE2 | ACTL6A | -0,499 | 3,61837E-13 |
| ACE2 | KSR1 | -0,499 | 3,62262E-13 |
| ACE2 | AC010524.4 | -0,499 | 3,63377E-13 |
| ACE2 | RP11-626E13.1 | -0,4991 | 3,57809E-13 |
| ACE2 | ARVCF | -0,4991 | 3,56508E-13 |
| ACE2 | RP11-511P7.6 | -0,4992 | 3,51587E-13 |
| ACE2 | UNC13C | -0,4992 | 3,52406E-13 |
| ACE2 | LINC01581 | -0,4992 | 3,52529E-13 |
| ACE2 | RP5-991G20.1 | -0,4992 | 3,52243E-13 |
| ACE2 | WDR45B | -0,4992 | 3,55651E-13 |
| ACE2 | TMA7 | -0,4993 | 3,48766E-13 |
| ACE2 | ZDHHC14 | -0,4993 | 3,49567E-13 |
| ACE2 | RP11-291L22.9 | -0,4993 | 3,49988E-13 |
| ACE2 | CTD-2002J20.1 | -0,4994 | 3,47185E-13 |
| ACE2 | STAT2 | -0,4994 | 3,45769E-13 |
| ACE2 | DIEXF | -0,4995 | 3,39704E-13 |
| ACE2 | LDHAP2 | -0,4995 | 3,4114E-13 |
| ACE2 | AC096574.5 | -0,4996 | 3,38078E-13 |
| ACE2 | WDR1 | -0,4996 | 3,35173E-13 |

| | | | |
|---|---|---|---|
| ACE2 | GRPEL2 | -0,4996 | 3,36291E-13 |
| ACE2 | MKS1 | -0,4996 | 3,37431E-13 |
| ACE2 | RP1-168L15.6 | -0,4997 | 3,32538E-13 |
| ACE2 | GNB2 | -0,4997 | 3,33685E-13 |
| ACE2 | ADAMTS13 | -0,4997 | 3,31033E-13 |
| ACE2 | AP000695.6 | -0,4997 | 3,3344E-13 |
| ACE2 | GAGE10 | -0,4997 | 3,31716E-13 |
| ACE2 | PQLC2L | -0,4998 | 3,26156E-13 |
| ACE2 | MAFK | -0,4999 | 3,23962E-13 |
| ACE2 | MADD | -0,4999 | 3,24222E-13 |
| ACE2 | CKMT2 | -0,5 | 3,2026E-13 |
| ACE2 | GRP | -0,5 | 3,19409E-13 |
| ACE2 | L3MBTL3 | -0,5001 | 3,165E-13 |
| ACE2 | METTL25 | -0,5001 | 3,1484E-13 |
| ACE2 | RP11-119B16.2 | -0,5001 | 3,17657E-13 |
| ACE2 | PTH2R | -0,5002 | 3,12503E-13 |
| ACE2 | RP11-20I20.4 | -0,5002 | 3,13909E-13 |
| ACE2 | LINC00539 | -0,5002 | 3,14035E-13 |
| ACE2 | BLMH | -0,5002 | 3,1115E-13 |
| ACE2 | SNRK | -0,5003 | 3,09357E-13 |
| ACE2 | RP11-696F12.1 | -0,5003 | 3,07109E-13 |
| ACE2 | CMB9-22P13.1 | -0,5003 | 3,07902E-13 |
| ACE2 | SLC16A11 | -0,5003 | 3,09117E-13 |
| ACE2 | AAR2 | -0,5003 | 3,06664E-13 |
| ACE2 | LYVE1 | -0,5004 | 3,05142E-13 |
| ACE2 | FAM104A | -0,5004 | 3,04519E-13 |
| ACE2 | PSTPIP2 | -0,5004 | 3,0632E-13 |
| ACE2 | CXXC1 | -0,5004 | 3,02815E-13 |
| ACE2 | DGKH | -0,5005 | 2,99358E-13 |
| ACE2 | ELMSAN1 | -0,5005 | 3,02441E-13 |
| ACE2 | TMEM14C | -0,5006 | 2,97768E-13 |
| ACE2 | TEAD4 | -0,5006 | 2,95395E-13 |
| ACE2 | PIF1 | -0,5006 | 2,98601E-13 |
| ACE2 | CTA-292E10.6 | -0,5006 | 2,95106E-13 |
| ACE2 | SCAMP1-AS1 | -0,5008 | 2,89983E-13 |
| ACE2 | INTS9 | -0,5008 | 2,91164E-13 |
| ACE2 | PARD3B | -0,501 | 2,82307E-13 |
| ACE2 | AZI2 | -0,501 | 2,82869E-13 |
| ACE2 | ZNF18 | -0,501 | 2,82894E-13 |
| ACE2 | DNMT1 | -0,501 | 2,83849E-13 |
| ACE2 | ZNF233 | -0,501 | 2,81697E-13 |
| ACE2 | NRSN1 | -0,5011 | 2,7792E-13 |
| ACE2 | RP3-375P9.2 | -0,5011 | 2,78708E-13 |
| ACE2 | MRPS24 | -0,5011 | 2,8053E-13 |
| ACE2 | SCARA5 | -0,5011 | 2,80264E-13 |
| ACE2 | NAA35 | -0,5011 | 2,78724E-13 |
| ACE2 | CLDND2 | -0,5012 | 2,77058E-13 |
| ACE2 | HMGXB3 | -0,5013 | 2,72846E-13 |
| ACE2 | TSPAN10 | -0,5013 | 2,70971E-13 |
| ACE2 | RPS15AP38 | -0,5013 | 2,73406E-13 |
| ACE2 | CAMKMT | -0,5014 | 2,68922E-13 |
| ACE2 | HIST1H2BE | -0,5014 | 2,6759E-13 |
| ACE2 | GATC | -0,5014 | 2,68651E-13 |
| ACE2 | HOXB3 | -0,5014 | 2,69972E-13 |

| | | | |
|---|---|---|---|
| ACE2 | RP11-399E6.4 | -0,5015 | 2,638E-13 |
| ACE2 | ZNF844 | -0,5015 | 2,64205E-13 |
| ACE2 | NEMP2 | -0,5016 | 2,63459E-13 |
| ACE2 | RP11-597D13.9 | -0,5016 | 2,60327E-13 |
| ACE2 | RGS7BP | -0,5016 | 2,62532E-13 |
| ACE2 | SNED1 | -0,5017 | 2,60094E-13 |
| ACE2 | ARID1B | -0,5017 | 2,58439E-13 |
| ACE2 | APLNR | -0,5017 | 2,58745E-13 |
| ACE2 | MEF2B | -0,5017 | 2,58267E-13 |
| ACE2 | COL24A1 | -0,5018 | 2,54864E-13 |
| ACE2 | ATP6V0CP3 | -0,5018 | 2,5577E-13 |
| ACE2 | EEF1DP4 | -0,5018 | 2,53829E-13 |
| ACE2 | RP11-849F2.9 | -0,5018 | 2,53739E-13 |
| ACE2 | SYT4 | -0,5018 | 2,54489E-13 |
| ACE2 | ADRM1 | -0,5018 | 2,55336E-13 |
| ACE2 | DPYSL5 | -0,5019 | 2,52007E-13 |
| ACE2 | ZNF654 | -0,5019 | 2,51876E-13 |
| ACE2 | GPR22 | -0,5019 | 2,52058E-13 |
| ACE2 | PENK | -0,5019 | 2,51705E-13 |
| ACE2 | RASGEF1A | -0,5019 | 2,51381E-13 |
| ACE2 | RP13-735L24.1 | -0,5019 | 2,52379E-13 |
| ACE2 | LA16c-312E8.4 | -0,5019 | 2,50982E-13 |
| ACE2 | BEND3P3 | -0,502 | 2,49159E-13 |
| ACE2 | XG | -0,502 | 2,49881E-13 |
| ACE2 | RP11-279O9.4 | -0,5021 | 2,45679E-13 |
| ACE2 | ZNF483 | -0,5021 | 2,44342E-13 |
| ACE2 | RP13-152O15.5 | -0,5021 | 2,45916E-13 |
| ACE2 | RPL10P16 | -0,5022 | 2,43417E-13 |
| ACE2 | UBE2D3 | -0,5023 | 2,39102E-13 |
| ACE2 | ZNF682 | -0,5023 | 2,38936E-13 |
| ACE2 | C2orf27A | -0,5024 | 2,37645E-13 |
| ACE2 | RP11-972P1.10 | -0,5024 | 2,3615E-13 |
| ACE2 | LINC00648 | -0,5024 | 2,36916E-13 |
| ACE2 | RPL7P3 | -0,5024 | 2,35344E-13 |
| ACE2 | AC006466.5 | -0,5025 | 2,33487E-13 |
| ACE2 | USPL1 | -0,5025 | 2,34664E-13 |
| ACE2 | DDX49 | -0,5025 | 2,34094E-13 |
| ACE2 | TMEM65 | -0,5026 | 2,29394E-13 |
| ACE2 | ATE1-AS1 | -0,5026 | 2,31075E-13 |
| ACE2 | HCFC1 | -0,5027 | 2,29055E-13 |
| ACE2 | SH3GL3 | -0,5028 | 2,24356E-13 |
| ACE2 | KCNJ15 | -0,5028 | 2,26034E-13 |
| ACE2 | AL592528.1 | -0,5028 | 2,2459E-13 |
| ACE2 | HS6ST3 | -0,5029 | 2,2249E-13 |
| ACE2 | SLC4A1AP | -0,503 | 2,19502E-13 |
| ACE2 | ELMOD1 | -0,503 | 2,20351E-13 |
| ACE2 | DYNC1H1 | -0,503 | 2,19375E-13 |
| ACE2 | CTC-435M10.12 | -0,503 | 2,19949E-13 |
| ACE2 | HIC2 | -0,503 | 2,20665E-13 |
| ACE2 | RP11-266J6.2 | -0,5031 | 2,17641E-13 |
| ACE2 | ABCA11P | -0,5031 | 2,17726E-13 |
| ACE2 | ASB6 | -0,5031 | 2,1765E-13 |
| ACE2 | DNAJC9-AS1 | -0,5031 | 2,17133E-13 |
| ACE2 | RAB3D | -0,5031 | 2,15563E-13 |

| | | | |
|---|---|---|---|
| ACE2 | GAP43 | -0,5032 | 2,15232E-13 |
| ACE2 | RP11-73K9.2 | -0,5032 | 2,14539E-13 |
| ACE2 | DAAM2 | -0,5032 | 2,14067E-13 |
| ACE2 | RP11-38L15.3 | -0,5032 | 2,14344E-13 |
| ACE2 | GTF2A1 | -0,5032 | 2,14141E-13 |
| ACE2 | TNFSF9 | -0,5032 | 2,1279E-13 |
| ACE2 | NBPF9 | -0,5033 | 2,11055E-13 |
| ACE2 | HARS2 | -0,5033 | 2,12395E-13 |
| ACE2 | RBM48 | -0,5033 | 2,10498E-13 |
| ACE2 | ZFHX4-AS1 | -0,5034 | 2,08273E-13 |
| ACE2 | RP11-131L12.3 | -0,5034 | 2,07269E-13 |
| ACE2 | RP11-3D4.3 | -0,5034 | 2,08398E-13 |
| ACE2 | RANBP6 | -0,5035 | 2,04904E-13 |
| ACE2 | MYO9B | -0,5035 | 2,07203E-13 |
| ACE2 | ZCCHC18 | -0,5035 | 2,04892E-13 |
| ACE2 | RP11-944L7.4 | -0,5036 | 2,02962E-13 |
| ACE2 | GPX1 | -0,5036 | 2,03971E-13 |
| ACE2 | KLHDC10 | -0,5036 | 2,04225E-13 |
| ACE2 | SART3 | -0,5036 | 2,02965E-13 |
| ACE2 | DLL4 | -0,5036 | 2,02582E-13 |
| ACE2 | CTD-3247H4.2 | -0,5036 | 2,03768E-13 |
| ACE2 | MED26 | -0,5036 | 2,02311E-13 |
| ACE2 | ILF2 | -0,5037 | 2,01003E-13 |
| ACE2 | AC008992.1 | -0,5037 | 2,01981E-13 |
| ACE2 | CDR1 | -0,5037 | 2,00194E-13 |
| ACE2 | CH17-408M7.1 | -0,5038 | 1,98822E-13 |
| ACE2 | SNX31 | -0,5038 | 1,97208E-13 |
| ACE2 | DISC1FP1 | -0,5038 | 1,97992E-13 |
| ACE2 | RSPH14 | -0,5038 | 1,98829E-13 |
| ACE2 | CSAD | -0,5039 | 1,94628E-13 |
| ACE2 | BRAP | -0,5039 | 1,97009E-13 |
| ACE2 | RPL22L1 | -0,504 | 1,93989E-13 |
| ACE2 | RP11-35G9.5 | -0,504 | 1,92431E-13 |
| ACE2 | RP11-586K2.1 | -0,5041 | 1,90942E-13 |
| ACE2 | RP11-319G9.3 | -0,5041 | 1,92017E-13 |
| ACE2 | TAF15 | -0,5041 | 1,91581E-13 |
| ACE2 | RP1-122P22.2 | -0,5041 | 1,90352E-13 |
| ACE2 | IL15 | -0,5042 | 1,88691E-13 |
| ACE2 | PPP1R17 | -0,5042 | 1,89423E-13 |
| ACE2 | STX17 | -0,5042 | 1,87436E-13 |
| ACE2 | ST6GALNAC4P1 | -0,5042 | 1,88613E-13 |
| ACE2 | FAM216B | -0,5042 | 1,8939E-13 |
| ACE2 | CCDC150P1 | -0,5044 | 1,83261E-13 |
| ACE2 | RPS21 | -0,5044 | 1,8408E-13 |
| ACE2 | SPIN2A | -0,5044 | 1,84E-13 |
| ACE2 | RPL7AP30 | -0,5045 | 1,81756E-13 |
| ACE2 | RP11-254F19.2 | -0,5045 | 1,81578E-13 |
| ACE2 | GART | -0,5045 | 1,82558E-13 |
| ACE2 | RGS1 | -0,5046 | 1,79407E-13 |
| ACE2 | ADAM17 | -0,5046 | 1,79015E-13 |
| ACE2 | GTF2A1L | -0,5046 | 1,80062E-13 |
| ACE2 | SDHAF3 | -0,5046 | 1,78265E-13 |
| ACE2 | RP11-363E7.4 | -0,5046 | 1,79386E-13 |
| ACE2 | HPS4 | -0,5046 | 1,78979E-13 |

| | | | |
|---|---|---|---|
| ACE2 | C2orf73 | -0,5047 | 1,75804E-13 |
| ACE2 | RP11-7F18.2 | -0,5047 | 1,77881E-13 |
| ACE2 | AC108448.2 | -0,5047 | 1,776E-13 |
| ACE2 | EMC8 | -0,5047 | 1,76057E-13 |
| ACE2 | PIRT | -0,5047 | 1,77932E-13 |
| ACE2 | POLH | -0,5048 | 1,74539E-13 |
| ACE2 | SF3A1 | -0,5048 | 1,7462E-13 |
| ACE2 | HMGN2 | -0,5049 | 1,71658E-13 |
| ACE2 | NID1 | -0,5049 | 1,72493E-13 |
| ACE2 | UNC80 | -0,5049 | 1,72906E-13 |
| ACE2 | GPR26 | -0,5049 | 1,73127E-13 |
| ACE2 | CLDN10 | -0,5049 | 1,72328E-13 |
| ACE2 | RP11-364B14.3 | -0,5049 | 1,72925E-13 |
| ACE2 | DUX4L51 | -0,505 | 1,71293E-13 |
| ACE2 | LINC01447 | -0,505 | 1,70788E-13 |
| ACE2 | WIPF3 | -0,5051 | 1,68973E-13 |
| ACE2 | NINJ1 | -0,5051 | 1,67093E-13 |
| ACE2 | RP11-248J18.2 | -0,5052 | 1,65789E-13 |
| ACE2 | ZFYVE26 | -0,5052 | 1,66513E-13 |
| ACE2 | PNN | -0,5053 | 1,63819E-13 |
| ACE2 | UBE2Z | -0,5053 | 1,63441E-13 |
| ACE2 | VSX1 | -0,5053 | 1,63217E-13 |
| ACE2 | PRG4 | -0,5054 | 1,61971E-13 |
| ACE2 | CACNA2D3-AS1 | -0,5054 | 1,60831E-13 |
| ACE2 | RPL7AP50 | -0,5054 | 1,61777E-13 |
| ACE2 | TMEM50B | -0,5054 | 1,62327E-13 |
| ACE2 | RP11-557H15.4 | -0,5055 | 1,5901E-13 |
| ACE2 | TMEM179 | -0,5055 | 1,59219E-13 |
| ACE2 | AC004156.3 | -0,5055 | 1,59592E-13 |
| ACE2 | GPR17 | -0,5056 | 1,5854E-13 |
| ACE2 | ATP6V1G1P2 | -0,5056 | 1,57276E-13 |
| ACE2 | MED14OS | -0,5056 | 1,57897E-13 |
| ACE2 | TMEM234 | -0,5057 | 1,55103E-13 |
| ACE2 | PRRC2B | -0,5057 | 1,54985E-13 |
| ACE2 | MSTO2P | -0,5058 | 1,53463E-13 |
| ACE2 | GTF2I | -0,5058 | 1,53823E-13 |
| ACE2 | RP11-503E24.2 | -0,5058 | 1,52983E-13 |
| ACE2 | PKN3 | -0,5058 | 1,54428E-13 |
| ACE2 | GTPBP8 | -0,5059 | 1,52326E-13 |
| ACE2 | RPS20P14 | -0,5059 | 1,51836E-13 |
| ACE2 | RPL18AP3 | -0,5059 | 1,52032E-13 |
| ACE2 | MSL2 | -0,506 | 1,49138E-13 |
| ACE2 | CYYR1-AS1 | -0,506 | 1,48924E-13 |
| ACE2 | EFCC1 | -0,5061 | 1,47508E-13 |
| ACE2 | ETFBKMT | -0,5061 | 1,47577E-13 |
| ACE2 | RP11-2C24.3 | -0,5061 | 1,48684E-13 |
| ACE2 | RP11-347L15.1 | -0,5061 | 1,48678E-13 |
| ACE2 | RP11-1246C19.1 | -0,5062 | 1,46964E-13 |
| ACE2 | RBM12B | -0,5062 | 1,4537E-13 |
| ACE2 | RP1-29C18.9 | -0,5062 | 1,46925E-13 |
| ACE2 | TEKT2 | -0,5063 | 1,44258E-13 |
| ACE2 | PTPN4 | -0,5063 | 1,45114E-13 |
| ACE2 | RP11-17G12.2 | -0,5063 | 1,43944E-13 |
| ACE2 | RP11-672A2.5 | -0,5064 | 1,41537E-13 |

| | | | |
|---|---|---|---|
| ACE2 | TBC1D4 | -0,5064 | 1,43283E-13 |
| ACE2 | RP11-351M8.1 | -0,5064 | 1,4167E-13 |
| ACE2 | UPF1 | -0,5064 | 1,42608E-13 |
| ACE2 | AC004985.12 | -0,5065 | 1,3968E-13 |
| ACE2 | LCNL1 | -0,5065 | 1,39775E-13 |
| ACE2 | C10orf90 | -0,5065 | 1,40644E-13 |
| ACE2 | SAMD1 | -0,5065 | 1,40899E-13 |
| ACE2 | MICALL1 | -0,5065 | 1,4135E-13 |
| ACE2 | ZFP69B | -0,5066 | 1,38614E-13 |
| ACE2 | NUP210L | -0,5066 | 1,391E-13 |
| ACE2 | UBN2 | -0,5066 | 1,38252E-13 |
| ACE2 | SLC9A7P1 | -0,5066 | 1,38732E-13 |
| ACE2 | KIAA1107 | -0,5067 | 1,37823E-13 |
| ACE2 | TYW1 | -0,5067 | 1,37673E-13 |
| ACE2 | CSPG4P12 | -0,5067 | 1,36305E-13 |
| ACE2 | BASP1 | -0,5068 | 1,3585E-13 |
| ACE2 | CTD-2341M24.1 | -0,5068 | 1,345E-13 |
| ACE2 | ANAPC11 | -0,5068 | 1,36113E-13 |
| ACE2 | RP11-481J13.1 | -0,5069 | 1,34066E-13 |
| ACE2 | SH3BP5-AS1 | -0,5069 | 1,34121E-13 |
| ACE2 | ACTR8 | -0,5069 | 1,34089E-13 |
| ACE2 | CFAP53 | -0,5069 | 1,32823E-13 |
| ACE2 | KDM4B | -0,5069 | 1,34345E-13 |
| ACE2 | RPS3AP5 | -0,507 | 1,31235E-13 |
| ACE2 | DHRS12 | -0,507 | 1,32608E-13 |
| ACE2 | CFD | -0,507 | 1,32028E-13 |
| ACE2 | TOMM34 | -0,507 | 1,32118E-13 |
| ACE2 | NFYC | -0,5071 | 1,29999E-13 |
| ACE2 | PPP6R3 | -0,5071 | 1,2975E-13 |
| ACE2 | BIRC7 | -0,5071 | 1,30226E-13 |
| ACE2 | PRKAG2-AS1 | -0,5072 | 1,27935E-13 |
| ACE2 | MAP3K20-AS1 | -0,5073 | 1,26923E-13 |
| ACE2 | CTC-451P13.1 | -0,5073 | 1,27076E-13 |
| ACE2 | ADCK2 | -0,5073 | 1,26282E-13 |
| ACE2 | PSD2 | -0,5074 | 1,2483E-13 |
| ACE2 | RPS18P9 | -0,5074 | 1,24856E-13 |
| ACE2 | MSS51 | -0,5074 | 1,2556E-13 |
| ACE2 | RNF138 | -0,5074 | 1,26011E-13 |
| ACE2 | LRRFIP1 | -0,5075 | 1,23907E-13 |
| ACE2 | RP11-616M22.11 | -0,5075 | 1,23414E-13 |
| ACE2 | SV2C | -0,5076 | 1,21788E-13 |
| ACE2 | SVIL | -0,5076 | 1,22035E-13 |
| ACE2 | ZNF32-AS2 | -0,5076 | 1,21734E-13 |
| ACE2 | RP11-112J1.1 | -0,5076 | 1,21415E-13 |
| ACE2 | MPPED1 | -0,5076 | 1,22294E-13 |
| ACE2 | RP3-402G11.27 | -0,5076 | 1,21834E-13 |
| ACE2 | CADPS2 | -0,5078 | 1,19555E-13 |
| ACE2 | ZNF79 | -0,5078 | 1,19049E-13 |
| ACE2 | VPS39 | -0,5078 | 1,18723E-13 |
| ACE2 | CPNE3 | -0,5079 | 1,17362E-13 |
| ACE2 | MTHFD1 | -0,5079 | 1,17314E-13 |
| ACE2 | NDUFV3 | -0,5079 | 1,17002E-13 |
| ACE2 | LHFPL1 | -0,5079 | 1,17104E-13 |
| ACE2 | TIMM29 | -0,508 | 1,15579E-13 |

| | | | |
|---|---|---|---|
| ACE2 | ZFYVE9 | -0,5081 | 1,14042E-13 |
| ACE2 | ZNF740 | -0,5081 | 1,13938E-13 |
| ACE2 | RP11-749I16.3 | -0,5082 | 1,12561E-13 |
| ACE2 | RC3H1 | -0,5083 | 1,11602E-13 |
| ACE2 | LRRC3B | -0,5083 | 1,11421E-13 |
| ACE2 | RPL7P23 | -0,5084 | 1,1008E-13 |
| ACE2 | PDE6C | -0,5084 | 1,09632E-13 |
| ACE2 | POLA2 | -0,5084 | 1,10005E-13 |
| ACE2 | CHRNB1 | -0,5084 | 1,10236E-13 |
| ACE2 | AC159540.1 | -0,5085 | 1,09277E-13 |
| ACE2 | RP11-255C15.3 | -0,5085 | 1,09106E-13 |
| ACE2 | MUSK | -0,5085 | 1,09132E-13 |
| ACE2 | RPL13AP5 | -0,5085 | 1,08505E-13 |
| ACE2 | STMN1 | -0,5086 | 1,07259E-13 |
| ACE2 | SLC10A6 | -0,5086 | 1,07961E-13 |
| ACE2 | LYL1 | -0,5086 | 1,06914E-13 |
| ACE2 | YES1P1 | -0,5086 | 1,07456E-13 |
| ACE2 | RP11-295G24.5 | -0,5087 | 1,05417E-13 |
| ACE2 | REXO4 | -0,5087 | 1,05351E-13 |
| ACE2 | TGIF1 | -0,5087 | 1,06547E-13 |
| ACE2 | NIPBL | -0,5088 | 1,04196E-13 |
| ACE2 | NUP43 | -0,5088 | 1,04671E-13 |
| ACE2 | ZNF562 | -0,5088 | 1,04353E-13 |
| ACE2 | HHEX | -0,5089 | 1,03208E-13 |
| ACE2 | RP11-548P2.2 | -0,5091 | 1,0004E-13 |
| ACE2 | ZNHIT2 | -0,5091 | 1,00651E-13 |
| ACE2 | AC006946.17 | -0,5092 | 9,99219E-14 |
| ACE2 | ELAVL4 | -0,5093 | 9,78219E-14 |
| ACE2 | TANK | -0,5093 | 9,85399E-14 |
| ACE2 | RP11-680A11.5 | -0,5093 | 9,85868E-14 |
| ACE2 | LINS1 | -0,5093 | 9,78073E-14 |
| ACE2 | DBR1 | -0,5094 | 9,72979E-14 |
| ACE2 | TVP23A | -0,5094 | 9,69622E-14 |
| ACE2 | RP11-285J16.1 | -0,5095 | 9,53155E-14 |
| ACE2 | RAD54L2 | -0,5095 | 9,56151E-14 |
| ACE2 | OSBPL5 | -0,5095 | 9,55076E-14 |
| ACE2 | CFAP61 | -0,5095 | 9,56279E-14 |
| ACE2 | AKT3-IT1 | -0,5096 | 9,49308E-14 |
| ACE2 | RP11-54G14.1 | -0,5096 | 9,42749E-14 |
| ACE2 | MCL1 | -0,5097 | 9,35323E-14 |
| ACE2 | FGF5 | -0,5097 | 9,32067E-14 |
| ACE2 | METAP1 | -0,5097 | 9,28825E-14 |
| ACE2 | ZNF786 | -0,5097 | 9,26932E-14 |
| ACE2 | RANGAP1 | -0,5097 | 9,27482E-14 |
| ACE2 | CDCP2 | -0,5098 | 9,14629E-14 |
| ACE2 | USP5 | -0,5098 | 9,20439E-14 |
| ACE2 | LYPD3 | -0,5098 | 9,16108E-14 |
| ACE2 | MMP14 | -0,51 | 8,95863E-14 |
| ACE2 | RP6-24A23.7 | -0,51 | 8,98053E-14 |
| ACE2 | ACADL | -0,5101 | 8,89397E-14 |
| ACE2 | PSMA3-AS1 | -0,5101 | 8,81588E-14 |
| ACE2 | TBX2-AS1 | -0,5101 | 8,80072E-14 |
| ACE2 | RP11-1100L3.8 | -0,5102 | 8,72602E-14 |
| ACE2 | H2AFJ | -0,5103 | 8,59707E-14 |

| | | | |
|---|---|---|---|
| ACE2 | RSPO4 | -0,5103 | 8,64552E-14 |
| ACE2 | AC108463.1 | -0,5104 | 8,558E-14 |
| ACE2 | PTAR1 | -0,5104 | 8,50025E-14 |
| ACE2 | KLF7 | -0,5105 | 8,45188E-14 |
| ACE2 | P3H2-AS1 | -0,5105 | 8,45051E-14 |
| ACE2 | RP11-218E20.5 | -0,5105 | 8,43959E-14 |
| ACE2 | GPR68 | -0,5105 | 8,43912E-14 |
| ACE2 | ZNF43 | -0,5105 | 8,40807E-14 |
| ACE2 | CTD-2245F17.3 | -0,5105 | 8,44164E-14 |
| ACE2 | C22orf46 | -0,5106 | 8,30837E-14 |
| ACE2 | LINC01778 | -0,5107 | 8,22563E-14 |
| ACE2 | LINC01125 | -0,5107 | 8,16504E-14 |
| ACE2 | ELN | -0,5107 | 8,14202E-14 |
| ACE2 | STAT1 | -0,5108 | 8,09202E-14 |
| ACE2 | ADAMTS6 | -0,5108 | 8,0385E-14 |
| ACE2 | RP11-162A12.2 | -0,5108 | 8,08647E-14 |
| ACE2 | ATIC | -0,5109 | 7,95779E-14 |
| ACE2 | PLAU | -0,5109 | 8,02246E-14 |
| ACE2 | IRAK3 | -0,5109 | 7,94429E-14 |
| ACE2 | RP11-526N18.1 | -0,5109 | 8,01844E-14 |
| ACE2 | RPL12P12 | -0,5109 | 8,01722E-14 |
| ACE2 | RABL3 | -0,511 | 7,87292E-14 |
| ACE2 | NRCAM | -0,511 | 7,90958E-14 |
| ACE2 | LINC01239 | -0,511 | 7,84133E-14 |
| ACE2 | RHOT1 | -0,511 | 7,90267E-14 |
| ACE2 | PLAT | -0,5111 | 7,81056E-14 |
| ACE2 | DDX21 | -0,5111 | 7,7436E-14 |
| ACE2 | VPS26A | -0,5111 | 7,73576E-14 |
| ACE2 | H2AFZ | -0,5112 | 7,6923E-14 |
| ACE2 | PCDHAC1 | -0,5112 | 7,67853E-14 |
| ACE2 | SNCB | -0,5112 | 7,63003E-14 |
| ACE2 | PILRA | -0,5113 | 7,60933E-14 |
| ACE2 | GPR12 | -0,5113 | 7,55831E-14 |
| ACE2 | RP3-449M8.6 | -0,5113 | 7,58375E-14 |
| ACE2 | BCL2L12 | -0,5115 | 7,35717E-14 |
| ACE2 | CENPL | -0,5117 | 7,18471E-14 |
| ACE2 | CSTF1 | -0,5117 | 7,218E-14 |
| ACE2 | BRPF1 | -0,5118 | 7,05353E-14 |
| ACE2 | CD36 | -0,5118 | 7,05905E-14 |
| ACE2 | RPL17P50 | -0,5118 | 7,1041E-14 |
| ACE2 | WDR11-AS1 | -0,5118 | 7,12642E-14 |
| ACE2 | CELF5 | -0,5118 | 7,12512E-14 |
| ACE2 | RP4-668J24.2 | -0,5119 | 7,01366E-14 |
| ACE2 | RP11-513I15.6 | -0,5119 | 7,01391E-14 |
| ACE2 | FABP4 | -0,5119 | 6,97294E-14 |
| ACE2 | MRPS30 | -0,512 | 6,89813E-14 |
| ACE2 | AC006116.22 | -0,512 | 6,93966E-14 |
| ACE2 | AC005498.4 | -0,512 | 6,94703E-14 |
| ACE2 | ITSN2 | -0,5121 | 6,83043E-14 |
| ACE2 | UBA6-AS1 | -0,5121 | 6,86031E-14 |
| ACE2 | PIGBOS1 | -0,5121 | 6,84839E-14 |
| ACE2 | TAGLN3 | -0,5122 | 6,73514E-14 |
| ACE2 | C9orf43 | -0,5122 | 6,77655E-14 |
| ACE2 | RP3-449M8.9 | -0,5122 | 6,77633E-14 |

| | | | |
|---|---|---|---|
| ACE2 | ZFX | -0,5122 | 6,73834E-14 |
| ACE2 | RP11-1212A22.1 | -0,5123 | 6,61103E-14 |
| ACE2 | RP11-270M14.5 | -0,5124 | 6,54279E-14 |
| ACE2 | CTD-3018O17.5 | -0,5124 | 6,5597E-14 |
| ACE2 | HMGN2P19 | -0,5125 | 6,49216E-14 |
| ACE2 | RNF4 | -0,5125 | 6,4421E-14 |
| ACE2 | LINC01094 | -0,5125 | 6,49613E-14 |
| ACE2 | NOP2 | -0,5125 | 6,50371E-14 |
| ACE2 | TAF4B | -0,5125 | 6,46355E-14 |
| ACE2 | NBPF10 | -0,5126 | 6,41498E-14 |
| ACE2 | BCL2L2 | -0,5126 | 6,38242E-14 |
| ACE2 | ACACA | -0,5126 | 6,4073E-14 |
| ACE2 | CTD-2561J22.5 | -0,5126 | 6,42787E-14 |
| ACE2 | AC115522.3 | -0,5126 | 6,42011E-14 |
| ACE2 | RP5-978I12.1 | -0,5127 | 6,29955E-14 |
| ACE2 | AC068831.10 | -0,5127 | 6,31606E-14 |
| ACE2 | ZNF730 | -0,5127 | 6,27961E-14 |
| ACE2 | KDM5B | -0,5128 | 6,2015E-14 |
| ACE2 | RP11-53O19.3 | -0,5128 | 6,24272E-14 |
| ACE2 | RP11-661A12.8 | -0,5128 | 6,19984E-14 |
| ACE2 | RP11-685G9.4 | -0,5128 | 6,22009E-14 |
| ACE2 | ATXN1L | -0,5128 | 6,19093E-14 |
| ACE2 | PLIN4 | -0,5128 | 6,19262E-14 |
| ACE2 | PRR12 | -0,5128 | 6,23727E-14 |
| ACE2 | EEF1A1P6 | -0,5129 | 6,16577E-14 |
| ACE2 | NAP1L1P1 | -0,5129 | 6,1305E-14 |
| ACE2 | RP11-139H14.5 | -0,5129 | 6,18201E-14 |
| ACE2 | BANP | -0,5129 | 6,16744E-14 |
| ACE2 | RP4-535B20.4 | -0,513 | 6,03796E-14 |
| ACE2 | PTK2 | -0,513 | 6,07778E-14 |
| ACE2 | ACIN1 | -0,513 | 6,05915E-14 |
| ACE2 | EFCAB11 | -0,513 | 6,04302E-14 |
| ACE2 | CHRNE | -0,513 | 6,0728E-14 |
| ACE2 | ADPRHL2 | -0,5131 | 6,02074E-14 |
| ACE2 | AP2A2 | -0,5131 | 6,01895E-14 |
| ACE2 | RP11-1C8.6 | -0,5132 | 5,94126E-14 |
| ACE2 | RP11-481J2.1 | -0,5132 | 5,909E-14 |
| ACE2 | ZNF121 | -0,5132 | 5,93828E-14 |
| ACE2 | EIF4BP6 | -0,5133 | 5,86781E-14 |
| ACE2 | ITPRIPL2 | -0,5133 | 5,79759E-14 |
| ACE2 | AC018766.6 | -0,5133 | 5,8678E-14 |
| ACE2 | NUDT2 | -0,5134 | 5,76015E-14 |
| ACE2 | ZBTB39 | -0,5134 | 5,77003E-14 |
| ACE2 | TAMM41 | -0,5135 | 5,65716E-14 |
| ACE2 | ZNRF1 | -0,5135 | 5,68712E-14 |
| ACE2 | NBPF11 | -0,5136 | 5,60171E-14 |
| ACE2 | SWT1 | -0,5136 | 5,61349E-14 |
| ACE2 | RP11-452G18.2 | -0,5136 | 5,61368E-14 |
| ACE2 | LUC7L2 | -0,5137 | 5,51157E-14 |
| ACE2 | A1BG | -0,5137 | 5,56199E-14 |
| ACE2 | COA1 | -0,5138 | 5,47016E-14 |
| ACE2 | PLAA | -0,5138 | 5,43291E-14 |
| ACE2 | ZNF385C | -0,5138 | 5,47788E-14 |
| ACE2 | SS18L1 | -0,5139 | 5,39495E-14 |

| | | | |
|---|---|---|---|
| ACE2 | LRP1B | -0,514 | 5,34041E-14 |
| ACE2 | RPL21P135 | -0,514 | 5,33296E-14 |
| ACE2 | RP11-89C3.4 | -0,514 | 5,31907E-14 |
| ACE2 | RP11-131H24.4 | -0,514 | 5,32733E-14 |
| ACE2 | RP11-156L14.1 | -0,514 | 5,35426E-14 |
| ACE2 | TTPAL | -0,514 | 5,29888E-14 |
| ACE2 | GPKOW | -0,514 | 5,33789E-14 |
| ACE2 | METTL18 | -0,5141 | 5,22324E-14 |
| ACE2 | RP11-269F20.1 | -0,5141 | 5,23548E-14 |
| ACE2 | SNHG1 | -0,5141 | 5,25224E-14 |
| ACE2 | ALOX15 | -0,5141 | 5,23536E-14 |
| ACE2 | RP11-159D12.11 | -0,5141 | 5,2202E-14 |
| ACE2 | TMEM185B | -0,5142 | 5,18314E-14 |
| ACE2 | MPLKIP | -0,5142 | 5,17101E-14 |
| ACE2 | SMARCC2 | -0,5142 | 5,16337E-14 |
| ACE2 | ZNF324B | -0,5142 | 5,17832E-14 |
| ACE2 | TSPEAR | -0,5142 | 5,211E-14 |
| ACE2 | ODF3L2 | -0,5143 | 5,13178E-14 |
| ACE2 | FGR | -0,5144 | 5,07294E-14 |
| ACE2 | SRSF5 | -0,5144 | 5,04689E-14 |
| ACE2 | KIAA0895L | -0,5144 | 5,08211E-14 |
| ACE2 | RP5-1116H23.3 | -0,5144 | 5,0224E-14 |
| ACE2 | PCBP3-OT1 | -0,5145 | 5,00135E-14 |
| ACE2 | GAR1 | -0,5146 | 4,919E-14 |
| ACE2 | RP11-49G2.3 | -0,5146 | 4,90474E-14 |
| ACE2 | C16orf70 | -0,5147 | 4,83066E-14 |
| ACE2 | IWS1 | -0,5149 | 4,72808E-14 |
| ACE2 | EDRF1 | -0,5149 | 4,71043E-14 |
| ACE2 | RP11-95O2.5 | -0,5149 | 4,73381E-14 |
| ACE2 | MIR663AHG | -0,5149 | 4,73962E-14 |
| ACE2 | AC009784.3 | -0,515 | 4,67434E-14 |
| ACE2 | UFC1 | -0,5151 | 4,61491E-14 |
| ACE2 | RP11-367H1.1 | -0,5151 | 4,60033E-14 |
| ACE2 | TMEM70 | -0,5151 | 4,59625E-14 |
| ACE2 | IRF9 | -0,5151 | 4,59448E-14 |
| ACE2 | GSN | -0,5152 | 4,53686E-14 |
| ACE2 | TEAD1 | -0,5152 | 4,52183E-14 |
| ACE2 | RP11-872J21.5 | -0,5152 | 4,52264E-14 |
| ACE2 | RASGRP4 | -0,5152 | 4,51786E-14 |
| ACE2 | AC006007.1 | -0,5153 | 4,50609E-14 |
| ACE2 | LINC01436 | -0,5153 | 4,49691E-14 |
| ACE2 | FAM46B | -0,5154 | 4,43861E-14 |
| ACE2 | HSPA8P11 | -0,5154 | 4,4456E-14 |
| ACE2 | RP11-477N3.1 | -0,5155 | 4,39018E-14 |
| ACE2 | FBXO38 | -0,5155 | 4,34818E-14 |
| ACE2 | SFTPD-AS1 | -0,5155 | 4,37376E-14 |
| ACE2 | P2RY11 | -0,5156 | 4,32889E-14 |
| ACE2 | AP000223.42 | -0,5156 | 4,33533E-14 |
| ACE2 | KCNH1 | -0,5157 | 4,26037E-14 |
| ACE2 | VPS13B | -0,5157 | 4,24736E-14 |
| ACE2 | OSBPL8 | -0,5157 | 4,22954E-14 |
| ACE2 | RALGDS | -0,5158 | 4,17425E-14 |
| ACE2 | CACNA1B | -0,5158 | 4,21143E-14 |
| ACE2 | PHF12 | -0,5158 | 4,22006E-14 |

| | | | |
|---|---|---|---|
| ACE2 | ABCB8 | -0,5159 | 4,1538E-14 |
| ACE2 | LINC00844 | -0,5159 | 4,12919E-14 |
| ACE2 | TARBP2 | -0,5159 | 4,15208E-14 |
| ACE2 | POSTN | -0,5159 | 4,1677E-14 |
| ACE2 | MTCYBP21 | -0,5159 | 4,14688E-14 |
| ACE2 | RP4-706A16.3 | -0,516 | 4,09096E-14 |
| ACE2 | CSF3 | -0,516 | 4,09008E-14 |
| ACE2 | LLNLR-285B5.1 | -0,516 | 4,10622E-14 |
| ACE2 | RP11-488C13.6 | -0,5161 | 4,04747E-14 |
| ACE2 | FLJ16779 | -0,5161 | 4,05955E-14 |
| ACE2 | CCDC160 | -0,5161 | 4,05601E-14 |
| ACE2 | CTD-2161E19.1 | -0,5162 | 3,97659E-14 |
| ACE2 | LRRC15 | -0,5163 | 3,92239E-14 |
| ACE2 | ATM | -0,5163 | 3,94556E-14 |
| ACE2 | COL4A2-AS1 | -0,5163 | 3,91649E-14 |
| ACE2 | RP11-861E21.3 | -0,5163 | 3,94025E-14 |
| ACE2 | PIH1D1 | -0,5164 | 3,87822E-14 |
| ACE2 | ZNF586 | -0,5165 | 3,80993E-14 |
| ACE2 | AC073072.5 | -0,5166 | 3,78425E-14 |
| ACE2 | KIAA0895 | -0,5166 | 3,76672E-14 |
| ACE2 | RP11-654A16.3 | -0,5166 | 3,77039E-14 |
| ACE2 | HIST2H3PS2 | -0,5167 | 3,71343E-14 |
| ACE2 | LINC01587 | -0,5167 | 3,73406E-14 |
| ACE2 | ACSM1 | -0,5167 | 3,71283E-14 |
| ACE2 | RP11-583F2.2 | -0,5167 | 3,73783E-14 |
| ACE2 | CTD-2201E18.5 | -0,5168 | 3,68153E-14 |
| ACE2 | SYNGR3 | -0,5168 | 3,68466E-14 |
| ACE2 | SLC25A42 | -0,5168 | 3,67649E-14 |
| ACE2 | ELK4 | -0,5169 | 3,63918E-14 |
| ACE2 | PARP1 | -0,5169 | 3,62694E-14 |
| ACE2 | SELENOH | -0,517 | 3,58141E-14 |
| ACE2 | DDX27 | -0,517 | 3,59144E-14 |
| ACE2 | THAP10 | -0,5171 | 3,53451E-14 |
| ACE2 | AC009228.1 | -0,5172 | 3,48994E-14 |
| ACE2 | UNC5A | -0,5172 | 3,47224E-14 |
| ACE2 | ZNF366 | -0,5173 | 3,45958E-14 |
| ACE2 | CCDC167 | -0,5173 | 3,43663E-14 |
| ACE2 | CYSLTR1 | -0,5173 | 3,44312E-14 |
| ACE2 | LYRM7 | -0,5174 | 3,41009E-14 |
| ACE2 | PHF23 | -0,5174 | 3,41674E-14 |
| ACE2 | ZNF257 | -0,5174 | 3,4042E-14 |
| ACE2 | DPF1 | -0,5174 | 3,40231E-14 |
| ACE2 | RP6-109B7.4 | -0,5174 | 3,37819E-14 |
| ACE2 | KDM3B | -0,5175 | 3,33785E-14 |
| ACE2 | MAFF | -0,5175 | 3,37594E-14 |
| ACE2 | BTN2A2 | -0,5176 | 3,32882E-14 |
| ACE2 | CBX3 | -0,5176 | 3,32526E-14 |
| ACE2 | PLXNC1 | -0,5176 | 3,32406E-14 |
| ACE2 | LRPPRC | -0,5177 | 3,25527E-14 |
| ACE2 | SGCB | -0,5177 | 3,24806E-14 |
| ACE2 | UTP15 | -0,5177 | 3,27227E-14 |
| ACE2 | RP11-247A12.7 | -0,5177 | 3,26628E-14 |
| ACE2 | RP11-118K6.3 | -0,5177 | 3,26003E-14 |
| ACE2 | SLK | -0,5178 | 3,22706E-14 |

| | | | |
|---|---|---|---|
| ACE2 | LAMA5-AS1 | -0,5178 | 3,22744E-14 |
| ACE2 | ADORA2A | -0,5178 | 3,24055E-14 |
| ACE2 | KLHL18 | -0,5179 | 3,18522E-14 |
| ACE2 | TSFM | -0,5179 | 3,17333E-14 |
| ACE2 | LL0XNC01-237H1.2 | -0,5179 | 3,16505E-14 |
| ACE2 | COG7 | -0,518 | 3,15375E-14 |
| ACE2 | CTD-2619J13.19 | -0,518 | 3,11871E-14 |
| ACE2 | PXDC1 | -0,5181 | 3,09373E-14 |
| ACE2 | HLA-E | -0,5181 | 3,0892E-14 |
| ACE2 | NDUFAF4P3 | -0,5182 | 3,07089E-14 |
| ACE2 | CD47 | -0,5182 | 3,04906E-14 |
| ACE2 | WISP2 | -0,5182 | 3,06118E-14 |
| ACE2 | PROK2 | -0,5183 | 3,01293E-14 |
| ACE2 | GIT2 | -0,5183 | 3,0015E-14 |
| ACE2 | RP5-837J1.4 | -0,5183 | 3,02129E-14 |
| ACE2 | LINC01521 | -0,5183 | 3,03423E-14 |
| ACE2 | CYB5R3 | -0,5183 | 3,00876E-14 |
| ACE2 | ZBTB11 | -0,5184 | 2,98073E-14 |
| ACE2 | MFN1 | -0,5184 | 2,98045E-14 |
| ACE2 | CTB-33O18.3 | -0,5184 | 2,95837E-14 |
| ACE2 | PDCL3P5 | -0,5184 | 2,97302E-14 |
| ACE2 | MT1L | -0,5184 | 2,95944E-14 |
| ACE2 | AC084219.4 | -0,5184 | 2,99478E-14 |
| ACE2 | TTC32 | -0,5185 | 2,91944E-14 |
| ACE2 | CTBP1 | -0,5185 | 2,94342E-14 |
| ACE2 | FPGS | -0,5185 | 2,94368E-14 |
| ACE2 | BHLHE41 | -0,5185 | 2,92881E-14 |
| ACE2 | CCDC22 | -0,5185 | 2,93839E-14 |
| ACE2 | LRP1 | -0,5186 | 2,9077E-14 |
| ACE2 | ZCCHC8 | -0,5186 | 2,91044E-14 |
| ACE2 | KLHL17 | -0,5187 | 2,85594E-14 |
| ACE2 | DGUOK | -0,5187 | 2,86486E-14 |
| ACE2 | ACSM2A | -0,5187 | 2,84385E-14 |
| ACE2 | KCNIP2 | -0,5188 | 2,83371E-14 |
| ACE2 | ODF3B | -0,5188 | 2,83093E-14 |
| ACE2 | ELOA | -0,5189 | 2,77848E-14 |
| ACE2 | FAM185BP | -0,5189 | 2,80315E-14 |
| ACE2 | INTS6L | -0,5189 | 2,78997E-14 |
| ACE2 | ZNF275 | -0,5189 | 2,79295E-14 |
| ACE2 | PLEKHG1 | -0,519 | 2,74289E-14 |
| ACE2 | FAM66A | -0,519 | 2,74194E-14 |
| ACE2 | RP11-285F16.1 | -0,519 | 2,74604E-14 |
| ACE2 | CCNT1 | -0,519 | 2,73283E-14 |
| ACE2 | RP11-318A15.8 | -0,519 | 2,75399E-14 |
| ACE2 | TP53BP2 | -0,5191 | 2,7194E-14 |
| ACE2 | EFHD1 | -0,5191 | 2,71502E-14 |
| ACE2 | SLC16A12 | -0,5191 | 2,71452E-14 |
| ACE2 | CTD-2231E14.2 | -0,5191 | 2,70307E-14 |
| ACE2 | JAK1 | -0,5192 | 2,65913E-14 |
| ACE2 | NARS2 | -0,5192 | 2,66123E-14 |
| ACE2 | ZNF833P | -0,5192 | 2,67643E-14 |
| ACE2 | FAM196B | -0,5193 | 2,64684E-14 |
| ACE2 | TAF1C | -0,5193 | 2,64824E-14 |
| ACE2 | MIS12 | -0,5193 | 2,64856E-14 |

| | | | |
|---|---|---|---|
| ACE2 | TOX2 | -0,5194 | 2,59617E-14 |
| ACE2 | NDP | -0,5194 | 2,59475E-14 |
| ACE2 | ART5 | -0,5195 | 2,57676E-14 |
| ACE2 | FKBP3 | -0,5195 | 2,57874E-14 |
| ACE2 | RP11-259O2.1 | -0,5196 | 2,53945E-14 |
| ACE2 | MGC16275 | -0,5196 | 2,52745E-14 |
| ACE2 | ZNF317 | -0,5196 | 2,52527E-14 |
| ACE2 | MMP11 | -0,5196 | 2,5387E-14 |
| ACE2 | RP11-176H8.1 | -0,5197 | 2,50962E-14 |
| ACE2 | SLC25A19 | -0,5197 | 2,4975E-14 |
| ACE2 | PABPC1P7 | -0,5198 | 2,47537E-14 |
| ACE2 | CDC20P1 | -0,5198 | 2,47072E-14 |
| ACE2 | MED13L | -0,5198 | 2,45741E-14 |
| ACE2 | RP11-338N10.1 | -0,5199 | 2,4268E-14 |
| ACE2 | ZNF674 | -0,5199 | 2,44898E-14 |
| ACE2 | CH17-140K24.6 | -0,52 | 2,39961E-14 |
| ACE2 | FGF14-IT1 | -0,52 | 2,41738E-14 |
| ACE2 | RP11-857B24.5 | -0,52 | 2,39865E-14 |
| ACE2 | RP11-305K5.1 | -0,5202 | 2,33341E-14 |
| ACE2 | HDLBP | -0,5203 | 2,29778E-14 |
| ACE2 | FGF17 | -0,5204 | 2,28717E-14 |
| ACE2 | AHCYP2 | -0,5204 | 2,27102E-14 |
| ACE2 | PTPRQ | -0,5204 | 2,26783E-14 |
| ACE2 | SRCIN1 | -0,5204 | 2,27853E-14 |
| ACE2 | SNAP29 | -0,5204 | 2,27677E-14 |
| ACE2 | AMOTL2 | -0,5205 | 2,23698E-14 |
| ACE2 | RP11-799M12.2 | -0,5205 | 2,26119E-14 |
| ACE2 | RP3-406P24.5 | -0,5205 | 2,25764E-14 |
| ACE2 | FOXO3 | -0,5205 | 2,24691E-14 |
| ACE2 | HMBOX1 | -0,5205 | 2,2539E-14 |
| ACE2 | METTL15 | -0,5205 | 2,24098E-14 |
| ACE2 | ETV1 | -0,5206 | 2,20505E-14 |
| ACE2 | TRIM5 | -0,5206 | 2,2058E-14 |
| ACE2 | PRPH | -0,5206 | 2,21212E-14 |
| ACE2 | ZSWIM4 | -0,5206 | 2,21777E-14 |
| ACE2 | RIMS3 | -0,5207 | 2,18709E-14 |
| ACE2 | LINC01625 | -0,5207 | 2,18579E-14 |
| ACE2 | APCDD1 | -0,5207 | 2,17798E-14 |
| ACE2 | RASGRP3 | -0,5208 | 2,15658E-14 |
| ACE2 | NDUFB4P11 | -0,5208 | 2,1515E-14 |
| ACE2 | FAM86C2P | -0,5209 | 2,12889E-14 |
| ACE2 | CCDC77 | -0,5209 | 2,14026E-14 |
| ACE2 | GSAP | -0,521 | 2,10281E-14 |
| ACE2 | RP11-506M13.3 | -0,521 | 2,10442E-14 |
| ACE2 | RP11-314N13.3 | -0,521 | 2,09447E-14 |
| ACE2 | RP4-564M11.2 | -0,5211 | 2,07765E-14 |
| ACE2 | AC004951.5 | -0,5211 | 2,07917E-14 |
| ACE2 | RP4-593H12.1 | -0,5211 | 2,08149E-14 |
| ACE2 | SURF2 | -0,5211 | 2,06278E-14 |
| ACE2 | RP11-326C3.13 | -0,5211 | 2,06639E-14 |
| ACE2 | FAM153A | -0,5212 | 2,04258E-14 |
| ACE2 | SNX19 | -0,5212 | 2,06087E-14 |
| ACE2 | ARID2 | -0,5212 | 2,04835E-14 |
| ACE2 | PHC3 | -0,5213 | 2,02342E-14 |

| | | | |
|---|---|---|---|
| ACE2 | RP11-54O7.1 | -0,5215 | 1,97616E-14 |
| ACE2 | CNTN4 | -0,5215 | 1,97148E-14 |
| ACE2 | AC004453.8 | -0,5215 | 1,96971E-14 |
| ACE2 | RP11-218M22.1 | -0,5215 | 1,97896E-14 |
| ACE2 | RP11-178H8.7 | -0,5217 | 1,90961E-14 |
| ACE2 | ZC3H18 | -0,5217 | 1,92689E-14 |
| ACE2 | HNRNPD | -0,5218 | 1,89539E-14 |
| ACE2 | CFAP46 | -0,5218 | 1,88916E-14 |
| ACE2 | NLRP2B | -0,5218 | 1,8933E-14 |
| ACE2 | WI2-85898F10.2 | -0,5219 | 1,86795E-14 |
| ACE2 | CLHC1 | -0,522 | 1,83554E-14 |
| ACE2 | EBAG9 | -0,522 | 1,84876E-14 |
| ACE2 | RP11-278C7.5 | -0,522 | 1,83038E-14 |
| ACE2 | FBXL16 | -0,522 | 1,83837E-14 |
| ACE2 | SMG1P2 | -0,522 | 1,84298E-14 |
| ACE2 | TMEM101 | -0,522 | 1,84012E-14 |
| ACE2 | FBXO6 | -0,5221 | 1,81923E-14 |
| ACE2 | RP11-337N6.3 | -0,5221 | 1,80854E-14 |
| ACE2 | EEF1A1P13 | -0,5221 | 1,80685E-14 |
| ACE2 | RP11-84G21.1 | -0,5221 | 1,82216E-14 |
| ACE2 | LINC00682 | -0,5222 | 1,79383E-14 |
| ACE2 | TKTL1 | -0,5222 | 1,79291E-14 |
| ACE2 | RP11-134K13.4 | -0,5223 | 1,75584E-14 |
| ACE2 | SETX | -0,5223 | 1,75425E-14 |
| ACE2 | ZNF518B | -0,5224 | 1,73674E-14 |
| ACE2 | FASTKD2 | -0,5225 | 1,72809E-14 |
| ACE2 | RUVBL2 | -0,5225 | 1,7148E-14 |
| ACE2 | RP5-1115A15.1 | -0,5226 | 1,69287E-14 |
| ACE2 | RP1-20B11.2 | -0,5226 | 1,6847E-14 |
| ACE2 | RP11-1109F11.3 | -0,5226 | 1,70317E-14 |
| ACE2 | CSPG4P11 | -0,5226 | 1,69242E-14 |
| ACE2 | RP11-58A12.3 | -0,5227 | 1,67959E-14 |
| ACE2 | RP11-617F23.1 | -0,5227 | 1,68221E-14 |
| ACE2 | GP6 | -0,5227 | 1,6801E-14 |
| ACE2 | DSCAM | -0,5227 | 1,66449E-14 |
| ACE2 | ZBTB37 | -0,5228 | 1,66205E-14 |
| ACE2 | RAMP2-AS1 | -0,5228 | 1,64898E-14 |
| ACE2 | KCNK12 | -0,5229 | 1,63192E-14 |
| ACE2 | AQP5 | -0,5229 | 1,62016E-14 |
| ACE2 | CTC-523E23.1 | -0,5229 | 1,62933E-14 |
| ACE2 | SP3 | -0,523 | 1,61192E-14 |
| ACE2 | APBA3 | -0,523 | 1,60475E-14 |
| ACE2 | ARFRP1 | -0,523 | 1,61446E-14 |
| ACE2 | RNF182 | -0,5231 | 1,5915E-14 |
| ACE2 | PHLDB3 | -0,5231 | 1,59479E-14 |
| ACE2 | RP5-858L17.1 | -0,5231 | 1,59376E-14 |
| ACE2 | MTOR | -0,5232 | 1,56384E-14 |
| ACE2 | TRIM60P18 | -0,5232 | 1,57164E-14 |
| ACE2 | IKBKB | -0,5233 | 1,54935E-14 |
| ACE2 | AC093110.3 | -0,5234 | 1,51343E-14 |
| ACE2 | KAT2B | -0,5234 | 1,52853E-14 |
| ACE2 | ZNF721 | -0,5234 | 1,52911E-14 |
| ACE2 | CHST11 | -0,5234 | 1,53257E-14 |
| ACE2 | CDCA4 | -0,5234 | 1,53238E-14 |

| | | | |
|---|---|---|---|
| ACE2 | RP11-661O13.1 | -0,5234 | 1,5248E-14 |
| ACE2 | SLC1A4 | -0,5235 | 1,50543E-14 |
| ACE2 | HGSNAT | -0,5235 | 1,50067E-14 |
| ACE2 | ZW10 | -0,5235 | 1,49253E-14 |
| ACE2 | RP11-295I5.3 | -0,5235 | 1,50296E-14 |
| ACE2 | RP3-473L9.4 | -0,5235 | 1,49198E-14 |
| ACE2 | ZRANB2-AS2 | -0,5236 | 1,48856E-14 |
| ACE2 | CSNK1A1 | -0,5236 | 1,48033E-14 |
| ACE2 | PHTF2 | -0,5236 | 1,48696E-14 |
| ACE2 | RP11-364L4.1 | -0,5237 | 1,46989E-14 |
| ACE2 | ZNF138 | -0,5237 | 1,46575E-14 |
| ACE2 | LRRC42 | -0,5238 | 1,44566E-14 |
| ACE2 | ACP1 | -0,5238 | 1,45135E-14 |
| ACE2 | IGSF10 | -0,5238 | 1,43316E-14 |
| ACE2 | RP11-373N22.3 | -0,5238 | 1,43588E-14 |
| ACE2 | CTD-3076O17.1 | -0,5238 | 1,4489E-14 |
| ACE2 | CTD-2013N17.6 | -0,5238 | 1,43356E-14 |
| ACE2 | NFE2L2 | -0,5239 | 1,4265E-14 |
| ACE2 | RP11-479G22.8 | -0,5239 | 1,41427E-14 |
| ACE2 | MTIF3 | -0,524 | 1,41278E-14 |
| ACE2 | CGRRF1 | -0,5241 | 1,37683E-14 |
| ACE2 | TCEANC2 | -0,5242 | 1,36809E-14 |
| ACE2 | CTD-2008E3.1 | -0,5242 | 1,3611E-14 |
| ACE2 | STK19B | -0,5242 | 1,37506E-14 |
| ACE2 | GLIS3 | -0,5242 | 1,35832E-14 |
| ACE2 | HERC4 | -0,5242 | 1,3597E-14 |
| ACE2 | CTB-152G17.6 | -0,5243 | 1,34104E-14 |
| ACE2 | CTD-2206N4.4 | -0,5243 | 1,34008E-14 |
| ACE2 | SDR9C7 | -0,5244 | 1,32511E-14 |
| ACE2 | SDCBP2-AS1 | -0,5244 | 1,3368E-14 |
| ACE2 | PCBP1-AS1 | -0,5245 | 1,31048E-14 |
| ACE2 | RERGL | -0,5245 | 1,31957E-14 |
| ACE2 | TPST2 | -0,5245 | 1,31232E-14 |
| ACE2 | MYT1L | -0,5246 | 1,29404E-14 |
| ACE2 | SLC25A21 | -0,5246 | 1,29676E-14 |
| ACE2 | RP11-1260E13.4 | -0,5246 | 1,29128E-14 |
| ACE2 | SCPEP1 | -0,5246 | 1,28951E-14 |
| ACE2 | RP11-190A12.8 | -0,5247 | 1,2688E-14 |
| ACE2 | POU3F3 | -0,5247 | 1,2745E-14 |
| ACE2 | GIMAP1 | -0,5247 | 1,27807E-14 |
| ACE2 | CTC-325H20.4 | -0,5247 | 1,27935E-14 |
| ACE2 | CTD-2553C6.1 | -0,5247 | 1,26967E-14 |
| ACE2 | ZNF114 | -0,5247 | 1,28436E-14 |
| ACE2 | FAM156A | -0,5247 | 1,27735E-14 |
| ACE2 | RIT1 | -0,5248 | 1,25952E-14 |
| ACE2 | TBP | -0,5248 | 1,25186E-14 |
| ACE2 | C1QTNF9 | -0,5248 | 1,26485E-14 |
| ACE2 | NUBP1 | -0,5248 | 1,26615E-14 |
| ACE2 | KLK14 | -0,5248 | 1,25739E-14 |
| ACE2 | TRAK2 | -0,5249 | 1,23754E-14 |
| ACE2 | C3orf67 | -0,5249 | 1,23551E-14 |
| ACE2 | TRAPPC3L | -0,5249 | 1,23402E-14 |
| ACE2 | RP11-4B16.3 | -0,5249 | 1,24914E-14 |
| ACE2 | ZNF804B | -0,525 | 1,22567E-14 |

| | | | |
|---|---|---|---|
| ACE2 | RP5-1042I8.7 | -0,5251 | 1,20275E-14 |
| ACE2 | CYTOR | -0,5251 | 1,20432E-14 |
| ACE2 | FCN1 | -0,5251 | 1,21315E-14 |
| ACE2 | GACAT2 | -0,5251 | 1,21119E-14 |
| ACE2 | CTTNBP2 | -0,5252 | 1,19266E-14 |
| ACE2 | INA | -0,5252 | 1,19225E-14 |
| ACE2 | MGMT | -0,5252 | 1,18778E-14 |
| ACE2 | HMGN2P15 | -0,5252 | 1,19799E-14 |
| ACE2 | RP4-639F20.1 | -0,5253 | 1,17401E-14 |
| ACE2 | CRTC2 | -0,5253 | 1,1814E-14 |
| ACE2 | FAM114A1 | -0,5254 | 1,1567E-14 |
| ACE2 | MORN5 | -0,5254 | 1,16115E-14 |
| ACE2 | RP11-21A7A.4 | -0,5254 | 1,16801E-14 |
| ACE2 | RP11-396B14.2 | -0,5254 | 1,15532E-14 |
| ACE2 | CTC-524C5.2 | -0,5254 | 1,15291E-14 |
| ACE2 | PPM1N | -0,5255 | 1,14857E-14 |
| ACE2 | DCAF1 | -0,5256 | 1,12643E-14 |
| ACE2 | AKIRIN2 | -0,5256 | 1,12992E-14 |
| ACE2 | RP11-334J6.6 | -0,5256 | 1,13505E-14 |
| ACE2 | RP11-149I23.3 | -0,5256 | 1,12301E-14 |
| ACE2 | RP11-230G5.2 | -0,5257 | 1,11867E-14 |
| ACE2 | AC090044.2 | -0,5258 | 1,10243E-14 |
| ACE2 | AQR | -0,5258 | 1,09563E-14 |
| ACE2 | GBP4 | -0,5259 | 1,0785E-14 |
| ACE2 | CEBPZ | -0,5259 | 1,07918E-14 |
| ACE2 | RP11-187C18.3 | -0,5259 | 1,0857E-14 |
| ACE2 | MAU2 | -0,5259 | 1,0891E-14 |
| ACE2 | SYNDIG1 | -0,5259 | 1,07857E-14 |
| ACE2 | POLR2C | -0,526 | 1,06626E-14 |
| ACE2 | RABAC1 | -0,526 | 1,0726E-14 |
| ACE2 | CAV3 | -0,5261 | 1,05132E-14 |
| ACE2 | CCDC39 | -0,5262 | 1,03671E-14 |
| ACE2 | RP11-109A6.5 | -0,5263 | 1,02479E-14 |
| ACE2 | GOLGA6L5P | -0,5263 | 1,02293E-14 |
| ACE2 | CCDC71L | -0,5265 | 9,98211E-15 |
| ACE2 | UBXN2B | -0,5265 | 9,94097E-15 |
| ACE2 | CLSTN3 | -0,5265 | 1,00302E-14 |
| ACE2 | RP11-396F22.1 | -0,5265 | 9,95603E-15 |
| ACE2 | MED1 | -0,5265 | 9,96027E-15 |
| ACE2 | WDR43 | -0,5266 | 9,91854E-15 |
| ACE2 | RP11-381K20.2 | -0,5267 | 9,77533E-15 |
| ACE2 | SPRED1 | -0,5267 | 9,74882E-15 |
| ACE2 | TUBGCP4 | -0,5268 | 9,60376E-15 |
| ACE2 | MCF2 | -0,5268 | 9,53715E-15 |
| ACE2 | PLXNA1 | -0,5269 | 9,49153E-15 |
| ACE2 | RP11-392A14.8 | -0,5269 | 9,48007E-15 |
| ACE2 | RP11-111M22.4 | -0,5269 | 9,41996E-15 |
| ACE2 | TTC9B | -0,527 | 9,37725E-15 |
| ACE2 | ZNF8 | -0,5271 | 9,18028E-15 |
| ACE2 | PLA2G16 | -0,5272 | 9,06656E-15 |
| ACE2 | FZD10 | -0,5272 | 9,13202E-15 |
| ACE2 | KL | -0,5272 | 9,13491E-15 |
| ACE2 | RP11-403A3.2 | -0,5272 | 9,04498E-15 |
| ACE2 | HDAC6 | -0,5272 | 9,07088E-15 |

| | | | |
|---|---|---|---|
| ACE2 | DAB2 | -0,5273 | 8,92475E-15 |
| ACE2 | RP3-468K18.6 | -0,5273 | 8,9686E-15 |
| ACE2 | TBX6 | -0,5273 | 8,92868E-15 |
| ACE2 | MIR503HG | -0,5273 | 9,0005E-15 |
| ACE2 | HS3ST5 | -0,5274 | 8,85296E-15 |
| ACE2 | SF3B3 | -0,5274 | 8,87265E-15 |
| ACE2 | LINC02091 | -0,5274 | 8,84771E-15 |
| ACE2 | RBM6 | -0,5275 | 8,75087E-15 |
| ACE2 | CTD-2366F13.1 | -0,5275 | 8,6966E-15 |
| ACE2 | GPR137 | -0,5275 | 8,76733E-15 |
| ACE2 | MAP2K7 | -0,5275 | 8,72063E-15 |
| ACE2 | DGCR5 | -0,5275 | 8,74922E-15 |
| ACE2 | RP11-7O11.3 | -0,5277 | 8,48752E-15 |
| ACE2 | RP11-426C22.4 | -0,5277 | 8,51533E-15 |
| ACE2 | SON | -0,5277 | 8,43875E-15 |
| ACE2 | RP11-755F10.1 | -0,5278 | 8,33426E-15 |
| ACE2 | RP11-197M22.2 | -0,5279 | 8,23131E-15 |
| ACE2 | ZNF770 | -0,5279 | 8,28899E-15 |
| ACE2 | RP11-977B10.2 | -0,528 | 8,18409E-15 |
| ACE2 | RPS3AP6 | -0,528 | 8,10097E-15 |
| ACE2 | FUS | -0,528 | 8,12829E-15 |
| ACE2 | SLC39A3 | -0,528 | 8,09995E-15 |
| ACE2 | PURA | -0,5281 | 8,06255E-15 |
| ACE2 | SMYD4 | -0,5281 | 8,03282E-15 |
| ACE2 | B3GALT6 | -0,5282 | 7,90011E-15 |
| ACE2 | AC005197.2 | -0,5282 | 7,96808E-15 |
| ACE2 | AC074212.6 | -0,5282 | 7,89387E-15 |
| ACE2 | RP11-1081M5.1 | -0,5283 | 7,78975E-15 |
| ACE2 | KCNC1 | -0,5283 | 7,79002E-15 |
| ACE2 | RP5-1057I20.6 | -0,5283 | 7,85214E-15 |
| ACE2 | LINC01376 | -0,5284 | 7,73434E-15 |
| ACE2 | AC009948.7 | -0,5285 | 7,59475E-15 |
| ACE2 | IQCA1 | -0,5285 | 7,64192E-15 |
| ACE2 | MSC | -0,5285 | 7,62894E-15 |
| ACE2 | GSTO1 | -0,5285 | 7,62145E-15 |
| ACE2 | DBNDD1 | -0,5285 | 7,56684E-15 |
| ACE2 | RP11-466H18.1 | -0,5286 | 7,4576E-15 |
| ACE2 | CTD-2262B20.1 | -0,5286 | 7,45056E-15 |
| ACE2 | LINC00674 | -0,5286 | 7,49512E-15 |
| ACE2 | C10orf10 | -0,5287 | 7,41189E-15 |
| ACE2 | RYR1 | -0,5287 | 7,42071E-15 |
| ACE2 | BNIP3P28 | -0,5288 | 7,32279E-15 |
| ACE2 | TMLHE | -0,5288 | 7,32138E-15 |
| ACE2 | ZSCAN31 | -0,529 | 7,06832E-15 |
| ACE2 | CTA-268H5.14 | -0,529 | 7,11216E-15 |
| ACE2 | MTHFD2 | -0,5291 | 6,99921E-15 |
| ACE2 | P2RX3 | -0,5291 | 7,03505E-15 |
| ACE2 | RP11-800A3.7 | -0,5291 | 6,95428E-15 |
| ACE2 | R3HDM2 | -0,5291 | 6,99567E-15 |
| ACE2 | SLC26A11 | -0,5291 | 7,0192E-15 |
| ACE2 | ZNF274 | -0,5291 | 6,98341E-15 |
| ACE2 | PWRN1 | -0,5292 | 6,89371E-15 |
| ACE2 | LGSN | -0,5293 | 6,8092E-15 |
| ACE2 | MIR181A2HG | -0,5293 | 6,80276E-15 |

| | | | |
|---|---|---|---|
| ACE2 | MMS19 | -0,5293 | 6,77203E-15 |
| ACE2 | MYCBPAP | -0,5293 | 6,8257E-15 |
| ACE2 | TRIM60P17 | -0,5294 | 6,74936E-15 |
| ACE2 | FAM131B | -0,5294 | 6,71289E-15 |
| ACE2 | DDX11-AS1 | -0,5294 | 6,74228E-15 |
| ACE2 | RP11-63G10.4 | -0,5295 | 6,59188E-15 |
| ACE2 | SHISA6 | -0,5295 | 6,65206E-15 |
| ACE2 | RP11-490M8.1 | -0,5296 | 6,49414E-15 |
| ACE2 | SNHG5 | -0,5297 | 6,40959E-15 |
| ACE2 | RP11-585P4.6 | -0,5297 | 6,40186E-15 |
| ACE2 | RP11-936I5.1 | -0,5297 | 6,48099E-15 |
| ACE2 | RIMS4 | -0,5297 | 6,43215E-15 |
| ACE2 | TRIP12 | -0,5298 | 6,36762E-15 |
| ACE2 | POMT1 | -0,5298 | 6,34623E-15 |
| ACE2 | SUZ12 | -0,5298 | 6,32362E-15 |
| ACE2 | FAM87B | -0,5299 | 6,26072E-15 |
| ACE2 | KIAA1191 | -0,5299 | 6,29467E-15 |
| ACE2 | URM1 | -0,5299 | 6,22486E-15 |
| ACE2 | RP11-282I1.1 | -0,5299 | 6,29373E-15 |
| ACE2 | DDX46 | -0,53 | 6,14855E-15 |
| ACE2 | SYCP2L | -0,53 | 6,18165E-15 |
| ACE2 | CH17-13I23.3 | -0,5301 | 6,06785E-15 |
| ACE2 | LINC02147 | -0,5301 | 6,07576E-15 |
| ACE2 | CKB | -0,5301 | 6,09696E-15 |
| ACE2 | RP1-266L20.2 | -0,5302 | 6,02837E-15 |
| ACE2 | TAC1 | -0,5302 | 6,01421E-15 |
| ACE2 | TOX4 | -0,5302 | 6,00797E-15 |
| ACE2 | GOLGA8K | -0,5302 | 6,04271E-15 |
| ACE2 | TK2 | -0,5303 | 5,89884E-15 |
| ACE2 | NATD1 | -0,5303 | 5,88865E-15 |
| ACE2 | ZNF611 | -0,5303 | 5,9565E-15 |
| ACE2 | SPON2 | -0,5304 | 5,80769E-15 |
| ACE2 | PCDHB17P | -0,5304 | 5,85594E-15 |
| ACE2 | RP11-546D6.3 | -0,5305 | 5,7836E-15 |
| ACE2 | FLYWCH1 | -0,5305 | 5,73261E-15 |
| ACE2 | SYS1 | -0,5305 | 5,80086E-15 |
| ACE2 | GM2A | -0,5306 | 5,70699E-15 |
| ACE2 | LAMTOR4 | -0,5306 | 5,6846E-15 |
| ACE2 | LINC00462 | -0,5306 | 5,64799E-15 |
| ACE2 | AC073410.1 | -0,5307 | 5,62514E-15 |
| ACE2 | BTBD1 | -0,5307 | 5,60889E-15 |
| ACE2 | TMPRSS5 | -0,5308 | 5,54003E-15 |
| ACE2 | USP12 | -0,5308 | 5,5117E-15 |
| ACE2 | GLIS2-AS1 | -0,5308 | 5,5597E-15 |
| ACE2 | CSPG4P13 | -0,5309 | 5,43051E-15 |
| ACE2 | AATF | -0,5309 | 5,44539E-15 |
| ACE2 | PRPF6 | -0,5309 | 5,49072E-15 |
| ACE2 | PES1 | -0,5309 | 5,42171E-15 |
| ACE2 | RP3-522J7.6 | -0,5309 | 5,48022E-15 |
| ACE2 | TIMM8A | -0,5309 | 5,48049E-15 |
| ACE2 | RPS2P46 | -0,531 | 5,3778E-15 |
| ACE2 | TMEM18 | -0,5311 | 5,30447E-15 |
| ACE2 | DNAJC9 | -0,5311 | 5,29588E-15 |
| ACE2 | MSTO1 | -0,5312 | 5,2016E-15 |

| | | | |
|---|---|---|---|
| ACE2 | INO80B | -0,5312 | 5,23432E-15 |
| ACE2 | RP11-159D12.6 | -0,5312 | 5,21304E-15 |
| ACE2 | RP1-151F17.1 | -0,5313 | 5,13121E-15 |
| ACE2 | SCFD2 | -0,5314 | 5,05729E-15 |
| ACE2 | AC015849.2 | -0,5314 | 5,05627E-15 |
| ACE2 | RP11-15A1.8 | -0,5314 | 5,10076E-15 |
| ACE2 | NCOA3 | -0,5314 | 5,09215E-15 |
| ACE2 | RHOBTB3 | -0,5315 | 4,99509E-15 |
| ACE2 | FBXO18 | -0,5315 | 4,99811E-15 |
| ACE2 | NUTM2A-AS1 | -0,5315 | 5,03445E-15 |
| ACE2 | ARHGAP19 | -0,5315 | 5,02932E-15 |
| ACE2 | RASL10B | -0,5315 | 5,03671E-15 |
| ACE2 | ZNF90 | -0,5315 | 4,99768E-15 |
| ACE2 | TAF8 | -0,5317 | 4,87301E-15 |
| ACE2 | GRIK4 | -0,5317 | 4,89772E-15 |
| ACE2 | CENPC | -0,5318 | 4,84105E-15 |
| ACE2 | SMC5 | -0,5318 | 4,78947E-15 |
| ACE2 | RP13-514E23.2 | -0,5321 | 4,60578E-15 |
| ACE2 | C3orf80 | -0,5322 | 4,53404E-15 |
| ACE2 | PHBP13 | -0,5322 | 4,52768E-15 |
| ACE2 | RP11-666A8.7 | -0,5322 | 4,56658E-15 |
| ACE2 | GYS1 | -0,5323 | 4,49797E-15 |
| ACE2 | RP11-195B17.1 | -0,5324 | 4,41169E-15 |
| ACE2 | FAM86HP | -0,5324 | 4,43225E-15 |
| ACE2 | RP11-64B16.4 | -0,5324 | 4,44915E-15 |
| ACE2 | SZRD1 | -0,5325 | 4,39095E-15 |
| ACE2 | RP11-285F7.2 | -0,5325 | 4,361E-15 |
| ACE2 | PCDHA5 | -0,5325 | 4,38787E-15 |
| ACE2 | NAP1L4 | -0,5325 | 4,38123E-15 |
| ACE2 | XXbac-B562F10.11 | -0,5325 | 4,36334E-15 |
| ACE2 | RP11-549J18.1 | -0,5326 | 4,30465E-15 |
| ACE2 | CDH8 | -0,5326 | 4,27904E-15 |
| ACE2 | HTR1B | -0,5327 | 4,25776E-15 |
| ACE2 | RICTOR | -0,5328 | 4,1888E-15 |
| ACE2 | MFAP1P1 | -0,5328 | 4,18199E-15 |
| ACE2 | CCNJ | -0,5329 | 4,14561E-15 |
| ACE2 | PAXBP1 | -0,5329 | 4,11811E-15 |
| ACE2 | CREB1 | -0,533 | 4,09938E-15 |
| ACE2 | VENTX | -0,533 | 4,09498E-15 |
| ACE2 | SLITRK1 | -0,533 | 4,07318E-15 |
| ACE2 | CTB-58E17.1 | -0,533 | 4,06897E-15 |
| ACE2 | ZNF358 | -0,533 | 4,067E-15 |
| ACE2 | CXorf56 | -0,533 | 4,0686E-15 |
| ACE2 | RP11-350J20.5 | -0,5331 | 4,04043E-15 |
| ACE2 | HSPB1P1 | -0,5331 | 4,03135E-15 |
| ACE2 | RP11-690D19.3 | -0,5331 | 4,01861E-15 |
| ACE2 | LINC01489 | -0,5331 | 4,00628E-15 |
| ACE2 | HACD3 | -0,5332 | 3,93785E-15 |
| ACE2 | LMF1-AS1 | -0,5332 | 3,94496E-15 |
| ACE2 | NOL4 | -0,5332 | 3,98346E-15 |
| ACE2 | EEF1A1P19 | -0,5333 | 3,90739E-15 |
| ACE2 | MYO3A | -0,5333 | 3,89416E-15 |
| ACE2 | CCDC144NL-AS1 | -0,5333 | 3,92187E-15 |
| ACE2 | RP11-517P14.2 | -0,5334 | 3,86112E-15 |

| | | | |
|---|---|---|---|
| ACE2 | H2AFY2 | -0,5334 | 3,87785E-15 |
| ACE2 | RP11-613D13.8 | -0,5334 | 3,84903E-15 |
| ACE2 | RNF217-AS1 | -0,5335 | 3,80886E-15 |
| ACE2 | ACVR2B | -0,5336 | 3,75682E-15 |
| ACE2 | NUDT1 | -0,5336 | 3,73749E-15 |
| ACE2 | SNORA66 | -0,5337 | 3,67022E-15 |
| ACE2 | STAG3L4 | -0,5337 | 3,69123E-15 |
| ACE2 | C9orf142 | -0,5337 | 3,69648E-15 |
| ACE2 | ATF7IP | -0,5337 | 3,6986E-15 |
| ACE2 | DYNLRB2 | -0,5337 | 3,6812E-15 |
| ACE2 | NOP16 | -0,5338 | 3,62378E-15 |
| ACE2 | DPY19L2P2 | -0,5338 | 3,64652E-15 |
| ACE2 | SNRNP35 | -0,5338 | 3,66359E-15 |
| ACE2 | MAP2K4P1 | -0,5338 | 3,62288E-15 |
| ACE2 | GPR62 | -0,5339 | 3,60175E-15 |
| ACE2 | PMS2P3 | -0,5339 | 3,60519E-15 |
| ACE2 | SNX6 | -0,5339 | 3,61096E-15 |
| ACE2 | C21orf2 | -0,5339 | 3,5773E-15 |
| ACE2 | THAP6 | -0,534 | 3,52291E-15 |
| ACE2 | WDR74 | -0,534 | 3,53037E-15 |
| ACE2 | PEX12 | -0,534 | 3,54686E-15 |
| ACE2 | RHBDD3 | -0,534 | 3,54215E-15 |
| ACE2 | ATP13A3 | -0,5341 | 3,48363E-15 |
| ACE2 | LTA4H | -0,5341 | 3,5036E-15 |
| ACE2 | RP11-227G15.11 | -0,5341 | 3,51321E-15 |
| ACE2 | ITIH3 | -0,5342 | 3,44855E-15 |
| ACE2 | FABP5 | -0,5342 | 3,46354E-15 |
| ACE2 | DIRAS1 | -0,5342 | 3,45577E-15 |
| ACE2 | ADAMTSL2 | -0,5344 | 3,34259E-15 |
| ACE2 | ST3GAL1 | -0,5345 | 3,30078E-15 |
| ACE2 | AMER2 | -0,5345 | 3,2896E-15 |
| ACE2 | EXOSC6 | -0,5345 | 3,28783E-15 |
| ACE2 | DNAJC7 | -0,5345 | 3,32251E-15 |
| ACE2 | JMJD4 | -0,5346 | 3,26103E-15 |
| ACE2 | MDGA2 | -0,5346 | 3,23731E-15 |
| ACE2 | MRPL45 | -0,5346 | 3,25594E-15 |
| ACE2 | SLC5A3 | -0,5346 | 3,25106E-15 |
| ACE2 | WDR36 | -0,5347 | 3,21791E-15 |
| ACE2 | SWI5 | -0,5347 | 3,22048E-15 |
| ACE2 | H3F3B | -0,5347 | 3,22206E-15 |
| ACE2 | PIK3CD-AS2 | -0,5348 | 3,15933E-15 |
| ACE2 | PRG2 | -0,5348 | 3,1733E-15 |
| ACE2 | RP11-315O6.1 | -0,5348 | 3,1495E-15 |
| ACE2 | SMS | -0,5348 | 3,16707E-15 |
| ACE2 | RP11-646I6.5 | -0,5349 | 3,12393E-15 |
| ACE2 | GOLGA2 | -0,5349 | 3,10915E-15 |
| ACE2 | LUZP2 | -0,5349 | 3,10678E-15 |
| ACE2 | RP5-940J5.8 | -0,5349 | 3,10958E-15 |
| ACE2 | RP11-175B9.3 | -0,535 | 3,05978E-15 |
| ACE2 | MYC | -0,535 | 3,07014E-15 |
| ACE2 | RP11-408P14.1 | -0,5351 | 3,05563E-15 |
| ACE2 | RP11-188P17.2 | -0,5351 | 3,03254E-15 |
| ACE2 | NPIPA2 | -0,5351 | 3,04844E-15 |
| ACE2 | ZSCAN1 | -0,5351 | 3,0554E-15 |

| | | | |
|---|---|---|---|
| ACE2 | HAUS7 | -0,5351 | 3,02945E-15 |
| ACE2 | LINC01715 | -0,5352 | 3,0036E-15 |
| ACE2 | MATN3 | -0,5352 | 2,98968E-15 |
| ACE2 | POC5 | -0,5352 | 2,9971E-15 |
| ACE2 | RP11-401P9.7 | -0,5352 | 3,01063E-15 |
| ACE2 | ZNF761 | -0,5353 | 2,97016E-15 |
| ACE2 | CTB-114C7.4 | -0,5354 | 2,92458E-15 |
| ACE2 | LRCOL1 | -0,5354 | 2,89945E-15 |
| ACE2 | TFDP1 | -0,5354 | 2,92594E-15 |
| ACE2 | RP11-252A24.2 | -0,5354 | 2,90647E-15 |
| ACE2 | CTA-984G1.5 | -0,5354 | 2,9119E-15 |
| ACE2 | SYNE2 | -0,5355 | 2,86445E-15 |
| ACE2 | HNRNPA1P16 | -0,5355 | 2,85217E-15 |
| ACE2 | NME5 | -0,5356 | 2,83624E-15 |
| ACE2 | TSPEAR-AS1 | -0,5356 | 2,82792E-15 |
| ACE2 | LINC00638 | -0,5357 | 2,78623E-15 |
| ACE2 | CIRBP | -0,5357 | 2,77628E-15 |
| ACE2 | THAP12 | -0,5358 | 2,75264E-15 |
| ACE2 | POLG2 | -0,5358 | 2,74946E-15 |
| ACE2 | CYP2T1P | -0,5358 | 2,73554E-15 |
| ACE2 | FADS2 | -0,5359 | 2,69806E-15 |
| ACE2 | RP4-671G15.2 | -0,536 | 2,67989E-15 |
| ACE2 | RPL29P11 | -0,536 | 2,67584E-15 |
| ACE2 | PDE6B | -0,536 | 2,65627E-15 |
| ACE2 | RP11-635N19.1 | -0,536 | 2,66236E-15 |
| ACE2 | EIF2B5 | -0,5361 | 2,64433E-15 |
| ACE2 | RP11-479O17.12 | -0,5361 | 2,63951E-15 |
| ACE2 | NFKB1 | -0,5362 | 2,60488E-15 |
| ACE2 | SNORD87 | -0,5362 | 2,59799E-15 |
| ACE2 | RP11-365P13.5 | -0,5362 | 2,60041E-15 |
| ACE2 | CERS3 | -0,5362 | 2,59922E-15 |
| ACE2 | RP4-635E18.8 | -0,5363 | 2,5793E-15 |
| ACE2 | C1GALT1C1L | -0,5363 | 2,5463E-15 |
| ACE2 | AC007750.5 | -0,5363 | 2,56546E-15 |
| ACE2 | WDR6 | -0,5363 | 2,56719E-15 |
| ACE2 | ADGRL3 | -0,5363 | 2,5811E-15 |
| ACE2 | MPO | -0,5363 | 2,54564E-15 |
| ACE2 | MTHFD2P1 | -0,5364 | 2,52614E-15 |
| ACE2 | ZNF12 | -0,5364 | 2,53492E-15 |
| ACE2 | RBFOX1 | -0,5364 | 2,54421E-15 |
| ACE2 | AC112715.2 | -0,5365 | 2,50072E-15 |
| ACE2 | RP11-109N23.4 | -0,5365 | 2,48351E-15 |
| ACE2 | RP11-262A16.1 | -0,5365 | 2,50192E-15 |
| ACE2 | RFC2 | -0,5366 | 2,47001E-15 |
| ACE2 | GOLGA8H | -0,5366 | 2,46582E-15 |
| ACE2 | UXT | -0,5366 | 2,46957E-15 |
| ACE2 | GAL | -0,5367 | 2,43192E-15 |
| ACE2 | CTA-276F8.1 | -0,5367 | 2,42871E-15 |
| ACE2 | INTS4 | -0,5368 | 2,39673E-15 |
| ACE2 | GABPB1 | -0,5368 | 2,37838E-15 |
| ACE2 | INTS12 | -0,5369 | 2,34279E-15 |
| ACE2 | MTX3 | -0,5369 | 2,35987E-15 |
| ACE2 | TTC9C | -0,5369 | 2,33829E-15 |
| ACE2 | KLHL9 | -0,537 | 2,32094E-15 |

| | | | |
|---|---|---|---|
| ACE2 | DIABLO | -0,537 | 2,30837E-15 |
| ACE2 | RP11-710C12.1 | -0,5371 | 2,28649E-15 |
| ACE2 | ZBTB34 | -0,5371 | 2,29964E-15 |
| ACE2 | GLDN | -0,5371 | 2,28975E-15 |
| ACE2 | WWTR1-AS1 | -0,5372 | 2,27197E-15 |
| ACE2 | NNMT | -0,5373 | 2,21087E-15 |
| ACE2 | RP11-1365D11.1 | -0,5374 | 2,18752E-15 |
| ACE2 | PNLDC1 | -0,5374 | 2,18793E-15 |
| ACE2 | MAP3K7CL | -0,5374 | 2,20291E-15 |
| ACE2 | CNOT9 | -0,5375 | 2,15425E-15 |
| ACE2 | ID4 | -0,5375 | 2,15205E-15 |
| ACE2 | CBWD1 | -0,5375 | 2,15101E-15 |
| ACE2 | COL4A3BP | -0,5377 | 2,09447E-15 |
| ACE2 | CHTOP | -0,5378 | 2,072E-15 |
| ACE2 | MX2 | -0,5378 | 2,07774E-15 |
| ACE2 | RP1-224A6.9 | -0,5379 | 2,0481E-15 |
| ACE2 | MARCKS | -0,5379 | 2,03349E-15 |
| ACE2 | RP11-147L13.12 | -0,5379 | 2,05076E-15 |
| ACE2 | NTSR1 | -0,5379 | 2,03659E-15 |
| ACE2 | RRP7A | -0,5379 | 2,05679E-15 |
| ACE2 | ARPC2 | -0,5381 | 1,99901E-15 |
| ACE2 | RP5-907C10.3 | -0,5381 | 1,97544E-15 |
| ACE2 | RP11-357H14.16 | -0,5382 | 1,96748E-15 |
| ACE2 | ACP6 | -0,5383 | 1,936E-15 |
| ACE2 | LINC01948 | -0,5383 | 1,92947E-15 |
| ACE2 | CCDC61 | -0,5383 | 1,92911E-15 |
| ACE2 | AP001596.6 | -0,5383 | 1,9296E-15 |
| ACE2 | NANP | -0,5384 | 1,91517E-15 |
| ACE2 | SPOPL | -0,5385 | 1,89019E-15 |
| ACE2 | CCP110 | -0,5385 | 1,87246E-15 |
| ACE2 | KCTD18 | -0,5386 | 1,85719E-15 |
| ACE2 | RP11-673E1.3 | -0,5386 | 1,86151E-15 |
| ACE2 | HAS2 | -0,5386 | 1,85879E-15 |
| ACE2 | ANKLE1 | -0,5386 | 1,85503E-15 |
| ACE2 | SRRT | -0,5387 | 1,81861E-15 |
| ACE2 | TBC1D15 | -0,5387 | 1,81634E-15 |
| ACE2 | ATP1A4 | -0,5388 | 1,79419E-15 |
| ACE2 | RP11-105N14.1 | -0,5388 | 1,79989E-15 |
| ACE2 | ZNF704 | -0,5388 | 1,79633E-15 |
| ACE2 | CACNA1C-AS1 | -0,5389 | 1,76771E-15 |
| ACE2 | GNG7 | -0,5389 | 1,78511E-15 |
| ACE2 | RSPO2 | -0,539 | 1,75248E-15 |
| ACE2 | CHRAC1 | -0,539 | 1,74224E-15 |
| ACE2 | RP11-574K11.24 | -0,539 | 1,73579E-15 |
| ACE2 | FOXN3-AS1 | -0,539 | 1,73827E-15 |
| ACE2 | UBE2Q2L | -0,539 | 1,73598E-15 |
| ACE2 | RP11-449L23.2 | -0,539 | 1,75273E-15 |
| ACE2 | OR7E12P | -0,5391 | 1,71377E-15 |
| ACE2 | LIPI | -0,5391 | 1,72464E-15 |
| ACE2 | EMC1 | -0,5392 | 1,70008E-15 |
| ACE2 | LINC00619 | -0,5392 | 1,70415E-15 |
| ACE2 | TFAM | -0,5392 | 1,70136E-15 |
| ACE2 | GCDH | -0,5392 | 1,69221E-15 |
| ACE2 | TMEM130 | -0,5393 | 1,681E-15 |

| | | | |
|---|---|---|---|
| ACE2 | RNF31 | -0,5393 | 1,6681E-15 |
| ACE2 | MRPS9 | -0,5394 | 1,65603E-15 |
| ACE2 | RP11-543P15.1 | -0,5394 | 1,63994E-15 |
| ACE2 | ECSIT | -0,5394 | 1,64688E-15 |
| ACE2 | PTPN1 | -0,5394 | 1,65385E-15 |
| ACE2 | VPS13D | -0,5395 | 1,63666E-15 |
| ACE2 | TMEM206 | -0,5395 | 1,63889E-15 |
| ACE2 | RP11-493E12.1 | -0,5395 | 1,63848E-15 |
| ACE2 | RP11-140I16.3 | -0,5396 | 1,61256E-15 |
| ACE2 | ABHD17AP6 | -0,5396 | 1,59507E-15 |
| ACE2 | MITD1 | -0,5397 | 1,57773E-15 |
| ACE2 | XXbac-BPGBPG55C: | -0,5397 | 1,57893E-15 |
| ACE2 | AC004980.7 | -0,5397 | 1,59055E-15 |
| ACE2 | RNF169 | -0,5397 | 1,57724E-15 |
| ACE2 | MDN1 | -0,5398 | 1,55184E-15 |
| ACE2 | GNRH1 | -0,5398 | 1,56689E-15 |
| ACE2 | WRN | -0,5398 | 1,56014E-15 |
| ACE2 | SOX2 | -0,5399 | 1,5405E-15 |
| ACE2 | RP11-148L24.1 | -0,5399 | 1,52697E-15 |
| ACE2 | RAB32 | -0,5399 | 1,53896E-15 |
| ACE2 | WDTC1 | -0,54 | 1,5076E-15 |
| ACE2 | LINC02019 | -0,54 | 1,51611E-15 |
| ACE2 | RP1-240B8.3 | -0,54 | 1,51963E-15 |
| ACE2 | PPP6C | -0,54 | 1,51537E-15 |
| ACE2 | ZNF747 | -0,54 | 1,51398E-15 |
| ACE2 | TRIM17 | -0,5401 | 1,49069E-15 |
| ACE2 | RIPPLY2 | -0,5401 | 1,48775E-15 |
| ACE2 | PTGDS | -0,5402 | 1,47748E-15 |
| ACE2 | DAZAP1 | -0,5402 | 1,47959E-15 |
| ACE2 | KB-1042C11.5 | -0,5403 | 1,45232E-15 |
| ACE2 | RP11-417L19.4 | -0,5403 | 1,44783E-15 |
| ACE2 | HECTD1 | -0,5403 | 1,44502E-15 |
| ACE2 | DLX4 | -0,5403 | 1,44792E-15 |
| ACE2 | RP11-517B11.7 | -0,5404 | 1,42977E-15 |
| ACE2 | HSD17B1 | -0,5404 | 1,43376E-15 |
| ACE2 | CHST7 | -0,5404 | 1,42685E-15 |
| ACE2 | TAF1A | -0,5405 | 1,40946E-15 |
| ACE2 | AMT | -0,5405 | 1,41407E-15 |
| ACE2 | ALG8 | -0,5405 | 1,4202E-15 |
| ACE2 | LINC01198 | -0,5405 | 1,40502E-15 |
| ACE2 | EDNRB | -0,5405 | 1,40336E-15 |
| ACE2 | RP11-640N20.4 | -0,5405 | 1,412E-15 |
| ACE2 | WFIKKN2 | -0,5405 | 1,40335E-15 |
| ACE2 | DPH2 | -0,5406 | 1,39334E-15 |
| ACE2 | SLC35B2 | -0,5406 | 1,39468E-15 |
| ACE2 | PIN4 | -0,5406 | 1,38837E-15 |
| ACE2 | ELP6 | -0,5407 | 1,37496E-15 |
| ACE2 | ZNF436-AS1 | -0,5408 | 1,34384E-15 |
| ACE2 | C4orf50 | -0,5408 | 1,34981E-15 |
| ACE2 | CUX2 | -0,5408 | 1,35121E-15 |
| ACE2 | PRCD | -0,5408 | 1,351E-15 |
| ACE2 | EED | -0,5409 | 1,33044E-15 |
| ACE2 | AC100830.5 | -0,5409 | 1,33371E-15 |
| ACE2 | TBX18 | -0,541 | 1,30902E-15 |

| | | | |
|---|---|---|---|
| ACE2 | KCNE2 | -0,541 | 1,32108E-15 |
| ACE2 | APOBEC3F | -0,541 | 1,31682E-15 |
| ACE2 | RP11-377G16.2 | -0,5411 | 1,30268E-15 |
| ACE2 | CTC-548K16.6 | -0,5411 | 1,2991E-15 |
| ACE2 | AC093609.1 | -0,5412 | 1,26945E-15 |
| ACE2 | SERTAD2 | -0,5412 | 1,28239E-15 |
| ACE2 | RP11-46C24.7 | -0,5412 | 1,28432E-15 |
| ACE2 | CNKSR2 | -0,5412 | 1,26937E-15 |
| ACE2 | GTF2IRD1 | -0,5413 | 1,25591E-15 |
| ACE2 | RP11-174J11.1 | -0,5413 | 1,25504E-15 |
| ACE2 | CCL14 | -0,5413 | 1,26707E-15 |
| ACE2 | SEPT7-AS1 | -0,5414 | 1,23421E-15 |
| ACE2 | DCUN1D2 | -0,5414 | 1,23331E-15 |
| ACE2 | SYMPK | -0,5414 | 1,23952E-15 |
| ACE2 | CEP250 | -0,5414 | 1,24121E-15 |
| ACE2 | ANKRD7 | -0,5415 | 1,23011E-15 |
| ACE2 | TRUB1 | -0,5415 | 1,22224E-15 |
| ACE2 | HMSD | -0,5415 | 1,21646E-15 |
| ACE2 | GAB3 | -0,5415 | 1,2195E-15 |
| ACE2 | PCDHA2 | -0,5416 | 1,2028E-15 |
| ACE2 | RP11-658F2.8 | -0,5416 | 1,2012E-15 |
| ACE2 | ENTPD3 | -0,5417 | 1,19647E-15 |
| ACE2 | CCNT2-AS1 | -0,5418 | 1,17948E-15 |
| ACE2 | ZNF613 | -0,5418 | 1,16317E-15 |
| ACE2 | CRYBB1 | -0,5418 | 1,17786E-15 |
| ACE2 | RP11-60A14.1 | -0,5419 | 1,15563E-15 |
| ACE2 | NDNF | -0,5419 | 1,16228E-15 |
| ACE2 | RP11-500C11.3 | -0,542 | 1,13067E-15 |
| ACE2 | KMT2E | -0,542 | 1,13391E-15 |
| ACE2 | HERC2 | -0,542 | 1,14561E-15 |
| ACE2 | AP000280.66 | -0,542 | 1,14129E-15 |
| ACE2 | RPL5P34 | -0,542 | 1,13724E-15 |
| ACE2 | SAP30 | -0,5421 | 1,12393E-15 |
| ACE2 | DNM1P46 | -0,5421 | 1,12543E-15 |
| ACE2 | RP11-390P24.1 | -0,5421 | 1,12957E-15 |
| ACE2 | C17orf67 | -0,5421 | 1,11444E-15 |
| ACE2 | NBPF20 | -0,5422 | 1,11196E-15 |
| ACE2 | SDHAF4 | -0,5422 | 1,10144E-15 |
| ACE2 | DNAJA3 | -0,5422 | 1,09976E-15 |
| ACE2 | JAKMIP1 | -0,5423 | 1,09491E-15 |
| ACE2 | CTB-113P19.1 | -0,5423 | 1,08449E-15 |
| ACE2 | KLHL33 | -0,5423 | 1,08746E-15 |
| ACE2 | PIAS4 | -0,5423 | 1,09204E-15 |
| ACE2 | ST3GAL5-AS1 | -0,5424 | 1,06701E-15 |
| ACE2 | HIVEP1 | -0,5425 | 1,05642E-15 |
| ACE2 | GPR20 | -0,5425 | 1,06338E-15 |
| ACE2 | RBM14 | -0,5425 | 1,05851E-15 |
| ACE2 | RBM23 | -0,5425 | 1,06622E-15 |
| ACE2 | CTC-542B22.2 | -0,5425 | 1,06275E-15 |
| ACE2 | AD001527.4 | -0,5425 | 1,05847E-15 |
| ACE2 | MTPAP | -0,5426 | 1,04821E-15 |
| ACE2 | FANCE | -0,5427 | 1,02926E-15 |
| ACE2 | RP11-382J24.2 | -0,5427 | 1,03339E-15 |
| ACE2 | SETD1A | -0,5427 | 1,02359E-15 |

| | | | |
|---|---|---|---|
| ACE2 | AC124789.1 | -0,5427 | 1,02654E-15 |
| ACE2 | PAXBP1-AS1 | -0,5427 | 1,02683E-15 |
| ACE2 | COX7A2L | -0,5428 | 1,02109E-15 |
| ACE2 | HSBP1 | -0,5428 | 1,00822E-15 |
| ACE2 | ISM1 | -0,5428 | 1,01632E-15 |
| ACE2 | RP11-9N20.3 | -0,5429 | 9,94337E-16 |
| ACE2 | LINC01314 | -0,5429 | 1,00306E-15 |
| ACE2 | CTD-3157E16.1 | -0,543 | 9,79189E-16 |
| ACE2 | TRMT11 | -0,5431 | 9,72315E-16 |
| ACE2 | LL22NC03-86G7.1 | -0,5431 | 9,71056E-16 |
| ACE2 | TGFBRAP1 | -0,5432 | 9,61202E-16 |
| ACE2 | NEU3 | -0,5432 | 9,57591E-16 |
| ACE2 | AC073316.1 | -0,5433 | 9,40291E-16 |
| ACE2 | CCNYL2 | -0,5433 | 9,38868E-16 |
| ACE2 | RP11-762L8.6 | -0,5433 | 9,46796E-16 |
| ACE2 | LMO7DN | -0,5433 | 9,48099E-16 |
| ACE2 | GNPTG | -0,5434 | 9,36109E-16 |
| ACE2 | RP11-615I2.6 | -0,5434 | 9,27622E-16 |
| ACE2 | HIVEP3 | -0,5435 | 9,17827E-16 |
| ACE2 | MANBA | -0,5435 | 9,13578E-16 |
| ACE2 | RP5-1142A6.2 | -0,5435 | 9,22739E-16 |
| ACE2 | SMG9 | -0,5435 | 9,10884E-16 |
| ACE2 | PLCG1-AS1 | -0,5435 | 9,23496E-16 |
| ACE2 | GNL3 | -0,5436 | 8,97942E-16 |
| ACE2 | PDLIM1P4 | -0,5436 | 9,02345E-16 |
| ACE2 | RP11-216B9.9 | -0,5436 | 9,09129E-16 |
| ACE2 | IL34 | -0,5436 | 9,03284E-16 |
| ACE2 | HRH4 | -0,5436 | 9,0695E-16 |
| ACE2 | IGF2BP3 | -0,5438 | 8,78304E-16 |
| ACE2 | RP11-662G23.1 | -0,5438 | 8,77348E-16 |
| ACE2 | HPD | -0,5438 | 8,76838E-16 |
| ACE2 | RP11-403A3.3 | -0,5438 | 8,8346E-16 |
| ACE2 | SETD6 | -0,5438 | 8,81165E-16 |
| ACE2 | SLC22A15 | -0,5439 | 8,60154E-16 |
| ACE2 | C19orf12 | -0,5439 | 8,63671E-16 |
| ACE2 | RP11-468H14.2 | -0,544 | 8,51016E-16 |
| ACE2 | ING1 | -0,544 | 8,53803E-16 |
| ACE2 | FBXO15 | -0,544 | 8,48214E-16 |
| ACE2 | BEX1 | -0,544 | 8,51322E-16 |
| ACE2 | MAP6D1 | -0,5441 | 8,39107E-16 |
| ACE2 | DAXX | -0,5441 | 8,41949E-16 |
| ACE2 | ITGAM | -0,5441 | 8,40781E-16 |
| ACE2 | RB1CC1 | -0,5442 | 8,23494E-16 |
| ACE2 | PPM1E | -0,5442 | 8,33391E-16 |
| ACE2 | FAM131A | -0,5443 | 8,22499E-16 |
| ACE2 | CDK19 | -0,5443 | 8,16336E-16 |
| ACE2 | RRM2P3 | -0,5443 | 8,14501E-16 |
| ACE2 | NPAP1 | -0,5444 | 8,00286E-16 |
| ACE2 | RP11-227D13.1 | -0,5444 | 8,00124E-16 |
| ACE2 | IGLON5 | -0,5444 | 8,05512E-16 |
| ACE2 | PPEF1 | -0,5444 | 8,07959E-16 |
| ACE2 | SACS-AS1 | -0,5445 | 7,96918E-16 |
| ACE2 | CCDC192 | -0,5446 | 7,81999E-16 |
| ACE2 | CFL1P1 | -0,5446 | 7,81034E-16 |

| | | | |
|---|---|---|---|
| ACE2 | RSRC2 | -0,5446 | 7,84538E-16 |
| ACE2 | PSMA7 | -0,5446 | 7,81517E-16 |
| ACE2 | PTP4A2P2 | -0,5447 | 7,69285E-16 |
| ACE2 | GPATCH11 | -0,5448 | 7,54713E-16 |
| ACE2 | LINC00877 | -0,5448 | 7,54165E-16 |
| ACE2 | RP11-153F10.2 | -0,5448 | 7,62455E-16 |
| ACE2 | NMT1 | -0,5448 | 7,62693E-16 |
| ACE2 | SNRPD1 | -0,5448 | 7,57166E-16 |
| ACE2 | UHRF2 | -0,5449 | 7,46706E-16 |
| ACE2 | CES1 | -0,5449 | 7,45206E-16 |
| ACE2 | MSI2 | -0,5449 | 7,47629E-16 |
| ACE2 | IRF2BP1 | -0,5449 | 7,5344E-16 |
| ACE2 | TMCO6 | -0,545 | 7,41516E-16 |
| ACE2 | SFR1 | -0,545 | 7,34214E-16 |
| ACE2 | RBM25 | -0,545 | 7,36245E-16 |
| ACE2 | IGFBP6 | -0,5451 | 7,31675E-16 |
| ACE2 | ANGEL2 | -0,5452 | 7,13326E-16 |
| ACE2 | RP11-486I11.2 | -0,5452 | 7,20275E-16 |
| ACE2 | GCSAML | -0,5453 | 7,06775E-16 |
| ACE2 | RASA4B | -0,5453 | 7,03646E-16 |
| ACE2 | RASA4 | -0,5453 | 7,04115E-16 |
| ACE2 | PSMB5 | -0,5453 | 7,04141E-16 |
| ACE2 | CTA-390C10.10 | -0,5453 | 7,01559E-16 |
| ACE2 | NUP107 | -0,5454 | 7,01278E-16 |
| ACE2 | HEXA | -0,5454 | 6,97106E-16 |
| ACE2 | CNGB1 | -0,5454 | 6,92816E-16 |
| ACE2 | RP11-73M18.8 | -0,5455 | 6,85618E-16 |
| ACE2 | ZNF776 | -0,5455 | 6,85044E-16 |
| ACE2 | TUBBP1 | -0,5456 | 6,72744E-16 |
| ACE2 | FAM69C | -0,5457 | 6,65871E-16 |
| ACE2 | AKAP8L | -0,5457 | 6,66571E-16 |
| ACE2 | PLS3-AS1 | -0,5457 | 6,63145E-16 |
| ACE2 | CTD-2311M21.2 | -0,5458 | 6,56877E-16 |
| ACE2 | MED12 | -0,5458 | 6,52337E-16 |
| ACE2 | MAPK14 | -0,5459 | 6,42717E-16 |
| ACE2 | TNNT3 | -0,5459 | 6,47404E-16 |
| ACE2 | RP11-426C22.1 | -0,5459 | 6,51063E-16 |
| ACE2 | RFPL1S | -0,546 | 6,37552E-16 |
| ACE2 | NDUFS5 | -0,5461 | 6,29827E-16 |
| ACE2 | RP11-203B9.4 | -0,5461 | 6,28306E-16 |
| ACE2 | SLC38A1 | -0,5461 | 6,30854E-16 |
| ACE2 | ZNF529-AS1 | -0,5461 | 6,24301E-16 |
| ACE2 | MYOZ3 | -0,5462 | 6,20766E-16 |
| ACE2 | ZCCHC2 | -0,5463 | 6,08832E-16 |
| ACE2 | RP4-575N6.5 | -0,5464 | 6,00598E-16 |
| ACE2 | RP11-385F5.5 | -0,5464 | 6,02312E-16 |
| ACE2 | LIPT1 | -0,5464 | 6,03809E-16 |
| ACE2 | SNX13 | -0,5464 | 5,9992E-16 |
| ACE2 | RP11-5C23.2 | -0,5464 | 5,97515E-16 |
| ACE2 | RP11-1134I14.4 | -0,5464 | 6,02821E-16 |
| ACE2 | MYCBP2-AS1 | -0,5464 | 6,00343E-16 |
| ACE2 | GPS2 | -0,5464 | 6,04617E-16 |
| ACE2 | PTRH2 | -0,5464 | 5,98658E-16 |
| ACE2 | JAG1 | -0,5464 | 6,03338E-16 |

| | | | |
|---|---|---|---|
| ACE2 | FGD5-AS1 | -0,5465 | 5,94045E-16 |
| ACE2 | CNTNAP3B | -0,5465 | 5,95802E-16 |
| ACE2 | PMPCA | -0,5465 | 5,95352E-16 |
| ACE2 | RP11-977G19.12 | -0,5465 | 5,96854E-16 |
| ACE2 | ANKIB1 | -0,5466 | 5,82362E-16 |
| ACE2 | PEMT | -0,5466 | 5,81696E-16 |
| ACE2 | RP11-234K19.1 | -0,5467 | 5,73872E-16 |
| ACE2 | ZNF221 | -0,5467 | 5,75874E-16 |
| ACE2 | RPL7P1 | -0,5468 | 5,71701E-16 |
| ACE2 | TSPAN31 | -0,5468 | 5,68831E-16 |
| ACE2 | SERPINA5 | -0,5468 | 5,67544E-16 |
| ACE2 | MEG9 | -0,5468 | 5,66266E-16 |
| ACE2 | Z97634.3 | -0,5468 | 5,67883E-16 |
| ACE2 | DLX6-AS1 | -0,5469 | 5,6117E-16 |
| ACE2 | RHEBL1 | -0,5469 | 5,6343E-16 |
| ACE2 | APEX1 | -0,5469 | 5,58309E-16 |
| ACE2 | ASXL2 | -0,547 | 5,54789E-16 |
| ACE2 | TLR4 | -0,547 | 5,47378E-16 |
| ACE2 | SNX2 | -0,5471 | 5,46337E-16 |
| ACE2 | RP11-418H16.1 | -0,5472 | 5,38318E-16 |
| ACE2 | BTN2A1 | -0,5472 | 5,37612E-16 |
| ACE2 | ZMYND10 | -0,5473 | 5,24144E-16 |
| ACE2 | PPP1R9A | -0,5473 | 5,26355E-16 |
| ACE2 | ZNF862 | -0,5473 | 5,26034E-16 |
| ACE2 | GFRA2 | -0,5474 | 5,19214E-16 |
| ACE2 | ZNF85 | -0,5474 | 5,20001E-16 |
| ACE2 | OR51E1 | -0,5475 | 5,11262E-16 |
| ACE2 | VDAC3 | -0,5476 | 5,04228E-16 |
| ACE2 | ACTN3 | -0,5476 | 5,04111E-16 |
| ACE2 | CTC-265F19.2 | -0,5476 | 5,04999E-16 |
| ACE2 | RCAN2 | -0,5477 | 4,95863E-16 |
| ACE2 | TRIM4 | -0,5477 | 4,96602E-16 |
| ACE2 | RBL2 | -0,5477 | 4,98165E-16 |
| ACE2 | LINC01535 | -0,5477 | 4,95578E-16 |
| ACE2 | GTF2IP1 | -0,5478 | 4,92743E-16 |
| ACE2 | RPS7P1 | -0,5478 | 4,93505E-16 |
| ACE2 | AADACL4 | -0,5479 | 4,85326E-16 |
| ACE2 | RP11-890B15.3 | -0,5479 | 4,85325E-16 |
| ACE2 | RP11-946P6.4 | -0,5479 | 4,84258E-16 |
| ACE2 | ADSSL1 | -0,5479 | 4,79826E-16 |
| ACE2 | TOP3A | -0,5479 | 4,85385E-16 |
| ACE2 | ZFP14 | -0,5479 | 4,83407E-16 |
| ACE2 | CDK5RAP1 | -0,5479 | 4,80554E-16 |
| ACE2 | SLMAP | -0,548 | 4,73105E-16 |
| ACE2 | CCND2-AS2 | -0,548 | 4,77027E-16 |
| ACE2 | MID1 | -0,5481 | 4,69478E-16 |
| ACE2 | RP4-751H13.7 | -0,5482 | 4,61084E-16 |
| ACE2 | FAM66B | -0,5482 | 4,62961E-16 |
| ACE2 | CEBPD | -0,5482 | 4,62044E-16 |
| ACE2 | GRWD1 | -0,5483 | 4,58971E-16 |
| ACE2 | ZNF581 | -0,5483 | 4,53691E-16 |
| ACE2 | B4GALT7 | -0,5484 | 4,47454E-16 |
| ACE2 | RP11-344B5.2 | -0,5484 | 4,46294E-16 |
| ACE2 | TATDN3 | -0,5486 | 4,37905E-16 |

| | | | |
|---|---|---|---|
| ACE2 | AC025171.1 | -0,5486 | 4,35478E-16 |
| ACE2 | PCYOX1L | -0,5486 | 4,33385E-16 |
| ACE2 | DMGDH | -0,5487 | 4,31852E-16 |
| ACE2 | PCDHA4 | -0,5487 | 4,28635E-16 |
| ACE2 | CTB-113P19.5 | -0,5487 | 4,30879E-16 |
| ACE2 | DNAJC30 | -0,5487 | 4,32801E-16 |
| ACE2 | SEC23IP | -0,5487 | 4,30746E-16 |
| ACE2 | IMPDH1P8 | -0,5488 | 4,2211E-16 |
| ACE2 | NRXN2 | -0,5488 | 4,20975E-16 |
| ACE2 | USP32 | -0,5488 | 4,24694E-16 |
| ACE2 | ZNF792 | -0,5489 | 4,19192E-16 |
| ACE2 | UBE2S | -0,5489 | 4,15923E-16 |
| ACE2 | BOLA1 | -0,549 | 4,13769E-16 |
| ACE2 | AC053503.12 | -0,549 | 4,08427E-16 |
| ACE2 | RP11-272P10.2 | -0,549 | 4,13519E-16 |
| ACE2 | CDC123 | -0,549 | 4,08531E-16 |
| ACE2 | CEP295NL | -0,549 | 4,13541E-16 |
| ACE2 | CHRM3 | -0,5491 | 4,073E-16 |
| ACE2 | RP11-69I8.3 | -0,5491 | 4,07784E-16 |
| ACE2 | RP11-463O9.1 | -0,5491 | 4,05066E-16 |
| ACE2 | ZNF534 | -0,5491 | 4,0619E-16 |
| ACE2 | OPRD1 | -0,5492 | 3,96672E-16 |
| ACE2 | LINC00466 | -0,5492 | 3,97229E-16 |
| ACE2 | FBXO3 | -0,5492 | 3,99326E-16 |
| ACE2 | ZNF407 | -0,5492 | 4,01116E-16 |
| ACE2 | FTX | -0,5492 | 3,97344E-16 |
| ACE2 | RP11-307L14.1 | -0,5493 | 3,92638E-16 |
| ACE2 | RP5-855D21.1 | -0,5493 | 3,96069E-16 |
| ACE2 | ACTN1-AS1 | -0,5493 | 3,9244E-16 |
| ACE2 | DIP2C | -0,5494 | 3,88601E-16 |
| ACE2 | ZNF573 | -0,5494 | 3,8587E-16 |
| ACE2 | ATOH8 | -0,5495 | 3,80401E-16 |
| ACE2 | IQCB1 | -0,5495 | 3,79368E-16 |
| ACE2 | SNRPF | -0,5495 | 3,79639E-16 |
| ACE2 | GCN1 | -0,5495 | 3,83203E-16 |
| ACE2 | RPS26P21 | -0,5496 | 3,7555E-16 |
| ACE2 | ZBED3-AS1 | -0,5496 | 3,74073E-16 |
| ACE2 | TRPC6 | -0,5496 | 3,77261E-16 |
| ACE2 | TSC22D1 | -0,5496 | 3,73719E-16 |
| ACE2 | RP11-342K6.1 | -0,5497 | 3,6846E-16 |
| ACE2 | APOLD1 | -0,5497 | 3,6826E-16 |
| ACE2 | GOLGA8A | -0,5498 | 3,64702E-16 |
| ACE2 | SCGB1B2P | -0,5498 | 3,66763E-16 |
| ACE2 | CTD-2104P17.1 | -0,5499 | 3,60057E-16 |
| ACE2 | RP11-90L20.2 | -0,5501 | 3,50128E-16 |
| ACE2 | SUSD4 | -0,5502 | 3,46344E-16 |
| ACE2 | NOP14 | -0,5502 | 3,42575E-16 |
| ACE2 | AFF4 | -0,5502 | 3,43291E-16 |
| ACE2 | AC022819.3 | -0,5502 | 3,45449E-16 |
| ACE2 | SNORD21 | -0,5503 | 3,39933E-16 |
| ACE2 | SRGAP3 | -0,5503 | 3,3806E-16 |
| ACE2 | GTPBP10 | -0,5503 | 3,39849E-16 |
| ACE2 | SCNM1 | -0,5504 | 3,35891E-16 |
| ACE2 | PCDHGB8P | -0,5504 | 3,33145E-16 |

| | | | |
|---|---|---|---|
| ACE2 | RP11-155G14.5 | -0,5504 | 3,34763E-16 |
| ACE2 | ZNF574 | -0,5504 | 3,32045E-16 |
| ACE2 | KCNE5 | -0,5504 | 3,34644E-16 |
| ACE2 | IL1R1 | -0,5505 | 3,27623E-16 |
| ACE2 | IMPAD1 | -0,5505 | 3,31178E-16 |
| ACE2 | MID1IP1 | -0,5505 | 3,30362E-16 |
| ACE2 | FKBP15 | -0,5506 | 3,26976E-16 |
| ACE2 | IPO8 | -0,5507 | 3,20206E-16 |
| ACE2 | UBE2L5P | -0,5507 | 3,18259E-16 |
| ACE2 | POLR1A | -0,5508 | 3,13681E-16 |
| ACE2 | SCN3A | -0,5508 | 3,14718E-16 |
| ACE2 | SPICE1 | -0,5508 | 3,16666E-16 |
| ACE2 | B9D2 | -0,5508 | 3,16256E-16 |
| ACE2 | RP11-10L12.4 | -0,5509 | 3,11265E-16 |
| ACE2 | MCHR1 | -0,5509 | 3,10912E-16 |
| ACE2 | RPA4 | -0,5509 | 3,10307E-16 |
| ACE2 | GPAT2 | -0,551 | 3,04224E-16 |
| ACE2 | WFDC2 | -0,551 | 3,06625E-16 |
| ACE2 | HSD17B10 | -0,551 | 3,04318E-16 |
| ACE2 | TCEAL6 | -0,551 | 3,0383E-16 |
| ACE2 | HIST2H3DP1 | -0,5511 | 3,02361E-16 |
| ACE2 | CTD-3128G10.7 | -0,5511 | 3,00094E-16 |
| ACE2 | STK3 | -0,5512 | 2,98389E-16 |
| ACE2 | CPB2-AS1 | -0,5512 | 2,98305E-16 |
| ACE2 | RP11-299L17.3 | -0,5513 | 2,93235E-16 |
| ACE2 | AP4E1 | -0,5513 | 2,94237E-16 |
| ACE2 | PSMB6 | -0,5513 | 2,93514E-16 |
| ACE2 | RP11-795F19.5 | -0,5513 | 2,91348E-16 |
| ACE2 | XRCC5 | -0,5514 | 2,87148E-16 |
| ACE2 | ALKBH1 | -0,5514 | 2,87231E-16 |
| ACE2 | PPP1R7 | -0,5515 | 2,84863E-16 |
| ACE2 | OTOGL | -0,5515 | 2,83183E-16 |
| ACE2 | MAPK4 | -0,5515 | 2,84395E-16 |
| ACE2 | CWF19L1 | -0,5516 | 2,80163E-16 |
| ACE2 | ACTR5 | -0,5516 | 2,81121E-16 |
| ACE2 | NPY1R | -0,5517 | 2,73973E-16 |
| ACE2 | KB-1507C5.4 | -0,5517 | 2,77704E-16 |
| ACE2 | MTBP | -0,5517 | 2,76863E-16 |
| ACE2 | SEH1L | -0,5517 | 2,76043E-16 |
| ACE2 | MRPL4 | -0,5517 | 2,76674E-16 |
| ACE2 | RP11-6O2.4 | -0,5519 | 2,69743E-16 |
| ACE2 | ABCA9 | -0,5519 | 2,65847E-16 |
| ACE2 | SRSF7 | -0,5521 | 2,58427E-16 |
| ACE2 | RP5-855D21.2 | -0,5521 | 2,58055E-16 |
| ACE2 | DPP8 | -0,5521 | 2,59927E-16 |
| ACE2 | LA16c-385E7.1 | -0,5522 | 2,54606E-16 |
| ACE2 | NCKAP1L | -0,5523 | 2,52053E-16 |
| ACE2 | TDRD9 | -0,5523 | 2,53392E-16 |
| ACE2 | TMPRSS9 | -0,5523 | 2,51476E-16 |
| ACE2 | RP11-69E11.4 | -0,5524 | 2,49938E-16 |
| ACE2 | DYNLL1P4 | -0,5524 | 2,49622E-16 |
| ACE2 | AC025335.1 | -0,5524 | 2,47637E-16 |
| ACE2 | ATP2B3 | -0,5525 | 2,44361E-16 |
| ACE2 | RP11-342D11.2 | -0,5526 | 2,39679E-16 |

| | | | |
|---|---|---|---|
| ACE2 | ACSM3 | -0,5526 | 2,42256E-16 |
| ACE2 | RP11-654E17.2 | -0,5526 | 2,40233E-16 |
| ACE2 | RAB29 | -0,5527 | 2,37437E-16 |
| ACE2 | NR1D2 | -0,5527 | 2,38573E-16 |
| ACE2 | DNAJC24 | -0,5527 | 2,37205E-16 |
| ACE2 | TESC | -0,5527 | 2,39164E-16 |
| ACE2 | AC061992.2 | -0,5527 | 2,38938E-16 |
| ACE2 | SLC18A2 | -0,5528 | 2,35094E-16 |
| ACE2 | KCNJ5 | -0,5528 | 2,32586E-16 |
| ACE2 | SMARCC1 | -0,5529 | 2,29811E-16 |
| ACE2 | MMRN1 | -0,5529 | 2,30257E-16 |
| ACE2 | RP5-942I16.1 | -0,5529 | 2,32231E-16 |
| ACE2 | TES | -0,553 | 2,29067E-16 |
| ACE2 | AKAP6 | -0,553 | 2,26348E-16 |
| ACE2 | MSL1 | -0,553 | 2,26813E-16 |
| ACE2 | RP11-496N12.6 | -0,5531 | 2,24724E-16 |
| ACE2 | RP11-360F5.1 | -0,5531 | 2,24268E-16 |
| ACE2 | API5 | -0,5531 | 2,24936E-16 |
| ACE2 | MSH4 | -0,5532 | 2,21795E-16 |
| ACE2 | INHA | -0,5532 | 2,21375E-16 |
| ACE2 | RP11-298J23.10 | -0,5532 | 2,19543E-16 |
| ACE2 | MAGED1 | -0,5532 | 2,19646E-16 |
| ACE2 | LPP | -0,5533 | 2,16368E-16 |
| ACE2 | TRAFD1 | -0,5533 | 2,1613E-16 |
| ACE2 | ASB7 | -0,5533 | 2,18784E-16 |
| ACE2 | EFCAB13 | -0,5533 | 2,17134E-16 |
| ACE2 | RPL28 | -0,5533 | 2,16926E-16 |
| ACE2 | NMD3P1 | -0,5534 | 2,14796E-16 |
| ACE2 | RP11-459F6.3 | -0,5534 | 2,12916E-16 |
| ACE2 | PEX5L | -0,5535 | 2,11686E-16 |
| ACE2 | BMPR1A | -0,5535 | 2,10442E-16 |
| ACE2 | TMEM132B | -0,5535 | 2,11557E-16 |
| ACE2 | TMEM132E | -0,5535 | 2,10632E-16 |
| ACE2 | CECR7 | -0,5536 | 2,09399E-16 |
| ACE2 | ANKRD45 | -0,5537 | 2,05712E-16 |
| ACE2 | TSPEAR-AS2 | -0,5537 | 2,03434E-16 |
| ACE2 | AC007969.5 | -0,5538 | 2,01506E-16 |
| ACE2 | RPL3P4 | -0,5538 | 2,03171E-16 |
| ACE2 | POLL | -0,5539 | 1,98575E-16 |
| ACE2 | CIR1 | -0,554 | 1,94744E-16 |
| ACE2 | MANEA-AS1 | -0,554 | 1,97095E-16 |
| ACE2 | RAPGEF2 | -0,5541 | 1,92321E-16 |
| ACE2 | RP11-1280N14.3 | -0,5541 | 1,93821E-16 |
| ACE2 | CPSF7 | -0,5541 | 1,91947E-16 |
| ACE2 | RP11-13A1.1 | -0,5541 | 1,91996E-16 |
| ACE2 | RAB35 | -0,5541 | 1,92304E-16 |
| ACE2 | TCAP | -0,5541 | 1,93818E-16 |
| ACE2 | CRB1 | -0,5542 | 1,89209E-16 |
| ACE2 | AC064875.2 | -0,5542 | 1,89537E-16 |
| ACE2 | RP11-446H18.6 | -0,5542 | 1,91039E-16 |
| ACE2 | IPO4 | -0,5542 | 1,90146E-16 |
| ACE2 | RP11-3D4.4 | -0,5542 | 1,89865E-16 |
| ACE2 | RBM39 | -0,5542 | 1,89512E-16 |
| ACE2 | MSANTD1 | -0,5543 | 1,865E-16 |

| | | | |
|---|---|---|---|
| ACE2 | A2MP1 | -0,5543 | 1,86885E-16 |
| ACE2 | ATP2C1 | -0,5544 | 1,83659E-16 |
| ACE2 | MNT | -0,5544 | 1,84266E-16 |
| ACE2 | ESCO1 | -0,5544 | 1,83878E-16 |
| ACE2 | CP | -0,5545 | 1,82074E-16 |
| ACE2 | TMEM129 | -0,5545 | 1,82561E-16 |
| ACE2 | HPSE2 | -0,5545 | 1,82042E-16 |
| ACE2 | NUP85 | -0,5545 | 1,81102E-16 |
| ACE2 | CTD-2561B21.10 | -0,5545 | 1,80862E-16 |
| ACE2 | OSGEP | -0,5546 | 1,79801E-16 |
| ACE2 | UBE2G1 | -0,5546 | 1,78743E-16 |
| ACE2 | TRAF5 | -0,5547 | 1,7681E-16 |
| ACE2 | LINC00887 | -0,5547 | 1,77354E-16 |
| ACE2 | HNRNPK | -0,5547 | 1,75021E-16 |
| ACE2 | RP11-973D8.5 | -0,5547 | 1,76761E-16 |
| ACE2 | DLGAP4-AS1 | -0,5547 | 1,75979E-16 |
| ACE2 | LL22NC03-80A10.6 | -0,5547 | 1,76473E-16 |
| ACE2 | OTUD5 | -0,5547 | 1,76966E-16 |
| ACE2 | CDC42SE1 | -0,5548 | 1,73418E-16 |
| ACE2 | RANBP10 | -0,5548 | 1,72793E-16 |
| ACE2 | TMEM222 | -0,5549 | 1,70034E-16 |
| ACE2 | NRROS | -0,5549 | 1,70295E-16 |
| ACE2 | RP11-802E16.3 | -0,5549 | 1,70527E-16 |
| ACE2 | RP11-775C24.5 | -0,555 | 1,69253E-16 |
| ACE2 | ACKR2 | -0,5551 | 1,6601E-16 |
| ACE2 | RCN1P2 | -0,5551 | 1,66599E-16 |
| ACE2 | DICER1 | -0,5551 | 1,67E-16 |
| ACE2 | SLC12A1 | -0,5551 | 1,66713E-16 |
| ACE2 | FAM180A | -0,5552 | 1,63084E-16 |
| ACE2 | LINC01158 | -0,5554 | 1,58886E-16 |
| ACE2 | POMZP3 | -0,5554 | 1,57632E-16 |
| ACE2 | NWD1 | -0,5554 | 1,58485E-16 |
| ACE2 | TMEM39A | -0,5555 | 1,56652E-16 |
| ACE2 | SNHG8 | -0,5555 | 1,56401E-16 |
| ACE2 | LRRIQ1 | -0,5555 | 1,55575E-16 |
| ACE2 | OST4 | -0,5556 | 1,53338E-16 |
| ACE2 | THUMPD3 | -0,5556 | 1,5418E-16 |
| ACE2 | SUPT20HL2 | -0,5556 | 1,54104E-16 |
| ACE2 | RPGR | -0,5556 | 1,52909E-16 |
| ACE2 | LRRC37B | -0,5557 | 1,5103E-16 |
| ACE2 | CTD-2010I16.1 | -0,5558 | 1,48979E-16 |
| ACE2 | MARCH10 | -0,5558 | 1,48543E-16 |
| ACE2 | RRP9 | -0,5559 | 1,4625E-16 |
| ACE2 | RP11-420L9.5 | -0,5559 | 1,47997E-16 |
| ACE2 | NSUN5 | -0,5559 | 1,48079E-16 |
| ACE2 | TPT1-AS1 | -0,5559 | 1,47089E-16 |
| ACE2 | SMURF2P1 | -0,5559 | 1,47318E-16 |
| ACE2 | SUN2 | -0,5559 | 1,47144E-16 |
| ACE2 | WDR77 | -0,556 | 1,44719E-16 |
| ACE2 | GRSF1 | -0,556 | 1,45084E-16 |
| ACE2 | ADRB3 | -0,556 | 1,45121E-16 |
| ACE2 | RP11-472N13.3 | -0,556 | 1,44091E-16 |
| ACE2 | RAD52 | -0,556 | 1,45898E-16 |
| ACE2 | C20orf166-AS1 | -0,556 | 1,45526E-16 |

| | | | |
|---|---|---|---|
| ACE2 | CIT | -0,5561 | 1,41925E-16 |
| ACE2 | MMP2 | -0,5561 | 1,42172E-16 |
| ACE2 | SPATA20 | -0,5561 | 1,43793E-16 |
| ACE2 | RP5-1068E13.7 | -0,5561 | 1,43068E-16 |
| ACE2 | RP11-430G17.3 | -0,5562 | 1,41633E-16 |
| ACE2 | LAPTM4A | -0,5562 | 1,40382E-16 |
| ACE2 | CLK4 | -0,5562 | 1,4083E-16 |
| ACE2 | DNAJC2 | -0,5562 | 1,39697E-16 |
| ACE2 | RP11-629O1.2 | -0,5562 | 1,40763E-16 |
| ACE2 | TPP2 | -0,5562 | 1,41059E-16 |
| ACE2 | AC005154.7 | -0,5563 | 1,38197E-16 |
| ACE2 | CTD-2368P22.1 | -0,5563 | 1,38726E-16 |
| ACE2 | RP11-98I9.4 | -0,5564 | 1,36452E-16 |
| ACE2 | ZNF625 | -0,5564 | 1,35771E-16 |
| ACE2 | PARP10 | -0,5565 | 1,34946E-16 |
| ACE2 | RP11-2B6.3 | -0,5565 | 1,35323E-16 |
| ACE2 | RP11-240G22.5 | -0,5565 | 1,34282E-16 |
| ACE2 | XXbac-B444P24.13 | -0,5565 | 1,34331E-16 |
| ACE2 | BRD9 | -0,5566 | 1,32605E-16 |
| ACE2 | ZMYM2 | -0,5566 | 1,32797E-16 |
| ACE2 | TNRC6A | -0,5566 | 1,31532E-16 |
| ACE2 | ASGR1 | -0,5566 | 1,3249E-16 |
| ACE2 | RP11-893F2.18 | -0,5566 | 1,31497E-16 |
| ACE2 | RP5-1116H23.4 | -0,5566 | 1,32045E-16 |
| ACE2 | LRRTM2 | -0,5567 | 1,30481E-16 |
| ACE2 | FLI1 | -0,5567 | 1,29575E-16 |
| ACE2 | TRIM13 | -0,5567 | 1,30976E-16 |
| ACE2 | SNORD10 | -0,5567 | 1,30605E-16 |
| ACE2 | USP16 | -0,5567 | 1,30434E-16 |
| ACE2 | OPTC | -0,5568 | 1,28037E-16 |
| ACE2 | RP11-174O3.3 | -0,5568 | 1,28898E-16 |
| ACE2 | UBXN8 | -0,5568 | 1,29338E-16 |
| ACE2 | C8orf44 | -0,5568 | 1,28182E-16 |
| ACE2 | C2CD4C | -0,5568 | 1,29022E-16 |
| ACE2 | SIRT2 | -0,5568 | 1,2836E-16 |
| ACE2 | AP000436.4 | -0,5569 | 1,26775E-16 |
| ACE2 | SNHG17 | -0,5569 | 1,26355E-16 |
| ACE2 | ZNRD1ASP | -0,557 | 1,24075E-16 |
| ACE2 | C7orf25 | -0,557 | 1,25306E-16 |
| ACE2 | JAK2 | -0,557 | 1,23823E-16 |
| ACE2 | BCAT1 | -0,557 | 1,25293E-16 |
| ACE2 | CDC34 | -0,557 | 1,2427E-16 |
| ACE2 | SLC44A2 | -0,557 | 1,23794E-16 |
| ACE2 | AP001258.4 | -0,5571 | 1,23482E-16 |
| ACE2 | TMED8 | -0,5571 | 1,22429E-16 |
| ACE2 | DNM1P47 | -0,5571 | 1,22867E-16 |
| ACE2 | GPR1 | -0,5572 | 1,20874E-16 |
| ACE2 | AC004069.2 | -0,5572 | 1,2046E-16 |
| ACE2 | STMN2 | -0,5572 | 1,21797E-16 |
| ACE2 | SUV39H2 | -0,5572 | 1,21484E-16 |
| ACE2 | RAB30-AS1 | -0,5572 | 1,20375E-16 |
| ACE2 | RP11-631N16.4 | -0,5572 | 1,20527E-16 |
| ACE2 | PTPRB | -0,5572 | 1,20635E-16 |
| ACE2 | CILP2 | -0,5572 | 1,20252E-16 |

| | | | |
|---|---|---|---|
| ACE2 | PDPN | -0,5573 | 1,18334E-16 |
| ACE2 | DNAH14 | -0,5573 | 1,1833E-16 |
| ACE2 | COL11A2 | -0,5573 | 1,19438E-16 |
| ACE2 | CAD | -0,5574 | 1,17572E-16 |
| ACE2 | ZNF658B | -0,5574 | 1,16922E-16 |
| ACE2 | RP11-87H9.4 | -0,5574 | 1,17801E-16 |
| ACE2 | RP11-20O24.4 | -0,5575 | 1,16233E-16 |
| ACE2 | NPRL2 | -0,5576 | 1,13127E-16 |
| ACE2 | LIMCH1 | -0,5576 | 1,14566E-16 |
| ACE2 | ZNF460 | -0,5576 | 1,14389E-16 |
| ACE2 | RP11-365O16.6 | -0,5577 | 1,12198E-16 |
| ACE2 | RP11-834C11.11 | -0,5577 | 1,1145E-16 |
| ACE2 | TRIP11 | -0,5577 | 1,11714E-16 |
| ACE2 | INTS11 | -0,5578 | 1,10685E-16 |
| ACE2 | RP11-545G3.1 | -0,5578 | 1,10673E-16 |
| ACE2 | CTD-2349P21.12 | -0,5578 | 1,10308E-16 |
| ACE2 | GDNF | -0,5579 | 1,08436E-16 |
| ACE2 | KLRA1P | -0,5579 | 1,08477E-16 |
| ACE2 | FLJ12825 | -0,5579 | 1,09247E-16 |
| ACE2 | CCNL2 | -0,558 | 1,0712E-16 |
| ACE2 | GTF2IRD2P1 | -0,558 | 1,07144E-16 |
| ACE2 | USP10 | -0,558 | 1,07911E-16 |
| ACE2 | PPP2R2C | -0,5581 | 1,05183E-16 |
| ACE2 | RP11-146N23.1 | -0,5581 | 1,05655E-16 |
| ACE2 | C10orf88 | -0,5581 | 1,05777E-16 |
| ACE2 | ALMS1-IT1 | -0,5583 | 1,02142E-16 |
| ACE2 | GPR37 | -0,5583 | 1,02201E-16 |
| ACE2 | RP11-159H10.3 | -0,5583 | 1,02296E-16 |
| ACE2 | RP11-355O1.11 | -0,5583 | 1,02362E-16 |
| ACE2 | ZNF365 | -0,5583 | 1,01771E-16 |
| ACE2 | MRE11 | -0,5583 | 1,0288E-16 |
| ACE2 | RP11-114M1.1 | -0,5584 | 1,00901E-16 |
| ACE2 | HIST2H2AC | -0,5586 | 9,80206E-17 |
| ACE2 | ELOVL3 | -0,5586 | 9,74849E-17 |
| ACE2 | ERH | -0,5586 | 9,8285E-17 |
| ACE2 | CTC-296K1.4 | -0,5586 | 9,85715E-17 |
| ACE2 | ATP6AP1L | -0,5587 | 9,67982E-17 |
| ACE2 | RP11-314C16.1 | -0,5587 | 9,70766E-17 |
| ACE2 | ZC3HAV1L | -0,5587 | 9,62844E-17 |
| ACE2 | MAP2K1 | -0,5588 | 9,4229E-17 |
| ACE2 | ELP5 | -0,5588 | 9,51177E-17 |
| ACE2 | RP11-630A13.4 | -0,5589 | 9,32516E-17 |
| ACE2 | SBF2 | -0,5589 | 9,28246E-17 |
| ACE2 | MYH9 | -0,5589 | 9,32797E-17 |
| ACE2 | TIA1 | -0,559 | 9,16712E-17 |
| ACE2 | RP11-260M2.1 | -0,559 | 9,21203E-17 |
| ACE2 | PMPCB | -0,559 | 9,2603E-17 |
| ACE2 | FBXO39 | -0,5591 | 9,10116E-17 |
| ACE2 | SCLT1 | -0,5592 | 8,92158E-17 |
| ACE2 | FAM86GP | -0,5592 | 8,9608E-17 |
| ACE2 | SCAPER | -0,5592 | 8,95239E-17 |
| ACE2 | PLCG1 | -0,5592 | 8,91869E-17 |
| ACE2 | TTC23L | -0,5593 | 8,78845E-17 |
| ACE2 | SOX2-OT | -0,5594 | 8,6519E-17 |

| | | | |
|---|---|---|---|
| ACE2 | XXbac-BPG252P9.9 | -0,5594 | 8,6059E-17 |
| ACE2 | CSNK2B | -0,5594 | 8,68733E-17 |
| ACE2 | EIF2S2P3 | -0,5595 | 8,48748E-17 |
| ACE2 | SORBS1 | -0,5595 | 8,51174E-17 |
| ACE2 | PSMD13 | -0,5595 | 8,48683E-17 |
| ACE2 | RP11-341G23.4 | -0,5595 | 8,56614E-17 |
| ACE2 | NFATC2 | -0,5595 | 8,47065E-17 |
| ACE2 | RP1-79C4.4 | -0,5596 | 8,38415E-17 |
| ACE2 | SRP9 | -0,5596 | 8,39553E-17 |
| ACE2 | MLLT6 | -0,5596 | 8,36552E-17 |
| ACE2 | GNB1L | -0,5596 | 8,45682E-17 |
| ACE2 | LINC01589 | -0,5596 | 8,37662E-17 |
| ACE2 | SYT2 | -0,5597 | 8,31218E-17 |
| ACE2 | RABGEF1 | -0,5597 | 8,33948E-17 |
| ACE2 | SCN4A | -0,5597 | 8,29623E-17 |
| ACE2 | USP48 | -0,5598 | 8,1644E-17 |
| ACE2 | KCNC4 | -0,5598 | 8,12279E-17 |
| ACE2 | RP11-422P24.11 | -0,5598 | 8,1545E-17 |
| ACE2 | EMG1 | -0,5598 | 8,15108E-17 |
| ACE2 | POLR2E | -0,5598 | 8,20093E-17 |
| ACE2 | ARHGEF26-AS1 | -0,5599 | 8,0315E-17 |
| ACE2 | RP11-119F19.5 | -0,56 | 7,93437E-17 |
| ACE2 | NSL1 | -0,5601 | 7,82777E-17 |
| ACE2 | GLYATL2 | -0,5601 | 7,73185E-17 |
| ACE2 | C11orf1 | -0,5601 | 7,76628E-17 |
| ACE2 | SHISA2 | -0,5601 | 7,76332E-17 |
| ACE2 | HTR1F | -0,5602 | 7,61425E-17 |
| ACE2 | MYL6 | -0,5602 | 7,61277E-17 |
| ACE2 | PAN3-AS1 | -0,5602 | 7,67414E-17 |
| ACE2 | RP11-696D21.2 | -0,5602 | 7,70915E-17 |
| ACE2 | DNAJA2 | -0,5602 | 7,64824E-17 |
| ACE2 | GANC | -0,5603 | 7,52363E-17 |
| ACE2 | IKZF2 | -0,5604 | 7,47286E-17 |
| ACE2 | FNDC11 | -0,5604 | 7,43789E-17 |
| ACE2 | VAMP3 | -0,5605 | 7,35489E-17 |
| ACE2 | DUS4L | -0,5605 | 7,31212E-17 |
| ACE2 | KDM2A | -0,5605 | 7,29675E-17 |
| ACE2 | FAM86DP | -0,5606 | 7,23918E-17 |
| ACE2 | TCP1 | -0,5606 | 7,24166E-17 |
| ACE2 | ZNF772 | -0,5606 | 7,26189E-17 |
| ACE2 | CCSAP | -0,5607 | 7,10999E-17 |
| ACE2 | BRE | -0,5607 | 7,14521E-17 |
| ACE2 | CEP72 | -0,5607 | 7,10057E-17 |
| ACE2 | CDK13 | -0,5607 | 7,07955E-17 |
| ACE2 | INTS8 | -0,5607 | 7,07899E-17 |
| ACE2 | UBIAD1 | -0,5608 | 6,96472E-17 |
| ACE2 | ZNF602P | -0,5608 | 6,99317E-17 |
| ACE2 | RP11-513M16.8 | -0,5608 | 7,03177E-17 |
| ACE2 | RP11-107E5.3 | -0,5609 | 6,8909E-17 |
| ACE2 | SLC8A3 | -0,5609 | 6,87334E-17 |
| ACE2 | ZNF737 | -0,5609 | 6,84182E-17 |
| ACE2 | HGFAC | -0,561 | 6,81851E-17 |
| ACE2 | HNRNPLP2 | -0,561 | 6,7724E-17 |
| ACE2 | AC006116.20 | -0,561 | 6,77606E-17 |

| | | | |
|---|---|---|---|
| ACE2 | KB-68A7.1 | -0,561 | 6,8344E-17 |
| ACE2 | ZNF141 | -0,5611 | 6,70918E-17 |
| ACE2 | SHPRH | -0,5611 | 6,7339E-17 |
| ACE2 | SPAG17 | -0,5612 | 6,62225E-17 |
| ACE2 | CAPN7 | -0,5612 | 6,57418E-17 |
| ACE2 | NOD2 | -0,5612 | 6,6151E-17 |
| ACE2 | SDK2 | -0,5612 | 6,54359E-17 |
| ACE2 | PCDH18 | -0,5613 | 6,48195E-17 |
| ACE2 | ZBTB12 | -0,5613 | 6,4783E-17 |
| ACE2 | PHYHIP | -0,5613 | 6,46737E-17 |
| ACE2 | SPESP1 | -0,5613 | 6,47127E-17 |
| ACE2 | ANGPTL1 | -0,5614 | 6,4187E-17 |
| ACE2 | SPCS2 | -0,5614 | 6,40737E-17 |
| ACE2 | SYNE3 | -0,5614 | 6,40063E-17 |
| ACE2 | RP11-416A14.1 | -0,5615 | 6,30947E-17 |
| ACE2 | P2RY14 | -0,5615 | 6,24639E-17 |
| ACE2 | IKZF5 | -0,5615 | 6,26969E-17 |
| ACE2 | CTD-2196E14.6 | -0,5615 | 6,25921E-17 |
| ACE2 | GPC4 | -0,5615 | 6,28234E-17 |
| ACE2 | LINC01273 | -0,5616 | 6,22347E-17 |
| ACE2 | PANK4 | -0,5617 | 6,10624E-17 |
| ACE2 | LRTOMT | -0,5617 | 6,0761E-17 |
| ACE2 | MRPL52 | -0,5617 | 6,11892E-17 |
| ACE2 | LINC00473 | -0,5618 | 6,00604E-17 |
| ACE2 | CNGB3 | -0,5618 | 5,97232E-17 |
| ACE2 | EEA1 | -0,5618 | 6,02614E-17 |
| ACE2 | RP11-266L9.4 | -0,5618 | 5,95761E-17 |
| ACE2 | DNAH9 | -0,5618 | 6,0035E-17 |
| ACE2 | ZNF254 | -0,5618 | 5,95892E-17 |
| ACE2 | ZNF432 | -0,5618 | 6,02686E-17 |
| ACE2 | GRPR | -0,5618 | 5,96768E-17 |
| ACE2 | PGS1 | -0,5619 | 5,91423E-17 |
| ACE2 | CTD-2528L19.6 | -0,5619 | 5,9449E-17 |
| ACE2 | HIST3H2A | -0,562 | 5,79232E-17 |
| ACE2 | RP11-324I22.4 | -0,562 | 5,80305E-17 |
| ACE2 | UBE2Q2P1 | -0,562 | 5,82966E-17 |
| ACE2 | GNA15 | -0,5621 | 5,68992E-17 |
| ACE2 | HSPA6 | -0,5622 | 5,62174E-17 |
| ACE2 | TOPORS | -0,5622 | 5,607E-17 |
| ACE2 | NCAM1-AS1 | -0,5622 | 5,66212E-17 |
| ACE2 | TTC12 | -0,5622 | 5,65733E-17 |
| ACE2 | ZNF93 | -0,5622 | 5,67241E-17 |
| ACE2 | NBPF26 | -0,5623 | 5,52054E-17 |
| ACE2 | POU2F1 | -0,5623 | 5,55055E-17 |
| ACE2 | NPY5R | -0,5623 | 5,51712E-17 |
| ACE2 | RP11-494K3.2 | -0,5624 | 5,454E-17 |
| ACE2 | SHQ1 | -0,5624 | 5,51508E-17 |
| ACE2 | RAB30 | -0,5624 | 5,44335E-17 |
| ACE2 | RP11-1090M7.1 | -0,5624 | 5,49767E-17 |
| ACE2 | CEP104 | -0,5625 | 5,42076E-17 |
| ACE2 | OSBPL10 | -0,5625 | 5,38326E-17 |
| ACE2 | LINC01801 | -0,5625 | 5,35519E-17 |
| ACE2 | BCAT2 | -0,5625 | 5,37763E-17 |
| ACE2 | NF2 | -0,5625 | 5,35888E-17 |

| | | | |
|---|---|---|---|
| ACE2 | RFX3-AS1 | -0,5626 | 5,30377E-17 |
| ACE2 | FBXW2 | -0,5627 | 5,18719E-17 |
| ACE2 | ADAMTS16 | -0,5628 | 5,11909E-17 |
| ACE2 | C5orf64 | -0,5628 | 5,11093E-17 |
| ACE2 | DPY19L4 | -0,5628 | 5,11567E-17 |
| ACE2 | ZNF485 | -0,5628 | 5,12302E-17 |
| ACE2 | RP11-366L20.2 | -0,5628 | 5,15971E-17 |
| ACE2 | NBPF19 | -0,5629 | 5,09268E-17 |
| ACE2 | RP11-758N13.3 | -0,563 | 4,98206E-17 |
| ACE2 | PRICKLE3 | -0,563 | 5,01902E-17 |
| ACE2 | PTP4A2 | -0,5631 | 4,92475E-17 |
| ACE2 | RP11-635L1.2 | -0,5631 | 4,9237E-17 |
| ACE2 | CFAP157 | -0,5631 | 4,92884E-17 |
| ACE2 | ZNF506 | -0,5631 | 4,92316E-17 |
| ACE2 | CTC-444N24.6 | -0,5631 | 4,91361E-17 |
| ACE2 | TRPC3 | -0,5632 | 4,83034E-17 |
| ACE2 | ATG10 | -0,5632 | 4,83115E-17 |
| ACE2 | RP11-307O13.1 | -0,5632 | 4,83561E-17 |
| ACE2 | ZSCAN2 | -0,5632 | 4,82733E-17 |
| ACE2 | RUNDC3A | -0,5632 | 4,82449E-17 |
| ACE2 | TGFB1 | -0,5632 | 4,8514E-17 |
| ACE2 | PLA2G6 | -0,5632 | 4,8374E-17 |
| ACE2 | ZNF157 | -0,5632 | 4,8213E-17 |
| ACE2 | IER5 | -0,5633 | 4,75915E-17 |
| ACE2 | WISP3 | -0,5633 | 4,75271E-17 |
| ACE2 | RP11-475D10.4 | -0,5633 | 4,74472E-17 |
| ACE2 | PGBD5 | -0,5634 | 4,71228E-17 |
| ACE2 | ZNF184 | -0,5634 | 4,68575E-17 |
| ACE2 | RP11-57H12.3 | -0,5635 | 4,61049E-17 |
| ACE2 | HMGB1P5 | -0,5635 | 4,62645E-17 |
| ACE2 | NEXN | -0,5636 | 4,53091E-17 |
| ACE2 | HCG23 | -0,5637 | 4,45819E-17 |
| ACE2 | RENBP | -0,5637 | 4,4526E-17 |
| ACE2 | CPSF2 | -0,5638 | 4,38632E-17 |
| ACE2 | ZNF878 | -0,5638 | 4,39307E-17 |
| ACE2 | RP11-324I22.3 | -0,5639 | 4,35627E-17 |
| ACE2 | RP11-677M14.8 | -0,5639 | 4,34853E-17 |
| ACE2 | TMCC1-AS1 | -0,564 | 4,29636E-17 |
| ACE2 | GPR63 | -0,564 | 4,30596E-17 |
| ACE2 | RP4-800J21.3 | -0,564 | 4,24413E-17 |
| ACE2 | KLHL13 | -0,564 | 4,25906E-17 |
| ACE2 | HRNR | -0,5641 | 4,23158E-17 |
| ACE2 | AFF3 | -0,5641 | 4,18131E-17 |
| ACE2 | MBIP | -0,5642 | 4,15881E-17 |
| ACE2 | RP11-1212A22.4 | -0,5642 | 4,1365E-17 |
| ACE2 | ZNF623 | -0,5643 | 4,07408E-17 |
| ACE2 | ATP5F1P5 | -0,5643 | 4,09772E-17 |
| ACE2 | GNPAT | -0,5644 | 4,04745E-17 |
| ACE2 | C10orf128 | -0,5644 | 4,016E-17 |
| ACE2 | PEX10 | -0,5645 | 3,9618E-17 |
| ACE2 | ZBTB17 | -0,5645 | 3,95337E-17 |
| ACE2 | RNF175 | -0,5645 | 3,94108E-17 |
| ACE2 | ATCAY | -0,5645 | 3,9446E-17 |
| ACE2 | DEAF1 | -0,5647 | 3,83682E-17 |

| | | | |
|---|---|---|---|
| ACE2 | TEX21P | -0,5647 | 3,8278E-17 |
| ACE2 | SPATA9 | -0,5648 | 3,75517E-17 |
| ACE2 | UFM1 | -0,5648 | 3,8005E-17 |
| ACE2 | ZACN | -0,5648 | 3,79501E-17 |
| ACE2 | SAFB | -0,5649 | 3,73476E-17 |
| ACE2 | LLNLR-268E12.1 | -0,5649 | 3,70796E-17 |
| ACE2 | OR2W3 | -0,565 | 3,66324E-17 |
| ACE2 | CCIN | -0,565 | 3,66786E-17 |
| ACE2 | SSRP1 | -0,565 | 3,67761E-17 |
| ACE2 | GREM2 | -0,5651 | 3,58529E-17 |
| ACE2 | PPAN | -0,5651 | 3,58084E-17 |
| ACE2 | ZNF548 | -0,5651 | 3,62241E-17 |
| ACE2 | NPHP4 | -0,5652 | 3,52864E-17 |
| ACE2 | RAB6C | -0,5652 | 3,53611E-17 |
| ACE2 | PPIP5K2 | -0,5652 | 3,55613E-17 |
| ACE2 | HYMAI | -0,5653 | 3,51108E-17 |
| ACE2 | CTD-2373N4.3 | -0,5653 | 3,49239E-17 |
| ACE2 | RP11-566K19.5 | -0,5653 | 3,47374E-17 |
| ACE2 | SLC6A17 | -0,5654 | 3,45657E-17 |
| ACE2 | SLC27A6 | -0,5654 | 3,46095E-17 |
| ACE2 | WDR91 | -0,5654 | 3,46271E-17 |
| ACE2 | AC100830.4 | -0,5654 | 3,43635E-17 |
| ACE2 | WDR83OS | -0,5654 | 3,4527E-17 |
| ACE2 | ADAMTS4 | -0,5655 | 3,40636E-17 |
| ACE2 | BAALC-AS1 | -0,5655 | 3,41255E-17 |
| ACE2 | PAFAH1B2 | -0,5655 | 3,36678E-17 |
| ACE2 | CTD-2623N2.3 | -0,5655 | 3,3655E-17 |
| ACE2 | ZNF561 | -0,5657 | 3,26644E-17 |
| ACE2 | C1orf112 | -0,5658 | 3,25539E-17 |
| ACE2 | RP11-467D6.1 | -0,5658 | 3,23605E-17 |
| ACE2 | TCF25 | -0,5658 | 3,21755E-17 |
| ACE2 | BX322557.10 | -0,5658 | 3,25101E-17 |
| ACE2 | DANCR | -0,5659 | 3,20411E-17 |
| ACE2 | NMT2 | -0,5659 | 3,16163E-17 |
| ACE2 | ADAM21 | -0,5659 | 3,19413E-17 |
| ACE2 | ZNF345 | -0,566 | 3,126E-17 |
| ACE2 | TMCO4 | -0,5661 | 3,10482E-17 |
| ACE2 | MRPL51 | -0,5661 | 3,09892E-17 |
| ACE2 | TMEM81 | -0,5662 | 3,03427E-17 |
| ACE2 | RNF13 | -0,5662 | 3,03074E-17 |
| ACE2 | CTB-131K11.1 | -0,5662 | 3,01795E-17 |
| ACE2 | RP4-681N20.5 | -0,5662 | 3,01653E-17 |
| ACE2 | TRIL | -0,5663 | 3,00146E-17 |
| ACE2 | GPAT4 | -0,5663 | 2,976E-17 |
| ACE2 | RP11-330O11.3 | -0,5663 | 2,97788E-17 |
| ACE2 | STX5 | -0,5663 | 2,96967E-17 |
| ACE2 | RP11-283I3.6 | -0,5663 | 3,00263E-17 |
| ACE2 | EIF3K | -0,5663 | 2,99835E-17 |
| ACE2 | AC142528.1 | -0,5664 | 2,9676E-17 |
| ACE2 | CTD-2527I21.14 | -0,5664 | 2,95767E-17 |
| ACE2 | PDCL | -0,5666 | 2,85054E-17 |
| ACE2 | AGBL4 | -0,5667 | 2,81359E-17 |
| ACE2 | RUBCN | -0,5667 | 2,82041E-17 |
| ACE2 | CSTF2T | -0,5667 | 2,82915E-17 |

| | | | |
|---|---|---|---|
| ACE2 | CCDC181 | -0,5668 | 2,76502E-17 |
| ACE2 | PCDHA7 | -0,5668 | 2,76381E-17 |
| ACE2 | AF186192.5 | -0,5668 | 2,75615E-17 |
| ACE2 | FBXO10 | -0,5668 | 2,76321E-17 |
| ACE2 | RPS3 | -0,5668 | 2,7482E-17 |
| ACE2 | UBXN6 | -0,5668 | 2,74908E-17 |
| ACE2 | EPB41L1 | -0,5668 | 2,7623E-17 |
| ACE2 | AGO4 | -0,5669 | 2,71522E-17 |
| ACE2 | PI4KB | -0,5669 | 2,72377E-17 |
| ACE2 | PSME4 | -0,5669 | 2,72026E-17 |
| ACE2 | CTC-325H20.7 | -0,5669 | 2,72281E-17 |
| ACE2 | LINC00891 | -0,567 | 2,67922E-17 |
| ACE2 | TENM2 | -0,5671 | 2,62659E-17 |
| ACE2 | TAF2 | -0,5671 | 2,65926E-17 |
| ACE2 | RP11-2C24.7 | -0,5671 | 2,65173E-17 |
| ACE2 | LINC01529 | -0,5671 | 2,62035E-17 |
| ACE2 | CYLD | -0,5674 | 2,53162E-17 |
| ACE2 | GAS8 | -0,5674 | 2,52835E-17 |
| ACE2 | SIPA1 | -0,5675 | 2,47908E-17 |
| ACE2 | S100A11 | -0,5676 | 2,4569E-17 |
| ACE2 | C15orf61 | -0,5676 | 2,43608E-17 |
| ACE2 | RP11-671M22.4 | -0,5676 | 2,43235E-17 |
| ACE2 | DCAF12L2 | -0,5676 | 2,45911E-17 |
| ACE2 | RP11-446H18.5 | -0,5677 | 2,38786E-17 |
| ACE2 | RPL10P7 | -0,5677 | 2,38768E-17 |
| ACE2 | HSPC324 | -0,5677 | 2,40173E-17 |
| ACE2 | RP11-57H14.4 | -0,5677 | 2,41221E-17 |
| ACE2 | SLC24A4 | -0,5677 | 2,41161E-17 |
| ACE2 | CTD-3252C9.4 | -0,5677 | 2,4085E-17 |
| ACE2 | RP11-552M11.8 | -0,5678 | 2,35058E-17 |
| ACE2 | LINC00623 | -0,5678 | 2,36788E-17 |
| ACE2 | LINC00504 | -0,5678 | 2,37142E-17 |
| ACE2 | PTCD1 | -0,5679 | 2,32693E-17 |
| ACE2 | IFT20 | -0,5679 | 2,32842E-17 |
| ACE2 | HSD17B1P1 | -0,5679 | 2,32101E-17 |
| ACE2 | AC141928.1 | -0,568 | 2,2761E-17 |
| ACE2 | TMEM14B | -0,568 | 2,29596E-17 |
| ACE2 | SRGN | -0,568 | 2,27552E-17 |
| ACE2 | CHRNA5 | -0,568 | 2,28419E-17 |
| ACE2 | RPL9P29 | -0,568 | 2,28701E-17 |
| ACE2 | AJ011931.1 | -0,568 | 2,28531E-17 |
| ACE2 | GS1-358P8.4 | -0,568 | 2,30333E-17 |
| ACE2 | TRMT13 | -0,5681 | 2,25646E-17 |
| ACE2 | HYDIN2 | -0,5681 | 2,27168E-17 |
| ACE2 | PPM1D | -0,5681 | 2,27154E-17 |
| ACE2 | OMA1 | -0,5682 | 2,22091E-17 |
| ACE2 | RP11-301L8.2 | -0,5682 | 2,22278E-17 |
| ACE2 | CTC-325H20.8 | -0,5682 | 2,2117E-17 |
| ACE2 | NFIA-AS2 | -0,5683 | 2,2028E-17 |
| ACE2 | LINC00607 | -0,5683 | 2,19363E-17 |
| ACE2 | LSM3 | -0,5683 | 2,19676E-17 |
| ACE2 | RPL29P19 | -0,5683 | 2,18465E-17 |
| ACE2 | YME1L1 | -0,5683 | 2,17338E-17 |
| ACE2 | CTNNA3 | -0,5683 | 2,19596E-17 |

| | | | |
|---|---|---|---|
| ACE2 | PPP1R8 | -0,5684 | 2,14539E-17 |
| ACE2 | IFITM4P | -0,5684 | 2,16073E-17 |
| ACE2 | UCN | -0,5686 | 2,09488E-17 |
| ACE2 | TSC22D4 | -0,5686 | 2,09868E-17 |
| ACE2 | DMTN | -0,5686 | 2,08829E-17 |
| ACE2 | SIAH3 | -0,5686 | 2,0735E-17 |
| ACE2 | DNM1P51 | -0,5686 | 2,08116E-17 |
| ACE2 | RP3-368A4.6 | -0,5686 | 2,08348E-17 |
| ACE2 | KAZN | -0,5687 | 2,05529E-17 |
| ACE2 | ATG9B | -0,5687 | 2,0547E-17 |
| ACE2 | NOL6 | -0,5687 | 2,0452E-17 |
| ACE2 | ERAL1 | -0,5687 | 2,06612E-17 |
| ACE2 | IER2 | -0,5687 | 2,06748E-17 |
| ACE2 | AP000318.2 | -0,5687 | 2,03994E-17 |
| ACE2 | EIF4A1 | -0,5688 | 2,03729E-17 |
| ACE2 | LINC00639 | -0,5689 | 1,99235E-17 |
| ACE2 | AC004237.1 | -0,569 | 1,95334E-17 |
| ACE2 | PHLDB1 | -0,569 | 1,9493E-17 |
| ACE2 | CCDC13-AS1 | -0,5691 | 1,93895E-17 |
| ACE2 | ZMAT2 | -0,5691 | 1,93496E-17 |
| ACE2 | RP11-399O19.9 | -0,5691 | 1,93708E-17 |
| ACE2 | RPL14P1 | -0,5691 | 1,9279E-17 |
| ACE2 | SOX1 | -0,5691 | 1,91564E-17 |
| ACE2 | SAXO2 | -0,5691 | 1,9346E-17 |
| ACE2 | CRTC1 | -0,5691 | 1,9388E-17 |
| ACE2 | CAMK1G | -0,5692 | 1,91059E-17 |
| ACE2 | RPS2P32 | -0,5693 | 1,88377E-17 |
| ACE2 | AP001189.4 | -0,5693 | 1,88217E-17 |
| ACE2 | RRN3 | -0,5693 | 1,86789E-17 |
| ACE2 | RP11-517I3.2 | -0,5694 | 1,85172E-17 |
| ACE2 | DEK | -0,5694 | 1,83831E-17 |
| ACE2 | AC130469.2 | -0,5694 | 1,82914E-17 |
| ACE2 | RP11-571L19.8 | -0,5695 | 1,81311E-17 |
| ACE2 | RBM7 | -0,5695 | 1,81803E-17 |
| ACE2 | SARM1 | -0,5695 | 1,82433E-17 |
| ACE2 | GRK3 | -0,5695 | 1,81206E-17 |
| ACE2 | MCCC1-AS1 | -0,5696 | 1,77067E-17 |
| ACE2 | CNTNAP4 | -0,5696 | 1,7827E-17 |
| ACE2 | OSR1 | -0,5697 | 1,7676E-17 |
| ACE2 | TIMM10B | -0,5697 | 1,74763E-17 |
| ACE2 | AC026271.5 | -0,5697 | 1,74902E-17 |
| ACE2 | TLK1 | -0,5699 | 1,69756E-17 |
| ACE2 | RP11-395I6.3 | -0,5699 | 1,7128E-17 |
| ACE2 | NFATC1 | -0,5699 | 1,71319E-17 |
| ACE2 | ZNF230 | -0,5699 | 1,69061E-17 |
| ACE2 | RP11-108M9.3 | -0,57 | 1,66282E-17 |
| ACE2 | NUP153 | -0,57 | 1,67925E-17 |
| ACE2 | CHID1 | -0,57 | 1,67356E-17 |
| ACE2 | PAN2 | -0,57 | 1,68541E-17 |
| ACE2 | MTUS2 | -0,5701 | 1,63659E-17 |
| ACE2 | RP11-1166P10.1 | -0,5701 | 1,64132E-17 |
| ACE2 | FN3KRP | -0,5702 | 1,62527E-17 |
| ACE2 | EMILIN2 | -0,5702 | 1,62E-17 |
| ACE2 | CRYBB2P1 | -0,5702 | 1,6282E-17 |

| | | | |
|---|---|---|---|
| ACE2 | OBSCN | -0,5703 | 1,59245E-17 |
| ACE2 | MYLIP | -0,5703 | 1,59476E-17 |
| ACE2 | HIST1H1E | -0,5703 | 1,59202E-17 |
| ACE2 | KB-1410C5.5 | -0,5703 | 1,60503E-17 |
| ACE2 | RPS3AP47 | -0,5703 | 1,59159E-17 |
| ACE2 | TCF12 | -0,5703 | 1,59029E-17 |
| ACE2 | ASH1L | -0,5704 | 1,55988E-17 |
| ACE2 | KCNK4 | -0,5704 | 1,55969E-17 |
| ACE2 | CDH26 | -0,5704 | 1,58189E-17 |
| ACE2 | LINC01352 | -0,5705 | 1,54702E-17 |
| ACE2 | ALDH8A1 | -0,5705 | 1,55862E-17 |
| ACE2 | U47924.31 | -0,5706 | 1,52256E-17 |
| ACE2 | URAHP | -0,5706 | 1,52373E-17 |
| ACE2 | RP11-430H10.1 | -0,5707 | 1,50766E-17 |
| ACE2 | RASSF2 | -0,5707 | 1,5097E-17 |
| ACE2 | GEMIN6 | -0,5708 | 1,48131E-17 |
| ACE2 | ZNF783 | -0,5708 | 1,46936E-17 |
| ACE2 | GOLGA8N | -0,5708 | 1,47162E-17 |
| ACE2 | APOBEC3G | -0,5708 | 1,47404E-17 |
| ACE2 | PHF21B | -0,5708 | 1,46656E-17 |
| ACE2 | SRP14-AS1 | -0,5709 | 1,45528E-17 |
| ACE2 | NOL9 | -0,571 | 1,42168E-17 |
| ACE2 | RERG-IT1 | -0,571 | 1,42008E-17 |
| ACE2 | DMKN | -0,5711 | 1,41558E-17 |
| ACE2 | HSPBP1 | -0,5711 | 1,41138E-17 |
| ACE2 | RP11-87H9.5 | -0,5712 | 1,37648E-17 |
| ACE2 | C19orf60 | -0,5712 | 1,38178E-17 |
| ACE2 | PODN | -0,5713 | 1,3667E-17 |
| ACE2 | CIDECP | -0,5713 | 1,35254E-17 |
| ACE2 | PRPF38A | -0,5714 | 1,34167E-17 |
| ACE2 | RP11-603J24.5 | -0,5714 | 1,34721E-17 |
| ACE2 | C21orf62 | -0,5714 | 1,34564E-17 |
| ACE2 | HPCAL4 | -0,5715 | 1,32239E-17 |
| ACE2 | LINC01096 | -0,5715 | 1,31596E-17 |
| ACE2 | RP11-839D17.3 | -0,5715 | 1,32734E-17 |
| ACE2 | RP11-171I2.1 | -0,5716 | 1,29031E-17 |
| ACE2 | ATG12 | -0,5716 | 1,29315E-17 |
| ACE2 | SLC2A4 | -0,5716 | 1,29999E-17 |
| ACE2 | FECH | -0,5716 | 1,29864E-17 |
| ACE2 | PCOLCE-AS1 | -0,5717 | 1,27411E-17 |
| ACE2 | ENDOU | -0,5717 | 1,28819E-17 |
| ACE2 | COL4A6 | -0,5717 | 1,28338E-17 |
| ACE2 | RP11-212I21.4 | -0,5718 | 1,2503E-17 |
| ACE2 | ME3 | -0,5719 | 1,23892E-17 |
| ACE2 | BMP4 | -0,5719 | 1,23737E-17 |
| ACE2 | LSM14B | -0,5719 | 1,23742E-17 |
| ACE2 | FZD7 | -0,572 | 1,22575E-17 |
| ACE2 | ANAPC4 | -0,572 | 1,21892E-17 |
| ACE2 | DDX31 | -0,572 | 1,22785E-17 |
| ACE2 | UBE3A | -0,572 | 1,22002E-17 |
| ACE2 | RP11-762H8.2 | -0,572 | 1,22748E-17 |
| ACE2 | FRMD6-AS2 | -0,5721 | 1,1956E-17 |
| ACE2 | ZNF791 | -0,5721 | 1,20651E-17 |
| ACE2 | PDHB | -0,5723 | 1,15941E-17 |

| | | | |
|---|---|---|---|
| ACE2 | SF1 | -0,5723 | 1,15424E-17 |
| ACE2 | INO80 | -0,5723 | 1,16456E-17 |
| ACE2 | HMG20A | -0,5723 | 1,15546E-17 |
| ACE2 | CHTF8 | -0,5723 | 1,16788E-17 |
| ACE2 | LYRM9 | -0,5723 | 1,15528E-17 |
| ACE2 | CTD-3099C6.9 | -0,5723 | 1,1566E-17 |
| ACE2 | BBIP1 | -0,5724 | 1,14126E-17 |
| ACE2 | FAM122B | -0,5724 | 1,13656E-17 |
| ACE2 | LLPH | -0,5725 | 1,11918E-17 |
| ACE2 | COCH | -0,5725 | 1,12866E-17 |
| ACE2 | ITGA11 | -0,5725 | 1,13322E-17 |
| ACE2 | SOCS7 | -0,5725 | 1,13374E-17 |
| ACE2 | SCAF1 | -0,5725 | 1,12526E-17 |
| ACE2 | NOTCH2NL | -0,5726 | 1,10027E-17 |
| ACE2 | GTF2E2 | -0,5726 | 1,11157E-17 |
| ACE2 | KB-1208A12.3 | -0,5726 | 1,11131E-17 |
| ACE2 | TBC1D10A | -0,5726 | 1,11064E-17 |
| ACE2 | PHLDA1 | -0,5727 | 1,09101E-17 |
| ACE2 | AC053503.6 | -0,5728 | 1,06666E-17 |
| ACE2 | PAXIP1-AS2 | -0,5728 | 1,07348E-17 |
| ACE2 | CCDC169 | -0,5728 | 1,07241E-17 |
| ACE2 | ARHGEF7 | -0,5728 | 1,07992E-17 |
| ACE2 | LINC02076 | -0,5728 | 1,07472E-17 |
| ACE2 | JUNB | -0,5728 | 1,07868E-17 |
| ACE2 | ZHX2 | -0,5729 | 1,06031E-17 |
| ACE2 | ITFG2 | -0,5729 | 1,05796E-17 |
| ACE2 | SOCS2 | -0,5729 | 1,05108E-17 |
| ACE2 | AL022476.2 | -0,5729 | 1,05998E-17 |
| ACE2 | NBPF14 | -0,573 | 1,03577E-17 |
| ACE2 | RP11-611L7.2 | -0,573 | 1,04103E-17 |
| ACE2 | FAM208B | -0,573 | 1,03657E-17 |
| ACE2 | NFE2 | -0,573 | 1,03473E-17 |
| ACE2 | EML5 | -0,573 | 1,03199E-17 |
| ACE2 | CCDC191 | -0,5731 | 1,02247E-17 |
| ACE2 | RP11-735G4.1 | -0,5731 | 1,02585E-17 |
| ACE2 | CELF1 | -0,5731 | 1,02815E-17 |
| ACE2 | SNHG21 | -0,5731 | 1,02721E-17 |
| ACE2 | CCBE1 | -0,5731 | 1,03037E-17 |
| ACE2 | RP11-362F19.1 | -0,5732 | 1,00418E-17 |
| ACE2 | RP11-148K1.12 | -0,5732 | 1,00784E-17 |
| ACE2 | AC006946.16 | -0,5732 | 1,00906E-17 |
| ACE2 | WFDC1 | -0,5733 | 9,84777E-18 |
| ACE2 | PI4KA | -0,5733 | 9,85106E-18 |
| ACE2 | CYP20A1 | -0,5734 | 9,79697E-18 |
| ACE2 | KIAA0040 | -0,5735 | 9,65442E-18 |
| ACE2 | MAP4K2 | -0,5735 | 9,66758E-18 |
| ACE2 | CTD-3185P2.1 | -0,5735 | 9,59636E-18 |
| ACE2 | RBM12 | -0,5735 | 9,62041E-18 |
| ACE2 | RAB3C | -0,5736 | 9,493E-18 |
| ACE2 | RBM27 | -0,5736 | 9,43767E-18 |
| ACE2 | PTRH1 | -0,5736 | 9,48902E-18 |
| ACE2 | MGEA5 | -0,5736 | 9,51869E-18 |
| ACE2 | RP11-390G14.1 | -0,5737 | 9,31169E-18 |
| ACE2 | NDUFA4 | -0,5737 | 9,2792E-18 |

| | | | |
|---|---|---|---|
| ACE2 | MEOX2-AS1 | -0,5737 | 9,34012E-18 |
| ACE2 | RP11-126K1.6 | -0,5738 | 9,17553E-18 |
| ACE2 | ARID4B | -0,5738 | 9,20481E-18 |
| ACE2 | UBAC2-AS1 | -0,5738 | 9,18361E-18 |
| ACE2 | WDR3 | -0,5739 | 9,05228E-18 |
| ACE2 | ELMO1 | -0,5739 | 9,04336E-18 |
| ACE2 | ZNF277 | -0,5739 | 9,05189E-18 |
| ACE2 | GOLGA3 | -0,5739 | 8,93494E-18 |
| ACE2 | B4GALT2 | -0,574 | 8,81429E-18 |
| ACE2 | LINC02062 | -0,574 | 8,84728E-18 |
| ACE2 | EEF1A1P5 | -0,574 | 8,82614E-18 |
| ACE2 | CTD-2047H16.4 | -0,574 | 8,79563E-18 |
| ACE2 | DNAJC27-AS1 | -0,5741 | 8,72907E-18 |
| ACE2 | KB-1460A1.5 | -0,5741 | 8,79195E-18 |
| ACE2 | NDUFB8 | -0,5741 | 8,68153E-18 |
| ACE2 | RP11-425L10.1 | -0,5742 | 8,6267E-18 |
| ACE2 | SAP18 | -0,5742 | 8,63812E-18 |
| ACE2 | MEDAG | -0,5742 | 8,59778E-18 |
| ACE2 | CYB5R2 | -0,5743 | 8,46975E-18 |
| ACE2 | CHST1 | -0,5743 | 8,46877E-18 |
| ACE2 | NAA30 | -0,5743 | 8,4041E-18 |
| ACE2 | PAPLN | -0,5744 | 8,31804E-18 |
| ACE2 | LAMP5 | -0,5745 | 8,15724E-18 |
| ACE2 | ZNF551 | -0,5746 | 8,06076E-18 |
| ACE2 | NSG1 | -0,5747 | 7,93097E-18 |
| ACE2 | RINT1 | -0,5747 | 7,98484E-18 |
| ACE2 | RP11-687F6.5 | -0,5747 | 7,85957E-18 |
| ACE2 | AC097359.2 | -0,5748 | 7,81462E-18 |
| ACE2 | ACSL1 | -0,5748 | 7,79642E-18 |
| ACE2 | PRPF4 | -0,5748 | 7,84106E-18 |
| ACE2 | TMEM132C | -0,5748 | 7,8172E-18 |
| ACE2 | DRG2 | -0,5748 | 7,8447E-18 |
| ACE2 | DDX20 | -0,5749 | 7,69334E-18 |
| ACE2 | AC005154.6 | -0,5749 | 7,72969E-18 |
| ACE2 | EPOP | -0,575 | 7,53945E-18 |
| ACE2 | WDR64 | -0,5752 | 7,28811E-18 |
| ACE2 | ZSCAN23 | -0,5752 | 7,2544E-18 |
| ACE2 | RPLP0 | -0,5752 | 7,26479E-18 |
| ACE2 | VPS29 | -0,5753 | 7,16052E-18 |
| ACE2 | SLFN13 | -0,5753 | 7,17518E-18 |
| ACE2 | ANAPC1 | -0,5754 | 7,03382E-18 |
| ACE2 | RP11-1336O20.2 | -0,5754 | 7,02986E-18 |
| ACE2 | ZNF431 | -0,5754 | 7,09316E-18 |
| ACE2 | RP11-23J9.6 | -0,5755 | 6,93551E-18 |
| ACE2 | NEMF | -0,5755 | 6,93397E-18 |
| ACE2 | C17orf107 | -0,5755 | 7,00452E-18 |
| ACE2 | ETF1 | -0,5756 | 6,85284E-18 |
| ACE2 | FMC1 | -0,5756 | 6,83188E-18 |
| ACE2 | BCS1L | -0,5757 | 6,79783E-18 |
| ACE2 | FAM86EP | -0,5757 | 6,75451E-18 |
| ACE2 | TP53RK | -0,5757 | 6,79955E-18 |
| ACE2 | YTHDC1 | -0,5758 | 6,63063E-18 |
| ACE2 | RP11-167B3.2 | -0,5758 | 6,59467E-18 |
| ACE2 | PLXNA4 | -0,5759 | 6,57127E-18 |

| | | | |
|---|---|---|---|
| ACE2 | FAM133A | -0,5759 | 6,55672E-18 |
| ACE2 | FAM102B | -0,576 | 6,45627E-18 |
| ACE2 | TOGARAM2 | -0,576 | 6,43749E-18 |
| ACE2 | RP11-351D16.3 | -0,576 | 6,40264E-18 |
| ACE2 | CDIPT | -0,576 | 6,4642E-18 |
| ACE2 | ARFGAP2 | -0,5761 | 6,28575E-18 |
| ACE2 | FBL | -0,5761 | 6,32785E-18 |
| ACE2 | WDR44 | -0,5761 | 6,28301E-18 |
| ACE2 | SLC2A1-AS1 | -0,5762 | 6,22109E-18 |
| ACE2 | FAM212A | -0,5762 | 6,17969E-18 |
| ACE2 | ZNF316 | -0,5762 | 6,2026E-18 |
| ACE2 | FAM86B1 | -0,5762 | 6,21033E-18 |
| ACE2 | NPAS4 | -0,5762 | 6,25818E-18 |
| ACE2 | RP11-930O11.3 | -0,5762 | 6,18321E-18 |
| ACE2 | HDHD2 | -0,5762 | 6,18163E-18 |
| ACE2 | AC004637.1 | -0,5762 | 6,21679E-18 |
| ACE2 | AP1M1 | -0,5762 | 6,219E-18 |
| ACE2 | KIAA1644 | -0,5762 | 6,17786E-18 |
| ACE2 | RP1-315G1.3 | -0,5762 | 6,24845E-18 |
| ACE2 | FALEC | -0,5763 | 6,12831E-18 |
| ACE2 | RP11-295M18.6 | -0,5763 | 6,13318E-18 |
| ACE2 | RP11-74J13.8 | -0,5763 | 6,14075E-18 |
| ACE2 | CTD-3116E22.7 | -0,5763 | 6,15675E-18 |
| ACE2 | PLK3 | -0,5764 | 5,99941E-18 |
| ACE2 | APBB3 | -0,5764 | 6,06435E-18 |
| ACE2 | SCAF8 | -0,5764 | 6,04411E-18 |
| ACE2 | POM121C | -0,5764 | 6,03684E-18 |
| ACE2 | AKR1E2 | -0,5764 | 6,01758E-18 |
| ACE2 | FOXI2 | -0,5765 | 5,91584E-18 |
| ACE2 | RP1-101A2.1 | -0,5765 | 5,9231E-18 |
| ACE2 | DPY30 | -0,5766 | 5,81011E-18 |
| ACE2 | ERI1 | -0,5766 | 5,85977E-18 |
| ACE2 | ANKRD46 | -0,5766 | 5,87405E-18 |
| ACE2 | AC012531.25 | -0,5766 | 5,79133E-18 |
| ACE2 | LMF1 | -0,5766 | 5,81269E-18 |
| ACE2 | RP11-676J12.9 | -0,5767 | 5,77259E-18 |
| ACE2 | RPSA | -0,5768 | 5,64004E-18 |
| ACE2 | PRSS46 | -0,5768 | 5,61026E-18 |
| ACE2 | CTD-2553L13.10 | -0,5768 | 5,62417E-18 |
| ACE2 | BRAT1 | -0,5769 | 5,52518E-18 |
| ACE2 | OR10AB1P | -0,5769 | 5,5848E-18 |
| ACE2 | AMMECR1L | -0,577 | 5,42929E-18 |
| ACE2 | GDF10 | -0,577 | 5,45828E-18 |
| ACE2 | TECPR2 | -0,577 | 5,45319E-18 |
| ACE2 | RP11-689P11.2 | -0,5771 | 5,34694E-18 |
| ACE2 | NUP98 | -0,5771 | 5,41815E-18 |
| ACE2 | PSMF1 | -0,5771 | 5,41057E-18 |
| ACE2 | RGAG1 | -0,5771 | 5,39158E-18 |
| ACE2 | RANBP2 | -0,5772 | 5,31099E-18 |
| ACE2 | HMCN2 | -0,5772 | 5,33709E-18 |
| ACE2 | TRMT10C | -0,5773 | 5,25151E-18 |
| ACE2 | PARK2 | -0,5773 | 5,22962E-18 |
| ACE2 | WBP11 | -0,5773 | 5,19456E-18 |
| ACE2 | ORAI1 | -0,5773 | 5,22629E-18 |

| | | | |
|---|---|---|---|
| ACE2 | AC005498.3 | -0,5773 | 5,20949E-18 |
| ACE2 | FAM138B | -0,5774 | 5,11383E-18 |
| ACE2 | RP11-550I24.2 | -0,5774 | 5,11916E-18 |
| ACE2 | SMARCAD1 | -0,5774 | 5,12007E-18 |
| ACE2 | HEPACAM | -0,5775 | 5,07864E-18 |
| ACE2 | RP11-182J1.18 | -0,5775 | 5,01727E-18 |
| ACE2 | IMP4 | -0,5776 | 4,9539E-18 |
| ACE2 | CEP95 | -0,5776 | 4,96147E-18 |
| ACE2 | HTR2B | -0,5777 | 4,91266E-18 |
| ACE2 | SEC31B | -0,5777 | 4,92279E-18 |
| ACE2 | PROCR | -0,5777 | 4,85941E-18 |
| ACE2 | RBM4B | -0,5778 | 4,83355E-18 |
| ACE2 | AKIRIN1 | -0,5779 | 4,69851E-18 |
| ACE2 | CAPN10 | -0,5779 | 4,7082E-18 |
| ACE2 | SCOC | -0,5779 | 4,73087E-18 |
| ACE2 | C16orf72 | -0,5779 | 4,74527E-18 |
| ACE2 | AIMP1 | -0,578 | 4,62537E-18 |
| ACE2 | TLL1 | -0,578 | 4,68952E-18 |
| ACE2 | MAPK15 | -0,578 | 4,65208E-18 |
| ACE2 | COMMD5 | -0,578 | 4,65662E-18 |
| ACE2 | RP11-247A12.2 | -0,578 | 4,65347E-18 |
| ACE2 | RP1-310O13.7 | -0,578 | 4,66923E-18 |
| ACE2 | ZC3H7B | -0,578 | 4,68066E-18 |
| ACE2 | NKAPP1 | -0,578 | 4,61532E-18 |
| ACE2 | KIF1BP | -0,5781 | 4,56969E-18 |
| ACE2 | LENG1 | -0,5781 | 4,60254E-18 |
| ACE2 | FMN2 | -0,5782 | 4,49934E-18 |
| ACE2 | GRB10 | -0,5782 | 4,4765E-18 |
| ACE2 | CEBPB | -0,5782 | 4,5274E-18 |
| ACE2 | YIPF4 | -0,5783 | 4,4557E-18 |
| ACE2 | TBCCD1 | -0,5783 | 4,45465E-18 |
| ACE2 | RP1-111C20.4 | -0,5783 | 4,44414E-18 |
| ACE2 | EML3 | -0,5783 | 4,45133E-18 |
| ACE2 | DNMT3B | -0,5783 | 4,46481E-18 |
| ACE2 | PPP1R32 | -0,5784 | 4,32732E-18 |
| ACE2 | ISLR2 | -0,5784 | 4,36583E-18 |
| ACE2 | SGCG | -0,5785 | 4,27978E-18 |
| ACE2 | RP11-861E21.1 | -0,5785 | 4,26891E-18 |
| ACE2 | EPG5 | -0,5785 | 4,29499E-18 |
| ACE2 | ILF3 | -0,5785 | 4,29078E-18 |
| ACE2 | RP11-181K3.4 | -0,5786 | 4,19566E-18 |
| ACE2 | ST6GAL2 | -0,5786 | 4,19986E-18 |
| ACE2 | TCF21 | -0,5786 | 4,25319E-18 |
| ACE2 | RP11-11N9.4 | -0,5786 | 4,21214E-18 |
| ACE2 | SARDH | -0,5786 | 4,24341E-18 |
| ACE2 | OVCH1 | -0,5786 | 4,23457E-18 |
| ACE2 | MNS1 | -0,5786 | 4,23456E-18 |
| ACE2 | RP11-676J12.6 | -0,5786 | 4,22976E-18 |
| ACE2 | CTD-2260A17.2 | -0,5787 | 4,14453E-18 |
| ACE2 | HNRNPH3 | -0,5787 | 4,17398E-18 |
| ACE2 | DIS3 | -0,5787 | 4,18458E-18 |
| ACE2 | HNRNPUL1 | -0,5787 | 4,18494E-18 |
| ACE2 | AURKC | -0,5787 | 4,13994E-18 |
| ACE2 | LINC00403 | -0,5788 | 4,11166E-18 |

| | | | |
|---|---|---|---|
| ACE2 | MIR133A1HG | -0,5788 | 4,0552E-18 |
| ACE2 | MYLK-AS1 | -0,5789 | 4,03811E-18 |
| ACE2 | TRIM52 | -0,5789 | 3,99946E-18 |
| ACE2 | WTAP | -0,5789 | 3,99005E-18 |
| ACE2 | MRPS2 | -0,5789 | 4,04103E-18 |
| ACE2 | LRFN1 | -0,5789 | 4,03113E-18 |
| ACE2 | WDR89 | -0,579 | 3,96833E-18 |
| ACE2 | CD276 | -0,579 | 3,96519E-18 |
| ACE2 | LINC02185 | -0,579 | 3,97087E-18 |
| ACE2 | TMSB15B | -0,579 | 3,94182E-18 |
| ACE2 | TRAP1 | -0,5791 | 3,85922E-18 |
| ACE2 | CTC-510F12.7 | -0,5791 | 3,8895E-18 |
| ACE2 | GDAP1L1 | -0,5791 | 3,91474E-18 |
| ACE2 | TSTD2 | -0,5793 | 3,79146E-18 |
| ACE2 | NDUFV2-AS1 | -0,5794 | 3,6934E-18 |
| ACE2 | FAM228A | -0,5796 | 3,56554E-18 |
| ACE2 | GGCX | -0,5796 | 3,59809E-18 |
| ACE2 | CLK1 | -0,5796 | 3,60672E-18 |
| ACE2 | NUS1 | -0,5796 | 3,59118E-18 |
| ACE2 | ZFAND5 | -0,5796 | 3,56325E-18 |
| ACE2 | RP11-98F14.12 | -0,5796 | 3,5848E-18 |
| ACE2 | HNRNPM | -0,5797 | 3,52239E-18 |
| ACE2 | AC006116.24 | -0,5797 | 3,51128E-18 |
| ACE2 | PCNT | -0,5797 | 3,5039E-18 |
| ACE2 | HUS1B | -0,5798 | 3,47707E-18 |
| ACE2 | NDEL1 | -0,5798 | 3,46431E-18 |
| ACE2 | IFT27 | -0,5798 | 3,46979E-18 |
| ACE2 | BMS1P21 | -0,5799 | 3,42851E-18 |
| ACE2 | IGF2-AS | -0,5799 | 3,40762E-18 |
| ACE2 | CREG1 | -0,58 | 3,35197E-18 |
| ACE2 | HDAC3 | -0,58 | 3,36254E-18 |
| ACE2 | ARGLU1 | -0,58 | 3,35311E-18 |
| ACE2 | MAP2K4 | -0,58 | 3,37644E-18 |
| ACE2 | PSMD10 | -0,58 | 3,36311E-18 |
| ACE2 | DHX9 | -0,5801 | 3,28974E-18 |
| ACE2 | TOMM70 | -0,5801 | 3,30753E-18 |
| ACE2 | FAM71F2 | -0,5801 | 3,31748E-18 |
| ACE2 | KCNH3 | -0,5801 | 3,28776E-18 |
| ACE2 | SNF8 | -0,5801 | 3,31504E-18 |
| ACE2 | STRN4 | -0,5801 | 3,29248E-18 |
| ACE2 | RP11-386I23.1 | -0,5802 | 3,25844E-18 |
| ACE2 | SYT6 | -0,5802 | 3,27116E-18 |
| ACE2 | LIX1L-AS1 | -0,5802 | 3,23902E-18 |
| ACE2 | HNRNPCP2 | -0,5802 | 3,23888E-18 |
| ACE2 | RPS19 | -0,5802 | 3,27644E-18 |
| ACE2 | ZNF620 | -0,5803 | 3,17739E-18 |
| ACE2 | KIAA0319 | -0,5803 | 3,22234E-18 |
| ACE2 | PLA2G4B | -0,5803 | 3,17626E-18 |
| ACE2 | CTD-2515H24.2 | -0,5803 | 3,19209E-18 |
| ACE2 | CTC-296K1.3 | -0,5803 | 3,18738E-18 |
| ACE2 | RP11-10N23.2 | -0,5804 | 3,16327E-18 |
| ACE2 | LINC00173 | -0,5804 | 3,13821E-18 |
| ACE2 | HES7 | -0,5804 | 3,12317E-18 |
| ACE2 | DMC1 | -0,5804 | 3,16131E-18 |

| | | | |
|---|---|---|---|
| ACE2 | U82695.9 | -0,5804 | 3,13088E-18 |
| ACE2 | RP11-389C8.2 | -0,5805 | 3,08171E-18 |
| ACE2 | SCAMP1 | -0,5805 | 3,11087E-18 |
| ACE2 | AP4M1 | -0,5805 | 3,0883E-18 |
| ACE2 | STKLD1 | -0,5805 | 3,12036E-18 |
| ACE2 | C17orf49 | -0,5805 | 3,10933E-18 |
| ACE2 | HIVEP2 | -0,5806 | 3,04445E-18 |
| ACE2 | KCTD21 | -0,5806 | 3,04673E-18 |
| ACE2 | S100PBP | -0,5807 | 2,97871E-18 |
| ACE2 | IGF1 | -0,5807 | 3,01318E-18 |
| ACE2 | RP11-580I16.2 | -0,5807 | 3,00582E-18 |
| ACE2 | TAF5L | -0,5808 | 2,94114E-18 |
| ACE2 | WEE1 | -0,5808 | 2,95349E-18 |
| ACE2 | UNC13A | -0,5808 | 2,92415E-18 |
| ACE2 | NOP56 | -0,5808 | 2,96717E-18 |
| ACE2 | PSPH | -0,5809 | 2,88757E-18 |
| ACE2 | RP11-111J6.2 | -0,581 | 2,85924E-18 |
| ACE2 | GMPR | -0,581 | 2,83392E-18 |
| ACE2 | MBL1P | -0,581 | 2,82929E-18 |
| ACE2 | VAMP1 | -0,581 | 2,83388E-18 |
| ACE2 | RP11-4B16.4 | -0,581 | 2,83873E-18 |
| ACE2 | RP1-309F20.3 | -0,581 | 2,86983E-18 |
| ACE2 | MORN2 | -0,5811 | 2,81964E-18 |
| ACE2 | SNURF | -0,5811 | 2,78896E-18 |
| ACE2 | NPM1P27 | -0,5813 | 2,71788E-18 |
| ACE2 | RP11-553A21.3 | -0,5813 | 2,72985E-18 |
| ACE2 | MRPL46 | -0,5813 | 2,72656E-18 |
| ACE2 | CTC-203F4.2 | -0,5814 | 2,65229E-18 |
| ACE2 | CTD-2278I10.4 | -0,5814 | 2,6838E-18 |
| ACE2 | RP11-155D18.13 | -0,5815 | 2,6477E-18 |
| ACE2 | RP11-573M3.3 | -0,5815 | 2,63743E-18 |
| ACE2 | FAM19A2 | -0,5815 | 2,62447E-18 |
| ACE2 | POP5 | -0,5815 | 2,63482E-18 |
| ACE2 | RP11-393I2.4 | -0,5816 | 2,56608E-18 |
| ACE2 | RP11-759A24.1 | -0,5816 | 2,56672E-18 |
| ACE2 | FXR2 | -0,5816 | 2,58788E-18 |
| ACE2 | RIBC1 | -0,5816 | 2,59558E-18 |
| ACE2 | SLC5A7 | -0,5817 | 2,54024E-18 |
| ACE2 | UBE2QL1 | -0,5817 | 2,54358E-18 |
| ACE2 | CRY2 | -0,5817 | 2,54474E-18 |
| ACE2 | PDCD11 | -0,5818 | 2,50571E-18 |
| ACE2 | PSPC1-AS2 | -0,5818 | 2,51763E-18 |
| ACE2 | TUBGCP2 | -0,5819 | 2,47518E-18 |
| ACE2 | EFNB3 | -0,5819 | 2,46708E-18 |
| ACE2 | TXNL1 | -0,5819 | 2,4486E-18 |
| ACE2 | BTG3 | -0,5819 | 2,47923E-18 |
| ACE2 | CDKN1B | -0,582 | 2,4189E-18 |
| ACE2 | DNAL4 | -0,582 | 2,4075E-18 |
| ACE2 | RP1-198K11.5 | -0,5821 | 2,36514E-18 |
| ACE2 | RP11-875O11.1 | -0,5823 | 2,31571E-18 |
| ACE2 | MYO7A | -0,5823 | 2,28616E-18 |
| ACE2 | HSP90AA1 | -0,5823 | 2,31824E-18 |
| ACE2 | METTL23 | -0,5823 | 2,30012E-18 |
| ACE2 | CTA-29F11.1 | -0,5823 | 2,30608E-18 |

| | | | |
|---|---|---|---|
| ACE2 | LINC01359 | -0,5824 | 2,27784E-18 |
| ACE2 | RP11-302B13.1 | -0,5824 | 2,27582E-18 |
| ACE2 | RPS2 | -0,5824 | 2,27843E-18 |
| ACE2 | KYAT3 | -0,5826 | 2,20485E-18 |
| ACE2 | RRP12 | -0,5826 | 2,19539E-18 |
| ACE2 | RAN | -0,5826 | 2,18828E-18 |
| ACE2 | CTD-2536I1.3 | -0,5827 | 2,15987E-18 |
| ACE2 | PFN1P2 | -0,5828 | 2,11977E-18 |
| ACE2 | RP11-646I6.6 | -0,5828 | 2,11184E-18 |
| ACE2 | RRN3P1 | -0,5828 | 2,12458E-18 |
| ACE2 | MIF4GD | -0,5828 | 2,13702E-18 |
| ACE2 | ALMS1P1 | -0,5829 | 2,07258E-18 |
| ACE2 | FAM186B | -0,5829 | 2,09511E-18 |
| ACE2 | RP11-651L5.3 | -0,5829 | 2,09873E-18 |
| ACE2 | TGFBR3 | -0,583 | 2,03723E-18 |
| ACE2 | RPL15P3 | -0,5831 | 2,01074E-18 |
| ACE2 | RAPSN | -0,5831 | 2,02655E-18 |
| ACE2 | MEX3C | -0,5831 | 2,00478E-18 |
| ACE2 | AP001469.9 | -0,5831 | 2,00508E-18 |
| ACE2 | RP11-350N15.6 | -0,5832 | 1,98833E-18 |
| ACE2 | ZNF572 | -0,5832 | 1,97517E-18 |
| ACE2 | RP11-392O17.2 | -0,5833 | 1,96099E-18 |
| ACE2 | HEBP2 | -0,5833 | 1,96768E-18 |
| ACE2 | FAM200A | -0,5833 | 1,96035E-18 |
| ACE2 | EPHB6 | -0,5833 | 1,96444E-18 |
| ACE2 | GPATCH2 | -0,5834 | 1,92267E-18 |
| ACE2 | RTEL1 | -0,5834 | 1,91797E-18 |
| ACE2 | SLC4A5 | -0,5835 | 1,89657E-18 |
| ACE2 | RP11-411B10.8 | -0,5835 | 1,89826E-18 |
| ACE2 | ATG7 | -0,5836 | 1,87385E-18 |
| ACE2 | EYA4 | -0,5836 | 1,84731E-18 |
| ACE2 | FAM90A1 | -0,5836 | 1,85304E-18 |
| ACE2 | TM6SF1 | -0,5836 | 1,87397E-18 |
| ACE2 | LCMT1-AS1 | -0,5836 | 1,85088E-18 |
| ACE2 | ADNP | -0,5836 | 1,85445E-18 |
| ACE2 | CCT3 | -0,5837 | 1,81778E-18 |
| ACE2 | FAM124B | -0,5839 | 1,75683E-18 |
| ACE2 | HUWE1 | -0,5839 | 1,76534E-18 |
| ACE2 | PPM1K | -0,584 | 1,74735E-18 |
| ACE2 | LINC01505 | -0,584 | 1,7264E-18 |
| ACE2 | LINC00342 | -0,5841 | 1,69857E-18 |
| ACE2 | HEIH | -0,5841 | 1,71786E-18 |
| ACE2 | RBFOX3 | -0,5841 | 1,7019E-18 |
| ACE2 | LINC00957 | -0,5842 | 1,67323E-18 |
| ACE2 | CTD-2006H14.2 | -0,5842 | 1,68414E-18 |
| ACE2 | PSMB7 | -0,5842 | 1,67083E-18 |
| ACE2 | RP11-230F18.5 | -0,5842 | 1,67563E-18 |
| ACE2 | GADD45B | -0,5842 | 1,68509E-18 |
| ACE2 | CBR3 | -0,5842 | 1,6823E-18 |
| ACE2 | ZMAT5 | -0,5842 | 1,6823E-18 |
| ACE2 | RFT1 | -0,5843 | 1,65572E-18 |
| ACE2 | COL28A1 | -0,5843 | 1,668E-18 |
| ACE2 | GPR158 | -0,5843 | 1,65109E-18 |
| ACE2 | RP11-218E20.3 | -0,5844 | 1,63258E-18 |

| | | | |
|---|---|---|---|
| ACE2 | ZNF571-AS1 | -0,5844 | 1,6304E-18 |
| ACE2 | LINC00323 | -0,5844 | 1,6387E-18 |
| ACE2 | SYF2 | -0,5845 | 1,59547E-18 |
| ACE2 | PTCD3 | -0,5845 | 1,60514E-18 |
| ACE2 | PPP1R2 | -0,5845 | 1,59415E-18 |
| ACE2 | FMNL3 | -0,5845 | 1,60045E-18 |
| ACE2 | SFSWAP | -0,5845 | 1,61091E-18 |
| ACE2 | ZGLP1 | -0,5845 | 1,60438E-18 |
| ACE2 | NTN5 | -0,5845 | 1,60386E-18 |
| ACE2 | AATBC | -0,5845 | 1,59589E-18 |
| ACE2 | LINC00235 | -0,5846 | 1,58107E-18 |
| ACE2 | KB-1125A3.11 | -0,5846 | 1,57586E-18 |
| ACE2 | AC002456.2 | -0,5847 | 1,55473E-18 |
| ACE2 | C8orf46 | -0,5847 | 1,55575E-18 |
| ACE2 | SLC29A3 | -0,5848 | 1,52673E-18 |
| ACE2 | YARS2 | -0,5848 | 1,52239E-18 |
| ACE2 | RP11-571M6.17 | -0,5848 | 1,52721E-18 |
| ACE2 | DZANK1 | -0,5848 | 1,52457E-18 |
| ACE2 | LRRC14 | -0,5849 | 1,49483E-18 |
| ACE2 | C19orf18 | -0,5849 | 1,50876E-18 |
| ACE2 | TNRC18 | -0,585 | 1,48451E-18 |
| ACE2 | LINC00652 | -0,585 | 1,46539E-18 |
| ACE2 | NAT6 | -0,5851 | 1,45207E-18 |
| ACE2 | TBL1XR1 | -0,5851 | 1,45464E-18 |
| ACE2 | SLAIN2 | -0,5852 | 1,41751E-18 |
| ACE2 | TSLP | -0,5852 | 1,43559E-18 |
| ACE2 | E2F5 | -0,5853 | 1,39413E-18 |
| ACE2 | RP11-713P17.4 | -0,5853 | 1,4045E-18 |
| ACE2 | CACTIN-AS1 | -0,5853 | 1,39133E-18 |
| ACE2 | PUM1 | -0,5854 | 1,38343E-18 |
| ACE2 | PDZRN4 | -0,5854 | 1,3822E-18 |
| ACE2 | RPL36AL | -0,5854 | 1,38283E-18 |
| ACE2 | SLC25A38 | -0,5855 | 1,35719E-18 |
| ACE2 | SOD2 | -0,5855 | 1,3471E-18 |
| ACE2 | TACR2 | -0,5855 | 1,36152E-18 |
| ACE2 | RPL21P16 | -0,5855 | 1,35293E-18 |
| ACE2 | MOK | -0,5855 | 1,36203E-18 |
| ACE2 | RP11-567M16.6 | -0,5855 | 1,36369E-18 |
| ACE2 | PDGFC | -0,5856 | 1,33189E-18 |
| ACE2 | MRS2 | -0,5856 | 1,34169E-18 |
| ACE2 | HMGN3-AS1 | -0,5857 | 1,30484E-18 |
| ACE2 | TSN | -0,5858 | 1,28549E-18 |
| ACE2 | PHC1P1 | -0,5858 | 1,27994E-18 |
| ACE2 | VPS13C | -0,5858 | 1,2817E-18 |
| ACE2 | C20orf96 | -0,5858 | 1,28431E-18 |
| ACE2 | LINC00694 | -0,5859 | 1,26652E-18 |
| ACE2 | TUBGCP3 | -0,5859 | 1,27717E-18 |
| ACE2 | TMEM88 | -0,5859 | 1,27491E-18 |
| ACE2 | ZNF624 | -0,5859 | 1,27461E-18 |
| ACE2 | NR4A1 | -0,586 | 1,24477E-18 |
| ACE2 | TMTC3 | -0,586 | 1,24211E-18 |
| ACE2 | AF001548.6 | -0,586 | 1,25115E-18 |
| ACE2 | CLCNKB | -0,5861 | 1,21755E-18 |
| ACE2 | ACTR1B | -0,5862 | 1,21343E-18 |

| | | | |
|---|---|---|---|
| ACE2 | POLR1C | -0,5862 | 1,20146E-18 |
| ACE2 | CH25H | -0,5862 | 1,20615E-18 |
| ACE2 | RP11-342A23.2 | -0,5862 | 1,19751E-18 |
| ACE2 | ACSBG2 | -0,5862 | 1,21324E-18 |
| ACE2 | ZNF547 | -0,5862 | 1,21436E-18 |
| ACE2 | WLS | -0,5863 | 1,18026E-18 |
| ACE2 | ZNF680 | -0,5864 | 1,17328E-18 |
| ACE2 | FBXO33 | -0,5864 | 1,17376E-18 |
| ACE2 | C2orf40 | -0,5865 | 1,14656E-18 |
| ACE2 | GRB14 | -0,5865 | 1,1433E-18 |
| ACE2 | CTD-2013N24.2 | -0,5865 | 1,15E-18 |
| ACE2 | CRYBG3 | -0,5865 | 1,14108E-18 |
| ACE2 | PCDHB19P | -0,5865 | 1,15121E-18 |
| ACE2 | CNTNAP3P2 | -0,5865 | 1,1546E-18 |
| ACE2 | LPIN1 | -0,5866 | 1,12337E-18 |
| ACE2 | MORN1 | -0,5867 | 1,10285E-18 |
| ACE2 | DCAF17 | -0,5867 | 1,11624E-18 |
| ACE2 | RP11-708J19.1 | -0,5867 | 1,11839E-18 |
| ACE2 | PAFAH1B1 | -0,5867 | 1,11547E-18 |
| ACE2 | RP11-723O4.6 | -0,5868 | 1,10069E-18 |
| ACE2 | RP11-329B9.3 | -0,5868 | 1,08822E-18 |
| ACE2 | PPWD1 | -0,5868 | 1,09323E-18 |
| ACE2 | MBD1 | -0,5868 | 1,08866E-18 |
| ACE2 | PRDM2 | -0,5869 | 1,06516E-18 |
| ACE2 | ZEB2-AS1 | -0,5869 | 1,07201E-18 |
| ACE2 | RPLP2 | -0,5869 | 1,07885E-18 |
| ACE2 | ZNF544 | -0,5869 | 1,0772E-18 |
| ACE2 | FUBP1 | -0,587 | 1,05956E-18 |
| ACE2 | RP11-520H14.7 | -0,587 | 1,05072E-18 |
| ACE2 | RP11-16P20.3 | -0,587 | 1,05565E-18 |
| ACE2 | CHL1-AS1 | -0,5871 | 1,03279E-18 |
| ACE2 | TAS2R19 | -0,5871 | 1,03985E-18 |
| ACE2 | RP11-261C10.5 | -0,5872 | 1,021E-18 |
| ACE2 | IQCH | -0,5872 | 1,02635E-18 |
| ACE2 | RAB40A | -0,5872 | 1,0196E-18 |
| ACE2 | TRMT1L | -0,5873 | 1,01197E-18 |
| ACE2 | UBR3 | -0,5873 | 1,00125E-18 |
| ACE2 | CPQ | -0,5873 | 1,00637E-18 |
| ACE2 | MTSS1 | -0,5873 | 1,00384E-18 |
| ACE2 | RPA1 | -0,5874 | 9,9099E-19 |
| ACE2 | AC099850.1 | -0,5874 | 9,79583E-19 |
| ACE2 | RANBP1 | -0,5874 | 9,91187E-19 |
| ACE2 | TAF1 | -0,5874 | 9,95383E-19 |
| ACE2 | ZNF131 | -0,5876 | 9,50575E-19 |
| ACE2 | TRIM68 | -0,5876 | 9,62409E-19 |
| ACE2 | RP11-161M6.2 | -0,5876 | 9,59921E-19 |
| ACE2 | TIAL1 | -0,5877 | 9,31745E-19 |
| ACE2 | LINC00900 | -0,5877 | 9,43708E-19 |
| ACE2 | FAM92A1P1 | -0,5877 | 9,36737E-19 |
| ACE2 | PGPEP1L | -0,5877 | 9,43161E-19 |
| ACE2 | FAM173B | -0,5879 | 9,0717E-19 |
| ACE2 | C8orf31 | -0,5879 | 9,00667E-19 |
| ACE2 | NELL1 | -0,5879 | 9,05589E-19 |
| ACE2 | IPO9-AS1 | -0,588 | 8,89948E-19 |

| | | | |
|---|---|---|---|
| ACE2 | RP11-299J3.8 | -0,588 | 8,86117E-19 |
| ACE2 | PMS2P4 | -0,588 | 8,95393E-19 |
| ACE2 | NUP205 | -0,588 | 8,8754E-19 |
| ACE2 | CDHR4 | -0,5881 | 8,75489E-19 |
| ACE2 | INPP5A | -0,5881 | 8,77126E-19 |
| ACE2 | RP11-715J22.6 | -0,5881 | 8,73507E-19 |
| ACE2 | UPB1 | -0,5881 | 8,79339E-19 |
| ACE2 | CYP1B1 | -0,5882 | 8,62229E-19 |
| ACE2 | CTC-471J1.11 | -0,5882 | 8,69275E-19 |
| ACE2 | DYRK1A | -0,5882 | 8,65229E-19 |
| ACE2 | RP3-414A15.12 | -0,5883 | 8,4385E-19 |
| ACE2 | NSFL1C | -0,5883 | 8,54342E-19 |
| ACE2 | SPRTN | -0,5884 | 8,28104E-19 |
| ACE2 | RERE | -0,5885 | 8,14515E-19 |
| ACE2 | CARF | -0,5885 | 8,16337E-19 |
| ACE2 | ANKRD19P | -0,5885 | 8,25256E-19 |
| ACE2 | DROSHA | -0,5886 | 8,05738E-19 |
| ACE2 | STARD13 | -0,5886 | 8,079E-19 |
| ACE2 | AC016995.3 | -0,5887 | 8,00143E-19 |
| ACE2 | AC010524.2 | -0,5887 | 7,91693E-19 |
| ACE2 | HCG18 | -0,5888 | 7,86444E-19 |
| ACE2 | ACVRL1 | -0,5888 | 7,76064E-19 |
| ACE2 | RP11-327J17.7 | -0,5888 | 7,80867E-19 |
| ACE2 | LA16c-380F5.1 | -0,5888 | 7,8688E-19 |
| ACE2 | TERF2 | -0,5888 | 7,84506E-19 |
| ACE2 | KANK4 | -0,5889 | 7,67595E-19 |
| ACE2 | C11orf88 | -0,5889 | 7,61824E-19 |
| ACE2 | RAB11B | -0,5889 | 7,62607E-19 |
| ACE2 | EXOSC5 | -0,589 | 7,50199E-19 |
| ACE2 | PPIG | -0,5891 | 7,3919E-19 |
| ACE2 | MCUB | -0,5891 | 7,36505E-19 |
| ACE2 | PRPF8 | -0,5891 | 7,4463E-19 |
| ACE2 | APPL1 | -0,5892 | 7,26486E-19 |
| ACE2 | RP11-386M24.4 | -0,5893 | 7,13827E-19 |
| ACE2 | HSD11B1L | -0,5893 | 7,18575E-19 |
| ACE2 | RP3-412A9.16 | -0,5893 | 7,17653E-19 |
| ACE2 | GPR182 | -0,5894 | 7,07364E-19 |
| ACE2 | CPSF6 | -0,5894 | 7,09021E-19 |
| ACE2 | ZC3H13 | -0,5894 | 7,08363E-19 |
| ACE2 | RP11-477I4.4 | -0,5894 | 7,05472E-19 |
| ACE2 | GPN1 | -0,5895 | 6,9133E-19 |
| ACE2 | CCL11 | -0,5895 | 6,93116E-19 |
| ACE2 | ZNF514 | -0,5896 | 6,86929E-19 |
| ACE2 | CUL4A | -0,5896 | 6,87442E-19 |
| ACE2 | RLBP1 | -0,5896 | 6,82689E-19 |
| ACE2 | PLXNB3 | -0,5896 | 6,77828E-19 |
| ACE2 | PDS5A | -0,5897 | 6,71307E-19 |
| ACE2 | NFKBIL1 | -0,5897 | 6,71356E-19 |
| ACE2 | MPIG6B | -0,5897 | 6,65512E-19 |
| ACE2 | BRD2 | -0,5897 | 6,66228E-19 |
| ACE2 | TARSL2 | -0,5897 | 6,75562E-19 |
| ACE2 | EMP2 | -0,5897 | 6,72563E-19 |
| ACE2 | DES | -0,5898 | 6,57951E-19 |
| ACE2 | NCOA7 | -0,5898 | 6,63071E-19 |

| | | | |
|---|---|---|---|
| ACE2 | THBS1 | -0,5898 | 6,6106E-19 |
| ACE2 | SMIM17 | -0,5898 | 6,55011E-19 |
| ACE2 | RBX1 | -0,5898 | 6,56668E-19 |
| ACE2 | CDH9 | -0,5899 | 6,53417E-19 |
| ACE2 | RP11-455I9.1 | -0,5899 | 6,4552E-19 |
| ACE2 | PSMC2 | -0,5899 | 6,47166E-19 |
| ACE2 | TSC1 | -0,5899 | 6,43896E-19 |
| ACE2 | TF | -0,59 | 6,38351E-19 |
| ACE2 | MOB4 | -0,5901 | 6,2959E-19 |
| ACE2 | OTUD1 | -0,5901 | 6,28448E-19 |
| ACE2 | GPATCH2L | -0,5901 | 6,28787E-19 |
| ACE2 | ABCA8 | -0,5901 | 6,25654E-19 |
| ACE2 | SLC23A2 | -0,5901 | 6,28465E-19 |
| ACE2 | RP13-516M14.1 | -0,5902 | 6,16946E-19 |
| ACE2 | SFPQ | -0,5903 | 6,04596E-19 |
| ACE2 | AC020594.5 | -0,5903 | 6,06426E-19 |
| ACE2 | CCL21 | -0,5903 | 6,06034E-19 |
| ACE2 | FAT3 | -0,5903 | 6,01291E-19 |
| ACE2 | FZD10-AS1 | -0,5903 | 6,08091E-19 |
| ACE2 | PITPNA-AS1 | -0,5903 | 6,10579E-19 |
| ACE2 | CYP4X1 | -0,5905 | 5,84136E-19 |
| ACE2 | WAC | -0,5905 | 5,82541E-19 |
| ACE2 | CSTF3 | -0,5905 | 5,83266E-19 |
| ACE2 | OPCML | -0,5905 | 5,88418E-19 |
| ACE2 | CPAMD8 | -0,5905 | 5,85526E-19 |
| ACE2 | RBL1 | -0,5905 | 5,84617E-19 |
| ACE2 | LINC01358 | -0,5906 | 5,79036E-19 |
| ACE2 | BCAS2 | -0,5906 | 5,77661E-19 |
| ACE2 | CASQ1 | -0,5906 | 5,7884E-19 |
| ACE2 | ACTG2 | -0,5907 | 5,61542E-19 |
| ACE2 | RPL23AP42 | -0,5907 | 5,64019E-19 |
| ACE2 | ASTN2 | -0,5907 | 5,70813E-19 |
| ACE2 | EPHA5 | -0,5908 | 5,55962E-19 |
| ACE2 | LINC01475 | -0,5908 | 5,52196E-19 |
| ACE2 | RP11-677M14.5 | -0,5908 | 5,56967E-19 |
| ACE2 | SLC16A7 | -0,5909 | 5,48709E-19 |
| ACE2 | ACTR10 | -0,5909 | 5,4883E-19 |
| ACE2 | TRAPPC2L | -0,5909 | 5,49865E-19 |
| ACE2 | BTBD2 | -0,5909 | 5,43117E-19 |
| ACE2 | DUSP15 | -0,5909 | 5,4888E-19 |
| ACE2 | IPO13 | -0,591 | 5,39473E-19 |
| ACE2 | RP4-575N6.4 | -0,591 | 5,4253E-19 |
| ACE2 | RP11-326I11.3 | -0,591 | 5,33715E-19 |
| ACE2 | ZBTB44 | -0,591 | 5,42064E-19 |
| ACE2 | CAND1 | -0,591 | 5,37378E-19 |
| ACE2 | LINC01018 | -0,5911 | 5,27457E-19 |
| ACE2 | FAM200B | -0,5912 | 5,17715E-19 |
| ACE2 | SRP19 | -0,5912 | 5,18916E-19 |
| ACE2 | VEZT | -0,5912 | 5,2449E-19 |
| ACE2 | RTN1 | -0,5912 | 5,17659E-19 |
| ACE2 | VSTM2L | -0,5912 | 5,19802E-19 |
| ACE2 | G3BP1 | -0,5913 | 5,11104E-19 |
| ACE2 | RP11-723D22.3 | -0,5914 | 5,00383E-19 |
| ACE2 | RP11-589M4.4 | -0,5914 | 5,02431E-19 |

| | | | |
|---|---|---|---|
| ACE2 | TMEM145 | -0,5914 | 4,99728E-19 |
| ACE2 | ZNF444 | -0,5914 | 5,05512E-19 |
| ACE2 | MAD1L1 | -0,5915 | 4,90531E-19 |
| ACE2 | RAB8B | -0,5915 | 4,94634E-19 |
| ACE2 | KRI1 | -0,5916 | 4,84719E-19 |
| ACE2 | CLIP1 | -0,5917 | 4,73722E-19 |
| ACE2 | CAPN3 | -0,5917 | 4,75343E-19 |
| ACE2 | RP11-503P10.1 | -0,5918 | 4,69048E-19 |
| ACE2 | RP11-679C8.2 | -0,5918 | 4,66548E-19 |
| ACE2 | KBTBD3 | -0,5918 | 4,68135E-19 |
| ACE2 | PCED1B | -0,5918 | 4,68974E-19 |
| ACE2 | CASKIN1 | -0,5918 | 4,71418E-19 |
| ACE2 | ZNF211 | -0,5918 | 4,71478E-19 |
| ACE2 | SLC10A4 | -0,5919 | 4,63348E-19 |
| ACE2 | GS1-57L11.1 | -0,5919 | 4,57994E-19 |
| ACE2 | LRRC63 | -0,5919 | 4,59783E-19 |
| ACE2 | PPP4R1L | -0,5919 | 4,62936E-19 |
| ACE2 | RPL36A | -0,5919 | 4,61695E-19 |
| ACE2 | SENP2 | -0,592 | 4,51019E-19 |
| ACE2 | NALCN | -0,592 | 4,57566E-19 |
| ACE2 | NCOA5 | -0,592 | 4,56247E-19 |
| ACE2 | WIPF1 | -0,5921 | 4,46968E-19 |
| ACE2 | PPP1R3B | -0,5921 | 4,43393E-19 |
| ACE2 | METTL12 | -0,5921 | 4,47906E-19 |
| ACE2 | RP13-1016M1.2 | -0,5922 | 4,3702E-19 |
| ACE2 | SIK2 | -0,5922 | 4,40761E-19 |
| ACE2 | RP11-69M1.6 | -0,5922 | 4,37841E-19 |
| ACE2 | HIST3H2BB | -0,5923 | 4,33919E-19 |
| ACE2 | FN1 | -0,5923 | 4,27958E-19 |
| ACE2 | RFC4 | -0,5923 | 4,29697E-19 |
| ACE2 | GRID1 | -0,5924 | 4,27057E-19 |
| ACE2 | GATS | -0,5925 | 4,15229E-19 |
| ACE2 | RAP1B | -0,5925 | 4,18236E-19 |
| ACE2 | TMEM72-AS1 | -0,5926 | 4,11438E-19 |
| ACE2 | ARHGAP24 | -0,5927 | 4,06343E-19 |
| ACE2 | LINC00028 | -0,5927 | 3,99839E-19 |
| ACE2 | METTL8 | -0,5928 | 3,95072E-19 |
| ACE2 | PP12613 | -0,5928 | 3,95491E-19 |
| ACE2 | OTUD4 | -0,5928 | 3,96251E-19 |
| ACE2 | N4BP3 | -0,5928 | 3,97979E-19 |
| ACE2 | DET1 | -0,5928 | 3,98547E-19 |
| ACE2 | PRPSAP1 | -0,5928 | 3,93424E-19 |
| ACE2 | PART1 | -0,5929 | 3,87442E-19 |
| ACE2 | ZCWPW2 | -0,593 | 3,8275E-19 |
| ACE2 | VKORC1L1 | -0,593 | 3,79933E-19 |
| ACE2 | RP11-359E10.1 | -0,5931 | 3,77036E-19 |
| ACE2 | RCVRN | -0,5931 | 3,75563E-19 |
| ACE2 | GPR4 | -0,5931 | 3,75027E-19 |
| ACE2 | PHC2 | -0,5932 | 3,72717E-19 |
| ACE2 | ABCC1 | -0,5932 | 3,67372E-19 |
| ACE2 | YIF1B | -0,5932 | 3,71089E-19 |
| ACE2 | TULP2 | -0,5932 | 3,71501E-19 |
| ACE2 | GIMAP8 | -0,5933 | 3,65124E-19 |
| ACE2 | GLULP4 | -0,5933 | 3,64291E-19 |

| | | | |
|---|---|---|---|
| ACE2 | TSR3 | -0,5933 | 3,63637E-19 |
| ACE2 | RP11-475B2.1 | -0,5934 | 3,59535E-19 |
| ACE2 | PWAR5 | -0,5934 | 3,60077E-19 |
| ACE2 | TNFSF14 | -0,5934 | 3,56774E-19 |
| ACE2 | UGDH-AS1 | -0,5935 | 3,53896E-19 |
| ACE2 | NSD3 | -0,5935 | 3,52665E-19 |
| ACE2 | DSCC1 | -0,5935 | 3,48897E-19 |
| ACE2 | TMEM268 | -0,5935 | 3,5082E-19 |
| ACE2 | AC012146.7 | -0,5935 | 3,54196E-19 |
| ACE2 | ARHGAP28 | -0,5935 | 3,486E-19 |
| ACE2 | ZNF446 | -0,5935 | 3,50689E-19 |
| ACE2 | AP001476.2 | -0,5935 | 3,494E-19 |
| ACE2 | GATAD1 | -0,5936 | 3,4341E-19 |
| ACE2 | BEST1 | -0,5936 | 3,45602E-19 |
| ACE2 | UCP3 | -0,5936 | 3,47813E-19 |
| ACE2 | SF3A2 | -0,5936 | 3,46059E-19 |
| ACE2 | ASXL1 | -0,5936 | 3,44247E-19 |
| ACE2 | CLDN5 | -0,5936 | 3,47141E-19 |
| ACE2 | TMEM42 | -0,5937 | 3,38152E-19 |
| ACE2 | TRAPPC13 | -0,5937 | 3,3901E-19 |
| ACE2 | EPHB4 | -0,5937 | 3,40964E-19 |
| ACE2 | E4F1 | -0,5937 | 3,40531E-19 |
| ACE2 | CHD5 | -0,5938 | 3,35452E-19 |
| ACE2 | AC007383.3 | -0,5938 | 3,35263E-19 |
| ACE2 | AC006262.6 | -0,5938 | 3,34016E-19 |
| ACE2 | EEF1A2 | -0,5938 | 3,3431E-19 |
| ACE2 | DUSP19 | -0,5939 | 3,2714E-19 |
| ACE2 | RP11-304C12.5 | -0,5939 | 3,27772E-19 |
| ACE2 | EEF2K | -0,5939 | 3,28616E-19 |
| ACE2 | ZIK1 | -0,5939 | 3,28061E-19 |
| ACE2 | RAB9A | -0,5939 | 3,26494E-19 |
| ACE2 | WDR78 | -0,594 | 3,20311E-19 |
| ACE2 | CTC-487M23.6 | -0,594 | 3,21189E-19 |
| ACE2 | ZNF7 | -0,594 | 3,21867E-19 |
| ACE2 | EGR2 | -0,594 | 3,24659E-19 |
| ACE2 | SEC11A | -0,594 | 3,22622E-19 |
| ACE2 | RPL23P2 | -0,594 | 3,20059E-19 |
| ACE2 | RP11-216B9.6 | -0,5941 | 3,14602E-19 |
| ACE2 | RP11-753H16.5 | -0,5941 | 3,18637E-19 |
| ACE2 | FAM168A | -0,5942 | 3,1282E-19 |
| ACE2 | GTF2E1 | -0,5943 | 3,08263E-19 |
| ACE2 | SMC2-AS1 | -0,5943 | 3,07482E-19 |
| ACE2 | GLIPR1L2 | -0,5943 | 3,0472E-19 |
| ACE2 | RAD51D | -0,5943 | 3,06549E-19 |
| ACE2 | ZNF99 | -0,5943 | 3,04452E-19 |
| ACE2 | RP11-802O23.3 | -0,5944 | 3,00384E-19 |
| ACE2 | KCTD11 | -0,5944 | 3,01907E-19 |
| ACE2 | STAG1 | -0,5945 | 2,9854E-19 |
| ACE2 | RP11-531A24.7 | -0,5945 | 2,93943E-19 |
| ACE2 | CASQ2 | -0,5947 | 2,87389E-19 |
| ACE2 | DNAJC10 | -0,5947 | 2,84492E-19 |
| ACE2 | TRIAP1 | -0,5947 | 2,85336E-19 |
| ACE2 | ATP10A | -0,5947 | 2,83961E-19 |
| ACE2 | RPL39 | -0,5947 | 2,87394E-19 |

| | | | |
|---|---|---|---|
| ACE2 | KCNF1 | -0,5949 | 2,76159E-19 |
| ACE2 | TLE1 | -0,5949 | 2,75807E-19 |
| ACE2 | NT5DC3 | -0,5949 | 2,76195E-19 |
| ACE2 | UNC50 | -0,595 | 2,72213E-19 |
| ACE2 | MYNN | -0,595 | 2,72781E-19 |
| ACE2 | AJ011932.1 | -0,595 | 2,70224E-19 |
| ACE2 | MRPS10 | -0,5952 | 2,61074E-19 |
| ACE2 | RP11-1105G2.3 | -0,5952 | 2,62903E-19 |
| ACE2 | HERC5 | -0,5953 | 2,57241E-19 |
| ACE2 | INPP4B | -0,5953 | 2,58169E-19 |
| ACE2 | PEBP4 | -0,5953 | 2,59942E-19 |
| ACE2 | RC3H2 | -0,5953 | 2,60111E-19 |
| ACE2 | XAF1 | -0,5953 | 2,58071E-19 |
| ACE2 | METTL21A | -0,5954 | 2,54547E-19 |
| ACE2 | GS1-124K5.10 | -0,5954 | 2,54053E-19 |
| ACE2 | ZNF607 | -0,5954 | 2,5224E-19 |
| ACE2 | AGPAT4-IT1 | -0,5955 | 2,4764E-19 |
| ACE2 | GGTA1P | -0,5955 | 2,48244E-19 |
| ACE2 | UBXN7 | -0,5956 | 2,44294E-19 |
| ACE2 | ELP3 | -0,5958 | 2,35462E-19 |
| ACE2 | CDR2 | -0,5958 | 2,35406E-19 |
| ACE2 | CD300LG | -0,5958 | 2,38549E-19 |
| ACE2 | OLA1 | -0,5959 | 2,32086E-19 |
| ACE2 | TMEM241 | -0,5959 | 2,30943E-19 |
| ACE2 | SNX29P2 | -0,596 | 2,26957E-19 |
| ACE2 | RP11-888D10.3 | -0,596 | 2,27596E-19 |
| ACE2 | SRPX2 | -0,596 | 2,28808E-19 |
| ACE2 | ANKRD34A | -0,5961 | 2,25687E-19 |
| ACE2 | BCR | -0,5961 | 2,24429E-19 |
| ACE2 | RASGRF2-AS1 | -0,5962 | 2,2141E-19 |
| ACE2 | RP11-326C3.15 | -0,5962 | 2,21404E-19 |
| ACE2 | PRKAG1 | -0,5962 | 2,20487E-19 |
| ACE2 | RP11-419I17.1 | -0,5963 | 2,17115E-19 |
| ACE2 | GPIHBP1 | -0,5963 | 2,15445E-19 |
| ACE2 | BCDIN3D | -0,5963 | 2,18792E-19 |
| ACE2 | FBXO2 | -0,5964 | 2,14814E-19 |
| ACE2 | TTN | -0,5964 | 2,12313E-19 |
| ACE2 | HECTD4 | -0,5964 | 2,14494E-19 |
| ACE2 | PPP4R4 | -0,5965 | 2,08947E-19 |
| ACE2 | PI4KAP2 | -0,5965 | 2,09068E-19 |
| ACE2 | SLCO5A1 | -0,5966 | 2,06991E-19 |
| ACE2 | SNAI3-AS1 | -0,5966 | 2,05423E-19 |
| ACE2 | C5orf34 | -0,5967 | 2,03971E-19 |
| ACE2 | CCT6P3 | -0,5967 | 2,01585E-19 |
| ACE2 | AKTIP | -0,5968 | 2,00547E-19 |
| ACE2 | DKKL1 | -0,5968 | 2,00819E-19 |
| ACE2 | HIST2H2BA | -0,5969 | 1,95128E-19 |
| ACE2 | PCDHGC4 | -0,5969 | 1,97377E-19 |
| ACE2 | PRKDC | -0,5969 | 1,96389E-19 |
| ACE2 | VIPAS39 | -0,5969 | 1,97256E-19 |
| ACE2 | RWDD3 | -0,597 | 1,91139E-19 |
| ACE2 | TPRA1 | -0,5972 | 1,86794E-19 |
| ACE2 | NEFH | -0,5972 | 1,86082E-19 |
| ACE2 | RP11-569A11.1 | -0,5973 | 1,83852E-19 |

| | | | |
|---|---|---|---|
| ACE2 | RP11-290O12.2 | -0,5973 | 1,83398E-19 |
| ACE2 | ZDHHC2 | -0,5973 | 1,8229E-19 |
| ACE2 | LINC01537 | -0,5974 | 1,79829E-19 |
| ACE2 | RP11-340F14.6 | -0,5974 | 1,79112E-19 |
| ACE2 | ZSCAN30 | -0,5974 | 1,80989E-19 |
| ACE2 | CTC-439O9.1 | -0,5974 | 1,78137E-19 |
| ACE2 | ZNF713 | -0,5975 | 1,76764E-19 |
| ACE2 | PTGES | -0,5975 | 1,77187E-19 |
| ACE2 | RP11-444D3.1 | -0,5975 | 1,76513E-19 |
| ACE2 | RP11-817O13.9 | -0,5975 | 1,75748E-19 |
| ACE2 | CAPS | -0,5975 | 1,76682E-19 |
| ACE2 | MIIP | -0,5976 | 1,74827E-19 |
| ACE2 | PTPRG-AS1 | -0,5976 | 1,7454E-19 |
| ACE2 | SKP1 | -0,5976 | 1,71881E-19 |
| ACE2 | PTENP1-AS | -0,5976 | 1,74117E-19 |
| ACE2 | GJA5 | -0,5977 | 1,69836E-19 |
| ACE2 | LMOD1 | -0,5977 | 1,71787E-19 |
| ACE2 | ART4 | -0,5977 | 1,69172E-19 |
| ACE2 | SELE | -0,5978 | 1,6855E-19 |
| ACE2 | LMCD1-AS1 | -0,5978 | 1,68347E-19 |
| ACE2 | CCSER2 | -0,5978 | 1,68515E-19 |
| ACE2 | PSPC1 | -0,5978 | 1,66041E-19 |
| ACE2 | ABHD12B | -0,5978 | 1,68763E-19 |
| ACE2 | FOXO6 | -0,5979 | 1,64323E-19 |
| ACE2 | DCP1A | -0,5979 | 1,63612E-19 |
| ACE2 | KIT | -0,5979 | 1,63725E-19 |
| ACE2 | LINC00404 | -0,5979 | 1,65938E-19 |
| ACE2 | NOL11 | -0,5979 | 1,6488E-19 |
| ACE2 | FXYD5 | -0,5979 | 1,64901E-19 |
| ACE2 | EPC2 | -0,598 | 1,63107E-19 |
| ACE2 | ITGB3 | -0,598 | 1,62577E-19 |
| ACE2 | RP11-552F3.9 | -0,598 | 1,61216E-19 |
| ACE2 | XRCC6 | -0,598 | 1,63071E-19 |
| ACE2 | CCNH | -0,5981 | 1,58136E-19 |
| ACE2 | EXOC7 | -0,5981 | 1,58194E-19 |
| ACE2 | SAMHD1 | -0,5981 | 1,59584E-19 |
| ACE2 | RP11-434H6.7 | -0,5982 | 1,55243E-19 |
| ACE2 | PCDHGB9P | -0,5982 | 1,57515E-19 |
| ACE2 | TRERF1 | -0,5982 | 1,5617E-19 |
| ACE2 | TXNL4A | -0,5982 | 1,56108E-19 |
| ACE2 | GBP1 | -0,5983 | 1,54321E-19 |
| ACE2 | HEATR1 | -0,5983 | 1,52825E-19 |
| ACE2 | NPFFR2 | -0,5983 | 1,52221E-19 |
| ACE2 | CTB-47B11.3 | -0,5983 | 1,52913E-19 |
| ACE2 | EIF2AK2 | -0,5984 | 1,52161E-19 |
| ACE2 | SWAP70 | -0,5985 | 1,47885E-19 |
| ACE2 | KLHL2 | -0,5986 | 1,44721E-19 |
| ACE2 | TCERG1 | -0,5986 | 1,46654E-19 |
| ACE2 | SLC4A2 | -0,5986 | 1,45447E-19 |
| ACE2 | ASPN | -0,5986 | 1,46427E-19 |
| ACE2 | RPAIN | -0,5986 | 1,44714E-19 |
| ACE2 | CTC-490E21.11 | -0,5986 | 1,44615E-19 |
| ACE2 | FUT1 | -0,5986 | 1,45601E-19 |
| ACE2 | MIER1 | -0,5987 | 1,42542E-19 |

| | | | |
|---|---|---|---|
| ACE2 | WNT9A | -0,5987 | 1,44158E-19 |
| ACE2 | RP11-77K12.9 | -0,5987 | 1,43379E-19 |
| ACE2 | SMG6 | -0,5987 | 1,42073E-19 |
| ACE2 | FAM58A | -0,5987 | 1,44031E-19 |
| ACE2 | USP22 | -0,5988 | 1,41647E-19 |
| ACE2 | HLF | -0,5988 | 1,41667E-19 |
| ACE2 | SNHG15 | -0,5989 | 1,37085E-19 |
| ACE2 | ARC | -0,5989 | 1,38723E-19 |
| ACE2 | NXF1 | -0,5989 | 1,37136E-19 |
| ACE2 | MUS81 | -0,5989 | 1,39397E-19 |
| ACE2 | METTL1 | -0,5989 | 1,3741E-19 |
| ACE2 | ZNF585B | -0,5989 | 1,38317E-19 |
| ACE2 | RAB3GAP2 | -0,599 | 1,35919E-19 |
| ACE2 | YY2 | -0,599 | 1,35441E-19 |
| ACE2 | MAGOH | -0,5991 | 1,33136E-19 |
| ACE2 | PLD5 | -0,5991 | 1,33456E-19 |
| ACE2 | PLEKHG3 | -0,5991 | 1,34527E-19 |
| ACE2 | ZNF720 | -0,5991 | 1,34382E-19 |
| ACE2 | FAM167B | -0,5992 | 1,302E-19 |
| ACE2 | HILS1 | -0,5992 | 1,30146E-19 |
| ACE2 | RGS17 | -0,5993 | 1,27829E-19 |
| ACE2 | BAIAP2 | -0,5993 | 1,29307E-19 |
| ACE2 | TRMT2A | -0,5993 | 1,28509E-19 |
| ACE2 | GATAD2B | -0,5994 | 1,27709E-19 |
| ACE2 | AL133243.1 | -0,5994 | 1,26407E-19 |
| ACE2 | ZNF148 | -0,5994 | 1,26695E-19 |
| ACE2 | ACTR1A | -0,5994 | 1,26061E-19 |
| ACE2 | RP11-843B15.4 | -0,5994 | 1,26953E-19 |
| ACE2 | ZBTB25 | -0,5994 | 1,26424E-19 |
| ACE2 | PLIN5 | -0,5994 | 1,26716E-19 |
| ACE2 | LY96 | -0,5995 | 1,24668E-19 |
| ACE2 | EXOSC2 | -0,5995 | 1,23615E-19 |
| ACE2 | IRF2BP2 | -0,5996 | 1,22866E-19 |
| ACE2 | RSAD2 | -0,5996 | 1,23303E-19 |
| ACE2 | NPHP3 | -0,5996 | 1,23314E-19 |
| ACE2 | P4HA1 | -0,5996 | 1,21945E-19 |
| ACE2 | RP11-73M18.6 | -0,5996 | 1,2202E-19 |
| ACE2 | HAUS3 | -0,5997 | 1,20022E-19 |
| ACE2 | ANKRD31 | -0,5997 | 1,193E-19 |
| ACE2 | SCN2B | -0,5997 | 1,19273E-19 |
| ACE2 | DHPS | -0,5997 | 1,20435E-19 |
| ACE2 | RAD1 | -0,5998 | 1,18162E-19 |
| ACE2 | AC005754.8 | -0,5999 | 1,15934E-19 |
| ACE2 | PTCHD4 | -0,5999 | 1,15536E-19 |
| ACE2 | KAT5 | -0,5999 | 1,17086E-19 |
| ACE2 | GTF2H3 | -0,5999 | 1,16666E-19 |
| ACE2 | RP11-758H9.2 | -0,5999 | 1,15162E-19 |
| ACE2 | POLR3C | -0,6 | 1,1429E-19 |
| ACE2 | PUM2 | -0,6 | 1,14171E-19 |
| ACE2 | DCAF16 | -0,6 | 1,1461E-19 |
| ACE2 | PABPC1 | -0,6 | 1,13878E-19 |
| ACE2 | HYAL1 | -0,6001 | 1,12931E-19 |
| ACE2 | STEAP1 | -0,6001 | 1,13018E-19 |
| ACE2 | SLC35E3 | -0,6001 | 1,12018E-19 |

| | | | |
|---|---|---|---|
| ACE2 | NEURL4 | -0,6001 | 1,12398E-19 |
| ACE2 | RBM5 | -0,6002 | 1,1022E-19 |
| ACE2 | ZC3H14 | -0,6002 | 1,10514E-19 |
| ACE2 | RASL11B | -0,6003 | 1,08122E-19 |
| ACE2 | DLX5 | -0,6003 | 1,07916E-19 |
| ACE2 | EIF4H | -0,6004 | 1,0712E-19 |
| ACE2 | CTD-2537I9.18 | -0,6004 | 1,06582E-19 |
| ACE2 | POLN | -0,6005 | 1,0443E-19 |
| ACE2 | CELF2 | -0,6005 | 1,04237E-19 |
| ACE2 | RP11-411A19.5 | -0,6005 | 1,03765E-19 |
| ACE2 | FRS2 | -0,6005 | 1,04644E-19 |
| ACE2 | FIZ1 | -0,6005 | 1,03633E-19 |
| ACE2 | DYNLT3 | -0,6005 | 1,05235E-19 |
| ACE2 | MRGPRE | -0,6006 | 1,02805E-19 |
| ACE2 | HSF2BP | -0,6006 | 1,029E-19 |
| ACE2 | ZNF596 | -0,6007 | 1,00367E-19 |
| ACE2 | RP11-588H23.3 | -0,6007 | 1,00922E-19 |
| ACE2 | MPRIP | -0,6007 | 1,00672E-19 |
| ACE2 | SLITRK4 | -0,6007 | 1,00591E-19 |
| ACE2 | MSTN | -0,6008 | 9,97744E-20 |
| ACE2 | LINC01579 | -0,6008 | 9,95128E-20 |
| ACE2 | IQCH-AS1 | -0,6009 | 9,75871E-20 |
| ACE2 | HSPB7 | -0,601 | 9,55115E-20 |
| ACE2 | IMPG2 | -0,601 | 9,5769E-20 |
| ACE2 | TNXA | -0,601 | 9,61355E-20 |
| ACE2 | ZNF319 | -0,601 | 9,60256E-20 |
| ACE2 | RP11-972P1.11 | -0,6011 | 9,39703E-20 |
| ACE2 | TIMM44 | -0,6011 | 9,36345E-20 |
| ACE2 | LL0XNC01-240C2.1 | -0,6011 | 9,48366E-20 |
| ACE2 | GSTM3 | -0,6012 | 9,23999E-20 |
| ACE2 | RFX8 | -0,6013 | 9,15305E-20 |
| ACE2 | DLX2 | -0,6013 | 9,03407E-20 |
| ACE2 | GEMIN4 | -0,6013 | 9,02938E-20 |
| ACE2 | COL25A1 | -0,6014 | 8,91111E-20 |
| ACE2 | FGF18 | -0,6014 | 8,87021E-20 |
| ACE2 | LUM | -0,6014 | 8,98332E-20 |
| ACE2 | SHISA7 | -0,6014 | 8,92839E-20 |
| ACE2 | ZNF416 | -0,6014 | 8,95494E-20 |
| ACE2 | RP11-403I13.8 | -0,6015 | 8,81923E-20 |
| ACE2 | FBXO45 | -0,6015 | 8,75482E-20 |
| ACE2 | CTHRC1 | -0,6015 | 8,74394E-20 |
| ACE2 | ZNF256 | -0,6015 | 8,84133E-20 |
| ACE2 | BCL2 | -0,6016 | 8,57375E-20 |
| ACE2 | YEATS2-AS1 | -0,6017 | 8,45653E-20 |
| ACE2 | C11orf49 | -0,6017 | 8,47048E-20 |
| ACE2 | RP11-425D17.1 | -0,6017 | 8,45775E-20 |
| ACE2 | PPFIA2 | -0,6017 | 8,45175E-20 |
| ACE2 | CCT8 | -0,6017 | 8,4672E-20 |
| ACE2 | LINC01679 | -0,6017 | 8,46829E-20 |
| ACE2 | FAM69B | -0,6018 | 8,36398E-20 |
| ACE2 | RP11-291I6.2 | -0,6018 | 8,3399E-20 |
| ACE2 | IGDCC4 | -0,6018 | 8,2858E-20 |
| ACE2 | NOB1 | -0,6018 | 8,33078E-20 |
| ACE2 | TAF7 | -0,6019 | 8,22813E-20 |

| | | | |
|---|---|---|---|
| ACE2 | MRPL16 | -0,6019 | 8,22534E-20 |
| ACE2 | RP11-102G14.1 | -0,6019 | 8,10742E-20 |
| ACE2 | RNF216-IT1 | -0,602 | 7,98061E-20 |
| ACE2 | DCLK3 | -0,6021 | 7,88775E-20 |
| ACE2 | NDST2 | -0,6021 | 7,93191E-20 |
| ACE2 | C1QTNF4 | -0,6021 | 7,85553E-20 |
| ACE2 | LINC00899 | -0,6021 | 7,94912E-20 |
| ACE2 | MAGEE2 | -0,6021 | 7,91191E-20 |
| ACE2 | LINC00622 | -0,6022 | 7,73955E-20 |
| ACE2 | ZNF692 | -0,6022 | 7,80332E-20 |
| ACE2 | CCDC7 | -0,6022 | 7,71204E-20 |
| ACE2 | PSMC6 | -0,6022 | 7,68636E-20 |
| ACE2 | SNRNP25 | -0,6022 | 7,7249E-20 |
| ACE2 | GSE1 | -0,6023 | 7,63562E-20 |
| ACE2 | TRAPPC2 | -0,6023 | 7,55433E-20 |
| ACE2 | NCL | -0,6024 | 7,51455E-20 |
| ACE2 | PLBD2 | -0,6024 | 7,53478E-20 |
| ACE2 | GPR179 | -0,6024 | 7,53967E-20 |
| ACE2 | ANKRD29 | -0,6024 | 7,5437E-20 |
| ACE2 | SNHG7 | -0,6026 | 7,27802E-20 |
| ACE2 | TAPT1 | -0,6027 | 7,08924E-20 |
| ACE2 | PPP3R1 | -0,6028 | 7,01697E-20 |
| ACE2 | CTD-2287O16.1 | -0,6028 | 6,97182E-20 |
| ACE2 | EEF1G | -0,6028 | 6,91011E-20 |
| ACE2 | RORA | -0,6028 | 7,00855E-20 |
| ACE2 | AL590762.7 | -0,6028 | 6,92672E-20 |
| ACE2 | LMO3 | -0,6029 | 6,79801E-20 |
| ACE2 | WDR76 | -0,6029 | 6,90653E-20 |
| ACE2 | ZNF493 | -0,6029 | 6,90104E-20 |
| ACE2 | NDUFB11 | -0,6029 | 6,90114E-20 |
| ACE2 | RAPGEF6 | -0,603 | 6,69616E-20 |
| ACE2 | KB-431C1.5 | -0,603 | 6,74778E-20 |
| ACE2 | B3GNTL1 | -0,603 | 6,68627E-20 |
| ACE2 | GFOD1 | -0,6031 | 6,65488E-20 |
| ACE2 | WDR27 | -0,6031 | 6,5614E-20 |
| ACE2 | SLC13A4 | -0,6031 | 6,58757E-20 |
| ACE2 | AKT1 | -0,6031 | 6,58425E-20 |
| ACE2 | ZNF280B | -0,6031 | 6,57432E-20 |
| ACE2 | ARL2BP | -0,6032 | 6,52751E-20 |
| ACE2 | FBXW4P1 | -0,6032 | 6,4706E-20 |
| ACE2 | RP11-722G7.1 | -0,6032 | 6,50036E-20 |
| ACE2 | FGFBP2 | -0,6033 | 6,38975E-20 |
| ACE2 | SRA1 | -0,6033 | 6,35567E-20 |
| ACE2 | FBXO24 | -0,6033 | 6,35733E-20 |
| ACE2 | CCDC148 | -0,6034 | 6,22926E-20 |
| ACE2 | PRIM2 | -0,6034 | 6,26878E-20 |
| ACE2 | PI15 | -0,6034 | 6,23392E-20 |
| ACE2 | PAX9 | -0,6034 | 6,2147E-20 |
| ACE2 | CTD-3014M21.1 | -0,6034 | 6,22663E-20 |
| ACE2 | ASB9 | -0,6034 | 6,27122E-20 |
| ACE2 | ZSCAN9 | -0,6035 | 6,16828E-20 |
| ACE2 | ADH5 | -0,6036 | 6,02816E-20 |
| ACE2 | STC1 | -0,6036 | 6,00467E-20 |
| ACE2 | FAM181B | -0,6036 | 6,09184E-20 |

| | | | |
|---|---|---|---|
| ACE2 | TMEM126A | -0,6036 | 6,0841E-20 |
| ACE2 | PARN | -0,6036 | 6,01522E-20 |
| ACE2 | RP4-737E23.6 | -0,6036 | 6,04366E-20 |
| ACE2 | LINC02102 | -0,6037 | 5,956E-20 |
| ACE2 | N4BP2L1 | -0,6037 | 5,95523E-20 |
| ACE2 | DDHD1 | -0,6037 | 5,9116E-20 |
| ACE2 | ZNF222 | -0,6037 | 5,96183E-20 |
| ACE2 | VGLL4 | -0,6038 | 5,85309E-20 |
| ACE2 | SCOC-AS1 | -0,6038 | 5,84596E-20 |
| ACE2 | C8orf48 | -0,6038 | 5,8532E-20 |
| ACE2 | PAM16 | -0,6038 | 5,86922E-20 |
| ACE2 | NONO | -0,6038 | 5,82107E-20 |
| ACE2 | ATP6V1F | -0,6039 | 5,73629E-20 |
| ACE2 | DDX55 | -0,6039 | 5,73939E-20 |
| ACE2 | CPNE1 | -0,6039 | 5,7644E-20 |
| ACE2 | SEMA3E | -0,604 | 5,63804E-20 |
| ACE2 | C9orf72 | -0,604 | 5,65966E-20 |
| ACE2 | NTRK3 | -0,604 | 5,61843E-20 |
| ACE2 | RNMT | -0,604 | 5,58655E-20 |
| ACE2 | CD3EAP | -0,604 | 5,67554E-20 |
| ACE2 | ARHGEF4 | -0,6041 | 5,52105E-20 |
| ACE2 | PCDHA10 | -0,6041 | 5,52024E-20 |
| ACE2 | LOH12CR2 | -0,6041 | 5,52285E-20 |
| ACE2 | MRM3 | -0,6041 | 5,57156E-20 |
| ACE2 | LGI4 | -0,6041 | 5,56232E-20 |
| ACE2 | BSN | -0,6042 | 5,39367E-20 |
| ACE2 | LINC00377 | -0,6042 | 5,48707E-20 |
| ACE2 | FANCM | -0,6042 | 5,40186E-20 |
| ACE2 | C21orf62-AS1 | -0,6042 | 5,46687E-20 |
| ACE2 | LRRD1 | -0,6043 | 5,34528E-20 |
| ACE2 | SPIN3 | -0,6043 | 5,36404E-20 |
| ACE2 | EHD3 | -0,6044 | 5,20604E-20 |
| ACE2 | WDR46 | -0,6044 | 5,26024E-20 |
| ACE2 | ITGA7 | -0,6044 | 5,28924E-20 |
| ACE2 | RP11-231C14.6 | -0,6044 | 5,26393E-20 |
| ACE2 | COQ8A | -0,6045 | 5,1912E-20 |
| ACE2 | ERBB4 | -0,6045 | 5,11551E-20 |
| ACE2 | RP11-472B18.1 | -0,6046 | 5,04519E-20 |
| ACE2 | STN1 | -0,6046 | 5,03358E-20 |
| ACE2 | SORCS3 | -0,6046 | 5,07551E-20 |
| ACE2 | C18orf25 | -0,6046 | 5,06317E-20 |
| ACE2 | UBE4B | -0,6047 | 4,97938E-20 |
| ACE2 | SPTBN1 | -0,6047 | 5,00091E-20 |
| ACE2 | AC016757.3 | -0,6047 | 4,97607E-20 |
| ACE2 | HNRNPCP7 | -0,6048 | 4,8897E-20 |
| ACE2 | CPSF4 | -0,6048 | 4,86976E-20 |
| ACE2 | SFRP1 | -0,6048 | 4,87336E-20 |
| ACE2 | CDH23 | -0,6048 | 4,86985E-20 |
| ACE2 | IFT80 | -0,6049 | 4,7696E-20 |
| ACE2 | MDGA1 | -0,6049 | 4,76664E-20 |
| ACE2 | RPUSD4 | -0,6049 | 4,83772E-20 |
| ACE2 | UBTF | -0,6049 | 4,83331E-20 |
| ACE2 | RP11-62F24.2 | -0,605 | 4,68247E-20 |
| ACE2 | TRPV2 | -0,605 | 4,73589E-20 |

| | | | |
|---|---|---|---|
| ACE2 | C17orf75 | -0,605 | 4,74822E-20 |
| ACE2 | MYH7B | -0,605 | 4,68437E-20 |
| ACE2 | KCNK15-AS1 | -0,605 | 4,75333E-20 |
| ACE2 | BCL2L13 | -0,605 | 4,71611E-20 |
| ACE2 | KLHL30-AS1 | -0,6051 | 4,67306E-20 |
| ACE2 | RP4-751H13.5 | -0,6051 | 4,59697E-20 |
| ACE2 | ZNF503 | -0,6051 | 4,66672E-20 |
| ACE2 | bP-21264C1.1 | -0,6051 | 4,67489E-20 |
| ACE2 | FKTN | -0,6052 | 4,53898E-20 |
| ACE2 | LONP1 | -0,6052 | 4,54366E-20 |
| ACE2 | CTXN1 | -0,6052 | 4,5209E-20 |
| ACE2 | LUC7L3 | -0,6053 | 4,47343E-20 |
| ACE2 | SNRPA | -0,6053 | 4,45E-20 |
| ACE2 | SUPT20HL1 | -0,6053 | 4,441E-20 |
| ACE2 | POLE3 | -0,6054 | 4,42415E-20 |
| ACE2 | PARL | -0,6055 | 4,33679E-20 |
| ACE2 | FUBP3 | -0,6055 | 4,32473E-20 |
| ACE2 | LEPROT | -0,6056 | 4,21993E-20 |
| ACE2 | CNR1 | -0,6056 | 4,25012E-20 |
| ACE2 | LRP12 | -0,6056 | 4,24546E-20 |
| ACE2 | NDUFA11 | -0,6056 | 4,2263E-20 |
| ACE2 | GRIK1-AS1 | -0,6056 | 4,23222E-20 |
| ACE2 | RP11-326C3.11 | -0,6057 | 4,13154E-20 |
| ACE2 | CPA6 | -0,6058 | 4,09633E-20 |
| ACE2 | SPIN1 | -0,6058 | 4,11963E-20 |
| ACE2 | BORCS7 | -0,6058 | 4,05418E-20 |
| ACE2 | PPP2R3C | -0,6059 | 3,98371E-20 |
| ACE2 | BPTF | -0,6059 | 4,01829E-20 |
| ACE2 | ASCC1 | -0,606 | 3,93701E-20 |
| ACE2 | HIPK4 | -0,606 | 3,94938E-20 |
| ACE2 | UNC119B | -0,6061 | 3,85493E-20 |
| ACE2 | PCSK2 | -0,6061 | 3,84661E-20 |
| ACE2 | SSBP3 | -0,6062 | 3,82005E-20 |
| ACE2 | RP11-170K4.2 | -0,6062 | 3,83652E-20 |
| ACE2 | ZNF264 | -0,6062 | 3,82583E-20 |
| ACE2 | FLJ20021 | -0,6063 | 3,76179E-20 |
| ACE2 | HILPDA | -0,6063 | 3,73055E-20 |
| ACE2 | MRPL11 | -0,6063 | 3,73962E-20 |
| ACE2 | ZNF641 | -0,6063 | 3,7417E-20 |
| ACE2 | GS1-259H13.2 | -0,6064 | 3,66338E-20 |
| ACE2 | RP11-70L8.4 | -0,6064 | 3,64969E-20 |
| ACE2 | SAP30BP | -0,6064 | 3,65184E-20 |
| ACE2 | LINC00334 | -0,6064 | 3,66681E-20 |
| ACE2 | GLTSCR1L | -0,6065 | 3,63347E-20 |
| ACE2 | SLC38A2 | -0,6065 | 3,6376E-20 |
| ACE2 | RP11-147L13.15 | -0,6065 | 3,62864E-20 |
| ACE2 | SMAD4 | -0,6065 | 3,6172E-20 |
| ACE2 | TXNRD2 | -0,6065 | 3,58289E-20 |
| ACE2 | FCF1P2 | -0,6066 | 3,54086E-20 |
| ACE2 | ADAM19 | -0,6066 | 3,56534E-20 |
| ACE2 | RNASEH1 | -0,6067 | 3,46494E-20 |
| ACE2 | RPL12P16 | -0,6067 | 3,46649E-20 |
| ACE2 | ADAL | -0,6067 | 3,45954E-20 |
| ACE2 | ZNF671 | -0,6067 | 3,49551E-20 |

| | | | |
|---|---|---|---|
| ACE2 | MAN2B2 | -0,6068 | 3,42008E-20 |
| ACE2 | UBQLNL | -0,6068 | 3,40551E-20 |
| ACE2 | ZBTB1 | -0,6068 | 3,38985E-20 |
| ACE2 | ZNF541 | -0,6068 | 3,40887E-20 |
| ACE2 | RP11-565F19.2 | -0,6069 | 3,37039E-20 |
| ACE2 | PIGK | -0,6071 | 3,21765E-20 |
| ACE2 | RP11-326G21.1 | -0,6071 | 3,21212E-20 |
| ACE2 | CLEC3B | -0,6071 | 3,25673E-20 |
| ACE2 | TMEM216 | -0,6071 | 3,24335E-20 |
| ACE2 | RP11-578F21.9 | -0,6071 | 3,26685E-20 |
| ACE2 | CACNA1H | -0,6071 | 3,24944E-20 |
| ACE2 | BCAS3 | -0,6072 | 3,15847E-20 |
| ACE2 | CNN1 | -0,6072 | 3,19911E-20 |
| ACE2 | AC002398.13 | -0,6072 | 3,17155E-20 |
| ACE2 | CD81-AS1 | -0,6073 | 3,11289E-20 |
| ACE2 | GABPA | -0,6073 | 3,11837E-20 |
| ACE2 | SH3RF3-AS1 | -0,6074 | 3,09152E-20 |
| ACE2 | ISCA1 | -0,6074 | 3,07454E-20 |
| ACE2 | CTC-1337H24.4 | -0,6074 | 3,07257E-20 |
| ACE2 | EEF2KMT | -0,6074 | 3,04634E-20 |
| ACE2 | BNIP2 | -0,6075 | 2,99032E-20 |
| ACE2 | CTC-241N9.1 | -0,6076 | 2,9423E-20 |
| ACE2 | DNAJB1 | -0,6076 | 2,94861E-20 |
| ACE2 | SPATC1L | -0,6076 | 2,94196E-20 |
| ACE2 | RP11-822E23.6 | -0,6077 | 2,90744E-20 |
| ACE2 | RASSF1 | -0,6078 | 2,86061E-20 |
| ACE2 | TESK1 | -0,6078 | 2,83896E-20 |
| ACE2 | NANOS1 | -0,6078 | 2,85038E-20 |
| ACE2 | IFITM1 | -0,6078 | 2,84149E-20 |
| ACE2 | SMAD9-IT1 | -0,6078 | 2,86125E-20 |
| ACE2 | RP11-378J18.8 | -0,6079 | 2,79201E-20 |
| ACE2 | C4orf46 | -0,6079 | 2,79574E-20 |
| ACE2 | RP11-247C2.2 | -0,6079 | 2,80944E-20 |
| ACE2 | VASH2 | -0,608 | 2,78018E-20 |
| ACE2 | RP11-154H23.3 | -0,608 | 2,76612E-20 |
| ACE2 | SEC24B | -0,608 | 2,73345E-20 |
| ACE2 | SHC3 | -0,608 | 2,73491E-20 |
| ACE2 | RP5-1039K5.12 | -0,608 | 2,78078E-20 |
| ACE2 | RPL41 | -0,6081 | 2,72394E-20 |
| ACE2 | YPEL1 | -0,6081 | 2,72935E-20 |
| ACE2 | PCDHGB5 | -0,6083 | 2,63044E-20 |
| ACE2 | EMC2 | -0,6083 | 2,61228E-20 |
| ACE2 | TMEM267 | -0,6084 | 2,5551E-20 |
| ACE2 | AC003104.1 | -0,6084 | 2,564E-20 |
| ACE2 | CCDC114 | -0,6084 | 2,57573E-20 |
| ACE2 | GSTT2B | -0,6084 | 2,54919E-20 |
| ACE2 | TULP1 | -0,6085 | 2,52616E-20 |
| ACE2 | PHF3 | -0,6085 | 2,51853E-20 |
| ACE2 | RBM42 | -0,6085 | 2,525E-20 |
| ACE2 | RP11-290L1.4 | -0,6086 | 2,48646E-20 |
| ACE2 | ZNF24 | -0,6086 | 2,46541E-20 |
| ACE2 | SLC25A33 | -0,6087 | 2,42773E-20 |
| ACE2 | RP4-621B10.8 | -0,6087 | 2,41631E-20 |
| ACE2 | RP4-597N16.4 | -0,6087 | 2,43266E-20 |

| | | | |
|---|---|---|---|
| ACE2 | TRAPPC12 | -0,6087 | 2,41706E-20 |
| ACE2 | TFAP2A-AS1 | -0,6087 | 2,42899E-20 |
| ACE2 | GAPVD1 | -0,6087 | 2,4173E-20 |
| ACE2 | SCFD1 | -0,6087 | 2,4507E-20 |
| ACE2 | CTC-543D15.8 | -0,6087 | 2,42745E-20 |
| ACE2 | ZNF528-AS1 | -0,6087 | 2,41047E-20 |
| ACE2 | TARDBP | -0,6088 | 2,37468E-20 |
| ACE2 | STX11 | -0,6088 | 2,38532E-20 |
| ACE2 | NELFCD | -0,6088 | 2,38508E-20 |
| ACE2 | RFX5 | -0,6089 | 2,33286E-20 |
| ACE2 | HINT3 | -0,6089 | 2,32799E-20 |
| ACE2 | VWC2 | -0,6089 | 2,36302E-20 |
| ACE2 | DENND3 | -0,6089 | 2,34302E-20 |
| ACE2 | TSEN34 | -0,6089 | 2,3415E-20 |
| ACE2 | RP11-171I2.3 | -0,609 | 2,29802E-20 |
| ACE2 | MRPL22 | -0,609 | 2,31744E-20 |
| ACE2 | RP11-1391J7.1 | -0,609 | 2,28597E-20 |
| ACE2 | FBLN5 | -0,609 | 2,30752E-20 |
| ACE2 | AC105760.2 | -0,6091 | 2,27927E-20 |
| ACE2 | TUSC2 | -0,6091 | 2,27733E-20 |
| ACE2 | CEP83-AS1 | -0,6091 | 2,2784E-20 |
| ACE2 | RP11-21C4.1 | -0,6092 | 2,21408E-20 |
| ACE2 | FBXL22 | -0,6092 | 2,23465E-20 |
| ACE2 | UBE2O | -0,6092 | 2,24025E-20 |
| ACE2 | PDRG1 | -0,6092 | 2,21755E-20 |
| ACE2 | RPA2 | -0,6093 | 2,17109E-20 |
| ACE2 | SNRNP40 | -0,6093 | 2,16804E-20 |
| ACE2 | RAB4A | -0,6093 | 2,19923E-20 |
| ACE2 | SDCCAG3 | -0,6093 | 2,19358E-20 |
| ACE2 | HMG20B | -0,6093 | 2,19569E-20 |
| ACE2 | RHOB | -0,6094 | 2,1229E-20 |
| ACE2 | TAS2R15P | -0,6095 | 2,09913E-20 |
| ACE2 | NPIPB6 | -0,6095 | 2,11187E-20 |
| ACE2 | PRR36 | -0,6095 | 2,10651E-20 |
| ACE2 | HAUS5 | -0,6095 | 2,08607E-20 |
| ACE2 | TMEM35A | -0,6095 | 2,1147E-20 |
| ACE2 | ZNF300 | -0,6096 | 2,05993E-20 |
| ACE2 | RP11-326C3.12 | -0,6096 | 2,05254E-20 |
| ACE2 | AVEN | -0,6096 | 2,07425E-20 |
| ACE2 | RP11-395L14.18 | -0,6097 | 2,0245E-20 |
| ACE2 | WDR48 | -0,6097 | 2,01529E-20 |
| ACE2 | CTC-487M23.5 | -0,6097 | 2,03278E-20 |
| ACE2 | LHX6 | -0,6097 | 2,04244E-20 |
| ACE2 | BCDIN3D-AS1 | -0,6097 | 2,01815E-20 |
| ACE2 | RP11-799B12.1 | -0,6097 | 2,03868E-20 |
| ACE2 | C20orf203 | -0,6097 | 2,01376E-20 |
| ACE2 | XXyac-YX65C7_A.2 | -0,6098 | 2,00965E-20 |
| ACE2 | VWF | -0,6098 | 1,99103E-20 |
| ACE2 | CTD-2545G14.4 | -0,6099 | 1,93917E-20 |
| ACE2 | CBL | -0,61 | 1,92445E-20 |
| ACE2 | SETDB2 | -0,61 | 1,919E-20 |
| ACE2 | RP11-72I8.1 | -0,61 | 1,93134E-20 |
| ACE2 | RRP1 | -0,61 | 1,91623E-20 |
| ACE2 | RBMS3-AS3 | -0,6101 | 1,89127E-20 |

| | | | |
|---|---|---|---|
| ACE2 | PCDHB3 | -0,6101 | 1,87133E-20 |
| ACE2 | SFTPD | -0,6101 | 1,89654E-20 |
| ACE2 | RP3-414A15.11 | -0,6101 | 1,89149E-20 |
| ACE2 | NDUFAF5 | -0,6101 | 1,88417E-20 |
| ACE2 | RTFDC1 | -0,6101 | 1,87932E-20 |
| ACE2 | DDX6 | -0,6102 | 1,86389E-20 |
| ACE2 | KTN1 | -0,6102 | 1,86831E-20 |
| ACE2 | THEM4 | -0,6103 | 1,80988E-20 |
| ACE2 | ABTB1 | -0,6103 | 1,81941E-20 |
| ACE2 | AKR1B1 | -0,6103 | 1,82295E-20 |
| ACE2 | ZNF865 | -0,6103 | 1,83455E-20 |
| ACE2 | H1FX | -0,6104 | 1,78455E-20 |
| ACE2 | MTRR | -0,6104 | 1,80218E-20 |
| ACE2 | RPS6KA2 | -0,6104 | 1,77199E-20 |
| ACE2 | DBIL5P | -0,6104 | 1,77146E-20 |
| ACE2 | SIN3B | -0,6104 | 1,79472E-20 |
| ACE2 | ERICH3 | -0,6105 | 1,74585E-20 |
| ACE2 | CEP120 | -0,6106 | 1,70776E-20 |
| ACE2 | SGTA | -0,6106 | 1,73112E-20 |
| ACE2 | RP11-267M23.4 | -0,6107 | 1,68391E-20 |
| ACE2 | TOMM5 | -0,6107 | 1,67964E-20 |
| ACE2 | CALCOCO2 | -0,6107 | 1,7019E-20 |
| ACE2 | PANK2 | -0,6107 | 1,68833E-20 |
| ACE2 | TTC33 | -0,6108 | 1,66006E-20 |
| ACE2 | MYOF | -0,6108 | 1,6466E-20 |
| ACE2 | RAB5B | -0,6108 | 1,66225E-20 |
| ACE2 | SNW1 | -0,6108 | 1,67283E-20 |
| ACE2 | MIR637 | -0,6108 | 1,64638E-20 |
| ACE2 | RP11-97C16.1 | -0,6109 | 1,61665E-20 |
| ACE2 | CXXC5 | -0,6109 | 1,62187E-20 |
| ACE2 | ZSWIM8-AS1 | -0,6109 | 1,63565E-20 |
| ACE2 | SLC1A2 | -0,6109 | 1,64396E-20 |
| ACE2 | ATP8A2 | -0,6109 | 1,63827E-20 |
| ACE2 | RP11-679B19.1 | -0,6109 | 1,63846E-20 |
| ACE2 | RPL18 | -0,6109 | 1,62162E-20 |
| ACE2 | AC097721.2 | -0,611 | 1,59633E-20 |
| ACE2 | RP1-142L7.5 | -0,611 | 1,59324E-20 |
| ACE2 | LAG3 | -0,611 | 1,60031E-20 |
| ACE2 | MYH11 | -0,611 | 1,59784E-20 |
| ACE2 | SNX11 | -0,611 | 1,61382E-20 |
| ACE2 | EFCAB14 | -0,6111 | 1,57206E-20 |
| ACE2 | AC008073.7 | -0,6111 | 1,56505E-20 |
| ACE2 | ZMAT4 | -0,6111 | 1,58607E-20 |
| ACE2 | PRKG1 | -0,6111 | 1,58334E-20 |
| ACE2 | LINC01355 | -0,6112 | 1,55218E-20 |
| ACE2 | ATL2 | -0,6112 | 1,54899E-20 |
| ACE2 | ARL6IP5 | -0,6112 | 1,53236E-20 |
| ACE2 | GAS6-AS2 | -0,6112 | 1,54606E-20 |
| ACE2 | CTD-2547L24.4 | -0,6112 | 1,54076E-20 |
| ACE2 | NRG3 | -0,6113 | 1,50292E-20 |
| ACE2 | PSMA1 | -0,6113 | 1,51755E-20 |
| ACE2 | UBE3B | -0,6113 | 1,51475E-20 |
| ACE2 | BBOF1 | -0,6113 | 1,51253E-20 |
| ACE2 | DHX33 | -0,6113 | 1,50413E-20 |

| | | | |
|---|---|---|---|
| ACE2 | FOXK1 | -0,6114 | 1,48694E-20 |
| ACE2 | AQP6 | -0,6114 | 1,48433E-20 |
| ACE2 | CLCN4 | -0,6114 | 1,48734E-20 |
| ACE2 | LINC00847 | -0,6115 | 1,47116E-20 |
| ACE2 | CTD-2311M21.3 | -0,6115 | 1,45413E-20 |
| ACE2 | CTD-2331H12.5 | -0,6115 | 1,44969E-20 |
| ACE2 | DDB2 | -0,6116 | 1,4273E-20 |
| ACE2 | ZXDA | -0,6116 | 1,44002E-20 |
| ACE2 | C1orf122 | -0,6117 | 1,40063E-20 |
| ACE2 | CRIM1 | -0,6117 | 1,39916E-20 |
| ACE2 | NUP155 | -0,6118 | 1,38615E-20 |
| ACE2 | CEP295 | -0,6118 | 1,3919E-20 |
| ACE2 | RP11-350N15.3 | -0,6119 | 1,36697E-20 |
| ACE2 | C9orf50 | -0,6119 | 1,36588E-20 |
| ACE2 | MRPS31 | -0,6119 | 1,35236E-20 |
| ACE2 | RPL6P27 | -0,6119 | 1,37071E-20 |
| ACE2 | CTC-360J11.4 | -0,6119 | 1,36327E-20 |
| ACE2 | ARHGAP1 | -0,612 | 1,33804E-20 |
| ACE2 | PRKRIP1 | -0,6121 | 1,30666E-20 |
| ACE2 | PRKAA2 | -0,6122 | 1,28376E-20 |
| ACE2 | MGARP | -0,6122 | 1,28442E-20 |
| ACE2 | RP11-352M15.2 | -0,6123 | 1,25946E-20 |
| ACE2 | SUSD5 | -0,6124 | 1,23816E-20 |
| ACE2 | KCND2 | -0,6124 | 1,24703E-20 |
| ACE2 | TNFRSF8 | -0,6126 | 1,19132E-20 |
| ACE2 | HP1BP3 | -0,6126 | 1,18894E-20 |
| ACE2 | RGS8 | -0,6126 | 1,19886E-20 |
| ACE2 | WDSUB1 | -0,6126 | 1,19106E-20 |
| ACE2 | CABP1 | -0,6126 | 1,19436E-20 |
| ACE2 | SCAND2P | -0,6126 | 1,19306E-20 |
| ACE2 | YWHAG | -0,6127 | 1,18482E-20 |
| ACE2 | CEP131 | -0,6127 | 1,17276E-20 |
| ACE2 | CTD-2001J20.1 | -0,6127 | 1,17708E-20 |
| ACE2 | TSKS | -0,6127 | 1,1847E-20 |
| ACE2 | ZNF549 | -0,6127 | 1,18404E-20 |
| ACE2 | HOXD10 | -0,6128 | 1,14943E-20 |
| ACE2 | SLC26A4-AS1 | -0,6128 | 1,15613E-20 |
| ACE2 | SPATA41 | -0,6128 | 1,1496E-20 |
| ACE2 | CD2BP2 | -0,6128 | 1,14484E-20 |
| ACE2 | DAPK3 | -0,6128 | 1,1632E-20 |
| ACE2 | TMEM158 | -0,6129 | 1,14121E-20 |
| ACE2 | FRG1HP | -0,6129 | 1,1298E-20 |
| ACE2 | SYT15 | -0,6129 | 1,13835E-20 |
| ACE2 | C19orf43 | -0,6129 | 1,13005E-20 |
| ACE2 | NAPB | -0,6129 | 1,13439E-20 |
| ACE2 | TMEM17 | -0,613 | 1,11778E-20 |
| ACE2 | KPNA1 | -0,613 | 1,1192E-20 |
| ACE2 | RPL21P75 | -0,613 | 1,11746E-20 |
| ACE2 | ALKBH6 | -0,613 | 1,10517E-20 |
| ACE2 | GOLGA7 | -0,6131 | 1,10149E-20 |
| ACE2 | WDFY2 | -0,6131 | 1,09071E-20 |
| ACE2 | RP1-137D17.2 | -0,6132 | 1,07499E-20 |
| ACE2 | RP11-91J19.2 | -0,6132 | 1,06247E-20 |
| ACE2 | C14orf28 | -0,6132 | 1,0697E-20 |

| | | | |
|---|---|---|---|
| ACE2 | ART3 | -0,6133 | 1,05568E-20 |
| ACE2 | CCL27 | -0,6133 | 1,04701E-20 |
| ACE2 | TRPM3 | -0,6133 | 1,05747E-20 |
| ACE2 | RP4-621F18.2 | -0,6134 | 1,02759E-20 |
| ACE2 | C2orf49 | -0,6134 | 1,03358E-20 |
| ACE2 | RP9 | -0,6134 | 1,03296E-20 |
| ACE2 | ZNF77 | -0,6134 | 1,02947E-20 |
| ACE2 | ATP1A2 | -0,6135 | 1,0206E-20 |
| ACE2 | TEX26-AS1 | -0,6135 | 1,01543E-20 |
| ACE2 | CABYR | -0,6135 | 1,00689E-20 |
| ACE2 | ZNF577 | -0,6135 | 1,01017E-20 |
| ACE2 | SOCS4 | -0,6136 | 9,96838E-21 |
| ACE2 | ESF1 | -0,6136 | 9,90437E-21 |
| ACE2 | SPOUT1 | -0,6137 | 9,80128E-21 |
| ACE2 | PLP2 | -0,6138 | 9,66183E-21 |
| ACE2 | NKRF | -0,6138 | 9,59775E-21 |
| ACE2 | GLIPR1 | -0,6139 | 9,36757E-21 |
| ACE2 | RP11-622C24.2 | -0,6139 | 9,41559E-21 |
| ACE2 | RP11-499E18.1 | -0,614 | 9,33601E-21 |
| ACE2 | SOCS2-AS1 | -0,614 | 9,30238E-21 |
| ACE2 | BRF1 | -0,614 | 9,17854E-21 |
| ACE2 | ZNF23 | -0,614 | 9,28935E-21 |
| ACE2 | SOCS3 | -0,614 | 9,2861E-21 |
| ACE2 | RPTOR | -0,614 | 9,30299E-21 |
| ACE2 | RFXANK | -0,614 | 9,21639E-21 |
| ACE2 | ZNF182 | -0,614 | 9,31386E-21 |
| ACE2 | EIF1B-AS1 | -0,6141 | 9,04996E-21 |
| ACE2 | WIPI2 | -0,6141 | 9,02203E-21 |
| ACE2 | WBP1L | -0,6141 | 9,05866E-21 |
| ACE2 | CTC-526N19.1 | -0,6141 | 9,14921E-21 |
| ACE2 | BCHE | -0,6142 | 8,85392E-21 |
| ACE2 | PHF20L1 | -0,6142 | 8,8922E-21 |
| ACE2 | SLC6A15 | -0,6142 | 8,90386E-21 |
| ACE2 | CARHSP1 | -0,6142 | 8,99045E-21 |
| ACE2 | BARD1 | -0,6143 | 8,70369E-21 |
| ACE2 | PHF5A | -0,6143 | 8,78134E-21 |
| ACE2 | HESX1 | -0,6144 | 8,59554E-21 |
| ACE2 | CCDC107 | -0,6145 | 8,50826E-21 |
| ACE2 | CTD-2547L24.3 | -0,6145 | 8,47955E-21 |
| ACE2 | KDSR | -0,6145 | 8,40917E-21 |
| ACE2 | FAM210B | -0,6145 | 8,47305E-21 |
| ACE2 | RP3-333A15.2 | -0,6146 | 8,33802E-21 |
| ACE2 | GINM1 | -0,6146 | 8,25978E-21 |
| ACE2 | RP11-274B21.9 | -0,6146 | 8,34173E-21 |
| ACE2 | RP11-613M10.6 | -0,6146 | 8,25935E-21 |
| ACE2 | TRIM65 | -0,6146 | 8,21295E-21 |
| ACE2 | WDR33 | -0,6147 | 8,08502E-21 |
| ACE2 | RP11-389G6.3 | -0,6147 | 8,19013E-21 |
| ACE2 | ANGPT2 | -0,6147 | 8,14601E-21 |
| ACE2 | APH1B | -0,6147 | 8,19395E-21 |
| ACE2 | CTB-25B13.5 | -0,6147 | 8,18841E-21 |
| ACE2 | EIF4E | -0,6148 | 7,93565E-21 |
| ACE2 | CTC-255N20.1 | -0,6148 | 7,92282E-21 |
| ACE2 | KRIT1 | -0,6149 | 7,85424E-21 |

| | | | |
|---|---|---|---|
| ACE2 | ZKSCAN2 | -0,6149 | 7,89315E-21 |
| ACE2 | C22orf23 | -0,6149 | 7,84351E-21 |
| ACE2 | B3GLCT | -0,615 | 7,72187E-21 |
| ACE2 | FRMD6-AS1 | -0,615 | 7,75059E-21 |
| ACE2 | RP11-650L12.2 | -0,615 | 7,68695E-21 |
| ACE2 | RAE1 | -0,615 | 7,64271E-21 |
| ACE2 | ECE1 | -0,6151 | 7,4854E-21 |
| ACE2 | ZKSCAN8 | -0,6152 | 7,4773E-21 |
| ACE2 | TRIM39 | -0,6152 | 7,47389E-21 |
| ACE2 | NUMA1 | -0,6152 | 7,48354E-21 |
| ACE2 | RP11-712B9.5 | -0,6152 | 7,46317E-21 |
| ACE2 | RP11-426C22.5 | -0,6152 | 7,41915E-21 |
| ACE2 | ZXDB | -0,6152 | 7,44278E-21 |
| ACE2 | GAS1 | -0,6153 | 7,25167E-21 |
| ACE2 | RN7SL417P | -0,6153 | 7,24184E-21 |
| ACE2 | ZNF594 | -0,6153 | 7,28433E-21 |
| ACE2 | LINC00315 | -0,6153 | 7,28818E-21 |
| ACE2 | GABPB2 | -0,6154 | 7,21046E-21 |
| ACE2 | ZFAT | -0,6154 | 7,16985E-21 |
| ACE2 | LINC01082 | -0,6154 | 7,14268E-21 |
| ACE2 | ZNF563 | -0,6154 | 7,18127E-21 |
| ACE2 | PRPF3 | -0,6155 | 7,08126E-21 |
| ACE2 | ATP2B2 | -0,6155 | 7,01266E-21 |
| ACE2 | DENND6A | -0,6155 | 7,04779E-21 |
| ACE2 | PPIL6 | -0,6155 | 7,03842E-21 |
| ACE2 | AC010980.2 | -0,6156 | 6,84471E-21 |
| ACE2 | HSPA1L | -0,6156 | 6,90674E-21 |
| ACE2 | C9orf78 | -0,6156 | 6,94901E-21 |
| ACE2 | RNASEK | -0,6156 | 6,87372E-21 |
| ACE2 | RP3-523C21.2 | -0,6157 | 6,77329E-21 |
| ACE2 | CNN2 | -0,6157 | 6,70536E-21 |
| ACE2 | DHX34 | -0,6157 | 6,72211E-21 |
| ACE2 | FAM208A | -0,6158 | 6,69032E-21 |
| ACE2 | ZNF232 | -0,6158 | 6,60754E-21 |
| ACE2 | ZNF236 | -0,6158 | 6,69045E-21 |
| ACE2 | XRN2 | -0,6158 | 6,64437E-21 |
| ACE2 | RRP7BP | -0,6158 | 6,59952E-21 |
| ACE2 | CREB3L2 | -0,6159 | 6,52295E-21 |
| ACE2 | ID2 | -0,616 | 6,39572E-21 |
| ACE2 | RP11-1020A11.2 | -0,616 | 6,45241E-21 |
| ACE2 | RP11-358D17.2 | -0,616 | 6,44255E-21 |
| ACE2 | GSTA4 | -0,616 | 6,38202E-21 |
| ACE2 | LINC00856 | -0,6161 | 6,25626E-21 |
| ACE2 | FAM234B | -0,6161 | 6,26658E-21 |
| ACE2 | DHX36 | -0,6162 | 6,18166E-21 |
| ACE2 | LRRC77P | -0,6162 | 6,12309E-21 |
| ACE2 | RP11-541N10.3 | -0,6162 | 6,18033E-21 |
| ACE2 | PIH1D2 | -0,6162 | 6,13124E-21 |
| ACE2 | YLPM1 | -0,6162 | 6,16426E-21 |
| ACE2 | CYP11A1 | -0,6162 | 6,16503E-21 |
| ACE2 | DACT3-AS1 | -0,6162 | 6,13092E-21 |
| ACE2 | TAF11 | -0,6163 | 6,02679E-21 |
| ACE2 | UBE2L6 | -0,6163 | 6,06763E-21 |
| ACE2 | CCDC86 | -0,6163 | 6,0509E-21 |

| | | | |
|---|---|---|---|
| ACE2 | MACROD2 | -0,6163 | 6,10107E-21 |
| ACE2 | C1orf162 | -0,6164 | 5,97233E-21 |
| ACE2 | SOX11 | -0,6164 | 5,96654E-21 |
| ACE2 | RP11-282O18.3 | -0,6164 | 5,9282E-21 |
| ACE2 | SNX17 | -0,6165 | 5,83585E-21 |
| ACE2 | SUPV3L1 | -0,6165 | 5,88127E-21 |
| ACE2 | TRPC4 | -0,6165 | 5,87819E-21 |
| ACE2 | EGR1 | -0,6166 | 5,70852E-21 |
| ACE2 | KIAA0141 | -0,6166 | 5,68896E-21 |
| ACE2 | IFT43 | -0,6166 | 5,76609E-21 |
| ACE2 | SOX8 | -0,6166 | 5,68754E-21 |
| ACE2 | AC009120.6 | -0,6167 | 5,58956E-21 |
| ACE2 | ATR | -0,6168 | 5,46428E-21 |
| ACE2 | HAND2 | -0,6168 | 5,51886E-21 |
| ACE2 | TWIST1 | -0,6168 | 5,54839E-21 |
| ACE2 | LSM5 | -0,6168 | 5,54853E-21 |
| ACE2 | B3GAT3 | -0,6168 | 5,47296E-21 |
| ACE2 | ZNF773 | -0,6168 | 5,47181E-21 |
| ACE2 | XPO1 | -0,6169 | 5,42096E-21 |
| ACE2 | RP11-307E17.8 | -0,6169 | 5,45068E-21 |
| ACE2 | CBLN3 | -0,6169 | 5,37585E-21 |
| ACE2 | HIST2H2BB | -0,617 | 5,33369E-21 |
| ACE2 | RNASEH1-AS1 | -0,617 | 5,26632E-21 |
| ACE2 | OXTR | -0,617 | 5,28675E-21 |
| ACE2 | PCDHA3 | -0,617 | 5,28252E-21 |
| ACE2 | PMS2 | -0,617 | 5,32085E-21 |
| ACE2 | SNHG9 | -0,617 | 5,34673E-21 |
| ACE2 | RP11-686D22.5 | -0,617 | 5,30736E-21 |
| ACE2 | EID2 | -0,617 | 5,2955E-21 |
| ACE2 | ZNF227 | -0,617 | 5,30669E-21 |
| ACE2 | HMGN4 | -0,6171 | 5,18287E-21 |
| ACE2 | TPD52L1 | -0,6172 | 5,1263E-21 |
| ACE2 | MRM2 | -0,6172 | 5,12468E-21 |
| ACE2 | ERICH1 | -0,6172 | 5,09136E-21 |
| ACE2 | BAD | -0,6172 | 5,12341E-21 |
| ACE2 | RP11-819C21.1 | -0,6172 | 5,09196E-21 |
| ACE2 | AF001548.3 | -0,6174 | 4,97287E-21 |
| ACE2 | LINC00950 | -0,6175 | 4,82366E-21 |
| ACE2 | SAMD8 | -0,6175 | 4,85694E-21 |
| ACE2 | QRICH2 | -0,6175 | 4,81296E-21 |
| ACE2 | KCNIP4 | -0,6177 | 4,63324E-21 |
| ACE2 | TNPO1 | -0,6177 | 4,66441E-21 |
| ACE2 | AC004947.2 | -0,6177 | 4,7015E-21 |
| ACE2 | TLK2 | -0,6177 | 4,68126E-21 |
| ACE2 | TIMP3 | -0,6177 | 4,65128E-21 |
| ACE2 | RAB21 | -0,6178 | 4,60208E-21 |
| ACE2 | CARS2 | -0,6178 | 4,56421E-21 |
| ACE2 | RP11-632P5.1 | -0,6179 | 4,4681E-21 |
| ACE2 | LIG3 | -0,6179 | 4,5018E-21 |
| ACE2 | ZNF841 | -0,6179 | 4,53201E-21 |
| ACE2 | SLC2A11 | -0,6179 | 4,48808E-21 |
| ACE2 | MXD4 | -0,618 | 4,4399E-21 |
| ACE2 | RP11-867G23.10 | -0,618 | 4,37E-21 |
| ACE2 | SDR39U1 | -0,618 | 4,3888E-21 |

| | | | |
|---|---|---|---|
| ACE2 | LVRN | -0,6181 | 4,34321E-21 |
| ACE2 | SYT10 | -0,6181 | 4,3658E-21 |
| ACE2 | WARS | -0,6181 | 4,34292E-21 |
| ACE2 | SFRP2 | -0,6182 | 4,24607E-21 |
| ACE2 | DCTN4 | -0,6182 | 4,27162E-21 |
| ACE2 | TTC23 | -0,6182 | 4,27847E-21 |
| ACE2 | PRIMA1 | -0,6183 | 4,16043E-21 |
| ACE2 | USP1 | -0,6184 | 4,07068E-21 |
| ACE2 | RP11-141M3.6 | -0,6184 | 4,11195E-21 |
| ACE2 | SYNGAP1 | -0,6184 | 4,10528E-21 |
| ACE2 | CLEC2L | -0,6184 | 4,13052E-21 |
| ACE2 | HAUS6 | -0,6184 | 4,05525E-21 |
| ACE2 | STK33 | -0,6184 | 4,10404E-21 |
| ACE2 | RP11-88I18.3 | -0,6185 | 4,00205E-21 |
| ACE2 | ATP5G2 | -0,6185 | 3,98924E-21 |
| ACE2 | CERS3-AS1 | -0,6185 | 4,01468E-21 |
| ACE2 | RP11-2E11.9 | -0,6186 | 3,91726E-21 |
| ACE2 | AF186192.1 | -0,6186 | 3,95414E-21 |
| ACE2 | LBH | -0,6187 | 3,88821E-21 |
| ACE2 | MTPN | -0,6187 | 3,83799E-21 |
| ACE2 | GBGT1 | -0,6187 | 3,89142E-21 |
| ACE2 | AP001471.1 | -0,6187 | 3,86103E-21 |
| ACE2 | IL4R | -0,6188 | 3,77615E-21 |
| ACE2 | INTS2 | -0,6188 | 3,82759E-21 |
| ACE2 | C5orf51 | -0,6189 | 3,70074E-21 |
| ACE2 | FRMPD1 | -0,6189 | 3,71461E-21 |
| ACE2 | CD40 | -0,6189 | 3,70785E-21 |
| ACE2 | IBA57-AS1 | -0,619 | 3,67799E-21 |
| ACE2 | BHLHE40 | -0,619 | 3,66034E-21 |
| ACE2 | OSM | -0,619 | 3,6744E-21 |
| ACE2 | PPP1CB | -0,6191 | 3,58223E-21 |
| ACE2 | TMEM30A | -0,6191 | 3,59139E-21 |
| ACE2 | SP4 | -0,6191 | 3,58237E-21 |
| ACE2 | HAUS2 | -0,6191 | 3,56687E-21 |
| ACE2 | RP11-327J17.2 | -0,6191 | 3,57288E-21 |
| ACE2 | TPO | -0,6192 | 3,50965E-21 |
| ACE2 | C8orf59 | -0,6192 | 3,53901E-21 |
| ACE2 | SVILP1 | -0,6192 | 3,52435E-21 |
| ACE2 | RP11-666A8.9 | -0,6192 | 3,53195E-21 |
| ACE2 | THAP8 | -0,6192 | 3,53285E-21 |
| ACE2 | NOC2L | -0,6194 | 3,37005E-21 |
| ACE2 | OPN1SW | -0,6194 | 3,38301E-21 |
| ACE2 | RP11-129J12.1 | -0,6194 | 3,42105E-21 |
| ACE2 | POLR3E | -0,6194 | 3,41734E-21 |
| ACE2 | RP11-380B4.3 | -0,6195 | 3,35411E-21 |
| ACE2 | FRMPD4 | -0,6195 | 3,31241E-21 |
| ACE2 | NEXN-AS1 | -0,6196 | 3,28382E-21 |
| ACE2 | PRH1 | -0,6196 | 3,25412E-21 |
| ACE2 | DCC | -0,6197 | 3,20281E-21 |
| ACE2 | NPM3 | -0,6198 | 3,14686E-21 |
| ACE2 | PRSS45 | -0,6199 | 3,08533E-21 |
| ACE2 | RP11-281O15.4 | -0,6199 | 3,09247E-21 |
| ACE2 | SUFU | -0,6199 | 3,09416E-21 |
| ACE2 | IRX3 | -0,6199 | 3,07694E-21 |

| | | | |
|---|---|---|---|
| ACE2 | FAM227A | -0,6199 | 3,10501E-21 |
| ACE2 | FKBP1A | -0,62 | 3,04713E-21 |
| ACE2 | MCFD2 | -0,6201 | 2,9575E-21 |
| ACE2 | RBM19 | -0,6201 | 2,9993E-21 |
| ACE2 | CHRM2 | -0,6202 | 2,93892E-21 |
| ACE2 | SH2D3C | -0,6202 | 2,92367E-21 |
| ACE2 | LTBP4 | -0,6202 | 2,90668E-21 |
| ACE2 | GYS2 | -0,6203 | 2,86643E-21 |
| ACE2 | RP11-66N11.8 | -0,6203 | 2,84458E-21 |
| ACE2 | ZNF879 | -0,6204 | 2,81934E-21 |
| ACE2 | HID1-AS1 | -0,6204 | 2,82317E-21 |
| ACE2 | ZYG11B | -0,6205 | 2,74911E-21 |
| ACE2 | WDR26 | -0,6205 | 2,76325E-21 |
| ACE2 | NEK11 | -0,6205 | 2,7603E-21 |
| ACE2 | TRAM2-AS1 | -0,6205 | 2,7803E-21 |
| ACE2 | C9orf170 | -0,6205 | 2,75345E-21 |
| ACE2 | NUP50-AS1 | -0,6205 | 2,78096E-21 |
| ACE2 | AMER1 | -0,6205 | 2,75533E-21 |
| ACE2 | RP11-177G23.1 | -0,6206 | 2,69216E-21 |
| ACE2 | DIS3L2 | -0,6207 | 2,66787E-21 |
| ACE2 | EPM2AIP1 | -0,6207 | 2,67515E-21 |
| ACE2 | TPTE2P3 | -0,6207 | 2,63696E-21 |
| ACE2 | AC006538.1 | -0,6207 | 2,67644E-21 |
| ACE2 | ADRA2C | -0,6208 | 2,61881E-21 |
| ACE2 | GHc-857G6.7 | -0,6208 | 2,62757E-21 |
| ACE2 | GRHL1 | -0,6209 | 2,55927E-21 |
| ACE2 | CTD-2287O16.5 | -0,6209 | 2,56547E-21 |
| ACE2 | LRRC27 | -0,6209 | 2,55043E-21 |
| ACE2 | PPM1L | -0,621 | 2,51534E-21 |
| ACE2 | F2RL2 | -0,621 | 2,50899E-21 |
| ACE2 | GTF2H1 | -0,621 | 2,51832E-21 |
| ACE2 | LINC01638 | -0,621 | 2,53613E-21 |
| ACE2 | SS18 | -0,6211 | 2,47436E-21 |
| ACE2 | PBX1 | -0,6212 | 2,40856E-21 |
| ACE2 | FYN | -0,6212 | 2,41817E-21 |
| ACE2 | DHX35 | -0,6212 | 2,42428E-21 |
| ACE2 | MMP23B | -0,6213 | 2,38738E-21 |
| ACE2 | RP11-159N11.4 | -0,6213 | 2,36692E-21 |
| ACE2 | RP11-421F16.3 | -0,6213 | 2,36191E-21 |
| ACE2 | ROPN1L | -0,6214 | 2,31125E-21 |
| ACE2 | GIMAP4 | -0,6214 | 2,33811E-21 |
| ACE2 | KCNA5 | -0,6214 | 2,32599E-21 |
| ACE2 | NUP93 | -0,6214 | 2,34123E-21 |
| ACE2 | BORCS8 | -0,6214 | 2,31419E-21 |
| ACE2 | WBP1LP2 | -0,6215 | 2,27656E-21 |
| ACE2 | ASIC3 | -0,6216 | 2,25403E-21 |
| ACE2 | RB1 | -0,6216 | 2,25291E-21 |
| ACE2 | AC068134.10 | -0,6217 | 2,22233E-21 |
| ACE2 | CAV2 | -0,6217 | 2,21812E-21 |
| ACE2 | RP11-92C4.6 | -0,6217 | 2,21221E-21 |
| ACE2 | LINC02094 | -0,6217 | 2,19052E-21 |
| ACE2 | ZNF826P | -0,6217 | 2,19132E-21 |
| ACE2 | NCOA1 | -0,6218 | 2,14162E-21 |
| ACE2 | SAAL1 | -0,6218 | 2,16558E-21 |

| | | | |
|---|---|---|---|
| ACE2 | RP11-49C9.2 | -0,6218 | 2,17353E-21 |
| ACE2 | HSD11B1 | -0,6219 | 2,13862E-21 |
| ACE2 | TRIM52-AS1 | -0,6219 | 2,10583E-21 |
| ACE2 | CAMK1D | -0,6219 | 2,12927E-21 |
| ACE2 | SLC10A3 | -0,622 | 2,07118E-21 |
| ACE2 | DCAF8 | -0,6221 | 2,05265E-21 |
| ACE2 | GPR156 | -0,6221 | 2,05169E-21 |
| ACE2 | ZNF474 | -0,6221 | 2,04567E-21 |
| ACE2 | PCDHB8 | -0,6221 | 2,03453E-21 |
| ACE2 | XPO5 | -0,6221 | 2,02927E-21 |
| ACE2 | ABC7-42404400C24. | -0,6221 | 2,03278E-21 |
| ACE2 | RP11-1180F24.1 | -0,6221 | 2,04121E-21 |
| ACE2 | ADAMTS9-AS1 | -0,6222 | 2,00982E-21 |
| ACE2 | IL21R-AS1 | -0,6222 | 1,99682E-21 |
| ACE2 | RP11-44F14.6 | -0,6222 | 1,99105E-21 |
| ACE2 | PTGIS | -0,6222 | 2,01986E-21 |
| ACE2 | BNIP3P30 | -0,6223 | 1,96007E-21 |
| ACE2 | TUG1 | -0,6223 | 1,96597E-21 |
| ACE2 | THRAP3 | -0,6224 | 1,93149E-21 |
| ACE2 | ZNF76 | -0,6224 | 1,9247E-21 |
| ACE2 | CSPP1 | -0,6224 | 1,94499E-21 |
| ACE2 | NEK5 | -0,6224 | 1,91259E-21 |
| ACE2 | TRNP1 | -0,6225 | 1,87501E-21 |
| ACE2 | RSPH9 | -0,6225 | 1,89757E-21 |
| ACE2 | PDE7B | -0,6225 | 1,87687E-21 |
| ACE2 | SSBP1 | -0,6225 | 1,88151E-21 |
| ACE2 | RP11-313J2.1 | -0,6225 | 1,89534E-21 |
| ACE2 | FCF1 | -0,6225 | 1,8774E-21 |
| ACE2 | MYLK3 | -0,6225 | 1,88503E-21 |
| ACE2 | CEP85L | -0,6226 | 1,85048E-21 |
| ACE2 | CENPBD1P1 | -0,6226 | 1,86294E-21 |
| ACE2 | MTG2 | -0,6226 | 1,86452E-21 |
| ACE2 | NEK7 | -0,6227 | 1,82463E-21 |
| ACE2 | THSD7B | -0,6227 | 1,81015E-21 |
| ACE2 | TRIM44 | -0,6227 | 1,81227E-21 |
| ACE2 | GABARAP | -0,6227 | 1,82281E-21 |
| ACE2 | C19orf47 | -0,6227 | 1,82162E-21 |
| ACE2 | TFPT | -0,6227 | 1,83098E-21 |
| ACE2 | DSTN | -0,6227 | 1,83415E-21 |
| ACE2 | P2RX6 | -0,6227 | 1,83235E-21 |
| ACE2 | ZNF143 | -0,6228 | 1,79207E-21 |
| ACE2 | SAMD14 | -0,6228 | 1,79111E-21 |
| ACE2 | ORC4 | -0,6229 | 1,74897E-21 |
| ACE2 | FNDC3B | -0,6229 | 1,74332E-21 |
| ACE2 | WDR31 | -0,6229 | 1,76226E-21 |
| ACE2 | DPH7 | -0,6229 | 1,74627E-21 |
| ACE2 | PAPOLG | -0,623 | 1,73099E-21 |
| ACE2 | HNRNPA0 | -0,6231 | 1,6795E-21 |
| ACE2 | TRAM2 | -0,6231 | 1,70259E-21 |
| ACE2 | INTS10 | -0,6231 | 1,67847E-21 |
| ACE2 | RP11-551L14.8 | -0,6231 | 1,67337E-21 |
| ACE2 | ANAPC5 | -0,6231 | 1,68908E-21 |
| ACE2 | ZBTB21 | -0,6231 | 1,67588E-21 |
| ACE2 | EXOSC9 | -0,6232 | 1,66692E-21 |

| | | | |
|---|---|---|---|
| ACE2 | TCAF1 | -0,6232 | 1,64441E-21 |
| ACE2 | ZNF605 | -0,6232 | 1,64393E-21 |
| ACE2 | RPE | -0,6233 | 1,62831E-21 |
| ACE2 | THRB | -0,6233 | 1,63099E-21 |
| ACE2 | ERC2 | -0,6233 | 1,62642E-21 |
| ACE2 | RP11-269C23.5 | -0,6233 | 1,61365E-21 |
| ACE2 | KLHDC8A | -0,6234 | 1,60556E-21 |
| ACE2 | KCNQ5 | -0,6234 | 1,58102E-21 |
| ACE2 | CASC2 | -0,6234 | 1,59124E-21 |
| ACE2 | AF146191.4 | -0,6235 | 1,56909E-21 |
| ACE2 | LINC00472 | -0,6235 | 1,55144E-21 |
| ACE2 | EMILIN3 | -0,6235 | 1,57274E-21 |
| ACE2 | GPR27 | -0,6236 | 1,52599E-21 |
| ACE2 | SNRNP70 | -0,6236 | 1,53953E-21 |
| ACE2 | TMEM150A | -0,6237 | 1,51372E-21 |
| ACE2 | PCOLCE2 | -0,6237 | 1,5074E-21 |
| ACE2 | FAR2P2 | -0,6238 | 1,49047E-21 |
| ACE2 | ITIH4 | -0,6238 | 1,47649E-21 |
| ACE2 | DCPS | -0,6238 | 1,46679E-21 |
| ACE2 | MOB3A | -0,6238 | 1,49174E-21 |
| ACE2 | ELK1 | -0,6238 | 1,48505E-21 |
| ACE2 | SETD1B | -0,624 | 1,43567E-21 |
| ACE2 | CHODL | -0,624 | 1,42939E-21 |
| ACE2 | ASIC4 | -0,6241 | 1,40827E-21 |
| ACE2 | EP400NL | -0,6241 | 1,40036E-21 |
| ACE2 | UBR5 | -0,6242 | 1,37769E-21 |
| ACE2 | TEPP | -0,6242 | 1,35998E-21 |
| ACE2 | LRRC8C | -0,6243 | 1,34001E-21 |
| ACE2 | PRSS42 | -0,6243 | 1,33184E-21 |
| ACE2 | RGS4 | -0,6244 | 1,31422E-21 |
| ACE2 | ZNF397 | -0,6244 | 1,32516E-21 |
| ACE2 | PATZ1 | -0,6244 | 1,30712E-21 |
| ACE2 | DESI2 | -0,6245 | 1,29741E-21 |
| ACE2 | MGA | -0,6245 | 1,29358E-21 |
| ACE2 | ZNF502 | -0,6246 | 1,26349E-21 |
| ACE2 | IGHMBP2 | -0,6246 | 1,26551E-21 |
| ACE2 | SUGP2 | -0,6246 | 1,25775E-21 |
| ACE2 | LUC7L | -0,6247 | 1,25195E-21 |
| ACE2 | CTD-3220F14.3 | -0,6247 | 1,24927E-21 |
| ACE2 | ELOVL2 | -0,6248 | 1,22344E-21 |
| ACE2 | PNMA6A | -0,6248 | 1,2193E-21 |
| ACE2 | CYP8B1 | -0,6249 | 1,18867E-21 |
| ACE2 | CTC-366B18.4 | -0,6249 | 1,19986E-21 |
| ACE2 | FAM120AOS | -0,6249 | 1,20095E-21 |
| ACE2 | UGGT2 | -0,6249 | 1,18728E-21 |
| ACE2 | LINC00339 | -0,625 | 1,1776E-21 |
| ACE2 | RPL5P4 | -0,625 | 1,18095E-21 |
| ACE2 | ETAA1 | -0,625 | 1,17692E-21 |
| ACE2 | DNM1 | -0,625 | 1,16666E-21 |
| ACE2 | RECQL | -0,625 | 1,16963E-21 |
| ACE2 | FEM1B | -0,625 | 1,18466E-21 |
| ACE2 | ELAC2 | -0,625 | 1,18256E-21 |
| ACE2 | ZNF283 | -0,625 | 1,16671E-21 |
| ACE2 | CGGBP1 | -0,6251 | 1,14693E-21 |

| | | | |
|---|---|---|---|
| ACE2 | FAM114A2 | -0,6251 | 1,1431E-21 |
| ACE2 | DNAJB2 | -0,6252 | 1,13796E-21 |
| ACE2 | ZNF891 | -0,6252 | 1,12334E-21 |
| ACE2 | RP11-411B10.2 | -0,6252 | 1,12183E-21 |
| ACE2 | ZNF322 | -0,6253 | 1,11117E-21 |
| ACE2 | EXOSC8 | -0,6253 | 1,11217E-21 |
| ACE2 | YY1 | -0,6253 | 1,10965E-21 |
| ACE2 | RBM10 | -0,6253 | 1,11298E-21 |
| ACE2 | UPF3B | -0,6253 | 1,11682E-21 |
| ACE2 | ZCCHC11 | -0,6254 | 1,07967E-21 |
| ACE2 | MB21D1 | -0,6254 | 1,09096E-21 |
| ACE2 | MGAT4C | -0,6254 | 1,08029E-21 |
| ACE2 | LYPLAL1-AS1 | -0,6255 | 1,07398E-21 |
| ACE2 | UNC79 | -0,6255 | 1,06002E-21 |
| ACE2 | RP11-848P1.9 | -0,6255 | 1,06338E-21 |
| ACE2 | CTD-2027I19.2 | -0,6255 | 1,06775E-21 |
| ACE2 | LCA5L | -0,6255 | 1,07069E-21 |
| ACE2 | SOS1-IT1 | -0,6256 | 1,04448E-21 |
| ACE2 | ANKRD37 | -0,6256 | 1,0438E-21 |
| ACE2 | SDCBP | -0,6256 | 1,05737E-21 |
| ACE2 | ZNF597 | -0,6256 | 1,04576E-21 |
| ACE2 | C18orf21 | -0,6256 | 1,04003E-21 |
| ACE2 | FAXC | -0,6257 | 1,0289E-21 |
| ACE2 | LRRC10B | -0,6257 | 1,03017E-21 |
| ACE2 | NOP58 | -0,6258 | 1,00139E-21 |
| ACE2 | RBM22P2 | -0,6259 | 9,9449E-22 |
| ACE2 | RP11-452L6.7 | -0,6259 | 9,80426E-22 |
| ACE2 | DNTTIP1 | -0,6259 | 9,89261E-22 |
| ACE2 | MYOZ2 | -0,626 | 9,73114E-22 |
| ACE2 | UBE2B | -0,626 | 9,67888E-22 |
| ACE2 | CTA-941F9.10 | -0,626 | 9,65824E-22 |
| ACE2 | RP11-365H22.2 | -0,6261 | 9,42847E-22 |
| ACE2 | NR4A3 | -0,6261 | 9,5533E-22 |
| ACE2 | CAPRIN1 | -0,6261 | 9,56467E-22 |
| ACE2 | LINC01578 | -0,6261 | 9,46563E-22 |
| ACE2 | RBMX | -0,6261 | 9,52588E-22 |
| ACE2 | ETV5 | -0,6263 | 9,23589E-22 |
| ACE2 | CMC2 | -0,6263 | 9,16147E-22 |
| ACE2 | RP11-417F21.1 | -0,6265 | 8,86564E-22 |
| ACE2 | PKD1L1 | -0,6265 | 8,8505E-22 |
| ACE2 | RCL1 | -0,6265 | 8,89276E-22 |
| ACE2 | NIN | -0,6265 | 8,8859E-22 |
| ACE2 | AC103563.9 | -0,6266 | 8,6039E-22 |
| ACE2 | PIP4K2A | -0,6266 | 8,72082E-22 |
| ACE2 | YAF2 | -0,6266 | 8,65621E-22 |
| ACE2 | C14orf79 | -0,6266 | 8,60146E-22 |
| ACE2 | AC006942.4 | -0,6266 | 8,70487E-22 |
| ACE2 | STX16 | -0,6266 | 8,57369E-22 |
| ACE2 | EXOG | -0,6267 | 8,4118E-22 |
| ACE2 | JPH3 | -0,6267 | 8,40568E-22 |
| ACE2 | RP11-552F3.10 | -0,6267 | 8,54998E-22 |
| ACE2 | VN1R85P | -0,6267 | 8,45435E-22 |
| ACE2 | LRRC41 | -0,6268 | 8,28313E-22 |
| ACE2 | SH2B3 | -0,627 | 8,02394E-22 |

| | | | |
|---|---|---|---|
| ACE2 | TXNDC16 | -0,627 | 8,06617E-22 |
| ACE2 | FRMPD3 | -0,627 | 8,02589E-22 |
| ACE2 | SRSF4 | -0,6272 | 7,72609E-22 |
| ACE2 | TAF1D | -0,6272 | 7,72566E-22 |
| ACE2 | RP11-680G24.4 | -0,6272 | 7,757E-22 |
| ACE2 | GSTCD | -0,6273 | 7,54545E-22 |
| ACE2 | LA16c-380F5.3 | -0,6273 | 7,55832E-22 |
| ACE2 | EDIL3 | -0,6274 | 7,41125E-22 |
| ACE2 | FTSJ3 | -0,6274 | 7,34941E-22 |
| ACE2 | LINC00890 | -0,6274 | 7,44291E-22 |
| ACE2 | KCNMB1 | -0,6275 | 7,2522E-22 |
| ACE2 | AC079630.4 | -0,6275 | 7,31517E-22 |
| ACE2 | STX1B | -0,6275 | 7,20498E-22 |
| ACE2 | HERPUD2 | -0,6276 | 7,12555E-22 |
| ACE2 | RCBTB1 | -0,6276 | 7,10829E-22 |
| ACE2 | TOB1-AS1 | -0,6277 | 6,93731E-22 |
| ACE2 | CTD-2666L21.1 | -0,6277 | 7,03132E-22 |
| ACE2 | SIK3 | -0,6278 | 6,8352E-22 |
| ACE2 | RP11-514D23.3 | -0,6278 | 6,89156E-22 |
| ACE2 | AF001548.5 | -0,6279 | 6,76676E-22 |
| ACE2 | ZC3H6 | -0,628 | 6,55953E-22 |
| ACE2 | MROH2B | -0,628 | 6,59778E-22 |
| ACE2 | TMEM100 | -0,628 | 6,64306E-22 |
| ACE2 | TXLNG | -0,628 | 6,56309E-22 |
| ACE2 | COL9A1 | -0,6281 | 6,51616E-22 |
| ACE2 | STRAP | -0,6281 | 6,45407E-22 |
| ACE2 | CCDC154 | -0,6281 | 6,52354E-22 |
| ACE2 | SLC6A1 | -0,6282 | 6,32439E-22 |
| ACE2 | RIOK2 | -0,6282 | 6,30008E-22 |
| ACE2 | RP11-863K10.4 | -0,6282 | 6,39823E-22 |
| ACE2 | DNAJC13 | -0,6283 | 6,23301E-22 |
| ACE2 | CRAMP1 | -0,6283 | 6,22637E-22 |
| ACE2 | PHF11 | -0,6284 | 6,05238E-22 |
| ACE2 | GPAT2P2 | -0,6286 | 5,8475E-22 |
| ACE2 | LINC01881 | -0,6286 | 5,85454E-22 |
| ACE2 | IL1RAP | -0,6286 | 5,83589E-22 |
| ACE2 | CCM2 | -0,6286 | 5,83103E-22 |
| ACE2 | NUDT9P1 | -0,6286 | 5,84245E-22 |
| ACE2 | APOPT1 | -0,6286 | 5,84863E-22 |
| ACE2 | CTD-3064M3.7 | -0,6287 | 5,76977E-22 |
| ACE2 | CACUL1 | -0,6287 | 5,81381E-22 |
| ACE2 | CDK2 | -0,6287 | 5,80619E-22 |
| ACE2 | ZNF684 | -0,6288 | 5,62166E-22 |
| ACE2 | RP11-211G3.2 | -0,6288 | 5,70242E-22 |
| ACE2 | DGKB | -0,6288 | 5,67206E-22 |
| ACE2 | RP4-584D14.7 | -0,6288 | 5,60551E-22 |
| ACE2 | CTIF | -0,6288 | 5,60719E-22 |
| ACE2 | AC083899.3 | -0,6289 | 5,51058E-22 |
| ACE2 | C4A | -0,6289 | 5,55417E-22 |
| ACE2 | AC092667.2 | -0,629 | 5,38904E-22 |
| ACE2 | CA5BP1 | -0,629 | 5,4049E-22 |
| ACE2 | ZFP92 | -0,629 | 5,3969E-22 |
| ACE2 | LINC02157 | -0,6291 | 5,38372E-22 |
| ACE2 | FLNC | -0,6292 | 5,24301E-22 |

| | | | |
|---|---|---|---|
| ACE2 | TUBB3 | -0,6292 | 5,26301E-22 |
| ACE2 | CENPB | -0,6292 | 5,24306E-22 |
| ACE2 | EIF4G3 | -0,6294 | 4,98418E-22 |
| ACE2 | ATPAF1 | -0,6295 | 4,91584E-22 |
| ACE2 | PSMC3 | -0,6295 | 4,97058E-22 |
| ACE2 | HAPLN3 | -0,6295 | 4,96969E-22 |
| ACE2 | AC005740.6 | -0,6296 | 4,79347E-22 |
| ACE2 | MED7 | -0,6296 | 4,79288E-22 |
| ACE2 | RPL4P5 | -0,6296 | 4,87311E-22 |
| ACE2 | RALGAPA1 | -0,6296 | 4,7954E-22 |
| ACE2 | DDX25 | -0,6297 | 4,72598E-22 |
| ACE2 | CBFB | -0,6297 | 4,7782E-22 |
| ACE2 | RABL2B | -0,6297 | 4,76128E-22 |
| ACE2 | S100A3 | -0,6298 | 4,66534E-22 |
| ACE2 | PDGFRA | -0,6298 | 4,6343E-22 |
| ACE2 | DMTF1 | -0,6298 | 4,6313E-22 |
| ACE2 | RP4-800G7.2 | -0,6298 | 4,64761E-22 |
| ACE2 | RP11-573G6.6 | -0,6298 | 4,64204E-22 |
| ACE2 | ABHD17A | -0,6298 | 4,64398E-22 |
| ACE2 | MED29 | -0,6298 | 4,63557E-22 |
| ACE2 | ERCC2 | -0,6298 | 4,65986E-22 |
| ACE2 | TRIM46 | -0,63 | 4,44719E-22 |
| ACE2 | GTPBP4 | -0,63 | 4,44594E-22 |
| ACE2 | METTL14 | -0,6302 | 4,29536E-22 |
| ACE2 | NTN1 | -0,6302 | 4,29067E-22 |
| ACE2 | TRMT1 | -0,6302 | 4,28123E-22 |
| ACE2 | CPXM2 | -0,6303 | 4,20281E-22 |
| ACE2 | RP5-1139B12.2 | -0,6304 | 4,10661E-22 |
| ACE2 | RHPN1-AS1 | -0,6305 | 4,08806E-22 |
| ACE2 | TIPRL | -0,6307 | 3,89162E-22 |
| ACE2 | PA2G4 | -0,6307 | 3,94022E-22 |
| ACE2 | PDE5A | -0,6308 | 3,82747E-22 |
| ACE2 | ZDHHC4 | -0,6308 | 3,79836E-22 |
| ACE2 | CYB5D1 | -0,6308 | 3,84444E-22 |
| ACE2 | LARP7 | -0,6309 | 3,74854E-22 |
| ACE2 | LY6G5C | -0,6309 | 3,75129E-22 |
| ACE2 | ACSM5 | -0,6309 | 3,74826E-22 |
| ACE2 | CASP8AP2 | -0,631 | 3,70611E-22 |
| ACE2 | NUAK1 | -0,631 | 3,69263E-22 |
| ACE2 | ZNF540 | -0,631 | 3,71831E-22 |
| ACE2 | DNTTIP2 | -0,6312 | 3,57537E-22 |
| ACE2 | OXCT1-AS1 | -0,6312 | 3,51079E-22 |
| ACE2 | STIM1 | -0,6312 | 3,5516E-22 |
| ACE2 | RP11-134G8.7 | -0,6313 | 3,48837E-22 |
| ACE2 | RP11-228B15.4 | -0,6313 | 3,47071E-22 |
| ACE2 | ZNF461 | -0,6313 | 3,466E-22 |
| ACE2 | CTB-109A12.1 | -0,6314 | 3,42816E-22 |
| ACE2 | RNF10 | -0,6314 | 3,41218E-22 |
| ACE2 | FLOT2 | -0,6314 | 3,41057E-22 |
| ACE2 | ZSCAN22 | -0,6314 | 3,38322E-22 |
| ACE2 | RP11-572M11.4 | -0,6315 | 3,37126E-22 |
| ACE2 | ASF1A | -0,6315 | 3,37158E-22 |
| ACE2 | METTL2B | -0,6315 | 3,33618E-22 |
| ACE2 | NRBP2 | -0,6315 | 3,3448E-22 |

| | | | |
|---|---|---|---|
| ACE2 | PAPOLA | -0,6315 | 3,35345E-22 |
| ACE2 | AP001347.6 | -0,6315 | 3,34082E-22 |
| ACE2 | FLG-AS1 | -0,6316 | 3,29269E-22 |
| ACE2 | SORBS2 | -0,6316 | 3,285E-22 |
| ACE2 | RBPMS2 | -0,6316 | 3,25816E-22 |
| ACE2 | ZCCHC5 | -0,6317 | 3,22444E-22 |
| ACE2 | MKNK1 | -0,6318 | 3,16556E-22 |
| ACE2 | ANKS1A | -0,6318 | 3,16442E-22 |
| ACE2 | FAM160B2 | -0,6318 | 3,12292E-22 |
| ACE2 | FMO3 | -0,632 | 2,99896E-22 |
| ACE2 | GRM6 | -0,632 | 3,01629E-22 |
| ACE2 | MED31 | -0,632 | 3,04952E-22 |
| ACE2 | IL15RA | -0,6321 | 2,9823E-22 |
| ACE2 | ALG9 | -0,6321 | 2,9506E-22 |
| ACE2 | NAA16 | -0,6321 | 2,9678E-22 |
| ACE2 | RP11-463O9.9 | -0,6321 | 2,96316E-22 |
| ACE2 | EIF1AX | -0,6321 | 2,99107E-22 |
| ACE2 | RP11-378A13.1 | -0,6322 | 2,93028E-22 |
| ACE2 | ERICH6-AS1 | -0,6322 | 2,89163E-22 |
| ACE2 | PPIL1 | -0,6322 | 2,92151E-22 |
| ACE2 | KRTCAP2 | -0,6323 | 2,846E-22 |
| ACE2 | CTD-2530N21.4 | -0,6323 | 2,85197E-22 |
| ACE2 | NUP160 | -0,6324 | 2,7792E-22 |
| ACE2 | ADAM20P1 | -0,6325 | 2,76452E-22 |
| ACE2 | ZNF185 | -0,6325 | 2,75245E-22 |
| ACE2 | WRNIP1 | -0,6326 | 2,69967E-22 |
| ACE2 | GPATCH4 | -0,6327 | 2,62218E-22 |
| ACE2 | LEFTY2 | -0,6327 | 2,64455E-22 |
| ACE2 | CCDC82 | -0,6327 | 2,62759E-22 |
| ACE2 | VAMP4 | -0,6328 | 2,59349E-22 |
| ACE2 | ZNF385B | -0,6329 | 2,55964E-22 |
| ACE2 | RPS18 | -0,6329 | 2,53617E-22 |
| ACE2 | RP11-305E6.4 | -0,6329 | 2,51263E-22 |
| ACE2 | TEX30 | -0,6329 | 2,5224E-22 |
| ACE2 | RBBP6 | -0,6329 | 2,54305E-22 |
| ACE2 | PAF1 | -0,6329 | 2,51593E-22 |
| ACE2 | KIDINS220 | -0,633 | 2,48615E-22 |
| ACE2 | ZNF616 | -0,633 | 2,49524E-22 |
| ACE2 | ANP32E | -0,6331 | 2,43246E-22 |
| ACE2 | HTR2A | -0,6331 | 2,44687E-22 |
| ACE2 | RTN4RL1 | -0,6331 | 2,42677E-22 |
| ACE2 | ZNF281 | -0,6332 | 2,40958E-22 |
| ACE2 | ADAMTS9 | -0,6332 | 2,41438E-22 |
| ACE2 | MKL2 | -0,6332 | 2,40479E-22 |
| ACE2 | UBXN11 | -0,6333 | 2,34961E-22 |
| ACE2 | ZNF300P1 | -0,6333 | 2,34536E-22 |
| ACE2 | EXOSC3 | -0,6333 | 2,3576E-22 |
| ACE2 | PGM5-AS1 | -0,6333 | 2,32282E-22 |
| ACE2 | GAS5 | -0,6334 | 2,30909E-22 |
| ACE2 | LRRTM4 | -0,6334 | 2,29832E-22 |
| ACE2 | DLX1 | -0,6334 | 2,31318E-22 |
| ACE2 | CD248 | -0,6334 | 2,29208E-22 |
| ACE2 | RP11-455O6.2 | -0,6335 | 2,235E-22 |
| ACE2 | MORF4L2 | -0,6335 | 2,24553E-22 |

| | | | |
|---|---|---|---|
| ACE2 | THAP9 | -0,6336 | 2,22777E-22 |
| ACE2 | TTC5 | -0,6336 | 2,2055E-22 |
| ACE2 | XAB2 | -0,6336 | 2,20512E-22 |
| ACE2 | RBBP4 | -0,6337 | 2,17037E-22 |
| ACE2 | FAM189B | -0,6337 | 2,17043E-22 |
| ACE2 | MEIS1-AS2 | -0,6337 | 2,16832E-22 |
| ACE2 | ZNF35 | -0,6337 | 2,16123E-22 |
| ACE2 | CDH10 | -0,6337 | 2,14631E-22 |
| ACE2 | AGER | -0,6337 | 2,17095E-22 |
| ACE2 | ING3 | -0,6337 | 2,15224E-22 |
| ACE2 | DGKI | -0,6337 | 2,18023E-22 |
| ACE2 | NKD1 | -0,6337 | 2,17653E-22 |
| ACE2 | RPL13A | -0,6337 | 2,16998E-22 |
| ACE2 | USP25 | -0,6337 | 2,18039E-22 |
| ACE2 | FBF1 | -0,6338 | 2,12151E-22 |
| ACE2 | ZNF337-AS1 | -0,6338 | 2,1066E-22 |
| ACE2 | RP3-449O17.1 | -0,6338 | 2,13566E-22 |
| ACE2 | ZMAT1 | -0,6338 | 2,11728E-22 |
| ACE2 | FBLN2 | -0,6339 | 2,10251E-22 |
| ACE2 | RNF144B | -0,6339 | 2,10299E-22 |
| ACE2 | ATXN7L2 | -0,634 | 2,03216E-22 |
| ACE2 | RP11-383I23.2 | -0,634 | 2,02543E-22 |
| ACE2 | PIK3R4 | -0,634 | 2,04824E-22 |
| ACE2 | SLITRK5 | -0,634 | 2,03522E-22 |
| ACE2 | SHISA9 | -0,634 | 2,02834E-22 |
| ACE2 | NAE1 | -0,634 | 2,03795E-22 |
| ACE2 | AC016582.2 | -0,634 | 2,04654E-22 |
| ACE2 | TSPYL2 | -0,634 | 2,02461E-22 |
| ACE2 | DCDC1 | -0,6342 | 1,96632E-22 |
| ACE2 | MED17 | -0,6342 | 1,97377E-22 |
| ACE2 | ETFRF1 | -0,6342 | 1,98214E-22 |
| ACE2 | DHH | -0,6342 | 1,95133E-22 |
| ACE2 | RP11-395N3.2 | -0,6343 | 1,91868E-22 |
| ACE2 | RGL3 | -0,6343 | 1,90824E-22 |
| ACE2 | NR1H2 | -0,6343 | 1,91148E-22 |
| ACE2 | TFPI2 | -0,6344 | 1,88208E-22 |
| ACE2 | NME3 | -0,6344 | 1,89754E-22 |
| ACE2 | WWOX | -0,6344 | 1,87834E-22 |
| ACE2 | L3MBTL1 | -0,6344 | 1,90363E-22 |
| ACE2 | RP11-167N5.5 | -0,6345 | 1,84629E-22 |
| ACE2 | PRMT5 | -0,6345 | 1,85003E-22 |
| ACE2 | DDX17 | -0,6345 | 1,85584E-22 |
| ACE2 | ATP6V1C1 | -0,6346 | 1,81248E-22 |
| ACE2 | FZD6 | -0,6346 | 1,82992E-22 |
| ACE2 | AIF1L | -0,6347 | 1,76331E-22 |
| ACE2 | SLC15A4 | -0,6347 | 1,77086E-22 |
| ACE2 | RP1-18D14.7 | -0,6348 | 1,7333E-22 |
| ACE2 | LINC01266 | -0,6348 | 1,74319E-22 |
| ACE2 | BZW2 | -0,6348 | 1,7288E-22 |
| ACE2 | METTL3 | -0,6348 | 1,74936E-22 |
| ACE2 | KCNN1 | -0,6348 | 1,7288E-22 |
| ACE2 | RPP40 | -0,6349 | 1,70401E-22 |
| ACE2 | PARP6 | -0,6349 | 1,6971E-22 |
| ACE2 | RP1-93H18.1 | -0,635 | 1,66966E-22 |

| | | | |
|---|---|---|---|
| ACE2 | CNOT7 | -0,635 | 1,68298E-22 |
| ACE2 | AARD | -0,635 | 1,66811E-22 |
| ACE2 | BUD13P1 | -0,635 | 1,67941E-22 |
| ACE2 | FUNDC1 | -0,635 | 1,66472E-22 |
| ACE2 | CTD-2591A1.1 | -0,6351 | 1,63331E-22 |
| ACE2 | PXDNL | -0,6351 | 1,6281E-22 |
| ACE2 | FLRT1 | -0,6351 | 1,64978E-22 |
| ACE2 | FAM57B | -0,6351 | 1,6506E-22 |
| ACE2 | RGN | -0,6351 | 1,6421E-22 |
| ACE2 | EDA2R | -0,6351 | 1,62654E-22 |
| ACE2 | AKNAD1 | -0,6353 | 1,57376E-22 |
| ACE2 | PXDN | -0,6353 | 1,59147E-22 |
| ACE2 | RYBP | -0,6353 | 1,58616E-22 |
| ACE2 | PHIP | -0,6353 | 1,58992E-22 |
| ACE2 | RBBP9 | -0,6353 | 1,57118E-22 |
| ACE2 | TUBB2B | -0,6355 | 1,50263E-22 |
| ACE2 | NARF | -0,6355 | 1,52131E-22 |
| ACE2 | NRSN2-AS1 | -0,6355 | 1,52185E-22 |
| ACE2 | RPL7P9 | -0,6356 | 1,49808E-22 |
| ACE2 | FAM120B | -0,6356 | 1,50138E-22 |
| ACE2 | TADA3 | -0,6357 | 1,46198E-22 |
| ACE2 | ZNF718 | -0,6357 | 1,45163E-22 |
| ACE2 | CCDC92 | -0,6357 | 1,4543E-22 |
| ACE2 | GPC5 | -0,6357 | 1,44504E-22 |
| ACE2 | TNFAIP2 | -0,6357 | 1,45488E-22 |
| ACE2 | SLC9A3R2 | -0,6357 | 1,46875E-22 |
| ACE2 | GDPD5 | -0,6358 | 1,4237E-22 |
| ACE2 | FCHSD2 | -0,6359 | 1,40384E-22 |
| ACE2 | MAML2 | -0,6359 | 1,40734E-22 |
| ACE2 | CTD-2619J13.17 | -0,6359 | 1,40554E-22 |
| ACE2 | N6AMT1 | -0,6359 | 1,41171E-22 |
| ACE2 | NKAIN3 | -0,636 | 1,38017E-22 |
| ACE2 | ZNF728 | -0,636 | 1,38622E-22 |
| ACE2 | PCMTD2 | -0,636 | 1,3812E-22 |
| ACE2 | GSTM2 | -0,6361 | 1,33488E-22 |
| ACE2 | PMF1 | -0,6361 | 1,35628E-22 |
| ACE2 | IFI16 | -0,6361 | 1,34082E-22 |
| ACE2 | BATF3 | -0,6361 | 1,33879E-22 |
| ACE2 | THBS4 | -0,6361 | 1,35462E-22 |
| ACE2 | RP11-225H22.4 | -0,6361 | 1,35501E-22 |
| ACE2 | ZNF268 | -0,6361 | 1,35969E-22 |
| ACE2 | SIN3A | -0,6361 | 1,35058E-22 |
| ACE2 | IQGAP1 | -0,6361 | 1,35734E-22 |
| ACE2 | MICAL3 | -0,6361 | 1,33806E-22 |
| ACE2 | SF3B2 | -0,6362 | 1,31593E-22 |
| ACE2 | HELQ | -0,6363 | 1,30289E-22 |
| ACE2 | PFDN1 | -0,6363 | 1,28639E-22 |
| ACE2 | SYT9 | -0,6363 | 1,28961E-22 |
| ACE2 | KANSL2 | -0,6363 | 1,28434E-22 |
| ACE2 | INTS6 | -0,6363 | 1,28225E-22 |
| ACE2 | TNK2 | -0,6365 | 1,2379E-22 |
| ACE2 | ESYT2 | -0,6365 | 1,24211E-22 |
| ACE2 | RP11-867G23.1 | -0,6365 | 1,24182E-22 |
| ACE2 | RP11-693M3.1 | -0,6365 | 1,24465E-22 |

| | | | |
|---|---|---|---|
| ACE2 | FBXO7 | -0,6365 | 1,25455E-22 |
| ACE2 | RP11-1109F11.5 | -0,6367 | 1,19374E-22 |
| ACE2 | SETD5 | -0,6368 | 1,15932E-22 |
| ACE2 | AMN1 | -0,6368 | 1,16507E-22 |
| ACE2 | TPRG1 | -0,6369 | 1,14259E-22 |
| ACE2 | RIC8A | -0,6369 | 1,14364E-22 |
| ACE2 | SMU1 | -0,637 | 1,11562E-22 |
| ACE2 | ALKBH3 | -0,637 | 1,12373E-22 |
| ACE2 | AC007228.11 | -0,637 | 1,12417E-22 |
| ACE2 | LINC00982 | -0,6371 | 1,10394E-22 |
| ACE2 | STUM | -0,6371 | 1,10827E-22 |
| ACE2 | CCDC9 | -0,6371 | 1,11282E-22 |
| ACE2 | ZBTB8A | -0,6373 | 1,05855E-22 |
| ACE2 | RP5-1139B12.3 | -0,6373 | 1,06644E-22 |
| ACE2 | ZSCAN12 | -0,6373 | 1,06761E-22 |
| ACE2 | ELFN1 | -0,6373 | 1,06638E-22 |
| ACE2 | RP11-6O2.2 | -0,6373 | 1,05624E-22 |
| ACE2 | LINC01354 | -0,6374 | 1,04269E-22 |
| ACE2 | BYSL | -0,6374 | 1,04519E-22 |
| ACE2 | OTUD6B-AS1 | -0,6374 | 1,04787E-22 |
| ACE2 | STAT5A | -0,6374 | 1,03551E-22 |
| ACE2 | CELF4 | -0,6374 | 1,03589E-22 |
| ACE2 | ZNF486 | -0,6374 | 1,03633E-22 |
| ACE2 | SMARCAL1 | -0,6375 | 1,01791E-22 |
| ACE2 | COQ3 | -0,6375 | 1,02693E-22 |
| ACE2 | UNK | -0,6376 | 1,00049E-22 |
| ACE2 | UCHL1 | -0,6377 | 9,86419E-23 |
| ACE2 | RP5-1057I20.5 | -0,6377 | 9,84994E-23 |
| ACE2 | ZNHIT3 | -0,6377 | 9,8571E-23 |
| ACE2 | SRRM1 | -0,6378 | 9,55177E-23 |
| ACE2 | PTN | -0,6378 | 9,63356E-23 |
| ACE2 | WISP1 | -0,6378 | 9,51446E-23 |
| ACE2 | GOLGA8VP | -0,6378 | 9,53043E-23 |
| ACE2 | STAC2 | -0,6378 | 9,50334E-23 |
| ACE2 | RPS27 | -0,6379 | 9,41231E-23 |
| ACE2 | RP11-346C20.4 | -0,6379 | 9,30664E-23 |
| ACE2 | AC106801.1 | -0,638 | 9,20342E-23 |
| ACE2 | GLT1D1 | -0,638 | 9,16671E-23 |
| ACE2 | SKINT1L | -0,6381 | 9,08326E-23 |
| ACE2 | CDC73 | -0,6381 | 8,94288E-23 |
| ACE2 | SIM2 | -0,6381 | 9,09444E-23 |
| ACE2 | FBXW7 | -0,6382 | 8,90016E-23 |
| ACE2 | LINC00840 | -0,6382 | 8,83176E-23 |
| ACE2 | METTL10 | -0,6382 | 8,85758E-23 |
| ACE2 | ZNF202 | -0,6382 | 8,79314E-23 |
| ACE2 | RAD51B | -0,6382 | 8,83165E-23 |
| ACE2 | EMC4 | -0,6382 | 8,78711E-23 |
| ACE2 | HSPG2 | -0,6383 | 8,70706E-23 |
| ACE2 | TCAIM | -0,6383 | 8,75803E-23 |
| ACE2 | C8orf58 | -0,6383 | 8,71034E-23 |
| ACE2 | LIN37 | -0,6383 | 8,73033E-23 |
| ACE2 | ELOF1 | -0,6384 | 8,42061E-23 |
| ACE2 | SYT3 | -0,6384 | 8,48373E-23 |
| ACE2 | AGBL5-IT1 | -0,6385 | 8,35448E-23 |

| | | | |
|---|---|---|---|
| ACE2 | IL13RA2 | -0,6385 | 8,26049E-23 |
| ACE2 | LAGE3 | -0,6385 | 8,37397E-23 |
| ACE2 | PNMAL1 | -0,6386 | 8,16602E-23 |
| ACE2 | RAB7B | -0,6387 | 8,07417E-23 |
| ACE2 | HOXD4 | -0,6387 | 8,0553E-23 |
| ACE2 | PRRX2 | -0,6387 | 7,92801E-23 |
| ACE2 | VN1R108P | -0,6387 | 7,97556E-23 |
| ACE2 | RP11-421L21.3 | -0,6388 | 7,89297E-23 |
| ACE2 | TGFBI | -0,6388 | 7,88715E-23 |
| ACE2 | HERC1 | -0,6388 | 7,83116E-23 |
| ACE2 | FBXL5 | -0,6389 | 7,65616E-23 |
| ACE2 | POLR1D | -0,6389 | 7,75731E-23 |
| ACE2 | RFX3 | -0,639 | 7,56245E-23 |
| ACE2 | RP11-6O2.3 | -0,639 | 7,58719E-23 |
| ACE2 | C10orf142 | -0,6391 | 7,33449E-23 |
| ACE2 | DSE | -0,6393 | 7,09798E-23 |
| ACE2 | C9orf16 | -0,6393 | 7,14472E-23 |
| ACE2 | NOS1 | -0,6393 | 7,06868E-23 |
| ACE2 | SRSF10 | -0,6394 | 6,96772E-23 |
| ACE2 | ATP5E | -0,6395 | 6,8258E-23 |
| ACE2 | CTSK | -0,6396 | 6,72921E-23 |
| ACE2 | BRSK2 | -0,6396 | 6,65242E-23 |
| ACE2 | DENR | -0,6396 | 6,68456E-23 |
| ACE2 | ZNF702P | -0,6396 | 6,66514E-23 |
| ACE2 | ZNF451 | -0,6397 | 6,54935E-23 |
| ACE2 | VPS50 | -0,6397 | 6,51229E-23 |
| ACE2 | PIK3C3 | -0,6397 | 6,54571E-23 |
| ACE2 | RFX2 | -0,6397 | 6,53075E-23 |
| ACE2 | WBSCR17 | -0,6398 | 6,40732E-23 |
| ACE2 | ST8SIA6 | -0,6398 | 6,45699E-23 |
| ACE2 | SSR4P1 | -0,6398 | 6,34941E-23 |
| ACE2 | RP11-436K8.1 | -0,6399 | 6,23762E-23 |
| ACE2 | LAT | -0,6399 | 6,24943E-23 |
| ACE2 | NPEPL1 | -0,6399 | 6,29206E-23 |
| ACE2 | EVA1C | -0,6399 | 6,24411E-23 |
| ACE2 | FLJ37453 | -0,64 | 6,11647E-23 |
| ACE2 | DNALI1 | -0,6401 | 5,97554E-23 |
| ACE2 | MARK1 | -0,6401 | 6,06456E-23 |
| ACE2 | BCLAF1 | -0,6401 | 5,98371E-23 |
| ACE2 | LINC01703 | -0,6402 | 5,88143E-23 |
| ACE2 | PLN | -0,6402 | 5,94696E-23 |
| ACE2 | AC003090.1 | -0,6402 | 5,87986E-23 |
| ACE2 | EYA1 | -0,6402 | 5,87589E-23 |
| ACE2 | SLITRK2 | -0,6402 | 5,90356E-23 |
| ACE2 | LYRM2 | -0,6403 | 5,77714E-23 |
| ACE2 | PCDHGB1 | -0,6404 | 5,70475E-23 |
| ACE2 | SRR | -0,6404 | 5,64214E-23 |
| ACE2 | PRKACA | -0,6404 | 5,66614E-23 |
| ACE2 | BCOR | -0,6404 | 5,65611E-23 |
| ACE2 | RPRD2 | -0,6405 | 5,52708E-23 |
| ACE2 | IP6K1 | -0,6405 | 5,59244E-23 |
| ACE2 | KCNIP1 | -0,6405 | 5,58954E-23 |
| ACE2 | ULK1 | -0,6405 | 5,55958E-23 |
| ACE2 | CSPG4 | -0,6406 | 5,41213E-23 |

| | | | |
|---|---|---|---|
| ACE2 | NSUN4 | -0,6407 | 5,29443E-23 |
| ACE2 | LINC00240 | -0,6407 | 5,33823E-23 |
| ACE2 | RP11-308D16.2 | -0,6407 | 5,33792E-23 |
| ACE2 | THAP5 | -0,6408 | 5,2543E-23 |
| ACE2 | MRPL50 | -0,6408 | 5,24512E-23 |
| ACE2 | BIVM | -0,6408 | 5,2807E-23 |
| ACE2 | MRPS23 | -0,6408 | 5,28456E-23 |
| ACE2 | TDRD6 | -0,6409 | 5,17501E-23 |
| ACE2 | RBPMS-AS1 | -0,6409 | 5,16955E-23 |
| ACE2 | COL13A1 | -0,6409 | 5,10901E-23 |
| ACE2 | CBARP | -0,6409 | 5,10677E-23 |
| ACE2 | MAP3K8 | -0,641 | 5,07521E-23 |
| ACE2 | SYT1 | -0,641 | 5,0573E-23 |
| ACE2 | CCNY | -0,6411 | 4,87855E-23 |
| ACE2 | ABHD10 | -0,6412 | 4,8644E-23 |
| ACE2 | JADE1 | -0,6412 | 4,8635E-23 |
| ACE2 | C6orf89 | -0,6412 | 4,85984E-23 |
| ACE2 | PNRC2 | -0,6413 | 4,70745E-23 |
| ACE2 | CLSTN2 | -0,6413 | 4,70833E-23 |
| ACE2 | IK | -0,6413 | 4,76455E-23 |
| ACE2 | BUD31 | -0,6413 | 4,711E-23 |
| ACE2 | ZNF746 | -0,6413 | 4,72443E-23 |
| ACE2 | YPEL2 | -0,6413 | 4,69085E-23 |
| ACE2 | ZNF836 | -0,6413 | 4,77235E-23 |
| ACE2 | OR7E94P | -0,6415 | 4,52527E-23 |
| ACE2 | CNTNAP3 | -0,6415 | 4,56547E-23 |
| ACE2 | RP11-175K6.2 | -0,6416 | 4,41448E-23 |
| ACE2 | C7orf31 | -0,6416 | 4,46252E-23 |
| ACE2 | BMS1 | -0,6416 | 4,44707E-23 |
| ACE2 | KDR | -0,6417 | 4,35645E-23 |
| ACE2 | TMEM165 | -0,6417 | 4,40147E-23 |
| ACE2 | PDE8B | -0,6417 | 4,37839E-23 |
| ACE2 | AC004448.5 | -0,6418 | 4,25847E-23 |
| ACE2 | DDX42 | -0,6418 | 4,29587E-23 |
| ACE2 | ZNF780B | -0,6419 | 4,19103E-23 |
| ACE2 | UBP1 | -0,642 | 4,10404E-23 |
| ACE2 | TRAM1L1 | -0,642 | 4,08059E-23 |
| ACE2 | SYNPO2 | -0,642 | 4,10327E-23 |
| ACE2 | NCOR2 | -0,642 | 4,07425E-23 |
| ACE2 | VTN | -0,642 | 4,13611E-23 |
| ACE2 | FAM19A5 | -0,642 | 4,13299E-23 |
| ACE2 | SKIV2L2 | -0,6421 | 4,05051E-23 |
| ACE2 | TOPORS-AS1 | -0,6421 | 4,04214E-23 |
| ACE2 | CASC1 | -0,6421 | 4,03564E-23 |
| ACE2 | CXorf67 | -0,6421 | 3,9931E-23 |
| ACE2 | BAP1 | -0,6422 | 3,93213E-23 |
| ACE2 | LPCAT1 | -0,6422 | 3,93139E-23 |
| ACE2 | TRAF1 | -0,6422 | 3,95044E-23 |
| ACE2 | CTD-3157E16.2 | -0,6422 | 3,97573E-23 |
| ACE2 | AC139100.3 | -0,6422 | 3,9407E-23 |
| ACE2 | RP11-752L20.3 | -0,6423 | 3,85641E-23 |
| ACE2 | RP11-398K22.12 | -0,6423 | 3,83937E-23 |
| ACE2 | SEC61A2 | -0,6423 | 3,86797E-23 |
| ACE2 | MYCBP2 | -0,6423 | 3,87327E-23 |

| | | | |
|---|---|---|---|
| ACE2 | SNX29 | -0,6423 | 3,88586E-23 |
| ACE2 | TNFAIP1 | -0,6423 | 3,87816E-23 |
| ACE2 | BPI | -0,6423 | 3,85853E-23 |
| ACE2 | RP11-1148L6.5 | -0,6423 | 3,85615E-23 |
| ACE2 | CSDE1 | -0,6424 | 3,74438E-23 |
| ACE2 | FBXO11 | -0,6424 | 3,75488E-23 |
| ACE2 | UBA3 | -0,6424 | 3,76541E-23 |
| ACE2 | NIPAL4 | -0,6424 | 3,7847E-23 |
| ACE2 | LINC01119 | -0,6425 | 3,71711E-23 |
| ACE2 | ING4 | -0,6425 | 3,68986E-23 |
| ACE2 | PPTC7 | -0,6425 | 3,74248E-23 |
| ACE2 | NMNAT2 | -0,6426 | 3,62064E-23 |
| ACE2 | RPL23AP1 | -0,6426 | 3,6667E-23 |
| ACE2 | NRBP1 | -0,6427 | 3,55604E-23 |
| ACE2 | ANKRD36C | -0,6427 | 3,56936E-23 |
| ACE2 | CTD-2349P21.9 | -0,6427 | 3,59004E-23 |
| ACE2 | NOL10 | -0,6428 | 3,46852E-23 |
| ACE2 | MAP3K7 | -0,6428 | 3,46425E-23 |
| ACE2 | EGR3 | -0,6428 | 3,49003E-23 |
| ACE2 | PDCD1LG2 | -0,6428 | 3,51235E-23 |
| ACE2 | PLEKHA2 | -0,6429 | 3,38758E-23 |
| ACE2 | TJAP1 | -0,643 | 3,31475E-23 |
| ACE2 | RP11-13P5.2 | -0,643 | 3,37747E-23 |
| ACE2 | ERV3-1 | -0,643 | 3,35766E-23 |
| ACE2 | MCMDC2 | -0,643 | 3,34304E-23 |
| ACE2 | HIGD1B | -0,643 | 3,32966E-23 |
| ACE2 | RTTN | -0,643 | 3,3516E-23 |
| ACE2 | NKAIN4 | -0,643 | 3,3183E-23 |
| ACE2 | CUL5 | -0,6431 | 3,25683E-23 |
| ACE2 | FBXO28 | -0,6432 | 3,18074E-23 |
| ACE2 | ZNF670 | -0,6432 | 3,20709E-23 |
| ACE2 | EFHC1 | -0,6432 | 3,19029E-23 |
| ACE2 | EP400 | -0,6432 | 3,22703E-23 |
| ACE2 | CCNA1 | -0,6432 | 3,22098E-23 |
| ACE2 | RARB | -0,6433 | 3,11836E-23 |
| ACE2 | GNAI1 | -0,6433 | 3,13036E-23 |
| ACE2 | TMEM132A | -0,6433 | 3,12332E-23 |
| ACE2 | C1orf35 | -0,6434 | 3,09515E-23 |
| ACE2 | TTC37 | -0,6434 | 3,11203E-23 |
| ACE2 | FXYD7 | -0,6434 | 3,07004E-23 |
| ACE2 | bP-2189O9.2 | -0,6434 | 3,08722E-23 |
| ACE2 | DRG1 | -0,6434 | 3,0793E-23 |
| ACE2 | KCNQ4 | -0,6435 | 3,01444E-23 |
| ACE2 | MORF4L1P1 | -0,6435 | 3,04251E-23 |
| ACE2 | SYCE2 | -0,6435 | 3,01066E-23 |
| ACE2 | RP11-242D8.1 | -0,6436 | 2,98009E-23 |
| ACE2 | RP11-348F1.2 | -0,6436 | 2,96287E-23 |
| ACE2 | PEX3 | -0,6437 | 2,8727E-23 |
| ACE2 | TYRO3 | -0,6437 | 2,87811E-23 |
| ACE2 | RLF | -0,6438 | 2,81917E-23 |
| ACE2 | SNTG2 | -0,6438 | 2,85653E-23 |
| ACE2 | AC011239.2 | -0,6438 | 2,85076E-23 |
| ACE2 | CDAN1 | -0,6438 | 2,81369E-23 |
| ACE2 | TIPIN | -0,6438 | 2,83602E-23 |

| | | | |
|---|---|---|---|
| ACE2 | KLHL26 | -0,6438 | 2,85614E-23 |
| ACE2 | MBTPS2 | -0,6438 | 2,81333E-23 |
| ACE2 | NDUFAF3 | -0,6439 | 2,80542E-23 |
| ACE2 | RP11-329B9.4 | -0,6439 | 2,78756E-23 |
| ACE2 | OR2H2 | -0,6439 | 2,77172E-23 |
| ACE2 | ZNF727 | -0,6439 | 2,80051E-23 |
| ACE2 | GNL2 | -0,644 | 2,73477E-23 |
| ACE2 | STK25 | -0,644 | 2,75105E-23 |
| ACE2 | RP11-286H14.8 | -0,644 | 2,73283E-23 |
| ACE2 | RRS1 | -0,644 | 2,7282E-23 |
| ACE2 | LINC00202-1 | -0,6441 | 2,65024E-23 |
| ACE2 | RP11-66B24.1 | -0,6441 | 2,65993E-23 |
| ACE2 | LINC01828 | -0,6442 | 2,62641E-23 |
| ACE2 | RP3-425C14.4 | -0,6442 | 2,61902E-23 |
| ACE2 | RPL4P4 | -0,6443 | 2,53886E-23 |
| ACE2 | CCDC50 | -0,6443 | 2,55115E-23 |
| ACE2 | RFESD | -0,6444 | 2,49357E-23 |
| ACE2 | TFAP2A | -0,6444 | 2,522E-23 |
| ACE2 | ZNF181 | -0,6444 | 2,51725E-23 |
| ACE2 | RRP1B | -0,6444 | 2,51451E-23 |
| ACE2 | SRPX | -0,6444 | 2,51615E-23 |
| ACE2 | PRELP | -0,6445 | 2,47213E-23 |
| ACE2 | PLSCR4 | -0,6445 | 2,45017E-23 |
| ACE2 | SPTY2D1-AS1 | -0,6445 | 2,4863E-23 |
| ACE2 | TMEM38A | -0,6446 | 2,43022E-23 |
| ACE2 | PCDHGA8 | -0,6447 | 2,35873E-23 |
| ACE2 | KIAA1549L | -0,6447 | 2,35826E-23 |
| ACE2 | DGKE | -0,6447 | 2,34644E-23 |
| ACE2 | TMSB15A | -0,6447 | 2,36721E-23 |
| ACE2 | TMEM167B | -0,6448 | 2,31705E-23 |
| ACE2 | CTB-55O6.12 | -0,6448 | 2,30026E-23 |
| ACE2 | RNF187 | -0,645 | 2,23309E-23 |
| ACE2 | CFAP70 | -0,645 | 2,22241E-23 |
| ACE2 | RP11-497E19.1 | -0,645 | 2,24209E-23 |
| ACE2 | LPCAT2 | -0,645 | 2,22339E-23 |
| ACE2 | JPH2 | -0,645 | 2,2104E-23 |
| ACE2 | RGL1 | -0,6451 | 2,19234E-23 |
| ACE2 | MRPL32 | -0,6451 | 2,17878E-23 |
| ACE2 | OMG | -0,6451 | 2,18054E-23 |
| ACE2 | IVD | -0,6452 | 2,13624E-23 |
| ACE2 | PITPNC1 | -0,6452 | 2,12508E-23 |
| ACE2 | KDM1A | -0,6453 | 2,10006E-23 |
| ACE2 | C1QTNF7 | -0,6453 | 2,08317E-23 |
| ACE2 | PQLC1 | -0,6453 | 2,099E-23 |
| ACE2 | RP11-1398P2.1 | -0,6454 | 2,05139E-23 |
| ACE2 | MTHFD2L | -0,6454 | 2,06033E-23 |
| ACE2 | RP11-714G18.1 | -0,6454 | 2,03068E-23 |
| ACE2 | GHR | -0,6454 | 2,06448E-23 |
| ACE2 | GPBP1 | -0,6454 | 2,0258E-23 |
| ACE2 | FNDC5 | -0,6455 | 2,00116E-23 |
| ACE2 | ZMYM1 | -0,6455 | 2,01104E-23 |
| ACE2 | PSMG3-AS1 | -0,6455 | 1,99213E-23 |
| ACE2 | LINC00346 | -0,6455 | 2,01904E-23 |
| ACE2 | LINC00632 | -0,6455 | 2,01156E-23 |

| | | | |
|---|---|---|---|
| ACE2 | RP11-307C12.12 | -0,6456 | 1,96689E-23 |
| ACE2 | RPUSD3 | -0,6456 | 1,96208E-23 |
| ACE2 | TXLNB | -0,6456 | 1,95733E-23 |
| ACE2 | ZNF484 | -0,6456 | 1,97434E-23 |
| ACE2 | CCDC178 | -0,6456 | 1,94466E-23 |
| ACE2 | MTHFD1L | -0,6457 | 1,93425E-23 |
| ACE2 | KIAA0586 | -0,6457 | 1,9228E-23 |
| ACE2 | HAUS1 | -0,6457 | 1,90587E-23 |
| ACE2 | ZNF30 | -0,6457 | 1,92098E-23 |
| ACE2 | HNRNPR | -0,6458 | 1,89235E-23 |
| ACE2 | PLEKHH2 | -0,6458 | 1,89163E-23 |
| ACE2 | NRF1 | -0,6458 | 1,88937E-23 |
| ACE2 | LDB3 | -0,6458 | 1,87202E-23 |
| ACE2 | ZRSR2 | -0,6458 | 1,86778E-23 |
| ACE2 | DIRAS3 | -0,6459 | 1,86046E-23 |
| ACE2 | NEK4 | -0,6459 | 1,83969E-23 |
| ACE2 | CTD-2325P2.4 | -0,6459 | 1,84002E-23 |
| ACE2 | PTPN11 | -0,646 | 1,81635E-23 |
| ACE2 | ABCC4 | -0,646 | 1,80228E-23 |
| ACE2 | RSAD1 | -0,646 | 1,79524E-23 |
| ACE2 | EIF3D | -0,646 | 1,79388E-23 |
| ACE2 | FARS2 | -0,6461 | 1,77653E-23 |
| ACE2 | MDFI | -0,6461 | 1,78302E-23 |
| ACE2 | KIAA0430 | -0,6461 | 1,78822E-23 |
| ACE2 | MAST1 | -0,6462 | 1,74808E-23 |
| ACE2 | RELN | -0,6464 | 1,64653E-23 |
| ACE2 | AGK | -0,6464 | 1,65805E-23 |
| ACE2 | KIAA2026 | -0,6464 | 1,65395E-23 |
| ACE2 | MFSD14C | -0,6464 | 1,64945E-23 |
| ACE2 | CHST8 | -0,6464 | 1,65394E-23 |
| ACE2 | PPP1R14A | -0,6464 | 1,66161E-23 |
| ACE2 | POMT2 | -0,6465 | 1,64498E-23 |
| ACE2 | RP1-39G22.7 | -0,6466 | 1,59664E-23 |
| ACE2 | PRSS27 | -0,6466 | 1,59724E-23 |
| ACE2 | JPX | -0,6466 | 1,60078E-23 |
| ACE2 | C4orf47 | -0,6467 | 1,56653E-23 |
| ACE2 | CHST9 | -0,6467 | 1,57304E-23 |
| ACE2 | HTRA3 | -0,6468 | 1,52128E-23 |
| ACE2 | PAMR1 | -0,6469 | 1,51387E-23 |
| ACE2 | MIA | -0,6469 | 1,5087E-23 |
| ACE2 | AC017104.6 | -0,647 | 1,46606E-23 |
| ACE2 | SCHIP1 | -0,647 | 1,48243E-23 |
| ACE2 | ANTXR2 | -0,647 | 1,48027E-23 |
| ACE2 | SASH1 | -0,647 | 1,45956E-23 |
| ACE2 | LINC02044 | -0,6471 | 1,43715E-23 |
| ACE2 | ACAP3 | -0,6472 | 1,40576E-23 |
| ACE2 | RPS14 | -0,6472 | 1,42254E-23 |
| ACE2 | DCT | -0,6472 | 1,4113E-23 |
| ACE2 | KLHDC1 | -0,6472 | 1,42037E-23 |
| ACE2 | RP11-805I24.3 | -0,6472 | 1,41713E-23 |
| ACE2 | TMEM169 | -0,6473 | 1,38046E-23 |
| ACE2 | NKAPL | -0,6473 | 1,3881E-23 |
| ACE2 | SNTB2 | -0,6473 | 1,38646E-23 |
| ACE2 | AC010136.2 | -0,6475 | 1,32237E-23 |

| | | | |
|---|---|---|---|
| ACE2 | SEPT6 | -0,6475 | 1,32252E-23 |
| ACE2 | BORCS5 | -0,6477 | 1,27157E-23 |
| ACE2 | C18orf54 | -0,6477 | 1,26858E-23 |
| ACE2 | OGN | -0,6478 | 1,23355E-23 |
| ACE2 | FBXO44 | -0,6479 | 1,22402E-23 |
| ACE2 | TTC14 | -0,6479 | 1,21579E-23 |
| ACE2 | CUTC | -0,6479 | 1,22917E-23 |
| ACE2 | HSF4 | -0,6479 | 1,22964E-23 |
| ACE2 | ASB3 | -0,648 | 1,20171E-23 |
| ACE2 | MYL3 | -0,648 | 1,2021E-23 |
| ACE2 | SSR3 | -0,648 | 1,19143E-23 |
| ACE2 | DHX29 | -0,648 | 1,19539E-23 |
| ACE2 | CTD-2521M24.13 | -0,648 | 1,18659E-23 |
| ACE2 | ZNF449 | -0,648 | 1,19888E-23 |
| ACE2 | PEX19 | -0,6481 | 1,16198E-23 |
| ACE2 | EBF2 | -0,6481 | 1,16486E-23 |
| ACE2 | GSPT1 | -0,6481 | 1,16926E-23 |
| ACE2 | EFCAB6 | -0,6481 | 1,17673E-23 |
| ACE2 | RP11-23J9.7 | -0,6482 | 1,15115E-23 |
| ACE2 | AGAP6 | -0,6482 | 1,14705E-23 |
| ACE2 | RBM4 | -0,6482 | 1,14866E-23 |
| ACE2 | OLFML2B | -0,6483 | 1,12842E-23 |
| ACE2 | TFP1 | -0,6483 | 1,12003E-23 |
| ACE2 | RP3-428L16.2 | -0,6483 | 1,11114E-23 |
| ACE2 | CTC-360G5.9 | -0,6483 | 1,12598E-23 |
| ACE2 | SUPT7L | -0,6484 | 1,09602E-23 |
| ACE2 | C5orf49 | -0,6484 | 1,09668E-23 |
| ACE2 | RP11-333E1.1 | -0,6484 | 1,09323E-23 |
| ACE2 | LINC00630 | -0,6484 | 1,08694E-23 |
| ACE2 | PBXIP1 | -0,6485 | 1,08579E-23 |
| ACE2 | ZNF578 | -0,6485 | 1,0646E-23 |
| ACE2 | KCND1 | -0,6485 | 1,07978E-23 |
| ACE2 | TFAP2E | -0,6486 | 1,05459E-23 |
| ACE2 | LMBR1 | -0,6486 | 1,05797E-23 |
| ACE2 | CTD-3064M3.3 | -0,6486 | 1,04461E-23 |
| ACE2 | LRRCC1 | -0,6488 | 1,01006E-23 |
| ACE2 | TMEM107 | -0,6488 | 1,00696E-23 |
| ACE2 | CHADL | -0,6488 | 1,00004E-23 |
| ACE2 | KANTR | -0,6488 | 1,00443E-23 |
| ACE2 | TIMM50 | -0,6489 | 9,95622E-24 |
| ACE2 | AC074141.3 | -0,6489 | 9,94435E-24 |
| ACE2 | TSC2 | -0,649 | 9,63974E-24 |
| ACE2 | CTB-108O6.2 | -0,6491 | 9,42537E-24 |
| ACE2 | RFNG | -0,6491 | 9,3885E-24 |
| ACE2 | TAF9 | -0,6492 | 9,38378E-24 |
| ACE2 | CTD-2269F5.1 | -0,6492 | 9,2686E-24 |
| ACE2 | MAPK9 | -0,6492 | 9,34445E-24 |
| ACE2 | CAV1 | -0,6492 | 9,29772E-24 |
| ACE2 | RHEB | -0,6492 | 9,31495E-24 |
| ACE2 | ABCA1 | -0,6492 | 9,20303E-24 |
| ACE2 | LINC01798 | -0,6493 | 9,07861E-24 |
| ACE2 | ANO1 | -0,6493 | 9,19358E-24 |
| ACE2 | DEDD | -0,6494 | 8,88033E-24 |
| ACE2 | SNRPE | -0,6494 | 8,8225E-24 |

| | | | |
|---|---|---|---|
| ACE2 | CFLAR | -0,6495 | 8,77181E-24 |
| ACE2 | SCN4B | -0,6495 | 8,75969E-24 |
| ACE2 | RP11-701H24.7 | -0,6495 | 8,67084E-24 |
| ACE2 | NPTX1 | -0,6495 | 8,76612E-24 |
| ACE2 | SLC6A16 | -0,6495 | 8,70685E-24 |
| ACE2 | FTCD | -0,6495 | 8,74372E-24 |
| ACE2 | SHF | -0,6496 | 8,60876E-24 |
| ACE2 | RP11-686D22.4 | -0,6497 | 8,34861E-24 |
| ACE2 | IGFBP5 | -0,6498 | 8,13989E-24 |
| ACE2 | TRAPPC11 | -0,6498 | 8,24497E-24 |
| ACE2 | RFLNB | -0,6498 | 8,26556E-24 |
| ACE2 | ZFAS1 | -0,6498 | 8,2059E-24 |
| ACE2 | AC007283.4 | -0,6499 | 8,06358E-24 |
| ACE2 | ATP6V1H | -0,6499 | 8,02289E-24 |
| ACE2 | SNUPN | -0,6499 | 8,09992E-24 |
| ACE2 | RP11-12M5.3 | -0,65 | 7,93745E-24 |
| ACE2 | HYAL3 | -0,65 | 7,8942E-24 |
| ACE2 | NKIRAS2 | -0,65 | 7,93073E-24 |
| ACE2 | MORC3 | -0,65 | 7,90103E-24 |
| ACE2 | ALDH4A1 | -0,6501 | 7,69128E-24 |
| ACE2 | SCN2A | -0,6501 | 7,70009E-24 |
| ACE2 | RP5-1136G13.2 | -0,6501 | 7,67796E-24 |
| ACE2 | RAB2B | -0,6501 | 7,66201E-24 |
| ACE2 | AZIN2 | -0,6503 | 7,35795E-24 |
| ACE2 | CH17-189H20.1 | -0,6503 | 7,30572E-24 |
| ACE2 | PPAT | -0,6503 | 7,33439E-24 |
| ACE2 | BLOC1S5 | -0,6503 | 7,44783E-24 |
| ACE2 | UBE2N | -0,6503 | 7,30995E-24 |
| ACE2 | PCDH19 | -0,6503 | 7,37499E-24 |
| ACE2 | ARHGAP10 | -0,6505 | 7,11808E-24 |
| ACE2 | CALY | -0,6505 | 7,07475E-24 |
| ACE2 | FIGNL2 | -0,6505 | 7,01589E-24 |
| ACE2 | MPHOSPH8 | -0,6505 | 7,01836E-24 |
| ACE2 | CTD-2033D15.2 | -0,6505 | 7,01766E-24 |
| ACE2 | FNTB | -0,6506 | 6,96655E-24 |
| ACE2 | ZNF678 | -0,6507 | 6,82901E-24 |
| ACE2 | RP11-290D2.6 | -0,6507 | 6,81756E-24 |
| ACE2 | CCDC106 | -0,6508 | 6,69502E-24 |
| ACE2 | SIAH2 | -0,651 | 6,317E-24 |
| ACE2 | ZNF516 | -0,651 | 6,39972E-24 |
| ACE2 | RPL21P28 | -0,6511 | 6,26271E-24 |
| ACE2 | MRPL36 | -0,6511 | 6,18101E-24 |
| ACE2 | ZNF473 | -0,6511 | 6,27364E-24 |
| ACE2 | ADCY1 | -0,6512 | 6,07905E-24 |
| ACE2 | REPS1 | -0,6513 | 5,98577E-24 |
| ACE2 | PAPPA | -0,6513 | 5,94537E-24 |
| ACE2 | MRPL55 | -0,6514 | 5,81645E-24 |
| ACE2 | EZH1 | -0,6514 | 5,81195E-24 |
| ACE2 | LRRC3 | -0,6514 | 5,86317E-24 |
| ACE2 | NOP14-AS1 | -0,6515 | 5,69321E-24 |
| ACE2 | CINP | -0,6516 | 5,63299E-24 |
| ACE2 | HMGCLL1 | -0,6517 | 5,52739E-24 |
| ACE2 | OGFOD3 | -0,6517 | 5,46499E-24 |
| ACE2 | PEX5 | -0,6518 | 5,42738E-24 |

| | | | |
|---|---|---|---|
| ACE2 | UBR7 | -0,6518 | 5,34701E-24 |
| ACE2 | ICE1 | -0,6519 | 5,32036E-24 |
| ACE2 | CARNS1 | -0,6519 | 5,24264E-24 |
| ACE2 | RP11-346E8.1 | -0,6519 | 5,26134E-24 |
| ACE2 | ATF6 | -0,6521 | 5,04754E-24 |
| ACE2 | LINC01117 | -0,6521 | 5,08576E-24 |
| ACE2 | FGF10-AS1 | -0,6521 | 5,01573E-24 |
| ACE2 | RP11-379F4.8 | -0,6522 | 4,9502E-24 |
| ACE2 | SPATS2 | -0,6522 | 4,91763E-24 |
| ACE2 | CALCOCO1 | -0,6522 | 4,99357E-24 |
| ACE2 | ADAM23 | -0,6523 | 4,85691E-24 |
| ACE2 | ZBTB20 | -0,6523 | 4,79603E-24 |
| ACE2 | KIF6 | -0,6523 | 4,86949E-24 |
| ACE2 | NIPSNAP3B | -0,6523 | 4,81749E-24 |
| ACE2 | ALDH1A3 | -0,6523 | 4,79974E-24 |
| ACE2 | WDR7 | -0,6523 | 4,84831E-24 |
| ACE2 | CTD-2006M22.2 | -0,6524 | 4,74817E-24 |
| ACE2 | NEFM | -0,6524 | 4,69042E-24 |
| ACE2 | GTF3C4 | -0,6524 | 4,74322E-24 |
| ACE2 | CFAP58-AS1 | -0,6524 | 4,71743E-24 |
| ACE2 | PAPSS1 | -0,6525 | 4,66409E-24 |
| ACE2 | HAS2-AS1 | -0,6525 | 4,5975E-24 |
| ACE2 | ITPR2 | -0,6525 | 4,65952E-24 |
| ACE2 | FOXC2 | -0,6525 | 4,63006E-24 |
| ACE2 | LINC00506 | -0,6526 | 4,52101E-24 |
| ACE2 | HCN2 | -0,6526 | 4,55748E-24 |
| ACE2 | CTB-36H16.2 | -0,6527 | 4,41906E-24 |
| ACE2 | MPP3 | -0,6527 | 4,40078E-24 |
| ACE2 | HIPK1 | -0,6528 | 4,31646E-24 |
| ACE2 | POLR2I | -0,6528 | 4,33761E-24 |
| ACE2 | ZNF546 | -0,6528 | 4,36017E-24 |
| ACE2 | PGM2 | -0,6529 | 4,22425E-24 |
| ACE2 | PNISR | -0,6529 | 4,24553E-24 |
| ACE2 | CCT6A | -0,6529 | 4,22464E-24 |
| ACE2 | RCC1L | -0,6529 | 4,24031E-24 |
| ACE2 | LINC00894 | -0,6529 | 4,27425E-24 |
| ACE2 | PDCL3 | -0,653 | 4,21563E-24 |
| ACE2 | ZBED5 | -0,653 | 4,16884E-24 |
| ACE2 | PGM5P2 | -0,6531 | 4,10552E-24 |
| ACE2 | ZNF32-AS1 | -0,6531 | 4,06888E-24 |
| ACE2 | RAP2A | -0,6531 | 4,09355E-24 |
| ACE2 | RPL36 | -0,6531 | 4,0906E-24 |
| ACE2 | PCDHGC5 | -0,6532 | 3,97232E-24 |
| ACE2 | SFRP4 | -0,6532 | 4,04215E-24 |
| ACE2 | SEC14L1 | -0,6532 | 3,99874E-24 |
| ACE2 | R3HCC1 | -0,6533 | 3,9415E-24 |
| ACE2 | PGM5 | -0,6533 | 3,87512E-24 |
| ACE2 | RBFOX2 | -0,6533 | 3,94225E-24 |
| ACE2 | RP11-34A14.3 | -0,6534 | 3,86018E-24 |
| ACE2 | RP11-707G18.1 | -0,6534 | 3,79499E-24 |
| ACE2 | ATG4C | -0,6535 | 3,76478E-24 |
| ACE2 | WBP1 | -0,6535 | 3,79303E-24 |
| ACE2 | VPS11 | -0,6535 | 3,76553E-24 |
| ACE2 | LPAR4 | -0,6535 | 3,75357E-24 |

| | | | |
|---|---|---|---|
| ACE2 | TRIM8 | -0,6536 | 3,666E-24 |
| ACE2 | TFCP2 | -0,6536 | 3,68972E-24 |
| ACE2 | AC092835.2 | -0,6537 | 3,63513E-24 |
| ACE2 | ENY2 | -0,6537 | 3,61609E-24 |
| ACE2 | RP11-296O14.3 | -0,6538 | 3,52063E-24 |
| ACE2 | RP11-666A8.13 | -0,6538 | 3,53555E-24 |
| ACE2 | CABIN1 | -0,6538 | 3,55541E-24 |
| ACE2 | CENPQ | -0,6539 | 3,46223E-24 |
| ACE2 | ZC3H7A | -0,6539 | 3,46859E-24 |
| ACE2 | CCR10 | -0,6539 | 3,46811E-24 |
| ACE2 | SP2-AS1 | -0,6539 | 3,44875E-24 |
| ACE2 | PPM1J | -0,654 | 3,34436E-24 |
| ACE2 | VANGL2 | -0,654 | 3,37829E-24 |
| ACE2 | RPAP1 | -0,654 | 3,39608E-24 |
| ACE2 | EFCAB2 | -0,6541 | 3,28467E-24 |
| ACE2 | NAT10 | -0,6541 | 3,29357E-24 |
| ACE2 | FHL5 | -0,6542 | 3,22477E-24 |
| ACE2 | ABCD2 | -0,6542 | 3,23866E-24 |
| ACE2 | HSPA2 | -0,6542 | 3,20402E-24 |
| ACE2 | YWHAH | -0,6542 | 3,21911E-24 |
| ACE2 | STX6 | -0,6543 | 3,15965E-24 |
| ACE2 | KRT8P12 | -0,6543 | 3,19817E-24 |
| ACE2 | SRFBP1 | -0,6543 | 3,19318E-24 |
| ACE2 | LBR | -0,6544 | 3,07909E-24 |
| ACE2 | WDR82 | -0,6544 | 3,10632E-24 |
| ACE2 | RP11-182J1.14 | -0,6544 | 3,08854E-24 |
| ACE2 | RP11-339B21.13 | -0,6545 | 3,03246E-24 |
| ACE2 | RBM38 | -0,6545 | 3,03482E-24 |
| ACE2 | PGM5P4-AS1 | -0,6546 | 2,99826E-24 |
| ACE2 | ZKSCAN3 | -0,6546 | 2,97511E-24 |
| ACE2 | CSGALNACT1 | -0,6546 | 2,97366E-24 |
| ACE2 | HECW2 | -0,6547 | 2,87684E-24 |
| ACE2 | PALM | -0,6547 | 2,8838E-24 |
| ACE2 | LINC01933 | -0,6548 | 2,87384E-24 |
| ACE2 | ZNF454 | -0,6548 | 2,85028E-24 |
| ACE2 | ZBTB2 | -0,6548 | 2,86897E-24 |
| ACE2 | APIP | -0,6548 | 2,84805E-24 |
| ACE2 | TP53I11 | -0,6548 | 2,82379E-24 |
| ACE2 | UBAP1L | -0,6548 | 2,87008E-24 |
| ACE2 | FOXS1 | -0,6548 | 2,86306E-24 |
| ACE2 | UHRF1BP1 | -0,6549 | 2,76121E-24 |
| ACE2 | PPP1R3C | -0,6549 | 2,80209E-24 |
| ACE2 | RHBDF1 | -0,6549 | 2,78477E-24 |
| ACE2 | EXTL3-AS1 | -0,655 | 2,70688E-24 |
| ACE2 | PLPP3 | -0,6551 | 2,65014E-24 |
| ACE2 | RP4-561L24.3 | -0,6551 | 2,67189E-24 |
| ACE2 | AC079584.3 | -0,6551 | 2,64756E-24 |
| ACE2 | CCDC149 | -0,6551 | 2,66003E-24 |
| ACE2 | MRPL1 | -0,6551 | 2,69384E-24 |
| ACE2 | C1QL1 | -0,6551 | 2,65404E-24 |
| ACE2 | SMIM7 | -0,6551 | 2,65754E-24 |
| ACE2 | PRRT4 | -0,6552 | 2,58828E-24 |
| ACE2 | DPH1 | -0,6552 | 2,58643E-24 |
| ACE2 | RP11-333I13.1 | -0,6553 | 2,53349E-24 |

| | | | |
|---|---|---|---|
| ACE2 | NKAIN2 | -0,6553 | 2,53376E-24 |
| ACE2 | KIAA1429 | -0,6553 | 2,56525E-24 |
| ACE2 | TIMM23B | -0,6553 | 2,54013E-24 |
| ACE2 | LINC01081 | -0,6553 | 2,53682E-24 |
| ACE2 | ADAMTS1 | -0,6553 | 2,58356E-24 |
| ACE2 | RP11-446H18.1 | -0,6554 | 2,48922E-24 |
| ACE2 | RPP21 | -0,6554 | 2,49907E-24 |
| ACE2 | RPL27A | -0,6554 | 2,48162E-24 |
| ACE2 | WDR66 | -0,6554 | 2,52637E-24 |
| ACE2 | VPS54 | -0,6555 | 2,45088E-24 |
| ACE2 | REST | -0,6555 | 2,44741E-24 |
| ACE2 | HGF | -0,6555 | 2,45945E-24 |
| ACE2 | SOGA3 | -0,6556 | 2,41145E-24 |
| ACE2 | PARP11 | -0,6556 | 2,38951E-24 |
| ACE2 | FAM98B | -0,6556 | 2,4219E-24 |
| ACE2 | FOXC1 | -0,6557 | 2,35091E-24 |
| ACE2 | RP4-758J18.7 | -0,6558 | 2,27416E-24 |
| ACE2 | FOXF2 | -0,6558 | 2,28662E-24 |
| ACE2 | CYP21A1P | -0,6558 | 2,27689E-24 |
| ACE2 | AK8 | -0,6558 | 2,31741E-24 |
| ACE2 | TMEM45A | -0,6559 | 2,27197E-24 |
| ACE2 | CTDSP2 | -0,6559 | 2,27244E-24 |
| ACE2 | CACTIN | -0,6559 | 2,22991E-24 |
| ACE2 | EEF2 | -0,6559 | 2,27035E-24 |
| ACE2 | DCLRE1B | -0,656 | 2,18859E-24 |
| ACE2 | NUCB2 | -0,656 | 2,18815E-24 |
| ACE2 | GBAP1 | -0,6561 | 2,15255E-24 |
| ACE2 | TAS2R14 | -0,6561 | 2,16829E-24 |
| ACE2 | UBL7 | -0,6561 | 2,14226E-24 |
| ACE2 | AKAP10 | -0,6561 | 2,13576E-24 |
| ACE2 | TBX19 | -0,6562 | 2,10296E-24 |
| ACE2 | C4B | -0,6562 | 2,10779E-24 |
| ACE2 | NSMCE1 | -0,6562 | 2,11142E-24 |
| ACE2 | SH3BGR | -0,6562 | 2,13076E-24 |
| ACE2 | CH17-437K3.1 | -0,6563 | 2,04467E-24 |
| ACE2 | MEF2D | -0,6563 | 2,0762E-24 |
| ACE2 | RXRG | -0,6563 | 2,07225E-24 |
| ACE2 | C1orf101 | -0,6563 | 2,0599E-24 |
| ACE2 | ST3GAL6-AS1 | -0,6563 | 2,04491E-24 |
| ACE2 | ANKRD33B | -0,6563 | 2,05733E-24 |
| ACE2 | ACTR3B | -0,6563 | 2,04332E-24 |
| ACE2 | MPG | -0,6563 | 2,0518E-24 |
| ACE2 | AP000473.5 | -0,6563 | 2,06309E-24 |
| ACE2 | RP5-1007F24.1 | -0,6564 | 2,01055E-24 |
| ACE2 | EEFSEC | -0,6565 | 1,99032E-24 |
| ACE2 | CTC-463A16.1 | -0,6565 | 1,97312E-24 |
| ACE2 | SEPHS1 | -0,6565 | 1,99675E-24 |
| ACE2 | GS1-590J6.3 | -0,6565 | 1,99238E-24 |
| ACE2 | LINC01797 | -0,6566 | 1,91413E-24 |
| ACE2 | KCNAB1 | -0,6566 | 1,93089E-24 |
| ACE2 | RPL37 | -0,6566 | 1,9286E-24 |
| ACE2 | C1QTNF2 | -0,6566 | 1,93941E-24 |
| ACE2 | MAP1LC3B | -0,6566 | 1,93533E-24 |
| ACE2 | PLD3 | -0,6566 | 1,91534E-24 |

| | | | |
|---|---|---|---|
| ACE2 | UQCRB | -0,6567 | 1,90616E-24 |
| ACE2 | CELF6 | -0,6567 | 1,89852E-24 |
| ACE2 | GINS3 | -0,6567 | 1,88756E-24 |
| ACE2 | IQSEC1 | -0,6568 | 1,8514E-24 |
| ACE2 | RP11-158I9.8 | -0,6568 | 1,84483E-24 |
| ACE2 | LEPR | -0,6569 | 1,80257E-24 |
| ACE2 | AC097662.2 | -0,6569 | 1,80882E-24 |
| ACE2 | DPYSL4 | -0,6569 | 1,82406E-24 |
| ACE2 | SUGT1 | -0,6569 | 1,80554E-24 |
| ACE2 | MBNL1 | -0,657 | 1,76428E-24 |
| ACE2 | COX20 | -0,6571 | 1,74613E-24 |
| ACE2 | PTENP1 | -0,6571 | 1,73266E-24 |
| ACE2 | SECISBP2L | -0,6571 | 1,74901E-24 |
| ACE2 | ZDHHC8 | -0,6571 | 1,73821E-24 |
| ACE2 | AGBL3 | -0,6572 | 1,70699E-24 |
| ACE2 | DTD2 | -0,6572 | 1,70992E-24 |
| ACE2 | FBN1 | -0,6572 | 1,70317E-24 |
| ACE2 | MEGF9 | -0,6573 | 1,65841E-24 |
| ACE2 | VLDLR-AS1 | -0,6574 | 1,61797E-24 |
| ACE2 | BTBD10 | -0,6574 | 1,63642E-24 |
| ACE2 | PPIEL | -0,6575 | 1,60704E-24 |
| ACE2 | RP11-3B7.1 | -0,6575 | 1,5887E-24 |
| ACE2 | NME2 | -0,6575 | 1,59059E-24 |
| ACE2 | IFT57 | -0,6576 | 1,56559E-24 |
| ACE2 | NSMAF | -0,6576 | 1,56353E-24 |
| ACE2 | CCPG1 | -0,6576 | 1,56787E-24 |
| ACE2 | VAMP2 | -0,6576 | 1,56085E-24 |
| ACE2 | SLC7A2 | -0,6577 | 1,53721E-24 |
| ACE2 | IQCD | -0,6577 | 1,51464E-24 |
| ACE2 | RP11-566E18.1 | -0,6578 | 1,4923E-24 |
| ACE2 | HNRNPC | -0,6578 | 1,50726E-24 |
| ACE2 | ANKRD23 | -0,6579 | 1,455E-24 |
| ACE2 | SRD5A3-AS1 | -0,6579 | 1,45568E-24 |
| ACE2 | UBE2V2 | -0,6579 | 1,46834E-24 |
| ACE2 | STK36 | -0,658 | 1,42145E-24 |
| ACE2 | ZCCHC7 | -0,658 | 1,42248E-24 |
| ACE2 | VASH1 | -0,658 | 1,44504E-24 |
| ACE2 | MOAP1 | -0,658 | 1,41994E-24 |
| ACE2 | ICAM1 | -0,658 | 1,43931E-24 |
| ACE2 | RP1-151F17.2 | -0,6581 | 1,39952E-24 |
| ACE2 | NOL8 | -0,6581 | 1,38975E-24 |
| ACE2 | CENPP | -0,6581 | 1,41273E-24 |
| ACE2 | HRC | -0,6581 | 1,39296E-24 |
| ACE2 | TIAM1 | -0,6581 | 1,41198E-24 |
| ACE2 | LAMB2 | -0,6582 | 1,37148E-24 |
| ACE2 | PTPA | -0,6582 | 1,36252E-24 |
| ACE2 | FGF13-AS1 | -0,6582 | 1,36622E-24 |
| ACE2 | PCDH15 | -0,6583 | 1,33343E-24 |
| ACE2 | MTMR6 | -0,6583 | 1,34547E-24 |
| ACE2 | SPTB | -0,6583 | 1,33529E-24 |
| ACE2 | ZNF383 | -0,6583 | 1,32903E-24 |
| ACE2 | SPEF1 | -0,6584 | 1,3225E-24 |
| ACE2 | RP11-395A13.2 | -0,6585 | 1,2949E-24 |
| ACE2 | NOX4 | -0,6585 | 1,29121E-24 |

| | | | |
|---|---|---|---|
| ACE2 | RPL23A | -0,6585 | 1,29696E-24 |
| ACE2 | DLGAP1-AS1 | -0,6585 | 1,27709E-24 |
| ACE2 | ANKRD27 | -0,6585 | 1,29173E-24 |
| ACE2 | PTMA | -0,6586 | 1,26656E-24 |
| ACE2 | KCNN2 | -0,6586 | 1,25262E-24 |
| ACE2 | RP11-479O9.4 | -0,6586 | 1,25204E-24 |
| ACE2 | RP11-956J14.1 | -0,6586 | 1,26908E-24 |
| ACE2 | CFAP20 | -0,6586 | 1,26413E-24 |
| ACE2 | KCNJ12 | -0,6586 | 1,26967E-24 |
| ACE2 | TLE6 | -0,6586 | 1,24468E-24 |
| ACE2 | FBXW11 | -0,6587 | 1,23245E-24 |
| ACE2 | GAL3ST4 | -0,6587 | 1,21669E-24 |
| ACE2 | CADM2 | -0,6588 | 1,21408E-24 |
| ACE2 | LINC00689 | -0,6588 | 1,21388E-24 |
| ACE2 | AK6 | -0,6589 | 1,17149E-24 |
| ACE2 | VCAN | -0,6589 | 1,18037E-24 |
| ACE2 | UBAP1 | -0,6589 | 1,18382E-24 |
| ACE2 | ANGEL1 | -0,6589 | 1,18595E-24 |
| ACE2 | ZNF558 | -0,6589 | 1,1728E-24 |
| ACE2 | CNN3 | -0,659 | 1,14694E-24 |
| ACE2 | MED28 | -0,659 | 1,15996E-24 |
| ACE2 | PNMA2 | -0,659 | 1,13988E-24 |
| ACE2 | ATP6AP2 | -0,659 | 1,15512E-24 |
| ACE2 | RPAP2 | -0,6591 | 1,12388E-24 |
| ACE2 | FAM218A | -0,6591 | 1,13724E-24 |
| ACE2 | CDKAL1 | -0,6591 | 1,12623E-24 |
| ACE2 | GTF3C1 | -0,6591 | 1,12172E-24 |
| ACE2 | CLN5 | -0,6592 | 1,09504E-24 |
| ACE2 | SUPT16H | -0,6592 | 1,10839E-24 |
| ACE2 | MYOCD | -0,6592 | 1,1089E-24 |
| ACE2 | B4GAT1 | -0,6593 | 1,08155E-24 |
| ACE2 | FNDC1 | -0,6594 | 1,05539E-24 |
| ACE2 | GPNMB | -0,6594 | 1,04913E-24 |
| ACE2 | CPEB1-AS1 | -0,6594 | 1,06542E-24 |
| ACE2 | CRABP2 | -0,6595 | 1,04391E-24 |
| ACE2 | RHOA | -0,6595 | 1,03156E-24 |
| ACE2 | SMNDC1 | -0,6595 | 1,03396E-24 |
| ACE2 | CTC-471J1.2 | -0,6595 | 1,03286E-24 |
| ACE2 | CDV3 | -0,6598 | 9,63876E-25 |
| ACE2 | SPP1 | -0,6598 | 9,62975E-25 |
| ACE2 | ZNF517 | -0,6598 | 9,73805E-25 |
| ACE2 | NXNL2 | -0,6598 | 9,58065E-25 |
| ACE2 | THCAT158 | -0,6598 | 9,77635E-25 |
| ACE2 | GPC1 | -0,6599 | 9,44843E-25 |
| ACE2 | CTD-2036P10.5 | -0,6599 | 9,5493E-25 |
| ACE2 | EXOSC10 | -0,66 | 9,35108E-25 |
| ACE2 | SERINC3 | -0,66 | 9,26303E-25 |
| ACE2 | ARHGEF9 | -0,66 | 9,21686E-25 |
| ACE2 | ITGB3BP | -0,6601 | 9,01938E-25 |
| ACE2 | HCFC2 | -0,6601 | 9,15747E-25 |
| ACE2 | HSPB6 | -0,6601 | 9,05432E-25 |
| ACE2 | SLC17A7 | -0,6601 | 9,03924E-25 |
| ACE2 | GRIA3 | -0,6601 | 8,99717E-25 |
| ACE2 | LYSMD2 | -0,6602 | 8,91184E-25 |

| | | | |
|---|---|---|---|
| ACE2 | UBALD1 | -0,6602 | 8,92464E-25 |
| ACE2 | BEX2 | -0,6602 | 8,93681E-25 |
| ACE2 | SERTAD3 | -0,6603 | 8,59547E-25 |
| ACE2 | KPNA4 | -0,6604 | 8,44257E-25 |
| ACE2 | GOPC | -0,6604 | 8,49116E-25 |
| ACE2 | CUL2 | -0,6604 | 8,56374E-25 |
| ACE2 | CHD4 | -0,6605 | 8,29678E-25 |
| ACE2 | RP11-322E11.5 | -0,6605 | 8,30952E-25 |
| ACE2 | CD320 | -0,6605 | 8,32563E-25 |
| ACE2 | RP11-573G6.4 | -0,6606 | 8,10623E-25 |
| ACE2 | NBPF8 | -0,6607 | 7,94383E-25 |
| ACE2 | PHAX | -0,6607 | 8,04207E-25 |
| ACE2 | CAPRIN2 | -0,6607 | 8,01633E-25 |
| ACE2 | FAM50B | -0,6608 | 7,70751E-25 |
| ACE2 | FADS3 | -0,6608 | 7,82164E-25 |
| ACE2 | TRIM47 | -0,6608 | 7,80441E-25 |
| ACE2 | CRELD1 | -0,6609 | 7,56813E-25 |
| ACE2 | ARL8B | -0,661 | 7,4936E-25 |
| ACE2 | ZNF37BP | -0,661 | 7,41783E-25 |
| ACE2 | TUBGCP5 | -0,661 | 7,44316E-25 |
| ACE2 | RBM8A | -0,6611 | 7,28069E-25 |
| ACE2 | RUFY1 | -0,6611 | 7,29775E-25 |
| ACE2 | UG0898H09 | -0,6611 | 7,32267E-25 |
| ACE2 | ERVK3-1 | -0,6611 | 7,34157E-25 |
| ACE2 | RP11-508N22.12 | -0,6612 | 7,16324E-25 |
| ACE2 | PDDC1 | -0,6612 | 7,06849E-25 |
| ACE2 | DNASE2 | -0,6612 | 7,19254E-25 |
| ACE2 | CTD-3220F14.1 | -0,6613 | 6,95393E-25 |
| ACE2 | HDDC2 | -0,6614 | 6,81154E-25 |
| ACE2 | FAM122A | -0,6614 | 6,82754E-25 |
| ACE2 | MARS | -0,6614 | 6,78466E-25 |
| ACE2 | PRR29 | -0,6614 | 6,77239E-25 |
| ACE2 | TNFAIP6 | -0,6615 | 6,67822E-25 |
| ACE2 | RNF168 | -0,6615 | 6,64655E-25 |
| ACE2 | UBE2D1 | -0,6615 | 6,7101E-25 |
| ACE2 | A2M | -0,6615 | 6,752E-25 |
| ACE2 | HIST2H2BE | -0,6616 | 6,48112E-25 |
| ACE2 | DPM1 | -0,6616 | 6,55367E-25 |
| ACE2 | MEGF6 | -0,6617 | 6,37596E-25 |
| ACE2 | ZNF691 | -0,6617 | 6,44171E-25 |
| ACE2 | PCDHGB3 | -0,6617 | 6,36612E-25 |
| ACE2 | EPC1 | -0,6617 | 6,38409E-25 |
| ACE2 | GLG1 | -0,6617 | 6,3393E-25 |
| ACE2 | MALAT1 | -0,6618 | 6,19053E-25 |
| ACE2 | LTV1 | -0,6619 | 6,11089E-25 |
| ACE2 | CC2D2B | -0,6619 | 6,13306E-25 |
| ACE2 | RAD9B | -0,6619 | 6,05738E-25 |
| ACE2 | NEDD8 | -0,6619 | 6,10002E-25 |
| ACE2 | ZNF142 | -0,662 | 5,9694E-25 |
| ACE2 | CCDC127 | -0,662 | 5,9722E-25 |
| ACE2 | LINC00641 | -0,662 | 5,95938E-25 |
| ACE2 | CTC-301O7.4 | -0,662 | 5,96418E-25 |
| ACE2 | CTNNBL1 | -0,662 | 6,04744E-25 |
| ACE2 | KPNA6 | -0,6621 | 5,90006E-25 |

| | | | |
|---|---|---|---|
| ACE2 | TSNAX | -0,6621 | 5,89096E-25 |
| ACE2 | CDO1 | -0,6621 | 5,84956E-25 |
| ACE2 | MDC1 | -0,6621 | 5,86215E-25 |
| ACE2 | HSPH1 | -0,6621 | 5,91871E-25 |
| ACE2 | TBC1D17 | -0,6621 | 5,83543E-25 |
| ACE2 | PANX2 | -0,6621 | 5,89016E-25 |
| ACE2 | NFU1 | -0,6622 | 5,68513E-25 |
| ACE2 | DNAJA1 | -0,6622 | 5,71556E-25 |
| ACE2 | RP11-535M15.2 | -0,6622 | 5,71633E-25 |
| ACE2 | MSANTD2 | -0,6623 | 5,56884E-25 |
| ACE2 | CCDC102B | -0,6623 | 5,64325E-25 |
| ACE2 | USP21 | -0,6624 | 5,50186E-25 |
| ACE2 | RRM2B | -0,6624 | 5,51185E-25 |
| ACE2 | PGM5P3-AS1 | -0,6624 | 5,47458E-25 |
| ACE2 | CAMKK1 | -0,6624 | 5,44336E-25 |
| ACE2 | ZRANB3 | -0,6625 | 5,33821E-25 |
| ACE2 | RP11-162D16.2 | -0,6625 | 5,40441E-25 |
| ACE2 | TLR5 | -0,6626 | 5,29529E-25 |
| ACE2 | PDGFRL | -0,6626 | 5,30206E-25 |
| ACE2 | CLEC2B | -0,6627 | 5,14911E-25 |
| ACE2 | HERC2P9 | -0,6627 | 5,18109E-25 |
| ACE2 | NADK2 | -0,6628 | 4,98104E-25 |
| ACE2 | RP11-478C6.6 | -0,6631 | 4,6933E-25 |
| ACE2 | CTSL | -0,6631 | 4,7158E-25 |
| ACE2 | ALDH1L2 | -0,6631 | 4,68854E-25 |
| ACE2 | RIF1 | -0,6632 | 4,59124E-25 |
| ACE2 | CCDC141 | -0,6632 | 4,60256E-25 |
| ACE2 | MED6 | -0,6632 | 4,59559E-25 |
| ACE2 | ZNF136 | -0,6632 | 4,58691E-25 |
| ACE2 | PCDHAC2 | -0,6633 | 4,54061E-25 |
| ACE2 | DPY19L2P4 | -0,6633 | 4,45479E-25 |
| ACE2 | XPO4 | -0,6633 | 4,45306E-25 |
| ACE2 | CCDC30 | -0,6634 | 4,37324E-25 |
| ACE2 | AC009501.4 | -0,6634 | 4,39899E-25 |
| ACE2 | LIX1 | -0,6634 | 4,44376E-25 |
| ACE2 | DDX39B | -0,6634 | 4,3759E-25 |
| ACE2 | RIMS1 | -0,6634 | 4,36822E-25 |
| ACE2 | CCDC151 | -0,6634 | 4,40821E-25 |
| ACE2 | HEYL | -0,6635 | 4,3362E-25 |
| ACE2 | CYB5RL | -0,6635 | 4,26852E-25 |
| ACE2 | AP3B2 | -0,6635 | 4,34939E-25 |
| ACE2 | ZNF287 | -0,6635 | 4,32538E-25 |
| ACE2 | DDX18 | -0,6636 | 4,24265E-25 |
| ACE2 | RPS10 | -0,6636 | 4,16936E-25 |
| ACE2 | VWCE | -0,6636 | 4,16598E-25 |
| ACE2 | LINC00987 | -0,6636 | 4,22778E-25 |
| ACE2 | ATP2B1-AS1 | -0,6636 | 4,22707E-25 |
| ACE2 | RP11-481J2.4 | -0,6636 | 4,22007E-25 |
| ACE2 | ZFHX3 | -0,6636 | 4,17781E-25 |
| ACE2 | MON2 | -0,6637 | 4,14548E-25 |
| ACE2 | CSMD2 | -0,6638 | 4,059E-25 |
| ACE2 | SIX2 | -0,6638 | 3,98487E-25 |
| ACE2 | TBCK | -0,6638 | 4,0453E-25 |
| ACE2 | POT1-AS1 | -0,6638 | 4,04312E-25 |

| | | | |
|---|---|---|---|
| ACE2 | FEM1A | -0,6638 | 4,05105E-25 |
| ACE2 | LRRFIP1P1 | -0,6639 | 3,94056E-25 |
| ACE2 | HACE1 | -0,6639 | 3,91374E-25 |
| ACE2 | CRYGN | -0,6639 | 3,92716E-25 |
| ACE2 | SPG21 | -0,6639 | 3,93537E-25 |
| ACE2 | MYADM | -0,6639 | 3,97186E-25 |
| ACE2 | MRPL30 | -0,664 | 3,86011E-25 |
| ACE2 | CPED1 | -0,664 | 3,88473E-25 |
| ACE2 | KANSL3 | -0,6641 | 3,75628E-25 |
| ACE2 | ZNF251 | -0,6641 | 3,75983E-25 |
| ACE2 | LINC01694 | -0,6641 | 3,80359E-25 |
| ACE2 | ID3 | -0,6642 | 3,71531E-25 |
| ACE2 | ANKRD13C | -0,6642 | 3,64851E-25 |
| ACE2 | CDCA7L | -0,6642 | 3,66187E-25 |
| ACE2 | ZFAND4 | -0,6642 | 3,70577E-25 |
| ACE2 | IFIT5 | -0,6642 | 3,68776E-25 |
| ACE2 | RP11-348P10.2 | -0,6643 | 3,58677E-25 |
| ACE2 | NICN1 | -0,6643 | 3,58082E-25 |
| ACE2 | RNF217 | -0,6643 | 3,64184E-25 |
| ACE2 | PACRG | -0,6643 | 3,59114E-25 |
| ACE2 | KHDRBS3 | -0,6643 | 3,63445E-25 |
| ACE2 | NAA38 | -0,6643 | 3,60821E-25 |
| ACE2 | C6orf226 | -0,6644 | 3,53874E-25 |
| ACE2 | CRCP | -0,6644 | 3,54427E-25 |
| ACE2 | KLRG1 | -0,6644 | 3,49561E-25 |
| ACE2 | HEMK1 | -0,6645 | 3,46119E-25 |
| ACE2 | NMB | -0,6645 | 3,46424E-25 |
| ACE2 | RGS9 | -0,6645 | 3,43968E-25 |
| ACE2 | C1QTNF1 | -0,6645 | 3,42618E-25 |
| ACE2 | RP5-1054A22.4 | -0,6645 | 3,41389E-25 |
| ACE2 | PTCHD1 | -0,6645 | 3,44434E-25 |
| ACE2 | ANGPTL7 | -0,6646 | 3,38393E-25 |
| ACE2 | FER1L5 | -0,6646 | 3,34515E-25 |
| ACE2 | STIP1 | -0,6646 | 3,38267E-25 |
| ACE2 | CEP128 | -0,6646 | 3,39485E-25 |
| ACE2 | HIRIP3 | -0,6646 | 3,40869E-25 |
| ACE2 | PRMT7 | -0,6646 | 3,36655E-25 |
| ACE2 | PLK2 | -0,6647 | 3,28991E-25 |
| ACE2 | ZNF529 | -0,6647 | 3,33473E-25 |
| ACE2 | C7orf61 | -0,6648 | 3,2233E-25 |
| ACE2 | LINC01503 | -0,6648 | 3,1965E-25 |
| ACE2 | RPL10AP6 | -0,6649 | 3,13526E-25 |
| ACE2 | HAND2-AS1 | -0,6649 | 3,1751E-25 |
| ACE2 | TNXB | -0,6649 | 3,17833E-25 |
| ACE2 | MRPS17 | -0,6649 | 3,12718E-25 |
| ACE2 | INPP5E | -0,665 | 3,11447E-25 |
| ACE2 | RAB6C-AS1 | -0,6651 | 3,0409E-25 |
| ACE2 | TTC41P | -0,6651 | 3,03759E-25 |
| ACE2 | XXbac-BPG283O16.9 | -0,6652 | 2,96339E-25 |
| ACE2 | EIF2A | -0,6653 | 2,88331E-25 |
| ACE2 | RP11-582E3.6 | -0,6653 | 2,88214E-25 |
| ACE2 | IQUB | -0,6654 | 2,80175E-25 |
| ACE2 | USP20 | -0,6654 | 2,84437E-25 |
| ACE2 | NAV3 | -0,6654 | 2,80899E-25 |

| | | | |
|---|---|---|---|
| ACE2 | TRAF3IP1 | -0,6655 | 2,75468E-25 |
| ACE2 | PSMB1 | -0,6655 | 2,77179E-25 |
| ACE2 | MLXIP | -0,6655 | 2,74719E-25 |
| ACE2 | COL3A1 | -0,6656 | 2,67541E-25 |
| ACE2 | RUVBL1 | -0,6656 | 2,72726E-25 |
| ACE2 | DLGAP2 | -0,6656 | 2,69057E-25 |
| ACE2 | UROS | -0,6656 | 2,6902E-25 |
| ACE2 | PRCC | -0,6658 | 2,56706E-25 |
| ACE2 | RABL2A | -0,6658 | 2,59874E-25 |
| ACE2 | ITGA9 | -0,6658 | 2,6108E-25 |
| ACE2 | C2CD3 | -0,6658 | 2,60352E-25 |
| ACE2 | FAM50A | -0,6658 | 2,57197E-25 |
| ACE2 | ARSG | -0,6659 | 2,52629E-25 |
| ACE2 | JMJD6 | -0,6659 | 2,51101E-25 |
| ACE2 | IDS | -0,6659 | 2,54443E-25 |
| ACE2 | KLHL20 | -0,666 | 2,49294E-25 |
| ACE2 | CSPG5 | -0,666 | 2,44597E-25 |
| ACE2 | MCMBP | -0,666 | 2,49787E-25 |
| ACE2 | SENP3 | -0,666 | 2,46741E-25 |
| ACE2 | LRRC37A7P | -0,666 | 2,4651E-25 |
| ACE2 | SHROOM4 | -0,666 | 2,4575E-25 |
| ACE2 | BAZ1B | -0,6661 | 2,4107E-25 |
| ACE2 | NOC3L | -0,6661 | 2,40528E-25 |
| ACE2 | CTD-2012K14.8 | -0,6661 | 2,42042E-25 |
| ACE2 | PSMB4 | -0,6662 | 2,33877E-25 |
| ACE2 | ANKRD36 | -0,6662 | 2,34789E-25 |
| ACE2 | EPHB1 | -0,6662 | 2,34356E-25 |
| ACE2 | C1orf229 | -0,6663 | 2,28622E-25 |
| ACE2 | FBXO25 | -0,6663 | 2,33558E-25 |
| ACE2 | CFAP54 | -0,6663 | 2,32065E-25 |
| ACE2 | ZNF14 | -0,6663 | 2,28981E-25 |
| ACE2 | KLHL30 | -0,6664 | 2,26725E-25 |
| ACE2 | NELFE | -0,6664 | 2,27128E-25 |
| ACE2 | TTC39C | -0,6664 | 2,24568E-25 |
| ACE2 | ANAPC13 | -0,6665 | 2,2269E-25 |
| ACE2 | PPME1 | -0,6665 | 2,20194E-25 |
| ACE2 | TMEM120B | -0,6665 | 2,19984E-25 |
| ACE2 | NSMCE3 | -0,6665 | 2,21265E-25 |
| ACE2 | CYR61 | -0,6666 | 2,16702E-25 |
| ACE2 | RPL30 | -0,6666 | 2,1417E-25 |
| ACE2 | RRAGA | -0,6666 | 2,18173E-25 |
| ACE2 | CDKN2D | -0,6666 | 2,14478E-25 |
| ACE2 | GDNF-AS1 | -0,6667 | 2,11653E-25 |
| ACE2 | CCDC90B | -0,6667 | 2,13177E-25 |
| ACE2 | THAP7 | -0,6667 | 2,09837E-25 |
| ACE2 | GSPT2 | -0,6667 | 2,13367E-25 |
| ACE2 | WDR54 | -0,6668 | 2,08137E-25 |
| ACE2 | RPLP1 | -0,6668 | 2,08251E-25 |
| ACE2 | UBR1 | -0,6669 | 2,0081E-25 |
| ACE2 | ZFR | -0,667 | 1,99045E-25 |
| ACE2 | IFT74 | -0,667 | 1,97701E-25 |
| ACE2 | FBXW8 | -0,667 | 1,97336E-25 |
| ACE2 | RP11-455F5.3 | -0,667 | 1,96494E-25 |
| ACE2 | RP11-441O15.3 | -0,6671 | 1,9135E-25 |

| | | | |
|---|---|---|---|
| ACE2 | SCN8A | -0,6671 | 1,92203E-25 |
| ACE2 | SGCA | -0,6671 | 1,9135E-25 |
| ACE2 | AP3M1 | -0,6672 | 1,88501E-25 |
| ACE2 | ADM5 | -0,6672 | 1,90552E-25 |
| ACE2 | PCNX4 | -0,6673 | 1,85161E-25 |
| ACE2 | ZBTB11-AS1 | -0,6674 | 1,81667E-25 |
| ACE2 | AC025165.8 | -0,6674 | 1,79747E-25 |
| ACE2 | SERP2 | -0,6674 | 1,80364E-25 |
| ACE2 | AGO2 | -0,6675 | 1,76729E-25 |
| ACE2 | NLE1 | -0,6675 | 1,75521E-25 |
| ACE2 | ZNF404 | -0,6675 | 1,77774E-25 |
| ACE2 | HS6ST1 | -0,6676 | 1,723E-25 |
| ACE2 | RP11-267M23.1 | -0,6676 | 1,72786E-25 |
| ACE2 | ATP9B | -0,6677 | 1,68802E-25 |
| ACE2 | SRP72 | -0,6678 | 1,66306E-25 |
| ACE2 | DONSON | -0,6678 | 1,65907E-25 |
| ACE2 | GBP2 | -0,6679 | 1,63275E-25 |
| ACE2 | RFTN1 | -0,6679 | 1,62715E-25 |
| ACE2 | ADAT2 | -0,6679 | 1,62548E-25 |
| ACE2 | MKL1 | -0,6679 | 1,63364E-25 |
| ACE2 | LRRC8A | -0,668 | 1,56274E-25 |
| ACE2 | INPP5K | -0,6681 | 1,53264E-25 |
| ACE2 | CDC25B | -0,6681 | 1,53053E-25 |
| ACE2 | RBM43 | -0,6682 | 1,51879E-25 |
| ACE2 | AASDH | -0,6682 | 1,51233E-25 |
| ACE2 | GTF2H4 | -0,6682 | 1,50073E-25 |
| ACE2 | NDRG2 | -0,6682 | 1,52392E-25 |
| ACE2 | SHC2 | -0,6682 | 1,51775E-25 |
| ACE2 | XXbac-BPGBPG55C: | -0,6683 | 1,48748E-25 |
| ACE2 | BAG5 | -0,6683 | 1,47069E-25 |
| ACE2 | KATNAL2 | -0,6683 | 1,48278E-25 |
| ACE2 | NEGR1 | -0,6684 | 1,44392E-25 |
| ACE2 | CHL1-AS2 | -0,6684 | 1,44761E-25 |
| ACE2 | SUPT4H1 | -0,6684 | 1,44214E-25 |
| ACE2 | ZSWIM7 | -0,6685 | 1,41588E-25 |
| ACE2 | AOC4P | -0,6685 | 1,39806E-25 |
| ACE2 | SAMD4B | -0,6685 | 1,4237E-25 |
| ACE2 | ZNF512B | -0,6685 | 1,42605E-25 |
| ACE2 | NOTCH2 | -0,6686 | 1,39019E-25 |
| ACE2 | DNM3OS | -0,6686 | 1,38933E-25 |
| ACE2 | RPS9 | -0,6686 | 1,3725E-25 |
| ACE2 | PRRG3 | -0,6686 | 1,36655E-25 |
| ACE2 | NBAT1 | -0,6687 | 1,35445E-25 |
| ACE2 | DRP2 | -0,6687 | 1,33704E-25 |
| ACE2 | RGS12 | -0,6688 | 1,32868E-25 |
| ACE2 | SNX16 | -0,6688 | 1,30481E-25 |
| ACE2 | JAKMIP3 | -0,6688 | 1,32183E-25 |
| ACE2 | CTD-2319I12.4 | -0,6688 | 1,33088E-25 |
| ACE2 | LAMC1 | -0,6689 | 1,29791E-25 |
| ACE2 | CACNA2D2 | -0,6689 | 1,28402E-25 |
| ACE2 | FAN1 | -0,6689 | 1,29424E-25 |
| ACE2 | HNRNPA1L2 | -0,669 | 1,26268E-25 |
| ACE2 | CDC16 | -0,669 | 1,2484E-25 |
| ACE2 | DBF4B | -0,669 | 1,26238E-25 |

| | | | |
|---|---|---|---|
| ACE2 | ZNF528 | -0,669 | 1,25314E-25 |
| ACE2 | CBX5 | -0,6691 | 1,22333E-25 |
| ACE2 | EVA1B | -0,6692 | 1,19273E-25 |
| ACE2 | DCTN1 | -0,6692 | 1,20215E-25 |
| ACE2 | ULK4 | -0,6692 | 1,19669E-25 |
| ACE2 | RP3-331H24.7 | -0,6692 | 1,19884E-25 |
| ACE2 | TMEM147 | -0,6692 | 1,21106E-25 |
| ACE2 | AHSA1 | -0,6693 | 1,16965E-25 |
| ACE2 | STAG2 | -0,6693 | 1,16989E-25 |
| ACE2 | AC016747.3 | -0,6694 | 1,16305E-25 |
| ACE2 | HRAS | -0,6694 | 1,15305E-25 |
| ACE2 | CCT2 | -0,6694 | 1,16457E-25 |
| ACE2 | SELENOW | -0,6694 | 1,15122E-25 |
| ACE2 | IL6ST | -0,6695 | 1,1344E-25 |
| ACE2 | RILPL2 | -0,6695 | 1,11643E-25 |
| ACE2 | DUOX1 | -0,6695 | 1,13762E-25 |
| ACE2 | TMC3 | -0,6695 | 1,13495E-25 |
| ACE2 | USP36 | -0,6695 | 1,12674E-25 |
| ACE2 | ATP6V1E1 | -0,6695 | 1,11659E-25 |
| ACE2 | POLDIP3 | -0,6695 | 1,11489E-25 |
| ACE2 | LRRC47 | -0,6696 | 1,1058E-25 |
| ACE2 | KLF3-AS1 | -0,6696 | 1,10438E-25 |
| ACE2 | LINC02104 | -0,6696 | 1,11294E-25 |
| ACE2 | ZDHHC1 | -0,6696 | 1,09541E-25 |
| ACE2 | HSD17B14 | -0,6696 | 1,10587E-25 |
| ACE2 | GNB5 | -0,6697 | 1,06892E-25 |
| ACE2 | DDX59 | -0,6698 | 1,06435E-25 |
| ACE2 | RP11-793H13.11 | -0,6698 | 1,05183E-25 |
| ACE2 | RP11-346C20.3 | -0,6698 | 1,0584E-25 |
| ACE2 | AP000473.8 | -0,6698 | 1,05339E-25 |
| ACE2 | FOXD3-AS1 | -0,6699 | 1,02801E-25 |
| ACE2 | HAT1 | -0,67 | 1,00702E-25 |
| ACE2 | LDB1 | -0,6701 | 9,73424E-26 |
| ACE2 | TMEM159 | -0,6701 | 9,84065E-26 |
| ACE2 | PRR3 | -0,6702 | 9,52669E-26 |
| ACE2 | PMP2 | -0,6702 | 9,52013E-26 |
| ACE2 | RP3-333H23.9 | -0,6702 | 9,53038E-26 |
| ACE2 | BTG2 | -0,6703 | 9,32355E-26 |
| ACE2 | CEP57L1 | -0,6703 | 9,44716E-26 |
| ACE2 | USP46 | -0,6704 | 9,14744E-26 |
| ACE2 | CYP2E1 | -0,6704 | 9,18876E-26 |
| ACE2 | DDIT3 | -0,6704 | 9,18048E-26 |
| ACE2 | ZNF567 | -0,6704 | 9,21111E-26 |
| ACE2 | CUTA | -0,6705 | 8,90387E-26 |
| ACE2 | CKAP5 | -0,6705 | 9,01767E-26 |
| ACE2 | TAL1 | -0,6707 | 8,51999E-26 |
| ACE2 | FAM129A | -0,6707 | 8,63074E-26 |
| ACE2 | GNA12 | -0,6707 | 8,52948E-26 |
| ACE2 | PRR14 | -0,6707 | 8,66134E-26 |
| ACE2 | PACS1 | -0,6708 | 8,41943E-26 |
| ACE2 | TRIM16L | -0,6708 | 8,44108E-26 |
| ACE2 | RPL18A | -0,6708 | 8,42586E-26 |
| ACE2 | RPL11 | -0,6709 | 8,20749E-26 |
| ACE2 | GLRB | -0,6709 | 8,11901E-26 |

| | | | |
|---|---|---|---|
| ACE2 | SLIT1 | -0,6709 | 8,22233E-26 |
| ACE2 | PTMS | -0,6709 | 8,29002E-26 |
| ACE2 | RP3-508I15.9 | -0,6709 | 8,27793E-26 |
| ACE2 | TM2D1 | -0,671 | 8,06976E-26 |
| ACE2 | RP11-864N7.4 | -0,671 | 7,93466E-26 |
| ACE2 | AGL | -0,6711 | 7,92882E-26 |
| ACE2 | C11orf74 | -0,6711 | 7,84222E-26 |
| ACE2 | PCP4L1 | -0,6712 | 7,68938E-26 |
| ACE2 | COMMD1 | -0,6712 | 7,73119E-26 |
| ACE2 | EXOC4 | -0,6712 | 7,67439E-26 |
| ACE2 | RNF139 | -0,6712 | 7,72368E-26 |
| ACE2 | TRDMT1 | -0,6712 | 7,68669E-26 |
| ACE2 | RIC3 | -0,6712 | 7,60849E-26 |
| ACE2 | POLR2G | -0,6712 | 7,61061E-26 |
| ACE2 | EARS2 | -0,6712 | 7,57733E-26 |
| ACE2 | ZNF304 | -0,6712 | 7,65202E-26 |
| ACE2 | C2orf42 | -0,6713 | 7,53423E-26 |
| ACE2 | KAT6B | -0,6713 | 7,55625E-26 |
| ACE2 | IL17D | -0,6713 | 7,45703E-26 |
| ACE2 | KPTN | -0,6713 | 7,51433E-26 |
| ACE2 | PTCD2 | -0,6714 | 7,27805E-26 |
| ACE2 | RPS4X | -0,6714 | 7,32737E-26 |
| ACE2 | BHMT2 | -0,6715 | 7,19691E-26 |
| ACE2 | KMT2A | -0,6715 | 7,11021E-26 |
| ACE2 | PLCXD3 | -0,6716 | 6,92736E-26 |
| ACE2 | OGFRL1 | -0,6716 | 7,0016E-26 |
| ACE2 | RAP2C-AS1 | -0,6716 | 6,99818E-26 |
| ACE2 | ABHD1 | -0,6718 | 6,65566E-26 |
| ACE2 | SLC9B1 | -0,6718 | 6,64547E-26 |
| ACE2 | INVS | -0,6718 | 6,69896E-26 |
| ACE2 | COL4A5 | -0,6718 | 6,65566E-26 |
| ACE2 | FAM110D | -0,6719 | 6,49289E-26 |
| ACE2 | UQCC2 | -0,6719 | 6,54928E-26 |
| ACE2 | SCRG1 | -0,672 | 6,32784E-26 |
| ACE2 | NUBPL | -0,672 | 6,39043E-26 |
| ACE2 | ZNF500 | -0,6721 | 6,27989E-26 |
| ACE2 | TEX41 | -0,6722 | 6,04298E-26 |
| ACE2 | USP35 | -0,6722 | 6,11121E-26 |
| ACE2 | CCDC121 | -0,6723 | 5,97198E-26 |
| ACE2 | MASP1 | -0,6723 | 5,91777E-26 |
| ACE2 | SYPL1 | -0,6723 | 5,96447E-26 |
| ACE2 | TCF15 | -0,6723 | 5,90221E-26 |
| ACE2 | MPHOSPH10 | -0,6724 | 5,85545E-26 |
| ACE2 | TTI1 | -0,6724 | 5,84521E-26 |
| ACE2 | TRABD2B | -0,6725 | 5,67255E-26 |
| ACE2 | METTL6 | -0,6725 | 5,71973E-26 |
| ACE2 | WBSCR22 | -0,6725 | 5,63339E-26 |
| ACE2 | PTH1R | -0,6726 | 5,51566E-26 |
| ACE2 | FBXO21 | -0,6726 | 5,52878E-26 |
| ACE2 | TMEM184B | -0,6726 | 5,61983E-26 |
| ACE2 | EPHA6 | -0,6727 | 5,39615E-26 |
| ACE2 | PFDN6 | -0,6727 | 5,46496E-26 |
| ACE2 | CDC5L | -0,6727 | 5,43749E-26 |
| ACE2 | GS1-124K5.12 | -0,6727 | 5,42952E-26 |

| | | | |
|---|---|---|---|
| ACE2 | TRMT12 | -0,6727 | 5,40413E-26 |
| ACE2 | BTBD19 | -0,6728 | 5,28404E-26 |
| ACE2 | GPR137B | -0,6729 | 5,25932E-26 |
| ACE2 | TBPL1 | -0,6729 | 5,18546E-26 |
| ACE2 | RPS15A | -0,6729 | 5,18182E-26 |
| ACE2 | ZIM2-AS1 | -0,6729 | 5,15275E-26 |
| ACE2 | CPNE9 | -0,673 | 5,09761E-26 |
| ACE2 | DNAAF1 | -0,673 | 5,07303E-26 |
| ACE2 | ASB1 | -0,6731 | 4,96748E-26 |
| ACE2 | TRIM61 | -0,6731 | 4,93969E-26 |
| ACE2 | ZNF337 | -0,6731 | 4,99928E-26 |
| ACE2 | PTTG2 | -0,6732 | 4,80731E-26 |
| ACE2 | MEA1 | -0,6733 | 4,77728E-26 |
| ACE2 | SCAI | -0,6733 | 4,70512E-26 |
| ACE2 | CREBL2 | -0,6733 | 4,71967E-26 |
| ACE2 | PALLD | -0,6734 | 4,62915E-26 |
| ACE2 | PLPP1 | -0,6734 | 4,59485E-26 |
| ACE2 | PI16 | -0,6734 | 4,63566E-26 |
| ACE2 | CTD-2561B21.11 | -0,6734 | 4,58969E-26 |
| ACE2 | ZFYVE16 | -0,6735 | 4,50669E-26 |
| ACE2 | RNF145 | -0,6735 | 4,58197E-26 |
| ACE2 | FBXO9 | -0,6735 | 4,57168E-26 |
| ACE2 | RP11-195F19.9 | -0,6735 | 4,49343E-26 |
| ACE2 | TRNT1 | -0,6736 | 4,46514E-26 |
| ACE2 | C3orf14 | -0,6736 | 4,39575E-26 |
| ACE2 | UBAP2 | -0,6736 | 4,42375E-26 |
| ACE2 | VCL | -0,6736 | 4,38589E-26 |
| ACE2 | C19orf68 | -0,6736 | 4,38743E-26 |
| ACE2 | PCMT1 | -0,6737 | 4,34115E-26 |
| ACE2 | RP11-834C11.8 | -0,6737 | 4,35013E-26 |
| ACE2 | ADAM11 | -0,6737 | 4,2842E-26 |
| ACE2 | PCIF1 | -0,6737 | 4,35397E-26 |
| ACE2 | PTPMT1 | -0,6738 | 4,20012E-26 |
| ACE2 | NAT9 | -0,6738 | 4,21624E-26 |
| ACE2 | EPS15 | -0,6739 | 4,13189E-26 |
| ACE2 | GIN1 | -0,6739 | 4,1804E-26 |
| ACE2 | EIF3J-AS1 | -0,6739 | 4,14587E-26 |
| ACE2 | ECM1 | -0,674 | 4,07163E-26 |
| ACE2 | ZKSCAN5 | -0,674 | 4,01194E-26 |
| ACE2 | ZNF491 | -0,674 | 4,03511E-26 |
| ACE2 | HS6ST2 | -0,674 | 4,02266E-26 |
| ACE2 | TRNAU1AP | -0,6741 | 3,94783E-26 |
| ACE2 | COMMD3 | -0,6741 | 3,9634E-26 |
| ACE2 | ST5 | -0,6741 | 3,93009E-26 |
| ACE2 | PLPPR4 | -0,6742 | 3,86075E-26 |
| ACE2 | ZNF2 | -0,6742 | 3,81429E-26 |
| ACE2 | CYP21A2 | -0,6742 | 3,83879E-26 |
| ACE2 | HOXA4 | -0,6742 | 3,88036E-26 |
| ACE2 | IGSF9B | -0,6742 | 3,8245E-26 |
| ACE2 | FOXO1 | -0,6742 | 3,89896E-26 |
| ACE2 | UTP6 | -0,6742 | 3,81614E-26 |
| ACE2 | DNAJC6 | -0,6743 | 3,80034E-26 |
| ACE2 | MAN1A2 | -0,6743 | 3,77314E-26 |
| ACE2 | NOS3 | -0,6743 | 3,7509E-26 |

| | | | |
|---|---|---|---|
| ACE2 | QTRT1 | -0,6743 | 3,77156E-26 |
| ACE2 | AC093323.3 | -0,6744 | 3,69999E-26 |
| ACE2 | GAB1 | -0,6744 | 3,69923E-26 |
| ACE2 | LINC00461 | -0,6744 | 3,6552E-26 |
| ACE2 | SIRT1 | -0,6744 | 3,68413E-26 |
| ACE2 | C7orf26 | -0,6745 | 3,61853E-26 |
| ACE2 | MIR4435-2HG | -0,6746 | 3,49843E-26 |
| ACE2 | SLC25A27 | -0,6746 | 3,48169E-26 |
| ACE2 | ARMCX4 | -0,6746 | 3,48194E-26 |
| ACE2 | RP11-506B6.6 | -0,6747 | 3,40846E-26 |
| ACE2 | NTM | -0,6747 | 3,42592E-26 |
| ACE2 | RP11-958N24.1 | -0,6747 | 3,45503E-26 |
| ACE2 | FOXK2 | -0,6747 | 3,43316E-26 |
| ACE2 | STARD8 | -0,6747 | 3,40904E-26 |
| ACE2 | KHDRBS1 | -0,6748 | 3,36486E-26 |
| ACE2 | RP11-664I21.6 | -0,6748 | 3,36863E-26 |
| ACE2 | CLTCL1 | -0,6749 | 3,26913E-26 |
| ACE2 | RP11-173C1.1 | -0,675 | 3,22864E-26 |
| ACE2 | MFGE8 | -0,675 | 3,23789E-26 |
| ACE2 | THAP11 | -0,675 | 3,21591E-26 |
| ACE2 | PIK3R3 | -0,6751 | 3,15596E-26 |
| ACE2 | AHCTF1 | -0,6751 | 3,16799E-26 |
| ACE2 | SENP1 | -0,6751 | 3,13308E-26 |
| ACE2 | CLEC14A | -0,6751 | 3,13772E-26 |
| ACE2 | FZD2 | -0,6751 | 3,15082E-26 |
| ACE2 | PCDH10 | -0,6752 | 3,05851E-26 |
| ACE2 | TP63 | -0,6753 | 2,98779E-26 |
| ACE2 | PCDH7 | -0,6753 | 2,968E-26 |
| ACE2 | BBS2 | -0,6753 | 2,9861E-26 |
| ACE2 | B9D1 | -0,6753 | 2,97852E-26 |
| ACE2 | PGR-AS1 | -0,6754 | 2,93503E-26 |
| ACE2 | RP11-128A17.2 | -0,6754 | 2,91879E-26 |
| ACE2 | CDH13 | -0,6754 | 2,90653E-26 |
| ACE2 | SLC25A14 | -0,6755 | 2,84107E-26 |
| ACE2 | RXRB | -0,6756 | 2,8067E-26 |
| ACE2 | AK9 | -0,6756 | 2,77058E-26 |
| ACE2 | TRAPPC3 | -0,6757 | 2,71888E-26 |
| ACE2 | ZNF543 | -0,6757 | 2,72045E-26 |
| ACE2 | TRIM6 | -0,6758 | 2,65704E-26 |
| ACE2 | PDS5B | -0,6758 | 2,65817E-26 |
| ACE2 | COL11A1 | -0,6759 | 2,61096E-26 |
| ACE2 | NUDT17 | -0,6759 | 2,57731E-26 |
| ACE2 | NEPRO | -0,6759 | 2,60467E-26 |
| ACE2 | CPE | -0,6759 | 2,59491E-26 |
| ACE2 | GRIK1 | -0,6759 | 2,62878E-26 |
| ACE2 | KCNK2 | -0,676 | 2,56456E-26 |
| ACE2 | ZNF250 | -0,676 | 2,54093E-26 |
| ACE2 | NXPH3 | -0,676 | 2,54807E-26 |
| ACE2 | PTEN | -0,6761 | 2,49038E-26 |
| ACE2 | MRPL43 | -0,6761 | 2,4677E-26 |
| ACE2 | BCL7A | -0,6761 | 2,48932E-26 |
| ACE2 | RNF112 | -0,6761 | 2,46457E-26 |
| ACE2 | BAG3 | -0,6762 | 2,44993E-26 |
| ACE2 | NYNRIN | -0,6762 | 2,43607E-26 |

| | | | |
|---|---|---|---|
| ACE2 | RP11-66B24.2 | -0,6762 | 2,42125E-26 |
| ACE2 | UROD | -0,6763 | 2,3647E-26 |
| ACE2 | SCYL1 | -0,6763 | 2,38239E-26 |
| ACE2 | ZNF263 | -0,6763 | 2,35181E-26 |
| ACE2 | STARD3NL | -0,6764 | 2,30381E-26 |
| ACE2 | WNT5B | -0,6764 | 2,33252E-26 |
| ACE2 | TTC1 | -0,6765 | 2,2595E-26 |
| ACE2 | BMPR1B | -0,6766 | 2,21764E-26 |
| ACE2 | C16orf71 | -0,6767 | 2,17137E-26 |
| ACE2 | PSMD6 | -0,6769 | 2,0627E-26 |
| ACE2 | MAK | -0,6769 | 2,05787E-26 |
| ACE2 | RPS19BP1 | -0,6769 | 2,08126E-26 |
| ACE2 | NAGK | -0,677 | 1,99885E-26 |
| ACE2 | PIK3C2A | -0,677 | 2,04063E-26 |
| ACE2 | MAPK8IP2 | -0,677 | 2,01857E-26 |
| ACE2 | PLXDC1 | -0,6771 | 1,97536E-26 |
| ACE2 | TRA2A | -0,6772 | 1,94771E-26 |
| ACE2 | RP11-46H11.3 | -0,6773 | 1,8645E-26 |
| ACE2 | RPSAP18 | -0,6774 | 1,83798E-26 |
| ACE2 | FGF12 | -0,6774 | 1,83982E-26 |
| ACE2 | CHST15 | -0,6774 | 1,86012E-26 |
| ACE2 | USE1 | -0,6774 | 1,85257E-26 |
| ACE2 | VN1R1 | -0,6774 | 1,84321E-26 |
| ACE2 | WDR45 | -0,6775 | 1,78834E-26 |
| ACE2 | RANP4 | -0,6775 | 1,80596E-26 |
| ACE2 | CCT7 | -0,6776 | 1,7702E-26 |
| ACE2 | WDFY3-AS2 | -0,6776 | 1,74301E-26 |
| ACE2 | ZNF442 | -0,6776 | 1,75737E-26 |
| ACE2 | AP001505.10 | -0,6776 | 1,77122E-26 |
| ACE2 | CH17-472G23.2 | -0,6777 | 1,72729E-26 |
| ACE2 | LYST | -0,6777 | 1,72139E-26 |
| ACE2 | DNMT3A | -0,6777 | 1,71659E-26 |
| ACE2 | ATP6V1E2 | -0,6777 | 1,70415E-26 |
| ACE2 | GFPT2 | -0,6777 | 1,71232E-26 |
| ACE2 | PMS2P1 | -0,6777 | 1,70781E-26 |
| ACE2 | KAT8 | -0,6777 | 1,73234E-26 |
| ACE2 | CIC | -0,6777 | 1,72593E-26 |
| ACE2 | CTA-445C9.14 | -0,6777 | 1,72175E-26 |
| ACE2 | ZNF630 | -0,6777 | 1,73346E-26 |
| ACE2 | PPP3CA | -0,6779 | 1,62895E-26 |
| ACE2 | PUS7 | -0,6779 | 1,63842E-26 |
| ACE2 | NTNG1 | -0,6781 | 1,57675E-26 |
| ACE2 | DTNB | -0,6781 | 1,56095E-26 |
| ACE2 | ALK | -0,6781 | 1,56173E-26 |
| ACE2 | FARSB | -0,6781 | 1,56049E-26 |
| ACE2 | ZNF425 | -0,6781 | 1,57265E-26 |
| ACE2 | NRGN | -0,6782 | 1,51857E-26 |
| ACE2 | RPL13P5 | -0,6782 | 1,53505E-26 |
| ACE2 | UTP20 | -0,6782 | 1,54297E-26 |
| ACE2 | SNAI1 | -0,6782 | 1,53111E-26 |
| ACE2 | FOXRED2 | -0,6782 | 1,53824E-26 |
| ACE2 | SHARPIN | -0,6783 | 1,49582E-26 |
| ACE2 | SMCO3 | -0,6783 | 1,47686E-26 |
| ACE2 | FAM117A | -0,6783 | 1,48825E-26 |

| | | | |
|---|---|---|---|
| ACE2 | MRPS31P4 | -0,6784 | 1,44079E-26 |
| ACE2 | SGF29 | -0,6784 | 1,44884E-26 |
| ACE2 | AC007405.6 | -0,6785 | 1,42836E-26 |
| ACE2 | ZRANB2 | -0,6786 | 1,39711E-26 |
| ACE2 | PPRC1 | -0,6786 | 1,37876E-26 |
| ACE2 | AP000648.5 | -0,6786 | 1,40072E-26 |
| ACE2 | ACAT1 | -0,6786 | 1,38519E-26 |
| ACE2 | SOX15 | -0,6786 | 1,39052E-26 |
| ACE2 | LZTFL1 | -0,6787 | 1,35979E-26 |
| ACE2 | CTD-2124B8.2 | -0,6787 | 1,35745E-26 |
| ACE2 | MZT1 | -0,6787 | 1,34823E-26 |
| ACE2 | RIN3 | -0,6787 | 1,35415E-26 |
| ACE2 | CDK9 | -0,6789 | 1,30959E-26 |
| ACE2 | STRADA | -0,6789 | 1,28797E-26 |
| ACE2 | BCORL1 | -0,6789 | 1,2878E-26 |
| ACE2 | MRPL57 | -0,679 | 1,25831E-26 |
| ACE2 | XYLT2 | -0,679 | 1,27465E-26 |
| ACE2 | ETNK2 | -0,6791 | 1,2228E-26 |
| ACE2 | RPL34 | -0,6791 | 1,22913E-26 |
| ACE2 | RP11-798M19.6 | -0,6791 | 1,22148E-26 |
| ACE2 | ZNF767P | -0,6791 | 1,24528E-26 |
| ACE2 | COL22A1 | -0,6791 | 1,23134E-26 |
| ACE2 | CALCB | -0,6791 | 1,22945E-26 |
| ACE2 | SRSF1 | -0,6791 | 1,22481E-26 |
| ACE2 | ERICH2 | -0,6792 | 1,19536E-26 |
| ACE2 | LIFR | -0,6792 | 1,20226E-26 |
| ACE2 | MAP1B | -0,6792 | 1,21467E-26 |
| ACE2 | JMY | -0,6792 | 1,21335E-26 |
| ACE2 | RP11-551L14.4 | -0,6792 | 1,19592E-26 |
| ACE2 | RP4-613B23.1 | -0,6793 | 1,16604E-26 |
| ACE2 | NLGN1 | -0,6793 | 1,19024E-26 |
| ACE2 | SBDSP1 | -0,6793 | 1,16536E-26 |
| ACE2 | RP11-817I4.2 | -0,6793 | 1,16688E-26 |
| ACE2 | NOG | -0,6793 | 1,17985E-26 |
| ACE2 | RNF24 | -0,6794 | 1,14676E-26 |
| ACE2 | RYR2 | -0,6795 | 1,12692E-26 |
| ACE2 | VAX2 | -0,6795 | 1,11579E-26 |
| ACE2 | KYAT1 | -0,6795 | 1,12805E-26 |
| ACE2 | FAM149B1 | -0,6795 | 1,11965E-26 |
| ACE2 | GPR83 | -0,6795 | 1,13578E-26 |
| ACE2 | CWC15 | -0,6795 | 1,11483E-26 |
| ACE2 | HNRNPA1 | -0,6795 | 1,11499E-26 |
| ACE2 | CEBPZOS | -0,6796 | 1,08852E-26 |
| ACE2 | TM4SF1 | -0,6796 | 1,10193E-26 |
| ACE2 | CMYA5 | -0,6796 | 1,09396E-26 |
| ACE2 | PSD | -0,6796 | 1,11155E-26 |
| ACE2 | ACADSB | -0,6796 | 1,09135E-26 |
| ACE2 | YPEL4 | -0,6796 | 1,09979E-26 |
| ACE2 | ZNF843 | -0,6796 | 1,09088E-26 |
| ACE2 | FRS3 | -0,6797 | 1,07748E-26 |
| ACE2 | LATS1 | -0,6797 | 1,06139E-26 |
| ACE2 | RP11-767C1.2 | -0,6798 | 1,04475E-26 |
| ACE2 | RP11-448G15.3 | -0,6798 | 1,04278E-26 |
| ACE2 | SH3TC2 | -0,6798 | 1,0362E-26 |

| | | | |
|---|---|---|---|
| ACE2 | SMARCA2 | -0,6798 | 1,0566E-26 |
| ACE2 | RP11-452L6.6 | -0,6798 | 1,05077E-26 |
| ACE2 | ADCY2 | -0,6799 | 1,03055E-26 |
| ACE2 | FBXO42 | -0,68 | 9,89382E-27 |
| ACE2 | RP11-168A11.4 | -0,68 | 1,01098E-26 |
| ACE2 | FAM172A | -0,68 | 1,00121E-26 |
| ACE2 | EXOC2 | -0,68 | 9,92453E-27 |
| ACE2 | EIF4G2 | -0,68 | 1,00961E-26 |
| ACE2 | NBPF15 | -0,6801 | 9,7425E-27 |
| ACE2 | SVEP1 | -0,6801 | 9,87968E-27 |
| ACE2 | AKAP11 | -0,6801 | 9,87396E-27 |
| ACE2 | TUBB | -0,6802 | 9,59828E-27 |
| ACE2 | ADCYAP1R1 | -0,6802 | 9,58248E-27 |
| ACE2 | NPIPA1 | -0,6802 | 9,5342E-27 |
| ACE2 | ENOX2 | -0,6803 | 9,26994E-27 |
| ACE2 | ARMC9 | -0,6804 | 9,06302E-27 |
| ACE2 | CBLL1 | -0,6804 | 9,04695E-27 |
| ACE2 | BBS10 | -0,6804 | 9,17418E-27 |
| ACE2 | LTBP2 | -0,6804 | 9,00632E-27 |
| ACE2 | PRDM16 | -0,6805 | 8,82997E-27 |
| ACE2 | VCAM1 | -0,6805 | 8,94223E-27 |
| ACE2 | SARS | -0,6805 | 8,9687E-27 |
| ACE2 | ZKSCAN7 | -0,6805 | 8,8294E-27 |
| ACE2 | BNIP3 | -0,6805 | 8,88046E-27 |
| ACE2 | FAM122C | -0,6805 | 8,99004E-27 |
| ACE2 | FAM49A | -0,6806 | 8,71608E-27 |
| ACE2 | THADA | -0,6806 | 8,67522E-27 |
| ACE2 | ZNF709 | -0,6806 | 8,62218E-27 |
| ACE2 | AC005301.9 | -0,6806 | 8,69659E-27 |
| ACE2 | AC007743.1 | -0,6807 | 8,44049E-27 |
| ACE2 | SPARCL1 | -0,6807 | 8,52494E-27 |
| ACE2 | MBLAC2 | -0,6807 | 8,45696E-27 |
| ACE2 | ZNF106 | -0,6807 | 8,47831E-27 |
| ACE2 | CALCRL | -0,6808 | 8,3719E-27 |
| ACE2 | DVL3 | -0,6808 | 8,30681E-27 |
| ACE2 | USP49 | -0,6808 | 8,33673E-27 |
| ACE2 | HCFC1R1 | -0,6808 | 8,3719E-27 |
| ACE2 | FAM65C | -0,6808 | 8,27271E-27 |
| ACE2 | GNRHR2 | -0,6809 | 8,062E-27 |
| ACE2 | IL11RA | -0,6809 | 8,10466E-27 |
| ACE2 | SUPT20H | -0,6809 | 8,17811E-27 |
| ACE2 | ZBTB49 | -0,681 | 7,98489E-27 |
| ACE2 | NAXD | -0,681 | 7,88559E-27 |
| ACE2 | PDAP1 | -0,6811 | 7,71362E-27 |
| ACE2 | NR2C2AP | -0,6812 | 7,44928E-27 |
| ACE2 | EIF2S2 | -0,6812 | 7,58086E-27 |
| ACE2 | ERO1B | -0,6813 | 7,31305E-27 |
| ACE2 | GUSBP4 | -0,6814 | 7,17871E-27 |
| ACE2 | ZCWPW1 | -0,6815 | 6,99653E-27 |
| ACE2 | RP11-667F14.1 | -0,6815 | 6,9611E-27 |
| ACE2 | PUM3 | -0,6815 | 7,05623E-27 |
| ACE2 | ANKRD13D | -0,6815 | 7,01224E-27 |
| ACE2 | CTD-2555O16.2 | -0,6815 | 6,95154E-27 |
| ACE2 | DOCK7 | -0,6816 | 6,84061E-27 |

| | | | |
|---|---|---|---|
| ACE2 | BEX5 | -0,6816 | 6,92693E-27 |
| ACE2 | COPS9 | -0,6817 | 6,73867E-27 |
| ACE2 | IQSEC3 | -0,6817 | 6,77102E-27 |
| ACE2 | COPS7A | -0,6817 | 6,71972E-27 |
| ACE2 | FHL1 | -0,6817 | 6,66105E-27 |
| ACE2 | EBNA1BP2 | -0,6818 | 6,50678E-27 |
| ACE2 | IQCJ-SCHIP1 | -0,6818 | 6,5632E-27 |
| ACE2 | COPS4 | -0,6818 | 6,46911E-27 |
| ACE2 | AGTR1 | -0,6819 | 6,33747E-27 |
| ACE2 | HCG11 | -0,6819 | 6,45651E-27 |
| ACE2 | PDCD2 | -0,6819 | 6,4125E-27 |
| ACE2 | KCNK7 | -0,6819 | 6,44857E-27 |
| ACE2 | LINC00310 | -0,6819 | 6,43499E-27 |
| ACE2 | SCUBE3 | -0,682 | 6,24637E-27 |
| ACE2 | ZNF652 | -0,682 | 6,19213E-27 |
| ACE2 | TGIF2 | -0,682 | 6,19983E-27 |
| ACE2 | MBD3 | -0,6821 | 6,04996E-27 |
| ACE2 | ZNF793-AS1 | -0,6821 | 6,13669E-27 |
| ACE2 | SHANK1 | -0,6821 | 6,08481E-27 |
| ACE2 | BRD8 | -0,6822 | 5,92689E-27 |
| ACE2 | IL6 | -0,6823 | 5,86044E-27 |
| ACE2 | HHLA3 | -0,6825 | 5,56977E-27 |
| ACE2 | NUPL2 | -0,6825 | 5,48727E-27 |
| ACE2 | RP11-64B16.2 | -0,6825 | 5,58761E-27 |
| ACE2 | ARTN | -0,6826 | 5,43411E-27 |
| ACE2 | POLR2D | -0,6826 | 5,37821E-27 |
| ACE2 | PAM | -0,6826 | 5,3491E-27 |
| ACE2 | ZRANB1 | -0,6826 | 5,37723E-27 |
| ACE2 | ZNF140 | -0,6826 | 5,4682E-27 |
| ACE2 | BNC1 | -0,6826 | 5,37474E-27 |
| ACE2 | ERCC4 | -0,6826 | 5,34931E-27 |
| ACE2 | SEC14L6 | -0,6826 | 5,41989E-27 |
| ACE2 | RACK1 | -0,6827 | 5,25212E-27 |
| ACE2 | RP11-789C17.1 | -0,6827 | 5,26209E-27 |
| ACE2 | NAA20 | -0,6827 | 5,2905E-27 |
| ACE2 | STAM | -0,6828 | 5,10565E-27 |
| ACE2 | CCS | -0,6828 | 5,12448E-27 |
| ACE2 | RNF219 | -0,6828 | 5,10023E-27 |
| ACE2 | MYO15A | -0,6828 | 5,14242E-27 |
| ACE2 | FMO2 | -0,6829 | 5,06178E-27 |
| ACE2 | THAP2 | -0,6829 | 5,00128E-27 |
| ACE2 | NPR1 | -0,683 | 4,89922E-27 |
| ACE2 | RP11-774O3.3 | -0,683 | 4,86469E-27 |
| ACE2 | RANBP9 | -0,683 | 4,86716E-27 |
| ACE2 | WDR83 | -0,683 | 4,96099E-27 |
| ACE2 | SMTN | -0,683 | 4,93776E-27 |
| ACE2 | FAM133B | -0,6831 | 4,83709E-27 |
| ACE2 | PTPRM | -0,6831 | 4,78514E-27 |
| ACE2 | CDH19 | -0,6831 | 4,81742E-27 |
| ACE2 | URB1 | -0,6831 | 4,84135E-27 |
| ACE2 | SENP5 | -0,6832 | 4,68505E-27 |
| ACE2 | WDR86-AS1 | -0,6832 | 4,65477E-27 |
| ACE2 | UBA52 | -0,6832 | 4,73448E-27 |
| ACE2 | SMN1 | -0,6833 | 4,57229E-27 |

| | | | |
|---|---|---|---|
| ACE2 | IGFBP7-AS1 | -0,6834 | 4,50537E-27 |
| ACE2 | UTRN | -0,6834 | 4,49584E-27 |
| ACE2 | CDON | -0,6834 | 4,49032E-27 |
| ACE2 | RPL7L1 | -0,6835 | 4,40071E-27 |
| ACE2 | LINC00933 | -0,6835 | 4,36332E-27 |
| ACE2 | ZNF177 | -0,6835 | 4,37287E-27 |
| ACE2 | AXL | -0,6835 | 4,3908E-27 |
| ACE2 | RPS27A | -0,6836 | 4,27712E-27 |
| ACE2 | SDPR | -0,6836 | 4,21698E-27 |
| ACE2 | CYP2U1 | -0,6836 | 4,21912E-27 |
| ACE2 | RPS12 | -0,6836 | 4,29751E-27 |
| ACE2 | PRMT3 | -0,6836 | 4,2219E-27 |
| ACE2 | ZNF609 | -0,6836 | 4,25222E-27 |
| ACE2 | SGSM2 | -0,6836 | 4,23797E-27 |
| ACE2 | UBE2M | -0,6836 | 4,30602E-27 |
| ACE2 | TBX1 | -0,6836 | 4,28343E-27 |
| ACE2 | SH3BP5 | -0,6837 | 4,14699E-27 |
| ACE2 | PCDHB11 | -0,6837 | 4,19454E-27 |
| ACE2 | ADO | -0,6837 | 4,17771E-27 |
| ACE2 | SRL | -0,6837 | 4,2059E-27 |
| ACE2 | AP2M1 | -0,6838 | 4,09932E-27 |
| ACE2 | ZBTB26 | -0,6838 | 4,01866E-27 |
| ACE2 | CRNDE | -0,6838 | 4,04299E-27 |
| ACE2 | TIMM22 | -0,6838 | 4,06891E-27 |
| ACE2 | LINC00598 | -0,6839 | 3,96265E-27 |
| ACE2 | RP11-139H15.5 | -0,6839 | 3,94646E-27 |
| ACE2 | TSR2 | -0,6839 | 4,00166E-27 |
| ACE2 | NCBP2 | -0,684 | 3,88784E-27 |
| ACE2 | ELP4 | -0,684 | 3,8814E-27 |
| ACE2 | C12orf76 | -0,684 | 3,89398E-27 |
| ACE2 | TPT1 | -0,684 | 3,89724E-27 |
| ACE2 | TAF1B | -0,6841 | 3,8248E-27 |
| ACE2 | PSMD5 | -0,6841 | 3,80127E-27 |
| ACE2 | WBP4 | -0,6841 | 3,74952E-27 |
| ACE2 | SYNM | -0,6841 | 3,75902E-27 |
| ACE2 | MAP3K4 | -0,6842 | 3,69188E-27 |
| ACE2 | UBXN1 | -0,6842 | 3,68483E-27 |
| ACE2 | GPS1 | -0,6842 | 3,6838E-27 |
| ACE2 | PACERR | -0,6843 | 3,57772E-27 |
| ACE2 | TMEM74 | -0,6843 | 3,63587E-27 |
| ACE2 | OSGEPL1 | -0,6845 | 3,47003E-27 |
| ACE2 | TMEM110 | -0,6845 | 3,44844E-27 |
| ACE2 | SENP7 | -0,6845 | 3,40002E-27 |
| ACE2 | CLGN | -0,6845 | 3,39804E-27 |
| ACE2 | WWC2-AS2 | -0,6845 | 3,4419E-27 |
| ACE2 | AMD1 | -0,6845 | 3,46064E-27 |
| ACE2 | HDAC2 | -0,6845 | 3,47149E-27 |
| ACE2 | KMT5A | -0,6845 | 3,43579E-27 |
| ACE2 | NORAD | -0,6845 | 3,44894E-27 |
| ACE2 | COLGALT1 | -0,6846 | 3,36528E-27 |
| ACE2 | CSDC2 | -0,6848 | 3,22719E-27 |
| ACE2 | CREB5 | -0,6849 | 3,1574E-27 |
| ACE2 | CHORDC1 | -0,6849 | 3,15208E-27 |
| ACE2 | METTL9 | -0,6849 | 3,08336E-27 |

| | | | |
|---|---|---|---|
| ACE2 | OXCT1 | -0,685 | 3,02195E-27 |
| ACE2 | TMEM233 | -0,685 | 3,07888E-27 |
| ACE2 | LINC00304 | -0,685 | 3,05769E-27 |
| ACE2 | ALKBH2 | -0,6851 | 3,00597E-27 |
| ACE2 | GTF3A | -0,6851 | 2,94348E-27 |
| ACE2 | CTB-12A17.2 | -0,6851 | 2,97E-27 |
| ACE2 | ZNF154 | -0,6851 | 2,95325E-27 |
| ACE2 | KCNT2 | -0,6853 | 2,83745E-27 |
| ACE2 | PPP1R21 | -0,6853 | 2,83491E-27 |
| ACE2 | NAA50 | -0,6853 | 2,80319E-27 |
| ACE2 | CLCNKA | -0,6855 | 2,72852E-27 |
| ACE2 | SNRPB2 | -0,6855 | 2,69876E-27 |
| ACE2 | TNC | -0,6856 | 2,64606E-27 |
| ACE2 | FLYWCH2 | -0,6856 | 2,60519E-27 |
| ACE2 | RP1-59D14.5 | -0,6856 | 2,64137E-27 |
| ACE2 | SPSB4 | -0,6857 | 2,56384E-27 |
| ACE2 | RP11-535M15.1 | -0,6857 | 2,60046E-27 |
| ACE2 | PIGQ | -0,6857 | 2,55653E-27 |
| ACE2 | PLP1 | -0,6857 | 2,56677E-27 |
| ACE2 | AC073343.13 | -0,6858 | 2,51863E-27 |
| ACE2 | PARVA | -0,6858 | 2,51825E-27 |
| ACE2 | ZNF627 | -0,6859 | 2,47812E-27 |
| ACE2 | ARNT | -0,686 | 2,37206E-27 |
| ACE2 | YPEL5 | -0,686 | 2,41374E-27 |
| ACE2 | APTX | -0,686 | 2,37362E-27 |
| ACE2 | PITRM1 | -0,6861 | 2,35291E-27 |
| ACE2 | HSDL1 | -0,6861 | 2,34839E-27 |
| ACE2 | NCAPH2 | -0,6861 | 2,33396E-27 |
| ACE2 | BDP1 | -0,6862 | 2,30294E-27 |
| ACE2 | ZNF732 | -0,6863 | 2,248E-27 |
| ACE2 | ZNF658 | -0,6863 | 2,22127E-27 |
| ACE2 | ZNF536 | -0,6863 | 2,23623E-27 |
| ACE2 | LINC00493 | -0,6863 | 2,23218E-27 |
| ACE2 | ALPK3 | -0,6864 | 2,16779E-27 |
| ACE2 | HEATR6 | -0,6864 | 2,1625E-27 |
| ACE2 | ZNF580 | -0,6864 | 2,16649E-27 |
| ACE2 | CACYBP | -0,6865 | 2,10042E-27 |
| ACE2 | WDR75 | -0,6865 | 2,10789E-27 |
| ACE2 | CYTL1 | -0,6865 | 2,14247E-27 |
| ACE2 | ZNF398 | -0,6866 | 2,07637E-27 |
| ACE2 | MCPH1 | -0,6866 | 2,06778E-27 |
| ACE2 | MAPK1 | -0,6866 | 2,06379E-27 |
| ACE2 | CTD-2135J3.3 | -0,6868 | 1,98477E-27 |
| ACE2 | SHISA3 | -0,6869 | 1,94464E-27 |
| ACE2 | TSTD3 | -0,6869 | 1,94824E-27 |
| ACE2 | ZFHX2 | -0,6869 | 1,9145E-27 |
| ACE2 | MAPK1IP1L | -0,6869 | 1,94621E-27 |
| ACE2 | CTC-444N24.11 | -0,6869 | 1,90636E-27 |
| ACE2 | SERPINF1 | -0,687 | 1,86242E-27 |
| ACE2 | RP11-400K9.4 | -0,6871 | 1,8167E-27 |
| ACE2 | RP11-1148L6.8 | -0,6871 | 1,83375E-27 |
| ACE2 | TROVE2 | -0,6872 | 1,78387E-27 |
| ACE2 | ABCC9 | -0,6872 | 1,79648E-27 |
| ACE2 | TTLL3 | -0,6873 | 1,76677E-27 |

| | | | |
|---|---|---|---|
| ACE2 | GPM6A | -0,6873 | 1,73315E-27 |
| ACE2 | SREK1IP1 | -0,6873 | 1,75429E-27 |
| ACE2 | EHMT2 | -0,6873 | 1,74308E-27 |
| ACE2 | RP11-531A24.5 | -0,6873 | 1,74426E-27 |
| ACE2 | TOP1MT | -0,6873 | 1,75424E-27 |
| ACE2 | LURAP1L | -0,6873 | 1,75723E-27 |
| ACE2 | PLA2G4C | -0,6873 | 1,76965E-27 |
| ACE2 | TDRKH | -0,6874 | 1,69755E-27 |
| ACE2 | RP11-273G15.2 | -0,6874 | 1,7032E-27 |
| ACE2 | REEP5 | -0,6875 | 1,68312E-27 |
| ACE2 | CACNA1G | -0,6875 | 1,65048E-27 |
| ACE2 | ADRA1A | -0,6876 | 1,6091E-27 |
| ACE2 | PDE2A | -0,6877 | 1,5761E-27 |
| ACE2 | MTA1 | -0,6877 | 1,5862E-27 |
| ACE2 | MECP2 | -0,6877 | 1,57014E-27 |
| ACE2 | MBNL1-AS1 | -0,6878 | 1,54297E-27 |
| ACE2 | CEP126 | -0,6878 | 1,55865E-27 |
| ACE2 | CAPZB | -0,6879 | 1,49587E-27 |
| ACE2 | HAX1 | -0,6879 | 1,50168E-27 |
| ACE2 | EXD2 | -0,6879 | 1,49445E-27 |
| ACE2 | NRXN1 | -0,688 | 1,47087E-27 |
| ACE2 | LRRTM1 | -0,688 | 1,46846E-27 |
| ACE2 | MDH1B | -0,688 | 1,45996E-27 |
| ACE2 | RP11-48B3.4 | -0,688 | 1,46192E-27 |
| ACE2 | ANP32B | -0,688 | 1,45884E-27 |
| ACE2 | LINC00959 | -0,688 | 1,46158E-27 |
| ACE2 | PTGES3 | -0,688 | 1,49139E-27 |
| ACE2 | TRMT5 | -0,688 | 1,47217E-27 |
| ACE2 | LITAF | -0,688 | 1,48985E-27 |
| ACE2 | TATDN1 | -0,6881 | 1,43026E-27 |
| ACE2 | LINC01415 | -0,6881 | 1,42718E-27 |
| ACE2 | SPEG | -0,6882 | 1,40128E-27 |
| ACE2 | PRRT3 | -0,6882 | 1,40153E-27 |
| ACE2 | TRA2B | -0,6882 | 1,40175E-27 |
| ACE2 | MYLK | -0,6883 | 1,37796E-27 |
| ACE2 | ANKRD53 | -0,6884 | 1,33931E-27 |
| ACE2 | NIFK | -0,6884 | 1,343E-27 |
| ACE2 | SMPD4 | -0,6884 | 1,35043E-27 |
| ACE2 | ZNF260 | -0,6885 | 1,3032E-27 |
| ACE2 | CTD-2245F17.6 | -0,6885 | 1,29063E-27 |
| ACE2 | ZNF197 | -0,6886 | 1,27743E-27 |
| ACE2 | RAET1G | -0,6886 | 1,2633E-27 |
| ACE2 | GNG10 | -0,6886 | 1,26333E-27 |
| ACE2 | TNPO2 | -0,6886 | 1,2653E-27 |
| ACE2 | MYL9 | -0,6886 | 1,26078E-27 |
| ACE2 | RMND5B | -0,6887 | 1,23067E-27 |
| ACE2 | MAML1 | -0,6887 | 1,24294E-27 |
| ACE2 | MTCL1 | -0,6887 | 1,24956E-27 |
| ACE2 | ZMYM3 | -0,6887 | 1,24591E-27 |
| ACE2 | UIMC1 | -0,6888 | 1,20097E-27 |
| ACE2 | TLE4 | -0,6888 | 1,2002E-27 |
| ACE2 | DNLZ | -0,6888 | 1,21603E-27 |
| ACE2 | COL4A1 | -0,6888 | 1,22109E-27 |
| ACE2 | TPD52L2 | -0,6888 | 1,2074E-27 |

| | | | |
|---|---|---|---|
| ACE2 | TCP10L | -0,6888 | 1,20238E-27 |
| ACE2 | EIF2B1 | -0,6889 | 1,18677E-27 |
| ACE2 | CREB3 | -0,689 | 1,15678E-27 |
| ACE2 | APOBEC3C | -0,689 | 1,16067E-27 |
| ACE2 | TNNI3K | -0,6891 | 1,1288E-27 |
| ACE2 | RP11-120J1.1 | -0,6891 | 1,12863E-27 |
| ACE2 | HNRNPLL | -0,6892 | 1,11228E-27 |
| ACE2 | RP11-977G19.5 | -0,6892 | 1,09121E-27 |
| ACE2 | SERPINE2 | -0,6893 | 1,07192E-27 |
| ACE2 | NOLC1 | -0,6893 | 1,08692E-27 |
| ACE2 | LINGO1 | -0,6893 | 1,07448E-27 |
| ACE2 | C16orf46 | -0,6893 | 1,07255E-27 |
| ACE2 | DYX1C1 | -0,6894 | 1,04197E-27 |
| ACE2 | PDE4D | -0,6895 | 1,0332E-27 |
| ACE2 | EPB41L4A-AS1 | -0,6895 | 1,03374E-27 |
| ACE2 | RPL8 | -0,6896 | 1,0095E-27 |
| ACE2 | ZNF561-AS1 | -0,6896 | 1,00093E-27 |
| ACE2 | TOGARAM1 | -0,6897 | 9,77319E-28 |
| ACE2 | RP11-20J15.3 | -0,6898 | 9,49783E-28 |
| ACE2 | TMEM221 | -0,6898 | 9,44518E-28 |
| ACE2 | TMEM39B | -0,69 | 9,07457E-28 |
| ACE2 | FLJ27354 | -0,69 | 9,03673E-28 |
| ACE2 | TICAM2 | -0,69 | 9,05497E-28 |
| ACE2 | TMEM126B | -0,69 | 9,12099E-28 |
| ACE2 | RP11-861E21.2 | -0,69 | 9,07409E-28 |
| ACE2 | SNX25 | -0,6901 | 8,94095E-28 |
| ACE2 | B4GALNT4 | -0,6901 | 8,86692E-28 |
| ACE2 | COQ10A | -0,6901 | 8,74993E-28 |
| ACE2 | ANKFY1 | -0,6901 | 8,7868E-28 |
| ACE2 | ARHGAP33 | -0,6901 | 8,73345E-28 |
| ACE2 | ZNF585A | -0,6901 | 8,80926E-28 |
| ACE2 | ABCF2 | -0,6902 | 8,7237E-28 |
| ACE2 | RP4-647C14.2 | -0,6902 | 8,67422E-28 |
| ACE2 | CECR2 | -0,6902 | 8,72316E-28 |
| ACE2 | ATP6V1G1 | -0,6903 | 8,49632E-28 |
| ACE2 | MTMR9LP | -0,6904 | 8,28797E-28 |
| ACE2 | ENSA | -0,6904 | 8,25774E-28 |
| ACE2 | C5 | -0,6904 | 8,28977E-28 |
| ACE2 | DPP7 | -0,6904 | 8,21364E-28 |
| ACE2 | RP11-170N16.3 | -0,6905 | 8,04392E-28 |
| ACE2 | RP11-557N21.1 | -0,6906 | 7,72531E-28 |
| ACE2 | MAP1A | -0,6906 | 7,88603E-28 |
| ACE2 | SRGAP2B | -0,6907 | 7,62725E-28 |
| ACE2 | RP11-254F7.2 | -0,6907 | 7,59286E-28 |
| ACE2 | ACTR6 | -0,6907 | 7,61794E-28 |
| ACE2 | HMGB1 | -0,6907 | 7,56499E-28 |
| ACE2 | KLF12 | -0,6907 | 7,60605E-28 |
| ACE2 | MR1 | -0,6908 | 7,39515E-28 |
| ACE2 | GPRC5B | -0,6908 | 7,41006E-28 |
| ACE2 | MRPS26 | -0,6908 | 7,52134E-28 |
| ACE2 | ARMCX5-GPRASP2 | -0,6908 | 7,42912E-28 |
| ACE2 | PRKCE | -0,6909 | 7,25454E-28 |
| ACE2 | PCDHB2 | -0,691 | 7,04693E-28 |
| ACE2 | THBS2 | -0,691 | 7,09963E-28 |

| | | | |
|---|---|---|---|
| ACE2 | DUSP10 | -0,6911 | 6,91505E-28 |
| ACE2 | ASNS | -0,6911 | 6,93635E-28 |
| ACE2 | RP11-728F11.3 | -0,6911 | 6,97198E-28 |
| ACE2 | SPECC1L | -0,6911 | 6,8419E-28 |
| ACE2 | CITED4 | -0,6912 | 6,66454E-28 |
| ACE2 | SEC22C | -0,6912 | 6,6609E-28 |
| ACE2 | ISPD | -0,6912 | 6,66953E-28 |
| ACE2 | RNF20 | -0,6912 | 6,69859E-28 |
| ACE2 | SLF2 | -0,6912 | 6,7958E-28 |
| ACE2 | DKK3 | -0,6912 | 6,77128E-28 |
| ACE2 | PRICKLE2 | -0,6913 | 6,63016E-28 |
| ACE2 | GRIA1 | -0,6913 | 6,57277E-28 |
| ACE2 | RP5-1142A6.9 | -0,6913 | 6,60697E-28 |
| ACE2 | SHISA5 | -0,6914 | 6,39098E-28 |
| ACE2 | COMMD2 | -0,6914 | 6,46406E-28 |
| ACE2 | FAM219B | -0,6915 | 6,3005E-28 |
| ACE2 | FAM198A | -0,6916 | 6,15081E-28 |
| ACE2 | LINC02210 | -0,6916 | 6,05139E-28 |
| ACE2 | BOK | -0,6917 | 5,94453E-28 |
| ACE2 | TSSC2 | -0,6917 | 5,90341E-28 |
| ACE2 | INPP5B | -0,6918 | 5,84177E-28 |
| ACE2 | PYCR2 | -0,6918 | 5,8066E-28 |
| ACE2 | PROS1 | -0,6918 | 5,75468E-28 |
| ACE2 | TAOK1 | -0,6918 | 5,79862E-28 |
| ACE2 | FSTL3 | -0,6918 | 5,87008E-28 |
| ACE2 | KLF2 | -0,6918 | 5,83388E-28 |
| ACE2 | ENOPH1 | -0,6919 | 5,61506E-28 |
| ACE2 | LNPEP | -0,6919 | 5,69451E-28 |
| ACE2 | TCEA1 | -0,6919 | 5,6966E-28 |
| ACE2 | RSF1 | -0,6919 | 5,66535E-28 |
| ACE2 | FAM182B | -0,6919 | 5,66456E-28 |
| ACE2 | MRPL20 | -0,692 | 5,58426E-28 |
| ACE2 | RP1-153G14.4 | -0,692 | 5,56608E-28 |
| ACE2 | GNA14 | -0,692 | 5,57939E-28 |
| ACE2 | GFRA3 | -0,6922 | 5,27647E-28 |
| ACE2 | TRIM7 | -0,6922 | 5,29441E-28 |
| ACE2 | USP28 | -0,6922 | 5,21171E-28 |
| ACE2 | TINF2 | -0,6922 | 5,31287E-28 |
| ACE2 | PARD6A | -0,6922 | 5,25652E-28 |
| ACE2 | MEGF8 | -0,6922 | 5,27518E-28 |
| ACE2 | RP11-88I18.2 | -0,6923 | 5,08404E-28 |
| ACE2 | PIEZO2 | -0,6923 | 5,11378E-28 |
| ACE2 | DNAJC21 | -0,6924 | 4,9706E-28 |
| ACE2 | PPP1R18 | -0,6924 | 5,06756E-28 |
| ACE2 | RP11-594N15.3 | -0,6924 | 4,95464E-28 |
| ACE2 | CEP78 | -0,6924 | 5,07637E-28 |
| ACE2 | LINC02001 | -0,6924 | 5,04482E-28 |
| ACE2 | KLF11 | -0,6925 | 4,94989E-28 |
| ACE2 | ARIH2 | -0,6925 | 4,84616E-28 |
| ACE2 | MED12L | -0,6925 | 4,84976E-28 |
| ACE2 | RP5-1142A6.10 | -0,6926 | 4,77544E-28 |
| ACE2 | MEMO1 | -0,6927 | 4,67834E-28 |
| ACE2 | GS1-124K5.4 | -0,6927 | 4,61418E-28 |
| ACE2 | DCTN6 | -0,6929 | 4,3789E-28 |

| | | | |
|---|---|---|---|
| ACE2 | ADAMTS8 | -0,6929 | 4,37893E-28 |
| ACE2 | KPNB1 | -0,6929 | 4,3998E-28 |
| ACE2 | ZNF285 | -0,6929 | 4,39041E-28 |
| ACE2 | RPL23AP7 | -0,693 | 4,31837E-28 |
| ACE2 | HDAC9 | -0,693 | 4,28057E-28 |
| ACE2 | RP11-410L14.2 | -0,693 | 4,34834E-28 |
| ACE2 | INPPL1 | -0,693 | 4,35558E-28 |
| ACE2 | WDR73 | -0,693 | 4,30599E-28 |
| ACE2 | FIG4 | -0,6931 | 4,20102E-28 |
| ACE2 | PALD1 | -0,6931 | 4,20839E-28 |
| ACE2 | CRTAC1 | -0,6931 | 4,19885E-28 |
| ACE2 | FAU | -0,6931 | 4,17692E-28 |
| ACE2 | EXOSC7 | -0,6932 | 4,12202E-28 |
| ACE2 | SMARCA5 | -0,6933 | 3,96873E-28 |
| ACE2 | CAP2 | -0,6934 | 3,90803E-28 |
| ACE2 | MTERF2 | -0,6934 | 3,88458E-28 |
| ACE2 | PALMD | -0,6935 | 3,85431E-28 |
| ACE2 | PAK1IP1 | -0,6935 | 3,77109E-28 |
| ACE2 | SLFN11 | -0,6935 | 3,82716E-28 |
| ACE2 | VEGFD | -0,6935 | 3,85844E-28 |
| ACE2 | DOCK11 | -0,6936 | 3,75851E-28 |
| ACE2 | DBT | -0,6937 | 3,66868E-28 |
| ACE2 | ANKDD1A | -0,6937 | 3,58891E-28 |
| ACE2 | SDAD1 | -0,6939 | 3,43564E-28 |
| ACE2 | PCDHGA1 | -0,6939 | 3,47252E-28 |
| ACE2 | CTA-204B4.2 | -0,6939 | 3,46198E-28 |
| ACE2 | GALNT18 | -0,6939 | 3,48971E-28 |
| ACE2 | RP11-266K4.13 | -0,6939 | 3,41322E-28 |
| ACE2 | CTD-2555O16.4 | -0,694 | 3,37192E-28 |
| ACE2 | DAP3 | -0,6941 | 3,28891E-28 |
| ACE2 | RP11-758N13.1 | -0,6941 | 3,30402E-28 |
| ACE2 | TBC1D25 | -0,6941 | 3,29516E-28 |
| ACE2 | LINC00578 | -0,6943 | 3,15534E-28 |
| ACE2 | KBTBD2 | -0,6943 | 3,10319E-28 |
| ACE2 | HTR3A | -0,6943 | 3,12131E-28 |
| ACE2 | TBCEL | -0,6943 | 3,11798E-28 |
| ACE2 | CTD-2267D19.3 | -0,6943 | 3,12062E-28 |
| ACE2 | ARMC1 | -0,6944 | 3,01913E-28 |
| ACE2 | FGD5 | -0,6946 | 2,91676E-28 |
| ACE2 | GPR107 | -0,6946 | 2,88719E-28 |
| ACE2 | C10orf25 | -0,6946 | 2,88562E-28 |
| ACE2 | RASSF9 | -0,6946 | 2,93153E-28 |
| ACE2 | RAB3GAP1 | -0,6948 | 2,73623E-28 |
| ACE2 | TTN-AS1 | -0,6948 | 2,7409E-28 |
| ACE2 | C12orf65 | -0,6948 | 2,79162E-28 |
| ACE2 | ARL4D | -0,6948 | 2,73691E-28 |
| ACE2 | SERBP1 | -0,695 | 2,60916E-28 |
| ACE2 | CCDC91 | -0,695 | 2,59879E-28 |
| ACE2 | MAPT | -0,695 | 2,62679E-28 |
| ACE2 | ATRAID | -0,6951 | 2,59011E-28 |
| ACE2 | RP11-360F5.3 | -0,6951 | 2,53887E-28 |
| ACE2 | PREX2 | -0,6951 | 2,55247E-28 |
| ACE2 | NPRL3 | -0,6951 | 2,54115E-28 |
| ACE2 | RPL13 | -0,6951 | 2,5881E-28 |

| | | | |
|---|---|---|---|
| ACE2 | THUMPD2 | -0,6952 | 2,50052E-28 |
| ACE2 | HTRA2 | -0,6952 | 2,46943E-28 |
| ACE2 | LINC00989 | -0,6952 | 2,47449E-28 |
| ACE2 | FAM66D | -0,6952 | 2,49465E-28 |
| ACE2 | AGTPBP1 | -0,6952 | 2,49595E-28 |
| ACE2 | PAGR1 | -0,6952 | 2,50488E-28 |
| ACE2 | THOC7 | -0,6953 | 2,46003E-28 |
| ACE2 | KIAA1324L | -0,6953 | 2,41983E-28 |
| ACE2 | NVL | -0,6954 | 2,36159E-28 |
| ACE2 | HNRNPU | -0,6954 | 2,3636E-28 |
| ACE2 | POPDC2 | -0,6954 | 2,37722E-28 |
| ACE2 | TM4SF18 | -0,6954 | 2,35097E-28 |
| ACE2 | CMTR1 | -0,6954 | 2,38922E-28 |
| ACE2 | AC005682.5 | -0,6954 | 2,34789E-28 |
| ACE2 | PTPRZ1 | -0,6954 | 2,35782E-28 |
| ACE2 | IRF2BPL | -0,6954 | 2,36178E-28 |
| ACE2 | RP11-394B2.1 | -0,6954 | 2,35727E-28 |
| ACE2 | SMIM10L2B | -0,6954 | 2,35132E-28 |
| ACE2 | LINC01140 | -0,6955 | 2,29046E-28 |
| ACE2 | TRMT10B | -0,6955 | 2,31623E-28 |
| ACE2 | PHACTR2 | -0,6956 | 2,24251E-28 |
| ACE2 | NTRK2 | -0,6956 | 2,23924E-28 |
| ACE2 | NALT1 | -0,6956 | 2,25252E-28 |
| ACE2 | CNNM1 | -0,6956 | 2,24215E-28 |
| ACE2 | TENM4 | -0,6956 | 2,27855E-28 |
| ACE2 | RP11-266O8.1 | -0,6956 | 2,26565E-28 |
| ACE2 | DMD | -0,6956 | 2,26794E-28 |
| ACE2 | EIF4B | -0,6957 | 2,20209E-28 |
| ACE2 | RP11-156P1.3 | -0,6957 | 2,1804E-28 |
| ACE2 | DTNBP1 | -0,6958 | 2,16325E-28 |
| ACE2 | FRY | -0,6958 | 2,17323E-28 |
| ACE2 | SLC16A1-AS1 | -0,6959 | 2,12017E-28 |
| ACE2 | RRP36 | -0,6959 | 2,12079E-28 |
| ACE2 | FRMD5 | -0,6959 | 2,07349E-28 |
| ACE2 | RPS5 | -0,6959 | 2,11272E-28 |
| ACE2 | MUM1 | -0,696 | 2,02095E-28 |
| ACE2 | PLEKHM2 | -0,6961 | 2,0018E-28 |
| ACE2 | RP11-73M7.6 | -0,6961 | 1,98894E-28 |
| ACE2 | CTD-2336O2.1 | -0,6961 | 1,97612E-28 |
| ACE2 | RP11-380L11.4 | -0,6961 | 2,0075E-28 |
| ACE2 | ZNF391 | -0,6962 | 1,93493E-28 |
| ACE2 | CEP97 | -0,6963 | 1,88879E-28 |
| ACE2 | SUN1 | -0,6963 | 1,88255E-28 |
| ACE2 | MALSU1 | -0,6963 | 1,90774E-28 |
| ACE2 | SPTBN4 | -0,6963 | 1,88444E-28 |
| ACE2 | TASP1 | -0,6963 | 1,91583E-28 |
| ACE2 | MAPKAP1 | -0,6964 | 1,83829E-28 |
| ACE2 | HRASLS5 | -0,6964 | 1,84172E-28 |
| ACE2 | RP11-983P16.4 | -0,6964 | 1,86368E-28 |
| ACE2 | TNFRSF19 | -0,6966 | 1,75844E-28 |
| ACE2 | CTD-2541J13.1 | -0,6967 | 1,72215E-28 |
| ACE2 | GNG8 | -0,6968 | 1,68158E-28 |
| ACE2 | USP27X | -0,6968 | 1,68306E-28 |
| ACE2 | RP11-5C23.1 | -0,6969 | 1,61244E-28 |

| | | | |
|---|---|---|---|
| ACE2 | DPCD | -0,6969 | 1,61643E-28 |
| ACE2 | RP11-156E6.1 | -0,6969 | 1,6273E-28 |
| ACE2 | EIF2S3 | -0,697 | 1,57726E-28 |
| ACE2 | IFI44L | -0,6971 | 1,55877E-28 |
| ACE2 | TNS1 | -0,6971 | 1,55026E-28 |
| ACE2 | TEX10 | -0,6971 | 1,54247E-28 |
| ACE2 | ZNF829 | -0,6971 | 1,53038E-28 |
| ACE2 | RP11-379F4.4 | -0,6972 | 1,5052E-28 |
| ACE2 | IMMP1L | -0,6972 | 1,50147E-28 |
| ACE2 | GIT1 | -0,6972 | 1,52375E-28 |
| ACE2 | CENPT | -0,6974 | 1,44686E-28 |
| ACE2 | HSPA4L | -0,6975 | 1,41119E-28 |
| ACE2 | MCTP1 | -0,6975 | 1,40553E-28 |
| ACE2 | MVB12B | -0,6975 | 1,40648E-28 |
| ACE2 | SCN11A | -0,6976 | 1,38152E-28 |
| ACE2 | JAZF1-AS1 | -0,6976 | 1,37737E-28 |
| ACE2 | ARMC4 | -0,6976 | 1,3584E-28 |
| ACE2 | TBC1D5 | -0,6977 | 1,33703E-28 |
| ACE2 | CEP70 | -0,6977 | 1,32102E-28 |
| ACE2 | CTD-3138B18.5 | -0,6977 | 1,33004E-28 |
| ACE2 | OSMR-AS1 | -0,6978 | 1,29832E-28 |
| ACE2 | NUBP2 | -0,6978 | 1,29787E-28 |
| ACE2 | RBP7 | -0,6979 | 1,26616E-28 |
| ACE2 | ROBO1 | -0,6979 | 1,27186E-28 |
| ACE2 | PHF14 | -0,6979 | 1,27866E-28 |
| ACE2 | FAM185A | -0,6979 | 1,2688E-28 |
| ACE2 | PEAR1 | -0,698 | 1,24324E-28 |
| ACE2 | MTMR12 | -0,698 | 1,24425E-28 |
| ACE2 | PANX1 | -0,698 | 1,23279E-28 |
| ACE2 | CAPS2 | -0,698 | 1,21957E-28 |
| ACE2 | AOC2 | -0,698 | 1,23095E-28 |
| ACE2 | CSRP1 | -0,6981 | 1,20297E-28 |
| ACE2 | LA16c-349E10.1 | -0,6981 | 1,19296E-28 |
| ACE2 | DTD1 | -0,6981 | 1,21712E-28 |
| ACE2 | ZNF70 | -0,6981 | 1,19443E-28 |
| ACE2 | FOXD3 | -0,6982 | 1,18559E-28 |
| ACE2 | RP11-977G19.11 | -0,6982 | 1,17961E-28 |
| ACE2 | GALNT9 | -0,6982 | 1,17515E-28 |
| ACE2 | C2orf81 | -0,6983 | 1,14236E-28 |
| ACE2 | CDC23 | -0,6983 | 1,15256E-28 |
| ACE2 | RP11-88E10.4 | -0,6983 | 1,13812E-28 |
| ACE2 | PCDHB13 | -0,6984 | 1,10198E-28 |
| ACE2 | RP11-124N14.3 | -0,6984 | 1,11133E-28 |
| ACE2 | SSC5D | -0,6984 | 1,12522E-28 |
| ACE2 | CPLX1 | -0,6985 | 1,08795E-28 |
| ACE2 | RBM24 | -0,6985 | 1,07675E-28 |
| ACE2 | LRRC55 | -0,6985 | 1,09075E-28 |
| ACE2 | CAPN6 | -0,6985 | 1,0978E-28 |
| ACE2 | MAP3K6 | -0,6987 | 1,03553E-28 |
| ACE2 | QSER1 | -0,6987 | 1,0357E-28 |
| ACE2 | CCDC113 | -0,6987 | 1,02244E-28 |
| ACE2 | DDX5 | -0,6987 | 1,03967E-28 |
| ACE2 | LINC01936 | -0,6988 | 1,01091E-28 |
| ACE2 | WDR17 | -0,6988 | 1,0124E-28 |

| | | | |
|---|---|---|---|
| ACE2 | PARP16 | -0,6988 | 1,00963E-28 |
| ACE2 | RP11-1024P17.1 | -0,6989 | 9,69939E-29 |
| ACE2 | APBB2 | -0,6989 | 9,82552E-29 |
| ACE2 | RP3-525N10.2 | -0,6989 | 9,86616E-29 |
| ACE2 | HOXC-AS1 | -0,6989 | 9,82403E-29 |
| ACE2 | GJA1 | -0,699 | 9,48622E-29 |
| ACE2 | PCDHGB2 | -0,6991 | 9,36885E-29 |
| ACE2 | C21orf91 | -0,6991 | 9,34341E-29 |
| ACE2 | AC012358.8 | -0,6992 | 9,142E-29 |
| ACE2 | MSC-AS1 | -0,6992 | 9,19767E-29 |
| ACE2 | ACTA2-AS1 | -0,6992 | 9,12985E-29 |
| ACE2 | LRRC32 | -0,6992 | 9,01471E-29 |
| ACE2 | SNX1 | -0,6992 | 9,13411E-29 |
| ACE2 | JMJD8 | -0,6992 | 9,142E-29 |
| ACE2 | PGM5P4 | -0,6993 | 8,8678E-29 |
| ACE2 | MZT2B | -0,6993 | 8,77978E-29 |
| ACE2 | EIF3EP1 | -0,6993 | 8,89082E-29 |
| ACE2 | ST6GALNAC2 | -0,6993 | 8,86448E-29 |
| ACE2 | DXO | -0,6994 | 8,7553E-29 |
| ACE2 | CTD-2525P14.5 | -0,6994 | 8,64289E-29 |
| ACE2 | CERS1 | -0,6994 | 8,55846E-29 |
| ACE2 | TCTN3 | -0,6995 | 8,46921E-29 |
| ACE2 | NUP58 | -0,6995 | 8,37636E-29 |
| ACE2 | CTC-359D24.3 | -0,6995 | 8,4801E-29 |
| ACE2 | L1CAM | -0,6995 | 8,52392E-29 |
| ACE2 | RIOK1 | -0,6996 | 8,1279E-29 |
| ACE2 | RPS15 | -0,6996 | 8,27774E-29 |
| ACE2 | SCN7A | -0,6997 | 7,95872E-29 |
| ACE2 | DIRC3 | -0,6997 | 7,94319E-29 |
| ACE2 | LINC02202 | -0,6997 | 8,01226E-29 |
| ACE2 | GTF2IRD2 | -0,6997 | 8,01592E-29 |
| ACE2 | SPIDR | -0,6997 | 7,97784E-29 |
| ACE2 | ANOS1 | -0,6997 | 8,06813E-29 |
| ACE2 | CD200 | -0,6998 | 7,88125E-29 |
| ACE2 | RWDD4 | -0,6998 | 7,89066E-29 |
| ACE2 | BNIP3L | -0,6998 | 7,80497E-29 |
| ACE2 | PLAC9 | -0,6998 | 7,78736E-29 |
| ACE2 | UVRAG | -0,6998 | 7,80059E-29 |
| ACE2 | SPG20-AS1 | -0,6998 | 7,74981E-29 |
| ACE2 | GBP1P1 | -0,6999 | 7,54858E-29 |
| ACE2 | MRPL24 | -0,6999 | 7,61847E-29 |
| ACE2 | ADGRB1 | -0,6999 | 7,64286E-29 |
| ACE2 | TSSC4 | -0,6999 | 7,57799E-29 |
| ACE2 | CHRDL2 | -0,6999 | 7,57338E-29 |
| ACE2 | DDX51 | -0,6999 | 7,53181E-29 |
| ACE2 | ACAD9 | -0,7 | 7,38938E-29 |
| ACE2 | RPS11 | -0,7 | 7,47977E-29 |
| ACE2 | ZNF84 | -0,7001 | 7,22153E-29 |
| ACE2 | TENM1 | -0,7001 | 7,26175E-29 |
| ACE2 | IGSF11 | -0,7002 | 6,98844E-29 |
| ACE2 | ACTA2 | -0,7002 | 7,10184E-29 |
| ACE2 | LRRC2 | -0,7003 | 6,88263E-29 |
| ACE2 | ANKRD42 | -0,7003 | 6,87029E-29 |
| ACE2 | TNS2 | -0,7003 | 6,90097E-29 |

| | | | |
|---|---|---|---|
| ACE2 | DDX24 | -0,7003 | 6,92432E-29 |
| ACE2 | PLD2 | -0,7003 | 6,96047E-29 |
| ACE2 | WSB1 | -0,7003 | 6,85693E-29 |
| ACE2 | TAF12 | -0,7004 | 6,70759E-29 |
| ACE2 | COL12A1 | -0,7004 | 6,78174E-29 |
| ACE2 | AR | -0,7004 | 6,76838E-29 |
| ACE2 | NISCH | -0,7005 | 6,54762E-29 |
| ACE2 | ZNF766 | -0,7005 | 6,60016E-29 |
| ACE2 | SNTA1 | -0,7005 | 6,57789E-29 |
| ACE2 | ATP7A | -0,7005 | 6,58461E-29 |
| ACE2 | RP11-355F16.1 | -0,7006 | 6,30973E-29 |
| ACE2 | GAS1RR | -0,7006 | 6,35427E-29 |
| ACE2 | ISL2 | -0,7006 | 6,43439E-29 |
| ACE2 | FOXO3B | -0,7006 | 6,28739E-29 |
| ACE2 | SYN3 | -0,7006 | 6,3265E-29 |
| ACE2 | STARD4-AS1 | -0,7007 | 6,24815E-29 |
| ACE2 | RTF1 | -0,7007 | 6,16894E-29 |
| ACE2 | TRIT1 | -0,7008 | 6,01935E-29 |
| ACE2 | RP11-263K19.4 | -0,7008 | 6,09789E-29 |
| ACE2 | CRBN | -0,7009 | 5,88376E-29 |
| ACE2 | C14orf159 | -0,7009 | 5,8765E-29 |
| ACE2 | HMGB1P6 | -0,7009 | 5,88501E-29 |
| ACE2 | MOV10L1 | -0,7009 | 5,94055E-29 |
| ACE2 | TTC27 | -0,7011 | 5,64428E-29 |
| ACE2 | GRIP1 | -0,7011 | 5,6643E-29 |
| ACE2 | ST3GAL2 | -0,7011 | 5,65179E-29 |
| ACE2 | PABPC4 | -0,7012 | 5,40974E-29 |
| ACE2 | CLDN11 | -0,7012 | 5,51046E-29 |
| ACE2 | FAM53A | -0,7012 | 5,42077E-29 |
| ACE2 | PPIL4 | -0,7012 | 5,48444E-29 |
| ACE2 | TBC1D26 | -0,7012 | 5,51895E-29 |
| ACE2 | CA5B | -0,7012 | 5,45534E-29 |
| ACE2 | C7orf50 | -0,7013 | 5,3317E-29 |
| ACE2 | RAD21 | -0,7013 | 5,35353E-29 |
| ACE2 | BTRC | -0,7013 | 5,32291E-29 |
| ACE2 | SART1 | -0,7013 | 5,30876E-29 |
| ACE2 | GLTSCR2 | -0,7013 | 5,30195E-29 |
| ACE2 | LINC00327 | -0,7014 | 5,14985E-29 |
| ACE2 | IRGQ | -0,7014 | 5,22557E-29 |
| ACE2 | IARS | -0,7015 | 5,08663E-29 |
| ACE2 | ARID5B | -0,7016 | 4,93972E-29 |
| ACE2 | KNDC1 | -0,7016 | 4,90224E-29 |
| ACE2 | MIPOL1 | -0,7016 | 4,96273E-29 |
| ACE2 | FAP | -0,7017 | 4,7456E-29 |
| ACE2 | PLEKHM3 | -0,7017 | 4,82531E-29 |
| ACE2 | NPFF | -0,7017 | 4,79241E-29 |
| ACE2 | HSPB11 | -0,7018 | 4,66812E-29 |
| ACE2 | CTNND2 | -0,7018 | 4,67921E-29 |
| ACE2 | CARMN | -0,7018 | 4,62867E-29 |
| ACE2 | RPS6 | -0,7018 | 4,72175E-29 |
| ACE2 | CCDC184 | -0,7018 | 4,62578E-29 |
| ACE2 | MYO16 | -0,7018 | 4,7251E-29 |
| ACE2 | TRIP10 | -0,7018 | 4,62681E-29 |
| ACE2 | PPP2R5C | -0,7019 | 4,56712E-29 |

| | | | |
|---|---|---|---|
| ACE2 | USP4 | -0,702 | 4,50106E-29 |
| ACE2 | RP11-95D17.1 | -0,7021 | 4,33921E-29 |
| ACE2 | UTP23 | -0,7021 | 4,38389E-29 |
| ACE2 | C11orf57 | -0,7021 | 4,30138E-29 |
| ACE2 | RP11-817I4.1 | -0,7021 | 4,30418E-29 |
| ACE2 | HSPA13 | -0,7021 | 4,31365E-29 |
| ACE2 | FLT1 | -0,7022 | 4,2762E-29 |
| ACE2 | DCAF5 | -0,7022 | 4,17351E-29 |
| ACE2 | AKT2 | -0,7022 | 4,2342E-29 |
| ACE2 | AC012074.2 | -0,7023 | 4,11808E-29 |
| ACE2 | RHBDD1 | -0,7023 | 4,14279E-29 |
| ACE2 | ARMC8 | -0,7023 | 4,16297E-29 |
| ACE2 | RBM18 | -0,7023 | 4,1229E-29 |
| ACE2 | PLEKHB1 | -0,7023 | 4,06701E-29 |
| ACE2 | RP11-644F5.11 | -0,7023 | 4,06774E-29 |
| ACE2 | CHRDL1 | -0,7024 | 4,05294E-29 |
| ACE2 | FAM161A | -0,7025 | 3,88207E-29 |
| ACE2 | CEP44 | -0,7025 | 3,86191E-29 |
| ACE2 | ZNF10 | -0,7025 | 3,94446E-29 |
| ACE2 | PWAR6 | -0,7025 | 3,86163E-29 |
| ACE2 | UBAP2L | -0,7026 | 3,78168E-29 |
| ACE2 | SNRNP27 | -0,7026 | 3,83646E-29 |
| ACE2 | NMNAT3 | -0,7026 | 3,80499E-29 |
| ACE2 | FXN | -0,7026 | 3,80027E-29 |
| ACE2 | INTS13 | -0,7026 | 3,8113E-29 |
| ACE2 | SPIN4 | -0,7026 | 3,82134E-29 |
| ACE2 | CALD1 | -0,7027 | 3,74607E-29 |
| ACE2 | PPP3CC | -0,7027 | 3,72037E-29 |
| ACE2 | GLRX3 | -0,7027 | 3,74584E-29 |
| ACE2 | COL4A2 | -0,7027 | 3,74695E-29 |
| ACE2 | CACNA1C | -0,7028 | 3,60188E-29 |
| ACE2 | SEC62 | -0,7029 | 3,50242E-29 |
| ACE2 | FNDC9 | -0,7029 | 3,55324E-29 |
| ACE2 | RABEP1 | -0,7029 | 3,52245E-29 |
| ACE2 | ZNF644 | -0,703 | 3,4691E-29 |
| ACE2 | FIGN | -0,7031 | 3,31631E-29 |
| ACE2 | ERCC6L2 | -0,7031 | 3,33406E-29 |
| ACE2 | ANKRD26 | -0,7031 | 3,37442E-29 |
| ACE2 | CCT6P1 | -0,7032 | 3,2963E-29 |
| ACE2 | MRPL17 | -0,7032 | 3,22339E-29 |
| ACE2 | SLC25A30 | -0,7032 | 3,25644E-29 |
| ACE2 | CMTM5 | -0,7032 | 3,28049E-29 |
| ACE2 | ANKAR | -0,7033 | 3,17353E-29 |
| ACE2 | TSC22D2 | -0,7033 | 3,15114E-29 |
| ACE2 | GDF6 | -0,7033 | 3,19588E-29 |
| ACE2 | TMEM261 | -0,7033 | 3,15485E-29 |
| ACE2 | RP11-736K20.5 | -0,7033 | 3,19235E-29 |
| ACE2 | CAB39L | -0,7033 | 3,15227E-29 |
| ACE2 | C14orf37 | -0,7033 | 3,18932E-29 |
| ACE2 | ZNF134 | -0,7033 | 3,21747E-29 |
| ACE2 | RPL10 | -0,7033 | 3,15635E-29 |
| ACE2 | COPS7B | -0,7034 | 3,06861E-29 |
| ACE2 | CCDC3 | -0,7035 | 3,05487E-29 |
| ACE2 | ATXN2 | -0,7035 | 3,04873E-29 |

| | | | |
|---|---|---|---|
| ACE2 | GATA2-AS1 | -0,7036 | 2,95719E-29 |
| ACE2 | IRAK1BP1 | -0,7036 | 2,94336E-29 |
| ACE2 | NUP188 | -0,7036 | 2,96228E-29 |
| ACE2 | GPR180 | -0,7036 | 2,95206E-29 |
| ACE2 | ARHGEF33 | -0,7037 | 2,85374E-29 |
| ACE2 | KIAA0930 | -0,7037 | 2,85894E-29 |
| ACE2 | C1orf204 | -0,7038 | 2,79484E-29 |
| ACE2 | BCL6 | -0,7039 | 2,69638E-29 |
| ACE2 | ALKBH8 | -0,7039 | 2,74977E-29 |
| ACE2 | RNPS1 | -0,7039 | 2,68993E-29 |
| ACE2 | SNN | -0,7039 | 2,7369E-29 |
| ACE2 | RPL38 | -0,7039 | 2,70561E-29 |
| ACE2 | MTCH1 | -0,704 | 2,63274E-29 |
| ACE2 | NOV | -0,704 | 2,64423E-29 |
| ACE2 | NEMP1 | -0,7041 | 2,55665E-29 |
| ACE2 | DNAH10OS | -0,7041 | 2,56196E-29 |
| ACE2 | SNTB1 | -0,7042 | 2,49306E-29 |
| ACE2 | BRD3 | -0,7042 | 2,51244E-29 |
| ACE2 | RAI1 | -0,7042 | 2,54627E-29 |
| ACE2 | EIF3I | -0,7043 | 2,48199E-29 |
| ACE2 | ENTPD1-AS1 | -0,7043 | 2,47293E-29 |
| ACE2 | CELSR2 | -0,7046 | 2,29192E-29 |
| ACE2 | TRIM58 | -0,7046 | 2,27024E-29 |
| ACE2 | MOCS2 | -0,7046 | 2,26528E-29 |
| ACE2 | DUSP22 | -0,7046 | 2,27602E-29 |
| ACE2 | RP11-629G13.1 | -0,7046 | 2,25398E-29 |
| ACE2 | CTD-2595P9.4 | -0,7046 | 2,2882E-29 |
| ACE2 | C10orf35 | -0,7047 | 2,21825E-29 |
| ACE2 | AHNAK2 | -0,7047 | 2,22872E-29 |
| ACE2 | KIAA2022 | -0,7047 | 2,22231E-29 |
| ACE2 | BMP7 | -0,7048 | 2,1456E-29 |
| ACE2 | FTSJ1 | -0,7048 | 2,15452E-29 |
| ACE2 | MRPS15 | -0,7049 | 2,08821E-29 |
| ACE2 | ADD2 | -0,7049 | 2,07416E-29 |
| ACE2 | RP11-834C11.6 | -0,705 | 2,01821E-29 |
| ACE2 | COL5A3 | -0,705 | 2,03611E-29 |
| ACE2 | PLA2G5 | -0,7051 | 1,97512E-29 |
| ACE2 | MLLT11 | -0,7051 | 1,9934E-29 |
| ACE2 | KLHDC3 | -0,7051 | 2,00822E-29 |
| ACE2 | CDC37 | -0,7051 | 1,98048E-29 |
| ACE2 | PIGP | -0,7051 | 2,00935E-29 |
| ACE2 | PCBP4 | -0,7052 | 1,91788E-29 |
| ACE2 | C6orf3 | -0,7052 | 1,95827E-29 |
| ACE2 | LINC00963 | -0,7052 | 1,93682E-29 |
| ACE2 | UCK1 | -0,7052 | 1,91585E-29 |
| ACE2 | BACE1-AS | -0,7053 | 1,8943E-29 |
| ACE2 | PRR4 | -0,7053 | 1,89839E-29 |
| ACE2 | RPS6KL1 | -0,7053 | 1,88615E-29 |
| ACE2 | ZNF419 | -0,7053 | 1,88799E-29 |
| ACE2 | UBE2D2 | -0,7054 | 1,83123E-29 |
| ACE2 | KPNA5 | -0,7054 | 1,85555E-29 |
| ACE2 | SIVA1 | -0,7054 | 1,86118E-29 |
| ACE2 | VWA3A | -0,7054 | 1,85479E-29 |
| ACE2 | BVES-AS1 | -0,7055 | 1,78615E-29 |

| | | | |
|---|---|---|---|
| ACE2 | ABL1 | -0,7055 | 1,789E-29 |
| ACE2 | ZNF37A | -0,7055 | 1,78673E-29 |
| ACE2 | LRFN5 | -0,7055 | 1,77903E-29 |
| ACE2 | CTC-444N24.8 | -0,7055 | 1,79977E-29 |
| ACE2 | PXK | -0,7056 | 1,73661E-29 |
| ACE2 | SENCR | -0,7056 | 1,74051E-29 |
| ACE2 | CLASP1 | -0,7057 | 1,71349E-29 |
| ACE2 | PRRC2A | -0,7057 | 1,71393E-29 |
| ACE2 | LRFN4 | -0,7057 | 1,70279E-29 |
| ACE2 | PLEKHF1 | -0,7057 | 1,7203E-29 |
| ACE2 | MAP3K10 | -0,7057 | 1,68745E-29 |
| ACE2 | LZTR1 | -0,7057 | 1,70084E-29 |
| ACE2 | INHBA | -0,7058 | 1,6711E-29 |
| ACE2 | GEMIN2 | -0,7058 | 1,67056E-29 |
| ACE2 | KPNA3 | -0,7059 | 1,62072E-29 |
| ACE2 | PRIMPOL | -0,706 | 1,56213E-29 |
| ACE2 | NXPE3 | -0,7061 | 1,52046E-29 |
| ACE2 | UBLCP1 | -0,7062 | 1,50817E-29 |
| ACE2 | KCNH2 | -0,7062 | 1,50397E-29 |
| ACE2 | ANKRD16 | -0,7062 | 1,4904E-29 |
| ACE2 | STUB1 | -0,7062 | 1,48836E-29 |
| ACE2 | TBXA2R | -0,7062 | 1,51184E-29 |
| ACE2 | FAM76A | -0,7063 | 1,47188E-29 |
| ACE2 | C3orf62 | -0,7063 | 1,4654E-29 |
| ACE2 | ORC5 | -0,7064 | 1,41167E-29 |
| ACE2 | PPP2R5B | -0,7064 | 1,4306E-29 |
| ACE2 | NFIX | -0,7064 | 1,41189E-29 |
| ACE2 | ISY1 | -0,7065 | 1,38881E-29 |
| ACE2 | MINDY3 | -0,7065 | 1,37683E-29 |
| ACE2 | CCDC36 | -0,7066 | 1,3436E-29 |
| ACE2 | PCDHB6 | -0,7066 | 1,34166E-29 |
| ACE2 | CDC37L1 | -0,7066 | 1,34775E-29 |
| ACE2 | DSCAML1 | -0,7066 | 1,35497E-29 |
| ACE2 | LRRC49 | -0,7066 | 1,33597E-29 |
| ACE2 | CERS4 | -0,7066 | 1,33285E-29 |
| ACE2 | FAM43B | -0,7067 | 1,29311E-29 |
| ACE2 | VRK2 | -0,7067 | 1,32294E-29 |
| ACE2 | TMEM155 | -0,7067 | 1,32378E-29 |
| ACE2 | SPX | -0,7067 | 1,32575E-29 |
| ACE2 | AGAP2-AS1 | -0,7067 | 1,32474E-29 |
| ACE2 | CHURC1 | -0,7067 | 1,32558E-29 |
| ACE2 | ARHGAP22 | -0,7069 | 1,24248E-29 |
| ACE2 | DFFA | -0,707 | 1,20293E-29 |
| ACE2 | ZFP69 | -0,707 | 1,22435E-29 |
| ACE2 | ZNF346 | -0,707 | 1,20056E-29 |
| ACE2 | COL21A1 | -0,707 | 1,21551E-29 |
| ACE2 | RPS3A | -0,7071 | 1,18204E-29 |
| ACE2 | EPHA7 | -0,7071 | 1,19057E-29 |
| ACE2 | PHF1 | -0,7072 | 1,13763E-29 |
| ACE2 | KCTD7 | -0,7073 | 1,1266E-29 |
| ACE2 | SMIM10L1 | -0,7073 | 1,10598E-29 |
| ACE2 | ITGA2B | -0,7073 | 1,12444E-29 |
| ACE2 | ROCK1 | -0,7074 | 1,10067E-29 |
| ACE2 | RP11-967K21.1 | -0,7075 | 1,06358E-29 |

| | | | |
|---|---|---|---|
| ACE2 | TCHP | -0,7075 | 1,05459E-29 |
| ACE2 | LINC00888 | -0,7076 | 1,03E-29 |
| ACE2 | EMSY | -0,7076 | 1,04134E-29 |
| ACE2 | SDCCAG8 | -0,7078 | 9,80633E-30 |
| ACE2 | MARCH7 | -0,7078 | 9,81805E-30 |
| ACE2 | RPL29 | -0,7078 | 9,75326E-30 |
| ACE2 | PBRM1 | -0,7078 | 9,76709E-30 |
| ACE2 | MTR | -0,7079 | 9,5315E-30 |
| ACE2 | ZNF132 | -0,7079 | 9,50892E-30 |
| ACE2 | VPS41 | -0,708 | 9,20594E-30 |
| ACE2 | RASA3 | -0,708 | 9,1812E-30 |
| ACE2 | CTD-2036P10.6 | -0,708 | 9,38651E-30 |
| ACE2 | IFI44 | -0,7081 | 9,0898E-30 |
| ACE2 | LONRF2 | -0,7081 | 9,05243E-30 |
| ACE2 | PLXND1 | -0,7081 | 8,9839E-30 |
| ACE2 | RRAS | -0,7081 | 9,15011E-30 |
| ACE2 | S1PR1 | -0,7082 | 8,86921E-30 |
| ACE2 | PNPT1 | -0,7082 | 8,77807E-30 |
| ACE2 | PRDM11 | -0,7082 | 8,77147E-30 |
| ACE2 | ZFP30 | -0,7082 | 8,82984E-30 |
| ACE2 | NKX3-2 | -0,7083 | 8,55166E-30 |
| ACE2 | RP11-736K20.4 | -0,7083 | 8,68408E-30 |
| ACE2 | AVPR1A | -0,7083 | 8,49244E-30 |
| ACE2 | KIF9-AS1 | -0,7085 | 8,13003E-30 |
| ACE2 | DKK2 | -0,7085 | 8,14306E-30 |
| ACE2 | RSBN1L | -0,7085 | 8,18395E-30 |
| ACE2 | POLB | -0,7085 | 8,17328E-30 |
| ACE2 | LRP3 | -0,7085 | 8,1385E-30 |
| ACE2 | LIPE-AS1 | -0,7085 | 8,12382E-30 |
| ACE2 | ZNF160 | -0,7085 | 8,19202E-30 |
| ACE2 | ITGA9-AS1 | -0,7086 | 7,85693E-30 |
| ACE2 | ZNF615 | -0,7086 | 7,93114E-30 |
| ACE2 | MURC | -0,7087 | 7,68525E-30 |
| ACE2 | ZNF846 | -0,7087 | 7,7196E-30 |
| ACE2 | DYNC1I1 | -0,7088 | 7,44325E-30 |
| ACE2 | SMAD2 | -0,7088 | 7,57585E-30 |
| ACE2 | TUBB4A | -0,7088 | 7,47279E-30 |
| ACE2 | PMS1 | -0,7089 | 7,24191E-30 |
| ACE2 | PDZRN3 | -0,7089 | 7,41912E-30 |
| ACE2 | CHMP4A | -0,7089 | 7,34196E-30 |
| ACE2 | DCUN1D3 | -0,7089 | 7,40729E-30 |
| ACE2 | IL1RAPL1 | -0,7089 | 7,28647E-30 |
| ACE2 | ST3GAL5 | -0,709 | 7,20451E-30 |
| ACE2 | CRYAB | -0,7091 | 6,93519E-30 |
| ACE2 | SUGT1P1 | -0,7092 | 6,74665E-30 |
| ACE2 | RPL21 | -0,7092 | 6,76161E-30 |
| ACE2 | USP31 | -0,7092 | 6,84797E-30 |
| ACE2 | MPHOSPH6 | -0,7092 | 6,8273E-30 |
| ACE2 | SMIM1 | -0,7093 | 6,5632E-30 |
| ACE2 | HOPX | -0,7093 | 6,56913E-30 |
| ACE2 | CAMLG | -0,7093 | 6,53402E-30 |
| ACE2 | PEG10 | -0,7093 | 6,59696E-30 |
| ACE2 | DSEL | -0,7093 | 6,67338E-30 |
| ACE2 | N4BP2L2 | -0,7094 | 6,4841E-30 |

| | | | |
|---|---|---|---|
| ACE2 | FBLN1 | -0,7094 | 6,35037E-30 |
| ACE2 | TSPAN2 | -0,7095 | 6,16879E-30 |
| ACE2 | ESR1 | -0,7095 | 6,29083E-30 |
| ACE2 | XPO7 | -0,7095 | 6,23048E-30 |
| ACE2 | ITPR1-AS1 | -0,7096 | 6,02362E-30 |
| ACE2 | ZNF777 | -0,7096 | 6,12479E-30 |
| ACE2 | DOCK9 | -0,7096 | 6,03634E-30 |
| ACE2 | TRIM9 | -0,7097 | 5,93993E-30 |
| ACE2 | FAM20A | -0,7097 | 5,93072E-30 |
| ACE2 | NCAM1 | -0,7098 | 5,72871E-30 |
| ACE2 | TECTA | -0,7098 | 5,80522E-30 |
| ACE2 | UBE2D4 | -0,7099 | 5,60331E-30 |
| ACE2 | MAP2K5 | -0,7099 | 5,55973E-30 |
| ACE2 | ZC3H12B | -0,7099 | 5,6683E-30 |
| ACE2 | TFB1M | -0,71 | 5,47297E-30 |
| ACE2 | FGF14-AS2 | -0,71 | 5,45087E-30 |
| ACE2 | MRPL40 | -0,71 | 5,44089E-30 |
| ACE2 | RPS8 | -0,7101 | 5,30481E-30 |
| ACE2 | MAGI2 | -0,7101 | 5,31092E-30 |
| ACE2 | CIZ1 | -0,7101 | 5,29944E-30 |
| ACE2 | CDH24 | -0,7101 | 5,28289E-30 |
| ACE2 | VAT1 | -0,7101 | 5,31008E-30 |
| ACE2 | PSMD4 | -0,7102 | 5,18846E-30 |
| ACE2 | AC005618.6 | -0,7102 | 5,15071E-30 |
| ACE2 | ZNF793 | -0,7102 | 5,15359E-30 |
| ACE2 | TMX4 | -0,7102 | 5,1491E-30 |
| ACE2 | EOGT | -0,7103 | 5,11592E-30 |
| ACE2 | ANXA7 | -0,7103 | 4,99632E-30 |
| ACE2 | NGFR | -0,7103 | 5,10673E-30 |
| ACE2 | ABCG4 | -0,7104 | 4,97279E-30 |
| ACE2 | IGDCC3 | -0,7104 | 4,94527E-30 |
| ACE2 | HDAC4 | -0,7105 | 4,7459E-30 |
| ACE2 | VIPR2 | -0,7105 | 4,78851E-30 |
| ACE2 | RAB3A | -0,7105 | 4,77563E-30 |
| ACE2 | C8orf37 | -0,7106 | 4,64033E-30 |
| ACE2 | NHP2 | -0,7107 | 4,56439E-30 |
| ACE2 | ZNF618 | -0,7107 | 4,5217E-30 |
| ACE2 | ZFYVE21 | -0,7107 | 4,57295E-30 |
| ACE2 | WDR61 | -0,7107 | 4,52835E-30 |
| ACE2 | FAM207A | -0,7107 | 4,56432E-30 |
| ACE2 | FAM163A | -0,7108 | 4,36274E-30 |
| ACE2 | SLC4A3 | -0,7108 | 4,40079E-30 |
| ACE2 | RP11-119F7.5 | -0,7108 | 4,48049E-30 |
| ACE2 | STYX | -0,7108 | 4,43212E-30 |
| ACE2 | PCDHB16 | -0,711 | 4,1925E-30 |
| ACE2 | PCDHGB7 | -0,711 | 4,21032E-30 |
| ACE2 | GOLT1B | -0,711 | 4,20118E-30 |
| ACE2 | SRGAP2 | -0,7111 | 4,09654E-30 |
| ACE2 | MIPEPP3 | -0,7111 | 4,03198E-30 |
| ACE2 | EBPL | -0,7111 | 4,04572E-30 |
| ACE2 | RP11-53O19.1 | -0,7112 | 3,93039E-30 |
| ACE2 | BHLHE22 | -0,7112 | 3,94683E-30 |
| ACE2 | POGK | -0,7113 | 3,8181E-30 |
| ACE2 | ARL8A | -0,7113 | 3,91491E-30 |

| | | | |
|---|---|---|---|
| ACE2 | C16orf62 | -0,7113 | 3,87602E-30 |
| ACE2 | FABP3 | -0,7114 | 3,79718E-30 |
| ACE2 | HACD1 | -0,7115 | 3,70148E-30 |
| ACE2 | PSMG2 | -0,7115 | 3,6603E-30 |
| ACE2 | L3MBTL2 | -0,7115 | 3,65549E-30 |
| ACE2 | TPBGL | -0,7116 | 3,58843E-30 |
| ACE2 | POLI | -0,7116 | 3,58931E-30 |
| ACE2 | SPACA6P-AS | -0,7116 | 3,54108E-30 |
| ACE2 | CBX2 | -0,7117 | 3,44307E-30 |
| ACE2 | MDFIC | -0,7118 | 3,38878E-30 |
| ACE2 | BRAF | -0,7118 | 3,35133E-30 |
| ACE2 | AOC3 | -0,7118 | 3,35876E-30 |
| ACE2 | PCDHGC3 | -0,7119 | 3,31346E-30 |
| ACE2 | ZBED8 | -0,7119 | 3,32514E-30 |
| ACE2 | C3 | -0,7119 | 3,28849E-30 |
| ACE2 | PCDHB10 | -0,712 | 3,24533E-30 |
| ACE2 | FJX1 | -0,712 | 3,24641E-30 |
| ACE2 | AC156455.1 | -0,712 | 3,23626E-30 |
| ACE2 | NUTF2 | -0,712 | 3,22875E-30 |
| ACE2 | DEFB124 | -0,712 | 3,19703E-30 |
| ACE2 | KLF8 | -0,712 | 3,18535E-30 |
| ACE2 | ATRIP | -0,7121 | 3,12239E-30 |
| ACE2 | U2SURP | -0,7121 | 3,16211E-30 |
| ACE2 | C4orf3 | -0,7121 | 3,12982E-30 |
| ACE2 | CTD-3035K23.3 | -0,7121 | 3,08213E-30 |
| ACE2 | ADH1B | -0,7122 | 3,04437E-30 |
| ACE2 | FAT4 | -0,7122 | 3,07075E-30 |
| ACE2 | CCDC59 | -0,7122 | 3,07168E-30 |
| ACE2 | RP11-89K11.1 | -0,7122 | 3,05188E-30 |
| ACE2 | MECR | -0,7123 | 2,91841E-30 |
| ACE2 | STON1 | -0,7123 | 2,95318E-30 |
| ACE2 | EXOSC1 | -0,7123 | 2,93254E-30 |
| ACE2 | LOXL1-AS1 | -0,7123 | 2,98878E-30 |
| ACE2 | ATL3 | -0,7124 | 2,84828E-30 |
| ACE2 | SAE1 | -0,7124 | 2,88024E-30 |
| ACE2 | ADAMTS9-AS2 | -0,7125 | 2,82277E-30 |
| ACE2 | SMARCE1 | -0,7125 | 2,8104E-30 |
| ACE2 | USHBP1 | -0,7125 | 2,80926E-30 |
| ACE2 | CSRNP3 | -0,7126 | 2,70872E-30 |
| ACE2 | LINC00882 | -0,7126 | 2,71049E-30 |
| ACE2 | ICAM2 | -0,7126 | 2,73914E-30 |
| ACE2 | WDCP | -0,7127 | 2,64517E-30 |
| ACE2 | LMCD1 | -0,7127 | 2,64868E-30 |
| ACE2 | ZNF174 | -0,7127 | 2,68819E-30 |
| ACE2 | ILF3-AS1 | -0,7127 | 2,67367E-30 |
| ACE2 | ATN1 | -0,7128 | 2,59013E-30 |
| ACE2 | ZNF771 | -0,7128 | 2,6004E-30 |
| ACE2 | ZNF83 | -0,7128 | 2,60407E-30 |
| ACE2 | RP11-696N14.1 | -0,7129 | 2,54906E-30 |
| ACE2 | RP1-142L7.9 | -0,713 | 2,44085E-30 |
| ACE2 | VAMP5 | -0,7131 | 2,41347E-30 |
| ACE2 | THOC1 | -0,7131 | 2,4056E-30 |
| ACE2 | BABAM1 | -0,7131 | 2,39312E-30 |
| ACE2 | TMEM91 | -0,7131 | 2,37491E-30 |

| | | | |
|---|---|---|---|
| ACE2 | STMN3 | -0,7131 | 2,39956E-30 |
| ACE2 | CCDC80 | -0,7132 | 2,33646E-30 |
| ACE2 | DCAF10 | -0,7132 | 2,29415E-30 |
| ACE2 | RIMKLB | -0,7132 | 2,3165E-30 |
| ACE2 | MAGOHB | -0,7132 | 2,30697E-30 |
| ACE2 | ARL9 | -0,7133 | 2,23734E-30 |
| ACE2 | CASC10 | -0,7133 | 2,28003E-30 |
| ACE2 | GPR162 | -0,7133 | 2,26491E-30 |
| ACE2 | RP11-454H19.2 | -0,7134 | 2,19551E-30 |
| ACE2 | PHKG1 | -0,7135 | 2,12787E-30 |
| ACE2 | SLC27A5 | -0,7135 | 2,15285E-30 |
| ACE2 | MRPS21 | -0,7136 | 2,11019E-30 |
| ACE2 | FOCAD | -0,7137 | 2,02788E-30 |
| ACE2 | KARS | -0,7137 | 2,00553E-30 |
| ACE2 | RP11-154D6.1 | -0,7138 | 1,98801E-30 |
| ACE2 | ACSS3 | -0,7138 | 1,95147E-30 |
| ACE2 | MIB1 | -0,7138 | 1,98079E-30 |
| ACE2 | ZNF649 | -0,7138 | 1,94974E-30 |
| ACE2 | RNF216P1 | -0,7139 | 1,90096E-30 |
| ACE2 | RPL19 | -0,7139 | 1,8956E-30 |
| ACE2 | PIFO | -0,714 | 1,88433E-30 |
| ACE2 | WDR70 | -0,714 | 1,86778E-30 |
| ACE2 | MRPL38 | -0,714 | 1,85549E-30 |
| ACE2 | CHM | -0,714 | 1,8731E-30 |
| ACE2 | MARCH2 | -0,7141 | 1,82261E-30 |
| ACE2 | CEBPB-AS1 | -0,7141 | 1,80275E-30 |
| ACE2 | CDKL2 | -0,7142 | 1,78706E-30 |
| ACE2 | RP11-366M4.11 | -0,7142 | 1,75039E-30 |
| ACE2 | GPR85 | -0,7142 | 1,76947E-30 |
| ACE2 | AIP | -0,7142 | 1,76959E-30 |
| ACE2 | PNO1 | -0,7143 | 1,70774E-30 |
| ACE2 | MRFAP1 | -0,7143 | 1,74533E-30 |
| ACE2 | KHDC1 | -0,7143 | 1,70874E-30 |
| ACE2 | RBM28 | -0,7143 | 1,71449E-30 |
| ACE2 | ATP6V0E2 | -0,7143 | 1,70271E-30 |
| ACE2 | RPS17 | -0,7143 | 1,71184E-30 |
| ACE2 | SF3A3 | -0,7144 | 1,67125E-30 |
| ACE2 | TOMM20 | -0,7144 | 1,6798E-30 |
| ACE2 | DHX30 | -0,7144 | 1,67309E-30 |
| ACE2 | VGLL3 | -0,7144 | 1,69269E-30 |
| ACE2 | CD27-AS1 | -0,7144 | 1,66576E-30 |
| ACE2 | DENND4A | -0,7144 | 1,68635E-30 |
| ACE2 | CHD6 | -0,7144 | 1,68776E-30 |
| ACE2 | FMNL2 | -0,7145 | 1,61063E-30 |
| ACE2 | HBS1L | -0,7145 | 1,65254E-30 |
| ACE2 | ZNF229 | -0,7145 | 1,6359E-30 |
| ACE2 | SMYD5 | -0,7146 | 1,58482E-30 |
| ACE2 | CTA-14H9.5 | -0,7146 | 1,57397E-30 |
| ACE2 | MKX | -0,7146 | 1,56885E-30 |
| ACE2 | FRA10AC1 | -0,7146 | 1,58352E-30 |
| ACE2 | NUFIP1 | -0,7146 | 1,59577E-30 |
| ACE2 | EIF2S1 | -0,7146 | 1,60063E-30 |
| ACE2 | RP11-210M15.2 | -0,7146 | 1,58722E-30 |
| ACE2 | ZNF280C | -0,7146 | 1,57405E-30 |

| | | | |
|---|---|---|---|
| ACE2 | ATXN1 | -0,7148 | 1,49814E-30 |
| ACE2 | NRXN3 | -0,7148 | 1,49939E-30 |
| ACE2 | MAPRE1 | -0,7148 | 1,5045E-30 |
| ACE2 | PCDHB9 | -0,7149 | 1,47355E-30 |
| ACE2 | TAGLN | -0,7149 | 1,44987E-30 |
| ACE2 | CORO1C | -0,7149 | 1,45941E-30 |
| ACE2 | ARIH1 | -0,7149 | 1,47166E-30 |
| ACE2 | CTD-2536I1.2 | -0,7149 | 1,48292E-30 |
| ACE2 | CTU2 | -0,7149 | 1,47515E-30 |
| ACE2 | FAM212B | -0,715 | 1,43215E-30 |
| ACE2 | TNFSF4 | -0,715 | 1,44037E-30 |
| ACE2 | CDYL | -0,715 | 1,43286E-30 |
| ACE2 | C12orf4 | -0,715 | 1,44338E-30 |
| ACE2 | KBTBD6 | -0,7151 | 1,37836E-30 |
| ACE2 | CWC22 | -0,7152 | 1,35455E-30 |
| ACE2 | CHSY3 | -0,7152 | 1,35109E-30 |
| ACE2 | ANAPC16 | -0,7152 | 1,35672E-30 |
| ACE2 | C1orf109 | -0,7153 | 1,29882E-30 |
| ACE2 | P3H2 | -0,7155 | 1,23595E-30 |
| ACE2 | CETN2 | -0,7155 | 1,23542E-30 |
| ACE2 | IAH1 | -0,7156 | 1,20023E-30 |
| ACE2 | KIF5A | -0,7156 | 1,20761E-30 |
| ACE2 | RYR3 | -0,7156 | 1,20824E-30 |
| ACE2 | LRCH2 | -0,7156 | 1,20681E-30 |
| ACE2 | EIF1B | -0,7157 | 1,19168E-30 |
| ACE2 | LZTS1 | -0,7157 | 1,1757E-30 |
| ACE2 | BICD1 | -0,7157 | 1,1762E-30 |
| ACE2 | KCTD2 | -0,7158 | 1,1379E-30 |
| ACE2 | SUZ12P1 | -0,7159 | 1,12386E-30 |
| ACE2 | PREPL | -0,716 | 1,07454E-30 |
| ACE2 | RSRC1 | -0,716 | 1,09764E-30 |
| ACE2 | TDRD10 | -0,7161 | 1,06636E-30 |
| ACE2 | ZNF662 | -0,7161 | 1,05298E-30 |
| ACE2 | RNH1 | -0,7161 | 1,04422E-30 |
| ACE2 | STRN3 | -0,7161 | 1,04241E-30 |
| ACE2 | RALBP1 | -0,7161 | 1,04904E-30 |
| ACE2 | NOTCH4 | -0,7162 | 1,01894E-30 |
| ACE2 | RPS20P22 | -0,7162 | 1,02803E-30 |
| ACE2 | P2RX7 | -0,7162 | 1,02805E-30 |
| ACE2 | TPGS2 | -0,7162 | 1,03119E-30 |
| ACE2 | ZNF445 | -0,7163 | 9,87807E-31 |
| ACE2 | RBP1 | -0,7163 | 1,0001E-30 |
| ACE2 | ZCRB1 | -0,7163 | 9,99671E-31 |
| ACE2 | RP11-324L17.1 | -0,7164 | 9,85807E-31 |
| ACE2 | MAB21L2 | -0,7164 | 9,77113E-31 |
| ACE2 | FBXW4 | -0,7164 | 9,62019E-31 |
| ACE2 | HSPA1A | -0,7165 | 9,37583E-31 |
| ACE2 | AGAP3 | -0,7165 | 9,49452E-31 |
| ACE2 | TMEM200C | -0,7165 | 9,34008E-31 |
| ACE2 | ANO2 | -0,7166 | 9,16027E-31 |
| ACE2 | ZMYM5 | -0,7166 | 9,24501E-31 |
| ACE2 | HS3ST3A1 | -0,7166 | 9,2412E-31 |
| ACE2 | LSM14A | -0,7166 | 9,22329E-31 |
| ACE2 | DPY19L1 | -0,7167 | 9,06034E-31 |

| | | | |
|---|---|---|---|
| ACE2 | ZNF331 | -0,7167 | 8,98839E-31 |
| ACE2 | ZBTB41 | -0,7168 | 8,72718E-31 |
| ACE2 | RP11-545I5.3 | -0,7168 | 8,74837E-31 |
| ACE2 | ZC3HC1 | -0,7168 | 8,8094E-31 |
| ACE2 | CCND2 | -0,7168 | 8,71135E-31 |
| ACE2 | TIGD7 | -0,7168 | 8,74617E-31 |
| ACE2 | FILIP1 | -0,7169 | 8,55361E-31 |
| ACE2 | LDLRAD3 | -0,7169 | 8,51359E-31 |
| ACE2 | AP000662.4 | -0,7169 | 8,53165E-31 |
| ACE2 | RPL26 | -0,7169 | 8,53319E-31 |
| ACE2 | PIAS2 | -0,717 | 8,24666E-31 |
| ACE2 | TIMMDC1 | -0,7171 | 8,11295E-31 |
| ACE2 | ZNF286B | -0,7171 | 7,98123E-31 |
| ACE2 | WIZ | -0,7171 | 7,94754E-31 |
| ACE2 | RP11-815I9.4 | -0,7172 | 7,79572E-31 |
| ACE2 | MAGEL2 | -0,7173 | 7,67735E-31 |
| ACE2 | RNF212 | -0,7174 | 7,40619E-31 |
| ACE2 | CNTROB | -0,7174 | 7,31575E-31 |
| ACE2 | GATA2 | -0,7175 | 7,10555E-31 |
| ACE2 | CEP290 | -0,7175 | 7,24676E-31 |
| ACE2 | PKD1P6 | -0,7175 | 7,22771E-31 |
| ACE2 | ST6GALNAC5 | -0,7177 | 6,88265E-31 |
| ACE2 | ZNF835 | -0,7177 | 6,8855E-31 |
| ACE2 | MAF1 | -0,7178 | 6,57108E-31 |
| ACE2 | SNAPC1 | -0,7178 | 6,65533E-31 |
| ACE2 | PRR16 | -0,7179 | 6,47607E-31 |
| ACE2 | NRM | -0,7179 | 6,51299E-31 |
| ACE2 | HAR1A | -0,7179 | 6,48262E-31 |
| ACE2 | SYNCRIP | -0,718 | 6,31741E-31 |
| ACE2 | IMPDH2 | -0,7181 | 6,03028E-31 |
| ACE2 | CCDC146 | -0,7181 | 6,03823E-31 |
| ACE2 | LRSAM1 | -0,7181 | 6,09734E-31 |
| ACE2 | COPS3 | -0,7181 | 6,13968E-31 |
| ACE2 | LYPD5 | -0,7182 | 6,00355E-31 |
| ACE2 | SYN1 | -0,7182 | 5,95008E-31 |
| ACE2 | NT5DC2 | -0,7183 | 5,82251E-31 |
| ACE2 | GPLD1 | -0,7183 | 5,80097E-31 |
| ACE2 | RPL7A | -0,7183 | 5,80862E-31 |
| ACE2 | SPAG7 | -0,7183 | 5,72943E-31 |
| ACE2 | THOC5 | -0,7183 | 5,70948E-31 |
| ACE2 | YWHAQ | -0,7184 | 5,65051E-31 |
| ACE2 | AHR | -0,7184 | 5,68938E-31 |
| ACE2 | SCUBE2 | -0,7184 | 5,63735E-31 |
| ACE2 | HSPA12B | -0,7184 | 5,54901E-31 |
| ACE2 | F8 | -0,7184 | 5,60359E-31 |
| ACE2 | CUL1 | -0,7185 | 5,42519E-31 |
| ACE2 | RP11-728F11.4 | -0,7185 | 5,45059E-31 |
| ACE2 | SOX18 | -0,7185 | 5,53647E-31 |
| ACE2 | GATSL3 | -0,7185 | 5,5075E-31 |
| ACE2 | CYP39A1 | -0,7186 | 5,34641E-31 |
| ACE2 | DHODH | -0,7186 | 5,35921E-31 |
| ACE2 | SMTNL2 | -0,7186 | 5,30294E-31 |
| ACE2 | RP11-147L13.11 | -0,7186 | 5,35639E-31 |
| ACE2 | AC005682.6 | -0,7187 | 5,11411E-31 |

| | | | |
|---|---|---|---|
| ACE2 | MYO5A | -0,7187 | 5,12319E-31 |
| ACE2 | TTYH1 | -0,7187 | 5,23748E-31 |
| ACE2 | ZBTB46 | -0,7187 | 5,21675E-31 |
| ACE2 | PDZD2 | -0,7188 | 5,07019E-31 |
| ACE2 | BHLHB9 | -0,7189 | 4,89553E-31 |
| ACE2 | CLCC1 | -0,719 | 4,81263E-31 |
| ACE2 | TOP2B | -0,719 | 4,70322E-31 |
| ACE2 | HNRNPDL | -0,719 | 4,7958E-31 |
| ACE2 | BTNL9 | -0,719 | 4,82129E-31 |
| ACE2 | ITGAV | -0,7191 | 4,6431E-31 |
| ACE2 | FAF1 | -0,7192 | 4,43896E-31 |
| ACE2 | RNF8 | -0,7192 | 4,44641E-31 |
| ACE2 | PNPLA8 | -0,7192 | 4,52811E-31 |
| ACE2 | DOK6 | -0,7192 | 4,54623E-31 |
| ACE2 | TPTEP1 | -0,7192 | 4,4824E-31 |
| ACE2 | ARHGEF6 | -0,7193 | 4,3787E-31 |
| ACE2 | SAMD4A | -0,7195 | 4,12357E-31 |
| ACE2 | SCMH1 | -0,7196 | 4,0219E-31 |
| ACE2 | TCOF1 | -0,7196 | 4,02198E-31 |
| ACE2 | MAPK8IP1 | -0,7196 | 4,0299E-31 |
| ACE2 | ITGA5 | -0,7196 | 3,99748E-31 |
| ACE2 | ARL6IP4 | -0,7196 | 4,02074E-31 |
| ACE2 | RCN1 | -0,7198 | 3,83164E-31 |
| ACE2 | EIF3M | -0,7198 | 3,83434E-31 |
| ACE2 | MAB21L1 | -0,7198 | 3,86273E-31 |
| ACE2 | ZNF880 | -0,7198 | 3,85585E-31 |
| ACE2 | CD93 | -0,7198 | 3,82746E-31 |
| ACE2 | BSCL2 | -0,7199 | 3,75719E-31 |
| ACE2 | C12orf73 | -0,7199 | 3,70496E-31 |
| ACE2 | BCL9 | -0,72 | 3,62605E-31 |
| ACE2 | ENG | -0,72 | 3,62646E-31 |
| ACE2 | UBQLN4 | -0,7201 | 3,46646E-31 |
| ACE2 | RP11-532F6.3 | -0,7201 | 3,46636E-31 |
| ACE2 | PLD6 | -0,7201 | 3,47521E-31 |
| ACE2 | CBWD2 | -0,7202 | 3,39176E-31 |
| ACE2 | DOC2GP | -0,7202 | 3,36915E-31 |
| ACE2 | PRSS23 | -0,7202 | 3,37783E-31 |
| ACE2 | NFIC | -0,7202 | 3,36375E-31 |
| ACE2 | PTGS2 | -0,7203 | 3,34095E-31 |
| ACE2 | TMEM161A | -0,7203 | 3,27111E-31 |
| ACE2 | GLT8D1 | -0,7205 | 3,1214E-31 |
| ACE2 | WWP1 | -0,7205 | 3,09388E-31 |
| ACE2 | OSER1-AS1 | -0,7205 | 3,10208E-31 |
| ACE2 | SNX3 | -0,7206 | 3,07125E-31 |
| ACE2 | GPR146 | -0,7207 | 2,95421E-31 |
| ACE2 | ZNF571 | -0,7207 | 2,96964E-31 |
| ACE2 | KXD1 | -0,7208 | 2,87436E-31 |
| ACE2 | ADA | -0,7208 | 2,86807E-31 |
| ACE2 | ADRA2B | -0,7209 | 2,78227E-31 |
| ACE2 | TRIM23 | -0,7209 | 2,79917E-31 |
| ACE2 | GABARAPL2 | -0,721 | 2,75541E-31 |
| ACE2 | KLHL10 | -0,721 | 2,73189E-31 |
| ACE2 | TIMM21 | -0,721 | 2,71028E-31 |
| ACE2 | REEP2 | -0,7211 | 2,65412E-31 |

| | | | |
|---|---|---|---|
| ACE2 | FBXL13 | -0,7211 | 2,63417E-31 |
| ACE2 | PODXL | -0,7211 | 2,65845E-31 |
| ACE2 | IFFO1 | -0,7211 | 2,64326E-31 |
| ACE2 | RP11-834C11.15 | -0,7211 | 2,63977E-31 |
| ACE2 | KLHL4 | -0,7211 | 2,63115E-31 |
| ACE2 | RP11-244O19.1 | -0,7212 | 2,55711E-31 |
| ACE2 | NAA25 | -0,7212 | 2,56559E-31 |
| ACE2 | POLR3H | -0,7212 | 2,57249E-31 |
| ACE2 | STOM | -0,7213 | 2,51373E-31 |
| ACE2 | MYO9A | -0,7213 | 2,48981E-31 |
| ACE2 | ARMCX6 | -0,7213 | 2,47752E-31 |
| ACE2 | EFEMP1 | -0,7214 | 2,46611E-31 |
| ACE2 | COL9A3 | -0,7215 | 2,34168E-31 |
| ACE2 | C21orf59 | -0,7215 | 2,35092E-31 |
| ACE2 | COL18A1 | -0,7215 | 2,3915E-31 |
| ACE2 | EEF1E1 | -0,7216 | 2,33165E-31 |
| ACE2 | AC093673.5 | -0,7217 | 2,26495E-31 |
| ACE2 | RALYL | -0,7217 | 2,24097E-31 |
| ACE2 | PTGS1 | -0,7217 | 2,26284E-31 |
| ACE2 | ENAH | -0,7218 | 2,17484E-31 |
| ACE2 | ADPRH | -0,7218 | 2,18513E-31 |
| ACE2 | RING1 | -0,7218 | 2,20807E-31 |
| ACE2 | UNC45A | -0,7218 | 2,18661E-31 |
| ACE2 | RNF216 | -0,7219 | 2,14967E-31 |
| ACE2 | PFN4 | -0,722 | 2,05271E-31 |
| ACE2 | AC093838.4 | -0,722 | 2,06958E-31 |
| ACE2 | STK38L | -0,722 | 2,05072E-31 |
| ACE2 | CHRNB4 | -0,722 | 2,08527E-31 |
| ACE2 | TMEM205 | -0,722 | 2,08503E-31 |
| ACE2 | RP11-284E5.1 | -0,7221 | 2,02927E-31 |
| ACE2 | MAP2 | -0,7221 | 2,01767E-31 |
| ACE2 | RPL32 | -0,7221 | 1,99089E-31 |
| ACE2 | LRRC34 | -0,7221 | 1,99752E-31 |
| ACE2 | SNCAIP | -0,7221 | 1,97815E-31 |
| ACE2 | NOD1 | -0,7221 | 1,98731E-31 |
| ACE2 | RP11-182L21.6 | -0,7221 | 1,98244E-31 |
| ACE2 | FAM155A | -0,7221 | 2,01371E-31 |
| ACE2 | TACR1 | -0,7222 | 1,96376E-31 |
| ACE2 | DYNC1LI1 | -0,7222 | 1,967E-31 |
| ACE2 | THG1L | -0,7222 | 1,95465E-31 |
| ACE2 | KLHL15 | -0,7222 | 1,92992E-31 |
| ACE2 | ZNF501 | -0,7223 | 1,88963E-31 |
| ACE2 | RP11-208N14.5 | -0,7223 | 1,91821E-31 |
| ACE2 | ZFHX4 | -0,7223 | 1,91627E-31 |
| ACE2 | UPF2 | -0,7223 | 1,91983E-31 |
| ACE2 | NKX2-3 | -0,7223 | 1,87684E-31 |
| ACE2 | RP11-158I9.5 | -0,7223 | 1,87776E-31 |
| ACE2 | AGTRAP | -0,7224 | 1,82361E-31 |
| ACE2 | PPFIA4 | -0,7224 | 1,8589E-31 |
| ACE2 | SLC16A14 | -0,7224 | 1,85711E-31 |
| ACE2 | CEND1 | -0,7224 | 1,81798E-31 |
| ACE2 | HOXC-AS2 | -0,7224 | 1,85886E-31 |
| ACE2 | BCL6B | -0,7225 | 1,78103E-31 |
| ACE2 | PRX | -0,7225 | 1,80089E-31 |

| | | | |
|---|---|---|---|
| ACE2 | NIT2 | -0,7226 | 1,72134E-31 |
| ACE2 | HMGB3P22 | -0,7226 | 1,74619E-31 |
| ACE2 | NOL7 | -0,7226 | 1,73744E-31 |
| ACE2 | DNAJB6 | -0,7226 | 1,74124E-31 |
| ACE2 | KIAA0355 | -0,7226 | 1,75016E-31 |
| ACE2 | HOXD-AS2 | -0,7227 | 1,69203E-31 |
| ACE2 | RANBP3L | -0,7227 | 1,69454E-31 |
| ACE2 | FZD4 | -0,7227 | 1,6859E-31 |
| ACE2 | COPS2 | -0,7227 | 1,71646E-31 |
| ACE2 | HDAC5 | -0,7227 | 1,67761E-31 |
| ACE2 | ZNF582-AS1 | -0,7227 | 1,69565E-31 |
| ACE2 | FAM20B | -0,7228 | 1,64797E-31 |
| ACE2 | ICA1L | -0,7228 | 1,64498E-31 |
| ACE2 | CLCF1 | -0,7228 | 1,6512E-31 |
| ACE2 | ARSJ | -0,7229 | 1,60553E-31 |
| ACE2 | SRRM3 | -0,7229 | 1,62049E-31 |
| ACE2 | TMTC1 | -0,7229 | 1,61532E-31 |
| ACE2 | ITPKB | -0,723 | 1,54701E-31 |
| ACE2 | SMIM13 | -0,723 | 1,54381E-31 |
| ACE2 | ODF2 | -0,723 | 1,57367E-31 |
| ACE2 | TSR1 | -0,723 | 1,56459E-31 |
| ACE2 | CCM2L | -0,723 | 1,54196E-31 |
| ACE2 | RBM11 | -0,7231 | 1,53481E-31 |
| ACE2 | FSTL1 | -0,7232 | 1,45168E-31 |
| ACE2 | RP11-177H2.1 | -0,7232 | 1,48974E-31 |
| ACE2 | TMEM204 | -0,7232 | 1,47057E-31 |
| ACE2 | RPS16 | -0,7232 | 1,46499E-31 |
| ACE2 | HTATSF1 | -0,7232 | 1,46978E-31 |
| ACE2 | FAM178B | -0,7233 | 1,45103E-31 |
| ACE2 | PIP5KL1 | -0,7233 | 1,45052E-31 |
| ACE2 | LINC00908 | -0,7233 | 1,43974E-31 |
| ACE2 | C12orf45 | -0,7234 | 1,39674E-31 |
| ACE2 | HNRNPA3 | -0,7235 | 1,3389E-31 |
| ACE2 | MAST4 | -0,7235 | 1,34104E-31 |
| ACE2 | E2F6 | -0,7236 | 1,32557E-31 |
| ACE2 | SLBP | -0,7236 | 1,30262E-31 |
| ACE2 | PHF24 | -0,7236 | 1,32606E-31 |
| ACE2 | SPOP | -0,7237 | 1,27482E-31 |
| ACE2 | POLR2F | -0,7237 | 1,27419E-31 |
| ACE2 | AFAP1 | -0,7238 | 1,22961E-31 |
| ACE2 | SPCS3 | -0,7238 | 1,25897E-31 |
| ACE2 | CDK5RAP2 | -0,7238 | 1,25036E-31 |
| ACE2 | MSRB3 | -0,7238 | 1,25888E-31 |
| ACE2 | ATXN3 | -0,7238 | 1,24328E-31 |
| ACE2 | EIF3J | -0,7238 | 1,23568E-31 |
| ACE2 | CTD-3035K23.7 | -0,7238 | 1,24566E-31 |
| ACE2 | RP11-1334A24.5 | -0,7239 | 1,19886E-31 |
| ACE2 | SLAIN1 | -0,7239 | 1,20999E-31 |
| ACE2 | ALKBH5 | -0,7239 | 1,20248E-31 |
| ACE2 | CCDC8 | -0,7239 | 1,20234E-31 |
| ACE2 | POLR3G | -0,724 | 1,18575E-31 |
| ACE2 | MRPL18 | -0,724 | 1,18075E-31 |
| ACE2 | IGF1R | -0,724 | 1,16864E-31 |
| ACE2 | RAB13 | -0,7241 | 1,13261E-31 |

| | | | |
|---|---|---|---|
| ACE2 | SPEF2 | -0,7241 | 1,13967E-31 |
| ACE2 | ATAD1 | -0,7241 | 1,15591E-31 |
| ACE2 | MRGPRF-AS1 | -0,7241 | 1,13114E-31 |
| ACE2 | ZNF471 | -0,7241 | 1,13977E-31 |
| ACE2 | NTNG2 | -0,7243 | 1,09397E-31 |
| ACE2 | MORC4 | -0,7243 | 1,09281E-31 |
| ACE2 | VPS72 | -0,7244 | 1,03624E-31 |
| ACE2 | NAA15 | -0,7244 | 1,05845E-31 |
| ACE2 | SHC4 | -0,7244 | 1,05416E-31 |
| ACE2 | HOMER2 | -0,7244 | 1,06138E-31 |
| ACE2 | ZNF48 | -0,7244 | 1,03446E-31 |
| ACE2 | SLFN12 | -0,7244 | 1,04469E-31 |
| ACE2 | ZNF382 | -0,7245 | 1,03102E-31 |
| ACE2 | CTA-212A2.1 | -0,7245 | 1,0191E-31 |
| ACE2 | CTD-3162L10.1 | -0,7246 | 9,82993E-32 |
| ACE2 | CCDC110 | -0,7247 | 9,64743E-32 |
| ACE2 | FRG1 | -0,7248 | 9,3396E-32 |
| ACE2 | IKBKAP | -0,7248 | 9,32489E-32 |
| ACE2 | SLC25A29 | -0,7248 | 9,29788E-32 |
| ACE2 | MEF2A | -0,7248 | 9,41129E-32 |
| ACE2 | FSIP1 | -0,7249 | 9,20564E-32 |
| ACE2 | COL6A3 | -0,725 | 8,86891E-32 |
| ACE2 | RFC1 | -0,725 | 8,85644E-32 |
| ACE2 | PCDHB5 | -0,725 | 8,85358E-32 |
| ACE2 | CALU | -0,725 | 8,93789E-32 |
| ACE2 | ABCA17P | -0,725 | 8,76091E-32 |
| ACE2 | RRAD | -0,7251 | 8,54209E-32 |
| ACE2 | APOL3 | -0,7251 | 8,7009E-32 |
| ACE2 | AGO3 | -0,7252 | 8,4725E-32 |
| ACE2 | LMLN | -0,7252 | 8,41602E-32 |
| ACE2 | TRMT44 | -0,7252 | 8,24892E-32 |
| ACE2 | NR3C1 | -0,7252 | 8,38375E-32 |
| ACE2 | PTGIR | -0,7252 | 8,34981E-32 |
| ACE2 | CCT4 | -0,7253 | 8,02726E-32 |
| ACE2 | FGF1 | -0,7253 | 8,0667E-32 |
| ACE2 | SGCD | -0,7253 | 8,11199E-32 |
| ACE2 | STEAP4 | -0,7253 | 8,04643E-32 |
| ACE2 | ZNF239 | -0,7253 | 8,03575E-32 |
| ACE2 | FAM216A | -0,7253 | 8,11912E-32 |
| ACE2 | BVES | -0,7254 | 7,79358E-32 |
| ACE2 | LMAN2L | -0,7256 | 7,37026E-32 |
| ACE2 | DNAJC19 | -0,7256 | 7,43579E-32 |
| ACE2 | RP11-350N15.4 | -0,7256 | 7,4059E-32 |
| ACE2 | IZUMO4 | -0,7256 | 7,4603E-32 |
| ACE2 | C1orf52 | -0,7257 | 7,25728E-32 |
| ACE2 | DPF2 | -0,7257 | 7,29481E-32 |
| ACE2 | RP11-162A12.4 | -0,7257 | 7,15194E-32 |
| ACE2 | RPL3 | -0,7257 | 7,18261E-32 |
| ACE2 | NTHL1 | -0,7258 | 6,9678E-32 |
| ACE2 | TTLL7 | -0,7259 | 6,80947E-32 |
| ACE2 | LRRC58 | -0,7259 | 6,75252E-32 |
| ACE2 | MED30 | -0,7259 | 6,90869E-32 |
| ACE2 | CD44 | -0,7259 | 6,74802E-32 |
| ACE2 | TRIB2 | -0,726 | 6,63313E-32 |

| | | | |
|---|---|---|---|
| ACE2 | NGDN | -0,726 | 6,62801E-32 |
| ACE2 | MRPS27 | -0,7261 | 6,41865E-32 |
| ACE2 | RASGEF1C | -0,7261 | 6,42706E-32 |
| ACE2 | OSCP1 | -0,7262 | 6,34236E-32 |
| ACE2 | RP11-31F15.2 | -0,7262 | 6,30756E-32 |
| ACE2 | THSD7A | -0,7262 | 6,2124E-32 |
| ACE2 | CEP41 | -0,7263 | 6,03639E-32 |
| ACE2 | BMPR2 | -0,7264 | 5,92743E-32 |
| ACE2 | OLFML1 | -0,7264 | 5,98066E-32 |
| ACE2 | BIN1 | -0,7265 | 5,68537E-32 |
| ACE2 | FHOD3 | -0,7265 | 5,80543E-32 |
| ACE2 | RP11-428J1.5 | -0,7266 | 5,66735E-32 |
| ACE2 | CC2D2A | -0,7267 | 5,50957E-32 |
| ACE2 | RP11-356J5.12 | -0,7267 | 5,3783E-32 |
| ACE2 | MED22 | -0,7268 | 5,32386E-32 |
| ACE2 | NSMCE4A | -0,7268 | 5,28458E-32 |
| ACE2 | HKR1 | -0,7268 | 5,31141E-32 |
| ACE2 | PTOV1-AS1 | -0,7268 | 5,30735E-32 |
| ACE2 | MAPK12 | -0,7268 | 5,25236E-32 |
| ACE2 | ARPP19 | -0,7269 | 5,07749E-32 |
| ACE2 | NDRG4 | -0,7269 | 5,10121E-32 |
| ACE2 | MARVELD1 | -0,727 | 5,00686E-32 |
| ACE2 | ATP1A3 | -0,727 | 5,0382E-32 |
| ACE2 | WDR53 | -0,7271 | 4,84722E-32 |
| ACE2 | PABPC4L | -0,7271 | 4,92635E-32 |
| ACE2 | RBMS2 | -0,7271 | 4,90378E-32 |
| ACE2 | DCN | -0,7271 | 4,86709E-32 |
| ACE2 | ZNF420 | -0,7271 | 4,85349E-32 |
| ACE2 | ZNF418 | -0,7271 | 4,86197E-32 |
| ACE2 | TTC3P1 | -0,7271 | 4,87525E-32 |
| ACE2 | STEAP1B | -0,7272 | 4,78043E-32 |
| ACE2 | AC004540.5 | -0,7272 | 4,65457E-32 |
| ACE2 | MINDY4 | -0,7272 | 4,74534E-32 |
| ACE2 | MXRA7 | -0,7272 | 4,6712E-32 |
| ACE2 | LINC00663 | -0,7272 | 4,74942E-32 |
| ACE2 | PCDH12 | -0,7273 | 4,62127E-32 |
| ACE2 | WASF3 | -0,7273 | 4,5949E-32 |
| ACE2 | EIF3B | -0,7274 | 4,44395E-32 |
| ACE2 | HSPB1 | -0,7274 | 4,50309E-32 |
| ACE2 | RP11-667K14.3 | -0,7274 | 4,44214E-32 |
| ACE2 | POGZ | -0,7275 | 4,37047E-32 |
| ACE2 | TSEN2 | -0,7276 | 4,24679E-32 |
| ACE2 | DOCK3 | -0,7276 | 4,18344E-32 |
| ACE2 | SKIDA1 | -0,7276 | 4,1966E-32 |
| ACE2 | ARL2 | -0,7276 | 4,15177E-32 |
| ACE2 | UXS1 | -0,7278 | 4,01141E-32 |
| ACE2 | AKAP12 | -0,7278 | 3,95863E-32 |
| ACE2 | TMOD1 | -0,7278 | 4,00352E-32 |
| ACE2 | AASDHPPT | -0,7278 | 3,94318E-32 |
| ACE2 | TCEAL8 | -0,7278 | 4,02143E-32 |
| ACE2 | ADD1 | -0,7279 | 3,89138E-32 |
| ACE2 | GPR137C | -0,7279 | 3,89821E-32 |
| ACE2 | ASTN1 | -0,728 | 3,79589E-32 |
| ACE2 | RP11-101E13.5 | -0,728 | 3,76946E-32 |

| | | | |
|---|---|---|---|
| ACE2 | ARID5A | -0,7281 | 3,66105E-32 |
| ACE2 | SOX5 | -0,7281 | 3,63808E-32 |
| ACE2 | THAP3 | -0,7282 | 3,52541E-32 |
| ACE2 | GIPC3 | -0,7282 | 3,57465E-32 |
| ACE2 | ADSL | -0,7282 | 3,52067E-32 |
| ACE2 | AP000892.6 | -0,7283 | 3,42318E-32 |
| ACE2 | EID3 | -0,7283 | 3,43315E-32 |
| ACE2 | ATAD3C | -0,7284 | 3,31356E-32 |
| ACE2 | ARRDC3 | -0,7284 | 3,30194E-32 |
| ACE2 | C6orf48 | -0,7284 | 3,33869E-32 |
| ACE2 | ACYP2 | -0,7285 | 3,23096E-32 |
| ACE2 | NOM1 | -0,7285 | 3,22296E-32 |
| ACE2 | ANKRD10 | -0,7286 | 3,12079E-32 |
| ACE2 | YY1AP1 | -0,7287 | 3,08157E-32 |
| ACE2 | NYAP1 | -0,7287 | 3,0535E-32 |
| ACE2 | TMEM218 | -0,7287 | 3,04754E-32 |
| ACE2 | PPP2R5E | -0,7287 | 3,09273E-32 |
| ACE2 | ACD | -0,7288 | 3,00373E-32 |
| ACE2 | MYOM1 | -0,7288 | 3,0205E-32 |
| ACE2 | IFT122 | -0,7289 | 2,85139E-32 |
| ACE2 | RPS23 | -0,7289 | 2,92401E-32 |
| ACE2 | LSM2 | -0,7289 | 2,91662E-32 |
| ACE2 | UBE2I | -0,7289 | 2,93051E-32 |
| ACE2 | PCDHGB4 | -0,729 | 2,83597E-32 |
| ACE2 | TBC1D20 | -0,729 | 2,7907E-32 |
| ACE2 | CADM3-AS1 | -0,7291 | 2,72872E-32 |
| ACE2 | COL5A1 | -0,7291 | 2,76804E-32 |
| ACE2 | NFKB2 | -0,7291 | 2,73616E-32 |
| ACE2 | KCNJ8 | -0,7292 | 2,68319E-32 |
| ACE2 | PACS2 | -0,7293 | 2,56337E-32 |
| ACE2 | ARHGEF40 | -0,7294 | 2,47033E-32 |
| ACE2 | ASB2 | -0,7294 | 2,49529E-32 |
| ACE2 | PPP1R3G | -0,7295 | 2,45515E-32 |
| ACE2 | FBXO30 | -0,7295 | 2,40307E-32 |
| ACE2 | TOMM7 | -0,7295 | 2,43771E-32 |
| ACE2 | LINC-PINT | -0,7295 | 2,46342E-32 |
| ACE2 | FUT11 | -0,7295 | 2,45355E-32 |
| ACE2 | USP14 | -0,7295 | 2,39935E-32 |
| ACE2 | IGFBP3 | -0,7296 | 2,35451E-32 |
| ACE2 | RPF1 | -0,7297 | 2,26939E-32 |
| ACE2 | PLEKHA3 | -0,7297 | 2,29874E-32 |
| ACE2 | RNF2 | -0,7298 | 2,21587E-32 |
| ACE2 | MRFAP1L1 | -0,7298 | 2,20676E-32 |
| ACE2 | MGP | -0,7299 | 2,13667E-32 |
| ACE2 | PARP2 | -0,7299 | 2,18703E-32 |
| ACE2 | NUP35 | -0,73 | 2,13149E-32 |
| ACE2 | SCN5A | -0,73 | 2,07438E-32 |
| ACE2 | WFS1 | -0,73 | 2,07715E-32 |
| ACE2 | ABCE1 | -0,73 | 2,0953E-32 |
| ACE2 | PAIP1 | -0,73 | 2,08022E-32 |
| ACE2 | PJA2 | -0,73 | 2,11396E-32 |
| ACE2 | CD109 | -0,73 | 2,07493E-32 |
| ACE2 | CYHR1 | -0,73 | 2,12454E-32 |
| ACE2 | MDP1 | -0,73 | 2,07747E-32 |

| | | | |
|---|---|---|---|
| ACE2 | DVL2 | -0,73 | 2,12073E-32 |
| ACE2 | PCDHB4 | -0,7301 | 2,0221E-32 |
| ACE2 | RPL12 | -0,7301 | 2,04746E-32 |
| ACE2 | ADAMTSL3 | -0,7301 | 2,07011E-32 |
| ACE2 | THUMPD1 | -0,7301 | 2,07151E-32 |
| ACE2 | C11orf96 | -0,7302 | 1,97428E-32 |
| ACE2 | DGKG | -0,7303 | 1,95271E-32 |
| ACE2 | MFAP3L | -0,7303 | 1,94538E-32 |
| ACE2 | DIXDC1 | -0,7303 | 1,93954E-32 |
| ACE2 | COX4I2 | -0,7303 | 1,92421E-32 |
| ACE2 | CETN3 | -0,7304 | 1,87785E-32 |
| ACE2 | KRR1 | -0,7304 | 1,86821E-32 |
| ACE2 | RP5-1085F17.3 | -0,7304 | 1,84998E-32 |
| ACE2 | ZNF354C | -0,7306 | 1,7906E-32 |
| ACE2 | GNB3 | -0,7306 | 1,74565E-32 |
| ACE2 | NHS | -0,7306 | 1,76449E-32 |
| ACE2 | CUL4B | -0,7306 | 1,74367E-32 |
| ACE2 | ACVR1 | -0,7307 | 1,72322E-32 |
| ACE2 | RP11-379H18.1 | -0,7307 | 1,73246E-32 |
| ACE2 | SNX21 | -0,7307 | 1,70071E-32 |
| ACE2 | DUSP12 | -0,7308 | 1,66123E-32 |
| ACE2 | POLD2 | -0,7308 | 1,65863E-32 |
| ACE2 | CAMTA1 | -0,7309 | 1,60362E-32 |
| ACE2 | PITHD1 | -0,7309 | 1,61215E-32 |
| ACE2 | PFDN2 | -0,7309 | 1,60047E-32 |
| ACE2 | UBE2W | -0,7309 | 1,597E-32 |
| ACE2 | EIF3F | -0,7309 | 1,6412E-32 |
| ACE2 | MRPL10 | -0,7309 | 1,60304E-32 |
| ACE2 | EVI5L | -0,7309 | 1,61494E-32 |
| ACE2 | BGLAP | -0,7311 | 1,54219E-32 |
| ACE2 | MARK3 | -0,7311 | 1,52375E-32 |
| ACE2 | VEZF1 | -0,7311 | 1,52279E-32 |
| ACE2 | PRPS1 | -0,7311 | 1,52926E-32 |
| ACE2 | QPCT | -0,7312 | 1,48059E-32 |
| ACE2 | PCDHGA4 | -0,7313 | 1,45814E-32 |
| ACE2 | SMIM8 | -0,7313 | 1,45551E-32 |
| ACE2 | DIS3L | -0,7313 | 1,4231E-32 |
| ACE2 | SVBP | -0,7314 | 1,41039E-32 |
| ACE2 | WDR92 | -0,7314 | 1,39831E-32 |
| ACE2 | NMD3 | -0,7314 | 1,39724E-32 |
| ACE2 | RPAP3 | -0,7314 | 1,38247E-32 |
| ACE2 | GRIP2 | -0,7315 | 1,36805E-32 |
| ACE2 | IQCK | -0,7315 | 1,36602E-32 |
| ACE2 | ZNF599 | -0,7315 | 1,36735E-32 |
| ACE2 | VEGFC | -0,7316 | 1,32855E-32 |
| ACE2 | ADAMTSL4 | -0,7317 | 1,27679E-32 |
| ACE2 | GCFC2 | -0,7317 | 1,29653E-32 |
| ACE2 | TPM2 | -0,7317 | 1,27179E-32 |
| ACE2 | CDS2 | -0,7317 | 1,29055E-32 |
| ACE2 | PWP1 | -0,7318 | 1,23783E-32 |
| ACE2 | PDE6D | -0,7319 | 1,19233E-32 |
| ACE2 | ZMAT3 | -0,7319 | 1,19939E-32 |
| ACE2 | RP11-536O18.1 | -0,7319 | 1,20186E-32 |
| ACE2 | RGS6 | -0,7319 | 1,20233E-32 |

| | | | |
|---|---|---|---|
| ACE2 | MGAT5B | -0,7319 | 1,19308E-32 |
| ACE2 | MED10 | -0,732 | 1,16341E-32 |
| ACE2 | FOXJ2 | -0,732 | 1,18717E-32 |
| ACE2 | IGSF1 | -0,732 | 1,1572E-32 |
| ACE2 | SYPL2 | -0,7321 | 1,1406E-32 |
| ACE2 | RGCC | -0,7321 | 1,13681E-32 |
| ACE2 | TRIM69 | -0,7321 | 1,12706E-32 |
| ACE2 | LRRC70 | -0,7322 | 1,09576E-32 |
| ACE2 | AC113189.5 | -0,7322 | 1,11756E-32 |
| ACE2 | USP51 | -0,7322 | 1,11819E-32 |
| ACE2 | PLCD4 | -0,7323 | 1,08278E-32 |
| ACE2 | WNT6 | -0,7323 | 1,05986E-32 |
| ACE2 | RPS20 | -0,7323 | 1,06543E-32 |
| ACE2 | RTN4RL2 | -0,7323 | 1,05975E-32 |
| ACE2 | WNK1 | -0,7323 | 1,07509E-32 |
| ACE2 | METTL21B | -0,7323 | 1,08663E-32 |
| ACE2 | APPBP2 | -0,7323 | 1,07423E-32 |
| ACE2 | AC106786.1 | -0,7324 | 1,04004E-32 |
| ACE2 | RP11-522B15.3 | -0,7324 | 1,04691E-32 |
| ACE2 | TMEM173 | -0,7325 | 1,0199E-32 |
| ACE2 | RRN3P2 | -0,7325 | 1,00598E-32 |
| ACE2 | TCEAL2 | -0,7325 | 1,02065E-32 |
| ACE2 | NRDC | -0,7326 | 9,97677E-33 |
| ACE2 | RNF144A | -0,7326 | 9,894E-33 |
| ACE2 | LOXL3 | -0,7326 | 9,71518E-33 |
| ACE2 | NOTCH1 | -0,7326 | 9,88112E-33 |
| ACE2 | CHST3 | -0,7326 | 9,82223E-33 |
| ACE2 | MORC2 | -0,7326 | 9,89347E-33 |
| ACE2 | VBP1 | -0,7326 | 9,85601E-33 |
| ACE2 | DNAJB5 | -0,7327 | 9,54281E-33 |
| ACE2 | SERTAD4-AS1 | -0,7328 | 9,21883E-33 |
| ACE2 | MUSTN1 | -0,7328 | 9,23759E-33 |
| ACE2 | LOXL2 | -0,7328 | 9,26939E-33 |
| ACE2 | SLC13A3 | -0,7328 | 9,33971E-33 |
| ACE2 | SEPT7P2 | -0,7329 | 8,94795E-33 |
| ACE2 | RLIM | -0,733 | 8,74668E-33 |
| ACE2 | LINC01750 | -0,7331 | 8,49382E-33 |
| ACE2 | LINC00839 | -0,7331 | 8,59898E-33 |
| ACE2 | SEMA4F | -0,7332 | 8,35184E-33 |
| ACE2 | SORCS2 | -0,7332 | 8,15245E-33 |
| ACE2 | RPL9 | -0,7332 | 8,36434E-33 |
| ACE2 | MEF2C | -0,7332 | 8,36826E-33 |
| ACE2 | RP11-37B2.1 | -0,7332 | 8,35387E-33 |
| ACE2 | PAAF1 | -0,7332 | 8,15525E-33 |
| ACE2 | ZDBF2 | -0,7333 | 8,11726E-33 |
| ACE2 | VPS8 | -0,7333 | 8,09847E-33 |
| ACE2 | RADIL | -0,7333 | 7,96422E-33 |
| ACE2 | IL17RD | -0,7334 | 7,67666E-33 |
| ACE2 | ZNF248 | -0,7334 | 7,66247E-33 |
| ACE2 | RPL27 | -0,7334 | 7,66061E-33 |
| ACE2 | TUBD1 | -0,7334 | 7,76134E-33 |
| ACE2 | SNRPD2 | -0,7334 | 7,816E-33 |
| ACE2 | EHD2 | -0,7334 | 7,78658E-33 |
| ACE2 | BGN | -0,7334 | 7,66759E-33 |

| | | | |
|---|---|---|---|
| ACE2 | PTGES3P1 | -0,7336 | 7,37232E-33 |
| ACE2 | DHX57 | -0,7336 | 7,39839E-33 |
| ACE2 | SCN3B | -0,7336 | 7,3423E-33 |
| ACE2 | CACHD1 | -0,7337 | 7,03721E-33 |
| ACE2 | TMA16 | -0,7337 | 7,05752E-33 |
| ACE2 | BTF3 | -0,7337 | 7,03169E-33 |
| ACE2 | XPA | -0,7337 | 7,12845E-33 |
| ACE2 | TANGO6 | -0,7339 | 6,78891E-33 |
| ACE2 | PTGFR | -0,734 | 6,55878E-33 |
| ACE2 | IL12A | -0,734 | 6,53988E-33 |
| ACE2 | RNF130 | -0,734 | 6,45647E-33 |
| ACE2 | TAB2 | -0,734 | 6,58374E-33 |
| ACE2 | TAF6 | -0,734 | 6,47068E-33 |
| ACE2 | ZFR2 | -0,734 | 6,52617E-33 |
| ACE2 | PCDHGA12 | -0,7341 | 6,40226E-33 |
| ACE2 | GPER1 | -0,7342 | 6,21007E-33 |
| ACE2 | MFHAS1 | -0,7342 | 6,20574E-33 |
| ACE2 | NEIL2 | -0,7342 | 6,1685E-33 |
| ACE2 | ZNF559 | -0,7342 | 6,17688E-33 |
| ACE2 | PNMAL2 | -0,7343 | 6,04635E-33 |
| ACE2 | UBE3D | -0,7344 | 5,8189E-33 |
| ACE2 | ANKS1B | -0,7344 | 5,77023E-33 |
| ACE2 | SNAP91 | -0,7345 | 5,65667E-33 |
| ACE2 | ERGIC2 | -0,7345 | 5,65918E-33 |
| ACE2 | RFXAP | -0,7345 | 5,61794E-33 |
| ACE2 | TMEM255B | -0,7345 | 5,6292E-33 |
| ACE2 | KITLG | -0,7346 | 5,48953E-33 |
| ACE2 | ORAI3 | -0,7346 | 5,48435E-33 |
| ACE2 | FAM184A | -0,7347 | 5,21623E-33 |
| ACE2 | ZNF22 | -0,7347 | 5,2906E-33 |
| ACE2 | DNM1P35 | -0,7347 | 5,30213E-33 |
| ACE2 | PTOV1 | -0,7347 | 5,24332E-33 |
| ACE2 | MN1 | -0,7347 | 5,34362E-33 |
| ACE2 | CCDC117 | -0,7347 | 5,22821E-33 |
| ACE2 | C15orf59 | -0,7348 | 5,16586E-33 |
| ACE2 | NSRP1 | -0,7348 | 5,11243E-33 |
| ACE2 | GCK | -0,7349 | 5,0415E-33 |
| ACE2 | RP11-177H13.2 | -0,7349 | 5,05612E-33 |
| ACE2 | HMGN1P3 | -0,7349 | 4,99007E-33 |
| ACE2 | NUDT3 | -0,735 | 4,79693E-33 |
| ACE2 | DGCR6L | -0,735 | 4,9005E-33 |
| ACE2 | GTDC1 | -0,7351 | 4,72492E-33 |
| ACE2 | OBSL1 | -0,7351 | 4,74128E-33 |
| ACE2 | SET | -0,7351 | 4,63237E-33 |
| ACE2 | OR51E2 | -0,7351 | 4,66145E-33 |
| ACE2 | TMEM230 | -0,7351 | 4,75894E-33 |
| ACE2 | RP11-428J1.4 | -0,7352 | 4,54174E-33 |
| ACE2 | YBX3 | -0,7352 | 4,49677E-33 |
| ACE2 | WTIP | -0,7352 | 4,61187E-33 |
| ACE2 | RP11-384P7.7 | -0,7353 | 4,41947E-33 |
| ACE2 | AAAS | -0,7353 | 4,39004E-33 |
| ACE2 | TMEM240 | -0,7354 | 4,26302E-33 |
| ACE2 | MRPL9 | -0,7354 | 4,26376E-33 |
| ACE2 | PPP1R12B | -0,7354 | 4,34161E-33 |

| | | | |
|---|---|---|---|
| ACE2 | RPL35A | -0,7354 | 4,24169E-33 |
| ACE2 | DUSP26 | -0,7354 | 4,2852E-33 |
| ACE2 | PCGF6 | -0,7354 | 4,30663E-33 |
| ACE2 | BPGM | -0,7355 | 4,20109E-33 |
| ACE2 | COL15A1 | -0,7355 | 4,15309E-33 |
| ACE2 | TIMP2 | -0,7355 | 4,12553E-33 |
| ACE2 | ATRX | -0,7355 | 4,17147E-33 |
| ACE2 | LINC00998 | -0,7356 | 4,03028E-33 |
| ACE2 | LYNX1 | -0,7356 | 4,0818E-33 |
| ACE2 | CTB-50L17.10 | -0,7356 | 3,99736E-33 |
| ACE2 | DGCR14 | -0,7356 | 4,05902E-33 |
| ACE2 | KAT7 | -0,7357 | 3,88624E-33 |
| ACE2 | LRRN4CL | -0,7358 | 3,786E-33 |
| ACE2 | TMEM47 | -0,7358 | 3,82909E-33 |
| ACE2 | OLFML2A | -0,7359 | 3,69665E-33 |
| ACE2 | S100A4 | -0,736 | 3,61793E-33 |
| ACE2 | COLGALT2 | -0,736 | 3,64178E-33 |
| ACE2 | FAM13B | -0,736 | 3,61908E-33 |
| ACE2 | PIANP | -0,736 | 3,58466E-33 |
| ACE2 | RGMA | -0,736 | 3,64514E-33 |
| ACE2 | PTPN14 | -0,7361 | 3,46186E-33 |
| ACE2 | ZNF354B | -0,7361 | 3,47805E-33 |
| ACE2 | SOX17 | -0,7361 | 3,48852E-33 |
| ACE2 | ZNF610 | -0,7361 | 3,45144E-33 |
| ACE2 | ANKRD54 | -0,7361 | 3,53775E-33 |
| ACE2 | COL16A1 | -0,7362 | 3,40109E-33 |
| ACE2 | GEMIN5 | -0,7362 | 3,39447E-33 |
| ACE2 | HOXC10 | -0,7363 | 3,25738E-33 |
| ACE2 | TCEAL5 | -0,7363 | 3,3106E-33 |
| ACE2 | FYCO1 | -0,7364 | 3,18843E-33 |
| ACE2 | EDNRA | -0,7364 | 3,2361E-33 |
| ACE2 | TRAPPC6B | -0,7364 | 3,16362E-33 |
| ACE2 | HMCN1 | -0,7365 | 3,0591E-33 |
| ACE2 | LOXL4 | -0,7365 | 3,05773E-33 |
| ACE2 | BCCIP | -0,7365 | 3,07399E-33 |
| ACE2 | SAMD11 | -0,7366 | 2,99975E-33 |
| ACE2 | AC058791.1 | -0,7366 | 3,04416E-33 |
| ACE2 | CDH2 | -0,7366 | 2,97617E-33 |
| ACE2 | SCARF2 | -0,7366 | 3,01078E-33 |
| ACE2 | GNAO1 | -0,7367 | 2,92731E-33 |
| ACE2 | NCK1 | -0,7368 | 2,8199E-33 |
| ACE2 | INMT | -0,7368 | 2,82587E-33 |
| ACE2 | ITGB1BP2 | -0,7368 | 2,80025E-33 |
| ACE2 | DPT | -0,7369 | 2,72094E-33 |
| ACE2 | GUK1 | -0,7369 | 2,70657E-33 |
| ACE2 | RDH14 | -0,7369 | 2,73258E-33 |
| ACE2 | SLC15A3 | -0,7369 | 2,7352E-33 |
| ACE2 | SPATA6 | -0,737 | 2,66119E-33 |
| ACE2 | ALMS1 | -0,737 | 2,67808E-33 |
| ACE2 | PDE1C | -0,737 | 2,65588E-33 |
| ACE2 | LATS2 | -0,737 | 2,69812E-33 |
| ACE2 | PTPRS | -0,737 | 2,67688E-33 |
| ACE2 | SULT1C4 | -0,7371 | 2,57378E-33 |
| ACE2 | RP11-548H3.1 | -0,7372 | 2,5014E-33 |

| | | | |
|---|---|---|---|
| ACE2 | POLR2M | -0,7372 | 2,50462E-33 |
| ACE2 | AARS | -0,7372 | 2,51641E-33 |
| ACE2 | C17orf80 | -0,7372 | 2,51937E-33 |
| ACE2 | FXYD1 | -0,7372 | 2,50165E-33 |
| ACE2 | POLA1 | -0,7372 | 2,48269E-33 |
| ACE2 | IGIP | -0,7373 | 2,45549E-33 |
| ACE2 | SPRYD3 | -0,7373 | 2,40736E-33 |
| ACE2 | NID2 | -0,7374 | 2,38178E-33 |
| ACE2 | SPHK1 | -0,7374 | 2,35461E-33 |
| ACE2 | PPP1R12C | -0,7374 | 2,36358E-33 |
| ACE2 | CNOT4 | -0,7375 | 2,30343E-33 |
| ACE2 | RPL23AP53 | -0,7375 | 2,2929E-33 |
| ACE2 | OTUD6B | -0,7375 | 2,3172E-33 |
| ACE2 | UQCC1 | -0,7375 | 2,32576E-33 |
| ACE2 | DLGAP4 | -0,7375 | 2,30592E-33 |
| ACE2 | FAM98A | -0,7377 | 2,18932E-33 |
| ACE2 | ANXA1 | -0,7377 | 2,15372E-33 |
| ACE2 | AJUBA | -0,7377 | 2,17652E-33 |
| ACE2 | AC022007.5 | -0,7378 | 2,10366E-33 |
| ACE2 | TSPAN5 | -0,7378 | 2,08658E-33 |
| ACE2 | PCDHB18P | -0,7378 | 2,09509E-33 |
| ACE2 | INPP5F | -0,7378 | 2,09198E-33 |
| ACE2 | RPL17 | -0,7378 | 2,1054E-33 |
| ACE2 | ERCC8 | -0,7379 | 2,02765E-33 |
| ACE2 | CCDC136 | -0,7379 | 2,05591E-33 |
| ACE2 | APBA1 | -0,7379 | 2,05988E-33 |
| ACE2 | FNDC4 | -0,738 | 2,00266E-33 |
| ACE2 | PIP5K1C | -0,738 | 1,97711E-33 |
| ACE2 | ZNF565 | -0,738 | 2,00178E-33 |
| ACE2 | SCD5 | -0,7381 | 1,92926E-33 |
| ACE2 | TUBB6 | -0,7381 | 1,93145E-33 |
| ACE2 | PIN1 | -0,7381 | 1,92394E-33 |
| ACE2 | ZNF790 | -0,7381 | 1,90752E-33 |
| ACE2 | BRIX1 | -0,7382 | 1,87933E-33 |
| ACE2 | DACT1 | -0,7382 | 1,83697E-33 |
| ACE2 | COX7A1 | -0,7382 | 1,88445E-33 |
| ACE2 | PLEKHG2 | -0,7382 | 1,85401E-33 |
| ACE2 | KLHL21 | -0,7383 | 1,81557E-33 |
| ACE2 | TEX14 | -0,7383 | 1,78702E-33 |
| ACE2 | TEK | -0,7384 | 1,76223E-33 |
| ACE2 | WASF1 | -0,7385 | 1,71969E-33 |
| ACE2 | KIAA1143 | -0,7386 | 1,65532E-33 |
| ACE2 | MOB2 | -0,7386 | 1,64121E-33 |
| ACE2 | FARP1 | -0,7386 | 1,64348E-33 |
| ACE2 | DLC1 | -0,7387 | 1,59745E-33 |
| ACE2 | STK40 | -0,7388 | 1,55389E-33 |
| ACE2 | RP11-356B19.11 | -0,7388 | 1,53901E-33 |
| ACE2 | ADK | -0,7388 | 1,56262E-33 |
| ACE2 | MBTPS1 | -0,7388 | 1,53822E-33 |
| ACE2 | LINC01686 | -0,7389 | 1,49578E-33 |
| ACE2 | LIMS2 | -0,7389 | 1,49343E-33 |
| ACE2 | COL5A2 | -0,7389 | 1,51372E-33 |
| ACE2 | WDYHV1 | -0,7389 | 1,51135E-33 |
| ACE2 | RP11-429G19.3 | -0,7389 | 1,5175E-33 |

| | | | |
|---|---|---|---|
| ACE2 | PCDH9 | -0,7389 | 1,51861E-33 |
| ACE2 | COA5 | -0,739 | 1,45185E-33 |
| ACE2 | PURG | -0,739 | 1,44807E-33 |
| ACE2 | TMEM108 | -0,7391 | 1,41719E-33 |
| ACE2 | CCNI | -0,7391 | 1,39586E-33 |
| ACE2 | RNF41 | -0,7391 | 1,40408E-33 |
| ACE2 | C14orf132 | -0,7391 | 1,42617E-33 |
| ACE2 | WWC3 | -0,7391 | 1,42811E-33 |
| ACE2 | TMEM35B | -0,7392 | 1,36588E-33 |
| ACE2 | SPIRE1 | -0,7392 | 1,3734E-33 |
| ACE2 | FKRP | -0,7392 | 1,38667E-33 |
| ACE2 | OSTM1 | -0,7393 | 1,33312E-33 |
| ACE2 | ASIC1 | -0,7393 | 1,35007E-33 |
| ACE2 | GPR88 | -0,7394 | 1,29939E-33 |
| ACE2 | RP11-461O7.1 | -0,7394 | 1,30419E-33 |
| ACE2 | RNF146 | -0,7396 | 1,21736E-33 |
| ACE2 | DACT3 | -0,7397 | 1,16769E-33 |
| ACE2 | ZNF17 | -0,7397 | 1,16432E-33 |
| ACE2 | TWIST2 | -0,7398 | 1,13905E-33 |
| ACE2 | CREM | -0,7398 | 1,15858E-33 |
| ACE2 | COL1A1 | -0,7398 | 1,14897E-33 |
| ACE2 | DOK5 | -0,7398 | 1,1568E-33 |
| ACE2 | KCTD20 | -0,7399 | 1,10075E-33 |
| ACE2 | AC000403.4 | -0,7399 | 1,10063E-33 |
| ACE2 | RP11-566K19.6 | -0,7399 | 1,10887E-33 |
| ACE2 | CTF1 | -0,7399 | 1,10385E-33 |
| ACE2 | TANC2 | -0,7399 | 1,11779E-33 |
| ACE2 | PCDHGA3 | -0,74 | 1,08929E-33 |
| ACE2 | ANGPT1 | -0,74 | 1,09123E-33 |
| ACE2 | AS3MT | -0,74 | 1,08463E-33 |
| ACE2 | ZNF527 | -0,74 | 1,08086E-33 |
| ACE2 | USP27X-AS1 | -0,74 | 1,09302E-33 |
| ACE2 | MLF1 | -0,7401 | 1,03631E-33 |
| ACE2 | REV3L | -0,7401 | 1,03267E-33 |
| ACE2 | TMEM185A | -0,7401 | 1,05178E-33 |
| ACE2 | ADAM33 | -0,7403 | 9,73828E-34 |
| ACE2 | PQLC3 | -0,7404 | 9,55747E-34 |
| ACE2 | DNAJC27 | -0,7404 | 9,54936E-34 |
| ACE2 | ZBTB47 | -0,7404 | 9,49585E-34 |
| ACE2 | MCF2L2 | -0,7404 | 9,42545E-34 |
| ACE2 | KLHL42 | -0,7404 | 9,43438E-34 |
| ACE2 | ZNF584 | -0,7404 | 9,46338E-34 |
| ACE2 | MRPS6 | -0,7404 | 9,54373E-34 |
| ACE2 | CROCC | -0,7405 | 9,23156E-34 |
| ACE2 | KIF26B | -0,7405 | 9,31546E-34 |
| ACE2 | TWISTNB | -0,7405 | 9,37816E-34 |
| ACE2 | IGF2 | -0,7405 | 9,26517E-34 |
| ACE2 | PACSIN3 | -0,7405 | 9,33139E-34 |
| ACE2 | RP11-999E24.3 | -0,7405 | 9,26804E-34 |
| ACE2 | DAAM1 | -0,7405 | 9,28975E-34 |
| ACE2 | SIX4 | -0,7405 | 9,13335E-34 |
| ACE2 | MMD | -0,7405 | 9,3145E-34 |
| ACE2 | DSTNP2 | -0,7406 | 9,0372E-34 |
| ACE2 | PPFIBP1 | -0,7406 | 9,05012E-34 |

| | | | |
|---|---|---|---|
| ACE2 | RAPGEF3 | -0,7406 | 8,8443E-34 |
| ACE2 | RAMP2 | -0,7406 | 9,05199E-34 |
| ACE2 | ZZZ3 | -0,7407 | 8,79286E-34 |
| ACE2 | IQCG | -0,7407 | 8,72055E-34 |
| ACE2 | BBS7 | -0,7407 | 8,78303E-34 |
| ACE2 | C1QTNF3 | -0,7407 | 8,76303E-34 |
| ACE2 | TLN1 | -0,7407 | 8,74498E-34 |
| ACE2 | EML1 | -0,7407 | 8,84182E-34 |
| ACE2 | SNX18 | -0,7408 | 8,52273E-34 |
| ACE2 | TAF3 | -0,7408 | 8,55051E-34 |
| ACE2 | GPR135 | -0,7408 | 8,41693E-34 |
| ACE2 | PECAM1 | -0,7409 | 8,14051E-34 |
| ACE2 | BCAM | -0,7409 | 8,1808E-34 |
| ACE2 | TAB1 | -0,7409 | 8,30518E-34 |
| ACE2 | UBQLN2 | -0,7409 | 8,25272E-34 |
| ACE2 | ZNF555 | -0,741 | 7,9218E-34 |
| ACE2 | MAP3K20 | -0,7411 | 7,68962E-34 |
| ACE2 | RAMP3 | -0,7411 | 7,81621E-34 |
| ACE2 | TNFAIP8L1 | -0,7411 | 7,68938E-34 |
| ACE2 | ZNF333 | -0,7411 | 7,74345E-34 |
| ACE2 | CORO2B | -0,7412 | 7,40999E-34 |
| ACE2 | DISC1 | -0,7413 | 7,18863E-34 |
| ACE2 | SUB1 | -0,7413 | 7,15041E-34 |
| ACE2 | RPP30 | -0,7413 | 7,18609E-34 |
| ACE2 | RBFADN | -0,7413 | 7,27611E-34 |
| ACE2 | ZNF426 | -0,7413 | 7,19947E-34 |
| ACE2 | ZNF665 | -0,7413 | 7,16045E-34 |
| ACE2 | PCOLCE | -0,7414 | 7,0046E-34 |
| ACE2 | TTC25 | -0,7414 | 7,09614E-34 |
| ACE2 | CUEDC1 | -0,7414 | 7,10588E-34 |
| ACE2 | PCDHGA11 | -0,7415 | 6,85866E-34 |
| ACE2 | SMOC2 | -0,7415 | 6,82961E-34 |
| ACE2 | COL1A2 | -0,7415 | 6,79278E-34 |
| ACE2 | GNAZ | -0,7415 | 6,89247E-34 |
| ACE2 | RSPO1 | -0,7416 | 6,55035E-34 |
| ACE2 | RPL31 | -0,7416 | 6,70656E-34 |
| ACE2 | NKIRAS1 | -0,7416 | 6,71151E-34 |
| ACE2 | NSA2 | -0,7416 | 6,6093E-34 |
| ACE2 | ARSK | -0,7416 | 6,51845E-34 |
| ACE2 | KIRREL3 | -0,7417 | 6,41507E-34 |
| ACE2 | MIR99AHG | -0,7417 | 6,47349E-34 |
| ACE2 | C7 | -0,7418 | 6,12923E-34 |
| ACE2 | POPDC3 | -0,7418 | 6,21822E-34 |
| ACE2 | RNF38 | -0,7418 | 6,28659E-34 |
| ACE2 | FAM189A2 | -0,7419 | 6,07259E-34 |
| ACE2 | SDAD1P1 | -0,742 | 5,93701E-34 |
| ACE2 | APLP1 | -0,742 | 5,8782E-34 |
| ACE2 | MTA3 | -0,7421 | 5,69017E-34 |
| ACE2 | SPTAN1 | -0,7421 | 5,60763E-34 |
| ACE2 | RASL12 | -0,7421 | 5,74347E-34 |
| ACE2 | PRKACB | -0,7422 | 5,47463E-34 |
| ACE2 | LANCL1 | -0,7422 | 5,47538E-34 |
| ACE2 | EPHA4 | -0,7422 | 5,49872E-34 |
| ACE2 | CCDC66 | -0,7422 | 5,50744E-34 |

| | | | |
|---|---|---|---|
| ACE2 | CASC4 | -0,7422 | 5,43635E-34 |
| ACE2 | SNX27 | -0,7423 | 5,27154E-34 |
| ACE2 | GNPDA2 | -0,7423 | 5,37879E-34 |
| ACE2 | NPR3 | -0,7423 | 5,31139E-34 |
| ACE2 | RP11-266K4.14 | -0,7423 | 5,29158E-34 |
| ACE2 | CBX7 | -0,7423 | 5,30579E-34 |
| ACE2 | ITM2A | -0,7424 | 5,24887E-34 |
| ACE2 | YBX1 | -0,7425 | 5,08085E-34 |
| ACE2 | MRVI1 | -0,7425 | 5,08856E-34 |
| ACE2 | MIR193BHG | -0,7425 | 5,02563E-34 |
| ACE2 | ELAVL1 | -0,7425 | 4,95383E-34 |
| ACE2 | UBE2V1 | -0,7425 | 4,95871E-34 |
| ACE2 | ZNF674-AS1 | -0,7425 | 5,00687E-34 |
| ACE2 | CHIC1 | -0,7425 | 5,05422E-34 |
| ACE2 | CYS1 | -0,7426 | 4,87958E-34 |
| ACE2 | MSN | -0,7426 | 4,82379E-34 |
| ACE2 | HEY1 | -0,7427 | 4,76205E-34 |
| ACE2 | NUP54 | -0,7428 | 4,5281E-34 |
| ACE2 | POLR3D | -0,7428 | 4,62161E-34 |
| ACE2 | VASN | -0,7429 | 4,38405E-34 |
| ACE2 | SERTAD4 | -0,743 | 4,36117E-34 |
| ACE2 | MITF | -0,743 | 4,31655E-34 |
| ACE2 | EIF4A2 | -0,743 | 4,28434E-34 |
| ACE2 | C1RL | -0,743 | 4,36628E-34 |
| ACE2 | SLFN5 | -0,743 | 4,35286E-34 |
| ACE2 | POU6F1 | -0,7431 | 4,16677E-34 |
| ACE2 | C16orf89 | -0,7431 | 4,23553E-34 |
| ACE2 | PEG3 | -0,7431 | 4,2276E-34 |
| ACE2 | SEC63 | -0,7432 | 4,02808E-34 |
| ACE2 | DMWD | -0,7432 | 4,09402E-34 |
| ACE2 | ST13 | -0,7432 | 4,05898E-34 |
| ACE2 | PLAGL1 | -0,7433 | 3,94722E-34 |
| ACE2 | CNTN1 | -0,7433 | 3,96697E-34 |
| ACE2 | ARHGAP6 | -0,7433 | 3,93578E-34 |
| ACE2 | NUP133 | -0,7434 | 3,74732E-34 |
| ACE2 | CACNA1E | -0,7435 | 3,63382E-34 |
| ACE2 | PDCD2L | -0,7435 | 3,64672E-34 |
| ACE2 | WDR35 | -0,7437 | 3,44881E-34 |
| ACE2 | FNTA | -0,7437 | 3,50367E-34 |
| ACE2 | WSCD2 | -0,7437 | 3,49609E-34 |
| ACE2 | ADAMTS3 | -0,7438 | 3,35422E-34 |
| ACE2 | SOX7 | -0,7438 | 3,32707E-34 |
| ACE2 | ROBO3 | -0,7438 | 3,39108E-34 |
| ACE2 | C16orf52 | -0,7438 | 3,4018E-34 |
| ACE2 | IPO9 | -0,7439 | 3,26393E-34 |
| ACE2 | RPS6KC1 | -0,7439 | 3,30872E-34 |
| ACE2 | ERCC3 | -0,7439 | 3,30168E-34 |
| ACE2 | SLC45A1 | -0,744 | 3,14059E-34 |
| ACE2 | PRDM5 | -0,744 | 3,15792E-34 |
| ACE2 | IL17B | -0,744 | 3,14013E-34 |
| ACE2 | RET | -0,744 | 3,13356E-34 |
| ACE2 | C11orf68 | -0,744 | 3,14996E-34 |
| ACE2 | NDUFAF8 | -0,7441 | 3,01821E-34 |
| ACE2 | ZNF271P | -0,7441 | 3,04632E-34 |

| | | | |
|---|---|---|---|
| ACE2 | ZNF566 | -0,7441 | 3,0213E-34 |
| ACE2 | FAM127B | -0,7441 | 3,04923E-34 |
| ACE2 | PRDX6 | -0,7442 | 2,95296E-34 |
| ACE2 | ORC3 | -0,7442 | 2,96917E-34 |
| ACE2 | GPALPP1 | -0,7442 | 2,95756E-34 |
| ACE2 | FERMT2 | -0,7442 | 3,0093E-34 |
| ACE2 | MOB3C | -0,7443 | 2,88724E-34 |
| ACE2 | MTSS1L | -0,7443 | 2,89339E-34 |
| ACE2 | RAB28P5 | -0,7443 | 2,86727E-34 |
| ACE2 | GRIK3 | -0,7444 | 2,77051E-34 |
| ACE2 | CCDC85A | -0,7444 | 2,82989E-34 |
| ACE2 | HOXD3 | -0,7445 | 2,70958E-34 |
| ACE2 | ADGRF5 | -0,7445 | 2,71556E-34 |
| ACE2 | TSPY26P | -0,7445 | 2,7063E-34 |
| ACE2 | NCAM2 | -0,7445 | 2,70071E-34 |
| ACE2 | DOCK4 | -0,7446 | 2,59458E-34 |
| ACE2 | FAM89A | -0,7448 | 2,5019E-34 |
| ACE2 | MKRN2 | -0,7448 | 2,43426E-34 |
| ACE2 | LRRK2 | -0,7448 | 2,47081E-34 |
| ACE2 | MAP4K4 | -0,7449 | 2,4112E-34 |
| ACE2 | XRCC1 | -0,7449 | 2,4262E-34 |
| ACE2 | ZNF510 | -0,745 | 2,33162E-34 |
| ACE2 | CDK4 | -0,745 | 2,29596E-34 |
| ACE2 | FAM175A | -0,7451 | 2,24516E-34 |
| ACE2 | NAF1 | -0,7451 | 2,25213E-34 |
| ACE2 | FLOT1 | -0,7451 | 2,23222E-34 |
| ACE2 | NR2C1 | -0,7451 | 2,25971E-34 |
| ACE2 | BLOC1S6 | -0,7451 | 2,26093E-34 |
| ACE2 | ARNT2 | -0,7451 | 2,22759E-34 |
| ACE2 | GBAS | -0,7452 | 2,18116E-34 |
| ACE2 | ATRNL1 | -0,7452 | 2,16541E-34 |
| ACE2 | HES4 | -0,7453 | 2,10184E-34 |
| ACE2 | LINC01770 | -0,7453 | 2,1151E-34 |
| ACE2 | CCDC74B | -0,7453 | 2,12336E-34 |
| ACE2 | THPO | -0,7453 | 2,10463E-34 |
| ACE2 | KLHL3 | -0,7453 | 2,13889E-34 |
| ACE2 | MAP6 | -0,7453 | 2,08434E-34 |
| ACE2 | SLC26A10 | -0,7453 | 2,12464E-34 |
| ACE2 | DUT | -0,7453 | 2,12799E-34 |
| ACE2 | SLC24A3 | -0,7453 | 2,11873E-34 |
| ACE2 | SEC22A | -0,7454 | 2,03089E-34 |
| ACE2 | RP11-16P20.4 | -0,7454 | 2,03649E-34 |
| ACE2 | PLEKHA8P1 | -0,7454 | 2,05822E-34 |
| ACE2 | CDH6 | -0,7455 | 1,97482E-34 |
| ACE2 | ATP2B4 | -0,7456 | 1,9492E-34 |
| ACE2 | ST6GALNAC3 | -0,7457 | 1,86688E-34 |
| ACE2 | LYAR | -0,7457 | 1,88548E-34 |
| ACE2 | ZNF660 | -0,7458 | 1,81723E-34 |
| ACE2 | LL21NC02-21A1.1 | -0,7458 | 1,81054E-34 |
| ACE2 | SGSM1 | -0,7458 | 1,81086E-34 |
| ACE2 | P3H1 | -0,7459 | 1,77593E-34 |
| ACE2 | UTP14A | -0,7459 | 1,77231E-34 |
| ACE2 | RPS25 | -0,746 | 1,69486E-34 |
| ACE2 | TCEAL9 | -0,746 | 1,6999E-34 |

| | | | |
|---|---|---|---|
| ACE2 | ITGA1 | -0,7461 | 1,63974E-34 |
| ACE2 | DCTN3 | -0,7461 | 1,65529E-34 |
| ACE2 | NASP | -0,7462 | 1,61436E-34 |
| ACE2 | MED4 | -0,7462 | 1,57692E-34 |
| ACE2 | PPP5D1 | -0,7462 | 1,60449E-34 |
| ACE2 | ZNF579 | -0,7462 | 1,60086E-34 |
| ACE2 | PCDHGA2 | -0,7464 | 1,50114E-34 |
| ACE2 | RP11-686D22.3 | -0,7465 | 1,43451E-34 |
| ACE2 | FMOD | -0,7466 | 1,41317E-34 |
| ACE2 | DPY19L2 | -0,7466 | 1,39184E-34 |
| ACE2 | MROH8 | -0,7467 | 1,36956E-34 |
| ACE2 | IKBIP | -0,7468 | 1,30173E-34 |
| ACE2 | PACRGL | -0,7469 | 1,26123E-34 |
| ACE2 | RP11-893F2.14 | -0,7469 | 1,26461E-34 |
| ACE2 | DCTD | -0,747 | 1,2489E-34 |
| ACE2 | SKA2 | -0,747 | 1,25566E-34 |
| ACE2 | RAP1A | -0,7471 | 1,18431E-34 |
| ACE2 | SEPT2 | -0,7471 | 1,19572E-34 |
| ACE2 | TBC1D24 | -0,7471 | 1,19671E-34 |
| ACE2 | CASC3 | -0,7471 | 1,19542E-34 |
| ACE2 | RPS7 | -0,7472 | 1,16459E-34 |
| ACE2 | LIMS1 | -0,7472 | 1,16812E-34 |
| ACE2 | MEG3 | -0,7472 | 1,16434E-34 |
| ACE2 | MEAF6 | -0,7473 | 1,1409E-34 |
| ACE2 | AC074286.1 | -0,7473 | 1,13189E-34 |
| ACE2 | FAM171A2 | -0,7473 | 1,1242E-34 |
| ACE2 | HOXB-AS1 | -0,7473 | 1,12769E-34 |
| ACE2 | ST7L | -0,7474 | 1,07719E-34 |
| ACE2 | NDUFAF7 | -0,7474 | 1,09506E-34 |
| ACE2 | RPL37A | -0,7475 | 1,04751E-34 |
| ACE2 | ZNF470 | -0,7475 | 1,04348E-34 |
| ACE2 | GPRASP1 | -0,7475 | 1,04997E-34 |
| ACE2 | XXYLT1 | -0,7476 | 1,01706E-34 |
| ACE2 | CEP19 | -0,7476 | 1,0179E-34 |
| ACE2 | PTGER1 | -0,7476 | 1,04121E-34 |
| ACE2 | TTC21B | -0,7477 | 9,83987E-35 |
| ACE2 | COL8A1 | -0,7477 | 1,00254E-34 |
| ACE2 | LARS | -0,7477 | 1,00359E-34 |
| ACE2 | AP2B1 | -0,7477 | 9,88657E-35 |
| ACE2 | PEF1 | -0,7479 | 9,29391E-35 |
| ACE2 | TNRC6C | -0,7479 | 9,28283E-35 |
| ACE2 | NCS1 | -0,748 | 9,08548E-35 |
| ACE2 | NAB2 | -0,748 | 8,95493E-35 |
| ACE2 | RND2 | -0,748 | 9,12864E-35 |
| ACE2 | CCDC124 | -0,748 | 9,00856E-35 |
| ACE2 | HSP90AB1 | -0,7481 | 8,73153E-35 |
| ACE2 | RP11-395G23.3 | -0,7481 | 8,76098E-35 |
| ACE2 | RP11-82L18.2 | -0,7481 | 8,71132E-35 |
| ACE2 | OLFM2 | -0,7481 | 8,83902E-35 |
| ACE2 | LINC00205 | -0,7481 | 8,85663E-35 |
| ACE2 | EGFL7 | -0,7482 | 8,51887E-35 |
| ACE2 | MYEF2 | -0,7482 | 8,43903E-35 |
| ACE2 | ZNF775 | -0,7483 | 8,1102E-35 |
| ACE2 | RP11-195F19.5 | -0,7483 | 8,35388E-35 |

| | | | |
|---|---|---|---|
| ACE2 | SIX5 | -0,7485 | 7,76325E-35 |
| ACE2 | SHANK3 | -0,7485 | 7,73262E-35 |
| ACE2 | FOPNL | -0,7486 | 7,42528E-35 |
| ACE2 | SNU13 | -0,7486 | 7,4818E-35 |
| ACE2 | LINC01160 | -0,7487 | 7,14629E-35 |
| ACE2 | SNHG14 | -0,7487 | 7,24794E-35 |
| ACE2 | EIF5B | -0,7488 | 6,93661E-35 |
| ACE2 | ADAMTSL1 | -0,7488 | 7,04401E-35 |
| ACE2 | IFITM3 | -0,7488 | 6,92755E-35 |
| ACE2 | THY1 | -0,7488 | 6,9773E-35 |
| ACE2 | XKR4 | -0,7489 | 6,7237E-35 |
| ACE2 | RP11-48B3.5 | -0,7489 | 6,82115E-35 |
| ACE2 | ST7 | -0,749 | 6,59038E-35 |
| ACE2 | SRM | -0,7491 | 6,35966E-35 |
| ACE2 | IMMP2L | -0,7491 | 6,42079E-35 |
| ACE2 | TTLL11 | -0,7491 | 6,3345E-35 |
| ACE2 | PINX1 | -0,7492 | 6,26871E-35 |
| ACE2 | CCDC28B | -0,7493 | 5,97098E-35 |
| ACE2 | RAB23 | -0,7493 | 6,04183E-35 |
| ACE2 | ESD | -0,7493 | 5,95427E-35 |
| ACE2 | SULF2 | -0,7493 | 6,05173E-35 |
| ACE2 | TCEAL3 | -0,7493 | 6,0842E-35 |
| ACE2 | ZNF511 | -0,7494 | 5,78036E-35 |
| ACE2 | C20orf194 | -0,7494 | 5,81054E-35 |
| ACE2 | TGFB2 | -0,7495 | 5,70816E-35 |
| ACE2 | RBPJ | -0,7495 | 5,69135E-35 |
| ACE2 | GTF2IRD2B | -0,7495 | 5,62865E-35 |
| ACE2 | PRKAR2B | -0,7495 | 5,55266E-35 |
| ACE2 | AMPH | -0,7496 | 5,43471E-35 |
| ACE2 | RP11-236L14.2 | -0,7496 | 5,4695E-35 |
| ACE2 | ZNF790-AS1 | -0,7496 | 5,38089E-35 |
| ACE2 | ELF2 | -0,7497 | 5,34335E-35 |
| ACE2 | RNF215 | -0,7497 | 5,25886E-35 |
| ACE2 | NUDT21 | -0,7498 | 5,14819E-35 |
| ACE2 | FAM104B | -0,7498 | 5,18022E-35 |
| ACE2 | DZIP3 | -0,7499 | 4,90209E-35 |
| ACE2 | KCNT1 | -0,7499 | 4,96658E-35 |
| ACE2 | P4HA3 | -0,7499 | 5,03265E-35 |
| ACE2 | ZNF788 | -0,7499 | 4,98675E-35 |
| ACE2 | ZSCAN21 | -0,75 | 4,75676E-35 |
| ACE2 | PCM1 | -0,75 | 4,79356E-35 |
| ACE2 | MPPED2 | -0,75 | 4,83012E-35 |
| ACE2 | AES | -0,75 | 4,74582E-35 |
| ACE2 | PARVB | -0,75 | 4,81058E-35 |
| ACE2 | KIAA0408 | -0,7502 | 4,5437E-35 |
| ACE2 | FIBIN | -0,7502 | 4,55651E-35 |
| ACE2 | FGFR1OP2 | -0,7502 | 4,56585E-35 |
| ACE2 | PELP1 | -0,7502 | 4,53259E-35 |
| ACE2 | ACOT9 | -0,7502 | 4,49759E-35 |
| ACE2 | COL8A2 | -0,7503 | 4,34323E-35 |
| ACE2 | ZC3H10 | -0,7503 | 4,34227E-35 |
| ACE2 | USP39 | -0,7504 | 4,17326E-35 |
| ACE2 | FAM26E | -0,7504 | 4,21131E-35 |
| ACE2 | PDLIM3 | -0,7505 | 4,08544E-35 |

| | | | |
|---|---|---|---|
| ACE2 | IER5L | -0,7505 | 4,13237E-35 |
| ACE2 | EHBP1L1 | -0,7505 | 4,15284E-35 |
| ACE2 | RP11-554A11.4 | -0,7505 | 4,03622E-35 |
| ACE2 | COLEC12 | -0,7505 | 4,07797E-35 |
| ACE2 | RAD23A | -0,7505 | 4,06857E-35 |
| ACE2 | RNF25 | -0,7506 | 3,91995E-35 |
| ACE2 | DCUN1D4 | -0,7506 | 4,00236E-35 |
| ACE2 | FAM162B | -0,7506 | 4,00706E-35 |
| ACE2 | USP47 | -0,7506 | 3,93982E-35 |
| ACE2 | CD59 | -0,7506 | 4,00677E-35 |
| ACE2 | HOXC8 | -0,7506 | 3,91938E-35 |
| ACE2 | FLNA | -0,7506 | 4,00838E-35 |
| ACE2 | CSF1 | -0,7507 | 3,87867E-35 |
| ACE2 | ZDHHC17 | -0,7507 | 3,81453E-35 |
| ACE2 | FENDRR | -0,7507 | 3,83415E-35 |
| ACE2 | SPAG16 | -0,7508 | 3,74841E-35 |
| ACE2 | RASSF3 | -0,7508 | 3,77088E-35 |
| ACE2 | LINC00924 | -0,7508 | 3,75901E-35 |
| ACE2 | TERF2IP | -0,7508 | 3,68167E-35 |
| ACE2 | SDC2 | -0,7509 | 3,62527E-35 |
| ACE2 | PIK3CA | -0,751 | 3,4957E-35 |
| ACE2 | LINC01772 | -0,7511 | 3,39853E-35 |
| ACE2 | EHBP1 | -0,7511 | 3,35132E-35 |
| ACE2 | MSANTD4 | -0,7511 | 3,40965E-35 |
| ACE2 | TXLNA | -0,7512 | 3,23675E-35 |
| ACE2 | KCNMB3 | -0,7512 | 3,25404E-35 |
| ACE2 | ZFP37 | -0,7512 | 3,30002E-35 |
| ACE2 | RPS13 | -0,7512 | 3,31131E-35 |
| ACE2 | UTP4 | -0,7512 | 3,25213E-35 |
| ACE2 | C19orf53 | -0,7512 | 3,2269E-35 |
| ACE2 | PAIP2B | -0,7513 | 3,14009E-35 |
| ACE2 | SLC25A26 | -0,7513 | 3,20397E-35 |
| ACE2 | GREM1 | -0,7513 | 3,12823E-35 |
| ACE2 | ISYNA1 | -0,7513 | 3,14079E-35 |
| ACE2 | TRIM28 | -0,7513 | 3,19105E-35 |
| ACE2 | AADAT | -0,7514 | 3,03812E-35 |
| ACE2 | TCTN2 | -0,7514 | 3,05818E-35 |
| ACE2 | ADAMTS7 | -0,7514 | 3,05279E-35 |
| ACE2 | BANF1 | -0,7515 | 3,00478E-35 |
| ACE2 | ACTN1 | -0,7516 | 2,88677E-35 |
| ACE2 | ZNF582 | -0,7517 | 2,81065E-35 |
| ACE2 | STAC | -0,7518 | 2,66252E-35 |
| ACE2 | PRKAR1B | -0,7518 | 2,67175E-35 |
| ACE2 | SHE | -0,7519 | 2,63431E-35 |
| ACE2 | RAB31 | -0,7519 | 2,5994E-35 |
| ACE2 | RRAGB | -0,7519 | 2,63907E-35 |
| ACE2 | MRPL33 | -0,752 | 2,5219E-35 |
| ACE2 | ZNF876P | -0,752 | 2,56627E-35 |
| ACE2 | PZP | -0,7521 | 2,47463E-35 |
| ACE2 | PCDHGA9 | -0,7522 | 2,36763E-35 |
| ACE2 | RP11-893F2.13 | -0,7522 | 2,34085E-35 |
| ACE2 | TIMP1 | -0,7522 | 2,34008E-35 |
| ACE2 | NGF | -0,7523 | 2,32931E-35 |
| ACE2 | C8orf88 | -0,7523 | 2,29626E-35 |

| | | | |
|---|---|---|---|
| ACE2 | FAM198B | -0,7524 | 2,20556E-35 |
| ACE2 | MEGF10 | -0,7524 | 2,19735E-35 |
| ACE2 | CASP12 | -0,7525 | 2,15234E-35 |
| ACE2 | LRRC37A17P | -0,7525 | 2,16847E-35 |
| ACE2 | PDGFB | -0,7525 | 2,1667E-35 |
| ACE2 | RAI2 | -0,7525 | 2,15719E-35 |
| ACE2 | PRRT3-AS1 | -0,7526 | 2,10777E-35 |
| ACE2 | KANK2 | -0,7526 | 2,10917E-35 |
| ACE2 | POLR3F | -0,7526 | 2,07582E-35 |
| ACE2 | TCEAL1 | -0,7526 | 2,07659E-35 |
| ACE2 | ROR2 | -0,7527 | 2,00815E-35 |
| ACE2 | BRD7 | -0,7527 | 2,01298E-35 |
| ACE2 | COLQ | -0,7528 | 1,98789E-35 |
| ACE2 | LSM11 | -0,7528 | 1,97594E-35 |
| ACE2 | TMEM71 | -0,7528 | 1,94128E-35 |
| ACE2 | SETMAR | -0,7529 | 1,91388E-35 |
| ACE2 | ARL14EP | -0,7529 | 1,88316E-35 |
| ACE2 | PCBP3 | -0,753 | 1,81004E-35 |
| ACE2 | PCDHB7 | -0,7531 | 1,80147E-35 |
| ACE2 | BACE1 | -0,7531 | 1,76289E-35 |
| ACE2 | MFSD14A | -0,7532 | 1,69853E-35 |
| ACE2 | TMEM43 | -0,7532 | 1,71747E-35 |
| ACE2 | PRMT1 | -0,7532 | 1,74191E-35 |
| ACE2 | NKAP | -0,7532 | 1,71963E-35 |
| ACE2 | ATP6V1G2 | -0,7533 | 1,6896E-35 |
| ACE2 | EXTL3 | -0,7533 | 1,64574E-35 |
| ACE2 | MCAM | -0,7533 | 1,65621E-35 |
| ACE2 | ATP11C | -0,7533 | 1,66255E-35 |
| ACE2 | CFAP58 | -0,7534 | 1,61764E-35 |
| ACE2 | AKIP1 | -0,7534 | 1,58812E-35 |
| ACE2 | LRRC37BP1 | -0,7534 | 1,59492E-35 |
| ACE2 | LOXL1 | -0,7535 | 1,56657E-35 |
| ACE2 | ZNF208 | -0,7535 | 1,54519E-35 |
| ACE2 | ACKR1 | -0,7537 | 1,47424E-35 |
| ACE2 | CSE1L | -0,7537 | 1,47238E-35 |
| ACE2 | ZC4H2 | -0,7537 | 1,47835E-35 |
| ACE2 | GGPS1 | -0,7538 | 1,40451E-35 |
| ACE2 | TTF1 | -0,7539 | 1,35957E-35 |
| ACE2 | LINC00869 | -0,754 | 1,3163E-35 |
| ACE2 | GRK5 | -0,754 | 1,325E-35 |
| ACE2 | EMCN | -0,7541 | 1,30202E-35 |
| ACE2 | BNC2 | -0,7541 | 1,29467E-35 |
| ACE2 | GLMN | -0,7542 | 1,22476E-35 |
| ACE2 | ZNF512 | -0,7542 | 1,24737E-35 |
| ACE2 | PCMTD1 | -0,7542 | 1,22626E-35 |
| ACE2 | MPP2 | -0,7542 | 1,26144E-35 |
| ACE2 | PPOX | -0,7543 | 1,19784E-35 |
| ACE2 | DEGS1 | -0,7543 | 1,19112E-35 |
| ACE2 | DHX40 | -0,7543 | 1,20196E-35 |
| ACE2 | LRP11 | -0,7544 | 1,15369E-35 |
| ACE2 | PPP1R12A | -0,7544 | 1,17318E-35 |
| ACE2 | MAGEE1 | -0,7544 | 1,15783E-35 |
| ACE2 | PHTF1 | -0,7545 | 1,14443E-35 |
| ACE2 | C3orf58 | -0,7545 | 1,13839E-35 |

| | | | |
|---|---|---|---|
| ACE2 | MRGPRF | -0,7545 | 1,138E-35 |
| ACE2 | ZFP90 | -0,7545 | 1,13537E-35 |
| ACE2 | TMEM231 | -0,7545 | 1,13448E-35 |
| ACE2 | FILIP1L | -0,7546 | 1,07716E-35 |
| ACE2 | SRSF3 | -0,7546 | 1,08123E-35 |
| ACE2 | RP11-54O7.3 | -0,7548 | 1,03693E-35 |
| ACE2 | GRK4 | -0,7548 | 1,01983E-35 |
| ACE2 | AHI1 | -0,7548 | 1,03033E-35 |
| ACE2 | TSPAN9 | -0,7548 | 1,02132E-35 |
| ACE2 | TSPYL5 | -0,7549 | 9,92895E-36 |
| ACE2 | VAT1L | -0,7549 | 9,87123E-36 |
| ACE2 | METTL5 | -0,7551 | 9,22694E-36 |
| ACE2 | IFI27L2 | -0,7552 | 8,97056E-36 |
| ACE2 | ARSB | -0,7553 | 8,78568E-36 |
| ACE2 | NUDCD3 | -0,7553 | 8,79745E-36 |
| ACE2 | RPL35 | -0,7553 | 8,74104E-36 |
| ACE2 | STX8 | -0,7553 | 8,71749E-36 |
| ACE2 | TCEAL4 | -0,7553 | 8,79294E-36 |
| ACE2 | SGIP1 | -0,7554 | 8,52757E-36 |
| ACE2 | SLC2A4RG | -0,7554 | 8,37772E-36 |
| ACE2 | ADGRL4 | -0,7555 | 8,05976E-36 |
| ACE2 | CSNK2A2 | -0,7555 | 8,02025E-36 |
| ACE2 | RP11-686D22.7 | -0,7556 | 7,83412E-36 |
| ACE2 | ZFP82 | -0,7557 | 7,66287E-36 |
| ACE2 | TRIOBP | -0,7557 | 7,55856E-36 |
| ACE2 | MOSPD1 | -0,7557 | 7,53534E-36 |
| ACE2 | SDK1 | -0,7558 | 7,40634E-36 |
| ACE2 | NRP1 | -0,7558 | 7,46441E-36 |
| ACE2 | TMEM5 | -0,7558 | 7,38467E-36 |
| ACE2 | DOC2B | -0,7558 | 7,30523E-36 |
| ACE2 | PBX2 | -0,7559 | 7,22094E-36 |
| ACE2 | MAK16 | -0,756 | 6,94443E-36 |
| ACE2 | SLC26A4 | -0,7561 | 6,71292E-36 |
| ACE2 | FAM65A | -0,7561 | 6,6842E-36 |
| ACE2 | ERCC1 | -0,7561 | 6,79927E-36 |
| ACE2 | C10orf11 | -0,7562 | 6,47679E-36 |
| ACE2 | ANTXR1 | -0,7563 | 6,26275E-36 |
| ACE2 | LIFR-AS1 | -0,7563 | 6,16729E-36 |
| ACE2 | RP11-1134I14.8 | -0,7563 | 6,25756E-36 |
| ACE2 | SNHG23 | -0,7563 | 6,23569E-36 |
| ACE2 | SRPK2 | -0,7564 | 6,04495E-36 |
| ACE2 | C9orf3 | -0,7564 | 6,1284E-36 |
| ACE2 | CEP164 | -0,7564 | 6,09844E-36 |
| ACE2 | ZNF330 | -0,7565 | 5,93663E-36 |
| ACE2 | KCNMB4 | -0,7565 | 5,8366E-36 |
| ACE2 | TMEM263 | -0,7565 | 5,93081E-36 |
| ACE2 | TMEM255A | -0,7565 | 5,94762E-36 |
| ACE2 | PTPN13 | -0,7566 | 5,75471E-36 |
| ACE2 | ANGPTL2 | -0,7566 | 5,6207E-36 |
| ACE2 | MFAP4 | -0,7566 | 5,6273E-36 |
| ACE2 | FBXL2 | -0,7567 | 5,41514E-36 |
| ACE2 | MOSPD3 | -0,7567 | 5,40735E-36 |
| ACE2 | OXR1 | -0,7567 | 5,54428E-36 |
| ACE2 | PIBF1 | -0,7567 | 5,45121E-36 |

| | | | |
|---|---|---|---|
| ACE2 | HIC1 | -0,7567 | 5,47201E-36 |
| ACE2 | LINC00665 | -0,7567 | 5,56126E-36 |
| ACE2 | GPR161 | -0,7568 | 5,354E-36 |
| ACE2 | RNF122 | -0,7568 | 5,37606E-36 |
| ACE2 | ZNF415 | -0,7568 | 5,27358E-36 |
| ACE2 | NDUFAF2 | -0,7569 | 5,14671E-36 |
| ACE2 | STARD9 | -0,7569 | 5,17168E-36 |
| ACE2 | STK32A | -0,757 | 4,9761E-36 |
| ACE2 | RNASEH2C | -0,757 | 5,00323E-36 |
| ACE2 | KAT2A | -0,757 | 5,02633E-36 |
| ACE2 | LTBP1 | -0,7571 | 4,8876E-36 |
| ACE2 | OSMR | -0,7571 | 4,88078E-36 |
| ACE2 | EBF3 | -0,7571 | 4,75894E-36 |
| ACE2 | TTLL5 | -0,7571 | 4,82338E-36 |
| ACE2 | SLC25A32 | -0,7572 | 4,65271E-36 |
| ACE2 | ZNF133 | -0,7572 | 4,69443E-36 |
| ACE2 | THBS3 | -0,7573 | 4,48991E-36 |
| ACE2 | NEDD1 | -0,7573 | 4,44427E-36 |
| ACE2 | ISCU | -0,7573 | 4,54336E-36 |
| ACE2 | MINDY2 | -0,7573 | 4,54046E-36 |
| ACE2 | NPM1 | -0,7575 | 4,19199E-36 |
| ACE2 | ADGRB3 | -0,7575 | 4,22324E-36 |
| ACE2 | CEP83 | -0,7575 | 4,17172E-36 |
| ACE2 | UBE2G2 | -0,7575 | 4,22593E-36 |
| ACE2 | TTLL1 | -0,7575 | 4,25872E-36 |
| ACE2 | HFM1 | -0,7576 | 4,04247E-36 |
| ACE2 | PHF10 | -0,7576 | 4,07707E-36 |
| ACE2 | DTNA | -0,7576 | 4,14795E-36 |
| ACE2 | GPX7 | -0,7578 | 3,82048E-36 |
| ACE2 | MTERF3 | -0,7578 | 3,7594E-36 |
| ACE2 | HSPB8 | -0,7578 | 3,79308E-36 |
| ACE2 | FKBP8 | -0,7578 | 3,76564E-36 |
| ACE2 | DYNC2H1 | -0,7579 | 3,69523E-36 |
| ACE2 | METTL26 | -0,7579 | 3,68864E-36 |
| ACE2 | NOTCH3 | -0,7579 | 3,75785E-36 |
| ACE2 | RTN2 | -0,7579 | 3,71504E-36 |
| ACE2 | NRP2 | -0,758 | 3,60714E-36 |
| ACE2 | FOXP1 | -0,758 | 3,54187E-36 |
| ACE2 | SERPING1 | -0,758 | 3,53287E-36 |
| ACE2 | DRAM1 | -0,758 | 3,52766E-36 |
| ACE2 | INAFM2 | -0,758 | 3,52899E-36 |
| ACE2 | GADD45GIP1 | -0,758 | 3,56494E-36 |
| ACE2 | EEF1D | -0,7581 | 3,43603E-36 |
| ACE2 | TBC1D1 | -0,7582 | 3,38346E-36 |
| ACE2 | KLHL7 | -0,7582 | 3,33003E-36 |
| ACE2 | TMEM8B | -0,7583 | 3,2521E-36 |
| ACE2 | METTL22 | -0,7583 | 3,27557E-36 |
| ACE2 | CACNA2D3 | -0,7584 | 3,17939E-36 |
| ACE2 | CADM4 | -0,7584 | 3,1713E-36 |
| ACE2 | NTRK1 | -0,7585 | 3,01671E-36 |
| ACE2 | MBNL2 | -0,7585 | 3,06853E-36 |
| ACE2 | EXOC5 | -0,7585 | 3,02026E-36 |
| ACE2 | EIF1 | -0,7585 | 3,0753E-36 |
| ACE2 | S100B | -0,7585 | 3,00684E-36 |

| | | | |
|---|---|---|---|
| ACE2 | OLFML3 | -0,7587 | 2,82851E-36 |
| ACE2 | SNAP47 | -0,7587 | 2,86723E-36 |
| ACE2 | OSGIN2 | -0,7587 | 2,84472E-36 |
| ACE2 | SLC29A1 | -0,7588 | 2,75903E-36 |
| ACE2 | ZNF676 | -0,7588 | 2,77811E-36 |
| ACE2 | RAB33B | -0,7589 | 2,65879E-36 |
| ACE2 | AC004540.4 | -0,7589 | 2,68619E-36 |
| ACE2 | INTS3 | -0,759 | 2,54512E-36 |
| ACE2 | KIF5C | -0,759 | 2,52779E-36 |
| ACE2 | FNBP1 | -0,759 | 2,58819E-36 |
| ACE2 | ZNF302 | -0,759 | 2,55981E-36 |
| ACE2 | STK19 | -0,7591 | 2,4527E-36 |
| ACE2 | NECAB1 | -0,7591 | 2,44862E-36 |
| ACE2 | SSBP4 | -0,7591 | 2,51365E-36 |
| ACE2 | PARK7 | -0,7592 | 2,41691E-36 |
| ACE2 | DSTYK | -0,7592 | 2,4325E-36 |
| ACE2 | DRC3 | -0,7592 | 2,37605E-36 |
| ACE2 | MAPRE2 | -0,7592 | 2,39527E-36 |
| ACE2 | NDUFAF4 | -0,7593 | 2,34602E-36 |
| ACE2 | NCALD | -0,7593 | 2,36078E-36 |
| ACE2 | IPO7 | -0,7593 | 2,35213E-36 |
| ACE2 | NUMBL | -0,7593 | 2,33779E-36 |
| ACE2 | TOMM6 | -0,7594 | 2,21764E-36 |
| ACE2 | KATNA1 | -0,7594 | 2,24493E-36 |
| ACE2 | RP11-755F10.3 | -0,7594 | 2,27082E-36 |
| ACE2 | PDGFD | -0,7594 | 2,27943E-36 |
| ACE2 | METAP2 | -0,7595 | 2,19715E-36 |
| ACE2 | EPS15L1 | -0,7596 | 2,10656E-36 |
| ACE2 | CA11 | -0,7597 | 2,03512E-36 |
| ACE2 | EBF1 | -0,7598 | 1,93831E-36 |
| ACE2 | GNL1 | -0,7598 | 1,93941E-36 |
| ACE2 | ZNF639 | -0,7599 | 1,87069E-36 |
| ACE2 | SPRY1 | -0,7599 | 1,91506E-36 |
| ACE2 | RELA | -0,7599 | 1,8783E-36 |
| ACE2 | INAFM1 | -0,7599 | 1,87929E-36 |
| ACE2 | PPT2 | -0,76 | 1,84959E-36 |
| ACE2 | PELI3 | -0,76 | 1,85257E-36 |
| ACE2 | KIAA1614 | -0,7601 | 1,76E-36 |
| ACE2 | CTD-2006C1.2 | -0,7601 | 1,76454E-36 |
| ACE2 | GSTM5 | -0,7602 | 1,69475E-36 |
| ACE2 | SH3RF3 | -0,7602 | 1,72804E-36 |
| ACE2 | KDELC1 | -0,7602 | 1,74255E-36 |
| ACE2 | CD99L2 | -0,7602 | 1,73447E-36 |
| ACE2 | MACF1 | -0,7603 | 1,68192E-36 |
| ACE2 | SELP | -0,7603 | 1,64533E-36 |
| ACE2 | ZNHIT6 | -0,7604 | 1,62686E-36 |
| ACE2 | MTMR9 | -0,7604 | 1,60293E-36 |
| ACE2 | ELOC | -0,7606 | 1,52425E-36 |
| ACE2 | USP44 | -0,7606 | 1,5105E-36 |
| ACE2 | CDK2AP1 | -0,7606 | 1,49529E-36 |
| ACE2 | DEF8 | -0,7606 | 1,50207E-36 |
| ACE2 | HOXC5 | -0,7607 | 1,47208E-36 |
| ACE2 | SLC35F1 | -0,7608 | 1,40265E-36 |
| ACE2 | NR2F2 | -0,7608 | 1,41554E-36 |

| | | | |
|---|---|---|---|
| ACE2 | PORCN | -0,7608 | 1,42648E-36 |
| ACE2 | RBSN | -0,7609 | 1,35062E-36 |
| ACE2 | ADGRL2 | -0,761 | 1,30869E-36 |
| ACE2 | LRP2BP | -0,761 | 1,30025E-36 |
| ACE2 | GDI1 | -0,761 | 1,32671E-36 |
| ACE2 | ARAP3 | -0,7611 | 1,26035E-36 |
| ACE2 | CAHM | -0,7611 | 1,2873E-36 |
| ACE2 | GRASP | -0,7611 | 1,26122E-36 |
| ACE2 | FAM227B | -0,7611 | 1,26854E-36 |
| ACE2 | POLR3GL | -0,7612 | 1,24985E-36 |
| ACE2 | ATP8B2 | -0,7612 | 1,23672E-36 |
| ACE2 | MSX1 | -0,7612 | 1,2314E-36 |
| ACE2 | CYYR1 | -0,7612 | 1,21316E-36 |
| ACE2 | TACC1 | -0,7613 | 1,19445E-36 |
| ACE2 | PHYHD1 | -0,7613 | 1,20398E-36 |
| ACE2 | UFSP2 | -0,7614 | 1,14636E-36 |
| ACE2 | PCDHB14 | -0,7614 | 1,1454E-36 |
| ACE2 | PKMP3 | -0,7616 | 1,06454E-36 |
| ACE2 | CLMP | -0,7616 | 1,07104E-36 |
| ACE2 | MDM1 | -0,7616 | 1,06799E-36 |
| ACE2 | TM2D3 | -0,7616 | 1,06094E-36 |
| ACE2 | VMA21 | -0,7616 | 1,08199E-36 |
| ACE2 | SH3PXD2A | -0,7617 | 1,03909E-36 |
| ACE2 | METTL16 | -0,7617 | 1,03772E-36 |
| ACE2 | ZNF570 | -0,7617 | 1,03997E-36 |
| ACE2 | ATF4 | -0,7617 | 1,03608E-36 |
| ACE2 | GLI2 | -0,7618 | 1,00998E-36 |
| ACE2 | SLIT2 | -0,7619 | 9,84685E-37 |
| ACE2 | HARS | -0,7619 | 9,71999E-37 |
| ACE2 | SSH1 | -0,7619 | 9,71741E-37 |
| ACE2 | ADAMTS12 | -0,762 | 9,36595E-37 |
| ACE2 | PCDHB12 | -0,762 | 9,45559E-37 |
| ACE2 | CASD1 | -0,762 | 9,29304E-37 |
| ACE2 | PLPP7 | -0,762 | 9,38738E-37 |
| ACE2 | ZNF595 | -0,7621 | 9,16375E-37 |
| ACE2 | FAM66C | -0,7621 | 9,17646E-37 |
| ACE2 | DCLK1 | -0,7621 | 9,1439E-37 |
| ACE2 | C14orf166 | -0,7621 | 9,1793E-37 |
| ACE2 | PRPF31 | -0,7621 | 9,03369E-37 |
| ACE2 | LAMA4 | -0,7622 | 8,77124E-37 |
| ACE2 | NDN | -0,7622 | 8,76555E-37 |
| ACE2 | NUDT10 | -0,7622 | 8,76995E-37 |
| ACE2 | LINC01278 | -0,7623 | 8,61063E-37 |
| ACE2 | DCHS2 | -0,7624 | 8,09809E-37 |
| ACE2 | RP11-1114A5.4 | -0,7625 | 7,9839E-37 |
| ACE2 | FSCN1 | -0,7626 | 7,75796E-37 |
| ACE2 | MLLT10 | -0,7626 | 7,55048E-37 |
| ACE2 | TWSG1 | -0,7626 | 7,69945E-37 |
| ACE2 | TXNRD3 | -0,7627 | 7,30846E-37 |
| ACE2 | C5orf24 | -0,7627 | 7,44253E-37 |
| ACE2 | CMAHP | -0,7627 | 7,52217E-37 |
| ACE2 | UBE2H | -0,7627 | 7,47204E-37 |
| ACE2 | SPAG8 | -0,7628 | 7,26942E-37 |
| ACE2 | CCND2-AS1 | -0,763 | 6,59433E-37 |

| | | | |
|---|---|---|---|
| ACE2 | SUMO2 | -0,763 | 6,63526E-37 |
| ACE2 | EGLN2 | -0,763 | 6,73536E-37 |
| ACE2 | RAPGEF4 | -0,7631 | 6,52414E-37 |
| ACE2 | KCTD15 | -0,7631 | 6,39695E-37 |
| ACE2 | EIF4ENIF1 | -0,7631 | 6,57042E-37 |
| ACE2 | WDR19 | -0,7632 | 6,22863E-37 |
| ACE2 | ROBO4 | -0,7632 | 6,17619E-37 |
| ACE2 | RNF115 | -0,7633 | 5,98758E-37 |
| ACE2 | SERPINH1 | -0,7634 | 5,84428E-37 |
| ACE2 | PRELID3A | -0,7634 | 5,89368E-37 |
| ACE2 | SOX12 | -0,7634 | 5,91182E-37 |
| ACE2 | SENP6 | -0,7635 | 5,64207E-37 |
| ACE2 | SLC2A3 | -0,7635 | 5,64015E-37 |
| ACE2 | MXRA8 | -0,7636 | 5,45471E-37 |
| ACE2 | CTSF | -0,7636 | 5,38016E-37 |
| ACE2 | RP11-713P17.3 | -0,7636 | 5,50864E-37 |
| ACE2 | KANK3 | -0,7636 | 5,42077E-37 |
| ACE2 | NES | -0,7637 | 5,31671E-37 |
| ACE2 | RUFY2 | -0,7637 | 5,2129E-37 |
| ACE2 | MIR4697HG | -0,7637 | 5,21516E-37 |
| ACE2 | PRKD3 | -0,7638 | 5,12374E-37 |
| ACE2 | PRMT9 | -0,7638 | 5,1514E-37 |
| ACE2 | CYP7B1 | -0,7638 | 5,02799E-37 |
| ACE2 | PABPC1P4 | -0,7638 | 5,02388E-37 |
| ACE2 | SNHG19 | -0,7638 | 5,069E-37 |
| ACE2 | CEP89 | -0,7638 | 5,05872E-37 |
| ACE2 | NXT1 | -0,7638 | 5,16206E-37 |
| ACE2 | SRF | -0,7639 | 4,89042E-37 |
| ACE2 | ECD | -0,7639 | 4,90434E-37 |
| ACE2 | TTC8 | -0,7639 | 4,86929E-37 |
| ACE2 | SLC38A6 | -0,764 | 4,82872E-37 |
| ACE2 | UTP11 | -0,7641 | 4,5756E-37 |
| ACE2 | CD34 | -0,7641 | 4,55208E-37 |
| ACE2 | ZSCAN16-AS1 | -0,7641 | 4,66342E-37 |
| ACE2 | ENTPD1 | -0,7641 | 4,54869E-37 |
| ACE2 | PHF21A | -0,7641 | 4,67274E-37 |
| ACE2 | ZNF569 | -0,7642 | 4,52947E-37 |
| ACE2 | METAP1D | -0,7643 | 4,32564E-37 |
| ACE2 | TDRD3 | -0,7643 | 4,27829E-37 |
| ACE2 | GZF1 | -0,7643 | 4,28468E-37 |
| ACE2 | JAKMIP2 | -0,7644 | 4,11626E-37 |
| ACE2 | EEF1A1 | -0,7644 | 4,19384E-37 |
| ACE2 | EPB41L2 | -0,7644 | 4,21332E-37 |
| ACE2 | POLR2K | -0,7644 | 4,10803E-37 |
| ACE2 | PCID2 | -0,7644 | 4,18025E-37 |
| ACE2 | CKMT2-AS1 | -0,7645 | 3,99693E-37 |
| ACE2 | SKI | -0,7646 | 3,83153E-37 |
| ACE2 | TMEM201 | -0,7646 | 3,87093E-37 |
| ACE2 | RP11-798K23.5 | -0,7646 | 3,90309E-37 |
| ACE2 | PTHLH | -0,7646 | 3,8764E-37 |
| ACE2 | LGI2 | -0,7647 | 3,7728E-37 |
| ACE2 | TMEM178B | -0,7647 | 3,79967E-37 |
| ACE2 | MED27 | -0,7648 | 3,59895E-37 |
| ACE2 | MORN4 | -0,7648 | 3,64517E-37 |

| | | | |
|---|---|---|---|
| ACE2 | XPOT | -0,7648 | 3,67562E-37 |
| ACE2 | C16orf86 | -0,7648 | 3,60321E-37 |
| ACE2 | NLGN4X | -0,7648 | 3,67535E-37 |
| ACE2 | PRRT1 | -0,7649 | 3,46532E-37 |
| ACE2 | CEP170 | -0,765 | 3,42204E-37 |
| ACE2 | EPM2A | -0,765 | 3,43996E-37 |
| ACE2 | RP11-213G2.3 | -0,765 | 3,38568E-37 |
| ACE2 | IMPDH1 | -0,7651 | 3,30145E-37 |
| ACE2 | RIC8B | -0,7651 | 3,22286E-37 |
| ACE2 | TBCA | -0,7652 | 3,20832E-37 |
| ACE2 | CPNE8 | -0,7652 | 3,1497E-37 |
| ACE2 | FGF14 | -0,7652 | 3,13206E-37 |
| ACE2 | SACS | -0,7653 | 3,11106E-37 |
| ACE2 | AHDC1 | -0,7654 | 2,95905E-37 |
| ACE2 | XPNPEP3 | -0,7654 | 2,91353E-37 |
| ACE2 | HDAC8 | -0,7654 | 2,96385E-37 |
| ACE2 | ZEB1-AS1 | -0,7655 | 2,86508E-37 |
| ACE2 | SNHG16 | -0,7655 | 2,87852E-37 |
| ACE2 | VPS45 | -0,7656 | 2,71237E-37 |
| ACE2 | TOR1AIP1 | -0,7656 | 2,74549E-37 |
| ACE2 | ADCY5 | -0,7656 | 2,71562E-37 |
| ACE2 | TMEM242 | -0,7656 | 2,80068E-37 |
| ACE2 | PMEPA1 | -0,7656 | 2,75677E-37 |
| ACE2 | ABL2 | -0,7657 | 2,71069E-37 |
| ACE2 | SOCS5 | -0,7657 | 2,62055E-37 |
| ACE2 | EVC | -0,7657 | 2,63286E-37 |
| ACE2 | BEND6 | -0,7657 | 2,65247E-37 |
| ACE2 | PLSCR3 | -0,7657 | 2,63069E-37 |
| ACE2 | FAIM | -0,7658 | 2,56898E-37 |
| ACE2 | SEPT7 | -0,7658 | 2,6011E-37 |
| ACE2 | TMEFF1 | -0,7658 | 2,60908E-37 |
| ACE2 | CTD-2054N24.2 | -0,7658 | 2,59883E-37 |
| ACE2 | ZNF71 | -0,7658 | 2,55109E-37 |
| ACE2 | CBY1 | -0,7658 | 2,6196E-37 |
| ACE2 | SORCS1 | -0,7659 | 2,48984E-37 |
| ACE2 | HSPB2 | -0,7659 | 2,47477E-37 |
| ACE2 | MTCP1 | -0,7659 | 2,47477E-37 |
| ACE2 | GUCY1A2 | -0,7661 | 2,30527E-37 |
| ACE2 | MED21 | -0,7661 | 2,31465E-37 |
| ACE2 | SNPH | -0,7661 | 2,30937E-37 |
| ACE2 | PRRX1 | -0,7662 | 2,25554E-37 |
| ACE2 | MPV17 | -0,7662 | 2,23581E-37 |
| ACE2 | NBN | -0,7662 | 2,208E-37 |
| ACE2 | LPAR1 | -0,7662 | 2,23654E-37 |
| ACE2 | TTC3 | -0,7662 | 2,2782E-37 |
| ACE2 | IKZF4 | -0,7663 | 2,17913E-37 |
| ACE2 | ADCY4 | -0,7663 | 2,20172E-37 |
| ACE2 | HOXB4 | -0,7663 | 2,16855E-37 |
| ACE2 | NKILA | -0,7663 | 2,1856E-37 |
| ACE2 | TTC28-AS1 | -0,7663 | 2,17722E-37 |
| ACE2 | PRKCDBP | -0,7664 | 2,06461E-37 |
| ACE2 | GTF2H5 | -0,7665 | 2,04856E-37 |
| ACE2 | KIF26A | -0,7665 | 1,99954E-37 |
| ACE2 | NHSL2 | -0,7665 | 2,02054E-37 |

| | | | |
|---|---|---|---|
| ACE2 | RRP8 | -0,7666 | 1,9398E-37 |
| ACE2 | ARHGEF25 | -0,7666 | 1,93479E-37 |
| ACE2 | AC003973.3 | -0,7666 | 1,95865E-37 |
| ACE2 | ADRB2 | -0,7668 | 1,80085E-37 |
| ACE2 | PHC1 | -0,7668 | 1,85041E-37 |
| ACE2 | DDX10 | -0,7669 | 1,7638E-37 |
| ACE2 | ACSL4 | -0,7669 | 1,74931E-37 |
| ACE2 | ELOVL4 | -0,767 | 1,69186E-37 |
| ACE2 | SLC48A1 | -0,767 | 1,69023E-37 |
| ACE2 | ZNF343 | -0,767 | 1,6889E-37 |
| ACE2 | SPIN2B | -0,767 | 1,7235E-37 |
| ACE2 | PPT1 | -0,7671 | 1,63918E-37 |
| ACE2 | MFF | -0,7671 | 1,62142E-37 |
| ACE2 | COMMD10 | -0,7671 | 1,64797E-37 |
| ACE2 | TGFBR1 | -0,7672 | 1,60796E-37 |
| ACE2 | BEND5 | -0,7673 | 1,54621E-37 |
| ACE2 | ANKRD39 | -0,7673 | 1,52142E-37 |
| ACE2 | MLH1 | -0,7673 | 1,51001E-37 |
| ACE2 | RABGAP1 | -0,7673 | 1,51104E-37 |
| ACE2 | TUBA1A | -0,7673 | 1,54973E-37 |
| ACE2 | ZNF614 | -0,7673 | 1,52941E-37 |
| ACE2 | FGF10 | -0,7674 | 1,47766E-37 |
| ACE2 | NR2F1 | -0,7674 | 1,46009E-37 |
| ACE2 | PYROXD2 | -0,7674 | 1,48107E-37 |
| ACE2 | PCAT19 | -0,7674 | 1,48804E-37 |
| ACE2 | SLC27A3 | -0,7675 | 1,40839E-37 |
| ACE2 | ITIH5 | -0,7675 | 1,43733E-37 |
| ACE2 | PRICKLE1 | -0,7675 | 1,41194E-37 |
| ACE2 | DMPK | -0,7675 | 1,44898E-37 |
| ACE2 | ZNF74 | -0,7675 | 1,41381E-37 |
| ACE2 | STX2 | -0,7676 | 1,37846E-37 |
| ACE2 | ZNF626 | -0,7677 | 1,3298E-37 |
| ACE2 | ABI3BP | -0,7678 | 1,28537E-37 |
| ACE2 | PM20D2 | -0,7678 | 1,28042E-37 |
| ACE2 | GPC6 | -0,7678 | 1,30962E-37 |
| ACE2 | LDLRAD2 | -0,768 | 1,22017E-37 |
| ACE2 | EXTL1 | -0,768 | 1,19782E-37 |
| ACE2 | BMT2 | -0,768 | 1,21975E-37 |
| ACE2 | SLC36A4 | -0,768 | 1,21498E-37 |
| ACE2 | COPZ2 | -0,768 | 1,20022E-37 |
| ACE2 | DDR2 | -0,7681 | 1,14361E-37 |
| ACE2 | MEF2C-AS1 | -0,7681 | 1,14676E-37 |
| ACE2 | ZNF329 | -0,7681 | 1,17815E-37 |
| ACE2 | RRP15 | -0,7682 | 1,1242E-37 |
| ACE2 | ASNSD1 | -0,7682 | 1,13524E-37 |
| ACE2 | LRRC23 | -0,7682 | 1,11499E-37 |
| ACE2 | GRAMD1A | -0,7682 | 1,11242E-37 |
| ACE2 | C9orf47 | -0,7683 | 1,08825E-37 |
| ACE2 | PYGM | -0,7683 | 1,08352E-37 |
| ACE2 | PHF2 | -0,7684 | 1,02979E-37 |
| ACE2 | TCF4 | -0,7684 | 1,05916E-37 |
| ACE2 | LMO2 | -0,7686 | 9,89234E-38 |
| ACE2 | MRC2 | -0,7686 | 9,90306E-38 |
| ACE2 | TSPYL4 | -0,7687 | 9,48434E-38 |

| | | | |
|---|---|---|---|
| ACE2 | TBCD | -0,7687 | 9,27141E-38 |
| ACE2 | EIF5A2 | -0,7688 | 8,96921E-38 |
| ACE2 | LRRC17 | -0,7688 | 8,97286E-38 |
| ACE2 | HOXC9 | -0,7689 | 8,83232E-38 |
| ACE2 | CCL2 | -0,7689 | 8,66347E-38 |
| ACE2 | BFSP1 | -0,7689 | 8,69857E-38 |
| ACE2 | NACAD | -0,769 | 8,59496E-38 |
| ACE2 | TMEM121 | -0,769 | 8,47185E-38 |
| ACE2 | RP11-456K23.1 | -0,769 | 8,44681E-38 |
| ACE2 | MTURN | -0,7691 | 8,12167E-38 |
| ACE2 | RAB6B | -0,7692 | 7,80193E-38 |
| ACE2 | SULF1 | -0,7692 | 7,89398E-38 |
| ACE2 | FUNDC2 | -0,7692 | 7,78586E-38 |
| ACE2 | PHF13 | -0,7693 | 7,77194E-38 |
| ACE2 | GFRA1 | -0,7693 | 7,62158E-38 |
| ACE2 | EYA2 | -0,7693 | 7,66822E-38 |
| ACE2 | SPACA9 | -0,7694 | 7,32162E-38 |
| ACE2 | UPF3A | -0,7694 | 7,39691E-38 |
| ACE2 | KIAA1462 | -0,7695 | 7,23276E-38 |
| ACE2 | PTPRA | -0,7696 | 6,79676E-38 |
| ACE2 | TTL | -0,7698 | 6,37184E-38 |
| ACE2 | RNF180 | -0,7698 | 6,47523E-38 |
| ACE2 | C6orf1 | -0,7698 | 6,39587E-38 |
| ACE2 | SALL2 | -0,7698 | 6,49634E-38 |
| ACE2 | MEIS3 | -0,7698 | 6,36979E-38 |
| ACE2 | SNHG6 | -0,7699 | 6,27349E-38 |
| ACE2 | NUDCD1 | -0,7699 | 6,16549E-38 |
| ACE2 | EPOR | -0,7699 | 6,23561E-38 |
| ACE2 | CLIC4 | -0,77 | 6,04162E-38 |
| ACE2 | ZFAND1 | -0,77 | 5,91161E-38 |
| ACE2 | NEK9 | -0,77 | 5,98186E-38 |
| ACE2 | MEX3B | -0,77 | 5,90891E-38 |
| ACE2 | TMIE | -0,7701 | 5,68129E-38 |
| ACE2 | PFDN5 | -0,7701 | 5,8705E-38 |
| ACE2 | IGFBP4 | -0,7702 | 5,49346E-38 |
| ACE2 | RPS6KB1 | -0,7702 | 5,67621E-38 |
| ACE2 | TSHZ3 | -0,7702 | 5,49101E-38 |
| ACE2 | RP4-756G23.5 | -0,7702 | 5,55267E-38 |
| ACE2 | MFAP2 | -0,7703 | 5,39347E-38 |
| ACE2 | C3orf33 | -0,7703 | 5,30872E-38 |
| ACE2 | ST8SIA1 | -0,7703 | 5,34389E-38 |
| ACE2 | ZSCAN26 | -0,7704 | 5,18789E-38 |
| ACE2 | BAALC | -0,7704 | 5,18937E-38 |
| ACE2 | AMOTL1 | -0,7704 | 5,18274E-38 |
| ACE2 | FAM53C | -0,7706 | 4,84387E-38 |
| ACE2 | PHF20 | -0,7706 | 4,7776E-38 |
| ACE2 | DCUN1D5 | -0,7707 | 4,67149E-38 |
| ACE2 | EIF2AK4 | -0,7707 | 4,61535E-38 |
| ACE2 | GCSH | -0,7708 | 4,51088E-38 |
| ACE2 | ZDHHC15 | -0,7708 | 4,44269E-38 |
| ACE2 | TP73-AS1 | -0,7709 | 4,37572E-38 |
| ACE2 | FLT4 | -0,771 | 4,194E-38 |
| ACE2 | RP11-359B12.2 | -0,771 | 4,25472E-38 |
| ACE2 | LSM10 | -0,7711 | 4,05656E-38 |

| | | | |
|---|---|---|---|
| ACE2 | SNCA | -0,7711 | 4,09537E-38 |
| ACE2 | DBN1 | -0,7711 | 4,04374E-38 |
| ACE2 | COL27A1 | -0,7711 | 4,03116E-38 |
| ACE2 | LARP6 | -0,7711 | 4,00122E-38 |
| ACE2 | CWC27 | -0,7712 | 3,8946E-38 |
| ACE2 | PPM1M | -0,7713 | 3,82538E-38 |
| ACE2 | C12orf10 | -0,7713 | 3,83795E-38 |
| ACE2 | GUSBP1 | -0,7714 | 3,70282E-38 |
| ACE2 | ERICD | -0,7714 | 3,70208E-38 |
| ACE2 | DCHS1 | -0,7714 | 3,66575E-38 |
| ACE2 | CCDC81 | -0,7714 | 3,718E-38 |
| ACE2 | SDC3 | -0,7715 | 3,52729E-38 |
| ACE2 | NUPR1 | -0,7715 | 3,5676E-38 |
| ACE2 | LONRF1 | -0,7716 | 3,46148E-38 |
| ACE2 | PGM2L1 | -0,7716 | 3,36022E-38 |
| ACE2 | MPDZ | -0,7719 | 3,01205E-38 |
| ACE2 | NRIP2 | -0,7719 | 3,05266E-38 |
| ACE2 | RP5-1057I20.4 | -0,7719 | 3,05365E-38 |
| ACE2 | WWTR1 | -0,772 | 2,93823E-38 |
| ACE2 | RPS24 | -0,772 | 3,00557E-38 |
| ACE2 | AVPR2 | -0,772 | 2,97481E-38 |
| ACE2 | ZNF385A | -0,7721 | 2,87083E-38 |
| ACE2 | FAM124A | -0,7721 | 2,82804E-38 |
| ACE2 | LINC01315 | -0,7721 | 2,90058E-38 |
| ACE2 | DENND2C | -0,7722 | 2,79304E-38 |
| ACE2 | RAI14 | -0,7722 | 2,76823E-38 |
| ACE2 | LLGL1 | -0,7722 | 2,72202E-38 |
| ACE2 | ARL10 | -0,7723 | 2,69293E-38 |
| ACE2 | PDLIM7 | -0,7723 | 2,68699E-38 |
| ACE2 | PRPH2 | -0,7723 | 2,6692E-38 |
| ACE2 | PLS3 | -0,7723 | 2,65907E-38 |
| ACE2 | FRMD6 | -0,7724 | 2,55039E-38 |
| ACE2 | CDK17 | -0,7725 | 2,45159E-38 |
| ACE2 | RCN2 | -0,7725 | 2,50246E-38 |
| ACE2 | PHLDA3 | -0,7726 | 2,3947E-38 |
| ACE2 | NSMCE2 | -0,7726 | 2,39831E-38 |
| ACE2 | AFAP1L1 | -0,7727 | 2,27322E-38 |
| ACE2 | NRIP3 | -0,7727 | 2,28198E-38 |
| ACE2 | PTRF | -0,7727 | 2,34029E-38 |
| ACE2 | MAL | -0,7728 | 2,22937E-38 |
| ACE2 | NNT-AS1 | -0,7728 | 2,22763E-38 |
| ACE2 | SGTB | -0,7728 | 2,22937E-38 |
| ACE2 | SPARC | -0,7728 | 2,25575E-38 |
| ACE2 | H2AFV | -0,7728 | 2,19538E-38 |
| ACE2 | CROCCP3 | -0,7729 | 2,12502E-38 |
| ACE2 | SLIT3 | -0,7729 | 2,14624E-38 |
| ACE2 | RANBP3 | -0,7729 | 2,14921E-38 |
| ACE2 | KCNA2 | -0,7731 | 1,99396E-38 |
| ACE2 | SERPINI1 | -0,7731 | 2,02589E-38 |
| ACE2 | KCNN3 | -0,7732 | 1,95239E-38 |
| ACE2 | SMYD3 | -0,7732 | 1,95002E-38 |
| ACE2 | ASAP1 | -0,7732 | 1,94407E-38 |
| ACE2 | LRCH1 | -0,7732 | 1,91433E-38 |
| ACE2 | ZNF117 | -0,7733 | 1,868E-38 |

| | | | |
|---|---|---|---|
| ACE2 | GID4 | -0,7733 | 1,88038E-38 |
| ACE2 | KCND3 | -0,7735 | 1,71792E-38 |
| ACE2 | PNRC1 | -0,7735 | 1,75325E-38 |
| ACE2 | FDX2 | -0,7735 | 1,70806E-38 |
| ACE2 | CLEC11A | -0,7735 | 1,72442E-38 |
| ACE2 | RWDD2B | -0,7735 | 1,76257E-38 |
| ACE2 | HOMER1 | -0,7736 | 1,67243E-38 |
| ACE2 | HSF2 | -0,7737 | 1,64034E-38 |
| ACE2 | PKNOX1 | -0,7737 | 1,58548E-38 |
| ACE2 | PTPDC1 | -0,7739 | 1,49552E-38 |
| ACE2 | ILK | -0,7739 | 1,4973E-38 |
| ACE2 | ZNF146 | -0,7739 | 1,48994E-38 |
| ACE2 | CLASP2 | -0,774 | 1,42599E-38 |
| ACE2 | RPF2 | -0,774 | 1,46238E-38 |
| ACE2 | FAM220A | -0,774 | 1,44448E-38 |
| ACE2 | RUFY3 | -0,7741 | 1,37883E-38 |
| ACE2 | TMEM150C | -0,7741 | 1,39121E-38 |
| ACE2 | RNF150 | -0,7741 | 1,40923E-38 |
| ACE2 | ICE2 | -0,7741 | 1,40869E-38 |
| ACE2 | GRIK5 | -0,7741 | 1,41525E-38 |
| ACE2 | IFFO2 | -0,7743 | 1,29223E-38 |
| ACE2 | DLG2 | -0,7743 | 1,29597E-38 |
| ACE2 | MSH6 | -0,7745 | 1,19728E-38 |
| ACE2 | PSIP1 | -0,7745 | 1,19625E-38 |
| ACE2 | SRP14 | -0,7745 | 1,23016E-38 |
| ACE2 | C12orf43 | -0,7746 | 1,18613E-38 |
| ACE2 | SAV1 | -0,7746 | 1,16694E-38 |
| ACE2 | RBMXL1 | -0,7747 | 1,10619E-38 |
| ACE2 | ARSI | -0,7747 | 1,11689E-38 |
| ACE2 | ZMYND11 | -0,7747 | 1,1342E-38 |
| ACE2 | LAMA5 | -0,7748 | 1,09281E-38 |
| ACE2 | AAED1 | -0,7749 | 1,03966E-38 |
| ACE2 | EBF4 | -0,7749 | 1,06537E-38 |
| ACE2 | PTGER3 | -0,775 | 1,02667E-38 |
| ACE2 | MEOX2 | -0,775 | 1,00265E-38 |
| ACE2 | CCDC180 | -0,775 | 1,01656E-38 |
| ACE2 | ATP1B2 | -0,775 | 1,01652E-38 |
| ACE2 | MRPS14 | -0,7751 | 9,88918E-39 |
| ACE2 | LRIG1 | -0,7751 | 9,62582E-39 |
| ACE2 | RP3-329A5.8 | -0,7751 | 9,79981E-39 |
| ACE2 | ZNF638 | -0,7752 | 9,44025E-39 |
| ACE2 | ERG | -0,7753 | 9,20154E-39 |
| ACE2 | ZYX | -0,7754 | 8,63948E-39 |
| ACE2 | LRRC20 | -0,7754 | 8,80803E-39 |
| ACE2 | MAP4K5 | -0,7754 | 8,8405E-39 |
| ACE2 | LURAP1 | -0,7755 | 8,31455E-39 |
| ACE2 | TTC4 | -0,7755 | 8,56126E-39 |
| ACE2 | YEATS2 | -0,7755 | 8,3534E-39 |
| ACE2 | RAB11FIP2 | -0,7755 | 8,35046E-39 |
| ACE2 | C1orf54 | -0,7756 | 8,07754E-39 |
| ACE2 | ACBD6 | -0,7756 | 8,10574E-39 |
| ACE2 | RP11-401P9.5 | -0,7757 | 7,8058E-39 |
| ACE2 | MEOX1 | -0,7757 | 7,80363E-39 |
| ACE2 | TNFAIP8L3 | -0,7758 | 7,62894E-39 |

| | | | |
|---|---|---|---|
| ACE2 | PABPC5 | -0,7758 | 7,63359E-39 |
| ACE2 | LINC01197 | -0,7759 | 7,34316E-39 |
| ACE2 | ZEB2 | -0,7761 | 6,78585E-39 |
| ACE2 | PIEZO1 | -0,7761 | 6,81121E-39 |
| ACE2 | TRAPPC2B | -0,7761 | 6,69656E-39 |
| ACE2 | PRDM8 | -0,7762 | 6,59709E-39 |
| ACE2 | PURB | -0,7762 | 6,4944E-39 |
| ACE2 | LINC00094 | -0,7762 | 6,66374E-39 |
| ACE2 | RBMS3 | -0,7763 | 6,39305E-39 |
| ACE2 | RAB36 | -0,7764 | 6,08507E-39 |
| ACE2 | FAIM2 | -0,7765 | 5,85011E-39 |
| ACE2 | EPN2 | -0,7765 | 5,96282E-39 |
| ACE2 | BICD2 | -0,7766 | 5,63006E-39 |
| ACE2 | LRCH3 | -0,7768 | 5,24447E-39 |
| ACE2 | ERC1 | -0,7768 | 5,31453E-39 |
| ACE2 | NAP1L2 | -0,7768 | 5,24204E-39 |
| ACE2 | RSPO3 | -0,7769 | 5,01842E-39 |
| ACE2 | ABHD14A | -0,777 | 4,98132E-39 |
| ACE2 | INTU | -0,7771 | 4,77318E-39 |
| ACE2 | LRRC37A3 | -0,7771 | 4,72321E-39 |
| ACE2 | RBFA | -0,7771 | 4,72241E-39 |
| ACE2 | AIDA | -0,7772 | 4,51043E-39 |
| ACE2 | SLC25A12 | -0,7773 | 4,42025E-39 |
| ACE2 | PPP2R2B | -0,7773 | 4,45538E-39 |
| ACE2 | PDLIM4 | -0,7774 | 4,23178E-39 |
| ACE2 | CFL2 | -0,7774 | 4,26901E-39 |
| ACE2 | PSMC5 | -0,7774 | 4,179E-39 |
| ACE2 | WDPCP | -0,7776 | 3,98159E-39 |
| ACE2 | DCBLD2 | -0,7776 | 3,98938E-39 |
| ACE2 | GYG1 | -0,7776 | 3,89169E-39 |
| ACE2 | CHSY1 | -0,7776 | 3,97256E-39 |
| ACE2 | PGR | -0,7777 | 3,77598E-39 |
| ACE2 | MAP3K3 | -0,7777 | 3,85735E-39 |
| ACE2 | POLK | -0,7778 | 3,64486E-39 |
| ACE2 | ILKAP | -0,778 | 3,39197E-39 |
| ACE2 | TCTEX1D2 | -0,778 | 3,45995E-39 |
| ACE2 | ELP2 | -0,778 | 3,36631E-39 |
| ACE2 | ATF2 | -0,7781 | 3,25658E-39 |
| ACE2 | MMRN2 | -0,7782 | 3,1708E-39 |
| ACE2 | DYRK4 | -0,7782 | 3,21737E-39 |
| ACE2 | RP11-834C11.7 | -0,7782 | 3,18939E-39 |
| ACE2 | IFT140 | -0,7782 | 3,10459E-39 |
| ACE2 | ZNF433 | -0,7782 | 3,12998E-39 |
| ACE2 | TMEM136 | -0,7783 | 3,07591E-39 |
| ACE2 | JPH4 | -0,7783 | 3,09978E-39 |
| ACE2 | PRDM6 | -0,7784 | 2,98359E-39 |
| ACE2 | CAMSAP2 | -0,7785 | 2,79388E-39 |
| ACE2 | KLHDC8B | -0,7785 | 2,82213E-39 |
| ACE2 | S1PR3 | -0,7785 | 2,78095E-39 |
| ACE2 | RCBTB2 | -0,7785 | 2,87717E-39 |
| ACE2 | NOVA2 | -0,7785 | 2,8022E-39 |
| ACE2 | SNCG | -0,7786 | 2,77253E-39 |
| ACE2 | ENOX1 | -0,7786 | 2,70647E-39 |
| ACE2 | CMTM3 | -0,7786 | 2,69212E-39 |

| | | | |
|---|---|---|---|
| ACE2 | TRMT61A | -0,7787 | 2,6119E-39 |
| ACE2 | GJC1 | -0,7787 | 2,65788E-39 |
| ACE2 | DYM | -0,7787 | 2,67244E-39 |
| ACE2 | RPL23 | -0,7789 | 2,48678E-39 |
| ACE2 | CLDND1 | -0,779 | 2,35881E-39 |
| ACE2 | SEMA3A | -0,779 | 2,36102E-39 |
| ACE2 | ASAP3 | -0,7791 | 2,24346E-39 |
| ACE2 | HSF1 | -0,7791 | 2,2325E-39 |
| ACE2 | DRAP1 | -0,7791 | 2,26091E-39 |
| ACE2 | TIE1 | -0,7792 | 2,21626E-39 |
| ACE2 | LSAMP | -0,7792 | 2,22781E-39 |
| ACE2 | TUSC3 | -0,7792 | 2,18931E-39 |
| ACE2 | HSPA12A | -0,7792 | 2,16695E-39 |
| ACE2 | RANGRF | -0,7792 | 2,15891E-39 |
| ACE2 | GPM6B | -0,7792 | 2,20746E-39 |
| ACE2 | PLCL1 | -0,7793 | 2,09962E-39 |
| ACE2 | RBPMS | -0,7794 | 2,0452E-39 |
| ACE2 | ITPR1 | -0,7795 | 1,97335E-39 |
| ACE2 | NRG2 | -0,7795 | 1,96509E-39 |
| ACE2 | TSPAN18 | -0,7795 | 1,96813E-39 |
| ACE2 | PTCH2 | -0,7797 | 1,81775E-39 |
| ACE2 | ROR1 | -0,7797 | 1,80547E-39 |
| ACE2 | TERF1 | -0,7797 | 1,79196E-39 |
| ACE2 | ARHGAP29 | -0,7798 | 1,77034E-39 |
| ACE2 | TET1 | -0,7798 | 1,73072E-39 |
| ACE2 | KIRREL | -0,78 | 1,60583E-39 |
| ACE2 | ZNF853 | -0,78 | 1,63564E-39 |
| ACE2 | ESYT1 | -0,78 | 1,63808E-39 |
| ACE2 | PKD1 | -0,78 | 1,61761E-39 |
| ACE2 | MAP4 | -0,7801 | 1,54878E-39 |
| ACE2 | BAIAP2-AS1 | -0,7801 | 1,59497E-39 |
| ACE2 | TGFB1I1 | -0,7802 | 1,50449E-39 |
| ACE2 | AK5 | -0,7804 | 1,40583E-39 |
| ACE2 | DPH5 | -0,7804 | 1,40396E-39 |
| ACE2 | WDR47 | -0,7804 | 1,3987E-39 |
| ACE2 | PCDHGA7 | -0,7804 | 1,38571E-39 |
| ACE2 | RPL10A | -0,7804 | 1,3827E-39 |
| ACE2 | COL14A1 | -0,7804 | 1,39024E-39 |
| ACE2 | NBEA | -0,7805 | 1,34513E-39 |
| ACE2 | ZNF677 | -0,7805 | 1,35127E-39 |
| ACE2 | RP11-486G15.2 | -0,7806 | 1,2849E-39 |
| ACE2 | AP3M2 | -0,7806 | 1,30022E-39 |
| ACE2 | AP4S1 | -0,7806 | 1,32441E-39 |
| ACE2 | MORF4L1 | -0,7806 | 1,32632E-39 |
| ACE2 | SNAPIN | -0,7808 | 1,21033E-39 |
| ACE2 | TBCE | -0,7808 | 1,21832E-39 |
| ACE2 | ZNF883 | -0,7808 | 1,19768E-39 |
| ACE2 | FXYD6 | -0,7808 | 1,23322E-39 |
| ACE2 | RIOX2 | -0,7809 | 1,18258E-39 |
| ACE2 | MID2 | -0,7809 | 1,18712E-39 |
| ACE2 | PKD2 | -0,781 | 1,14249E-39 |
| ACE2 | SAR1A | -0,781 | 1,13018E-39 |
| ACE2 | BEX3 | -0,781 | 1,14187E-39 |
| ACE2 | PTPN12 | -0,7811 | 1,07158E-39 |

| | | | |
|---|---|---|---|
| ACE2 | HK1 | -0,7811 | 1,08982E-39 |
| ACE2 | GDF11 | -0,7811 | 1,07378E-39 |
| ACE2 | PLCB1 | -0,7811 | 1,0852E-39 |
| ACE2 | GNAL | -0,7812 | 1,03808E-39 |
| ACE2 | MAATS1 | -0,7813 | 1,00807E-39 |
| ACE2 | HTRA1 | -0,7814 | 9,62022E-40 |
| ACE2 | PYGL | -0,7814 | 9,61765E-40 |
| ACE2 | DNAJC8 | -0,7815 | 9,32179E-40 |
| ACE2 | RWDD1 | -0,7815 | 9,38716E-40 |
| ACE2 | EGFL8 | -0,7816 | 9,0745E-40 |
| ACE2 | POT1 | -0,7816 | 8,85789E-40 |
| ACE2 | ANKRD40 | -0,7816 | 8,87676E-40 |
| ACE2 | CNBP | -0,7817 | 8,77707E-40 |
| ACE2 | FKBP14 | -0,7817 | 8,55625E-40 |
| ACE2 | C8orf33 | -0,7817 | 8,70229E-40 |
| ACE2 | MAP7D1 | -0,7818 | 8,23489E-40 |
| ACE2 | CHL1 | -0,7818 | 8,21174E-40 |
| ACE2 | CADM1 | -0,7818 | 8,30287E-40 |
| ACE2 | OGFOD1 | -0,7818 | 8,29416E-40 |
| ACE2 | ZNF354A | -0,7819 | 8,17476E-40 |
| ACE2 | C12orf60 | -0,7819 | 8,07948E-40 |
| ACE2 | RP11-412D9.4 | -0,7819 | 8,02215E-40 |
| ACE2 | PRTG | -0,7819 | 8,06673E-40 |
| ACE2 | RPL23AP82 | -0,7819 | 8,01116E-40 |
| ACE2 | DZIP1L | -0,782 | 7,64509E-40 |
| ACE2 | NIM1K | -0,782 | 7,70562E-40 |
| ACE2 | DNAJB4 | -0,7821 | 7,42907E-40 |
| ACE2 | WWC2 | -0,7821 | 7,52267E-40 |
| ACE2 | CSGALNACT2 | -0,7821 | 7,51309E-40 |
| ACE2 | DCAKD | -0,7821 | 7,35848E-40 |
| ACE2 | RASGRF2 | -0,7822 | 7,242E-40 |
| ACE2 | GPR176 | -0,7822 | 7,31219E-40 |
| ACE2 | CDC27 | -0,7822 | 7,148E-40 |
| ACE2 | AC018647.3 | -0,7823 | 6,88965E-40 |
| ACE2 | ZNF273 | -0,7823 | 7,01326E-40 |
| ACE2 | C22orf39 | -0,7823 | 6,93631E-40 |
| ACE2 | POMGNT2 | -0,7824 | 6,75234E-40 |
| ACE2 | PCDHGA5 | -0,7824 | 6,69485E-40 |
| ACE2 | WNK3 | -0,7824 | 6,55662E-40 |
| ACE2 | NAP1L5 | -0,7825 | 6,39386E-40 |
| ACE2 | DDIT4L | -0,7825 | 6,4396E-40 |
| ACE2 | MAP9 | -0,7825 | 6,52963E-40 |
| ACE2 | CAMSAP1 | -0,7825 | 6,43037E-40 |
| ACE2 | FADS1 | -0,7825 | 6,53412E-40 |
| ACE2 | CWF19L2 | -0,7825 | 6,43652E-40 |
| ACE2 | TMCC1 | -0,7826 | 6,23396E-40 |
| ACE2 | RRAGD | -0,7827 | 5,89883E-40 |
| ACE2 | ATF7 | -0,7828 | 5,76408E-40 |
| ACE2 | FLRT2 | -0,7828 | 5,67299E-40 |
| ACE2 | MBD5 | -0,7829 | 5,62284E-40 |
| ACE2 | FBXL7 | -0,783 | 5,36288E-40 |
| ACE2 | GKAP1 | -0,783 | 5,41739E-40 |
| ACE2 | NGRN | -0,783 | 5,30568E-40 |
| ACE2 | CLU | -0,7833 | 4,69067E-40 |

| | | | |
|---|---|---|---|
| ACE2 | MIER2 | -0,7833 | 4,71793E-40 |
| ACE2 | DCP1B | -0,7834 | 4,58717E-40 |
| ACE2 | GALNT16 | -0,7834 | 4,584E-40 |
| ACE2 | SETBP1 | -0,7834 | 4,641E-40 |
| ACE2 | CACNB4 | -0,7835 | 4,35612E-40 |
| ACE2 | CDH5 | -0,7835 | 4,43506E-40 |
| ACE2 | SOD3 | -0,7837 | 4,13532E-40 |
| ACE2 | KIAA1328 | -0,7838 | 3,90422E-40 |
| ACE2 | S1PR2 | -0,7838 | 3,97837E-40 |
| ACE2 | RHBDL3 | -0,7839 | 3,78751E-40 |
| ACE2 | PRKAR1A | -0,7839 | 3,80517E-40 |
| ACE2 | FAAP20 | -0,784 | 3,61351E-40 |
| ACE2 | AKT3 | -0,784 | 3,63936E-40 |
| ACE2 | EMILIN1 | -0,7841 | 3,4884E-40 |
| ACE2 | MAMSTR | -0,7842 | 3,35316E-40 |
| ACE2 | CRIP2 | -0,7844 | 3,11441E-40 |
| ACE2 | FUZ | -0,7844 | 3,10558E-40 |
| ACE2 | ZNF286A | -0,7845 | 3,07808E-40 |
| ACE2 | PHGDH | -0,7846 | 2,86421E-40 |
| ACE2 | WDR86 | -0,7846 | 2,94067E-40 |
| ACE2 | RP11-91J19.3 | -0,7846 | 2,86062E-40 |
| ACE2 | HEG1 | -0,7847 | 2,77498E-40 |
| ACE2 | ZFP28 | -0,7847 | 2,75399E-40 |
| ACE2 | FAM127A | -0,7847 | 2,81071E-40 |
| ACE2 | CLIC2 | -0,7847 | 2,80977E-40 |
| ACE2 | PDGFRB | -0,7848 | 2,71424E-40 |
| ACE2 | TSPAN4 | -0,7848 | 2,65522E-40 |
| ACE2 | ELMO2 | -0,7848 | 2,70297E-40 |
| ACE2 | ZNF362 | -0,7849 | 2,61326E-40 |
| ACE2 | INPP4A | -0,7849 | 2,57943E-40 |
| ACE2 | HEY2 | -0,7849 | 2,5613E-40 |
| ACE2 | KIAA1755 | -0,7849 | 2,64651E-40 |
| ACE2 | EDA | -0,7849 | 2,56576E-40 |
| ACE2 | SH3BGRL | -0,7849 | 2,58029E-40 |
| ACE2 | EIF2B3 | -0,785 | 2,50521E-40 |
| ACE2 | RYK | -0,7851 | 2,41698E-40 |
| ACE2 | MAGEF1 | -0,7851 | 2,43107E-40 |
| ACE2 | KCTD1 | -0,7851 | 2,4561E-40 |
| ACE2 | MAD2L2 | -0,7852 | 2,3246E-40 |
| ACE2 | C1R | -0,7853 | 2,22655E-40 |
| ACE2 | IPO11 | -0,7855 | 2,07494E-40 |
| ACE2 | EIF3L | -0,7856 | 1,96253E-40 |
| ACE2 | CAMK1 | -0,7857 | 1,94029E-40 |
| ACE2 | CCDC85B | -0,7857 | 1,89882E-40 |
| ACE2 | BBX | -0,7858 | 1,8534E-40 |
| ACE2 | PRPS2 | -0,7858 | 1,87269E-40 |
| ACE2 | HIP1 | -0,7859 | 1,78147E-40 |
| ACE2 | CALHM2 | -0,7859 | 1,7618E-40 |
| ACE2 | ARHGAP20 | -0,7859 | 1,76231E-40 |
| ACE2 | ZFP1 | -0,7859 | 1,75332E-40 |
| ACE2 | TGFBR2 | -0,786 | 1,69251E-40 |
| ACE2 | CEP162 | -0,786 | 1,6905E-40 |
| ACE2 | ENKD1 | -0,786 | 1,71112E-40 |
| ACE2 | C1orf56 | -0,7861 | 1,62043E-40 |

| | | | |
|---|---|---|---|
| ACE2 | IFT172 | -0,7861 | 1,6378E-40 |
| ACE2 | ZNF385D | -0,7861 | 1,67036E-40 |
| ACE2 | GAS7 | -0,7861 | 1,64088E-40 |
| ACE2 | FYTTD1 | -0,7864 | 1,44402E-40 |
| ACE2 | SLC9B2 | -0,7864 | 1,48555E-40 |
| ACE2 | GPX8 | -0,7864 | 1,47579E-40 |
| ACE2 | MTMR2 | -0,7864 | 1,46764E-40 |
| ACE2 | SPATA7 | -0,7864 | 1,49328E-40 |
| ACE2 | TPM4 | -0,7864 | 1,4598E-40 |
| ACE2 | RBBP7 | -0,7864 | 1,47796E-40 |
| ACE2 | RORB | -0,7866 | 1,38452E-40 |
| ACE2 | EIF3G | -0,7866 | 1,38497E-40 |
| ACE2 | FES | -0,7867 | 1,32437E-40 |
| ACE2 | R3HDM1 | -0,7869 | 1,22905E-40 |
| ACE2 | CYB5R1 | -0,787 | 1,14646E-40 |
| ACE2 | SNAPC2 | -0,787 | 1,16962E-40 |
| ACE2 | PRMT2 | -0,787 | 1,18704E-40 |
| ACE2 | STK32B | -0,7871 | 1,14226E-40 |
| ACE2 | NCDN | -0,7872 | 1,07457E-40 |
| ACE2 | KTN1-AS1 | -0,7872 | 1,07213E-40 |
| ACE2 | SRP68 | -0,7872 | 1,07085E-40 |
| ACE2 | PFDN4 | -0,7872 | 1,08577E-40 |
| ACE2 | RSU1 | -0,7873 | 1,04378E-40 |
| ACE2 | RP11-327J17.9 | -0,7873 | 1,05828E-40 |
| ACE2 | PAK3 | -0,7873 | 1,05861E-40 |
| ACE2 | SHISA4 | -0,7875 | 9,75205E-41 |
| ACE2 | HACD4 | -0,7875 | 9,48917E-41 |
| ACE2 | FOXF1 | -0,7875 | 9,57465E-41 |
| ACE2 | SYNGR1 | -0,7875 | 9,48187E-41 |
| ACE2 | SETD7 | -0,7876 | 9,41039E-41 |
| ACE2 | TTC26 | -0,7877 | 9,00077E-41 |
| ACE2 | ESAM | -0,7877 | 8,87127E-41 |
| ACE2 | RPL14 | -0,7878 | 8,47626E-41 |
| ACE2 | CRISPLD1 | -0,7878 | 8,5497E-41 |
| ACE2 | LHFP | -0,7878 | 8,53341E-41 |
| ACE2 | RP11-157P1.4 | -0,7878 | 8,71332E-41 |
| ACE2 | GJC2 | -0,7879 | 8,26538E-41 |
| ACE2 | KATNBL1 | -0,7879 | 8,28011E-41 |
| ACE2 | DDX19A | -0,7879 | 8,22701E-41 |
| ACE2 | COL6A1 | -0,7879 | 8,37842E-41 |
| ACE2 | FAM110B | -0,788 | 7,80199E-41 |
| ACE2 | ANAPC7 | -0,788 | 8,07864E-41 |
| ACE2 | FXR1 | -0,7881 | 7,53905E-41 |
| ACE2 | CNTLN | -0,7881 | 7,59534E-41 |
| ACE2 | CAND2 | -0,7882 | 7,29824E-41 |
| ACE2 | PPM1F | -0,7882 | 7,21755E-41 |
| ACE2 | NME7 | -0,7883 | 7,05304E-41 |
| ACE2 | GMPS | -0,7883 | 7,17464E-41 |
| ACE2 | EIF2B4 | -0,7884 | 6,8068E-41 |
| ACE2 | FAM126A | -0,7884 | 6,73756E-41 |
| ACE2 | URI1 | -0,7884 | 6,71665E-41 |
| ACE2 | MARK4 | -0,7884 | 6,9345E-41 |
| ACE2 | PTBP2 | -0,7885 | 6,45841E-41 |
| ACE2 | HLX | -0,7885 | 6,66098E-41 |

| | | | |
|---|---|---|---|
| ACE2 | CHST14 | -0,7885 | 6,45751E-41 |
| ACE2 | CX3CL1 | -0,7885 | 6,64638E-41 |
| ACE2 | BOC | -0,7886 | 6,37322E-41 |
| ACE2 | LDB2 | -0,7886 | 6,37912E-41 |
| ACE2 | KDELC2 | -0,7887 | 6,05409E-41 |
| ACE2 | ST3GAL4-AS1 | -0,7887 | 5,99925E-41 |
| ACE2 | HDX | -0,7887 | 6,0219E-41 |
| ACE2 | FAM127C | -0,7888 | 5,79302E-41 |
| ACE2 | ANAPC10 | -0,7889 | 5,50024E-41 |
| ACE2 | DCLK2 | -0,7889 | 5,59578E-41 |
| ACE2 | RSL1D1 | -0,7889 | 5,59897E-41 |
| ACE2 | GPRASP2 | -0,7889 | 5,58981E-41 |
| ACE2 | KCTD10 | -0,789 | 5,44729E-41 |
| ACE2 | LINC00662 | -0,789 | 5,38595E-41 |
| ACE2 | EFCAB7 | -0,7891 | 5,15883E-41 |
| ACE2 | CYP27C1 | -0,7891 | 5,23285E-41 |
| ACE2 | IFT22 | -0,7892 | 5,07608E-41 |
| ACE2 | ABCA3 | -0,7892 | 5,06847E-41 |
| ACE2 | TFE3 | -0,7892 | 5,08184E-41 |
| ACE2 | MATR3 | -0,7893 | 4,81664E-41 |
| ACE2 | RASIP1 | -0,7893 | 4,83488E-41 |
| ACE2 | RP5-1039K5.19 | -0,7893 | 4,76423E-41 |
| ACE2 | PLEKHO1 | -0,7894 | 4,64193E-41 |
| ACE2 | ZNF496 | -0,7894 | 4,59847E-41 |
| ACE2 | GJA4 | -0,7895 | 4,38997E-41 |
| ACE2 | ZMYM4 | -0,7895 | 4,42873E-41 |
| ACE2 | STOX2 | -0,7896 | 4,25325E-41 |
| ACE2 | FGF7 | -0,7897 | 4,05837E-41 |
| ACE2 | C2CD2 | -0,7897 | 4,11722E-41 |
| ACE2 | SRRD | -0,7897 | 4,15123E-41 |
| ACE2 | ATP10D | -0,7898 | 3,98547E-41 |
| ACE2 | DIMT1 | -0,7898 | 3,95176E-41 |
| ACE2 | KIAA1586 | -0,7898 | 3,94975E-41 |
| ACE2 | ZC2HC1A | -0,7898 | 3,97267E-41 |
| ACE2 | BBS1 | -0,7898 | 4,02904E-41 |
| ACE2 | PRPF40B | -0,7898 | 3,87907E-41 |
| ACE2 | SMAD5 | -0,7899 | 3,87261E-41 |
| ACE2 | ARHGEF15 | -0,7899 | 3,7369E-41 |
| ACE2 | GAMT | -0,7899 | 3,8252E-41 |
| ACE2 | DUSP7 | -0,7901 | 3,49831E-41 |
| ACE2 | COPS6 | -0,7901 | 3,57381E-41 |
| ACE2 | GALNT11 | -0,7902 | 3,40805E-41 |
| ACE2 | SSSCA1 | -0,7902 | 3,3721E-41 |
| ACE2 | RERG | -0,7902 | 3,34367E-41 |
| ACE2 | COL6A2 | -0,7902 | 3,42209E-41 |
| ACE2 | GNG11 | -0,7904 | 3,12151E-41 |
| ACE2 | ATP6V0E2-AS1 | -0,7904 | 3,08277E-41 |
| ACE2 | FAM219A | -0,7904 | 3,08875E-41 |
| ACE2 | CACNA2D1 | -0,7906 | 2,92834E-41 |
| ACE2 | NKX3-1 | -0,7906 | 2,83706E-41 |
| ACE2 | C20orf27 | -0,7906 | 2,85895E-41 |
| ACE2 | TMEM9 | -0,7907 | 2,74581E-41 |
| ACE2 | NPTXR | -0,7907 | 2,76183E-41 |
| ACE2 | RFTN2 | -0,7909 | 2,57913E-41 |

| | | | |
|---|---|---|---|
| ACE2 | AC010226.4 | -0,791 | 2,42436E-41 |
| ACE2 | CPEB1 | -0,791 | 2,50851E-41 |
| ACE2 | KRBOX4 | -0,791 | 2,49499E-41 |
| ACE2 | TENM3 | -0,7911 | 2,37699E-41 |
| ACE2 | SRSF12 | -0,7911 | 2,334E-41 |
| ACE2 | RNASEH2B | -0,7911 | 2,36504E-41 |
| ACE2 | PLEKHO2 | -0,7911 | 2,37085E-41 |
| ACE2 | SMARCA1 | -0,7911 | 2,38008E-41 |
| ACE2 | C2orf76 | -0,7912 | 2,31534E-41 |
| ACE2 | IFITM2 | -0,7912 | 2,24696E-41 |
| ACE2 | SLC43A3 | -0,7912 | 2,26289E-41 |
| ACE2 | CHST12 | -0,7914 | 2,09413E-41 |
| ACE2 | WASHC3 | -0,7914 | 2,14086E-41 |
| ACE2 | MEIS2 | -0,7914 | 2,11324E-41 |
| ACE2 | REM1 | -0,7914 | 2,09095E-41 |
| ACE2 | RASSF8 | -0,7915 | 2,02898E-41 |
| ACE2 | TMEM123 | -0,7918 | 1,82341E-41 |
| ACE2 | SCML2 | -0,7918 | 1,79466E-41 |
| ACE2 | AEBP1 | -0,7919 | 1,76174E-41 |
| ACE2 | ANKRD13B | -0,7919 | 1,76454E-41 |
| ACE2 | LRRC4B | -0,7919 | 1,73506E-41 |
| ACE2 | BRK1 | -0,792 | 1,65172E-41 |
| ACE2 | ANKRD6 | -0,792 | 1,69243E-41 |
| ACE2 | RPRD1A | -0,792 | 1,66594E-41 |
| ACE2 | PDZD4 | -0,792 | 1,66641E-41 |
| ACE2 | PHF19 | -0,7921 | 1,5772E-41 |
| ACE2 | SSPN | -0,7921 | 1,59558E-41 |
| ACE2 | AMIGO2 | -0,7921 | 1,58384E-41 |
| ACE2 | GNAS | -0,7921 | 1,6353E-41 |
| ACE2 | BAG2 | -0,7922 | 1,5123E-41 |
| ACE2 | DENND2A | -0,7922 | 1,52087E-41 |
| ACE2 | HECTD2 | -0,7923 | 1,46272E-41 |
| ACE2 | NUDT11 | -0,7924 | 1,43591E-41 |
| ACE2 | SPATA5 | -0,7925 | 1,34672E-41 |
| ACE2 | POLR1E | -0,7926 | 1,3177E-41 |
| ACE2 | ZNF568 | -0,7926 | 1,34213E-41 |
| ACE2 | SLC43A1 | -0,7927 | 1,2607E-41 |
| ACE2 | SCRN1 | -0,7928 | 1,2256E-41 |
| ACE2 | BCAP29 | -0,7928 | 1,20683E-41 |
| ACE2 | FGD1 | -0,7928 | 1,23593E-41 |
| ACE2 | ECSCR | -0,7929 | 1,18366E-41 |
| ACE2 | VWA1 | -0,793 | 1,10861E-41 |
| ACE2 | NAB1 | -0,7931 | 1,07562E-41 |
| ACE2 | RPGRIP1L | -0,7931 | 1,091E-41 |
| ACE2 | STAT5B | -0,7932 | 1,03542E-41 |
| ACE2 | C11orf70 | -0,7935 | 9,11653E-42 |
| ACE2 | PARD6G | -0,7935 | 9,38309E-42 |
| ACE2 | EFR3B | -0,7936 | 9,00146E-42 |
| ACE2 | SNAPC5 | -0,7936 | 8,69853E-42 |
| ACE2 | EPHA3 | -0,7937 | 8,3648E-42 |
| ACE2 | FAM13C | -0,7937 | 8,42767E-42 |
| ACE2 | TBCB | -0,7937 | 8,66929E-42 |
| ACE2 | TMTC2 | -0,7938 | 8,20406E-42 |
| ACE2 | JAM2 | -0,7938 | 8,15325E-42 |

| | | | |
|---|---|---|---|
| ACE2 | SMIM10 | -0,7939 | 7,99507E-42 |
| ACE2 | GARNL3 | -0,794 | 7,49176E-42 |
| ACE2 | DNM1L | -0,794 | 7,5308E-42 |
| ACE2 | AMZ2 | -0,794 | 7,4116E-42 |
| ACE2 | SSB | -0,7941 | 7,15542E-42 |
| ACE2 | ST3GAL6 | -0,7941 | 7,13738E-42 |
| ACE2 | PGBD1 | -0,7941 | 7,11183E-42 |
| ACE2 | EVC2 | -0,7942 | 7,09672E-42 |
| ACE2 | TRAF3IP2-AS1 | -0,7942 | 7,07855E-42 |
| ACE2 | PKIA | -0,7942 | 6,94959E-42 |
| ACE2 | PCDH17 | -0,7942 | 7,0238E-42 |
| ACE2 | SMIM10L2A | -0,7942 | 7,0408E-42 |
| ACE2 | NFASC | -0,7943 | 6,69422E-42 |
| ACE2 | DFNB59 | -0,7943 | 6,67168E-42 |
| ACE2 | GPSM1 | -0,7943 | 6,72372E-42 |
| ACE2 | ZCCHC24 | -0,7943 | 6,64387E-42 |
| ACE2 | ARHGEF17 | -0,7944 | 6,34395E-42 |
| ACE2 | TTC28 | -0,7944 | 6,41599E-42 |
| ACE2 | PAIP2 | -0,7945 | 6,12848E-42 |
| ACE2 | IFT46 | -0,7945 | 6,29392E-42 |
| ACE2 | UBFD1 | -0,7945 | 6,23486E-42 |
| ACE2 | CNTN2 | -0,7946 | 5,8739E-42 |
| ACE2 | RPL6 | -0,7946 | 6,05873E-42 |
| ACE2 | KLHL22 | -0,7946 | 6,03654E-42 |
| ACE2 | AGBL5 | -0,7947 | 5,78829E-42 |
| ACE2 | AK1 | -0,7947 | 5,77768E-42 |
| ACE2 | COMMD6 | -0,7947 | 5,71351E-42 |
| ACE2 | MAGI2-AS3 | -0,7948 | 5,57461E-42 |
| ACE2 | RPL7 | -0,7948 | 5,50419E-42 |
| ACE2 | SLC25A28 | -0,7948 | 5,56072E-42 |
| ACE2 | NLGN3 | -0,7948 | 5,37804E-42 |
| ACE2 | DPYSL2 | -0,7949 | 5,27584E-42 |
| ACE2 | MMP17 | -0,7949 | 5,16504E-42 |
| ACE2 | FGFR1 | -0,795 | 5,09674E-42 |
| ACE2 | SCN1B | -0,795 | 5,13437E-42 |
| ACE2 | TMEM237 | -0,7951 | 4,86118E-42 |
| ACE2 | GAREM2 | -0,7952 | 4,65861E-42 |
| ACE2 | C2orf74 | -0,7952 | 4,63808E-42 |
| ACE2 | TTYH2 | -0,7952 | 4,62885E-42 |
| ACE2 | ZSCAN18 | -0,7952 | 4,6655E-42 |
| ACE2 | CDKL5 | -0,7952 | 4,74973E-42 |
| ACE2 | USP11 | -0,7952 | 4,7396E-42 |
| ACE2 | CCDC69 | -0,7953 | 4,50039E-42 |
| ACE2 | C19orf44 | -0,7953 | 4,41528E-42 |
| ACE2 | TMEM200B | -0,7954 | 4,28855E-42 |
| ACE2 | SPG20 | -0,7954 | 4,28445E-42 |
| ACE2 | GTF2F2 | -0,7954 | 4,30758E-42 |
| ACE2 | HOXD8 | -0,7955 | 4,07874E-42 |
| ACE2 | AC009948.5 | -0,7955 | 4,15743E-42 |
| ACE2 | NFIA | -0,7956 | 3,99855E-42 |
| ACE2 | CXorf36 | -0,7956 | 4,0386E-42 |
| ACE2 | NPIPP1 | -0,7957 | 3,77922E-42 |
| ACE2 | C1orf198 | -0,7958 | 3,66184E-42 |
| ACE2 | SLC39A6 | -0,7958 | 3,74005E-42 |

| | | | |
|---|---|---|---|
| ACE2 | ERI3 | -0,7959 | 3,45971E-42 |
| ACE2 | CHCHD6 | -0,7959 | 3,46048E-42 |
| ACE2 | HLTF | -0,7959 | 3,51802E-42 |
| ACE2 | RASSF8-AS1 | -0,7959 | 3,54098E-42 |
| ACE2 | HOXC6 | -0,7959 | 3,48822E-42 |
| ACE2 | UBA2 | -0,7959 | 3,55305E-42 |
| ACE2 | ANXA6 | -0,796 | 3,41976E-42 |
| ACE2 | MYH10 | -0,796 | 3,34989E-42 |
| ACE2 | RPL39L | -0,7962 | 3,12192E-42 |
| ACE2 | FKBP7 | -0,7964 | 2,86516E-42 |
| ACE2 | NBDY | -0,7964 | 2,83011E-42 |
| ACE2 | TUBE1 | -0,7965 | 2,78947E-42 |
| ACE2 | TTBK2 | -0,7966 | 2,65278E-42 |
| ACE2 | NAP1L3 | -0,7966 | 2,62691E-42 |
| ACE2 | UST | -0,7967 | 2,59897E-42 |
| ACE2 | MICU3 | -0,7967 | 2,54719E-42 |
| ACE2 | PYGO1 | -0,7967 | 2,58761E-42 |
| ACE2 | NFYB | -0,7969 | 2,38117E-42 |
| ACE2 | PLEKHA4 | -0,7969 | 2,31895E-42 |
| ACE2 | C5orf42 | -0,797 | 2,27623E-42 |
| ACE2 | TSPOAP1 | -0,797 | 2,2671E-42 |
| ACE2 | NAV1 | -0,7971 | 2,14665E-42 |
| ACE2 | RECK | -0,7971 | 2,16987E-42 |
| ACE2 | ISLR | -0,7971 | 2,18698E-42 |
| ACE2 | SOGA1 | -0,7971 | 2,13065E-42 |
| ACE2 | MAMLD1 | -0,7971 | 2,15137E-42 |
| ACE2 | PRKD1 | -0,7972 | 2,06534E-42 |
| ACE2 | LDOC1 | -0,7972 | 2,07645E-42 |
| ACE2 | TMEM170B | -0,7973 | 1,96652E-42 |
| ACE2 | GLI3 | -0,7974 | 1,92202E-42 |
| ACE2 | CDH11 | -0,7974 | 1,95074E-42 |
| ACE2 | EMD | -0,7974 | 1,9406E-42 |
| ACE2 | HOXD9 | -0,7975 | 1,82056E-42 |
| ACE2 | IFT52 | -0,7975 | 1,86309E-42 |
| ACE2 | GXYLT2 | -0,7976 | 1,75308E-42 |
| ACE2 | GLT8D2 | -0,7976 | 1,78184E-42 |
| ACE2 | BRMS1L | -0,7976 | 1,77733E-42 |
| ACE2 | FGF2 | -0,7977 | 1,72503E-42 |
| ACE2 | ZNF438 | -0,7977 | 1,7342E-42 |
| ACE2 | NFATC4 | -0,7977 | 1,69072E-42 |
| ACE2 | B3GNT9 | -0,7977 | 1,69654E-42 |
| ACE2 | SEPT5 | -0,7978 | 1,64856E-42 |
| ACE2 | CDC42EP3 | -0,7979 | 1,5831E-42 |
| ACE2 | RP11-454E5.4 | -0,798 | 1,5306E-42 |
| ACE2 | FGF13 | -0,798 | 1,52056E-42 |
| ACE2 | ITPRIP | -0,7981 | 1,45369E-42 |
| ACE2 | EEF1AKMT1 | -0,7981 | 1,4488E-42 |
| ACE2 | SELENON | -0,7982 | 1,38951E-42 |
| ACE2 | TCEAL7 | -0,7982 | 1,37419E-42 |
| ACE2 | KANSL1L | -0,7983 | 1,32375E-42 |
| ACE2 | SYT11 | -0,7984 | 1,29228E-42 |
| ACE2 | ABLIM3 | -0,7984 | 1,25531E-42 |
| ACE2 | KCTD12 | -0,7988 | 1,07841E-42 |
| ACE2 | KCNE4 | -0,7989 | 1,04615E-42 |

| | | | |
|---|---|---|---|
| ACE2 | AEBP2 | -0,7989 | 1,03527E-42 |
| ACE2 | BICC1 | -0,799 | 1,0111E-42 |
| ACE2 | PTDSS2 | -0,799 | 9,8653E-43 |
| ACE2 | CHRNA3 | -0,799 | 9,86603E-43 |
| ACE2 | C16orf45 | -0,799 | 1,01446E-42 |
| ACE2 | COPS8 | -0,7991 | 9,63378E-43 |
| ACE2 | AMZ2P1 | -0,7991 | 9,50124E-43 |
| ACE2 | NINL | -0,7992 | 9,37368E-43 |
| ACE2 | RGL2 | -0,7993 | 8,98511E-43 |
| ACE2 | SNRPN | -0,7993 | 8,92034E-43 |
| ACE2 | LY6E | -0,7997 | 7,38171E-43 |
| ACE2 | MRAS | -0,7998 | 7,21243E-43 |
| ACE2 | CERCAM | -0,7998 | 7,28254E-43 |
| ACE2 | SYDE1 | -0,7998 | 7,08276E-43 |
| ACE2 | FER | -0,7999 | 6,93596E-43 |
| ACE2 | CCDC88A | -0,8 | 6,58021E-43 |
| ACE2 | SMAD9 | -0,8 | 6,70739E-43 |
| ACE2 | SV2B | -0,8 | 6,49756E-43 |
| ACE2 | ZNF347 | -0,8 | 6,56877E-43 |
| ACE2 | ARMCX1 | -0,8 | 6,72569E-43 |
| ACE2 | ZNF204P | -0,8001 | 6,40139E-43 |
| ACE2 | EIF3H | -0,8001 | 6,28564E-43 |
| ACE2 | PKN1 | -0,8001 | 6,33533E-43 |
| ACE2 | MPZ | -0,8002 | 6,11749E-43 |
| ACE2 | PHLDB2 | -0,8002 | 6,02687E-43 |
| ACE2 | MYCT1 | -0,8002 | 6,15887E-43 |
| ACE2 | LRRC4C | -0,8002 | 6,12955E-43 |
| ACE2 | NAALAD2 | -0,8002 | 6,15514E-43 |
| ACE2 | FAM171B | -0,8003 | 5,84735E-43 |
| ACE2 | AGPAT4 | -0,8003 | 5,92633E-43 |
| ACE2 | PIP4K2B | -0,8005 | 5,37034E-43 |
| ACE2 | CDK14 | -0,8006 | 5,25182E-43 |
| ACE2 | RAB9B | -0,8006 | 5,17779E-43 |
| ACE2 | ARHGEF3 | -0,8007 | 5,02134E-43 |
| ACE2 | NOVA1 | -0,8007 | 4,91752E-43 |
| ACE2 | CNTNAP1 | -0,8007 | 5,03242E-43 |
| ACE2 | CLIP3 | -0,8007 | 5,06616E-43 |
| ACE2 | RP4-665J23.1 | -0,8008 | 4,69331E-43 |
| ACE2 | GPN3 | -0,8009 | 4,60844E-43 |
| ACE2 | TP53BP1 | -0,8009 | 4,63938E-43 |
| ACE2 | PRCP | -0,801 | 4,32181E-43 |
| ACE2 | CCDC74A | -0,8011 | 4,14836E-43 |
| ACE2 | PCDHGB6 | -0,8011 | 4,20753E-43 |
| ACE2 | MTAP | -0,8012 | 3,97843E-43 |
| ACE2 | PPP1R3F | -0,8012 | 4,01315E-43 |
| ACE2 | FAM229B | -0,8013 | 3,88969E-43 |
| ACE2 | TSNAXIP1 | -0,8014 | 3,68694E-43 |
| ACE2 | GNAI2 | -0,8017 | 3,23048E-43 |
| ACE2 | KIF3A | -0,8017 | 3,30944E-43 |
| ACE2 | RUSC2 | -0,8017 | 3,23815E-43 |
| ACE2 | NAP1L1 | -0,8017 | 3,33507E-43 |
| ACE2 | ABHD8 | -0,8017 | 3,34017E-43 |
| ACE2 | SLC9A9 | -0,8018 | 3,10174E-43 |
| ACE2 | NAA10 | -0,8018 | 3,18537E-43 |

| | | | |
|---|---|---|---|
| ACE2 | FEZ1 | -0,8019 | 3,04231E-43 |
| ACE2 | FAM69A | -0,802 | 2,94791E-43 |
| ACE2 | MEIS1 | -0,802 | 2,94612E-43 |
| ACE2 | RAB3IL1 | -0,802 | 2,8936E-43 |
| ACE2 | CHD3 | -0,802 | 2,8442E-43 |
| ACE2 | DARS | -0,8022 | 2,66371E-43 |
| ACE2 | CCNB1IP1 | -0,8023 | 2,5174E-43 |
| ACE2 | CPT1C | -0,8023 | 2,56415E-43 |
| ACE2 | RBMX2 | -0,8023 | 2,57873E-43 |
| ACE2 | PCDHGA6 | -0,8024 | 2,47092E-43 |
| ACE2 | TPST1 | -0,8024 | 2,42566E-43 |
| ACE2 | TBL1X | -0,8024 | 2,4494E-43 |
| ACE2 | ADAMTS10 | -0,8026 | 2,21218E-43 |
| ACE2 | LAYN | -0,8027 | 2,16443E-43 |
| ACE2 | BBS4 | -0,8027 | 2,19036E-43 |
| ACE2 | SLCO3A1 | -0,8027 | 2,12324E-43 |
| ACE2 | BLVRA | -0,8028 | 2,09189E-43 |
| ACE2 | TUB | -0,8028 | 2,08555E-43 |
| ACE2 | EFS | -0,8028 | 2,06386E-43 |
| ACE2 | FLT3LG | -0,8028 | 2,08948E-43 |
| ACE2 | WDR41 | -0,8029 | 1,97062E-43 |
| ACE2 | KLC2 | -0,8029 | 2,01399E-43 |
| ACE2 | PPP3CB | -0,8032 | 1,77361E-43 |
| ACE2 | FXYD2 | -0,8032 | 1,73228E-43 |
| ACE2 | RNF165 | -0,8034 | 1,61401E-43 |
| ACE2 | C11orf63 | -0,8036 | 1,48335E-43 |
| ACE2 | NT5M | -0,8036 | 1,48628E-43 |
| ACE2 | CTD-2554C21.2 | -0,8037 | 1,42071E-43 |
| ACE2 | KIF3C | -0,8039 | 1,32336E-43 |
| ACE2 | ADGRA2 | -0,8039 | 1,30143E-43 |
| ACE2 | SLC35G2 | -0,804 | 1,25135E-43 |
| ACE2 | CFAP97 | -0,804 | 1,25853E-43 |
| ACE2 | DUSP14 | -0,804 | 1,2415E-43 |
| ACE2 | FZD8 | -0,8041 | 1,20971E-43 |
| ACE2 | ULK2 | -0,8041 | 1,18283E-43 |
| ACE2 | RUNX1T1 | -0,8042 | 1,13268E-43 |
| ACE2 | CEP63 | -0,8043 | 1,10449E-43 |
| ACE2 | CGNL1 | -0,8043 | 1,09817E-43 |
| ACE2 | RGS10 | -0,8044 | 1,078E-43 |
| ACE2 | KATNAL1 | -0,8044 | 1,03553E-43 |
| ACE2 | MOXD1 | -0,8045 | 9,94091E-44 |
| ACE2 | SYNE1 | -0,8045 | 1,0013E-43 |
| ACE2 | CTNNAL1 | -0,8045 | 1,02553E-43 |
| ACE2 | C1S | -0,8045 | 1,00114E-43 |
| ACE2 | KLHL5 | -0,8046 | 9,54584E-44 |
| ACE2 | PELO | -0,8047 | 9,27445E-44 |
| ACE2 | BOLA3-AS1 | -0,8048 | 9,05787E-44 |
| ACE2 | ZNF781 | -0,8048 | 8,87722E-44 |
| ACE2 | RGS16 | -0,8049 | 8,72116E-44 |
| ACE2 | MYL6B | -0,8049 | 8,72253E-44 |
| ACE2 | NDE1 | -0,8049 | 8,44336E-44 |
| ACE2 | IL33 | -0,805 | 8,04731E-44 |
| ACE2 | FTO | -0,805 | 8,22983E-44 |
| ACE2 | DPYSL3 | -0,8051 | 7,96248E-44 |

| | | | |
|---|---|---|---|
| ACE2 | NIFK-AS1 | -0,8052 | 7,56568E-44 |
| ACE2 | ZNF280D | -0,8053 | 7,18053E-44 |
| ACE2 | RGAG4 | -0,8054 | 6,90091E-44 |
| ACE2 | CSNK1E | -0,8057 | 5,96333E-44 |
| ACE2 | KCTD6 | -0,8058 | 5,92797E-44 |
| ACE2 | SBSPON | -0,8058 | 5,75884E-44 |
| ACE2 | DYSF | -0,806 | 5,33731E-44 |
| ACE2 | LNP1 | -0,806 | 5,2387E-44 |
| ACE2 | ANKRD35 | -0,8061 | 5,15526E-44 |
| ACE2 | AKAP2 | -0,8061 | 5,21087E-44 |
| ACE2 | NXN | -0,8063 | 4,71702E-44 |
| ACE2 | CRTAP | -0,8064 | 4,57267E-44 |
| ACE2 | BOD1 | -0,8064 | 4,46343E-44 |
| ACE2 | NREP | -0,8065 | 4,37564E-44 |
| ACE2 | PDCD7 | -0,8065 | 4,34718E-44 |
| ACE2 | ARMC2 | -0,8067 | 4,04429E-44 |
| ACE2 | A4GALT | -0,8069 | 3,56783E-44 |
| ACE2 | ARMCX3 | -0,8069 | 3,63359E-44 |
| ACE2 | SV2A | -0,807 | 3,56108E-44 |
| ACE2 | CADM3 | -0,8071 | 3,37107E-44 |
| ACE2 | GABBR1 | -0,8071 | 3,3317E-44 |
| ACE2 | TRPS1 | -0,8072 | 3,19131E-44 |
| ACE2 | FAM204A | -0,8073 | 3,06979E-44 |
| ACE2 | BRSK1 | -0,8073 | 3,03894E-44 |
| ACE2 | GEMIN8 | -0,8073 | 3,12059E-44 |
| ACE2 | DIAPH2 | -0,8073 | 3,09144E-44 |
| ACE2 | LDHB | -0,8074 | 2,96944E-44 |
| ACE2 | CRYZL1 | -0,8074 | 2,91369E-44 |
| ACE2 | SOBP | -0,8075 | 2,76589E-44 |
| ACE2 | LINC01116 | -0,8076 | 2,74368E-44 |
| ACE2 | CCDC102A | -0,8076 | 2,72877E-44 |
| ACE2 | MMP16 | -0,8077 | 2,53026E-44 |
| ACE2 | EXTL2 | -0,8078 | 2,46308E-44 |
| ACE2 | BBS5 | -0,8078 | 2,52105E-44 |
| ACE2 | PDE4A | -0,8078 | 2,44277E-44 |
| ACE2 | HSCB | -0,8078 | 2,43937E-44 |
| ACE2 | CASC15 | -0,808 | 2,26169E-44 |
| ACE2 | ZNF532 | -0,8081 | 2,15014E-44 |
| ACE2 | PDCD5 | -0,8082 | 2,05736E-44 |
| ACE2 | BBS9 | -0,8084 | 1,9443E-44 |
| ACE2 | PFKM | -0,8084 | 1,87887E-44 |
| ACE2 | ZFP36L1 | -0,8084 | 1,89777E-44 |
| ACE2 | TMEM109 | -0,8085 | 1,80568E-44 |
| ACE2 | REXO2 | -0,8085 | 1,84617E-44 |
| ACE2 | PDE1A | -0,8086 | 1,71185E-44 |
| ACE2 | C12orf29 | -0,8086 | 1,7707E-44 |
| ACE2 | BCL7C | -0,8087 | 1,66457E-44 |
| ACE2 | ZNF519 | -0,8087 | 1,6991E-44 |
| ACE2 | ZNF667 | -0,8087 | 1,64548E-44 |
| ACE2 | ZBTB4 | -0,8089 | 1,54553E-44 |
| ACE2 | LAS1L | -0,8089 | 1,51371E-44 |
| ACE2 | NDUFA4L2 | -0,809 | 1,46027E-44 |
| ACE2 | DNM3 | -0,8091 | 1,43065E-44 |
| ACE2 | EIF2D | -0,8092 | 1,3733E-44 |

| | | | |
|---|---|---|---|
| ACE2 | USP13 | -0,8092 | 1,35263E-44 |
| ACE2 | EID2B | -0,8093 | 1,31369E-44 |
| ACE2 | TRPC1 | -0,8094 | 1,23218E-44 |
| ACE2 | MOB1B | -0,8095 | 1,1958E-44 |
| ACE2 | EIF3E | -0,8095 | 1,17757E-44 |
| ACE2 | RAB12 | -0,8095 | 1,1914E-44 |
| ACE2 | MAGED2 | -0,8095 | 1,1823E-44 |
| ACE2 | TSPAN11 | -0,8096 | 1,14042E-44 |
| ACE2 | EID1 | -0,8096 | 1,14462E-44 |
| ACE2 | PML | -0,8096 | 1,14452E-44 |
| ACE2 | ANXA5 | -0,8098 | 1,03059E-44 |
| ACE2 | PCDHB15 | -0,8098 | 1,02238E-44 |
| ACE2 | LINC01128 | -0,8099 | 1,00298E-44 |
| ACE2 | NPAS3 | -0,8099 | 9,90924E-45 |
| ACE2 | ARHGAP23 | -0,8099 | 9,76805E-45 |
| ACE2 | FAHD2CP | -0,8101 | 9,24898E-45 |
| ACE2 | FAHD2B | -0,8101 | 8,97758E-45 |
| ACE2 | VLDLR | -0,8101 | 8,99863E-45 |
| ACE2 | NME4 | -0,8102 | 8,79933E-45 |
| ACE2 | EGFLAM | -0,8104 | 8,08575E-45 |
| ACE2 | RABEPK | -0,8104 | 7,91648E-45 |
| ACE2 | FAM78B | -0,8105 | 7,53588E-45 |
| ACE2 | GUCY1A3 | -0,8105 | 7,53711E-45 |
| ACE2 | FBXO17 | -0,8105 | 7,73361E-45 |
| ACE2 | LGALS1 | -0,8105 | 7,48689E-45 |
| ACE2 | NUDC | -0,8108 | 6,56495E-45 |
| ACE2 | PDE10A | -0,8108 | 6,55916E-45 |
| ACE2 | TGFB3 | -0,8108 | 6,63734E-45 |
| ACE2 | SLC16A2 | -0,8108 | 6,57239E-45 |
| ACE2 | APBB1 | -0,811 | 6,23631E-45 |
| ACE2 | C15orf41 | -0,811 | 6,08185E-45 |
| ACE2 | NACA | -0,8111 | 5,91914E-45 |
| ACE2 | ZNF521 | -0,8111 | 5,81223E-45 |
| ACE2 | ZNF583 | -0,8113 | 5,37338E-45 |
| ACE2 | RPL24 | -0,8114 | 5,05478E-45 |
| ACE2 | CDPF1 | -0,8114 | 5,121E-45 |
| ACE2 | CBX6 | -0,8115 | 4,93559E-45 |
| ACE2 | S100A13 | -0,8116 | 4,69672E-45 |
| ACE2 | CDKL3 | -0,8119 | 4,06059E-45 |
| ACE2 | ZNF467 | -0,8119 | 4,18089E-45 |
| ACE2 | NTAN1 | -0,8119 | 4,1819E-45 |
| ACE2 | ROM1 | -0,812 | 3,86651E-45 |
| ACE2 | RAD51C | -0,812 | 3,9708E-45 |
| ACE2 | SELENOM | -0,812 | 4,00985E-45 |
| ACE2 | TYW3 | -0,8125 | 3,16052E-45 |
| ACE2 | TXNDC15 | -0,8125 | 3,17107E-45 |
| ACE2 | PLEKHH3 | -0,8125 | 3,21999E-45 |
| ACE2 | SPACA6 | -0,8125 | 3,1367E-45 |
| ACE2 | PNMA1 | -0,8127 | 2,84298E-45 |
| ACE2 | CRMP1 | -0,813 | 2,58657E-45 |
| ACE2 | C11orf84 | -0,8132 | 2,33839E-45 |
| ACE2 | CNP | -0,8132 | 2,33743E-45 |
| ACE2 | HPF1 | -0,8133 | 2,23578E-45 |
| ACE2 | FAM228B | -0,8134 | 2,10962E-45 |

| | | | |
|---|---|---|---|
| ACE2 | AFAP1L2 | -0,8136 | 1,97309E-45 |
| ACE2 | RPL5 | -0,8139 | 1,70249E-45 |
| ACE2 | ANK2 | -0,8139 | 1,67892E-45 |
| ACE2 | GPR173 | -0,8139 | 1,67561E-45 |
| ACE2 | ANKRD65 | -0,814 | 1,58815E-45 |
| ACE2 | MIR100HG | -0,814 | 1,65636E-45 |
| ACE2 | AC108488.4 | -0,8141 | 1,5222E-45 |
| ACE2 | LINC01184 | -0,8142 | 1,45001E-45 |
| ACE2 | SARNP | -0,8142 | 1,46727E-45 |
| ACE2 | MSANTD3 | -0,8144 | 1,38448E-45 |
| ACE2 | CLIP4 | -0,8145 | 1,3144E-45 |
| ACE2 | CSRP2 | -0,8145 | 1,2674E-45 |
| ACE2 | UBE2E2 | -0,8146 | 1,24514E-45 |
| ACE2 | FAM120C | -0,8147 | 1,20543E-45 |
| ACE2 | APLF | -0,8148 | 1,11844E-45 |
| ACE2 | BMI1 | -0,8148 | 1,11448E-45 |
| ACE2 | NPHP1 | -0,8149 | 1,07339E-45 |
| ACE2 | C3orf18 | -0,8149 | 1,09532E-45 |
| ACE2 | C11orf58 | -0,8149 | 1,06466E-45 |
| ACE2 | VEGFB | -0,8149 | 1,0664E-45 |
| ACE2 | RP11-588K22.2 | -0,815 | 1,04272E-45 |
| ACE2 | TRIM37 | -0,8151 | 9,99347E-46 |
| ACE2 | TTC7B | -0,8152 | 9,28287E-46 |
| ACE2 | ZNF428 | -0,8152 | 9,4158E-46 |
| ACE2 | SRGAP2C | -0,8153 | 9,07468E-46 |
| ACE2 | DYRK3 | -0,8153 | 9,19326E-46 |
| ACE2 | ZFPM2 | -0,8153 | 8,97535E-46 |
| ACE2 | JAM3 | -0,8153 | 9,09759E-46 |
| ACE2 | CXCL12 | -0,8155 | 8,12068E-46 |
| ACE2 | SEPT4 | -0,8155 | 8,22726E-46 |
| ACE2 | CHST10 | -0,8156 | 8,00433E-46 |
| ACE2 | CLEC1A | -0,8156 | 7,90673E-46 |
| ACE2 | CDADC1 | -0,8156 | 8,03758E-46 |
| ACE2 | METRN | -0,8156 | 7,85963E-46 |
| ACE2 | KCTD17 | -0,8156 | 7,78825E-46 |
| ACE2 | ZFP2 | -0,8157 | 7,52837E-46 |
| ACE2 | KIAA0753 | -0,8157 | 7,55596E-46 |
| ACE2 | ZEB1 | -0,8158 | 7,27717E-46 |
| ACE2 | A2M-AS1 | -0,8158 | 7,32746E-46 |
| ACE2 | CRTC3 | -0,8158 | 7,12997E-46 |
| ACE2 | RPL22 | -0,8159 | 7,01888E-46 |
| ACE2 | VSTM4 | -0,8159 | 6,76705E-46 |
| ACE2 | CNPY4 | -0,816 | 6,50557E-46 |
| ACE2 | STK32C | -0,816 | 6,49156E-46 |
| ACE2 | ZNF423 | -0,816 | 6,50866E-46 |
| ACE2 | KRBA1 | -0,8161 | 6,33318E-46 |
| ACE2 | KLHL29 | -0,8163 | 5,6993E-46 |
| ACE2 | KLC1 | -0,8164 | 5,55463E-46 |
| ACE2 | GPR155 | -0,8167 | 4,69667E-46 |
| ACE2 | B3GALNT1 | -0,8167 | 4,87611E-46 |
| ACE2 | DNAJC18 | -0,8167 | 4,71086E-46 |
| ACE2 | RAB11FIP5 | -0,817 | 4,22905E-46 |
| ACE2 | CACNB1 | -0,8171 | 3,9942E-46 |
| ACE2 | PCNP | -0,8172 | 3,84456E-46 |

| | | | |
|---|---|---|---|
| ACE2 | NEK1 | -0,8172 | 3,8202E-46 |
| ACE2 | TCEA2 | -0,8174 | 3,49466E-46 |
| ACE2 | ARL6 | -0,8177 | 3,09976E-46 |
| ACE2 | CHN1 | -0,8179 | 2,80763E-46 |
| ACE2 | UBTD1 | -0,818 | 2,62911E-46 |
| ACE2 | SERGEF | -0,8181 | 2,49034E-46 |
| ACE2 | TRMT112 | -0,8182 | 2,40395E-46 |
| ACE2 | CSRNP2 | -0,8182 | 2,43924E-46 |
| ACE2 | C1orf123 | -0,8183 | 2,33981E-46 |
| ACE2 | WHAMMP2 | -0,8185 | 2,09788E-46 |
| ACE2 | PINLYP | -0,8185 | 2,0878E-46 |
| ACE2 | TCFL5 | -0,8185 | 2,12461E-46 |
| ACE2 | PJA1 | -0,8185 | 2,05903E-46 |
| ACE2 | C4orf32 | -0,8187 | 1,90495E-46 |
| ACE2 | EFNA5 | -0,8187 | 1,92145E-46 |
| ACE2 | TSHZ2 | -0,8187 | 1,91026E-46 |
| ACE2 | LETM2 | -0,8188 | 1,79851E-46 |
| ACE2 | C17orf51 | -0,8188 | 1,85674E-46 |
| ACE2 | MLLT1 | -0,8188 | 1,84674E-46 |
| ACE2 | ZNF667-AS1 | -0,8188 | 1,79833E-46 |
| ACE2 | SLC41A1 | -0,8189 | 1,76398E-46 |
| ACE2 | ARL3 | -0,819 | 1,69359E-46 |
| ACE2 | HOXC4 | -0,819 | 1,6866E-46 |
| ACE2 | RPL4 | -0,819 | 1,67722E-46 |
| ACE2 | PTPRU | -0,8191 | 1,60196E-46 |
| ACE2 | FHL3 | -0,8191 | 1,56772E-46 |
| ACE2 | AARSD1 | -0,8191 | 1,62795E-46 |
| ACE2 | RIPK2 | -0,8192 | 1,50672E-46 |
| ACE2 | LZTS2 | -0,8192 | 1,53316E-46 |
| ACE2 | ZNF542P | -0,8192 | 1,52415E-46 |
| ACE2 | MAPK11 | -0,8192 | 1,54749E-46 |
| ACE2 | C12orf57 | -0,8193 | 1,42474E-46 |
| ACE2 | TLE2 | -0,8194 | 1,4019E-46 |
| ACE2 | HOXB2 | -0,8195 | 1,348E-46 |
| ACE2 | BTF3L4 | -0,8196 | 1,27002E-46 |
| ACE2 | PRNP | -0,8198 | 1,18031E-46 |
| ACE2 | PRKRA | -0,8199 | 1,12432E-46 |
| ACE2 | ST3GAL3 | -0,8203 | 9,28443E-47 |
| ACE2 | TRIP6 | -0,8203 | 9,1852E-47 |
| ACE2 | ARHGAP31 | -0,8205 | 8,47583E-47 |
| ACE2 | RP11-382A20.3 | -0,8205 | 8,20148E-47 |
| ACE2 | HHIPL1 | -0,8206 | 7,78267E-47 |
| ACE2 | TBKBP1 | -0,8209 | 6,81424E-47 |
| ACE2 | BEX4 | -0,821 | 6,72668E-47 |
| ACE2 | TBC1D19 | -0,8212 | 6,13843E-47 |
| ACE2 | DDX50 | -0,8212 | 6,1527E-47 |
| ACE2 | CERS5 | -0,8212 | 6,06512E-47 |
| ACE2 | NAT14 | -0,8212 | 6,09905E-47 |
| ACE2 | FZD3 | -0,8215 | 5,22163E-47 |
| ACE2 | ANKS3 | -0,8217 | 4,6961E-47 |
| ACE2 | NRN1 | -0,8218 | 4,62161E-47 |
| ACE2 | SCARA3 | -0,8219 | 4,39787E-47 |
| ACE2 | LYRM1 | -0,822 | 4,05706E-47 |
| ACE2 | B4GALNT1 | -0,8223 | 3,51952E-47 |

| | | | |
|---|---|---|---|
| ACE2 | THSD1 | -0,8223 | 3,58017E-47 |
| ACE2 | HOMER3 | -0,8225 | 3,28556E-47 |
| ACE2 | TAF9B | -0,8225 | 3,27429E-47 |
| ACE2 | NENF | -0,8226 | 3,08836E-47 |
| ACE2 | GEM | -0,8226 | 3,086E-47 |
| ACE2 | ZNF34 | -0,8226 | 3,12905E-47 |
| ACE2 | AP1S2 | -0,8228 | 2,87039E-47 |
| ACE2 | SUPT3H | -0,8229 | 2,67171E-47 |
| ACE2 | RILPL1 | -0,8229 | 2,65919E-47 |
| ACE2 | PDE1B | -0,823 | 2,54807E-47 |
| ACE2 | TMCC2 | -0,8231 | 2,44209E-47 |
| ACE2 | FKBP10 | -0,8233 | 2,27352E-47 |
| ACE2 | ARL13B | -0,8234 | 2,14254E-47 |
| ACE2 | RP9P | -0,8234 | 2,16426E-47 |
| ACE2 | SGCE | -0,8234 | 2,1236E-47 |
| ACE2 | PLPPR2 | -0,8234 | 2,11961E-47 |
| ACE2 | COPRS | -0,8235 | 2,04594E-47 |
| ACE2 | UBE2E1 | -0,8236 | 1,90903E-47 |
| ACE2 | LYRM4 | -0,8236 | 1,90737E-47 |
| ACE2 | CYP46A1 | -0,8237 | 1,84824E-47 |
| ACE2 | LTBP3 | -0,8238 | 1,76846E-47 |
| ACE2 | ZBTB10 | -0,8239 | 1,68E-47 |
| ACE2 | HDAC7 | -0,8239 | 1,66032E-47 |
| ACE2 | FRMD4A | -0,8244 | 1,32611E-47 |
| ACE2 | ZNF25 | -0,8244 | 1,34013E-47 |
| ACE2 | SLC22A17 | -0,8246 | 1,22116E-47 |
| ACE2 | GYPC | -0,8247 | 1,12351E-47 |
| ACE2 | RDX | -0,8247 | 1,13201E-47 |
| ACE2 | MAP7D3 | -0,8249 | 1,02379E-47 |
| ACE2 | IPO5 | -0,825 | 9,94625E-48 |
| ACE2 | RAB34 | -0,825 | 1,01272E-47 |
| ACE2 | DDAH2 | -0,8251 | 9,67703E-48 |
| ACE2 | OLFM1 | -0,8252 | 9,08629E-48 |
| ACE2 | THYN1 | -0,8252 | 9,16566E-48 |
| ACE2 | ARMC10 | -0,8253 | 8,78707E-48 |
| ACE2 | CUEDC2 | -0,8253 | 8,56481E-48 |
| ACE2 | RP11-620J15.3 | -0,8253 | 8,82289E-48 |
| ACE2 | PITPNB | -0,8255 | 7,93899E-48 |
| ACE2 | DTX3 | -0,8257 | 6,96505E-48 |
| ACE2 | ATXN10 | -0,8257 | 7,2857E-48 |
| ACE2 | MAPK10 | -0,8259 | 6,57025E-48 |
| ACE2 | WHAMMP3 | -0,8259 | 6,56734E-48 |
| ACE2 | STXBP1 | -0,8261 | 5,77871E-48 |
| ACE2 | SLC12A4 | -0,8263 | 5,24859E-48 |
| ACE2 | PIAS3 | -0,8267 | 4,41665E-48 |
| ACE2 | IGFBP7 | -0,8267 | 4,3001E-48 |
| ACE2 | EFEMP2 | -0,8269 | 3,89536E-48 |
| ACE2 | PKIG | -0,827 | 3,83143E-48 |
| ACE2 | CNIH3 | -0,8271 | 3,57201E-48 |
| ACE2 | NLGN2 | -0,8272 | 3,4069E-48 |
| ACE2 | TCF7L1 | -0,8274 | 3,0776E-48 |
| ACE2 | CTB-193M12.5 | -0,8274 | 3,11386E-48 |
| ACE2 | MNAT1 | -0,8275 | 2,98398E-48 |
| ACE2 | KIF7 | -0,8275 | 2,94635E-48 |

| | | | |
|---|---|---|---|
| ACE2 | TMEM67 | -0,8278 | 2,63145E-48 |
| ACE2 | RARRES2 | -0,8279 | 2,39773E-48 |
| ACE2 | MAP3K12 | -0,828 | 2,37209E-48 |
| ACE2 | LCA5 | -0,8282 | 2,10367E-48 |
| ACE2 | PKNOX2 | -0,8282 | 2,11313E-48 |
| ACE2 | GNB4 | -0,8284 | 1,93984E-48 |
| ACE2 | SLC39A13 | -0,8286 | 1,75699E-48 |
| ACE2 | CLUAP1 | -0,8287 | 1,6817E-48 |
| ACE2 | ENO2 | -0,829 | 1,41387E-48 |
| ACE2 | QKI | -0,8294 | 1,20052E-48 |
| ACE2 | RSL24D1 | -0,8294 | 1,15756E-48 |
| ACE2 | ZCCHC17 | -0,8295 | 1,11599E-48 |
| ACE2 | GNG2 | -0,8296 | 1,06238E-48 |
| ACE2 | GUCY1B3 | -0,8298 | 9,4731E-49 |
| ACE2 | HMGN3 | -0,83 | 8,92484E-49 |
| ACE2 | VIM | -0,8301 | 8,48013E-49 |
| ACE2 | PTP4A3 | -0,8302 | 7,91363E-49 |
| ACE2 | NUCKS1 | -0,8303 | 7,5653E-49 |
| ACE2 | TDRP | -0,8303 | 7,61133E-49 |
| ACE2 | GLIS2 | -0,8305 | 6,82762E-49 |
| ACE2 | DLG4 | -0,8306 | 6,37464E-49 |
| ACE2 | NR2F1-AS1 | -0,831 | 5,4328E-49 |
| ACE2 | SMO | -0,831 | 5,31921E-49 |
| ACE2 | ARHGEF10 | -0,831 | 5,21769E-49 |
| ACE2 | ADARB1 | -0,831 | 5,22227E-49 |
| ACE2 | MIR497HG | -0,8313 | 4,51002E-49 |
| ACE2 | MAN1C1 | -0,8315 | 4,21388E-49 |
| ACE2 | CMSS1 | -0,8315 | 4,14649E-49 |
| ACE2 | PBX3 | -0,8315 | 4,24067E-49 |
| ACE2 | TMOD2 | -0,8315 | 4,17286E-49 |
| ACE2 | CD81 | -0,8319 | 3,39996E-49 |
| ACE2 | ZNF32 | -0,8322 | 2,98114E-49 |
| ACE2 | NR2F2-AS1 | -0,8322 | 2,86482E-49 |
| ACE2 | FAM89B | -0,8323 | 2,79121E-49 |
| ACE2 | ELK3 | -0,8323 | 2,78585E-49 |
| ACE2 | ZNF135 | -0,8327 | 2,23667E-49 |
| ACE2 | CNRIP1 | -0,8328 | 2,11832E-49 |
| ACE2 | ST3GAL4 | -0,8328 | 2,13833E-49 |
| ACE2 | P3H3 | -0,8328 | 2,18371E-49 |
| ACE2 | CEP68 | -0,8331 | 1,85693E-49 |
| ACE2 | CEP112 | -0,8331 | 1,87046E-49 |
| ACE2 | RPL15 | -0,8333 | 1,66421E-49 |
| ACE2 | DZIP1 | -0,8336 | 1,4639E-49 |
| ACE2 | CBX1 | -0,8337 | 1,36984E-49 |
| ACE2 | TEAD2 | -0,8344 | 9,66152E-50 |
| ACE2 | IFT81 | -0,8345 | 9,19913E-50 |
| ACE2 | DENND5A | -0,8347 | 8,37244E-50 |
| ACE2 | CHRD | -0,8348 | 7,93189E-50 |
| ACE2 | RHOJ | -0,8348 | 7,91178E-50 |
| ACE2 | EMP3 | -0,8351 | 6,63615E-50 |
| ACE2 | UBE2Q2 | -0,8353 | 6,00855E-50 |
| ACE2 | RAB28 | -0,8357 | 5,03977E-50 |
| ACE2 | HABP4 | -0,8359 | 4,51103E-50 |
| ACE2 | NPR2 | -0,8361 | 4,00511E-50 |

| | | | |
|---|---|---|---|
| ACE2 | TNFSF12 | -0,8362 | 3,90677E-50 |
| ACE2 | TIMM9 | -0,8366 | 3,09143E-50 |
| ACE2 | PEA15 | -0,8367 | 2,91929E-50 |
| ACE2 | NRSN2 | -0,837 | 2,49681E-50 |
| ACE2 | RBMS1 | -0,8375 | 1,93411E-50 |
| ACE2 | DNAL1 | -0,8377 | 1,73374E-50 |
| ACE2 | TSEN15 | -0,8378 | 1,68036E-50 |
| ACE2 | MCC | -0,8382 | 1,31599E-50 |
| ACE2 | MAPRE3 | -0,8383 | 1,26826E-50 |
| ACE2 | C11orf95 | -0,8386 | 1,09996E-50 |
| ACE2 | TMEM55A | -0,8387 | 1,05625E-50 |
| ACE2 | DUBR | -0,8388 | 1,00982E-50 |
| ACE2 | SEMA4C | -0,8393 | 7,64517E-51 |
| ACE2 | TULP3 | -0,8393 | 7,768E-51 |
| ACE2 | TRO | -0,8394 | 7,23395E-51 |
| ACE2 | SSBP2 | -0,8398 | 5,70856E-51 |
| ACE2 | COMMD7 | -0,8401 | 5,10093E-51 |
| ACE2 | SLC27A1 | -0,8402 | 4,68478E-51 |
| ACE2 | ST6GALNAC6 | -0,8407 | 3,54794E-51 |
| ACE2 | BST2 | -0,8407 | 3,58025E-51 |
| ACE2 | JAZF1 | -0,8415 | 2,39267E-51 |
| ACE2 | RP11-834C11.4 | -0,8418 | 2,05897E-51 |
| ACE2 | C1orf216 | -0,8422 | 1,61462E-51 |
| ACE2 | ITGB1BP1 | -0,8425 | 1,41407E-51 |
| ACE2 | MCRIP1 | -0,8425 | 1,37715E-51 |
| ACE2 | HIKESHI | -0,8428 | 1,17621E-51 |
| ACE2 | LCAT | -0,8434 | 8,59309E-52 |
| ACE2 | CFAP36 | -0,8444 | 5,04393E-52 |
| ACE2 | ARMCX2 | -0,8455 | 2,72074E-52 |
| ACE2 | TUBG2 | -0,8465 | 1,59039E-52 |
| ACE2 | NCKAP5L | -0,8467 | 1,38572E-52 |
| ACE2 | FAM92A | -0,8477 | 8,15636E-53 |
| ACE2 | RHOQ | -0,8489 | 4,06779E-53 |
| ACE2 | LIX1L | -0,8508 | 1,36489E-53 |
| ACE2 | PRAF2 | -0,8509 | 1,32998E-53 |
| ACE2 | MAGEH1 | -0,8514 | 9,6578E-54 |
| ACE2 | GLIPR2 | -0,8516 | 8,73063E-54 |
| ACE2 | ABI2 | -0,853 | 3,85204E-54 |
| ACE2 | OAZ2 | -0,8531 | 3,66608E-54 |
| ACE2 | NT5C3B | -0,8549 | 1,27423E-54 |
| ACE2 | DTWD1 | -0,8552 | 1,0468E-54 |
| ACE2 | SMARCD3 | -0,8587 | 1,31809E-55 |
| ACE2 | RARG | -0,8587 | 1,34186E-55 |
| ACE2 | TCTN1 | -0,8598 | 6,86955E-56 |
