## Supplementary table 5 colon for "Bioinformatic characterization of angiotensin-converting enzyme 2, the entry receptor for SARS-CoV-2"

| target_gene | correlated_gene | correlation | pval |
|---|---|---|---|
| ACE2 | ACE2 | 1 | 0 |
| ACE2 | MEP1A | 0,9056 | 1,5248E-152 |
| ACE2 | GDA | 0,8964 | 8,1914E-145 |
| ACE2 | MGAM2 | 0,8963 | 8,9961E-145 |
| ACE2 | TMEM236 | 0,894 | 6,663E-143 |
| ACE2 | GDPD2 | 0,8935 | 1,5818E-142 |
| ACE2 | PLS1 | 0,8932 | 2,8116E-142 |
| ACE2 | HHLA2 | 0,893 | 3,5864E-142 |
| ACE2 | SLC3A1 | 0,8928 | 5,268E-142 |
| ACE2 | COL17A1 | 0,8914 | 6,1163E-141 |
| ACE2 | TINAG | 0,8905 | 2,9425E-140 |
| ACE2 | CDHR2 | 0,8897 | 1,2368E-139 |
| ACE2 | RETSAT | 0,889 | 4,278E-139 |
| ACE2 | TUBAL3 | 0,8881 | 1,915E-138 |
| ACE2 | MUC13 | 0,8878 | 2,9654E-138 |
| ACE2 | GUCA2A | 0,8871 | 1,0164E-137 |
| ACE2 | MYH14 | 0,8847 | 5,3493E-136 |
| ACE2 | TMEM45B | 0,8845 | 7,2732E-136 |
| ACE2 | PLA2G12B | 0,8833 | 5,3966E-135 |
| ACE2 | MARVELD3 | 0,8831 | 7,1143E-135 |
| ACE2 | DSG2 | 0,8819 | 5,5172E-134 |
| ACE2 | TMIGD1 | 0,8813 | 1,4578E-133 |
| ACE2 | BTNL8 | 0,8812 | 1,617E-133 |
| ACE2 | SLC26A3 | 0,881 | 2,1512E-133 |
| ACE2 | LYPD8 | 0,8808 | 2,9997E-133 |
| ACE2 | SCIN | 0,8804 | 5,8302E-133 |
| ACE2 | CDHR5 | 0,8803 | 6,3151E-133 |
| ACE2 | CES2 | 0,8799 | 1,1641E-132 |
| ACE2 | TSPAN1 | 0,8789 | 6,3398E-132 |
| ACE2 | AOC1 | 0,8775 | 5,511E-131 |
| ACE2 | TMEM164 | 0,8762 | 4,0192E-130 |
| ACE2 | ANPEP | 0,8755 | 1,0864E-129 |
| ACE2 | SIPA1L3 | 0,8753 | 1,4826E-129 |
| ACE2 | NR1I2 | 0,8748 | 3,1954E-129 |
| ACE2 | CDH1 | 0,8747 | 3,6297E-129 |
| ACE2 | PLA2G4F | 0,8741 | 9,8067E-129 |
| ACE2 | PRAP1 | 0,874 | 1,1034E-128 |
| ACE2 | SGPP2 | 0,8739 | 1,2709E-128 |
| ACE2 | TMEM37 | 0,8736 | 1,8838E-128 |
| ACE2 | SLC17A4 | 0,8736 | 1,8852E-128 |
| ACE2 | CLRN3 | 0,8735 | 2,3521E-128 |
| ACE2 | IL22RA1 | 0,8734 | 2,7094E-128 |
| ACE2 | LIPH | 0,8734 | 2,4679E-128 |
| ACE2 | NDFIP2 | 0,8732 | 3,7524E-128 |
| ACE2 | ALPI | 0,8726 | 8,5275E-128 |
| ACE2 | TMEM82 | 0,8724 | 1,1005E-127 |
| ACE2 | STYK1 | 0,8723 | 1,4255E-127 |
| ACE2 | SDCBP2 | 0,8721 | 1,9333E-127 |
| ACE2 | CXADR | 0,8721 | 1,9043E-127 |
| ACE2 | F2RL1 | 0,872 | 2,1513E-127 |
| ACE2 | CGN | 0,8718 | 2,7491E-127 |
| ACE2 | DSC2 | 0,8717 | 3,0556E-127 |
| ACE2 | EPCAM | 0,8713 | 5,8133E-127 |

| | | | |
|---|---|---|---|
| ACE2 | PRSS8 | 0,8709 | 1,0201E-126 |
| ACE2 | AC007182.6 | 0,8704 | 2,3239E-126 |
| ACE2 | GPA33 | 0,8703 | 2,5295E-126 |
| ACE2 | LRRC19 | 0,8703 | 2,4813E-126 |
| ACE2 | C1orf106 | 0,8702 | 2,7941E-126 |
| ACE2 | SLC5A1 | 0,8692 | 1,2823E-125 |
| ACE2 | HNF4G | 0,8691 | 1,327E-125 |
| ACE2 | PBLD | 0,8691 | 1,3417E-125 |
| ACE2 | ENTPD8 | 0,8685 | 3,1896E-125 |
| ACE2 | MISP | 0,8685 | 3,1361E-125 |
| ACE2 | ABCC13 | 0,8685 | 3,5149E-125 |
| ACE2 | LSR | 0,8679 | 7,5888E-125 |
| ACE2 | TTC22 | 0,8673 | 1,8611E-124 |
| ACE2 | PTPRH | 0,8673 | 1,7522E-124 |
| ACE2 | SERINC2 | 0,867 | 2,6564E-124 |
| ACE2 | HGD | 0,867 | 2,9384E-124 |
| ACE2 | SPINT1 | 0,867 | 2,7582E-124 |
| ACE2 | ANKS4B | 0,8669 | 3,4008E-124 |
| ACE2 | SLC51B | 0,8666 | 4,6165E-124 |
| ACE2 | CLCA4 | 0,8663 | 7,1794E-124 |
| ACE2 | TJP3 | 0,8659 | 1,3566E-123 |
| ACE2 | HKDC1 | 0,8656 | 1,9709E-123 |
| ACE2 | SLC17A8 | 0,8656 | 2,0517E-123 |
| ACE2 | RBM47 | 0,8644 | 1,1303E-122 |
| ACE2 | LINC01207 | 0,8636 | 3,2712E-122 |
| ACE2 | ITGB6 | 0,8627 | 1,0394E-121 |
| ACE2 | DSP | 0,8627 | 1,1275E-121 |
| ACE2 | LINC00483 | 0,8627 | 1,145E-121 |
| ACE2 | VDR | 0,8624 | 1,5742E-121 |
| ACE2 | ADGRG7 | 0,8622 | 2,1292E-121 |
| ACE2 | SOWAHA | 0,8622 | 2,3081E-121 |
| ACE2 | EDN3 | 0,8621 | 2,4493E-121 |
| ACE2 | IL17RE | 0,862 | 3,0267E-121 |
| ACE2 | TMEM171 | 0,8618 | 3,7092E-121 |
| ACE2 | MYO7B | 0,8615 | 5,353E-121 |
| ACE2 | CLCN2 | 0,8614 | 6,106E-121 |
| ACE2 | FLJ22763 | 0,8613 | 6,8858E-121 |
| ACE2 | XDH | 0,8609 | 1,1917E-120 |
| ACE2 | PRKCD | 0,8608 | 1,473E-120 |
| ACE2 | SULT1B1 | 0,8608 | 1,4548E-120 |
| ACE2 | ATP1A1 | 0,8607 | 1,7537E-120 |
| ACE2 | ACY1 | 0,8607 | 1,7402E-120 |
| ACE2 | SPINT2 | 0,8602 | 3,2281E-120 |
| ACE2 | GLOD5 | 0,86 | 4,0882E-120 |
| ACE2 | CYP2C18 | 0,8597 | 5,9213E-120 |
| ACE2 | LAMB3 | 0,8595 | 7,8987E-120 |
| ACE2 | MALL | 0,8595 | 7,6735E-120 |
| ACE2 | GUCY2C | 0,8594 | 9,4391E-120 |
| ACE2 | SCNN1A | 0,8593 | 1,0076E-119 |
| ACE2 | EPB41L4B | 0,8592 | 1,2659E-119 |
| ACE2 | NAT2 | 0,8591 | 1,3494E-119 |
| ACE2 | HNF4A | 0,8588 | 1,9614E-119 |
| ACE2 | CDKN2B | 0,8587 | 2,4686E-119 |
| ACE2 | CLDN7 | 0,8587 | 2,3837E-119 |

| | | | |
|---|---|---|---|
| ACE2 | TSPAN3 | 0,8586 | 2,6184E-119 |
| ACE2 | B3GNT3 | 0,8583 | 3,9633E-119 |
| ACE2 | TMEM253 | 0,8582 | 4,4547E-119 |
| ACE2 | STX19 | 0,8579 | 7,0619E-119 |
| ACE2 | CORO2A | 0,8578 | 7,3326E-119 |
| ACE2 | SCAMP2 | 0,8577 | 9,3728E-119 |
| ACE2 | IYD | 0,8576 | 1,0099E-118 |
| ACE2 | PLEKHG6 | 0,8573 | 1,5892E-118 |
| ACE2 | SLC9A3R1 | 0,8572 | 1,7278E-118 |
| ACE2 | POF1B | 0,8572 | 1,705E-118 |
| ACE2 | C5orf30 | 0,857 | 2,2214E-118 |
| ACE2 | CDH17 | 0,8566 | 3,8075E-118 |
| ACE2 | MYO1A | 0,8564 | 5,0219E-118 |
| ACE2 | FMO5 | 0,8562 | 5,8925E-118 |
| ACE2 | MAB21L3 | 0,8559 | 9,6984E-118 |
| ACE2 | ST14 | 0,8557 | 1,2406E-117 |
| ACE2 | AIM1L | 0,8554 | 1,6551E-117 |
| ACE2 | SLAMF6P1 | 0,8553 | 1,9808E-117 |
| ACE2 | CCL15 | 0,8548 | 4,0309E-117 |
| ACE2 | FLVCR2 | 0,8545 | 5,4376E-117 |
| ACE2 | CEACAM7 | 0,8545 | 5,3738E-117 |
| ACE2 | ACSL5 | 0,8542 | 8,6715E-117 |
| ACE2 | LMO7 | 0,8542 | 7,9587E-117 |
| ACE2 | RAB11FIP1 | 0,8541 | 9,0798E-117 |
| ACE2 | BTNL3 | 0,8539 | 1,225E-116 |
| ACE2 | SLC13A2 | 0,8534 | 2,2748E-116 |
| ACE2 | SLC44A4 | 0,8531 | 3,2282E-116 |
| ACE2 | MUC12 | 0,853 | 3,707E-116 |
| ACE2 | KRT19 | 0,8528 | 4,5889E-116 |
| ACE2 | FRK | 0,8524 | 8,2181E-116 |
| ACE2 | SLC6A19 | 0,8523 | 9,6008E-116 |
| ACE2 | TMPRSS2 | 0,852 | 1,2857E-115 |
| ACE2 | MICALCL | 0,8519 | 1,5732E-115 |
| ACE2 | MYO1D | 0,8518 | 1,7118E-115 |
| ACE2 | FER1L6 | 0,8515 | 2,3738E-115 |
| ACE2 | CCL28 | 0,8514 | 2,9107E-115 |
| ACE2 | FXYD3 | 0,8509 | 5,5328E-115 |
| ACE2 | C1orf210 | 0,8508 | 6,1619E-115 |
| ACE2 | CLDN4 | 0,8507 | 6,5832E-115 |
| ACE2 | CTAGE5 | 0,8505 | 8,8437E-115 |
| ACE2 | SLC22A18 | 0,8504 | 1,0122E-114 |
| ACE2 | CYP2C19 | 0,8503 | 1,1563E-114 |
| ACE2 | C1orf116 | 0,8502 | 1,2856E-114 |
| ACE2 | MIA2 | 0,8495 | 3,0632E-114 |
| ACE2 | C4orf19 | 0,8494 | 3,1881E-114 |
| ACE2 | A1CF | 0,8494 | 3,2532E-114 |
| ACE2 | TMEM139 | 0,8491 | 4,8227E-114 |
| ACE2 | CLDN23 | 0,8488 | 7,1114E-114 |
| ACE2 | VIPR1 | 0,8487 | 7,6971E-114 |
| ACE2 | SMIM6 | 0,8484 | 1,1648E-113 |
| ACE2 | GCNT3 | 0,8483 | 1,3619E-113 |
| ACE2 | BCAS1 | 0,8483 | 1,3201E-113 |
| ACE2 | SMPDL3A | 0,848 | 1,997E-113 |
| ACE2 | KRT20 | 0,848 | 1,9324E-113 |

| | | | |
|---|---|---|---|
| ACE2 | KCNK1 | 0,8478 | 2,3631E-113 |
| ACE2 | MARVELD2 | 0,8472 | 4,8694E-113 |
| ACE2 | CDKN2B-AS1 | 0,8472 | 4,7459E-113 |
| ACE2 | MYO5B | 0,847 | 6,7493E-113 |
| ACE2 | TMC5 | 0,8468 | 8,2646E-113 |
| ACE2 | IRF6 | 0,8465 | 1,1478E-112 |
| ACE2 | TRIM36 | 0,8463 | 1,4228E-112 |
| ACE2 | TSPAN8 | 0,846 | 2,0789E-112 |
| ACE2 | LETM1 | 0,8459 | 2,5178E-112 |
| ACE2 | SOWAHB | 0,8459 | 2,4878E-112 |
| ACE2 | CARMIL1 | 0,8459 | 2,3314E-112 |
| ACE2 | GGT6 | 0,8459 | 2,2663E-112 |
| ACE2 | DOLPP1 | 0,8458 | 2,6363E-112 |
| ACE2 | TMEM30B | 0,8458 | 2,869E-112 |
| ACE2 | GALNT7 | 0,8457 | 3,1251E-112 |
| ACE2 | OCLN | 0,8456 | 3,585E-112 |
| ACE2 | TRIM31 | 0,8456 | 3,4016E-112 |
| ACE2 | TGFA | 0,8454 | 4,5963E-112 |
| ACE2 | FAM151A | 0,8453 | 4,8291E-112 |
| ACE2 | PRSS3 | 0,8449 | 7,7564E-112 |
| ACE2 | METTL7B | 0,8449 | 7,8477E-112 |
| ACE2 | RPL12P14 | 0,8448 | 8,6494E-112 |
| ACE2 | USH1C | 0,8447 | 1,0005E-111 |
| ACE2 | VILL | 0,8442 | 1,7716E-111 |
| ACE2 | GPRIN2 | 0,8442 | 1,7561E-111 |
| ACE2 | RASSF6 | 0,8441 | 2,116E-111 |
| ACE2 | PKIB | 0,8441 | 1,9571E-111 |
| ACE2 | SLC37A1 | 0,8441 | 2,1701E-111 |
| ACE2 | PRRG2 | 0,844 | 2,2453E-111 |
| ACE2 | DHRS9 | 0,8438 | 2,9932E-111 |
| ACE2 | EZR | 0,8438 | 2,9682E-111 |
| ACE2 | DNM2 | 0,8438 | 3,0177E-111 |
| ACE2 | LINC01843 | 0,8436 | 3,7204E-111 |
| ACE2 | TFCP2L1 | 0,8434 | 4,7268E-111 |
| ACE2 | CRB3 | 0,8434 | 4,9313E-111 |
| ACE2 | ABHD17C | 0,8432 | 5,8054E-111 |
| ACE2 | ABHD11-AS1 | 0,8428 | 9,77E-111 |
| ACE2 | RP11-680F8.1 | 0,8427 | 1,0632E-110 |
| ACE2 | CLIC5 | 0,8423 | 1,6772E-110 |
| ACE2 | LRRC75A | 0,8419 | 2,8729E-110 |
| ACE2 | XXbac-B476C20.9 | 0,8419 | 2,876E-110 |
| ACE2 | C6orf222 | 0,8409 | 8,9453E-110 |
| ACE2 | ZBTB7B | 0,8408 | 9,8145E-110 |
| ACE2 | ZG16 | 0,8406 | 1,1851E-109 |
| ACE2 | GPR39 | 0,8404 | 1,6181E-109 |
| ACE2 | ENTPD5 | 0,8404 | 1,5675E-109 |
| ACE2 | RP11-297L17.2 | 0,8404 | 1,6528E-109 |
| ACE2 | RP11-486A14.1 | 0,8403 | 1,8551E-109 |
| ACE2 | CLDN3 | 0,8402 | 2,073E-109 |
| ACE2 | APOBR | 0,8401 | 2,3408E-109 |
| ACE2 | TRIM10 | 0,84 | 2,4941E-109 |
| ACE2 | PAPSS2 | 0,84 | 2,4421E-109 |
| ACE2 | GATA6 | 0,84 | 2,5987E-109 |
| ACE2 | ESPN | 0,8399 | 2,7725E-109 |

| | | | |
|---|---|---|---|
| ACE2 | KIF13B | 0,8399 | 2,8785E-109 |
| ACE2 | ACVR1B | 0,8399 | 2,8763E-109 |
| ACE2 | TRPM6 | 0,8396 | 4,1083E-109 |
| ACE2 | GUCA2B | 0,8395 | 4,2995E-109 |
| ACE2 | PRR15 | 0,8394 | 4,9626E-109 |
| ACE2 | CALML4 | 0,8391 | 6,6953E-109 |
| ACE2 | DNMBP-AS1 | 0,839 | 7,834E-109 |
| ACE2 | NAT1 | 0,8389 | 9,3125E-109 |
| ACE2 | MAPK13 | 0,8387 | 1,0752E-108 |
| ACE2 | CEACAM1 | 0,8386 | 1,2388E-108 |
| ACE2 | DHRS11 | 0,8384 | 1,5624E-108 |
| ACE2 | KIAA1522 | 0,8383 | 1,7047E-108 |
| ACE2 | CTSS | 0,8383 | 1,7531E-108 |
| ACE2 | SEPHS2 | 0,8383 | 1,7157E-108 |
| ACE2 | BARX2 | 0,8382 | 1,9101E-108 |
| ACE2 | PLEKHA7 | 0,8381 | 2,2865E-108 |
| ACE2 | SLC41A2 | 0,838 | 2,4858E-108 |
| ACE2 | JUP | 0,838 | 2,41E-108 |
| ACE2 | SP6 | 0,838 | 2,5088E-108 |
| ACE2 | LINC02038 | 0,8378 | 3,119E-108 |
| ACE2 | SLC35A3 | 0,8377 | 3,3373E-108 |
| ACE2 | CDCP1 | 0,8375 | 4,6E-108 |
| ACE2 | PPARG | 0,8374 | 4,6484E-108 |
| ACE2 | CEACAM6 | 0,8374 | 4,6745E-108 |
| ACE2 | CDC42BPG | 0,8372 | 6,0255E-108 |
| ACE2 | PCK2 | 0,8372 | 5,8145E-108 |
| ACE2 | ABHD3 | 0,8372 | 6,0214E-108 |
| ACE2 | GRAMD4 | 0,8372 | 6,0793E-108 |
| ACE2 | P2RX4 | 0,8371 | 7,0397E-108 |
| ACE2 | SI | 0,837 | 7,3419E-108 |
| ACE2 | CDS1 | 0,8366 | 1,2288E-107 |
| ACE2 | SLC26A2 | 0,8364 | 1,5733E-107 |
| ACE2 | SYTL5 | 0,8362 | 1,8285E-107 |
| ACE2 | LY75 | 0,8361 | 2,2168E-107 |
| ACE2 | MROH3P | 0,8359 | 2,8054E-107 |
| ACE2 | AKR1C3 | 0,8357 | 3,3028E-107 |
| ACE2 | RPS6KA1 | 0,8349 | 8,3638E-107 |
| ACE2 | CTD-2396E7.11 | 0,8349 | 7,764E-107 |
| ACE2 | AMN | 0,8348 | 9,5544E-107 |
| ACE2 | CEACAM5 | 0,8348 | 8,9234E-107 |
| ACE2 | HNRNPA1P17 | 0,8347 | 9,8483E-107 |
| ACE2 | PTK6 | 0,8347 | 1,0659E-106 |
| ACE2 | FSIP2 | 0,8346 | 1,0893E-106 |
| ACE2 | VAMP8 | 0,8345 | 1,2447E-106 |
| ACE2 | SLC4A4 | 0,8345 | 1,2898E-106 |
| ACE2 | PLAC8 | 0,8344 | 1,3596E-106 |
| ACE2 | TM4SF5 | 0,8343 | 1,5703E-106 |
| ACE2 | PRSS12 | 0,8342 | 1,8398E-106 |
| ACE2 | NHSL1 | 0,8341 | 2,0356E-106 |
| ACE2 | CHMP4C | 0,834 | 2,1746E-106 |
| ACE2 | GBA3 | 0,8338 | 2,8322E-106 |
| ACE2 | CYP2J2 | 0,8335 | 3,8921E-106 |
| ACE2 | SMIM24 | 0,8333 | 4,9023E-106 |
| ACE2 | TMEM150B | 0,8333 | 4,8395E-106 |

| | | | |
|---|---|---|---|
| ACE2 | IFNLR1 | 0,8331 | 5,9567E-106 |
| ACE2 | RP11-800A3.4 | 0,8331 | 5,961E-106 |
| ACE2 | MFSD9 | 0,833 | 6,3588E-106 |
| ACE2 | PPP1R14C | 0,833 | 6,4349E-106 |
| ACE2 | B3GNT5 | 0,8326 | 1,0129E-105 |
| ACE2 | SLC39A4 | 0,8326 | 1,076E-105 |
| ACE2 | TM4SF20 | 0,8325 | 1,1073E-105 |
| ACE2 | NGEF | 0,8324 | 1,2804E-105 |
| ACE2 | SMIM2-AS1 | 0,8321 | 1,8919E-105 |
| ACE2 | FCGBP | 0,8319 | 2,3108E-105 |
| ACE2 | TPRN | 0,8317 | 2,8574E-105 |
| ACE2 | ACSS2 | 0,8317 | 2,9033E-105 |
| ACE2 | CYP3A5 | 0,8316 | 3,1742E-105 |
| ACE2 | MUC17 | 0,8313 | 4,4265E-105 |
| ACE2 | ITM2C | 0,8312 | 4,9299E-105 |
| ACE2 | P2RY2 | 0,8311 | 5,3459E-105 |
| ACE2 | SGK2 | 0,8311 | 5,3272E-105 |
| ACE2 | CD2AP | 0,831 | 5,9275E-105 |
| ACE2 | PTPN3 | 0,831 | 6,2762E-105 |
| ACE2 | DNAJC22 | 0,8307 | 8,203E-105 |
| ACE2 | IGSF9 | 0,8305 | 1,0387E-104 |
| ACE2 | CHP1 | 0,8304 | 1,1712E-104 |
| ACE2 | MEP1B | 0,8303 | 1,2273E-104 |
| ACE2 | RP11-27G14.4 | 0,8302 | 1,3553E-104 |
| ACE2 | RAB17 | 0,83 | 1,8171E-104 |
| ACE2 | KRT8 | 0,83 | 1,6808E-104 |
| ACE2 | LINC00974 | 0,83 | 1,6719E-104 |
| ACE2 | FAM110C | 0,8299 | 1,9771E-104 |
| ACE2 | MOGAT2 | 0,8299 | 1,863E-104 |
| ACE2 | HK2 | 0,8298 | 2,1202E-104 |
| ACE2 | UGT2A3 | 0,8298 | 2,2569E-104 |
| ACE2 | PIP5K1B | 0,8298 | 2,2846E-104 |
| ACE2 | MUC3A | 0,8295 | 3,0962E-104 |
| ACE2 | ADTRP | 0,8293 | 3,7923E-104 |
| ACE2 | TMEM102 | 0,8293 | 3,6042E-104 |
| ACE2 | PDZD3 | 0,8291 | 4,6485E-104 |
| ACE2 | LINC02177 | 0,829 | 5,2219E-104 |
| ACE2 | TMC4 | 0,829 | 5,4202E-104 |
| ACE2 | SORL1 | 0,8289 | 5,591E-104 |
| ACE2 | PDZK1IP1 | 0,8287 | 7,3739E-104 |
| ACE2 | CTSE | 0,8287 | 6,7833E-104 |
| ACE2 | LINC01559 | 0,8287 | 7,1183E-104 |
| ACE2 | FAM83F | 0,8287 | 7,0749E-104 |
| ACE2 | OR7E14P | 0,8286 | 7,5758E-104 |
| ACE2 | NR1H4 | 0,8285 | 8,844E-104 |
| ACE2 | HSD11B2 | 0,8285 | 9,0398E-104 |
| ACE2 | C19orf33 | 0,8285 | 9,1726E-104 |
| ACE2 | CAPN13 | 0,8283 | 1,0699E-103 |
| ACE2 | ISX | 0,8282 | 1,2459E-103 |
| ACE2 | HTATIP2 | 0,8281 | 1,3816E-103 |
| ACE2 | SLC30A10 | 0,8278 | 1,9277E-103 |
| ACE2 | FAM83H | 0,8276 | 2,3337E-103 |
| ACE2 | C11orf86 | 0,8274 | 2,7764E-103 |
| ACE2 | MYH15 | 0,8272 | 3,6015E-103 |

| | | | |
|---|---|---|---|
| ACE2 | RP11-35P15.1 | 0,8272 | 3,4285E-103 |
| ACE2 | PRR15L | 0,8272 | 3,4518E-103 |
| ACE2 | VIPR1-AS1 | 0,8271 | 3,8122E-103 |
| ACE2 | SCNN1B | 0,8271 | 4,0884E-103 |
| ACE2 | ILDR1 | 0,8269 | 4,822E-103 |
| ACE2 | LRRC31 | 0,8267 | 5,9677E-103 |
| ACE2 | CA4 | 0,8264 | 8,185E-103 |
| ACE2 | PTPRF | 0,8263 | 8,6447E-103 |
| ACE2 | CRAT8 | 0,8263 | 8,9371E-103 |
| ACE2 | RP11-191G24.1 | 0,8262 | 1,0343E-102 |
| ACE2 | SPECC1 | 0,8261 | 1,0863E-102 |
| ACE2 | MKRN2OS | 0,826 | 1,265E-102 |
| ACE2 | LINC02023 | 0,826 | 1,1823E-102 |
| ACE2 | AKR1B10 | 0,826 | 1,2035E-102 |
| ACE2 | PRELID2 | 0,8259 | 1,3266E-102 |
| ACE2 | AHCYL2 | 0,8259 | 1,4544E-102 |
| ACE2 | BEST4 | 0,8257 | 1,6978E-102 |
| ACE2 | TMPRSS4 | 0,8256 | 1,8798E-102 |
| ACE2 | OIT3 | 0,8253 | 2,5497E-102 |
| ACE2 | RNF128 | 0,8252 | 2,7515E-102 |
| ACE2 | RP11-63P12.7 | 0,8251 | 3,0848E-102 |
| ACE2 | ATP8B1 | 0,8248 | 4,4482E-102 |
| ACE2 | SIAE | 0,8247 | 4,7751E-102 |
| ACE2 | LEXM | 0,8246 | 5,6364E-102 |
| ACE2 | MGAT4B | 0,8245 | 5,9368E-102 |
| ACE2 | GSKIP | 0,8244 | 6,7917E-102 |
| ACE2 | SLC35D1 | 0,8243 | 7,2848E-102 |
| ACE2 | HRASLS2 | 0,8242 | 8,542E-102 |
| ACE2 | APOBEC1 | 0,8241 | 9,4519E-102 |
| ACE2 | CYP4F3 | 0,8241 | 8,6863E-102 |
| ACE2 | C6orf132 | 0,8238 | 1,29E-101 |
| ACE2 | KDF1 | 0,8235 | 1,6464E-101 |
| ACE2 | SMPDL3B | 0,8234 | 1,837E-101 |
| ACE2 | CNNM4 | 0,8234 | 1,9372E-101 |
| ACE2 | ESRRA | 0,8234 | 1,8095E-101 |
| ACE2 | CLMN | 0,8232 | 2,312E-101 |
| ACE2 | GALNT3 | 0,8229 | 3,1522E-101 |
| ACE2 | RP5-1185I7.1 | 0,8229 | 3,0048E-101 |
| ACE2 | NR5A2 | 0,8228 | 3,6152E-101 |
| ACE2 | ARL14 | 0,8228 | 3,3418E-101 |
| ACE2 | TRIM15 | 0,8228 | 3,3634E-101 |
| ACE2 | HOOK1 | 0,8225 | 4,5186E-101 |
| ACE2 | NAAA | 0,8224 | 5,3167E-101 |
| ACE2 | UGT2B7 | 0,8221 | 7,1505E-101 |
| ACE2 | PLEKHB2 | 0,8217 | 1,0637E-100 |
| ACE2 | NCEH1 | 0,8217 | 1,0328E-100 |
| ACE2 | LRRC66 | 0,8217 | 1,0238E-100 |
| ACE2 | EPS8L2 | 0,8217 | 1,043E-100 |
| ACE2 | VIL1 | 0,8216 | 1,1858E-100 |
| ACE2 | PFKFB2 | 0,8215 | 1,3627E-100 |
| ACE2 | LINC02000 | 0,8212 | 1,7274E-100 |
| ACE2 | PPP1R14D | 0,821 | 2,1309E-100 |
| ACE2 | FAM160A1 | 0,8208 | 2,7957E-100 |
| ACE2 | TBX10 | 0,8207 | 3,0053E-100 |

| | | | |
|---|---|---|---|
| ACE2 | DOK4 | 0,8203 | 4,5728E-100 |
| ACE2 | MTND4P20 | 0,8201 | 5,4664E-100 |
| ACE2 | OTOP2 | 0,8201 | 5,4579E-100 |
| ACE2 | FUT3 | 0,82 | 6,2995E-100 |
| ACE2 | CASP16P | 0,8199 | 6,9755E-100 |
| ACE2 | STXBP2 | 0,8199 | 6,5171E-100 |
| ACE2 | RP11-10C24.3 | 0,8197 | 8,6353E-100 |
| ACE2 | RHPN2 | 0,8197 | 8,3478E-100 |
| ACE2 | AGR3 | 0,8195 | 1,0039E-99 |
| ACE2 | CHST5 | 0,8194 | 1,0921E-99 |
| ACE2 | ITPKA | 0,8191 | 1,4877E-99 |
| ACE2 | GPRC5C | 0,8188 | 1,9939E-99 |
| ACE2 | FRYL | 0,8187 | 2,2276E-99 |
| ACE2 | SLC22A18AS | 0,8187 | 2,1743E-99 |
| ACE2 | MS4A12 | 0,8187 | 2,3319E-99 |
| ACE2 | PAQR8 | 0,8186 | 2,4186E-99 |
| ACE2 | SYT8 | 0,8186 | 2,4401E-99 |
| ACE2 | STAP2 | 0,8186 | 2,549E-99 |
| ACE2 | ENPP3 | 0,8184 | 2,92E-99 |
| ACE2 | TDP2 | 0,8179 | 5,0814E-99 |
| ACE2 | AIM1 | 0,8177 | 6,1091E-99 |
| ACE2 | CES3 | 0,8176 | 6,7373E-99 |
| ACE2 | GRAMD1C | 0,8174 | 8,5077E-99 |
| ACE2 | FFAR4 | 0,8174 | 8,0472E-99 |
| ACE2 | ZBTB7C | 0,8174 | 8,1049E-99 |
| ACE2 | C1QTNF12 | 0,8173 | 9,2805E-99 |
| ACE2 | PLA2G10 | 0,8172 | 1,05377E-98 |
| ACE2 | B3GALT5 | 0,8172 | 1,00757E-98 |
| ACE2 | PRR13 | 0,8171 | 1,08572E-98 |
| ACE2 | APPL2 | 0,8171 | 1,10725E-98 |
| ACE2 | ARHGEF16 | 0,817 | 1,19083E-98 |
| ACE2 | NIPAL1 | 0,817 | 1,25684E-98 |
| ACE2 | TPMT | 0,817 | 1,2374E-98 |
| ACE2 | MOGAT3 | 0,817 | 1,20242E-98 |
| ACE2 | KLF5 | 0,8169 | 1,40213E-98 |
| ACE2 | PKP3 | 0,8168 | 1,55461E-98 |
| ACE2 | ACOX1 | 0,8167 | 1,6722E-98 |
| ACE2 | RP11-532F12.5 | 0,8166 | 1,85785E-98 |
| ACE2 | PRKCZ | 0,8165 | 2,06706E-98 |
| ACE2 | RP11-96D1.6 | 0,8163 | 2,56579E-98 |
| ACE2 | IMPA1 | 0,8162 | 2,76602E-98 |
| ACE2 | PXMP2 | 0,8159 | 3,78958E-98 |
| ACE2 | EPS8L3 | 0,8157 | 4,62089E-98 |
| ACE2 | FUT6 | 0,8156 | 4,85072E-98 |
| ACE2 | TST | 0,8152 | 7,31278E-98 |
| ACE2 | B4GALNT3 | 0,8148 | 1,11812E-97 |
| ACE2 | ABCC3 | 0,8148 | 1,10986E-97 |
| ACE2 | LGALS3 | 0,8145 | 1,46019E-97 |
| ACE2 | ZNF552 | 0,8145 | 1,413E-97 |
| ACE2 | FAM83G | 0,8143 | 1,71055E-97 |
| ACE2 | KRT8P3 | 0,8142 | 1,86854E-97 |
| ACE2 | CAMSAP3 | 0,8139 | 2,51211E-97 |
| ACE2 | MPST | 0,8138 | 2,74028E-97 |
| ACE2 | ALDH18A1 | 0,8137 | 3,27202E-97 |

| | | | |
|---|---|---|---|
| ACE2 | DENND1B | 0,8136 | 3,39418E-97 |
| ACE2 | NUDT16P1 | 0,8135 | 3,97886E-97 |
| ACE2 | DHCR24 | 0,8133 | 4,80638E-97 |
| ACE2 | RAB19 | 0,8131 | 5,48065E-97 |
| ACE2 | SMIM22 | 0,8131 | 5,43255E-97 |
| ACE2 | BCL2L14 | 0,813 | 6,4112E-97 |
| ACE2 | F11R | 0,8128 | 7,5632E-97 |
| ACE2 | LAD1 | 0,8127 | 8,49285E-97 |
| ACE2 | RAPGEFL1 | 0,8127 | 8,00841E-97 |
| ACE2 | SYTL2 | 0,8126 | 9,17271E-97 |
| ACE2 | VAV3 | 0,8125 | 9,86229E-97 |
| ACE2 | PARD6B | 0,8123 | 1,2556E-96 |
| ACE2 | GALNT5 | 0,812 | 1,5914E-96 |
| ACE2 | RASEF | 0,8119 | 1,86818E-96 |
| ACE2 | CAPN9 | 0,8118 | 2,01866E-96 |
| ACE2 | EHHADH | 0,8118 | 1,98701E-96 |
| ACE2 | HPSE | 0,8118 | 2,07311E-96 |
| ACE2 | FABP2 | 0,8118 | 1,96985E-96 |
| ACE2 | OVOL1 | 0,8115 | 2,57094E-96 |
| ACE2 | CDKN2B-AS | 0,8113 | 3,32639E-96 |
| ACE2 | PIGR | 0,8112 | 3,48744E-96 |
| ACE2 | SHROOM3 | 0,8112 | 3,44748E-96 |
| ACE2 | EFNA2 | 0,8109 | 4,5464E-96 |
| ACE2 | SLC9A2 | 0,8108 | 5,41852E-96 |
| ACE2 | PRSS16 | 0,8107 | 5,69276E-96 |
| ACE2 | SATB2 | 0,8106 | 6,19159E-96 |
| ACE2 | NMRAL2P | 0,8106 | 6,15557E-96 |
| ACE2 | SMPD3 | 0,8106 | 6,21364E-96 |
| ACE2 | TMEM184A | 0,8101 | 9,73654E-96 |
| ACE2 | HNF1A | 0,8101 | 1,0246E-95 |
| ACE2 | SULT1A2 | 0,81 | 1,16982E-95 |
| ACE2 | RP11-71L14.3 | 0,8098 | 1,38654E-95 |
| ACE2 | LLGL2 | 0,8097 | 1,51072E-95 |
| ACE2 | TP53I3 | 0,8096 | 1,71224E-95 |
| ACE2 | CCDC162P | 0,8096 | 1,6916E-95 |
| ACE2 | MGAT4A | 0,8094 | 1,94974E-95 |
| ACE2 | ABCC6 | 0,8094 | 2,01131E-95 |
| ACE2 | ARHGEF5 | 0,8092 | 2,45195E-95 |
| ACE2 | SLC27A2 | 0,8091 | 2,6986E-95 |
| ACE2 | RILP | 0,8091 | 2,55155E-95 |
| ACE2 | UGT1A10 | 0,8087 | 3,84378E-95 |
| ACE2 | PARP4 | 0,8087 | 3,76604E-95 |
| ACE2 | GALNT4 | 0,8084 | 5,04792E-95 |
| ACE2 | AC106876.2 | 0,8083 | 5,70385E-95 |
| ACE2 | HSD17B2 | 0,8082 | 6,386E-95 |
| ACE2 | LINC01133 | 0,8081 | 6,57722E-95 |
| ACE2 | BPNT1 | 0,8079 | 8,50981E-95 |
| ACE2 | MS4A8 | 0,8078 | 9,14007E-95 |
| ACE2 | CPT2 | 0,8077 | 9,66109E-95 |
| ACE2 | PRAG1 | 0,8077 | 1,01818E-94 |
| ACE2 | SH2D3A | 0,8077 | 1,0054E-94 |
| ACE2 | PHGR1 | 0,8076 | 1,06048E-94 |
| ACE2 | CLDN8 | 0,8075 | 1,18712E-94 |
| ACE2 | SLC51A | 0,8073 | 1,45386E-94 |

| | | | |
|---|---|---|---|
| ACE2 | CTB-186H2.3 | 0,807 | 1,83864E-94 |
| ACE2 | ZDHHC12 | 0,8069 | 1,99591E-94 |
| ACE2 | NAT8B | 0,8059 | 5,38138E-94 |
| ACE2 | MTM1 | 0,8057 | 6,4754E-94 |
| ACE2 | TMEM9B | 0,8055 | 8,01774E-94 |
| ACE2 | TAT | 0,8055 | 7,49877E-94 |
| ACE2 | ARHGAP27 | 0,8054 | 8,14126E-94 |
| ACE2 | ABRACL | 0,8052 | 9,89776E-94 |
| ACE2 | CANT1 | 0,8048 | 1,51025E-93 |
| ACE2 | SRI | 0,8047 | 1,59612E-93 |
| ACE2 | RP5-1148A21.3 | 0,8045 | 2,01974E-93 |
| ACE2 | OPN3 | 0,8044 | 2,12935E-93 |
| ACE2 | BAIAP2L2 | 0,8042 | 2,51683E-93 |
| ACE2 | CDC42SE2 | 0,8039 | 3,32295E-93 |
| ACE2 | MUC2 | 0,8039 | 3,33231E-93 |
| ACE2 | RP11-395B7.2 | 0,8033 | 5,69716E-93 |
| ACE2 | PEX26 | 0,8033 | 5,65067E-93 |
| ACE2 | CASP5 | 0,8031 | 7,07795E-93 |
| ACE2 | TRIM31-AS1 | 0,8028 | 9,62837E-93 |
| ACE2 | MAP7 | 0,8028 | 9,28691E-93 |
| ACE2 | IL18 | 0,8026 | 1,124E-92 |
| ACE2 | USP43 | 0,8022 | 1,65684E-92 |
| ACE2 | HSD3B2 | 0,8021 | 1,6937E-92 |
| ACE2 | CMAS | 0,8021 | 1,82661E-92 |
| ACE2 | HEPACAM2 | 0,802 | 1,9821E-92 |
| ACE2 | NAGS | 0,8019 | 2,03124E-92 |
| ACE2 | TOX3 | 0,8018 | 2,39349E-92 |
| ACE2 | TMEM61 | 0,8016 | 2,86603E-92 |
| ACE2 | SLC35D2 | 0,8014 | 3,23603E-92 |
| ACE2 | B3GNT6 | 0,8014 | 3,39817E-92 |
| ACE2 | DAO | 0,8013 | 3,51193E-92 |
| ACE2 | CDX1 | 0,8007 | 6,44413E-92 |
| ACE2 | AGPAT2 | 0,8007 | 6,37645E-92 |
| ACE2 | FA2H | 0,8007 | 6,33198E-92 |
| ACE2 | ACAA2 | 0,8006 | 6,51395E-92 |
| ACE2 | EPHA10 | 0,8005 | 7,53395E-92 |
| ACE2 | MIR194-2HG | 0,8004 | 7,89637E-92 |
| ACE2 | RP4-651E10.4 | 0,8003 | 9,15414E-92 |
| ACE2 | RP11-276H7.3 | 0,8002 | 9,53789E-92 |
| ACE2 | IQGAP2 | 0,8 | 1,18641E-91 |
| ACE2 | EHF | 0,7999 | 1,2666E-91 |
| ACE2 | RP1-117B12.4 | 0,7999 | 1,27135E-91 |
| ACE2 | AKR7A3 | 0,7994 | 1,94399E-91 |
| ACE2 | LINC01752 | 0,7994 | 2,01031E-91 |
| ACE2 | GPD2 | 0,7993 | 2,14322E-91 |
| ACE2 | RP11-102C16.3 | 0,7992 | 2,47152E-91 |
| ACE2 | ACADS | 0,7992 | 2,50271E-91 |
| ACE2 | CTD-2021H9.3 | 0,7991 | 2,57777E-91 |
| ACE2 | UGDH | 0,799 | 2,76981E-91 |
| ACE2 | DGAT1 | 0,799 | 2,93367E-91 |
| ACE2 | COX5A | 0,7989 | 3,20889E-91 |
| ACE2 | UGT1A1 | 0,7988 | 3,44862E-91 |
| ACE2 | CCRL2 | 0,7987 | 3,78293E-91 |
| ACE2 | DSG2-AS1 | 0,7987 | 3,72894E-91 |

| | | | |
|---|---|---|---|
| ACE2 | WTH3DI | 0,7986 | 4,10808E-91 |
| ACE2 | CNGA1 | 0,7986 | 4,2105E-91 |
| ACE2 | CA12 | 0,7986 | 3,96423E-91 |
| ACE2 | CHRNA1 | 0,7985 | 4,63408E-91 |
| ACE2 | PAFAH2 | 0,7984 | 4,83544E-91 |
| ACE2 | TMEM125 | 0,7983 | 5,31936E-91 |
| ACE2 | FAM3D | 0,7983 | 5,4439E-91 |
| ACE2 | B4GALT4 | 0,7983 | 5,13262E-91 |
| ACE2 | SLC25A5 | 0,7983 | 5,27301E-91 |
| ACE2 | HDHD3 | 0,7981 | 6,32672E-91 |
| ACE2 | OTOP3 | 0,798 | 7,23715E-91 |
| ACE2 | NEU4 | 0,7978 | 8,34934E-91 |
| ACE2 | DDR1 | 0,7978 | 8,13815E-91 |
| ACE2 | AP000439.3 | 0,7978 | 7,98824E-91 |
| ACE2 | CCDC68 | 0,7977 | 8,79742E-91 |
| ACE2 | TSTD1 | 0,7975 | 1,04366E-90 |
| ACE2 | ATP10B | 0,7975 | 1,12559E-90 |
| ACE2 | RAB11FIP4 | 0,7974 | 1,23382E-90 |
| ACE2 | PADI2 | 0,7973 | 1,25096E-90 |
| ACE2 | CAPN5 | 0,7972 | 1,47073E-90 |
| ACE2 | WSCD1 | 0,7972 | 1,37729E-90 |
| ACE2 | ZNF57 | 0,797 | 1,69476E-90 |
| ACE2 | AP1M2 | 0,797 | 1,67263E-90 |
| ACE2 | ATP1B1 | 0,7968 | 2,07653E-90 |
| ACE2 | FHDC1 | 0,7968 | 1,93753E-90 |
| ACE2 | PDCD4 | 0,7967 | 2,24298E-90 |
| ACE2 | KB-1247B1.1 | 0,7966 | 2,33662E-90 |
| ACE2 | DHDDS | 0,7965 | 2,69248E-90 |
| ACE2 | C1orf115 | 0,7965 | 2,67282E-90 |
| ACE2 | UBE2FP1 | 0,796 | 3,95076E-90 |
| ACE2 | NXPE4 | 0,7959 | 4,46041E-90 |
| ACE2 | LGALS4 | 0,7958 | 4,86338E-90 |
| ACE2 | SHD | 0,7957 | 5,16504E-90 |
| ACE2 | SLC36A1 | 0,7956 | 5,65785E-90 |
| ACE2 | SFN | 0,7954 | 7,02942E-90 |
| ACE2 | RMDN2 | 0,7954 | 6,68311E-90 |
| ACE2 | CTSA | 0,7954 | 6,7936E-90 |
| ACE2 | SLC15A1 | 0,795 | 9,96427E-90 |
| ACE2 | FUT2 | 0,7949 | 1,03856E-89 |
| ACE2 | KIF16B | 0,7949 | 1,02854E-89 |
| ACE2 | RP11-94C24.13 | 0,7948 | 1,20531E-89 |
| ACE2 | TMEM53 | 0,7947 | 1,32444E-89 |
| ACE2 | BRI3BP | 0,7944 | 1,68885E-89 |
| ACE2 | LDHD | 0,7944 | 1,62206E-89 |
| ACE2 | DQX1 | 0,7943 | 1,80626E-89 |
| ACE2 | C15orf48 | 0,7943 | 1,88017E-89 |
| ACE2 | RP11-433O3.1 | 0,7942 | 2,01244E-89 |
| ACE2 | ETHE1 | 0,7941 | 2,15072E-89 |
| ACE2 | TSPAN6 | 0,7941 | 2,22096E-89 |
| ACE2 | CTNND1 | 0,794 | 2,33992E-89 |
| ACE2 | CDX2 | 0,794 | 2,34624E-89 |
| ACE2 | HECTD3 | 0,7939 | 2,46791E-89 |
| ACE2 | FAR2 | 0,7939 | 2,58279E-89 |
| ACE2 | IHH | 0,7937 | 3,16036E-89 |

| | | | |
|---|---|---|---|
| ACE2 | SLC52A3 | 0,7935 | 3,75969E-89 |
| ACE2 | LPAR5 | 0,7933 | 4,21895E-89 |
| ACE2 | LMTK2 | 0,793 | 5,40872E-89 |
| ACE2 | LGALS9 | 0,793 | 5,69891E-89 |
| ACE2 | IDH1 | 0,7923 | 9,91308E-89 |
| ACE2 | SLC25A24 | 0,7922 | 1,14148E-88 |
| ACE2 | UGT8 | 0,792 | 1,31769E-88 |
| ACE2 | CEBPA | 0,792 | 1,35928E-88 |
| ACE2 | ZDHHC23 | 0,7919 | 1,48276E-88 |
| ACE2 | FUCA1 | 0,7918 | 1,53699E-88 |
| ACE2 | PRR5L | 0,7915 | 1,97634E-88 |
| ACE2 | CMTM4 | 0,7915 | 2,03885E-88 |
| ACE2 | SLC39A11 | 0,7915 | 2,03465E-88 |
| ACE2 | RP11-542M13.2 | 0,7914 | 2,14271E-88 |
| ACE2 | PLPP2 | 0,7914 | 2,28942E-88 |
| ACE2 | ADH6 | 0,7912 | 2,56378E-88 |
| ACE2 | STX3 | 0,7912 | 2,57091E-88 |
| ACE2 | GALE | 0,7911 | 2,76266E-88 |
| ACE2 | CD177 | 0,7911 | 2,80015E-88 |
| ACE2 | GJB2 | 0,7909 | 3,44702E-88 |
| ACE2 | FAM83E | 0,7909 | 3,54302E-88 |
| ACE2 | B3GNT8 | 0,7908 | 3,58352E-88 |
| ACE2 | SH3BGRL2 | 0,7906 | 4,52538E-88 |
| ACE2 | SLC26A6 | 0,7905 | 4,65212E-88 |
| ACE2 | FBXO34 | 0,7904 | 5,36469E-88 |
| ACE2 | BICDL2 | 0,7899 | 7,82095E-88 |
| ACE2 | CYP2C9 | 0,7897 | 9,3678E-88 |
| ACE2 | HMGCR | 0,7896 | 9,92524E-88 |
| ACE2 | NET1 | 0,7894 | 1,17022E-87 |
| ACE2 | DAPK2 | 0,7894 | 1,1782E-87 |
| ACE2 | RP4-811M8.1 | 0,7894 | 1,26629E-87 |
| ACE2 | SDC1 | 0,7892 | 1,48385E-87 |
| ACE2 | SERINC5 | 0,7887 | 2,18688E-87 |
| ACE2 | AP006621.9 | 0,7886 | 2,3695E-87 |
| ACE2 | LINC00675 | 0,7886 | 2,44888E-87 |
| ACE2 | AQP8 | 0,7885 | 2,57375E-87 |
| ACE2 | CASC16 | 0,7885 | 2,52594E-87 |
| ACE2 | SLC52A1 | 0,7883 | 3,11388E-87 |
| ACE2 | RP11-336A10.5 | 0,7882 | 3,41117E-87 |
| ACE2 | DOPEY2 | 0,7882 | 3,34571E-87 |
| ACE2 | SLC13A1 | 0,7881 | 3,73446E-87 |
| ACE2 | RIPK3 | 0,788 | 3,87523E-87 |
| ACE2 | GPRC5A | 0,7879 | 4,23794E-87 |
| ACE2 | BCL2L10 | 0,7879 | 4,46647E-87 |
| ACE2 | HNF1B | 0,7879 | 4,3112E-87 |
| ACE2 | ATG4D | 0,7878 | 4,72628E-87 |
| ACE2 | ZDHHC13 | 0,7876 | 5,48367E-87 |
| ACE2 | CA7 | 0,7876 | 5,34123E-87 |
| ACE2 | ACVR1C | 0,7875 | 6,15103E-87 |
| ACE2 | GEMIN8P4 | 0,7874 | 6,50068E-87 |
| ACE2 | BAIAP2L1 | 0,7874 | 6,83218E-87 |
| ACE2 | FGFR3 | 0,7872 | 8,06551E-87 |
| ACE2 | HN1 | 0,7872 | 7,72892E-87 |
| ACE2 | TTC38 | 0,7872 | 7,98724E-87 |

| | | | |
|---|---|---|---|
| ACE2 | GPR35 | 0,787 | 9,48167E-87 |
| ACE2 | LINC00543 | 0,787 | 9,51773E-87 |
| ACE2 | GJB1 | 0,787 | 9,03473E-87 |
| ACE2 | C11orf52 | 0,7865 | 1,42686E-86 |
| ACE2 | LASP1 | 0,7864 | 1,4918E-86 |
| ACE2 | NANS | 0,7862 | 1,72245E-86 |
| ACE2 | PRRG4 | 0,7862 | 1,78631E-86 |
| ACE2 | ACY3 | 0,7862 | 1,75615E-86 |
| ACE2 | ERVH-1 | 0,7861 | 1,90183E-86 |
| ACE2 | TRAF4 | 0,7861 | 1,92599E-86 |
| ACE2 | CLN3 | 0,786 | 2,03928E-86 |
| ACE2 | RP11-164P12.5 | 0,7857 | 2,81355E-86 |
| ACE2 | SIRT6 | 0,7856 | 2,9405E-86 |
| ACE2 | TET3 | 0,7854 | 3,58498E-86 |
| ACE2 | PLEKHJ1 | 0,7854 | 3,51902E-86 |
| ACE2 | TFF3 | 0,7853 | 3,7771E-86 |
| ACE2 | BDH1 | 0,7852 | 4,2788E-86 |
| ACE2 | RP11-2I17.4 | 0,7852 | 3,97465E-86 |
| ACE2 | RHBDL2 | 0,7849 | 5,10056E-86 |
| ACE2 | RP11-276H7.2 | 0,7848 | 5,68494E-86 |
| ACE2 | LAMC2 | 0,7848 | 5,85733E-86 |
| ACE2 | TMEM54 | 0,7847 | 6,46269E-86 |
| ACE2 | SPAG1 | 0,7847 | 6,05454E-86 |
| ACE2 | PI3 | 0,7847 | 6,01409E-86 |
| ACE2 | RAB3IP | 0,7846 | 6,8389E-86 |
| ACE2 | CHAD | 0,7846 | 6,81963E-86 |
| ACE2 | BEST2 | 0,7844 | 7,91309E-86 |
| ACE2 | RP4-568C11.4 | 0,7842 | 9,40484E-86 |
| ACE2 | ERBB3 | 0,7839 | 1,23593E-85 |
| ACE2 | RP11-187E13.1 | 0,7838 | 1,28016E-85 |
| ACE2 | SEMA4G | 0,7837 | 1,44965E-85 |
| ACE2 | GRTP1 | 0,7837 | 1,4058E-85 |
| ACE2 | ST6GALNAC1 | 0,7837 | 1,40649E-85 |
| ACE2 | SLC39A5 | 0,7836 | 1,60518E-85 |
| ACE2 | RAB20 | 0,7835 | 1,66677E-85 |
| ACE2 | FRMD1 | 0,7834 | 1,79591E-85 |
| ACE2 | LRRC8D | 0,7831 | 2,38646E-85 |
| ACE2 | ESRP1 | 0,7831 | 2,30082E-85 |
| ACE2 | ABCB11 | 0,783 | 2,4697E-85 |
| ACE2 | RP3-523K23.2 | 0,783 | 2,55673E-85 |
| ACE2 | RP11-473M20.5 | 0,783 | 2,46373E-85 |
| ACE2 | RNF186 | 0,7828 | 3,08971E-85 |
| ACE2 | CASP3 | 0,7828 | 2,93983E-85 |
| ACE2 | LINC00704 | 0,7826 | 3,61909E-85 |
| ACE2 | CAPN8 | 0,7825 | 3,82116E-85 |
| ACE2 | TMEM63B | 0,7825 | 3,72314E-85 |
| ACE2 | GIPC2 | 0,7821 | 5,25336E-85 |
| ACE2 | ABO | 0,7821 | 5,45776E-85 |
| ACE2 | EXPH5 | 0,7821 | 5,09947E-85 |
| ACE2 | STRADB | 0,782 | 5,52241E-85 |
| ACE2 | RP11-209E8.1 | 0,7819 | 6,21188E-85 |
| ACE2 | CYP2B6 | 0,7817 | 7,49644E-85 |
| ACE2 | B3GNT7 | 0,7815 | 8,36968E-85 |
| ACE2 | AKR7L | 0,7814 | 9,59104E-85 |

| | | | |
|---|---|---|---|
| ACE2 | JHDM1D-AS1 | 0,7814 | 9,17813E-85 |
| ACE2 | B4GALNT2 | 0,7814 | 9,62054E-85 |
| ACE2 | FTLP14 | 0,7813 | 1,01812E-84 |
| ACE2 | CNKSR1 | 0,781 | 1,33787E-84 |
| ACE2 | SLC16A9 | 0,7809 | 1,3413E-84 |
| ACE2 | CMBL | 0,7808 | 1,49646E-84 |
| ACE2 | ACSF2 | 0,7808 | 1,56879E-84 |
| ACE2 | TTC39A | 0,7804 | 2,09085E-84 |
| ACE2 | ATOH1 | 0,7803 | 2,31367E-84 |
| ACE2 | RP11-284F21.7 | 0,7801 | 2,69622E-84 |
| ACE2 | TLCD1 | 0,7801 | 2,66939E-84 |
| ACE2 | KRT15 | 0,7801 | 2,63489E-84 |
| ACE2 | RP11-478K15.6 | 0,78 | 2,78133E-84 |
| ACE2 | TRPM4 | 0,78 | 2,89777E-84 |
| ACE2 | DDC | 0,7799 | 3,01621E-84 |
| ACE2 | GFPT1 | 0,7797 | 3,71694E-84 |
| ACE2 | PLPP6 | 0,7797 | 3,73847E-84 |
| ACE2 | TLCD2 | 0,7797 | 3,6219E-84 |
| ACE2 | MUC20 | 0,7796 | 3,85208E-84 |
| ACE2 | PHLPP2 | 0,7793 | 5,19E-84 |
| ACE2 | GRHL2 | 0,7792 | 5,27762E-84 |
| ACE2 | PANK1 | 0,7792 | 5,40869E-84 |
| ACE2 | TBC1D2 | 0,7791 | 5,71978E-84 |
| ACE2 | GPT | 0,7789 | 6,8941E-84 |
| ACE2 | PCDH1 | 0,7787 | 8,50517E-84 |
| ACE2 | MMP15 | 0,7786 | 8,95617E-84 |
| ACE2 | CA2 | 0,7785 | 9,85439E-84 |
| ACE2 | PPARGC1A | 0,7784 | 1,00099E-83 |
| ACE2 | FBP1 | 0,7784 | 1,04708E-83 |
| ACE2 | SLC35G1 | 0,7779 | 1,55447E-83 |
| ACE2 | PDE6A | 0,7777 | 1,86011E-83 |
| ACE2 | PKLR | 0,7776 | 1,9599E-83 |
| ACE2 | KRT8P10 | 0,7775 | 2,07688E-83 |
| ACE2 | AFG3L2 | 0,7775 | 2,0849E-83 |
| ACE2 | TRIM14 | 0,7773 | 2,56284E-83 |
| ACE2 | WDR78 | 0,7771 | 3,01812E-83 |
| ACE2 | ERMP1 | 0,7771 | 2,99948E-83 |
| ACE2 | S100A14 | 0,777 | 3,17613E-83 |
| ACE2 | BCL2L15 | 0,7769 | 3,56111E-83 |
| ACE2 | AC007556.3 | 0,7768 | 3,70264E-83 |
| ACE2 | SLC22A23 | 0,7768 | 3,79257E-83 |
| ACE2 | EMX1 | 0,7767 | 3,88009E-83 |
| ACE2 | IGF2BP2 | 0,7765 | 4,87415E-83 |
| ACE2 | HSD17B11 | 0,7765 | 4,88965E-83 |
| ACE2 | TOR4A | 0,7765 | 4,76887E-83 |
| ACE2 | RP11-284F21.10 | 0,7764 | 5,29469E-83 |
| ACE2 | ELMO3 | 0,7764 | 5,15389E-83 |
| ACE2 | CBLC | 0,7763 | 5,52467E-83 |
| ACE2 | CACFD1 | 0,7762 | 6,09985E-83 |
| ACE2 | FABP1 | 0,7759 | 7,32594E-83 |
| ACE2 | C3orf52 | 0,7759 | 7,75233E-83 |
| ACE2 | CHP2 | 0,7759 | 7,30355E-83 |
| ACE2 | SELENBP1 | 0,7756 | 9,6989E-83 |
| ACE2 | KRT12 | 0,7756 | 9,35721E-83 |

| | | | |
|---|---|---|---|
| ACE2 | ERBB2 | 0,7752 | 1,34603E-82 |
| ACE2 | CNDP2 | 0,7752 | 1,26852E-82 |
| ACE2 | AC083900.1 | 0,7751 | 1,46681E-82 |
| ACE2 | RP11-396O20.2 | 0,7748 | 1,85022E-82 |
| ACE2 | ZNF440 | 0,7746 | 2,09865E-82 |
| ACE2 | ELF4 | 0,7744 | 2,56239E-82 |
| ACE2 | SLCO4C1 | 0,7743 | 2,70514E-82 |
| ACE2 | SERPINA1 | 0,7743 | 2,59976E-82 |
| ACE2 | CASP7 | 0,7742 | 2,81893E-82 |
| ACE2 | NUAK2 | 0,774 | 3,38182E-82 |
| ACE2 | AC009014.3 | 0,774 | 3,3286E-82 |
| ACE2 | PLD1 | 0,7739 | 3,51695E-82 |
| ACE2 | PROM1 | 0,7739 | 3,58673E-82 |
| ACE2 | GOLM1 | 0,7738 | 4,03253E-82 |
| ACE2 | C12orf75 | 0,7738 | 3,90668E-82 |
| ACE2 | GGT8P | 0,7737 | 4,2318E-82 |
| ACE2 | LRRIQ4 | 0,7737 | 4,13511E-82 |
| ACE2 | NEURL1 | 0,7732 | 6,11015E-82 |
| ACE2 | MST1R | 0,7731 | 7,05028E-82 |
| ACE2 | RP11-588L15.2 | 0,7729 | 8,08025E-82 |
| ACE2 | PKP2 | 0,7727 | 9,00641E-82 |
| ACE2 | BAK1 | 0,7726 | 9,9538E-82 |
| ACE2 | DPP3 | 0,7724 | 1,21261E-81 |
| ACE2 | KRT18 | 0,7724 | 1,14457E-81 |
| ACE2 | VPS37B | 0,7724 | 1,13172E-81 |
| ACE2 | SPIRE2 | 0,7724 | 1,14245E-81 |
| ACE2 | SLC9A3 | 0,7723 | 1,24109E-81 |
| ACE2 | RP11-380P13.1 | 0,7722 | 1,39015E-81 |
| ACE2 | BCL10 | 0,772 | 1,56114E-81 |
| ACE2 | CCT8P1 | 0,772 | 1,63181E-81 |
| ACE2 | CTB-39G8.3 | 0,772 | 1,55305E-81 |
| ACE2 | FAM105A | 0,7719 | 1,69112E-81 |
| ACE2 | FAM83B | 0,7717 | 1,97403E-81 |
| ACE2 | KBTBD11-OT1 | 0,7717 | 2,05918E-81 |
| ACE2 | TMEM87B | 0,7713 | 2,66774E-81 |
| ACE2 | GPR160 | 0,7713 | 2,68628E-81 |
| ACE2 | RP11-545A16.1 | 0,7712 | 2,93946E-81 |
| ACE2 | SPPL2A | 0,7712 | 2,97155E-81 |
| ACE2 | POLD4 | 0,7711 | 3,1583E-81 |
| ACE2 | LINC01080 | 0,7711 | 3,15741E-81 |
| ACE2 | SLC16A5 | 0,771 | 3,48964E-81 |
| ACE2 | CKMT1B | 0,7709 | 3,7566E-81 |
| ACE2 | CYP4F2 | 0,7709 | 3,5877E-81 |
| ACE2 | OVOL2 | 0,7709 | 3,61054E-81 |
| ACE2 | RP11-356M20.1 | 0,7708 | 3,91253E-81 |
| ACE2 | RP11-356O9.2 | 0,7707 | 4,48124E-81 |
| ACE2 | CFTR | 0,7705 | 4,97364E-81 |
| ACE2 | MLPH | 0,7704 | 5,50005E-81 |
| ACE2 | SCGB2A1 | 0,7703 | 6,04461E-81 |
| ACE2 | RAB25 | 0,77 | 7,29786E-81 |
| ACE2 | AC073283.4 | 0,77 | 7,67955E-81 |
| ACE2 | NETO2 | 0,77 | 7,32221E-81 |
| ACE2 | KRTAP13-2 | 0,7699 | 7,99278E-81 |
| ACE2 | HNF1A-AS1 | 0,7695 | 1,05797E-80 |

| | | | |
|---|---|---|---|
| ACE2 | C8G | 0,7694 | 1,13999E-80 |
| ACE2 | TMEM238 | 0,7694 | 1,16783E-80 |
| ACE2 | AMACR | 0,7693 | 1,24275E-80 |
| ACE2 | GSR | 0,7692 | 1,32395E-80 |
| ACE2 | WWC1 | 0,7691 | 1,42456E-80 |
| ACE2 | PTK2B | 0,7691 | 1,43287E-80 |
| ACE2 | TPD52 | 0,7691 | 1,4562E-80 |
| ACE2 | PLEKHS1 | 0,7691 | 1,50994E-80 |
| ACE2 | CTSV | 0,769 | 1,56613E-80 |
| ACE2 | CTD-2385L22.2 | 0,7689 | 1,70464E-80 |
| ACE2 | LCN15 | 0,7688 | 1,80646E-80 |
| ACE2 | MCU | 0,7688 | 1,87685E-80 |
| ACE2 | MYO5C | 0,7688 | 1,89557E-80 |
| ACE2 | LINC01687 | 0,7688 | 1,91687E-80 |
| ACE2 | DNAH3 | 0,7687 | 2,01412E-80 |
| ACE2 | ROS1 | 0,7686 | 2,13564E-80 |
| ACE2 | CKMT1A | 0,7686 | 2,20847E-80 |
| ACE2 | RP13-122B23.8 | 0,7685 | 2,26241E-80 |
| ACE2 | PAQR5 | 0,7684 | 2,4788E-80 |
| ACE2 | PRELID3B | 0,7684 | 2,43142E-80 |
| ACE2 | KLHL34 | 0,7683 | 2,74384E-80 |
| ACE2 | SLC35C1 | 0,7679 | 3,81601E-80 |
| ACE2 | MCTP2 | 0,7678 | 3,95463E-80 |
| ACE2 | C6orf136 | 0,7677 | 4,25253E-80 |
| ACE2 | FOXA1 | 0,7677 | 4,31049E-80 |
| ACE2 | RPS29P12 | 0,7676 | 4,5295E-80 |
| ACE2 | TREH | 0,7676 | 4,56595E-80 |
| ACE2 | TNFRSF11A | 0,7676 | 4,56498E-80 |
| ACE2 | RASSF7 | 0,7675 | 5,12285E-80 |
| ACE2 | CALR4P | 0,7674 | 5,47908E-80 |
| ACE2 | MTMR11 | 0,7674 | 5,45903E-80 |
| ACE2 | TRPC7-AS1 | 0,7674 | 5,34028E-80 |
| ACE2 | SERPINB1 | 0,7672 | 6,41988E-80 |
| ACE2 | BRINP3 | 0,7669 | 7,99077E-80 |
| ACE2 | NEO1 | 0,7669 | 7,78881E-80 |
| ACE2 | RP11-357H14.17 | 0,7668 | 8,23387E-80 |
| ACE2 | HES2 | 0,7667 | 9,05919E-80 |
| ACE2 | GUCY2GP | 0,7663 | 1,25169E-79 |
| ACE2 | C1orf168 | 0,7662 | 1,365E-79 |
| ACE2 | CIB1 | 0,7662 | 1,27959E-79 |
| ACE2 | LA16c-380H5.5 | 0,7661 | 1,45159E-79 |
| ACE2 | SPATA2 | 0,766 | 1,59386E-79 |
| ACE2 | MIER3 | 0,7659 | 1,68019E-79 |
| ACE2 | RFLNA | 0,7659 | 1,60126E-79 |
| ACE2 | CPTP | 0,7656 | 2,04233E-79 |
| ACE2 | SLC37A4 | 0,7655 | 2,26209E-79 |
| ACE2 | ANO9 | 0,7654 | 2,47473E-79 |
| ACE2 | CTSD | 0,7652 | 2,75527E-79 |
| ACE2 | PPP1R16A | 0,7651 | 3,1173E-79 |
| ACE2 | ADORA2B | 0,7651 | 3,12378E-79 |
| ACE2 | RPS4XP7 | 0,7649 | 3,38957E-79 |
| ACE2 | PC | 0,7646 | 4,47505E-79 |
| ACE2 | PRKAB1 | 0,7646 | 4,32977E-79 |
| ACE2 | OCIAD2 | 0,7645 | 4,58142E-79 |

| | | | |
|---|---|---|---|
| ACE2 | KPNA7 | 0,7643 | 5,60305E-79 |
| ACE2 | DEFB1 | 0,7643 | 5,62108E-79 |
| ACE2 | UQCRC1 | 0,764 | 6,73992E-79 |
| ACE2 | KLB | 0,764 | 7,0044E-79 |
| ACE2 | DECR1 | 0,764 | 6,96257E-79 |
| ACE2 | SMAGP | 0,7639 | 7,30332E-79 |
| ACE2 | FOXH1 | 0,7635 | 9,9129E-79 |
| ACE2 | AC009133.12 | 0,7635 | 9,71853E-79 |
| ACE2 | RP11-61A14.1 | 0,7635 | 9,94741E-79 |
| ACE2 | BSPRY | 0,7633 | 1,18469E-78 |
| ACE2 | RP11-844P9.6 | 0,7631 | 1,34992E-78 |
| ACE2 | MPDU1 | 0,7628 | 1,69969E-78 |
| ACE2 | MVP | 0,7624 | 2,25683E-78 |
| ACE2 | TRIM40 | 0,762 | 3,05886E-78 |
| ACE2 | KIAA1211 | 0,7619 | 3,15131E-78 |
| ACE2 | CCNJL | 0,7618 | 3,47425E-78 |
| ACE2 | EBP | 0,7618 | 3,52825E-78 |
| ACE2 | DLG3 | 0,7618 | 3,3605E-78 |
| ACE2 | TMEM63C | 0,7617 | 3,7749E-78 |
| ACE2 | RP11-99J16__A.2 | 0,7616 | 3,97354E-78 |
| ACE2 | SOX9 | 0,7616 | 3,90538E-78 |
| ACE2 | SQRDL | 0,7614 | 4,66508E-78 |
| ACE2 | AXDND1 | 0,7613 | 4,88549E-78 |
| ACE2 | SLC22A5 | 0,7613 | 4,98291E-78 |
| ACE2 | MLX | 0,7611 | 5,94989E-78 |
| ACE2 | CTD-3010D24.3 | 0,7611 | 5,71765E-78 |
| ACE2 | DGKQ | 0,761 | 6,0503E-78 |
| ACE2 | CTB-167B5.1 | 0,761 | 6,18454E-78 |
| ACE2 | SNX30 | 0,761 | 6,43574E-78 |
| ACE2 | SAMD13 | 0,7606 | 8,35613E-78 |
| ACE2 | SLC35A2 | 0,7606 | 8,23705E-78 |
| ACE2 | ASL | 0,7604 | 9,79233E-78 |
| ACE2 | KLF4 | 0,7601 | 1,2336E-77 |
| ACE2 | OVOL1-AS1 | 0,7601 | 1,17247E-77 |
| ACE2 | UGT1A8 | 0,7599 | 1,40958E-77 |
| ACE2 | VSIG10 | 0,7597 | 1,62033E-77 |
| ACE2 | CREB3L3 | 0,7595 | 1,86614E-77 |
| ACE2 | RP11-297D21.2 | 0,7589 | 2,98344E-77 |
| ACE2 | STXBP5-AS1 | 0,7588 | 3,1E-77 |
| ACE2 | C10orf99 | 0,7587 | 3,4428E-77 |
| ACE2 | EPS8L1 | 0,7586 | 3,63243E-77 |
| ACE2 | OMP | 0,7585 | 3,96434E-77 |
| ACE2 | SH3BP1 | 0,7585 | 3,94858E-77 |
| ACE2 | MYO6 | 0,7583 | 4,59719E-77 |
| ACE2 | ABCC6P1 | 0,7581 | 5,28539E-77 |
| ACE2 | ESRP2 | 0,7579 | 6,00269E-77 |
| ACE2 | CRYL1 | 0,7576 | 7,25547E-77 |
| ACE2 | MYO1E | 0,7576 | 7,14582E-77 |
| ACE2 | TM6SF2 | 0,7576 | 7,56191E-77 |
| ACE2 | LINC00501 | 0,7573 | 9,32649E-77 |
| ACE2 | RP5-1180D12.1 | 0,7573 | 8,97679E-77 |
| ACE2 | ARHGEF38 | 0,7572 | 9,72269E-77 |
| ACE2 | PAK1 | 0,7572 | 9,58303E-77 |
| ACE2 | LGALS9B | 0,757 | 1,17442E-76 |

| | | | |
|---|---|---|---|
| ACE2 | RNPEP | 0,7568 | 1,26478E-76 |
| ACE2 | GOT1 | 0,7568 | 1,33621E-76 |
| ACE2 | RP11-30P6.6 | 0,7566 | 1,5089E-76 |
| ACE2 | TM9SF2 | 0,7565 | 1,64546E-76 |
| ACE2 | LINC01637 | 0,7564 | 1,74802E-76 |
| ACE2 | REP15 | 0,7563 | 1,82896E-76 |
| ACE2 | RP5-1033H22.2 | 0,756 | 2,38227E-76 |
| ACE2 | GEN1 | 0,756 | 2,33967E-76 |
| ACE2 | SLC25A13 | 0,756 | 2,29241E-76 |
| ACE2 | TMEM72 | 0,756 | 2,36213E-76 |
| ACE2 | RP11-384L8.1 | 0,7559 | 2,45191E-76 |
| ACE2 | CASP6 | 0,7559 | 2,53625E-76 |
| ACE2 | SULT1A1 | 0,7558 | 2,72225E-76 |
| ACE2 | EFNA4 | 0,7555 | 3,4173E-76 |
| ACE2 | HMGN1P7 | 0,7555 | 3,31174E-76 |
| ACE2 | TFF1 | 0,7555 | 3,35051E-76 |
| ACE2 | ADGRF4 | 0,7554 | 3,55123E-76 |
| ACE2 | SHTN1 | 0,7554 | 3,44768E-76 |
| ACE2 | CTD-2288F12.1 | 0,7554 | 3,49319E-76 |
| ACE2 | MCCC2 | 0,7553 | 3,70317E-76 |
| ACE2 | CYP2S1 | 0,7553 | 3,69748E-76 |
| ACE2 | SH3RF2 | 0,7552 | 4,10822E-76 |
| ACE2 | PCK1 | 0,7552 | 4,04588E-76 |
| ACE2 | PANK3 | 0,7549 | 5,25545E-76 |
| ACE2 | SHROOM1 | 0,7548 | 5,46754E-76 |
| ACE2 | NR2F6 | 0,7548 | 5,34736E-76 |
| ACE2 | LINC00992 | 0,7546 | 6,15398E-76 |
| ACE2 | ENDOG | 0,7543 | 7,52095E-76 |
| ACE2 | LPCAT4 | 0,7542 | 8,13414E-76 |
| ACE2 | MAP2K6 | 0,7542 | 8,07772E-76 |
| ACE2 | KLC4 | 0,7541 | 8,91479E-76 |
| ACE2 | CD24 | 0,754 | 9,50712E-76 |
| ACE2 | PRLR | 0,7538 | 1,12387E-75 |
| ACE2 | CDC42EP5 | 0,7535 | 1,3412E-75 |
| ACE2 | COMTD1 | 0,7534 | 1,51677E-75 |
| ACE2 | SIRT7 | 0,7534 | 1,44006E-75 |
| ACE2 | TNNI1 | 0,7533 | 1,56523E-75 |
| ACE2 | LINC01134 | 0,7532 | 1,6362E-75 |
| ACE2 | C4BPB | 0,7531 | 1,76349E-75 |
| ACE2 | TNK1 | 0,7531 | 1,85252E-75 |
| ACE2 | URAD | 0,753 | 1,92496E-75 |
| ACE2 | C2orf72 | 0,7529 | 2,09623E-75 |
| ACE2 | SDHA | 0,7529 | 2,01832E-75 |
| ACE2 | SSTR1 | 0,7528 | 2,18321E-75 |
| ACE2 | ELF3 | 0,7527 | 2,47087E-75 |
| ACE2 | PIGZ | 0,7527 | 2,43743E-75 |
| ACE2 | SLC25A10 | 0,7527 | 2,47953E-75 |
| ACE2 | PPP1CA | 0,7526 | 2,51389E-75 |
| ACE2 | MYO10 | 0,7525 | 2,76985E-75 |
| ACE2 | MREG | 0,7524 | 2,96672E-75 |
| ACE2 | AP001610.9 | 0,7524 | 2,86962E-75 |
| ACE2 | FZD5 | 0,752 | 4,0539E-75 |
| ACE2 | PLCD3 | 0,752 | 3,8269E-75 |
| ACE2 | SLC44A3 | 0,7517 | 4,74763E-75 |

| | | | |
|---|---|---|---|
| ACE2 | TOR2A | 0,7516 | 5,1683E-75 |
| ACE2 | CYP4F12 | 0,7512 | 7,03411E-75 |
| ACE2 | RNF223 | 0,751 | 8,16955E-75 |
| ACE2 | TMEM141 | 0,751 | 8,17132E-75 |
| ACE2 | KIAA1211L | 0,7509 | 8,48555E-75 |
| ACE2 | C9orf152 | 0,7509 | 8,28812E-75 |
| ACE2 | ZSWIM5 | 0,7508 | 9,37586E-75 |
| ACE2 | SLC31A1 | 0,7508 | 8,90798E-75 |
| ACE2 | AC007255.8 | 0,7506 | 1,07936E-74 |
| ACE2 | SPIB | 0,7506 | 1,02908E-74 |
| ACE2 | CTNNA1P1 | 0,7505 | 1,10684E-74 |
| ACE2 | NAPRT | 0,7504 | 1,20216E-74 |
| ACE2 | CTB-78O21.1 | 0,7503 | 1,27635E-74 |
| ACE2 | ARHGAP32 | 0,7503 | 1,2998E-74 |
| ACE2 | ACOT11 | 0,7502 | 1,386E-74 |
| ACE2 | KRT8P45 | 0,7501 | 1,43658E-74 |
| ACE2 | ANKEF1 | 0,7501 | 1,45629E-74 |
| ACE2 | VSIG2 | 0,7499 | 1,75795E-74 |
| ACE2 | ELOVL6 | 0,7498 | 1,82705E-74 |
| ACE2 | PSMD10P2 | 0,7494 | 2,46874E-74 |
| ACE2 | MXD1 | 0,7493 | 2,55388E-74 |
| ACE2 | CLCN3 | 0,7493 | 2,52182E-74 |
| ACE2 | SPACA4 | 0,7493 | 2,61048E-74 |
| ACE2 | PLGRKT | 0,7491 | 2,90293E-74 |
| ACE2 | RP1-172H20.4 | 0,7491 | 2,92106E-74 |
| ACE2 | ARRDC1 | 0,749 | 3,08153E-74 |
| ACE2 | TPSG1 | 0,749 | 3,09025E-74 |
| ACE2 | PLEKHH1 | 0,7489 | 3,50621E-74 |
| ACE2 | PTMAP3 | 0,7489 | 3,40664E-74 |
| ACE2 | RP11-319C21.1 | 0,7488 | 3,65946E-74 |
| ACE2 | RP11-519G16.3 | 0,7485 | 4,43266E-74 |
| ACE2 | INPP5J | 0,7483 | 5,10819E-74 |
| ACE2 | RP11-255H23.2 | 0,7482 | 5,58816E-74 |
| ACE2 | ARSD | 0,7481 | 5,89288E-74 |
| ACE2 | LINC00507 | 0,748 | 6,35516E-74 |
| ACE2 | ATP2C2 | 0,748 | 6,41322E-74 |
| ACE2 | RP13-726E6.2 | 0,7477 | 7,98064E-74 |
| ACE2 | PYY | 0,7477 | 8,02281E-74 |
| ACE2 | ZNRF2 | 0,7475 | 8,83883E-74 |
| ACE2 | MGST1 | 0,7474 | 9,54338E-74 |
| ACE2 | GALM | 0,7472 | 1,07049E-73 |
| ACE2 | KCNG3 | 0,7472 | 1,10722E-73 |
| ACE2 | TSPAN13 | 0,7472 | 1,06811E-73 |
| ACE2 | ADH1C | 0,7471 | 1,17281E-73 |
| ACE2 | KNG1 | 0,747 | 1,29381E-73 |
| ACE2 | RP11-22C11.2 | 0,747 | 1,25407E-73 |
| ACE2 | LINC00705 | 0,747 | 1,27907E-73 |
| ACE2 | RP11-103J17.2 | 0,7469 | 1,35952E-73 |
| ACE2 | RP11-867G23.8 | 0,7469 | 1,32186E-73 |
| ACE2 | RP11-48O20.5 | 0,7465 | 1,81193E-73 |
| ACE2 | SH2D4A | 0,7465 | 1,72621E-73 |
| ACE2 | EPHX2 | 0,7465 | 1,82336E-73 |
| ACE2 | AC002128.5 | 0,7464 | 1,91611E-73 |
| ACE2 | RP4-659I19.1 | 0,7463 | 2,06523E-73 |

| | | | |
|---|---|---|---|
| ACE2 | ACPP | 0,7463 | 1,97699E-73 |
| ACE2 | HMGCS2 | 0,7462 | 2,18196E-73 |
| ACE2 | EFNB1 | 0,7461 | 2,31451E-73 |
| ACE2 | GK | 0,746 | 2,40871E-73 |
| ACE2 | BNIP3P39 | 0,7458 | 2,91687E-73 |
| ACE2 | FAM109A | 0,7455 | 3,50981E-73 |
| ACE2 | CORO1B | 0,7452 | 4,26736E-73 |
| ACE2 | LINC00511 | 0,7452 | 4,28163E-73 |
| ACE2 | MAP3K21 | 0,7451 | 4,5889E-73 |
| ACE2 | FOXD2 | 0,745 | 4,77782E-73 |
| ACE2 | MYCL | 0,7449 | 5,38953E-73 |
| ACE2 | MAP3K9 | 0,7449 | 5,25413E-73 |
| ACE2 | RP11-336K24.5 | 0,7447 | 5,9109E-73 |
| ACE2 | PSD4 | 0,7447 | 6,05941E-73 |
| ACE2 | LGALS9C | 0,7447 | 6,03883E-73 |
| ACE2 | RP11-647P12.2 | 0,7443 | 7,72082E-73 |
| ACE2 | KRTCAP3 | 0,7442 | 8,16708E-73 |
| ACE2 | MOB3B | 0,7442 | 8,36003E-73 |
| ACE2 | EPN3 | 0,7442 | 8,43198E-73 |
| ACE2 | RAVER2 | 0,7441 | 8,85339E-73 |
| ACE2 | RP11-311D14.1 | 0,7441 | 8,79604E-73 |
| ACE2 | SCNN1G | 0,744 | 9,69321E-73 |
| ACE2 | ARHGAP44 | 0,744 | 9,46342E-73 |
| ACE2 | COX6A1 | 0,7439 | 1,06176E-72 |
| ACE2 | Z83844.1 | 0,7439 | 1,01517E-72 |
| ACE2 | SLC6A7 | 0,7437 | 1,21204E-72 |
| ACE2 | CTB-171A8.1 | 0,7435 | 1,31014E-72 |
| ACE2 | CWH43 | 0,7434 | 1,39377E-72 |
| ACE2 | MYOM3 | 0,7433 | 1,55073E-72 |
| ACE2 | SIDT1 | 0,7432 | 1,65539E-72 |
| ACE2 | SLC25A25-AS1 | 0,7431 | 1,80756E-72 |
| ACE2 | PREP | 0,7428 | 2,22342E-72 |
| ACE2 | CD164 | 0,7427 | 2,29116E-72 |
| ACE2 | EVX1 | 0,7427 | 2,29286E-72 |
| ACE2 | RP11-320N21.1 | 0,7426 | 2,38851E-72 |
| ACE2 | RP3-400B16.1 | 0,7425 | 2,70884E-72 |
| ACE2 | TMEM2 | 0,7425 | 2,58799E-72 |
| ACE2 | ARF6 | 0,7423 | 2,95551E-72 |
| ACE2 | TTLL12 | 0,7423 | 2,97941E-72 |
| ACE2 | SAMD12 | 0,7422 | 3,19915E-72 |
| ACE2 | ZZEF1 | 0,7419 | 3,93627E-72 |
| ACE2 | RP11-10C24.1 | 0,7418 | 4,24254E-72 |
| ACE2 | E2F2 | 0,7417 | 4,48133E-72 |
| ACE2 | SUCLG1 | 0,7416 | 4,69E-72 |
| ACE2 | GSDMB | 0,7415 | 4,98252E-72 |
| ACE2 | FAM83H-AS1 | 0,7414 | 5,49109E-72 |
| ACE2 | OAS1 | 0,7413 | 5,85153E-72 |
| ACE2 | CLCA1 | 0,7412 | 6,15618E-72 |
| ACE2 | VWA7 | 0,7412 | 6,39795E-72 |
| ACE2 | KCTD5 | 0,7412 | 6,18431E-72 |
| ACE2 | KLK1 | 0,7409 | 7,88561E-72 |
| ACE2 | UGT1A9 | 0,7408 | 8,13857E-72 |
| ACE2 | RP11-4O1.2 | 0,7408 | 8,25567E-72 |
| ACE2 | RP11-40A7.2 | 0,7407 | 8,50077E-72 |

| | | | |
|---|---|---|---|
| ACE2 | PGPEP1 | 0,7405 | 1,02469E-71 |
| ACE2 | FGFBP1 | 0,7404 | 1,10067E-71 |
| ACE2 | SELENOS | 0,7404 | 1,09849E-71 |
| ACE2 | TMC7 | 0,7403 | 1,14265E-71 |
| ACE2 | CTD-2231E14.8 | 0,7402 | 1,18793E-71 |
| ACE2 | FMN1 | 0,7401 | 1,30084E-71 |
| ACE2 | ZFYVE28 | 0,7398 | 1,54433E-71 |
| ACE2 | RP5-903G2.2 | 0,7398 | 1,56309E-71 |
| ACE2 | RPL15P21 | 0,7398 | 1,60405E-71 |
| ACE2 | ICA1 | 0,7397 | 1,66056E-71 |
| ACE2 | SHH | 0,7397 | 1,65694E-71 |
| ACE2 | MBNL3 | 0,7397 | 1,69177E-71 |
| ACE2 | LMNTD2 | 0,7396 | 1,76073E-71 |
| ACE2 | LINC00336 | 0,7391 | 2,59999E-71 |
| ACE2 | RXFP4 | 0,739 | 2,65515E-71 |
| ACE2 | RP11-479A21.1 | 0,7389 | 2,82482E-71 |
| ACE2 | PSEN1 | 0,7388 | 3,09556E-71 |
| ACE2 | PLCXD2 | 0,7387 | 3,1798E-71 |
| ACE2 | KRT8P32 | 0,7387 | 3,23351E-71 |
| ACE2 | UGT1A6 | 0,7386 | 3,46537E-71 |
| ACE2 | MAL2 | 0,7381 | 4,85388E-71 |
| ACE2 | SLC46A3 | 0,7381 | 4,81254E-71 |
| ACE2 | CTD-2385L22.1 | 0,7381 | 4,905E-71 |
| ACE2 | RP13-497K6.1 | 0,7378 | 5,86668E-71 |
| ACE2 | GGH | 0,7378 | 5,87024E-71 |
| ACE2 | PDSS1 | 0,7378 | 6,09492E-71 |
| ACE2 | ATP5B | 0,7378 | 5,82454E-71 |
| ACE2 | MESP2 | 0,7378 | 6,04766E-71 |
| ACE2 | CA1 | 0,7377 | 6,1384E-71 |
| ACE2 | B4GALT1-AS1 | 0,7374 | 7,57085E-71 |
| ACE2 | GRB7 | 0,7374 | 7,91245E-71 |
| ACE2 | RP11-27K13.3 | 0,7373 | 8,30414E-71 |
| ACE2 | RP11-757F18.5 | 0,7373 | 8,21427E-71 |
| ACE2 | CIR1P2 | 0,7373 | 8,0711E-71 |
| ACE2 | RHOF | 0,7373 | 8,45042E-71 |
| ACE2 | AC106869.2 | 0,7372 | 8,76217E-71 |
| ACE2 | KALRN | 0,7372 | 8,5543E-71 |
| ACE2 | STRBP | 0,7372 | 8,99661E-71 |
| ACE2 | DNASE2B | 0,737 | 9,99816E-71 |
| ACE2 | SPINK5 | 0,737 | 9,79452E-71 |
| ACE2 | KIAA1324 | 0,7367 | 1,24763E-70 |
| ACE2 | RP11-1149M10.2 | 0,7366 | 1,27282E-70 |
| ACE2 | EPHA1 | 0,7365 | 1,36329E-70 |
| ACE2 | AGMO | 0,7363 | 1,57455E-70 |
| ACE2 | LINC00482 | 0,7363 | 1,53414E-70 |
| ACE2 | RP11-284F21.9 | 0,7361 | 1,81871E-70 |
| ACE2 | SLC1A1 | 0,7361 | 1,79566E-70 |
| ACE2 | LINC01814 | 0,7359 | 2,11716E-70 |
| ACE2 | PLCB3 | 0,7359 | 2,11549E-70 |
| ACE2 | GCNT1 | 0,7358 | 2,12172E-70 |
| ACE2 | XYLB | 0,7356 | 2,46055E-70 |
| ACE2 | RP11-731D1.4 | 0,7356 | 2,50575E-70 |
| ACE2 | CTA-363E6.6 | 0,7355 | 2,69203E-70 |
| ACE2 | ERN2 | 0,7355 | 2,65785E-70 |

| | | | |
|---|---|---|---|
| ACE2 | HOOK2 | 0,7354 | 2,91919E-70 |
| ACE2 | MUC1 | 0,7352 | 3,27261E-70 |
| ACE2 | RSPH1 | 0,7352 | 3,19398E-70 |
| ACE2 | RAD17P1 | 0,7349 | 3,95586E-70 |
| ACE2 | RORC | 0,7348 | 4,29635E-70 |
| ACE2 | PPP1R1B | 0,7347 | 4,46601E-70 |
| ACE2 | CRACR2A | 0,7344 | 5,51934E-70 |
| ACE2 | INSL5 | 0,7342 | 6,09513E-70 |
| ACE2 | VWA3B | 0,7342 | 6,09727E-70 |
| ACE2 | FOXA3 | 0,7342 | 6,30639E-70 |
| ACE2 | SLC44A1 | 0,7341 | 6,66534E-70 |
| ACE2 | ADAT3 | 0,7341 | 6,72651E-70 |
| ACE2 | RIPK4 | 0,734 | 7,08674E-70 |
| ACE2 | ANO7 | 0,7338 | 8,06502E-70 |
| ACE2 | BIK | 0,7338 | 7,78207E-70 |
| ACE2 | HSD3B1 | 0,7336 | 9,14596E-70 |
| ACE2 | AGMAT | 0,7335 | 9,38475E-70 |
| ACE2 | RBBP8NL | 0,7335 | 9,48351E-70 |
| ACE2 | MARK2 | 0,7333 | 1,11947E-69 |
| ACE2 | FAM96A | 0,7331 | 1,25364E-69 |
| ACE2 | RP11-844P9.5 | 0,7329 | 1,40795E-69 |
| ACE2 | ELL3 | 0,7329 | 1,40115E-69 |
| ACE2 | GLYCTK | 0,7327 | 1,56655E-69 |
| ACE2 | FRAT2 | 0,7326 | 1,72025E-69 |
| ACE2 | FAHD2P1 | 0,7326 | 1,7654E-69 |
| ACE2 | DEGS2 | 0,7326 | 1,69828E-69 |
| ACE2 | RP11-227H15.4 | 0,7325 | 1,81681E-69 |
| ACE2 | RP11-1260E13.3 | 0,732 | 2,49295E-69 |
| ACE2 | CA13 | 0,7318 | 2,79905E-69 |
| ACE2 | C5orf52 | 0,7315 | 3,53892E-69 |
| ACE2 | ARHGEF34P | 0,7313 | 3,88948E-69 |
| ACE2 | NXPE2 | 0,7313 | 3,99255E-69 |
| ACE2 | LRG1 | 0,7313 | 4,06865E-69 |
| ACE2 | UGT1A7 | 0,7311 | 4,41923E-69 |
| ACE2 | CHDH | 0,7311 | 4,35774E-69 |
| ACE2 | AC012485.2 | 0,731 | 4,77693E-69 |
| ACE2 | LRRC26 | 0,731 | 4,70502E-69 |
| ACE2 | RP11-77K12.5 | 0,731 | 4,78228E-69 |
| ACE2 | GMDS | 0,7307 | 5,69503E-69 |
| ACE2 | RP11-844P9.2 | 0,7306 | 6,27615E-69 |
| ACE2 | SPDEF | 0,7305 | 6,51488E-69 |
| ACE2 | RP11-677M14.3 | 0,7304 | 7,19052E-69 |
| ACE2 | U47924.27 | 0,7304 | 6,87764E-69 |
| ACE2 | GIPC1 | 0,7303 | 7,63458E-69 |
| ACE2 | MS4A10 | 0,7301 | 8,31215E-69 |
| ACE2 | CDH16 | 0,7301 | 8,2533E-69 |
| ACE2 | LEFTY1 | 0,73 | 8,78052E-69 |
| ACE2 | CTD-2547H18.1 | 0,73 | 9,13817E-69 |
| ACE2 | ABHD11 | 0,7299 | 9,61234E-69 |
| ACE2 | NR3C2 | 0,7298 | 1,02432E-68 |
| ACE2 | NUDT8 | 0,7298 | 1,05598E-68 |
| ACE2 | NXPE1 | 0,7296 | 1,13669E-68 |
| ACE2 | AC084809.2 | 0,7294 | 1,30692E-68 |
| ACE2 | TUSC8 | 0,7292 | 1,47233E-68 |

| | | | |
|---|---|---|---|
| ACE2 | SLC25A1 | 0,729 | 1,74132E-68 |
| ACE2 | CA3-AS1 | 0,7289 | 1,76843E-68 |
| ACE2 | SYT7 | 0,7288 | 1,9223E-68 |
| ACE2 | RP11-95M15.1 | 0,7284 | 2,51842E-68 |
| ACE2 | RP11-502I4.3 | 0,7284 | 2,51961E-68 |
| ACE2 | UGT2B15 | 0,7282 | 2,72832E-68 |
| ACE2 | TRBV29OR9-2 | 0,7282 | 2,77549E-68 |
| ACE2 | RP11-855O10.2 | 0,7281 | 3,04059E-68 |
| ACE2 | MFSD6L | 0,7281 | 2,89548E-68 |
| ACE2 | CERS6 | 0,7274 | 4,54845E-68 |
| ACE2 | TTC39A-AS1 | 0,7273 | 5,04989E-68 |
| ACE2 | ZNF321P | 0,7272 | 5,09059E-68 |
| ACE2 | FAAH2 | 0,7272 | 5,29526E-68 |
| ACE2 | AP1S1 | 0,7271 | 5,57859E-68 |
| ACE2 | APOBEC3B | 0,7271 | 5,44686E-68 |
| ACE2 | NRAS | 0,7268 | 6,72632E-68 |
| ACE2 | RP11-757F18.3 | 0,7268 | 6,91454E-68 |
| ACE2 | FERMT1 | 0,7264 | 8,47792E-68 |
| ACE2 | RP11-1036E20.9 | 0,7263 | 9,05556E-68 |
| ACE2 | MAGIX | 0,7263 | 8,93257E-68 |
| ACE2 | ATP13A4 | 0,7262 | 9,59293E-68 |
| ACE2 | ATP6V0D1 | 0,7262 | 9,87029E-68 |
| ACE2 | RP11-104J23.1 | 0,7262 | 9,94488E-68 |
| ACE2 | UOX | 0,7261 | 1,07313E-67 |
| ACE2 | RP11-610P16.1 | 0,7261 | 1,05274E-67 |
| ACE2 | SREBF2 | 0,7261 | 1,06266E-67 |
| ACE2 | KCNJ2 | 0,7259 | 1,1832E-67 |
| ACE2 | PLEKHA6 | 0,7257 | 1,30011E-67 |
| ACE2 | CDCA4P2 | 0,7256 | 1,42099E-67 |
| ACE2 | XXyac-YM21GA2.7 | 0,7256 | 1,44151E-67 |
| ACE2 | CREB3L1 | 0,7255 | 1,51573E-67 |
| ACE2 | CTD-2196E14.8 | 0,7255 | 1,53246E-67 |
| ACE2 | HDAC1 | 0,7251 | 1,89694E-67 |
| ACE2 | FAM118B | 0,7249 | 2,16095E-67 |
| ACE2 | RP11-432I5.2 | 0,7249 | 2,24049E-67 |
| ACE2 | GSS | 0,7248 | 2,32289E-67 |
| ACE2 | RP11-579D7.4 | 0,7247 | 2,51412E-67 |
| ACE2 | RP11-83M16.4 | 0,7246 | 2,68293E-67 |
| ACE2 | RP11-466A19.1 | 0,7246 | 2,59245E-67 |
| ACE2 | RP11-1060J15.5 | 0,7245 | 2,84096E-67 |
| ACE2 | PGAP3 | 0,7245 | 2,78941E-67 |
| ACE2 | DBI | 0,7243 | 3,08237E-67 |
| ACE2 | HOTTIP | 0,7243 | 3,09874E-67 |
| ACE2 | RP11-321G12.1 | 0,7242 | 3,4056E-67 |
| ACE2 | FAM84A | 0,7241 | 3,67016E-67 |
| ACE2 | RP11-725G5.2 | 0,7241 | 3,63991E-67 |
| ACE2 | STOML3 | 0,7239 | 4,04604E-67 |
| ACE2 | CTD-3094K11.1 | 0,7239 | 4,18397E-67 |
| ACE2 | SH3RF1 | 0,7236 | 4,90434E-67 |
| ACE2 | LINC01224 | 0,7236 | 4,99454E-67 |
| ACE2 | SVOPL | 0,7235 | 5,13318E-67 |
| ACE2 | MNX1 | 0,7235 | 5,17232E-67 |
| ACE2 | AC046143.3 | 0,7233 | 5,89841E-67 |
| ACE2 | RN7SL541P | 0,7232 | 6,34778E-67 |

| | | | |
|---|---|---|---|
| ACE2 | ZG16B | 0,7231 | 6,47918E-67 |
| ACE2 | LINC01389 | 0,723 | 7,00461E-67 |
| ACE2 | SUSD1 | 0,723 | 7,23079E-67 |
| ACE2 | SOWAHD | 0,723 | 7,11389E-67 |
| ACE2 | PP7080 | 0,7228 | 7,88358E-67 |
| ACE2 | FADS6 | 0,7228 | 7,99793E-67 |
| ACE2 | GMIP | 0,7228 | 7,9212E-67 |
| ACE2 | RP11-266I3.9 | 0,7228 | 8,15888E-67 |
| ACE2 | RP1-80N2.3 | 0,7225 | 9,33583E-67 |
| ACE2 | PROB1 | 0,7222 | 1,1535E-66 |
| ACE2 | NBPF7 | 0,722 | 1,27981E-66 |
| ACE2 | NEURL3 | 0,722 | 1,26762E-66 |
| ACE2 | SH3D21 | 0,7219 | 1,35699E-66 |
| ACE2 | RP11-307N16.7 | 0,7219 | 1,36359E-66 |
| ACE2 | RP11-627G23.1 | 0,7216 | 1,63977E-66 |
| ACE2 | HOXB7 | 0,7215 | 1,73894E-66 |
| ACE2 | AC019330.1 | 0,7214 | 1,83029E-66 |
| ACE2 | MPND | 0,7214 | 1,9036E-66 |
| ACE2 | PATJ | 0,7212 | 2,13364E-66 |
| ACE2 | RRS1-AS1 | 0,7212 | 2,15706E-66 |
| ACE2 | SLC16A13 | 0,7211 | 2,25407E-66 |
| ACE2 | NUDT22 | 0,721 | 2,41864E-66 |
| ACE2 | PPP1R36 | 0,721 | 2,37735E-66 |
| ACE2 | AC024592.9 | 0,7207 | 2,92851E-66 |
| ACE2 | LINC01123 | 0,7206 | 2,99024E-66 |
| ACE2 | ATP5G3 | 0,7206 | 2,97556E-66 |
| ACE2 | APH1A | 0,7205 | 3,29399E-66 |
| ACE2 | ANO10 | 0,7205 | 3,21588E-66 |
| ACE2 | HMGB3 | 0,7203 | 3,67476E-66 |
| ACE2 | RDH13 | 0,7202 | 3,92093E-66 |
| ACE2 | DCST1-AS1 | 0,7199 | 4,77789E-66 |
| ACE2 | NUTM1 | 0,7197 | 5,15712E-66 |
| ACE2 | LGR4 | 0,7196 | 5,50041E-66 |
| ACE2 | ANKRD9 | 0,7195 | 5,84683E-66 |
| ACE2 | RAP1GAP | 0,7194 | 6,4139E-66 |
| ACE2 | RP11-255H23.4 | 0,7192 | 6,95163E-66 |
| ACE2 | WNK4 | 0,7191 | 7,66094E-66 |
| ACE2 | MCOLN2 | 0,719 | 8,2664E-66 |
| ACE2 | LINC01588 | 0,7188 | 8,9779E-66 |
| ACE2 | AC114730.3 | 0,7186 | 1,0013E-65 |
| ACE2 | SLC23A1 | 0,7185 | 1,11648E-65 |
| ACE2 | C4orf33 | 0,7184 | 1,15041E-65 |
| ACE2 | C4BPA | 0,7183 | 1,20388E-65 |
| ACE2 | PTPRD | 0,7183 | 1,20504E-65 |
| ACE2 | GALNT12 | 0,7182 | 1,30414E-65 |
| ACE2 | RP3-395M20.8 | 0,718 | 1,4655E-65 |
| ACE2 | LINC02012 | 0,7179 | 1,53506E-65 |
| ACE2 | LINC01842 | 0,7179 | 1,55568E-65 |
| ACE2 | RP11-496I9.1 | 0,7178 | 1,70247E-65 |
| ACE2 | RN7SKP127 | 0,7178 | 1,62794E-65 |
| ACE2 | LINC01106 | 0,7177 | 1,72558E-65 |
| ACE2 | C7orf43 | 0,7176 | 1,84075E-65 |
| ACE2 | SDSL | 0,7176 | 1,89149E-65 |
| ACE2 | RP11-519G16.5 | 0,7176 | 1,86151E-65 |

| | | | |
|---|---|---|---|
| ACE2 | GOLT1A | 0,717 | 2,61642E-65 |
| ACE2 | GNE | 0,717 | 2,66059E-65 |
| ACE2 | ACAA1 | 0,7169 | 2,84704E-65 |
| ACE2 | TLN2 | 0,7169 | 2,76838E-65 |
| ACE2 | KRT19P1 | 0,7168 | 3,05752E-65 |
| ACE2 | RP5-881L22.5 | 0,7167 | 3,10619E-65 |
| ACE2 | RP11-480I12.5 | 0,7166 | 3,41237E-65 |
| ACE2 | DHX32 | 0,7166 | 3,462E-65 |
| ACE2 | LINC02057 | 0,7162 | 4,34307E-65 |
| ACE2 | LENG9 | 0,7162 | 4,32002E-65 |
| ACE2 | KB-1991G8.1 | 0,7161 | 4,47152E-65 |
| ACE2 | RP5-965G21.4 | 0,7161 | 4,51249E-65 |
| ACE2 | RP4-695O20.1 | 0,7158 | 5,52146E-65 |
| ACE2 | CTD-2008P7.9 | 0,7157 | 5,90852E-65 |
| ACE2 | ATP7B | 0,7155 | 6,52968E-65 |
| ACE2 | C12orf49 | 0,7152 | 7,98357E-65 |
| ACE2 | FAM162A | 0,7151 | 8,36095E-65 |
| ACE2 | RP1-80N2.4 | 0,7151 | 8,17098E-65 |
| ACE2 | SUCLG2 | 0,7148 | 9,81536E-65 |
| ACE2 | PAG1 | 0,7147 | 1,05247E-64 |
| ACE2 | CCNYL1 | 0,7146 | 1,13678E-64 |
| ACE2 | RP11-227H15.5 | 0,7146 | 1,12184E-64 |
| ACE2 | TMEM92 | 0,7146 | 1,1403E-64 |
| ACE2 | MACC1 | 0,7145 | 1,18646E-64 |
| ACE2 | RP11-467J12.4 | 0,7144 | 1,28585E-64 |
| ACE2 | AC010745.1 | 0,7143 | 1,31757E-64 |
| ACE2 | HRCT1 | 0,7141 | 1,46602E-64 |
| ACE2 | KRT18P28 | 0,714 | 1,53937E-64 |
| ACE2 | FAAH | 0,7138 | 1,8341E-64 |
| ACE2 | FASTKD1 | 0,7138 | 1,80643E-64 |
| ACE2 | AC097468.4 | 0,7137 | 1,86865E-64 |
| ACE2 | TMEM56 | 0,7136 | 2,00648E-64 |
| ACE2 | NBEAL1 | 0,7136 | 2,0036E-64 |
| ACE2 | UCA1 | 0,7134 | 2,31979E-64 |
| ACE2 | NOX1 | 0,7133 | 2,40385E-64 |
| ACE2 | XKRX | 0,7132 | 2,53925E-64 |
| ACE2 | RP11-234B24.2 | 0,7131 | 2,71049E-64 |
| ACE2 | CARD10 | 0,7131 | 2,62223E-64 |
| ACE2 | EDN2 | 0,713 | 2,90209E-64 |
| ACE2 | KCNQ1 | 0,7128 | 3,15263E-64 |
| ACE2 | MED18 | 0,7127 | 3,34952E-64 |
| ACE2 | LINC01645 | 0,7127 | 3,44344E-64 |
| ACE2 | GIPR | 0,7127 | 3,3882E-64 |
| ACE2 | GSTZ1 | 0,7126 | 3,53482E-64 |
| ACE2 | SEL1L3 | 0,7121 | 4,93377E-64 |
| ACE2 | PEX11A | 0,7121 | 4,96122E-64 |
| ACE2 | RP11-132F7.2 | 0,7121 | 4,88048E-64 |
| ACE2 | AC011523.2 | 0,7121 | 4,75E-64 |
| ACE2 | CRYBA2 | 0,712 | 5,25948E-64 |
| ACE2 | SCTR | 0,7117 | 6,26229E-64 |
| ACE2 | TMEM8A | 0,7117 | 5,98554E-64 |
| ACE2 | MFSD4A | 0,7116 | 6,5211E-64 |
| ACE2 | HOXB-AS4 | 0,7114 | 7,24906E-64 |
| ACE2 | UGT2B10 | 0,7113 | 7,56808E-64 |

| | | | |
|---|---|---|---|
| ACE2 | RP5-943J3.2 | 0,7112 | 8,3699E-64 |
| ACE2 | RP11-569G9.7 | 0,7111 | 8,61176E-64 |
| ACE2 | RP11-296E3.2 | 0,7111 | 8,63878E-64 |
| ACE2 | HSBP1L1 | 0,7111 | 8,71428E-64 |
| ACE2 | RP11-44F21.5 | 0,7109 | 9,95056E-64 |
| ACE2 | AC114812.10 | 0,7108 | 1,04137E-63 |
| ACE2 | SYT13 | 0,7106 | 1,16375E-63 |
| ACE2 | RPL10L | 0,7106 | 1,14259E-63 |
| ACE2 | STARD5 | 0,7099 | 1,71411E-63 |
| ACE2 | PARM1 | 0,7096 | 2,10881E-63 |
| ACE2 | RP11-89H19.2 | 0,7095 | 2,19719E-63 |
| ACE2 | TJP2 | 0,7094 | 2,27561E-63 |
| ACE2 | FAM160A2 | 0,7094 | 2,36139E-63 |
| ACE2 | KBTBD11 | 0,7093 | 2,51941E-63 |
| ACE2 | CC2D1A | 0,7092 | 2,63831E-63 |
| ACE2 | FOXA2 | 0,7092 | 2,68756E-63 |
| ACE2 | RP11-526A4.1 | 0,7087 | 3,54987E-63 |
| ACE2 | AURKA | 0,7087 | 3,53284E-63 |
| ACE2 | LPIN3 | 0,7086 | 3,6942E-63 |
| ACE2 | YBX2 | 0,7084 | 4,12738E-63 |
| ACE2 | RP11-797A18.5 | 0,7083 | 4,42159E-63 |
| ACE2 | PAK4 | 0,7082 | 4,60686E-63 |
| ACE2 | MISP3 | 0,708 | 5,12455E-63 |
| ACE2 | RP11-369K16.1 | 0,7079 | 5,63038E-63 |
| ACE2 | BOLA3P3 | 0,7078 | 5,77831E-63 |
| ACE2 | ANXA13 | 0,7078 | 5,91856E-63 |
| ACE2 | MSLN | 0,7078 | 5,95123E-63 |
| ACE2 | MTNR1A | 0,7074 | 7,3442E-63 |
| ACE2 | RP11-168F9.2 | 0,7072 | 8,26198E-63 |
| ACE2 | DLG1-AS1 | 0,7071 | 8,63671E-63 |
| ACE2 | MINK1 | 0,7071 | 8,95973E-63 |
| ACE2 | PLCD1 | 0,707 | 9,46504E-63 |
| ACE2 | ALG14 | 0,7068 | 1,04819E-62 |
| ACE2 | TMEM89 | 0,7067 | 1,12828E-62 |
| ACE2 | PLXNA2 | 0,7066 | 1,15196E-62 |
| ACE2 | RREB1 | 0,7066 | 1,15948E-62 |
| ACE2 | RP11-259N19.1 | 0,7065 | 1,24866E-62 |
| ACE2 | ABCB1 | 0,7065 | 1,23689E-62 |
| ACE2 | C2orf88 | 0,7064 | 1,34189E-62 |
| ACE2 | OXNAD1 | 0,7064 | 1,30219E-62 |
| ACE2 | RP11-1260E13.2 | 0,7063 | 1,39597E-62 |
| ACE2 | ELF3-AS1 | 0,7062 | 1,45321E-62 |
| ACE2 | ARFGEF3 | 0,706 | 1,62452E-62 |
| ACE2 | TCP10 | 0,7058 | 1,80784E-62 |
| ACE2 | ABR | 0,7057 | 1,93861E-62 |
| ACE2 | ATP2A3 | 0,7056 | 2,02969E-62 |
| ACE2 | POU5F1 | 0,7055 | 2,22594E-62 |
| ACE2 | C2orf54 | 0,7051 | 2,7165E-62 |
| ACE2 | PCYT2 | 0,7051 | 2,80864E-62 |
| ACE2 | ARHGEF35 | 0,705 | 2,86126E-62 |
| ACE2 | CDA | 0,7049 | 3,04397E-62 |
| ACE2 | SLC35F5 | 0,7049 | 3,11364E-62 |
| ACE2 | ABCD3 | 0,7047 | 3,46396E-62 |
| ACE2 | PFKL | 0,7047 | 3,49665E-62 |

| | | | |
|---|---|---|---|
| ACE2 | RP11-14C10.3 | 0,7046 | 3,7173E-62 |
| ACE2 | RNU6-1065P | 0,7046 | 3,69122E-62 |
| ACE2 | TTC6 | 0,7044 | 4,03629E-62 |
| ACE2 | TDRD5 | 0,7041 | 4,71233E-62 |
| ACE2 | HMGCS1 | 0,7041 | 4,93634E-62 |
| ACE2 | CYCS | 0,7041 | 4,95214E-62 |
| ACE2 | AP001625.6 | 0,7039 | 5,48389E-62 |
| ACE2 | ANO7L1 | 0,7038 | 5,59354E-62 |
| ACE2 | SLC35F2 | 0,7037 | 6,14384E-62 |
| ACE2 | LFNG | 0,7036 | 6,23456E-62 |
| ACE2 | PDXDC1 | 0,7036 | 6,48707E-62 |
| ACE2 | GAL3ST2 | 0,7035 | 6,87539E-62 |
| ACE2 | TEX43 | 0,7035 | 6,66796E-62 |
| ACE2 | DARS2 | 0,7033 | 7,62952E-62 |
| ACE2 | LINC00668 | 0,7033 | 7,67686E-62 |
| ACE2 | AC114730.5 | 0,7029 | 9,73103E-62 |
| ACE2 | SLC39A8 | 0,7027 | 1,05591E-61 |
| ACE2 | NDUFA9 | 0,7026 | 1,10397E-61 |
| ACE2 | ADGRF1 | 0,7025 | 1,17176E-61 |
| ACE2 | SGPL1 | 0,7025 | 1,22153E-61 |
| ACE2 | MNX1-AS2 | 0,7024 | 1,22597E-61 |
| ACE2 | SERPINB5 | 0,7024 | 1,25135E-61 |
| ACE2 | MUC4 | 0,7023 | 1,31786E-61 |
| ACE2 | SLC38A4 | 0,7023 | 1,30006E-61 |
| ACE2 | HMGN2P46 | 0,7023 | 1,30299E-61 |
| ACE2 | GNA11 | 0,7021 | 1,46209E-61 |
| ACE2 | LGALS2 | 0,702 | 1,55317E-61 |
| ACE2 | MTND4P19 | 0,7019 | 1,67768E-61 |
| ACE2 | CEP170B | 0,7018 | 1,76544E-61 |
| ACE2 | RP11-482M8.1 | 0,7017 | 1,81558E-61 |
| ACE2 | PDXK | 0,7017 | 1,8276E-61 |
| ACE2 | PLCH1 | 0,7016 | 1,97794E-61 |
| ACE2 | ZBED9 | 0,7016 | 1,96728E-61 |
| ACE2 | ZDHHC3 | 0,7013 | 2,38084E-61 |
| ACE2 | NECTIN1 | 0,7011 | 2,57448E-61 |
| ACE2 | RNF43 | 0,7011 | 2,54365E-61 |
| ACE2 | SLC4A10 | 0,7007 | 3,21263E-61 |
| ACE2 | MYB | 0,7007 | 3,2362E-61 |
| ACE2 | SLC25A48 | 0,7005 | 3,65487E-61 |
| ACE2 | MTCH2 | 0,7005 | 3,57872E-61 |
| ACE2 | RP11-172C16.5 | 0,7005 | 3,62868E-61 |
| ACE2 | CTD-3051D23.4 | 0,7005 | 3,62103E-61 |
| ACE2 | SPTLC2 | 0,7004 | 3,74561E-61 |
| ACE2 | ENTPD6 | 0,7004 | 3,87617E-61 |
| ACE2 | RP11-462G2.1 | 0,7003 | 4,11788E-61 |
| ACE2 | SMCO4 | 0,7003 | 4,17041E-61 |
| ACE2 | AIFM3 | 0,7002 | 4,33088E-61 |
| ACE2 | RP11-524H19.2 | 0,6997 | 5,66566E-61 |
| ACE2 | VDAC1 | 0,6996 | 6,08097E-61 |
| ACE2 | MAD2L1BP | 0,6995 | 6,44004E-61 |
| ACE2 | AACS | 0,6995 | 6,39137E-61 |
| ACE2 | NECTIN4 | 0,6994 | 6,71701E-61 |
| ACE2 | FMO4 | 0,6994 | 6,78494E-61 |
| ACE2 | SPINK1 | 0,6994 | 6,60465E-61 |

| | | | |
|---|---|---|---|
| ACE2 | SGK1 | 0,6994 | 6,68545E-61 |
| ACE2 | STEAP3 | 0,6993 | 7,151E-61 |
| ACE2 | MFSD3 | 0,699 | 8,58497E-61 |
| ACE2 | RP11-640L9.2 | 0,6988 | 9,48225E-61 |
| ACE2 | GSTK1 | 0,6988 | 9,31106E-61 |
| ACE2 | CEACAM3 | 0,6987 | 9,65013E-61 |
| ACE2 | CRACR2B | 0,6986 | 1,06325E-60 |
| ACE2 | AQP11 | 0,6985 | 1,10589E-60 |
| ACE2 | ATP8A1 | 0,6982 | 1,28562E-60 |
| ACE2 | MCRIP2 | 0,6982 | 1,29732E-60 |
| ACE2 | STIL | 0,6981 | 1,35475E-60 |
| ACE2 | GS1-204I12.4 | 0,6981 | 1,34101E-60 |
| ACE2 | PLEK2 | 0,698 | 1,46404E-60 |
| ACE2 | ZNF888 | 0,6979 | 1,57038E-60 |
| ACE2 | GPAT3 | 0,6975 | 1,96092E-60 |
| ACE2 | AC009495.2 | 0,6973 | 2,17995E-60 |
| ACE2 | GJB3 | 0,697 | 2,49798E-60 |
| ACE2 | SDR16C5 | 0,697 | 2,47814E-60 |
| ACE2 | OR5BA1P | 0,697 | 2,55357E-60 |
| ACE2 | SPATA32 | 0,697 | 2,58095E-60 |
| ACE2 | IL1R2 | 0,6967 | 3,01234E-60 |
| ACE2 | MPZL3 | 0,6967 | 2,98258E-60 |
| ACE2 | RP11-486B10.4 | 0,6963 | 3,60989E-60 |
| ACE2 | NQO1 | 0,6963 | 3,61967E-60 |
| ACE2 | AGR2 | 0,6962 | 4,00442E-60 |
| ACE2 | NUDT14 | 0,6961 | 4,08745E-60 |
| ACE2 | MAPK3 | 0,696 | 4,36367E-60 |
| ACE2 | RP11-285E9.6 | 0,6958 | 4,80802E-60 |
| ACE2 | TMEM170A | 0,6957 | 5,13418E-60 |
| ACE2 | TXNDC17 | 0,6953 | 6,40165E-60 |
| ACE2 | RP11-627G18.1 | 0,6953 | 6,23203E-60 |
| ACE2 | TFRC | 0,6951 | 7,30149E-60 |
| ACE2 | PNKD | 0,6948 | 8,27508E-60 |
| ACE2 | ETFA | 0,6948 | 8,29991E-60 |
| ACE2 | RRBP1 | 0,6948 | 8,46507E-60 |
| ACE2 | RP11-320N21.2 | 0,6946 | 9,34321E-60 |
| ACE2 | RP11-448G15.1 | 0,6945 | 1,01093E-59 |
| ACE2 | AGFG2 | 0,6945 | 9,64127E-60 |
| ACE2 | REEP4 | 0,6944 | 1,02475E-59 |
| ACE2 | ZNF823 | 0,6944 | 1,03471E-59 |
| ACE2 | CEBPA-AS1 | 0,6944 | 1,06968E-59 |
| ACE2 | HPDL | 0,6943 | 1,08076E-59 |
| ACE2 | PDHA1 | 0,6942 | 1,13665E-59 |
| ACE2 | RP11-92K15.3 | 0,6941 | 1,23691E-59 |
| ACE2 | RP11-465B22.8 | 0,6939 | 1,39448E-59 |
| ACE2 | SLC10A5 | 0,6937 | 1,51532E-59 |
| ACE2 | LIMK2 | 0,6937 | 1,54085E-59 |
| ACE2 | FGD4 | 0,6935 | 1,66185E-59 |
| ACE2 | TCEA3 | 0,6934 | 1,74529E-59 |
| ACE2 | RPL7P28 | 0,6934 | 1,83181E-59 |
| ACE2 | RPSAP75 | 0,6934 | 1,79617E-59 |
| ACE2 | CENPV | 0,6934 | 1,75604E-59 |
| ACE2 | MBOAT7 | 0,6934 | 1,82641E-59 |
| ACE2 | RP11-486F17.1 | 0,6928 | 2,49805E-59 |

| | | | |
|---|---|---|---|
| ACE2 | ARSE | 0,6928 | 2,51656E-59 |
| ACE2 | MIR429 | 0,6921 | 3,67621E-59 |
| ACE2 | RFX6 | 0,692 | 3,74084E-59 |
| ACE2 | EPN1 | 0,6918 | 4,268E-59 |
| ACE2 | CAPN15 | 0,6917 | 4,54442E-59 |
| ACE2 | C4orf48 | 0,6914 | 5,13217E-59 |
| ACE2 | TMED6 | 0,6914 | 5,27995E-59 |
| ACE2 | CD24P4 | 0,6914 | 5,28498E-59 |
| ACE2 | FAM177B | 0,691 | 6,36833E-59 |
| ACE2 | MROH2A | 0,6909 | 7,02939E-59 |
| ACE2 | IL32 | 0,6907 | 7,64999E-59 |
| ACE2 | DTX4 | 0,6905 | 8,5926E-59 |
| ACE2 | MYRF | 0,6905 | 8,55114E-59 |
| ACE2 | RP11-467L13.4 | 0,6905 | 8,63472E-59 |
| ACE2 | SNORA44 | 0,6904 | 8,78998E-59 |
| ACE2 | CISD3 | 0,6904 | 8,86881E-59 |
| ACE2 | MGC32805 | 0,6902 | 9,97917E-59 |
| ACE2 | ACOT4 | 0,6902 | 1,02397E-58 |
| ACE2 | AC012507.3 | 0,69 | 1,08618E-58 |
| ACE2 | PWWP2B | 0,6899 | 1,14349E-58 |
| ACE2 | RP11-354M1.2 | 0,6899 | 1,18336E-58 |
| ACE2 | RAB27B | 0,6899 | 1,16807E-58 |
| ACE2 | ALAS1 | 0,6897 | 1,28104E-58 |
| ACE2 | RP11-794G24.1 | 0,6897 | 1,29276E-58 |
| ACE2 | AGPAT5 | 0,6896 | 1,38219E-58 |
| ACE2 | TUBA1C | 0,6896 | 1,36551E-58 |
| ACE2 | RP13-580F15.2 | 0,6896 | 1,36748E-58 |
| ACE2 | AC084809.3 | 0,6896 | 1,40183E-58 |
| ACE2 | ACAT2 | 0,6895 | 1,48623E-58 |
| ACE2 | ITLN1 | 0,6893 | 1,62005E-58 |
| ACE2 | RP11-390F4.3 | 0,6892 | 1,70001E-58 |
| ACE2 | AQP7 | 0,6892 | 1,67456E-58 |
| ACE2 | RPSAP9 | 0,6892 | 1,69864E-58 |
| ACE2 | STARD10 | 0,6891 | 1,78351E-58 |
| ACE2 | ESPL1 | 0,689 | 1,92923E-58 |
| ACE2 | RNU6-140P | 0,6889 | 1,96048E-58 |
| ACE2 | GSTO2 | 0,6887 | 2,21888E-58 |
| ACE2 | LINC01348 | 0,6886 | 2,40592E-58 |
| ACE2 | RP3-406A7.7 | 0,6885 | 2,53377E-58 |
| ACE2 | HOXB9 | 0,6885 | 2,4743E-58 |
| ACE2 | NOXA1 | 0,6883 | 2,74343E-58 |
| ACE2 | AC005077.14 | 0,6882 | 2,84459E-58 |
| ACE2 | HNF4A-AS1 | 0,6881 | 3,03504E-58 |
| ACE2 | LINC00964 | 0,688 | 3,19733E-58 |
| ACE2 | HACD2 | 0,6877 | 3,86136E-58 |
| ACE2 | TXN | 0,6877 | 3,69824E-58 |
| ACE2 | PYCARD | 0,6877 | 3,82905E-58 |
| ACE2 | TSPO | 0,6877 | 3,86515E-58 |
| ACE2 | ZC3H12D | 0,6876 | 3,93437E-58 |
| ACE2 | PLBD1 | 0,6876 | 3,97415E-58 |
| ACE2 | KCNH6 | 0,6875 | 4,1078E-58 |
| ACE2 | LINC00628 | 0,6873 | 4,77322E-58 |
| ACE2 | KRT9 | 0,6873 | 4,68201E-58 |
| ACE2 | NOVA1-AS1 | 0,6871 | 5,09519E-58 |

| | | | |
|---|---|---|---|
| ACE2 | SSTR5-AS1 | 0,687 | 5,35482E-58 |
| ACE2 | SLC7A4 | 0,6866 | 6,8349E-58 |
| ACE2 | SLC17A5 | 0,6864 | 7,49946E-58 |
| ACE2 | RP11-797A18.6 | 0,6862 | 8,18089E-58 |
| ACE2 | UBALD2 | 0,6858 | 1,03382E-57 |
| ACE2 | MLEC | 0,6857 | 1,06258E-57 |
| ACE2 | HCN3 | 0,6854 | 1,27834E-57 |
| ACE2 | PPARA | 0,6853 | 1,34952E-57 |
| ACE2 | RP11-67L3.2 | 0,6851 | 1,49765E-57 |
| ACE2 | RP11-704M14.1 | 0,6849 | 1,656E-57 |
| ACE2 | SDHB | 0,6848 | 1,72769E-57 |
| ACE2 | AC017074.2 | 0,6848 | 1,77216E-57 |
| ACE2 | AC138517.6 | 0,6848 | 1,72896E-57 |
| ACE2 | AKR1B10P1 | 0,6847 | 1,80538E-57 |
| ACE2 | RP11-730K11.1 | 0,6847 | 1,87223E-57 |
| ACE2 | CTD-2134A5.4 | 0,6847 | 1,78077E-57 |
| ACE2 | BUB1P1 | 0,6846 | 1,9428E-57 |
| ACE2 | RP11-8L2.1 | 0,6845 | 2,03127E-57 |
| ACE2 | PCLO | 0,6841 | 2,45821E-57 |
| ACE2 | DMBT1 | 0,684 | 2,67768E-57 |
| ACE2 | RP11-595B24.2 | 0,684 | 2,60924E-57 |
| ACE2 | CYP51A1 | 0,6839 | 2,78087E-57 |
| ACE2 | TESMIN | 0,6839 | 2,82631E-57 |
| ACE2 | KRT18P11 | 0,6837 | 3,05479E-57 |
| ACE2 | CYP3A4 | 0,6833 | 3,87757E-57 |
| ACE2 | OASL | 0,6832 | 3,93693E-57 |
| ACE2 | MT-CO1 | 0,6831 | 4,12775E-57 |
| ACE2 | RP11-10C24.2 | 0,6829 | 4,71663E-57 |
| ACE2 | KLK15 | 0,6829 | 4,57505E-57 |
| ACE2 | RP5-984P4.6 | 0,6829 | 4,5782E-57 |
| ACE2 | KLHDC7A | 0,6827 | 5,23858E-57 |
| ACE2 | LINC01975 | 0,6827 | 5,18158E-57 |
| ACE2 | MUC5B | 0,6826 | 5,48081E-57 |
| ACE2 | AC004870.4 | 0,6824 | 6,1329E-57 |
| ACE2 | C10orf95 | 0,6824 | 5,9027E-57 |
| ACE2 | HIST1H3E | 0,6823 | 6,37771E-57 |
| ACE2 | SLC18A1 | 0,6823 | 6,3491E-57 |
| ACE2 | RP5-1125A11.7 | 0,6822 | 6,74404E-57 |
| ACE2 | OXSM | 0,6818 | 8,15601E-57 |
| ACE2 | CFAP99 | 0,6818 | 8,16153E-57 |
| ACE2 | MRAP2 | 0,6817 | 8,55931E-57 |
| ACE2 | NRARP | 0,6817 | 8,83476E-57 |
| ACE2 | KCNV1 | 0,6813 | 1,07897E-56 |
| ACE2 | CEBPG | 0,6812 | 1,14832E-56 |
| ACE2 | RP11-649A18.4 | 0,6811 | 1,18764E-56 |
| ACE2 | LINC01806 | 0,681 | 1,27145E-56 |
| ACE2 | RP5-1056H1.2 | 0,681 | 1,23575E-56 |
| ACE2 | UGT1A13P | 0,6808 | 1,35739E-56 |
| ACE2 | STPG3 | 0,6807 | 1,444E-56 |
| ACE2 | RNF152 | 0,6807 | 1,48091E-56 |
| ACE2 | OR2A7 | 0,6805 | 1,593E-56 |
| ACE2 | SLC52A2 | 0,6803 | 1,74853E-56 |
| ACE2 | RP11-627G18.2 | 0,6803 | 1,80264E-56 |
| ACE2 | CCNI2 | 0,6802 | 1,8308E-56 |

| | | | |
|---|---|---|---|
| ACE2 | RP11-465N4.5 | 0,6801 | 2,00071E-56 |
| ACE2 | KRT8P35 | 0,6799 | 2,24087E-56 |
| ACE2 | GMNN | 0,6799 | 2,21834E-56 |
| ACE2 | RP11-77K12.10 | 0,6799 | 2,17425E-56 |
| ACE2 | C2orf82 | 0,6797 | 2,45104E-56 |
| ACE2 | GALNT6 | 0,6796 | 2,49851E-56 |
| ACE2 | GCG | 0,6794 | 2,77908E-56 |
| ACE2 | R3HDML | 0,6794 | 2,77421E-56 |
| ACE2 | KRTAP5-AS1 | 0,6793 | 3,04775E-56 |
| ACE2 | IGHVII-78-1 | 0,6792 | 3,14339E-56 |
| ACE2 | RP11-93B14.9 | 0,679 | 3,52285E-56 |
| ACE2 | RP11-649A18.5 | 0,6789 | 3,71696E-56 |
| ACE2 | ATP5F1 | 0,6785 | 4,48237E-56 |
| ACE2 | RP11-391H12.8 | 0,6785 | 4,554E-56 |
| ACE2 | LAMA1 | 0,6784 | 4,63695E-56 |
| ACE2 | FARP2 | 0,6783 | 4,87279E-56 |
| ACE2 | GRN | 0,6783 | 4,97177E-56 |
| ACE2 | RN7SL272P | 0,6782 | 5,14593E-56 |
| ACE2 | FCAMR | 0,678 | 5,83756E-56 |
| ACE2 | ERICH5 | 0,678 | 5,71536E-56 |
| ACE2 | RP11-514F3.5 | 0,6777 | 6,62431E-56 |
| ACE2 | AC112229.1 | 0,6776 | 6,89738E-56 |
| ACE2 | ANKRD22 | 0,6772 | 8,45428E-56 |
| ACE2 | RP11-10N16.2 | 0,677 | 9,63108E-56 |
| ACE2 | ADGB | 0,677 | 9,36964E-56 |
| ACE2 | RP13-870H17.3 | 0,6769 | 1,00292E-55 |
| ACE2 | CD177P1 | 0,6768 | 1,06537E-55 |
| ACE2 | TPM3 | 0,6767 | 1,13314E-55 |
| ACE2 | RBP2 | 0,6767 | 1,12506E-55 |
| ACE2 | ADCY6 | 0,6767 | 1,09129E-55 |
| ACE2 | CASK | 0,6765 | 1,24997E-55 |
| ACE2 | RP11-517C16.2 | 0,6761 | 1,47706E-55 |
| ACE2 | AC092569.2 | 0,676 | 1,61586E-55 |
| ACE2 | EHMT2-AS1 | 0,676 | 1,57475E-55 |
| ACE2 | TMEM44 | 0,6759 | 1,6616E-55 |
| ACE2 | TTLL6 | 0,6759 | 1,6811E-55 |
| ACE2 | SLC45A2 | 0,6753 | 2,23741E-55 |
| ACE2 | SERPINA6 | 0,6753 | 2,25521E-55 |
| ACE2 | LINC01226 | 0,6752 | 2,39072E-55 |
| ACE2 | HMGA2 | 0,6749 | 2,70158E-55 |
| ACE2 | AP006748.1 | 0,6749 | 2,78705E-55 |
| ACE2 | RP11-386E5.1 | 0,6747 | 3,05057E-55 |
| ACE2 | AC005083.1 | 0,6747 | 2,97945E-55 |
| ACE2 | RP11-211N11.5 | 0,6747 | 3,07067E-55 |
| ACE2 | ECHS1 | 0,6747 | 3,01307E-55 |
| ACE2 | CTD-2566J3.1 | 0,6745 | 3,4195E-55 |
| ACE2 | AK7 | 0,6745 | 3,42252E-55 |
| ACE2 | PITPNM1 | 0,6742 | 4,00965E-55 |
| ACE2 | ITGA3 | 0,6741 | 4,0897E-55 |
| ACE2 | ABCG2 | 0,674 | 4,31266E-55 |
| ACE2 | TLX1 | 0,6739 | 4,63663E-55 |
| ACE2 | RP11-93B14.10 | 0,6739 | 4,48638E-55 |
| ACE2 | IL10RB | 0,6739 | 4,65377E-55 |
| ACE2 | AKR1B15 | 0,6738 | 4,75998E-55 |

| | | | |
|---|---|---|---|
| ACE2 | CASC9 | 0,6736 | 5,16871E-55 |
| ACE2 | AC011747.3 | 0,6734 | 5,98761E-55 |
| ACE2 | MAOA | 0,6734 | 5,96919E-55 |
| ACE2 | MARS2 | 0,6732 | 6,32915E-55 |
| ACE2 | GPX2 | 0,6732 | 6,39776E-55 |
| ACE2 | RP11-963H4.3 | 0,6732 | 6,37004E-55 |
| ACE2 | SLC29A2 | 0,6731 | 6,71543E-55 |
| ACE2 | PAX6 | 0,673 | 7,16601E-55 |
| ACE2 | GATA6-AS1 | 0,6729 | 7,62788E-55 |
| ACE2 | GLTP | 0,6725 | 8,98682E-55 |
| ACE2 | AC087491.2 | 0,6724 | 9,55191E-55 |
| ACE2 | CHGA | 0,6722 | 1,04646E-54 |
| ACE2 | MIR559 | 0,672 | 1,19354E-54 |
| ACE2 | SPATA24 | 0,6715 | 1,49748E-54 |
| ACE2 | GFI1B | 0,6713 | 1,62724E-54 |
| ACE2 | CTD-2020K17.4 | 0,6712 | 1,78718E-54 |
| ACE2 | ENPP7P11 | 0,6709 | 1,99631E-54 |
| ACE2 | CEACAM8 | 0,6709 | 2,04975E-54 |
| ACE2 | COX6B2 | 0,6707 | 2,19373E-54 |
| ACE2 | RP11-807H22.5 | 0,6706 | 2,39208E-54 |
| ACE2 | SCML2P1 | 0,6705 | 2,44384E-54 |
| ACE2 | TEX11 | 0,6705 | 2,52971E-54 |
| ACE2 | COPG1 | 0,6704 | 2,63498E-54 |
| ACE2 | RP11-172C16.4 | 0,6704 | 2,62618E-54 |
| ACE2 | EYA2 | 0,6704 | 2,55939E-54 |
| ACE2 | AC027612.4 | 0,6702 | 2,92895E-54 |
| ACE2 | DGAT2L7P | 0,6702 | 2,79513E-54 |
| ACE2 | CTA-363E6.1 | 0,6702 | 2,84032E-54 |
| ACE2 | REG4 | 0,67 | 3,15594E-54 |
| ACE2 | HNRNPA1P26 | 0,6699 | 3,34087E-54 |
| ACE2 | RP1-80N2.2 | 0,6698 | 3,44728E-54 |
| ACE2 | SEC23B | 0,6698 | 3,50231E-54 |
| ACE2 | CTD-2270L9.4 | 0,6697 | 3,73188E-54 |
| ACE2 | SLC50A1 | 0,6696 | 3,92349E-54 |
| ACE2 | SUGCT | 0,6694 | 4,27459E-54 |
| ACE2 | PMM2 | 0,6693 | 4,386E-54 |
| ACE2 | FEV | 0,669 | 5,27991E-54 |
| ACE2 | EFNB2 | 0,6689 | 5,3729E-54 |
| ACE2 | FAHD1 | 0,6688 | 5,60363E-54 |
| ACE2 | DEPDC1B | 0,6684 | 7,06725E-54 |
| ACE2 | HIP1R | 0,6684 | 6,81083E-54 |
| ACE2 | DDX60 | 0,6683 | 7,36118E-54 |
| ACE2 | PLCL2 | 0,6681 | 7,9647E-54 |
| ACE2 | RP11-616M22.3 | 0,668 | 8,37988E-54 |
| ACE2 | LINC00570 | 0,6679 | 8,84515E-54 |
| ACE2 | RP11-798K3.2 | 0,6679 | 8,76277E-54 |
| ACE2 | DCAF11 | 0,6677 | 9,6048E-54 |
| ACE2 | KIAA1671 | 0,6675 | 1,09438E-53 |
| ACE2 | ALPPL2 | 0,6674 | 1,12542E-53 |
| ACE2 | TRPC7 | 0,6674 | 1,12835E-53 |
| ACE2 | BEND3 | 0,6673 | 1,2128E-53 |
| ACE2 | DCTPP1 | 0,6672 | 1,24269E-53 |
| ACE2 | ARR3 | 0,6672 | 1,23748E-53 |
| ACE2 | RNF157-AS1 | 0,667 | 1,36583E-53 |

| | | | |
|---|---|---|---|
| ACE2 | ACTR3C | 0,6668 | 1,49287E-53 |
| ACE2 | RP5-1142J19.1 | 0,6667 | 1,56267E-53 |
| ACE2 | VPS25 | 0,6666 | 1,6954E-53 |
| ACE2 | GLCE | 0,6662 | 2,05615E-53 |
| ACE2 | C11orf53 | 0,6661 | 2,16094E-53 |
| ACE2 | FOXJ1 | 0,6659 | 2,32433E-53 |
| ACE2 | ZP2 | 0,6658 | 2,51812E-53 |
| ACE2 | HAS3 | 0,6658 | 2,43249E-53 |
| ACE2 | FBLIM1 | 0,6657 | 2,61877E-53 |
| ACE2 | RP11-395L14.13 | 0,6657 | 2,56315E-53 |
| ACE2 | CYB561 | 0,6657 | 2,60774E-53 |
| ACE2 | STPG3-AS1 | 0,6654 | 3,02351E-53 |
| ACE2 | RP11-281P23.2 | 0,6652 | 3,2896E-53 |
| ACE2 | SMPD2 | 0,6652 | 3,24729E-53 |
| ACE2 | RP4-724E13.2 | 0,6652 | 3,32771E-53 |
| ACE2 | OPLAH | 0,6652 | 3,31205E-53 |
| ACE2 | CCDC183-AS1 | 0,6651 | 3,46205E-53 |
| ACE2 | PTRHD1 | 0,665 | 3,57194E-53 |
| ACE2 | RHOV | 0,6648 | 3,91079E-53 |
| ACE2 | NCLP1 | 0,6645 | 4,6649E-53 |
| ACE2 | KRTAP4-1 | 0,6645 | 4,6377E-53 |
| ACE2 | EFHD2 | 0,6644 | 4,92107E-53 |
| ACE2 | HEXA-AS1 | 0,6641 | 5,6482E-53 |
| ACE2 | B3GALT1 | 0,6638 | 6,38673E-53 |
| ACE2 | RP11-44F14.8 | 0,6636 | 7,27468E-53 |
| ACE2 | SLC25A34 | 0,6635 | 7,60569E-53 |
| ACE2 | XXyac-YM21GA2.4 | 0,6633 | 8,08562E-53 |
| ACE2 | GCSAM | 0,6632 | 8,62015E-53 |
| ACE2 | MNX1-AS1 | 0,6631 | 8,89357E-53 |
| ACE2 | ENPP1 | 0,663 | 9,49474E-53 |
| ACE2 | RP11-231E6.1 | 0,6629 | 9,71782E-53 |
| ACE2 | AC010761.9 | 0,6629 | 9,84735E-53 |
| ACE2 | RP11-314A15.2 | 0,6629 | 9,77785E-53 |
| ACE2 | UCK2 | 0,6628 | 1,03795E-52 |
| ACE2 | HNRNPA1P59 | 0,6627 | 1,09699E-52 |
| ACE2 | AIFM1 | 0,6626 | 1,16188E-52 |
| ACE2 | RP3-325F22.5 | 0,6625 | 1,22729E-52 |
| ACE2 | RP11-626P14.2 | 0,6625 | 1,22173E-52 |
| ACE2 | ACO2 | 0,6625 | 1,22148E-52 |
| ACE2 | LCN12 | 0,6624 | 1,27756E-52 |
| ACE2 | RP5-908M14.5 | 0,6623 | 1,35542E-52 |
| ACE2 | AGXT | 0,6622 | 1,41857E-52 |
| ACE2 | MYD88 | 0,6622 | 1,38826E-52 |
| ACE2 | LINC00955 | 0,6622 | 1,4224E-52 |
| ACE2 | NMNAT1 | 0,662 | 1,49719E-52 |
| ACE2 | LINC02086 | 0,6619 | 1,63196E-52 |
| ACE2 | LXN | 0,6618 | 1,66266E-52 |
| ACE2 | RP11-150O12.3 | 0,6617 | 1,76501E-52 |
| ACE2 | LIPM | 0,6617 | 1,77272E-52 |
| ACE2 | ARHGAP8 | 0,6617 | 1,75161E-52 |
| ACE2 | DCST1 | 0,6615 | 1,90432E-52 |
| ACE2 | GOLIM4 | 0,6615 | 1,95401E-52 |
| ACE2 | PIGO | 0,6615 | 1,9881E-52 |
| ACE2 | RP1-193H18.2 | 0,6615 | 1,978E-52 |

| | | | |
|---|---|---|---|
| ACE2 | RP11-44F14.10 | 0,6611 | 2,37184E-52 |
| ACE2 | RP11-385F7.1 | 0,661 | 2,50514E-52 |
| ACE2 | ARFGEF2 | 0,6609 | 2,55318E-52 |
| ACE2 | DCDC2 | 0,6608 | 2,71019E-52 |
| ACE2 | PLA2G4D | 0,6608 | 2,65392E-52 |
| ACE2 | RP13-128O4.3 | 0,6608 | 2,72446E-52 |
| ACE2 | CDCA8 | 0,6607 | 2,87562E-52 |
| ACE2 | SLC27A4 | 0,6607 | 2,80325E-52 |
| ACE2 | AC022182.2 | 0,6606 | 3,0468E-52 |
| ACE2 | LYPLA2 | 0,6605 | 3,09266E-52 |
| ACE2 | PROSER1 | 0,6605 | 3,15403E-52 |
| ACE2 | RP1-60O19.1 | 0,6604 | 3,22925E-52 |
| ACE2 | PRSS22 | 0,6602 | 3,57161E-52 |
| ACE2 | RP11-395C3.1 | 0,6601 | 3,87109E-52 |
| ACE2 | ADGRV1 | 0,66 | 4,02119E-52 |
| ACE2 | RP5-1057J7.7 | 0,6598 | 4,30616E-52 |
| ACE2 | NHEJ1 | 0,6596 | 4,88924E-52 |
| ACE2 | FUT4 | 0,6596 | 4,84631E-52 |
| ACE2 | SLC6A20 | 0,6592 | 5,83462E-52 |
| ACE2 | SRSF9P1 | 0,6589 | 6,63322E-52 |
| ACE2 | LINC01612 | 0,6587 | 7,35041E-52 |
| ACE2 | TNFSF13 | 0,6587 | 7,31278E-52 |
| ACE2 | EGF | 0,6586 | 7,61152E-52 |
| ACE2 | RN7SL208P | 0,6582 | 9,37673E-52 |
| ACE2 | KRT8P49 | 0,6582 | 9,09383E-52 |
| ACE2 | AC073046.25 | 0,6581 | 9,93026E-52 |
| ACE2 | CCL20 | 0,6581 | 9,51828E-52 |
| ACE2 | AOAH | 0,6581 | 9,73345E-52 |
| ACE2 | IMPA2 | 0,6581 | 9,51791E-52 |
| ACE2 | UNC5B | 0,658 | 1,0155E-51 |
| ACE2 | ITPR3 | 0,6578 | 1,09577E-51 |
| ACE2 | DISP2 | 0,6577 | 1,15612E-51 |
| ACE2 | CXCR3 | 0,6577 | 1,19542E-51 |
| ACE2 | KIR2DL4 | 0,6576 | 1,22639E-51 |
| ACE2 | NEDD4L | 0,6575 | 1,28527E-51 |
| ACE2 | ABHD15 | 0,6573 | 1,44693E-51 |
| ACE2 | SLC28A2 | 0,6572 | 1,45393E-51 |
| ACE2 | MPI | 0,6571 | 1,58777E-51 |
| ACE2 | TRABD2A | 0,6568 | 1,78328E-51 |
| ACE2 | KCNJ2-AS1 | 0,6567 | 1,86619E-51 |
| ACE2 | STK17B | 0,6564 | 2,16947E-51 |
| ACE2 | PPFIBP2 | 0,6564 | 2,1711E-51 |
| ACE2 | RP11-1100L3.4 | 0,6562 | 2,40307E-51 |
| ACE2 | HSD3BP2 | 0,6558 | 2,8696E-51 |
| ACE2 | PROM2 | 0,6558 | 2,85929E-51 |
| ACE2 | RP11-286N22.16 | 0,6557 | 2,99394E-51 |
| ACE2 | RP11-21L23.4 | 0,6554 | 3,37303E-51 |
| ACE2 | EFNA3 | 0,6551 | 3,98487E-51 |
| ACE2 | C1orf53 | 0,6551 | 4,04417E-51 |
| ACE2 | CDHR1 | 0,655 | 4,23159E-51 |
| ACE2 | NUDT15 | 0,655 | 4,14135E-51 |
| ACE2 | RP11-123J14.4 | 0,6549 | 4,30674E-51 |
| ACE2 | RMI1 | 0,6549 | 4,35332E-51 |
| ACE2 | LRIT2 | 0,6549 | 4,34364E-51 |

| | | | |
|---|---|---|---|
| ACE2 | PNLIPRP2 | 0,6549 | 4,37284E-51 |
| ACE2 | USP30-AS1 | 0,6549 | 4,39696E-51 |
| ACE2 | RP11-708H21.1 | 0,6546 | 5,05332E-51 |
| ACE2 | RP11-465B22.3 | 0,6545 | 5,23698E-51 |
| ACE2 | RETNLB | 0,6545 | 5,31122E-51 |
| ACE2 | NT5C3A | 0,6545 | 5,24521E-51 |
| ACE2 | MBOAT1 | 0,6542 | 6,09714E-51 |
| ACE2 | BMP2 | 0,654 | 6,56169E-51 |
| ACE2 | KRT8P8 | 0,654 | 6,69507E-51 |
| ACE2 | RP3-395M20.9 | 0,6538 | 7,32149E-51 |
| ACE2 | CNNM2 | 0,6538 | 7,39347E-51 |
| ACE2 | CD164L2 | 0,6537 | 7,50584E-51 |
| ACE2 | ETNK1 | 0,6537 | 7,41025E-51 |
| ACE2 | SF3B4 | 0,6534 | 8,71852E-51 |
| ACE2 | ZNF165 | 0,6534 | 8,77432E-51 |
| ACE2 | ZFYVE27 | 0,6533 | 9,03094E-51 |
| ACE2 | DPP10-AS1 | 0,6529 | 1,09861E-50 |
| ACE2 | CAPN1 | 0,6529 | 1,12011E-50 |
| ACE2 | SNX18P3 | 0,6527 | 1,22455E-50 |
| ACE2 | ABCA5 | 0,6527 | 1,18032E-50 |
| ACE2 | RP11-439C15.5 | 0,6526 | 1,28658E-50 |
| ACE2 | BCL2L11 | 0,6525 | 1,32607E-50 |
| ACE2 | RP11-57H12.2 | 0,6523 | 1,43919E-50 |
| ACE2 | STMND1 | 0,6523 | 1,42134E-50 |
| ACE2 | MCF2L-AS1 | 0,652 | 1,68544E-50 |
| ACE2 | RPS6KA6 | 0,6519 | 1,71051E-50 |
| ACE2 | NEUROD1 | 0,6515 | 2,06004E-50 |
| ACE2 | TMCC3 | 0,6515 | 2,06585E-50 |
| ACE2 | ZNF774 | 0,6513 | 2,25749E-50 |
| ACE2 | CLCA2 | 0,6512 | 2,36617E-50 |
| ACE2 | LCN2 | 0,6512 | 2,35618E-50 |
| ACE2 | PDE8A | 0,6512 | 2,4482E-50 |
| ACE2 | SATB2-AS1 | 0,6511 | 2,52217E-50 |
| ACE2 | OTUD7A | 0,6508 | 2,89562E-50 |
| ACE2 | RP11-13N13.5 | 0,6508 | 2,90686E-50 |
| ACE2 | SCO2 | 0,6508 | 2,81992E-50 |
| ACE2 | RP11-1267H10.4 | 0,6507 | 3,05536E-50 |
| ACE2 | TOM1L1 | 0,6507 | 2,98514E-50 |
| ACE2 | FAM177A1 | 0,6506 | 3,08146E-50 |
| ACE2 | CRYM | 0,6506 | 3,18947E-50 |
| ACE2 | CTB-175E5.7 | 0,6504 | 3,38325E-50 |
| ACE2 | EPHB2 | 0,6503 | 3,57985E-50 |
| ACE2 | AC114812.8 | 0,6503 | 3,69682E-50 |
| ACE2 | PDX1 | 0,6502 | 3,72468E-50 |
| ACE2 | RP11-223I10.1 | 0,6501 | 3,87246E-50 |
| ACE2 | TRAF3IP2 | 0,6501 | 3,96107E-50 |
| ACE2 | MECOM | 0,65 | 4,15542E-50 |
| ACE2 | ANXA4 | 0,6499 | 4,24256E-50 |
| ACE2 | CTA-363E6.2 | 0,6499 | 4,30599E-50 |
| ACE2 | RBKS | 0,6498 | 4,47041E-50 |
| ACE2 | RNF157 | 0,6497 | 4,74443E-50 |
| ACE2 | LCOR | 0,6493 | 5,64605E-50 |
| ACE2 | LINC00239 | 0,6492 | 5,85485E-50 |
| ACE2 | LA16c-380H5.2 | 0,6492 | 5,84197E-50 |

| | | | |
|---|---|---|---|
| ACE2 | RP11-503C24.6 | 0,6491 | 6,23008E-50 |
| ACE2 | KCNK10 | 0,6491 | 6,33572E-50 |
| ACE2 | ELOVL1 | 0,649 | 6,68474E-50 |
| ACE2 | OSBPL7 | 0,649 | 6,50612E-50 |
| ACE2 | TRIM36-IT1 | 0,6489 | 6,8796E-50 |
| ACE2 | LINC01351 | 0,6488 | 7,16027E-50 |
| ACE2 | TPH1 | 0,6488 | 7,21489E-50 |
| ACE2 | TMEM177 | 0,6486 | 7,77821E-50 |
| ACE2 | RP11-33O4.1 | 0,6486 | 7,68944E-50 |
| ACE2 | LINC00261 | 0,6486 | 7,72379E-50 |
| ACE2 | CTA-243E7.1 | 0,6483 | 8,85844E-50 |
| ACE2 | C2orf15 | 0,6481 | 9,75516E-50 |
| ACE2 | TMEM246 | 0,6481 | 1,00364E-49 |
| ACE2 | RAB26 | 0,6481 | 9,66435E-50 |
| ACE2 | RP11-466A19.4 | 0,648 | 1,04533E-49 |
| ACE2 | LINC00479 | 0,648 | 1,02823E-49 |
| ACE2 | MARCKSL1 | 0,6479 | 1,10105E-49 |
| ACE2 | ROPN1 | 0,6478 | 1,12989E-49 |
| ACE2 | NUP62CL | 0,6478 | 1,10881E-49 |
| ACE2 | AC114812.5 | 0,6477 | 1,20351E-49 |
| ACE2 | CTC-558O2.1 | 0,6476 | 1,21467E-49 |
| ACE2 | MYB-AS1 | 0,6473 | 1,41338E-49 |
| ACE2 | GORAB | 0,6472 | 1,4941E-49 |
| ACE2 | PAX4 | 0,6469 | 1,65674E-49 |
| ACE2 | TTPA | 0,6466 | 1,94392E-49 |
| ACE2 | LPCAT3 | 0,6465 | 1,98993E-49 |
| ACE2 | LNX2 | 0,6464 | 2,09306E-49 |
| ACE2 | AP1G2 | 0,6464 | 2,16032E-49 |
| ACE2 | ZNF69 | 0,6461 | 2,39339E-49 |
| ACE2 | RP11-44F14.2 | 0,6459 | 2,66529E-49 |
| ACE2 | ATP1B3 | 0,6457 | 2,84876E-49 |
| ACE2 | KRT8P37 | 0,6457 | 2,90068E-49 |
| ACE2 | AFDN-AS1 | 0,6456 | 3,06926E-49 |
| ACE2 | PCCA | 0,6456 | 3,07176E-49 |
| ACE2 | FRRS1 | 0,6454 | 3,34443E-49 |
| ACE2 | RP11-350E12.4 | 0,6453 | 3,44255E-49 |
| ACE2 | FAM201A | 0,6451 | 3,74778E-49 |
| ACE2 | RP11-342A1.1 | 0,6449 | 4,18016E-49 |
| ACE2 | SLC2A1 | 0,6447 | 4,515E-49 |
| ACE2 | RP11-832A4.7 | 0,6446 | 4,82822E-49 |
| ACE2 | GDPGP1 | 0,6446 | 4,65057E-49 |
| ACE2 | DPF3 | 0,6444 | 5,25586E-49 |
| ACE2 | PNPLA1 | 0,6442 | 5,73499E-49 |
| ACE2 | CMTM8 | 0,6441 | 6,01059E-49 |
| ACE2 | CLYBL | 0,6441 | 5,90429E-49 |
| ACE2 | CTC-304I17.6 | 0,6439 | 6,59669E-49 |
| ACE2 | TACO1 | 0,6439 | 6,6127E-49 |
| ACE2 | TMCO1 | 0,6438 | 6,91844E-49 |
| ACE2 | NIPAL2 | 0,6438 | 6,83164E-49 |
| ACE2 | S100P | 0,6436 | 7,33767E-49 |
| ACE2 | CTD-2005D20.1 | 0,6431 | 9,18517E-49 |
| ACE2 | SFXN1 | 0,6431 | 9,07023E-49 |
| ACE2 | AC034220.3 | 0,643 | 9,51609E-49 |
| ACE2 | AC073133.1 | 0,6428 | 1,03682E-48 |

| | | | |
|---|---|---|---|
| ACE2 | ZNF296 | 0,6428 | 1,07419E-48 |
| ACE2 | TMEM106C | 0,6427 | 1,12852E-48 |
| ACE2 | COA3 | 0,6427 | 1,12903E-48 |
| ACE2 | WI2-2118C23.2 | 0,6426 | 1,13901E-48 |
| ACE2 | MEP1AP4 | 0,6425 | 1,2023E-48 |
| ACE2 | CCDC129 | 0,6423 | 1,29894E-48 |
| ACE2 | RP11-120K24.3 | 0,6422 | 1,41043E-48 |
| ACE2 | SESN2 | 0,6421 | 1,44781E-48 |
| ACE2 | RP11-670E13.6 | 0,6421 | 1,43332E-48 |
| ACE2 | REPS2 | 0,6421 | 1,42362E-48 |
| ACE2 | NBPF4 | 0,642 | 1,52315E-48 |
| ACE2 | KRT8P7 | 0,642 | 1,53296E-48 |
| ACE2 | HAL | 0,6419 | 1,55207E-48 |
| ACE2 | LINC00513 | 0,6418 | 1,63541E-48 |
| ACE2 | FAM213B | 0,6416 | 1,80819E-48 |
| ACE2 | TMEM59 | 0,6413 | 2,0868E-48 |
| ACE2 | AREL1 | 0,6413 | 2,01876E-48 |
| ACE2 | TUFM | 0,6412 | 2,13825E-48 |
| ACE2 | MKRN1 | 0,6411 | 2,27687E-48 |
| ACE2 | BCO1 | 0,6407 | 2,73984E-48 |
| ACE2 | CTA-363E6.3 | 0,6405 | 2,92062E-48 |
| ACE2 | HADH | 0,6404 | 3,0947E-48 |
| ACE2 | RP11-452D12.1 | 0,6404 | 3,02887E-48 |
| ACE2 | SYK | 0,6403 | 3,20867E-48 |
| ACE2 | MTFP1 | 0,64 | 3,63458E-48 |
| ACE2 | SIGIRR | 0,6398 | 3,95384E-48 |
| ACE2 | NCR3LG1 | 0,6397 | 4,1598E-48 |
| ACE2 | CTD-2284J15.1 | 0,6395 | 4,47728E-48 |
| ACE2 | CTC-428G20.6 | 0,6393 | 5,02479E-48 |
| ACE2 | LINC00637 | 0,6393 | 4,88216E-48 |
| ACE2 | MLIP-AS1 | 0,6391 | 5,34515E-48 |
| ACE2 | RP3-512B11.3 | 0,6389 | 5,8568E-48 |
| ACE2 | PLCE1 | 0,6389 | 6,02505E-48 |
| ACE2 | OR5G5P | 0,6389 | 5,91022E-48 |
| ACE2 | AC104534.2 | 0,6389 | 5,84909E-48 |
| ACE2 | SLC23A3 | 0,6386 | 6,59813E-48 |
| ACE2 | RP11-351J23.1 | 0,6382 | 7,94045E-48 |
| ACE2 | LRRK1 | 0,6382 | 7,87907E-48 |
| ACE2 | RP11-719K4.3 | 0,6382 | 8,03274E-48 |
| ACE2 | MSMO1 | 0,6379 | 9,24322E-48 |
| ACE2 | GP9 | 0,6378 | 9,76459E-48 |
| ACE2 | RP11-91K11.2 | 0,6377 | 9,8879E-48 |
| ACE2 | CACNA1F | 0,6377 | 1,01013E-47 |
| ACE2 | ANKDD1B | 0,6374 | 1,12236E-47 |
| ACE2 | MGST2 | 0,6372 | 1,21722E-47 |
| ACE2 | RP11-64C12.3 | 0,6369 | 1,42732E-47 |
| ACE2 | PPFIA3 | 0,6369 | 1,41815E-47 |
| ACE2 | CCDC125 | 0,6367 | 1,56566E-47 |
| ACE2 | HAVCR1 | 0,6367 | 1,53841E-47 |
| ACE2 | RP11-7K24.3 | 0,6364 | 1,76474E-47 |
| ACE2 | THSD4 | 0,6364 | 1,78442E-47 |
| ACE2 | EML2 | 0,6364 | 1,78313E-47 |
| ACE2 | ZRANB2-AS1 | 0,6363 | 1,84795E-47 |
| ACE2 | UGT2B17 | 0,6363 | 1,85681E-47 |

| | | | |
|---|---|---|---|
| ACE2 | MMAA | 0,636 | 2,07883E-47 |
| ACE2 | PTPN18 | 0,6359 | 2,21829E-47 |
| ACE2 | RNASET2 | 0,6359 | 2,20228E-47 |
| ACE2 | ALOX12B | 0,6356 | 2,49743E-47 |
| ACE2 | CAPG | 0,6352 | 2,97441E-47 |
| ACE2 | ASPG | 0,6345 | 4,04321E-47 |
| ACE2 | MGAM | 0,6343 | 4,326E-47 |
| ACE2 | SYNGR2 | 0,6342 | 4,55317E-47 |
| ACE2 | ZNF711 | 0,6342 | 4,51491E-47 |
| ACE2 | IVNS1ABP | 0,634 | 4,92346E-47 |
| ACE2 | C11orf42 | 0,634 | 4,98722E-47 |
| ACE2 | SRMS | 0,634 | 4,86263E-47 |
| ACE2 | RP11-661A12.9 | 0,6339 | 5,1616E-47 |
| ACE2 | RP11-95I16.6 | 0,6339 | 5,18028E-47 |
| ACE2 | RPS15AP11 | 0,6338 | 5,39809E-47 |
| ACE2 | LINC01012 | 0,6338 | 5,37115E-47 |
| ACE2 | EIF6 | 0,6335 | 6,02939E-47 |
| ACE2 | RP3-407E4.4 | 0,6331 | 7,22552E-47 |
| ACE2 | HTR4 | 0,633 | 7,73844E-47 |
| ACE2 | RP11-783K16.5 | 0,633 | 7,75101E-47 |
| ACE2 | DGKA | 0,633 | 7,46859E-47 |
| ACE2 | MGAT2 | 0,6329 | 7,8598E-47 |
| ACE2 | KLK3 | 0,6329 | 7,83739E-47 |
| ACE2 | MIR3131 | 0,6327 | 8,5138E-47 |
| ACE2 | AC022182.1 | 0,6327 | 8,61837E-47 |
| ACE2 | RP11-378I13.1 | 0,6325 | 9,29519E-47 |
| ACE2 | ARX | 0,6323 | 1,01896E-46 |
| ACE2 | FOXD2-AS1 | 0,6321 | 1,12127E-46 |
| ACE2 | GCH1 | 0,6317 | 1,33515E-46 |
| ACE2 | RN7SL684P | 0,6316 | 1,3814E-46 |
| ACE2 | LINC01342 | 0,6315 | 1,46573E-46 |
| ACE2 | KNL1 | 0,6314 | 1,49173E-46 |
| ACE2 | CPM | 0,6311 | 1,74771E-46 |
| ACE2 | APEH | 0,631 | 1,75978E-46 |
| ACE2 | ALDH3A1 | 0,631 | 1,77417E-46 |
| ACE2 | NKX2-2 | 0,631 | 1,78064E-46 |
| ACE2 | PLAGL2 | 0,6309 | 1,83634E-46 |
| ACE2 | CTD-2196E14.5 | 0,6308 | 1,95966E-46 |
| ACE2 | RP1-170O19.24 | 0,6307 | 2,04301E-46 |
| ACE2 | RASAL1 | 0,6306 | 2,09293E-46 |
| ACE2 | ACSF3 | 0,6305 | 2,2417E-46 |
| ACE2 | HMGCL | 0,6303 | 2,42384E-46 |
| ACE2 | KCNE3 | 0,63 | 2,69481E-46 |
| ACE2 | RP5-908M14.10 | 0,63 | 2,69406E-46 |
| ACE2 | KRT8P33 | 0,6299 | 2,87817E-46 |
| ACE2 | UGT2B29P | 0,6297 | 3,0662E-46 |
| ACE2 | RP4-584D14.5 | 0,6297 | 3,04176E-46 |
| ACE2 | P2RY4 | 0,6297 | 3,06988E-46 |
| ACE2 | CHST13 | 0,6296 | 3,2091E-46 |
| ACE2 | CCDC88C | 0,6296 | 3,17118E-46 |
| ACE2 | RPIA | 0,6294 | 3,58791E-46 |
| ACE2 | HK2P1 | 0,6294 | 3,51222E-46 |
| ACE2 | GRAMD2 | 0,6293 | 3,73571E-46 |
| ACE2 | RP11-368N21.5 | 0,6293 | 3,62893E-46 |

| | | | |
|---|---|---|---|
| ACE2 | NCMAP | 0,6291 | 3,99368E-46 |
| ACE2 | BATF2 | 0,6291 | 3,91192E-46 |
| ACE2 | CFAP65 | 0,6288 | 4,45427E-46 |
| ACE2 | IFI27 | 0,6288 | 4,50115E-46 |
| ACE2 | NANOGP9 | 0,6288 | 4,58883E-46 |
| ACE2 | IL2RG | 0,6288 | 4,5959E-46 |
| ACE2 | MYBPC1 | 0,6287 | 4,68463E-46 |
| ACE2 | CYC1 | 0,6285 | 5,0848E-46 |
| ACE2 | FUT5 | 0,6283 | 5,61828E-46 |
| ACE2 | AC013268.3 | 0,6279 | 6,5599E-46 |
| ACE2 | FOXN1 | 0,6279 | 6,75261E-46 |
| ACE2 | RP11-376N17.4 | 0,6278 | 6,97809E-46 |
| ACE2 | TMEM191A | 0,6277 | 7,11822E-46 |
| ACE2 | GSTA1 | 0,6275 | 7,6969E-46 |
| ACE2 | RP11-789C1.2 | 0,6273 | 8,35207E-46 |
| ACE2 | SPATA31C1 | 0,6273 | 8,44667E-46 |
| ACE2 | RP11-567C2.1 | 0,6271 | 9,28086E-46 |
| ACE2 | FAM95C | 0,6269 | 1,00354E-45 |
| ACE2 | NPM1P25 | 0,6268 | 1,06003E-45 |
| ACE2 | LLNLR-271C9.1 | 0,6268 | 1,02892E-45 |
| ACE2 | RP11-470P21.2 | 0,6265 | 1,18323E-45 |
| ACE2 | TMBIM6 | 0,6265 | 1,19302E-45 |
| ACE2 | TMPPE | 0,6263 | 1,27408E-45 |
| ACE2 | TNFRSF11B | 0,6263 | 1,2846E-45 |
| ACE2 | CSK | 0,6262 | 1,35148E-45 |
| ACE2 | ERLIN1 | 0,6259 | 1,51407E-45 |
| ACE2 | SRPK1 | 0,6258 | 1,60117E-45 |
| ACE2 | LRRC56 | 0,6258 | 1,60235E-45 |
| ACE2 | GPR143 | 0,6257 | 1,69335E-45 |
| ACE2 | KRTAP5-5 | 0,6256 | 1,71779E-45 |
| ACE2 | RP11-188C12.2 | 0,6253 | 1,94926E-45 |
| ACE2 | CCL24 | 0,6252 | 2,05893E-45 |
| ACE2 | REN | 0,625 | 2,21297E-45 |
| ACE2 | RP11-407N17.5 | 0,625 | 2,19335E-45 |
| ACE2 | RP11-93B14.6 | 0,625 | 2,24116E-45 |
| ACE2 | RP11-44F14.9 | 0,6248 | 2,44413E-45 |
| ACE2 | NPM1P32 | 0,6247 | 2,56744E-45 |
| ACE2 | SPHK2 | 0,6247 | 2,48192E-45 |
| ACE2 | SLC19A3 | 0,6246 | 2,62634E-45 |
| ACE2 | DAGLA | 0,6246 | 2,61028E-45 |
| ACE2 | FBXL14 | 0,6245 | 2,73889E-45 |
| ACE2 | U47924.29 | 0,6245 | 2,76808E-45 |
| ACE2 | DNAJB3 | 0,6244 | 2,85835E-45 |
| ACE2 | RP5-1125A11.6 | 0,624 | 3,31001E-45 |
| ACE2 | PLSCR2 | 0,6239 | 3,49211E-45 |
| ACE2 | OLFM4 | 0,6239 | 3,46739E-45 |
| ACE2 | HOXD1 | 0,6238 | 3,64746E-45 |
| ACE2 | KIF12 | 0,6234 | 4,37859E-45 |
| ACE2 | ARHGEF10L | 0,6232 | 4,69757E-45 |
| ACE2 | LINC00526 | 0,6229 | 5,32131E-45 |
| ACE2 | ATP5G1 | 0,6227 | 5,81666E-45 |
| ACE2 | CTA-929C8.6 | 0,6225 | 6,2441E-45 |
| ACE2 | NDUFS3 | 0,6224 | 6,47721E-45 |
| ACE2 | ACOT13 | 0,6223 | 6,72421E-45 |

| | | | |
|---|---|---|---|
| ACE2 | RUBCNL | 0,6222 | 6,95179E-45 |
| ACE2 | SLC1A5 | 0,6218 | 8,25832E-45 |
| ACE2 | RP11-104J23.2 | 0,6217 | 8,53996E-45 |
| ACE2 | LPAR2 | 0,6217 | 8,76022E-45 |
| ACE2 | SLC6A14 | 0,6216 | 9,14146E-45 |
| ACE2 | LRRC1 | 0,6215 | 9,37272E-45 |
| ACE2 | GHITM | 0,6213 | 1,01208E-44 |
| ACE2 | GJA9 | 0,6212 | 1,04901E-44 |
| ACE2 | RP11-523J2.1 | 0,6212 | 1,07318E-44 |
| ACE2 | TRIM2 | 0,6212 | 1,05131E-44 |
| ACE2 | AC000111.4 | 0,6211 | 1,11246E-44 |
| ACE2 | RP11-470M1.2 | 0,6211 | 1,10043E-44 |
| ACE2 | KRT8P24 | 0,6211 | 1,10967E-44 |
| ACE2 | LINC00940 | 0,6205 | 1,42738E-44 |
| ACE2 | TSGA10 | 0,6202 | 1,60218E-44 |
| ACE2 | DPEP1 | 0,6201 | 1,68399E-44 |
| ACE2 | SH3BGRL3 | 0,6199 | 1,83949E-44 |
| ACE2 | DENND2D | 0,6198 | 1,93045E-44 |
| ACE2 | UQCRFS1 | 0,6198 | 1,88285E-44 |
| ACE2 | RP11-396O20.1 | 0,6196 | 2,08154E-44 |
| ACE2 | RP11-314N13.9 | 0,6196 | 2,04848E-44 |
| ACE2 | RP5-1018K9.1 | 0,6193 | 2,36359E-44 |
| ACE2 | ABHD17B | 0,6192 | 2,45508E-44 |
| ACE2 | ONECUT2 | 0,6192 | 2,44688E-44 |
| ACE2 | RP11-290F24.6 | 0,6189 | 2,70411E-44 |
| ACE2 | KIF22 | 0,6189 | 2,7331E-44 |
| ACE2 | RP1-239B22.5 | 0,6188 | 2,80766E-44 |
| ACE2 | WWP2 | 0,6188 | 2,90881E-44 |
| ACE2 | GAK | 0,6186 | 3,08577E-44 |
| ACE2 | KB-1184D12.1 | 0,6186 | 3,14665E-44 |
| ACE2 | LINC01451 | 0,6183 | 3,5545E-44 |
| ACE2 | E2F8 | 0,6183 | 3,53695E-44 |
| ACE2 | PADI1 | 0,6181 | 3,78114E-44 |
| ACE2 | UMODL1 | 0,6178 | 4,21677E-44 |
| ACE2 | CTD-2574D22.3 | 0,6174 | 5,1127E-44 |
| ACE2 | TEX101 | 0,6174 | 4,94872E-44 |
| ACE2 | LINC01644 | 0,6174 | 4,95797E-44 |
| ACE2 | SLC20A1 | 0,6173 | 5,13851E-44 |
| ACE2 | RP11-151G14.2 | 0,6173 | 5,16203E-44 |
| ACE2 | RP11-83M16.6 | 0,6172 | 5,37771E-44 |
| ACE2 | RP11-510N19.5 | 0,6171 | 5,59567E-44 |
| ACE2 | SLC25A47P1 | 0,617 | 5,9981E-44 |
| ACE2 | CEACAM16 | 0,617 | 5,99988E-44 |
| ACE2 | MGLL | 0,6169 | 6,089E-44 |
| ACE2 | TMPRSS3 | 0,6165 | 7,09959E-44 |
| ACE2 | LMNB1 | 0,6164 | 7,61168E-44 |
| ACE2 | TMEM144 | 0,6163 | 7,79512E-44 |
| ACE2 | CROT | 0,6161 | 8,40303E-44 |
| ACE2 | CLUH | 0,6161 | 8,4503E-44 |
| ACE2 | ADM2 | 0,6161 | 8,4984E-44 |
| ACE2 | CIDEC | 0,616 | 8,93677E-44 |
| ACE2 | SLC17A1 | 0,6155 | 1,06869E-43 |
| ACE2 | HIGD1A | 0,6153 | 1,15617E-43 |
| ACE2 | RP11-255E6.5 | 0,615 | 1,29797E-43 |

| | | | |
|---|---|---|---|
| ACE2 | FAM135A | 0,615 | 1,31853E-43 |
| ACE2 | SH2D1B | 0,6148 | 1,41228E-43 |
| ACE2 | IL20RA | 0,6147 | 1,51361E-43 |
| ACE2 | ZNF503-AS1 | 0,6143 | 1,73991E-43 |
| ACE2 | RP3-323A16.1 | 0,6143 | 1,75396E-43 |
| ACE2 | SLC35B3 | 0,6138 | 2,16799E-43 |
| ACE2 | CTD-2280E9.1 | 0,6137 | 2,21943E-43 |
| ACE2 | CFL1 | 0,6134 | 2,47517E-43 |
| ACE2 | ZNF124 | 0,6133 | 2,62312E-43 |
| ACE2 | ATPIF1 | 0,613 | 2,98825E-43 |
| ACE2 | CLCA3P | 0,6129 | 3,11114E-43 |
| ACE2 | FANCF | 0,6128 | 3,15435E-43 |
| ACE2 | CTD-2555C10.3 | 0,6128 | 3,20049E-43 |
| ACE2 | IL23R | 0,6127 | 3,31299E-43 |
| ACE2 | HNRNPAB | 0,6125 | 3,54901E-43 |
| ACE2 | AC006042.6 | 0,6125 | 3,57326E-43 |
| ACE2 | DTX2 | 0,6124 | 3,7917E-43 |
| ACE2 | ADAM22 | 0,6123 | 3,95253E-43 |
| ACE2 | RIOK3 | 0,6123 | 3,89437E-43 |
| ACE2 | C5orf63 | 0,612 | 4,33568E-43 |
| ACE2 | C12orf74 | 0,612 | 4,42224E-43 |
| ACE2 | SLC12A8 | 0,6119 | 4,56261E-43 |
| ACE2 | FBXL6 | 0,6116 | 5,22328E-43 |
| ACE2 | UGT1A4 | 0,6115 | 5,40361E-43 |
| ACE2 | HCP5 | 0,6115 | 5,32057E-43 |
| ACE2 | KRT8P2 | 0,6115 | 5,39322E-43 |
| ACE2 | AKAP5 | 0,6112 | 6,0007E-43 |
| ACE2 | SLC9A1 | 0,6109 | 6,7203E-43 |
| ACE2 | HINT1 | 0,6109 | 6,74704E-43 |
| ACE2 | RP11-466A19.8 | 0,6106 | 7,50081E-43 |
| ACE2 | SLC31A2 | 0,6105 | 7,89496E-43 |
| ACE2 | FIBCD1 | 0,6104 | 8,10468E-43 |
| ACE2 | LINC02159 | 0,6103 | 8,54111E-43 |
| ACE2 | CCDC183 | 0,6099 | 1,0161E-42 |
| ACE2 | AADACP1 | 0,6095 | 1,18305E-42 |
| ACE2 | ESCO2 | 0,6095 | 1,17453E-42 |
| ACE2 | THEM6 | 0,6094 | 1,24646E-42 |
| ACE2 | SAPCD2 | 0,6094 | 1,212E-42 |
| ACE2 | RP4-612B15.3 | 0,6092 | 1,32966E-42 |
| ACE2 | RPH3AL | 0,6089 | 1,4915E-42 |
| ACE2 | NAPEPLD | 0,6086 | 1,68484E-42 |
| ACE2 | DECR2 | 0,6084 | 1,81615E-42 |
| ACE2 | RP11-597M12.2 | 0,6084 | 1,806E-42 |
| ACE2 | RP11-568K15.1 | 0,6083 | 1,92021E-42 |
| ACE2 | TFF2 | 0,6083 | 1,91093E-42 |
| ACE2 | DHCR7 | 0,6082 | 1,92949E-42 |
| ACE2 | MOCOS | 0,6082 | 1,96756E-42 |
| ACE2 | SDR42E1 | 0,6079 | 2,23371E-42 |
| ACE2 | RP11-429J17.7 | 0,6076 | 2,47312E-42 |
| ACE2 | ABCG8 | 0,6074 | 2,63515E-42 |
| ACE2 | EFCAB5 | 0,6073 | 2,83514E-42 |
| ACE2 | CYSTM1 | 0,6069 | 3,23354E-42 |
| ACE2 | MYRFL | 0,6069 | 3,27083E-42 |
| ACE2 | RP11-164P12.4 | 0,6065 | 3,80789E-42 |

| | | | |
|---|---|---|---|
| ACE2 | HTR3E | 0,6063 | 4,0906E-42 |
| ACE2 | IGHV3-79 | 0,6063 | 4,07142E-42 |
| ACE2 | RPSAP69 | 0,6062 | 4,29704E-42 |
| ACE2 | BORA | 0,6062 | 4,272E-42 |
| ACE2 | GOLGA2P5 | 0,6061 | 4,42537E-42 |
| ACE2 | DYRK2 | 0,606 | 4,70493E-42 |
| ACE2 | TNS4 | 0,606 | 4,56E-42 |
| ACE2 | TMEM41B | 0,6059 | 4,89513E-42 |
| ACE2 | MIR200B | 0,6058 | 4,98343E-42 |
| ACE2 | ORMDL2 | 0,6058 | 4,95819E-42 |
| ACE2 | CBR3-AS1 | 0,6058 | 4,97625E-42 |
| ACE2 | CAMK2N1 | 0,6057 | 5,27774E-42 |
| ACE2 | FAM171A1 | 0,6053 | 6,06728E-42 |
| ACE2 | APRT | 0,6053 | 6,10643E-42 |
| ACE2 | RP5-892K4.1 | 0,605 | 6,69684E-42 |
| ACE2 | GOLPH3L | 0,6046 | 8,0335E-42 |
| ACE2 | RP11-15F12.6 | 0,6045 | 8,36269E-42 |
| ACE2 | ASB13 | 0,6044 | 8,46415E-42 |
| ACE2 | ECI2 | 0,6043 | 8,7929E-42 |
| ACE2 | SPR | 0,604 | 1,0159E-41 |
| ACE2 | RP3-407E4.3 | 0,6038 | 1,09676E-41 |
| ACE2 | IL17RB | 0,6035 | 1,2108E-41 |
| ACE2 | PPP2R1B | 0,6035 | 1,20747E-41 |
| ACE2 | AP5M1 | 0,6035 | 1,21446E-41 |
| ACE2 | CAAP1 | 0,6034 | 1,27485E-41 |
| ACE2 | RP11-736N17.8 | 0,603 | 1,50026E-41 |
| ACE2 | MROH1 | 0,6027 | 1,63406E-41 |
| ACE2 | BLNK | 0,6026 | 1,73023E-41 |
| ACE2 | NR0B2 | 0,6025 | 1,81173E-41 |
| ACE2 | RP11-392P7.6 | 0,6023 | 1,96513E-41 |
| ACE2 | RP11-575A19.2 | 0,602 | 2,12341E-41 |
| ACE2 | C19orf57 | 0,6019 | 2,28313E-41 |
| ACE2 | PECR | 0,6017 | 2,38757E-41 |
| ACE2 | CTD-2184D3.3 | 0,6017 | 2,45164E-41 |
| ACE2 | C2CD4D | 0,6016 | 2,51378E-41 |
| ACE2 | RP11-119D9.1 | 0,6013 | 2,84976E-41 |
| ACE2 | RP11-226B15.1 | 0,6013 | 2,86439E-41 |
| ACE2 | KREMEN2 | 0,6012 | 2,89545E-41 |
| ACE2 | RP11-109E12.5 | 0,6009 | 3,33061E-41 |
| ACE2 | RPL3P13 | 0,6009 | 3,32082E-41 |
| ACE2 | TMEM98 | 0,6008 | 3,41819E-41 |
| ACE2 | ETF1P3 | 0,6006 | 3,7067E-41 |
| ACE2 | SCEL | 0,6005 | 3,80946E-41 |
| ACE2 | PP14571 | 0,6003 | 4,08558E-41 |
| ACE2 | TMEM210 | 0,6002 | 4,3755E-41 |
| ACE2 | RP11-43N5.1 | 0,6002 | 4,33889E-41 |
| ACE2 | NEK3 | 0,6 | 4,63014E-41 |
| ACE2 | YDJC | 0,6 | 4,71804E-41 |
| ACE2 | ZBED4 | 0,5996 | 5,48933E-41 |
| ACE2 | GNG13 | 0,5994 | 5,90933E-41 |
| ACE2 | PODXL2 | 0,5993 | 6,12542E-41 |
| ACE2 | SULT1E1 | 0,5993 | 6,07926E-41 |
| ACE2 | IKBKE | 0,5992 | 6,30775E-41 |
| ACE2 | SLC7A11 | 0,5991 | 6,42556E-41 |

| | | | |
|---|---|---|---|
| ACE2 | RP11-1143G9.5 | 0,5989 | 7,06185E-41 |
| ACE2 | ALS2CL | 0,5988 | 7,27457E-41 |
| ACE2 | IGSF23 | 0,5988 | 7,24879E-41 |
| ACE2 | DEPDC1 | 0,5986 | 7,9981E-41 |
| ACE2 | PITX2 | 0,5986 | 7,75932E-41 |
| ACE2 | CTD-2330K9.3 | 0,5985 | 8,20822E-41 |
| ACE2 | CCNB2 | 0,5985 | 8,23148E-41 |
| ACE2 | AP001627.1 | 0,5985 | 8,26657E-41 |
| ACE2 | RP11-293P20.2 | 0,5984 | 8,49649E-41 |
| ACE2 | CCDC175 | 0,5983 | 8,98015E-41 |
| ACE2 | HIST1H3G | 0,598 | 9,91183E-41 |
| ACE2 | KIF18A | 0,5979 | 1,02027E-40 |
| ACE2 | WIPI1 | 0,5978 | 1,06521E-40 |
| ACE2 | NEK2 | 0,5976 | 1,1583E-40 |
| ACE2 | PROC | 0,5976 | 1,17384E-40 |
| ACE2 | RP11-108L7.15 | 0,5976 | 1,15609E-40 |
| ACE2 | PLEKHG7 | 0,5974 | 1,26067E-40 |
| ACE2 | JPH1 | 0,5973 | 1,27718E-40 |
| ACE2 | MIR210HG | 0,5972 | 1,34149E-40 |
| ACE2 | CCNO | 0,5971 | 1,40487E-40 |
| ACE2 | GFER | 0,5971 | 1,39165E-40 |
| ACE2 | MEP1AP1 | 0,597 | 1,45919E-40 |
| ACE2 | SLC10A2 | 0,597 | 1,44534E-40 |
| ACE2 | ZBTB42 | 0,597 | 1,43966E-40 |
| ACE2 | TMEM87A | 0,597 | 1,45353E-40 |
| ACE2 | AC007009.1 | 0,5969 | 1,48211E-40 |
| ACE2 | CDC25A | 0,5968 | 1,55349E-40 |
| ACE2 | IL37 | 0,5967 | 1,59958E-40 |
| ACE2 | RP11-576I22.2 | 0,5966 | 1,67777E-40 |
| ACE2 | MKI67 | 0,5966 | 1,69039E-40 |
| ACE2 | CDC25C | 0,5965 | 1,75871E-40 |
| ACE2 | LINC00589 | 0,5965 | 1,71589E-40 |
| ACE2 | KIF21B | 0,5962 | 1,95442E-40 |
| ACE2 | RAB15 | 0,5962 | 1,96591E-40 |
| ACE2 | TPX2 | 0,5962 | 1,92929E-40 |
| ACE2 | KRT8P5 | 0,5959 | 2,14453E-40 |
| ACE2 | DHRS4L1 | 0,5958 | 2,30245E-40 |
| ACE2 | FDX1 | 0,5957 | 2,32239E-40 |
| ACE2 | SLC45A3 | 0,5956 | 2,48595E-40 |
| ACE2 | RP11-341G23.3 | 0,5956 | 2,47313E-40 |
| ACE2 | NDUFA10 | 0,5953 | 2,76984E-40 |
| ACE2 | COX8A | 0,5951 | 2,90476E-40 |
| ACE2 | AURKB | 0,5951 | 2,91828E-40 |
| ACE2 | RP11-20G13.1 | 0,595 | 3,0239E-40 |
| ACE2 | IP6K2 | 0,5943 | 3,94145E-40 |
| ACE2 | RP1-92O14.3 | 0,5939 | 4,65874E-40 |
| ACE2 | LINC01923 | 0,5938 | 4,72187E-40 |
| ACE2 | AGBL2 | 0,5938 | 4,80388E-40 |
| ACE2 | ASPM | 0,5937 | 5,00131E-40 |
| ACE2 | NR6A1 | 0,5937 | 4,8869E-40 |
| ACE2 | UNC93B1 | 0,5937 | 4,96011E-40 |
| ACE2 | RP11-223C24.1 | 0,5936 | 5,14736E-40 |
| ACE2 | RP11-83M16.3 | 0,5934 | 5,47147E-40 |
| ACE2 | XXyac-YR21CG7.1 | 0,5932 | 5,87348E-40 |

| | | | |
|---|---|---|---|
| ACE2 | ULK3 | 0,5932 | 5,91117E-40 |
| ACE2 | APEX2 | 0,5932 | 5,89531E-40 |
| ACE2 | LYPD6B | 0,5931 | 6,20229E-40 |
| ACE2 | NUDT16L1 | 0,5929 | 6,63407E-40 |
| ACE2 | RP4-657E11.10 | 0,5928 | 7,0384E-40 |
| ACE2 | RP6-65G23.3 | 0,5928 | 6,89457E-40 |
| ACE2 | TOP2A | 0,5928 | 6,90023E-40 |
| ACE2 | PLK1 | 0,5926 | 7,54549E-40 |
| ACE2 | C11orf54 | 0,5924 | 8,10691E-40 |
| ACE2 | PTBP3 | 0,5923 | 8,36957E-40 |
| ACE2 | NCLN | 0,5923 | 8,24313E-40 |
| ACE2 | PDCD6IP | 0,5922 | 8,77843E-40 |
| ACE2 | ASS1 | 0,5922 | 8,62996E-40 |
| ACE2 | OR2A1-AS1 | 0,592 | 9,44481E-40 |
| ACE2 | PRR13P5 | 0,592 | 9,20922E-40 |
| ACE2 | CTA-398F10.2 | 0,5918 | 1,02303E-39 |
| ACE2 | TMPRSS7 | 0,5916 | 1,09042E-39 |
| ACE2 | RAB8A | 0,5916 | 1,06657E-39 |
| ACE2 | MAGI3 | 0,5913 | 1,19345E-39 |
| ACE2 | CDC20 | 0,5912 | 1,24024E-39 |
| ACE2 | AP003774.1 | 0,5908 | 1,46249E-39 |
| ACE2 | AC009133.17 | 0,5908 | 1,46718E-39 |
| ACE2 | PTGDR | 0,5907 | 1,48329E-39 |
| ACE2 | ZNF468 | 0,5907 | 1,50541E-39 |
| ACE2 | QARS | 0,5905 | 1,62514E-39 |
| ACE2 | IRF8 | 0,5905 | 1,62164E-39 |
| ACE2 | HAGLR | 0,5901 | 1,9076E-39 |
| ACE2 | AP006621.8 | 0,5897 | 2,15579E-39 |
| ACE2 | PYM1 | 0,5897 | 2,14175E-39 |
| ACE2 | FSIP1 | 0,5897 | 2,18343E-39 |
| ACE2 | MT-TS1 | 0,5897 | 2,16664E-39 |
| ACE2 | HMGA1 | 0,5895 | 2,35672E-39 |
| ACE2 | HENMT1 | 0,5894 | 2,42477E-39 |
| ACE2 | PRDX5 | 0,5888 | 3,00815E-39 |
| ACE2 | ABCC6P2 | 0,5888 | 2,9814E-39 |
| ACE2 | AC005281.2 | 0,5885 | 3,42183E-39 |
| ACE2 | LINC00488 | 0,5882 | 3,80042E-39 |
| ACE2 | CELP | 0,5881 | 3,93254E-39 |
| ACE2 | CTD-2331H12.7 | 0,5881 | 3,92788E-39 |
| ACE2 | DDAH1 | 0,5874 | 4,96053E-39 |
| ACE2 | KLHL31 | 0,587 | 5,89564E-39 |
| ACE2 | ADGRG1 | 0,5867 | 6,48572E-39 |
| ACE2 | FAM71E2 | 0,5867 | 6,49299E-39 |
| ACE2 | MT-CO2 | 0,5864 | 7,30678E-39 |
| ACE2 | SPC25 | 0,5862 | 7,6845E-39 |
| ACE2 | KIFC1 | 0,5862 | 7,88781E-39 |
| ACE2 | RTN4IP1 | 0,5861 | 8,06103E-39 |
| ACE2 | TMEM52 | 0,586 | 8,32635E-39 |
| ACE2 | F12 | 0,5859 | 8,78608E-39 |
| ACE2 | TCP10L2 | 0,5859 | 8,68701E-39 |
| ACE2 | CTAGE4 | 0,5859 | 8,69971E-39 |
| ACE2 | MAIP1 | 0,5857 | 9,51513E-39 |
| ACE2 | POC1A | 0,5855 | 9,96591E-39 |
| ACE2 | BCL2L2-PABPN1 | 0,5855 | 1,01469E-38 |

| | | | |
|---|---|---|---|
| ACE2 | CPHL1P | 0,5851 | 1,15279E-38 |
| ACE2 | RP11-545E17.3 | 0,585 | 1,19565E-38 |
| ACE2 | MCM4 | 0,5849 | 1,2712E-38 |
| ACE2 | RP11-67L14.1 | 0,5847 | 1,35308E-38 |
| ACE2 | MYL12B | 0,5846 | 1,39139E-38 |
| ACE2 | UGT2B11 | 0,5845 | 1,44259E-38 |
| ACE2 | RP11-542B15.1 | 0,5845 | 1,42635E-38 |
| ACE2 | RP11-22C11.1 | 0,5843 | 1,53211E-38 |
| ACE2 | TEP1 | 0,5839 | 1,80554E-38 |
| ACE2 | PCSK5 | 0,5838 | 1,86628E-38 |
| ACE2 | DMRT2 | 0,5831 | 2,36522E-38 |
| ACE2 | VWA5A | 0,583 | 2,4945E-38 |
| ACE2 | TAS2R38 | 0,5828 | 2,62749E-38 |
| ACE2 | RP11-217B1.2 | 0,5828 | 2,62126E-38 |
| ACE2 | NOL4L | 0,5828 | 2,64756E-38 |
| ACE2 | PCSK9 | 0,5826 | 2,89642E-38 |
| ACE2 | RP11-324O2.3 | 0,5826 | 2,83292E-38 |
| ACE2 | AC093162.5 | 0,5825 | 2,99465E-38 |
| ACE2 | CHKA | 0,5825 | 2,91984E-38 |
| ACE2 | KRT40 | 0,5825 | 2,97807E-38 |
| ACE2 | HOXB8 | 0,5825 | 2,96626E-38 |
| ACE2 | OIP5 | 0,5824 | 3,03687E-38 |
| ACE2 | RP11-232L2.2 | 0,5823 | 3,14008E-38 |
| ACE2 | RP3-395M20.12 | 0,5822 | 3,3043E-38 |
| ACE2 | C20orf24 | 0,5822 | 3,30604E-38 |
| ACE2 | GS1-393G12.14 | 0,5821 | 3,46929E-38 |
| ACE2 | RP11-884K10.6 | 0,582 | 3,51504E-38 |
| ACE2 | LHFPL3-AS2 | 0,582 | 3,53836E-38 |
| ACE2 | RAD51 | 0,5819 | 3,69961E-38 |
| ACE2 | PGM1 | 0,5817 | 3,89539E-38 |
| ACE2 | MTIF2 | 0,5817 | 3,97314E-38 |
| ACE2 | DENND1C | 0,5817 | 3,87556E-38 |
| ACE2 | CSTF2 | 0,5816 | 4,0106E-38 |
| ACE2 | MRPL12 | 0,5815 | 4,16194E-38 |
| ACE2 | PDXP | 0,5815 | 4,23754E-38 |
| ACE2 | MIR135A1 | 0,5812 | 4,7239E-38 |
| ACE2 | FLNB | 0,5812 | 4,62909E-38 |
| ACE2 | KRT18P10 | 0,581 | 4,98291E-38 |
| ACE2 | SEMA5A | 0,5804 | 6,17848E-38 |
| ACE2 | BRPF3 | 0,5804 | 6,17497E-38 |
| ACE2 | RPL7P16 | 0,5801 | 6,97329E-38 |
| ACE2 | MIR200A | 0,58 | 7,32616E-38 |
| ACE2 | MELK | 0,5799 | 7,51037E-38 |
| ACE2 | NADSYN1 | 0,5798 | 7,75275E-38 |
| ACE2 | PDXDC2P | 0,5798 | 7,63174E-38 |
| ACE2 | SH3KBP1 | 0,5797 | 7,93682E-38 |
| ACE2 | SLC9C1 | 0,5796 | 8,25465E-38 |
| ACE2 | AP000619.5 | 0,5796 | 8,28016E-38 |
| ACE2 | SQLE | 0,5795 | 8,56947E-38 |
| ACE2 | TKFC | 0,5794 | 8,87628E-38 |
| ACE2 | PPP1R26-AS1 | 0,5792 | 9,65438E-38 |
| ACE2 | LINC02014 | 0,5791 | 9,82568E-38 |
| ACE2 | ARRDC3-AS1 | 0,5787 | 1,13533E-37 |
| ACE2 | PARP12 | 0,5787 | 1,12962E-37 |

| | | | |
|---|---|---|---|
| ACE2 | RP11-649A18.3 | 0,5787 | 1,12845E-37 |
| ACE2 | CTA-228A9.3 | 0,5785 | 1,21205E-37 |
| ACE2 | RFK | 0,5784 | 1,27226E-37 |
| ACE2 | LGR5 | 0,5781 | 1,42519E-37 |
| ACE2 | RN7SL619P | 0,5781 | 1,39076E-37 |
| ACE2 | ATP6V0D2 | 0,578 | 1,46044E-37 |
| ACE2 | INIP | 0,578 | 1,48278E-37 |
| ACE2 | SNHG4 | 0,5779 | 1,50346E-37 |
| ACE2 | SH2D7 | 0,5779 | 1,52085E-37 |
| ACE2 | AC098828.3 | 0,5775 | 1,73102E-37 |
| ACE2 | ENPP5 | 0,5774 | 1,77717E-37 |
| ACE2 | TTK | 0,5774 | 1,82663E-37 |
| ACE2 | CNPPD1 | 0,5773 | 1,8645E-37 |
| ACE2 | PF4 | 0,5772 | 1,9529E-37 |
| ACE2 | GLRX | 0,5771 | 2,01346E-37 |
| ACE2 | CHCHD10 | 0,577 | 2,09769E-37 |
| ACE2 | EXOC3L4 | 0,5767 | 2,34781E-37 |
| ACE2 | CENPF | 0,5765 | 2,49224E-37 |
| ACE2 | ELOVL7 | 0,5765 | 2,4486E-37 |
| ACE2 | CTD-2235C13.3 | 0,5764 | 2,5703E-37 |
| ACE2 | OR7E91P | 0,5762 | 2,7473E-37 |
| ACE2 | RALGPS1 | 0,5761 | 2,84647E-37 |
| ACE2 | AC073900.4 | 0,576 | 2,99023E-37 |
| ACE2 | FAM189A1 | 0,576 | 2,98934E-37 |
| ACE2 | TINCR | 0,5759 | 3,0569E-37 |
| ACE2 | RP11-538D16.3 | 0,5757 | 3,31423E-37 |
| ACE2 | NPHS2 | 0,5754 | 3,68171E-37 |
| ACE2 | FDFT1 | 0,5754 | 3,6646E-37 |
| ACE2 | HMMR | 0,5753 | 3,73614E-37 |
| ACE2 | DUSP16 | 0,5753 | 3,79717E-37 |
| ACE2 | LGALS3BP | 0,5751 | 4,07805E-37 |
| ACE2 | S100A16 | 0,5745 | 4,99475E-37 |
| ACE2 | MYEOV | 0,574 | 5,95975E-37 |
| ACE2 | LHFPL3 | 0,5738 | 6,29848E-37 |
| ACE2 | LDHAL6B | 0,5729 | 8,58056E-37 |
| ACE2 | RN7SL728P | 0,5724 | 1,03941E-36 |
| ACE2 | CD1D | 0,572 | 1,19766E-36 |
| ACE2 | TRIM25 | 0,5719 | 1,2145E-36 |
| ACE2 | NIPSNAP1 | 0,5719 | 1,22985E-36 |
| ACE2 | RP11-48B14.1 | 0,5716 | 1,37491E-36 |
| ACE2 | RALGAPA2 | 0,5715 | 1,4093E-36 |
| ACE2 | RP11-24F11.2 | 0,5714 | 1,44753E-36 |
| ACE2 | NEIL3 | 0,5713 | 1,51797E-36 |
| ACE2 | UGP2 | 0,5712 | 1,54253E-36 |
| ACE2 | CEP85 | 0,5709 | 1,73442E-36 |
| ACE2 | PKD2L1 | 0,5709 | 1,75731E-36 |
| ACE2 | PBK | 0,5708 | 1,7904E-36 |
| ACE2 | PTPRO | 0,5708 | 1,77866E-36 |
| ACE2 | AC004066.3 | 0,5707 | 1,83683E-36 |
| ACE2 | RP11-26J3.3 | 0,5707 | 1,8864E-36 |
| ACE2 | CTD-2540B15.9 | 0,5707 | 1,86955E-36 |
| ACE2 | KIAA1456 | 0,5706 | 1,91562E-36 |
| ACE2 | IMPA1P1 | 0,5706 | 1,90443E-36 |
| ACE2 | LINC01978 | 0,5702 | 2,20042E-36 |

| | | | |
|---|---|---|---|
| ACE2 | KB-1460A1.1 | 0,5701 | 2,25408E-36 |
| ACE2 | DNAJC11 | 0,5698 | 2,56912E-36 |
| ACE2 | RP11-796E10.1 | 0,5698 | 2,50122E-36 |
| ACE2 | KRT18P34 | 0,5698 | 2,49909E-36 |
| ACE2 | AP1AR | 0,5695 | 2,79615E-36 |
| ACE2 | AC129492.6 | 0,5695 | 2,80035E-36 |
| ACE2 | FCGRT | 0,5695 | 2,83758E-36 |
| ACE2 | LINC01659 | 0,5695 | 2,82582E-36 |
| ACE2 | SYNPR-AS1 | 0,5693 | 3,04324E-36 |
| ACE2 | RP11-452I5.2 | 0,5693 | 3,04494E-36 |
| ACE2 | CTC-490G23.2 | 0,5693 | 3,03183E-36 |
| ACE2 | GLB1 | 0,5691 | 3,19595E-36 |
| ACE2 | LINC01443 | 0,5691 | 3,26315E-36 |
| ACE2 | TMIGD2 | 0,5689 | 3,40094E-36 |
| ACE2 | FAM95B1 | 0,5687 | 3,64524E-36 |
| ACE2 | PMFBP1 | 0,5686 | 3,78506E-36 |
| ACE2 | RP11-739N20.2 | 0,5685 | 3,99182E-36 |
| ACE2 | NCAPH | 0,5685 | 3,91526E-36 |
| ACE2 | AC009133.21 | 0,5684 | 4,08989E-36 |
| ACE2 | RP11-807H22.6 | 0,5683 | 4,21832E-36 |
| ACE2 | RP11-626G11.1 | 0,5682 | 4,3824E-36 |
| ACE2 | PDP2 | 0,5682 | 4,41349E-36 |
| ACE2 | LINC01344 | 0,5678 | 4,93272E-36 |
| ACE2 | CTC-550B14.7 | 0,5678 | 5,07584E-36 |
| ACE2 | CNFN | 0,5677 | 5,10704E-36 |
| ACE2 | HOXB13 | 0,5676 | 5,30471E-36 |
| ACE2 | RP11-734K21.3 | 0,5673 | 5,83529E-36 |
| ACE2 | PHYKPL | 0,5672 | 6,16587E-36 |
| ACE2 | GTF2IP7 | 0,5669 | 6,78621E-36 |
| ACE2 | EZH2 | 0,5669 | 6,73491E-36 |
| ACE2 | PTPRR | 0,5666 | 7,52328E-36 |
| ACE2 | BUB1 | 0,5665 | 7,9065E-36 |
| ACE2 | GRID2IP | 0,5664 | 8,11225E-36 |
| ACE2 | KLRC1 | 0,5663 | 8,30714E-36 |
| ACE2 | DGUOK-AS1 | 0,5662 | 8,57159E-36 |
| ACE2 | YAP1P1 | 0,5662 | 8,72864E-36 |
| ACE2 | POR | 0,5662 | 8,52861E-36 |
| ACE2 | GON7 | 0,5662 | 8,61696E-36 |
| ACE2 | SLC44A3-AS1 | 0,566 | 9,35516E-36 |
| ACE2 | RP11-44F21.3 | 0,566 | 9,27702E-36 |
| ACE2 | AC098828.2 | 0,5659 | 9,51282E-36 |
| ACE2 | RP1-8B1.4 | 0,5659 | 9,58349E-36 |
| ACE2 | NOS2 | 0,5659 | 9,56927E-36 |
| ACE2 | RALY-AS1 | 0,5659 | 9,5317E-36 |
| ACE2 | BUB1B | 0,5658 | 9,6911E-36 |
| ACE2 | PAK6 | 0,5658 | 9,9854E-36 |
| ACE2 | RP11-347C12.10 | 0,5657 | 1,01183E-35 |
| ACE2 | GLRA4 | 0,5655 | 1,08526E-35 |
| ACE2 | TMEM131 | 0,5654 | 1,14245E-35 |
| ACE2 | NPIPB14P | 0,5653 | 1,1526E-35 |
| ACE2 | PRDX6 | 0,565 | 1,27267E-35 |
| ACE2 | NADK | 0,5649 | 1,31509E-35 |
| ACE2 | AP1S3 | 0,5649 | 1,32242E-35 |
| ACE2 | ENC1 | 0,5649 | 1,33621E-35 |

| | | | |
|---|---|---|---|
| ACE2 | ZWINT | 0,5648 | 1,4039E-35 |
| ACE2 | RP11-416N2.3 | 0,5646 | 1,49138E-35 |
| ACE2 | RP1-247F3.1 | 0,5645 | 1,54655E-35 |
| ACE2 | MRPS35 | 0,5644 | 1,59518E-35 |
| ACE2 | CYP2D6 | 0,5642 | 1,71327E-35 |
| ACE2 | CCNB1 | 0,5638 | 1,94231E-35 |
| ACE2 | SLC39A2 | 0,5638 | 1,955E-35 |
| ACE2 | RP11-172H24.4 | 0,5636 | 2,10087E-35 |
| ACE2 | TIGD6 | 0,5635 | 2,15639E-35 |
| ACE2 | GSG2 | 0,5635 | 2,1383E-35 |
| ACE2 | ASF1B | 0,5635 | 2,15769E-35 |
| ACE2 | PCSK6 | 0,5633 | 2,29976E-35 |
| ACE2 | COQ2 | 0,563 | 2,54669E-35 |
| ACE2 | RP11-5O24.1 | 0,5628 | 2,68654E-35 |
| ACE2 | CTC-428G20.1 | 0,5628 | 2,66335E-35 |
| ACE2 | RP11-867O8.5 | 0,5627 | 2,75617E-35 |
| ACE2 | CCDC24 | 0,5625 | 2,97432E-35 |
| ACE2 | KIF14 | 0,5624 | 3,06672E-35 |
| ACE2 | PMEL | 0,5624 | 3,06043E-35 |
| ACE2 | MICU2 | 0,5624 | 3,13049E-35 |
| ACE2 | STARD4 | 0,5623 | 3,19241E-35 |
| ACE2 | HOXA-AS3 | 0,5622 | 3,2686E-35 |
| ACE2 | PEX11G | 0,5621 | 3,40039E-35 |
| ACE2 | RP11-460I13.2 | 0,5618 | 3,73724E-35 |
| ACE2 | S100A10 | 0,5618 | 3,81156E-35 |
| ACE2 | APAF1 | 0,5618 | 3,74138E-35 |
| ACE2 | RP11-111E14.1 | 0,5617 | 3,84364E-35 |
| ACE2 | C19orf73 | 0,5617 | 3,89161E-35 |
| ACE2 | RNFT1 | 0,5616 | 4,09593E-35 |
| ACE2 | RP11-44N21.1 | 0,5611 | 4,77454E-35 |
| ACE2 | SPAG5 | 0,5611 | 4,75472E-35 |
| ACE2 | RP11-138I1.3 | 0,561 | 5,00168E-35 |
| ACE2 | SLC6A18 | 0,5605 | 5,89236E-35 |
| ACE2 | AC008937.3 | 0,5605 | 5,8397E-35 |
| ACE2 | BSG | 0,5605 | 5,7951E-35 |
| ACE2 | ANKRD62 | 0,5603 | 6,15011E-35 |
| ACE2 | MOBP | 0,5602 | 6,48656E-35 |
| ACE2 | NSF | 0,5602 | 6,392E-35 |
| ACE2 | RPL29P30 | 0,5599 | 7,09196E-35 |
| ACE2 | LGALSL | 0,5598 | 7,21449E-35 |
| ACE2 | RP3-355L5.4 | 0,5597 | 7,63981E-35 |
| ACE2 | LGALS9DP | 0,5597 | 7,63489E-35 |
| ACE2 | MAST3 | 0,5596 | 7,87732E-35 |
| ACE2 | LPAR3 | 0,5594 | 8,50255E-35 |
| ACE2 | GBP3 | 0,5592 | 8,85545E-35 |
| ACE2 | IGHV7-81 | 0,5592 | 8,85096E-35 |
| ACE2 | GRIN1 | 0,5591 | 9,17803E-35 |
| ACE2 | RP5-823G15.5 | 0,559 | 9,46486E-35 |
| ACE2 | HMGA1P5 | 0,5589 | 9,81991E-35 |
| ACE2 | SPPL2B | 0,5587 | 1,05503E-34 |
| ACE2 | RP11-1149O23.3 | 0,5585 | 1,1356E-34 |
| ACE2 | USP2 | 0,5585 | 1,11783E-34 |
| ACE2 | ANKRD18A | 0,5584 | 1,14917E-34 |
| ACE2 | UGT1A5 | 0,5581 | 1,2824E-34 |

| | | | |
|---|---|---|---|
| ACE2 | RGP1 | 0,5581 | 1,28841E-34 |
| ACE2 | RP11-28H5.2 | 0,558 | 1,31545E-34 |
| ACE2 | RP1-228P16.5 | 0,5578 | 1,43257E-34 |
| ACE2 | RP11-104N10.2 | 0,5578 | 1,4265E-34 |
| ACE2 | PHLDA2 | 0,5577 | 1,45875E-34 |
| ACE2 | SPATA2L | 0,5577 | 1,47447E-34 |
| ACE2 | CTA-221G9.7 | 0,5577 | 1,4891E-34 |
| ACE2 | IL1RN | 0,5573 | 1,69106E-34 |
| ACE2 | NIPA2 | 0,5573 | 1,67823E-34 |
| ACE2 | DOT1L | 0,5573 | 1,65902E-34 |
| ACE2 | ABHD12B | 0,5566 | 2,11063E-34 |
| ACE2 | DLGAP5 | 0,5564 | 2,23455E-34 |
| ACE2 | RP11-103J8.1 | 0,5561 | 2,47643E-34 |
| ACE2 | CT62 | 0,5558 | 2,71575E-34 |
| ACE2 | RP11-700H6.1 | 0,5556 | 2,90168E-34 |
| ACE2 | LINC01648 | 0,5555 | 2,99188E-34 |
| ACE2 | UCN3 | 0,5554 | 3,10619E-34 |
| ACE2 | RP11-290F5.2 | 0,5553 | 3,20576E-34 |
| ACE2 | KRT8P14 | 0,5553 | 3,25889E-34 |
| ACE2 | AKR1C2 | 0,5552 | 3,31164E-34 |
| ACE2 | KIF11 | 0,5552 | 3,32881E-34 |
| ACE2 | COASY | 0,5546 | 3,97767E-34 |
| ACE2 | AC114812.9 | 0,5542 | 4,53231E-34 |
| ACE2 | MROH6 | 0,5542 | 4,58791E-34 |
| ACE2 | DLG3-AS1 | 0,5542 | 4,63619E-34 |
| ACE2 | CASP8 | 0,5541 | 4,81904E-34 |
| ACE2 | TEN1 | 0,554 | 4,85474E-34 |
| ACE2 | CLEC19A | 0,5539 | 5,15507E-34 |
| ACE2 | ZNF648 | 0,5537 | 5,32942E-34 |
| ACE2 | HSD17B4 | 0,5537 | 5,36714E-34 |
| ACE2 | RP11-677M14.2 | 0,5534 | 5,94694E-34 |
| ACE2 | EAPP | 0,5534 | 5,93744E-34 |
| ACE2 | TIMM13 | 0,5533 | 6,25988E-34 |
| ACE2 | ACTL10 | 0,5533 | 6,21344E-34 |
| ACE2 | HEPH | 0,5533 | 6,22123E-34 |
| ACE2 | RTKN2 | 0,5531 | 6,6176E-34 |
| ACE2 | KCNK5 | 0,5529 | 7,02837E-34 |
| ACE2 | ACTG1P22 | 0,5528 | 7,21474E-34 |
| ACE2 | ACOXL | 0,5528 | 7,19642E-34 |
| ACE2 | PAUPAR | 0,5528 | 7,24105E-34 |
| ACE2 | BRIP1 | 0,5528 | 7,34394E-34 |
| ACE2 | AKAP9 | 0,5527 | 7,59051E-34 |
| ACE2 | CTC-391G2.1 | 0,5526 | 7,63904E-34 |
| ACE2 | RPSAP46 | 0,5524 | 8,35209E-34 |
| ACE2 | HORMAD1 | 0,5523 | 8,50504E-34 |
| ACE2 | TRPM5 | 0,5523 | 8,58294E-34 |
| ACE2 | SHPK | 0,5522 | 8,78183E-34 |
| ACE2 | RP11-443B7.1 | 0,5521 | 9,0175E-34 |
| ACE2 | UQCRQ | 0,552 | 9,46727E-34 |
| ACE2 | WFDC12 | 0,552 | 9,45839E-34 |
| ACE2 | ARL4A | 0,5519 | 9,68939E-34 |
| ACE2 | PUDP | 0,5516 | 1,05782E-33 |
| ACE2 | KDM4A | 0,5515 | 1,09192E-33 |
| ACE2 | NEK8 | 0,5515 | 1,11893E-33 |

| | | | |
|---|---|---|---|
| ACE2 | SPATA31C2 | 0,5513 | 1,17431E-33 |
| ACE2 | GTSE1 | 0,5513 | 1,15937E-33 |
| ACE2 | CACNA2D4 | 0,5511 | 1,26578E-33 |
| ACE2 | RP11-671J11.4 | 0,5507 | 1,40458E-33 |
| ACE2 | TAT-AS1 | 0,5507 | 1,41963E-33 |
| ACE2 | SH2D6 | 0,5503 | 1,63719E-33 |
| ACE2 | CDCA7 | 0,5503 | 1,61406E-33 |
| ACE2 | RP4-694A7.2 | 0,5502 | 1,68131E-33 |
| ACE2 | MIGA2 | 0,5502 | 1,65206E-33 |
| ACE2 | FANK1 | 0,5501 | 1,71004E-33 |
| ACE2 | COX5B | 0,5495 | 2,06833E-33 |
| ACE2 | KRT8P9 | 0,5495 | 2,10718E-33 |
| ACE2 | AC000068.5 | 0,5495 | 2,09706E-33 |
| ACE2 | SOX9-AS1 | 0,5492 | 2,31988E-33 |
| ACE2 | AC104667.3 | 0,5491 | 2,37444E-33 |
| ACE2 | RP1-118J21.25 | 0,549 | 2,43101E-33 |
| ACE2 | RP11-304L19.1 | 0,5489 | 2,51502E-33 |
| ACE2 | AL121578.7 | 0,5489 | 2,56107E-33 |
| ACE2 | CTA-929C8.7 | 0,5488 | 2,58573E-33 |
| ACE2 | RP11-771K4.1 | 0,5486 | 2,81009E-33 |
| ACE2 | ID1 | 0,5486 | 2,788E-33 |
| ACE2 | MAD2L1 | 0,5484 | 2,99356E-33 |
| ACE2 | EVX1-AS | 0,5483 | 3,08028E-33 |
| ACE2 | CHRM5 | 0,5482 | 3,21002E-33 |
| ACE2 | CDT1 | 0,5481 | 3,23924E-33 |
| ACE2 | KLRC3 | 0,548 | 3,42168E-33 |
| ACE2 | GPR55 | 0,5477 | 3,70043E-33 |
| ACE2 | RP11-599B13.9 | 0,5477 | 3,66244E-33 |
| ACE2 | MIR3142HG | 0,5474 | 4,14231E-33 |
| ACE2 | MT-TC | 0,5474 | 4,07136E-33 |
| ACE2 | AGAP1-IT1 | 0,5472 | 4,32694E-33 |
| ACE2 | ACP2 | 0,5472 | 4,39114E-33 |
| ACE2 | RP11-431M3.1 | 0,547 | 4,69448E-33 |
| ACE2 | AC107072.2 | 0,5468 | 4,96165E-33 |
| ACE2 | ANKRD20A21P | 0,5468 | 4,92984E-33 |
| ACE2 | AC094019.4 | 0,5466 | 5,30308E-33 |
| ACE2 | LACTB | 0,5466 | 5,29015E-33 |
| ACE2 | LINC00114 | 0,5466 | 5,23186E-33 |
| ACE2 | RP11-14C10.4 | 0,5463 | 5,84068E-33 |
| ACE2 | MGST3 | 0,5462 | 5,98037E-33 |
| ACE2 | H2AFY | 0,5462 | 5,91529E-33 |
| ACE2 | FAM81A | 0,5462 | 5,99635E-33 |
| ACE2 | ACER3 | 0,5461 | 6,13883E-33 |
| ACE2 | AC017060.1 | 0,5454 | 7,78611E-33 |
| ACE2 | RP11-285E9.5 | 0,5454 | 7,64117E-33 |
| ACE2 | TLDC2 | 0,5454 | 7,67098E-33 |
| ACE2 | TSKU | 0,5452 | 8,15619E-33 |
| ACE2 | RP1-47M23.3 | 0,5451 | 8,5241E-33 |
| ACE2 | PGD | 0,5448 | 9,34949E-33 |
| ACE2 | CDCA2 | 0,5448 | 9,24776E-33 |
| ACE2 | BIRC5 | 0,5448 | 9,33337E-33 |
| ACE2 | CYP4F11 | 0,5448 | 9,34618E-33 |
| ACE2 | MICAL1 | 0,5445 | 1,01326E-32 |
| ACE2 | ABCA13 | 0,5444 | 1,06596E-32 |

| | | | |
|---|---|---|---|
| ACE2 | SEZ6L2 | 0,5442 | 1,12013E-32 |
| ACE2 | LIMA1 | 0,544 | 1,18153E-32 |
| ACE2 | PKMYT1 | 0,5439 | 1,22535E-32 |
| ACE2 | RP11-55L4.2 | 0,5439 | 1,21777E-32 |
| ACE2 | PTPN22 | 0,5438 | 1,29199E-32 |
| ACE2 | HMGA1P4 | 0,5438 | 1,2809E-32 |
| ACE2 | KIF4A | 0,5436 | 1,35972E-32 |
| ACE2 | SLC25A15 | 0,5434 | 1,45191E-32 |
| ACE2 | C19orf54 | 0,5431 | 1,56404E-32 |
| ACE2 | KLK4 | 0,5429 | 1,70218E-32 |
| ACE2 | NLRP2 | 0,5429 | 1,70774E-32 |
| ACE2 | GPR82 | 0,5428 | 1,74036E-32 |
| ACE2 | C7orf31 | 0,5427 | 1,81212E-32 |
| ACE2 | NPAS2 | 0,5426 | 1,863E-32 |
| ACE2 | HOXA7 | 0,5426 | 1,83068E-32 |
| ACE2 | PPIP5K1 | 0,5426 | 1,83643E-32 |
| ACE2 | LINC01079 | 0,5422 | 2,09416E-32 |
| ACE2 | IPMK | 0,542 | 2,24637E-32 |
| ACE2 | CPA2 | 0,5419 | 2,2811E-32 |
| ACE2 | NUDT19 | 0,5418 | 2,3607E-32 |
| ACE2 | CXCR2P1 | 0,5417 | 2,48321E-32 |
| ACE2 | RP1-278O22.2 | 0,5416 | 2,56515E-32 |
| ACE2 | ACOT8 | 0,5416 | 2,54193E-32 |
| ACE2 | RP11-115J16.2 | 0,5415 | 2,59642E-32 |
| ACE2 | SYT17 | 0,5415 | 2,63875E-32 |
| ACE2 | RP11-432I5.8 | 0,5415 | 2,6431E-32 |
| ACE2 | CENPA | 0,5413 | 2,74524E-32 |
| ACE2 | FBXL19-AS1 | 0,541 | 3,07653E-32 |
| ACE2 | ARMC7 | 0,541 | 3,00964E-32 |
| ACE2 | CRYBB3 | 0,541 | 3,01158E-32 |
| ACE2 | KIF23 | 0,5409 | 3,16789E-32 |
| ACE2 | POLQ | 0,5408 | 3,21468E-32 |
| ACE2 | RP4-680D5.8 | 0,5406 | 3,48592E-32 |
| ACE2 | RP11-95G17.2 | 0,5406 | 3,46618E-32 |
| ACE2 | CTB-134H23.3 | 0,5405 | 3,62395E-32 |
| ACE2 | CCAT1 | 0,5404 | 3,72452E-32 |
| ACE2 | RP11-10J21.5 | 0,5403 | 3,77033E-32 |
| ACE2 | MET | 0,5399 | 4,23587E-32 |
| ACE2 | UGT1A2P | 0,5398 | 4,46562E-32 |
| ACE2 | BAX | 0,5396 | 4,75131E-32 |
| ACE2 | DUSP21 | 0,5396 | 4,74312E-32 |
| ACE2 | ORC1 | 0,5395 | 4,82656E-32 |
| ACE2 | RP11-64C12.4 | 0,5393 | 5,15046E-32 |
| ACE2 | PARS2 | 0,5392 | 5,30449E-32 |
| ACE2 | PRELID1 | 0,5392 | 5,4E-32 |
| ACE2 | RP11-14C10.6 | 0,5392 | 5,33216E-32 |
| ACE2 | GPRC5D | 0,5391 | 5,56585E-32 |
| ACE2 | PCP2 | 0,5391 | 5,49026E-32 |
| ACE2 | KLRC2 | 0,539 | 5,70748E-32 |
| ACE2 | DPP9-AS1 | 0,5389 | 5,76571E-32 |
| ACE2 | FBP2 | 0,5388 | 6,05125E-32 |
| ACE2 | SFTPA2 | 0,5388 | 6,0135E-32 |
| ACE2 | PTPN6 | 0,5388 | 6,04312E-32 |
| ACE2 | AP003068.23 | 0,5387 | 6,27726E-32 |

| | | | |
|---|---|---|---|
| ACE2 | PSMB10 | 0,5387 | 6,27417E-32 |
| ACE2 | CDK1 | 0,5385 | 6,72027E-32 |
| ACE2 | LINC01909 | 0,5385 | 6,58673E-32 |
| ACE2 | RN7SL663P | 0,5385 | 6,63139E-32 |
| ACE2 | KRT18P38 | 0,5384 | 6,86392E-32 |
| ACE2 | SFT2D1 | 0,5384 | 6,74952E-32 |
| ACE2 | EFL1 | 0,5383 | 7,13736E-32 |
| ACE2 | TMEM105 | 0,5383 | 7,03136E-32 |
| ACE2 | GCHFR | 0,5381 | 7,51746E-32 |
| ACE2 | AF064858.6 | 0,538 | 7,61065E-32 |
| ACE2 | AC016738.4 | 0,5379 | 7,98569E-32 |
| ACE2 | SLC25A20 | 0,5379 | 7,90612E-32 |
| ACE2 | SCN8A | 0,5379 | 8,05647E-32 |
| ACE2 | AC005336.4 | 0,5378 | 8,21975E-32 |
| ACE2 | SLC26A8 | 0,5375 | 8,92852E-32 |
| ACE2 | LINC01214 | 0,5373 | 9,45777E-32 |
| ACE2 | KIF20A | 0,5373 | 9,72773E-32 |
| ACE2 | SLC32A1 | 0,5373 | 9,51447E-32 |
| ACE2 | RP11-708J19.3 | 0,5372 | 9,7337E-32 |
| ACE2 | CYBA | 0,5371 | 1,01039E-31 |
| ACE2 | MRPL44 | 0,537 | 1,05458E-31 |
| ACE2 | CYP2C8 | 0,5369 | 1,09044E-31 |
| ACE2 | CENPM | 0,5368 | 1,11568E-31 |
| ACE2 | RP3-467K16.4 | 0,5367 | 1,13789E-31 |
| ACE2 | LINC02031 | 0,5367 | 1,14813E-31 |
| ACE2 | ZNF91 | 0,5367 | 1,14458E-31 |
| ACE2 | RP11-1149O23.2 | 0,5365 | 1,22016E-31 |
| ACE2 | DSG4 | 0,5363 | 1,30638E-31 |
| ACE2 | KIFC2 | 0,5362 | 1,36216E-31 |
| ACE2 | TONSL-AS1 | 0,5361 | 1,37725E-31 |
| ACE2 | DOC2A | 0,5358 | 1,52265E-31 |
| ACE2 | TOR1B | 0,5357 | 1,57998E-31 |
| ACE2 | KCNN4 | 0,5357 | 1,56923E-31 |
| ACE2 | TUBB4B | 0,5354 | 1,68814E-31 |
| ACE2 | UNG | 0,5354 | 1,69158E-31 |
| ACE2 | CYP27B1 | 0,5352 | 1,83535E-31 |
| ACE2 | OTUB2 | 0,5351 | 1,8966E-31 |
| ACE2 | AP000344.4 | 0,5351 | 1,8757E-31 |
| ACE2 | MTCO1P12 | 0,5348 | 2,08778E-31 |
| ACE2 | RP11-45M22.2 | 0,5348 | 2,03473E-31 |
| ACE2 | TLX1NB | 0,5346 | 2,16132E-31 |
| ACE2 | SNRPGP4 | 0,5344 | 2,3027E-31 |
| ACE2 | BCL11B | 0,5343 | 2,43099E-31 |
| ACE2 | GFI1 | 0,5342 | 2,44374E-31 |
| ACE2 | HOXA10-AS | 0,5342 | 2,48696E-31 |
| ACE2 | NRG4 | 0,5341 | 2,56485E-31 |
| ACE2 | LACTB2 | 0,5338 | 2,74975E-31 |
| ACE2 | REPIN1 | 0,5336 | 2,98136E-31 |
| ACE2 | IDI1 | 0,5336 | 2,95909E-31 |
| ACE2 | CKS2 | 0,5334 | 3,16348E-31 |
| ACE2 | MAPK6 | 0,5334 | 3,13319E-31 |
| ACE2 | SNHG18 | 0,5332 | 3,32594E-31 |
| ACE2 | NAGPA | 0,5332 | 3,29998E-31 |
| ACE2 | RP11-77E14.2 | 0,533 | 3,53248E-31 |

| | | | |
|---|---|---|---|
| ACE2 | PYCR1 | 0,533 | 3,60081E-31 |
| ACE2 | ZNF587 | 0,5329 | 3,69823E-31 |
| ACE2 | NPSR1 | 0,5324 | 4,32202E-31 |
| ACE2 | ZNF195 | 0,5324 | 4,26385E-31 |
| ACE2 | RP11-345K9.2 | 0,5321 | 4,61345E-31 |
| ACE2 | UNC5CL | 0,5319 | 4,96998E-31 |
| ACE2 | ARHGAP11B | 0,5317 | 5,21597E-31 |
| ACE2 | NECTIN3 | 0,5316 | 5,44855E-31 |
| ACE2 | RNFT1P2 | 0,5313 | 6,00972E-31 |
| ACE2 | RP11-491F9.1 | 0,5312 | 6,14327E-31 |
| ACE2 | LILRP2 | 0,5307 | 7,11583E-31 |
| ACE2 | PRC1 | 0,5304 | 7,74735E-31 |
| ACE2 | LINC00896 | 0,5303 | 7,98008E-31 |
| ACE2 | KRTAP5-9 | 0,5301 | 8,62662E-31 |
| ACE2 | MT-TY | 0,5301 | 8,54323E-31 |
| ACE2 | HIBCH | 0,53 | 8,71978E-31 |
| ACE2 | SIDT1-AS1 | 0,5297 | 9,60026E-31 |
| ACE2 | SMIM15 | 0,5296 | 9,87884E-31 |
| ACE2 | PSCA | 0,5296 | 9,92427E-31 |
| ACE2 | GMPPB | 0,5295 | 1,02768E-30 |
| ACE2 | SAMD10 | 0,5295 | 1,03373E-30 |
| ACE2 | NCAPG | 0,5294 | 1,05217E-30 |
| ACE2 | RNF19B | 0,5291 | 1,16429E-30 |
| ACE2 | AC003991.3 | 0,5291 | 1,16141E-30 |
| ACE2 | KIF18B | 0,5291 | 1,13248E-30 |
| ACE2 | RP11-386I14.2 | 0,529 | 1,17671E-30 |
| ACE2 | SNTN | 0,5289 | 1,23474E-30 |
| ACE2 | DTWD2 | 0,5284 | 1,42507E-30 |
| ACE2 | FAM221B | 0,5284 | 1,41035E-30 |
| ACE2 | UBE2C | 0,5284 | 1,40841E-30 |
| ACE2 | ITGA6 | 0,5281 | 1,55837E-30 |
| ACE2 | FOXM1 | 0,5279 | 1,6541E-30 |
| ACE2 | MIR600HG | 0,5278 | 1,71009E-30 |
| ACE2 | TMEM19 | 0,5274 | 1,92007E-30 |
| ACE2 | NOP56P3 | 0,5274 | 1,89967E-30 |
| ACE2 | FGFR4 | 0,5273 | 1,9526E-30 |
| ACE2 | NUF2 | 0,5272 | 2,00846E-30 |
| ACE2 | NIPSNAP3A | 0,5272 | 2,0489E-30 |
| ACE2 | CTD-3214H19.6 | 0,5272 | 2,04262E-30 |
| ACE2 | RP11-158H5.7 | 0,527 | 2,1679E-30 |
| ACE2 | RP5-965G21.3 | 0,5269 | 2,22176E-30 |
| ACE2 | MTCO1P2 | 0,5267 | 2,33228E-30 |
| ACE2 | CASR | 0,5266 | 2,4436E-30 |
| ACE2 | KPNA2P2 | 0,5266 | 2,4469E-30 |
| ACE2 | NBPF6 | 0,5263 | 2,64368E-30 |
| ACE2 | C10orf91 | 0,5263 | 2,62256E-30 |
| ACE2 | KRT8P13 | 0,5262 | 2,73167E-30 |
| ACE2 | RPL39P40 | 0,5262 | 2,69978E-30 |
| ACE2 | KLKP1 | 0,5258 | 3,07916E-30 |
| ACE2 | FBXL8 | 0,5256 | 3,26076E-30 |
| ACE2 | SLC9A4 | 0,5255 | 3,3081E-30 |
| ACE2 | CLIC1 | 0,5253 | 3,54776E-30 |
| ACE2 | DNAAF2 | 0,5253 | 3,49517E-30 |
| ACE2 | LBHD1 | 0,5251 | 3,73196E-30 |

| | | | |
|---|---|---|---|
| ACE2 | RP11-285G1.9 | 0,525 | 3,87774E-30 |
| ACE2 | SAMD9 | 0,5248 | 4,08543E-30 |
| ACE2 | UQCRC2 | 0,5245 | 4,47367E-30 |
| ACE2 | HCCS | 0,5245 | 4,53231E-30 |
| ACE2 | CCNE2 | 0,5244 | 4,57475E-30 |
| ACE2 | MRPL13 | 0,524 | 5,12986E-30 |
| ACE2 | GBA | 0,5239 | 5,34597E-30 |
| ACE2 | CTA-243E7.2 | 0,5239 | 5,34282E-30 |
| ACE2 | LTK | 0,5238 | 5,58141E-30 |
| ACE2 | AC004067.5 | 0,5236 | 5,92235E-30 |
| ACE2 | IQGAP3 | 0,5234 | 6,20158E-30 |
| ACE2 | EML4 | 0,5231 | 6,86638E-30 |
| ACE2 | LINC00870 | 0,5229 | 7,08557E-30 |
| ACE2 | SPATA31E1 | 0,5229 | 7,27042E-30 |
| ACE2 | HJURP | 0,5228 | 7,44597E-30 |
| ACE2 | FAM64A | 0,5226 | 7,9416E-30 |
| ACE2 | SRD5A1 | 0,5223 | 8,45279E-30 |
| ACE2 | RAB43 | 0,5222 | 8,78209E-30 |
| ACE2 | LINC02033 | 0,5221 | 9,02515E-30 |
| ACE2 | RP11-43P8.2 | 0,5221 | 8,97116E-30 |
| ACE2 | RP4-594I10.3 | 0,522 | 9,398E-30 |
| ACE2 | POU2F3 | 0,5219 | 9,73896E-30 |
| ACE2 | BOLA2 | 0,5219 | 9,70698E-30 |
| ACE2 | DNMBP | 0,5218 | 9,95751E-30 |
| ACE2 | ARL6IP1 | 0,5218 | 9,81043E-30 |
| ACE2 | PLK4 | 0,5217 | 1,00545E-29 |
| ACE2 | LUZP2 | 0,5217 | 1,01218E-29 |
| ACE2 | ZNF675 | 0,5216 | 1,03573E-29 |
| ACE2 | RP11-112J1.3 | 0,5215 | 1,08211E-29 |
| ACE2 | MED14 | 0,5213 | 1,15817E-29 |
| ACE2 | COX4I1 | 0,5212 | 1,18683E-29 |
| ACE2 | DERA | 0,5209 | 1,27616E-29 |
| ACE2 | FAM9C | 0,5209 | 1,28092E-29 |
| ACE2 | SMIM5 | 0,5206 | 1,39759E-29 |
| ACE2 | SLC8B1 | 0,5205 | 1,45755E-29 |
| ACE2 | RP11-483I13.6 | 0,5204 | 1,46899E-29 |
| ACE2 | KRT8P39 | 0,5204 | 1,50468E-29 |
| ACE2 | FAM134B | 0,5203 | 1,54257E-29 |
| ACE2 | TIPARP-AS1 | 0,5201 | 1,63673E-29 |
| ACE2 | LINC02183 | 0,5198 | 1,79144E-29 |
| ACE2 | RHOT1P1 | 0,5198 | 1,769E-29 |
| ACE2 | EVPLL | 0,5196 | 1,86371E-29 |
| ACE2 | RP3-368B9.2 | 0,5195 | 1,91641E-29 |
| ACE2 | LIPT2 | 0,5195 | 1,90851E-29 |
| ACE2 | DIAPH1 | 0,5194 | 2,0121E-29 |
| ACE2 | ANKRD20A5P | 0,5194 | 1,99351E-29 |
| ACE2 | ASTE1 | 0,5192 | 2,10596E-29 |
| ACE2 | RP11-9M16.2 | 0,5192 | 2,08042E-29 |
| ACE2 | SLC40A1 | 0,5189 | 2,31451E-29 |
| ACE2 | RP5-1132H15.3 | 0,5187 | 2,44346E-29 |
| ACE2 | RP11-386M24.3 | 0,5186 | 2,53462E-29 |
| ACE2 | PSORS1C3 | 0,5184 | 2,68522E-29 |
| ACE2 | RP11-510J16.5 | 0,5184 | 2,66757E-29 |
| ACE2 | CCDC47 | 0,5183 | 2,69117E-29 |

| | | | |
|---|---|---|---|
| ACE2 | ZNF217 | 0,5181 | 2,87539E-29 |
| ACE2 | PNPLA4 | 0,518 | 3,01741E-29 |
| ACE2 | KIAA1468 | 0,5179 | 3,09094E-29 |
| ACE2 | DEPDC5 | 0,5179 | 3,02684E-29 |
| ACE2 | CD300LF | 0,5178 | 3,13052E-29 |
| ACE2 | COG2 | 0,5177 | 3,27228E-29 |
| ACE2 | RP11-723C11.2 | 0,5176 | 3,2962E-29 |
| ACE2 | BICDL1 | 0,5176 | 3,36431E-29 |
| ACE2 | RAVER1 | 0,5176 | 3,37799E-29 |
| ACE2 | MIR210 | 0,5175 | 3,47818E-29 |
| ACE2 | RP1-257I20.14 | 0,5175 | 3,42005E-29 |
| ACE2 | LINC01016 | 0,5174 | 3,5113E-29 |
| ACE2 | HLA-A | 0,517 | 3,9831E-29 |
| ACE2 | RP11-944C7.1 | 0,5167 | 4,329E-29 |
| ACE2 | RP11-33I11.3 | 0,5166 | 4,41953E-29 |
| ACE2 | RP11-242G20.1 | 0,5164 | 4,70873E-29 |
| ACE2 | NDUFV1 | 0,5163 | 4,83019E-29 |
| ACE2 | HPGD | 0,516 | 5,28215E-29 |
| ACE2 | OR8D1 | 0,516 | 5,2396E-29 |
| ACE2 | MPC2 | 0,5157 | 5,76961E-29 |
| ACE2 | RP3-476K8.4 | 0,5156 | 5,90779E-29 |
| ACE2 | CEACAM22P | 0,5156 | 6,00077E-29 |
| ACE2 | RP11-366L5.1 | 0,5155 | 6,09939E-29 |
| ACE2 | LPIN2 | 0,5155 | 6,11495E-29 |
| ACE2 | STK38 | 0,5154 | 6,35417E-29 |
| ACE2 | CYP4A22-AS1 | 0,5153 | 6,49457E-29 |
| ACE2 | STBD1 | 0,5153 | 6,39099E-29 |
| ACE2 | CTA-989H11.1 | 0,5153 | 6,47418E-29 |
| ACE2 | RP11-46I1.2 | 0,5147 | 7,72607E-29 |
| ACE2 | RP11-61A14.4 | 0,5145 | 8,07797E-29 |
| ACE2 | MMP28 | 0,5143 | 8,63704E-29 |
| ACE2 | TUFT1 | 0,5142 | 8,82611E-29 |
| ACE2 | SELENOP | 0,5142 | 8,72871E-29 |
| ACE2 | STXBP5 | 0,514 | 9,40597E-29 |
| ACE2 | MESP1 | 0,514 | 9,29348E-29 |
| ACE2 | CH17-78J1.1 | 0,5137 | 1,02733E-28 |
| ACE2 | SLC16A10 | 0,5137 | 1,03025E-28 |
| ACE2 | TAX1BP3 | 0,5137 | 1,0244E-28 |
| ACE2 | RP11-416N2.4 | 0,5133 | 1,14379E-28 |
| ACE2 | ANXA10 | 0,5132 | 1,18757E-28 |
| ACE2 | TTC39B | 0,5132 | 1,16602E-28 |
| ACE2 | NDST3 | 0,513 | 1,23146E-28 |
| ACE2 | LMAN2 | 0,513 | 1,24776E-28 |
| ACE2 | DDT | 0,513 | 1,2495E-28 |
| ACE2 | RRM2 | 0,5129 | 1,26204E-28 |
| ACE2 | ZNF215 | 0,5129 | 1,28658E-28 |
| ACE2 | CYB561A3 | 0,5128 | 1,31252E-28 |
| ACE2 | DLEC1 | 0,5127 | 1,34754E-28 |
| ACE2 | ANKH | 0,5127 | 1,35215E-28 |
| ACE2 | LINC00337 | 0,5126 | 1,38271E-28 |
| ACE2 | LNX1 | 0,5124 | 1,452E-28 |
| ACE2 | SLC25A15P3 | 0,5122 | 1,55264E-28 |
| ACE2 | KLK13 | 0,5122 | 1,56982E-28 |
| ACE2 | SEPSECS | 0,5121 | 1,59001E-28 |

| | | | |
|---|---|---|---|
| ACE2 | TROAP | 0,512 | 1,6577E-28 |
| ACE2 | PCLAF | 0,5119 | 1,68402E-28 |
| ACE2 | SELENOI | 0,5118 | 1,72137E-28 |
| ACE2 | CKAP2L | 0,5118 | 1,73345E-28 |
| ACE2 | PCCB | 0,5118 | 1,72288E-28 |
| ACE2 | LA16c-380H5.3 | 0,5117 | 1,7807E-28 |
| ACE2 | AC074363.1 | 0,5116 | 1,84202E-28 |
| ACE2 | ADCK5 | 0,5116 | 1,84312E-28 |
| ACE2 | FOXE3 | 0,5113 | 1,9818E-28 |
| ACE2 | RP11-478J18.2 | 0,5111 | 2,11204E-28 |
| ACE2 | MTCO1P40 | 0,5111 | 2,14492E-28 |
| ACE2 | RP11-707F2.1 | 0,5108 | 2,33186E-28 |
| ACE2 | MPC1 | 0,5106 | 2,41623E-28 |
| ACE2 | SKA1 | 0,5105 | 2,53393E-28 |
| ACE2 | RAB4B | 0,5105 | 2,52061E-28 |
| ACE2 | RP11-150O12.5 | 0,5104 | 2,61883E-28 |
| ACE2 | TONSL | 0,5097 | 3,16236E-28 |
| ACE2 | MTCO1P53 | 0,5097 | 3,13892E-28 |
| ACE2 | PTP4A1 | 0,5096 | 3,24788E-28 |
| ACE2 | SLC6A4 | 0,5094 | 3,39722E-28 |
| ACE2 | HIST1H2AI | 0,5092 | 3,61399E-28 |
| ACE2 | SLC28A3 | 0,5091 | 3,73303E-28 |
| ACE2 | SWSAP1 | 0,5091 | 3,67891E-28 |
| ACE2 | NUSAP1 | 0,5089 | 3,96894E-28 |
| ACE2 | NDUFS2 | 0,5087 | 4,10134E-28 |
| ACE2 | C8orf82 | 0,5086 | 4,32124E-28 |
| ACE2 | RP11-646E18.4 | 0,5085 | 4,44023E-28 |
| ACE2 | CTD-2022H16.3 | 0,5085 | 4,4123E-28 |
| ACE2 | MAVS | 0,5084 | 4,53027E-28 |
| ACE2 | MTFR2 | 0,508 | 5,06158E-28 |
| ACE2 | TSEN54 | 0,508 | 5,07607E-28 |
| ACE2 | PIGF | 0,5075 | 5,88371E-28 |
| ACE2 | LINC01225 | 0,5074 | 6,05041E-28 |
| ACE2 | ATP23 | 0,5074 | 5,90533E-28 |
| ACE2 | USP53 | 0,5073 | 6,10699E-28 |
| ACE2 | RP11-57A1.1 | 0,5073 | 6,06456E-28 |
| ACE2 | MIPEP | 0,5071 | 6,44119E-28 |
| ACE2 | WNT3A | 0,5066 | 7,50939E-28 |
| ACE2 | WI2-2373I1.1 | 0,5065 | 7,63073E-28 |
| ACE2 | RPLP0P2 | 0,5065 | 7,68108E-28 |
| ACE2 | SFXN5 | 0,5063 | 8,03336E-28 |
| ACE2 | AC073043.1 | 0,5063 | 8,15599E-28 |
| ACE2 | KLK12 | 0,5058 | 9,23456E-28 |
| ACE2 | RP13-415G19.2 | 0,5057 | 9,47983E-28 |
| ACE2 | SLC39A7 | 0,5055 | 1,00964E-27 |
| ACE2 | PLA2G4E | 0,5054 | 1,05366E-27 |
| ACE2 | BHLHA15 | 0,5053 | 1,06587E-27 |
| ACE2 | UNC13B | 0,5051 | 1,12686E-27 |
| ACE2 | ECEL1P2 | 0,5049 | 1,17933E-27 |
| ACE2 | SCAP | 0,5049 | 1,20646E-27 |
| ACE2 | RABEP2 | 0,5049 | 1,18862E-27 |
| ACE2 | AKR1C4 | 0,5048 | 1,23439E-27 |
| ACE2 | CYP4F23P | 0,5048 | 1,23032E-27 |
| ACE2 | LINC02003 | 0,5046 | 1,2956E-27 |

| | | | |
|---|---|---|---|
| ACE2 | PRAC2 | 0,5043 | 1,3887E-27 |
| ACE2 | SEMA6A | 0,5041 | 1,5037E-27 |
| ACE2 | MFSD5 | 0,5041 | 1,49285E-27 |
| ACE2 | RP11-64C12.5 | 0,5039 | 1,58918E-27 |
| ACE2 | RN7SKP116 | 0,5038 | 1,63167E-27 |
| ACE2 | PIP4K2C | 0,5038 | 1,61028E-27 |
| ACE2 | RP11-175P13.3 | 0,5038 | 1,60746E-27 |
| ACE2 | HDDC3 | 0,5037 | 1,64553E-27 |
| ACE2 | PLA2G7 | 0,5034 | 1,81352E-27 |
| ACE2 | FAM84B | 0,5032 | 1,88596E-27 |
| ACE2 | SLC10A1 | 0,5032 | 1,90568E-27 |
| ACE2 | RRN3P3 | 0,5032 | 1,91614E-27 |
| ACE2 | PHBP8 | 0,503 | 1,99862E-27 |
| ACE2 | GSTA2 | 0,503 | 2,02397E-27 |
| ACE2 | FKBP4P1 | 0,5026 | 2,26696E-27 |
| ACE2 | UGT2B25P | 0,5023 | 2,45759E-27 |
| ACE2 | PDZK1 | 0,5021 | 2,60119E-27 |
| ACE2 | C11orf80 | 0,5021 | 2,54923E-27 |
| ACE2 | TRAK1 | 0,502 | 2,65862E-27 |
| ACE2 | RP11-963H4.5 | 0,502 | 2,6358E-27 |
| ACE2 | GRTP1-AS1 | 0,5014 | 3,11114E-27 |
| ACE2 | PIGN | 0,5013 | 3,18603E-27 |
| ACE2 | MT1H | 0,5012 | 3,3303E-27 |
| ACE2 | Z83851.4 | 0,5012 | 3,30758E-27 |
| ACE2 | LEO1 | 0,5011 | 3,35184E-27 |
| ACE2 | AAGAB | 0,5011 | 3,38477E-27 |
| ACE2 | DNASE1L3 | 0,5006 | 3,87921E-27 |
| ACE2 | RP5-958B11.2 | 0,5002 | 4,33645E-27 |
| ACE2 | LINC00365 | 0,5001 | 4,3861E-27 |
| ACE2 | MYBL2 | 0,5 | 4,55522E-27 |
| ACE2 | C6orf99 | 0,4999 | 4,69724E-27 |
| ACE2 | KCCAT198 | 0,4997 | 4,93967E-27 |
| ACE2 | NSDHL | 0,4996 | 5,08777E-27 |
| ACE2 | GRM7-AS2 | 0,4995 | 5,2675E-27 |
| ACE2 | RP5-875H18.10 | 0,4994 | 5,42713E-27 |
| ACE2 | STARD7 | 0,4993 | 5,56661E-27 |
| ACE2 | ISCA2 | 0,4992 | 5,63698E-27 |
| ACE2 | TMX2 | 0,4988 | 6,26218E-27 |
| ACE2 | RP11-747H7.3 | 0,4987 | 6,53457E-27 |
| ACE2 | SSFA2 | 0,4986 | 6,57978E-27 |
| ACE2 | PSMB8 | 0,4986 | 6,65412E-27 |
| ACE2 | ATP5C1 | 0,4986 | 6,6181E-27 |
| ACE2 | PLS1-AS1 | 0,4985 | 6,7878E-27 |
| ACE2 | STPG1 | 0,4983 | 7,16532E-27 |
| ACE2 | AC021218.2 | 0,4983 | 7,21505E-27 |
| ACE2 | RP11-498C9.15 | 0,4981 | 7,5288E-27 |
| ACE2 | RP11-57G10.8 | 0,498 | 7,84321E-27 |
| ACE2 | RAD51AP1 | 0,4979 | 7,9997E-27 |
| ACE2 | PRSS36 | 0,4979 | 8,03694E-27 |
| ACE2 | OTC | 0,4979 | 8,03841E-27 |
| ACE2 | RHOQP1 | 0,4978 | 8,2585E-27 |
| ACE2 | RP11-66N24.6 | 0,4977 | 8,58136E-27 |
| ACE2 | CECR3 | 0,4977 | 8,48367E-27 |
| ACE2 | MT-TL1 | 0,4977 | 8,48952E-27 |

| | | | |
|---|---|---|---|
| ACE2 | XX-CR54.1 | 0,4974 | 9,16713E-27 |
| ACE2 | SAMD12-AS1 | 0,4974 | 9,14589E-27 |
| ACE2 | AC128709.4 | 0,4973 | 9,55494E-27 |
| ACE2 | RCE1 | 0,4973 | 9,53064E-27 |
| ACE2 | NDC80 | 0,4973 | 9,4115E-27 |
| ACE2 | LINC00520 | 0,497 | 1,03755E-26 |
| ACE2 | TYMSOS | 0,497 | 1,01116E-26 |
| ACE2 | HMGN2P28 | 0,4969 | 1,06167E-26 |
| ACE2 | COX7A2 | 0,4969 | 1,04989E-26 |
| ACE2 | RP11-49I11.4 | 0,4966 | 1,1328E-26 |
| ACE2 | MTCO1P22 | 0,4963 | 1,24492E-26 |
| ACE2 | AVL9 | 0,496 | 1,32728E-26 |
| ACE2 | AC093642.1 | 0,4957 | 1,45409E-26 |
| ACE2 | HLA-C | 0,4957 | 1,44261E-26 |
| ACE2 | RP11-440I14.3 | 0,4955 | 1,53785E-26 |
| ACE2 | RP1-32B1.4 | 0,4954 | 1,57188E-26 |
| ACE2 | MAPKAPK5 | 0,4954 | 1,5621E-26 |
| ACE2 | SH3YL1 | 0,4953 | 1,59975E-26 |
| ACE2 | OR6M1 | 0,4951 | 1,70107E-26 |
| ACE2 | AC008694.3 | 0,4949 | 1,81189E-26 |
| ACE2 | CLDN15 | 0,4949 | 1,78075E-26 |
| ACE2 | CASP10 | 0,4948 | 1,83759E-26 |
| ACE2 | NEUROG3 | 0,4946 | 1,94297E-26 |
| ACE2 | CDCA5 | 0,4945 | 1,98741E-26 |
| ACE2 | AP004550.1 | 0,4944 | 2,02692E-26 |
| ACE2 | SPRYD4 | 0,4942 | 2,16391E-26 |
| ACE2 | RP4-799G3.2 | 0,494 | 2,29176E-26 |
| ACE2 | UGT2B27P | 0,494 | 2,31229E-26 |
| ACE2 | SLC26A1 | 0,4939 | 2,32776E-26 |
| ACE2 | ZSCAN16 | 0,4939 | 2,37705E-26 |
| ACE2 | VPS4B | 0,4939 | 2,363E-26 |
| ACE2 | TATDN2P2 | 0,4934 | 2,67508E-26 |
| ACE2 | ACAD10 | 0,4934 | 2,6863E-26 |
| ACE2 | QPRT | 0,4932 | 2,8326E-26 |
| ACE2 | ALG1 | 0,4931 | 2,88087E-26 |
| ACE2 | CEACAM20 | 0,4931 | 2,86293E-26 |
| ACE2 | SGO1 | 0,493 | 3,01464E-26 |
| ACE2 | RP4-587D13.1 | 0,493 | 3,00776E-26 |
| ACE2 | NLRP6 | 0,493 | 2,99759E-26 |
| ACE2 | RN7SL176P | 0,4929 | 3,0857E-26 |
| ACE2 | FAM47DP | 0,4927 | 3,20993E-26 |
| ACE2 | KIF2C | 0,4926 | 3,32069E-26 |
| ACE2 | RP11-734K21.5 | 0,4926 | 3,33589E-26 |
| ACE2 | KRT8P31 | 0,4926 | 3,34877E-26 |
| ACE2 | LDLRAP1 | 0,4924 | 3,49283E-26 |
| ACE2 | LRRC59 | 0,4924 | 3,46663E-26 |
| ACE2 | OR2S1P | 0,4922 | 3,72829E-26 |
| ACE2 | SEMA6D | 0,4922 | 3,69336E-26 |
| ACE2 | CTB-127C13.1 | 0,4921 | 3,79411E-26 |
| ACE2 | RP11-468N14.09 | 0,492 | 3,84611E-26 |
| ACE2 | RP11-575I8.1 | 0,492 | 3,90458E-26 |
| ACE2 | PACSIN2 | 0,4918 | 4,14229E-26 |
| ACE2 | RP11-463O12.5 | 0,4917 | 4,17219E-26 |
| ACE2 | RNU2-33P | 0,4917 | 4,24911E-26 |

| | | | |
|---|---|---|---|
| ACE2 | CATSPERB | 0,4916 | 4,3467E-26 |
| ACE2 | GINS1 | 0,4914 | 4,50955E-26 |
| ACE2 | SH3PXD2A-AS1 | 0,4911 | 4,90068E-26 |
| ACE2 | FITM1 | 0,4909 | 5,15348E-26 |
| ACE2 | AC005281.1 | 0,4907 | 5,50459E-26 |
| ACE2 | RP1-240K6.3 | 0,4907 | 5,53917E-26 |
| ACE2 | ADGRE5 | 0,4904 | 5,98892E-26 |
| ACE2 | OXGR1 | 0,4903 | 6,05486E-26 |
| ACE2 | HMOX1 | 0,4901 | 6,43103E-26 |
| ACE2 | LINC01022 | 0,49 | 6,62759E-26 |
| ACE2 | SOCS5P4 | 0,49 | 6,50747E-26 |
| ACE2 | IGSF3 | 0,4898 | 6,89223E-26 |
| ACE2 | GPRIN3 | 0,4897 | 7,04595E-26 |
| ACE2 | RP11-178C3.2 | 0,4897 | 7,14408E-26 |
| ACE2 | CTC-429L19.3 | 0,4897 | 7,1303E-26 |
| ACE2 | E2F3P2 | 0,4896 | 7,29266E-26 |
| ACE2 | USP30 | 0,4894 | 7,78789E-26 |
| ACE2 | RN7SL8P | 0,4893 | 7,79863E-26 |
| ACE2 | DNAJC28 | 0,4893 | 7,83486E-26 |
| ACE2 | RP11-400N9.1 | 0,4891 | 8,27641E-26 |
| ACE2 | ALG1L5P | 0,4891 | 8,42015E-26 |
| ACE2 | E2F7 | 0,4891 | 8,35893E-26 |
| ACE2 | PICK1 | 0,489 | 8,63845E-26 |
| ACE2 | LHX4 | 0,4888 | 9,0815E-26 |
| ACE2 | RPSAP11 | 0,4887 | 9,32061E-26 |
| ACE2 | RAD51AP2 | 0,4886 | 9,46124E-26 |
| ACE2 | FADD | 0,4886 | 9,55247E-26 |
| ACE2 | AC090616.2 | 0,488 | 1,09503E-25 |
| ACE2 | CEP55 | 0,4879 | 1,14584E-25 |
| ACE2 | AC007387.2 | 0,4877 | 1,2027E-25 |
| ACE2 | TCF7L2 | 0,4876 | 1,23178E-25 |
| ACE2 | KIAA1161 | 0,4875 | 1,25007E-25 |
| ACE2 | AC144836.1 | 0,4873 | 1,32954E-25 |
| ACE2 | CTD-2165H16.3 | 0,4872 | 1,38003E-25 |
| ACE2 | CTH | 0,4871 | 1,412E-25 |
| ACE2 | CLN8 | 0,4871 | 1,41541E-25 |
| ACE2 | SLAMF6 | 0,487 | 1,45179E-25 |
| ACE2 | FAM129B | 0,4868 | 1,52959E-25 |
| ACE2 | RP13-16H11.8 | 0,4867 | 1,55543E-25 |
| ACE2 | FUK | 0,4867 | 1,5597E-25 |
| ACE2 | C2CD2L | 0,4866 | 1,61627E-25 |
| ACE2 | RPP25 | 0,4865 | 1,62612E-25 |
| ACE2 | RP1-13D10.2 | 0,4863 | 1,7247E-25 |
| ACE2 | MEST | 0,4863 | 1,70704E-25 |
| ACE2 | TECTB | 0,4863 | 1,7466E-25 |
| ACE2 | GDPD3 | 0,4863 | 1,70968E-25 |
| ACE2 | SLC47A2 | 0,4861 | 1,80802E-25 |
| ACE2 | RPL7AP52 | 0,4859 | 1,8984E-25 |
| ACE2 | P4HB | 0,4859 | 1,92885E-25 |
| ACE2 | AREG | 0,4857 | 2,03486E-25 |
| ACE2 | KDM1B | 0,4857 | 2,00421E-25 |
| ACE2 | KRT8P11 | 0,4857 | 2,02089E-25 |
| ACE2 | RDM1 | 0,4856 | 2,08159E-25 |
| ACE2 | IL1RL2 | 0,4855 | 2,13066E-25 |

| | | | |
|---|---|---|---|
| ACE2 | ACTL8 | 0,4853 | 2,25859E-25 |
| ACE2 | NSD2 | 0,485 | 2,39765E-25 |
| ACE2 | RP13-714J12.1 | 0,485 | 2,38646E-25 |
| ACE2 | KCNJ11 | 0,4849 | 2,46109E-25 |
| ACE2 | GGT4P | 0,4848 | 2,52983E-25 |
| ACE2 | AD000684.2 | 0,4846 | 2,64978E-25 |
| ACE2 | MTG1 | 0,4843 | 2,91678E-25 |
| ACE2 | AL163953.2 | 0,4843 | 2,85923E-25 |
| ACE2 | LACE1 | 0,4842 | 2,99174E-25 |
| ACE2 | AC011298.2 | 0,4839 | 3,18578E-25 |
| ACE2 | RP11-253M7.6 | 0,4839 | 3,18095E-25 |
| ACE2 | MYCBP | 0,4838 | 3,3208E-25 |
| ACE2 | CAPN2 | 0,4838 | 3,30745E-25 |
| ACE2 | RP11-933H2.4 | 0,4838 | 3,32498E-25 |
| ACE2 | AC016735.1 | 0,4834 | 3,69465E-25 |
| ACE2 | RP11-408O19.5 | 0,4833 | 3,70453E-25 |
| ACE2 | RP11-386M24.6 | 0,4833 | 3,71465E-25 |
| ACE2 | CTD-3128G10.6 | 0,483 | 4,0893E-25 |
| ACE2 | AMBP | 0,4829 | 4,16673E-25 |
| ACE2 | RP11-8P13.5 | 0,4829 | 4,17359E-25 |
| ACE2 | DRAIC | 0,4829 | 4,17483E-25 |
| ACE2 | NAA60 | 0,4827 | 4,36381E-25 |
| ACE2 | AP1B1 | 0,4827 | 4,36533E-25 |
| ACE2 | RP11-685B14.3 | 0,4826 | 4,43396E-25 |
| ACE2 | KRT18P1 | 0,4825 | 4,54756E-25 |
| ACE2 | CCDC58 | 0,4823 | 4,88899E-25 |
| ACE2 | RP11-644A7.2 | 0,4823 | 4,86259E-25 |
| ACE2 | KB-1460A1.3 | 0,4822 | 5,00175E-25 |
| ACE2 | LINC01137 | 0,4821 | 5,13768E-25 |
| ACE2 | RP11-58O9.2 | 0,4819 | 5,3387E-25 |
| ACE2 | TSTA3 | 0,4818 | 5,53477E-25 |
| ACE2 | MT-CO3 | 0,4818 | 5,51168E-25 |
| ACE2 | FNBP1L | 0,4817 | 5,60028E-25 |
| ACE2 | TEX264 | 0,4816 | 5,81145E-25 |
| ACE2 | RP11-489O18.1 | 0,4814 | 6,03751E-25 |
| ACE2 | RP5-1171I10.5 | 0,4812 | 6,47551E-25 |
| ACE2 | RP11-49I11.1 | 0,481 | 6,81998E-25 |
| ACE2 | HADHA | 0,4809 | 6,86093E-25 |
| ACE2 | ITGB4 | 0,4808 | 7,0071E-25 |
| ACE2 | FASTKD3 | 0,4806 | 7,41946E-25 |
| ACE2 | CTC-348L5.1 | 0,4805 | 7,75091E-25 |
| ACE2 | GZMA | 0,4804 | 7,9199E-25 |
| ACE2 | TNFSF10 | 0,4802 | 8,36847E-25 |
| ACE2 | LINC00970 | 0,48 | 8,73225E-25 |
| ACE2 | LRFN4 | 0,4798 | 9,25563E-25 |
| ACE2 | RP11-24I21.1 | 0,4796 | 9,63084E-25 |
| ACE2 | ST3GAL4 | 0,479 | 1,11823E-24 |
| ACE2 | F11-AS1 | 0,4789 | 1,1495E-24 |
| ACE2 | CTD-2134A5.3 | 0,4789 | 1,1361E-24 |
| ACE2 | AC073254.1 | 0,4788 | 1,18819E-24 |
| ACE2 | NSUN7 | 0,4786 | 1,22431E-24 |
| ACE2 | LRR1 | 0,4781 | 1,40759E-24 |
| ACE2 | HIST1H2APS4 | 0,4778 | 1,53054E-24 |
| ACE2 | PDLIM1 | 0,4778 | 1,52621E-24 |

| | | | |
|---|---|---|---|
| ACE2 | HIST1H3F | 0,4777 | 1,54137E-24 |
| ACE2 | VPS9D1-AS1 | 0,4777 | 1,53933E-24 |
| ACE2 | FUCA2 | 0,4776 | 1,58858E-24 |
| ACE2 | TSC22D1-AS1 | 0,4776 | 1,57604E-24 |
| ACE2 | TTC19 | 0,4774 | 1,67479E-24 |
| ACE2 | RP11-480D4.7 | 0,4773 | 1,74213E-24 |
| ACE2 | RP11-359M6.1 | 0,4773 | 1,71383E-24 |
| ACE2 | ARL11 | 0,4773 | 1,73125E-24 |
| ACE2 | RP11-244M2.1 | 0,4772 | 1,77463E-24 |
| ACE2 | RP11-800A18.4 | 0,4772 | 1,76389E-24 |
| ACE2 | ANKRD26P2 | 0,4771 | 1,80622E-24 |
| ACE2 | SKA3 | 0,4768 | 1,96051E-24 |
| ACE2 | MMP23A | 0,4767 | 1,98838E-24 |
| ACE2 | RTCB | 0,4767 | 1,97934E-24 |
| ACE2 | RGMB-AS1 | 0,4766 | 2,06061E-24 |
| ACE2 | KB-1410C5.3 | 0,4766 | 2,0398E-24 |
| ACE2 | LINC02041 | 0,4761 | 2,33851E-24 |
| ACE2 | SRRM2-AS1 | 0,4761 | 2,3425E-24 |
| ACE2 | OR13Z2P | 0,476 | 2,40977E-24 |
| ACE2 | RP11-88L20.1 | 0,476 | 2,40389E-24 |
| ACE2 | IGHV5-78 | 0,4759 | 2,4597E-24 |
| ACE2 | PTGR2 | 0,4758 | 2,47985E-24 |
| ACE2 | FER1L6-AS1 | 0,4755 | 2,69907E-24 |
| ACE2 | FTLP12 | 0,4755 | 2,70434E-24 |
| ACE2 | PRR22 | 0,4754 | 2,80348E-24 |
| ACE2 | AC073133.2 | 0,4753 | 2,87874E-24 |
| ACE2 | PPIF | 0,4753 | 2,85719E-24 |
| ACE2 | AK2 | 0,4752 | 2,89551E-24 |
| ACE2 | TERT | 0,4752 | 2,89368E-24 |
| ACE2 | PIK3C2B | 0,4751 | 2,96257E-24 |
| ACE2 | IL7 | 0,4751 | 3,00049E-24 |
| ACE2 | PPP1R13B | 0,4749 | 3,13876E-24 |
| ACE2 | CTD-2616J11.10 | 0,4749 | 3,15987E-24 |
| ACE2 | RP11-205A8.3 | 0,4748 | 3,22423E-24 |
| ACE2 | RP11-138A9.1 | 0,4747 | 3,31598E-24 |
| ACE2 | C16orf54 | 0,4746 | 3,37193E-24 |
| ACE2 | CTD-2575K13.6 | 0,4746 | 3,42312E-24 |
| ACE2 | TC2N | 0,4744 | 3,5861E-24 |
| ACE2 | TMEM33 | 0,4741 | 3,82059E-24 |
| ACE2 | ADAP1 | 0,4741 | 3,8727E-24 |
| ACE2 | SHROOM2 | 0,4741 | 3,8312E-24 |
| ACE2 | KRT18P23 | 0,474 | 3,96809E-24 |
| ACE2 | COL2A1 | 0,4739 | 4,08157E-24 |
| ACE2 | QTRT1P1 | 0,4739 | 4,02079E-24 |
| ACE2 | RP11-420N3.2 | 0,4738 | 4,11622E-24 |
| ACE2 | CDC6 | 0,4738 | 4,10609E-24 |
| ACE2 | MYL12A | 0,4737 | 4,25153E-24 |
| ACE2 | ZNF681 | 0,4736 | 4,30815E-24 |
| ACE2 | RP3-425C14.5 | 0,4732 | 4,85465E-24 |
| ACE2 | RNF44 | 0,473 | 5,08452E-24 |
| ACE2 | RP11-326K13.4 | 0,4729 | 5,13097E-24 |
| ACE2 | MRPS12 | 0,4728 | 5,25813E-24 |
| ACE2 | MCIDAS | 0,4726 | 5,50096E-24 |
| ACE2 | RP11-219E7.4 | 0,4726 | 5,5966E-24 |

| | | | |
|---|---|---|---|
| ACE2 | ZNF101 | 0,4725 | 5,68946E-24 |
| ACE2 | L1TD1 | 0,4724 | 5,83727E-24 |
| ACE2 | ARHGAP42 | 0,4724 | 5,80871E-24 |
| ACE2 | RNU6-652P | 0,472 | 6,49133E-24 |
| ACE2 | BCAR1 | 0,472 | 6,41727E-24 |
| ACE2 | MSGN1 | 0,4719 | 6,66025E-24 |
| ACE2 | PIGU | 0,4718 | 6,73946E-24 |
| ACE2 | DEFA6 | 0,4716 | 7,16127E-24 |
| ACE2 | ARSEP1 | 0,4716 | 7,06768E-24 |
| ACE2 | RP11-84C10.4 | 0,4715 | 7,32587E-24 |
| ACE2 | LINC02097 | 0,4714 | 7,55913E-24 |
| ACE2 | LYZ | 0,4713 | 7,60406E-24 |
| ACE2 | CTD-2330K9.2 | 0,4712 | 7,81272E-24 |
| ACE2 | RP11-103J8.2 | 0,4711 | 7,99818E-24 |
| ACE2 | AKR1A1 | 0,4709 | 8,4564E-24 |
| ACE2 | KLHL32 | 0,4708 | 8,60105E-24 |
| ACE2 | CYP2W1 | 0,4705 | 9,42777E-24 |
| ACE2 | OGFRP1 | 0,4704 | 9,63587E-24 |
| ACE2 | ISG20 | 0,4703 | 9,93062E-24 |
| ACE2 | CTB-102L5.8 | 0,4703 | 9,92125E-24 |
| ACE2 | RP11-406A20.1 | 0,4699 | 1,07765E-23 |
| ACE2 | AC128709.3 | 0,4698 | 1,10891E-23 |
| ACE2 | SUCLG2P2 | 0,4698 | 1,098E-23 |
| ACE2 | RP3-400B16.4 | 0,4697 | 1,14031E-23 |
| ACE2 | NCBP2-AS2 | 0,4695 | 1,18903E-23 |
| ACE2 | RP5-894D12.5 | 0,4695 | 1,18217E-23 |
| ACE2 | RP5-881L22.6 | 0,4691 | 1,3198E-23 |
| ACE2 | PAWR | 0,4688 | 1,42526E-23 |
| ACE2 | LINC00244 | 0,4686 | 1,49818E-23 |
| ACE2 | CTC-499B15.6 | 0,4686 | 1,5001E-23 |
| ACE2 | ALOX15B | 0,4685 | 1,52068E-23 |
| ACE2 | PTGDR2 | 0,4683 | 1,59067E-23 |
| ACE2 | CTC-542B22.1 | 0,4683 | 1,60879E-23 |
| ACE2 | RP13-15M17.1 | 0,4682 | 1,62673E-23 |
| ACE2 | CTAGE8 | 0,4677 | 1,83869E-23 |
| ACE2 | RP11-345K9.3 | 0,4676 | 1,91458E-23 |
| ACE2 | SLC12A9-AS1 | 0,4675 | 1,93583E-23 |
| ACE2 | KRT8P22 | 0,4674 | 1,99129E-23 |
| ACE2 | RP11-525G13.2 | 0,4668 | 2,32997E-23 |
| ACE2 | MCM10 | 0,4668 | 2,3122E-23 |
| ACE2 | TYMS | 0,4668 | 2,33913E-23 |
| ACE2 | SGO2 | 0,4667 | 2,3874E-23 |
| ACE2 | RBM22P5 | 0,4666 | 2,45758E-23 |
| ACE2 | RNLS | 0,4666 | 2,45869E-23 |
| ACE2 | RP11-449J21.3 | 0,4666 | 2,43782E-23 |
| ACE2 | GGT1 | 0,4666 | 2,42116E-23 |
| ACE2 | PAX8 | 0,4665 | 2,51557E-23 |
| ACE2 | KRT8P18 | 0,4665 | 2,49066E-23 |
| ACE2 | CENPK | 0,4662 | 2,6875E-23 |
| ACE2 | RP11-407G23.2 | 0,466 | 2,84069E-23 |
| ACE2 | PLIN3 | 0,4657 | 3,03873E-23 |
| ACE2 | CAPZA1 | 0,4655 | 3,20196E-23 |
| ACE2 | INTS6L | 0,4654 | 3,21381E-23 |
| ACE2 | AC022201.4 | 0,465 | 3,60638E-23 |

| | | | |
|---|---|---|---|
| ACE2 | AC072052.7 | 0,465 | 3,60768E-23 |
| ACE2 | TUBBP5 | 0,465 | 3,61785E-23 |
| ACE2 | NUDT12 | 0,4649 | 3,69783E-23 |
| ACE2 | AC073842.19 | 0,4649 | 3,70309E-23 |
| ACE2 | ARHGAP45 | 0,4647 | 3,88363E-23 |
| ACE2 | HSH2D | 0,4647 | 3,85153E-23 |
| ACE2 | RP11-63M22.2 | 0,4644 | 4,12793E-23 |
| ACE2 | C17orf50 | 0,4644 | 4,1238E-23 |
| ACE2 | SLC35A1 | 0,4642 | 4,37247E-23 |
| ACE2 | ETFDH | 0,4641 | 4,4708E-23 |
| ACE2 | ARF3 | 0,4641 | 4,44787E-23 |
| ACE2 | MON1B | 0,4641 | 4,46506E-23 |
| ACE2 | LILRA4 | 0,464 | 4,58689E-23 |
| ACE2 | KRT8P36 | 0,4639 | 4,71063E-23 |
| ACE2 | SLC46A2 | 0,4637 | 4,85138E-23 |
| ACE2 | RP11-336A10.4 | 0,4635 | 5,14304E-23 |
| ACE2 | MT-TN | 0,4634 | 5,30652E-23 |
| ACE2 | AC147651.5 | 0,4633 | 5,3794E-23 |
| ACE2 | RP11-485G7.6 | 0,4633 | 5,34927E-23 |
| ACE2 | LBX2-AS1 | 0,4632 | 5,51023E-23 |
| ACE2 | PFKFB4 | 0,4632 | 5,58515E-23 |
| ACE2 | CNOT11 | 0,4631 | 5,62161E-23 |
| ACE2 | RP11-706O15.1 | 0,4631 | 5,6362E-23 |
| ACE2 | KRT8P43 | 0,463 | 5,81271E-23 |
| ACE2 | CACNA1I | 0,463 | 5,79139E-23 |
| ACE2 | DNAJC19P3 | 0,4628 | 6,04128E-23 |
| ACE2 | COBL | 0,4627 | 6,20573E-23 |
| ACE2 | CYP4F35P | 0,4626 | 6,42277E-23 |
| ACE2 | SUSD6 | 0,4624 | 6,67117E-23 |
| ACE2 | RP11-445K13.2 | 0,4622 | 6,94238E-23 |
| ACE2 | PDCD4-AS1 | 0,4622 | 7,09724E-23 |
| ACE2 | MIP | 0,4622 | 7,08187E-23 |
| ACE2 | PTGES2 | 0,4618 | 7,77797E-23 |
| ACE2 | RN7SKP281 | 0,4614 | 8,54012E-23 |
| ACE2 | KLF3 | 0,4613 | 8,64309E-23 |
| ACE2 | TRIM11 | 0,4612 | 8,81684E-23 |
| ACE2 | CASC8 | 0,4611 | 9,0973E-23 |
| ACE2 | RP5-1061H20.4 | 0,461 | 9,326E-23 |
| ACE2 | RP11-425D10.10 | 0,4607 | 1,00802E-22 |
| ACE2 | RP11-486M23.3 | 0,4607 | 9,9733E-23 |
| ACE2 | PIK3AP1 | 0,4606 | 1,02937E-22 |
| ACE2 | RP11-309J13.1 | 0,4606 | 1,04115E-22 |
| ACE2 | IGHV4-80 | 0,4606 | 1,02774E-22 |
| ACE2 | GBA2 | 0,4605 | 1,0533E-22 |
| ACE2 | AC002059.10 | 0,4605 | 1,06084E-22 |
| ACE2 | FAM46C | 0,4604 | 1,06902E-22 |
| ACE2 | AP000251.3 | 0,4603 | 1,10093E-22 |
| ACE2 | C9orf163 | 0,4601 | 1,17037E-22 |
| ACE2 | FANCD2 | 0,46 | 1,18863E-22 |
| ACE2 | MRPS18B | 0,4599 | 1,2168E-22 |
| ACE2 | DIAPH3 | 0,4599 | 1,20717E-22 |
| ACE2 | RP11-100K18.1 | 0,4598 | 1,25419E-22 |
| ACE2 | HAGLROS | 0,4596 | 1,30511E-22 |
| ACE2 | TAPBPL | 0,4595 | 1,33111E-22 |

| | | | |
|---|---|---|---|
| ACE2 | AC010148.1 | 0,4592 | 1,4403E-22 |
| ACE2 | NT5C | 0,4592 | 1,45015E-22 |
| ACE2 | MYL5 | 0,4591 | 1,48661E-22 |
| ACE2 | LINC01444 | 0,4591 | 1,47405E-22 |
| ACE2 | ZNF326 | 0,4587 | 1,61584E-22 |
| ACE2 | ECT2 | 0,4584 | 1,73209E-22 |
| ACE2 | SLC12A2 | 0,4583 | 1,77273E-22 |
| ACE2 | GINS2 | 0,4582 | 1,80395E-22 |
| ACE2 | RP11-160H22.5 | 0,4581 | 1,87408E-22 |
| ACE2 | ST3GAL1P1 | 0,4581 | 1,86181E-22 |
| ACE2 | CDC45 | 0,4577 | 2,0635E-22 |
| ACE2 | MAMDC4 | 0,4576 | 2,09368E-22 |
| ACE2 | BTBD7P1 | 0,4576 | 2,105E-22 |
| ACE2 | CTD-2260A17.1 | 0,4575 | 2,15018E-22 |
| ACE2 | LINC01572 | 0,4573 | 2,25029E-22 |
| ACE2 | RP11-230B22.1 | 0,4572 | 2,31555E-22 |
| ACE2 | TTLL10 | 0,457 | 2,39803E-22 |
| ACE2 | MND1 | 0,457 | 2,42603E-22 |
| ACE2 | AIMP2 | 0,457 | 2,39713E-22 |
| ACE2 | KIAA2013 | 0,4569 | 2,45125E-22 |
| ACE2 | GP2 | 0,4567 | 2,59786E-22 |
| ACE2 | ST6GAL1 | 0,4565 | 2,69696E-22 |
| ACE2 | ZBTB45P2 | 0,4563 | 2,83478E-22 |
| ACE2 | HTR3C | 0,4563 | 2,81981E-22 |
| ACE2 | RPSAP1 | 0,4563 | 2,81715E-22 |
| ACE2 | RP11-44F21.4 | 0,4561 | 2,99973E-22 |
| ACE2 | AZIN2 | 0,4558 | 3,21112E-22 |
| ACE2 | PKHD1 | 0,4557 | 3,25971E-22 |
| ACE2 | ARFIP2 | 0,4555 | 3,39705E-22 |
| ACE2 | CCDC120 | 0,4554 | 3,52948E-22 |
| ACE2 | TMEM179B | 0,4553 | 3,59386E-22 |
| ACE2 | AC000111.6 | 0,4552 | 3,69278E-22 |
| ACE2 | RP4-697K14.15 | 0,4551 | 3,77572E-22 |
| ACE2 | SLC25A43 | 0,4547 | 4,18613E-22 |
| ACE2 | EML6 | 0,4546 | 4,23041E-22 |
| ACE2 | PIGS | 0,4546 | 4,28334E-22 |
| ACE2 | LSM4 | 0,4546 | 4,22577E-22 |
| ACE2 | CTA-31J9.2 | 0,4545 | 4,30933E-22 |
| ACE2 | LINC00238 | 0,4544 | 4,44748E-22 |
| ACE2 | FRMD4B | 0,4543 | 4,50447E-22 |
| ACE2 | AC007386.4 | 0,4542 | 4,65632E-22 |
| ACE2 | RP11-159H22.2 | 0,4542 | 4,67815E-22 |
| ACE2 | AC073465.2 | 0,4537 | 5,22718E-22 |
| ACE2 | SDS | 0,4536 | 5,30662E-22 |
| ACE2 | CHN2 | 0,4535 | 5,42196E-22 |
| ACE2 | CENPE | 0,4534 | 5,67383E-22 |
| ACE2 | ORC6 | 0,4533 | 5,73829E-22 |
| ACE2 | KIF15 | 0,4532 | 5,91246E-22 |
| ACE2 | RN7SL125P | 0,4532 | 5,87306E-22 |
| ACE2 | RP11-108K3.2 | 0,4531 | 6,02141E-22 |
| ACE2 | RN7SL535P | 0,453 | 6,13757E-22 |
| ACE2 | PCAT18 | 0,453 | 6,09711E-22 |
| ACE2 | NOSTRIN | 0,4527 | 6,53669E-22 |
| ACE2 | NHLRC1 | 0,4527 | 6,61911E-22 |

| | | | |
|---|---|---|---|
| ACE2 | NCAPGP2 | 0,4527 | 6,60301E-22 |
| ACE2 | MRM1 | 0,4524 | 7,15753E-22 |
| ACE2 | RP11-511H23.2 | 0,4523 | 7,20449E-22 |
| ACE2 | RP11-79H23.3 | 0,4521 | 7,51999E-22 |
| ACE2 | AC005355.3 | 0,452 | 7,8007E-22 |
| ACE2 | LINC01268 | 0,452 | 7,83645E-22 |
| ACE2 | MT-ND6 | 0,452 | 7,68452E-22 |
| ACE2 | RP11-143J24.1 | 0,4519 | 7,96341E-22 |
| ACE2 | RP1-170O19.17 | 0,4517 | 8,39128E-22 |
| ACE2 | RP11-404P21.3 | 0,4517 | 8,24004E-22 |
| ACE2 | RER1 | 0,4515 | 8,77379E-22 |
| ACE2 | RP11-738O11.13 | 0,4515 | 8,79016E-22 |
| ACE2 | ATP6V0B | 0,4513 | 9,07822E-22 |
| ACE2 | TXK | 0,4512 | 9,27198E-22 |
| ACE2 | ENPP7P7 | 0,4511 | 9,50515E-22 |
| ACE2 | RP11-758I14.3 | 0,451 | 9,84525E-22 |
| ACE2 | CTAGE15 | 0,451 | 9,71393E-22 |
| ACE2 | RP11-809N8.5 | 0,451 | 9,89583E-22 |
| ACE2 | RP1-7G5.5 | 0,451 | 9,7458E-22 |
| ACE2 | DHRS1 | 0,4509 | 9,93872E-22 |
| ACE2 | DOCK5 | 0,4508 | 1,01317E-21 |
| ACE2 | RP11-465C12.1 | 0,4506 | 1,08447E-21 |
| ACE2 | FAT1 | 0,4505 | 1,09423E-21 |
| ACE2 | PLCE1-AS2 | 0,4502 | 1,17571E-21 |
| ACE2 | CCDC112 | 0,4501 | 1,21E-21 |
| ACE2 | HLA-B | 0,45 | 1,22227E-21 |
| ACE2 | CCR9 | 0,4499 | 1,27E-21 |
| ACE2 | DIRC2 | 0,4498 | 1,28633E-21 |
| ACE2 | RP11-541P9.3 | 0,4498 | 1,30006E-21 |
| ACE2 | BLM | 0,4498 | 1,30357E-21 |
| ACE2 | SPINK4 | 0,4496 | 1,35196E-21 |
| ACE2 | MYZAP | 0,4495 | 1,38084E-21 |
| ACE2 | VSNL1 | 0,4494 | 1,40465E-21 |
| ACE2 | TK1 | 0,4494 | 1,42217E-21 |
| ACE2 | RP11-650P15.1 | 0,4493 | 1,45528E-21 |
| ACE2 | RP11-521D12.4 | 0,4492 | 1,47345E-21 |
| ACE2 | TMSB4XP3 | 0,4491 | 1,52693E-21 |
| ACE2 | RP11-232A1.2 | 0,4491 | 1,51865E-21 |
| ACE2 | KRT8P17 | 0,4489 | 1,58499E-21 |
| ACE2 | B4GALT4-AS1 | 0,4488 | 1,60884E-21 |
| ACE2 | LDHAL6EP | 0,4487 | 1,66135E-21 |
| ACE2 | IGF2BP2-AS1 | 0,4485 | 1,74351E-21 |
| ACE2 | KCTD14 | 0,4485 | 1,72227E-21 |
| ACE2 | NOCT | 0,4483 | 1,82312E-21 |
| ACE2 | IFI30 | 0,4482 | 1,8761E-21 |
| ACE2 | RP11-250H24.2 | 0,4481 | 1,8898E-21 |
| ACE2 | RP11-781P6.1 | 0,4481 | 1,90343E-21 |
| ACE2 | RP11-670E13.3 | 0,4479 | 2,0091E-21 |
| ACE2 | CTC-492K19.4 | 0,4477 | 2,09854E-21 |
| ACE2 | RP11-80H5.2 | 0,4475 | 2,16218E-21 |
| ACE2 | GPRIN1 | 0,4474 | 2,22048E-21 |
| ACE2 | CD58 | 0,4473 | 2,26598E-21 |
| ACE2 | CD8A | 0,4473 | 2,26508E-21 |
| ACE2 | RPP25L | 0,4472 | 2,3311E-21 |

| | | | |
|---|---|---|---|
| ACE2 | SLPI | 0,447 | 2,45167E-21 |
| ACE2 | WDR55 | 0,4466 | 2,70946E-21 |
| ACE2 | TIGD3 | 0,4466 | 2,68399E-21 |
| ACE2 | LINC00032 | 0,4465 | 2,75504E-21 |
| ACE2 | ASTL | 0,4463 | 2,85842E-21 |
| ACE2 | CLEC18A | 0,4462 | 2,94284E-21 |
| ACE2 | RP11-1060J15.9 | 0,4459 | 3,11707E-21 |
| ACE2 | C2orf48 | 0,4458 | 3,23474E-21 |
| ACE2 | RP3-434P1.6 | 0,4457 | 3,285E-21 |
| ACE2 | GPT2 | 0,4454 | 3,5378E-21 |
| ACE2 | RP11-142I2.1 | 0,4453 | 3,61052E-21 |
| ACE2 | TYSND1 | 0,4452 | 3,66968E-21 |
| ACE2 | SYNGR4 | 0,445 | 3,86361E-21 |
| ACE2 | CXCR6 | 0,4449 | 3,96573E-21 |
| ACE2 | TRPA1 | 0,4449 | 3,97953E-21 |
| ACE2 | RP11-550A18.1 | 0,4448 | 4,07024E-21 |
| ACE2 | RP11-493E3.1 | 0,4447 | 4,12355E-21 |
| ACE2 | PLEKHF2 | 0,4445 | 4,28903E-21 |
| ACE2 | RP11-659E9.4 | 0,4443 | 4,54462E-21 |
| ACE2 | ACTN4 | 0,4443 | 4,49943E-21 |
| ACE2 | KRT18P15 | 0,4441 | 4,69266E-21 |
| ACE2 | RP11-471M2.3 | 0,4439 | 4,9722E-21 |
| ACE2 | AC012499.1 | 0,4437 | 5,22549E-21 |
| ACE2 | CTD-3199J23.6 | 0,4437 | 5,12659E-21 |
| ACE2 | RP11-480I12.9 | 0,4436 | 5,27541E-21 |
| ACE2 | LAMA3 | 0,4436 | 5,25911E-21 |
| ACE2 | RP11-303E16.9 | 0,4435 | 5,37959E-21 |
| ACE2 | MUT | 0,4433 | 5,62426E-21 |
| ACE2 | SLC25A39 | 0,4433 | 5,72286E-21 |
| ACE2 | ZNF575 | 0,4433 | 5,71773E-21 |
| ACE2 | RP11-245A18.1 | 0,443 | 6,04954E-21 |
| ACE2 | LINC01725 | 0,4429 | 6,20625E-21 |
| ACE2 | RP11-182N22.8 | 0,4427 | 6,50207E-21 |
| ACE2 | FDPS | 0,4426 | 6,5785E-21 |
| ACE2 | VEPH1 | 0,4426 | 6,56316E-21 |
| ACE2 | RP11-296O14.3 | 0,4423 | 7,15955E-21 |
| ACE2 | ZFP3 | 0,4423 | 7,07398E-21 |
| ACE2 | CTD-2377D24.2 | 0,4423 | 7,10266E-21 |
| ACE2 | MTCO2P12 | 0,4422 | 7,31226E-21 |
| ACE2 | IFI27L1 | 0,4422 | 7,2043E-21 |
| ACE2 | PRR5 | 0,4422 | 7,32728E-21 |
| ACE2 | ABCC11 | 0,4421 | 7,44901E-21 |
| ACE2 | NDRG1 | 0,4419 | 7,80485E-21 |
| ACE2 | PTER | 0,4418 | 7,84869E-21 |
| ACE2 | LINC00390 | 0,4418 | 7,84755E-21 |
| ACE2 | PRKG2 | 0,4417 | 8,17754E-21 |
| ACE2 | MT-TE | 0,4416 | 8,2202E-21 |
| ACE2 | HIST1H2AG | 0,4415 | 8,46168E-21 |
| ACE2 | UQCR10 | 0,4414 | 8,72695E-21 |
| ACE2 | CTD-2256P15.5 | 0,4412 | 9,06238E-21 |
| ACE2 | COX6A1P2 | 0,4412 | 8,99228E-21 |
| ACE2 | PARPBP | 0,4412 | 9,06468E-21 |
| ACE2 | C19orf45 | 0,4412 | 8,9985E-21 |
| ACE2 | RP11-352D13.6 | 0,4411 | 9,24725E-21 |

| | | | |
|---|---|---|---|
| ACE2 | SDHD | 0,441 | 9,39413E-21 |
| ACE2 | AC005522.7 | 0,4408 | 1,00235E-20 |
| ACE2 | RP11-428C19.4 | 0,4408 | 9,98833E-21 |
| ACE2 | RP11-509E10.1 | 0,4408 | 9,83224E-21 |
| ACE2 | RP11-1157N2__B.2 | 0,4408 | 9,9383E-21 |
| ACE2 | RP1-120G22.11 | 0,4407 | 1,00941E-20 |
| ACE2 | PTTG1 | 0,4405 | 1,05689E-20 |
| ACE2 | PLBD1-AS1 | 0,4405 | 1,06367E-20 |
| ACE2 | EPPK1 | 0,4404 | 1,07707E-20 |
| ACE2 | ASCL2 | 0,4404 | 1,0739E-20 |
| ACE2 | RP5-875H18.4 | 0,4404 | 1,0909E-20 |
| ACE2 | EXO1 | 0,4403 | 1,11093E-20 |
| ACE2 | RPL17P11 | 0,4401 | 1,14795E-20 |
| ACE2 | RORA-AS1 | 0,4401 | 1,1493E-20 |
| ACE2 | SSH3 | 0,44 | 1,19292E-20 |
| ACE2 | ATP5A1 | 0,4399 | 1,22256E-20 |
| ACE2 | RP11-417L19.6 | 0,4398 | 1,25015E-20 |
| ACE2 | EHD4 | 0,4398 | 1,23959E-20 |
| ACE2 | RP11-255E6.6 | 0,4396 | 1,29347E-20 |
| ACE2 | GULOP | 0,4396 | 1,29761E-20 |
| ACE2 | DMRTA1 | 0,4396 | 1,30352E-20 |
| ACE2 | AC018717.1 | 0,4395 | 1,318E-20 |
| ACE2 | CTC-251D13.1 | 0,4395 | 1,31844E-20 |
| ACE2 | CTDP1 | 0,4395 | 1,32117E-20 |
| ACE2 | RP4-669P10.19 | 0,4395 | 1,32999E-20 |
| ACE2 | RP11-67L3.4 | 0,4394 | 1,34878E-20 |
| ACE2 | RP11-379K17.9 | 0,4392 | 1,40449E-20 |
| ACE2 | TRDV1 | 0,439 | 1,4918E-20 |
| ACE2 | NEBL | 0,4388 | 1,5403E-20 |
| ACE2 | RP11-575F12.1 | 0,4385 | 1,66141E-20 |
| ACE2 | NARS | 0,4383 | 1,71708E-20 |
| ACE2 | TNIP3 | 0,4382 | 1,77932E-20 |
| ACE2 | BFSP2-AS1 | 0,4381 | 1,82185E-20 |
| ACE2 | FAM111B | 0,4381 | 1,82657E-20 |
| ACE2 | PNP | 0,4381 | 1,80623E-20 |
| ACE2 | AC005224.2 | 0,4381 | 1,79873E-20 |
| ACE2 | FAM110A | 0,4377 | 1,96369E-20 |
| ACE2 | IGSF8 | 0,4376 | 1,99644E-20 |
| ACE2 | CENPU | 0,4374 | 2,11337E-20 |
| ACE2 | SPATA3-AS1 | 0,4371 | 2,26393E-20 |
| ACE2 | IL36G | 0,437 | 2,28727E-20 |
| ACE2 | ATG4A | 0,437 | 2,29612E-20 |
| ACE2 | RP11-388N2.1 | 0,4369 | 2,33103E-20 |
| ACE2 | RP11-15F12.1 | 0,4367 | 2,48625E-20 |
| ACE2 | KLLN | 0,4363 | 2,67918E-20 |
| ACE2 | NUS1P3 | 0,4363 | 2,6819E-20 |
| ACE2 | RP11-47A17.2 | 0,4362 | 2,77322E-20 |
| ACE2 | MRPS36 | 0,4362 | 2,74845E-20 |
| ACE2 | AC015849.16 | 0,4362 | 2,75332E-20 |
| ACE2 | CTD-2116N20.1 | 0,4361 | 2,80267E-20 |
| ACE2 | ACAT1 | 0,4361 | 2,83321E-20 |
| ACE2 | TBPL2 | 0,4361 | 2,80107E-20 |
| ACE2 | CREB3L4 | 0,436 | 2,84479E-20 |
| ACE2 | KRT18P29 | 0,436 | 2,89726E-20 |

| | | | |
|---|---|---|---|
| ACE2 | SLX4IP | 0,4359 | 2,92787E-20 |
| ACE2 | RP11-379B8.1 | 0,4358 | 3,00688E-20 |
| ACE2 | KMO | 0,4357 | 3,06387E-20 |
| ACE2 | SEMG1 | 0,4357 | 3,05854E-20 |
| ACE2 | RP11-555M1.3 | 0,4356 | 3,11653E-20 |
| ACE2 | BRCC3 | 0,4356 | 3,15421E-20 |
| ACE2 | MYO19 | 0,4354 | 3,30774E-20 |
| ACE2 | AC069277.2 | 0,4353 | 3,3356E-20 |
| ACE2 | HSD17B8 | 0,4352 | 3,44354E-20 |
| ACE2 | CTD-2008A1.3 | 0,4352 | 3,40698E-20 |
| ACE2 | TMEM220-AS1 | 0,4352 | 3,45993E-20 |
| ACE2 | NT5DC4 | 0,4351 | 3,46939E-20 |
| ACE2 | AC024937.2 | 0,435 | 3,5658E-20 |
| ACE2 | NUDT16 | 0,4349 | 3,67063E-20 |
| ACE2 | ARID3B | 0,4348 | 3,7376E-20 |
| ACE2 | PI4K2B | 0,4345 | 3,97691E-20 |
| ACE2 | MTND2P11 | 0,4343 | 4,19655E-20 |
| ACE2 | C1GALT1C1 | 0,4343 | 4,12899E-20 |
| ACE2 | MT-ND4 | 0,4343 | 4,21051E-20 |
| ACE2 | DNPH1 | 0,4339 | 4,51037E-20 |
| ACE2 | RP11-539I5.1 | 0,4339 | 4,52818E-20 |
| ACE2 | ADGRD2 | 0,4338 | 4,601E-20 |
| ACE2 | WDR62 | 0,4337 | 4,70773E-20 |
| ACE2 | UGT1A3 | 0,4336 | 4,80465E-20 |
| ACE2 | RN7SL834P | 0,4335 | 4,99581E-20 |
| ACE2 | CASP2 | 0,4335 | 4,94192E-20 |
| ACE2 | XIAPP3 | 0,4334 | 5,12628E-20 |
| ACE2 | IGHV3-73 | 0,4334 | 5,05146E-20 |
| ACE2 | CHAF1A | 0,4334 | 5,07544E-20 |
| ACE2 | RAD54L | 0,4332 | 5,34316E-20 |
| ACE2 | SIGLEC10 | 0,4331 | 5,38254E-20 |
| ACE2 | RPARP-AS1 | 0,433 | 5,57378E-20 |
| ACE2 | NIT1 | 0,4325 | 6,22571E-20 |
| ACE2 | LINC00307 | 0,4324 | 6,26288E-20 |
| ACE2 | RP11-564D11.3 | 0,4321 | 6,71692E-20 |
| ACE2 | RP11-234B24.4 | 0,4321 | 6,77075E-20 |
| ACE2 | RP4-803J11.2 | 0,432 | 6,87152E-20 |
| ACE2 | RP11-241M13.2 | 0,432 | 6,82801E-20 |
| ACE2 | TMEM120A | 0,432 | 6,86426E-20 |
| ACE2 | LINC01977 | 0,432 | 6,81515E-20 |
| ACE2 | RP11-1399P15.1 | 0,4319 | 7,06873E-20 |
| ACE2 | SFXN2 | 0,4319 | 6,96453E-20 |
| ACE2 | IGHV3OR16-13 | 0,4318 | 7,21282E-20 |
| ACE2 | RN7SKP25 | 0,4317 | 7,27041E-20 |
| ACE2 | ALDOB | 0,4317 | 7,35162E-20 |
| ACE2 | RP11-490O6.2 | 0,4316 | 7,42562E-20 |
| ACE2 | SCML4 | 0,4315 | 7,69115E-20 |
| ACE2 | CCDC187 | 0,4315 | 7,65793E-20 |
| ACE2 | RP11-900F13.3 | 0,4313 | 8,07414E-20 |
| ACE2 | KDM7A | 0,4312 | 8,26291E-20 |
| ACE2 | FUOM | 0,4312 | 8,22265E-20 |
| ACE2 | RP1-170O19.14 | 0,431 | 8,54757E-20 |
| ACE2 | RMI2 | 0,431 | 8,56024E-20 |
| ACE2 | RP13-225O21.2 | 0,4309 | 8,75834E-20 |

| | | | |
|---|---|---|---|
| ACE2 | AC011513.4 | 0,4309 | 8,65745E-20 |
| ACE2 | GCLM | 0,4308 | 8,83255E-20 |
| ACE2 | SST | 0,4308 | 8,91519E-20 |
| ACE2 | RP11-800A3.2 | 0,4308 | 8,84149E-20 |
| ACE2 | GK-AS1 | 0,4308 | 8,93746E-20 |
| ACE2 | AC016644.1 | 0,4307 | 9,07917E-20 |
| ACE2 | SLC5A8 | 0,4307 | 9,02884E-20 |
| ACE2 | LINC01658 | 0,4306 | 9,35725E-20 |
| ACE2 | RP11-395B7.4 | 0,4303 | 9,95305E-20 |
| ACE2 | LINC00871 | 0,4302 | 1,0254E-19 |
| ACE2 | FADS2P1 | 0,4301 | 1,03352E-19 |
| ACE2 | AC006978.6 | 0,43 | 1,06461E-19 |
| ACE2 | RP11-338I21.1 | 0,43 | 1,0519E-19 |
| ACE2 | SLC22A4 | 0,4299 | 1,09356E-19 |
| ACE2 | CCDC14 | 0,4298 | 1,10755E-19 |
| ACE2 | DNM1P17 | 0,4297 | 1,1299E-19 |
| ACE2 | RP11-314A15.3 | 0,4297 | 1,13813E-19 |
| ACE2 | PGAP2 | 0,4295 | 1,17758E-19 |
| ACE2 | OR6T1 | 0,4295 | 1,17746E-19 |
| ACE2 | RP11-453E17.3 | 0,4292 | 1,24639E-19 |
| ACE2 | RP6-149D17.1 | 0,4292 | 1,2686E-19 |
| ACE2 | MED28P3 | 0,4291 | 1,2934E-19 |
| ACE2 | EIF2AK1 | 0,429 | 1,31159E-19 |
| ACE2 | RP1-313I6.12 | 0,4289 | 1,33423E-19 |
| ACE2 | CLSPN | 0,4284 | 1,50078E-19 |
| ACE2 | HES1 | 0,4284 | 1,47888E-19 |
| ACE2 | MB | 0,4284 | 1,49855E-19 |
| ACE2 | COQ4 | 0,4282 | 1,5574E-19 |
| ACE2 | RP1-150O5.3 | 0,4281 | 1,59827E-19 |
| ACE2 | PNLIPRP1 | 0,4281 | 1,58307E-19 |
| ACE2 | CD3G | 0,4281 | 1,58541E-19 |
| ACE2 | AC009403.2 | 0,428 | 1,64375E-19 |
| ACE2 | IGHV3-66 | 0,428 | 1,61399E-19 |
| ACE2 | MT1G | 0,428 | 1,64305E-19 |
| ACE2 | RP11-446N19.1 | 0,4279 | 1,66585E-19 |
| ACE2 | CTD-2184C24.2 | 0,4278 | 1,69737E-19 |
| ACE2 | EFNA1 | 0,4276 | 1,76504E-19 |
| ACE2 | NPW | 0,4275 | 1,80569E-19 |
| ACE2 | RP11-84C10.1 | 0,4274 | 1,85188E-19 |
| ACE2 | IGHV6-1 | 0,4272 | 1,93295E-19 |
| ACE2 | ARHGAP11A | 0,4271 | 1,97348E-19 |
| ACE2 | CTD-2227E11.1 | 0,427 | 2,00423E-19 |
| ACE2 | IGHV4-4 | 0,427 | 1,99674E-19 |
| ACE2 | RBMY2TP | 0,427 | 2,03445E-19 |
| ACE2 | MUC6 | 0,4269 | 2,04718E-19 |
| ACE2 | MARC1 | 0,4265 | 2,21808E-19 |
| ACE2 | RP11-697K23.3 | 0,4265 | 2,26477E-19 |
| ACE2 | TNFRSF10A | 0,4264 | 2,274E-19 |
| ACE2 | RP3-405J10.4 | 0,4264 | 2,27386E-19 |
| ACE2 | IGHV3-64 | 0,4263 | 2,32469E-19 |
| ACE2 | RP11-647P12.1 | 0,4262 | 2,40187E-19 |
| ACE2 | HOXA1 | 0,4262 | 2,40222E-19 |
| ACE2 | KRTAP5-10 | 0,4262 | 2,39232E-19 |
| ACE2 | IGHV1-3 | 0,4261 | 2,43639E-19 |

| | | | |
|---|---|---|---|
| ACE2 | DCST2 | 0,426 | 2,48799E-19 |
| ACE2 | DBF4 | 0,426 | 2,51311E-19 |
| ACE2 | G6PC | 0,426 | 2,49751E-19 |
| ACE2 | RECQL5 | 0,426 | 2,49445E-19 |
| ACE2 | CNTD1 | 0,4257 | 2,68369E-19 |
| ACE2 | OSBPL2 | 0,4257 | 2,67751E-19 |
| ACE2 | AC006486.10 | 0,4256 | 2,69747E-19 |
| ACE2 | RP6-91H8.5 | 0,4255 | 2,7522E-19 |
| ACE2 | KIAA1217 | 0,4254 | 2,83503E-19 |
| ACE2 | RP11-418I22.2 | 0,4251 | 2,99161E-19 |
| ACE2 | MTND5P11 | 0,425 | 3,05854E-19 |
| ACE2 | RP11-267D19.2 | 0,425 | 3,08423E-19 |
| ACE2 | RP11-362L22.1 | 0,4248 | 3,21846E-19 |
| ACE2 | IGHV1-69 | 0,4248 | 3,24894E-19 |
| ACE2 | C1orf127 | 0,4245 | 3,41378E-19 |
| ACE2 | CAPN12 | 0,4245 | 3,43298E-19 |
| ACE2 | DDOST | 0,4244 | 3,51337E-19 |
| ACE2 | RP11-982M15.5 | 0,4244 | 3,48815E-19 |
| ACE2 | AC068580.6 | 0,4241 | 3,69652E-19 |
| ACE2 | PGBD2 | 0,424 | 3,77133E-19 |
| ACE2 | SLC25A5P5 | 0,4239 | 3,87297E-19 |
| ACE2 | SIGLEC12 | 0,4239 | 3,86804E-19 |
| ACE2 | ARSA | 0,4239 | 3,86752E-19 |
| ACE2 | MT1F | 0,4238 | 3,93594E-19 |
| ACE2 | IGLV3-9 | 0,4238 | 3,9342E-19 |
| ACE2 | PRR11 | 0,4237 | 4,03749E-19 |
| ACE2 | RPL13AP20 | 0,4233 | 4,40241E-19 |
| ACE2 | RP11-812E19.9 | 0,4233 | 4,37068E-19 |
| ACE2 | IGLV5-45 | 0,4233 | 4,42605E-19 |
| ACE2 | IGHV4-61 | 0,4232 | 4,47558E-19 |
| ACE2 | CATSPERG | 0,4232 | 4,54979E-19 |
| ACE2 | CXorf40A | 0,4232 | 4,54536E-19 |
| ACE2 | PLSCR1 | 0,423 | 4,66751E-19 |
| ACE2 | SMC6 | 0,4229 | 4,82629E-19 |
| ACE2 | CPT1B | 0,4229 | 4,75521E-19 |
| ACE2 | AC004066.2 | 0,4228 | 4,91077E-19 |
| ACE2 | RP4-777L9.2 | 0,4227 | 5,01358E-19 |
| ACE2 | RP3-446N13.5 | 0,4226 | 5,06392E-19 |
| ACE2 | MPZL2 | 0,4225 | 5,17831E-19 |
| ACE2 | IL17REL | 0,4223 | 5,47357E-19 |
| ACE2 | IGHV3-74 | 0,4221 | 5,73698E-19 |
| ACE2 | TTC13 | 0,422 | 5,75381E-19 |
| ACE2 | RP11-710F7.2 | 0,422 | 5,79979E-19 |
| ACE2 | TRIB1 | 0,4219 | 5,95034E-19 |
| ACE2 | C14orf80 | 0,4218 | 6,08116E-19 |
| ACE2 | KRTAP9-1 | 0,4218 | 6,09151E-19 |
| ACE2 | NUGGC | 0,4215 | 6,49018E-19 |
| ACE2 | INF2 | 0,4211 | 6,94485E-19 |
| ACE2 | VWDE | 0,421 | 7,12598E-19 |
| ACE2 | SETD9 | 0,4209 | 7,31187E-19 |
| ACE2 | RP11-639F1.2 | 0,4208 | 7,40267E-19 |
| ACE2 | RP11-91K8.4 | 0,4207 | 7,65936E-19 |
| ACE2 | PRF1 | 0,4207 | 7,69184E-19 |
| ACE2 | SYNE4 | 0,4207 | 7,57009E-19 |

| | | | |
|---|---|---|---|
| ACE2 | MYO15B | 0,4206 | 7,83471E-19 |
| ACE2 | SAT1 | 0,4204 | 8,06254E-19 |
| ACE2 | ACADM | 0,4202 | 8,40718E-19 |
| ACE2 | APOBEC4 | 0,4202 | 8,41223E-19 |
| ACE2 | RP11-478K15.7 | 0,4202 | 8,48135E-19 |
| ACE2 | RP11-254F7.4 | 0,4201 | 8,71391E-19 |
| ACE2 | IGFBP2 | 0,4201 | 8,71055E-19 |
| ACE2 | LINC00330 | 0,42 | 8,74705E-19 |
| ACE2 | IGHV3-23 | 0,4194 | 9,96608E-19 |
| ACE2 | RP11-190C22.8 | 0,4192 | 1,04576E-18 |
| ACE2 | RP11-496H1.1 | 0,4191 | 1,06693E-18 |
| ACE2 | PHPT1 | 0,4191 | 1,06282E-18 |
| ACE2 | C17orf77 | 0,419 | 1,08762E-18 |
| ACE2 | SLC5A11 | 0,4189 | 1,10231E-18 |
| ACE2 | RP13-210D15.8 | 0,4189 | 1,10122E-18 |
| ACE2 | SECTM1 | 0,4188 | 1,13102E-18 |
| ACE2 | RP11-46D6.1 | 0,4187 | 1,16612E-18 |
| ACE2 | IGLV2-11 | 0,4185 | 1,21433E-18 |
| ACE2 | RP1-317E23.3 | 0,4184 | 1,22245E-18 |
| ACE2 | CBR1 | 0,4183 | 1,26227E-18 |
| ACE2 | RP11-153K11.3 | 0,4182 | 1,27689E-18 |
| ACE2 | IGHV3-13 | 0,4182 | 1,27963E-18 |
| ACE2 | RP1-10C16.1 | 0,418 | 1,34246E-18 |
| ACE2 | RP4-803A2.2 | 0,4176 | 1,44125E-18 |
| ACE2 | CTB-31N19.4 | 0,4175 | 1,47982E-18 |
| ACE2 | RP11-165F24.5 | 0,4172 | 1,57658E-18 |
| ACE2 | RP11-539L10.2 | 0,4169 | 1,68908E-18 |
| ACE2 | TECR | 0,4168 | 1,70132E-18 |
| ACE2 | IGKV1D-8 | 0,4166 | 1,78634E-18 |
| ACE2 | MRPL42P2 | 0,4166 | 1,7685E-18 |
| ACE2 | SUMF1 | 0,4165 | 1,809E-18 |
| ACE2 | CTC-756D1.3 | 0,4164 | 1,85531E-18 |
| ACE2 | RP11-108K3.1 | 0,4164 | 1,86034E-18 |
| ACE2 | UHRF1 | 0,4163 | 1,8828E-18 |
| ACE2 | XCR1 | 0,4162 | 1,94438E-18 |
| ACE2 | ATP6V0CP1 | 0,4161 | 1,96435E-18 |
| ACE2 | RP11-386M24.9 | 0,4159 | 2,06285E-18 |
| ACE2 | RP11-757O6.6 | 0,4159 | 2,04413E-18 |
| ACE2 | AC011899.10 | 0,4156 | 2,19923E-18 |
| ACE2 | SPACA3 | 0,4155 | 2,24759E-18 |
| ACE2 | RP13-254B10.1 | 0,4154 | 2,27014E-18 |
| ACE2 | IGHV3-15 | 0,4153 | 2,31296E-18 |
| ACE2 | AC007078.4 | 0,4151 | 2,43861E-18 |
| ACE2 | TRIP13 | 0,415 | 2,49338E-18 |
| ACE2 | CTAGE9 | 0,4149 | 2,52489E-18 |
| ACE2 | AC005546.2 | 0,4149 | 2,54725E-18 |
| ACE2 | RP11-387A1.6 | 0,4147 | 2,64562E-18 |
| ACE2 | PHF7 | 0,4147 | 2,61205E-18 |
| ACE2 | RP11-150O12.6 | 0,4146 | 2,71427E-18 |
| ACE2 | ZNF695 | 0,4145 | 2,72953E-18 |
| ACE2 | B3GNT2 | 0,4143 | 2,86211E-18 |
| ACE2 | CTD-2583A14.8 | 0,4142 | 2,93794E-18 |
| ACE2 | RP3-337O18.9 | 0,4142 | 2,92468E-18 |
| ACE2 | ANXA11 | 0,414 | 3,01377E-18 |

| | | | |
|---|---|---|---|
| ACE2 | SLC38A5 | 0,414 | 3,04587E-18 |
| ACE2 | TNFRSF17 | 0,4139 | 3,07601E-18 |
| ACE2 | DEPDC4 | 0,4138 | 3,14196E-18 |
| ACE2 | MOXD2P | 0,4137 | 3,24177E-18 |
| ACE2 | MYPN | 0,4137 | 3,2592E-18 |
| ACE2 | SLC7A11-AS1 | 0,4136 | 3,27923E-18 |
| ACE2 | POLR2H | 0,4134 | 3,40599E-18 |
| ACE2 | CTB-31O20.8 | 0,4134 | 3,41218E-18 |
| ACE2 | SMIM14 | 0,4133 | 3,50801E-18 |
| ACE2 | B2M | 0,4133 | 3,47909E-18 |
| ACE2 | LRRC8E | 0,4133 | 3,52297E-18 |
| ACE2 | ITGA2 | 0,4132 | 3,60026E-18 |
| ACE2 | RP11-542G1.1 | 0,4131 | 3,62862E-18 |
| ACE2 | IGHV3-49 | 0,4131 | 3,65931E-18 |
| ACE2 | KSR2 | 0,413 | 3,72124E-18 |
| ACE2 | IGHV3OR16-8 | 0,413 | 3,73214E-18 |
| ACE2 | XXbac-B476C20.11 | 0,4127 | 3,97668E-18 |
| ACE2 | IGHJ6 | 0,4125 | 4,09414E-18 |
| ACE2 | RP11-115A14.1 | 0,4124 | 4,16757E-18 |
| ACE2 | SNORA5B | 0,4124 | 4,21321E-18 |
| ACE2 | AC009961.5 | 0,4122 | 4,34097E-18 |
| ACE2 | CACNA1D | 0,4122 | 4,41779E-18 |
| ACE2 | RP11-864J10.4 | 0,4122 | 4,36777E-18 |
| ACE2 | IGHV3-53 | 0,4122 | 4,40026E-18 |
| ACE2 | LNX1-AS2 | 0,412 | 4,60489E-18 |
| ACE2 | IGHV3-72 | 0,412 | 4,60298E-18 |
| ACE2 | IGHV3-43 | 0,4119 | 4,64119E-18 |
| ACE2 | MYH13 | 0,4119 | 4,65254E-18 |
| ACE2 | C6orf203 | 0,4117 | 4,88716E-18 |
| ACE2 | PRDM1 | 0,4116 | 4,94817E-18 |
| ACE2 | DDX60L | 0,4115 | 5,01207E-18 |
| ACE2 | TEKT5 | 0,4113 | 5,24801E-18 |
| ACE2 | TMED1 | 0,4113 | 5,2837E-18 |
| ACE2 | RPL9P28 | 0,4112 | 5,32218E-18 |
| ACE2 | IGLV2-18 | 0,4111 | 5,48761E-18 |
| ACE2 | RP11-429J17.5 | 0,411 | 5,56095E-18 |
| ACE2 | DHRS4 | 0,411 | 5,54171E-18 |
| ACE2 | HPCA | 0,4109 | 5,68388E-18 |
| ACE2 | AC005532.5 | 0,4109 | 5,74981E-18 |
| ACE2 | RNA5SP434 | 0,4108 | 5,81109E-18 |
| ACE2 | MT-ND1 | 0,4107 | 5,91395E-18 |
| ACE2 | SLC39A14 | 0,4105 | 6,15247E-18 |
| ACE2 | LL0XNC01-131B10.2 | 0,4105 | 6,18135E-18 |
| ACE2 | SLC35A4 | 0,4104 | 6,3509E-18 |
| ACE2 | IGHV2-5 | 0,4104 | 6,29867E-18 |
| ACE2 | AC142119.1 | 0,4102 | 6,5307E-18 |
| ACE2 | PLA2G4E-AS1 | 0,41 | 6,82473E-18 |
| ACE2 | AMMECR1 | 0,41 | 6,89511E-18 |
| ACE2 | MCUR1 | 0,4099 | 7,01787E-18 |
| ACE2 | RBMY2EP | 0,4099 | 7,00103E-18 |
| ACE2 | RP11-649G15.2 | 0,4098 | 7,08157E-18 |
| ACE2 | ERCC6L | 0,4098 | 7,18437E-18 |
| ACE2 | RAB42 | 0,4097 | 7,28919E-18 |
| ACE2 | AC008592.3 | 0,4097 | 7,32048E-18 |

| | | | |
|---|---|---|---|
| ACE2 | RP11-166P13.4 | 0,4097 | 7,31823E-18 |
| ACE2 | KRT18P16 | 0,4096 | 7,46697E-18 |
| ACE2 | RP11-320N7.1 | 0,4096 | 7,4297E-18 |
| ACE2 | TARDBPP2 | 0,4096 | 7,41036E-18 |
| ACE2 | IGLV2-8 | 0,4096 | 7,40999E-18 |
| ACE2 | TRAPPC6A | 0,4095 | 7,62941E-18 |
| ACE2 | KRT8P42 | 0,4094 | 7,76438E-18 |
| ACE2 | MRPL2 | 0,4093 | 7,89939E-18 |
| ACE2 | IGHV3-11 | 0,4093 | 7,9089E-18 |
| ACE2 | RP11-489M13.3 | 0,4092 | 7,96098E-18 |
| ACE2 | PDCD6 | 0,4091 | 8,22754E-18 |
| ACE2 | IGHV3-25 | 0,4091 | 8,22742E-18 |
| ACE2 | FAR1 | 0,409 | 8,388E-18 |
| ACE2 | CDC37P2 | 0,409 | 8,32196E-18 |
| ACE2 | COG3 | 0,4089 | 8,47913E-18 |
| ACE2 | IGLV3-1 | 0,4089 | 8,5629E-18 |
| ACE2 | CDK3 | 0,4088 | 8,7732E-18 |
| ACE2 | CASC19 | 0,4087 | 8,87445E-18 |
| ACE2 | IGHJ2 | 0,4086 | 9,02302E-18 |
| ACE2 | CYTH1 | 0,4086 | 9,1091E-18 |
| ACE2 | DRGX | 0,4085 | 9,29939E-18 |
| ACE2 | RP11-148O7.2 | 0,4084 | 9,44633E-18 |
| ACE2 | IGHV3-7 | 0,4084 | 9,52201E-18 |
| ACE2 | RP11-329A14.1 | 0,4083 | 9,67515E-18 |
| ACE2 | RP11-480A16.1 | 0,4082 | 9,78448E-18 |
| ACE2 | RP11-484D2.2 | 0,4081 | 9,99696E-18 |
| ACE2 | B4GALT1 | 0,4077 | 1,08379E-17 |
| ACE2 | IGHV3-65 | 0,4077 | 1,08804E-17 |
| ACE2 | RP11-34D15.2 | 0,4074 | 1,14483E-17 |
| ACE2 | B3GALT4 | 0,4073 | 1,18419E-17 |
| ACE2 | RP11-466A19.7 | 0,4073 | 1,16476E-17 |
| ACE2 | RP11-632K5.2 | 0,4071 | 1,2351E-17 |
| ACE2 | CTD-2666L21.2 | 0,4071 | 1,21729E-17 |
| ACE2 | RP11-12G12.7 | 0,4068 | 1,29444E-17 |
| ACE2 | RP11-474P2.7 | 0,4067 | 1,31326E-17 |
| ACE2 | MIR17HG | 0,4067 | 1,31462E-17 |
| ACE2 | TSSK6 | 0,4067 | 1,32052E-17 |
| ACE2 | FXYD4 | 0,4065 | 1,39164E-17 |
| ACE2 | NDUFS1 | 0,4064 | 1,41779E-17 |
| ACE2 | RP11-20I23.2 | 0,4064 | 1,41233E-17 |
| ACE2 | ACSM3 | 0,4064 | 1,39617E-17 |
| ACE2 | IGHV3-64D | 0,406 | 1,51136E-17 |
| ACE2 | LINC01564 | 0,4059 | 1,55046E-17 |
| ACE2 | C16orf59 | 0,4059 | 1,54497E-17 |
| ACE2 | IGHV3-75 | 0,4057 | 1,61115E-17 |
| ACE2 | IGHV3-33 | 0,4056 | 1,63529E-17 |
| ACE2 | ZNF726 | 0,4056 | 1,64892E-17 |
| ACE2 | BRCA2 | 0,4055 | 1,69724E-17 |
| ACE2 | LINC00324 | 0,4055 | 1,6771E-17 |
| ACE2 | TPI1P2 | 0,4054 | 1,70532E-17 |
| ACE2 | TBC1D30 | 0,4054 | 1,70852E-17 |
| ACE2 | USP32P3 | 0,4054 | 1,72019E-17 |
| ACE2 | RP11-219O3.2 | 0,4053 | 1,76133E-17 |
| ACE2 | RP11-1223D19.5 | 0,4052 | 1,78203E-17 |

| | | | |
|---|---|---|---|
| ACE2 | UQCC3 | 0,4052 | 1,78353E-17 |
| ACE2 | NFE2L3 | 0,4051 | 1,8278E-17 |
| ACE2 | RP11-45P22.2 | 0,4051 | 1,81185E-17 |
| ACE2 | GALNT8 | 0,4051 | 1,83001E-17 |
| ACE2 | RP11-170L3.7 | 0,4051 | 1,82556E-17 |
| ACE2 | IGKV3D-7 | 0,405 | 1,85501E-17 |
| ACE2 | CRYBB2 | 0,4049 | 1,89025E-17 |
| ACE2 | AC112229.6 | 0,4048 | 1,93789E-17 |
| ACE2 | ARHGAP36 | 0,4047 | 1,98845E-17 |
| ACE2 | KLRC4 | 0,4046 | 1,99933E-17 |
| ACE2 | ABTB2 | 0,4045 | 2,0305E-17 |
| ACE2 | AP000479.1 | 0,4043 | 2,11329E-17 |
| ACE2 | OR4K7P | 0,4042 | 2,18364E-17 |
| ACE2 | NCR1 | 0,4042 | 2,16879E-17 |
| ACE2 | AC144835.1 | 0,4041 | 2,22685E-17 |
| ACE2 | TSPAN7 | 0,404 | 2,25523E-17 |
| ACE2 | SLCO2B1 | 0,4038 | 2,33137E-17 |
| ACE2 | EIF2S3L | 0,4036 | 2,45306E-17 |
| ACE2 | IGHV3-48 | 0,4036 | 2,42803E-17 |
| ACE2 | SMIM2-IT1 | 0,4034 | 2,56669E-17 |
| ACE2 | SHMT1 | 0,4031 | 2,72317E-17 |
| ACE2 | RP11-443B20.1 | 0,403 | 2,76924E-17 |
| ACE2 | RP11-27I1.4 | 0,403 | 2,73869E-17 |
| ACE2 | IGHV1-14 | 0,403 | 2,74396E-17 |
| ACE2 | IGHV3-69-1 | 0,403 | 2,7481E-17 |
| ACE2 | FANCA | 0,403 | 2,7371E-17 |
| ACE2 | AC109815.2 | 0,4029 | 2,79897E-17 |
| ACE2 | IGHV1-18 | 0,4029 | 2,79154E-17 |
| ACE2 | IGHV1-46 | 0,4029 | 2,8043E-17 |
| ACE2 | NSUN3 | 0,4028 | 2,87983E-17 |
| ACE2 | RP11-872J21.3 | 0,4028 | 2,85898E-17 |
| ACE2 | RP11-100N20.1 | 0,4027 | 2,90203E-17 |
| ACE2 | ENTPD7 | 0,4027 | 2,92319E-17 |
| ACE2 | RP11-158L12.4 | 0,4027 | 2,89982E-17 |
| ACE2 | AC104088.1 | 0,4024 | 3,09713E-17 |
| ACE2 | AADAC | 0,4024 | 3,13081E-17 |
| ACE2 | RUNX3 | 0,4022 | 3,25066E-17 |
| ACE2 | RP11-268P4.4 | 0,4022 | 3,21487E-17 |
| ACE2 | RP11-370B11.4 | 0,402 | 3,3439E-17 |
| ACE2 | RP11-1166P10.8 | 0,402 | 3,3361E-17 |
| ACE2 | SLC16A3 | 0,4019 | 3,41371E-17 |
| ACE2 | ETV2 | 0,4019 | 3,45112E-17 |
| ACE2 | RPS3AP40 | 0,4018 | 3,49537E-17 |
| ACE2 | AC093818.1 | 0,4015 | 3,70834E-17 |
| ACE2 | IGHV4-34 | 0,4015 | 3,68393E-17 |
| ACE2 | IKZF3 | 0,4014 | 3,78203E-17 |
| ACE2 | CXCL16 | 0,4012 | 3,95358E-17 |
| ACE2 | RP11-415C15.2 | 0,4011 | 4,0172E-17 |
| ACE2 | IGLV5-37 | 0,401 | 4,11102E-17 |
| ACE2 | RP11-540A21.2 | 0,4008 | 4,22031E-17 |
| ACE2 | RP11-568N6.1 | 0,4007 | 4,30526E-17 |
| ACE2 | IGHJ3 | 0,4007 | 4,30707E-17 |
| ACE2 | IGHV4-59 | 0,4007 | 4,34125E-17 |
| ACE2 | RMDN2-AS1 | 0,4006 | 4,43815E-17 |

| | | | |
|---|---|---|---|
| ACE2 | FEM1C | 0,4006 | 4,38135E-17 |
| ACE2 | PRSS21 | 0,4004 | 4,59056E-17 |
| ACE2 | KISS1 | 0,4003 | 4,67854E-17 |
| ACE2 | C10orf111 | 0,4003 | 4,67137E-17 |
| ACE2 | KRT8P41 | 0,4003 | 4,64549E-17 |
| ACE2 | AC004490.1 | 0,4003 | 4,70494E-17 |
| ACE2 | AKR7A2 | 0,4002 | 4,78301E-17 |
| ACE2 | AC114973.1 | 0,4002 | 4,75657E-17 |
| ACE2 | IGHV2-26 | 0,4002 | 4,74889E-17 |
| ACE2 | TBRG4 | 0,4 | 4,9928E-17 |
| ACE2 | XBP1 | 0,3998 | 5,18507E-17 |
| ACE2 | IGHM | 0,3995 | 5,51898E-17 |
| ACE2 | IGHV5-51 | 0,3995 | 5,4704E-17 |
| ACE2 | RP11-109E24.2 | 0,3994 | 5,52457E-17 |
| ACE2 | RP3-355L5.5 | 0,3994 | 5,60893E-17 |
| ACE2 | LARS2 | 0,3993 | 5,68237E-17 |
| ACE2 | RP11-783K16.14 | 0,3993 | 5,67353E-17 |
| ACE2 | MT-TI | 0,3993 | 5,73639E-17 |
| ACE2 | HIST1H4PS1 | 0,3992 | 5,82084E-17 |
| ACE2 | RP11-84A14.4 | 0,3991 | 5,95337E-17 |
| ACE2 | IGHD2-15 | 0,3991 | 5,9151E-17 |
| ACE2 | SLC25A40 | 0,399 | 5,99294E-17 |
| ACE2 | IGLV3-25 | 0,399 | 6,04301E-17 |
| ACE2 | RBP4 | 0,3989 | 6,11021E-17 |
| ACE2 | GRM8 | 0,3988 | 6,30193E-17 |
| ACE2 | VWA5B2 | 0,3987 | 6,43231E-17 |
| ACE2 | RP11-360A10.1 | 0,3987 | 6,35292E-17 |
| ACE2 | MRPL14 | 0,3986 | 6,46607E-17 |
| ACE2 | SERPINA10 | 0,3985 | 6,65219E-17 |
| ACE2 | GALNT1 | 0,3985 | 6,61026E-17 |
| ACE2 | MT-ND5 | 0,3985 | 6,60114E-17 |
| ACE2 | Six3os1_2 | 0,3984 | 6,80458E-17 |
| ACE2 | AC073869.2 | 0,3984 | 6,83139E-17 |
| ACE2 | RP11-124I4.2 | 0,3983 | 6,8606E-17 |
| ACE2 | RN7SL130P | 0,3982 | 7,0319E-17 |
| ACE2 | RUSC1-AS1 | 0,3982 | 7,05467E-17 |
| ACE2 | RP11-544M22.3 | 0,3982 | 6,98891E-17 |
| ACE2 | AC009005.2 | 0,3982 | 7,04552E-17 |
| ACE2 | GAL3ST1 | 0,3982 | 7,04123E-17 |
| ACE2 | OR2B6 | 0,398 | 7,35808E-17 |
| ACE2 | CTD-2522E6.4 | 0,398 | 7,32642E-17 |
| ACE2 | GNG12 | 0,3979 | 7,41119E-17 |
| ACE2 | IGHV3OR15-7 | 0,3979 | 7,45199E-17 |
| ACE2 | ADGRG5 | 0,3977 | 7,71823E-17 |
| ACE2 | LCT | 0,3976 | 7,8507E-17 |
| ACE2 | RP11-545M8.4 | 0,3976 | 7,9089E-17 |
| ACE2 | IGHV3-21 | 0,3976 | 7,93564E-17 |
| ACE2 | DEFB131 | 0,3975 | 8,09442E-17 |
| ACE2 | AF064860.7 | 0,3975 | 8,04141E-17 |
| ACE2 | SERPINA7 | 0,3975 | 8,00687E-17 |
| ACE2 | TMEM168 | 0,3974 | 8,21139E-17 |
| ACE2 | INSM1 | 0,3973 | 8,39722E-17 |
| ACE2 | HR | 0,3972 | 8,60785E-17 |
| ACE2 | SCARB1 | 0,3972 | 8,53361E-17 |

| | | | |
|---|---|---|---|
| ACE2 | BCKDHB | 0,3971 | 8,63984E-17 |
| ACE2 | KRTAP5-1 | 0,397 | 8,88202E-17 |
| ACE2 | IGHV1OR15-2 | 0,397 | 8,90485E-17 |
| ACE2 | IL17RA | 0,397 | 8,83277E-17 |
| ACE2 | IGHV3-20 | 0,3969 | 9,12764E-17 |
| ACE2 | IGLV1-51 | 0,3966 | 9,56402E-17 |
| ACE2 | RP11-783K16.13 | 0,3965 | 9,75784E-17 |
| ACE2 | IGHV1-67 | 0,3965 | 9,72587E-17 |
| ACE2 | ASB11 | 0,3965 | 9,81974E-17 |
| ACE2 | IGHV7-4-1 | 0,3964 | 1,00394E-16 |
| ACE2 | REEP6 | 0,3964 | 9,96629E-17 |
| ACE2 | PASK | 0,3963 | 1,00566E-16 |
| ACE2 | RP11-415F23.3 | 0,3963 | 1,00837E-16 |
| ACE2 | RP11-132A1.4 | 0,3963 | 1,01852E-16 |
| ACE2 | RP11-375N15.2 | 0,396 | 1,0673E-16 |
| ACE2 | RP11-123J14.1 | 0,3959 | 1,0894E-16 |
| ACE2 | CTD-2240J17.3 | 0,3959 | 1,0881E-16 |
| ACE2 | CTD-2576D5.4 | 0,3957 | 1,13216E-16 |
| ACE2 | TNIP1 | 0,3956 | 1,15372E-16 |
| ACE2 | ABAT | 0,3956 | 1,16337E-16 |
| ACE2 | RP11-839G9.1 | 0,3956 | 1,16828E-16 |
| ACE2 | CERS6-AS1 | 0,3955 | 1,18387E-16 |
| ACE2 | PTCHD3P3 | 0,3955 | 1,18843E-16 |
| ACE2 | LINC01620 | 0,3955 | 1,18419E-16 |
| ACE2 | IGLV2-23 | 0,3955 | 1,18288E-16 |
| ACE2 | IGLV3-19 | 0,3955 | 1,18201E-16 |
| ACE2 | TM7SF2 | 0,3954 | 1,19549E-16 |
| ACE2 | IGHV3-36 | 0,3954 | 1,19887E-16 |
| ACE2 | MTMR4 | 0,3954 | 1,21064E-16 |
| ACE2 | EPB41 | 0,3953 | 1,22262E-16 |
| ACE2 | IGHV3-62 | 0,3952 | 1,26441E-16 |
| ACE2 | TRAPPC10 | 0,3952 | 1,2504E-16 |
| ACE2 | IGHV4-55 | 0,3951 | 1,2795E-16 |
| ACE2 | IGHV3-63 | 0,3948 | 1,36437E-16 |
| ACE2 | TPI1 | 0,3946 | 1,40699E-16 |
| ACE2 | RP11-146F11.5 | 0,3945 | 1,42253E-16 |
| ACE2 | GPS2P2 | 0,3944 | 1,46488E-16 |
| ACE2 | MDH2 | 0,3944 | 1,45845E-16 |
| ACE2 | RP1-27K12.2 | 0,3941 | 1,55566E-16 |
| ACE2 | RP11-438P9.2 | 0,3941 | 1,53674E-16 |
| ACE2 | ZP1 | 0,3941 | 1,53843E-16 |
| ACE2 | IGHV3-38 | 0,3939 | 1,5928E-16 |
| ACE2 | IGHV4-39 | 0,3939 | 1,6158E-16 |
| ACE2 | FAM183A | 0,3938 | 1,65374E-16 |
| ACE2 | PTGES2-AS1 | 0,3938 | 1,62754E-16 |
| ACE2 | GLOD4 | 0,3938 | 1,62383E-16 |
| ACE2 | COX10 | 0,3938 | 1,63527E-16 |
| ACE2 | FH | 0,3937 | 1,68488E-16 |
| ACE2 | CTAGE12P | 0,3937 | 1,6778E-16 |
| ACE2 | IGHV3-30 | 0,3937 | 1,67141E-16 |
| ACE2 | OACYLP | 0,3936 | 1,70546E-16 |
| ACE2 | RP11-556I14.2 | 0,3935 | 1,73489E-16 |
| ACE2 | IGKV1-17 | 0,3932 | 1,82959E-16 |
| ACE2 | ACTBP13 | 0,3932 | 1,83137E-16 |

| | | | |
|---|---|---|---|
| ACE2 | RP11-485G7.5 | 0,3931 | 1,88754E-16 |
| ACE2 | RP11-138A9.2 | 0,393 | 1,92147E-16 |
| ACE2 | MT-TT | 0,393 | 1,89469E-16 |
| ACE2 | BCAR3 | 0,3928 | 1,96561E-16 |
| ACE2 | AP001171.1 | 0,3926 | 2,06021E-16 |
| ACE2 | RP11-499O7.7 | 0,3924 | 2,14732E-16 |
| ACE2 | RP11-157G21.2 | 0,3924 | 2,13477E-16 |
| ACE2 | IGHJ5 | 0,3924 | 2,13709E-16 |
| ACE2 | HIST1H2BN | 0,3923 | 2,17789E-16 |
| ACE2 | RP11-526F3.1 | 0,3922 | 2,23065E-16 |
| ACE2 | RP3-425P12.4 | 0,3922 | 2,20747E-16 |
| ACE2 | RIMKLA | 0,3921 | 2,2505E-16 |
| ACE2 | HLA-F | 0,3921 | 2,27984E-16 |
| ACE2 | AC073342.12 | 0,3921 | 2,26444E-16 |
| ACE2 | CHPF2 | 0,3921 | 2,28295E-16 |
| ACE2 | HIST1H3C | 0,392 | 2,28996E-16 |
| ACE2 | AC004009.2 | 0,392 | 2,28648E-16 |
| ACE2 | IGHV3-19 | 0,392 | 2,29967E-16 |
| ACE2 | PHB | 0,392 | 2,296E-16 |
| ACE2 | IGLV2-14 | 0,392 | 2,32676E-16 |
| ACE2 | C1orf74 | 0,3919 | 2,33317E-16 |
| ACE2 | RP11-618I10.1 | 0,3919 | 2,36683E-16 |
| ACE2 | AIFM2 | 0,3919 | 2,37265E-16 |
| ACE2 | IGLV3-21 | 0,3919 | 2,34597E-16 |
| ACE2 | RP11-455F5.4 | 0,3918 | 2,41108E-16 |
| ACE2 | MTND6P4 | 0,3916 | 2,50017E-16 |
| ACE2 | PPARGC1B | 0,3914 | 2,60648E-16 |
| ACE2 | TRHDE | 0,3914 | 2,57994E-16 |
| ACE2 | IGHJ4 | 0,3913 | 2,6306E-16 |
| ACE2 | PDCD10 | 0,3911 | 2,74198E-16 |
| ACE2 | AP000679.2 | 0,391 | 2,78208E-16 |
| ACE2 | SMAP1 | 0,3909 | 2,84182E-16 |
| ACE2 | WDR72 | 0,3907 | 2,97822E-16 |
| ACE2 | IGHV3OR16-6 | 0,3907 | 2,93724E-16 |
| ACE2 | IGLV7-46 | 0,3906 | 3,00558E-16 |
| ACE2 | MT-TA | 0,3906 | 3,03042E-16 |
| ACE2 | ATP5J2 | 0,3904 | 3,11781E-16 |
| ACE2 | TMC6 | 0,3904 | 3,1181E-16 |
| ACE2 | POLE2 | 0,3903 | 3,1713E-16 |
| ACE2 | UPP1 | 0,3901 | 3,33494E-16 |
| ACE2 | ZNF488 | 0,3901 | 3,31083E-16 |
| ACE2 | IGHJ1 | 0,3901 | 3,31909E-16 |
| ACE2 | MIR1227 | 0,3901 | 3,29431E-16 |
| ACE2 | IGLV1-44 | 0,3901 | 3,29969E-16 |
| ACE2 | OSBPL3 | 0,3898 | 3,51278E-16 |
| ACE2 | RP11-805L22.3 | 0,3896 | 3,63207E-16 |
| ACE2 | BMP8B | 0,3895 | 3,72474E-16 |
| ACE2 | FAM117B | 0,3895 | 3,71267E-16 |
| ACE2 | ARHGAP18 | 0,3895 | 3,67982E-16 |
| ACE2 | PYCARD-AS1 | 0,3895 | 3,67844E-16 |
| ACE2 | RP11-135A1.2 | 0,3894 | 3,80104E-16 |
| ACE2 | OAF | 0,3894 | 3,74236E-16 |
| ACE2 | MIR5195 | 0,3894 | 3,77907E-16 |
| ACE2 | IGHV1OR15-9 | 0,3894 | 3,78377E-16 |

| | | | |
|---|---|---|---|
| ACE2 | PDIK1L | 0,3893 | 3,84257E-16 |
| ACE2 | IL7R | 0,3893 | 3,82868E-16 |
| ACE2 | KDELR2 | 0,3893 | 3,80961E-16 |
| ACE2 | CABP2 | 0,389 | 4,04455E-16 |
| ACE2 | IGHV3-71 | 0,3889 | 4,12524E-16 |
| ACE2 | RP11-403F21.4 | 0,3888 | 4,22955E-16 |
| ACE2 | RP11-379C10.4 | 0,3888 | 4,20095E-16 |
| ACE2 | CPT1A | 0,3887 | 4,26805E-16 |
| ACE2 | IGHV4-28 | 0,3887 | 4,26681E-16 |
| ACE2 | COX6B1 | 0,3884 | 4,57177E-16 |
| ACE2 | RP5-1092A3.4 | 0,3883 | 4,63565E-16 |
| ACE2 | IGKV1D-39 | 0,3882 | 4,75191E-16 |
| ACE2 | BPHL | 0,3882 | 4,75072E-16 |
| ACE2 | RNU1-58P | 0,3882 | 4,74613E-16 |
| ACE2 | RP11-297M9.1 | 0,3882 | 4,7061E-16 |
| ACE2 | LINC00426 | 0,3881 | 4,83703E-16 |
| ACE2 | RN7SKP204 | 0,388 | 4,87577E-16 |
| ACE2 | RP11-16E12.2 | 0,3879 | 5,03231E-16 |
| ACE2 | AP3S2 | 0,3879 | 4,98299E-16 |
| ACE2 | AC114271.2 | 0,3878 | 5,09276E-16 |
| ACE2 | PAFAH1B3 | 0,3878 | 5,05502E-16 |
| ACE2 | IGHV3OR16-9 | 0,3877 | 5,19992E-16 |
| ACE2 | AC128709.2 | 0,3876 | 5,29036E-16 |
| ACE2 | FAM106DP | 0,3876 | 5,24306E-16 |
| ACE2 | RP11-150C16.1 | 0,3875 | 5,33812E-16 |
| ACE2 | DHRS7B | 0,3875 | 5,34997E-16 |
| ACE2 | MFSD1 | 0,3874 | 5,47125E-16 |
| ACE2 | RP11-70C1.3 | 0,387 | 5,86015E-16 |
| ACE2 | PDIA3P2 | 0,387 | 5,85517E-16 |
| ACE2 | TXNDC11 | 0,387 | 5,87057E-16 |
| ACE2 | RP11-1148O4.1 | 0,387 | 5,94916E-16 |
| ACE2 | RP11-511B23.2 | 0,3867 | 6,26784E-16 |
| ACE2 | IGHV3-6 | 0,3867 | 6,27648E-16 |
| ACE2 | IGHV3-41 | 0,3867 | 6,27059E-16 |
| ACE2 | AC005626.3 | 0,3867 | 6,22956E-16 |
| ACE2 | AC103563.5 | 0,3865 | 6,53375E-16 |
| ACE2 | CCSER1 | 0,3865 | 6,51671E-16 |
| ACE2 | MAN2A1 | 0,3863 | 6,68688E-16 |
| ACE2 | STK39 | 0,3862 | 6,79738E-16 |
| ACE2 | MOGAT1 | 0,3862 | 6,8295E-16 |
| ACE2 | FAM134A | 0,386 | 7,17734E-16 |
| ACE2 | CYP2R1 | 0,3858 | 7,42282E-16 |
| ACE2 | RNASE3 | 0,3858 | 7,3841E-16 |
| ACE2 | CPNE2 | 0,3858 | 7,38645E-16 |
| ACE2 | VDAC1P13 | 0,3857 | 7,54316E-16 |
| ACE2 | HLA-G | 0,3857 | 7,51679E-16 |
| ACE2 | RP11-407G23.5 | 0,3853 | 8,14094E-16 |
| ACE2 | PATL2 | 0,3852 | 8,21446E-16 |
| ACE2 | LYPLA1 | 0,3851 | 8,37554E-16 |
| ACE2 | RP11-69G7.1 | 0,385 | 8,51873E-16 |
| ACE2 | RP5-882C2.2 | 0,385 | 8,64237E-16 |
| ACE2 | IGLV3-10 | 0,385 | 8,54195E-16 |
| ACE2 | RP1-273G13.2 | 0,3849 | 8,75547E-16 |
| ACE2 | RP11-360K13.1 | 0,3849 | 8,71476E-16 |

| | | | |
|---|---|---|---|
| ACE2 | NBEAL2 | 0,3848 | 8,82122E-16 |
| ACE2 | ARHGEF38-IT1 | 0,3848 | 8,96593E-16 |
| ACE2 | CTD-2270P14.2 | 0,3847 | 9,12398E-16 |
| ACE2 | CLCN5 | 0,3847 | 9,00839E-16 |
| ACE2 | LARGE2 | 0,3845 | 9,45256E-16 |
| ACE2 | RP11-369C8.1 | 0,3845 | 9,4221E-16 |
| ACE2 | RN7SL706P | 0,3845 | 9,38715E-16 |
| ACE2 | PRKAR2A-AS1 | 0,3844 | 9,49914E-16 |
| ACE2 | HMBS | 0,3844 | 9,50165E-16 |
| ACE2 | RP11-305A4.5 | 0,3844 | 9,49656E-16 |
| ACE2 | IGKV1OR2-108 | 0,3842 | 9,91653E-16 |
| ACE2 | NPY1R | 0,3842 | 1,00147E-15 |
| ACE2 | IGHV1-2 | 0,3841 | 1,01498E-15 |
| ACE2 | CTD-2545G14.6 | 0,3841 | 1,01371E-15 |
| ACE2 | CISH | 0,3839 | 1,04233E-15 |
| ACE2 | IGLV8-61 | 0,3839 | 1,04032E-15 |
| ACE2 | IGHV3-52 | 0,3838 | 1,065E-15 |
| ACE2 | IGLV4-69 | 0,3838 | 1,06695E-15 |
| ACE2 | LINC00853 | 0,3837 | 1,08783E-15 |
| ACE2 | RP11-576N17.2 | 0,3837 | 1,09059E-15 |
| ACE2 | SSTR5 | 0,3837 | 1,08536E-15 |
| ACE2 | CES4A | 0,3836 | 1,10008E-15 |
| ACE2 | IGHV5-10-1 | 0,3834 | 1,16028E-15 |
| ACE2 | ENGASE | 0,3833 | 1,16278E-15 |
| ACE2 | IGLC6 | 0,3833 | 1,17169E-15 |
| ACE2 | NDUFB3 | 0,3832 | 1,19743E-15 |
| ACE2 | CTD-2537I9.13 | 0,3832 | 1,1983E-15 |
| ACE2 | RN7SL786P | 0,3831 | 1,22699E-15 |
| ACE2 | IGKV1-9 | 0,383 | 1,22761E-15 |
| ACE2 | RP11-739B23.1 | 0,383 | 1,23881E-15 |
| ACE2 | IGLV1-40 | 0,3829 | 1,2515E-15 |
| ACE2 | RP5-1184F4.5 | 0,3828 | 1,28684E-15 |
| ACE2 | UNQ6494 | 0,3827 | 1,31296E-15 |
| ACE2 | LARP4B | 0,3827 | 1,30946E-15 |
| ACE2 | KIR3DL2 | 0,3826 | 1,34254E-15 |
| ACE2 | RP11-248B24.1 | 0,3823 | 1,39847E-15 |
| ACE2 | RP11-294C11.2 | 0,3823 | 1,40252E-15 |
| ACE2 | RUNDC3B | 0,3822 | 1,44447E-15 |
| ACE2 | ENKUR | 0,3822 | 1,42707E-15 |
| ACE2 | CTD-2020K17.3 | 0,3822 | 1,44489E-15 |
| ACE2 | HLA-J | 0,3819 | 1,51245E-15 |
| ACE2 | VRTN | 0,3819 | 1,50608E-15 |
| ACE2 | IGHV1-24 | 0,3819 | 1,50253E-15 |
| ACE2 | LA16c-431H6.7 | 0,3819 | 1,50377E-15 |
| ACE2 | RP11-164P12.3 | 0,3817 | 1,56745E-15 |
| ACE2 | FRAT1 | 0,3817 | 1,57372E-15 |
| ACE2 | IGHV1-69D | 0,3817 | 1,57415E-15 |
| ACE2 | NRIP1 | 0,3817 | 1,56685E-15 |
| ACE2 | IGLV7-43 | 0,3817 | 1,57353E-15 |
| ACE2 | RP11-521M14.1 | 0,3816 | 1,60735E-15 |
| ACE2 | MTMR8 | 0,3816 | 1,59014E-15 |
| ACE2 | TNFRSF25 | 0,3814 | 1,65743E-15 |
| ACE2 | OSTCP2 | 0,3814 | 1,64774E-15 |
| ACE2 | FDPSP2 | 0,3813 | 1,70146E-15 |

| | | | |
|---|---|---|---|
| ACE2 | RP11-434H6.2 | 0,3812 | 1,73162E-15 |
| ACE2 | RP11-83J21.3 | 0,3812 | 1,73322E-15 |
| ACE2 | RPL23AP10 | 0,3812 | 1,73242E-15 |
| ACE2 | PTPRK | 0,3808 | 1,86869E-15 |
| ACE2 | CTD-2008P7.10 | 0,3808 | 1,85243E-15 |
| ACE2 | SEMA5A-AS1 | 0,3807 | 1,89144E-15 |
| ACE2 | IGLV1-41 | 0,3807 | 1,89865E-15 |
| ACE2 | MGC27382 | 0,3803 | 2,03332E-15 |
| ACE2 | FAM43A | 0,3803 | 2,03807E-15 |
| ACE2 | EGLN3-AS1 | 0,3803 | 2,03786E-15 |
| ACE2 | RP6-114E22.1 | 0,3803 | 2,02598E-15 |
| ACE2 | LINC01876 | 0,3802 | 2,08626E-15 |
| ACE2 | FN3K | 0,3802 | 2,07837E-15 |
| ACE2 | NAGA | 0,3802 | 2,06574E-15 |
| ACE2 | IGKV3D-11 | 0,38 | 2,13471E-15 |
| ACE2 | SRIP3 | 0,3799 | 2,19412E-15 |
| ACE2 | MMACHC | 0,3798 | 2,23222E-15 |
| ACE2 | PCBD2 | 0,3798 | 2,22617E-15 |
| ACE2 | OR5G3 | 0,3797 | 2,28376E-15 |
| ACE2 | NAA40 | 0,3797 | 2,24609E-15 |
| ACE2 | SLC2A5 | 0,3795 | 2,33573E-15 |
| ACE2 | IGKV1-5 | 0,3794 | 2,37827E-15 |
| ACE2 | CCNF | 0,3794 | 2,37968E-15 |
| ACE2 | LINC01450 | 0,3793 | 2,41713E-15 |
| ACE2 | IGKV2-30 | 0,3792 | 2,49356E-15 |
| ACE2 | LINC01855 | 0,379 | 2,55096E-15 |
| ACE2 | IGHV3OR16-15 | 0,3789 | 2,61324E-15 |
| ACE2 | IGHV1-45 | 0,3788 | 2,64935E-15 |
| ACE2 | NOXO1 | 0,3788 | 2,66149E-15 |
| ACE2 | RTN4R | 0,3788 | 2,67441E-15 |
| ACE2 | CR2 | 0,3787 | 2,73693E-15 |
| ACE2 | TMEM254 | 0,3787 | 2,72323E-15 |
| ACE2 | RP11-982M15.8 | 0,3787 | 2,69313E-15 |
| ACE2 | IGHV3-60 | 0,3787 | 2,72921E-15 |
| ACE2 | NPAS1 | 0,3787 | 2,73809E-15 |
| ACE2 | APOO | 0,3787 | 2,69598E-15 |
| ACE2 | CTB-31N19.2 | 0,3786 | 2,78298E-15 |
| ACE2 | IGLV3-12 | 0,3783 | 2,93374E-15 |
| ACE2 | MT-RNR1 | 0,3782 | 2,99356E-15 |
| ACE2 | C1QTNF1-AS1 | 0,378 | 3,07983E-15 |
| ACE2 | MARCH3 | 0,3779 | 3,13491E-15 |
| ACE2 | CTD-2203K17.1 | 0,3778 | 3,217E-15 |
| ACE2 | TMEM223 | 0,3778 | 3,17529E-15 |
| ACE2 | IGHV1OR16-1 | 0,3778 | 3,21805E-15 |
| ACE2 | OR8G5 | 0,3777 | 3,23523E-15 |
| ACE2 | DUSP23 | 0,3775 | 3,34955E-15 |
| ACE2 | CTC-343N3.1 | 0,3775 | 3,3932E-15 |
| ACE2 | MGME1 | 0,3775 | 3,39334E-15 |
| ACE2 | IGKV5-2 | 0,3774 | 3,43749E-15 |
| ACE2 | IGKV1D-16 | 0,3774 | 3,42972E-15 |
| ACE2 | HRK | 0,3774 | 3,46356E-15 |
| ACE2 | NXPE2P1 | 0,3773 | 3,52548E-15 |
| ACE2 | GNPTAB | 0,3772 | 3,59255E-15 |
| ACE2 | IGHV3-47 | 0,3772 | 3,54812E-15 |

| | | | |
|---|---|---|---|
| ACE2 | KLF13 | 0,3772 | 3,5716E-15 |
| ACE2 | IGLV6-57 | 0,3772 | 3,54947E-15 |
| ACE2 | IARS2 | 0,3771 | 3,61414E-15 |
| ACE2 | OR10G4 | 0,3769 | 3,75494E-15 |
| ACE2 | IGLV1-47 | 0,3769 | 3,76076E-15 |
| ACE2 | NAT8 | 0,3768 | 3,82986E-15 |
| ACE2 | IGKV1-8 | 0,3768 | 3,814E-15 |
| ACE2 | LIPC | 0,3768 | 3,81631E-15 |
| ACE2 | HOXB5 | 0,3768 | 3,85222E-15 |
| ACE2 | SLC30A1 | 0,3767 | 3,86644E-15 |
| ACE2 | RP11-756A22.7 | 0,3767 | 3,88844E-15 |
| ACE2 | RP11-573D15.8 | 0,3766 | 3,96871E-15 |
| ACE2 | RP11-535A5.1 | 0,3766 | 3,95478E-15 |
| ACE2 | KB-1460A1.2 | 0,3765 | 4,01977E-15 |
| ACE2 | HPS6 | 0,3764 | 4,0897E-15 |
| ACE2 | AC004009.3 | 0,3761 | 4,32023E-15 |
| ACE2 | RP11-740N7.3 | 0,376 | 4,3908E-15 |
| ACE2 | SLC10A5P1 | 0,3759 | 4,51395E-15 |
| ACE2 | RP3-400B16.2 | 0,3758 | 4,54907E-15 |
| ACE2 | RP11-21L23.2 | 0,3757 | 4,6874E-15 |
| ACE2 | IGKV1D-12 | 0,3756 | 4,74582E-15 |
| ACE2 | MINPP1 | 0,3756 | 4,75516E-15 |
| ACE2 | IGHV2-70 | 0,3756 | 4,78663E-15 |
| ACE2 | RP11-434D2.11 | 0,3756 | 4,78649E-15 |
| ACE2 | IGHV3-42 | 0,3754 | 4,9411E-15 |
| ACE2 | GPR171 | 0,3752 | 5,07082E-15 |
| ACE2 | RP11-295P22.2 | 0,3751 | 5,20403E-15 |
| ACE2 | RP11-361F15.5 | 0,3748 | 5,435E-15 |
| ACE2 | KIF24 | 0,3748 | 5,51962E-15 |
| ACE2 | IGHV1-69-2 | 0,3747 | 5,59548E-15 |
| ACE2 | CTD-2047H16.2 | 0,3747 | 5,58523E-15 |
| ACE2 | CD3E | 0,3746 | 5,64414E-15 |
| ACE2 | IGHV7-56 | 0,3746 | 5,70233E-15 |
| ACE2 | TAS1R3 | 0,3745 | 5,81785E-15 |
| ACE2 | LINC01833 | 0,3743 | 6,0337E-15 |
| ACE2 | IGKV3D-20 | 0,3743 | 5,95465E-15 |
| ACE2 | RP11-240E2.2 | 0,3742 | 6,12409E-15 |
| ACE2 | CHCHD4P2 | 0,3741 | 6,19577E-15 |
| ACE2 | CTD-2240J17.2 | 0,3741 | 6,2251E-15 |
| ACE2 | CTB-179K24.3 | 0,3741 | 6,18094E-15 |
| ACE2 | IGKV3-11 | 0,374 | 6,28973E-15 |
| ACE2 | IGKV2D-29 | 0,374 | 6,26736E-15 |
| ACE2 | RP11-366L18.1 | 0,3737 | 6,67829E-15 |
| ACE2 | RP11-476C8.2 | 0,3737 | 6,65258E-15 |
| ACE2 | PAX5 | 0,3737 | 6,69206E-15 |
| ACE2 | OR52W1 | 0,3735 | 6,84992E-15 |
| ACE2 | IGHD3-3 | 0,3735 | 6,91847E-15 |
| ACE2 | RP1-202O8.2 | 0,3734 | 7,0391E-15 |
| ACE2 | H2AFX | 0,3734 | 7,05604E-15 |
| ACE2 | IGLC7 | 0,3734 | 6,99698E-15 |
| ACE2 | IL2RB | 0,3734 | 6,99499E-15 |
| ACE2 | TP53AIP1 | 0,3733 | 7,18089E-15 |
| ACE2 | RP11-654D12.2 | 0,3731 | 7,4559E-15 |
| ACE2 | SIRPG | 0,3731 | 7,36253E-15 |

| | | | |
|---|---|---|---|
| ACE2 | COA7 | 0,373 | 7,54535E-15 |
| ACE2 | MRPS33 | 0,373 | 7,50704E-15 |
| ACE2 | RAD9A | 0,373 | 7,53194E-15 |
| ACE2 | MIEN1 | 0,3729 | 7,69807E-15 |
| ACE2 | CTB-176F20.3 | 0,3729 | 7,68763E-15 |
| ACE2 | IGLV2-33 | 0,3729 | 7,719E-15 |
| ACE2 | RP11-626P14.1 | 0,3728 | 7,75775E-15 |
| ACE2 | EIF4EBP3 | 0,3726 | 8,10493E-15 |
| ACE2 | RNU6-353P | 0,3726 | 8,05911E-15 |
| ACE2 | RP11-592N21.2 | 0,3726 | 8,09906E-15 |
| ACE2 | ACTG1P15 | 0,3725 | 8,22156E-15 |
| ACE2 | ALKBH7 | 0,3725 | 8,18782E-15 |
| ACE2 | PDSS2 | 0,3723 | 8,54864E-15 |
| ACE2 | IGHV3OR16-10 | 0,3723 | 8,48365E-15 |
| ACE2 | IGLV10-54 | 0,3723 | 8,59132E-15 |
| ACE2 | RPS7P7 | 0,3722 | 8,76925E-15 |
| ACE2 | IGHV3-35 | 0,3722 | 8,70892E-15 |
| ACE2 | CXCL3 | 0,3721 | 8,83549E-15 |
| ACE2 | F3 | 0,372 | 8,96862E-15 |
| ACE2 | CHST4 | 0,3719 | 9,17012E-15 |
| ACE2 | HIST1H4K | 0,3715 | 9,8205E-15 |
| ACE2 | YRDCP1 | 0,3715 | 9,84717E-15 |
| ACE2 | RP5-905G11.3 | 0,3715 | 9,90947E-15 |
| ACE2 | IGHV3-37 | 0,3714 | 1,00876E-14 |
| ACE2 | RP4-635A23.6 | 0,3713 | 1,01913E-14 |
| ACE2 | RP11-49K24.5 | 0,3713 | 1,01231E-14 |
| ACE2 | CDK8P2 | 0,3712 | 1,03718E-14 |
| ACE2 | CNN2P1 | 0,3712 | 1,03668E-14 |
| ACE2 | FDPSP8 | 0,3711 | 1,05422E-14 |
| ACE2 | IGHV4-31 | 0,3711 | 1,05959E-14 |
| ACE2 | IGLV4-60 | 0,3711 | 1,05755E-14 |
| ACE2 | IGKV1D-42 | 0,3709 | 1,10401E-14 |
| ACE2 | LINC01504 | 0,3709 | 1,10245E-14 |
| ACE2 | PRR14L | 0,3708 | 1,1184E-14 |
| ACE2 | IGKV2-24 | 0,3706 | 1,14569E-14 |
| ACE2 | RP11-361C13.1 | 0,3705 | 1,18062E-14 |
| ACE2 | EGLN3 | 0,3704 | 1,20529E-14 |
| ACE2 | RP11-600F24.7 | 0,3704 | 1,19578E-14 |
| ACE2 | RP11-734K21.2 | 0,3703 | 1,20956E-14 |
| ACE2 | GS1-24F4.2 | 0,3703 | 1,20896E-14 |
| ACE2 | IGLV4-3 | 0,3703 | 1,22332E-14 |
| ACE2 | ATP13A2 | 0,3702 | 1,24459E-14 |
| ACE2 | CHPT1 | 0,37 | 1,2829E-14 |
| ACE2 | RP11-297L17.6 | 0,3699 | 1,30035E-14 |
| ACE2 | TRPV6 | 0,3698 | 1,31943E-14 |
| ACE2 | SYNE2 | 0,3698 | 1,32133E-14 |
| ACE2 | SCP2 | 0,3697 | 1,3507E-14 |
| ACE2 | GGNBP1 | 0,3697 | 1,35665E-14 |
| ACE2 | RP11-575F12.3 | 0,3696 | 1,37662E-14 |
| ACE2 | XK | 0,3696 | 1,38644E-14 |
| ACE2 | TACC3 | 0,3695 | 1,41149E-14 |
| ACE2 | IGLJ2 | 0,3695 | 1,39693E-14 |
| ACE2 | RP11-73M7.1 | 0,3694 | 1,42E-14 |
| ACE2 | IGKV4-1 | 0,3694 | 1,4351E-14 |

| | | | |
|---|---|---|---|
| ACE2 | IGKV1-12 | 0,3694 | 1,41941E-14 |
| ACE2 | LINC00867 | 0,3694 | 1,41574E-14 |
| ACE2 | PNPO | 0,3693 | 1,46243E-14 |
| ACE2 | LLNLR-304A6.2 | 0,3691 | 1,50061E-14 |
| ACE2 | CASP1 | 0,369 | 1,54049E-14 |
| ACE2 | KRT23 | 0,3689 | 1,54601E-14 |
| ACE2 | CH17-212P11.8 | 0,3686 | 1,65005E-14 |
| ACE2 | RP3-347M6.3 | 0,3686 | 1,65244E-14 |
| ACE2 | ARSDP1 | 0,3685 | 1,6823E-14 |
| ACE2 | PRORY | 0,3684 | 1,69803E-14 |
| ACE2 | CHAC2 | 0,3683 | 1,74167E-14 |
| ACE2 | IGKV3-20 | 0,3683 | 1,7242E-14 |
| ACE2 | ELOCP19 | 0,3682 | 1,7474E-14 |
| ACE2 | LA16c-325D7.1 | 0,3682 | 1,75604E-14 |
| ACE2 | RP1-232L22__B.1 | 0,3682 | 1,75975E-14 |
| ACE2 | IGKV1OR22-5 | 0,3681 | 1,7784E-14 |
| ACE2 | IGLJ1 | 0,3681 | 1,78244E-14 |
| ACE2 | SPINK14 | 0,3679 | 1,86125E-14 |
| ACE2 | KLF16 | 0,3677 | 1,92387E-14 |
| ACE2 | NPY6R | 0,3676 | 1,96189E-14 |
| ACE2 | IGHV1-68 | 0,3676 | 1,96582E-14 |
| ACE2 | RP11-497D6.3 | 0,3674 | 2,01399E-14 |
| ACE2 | DCXR | 0,3673 | 2,07027E-14 |
| ACE2 | CTB-102L5.7 | 0,3673 | 2,05176E-14 |
| ACE2 | RP1-13D10.3 | 0,3672 | 2,07948E-14 |
| ACE2 | ARHGAP39 | 0,3672 | 2,10576E-14 |
| ACE2 | KRT18P18 | 0,3672 | 2,10462E-14 |
| ACE2 | PCNA | 0,3672 | 2,08061E-14 |
| ACE2 | STEAP1 | 0,3669 | 2,20056E-14 |
| ACE2 | RP11-677M24.1 | 0,3669 | 2,21975E-14 |
| ACE2 | LINC00566 | 0,3669 | 2,18962E-14 |
| ACE2 | CD79A | 0,3669 | 2,21022E-14 |
| ACE2 | IGLV3-16 | 0,3669 | 2,22366E-14 |
| ACE2 | RP4-539M6.22 | 0,3667 | 2,27662E-14 |
| ACE2 | GLYATL3 | 0,3665 | 2,36218E-14 |
| ACE2 | RP11-439C15.4 | 0,3663 | 2,4279E-14 |
| ACE2 | IGHVII-49-1 | 0,3663 | 2,44363E-14 |
| ACE2 | TBC1D22A-AS1 | 0,3662 | 2,50554E-14 |
| ACE2 | AP001439.2 | 0,366 | 2,56744E-14 |
| ACE2 | CTA-243E7.4 | 0,366 | 2,60071E-14 |
| ACE2 | RP5-968D22.3 | 0,3659 | 2,64521E-14 |
| ACE2 | IGLV1-36 | 0,3658 | 2,68546E-14 |
| ACE2 | IGLV3-29 | 0,3657 | 2,73984E-14 |
| ACE2 | PTPRJ | 0,3656 | 2,74042E-14 |
| ACE2 | MYO1B | 0,3655 | 2,81867E-14 |
| ACE2 | IGKV1-16 | 0,3654 | 2,8398E-14 |
| ACE2 | KLRF2 | 0,3654 | 2,8753E-14 |
| ACE2 | BBC3 | 0,3654 | 2,84178E-14 |
| ACE2 | SIRT5 | 0,3652 | 2,96673E-14 |
| ACE2 | UNC93A | 0,3651 | 2,99317E-14 |
| ACE2 | IGKV1-6 | 0,365 | 3,07769E-14 |
| ACE2 | KAZALD1 | 0,365 | 3,06489E-14 |
| ACE2 | RP11-426C22.7 | 0,365 | 3,07502E-14 |
| ACE2 | ANLN | 0,3649 | 3,12011E-14 |

| | | | |
|---|---|---|---|
| ACE2 | TXNP4 | 0,3649 | 3,09659E-14 |
| ACE2 | RP11-523H20.3 | 0,3648 | 3,1476E-14 |
| ACE2 | PFKFB1 | 0,3648 | 3,16642E-14 |
| ACE2 | CD8B | 0,3647 | 3,21346E-14 |
| ACE2 | IGKV1-39 | 0,3647 | 3,22362E-14 |
| ACE2 | LAMC3 | 0,3647 | 3,24981E-14 |
| ACE2 | RP11-384J4.2 | 0,3647 | 3,25447E-14 |
| ACE2 | CTA-243E7.3 | 0,3645 | 3,36181E-14 |
| ACE2 | NUP50 | 0,3643 | 3,48446E-14 |
| ACE2 | IGKV3OR2-268 | 0,3642 | 3,54062E-14 |
| ACE2 | RNF125 | 0,3642 | 3,52036E-14 |
| ACE2 | SUMO1P2 | 0,364 | 3,64356E-14 |
| ACE2 | SPINK2 | 0,364 | 3,64408E-14 |
| ACE2 | RP3-510L9.1 | 0,3639 | 3,70785E-14 |
| ACE2 | IGHV1-12 | 0,3639 | 3,71509E-14 |
| ACE2 | FOXI1 | 0,3637 | 3,81956E-14 |
| ACE2 | AC005220.3 | 0,3637 | 3,85915E-14 |
| ACE2 | MAGI1-IT1 | 0,3636 | 3,92539E-14 |
| ACE2 | TDRD7 | 0,3636 | 3,9185E-14 |
| ACE2 | ATP5J2P5 | 0,3636 | 3,90359E-14 |
| ACE2 | DGAT2 | 0,3634 | 4,03218E-14 |
| ACE2 | IGKV2D-30 | 0,3633 | 4,10978E-14 |
| ACE2 | IGHV2-70D | 0,3633 | 4,14367E-14 |
| ACE2 | RP11-744O11.2 | 0,3631 | 4,24448E-14 |
| ACE2 | RP5-1031D4.3 | 0,3631 | 4,22103E-14 |
| ACE2 | RP11-192M23.1 | 0,3631 | 4,21805E-14 |
| ACE2 | AC006077.3 | 0,363 | 4,2909E-14 |
| ACE2 | TAP1 | 0,363 | 4,31944E-14 |
| ACE2 | CTD-2006C1.12 | 0,363 | 4,32788E-14 |
| ACE2 | CEACAMP1 | 0,363 | 4,32596E-14 |
| ACE2 | IGHV1-58 | 0,3629 | 4,42562E-14 |
| ACE2 | JSRP1 | 0,3625 | 4,74522E-14 |
| ACE2 | AC005795.1 | 0,3625 | 4,70903E-14 |
| ACE2 | DGKD | 0,3624 | 4,81605E-14 |
| ACE2 | C6orf141 | 0,3624 | 4,77552E-14 |
| ACE2 | RP11-770J1.5 | 0,3624 | 4,83309E-14 |
| ACE2 | RP11-420A23.1 | 0,3623 | 4,89893E-14 |
| ACE2 | RP11-297C4.1 | 0,3623 | 4,89974E-14 |
| ACE2 | DDA1 | 0,3623 | 4,84154E-14 |
| ACE2 | C6orf223 | 0,3622 | 4,92963E-14 |
| ACE2 | APOC2 | 0,3622 | 4,95963E-14 |
| ACE2 | OS9 | 0,3621 | 5,05437E-14 |
| ACE2 | RP11-466A19.5 | 0,3621 | 5,05264E-14 |
| ACE2 | IGHV3-22 | 0,362 | 5,12296E-14 |
| ACE2 | MTCO2P2 | 0,3619 | 5,22797E-14 |
| ACE2 | SNRPGP15 | 0,3619 | 5,26384E-14 |
| ACE2 | IGKV2D-26 | 0,3618 | 5,30747E-14 |
| ACE2 | LINC00402 | 0,3615 | 5,55112E-14 |
| ACE2 | RP11-544A12.4 | 0,3614 | 5,69808E-14 |
| ACE2 | IGKV6-21 | 0,3613 | 5,74864E-14 |
| ACE2 | DUSP18 | 0,3612 | 5,8428E-14 |
| ACE2 | SOCS6 | 0,3611 | 5,99544E-14 |
| ACE2 | MT-ND4L | 0,3611 | 5,94357E-14 |
| ACE2 | RP11-329N15.3 | 0,361 | 6,0446E-14 |

| | | | |
|---|---|---|---|
| ACE2 | RP11-77K12.3 | 0,361 | 6,06503E-14 |
| ACE2 | ERAP1 | 0,3609 | 6,15509E-14 |
| ACE2 | MTCO1P24 | 0,3608 | 6,30584E-14 |
| ACE2 | CTD-2516F10.4 | 0,3608 | 6,35005E-14 |
| ACE2 | LRP4-AS1 | 0,3608 | 6,32247E-14 |
| ACE2 | RP11-316K19.3 | 0,3608 | 6,34841E-14 |
| ACE2 | VPREB1 | 0,3607 | 6,40377E-14 |
| ACE2 | DCAF8L1 | 0,3606 | 6,4657E-14 |
| ACE2 | DTX2P1 | 0,3605 | 6,66091E-14 |
| ACE2 | RP11-5G9.5 | 0,3604 | 6,75722E-14 |
| ACE2 | APMAP | 0,3603 | 6,88718E-14 |
| ACE2 | RP11-129M16.4 | 0,3602 | 6,94398E-14 |
| ACE2 | GMFB | 0,3602 | 6,96448E-14 |
| ACE2 | CRB2 | 0,3601 | 7,13672E-14 |
| ACE2 | RP11-277P12.20 | 0,3601 | 7,1388E-14 |
| ACE2 | IGHV3-16 | 0,3601 | 7,10076E-14 |
| ACE2 | LINC00852 | 0,36 | 7,16623E-14 |
| ACE2 | AP1G1 | 0,3599 | 7,32166E-14 |
| ACE2 | TNMD | 0,3598 | 7,51602E-14 |
| ACE2 | PAQR4 | 0,3597 | 7,65007E-14 |
| ACE2 | RP11-260M19.2 | 0,3596 | 7,7375E-14 |
| ACE2 | RP11-387M24.5 | 0,3595 | 7,91047E-14 |
| ACE2 | POU4F1 | 0,3595 | 7,82118E-14 |
| ACE2 | METRNL | 0,3595 | 7,87198E-14 |
| ACE2 | PANCR | 0,3594 | 7,97236E-14 |
| ACE2 | MSLNL | 0,3594 | 8,00073E-14 |
| ACE2 | IGKV3D-15 | 0,3593 | 8,18538E-14 |
| ACE2 | PIK3CB | 0,3592 | 8,28585E-14 |
| ACE2 | MAEA | 0,3592 | 8,32432E-14 |
| ACE2 | TKTL2 | 0,3592 | 8,22509E-14 |
| ACE2 | RP11-534L20.5 | 0,3591 | 8,44933E-14 |
| ACE2 | CTD-2310F14.1 | 0,3591 | 8,35131E-14 |
| ACE2 | RP11-1038A11.3 | 0,3591 | 8,40101E-14 |
| ACE2 | RP1-102K2.8 | 0,3588 | 8,85257E-14 |
| ACE2 | IGKJ1 | 0,3587 | 9,04521E-14 |
| ACE2 | RPL12P18 | 0,3585 | 9,36525E-14 |
| ACE2 | CTD-2589M5.4 | 0,3585 | 9,24723E-14 |
| ACE2 | NLRP7 | 0,3584 | 9,54253E-14 |
| ACE2 | RPS10P5 | 0,3583 | 9,60214E-14 |
| ACE2 | RP11-1038A11.1 | 0,3582 | 9,86905E-14 |
| ACE2 | PBOV1 | 0,3581 | 1,00104E-13 |
| ACE2 | FDPSP7 | 0,3581 | 9,96227E-14 |
| ACE2 | IGKV1OR10-1 | 0,3581 | 1,00173E-13 |
| ACE2 | RP11-400F19.12 | 0,3581 | 9,92021E-14 |
| ACE2 | IGKV1OR2-6 | 0,3578 | 1,0546E-13 |
| ACE2 | EMC10 | 0,3577 | 1,07343E-13 |
| ACE2 | IGHV3OR16-16 | 0,3574 | 1,11672E-13 |
| ACE2 | CASD1 | 0,3573 | 1,1476E-13 |
| ACE2 | NXT2 | 0,3573 | 1,13255E-13 |
| ACE2 | FCMR | 0,3571 | 1,17345E-13 |
| ACE2 | CYB561D1 | 0,3569 | 1,22331E-13 |
| ACE2 | CTD-2510F5.4 | 0,3569 | 1,22459E-13 |
| ACE2 | LINC00930 | 0,3565 | 1,30929E-13 |
| ACE2 | ABCA7 | 0,3564 | 1,33529E-13 |

| | | | |
|---|---|---|---|
| ACE2 | FER1L6-AS2 | 0,3563 | 1,35193E-13 |
| ACE2 | AC008074.3 | 0,3561 | 1,38601E-13 |
| ACE2 | YIPF2 | 0,3561 | 1,38812E-13 |
| ACE2 | IGLC5 | 0,3561 | 1,40431E-13 |
| ACE2 | SMIM2 | 0,356 | 1,41757E-13 |
| ACE2 | MBOAT2 | 0,3558 | 1,47541E-13 |
| ACE2 | CTC-756D1.2 | 0,3558 | 1,47008E-13 |
| ACE2 | IGHV3-76 | 0,3558 | 1,47466E-13 |
| ACE2 | IGLV5-48 | 0,3558 | 1,4562E-13 |
| ACE2 | LL22NC03-84E4.8 | 0,3558 | 1,45689E-13 |
| ACE2 | UPK3A | 0,3558 | 1,45439E-13 |
| ACE2 | KCNJ4 | 0,3557 | 1,4806E-13 |
| ACE2 | GUCD1 | 0,3556 | 1,51377E-13 |
| ACE2 | IGHV3OR16-7 | 0,3555 | 1,53493E-13 |
| ACE2 | LINC01431 | 0,3554 | 1,56219E-13 |
| ACE2 | ARPC3 | 0,3553 | 1,60353E-13 |
| ACE2 | IGHVII-51-2 | 0,3552 | 1,62474E-13 |
| ACE2 | SEC1P | 0,3552 | 1,60701E-13 |
| ACE2 | HID1 | 0,355 | 1,67668E-13 |
| ACE2 | CNR2 | 0,3549 | 1,68907E-13 |
| ACE2 | IGHD3-22 | 0,3547 | 1,77417E-13 |
| ACE2 | IGKV1-27 | 0,3546 | 1,80382E-13 |
| ACE2 | SLAMF7 | 0,3545 | 1,83533E-13 |
| ACE2 | GPD1 | 0,3545 | 1,83165E-13 |
| ACE2 | CD9 | 0,3542 | 1,92017E-13 |
| ACE2 | RP11-99A1.2 | 0,3542 | 1,91363E-13 |
| ACE2 | RP11-108P20.2 | 0,3542 | 1,90671E-13 |
| ACE2 | MRPS18A | 0,3541 | 1,9294E-13 |
| ACE2 | AZGP1P1 | 0,354 | 1,98895E-13 |
| ACE2 | PDIA4 | 0,354 | 1,96742E-13 |
| ACE2 | CTD-2033D15.1 | 0,354 | 1,98039E-13 |
| ACE2 | ARHGEF28 | 0,3539 | 1,99799E-13 |
| ACE2 | DLAT | 0,3539 | 2,02291E-13 |
| ACE2 | IGHD3-16 | 0,3539 | 2,00694E-13 |
| ACE2 | LDLR | 0,3539 | 2,00998E-13 |
| ACE2 | ZDHHC9 | 0,3539 | 2,00342E-13 |
| ACE2 | RP11-285G1.2 | 0,3538 | 2,05718E-13 |
| ACE2 | HIST1H1C | 0,3537 | 2,08318E-13 |
| ACE2 | RP11-618I10.2 | 0,3535 | 2,15056E-13 |
| ACE2 | SCO1 | 0,3535 | 2,16257E-13 |
| ACE2 | ABHD14B | 0,3534 | 2,18981E-13 |
| ACE2 | LINC01004 | 0,3534 | 2,17299E-13 |
| ACE2 | RP11-672A2.7 | 0,3534 | 2,18293E-13 |
| ACE2 | ZBTB3 | 0,3533 | 2,23615E-13 |
| ACE2 | FBXW11P1 | 0,3533 | 2,23032E-13 |
| ACE2 | IGKJ4 | 0,3532 | 2,26439E-13 |
| ACE2 | ALPP | 0,3531 | 2,28516E-13 |
| ACE2 | MIR3646 | 0,3531 | 2,31507E-13 |
| ACE2 | MANSC1 | 0,353 | 2,31627E-13 |
| ACE2 | IGHVIII-51-1 | 0,353 | 2,33441E-13 |
| ACE2 | UQCRFS1P1 | 0,353 | 2,3273E-13 |
| ACE2 | HSPA8P3 | 0,3529 | 2,38828E-13 |
| ACE2 | NDC1 | 0,3528 | 2,40493E-13 |
| ACE2 | KRT18P68 | 0,3528 | 2,42741E-13 |

| | | | |
|---|---|---|---|
| ACE2 | IGKV6D-41 | 0,3527 | 2,45093E-13 |
| ACE2 | IGKV2D-24 | 0,3523 | 2,61347E-13 |
| ACE2 | IGLV3-32 | 0,3523 | 2,63991E-13 |
| ACE2 | BOLA2B | 0,3522 | 2,64616E-13 |
| ACE2 | MIEF1 | 0,3522 | 2,64527E-13 |
| ACE2 | RN7SKP47 | 0,3521 | 2,70138E-13 |
| ACE2 | AC005915.1 | 0,352 | 2,77396E-13 |
| ACE2 | RPS29P14 | 0,352 | 2,73712E-13 |
| ACE2 | ZC3H12A | 0,3519 | 2,82234E-13 |
| ACE2 | RABGGTA | 0,3517 | 2,88523E-13 |
| ACE2 | IGHVIV-44-1 | 0,3517 | 2,88895E-13 |
| ACE2 | POLR2A | 0,3517 | 2,87352E-13 |
| ACE2 | POU5F1B | 0,3516 | 2,95378E-13 |
| ACE2 | RPSAP52 | 0,3516 | 2,96513E-13 |
| ACE2 | IGHV7-34-1 | 0,3515 | 3,01307E-13 |
| ACE2 | RP11-269G24.4 | 0,3514 | 3,05535E-13 |
| ACE2 | RP11-120E11.2 | 0,3512 | 3,13702E-13 |
| ACE2 | KRT8P20 | 0,3511 | 3,19706E-13 |
| ACE2 | CTD-2306A12.1 | 0,3509 | 3,32809E-13 |
| ACE2 | RP11-713M15.2 | 0,3508 | 3,37759E-13 |
| ACE2 | RP11-294C11.5 | 0,3508 | 3,35828E-13 |
| ACE2 | RP11-466A19.6 | 0,3506 | 3,46639E-13 |
| ACE2 | IGKV2D-28 | 0,3505 | 3,49813E-13 |
| ACE2 | APOC1 | 0,3505 | 3,52033E-13 |
| ACE2 | RP11-249C24.12 | 0,3504 | 3,57848E-13 |
| ACE2 | HOXA6 | 0,3503 | 3,62996E-13 |
| ACE2 | IGLV3-13 | 0,3502 | 3,70677E-13 |
| ACE2 | RP11-439E19.7 | 0,3501 | 3,77832E-13 |
| ACE2 | AC062028.1 | 0,35 | 3,82962E-13 |
| ACE2 | RNU6-554P | 0,35 | 3,82996E-13 |
| ACE2 | ODCP | 0,3499 | 3,90305E-13 |
| ACE2 | STK4-AS1 | 0,3499 | 3,87294E-13 |
| ACE2 | RP11-1084J3.3 | 0,3498 | 3,96711E-13 |
| ACE2 | ARSH | 0,3498 | 3,92961E-13 |
| ACE2 | IGKJ2 | 0,3496 | 4,06736E-13 |
| ACE2 | FAM120A | 0,3496 | 4,10663E-13 |
| ACE2 | RP11-196G11.4 | 0,3496 | 4,09788E-13 |
| ACE2 | MRPS21P9 | 0,3496 | 4,08296E-13 |
| ACE2 | IGLV3-30 | 0,3496 | 4,09305E-13 |
| ACE2 | EPS8 | 0,3494 | 4,21539E-13 |
| ACE2 | CTD-3105H18.7 | 0,3493 | 4,32634E-13 |
| ACE2 | SNRPCP10 | 0,3492 | 4,36507E-13 |
| ACE2 | JCHAIN | 0,3492 | 4,39626E-13 |
| ACE2 | OOSP1 | 0,3492 | 4,33238E-13 |
| ACE2 | GATAD2A | 0,3492 | 4,37812E-13 |
| ACE2 | ABHD5 | 0,3491 | 4,44833E-13 |
| ACE2 | POLR3B | 0,3491 | 4,46819E-13 |
| ACE2 | CYCSP33 | 0,3491 | 4,42815E-13 |
| ACE2 | PRADC1 | 0,349 | 4,53787E-13 |
| ACE2 | MCM2 | 0,3489 | 4,59179E-13 |
| ACE2 | RP4-547N15.3 | 0,3486 | 4,79492E-13 |
| ACE2 | MIR4653 | 0,3486 | 4,78648E-13 |
| ACE2 | KRAS | 0,3486 | 4,80644E-13 |
| ACE2 | IGHEP1 | 0,3486 | 4,78135E-13 |

| | | | |
|---|---|---|---|
| ACE2 | TICAM1 | 0,3486 | 4,84322E-13 |
| ACE2 | IGHV1OR21-1 | 0,3485 | 4,88675E-13 |
| ACE2 | CD2 | 0,3484 | 5,00776E-13 |
| ACE2 | RP11-343L5.2 | 0,3483 | 5,07127E-13 |
| ACE2 | IGKJ5 | 0,3482 | 5,17082E-13 |
| ACE2 | RP11-133K1.9 | 0,348 | 5,31897E-13 |
| ACE2 | SOWAHC | 0,3479 | 5,35016E-13 |
| ACE2 | FAM8A4P | 0,3479 | 5,4375E-13 |
| ACE2 | IGHA1 | 0,3478 | 5,4857E-13 |
| ACE2 | DTL | 0,3477 | 5,60638E-13 |
| ACE2 | RP5-1087E8.2 | 0,3477 | 5,56367E-13 |
| ACE2 | RP11-575F12.2 | 0,3477 | 5,53055E-13 |
| ACE2 | RP11-264B17.2 | 0,3477 | 5,61276E-13 |
| ACE2 | TIGD2 | 0,3476 | 5,70586E-13 |
| ACE2 | RP3-355L5.3 | 0,3476 | 5,65584E-13 |
| ACE2 | KIAA0391 | 0,3476 | 5,66939E-13 |
| ACE2 | IGLV2-28 | 0,3476 | 5,67186E-13 |
| ACE2 | LINC00397 | 0,3475 | 5,79055E-13 |
| ACE2 | IGLC1 | 0,3475 | 5,74356E-13 |
| ACE2 | RCC1 | 0,3474 | 5,84166E-13 |
| ACE2 | AC002511.3 | 0,3474 | 5,82635E-13 |
| ACE2 | ZFAND2B | 0,3472 | 6,03756E-13 |
| ACE2 | APOBEC3A | 0,3472 | 6,03178E-13 |
| ACE2 | LINC02150 | 0,347 | 6,24881E-13 |
| ACE2 | KIF3B | 0,347 | 6,2021E-13 |
| ACE2 | RP3-395M20.7 | 0,3469 | 6,37051E-13 |
| ACE2 | TREML2 | 0,3469 | 6,37301E-13 |
| ACE2 | ANG | 0,3469 | 6,37075E-13 |
| ACE2 | CTD-3193O13.12 | 0,3469 | 6,31446E-13 |
| ACE2 | RP11-810P12.1 | 0,3468 | 6,4488E-13 |
| ACE2 | SPTSSB | 0,3467 | 6,53103E-13 |
| ACE2 | CYP51A1P1 | 0,3466 | 6,65596E-13 |
| ACE2 | RP11-108M12.3 | 0,3466 | 6,67746E-13 |
| ACE2 | RP11-432I5.6 | 0,3466 | 6,71446E-13 |
| ACE2 | IGLV2-34 | 0,3466 | 6,67839E-13 |
| ACE2 | KHDRBS2 | 0,3465 | 6,82262E-13 |
| ACE2 | C19orf67 | 0,3464 | 6,90563E-13 |
| ACE2 | KRT18P21 | 0,3463 | 6,94023E-13 |
| ACE2 | MT-ATP6 | 0,3463 | 7,00088E-13 |
| ACE2 | MTATP6P1 | 0,3462 | 7,12533E-13 |
| ACE2 | RPL23AP66 | 0,3462 | 7,11693E-13 |
| ACE2 | MAZ | 0,3462 | 7,14589E-13 |
| ACE2 | BIN2 | 0,3461 | 7,24082E-13 |
| ACE2 | ZNF789 | 0,346 | 7,30579E-13 |
| ACE2 | JOSD1 | 0,346 | 7,28194E-13 |
| ACE2 | MUC5AC | 0,3459 | 7,48866E-13 |
| ACE2 | ERP27 | 0,3459 | 7,46308E-13 |
| ACE2 | DDTL | 0,3459 | 7,4619E-13 |
| ACE2 | HPCAL1 | 0,3458 | 7,60988E-13 |
| ACE2 | IGKV1-13 | 0,3458 | 7,53606E-13 |
| ACE2 | IGKV6D-21 | 0,3458 | 7,52819E-13 |
| ACE2 | CTC-445B21.1 | 0,3458 | 7,62596E-13 |
| ACE2 | OR4D5 | 0,3456 | 7,8354E-13 |
| ACE2 | ATP2A1 | 0,3456 | 7,77327E-13 |

| | | | |
|---|---|---|---|
| ACE2 | TNFRSF13C | 0,3456 | 7,84464E-13 |
| ACE2 | CS | 0,3455 | 7,95202E-13 |
| ACE2 | RP11-835E18.5 | 0,3455 | 7,98294E-13 |
| ACE2 | HUNK | 0,3454 | 8,06649E-13 |
| ACE2 | CLUHP3 | 0,3452 | 8,32257E-13 |
| ACE2 | IGKV2-28 | 0,345 | 8,68326E-13 |
| ACE2 | RP11-433A10.3 | 0,345 | 8,61925E-13 |
| ACE2 | IGKV1D-35 | 0,3449 | 8,76688E-13 |
| ACE2 | ELFN1-AS1 | 0,3449 | 8,79993E-13 |
| ACE2 | IGHVII-43-1 | 0,3449 | 8,75446E-13 |
| ACE2 | IGHV1OR16-3 | 0,3448 | 8,96302E-13 |
| ACE2 | CTD-2021H9.1 | 0,3447 | 9,01493E-13 |
| ACE2 | STK26 | 0,3446 | 9,22633E-13 |
| ACE2 | ZNF736P9Y | 0,3446 | 9,14623E-13 |
| ACE2 | LIPG | 0,3445 | 9,41864E-13 |
| ACE2 | D2HGDH | 0,3444 | 9,42665E-13 |
| ACE2 | ACBD5 | 0,3444 | 9,44367E-13 |
| ACE2 | GPR119 | 0,3444 | 9,49626E-13 |
| ACE2 | MUC20P1 | 0,3443 | 9,58265E-13 |
| ACE2 | RPL12P25 | 0,3442 | 9,78211E-13 |
| ACE2 | PPIAP30 | 0,3442 | 9,79334E-13 |
| ACE2 | KRT18P20 | 0,3441 | 9,91669E-13 |
| ACE2 | RP11-576N17.5 | 0,3439 | 1,0255E-12 |
| ACE2 | PGLS | 0,3439 | 1,03074E-12 |
| ACE2 | IGLJ7 | 0,3438 | 1,04539E-12 |
| ACE2 | ALDH1L1 | 0,3437 | 1,06295E-12 |
| ACE2 | IGLVI-63 | 0,3436 | 1,08737E-12 |
| ACE2 | RP1-122P22.4 | 0,3435 | 1,10363E-12 |
| ACE2 | AP001258.5 | 0,3434 | 1,1116E-12 |
| ACE2 | IGHVIII-5-1 | 0,3434 | 1,1103E-12 |
| ACE2 | BBIP1P1 | 0,3433 | 1,13561E-12 |
| ACE2 | LARP7P1 | 0,3432 | 1,14705E-12 |
| ACE2 | OR14I1 | 0,3432 | 1,15742E-12 |
| ACE2 | RP1-56K13.2 | 0,3431 | 1,16549E-12 |
| ACE2 | IGKV3-15 | 0,343 | 1,18232E-12 |
| ACE2 | SULT1C3 | 0,3428 | 1,23637E-12 |
| ACE2 | ATP5EP1 | 0,3427 | 1,25116E-12 |
| ACE2 | RP11-11M20.4 | 0,3427 | 1,24121E-12 |
| ACE2 | MEI1 | 0,3427 | 1,24714E-12 |
| ACE2 | FTH1P22 | 0,3426 | 1,26431E-12 |
| ACE2 | IGLJ3 | 0,3426 | 1,273E-12 |
| ACE2 | RP11-418I22.3 | 0,3425 | 1,2937E-12 |
| ACE2 | ESPNP | 0,3424 | 1,31084E-12 |
| ACE2 | RP11-277B15.1 | 0,3424 | 1,30089E-12 |
| ACE2 | CCR6 | 0,3423 | 1,33501E-12 |
| ACE2 | PTGR1 | 0,3423 | 1,32755E-12 |
| ACE2 | IGKV1D-17 | 0,3422 | 1,35729E-12 |
| ACE2 | RNF39 | 0,3422 | 1,34474E-12 |
| ACE2 | IGHV7-27 | 0,3422 | 1,34227E-12 |
| ACE2 | IGLC2 | 0,3422 | 1,35646E-12 |
| ACE2 | IGKJ3 | 0,3421 | 1,38042E-12 |
| ACE2 | MTND3P25 | 0,3421 | 1,38193E-12 |
| ACE2 | RP11-723J4.3 | 0,3421 | 1,37664E-12 |
| ACE2 | LRRFIP2 | 0,342 | 1,39173E-12 |

| | | | |
|---|---|---|---|
| ACE2 | RPL18P10 | 0,342 | 1,40432E-12 |
| ACE2 | AC093642.4 | 0,3419 | 1,42304E-12 |
| ACE2 | CDKN3 | 0,3419 | 1,41793E-12 |
| ACE2 | POU6F2-AS2 | 0,3417 | 1,47267E-12 |
| ACE2 | ADAMTS17 | 0,3416 | 1,49638E-12 |
| ACE2 | IGKC | 0,3414 | 1,54112E-12 |
| ACE2 | TMEM38B | 0,3412 | 1,59405E-12 |
| ACE2 | ZNF700 | 0,3412 | 1,58345E-12 |
| ACE2 | IGKV1D-33 | 0,341 | 1,63834E-12 |
| ACE2 | PTMAP1 | 0,3409 | 1,66752E-12 |
| ACE2 | RP11-203M5.6 | 0,3409 | 1,65581E-12 |
| ACE2 | RP1-140A9.1 | 0,3408 | 1,69428E-12 |
| ACE2 | AL450992.2 | 0,3407 | 1,71396E-12 |
| ACE2 | AC015971.2 | 0,3407 | 1,70763E-12 |
| ACE2 | RN7SL346P | 0,3407 | 1,70122E-12 |
| ACE2 | PRSS41 | 0,3407 | 1,71427E-12 |
| ACE2 | TRDJ1 | 0,3406 | 1,74899E-12 |
| ACE2 | IGHD3-9 | 0,3404 | 1,79949E-12 |
| ACE2 | CTC-458I2.2 | 0,3402 | 1,85698E-12 |
| ACE2 | XPR1 | 0,3399 | 1,9414E-12 |
| ACE2 | GPD1L | 0,3399 | 1,94925E-12 |
| ACE2 | RNA5SP151 | 0,3399 | 1,95386E-12 |
| ACE2 | NCR2 | 0,3398 | 1,96757E-12 |
| ACE2 | KRT18P65 | 0,3396 | 2,04836E-12 |
| ACE2 | IGHV1-17 | 0,3396 | 2,04025E-12 |
| ACE2 | RP11-325K4.3 | 0,3396 | 2,03016E-12 |
| ACE2 | NIF3L1 | 0,3395 | 2,08692E-12 |
| ACE2 | LL22NC03-N95F10.1 | 0,3395 | 2,0872E-12 |
| ACE2 | RNA5SP65 | 0,3394 | 2,10498E-12 |
| ACE2 | RP11-61F12.1 | 0,3394 | 2,09758E-12 |
| ACE2 | EIF2AK3 | 0,3393 | 2,13583E-12 |
| ACE2 | IGKV1D-27 | 0,3393 | 2,14007E-12 |
| ACE2 | IGLV3-22 | 0,3393 | 2,12816E-12 |
| ACE2 | KB-1517D11.4 | 0,3392 | 2,1647E-12 |
| ACE2 | SOX30 | 0,3391 | 2,19721E-12 |
| ACE2 | TSPO2 | 0,3391 | 2,21087E-12 |
| ACE2 | VRK3 | 0,3391 | 2,20166E-12 |
| ACE2 | IGHV4OR15-8 | 0,339 | 2,24228E-12 |
| ACE2 | PLCH1-AS1 | 0,3389 | 2,2629E-12 |
| ACE2 | AP001625.4 | 0,3389 | 2,27715E-12 |
| ACE2 | MANEAL | 0,3388 | 2,30616E-12 |
| ACE2 | IPO5P1 | 0,3388 | 2,30204E-12 |
| ACE2 | MT-TH | 0,3388 | 2,30203E-12 |
| ACE2 | CD3D | 0,3386 | 2,40714E-12 |
| ACE2 | RP3-337H4.8 | 0,3385 | 2,4408E-12 |
| ACE2 | XKR9 | 0,3384 | 2,46624E-12 |
| ACE2 | PRELID3BP11 | 0,3384 | 2,4594E-12 |
| ACE2 | EPHB3 | 0,3383 | 2,5095E-12 |
| ACE2 | RP11-588G21.2 | 0,3382 | 2,53294E-12 |
| ACE2 | HOXB6 | 0,3382 | 2,56312E-12 |
| ACE2 | BTLA | 0,3381 | 2,57693E-12 |
| ACE2 | TTF2 | 0,338 | 2,63185E-12 |
| ACE2 | RP11-379B18.5 | 0,338 | 2,63545E-12 |
| ACE2 | MIR22HG | 0,3379 | 2,66933E-12 |

| | | | |
|---|---|---|---|
| ACE2 | CTC-490G23.4 | 0,3379 | 2,65322E-12 |
| ACE2 | RP11-219B4.5 | 0,3378 | 2,70277E-12 |
| ACE2 | EQTN | 0,3378 | 2,72519E-12 |
| ACE2 | RP11-21K12.3 | 0,3377 | 2,75274E-12 |
| ACE2 | IGHA2 | 0,3376 | 2,80648E-12 |
| ACE2 | IGKV3-7 | 0,3375 | 2,82061E-12 |
| ACE2 | CPN1 | 0,3375 | 2,8519E-12 |
| ACE2 | CYP2T3P | 0,3374 | 2,87401E-12 |
| ACE2 | RP11-184A2.2 | 0,3373 | 2,91332E-12 |
| ACE2 | CAB39P1 | 0,3372 | 2,98482E-12 |
| ACE2 | KRT18P3 | 0,3372 | 2,9562E-12 |
| ACE2 | IGLV3-27 | 0,3372 | 2,97147E-12 |
| ACE2 | MT-TL2 | 0,3372 | 2,95625E-12 |
| ACE2 | AQP12A | 0,3371 | 3,02842E-12 |
| ACE2 | CTC-329D1.2 | 0,337 | 3,06252E-12 |
| ACE2 | ARL4AP2 | 0,3369 | 3,13397E-12 |
| ACE2 | DSCAS | 0,3368 | 3,16701E-12 |
| ACE2 | CCDC186 | 0,3367 | 3,24208E-12 |
| ACE2 | LRRC9 | 0,3367 | 3,2068E-12 |
| ACE2 | AP001469.5 | 0,3367 | 3,23695E-12 |
| ACE2 | GRAMD3 | 0,3364 | 3,37091E-12 |
| ACE2 | NEBL-AS1 | 0,3362 | 3,45187E-12 |
| ACE2 | RP11-21L23.3 | 0,3362 | 3,45304E-12 |
| ACE2 | RP11-21K12.2 | 0,3362 | 3,49105E-12 |
| ACE2 | LINC01218 | 0,336 | 3,60515E-12 |
| ACE2 | FRMD8 | 0,336 | 3,5967E-12 |
| ACE2 | RP11-769O8.3 | 0,336 | 3,60505E-12 |
| ACE2 | KRT8P48 | 0,3359 | 3,66777E-12 |
| ACE2 | AKR1C5P | 0,3359 | 3,61989E-12 |
| ACE2 | ATP1B3P1 | 0,3358 | 3,70864E-12 |
| ACE2 | RPL7P60 | 0,3358 | 3,7093E-12 |
| ACE2 | COLCA2 | 0,3358 | 3,72024E-12 |
| ACE2 | AC026471.6 | 0,3358 | 3,71026E-12 |
| ACE2 | FAM136A | 0,3357 | 3,75218E-12 |
| ACE2 | XPNPEP1 | 0,3355 | 3,89404E-12 |
| ACE2 | RN7SL363P | 0,3354 | 3,93806E-12 |
| ACE2 | BNIP3P4 | 0,3353 | 4,00197E-12 |
| ACE2 | IGHD2-21 | 0,3353 | 4,02701E-12 |
| ACE2 | HMGB3P21 | 0,3352 | 4,0954E-12 |
| ACE2 | CLYBL-AS2 | 0,3352 | 4,09213E-12 |
| ACE2 | RP11-288K12.1 | 0,3349 | 4,24209E-12 |
| ACE2 | IGHVII-22-1 | 0,3349 | 4,28122E-12 |
| ACE2 | KRT18P52 | 0,3347 | 4,41323E-12 |
| ACE2 | RP11-588L15.1 | 0,3347 | 4,41337E-12 |
| ACE2 | C11orf97 | 0,3347 | 4,40454E-12 |
| ACE2 | AC010525.4 | 0,3347 | 4,36598E-12 |
| ACE2 | RP11-126O1.6 | 0,3346 | 4,45042E-12 |
| ACE2 | TIGIT | 0,3345 | 4,51384E-12 |
| ACE2 | STAC3 | 0,3344 | 4,63575E-12 |
| ACE2 | UBE2T | 0,3343 | 4,66489E-12 |
| ACE2 | SIRPG-AS1 | 0,3343 | 4,69384E-12 |
| ACE2 | SIT1 | 0,3342 | 4,77551E-12 |
| ACE2 | IGHV3OR16-12 | 0,3342 | 4,74105E-12 |
| ACE2 | RP11-375H19.2 | 0,3341 | 4,82459E-12 |

| | | | |
|---|---|---|---|
| ACE2 | UGT2B26P | 0,3341 | 4,81935E-12 |
| ACE2 | RP11-109N23.6 | 0,334 | 4,89711E-12 |
| ACE2 | FAM217A | 0,3339 | 5,00183E-12 |
| ACE2 | AC005550.3 | 0,3339 | 4,95296E-12 |
| ACE2 | SLC25A5P6 | 0,3338 | 5,03193E-12 |
| ACE2 | MIR3936 | 0,3338 | 5,04036E-12 |
| ACE2 | ACVRL1 | 0,3338 | 5,04499E-12 |
| ACE2 | GGACT | 0,3338 | 5,0848E-12 |
| ACE2 | AGPAT3 | 0,3338 | 5,07127E-12 |
| ACE2 | ATOH7 | 0,3336 | 5,21356E-12 |
| ACE2 | RAB37 | 0,3336 | 5,21559E-12 |
| ACE2 | RP11-7G12.2 | 0,3335 | 5,32342E-12 |
| ACE2 | RP11-96H19.1 | 0,3335 | 5,3265E-12 |
| ACE2 | SUGT1P3 | 0,3335 | 5,27359E-12 |
| ACE2 | RP11-297C4.3 | 0,3335 | 5,27557E-12 |
| ACE2 | DGCR12 | 0,3335 | 5,30414E-12 |
| ACE2 | IGHD | 0,333 | 5,69175E-12 |
| ACE2 | RP11-74E22.3 | 0,333 | 5,69427E-12 |
| ACE2 | AC007349.7 | 0,3329 | 5,80638E-12 |
| ACE2 | UPK3B | 0,3329 | 5,81369E-12 |
| ACE2 | DKFZP434K028 | 0,3329 | 5,82557E-12 |
| ACE2 | RNF207 | 0,3328 | 5,89748E-12 |
| ACE2 | RP11-576N17.3 | 0,3327 | 6,01587E-12 |
| ACE2 | CTC-287O8.1 | 0,3327 | 5,99713E-12 |
| ACE2 | IGHV3-50 | 0,3327 | 6,00993E-12 |
| ACE2 | IGKV1D-13 | 0,3326 | 6,08713E-12 |
| ACE2 | RP11-65E22.2 | 0,3326 | 6,0732E-12 |
| ACE2 | MGC15885 | 0,3326 | 6,06801E-12 |
| ACE2 | CTA-440B3.1 | 0,3325 | 6,17538E-12 |
| ACE2 | AK3P4 | 0,3324 | 6,30304E-12 |
| ACE2 | SNORA5A | 0,3324 | 6,31557E-12 |
| ACE2 | RFPL4A | 0,3324 | 6,23421E-12 |
| ACE2 | CTD-2540F13.2 | 0,3323 | 6,38821E-12 |
| ACE2 | CHRM3-AS2 | 0,332 | 6,67956E-12 |
| ACE2 | SERPINA11 | 0,332 | 6,65543E-12 |
| ACE2 | ATP12A | 0,3318 | 6,83945E-12 |
| ACE2 | RP11-64D22.5 | 0,3317 | 6,96675E-12 |
| ACE2 | MARK3P1 | 0,3316 | 7,06303E-12 |
| ACE2 | RP13-16H11.5 | 0,3315 | 7,22038E-12 |
| ACE2 | RP11-174G6.5 | 0,3314 | 7,2696E-12 |
| ACE2 | LINC02154 | 0,3314 | 7,32596E-12 |
| ACE2 | RN7SL441P | 0,3313 | 7,40955E-12 |
| ACE2 | IGLV2-5 | 0,3313 | 7,45719E-12 |
| ACE2 | RP11-316J7.4 | 0,3312 | 7,54001E-12 |
| ACE2 | MAP3K11 | 0,3312 | 7,48248E-12 |
| ACE2 | RP11-1E11.1 | 0,3309 | 7,94039E-12 |
| ACE2 | PRAC1 | 0,3308 | 8,00776E-12 |
| ACE2 | RP11-734C14.2 | 0,3307 | 8,15722E-12 |
| ACE2 | LINC01555 | 0,3306 | 8,2419E-12 |
| ACE2 | HIST1H1B | 0,3306 | 8,20264E-12 |
| ACE2 | IGKV1-22 | 0,3305 | 8,42986E-12 |
| ACE2 | HIST1H2AC | 0,3304 | 8,56745E-12 |
| ACE2 | FABP12 | 0,3303 | 8,67768E-12 |
| ACE2 | RP11-187E13.3 | 0,3303 | 8,67935E-12 |

| | | | |
|---|---|---|---|
| ACE2 | P2RX5 | 0,3303 | 8,67673E-12 |
| ACE2 | LDLRAD1 | 0,3302 | 8,75114E-12 |
| ACE2 | RP11-468N14.5 | 0,3301 | 8,91706E-12 |
| ACE2 | IRF4 | 0,3301 | 8,85595E-12 |
| ACE2 | C1GALT1 | 0,3299 | 9,23756E-12 |
| ACE2 | XRCC2 | 0,3299 | 9,15535E-12 |
| ACE2 | PADI3 | 0,3298 | 9,28109E-12 |
| ACE2 | PTRH2 | 0,3298 | 9,37526E-12 |
| ACE2 | MAPKAPK5-AS1 | 0,3297 | 9,49995E-12 |
| ACE2 | AC005262.3 | 0,3297 | 9,48946E-12 |
| ACE2 | IGHD2-2 | 0,3295 | 9,71978E-12 |
| ACE2 | EME1 | 0,3294 | 9,89412E-12 |
| ACE2 | RP1-102E24.10 | 0,3291 | 1,04375E-11 |
| ACE2 | ADAM6 | 0,329 | 1,05429E-11 |
| ACE2 | IGLV1-50 | 0,3289 | 1,06283E-11 |
| ACE2 | TMC3-AS1 | 0,3288 | 1,08523E-11 |
| ACE2 | RP11-28B23.1 | 0,3287 | 1,10402E-11 |
| ACE2 | RP3-400B16.3 | 0,3285 | 1,12871E-11 |
| ACE2 | CTSZ | 0,3285 | 1,1418E-11 |
| ACE2 | LINC01395 | 0,3284 | 1,16118E-11 |
| ACE2 | OSTF1 | 0,3282 | 1,19281E-11 |
| ACE2 | CENPO | 0,3281 | 1,20945E-11 |
| ACE2 | TBC1D14 | 0,3281 | 1,20695E-11 |
| ACE2 | SERPINB8 | 0,328 | 1,22885E-11 |
| ACE2 | AC008079.10 | 0,328 | 1,21924E-11 |
| ACE2 | VDAC1P1 | 0,328 | 1,22743E-11 |
| ACE2 | SLC6A9 | 0,3279 | 1,23763E-11 |
| ACE2 | SLC25A47 | 0,3277 | 1,27526E-11 |
| ACE2 | IGKV1D-37 | 0,3275 | 1,32315E-11 |
| ACE2 | CLDN34 | 0,3275 | 1,31598E-11 |
| ACE2 | BMP5 | 0,3274 | 1,34832E-11 |
| ACE2 | RN7SL189P | 0,3274 | 1,33949E-11 |
| ACE2 | MT-TP | 0,3274 | 1,3382E-11 |
| ACE2 | RP11-407N17.4 | 0,327 | 1,4176E-11 |
| ACE2 | IGHV1OR15-1 | 0,327 | 1,43283E-11 |
| ACE2 | GABRB2 | 0,3269 | 1,45521E-11 |
| ACE2 | GINS4 | 0,3269 | 1,4447E-11 |
| ACE2 | RP13-20L14.6 | 0,3268 | 1,46925E-11 |
| ACE2 | IGKV2-29 | 0,3266 | 1,51768E-11 |
| ACE2 | PLOD2 | 0,3266 | 1,52635E-11 |
| ACE2 | MAPKAPK3 | 0,3264 | 1,55975E-11 |
| ACE2 | TMEM200A | 0,3263 | 1,59684E-11 |
| ACE2 | AC013410.2 | 0,3262 | 1,60648E-11 |
| ACE2 | SNX4 | 0,3262 | 1,61526E-11 |
| ACE2 | LINC01558 | 0,3262 | 1,61798E-11 |
| ACE2 | RP11-402J7.2 | 0,3261 | 1,62719E-11 |
| ACE2 | CYP4F25P | 0,3261 | 1,62531E-11 |
| ACE2 | FDXACB1 | 0,3261 | 1,62999E-11 |
| ACE2 | RP11-285E23.2 | 0,3261 | 1,64385E-11 |
| ACE2 | MFSD13B | 0,3259 | 1,68851E-11 |
| ACE2 | ICAM3 | 0,3259 | 1,68204E-11 |
| ACE2 | RP11-242C19.2 | 0,3259 | 1,69297E-11 |
| ACE2 | RP11-350N15.5 | 0,3257 | 1,74398E-11 |
| ACE2 | RP11-844P9.4 | 0,3256 | 1,75996E-11 |

| | | | |
|---|---|---|---|
| ACE2 | IGKV1-33 | 0,3255 | 1,78939E-11 |
| ACE2 | IGKV2-40 | 0,3255 | 1,78947E-11 |
| ACE2 | AC004775.5 | 0,3253 | 1,8533E-11 |
| ACE2 | RP11-538D16.2 | 0,3249 | 1,96993E-11 |
| ACE2 | IGKV7-3 | 0,3249 | 1,96504E-11 |
| ACE2 | REG1A | 0,3248 | 1,9762E-11 |
| ACE2 | RN7SL301P | 0,3248 | 1,97651E-11 |
| ACE2 | RP11-234K24.6 | 0,3248 | 1,97486E-11 |
| ACE2 | OR13C7 | 0,3247 | 2,03038E-11 |
| ACE2 | GAS8-AS1 | 0,3247 | 2,01761E-11 |
| ACE2 | ANXA2P1 | 0,3246 | 2,05727E-11 |
| ACE2 | CTB-73N10.1 | 0,3245 | 2,0846E-11 |
| ACE2 | CTD-3060P21.1 | 0,3245 | 2,06279E-11 |
| ACE2 | DERL3 | 0,3245 | 2,09149E-11 |
| ACE2 | TMEM176B | 0,3244 | 2,10211E-11 |
| ACE2 | ANXA2 | 0,3244 | 2,10967E-11 |
| ACE2 | MT-TD | 0,3244 | 2,11174E-11 |
| ACE2 | HES5 | 0,3243 | 2,12778E-11 |
| ACE2 | RN7SL386P | 0,3243 | 2,13914E-11 |
| ACE2 | CHAT | 0,3243 | 2,14842E-11 |
| ACE2 | RP11-982M15.7 | 0,3242 | 2,1682E-11 |
| ACE2 | LINC00467 | 0,3241 | 2,20596E-11 |
| ACE2 | ZBTB12BP | 0,3241 | 2,21938E-11 |
| ACE2 | C1QBP | 0,324 | 2,23255E-11 |
| ACE2 | AC010761.13 | 0,324 | 2,23425E-11 |
| ACE2 | RP11-89H19.1 | 0,3239 | 2,26631E-11 |
| ACE2 | IGHV3-33-2 | 0,3239 | 2,27381E-11 |
| ACE2 | GPR25 | 0,3238 | 2,30178E-11 |
| ACE2 | IGKV1OR2-118 | 0,3238 | 2,29665E-11 |
| ACE2 | AC103563.2 | 0,3238 | 2,29476E-11 |
| ACE2 | SLC25A5P1 | 0,3236 | 2,39013E-11 |
| ACE2 | RP11-486M3.3 | 0,3235 | 2,42616E-11 |
| ACE2 | RP11-77H9.2 | 0,3235 | 2,40928E-11 |
| ACE2 | SAMD5 | 0,3233 | 2,48508E-11 |
| ACE2 | LINC01900 | 0,3233 | 2,49409E-11 |
| ACE2 | MIR3944 | 0,3232 | 2,50494E-11 |
| ACE2 | CTC-537E7.2 | 0,3231 | 2,56848E-11 |
| ACE2 | KNOP1 | 0,3231 | 2,55823E-11 |
| ACE2 | RP13-644M16.4 | 0,323 | 2,59868E-11 |
| ACE2 | NLN | 0,323 | 2,59936E-11 |
| ACE2 | RP11-13N13.6 | 0,323 | 2,58969E-11 |
| ACE2 | ZSWIM3 | 0,323 | 2,61005E-11 |
| ACE2 | RPL32P33 | 0,3228 | 2,67323E-11 |
| ACE2 | KIAA0319L | 0,3227 | 2,73052E-11 |
| ACE2 | HIST1H4F | 0,3227 | 2,72803E-11 |
| ACE2 | RP5-890O15.3 | 0,3227 | 2,72651E-11 |
| ACE2 | NIP7 | 0,3225 | 2,78619E-11 |
| ACE2 | CYP2C58P | 0,3224 | 2,82882E-11 |
| ACE2 | RP13-122B23.9 | 0,3223 | 2,882E-11 |
| ACE2 | CCDC189 | 0,3223 | 2,90028E-11 |
| ACE2 | CMTR2 | 0,3223 | 2,89967E-11 |
| ACE2 | IGKV1-35 | 0,3222 | 2,92773E-11 |
| ACE2 | C6orf163 | 0,3222 | 2,92381E-11 |
| ACE2 | MTCO3P12 | 0,3221 | 2,98927E-11 |

| | | | |
|---|---|---|---|
| ACE2 | IGKV1D-43 | 0,3221 | 2,96753E-11 |
| ACE2 | INSL3 | 0,3221 | 2,96816E-11 |
| ACE2 | IGKV2D-40 | 0,322 | 3,01071E-11 |
| ACE2 | TMEM60 | 0,322 | 3,00738E-11 |
| ACE2 | RP11-1029J19.5 | 0,322 | 3,00718E-11 |
| ACE2 | RP5-991G20.2 | 0,322 | 3,02861E-11 |
| ACE2 | RNU6-418P | 0,3218 | 3,12368E-11 |
| ACE2 | LITAF | 0,3218 | 3,09066E-11 |
| ACE2 | WIPF2 | 0,3218 | 3,08719E-11 |
| ACE2 | IGLJ6 | 0,3218 | 3,08649E-11 |
| ACE2 | LAMP3 | 0,3217 | 3,13097E-11 |
| ACE2 | NT5C3AP1 | 0,3215 | 3,24434E-11 |
| ACE2 | RP11-1151B14.4 | 0,3215 | 3,26479E-11 |
| ACE2 | PPIAP28 | 0,3214 | 3,28453E-11 |
| ACE2 | LRRC37A9P | 0,3214 | 3,29776E-11 |
| ACE2 | AC006070.12 | 0,3214 | 3,28894E-11 |
| ACE2 | CTB-31O20.4 | 0,3214 | 3,29967E-11 |
| ACE2 | TP53TG1 | 0,3213 | 3,35767E-11 |
| ACE2 | POU2AF1 | 0,3213 | 3,33163E-11 |
| ACE2 | DIP2B | 0,3211 | 3,45631E-11 |
| ACE2 | ZNF622 | 0,321 | 3,50824E-11 |
| ACE2 | RP11-53B2.6 | 0,3209 | 3,56287E-11 |
| ACE2 | IGLV9-49 | 0,3209 | 3,54588E-11 |
| ACE2 | SULT1A3 | 0,3208 | 3,61776E-11 |
| ACE2 | ITGB7 | 0,3207 | 3,63655E-11 |
| ACE2 | SLA2 | 0,3207 | 3,67057E-11 |
| ACE2 | CD7 | 0,3206 | 3,67908E-11 |
| ACE2 | IGLC3 | 0,3206 | 3,68261E-11 |
| ACE2 | RP11-354K1.2 | 0,3204 | 3,79119E-11 |
| ACE2 | RP11-326C3.2 | 0,3202 | 3,91744E-11 |
| ACE2 | BLK | 0,3201 | 4,00502E-11 |
| ACE2 | RP11-503C24.4 | 0,32 | 4,02378E-11 |
| ACE2 | AJ003147.9 | 0,32 | 4,04552E-11 |
| ACE2 | AC005307.3 | 0,3198 | 4,18969E-11 |
| ACE2 | CTD-2583A14.11 | 0,3197 | 4,22061E-11 |
| ACE2 | GKN2 | 0,3196 | 4,30922E-11 |
| ACE2 | RP11-539L10.5 | 0,3195 | 4,38188E-11 |
| ACE2 | LRP4 | 0,3195 | 4,33757E-11 |
| ACE2 | CCDC88B | 0,3194 | 4,44924E-11 |
| ACE2 | AC013275.2 | 0,3193 | 4,47444E-11 |
| ACE2 | TMEM133 | 0,3191 | 4,64914E-11 |
| ACE2 | RP11-1029J19.4 | 0,319 | 4,65665E-11 |
| ACE2 | MAT1A | 0,3188 | 4,82831E-11 |
| ACE2 | RP11-347I19.7 | 0,3188 | 4,83984E-11 |
| ACE2 | LACC1 | 0,3188 | 4,83881E-11 |
| ACE2 | HS3ST6 | 0,3188 | 4,83259E-11 |
| ACE2 | ERMN | 0,3187 | 4,9071E-11 |
| ACE2 | BLOC1S1 | 0,3187 | 4,9095E-11 |
| ACE2 | ULBP3 | 0,3186 | 4,95014E-11 |
| ACE2 | MRPL15 | 0,3186 | 4,93942E-11 |
| ACE2 | RP4-706G24.1 | 0,3185 | 5,06494E-11 |
| ACE2 | RP11-489M13.1 | 0,3185 | 5,05314E-11 |
| ACE2 | C1orf226 | 0,3184 | 5,11841E-11 |
| ACE2 | MIXL1 | 0,3184 | 5,1025E-11 |

| | | | |
|---|---|---|---|
| ACE2 | CTC-806A22.1 | 0,3184 | 5,1406E-11 |
| ACE2 | RP4-742J24.2 | 0,3184 | 5,13535E-11 |
| ACE2 | PUS10 | 0,3183 | 5,22484E-11 |
| ACE2 | TRG-AS1 | 0,3181 | 5,37422E-11 |
| ACE2 | RP11-481L19.2 | 0,3181 | 5,32573E-11 |
| ACE2 | NONOP2 | 0,3179 | 5,49534E-11 |
| ACE2 | IGLV3-17 | 0,3179 | 5,50665E-11 |
| ACE2 | RP4-594A5.1 | 0,3176 | 5,7367E-11 |
| ACE2 | RP11-44D19.1 | 0,3176 | 5,73546E-11 |
| ACE2 | CCNE1 | 0,3176 | 5,73591E-11 |
| ACE2 | RPL10P1 | 0,3175 | 5,86782E-11 |
| ACE2 | RP5-1155K23.4 | 0,3173 | 5,99346E-11 |
| ACE2 | PHKG2 | 0,3173 | 6,01709E-11 |
| ACE2 | FCRL1 | 0,3172 | 6,11735E-11 |
| ACE2 | SPN | 0,317 | 6,30155E-11 |
| ACE2 | ZSCAN5DP | 0,317 | 6,31995E-11 |
| ACE2 | RP5-1092A11.5 | 0,3169 | 6,37129E-11 |
| ACE2 | FAM83A | 0,3169 | 6,32761E-11 |
| ACE2 | EME2 | 0,3169 | 6,40968E-11 |
| ACE2 | MLLT10P1 | 0,3169 | 6,36565E-11 |
| ACE2 | MTCYBP18 | 0,3168 | 6,50045E-11 |
| ACE2 | AC092573.2 | 0,3167 | 6,58613E-11 |
| ACE2 | RP1-179N16.3 | 0,3167 | 6,50984E-11 |
| ACE2 | DNAJC5B | 0,3165 | 6,72796E-11 |
| ACE2 | PCAT29 | 0,3164 | 6,88966E-11 |
| ACE2 | CTD-2132N18.2 | 0,3164 | 6,84309E-11 |
| ACE2 | REG1B | 0,3163 | 6,90724E-11 |
| ACE2 | RBBP4P2 | 0,3163 | 6,90782E-11 |
| ACE2 | RP11-358M14.2 | 0,3163 | 6,98722E-11 |
| ACE2 | IGHD3-10 | 0,3163 | 6,97485E-11 |
| ACE2 | MIR635 | 0,3162 | 7,0613E-11 |
| ACE2 | RP11-175D17.3 | 0,316 | 7,24046E-11 |
| ACE2 | MTND4P23 | 0,3159 | 7,38834E-11 |
| ACE2 | CLDN9 | 0,3159 | 7,3113E-11 |
| ACE2 | IGKV2-26 | 0,3156 | 7,66018E-11 |
| ACE2 | TIMD4 | 0,3155 | 7,78088E-11 |
| ACE2 | FAM98C | 0,3155 | 7,7936E-11 |
| ACE2 | CTB-88F18.5 | 0,3154 | 7,93814E-11 |
| ACE2 | RP11-166O4.4 | 0,3154 | 7,93172E-11 |
| ACE2 | SYTL1 | 0,3153 | 8,04863E-11 |
| ACE2 | ECEL1P1 | 0,315 | 8,38146E-11 |
| ACE2 | ACP5 | 0,3148 | 8,64524E-11 |
| ACE2 | NIP7P3 | 0,3147 | 8,72911E-11 |
| ACE2 | RP11-728K20.2 | 0,3147 | 8,76442E-11 |
| ACE2 | HELLS | 0,3147 | 8,71517E-11 |
| ACE2 | MTHFD1P1 | 0,3143 | 9,28592E-11 |
| ACE2 | HIST1H2BD | 0,3141 | 9,50471E-11 |
| ACE2 | LINC01619 | 0,3139 | 9,78202E-11 |
| ACE2 | IGLV11-55 | 0,3139 | 9,77433E-11 |
| ACE2 | IGHVIII-67-2 | 0,3137 | 1,00869E-10 |
| ACE2 | CELA3B | 0,3136 | 1,02476E-10 |
| ACE2 | RP11-575G13.2 | 0,3136 | 1,03107E-10 |
| ACE2 | AC096579.13 | 0,3135 | 1,03868E-10 |
| ACE2 | RNASE7 | 0,3135 | 1,0356E-10 |

| | | | |
|---|---|---|---|
| ACE2 | MMP25 | 0,3135 | 1,03446E-10 |
| ACE2 | LL22NC03-84E4.13 | 0,3135 | 1,0355E-10 |
| ACE2 | GPR31 | 0,3132 | 1,08418E-10 |
| ACE2 | LINC00659 | 0,3131 | 1,10153E-10 |
| ACE2 | TESK2 | 0,313 | 1,12086E-10 |
| ACE2 | LINC01889 | 0,313 | 1,11534E-10 |
| ACE2 | TRGV4 | 0,313 | 1,12457E-10 |
| ACE2 | RP11-74E22.8 | 0,313 | 1,1112E-10 |
| ACE2 | RP11-802D6.1 | 0,313 | 1,11182E-10 |
| ACE2 | RP11-288G3.4 | 0,3127 | 1,16934E-10 |
| ACE2 | GCA | 0,3126 | 1,18513E-10 |
| ACE2 | LRMP | 0,3123 | 1,23426E-10 |
| ACE2 | IGHD4-17 | 0,3123 | 1,23325E-10 |
| ACE2 | RP11-568A7.2 | 0,3122 | 1,2612E-10 |
| ACE2 | RP11-304L19.13 | 0,3122 | 1,25425E-10 |
| ACE2 | SLC35B1 | 0,3122 | 1,25283E-10 |
| ACE2 | RP11-61L19.1 | 0,3122 | 1,25131E-10 |
| ACE2 | GJB4 | 0,312 | 1,29676E-10 |
| ACE2 | SLC7A7 | 0,312 | 1,29823E-10 |
| ACE2 | BAIAP3 | 0,312 | 1,28642E-10 |
| ACE2 | PEX7 | 0,3118 | 1,32921E-10 |
| ACE2 | MIPOL1 | 0,3118 | 1,33394E-10 |
| ACE2 | CYP4F8 | 0,3117 | 1,33867E-10 |
| ACE2 | FLVCR1 | 0,3116 | 1,35805E-10 |
| ACE2 | LINC00996 | 0,3116 | 1,37272E-10 |
| ACE2 | AC017074.1 | 0,3115 | 1,38221E-10 |
| ACE2 | MIR25 | 0,3115 | 1,39114E-10 |
| ACE2 | ATP2B1 | 0,3115 | 1,39267E-10 |
| ACE2 | CLDN14 | 0,3115 | 1,3851E-10 |
| ACE2 | RP11-649E7.7 | 0,3114 | 1,41151E-10 |
| ACE2 | E2F3 | 0,3113 | 1,42106E-10 |
| ACE2 | BBOX1-AS1 | 0,3112 | 1,44604E-10 |
| ACE2 | PGGT1B | 0,3111 | 1,45756E-10 |
| ACE2 | AC068580.5 | 0,311 | 1,49277E-10 |
| ACE2 | RP11-626H12.2 | 0,311 | 1,47664E-10 |
| ACE2 | HCK | 0,311 | 1,49607E-10 |
| ACE2 | TMEM181 | 0,3109 | 1,49818E-10 |
| ACE2 | CDC14A | 0,3108 | 1,52719E-10 |
| ACE2 | MT-TR | 0,3108 | 1,52568E-10 |
| ACE2 | MS4A15 | 0,3107 | 1,54961E-10 |
| ACE2 | SLC39A9 | 0,3106 | 1,56607E-10 |
| ACE2 | MT-RNR2 | 0,3105 | 1,5919E-10 |
| ACE2 | AKR1C7P | 0,3104 | 1,61982E-10 |
| ACE2 | RP11-678G14.4 | 0,3104 | 1,6251E-10 |
| ACE2 | AC008982.2 | 0,3102 | 1,6703E-10 |
| ACE2 | TMEM44-AS1 | 0,3101 | 1,69205E-10 |
| ACE2 | TMOD3 | 0,3101 | 1,69556E-10 |
| ACE2 | HMGB1P23 | 0,31 | 1,71225E-10 |
| ACE2 | GDPD1 | 0,31 | 1,70648E-10 |
| ACE2 | CCDC60 | 0,3099 | 1,73646E-10 |
| ACE2 | TMEM208 | 0,3099 | 1,74866E-10 |
| ACE2 | ZBP1 | 0,3099 | 1,74499E-10 |
| ACE2 | OR56B4 | 0,3098 | 1,76888E-10 |
| ACE2 | STK24 | 0,3097 | 1,77899E-10 |

| | | | |
|---|---|---|---|
| ACE2 | BAZ1A | 0,3097 | 1,7921E-10 |
| ACE2 | NRAP | 0,3096 | 1,80911E-10 |
| ACE2 | MTCO2P22 | 0,3095 | 1,83664E-10 |
| ACE2 | EP300-AS1 | 0,3095 | 1,83048E-10 |
| ACE2 | IGKV3-25 | 0,3094 | 1,85945E-10 |
| ACE2 | RNU6-668P | 0,3094 | 1,87661E-10 |
| ACE2 | RP11-390E23.6 | 0,3094 | 1,85563E-10 |
| ACE2 | SRSF9 | 0,3094 | 1,87806E-10 |
| ACE2 | EBI3 | 0,3094 | 1,86787E-10 |
| ACE2 | IL1B | 0,3093 | 1,89338E-10 |
| ACE2 | SPRN | 0,3093 | 1,89915E-10 |
| ACE2 | LINC01215 | 0,3092 | 1,90887E-10 |
| ACE2 | LYN | 0,3092 | 1,90792E-10 |
| ACE2 | EPSTI1 | 0,3092 | 1,90614E-10 |
| ACE2 | CHRNG | 0,3091 | 1,94094E-10 |
| ACE2 | DSG3 | 0,3091 | 1,95092E-10 |
| ACE2 | RP11-545D22.1 | 0,309 | 1,97948E-10 |
| ACE2 | RP11-77H9.8 | 0,309 | 1,9695E-10 |
| ACE2 | RNA5SP187 | 0,3089 | 1,99027E-10 |
| ACE2 | RP1-256G22.2 | 0,3086 | 2,07471E-10 |
| ACE2 | MTND4P9 | 0,3084 | 2,14626E-10 |
| ACE2 | MGAT3-AS1 | 0,3084 | 2,15708E-10 |
| ACE2 | FANCI | 0,3082 | 2,19663E-10 |
| ACE2 | IGKV2-18 | 0,3078 | 2,32077E-10 |
| ACE2 | CPD | 0,3078 | 2,32888E-10 |
| ACE2 | NAPSB | 0,3076 | 2,39376E-10 |
| ACE2 | SLC25A5P3 | 0,3074 | 2,46242E-10 |
| ACE2 | PRKACG | 0,3074 | 2,4757E-10 |
| ACE2 | RP11-380G5.2 | 0,3074 | 2,4567E-10 |
| ACE2 | IGKV1-37 | 0,3073 | 2,51564E-10 |
| ACE2 | LINC01781 | 0,307 | 2,6156E-10 |
| ACE2 | AC226118.1 | 0,3069 | 2,66098E-10 |
| ACE2 | CTD-2562G15.3 | 0,3069 | 2,63385E-10 |
| ACE2 | RP11-128M1.1 | 0,3069 | 2,66029E-10 |
| ACE2 | IGLV3-6 | 0,3069 | 2,66106E-10 |
| ACE2 | RP4-736L20.3 | 0,3068 | 2,67501E-10 |
| ACE2 | RP11-1084J3.1 | 0,3067 | 2,72631E-10 |
| ACE2 | BCAT2 | 0,3067 | 2,73435E-10 |
| ACE2 | RP11-203P23.2 | 0,3066 | 2,74927E-10 |
| ACE2 | CTC-529I10.1 | 0,3066 | 2,7828E-10 |
| ACE2 | RNU6-850P | 0,3065 | 2,81694E-10 |
| ACE2 | RP11-570H19.2 | 0,3065 | 2,7982E-10 |
| ACE2 | CD163L1 | 0,3065 | 2,80884E-10 |
| ACE2 | RP11-133K1.11 | 0,3065 | 2,7869E-10 |
| ACE2 | CD247 | 0,3063 | 2,90248E-10 |
| ACE2 | MIR608 | 0,3063 | 2,87788E-10 |
| ACE2 | PRB3 | 0,3063 | 2,89477E-10 |
| ACE2 | RP11-1J11.1 | 0,3062 | 2,93692E-10 |
| ACE2 | LCN8 | 0,3062 | 2,93481E-10 |
| ACE2 | GIF | 0,3062 | 2,92854E-10 |
| ACE2 | RNU4-38P | 0,3061 | 2,94739E-10 |
| ACE2 | TBXAS1 | 0,3061 | 2,94826E-10 |
| ACE2 | CTC-450M9.1 | 0,3061 | 2,96735E-10 |
| ACE2 | KB-1090H4.2 | 0,306 | 3,02444E-10 |

| | | | |
|---|---|---|---|
| ACE2 | ADIPOR2 | 0,3059 | 3,06743E-10 |
| ACE2 | IGKV2D-18 | 0,3058 | 3,0867E-10 |
| ACE2 | AC005351.1 | 0,3058 | 3,08559E-10 |
| ACE2 | INCENP | 0,3057 | 3,13027E-10 |
| ACE2 | TRAF3 | 0,3056 | 3,15894E-10 |
| ACE2 | SSTR3 | 0,3056 | 3,18125E-10 |
| ACE2 | KB-1269D1.8 | 0,3055 | 3,2237E-10 |
| ACE2 | SLC5A12 | 0,3054 | 3,24809E-10 |
| ACE2 | RP11-337L12.1 | 0,3054 | 3,28682E-10 |
| ACE2 | AC090952.5 | 0,3053 | 3,33296E-10 |
| ACE2 | CD96 | 0,3052 | 3,38035E-10 |
| ACE2 | KB-1562D12.1 | 0,3052 | 3,35807E-10 |
| ACE2 | TRMT2B | 0,3052 | 3,36011E-10 |
| ACE2 | YWHAQP5 | 0,3051 | 3,41721E-10 |
| ACE2 | CCDC96 | 0,3049 | 3,50176E-10 |
| ACE2 | HIST1H2BH | 0,3048 | 3,56362E-10 |
| ACE2 | LLPH-AS1 | 0,3048 | 3,54051E-10 |
| ACE2 | IGLVVII-41-1 | 0,3048 | 3,54848E-10 |
| ACE2 | GLDC | 0,3047 | 3,61272E-10 |
| ACE2 | OR55B1P | 0,3047 | 3,62069E-10 |
| ACE2 | CFAP52 | 0,3047 | 3,62793E-10 |
| ACE2 | AC011816.1 | 0,3046 | 3,65128E-10 |
| ACE2 | CTC-458A3.1 | 0,3046 | 3,67509E-10 |
| ACE2 | RP5-928E24.2 | 0,3046 | 3,6317E-10 |
| ACE2 | RN7SL45P | 0,3045 | 3,71765E-10 |
| ACE2 | TNNC2 | 0,3045 | 3,68133E-10 |
| ACE2 | IGKV2OR22-4 | 0,3045 | 3,68249E-10 |
| ACE2 | IMMT | 0,3044 | 3,74052E-10 |
| ACE2 | NDUFA1 | 0,3042 | 3,8846E-10 |
| ACE2 | RP11-415F23.4 | 0,3041 | 3,92734E-10 |
| ACE2 | DNAH7 | 0,304 | 3,96232E-10 |
| ACE2 | VEGFA | 0,304 | 3,95008E-10 |
| ACE2 | LINC01624 | 0,304 | 3,96058E-10 |
| ACE2 | FAM60A | 0,304 | 3,971E-10 |
| ACE2 | RP13-942N8.1 | 0,304 | 3,95392E-10 |
| ACE2 | KRT8P25 | 0,3038 | 4,08612E-10 |
| ACE2 | CARD11 | 0,3035 | 4,23535E-10 |
| ACE2 | IGHV3OR16-11 | 0,3035 | 4,25646E-10 |
| ACE2 | HLA-H | 0,3033 | 4,36646E-10 |
| ACE2 | RP11-326K13.3 | 0,3031 | 4,51573E-10 |
| ACE2 | RN7SL192P | 0,3031 | 4,47658E-10 |
| ACE2 | AC096574.4 | 0,3029 | 4,6048E-10 |
| ACE2 | NANOGP7 | 0,3029 | 4,65285E-10 |
| ACE2 | IGLVIVOR22-1 | 0,3029 | 4,63179E-10 |
| ACE2 | COL9A2 | 0,3028 | 4,67785E-10 |
| ACE2 | PPY | 0,3027 | 4,73789E-10 |
| ACE2 | RP3-468B3.2 | 0,3026 | 4,78814E-10 |
| ACE2 | SNORA5C | 0,3026 | 4,83607E-10 |
| ACE2 | PHBP21 | 0,3026 | 4,78926E-10 |
| ACE2 | OR7E85P | 0,3025 | 4,90875E-10 |
| ACE2 | RP11-318A15.2 | 0,3025 | 4,88277E-10 |
| ACE2 | AC016734.2 | 0,3024 | 4,9468E-10 |
| ACE2 | RP11-424C20.2 | 0,3024 | 4,97778E-10 |
| ACE2 | RP6-91H8.1 | 0,3023 | 5,0511E-10 |

| | | | |
|---|---|---|---|
| ACE2 | TMEM62 | 0,3023 | 5,05145E-10 |
| ACE2 | RP11-186F10.2 | 0,3021 | 5,15362E-10 |
| ACE2 | CFAP73 | 0,3021 | 5,13639E-10 |
| ACE2 | HAUS8 | 0,3018 | 5,38935E-10 |
| ACE2 | AP5S1 | 0,3018 | 5,37627E-10 |
| ACE2 | MFSD2A | 0,3017 | 5,4384E-10 |
| ACE2 | RP11-18H21.1 | 0,3017 | 5,41826E-10 |
| ACE2 | CRYZP1 | 0,3017 | 5,44065E-10 |
| ACE2 | RP11-806L2.2 | 0,3016 | 5,49499E-10 |
| ACE2 | CTD-2616J11.2 | 0,3016 | 5,52704E-10 |
| ACE2 | FCRL4 | 0,3015 | 5,56585E-10 |
| ACE2 | HIST1H3H | 0,3015 | 5,56999E-10 |
| ACE2 | RP13-81N3.2 | 0,3015 | 5,60996E-10 |
| ACE2 | SSUH2 | 0,3013 | 5,78833E-10 |
| ACE2 | MTND4LP30 | 0,3012 | 5,81293E-10 |
| ACE2 | RP11-423H2.3 | 0,3012 | 5,83292E-10 |
| ACE2 | LUADT1 | 0,3012 | 5,81303E-10 |
| ACE2 | IGKV1OR22-1 | 0,3012 | 5,80595E-10 |
| ACE2 | RP11-394I13.2 | 0,3011 | 5,91361E-10 |
| ACE2 | SORD | 0,301 | 6,02597E-10 |
| ACE2 | HSD3BP1 | 0,3009 | 6,12248E-10 |
| ACE2 | SLC22A13 | 0,3009 | 6,08171E-10 |
| ACE2 | RP11-74J13.9 | 0,3009 | 6,12366E-10 |
| ACE2 | IGHV1OR15-3 | 0,3009 | 6,09317E-10 |
| ACE2 | RP11-367J11.2 | 0,3008 | 6,18406E-10 |
| ACE2 | CTA-363E6.5 | 0,3008 | 6,13475E-10 |
| ACE2 | CTD-2562J15.4 | 0,3008 | 6,16319E-10 |
| ACE2 | AC002511.2 | 0,3008 | 6,13019E-10 |
| ACE2 | DHTKD1 | 0,3007 | 6,23458E-10 |
| ACE2 | RP11-261P9.4 | 0,3007 | 6,27447E-10 |
| ACE2 | LL22NC03-24A12.9 | 0,3007 | 6,21509E-10 |
| ACE2 | AC016683.5 | 0,3006 | 6,34346E-10 |
| ACE2 | MTND3P5 | 0,3006 | 6,2967E-10 |
| ACE2 | IGSF6 | 0,3006 | 6,32296E-10 |
| ACE2 | CTD-2649C14.2 | 0,3006 | 6,31872E-10 |
| ACE2 | HMGB3P32 | 0,3006 | 6,31394E-10 |
| ACE2 | LINC01337 | 0,3005 | 6,43466E-10 |
| ACE2 | RP11-338L22.3 | 0,3005 | 6,42779E-10 |
| ACE2 | RP11-20B24.5 | 0,3003 | 6,61173E-10 |
| ACE2 | ATP5G1P4 | 0,3002 | 6,72921E-10 |
| ACE2 | RP4-676J13.2 | 0,3002 | 6,72937E-10 |
| ACE2 | C2CD4A | 0,3001 | 6,74213E-10 |
| ACE2 | SIGLEC8 | 0,3001 | 6,75214E-10 |
| ACE2 | MRPS34 | 0,2998 | 7,07475E-10 |
| ACE2 | NOS2P3 | 0,2998 | 7,02928E-10 |
| ACE2 | AP000350.5 | 0,2998 | 7,02515E-10 |
| ACE2 | GAPDHP74 | 0,2996 | 7,29487E-10 |
| ACE2 | NDUFB9 | 0,2996 | 7,3069E-10 |
| ACE2 | CTA-221G9.12 | 0,2996 | 7,30929E-10 |
| ACE2 | ENHO | 0,2995 | 7,41105E-10 |
| ACE2 | SDF4 | 0,2994 | 7,47741E-10 |
| ACE2 | ATP5LP5 | 0,2994 | 7,44681E-10 |
| ACE2 | RP11-379B18.6 | 0,2993 | 7,53292E-10 |
| ACE2 | RP11-1223D19.6 | 0,2991 | 7,78011E-10 |

| | | | |
|---|---|---|---|
| ACE2 | CTB-160O22.1 | 0,2991 | 7,76292E-10 |
| ACE2 | SLC6A12 | 0,299 | 7,90662E-10 |
| ACE2 | RP11-90P13.1 | 0,2989 | 7,98348E-10 |
| ACE2 | RP11-550A5.2 | 0,2989 | 8,01381E-10 |
| ACE2 | IGLV1-62 | 0,2989 | 7,93562E-10 |
| ACE2 | HUS1 | 0,2988 | 8,04863E-10 |
| ACE2 | MPP5 | 0,2988 | 8,09112E-10 |
| ACE2 | RP11-788H18.1 | 0,2987 | 8,15532E-10 |
| ACE2 | RP11-707P17.2 | 0,2987 | 8,25541E-10 |
| ACE2 | MAP2K2 | 0,2987 | 8,18799E-10 |
| ACE2 | CTD-2528A14.3 | 0,2986 | 8,342E-10 |
| ACE2 | MT-ATP8 | 0,2986 | 8,29805E-10 |
| ACE2 | P2RY13 | 0,2985 | 8,47231E-10 |
| ACE2 | LINC00525 | 0,2981 | 8,9385E-10 |
| ACE2 | METAP1 | 0,298 | 8,98855E-10 |
| ACE2 | PROSER2 | 0,2979 | 9,10553E-10 |
| ACE2 | CARNMT1 | 0,2978 | 9,2628E-10 |
| ACE2 | AUH | 0,2978 | 9,24531E-10 |
| ACE2 | RP11-53B2.3 | 0,2978 | 9,33317E-10 |
| ACE2 | MTND6P22 | 0,2977 | 9,40287E-10 |
| ACE2 | LINC01962 | 0,2977 | 9,37929E-10 |
| ACE2 | RP11-69E11.8 | 0,2976 | 9,46968E-10 |
| ACE2 | KCNK13 | 0,2976 | 9,57081E-10 |
| ACE2 | MZB1 | 0,2975 | 9,67168E-10 |
| ACE2 | RP11-328J14.1 | 0,2975 | 9,6723E-10 |
| ACE2 | DARS-AS1 | 0,2974 | 9,73688E-10 |
| ACE2 | GUSB | 0,2974 | 9,7631E-10 |
| ACE2 | NUDT18 | 0,2974 | 9,84233E-10 |
| ACE2 | MYO5BP3 | 0,2974 | 9,76565E-10 |
| ACE2 | RECQL4 | 0,2973 | 9,94751E-10 |
| ACE2 | PPP4C | 0,2973 | 9,97877E-10 |
| ACE2 | PKDREJ | 0,2973 | 9,91276E-10 |
| ACE2 | SALL1 | 0,2972 | 1,00265E-09 |
| ACE2 | TP53I11 | 0,2971 | 1,01612E-09 |
| ACE2 | IGHD4-23 | 0,2971 | 1,02015E-09 |
| ACE2 | RP11-131L12.2 | 0,297 | 1,02785E-09 |
| ACE2 | AC012314.8 | 0,297 | 1,03671E-09 |
| ACE2 | XXbac-BPG248L24.12 | 0,2969 | 1,04825E-09 |
| ACE2 | RP11-111M22.3 | 0,2969 | 1,04313E-09 |
| ACE2 | AC009970.1 | 0,2968 | 1,05955E-09 |
| ACE2 | AC000111.5 | 0,2968 | 1,06566E-09 |
| ACE2 | RP11-290F24.3 | 0,2967 | 1,07037E-09 |
| ACE2 | AC019097.7 | 0,2966 | 1,08827E-09 |
| ACE2 | LINC01967 | 0,2965 | 1,09805E-09 |
| ACE2 | RP11-575G13.3 | 0,2965 | 1,10059E-09 |
| ACE2 | LINC01979 | 0,2964 | 1,11585E-09 |
| ACE2 | RP5-997D16.2 | 0,2962 | 1,15135E-09 |
| ACE2 | RP1-118J21.5 | 0,2962 | 1,15014E-09 |
| ACE2 | NABP1 | 0,2962 | 1,1442E-09 |
| ACE2 | KLHL6 | 0,2961 | 1,16183E-09 |
| ACE2 | MYL12BP2 | 0,296 | 1,18647E-09 |
| ACE2 | CTC-218B8.3 | 0,2959 | 1,19905E-09 |
| ACE2 | CATSPER4 | 0,2958 | 1,22085E-09 |
| ACE2 | TPMTP1 | 0,2958 | 1,21357E-09 |

| | | | |
|---|---|---|---|
| ACE2 | UPK1B | 0,2957 | 1,23379E-09 |
| ACE2 | RP11-533K9.3 | 0,2957 | 1,22721E-09 |
| ACE2 | OR7E128P | 0,2957 | 1,22492E-09 |
| ACE2 | AC020951.1 | 0,2957 | 1,22816E-09 |
| ACE2 | AP001412.1 | 0,2957 | 1,22484E-09 |
| ACE2 | RPS6P12 | 0,2955 | 1,26728E-09 |
| ACE2 | DCBLD1 | 0,2954 | 1,28777E-09 |
| ACE2 | EDEM2 | 0,2954 | 1,28684E-09 |
| ACE2 | RP11-34P1.1 | 0,2953 | 1,29108E-09 |
| ACE2 | PNOC | 0,2953 | 1,30532E-09 |
| ACE2 | RP11-226L15.1 | 0,2952 | 1,30955E-09 |
| ACE2 | EAF2 | 0,2949 | 1,36112E-09 |
| ACE2 | UGT2B28 | 0,2948 | 1,39611E-09 |
| ACE2 | RP11-403H13.1 | 0,2948 | 1,38446E-09 |
| ACE2 | RP11-211G23.2 | 0,2947 | 1,40449E-09 |
| ACE2 | HIGD1AP1 | 0,2947 | 1,41476E-09 |
| ACE2 | FCRLA | 0,2946 | 1,41887E-09 |
| ACE2 | AP000647.3 | 0,2946 | 1,42394E-09 |
| ACE2 | JAML | 0,2946 | 1,42512E-09 |
| ACE2 | RP11-297C4.2 | 0,2946 | 1,42555E-09 |
| ACE2 | RP11-432I5.4 | 0,2946 | 1,42077E-09 |
| ACE2 | AC005102.1 | 0,2945 | 1,44553E-09 |
| ACE2 | CLN6 | 0,2945 | 1,4501E-09 |
| ACE2 | RP11-165N19.2 | 0,2943 | 1,48705E-09 |
| ACE2 | RP11-66N24.3 | 0,2943 | 1,49218E-09 |
| ACE2 | KB-173C10.2 | 0,294 | 1,53599E-09 |
| ACE2 | TRAV6 | 0,2939 | 1,56673E-09 |
| ACE2 | MYH4 | 0,2939 | 1,56537E-09 |
| ACE2 | TTC30B | 0,2938 | 1,58322E-09 |
| ACE2 | RN7SL445P | 0,2938 | 1,58357E-09 |
| ACE2 | RP11-126O1.5 | 0,2938 | 1,5854E-09 |
| ACE2 | RP5-827C21.2 | 0,2936 | 1,61811E-09 |
| ACE2 | RP11-10O17.1 | 0,2934 | 1,66273E-09 |
| ACE2 | RP11-338L22.2 | 0,2934 | 1,67908E-09 |
| ACE2 | RP11-456N14.6 | 0,2933 | 1,70496E-09 |
| ACE2 | ALG1L3P | 0,2931 | 1,7307E-09 |
| ACE2 | HNRNPA1P70 | 0,2931 | 1,73385E-09 |
| ACE2 | BMP8A | 0,293 | 1,76914E-09 |
| ACE2 | RP11-14N9.1 | 0,293 | 1,77138E-09 |
| ACE2 | RP11-235E17.5 | 0,2927 | 1,8344E-09 |
| ACE2 | CTAGE1 | 0,2927 | 1,84393E-09 |
| ACE2 | CTD-2201E9.2 | 0,2925 | 1,89293E-09 |
| ACE2 | RP1-128M12.3 | 0,2925 | 1,89204E-09 |
| ACE2 | RP11-599B13.8 | 0,2925 | 1,89435E-09 |
| ACE2 | RP11-135A1.3 | 0,2924 | 1,90974E-09 |
| ACE2 | CCDC134 | 0,2921 | 1,98254E-09 |
| ACE2 | LINC01251 | 0,292 | 2,02362E-09 |
| ACE2 | AC104024.1 | 0,292 | 2,02148E-09 |
| ACE2 | LINC00167 | 0,2919 | 2,04904E-09 |
| ACE2 | RP11-380O24.1 | 0,2918 | 2,06061E-09 |
| ACE2 | LINC01378 | 0,2917 | 2,10784E-09 |
| ACE2 | OR5M6P | 0,2917 | 2,1046E-09 |
| ACE2 | FOXI3 | 0,2916 | 2,12532E-09 |
| ACE2 | LINC00866 | 0,2915 | 2,15359E-09 |

| | | | |
|---|---|---|---|
| ACE2 | RP11-547D24.3 | 0,2914 | 2,17815E-09 |
| ACE2 | RP11-36B15.1 | 0,2912 | 2,24016E-09 |
| ACE2 | IGHVIII-25-1 | 0,2912 | 2,24121E-09 |
| ACE2 | RP11-255H23.5 | 0,2912 | 2,23816E-09 |
| ACE2 | AC004012.1 | 0,2911 | 2,27227E-09 |
| ACE2 | KRT89P | 0,291 | 2,29312E-09 |
| ACE2 | DNASE1L1 | 0,2909 | 2,33366E-09 |
| ACE2 | POLRMTP1 | 0,2908 | 2,37363E-09 |
| ACE2 | SPDYC | 0,2907 | 2,39549E-09 |
| ACE2 | NLRC5 | 0,2907 | 2,37551E-09 |
| ACE2 | RP11-203H2.2 | 0,2906 | 2,432E-09 |
| ACE2 | IRS3P | 0,2906 | 2,43685E-09 |
| ACE2 | FERMT3 | 0,2906 | 2,41564E-09 |
| ACE2 | LINC00449 | 0,2905 | 2,44691E-09 |
| ACE2 | HEATR5A | 0,2905 | 2,46473E-09 |
| ACE2 | RCC2P6 | 0,2904 | 2,47551E-09 |
| ACE2 | RP11-229D13.3 | 0,2902 | 2,56454E-09 |
| ACE2 | FASN | 0,2902 | 2,5509E-09 |
| ACE2 | RP11-156K23.3 | 0,2901 | 2,5924E-09 |
| ACE2 | AL161668.5 | 0,29 | 2,60805E-09 |
| ACE2 | AC010086.5 | 0,2899 | 2,66929E-09 |
| ACE2 | CD5 | 0,2898 | 2,70154E-09 |
| ACE2 | RP11-15E18.1 | 0,2897 | 2,72378E-09 |
| ACE2 | RP11-826F13.1 | 0,2895 | 2,78362E-09 |
| ACE2 | RP11-44F14.1 | 0,2895 | 2,78032E-09 |
| ACE2 | CTD-2561J22.2 | 0,2894 | 2,82282E-09 |
| ACE2 | SPTLC3 | 0,2893 | 2,87655E-09 |
| ACE2 | KIAA1958 | 0,2892 | 2,89998E-09 |
| ACE2 | IGHD2-8 | 0,2891 | 2,94219E-09 |
| ACE2 | RP2 | 0,289 | 2,98922E-09 |
| ACE2 | CIRBP-AS1 | 0,2889 | 3,01363E-09 |
| ACE2 | TUBB1 | 0,2889 | 3,01064E-09 |
| ACE2 | RP11-305O6.4 | 0,2887 | 3,11144E-09 |
| ACE2 | KDELR1 | 0,2887 | 3,12228E-09 |
| ACE2 | CCDC17 | 0,2886 | 3,15424E-09 |
| ACE2 | SMG1P6 | 0,2885 | 3,18897E-09 |
| ACE2 | ART1 | 0,2884 | 3,24756E-09 |
| ACE2 | CKS1B | 0,2883 | 3,25252E-09 |
| ACE2 | OVAAL | 0,2881 | 3,33847E-09 |
| ACE2 | CTC-277H1.6 | 0,288 | 3,38668E-09 |
| ACE2 | APOC1P1 | 0,288 | 3,39774E-09 |
| ACE2 | RP11-619A14.3 | 0,2879 | 3,43681E-09 |
| ACE2 | RIN1 | 0,2878 | 3,47772E-09 |
| ACE2 | LINC00221 | 0,2878 | 3,48283E-09 |
| ACE2 | RP11-51F16.9 | 0,2878 | 3,50959E-09 |
| ACE2 | RP5-1031D4.2 | 0,2877 | 3,54095E-09 |
| ACE2 | NPSR1-AS1 | 0,2875 | 3,62018E-09 |
| ACE2 | RP11-295M3.2 | 0,2875 | 3,63756E-09 |
| ACE2 | AC092168.2 | 0,2874 | 3,68102E-09 |
| ACE2 | CXCL13 | 0,2874 | 3,68436E-09 |
| ACE2 | RP11-81H14.1 | 0,2874 | 3,67501E-09 |
| ACE2 | CYP51A1P2 | 0,2874 | 3,66137E-09 |
| ACE2 | RPS15AP30 | 0,2873 | 3,72523E-09 |
| ACE2 | RP11-539L10.4 | 0,2872 | 3,78114E-09 |

| | | | |
|---|---|---|---|
| ACE2 | CXorf65 | 0,2871 | 3,823E-09 |
| ACE2 | RP11-95M15.3 | 0,287 | 3,85858E-09 |
| ACE2 | PIM3 | 0,2869 | 3,93638E-09 |
| ACE2 | RNU6-509P | 0,2868 | 3,95971E-09 |
| ACE2 | CYP2B7P | 0,2868 | 3,96448E-09 |
| ACE2 | RPL36P16 | 0,2868 | 3,97807E-09 |
| ACE2 | LINC01762 | 0,2867 | 4,0539E-09 |
| ACE2 | TTLL10-AS1 | 0,2866 | 4,10223E-09 |
| ACE2 | CTD-2325A15.5 | 0,2866 | 4,07778E-09 |
| ACE2 | LCMT1 | 0,2866 | 4,09478E-09 |
| ACE2 | GP1BA | 0,2865 | 4,13226E-09 |
| ACE2 | DLEU1 | 0,2864 | 4,20593E-09 |
| ACE2 | UTS2R | 0,2862 | 4,28044E-09 |
| ACE2 | TBC1D4 | 0,2861 | 4,34158E-09 |
| ACE2 | RP5-828H9.1 | 0,2861 | 4,3638E-09 |
| ACE2 | TMEM176A | 0,286 | 4,40521E-09 |
| ACE2 | CD27 | 0,286 | 4,39214E-09 |
| ACE2 | RALY | 0,286 | 4,41211E-09 |
| ACE2 | KCNJ18 | 0,2859 | 4,44941E-09 |
| ACE2 | DNAJC5 | 0,2859 | 4,44226E-09 |
| ACE2 | CSTB | 0,2859 | 4,48087E-09 |
| ACE2 | CYCSP52 | 0,2858 | 4,53352E-09 |
| ACE2 | AC016999.2 | 0,2858 | 4,50455E-09 |
| ACE2 | SLC9A8 | 0,2858 | 4,50349E-09 |
| ACE2 | RN7SL186P | 0,2857 | 4,58086E-09 |
| ACE2 | VDAC1P2 | 0,2855 | 4,71665E-09 |
| ACE2 | RSL24D1P1 | 0,2854 | 4,75332E-09 |
| ACE2 | RP11-175P13.2 | 0,2854 | 4,74037E-09 |
| ACE2 | LINC00935 | 0,2853 | 4,85805E-09 |
| ACE2 | ATP9A | 0,2853 | 4,81109E-09 |
| ACE2 | AKNA | 0,2852 | 4,88861E-09 |
| ACE2 | ETFB | 0,2852 | 4,88258E-09 |
| ACE2 | IGLVI-70 | 0,2852 | 4,89631E-09 |
| ACE2 | RARRES1 | 0,2851 | 4,97693E-09 |
| ACE2 | IGHV3-29 | 0,2851 | 4,94549E-09 |
| ACE2 | RP4-669H2.1 | 0,2851 | 4,93275E-09 |
| ACE2 | HCG4P11 | 0,285 | 5,02762E-09 |
| ACE2 | SAA1 | 0,2849 | 5,09736E-09 |
| ACE2 | RP11-747H7.1 | 0,2849 | 5,0596E-09 |
| ACE2 | FASLG | 0,2848 | 5,12376E-09 |
| ACE2 | RP11-497G19.1 | 0,2847 | 5,18602E-09 |
| ACE2 | RNU2-6P | 0,2846 | 5,31215E-09 |
| ACE2 | CMPK1 | 0,2845 | 5,37683E-09 |
| ACE2 | LEKR1 | 0,2845 | 5,35583E-09 |
| ACE2 | TPI1P1 | 0,2844 | 5,42667E-09 |
| ACE2 | TIFAB | 0,2844 | 5,3916E-09 |
| ACE2 | DPRXP2 | 0,2842 | 5,58358E-09 |
| ACE2 | SYAP1 | 0,2842 | 5,59246E-09 |
| ACE2 | PHYH | 0,2841 | 5,6169E-09 |
| ACE2 | RP11-298I3.3 | 0,2841 | 5,66563E-09 |
| ACE2 | RP11-141C7.3 | 0,284 | 5,70064E-09 |
| ACE2 | GHET1 | 0,284 | 5,72437E-09 |
| ACE2 | DAPP1 | 0,2839 | 5,76301E-09 |
| ACE2 | OXCT2 | 0,2838 | 5,85785E-09 |

| | | | |
|---|---|---|---|
| ACE2 | CTD-2410N18.3 | 0,2837 | 5,93093E-09 |
| ACE2 | ABHD12 | 0,2837 | 5,95696E-09 |
| ACE2 | FCRL5 | 0,2836 | 6,03235E-09 |
| ACE2 | MRPL53 | 0,2836 | 5,98657E-09 |
| ACE2 | RAB11A | 0,2836 | 6,00573E-09 |
| ACE2 | CLP1 | 0,2835 | 6,1198E-09 |
| ACE2 | L3MBTL4-AS1 | 0,2835 | 6,08671E-09 |
| ACE2 | PLEKHG4 | 0,2833 | 6,21381E-09 |
| ACE2 | MVD | 0,2833 | 6,25005E-09 |
| ACE2 | CTD-2228K2.7 | 0,2832 | 6,30555E-09 |
| ACE2 | RPL26P27 | 0,2832 | 6,34583E-09 |
| ACE2 | XACT | 0,2832 | 6,30263E-09 |
| ACE2 | RP11-37C7.3 | 0,2831 | 6,4218E-09 |
| ACE2 | RP11-818O24.3 | 0,2831 | 6,44107E-09 |
| ACE2 | RP11-697M17.2 | 0,283 | 6,47194E-09 |
| ACE2 | ASS1P11 | 0,2829 | 6,53451E-09 |
| ACE2 | TRAV9-2 | 0,2829 | 6,6033E-09 |
| ACE2 | RP11-421N8.1 | 0,2828 | 6,64901E-09 |
| ACE2 | RP4-803A2.1 | 0,2826 | 6,83842E-09 |
| ACE2 | CTC-756D1.1 | 0,2826 | 6,80303E-09 |
| ACE2 | EDAR | 0,2825 | 6,9006E-09 |
| ACE2 | VOPP1 | 0,2825 | 6,93731E-09 |
| ACE2 | IGKV2-14 | 0,2823 | 7,10356E-09 |
| ACE2 | CMC1 | 0,2823 | 7,1199E-09 |
| ACE2 | IGHE | 0,2823 | 7,11745E-09 |
| ACE2 | HOMER2P2 | 0,2823 | 7,05375E-09 |
| ACE2 | MT-TV | 0,2823 | 7,08598E-09 |
| ACE2 | AL928768.3 | 0,2822 | 7,16838E-09 |
| ACE2 | RP11-680E19.1 | 0,2821 | 7,27521E-09 |
| ACE2 | AC006380.3 | 0,282 | 7,39985E-09 |
| ACE2 | RP11-555G19.1 | 0,282 | 7,40995E-09 |
| ACE2 | CTC-436P18.5 | 0,2819 | 7,47253E-09 |
| ACE2 | AZGP1 | 0,2819 | 7,49978E-09 |
| ACE2 | POLR3GP1 | 0,2816 | 7,70683E-09 |
| ACE2 | AC002306.1 | 0,2815 | 7,80286E-09 |
| ACE2 | KBTBD8 | 0,2814 | 7,97346E-09 |
| ACE2 | SC5D | 0,2814 | 7,94541E-09 |
| ACE2 | RP11-25H12.1 | 0,2813 | 8,0419E-09 |
| ACE2 | HMMR-AS1 | 0,2813 | 8,05134E-09 |
| ACE2 | CTD-2555A7.3 | 0,2813 | 8,05728E-09 |
| ACE2 | IGHV7-40 | 0,2812 | 8,14399E-09 |
| ACE2 | PPCS | 0,2811 | 8,22131E-09 |
| ACE2 | RP11-227L6.1 | 0,2811 | 8,29926E-09 |
| ACE2 | MTCO3P44 | 0,281 | 8,3488E-09 |
| ACE2 | WARSP1 | 0,2809 | 8,45895E-09 |
| ACE2 | RN7SL351P | 0,2809 | 8,42774E-09 |
| ACE2 | ELK2AP | 0,2809 | 8,43853E-09 |
| ACE2 | RAB5CP2 | 0,2808 | 8,61209E-09 |
| ACE2 | YWHAZP5 | 0,2808 | 8,53884E-09 |
| ACE2 | HOMER2P1 | 0,2808 | 8,5343E-09 |
| ACE2 | RP1-151B14.6 | 0,2808 | 8,59535E-09 |
| ACE2 | MRPL22P1 | 0,2807 | 8,73454E-09 |
| ACE2 | RP11-74D7.3 | 0,2807 | 8,73294E-09 |
| ACE2 | HIST1H2AJ | 0,2806 | 8,827E-09 |

| | | | |
|---|---|---|---|
| ACE2 | RP11-791M20.1 | 0,2805 | 8,90553E-09 |
| ACE2 | CTC-215O4.4 | 0,2803 | 9,19132E-09 |
| ACE2 | RNF103 | 0,2802 | 9,24598E-09 |
| ACE2 | ATP13A5 | 0,2802 | 9,23529E-09 |
| ACE2 | RP11-20B24.2 | 0,2801 | 9,35497E-09 |
| ACE2 | AC027601.1 | 0,2801 | 9,41504E-09 |
| ACE2 | MRPL35 | 0,28 | 9,51557E-09 |
| ACE2 | RP11-435O5.7 | 0,28 | 9,46047E-09 |
| ACE2 | MTRNR2L12 | 0,2799 | 9,64139E-09 |
| ACE2 | RP11-496D24.2 | 0,2799 | 9,661E-09 |
| ACE2 | KRT18P13 | 0,2799 | 9,62294E-09 |
| ACE2 | CTD-2313F11.1 | 0,2797 | 9,82371E-09 |
| ACE2 | WBSCR28 | 0,2795 | 1,01568E-08 |
| ACE2 | THUMPD3P1 | 0,2794 | 1,02262E-08 |
| ACE2 | CCHCR1 | 0,2791 | 1,06887E-08 |
| ACE2 | IGHV3-30-2 | 0,2791 | 1,06338E-08 |
| ACE2 | IGLV3-4 | 0,279 | 1,07965E-08 |
| ACE2 | TPI1P4 | 0,2789 | 1,0959E-08 |
| ACE2 | KRT18P39 | 0,2788 | 1,0995E-08 |
| ACE2 | CTD-2521M24.6 | 0,2788 | 1,0986E-08 |
| ACE2 | RP11-661A12.5 | 0,2787 | 1,12274E-08 |
| ACE2 | PDE9A | 0,2787 | 1,11638E-08 |
| ACE2 | CCDC142 | 0,2785 | 1,14844E-08 |
| ACE2 | AVPI1 | 0,2785 | 1,15077E-08 |
| ACE2 | ZNF725P | 0,2785 | 1,14333E-08 |
| ACE2 | PLAUR | 0,2785 | 1,14399E-08 |
| ACE2 | RN7SL431P | 0,2784 | 1,16516E-08 |
| ACE2 | UMPS | 0,2784 | 1,158E-08 |
| ACE2 | VDAC1P11 | 0,2784 | 1,15529E-08 |
| ACE2 | RP11-152H18.4 | 0,2784 | 1,15979E-08 |
| ACE2 | RP11-755O11.2 | 0,2784 | 1,15467E-08 |
| ACE2 | CA3 | 0,2783 | 1,17863E-08 |
| ACE2 | HMGN1P15 | 0,2782 | 1,19081E-08 |
| ACE2 | MIR621 | 0,2782 | 1,19401E-08 |
| ACE2 | IGHV3-54 | 0,278 | 1,21924E-08 |
| ACE2 | RP6-159A1.3 | 0,2779 | 1,23325E-08 |
| ACE2 | AC073641.2 | 0,2778 | 1,24502E-08 |
| ACE2 | HIST1H2BL | 0,2777 | 1,27377E-08 |
| ACE2 | RP11-279F6.2 | 0,2776 | 1,27648E-08 |
| ACE2 | CTC-499B15.8 | 0,2776 | 1,28938E-08 |
| ACE2 | RAET1L | 0,2774 | 1,32101E-08 |
| ACE2 | GPR157 | 0,2773 | 1,33774E-08 |
| ACE2 | PRO1804 | 0,2773 | 1,3346E-08 |
| ACE2 | RP11-253M7.3 | 0,2772 | 1,3525E-08 |
| ACE2 | CD6 | 0,2771 | 1,37161E-08 |
| ACE2 | RN7SL492P | 0,2769 | 1,39511E-08 |
| ACE2 | YWHAEP7 | 0,2769 | 1,40328E-08 |
| ACE2 | RP11-126K1.2 | 0,2768 | 1,41257E-08 |
| ACE2 | RP11-79N23.1 | 0,2768 | 1,40797E-08 |
| ACE2 | MIR3176 | 0,2768 | 1,41874E-08 |
| ACE2 | NLRP11 | 0,2768 | 1,41623E-08 |
| ACE2 | RP11-30O15.1 | 0,2767 | 1,43777E-08 |
| ACE2 | RP1-287H17.1 | 0,2765 | 1,47254E-08 |
| ACE2 | RNA5SP38 | 0,2765 | 1,46193E-08 |

| | | | |
|---|---|---|---|
| ACE2 | RP5-1086K13.1 | 0,2764 | 1,49469E-08 |
| ACE2 | TMEM132D | 0,2764 | 1,48628E-08 |
| ACE2 | RP11-676J12.4 | 0,2764 | 1,48046E-08 |
| ACE2 | TP53 | 0,2764 | 1,4981E-08 |
| ACE2 | KLK2 | 0,2764 | 1,48993E-08 |
| ACE2 | ARHGAP26 | 0,2763 | 1,5032E-08 |
| ACE2 | RP11-341D18.5 | 0,2763 | 1,50705E-08 |
| ACE2 | POTEC | 0,2762 | 1,53531E-08 |
| ACE2 | CCDC170 | 0,2761 | 1,54827E-08 |
| ACE2 | WDR34 | 0,2761 | 1,53677E-08 |
| ACE2 | PRDX3P2 | 0,276 | 1,5724E-08 |
| ACE2 | TNFRSF21 | 0,276 | 1,56156E-08 |
| ACE2 | NPM1P43 | 0,276 | 1,56292E-08 |
| ACE2 | LINC01055 | 0,2759 | 1,5802E-08 |
| ACE2 | LL22NC03-24A12.8 | 0,2759 | 1,5755E-08 |
| ACE2 | LARP1P1 | 0,2758 | 1,59967E-08 |
| ACE2 | RP11-799B12.4 | 0,2758 | 1,6063E-08 |
| ACE2 | LL22NC03-63E9.3 | 0,2758 | 1,60814E-08 |
| ACE2 | RP11-71G12.1 | 0,2757 | 1,62923E-08 |
| ACE2 | TNFRSF18 | 0,2756 | 1,64222E-08 |
| ACE2 | GRHL3 | 0,2756 | 1,64785E-08 |
| ACE2 | CFC1 | 0,2755 | 1,66409E-08 |
| ACE2 | SAA2 | 0,2755 | 1,67263E-08 |
| ACE2 | TMSB4X | 0,2755 | 1,66935E-08 |
| ACE2 | SNRPFP4 | 0,2754 | 1,68298E-08 |
| ACE2 | CTD-2383M3.1 | 0,2754 | 1,68667E-08 |
| ACE2 | TOM1 | 0,2754 | 1,69607E-08 |
| ACE2 | SLC5A9 | 0,2753 | 1,70634E-08 |
| ACE2 | RP11-11M20.2 | 0,2753 | 1,70537E-08 |
| ACE2 | MRPL41 | 0,2752 | 1,73585E-08 |
| ACE2 | RASSF10 | 0,2752 | 1,71749E-08 |
| ACE2 | GRK2 | 0,2751 | 1,74137E-08 |
| ACE2 | AMPD1 | 0,2749 | 1,79363E-08 |
| ACE2 | RP11-845C23.2 | 0,2749 | 1,78371E-08 |
| ACE2 | TMSB10 | 0,2748 | 1,81979E-08 |
| ACE2 | KLRK1 | 0,2748 | 1,80977E-08 |
| ACE2 | IGHV2OR16-5 | 0,2747 | 1,82869E-08 |
| ACE2 | AC138472.4 | 0,2745 | 1,88E-08 |
| ACE2 | RP1-136B1.1 | 0,2744 | 1,9053E-08 |
| ACE2 | RP11-511H9.3 | 0,2744 | 1,89669E-08 |
| ACE2 | ASB7 | 0,2744 | 1,90507E-08 |
| ACE2 | RN7SL368P | 0,2744 | 1,89725E-08 |
| ACE2 | AC091492.2 | 0,2743 | 1,92032E-08 |
| ACE2 | LINC00861 | 0,2743 | 1,93341E-08 |
| ACE2 | LA16c-313F4.1 | 0,2743 | 1,93745E-08 |
| ACE2 | ARFIP1 | 0,2741 | 1,96839E-08 |
| ACE2 | SSR1P1 | 0,2741 | 1,9821E-08 |
| ACE2 | RP5-1142J19.2 | 0,2741 | 1,97805E-08 |
| ACE2 | CDRT15 | 0,274 | 2,01086E-08 |
| ACE2 | RHOH | 0,2739 | 2,03377E-08 |
| ACE2 | RP11-805F19.4 | 0,2739 | 2,028E-08 |
| ACE2 | FAM83C | 0,2738 | 2,04737E-08 |
| ACE2 | IGBP1-AS1 | 0,2738 | 2,06427E-08 |
| ACE2 | TXNP6 | 0,2737 | 2,07354E-08 |

| | | | |
|---|---|---|---|
| ACE2 | AP000437.3 | 0,2737 | 2,08801E-08 |
| ACE2 | RP11-651L5.2 | 0,2736 | 2,10194E-08 |
| ACE2 | ID2-AS1 | 0,2735 | 2,14256E-08 |
| ACE2 | RP11-63A11.1 | 0,2735 | 2,13996E-08 |
| ACE2 | RP11-372E1.4 | 0,2734 | 2,16934E-08 |
| ACE2 | RP11-182L7.1 | 0,2734 | 2,16269E-08 |
| ACE2 | CNTN3 | 0,2732 | 2,19813E-08 |
| ACE2 | TMOD4 | 0,2729 | 2,28021E-08 |
| ACE2 | IFNGR1 | 0,2729 | 2,29813E-08 |
| ACE2 | CLRN1 | 0,2727 | 2,34384E-08 |
| ACE2 | CTD-2313J17.5 | 0,2727 | 2,33661E-08 |
| ACE2 | TRAV12-2 | 0,2726 | 2,36688E-08 |
| ACE2 | RHCG | 0,2726 | 2,36751E-08 |
| ACE2 | DDIAS | 0,2725 | 2,41561E-08 |
| ACE2 | GNG5 | 0,2724 | 2,44682E-08 |
| ACE2 | RP11-148O21.2 | 0,2724 | 2,44675E-08 |
| ACE2 | RP11-511B23.1 | 0,2724 | 2,42459E-08 |
| ACE2 | LINC01361 | 0,2722 | 2,48534E-08 |
| ACE2 | ATP1A1-AS1 | 0,2722 | 2,50586E-08 |
| ACE2 | RP11-428K3.1 | 0,2722 | 2,502E-08 |
| ACE2 | TGM3 | 0,272 | 2,56746E-08 |
| ACE2 | PSD3 | 0,2717 | 2,65753E-08 |
| ACE2 | DNAH1 | 0,2716 | 2,6874E-08 |
| ACE2 | KDM4C | 0,2716 | 2,69079E-08 |
| ACE2 | HMGB3P6 | 0,2715 | 2,70533E-08 |
| ACE2 | LINC00593 | 0,2715 | 2,72355E-08 |
| ACE2 | RP11-500B12.1 | 0,2714 | 2,75781E-08 |
| ACE2 | OR7E115P | 0,2713 | 2,80065E-08 |
| ACE2 | TRAV3 | 0,2713 | 2,78593E-08 |
| ACE2 | RN7SL589P | 0,2712 | 2,81267E-08 |
| ACE2 | RP11-768B22.2 | 0,2712 | 2,83475E-08 |
| ACE2 | FLJ31104 | 0,2712 | 2,81548E-08 |
| ACE2 | WBSCR27 | 0,2712 | 2,81058E-08 |
| ACE2 | FAM30A | 0,2711 | 2,85756E-08 |
| ACE2 | AC002310.7 | 0,271 | 2,90558E-08 |
| ACE2 | ECH1 | 0,271 | 2,89009E-08 |
| ACE2 | RP4-654C18.1 | 0,2709 | 2,9167E-08 |
| ACE2 | FAM172BP | 0,2707 | 2,98348E-08 |
| ACE2 | RP11-895M11.2 | 0,2707 | 3,01054E-08 |
| ACE2 | DUSP5P2 | 0,2706 | 3,05004E-08 |
| ACE2 | CTD-2560E9.5 | 0,2706 | 3,03969E-08 |
| ACE2 | RP11-302F12.10 | 0,2703 | 3,14569E-08 |
| ACE2 | RP11-351C21.2 | 0,2703 | 3,16002E-08 |
| ACE2 | RP11-455O6.8 | 0,2703 | 3,16002E-08 |
| ACE2 | RP11-713M15.1 | 0,2702 | 3,17502E-08 |
| ACE2 | OR4E1 | 0,2702 | 3,19796E-08 |
| ACE2 | MMEL1 | 0,2701 | 3,24081E-08 |
| ACE2 | RP11-42O15.3 | 0,2701 | 3,21882E-08 |
| ACE2 | EDEM3 | 0,27 | 3,28431E-08 |
| ACE2 | CELA1 | 0,27 | 3,26021E-08 |
| ACE2 | RPL36AP43 | 0,27 | 3,26526E-08 |
| ACE2 | AGGF1P3 | 0,2699 | 3,30731E-08 |
| ACE2 | PTP4A2P1 | 0,2699 | 3,32278E-08 |
| ACE2 | ZNF587B | 0,2699 | 3,31177E-08 |

| | | | |
|---|---|---|---|
| ACE2 | DEF6 | 0,2698 | 3,35672E-08 |
| ACE2 | CTC-244M17.1 | 0,2697 | 3,37544E-08 |
| ACE2 | AC010761.14 | 0,2696 | 3,41485E-08 |
| ACE2 | MIR570 | 0,2695 | 3,46151E-08 |
| ACE2 | OR2AM1P | 0,2695 | 3,4535E-08 |
| ACE2 | SHCBP1 | 0,2694 | 3,49926E-08 |
| ACE2 | PPIAP11 | 0,2693 | 3,53407E-08 |
| ACE2 | KRT8P40 | 0,2692 | 3,58038E-08 |
| ACE2 | RP11-806L2.5 | 0,2692 | 3,60724E-08 |
| ACE2 | SMPD5 | 0,2691 | 3,62555E-08 |
| ACE2 | CKAP2 | 0,2691 | 3,64845E-08 |
| ACE2 | GAPDHP49 | 0,2688 | 3,77002E-08 |
| ACE2 | FAM86LP | 0,2688 | 3,78275E-08 |
| ACE2 | AC005794.1 | 0,2687 | 3,83145E-08 |
| ACE2 | TMEM63A | 0,2686 | 3,87265E-08 |
| ACE2 | FAM92B | 0,2686 | 3,86496E-08 |
| ACE2 | RP11-700H6.2 | 0,2686 | 3,86623E-08 |
| ACE2 | MIDN | 0,2686 | 3,86023E-08 |
| ACE2 | OCEL1 | 0,2686 | 3,86744E-08 |
| ACE2 | C22orf42 | 0,2686 | 3,88784E-08 |
| ACE2 | RP11-573D15.9 | 0,2685 | 3,90922E-08 |
| ACE2 | RP11-420K14.1 | 0,2685 | 3,89932E-08 |
| ACE2 | WNT4 | 0,2684 | 3,97208E-08 |
| ACE2 | RP11-903H12.3 | 0,2684 | 3,97254E-08 |
| ACE2 | RP11-705O1.8 | 0,2684 | 3,95249E-08 |
| ACE2 | TBC1D32 | 0,2683 | 4,00349E-08 |
| ACE2 | CDC42EP3P1 | 0,2683 | 4,02924E-08 |
| ACE2 | FAM25C | 0,2682 | 4,07447E-08 |
| ACE2 | AC011330.13 | 0,2681 | 4,08424E-08 |
| ACE2 | SORD2P | 0,268 | 4,15608E-08 |
| ACE2 | TNFRSF13B | 0,268 | 4,14422E-08 |
| ACE2 | RP11-466A19.3 | 0,268 | 4,15939E-08 |
| ACE2 | ANXA2P2 | 0,2679 | 4,2314E-08 |
| ACE2 | CTA-390C10.9 | 0,2679 | 4,22908E-08 |
| ACE2 | RN7SL559P | 0,2678 | 4,26835E-08 |
| ACE2 | CCNG2 | 0,2678 | 4,26462E-08 |
| ACE2 | RP11-290F5.1 | 0,2678 | 4,25037E-08 |
| ACE2 | RP11-195C7.1 | 0,2677 | 4,29476E-08 |
| ACE2 | KLRD1 | 0,2677 | 4,30092E-08 |
| ACE2 | LYPLA2P1 | 0,2676 | 4,35054E-08 |
| ACE2 | RP11-611E13.3 | 0,2676 | 4,36836E-08 |
| ACE2 | SPIC | 0,2676 | 4,35219E-08 |
| ACE2 | STRCP1 | 0,2676 | 4,35848E-08 |
| ACE2 | GUCY2D | 0,2676 | 4,36298E-08 |
| ACE2 | RP11-227D13.4 | 0,2675 | 4,40749E-08 |
| ACE2 | RP11-46H11.11 | 0,2673 | 4,54579E-08 |
| ACE2 | RP11-466M21.1 | 0,2672 | 4,55638E-08 |
| ACE2 | IGLV3-24 | 0,2672 | 4,58441E-08 |
| ACE2 | LINC01362 | 0,2671 | 4,61243E-08 |
| ACE2 | FCRL2 | 0,2669 | 4,7296E-08 |
| ACE2 | FETUB | 0,2669 | 4,74684E-08 |
| ACE2 | OR8D4 | 0,2668 | 4,77982E-08 |
| ACE2 | LINC01269 | 0,2668 | 4,80445E-08 |
| ACE2 | LINC01570 | 0,2668 | 4,77343E-08 |

| | | | |
|---|---|---|---|
| ACE2 | MTND4P12 | 0,2667 | 4,85433E-08 |
| ACE2 | APOC3 | 0,2667 | 4,88303E-08 |
| ACE2 | PYROXD1 | 0,2667 | 4,87213E-08 |
| ACE2 | RP11-367G18.1 | 0,2666 | 4,93947E-08 |
| ACE2 | RP11-356K23.1 | 0,2666 | 4,92961E-08 |
| ACE2 | RP11-299G20.5 | 0,2666 | 4,89969E-08 |
| ACE2 | RP11-220C2.1 | 0,2666 | 4,91014E-08 |
| ACE2 | RP11-557H15.5 | 0,2665 | 4,96544E-08 |
| ACE2 | RPL9P2 | 0,2665 | 4,98063E-08 |
| ACE2 | OFD1P16Y | 0,2665 | 5,00448E-08 |
| ACE2 | CTC-228N24.2 | 0,2664 | 5,02929E-08 |
| ACE2 | RP11-430B1.1 | 0,2664 | 5,01283E-08 |
| ACE2 | ME2 | 0,2664 | 5,03674E-08 |
| ACE2 | CENPX | 0,2663 | 5,11766E-08 |
| ACE2 | RNU6-704P | 0,2661 | 5,19346E-08 |
| ACE2 | RP3-405J10.3 | 0,2661 | 5,22116E-08 |
| ACE2 | IGHG2 | 0,2661 | 5,24771E-08 |
| ACE2 | CCDC152 | 0,266 | 5,29186E-08 |
| ACE2 | STK31 | 0,266 | 5,2748E-08 |
| ACE2 | LINC01671 | 0,266 | 5,2812E-08 |
| ACE2 | GYG2 | 0,266 | 5,25609E-08 |
| ACE2 | KRT8P1 | 0,2659 | 5,34163E-08 |
| ACE2 | GPBP1L1 | 0,2658 | 5,41933E-08 |
| ACE2 | IGHVIII-5-2 | 0,2658 | 5,41257E-08 |
| ACE2 | LINC01363 | 0,2657 | 5,48333E-08 |
| ACE2 | NCBP2-AS1 | 0,2657 | 5,4503E-08 |
| ACE2 | SPON1 | 0,2657 | 5,48986E-08 |
| ACE2 | RP3-522D1.1 | 0,2656 | 5,52042E-08 |
| ACE2 | SPC24 | 0,2655 | 5,59287E-08 |
| ACE2 | RP11-80H18.3 | 0,2654 | 5,65023E-08 |
| ACE2 | RP11-432I13.5 | 0,2653 | 5,74969E-08 |
| ACE2 | RP5-1092A3.5 | 0,2652 | 5,79521E-08 |
| ACE2 | EEF1E1P1 | 0,2652 | 5,79749E-08 |
| ACE2 | RP11-665C16.5 | 0,265 | 5,96117E-08 |
| ACE2 | RP11-470C13.4 | 0,265 | 5,98151E-08 |
| ACE2 | PARP4P2 | 0,2648 | 6,11026E-08 |
| ACE2 | RP13-766D20.2 | 0,2648 | 6,11861E-08 |
| ACE2 | TMEM265 | 0,2647 | 6,17987E-08 |
| ACE2 | ATP6V1A | 0,2646 | 6,20752E-08 |
| ACE2 | CTD-2233K9.1 | 0,2646 | 6,24307E-08 |
| ACE2 | SULT2A1 | 0,2646 | 6,21303E-08 |
| ACE2 | TDGF1 | 0,2643 | 6,47583E-08 |
| ACE2 | NOXRED1 | 0,2642 | 6,54916E-08 |
| ACE2 | S1PR4 | 0,2642 | 6,51437E-08 |
| ACE2 | RP11-278H7.4 | 0,264 | 6,67575E-08 |
| ACE2 | AC068134.6 | 0,264 | 6,71295E-08 |
| ACE2 | DANCR | 0,264 | 6,70225E-08 |
| ACE2 | PHACTR4 | 0,2636 | 7,03183E-08 |
| ACE2 | IGKV2-4 | 0,2636 | 7,02104E-08 |
| ACE2 | AC053503.4 | 0,2636 | 7,04521E-08 |
| ACE2 | RP11-473O4.4 | 0,2636 | 7,03792E-08 |
| ACE2 | RP11-168K11.2 | 0,2635 | 7,11505E-08 |
| ACE2 | RP11-807H22.7 | 0,2635 | 7,11923E-08 |
| ACE2 | COLCA1 | 0,2635 | 7,14093E-08 |

| | | | |
|---|---|---|---|
| ACE2 | LAX1 | 0,2634 | 7,20119E-08 |
| ACE2 | RNU6-230P | 0,2634 | 7,19924E-08 |
| ACE2 | LINC02153 | 0,2634 | 7,15806E-08 |
| ACE2 | MDK | 0,2634 | 7,18895E-08 |
| ACE2 | TRPV3 | 0,2634 | 7,14754E-08 |
| ACE2 | TGM4 | 0,2633 | 7,2373E-08 |
| ACE2 | PTGER4 | 0,2631 | 7,43859E-08 |
| ACE2 | LINC01435 | 0,2631 | 7,42373E-08 |
| ACE2 | LINC01942 | 0,263 | 7,4977E-08 |
| ACE2 | AKR1D1 | 0,263 | 7,53685E-08 |
| ACE2 | RP11-813F20.4 | 0,263 | 7,57626E-08 |
| ACE2 | TRUB2 | 0,2629 | 7,65265E-08 |
| ACE2 | UBASH3A | 0,2629 | 7,61935E-08 |
| ACE2 | RP5-1073O3.2 | 0,2628 | 7,72246E-08 |
| ACE2 | PIM2 | 0,2628 | 7,74298E-08 |
| ACE2 | RP11-478C1.7 | 0,2627 | 7,83608E-08 |
| ACE2 | RP11-855O10.3 | 0,2626 | 7,86027E-08 |
| ACE2 | RP11-684B2.3 | 0,2625 | 7,96587E-08 |
| ACE2 | RP11-467L13.7 | 0,2625 | 7,97866E-08 |
| ACE2 | KIF20B | 0,2624 | 8,1232E-08 |
| ACE2 | CTD-2063L20.1 | 0,2624 | 8,10834E-08 |
| ACE2 | PLUT | 0,2624 | 8,09534E-08 |
| ACE2 | RP4-550H1.5 | 0,2624 | 8,04838E-08 |
| ACE2 | C1orf147 | 0,2623 | 8,21814E-08 |
| ACE2 | LINC01293 | 0,2623 | 8,17156E-08 |
| ACE2 | MS4A1 | 0,2623 | 8,16647E-08 |
| ACE2 | GS1-594A7.3 | 0,2623 | 8,13688E-08 |
| ACE2 | CCR4 | 0,2622 | 8,26306E-08 |
| ACE2 | NOC2LP1 | 0,2621 | 8,38671E-08 |
| ACE2 | RP11-84D1.1 | 0,262 | 8,51257E-08 |
| ACE2 | REG3A | 0,262 | 8,51599E-08 |
| ACE2 | RP11-435F17.3 | 0,262 | 8,45669E-08 |
| ACE2 | RP11-148O21.4 | 0,262 | 8,46845E-08 |
| ACE2 | FKBP11 | 0,262 | 8,47043E-08 |
| ACE2 | ELOCP5 | 0,262 | 8,47941E-08 |
| ACE2 | RP11-282K24.3 | 0,2618 | 8,72306E-08 |
| ACE2 | TRAV29DV5 | 0,2618 | 8,68191E-08 |
| ACE2 | LLNLR-276H7.1 | 0,2617 | 8,77359E-08 |
| ACE2 | SNRPGP5 | 0,2616 | 8,86724E-08 |
| ACE2 | RP11-455F5.6 | 0,2616 | 8,92163E-08 |
| ACE2 | RP1-37N7.1 | 0,2616 | 8,83757E-08 |
| ACE2 | RP11-2B6.2 | 0,2615 | 9,02065E-08 |
| ACE2 | TNFSF11 | 0,2615 | 8,95555E-08 |
| ACE2 | AC104984.4 | 0,2615 | 8,93935E-08 |
| ACE2 | SNORA35 | 0,2614 | 9,08394E-08 |
| ACE2 | AC000111.3 | 0,2613 | 9,1542E-08 |
| ACE2 | RP11-344H11.5 | 0,2611 | 9,39955E-08 |
| ACE2 | MIR6740 | 0,2611 | 9,45671E-08 |
| ACE2 | CHRNA9 | 0,2611 | 9,46629E-08 |
| ACE2 | CD180 | 0,261 | 9,55427E-08 |
| ACE2 | OR52B2 | 0,2609 | 9,64397E-08 |
| ACE2 | CARMIL3 | 0,2609 | 9,60061E-08 |
| ACE2 | RP11-540B6.2 | 0,2609 | 9,59632E-08 |
| ACE2 | PGK1 | 0,2609 | 9,65916E-08 |

| | | | |
|---|---|---|---|
| ACE2 | FDXR | 0,2608 | 9,76333E-08 |
| ACE2 | RP11-597D13.7 | 0,2607 | 9,81128E-08 |
| ACE2 | RP11-1101H11.1 | 0,2607 | 9,90146E-08 |
| ACE2 | HNRNPA1P49 | 0,2607 | 9,8919E-08 |
| ACE2 | RP11-372B4.3 | 0,2606 | 9,96931E-08 |
| ACE2 | ZNF600 | 0,2606 | 9,9279E-08 |
| ACE2 | HIST1H3B | 0,2605 | 1,00628E-07 |
| ACE2 | RP11-401F2.3 | 0,2603 | 1,03329E-07 |
| ACE2 | HYAL4 | 0,2602 | 1,04641E-07 |
| ACE2 | CTD-3222D19.12 | 0,2602 | 1,04978E-07 |
| ACE2 | SNX25P1 | 0,2601 | 1,06377E-07 |
| ACE2 | IGKV2-10 | 0,26 | 1,06592E-07 |
| ACE2 | NRG1 | 0,26 | 1,06474E-07 |
| ACE2 | OSGIN1 | 0,26 | 1,06672E-07 |
| ACE2 | NWD1 | 0,26 | 1,07391E-07 |
| ACE2 | TFB2M | 0,2599 | 1,07675E-07 |
| ACE2 | FAM184B | 0,2599 | 1,0799E-07 |
| ACE2 | HOXA13 | 0,2599 | 1,07795E-07 |
| ACE2 | NPIPB7 | 0,2599 | 1,08322E-07 |
| ACE2 | TMEM97 | 0,2597 | 1,10729E-07 |
| ACE2 | LIF | 0,2597 | 1,10796E-07 |
| ACE2 | TRBV27 | 0,2595 | 1,12926E-07 |
| ACE2 | RP11-408H20.3 | 0,2594 | 1,14443E-07 |
| ACE2 | RP11-471G13.5 | 0,2592 | 1,17561E-07 |
| ACE2 | RP11-61D1.3 | 0,2591 | 1,18432E-07 |
| ACE2 | IDH3A | 0,259 | 1,19688E-07 |
| ACE2 | GOLGA2P10 | 0,259 | 1,19989E-07 |
| ACE2 | CTB-92J24.2 | 0,2589 | 1,22088E-07 |
| ACE2 | RP11-706D8.3 | 0,2588 | 1,23661E-07 |
| ACE2 | RP11-932O9.10 | 0,2588 | 1,22895E-07 |
| ACE2 | RP13-991F5.2 | 0,2588 | 1,22669E-07 |
| ACE2 | PDE11A | 0,2587 | 1,24148E-07 |
| ACE2 | WDFY1 | 0,2587 | 1,24334E-07 |
| ACE2 | RP11-485M7.1 | 0,2587 | 1,24758E-07 |
| ACE2 | SNORA71B | 0,2587 | 1,24605E-07 |
| ACE2 | RP11-74C1.2 | 0,2586 | 1,26314E-07 |
| ACE2 | RP11-57A19.2 | 0,2586 | 1,26E-07 |
| ACE2 | RP11-403P17.2 | 0,2586 | 1,25624E-07 |
| ACE2 | LINC01230 | 0,2585 | 1,27059E-07 |
| ACE2 | DDX28 | 0,2585 | 1,28082E-07 |
| ACE2 | RP11-178L8.6 | 0,2585 | 1,27231E-07 |
| ACE2 | RP11-326K13.5 | 0,2585 | 1,2785E-07 |
| ACE2 | ETV7 | 0,2584 | 1,29572E-07 |
| ACE2 | RP11-343C2.10 | 0,2583 | 1,30483E-07 |
| ACE2 | TBC1D22A | 0,2583 | 1,29753E-07 |
| ACE2 | PPA2 | 0,2582 | 1,3252E-07 |
| ACE2 | AGGF1P1 | 0,2582 | 1,32342E-07 |
| ACE2 | SNAI3 | 0,2582 | 1,31936E-07 |
| ACE2 | COX6B1P2 | 0,2581 | 1,3362E-07 |
| ACE2 | CACNA1A | 0,2581 | 1,33735E-07 |
| ACE2 | GLIS3-AS1 | 0,258 | 1,35016E-07 |
| ACE2 | RP4-569M23.4 | 0,258 | 1,35109E-07 |
| ACE2 | SLX1B | 0,2578 | 1,37292E-07 |
| ACE2 | RP11-568K15.2 | 0,2577 | 1,40412E-07 |

| | | | |
|---|---|---|---|
| ACE2 | SNRPCP5 | 0,2577 | 1,40005E-07 |
| ACE2 | RP11-295B17.6 | 0,2577 | 1,39829E-07 |
| ACE2 | LINC01484 | 0,2576 | 1,41734E-07 |
| ACE2 | IGHVII-60-1 | 0,2576 | 1,41453E-07 |
| ACE2 | RP11-473M20.16 | 0,2575 | 1,43171E-07 |
| ACE2 | NCSTN | 0,2574 | 1,44084E-07 |
| ACE2 | ECI1 | 0,2574 | 1,43912E-07 |
| ACE2 | C1orf159 | 0,2573 | 1,46018E-07 |
| ACE2 | MVK | 0,2573 | 1,46965E-07 |
| ACE2 | RP11-16D22.2 | 0,2573 | 1,46744E-07 |
| ACE2 | KRT18P8 | 0,2573 | 1,46254E-07 |
| ACE2 | RPL17P44 | 0,2573 | 1,45827E-07 |
| ACE2 | NPPA-AS1 | 0,2571 | 1,49217E-07 |
| ACE2 | TRAV26-1 | 0,2571 | 1,49532E-07 |
| ACE2 | RAC2 | 0,2571 | 1,50289E-07 |
| ACE2 | MT-TG | 0,2571 | 1,50381E-07 |
| ACE2 | RP11-225N10.1 | 0,257 | 1,52087E-07 |
| ACE2 | LINC00556 | 0,2569 | 1,52472E-07 |
| ACE2 | CNOT1 | 0,2569 | 1,52381E-07 |
| ACE2 | RNU6-570P | 0,2568 | 1,55485E-07 |
| ACE2 | DSG1-AS1 | 0,2568 | 1,55335E-07 |
| ACE2 | COX7B | 0,2568 | 1,54113E-07 |
| ACE2 | RP11-557J10.4 | 0,2567 | 1,56351E-07 |
| ACE2 | OR10D1P | 0,2567 | 1,56531E-07 |
| ACE2 | TRAV13-1 | 0,2567 | 1,56995E-07 |
| ACE2 | MTND6P3 | 0,2566 | 1,57767E-07 |
| ACE2 | CLTC | 0,2566 | 1,59289E-07 |
| ACE2 | CTA-714B7.7 | 0,2566 | 1,58729E-07 |
| ACE2 | IGHJ2P | 0,2565 | 1,60554E-07 |
| ACE2 | CDKN1A | 0,2564 | 1,61538E-07 |
| ACE2 | RP11-248G5.9 | 0,2564 | 1,62935E-07 |
| ACE2 | TMEM211 | 0,2564 | 1,62627E-07 |
| ACE2 | CYP4F62P | 0,2563 | 1,63932E-07 |
| ACE2 | PNPLA2 | 0,2563 | 1,64583E-07 |
| ACE2 | TCL6 | 0,2563 | 1,63625E-07 |
| ACE2 | RP11-388C12.5 | 0,2563 | 1,63917E-07 |
| ACE2 | KCNS2 | 0,2562 | 1,66411E-07 |
| ACE2 | FAM49B | 0,2561 | 1,67893E-07 |
| ACE2 | MIR3180-1 | 0,2561 | 1,6758E-07 |
| ACE2 | CELSR3-AS1 | 0,256 | 1,68969E-07 |
| ACE2 | CTD-2267D19.3 | 0,256 | 1,69611E-07 |
| ACE2 | IGLC4 | 0,256 | 1,70587E-07 |
| ACE2 | RP11-96K19.2 | 0,2559 | 1,72274E-07 |
| ACE2 | RP11-380N8.7 | 0,2559 | 1,72291E-07 |
| ACE2 | LY9 | 0,2558 | 1,74589E-07 |
| ACE2 | TLR10 | 0,2557 | 1,75712E-07 |
| ACE2 | NBPF5P | 0,2556 | 1,78315E-07 |
| ACE2 | UPK3BP1 | 0,2554 | 1,81651E-07 |
| ACE2 | RP11-1I2.1 | 0,2554 | 1,81917E-07 |
| ACE2 | FTH1 | 0,2553 | 1,83397E-07 |
| ACE2 | RP11-703I16.1 | 0,2552 | 1,85043E-07 |
| ACE2 | RP11-536C5.2 | 0,2551 | 1,88879E-07 |
| ACE2 | SLC12A7 | 0,2551 | 1,88157E-07 |
| ACE2 | LINC00582 | 0,255 | 1,89363E-07 |

| | | | |
|---|---|---|---|
| ACE2 | LINC01120 | 0,255 | 1,89251E-07 |
| ACE2 | PCNX3 | 0,255 | 1,90616E-07 |
| ACE2 | BSND | 0,2549 | 1,92624E-07 |
| ACE2 | RP11-6D1.3 | 0,2549 | 1,91828E-07 |
| ACE2 | VAV1 | 0,2548 | 1,9413E-07 |
| ACE2 | MT-TS2 | 0,2548 | 1,93712E-07 |
| ACE2 | C1orf141 | 0,2547 | 1,97617E-07 |
| ACE2 | RP11-93H24.3 | 0,2547 | 1,96139E-07 |
| ACE2 | RP11-510M2.6 | 0,2546 | 1,98325E-07 |
| ACE2 | RP1-161P9.5 | 0,2546 | 1,98198E-07 |
| ACE2 | RP11-645C24.4 | 0,2545 | 2,00577E-07 |
| ACE2 | DSCR3 | 0,2545 | 2,02116E-07 |
| ACE2 | LL22NC03-102D1.18 | 0,2545 | 2,01367E-07 |
| ACE2 | AC013472.3 | 0,2544 | 2,04745E-07 |
| ACE2 | RP11-259K5.2 | 0,2543 | 2,06923E-07 |
| ACE2 | SEMA6A-AS2 | 0,2543 | 2,04946E-07 |
| ACE2 | FAM8A6P | 0,2543 | 2,05096E-07 |
| ACE2 | IGHG1 | 0,2543 | 2,07161E-07 |
| ACE2 | IGHV1OR15-4 | 0,2543 | 2,06755E-07 |
| ACE2 | RP11-399F4.2 | 0,2542 | 2,08607E-07 |
| ACE2 | RP11-354B3.1 | 0,2542 | 2,08406E-07 |
| ACE2 | AP000230.1 | 0,2542 | 2,09431E-07 |
| ACE2 | BEND3P1 | 0,2541 | 2,11051E-07 |
| ACE2 | IGHG3 | 0,2541 | 2,10581E-07 |
| ACE2 | STK16 | 0,254 | 2,1319E-07 |
| ACE2 | KIAA0556 | 0,254 | 2,1356E-07 |
| ACE2 | TBC1D10C | 0,2539 | 2,14535E-07 |
| ACE2 | RP11-491F9.8 | 0,2539 | 2,14693E-07 |
| ACE2 | RP11-114J13.1 | 0,2538 | 2,17182E-07 |
| ACE2 | MSH5 | 0,2538 | 2,18471E-07 |
| ACE2 | TRGC2 | 0,2537 | 2,19748E-07 |
| ACE2 | ZDHHC5 | 0,2537 | 2,21484E-07 |
| ACE2 | RP11-291B21.2 | 0,2537 | 2,20803E-07 |
| ACE2 | FAM169A | 0,2536 | 2,22093E-07 |
| ACE2 | MGAT3 | 0,2536 | 2,22102E-07 |
| ACE2 | RP11-745A24.1 | 0,2535 | 2,25985E-07 |
| ACE2 | AP000253.1 | 0,2535 | 2,26215E-07 |
| ACE2 | ILVBL | 0,2534 | 2,27993E-07 |
| ACE2 | TRAV12-1 | 0,2533 | 2,30616E-07 |
| ACE2 | ACTN4P1 | 0,2532 | 2,32458E-07 |
| ACE2 | RP11-64C12.7 | 0,2531 | 2,35966E-07 |
| ACE2 | KB-1615E4.2 | 0,253 | 2,38446E-07 |
| ACE2 | RN7SKP249 | 0,2529 | 2,40196E-07 |
| ACE2 | RP11-425M5.7 | 0,2529 | 2,41175E-07 |
| ACE2 | TSPAN17 | 0,2528 | 2,4369E-07 |
| ACE2 | TRBV20-1 | 0,2528 | 2,45168E-07 |
| ACE2 | CH507-9B2.3 | 0,2527 | 2,4804E-07 |
| ACE2 | RP11-342H21.2 | 0,2526 | 2,50833E-07 |
| ACE2 | ARHGEF2 | 0,2526 | 2,49247E-07 |
| ACE2 | PCNAP4 | 0,2526 | 2,5103E-07 |
| ACE2 | MARCKSL1P1 | 0,2524 | 2,55948E-07 |
| ACE2 | CTD-2516F10.2 | 0,2524 | 2,55364E-07 |
| ACE2 | RASD1 | 0,2524 | 2,56583E-07 |
| ACE2 | GRK1 | 0,2523 | 2,58724E-07 |

| | | | |
|---|---|---|---|
| ACE2 | CLDN17 | 0,2522 | 2,62575E-07 |
| ACE2 | ELOCP7 | 0,2522 | 2,61113E-07 |
| ACE2 | RPS6P16 | 0,2521 | 2,63432E-07 |
| ACE2 | TRGJP2 | 0,252 | 2,67024E-07 |
| ACE2 | RP11-731F5.2 | 0,2519 | 2,71248E-07 |
| ACE2 | THEM5 | 0,2517 | 2,76598E-07 |
| ACE2 | RPL21P110 | 0,2517 | 2,77456E-07 |
| ACE2 | AC005037.3 | 0,2515 | 2,81346E-07 |
| ACE2 | VPREB3 | 0,2515 | 2,81333E-07 |
| ACE2 | GPR89P | 0,2514 | 2,84992E-07 |
| ACE2 | CD68 | 0,2514 | 2,86242E-07 |
| ACE2 | RP11-560I19.2 | 0,2512 | 2,90787E-07 |
| ACE2 | VTRNA1-2 | 0,2511 | 2,96405E-07 |
| ACE2 | RNF32 | 0,2509 | 3,01074E-07 |
| ACE2 | RP11-442J21.2 | 0,2509 | 3,03531E-07 |
| ACE2 | NANOGNBP3 | 0,2509 | 3,03681E-07 |
| ACE2 | RP11-27M9.1 | 0,2508 | 3,07046E-07 |
| ACE2 | RP11-82O19.1 | 0,2507 | 3,09549E-07 |
| ACE2 | RN7SL385P | 0,2507 | 3,09347E-07 |
| ACE2 | GRXCR2 | 0,2506 | 3,1163E-07 |
| ACE2 | HNRNPA1P9 | 0,2506 | 3,14093E-07 |
| ACE2 | RP4-550H1.4 | 0,2506 | 3,1126E-07 |
| ACE2 | LINC01709 | 0,2505 | 3,14342E-07 |
| ACE2 | RCC2 | 0,2504 | 3,18504E-07 |
| ACE2 | PRSS48 | 0,2503 | 3,23555E-07 |
| ACE2 | YWHAZ | 0,2501 | 3,29797E-07 |
| ACE2 | RP11-667M19.2 | 0,2501 | 3,29699E-07 |
| ACE2 | GTF2B | -0,2501 | 3,32212E-07 |
| ACE2 | RN7SKP275 | -0,2501 | 3,28966E-07 |
| ACE2 | RP11-12D24.10 | -0,2501 | 3,29468E-07 |
| ACE2 | RP11-253E3.3 | -0,2501 | 3,31157E-07 |
| ACE2 | COX7A2P1 | -0,2501 | 3,28763E-07 |
| ACE2 | RP11-24M17.4 | -0,2501 | 3,2988E-07 |
| ACE2 | AL022341.3 | -0,2501 | 3,28741E-07 |
| ACE2 | RP11-454P7.3 | -0,2501 | 3,28779E-07 |
| ACE2 | ZNF845 | -0,2501 | 3,31337E-07 |
| ACE2 | RP11-5P18.10 | -0,2502 | 3,27124E-07 |
| ACE2 | WDR3 | -0,2502 | 3,25428E-07 |
| ACE2 | CHST2 | -0,2502 | 3,27149E-07 |
| ACE2 | CUZD1 | -0,2502 | 3,25147E-07 |
| ACE2 | ZNF554 | -0,2502 | 3,26909E-07 |
| ACE2 | ZBBX | -0,2503 | 3,24516E-07 |
| ACE2 | RPS27AP11 | -0,2503 | 3,23784E-07 |
| ACE2 | UTY | -0,2503 | 3,24144E-07 |
| ACE2 | RP11-140K17.3 | -0,2504 | 3,19434E-07 |
| ACE2 | RPL3P7 | -0,2504 | 3,19481E-07 |
| ACE2 | PHBP20 | -0,2504 | 3,20185E-07 |
| ACE2 | CTB-61M7.2 | -0,2504 | 3,20916E-07 |
| ACE2 | CTA-992D9.11 | -0,2504 | 3,1926E-07 |
| ACE2 | RASGRP3 | -0,2505 | 3,17398E-07 |
| ACE2 | SNX18P7 | -0,2505 | 3,17621E-07 |
| ACE2 | LINC02175 | -0,2505 | 3,14451E-07 |
| ACE2 | CNOT8 | -0,2506 | 3,1237E-07 |
| ACE2 | AC068831.6 | -0,2506 | 3,11653E-07 |

| | | | |
|---|---|---|---|
| ACE2 | RP5-1024G6.7 | -0,2507 | 3,08743E-07 |
| ACE2 | RP11-356N1.2 | -0,2507 | 3,07903E-07 |
| ACE2 | RBM45 | -0,2507 | 3,08621E-07 |
| ACE2 | HIST1H2BJ | -0,2507 | 3,09721E-07 |
| ACE2 | RP11-996F15.2 | -0,2507 | 3,10211E-07 |
| ACE2 | ZNF814 | -0,2507 | 3,09494E-07 |
| ACE2 | CAPN10-AS1 | -0,2508 | 3,06801E-07 |
| ACE2 | LYSMD3 | -0,2508 | 3,06581E-07 |
| ACE2 | RALGAPA1P1 | -0,2508 | 3,05936E-07 |
| ACE2 | CCT8 | -0,2508 | 3,06731E-07 |
| ACE2 | CHCHD2P2 | -0,2509 | 3,02219E-07 |
| ACE2 | PLOD3 | -0,2509 | 3,02703E-07 |
| ACE2 | TEX15 | -0,2509 | 3,01323E-07 |
| ACE2 | FAM209B | -0,2509 | 3,02637E-07 |
| ACE2 | PPIH | -0,251 | 2,98949E-07 |
| ACE2 | AC005306.3 | -0,251 | 2,98312E-07 |
| ACE2 | THOP1 | -0,251 | 2,97672E-07 |
| ACE2 | XX-FW83128A1.2 | -0,251 | 2,98182E-07 |
| ACE2 | RP11-731K22.1 | -0,2511 | 2,96457E-07 |
| ACE2 | RP11-80H5.9 | -0,2511 | 2,94324E-07 |
| ACE2 | WASHC1 | -0,2512 | 2,92096E-07 |
| ACE2 | RP4-816N1.6 | -0,2512 | 2,90996E-07 |
| ACE2 | LINC02058 | -0,2513 | 2,88115E-07 |
| ACE2 | C17orf58 | -0,2513 | 2,88615E-07 |
| ACE2 | LINC02066 | -0,2514 | 2,86227E-07 |
| ACE2 | HCG9P5 | -0,2514 | 2,86137E-07 |
| ACE2 | RP11-90J7.2 | -0,2514 | 2,86376E-07 |
| ACE2 | MMADHC | -0,2515 | 2,82093E-07 |
| ACE2 | PTPN20 | -0,2515 | 2,81211E-07 |
| ACE2 | OR52V1P | -0,2515 | 2,84126E-07 |
| ACE2 | BRCA1 | -0,2515 | 2,81911E-07 |
| ACE2 | RP11-35G9.3 | -0,2515 | 2,82009E-07 |
| ACE2 | KB-1836B5.1 | -0,2516 | 2,79278E-07 |
| ACE2 | VPS50 | -0,2517 | 2,77047E-07 |
| ACE2 | AQP10 | -0,2518 | 2,72418E-07 |
| ACE2 | RP11-214K3.24 | -0,2518 | 2,74025E-07 |
| ACE2 | RP11-196B3.3 | -0,2518 | 2,74446E-07 |
| ACE2 | NCOA7-AS1 | -0,2519 | 2,68926E-07 |
| ACE2 | NPIPA8 | -0,2519 | 2,70708E-07 |
| ACE2 | RN7SL477P | -0,2519 | 2,69413E-07 |
| ACE2 | RP11-66N11.6 | -0,2519 | 2,71274E-07 |
| ACE2 | HCG2040054 | -0,252 | 2,67478E-07 |
| ACE2 | AC092684.1 | -0,252 | 2,66449E-07 |
| ACE2 | RP11-266N13.2 | -0,252 | 2,67158E-07 |
| ACE2 | RP11-293D9.2 | -0,252 | 2,68129E-07 |
| ACE2 | RP11-628E19.2 | -0,2521 | 2,6524E-07 |
| ACE2 | RP11-417O11.5 | -0,2521 | 2,64693E-07 |
| ACE2 | RP6-191P20.4 | -0,2521 | 2,64895E-07 |
| ACE2 | RP11-277A4.4 | -0,2522 | 2,62532E-07 |
| ACE2 | RP11-301M17.1 | -0,2522 | 2,61056E-07 |
| ACE2 | RP11-480D4.2 | -0,2523 | 2,5888E-07 |
| ACE2 | OR1B1 | -0,2523 | 2,57085E-07 |
| ACE2 | RP11-361H10.3 | -0,2523 | 2,57728E-07 |
| ACE2 | RP11-378J18.6 | -0,2524 | 2,54435E-07 |

| | | | |
|---|---|---|---|
| ACE2 | RP11-493E12.2 | -0,2524 | 2,55905E-07 |
| ACE2 | RN7SL68P | -0,2525 | 2,52766E-07 |
| ACE2 | LRRC73 | -0,2525 | 2,52145E-07 |
| ACE2 | RP11-278J20.2 | -0,2525 | 2,51234E-07 |
| ACE2 | RP11-464F9.21 | -0,2525 | 2,52887E-07 |
| ACE2 | CDC42EP1 | -0,2525 | 2,51713E-07 |
| ACE2 | RP11-688I9.5 | -0,2526 | 2,4845E-07 |
| ACE2 | ELOCP21 | -0,2527 | 2,4615E-07 |
| ACE2 | RP11-475C16.1 | -0,2527 | 2,47264E-07 |
| ACE2 | FOSL1 | -0,2527 | 2,47135E-07 |
| ACE2 | OR7E101P | -0,2527 | 2,46865E-07 |
| ACE2 | LINC01545 | -0,2527 | 2,4579E-07 |
| ACE2 | RP11-1338A24.1 | -0,2528 | 2,44996E-07 |
| ACE2 | LAMTOR3P2 | -0,2528 | 2,43736E-07 |
| ACE2 | OTUB1 | -0,2528 | 2,42913E-07 |
| ACE2 | RP5-971N18.3 | -0,2528 | 2,44188E-07 |
| ACE2 | AP000282.3 | -0,2528 | 2,43614E-07 |
| ACE2 | KCNMB3P1 | -0,2528 | 2,44341E-07 |
| ACE2 | RP11-407P15.2 | -0,2529 | 2,4129E-07 |
| ACE2 | RP11-170M17.1 | -0,2529 | 2,40497E-07 |
| ACE2 | RP11-22B23.1 | -0,2529 | 2,42022E-07 |
| ACE2 | CALML6 | -0,253 | 2,39869E-07 |
| ACE2 | TAS2R5 | -0,253 | 2,38351E-07 |
| ACE2 | RP11-90J7.3 | -0,253 | 2,3748E-07 |
| ACE2 | LINC01929 | -0,253 | 2,39846E-07 |
| ACE2 | AP001619.2 | -0,253 | 2,39543E-07 |
| ACE2 | RP11-415C15.3 | -0,2531 | 2,35329E-07 |
| ACE2 | SIMC1 | -0,2531 | 2,37009E-07 |
| ACE2 | LINC01372 | -0,2531 | 2,34948E-07 |
| ACE2 | LINC01996 | -0,2531 | 2,36289E-07 |
| ACE2 | GBP7 | -0,2532 | 2,33633E-07 |
| ACE2 | ANXA3 | -0,2532 | 2,34056E-07 |
| ACE2 | RP11-501J20.5 | -0,2532 | 2,32206E-07 |
| ACE2 | TMLHE | -0,2532 | 2,33501E-07 |
| ACE2 | RP13-585F24.1 | -0,2533 | 2,29549E-07 |
| ACE2 | RP11-259O2.2 | -0,2533 | 2,30387E-07 |
| ACE2 | SUPV3L1 | -0,2533 | 2,29455E-07 |
| ACE2 | CTD-2223O18.1 | -0,2534 | 2,2927E-07 |
| ACE2 | RP11-752G15.8 | -0,2534 | 2,27682E-07 |
| ACE2 | RP4-737E23.2 | -0,2534 | 2,28449E-07 |
| ACE2 | SLC1A3 | -0,2535 | 2,24609E-07 |
| ACE2 | RP11-440G5.2 | -0,2536 | 2,23101E-07 |
| ACE2 | HLA-DOA | -0,2537 | 2,20009E-07 |
| ACE2 | CTB-119C2.1 | -0,2538 | 2,19114E-07 |
| ACE2 | RP11-13K12.1 | -0,2538 | 2,1704E-07 |
| ACE2 | RP11-884K10.7 | -0,2539 | 2,16018E-07 |
| ACE2 | RP11-577H5.1 | -0,2539 | 2,14462E-07 |
| ACE2 | LL0XNC01-116E7.2 | -0,2539 | 2,15683E-07 |
| ACE2 | RHOQP3 | -0,254 | 2,12802E-07 |
| ACE2 | CMTM7 | -0,254 | 2,12214E-07 |
| ACE2 | CTC-482H14.5 | -0,254 | 2,13954E-07 |
| ACE2 | RPS27AP6 | -0,2541 | 2,10277E-07 |
| ACE2 | MEX3A | -0,2541 | 2,11888E-07 |
| ACE2 | XIRP1 | -0,2541 | 2,10911E-07 |

| | | | |
|---|---|---|---|
| ACE2 | RP11-234A1.1 | -0,2542 | 2,0845E-07 |
| ACE2 | LNCSRLR | -0,2542 | 2,08578E-07 |
| ACE2 | AP002530.2 | -0,2542 | 2,07563E-07 |
| ACE2 | LINC01895 | -0,2542 | 2,09361E-07 |
| ACE2 | HCST | -0,2542 | 2,09402E-07 |
| ACE2 | LINC00471 | -0,2543 | 2,06909E-07 |
| ACE2 | LINC01011 | -0,2543 | 2,06138E-07 |
| ACE2 | RP11-25E2.1 | -0,2543 | 2,06674E-07 |
| ACE2 | AC034228.3 | -0,2544 | 2,02734E-07 |
| ACE2 | POLM | -0,2544 | 2,04563E-07 |
| ACE2 | AC009542.2 | -0,2544 | 2,04005E-07 |
| ACE2 | PRKG1-AS1 | -0,2544 | 2,02862E-07 |
| ACE2 | AC090587.4 | -0,2544 | 2,03433E-07 |
| ACE2 | HOXC12 | -0,2544 | 2,03235E-07 |
| ACE2 | LINC02182 | -0,2544 | 2,03441E-07 |
| ACE2 | ZNF100 | -0,2544 | 2,03709E-07 |
| ACE2 | LIPE | -0,2544 | 2,02796E-07 |
| ACE2 | VPS16 | -0,2544 | 2,03575E-07 |
| ACE2 | OR56B1 | -0,2545 | 2,01303E-07 |
| ACE2 | RP11-397O8.7 | -0,2545 | 2,02285E-07 |
| ACE2 | CTD-2047H16.3 | -0,2545 | 2,02476E-07 |
| ACE2 | CDH22 | -0,2545 | 2,01943E-07 |
| ACE2 | UAP1L1 | -0,2546 | 1,99688E-07 |
| ACE2 | OR1E2 | -0,2546 | 1,98257E-07 |
| ACE2 | RP11-145O15.2 | -0,2548 | 1,94968E-07 |
| ACE2 | CTD-2210P15.3 | -0,2549 | 1,91812E-07 |
| ACE2 | IFNA5 | -0,255 | 1,90193E-07 |
| ACE2 | FGD6 | -0,255 | 1,91032E-07 |
| ACE2 | KHNYN | -0,255 | 1,91103E-07 |
| ACE2 | LL0XNC01-221F2.2 | -0,255 | 1,89566E-07 |
| ACE2 | RP11-196G18.23 | -0,2551 | 1,8893E-07 |
| ACE2 | AC004941.3 | -0,2551 | 1,87974E-07 |
| ACE2 | EBAG9 | -0,2551 | 1,88432E-07 |
| ACE2 | M6PR | -0,2551 | 1,87755E-07 |
| ACE2 | RP11-554D20.1 | -0,2551 | 1,88953E-07 |
| ACE2 | ARL4AP5 | -0,2552 | 1,86769E-07 |
| ACE2 | GLCCI1 | -0,2552 | 1,85303E-07 |
| ACE2 | SEPT7P6 | -0,2552 | 1,85416E-07 |
| ACE2 | GVQW1 | -0,2552 | 1,85306E-07 |
| ACE2 | LINC01583 | -0,2552 | 1,86764E-07 |
| ACE2 | LINC02120 | -0,2553 | 1,84493E-07 |
| ACE2 | RP11-862G15.1 | -0,2553 | 1,84697E-07 |
| ACE2 | RPS27L | -0,2553 | 1,84447E-07 |
| ACE2 | DNM1P41 | -0,2553 | 1,84637E-07 |
| ACE2 | LA16c-313D11.12 | -0,2553 | 1,83188E-07 |
| ACE2 | IFNW1 | -0,2554 | 1,82124E-07 |
| ACE2 | RP11-488C13.1 | -0,2554 | 1,82654E-07 |
| ACE2 | TRAPPC8 | -0,2555 | 1,80571E-07 |
| ACE2 | RP1-12G14.7 | -0,2556 | 1,77993E-07 |
| ACE2 | ANKRD33 | -0,2556 | 1,78549E-07 |
| ACE2 | C16orf74 | -0,2556 | 1,77075E-07 |
| ACE2 | RP11-473L15.3 | -0,2557 | 1,75015E-07 |
| ACE2 | IDNK | -0,2557 | 1,74998E-07 |
| ACE2 | DYNC1I2P1 | -0,2557 | 1,74981E-07 |

| | | | |
|---|---|---|---|
| ACE2 | AC022154.7 | -0,2557 | 1,7525E-07 |
| ACE2 | RUNX2 | -0,2558 | 1,73714E-07 |
| ACE2 | RP11-419C23.1 | -0,2558 | 1,73239E-07 |
| ACE2 | RPS26P45 | -0,2558 | 1,73717E-07 |
| ACE2 | KIAA0100 | -0,2558 | 1,72949E-07 |
| ACE2 | NOL11 | -0,2558 | 1,72715E-07 |
| ACE2 | SRSF2 | -0,2558 | 1,73559E-07 |
| ACE2 | ODC1 | -0,2559 | 1,70755E-07 |
| ACE2 | PROX1 | -0,2561 | 1,67015E-07 |
| ACE2 | CORIN | -0,2561 | 1,67808E-07 |
| ACE2 | G3BP2 | -0,2561 | 1,67331E-07 |
| ACE2 | CTB-37A13.1 | -0,2561 | 1,68403E-07 |
| ACE2 | RP11-310H4.3 | -0,2562 | 1,66095E-07 |
| ACE2 | RPL17P36 | -0,2562 | 1,6567E-07 |
| ACE2 | UPK1A | -0,2562 | 1,65983E-07 |
| ACE2 | RP11-298C3.2 | -0,2562 | 1,65018E-07 |
| ACE2 | AC144449.1 | -0,2563 | 1,6358E-07 |
| ACE2 | CTB-114C7.4 | -0,2563 | 1,63157E-07 |
| ACE2 | SNRPEP4 | -0,2563 | 1,6405E-07 |
| ACE2 | CBX3P3 | -0,2564 | 1,62179E-07 |
| ACE2 | LINC01287 | -0,2564 | 1,62899E-07 |
| ACE2 | RP11-51J9.6 | -0,2564 | 1,63059E-07 |
| ACE2 | HSPE1P3 | -0,2564 | 1,61528E-07 |
| ACE2 | RP11-2N1.1 | -0,2564 | 1,61852E-07 |
| ACE2 | SMIM11B | -0,2564 | 1,61747E-07 |
| ACE2 | PPP4R2 | -0,2565 | 1,59398E-07 |
| ACE2 | KIAA1024L | -0,2565 | 1,5968E-07 |
| ACE2 | CTB-174D11.2 | -0,2565 | 1,59623E-07 |
| ACE2 | RAD51-AS1 | -0,2565 | 1,6017E-07 |
| ACE2 | LILRB2 | -0,2565 | 1,611E-07 |
| ACE2 | NSUN5P1 | -0,2567 | 1,56045E-07 |
| ACE2 | RP11-510M2.2 | -0,2567 | 1,56071E-07 |
| ACE2 | RP11-471L13.3 | -0,2567 | 1,55925E-07 |
| ACE2 | CTD-2013N17.4 | -0,2567 | 1,56627E-07 |
| ACE2 | RP11-260N14.1 | -0,2568 | 1,54362E-07 |
| ACE2 | ADAMTS13 | -0,2568 | 1,54523E-07 |
| ACE2 | COG8 | -0,2568 | 1,54509E-07 |
| ACE2 | UNC13D | -0,2568 | 1,54256E-07 |
| ACE2 | RPL34P34 | -0,2568 | 1,54042E-07 |
| ACE2 | CYCSP10 | -0,2569 | 1,53584E-07 |
| ACE2 | RP11-958N24.2 | -0,2569 | 1,53573E-07 |
| ACE2 | VPS35 | -0,2569 | 1,53047E-07 |
| ACE2 | TMEM104 | -0,2569 | 1,52599E-07 |
| ACE2 | ABCA2 | -0,257 | 1,51612E-07 |
| ACE2 | EDC3 | -0,257 | 1,50625E-07 |
| ACE2 | CTC-454I21.4 | -0,257 | 1,51774E-07 |
| ACE2 | KIAA1324L | -0,2571 | 1,49743E-07 |
| ACE2 | C9orf131 | -0,2571 | 1,48798E-07 |
| ACE2 | RP11-319G9.5 | -0,2571 | 1,49577E-07 |
| ACE2 | CBX1P1 | -0,2571 | 1,49487E-07 |
| ACE2 | MIR3681HG | -0,2572 | 1,482E-07 |
| ACE2 | AC005358.3 | -0,2572 | 1,47523E-07 |
| ACE2 | LL22NC03-2H8.5 | -0,2572 | 1,47378E-07 |
| ACE2 | CTD-2647L4.4 | -0,2573 | 1,46408E-07 |

| | | | |
|---|---|---|---|
| ACE2 | RCAN3 | -0,2574 | 1,44819E-07 |
| ACE2 | CTB-17P3.4 | -0,2574 | 1,44618E-07 |
| ACE2 | RP11-544A12.8 | -0,2574 | 1,44014E-07 |
| ACE2 | RP11-523O18.1 | -0,2574 | 1,45043E-07 |
| ACE2 | RP11-357K6.1 | -0,2574 | 1,45304E-07 |
| ACE2 | ODF2L | -0,2575 | 1,43392E-07 |
| ACE2 | MUC19 | -0,2575 | 1,42236E-07 |
| ACE2 | RP11-415I12.2 | -0,2575 | 1,42347E-07 |
| ACE2 | RP11-13N12.1 | -0,2576 | 1,4084E-07 |
| ACE2 | ZC3H12C | -0,2576 | 1,40772E-07 |
| ACE2 | UTP14C | -0,2576 | 1,40831E-07 |
| ACE2 | CH17-258A22.4 | -0,2576 | 1,40683E-07 |
| ACE2 | ZNF415P1 | -0,2576 | 1,41738E-07 |
| ACE2 | RP11-338C15.3 | -0,2578 | 1,38054E-07 |
| ACE2 | LINC01614 | -0,2578 | 1,38177E-07 |
| ACE2 | MRS2 | -0,2578 | 1,38323E-07 |
| ACE2 | PRR33 | -0,2578 | 1,37911E-07 |
| ACE2 | CTD-2587H24.1 | -0,2578 | 1,37617E-07 |
| ACE2 | LINC01107 | -0,2579 | 1,3645E-07 |
| ACE2 | FAM13A | -0,2579 | 1,36384E-07 |
| ACE2 | MIR3907 | -0,2579 | 1,35792E-07 |
| ACE2 | AC105339.1 | -0,2579 | 1,36954E-07 |
| ACE2 | RP4-728D4.2 | -0,258 | 1,34362E-07 |
| ACE2 | NAA11 | -0,258 | 1,34543E-07 |
| ACE2 | OR2K2 | -0,258 | 1,34826E-07 |
| ACE2 | RP11-81A1.6 | -0,258 | 1,35591E-07 |
| ACE2 | RP11-583F2.6 | -0,258 | 1,35152E-07 |
| ACE2 | MIR222HG | -0,258 | 1,35033E-07 |
| ACE2 | CCDC190 | -0,2581 | 1,33028E-07 |
| ACE2 | RP11-443B7.3 | -0,2581 | 1,33767E-07 |
| ACE2 | TADA2B | -0,2581 | 1,33611E-07 |
| ACE2 | ST13P19 | -0,2582 | 1,32563E-07 |
| ACE2 | RP11-14I17.3 | -0,2584 | 1,28284E-07 |
| ACE2 | SIK1 | -0,2584 | 1,29085E-07 |
| ACE2 | ODF2-AS1 | -0,2585 | 1,26975E-07 |
| ACE2 | WASIR2 | -0,2585 | 1,27106E-07 |
| ACE2 | CTD-2008L17.2 | -0,2585 | 1,26958E-07 |
| ACE2 | LINC01374 | -0,2586 | 1,25944E-07 |
| ACE2 | SRP54-AS1 | -0,2586 | 1,25524E-07 |
| ACE2 | CDK5R1 | -0,2586 | 1,26253E-07 |
| ACE2 | GNA13 | -0,2586 | 1,25532E-07 |
| ACE2 | RP11-177C12.4 | -0,2588 | 1,23134E-07 |
| ACE2 | GS1-166A23.1 | -0,2588 | 1,22915E-07 |
| ACE2 | RP11-579D7.8 | -0,2588 | 1,2336E-07 |
| ACE2 | CTB-52I2.7 | -0,2588 | 1,22888E-07 |
| ACE2 | LINC00877 | -0,2589 | 1,21824E-07 |
| ACE2 | MBD6 | -0,2589 | 1,21222E-07 |
| ACE2 | AC093627.12 | -0,259 | 1,19652E-07 |
| ACE2 | RP11-157K17.5 | -0,259 | 1,20595E-07 |
| ACE2 | POM121L9P | -0,259 | 1,19886E-07 |
| ACE2 | NUP210L | -0,2591 | 1,18849E-07 |
| ACE2 | LINC00973 | -0,2591 | 1,18209E-07 |
| ACE2 | RP11-616K6.1 | -0,2591 | 1,19084E-07 |
| ACE2 | EARS2 | -0,2591 | 1,18348E-07 |

| | | | |
|---|---|---|---|
| ACE2 | HSFY3P | -0,2592 | 1,17465E-07 |
| ACE2 | AC141930.3 | -0,2593 | 1,15451E-07 |
| ACE2 | DRC7 | -0,2593 | 1,15484E-07 |
| ACE2 | LRP5L | -0,2593 | 1,16084E-07 |
| ACE2 | LINC00694 | -0,2594 | 1,15282E-07 |
| ACE2 | TVP23CP2 | -0,2594 | 1,14145E-07 |
| ACE2 | RBM17P1 | -0,2595 | 1,12995E-07 |
| ACE2 | AC005775.2 | -0,2595 | 1,13282E-07 |
| ACE2 | HIST2H3D | -0,2596 | 1,12498E-07 |
| ACE2 | MTHFD2 | -0,2596 | 1,12682E-07 |
| ACE2 | AC006373.1 | -0,2596 | 1,12456E-07 |
| ACE2 | CYP4F26P | -0,2596 | 1,1267E-07 |
| ACE2 | KDM5C | -0,2596 | 1,11883E-07 |
| ACE2 | RBM17P2 | -0,2597 | 1,10238E-07 |
| ACE2 | RP11-517O13.1 | -0,2597 | 1,10512E-07 |
| ACE2 | FABP6 | -0,2598 | 1,09732E-07 |
| ACE2 | UTP18 | -0,2598 | 1,09405E-07 |
| ACE2 | AC008991.1 | -0,2598 | 1,09724E-07 |
| ACE2 | RN7SL862P | -0,26 | 1,06924E-07 |
| ACE2 | CYTOR | -0,2601 | 1,0571E-07 |
| ACE2 | LINC01611 | -0,2601 | 1,05743E-07 |
| ACE2 | WEE2-AS1 | -0,2601 | 1,06261E-07 |
| ACE2 | IRAK4 | -0,2601 | 1,05574E-07 |
| ACE2 | RP11-736N17.11 | -0,2601 | 1,06233E-07 |
| ACE2 | RPL5P3 | -0,2601 | 1,06009E-07 |
| ACE2 | AF230666.2 | -0,2602 | 1,04546E-07 |
| ACE2 | RNU6-94P | -0,2602 | 1,04866E-07 |
| ACE2 | RP11-494H4.3 | -0,2603 | 1,03409E-07 |
| ACE2 | TFDP1 | -0,2603 | 1,03648E-07 |
| ACE2 | RP11-357N13.6 | -0,2603 | 1,02817E-07 |
| ACE2 | RP11-94B19.7 | -0,2603 | 1,0349E-07 |
| ACE2 | DUTP6 | -0,2604 | 1,02329E-07 |
| ACE2 | NOC4L | -0,2604 | 1,0253E-07 |
| ACE2 | LETMD1 | -0,2605 | 1,0097E-07 |
| ACE2 | LINC00165 | -0,2605 | 1,01398E-07 |
| ACE2 | RP5-1112D6.7 | -0,2606 | 9,93782E-08 |
| ACE2 | AC093690.1 | -0,2607 | 9,87113E-08 |
| ACE2 | RN7SL124P | -0,2607 | 9,83146E-08 |
| ACE2 | RP3-523C21.3 | -0,2607 | 9,87557E-08 |
| ACE2 | GID8 | -0,2607 | 9,89861E-08 |
| ACE2 | GYG1P1 | -0,2608 | 9,79469E-08 |
| ACE2 | RNU6-1010P | -0,2609 | 9,59965E-08 |
| ACE2 | SNORD14E | -0,2609 | 9,62344E-08 |
| ACE2 | RP11-592H6.3 | -0,2609 | 9,69378E-08 |
| ACE2 | RP11-152L20.3 | -0,2609 | 9,645E-08 |
| ACE2 | POP4 | -0,2609 | 9,59136E-08 |
| ACE2 | NPM1P39 | -0,261 | 9,54008E-08 |
| ACE2 | ARHGEF11 | -0,261 | 9,56614E-08 |
| ACE2 | LINC00843 | -0,261 | 9,53603E-08 |
| ACE2 | ATP2A1-AS1 | -0,261 | 9,57811E-08 |
| ACE2 | CTD-2619J13.9 | -0,2611 | 9,38125E-08 |
| ACE2 | DNAJC5G | -0,2612 | 9,27969E-08 |
| ACE2 | SBF2-AS1 | -0,2612 | 9,29671E-08 |
| ACE2 | TCAF2P1 | -0,2613 | 9,24349E-08 |

| | | | |
|---|---|---|---|
| ACE2 | CTD-2058B24.2 | -0,2613 | 9,22062E-08 |
| ACE2 | TBCAP1 | -0,2613 | 9,205E-08 |
| ACE2 | PINK1 | -0,2614 | 9,11903E-08 |
| ACE2 | COL4A3 | -0,2615 | 8,98314E-08 |
| ACE2 | RP11-177J6.1 | -0,2615 | 9,00673E-08 |
| ACE2 | RP11-557H15.4 | -0,2615 | 8,98502E-08 |
| ACE2 | CNTNAP3P1 | -0,2615 | 8,93689E-08 |
| ACE2 | CTD-2102P23.1 | -0,2615 | 8,96784E-08 |
| ACE2 | DOPEY1 | -0,2616 | 8,84725E-08 |
| ACE2 | NRN1L | -0,2616 | 8,83917E-08 |
| ACE2 | VN1R110P | -0,2616 | 8,92877E-08 |
| ACE2 | ARL4C | -0,2617 | 8,75546E-08 |
| ACE2 | LA16c-OS12.2 | -0,2617 | 8,78063E-08 |
| ACE2 | RESP18 | -0,2618 | 8,71965E-08 |
| ACE2 | CTD-3064M3.4 | -0,2618 | 8,7179E-08 |
| ACE2 | RP11-44F14.7 | -0,2618 | 8,72127E-08 |
| ACE2 | MFSD1P1 | -0,2618 | 8,68891E-08 |
| ACE2 | C20orf197 | -0,2618 | 8,68732E-08 |
| ACE2 | RP11-20E24.1 | -0,2619 | 8,54769E-08 |
| ACE2 | LINC01630 | -0,2619 | 8,53994E-08 |
| ACE2 | S100A7 | -0,262 | 8,4748E-08 |
| ACE2 | MIR765 | -0,262 | 8,51356E-08 |
| ACE2 | RP11-65L3.2 | -0,262 | 8,43773E-08 |
| ACE2 | SNRPCP3 | -0,262 | 8,43149E-08 |
| ACE2 | SSNA1 | -0,262 | 8,43273E-08 |
| ACE2 | RP11-464F9.22 | -0,262 | 8,48973E-08 |
| ACE2 | LINC01001 | -0,262 | 8,46262E-08 |
| ACE2 | TFAP4 | -0,262 | 8,46771E-08 |
| ACE2 | RP11-580J4.1 | -0,2621 | 8,40152E-08 |
| ACE2 | G0S2 | -0,2622 | 8,2931E-08 |
| ACE2 | RP11-1281K21.1 | -0,2622 | 8,26245E-08 |
| ACE2 | AF213884.2 | -0,2622 | 8,2975E-08 |
| ACE2 | SLC25A46 | -0,2622 | 8,2987E-08 |
| ACE2 | FAM133CP | -0,2623 | 8,16086E-08 |
| ACE2 | CHFR | -0,2623 | 8,20799E-08 |
| ACE2 | SMG1P1 | -0,2623 | 8,16793E-08 |
| ACE2 | RP11-465I4.2 | -0,2623 | 8,15564E-08 |
| ACE2 | UHRF2P1 | -0,2623 | 8,19773E-08 |
| ACE2 | CA8 | -0,2624 | 8,05114E-08 |
| ACE2 | RP11-712C19.1 | -0,2625 | 7,98732E-08 |
| ACE2 | FOXD1 | -0,2626 | 7,92064E-08 |
| ACE2 | RP11-313M3.1 | -0,2626 | 7,85894E-08 |
| ACE2 | PROSER2-AS1 | -0,2627 | 7,79153E-08 |
| ACE2 | ZNHIT2 | -0,2627 | 7,84503E-08 |
| ACE2 | LINC00158 | -0,2627 | 7,76777E-08 |
| ACE2 | RP11-343J18.1 | -0,2628 | 7,69502E-08 |
| ACE2 | ZNF85 | -0,2628 | 7,66915E-08 |
| ACE2 | HIRA | -0,2628 | 7,72944E-08 |
| ACE2 | PM20D1 | -0,2629 | 7,59728E-08 |
| ACE2 | LINC00857 | -0,2629 | 7,63739E-08 |
| ACE2 | CPPED1 | -0,2629 | 7,61603E-08 |
| ACE2 | RP11-329J18.4 | -0,2629 | 7,63324E-08 |
| ACE2 | CTD-2240E14.4 | -0,2629 | 7,62134E-08 |
| ACE2 | KNCN | -0,2631 | 7,48578E-08 |

| | | | |
|---|---|---|---|
| ACE2 | RP4-800O15.2 | -0,2631 | 7,47888E-08 |
| ACE2 | CCDC177 | -0,2631 | 7,41297E-08 |
| ACE2 | RP11-701H24.10 | -0,2631 | 7,43671E-08 |
| ACE2 | FHAD1 | -0,2632 | 7,35647E-08 |
| ACE2 | RP4-575N6.2 | -0,2633 | 7,27038E-08 |
| ACE2 | GUSBP2 | -0,2633 | 7,23087E-08 |
| ACE2 | OR4A40P | -0,2633 | 7,29666E-08 |
| ACE2 | FMO6P | -0,2634 | 7,15327E-08 |
| ACE2 | ZDHHC21 | -0,2634 | 7,14691E-08 |
| ACE2 | CCT2 | -0,2634 | 7,1525E-08 |
| ACE2 | TUBAP4 | -0,2634 | 7,16498E-08 |
| ACE2 | GPAM | -0,2635 | 7,12514E-08 |
| ACE2 | CTC-457E21.9 | -0,2635 | 7,12174E-08 |
| ACE2 | RP4-737E23.5 | -0,2635 | 7,10585E-08 |
| ACE2 | B3GALT2 | -0,2636 | 7,06064E-08 |
| ACE2 | EED | -0,2636 | 7,02493E-08 |
| ACE2 | RP11-31I22.2 | -0,2636 | 6,9914E-08 |
| ACE2 | SIGLEC16 | -0,2636 | 7,00238E-08 |
| ACE2 | RP11-46F15.2 | -0,2637 | 6,9717E-08 |
| ACE2 | RSG1 | -0,2637 | 6,93646E-08 |
| ACE2 | CPEB3 | -0,2637 | 6,95861E-08 |
| ACE2 | GRIN2A | -0,2637 | 6,91155E-08 |
| ACE2 | SNORD35A | -0,2637 | 6,95157E-08 |
| ACE2 | KIAA0907 | -0,2638 | 6,86778E-08 |
| ACE2 | SCFD2 | -0,2638 | 6,83734E-08 |
| ACE2 | RP11-159F24.5 | -0,2638 | 6,82219E-08 |
| ACE2 | RP11-561O23.5 | -0,2638 | 6,844E-08 |
| ACE2 | RP11-996F15.4 | -0,2638 | 6,86186E-08 |
| ACE2 | THBD | -0,2638 | 6,83995E-08 |
| ACE2 | AP001626.2 | -0,2638 | 6,86562E-08 |
| ACE2 | RP11-506O24.1 | -0,2639 | 6,79876E-08 |
| ACE2 | AGFG1 | -0,2639 | 6,77647E-08 |
| ACE2 | DLEU7 | -0,2639 | 6,77814E-08 |
| ACE2 | EIF4A2P4 | -0,2639 | 6,74732E-08 |
| ACE2 | GOLGA8M | -0,264 | 6,70644E-08 |
| ACE2 | CPSF1P1 | -0,264 | 6,70172E-08 |
| ACE2 | RP1-78B3.1 | -0,264 | 6,72639E-08 |
| ACE2 | BMX | -0,264 | 6,70131E-08 |
| ACE2 | RP11-553D4.2 | -0,2641 | 6,61775E-08 |
| ACE2 | LINC00311 | -0,2641 | 6,63058E-08 |
| ACE2 | CTC-575N7.1 | -0,2642 | 6,56119E-08 |
| ACE2 | UMAD1 | -0,2642 | 6,52598E-08 |
| ACE2 | TAX1BP1 | -0,2642 | 6,52894E-08 |
| ACE2 | EIF3A | -0,2642 | 6,5067E-08 |
| ACE2 | RP11-159D12.3 | -0,2642 | 6,52265E-08 |
| ACE2 | MSC | -0,2643 | 6,43708E-08 |
| ACE2 | RNA5SP298 | -0,2643 | 6,43705E-08 |
| ACE2 | MCOLN1 | -0,2643 | 6,44529E-08 |
| ACE2 | RP1-241P17.4 | -0,2643 | 6,48598E-08 |
| ACE2 | HSFY2 | -0,2643 | 6,46948E-08 |
| ACE2 | SLC25A51 | -0,2644 | 6,35441E-08 |
| ACE2 | LINC01785 | -0,2644 | 6,3997E-08 |
| ACE2 | THAP12P2 | -0,2645 | 6,32343E-08 |
| ACE2 | CTD-2339M3.1 | -0,2646 | 6,2578E-08 |

| | | | |
|---|---|---|---|
| ACE2 | RP11-521C20.5 | -0,2646 | 6,24235E-08 |
| ACE2 | PSMD8 | -0,2647 | 6,19221E-08 |
| ACE2 | APOOL | -0,2647 | 6,15311E-08 |
| ACE2 | FMC1 | -0,2648 | 6,0993E-08 |
| ACE2 | AC005954.3 | -0,2648 | 6,12147E-08 |
| ACE2 | NPM1P46 | -0,2649 | 6,05169E-08 |
| ACE2 | RP4-683L5.1 | -0,2649 | 6,01058E-08 |
| ACE2 | CTD-2562J17.4 | -0,2649 | 6,04931E-08 |
| ACE2 | RP3-461F17.1 | -0,265 | 5,9817E-08 |
| ACE2 | RP11-159L20.2 | -0,265 | 5,96559E-08 |
| ACE2 | RP11-16E23.4 | -0,265 | 5,9832E-08 |
| ACE2 | AC004019.10 | -0,265 | 5,97981E-08 |
| ACE2 | RP3-417G15.1 | -0,265 | 5,93249E-08 |
| ACE2 | AC005237.4 | -0,2651 | 5,90853E-08 |
| ACE2 | RP11-496I2.5 | -0,2651 | 5,85328E-08 |
| ACE2 | NPPA | -0,2652 | 5,81215E-08 |
| ACE2 | ITPKB-IT1 | -0,2652 | 5,80256E-08 |
| ACE2 | RP11-379I19.2 | -0,2652 | 5,79341E-08 |
| ACE2 | RP11-881M11.2 | -0,2652 | 5,82698E-08 |
| ACE2 | RP3-468K18.6 | -0,2653 | 5,74746E-08 |
| ACE2 | RP11-223P11.3 | -0,2653 | 5,74908E-08 |
| ACE2 | DLGAP1-AS5 | -0,2653 | 5,72185E-08 |
| ACE2 | RP11-65L3.4 | -0,2654 | 5,67896E-08 |
| ACE2 | LINC01238 | -0,2654 | 5,6532E-08 |
| ACE2 | GS1-124K5.7 | -0,2654 | 5,67402E-08 |
| ACE2 | RP11-620J15.4 | -0,2654 | 5,69942E-08 |
| ACE2 | RP11-650K20.2 | -0,2654 | 5,67936E-08 |
| ACE2 | EDARADD | -0,2656 | 5,51319E-08 |
| ACE2 | WNT11 | -0,2656 | 5,53742E-08 |
| ACE2 | KRT7-AS | -0,2656 | 5,54172E-08 |
| ACE2 | RPS2P5 | -0,2656 | 5,52758E-08 |
| ACE2 | LINC01402 | -0,2656 | 5,50852E-08 |
| ACE2 | HIST2H2BD | -0,2657 | 5,49394E-08 |
| ACE2 | CTD-2035E11.4 | -0,2657 | 5,45012E-08 |
| ACE2 | UTP15 | -0,2657 | 5,49167E-08 |
| ACE2 | RP11-378A12.1 | -0,2657 | 5,50357E-08 |
| ACE2 | RAB40AL | -0,2657 | 5,46627E-08 |
| ACE2 | RP11-404O13.4 | -0,2658 | 5,38485E-08 |
| ACE2 | PCDHA4 | -0,2658 | 5,3944E-08 |
| ACE2 | C17orf82 | -0,2658 | 5,42858E-08 |
| ACE2 | AC010504.2 | -0,2658 | 5,4373E-08 |
| ACE2 | GOLGA6L3 | -0,2659 | 5,34843E-08 |
| ACE2 | RP11-158M2.2 | -0,2659 | 5,35557E-08 |
| ACE2 | RPL17P43 | -0,2659 | 5,31691E-08 |
| ACE2 | RPL31P12 | -0,266 | 5,29118E-08 |
| ACE2 | EPRS | -0,266 | 5,26043E-08 |
| ACE2 | SLC6A13 | -0,266 | 5,31088E-08 |
| ACE2 | NMUR1 | -0,2661 | 5,2461E-08 |
| ACE2 | AC008753.6 | -0,2661 | 5,19574E-08 |
| ACE2 | TPT1P9 | -0,2662 | 5,14524E-08 |
| ACE2 | ACRV1 | -0,2662 | 5,15337E-08 |
| ACE2 | AC012363.4 | -0,2663 | 5,07097E-08 |
| ACE2 | KCNMB2-AS1 | -0,2663 | 5,09868E-08 |
| ACE2 | CTD-3092A11.3 | -0,2663 | 5,07829E-08 |

| | | | |
|---|---|---|---|
| ACE2 | RP11-505E24.3 | -0,2663 | 5,1093E-08 |
| ACE2 | RP11-93O14.2 | -0,2663 | 5,10741E-08 |
| ACE2 | CCDC12 | -0,2664 | 5,05794E-08 |
| ACE2 | HADHAP1 | -0,2664 | 5,05318E-08 |
| ACE2 | NTF4 | -0,2664 | 5,04371E-08 |
| ACE2 | AC012462.1 | -0,2665 | 4,99451E-08 |
| ACE2 | RP11-373E16.3 | -0,2665 | 4,99327E-08 |
| ACE2 | HOTAIRM1 | -0,2665 | 4,99204E-08 |
| ACE2 | LINC00265 | -0,2667 | 4,87353E-08 |
| ACE2 | AL359314.1 | -0,2667 | 4,8416E-08 |
| ACE2 | MKRN5P | -0,2667 | 4,86595E-08 |
| ACE2 | LA16c-60H5.7 | -0,2668 | 4,77411E-08 |
| ACE2 | RP11-977B10.2 | -0,2669 | 4,73105E-08 |
| ACE2 | RP11-305P14.1 | -0,267 | 4,66312E-08 |
| ACE2 | MCM3 | -0,267 | 4,71246E-08 |
| ACE2 | RP11-553A10.1 | -0,2671 | 4,62827E-08 |
| ACE2 | BCKDHA | -0,2671 | 4,61044E-08 |
| ACE2 | LL0XNC01-7P3.1 | -0,2671 | 4,61098E-08 |
| ACE2 | ATF4P1 | -0,2671 | 4,63638E-08 |
| ACE2 | AC026188.1 | -0,2672 | 4,55244E-08 |
| ACE2 | CTD-2232E5.2 | -0,2672 | 4,5755E-08 |
| ACE2 | RP11-407N8.5 | -0,2672 | 4,58953E-08 |
| ACE2 | GPAT2P1 | -0,2673 | 4,5386E-08 |
| ACE2 | NCOA6 | -0,2673 | 4,50108E-08 |
| ACE2 | STAB1 | -0,2674 | 4,48469E-08 |
| ACE2 | RN7SKP296 | -0,2674 | 4,45672E-08 |
| ACE2 | RN7SKP240 | -0,2674 | 4,44811E-08 |
| ACE2 | ASAH1 | -0,2674 | 4,49049E-08 |
| ACE2 | LIG4 | -0,2674 | 4,46524E-08 |
| ACE2 | PKP1 | -0,2675 | 4,4171E-08 |
| ACE2 | ADPGK | -0,2675 | 4,43829E-08 |
| ACE2 | RP11-5P22.1 | -0,2676 | 4,34736E-08 |
| ACE2 | RP11-427L15.2 | -0,2676 | 4,38181E-08 |
| ACE2 | LINC01877 | -0,2677 | 4,32721E-08 |
| ACE2 | LINC02151 | -0,2677 | 4,28992E-08 |
| ACE2 | CACNA1G-AS1 | -0,2677 | 4,30573E-08 |
| ACE2 | TTLL11-IT1 | -0,2678 | 4,27695E-08 |
| ACE2 | GOLGA2P11 | -0,2678 | 4,26046E-08 |
| ACE2 | NLRP3 | -0,268 | 4,13739E-08 |
| ACE2 | AMER3 | -0,268 | 4,17344E-08 |
| ACE2 | HSP90AB2P | -0,268 | 4,13516E-08 |
| ACE2 | RP11-157L3.10 | -0,268 | 4,14408E-08 |
| ACE2 | TSPYL6 | -0,2681 | 4,10269E-08 |
| ACE2 | CTD-2320G14.2 | -0,2681 | 4,10985E-08 |
| ACE2 | HTR1B | -0,2682 | 4,08003E-08 |
| ACE2 | RP11-775B15.3 | -0,2682 | 4,08161E-08 |
| ACE2 | RP11-478C6.2 | -0,2683 | 4,01474E-08 |
| ACE2 | CAPSL | -0,2683 | 3,98706E-08 |
| ACE2 | LINC00051 | -0,2683 | 3,99215E-08 |
| ACE2 | RP11-1H15.1 | -0,2683 | 4,02177E-08 |
| ACE2 | ATF4P2 | -0,2683 | 4,02835E-08 |
| ACE2 | RPS4XP16 | -0,2684 | 3,94897E-08 |
| ACE2 | C14orf119 | -0,2684 | 3,96482E-08 |
| ACE2 | GNAI3 | -0,2685 | 3,93214E-08 |

| | | | |
|---|---|---|---|
| ACE2 | C9orf139 | -0,2685 | 3,89367E-08 |
| ACE2 | RP11-97N19.3 | -0,2685 | 3,90915E-08 |
| ACE2 | KLF2P2 | -0,2686 | 3,86353E-08 |
| ACE2 | RP11-109I13.2 | -0,2686 | 3,86013E-08 |
| ACE2 | DOK1 | -0,2687 | 3,8105E-08 |
| ACE2 | LRRC37A11P | -0,2687 | 3,81409E-08 |
| ACE2 | MICF | -0,2688 | 3,79056E-08 |
| ACE2 | AC139099.4 | -0,2688 | 3,79075E-08 |
| ACE2 | RP11-405M12.3 | -0,2688 | 3,77499E-08 |
| ACE2 | SCML1 | -0,2688 | 3,78674E-08 |
| ACE2 | CYP51A1P3 | -0,2689 | 3,70794E-08 |
| ACE2 | OVCH2 | -0,2689 | 3,72383E-08 |
| ACE2 | RP11-116D17.5 | -0,2689 | 3,71018E-08 |
| ACE2 | RP5-994D16.3 | -0,269 | 3,66858E-08 |
| ACE2 | CTD-2015C24.1 | -0,2691 | 3,62232E-08 |
| ACE2 | RP11-212E8.1 | -0,2691 | 3,63512E-08 |
| ACE2 | RP11-126O22.1 | -0,2691 | 3,64254E-08 |
| ACE2 | RP4-713A8.1 | -0,2692 | 3,61625E-08 |
| ACE2 | U73166.2 | -0,2692 | 3,58033E-08 |
| ACE2 | RPL10P9 | -0,2692 | 3,57671E-08 |
| ACE2 | RP11-458K10.1 | -0,2692 | 3,59247E-08 |
| ACE2 | RP6-65G23.5 | -0,2692 | 3,61151E-08 |
| ACE2 | UNC45B | -0,2692 | 3,60925E-08 |
| ACE2 | FCAR | -0,2692 | 3,60822E-08 |
| ACE2 | AC092162.2 | -0,2693 | 3,56929E-08 |
| ACE2 | RPL12P6 | -0,2693 | 3,55942E-08 |
| ACE2 | RALGPS2 | -0,2694 | 3,5203E-08 |
| ACE2 | RP11-418J17.3 | -0,2695 | 3,48961E-08 |
| ACE2 | GSK3B | -0,2695 | 3,4661E-08 |
| ACE2 | C4orf36 | -0,2695 | 3,45896E-08 |
| ACE2 | FAM85B | -0,2695 | 3,47908E-08 |
| ACE2 | ZNF30-AS1 | -0,2695 | 3,46773E-08 |
| ACE2 | TMLHE-AS1 | -0,2695 | 3,45841E-08 |
| ACE2 | CTNNBIP1 | -0,2696 | 3,43454E-08 |
| ACE2 | GAPDHP47 | -0,2696 | 3,43111E-08 |
| ACE2 | AC091878.1 | -0,2696 | 3,44804E-08 |
| ACE2 | CTD-2278I10.4 | -0,2696 | 3,44773E-08 |
| ACE2 | GNAS-AS1 | -0,2696 | 3,42991E-08 |
| ACE2 | ADIPOR1 | -0,2697 | 3,3828E-08 |
| ACE2 | SLC25A31 | -0,2697 | 3,37018E-08 |
| ACE2 | RP5-872K7.7 | -0,2697 | 3,38022E-08 |
| ACE2 | CAMK4 | -0,2698 | 3,33422E-08 |
| ACE2 | RP11-347E10.1 | -0,2698 | 3,34044E-08 |
| ACE2 | RP3-388M5.9 | -0,2698 | 3,34578E-08 |
| ACE2 | RP11-89N17.4 | -0,2699 | 3,31021E-08 |
| ACE2 | GOLGA6A | -0,2699 | 3,30363E-08 |
| ACE2 | RP11-638L3.1 | -0,2699 | 3,29083E-08 |
| ACE2 | AC003986.6 | -0,2701 | 3,22519E-08 |
| ACE2 | RP11-217B7.2 | -0,2703 | 3,13027E-08 |
| ACE2 | WNT9B | -0,2703 | 3,14833E-08 |
| ACE2 | IFIT1B | -0,2704 | 3,12345E-08 |
| ACE2 | RP5-1065J22.2 | -0,2705 | 3,0866E-08 |
| ACE2 | VSX2 | -0,2705 | 3,07279E-08 |
| ACE2 | CTD-2526A2.4 | -0,2705 | 3,06808E-08 |

| | | | |
|---|---|---|---|
| ACE2 | CTD-2550O8.7 | -0,2706 | 3,03758E-08 |
| ACE2 | CTD-2528L19.3 | -0,2706 | 3,04226E-08 |
| ACE2 | MIR1231 | -0,2707 | 3,01713E-08 |
| ACE2 | LRRN1 | -0,2707 | 2,99611E-08 |
| ACE2 | RAB44 | -0,2707 | 2,99653E-08 |
| ACE2 | LL0XNC01-36H8.1 | -0,2707 | 3,01514E-08 |
| ACE2 | RNU1-16P | -0,2708 | 2,96528E-08 |
| ACE2 | NFS1 | -0,2708 | 2,94779E-08 |
| ACE2 | ZFX-AS1 | -0,2708 | 2,95631E-08 |
| ACE2 | NMD3P1 | -0,271 | 2,88067E-08 |
| ACE2 | AC002429.5 | -0,271 | 2,88418E-08 |
| ACE2 | LINC00354 | -0,271 | 2,8842E-08 |
| ACE2 | RP11-638I2.9 | -0,271 | 2,89602E-08 |
| ACE2 | RP11-264M12.2 | -0,271 | 2,90173E-08 |
| ACE2 | RP11-3L21.2 | -0,2711 | 2,85975E-08 |
| ACE2 | KLK5 | -0,2711 | 2,84355E-08 |
| ACE2 | MYO18B | -0,2711 | 2,84306E-08 |
| ACE2 | MAGEA8-AS1 | -0,2711 | 2,85987E-08 |
| ACE2 | RP11-673E1.1 | -0,2712 | 2,83411E-08 |
| ACE2 | SLC30A5 | -0,2712 | 2,81811E-08 |
| ACE2 | RPS20P33 | -0,2712 | 2,81402E-08 |
| ACE2 | RP11-477H21.2 | -0,2713 | 2,78173E-08 |
| ACE2 | RP11-331K21.1 | -0,2714 | 2,75447E-08 |
| ACE2 | AMD1P1 | -0,2714 | 2,76358E-08 |
| ACE2 | OR9Q1 | -0,2714 | 2,77194E-08 |
| ACE2 | ALOX12-AS1 | -0,2714 | 2,75762E-08 |
| ACE2 | RP4-718D20.3 | -0,2714 | 2,75146E-08 |
| ACE2 | OR51J1 | -0,2715 | 2,73834E-08 |
| ACE2 | RPL7AP11 | -0,2716 | 2,7041E-08 |
| ACE2 | CFDP1 | -0,2716 | 2,69072E-08 |
| ACE2 | LINC01423 | -0,2716 | 2,68823E-08 |
| ACE2 | RP11-439A17.4 | -0,2717 | 2,65709E-08 |
| ACE2 | COMMD9 | -0,2717 | 2,64014E-08 |
| ACE2 | LINC00561 | -0,2717 | 2,64504E-08 |
| ACE2 | LINC01028 | -0,2717 | 2,64373E-08 |
| ACE2 | MEPE | -0,2718 | 2,63705E-08 |
| ACE2 | DDX21 | -0,2718 | 2,6149E-08 |
| ACE2 | ANGPT4 | -0,2718 | 2,60874E-08 |
| ACE2 | PTGES3P3 | -0,2719 | 2,58446E-08 |
| ACE2 | GTF2IP9 | -0,2719 | 2,58515E-08 |
| ACE2 | CBLN1 | -0,2719 | 2,59134E-08 |
| ACE2 | HMGN1P34 | -0,2719 | 2,59124E-08 |
| ACE2 | PGRMC1 | -0,2719 | 2,59195E-08 |
| ACE2 | KBTBD12 | -0,272 | 2,55288E-08 |
| ACE2 | PUM3 | -0,272 | 2,55473E-08 |
| ACE2 | HSFX1 | -0,272 | 2,54536E-08 |
| ACE2 | LINC01780 | -0,2721 | 2,51932E-08 |
| ACE2 | RP11-2E11.5 | -0,2721 | 2,53091E-08 |
| ACE2 | CDC42P3 | -0,2721 | 2,53298E-08 |
| ACE2 | TRUB1 | -0,2721 | 2,52706E-08 |
| ACE2 | RP4-710M3.2 | -0,2721 | 2,52107E-08 |
| ACE2 | C11orf65 | -0,2721 | 2,5444E-08 |
| ACE2 | CTD-2293H3.2 | -0,2721 | 2,52078E-08 |
| ACE2 | PADI4 | -0,2722 | 2,49234E-08 |

| | | | |
|---|---|---|---|
| ACE2 | RP13-977J11.8 | -0,2722 | 2,50027E-08 |
| ACE2 | NCBP3 | -0,2722 | 2,50603E-08 |
| ACE2 | GGA3 | -0,2722 | 2,50922E-08 |
| ACE2 | JAKMIP2-AS1 | -0,2723 | 2,47038E-08 |
| ACE2 | GOLGA5 | -0,2723 | 2,45701E-08 |
| ACE2 | RP11-315D13.1 | -0,2724 | 2,4244E-08 |
| ACE2 | RP13-895J2.8 | -0,2724 | 2,44941E-08 |
| ACE2 | ZNF429 | -0,2725 | 2,39598E-08 |
| ACE2 | RPL21P5 | -0,2726 | 2,3745E-08 |
| ACE2 | RP11-143N13.2 | -0,2726 | 2,37338E-08 |
| ACE2 | KLHDC9 | -0,2727 | 2,35741E-08 |
| ACE2 | AC012363.13 | -0,2727 | 2,34221E-08 |
| ACE2 | BRD7P4 | -0,2727 | 2,35796E-08 |
| ACE2 | RP11-644C3.1 | -0,2728 | 2,31025E-08 |
| ACE2 | RPS7P6 | -0,2728 | 2,33219E-08 |
| ACE2 | RP11-118B22.4 | -0,2728 | 2,33383E-08 |
| ACE2 | ELOB | -0,2728 | 2,31174E-08 |
| ACE2 | DSC3 | -0,2728 | 2,33275E-08 |
| ACE2 | PROK1 | -0,2729 | 2,27987E-08 |
| ACE2 | RP11-506O24.2 | -0,2729 | 2,30172E-08 |
| ACE2 | SUMO2P1 | -0,2729 | 2,2867E-08 |
| ACE2 | AK3 | -0,2729 | 2,28747E-08 |
| ACE2 | INTS5 | -0,2729 | 2,29621E-08 |
| ACE2 | CAPNS2 | -0,2729 | 2,29305E-08 |
| ACE2 | USP9X | -0,2729 | 2,28969E-08 |
| ACE2 | RP11-716H6.1 | -0,273 | 2,2763E-08 |
| ACE2 | RPL34P33 | -0,273 | 2,26281E-08 |
| ACE2 | RP11-710E1.2 | -0,2732 | 2,20697E-08 |
| ACE2 | AC079610.2 | -0,2733 | 2,17635E-08 |
| ACE2 | RGS14 | -0,2733 | 2,19126E-08 |
| ACE2 | PAXIP1 | -0,2733 | 2,19248E-08 |
| ACE2 | RP11-613F22.8 | -0,2733 | 2,19141E-08 |
| ACE2 | MLXIP | -0,2733 | 2,19002E-08 |
| ACE2 | FAM167A-AS1 | -0,2734 | 2,15565E-08 |
| ACE2 | NADK2-AS1 | -0,2735 | 2,12814E-08 |
| ACE2 | RP3-380B8.4 | -0,2735 | 2,12966E-08 |
| ACE2 | AC008746.12 | -0,2735 | 2,13638E-08 |
| ACE2 | RP11-981G7.3 | -0,2736 | 2,1148E-08 |
| ACE2 | DLEU2 | -0,2736 | 2,10157E-08 |
| ACE2 | RP11-930O11.2 | -0,2736 | 2,1001E-08 |
| ACE2 | RP11-454F8.3 | -0,2736 | 2,10403E-08 |
| ACE2 | MCTS2P | -0,2736 | 2,10001E-08 |
| ACE2 | BCRP3 | -0,2736 | 2,09618E-08 |
| ACE2 | TBC1D10A | -0,2736 | 2,11651E-08 |
| ACE2 | JRK | -0,2737 | 2,07704E-08 |
| ACE2 | AGAP14 | -0,2737 | 2,08009E-08 |
| ACE2 | RP11-815J4.7 | -0,2737 | 2,07051E-08 |
| ACE2 | C1orf234 | -0,2738 | 2,05078E-08 |
| ACE2 | CC2D1B | -0,2738 | 2,06357E-08 |
| ACE2 | DTX3L | -0,2738 | 2,06565E-08 |
| ACE2 | RP11-91K8.5 | -0,2738 | 2,04861E-08 |
| ACE2 | ZNF267 | -0,2738 | 2,06481E-08 |
| ACE2 | CHERP | -0,2738 | 2,05535E-08 |
| ACE2 | SRGAP2-AS1 | -0,2739 | 2,0317E-08 |

| | | | |
|---|---|---|---|
| ACE2 | NUDT9 | -0,2739 | 2,01817E-08 |
| ACE2 | AP000355.2 | -0,2739 | 2,02757E-08 |
| ACE2 | COQ8A | -0,2741 | 1,98607E-08 |
| ACE2 | MYLK4 | -0,2741 | 1,96895E-08 |
| ACE2 | CALCOCO2 | -0,2741 | 1,96786E-08 |
| ACE2 | CTD-2561B21.3 | -0,2741 | 1,99066E-08 |
| ACE2 | RP11-34P13.13 | -0,2743 | 1,92628E-08 |
| ACE2 | RP11-153F1.1 | -0,2743 | 1,92863E-08 |
| ACE2 | RP11-446J8.1 | -0,2743 | 1,93513E-08 |
| ACE2 | ZNF608 | -0,2743 | 1,92454E-08 |
| ACE2 | RP3-394A18.1 | -0,2743 | 1,92904E-08 |
| ACE2 | SGO1-AS1 | -0,2744 | 1,9063E-08 |
| ACE2 | AC024704.2 | -0,2745 | 1,89415E-08 |
| ACE2 | TMEM183AP1 | -0,2745 | 1,8862E-08 |
| ACE2 | GOPC | -0,2745 | 1,87522E-08 |
| ACE2 | CACNA1C-AS3 | -0,2745 | 1,88521E-08 |
| ACE2 | RP11-188D8.1 | -0,2746 | 1,85219E-08 |
| ACE2 | RP3-410C9.2 | -0,2746 | 1,85909E-08 |
| ACE2 | RP11-467H10.1 | -0,2747 | 1,83675E-08 |
| ACE2 | GOLGA6L17P | -0,2747 | 1,84353E-08 |
| ACE2 | RNU1-123P | -0,2747 | 1,828E-08 |
| ACE2 | MYLK-AS2 | -0,2748 | 1,82481E-08 |
| ACE2 | ZNF121 | -0,2748 | 1,80455E-08 |
| ACE2 | ZNF341 | -0,2748 | 1,81667E-08 |
| ACE2 | RP11-241F15.10 | -0,275 | 1,78014E-08 |
| ACE2 | DDX4 | -0,275 | 1,77097E-08 |
| ACE2 | AC092171.4 | -0,275 | 1,77872E-08 |
| ACE2 | PABPN1L | -0,275 | 1,76905E-08 |
| ACE2 | RP11-543H23.1 | -0,275 | 1,77246E-08 |
| ACE2 | ZNF257 | -0,275 | 1,77951E-08 |
| ACE2 | FOLR2 | -0,2751 | 1,74773E-08 |
| ACE2 | SIPA1L1 | -0,2752 | 1,73332E-08 |
| ACE2 | CTA-299D3.8 | -0,2752 | 1,73023E-08 |
| ACE2 | C2 | -0,2753 | 1,71292E-08 |
| ACE2 | MTND3P10 | -0,2754 | 1,69019E-08 |
| ACE2 | LINC01149 | -0,2754 | 1,68147E-08 |
| ACE2 | WDR18 | -0,2754 | 1,6894E-08 |
| ACE2 | GART | -0,2754 | 1,68047E-08 |
| ACE2 | C17orf102 | -0,2755 | 1,65629E-08 |
| ACE2 | HMGN2P17 | -0,2756 | 1,64145E-08 |
| ACE2 | GRID2 | -0,2756 | 1,64418E-08 |
| ACE2 | RP11-430K21.2 | -0,2756 | 1,63938E-08 |
| ACE2 | DOCK11P1 | -0,2756 | 1,65345E-08 |
| ACE2 | RP11-544L8__B.4 | -0,2757 | 1,62681E-08 |
| ACE2 | RP11-47G11.2 | -0,2757 | 1,62377E-08 |
| ACE2 | RP11-111F16.2 | -0,2757 | 1,62033E-08 |
| ACE2 | RP11-700P18.1 | -0,2758 | 1,60172E-08 |
| ACE2 | CCT8L1P | -0,2758 | 1,60306E-08 |
| ACE2 | FAM135B | -0,2758 | 1,59507E-08 |
| ACE2 | MED13 | -0,2758 | 1,60016E-08 |
| ACE2 | RP11-314N13.3 | -0,2758 | 1,60331E-08 |
| ACE2 | SCGB1D2 | -0,2759 | 1,59131E-08 |
| ACE2 | WDR63 | -0,276 | 1,56899E-08 |
| ACE2 | RP11-439E19.6 | -0,276 | 1,56638E-08 |

| | | | |
|---|---|---|---|
| ACE2 | COL6A4P2 | -0,276 | 1,57273E-08 |
| ACE2 | AC007773.2 | -0,276 | 1,57371E-08 |
| ACE2 | CD33 | -0,276 | 1,56327E-08 |
| ACE2 | RN7SL724P | -0,2761 | 1,53732E-08 |
| ACE2 | WDR1 | -0,2761 | 1,55255E-08 |
| ACE2 | RP11-416A17.1 | -0,2761 | 1,55059E-08 |
| ACE2 | DYNLT3P1 | -0,2761 | 1,54038E-08 |
| ACE2 | RP13-228J13.1 | -0,2761 | 1,54591E-08 |
| ACE2 | RP11-521D12.1 | -0,2762 | 1,53196E-08 |
| ACE2 | CH17-140K24.2 | -0,2762 | 1,53523E-08 |
| ACE2 | RCOR2 | -0,2762 | 1,53213E-08 |
| ACE2 | RP11-32B5.1 | -0,2762 | 1,52397E-08 |
| ACE2 | LINC01142 | -0,2763 | 1,51551E-08 |
| ACE2 | SUV39H2 | -0,2763 | 1,51349E-08 |
| ACE2 | COX6CP10 | -0,2765 | 1,46941E-08 |
| ACE2 | RP11-651P23.4 | -0,2765 | 1,4789E-08 |
| ACE2 | INTS9 | -0,2765 | 1,46995E-08 |
| ACE2 | PPP1R26P1 | -0,2765 | 1,46974E-08 |
| ACE2 | RP11-252A24.3 | -0,2765 | 1,4637E-08 |
| ACE2 | WI2-80269A6.1 | -0,2765 | 1,47565E-08 |
| ACE2 | RP11-1319K7.1 | -0,2766 | 1,45994E-08 |
| ACE2 | RPL12P38 | -0,2766 | 1,46082E-08 |
| ACE2 | C21orf62 | -0,2766 | 1,4594E-08 |
| ACE2 | RP11-254F19.4 | -0,2767 | 1,43888E-08 |
| ACE2 | IBSP | -0,2768 | 1,41114E-08 |
| ACE2 | BAG6 | -0,2768 | 1,41951E-08 |
| ACE2 | APEX1 | -0,2768 | 1,42504E-08 |
| ACE2 | SPG7 | -0,2768 | 1,40935E-08 |
| ACE2 | CHRNB1 | -0,2768 | 1,41133E-08 |
| ACE2 | RPL23AP67 | -0,2769 | 1,39691E-08 |
| ACE2 | RP11-50O21.1 | -0,2769 | 1,39283E-08 |
| ACE2 | MIR325HG | -0,2769 | 1,39458E-08 |
| ACE2 | NDUFAF4P3 | -0,277 | 1,38701E-08 |
| ACE2 | AC003986.7 | -0,277 | 1,38685E-08 |
| ACE2 | MPRIP-AS1 | -0,277 | 1,38617E-08 |
| ACE2 | RP11-162A12.2 | -0,277 | 1,37734E-08 |
| ACE2 | KB-1000E4.2 | -0,2771 | 1,36631E-08 |
| ACE2 | RP13-554M15.8 | -0,2771 | 1,36248E-08 |
| ACE2 | GOLGA6L10 | -0,2771 | 1,36375E-08 |
| ACE2 | F13A1 | -0,2772 | 1,35298E-08 |
| ACE2 | RNU6-529P | -0,2773 | 1,32987E-08 |
| ACE2 | SERPINF2 | -0,2773 | 1,33043E-08 |
| ACE2 | LCE5A | -0,2774 | 1,31575E-08 |
| ACE2 | RP1-60O19.2 | -0,2774 | 1,31403E-08 |
| ACE2 | SIDT2 | -0,2774 | 1,31411E-08 |
| ACE2 | HMGB3P24 | -0,2775 | 1,29674E-08 |
| ACE2 | AC001226.7 | -0,2775 | 1,29339E-08 |
| ACE2 | TRAF7 | -0,2775 | 1,29389E-08 |
| ACE2 | CTD-3116E22.7 | -0,2775 | 1,30074E-08 |
| ACE2 | RP11-314B1.2 | -0,2776 | 1,28807E-08 |
| ACE2 | AF129075.5 | -0,2776 | 1,28546E-08 |
| ACE2 | TIFA | -0,2777 | 1,26083E-08 |
| ACE2 | RP11-458J1.1 | -0,2777 | 1,26601E-08 |
| ACE2 | EP300 | -0,2777 | 1,27048E-08 |

| | | | |
|---|---|---|---|
| ACE2 | CLEC5A | -0,2778 | 1,24595E-08 |
| ACE2 | DMTN | -0,2779 | 1,23326E-08 |
| ACE2 | INPP1 | -0,278 | 1,22255E-08 |
| ACE2 | RP11-787I22.3 | -0,278 | 1,21823E-08 |
| ACE2 | LINC02067 | -0,2781 | 1,20037E-08 |
| ACE2 | CTB-78F1.2 | -0,2781 | 1,2015E-08 |
| ACE2 | AC007036.4 | -0,2781 | 1,20329E-08 |
| ACE2 | RP11-452N17.1 | -0,2782 | 1,18847E-08 |
| ACE2 | KSR1P1 | -0,2782 | 1,19062E-08 |
| ACE2 | LAMTOR1 | -0,2783 | 1,17788E-08 |
| ACE2 | CTD-2524L6.3 | -0,2783 | 1,18155E-08 |
| ACE2 | RP11-148B18.1 | -0,2784 | 1,16604E-08 |
| ACE2 | RNU6-26P | -0,2784 | 1,16444E-08 |
| ACE2 | RP11-467I20.2 | -0,2785 | 1,14201E-08 |
| ACE2 | RND1 | -0,2785 | 1,13857E-08 |
| ACE2 | RPL32P34 | -0,2785 | 1,1479E-08 |
| ACE2 | SMG5 | -0,2786 | 1,12862E-08 |
| ACE2 | RP11-3D4.2 | -0,2786 | 1,12522E-08 |
| ACE2 | RP11-420C9.1 | -0,2787 | 1,11259E-08 |
| ACE2 | HMGN2P4 | -0,2787 | 1,1156E-08 |
| ACE2 | CTD-2587M2.1 | -0,2787 | 1,12064E-08 |
| ACE2 | RP3-337H4.6 | -0,2787 | 1,1135E-08 |
| ACE2 | RP11-381O7.3 | -0,2787 | 1,11349E-08 |
| ACE2 | MST1L | -0,2788 | 1,10963E-08 |
| ACE2 | CTA-253N17.1 | -0,2788 | 1,10624E-08 |
| ACE2 | GABRG1 | -0,2788 | 1,10725E-08 |
| ACE2 | NFKB1 | -0,2788 | 1,10085E-08 |
| ACE2 | RBP5 | -0,2788 | 1,09801E-08 |
| ACE2 | CERK | -0,2788 | 1,1017E-08 |
| ACE2 | MIR1244-2 | -0,2789 | 1,09583E-08 |
| ACE2 | RP1-278E11.3 | -0,2789 | 1,0881E-08 |
| ACE2 | RPS3AP29 | -0,2789 | 1,09124E-08 |
| ACE2 | LINC00664 | -0,2789 | 1,09364E-08 |
| ACE2 | KB-176G8.1 | -0,2789 | 1,09443E-08 |
| ACE2 | PLAC4 | -0,2789 | 1,08484E-08 |
| ACE2 | XXbac-B444P24.14 | -0,2789 | 1,09569E-08 |
| ACE2 | RNU1-47P | -0,279 | 1,07554E-08 |
| ACE2 | PTGER4P2 | -0,2791 | 1,06205E-08 |
| ACE2 | FCN2 | -0,2791 | 1,06762E-08 |
| ACE2 | MAGEC3 | -0,2792 | 1,04692E-08 |
| ACE2 | MIOS | -0,2794 | 1,02602E-08 |
| ACE2 | KCTD19 | -0,2794 | 1,02723E-08 |
| ACE2 | ZNF480 | -0,2794 | 1,02709E-08 |
| ACE2 | RP1-45I4.3 | -0,2795 | 1,00903E-08 |
| ACE2 | DDB1 | -0,2795 | 1,0115E-08 |
| ACE2 | C15orf59-AS1 | -0,2795 | 1,00778E-08 |
| ACE2 | RP11-775C24.5 | -0,2795 | 1,00971E-08 |
| ACE2 | RP11-775C24.4 | -0,2795 | 1,00876E-08 |
| ACE2 | TTR | -0,2795 | 1,00665E-08 |
| ACE2 | CTC-559E9.9 | -0,2795 | 1,01155E-08 |
| ACE2 | KCNG1 | -0,2795 | 1,01273E-08 |
| ACE2 | SPSB1 | -0,2796 | 9,99451E-09 |
| ACE2 | APOB | -0,2796 | 9,92271E-09 |
| ACE2 | FAM26D | -0,2796 | 9,93327E-09 |

| | | | |
|---|---|---|---|
| ACE2 | CDK20 | -0,2796 | 9,94129E-09 |
| ACE2 | DGCR8 | -0,2796 | 9,95909E-09 |
| ACE2 | AC068196.1 | -0,2797 | 9,89122E-09 |
| ACE2 | RP5-1049N15.2 | -0,2797 | 9,79619E-09 |
| ACE2 | TAF5 | -0,2797 | 9,81747E-09 |
| ACE2 | CTD-2526A2.2 | -0,2797 | 9,81679E-09 |
| ACE2 | IER3IP1 | -0,2797 | 9,85931E-09 |
| ACE2 | TRAPPC12-AS1 | -0,2798 | 9,7486E-09 |
| ACE2 | AC107079.1 | -0,2798 | 9,75466E-09 |
| ACE2 | USP19 | -0,2798 | 9,68256E-09 |
| ACE2 | C5orf22 | -0,2798 | 9,77835E-09 |
| ACE2 | CTC-448F2.6 | -0,2798 | 9,75503E-09 |
| ACE2 | KCNV2 | -0,2799 | 9,64317E-09 |
| ACE2 | RP11-383M4.6 | -0,2799 | 9,6526E-09 |
| ACE2 | ANP32D | -0,2799 | 9,58449E-09 |
| ACE2 | AC079250.1 | -0,28 | 9,42976E-09 |
| ACE2 | RP11-651P23.5 | -0,28 | 9,47397E-09 |
| ACE2 | RP11-760D2.5 | -0,28 | 9,53162E-09 |
| ACE2 | DNAH10 | -0,28 | 9,52816E-09 |
| ACE2 | RP11-325D8.1 | -0,28 | 9,5267E-09 |
| ACE2 | RP11-680G24.6 | -0,28 | 9,4944E-09 |
| ACE2 | CLEC10A | -0,28 | 9,44326E-09 |
| ACE2 | RP11-94D20.1 | -0,2801 | 9,36056E-09 |
| ACE2 | RP11-274B21.12 | -0,2801 | 9,40588E-09 |
| ACE2 | RP11-320A16.1 | -0,2801 | 9,35471E-09 |
| ACE2 | SEMA4B | -0,2801 | 9,40999E-09 |
| ACE2 | RP11-578O24.2 | -0,2802 | 9,26892E-09 |
| ACE2 | RN7SL405P | -0,2802 | 9,28081E-09 |
| ACE2 | KMT5C | -0,2802 | 9,22225E-09 |
| ACE2 | KLHL9 | -0,2803 | 9,11978E-09 |
| ACE2 | RP11-558A11.2 | -0,2803 | 9,10108E-09 |
| ACE2 | DDX49 | -0,2803 | 9,14984E-09 |
| ACE2 | ANKRD62P1-PARP4F | -0,2803 | 9,18601E-09 |
| ACE2 | AC138035.2 | -0,2804 | 9,03926E-09 |
| ACE2 | BAZ2A | -0,2804 | 8,9685E-09 |
| ACE2 | AC079781.8 | -0,2805 | 8,9133E-09 |
| ACE2 | MDM2 | -0,2805 | 8,86397E-09 |
| ACE2 | ZSCAN29 | -0,2805 | 8,94471E-09 |
| ACE2 | RP5-967N21.11 | -0,2805 | 8,86022E-09 |
| ACE2 | RP11-461L13.3 | -0,2806 | 8,83527E-09 |
| ACE2 | RP11-292B8.2 | -0,2806 | 8,74652E-09 |
| ACE2 | SIGLEC6 | -0,2806 | 8,76182E-09 |
| ACE2 | ZNF672 | -0,2807 | 8,63173E-09 |
| ACE2 | RP11-9N12.3 | -0,2807 | 8,63964E-09 |
| ACE2 | PKD1P5 | -0,2807 | 8,72871E-09 |
| ACE2 | MRPL45 | -0,2807 | 8,64593E-09 |
| ACE2 | NUMB | -0,2808 | 8,54499E-09 |
| ACE2 | L3MBTL3 | -0,2809 | 8,45055E-09 |
| ACE2 | AC104532.4 | -0,2809 | 8,46559E-09 |
| ACE2 | EEF1DP1 | -0,281 | 8,36411E-09 |
| ACE2 | PSAT1 | -0,2811 | 8,25955E-09 |
| ACE2 | RPL10P6 | -0,2812 | 8,11348E-09 |
| ACE2 | DUX4L26 | -0,2812 | 8,20017E-09 |
| ACE2 | RP11-144C15.2 | -0,2812 | 8,14372E-09 |

| | | | |
|---|---|---|---|
| ACE2 | RP11-622C24.1 | -0,2812 | 8,11977E-09 |
| ACE2 | LHFPL1 | -0,2812 | 8,10443E-09 |
| ACE2 | TMEM128 | -0,2813 | 8,03444E-09 |
| ACE2 | RP11-415D17.4 | -0,2813 | 8,06543E-09 |
| ACE2 | CTA-414D7.1 | -0,2813 | 7,99822E-09 |
| ACE2 | PSMC1P1 | -0,2814 | 7,98573E-09 |
| ACE2 | RP11-503C24.2 | -0,2814 | 7,91213E-09 |
| ACE2 | RP11-54O7.17 | -0,2815 | 7,85797E-09 |
| ACE2 | LINC02148 | -0,2815 | 7,83347E-09 |
| ACE2 | RP11-460I13.6 | -0,2816 | 7,75036E-09 |
| ACE2 | RP11-141M3.6 | -0,2816 | 7,79434E-09 |
| ACE2 | RPL36P19 | -0,2816 | 7,79562E-09 |
| ACE2 | COG1 | -0,2816 | 7,75171E-09 |
| ACE2 | WDR43 | -0,2817 | 7,68891E-09 |
| ACE2 | RP11-646I6.5 | -0,2817 | 7,62746E-09 |
| ACE2 | RP11-357D18.1 | -0,2817 | 7,63818E-09 |
| ACE2 | RP11-386G21.2 | -0,2817 | 7,68799E-09 |
| ACE2 | RPS2P7 | -0,2817 | 7,66749E-09 |
| ACE2 | CBLB | -0,2818 | 7,5082E-09 |
| ACE2 | AC006116.21 | -0,2818 | 7,51942E-09 |
| ACE2 | TMEM27 | -0,2818 | 7,5132E-09 |
| ACE2 | KCNJ10 | -0,2819 | 7,44677E-09 |
| ACE2 | RP11-394O4.3 | -0,2819 | 7,42047E-09 |
| ACE2 | KLHDC10 | -0,2819 | 7,46E-09 |
| ACE2 | SSR1 | -0,282 | 7,38615E-09 |
| ACE2 | FAAHP1 | -0,2821 | 7,29407E-09 |
| ACE2 | RAB4A | -0,2821 | 7,25577E-09 |
| ACE2 | RP11-744I24.2 | -0,2821 | 7,2356E-09 |
| ACE2 | ZNF705E | -0,2822 | 7,19526E-09 |
| ACE2 | HMGN2P3 | -0,2822 | 7,2143E-09 |
| ACE2 | AC009120.5 | -0,2822 | 7,19983E-09 |
| ACE2 | RPSAP44 | -0,2823 | 7,08267E-09 |
| ACE2 | MYCBP2-AS2 | -0,2823 | 7,12252E-09 |
| ACE2 | LA16c-3G11.7 | -0,2824 | 6,98644E-09 |
| ACE2 | THAP7-AS1 | -0,2824 | 6,95748E-09 |
| ACE2 | RP11-763B22.4 | -0,2825 | 6,93227E-09 |
| ACE2 | PRSS38 | -0,2825 | 6,90727E-09 |
| ACE2 | PSMD6-AS2 | -0,2825 | 6,94845E-09 |
| ACE2 | RP11-344F13.1 | -0,2827 | 6,76493E-09 |
| ACE2 | RP11-521B24.4 | -0,2827 | 6,76038E-09 |
| ACE2 | KB-208E9.1 | -0,2827 | 6,70321E-09 |
| ACE2 | NFIA-AS1 | -0,2828 | 6,66365E-09 |
| ACE2 | STRIP1 | -0,2828 | 6,63002E-09 |
| ACE2 | METTL18 | -0,2828 | 6,65201E-09 |
| ACE2 | SLC33A1 | -0,2828 | 6,64476E-09 |
| ACE2 | AC073626.2 | -0,2828 | 6,61545E-09 |
| ACE2 | CTD-2319I12.3 | -0,2828 | 6,6413E-09 |
| ACE2 | YWHAZP4 | -0,2829 | 6,55254E-09 |
| ACE2 | CMB9-14B22.1 | -0,2829 | 6,59677E-09 |
| ACE2 | RP11-474C8.7 | -0,2829 | 6,55154E-09 |
| ACE2 | RPS15AP36 | -0,2829 | 6,55027E-09 |
| ACE2 | MYOSLID | -0,2831 | 6,36923E-09 |
| ACE2 | RAD54B | -0,2831 | 6,37565E-09 |
| ACE2 | MTDHP2 | -0,2831 | 6,41929E-09 |

| | | | |
|---|---|---|---|
| ACE2 | ASIP | -0,2831 | 6,37129E-09 |
| ACE2 | RP11-486G15.1 | -0,2832 | 6,30839E-09 |
| ACE2 | MYCN | -0,2832 | 6,35904E-09 |
| ACE2 | SLC13A5 | -0,2832 | 6,33637E-09 |
| ACE2 | AC025811.3 | -0,2832 | 6,34262E-09 |
| ACE2 | RP5-856G1.2 | -0,2834 | 6,13316E-09 |
| ACE2 | RN7SL22P | -0,2834 | 6,15446E-09 |
| ACE2 | RP3-454B23.1 | -0,2834 | 6,13898E-09 |
| ACE2 | KB-1440D3.14 | -0,2835 | 6,06661E-09 |
| ACE2 | GK5 | -0,2836 | 6,00654E-09 |
| ACE2 | RP11-467I17.1 | -0,2836 | 6,00255E-09 |
| ACE2 | PRPF38AP2 | -0,2837 | 5,94954E-09 |
| ACE2 | PGGHG | -0,2837 | 5,94084E-09 |
| ACE2 | FAM32A | -0,2838 | 5,86743E-09 |
| ACE2 | SLC2A7 | -0,284 | 5,69817E-09 |
| ACE2 | LINC01819 | -0,284 | 5,69957E-09 |
| ACE2 | RN7SKP75 | -0,284 | 5,6925E-09 |
| ACE2 | RP11-563N4.1 | -0,2841 | 5,66199E-09 |
| ACE2 | RNF133 | -0,2841 | 5,65174E-09 |
| ACE2 | NUP214 | -0,2841 | 5,60137E-09 |
| ACE2 | CTD-2349P21.5 | -0,2841 | 5,65776E-09 |
| ACE2 | RP1-149C7.1 | -0,2842 | 5,57675E-09 |
| ACE2 | NACAP1 | -0,2842 | 5,5298E-09 |
| ACE2 | RP11-285G1.14 | -0,2842 | 5,54061E-09 |
| ACE2 | ACOT1 | -0,2842 | 5,57085E-09 |
| ACE2 | EFCAB6-AS1 | -0,2842 | 5,56447E-09 |
| ACE2 | NPM1P6 | -0,2843 | 5,47187E-09 |
| ACE2 | FOXD4 | -0,2843 | 5,48695E-09 |
| ACE2 | RP5-872K7.8 | -0,2843 | 5,52174E-09 |
| ACE2 | SDF2 | -0,2844 | 5,43689E-09 |
| ACE2 | PTTG1IP | -0,2844 | 5,40289E-09 |
| ACE2 | RP11-642D21.1 | -0,2845 | 5,37438E-09 |
| ACE2 | ST13P3 | -0,2845 | 5,35305E-09 |
| ACE2 | FGF7P8 | -0,2846 | 5,29175E-09 |
| ACE2 | RP11-1006G14.1 | -0,2846 | 5,30621E-09 |
| ACE2 | AC016629.3 | -0,2846 | 5,26558E-09 |
| ACE2 | PLXNA3 | -0,2847 | 5,18222E-09 |
| ACE2 | CDKL4 | -0,2848 | 5,17468E-09 |
| ACE2 | RP11-402G3.5 | -0,2848 | 5,12844E-09 |
| ACE2 | RP11-420A6.2 | -0,2848 | 5,15181E-09 |
| ACE2 | RP11-372E1.1 | -0,2849 | 5,0768E-09 |
| ACE2 | HBM | -0,2849 | 5,08302E-09 |
| ACE2 | RN7SKP44 | -0,2849 | 5,08327E-09 |
| ACE2 | IL13RA2 | -0,2849 | 5,11393E-09 |
| ACE2 | RP11-206L10.2 | -0,285 | 5,00678E-09 |
| ACE2 | TMED10 | -0,285 | 4,98634E-09 |
| ACE2 | RP11-204L24.2 | -0,2851 | 4,93373E-09 |
| ACE2 | RN7SL75P | -0,2851 | 4,93435E-09 |
| ACE2 | MS4A2 | -0,2851 | 4,97784E-09 |
| ACE2 | RP11-175B9.2 | -0,2852 | 4,91442E-09 |
| ACE2 | RPL29P14 | -0,2852 | 4,88724E-09 |
| ACE2 | PLEC | -0,2852 | 4,88704E-09 |
| ACE2 | ATE1-AS1 | -0,2852 | 4,87159E-09 |
| ACE2 | RP11-407N8.6 | -0,2852 | 4,89143E-09 |

| | | | |
|---|---|---|---|
| ACE2 | RP11-805I24.4 | -0,2852 | 4,91049E-09 |
| ACE2 | SDK2 | -0,2852 | 4,87415E-09 |
| ACE2 | RIN2 | -0,2852 | 4,89656E-09 |
| ACE2 | RP11-546B8.3 | -0,2853 | 4,8376E-09 |
| ACE2 | LY6G5B | -0,2853 | 4,82375E-09 |
| ACE2 | HINFP | -0,2853 | 4,82965E-09 |
| ACE2 | U47924.31 | -0,2853 | 4,8554E-09 |
| ACE2 | RP11-544I20.2 | -0,2853 | 4,83995E-09 |
| ACE2 | CCER2 | -0,2853 | 4,80641E-09 |
| ACE2 | PPP2R1A | -0,2853 | 4,813E-09 |
| ACE2 | GATA1 | -0,2853 | 4,83471E-09 |
| ACE2 | SIKE1 | -0,2855 | 4,69935E-09 |
| ACE2 | LTF | -0,2855 | 4,70973E-09 |
| ACE2 | RP11-355O1.11 | -0,2855 | 4,70252E-09 |
| ACE2 | RP11-521C20.2 | -0,2855 | 4,69708E-09 |
| ACE2 | RP11-62J1.4 | -0,2856 | 4,63796E-09 |
| ACE2 | RP11-96B5.3 | -0,2856 | 4,65109E-09 |
| ACE2 | CTD-2620I22.3 | -0,2857 | 4,57523E-09 |
| ACE2 | KCNS3 | -0,2858 | 4,50234E-09 |
| ACE2 | SAG | -0,2859 | 4,44765E-09 |
| ACE2 | RP11-464D20.2 | -0,2859 | 4,44772E-09 |
| ACE2 | ASAP2 | -0,286 | 4,41208E-09 |
| ACE2 | RP11-756P10.6 | -0,286 | 4,42059E-09 |
| ACE2 | ADD3 | -0,286 | 4,41362E-09 |
| ACE2 | AC006116.15 | -0,286 | 4,41564E-09 |
| ACE2 | RP11-385N23.1 | -0,2862 | 4,32504E-09 |
| ACE2 | IFITM9P | -0,2862 | 4,28205E-09 |
| ACE2 | LINC01550 | -0,2862 | 4,28069E-09 |
| ACE2 | BCS1L | -0,2863 | 4,23297E-09 |
| ACE2 | PEX6 | -0,2863 | 4,22386E-09 |
| ACE2 | PPP6C | -0,2863 | 4,22847E-09 |
| ACE2 | RP11-585P4.5 | -0,2863 | 4,24779E-09 |
| ACE2 | RP11-255G12.2 | -0,2863 | 4,21707E-09 |
| ACE2 | SERPINB11 | -0,2863 | 4,23534E-09 |
| ACE2 | RPL7AP10 | -0,2863 | 4,23892E-09 |
| ACE2 | RP4-794I6.4 | -0,2863 | 4,26474E-09 |
| ACE2 | RP11-274B21.10 | -0,2864 | 4,19052E-09 |
| ACE2 | IFNA21 | -0,2864 | 4,16704E-09 |
| ACE2 | RP11-73M11.3 | -0,2864 | 4,18848E-09 |
| ACE2 | CAPN3 | -0,2864 | 4,19207E-09 |
| ACE2 | OXTR | -0,2865 | 4,1559E-09 |
| ACE2 | RP11-120B7.1 | -0,2865 | 4,13004E-09 |
| ACE2 | KIAA0040 | -0,2866 | 4,09949E-09 |
| ACE2 | ABCB10 | -0,2866 | 4,07399E-09 |
| ACE2 | KIRREL3-AS1 | -0,2866 | 4,0966E-09 |
| ACE2 | ARRDC4 | -0,2866 | 4,07708E-09 |
| ACE2 | UST-AS1 | -0,2867 | 4,01739E-09 |
| ACE2 | RP11-1K3.1 | -0,2867 | 4,0231E-09 |
| ACE2 | CTB-25B13.9 | -0,2867 | 4,01412E-09 |
| ACE2 | NAPA-AS1 | -0,2867 | 4,02856E-09 |
| ACE2 | CTD-2619J13.5 | -0,2867 | 4,01807E-09 |
| ACE2 | HSPA8 | -0,2868 | 3,98494E-09 |
| ACE2 | FAM214A | -0,2868 | 3,95356E-09 |
| ACE2 | ZNF885P | -0,2868 | 3,99832E-09 |

| | | | |
|---|---|---|---|
| ACE2 | NEAT1 | -0,287 | 3,87714E-09 |
| ACE2 | NKPD1 | -0,287 | 3,87933E-09 |
| ACE2 | LINC01284 | -0,287 | 3,8965E-09 |
| ACE2 | PSAPL1 | -0,2871 | 3,8351E-09 |
| ACE2 | PATL1 | -0,2872 | 3,79966E-09 |
| ACE2 | AC137932.6 | -0,2872 | 3,76287E-09 |
| ACE2 | ALPK2 | -0,2872 | 3,78696E-09 |
| ACE2 | PFN1P6 | -0,2873 | 3,74835E-09 |
| ACE2 | P4HA2 | -0,2873 | 3,70709E-09 |
| ACE2 | RBM12B-AS1 | -0,2873 | 3,75187E-09 |
| ACE2 | LINC00539 | -0,2873 | 3,74396E-09 |
| ACE2 | RP11-182J1.5 | -0,2873 | 3,74941E-09 |
| ACE2 | MAP3K2 | -0,2874 | 3,67332E-09 |
| ACE2 | DMXL1 | -0,2874 | 3,66871E-09 |
| ACE2 | RP4-725G10.4 | -0,2874 | 3,69409E-09 |
| ACE2 | WASHC5 | -0,2874 | 3,68899E-09 |
| ACE2 | HEXIM2 | -0,2874 | 3,65821E-09 |
| ACE2 | CSNK1D | -0,2874 | 3,68672E-09 |
| ACE2 | AC010468.1 | -0,2875 | 3,61901E-09 |
| ACE2 | RP11-1167A19.6 | -0,2875 | 3,62888E-09 |
| ACE2 | GSPT1 | -0,2875 | 3,64944E-09 |
| ACE2 | RP11-196G11.3 | -0,2875 | 3,63481E-09 |
| ACE2 | RP11-927P21.1 | -0,2875 | 3,63944E-09 |
| ACE2 | KIRREL2 | -0,2876 | 3,58398E-09 |
| ACE2 | CYP4A22 | -0,2877 | 3,54541E-09 |
| ACE2 | RP4-537K23.4 | -0,2877 | 3,54596E-09 |
| ACE2 | RP11-803D5.1 | -0,2878 | 3,49041E-09 |
| ACE2 | OR7E89P | -0,2878 | 3,4835E-09 |
| ACE2 | LA16c-380A1.1 | -0,2878 | 3,50293E-09 |
| ACE2 | ENPP6 | -0,2879 | 3,46827E-09 |
| ACE2 | TUNAR | -0,2879 | 3,46685E-09 |
| ACE2 | MYH2 | -0,2879 | 3,42965E-09 |
| ACE2 | RP11-613M5.2 | -0,288 | 3,41519E-09 |
| ACE2 | PHF23 | -0,288 | 3,4132E-09 |
| ACE2 | RPL7AP26 | -0,2881 | 3,36022E-09 |
| ACE2 | POU3F2 | -0,2881 | 3,37886E-09 |
| ACE2 | RP11-49O14.2 | -0,2881 | 3,37787E-09 |
| ACE2 | KRBA2 | -0,2881 | 3,38117E-09 |
| ACE2 | SLC25A23 | -0,2881 | 3,355E-09 |
| ACE2 | COX10-AS1 | -0,2882 | 3,31267E-09 |
| ACE2 | RPL3P2 | -0,2883 | 3,25645E-09 |
| ACE2 | PAPPA-AS2 | -0,2883 | 3,25724E-09 |
| ACE2 | SPTLC1P1 | -0,2883 | 3,25459E-09 |
| ACE2 | RP11-672A2.6 | -0,2883 | 3,28895E-09 |
| ACE2 | UBQLN4P1 | -0,2884 | 3,24713E-09 |
| ACE2 | RP11-103C16.2 | -0,2884 | 3,23817E-09 |
| ACE2 | PIGCP1 | -0,2884 | 3,23396E-09 |
| ACE2 | FMNL1 | -0,2884 | 3,21662E-09 |
| ACE2 | C5AR1 | -0,2884 | 3,2444E-09 |
| ACE2 | RP11-454F8.2 | -0,2885 | 3,19577E-09 |
| ACE2 | IER2 | -0,2885 | 3,17267E-09 |
| ACE2 | RHD | -0,2886 | 3,15499E-09 |
| ACE2 | ASB6 | -0,2886 | 3,15074E-09 |
| ACE2 | RP5-1096D14.6 | -0,2886 | 3,12979E-09 |

| | | | |
|---|---|---|---|
| ACE2 | RERG-AS1 | -0,2886 | 3,16161E-09 |
| ACE2 | SDC4 | -0,2886 | 3,12863E-09 |
| ACE2 | TCF7L1-IT1 | -0,2888 | 3,08114E-09 |
| ACE2 | RP11-632C17__A.1 | -0,2888 | 3,07422E-09 |
| ACE2 | AC000123.3 | -0,2888 | 3,05963E-09 |
| ACE2 | LINC00865 | -0,2888 | 3,06451E-09 |
| ACE2 | RP11-360N9.2 | -0,2888 | 3,08156E-09 |
| ACE2 | RP11-437J2.3 | -0,2889 | 3,02669E-09 |
| ACE2 | RALGAPA1 | -0,2889 | 3,03502E-09 |
| ACE2 | ZNF688 | -0,289 | 2,98456E-09 |
| ACE2 | RP5-1027G4.3 | -0,289 | 2,98845E-09 |
| ACE2 | ARF4P2 | -0,289 | 3,0024E-09 |
| ACE2 | AC114877.3 | -0,2891 | 2,9578E-09 |
| ACE2 | RPL7AP49 | -0,2891 | 2,9522E-09 |
| ACE2 | RP11-326C3.14 | -0,2891 | 2,93599E-09 |
| ACE2 | RP1-29C18.8 | -0,2891 | 2,96391E-09 |
| ACE2 | RP11-650J17.1 | -0,2892 | 2,92329E-09 |
| ACE2 | RP11-218M11.1 | -0,2892 | 2,91074E-09 |
| ACE2 | LINC01634 | -0,2892 | 2,92168E-09 |
| ACE2 | RP11-452L6.1 | -0,2893 | 2,87476E-09 |
| ACE2 | CTD-2062F14.3 | -0,2894 | 2,82785E-09 |
| ACE2 | CTD-2026J24.1 | -0,2895 | 2,8143E-09 |
| ACE2 | LINC01159 | -0,2896 | 2,76426E-09 |
| ACE2 | CTD-3224K15.3 | -0,2896 | 2,76634E-09 |
| ACE2 | DBNDD1 | -0,2896 | 2,77931E-09 |
| ACE2 | LINC01678 | -0,2896 | 2,77628E-09 |
| ACE2 | TBL1XR1-AS1 | -0,2897 | 2,71849E-09 |
| ACE2 | RP11-1101K5.1 | -0,2897 | 2,72401E-09 |
| ACE2 | LINC01141 | -0,2898 | 2,69811E-09 |
| ACE2 | AC062017.1 | -0,2898 | 2,69263E-09 |
| ACE2 | ATP2C1 | -0,2898 | 2,70686E-09 |
| ACE2 | RP3-476K8.3 | -0,2899 | 2,65848E-09 |
| ACE2 | RP5-1198O20.5 | -0,2899 | 2,64227E-09 |
| ACE2 | RP11-807H7.2 | -0,2899 | 2,64808E-09 |
| ACE2 | CH17-302M23.1 | -0,2899 | 2,65553E-09 |
| ACE2 | RNVU1-3 | -0,29 | 2,6075E-09 |
| ACE2 | FLG2 | -0,29 | 2,63204E-09 |
| ACE2 | KHDC1L | -0,29 | 2,61098E-09 |
| ACE2 | RP11-441F2.4 | -0,29 | 2,63157E-09 |
| ACE2 | RP11-1055B8.9 | -0,29 | 2,63019E-09 |
| ACE2 | CIAO1 | -0,2901 | 2,5779E-09 |
| ACE2 | CATIP | -0,2901 | 2,5793E-09 |
| ACE2 | LINC01094 | -0,2901 | 2,59756E-09 |
| ACE2 | RP11-452G18.2 | -0,2901 | 2,57335E-09 |
| ACE2 | SOX21-AS1 | -0,2901 | 2,56979E-09 |
| ACE2 | RP11-932O9.8 | -0,2901 | 2,57384E-09 |
| ACE2 | EEF1A1P22 | -0,2901 | 2,59614E-09 |
| ACE2 | APC | -0,2902 | 2,55427E-09 |
| ACE2 | EIF3LP2 | -0,2902 | 2,53905E-09 |
| ACE2 | CTB-96E2.6 | -0,2902 | 2,54348E-09 |
| ACE2 | NHLH2 | -0,2903 | 2,50824E-09 |
| ACE2 | RP11-707M1.1 | -0,2903 | 2,53432E-09 |
| ACE2 | RP11-629N8.3 | -0,2903 | 2,50828E-09 |
| ACE2 | RP11-171I2.4 | -0,2904 | 2,49127E-09 |

| | | | |
|---|---|---|---|
| ACE2 | MAFIP | -0,2904 | 2,48815E-09 |
| ACE2 | RPE65 | -0,2905 | 2,46025E-09 |
| ACE2 | HLA-DRB1 | -0,2905 | 2,46441E-09 |
| ACE2 | UBAC2-AS1 | -0,2906 | 2,4378E-09 |
| ACE2 | RP5-1024N4.6 | -0,2907 | 2,39093E-09 |
| ACE2 | HNRNPA1P10 | -0,2907 | 2,38869E-09 |
| ACE2 | RABGAP1L | -0,2908 | 2,35712E-09 |
| ACE2 | RPL35P1 | -0,2908 | 2,35881E-09 |
| ACE2 | ZNF385D-AS1 | -0,2908 | 2,3711E-09 |
| ACE2 | LINC00707 | -0,2908 | 2,358E-09 |
| ACE2 | RN7SL399P | -0,2908 | 2,34635E-09 |
| ACE2 | KLK14 | -0,2908 | 2,36817E-09 |
| ACE2 | RN7SKP80 | -0,2909 | 2,33037E-09 |
| ACE2 | ZC3H11A | -0,291 | 2,30651E-09 |
| ACE2 | AC090018.1 | -0,291 | 2,28543E-09 |
| ACE2 | RP5-1186P10.2 | -0,291 | 2,29334E-09 |
| ACE2 | TRPV5 | -0,291 | 2,29779E-09 |
| ACE2 | RP11-110G21.1 | -0,291 | 2,30772E-09 |
| ACE2 | PTF1A | -0,2911 | 2,26446E-09 |
| ACE2 | NTM-AS1 | -0,2911 | 2,27015E-09 |
| ACE2 | RP11-923I11.5 | -0,2911 | 2,2776E-09 |
| ACE2 | RN7SL213P | -0,2911 | 2,2718E-09 |
| ACE2 | RP1 | -0,2912 | 2,2382E-09 |
| ACE2 | RP11-264B14.1 | -0,2912 | 2,23422E-09 |
| ACE2 | AC098820.3 | -0,2913 | 2,2009E-09 |
| ACE2 | ULBP1 | -0,2913 | 2,20547E-09 |
| ACE2 | RP11-642D21.2 | -0,2913 | 2,20364E-09 |
| ACE2 | RAET1E | -0,2914 | 2,18167E-09 |
| ACE2 | AC010976.2 | -0,2915 | 2,1576E-09 |
| ACE2 | RPL10AP2 | -0,2915 | 2,15617E-09 |
| ACE2 | GSPT2 | -0,2915 | 2,14226E-09 |
| ACE2 | CT45A11P | -0,2915 | 2,16517E-09 |
| ACE2 | SNRPF | -0,2916 | 2,12548E-09 |
| ACE2 | RP11-47I22.1 | -0,2916 | 2,11476E-09 |
| ACE2 | AC139099.5 | -0,2916 | 2,11592E-09 |
| ACE2 | CTD-3105H18.4 | -0,2916 | 2,1155E-09 |
| ACE2 | ZNF765 | -0,2916 | 2,12141E-09 |
| ACE2 | TMA7 | -0,2918 | 2,07978E-09 |
| ACE2 | RP11-115D19.2 | -0,2918 | 2,06993E-09 |
| ACE2 | ARMC4P1 | -0,2918 | 2,06696E-09 |
| ACE2 | NINJ2 | -0,2918 | 2,08015E-09 |
| ACE2 | TBX5 | -0,2918 | 2,06632E-09 |
| ACE2 | TPTE2 | -0,2918 | 2,06442E-09 |
| ACE2 | RP11-305O4.3 | -0,2919 | 2,04186E-09 |
| ACE2 | XXbac-BPG55C20.7 | -0,2919 | 2,05016E-09 |
| ACE2 | RORB-AS1 | -0,2919 | 2,03763E-09 |
| ACE2 | RP11-327F22.4 | -0,2919 | 2,03512E-09 |
| ACE2 | EIF1AY | -0,2919 | 2,04714E-09 |
| ACE2 | RP3-426I6.5 | -0,292 | 2,00608E-09 |
| ACE2 | CTB-33O18.1 | -0,292 | 2,02119E-09 |
| ACE2 | OPRM1 | -0,292 | 2,01509E-09 |
| ACE2 | TNK2-AS1 | -0,2921 | 1,99092E-09 |
| ACE2 | NACA3P | -0,2921 | 1,99456E-09 |
| ACE2 | RP3-393E18.2 | -0,2921 | 1,97787E-09 |

| | | | |
|---|---|---|---|
| ACE2 | CAMKK2 | -0,2921 | 1,98684E-09 |
| ACE2 | RP11-90O23.1 | -0,2922 | 1,9607E-09 |
| ACE2 | RP11-64D22.1 | -0,2922 | 1,9485E-09 |
| ACE2 | CTD-2270L9.2 | -0,2922 | 1,95009E-09 |
| ACE2 | RP11-116O18.1 | -0,2922 | 1,95754E-09 |
| ACE2 | RP11-269M20.3 | -0,2923 | 1,93909E-09 |
| ACE2 | RP11-532F6.5 | -0,2923 | 1,94242E-09 |
| ACE2 | TARBP2 | -0,2923 | 1,92997E-09 |
| ACE2 | UBE2J2 | -0,2924 | 1,90513E-09 |
| ACE2 | WARS2-IT1 | -0,2924 | 1,91439E-09 |
| ACE2 | AC079922.2 | -0,2924 | 1,90108E-09 |
| ACE2 | RP11-109A6.3 | -0,2924 | 1,91281E-09 |
| ACE2 | RPL23AP65 | -0,2924 | 1,90155E-09 |
| ACE2 | RPL12P7 | -0,2924 | 1,8994E-09 |
| ACE2 | SMAD6 | -0,2924 | 1,90009E-09 |
| ACE2 | SMG1 | -0,2924 | 1,90974E-09 |
| ACE2 | GMEB1 | -0,2925 | 1,88647E-09 |
| ACE2 | NFE4 | -0,2925 | 1,88751E-09 |
| ACE2 | BRF2 | -0,2925 | 1,89628E-09 |
| ACE2 | KRT8P15 | -0,2926 | 1,85241E-09 |
| ACE2 | MRPL1 | -0,2926 | 1,8714E-09 |
| ACE2 | RP11-477D19.2 | -0,2926 | 1,84809E-09 |
| ACE2 | RP11-33N14.3 | -0,2926 | 1,86686E-09 |
| ACE2 | RP1-257A7.4 | -0,2927 | 1,83732E-09 |
| ACE2 | LPAR6 | -0,2927 | 1,83504E-09 |
| ACE2 | GUCY2F | -0,2927 | 1,84057E-09 |
| ACE2 | CXorf40B | -0,2927 | 1,84418E-09 |
| ACE2 | RP11-328C8.4 | -0,2928 | 1,81971E-09 |
| ACE2 | SERTAD1 | -0,2928 | 1,81648E-09 |
| ACE2 | PTCHD3P2 | -0,2929 | 1,79732E-09 |
| ACE2 | FASTKD2 | -0,2929 | 1,79324E-09 |
| ACE2 | RANBP6 | -0,2929 | 1,79671E-09 |
| ACE2 | PSMA3 | -0,2929 | 1,78786E-09 |
| ACE2 | RP13-395E19.2 | -0,2929 | 1,78336E-09 |
| ACE2 | SNORD91A | -0,2929 | 1,79449E-09 |
| ACE2 | DDRGK1 | -0,2929 | 1,77786E-09 |
| ACE2 | RP11-453E17.4 | -0,293 | 1,77143E-09 |
| ACE2 | RP11-317B17.4 | -0,293 | 1,7727E-09 |
| ACE2 | LRRC37A6P | -0,293 | 1,75424E-09 |
| ACE2 | RP11-578F21.2 | -0,293 | 1,77052E-09 |
| ACE2 | SCAMP1-AS1 | -0,2932 | 1,71115E-09 |
| ACE2 | AC005324.6 | -0,2932 | 1,70747E-09 |
| ACE2 | EIF4EBP2 | -0,2933 | 1,69276E-09 |
| ACE2 | TNNI2 | -0,2933 | 1,70225E-09 |
| ACE2 | RP11-135J2.3 | -0,2934 | 1,68145E-09 |
| ACE2 | EPB41L5 | -0,2934 | 1,67849E-09 |
| ACE2 | AC010900.2 | -0,2934 | 1,68341E-09 |
| ACE2 | RP11-404G16.2 | -0,2934 | 1,67078E-09 |
| ACE2 | ANKRA2 | -0,2934 | 1,68043E-09 |
| ACE2 | ANKHD1 | -0,2934 | 1,68127E-09 |
| ACE2 | RP11-718B12.2 | -0,2934 | 1,6822E-09 |
| ACE2 | KRR1P1 | -0,2934 | 1,6822E-09 |
| ACE2 | GFAP | -0,2934 | 1,67976E-09 |
| ACE2 | RP11-120M18.5 | -0,2934 | 1,67074E-09 |

| | | | |
|---|---|---|---|
| ACE2 | CTD-2332E11.2 | -0,2935 | 1,65362E-09 |
| ACE2 | ZC3H4 | -0,2935 | 1,64589E-09 |
| ACE2 | LINC00284 | -0,2936 | 1,63543E-09 |
| ACE2 | RP11-403A3.1 | -0,2936 | 1,63331E-09 |
| ACE2 | VWFP1 | -0,2936 | 1,63931E-09 |
| ACE2 | RP11-364L4.1 | -0,2937 | 1,60689E-09 |
| ACE2 | HAUS4 | -0,2937 | 1,59899E-09 |
| ACE2 | RPS3AP33 | -0,2938 | 1,59021E-09 |
| ACE2 | RP11-53I6.4 | -0,2938 | 1,57754E-09 |
| ACE2 | AC073464.4 | -0,2939 | 1,55672E-09 |
| ACE2 | RP13-884E18.2 | -0,294 | 1,53874E-09 |
| ACE2 | RP11-289F5.1 | -0,294 | 1,5534E-09 |
| ACE2 | KB-68A7.2 | -0,294 | 1,54653E-09 |
| ACE2 | AGT | -0,2941 | 1,51441E-09 |
| ACE2 | DPYD-IT1 | -0,2942 | 1,51156E-09 |
| ACE2 | C5orf67 | -0,2942 | 1,49713E-09 |
| ACE2 | RP11-354E11.2 | -0,2942 | 1,50979E-09 |
| ACE2 | RP11-493L12.7 | -0,2943 | 1,47515E-09 |
| ACE2 | RP11-352G18.2 | -0,2943 | 1,48504E-09 |
| ACE2 | CSTF1 | -0,2943 | 1,4902E-09 |
| ACE2 | RPSA | -0,2944 | 1,47335E-09 |
| ACE2 | HECW1 | -0,2945 | 1,45275E-09 |
| ACE2 | RNA5SP449 | -0,2945 | 1,4499E-09 |
| ACE2 | HERC3 | -0,2946 | 1,43361E-09 |
| ACE2 | ACTL6A | -0,2947 | 1,4015E-09 |
| ACE2 | SLC18A2 | -0,2947 | 1,40961E-09 |
| ACE2 | RP11-617B3.2 | -0,2947 | 1,40178E-09 |
| ACE2 | RNF34 | -0,2947 | 1,41474E-09 |
| ACE2 | KLHDC4 | -0,2947 | 1,40735E-09 |
| ACE2 | RP11-315D16.4 | -0,2948 | 1,38746E-09 |
| ACE2 | IL10RB-AS1 | -0,2948 | 1,38261E-09 |
| ACE2 | C1QL2 | -0,295 | 1,35354E-09 |
| ACE2 | TBC1D31 | -0,295 | 1,35111E-09 |
| ACE2 | RSPRY1 | -0,2951 | 1,33975E-09 |
| ACE2 | RNU6-510P | -0,2952 | 1,31468E-09 |
| ACE2 | RP11-810P8.1 | -0,2952 | 1,31902E-09 |
| ACE2 | APIP | -0,2952 | 1,31611E-09 |
| ACE2 | ZNF682 | -0,2952 | 1,32212E-09 |
| ACE2 | SLC16A8 | -0,2952 | 1,30796E-09 |
| ACE2 | CCDC163 | -0,2953 | 1,29861E-09 |
| ACE2 | PSTK | -0,2953 | 1,30639E-09 |
| ACE2 | IPO4 | -0,2953 | 1,29722E-09 |
| ACE2 | RP11-288I21.1 | -0,2954 | 1,28208E-09 |
| ACE2 | RP11-71E19.2 | -0,2954 | 1,27317E-09 |
| ACE2 | IL5 | -0,2954 | 1,28519E-09 |
| ACE2 | PARG | -0,2954 | 1,28049E-09 |
| ACE2 | ISPD-AS1 | -0,2955 | 1,26421E-09 |
| ACE2 | AC093702.1 | -0,2956 | 1,24393E-09 |
| ACE2 | CTC-501O10.1 | -0,2956 | 1,23919E-09 |
| ACE2 | ARL17A | -0,2956 | 1,23917E-09 |
| ACE2 | LL22NC03-2H8.4 | -0,2956 | 1,24307E-09 |
| ACE2 | MIR4458HG | -0,2957 | 1,22791E-09 |
| ACE2 | RP11-358B23.6 | -0,2957 | 1,23081E-09 |
| ACE2 | RP11-744I24.3 | -0,2958 | 1,21555E-09 |

| | | | |
|---|---|---|---|
| ACE2 | FAM86B3P | -0,2958 | 1,21845E-09 |
| ACE2 | KBTBD4 | -0,2958 | 1,21121E-09 |
| ACE2 | RPL12P10 | -0,2959 | 1,19965E-09 |
| ACE2 | CCL26 | -0,2959 | 1,20065E-09 |
| ACE2 | TMEM243 | -0,2959 | 1,20181E-09 |
| ACE2 | RNU6-301P | -0,2959 | 1,20287E-09 |
| ACE2 | DGUOK | -0,296 | 1,17406E-09 |
| ACE2 | SPDL1 | -0,296 | 1,18001E-09 |
| ACE2 | AK3P2 | -0,296 | 1,18537E-09 |
| ACE2 | CTD-2562J15.6 | -0,296 | 1,18066E-09 |
| ACE2 | UQCRBP1 | -0,2961 | 1,1665E-09 |
| ACE2 | CES5AP1 | -0,2962 | 1,14471E-09 |
| ACE2 | SLC35B2 | -0,2963 | 1,13006E-09 |
| ACE2 | AJ011931.1 | -0,2963 | 1,13447E-09 |
| ACE2 | RP11-236P24.3 | -0,2965 | 1,10727E-09 |
| ACE2 | PBDC1 | -0,2965 | 1,10698E-09 |
| ACE2 | RP11-724N1.1 | -0,2966 | 1,08499E-09 |
| ACE2 | RNASEH2B-AS1 | -0,2966 | 1,08646E-09 |
| ACE2 | AC097724.3 | -0,2967 | 1,07273E-09 |
| ACE2 | AC026150.8 | -0,2967 | 1,07301E-09 |
| ACE2 | NUTF2P6 | -0,2967 | 1,08048E-09 |
| ACE2 | NAA35 | -0,2968 | 1,06661E-09 |
| ACE2 | RP3-323P24.3 | -0,2968 | 1,06056E-09 |
| ACE2 | ACSL1 | -0,2969 | 1,05403E-09 |
| ACE2 | TRIM17 | -0,2971 | 1,01898E-09 |
| ACE2 | RRP9 | -0,2972 | 1,00945E-09 |
| ACE2 | HSP90AA5P | -0,2972 | 1,0081E-09 |
| ACE2 | RP11-23F23.2 | -0,2973 | 9,93756E-10 |
| ACE2 | RP11-111M22.5 | -0,2973 | 9,96343E-10 |
| ACE2 | CLEC12B | -0,2973 | 9,86413E-10 |
| ACE2 | RN7SL15P | -0,2973 | 9,98233E-10 |
| ACE2 | RP11-73M7.9 | -0,2974 | 9,75806E-10 |
| ACE2 | LLNLR-245B6.1 | -0,2974 | 9,79641E-10 |
| ACE2 | MIR34AHG | -0,2975 | 9,5933E-10 |
| ACE2 | ADARB2-AS1 | -0,2975 | 9,64677E-10 |
| ACE2 | FAM222A-AS1 | -0,2975 | 9,59588E-10 |
| ACE2 | ARMC5 | -0,2975 | 9,71732E-10 |
| ACE2 | CGB7 | -0,2975 | 9,59389E-10 |
| ACE2 | RP3-426I6.6 | -0,2976 | 9,53094E-10 |
| ACE2 | EIF4G1 | -0,2976 | 9,48703E-10 |
| ACE2 | CCNC | -0,2976 | 9,51053E-10 |
| ACE2 | INPP4B | -0,2977 | 9,36209E-10 |
| ACE2 | GIMAP6 | -0,2977 | 9,37644E-10 |
| ACE2 | DOCK1 | -0,2977 | 9,40165E-10 |
| ACE2 | DPPA3P3 | -0,2977 | 9,41631E-10 |
| ACE2 | RP11-96D1.8 | -0,2977 | 9,3853E-10 |
| ACE2 | RP11-704J17.5 | -0,2978 | 9,23485E-10 |
| ACE2 | SPECC1L | -0,2978 | 9,32295E-10 |
| ACE2 | RP11-60A8.1 | -0,298 | 9,0238E-10 |
| ACE2 | RNF213 | -0,298 | 8,98418E-10 |
| ACE2 | LINC01730 | -0,298 | 9,07367E-10 |
| ACE2 | ZNF800 | -0,2981 | 8,93388E-10 |
| ACE2 | LA16c-425C2.1 | -0,2981 | 8,95911E-10 |
| ACE2 | FABP7P1 | -0,2982 | 8,78655E-10 |

| | | | |
|---|---|---|---|
| ACE2 | COX7C | -0,2982 | 8,73336E-10 |
| ACE2 | RP11-295H24.5 | -0,2982 | 8,7942E-10 |
| ACE2 | VSIR | -0,2983 | 8,71568E-10 |
| ACE2 | RP11-173M1.5 | -0,2983 | 8,69699E-10 |
| ACE2 | CCDC97 | -0,2983 | 8,61782E-10 |
| ACE2 | RP11-2L8.1 | -0,2984 | 8,50879E-10 |
| ACE2 | RP1-178F10.1 | -0,2984 | 8,50195E-10 |
| ACE2 | RP1-199J3.7 | -0,2985 | 8,3984E-10 |
| ACE2 | MFAP1 | -0,2985 | 8,40371E-10 |
| ACE2 | AC012506.4 | -0,2986 | 8,27826E-10 |
| ACE2 | LURAP1L-AS1 | -0,2986 | 8,31644E-10 |
| ACE2 | AC006019.3 | -0,2987 | 8,15542E-10 |
| ACE2 | RP11-261P13.5 | -0,2987 | 8,20538E-10 |
| ACE2 | TMCC1-AS1 | -0,2988 | 8,14743E-10 |
| ACE2 | NUDT19P5 | -0,2988 | 8,14991E-10 |
| ACE2 | MKNK2 | -0,2988 | 8,1322E-10 |
| ACE2 | RP11-517B11.4 | -0,2989 | 7,95978E-10 |
| ACE2 | THEM7P | -0,2989 | 7,94541E-10 |
| ACE2 | PYDC1 | -0,299 | 7,8678E-10 |
| ACE2 | PRPF40A | -0,2991 | 7,81656E-10 |
| ACE2 | RP11-157J24.1 | -0,2992 | 7,67026E-10 |
| ACE2 | RP11-133N21.7 | -0,2992 | 7,65686E-10 |
| ACE2 | RN7SL568P | -0,2992 | 7,70427E-10 |
| ACE2 | DFFB | -0,2994 | 7,43399E-10 |
| ACE2 | VARS2 | -0,2994 | 7,44812E-10 |
| ACE2 | SMARCE1P1 | -0,2994 | 7,45779E-10 |
| ACE2 | PHBP13 | -0,2994 | 7,41624E-10 |
| ACE2 | DKFZP434H168 | -0,2994 | 7,47913E-10 |
| ACE2 | AC015922.7 | -0,2994 | 7,50317E-10 |
| ACE2 | AMD1P3 | -0,2996 | 7,23247E-10 |
| ACE2 | PKIA-AS1 | -0,2996 | 7,29399E-10 |
| ACE2 | RP11-1110F20.1 | -0,2996 | 7,226E-10 |
| ACE2 | RP4-761J14.8 | -0,2996 | 7,2887E-10 |
| ACE2 | RP11-553P9.1 | -0,2997 | 7,15378E-10 |
| ACE2 | TNFRSF10C | -0,2997 | 7,17498E-10 |
| ACE2 | CHUK | -0,2997 | 7,16994E-10 |
| ACE2 | RP11-175G14.2 | -0,2998 | 7,05185E-10 |
| ACE2 | RP11-157E16.1 | -0,2998 | 7,0241E-10 |
| ACE2 | SLC24A1 | -0,2998 | 7,02437E-10 |
| ACE2 | RP5-858B6.3 | -0,2999 | 6,99584E-10 |
| ACE2 | WDR89 | -0,2999 | 6,95241E-10 |
| ACE2 | CTD-2012K14.7 | -0,2999 | 6,97162E-10 |
| ACE2 | RPS7P11 | -0,2999 | 6,92692E-10 |
| ACE2 | RP11-27I1.6 | -0,3 | 6,92107E-10 |
| ACE2 | DAB2IP | -0,3 | 6,85735E-10 |
| ACE2 | RP11-85A1.3 | -0,3 | 6,87639E-10 |
| ACE2 | ISG20L2 | -0,3001 | 6,76773E-10 |
| ACE2 | BUB3 | -0,3001 | 6,75755E-10 |
| ACE2 | TRAM1 | -0,3002 | 6,68747E-10 |
| ACE2 | ST13P18 | -0,3002 | 6,72566E-10 |
| ACE2 | RP11-342M1.3 | -0,3003 | 6,60634E-10 |
| ACE2 | AC091801.1 | -0,3003 | 6,64676E-10 |
| ACE2 | RP11-290L7.5 | -0,3003 | 6,61346E-10 |
| ACE2 | OR3A2 | -0,3003 | 6,62014E-10 |

| | | | |
|---|---|---|---|
| ACE2 | STS | -0,3003 | 6,61046E-10 |
| ACE2 | NTAN1P2 | -0,3004 | 6,54247E-10 |
| ACE2 | RP11-84G21.1 | -0,3004 | 6,55184E-10 |
| ACE2 | RP1L1 | -0,3005 | 6,42695E-10 |
| ACE2 | LINC00601 | -0,3005 | 6,44088E-10 |
| ACE2 | ARL5C | -0,3005 | 6,40456E-10 |
| ACE2 | ENTPD3-AS1 | -0,3006 | 6,31796E-10 |
| ACE2 | RP11-33M22.2 | -0,3007 | 6,25918E-10 |
| ACE2 | RP11-662I13.3 | -0,3007 | 6,22394E-10 |
| ACE2 | GPI | -0,3007 | 6,23018E-10 |
| ACE2 | RP11-657O9.1 | -0,3008 | 6,17786E-10 |
| ACE2 | AC012358.4 | -0,3009 | 6,06151E-10 |
| ACE2 | RP11-136C24.1 | -0,301 | 5,95956E-10 |
| ACE2 | OCIAD1 | -0,301 | 5,97167E-10 |
| ACE2 | MKRN3-AS1 | -0,301 | 5,99831E-10 |
| ACE2 | KB-1958F4.2 | -0,3011 | 5,9573E-10 |
| ACE2 | CTC-297N7.7 | -0,3011 | 5,90135E-10 |
| ACE2 | ITPK1 | -0,3012 | 5,83834E-10 |
| ACE2 | NT5C1A | -0,3013 | 5,73135E-10 |
| ACE2 | GLUD1P8 | -0,3013 | 5,76654E-10 |
| ACE2 | C11orf21 | -0,3013 | 5,74344E-10 |
| ACE2 | TCIRG1 | -0,3013 | 5,75628E-10 |
| ACE2 | ZBTB16 | -0,3013 | 5,77247E-10 |
| ACE2 | GOLGA8O | -0,3013 | 5,7385E-10 |
| ACE2 | CPAMD8 | -0,3013 | 5,77626E-10 |
| ACE2 | EPYC | -0,3014 | 5,69402E-10 |
| ACE2 | SPATA33 | -0,3014 | 5,71916E-10 |
| ACE2 | EMC6 | -0,3014 | 5,69689E-10 |
| ACE2 | VPS36 | -0,3016 | 5,53482E-10 |
| ACE2 | RP11-349H17.2 | -0,3016 | 5,55846E-10 |
| ACE2 | RPL18AP7 | -0,3017 | 5,48468E-10 |
| ACE2 | CTC-339F2.2 | -0,3017 | 5,48803E-10 |
| ACE2 | RP11-524F11.1 | -0,3017 | 5,48001E-10 |
| ACE2 | MYLKP1 | -0,3018 | 5,3685E-10 |
| ACE2 | CTD-2007L18.5 | -0,3018 | 5,40307E-10 |
| ACE2 | PXN-AS1 | -0,3018 | 5,38821E-10 |
| ACE2 | AC006145.4 | -0,3019 | 5,27844E-10 |
| ACE2 | ACTBP7 | -0,3019 | 5,27346E-10 |
| ACE2 | RNU6-30P | -0,3019 | 5,29443E-10 |
| ACE2 | BRINP2 | -0,302 | 5,20557E-10 |
| ACE2 | FBXO41 | -0,302 | 5,25155E-10 |
| ACE2 | SNX15 | -0,302 | 5,26025E-10 |
| ACE2 | RP1-41C23.1 | -0,302 | 5,23313E-10 |
| ACE2 | MIR4257 | -0,3021 | 5,13319E-10 |
| ACE2 | RBAKDN | -0,3021 | 5,19058E-10 |
| ACE2 | CD300E | -0,3021 | 5,14629E-10 |
| ACE2 | RP11-343J3.2 | -0,3022 | 5,09344E-10 |
| ACE2 | SIX1 | -0,3022 | 5,11027E-10 |
| ACE2 | RP11-4B16.4 | -0,3022 | 5,06743E-10 |
| ACE2 | KB-1995A5.5 | -0,3022 | 5,108E-10 |
| ACE2 | CTD-2287O16.4 | -0,3023 | 5,02706E-10 |
| ACE2 | DNAJC15 | -0,3023 | 5,03995E-10 |
| ACE2 | LA16c-361A3.3 | -0,3024 | 4,96894E-10 |
| ACE2 | CTD-2619J13.16 | -0,3024 | 4,9244E-10 |

| | | | |
|---|---|---|---|
| ACE2 | RP11-335E6.3 | -0,3025 | 4,87844E-10 |
| ACE2 | UBL4B | -0,3025 | 4,90418E-10 |
| ACE2 | AC007563.3 | -0,3025 | 4,89494E-10 |
| ACE2 | AC002064.4 | -0,3025 | 4,88675E-10 |
| ACE2 | AC078842.3 | -0,3025 | 4,86139E-10 |
| ACE2 | HNRNPCP1 | -0,3025 | 4,9122E-10 |
| ACE2 | RP11-849F2.9 | -0,3025 | 4,85659E-10 |
| ACE2 | NPM1P8 | -0,3025 | 4,90789E-10 |
| ACE2 | SPAST | -0,3026 | 4,80963E-10 |
| ACE2 | RP11-65J3.6 | -0,3026 | 4,79032E-10 |
| ACE2 | LINC00863 | -0,3026 | 4,84117E-10 |
| ACE2 | PSMA4 | -0,3026 | 4,81945E-10 |
| ACE2 | C17orf105 | -0,3026 | 4,82861E-10 |
| ACE2 | RP11-452J21.2 | -0,3027 | 4,77153E-10 |
| ACE2 | RP11-1415C14.3 | -0,3027 | 4,78128E-10 |
| ACE2 | SEPT7P3 | -0,3027 | 4,76975E-10 |
| ACE2 | AJ003147.11 | -0,3027 | 4,76193E-10 |
| ACE2 | OR3A3 | -0,3027 | 4,71956E-10 |
| ACE2 | RP1-124C6.1 | -0,3028 | 4,70631E-10 |
| ACE2 | RP11-350J20.9 | -0,3028 | 4,65924E-10 |
| ACE2 | HSP90B1 | -0,3028 | 4,69802E-10 |
| ACE2 | CKS1BP1 | -0,3028 | 4,71529E-10 |
| ACE2 | bP-21201H5.1 | -0,3028 | 4,7115E-10 |
| ACE2 | FUT10 | -0,3029 | 4,63188E-10 |
| ACE2 | LINC01219 | -0,3029 | 4,63379E-10 |
| ACE2 | SDCBP2-AS1 | -0,3029 | 4,62283E-10 |
| ACE2 | ELOVL2-AS1 | -0,303 | 4,57901E-10 |
| ACE2 | CTBP2P7 | -0,303 | 4,53733E-10 |
| ACE2 | SETP20 | -0,3031 | 4,48648E-10 |
| ACE2 | CYP3A43 | -0,3031 | 4,51916E-10 |
| ACE2 | GADD45G | -0,3031 | 4,47623E-10 |
| ACE2 | KHK | -0,3032 | 4,46275E-10 |
| ACE2 | RP11-778J15.1 | -0,3032 | 4,45477E-10 |
| ACE2 | RP11-513O13.1 | -0,3032 | 4,45213E-10 |
| ACE2 | RP11-271K11.1 | -0,3032 | 4,45118E-10 |
| ACE2 | SIRPAP1 | -0,3032 | 4,45818E-10 |
| ACE2 | CALHM3 | -0,3033 | 4,39053E-10 |
| ACE2 | RP1-228P16.1 | -0,3033 | 4,38362E-10 |
| ACE2 | RP3-414A15.2 | -0,3033 | 4,34682E-10 |
| ACE2 | CTD-2017C7.1 | -0,3033 | 4,40328E-10 |
| ACE2 | RP11-830F9.6 | -0,3033 | 4,38492E-10 |
| ACE2 | AGRN | -0,3034 | 4,33213E-10 |
| ACE2 | RP11-540B6.6 | -0,3034 | 4,33066E-10 |
| ACE2 | RP11-759A24.3 | -0,3034 | 4,30094E-10 |
| ACE2 | MCRS1 | -0,3035 | 4,22671E-10 |
| ACE2 | AC109631.1 | -0,3035 | 4,27934E-10 |
| ACE2 | SELENOV | -0,3035 | 4,2363E-10 |
| ACE2 | PRKAG3 | -0,3036 | 4,21489E-10 |
| ACE2 | RP11-797H7.5 | -0,3036 | 4,20661E-10 |
| ACE2 | CLEC4GP1 | -0,3036 | 4,21886E-10 |
| ACE2 | RN7SL704P | -0,3037 | 4,11126E-10 |
| ACE2 | RP11-44B19.1 | -0,3038 | 4,08624E-10 |
| ACE2 | IFT88 | -0,3038 | 4,06199E-10 |
| ACE2 | RP11-74E22.6 | -0,3038 | 4,09899E-10 |

| | | | |
|---|---|---|---|
| ACE2 | ZBTB18 | -0,304 | 3,95139E-10 |
| ACE2 | RP11-535A19.2 | -0,304 | 3,95028E-10 |
| ACE2 | CCDC77 | -0,304 | 3,96315E-10 |
| ACE2 | CTD-2303H24.2 | -0,304 | 3,94783E-10 |
| ACE2 | BPIFB9P | -0,304 | 3,95713E-10 |
| ACE2 | ROCK2 | -0,3041 | 3,93922E-10 |
| ACE2 | COPG2 | -0,3041 | 3,93849E-10 |
| ACE2 | RP11-180M15.6 | -0,3041 | 3,89409E-10 |
| ACE2 | NAP1L4P3 | -0,3041 | 3,89273E-10 |
| ACE2 | RP11-218M11.7 | -0,3041 | 3,90495E-10 |
| ACE2 | LAMTOR5 | -0,3042 | 3,84686E-10 |
| ACE2 | RP11-365O16.3 | -0,3042 | 3,8518E-10 |
| ACE2 | HOXD10 | -0,3042 | 3,85642E-10 |
| ACE2 | CTD-3162L10.5 | -0,3042 | 3,8695E-10 |
| ACE2 | AP000343.2 | -0,3042 | 3,86312E-10 |
| ACE2 | ZXDB | -0,3042 | 3,88446E-10 |
| ACE2 | IFT74 | -0,3043 | 3,83443E-10 |
| ACE2 | RP11-27G24.3 | -0,3043 | 3,82829E-10 |
| ACE2 | HAS1 | -0,3043 | 3,80961E-10 |
| ACE2 | ACTBP11 | -0,3044 | 3,77414E-10 |
| ACE2 | RP11-423E7.2 | -0,3044 | 3,77657E-10 |
| ACE2 | DIEXF | -0,3045 | 3,71635E-10 |
| ACE2 | HDAC1P2 | -0,3045 | 3,68633E-10 |
| ACE2 | RP11-16F15.2 | -0,3045 | 3,71945E-10 |
| ACE2 | CTC-525D6.2 | -0,3045 | 3,69463E-10 |
| ACE2 | SND1 | -0,3046 | 3,66742E-10 |
| ACE2 | LINC01776 | -0,3047 | 3,58092E-10 |
| ACE2 | LINC01164 | -0,3047 | 3,61979E-10 |
| ACE2 | SPPL3 | -0,3047 | 3,60457E-10 |
| ACE2 | RN7SL812P | -0,3047 | 3,57956E-10 |
| ACE2 | HS6ST2 | -0,3047 | 3,58133E-10 |
| ACE2 | RNASEH1P1 | -0,3048 | 3,5511E-10 |
| ACE2 | RP11-312J18.5 | -0,3049 | 3,49971E-10 |
| ACE2 | AC004945.1 | -0,3049 | 3,49097E-10 |
| ACE2 | AC005009.2 | -0,3049 | 3,52232E-10 |
| ACE2 | RP11-736E3.3 | -0,3049 | 3,52659E-10 |
| ACE2 | RP11-370I10.12 | -0,3049 | 3,51412E-10 |
| ACE2 | FANCG | -0,305 | 3,47021E-10 |
| ACE2 | RP11-370I10.10 | -0,305 | 3,468E-10 |
| ACE2 | CRNKL1 | -0,305 | 3,47071E-10 |
| ACE2 | ORMDL1 | -0,3051 | 3,41859E-10 |
| ACE2 | SPDYE3 | -0,3051 | 3,38657E-10 |
| ACE2 | RP11-116D17.3 | -0,3051 | 3,42043E-10 |
| ACE2 | DLL4 | -0,3051 | 3,42857E-10 |
| ACE2 | LINC02135 | -0,3051 | 3,42256E-10 |
| ACE2 | AC055764.1 | -0,3052 | 3,36884E-10 |
| ACE2 | GIMAP2 | -0,3052 | 3,35565E-10 |
| ACE2 | RP11-466H18.1 | -0,3052 | 3,37502E-10 |
| ACE2 | RP11-128N14.5 | -0,3052 | 3,35552E-10 |
| ACE2 | NPIPB3 | -0,3052 | 3,3698E-10 |
| ACE2 | RP11-849N15.1 | -0,3052 | 3,35891E-10 |
| ACE2 | TRMT6 | -0,3052 | 3,35754E-10 |
| ACE2 | DNAL4 | -0,3052 | 3,36122E-10 |
| ACE2 | DENND6A-AS1 | -0,3053 | 3,32074E-10 |

| | | | |
|---|---|---|---|
| ACE2 | CTA-503F6.2 | -0,3053 | 3,31355E-10 |
| ACE2 | RP11-1102P16.1 | -0,3054 | 3,26017E-10 |
| ACE2 | RP11-536K7.3 | -0,3055 | 3,22154E-10 |
| ACE2 | RP11-218E20.5 | -0,3055 | 3,2347E-10 |
| ACE2 | CCDC157 | -0,3055 | 3,20575E-10 |
| ACE2 | GTSE1-AS1 | -0,3056 | 3,18569E-10 |
| ACE2 | EIF4BP7 | -0,3056 | 3,18801E-10 |
| ACE2 | CLPX | -0,3057 | 3,12636E-10 |
| ACE2 | CYP4A11 | -0,3058 | 3,10344E-10 |
| ACE2 | RPL23AP88 | -0,3058 | 3,07032E-10 |
| ACE2 | RP4-625H18.2 | -0,3058 | 3,1001E-10 |
| ACE2 | RP11-212P7.1 | -0,3058 | 3,0992E-10 |
| ACE2 | LRRC46 | -0,3058 | 3,07891E-10 |
| ACE2 | RP11-53O19.2 | -0,3059 | 3,04786E-10 |
| ACE2 | RP11-390K5.6 | -0,306 | 3,00043E-10 |
| ACE2 | AP000688.29 | -0,3061 | 2,96353E-10 |
| ACE2 | RP11-409I10.2 | -0,3062 | 2,9143E-10 |
| ACE2 | ZNF43 | -0,3062 | 2,90876E-10 |
| ACE2 | ADORA1 | -0,3063 | 2,89058E-10 |
| ACE2 | RIMKLBP2 | -0,3063 | 2,88828E-10 |
| ACE2 | CEP70 | -0,3063 | 2,8685E-10 |
| ACE2 | DCAF13P3 | -0,3063 | 2,86847E-10 |
| ACE2 | EEF2KMT | -0,3063 | 2,87144E-10 |
| ACE2 | ZKSCAN4 | -0,3064 | 2,85468E-10 |
| ACE2 | SARAF | -0,3064 | 2,85458E-10 |
| ACE2 | RP4-657D16.6 | -0,3065 | 2,787E-10 |
| ACE2 | RP11-278C7.5 | -0,3065 | 2,81012E-10 |
| ACE2 | DLL3 | -0,3065 | 2,82255E-10 |
| ACE2 | AGGF1 | -0,3066 | 2,77476E-10 |
| ACE2 | AP000577.2 | -0,3066 | 2,78085E-10 |
| ACE2 | RPS4XP2 | -0,3066 | 2,76701E-10 |
| ACE2 | PPP1R35 | -0,3067 | 2,7326E-10 |
| ACE2 | RP11-756G20.1 | -0,3068 | 2,68649E-10 |
| ACE2 | U2AF2 | -0,3068 | 2,69304E-10 |
| ACE2 | BHLHE41 | -0,3069 | 2,66727E-10 |
| ACE2 | DUXAP9 | -0,3069 | 2,64335E-10 |
| ACE2 | MTFMT | -0,3069 | 2,65714E-10 |
| ACE2 | CDIPT-AS1 | -0,3069 | 2,64595E-10 |
| ACE2 | CARHSP1 | -0,307 | 2,60703E-10 |
| ACE2 | ATAD2B | -0,3071 | 2,56876E-10 |
| ACE2 | OR52B3P | -0,3071 | 2,59079E-10 |
| ACE2 | GTPBP3 | -0,3071 | 2,59053E-10 |
| ACE2 | PPP1R14BP3 | -0,3072 | 2,53362E-10 |
| ACE2 | USP6NL | -0,3072 | 2,5498E-10 |
| ACE2 | RP11-513D5.5 | -0,3073 | 2,50528E-10 |
| ACE2 | PTDSS1 | -0,3073 | 2,50456E-10 |
| ACE2 | RPL21P42 | -0,3074 | 2,46961E-10 |
| ACE2 | CTD-2184D3.5 | -0,3074 | 2,46585E-10 |
| ACE2 | CTD-2319I12.2 | -0,3074 | 2,48301E-10 |
| ACE2 | RP11-700N1.1 | -0,3075 | 2,43467E-10 |
| ACE2 | PPP1R17 | -0,3075 | 2,42925E-10 |
| ACE2 | MAMDC2-AS1 | -0,3075 | 2,44857E-10 |
| ACE2 | UBE2Q2P2 | -0,3075 | 2,4237E-10 |
| ACE2 | LINC01615 | -0,3076 | 2,40625E-10 |

| | | | |
|---|---|---|---|
| ACE2 | LDAH | -0,3077 | 2,37498E-10 |
| ACE2 | PARP3 | -0,3077 | 2,38478E-10 |
| ACE2 | CES5A | -0,3077 | 2,38378E-10 |
| ACE2 | NUTM2D | -0,3078 | 2,34095E-10 |
| ACE2 | MAPK7 | -0,3078 | 2,33968E-10 |
| ACE2 | KLK6 | -0,3078 | 2,32603E-10 |
| ACE2 | IFRD2 | -0,3079 | 2,29928E-10 |
| ACE2 | MRE11P1 | -0,3079 | 2,31818E-10 |
| ACE2 | GSTT2 | -0,3079 | 2,31023E-10 |
| ACE2 | GPR137 | -0,308 | 2,26366E-10 |
| ACE2 | PKD1P1 | -0,308 | 2,26092E-10 |
| ACE2 | RPL7P23 | -0,3081 | 2,23174E-10 |
| ACE2 | EEF1A1P4 | -0,3081 | 2,25562E-10 |
| ACE2 | RP11-85K15.3 | -0,3081 | 2,24521E-10 |
| ACE2 | TLR2 | -0,3082 | 2,20222E-10 |
| ACE2 | RPL21P120 | -0,3082 | 2,20719E-10 |
| ACE2 | MIR8083 | -0,3083 | 2,17092E-10 |
| ACE2 | CTD-3080F16.3 | -0,3083 | 2,16328E-10 |
| ACE2 | RP11-203L2.3 | -0,3083 | 2,18462E-10 |
| ACE2 | CDK2AP2P2 | -0,3084 | 2,1589E-10 |
| ACE2 | RP5-1163L11.2 | -0,3084 | 2,14986E-10 |
| ACE2 | HSD3B7 | -0,3084 | 2,15334E-10 |
| ACE2 | CTB-96E2.10 | -0,3084 | 2,14326E-10 |
| ACE2 | PDHB | -0,3085 | 2,10861E-10 |
| ACE2 | PRICKLE2-AS3 | -0,3085 | 2,11932E-10 |
| ACE2 | RP1-168L15.6 | -0,3085 | 2,10493E-10 |
| ACE2 | SHOC2 | -0,3085 | 2,12752E-10 |
| ACE2 | ENPP7P2 | -0,3086 | 2,08493E-10 |
| ACE2 | ACSM5P1 | -0,3086 | 2,07746E-10 |
| ACE2 | RN7SL608P | -0,3086 | 2,08608E-10 |
| ACE2 | RP11-332H14.2 | -0,3087 | 2,07162E-10 |
| ACE2 | ALAS2 | -0,3087 | 2,05866E-10 |
| ACE2 | RNF141 | -0,3088 | 2,04408E-10 |
| ACE2 | EMG1 | -0,3088 | 2,01922E-10 |
| ACE2 | COL4A4 | -0,3089 | 2,00185E-10 |
| ACE2 | RN7SKP56 | -0,3089 | 2,01235E-10 |
| ACE2 | AC079741.2 | -0,3089 | 1,98973E-10 |
| ACE2 | LSM12 | -0,309 | 1,98593E-10 |
| ACE2 | LA16c-359F1.1 | -0,3091 | 1,94551E-10 |
| ACE2 | RPL23AP87 | -0,3091 | 1,94717E-10 |
| ACE2 | RP11-615I2.2 | -0,3092 | 1,90741E-10 |
| ACE2 | SELE | -0,3093 | 1,88605E-10 |
| ACE2 | PPIAP16 | -0,3093 | 1,89195E-10 |
| ACE2 | RP11-313E19.2 | -0,3093 | 1,9014E-10 |
| ACE2 | RP11-115N4.1 | -0,3093 | 1,89957E-10 |
| ACE2 | POLR2J | -0,3093 | 1,886E-10 |
| ACE2 | RP11-530C5.1 | -0,3093 | 1,89519E-10 |
| ACE2 | RP1-265C24.8 | -0,3095 | 1,82782E-10 |
| ACE2 | FBXL3 | -0,3095 | 1,84459E-10 |
| ACE2 | PCDHGB9P | -0,3096 | 1,81718E-10 |
| ACE2 | DGCR9 | -0,3096 | 1,8169E-10 |
| ACE2 | CEP83-AS1 | -0,3097 | 1,77572E-10 |
| ACE2 | ACOT2 | -0,3097 | 1,78767E-10 |
| ACE2 | RP3-333A15.1 | -0,3098 | 1,77375E-10 |

| | | | |
|---|---|---|---|
| ACE2 | RP11-350G8.5 | -0,3098 | 1,75649E-10 |
| ACE2 | AHSP | -0,3098 | 1,75888E-10 |
| ACE2 | AC012501.3 | -0,3099 | 1,74992E-10 |
| ACE2 | C4orf26 | -0,3099 | 1,73075E-10 |
| ACE2 | DNAJC7 | -0,3099 | 1,74018E-10 |
| ACE2 | NACA2 | -0,3099 | 1,72819E-10 |
| ACE2 | SMARCD2 | -0,3099 | 1,74682E-10 |
| ACE2 | RP11-380M21.1 | -0,3099 | 1,73723E-10 |
| ACE2 | NXNL1 | -0,3099 | 1,74826E-10 |
| ACE2 | ZNF180 | -0,3099 | 1,73611E-10 |
| ACE2 | MED7 | -0,31 | 1,7052E-10 |
| ACE2 | AC000123.4 | -0,31 | 1,70739E-10 |
| ACE2 | RP11-849H4.2 | -0,31 | 1,71945E-10 |
| ACE2 | ZNF687 | -0,3101 | 1,67872E-10 |
| ACE2 | LINC01506 | -0,3101 | 1,69838E-10 |
| ACE2 | RPS6KA5 | -0,3101 | 1,69307E-10 |
| ACE2 | AC004004.2 | -0,3101 | 1,68368E-10 |
| ACE2 | ATP6V1B2 | -0,3102 | 1,67347E-10 |
| ACE2 | RP1-257I9.2 | -0,3103 | 1,64589E-10 |
| ACE2 | RP11-119H12.6 | -0,3104 | 1,61337E-10 |
| ACE2 | CSTF2T | -0,3104 | 1,61532E-10 |
| ACE2 | RP11-701H16.4 | -0,3104 | 1,62299E-10 |
| ACE2 | ZC2HC1B | -0,3105 | 1,59854E-10 |
| ACE2 | RP11-6I2.3 | -0,3105 | 1,59187E-10 |
| ACE2 | RPL13P12 | -0,3105 | 1,59054E-10 |
| ACE2 | BOP1 | -0,3106 | 1,57039E-10 |
| ACE2 | KRT1 | -0,3106 | 1,57289E-10 |
| ACE2 | RP11-61E11.2 | -0,3106 | 1,58294E-10 |
| ACE2 | RP11-218M22.1 | -0,3107 | 1,55565E-10 |
| ACE2 | CTD-2540B15.10 | -0,3107 | 1,54989E-10 |
| ACE2 | UBXN8 | -0,3108 | 1,53347E-10 |
| ACE2 | OR51AB1P | -0,3108 | 1,52712E-10 |
| ACE2 | AP000442.1 | -0,3108 | 1,52879E-10 |
| ACE2 | RP11-677M14.8 | -0,3108 | 1,53853E-10 |
| ACE2 | USP8 | -0,3108 | 1,53227E-10 |
| ACE2 | TDRD12 | -0,3108 | 1,53613E-10 |
| ACE2 | RP11-384C4.2 | -0,3109 | 1,50047E-10 |
| ACE2 | RP11-705C15.3 | -0,3109 | 1,50553E-10 |
| ACE2 | RP11-863P13.5 | -0,3109 | 1,50562E-10 |
| ACE2 | KRTDAP | -0,3109 | 1,51484E-10 |
| ACE2 | MIR4269 | -0,311 | 1,48751E-10 |
| ACE2 | AC133680.1 | -0,311 | 1,49491E-10 |
| ACE2 | CADM2-AS2 | -0,311 | 1,47758E-10 |
| ACE2 | RP11-259O2.3 | -0,311 | 1,4863E-10 |
| ACE2 | MACROD1 | -0,311 | 1,49584E-10 |
| ACE2 | ATP5H | -0,311 | 1,4927E-10 |
| ACE2 | RP11-122G18.7 | -0,3111 | 1,47506E-10 |
| ACE2 | BTF3L4P2 | -0,3111 | 1,47399E-10 |
| ACE2 | RNF224 | -0,3111 | 1,46549E-10 |
| ACE2 | HOXC-AS3 | -0,3111 | 1,46658E-10 |
| ACE2 | RP11-90L1.8 | -0,3111 | 1,45689E-10 |
| ACE2 | ALG1L12P | -0,3112 | 1,44802E-10 |
| ACE2 | FAM120AOS | -0,3112 | 1,43708E-10 |
| ACE2 | AB019441.29 | -0,3112 | 1,43976E-10 |

| | | | |
|---|---|---|---|
| ACE2 | SEC11A | -0,3112 | 1,45238E-10 |
| ACE2 | PAGE4 | -0,3112 | 1,43622E-10 |
| ACE2 | RP11-861A13.3 | -0,3113 | 1,415E-10 |
| ACE2 | SLC2A3P2 | -0,3114 | 1,40967E-10 |
| ACE2 | CTC-338M12.1 | -0,3114 | 1,39986E-10 |
| ACE2 | FIGNL1 | -0,3114 | 1,39858E-10 |
| ACE2 | KRT87P | -0,3114 | 1,40965E-10 |
| ACE2 | AL672294.1 | -0,3115 | 1,39094E-10 |
| ACE2 | SEMA5B | -0,3115 | 1,38241E-10 |
| ACE2 | FAM86B2 | -0,3115 | 1,37826E-10 |
| ACE2 | ZDHHC2 | -0,3115 | 1,38733E-10 |
| ACE2 | KANK4 | -0,3116 | 1,35695E-10 |
| ACE2 | RP11-12A20.7 | -0,3116 | 1,36205E-10 |
| ACE2 | CFAP157 | -0,3116 | 1,36471E-10 |
| ACE2 | CTD-2195B23.3 | -0,3116 | 1,37447E-10 |
| ACE2 | ARFRP1 | -0,3116 | 1,36922E-10 |
| ACE2 | RP1-251M9.2 | -0,3117 | 1,34499E-10 |
| ACE2 | GAPDHP15 | -0,3118 | 1,32028E-10 |
| ACE2 | SETD4 | -0,3118 | 1,33314E-10 |
| ACE2 | GAPDHP33 | -0,3119 | 1,30884E-10 |
| ACE2 | FAM96AP2 | -0,3119 | 1,30511E-10 |
| ACE2 | TMED7 | -0,3119 | 1,30786E-10 |
| ACE2 | LINC01411 | -0,3119 | 1,3124E-10 |
| ACE2 | SFMBT2 | -0,3119 | 1,31606E-10 |
| ACE2 | BAMBI | -0,3119 | 1,30404E-10 |
| ACE2 | SYNPO2L | -0,312 | 1,29636E-10 |
| ACE2 | RP11-359J14.2 | -0,312 | 1,2835E-10 |
| ACE2 | TTI2 | -0,3122 | 1,25884E-10 |
| ACE2 | RN7SL239P | -0,3122 | 1,25312E-10 |
| ACE2 | CTD-2358C21.5 | -0,3122 | 1,26051E-10 |
| ACE2 | EPB41L1 | -0,3122 | 1,25339E-10 |
| ACE2 | GDF9 | -0,3123 | 1,2426E-10 |
| ACE2 | OR10AD1 | -0,3123 | 1,24157E-10 |
| ACE2 | FAM199X | -0,3123 | 1,2279E-10 |
| ACE2 | LINC00906 | -0,3124 | 1,21359E-10 |
| ACE2 | SERPIND1 | -0,3124 | 1,22385E-10 |
| ACE2 | AC142293.3 | -0,3125 | 1,19638E-10 |
| ACE2 | KDM6B | -0,3125 | 1,20884E-10 |
| ACE2 | CHML | -0,3126 | 1,1795E-10 |
| ACE2 | RP11-439M11.1 | -0,3126 | 1,18761E-10 |
| ACE2 | PTGES3P2 | -0,3126 | 1,18953E-10 |
| ACE2 | RPL26P19 | -0,3126 | 1,18156E-10 |
| ACE2 | LIPA | -0,3126 | 1,19045E-10 |
| ACE2 | RP11-450H6.3 | -0,3126 | 1,18859E-10 |
| ACE2 | GOLGA6L7P | -0,3126 | 1,18157E-10 |
| ACE2 | POLDIP2 | -0,3126 | 1,17938E-10 |
| ACE2 | BX322557.13 | -0,3126 | 1,18468E-10 |
| ACE2 | LINC02172 | -0,3127 | 1,17149E-10 |
| ACE2 | GLUD1P3 | -0,3127 | 1,16404E-10 |
| ACE2 | LINC01143 | -0,3129 | 1,14103E-10 |
| ACE2 | TOR1A | -0,3129 | 1,13269E-10 |
| ACE2 | HNRNPA3P9 | -0,3129 | 1,12655E-10 |
| ACE2 | KIF1C | -0,3129 | 1,1347E-10 |
| ACE2 | CTC-563A5.2 | -0,313 | 1,12496E-10 |

| | | | |
|---|---|---|---|
| ACE2 | CLK3P2 | -0,3131 | 1,10305E-10 |
| ACE2 | DCLRE1A | -0,3131 | 1,1031E-10 |
| ACE2 | ZNRF1 | -0,3131 | 1,10925E-10 |
| ACE2 | GAS2L2 | -0,3131 | 1,09743E-10 |
| ACE2 | AC079776.1 | -0,3132 | 1,08863E-10 |
| ACE2 | IQCF1 | -0,3132 | 1,08099E-10 |
| ACE2 | ZNF696 | -0,3132 | 1,08495E-10 |
| ACE2 | CTD-3030D20.1 | -0,3132 | 1,08998E-10 |
| ACE2 | TRAPPC13P1 | -0,3132 | 1,0904E-10 |
| ACE2 | ZNF629 | -0,3133 | 1,06731E-10 |
| ACE2 | TIGD1 | -0,3134 | 1,04851E-10 |
| ACE2 | METTL12 | -0,3134 | 1,04812E-10 |
| ACE2 | NELFB | -0,3136 | 1,02614E-10 |
| ACE2 | EBAG9P1 | -0,3136 | 1,02912E-10 |
| ACE2 | RP11-484D2.4 | -0,3136 | 1,02462E-10 |
| ACE2 | ST13P15 | -0,3137 | 1,0092E-10 |
| ACE2 | RNU6-593P | -0,3137 | 1,00852E-10 |
| ACE2 | DDX18P1 | -0,3137 | 1,0091E-10 |
| ACE2 | SMAD3 | -0,3137 | 1,00853E-10 |
| ACE2 | RP11-1055B8.10 | -0,3137 | 1,01115E-10 |
| ACE2 | NLRP12 | -0,3137 | 1,01038E-10 |
| ACE2 | RP6-201G10.2 | -0,3137 | 1,0078E-10 |
| ACE2 | RP11-259P15.4 | -0,3138 | 9,89666E-11 |
| ACE2 | ZDHHC18 | -0,3139 | 9,8189E-11 |
| ACE2 | INHBA-AS1 | -0,3139 | 9,82416E-11 |
| ACE2 | RP11-597M17.3 | -0,3139 | 9,80131E-11 |
| ACE2 | IQCC | -0,314 | 9,73071E-11 |
| ACE2 | RP11-155O18.6 | -0,314 | 9,65613E-11 |
| ACE2 | GVINP2 | -0,314 | 9,70873E-11 |
| ACE2 | EEF1A1P16 | -0,314 | 9,73936E-11 |
| ACE2 | RP11-461F11.2 | -0,314 | 9,61735E-11 |
| ACE2 | RP11-109P11.1 | -0,314 | 9,73818E-11 |
| ACE2 | AC118754.4 | -0,314 | 9,62518E-11 |
| ACE2 | AC073326.3 | -0,3141 | 9,60986E-11 |
| ACE2 | RP11-395I14.2 | -0,3141 | 9,57817E-11 |
| ACE2 | C9orf64 | -0,3141 | 9,55509E-11 |
| ACE2 | RGS3 | -0,3141 | 9,55306E-11 |
| ACE2 | PAPD4 | -0,3142 | 9,34423E-11 |
| ACE2 | RP11-386G11.11 | -0,3142 | 9,39489E-11 |
| ACE2 | RP11-616M22.12 | -0,3142 | 9,38989E-11 |
| ACE2 | RP11-599J14.2 | -0,3143 | 9,2621E-11 |
| ACE2 | RP11-327P2.5 | -0,3143 | 9,238E-11 |
| ACE2 | FOLH1B | -0,3144 | 9,14929E-11 |
| ACE2 | CTD-2293H3.1 | -0,3144 | 9,15502E-11 |
| ACE2 | AC112497.1 | -0,3144 | 9,09973E-11 |
| ACE2 | ALOX12P2 | -0,3145 | 8,97015E-11 |
| ACE2 | NXF4 | -0,3145 | 9,06938E-11 |
| ACE2 | TBC1D9B | -0,3146 | 8,87488E-11 |
| ACE2 | RP11-598F7.4 | -0,3146 | 8,89752E-11 |
| ACE2 | PES1 | -0,3146 | 8,89541E-11 |
| ACE2 | GTF2E1 | -0,3147 | 8,77168E-11 |
| ACE2 | XYLT1 | -0,3147 | 8,81322E-11 |
| ACE2 | ZBTB39 | -0,3148 | 8,60979E-11 |
| ACE2 | RP11-123O10.4 | -0,3149 | 8,49812E-11 |

| | | | |
|---|---|---|---|
| ACE2 | RP11-218I21.2 | -0,3149 | 8,53285E-11 |
| ACE2 | AC016910.1 | -0,315 | 8,40138E-11 |
| ACE2 | ROPN1L | -0,315 | 8,42131E-11 |
| ACE2 | AGAP12P | -0,315 | 8,4057E-11 |
| ACE2 | DNAJC9-AS1 | -0,315 | 8,37483E-11 |
| ACE2 | RP5-1181K21.4 | -0,3152 | 8,11374E-11 |
| ACE2 | P2RX6P | -0,3152 | 8,09052E-11 |
| ACE2 | UBE2CP3 | -0,3153 | 8,03251E-11 |
| ACE2 | AC012531.2 | -0,3153 | 8,01179E-11 |
| ACE2 | SMPD4P1 | -0,3153 | 8,0116E-11 |
| ACE2 | GAPDHP32 | -0,3154 | 7,92388E-11 |
| ACE2 | RBM34 | -0,3154 | 7,88567E-11 |
| ACE2 | RP13-578N3.3 | -0,3154 | 7,96575E-11 |
| ACE2 | OR5AK4P | -0,3154 | 7,96066E-11 |
| ACE2 | ARNTL | -0,3155 | 7,75047E-11 |
| ACE2 | GS1-531I17.3 | -0,3155 | 7,80279E-11 |
| ACE2 | PMPCB | -0,3157 | 7,56538E-11 |
| ACE2 | CCT6B | -0,3157 | 7,53779E-11 |
| ACE2 | ARID1A | -0,3158 | 7,45201E-11 |
| ACE2 | RPLP0 | -0,3158 | 7,51119E-11 |
| ACE2 | RP11-358B23.5 | -0,316 | 7,23822E-11 |
| ACE2 | NPLOC4 | -0,316 | 7,30124E-11 |
| ACE2 | EXOSC3P1 | -0,316 | 7,28767E-11 |
| ACE2 | LINC02197 | -0,3161 | 7,13288E-11 |
| ACE2 | LINC00174 | -0,3161 | 7,17928E-11 |
| ACE2 | RP11-423G4.10 | -0,3161 | 7,16447E-11 |
| ACE2 | RP11-44N12.5 | -0,3162 | 7,00593E-11 |
| ACE2 | RP11-543C4.1 | -0,3162 | 7,01169E-11 |
| ACE2 | C2orf91 | -0,3163 | 6,96169E-11 |
| ACE2 | PROCA1 | -0,3163 | 6,97762E-11 |
| ACE2 | FOXN2 | -0,3165 | 6,77813E-11 |
| ACE2 | NDST4 | -0,3165 | 6,71698E-11 |
| ACE2 | RPS3AP37 | -0,3165 | 6,73917E-11 |
| ACE2 | GRWD1 | -0,3165 | 6,73222E-11 |
| ACE2 | PEX10 | -0,3166 | 6,6796E-11 |
| ACE2 | USP37 | -0,3166 | 6,67576E-11 |
| ACE2 | RP11-119B16.2 | -0,3166 | 6,68842E-11 |
| ACE2 | RP11-424G14.1 | -0,3166 | 6,68056E-11 |
| ACE2 | SHCBP1L | -0,3167 | 6,58492E-11 |
| ACE2 | CTB-46B19.2 | -0,3167 | 6,51154E-11 |
| ACE2 | SCGB2B2 | -0,3167 | 6,58369E-11 |
| ACE2 | RP11-332J15.2 | -0,3168 | 6,4526E-11 |
| ACE2 | AC005754.7 | -0,3168 | 6,43466E-11 |
| ACE2 | RPL5P24 | -0,3168 | 6,49785E-11 |
| ACE2 | RP11-567C20.2 | -0,3169 | 6,35572E-11 |
| ACE2 | GLRX2 | -0,3169 | 6,36779E-11 |
| ACE2 | RP11-115C10.1 | -0,3169 | 6,39177E-11 |
| ACE2 | RP11-7O11.3 | -0,317 | 6,25361E-11 |
| ACE2 | GRM1 | -0,317 | 6,24886E-11 |
| ACE2 | PSPH | -0,317 | 6,24704E-11 |
| ACE2 | RP11-429J17.2 | -0,317 | 6,25167E-11 |
| ACE2 | RP11-332H18.5 | -0,317 | 6,30129E-11 |
| ACE2 | SNORA11 | -0,3171 | 6,14104E-11 |
| ACE2 | MIA3 | -0,3172 | 6,06023E-11 |

| | | | |
|---|---|---|---|
| ACE2 | HS1BP3-IT1 | -0,3172 | 6,10417E-11 |
| ACE2 | POLR2J3 | -0,3172 | 6,10454E-11 |
| ACE2 | LINC00937 | -0,3172 | 6,11363E-11 |
| ACE2 | GGNBP2 | -0,3172 | 6,0583E-11 |
| ACE2 | RAP2C | -0,3172 | 6,12243E-11 |
| ACE2 | RP11-847H18.3 | -0,3173 | 6,033E-11 |
| ACE2 | RP11-247I13.11 | -0,3173 | 5,98403E-11 |
| ACE2 | AC002366.3 | -0,3173 | 6,03934E-11 |
| ACE2 | GXYLT1P6 | -0,3174 | 5,88399E-11 |
| ACE2 | THUMPD3 | -0,3175 | 5,82951E-11 |
| ACE2 | C9orf153 | -0,3175 | 5,84159E-11 |
| ACE2 | CRNDE | -0,3175 | 5,87635E-11 |
| ACE2 | RP11-400F19.6 | -0,3175 | 5,8133E-11 |
| ACE2 | RP13-43E11.1 | -0,3175 | 5,81816E-11 |
| ACE2 | ACTG1P23 | -0,3176 | 5,73882E-11 |
| ACE2 | HSPE1P11 | -0,3176 | 5,71443E-11 |
| ACE2 | CYSRT1 | -0,3176 | 5,79098E-11 |
| ACE2 | ULK4P1 | -0,3176 | 5,7735E-11 |
| ACE2 | RP11-669E14.6 | -0,3176 | 5,77582E-11 |
| ACE2 | TMF1 | -0,3177 | 5,67291E-11 |
| ACE2 | CTC-498M16.4 | -0,3177 | 5,66946E-11 |
| ACE2 | SRRM2 | -0,3177 | 5,66832E-11 |
| ACE2 | RP11-481F24.3 | -0,3177 | 5,64331E-11 |
| ACE2 | USP32P1 | -0,3177 | 5,68659E-11 |
| ACE2 | CTC-457E21.1 | -0,3177 | 5,62763E-11 |
| ACE2 | RPS2P46 | -0,3178 | 5,61868E-11 |
| ACE2 | RP13-52K8.1 | -0,3178 | 5,57644E-11 |
| ACE2 | RP11-11N5.3 | -0,3181 | 5,36303E-11 |
| ACE2 | LIN7B | -0,3181 | 5,34255E-11 |
| ACE2 | SNHG12 | -0,3182 | 5,25433E-11 |
| ACE2 | LINC02048 | -0,3182 | 5,28955E-11 |
| ACE2 | RP11-509J21.2 | -0,3182 | 5,26679E-11 |
| ACE2 | TMEM106A | -0,3182 | 5,25674E-11 |
| ACE2 | LINC01324 | -0,3183 | 5,22086E-11 |
| ACE2 | RP11-1415C14.4 | -0,3183 | 5,22131E-11 |
| ACE2 | SAAL1 | -0,3183 | 5,22476E-11 |
| ACE2 | CTC-575I10.1 | -0,3184 | 5,14035E-11 |
| ACE2 | NME8 | -0,3184 | 5,09699E-11 |
| ACE2 | C9orf84 | -0,3184 | 5,12064E-11 |
| ACE2 | SAMD15 | -0,3184 | 5,09085E-11 |
| ACE2 | RP11-498P14.5 | -0,3185 | 5,05232E-11 |
| ACE2 | RP11-46A10.5 | -0,3186 | 4,98101E-11 |
| ACE2 | RP11-413B19.2 | -0,3186 | 4,98879E-11 |
| ACE2 | EEF1A1P31 | -0,3186 | 4,98495E-11 |
| ACE2 | MROH5 | -0,3187 | 4,87213E-11 |
| ACE2 | SEC16A | -0,3187 | 4,92789E-11 |
| ACE2 | CERNA1 | -0,3187 | 4,86554E-11 |
| ACE2 | LA16c-312E8.5 | -0,3187 | 4,91197E-11 |
| ACE2 | RP11-266L9.6 | -0,3187 | 4,89449E-11 |
| ACE2 | RP11-739N10.1 | -0,3187 | 4,91763E-11 |
| ACE2 | UGCG | -0,3188 | 4,80897E-11 |
| ACE2 | GTF3C3 | -0,3189 | 4,74717E-11 |
| ACE2 | TARS | -0,3189 | 4,75052E-11 |
| ACE2 | RP11-10E18.7 | -0,3189 | 4,72139E-11 |

| | | | |
|---|---|---|---|
| ACE2 | LINC02193 | -0,3189 | 4,75687E-11 |
| ACE2 | RHOXF1P1 | -0,319 | 4,71616E-11 |
| ACE2 | RP11-508N22.6 | -0,3191 | 4,61917E-11 |
| ACE2 | HEATR4 | -0,3191 | 4,62969E-11 |
| ACE2 | CTD-2086O20.1 | -0,3191 | 4,59661E-11 |
| ACE2 | RP11-472M19.2 | -0,3192 | 4,57448E-11 |
| ACE2 | RP11-361M10.3 | -0,3192 | 4,5688E-11 |
| ACE2 | LINC00476 | -0,3193 | 4,49819E-11 |
| ACE2 | RP11-307C19.2 | -0,3193 | 4,51642E-11 |
| ACE2 | FIRRE | -0,3193 | 4,45867E-11 |
| ACE2 | TFAMP1 | -0,3194 | 4,42544E-11 |
| ACE2 | RP11-638I2.8 | -0,3194 | 4,44684E-11 |
| ACE2 | EIF4E1B | -0,3195 | 4,3828E-11 |
| ACE2 | AP000892.4 | -0,3195 | 4,35661E-11 |
| ACE2 | UBE2D3P3 | -0,3196 | 4,27155E-11 |
| ACE2 | EVA1A | -0,3197 | 4,24649E-11 |
| ACE2 | RPSAP3 | -0,3197 | 4,22499E-11 |
| ACE2 | AC002407.3 | -0,3197 | 4,20084E-11 |
| ACE2 | AKR1C1 | -0,3198 | 4,19141E-11 |
| ACE2 | RP11-17E3.1 | -0,3198 | 4,18114E-11 |
| ACE2 | HIST2H2BF | -0,3199 | 4,11853E-11 |
| ACE2 | SAPCD2P3 | -0,3199 | 4,09472E-11 |
| ACE2 | RP11-266A24.1 | -0,3199 | 4,12637E-11 |
| ACE2 | CYP4F24P | -0,3199 | 4,12376E-11 |
| ACE2 | RP1-101D8.1 | -0,3199 | 4,07799E-11 |
| ACE2 | KRT8P50 | -0,32 | 4,06436E-11 |
| ACE2 | RP11-505K9.1 | -0,32 | 4,05154E-11 |
| ACE2 | FAM19A2 | -0,3201 | 3,97246E-11 |
| ACE2 | PLGLB1 | -0,3202 | 3,92629E-11 |
| ACE2 | TLR1 | -0,3202 | 3,95315E-11 |
| ACE2 | TSPYL1 | -0,3202 | 3,93111E-11 |
| ACE2 | VCP | -0,3202 | 3,94046E-11 |
| ACE2 | CATSPERD | -0,3202 | 3,92269E-11 |
| ACE2 | RP11-54D18.4 | -0,3203 | 3,88043E-11 |
| ACE2 | RNF219-AS1 | -0,3203 | 3,84969E-11 |
| ACE2 | LRRC37A2 | -0,3203 | 3,87901E-11 |
| ACE2 | RP11-446E9.1 | -0,3204 | 3,8143E-11 |
| ACE2 | RP1-102D24.5 | -0,3204 | 3,78791E-11 |
| ACE2 | RP11-66N24.7 | -0,3205 | 3,77165E-11 |
| ACE2 | AC068499.10 | -0,3205 | 3,77964E-11 |
| ACE2 | MIATNB | -0,3205 | 3,75131E-11 |
| ACE2 | RP11-310H4.2 | -0,3206 | 3,71085E-11 |
| ACE2 | MED25 | -0,3206 | 3,71988E-11 |
| ACE2 | RP11-848P1.7 | -0,3207 | 3,66843E-11 |
| ACE2 | RP6-109B7.4 | -0,3207 | 3,67186E-11 |
| ACE2 | RP11-174G17.3 | -0,3208 | 3,60179E-11 |
| ACE2 | GNRHR | -0,3208 | 3,58959E-11 |
| ACE2 | RP11-44N11.2 | -0,3208 | 3,57684E-11 |
| ACE2 | ZNF624 | -0,3208 | 3,58285E-11 |
| ACE2 | RP11-68I3.11 | -0,3208 | 3,61518E-11 |
| ACE2 | AC104634.3 | -0,3209 | 3,55912E-11 |
| ACE2 | RP11-227F19.5 | -0,3209 | 3,55443E-11 |
| ACE2 | SMIM19 | -0,3209 | 3,54142E-11 |
| ACE2 | TACSTD2 | -0,321 | 3,51475E-11 |

| | | | |
|---|---|---|---|
| ACE2 | RP11-105N14.1 | -0,321 | 3,47043E-11 |
| ACE2 | MFN1 | -0,321 | 3,5067E-11 |
| ACE2 | RP11-434C1.2 | -0,321 | 3,47027E-11 |
| ACE2 | BTN3A3 | -0,3211 | 3,43364E-11 |
| ACE2 | SNAI2 | -0,3211 | 3,44061E-11 |
| ACE2 | FOS | -0,3211 | 3,4611E-11 |
| ACE2 | IFI35 | -0,3211 | 3,44462E-11 |
| ACE2 | RPL9P7 | -0,3211 | 3,45076E-11 |
| ACE2 | NAXE | -0,3213 | 3,32655E-11 |
| ACE2 | EEF1B2 | -0,3213 | 3,33435E-11 |
| ACE2 | CRHBP | -0,3213 | 3,34199E-11 |
| ACE2 | AP000560.3 | -0,3213 | 3,32064E-11 |
| ACE2 | DDX23 | -0,3214 | 3,27228E-11 |
| ACE2 | ATF5 | -0,3214 | 3,30442E-11 |
| ACE2 | CECR5-AS1 | -0,3214 | 3,28594E-11 |
| ACE2 | DLGAP3 | -0,3215 | 3,25631E-11 |
| ACE2 | UROC1 | -0,3216 | 3,1768E-11 |
| ACE2 | RP11-354M20.3 | -0,3216 | 3,19963E-11 |
| ACE2 | GCSAML | -0,3217 | 3,1449E-11 |
| ACE2 | SCRN3 | -0,3217 | 3,12793E-11 |
| ACE2 | AC002550.6 | -0,3217 | 3,12817E-11 |
| ACE2 | RP11-298D21.1 | -0,3217 | 3,13219E-11 |
| ACE2 | RP1-274L7.4 | -0,3217 | 3,16405E-11 |
| ACE2 | REXO4 | -0,3218 | 3,09574E-11 |
| ACE2 | RN7SKP287 | -0,3218 | 3,12576E-11 |
| ACE2 | ZNF384 | -0,3218 | 3,08898E-11 |
| ACE2 | SERPINA3 | -0,3218 | 3,08337E-11 |
| ACE2 | NDUFAF6 | -0,3219 | 3,05178E-11 |
| ACE2 | PTP4A2 | -0,322 | 2,99943E-11 |
| ACE2 | RP11-462D18.2 | -0,322 | 3,03521E-11 |
| ACE2 | SLIRP | -0,3221 | 2,95268E-11 |
| ACE2 | AP000347.4 | -0,3222 | 2,90492E-11 |
| ACE2 | RP11-371A22.1 | -0,3224 | 2,84746E-11 |
| ACE2 | HERC2P3 | -0,3224 | 2,83824E-11 |
| ACE2 | CTA-292E10.6 | -0,3225 | 2,79176E-11 |
| ACE2 | RP1-149A16.17 | -0,3225 | 2,80583E-11 |
| ACE2 | AC125232.1 | -0,3226 | 2,74059E-11 |
| ACE2 | POP7 | -0,3226 | 2,76016E-11 |
| ACE2 | RP11-524G24.2 | -0,3226 | 2,7674E-11 |
| ACE2 | ARNTL2-AS1 | -0,3226 | 2,74328E-11 |
| ACE2 | AC006116.13 | -0,3226 | 2,76345E-11 |
| ACE2 | RP11-218C14.8 | -0,3226 | 2,74917E-11 |
| ACE2 | MIR663AHG | -0,3226 | 2,76378E-11 |
| ACE2 | RP11-190A12.9 | -0,3227 | 2,71419E-11 |
| ACE2 | RP11-231L11.3 | -0,3227 | 2,69766E-11 |
| ACE2 | ANKRD17 | -0,3227 | 2,71448E-11 |
| ACE2 | TNKS2-AS1 | -0,3227 | 2,70976E-11 |
| ACE2 | RP11-282A11.4 | -0,3227 | 2,71046E-11 |
| ACE2 | GRAP | -0,3227 | 2,73381E-11 |
| ACE2 | LINC01841 | -0,3227 | 2,72355E-11 |
| ACE2 | ITFG2 | -0,3229 | 2,65395E-11 |
| ACE2 | RP11-362A9.3 | -0,323 | 2,6156E-11 |
| ACE2 | RN7SL494P | -0,323 | 2,59717E-11 |
| ACE2 | ACP6 | -0,3231 | 2,55382E-11 |

| | | | |
|---|---|---|---|
| ACE2 | RP11-71N10.1 | -0,3231 | 2,54658E-11 |
| ACE2 | RP13-487K5.1 | -0,3231 | 2,55154E-11 |
| ACE2 | RP11-252A24.8 | -0,3231 | 2,56926E-11 |
| ACE2 | HTR6 | -0,3232 | 2,51531E-11 |
| ACE2 | MERTK | -0,3232 | 2,51238E-11 |
| ACE2 | RP11-761N21.2 | -0,3232 | 2,51291E-11 |
| ACE2 | RASSF1-AS1 | -0,3232 | 2,50744E-11 |
| ACE2 | RBMXP4 | -0,3232 | 2,51178E-11 |
| ACE2 | AC098795.1 | -0,3233 | 2,48764E-11 |
| ACE2 | RP11-12A2.1 | -0,3234 | 2,45334E-11 |
| ACE2 | PCNPP5 | -0,3234 | 2,44216E-11 |
| ACE2 | AQR | -0,3234 | 2,43943E-11 |
| ACE2 | LINC01473 | -0,3235 | 2,42243E-11 |
| ACE2 | POLD2P1 | -0,3235 | 2,40426E-11 |
| ACE2 | SLC25A41 | -0,3235 | 2,39959E-11 |
| ACE2 | MKKS | -0,3235 | 2,40246E-11 |
| ACE2 | AC000068.9 | -0,3235 | 2,41797E-11 |
| ACE2 | RP11-46A10.8 | -0,3236 | 2,39082E-11 |
| ACE2 | RP11-480D4.6 | -0,3236 | 2,38408E-11 |
| ACE2 | BYSL | -0,3236 | 2,38116E-11 |
| ACE2 | HS6ST1P1 | -0,3237 | 2,32802E-11 |
| ACE2 | LIN28B | -0,3237 | 2,34296E-11 |
| ACE2 | MAPRE1P1 | -0,3237 | 2,3571E-11 |
| ACE2 | FAM175B | -0,3237 | 2,34766E-11 |
| ACE2 | IGFBPL1 | -0,3238 | 2,2898E-11 |
| ACE2 | CTD-3203P2.1 | -0,3239 | 2,2779E-11 |
| ACE2 | AC009474.1 | -0,3241 | 2,21989E-11 |
| ACE2 | UNC119B | -0,3241 | 2,21343E-11 |
| ACE2 | RP4-539M6.20 | -0,3241 | 2,21663E-11 |
| ACE2 | CYP4Z2P | -0,3242 | 2,16582E-11 |
| ACE2 | RP11-598F7.5 | -0,3242 | 2,17395E-11 |
| ACE2 | GABRA5 | -0,3242 | 2,16645E-11 |
| ACE2 | PSMD10P1 | -0,3242 | 2,15968E-11 |
| ACE2 | TTTY15 | -0,3242 | 2,15781E-11 |
| ACE2 | RP11-597D13.2 | -0,3243 | 2,12714E-11 |
| ACE2 | CTD-2371O3.2 | -0,3243 | 2,12719E-11 |
| ACE2 | RP11-507K2.2 | -0,3243 | 2,15105E-11 |
| ACE2 | RP11-196G11.2 | -0,3243 | 2,137E-11 |
| ACE2 | LINC02137 | -0,3243 | 2,13933E-11 |
| ACE2 | FTCDNL1 | -0,3244 | 2,09763E-11 |
| ACE2 | AC011247.3 | -0,3245 | 2,06801E-11 |
| ACE2 | RP11-399O19.9 | -0,3245 | 2,06677E-11 |
| ACE2 | PNPLA6 | -0,3245 | 2,08099E-11 |
| ACE2 | UBE3C | -0,3246 | 2,0341E-11 |
| ACE2 | RP11-403D15.2 | -0,3246 | 2,0479E-11 |
| ACE2 | RP4-657D16.3 | -0,3247 | 2,02352E-11 |
| ACE2 | CTA-254O6.1 | -0,3247 | 2,01468E-11 |
| ACE2 | CACNA1C-IT3 | -0,3247 | 2,01399E-11 |
| ACE2 | DCUN1D2-AS | -0,3247 | 2,02983E-11 |
| ACE2 | HSD17B7 | -0,3248 | 2,00043E-11 |
| ACE2 | VDAC3 | -0,3248 | 1,99065E-11 |
| ACE2 | LINC01567 | -0,3248 | 1,97164E-11 |
| ACE2 | OR7E99P | -0,3249 | 1,96247E-11 |
| ACE2 | CSF3 | -0,3249 | 1,96575E-11 |

| | | | |
|---|---|---|---|
| ACE2 | HSPB1P2 | -0,3249 | 1,95861E-11 |
| ACE2 | RP3-510D11.4 | -0,325 | 1,93407E-11 |
| ACE2 | RPL5P9 | -0,325 | 1,91938E-11 |
| ACE2 | PDZD9 | -0,325 | 1,92281E-11 |
| ACE2 | MRPL54 | -0,325 | 1,93583E-11 |
| ACE2 | KCNE1B | -0,325 | 1,92015E-11 |
| ACE2 | AC000089.3 | -0,325 | 1,9385E-11 |
| ACE2 | AC093724.2 | -0,3251 | 1,8866E-11 |
| ACE2 | MTBP | -0,3251 | 1,88451E-11 |
| ACE2 | RN7SKP161 | -0,3251 | 1,89436E-11 |
| ACE2 | FAUP1 | -0,3251 | 1,90607E-11 |
| ACE2 | AP001432.14 | -0,3251 | 1,89154E-11 |
| ACE2 | LRRC53 | -0,3252 | 1,87807E-11 |
| ACE2 | RP11-459I19.1 | -0,3253 | 1,84036E-11 |
| ACE2 | MTHFD2L | -0,3253 | 1,83829E-11 |
| ACE2 | RP5-844F9.1 | -0,3254 | 1,82648E-11 |
| ACE2 | CBS | -0,3254 | 1,81831E-11 |
| ACE2 | ELL2 | -0,3255 | 1,79878E-11 |
| ACE2 | SDHAF2 | -0,3255 | 1,78024E-11 |
| ACE2 | RP11-569D9.5 | -0,3255 | 1,77495E-11 |
| ACE2 | RP11-576D8.4 | -0,3256 | 1,75049E-11 |
| ACE2 | LINC01410 | -0,3256 | 1,76281E-11 |
| ACE2 | ESRRAP2 | -0,3256 | 1,76445E-11 |
| ACE2 | CTC-507E2.2 | -0,3256 | 1,76552E-11 |
| ACE2 | RP5-1070G24.2 | -0,3257 | 1,72533E-11 |
| ACE2 | KB-1043D8.6 | -0,3257 | 1,73753E-11 |
| ACE2 | MRPL58 | -0,3257 | 1,74265E-11 |
| ACE2 | TUBA1B | -0,3258 | 1,70229E-11 |
| ACE2 | TMX2P1 | -0,3259 | 1,69581E-11 |
| ACE2 | RP11-255M2.3 | -0,3259 | 1,68085E-11 |
| ACE2 | ELK1 | -0,3259 | 1,69377E-11 |
| ACE2 | RP11-400L8.2 | -0,326 | 1,65409E-11 |
| ACE2 | RP11-285J16.1 | -0,326 | 1,66598E-11 |
| ACE2 | OFD1P17 | -0,326 | 1,66875E-11 |
| ACE2 | TSC22D1 | -0,326 | 1,65012E-11 |
| ACE2 | NDUFB1 | -0,326 | 1,65346E-11 |
| ACE2 | RP11-323F24.4 | -0,326 | 1,65513E-11 |
| ACE2 | COQ7 | -0,326 | 1,66677E-11 |
| ACE2 | RP13-638C3.6 | -0,326 | 1,64865E-11 |
| ACE2 | RP11-399K21.10 | -0,3261 | 1,64477E-11 |
| ACE2 | ETV4 | -0,3261 | 1,6234E-11 |
| ACE2 | RP11-108M9.4 | -0,3262 | 1,62156E-11 |
| ACE2 | RP4-791M13.3 | -0,3262 | 1,60505E-11 |
| ACE2 | DCTN5 | -0,3262 | 1,62009E-11 |
| ACE2 | RP11-46J23.1 | -0,3263 | 1,59385E-11 |
| ACE2 | KLHL40 | -0,3264 | 1,5686E-11 |
| ACE2 | RP11-9N12.2 | -0,3264 | 1,56449E-11 |
| ACE2 | LINC00463 | -0,3264 | 1,55707E-11 |
| ACE2 | RP13-143G15.4 | -0,3265 | 1,53746E-11 |
| ACE2 | RP11-491F9.5 | -0,3265 | 1,53828E-11 |
| ACE2 | CCRL1P1 | -0,3266 | 1,51069E-11 |
| ACE2 | TNKS1BP1 | -0,3266 | 1,51468E-11 |
| ACE2 | OAS2 | -0,3268 | 1,46853E-11 |
| ACE2 | ZNF295-AS1 | -0,3268 | 1,46656E-11 |

| | | | |
|---|---|---|---|
| ACE2 | EPDR1 | -0,3269 | 1,44278E-11 |
| ACE2 | C12orf45 | -0,3269 | 1,45625E-11 |
| ACE2 | TRIM7 | -0,327 | 1,4292E-11 |
| ACE2 | CTD-2024I7.18 | -0,3271 | 1,41359E-11 |
| ACE2 | INTS4P1 | -0,3271 | 1,41258E-11 |
| ACE2 | AC006378.2 | -0,3272 | 1,37846E-11 |
| ACE2 | SMCR5 | -0,3272 | 1,38614E-11 |
| ACE2 | RP11-96L14.7 | -0,3273 | 1,36088E-11 |
| ACE2 | ABLIM1 | -0,3273 | 1,36802E-11 |
| ACE2 | STAG1 | -0,3274 | 1,34162E-11 |
| ACE2 | GS1-44D20.1 | -0,3274 | 1,34984E-11 |
| ACE2 | URM1 | -0,3274 | 1,3468E-11 |
| ACE2 | RP11-51J9.4 | -0,3275 | 1,31837E-11 |
| ACE2 | RP11-909N17.2 | -0,3275 | 1,3195E-11 |
| ACE2 | LMO7-AS1 | -0,3275 | 1,3179E-11 |
| ACE2 | ALDOC | -0,3275 | 1,32273E-11 |
| ACE2 | NCBP2L | -0,3275 | 1,31322E-11 |
| ACE2 | RP13-228J13.6 | -0,3275 | 1,32546E-11 |
| ACE2 | AAK1 | -0,3276 | 1,29872E-11 |
| ACE2 | AC073130.1 | -0,3277 | 1,2787E-11 |
| ACE2 | AP001062.7 | -0,3277 | 1,2778E-11 |
| ACE2 | FAM225B | -0,3278 | 1,25862E-11 |
| ACE2 | MIR770 | -0,3278 | 1,26675E-11 |
| ACE2 | AC026954.6 | -0,3278 | 1,25722E-11 |
| ACE2 | RP11-513M1.1 | -0,3278 | 1,26461E-11 |
| ACE2 | ATP5J | -0,328 | 1,21869E-11 |
| ACE2 | MYL12BP1 | -0,3282 | 1,1899E-11 |
| ACE2 | MAP3K15 | -0,3282 | 1,18168E-11 |
| ACE2 | GORASP2 | -0,3283 | 1,1698E-11 |
| ACE2 | PPTC7 | -0,3283 | 1,17772E-11 |
| ACE2 | RRP12 | -0,3284 | 1,15072E-11 |
| ACE2 | BACH1-IT2 | -0,3284 | 1,15018E-11 |
| ACE2 | C9orf43 | -0,3285 | 1,14113E-11 |
| ACE2 | PLAU | -0,3285 | 1,14162E-11 |
| ACE2 | CTD-2525I3.2 | -0,3285 | 1,14522E-11 |
| ACE2 | DZANK1 | -0,3285 | 1,138E-11 |
| ACE2 | IQCA1L | -0,3286 | 1,12395E-11 |
| ACE2 | CDKN2AIP | -0,3287 | 1,1006E-11 |
| ACE2 | DRD1 | -0,3287 | 1,10936E-11 |
| ACE2 | MXI1 | -0,3287 | 1,10103E-11 |
| ACE2 | CBWD1 | -0,3289 | 1,06655E-11 |
| ACE2 | RP11-165D7.5 | -0,3289 | 1,07755E-11 |
| ACE2 | CDRT1 | -0,3289 | 1,06614E-11 |
| ACE2 | AP001372.2 | -0,329 | 1,04806E-11 |
| ACE2 | RP11-530C5.2 | -0,3291 | 1,0332E-11 |
| ACE2 | SMIM20 | -0,3292 | 1,02587E-11 |
| ACE2 | RPL5P18 | -0,3293 | 1,00268E-11 |
| ACE2 | LOC440461 | -0,3293 | 1,01182E-11 |
| ACE2 | TMC2 | -0,3293 | 1,00335E-11 |
| ACE2 | RPL7P18 | -0,3294 | 9,9943E-12 |
| ACE2 | ALG1L11P | -0,3294 | 9,84885E-12 |
| ACE2 | LRRC69 | -0,3294 | 9,92872E-12 |
| ACE2 | MRPS31P5 | -0,3295 | 9,80815E-12 |
| ACE2 | LINC02170 | -0,3295 | 9,84089E-12 |

| | | | |
|---|---|---|---|
| ACE2 | GS1-304P7.3 | -0,3296 | 9,61669E-12 |
| ACE2 | MIR641 | -0,3296 | 9,68711E-12 |
| ACE2 | SELENOO | -0,3296 | 9,6474E-12 |
| ACE2 | AP3D1 | -0,3297 | 9,43477E-12 |
| ACE2 | TIMP4 | -0,3298 | 9,34053E-12 |
| ACE2 | RP11-48G14.1 | -0,33 | 9,06396E-12 |
| ACE2 | HMGB1P37 | -0,3301 | 8,85213E-12 |
| ACE2 | THBS1 | -0,3301 | 8,88433E-12 |
| ACE2 | ZNF33CP | -0,3302 | 8,73953E-12 |
| ACE2 | CEMIP | -0,3302 | 8,79652E-12 |
| ACE2 | RP11-362K14.7 | -0,3303 | 8,59257E-12 |
| ACE2 | RP11-359P5.1 | -0,3303 | 8,67021E-12 |
| ACE2 | RP11-467D6.1 | -0,3304 | 8,58035E-12 |
| ACE2 | CSDAP1 | -0,3304 | 8,56171E-12 |
| ACE2 | UBE2S | -0,3304 | 8,51592E-12 |
| ACE2 | RP11-123O22.1 | -0,3305 | 8,39097E-12 |
| ACE2 | RP11-651L5.3 | -0,3305 | 8,32589E-12 |
| ACE2 | RP11-455O6.9 | -0,3305 | 8,40049E-12 |
| ACE2 | SLC25A18 | -0,3305 | 8,44715E-12 |
| ACE2 | UBE2Q2P11 | -0,3306 | 8,32068E-12 |
| ACE2 | EPHX3 | -0,3307 | 8,07656E-12 |
| ACE2 | ZNF586 | -0,3307 | 8,1664E-12 |
| ACE2 | RP11-452C8.1 | -0,3308 | 8,03293E-12 |
| ACE2 | ALKBH4 | -0,3308 | 8,00576E-12 |
| ACE2 | RNF26 | -0,3308 | 7,95321E-12 |
| ACE2 | SPTA1 | -0,3309 | 7,86479E-12 |
| ACE2 | SH3BP5L | -0,3309 | 7,91537E-12 |
| ACE2 | SLC22A14 | -0,3309 | 7,88253E-12 |
| ACE2 | METTL15P3 | -0,3309 | 7,89384E-12 |
| ACE2 | PET117 | -0,331 | 7,76862E-12 |
| ACE2 | HMGB3P10 | -0,3311 | 7,6113E-12 |
| ACE2 | IPO8P1 | -0,3311 | 7,70468E-12 |
| ACE2 | AOX3P | -0,3311 | 7,6411E-12 |
| ACE2 | CTD-2083E4.6 | -0,3311 | 7,64695E-12 |
| ACE2 | METTL2B | -0,3311 | 7,68486E-12 |
| ACE2 | RP11-121G22.3 | -0,3311 | 7,68914E-12 |
| ACE2 | CCDC171 | -0,3312 | 7,5207E-12 |
| ACE2 | ADAM1B | -0,3312 | 7,5274E-12 |
| ACE2 | RP11-631N16.2 | -0,3314 | 7,25692E-12 |
| ACE2 | TBX5-AS1 | -0,3314 | 7,35699E-12 |
| ACE2 | NRAV | -0,3314 | 7,3103E-12 |
| ACE2 | RP1-302D9.1 | -0,3314 | 7,33294E-12 |
| ACE2 | NBPF10 | -0,3315 | 7,24159E-12 |
| ACE2 | TNRC18P1 | -0,3315 | 7,15579E-12 |
| ACE2 | BX255923.3 | -0,3315 | 7,17481E-12 |
| ACE2 | NKX6-2 | -0,3315 | 7,14708E-12 |
| ACE2 | ATP5L | -0,3316 | 7,03998E-12 |
| ACE2 | PPP1R3E | -0,3316 | 7,05591E-12 |
| ACE2 | RPL39P5 | -0,3317 | 6,98306E-12 |
| ACE2 | AC004987.9 | -0,3318 | 6,8514E-12 |
| ACE2 | RP11-409C19.2 | -0,3318 | 6,87947E-12 |
| ACE2 | NME2P1 | -0,3318 | 6,87588E-12 |
| ACE2 | WASH3P | -0,3318 | 6,82753E-12 |
| ACE2 | VMAC | -0,3318 | 6,89358E-12 |

| | | | |
|---|---|---|---|
| ACE2 | UACA | -0,3319 | 6,74084E-12 |
| ACE2 | HMGB1P24 | -0,332 | 6,61617E-12 |
| ACE2 | CTU1 | -0,332 | 6,67402E-12 |
| ACE2 | SNRPG | -0,3321 | 6,53364E-12 |
| ACE2 | GS1-124K5.9 | -0,3321 | 6,5274E-12 |
| ACE2 | MAML2 | -0,3321 | 6,57043E-12 |
| ACE2 | OR7M1P | -0,3322 | 6,48187E-12 |
| ACE2 | AC116366.6 | -0,3324 | 6,24295E-12 |
| ACE2 | UBAP2 | -0,3324 | 6,27117E-12 |
| ACE2 | PDCD11 | -0,3324 | 6,26966E-12 |
| ACE2 | RP11-122K13.12 | -0,3324 | 6,23183E-12 |
| ACE2 | RP11-108K14.12 | -0,3324 | 6,22574E-12 |
| ACE2 | VMO1 | -0,3324 | 6,28428E-12 |
| ACE2 | RP11-63M22.1 | -0,3326 | 6,04433E-12 |
| ACE2 | CD70 | -0,3326 | 6,08346E-12 |
| ACE2 | RP11-326I11.4 | -0,3327 | 5,95406E-12 |
| ACE2 | CCDC27 | -0,3328 | 5,92258E-12 |
| ACE2 | RP11-96D1.5 | -0,3328 | 5,85361E-12 |
| ACE2 | AC114808.2 | -0,3329 | 5,80293E-12 |
| ACE2 | FIP1L1 | -0,3329 | 5,79839E-12 |
| ACE2 | RIMBP2 | -0,3329 | 5,81784E-12 |
| ACE2 | ZBTB24 | -0,333 | 5,72214E-12 |
| ACE2 | RP11-320L11.2 | -0,333 | 5,70247E-12 |
| ACE2 | RP11-415J8.7 | -0,3331 | 5,59903E-12 |
| ACE2 | SIPA1L2 | -0,3331 | 5,61689E-12 |
| ACE2 | SLC30A3 | -0,3331 | 5,62379E-12 |
| ACE2 | RP1-53O8.2 | -0,3331 | 5,60128E-12 |
| ACE2 | RNF113A | -0,3332 | 5,50855E-12 |
| ACE2 | CYB5R2 | -0,3333 | 5,4653E-12 |
| ACE2 | CTD-2315E11.1 | -0,3335 | 5,2695E-12 |
| ACE2 | RP11-181G12.4 | -0,3336 | 5,2488E-12 |
| ACE2 | LINC00928 | -0,3336 | 5,24574E-12 |
| ACE2 | CTD-2095E4.3 | -0,3336 | 5,17361E-12 |
| ACE2 | LAMTOR3 | -0,3337 | 5,12308E-12 |
| ACE2 | CTD-3076O17.1 | -0,3337 | 5,11627E-12 |
| ACE2 | CLDND2 | -0,3337 | 5,16629E-12 |
| ACE2 | CTD-2337A12.1 | -0,3339 | 4,96176E-12 |
| ACE2 | AC010731.2 | -0,334 | 4,89706E-12 |
| ACE2 | MCM9 | -0,334 | 4,92326E-12 |
| ACE2 | CTD-2248H3.1 | -0,3341 | 4,83158E-12 |
| ACE2 | SLC26A4-AS1 | -0,3341 | 4,85368E-12 |
| ACE2 | LRRC37A | -0,3341 | 4,82669E-12 |
| ACE2 | RP11-51F16.1 | -0,3341 | 4,81866E-12 |
| ACE2 | RP11-234K24.3 | -0,3341 | 4,79125E-12 |
| ACE2 | ALLC | -0,3342 | 4,72869E-12 |
| ACE2 | RP11-337N6.3 | -0,3342 | 4,72238E-12 |
| ACE2 | RPL35P5 | -0,3342 | 4,74407E-12 |
| ACE2 | BDNF | -0,3342 | 4,76533E-12 |
| ACE2 | DUX4L27 | -0,3342 | 4,76616E-12 |
| ACE2 | RP11-1072A3.4 | -0,3342 | 4,74937E-12 |
| ACE2 | ZBTB8OS | -0,3344 | 4,57941E-12 |
| ACE2 | RP11-397E7.4 | -0,3344 | 4,58433E-12 |
| ACE2 | RP11-1280N14.3 | -0,3344 | 4,6228E-12 |
| ACE2 | GLYR1 | -0,3344 | 4,61811E-12 |

| | | | |
|---|---|---|---|
| ACE2 | EMC1 | -0,3345 | 4,5231E-12 |
| ACE2 | VPS28 | -0,3345 | 4,53438E-12 |
| ACE2 | PAICSP1 | -0,3345 | 4,54143E-12 |
| ACE2 | SBNO1 | -0,3346 | 4,45033E-12 |
| ACE2 | PRICKLE3 | -0,3346 | 4,49164E-12 |
| ACE2 | SDR42E2 | -0,3347 | 4,37508E-12 |
| ACE2 | TCAF1P1 | -0,3348 | 4,33224E-12 |
| ACE2 | RPL5P23 | -0,3348 | 4,33653E-12 |
| ACE2 | RP11-371F15.3 | -0,3349 | 4,28762E-12 |
| ACE2 | AF131215.8 | -0,3349 | 4,2483E-12 |
| ACE2 | HTR7 | -0,3349 | 4,23296E-12 |
| ACE2 | MIR3180-4 | -0,3349 | 4,28796E-12 |
| ACE2 | MINOS1 | -0,335 | 4,18793E-12 |
| ACE2 | SLC37A3 | -0,335 | 4,19764E-12 |
| ACE2 | NPM3 | -0,335 | 4,16294E-12 |
| ACE2 | EIF3FP3 | -0,3351 | 4,12326E-12 |
| ACE2 | PILRA | -0,3351 | 4,11871E-12 |
| ACE2 | CTTN | -0,3351 | 4,0991E-12 |
| ACE2 | RP11-69H7.2 | -0,3351 | 4,12821E-12 |
| ACE2 | SERHL | -0,3351 | 4,13946E-12 |
| ACE2 | CX3CR1 | -0,3352 | 4,08484E-12 |
| ACE2 | CYP1A1 | -0,3352 | 4,04492E-12 |
| ACE2 | XXbac-BPG252P9.10 | -0,3353 | 4,00622E-12 |
| ACE2 | RP11-85I17.2 | -0,3353 | 3,98172E-12 |
| ACE2 | SSU72 | -0,3354 | 3,94589E-12 |
| ACE2 | ERLEC1 | -0,3354 | 3,94764E-12 |
| ACE2 | SH2D4B | -0,3354 | 3,95449E-12 |
| ACE2 | RNF126P1 | -0,3354 | 3,95476E-12 |
| ACE2 | AKT3-IT1 | -0,3355 | 3,88308E-12 |
| ACE2 | DNTTIP1 | -0,3356 | 3,84165E-12 |
| ACE2 | DDX50P1 | -0,3357 | 3,75571E-12 |
| ACE2 | RP5-891H21.5 | -0,3358 | 3,69381E-12 |
| ACE2 | RNA5SP290 | -0,3358 | 3,70729E-12 |
| ACE2 | ZNF574 | -0,3358 | 3,72495E-12 |
| ACE2 | CTC-484P3.3 | -0,3359 | 3,6397E-12 |
| ACE2 | RP11-207C16.4 | -0,3359 | 3,63204E-12 |
| ACE2 | RP11-80H5.7 | -0,3359 | 3,62524E-12 |
| ACE2 | COIL | -0,3359 | 3,62031E-12 |
| ACE2 | RP11-93I21.3 | -0,336 | 3,56122E-12 |
| ACE2 | ATP8B5P | -0,336 | 3,56525E-12 |
| ACE2 | RP11-286N22.10 | -0,336 | 3,5775E-12 |
| ACE2 | RP1-63M2.5 | -0,336 | 3,59772E-12 |
| ACE2 | LINC00605 | -0,3361 | 3,51991E-12 |
| ACE2 | ZBTB20-AS1 | -0,3362 | 3,49787E-12 |
| ACE2 | AC009245.3 | -0,3362 | 3,50093E-12 |
| ACE2 | SLC8A3 | -0,3362 | 3,49648E-12 |
| ACE2 | RP4-753F5.1 | -0,3363 | 3,40341E-12 |
| ACE2 | MFSD10 | -0,3363 | 3,41715E-12 |
| ACE2 | RP11-61I13.3 | -0,3363 | 3,41413E-12 |
| ACE2 | RP11-252P19.1 | -0,3363 | 3,44234E-12 |
| ACE2 | NFXL1 | -0,3364 | 3,39443E-12 |
| ACE2 | RWDD4P2 | -0,3364 | 3,39276E-12 |
| ACE2 | RP11-45M22.3 | -0,3364 | 3,34785E-12 |
| ACE2 | CH17-172B3.1 | -0,3365 | 3,31166E-12 |

| | | | |
|---|---|---|---|
| ACE2 | KDM4E | -0,3365 | 3,33937E-12 |
| ACE2 | RP11-589P10.5 | -0,3365 | 3,34313E-12 |
| ACE2 | GRIN3B | -0,3365 | 3,33513E-12 |
| ACE2 | SSTR4 | -0,3365 | 3,34146E-12 |
| ACE2 | RP11-54O7.16 | -0,3366 | 3,29202E-12 |
| ACE2 | AC007365.3 | -0,3366 | 3,26104E-12 |
| ACE2 | RP11-446F17.3 | -0,3366 | 3,24415E-12 |
| ACE2 | RP5-832C2.5 | -0,3367 | 3,20798E-12 |
| ACE2 | MARCH11 | -0,3367 | 3,20025E-12 |
| ACE2 | AC010729.1 | -0,3368 | 3,14482E-12 |
| ACE2 | CTD-2201G3.1 | -0,3368 | 3,14961E-12 |
| ACE2 | RP11-342M3.5 | -0,3368 | 3,18535E-12 |
| ACE2 | CTA-212D2.2 | -0,3369 | 3,12717E-12 |
| ACE2 | MYL7 | -0,3369 | 3,10221E-12 |
| ACE2 | RP11-236J17.7 | -0,3369 | 3,1266E-12 |
| ACE2 | RP13-349O20.2 | -0,3369 | 3,12679E-12 |
| ACE2 | LPAL2 | -0,337 | 3,07769E-12 |
| ACE2 | VPS26A | -0,337 | 3,07656E-12 |
| ACE2 | NME2 | -0,337 | 3,08367E-12 |
| ACE2 | KCNG2 | -0,337 | 3,05765E-12 |
| ACE2 | ADNP2 | -0,337 | 3,0877E-12 |
| ACE2 | ST3GAL5-AS1 | -0,3371 | 3,03814E-12 |
| ACE2 | RP11-445L6.3 | -0,3371 | 3,02594E-12 |
| ACE2 | DNAJB1 | -0,3371 | 3,01025E-12 |
| ACE2 | XXyac-YR38GF2.1 | -0,3372 | 2,95437E-12 |
| ACE2 | CTD-2003C8.2 | -0,3372 | 2,99355E-12 |
| ACE2 | RP11-893F2.15 | -0,3372 | 2,98825E-12 |
| ACE2 | RP11-430H12.2 | -0,3373 | 2,95132E-12 |
| ACE2 | RP11-163E9.1 | -0,3373 | 2,92092E-12 |
| ACE2 | HPR | -0,3373 | 2,94808E-12 |
| ACE2 | COX11 | -0,3373 | 2,92195E-12 |
| ACE2 | AC010524.4 | -0,3373 | 2,95118E-12 |
| ACE2 | LINC00686 | -0,3373 | 2,93935E-12 |
| ACE2 | C1orf21 | -0,3374 | 2,89072E-12 |
| ACE2 | AC004453.8 | -0,3374 | 2,89879E-12 |
| ACE2 | PTS | -0,3374 | 2,86854E-12 |
| ACE2 | RP11-287D1.4 | -0,3375 | 2,82152E-12 |
| ACE2 | RTN4RL1 | -0,3375 | 2,85855E-12 |
| ACE2 | RN7SL263P | -0,3375 | 2,82676E-12 |
| ACE2 | PABPC1P3 | -0,3375 | 2,81769E-12 |
| ACE2 | RP11-147G7.3 | -0,3376 | 2,81352E-12 |
| ACE2 | OR52K3P | -0,3376 | 2,78136E-12 |
| ACE2 | RPS26P21 | -0,3377 | 2,75039E-12 |
| ACE2 | RP11-401O9.4 | -0,3377 | 2,74405E-12 |
| ACE2 | CNBD2 | -0,3377 | 2,75415E-12 |
| ACE2 | AC133106.2 | -0,3378 | 2,69072E-12 |
| ACE2 | XXbac-BPG294E21.9 | -0,3378 | 2,71604E-12 |
| ACE2 | ATXN8OS | -0,3378 | 2,72953E-12 |
| ACE2 | TKTL1 | -0,3378 | 2,71131E-12 |
| ACE2 | AC011290.5 | -0,338 | 2,60805E-12 |
| ACE2 | UQCR11 | -0,338 | 2,6113E-12 |
| ACE2 | RP11-561N12.5 | -0,3382 | 2,54281E-12 |
| ACE2 | MEPCE | -0,3382 | 2,54778E-12 |
| ACE2 | MED8 | -0,3383 | 2,52266E-12 |

| | | | |
|---|---|---|---|
| ACE2 | NSUN6 | -0,3383 | 2,51419E-12 |
| ACE2 | POLA2 | -0,3383 | 2,49849E-12 |
| ACE2 | RP11-342K6.1 | -0,3384 | 2,46585E-12 |
| ACE2 | RP11-344A5.1 | -0,3384 | 2,48233E-12 |
| ACE2 | N4BP2L2-IT2 | -0,3384 | 2,44699E-12 |
| ACE2 | MKRN7P | -0,3384 | 2,45936E-12 |
| ACE2 | LINC01426 | -0,3384 | 2,46081E-12 |
| ACE2 | CTC-529L17.2 | -0,3385 | 2,41264E-12 |
| ACE2 | RP11-498P14.3 | -0,3385 | 2,42553E-12 |
| ACE2 | PGAM1 | -0,3385 | 2,41862E-12 |
| ACE2 | GRK5-IT1 | -0,3385 | 2,4102E-12 |
| ACE2 | ATXN7L3B | -0,3385 | 2,41401E-12 |
| ACE2 | TNFSF13B | -0,3385 | 2,43907E-12 |
| ACE2 | RP1-67K17.4 | -0,3386 | 2,39229E-12 |
| ACE2 | SMARCE1P5 | -0,3386 | 2,40244E-12 |
| ACE2 | RPL7AP66 | -0,3386 | 2,3802E-12 |
| ACE2 | DDX50P2 | -0,3387 | 2,34917E-12 |
| ACE2 | ZSCAN25 | -0,3387 | 2,35618E-12 |
| ACE2 | RP11-677M14.7 | -0,3387 | 2,34604E-12 |
| ACE2 | RP4-614O4.13 | -0,3387 | 2,35764E-12 |
| ACE2 | CH17-140K24.5 | -0,3388 | 2,32097E-12 |
| ACE2 | NOX3 | -0,3388 | 2,32784E-12 |
| ACE2 | AF038458.5 | -0,3388 | 2,30952E-12 |
| ACE2 | OSGEPL1-AS1 | -0,3389 | 2,27268E-12 |
| ACE2 | ADH1A | -0,3389 | 2,28941E-12 |
| ACE2 | RP11-679B19.2 | -0,339 | 2,24762E-12 |
| ACE2 | AC108463.1 | -0,3391 | 2,19868E-12 |
| ACE2 | RP3-466P17.3 | -0,3391 | 2,20659E-12 |
| ACE2 | TRPV4 | -0,3391 | 2,22285E-12 |
| ACE2 | ACTG1P17 | -0,3391 | 2,21915E-12 |
| ACE2 | NFYC | -0,3392 | 2,16688E-12 |
| ACE2 | RP11-365F18.1 | -0,3392 | 2,15692E-12 |
| ACE2 | DUSP4 | -0,3392 | 2,15762E-12 |
| ACE2 | RP11-111M22.2 | -0,3393 | 2,14637E-12 |
| ACE2 | CTD-2162K18.4 | -0,3393 | 2,13008E-12 |
| ACE2 | RP13-216E22.4 | -0,3393 | 2,14398E-12 |
| ACE2 | AC092614.2 | -0,3394 | 2,09796E-12 |
| ACE2 | FAM104A | -0,3394 | 2,0926E-12 |
| ACE2 | RP11-986G18.3 | -0,3395 | 2,08475E-12 |
| ACE2 | APLNR | -0,3396 | 2,0472E-12 |
| ACE2 | RP11-50B3.2 | -0,3396 | 2,0545E-12 |
| ACE2 | PABPC5-AS1 | -0,3396 | 2,05023E-12 |
| ACE2 | UPP2 | -0,3397 | 1,99817E-12 |
| ACE2 | RP11-390P2.4 | -0,3397 | 1,99592E-12 |
| ACE2 | RP11-24B13.2 | -0,3398 | 1,98335E-12 |
| ACE2 | FAF2 | -0,3399 | 1,95094E-12 |
| ACE2 | VAPA | -0,3399 | 1,95171E-12 |
| ACE2 | SNORD23 | -0,3399 | 1,94024E-12 |
| ACE2 | VN1R87P | -0,3399 | 1,95325E-12 |
| ACE2 | TCERG1L | -0,34 | 1,91481E-12 |
| ACE2 | RP11-15A1.7 | -0,34 | 1,90105E-12 |
| ACE2 | PPP1R37 | -0,34 | 1,91116E-12 |
| ACE2 | AC093787.1 | -0,3401 | 1,88557E-12 |
| ACE2 | NALCN-AS1 | -0,3401 | 1,88438E-12 |

| | | | |
|---|---|---|---|
| ACE2 | CES1P2 | -0,3401 | 1,88803E-12 |
| ACE2 | RP11-1072C15.1 | -0,3401 | 1,87337E-12 |
| ACE2 | AC004691.5 | -0,3402 | 1,84907E-12 |
| ACE2 | RP11-179B2.2 | -0,3402 | 1,85039E-12 |
| ACE2 | RP1-4G17.2 | -0,3402 | 1,84159E-12 |
| ACE2 | RAB33A | -0,3402 | 1,86183E-12 |
| ACE2 | RP11-180O5.2 | -0,3403 | 1,82531E-12 |
| ACE2 | AC023481.1 | -0,3403 | 1,83252E-12 |
| ACE2 | EPHB4 | -0,3403 | 1,81518E-12 |
| ACE2 | ST13P6 | -0,3403 | 1,82207E-12 |
| ACE2 | RP11-91J19.2 | -0,3404 | 1,79025E-12 |
| ACE2 | CPSF3 | -0,3405 | 1,77573E-12 |
| ACE2 | TSN | -0,3406 | 1,74891E-12 |
| ACE2 | ATP6V1G1P2 | -0,3406 | 1,74932E-12 |
| ACE2 | RP11-100L22.1 | -0,3407 | 1,70528E-12 |
| ACE2 | SLC4A1 | -0,3407 | 1,703E-12 |
| ACE2 | RP11-400F19.18 | -0,3408 | 1,69019E-12 |
| ACE2 | RP11-396C23.4 | -0,3409 | 1,66782E-12 |
| ACE2 | COL6A6 | -0,3409 | 1,66638E-12 |
| ACE2 | TRAV8-5 | -0,3409 | 1,65316E-12 |
| ACE2 | CTD-2270N23.1 | -0,3409 | 1,66193E-12 |
| ACE2 | RP11-114H24.2 | -0,3409 | 1,67205E-12 |
| ACE2 | RBCK1 | -0,3409 | 1,66171E-12 |
| ACE2 | AC008277.1 | -0,341 | 1,64113E-12 |
| ACE2 | ZFP62 | -0,341 | 1,62756E-12 |
| ACE2 | AC005481.5 | -0,3411 | 1,60938E-12 |
| ACE2 | CCDC13 | -0,3412 | 1,5802E-12 |
| ACE2 | RPS4XP6 | -0,3412 | 1,58055E-12 |
| ACE2 | HLA-DQB1-AS1 | -0,3412 | 1,57705E-12 |
| ACE2 | CTD-2649C14.3 | -0,3412 | 1,58615E-12 |
| ACE2 | CLPP | -0,3412 | 1,57964E-12 |
| ACE2 | AC002984.2 | -0,3412 | 1,58849E-12 |
| ACE2 | RP11-506F3.1 | -0,3413 | 1,55583E-12 |
| ACE2 | RP11-977G19.14 | -0,3413 | 1,56557E-12 |
| ACE2 | RP11-160C18.2 | -0,3413 | 1,5608E-12 |
| ACE2 | SLC5A2 | -0,3413 | 1,55228E-12 |
| ACE2 | CTA-390C10.10 | -0,3413 | 1,56524E-12 |
| ACE2 | SOAT1 | -0,3414 | 1,54526E-12 |
| ACE2 | RP11-294N21.3 | -0,3415 | 1,50891E-12 |
| ACE2 | USP45 | -0,3416 | 1,49727E-12 |
| ACE2 | OR51B4 | -0,3416 | 1,48087E-12 |
| ACE2 | GP5 | -0,3417 | 1,46636E-12 |
| ACE2 | U2AF1L4 | -0,3417 | 1,46906E-12 |
| ACE2 | CTB-113P19.3 | -0,3418 | 1,44244E-12 |
| ACE2 | RP13-1032I1.11 | -0,3418 | 1,43533E-12 |
| ACE2 | UPK1A-AS1 | -0,3418 | 1,43237E-12 |
| ACE2 | ECE2 | -0,3419 | 1,40718E-12 |
| ACE2 | C1RL-AS1 | -0,3419 | 1,41562E-12 |
| ACE2 | LINC01684 | -0,3419 | 1,41158E-12 |
| ACE2 | RBBP5 | -0,342 | 1,38801E-12 |
| ACE2 | CCDC73 | -0,342 | 1,40413E-12 |
| ACE2 | ACCS | -0,342 | 1,38482E-12 |
| ACE2 | SPTY2D1-AS1 | -0,3421 | 1,37041E-12 |
| ACE2 | IDH2 | -0,3421 | 1,36236E-12 |

| | | | |
|---|---|---|---|
| ACE2 | LRRC28 | -0,3421 | 1,36957E-12 |
| ACE2 | ZNF560 | -0,3421 | 1,36968E-12 |
| ACE2 | AP000280.66 | -0,3421 | 1,38175E-12 |
| ACE2 | TMEM214 | -0,3422 | 1,34933E-12 |
| ACE2 | GABRA1 | -0,3422 | 1,35165E-12 |
| ACE2 | KRT18P59 | -0,3422 | 1,35686E-12 |
| ACE2 | RP11-490B18.5 | -0,3422 | 1,35688E-12 |
| ACE2 | ZNF155 | -0,3422 | 1,34919E-12 |
| ACE2 | YBX1P2 | -0,3423 | 1,32926E-12 |
| ACE2 | ANKRD30BL | -0,3424 | 1,30792E-12 |
| ACE2 | EEF1A1P8 | -0,3424 | 1,3044E-12 |
| ACE2 | CTD-2568A17.1 | -0,3425 | 1,28245E-12 |
| ACE2 | FCN1 | -0,3426 | 1,25817E-12 |
| ACE2 | RP11-504P24.9 | -0,3427 | 1,24644E-12 |
| ACE2 | RUVBL1 | -0,3427 | 1,24881E-12 |
| ACE2 | RP11-146F11.1 | -0,3427 | 1,24316E-12 |
| ACE2 | RP11-497H17.1 | -0,3427 | 1,25113E-12 |
| ACE2 | FTL | -0,3427 | 1,23942E-12 |
| ACE2 | CCNL1 | -0,3428 | 1,22188E-12 |
| ACE2 | CHTF8 | -0,3428 | 1,22597E-12 |
| ACE2 | RP11-191L17.1 | -0,3429 | 1,21652E-12 |
| ACE2 | RP11-521O16.1 | -0,3429 | 1,21508E-12 |
| ACE2 | CTSL3P | -0,3429 | 1,21341E-12 |
| ACE2 | AP000487.6 | -0,3429 | 1,21618E-12 |
| ACE2 | SLC24A5 | -0,3429 | 1,21431E-12 |
| ACE2 | NAP1L1P3 | -0,343 | 1,17962E-12 |
| ACE2 | RP13-192B19.2 | -0,343 | 1,19313E-12 |
| ACE2 | THG1L | -0,3431 | 1,17819E-12 |
| ACE2 | RPL26P20 | -0,3432 | 1,15135E-12 |
| ACE2 | ASB4 | -0,3432 | 1,15526E-12 |
| ACE2 | SNORD116-24 | -0,3432 | 1,16046E-12 |
| ACE2 | FAM201B | -0,3433 | 1,13139E-12 |
| ACE2 | DOCK6 | -0,3433 | 1,13627E-12 |
| ACE2 | EIF4A1 | -0,3434 | 1,11745E-12 |
| ACE2 | RP11-365O16.6 | -0,3435 | 1,09305E-12 |
| ACE2 | NPM1P37 | -0,3435 | 1,10177E-12 |
| ACE2 | SMLR1 | -0,3435 | 1,10255E-12 |
| ACE2 | BARX1 | -0,3435 | 1,10064E-12 |
| ACE2 | RP11-390B4.3 | -0,3435 | 1,09226E-12 |
| ACE2 | SHANK2 | -0,3436 | 1,07211E-12 |
| ACE2 | RP11-321P16.1 | -0,3436 | 1,08543E-12 |
| ACE2 | RP11-295P9.12 | -0,3437 | 1,058E-12 |
| ACE2 | RP11-22A3.2 | -0,3438 | 1,04337E-12 |
| ACE2 | C8orf34 | -0,3438 | 1,03836E-12 |
| ACE2 | CTD-2189E23.1 | -0,3438 | 1,05169E-12 |
| ACE2 | CASZ1 | -0,3439 | 1,03543E-12 |
| ACE2 | RP11-478C6.5 | -0,3439 | 1,03335E-12 |
| ACE2 | RP11-306G20.1 | -0,3439 | 1,03046E-12 |
| ACE2 | CYB5A | -0,3439 | 1,02293E-12 |
| ACE2 | CORT | -0,344 | 1,01772E-12 |
| ACE2 | VPS18 | -0,344 | 1,0047E-12 |
| ACE2 | PRSS57 | -0,344 | 1,01509E-12 |
| ACE2 | CBX1P3 | -0,3441 | 9,92863E-13 |
| ACE2 | RP11-371M22.1 | -0,3443 | 9,62779E-13 |

| | | | |
|---|---|---|---|
| ACE2 | RP11-63E9.1 | -0,3443 | 9,59093E-13 |
| ACE2 | AC092610.12 | -0,3443 | 9,69768E-13 |
| ACE2 | AGBL1 | -0,3443 | 9,64163E-13 |
| ACE2 | RP11-358N2.2 | -0,3443 | 9,59712E-13 |
| ACE2 | EPAS1 | -0,3444 | 9,44126E-13 |
| ACE2 | CDC123 | -0,3444 | 9,56283E-13 |
| ACE2 | MS4A14 | -0,3444 | 9,43408E-13 |
| ACE2 | RP11-789C17.3 | -0,3445 | 9,30365E-13 |
| ACE2 | RCAN1 | -0,3446 | 9,22269E-13 |
| ACE2 | RBMS3-AS1 | -0,3447 | 8,99403E-13 |
| ACE2 | RPL5P11 | -0,3447 | 9,01835E-13 |
| ACE2 | SLC7A5 | -0,3447 | 9,06477E-13 |
| ACE2 | ZNF703 | -0,3448 | 8,90715E-13 |
| ACE2 | JMJD1C-AS1 | -0,3448 | 8,88032E-13 |
| ACE2 | TRMU | -0,3448 | 8,83819E-13 |
| ACE2 | RP11-109L13.1 | -0,3449 | 8,81939E-13 |
| ACE2 | CTD-2382E5.6 | -0,3449 | 8,71733E-13 |
| ACE2 | OCRL | -0,3449 | 8,82503E-13 |
| ACE2 | DRAXIN | -0,345 | 8,58166E-13 |
| ACE2 | ZNF736 | -0,345 | 8,68487E-13 |
| ACE2 | GPR19 | -0,345 | 8,56319E-13 |
| ACE2 | RP11-70D24.3 | -0,345 | 8,57104E-13 |
| ACE2 | DDI2 | -0,3451 | 8,4291E-13 |
| ACE2 | AC108463.2 | -0,3451 | 8,43078E-13 |
| ACE2 | XXcos-LUCA16.1 | -0,3451 | 8,53182E-13 |
| ACE2 | ZFP91 | -0,3451 | 8,43728E-13 |
| ACE2 | UBE2N | -0,3451 | 8,46188E-13 |
| ACE2 | AC005954.4 | -0,3451 | 8,50972E-13 |
| ACE2 | AP000654.4 | -0,3452 | 8,35278E-13 |
| ACE2 | RP11-664H17.1 | -0,3452 | 8,40189E-13 |
| ACE2 | PHRF1 | -0,3453 | 8,21623E-13 |
| ACE2 | CLEC4M | -0,3453 | 8,26746E-13 |
| ACE2 | UXS1 | -0,3454 | 8,14372E-13 |
| ACE2 | RP11-857B24.1 | -0,3454 | 8,13879E-13 |
| ACE2 | RP11-732A19.2 | -0,3455 | 7,92399E-13 |
| ACE2 | RP11-219B17.3 | -0,3456 | 7,82904E-13 |
| ACE2 | TTTY14 | -0,3456 | 7,78861E-13 |
| ACE2 | RP11-50D9.4 | -0,3457 | 7,69198E-13 |
| ACE2 | LY96 | -0,3457 | 7,76556E-13 |
| ACE2 | TRIM16 | -0,3457 | 7,74465E-13 |
| ACE2 | RP11-430E17.1 | -0,3457 | 7,6421E-13 |
| ACE2 | RP5-874C20.6 | -0,3458 | 7,52232E-13 |
| ACE2 | CTD-2291D10.3 | -0,3458 | 7,6042E-13 |
| ACE2 | RPP40 | -0,3459 | 7,47349E-13 |
| ACE2 | SEPT10P1 | -0,3459 | 7,48127E-13 |
| ACE2 | DNAJC1 | -0,346 | 7,33086E-13 |
| ACE2 | CTB-107G13.1 | -0,3461 | 7,17758E-13 |
| ACE2 | LINC00535 | -0,3461 | 7,20511E-13 |
| ACE2 | TMEM9B-AS1 | -0,3461 | 7,21191E-13 |
| ACE2 | ASB16 | -0,3461 | 7,16915E-13 |
| ACE2 | A4GNT | -0,3462 | 7,14704E-13 |
| ACE2 | RP11-146N23.1 | -0,3462 | 7,11737E-13 |
| ACE2 | CDH15 | -0,3462 | 7,13707E-13 |
| ACE2 | RP11-186B7.7 | -0,3462 | 7,15363E-13 |

| | | | |
|---|---|---|---|
| ACE2 | AC006195.2 | -0,3463 | 6,97196E-13 |
| ACE2 | RP11-560I19.1 | -0,3463 | 6,97222E-13 |
| ACE2 | ERCC6 | -0,3464 | 6,90098E-13 |
| ACE2 | RP5-1014D13.2 | -0,3464 | 6,83591E-13 |
| ACE2 | RP11-710C12.1 | -0,3465 | 6,7342E-13 |
| ACE2 | CTD-2319I12.4 | -0,3466 | 6,63565E-13 |
| ACE2 | ANKRD36B | -0,3467 | 6,5992E-13 |
| ACE2 | SNHG15 | -0,3467 | 6,57059E-13 |
| ACE2 | RP11-281B1.2 | -0,3467 | 6,56605E-13 |
| ACE2 | CD55 | -0,3468 | 6,4783E-13 |
| ACE2 | RP11-567F11.2 | -0,3468 | 6,43635E-13 |
| ACE2 | VWC2L | -0,3468 | 6,4591E-13 |
| ACE2 | CTD-2373H9.3 | -0,3468 | 6,45452E-13 |
| ACE2 | RP11-548P2.2 | -0,3469 | 6,29609E-13 |
| ACE2 | UNC119 | -0,3469 | 6,37363E-13 |
| ACE2 | LINC01111 | -0,347 | 6,2626E-13 |
| ACE2 | RP4-753D10.3 | -0,347 | 6,27437E-13 |
| ACE2 | RP4-740C4.7 | -0,3471 | 6,10693E-13 |
| ACE2 | SDHDP6 | -0,3471 | 6,18167E-13 |
| ACE2 | RP11-44D5.2 | -0,3471 | 6,1684E-13 |
| ACE2 | TREM1 | -0,3471 | 6,15569E-13 |
| ACE2 | ZNF19 | -0,3471 | 6,09969E-13 |
| ACE2 | HSPB9 | -0,3471 | 6,17738E-13 |
| ACE2 | GUCA1A | -0,3472 | 6,01494E-13 |
| ACE2 | PMS2P5 | -0,3472 | 6,00595E-13 |
| ACE2 | CTC-137K3.1 | -0,3472 | 6,08319E-13 |
| ACE2 | RP4-773N10.4 | -0,3473 | 5,96856E-13 |
| ACE2 | ZNF259P1 | -0,3473 | 5,95073E-13 |
| ACE2 | KMT2E | -0,3473 | 5,93048E-13 |
| ACE2 | KLF10 | -0,3473 | 5,9545E-13 |
| ACE2 | TGDS | -0,3473 | 5,91337E-13 |
| ACE2 | ALG1L10P | -0,3474 | 5,87393E-13 |
| ACE2 | RP13-895J2.7 | -0,3474 | 5,87335E-13 |
| ACE2 | RP5-968J1.1 | -0,3474 | 5,82807E-13 |
| ACE2 | ATMIN | -0,3475 | 5,79541E-13 |
| ACE2 | CTD-2619J13.19 | -0,3475 | 5,77473E-13 |
| ACE2 | RP11-190P13.2 | -0,3476 | 5,68608E-13 |
| ACE2 | ADCY8 | -0,3476 | 5,67321E-13 |
| ACE2 | DUX4L50 | -0,3476 | 5,64783E-13 |
| ACE2 | ETV3 | -0,3477 | 5,57095E-13 |
| ACE2 | LINC01447 | -0,3477 | 5,6083E-13 |
| ACE2 | CHRFAM7A | -0,3478 | 5,46464E-13 |
| ACE2 | RP11-325L12.7 | -0,3478 | 5,44588E-13 |
| ACE2 | RP13-977J11.2 | -0,3479 | 5,37532E-13 |
| ACE2 | RPS6P25 | -0,3479 | 5,40425E-13 |
| ACE2 | AC004019.13 | -0,3479 | 5,35038E-13 |
| ACE2 | MOB1A | -0,348 | 5,28925E-13 |
| ACE2 | NDUFB4 | -0,3481 | 5,23563E-13 |
| ACE2 | KRT81 | -0,3481 | 5,21626E-13 |
| ACE2 | INO80E | -0,3481 | 5,22098E-13 |
| ACE2 | SPATA1 | -0,3482 | 5,12626E-13 |
| ACE2 | RP11-87F15.2 | -0,3482 | 5,11237E-13 |
| ACE2 | TCP11 | -0,3482 | 5,15787E-13 |
| ACE2 | MTHFSD | -0,3482 | 5,10627E-13 |

| | | | |
|---|---|---|---|
| ACE2 | IL11 | -0,3482 | 5,16858E-13 |
| ACE2 | OAZ3 | -0,3483 | 5,05224E-13 |
| ACE2 | FAM8A1 | -0,3483 | 5,04837E-13 |
| ACE2 | AC145676.2 | -0,3483 | 5,03824E-13 |
| ACE2 | NFX1 | -0,3483 | 5,02861E-13 |
| ACE2 | FAM71C | -0,3483 | 5,06272E-13 |
| ACE2 | RP11-407P15.1 | -0,3485 | 4,8706E-13 |
| ACE2 | ZNF806 | -0,3485 | 4,87509E-13 |
| ACE2 | SPDYE18 | -0,3485 | 4,89763E-13 |
| ACE2 | TMEM258 | -0,3487 | 4,76989E-13 |
| ACE2 | KRT27 | -0,3487 | 4,76446E-13 |
| ACE2 | RP11-106M3.3 | -0,3488 | 4,63699E-13 |
| ACE2 | SGSH | -0,3489 | 4,61524E-13 |
| ACE2 | RP11-396M11.1 | -0,349 | 4,48551E-13 |
| ACE2 | RP11-618M23.5 | -0,349 | 4,53376E-13 |
| ACE2 | RP11-162G10.5 | -0,349 | 4,52231E-13 |
| ACE2 | TAMM41 | -0,3491 | 4,44473E-13 |
| ACE2 | SYNE1-AS1 | -0,3491 | 4,44392E-13 |
| ACE2 | AC011242.5 | -0,3492 | 4,33507E-13 |
| ACE2 | AC007285.6 | -0,3492 | 4,37986E-13 |
| ACE2 | NOB1 | -0,3492 | 4,39387E-13 |
| ACE2 | AC061992.2 | -0,3492 | 4,39247E-13 |
| ACE2 | SASS6 | -0,3493 | 4,27897E-13 |
| ACE2 | AC092162.1 | -0,3493 | 4,30229E-13 |
| ACE2 | RP11-89B16.1 | -0,3493 | 4,3164E-13 |
| ACE2 | RPL13AP5 | -0,3493 | 4,26788E-13 |
| ACE2 | IGLV5-52 | -0,3493 | 4,31832E-13 |
| ACE2 | DNAJC30 | -0,3494 | 4,20019E-13 |
| ACE2 | FAM27E3 | -0,3494 | 4,22028E-13 |
| ACE2 | TBX18-AS1 | -0,3497 | 3,99873E-13 |
| ACE2 | TAF5L | -0,3498 | 3,9615E-13 |
| ACE2 | GALNT15 | -0,3498 | 3,96127E-13 |
| ACE2 | CSRNP1 | -0,3498 | 3,93588E-13 |
| ACE2 | RP11-511P7.6 | -0,3499 | 3,90482E-13 |
| ACE2 | RBM7 | -0,3499 | 3,88215E-13 |
| ACE2 | IQCH | -0,3499 | 3,89152E-13 |
| ACE2 | PISD | -0,3499 | 3,90729E-13 |
| ACE2 | G3BP1 | -0,35 | 3,81154E-13 |
| ACE2 | RP3-483K16.4 | -0,35 | 3,79971E-13 |
| ACE2 | PSMG3-AS1 | -0,35 | 3,81366E-13 |
| ACE2 | CBX3P2 | -0,35 | 3,85321E-13 |
| ACE2 | BMS1P2 | -0,3501 | 3,77545E-13 |
| ACE2 | RP11-757A13.1 | -0,3502 | 3,72964E-13 |
| ACE2 | PLAA | -0,3502 | 3,70917E-13 |
| ACE2 | LINC00159 | -0,3502 | 3,68439E-13 |
| ACE2 | HERC4 | -0,3504 | 3,5789E-13 |
| ACE2 | MTUS2-AS1 | -0,3504 | 3,57235E-13 |
| ACE2 | PSMC1 | -0,3504 | 3,57653E-13 |
| ACE2 | RP11-288L9.1 | -0,3505 | 3,52899E-13 |
| ACE2 | ABHD17AP3 | -0,3505 | 3,53818E-13 |
| ACE2 | CNN2P8 | -0,3505 | 3,51124E-13 |
| ACE2 | LIMD1-AS1 | -0,3506 | 3,4424E-13 |
| ACE2 | RP11-583F2.2 | -0,3506 | 3,48944E-13 |
| ACE2 | SLC35E1 | -0,3508 | 3,37024E-13 |

| | | | |
|---|---|---|---|
| ACE2 | RBM15B | -0,3509 | 3,27869E-13 |
| ACE2 | RP11-1398P2.1 | -0,3509 | 3,29343E-13 |
| ACE2 | ZNF200 | -0,3509 | 3,31895E-13 |
| ACE2 | ZNF556 | -0,3509 | 3,3152E-13 |
| ACE2 | PLVAP | -0,3509 | 3,27594E-13 |
| ACE2 | RS1 | -0,3509 | 3,31321E-13 |
| ACE2 | RBMXL3 | -0,3509 | 3,32694E-13 |
| ACE2 | ZNF784 | -0,351 | 3,25831E-13 |
| ACE2 | HAMP | -0,3511 | 3,21197E-13 |
| ACE2 | CFAP100 | -0,3512 | 3,15794E-13 |
| ACE2 | ACAN | -0,3512 | 3,14843E-13 |
| ACE2 | RP5-827E24.1 | -0,3512 | 3,16458E-13 |
| ACE2 | ANAPC1 | -0,3513 | 3,08976E-13 |
| ACE2 | ATAD3B | -0,3514 | 3,06288E-13 |
| ACE2 | NDST1-AS1 | -0,3514 | 3,03039E-13 |
| ACE2 | RP11-158I3.1 | -0,3514 | 3,02434E-13 |
| ACE2 | ZNF207 | -0,3514 | 3,06114E-13 |
| ACE2 | LINC01345 | -0,3515 | 2,9911E-13 |
| ACE2 | LINC01940 | -0,3515 | 2,99335E-13 |
| ACE2 | SMC5 | -0,3515 | 3,01411E-13 |
| ACE2 | PATE2 | -0,3515 | 2,98512E-13 |
| ACE2 | RP11-961A15.3 | -0,3515 | 3,01073E-13 |
| ACE2 | CTD-3014M21.1 | -0,3515 | 2,98703E-13 |
| ACE2 | CXXC4-AS1 | -0,3516 | 2,95303E-13 |
| ACE2 | FAM66E | -0,3516 | 2,91883E-13 |
| ACE2 | RNU6-1039P | -0,3516 | 2,93784E-13 |
| ACE2 | NEU3 | -0,3516 | 2,9441E-13 |
| ACE2 | ZNF689 | -0,3516 | 2,92867E-13 |
| ACE2 | CSNK1G2 | -0,3516 | 2,92517E-13 |
| ACE2 | CD3EAP | -0,3516 | 2,92253E-13 |
| ACE2 | RP5-894A10.2 | -0,3517 | 2,87544E-13 |
| ACE2 | C5orf58 | -0,3518 | 2,86298E-13 |
| ACE2 | RP11-157H4.1 | -0,3518 | 2,84972E-13 |
| ACE2 | TOX2 | -0,3518 | 2,83963E-13 |
| ACE2 | EMC3-AS1 | -0,3519 | 2,79336E-13 |
| ACE2 | IFITM1 | -0,3519 | 2,78457E-13 |
| ACE2 | RP11-688I9.4 | -0,3519 | 2,82174E-13 |
| ACE2 | LINC00441 | -0,3519 | 2,7925E-13 |
| ACE2 | RPL31P61 | -0,3519 | 2,80899E-13 |
| ACE2 | MPL | -0,352 | 2,73678E-13 |
| ACE2 | TMEM99 | -0,352 | 2,77122E-13 |
| ACE2 | CTD-2639E6.4 | -0,352 | 2,74641E-13 |
| ACE2 | SRGAP3 | -0,3521 | 2,6944E-13 |
| ACE2 | ACO1 | -0,3521 | 2,69293E-13 |
| ACE2 | KARSP2 | -0,3522 | 2,67512E-13 |
| ACE2 | ZNF525 | -0,3522 | 2,65794E-13 |
| ACE2 | RPL24P4 | -0,3523 | 2,61146E-13 |
| ACE2 | SMKR1 | -0,3523 | 2,62356E-13 |
| ACE2 | JUNB | -0,3523 | 2,63719E-13 |
| ACE2 | RP11-1C8.6 | -0,3524 | 2,56007E-13 |
| ACE2 | RP11-680F8.3 | -0,3525 | 2,54156E-13 |
| ACE2 | C1orf43 | -0,3526 | 2,48471E-13 |
| ACE2 | RP11-245J9.5 | -0,3526 | 2,47459E-13 |
| ACE2 | STIM2 | -0,3526 | 2,51523E-13 |

| | | | |
|---|---|---|---|
| ACE2 | MTFR1 | -0,3526 | 2,49679E-13 |
| ACE2 | RP11-613D13.4 | -0,3526 | 2,51339E-13 |
| ACE2 | MIR6745 | -0,3526 | 2,51362E-13 |
| ACE2 | RNASE4 | -0,3526 | 2,47802E-13 |
| ACE2 | RP3-453C12.14 | -0,3526 | 2,49955E-13 |
| ACE2 | DDX20 | -0,3527 | 2,44732E-13 |
| ACE2 | KIF6 | -0,3527 | 2,47385E-13 |
| ACE2 | RP11-109A6.5 | -0,3527 | 2,45696E-13 |
| ACE2 | CSNK1A1L | -0,3527 | 2,45578E-13 |
| ACE2 | SHBG | -0,3527 | 2,46921E-13 |
| ACE2 | RP11-234N17.1 | -0,3528 | 2,40626E-13 |
| ACE2 | RP4-593H12.1 | -0,3528 | 2,40534E-13 |
| ACE2 | CTA-992D9.8 | -0,3528 | 2,41832E-13 |
| ACE2 | PTK7 | -0,3529 | 2,39126E-13 |
| ACE2 | CTD-2014D20.1 | -0,3529 | 2,35545E-13 |
| ACE2 | AK1 | -0,3529 | 2,37082E-13 |
| ACE2 | ATF7IP2 | -0,3529 | 2,37277E-13 |
| ACE2 | RP11-477J21.6 | -0,353 | 2,3302E-13 |
| ACE2 | PDCL | -0,353 | 2,33992E-13 |
| ACE2 | TBX4 | -0,353 | 2,34235E-13 |
| ACE2 | ARSF | -0,353 | 2,34904E-13 |
| ACE2 | AC004221.2 | -0,3531 | 2,31166E-13 |
| ACE2 | PLB1 | -0,3532 | 2,25801E-13 |
| ACE2 | ZGRF1 | -0,3532 | 2,27134E-13 |
| ACE2 | SNORA65 | -0,3532 | 2,24543E-13 |
| ACE2 | POGLUT1 | -0,3533 | 2,20967E-13 |
| ACE2 | MRM3 | -0,3533 | 2,23598E-13 |
| ACE2 | CTB-133G6.2 | -0,3533 | 2,23243E-13 |
| ACE2 | RPL31P52 | -0,3534 | 2,19761E-13 |
| ACE2 | RP11-159D12.2 | -0,3535 | 2,13228E-13 |
| ACE2 | TBL3 | -0,3536 | 2,12865E-13 |
| ACE2 | RP11-313A24.1 | -0,3537 | 2,06462E-13 |
| ACE2 | FTCD-AS1 | -0,3537 | 2,07267E-13 |
| ACE2 | RP11-308N19.3 | -0,3538 | 2,02999E-13 |
| ACE2 | RP11-680F20.9 | -0,3538 | 2,0527E-13 |
| ACE2 | RP11-66D17.5 | -0,3539 | 2,02118E-13 |
| ACE2 | SNRK-AS1 | -0,3539 | 2,01081E-13 |
| ACE2 | LRRC15 | -0,3539 | 2,00756E-13 |
| ACE2 | PROK2 | -0,354 | 1,97986E-13 |
| ACE2 | LINC01063 | -0,3541 | 1,95255E-13 |
| ACE2 | ZNF706 | -0,3541 | 1,93912E-13 |
| ACE2 | CD274 | -0,3541 | 1,94775E-13 |
| ACE2 | INTS6-AS1 | -0,3541 | 1,94666E-13 |
| ACE2 | RP11-45A16.4 | -0,3542 | 1,92616E-13 |
| ACE2 | RP11-768F21.1 | -0,3542 | 1,90193E-13 |
| ACE2 | RPL7AP6 | -0,3542 | 1,91877E-13 |
| ACE2 | AC067959.1 | -0,3543 | 1,87425E-13 |
| ACE2 | XXbac-BPG13B8.10 | -0,3543 | 1,89228E-13 |
| ACE2 | CTC-543D15.3 | -0,3543 | 1,8884E-13 |
| ACE2 | PICSAR | -0,3543 | 1,87435E-13 |
| ACE2 | GNRH1 | -0,3544 | 1,84047E-13 |
| ACE2 | KLHL7-AS1 | -0,3545 | 1,82384E-13 |
| ACE2 | RP11-99L13.2 | -0,3545 | 1,83216E-13 |
| ACE2 | LRRC37A15P | -0,3546 | 1,79437E-13 |

| | | | |
|---|---|---|---|
| ACE2 | LINC01135 | -0,3547 | 1,75344E-13 |
| ACE2 | RP11-1299A16.3 | -0,3547 | 1,76111E-13 |
| ACE2 | RP11-697E2.11 | -0,3547 | 1,76755E-13 |
| ACE2 | CDK15 | -0,3548 | 1,73156E-13 |
| ACE2 | KLKB1 | -0,3548 | 1,71774E-13 |
| ACE2 | ERVFRD-1 | -0,3548 | 1,7277E-13 |
| ACE2 | RP11-399C16.3 | -0,3548 | 1,7317E-13 |
| ACE2 | IRX6 | -0,3548 | 1,7188E-13 |
| ACE2 | RP5-1017F8.2 | -0,3548 | 1,7416E-13 |
| ACE2 | PCA3 | -0,3549 | 1,69385E-13 |
| ACE2 | GOLGA6L9 | -0,3549 | 1,70624E-13 |
| ACE2 | RP1-101K10.6 | -0,355 | 1,6872E-13 |
| ACE2 | CTD-2145A24.5 | -0,355 | 1,66654E-13 |
| ACE2 | NDUFB1P2 | -0,3551 | 1,63741E-13 |
| ACE2 | CCNT2 | -0,3551 | 1,65731E-13 |
| ACE2 | CYP8B1 | -0,3551 | 1,64154E-13 |
| ACE2 | SECISBP2 | -0,3551 | 1,65919E-13 |
| ACE2 | CTD-2507G9.1 | -0,3551 | 1,65064E-13 |
| ACE2 | RP11-244F12.3 | -0,3551 | 1,63856E-13 |
| ACE2 | CH17-195P21.2 | -0,3554 | 1,57374E-13 |
| ACE2 | RP11-88H12.2 | -0,3554 | 1,55553E-13 |
| ACE2 | CTB-147C13.1 | -0,3554 | 1,57733E-13 |
| ACE2 | C16orf96 | -0,3554 | 1,57557E-13 |
| ACE2 | CHCHD4 | -0,3555 | 1,53015E-13 |
| ACE2 | HERC6 | -0,3555 | 1,53599E-13 |
| ACE2 | CANX | -0,3555 | 1,53846E-13 |
| ACE2 | CTB-13F3.1 | -0,3556 | 1,52684E-13 |
| ACE2 | DCLRE1C | -0,3556 | 1,51037E-13 |
| ACE2 | ANKRD12 | -0,3556 | 1,52761E-13 |
| ACE2 | C1orf87 | -0,3557 | 1,49486E-13 |
| ACE2 | RP11-13A1.1 | -0,3557 | 1,48671E-13 |
| ACE2 | SSBP3-AS1 | -0,3558 | 1,45677E-13 |
| ACE2 | SFT2D2 | -0,3558 | 1,45998E-13 |
| ACE2 | GIMAP7 | -0,3558 | 1,4766E-13 |
| ACE2 | OR51C1P | -0,3558 | 1,47368E-13 |
| ACE2 | AK4P1 | -0,3558 | 1,47332E-13 |
| ACE2 | CTD-2561J22.5 | -0,3558 | 1,45745E-13 |
| ACE2 | RP11-242D8.3 | -0,3559 | 1,43758E-13 |
| ACE2 | CTD-2561B21.10 | -0,3559 | 1,43164E-13 |
| ACE2 | PSME1 | -0,356 | 1,41676E-13 |
| ACE2 | MCL1 | -0,3563 | 1,3501E-13 |
| ACE2 | CTD-2531D15.4 | -0,3563 | 1,34041E-13 |
| ACE2 | RP11-271M24.2 | -0,3565 | 1,30957E-13 |
| ACE2 | RP11-382A20.6 | -0,3565 | 1,29697E-13 |
| ACE2 | HRAT17 | -0,3566 | 1,27267E-13 |
| ACE2 | AC009784.3 | -0,3566 | 1,28072E-13 |
| ACE2 | KLHL35 | -0,3566 | 1,27815E-13 |
| ACE2 | REM2 | -0,3566 | 1,29177E-13 |
| ACE2 | PRPF18 | -0,3567 | 1,25569E-13 |
| ACE2 | RP11-696N14.1 | -0,3568 | 1,23054E-13 |
| ACE2 | LYPD6 | -0,3569 | 1,21134E-13 |
| ACE2 | RP11-930O11.1 | -0,3569 | 1,22836E-13 |
| ACE2 | RP1-56K13.5 | -0,3569 | 1,21234E-13 |
| ACE2 | RP11-171I2.2 | -0,357 | 1,19818E-13 |

| | | | |
|---|---|---|---|
| ACE2 | LIMK1 | -0,357 | 1,20796E-13 |
| ACE2 | C4A-AS1 | -0,3571 | 1,17073E-13 |
| ACE2 | RP11-513D5.2 | -0,3571 | 1,18058E-13 |
| ACE2 | CYCSP23 | -0,3571 | 1,18087E-13 |
| ACE2 | FAM35BP | -0,3571 | 1,17087E-13 |
| ACE2 | RP11-653J6.1 | -0,3571 | 1,17826E-13 |
| ACE2 | TMED10P2 | -0,3572 | 1,15239E-13 |
| ACE2 | LILRB3 | -0,3572 | 1,1656E-13 |
| ACE2 | ZC3HAV1 | -0,3573 | 1,13211E-13 |
| ACE2 | RP11-382D8.5 | -0,3574 | 1,12443E-13 |
| ACE2 | FGF7P6 | -0,3574 | 1,12688E-13 |
| ACE2 | KEAP1 | -0,3574 | 1,11725E-13 |
| ACE2 | METTL21A | -0,3575 | 1,09793E-13 |
| ACE2 | OR2A1 | -0,3575 | 1,09273E-13 |
| ACE2 | RBM12 | -0,3575 | 1,09239E-13 |
| ACE2 | FBXO45 | -0,3576 | 1,09166E-13 |
| ACE2 | RP4-529N6.1 | -0,3576 | 1,09198E-13 |
| ACE2 | RP11-561N12.6 | -0,3576 | 1,08307E-13 |
| ACE2 | ARMCX7P | -0,3576 | 1,08496E-13 |
| ACE2 | IFT57 | -0,3577 | 1,07143E-13 |
| ACE2 | RP1-142L7.8 | -0,3578 | 1,04621E-13 |
| ACE2 | KDM5D | -0,3578 | 1,03949E-13 |
| ACE2 | BTNL2 | -0,3579 | 1,03465E-13 |
| ACE2 | RP11-730G20.2 | -0,3579 | 1,02376E-13 |
| ACE2 | DYNLL1 | -0,3579 | 1,02802E-13 |
| ACE2 | HM13 | -0,3579 | 1,03422E-13 |
| ACE2 | TTTY10 | -0,3579 | 1,02385E-13 |
| ACE2 | SBF1P1 | -0,358 | 1,01303E-13 |
| ACE2 | ANKRD10-IT1 | -0,358 | 1,00579E-13 |
| ACE2 | AC142472.6 | -0,358 | 1,01612E-13 |
| ACE2 | RP11-80P20.3 | -0,3582 | 9,71862E-14 |
| ACE2 | RP11-353K11.1 | -0,3583 | 9,70104E-14 |
| ACE2 | AC013271.3 | -0,3583 | 9,6652E-14 |
| ACE2 | TINAGL1 | -0,3584 | 9,42138E-14 |
| ACE2 | LINC01562 | -0,3584 | 9,41801E-14 |
| ACE2 | GOLGA6L4 | -0,3584 | 9,51329E-14 |
| ACE2 | MFSD8 | -0,3585 | 9,33396E-14 |
| ACE2 | GNA14 | -0,3585 | 9,3593E-14 |
| ACE2 | FAM181A | -0,3585 | 9,29245E-14 |
| ACE2 | RPS23P8 | -0,3586 | 9,13098E-14 |
| ACE2 | RDH10 | -0,3587 | 9,04613E-14 |
| ACE2 | ZBTB5 | -0,3587 | 8,95817E-14 |
| ACE2 | NELFA | -0,3588 | 8,84164E-14 |
| ACE2 | DMP1 | -0,3588 | 8,90348E-14 |
| ACE2 | UFL1 | -0,3588 | 8,87884E-14 |
| ACE2 | RP11-264E20.2 | -0,3588 | 8,81278E-14 |
| ACE2 | RP11-196B3.1 | -0,3588 | 8,86117E-14 |
| ACE2 | ACP1 | -0,359 | 8,49603E-14 |
| ACE2 | PPEF2 | -0,359 | 8,51138E-14 |
| ACE2 | C5orf64-AS1 | -0,359 | 8,49144E-14 |
| ACE2 | RN7SL749P | -0,359 | 8,52019E-14 |
| ACE2 | RP11-1078H9.5 | -0,359 | 8,62104E-14 |
| ACE2 | RP11-112N19.2 | -0,359 | 8,49976E-14 |
| ACE2 | CTD-2576F9.1 | -0,359 | 8,51409E-14 |

| | | | |
|---|---|---|---|
| ACE2 | DYNC1LI2 | -0,3591 | 8,4141E-14 |
| ACE2 | RP11-308D16.2 | -0,3591 | 8,47216E-14 |
| ACE2 | KRT18P60 | -0,3592 | 8,30702E-14 |
| ACE2 | IMP4 | -0,3594 | 8,0503E-14 |
| ACE2 | RP11-114G11.5 | -0,3594 | 8,00909E-14 |
| ACE2 | RP11-231I16.1 | -0,3594 | 7,98794E-14 |
| ACE2 | ABALON | -0,3594 | 7,94941E-14 |
| ACE2 | MED14OS | -0,3594 | 8,05897E-14 |
| ACE2 | NPIPB13 | -0,3595 | 7,83039E-14 |
| ACE2 | RP5-908M14.9 | -0,3595 | 7,81819E-14 |
| ACE2 | AC019181.3 | -0,3596 | 7,70089E-14 |
| ACE2 | LINC01114 | -0,3597 | 7,61259E-14 |
| ACE2 | BRD7P2 | -0,3597 | 7,63319E-14 |
| ACE2 | ALDH3B1 | -0,3597 | 7,55754E-14 |
| ACE2 | RP11-872J21.5 | -0,3597 | 7,55231E-14 |
| ACE2 | LINC01703 | -0,3598 | 7,44492E-14 |
| ACE2 | CTD-2561B21.4 | -0,3598 | 7,48934E-14 |
| ACE2 | FAM109B | -0,3598 | 7,41706E-14 |
| ACE2 | RP11-145P16.2 | -0,3599 | 7,32396E-14 |
| ACE2 | AC116614.1 | -0,36 | 7,15903E-14 |
| ACE2 | AC073257.2 | -0,36 | 7,20917E-14 |
| ACE2 | RP11-770J1.3 | -0,36 | 7,23933E-14 |
| ACE2 | HEBP1 | -0,36 | 7,17931E-14 |
| ACE2 | SPATA17 | -0,3601 | 7,12318E-14 |
| ACE2 | SYNE3 | -0,3601 | 7,06818E-14 |
| ACE2 | TIMM23 | -0,3602 | 6,96979E-14 |
| ACE2 | CACNA1S | -0,3603 | 6,81977E-14 |
| ACE2 | AC015977.6 | -0,3603 | 6,8766E-14 |
| ACE2 | INO80D | -0,3603 | 6,91353E-14 |
| ACE2 | CRYGS | -0,3603 | 6,91762E-14 |
| ACE2 | RP11-1046B16.3 | -0,3603 | 6,82728E-14 |
| ACE2 | RP11-87N3.6 | -0,3603 | 6,89642E-14 |
| ACE2 | ANKRD45 | -0,3604 | 6,76921E-14 |
| ACE2 | AC005592.3 | -0,3604 | 6,77009E-14 |
| ACE2 | SCFD1 | -0,3604 | 6,77062E-14 |
| ACE2 | APOL4 | -0,3604 | 6,69731E-14 |
| ACE2 | RP11-419I17.1 | -0,3605 | 6,6339E-14 |
| ACE2 | RP11-732A21.2 | -0,3606 | 6,56603E-14 |
| ACE2 | RP11-598F7.6 | -0,3606 | 6,54432E-14 |
| ACE2 | RP11-818F20.5 | -0,3609 | 6,21269E-14 |
| ACE2 | CTD-3096M3.2 | -0,3609 | 6,17073E-14 |
| ACE2 | PKD1L2 | -0,361 | 6,05025E-14 |
| ACE2 | FGR | -0,3612 | 5,92878E-14 |
| ACE2 | RPL23AP2 | -0,3612 | 5,88753E-14 |
| ACE2 | DNAH12 | -0,3613 | 5,76082E-14 |
| ACE2 | WRNIP1 | -0,3613 | 5,75015E-14 |
| ACE2 | CTD-2525I3.5 | -0,3613 | 5,73768E-14 |
| ACE2 | SF3B6 | -0,3614 | 5,67898E-14 |
| ACE2 | MIR6892 | -0,3614 | 5,64987E-14 |
| ACE2 | VIM-AS1 | -0,3614 | 5,70165E-14 |
| ACE2 | RP11-513G19.1 | -0,3614 | 5,70736E-14 |
| ACE2 | PITRM1-AS1 | -0,3615 | 5,55943E-14 |
| ACE2 | SLC38A2 | -0,3615 | 5,57738E-14 |
| ACE2 | RP11-855A2.2 | -0,3615 | 5,56671E-14 |

| | | | |
|---|---|---|---|
| ACE2 | TMEM138 | -0,3616 | 5,48651E-14 |
| ACE2 | LYG1 | -0,3618 | 5,32218E-14 |
| ACE2 | RP1-179N16.6 | -0,3618 | 5,30851E-14 |
| ACE2 | RP11-543P15.3 | -0,3618 | 5,32603E-14 |
| ACE2 | RP11-993B23.3 | -0,3618 | 5,27523E-14 |
| ACE2 | SNAPC3 | -0,3619 | 5,26335E-14 |
| ACE2 | RP11-76C10.2 | -0,3619 | 5,25193E-14 |
| ACE2 | SMURF1 | -0,362 | 5,09903E-14 |
| ACE2 | AC009487.6 | -0,3621 | 5,04826E-14 |
| ACE2 | AF131215.9 | -0,3621 | 5,08595E-14 |
| ACE2 | ASB8 | -0,3622 | 5,00107E-14 |
| ACE2 | GABRA2 | -0,3623 | 4,87596E-14 |
| ACE2 | ZNF658B | -0,3623 | 4,83824E-14 |
| ACE2 | RP11-693N9.2 | -0,3623 | 4,84265E-14 |
| ACE2 | RP11-508N22.9 | -0,3624 | 4,78817E-14 |
| ACE2 | RP11-59H7.3 | -0,3624 | 4,78742E-14 |
| ACE2 | C2CD4B | -0,3624 | 4,82154E-14 |
| ACE2 | RP11-441F2.2 | -0,3624 | 4,75725E-14 |
| ACE2 | RHO | -0,3625 | 4,72695E-14 |
| ACE2 | KLHL25 | -0,3625 | 4,71773E-14 |
| ACE2 | RP11-449L23.2 | -0,3625 | 4,70926E-14 |
| ACE2 | PHLDB3 | -0,3625 | 4,68063E-14 |
| ACE2 | IRF2 | -0,3626 | 4,59719E-14 |
| ACE2 | ABCC10 | -0,3626 | 4,5974E-14 |
| ACE2 | DTX1 | -0,3626 | 4,61243E-14 |
| ACE2 | UPF1 | -0,3626 | 4,65824E-14 |
| ACE2 | AC006372.6 | -0,3627 | 4,56216E-14 |
| ACE2 | TIMM29 | -0,3627 | 4,58148E-14 |
| ACE2 | ADGRA1-AS1 | -0,3628 | 4,46348E-14 |
| ACE2 | NFE2 | -0,3628 | 4,49805E-14 |
| ACE2 | RP11-807C20.2 | -0,3629 | 4,43614E-14 |
| ACE2 | RPL36AP33 | -0,363 | 4,30817E-14 |
| ACE2 | MIR6883 | -0,363 | 4,32988E-14 |
| ACE2 | DUSP5P1 | -0,3631 | 4,2611E-14 |
| ACE2 | CADM2-AS1 | -0,3631 | 4,28136E-14 |
| ACE2 | LSM6 | -0,3631 | 4,2314E-14 |
| ACE2 | RP11-363J20.2 | -0,3631 | 4,27684E-14 |
| ACE2 | HSD3BP4 | -0,3632 | 4,17035E-14 |
| ACE2 | SKP2 | -0,3632 | 4,21419E-14 |
| ACE2 | NKX2-5 | -0,3632 | 4,20261E-14 |
| ACE2 | RP11-346C4.3 | -0,3632 | 4,21242E-14 |
| ACE2 | PSTPIP1 | -0,3632 | 4,15763E-14 |
| ACE2 | CTD-3107M8.2 | -0,3633 | 4,0838E-14 |
| ACE2 | RP11-340L3.1 | -0,3633 | 4,0842E-14 |
| ACE2 | TPRXL | -0,3634 | 4,009E-14 |
| ACE2 | HSPD1P10 | -0,3634 | 4,02297E-14 |
| ACE2 | RP11-157J24.2 | -0,3635 | 3,95694E-14 |
| ACE2 | RP11-425A6.5 | -0,3635 | 3,97157E-14 |
| ACE2 | RP11-264E20.1 | -0,3635 | 3,97795E-14 |
| ACE2 | RP11-95G6.1 | -0,3635 | 3,96093E-14 |
| ACE2 | LINC01532 | -0,3635 | 3,9399E-14 |
| ACE2 | ZNF780A | -0,3635 | 3,94285E-14 |
| ACE2 | ACTG1P10 | -0,3636 | 3,90871E-14 |
| ACE2 | RPL5P12 | -0,3637 | 3,84072E-14 |

| | | | |
|---|---|---|---|
| ACE2 | ZNF807 | -0,3637 | 3,83258E-14 |
| ACE2 | BDNF-AS | -0,3638 | 3,75264E-14 |
| ACE2 | KCNK6 | -0,3638 | 3,73868E-14 |
| ACE2 | FAM19A1 | -0,364 | 3,63539E-14 |
| ACE2 | SLC7A9 | -0,3641 | 3,60249E-14 |
| ACE2 | RNGTTP1 | -0,3641 | 3,57004E-14 |
| ACE2 | RP11-300E4.2 | -0,3642 | 3,52752E-14 |
| ACE2 | RP11-324I22.2 | -0,3642 | 3,51492E-14 |
| ACE2 | ADD3-AS1 | -0,3642 | 3,49026E-14 |
| ACE2 | RNU6-343P | -0,3642 | 3,49908E-14 |
| ACE2 | RP11-180P8.1 | -0,3642 | 3,49761E-14 |
| ACE2 | TDO2 | -0,3643 | 3,46716E-14 |
| ACE2 | ZNF253 | -0,3643 | 3,4862E-14 |
| ACE2 | RP11-288L9.4 | -0,3644 | 3,39977E-14 |
| ACE2 | POLR1A | -0,3644 | 3,41759E-14 |
| ACE2 | ADAMTS15 | -0,3644 | 3,39381E-14 |
| ACE2 | SLC16A14 | -0,3645 | 3,36323E-14 |
| ACE2 | SLC6A1-AS1 | -0,3645 | 3,31799E-14 |
| ACE2 | HLA-E | -0,3645 | 3,32249E-14 |
| ACE2 | RP11-777F6.3 | -0,3645 | 3,32693E-14 |
| ACE2 | C15orf61 | -0,3646 | 3,26981E-14 |
| ACE2 | ZNF557 | -0,3646 | 3,28659E-14 |
| ACE2 | GAN | -0,3647 | 3,2207E-14 |
| ACE2 | ARID3A | -0,3647 | 3,20225E-14 |
| ACE2 | RPS18 | -0,3648 | 3,18535E-14 |
| ACE2 | LINC00235 | -0,3649 | 3,13018E-14 |
| ACE2 | PLA2R1 | -0,365 | 3,06486E-14 |
| ACE2 | ZSCAN32 | -0,365 | 3,08836E-14 |
| ACE2 | NUP50-AS1 | -0,365 | 3,07304E-14 |
| ACE2 | RP11-545G3.2 | -0,3651 | 3,0145E-14 |
| ACE2 | MAG | -0,3651 | 2,99701E-14 |
| ACE2 | FARS2 | -0,3652 | 2,97577E-14 |
| ACE2 | GTPBP4 | -0,3652 | 2,94113E-14 |
| ACE2 | FICD | -0,3652 | 2,94321E-14 |
| ACE2 | AC013460.1 | -0,3653 | 2,90127E-14 |
| ACE2 | RP11-885N19.6 | -0,3653 | 2,88815E-14 |
| ACE2 | RP11-392O17.1 | -0,3654 | 2,87739E-14 |
| ACE2 | MRPS25 | -0,3655 | 2,80348E-14 |
| ACE2 | GARS | -0,3655 | 2,82773E-14 |
| ACE2 | FSCN3 | -0,3655 | 2,79185E-14 |
| ACE2 | RP11-326A13.1 | -0,3655 | 2,79515E-14 |
| ACE2 | AC015933.2 | -0,3655 | 2,81638E-14 |
| ACE2 | FRG1GP | -0,3655 | 2,83292E-14 |
| ACE2 | VN1R51P | -0,3656 | 2,74693E-14 |
| ACE2 | RPL21P99 | -0,3656 | 2,78196E-14 |
| ACE2 | PCDHGB8P | -0,3657 | 2,72718E-14 |
| ACE2 | SUB1P1 | -0,3658 | 2,68684E-14 |
| ACE2 | RP11-91I8.2 | -0,3658 | 2,65412E-14 |
| ACE2 | CXCL2 | -0,3659 | 2,60564E-14 |
| ACE2 | GS1-72M22.1 | -0,3659 | 2,62918E-14 |
| ACE2 | RP11-329B9.5 | -0,366 | 2,57828E-14 |
| ACE2 | RP11-1277A3.2 | -0,366 | 2,58452E-14 |
| ACE2 | PA2G4P4 | -0,3661 | 2,52227E-14 |
| ACE2 | RP11-360I2.1 | -0,3661 | 2,55257E-14 |

| | | | |
|---|---|---|---|
| ACE2 | PTCHD3 | -0,3661 | 2,53443E-14 |
| ACE2 | ADCY9 | -0,3661 | 2,55076E-14 |
| ACE2 | SCML2P2 | -0,3661 | 2,55543E-14 |
| ACE2 | IZUMO1 | -0,3661 | 2,53196E-14 |
| ACE2 | SCAF4 | -0,3661 | 2,54974E-14 |
| ACE2 | RP1-20B11.2 | -0,3662 | 2,49614E-14 |
| ACE2 | AC011239.1 | -0,3663 | 2,43431E-14 |
| ACE2 | BNIP3P1 | -0,3663 | 2,44455E-14 |
| ACE2 | AC009227.2 | -0,3664 | 2,39132E-14 |
| ACE2 | RP5-907C10.3 | -0,3664 | 2,41465E-14 |
| ACE2 | DUSP8P3 | -0,3664 | 2,41829E-14 |
| ACE2 | TP53TG3D | -0,3664 | 2,38746E-14 |
| ACE2 | NOC2L | -0,3665 | 2,37845E-14 |
| ACE2 | RSBN1 | -0,3665 | 2,37156E-14 |
| ACE2 | RP11-348H3.2 | -0,3665 | 2,37084E-14 |
| ACE2 | RP11-575L7.8 | -0,3665 | 2,35698E-14 |
| ACE2 | CEP164P1 | -0,3666 | 2,30458E-14 |
| ACE2 | FRG1EP | -0,3666 | 2,33786E-14 |
| ACE2 | CTB-167G5.6 | -0,3667 | 2,26497E-14 |
| ACE2 | CTC-428H11.2 | -0,3668 | 2,24308E-14 |
| ACE2 | WRAP73 | -0,3669 | 2,22283E-14 |
| ACE2 | RP11-349F21.5 | -0,3669 | 2,20011E-14 |
| ACE2 | ZNF224 | -0,3669 | 2,20005E-14 |
| ACE2 | DDX3Y | -0,3669 | 2,18947E-14 |
| ACE2 | CYP20A1 | -0,367 | 2,16459E-14 |
| ACE2 | CTD-2588E21.1 | -0,367 | 2,18381E-14 |
| ACE2 | DUSP11 | -0,3671 | 2,1481E-14 |
| ACE2 | SNORD94 | -0,3672 | 2,11095E-14 |
| ACE2 | GHRLOS | -0,3672 | 2,11071E-14 |
| ACE2 | LAGE3P1 | -0,3672 | 2,07969E-14 |
| ACE2 | RP11-732A21.3 | -0,3672 | 2,07575E-14 |
| ACE2 | TCEAL3-AS1 | -0,3672 | 2,0882E-14 |
| ACE2 | RP11-166B2.8 | -0,3673 | 2,06758E-14 |
| ACE2 | TYW1 | -0,3674 | 2,03549E-14 |
| ACE2 | LINC02139 | -0,3674 | 2,02718E-14 |
| ACE2 | BCR | -0,3674 | 2,03066E-14 |
| ACE2 | BSDC1 | -0,3675 | 1,9723E-14 |
| ACE2 | PSMD2 | -0,3675 | 1,98899E-14 |
| ACE2 | N6AMT1 | -0,3675 | 1,98457E-14 |
| ACE2 | ANKRD28 | -0,3676 | 1,95219E-14 |
| ACE2 | RAN | -0,3676 | 1,95445E-14 |
| ACE2 | RP11-73K9.2 | -0,3678 | 1,87881E-14 |
| ACE2 | RP11-727A23.4 | -0,3678 | 1,89406E-14 |
| ACE2 | SLC28A1 | -0,3678 | 1,8764E-14 |
| ACE2 | SPATS1 | -0,3679 | 1,85518E-14 |
| ACE2 | NTN3 | -0,3679 | 1,869E-14 |
| ACE2 | RP3-500L14.2 | -0,368 | 1,81135E-14 |
| ACE2 | CTD-3051D23.1 | -0,368 | 1,83287E-14 |
| ACE2 | C15orf65 | -0,368 | 1,82283E-14 |
| ACE2 | CTD-2349P21.10 | -0,3681 | 1,80227E-14 |
| ACE2 | RP11-481C4.2 | -0,3681 | 1,79427E-14 |
| ACE2 | TRIM63 | -0,3682 | 1,75456E-14 |
| ACE2 | RP11-991C1.1 | -0,3682 | 1,75469E-14 |
| ACE2 | FCGR2B | -0,3683 | 1,73187E-14 |

| | | | |
|---|---|---|---|
| ACE2 | RP11-728K20.3 | -0,3683 | 1,74131E-14 |
| ACE2 | HNRNPMP1 | -0,3683 | 1,72409E-14 |
| ACE2 | SNORA2 | -0,3684 | 1,70439E-14 |
| ACE2 | SLC12A9 | -0,3684 | 1,7028E-14 |
| ACE2 | ARHGAP28 | -0,3684 | 1,68576E-14 |
| ACE2 | CCDC18-AS1 | -0,3685 | 1,66359E-14 |
| ACE2 | RP4-725G10.3 | -0,3685 | 1,67293E-14 |
| ACE2 | RP11-573M3.3 | -0,3685 | 1,66414E-14 |
| ACE2 | AC007618.3 | -0,3685 | 1,66522E-14 |
| ACE2 | RPL39 | -0,3685 | 1,65676E-14 |
| ACE2 | SLC19A2 | -0,3686 | 1,64478E-14 |
| ACE2 | SCARB2 | -0,3686 | 1,64746E-14 |
| ACE2 | RP11-179B15.6 | -0,3686 | 1,65325E-14 |
| ACE2 | FAM86HP | -0,3687 | 1,62457E-14 |
| ACE2 | RP11-57B24.1 | -0,3687 | 1,61806E-14 |
| ACE2 | CTB-31O20.2 | -0,3687 | 1,61443E-14 |
| ACE2 | ZNF324B | -0,3687 | 1,62101E-14 |
| ACE2 | LINC01412 | -0,3688 | 1,58496E-14 |
| ACE2 | RP11-70D24.2 | -0,3688 | 1,58008E-14 |
| ACE2 | IGBP1 | -0,3688 | 1,58382E-14 |
| ACE2 | TIPRL | -0,3689 | 1,56882E-14 |
| ACE2 | FABP5P3 | -0,3689 | 1,552E-14 |
| ACE2 | AC006372.5 | -0,3689 | 1,55675E-14 |
| ACE2 | COX6A2 | -0,369 | 1,52689E-14 |
| ACE2 | RP11-247I13.3 | -0,369 | 1,51752E-14 |
| ACE2 | RP5-1029F21.2 | -0,3691 | 1,50686E-14 |
| ACE2 | RP11-358D17.1 | -0,3692 | 1,48341E-14 |
| ACE2 | RP11-124D2.7 | -0,3692 | 1,46981E-14 |
| ACE2 | TMEM269 | -0,3693 | 1,45939E-14 |
| ACE2 | AC091633.2 | -0,3693 | 1,45802E-14 |
| ACE2 | CASP9 | -0,3694 | 1,41482E-14 |
| ACE2 | BLID | -0,3694 | 1,42433E-14 |
| ACE2 | FBXL15 | -0,3695 | 1,39468E-14 |
| ACE2 | RP11-876N24.3 | -0,3695 | 1,40394E-14 |
| ACE2 | C19orf70 | -0,3695 | 1,4067E-14 |
| ACE2 | GALNT14 | -0,3696 | 1,37859E-14 |
| ACE2 | TMEM14EP | -0,3696 | 1,38387E-14 |
| ACE2 | PWRN1 | -0,3696 | 1,3767E-14 |
| ACE2 | SEPHS1P6 | -0,3697 | 1,35397E-14 |
| ACE2 | PPP2R5D | -0,3697 | 1,35476E-14 |
| ACE2 | RP13-317D12.3 | -0,3697 | 1,34636E-14 |
| ACE2 | EEF1A1P12 | -0,3698 | 1,32557E-14 |
| ACE2 | FAM87A | -0,3698 | 1,32856E-14 |
| ACE2 | GXYLT1 | -0,3698 | 1,31966E-14 |
| ACE2 | CCDC43 | -0,3698 | 1,3385E-14 |
| ACE2 | RPL36A | -0,3698 | 1,31753E-14 |
| ACE2 | LYRM2 | -0,3699 | 1,31108E-14 |
| ACE2 | SCX | -0,3699 | 1,31597E-14 |
| ACE2 | RP11-541N10.3 | -0,3699 | 1,30441E-14 |
| ACE2 | ZNF18 | -0,3699 | 1,29637E-14 |
| ACE2 | TMEM259 | -0,3699 | 1,31458E-14 |
| ACE2 | HBA1 | -0,37 | 1,28308E-14 |
| ACE2 | SPCS2 | -0,3701 | 1,26714E-14 |
| ACE2 | GAL3ST4 | -0,3702 | 1,24407E-14 |

| | | | |
|---|---|---|---|
| ACE2 | AGAP5 | -0,3702 | 1,24504E-14 |
| ACE2 | RP11-98D18.15 | -0,3703 | 1,21286E-14 |
| ACE2 | ANKZF1 | -0,3703 | 1,21082E-14 |
| ACE2 | POU5F2 | -0,3703 | 1,22537E-14 |
| ACE2 | SLC7A6 | -0,3703 | 1,22605E-14 |
| ACE2 | DDX19B | -0,3703 | 1,207E-14 |
| ACE2 | AC006548.28 | -0,3703 | 1,21998E-14 |
| ACE2 | RPL21P134 | -0,3703 | 1,22031E-14 |
| ACE2 | DYRK1B | -0,3704 | 1,20458E-14 |
| ACE2 | RP11-337N6.2 | -0,3705 | 1,17972E-14 |
| ACE2 | LINC01208 | -0,3705 | 1,18009E-14 |
| ACE2 | RP11-713H12.2 | -0,3705 | 1,17352E-14 |
| ACE2 | LINC01497 | -0,3705 | 1,18479E-14 |
| ACE2 | RPS15AP1 | -0,3705 | 1,17879E-14 |
| ACE2 | LAMP2 | -0,3705 | 1,1689E-14 |
| ACE2 | LINC01763 | -0,3706 | 1,15178E-14 |
| ACE2 | RP13-941N14.1 | -0,3706 | 1,14675E-14 |
| ACE2 | CDYL2 | -0,3706 | 1,14794E-14 |
| ACE2 | DNMT3B | -0,3706 | 1,15003E-14 |
| ACE2 | CAV3 | -0,3707 | 1,12723E-14 |
| ACE2 | RP11-332H18.3 | -0,3707 | 1,14019E-14 |
| ACE2 | RP11-353N4.5 | -0,3708 | 1,10935E-14 |
| ACE2 | AC019186.1 | -0,3709 | 1,09277E-14 |
| ACE2 | CPNE3 | -0,3709 | 1,08726E-14 |
| ACE2 | RP11-888D10.3 | -0,3709 | 1,10216E-14 |
| ACE2 | GTF2F1 | -0,3709 | 1,0938E-14 |
| ACE2 | LL0XNC01-16G2.1 | -0,3709 | 1,10091E-14 |
| ACE2 | RP11-972P1.10 | -0,371 | 1,07509E-14 |
| ACE2 | RPL21P119 | -0,371 | 1,06683E-14 |
| ACE2 | MYCBPAP | -0,371 | 1,08329E-14 |
| ACE2 | SLC30A6 | -0,3713 | 1,01584E-14 |
| ACE2 | RP11-90J7.4 | -0,3713 | 1,02049E-14 |
| ACE2 | ULK4P3 | -0,3713 | 1,02732E-14 |
| ACE2 | CNOT6 | -0,3714 | 9,96821E-15 |
| ACE2 | ABCB8 | -0,3714 | 9,98333E-15 |
| ACE2 | LCTL | -0,3714 | 9,9969E-15 |
| ACE2 | NPIPA5 | -0,3714 | 1,00762E-14 |
| ACE2 | CTD-2630F21.1 | -0,3714 | 1,00539E-14 |
| ACE2 | DDX3X | -0,3714 | 1,00329E-14 |
| ACE2 | SRP9 | -0,3715 | 9,77152E-15 |
| ACE2 | RFPL4B | -0,3715 | 9,89193E-15 |
| ACE2 | RP13-735L24.1 | -0,3715 | 9,77863E-15 |
| ACE2 | UBR7 | -0,3715 | 9,78596E-15 |
| ACE2 | RP3-425C14.6 | -0,3716 | 9,71277E-15 |
| ACE2 | LINC01165 | -0,3716 | 9,74305E-15 |
| ACE2 | RP11-227G15.8 | -0,3716 | 9,6817E-15 |
| ACE2 | CTD-2020K17.1 | -0,3716 | 9,59331E-15 |
| ACE2 | CTD-2245F17.3 | -0,3716 | 9,5967E-15 |
| ACE2 | SLC25A36P1 | -0,3716 | 9,65817E-15 |
| ACE2 | RASAL2-AS1 | -0,3717 | 9,53077E-15 |
| ACE2 | FCHO2 | -0,3717 | 9,42989E-15 |
| ACE2 | AC051649.12 | -0,3717 | 9,43232E-15 |
| ACE2 | HNRNPUL2 | -0,3717 | 9,56165E-15 |
| ACE2 | LRP1-AS | -0,3717 | 9,48255E-15 |

| | | | |
|---|---|---|---|
| ACE2 | RP11-475D10.4 | -0,3717 | 9,57029E-15 |
| ACE2 | TBX2-AS1 | -0,3717 | 9,47939E-15 |
| ACE2 | CTB-167G5.7 | -0,3717 | 9,49376E-15 |
| ACE2 | TMEM31 | -0,3717 | 9,50529E-15 |
| ACE2 | ZFP36L2 | -0,3718 | 9,4205E-15 |
| ACE2 | AC007405.4 | -0,3718 | 9,40286E-15 |
| ACE2 | RMDN3 | -0,3718 | 9,30412E-15 |
| ACE2 | COL9A1 | -0,3719 | 9,10637E-15 |
| ACE2 | VN1R42P | -0,3719 | 9,15301E-15 |
| ACE2 | RP11-229P13.25 | -0,3719 | 9,18635E-15 |
| ACE2 | RP3-467K16.7 | -0,372 | 8,98556E-15 |
| ACE2 | RP11-342K6.3 | -0,372 | 8,94028E-15 |
| ACE2 | AQP7P1 | -0,372 | 8,95448E-15 |
| ACE2 | AC008088.4 | -0,372 | 9,05013E-15 |
| ACE2 | LINC01927 | -0,372 | 9,02714E-15 |
| ACE2 | RP11-863P13.4 | -0,3721 | 8,88899E-15 |
| ACE2 | RP11-474I16.8 | -0,3722 | 8,63321E-15 |
| ACE2 | ING5 | -0,3722 | 8,76674E-15 |
| ACE2 | ELOVL3 | -0,3722 | 8,72667E-15 |
| ACE2 | RN7SL665P | -0,3723 | 8,52423E-15 |
| ACE2 | CTD-2126E3.4 | -0,3723 | 8,55199E-15 |
| ACE2 | MCFD2P1 | -0,3724 | 8,42597E-15 |
| ACE2 | PRDX3 | -0,3724 | 8,32682E-15 |
| ACE2 | AC083899.3 | -0,3725 | 8,28455E-15 |
| ACE2 | HDAC11 | -0,3725 | 8,26309E-15 |
| ACE2 | SKIL | -0,3725 | 8,28165E-15 |
| ACE2 | LINC01600 | -0,3725 | 8,23201E-15 |
| ACE2 | HNRNPA1P14 | -0,3725 | 8,27096E-15 |
| ACE2 | KRT24 | -0,3725 | 8,21579E-15 |
| ACE2 | GIPC3 | -0,3725 | 8,28412E-15 |
| ACE2 | ST7-AS1 | -0,3726 | 8,09145E-15 |
| ACE2 | RP11-12J10.4 | -0,3727 | 7,95239E-15 |
| ACE2 | RP11-424M22.3 | -0,3727 | 7,96399E-15 |
| ACE2 | ELL | -0,3727 | 7,9256E-15 |
| ACE2 | FDX1P2 | -0,3727 | 7,92231E-15 |
| ACE2 | RP11-274B21.3 | -0,3728 | 7,80112E-15 |
| ACE2 | AF131216.7 | -0,3728 | 7,80324E-15 |
| ACE2 | RYKP1 | -0,3728 | 7,82848E-15 |
| ACE2 | LINC02208 | -0,3729 | 7,75021E-15 |
| ACE2 | AC006128.2 | -0,3729 | 7,64677E-15 |
| ACE2 | MORN1 | -0,373 | 7,48939E-15 |
| ACE2 | RP11-1072C15.4 | -0,373 | 7,50516E-15 |
| ACE2 | FAM174A | -0,3731 | 7,41389E-15 |
| ACE2 | C8orf89 | -0,3732 | 7,30388E-15 |
| ACE2 | Z69720.2 | -0,3732 | 7,3043E-15 |
| ACE2 | RP11-715H19.2 | -0,3733 | 7,14073E-15 |
| ACE2 | TMEM51 | -0,3734 | 7,03576E-15 |
| ACE2 | HSPA6 | -0,3734 | 7,06953E-15 |
| ACE2 | SLC39A10 | -0,3734 | 7,01569E-15 |
| ACE2 | CTD-2196P11.2 | -0,3735 | 6,90806E-15 |
| ACE2 | PDCL3P5 | -0,3735 | 6,8962E-15 |
| ACE2 | LINC02188 | -0,3735 | 6,94352E-15 |
| ACE2 | PTX3 | -0,3736 | 6,77966E-15 |
| ACE2 | LINC00706 | -0,3736 | 6,81865E-15 |

| | | | |
|---|---|---|---|
| ACE2 | TPTE2P2 | -0,3737 | 6,62638E-15 |
| ACE2 | RN7SKP23 | -0,3737 | 6,70871E-15 |
| ACE2 | RP11-822E23.7 | -0,3737 | 6,70743E-15 |
| ACE2 | INTS8 | -0,3738 | 6,60454E-15 |
| ACE2 | TMEM55B | -0,3738 | 6,59001E-15 |
| ACE2 | RP11-701H24.3 | -0,3738 | 6,53968E-15 |
| ACE2 | RP11-250B2.5 | -0,3739 | 6,46159E-15 |
| ACE2 | RP11-166N17.1 | -0,3739 | 6,45131E-15 |
| ACE2 | CLEC2D | -0,3739 | 6,43724E-15 |
| ACE2 | RP11-230F18.6 | -0,3739 | 6,44596E-15 |
| ACE2 | GCC2-AS1 | -0,374 | 6,31312E-15 |
| ACE2 | AC006160.5 | -0,374 | 6,31563E-15 |
| ACE2 | CARD19 | -0,3741 | 6,16595E-15 |
| ACE2 | NDUFB11 | -0,3741 | 6,19918E-15 |
| ACE2 | RP11-641D5.1 | -0,3742 | 6,06313E-15 |
| ACE2 | CH17-385C13.2 | -0,3743 | 5,95807E-15 |
| ACE2 | PPP1R10 | -0,3743 | 5,96229E-15 |
| ACE2 | HAS2 | -0,3743 | 5,9409E-15 |
| ACE2 | CTC-412M14.5 | -0,3744 | 5,88519E-15 |
| ACE2 | AC007879.4 | -0,3745 | 5,82884E-15 |
| ACE2 | RP11-177C12.1 | -0,3745 | 5,73126E-15 |
| ACE2 | KLHL28 | -0,3745 | 5,74738E-15 |
| ACE2 | SLC9C2 | -0,3746 | 5,65133E-15 |
| ACE2 | NOP14 | -0,3746 | 5,6656E-15 |
| ACE2 | LINC01422 | -0,3746 | 5,71783E-15 |
| ACE2 | SYNC | -0,3747 | 5,61596E-15 |
| ACE2 | TTLL7-IT1 | -0,3747 | 5,5714E-15 |
| ACE2 | MZF1-AS1 | -0,3749 | 5,34695E-15 |
| ACE2 | SLC25A15P5 | -0,3749 | 5,36271E-15 |
| ACE2 | CTD-2308G16.1 | -0,3751 | 5,22734E-15 |
| ACE2 | CHMP1B2P | -0,3751 | 5,21423E-15 |
| ACE2 | SNX10 | -0,3752 | 5,06475E-15 |
| ACE2 | RTL1 | -0,3753 | 4,97311E-15 |
| ACE2 | HMSD | -0,3753 | 4,99289E-15 |
| ACE2 | PRTN3 | -0,3753 | 4,97248E-15 |
| ACE2 | PGRMC2 | -0,3754 | 4,94955E-15 |
| ACE2 | RASGEF1A | -0,3754 | 4,9296E-15 |
| ACE2 | RP1-197B17.7 | -0,3754 | 4,91114E-15 |
| ACE2 | RP11-118F19.1 | -0,3754 | 4,95407E-15 |
| ACE2 | LINC00517 | -0,3755 | 4,79124E-15 |
| ACE2 | STMN1 | -0,3756 | 4,77018E-15 |
| ACE2 | CACNA1C-AS4 | -0,3756 | 4,72591E-15 |
| ACE2 | RP4-533D7.4 | -0,3757 | 4,63715E-15 |
| ACE2 | MADD | -0,3757 | 4,67141E-15 |
| ACE2 | RP5-901A4.1 | -0,3757 | 4,68203E-15 |
| ACE2 | LINC02065 | -0,3758 | 4,54108E-15 |
| ACE2 | VN2R19P | -0,3758 | 4,5999E-15 |
| ACE2 | SH3GLB1 | -0,3759 | 4,48261E-15 |
| ACE2 | SLC35F6 | -0,3759 | 4,48316E-15 |
| ACE2 | HOXA-AS2 | -0,3759 | 4,52321E-15 |
| ACE2 | RP11-236B18.5 | -0,3759 | 4,46251E-15 |
| ACE2 | RP11-876N24.4 | -0,3759 | 4,46066E-15 |
| ACE2 | RP11-350J20.13 | -0,376 | 4,43048E-15 |
| ACE2 | ITPRIPL2 | -0,376 | 4,39355E-15 |

| | | | |
|---|---|---|---|
| ACE2 | C4B-AS1 | -0,3761 | 4,34018E-15 |
| ACE2 | RP11-626G11.6 | -0,3761 | 4,35753E-15 |
| ACE2 | TSSK1A | -0,3762 | 4,28672E-15 |
| ACE2 | NDUFB6 | -0,3764 | 4,08519E-15 |
| ACE2 | KRT223P | -0,3764 | 4,0994E-15 |
| ACE2 | PPAN | -0,3764 | 4,10461E-15 |
| ACE2 | DDX3P1 | -0,3764 | 4,10694E-15 |
| ACE2 | SLC35D3 | -0,3765 | 4,06287E-15 |
| ACE2 | OR7E2P | -0,3765 | 4,01584E-15 |
| ACE2 | RBM22 | -0,3766 | 3,99326E-15 |
| ACE2 | GTF3C4 | -0,3766 | 3,94068E-15 |
| ACE2 | LINC01229 | -0,3766 | 3,96495E-15 |
| ACE2 | GTPBP1 | -0,3767 | 3,88397E-15 |
| ACE2 | CLK2 | -0,3768 | 3,82184E-15 |
| ACE2 | AC006019.4 | -0,3768 | 3,81274E-15 |
| ACE2 | AQP7P2 | -0,3768 | 3,79777E-15 |
| ACE2 | RPL12P12 | -0,3768 | 3,80736E-15 |
| ACE2 | USP9Y | -0,3768 | 3,82733E-15 |
| ACE2 | RP5-1011O1.2 | -0,3769 | 3,73211E-15 |
| ACE2 | HSP90AB3P | -0,3769 | 3,74514E-15 |
| ACE2 | LAPTM4B | -0,3769 | 3,79096E-15 |
| ACE2 | C3orf49 | -0,377 | 3,71528E-15 |
| ACE2 | CNPY1 | -0,377 | 3,71029E-15 |
| ACE2 | RBMS2P1 | -0,377 | 3,66471E-15 |
| ACE2 | RAPGEF6 | -0,3771 | 3,61402E-15 |
| ACE2 | RP11-205K6.3 | -0,3771 | 3,64079E-15 |
| ACE2 | USP15 | -0,3771 | 3,6306E-15 |
| ACE2 | PLEKHG3 | -0,3771 | 3,59742E-15 |
| ACE2 | ASH2L | -0,3772 | 3,53923E-15 |
| ACE2 | LINC00427 | -0,3772 | 3,56456E-15 |
| ACE2 | AP000962.2 | -0,3772 | 3,53336E-15 |
| ACE2 | RP11-726G1.1 | -0,3773 | 3,48416E-15 |
| ACE2 | RP11-401O9.3 | -0,3773 | 3,4872E-15 |
| ACE2 | ZCCHC3 | -0,3774 | 3,46029E-15 |
| ACE2 | AEN | -0,3776 | 3,30548E-15 |
| ACE2 | TSPAN33 | -0,3777 | 3,23716E-15 |
| ACE2 | SMG1P7 | -0,3777 | 3,25582E-15 |
| ACE2 | LINC00266-1 | -0,3777 | 3,24438E-15 |
| ACE2 | COX7A2L | -0,3778 | 3,2168E-15 |
| ACE2 | RP11-112J1.1 | -0,3778 | 3,2125E-15 |
| ACE2 | IQCJ | -0,3779 | 3,13983E-15 |
| ACE2 | IL4 | -0,3779 | 3,16602E-15 |
| ACE2 | ANKK1 | -0,378 | 3,05759E-15 |
| ACE2 | SP7 | -0,378 | 3,07736E-15 |
| ACE2 | HS3ST2 | -0,378 | 3,08598E-15 |
| ACE2 | CCDC54 | -0,3781 | 3,04047E-15 |
| ACE2 | PSMA2 | -0,3781 | 3,03068E-15 |
| ACE2 | RP11-298I3.4 | -0,3781 | 3,02837E-15 |
| ACE2 | LINC01987 | -0,3782 | 2,98576E-15 |
| ACE2 | OXCT2P1 | -0,3783 | 2,90647E-15 |
| ACE2 | Vax2os1_3 | -0,3783 | 2,9235E-15 |
| ACE2 | RP11-284M14.1 | -0,3783 | 2,938E-15 |
| ACE2 | FAM173B | -0,3783 | 2,91193E-15 |
| ACE2 | RP1-199J3.5 | -0,3783 | 2,91487E-15 |

| | | | |
|---|---|---|---|
| ACE2 | CYP7A1 | -0,3783 | 2,94191E-15 |
| ACE2 | RP11-18B16.2 | -0,3783 | 2,89452E-15 |
| ACE2 | PPP4R3A | -0,3783 | 2,91432E-15 |
| ACE2 | SWT1 | -0,3784 | 2,88873E-15 |
| ACE2 | HOXA2 | -0,3784 | 2,86877E-15 |
| ACE2 | ANAPC2 | -0,3784 | 2,86267E-15 |
| ACE2 | RP11-64B16.3 | -0,3784 | 2,86828E-15 |
| ACE2 | COMMD4 | -0,3785 | 2,79811E-15 |
| ACE2 | SLC27A6 | -0,3786 | 2,75798E-15 |
| ACE2 | RPS3AP38 | -0,3786 | 2,78593E-15 |
| ACE2 | LY6G5C | -0,3787 | 2,71288E-15 |
| ACE2 | CTC-490E21.10 | -0,3787 | 2,72254E-15 |
| ACE2 | KLK7 | -0,3787 | 2,70081E-15 |
| ACE2 | FBXO4 | -0,3788 | 2,68584E-15 |
| ACE2 | WDR90 | -0,3788 | 2,68117E-15 |
| ACE2 | RP11-350J20.5 | -0,3789 | 2,5966E-15 |
| ACE2 | CASP4 | -0,3789 | 2,61966E-15 |
| ACE2 | RP11-592N21.1 | -0,3789 | 2,59794E-15 |
| ACE2 | SNORD124 | -0,3789 | 2,63576E-15 |
| ACE2 | RP11-122G18.11 | -0,379 | 2,5697E-15 |
| ACE2 | B4GALT7 | -0,379 | 2,55772E-15 |
| ACE2 | NMT2 | -0,379 | 2,57701E-15 |
| ACE2 | DOHH | -0,379 | 2,57655E-15 |
| ACE2 | Z95704.3 | -0,3791 | 2,53482E-15 |
| ACE2 | RP11-30K9.1 | -0,3791 | 2,54224E-15 |
| ACE2 | AC091153.4 | -0,3791 | 2,50644E-15 |
| ACE2 | AL590762.10 | -0,3791 | 2,51671E-15 |
| ACE2 | LINC01118 | -0,3792 | 2,48521E-15 |
| ACE2 | RP11-15L13.4 | -0,3792 | 2,46864E-15 |
| ACE2 | CCDC62 | -0,3792 | 2,48649E-15 |
| ACE2 | CNGA4 | -0,3793 | 2,43179E-15 |
| ACE2 | RP11-789C17.5 | -0,3793 | 2,41452E-15 |
| ACE2 | DSTNP1 | -0,3793 | 2,42295E-15 |
| ACE2 | WDR11 | -0,3794 | 2,37316E-15 |
| ACE2 | BTBD7 | -0,3794 | 2,381E-15 |
| ACE2 | FAM96B | -0,3794 | 2,39481E-15 |
| ACE2 | ZNF77 | -0,3794 | 2,38107E-15 |
| ACE2 | RPP14 | -0,3795 | 2,36106E-15 |
| ACE2 | RP11-727F15.11 | -0,3795 | 2,36463E-15 |
| ACE2 | ABCB9 | -0,3795 | 2,34655E-15 |
| ACE2 | HIST2H3PS2 | -0,3796 | 2,30451E-15 |
| ACE2 | GPR182 | -0,3796 | 2,30843E-15 |
| ACE2 | CCDC188 | -0,3796 | 2,30172E-15 |
| ACE2 | RP11-262A16.1 | -0,3797 | 2,26527E-15 |
| ACE2 | RP11-461F11.3 | -0,3797 | 2,27806E-15 |
| ACE2 | AC004383.3 | -0,3797 | 2,26179E-15 |
| ACE2 | ZNF138 | -0,3798 | 2,22599E-15 |
| ACE2 | TTC31 | -0,3802 | 2,0672E-15 |
| ACE2 | AC016739.2 | -0,3802 | 2,0574E-15 |
| ACE2 | ZNF350 | -0,3802 | 2,07252E-15 |
| ACE2 | MYLK2 | -0,3802 | 2,06607E-15 |
| ACE2 | HNRNPA2B1 | -0,3803 | 2,02197E-15 |
| ACE2 | CH17-351M24.1 | -0,3803 | 2,02208E-15 |
| ACE2 | ZNF620 | -0,3804 | 2,00128E-15 |

| | | | |
|---|---|---|---|
| ACE2 | ULK4P2 | -0,3804 | 2,00735E-15 |
| ACE2 | RP11-147K16.3 | -0,3804 | 2,00158E-15 |
| ACE2 | AKAP8 | -0,3804 | 1,9824E-15 |
| ACE2 | LMNA | -0,3805 | 1,94631E-15 |
| ACE2 | TPTE2P6 | -0,3805 | 1,94034E-15 |
| ACE2 | RP5-1102E8.3 | -0,3806 | 1,92781E-15 |
| ACE2 | DBR1 | -0,3806 | 1,9292E-15 |
| ACE2 | RP11-16E12.1 | -0,3806 | 1,90837E-15 |
| ACE2 | WDR38 | -0,3807 | 1,87825E-15 |
| ACE2 | RPL7AP50 | -0,3807 | 1,87272E-15 |
| ACE2 | RP11-342L8.2 | -0,3808 | 1,84935E-15 |
| ACE2 | CEBPZ | -0,381 | 1,79571E-15 |
| ACE2 | ABC7-42389800N19.1 | -0,381 | 1,7761E-15 |
| ACE2 | RP11-218C14.5 | -0,3811 | 1,76547E-15 |
| ACE2 | RP11-80I15.4 | -0,3812 | 1,73584E-15 |
| ACE2 | RP11-109N23.4 | -0,3812 | 1,7164E-15 |
| ACE2 | RP11-34P13.18 | -0,3813 | 1,68658E-15 |
| ACE2 | DDX1 | -0,3813 | 1,67969E-15 |
| ACE2 | LINC01301 | -0,3813 | 1,70058E-15 |
| ACE2 | RP11-38L15.8 | -0,3813 | 1,69023E-15 |
| ACE2 | TEX13D | -0,3813 | 1,70114E-15 |
| ACE2 | FCER1A | -0,3814 | 1,67273E-15 |
| ACE2 | RP11-222K16.1 | -0,3814 | 1,65733E-15 |
| ACE2 | RP11-803P9.1 | -0,3815 | 1,63347E-15 |
| ACE2 | SLC46A1 | -0,3815 | 1,642E-15 |
| ACE2 | AP000487.5 | -0,3816 | 1,6143E-15 |
| ACE2 | PI4KAP2 | -0,3816 | 1,59939E-15 |
| ACE2 | ELOA-AS1 | -0,3817 | 1,56625E-15 |
| ACE2 | PTPRB | -0,3818 | 1,54587E-15 |
| ACE2 | RP3-466P17.1 | -0,3819 | 1,50268E-15 |
| ACE2 | AC093627.8 | -0,3819 | 1,51604E-15 |
| ACE2 | TMED4 | -0,3819 | 1,52346E-15 |
| ACE2 | FLII | -0,3819 | 1,5082E-15 |
| ACE2 | RP11-415J8.3 | -0,382 | 1,48832E-15 |
| ACE2 | ADAMTS19-AS1 | -0,382 | 1,50066E-15 |
| ACE2 | PSMD11 | -0,382 | 1,48069E-15 |
| ACE2 | RP11-10L7.1 | -0,3821 | 1,45941E-15 |
| ACE2 | GS1-25M2.1 | -0,3821 | 1,45762E-15 |
| ACE2 | RP11-956E11.1 | -0,3821 | 1,45796E-15 |
| ACE2 | SNORA66 | -0,3823 | 1,40597E-15 |
| ACE2 | RP11-79M19.2 | -0,3823 | 1,41453E-15 |
| ACE2 | BBOX1 | -0,3823 | 1,42002E-15 |
| ACE2 | CTD-2595P9.4 | -0,3823 | 1,41136E-15 |
| ACE2 | EMC8 | -0,3823 | 1,40765E-15 |
| ACE2 | THAP12 | -0,3824 | 1,39078E-15 |
| ACE2 | RP11-762H8.1 | -0,3824 | 1,37325E-15 |
| ACE2 | RRS1 | -0,3825 | 1,36469E-15 |
| ACE2 | STX17-AS1 | -0,3825 | 1,36028E-15 |
| ACE2 | RP11-976B16.1 | -0,3825 | 1,34865E-15 |
| ACE2 | RP11-266K4.1 | -0,3826 | 1,32841E-15 |
| ACE2 | RP13-977J11.3 | -0,3826 | 1,33823E-15 |
| ACE2 | C22orf29 | -0,3826 | 1,32738E-15 |
| ACE2 | RP11-174N3.4 | -0,3827 | 1,30707E-15 |
| ACE2 | TPH2 | -0,3828 | 1,28712E-15 |

| | | | |
|---|---|---|---|
| ACE2 | AP000692.9 | -0,3828 | 1,27972E-15 |
| ACE2 | WASH7P | -0,3829 | 1,27029E-15 |
| ACE2 | MYRIP | -0,3829 | 1,25145E-15 |
| ACE2 | P2RY14 | -0,3829 | 1,25557E-15 |
| ACE2 | RP11-244G12.1 | -0,3829 | 1,26125E-15 |
| ACE2 | NUTM2A-AS1 | -0,3829 | 1,26056E-15 |
| ACE2 | HTR3B | -0,3829 | 1,25081E-15 |
| ACE2 | RP11-442G21.2 | -0,3829 | 1,25332E-15 |
| ACE2 | LMF1-AS1 | -0,3829 | 1,27173E-15 |
| ACE2 | RP11-446E24.3 | -0,383 | 1,24504E-15 |
| ACE2 | RNF144A-AS1 | -0,383 | 1,24082E-15 |
| ACE2 | FBXL17 | -0,383 | 1,24856E-15 |
| ACE2 | RP11-462B18.2 | -0,383 | 1,23943E-15 |
| ACE2 | LY6K | -0,3831 | 1,2183E-15 |
| ACE2 | RP11-680C21.1 | -0,3831 | 1,20665E-15 |
| ACE2 | TRAPPC3L | -0,3832 | 1,18782E-15 |
| ACE2 | AC002480.3 | -0,3832 | 1,18768E-15 |
| ACE2 | ANO5 | -0,3832 | 1,19148E-15 |
| ACE2 | PCAT6 | -0,3833 | 1,1815E-15 |
| ACE2 | CTD-2547L16.1 | -0,3833 | 1,16592E-15 |
| ACE2 | RP11-405A12.2 | -0,3833 | 1,17997E-15 |
| ACE2 | RP11-368J22.2 | -0,3834 | 1,1597E-15 |
| ACE2 | PROSER3 | -0,3834 | 1,14981E-15 |
| ACE2 | RIOK2 | -0,3835 | 1,12197E-15 |
| ACE2 | RP11-243M5.4 | -0,3836 | 1,10162E-15 |
| ACE2 | RP3-326L13.2 | -0,3836 | 1,10236E-15 |
| ACE2 | RAB7A | -0,3837 | 1,09044E-15 |
| ACE2 | ABHD16A | -0,3837 | 1,09206E-15 |
| ACE2 | TRIM35 | -0,3837 | 1,07983E-15 |
| ACE2 | SH2B3 | -0,3837 | 1,09007E-15 |
| ACE2 | RP3-509L4.3 | -0,3838 | 1,0746E-15 |
| ACE2 | CTD-2653M23.3 | -0,3839 | 1,04173E-15 |
| ACE2 | RP1-140K8.5 | -0,3839 | 1,0552E-15 |
| ACE2 | RCL1 | -0,3839 | 1,04936E-15 |
| ACE2 | ARHGEF1 | -0,3839 | 1,0467E-15 |
| ACE2 | MCM8 | -0,3839 | 1,04836E-15 |
| ACE2 | USH2A | -0,384 | 1,03481E-15 |
| ACE2 | CTD-2005H7.2 | -0,384 | 1,03575E-15 |
| ACE2 | GS1-57L11.1 | -0,3842 | 9,87367E-16 |
| ACE2 | MCMDC2 | -0,3842 | 9,84571E-16 |
| ACE2 | RP11-605F22.1 | -0,3842 | 9,95038E-16 |
| ACE2 | RP11-12M9.4 | -0,3842 | 9,8835E-16 |
| ACE2 | ZNF212 | -0,3843 | 9,77E-16 |
| ACE2 | RP11-468O2.1 | -0,3843 | 9,71922E-16 |
| ACE2 | RP11-456P18.2 | -0,3845 | 9,42295E-16 |
| ACE2 | SEMA3B | -0,3845 | 9,44075E-16 |
| ACE2 | PAK2 | -0,3845 | 9,31557E-16 |
| ACE2 | CTSB | -0,3845 | 9,37585E-16 |
| ACE2 | AC004076.5 | -0,3845 | 9,35425E-16 |
| ACE2 | RPS4Y1 | -0,3846 | 9,24479E-16 |
| ACE2 | AC079145.4 | -0,3847 | 9,11617E-16 |
| ACE2 | PIGX | -0,3847 | 9,05488E-16 |
| ACE2 | ARAP1-AS2 | -0,3847 | 9,10622E-16 |
| ACE2 | RP11-315O6.1 | -0,3847 | 9,09913E-16 |

| | | | |
|---|---|---|---|
| ACE2 | TELO2 | -0,3847 | 9,13676E-16 |
| ACE2 | CTA-276F8.1 | -0,3847 | 9,02548E-16 |
| ACE2 | RP11-799B12.2 | -0,3847 | 8,99701E-16 |
| ACE2 | AC006116.20 | -0,3847 | 9,03715E-16 |
| ACE2 | ENTPD4 | -0,3849 | 8,79001E-16 |
| ACE2 | TH | -0,3849 | 8,66909E-16 |
| ACE2 | EFNB3 | -0,3849 | 8,68177E-16 |
| ACE2 | RP11-798G7.6 | -0,3849 | 8,73543E-16 |
| ACE2 | XRN2 | -0,3849 | 8,75802E-16 |
| ACE2 | MED20 | -0,385 | 8,63765E-16 |
| ACE2 | RP11-121C2.2 | -0,3851 | 8,39438E-16 |
| ACE2 | DUS4L | -0,3852 | 8,18092E-16 |
| ACE2 | AP006547.3 | -0,3852 | 8,20825E-16 |
| ACE2 | RP6-74O6.6 | -0,3853 | 8,11718E-16 |
| ACE2 | RAD50 | -0,3853 | 8,0979E-16 |
| ACE2 | ASCC3 | -0,3853 | 8,08341E-16 |
| ACE2 | RP11-864I4.4 | -0,3853 | 8,08335E-16 |
| ACE2 | TM2D1 | -0,3854 | 7,93389E-16 |
| ACE2 | RP11-10A14.3 | -0,3854 | 8,01653E-16 |
| ACE2 | CRK | -0,3854 | 7,91414E-16 |
| ACE2 | TFAP2B | -0,3856 | 7,63468E-16 |
| ACE2 | KHSRP | -0,3856 | 7,65978E-16 |
| ACE2 | CTC-523E23.1 | -0,3856 | 7,59488E-16 |
| ACE2 | RP11-245J9.6 | -0,3857 | 7,49998E-16 |
| ACE2 | PCDHA13 | -0,3858 | 7,38615E-16 |
| ACE2 | RP11-196E1.3 | -0,3858 | 7,34284E-16 |
| ACE2 | ANXA2R | -0,3859 | 7,19678E-16 |
| ACE2 | TCP11L2 | -0,3859 | 7,20537E-16 |
| ACE2 | LINC00452 | -0,3859 | 7,26465E-16 |
| ACE2 | ZNF430 | -0,3859 | 7,21078E-16 |
| ACE2 | WDR5 | -0,386 | 7,13205E-16 |
| ACE2 | NFYC-AS1 | -0,3861 | 6,97789E-16 |
| ACE2 | RP1-43E13.2 | -0,3862 | 6,81378E-16 |
| ACE2 | ARCN1 | -0,3862 | 6,80752E-16 |
| ACE2 | USP32P2 | -0,3862 | 6,793E-16 |
| ACE2 | ZNF678 | -0,3863 | 6,78976E-16 |
| ACE2 | RP11-1246C19.1 | -0,3863 | 6,77705E-16 |
| ACE2 | CTD-2308N23.2 | -0,3864 | 6,5891E-16 |
| ACE2 | STAT6 | -0,3865 | 6,52916E-16 |
| ACE2 | LLPH | -0,3865 | 6,52232E-16 |
| ACE2 | DTD2 | -0,3865 | 6,44564E-16 |
| ACE2 | PARD3-AS1 | -0,3866 | 6,34268E-16 |
| ACE2 | SLC29A3 | -0,3866 | 6,3137E-16 |
| ACE2 | GTF2A1 | -0,3866 | 6,3342E-16 |
| ACE2 | RP11-544O24.2 | -0,3867 | 6,26264E-16 |
| ACE2 | SELENOT | -0,3867 | 6,2851E-16 |
| ACE2 | KIAA1551 | -0,3867 | 6,28618E-16 |
| ACE2 | HCG4 | -0,3868 | 6,14746E-16 |
| ACE2 | LARP4 | -0,3868 | 6,13174E-16 |
| ACE2 | TUBB2BP1 | -0,3869 | 6,03617E-16 |
| ACE2 | RP11-203M5.7 | -0,3869 | 5,97383E-16 |
| ACE2 | QTRT1 | -0,3869 | 6,0561E-16 |
| ACE2 | SPIN2A | -0,3871 | 5,83725E-16 |
| ACE2 | DNAJC13 | -0,3872 | 5,68228E-16 |

| | | | |
|---|---|---|---|
| ACE2 | ZNF252P | -0,3872 | 5,73329E-16 |
| ACE2 | DCDC2C | -0,3873 | 5,63071E-16 |
| ACE2 | COQ10B | -0,3873 | 5,53845E-16 |
| ACE2 | RABL3 | -0,3873 | 5,59498E-16 |
| ACE2 | SMIM18 | -0,3874 | 5,5172E-16 |
| ACE2 | GMPR2 | -0,3875 | 5,4281E-16 |
| ACE2 | MIPEPP1 | -0,3876 | 5,27324E-16 |
| ACE2 | ZGLP1 | -0,3876 | 5,24859E-16 |
| ACE2 | PRPF38A | -0,3877 | 5,18458E-16 |
| ACE2 | CTB-35F21.4 | -0,3877 | 5,16923E-16 |
| ACE2 | RP11-168K11.3 | -0,3877 | 5,22862E-16 |
| ACE2 | ELMSAN1 | -0,3877 | 5,15908E-16 |
| ACE2 | AP001476.2 | -0,3877 | 5,19776E-16 |
| ACE2 | FAM86B1 | -0,3878 | 5,04493E-16 |
| ACE2 | EWSAT1 | -0,3878 | 5,13181E-16 |
| ACE2 | LINC01894 | -0,3878 | 5,07144E-16 |
| ACE2 | ABHD2 | -0,3879 | 4,99631E-16 |
| ACE2 | RP11-20G13.5 | -0,3879 | 5,02277E-16 |
| ACE2 | CARM1 | -0,3879 | 4,99554E-16 |
| ACE2 | RPL36 | -0,388 | 4,88544E-16 |
| ACE2 | RP11-326I11.3 | -0,3881 | 4,84195E-16 |
| ACE2 | AC006372.4 | -0,3881 | 4,77201E-16 |
| ACE2 | C9orf92 | -0,3882 | 4,70937E-16 |
| ACE2 | GPR149 | -0,3883 | 4,6325E-16 |
| ACE2 | CST3 | -0,3883 | 4,67404E-16 |
| ACE2 | RP11-408P14.1 | -0,3884 | 4,5285E-16 |
| ACE2 | PDK4 | -0,3884 | 4,56442E-16 |
| ACE2 | ZRANB2-AS2 | -0,3885 | 4,48829E-16 |
| ACE2 | AC008063.2 | -0,3885 | 4,44843E-16 |
| ACE2 | RP11-241F15.1 | -0,3885 | 4,47185E-16 |
| ACE2 | OR11H7 | -0,3885 | 4,43386E-16 |
| ACE2 | RP11-54O7.1 | -0,3886 | 4,3512E-16 |
| ACE2 | AC005082.12 | -0,3886 | 4,34336E-16 |
| ACE2 | TNPO3 | -0,3886 | 4,36301E-16 |
| ACE2 | ZNF830 | -0,3886 | 4,38563E-16 |
| ACE2 | G6PC3 | -0,3886 | 4,35563E-16 |
| ACE2 | TMEM143 | -0,3886 | 4,40383E-16 |
| ACE2 | MFN2 | -0,3887 | 4,29182E-16 |
| ACE2 | RP11-80H18.4 | -0,3887 | 4,3131E-16 |
| ACE2 | RERGL | -0,3887 | 4,31563E-16 |
| ACE2 | FN3KRP | -0,3887 | 4,32161E-16 |
| ACE2 | RP4-665N4.4 | -0,3888 | 4,23265E-16 |
| ACE2 | LINC01252 | -0,3888 | 4,22342E-16 |
| ACE2 | RP11-2C24.3 | -0,3888 | 4,24002E-16 |
| ACE2 | SNIP1 | -0,3889 | 4,16641E-16 |
| ACE2 | GCKR | -0,3889 | 4,13215E-16 |
| ACE2 | RP11-756G12.1 | -0,3889 | 4,16004E-16 |
| ACE2 | SEL1L2 | -0,3889 | 4,16199E-16 |
| ACE2 | CTB-174D11.1 | -0,389 | 4,08508E-16 |
| ACE2 | TRIM27 | -0,389 | 4,06427E-16 |
| ACE2 | TSC22D3 | -0,389 | 4,09406E-16 |
| ACE2 | RP11-562I5.1 | -0,3891 | 3,95292E-16 |
| ACE2 | RP11-359I18.5 | -0,3891 | 3,96307E-16 |
| ACE2 | SNORD68 | -0,3891 | 3,95674E-16 |

| | | | |
|---|---|---|---|
| ACE2 | RP11-687M24.8 | -0,3892 | 3,8833E-16 |
| ACE2 | CDCA4 | -0,3892 | 3,90374E-16 |
| ACE2 | CDH3 | -0,3892 | 3,92353E-16 |
| ACE2 | BACH1-IT1 | -0,3892 | 3,90047E-16 |
| ACE2 | EEF1A1P9 | -0,3893 | 3,82506E-16 |
| ACE2 | PAPOLB | -0,3894 | 3,77484E-16 |
| ACE2 | RP11-734J24.1 | -0,3894 | 3,77387E-16 |
| ACE2 | SMCHD1 | -0,3894 | 3,74068E-16 |
| ACE2 | RP4-569M23.2 | -0,3894 | 3,79802E-16 |
| ACE2 | RP5-1024N4.2 | -0,3895 | 3,72584E-16 |
| ACE2 | ZNF786 | -0,3895 | 3,67466E-16 |
| ACE2 | RP11-473M20.9 | -0,3895 | 3,72154E-16 |
| ACE2 | ARRDC1-AS1 | -0,3896 | 3,65236E-16 |
| ACE2 | RP11-142M10.2 | -0,3896 | 3,60067E-16 |
| ACE2 | RP11-417L19.4 | -0,3896 | 3,66295E-16 |
| ACE2 | CD82 | -0,3897 | 3,54862E-16 |
| ACE2 | CACNG2 | -0,3897 | 3,5341E-16 |
| ACE2 | RFT1 | -0,3898 | 3,46875E-16 |
| ACE2 | CCZ1B | -0,3898 | 3,47195E-16 |
| ACE2 | RP11-165D6.1 | -0,3898 | 3,49673E-16 |
| ACE2 | GLUD2 | -0,3898 | 3,49366E-16 |
| ACE2 | ZFAND6 | -0,3899 | 3,41189E-16 |
| ACE2 | RP11-570L15.1 | -0,39 | 3,34211E-16 |
| ACE2 | RP11-522I20.3 | -0,39 | 3,34412E-16 |
| ACE2 | RP11-104L21.2 | -0,3901 | 3,29135E-16 |
| ACE2 | HTR5A | -0,3901 | 3,27663E-16 |
| ACE2 | RP11-30K9.5 | -0,3901 | 3,32625E-16 |
| ACE2 | NEK11 | -0,3902 | 3,21896E-16 |
| ACE2 | AC132008.1 | -0,3902 | 3,22346E-16 |
| ACE2 | PFDN6 | -0,3902 | 3,22364E-16 |
| ACE2 | CTC-559E9.5 | -0,3902 | 3,23257E-16 |
| ACE2 | ECSIT | -0,3903 | 3,15763E-16 |
| ACE2 | RP11-416H1.2 | -0,3904 | 3,1363E-16 |
| ACE2 | ZNF90P1 | -0,3904 | 3,10562E-16 |
| ACE2 | RASA2 | -0,3904 | 3,10432E-16 |
| ACE2 | EXD2 | -0,3905 | 3,0817E-16 |
| ACE2 | TCEA1P2 | -0,3906 | 2,99392E-16 |
| ACE2 | AC090283.3 | -0,3906 | 3,01437E-16 |
| ACE2 | AC007228.9 | -0,3906 | 3,03301E-16 |
| ACE2 | MBLAC1 | -0,3907 | 2,94771E-16 |
| ACE2 | CTC-246B18.8 | -0,3907 | 2,93005E-16 |
| ACE2 | RP3-395P12.2 | -0,3908 | 2,87603E-16 |
| ACE2 | DYNLT1 | -0,3908 | 2,91589E-16 |
| ACE2 | DUXAP4 | -0,3908 | 2,87258E-16 |
| ACE2 | FKBP3 | -0,3908 | 2,88397E-16 |
| ACE2 | CPLX3 | -0,3908 | 2,89337E-16 |
| ACE2 | HNRNPA1P48 | -0,3908 | 2,87563E-16 |
| ACE2 | SEC23IP | -0,3909 | 2,82459E-16 |
| ACE2 | RP5-867C24.1 | -0,3909 | 2,84613E-16 |
| ACE2 | TMEM147-AS1 | -0,3909 | 2,84674E-16 |
| ACE2 | AL022393.9 | -0,391 | 2,81384E-16 |
| ACE2 | NPM2 | -0,391 | 2,81315E-16 |
| ACE2 | CTD-3051D23.3 | -0,391 | 2,80819E-16 |
| ACE2 | CISD1 | -0,3911 | 2,73444E-16 |

| | | | |
|---|---|---|---|
| ACE2 | PPIL3 | -0,3912 | 2,69138E-16 |
| ACE2 | CTC-295J13.3 | -0,3912 | 2,66196E-16 |
| ACE2 | STK4 | -0,3912 | 2,65985E-16 |
| ACE2 | CTCF | -0,3913 | 2,64911E-16 |
| ACE2 | POTEE | -0,3915 | 2,54866E-16 |
| ACE2 | ZMAT2 | -0,3915 | 2,55282E-16 |
| ACE2 | RPS3AP5 | -0,3915 | 2,52217E-16 |
| ACE2 | CTC-304I17.5 | -0,3915 | 2,55777E-16 |
| ACE2 | UFC1 | -0,3916 | 2,49258E-16 |
| ACE2 | RP11-365F18.3 | -0,3916 | 2,47789E-16 |
| ACE2 | VN1R107P | -0,3916 | 2,50994E-16 |
| ACE2 | RFC2 | -0,3917 | 2,44803E-16 |
| ACE2 | PIAS1 | -0,3917 | 2,42063E-16 |
| ACE2 | TADA1 | -0,3918 | 2,40785E-16 |
| ACE2 | C2orf69 | -0,3918 | 2,41578E-16 |
| ACE2 | DCUN1D1 | -0,3918 | 2,37477E-16 |
| ACE2 | ANKRD52 | -0,3918 | 2,40913E-16 |
| ACE2 | RP11-221N13.3 | -0,3919 | 2,33749E-16 |
| ACE2 | LILRA6 | -0,3919 | 2,34647E-16 |
| ACE2 | MRPL22 | -0,392 | 2,3275E-16 |
| ACE2 | RP11-553K23.2 | -0,3921 | 2,2736E-16 |
| ACE2 | TCTN3 | -0,3921 | 2,25733E-16 |
| ACE2 | SEMA3B-AS1 | -0,3922 | 2,20081E-16 |
| ACE2 | MMP3 | -0,3922 | 2,21682E-16 |
| ACE2 | CTC-232P5.1 | -0,3922 | 2,20496E-16 |
| ACE2 | KCNC4 | -0,3923 | 2,15926E-16 |
| ACE2 | RP11-889L3.1 | -0,3923 | 2,18071E-16 |
| ACE2 | CTB-16J4.1 | -0,3923 | 2,16441E-16 |
| ACE2 | RP11-2E11.9 | -0,3923 | 2,19899E-16 |
| ACE2 | RBM14 | -0,3923 | 2,18261E-16 |
| ACE2 | FZD10 | -0,3923 | 2,18448E-16 |
| ACE2 | EPOR | -0,3923 | 2,19473E-16 |
| ACE2 | ESF1 | -0,3923 | 2,167E-16 |
| ACE2 | TAF9 | -0,3924 | 2,12304E-16 |
| ACE2 | SVIP | -0,3924 | 2,12975E-16 |
| ACE2 | EIF4E | -0,3925 | 2,0943E-16 |
| ACE2 | RHBDD2 | -0,3925 | 2,08873E-16 |
| ACE2 | C21orf62-AS1 | -0,3925 | 2,10149E-16 |
| ACE2 | BEGAIN | -0,3926 | 2,06423E-16 |
| ACE2 | HMGN3-AS1 | -0,3927 | 2,02923E-16 |
| ACE2 | KCNAB2 | -0,3928 | 1,96659E-16 |
| ACE2 | DBIL5P2 | -0,3928 | 1,98397E-16 |
| ACE2 | CIPC | -0,3928 | 1,97482E-16 |
| ACE2 | RP11-119H12.4 | -0,3929 | 1,93695E-16 |
| ACE2 | SPRED1 | -0,3929 | 1,95437E-16 |
| ACE2 | MTND4P26 | -0,393 | 1,90592E-16 |
| ACE2 | ZNF805 | -0,393 | 1,89729E-16 |
| ACE2 | RP5-1116H23.5 | -0,393 | 1,92234E-16 |
| ACE2 | TRAK2 | -0,3931 | 1,86669E-16 |
| ACE2 | MALT1 | -0,3931 | 1,88766E-16 |
| ACE2 | RPL28 | -0,3931 | 1,86856E-16 |
| ACE2 | TRMT1L | -0,3932 | 1,83732E-16 |
| ACE2 | HPS3 | -0,3932 | 1,83575E-16 |
| ACE2 | RP11-1112C15.2 | -0,3932 | 1,83449E-16 |

| | | | |
|---|---|---|---|
| ACE2 | C16orf70 | -0,3932 | 1,84409E-16 |
| ACE2 | RP1-182O16.2 | -0,3933 | 1,79628E-16 |
| ACE2 | GOLGA8T | -0,3933 | 1,78963E-16 |
| ACE2 | GPR89A | -0,3934 | 1,75665E-16 |
| ACE2 | RP11-276H19.2 | -0,3935 | 1,73939E-16 |
| ACE2 | RP11-493L12.4 | -0,3936 | 1,69761E-16 |
| ACE2 | RP11-445F6.2 | -0,3937 | 1,66872E-16 |
| ACE2 | RP11-414H17.5 | -0,3937 | 1,66346E-16 |
| ACE2 | FSHR | -0,3938 | 1,6315E-16 |
| ACE2 | MITD1 | -0,3938 | 1,63462E-16 |
| ACE2 | AC004540.4 | -0,3938 | 1,63926E-16 |
| ACE2 | TXNL4B | -0,3938 | 1,64157E-16 |
| ACE2 | OSR2 | -0,3939 | 1,59517E-16 |
| ACE2 | KCTD21-AS1 | -0,3939 | 1,62291E-16 |
| ACE2 | MAP1LC3B2 | -0,3939 | 1,59811E-16 |
| ACE2 | RP11-995C19.2 | -0,3939 | 1,5947E-16 |
| ACE2 | AP000962.1 | -0,3939 | 1,60007E-16 |
| ACE2 | KLF7-IT1 | -0,394 | 1,59095E-16 |
| ACE2 | APOA1-AS | -0,394 | 1,58156E-16 |
| ACE2 | RP5-1021I20.1 | -0,3941 | 1,5414E-16 |
| ACE2 | AC083873.4 | -0,3942 | 1,52511E-16 |
| ACE2 | C11orf87 | -0,3942 | 1,50946E-16 |
| ACE2 | NXF5 | -0,3942 | 1,52622E-16 |
| ACE2 | COPS5 | -0,3943 | 1,49276E-16 |
| ACE2 | RP11-83N9.6 | -0,3943 | 1,49648E-16 |
| ACE2 | RP11-310I24.1 | -0,3943 | 1,50109E-16 |
| ACE2 | PRKAG2 | -0,3944 | 1,45458E-16 |
| ACE2 | PCBP2-OT1 | -0,3944 | 1,47439E-16 |
| ACE2 | CNOT2 | -0,3944 | 1,45346E-16 |
| ACE2 | CTD-2553L13.10 | -0,3944 | 1,46737E-16 |
| ACE2 | ASAP3 | -0,3945 | 1,43284E-16 |
| ACE2 | CNGB3 | -0,3945 | 1,44241E-16 |
| ACE2 | PCBP1 | -0,3946 | 1,39684E-16 |
| ACE2 | EGFEM1P | -0,3946 | 1,39459E-16 |
| ACE2 | ACSM2B | -0,3946 | 1,41848E-16 |
| ACE2 | FANCL | -0,3947 | 1,39209E-16 |
| ACE2 | RP11-326A19.4 | -0,3947 | 1,3882E-16 |
| ACE2 | CTA-212A2.3 | -0,3947 | 1,3738E-16 |
| ACE2 | HSD17B10 | -0,3947 | 1,37585E-16 |
| ACE2 | RP5-1071N3.1 | -0,3948 | 1,35746E-16 |
| ACE2 | SLC6A6 | -0,3948 | 1,34401E-16 |
| ACE2 | MCHR2 | -0,3948 | 1,35671E-16 |
| ACE2 | RPL13AP25 | -0,3948 | 1,36254E-16 |
| ACE2 | LINC02091 | -0,3951 | 1,2791E-16 |
| ACE2 | PAX1 | -0,3951 | 1,28937E-16 |
| ACE2 | CLCNKB | -0,3952 | 1,26403E-16 |
| ACE2 | RP3-442L6.4 | -0,3952 | 1,24797E-16 |
| ACE2 | LINCR-0001 | -0,3952 | 1,26339E-16 |
| ACE2 | MMP21 | -0,3952 | 1,24589E-16 |
| ACE2 | SLC25A37 | -0,3954 | 1,21492E-16 |
| ACE2 | KIAA0513 | -0,3954 | 1,2033E-16 |
| ACE2 | LINC01121 | -0,3955 | 1,18705E-16 |
| ACE2 | CDH9 | -0,3955 | 1,18369E-16 |
| ACE2 | CTXN2 | -0,3955 | 1,18577E-16 |

| | | | |
|---|---|---|---|
| ACE2 | AC104389.28 | -0,3956 | 1,16696E-16 |
| ACE2 | TMCO3 | -0,3956 | 1,15965E-16 |
| ACE2 | RP11-99E15.2 | -0,3956 | 1,15767E-16 |
| ACE2 | DEFB109P3 | -0,3957 | 1,14478E-16 |
| ACE2 | ZBTB10 | -0,3957 | 1,14135E-16 |
| ACE2 | RRAS2 | -0,3957 | 1,14986E-16 |
| ACE2 | NEK4P2 | -0,3957 | 1,13682E-16 |
| ACE2 | AOX1 | -0,3958 | 1,11705E-16 |
| ACE2 | EXD3 | -0,3958 | 1,11467E-16 |
| ACE2 | MRPL11 | -0,3958 | 1,11487E-16 |
| ACE2 | RP11-687E1.2 | -0,3958 | 1,11791E-16 |
| ACE2 | MAT2A | -0,3959 | 1,09055E-16 |
| ACE2 | OR10AB1P | -0,3959 | 1,09957E-16 |
| ACE2 | MCM3AP | -0,3959 | 1,10195E-16 |
| ACE2 | AC007391.2 | -0,396 | 1,08507E-16 |
| ACE2 | EGR4 | -0,396 | 1,0751E-16 |
| ACE2 | UNC5B-AS1 | -0,396 | 1,06639E-16 |
| ACE2 | RAD17 | -0,3961 | 1,05876E-16 |
| ACE2 | STEAP2 | -0,3961 | 1,06406E-16 |
| ACE2 | HSD17B3 | -0,3961 | 1,06267E-16 |
| ACE2 | NMRAL1 | -0,3961 | 1,05318E-16 |
| ACE2 | RP11-71H17.8 | -0,3962 | 1,03009E-16 |
| ACE2 | RP11-493E12.1 | -0,3963 | 1,01837E-16 |
| ACE2 | RP4-555D20.4 | -0,3963 | 1,00629E-16 |
| ACE2 | RP11-347C18.3 | -0,3963 | 1,02342E-16 |
| ACE2 | LINC00115 | -0,3964 | 9,93247E-17 |
| ACE2 | AC093732.1 | -0,3964 | 1,00267E-16 |
| ACE2 | XKR7 | -0,3964 | 1,00362E-16 |
| ACE2 | C1orf64 | -0,3965 | 9,71532E-17 |
| ACE2 | RP4-640E24.1 | -0,3965 | 9,86075E-17 |
| ACE2 | RPS23P6 | -0,3966 | 9,59825E-17 |
| ACE2 | ITGB1P1 | -0,3966 | 9,63644E-17 |
| ACE2 | MYLK-AS1 | -0,3967 | 9,36961E-17 |
| ACE2 | LL22NC03-86G7.1 | -0,3968 | 9,18687E-17 |
| ACE2 | MPP7 | -0,3969 | 8,99552E-17 |
| ACE2 | OSM | -0,3969 | 8,96483E-17 |
| ACE2 | RP11-307B6.3 | -0,397 | 8,78679E-17 |
| ACE2 | RPS17P1 | -0,397 | 8,80143E-17 |
| ACE2 | RP5-1116H23.6 | -0,397 | 8,7897E-17 |
| ACE2 | RP11-149I23.3 | -0,3971 | 8,65774E-17 |
| ACE2 | LINC02080 | -0,3971 | 8,77151E-17 |
| ACE2 | RP11-455J20.3 | -0,3973 | 8,33049E-17 |
| ACE2 | DNAJB5-AS1 | -0,3973 | 8,36824E-17 |
| ACE2 | CTD-2331H12.5 | -0,3973 | 8,4149E-17 |
| ACE2 | RP4-530I15.9 | -0,3973 | 8,39792E-17 |
| ACE2 | SMDT1 | -0,3973 | 8,44238E-17 |
| ACE2 | RP11-297D21.4 | -0,3975 | 8,08013E-17 |
| ACE2 | RP13-20L14.10 | -0,3975 | 8,09473E-17 |
| ACE2 | RBBP8P1 | -0,3977 | 7,75745E-17 |
| ACE2 | RP11-894J14.2 | -0,3978 | 7,54937E-17 |
| ACE2 | PDZRN3-AS1 | -0,3978 | 7,57285E-17 |
| ACE2 | RP11-646I6.3 | -0,3978 | 7,61696E-17 |
| ACE2 | SLC2A9 | -0,3979 | 7,51145E-17 |
| ACE2 | TNFAIP8 | -0,3979 | 7,41492E-17 |

| | | | |
|---|---|---|---|
| ACE2 | RP11-17G12.2 | -0,3979 | 7,44653E-17 |
| ACE2 | TLE6 | -0,3979 | 7,48277E-17 |
| ACE2 | UCKL1 | -0,398 | 7,31988E-17 |
| ACE2 | C1D | -0,3982 | 7,00969E-17 |
| ACE2 | PSMD10 | -0,3982 | 7,05702E-17 |
| ACE2 | RP11-499F19.3 | -0,3984 | 6,81882E-17 |
| ACE2 | RRN3P1 | -0,3985 | 6,63847E-17 |
| ACE2 | PFN1 | -0,3985 | 6,66756E-17 |
| ACE2 | RP11-147L13.13 | -0,3985 | 6,6879E-17 |
| ACE2 | RP11-13K12.5 | -0,3985 | 6,63487E-17 |
| ACE2 | FAM90A20P | -0,3986 | 6,56399E-17 |
| ACE2 | SELENOH | -0,3986 | 6,52874E-17 |
| ACE2 | RP5-1057I20.6 | -0,3986 | 6,497E-17 |
| ACE2 | RP11-311P8.2 | -0,3986 | 6,51394E-17 |
| ACE2 | FIGLA | -0,3987 | 6,39593E-17 |
| ACE2 | LINC02147 | -0,3987 | 6,33188E-17 |
| ACE2 | HSPH1 | -0,3987 | 6,39363E-17 |
| ACE2 | RP11-1252D15.1 | -0,3989 | 6,10184E-17 |
| ACE2 | DCP2 | -0,3989 | 6,18282E-17 |
| ACE2 | RAB2A | -0,3989 | 6,17368E-17 |
| ACE2 | LINC00092 | -0,3989 | 6,17663E-17 |
| ACE2 | ANKRD18CP | -0,3989 | 6,09007E-17 |
| ACE2 | LA16c-313D11.9 | -0,3989 | 6,10391E-17 |
| ACE2 | RRN3 | -0,3989 | 6,19194E-17 |
| ACE2 | AC007792.1 | -0,3989 | 6,14669E-17 |
| ACE2 | MAGOH | -0,399 | 5,99337E-17 |
| ACE2 | WDSUB1 | -0,399 | 6,07439E-17 |
| ACE2 | ABCF1 | -0,399 | 6,03005E-17 |
| ACE2 | CH17-340M24.3 | -0,399 | 6,06486E-17 |
| ACE2 | RP11-332L8.1 | -0,3991 | 5,91144E-17 |
| ACE2 | TIGD4 | -0,3991 | 5,92835E-17 |
| ACE2 | CMB9-22P13.1 | -0,3991 | 5,96229E-17 |
| ACE2 | SLC4A1AP | -0,3992 | 5,80233E-17 |
| ACE2 | DNAJB14 | -0,3992 | 5,8295E-17 |
| ACE2 | ACAD8 | -0,3992 | 5,77756E-17 |
| ACE2 | RP4-791K14.2 | -0,3992 | 5,76715E-17 |
| ACE2 | CHPF | -0,3993 | 5,71656E-17 |
| ACE2 | FAM151B | -0,3993 | 5,70712E-17 |
| ACE2 | RP11-348J24.2 | -0,3993 | 5,66711E-17 |
| ACE2 | CCDC28A | -0,3993 | 5,69024E-17 |
| ACE2 | CCDC168 | -0,3993 | 5,70507E-17 |
| ACE2 | GLUD1P2 | -0,3995 | 5,4252E-17 |
| ACE2 | RP11-571M6.18 | -0,3995 | 5,46195E-17 |
| ACE2 | CTA-941F9.10 | -0,3995 | 5,49104E-17 |
| ACE2 | RPL4P2 | -0,3996 | 5,36373E-17 |
| ACE2 | FBXO18 | -0,3996 | 5,32569E-17 |
| ACE2 | AK4P3 | -0,3996 | 5,35694E-17 |
| ACE2 | STK19B | -0,3997 | 5,22535E-17 |
| ACE2 | CLPB | -0,3997 | 5,29725E-17 |
| ACE2 | STAB2 | -0,3997 | 5,30296E-17 |
| ACE2 | RP11-295K2.3 | -0,3998 | 5,13622E-17 |
| ACE2 | DKK1 | -0,3998 | 5,14563E-17 |
| ACE2 | RP1-286D6.5 | -0,3999 | 5,05802E-17 |
| ACE2 | CCDC38 | -0,3999 | 5,10538E-17 |

| | | | |
|---|---|---|---|
| ACE2 | AIMP1 | -0,4 | 4,97835E-17 |
| ACE2 | RP11-513M16.7 | -0,4001 | 4,85979E-17 |
| ACE2 | GAS2L1P2 | -0,4001 | 4,87365E-17 |
| ACE2 | ZNF844 | -0,4001 | 4,8567E-17 |
| ACE2 | ITIH2 | -0,4002 | 4,74919E-17 |
| ACE2 | HNRNPL | -0,4002 | 4,75947E-17 |
| ACE2 | RP3-375P9.2 | -0,4003 | 4,63579E-17 |
| ACE2 | KCNH1-IT1 | -0,4004 | 4,62325E-17 |
| ACE2 | FAM205C | -0,4004 | 4,5987E-17 |
| ACE2 | CTXN3 | -0,4005 | 4,47606E-17 |
| ACE2 | HMGN5 | -0,4005 | 4,49634E-17 |
| ACE2 | ANKRD11 | -0,4007 | 4,32676E-17 |
| ACE2 | ZNF486 | -0,4007 | 4,35369E-17 |
| ACE2 | PLP2 | -0,4007 | 4,29441E-17 |
| ACE2 | HYMAI | -0,4009 | 4,12095E-17 |
| ACE2 | DPYD-AS1 | -0,401 | 4,0502E-17 |
| ACE2 | CCDC86 | -0,401 | 4,05104E-17 |
| ACE2 | SLC22A15 | -0,4011 | 4,02132E-17 |
| ACE2 | RP11-145M9.5 | -0,4011 | 3,98554E-17 |
| ACE2 | C10orf76 | -0,4011 | 3,96675E-17 |
| ACE2 | GS1-174L6.4 | -0,4013 | 3,83101E-17 |
| ACE2 | RP11-333B11.1 | -0,4014 | 3,79903E-17 |
| ACE2 | SLC1A6 | -0,4014 | 3,77801E-17 |
| ACE2 | RP11-2H3.6 | -0,4015 | 3,71551E-17 |
| ACE2 | RHPN1-AS1 | -0,4015 | 3,68779E-17 |
| ACE2 | RP11-406H21.2 | -0,4015 | 3,68568E-17 |
| ACE2 | SMC3 | -0,4016 | 3,63278E-17 |
| ACE2 | PEAK1 | -0,4016 | 3,62699E-17 |
| ACE2 | PPP1R8P1 | -0,4017 | 3,57527E-17 |
| ACE2 | RP11-521B24.3 | -0,4017 | 3,55046E-17 |
| ACE2 | HOXB1 | -0,4017 | 3,58463E-17 |
| ACE2 | TRAV18 | -0,4018 | 3,50963E-17 |
| ACE2 | RRP1 | -0,4018 | 3,50091E-17 |
| ACE2 | CTC-559E9.8 | -0,402 | 3,34478E-17 |
| ACE2 | GSK3A | -0,402 | 3,32294E-17 |
| ACE2 | OSBP2 | -0,402 | 3,32988E-17 |
| ACE2 | ATE1 | -0,4021 | 3,2633E-17 |
| ACE2 | ATPAF1 | -0,4022 | 3,24974E-17 |
| ACE2 | KB-1958F4.1 | -0,4022 | 3,22171E-17 |
| ACE2 | ELP5 | -0,4022 | 3,20002E-17 |
| ACE2 | RP11-196G18.22 | -0,4023 | 3,16035E-17 |
| ACE2 | KDM3B | -0,4023 | 3,1355E-17 |
| ACE2 | RP11-370F5.4 | -0,4023 | 3,15954E-17 |
| ACE2 | FAAP100 | -0,4023 | 3,17983E-17 |
| ACE2 | ULBP2 | -0,4024 | 3,11087E-17 |
| ACE2 | P2RX2 | -0,4024 | 3,10805E-17 |
| ACE2 | CROCCP2 | -0,4025 | 3,03554E-17 |
| ACE2 | MYBPHL | -0,4025 | 3,02312E-17 |
| ACE2 | UFD1L | -0,4025 | 3,01729E-17 |
| ACE2 | LINC00621 | -0,4026 | 2,98167E-17 |
| ACE2 | ZMYM6 | -0,4027 | 2,91518E-17 |
| ACE2 | RP11-92K2.2 | -0,4027 | 2,91306E-17 |
| ACE2 | RPL7AP30 | -0,4027 | 2,94638E-17 |
| ACE2 | ATF3 | -0,4029 | 2,83341E-17 |

| | | | |
|---|---|---|---|
| ACE2 | PTPN21 | -0,403 | 2,73535E-17 |
| ACE2 | AC005932.1 | -0,403 | 2,74145E-17 |
| ACE2 | LINC01963 | -0,4031 | 2,71259E-17 |
| ACE2 | LINC02054 | -0,4031 | 2,72272E-17 |
| ACE2 | FRG1JP | -0,4031 | 2,72612E-17 |
| ACE2 | TAS2R31 | -0,4031 | 2,68612E-17 |
| ACE2 | RP11-114F10.3 | -0,4031 | 2,69863E-17 |
| ACE2 | RP11-406H23.2 | -0,4031 | 2,68784E-17 |
| ACE2 | AP000704.5 | -0,4031 | 2,72738E-17 |
| ACE2 | ANTXRL | -0,4032 | 2,62843E-17 |
| ACE2 | LINC00642 | -0,4032 | 2,66714E-17 |
| ACE2 | ABCA10 | -0,4032 | 2,66839E-17 |
| ACE2 | DAZAP1 | -0,4033 | 2,57507E-17 |
| ACE2 | CTB-50L17.8 | -0,4033 | 2,58578E-17 |
| ACE2 | DNMT1 | -0,4033 | 2,59045E-17 |
| ACE2 | PRPS1P2 | -0,4034 | 2,54746E-17 |
| ACE2 | CRKL | -0,4034 | 2,56221E-17 |
| ACE2 | TAF13 | -0,4035 | 2,48012E-17 |
| ACE2 | AF146191.4 | -0,4035 | 2,50328E-17 |
| ACE2 | RP11-266K4.9 | -0,4035 | 2,51114E-17 |
| ACE2 | SEMA7A | -0,4035 | 2,50166E-17 |
| ACE2 | GSTCD | -0,4037 | 2,39857E-17 |
| ACE2 | CTD-2184D3.7 | -0,4037 | 2,41399E-17 |
| ACE2 | LANCL1-AS1 | -0,4038 | 2,34854E-17 |
| ACE2 | ENPP2 | -0,4038 | 2,34947E-17 |
| ACE2 | KCNH4 | -0,4038 | 2,34332E-17 |
| ACE2 | ADAM17 | -0,4039 | 2,29085E-17 |
| ACE2 | C7orf49 | -0,4039 | 2,31725E-17 |
| ACE2 | BAALC-AS2 | -0,4039 | 2,32265E-17 |
| ACE2 | CETN3 | -0,404 | 2,24596E-17 |
| ACE2 | MICE | -0,404 | 2,24414E-17 |
| ACE2 | HEXIM1 | -0,404 | 2,27191E-17 |
| ACE2 | LL21NC02-1C16.1 | -0,404 | 2,24801E-17 |
| ACE2 | IGSF21 | -0,4041 | 2,21643E-17 |
| ACE2 | SERPINE1 | -0,4041 | 2,22097E-17 |
| ACE2 | SERPINE3 | -0,4041 | 2,20673E-17 |
| ACE2 | SPAAR | -0,4042 | 2,18765E-17 |
| ACE2 | RP11-554E23.2 | -0,4042 | 2,15536E-17 |
| ACE2 | C20orf96 | -0,4042 | 2,18449E-17 |
| ACE2 | PCGF1 | -0,4043 | 2,13808E-17 |
| ACE2 | ZC3H8 | -0,4043 | 2,11555E-17 |
| ACE2 | FAM186B | -0,4043 | 2,1203E-17 |
| ACE2 | NLGN4Y | -0,4043 | 2,11846E-17 |
| ACE2 | C18orf65 | -0,4044 | 2,0819E-17 |
| ACE2 | KCNJ14 | -0,4044 | 2,09813E-17 |
| ACE2 | LSM5 | -0,4045 | 2,03599E-17 |
| ACE2 | RP11-280K24.4 | -0,4045 | 2,06932E-17 |
| ACE2 | GEMIN4 | -0,4045 | 2,05258E-17 |
| ACE2 | AC005076.5 | -0,4047 | 1,98131E-17 |
| ACE2 | SGK494 | -0,4047 | 1,96833E-17 |
| ACE2 | RP11-890B15.2 | -0,4048 | 1,9447E-17 |
| ACE2 | NDUFB4P11 | -0,4048 | 1,92877E-17 |
| ACE2 | RP11-567J20.2 | -0,4049 | 1,89918E-17 |
| ACE2 | LINC00484 | -0,4049 | 1,90896E-17 |

| | | | |
|---|---|---|---|
| ACE2 | TXNDC2 | -0,4049 | 1,90958E-17 |
| ACE2 | CTC-453G23.8 | -0,4049 | 1,87974E-17 |
| ACE2 | RPS11P6 | -0,405 | 1,86541E-17 |
| ACE2 | RNF144A | -0,4051 | 1,83727E-17 |
| ACE2 | INHBB | -0,4052 | 1,78512E-17 |
| ACE2 | PTH2R | -0,4052 | 1,77374E-17 |
| ACE2 | RP11-111F5.2 | -0,4052 | 1,78304E-17 |
| ACE2 | CCNT1 | -0,4052 | 1,78946E-17 |
| ACE2 | PAK1IP1 | -0,4053 | 1,75537E-17 |
| ACE2 | RP11-3D4.3 | -0,4053 | 1,76646E-17 |
| ACE2 | RP11-76E17.2 | -0,4053 | 1,76514E-17 |
| ACE2 | RP11-333E1.2 | -0,4053 | 1,74179E-17 |
| ACE2 | RSPH14 | -0,4053 | 1,76019E-17 |
| ACE2 | MIR4435-2HG | -0,4054 | 1,7073E-17 |
| ACE2 | ZNRD1ASP | -0,4054 | 1,71523E-17 |
| ACE2 | RP11-461A8.5 | -0,4054 | 1,72327E-17 |
| ACE2 | SNORD89 | -0,4055 | 1,68569E-17 |
| ACE2 | YAE1D1 | -0,4055 | 1,69466E-17 |
| ACE2 | RAB22A | -0,4055 | 1,6916E-17 |
| ACE2 | AC008592.8 | -0,4056 | 1,63678E-17 |
| ACE2 | CTC-429P9.3 | -0,4056 | 1,64955E-17 |
| ACE2 | BCL2L12 | -0,4056 | 1,66294E-17 |
| ACE2 | PSMD7 | -0,4057 | 1,61069E-17 |
| ACE2 | RP11-376P6.3 | -0,4057 | 1,60955E-17 |
| ACE2 | EGOT | -0,4058 | 1,59865E-17 |
| ACE2 | RPSAP36 | -0,4058 | 1,57345E-17 |
| ACE2 | FAM222A | -0,4058 | 1,57565E-17 |
| ACE2 | RDH11 | -0,4058 | 1,58726E-17 |
| ACE2 | RP11-15A1.2 | -0,4058 | 1,58703E-17 |
| ACE2 | ICMT | -0,406 | 1,51838E-17 |
| ACE2 | CENPS | -0,406 | 1,51344E-17 |
| ACE2 | IKZF2 | -0,406 | 1,50988E-17 |
| ACE2 | LINC01366 | -0,406 | 1,53461E-17 |
| ACE2 | ADAMTS7P4 | -0,4061 | 1,4999E-17 |
| ACE2 | FAM131C | -0,4062 | 1,47521E-17 |
| ACE2 | RPS13P2 | -0,4062 | 1,47388E-17 |
| ACE2 | DHX38 | -0,4062 | 1,47612E-17 |
| ACE2 | RP11-206P5.2 | -0,4063 | 1,44562E-17 |
| ACE2 | TWF2 | -0,4064 | 1,40255E-17 |
| ACE2 | RTP1 | -0,4064 | 1,39595E-17 |
| ACE2 | YIPF7 | -0,4064 | 1,39397E-17 |
| ACE2 | LINC01154 | -0,4064 | 1,40087E-17 |
| ACE2 | AC008073.9 | -0,4065 | 1,3701E-17 |
| ACE2 | ORC4 | -0,4065 | 1,38804E-17 |
| ACE2 | RBM23 | -0,4065 | 1,37152E-17 |
| ACE2 | SNHG17 | -0,4065 | 1,3847E-17 |
| ACE2 | TNN | -0,4066 | 1,36032E-17 |
| ACE2 | RP11-181E10.3 | -0,4066 | 1,35893E-17 |
| ACE2 | AC004980.7 | -0,4066 | 1,35736E-17 |
| ACE2 | NFE2L1 | -0,4066 | 1,35495E-17 |
| ACE2 | BRWD1 | -0,4066 | 1,36262E-17 |
| ACE2 | CEP72 | -0,4067 | 1,31325E-17 |
| ACE2 | RGS20 | -0,4067 | 1,33352E-17 |
| ACE2 | CHRM4 | -0,4067 | 1,32315E-17 |

| | | | |
|---|---|---|---|
| ACE2 | TVP23C | -0,4067 | 1,32604E-17 |
| ACE2 | RAD1P2 | -0,4068 | 1,28671E-17 |
| ACE2 | GUSBP6 | -0,4068 | 1,28847E-17 |
| ACE2 | RP11-506M13.3 | -0,4068 | 1,29442E-17 |
| ACE2 | RP11-666O2.2 | -0,4068 | 1,29812E-17 |
| ACE2 | CTC-471J1.9 | -0,4068 | 1,2909E-17 |
| ACE2 | ALS2CR11 | -0,4069 | 1,26442E-17 |
| ACE2 | GPR3 | -0,407 | 1,2373E-17 |
| ACE2 | RPS6KA2-IT1 | -0,407 | 1,24976E-17 |
| ACE2 | AC005253.2 | -0,407 | 1,25317E-17 |
| ACE2 | DROSHA | -0,4071 | 1,21553E-17 |
| ACE2 | MAFA-AS1 | -0,4071 | 1,22824E-17 |
| ACE2 | SNHG8 | -0,4073 | 1,17547E-17 |
| ACE2 | APCDD1 | -0,4073 | 1,1862E-17 |
| ACE2 | HSPBP1 | -0,4073 | 1,17771E-17 |
| ACE2 | LL22NC03-N14H11.1 | -0,4073 | 1,18231E-17 |
| ACE2 | DNAJC9 | -0,4074 | 1,15084E-17 |
| ACE2 | FLJ12825 | -0,4074 | 1,15775E-17 |
| ACE2 | RP11-242P2.2 | -0,4074 | 1,14535E-17 |
| ACE2 | RPL23AP3 | -0,4074 | 1,15165E-17 |
| ACE2 | KCNJ15 | -0,4074 | 1,16207E-17 |
| ACE2 | RP11-335E6.2 | -0,4075 | 1,12336E-17 |
| ACE2 | AC069542.1 | -0,4075 | 1,13921E-17 |
| ACE2 | CD63 | -0,4076 | 1,11214E-17 |
| ACE2 | NTN4 | -0,4078 | 1,06119E-17 |
| ACE2 | AP001172.2 | -0,4078 | 1,06157E-17 |
| ACE2 | WRN | -0,4079 | 1,03749E-17 |
| ACE2 | ERRFI1 | -0,408 | 1,02454E-17 |
| ACE2 | RP4-673D20.3 | -0,408 | 1,01858E-17 |
| ACE2 | ZNF717 | -0,4081 | 1,00475E-17 |
| ACE2 | RP11-258C19.4 | -0,4081 | 1,0062E-17 |
| ACE2 | RPS3AP26 | -0,4082 | 9,85722E-18 |
| ACE2 | ATP8B4 | -0,4082 | 9,89835E-18 |
| ACE2 | CSTF3 | -0,4083 | 9,66831E-18 |
| ACE2 | LINC00941 | -0,4084 | 9,50195E-18 |
| ACE2 | RBMS1P1 | -0,4084 | 9,37086E-18 |
| ACE2 | SRP14-AS1 | -0,4084 | 9,47937E-18 |
| ACE2 | DEXI | -0,4084 | 9,44598E-18 |
| ACE2 | BSNDP2 | -0,4084 | 9,34573E-18 |
| ACE2 | RP4-620E11.8 | -0,4084 | 9,39378E-18 |
| ACE2 | KLHL11 | -0,4085 | 9,23509E-18 |
| ACE2 | RP1-102K2.9 | -0,4085 | 9,24842E-18 |
| ACE2 | FAM107A | -0,4086 | 8,98644E-18 |
| ACE2 | C6orf118 | -0,4086 | 9,00371E-18 |
| ACE2 | MUSK | -0,4086 | 9,04674E-18 |
| ACE2 | IL15RA | -0,4086 | 8,99716E-18 |
| ACE2 | RP11-69M1.6 | -0,4086 | 9,12673E-18 |
| ACE2 | TRNT1 | -0,4088 | 8,74975E-18 |
| ACE2 | PRMT8 | -0,4088 | 8,679E-18 |
| ACE2 | RP11-122D10.1 | -0,4088 | 8,69802E-18 |
| ACE2 | AC091729.7 | -0,4089 | 8,58199E-18 |
| ACE2 | RSPO2 | -0,4089 | 8,45209E-18 |
| ACE2 | RP11-488P3.1 | -0,409 | 8,44005E-18 |
| ACE2 | RP11-397P13.7 | -0,409 | 8,33753E-18 |

| | | | |
|---|---|---|---|
| ACE2 | CTHRC1 | -0,409 | 8,29656E-18 |
| ACE2 | CYSLTR2 | -0,409 | 8,3965E-18 |
| ACE2 | DCAF1 | -0,4091 | 8,18157E-18 |
| ACE2 | CTA-363E19.2 | -0,4091 | 8,13575E-18 |
| ACE2 | DPYS | -0,4091 | 8,15166E-18 |
| ACE2 | C16orf87 | -0,4091 | 8,1561E-18 |
| ACE2 | RP11-1079K10.2 | -0,4091 | 8,12141E-18 |
| ACE2 | WTAPP1 | -0,4092 | 7,97206E-18 |
| ACE2 | ST13P11 | -0,4094 | 7,70717E-18 |
| ACE2 | FBL | -0,4094 | 7,64803E-18 |
| ACE2 | TTC32 | -0,4095 | 7,5675E-18 |
| ACE2 | HYAL2 | -0,4097 | 7,23925E-18 |
| ACE2 | RP11-490K7.4 | -0,4098 | 7,18761E-18 |
| ACE2 | TTLL4 | -0,4098 | 7,12739E-18 |
| ACE2 | LRRC14 | -0,4098 | 7,11796E-18 |
| ACE2 | FBXL20 | -0,4098 | 7,07004E-18 |
| ACE2 | RP11-677I18.3 | -0,4099 | 7,03792E-18 |
| ACE2 | RP11-1263C18.1 | -0,41 | 6,81405E-18 |
| ACE2 | OXR1 | -0,41 | 6,83188E-18 |
| ACE2 | CTD-3252C9.4 | -0,41 | 6,81955E-18 |
| ACE2 | RP11-13J10.1 | -0,4101 | 6,75192E-18 |
| ACE2 | AFF2 | -0,4101 | 6,65693E-18 |
| ACE2 | FLRT3 | -0,4102 | 6,61149E-18 |
| ACE2 | NUDCD2 | -0,4104 | 6,3273E-18 |
| ACE2 | IL18BP | -0,4104 | 6,30456E-18 |
| ACE2 | DAD1 | -0,4104 | 6,26275E-18 |
| ACE2 | RP11-167B3.3 | -0,4104 | 6,31295E-18 |
| ACE2 | CCDC173 | -0,4105 | 6,17291E-18 |
| ACE2 | ISPD | -0,4105 | 6,2304E-18 |
| ACE2 | MRPS31 | -0,4105 | 6,12173E-18 |
| ACE2 | RP11-1212A22.4 | -0,4105 | 6,12327E-18 |
| ACE2 | AC093326.3 | -0,4106 | 6,08368E-18 |
| ACE2 | GAD1 | -0,4106 | 6,04277E-18 |
| ACE2 | RP11-637O19.2 | -0,4107 | 5,93869E-18 |
| ACE2 | PTGER1 | -0,4107 | 5,94324E-18 |
| ACE2 | RP11-333O1.1 | -0,4108 | 5,79149E-18 |
| ACE2 | EMC9 | -0,4108 | 5,82965E-18 |
| ACE2 | TEKT4P2 | -0,4108 | 5,84468E-18 |
| ACE2 | LINC01124 | -0,4109 | 5,74455E-18 |
| ACE2 | RN7SL771P | -0,4109 | 5,70448E-18 |
| ACE2 | LINC01759 | -0,411 | 5,59526E-18 |
| ACE2 | RPL7AP14 | -0,411 | 5,62145E-18 |
| ACE2 | ITIH3 | -0,4111 | 5,42855E-18 |
| ACE2 | TRIM74 | -0,4111 | 5,4328E-18 |
| ACE2 | RP11-863K10.4 | -0,4111 | 5,52025E-18 |
| ACE2 | TTC9C | -0,4111 | 5,46953E-18 |
| ACE2 | MBD2 | -0,4111 | 5,49302E-18 |
| ACE2 | RHOXF1 | -0,4111 | 5,43634E-18 |
| ACE2 | RP4-575N6.5 | -0,4112 | 5,36369E-18 |
| ACE2 | DPY19L1 | -0,4112 | 5,38276E-18 |
| ACE2 | FAM205A | -0,4112 | 5,4063E-18 |
| ACE2 | SNHG21 | -0,4112 | 5,34031E-18 |
| ACE2 | UBE2E1-AS1 | -0,4113 | 5,25662E-18 |
| ACE2 | RP11-700A24.1 | -0,4113 | 5,26571E-18 |

| | | | |
|---|---|---|---|
| ACE2 | AC000078.5 | -0,4113 | 5,21705E-18 |
| ACE2 | RP1-302G2.5 | -0,4114 | 5,19692E-18 |
| ACE2 | DGKH | -0,4114 | 5,20475E-18 |
| ACE2 | AC023137.2 | -0,4115 | 5,06964E-18 |
| ACE2 | AC012066.1 | -0,4115 | 5,03298E-18 |
| ACE2 | CEP76 | -0,4115 | 5,03522E-18 |
| ACE2 | RNU6-1098P | -0,4116 | 4,9613E-18 |
| ACE2 | RPS3 | -0,4116 | 4,98964E-18 |
| ACE2 | CMC2 | -0,4117 | 4,82771E-18 |
| ACE2 | AL592528.1 | -0,4117 | 4,86134E-18 |
| ACE2 | MTMR3 | -0,4117 | 4,86409E-18 |
| ACE2 | ZNF184 | -0,4118 | 4,76367E-18 |
| ACE2 | AC069257.6 | -0,4119 | 4,63062E-18 |
| ACE2 | FAM27E2 | -0,4119 | 4,65742E-18 |
| ACE2 | GTF2H3 | -0,4119 | 4,6678E-18 |
| ACE2 | BACE2 | -0,4119 | 4,66615E-18 |
| ACE2 | SLC26A5 | -0,412 | 4,54936E-18 |
| ACE2 | LINC01629 | -0,412 | 4,53935E-18 |
| ACE2 | RP11-215P8.2 | -0,4121 | 4,49597E-18 |
| ACE2 | RP11-432J24.5 | -0,4121 | 4,49107E-18 |
| ACE2 | ANKRD26P1 | -0,4121 | 4,44308E-18 |
| ACE2 | OGT | -0,4121 | 4,49208E-18 |
| ACE2 | RP4-635E18.7 | -0,4122 | 4,35342E-18 |
| ACE2 | FAM208B | -0,4122 | 4,37204E-18 |
| ACE2 | GPX1 | -0,4123 | 4,25636E-18 |
| ACE2 | U91328.22 | -0,4123 | 4,33533E-18 |
| ACE2 | OVGP1 | -0,4124 | 4,17051E-18 |
| ACE2 | CP | -0,4124 | 4,22879E-18 |
| ACE2 | RP11-11N5.1 | -0,4124 | 4,19531E-18 |
| ACE2 | CTDSP1 | -0,4125 | 4,14216E-18 |
| ACE2 | PACRG-AS1 | -0,4125 | 4,14856E-18 |
| ACE2 | PPP2R2A | -0,4125 | 4,15117E-18 |
| ACE2 | RP11-802E16.3 | -0,4125 | 4,08744E-18 |
| ACE2 | CLEC4A | -0,4125 | 4,12157E-18 |
| ACE2 | ETV6 | -0,4125 | 4,08188E-18 |
| ACE2 | ABHD13 | -0,4125 | 4,12684E-18 |
| ACE2 | ZNF653 | -0,4125 | 4,13733E-18 |
| ACE2 | RP11-33A14.1 | -0,4126 | 4,0179E-18 |
| ACE2 | PTX4 | -0,4126 | 4,07481E-18 |
| ACE2 | STRIP2 | -0,4127 | 3,98467E-18 |
| ACE2 | RPL18AP3 | -0,4127 | 3,92178E-18 |
| ACE2 | SZT2 | -0,4128 | 3,90152E-18 |
| ACE2 | VSIG8 | -0,4128 | 3,89588E-18 |
| ACE2 | RP1-199J3.6 | -0,4128 | 3,88119E-18 |
| ACE2 | RP11-473E2.4 | -0,4128 | 3,89705E-18 |
| ACE2 | RP5-827C21.4 | -0,4129 | 3,81951E-18 |
| ACE2 | RP1-292B18.1 | -0,4129 | 3,82954E-18 |
| ACE2 | MAGED1 | -0,4129 | 3,80003E-18 |
| ACE2 | CTC-340A15.2 | -0,413 | 3,7554E-18 |
| ACE2 | GPR37 | -0,413 | 3,71476E-18 |
| ACE2 | CTC-336P14.1 | -0,4131 | 3,64009E-18 |
| ACE2 | HEBP2 | -0,4131 | 3,68283E-18 |
| ACE2 | FAM71F1 | -0,4131 | 3,64282E-18 |
| ACE2 | RP11-864N7.2 | -0,4131 | 3,65774E-18 |

| | | | |
|---|---|---|---|
| ACE2 | RP11-54O7.11 | -0,4132 | 3,58186E-18 |
| ACE2 | TUBG1 | -0,4132 | 3,59579E-18 |
| ACE2 | RN7SL583P | -0,4134 | 3,41579E-18 |
| ACE2 | NXPH2 | -0,4134 | 3,42847E-18 |
| ACE2 | CTD-2201E18.3 | -0,4134 | 3,45412E-18 |
| ACE2 | XXbac-BPG154L12.5 | -0,4134 | 3,4003E-18 |
| ACE2 | SNRPC | -0,4134 | 3,41485E-18 |
| ACE2 | PGM2L1 | -0,4134 | 3,45118E-18 |
| ACE2 | P3H4 | -0,4134 | 3,4484E-18 |
| ACE2 | NUBPL | -0,4135 | 3,39139E-18 |
| ACE2 | ZNF114 | -0,4135 | 3,3437E-18 |
| ACE2 | RP11-134L10.1 | -0,4136 | 3,2976E-18 |
| ACE2 | GSTP1 | -0,4136 | 3,32036E-18 |
| ACE2 | MLNR | -0,4136 | 3,27334E-18 |
| ACE2 | TIPIN | -0,4136 | 3,30325E-18 |
| ACE2 | RP11-274H2.2 | -0,4137 | 3,20193E-18 |
| ACE2 | SOD2 | -0,4137 | 3,233E-18 |
| ACE2 | OR7E158P | -0,4137 | 3,22013E-18 |
| ACE2 | SOCS3 | -0,4137 | 3,24809E-18 |
| ACE2 | KIAA1107 | -0,4138 | 3,19197E-18 |
| ACE2 | U47924.32 | -0,4138 | 3,16048E-18 |
| ACE2 | CALM1 | -0,4138 | 3,14413E-18 |
| ACE2 | HSPA8P4 | -0,4139 | 3,1057E-18 |
| ACE2 | CCNG1 | -0,4139 | 3,10162E-18 |
| ACE2 | RP11-180C16.1 | -0,4139 | 3,07458E-18 |
| ACE2 | RPS3AP6 | -0,4139 | 3,1095E-18 |
| ACE2 | ACSM2A | -0,4139 | 3,1184E-18 |
| ACE2 | CTD-2049O4.1 | -0,4139 | 3,1242E-18 |
| ACE2 | RP13-638C3.2 | -0,4139 | 3,08822E-18 |
| ACE2 | ZNF572 | -0,4141 | 2,96306E-18 |
| ACE2 | AC142528.1 | -0,4142 | 2,9144E-18 |
| ACE2 | RP11-796E2.4 | -0,4142 | 2,90984E-18 |
| ACE2 | PHAX | -0,4143 | 2,88222E-18 |
| ACE2 | RP11-182N22.9 | -0,4143 | 2,83444E-18 |
| ACE2 | OTUD1 | -0,4143 | 2,88592E-18 |
| ACE2 | RP11-666F17.1 | -0,4144 | 2,80831E-18 |
| ACE2 | GPS2 | -0,4144 | 2,7723E-18 |
| ACE2 | NCOA5 | -0,4144 | 2,79062E-18 |
| ACE2 | AC092675.4 | -0,4146 | 2,68743E-18 |
| ACE2 | RP11-158K1.3 | -0,4146 | 2,69173E-18 |
| ACE2 | CMKLR1 | -0,4146 | 2,70597E-18 |
| ACE2 | RAB35 | -0,4146 | 2,69006E-18 |
| ACE2 | CYCSP40 | -0,4146 | 2,70769E-18 |
| ACE2 | SLC9A6 | -0,4146 | 2,67568E-18 |
| ACE2 | LINC01126 | -0,4148 | 2,56984E-18 |
| ACE2 | AC010886.2 | -0,4148 | 2,56516E-18 |
| ACE2 | CST6 | -0,4148 | 2,58098E-18 |
| ACE2 | PIGG | -0,4149 | 2,5131E-18 |
| ACE2 | RP11-4L24.4 | -0,4149 | 2,50583E-18 |
| ACE2 | C1orf145 | -0,4151 | 2,43486E-18 |
| ACE2 | LEMD2 | -0,4151 | 2,40381E-18 |
| ACE2 | SESN3 | -0,4151 | 2,44266E-18 |
| ACE2 | PRPSAP2 | -0,4151 | 2,44181E-18 |
| ACE2 | RFPL3S | -0,4151 | 2,42205E-18 |

| | | | |
|---|---|---|---|
| ACE2 | ASS1P2 | -0,4152 | 2,3957E-18 |
| ACE2 | NDUFA8 | -0,4152 | 2,39417E-18 |
| ACE2 | NUP107 | -0,4152 | 2,36391E-18 |
| ACE2 | PSAT1P3 | -0,4153 | 2,31329E-18 |
| ACE2 | RSAD2 | -0,4153 | 2,35126E-18 |
| ACE2 | ABHD10 | -0,4153 | 2,3293E-18 |
| ACE2 | XXbac-BPG252P9.9 | -0,4153 | 2,32102E-18 |
| ACE2 | DCTN2 | -0,4154 | 2,28188E-18 |
| ACE2 | AKT1S1 | -0,4154 | 2,29426E-18 |
| ACE2 | FCGR2C | -0,4155 | 2,23119E-18 |
| ACE2 | TLR9 | -0,4155 | 2,22196E-18 |
| ACE2 | ZNF878 | -0,4155 | 2,25545E-18 |
| ACE2 | OXA1L | -0,4156 | 2,17882E-18 |
| ACE2 | PPP1R2P1 | -0,4157 | 2,13784E-18 |
| ACE2 | EPO | -0,4157 | 2,15919E-18 |
| ACE2 | AGL | -0,4158 | 2,09161E-18 |
| ACE2 | SRFBP1 | -0,4159 | 2,07176E-18 |
| ACE2 | TRAPPC9 | -0,4159 | 2,05428E-18 |
| ACE2 | MXRA5 | -0,4159 | 2,06736E-18 |
| ACE2 | EDA | -0,4159 | 2,05614E-18 |
| ACE2 | RP4-771M4.3 | -0,4161 | 1,96211E-18 |
| ACE2 | PREB | -0,4161 | 1,96962E-18 |
| ACE2 | ITM2B | -0,4161 | 1,98324E-18 |
| ACE2 | HMGN2 | -0,4162 | 1,9235E-18 |
| ACE2 | RP11-363D14.1 | -0,4162 | 1,94486E-18 |
| ACE2 | HOXD4 | -0,4162 | 1,92073E-18 |
| ACE2 | RP11-989F5.3 | -0,4162 | 1,92913E-18 |
| ACE2 | TNR | -0,4163 | 1,9051E-18 |
| ACE2 | POTEF | -0,4163 | 1,87678E-18 |
| ACE2 | RP11-89C3.3 | -0,4163 | 1,89748E-18 |
| ACE2 | CTB-32O4.2 | -0,4163 | 1,88596E-18 |
| ACE2 | CETN4P | -0,4164 | 1,86256E-18 |
| ACE2 | AC140912.1 | -0,4164 | 1,85372E-18 |
| ACE2 | ARL16 | -0,4164 | 1,86775E-18 |
| ACE2 | SNRPD3 | -0,4164 | 1,87365E-18 |
| ACE2 | HBB | -0,4165 | 1,82136E-18 |
| ACE2 | NSRP1P1 | -0,4167 | 1,74094E-18 |
| ACE2 | RNU1-36P | -0,4167 | 1,76265E-18 |
| ACE2 | RP11-468E2.11 | -0,4168 | 1,70936E-18 |
| ACE2 | CTD-2561B21.5 | -0,4168 | 1,69306E-18 |
| ACE2 | CCT3 | -0,4169 | 1,68844E-18 |
| ACE2 | RPL10P16 | -0,4169 | 1,6818E-18 |
| ACE2 | SCLT1 | -0,4172 | 1,57704E-18 |
| ACE2 | CHCHD2 | -0,4172 | 1,58513E-18 |
| ACE2 | AC012531.25 | -0,4173 | 1,5559E-18 |
| ACE2 | GABRA4 | -0,4174 | 1,49734E-18 |
| ACE2 | CYB5D1 | -0,4174 | 1,51578E-18 |
| ACE2 | EFTUD1P1 | -0,4176 | 1,45303E-18 |
| ACE2 | TIMM17B | -0,4176 | 1,45522E-18 |
| ACE2 | RP11-973D8.5 | -0,4177 | 1,42151E-18 |
| ACE2 | SPSB3 | -0,4177 | 1,42337E-18 |
| ACE2 | RP11-347L15.1 | -0,4177 | 1,4074E-18 |
| ACE2 | PCDHA8 | -0,4178 | 1,40396E-18 |
| ACE2 | CKM | -0,4178 | 1,3889E-18 |

| | | | |
|---|---|---|---|
| ACE2 | LINC00634 | -0,4178 | 1,38152E-18 |
| ACE2 | P2RX3 | -0,4179 | 1,3493E-18 |
| ACE2 | H2AFZ | -0,418 | 1,32274E-18 |
| ACE2 | FUT8 | -0,418 | 1,34294E-18 |
| ACE2 | TMSB15B | -0,418 | 1,33521E-18 |
| ACE2 | RP11-286N22.14 | -0,4181 | 1,30908E-18 |
| ACE2 | RP11-986G18.2 | -0,4181 | 1,29421E-18 |
| ACE2 | CTD-2623N2.3 | -0,4181 | 1,30044E-18 |
| ACE2 | BMP4 | -0,4182 | 1,27674E-18 |
| ACE2 | RP4-738P15.1 | -0,4183 | 1,25232E-18 |
| ACE2 | LINC00954 | -0,4185 | 1,192E-18 |
| ACE2 | PCSK1 | -0,4185 | 1,21554E-18 |
| ACE2 | AC006159.5 | -0,4185 | 1,21297E-18 |
| ACE2 | F2R | -0,4186 | 1,18856E-18 |
| ACE2 | ATRN | -0,4186 | 1,16963E-18 |
| ACE2 | RP13-152O15.5 | -0,4187 | 1,15964E-18 |
| ACE2 | SETP2 | -0,4188 | 1,13926E-18 |
| ACE2 | ZNF213-AS1 | -0,4188 | 1,1278E-18 |
| ACE2 | GPKOW | -0,4188 | 1,12111E-18 |
| ACE2 | ANKRD2 | -0,4189 | 1,10112E-18 |
| ACE2 | RPL36AL | -0,4189 | 1,10715E-18 |
| ACE2 | ZNF668 | -0,4189 | 1,11563E-18 |
| ACE2 | RP13-1016M1.2 | -0,419 | 1,07751E-18 |
| ACE2 | ZNRF2P1 | -0,419 | 1,07732E-18 |
| ACE2 | ODF3L2 | -0,419 | 1,07431E-18 |
| ACE2 | RP11-466G12.2 | -0,4191 | 1,05691E-18 |
| ACE2 | AP2B1 | -0,4191 | 1,05288E-18 |
| ACE2 | MAGED4 | -0,4192 | 1,04357E-18 |
| ACE2 | RP11-462G22.2 | -0,4193 | 1,01079E-18 |
| ACE2 | BTBD10 | -0,4193 | 1,02535E-18 |
| ACE2 | PPP2R3A | -0,4194 | 1,00149E-18 |
| ACE2 | PRPF4B | -0,4194 | 1,00686E-18 |
| ACE2 | CYP4Z1 | -0,4195 | 9,69453E-19 |
| ACE2 | RP11-580I16.2 | -0,4195 | 9,75617E-19 |
| ACE2 | UBIAD1 | -0,4196 | 9,50221E-19 |
| ACE2 | RP11-797D24.3 | -0,4196 | 9,50692E-19 |
| ACE2 | AC091133.1 | -0,4196 | 9,64655E-19 |
| ACE2 | CTC-559E9.12 | -0,4196 | 9,53406E-19 |
| ACE2 | AC007246.3 | -0,4197 | 9,29545E-19 |
| ACE2 | POLE4 | -0,4197 | 9,38951E-19 |
| ACE2 | ROPN1B | -0,4197 | 9,45058E-19 |
| ACE2 | AC005592.2 | -0,4197 | 9,38271E-19 |
| ACE2 | RAB24 | -0,4197 | 9,35409E-19 |
| ACE2 | TRPM7 | -0,4197 | 9,37547E-19 |
| ACE2 | RP11-320M16.2 | -0,4198 | 9,19612E-19 |
| ACE2 | GABRG3 | -0,4198 | 9,14443E-19 |
| ACE2 | RINL | -0,4198 | 9,26298E-19 |
| ACE2 | MIGA1 | -0,4199 | 9,05195E-19 |
| ACE2 | LINC02110 | -0,4199 | 8,95672E-19 |
| ACE2 | CTD-2571L23.8 | -0,4199 | 9,01121E-19 |
| ACE2 | RP11-98D18.3 | -0,42 | 8,77731E-19 |
| ACE2 | ALS2CR12 | -0,42 | 8,8996E-19 |
| ACE2 | RPL5P1 | -0,42 | 8,75073E-19 |
| ACE2 | GATM-AS1 | -0,42 | 8,76921E-19 |

| | | | |
|---|---|---|---|
| ACE2 | RHOXF1P3 | -0,42 | 8,83482E-19 |
| ACE2 | AKIRIN1 | -0,4201 | 8,61941E-19 |
| ACE2 | THNSL2 | -0,4201 | 8,55931E-19 |
| ACE2 | ABCF2 | -0,4201 | 8,58281E-19 |
| ACE2 | DPY19L2P1 | -0,4202 | 8,52734E-19 |
| ACE2 | ARHGEF39 | -0,4202 | 8,5308E-19 |
| ACE2 | TOP1MT | -0,4203 | 8,21219E-19 |
| ACE2 | TMEM203 | -0,4204 | 8,14004E-19 |
| ACE2 | TPP2 | -0,4204 | 8,08134E-19 |
| ACE2 | STXBP6 | -0,4204 | 8,18325E-19 |
| ACE2 | AC004158.3 | -0,4204 | 8,03559E-19 |
| ACE2 | USP6 | -0,4204 | 8,16737E-19 |
| ACE2 | GCNA | -0,4204 | 8,11145E-19 |
| ACE2 | RP11-363N22.3 | -0,4205 | 7,90141E-19 |
| ACE2 | RP11-637A17.2 | -0,4205 | 7,99877E-19 |
| ACE2 | STK35 | -0,4205 | 7,94127E-19 |
| ACE2 | PKP4-AS1 | -0,4206 | 7,83529E-19 |
| ACE2 | HYAL3 | -0,4206 | 7,79602E-19 |
| ACE2 | RP11-225N10.3 | -0,4206 | 7,83484E-19 |
| ACE2 | PCDHA12 | -0,4206 | 7,80433E-19 |
| ACE2 | RP11-831F12.2 | -0,4206 | 7,82041E-19 |
| ACE2 | NSMF | -0,4207 | 7,57646E-19 |
| ACE2 | RP11-164O23.8 | -0,4208 | 7,38255E-19 |
| ACE2 | KRT8P12 | -0,4208 | 7,46831E-19 |
| ACE2 | RP11-135F9.3 | -0,4208 | 7,38646E-19 |
| ACE2 | RP11-81K2.1 | -0,4208 | 7,53534E-19 |
| ACE2 | CTD-2538G9.5 | -0,4208 | 7,49467E-19 |
| ACE2 | U82695.5 | -0,4208 | 7,43196E-19 |
| ACE2 | CCDC110 | -0,4209 | 7,33994E-19 |
| ACE2 | HVCN1 | -0,4209 | 7,3335E-19 |
| ACE2 | GIGYF2 | -0,421 | 7,13402E-19 |
| ACE2 | CFAP43 | -0,421 | 7,10315E-19 |
| ACE2 | SEC23A | -0,421 | 7,1886E-19 |
| ACE2 | SEPHS1P1 | -0,4211 | 7,04143E-19 |
| ACE2 | HIF1AN | -0,4212 | 6,91883E-19 |
| ACE2 | HS2ST1 | -0,4213 | 6,69978E-19 |
| ACE2 | AC093802.1 | -0,4213 | 6,73632E-19 |
| ACE2 | LHFPL5 | -0,4213 | 6,76792E-19 |
| ACE2 | FAM206A | -0,4213 | 6,67099E-19 |
| ACE2 | LINC00216 | -0,4213 | 6,70413E-19 |
| ACE2 | EMC7 | -0,4213 | 6,65263E-19 |
| ACE2 | UCHL5 | -0,4214 | 6,63032E-19 |
| ACE2 | RP3-467D16.3 | -0,4214 | 6,5567E-19 |
| ACE2 | C1QTNF5 | -0,4214 | 6,53773E-19 |
| ACE2 | HBA2 | -0,4214 | 6,63274E-19 |
| ACE2 | ANO8 | -0,4214 | 6,6168E-19 |
| ACE2 | PLTP | -0,4214 | 6,55268E-19 |
| ACE2 | BIRC7 | -0,4214 | 6,52973E-19 |
| ACE2 | CRHR2 | -0,4215 | 6,38047E-19 |
| ACE2 | TYW1B | -0,4215 | 6,42466E-19 |
| ACE2 | DPH6-AS1 | -0,4215 | 6,45682E-19 |
| ACE2 | ZNF592 | -0,4215 | 6,41891E-19 |
| ACE2 | RPL22L1 | -0,4216 | 6,25735E-19 |
| ACE2 | XPO5 | -0,4216 | 6,25035E-19 |

| | | | |
|---|---|---|---|
| ACE2 | LINC01985 | -0,4217 | 6,14005E-19 |
| ACE2 | USP10 | -0,4217 | 6,2326E-19 |
| ACE2 | CH507-42P11.8 | -0,4218 | 6,02496E-19 |
| ACE2 | HOXD11 | -0,4219 | 5,87107E-19 |
| ACE2 | MPHOSPH6 | -0,4219 | 5,9341E-19 |
| ACE2 | DPRX | -0,4219 | 5,88105E-19 |
| ACE2 | RP11-111K18.2 | -0,422 | 5,81529E-19 |
| ACE2 | YME1L1 | -0,422 | 5,79625E-19 |
| ACE2 | RANGAP1 | -0,4221 | 5,70424E-19 |
| ACE2 | PTPN23 | -0,4222 | 5,54706E-19 |
| ACE2 | TMEM18 | -0,4223 | 5,45824E-19 |
| ACE2 | AC013474.4 | -0,4224 | 5,37797E-19 |
| ACE2 | RP11-255G12.3 | -0,4224 | 5,31157E-19 |
| ACE2 | MTX1P1 | -0,4225 | 5,20116E-19 |
| ACE2 | RP11-526D8.11 | -0,4225 | 5,25054E-19 |
| ACE2 | RP11-74M13.4 | -0,4225 | 5,25383E-19 |
| ACE2 | DNAH17 | -0,4225 | 5,2091E-19 |
| ACE2 | EIF3K | -0,4226 | 5,14969E-19 |
| ACE2 | BMPR1B-AS1 | -0,4227 | 5,05513E-19 |
| ACE2 | ZHX1 | -0,4227 | 5,0015E-19 |
| ACE2 | C1orf27 | -0,4228 | 4,87233E-19 |
| ACE2 | PUF60 | -0,4228 | 4,89009E-19 |
| ACE2 | NPFFR2 | -0,4229 | 4,81394E-19 |
| ACE2 | DIABLO | -0,4229 | 4,7875E-19 |
| ACE2 | NMRK2 | -0,4229 | 4,8155E-19 |
| ACE2 | CTD-2126E3.1 | -0,4229 | 4,80066E-19 |
| ACE2 | AC006116.24 | -0,4229 | 4,81467E-19 |
| ACE2 | KIF9 | -0,423 | 4,72317E-19 |
| ACE2 | RP11-678G15.1 | -0,423 | 4,71524E-19 |
| ACE2 | RP4-806M20.4 | -0,423 | 4,71686E-19 |
| ACE2 | KB-1448A5.1 | -0,4231 | 4,63345E-19 |
| ACE2 | ZW10 | -0,4231 | 4,596E-19 |
| ACE2 | RP11-353N4.6 | -0,4232 | 4,5189E-19 |
| ACE2 | AC004381.8 | -0,4232 | 4,47915E-19 |
| ACE2 | NARF-IT1 | -0,4232 | 4,54987E-19 |
| ACE2 | AC009948.7 | -0,4233 | 4,44572E-19 |
| ACE2 | OTULIN | -0,4233 | 4,41583E-19 |
| ACE2 | BMPR1APS1 | -0,4233 | 4,37073E-19 |
| ACE2 | RP11-323P17.2 | -0,4233 | 4,40199E-19 |
| ACE2 | NPC1 | -0,4233 | 4,41507E-19 |
| ACE2 | CDH7 | -0,4233 | 4,37137E-19 |
| ACE2 | SLC25A6P4 | -0,4233 | 4,45033E-19 |
| ACE2 | OR51A9P | -0,4234 | 4,33367E-19 |
| ACE2 | FGF22 | -0,4234 | 4,36383E-19 |
| ACE2 | CTC-444N24.7 | -0,4234 | 4,33638E-19 |
| ACE2 | NPDC1 | -0,4235 | 4,24241E-19 |
| ACE2 | CSPG4P13 | -0,4235 | 4,27098E-19 |
| ACE2 | BANP | -0,4235 | 4,26545E-19 |
| ACE2 | ZNF737 | -0,4235 | 4,27437E-19 |
| ACE2 | BLCAP | -0,4235 | 4,24103E-19 |
| ACE2 | AC007563.5 | -0,4236 | 4,18744E-19 |
| ACE2 | KLF15 | -0,4236 | 4,17066E-19 |
| ACE2 | AGAP9 | -0,4236 | 4,17625E-19 |
| ACE2 | CTD-2012K14.5 | -0,4236 | 4,15907E-19 |

| | | | |
|---|---|---|---|
| ACE2 | CCDC114 | -0,4236 | 4,11523E-19 |
| ACE2 | LINC00342 | -0,4239 | 3,92547E-19 |
| ACE2 | LINC01088 | -0,4239 | 3,8695E-19 |
| ACE2 | RP11-121E16.1 | -0,4239 | 3,85301E-19 |
| ACE2 | AC009120.3 | -0,4239 | 3,925E-19 |
| ACE2 | SLC16A11 | -0,4239 | 3,90921E-19 |
| ACE2 | EWSR1 | -0,4239 | 3,91395E-19 |
| ACE2 | SLIT2-IT1 | -0,424 | 3,84541E-19 |
| ACE2 | CER1 | -0,4241 | 3,7614E-19 |
| ACE2 | PPP4R1-AS1 | -0,4241 | 3,75161E-19 |
| ACE2 | KCNK17 | -0,4242 | 3,66672E-19 |
| ACE2 | MESTP1 | -0,4242 | 3,62096E-19 |
| ACE2 | CCDC7 | -0,4242 | 3,61335E-19 |
| ACE2 | UBE3B | -0,4242 | 3,64677E-19 |
| ACE2 | RP13-204A15.5 | -0,4244 | 3,53185E-19 |
| ACE2 | SPATA21 | -0,4245 | 3,44568E-19 |
| ACE2 | PRDX3P4 | -0,4245 | 3,43865E-19 |
| ACE2 | RP11-457P14.5 | -0,4245 | 3,40812E-19 |
| ACE2 | GLS2 | -0,4245 | 3,40091E-19 |
| ACE2 | CCDC22 | -0,4245 | 3,43972E-19 |
| ACE2 | SLC39A3 | -0,4246 | 3,38287E-19 |
| ACE2 | RP11-125O18.1 | -0,4247 | 3,2714E-19 |
| ACE2 | TMSB15A | -0,4247 | 3,27453E-19 |
| ACE2 | VAC14 | -0,4248 | 3,18537E-19 |
| ACE2 | RAB5A | -0,4249 | 3,12273E-19 |
| ACE2 | SF3B3 | -0,4249 | 3,14261E-19 |
| ACE2 | RWDD3 | -0,425 | 3,10563E-19 |
| ACE2 | RP11-676J12.7 | -0,425 | 3,06808E-19 |
| ACE2 | CTD-3099C6.5 | -0,425 | 3,07308E-19 |
| ACE2 | AP006222.2 | -0,4252 | 2,93743E-19 |
| ACE2 | BTBD10P2 | -0,4252 | 2,95569E-19 |
| ACE2 | NPIPA7 | -0,4252 | 2,9774E-19 |
| ACE2 | S100A5 | -0,4253 | 2,87608E-19 |
| ACE2 | FAM87B | -0,4254 | 2,81271E-19 |
| ACE2 | MAS1L | -0,4254 | 2,82017E-19 |
| ACE2 | RPS27AP16 | -0,4254 | 2,81723E-19 |
| ACE2 | RP11-244H3.1 | -0,4256 | 2,71075E-19 |
| ACE2 | VLDLR | -0,4256 | 2,72142E-19 |
| ACE2 | COL18A1-AS1 | -0,4256 | 2,68776E-19 |
| ACE2 | MAPK14 | -0,4257 | 2,64508E-19 |
| ACE2 | DDX43 | -0,4257 | 2,63913E-19 |
| ACE2 | HSD17B7P2 | -0,4257 | 2,64981E-19 |
| ACE2 | HYKK | -0,4257 | 2,6704E-19 |
| ACE2 | CTB-50L17.7 | -0,4257 | 2,63405E-19 |
| ACE2 | TCEANC | -0,4258 | 2,60437E-19 |
| ACE2 | RP11-531A24.7 | -0,4259 | 2,53944E-19 |
| ACE2 | AP002954.3 | -0,4259 | 2,55287E-19 |
| ACE2 | RP11-677M14.5 | -0,4259 | 2,56533E-19 |
| ACE2 | AC104653.1 | -0,426 | 2,4699E-19 |
| ACE2 | DUX4L51 | -0,426 | 2,51729E-19 |
| ACE2 | LINC01801 | -0,426 | 2,47627E-19 |
| ACE2 | SHB | -0,4261 | 2,44378E-19 |
| ACE2 | RP11-20D14.6 | -0,4261 | 2,46331E-19 |
| ACE2 | RP11-427M20.1 | -0,4262 | 2,40542E-19 |

| | | | |
|---|---|---|---|
| ACE2 | DLX4 | -0,4262 | 2,37526E-19 |
| ACE2 | HNRNPKP3 | -0,4263 | 2,3548E-19 |
| ACE2 | RP11-676J12.8 | -0,4263 | 2,33817E-19 |
| ACE2 | FOCAD | -0,4264 | 2,31383E-19 |
| ACE2 | RP11-73M18.7 | -0,4265 | 2,22963E-19 |
| ACE2 | KRT18P5 | -0,4265 | 2,22746E-19 |
| ACE2 | TBC1D23 | -0,4266 | 2,17622E-19 |
| ACE2 | NRGN | -0,4266 | 2,18365E-19 |
| ACE2 | RP11-141M1.4 | -0,4266 | 2,19008E-19 |
| ACE2 | CCDC144B | -0,4266 | 2,18044E-19 |
| ACE2 | TEX261 | -0,4267 | 2,13169E-19 |
| ACE2 | RP13-895J2.6 | -0,4267 | 2,12791E-19 |
| ACE2 | HECTD1 | -0,4267 | 2,13941E-19 |
| ACE2 | SMN2 | -0,4268 | 2,08422E-19 |
| ACE2 | TRIAP1 | -0,4268 | 2,08838E-19 |
| ACE2 | FOXP4 | -0,427 | 2,02949E-19 |
| ACE2 | DHODH | -0,427 | 1,99444E-19 |
| ACE2 | MAGEA8 | -0,427 | 2,01285E-19 |
| ACE2 | CH17-296N19.1 | -0,4271 | 1,96968E-19 |
| ACE2 | ELP4 | -0,4272 | 1,94026E-19 |
| ACE2 | TRIM72 | -0,4272 | 1,92947E-19 |
| ACE2 | RP11-227G15.12 | -0,4272 | 1,91448E-19 |
| ACE2 | FUNDC2P1 | -0,4273 | 1,90118E-19 |
| ACE2 | EN2 | -0,4273 | 1,89786E-19 |
| ACE2 | HNRNPK | -0,4273 | 1,89764E-19 |
| ACE2 | FGFR2 | -0,4273 | 1,90924E-19 |
| ACE2 | TEAD4 | -0,4273 | 1,88448E-19 |
| ACE2 | NFAT5 | -0,4273 | 1,88873E-19 |
| ACE2 | MYL4 | -0,4273 | 1,88449E-19 |
| ACE2 | THOC2 | -0,4273 | 1,89416E-19 |
| ACE2 | KIAA1109 | -0,4274 | 1,83467E-19 |
| ACE2 | PITPNA-AS1 | -0,4274 | 1,86602E-19 |
| ACE2 | RP11-729I10.3 | -0,4275 | 1,82026E-19 |
| ACE2 | RP11-154J22.1 | -0,4275 | 1,80926E-19 |
| ACE2 | CYP51A1-AS1 | -0,4277 | 1,71892E-19 |
| ACE2 | RPRML | -0,4278 | 1,702E-19 |
| ACE2 | RSL24D1P11 | -0,4278 | 1,69111E-19 |
| ACE2 | CTD-2224J9.8 | -0,4278 | 1,68074E-19 |
| ACE2 | ARRDC2 | -0,4279 | 1,66948E-19 |
| ACE2 | RP13-228J13.8 | -0,4279 | 1,65942E-19 |
| ACE2 | SNRNP200 | -0,428 | 1,6313E-19 |
| ACE2 | C4orf22 | -0,428 | 1,61956E-19 |
| ACE2 | LINC01547 | -0,428 | 1,61154E-19 |
| ACE2 | RP13-1039J1.2 | -0,4281 | 1,58847E-19 |
| ACE2 | TEX26 | -0,4281 | 1,57637E-19 |
| ACE2 | CFAP161 | -0,4281 | 1,58741E-19 |
| ACE2 | RP11-342D14.1 | -0,4281 | 1,57989E-19 |
| ACE2 | RP11-259O18.5 | -0,4282 | 1,55103E-19 |
| ACE2 | MTND1P8 | -0,4282 | 1,56725E-19 |
| ACE2 | CH17-140K24.3 | -0,4283 | 1,53445E-19 |
| ACE2 | TM4SF1 | -0,4284 | 1,50323E-19 |
| ACE2 | LINC00638 | -0,4284 | 1,50918E-19 |
| ACE2 | RP11-729I10.2 | -0,4285 | 1,46033E-19 |
| ACE2 | ADPRHL2 | -0,4286 | 1,43076E-19 |

| | | | |
|---|---|---|---|
| ACE2 | CDCP2 | -0,4286 | 1,44498E-19 |
| ACE2 | XRCC5 | -0,4286 | 1,42277E-19 |
| ACE2 | AC073052.1 | -0,4286 | 1,42563E-19 |
| ACE2 | INTS1 | -0,4286 | 1,44108E-19 |
| ACE2 | AC004947.2 | -0,4286 | 1,42932E-19 |
| ACE2 | AC026904.1 | -0,4286 | 1,42492E-19 |
| ACE2 | RP11-977G19.12 | -0,4286 | 1,44437E-19 |
| ACE2 | TBL1XR1 | -0,4287 | 1,40266E-19 |
| ACE2 | MIR181A2HG | -0,4287 | 1,40306E-19 |
| ACE2 | HSP90AA1 | -0,4287 | 1,40517E-19 |
| ACE2 | FAM153B | -0,4288 | 1,38119E-19 |
| ACE2 | HIST3H2BB | -0,4289 | 1,35037E-19 |
| ACE2 | RP11-379F4.1 | -0,429 | 1,31949E-19 |
| ACE2 | GJA1P1 | -0,429 | 1,32467E-19 |
| ACE2 | AHSA1 | -0,429 | 1,31481E-19 |
| ACE2 | RP11-455O6.2 | -0,4291 | 1,2723E-19 |
| ACE2 | HIVEP3 | -0,4292 | 1,25572E-19 |
| ACE2 | WBP1LP2 | -0,4292 | 1,27015E-19 |
| ACE2 | SKOR1 | -0,4292 | 1,24857E-19 |
| ACE2 | TRIM56 | -0,4293 | 1,22107E-19 |
| ACE2 | SH2B2 | -0,4294 | 1,20048E-19 |
| ACE2 | RHEBL1 | -0,4294 | 1,20741E-19 |
| ACE2 | AC100830.5 | -0,4294 | 1,20304E-19 |
| ACE2 | DNAJC12 | -0,4295 | 1,1816E-19 |
| ACE2 | CTA-984G1.5 | -0,4295 | 1,17788E-19 |
| ACE2 | PPAT | -0,4296 | 1,14315E-19 |
| ACE2 | RGS17 | -0,4296 | 1,15523E-19 |
| ACE2 | RP11-344B5.2 | -0,4297 | 1,11805E-19 |
| ACE2 | BFAR | -0,4297 | 1,13588E-19 |
| ACE2 | CTD-2012K14.6 | -0,4297 | 1,13856E-19 |
| ACE2 | ZNF750 | -0,4297 | 1,12962E-19 |
| ACE2 | TTC21A | -0,4298 | 1,1118E-19 |
| ACE2 | RNU6-1110P | -0,4298 | 1,11402E-19 |
| ACE2 | LINC00028 | -0,4299 | 1,07986E-19 |
| ACE2 | BRAFP1 | -0,4299 | 1,08939E-19 |
| ACE2 | RP11-307L14.2 | -0,4301 | 1,02607E-19 |
| ACE2 | CNIH1 | -0,4301 | 1,04525E-19 |
| ACE2 | GABRD | -0,4302 | 1,01825E-19 |
| ACE2 | MOB4 | -0,4302 | 1,00491E-19 |
| ACE2 | ADAMTS4 | -0,4303 | 9,90009E-20 |
| ACE2 | POTEKP | -0,4303 | 1,00302E-19 |
| ACE2 | PUS7 | -0,4304 | 9,7565E-20 |
| ACE2 | CTD-3001H11.2 | -0,4304 | 9,67762E-20 |
| ACE2 | NODAL | -0,4305 | 9,42463E-20 |
| ACE2 | OIP5-AS1 | -0,4306 | 9,34613E-20 |
| ACE2 | RP11-158I9.8 | -0,4307 | 9,15639E-20 |
| ACE2 | COX8C | -0,4307 | 9,10746E-20 |
| ACE2 | RP4-813D12.3 | -0,4308 | 8,83641E-20 |
| ACE2 | RP11-431K24.1 | -0,4309 | 8,69815E-20 |
| ACE2 | HOXB-AS2 | -0,4309 | 8,72318E-20 |
| ACE2 | WISP2 | -0,4309 | 8,72952E-20 |
| ACE2 | KCNK12 | -0,431 | 8,51018E-20 |
| ACE2 | CTD-2553C6.1 | -0,431 | 8,57975E-20 |
| ACE2 | SIAH1 | -0,4312 | 8,12953E-20 |

| | | | |
|---|---|---|---|
| ACE2 | C17orf49 | -0,4312 | 8,19946E-20 |
| ACE2 | RP4-555D20.2 | -0,4313 | 8,06714E-20 |
| ACE2 | RP11-415D17.3 | -0,4313 | 8,02098E-20 |
| ACE2 | RP11-208N14.4 | -0,4314 | 7,88043E-20 |
| ACE2 | NAPG | -0,4314 | 7,7893E-20 |
| ACE2 | PDCL3 | -0,4315 | 7,66225E-20 |
| ACE2 | UBE2E3 | -0,4315 | 7,62558E-20 |
| ACE2 | ANXA7 | -0,4315 | 7,64713E-20 |
| ACE2 | RP11-295I5.4 | -0,4315 | 7,70906E-20 |
| ACE2 | ABCD4 | -0,4315 | 7,66129E-20 |
| ACE2 | C15orf40 | -0,4315 | 7,69142E-20 |
| ACE2 | CRLF1 | -0,4315 | 7,68068E-20 |
| ACE2 | VRK2 | -0,4316 | 7,48289E-20 |
| ACE2 | HRH2 | -0,4316 | 7,53039E-20 |
| ACE2 | CTD-3224I3.3 | -0,4316 | 7,47737E-20 |
| ACE2 | GAL3ST3 | -0,4316 | 7,45858E-20 |
| ACE2 | SP3 | -0,4317 | 7,28178E-20 |
| ACE2 | OR7E7P | -0,4317 | 7,33717E-20 |
| ACE2 | RP11-536C5.7 | -0,4318 | 7,24208E-20 |
| ACE2 | CTB-113P19.1 | -0,4318 | 7,18224E-20 |
| ACE2 | SCUBE2 | -0,4318 | 7,18483E-20 |
| ACE2 | RP11-196B3.4 | -0,4318 | 7,24901E-20 |
| ACE2 | MAP1S | -0,4318 | 7,18134E-20 |
| ACE2 | C22orf34 | -0,4319 | 6,9776E-20 |
| ACE2 | RP11-317B7.2 | -0,432 | 6,88785E-20 |
| ACE2 | C8orf86 | -0,432 | 6,84329E-20 |
| ACE2 | RP1-41C23.4 | -0,432 | 6,88637E-20 |
| ACE2 | ZNF761 | -0,432 | 6,90155E-20 |
| ACE2 | IQCJ-SCHIP1-AS1 | -0,4321 | 6,69182E-20 |
| ACE2 | TLR4 | -0,4321 | 6,72653E-20 |
| ACE2 | LINC00640 | -0,4321 | 6,67909E-20 |
| ACE2 | RP11-92G12.3 | -0,4322 | 6,59159E-20 |
| ACE2 | AC000036.4 | -0,4322 | 6,52012E-20 |
| ACE2 | HDAC6 | -0,4324 | 6,34288E-20 |
| ACE2 | RP11-107E5.3 | -0,4325 | 6,20335E-20 |
| ACE2 | GABRR2 | -0,4325 | 6,15413E-20 |
| ACE2 | RRM2B | -0,4325 | 6,19202E-20 |
| ACE2 | SMG1P3 | -0,4325 | 6,16233E-20 |
| ACE2 | SMARCAD1 | -0,4326 | 6,03139E-20 |
| ACE2 | TNFRSF1B | -0,4327 | 5,94092E-20 |
| ACE2 | MARCH4 | -0,4327 | 5,86489E-20 |
| ACE2 | ZNF12 | -0,4327 | 5,84301E-20 |
| ACE2 | CCDC126 | -0,4327 | 5,88987E-20 |
| ACE2 | SNTG1 | -0,4327 | 5,93584E-20 |
| ACE2 | HNRNPA1P33 | -0,4327 | 5,86547E-20 |
| ACE2 | GPN1 | -0,4328 | 5,76874E-20 |
| ACE2 | XX-C00717C00720L.1 | -0,4328 | 5,79438E-20 |
| ACE2 | MPP1 | -0,4328 | 5,75707E-20 |
| ACE2 | RP11-133F8.2 | -0,4329 | 5,61326E-20 |
| ACE2 | CXorf56 | -0,4329 | 5,6766E-20 |
| ACE2 | ADGRG4 | -0,4329 | 5,59673E-20 |
| ACE2 | RP11-343N15.2 | -0,433 | 5,56793E-20 |
| ACE2 | NDNF | -0,433 | 5,58991E-20 |
| ACE2 | NOMO1 | -0,433 | 5,5542E-20 |

| | | | |
|---|---|---|---|
| ACE2 | WDR36 | -0,4331 | 5,45416E-20 |
| ACE2 | AF131216.5 | -0,4331 | 5,37218E-20 |
| ACE2 | G2E3 | -0,4331 | 5,40858E-20 |
| ACE2 | SEL1L | -0,4331 | 5,43294E-20 |
| ACE2 | RP3-340N1.2 | -0,4332 | 5,29623E-20 |
| ACE2 | ARHGAP15 | -0,4332 | 5,26305E-20 |
| ACE2 | FAM181A-AS1 | -0,4333 | 5,18885E-20 |
| ACE2 | NOD2 | -0,4333 | 5,13054E-20 |
| ACE2 | ITCH | -0,4334 | 5,03154E-20 |
| ACE2 | SSXP10 | -0,4335 | 4,93536E-20 |
| ACE2 | RP11-370I10.6 | -0,4336 | 4,85671E-20 |
| ACE2 | MBTD1 | -0,4336 | 4,84017E-20 |
| ACE2 | SLU7 | -0,4338 | 4,63542E-20 |
| ACE2 | RP1-283K11.3 | -0,4338 | 4,69459E-20 |
| ACE2 | KDM5A | -0,4338 | 4,68055E-20 |
| ACE2 | RPS7P1 | -0,4338 | 4,61495E-20 |
| ACE2 | HBE1 | -0,4339 | 4,57052E-20 |
| ACE2 | RP11-755H23.1 | -0,434 | 4,49745E-20 |
| ACE2 | PKHD1L1 | -0,434 | 4,42716E-20 |
| ACE2 | ANKRD18B | -0,434 | 4,4089E-20 |
| ACE2 | RP11-130L8.3 | -0,4341 | 4,37196E-20 |
| ACE2 | RN7SL868P | -0,4341 | 4,39558E-20 |
| ACE2 | TMEM5-AS1 | -0,4341 | 4,37352E-20 |
| ACE2 | LINC00211 | -0,4342 | 4,25264E-20 |
| ACE2 | MAGI1 | -0,4342 | 4,23698E-20 |
| ACE2 | RP4-777O23.1 | -0,4342 | 4,27986E-20 |
| ACE2 | DISC1FP1 | -0,4342 | 4,27684E-20 |
| ACE2 | HCCAT5 | -0,4343 | 4,18178E-20 |
| ACE2 | RGPD3 | -0,4344 | 4,10907E-20 |
| ACE2 | PVRL3-AS1 | -0,4344 | 4,07492E-20 |
| ACE2 | RPRD1B | -0,4344 | 4,11548E-20 |
| ACE2 | ZNF141 | -0,4345 | 3,94757E-20 |
| ACE2 | SSRP1 | -0,4345 | 3,95352E-20 |
| ACE2 | DAZL | -0,4346 | 3,89252E-20 |
| ACE2 | EIF2B5 | -0,4346 | 3,87738E-20 |
| ACE2 | RP11-483L5.1 | -0,4346 | 3,93193E-20 |
| ACE2 | C1orf174 | -0,4347 | 3,8083E-20 |
| ACE2 | ZNF93 | -0,4347 | 3,81116E-20 |
| ACE2 | CALM3 | -0,4348 | 3,71747E-20 |
| ACE2 | XXbac-B444P24.8 | -0,4348 | 3,72011E-20 |
| ACE2 | CTC-455F18.1 | -0,4349 | 3,63257E-20 |
| ACE2 | RP3-402G11.26 | -0,4349 | 3,68259E-20 |
| ACE2 | TOLLIP-AS1 | -0,435 | 3,56285E-20 |
| ACE2 | SH3GL1P1 | -0,435 | 3,56218E-20 |
| ACE2 | MOSPD2 | -0,435 | 3,61271E-20 |
| ACE2 | SREK1 | -0,4352 | 3,45902E-20 |
| ACE2 | MPV17L2 | -0,4352 | 3,44012E-20 |
| ACE2 | CTC-487M23.7 | -0,4353 | 3,37811E-20 |
| ACE2 | RP11-837J7.4 | -0,4353 | 3,33035E-20 |
| ACE2 | RP11-989F5.1 | -0,4353 | 3,36963E-20 |
| ACE2 | CMA1 | -0,4353 | 3,3201E-20 |
| ACE2 | NLE1 | -0,4353 | 3,33909E-20 |
| ACE2 | RP11-190A12.8 | -0,4354 | 3,30391E-20 |
| ACE2 | RP5-981O7.2 | -0,4354 | 3,26741E-20 |

| | | | |
|---|---|---|---|
| ACE2 | ALKBH3-AS1 | -0,4354 | 3,29933E-20 |
| ACE2 | AC005523.3 | -0,4354 | 3,2736E-20 |
| ACE2 | SNTB1 | -0,4355 | 3,23938E-20 |
| ACE2 | TTC12 | -0,4355 | 3,18E-20 |
| ACE2 | TNFSF4 | -0,4356 | 3,131E-20 |
| ACE2 | OR1L8 | -0,4356 | 3,15225E-20 |
| ACE2 | MRI1 | -0,4356 | 3,10869E-20 |
| ACE2 | A1BG-AS1 | -0,4356 | 3,15258E-20 |
| ACE2 | SNX12 | -0,4356 | 3,11562E-20 |
| ACE2 | RP11-367E12.4 | -0,4357 | 3,09777E-20 |
| ACE2 | RP11-390K5.1 | -0,4357 | 3,03765E-20 |
| ACE2 | GMNC | -0,4358 | 2,9814E-20 |
| ACE2 | RFC4 | -0,4359 | 2,92139E-20 |
| ACE2 | CBY3 | -0,4359 | 2,91007E-20 |
| ACE2 | RP11-230C9.4 | -0,4359 | 2,96562E-20 |
| ACE2 | DPY19L1P2 | -0,4359 | 2,96612E-20 |
| ACE2 | RP11-382J24.2 | -0,4359 | 2,93415E-20 |
| ACE2 | RP11-147L13.2 | -0,4359 | 2,94293E-20 |
| ACE2 | ABCA9-AS1 | -0,4359 | 2,94143E-20 |
| ACE2 | RP11-666A8.8 | -0,4359 | 2,93146E-20 |
| ACE2 | AC007787.3 | -0,4359 | 2,96323E-20 |
| ACE2 | EDNRB | -0,436 | 2,89136E-20 |
| ACE2 | UPK3BL | -0,4361 | 2,80904E-20 |
| ACE2 | ORAI2 | -0,4362 | 2,72451E-20 |
| ACE2 | GLULP4 | -0,4362 | 2,74513E-20 |
| ACE2 | ETS1 | -0,4362 | 2,73938E-20 |
| ACE2 | LINC01460 | -0,4363 | 2,68592E-20 |
| ACE2 | RP11-4K3__A.3 | -0,4364 | 2,64508E-20 |
| ACE2 | NR4A2 | -0,4364 | 2,63098E-20 |
| ACE2 | PDGFA | -0,4364 | 2,60526E-20 |
| ACE2 | ADAT1 | -0,4364 | 2,63945E-20 |
| ACE2 | DNAJB9 | -0,4365 | 2,55359E-20 |
| ACE2 | OR7E94P | -0,4366 | 2,52427E-20 |
| ACE2 | ADGRG3 | -0,4366 | 2,5236E-20 |
| ACE2 | SPIN4-AS1 | -0,4367 | 2,44047E-20 |
| ACE2 | ZBTB34 | -0,4368 | 2,40104E-20 |
| ACE2 | SQSTM1 | -0,4369 | 2,34375E-20 |
| ACE2 | TEX29 | -0,4369 | 2,35647E-20 |
| ACE2 | ZNF663P | -0,4369 | 2,34484E-20 |
| ACE2 | AP001604.3 | -0,4369 | 2,34433E-20 |
| ACE2 | EI24 | -0,437 | 2,28421E-20 |
| ACE2 | A1BG | -0,4371 | 2,234E-20 |
| ACE2 | ELOBP3 | -0,4372 | 2,18969E-20 |
| ACE2 | CES1P1 | -0,4372 | 2,20087E-20 |
| ACE2 | JUN | -0,4373 | 2,16567E-20 |
| ACE2 | CTD-2308L22.1 | -0,4374 | 2,13168E-20 |
| ACE2 | FBXO22 | -0,4374 | 2,11303E-20 |
| ACE2 | CCDC167 | -0,4375 | 2,07164E-20 |
| ACE2 | RP11-192P3.4 | -0,4375 | 2,0469E-20 |
| ACE2 | MAGED4B | -0,4375 | 2,04963E-20 |
| ACE2 | CTD-2263F21.1 | -0,4376 | 2,03566E-20 |
| ACE2 | P4HA1 | -0,4376 | 2,02323E-20 |
| ACE2 | PDE6C | -0,4377 | 1,96809E-20 |
| ACE2 | MESTIT1 | -0,4378 | 1,94018E-20 |

| | | | |
|---|---|---|---|
| ACE2 | GLA | -0,4378 | 1,9164E-20 |
| ACE2 | NBPF26 | -0,438 | 1,85613E-20 |
| ACE2 | RP11-762B21.5 | -0,438 | 1,8538E-20 |
| ACE2 | RP11-701H24.5 | -0,438 | 1,85976E-20 |
| ACE2 | BTBD1 | -0,438 | 1,86211E-20 |
| ACE2 | IL21R-AS1 | -0,438 | 1,8678E-20 |
| ACE2 | RP11-838N2.5 | -0,438 | 1,85766E-20 |
| ACE2 | RP11-327J17.8 | -0,4381 | 1,80002E-20 |
| ACE2 | AP000254.8 | -0,4381 | 1,80637E-20 |
| ACE2 | GADL1 | -0,4382 | 1,75808E-20 |
| ACE2 | CYP26B1 | -0,4383 | 1,72383E-20 |
| ACE2 | RP11-247A12.2 | -0,4383 | 1,72598E-20 |
| ACE2 | RP11-463O9.1 | -0,4383 | 1,71489E-20 |
| ACE2 | CTD-3193K9.11 | -0,4383 | 1,7329E-20 |
| ACE2 | PPM1J | -0,4384 | 1,6822E-20 |
| ACE2 | MTND5P28 | -0,4384 | 1,70635E-20 |
| ACE2 | CTD-2525I3.3 | -0,4384 | 1,70319E-20 |
| ACE2 | RP11-523O18.5 | -0,4385 | 1,64518E-20 |
| ACE2 | C19orf81 | -0,4386 | 1,61168E-20 |
| ACE2 | SEC14L4 | -0,4386 | 1,6145E-20 |
| ACE2 | UBR5-AS1 | -0,4387 | 1,59896E-20 |
| ACE2 | FBXO31 | -0,4387 | 1,56646E-20 |
| ACE2 | HSD3BP5 | -0,4388 | 1,56264E-20 |
| ACE2 | AMT | -0,4388 | 1,55992E-20 |
| ACE2 | ZMIZ2 | -0,4388 | 1,53432E-20 |
| ACE2 | TUT1 | -0,4388 | 1,53346E-20 |
| ACE2 | PAH | -0,4388 | 1,56381E-20 |
| ACE2 | AC010883.5 | -0,4389 | 1,50666E-20 |
| ACE2 | RP4-706A16.3 | -0,439 | 1,48901E-20 |
| ACE2 | RP11-665G4.1 | -0,439 | 1,47427E-20 |
| ACE2 | RP1-27K12.4 | -0,439 | 1,47596E-20 |
| ACE2 | RP11-244N9.4 | -0,439 | 1,47319E-20 |
| ACE2 | RP11-65J3.3 | -0,439 | 1,48335E-20 |
| ACE2 | GPATCH4 | -0,4391 | 1,44107E-20 |
| ACE2 | FAM27D1 | -0,4391 | 1,45859E-20 |
| ACE2 | POU3F1 | -0,4392 | 1,40635E-20 |
| ACE2 | PDE4B | -0,4392 | 1,42853E-20 |
| ACE2 | GCSAML-AS1 | -0,4392 | 1,42216E-20 |
| ACE2 | ASAP1-IT2 | -0,4392 | 1,41997E-20 |
| ACE2 | UBE2D1 | -0,4393 | 1,38647E-20 |
| ACE2 | RP11-973H7.3 | -0,4393 | 1,38269E-20 |
| ACE2 | NDUFA5 | -0,4395 | 1,32846E-20 |
| ACE2 | ROGDI | -0,4395 | 1,32008E-20 |
| ACE2 | ZNF571-AS1 | -0,4395 | 1,3103E-20 |
| ACE2 | SPATA41 | -0,4396 | 1,2914E-20 |
| ACE2 | PPM1D | -0,4396 | 1,29232E-20 |
| ACE2 | ODF3B | -0,4396 | 1,30539E-20 |
| ACE2 | GTF2I | -0,4397 | 1,28144E-20 |
| ACE2 | CTD-2576N18.1 | -0,4397 | 1,2789E-20 |
| ACE2 | TREX2 | -0,4397 | 1,26957E-20 |
| ACE2 | RP11-384F7.2 | -0,4398 | 1,25207E-20 |
| ACE2 | SLTM | -0,4398 | 1,23791E-20 |
| ACE2 | CTD-3193K9.4 | -0,4398 | 1,25309E-20 |
| ACE2 | RP11-204M4.3 | -0,4399 | 1,22399E-20 |

| | | | |
|---|---|---|---|
| ACE2 | BLOC1S2 | -0,4399 | 1,21044E-20 |
| ACE2 | RP11-582E3.4 | -0,4399 | 1,21033E-20 |
| ACE2 | SPIN4 | -0,4399 | 1,21085E-20 |
| ACE2 | AC008269.2 | -0,44 | 1,17887E-20 |
| ACE2 | NPY | -0,44 | 1,19278E-20 |
| ACE2 | PSMD9 | -0,4401 | 1,15416E-20 |
| ACE2 | ABCB6 | -0,4402 | 1,13218E-20 |
| ACE2 | NDUFAF3 | -0,4402 | 1,12297E-20 |
| ACE2 | HLX-AS1 | -0,4403 | 1,09781E-20 |
| ACE2 | SCOC-AS1 | -0,4403 | 1,1012E-20 |
| ACE2 | ZNF346-IT1 | -0,4403 | 1,12046E-20 |
| ACE2 | ACSBG2 | -0,4403 | 1,11461E-20 |
| ACE2 | LRRC71 | -0,4404 | 1,07913E-20 |
| ACE2 | PVT1 | -0,4404 | 1,08269E-20 |
| ACE2 | RPL7P1 | -0,4405 | 1,05513E-20 |
| ACE2 | LRTM2 | -0,4405 | 1,05071E-20 |
| ACE2 | AC018816.3 | -0,4406 | 1,03549E-20 |
| ACE2 | MSNP1 | -0,4406 | 1,03105E-20 |
| ACE2 | NOP56 | -0,4406 | 1,041E-20 |
| ACE2 | RP13-33H18.1 | -0,4406 | 1,03299E-20 |
| ACE2 | RP5-916L7.2 | -0,4407 | 1,02249E-20 |
| ACE2 | EIF2B2 | -0,4407 | 1,01375E-20 |
| ACE2 | GOSR1 | -0,4407 | 1,01925E-20 |
| ACE2 | RP11-157L3.6 | -0,4408 | 9,92891E-21 |
| ACE2 | CTC-548K16.6 | -0,4408 | 9,86779E-21 |
| ACE2 | CPVL | -0,4409 | 9,64405E-21 |
| ACE2 | TMEM248 | -0,4409 | 9,79544E-21 |
| ACE2 | KCNIP2-AS1 | -0,441 | 9,52489E-21 |
| ACE2 | CTD-2378E12.1 | -0,441 | 9,45281E-21 |
| ACE2 | HNRNPCP2 | -0,4411 | 9,31891E-21 |
| ACE2 | RP11-38M8.1 | -0,4412 | 9,16696E-21 |
| ACE2 | VPS29 | -0,4412 | 8,97953E-21 |
| ACE2 | INTS4P2 | -0,4414 | 8,64453E-21 |
| ACE2 | TGS1 | -0,4414 | 8,64566E-21 |
| ACE2 | GALT | -0,4414 | 8,64133E-21 |
| ACE2 | SPOCK2 | -0,4414 | 8,75034E-21 |
| ACE2 | RP11-106M7.1 | -0,4414 | 8,57738E-21 |
| ACE2 | PSMD6-AS1 | -0,4415 | 8,45175E-21 |
| ACE2 | DHFR2 | -0,4415 | 8,41455E-21 |
| ACE2 | CLNS1A | -0,4415 | 8,53921E-21 |
| ACE2 | C1orf112 | -0,4416 | 8,21089E-21 |
| ACE2 | RP11-620J15.2 | -0,4416 | 8,3814E-21 |
| ACE2 | CITED2 | -0,4417 | 8,10769E-21 |
| ACE2 | MYOZ1 | -0,4417 | 8,05587E-21 |
| ACE2 | KRT18P4 | -0,4418 | 7,95585E-21 |
| ACE2 | PTPRVP | -0,4419 | 7,7332E-21 |
| ACE2 | RP11-902B17.1 | -0,442 | 7,56746E-21 |
| ACE2 | SLC7A6OS | -0,442 | 7,54778E-21 |
| ACE2 | MTCO1P28 | -0,4421 | 7,48819E-21 |
| ACE2 | AC006994.2 | -0,4422 | 7,25163E-21 |
| ACE2 | HECA | -0,4422 | 7,27808E-21 |
| ACE2 | GIMAP5 | -0,4422 | 7,3195E-21 |
| ACE2 | AP000654.5 | -0,4422 | 7,21815E-21 |
| ACE2 | RP11-385G16.1 | -0,4422 | 7,2608E-21 |

| | | | |
|---|---|---|---|
| ACE2 | POTEI | -0,4423 | 7,04299E-21 |
| ACE2 | AP000442.4 | -0,4423 | 7,017E-21 |
| ACE2 | PPP2CA | -0,4424 | 6,85721E-21 |
| ACE2 | CTD-2339F6.1 | -0,4424 | 6,87902E-21 |
| ACE2 | RP5-837J1.4 | -0,4424 | 6,94597E-21 |
| ACE2 | ROR1-AS1 | -0,4425 | 6,71878E-21 |
| ACE2 | RP11-759A24.2 | -0,4425 | 6,83117E-21 |
| ACE2 | DKFZp779M0652 | -0,4427 | 6,44827E-21 |
| ACE2 | RP11-178H8.7 | -0,4427 | 6,52867E-21 |
| ACE2 | RP11-278C7.1 | -0,4427 | 6,46614E-21 |
| ACE2 | COQ6 | -0,4427 | 6,54085E-21 |
| ACE2 | TUBGCP4 | -0,4427 | 6,44852E-21 |
| ACE2 | CTD-2536I1.3 | -0,4427 | 6,5088E-21 |
| ACE2 | OR1E1 | -0,4427 | 6,49426E-21 |
| ACE2 | TTLL9 | -0,4427 | 6,47693E-21 |
| ACE2 | PKN2-AS1 | -0,4428 | 6,32637E-21 |
| ACE2 | SKIV2L | -0,4428 | 6,33722E-21 |
| ACE2 | BBS12 | -0,4429 | 6,19561E-21 |
| ACE2 | XRCC6 | -0,4429 | 6,1825E-21 |
| ACE2 | ADGRG2 | -0,4429 | 6,18044E-21 |
| ACE2 | RP4-535B20.1 | -0,443 | 6,08533E-21 |
| ACE2 | RSPO4 | -0,443 | 6,07358E-21 |
| ACE2 | C16orf58 | -0,4431 | 5,94182E-21 |
| ACE2 | ENPEP | -0,4432 | 5,73034E-21 |
| ACE2 | RP11-973H7.1 | -0,4432 | 5,79386E-21 |
| ACE2 | RP11-51J9.5 | -0,4433 | 5,69781E-21 |
| ACE2 | ZBTB11-AS1 | -0,4435 | 5,43102E-21 |
| ACE2 | RP11-358B23.7 | -0,4435 | 5,44654E-21 |
| ACE2 | PCF11 | -0,4436 | 5,27949E-21 |
| ACE2 | DPH6 | -0,4437 | 5,19114E-21 |
| ACE2 | PPL | -0,4437 | 5,1247E-21 |
| ACE2 | JAK1 | -0,4438 | 5,03488E-21 |
| ACE2 | RP11-438L19.1 | -0,4438 | 5,07462E-21 |
| ACE2 | NCK1 | -0,4438 | 5,05104E-21 |
| ACE2 | RP11-722E23.2 | -0,4438 | 5,04526E-21 |
| ACE2 | WDR74 | -0,4438 | 5,07821E-21 |
| ACE2 | THAP10 | -0,4438 | 5,00849E-21 |
| ACE2 | CTD-2311M21.2 | -0,4438 | 5,01197E-21 |
| ACE2 | AMPD2 | -0,4439 | 4,91925E-21 |
| ACE2 | CAPRIN1 | -0,4439 | 4,97009E-21 |
| ACE2 | RP11-154D6.2 | -0,444 | 4,8046E-21 |
| ACE2 | RP11-104L21.3 | -0,4441 | 4,74893E-21 |
| ACE2 | TACC2 | -0,4442 | 4,63064E-21 |
| ACE2 | RP11-932O9.7 | -0,4442 | 4,61192E-21 |
| ACE2 | RP11-1090M7.1 | -0,4442 | 4,67308E-21 |
| ACE2 | RP11-564C24.1 | -0,4444 | 4,43325E-21 |
| ACE2 | RP13-895J2.2 | -0,4444 | 4,40091E-21 |
| ACE2 | CTD-2311M21.3 | -0,4444 | 4,39688E-21 |
| ACE2 | TPGS1 | -0,4444 | 4,37605E-21 |
| ACE2 | PUM1 | -0,4445 | 4,33313E-21 |
| ACE2 | TTC41P | -0,4445 | 4,30862E-21 |
| ACE2 | FAM167B | -0,4447 | 4,113E-21 |
| ACE2 | HACD3 | -0,4447 | 4,09338E-21 |
| ACE2 | SNHG1 | -0,4448 | 4,05061E-21 |

| | | | |
|---|---|---|---|
| ACE2 | CTC-435M10.12 | -0,4448 | 4,08057E-21 |
| ACE2 | STX12 | -0,4449 | 3,95171E-21 |
| ACE2 | PBX2P1 | -0,4449 | 3,94325E-21 |
| ACE2 | BEND3P3 | -0,4449 | 3,94879E-21 |
| ACE2 | ANKRD20A18P | -0,4449 | 3,96723E-21 |
| ACE2 | MANEA | -0,445 | 3,87463E-21 |
| ACE2 | SNHG22 | -0,445 | 3,85211E-21 |
| ACE2 | CENPQ | -0,4451 | 3,76986E-21 |
| ACE2 | NEMF | -0,4451 | 3,75977E-21 |
| ACE2 | GLDN | -0,4451 | 3,81037E-21 |
| ACE2 | IGDCC4 | -0,4451 | 3,81117E-21 |
| ACE2 | RPL9P29 | -0,4451 | 3,7447E-21 |
| ACE2 | UBA6 | -0,4452 | 3,68322E-21 |
| ACE2 | RP11-1007O24.2 | -0,4452 | 3,6719E-21 |
| ACE2 | ZNF812P | -0,4452 | 3,64612E-21 |
| ACE2 | NOTCH2NL | -0,4453 | 3,61749E-21 |
| ACE2 | RP11-141M1.3 | -0,4453 | 3,6029E-21 |
| ACE2 | VPS13D | -0,4455 | 3,41843E-21 |
| ACE2 | NSG1 | -0,4455 | 3,41934E-21 |
| ACE2 | UQCC2 | -0,4455 | 3,44483E-21 |
| ACE2 | SLK | -0,4455 | 3,45225E-21 |
| ACE2 | RP11-71L14.4 | -0,4455 | 3,40815E-21 |
| ACE2 | RP11-284F21.11 | -0,4456 | 3,38197E-21 |
| ACE2 | RP11-644N4.1 | -0,4456 | 3,40114E-21 |
| ACE2 | WI2-85898F10.2 | -0,4456 | 3,35901E-21 |
| ACE2 | RNF14 | -0,4457 | 3,28482E-21 |
| ACE2 | KMT2C | -0,4457 | 3,29359E-21 |
| ACE2 | MKS1 | -0,4457 | 3,31467E-21 |
| ACE2 | FLJ38576 | -0,4458 | 3,23851E-21 |
| ACE2 | RP4-761J14.10 | -0,4458 | 3,21904E-21 |
| ACE2 | MAPRE3 | -0,4459 | 3,17373E-21 |
| ACE2 | ESPNL | -0,4459 | 3,13118E-21 |
| ACE2 | RP3-495K2.2 | -0,4459 | 3,15497E-21 |
| ACE2 | RP11-76C10.3 | -0,4459 | 3,12088E-21 |
| ACE2 | RP11-188P17.2 | -0,446 | 3,09133E-21 |
| ACE2 | XXbac-BPGBPG55C2( | -0,446 | 3,04722E-21 |
| ACE2 | RP11-410L14.1 | -0,4461 | 2,97911E-21 |
| ACE2 | CBX3P4 | -0,4461 | 3,00557E-21 |
| ACE2 | RFXANK | -0,4461 | 2,99096E-21 |
| ACE2 | RP11-1023L17.1 | -0,4463 | 2,87232E-21 |
| ACE2 | CTD-2206N4.4 | -0,4463 | 2,84632E-21 |
| ACE2 | ELMOD3 | -0,4464 | 2,79447E-21 |
| ACE2 | DSCC1 | -0,4464 | 2,81003E-21 |
| ACE2 | MAPK15 | -0,4464 | 2,81918E-21 |
| ACE2 | RP11-298I3.6 | -0,4464 | 2,78263E-21 |
| ACE2 | PIR | -0,4464 | 2,78626E-21 |
| ACE2 | GIMAP1 | -0,4465 | 2,75175E-21 |
| ACE2 | SPATA22 | -0,4465 | 2,71637E-21 |
| ACE2 | RP11-256I23.3 | -0,4466 | 2,68872E-21 |
| ACE2 | CTD-2245E15.3 | -0,4466 | 2,69784E-21 |
| ACE2 | ZXDA | -0,4466 | 2,65946E-21 |
| ACE2 | ZBTB12 | -0,4467 | 2,61968E-21 |
| ACE2 | TMEM117 | -0,4467 | 2,64564E-21 |
| ACE2 | STARD13-IT1 | -0,4467 | 2,64999E-21 |

| | | | |
|---|---|---|---|
| ACE2 | CYB5D2 | -0,4467 | 2,60814E-21 |
| ACE2 | DR1 | -0,4468 | 2,55401E-21 |
| ACE2 | LINC01336 | -0,4469 | 2,50177E-21 |
| ACE2 | WAPL | -0,4469 | 2,51761E-21 |
| ACE2 | WIF1 | -0,4469 | 2,50434E-21 |
| ACE2 | CDCA4P4 | -0,447 | 2,43873E-21 |
| ACE2 | SPRTN | -0,447 | 2,44449E-21 |
| ACE2 | TMEM185B | -0,4471 | 2,38111E-21 |
| ACE2 | RP5-1172N10.2 | -0,4471 | 2,42086E-21 |
| ACE2 | YARS | -0,4472 | 2,36E-21 |
| ACE2 | CFH | -0,4472 | 2,33571E-21 |
| ACE2 | KIAA0141 | -0,4472 | 2,32529E-21 |
| ACE2 | PSMD13 | -0,4472 | 2,34339E-21 |
| ACE2 | MAP2K1 | -0,4472 | 2,34413E-21 |
| ACE2 | HMGB1P10 | -0,4472 | 2,35285E-21 |
| ACE2 | MPIG6B | -0,4473 | 2,29395E-21 |
| ACE2 | LINC01844 | -0,4474 | 2,25504E-21 |
| ACE2 | PNLDC1 | -0,4474 | 2,24697E-21 |
| ACE2 | RP11-91P17.1 | -0,4474 | 2,21802E-21 |
| ACE2 | MAP3K8 | -0,4474 | 2,2348E-21 |
| ACE2 | OR5BT1P | -0,4474 | 2,24769E-21 |
| ACE2 | EEA1 | -0,4474 | 2,25514E-21 |
| ACE2 | ZBTB14 | -0,4475 | 2,19144E-21 |
| ACE2 | RPL32P3 | -0,4476 | 2,12153E-21 |
| ACE2 | CTD-2651B20.3 | -0,4476 | 2,15784E-21 |
| ACE2 | OXT | -0,4476 | 2,13839E-21 |
| ACE2 | CHRNA4 | -0,4476 | 2,14306E-21 |
| ACE2 | RP11-108M9.5 | -0,4477 | 2,0691E-21 |
| ACE2 | B3GNTL1P1 | -0,4477 | 2,07053E-21 |
| ACE2 | P3H2 | -0,4478 | 2,02017E-21 |
| ACE2 | SLC6A2 | -0,4478 | 2,06118E-21 |
| ACE2 | RP11-646I19.1 | -0,448 | 1,92954E-21 |
| ACE2 | RP11-894P9.2 | -0,448 | 1,96743E-21 |
| ACE2 | RP5-933K21.3 | -0,4481 | 1,88747E-21 |
| ACE2 | SH3GL2 | -0,4481 | 1,8847E-21 |
| ACE2 | ZC3H15 | -0,4482 | 1,85761E-21 |
| ACE2 | ARHGEF7 | -0,4482 | 1,87719E-21 |
| ACE2 | CTSG | -0,4482 | 1,87846E-21 |
| ACE2 | RP11-477I4.4 | -0,4482 | 1,84716E-21 |
| ACE2 | OGFOD1P1 | -0,4483 | 1,83759E-21 |
| ACE2 | CLK3 | -0,4483 | 1,80232E-21 |
| ACE2 | NNMT | -0,4484 | 1,76334E-21 |
| ACE2 | RP4-539M6.21 | -0,4485 | 1,74316E-21 |
| ACE2 | RPS20P14 | -0,4486 | 1,68407E-21 |
| ACE2 | ARMC12 | -0,4486 | 1,71204E-21 |
| ACE2 | AC006466.5 | -0,4486 | 1,71753E-21 |
| ACE2 | CLDN10 | -0,4487 | 1,65105E-21 |
| ACE2 | LENG1 | -0,4487 | 1,64872E-21 |
| ACE2 | ZBTB45 | -0,4487 | 1,6643E-21 |
| ACE2 | GMEB2 | -0,4487 | 1,67433E-21 |
| ACE2 | CCT6A | -0,4488 | 1,63854E-21 |
| ACE2 | PKDCC | -0,4489 | 1,59168E-21 |
| ACE2 | RP4-555L14.4 | -0,4489 | 1,585E-21 |
| ACE2 | AC145124.2 | -0,4489 | 1,59243E-21 |

| | | | |
|---|---|---|---|
| ACE2 | LINC01239 | -0,4489 | 1,58607E-21 |
| ACE2 | BANF1P1 | -0,4489 | 1,5943E-21 |
| ACE2 | RP11-159D12.6 | -0,4489 | 1,56873E-21 |
| ACE2 | TRIM38 | -0,449 | 1,55793E-21 |
| ACE2 | POM121 | -0,449 | 1,55601E-21 |
| ACE2 | ZNF704 | -0,449 | 1,54165E-21 |
| ACE2 | STX4 | -0,449 | 1,56732E-21 |
| ACE2 | MPP4 | -0,4491 | 1,52282E-21 |
| ACE2 | FKTN | -0,4491 | 1,51796E-21 |
| ACE2 | RP11-631N16.4 | -0,4492 | 1,47182E-21 |
| ACE2 | GPR107 | -0,4493 | 1,44952E-21 |
| ACE2 | RP11-371A19.2 | -0,4493 | 1,46063E-21 |
| ACE2 | ANAPC15 | -0,4493 | 1,44382E-21 |
| ACE2 | ZBED3 | -0,4494 | 1,40153E-21 |
| ACE2 | NVL | -0,4495 | 1,36923E-21 |
| ACE2 | PLXNB1 | -0,4495 | 1,37253E-21 |
| ACE2 | RP11-216B9.9 | -0,4495 | 1,39735E-21 |
| ACE2 | BRAP | -0,4495 | 1,37382E-21 |
| ACE2 | MRPL52 | -0,4495 | 1,37295E-21 |
| ACE2 | CTB-58E17.3 | -0,4495 | 1,39651E-21 |
| ACE2 | FZR1 | -0,4495 | 1,37462E-21 |
| ACE2 | FCRLB | -0,4496 | 1,33619E-21 |
| ACE2 | HHATL | -0,4496 | 1,33587E-21 |
| ACE2 | RP11-265B7.1 | -0,4497 | 1,33363E-21 |
| ACE2 | RP11-820I16.1 | -0,4497 | 1,30929E-21 |
| ACE2 | HSPA1B | -0,4498 | 1,30512E-21 |
| ACE2 | RP11-21A7A.2 | -0,4498 | 1,29085E-21 |
| ACE2 | SYNDIG1 | -0,4498 | 1,27923E-21 |
| ACE2 | TCEANC2 | -0,4499 | 1,25448E-21 |
| ACE2 | LINC01091 | -0,4499 | 1,269E-21 |
| ACE2 | SPDYE12P | -0,45 | 1,21975E-21 |
| ACE2 | RP11-403B2.7 | -0,45 | 1,24186E-21 |
| ACE2 | MZF1 | -0,45 | 1,23956E-21 |
| ACE2 | C1orf61 | -0,4501 | 1,2108E-21 |
| ACE2 | TNFSF14 | -0,4501 | 1,21043E-21 |
| ACE2 | LTBR | -0,4502 | 1,17603E-21 |
| ACE2 | LDHAP2 | -0,4504 | 1,12665E-21 |
| ACE2 | RNF167 | -0,4504 | 1,11313E-21 |
| ACE2 | CTD-2286N8.2 | -0,4504 | 1,13334E-21 |
| ACE2 | RAB11B-AS1 | -0,4504 | 1,13525E-21 |
| ACE2 | BCL2L13 | -0,4504 | 1,1232E-21 |
| ACE2 | GOLGA8IP | -0,4505 | 1,08775E-21 |
| ACE2 | SPEN | -0,4506 | 1,08378E-21 |
| ACE2 | AC097359.2 | -0,4506 | 1,0644E-21 |
| ACE2 | RP5-1007F24.1 | -0,4506 | 1,08259E-21 |
| ACE2 | HBP1 | -0,4506 | 1,07224E-21 |
| ACE2 | RP5-855D21.3 | -0,4506 | 1,0722E-21 |
| ACE2 | RP11-433J8.1 | -0,4506 | 1,07849E-21 |
| ACE2 | NPM1P27 | -0,4507 | 1,049E-21 |
| ACE2 | TBC1D29 | -0,4507 | 1,05631E-21 |
| ACE2 | RP11-391M7.3 | -0,4508 | 1,0293E-21 |
| ACE2 | LINC00612 | -0,4508 | 1,01717E-21 |
| ACE2 | ALDH1B1 | -0,4509 | 9,97162E-22 |
| ACE2 | RP11-114H24.6 | -0,4509 | 1,01201E-21 |

| | | | |
|---|---|---|---|
| ACE2 | RP5-1039K5.18 | -0,4509 | 1,01242E-21 |
| ACE2 | SPDYA | -0,451 | 9,78318E-22 |
| ACE2 | RP11-749I16.3 | -0,451 | 9,88636E-22 |
| ACE2 | TEF | -0,451 | 9,74716E-22 |
| ACE2 | RP5-1042I8.7 | -0,4511 | 9,50344E-22 |
| ACE2 | ATG16L1 | -0,4511 | 9,53347E-22 |
| ACE2 | HCG17 | -0,4511 | 9,66381E-22 |
| ACE2 | MIB2 | -0,4512 | 9,34061E-22 |
| ACE2 | RP5-884C9.2 | -0,4512 | 9,38519E-22 |
| ACE2 | IRS2 | -0,4512 | 9,41506E-22 |
| ACE2 | HSPC324 | -0,4513 | 9,23718E-22 |
| ACE2 | RP11-815J4.6 | -0,4513 | 9,04077E-22 |
| ACE2 | LYPD3 | -0,4513 | 9,23442E-22 |
| ACE2 | C22orf15 | -0,4513 | 9,14587E-22 |
| ACE2 | TMEM106B | -0,4514 | 9,00979E-22 |
| ACE2 | RP3-449M8.9 | -0,4514 | 9,02139E-22 |
| ACE2 | NUP62 | -0,4514 | 8,83246E-22 |
| ACE2 | PPP6R1 | -0,4514 | 8,98315E-22 |
| ACE2 | ELP6 | -0,4515 | 8,66648E-22 |
| ACE2 | COMMD5 | -0,4515 | 8,69643E-22 |
| ACE2 | RP11-395N3.1 | -0,4516 | 8,43156E-22 |
| ACE2 | SSC4D | -0,4516 | 8,59168E-22 |
| ACE2 | NOLC1 | -0,4516 | 8,43106E-22 |
| ACE2 | RP1-290F12.3 | -0,4516 | 8,60719E-22 |
| ACE2 | LINC01393 | -0,4517 | 8,40868E-22 |
| ACE2 | POMP | -0,4517 | 8,35081E-22 |
| ACE2 | RN7SL526P | -0,4517 | 8,2534E-22 |
| ACE2 | CHD9 | -0,4518 | 8,18756E-22 |
| ACE2 | RP3-495K2.4 | -0,4519 | 7,94317E-22 |
| ACE2 | PITPNA | -0,452 | 7,80596E-22 |
| ACE2 | SNAP29 | -0,452 | 7,82381E-22 |
| ACE2 | RP11-575H3.1 | -0,4521 | 7,65871E-22 |
| ACE2 | ZNF442 | -0,4521 | 7,55833E-22 |
| ACE2 | RP11-211G3.3 | -0,4522 | 7,33156E-22 |
| ACE2 | RP13-514E23.2 | -0,4522 | 7,46806E-22 |
| ACE2 | RP11-383B4.4 | -0,4522 | 7,493E-22 |
| ACE2 | FIBP | -0,4522 | 7,37908E-22 |
| ACE2 | SIAH2 | -0,4523 | 7,27685E-22 |
| ACE2 | MIR145 | -0,4523 | 7,28151E-22 |
| ACE2 | PKN3 | -0,4523 | 7,29948E-22 |
| ACE2 | RP11-279O9.4 | -0,4524 | 7,07541E-22 |
| ACE2 | RP11-30L15.4 | -0,4524 | 7,07728E-22 |
| ACE2 | CITF22-92A6.2 | -0,4524 | 7,1404E-22 |
| ACE2 | RP11-758M4.4 | -0,4525 | 6,87351E-22 |
| ACE2 | RP11-490B18.6 | -0,4525 | 6,84066E-22 |
| ACE2 | RNF7 | -0,4526 | 6,76966E-22 |
| ACE2 | LINC02062 | -0,4526 | 6,69017E-22 |
| ACE2 | RP11-856M7.2 | -0,4526 | 6,77298E-22 |
| ACE2 | FOXI2 | -0,4527 | 6,57907E-22 |
| ACE2 | AC004069.2 | -0,4528 | 6,44601E-22 |
| ACE2 | MYBL1 | -0,4528 | 6,49574E-22 |
| ACE2 | RP11-370I10.11 | -0,4528 | 6,43335E-22 |
| ACE2 | RPSAP14 | -0,4528 | 6,47485E-22 |
| ACE2 | HSPA8P11 | -0,4529 | 6,23555E-22 |

| | | | |
|---|---|---|---|
| ACE2 | MBL1P | -0,4529 | 6,32799E-22 |
| ACE2 | ZSCAN20 | -0,453 | 6,18106E-22 |
| ACE2 | JMJD4 | -0,4531 | 6,00724E-22 |
| ACE2 | AC159540.1 | -0,4531 | 6,02479E-22 |
| ACE2 | ZNF827 | -0,4531 | 6,03136E-22 |
| ACE2 | WDR46 | -0,4531 | 6,02621E-22 |
| ACE2 | CTD-2005H7.1 | -0,4531 | 5,95354E-22 |
| ACE2 | CTD-2325M2.1 | -0,4531 | 6,04179E-22 |
| ACE2 | AC073850.6 | -0,4532 | 5,89666E-22 |
| ACE2 | LA16c-358B7.3 | -0,4532 | 5,82887E-22 |
| ACE2 | TMEM241 | -0,4532 | 5,81365E-22 |
| ACE2 | BTG2 | -0,4533 | 5,79885E-22 |
| ACE2 | APOD | -0,4533 | 5,78697E-22 |
| ACE2 | CTD-2013N17.6 | -0,4533 | 5,79471E-22 |
| ACE2 | PDK3 | -0,4533 | 5,79205E-22 |
| ACE2 | NIFK | -0,4534 | 5,6342E-22 |
| ACE2 | AKIRIN2 | -0,4534 | 5,61685E-22 |
| ACE2 | IL31RA | -0,4535 | 5,48877E-22 |
| ACE2 | RP11-20I20.1 | -0,4536 | 5,37192E-22 |
| ACE2 | PDE7A | -0,4536 | 5,35955E-22 |
| ACE2 | GRIK4 | -0,4536 | 5,40081E-22 |
| ACE2 | STAT3 | -0,4536 | 5,32367E-22 |
| ACE2 | CCDC13-AS1 | -0,4537 | 5,21527E-22 |
| ACE2 | FCF1P2 | -0,4537 | 5,20051E-22 |
| ACE2 | RP11-29A19.2 | -0,4537 | 5,1846E-22 |
| ACE2 | NFRKB | -0,4537 | 5,25586E-22 |
| ACE2 | MYLIP | -0,4538 | 5,10552E-22 |
| ACE2 | KAT5 | -0,4538 | 5,13363E-22 |
| ACE2 | LTA4H | -0,4538 | 5,10447E-22 |
| ACE2 | RP11-284F21.8 | -0,4539 | 5,0299E-22 |
| ACE2 | LARP7 | -0,4539 | 5,01945E-22 |
| ACE2 | RNF31 | -0,4539 | 4,98367E-22 |
| ACE2 | MED31 | -0,4539 | 5,03892E-22 |
| ACE2 | AP001092.4 | -0,454 | 4,92925E-22 |
| ACE2 | GPATCH1 | -0,454 | 4,87238E-22 |
| ACE2 | TRIP12 | -0,4541 | 4,77506E-22 |
| ACE2 | RP1-78O14.1 | -0,4541 | 4,81673E-22 |
| ACE2 | TPR | -0,4542 | 4,65536E-22 |
| ACE2 | CRCP | -0,4542 | 4,64626E-22 |
| ACE2 | USP22 | -0,4542 | 4,67611E-22 |
| ACE2 | CBLL1 | -0,4543 | 4,51346E-22 |
| ACE2 | EFCAB8 | -0,4543 | 4,49739E-22 |
| ACE2 | AATK | -0,4544 | 4,39801E-22 |
| ACE2 | NOL10 | -0,4545 | 4,34908E-22 |
| ACE2 | ZNF718 | -0,4545 | 4,3421E-22 |
| ACE2 | RPL23AP32 | -0,4546 | 4,26761E-22 |
| ACE2 | UTS2B | -0,4548 | 4,06318E-22 |
| ACE2 | HCG22 | -0,4549 | 3,99176E-22 |
| ACE2 | TMEM5 | -0,4549 | 3,99098E-22 |
| ACE2 | HMG20A | -0,4549 | 3,99649E-22 |
| ACE2 | RP11-936I5.1 | -0,4549 | 3,92289E-22 |
| ACE2 | RP11-426J5.3 | -0,4549 | 3,90977E-22 |
| ACE2 | HOXD13 | -0,455 | 3,90675E-22 |
| ACE2 | RP11-690I21.1 | -0,455 | 3,88022E-22 |

| | | | |
|---|---|---|---|
| ACE2 | IRAK2 | -0,4551 | 3,79349E-22 |
| ACE2 | FNDC3B | -0,4551 | 3,74339E-22 |
| ACE2 | TRIM60P18 | -0,4551 | 3,75002E-22 |
| ACE2 | LPCAT2 | -0,4551 | 3,74196E-22 |
| ACE2 | MCF2 | -0,4552 | 3,64583E-22 |
| ACE2 | ALMS1P1 | -0,4553 | 3,59508E-22 |
| ACE2 | RP5-1000K24.2 | -0,4553 | 3,62636E-22 |
| ACE2 | RP11-326I19.2 | -0,4554 | 3,50545E-22 |
| ACE2 | TSPAN31 | -0,4554 | 3,54185E-22 |
| ACE2 | RP11-330M19.1 | -0,4555 | 3,43022E-22 |
| ACE2 | MOG | -0,4555 | 3,45013E-22 |
| ACE2 | IMMP1LP1 | -0,4555 | 3,40803E-22 |
| ACE2 | DIS3 | -0,4555 | 3,46805E-22 |
| ACE2 | RP11-66B24.9 | -0,4555 | 3,4199E-22 |
| ACE2 | NCL | -0,4556 | 3,33968E-22 |
| ACE2 | RP11-134K13.4 | -0,4556 | 3,38677E-22 |
| ACE2 | ATF4P4 | -0,4556 | 3,36552E-22 |
| ACE2 | SELENOK | -0,4557 | 3,30643E-22 |
| ACE2 | RP1-101A2.1 | -0,4557 | 3,26534E-22 |
| ACE2 | DIAPH2 | -0,4557 | 3,2794E-22 |
| ACE2 | SNRPE | -0,4558 | 3,21026E-22 |
| ACE2 | FBXO38 | -0,4558 | 3,19169E-22 |
| ACE2 | SNRNP35 | -0,4558 | 3,19314E-22 |
| ACE2 | LRCOL1 | -0,4558 | 3,20343E-22 |
| ACE2 | AP000695.4 | -0,4558 | 3,22328E-22 |
| ACE2 | IMMP1L | -0,4559 | 3,14589E-22 |
| ACE2 | RP11-460M2.1 | -0,456 | 3,07692E-22 |
| ACE2 | MTATP6P26 | -0,456 | 3,0537E-22 |
| ACE2 | HOXC-AS1 | -0,456 | 3,07998E-22 |
| ACE2 | NUFIP2 | -0,456 | 3,07433E-22 |
| ACE2 | XXbac-B444P24.10 | -0,456 | 3,0632E-22 |
| ACE2 | SPNS2 | -0,4561 | 2,95474E-22 |
| ACE2 | ARL6IP5 | -0,4562 | 2,92256E-22 |
| ACE2 | RP11-411K7.1 | -0,4562 | 2,93183E-22 |
| ACE2 | AF067845.1 | -0,4563 | 2,85332E-22 |
| ACE2 | RP11-15A1.9 | -0,4563 | 2,83112E-22 |
| ACE2 | ZBTB40 | -0,4564 | 2,77908E-22 |
| ACE2 | SUSD5 | -0,4564 | 2,80119E-22 |
| ACE2 | RPS4XP22 | -0,4564 | 2,78716E-22 |
| ACE2 | RN7SKP16 | -0,4565 | 2,73956E-22 |
| ACE2 | TSHB | -0,4565 | 2,73496E-22 |
| ACE2 | AC022007.5 | -0,4565 | 2,69845E-22 |
| ACE2 | IL4R | -0,4565 | 2,70923E-22 |
| ACE2 | RP11-96D1.10 | -0,4565 | 2,70012E-22 |
| ACE2 | PDPR | -0,4565 | 2,71092E-22 |
| ACE2 | ZADH2 | -0,4565 | 2,68995E-22 |
| ACE2 | AC092155.4 | -0,4566 | 2,64656E-22 |
| ACE2 | PPP2CB | -0,4566 | 2,62804E-22 |
| ACE2 | NPIPB4 | -0,4566 | 2,63423E-22 |
| ACE2 | CDK12 | -0,4566 | 2,63785E-22 |
| ACE2 | ZNF112 | -0,4566 | 2,6393E-22 |
| ACE2 | RP11-445H22.3 | -0,4566 | 2,64459E-22 |
| ACE2 | RP11-392O1.4 | -0,4567 | 2,57731E-22 |
| ACE2 | RPL12P4 | -0,4567 | 2,61906E-22 |

| | | | |
|---|---|---|---|
| ACE2 | ABCB7 | -0,4567 | 2,57086E-22 |
| ACE2 | NPIPB5 | -0,4568 | 2,52923E-22 |
| ACE2 | TRMT61B | -0,4569 | 2,47764E-22 |
| ACE2 | LINC00881 | -0,4569 | 2,4605E-22 |
| ACE2 | TRRAP | -0,457 | 2,41678E-22 |
| ACE2 | RP11-708L7.6 | -0,457 | 2,41189E-22 |
| ACE2 | NBPF11 | -0,4571 | 2,36093E-22 |
| ACE2 | NAT8L | -0,4571 | 2,37768E-22 |
| ACE2 | RP11-91P24.7 | -0,4571 | 2,34958E-22 |
| ACE2 | AC138969.4 | -0,4571 | 2,33273E-22 |
| ACE2 | LENG8-AS1 | -0,4571 | 2,38173E-22 |
| ACE2 | OGG1 | -0,4572 | 2,28908E-22 |
| ACE2 | ARIH2OS | -0,4572 | 2,30448E-22 |
| ACE2 | AC138649.1 | -0,4572 | 2,28361E-22 |
| ACE2 | RP11-299H22.3 | -0,4573 | 2,22777E-22 |
| ACE2 | FAM3C2 | -0,4573 | 2,25902E-22 |
| ACE2 | SCARA5 | -0,4574 | 2,18712E-22 |
| ACE2 | MSH4 | -0,4575 | 2,1697E-22 |
| ACE2 | MRPS24 | -0,4575 | 2,12866E-22 |
| ACE2 | SHISA2 | -0,4575 | 2,12718E-22 |
| ACE2 | PANK4 | -0,4576 | 2,08967E-22 |
| ACE2 | GPAT2 | -0,4577 | 2,02856E-22 |
| ACE2 | UBE2L3 | -0,4577 | 2,03864E-22 |
| ACE2 | RP4-683M8.2 | -0,4578 | 1,9917E-22 |
| ACE2 | EPHA5-AS1 | -0,4578 | 2,01566E-22 |
| ACE2 | CTD-2035E11.5 | -0,4578 | 2,01567E-22 |
| ACE2 | RBM12B | -0,4578 | 1,97929E-22 |
| ACE2 | FRY-AS1 | -0,4578 | 2,00664E-22 |
| ACE2 | SIAH3 | -0,4578 | 2,00489E-22 |
| ACE2 | GGT5 | -0,4578 | 1,97895E-22 |
| ACE2 | FBXO28 | -0,4579 | 1,94261E-22 |
| ACE2 | MRPS17 | -0,4579 | 1,95023E-22 |
| ACE2 | YBX1P1 | -0,4579 | 1,95563E-22 |
| ACE2 | LETM1P2 | -0,4579 | 1,97419E-22 |
| ACE2 | CH17-132F21.5 | -0,458 | 1,92744E-22 |
| ACE2 | AP000240.9 | -0,458 | 1,9181E-22 |
| ACE2 | RP11-365H22.2 | -0,4581 | 1,86385E-22 |
| ACE2 | RP11-348F1.3 | -0,4581 | 1,8431E-22 |
| ACE2 | RP11-243M5.3 | -0,4582 | 1,8104E-22 |
| ACE2 | EIF3D | -0,4582 | 1,81953E-22 |
| ACE2 | RP11-373D23.3 | -0,4583 | 1,75659E-22 |
| ACE2 | RP11-707A18.1 | -0,4583 | 1,76789E-22 |
| ACE2 | RP11-290H9.5 | -0,4583 | 1,7573E-22 |
| ACE2 | ID2 | -0,4584 | 1,75088E-22 |
| ACE2 | TMEM175 | -0,4584 | 1,75093E-22 |
| ACE2 | RP11-239L20.6 | -0,4584 | 1,74778E-22 |
| ACE2 | MKLN1-AS | -0,4586 | 1,67341E-22 |
| ACE2 | RPL41 | -0,4586 | 1,63831E-22 |
| ACE2 | FBXW10 | -0,4586 | 1,65487E-22 |
| ACE2 | IRF2BP1 | -0,4586 | 1,65956E-22 |
| ACE2 | RANBP17 | -0,4587 | 1,616E-22 |
| ACE2 | ALG1L9P | -0,4587 | 1,59909E-22 |
| ACE2 | RP11-632F7.3 | -0,4589 | 1,53519E-22 |
| ACE2 | PIGBOS1 | -0,4589 | 1,54544E-22 |

| | | | |
|---|---|---|---|
| ACE2 | ZBTB8B | -0,459 | 1,49737E-22 |
| ACE2 | LDLRAD4-AS1 | -0,459 | 1,49602E-22 |
| ACE2 | PCDHA9 | -0,4591 | 1,46621E-22 |
| ACE2 | RP11-661A12.8 | -0,4591 | 1,48509E-22 |
| ACE2 | ADPRHL1 | -0,4591 | 1,47455E-22 |
| ACE2 | ELANE | -0,4591 | 1,48573E-22 |
| ACE2 | FCF1 | -0,4592 | 1,42014E-22 |
| ACE2 | GTPBP8 | -0,4593 | 1,41593E-22 |
| ACE2 | AGAP4 | -0,4593 | 1,38813E-22 |
| ACE2 | MICU1 | -0,4593 | 1,41197E-22 |
| ACE2 | RP11-344N10.5 | -0,4593 | 1,40108E-22 |
| ACE2 | EPB41L3 | -0,4593 | 1,38747E-22 |
| ACE2 | TM4SF18 | -0,4594 | 1,38428E-22 |
| ACE2 | ABCC4 | -0,4594 | 1,3578E-22 |
| ACE2 | AP000695.6 | -0,4594 | 1,3841E-22 |
| ACE2 | RP11-138E2.1 | -0,4595 | 1,34096E-22 |
| ACE2 | ZSCAN4 | -0,4595 | 1,33884E-22 |
| ACE2 | GTPBP10 | -0,4597 | 1,26993E-22 |
| ACE2 | SETD6 | -0,4597 | 1,27933E-22 |
| ACE2 | RP11-521C22.2 | -0,4598 | 1,24333E-22 |
| ACE2 | RP11-94C24.6 | -0,4598 | 1,25866E-22 |
| ACE2 | ZNF708 | -0,4598 | 1,24551E-22 |
| ACE2 | PTPN4 | -0,4599 | 1,20766E-22 |
| ACE2 | RP11-766F14.2 | -0,4599 | 1,2113E-22 |
| ACE2 | GALNT10 | -0,4599 | 1,20443E-22 |
| ACE2 | SOAT2 | -0,4599 | 1,22399E-22 |
| ACE2 | ZNF776 | -0,4599 | 1,21017E-22 |
| ACE2 | RP11-394I13.3 | -0,46 | 1,19741E-22 |
| ACE2 | C9orf142 | -0,46 | 1,18029E-22 |
| ACE2 | TATDN2P3 | -0,4601 | 1,15076E-22 |
| ACE2 | LINC02102 | -0,4602 | 1,12559E-22 |
| ACE2 | ELOA | -0,4604 | 1,08997E-22 |
| ACE2 | RP11-571L19.8 | -0,4604 | 1,09051E-22 |
| ACE2 | ASF1A | -0,4604 | 1,06931E-22 |
| ACE2 | ATP6V0A1 | -0,4604 | 1,07281E-22 |
| ACE2 | TNFRSF6B | -0,4604 | 1,07272E-22 |
| ACE2 | ZNF436 | -0,4605 | 1,06532E-22 |
| ACE2 | IFIT2 | -0,4605 | 1,04789E-22 |
| ACE2 | RP11-140A10.3 | -0,4605 | 1,05323E-22 |
| ACE2 | ITGAM | -0,4605 | 1,06335E-22 |
| ACE2 | AFAP1-AS1 | -0,4606 | 1,03307E-22 |
| ACE2 | TRAF6 | -0,4606 | 1,03358E-22 |
| ACE2 | EEF1A1P11 | -0,4607 | 9,95897E-23 |
| ACE2 | PIM1 | -0,4607 | 1,00129E-22 |
| ACE2 | FAM166B | -0,4607 | 1,01505E-22 |
| ACE2 | RP1-40E16.9 | -0,4609 | 9,64704E-23 |
| ACE2 | HOGA1 | -0,4609 | 9,53752E-23 |
| ACE2 | HAGH | -0,4609 | 9,59475E-23 |
| ACE2 | ZNF785 | -0,4609 | 9,6957E-23 |
| ACE2 | CTC-471F3.5 | -0,4609 | 9,47656E-23 |
| ACE2 | PARGP1 | -0,461 | 9,36909E-23 |
| ACE2 | RP11-147L13.7 | -0,461 | 9,25072E-23 |
| ACE2 | AC016907.3 | -0,4611 | 9,04801E-23 |
| ACE2 | IL1R1 | -0,4611 | 9,20277E-23 |

| | | | |
|---|---|---|---|
| ACE2 | RP11-629O1.2 | -0,4611 | 9,11912E-23 |
| ACE2 | BEST1 | -0,4611 | 9,12641E-23 |
| ACE2 | USP5 | -0,4611 | 9,19318E-23 |
| ACE2 | RP1-257C22.2 | -0,4611 | 9,20831E-23 |
| ACE2 | MEI4 | -0,4612 | 8,96163E-23 |
| ACE2 | RNF139 | -0,4612 | 8,95165E-23 |
| ACE2 | RP11-709A23.2 | -0,4612 | 8,96955E-23 |
| ACE2 | IGFL4 | -0,4612 | 8,95264E-23 |
| ACE2 | FALEC | -0,4613 | 8,72908E-23 |
| ACE2 | PRIM2 | -0,4613 | 8,61564E-23 |
| ACE2 | SUMF2 | -0,4613 | 8,80228E-23 |
| ACE2 | CTD-2297D10.2 | -0,4615 | 8,39876E-23 |
| ACE2 | ORAI1 | -0,4615 | 8,392E-23 |
| ACE2 | AC068831.10 | -0,4615 | 8,33215E-23 |
| ACE2 | FHL2 | -0,4616 | 8,06883E-23 |
| ACE2 | BRD9P2 | -0,4616 | 8,12116E-23 |
| ACE2 | GPCPD1 | -0,4616 | 8,14028E-23 |
| ACE2 | C21orf2 | -0,4616 | 8,10885E-23 |
| ACE2 | TXNIP | -0,4618 | 7,81195E-23 |
| ACE2 | FMO3 | -0,4618 | 7,66677E-23 |
| ACE2 | PSME4 | -0,4618 | 7,81585E-23 |
| ACE2 | UBTD2 | -0,4618 | 7,71145E-23 |
| ACE2 | RP11-212E4.1 | -0,4618 | 7,64421E-23 |
| ACE2 | MSI2 | -0,4618 | 7,68909E-23 |
| ACE2 | ZC3H18 | -0,4619 | 7,56071E-23 |
| ACE2 | RP1-122P22.2 | -0,4619 | 7,49261E-23 |
| ACE2 | AC007362.3 | -0,462 | 7,30881E-23 |
| ACE2 | CDC40 | -0,462 | 7,28999E-23 |
| ACE2 | C8orf33 | -0,462 | 7,32343E-23 |
| ACE2 | RP11-472N13.2 | -0,462 | 7,43623E-23 |
| ACE2 | NCKAP1L | -0,462 | 7,4058E-23 |
| ACE2 | LINC00565 | -0,4621 | 7,18205E-23 |
| ACE2 | JAG2 | -0,4621 | 7,15125E-23 |
| ACE2 | MAFF | -0,4621 | 7,15519E-23 |
| ACE2 | GGTA1P | -0,4622 | 6,97299E-23 |
| ACE2 | MIR3164 | -0,4622 | 7,03092E-23 |
| ACE2 | SF3A1 | -0,4622 | 7,01402E-23 |
| ACE2 | RP11-3D4.4 | -0,4623 | 6,80399E-23 |
| ACE2 | MAF | -0,4623 | 6,78933E-23 |
| ACE2 | RP11-68L18.1 | -0,4624 | 6,64451E-23 |
| ACE2 | CTD-2589M5.5 | -0,4624 | 6,73646E-23 |
| ACE2 | CNOT6LP1 | -0,4624 | 6,64347E-23 |
| ACE2 | RP11-15A1.3 | -0,4624 | 6,62444E-23 |
| ACE2 | TPRG1LP1 | -0,4624 | 6,68831E-23 |
| ACE2 | DCLRE1B | -0,4625 | 6,47343E-23 |
| ACE2 | HDC | -0,4625 | 6,54211E-23 |
| ACE2 | ARHGAP23P1 | -0,4625 | 6,50075E-23 |
| ACE2 | PGF | -0,4626 | 6,42833E-23 |
| ACE2 | NDUFS4 | -0,4627 | 6,15803E-23 |
| ACE2 | FBRS | -0,4628 | 6,11568E-23 |
| ACE2 | PLK3 | -0,4629 | 5,88461E-23 |
| ACE2 | DCAF4L1 | -0,4629 | 5,90409E-23 |
| ACE2 | WAC-AS1 | -0,4629 | 5,8818E-23 |
| ACE2 | OGFOD3 | -0,4629 | 5,91237E-23 |

| | | | |
|---|---|---|---|
| ACE2 | RP11-403A21.1 | -0,4629 | 5,91491E-23 |
| ACE2 | RP11-679C8.2 | -0,463 | 5,8278E-23 |
| ACE2 | RP11-554A11.7 | -0,463 | 5,79135E-23 |
| ACE2 | E4F1 | -0,463 | 5,79069E-23 |
| ACE2 | BAG4 | -0,4631 | 5,59709E-23 |
| ACE2 | TUSC1 | -0,4631 | 5,69282E-23 |
| ACE2 | ATG14 | -0,4631 | 5,71711E-23 |
| ACE2 | CTD-2341M24.1 | -0,4631 | 5,611E-23 |
| ACE2 | MAN2C1 | -0,4631 | 5,66189E-23 |
| ACE2 | IL15 | -0,4632 | 5,57832E-23 |
| ACE2 | ARAP1 | -0,4632 | 5,56331E-23 |
| ACE2 | FUNDC1 | -0,4632 | 5,47715E-23 |
| ACE2 | XXbac-BPG154L12.4 | -0,4633 | 5,38199E-23 |
| ACE2 | FAM153C | -0,4634 | 5,24921E-23 |
| ACE2 | PSMG4 | -0,4634 | 5,27986E-23 |
| ACE2 | RP11-624M8.1 | -0,4634 | 5,26399E-23 |
| ACE2 | EIF4BP6 | -0,4634 | 5,26997E-23 |
| ACE2 | QRFP | -0,4634 | 5,2298E-23 |
| ACE2 | RP11-666A8.12 | -0,4634 | 5,31819E-23 |
| ACE2 | QSOX2 | -0,4635 | 5,18502E-23 |
| ACE2 | RP4-675C20.4 | -0,4636 | 4,99045E-23 |
| ACE2 | PLSCR5 | -0,4636 | 4,98898E-23 |
| ACE2 | EIF4H | -0,4636 | 5,01721E-23 |
| ACE2 | PALB2 | -0,4637 | 4,95137E-23 |
| ACE2 | YIF1B | -0,4637 | 4,93583E-23 |
| ACE2 | CTPS2 | -0,4637 | 4,93733E-23 |
| ACE2 | TOMM70 | -0,4638 | 4,79336E-23 |
| ACE2 | ZBTB38 | -0,4638 | 4,82975E-23 |
| ACE2 | RP11-17A4.2 | -0,4638 | 4,82427E-23 |
| ACE2 | HBG2 | -0,4638 | 4,83029E-23 |
| ACE2 | FANCE | -0,4639 | 4,68776E-23 |
| ACE2 | LINC00846 | -0,4639 | 4,67152E-23 |
| ACE2 | RP11-138I17.1 | -0,464 | 4,53277E-23 |
| ACE2 | ZNF561 | -0,464 | 4,57113E-23 |
| ACE2 | LINC00602 | -0,4641 | 4,47675E-23 |
| ACE2 | USP42 | -0,4641 | 4,4053E-23 |
| ACE2 | POTEJ | -0,4642 | 4,38009E-23 |
| ACE2 | DOCK4-AS1 | -0,4642 | 4,37847E-23 |
| ACE2 | TRMT5 | -0,4642 | 4,37204E-23 |
| ACE2 | OR2L13 | -0,4643 | 4,23752E-23 |
| ACE2 | CCKAR | -0,4643 | 4,28651E-23 |
| ACE2 | SCGB3A1 | -0,4643 | 4,27533E-23 |
| ACE2 | AC002066.1 | -0,4643 | 4,20109E-23 |
| ACE2 | WBP11 | -0,4643 | 4,26641E-23 |
| ACE2 | PABPC1 | -0,4644 | 4,11263E-23 |
| ACE2 | CTD-3131K8.2 | -0,4644 | 4,14203E-23 |
| ACE2 | RP1-187B23.1 | -0,4645 | 4,03356E-23 |
| ACE2 | CARS | -0,4645 | 4,08664E-23 |
| ACE2 | HDHD2 | -0,4645 | 4,01102E-23 |
| ACE2 | CCDC151 | -0,4645 | 4,08876E-23 |
| ACE2 | AL133243.2 | -0,4646 | 3,9798E-23 |
| ACE2 | RP11-539L10.3 | -0,4646 | 3,97992E-23 |
| ACE2 | NAF1 | -0,4646 | 3,97413E-23 |
| ACE2 | MLH3 | -0,4646 | 3,92304E-23 |

| | | | |
|---|---|---|---|
| ACE2 | SERAC1 | -0,4647 | 3,86063E-23 |
| ACE2 | FRMPD3 | -0,4647 | 3,80922E-23 |
| ACE2 | RP11-197M22.2 | -0,4648 | 3,78614E-23 |
| ACE2 | IRX3 | -0,4648 | 3,78487E-23 |
| ACE2 | FGF17 | -0,4649 | 3,66926E-23 |
| ACE2 | ZCCHC14 | -0,4649 | 3,65228E-23 |
| ACE2 | TLE1 | -0,465 | 3,59718E-23 |
| ACE2 | PHLDA1 | -0,465 | 3,59135E-23 |
| ACE2 | GCN1 | -0,4651 | 3,45617E-23 |
| ACE2 | ERFE | -0,4652 | 3,3715E-23 |
| ACE2 | ACTR5 | -0,4652 | 3,44379E-23 |
| ACE2 | CTSLP2 | -0,4653 | 3,29614E-23 |
| ACE2 | IDE | -0,4653 | 3,34979E-23 |
| ACE2 | NLRP9 | -0,4655 | 3,14691E-23 |
| ACE2 | EDIL3 | -0,4656 | 3,10632E-23 |
| ACE2 | AGPAT4-IT1 | -0,4656 | 3,10445E-23 |
| ACE2 | MRPL32 | -0,4656 | 3,10819E-23 |
| ACE2 | RP11-499P20.2 | -0,4656 | 3,08001E-23 |
| ACE2 | RP11-7F17.1 | -0,4656 | 3,1224E-23 |
| ACE2 | HNRNPH1 | -0,4657 | 3,00922E-23 |
| ACE2 | BAALC-AS1 | -0,4657 | 3,04952E-23 |
| ACE2 | RNF19A | -0,4658 | 2,96769E-23 |
| ACE2 | PCGF5 | -0,4658 | 2,94646E-23 |
| ACE2 | SMTNL1 | -0,4658 | 2,9497E-23 |
| ACE2 | CTC-529P8.1 | -0,4659 | 2,89674E-23 |
| ACE2 | METTL10 | -0,4659 | 2,8492E-23 |
| ACE2 | AC004237.1 | -0,466 | 2,79844E-23 |
| ACE2 | GOLGA8Q | -0,466 | 2,81564E-23 |
| ACE2 | COQ3 | -0,4661 | 2,7228E-23 |
| ACE2 | HDAC2 | -0,4661 | 2,73206E-23 |
| ACE2 | AC005154.7 | -0,4662 | 2,66703E-23 |
| ACE2 | SUMO3 | -0,4662 | 2,64854E-23 |
| ACE2 | RP11-692D12.1 | -0,4664 | 2,57561E-23 |
| ACE2 | UBE2D3 | -0,4664 | 2,53731E-23 |
| ACE2 | MED26 | -0,4664 | 2,53902E-23 |
| ACE2 | RP11-285F16.1 | -0,4665 | 2,47118E-23 |
| ACE2 | NBEAP1 | -0,4666 | 2,43899E-23 |
| ACE2 | HEATR1 | -0,4667 | 2,36678E-23 |
| ACE2 | CD47 | -0,4667 | 2,39532E-23 |
| ACE2 | MAP2K4P1 | -0,4667 | 2,39973E-23 |
| ACE2 | NAA38 | -0,4669 | 2,26019E-23 |
| ACE2 | SPATC1L | -0,4669 | 2,25939E-23 |
| ACE2 | SLC5A4 | -0,4669 | 2,24566E-23 |
| ACE2 | AP000255.6 | -0,467 | 2,18815E-23 |
| ACE2 | LONP1 | -0,4671 | 2,16015E-23 |
| ACE2 | VPS8 | -0,4673 | 2,06189E-23 |
| ACE2 | LINC01277 | -0,4673 | 2,03364E-23 |
| ACE2 | AC004623.3 | -0,4673 | 2,07248E-23 |
| ACE2 | RP11-316O14.1 | -0,4674 | 1,99045E-23 |
| ACE2 | ACAD9 | -0,4674 | 1,99693E-23 |
| ACE2 | C9orf24 | -0,4674 | 1,97944E-23 |
| ACE2 | RP11-47L3.1 | -0,4674 | 1,9889E-23 |
| ACE2 | GGN | -0,4674 | 2,01093E-23 |
| ACE2 | EXOSC6 | -0,4675 | 1,94557E-23 |

| | | | |
|---|---|---|---|
| ACE2 | ALG13 | -0,4675 | 1,95082E-23 |
| ACE2 | FUCA1P1 | -0,4676 | 1,90618E-23 |
| ACE2 | PAN3 | -0,4676 | 1,90059E-23 |
| ACE2 | SEMA6B | -0,4676 | 1,91053E-23 |
| ACE2 | MYPOP | -0,4676 | 1,92406E-23 |
| ACE2 | DDX27 | -0,4676 | 1,92642E-23 |
| ACE2 | USP1 | -0,4677 | 1,86624E-23 |
| ACE2 | AC005104.3 | -0,4677 | 1,84039E-23 |
| ACE2 | EXOC6 | -0,4677 | 1,84883E-23 |
| ACE2 | MZT1 | -0,4677 | 1,87169E-23 |
| ACE2 | DIDO1 | -0,4677 | 1,84882E-23 |
| ACE2 | LINC00384 | -0,4678 | 1,83525E-23 |
| ACE2 | CTD-2651B20.1 | -0,4678 | 1,79929E-23 |
| ACE2 | CYP2G1P | -0,4678 | 1,81857E-23 |
| ACE2 | PABPC1L2B | -0,4678 | 1,80387E-23 |
| ACE2 | RARA | -0,4679 | 1,79135E-23 |
| ACE2 | GYPE | -0,468 | 1,70996E-23 |
| ACE2 | RBM33 | -0,468 | 1,72907E-23 |
| ACE2 | SRGN | -0,468 | 1,73237E-23 |
| ACE2 | RP11-295I5.3 | -0,468 | 1,7202E-23 |
| ACE2 | QRICH2 | -0,468 | 1,7286E-23 |
| ACE2 | PTGES3 | -0,4681 | 1,68594E-23 |
| ACE2 | RP11-81A22.4 | -0,4681 | 1,68745E-23 |
| ACE2 | MATN1-AS1 | -0,4682 | 1,66245E-23 |
| ACE2 | C1orf162 | -0,4682 | 1,63565E-23 |
| ACE2 | ASXL2 | -0,4682 | 1,6512E-23 |
| ACE2 | RP11-246A10.1 | -0,4682 | 1,65161E-23 |
| ACE2 | PRR29-AS1 | -0,4682 | 1,65197E-23 |
| ACE2 | RP4-781B1.5 | -0,4682 | 1,66261E-23 |
| ACE2 | SNRNP40 | -0,4683 | 1,62416E-23 |
| ACE2 | TNFRSF1A | -0,4683 | 1,62007E-23 |
| ACE2 | IPO8 | -0,4683 | 1,62229E-23 |
| ACE2 | HSBP1 | -0,4683 | 1,60002E-23 |
| ACE2 | UBE2Z | -0,4683 | 1,62099E-23 |
| ACE2 | DPM1 | -0,4683 | 1,59765E-23 |
| ACE2 | IGSF5 | -0,4683 | 1,58918E-23 |
| ACE2 | GLYATL2 | -0,4684 | 1,55568E-23 |
| ACE2 | SNX11 | -0,4684 | 1,58296E-23 |
| ACE2 | EBNA1BP2 | -0,4685 | 1,51398E-23 |
| ACE2 | IGFN1 | -0,4685 | 1,51327E-23 |
| ACE2 | FGD5-AS1 | -0,4685 | 1,51686E-23 |
| ACE2 | SEC24D | -0,4685 | 1,51876E-23 |
| ACE2 | NDUFAF4 | -0,4685 | 1,51304E-23 |
| ACE2 | TOLLIP | -0,4685 | 1,53689E-23 |
| ACE2 | CGREF1 | -0,4686 | 1,48266E-23 |
| ACE2 | FZD8 | -0,4686 | 1,47991E-23 |
| ACE2 | RP11-554F20.1 | -0,4687 | 1,46403E-23 |
| ACE2 | CCDC25 | -0,4688 | 1,41918E-23 |
| ACE2 | RP11-1000B6.8 | -0,4688 | 1,40973E-23 |
| ACE2 | ACVR2B-AS1 | -0,4689 | 1,39869E-23 |
| ACE2 | SUDS3 | -0,4689 | 1,39781E-23 |
| ACE2 | RP11-442O1.3 | -0,4689 | 1,39226E-23 |
| ACE2 | DHX9 | -0,469 | 1,36724E-23 |
| ACE2 | RP11-981G7.1 | -0,469 | 1,3468E-23 |

| | | | |
|---|---|---|---|
| ACE2 | HSF2BP | -0,469 | 1,35562E-23 |
| ACE2 | PABPC1P7 | -0,4691 | 1,30428E-23 |
| ACE2 | GOLGA1 | -0,4691 | 1,33129E-23 |
| ACE2 | RP11-65J3.2 | -0,4691 | 1,32345E-23 |
| ACE2 | PA2G4 | -0,4691 | 1,33536E-23 |
| ACE2 | TOB2 | -0,4691 | 1,32755E-23 |
| ACE2 | GBP4 | -0,4692 | 1,29033E-23 |
| ACE2 | AP000708.1 | -0,4693 | 1,24376E-23 |
| ACE2 | RP11-118A3.1 | -0,4694 | 1,23142E-23 |
| ACE2 | SMIM7 | -0,4694 | 1,22449E-23 |
| ACE2 | GNL2 | -0,4695 | 1,18821E-23 |
| ACE2 | MAPKAPK2 | -0,4695 | 1,20058E-23 |
| ACE2 | AC091729.9 | -0,4695 | 1,20895E-23 |
| ACE2 | LINC02076 | -0,4695 | 1,1852E-23 |
| ACE2 | METTL4 | -0,4695 | 1,18276E-23 |
| ACE2 | IFNAR1 | -0,4695 | 1,18253E-23 |
| ACE2 | PHBP9 | -0,4696 | 1,18113E-23 |
| ACE2 | POU5F1P3 | -0,4696 | 1,16281E-23 |
| ACE2 | ACACA | -0,4696 | 1,15394E-23 |
| ACE2 | GPER1 | -0,4697 | 1,12554E-23 |
| ACE2 | RP11-554A11.8 | -0,4697 | 1,13975E-23 |
| ACE2 | KLHL17 | -0,4698 | 1,11344E-23 |
| ACE2 | MFHAS1 | -0,4698 | 1,10189E-23 |
| ACE2 | TRMT12 | -0,4698 | 1,10337E-23 |
| ACE2 | CCL14 | -0,4698 | 1,10928E-23 |
| ACE2 | ZNF792 | -0,4698 | 1,10509E-23 |
| ACE2 | PRR12 | -0,4698 | 1,11932E-23 |
| ACE2 | CALM2 | -0,4699 | 1,09572E-23 |
| ACE2 | RP4-671O14.7 | -0,4699 | 1,08359E-23 |
| ACE2 | RP11-181C21.4 | -0,47 | 1,06593E-23 |
| ACE2 | TAF15 | -0,47 | 1,06088E-23 |
| ACE2 | AC006116.12 | -0,47 | 1,04679E-23 |
| ACE2 | CADPS2 | -0,4701 | 1,04274E-23 |
| ACE2 | GPAT4 | -0,4701 | 1,0253E-23 |
| ACE2 | CDK13 | -0,4702 | 1,00717E-23 |
| ACE2 | DLGAP1 | -0,4702 | 1,01137E-23 |
| ACE2 | AC002398.11 | -0,4702 | 9,99373E-24 |
| ACE2 | CYP4V2 | -0,4703 | 9,8203E-24 |
| ACE2 | ASPSCR1 | -0,4703 | 9,94247E-24 |
| ACE2 | RP11-25K19.1 | -0,4704 | 9,56514E-24 |
| ACE2 | RP11-727F15.13 | -0,4704 | 9,59994E-24 |
| ACE2 | MMP24 | -0,4704 | 9,59627E-24 |
| ACE2 | CHID1 | -0,4705 | 9,42131E-24 |
| ACE2 | RP11-680A11.5 | -0,4705 | 9,33702E-24 |
| ACE2 | ZNF738 | -0,4705 | 9,38692E-24 |
| ACE2 | TIMM50 | -0,4705 | 9,37705E-24 |
| ACE2 | RRAGC | -0,4706 | 9,16507E-24 |
| ACE2 | LINC01930 | -0,4706 | 9,21114E-24 |
| ACE2 | DOCK9-AS2 | -0,4706 | 9,12396E-24 |
| ACE2 | PSMD12 | -0,4706 | 9,07788E-24 |
| ACE2 | CEP295NL | -0,4706 | 9,20175E-24 |
| ACE2 | RP11-187O7.3 | -0,4707 | 8,95361E-24 |
| ACE2 | FRG1DP | -0,4707 | 8,88075E-24 |
| ACE2 | RP11-686O6.2 | -0,4708 | 8,74304E-24 |

| | | | |
|---|---|---|---|
| ACE2 | AC005752.10 | -0,4708 | 8,75738E-24 |
| ACE2 | HFE | -0,4708 | 8,69008E-24 |
| ACE2 | SCAF8 | -0,4708 | 8,77213E-24 |
| ACE2 | ANKRD34C | -0,4708 | 8,65754E-24 |
| ACE2 | RP5-965F6.2 | -0,4709 | 8,43064E-24 |
| ACE2 | RP11-616M22.2 | -0,4709 | 8,57001E-24 |
| ACE2 | RP11-1212A22.1 | -0,4709 | 8,41119E-24 |
| ACE2 | NR2C2 | -0,471 | 8,27626E-24 |
| ACE2 | IRF2BP2 | -0,4711 | 8,09403E-24 |
| ACE2 | KB-1410C5.5 | -0,4711 | 7,97943E-24 |
| ACE2 | AP001258.4 | -0,4712 | 7,80356E-24 |
| ACE2 | MRM2 | -0,4713 | 7,62087E-24 |
| ACE2 | ADK | -0,4713 | 7,6403E-24 |
| ACE2 | TRIM62 | -0,4714 | 7,54448E-24 |
| ACE2 | MSC-AS1 | -0,4714 | 7,55845E-24 |
| ACE2 | ZEB2-AS1 | -0,4715 | 7,35769E-24 |
| ACE2 | ATG7 | -0,4715 | 7,31551E-24 |
| ACE2 | RP11-1079K10.3 | -0,4715 | 7,2693E-24 |
| ACE2 | RP11-95O2.1 | -0,4715 | 7,37705E-24 |
| ACE2 | ATP5E | -0,4715 | 7,29317E-24 |
| ACE2 | S100A11 | -0,4716 | 7,21379E-24 |
| ACE2 | RP11-484O2.1 | -0,4716 | 7,10057E-24 |
| ACE2 | AC100830.3 | -0,4716 | 7,09238E-24 |
| ACE2 | AMFR | -0,4716 | 7,2072E-24 |
| ACE2 | RP5-1159O4.2 | -0,4717 | 7,02243E-24 |
| ACE2 | C5orf34 | -0,4718 | 6,79446E-24 |
| ACE2 | RP11-121A14.2 | -0,4718 | 6,83714E-24 |
| ACE2 | TMEM216 | -0,4718 | 6,78036E-24 |
| ACE2 | TM2D2 | -0,4719 | 6,67813E-24 |
| ACE2 | BET1L | -0,4719 | 6,70459E-24 |
| ACE2 | LINC01085 | -0,472 | 6,41585E-24 |
| ACE2 | RPL9P9 | -0,472 | 6,4549E-24 |
| ACE2 | RPL23AP42 | -0,4722 | 6,08066E-24 |
| ACE2 | INTS10 | -0,4722 | 6,18376E-24 |
| ACE2 | IQCK | -0,4722 | 6,17213E-24 |
| ACE2 | RP11-97C16.1 | -0,4723 | 5,94352E-24 |
| ACE2 | BRI3 | -0,4723 | 5,96434E-24 |
| ACE2 | BAD | -0,4723 | 6,03686E-24 |
| ACE2 | CNNM3 | -0,4724 | 5,87189E-24 |
| ACE2 | MSX2 | -0,4724 | 5,87564E-24 |
| ACE2 | RP4-621B10.8 | -0,4725 | 5,72163E-24 |
| ACE2 | TMEM39A | -0,4725 | 5,77669E-24 |
| ACE2 | ABCA11P | -0,4725 | 5,75193E-24 |
| ACE2 | CKAP5 | -0,4725 | 5,64695E-24 |
| ACE2 | RP11-35O15.1 | -0,4725 | 5,68462E-24 |
| ACE2 | RP11-1102P22.1 | -0,4725 | 5,74828E-24 |
| ACE2 | CTDSPL | -0,4726 | 5,62184E-24 |
| ACE2 | LINC00519 | -0,4726 | 5,52084E-24 |
| ACE2 | PPM1N | -0,4726 | 5,63449E-24 |
| ACE2 | EGR1 | -0,4727 | 5,37351E-24 |
| ACE2 | ZNF282 | -0,4727 | 5,37461E-24 |
| ACE2 | LINC01122 | -0,4728 | 5,25155E-24 |
| ACE2 | CACNB3 | -0,4728 | 5,32675E-24 |
| ACE2 | MYOZ3 | -0,4729 | 5,11706E-24 |

| | | | |
|---|---|---|---|
| ACE2 | MAPK8 | -0,4729 | 5,23105E-24 |
| ACE2 | RBM41 | -0,473 | 5,03971E-24 |
| ACE2 | KLHL38 | -0,4731 | 4,92726E-24 |
| ACE2 | HCRTR1 | -0,4732 | 4,74948E-24 |
| ACE2 | RP5-1142A6.2 | -0,4734 | 4,56272E-24 |
| ACE2 | TIMM21 | -0,4734 | 4,58241E-24 |
| ACE2 | SDHAF1 | -0,4734 | 4,60202E-24 |
| ACE2 | NOX5 | -0,4735 | 4,5015E-24 |
| ACE2 | RP11-104F15.9 | -0,4736 | 4,30236E-24 |
| ACE2 | CFAP70 | -0,4737 | 4,2426E-24 |
| ACE2 | RP11-762H8.2 | -0,4737 | 4,28676E-24 |
| ACE2 | RALBP1 | -0,4737 | 4,26268E-24 |
| ACE2 | BASP1 | -0,4738 | 4,17824E-24 |
| ACE2 | SNRNP48 | -0,4738 | 4,08891E-24 |
| ACE2 | HSPA7 | -0,4739 | 4,04498E-24 |
| ACE2 | CELF2-AS1 | -0,4739 | 4,02073E-24 |
| ACE2 | PTCHD1-AS | -0,4739 | 4,02787E-24 |
| ACE2 | RP11-161D15.2 | -0,474 | 3,90602E-24 |
| ACE2 | RP11-266L9.4 | -0,474 | 3,93857E-24 |
| ACE2 | GLIPR1L1 | -0,4741 | 3,85758E-24 |
| ACE2 | RP11-45P15.4 | -0,4741 | 3,80667E-24 |
| ACE2 | RP13-270P17.2 | -0,4741 | 3,87292E-24 |
| ACE2 | ATG9B | -0,4742 | 3,72819E-24 |
| ACE2 | RAD21 | -0,4742 | 3,75649E-24 |
| ACE2 | RP11-293M10.6 | -0,4742 | 3,78826E-24 |
| ACE2 | RP11-676J12.6 | -0,4742 | 3,73847E-24 |
| ACE2 | ACSS1 | -0,4742 | 3,7495E-24 |
| ACE2 | AC007383.3 | -0,4743 | 3,60448E-24 |
| ACE2 | RP11-212I21.4 | -0,4743 | 3,62004E-24 |
| ACE2 | ANKRD62P1 | -0,4743 | 3,66033E-24 |
| ACE2 | TIPARP | -0,4744 | 3,52071E-24 |
| ACE2 | CRHR1 | -0,4744 | 3,53845E-24 |
| ACE2 | LINC01358 | -0,4745 | 3,47203E-24 |
| ACE2 | RP11-216N14.5 | -0,4745 | 3,49975E-24 |
| ACE2 | HERC2P2 | -0,4745 | 3,46915E-24 |
| ACE2 | FBXO39 | -0,4746 | 3,35471E-24 |
| ACE2 | PDLIM5 | -0,4747 | 3,31838E-24 |
| ACE2 | ZC3HAV1L | -0,4747 | 3,29109E-24 |
| ACE2 | ATL1 | -0,4747 | 3,2841E-24 |
| ACE2 | RP11-4B16.3 | -0,4747 | 3,31505E-24 |
| ACE2 | CTD-2027I19.2 | -0,4747 | 3,31183E-24 |
| ACE2 | APOL2 | -0,4747 | 3,28547E-24 |
| ACE2 | CTC-559E9.6 | -0,4748 | 3,25244E-24 |
| ACE2 | RPL7AP28 | -0,4749 | 3,12886E-24 |
| ACE2 | FAM76B | -0,4749 | 3,13591E-24 |
| ACE2 | RP11-687F6.5 | -0,4749 | 3,1564E-24 |
| ACE2 | PRND | -0,4749 | 3,1415E-24 |
| ACE2 | POLR3A | -0,475 | 3,07246E-24 |
| ACE2 | PRKCH | -0,475 | 3,06626E-24 |
| ACE2 | RP11-385H1.1 | -0,475 | 3,07236E-24 |
| ACE2 | OR7E28P | -0,4751 | 3,01662E-24 |
| ACE2 | MMP27 | -0,4751 | 3,00509E-24 |
| ACE2 | PRAL | -0,4751 | 2,96462E-24 |
| ACE2 | TTC23L | -0,4753 | 2,82534E-24 |

| | | | |
|---|---|---|---|
| ACE2 | XXbac-BPGBPG55C2( | -0,4753 | 2,80776E-24 |
| ACE2 | EHD1 | -0,4755 | 2,67083E-24 |
| ACE2 | RP11-944L7.4 | -0,4756 | 2,62902E-24 |
| ACE2 | C15orf57 | -0,4756 | 2,62541E-24 |
| ACE2 | USB1 | -0,4756 | 2,61861E-24 |
| ACE2 | RP11-1376P16.2 | -0,4756 | 2,63992E-24 |
| ACE2 | RP11-155G14.5 | -0,4757 | 2,59305E-24 |
| ACE2 | RP11-319G9.3 | -0,4757 | 2,54566E-24 |
| ACE2 | MYO1C | -0,4757 | 2,56326E-24 |
| ACE2 | RCAN2 | -0,4758 | 2,49908E-24 |
| ACE2 | PSMC6 | -0,4758 | 2,48005E-24 |
| ACE2 | ZFAND4 | -0,4759 | 2,47067E-24 |
| ACE2 | SYT16 | -0,4759 | 2,44713E-24 |
| ACE2 | HPN | -0,4759 | 2,41994E-24 |
| ACE2 | GHDC | -0,476 | 2,39033E-24 |
| ACE2 | DNAJC2 | -0,4761 | 2,34977E-24 |
| ACE2 | SCAF11 | -0,4761 | 2,33016E-24 |
| ACE2 | RP11-385F5.4 | -0,4762 | 2,25463E-24 |
| ACE2 | PTMA | -0,4762 | 2,29149E-24 |
| ACE2 | RAB40A | -0,4762 | 2,29606E-24 |
| ACE2 | RP1-67A8.3 | -0,4763 | 2,22961E-24 |
| ACE2 | NUS1 | -0,4763 | 2,23026E-24 |
| ACE2 | FBRSL1 | -0,4763 | 2,20683E-24 |
| ACE2 | ZNF573 | -0,4763 | 2,19212E-24 |
| ACE2 | CCT7 | -0,4764 | 2,13551E-24 |
| ACE2 | PIK3R4 | -0,4764 | 2,14369E-24 |
| ACE2 | PDLIM2 | -0,4764 | 2,15078E-24 |
| ACE2 | STEAP3-AS1 | -0,4765 | 2,1256E-24 |
| ACE2 | RP11-571M6.17 | -0,4765 | 2,12691E-24 |
| ACE2 | NTHL1 | -0,4765 | 2,08706E-24 |
| ACE2 | ERI1 | -0,4766 | 2,06838E-24 |
| ACE2 | RP11-1100L3.8 | -0,4766 | 2,02927E-24 |
| ACE2 | RNPC3 | -0,4767 | 1,98389E-24 |
| ACE2 | TMEM173 | -0,4767 | 1,98412E-24 |
| ACE2 | RP11-800A3.7 | -0,4767 | 2,00044E-24 |
| ACE2 | P2RY11 | -0,4767 | 1,97948E-24 |
| ACE2 | CHGB | -0,4767 | 1,99444E-24 |
| ACE2 | P2RX6 | -0,4767 | 2,00372E-24 |
| ACE2 | MTDHP1 | -0,4768 | 1,95025E-24 |
| ACE2 | COCH | -0,4768 | 1,93429E-24 |
| ACE2 | AP000347.2 | -0,4768 | 1,97624E-24 |
| ACE2 | ADH5 | -0,4769 | 1,8988E-24 |
| ACE2 | LINC01105 | -0,477 | 1,87656E-24 |
| ACE2 | AC011380.8 | -0,477 | 1,87259E-24 |
| ACE2 | XPO1 | -0,4771 | 1,81286E-24 |
| ACE2 | RP11-989E6.13 | -0,4771 | 1,81741E-24 |
| ACE2 | CTD-2231E14.2 | -0,4771 | 1,81764E-24 |
| ACE2 | RNF168 | -0,4772 | 1,7453E-24 |
| ACE2 | CD83 | -0,4772 | 1,78427E-24 |
| ACE2 | LINC01719 | -0,4773 | 1,72852E-24 |
| ACE2 | ECT2L | -0,4774 | 1,66621E-24 |
| ACE2 | NUBP1 | -0,4774 | 1,66355E-24 |
| ACE2 | B3GNTL1 | -0,4774 | 1,67664E-24 |
| ACE2 | HACL1 | -0,4775 | 1,64153E-24 |

| | | | |
|---|---|---|---|
| ACE2 | AC004854.5 | -0,4775 | 1,63778E-24 |
| ACE2 | CERS3 | -0,4775 | 1,64544E-24 |
| ACE2 | THAP9 | -0,4776 | 1,61249E-24 |
| ACE2 | PARP8 | -0,4776 | 1,5834E-24 |
| ACE2 | ENO4 | -0,4776 | 1,60279E-24 |
| ACE2 | TRIM5 | -0,4776 | 1,58963E-24 |
| ACE2 | RP11-479J7.2 | -0,4777 | 1,53992E-24 |
| ACE2 | EGLN1 | -0,4777 | 1,54063E-24 |
| ACE2 | RAPGEF2 | -0,4777 | 1,57097E-24 |
| ACE2 | RP11-504A18.1 | -0,4778 | 1,51449E-24 |
| ACE2 | OTOG | -0,4778 | 1,50315E-24 |
| ACE2 | RP11-642A1.2 | -0,4779 | 1,47467E-24 |
| ACE2 | OLFM5P | -0,4779 | 1,47292E-24 |
| ACE2 | HERC2 | -0,4779 | 1,49305E-24 |
| ACE2 | CYP11A1 | -0,4779 | 1,48549E-24 |
| ACE2 | SUPT5H | -0,4779 | 1,46736E-24 |
| ACE2 | NAA20 | -0,4779 | 1,4816E-24 |
| ACE2 | MMGT1 | -0,4779 | 1,46936E-24 |
| ACE2 | SLC2A6 | -0,478 | 1,44141E-24 |
| ACE2 | AGAP1 | -0,4781 | 1,42033E-24 |
| ACE2 | THAP1 | -0,4781 | 1,39115E-24 |
| ACE2 | ADAM12 | -0,4781 | 1,40167E-24 |
| ACE2 | NPPC | -0,4782 | 1,36586E-24 |
| ACE2 | EML5 | -0,4782 | 1,37709E-24 |
| ACE2 | RP11-862G15.2 | -0,4782 | 1,37226E-24 |
| ACE2 | CTA-212A2.4 | -0,4782 | 1,35766E-24 |
| ACE2 | XG | -0,4782 | 1,35757E-24 |
| ACE2 | FOXJ3 | -0,4783 | 1,32157E-24 |
| ACE2 | CUL7 | -0,4783 | 1,33469E-24 |
| ACE2 | AC107081.5 | -0,4784 | 1,2945E-24 |
| ACE2 | DDX10P1 | -0,4784 | 1,30475E-24 |
| ACE2 | MAP2K4 | -0,4784 | 1,29214E-24 |
| ACE2 | HCFC1 | -0,4784 | 1,2903E-24 |
| ACE2 | KLF14 | -0,4785 | 1,27831E-24 |
| ACE2 | RPL29P11 | -0,4786 | 1,23278E-24 |
| ACE2 | MRPL45P2 | -0,4788 | 1,17779E-24 |
| ACE2 | RPS7P14 | -0,4788 | 1,18616E-24 |
| ACE2 | QRICH1 | -0,4789 | 1,15299E-24 |
| ACE2 | CTD-2521M24.13 | -0,479 | 1,123E-24 |
| ACE2 | PRSS35 | -0,4791 | 1,10267E-24 |
| ACE2 | DDX31 | -0,4791 | 1,09126E-24 |
| ACE2 | RP5-1029K10.4 | -0,4791 | 1,09431E-24 |
| ACE2 | KMT2B | -0,4791 | 1,10114E-24 |
| ACE2 | ADRM1 | -0,4791 | 1,09703E-24 |
| ACE2 | F2 | -0,4792 | 1,05519E-24 |
| ACE2 | RP11-395I6.3 | -0,4793 | 1,04815E-24 |
| ACE2 | RPL15P3 | -0,4793 | 1,03489E-24 |
| ACE2 | EEF1E1 | -0,4795 | 9,91358E-25 |
| ACE2 | HRNR | -0,4796 | 9,51114E-25 |
| ACE2 | CTC-338M12.13 | -0,4796 | 9,74195E-25 |
| ACE2 | TRIM60P17 | -0,4796 | 9,70176E-25 |
| ACE2 | RP11-270M14.5 | -0,4796 | 9,50764E-25 |
| ACE2 | RP11-338N10.2 | -0,4797 | 9,29544E-25 |
| ACE2 | MTOR | -0,4797 | 9,37464E-25 |

| | | | |
|---|---|---|---|
| ACE2 | NUP155 | -0,4797 | 9,35215E-25 |
| ACE2 | RP11-374M1.5 | -0,4797 | 9,4913E-25 |
| ACE2 | OR7E13P | -0,4797 | 9,36648E-25 |
| ACE2 | RP11-467L19.16 | -0,4797 | 9,3139E-25 |
| ACE2 | RP11-226L15.5 | -0,4798 | 9,08169E-25 |
| ACE2 | SSBP1 | -0,4798 | 9,1181E-25 |
| ACE2 | RP11-498B4.5 | -0,4798 | 9,1617E-25 |
| ACE2 | RP11-305O6.3 | -0,4798 | 9,11604E-25 |
| ACE2 | RP11-209D14.2 | -0,4798 | 9,1577E-25 |
| ACE2 | TMEM232 | -0,4799 | 8,82134E-25 |
| ACE2 | CACYBP | -0,48 | 8,64793E-25 |
| ACE2 | VKORC1L1 | -0,48 | 8,70612E-25 |
| ACE2 | AQP5 | -0,4801 | 8,55448E-25 |
| ACE2 | DUOXA1 | -0,4801 | 8,54765E-25 |
| ACE2 | CIB2 | -0,4801 | 8,52846E-25 |
| ACE2 | LDOC1L | -0,4802 | 8,20643E-25 |
| ACE2 | CTA-29F11.1 | -0,4802 | 8,27076E-25 |
| ACE2 | ST6GALNAC4P1 | -0,4803 | 8,05744E-25 |
| ACE2 | RPL5P34 | -0,4803 | 8,14184E-25 |
| ACE2 | IP6K3 | -0,4804 | 7,93897E-25 |
| ACE2 | RP11-1000B6.3 | -0,4804 | 7,91345E-25 |
| ACE2 | KCTD13 | -0,4804 | 7,80242E-25 |
| ACE2 | LRRC42 | -0,4805 | 7,69543E-25 |
| ACE2 | PDS5A | -0,4805 | 7,61412E-25 |
| ACE2 | RP4-671O14.6 | -0,4805 | 7,58863E-25 |
| ACE2 | LRRD1 | -0,4806 | 7,41185E-25 |
| ACE2 | ZNF446 | -0,4806 | 7,45681E-25 |
| ACE2 | STAT1 | -0,4807 | 7,28665E-25 |
| ACE2 | LLNLR-285B5.1 | -0,4807 | 7,28544E-25 |
| ACE2 | RP11-1277A3.1 | -0,4808 | 7,00612E-25 |
| ACE2 | OPTN | -0,4809 | 6,90408E-25 |
| ACE2 | PRDM15 | -0,4809 | 6,83382E-25 |
| ACE2 | TBX15 | -0,481 | 6,7796E-25 |
| ACE2 | RPL23AP80 | -0,481 | 6,69183E-25 |
| ACE2 | RPUSD3 | -0,4811 | 6,5709E-25 |
| ACE2 | WDR27 | -0,4811 | 6,64829E-25 |
| ACE2 | CLEC4F | -0,4812 | 6,3621E-25 |
| ACE2 | GPX3 | -0,4812 | 6,42705E-25 |
| ACE2 | AMD1 | -0,4812 | 6,41898E-25 |
| ACE2 | FAM234A | -0,4812 | 6,47912E-25 |
| ACE2 | ANO4 | -0,4813 | 6,22648E-25 |
| ACE2 | NOP58 | -0,4814 | 6,08758E-25 |
| ACE2 | CDC14B | -0,4814 | 6,15226E-25 |
| ACE2 | CUL4A | -0,4814 | 6,15471E-25 |
| ACE2 | LINC02078 | -0,4814 | 6,15965E-25 |
| ACE2 | NUB1 | -0,4815 | 5,92236E-25 |
| ACE2 | ZNF613 | -0,4815 | 5,90416E-25 |
| ACE2 | MAK | -0,4816 | 5,84428E-25 |
| ACE2 | SCAND3P1 | -0,4816 | 5,81641E-25 |
| ACE2 | EPHX1 | -0,4817 | 5,67173E-25 |
| ACE2 | POMC | -0,4817 | 5,5964E-25 |
| ACE2 | CTB-32O4.3 | -0,4819 | 5,35115E-25 |
| ACE2 | ABCE1 | -0,482 | 5,23955E-25 |
| ACE2 | CTD-2562J17.6 | -0,482 | 5,25827E-25 |

| | | | |
|---|---|---|---|
| ACE2 | ZNF526 | -0,482 | 5,16546E-25 |
| ACE2 | RP11-74J13.8 | -0,4821 | 5,10185E-25 |
| ACE2 | FGF14-IT1 | -0,4821 | 5,09457E-25 |
| ACE2 | ME1 | -0,4822 | 4,90074E-25 |
| ACE2 | CTB-58E17.1 | -0,4823 | 4,82344E-25 |
| ACE2 | RP11-54C4.3 | -0,4824 | 4,67377E-25 |
| ACE2 | OR7E38P | -0,4824 | 4,69667E-25 |
| ACE2 | TMEM14B | -0,4825 | 4,60186E-25 |
| ACE2 | KEL | -0,4825 | 4,61253E-25 |
| ACE2 | EMP2 | -0,4825 | 4,58987E-25 |
| ACE2 | RP4-671G15.2 | -0,4826 | 4,46114E-25 |
| ACE2 | SESN1 | -0,4826 | 4,49915E-25 |
| ACE2 | TNRC6B | -0,4826 | 4,44379E-25 |
| ACE2 | TBCK | -0,4827 | 4,41378E-25 |
| ACE2 | SUMO4 | -0,4827 | 4,34586E-25 |
| ACE2 | SLC30A8 | -0,4827 | 4,35374E-25 |
| ACE2 | TOPORS | -0,4827 | 4,34474E-25 |
| ACE2 | CHST1 | -0,4827 | 4,32815E-25 |
| ACE2 | SLC38A7 | -0,4828 | 4,24156E-25 |
| ACE2 | SNX2 | -0,4829 | 4,1616E-25 |
| ACE2 | PROX2 | -0,483 | 4,05461E-25 |
| ACE2 | ZNF747 | -0,483 | 4,08494E-25 |
| ACE2 | LINC01902 | -0,483 | 4,04326E-25 |
| ACE2 | RAE1 | -0,483 | 4,05981E-25 |
| ACE2 | RP11-788M5.3 | -0,4831 | 3,92609E-25 |
| ACE2 | ARHGEF7-AS2 | -0,4831 | 3,90354E-25 |
| ACE2 | LOXHD1 | -0,4831 | 3,92755E-25 |
| ACE2 | HNRNPH2 | -0,4831 | 3,98237E-25 |
| ACE2 | LRP12 | -0,4832 | 3,8172E-25 |
| ACE2 | APTX | -0,4832 | 3,84203E-25 |
| ACE2 | FBXL4 | -0,4834 | 3,62541E-25 |
| ACE2 | MON2 | -0,4834 | 3,68529E-25 |
| ACE2 | ATF4P3 | -0,4834 | 3,69303E-25 |
| ACE2 | ARL5AP3 | -0,4835 | 3,5362E-25 |
| ACE2 | LINC01616 | -0,4835 | 3,5693E-25 |
| ACE2 | ETFBKMT | -0,4835 | 3,51232E-25 |
| ACE2 | GAA | -0,4836 | 3,48693E-25 |
| ACE2 | SCN9A | -0,4837 | 3,34202E-25 |
| ACE2 | COX17 | -0,4837 | 3,36692E-25 |
| ACE2 | PTP4A2P2 | -0,4837 | 3,35107E-25 |
| ACE2 | RC3H1 | -0,4838 | 3,29626E-25 |
| ACE2 | KIAA0825 | -0,4839 | 3,18828E-25 |
| ACE2 | STAG3 | -0,4839 | 3,22074E-25 |
| ACE2 | SEC31B | -0,4839 | 3,20551E-25 |
| ACE2 | FAM134C | -0,4839 | 3,24351E-25 |
| ACE2 | A2MP1 | -0,484 | 3,09514E-25 |
| ACE2 | RP11-963H4.6 | -0,484 | 3,14804E-25 |
| ACE2 | AC005785.2 | -0,484 | 3,10029E-25 |
| ACE2 | GTDC1 | -0,4841 | 3,05227E-25 |
| ACE2 | KCNK9 | -0,4841 | 3,07663E-25 |
| ACE2 | ORAOV1 | -0,4841 | 3,06773E-25 |
| ACE2 | RBM26 | -0,4841 | 3,08409E-25 |
| ACE2 | LINC02157 | -0,4841 | 3,04204E-25 |
| ACE2 | TCTE1 | -0,4842 | 2,96614E-25 |

| | | | |
|---|---|---|---|
| ACE2 | CPSF2 | -0,4842 | 2,97275E-25 |
| ACE2 | RP11-736N17.10 | -0,4842 | 2,97993E-25 |
| ACE2 | RP11-140I16.3 | -0,4844 | 2,84066E-25 |
| ACE2 | CTD-3092A11.1 | -0,4844 | 2,83314E-25 |
| ACE2 | MFSD12 | -0,4844 | 2,81614E-25 |
| ACE2 | AADACL2 | -0,4845 | 2,77563E-25 |
| ACE2 | FAM153A | -0,4845 | 2,76073E-25 |
| ACE2 | ZNF266 | -0,4847 | 2,63302E-25 |
| ACE2 | EXOC3-AS1 | -0,4848 | 2,54229E-25 |
| ACE2 | LINC01271 | -0,4848 | 2,52329E-25 |
| ACE2 | MTMR7 | -0,4849 | 2,48068E-25 |
| ACE2 | CTD-2325P2.3 | -0,4849 | 2,48725E-25 |
| ACE2 | B4GALT6 | -0,4849 | 2,45664E-25 |
| ACE2 | LRFN3 | -0,4849 | 2,48879E-25 |
| ACE2 | AC063976.7 | -0,485 | 2,44589E-25 |
| ACE2 | TPP1 | -0,485 | 2,42158E-25 |
| ACE2 | EEF1DP3 | -0,485 | 2,39892E-25 |
| ACE2 | LINC02060 | -0,4851 | 2,33126E-25 |
| ACE2 | SERPINB6 | -0,4851 | 2,32577E-25 |
| ACE2 | TP53BP2 | -0,4852 | 2,31678E-25 |
| ACE2 | NCK2 | -0,4852 | 2,30723E-25 |
| ACE2 | EIF3KP1 | -0,4852 | 2,31906E-25 |
| ACE2 | RP11-515O17.2 | -0,4852 | 2,26693E-25 |
| ACE2 | RP13-20L14.1 | -0,4852 | 2,29223E-25 |
| ACE2 | CTC-325H20.7 | -0,4853 | 2,21862E-25 |
| ACE2 | GLRA3 | -0,4855 | 2,10993E-25 |
| ACE2 | RP11-403A3.2 | -0,4855 | 2,14787E-25 |
| ACE2 | LINC02185 | -0,4855 | 2,14068E-25 |
| ACE2 | TP53RK | -0,4855 | 2,13983E-25 |
| ACE2 | BMS1 | -0,4856 | 2,06423E-25 |
| ACE2 | SKIV2L2 | -0,4857 | 2,04184E-25 |
| ACE2 | LRRC40 | -0,4858 | 1,98465E-25 |
| ACE2 | LL09NC01-139C3.1 | -0,4858 | 1,98589E-25 |
| ACE2 | ATAD1 | -0,4858 | 1,98749E-25 |
| ACE2 | RP4-607I7.1 | -0,4858 | 1,95868E-25 |
| ACE2 | RP11-227G15.9 | -0,4858 | 1,97877E-25 |
| ACE2 | BX322557.10 | -0,4858 | 1,98208E-25 |
| ACE2 | PPP3CA | -0,4859 | 1,91685E-25 |
| ACE2 | POLD2 | -0,4859 | 1,93024E-25 |
| ACE2 | RAB39B | -0,4859 | 1,92594E-25 |
| ACE2 | ZNF281 | -0,486 | 1,87899E-25 |
| ACE2 | IMPDH2 | -0,486 | 1,84865E-25 |
| ACE2 | RP11-290M5.4 | -0,4861 | 1,80079E-25 |
| ACE2 | RP11-20O24.4 | -0,4861 | 1,82212E-25 |
| ACE2 | PHACTR1 | -0,4861 | 1,79603E-25 |
| ACE2 | ITGA11 | -0,4861 | 1,80594E-25 |
| ACE2 | RP11-758H9.2 | -0,4861 | 1,80224E-25 |
| ACE2 | RP11-495P10.1 | -0,4862 | 1,75938E-25 |
| ACE2 | SURF1 | -0,4862 | 1,78361E-25 |
| ACE2 | NT5E | -0,4863 | 1,73412E-25 |
| ACE2 | BORCS5 | -0,4863 | 1,71667E-25 |
| ACE2 | RP11-93B14.4 | -0,4863 | 1,71346E-25 |
| ACE2 | CTD-2373J6.1 | -0,4864 | 1,69488E-25 |
| ACE2 | ZNF334 | -0,4864 | 1,68835E-25 |

| | | | |
|---|---|---|---|
| ACE2 | MST1P2 | -0,4865 | 1,63727E-25 |
| ACE2 | CEP83 | -0,4865 | 1,63624E-25 |
| ACE2 | TLX2 | -0,4867 | 1,54465E-25 |
| ACE2 | EEF1A1P19 | -0,4867 | 1,56293E-25 |
| ACE2 | ZNF277 | -0,4867 | 1,5529E-25 |
| ACE2 | PHF5A | -0,4867 | 1,55947E-25 |
| ACE2 | HMGB1P31 | -0,4868 | 1,51848E-25 |
| ACE2 | NPHP3 | -0,4868 | 1,535E-25 |
| ACE2 | RP11-543P15.1 | -0,4868 | 1,51092E-25 |
| ACE2 | PSMB2 | -0,4869 | 1,48369E-25 |
| ACE2 | CAND1 | -0,4869 | 1,46048E-25 |
| ACE2 | AC079613.1 | -0,487 | 1,44541E-25 |
| ACE2 | SLC11A2 | -0,487 | 1,42642E-25 |
| ACE2 | PLXNA1 | -0,4871 | 1,41179E-25 |
| ACE2 | GCOM2 | -0,4871 | 1,40558E-25 |
| ACE2 | L3HYPDH | -0,4871 | 1,39357E-25 |
| ACE2 | RP11-89C3.4 | -0,4872 | 1,37208E-25 |
| ACE2 | RPL24P2 | -0,4872 | 1,37996E-25 |
| ACE2 | LINC00176 | -0,4873 | 1,34484E-25 |
| ACE2 | NBPF14 | -0,4874 | 1,28586E-25 |
| ACE2 | RP11-399B17.1 | -0,4874 | 1,28847E-25 |
| ACE2 | AP001596.6 | -0,4874 | 1,30685E-25 |
| ACE2 | GAS2L1 | -0,4875 | 1,2652E-25 |
| ACE2 | HMGB1P5 | -0,4876 | 1,23685E-25 |
| ACE2 | FAM196B | -0,4876 | 1,21775E-25 |
| ACE2 | CTD-2292P10.4 | -0,4876 | 1,24017E-25 |
| ACE2 | RP11-182J1.13 | -0,4876 | 1,24708E-25 |
| ACE2 | KDM4B | -0,4876 | 1,21767E-25 |
| ACE2 | DDX18 | -0,4878 | 1,16211E-25 |
| ACE2 | PCDHA5 | -0,4878 | 1,15743E-25 |
| ACE2 | COL26A1 | -0,4878 | 1,18145E-25 |
| ACE2 | RP11-386M24.4 | -0,4878 | 1,17679E-25 |
| ACE2 | VPS53 | -0,4878 | 1,17251E-25 |
| ACE2 | RPS19 | -0,4878 | 1,16907E-25 |
| ACE2 | PQLC2L | -0,4879 | 1,13838E-25 |
| ACE2 | ARHGAP21 | -0,4881 | 1,07898E-25 |
| ACE2 | ZMIZ1-AS1 | -0,4881 | 1,08334E-25 |
| ACE2 | NRDE2 | -0,4882 | 1,04041E-25 |
| ACE2 | ELAC2 | -0,4882 | 1,06461E-25 |
| ACE2 | RP1-164F3.8 | -0,4883 | 1,01631E-25 |
| ACE2 | RP1-283K11.2 | -0,4885 | 9,6566E-26 |
| ACE2 | RP11-530C5.4 | -0,4885 | 9,79968E-26 |
| ACE2 | ZNF770 | -0,4885 | 9,74266E-26 |
| ACE2 | YIPF6 | -0,4885 | 9,65298E-26 |
| ACE2 | RP11-1134I14.4 | -0,4886 | 9,55064E-26 |
| ACE2 | ZNF202 | -0,4887 | 9,35223E-26 |
| ACE2 | NDUFAF1 | -0,4887 | 9,25043E-26 |
| ACE2 | UBL4A | -0,4887 | 9,29598E-26 |
| ACE2 | COG5 | -0,4888 | 9,04182E-26 |
| ACE2 | RP11-574K11.24 | -0,4888 | 8,94858E-26 |
| ACE2 | PXYLP1 | -0,489 | 8,57767E-26 |
| ACE2 | PIWIL2 | -0,489 | 8,4574E-26 |
| ACE2 | RP5-1021I20.5 | -0,489 | 8,57572E-26 |
| ACE2 | RP11-461M2.2 | -0,4891 | 8,38382E-26 |

| | | | |
|---|---|---|---|
| ACE2 | EGR3 | -0,4891 | 8,33383E-26 |
| ACE2 | KIF1BP | -0,4891 | 8,25023E-26 |
| ACE2 | MRPL18 | -0,4892 | 8,15626E-26 |
| ACE2 | FAN1 | -0,4892 | 8,02302E-26 |
| ACE2 | C19orf60 | -0,4892 | 8,20585E-26 |
| ACE2 | NUP205 | -0,4893 | 7,85741E-26 |
| ACE2 | LAMP1 | -0,4893 | 7,93415E-26 |
| ACE2 | RP11-462G12.1 | -0,4894 | 7,60633E-26 |
| ACE2 | RP11-665C14.2 | -0,4896 | 7,27456E-26 |
| ACE2 | CYP39A1 | -0,4896 | 7,36979E-26 |
| ACE2 | DENND5B-AS1 | -0,4896 | 7,27217E-26 |
| ACE2 | CTD-2651B20.4 | -0,4896 | 7,26673E-26 |
| ACE2 | RP11-486I11.2 | -0,4896 | 7,34747E-26 |
| ACE2 | NANP | -0,4896 | 7,21714E-26 |
| ACE2 | RP11-757G1.5 | -0,4897 | 7,14166E-26 |
| ACE2 | RP11-507K2.3 | -0,4897 | 7,1734E-26 |
| ACE2 | AHCYL1 | -0,4898 | 6,84165E-26 |
| ACE2 | RP11-357H14.16 | -0,4901 | 6,43723E-26 |
| ACE2 | RP4-545K15.5 | -0,4901 | 6,46267E-26 |
| ACE2 | UNC50 | -0,4902 | 6,31216E-26 |
| ACE2 | ATP6V0E1 | -0,4902 | 6,29847E-26 |
| ACE2 | RP11-131L12.4 | -0,4903 | 6,02523E-26 |
| ACE2 | TOP3A | -0,4903 | 6,07917E-26 |
| ACE2 | GJA5 | -0,4904 | 5,94089E-26 |
| ACE2 | CISTR | -0,4904 | 5,86408E-26 |
| ACE2 | GLTSCR2-AS1 | -0,4904 | 5,90449E-26 |
| ACE2 | RP11-98I9.4 | -0,4905 | 5,71919E-26 |
| ACE2 | PGS1 | -0,4906 | 5,5913E-26 |
| ACE2 | UBL5 | -0,4906 | 5,61002E-26 |
| ACE2 | RP11-281O15.8 | -0,4907 | 5,41476E-26 |
| ACE2 | RP11-3B12.4 | -0,4907 | 5,54085E-26 |
| ACE2 | ZNF616 | -0,4907 | 5,48636E-26 |
| ACE2 | LINC01102 | -0,4909 | 5,20478E-26 |
| ACE2 | RP11-147L13.12 | -0,4909 | 5,19816E-26 |
| ACE2 | RP1-161N10.1 | -0,4909 | 5,23085E-26 |
| ACE2 | CDK11B | -0,491 | 4,98844E-26 |
| ACE2 | MEIS3P2 | -0,491 | 5,05788E-26 |
| ACE2 | RNF138P1 | -0,4911 | 4,9783E-26 |
| ACE2 | RP11-335K5.2 | -0,4911 | 4,93815E-26 |
| ACE2 | KCNF1 | -0,4912 | 4,85477E-26 |
| ACE2 | KCTD4 | -0,4912 | 4,756E-26 |
| ACE2 | HEMK1 | -0,4913 | 4,6783E-26 |
| ACE2 | ACTBP2 | -0,4913 | 4,67957E-26 |
| ACE2 | MPRIPP1 | -0,4914 | 4,60164E-26 |
| ACE2 | FOXO4 | -0,4915 | 4,37907E-26 |
| ACE2 | RNF145 | -0,4917 | 4,23411E-26 |
| ACE2 | CILP | -0,4917 | 4,19585E-26 |
| ACE2 | DBNL | -0,4918 | 4,06E-26 |
| ACE2 | MTDH | -0,4918 | 4,1078E-26 |
| ACE2 | AC100830.4 | -0,4918 | 4,06757E-26 |
| ACE2 | WASH4P | -0,4918 | 4,07157E-26 |
| ACE2 | AC097721.2 | -0,4919 | 3,93696E-26 |
| ACE2 | ZDHHC14 | -0,4919 | 3,95285E-26 |
| ACE2 | LINC00994 | -0,492 | 3,89003E-26 |

| | | | |
|---|---|---|---|
| ACE2 | CFAP221 | -0,4921 | 3,7563E-26 |
| ACE2 | DAB2 | -0,4921 | 3,76601E-26 |
| ACE2 | CCNYL2 | -0,4921 | 3,79248E-26 |
| ACE2 | GS1-304P7.1 | -0,4922 | 3,6816E-26 |
| ACE2 | C2orf16 | -0,4922 | 3,69538E-26 |
| ACE2 | MAPT-AS1 | -0,4922 | 3,65348E-26 |
| ACE2 | SUN2 | -0,4922 | 3,70209E-26 |
| ACE2 | RP6-24A23.7 | -0,4922 | 3,71192E-26 |
| ACE2 | PUM2 | -0,4923 | 3,57018E-26 |
| ACE2 | FBLN7 | -0,4923 | 3,56814E-26 |
| ACE2 | NFIL3 | -0,4923 | 3,54339E-26 |
| ACE2 | RP11-262I2.2 | -0,4923 | 3,56077E-26 |
| ACE2 | RP11-326C3.13 | -0,4923 | 3,56156E-26 |
| ACE2 | RP11-646I6.6 | -0,4924 | 3,53029E-26 |
| ACE2 | RP11-45A17.4 | -0,4924 | 3,44859E-26 |
| ACE2 | RP11-417L19.5 | -0,4924 | 3,52083E-26 |
| ACE2 | ANKLE2 | -0,4924 | 3,49735E-26 |
| ACE2 | PNRC2 | -0,4925 | 3,3581E-26 |
| ACE2 | RP4-614C10.2 | -0,4925 | 3,43019E-26 |
| ACE2 | WDR5B | -0,4925 | 3,3872E-26 |
| ACE2 | LA16c-390E6.4 | -0,4925 | 3,36572E-26 |
| ACE2 | STON1-GTF2A1L | -0,4926 | 3,34205E-26 |
| ACE2 | RP11-696F12.1 | -0,4926 | 3,30937E-26 |
| ACE2 | RP11-158H5.8 | -0,4926 | 3,27892E-26 |
| ACE2 | HTR1E | -0,4927 | 3,2257E-26 |
| ACE2 | RPS15AP38 | -0,4927 | 3,19065E-26 |
| ACE2 | PRPS2 | -0,4927 | 3,25907E-26 |
| ACE2 | USP34 | -0,4929 | 3,08496E-26 |
| ACE2 | EEPD1 | -0,4929 | 3,02694E-26 |
| ACE2 | AC022153.1 | -0,4929 | 3,07765E-26 |
| ACE2 | LINC01285 | -0,4929 | 3,04745E-26 |
| ACE2 | PLAC9P1 | -0,493 | 2,959E-26 |
| ACE2 | CDH18 | -0,493 | 3,00129E-26 |
| ACE2 | XRRA1 | -0,493 | 2,96464E-26 |
| ACE2 | DGCR6 | -0,493 | 2,97208E-26 |
| ACE2 | PPIE | -0,4931 | 2,86945E-26 |
| ACE2 | C1orf194 | -0,4931 | 2,92417E-26 |
| ACE2 | ICAM1 | -0,4931 | 2,87182E-26 |
| ACE2 | RHOQP2 | -0,4932 | 2,85353E-26 |
| ACE2 | EIF4E3 | -0,4932 | 2,84122E-26 |
| ACE2 | TBRG1 | -0,4932 | 2,82216E-26 |
| ACE2 | AP000345.4 | -0,4932 | 2,82641E-26 |
| ACE2 | CHRNE | -0,4933 | 2,74905E-26 |
| ACE2 | PTGES | -0,4934 | 2,67532E-26 |
| ACE2 | RP11-325L12.6 | -0,4934 | 2,66855E-26 |
| ACE2 | PEMT | -0,4934 | 2,68908E-26 |
| ACE2 | FOXS1 | -0,4934 | 2,65148E-26 |
| ACE2 | SLC12A5 | -0,4934 | 2,69462E-26 |
| ACE2 | ESRRB | -0,4935 | 2,63485E-26 |
| ACE2 | AC007038.7 | -0,4936 | 2,52488E-26 |
| ACE2 | ZNRF2P2 | -0,4936 | 2,51584E-26 |
| ACE2 | RP11-626H12.1 | -0,4936 | 2,51489E-26 |
| ACE2 | ACTR6 | -0,4936 | 2,51837E-26 |
| ACE2 | CABP1 | -0,4936 | 2,52238E-26 |

| | | | |
|---|---|---|---|
| ACE2 | FANCM | -0,4936 | 2,55522E-26 |
| ACE2 | CTD-2561B21.11 | -0,4936 | 2,55191E-26 |
| ACE2 | EEF1A1P13 | -0,4937 | 2,50401E-26 |
| ACE2 | ZBED3-AS1 | -0,4938 | 2,38823E-26 |
| ACE2 | RP11-708J19.1 | -0,4939 | 2,35635E-26 |
| ACE2 | RP11-77K12.9 | -0,4939 | 2,32143E-26 |
| ACE2 | TSKS | -0,4939 | 2,34289E-26 |
| ACE2 | KCMF1 | -0,494 | 2,25431E-26 |
| ACE2 | MTTP | -0,494 | 2,30684E-26 |
| ACE2 | LINC01948 | -0,494 | 2,2578E-26 |
| ACE2 | PSPC1 | -0,494 | 2,31013E-26 |
| ACE2 | RP11-888D10.4 | -0,494 | 2,27782E-26 |
| ACE2 | RPL10P15 | -0,494 | 2,31132E-26 |
| ACE2 | RP11-136C24.2 | -0,4942 | 2,14122E-26 |
| ACE2 | NR4A3 | -0,4942 | 2,14357E-26 |
| ACE2 | BMP1 | -0,4943 | 2,13465E-26 |
| ACE2 | C22orf46 | -0,4943 | 2,12579E-26 |
| ACE2 | AC093616.4 | -0,4944 | 2,05611E-26 |
| ACE2 | DCPS | -0,4944 | 2,07249E-26 |
| ACE2 | CTDNEP1 | -0,4944 | 2,02651E-26 |
| ACE2 | IMPACT | -0,4944 | 2,07494E-26 |
| ACE2 | RANBP1 | -0,4944 | 2,05695E-26 |
| ACE2 | STIP1 | -0,4945 | 2,00898E-26 |
| ACE2 | CSPG4P12 | -0,4945 | 1,98086E-26 |
| ACE2 | ZNF713 | -0,4946 | 1,97082E-26 |
| ACE2 | RP11-251G23.5 | -0,4946 | 1,92815E-26 |
| ACE2 | SVILP1 | -0,4947 | 1,91978E-26 |
| ACE2 | BTBD16 | -0,4947 | 1,91749E-26 |
| ACE2 | RP11-793H13.11 | -0,4947 | 1,88948E-26 |
| ACE2 | MFSD2B | -0,4948 | 1,85747E-26 |
| ACE2 | ATG10 | -0,4948 | 1,86608E-26 |
| ACE2 | ZNF603P | -0,4948 | 1,82827E-26 |
| ACE2 | LINC01587 | -0,4949 | 1,80169E-26 |
| ACE2 | CTB-51J22.1 | -0,4949 | 1,79948E-26 |
| ACE2 | RPL17P50 | -0,4949 | 1,81642E-26 |
| ACE2 | CBX4 | -0,4949 | 1,81375E-26 |
| ACE2 | CTD-2194A8.2 | -0,4951 | 1,7068E-26 |
| ACE2 | NUP93 | -0,4951 | 1,72487E-26 |
| ACE2 | AC009404.2 | -0,4952 | 1,65115E-26 |
| ACE2 | TAOK2 | -0,4952 | 1,66925E-26 |
| ACE2 | RP5-890E16.5 | -0,4952 | 1,65578E-26 |
| ACE2 | COL4A5 | -0,4952 | 1,66599E-26 |
| ACE2 | RP4-535B20.4 | -0,4953 | 1,60901E-26 |
| ACE2 | RP11-111H13.1 | -0,4953 | 1,61583E-26 |
| ACE2 | AC004967.7 | -0,4953 | 1,62297E-26 |
| ACE2 | RPL14P1 | -0,4953 | 1,61399E-26 |
| ACE2 | MRPL28 | -0,4953 | 1,60443E-26 |
| ACE2 | B3GALT6 | -0,4954 | 1,57812E-26 |
| ACE2 | FAM98B | -0,4954 | 1,59216E-26 |
| ACE2 | ZSCAN2 | -0,4954 | 1,56672E-26 |
| ACE2 | RP11-643C9.2 | -0,4955 | 1,51214E-26 |
| ACE2 | HOXC11 | -0,4955 | 1,53289E-26 |
| ACE2 | HSD17B1 | -0,4955 | 1,54862E-26 |
| ACE2 | ZNF628 | -0,4956 | 1,5028E-26 |

| | | | |
|---|---|---|---|
| ACE2 | LHFPL4 | -0,4957 | 1,45536E-26 |
| ACE2 | EGR2 | -0,4957 | 1,44364E-26 |
| ACE2 | DAXX | -0,4958 | 1,39924E-26 |
| ACE2 | RP11-21B21.4 | -0,4958 | 1,42098E-26 |
| ACE2 | GATSL2 | -0,4959 | 1,36606E-26 |
| ACE2 | ZNF544 | -0,4959 | 1,35932E-26 |
| ACE2 | RP5-1024N4.5 | -0,4961 | 1,31391E-26 |
| ACE2 | ZFC3H1 | -0,4961 | 1,29089E-26 |
| ACE2 | SSR2 | -0,4962 | 1,26E-26 |
| ACE2 | SUCO | -0,4962 | 1,26795E-26 |
| ACE2 | CTD-2530N21.4 | -0,4962 | 1,2618E-26 |
| ACE2 | PAPD5 | -0,4962 | 1,27644E-26 |
| ACE2 | CTD-3113P16.11 | -0,4962 | 1,28508E-26 |
| ACE2 | LZTS3 | -0,4962 | 1,27769E-26 |
| ACE2 | FGD5P1 | -0,4963 | 1,22085E-26 |
| ACE2 | YTHDC2 | -0,4963 | 1,22992E-26 |
| ACE2 | AC007193.10 | -0,4963 | 1,23006E-26 |
| ACE2 | GAS5 | -0,4964 | 1,19147E-26 |
| ACE2 | RP11-131M11.2 | -0,4965 | 1,17081E-26 |
| ACE2 | RP11-90C4.1 | -0,4965 | 1,17957E-26 |
| ACE2 | LINC01139 | -0,4965 | 1,16812E-26 |
| ACE2 | FXN | -0,4965 | 1,16666E-26 |
| ACE2 | FIGNL2 | -0,4965 | 1,16132E-26 |
| ACE2 | C5orf51 | -0,4966 | 1,1355E-26 |
| ACE2 | STIM1 | -0,4966 | 1,14003E-26 |
| ACE2 | CCND2-AS2 | -0,4966 | 1,12698E-26 |
| ACE2 | TIA1 | -0,4967 | 1,1029E-26 |
| ACE2 | LARP1 | -0,4967 | 1,10012E-26 |
| ACE2 | MORN3 | -0,4967 | 1,12438E-26 |
| ACE2 | NEK5 | -0,4967 | 1,10459E-26 |
| ACE2 | RNF111 | -0,4967 | 1,11337E-26 |
| ACE2 | C1orf50 | -0,4968 | 1,08001E-26 |
| ACE2 | SRA1 | -0,4968 | 1,06876E-26 |
| ACE2 | CPEB4 | -0,4968 | 1,08514E-26 |
| ACE2 | NRROS | -0,4969 | 1,04553E-26 |
| ACE2 | RP11-53O19.3 | -0,4969 | 1,05935E-26 |
| ACE2 | MARCH5 | -0,4969 | 1,05912E-26 |
| ACE2 | SERPINA5 | -0,4969 | 1,04346E-26 |
| ACE2 | WI2-1896O14.1 | -0,497 | 1,03021E-26 |
| ACE2 | RP11-255C15.3 | -0,497 | 1,01251E-26 |
| ACE2 | ZNF815P | -0,497 | 1,0241E-26 |
| ACE2 | KMT5A | -0,497 | 1,02073E-26 |
| ACE2 | SETDB1 | -0,4971 | 1,00456E-26 |
| ACE2 | PDCD2 | -0,4971 | 1,00522E-26 |
| ACE2 | TTPAL | -0,4971 | 9,92399E-27 |
| ACE2 | DPH3 | -0,4972 | 9,74432E-27 |
| ACE2 | PSTPIP2 | -0,4972 | 9,62905E-27 |
| ACE2 | MTCO3P43 | -0,4974 | 9,28121E-27 |
| ACE2 | MATN1 | -0,4975 | 8,85136E-27 |
| ACE2 | STRA6 | -0,4975 | 8,84377E-27 |
| ACE2 | RP11-514D23.3 | -0,4975 | 9,01748E-27 |
| ACE2 | RPGR | -0,4975 | 9,06261E-27 |
| ACE2 | GDF7 | -0,4976 | 8,70295E-27 |
| ACE2 | VTI1B | -0,4976 | 8,80668E-27 |

| | | | |
|---|---|---|---|
| ACE2 | TMEM165 | -0,4977 | 8,40368E-27 |
| ACE2 | HBEGF | -0,4977 | 8,45807E-27 |
| ACE2 | MRPL51 | -0,4977 | 8,37755E-27 |
| ACE2 | CTC-591M7.1 | -0,4977 | 8,54452E-27 |
| ACE2 | RP11-588H23.3 | -0,4978 | 8,2344E-27 |
| ACE2 | CTD-2236F14.1 | -0,4979 | 8,12284E-27 |
| ACE2 | CSNK1A1P1 | -0,4979 | 7,97799E-27 |
| ACE2 | SNRNP25 | -0,498 | 7,90189E-27 |
| ACE2 | RP11-797H7.1 | -0,4981 | 7,71859E-27 |
| ACE2 | MGC16275 | -0,4981 | 7,68295E-27 |
| ACE2 | RP11-932O9.9 | -0,4982 | 7,36486E-27 |
| ACE2 | RP11-327J17.2 | -0,4982 | 7,49633E-27 |
| ACE2 | CTB-33G10.1 | -0,4982 | 7,49176E-27 |
| ACE2 | TCERG1 | -0,4983 | 7,30678E-27 |
| ACE2 | CTGF | -0,4983 | 7,24072E-27 |
| ACE2 | RP1-310O13.7 | -0,4983 | 7,20656E-27 |
| ACE2 | NDRG2 | -0,4984 | 6,97804E-27 |
| ACE2 | RP11-643M14.1 | -0,4984 | 7,06327E-27 |
| ACE2 | MRPS23 | -0,4984 | 6,98598E-27 |
| ACE2 | TLK2 | -0,4984 | 7,10071E-27 |
| ACE2 | ZMYND10 | -0,4985 | 6,85145E-27 |
| ACE2 | RP11-48B3.5 | -0,4985 | 6,75684E-27 |
| ACE2 | NOL6 | -0,4985 | 6,87774E-27 |
| ACE2 | PPP1R8 | -0,4986 | 6,65082E-27 |
| ACE2 | HIST2H2BE | -0,4986 | 6,59641E-27 |
| ACE2 | GAREM2 | -0,4986 | 6,58325E-27 |
| ACE2 | XPO4 | -0,4986 | 6,64675E-27 |
| ACE2 | OSER1 | -0,4986 | 6,66638E-27 |
| ACE2 | RBM48 | -0,4988 | 6,37067E-27 |
| ACE2 | LMNTD1 | -0,4988 | 6,33733E-27 |
| ACE2 | LRRC52 | -0,4989 | 6,17822E-27 |
| ACE2 | LINC01695 | -0,4989 | 6,19449E-27 |
| ACE2 | RP11-14N7.2 | -0,499 | 5,94544E-27 |
| ACE2 | SPICE1 | -0,499 | 6,02536E-27 |
| ACE2 | PDCD6IPP2 | -0,499 | 6,0421E-27 |
| ACE2 | PPP4R3B | -0,4991 | 5,83611E-27 |
| ACE2 | KCNA4 | -0,4991 | 5,78733E-27 |
| ACE2 | XRCC3 | -0,4991 | 5,8313E-27 |
| ACE2 | CTD-3065B20.3 | -0,4991 | 5,76077E-27 |
| ACE2 | PRKDC | -0,4992 | 5,62063E-27 |
| ACE2 | RP11-834C11.11 | -0,4992 | 5,6897E-27 |
| ACE2 | RP11-76C10.4 | -0,4993 | 5,49122E-27 |
| ACE2 | GSC | -0,4993 | 5,50269E-27 |
| ACE2 | RUNDC3A-AS1 | -0,4993 | 5,5357E-27 |
| ACE2 | C6 | -0,4994 | 5,29802E-27 |
| ACE2 | UBE2D2 | -0,4994 | 5,35028E-27 |
| ACE2 | ZNF26 | -0,4994 | 5,34041E-27 |
| ACE2 | RBL1 | -0,4994 | 5,40917E-27 |
| ACE2 | CFAP45 | -0,4995 | 5,26261E-27 |
| ACE2 | ADORA2A | -0,4995 | 5,19216E-27 |
| ACE2 | SNED1 | -0,4996 | 5,0458E-27 |
| ACE2 | FMR1 | -0,4996 | 5,1159E-27 |
| ACE2 | BAG5 | -0,4997 | 4,92606E-27 |
| ACE2 | TMEM147 | -0,4998 | 4,7824E-27 |

| | | | |
|---|---|---|---|
| ACE2 | RP11-791G15.2 | -0,4999 | 4,63008E-27 |
| ACE2 | DNAJB12 | -0,4999 | 4,67414E-27 |
| ACE2 | RP11-179A10.1 | -0,5 | 4,49716E-27 |
| ACE2 | RBM10 | -0,5 | 4,57893E-27 |
| ACE2 | RP11-425L10.1 | -0,5001 | 4,45061E-27 |
| ACE2 | HNRNPM | -0,5001 | 4,40025E-27 |
| ACE2 | CTC-451P13.1 | -0,5002 | 4,35169E-27 |
| ACE2 | CD36 | -0,5002 | 4,33243E-27 |
| ACE2 | TSTD2 | -0,5002 | 4,34833E-27 |
| ACE2 | C4orf46 | -0,5003 | 4,19528E-27 |
| ACE2 | RP11-685M7.3 | -0,5003 | 4,20845E-27 |
| ACE2 | MANEA-AS1 | -0,5004 | 4,09737E-27 |
| ACE2 | TCEA1P4 | -0,5004 | 4,06306E-27 |
| ACE2 | RP11-686D22.5 | -0,5004 | 4,09265E-27 |
| ACE2 | CTD-2319I12.10 | -0,5004 | 4,13885E-27 |
| ACE2 | CCDC178 | -0,5004 | 4,1239E-27 |
| ACE2 | PCNX2 | -0,5005 | 3,94759E-27 |
| ACE2 | GPBAR1 | -0,5005 | 4,0254E-27 |
| ACE2 | MAP1LC3B | -0,5005 | 4,01732E-27 |
| ACE2 | MZT2A | -0,5006 | 3,89764E-27 |
| ACE2 | ITGAE | -0,5006 | 3,86852E-27 |
| ACE2 | ABCA4 | -0,5007 | 3,79932E-27 |
| ACE2 | GS1-304P7.2 | -0,5008 | 3,6663E-27 |
| ACE2 | NOD1 | -0,5009 | 3,56987E-27 |
| ACE2 | UBE2V1P1 | -0,5009 | 3,52356E-27 |
| ACE2 | RP5-1119A7.17 | -0,5009 | 3,54044E-27 |
| ACE2 | XXbac-B444P24.13 | -0,501 | 3,49161E-27 |
| ACE2 | PCDH19 | -0,501 | 3,43042E-27 |
| ACE2 | ADAMTS2 | -0,5011 | 3,39651E-27 |
| ACE2 | CTC-338M12.9 | -0,5011 | 3,39551E-27 |
| ACE2 | CDYL | -0,5011 | 3,38374E-27 |
| ACE2 | XXbac-BPG170G13.32 | -0,5011 | 3,37365E-27 |
| ACE2 | CTD-2207L17.1 | -0,5013 | 3,18928E-27 |
| ACE2 | RP11-147I3.1 | -0,5013 | 3,2318E-27 |
| ACE2 | ZNF644 | -0,5014 | 3,07759E-27 |
| ACE2 | PPP1R7 | -0,5014 | 3,08793E-27 |
| ACE2 | RP11-329B9.3 | -0,5014 | 3,13523E-27 |
| ACE2 | UGDH-AS1 | -0,5016 | 2,92025E-27 |
| ACE2 | INTS2 | -0,5016 | 2,96173E-27 |
| ACE2 | RP11-60A14.1 | -0,5017 | 2,83975E-27 |
| ACE2 | NUP153 | -0,5017 | 2,85336E-27 |
| ACE2 | MDN1 | -0,5017 | 2,85489E-27 |
| ACE2 | ANXA8 | -0,5017 | 2,88631E-27 |
| ACE2 | PIK3IP1 | -0,5017 | 2,90512E-27 |
| ACE2 | ZNF131 | -0,5018 | 2,81207E-27 |
| ACE2 | ISM2 | -0,5018 | 2,76437E-27 |
| ACE2 | DYNC1H1 | -0,5018 | 2,76385E-27 |
| ACE2 | FBXL16 | -0,5018 | 2,77186E-27 |
| ACE2 | AC007277.3 | -0,5019 | 2,75445E-27 |
| ACE2 | AC005387.2 | -0,5019 | 2,72901E-27 |
| ACE2 | ZNF670 | -0,502 | 2,67235E-27 |
| ACE2 | RP11-185E8.2 | -0,502 | 2,62875E-27 |
| ACE2 | LINC02015 | -0,502 | 2,66908E-27 |
| ACE2 | EXOC3 | -0,502 | 2,64135E-27 |

| | | | |
|---|---|---|---|
| ACE2 | AC009264.1 | -0,502 | 2,65452E-27 |
| ACE2 | MIS12 | -0,502 | 2,62356E-27 |
| ACE2 | RP11-27G24.1 | -0,502 | 2,635E-27 |
| ACE2 | COX14 | -0,5021 | 2,53894E-27 |
| ACE2 | AC109309.4 | -0,5022 | 2,53599E-27 |
| ACE2 | RP1-228P16.8 | -0,5022 | 2,47308E-27 |
| ACE2 | KCNB2 | -0,5023 | 2,45593E-27 |
| ACE2 | DDX58 | -0,5023 | 2,41686E-27 |
| ACE2 | RP11-467K18.2 | -0,5023 | 2,41265E-27 |
| ACE2 | RARRES3 | -0,5023 | 2,43804E-27 |
| ACE2 | BRICD5 | -0,5023 | 2,41887E-27 |
| ACE2 | PCBD1 | -0,5024 | 2,35011E-27 |
| ACE2 | CCSAP | -0,5025 | 2,28144E-27 |
| ACE2 | RPE | -0,5025 | 2,3272E-27 |
| ACE2 | RP11-799D4.4 | -0,5025 | 2,28915E-27 |
| ACE2 | RP11-481H12.1 | -0,5026 | 2,22853E-27 |
| ACE2 | RAB30-AS1 | -0,5026 | 2,25469E-27 |
| ACE2 | RAB38 | -0,5026 | 2,23461E-27 |
| ACE2 | EFCAB14 | -0,5027 | 2,1667E-27 |
| ACE2 | PNPLA8 | -0,5027 | 2,19326E-27 |
| ACE2 | MROH7 | -0,5028 | 2,10071E-27 |
| ACE2 | GEMIN6 | -0,5029 | 2,07533E-27 |
| ACE2 | REST | -0,5029 | 2,05127E-27 |
| ACE2 | LSM1 | -0,5029 | 2,09402E-27 |
| ACE2 | NUP98 | -0,5029 | 2,06574E-27 |
| ACE2 | RP4-734G22.3 | -0,5031 | 1,93665E-27 |
| ACE2 | RP13-554M15.2 | -0,5031 | 1,95816E-27 |
| ACE2 | RP11-290D2.6 | -0,5031 | 1,93229E-27 |
| ACE2 | RP11-297J22.1 | -0,5032 | 1,91547E-27 |
| ACE2 | ZNF395 | -0,5032 | 1,91829E-27 |
| ACE2 | ZER1 | -0,5032 | 1,9065E-27 |
| ACE2 | MRPL17 | -0,5032 | 1,88531E-27 |
| ACE2 | TTC9 | -0,5032 | 1,87924E-27 |
| ACE2 | C14orf79 | -0,5032 | 1,8868E-27 |
| ACE2 | KCNJ12 | -0,5032 | 1,90531E-27 |
| ACE2 | RP1-39G22.7 | -0,5033 | 1,85334E-27 |
| ACE2 | SMARCAL1 | -0,5033 | 1,8452E-27 |
| ACE2 | GS1-5L10.1 | -0,5033 | 1,8365E-27 |
| ACE2 | LDLRAD4 | -0,5033 | 1,82882E-27 |
| ACE2 | DBX2 | -0,5034 | 1,78666E-27 |
| ACE2 | LA16c-360H6.1 | -0,5034 | 1,80459E-27 |
| ACE2 | AC135178.7 | -0,5034 | 1,80238E-27 |
| ACE2 | EFHC2 | -0,5034 | 1,81831E-27 |
| ACE2 | TLK2P2 | -0,5035 | 1,75975E-27 |
| ACE2 | ATP8B3 | -0,5035 | 1,73153E-27 |
| ACE2 | UBR4 | -0,5036 | 1,7291E-27 |
| ACE2 | RP11-43F13.1 | -0,5036 | 1,71389E-27 |
| ACE2 | MBLAC2 | -0,5036 | 1,68315E-27 |
| ACE2 | CTB-33O18.2 | -0,5036 | 1,72326E-27 |
| ACE2 | AC005387.3 | -0,5036 | 1,71188E-27 |
| ACE2 | GBAP1 | -0,5037 | 1,67424E-27 |
| ACE2 | CIT | -0,5037 | 1,65304E-27 |
| ACE2 | HAT1 | -0,5038 | 1,59818E-27 |
| ACE2 | SYNPO | -0,5038 | 1,59478E-27 |

| | | | |
|---|---|---|---|
| ACE2 | GATC | -0,5038 | 1,59738E-27 |
| ACE2 | ATP5SL | -0,5038 | 1,60854E-27 |
| ACE2 | FAM122B | -0,5038 | 1,60464E-27 |
| ACE2 | CTD-2260A17.2 | -0,5039 | 1,58051E-27 |
| ACE2 | AC092675.3 | -0,504 | 1,54865E-27 |
| ACE2 | RGS22 | -0,504 | 1,5238E-27 |
| ACE2 | RP11-426C22.1 | -0,504 | 1,5407E-27 |
| ACE2 | SHPRH | -0,5041 | 1,47194E-27 |
| ACE2 | WDR11-AS1 | -0,5041 | 1,50063E-27 |
| ACE2 | SNF8 | -0,5041 | 1,48469E-27 |
| ACE2 | TASP1 | -0,5041 | 1,49596E-27 |
| ACE2 | AC093390.1 | -0,5042 | 1,42968E-27 |
| ACE2 | LINC00339 | -0,5043 | 1,39462E-27 |
| ACE2 | PCGF2 | -0,5043 | 1,41065E-27 |
| ACE2 | TLK1 | -0,5045 | 1,32086E-27 |
| ACE2 | RP11-340E6.1 | -0,5045 | 1,32178E-27 |
| ACE2 | ZNF33BP1 | -0,5046 | 1,30934E-27 |
| ACE2 | HMGN2P15 | -0,5046 | 1,30287E-27 |
| ACE2 | ERCC2 | -0,5046 | 1,30539E-27 |
| ACE2 | GABRB1 | -0,5047 | 1,26743E-27 |
| ACE2 | RPL3P4 | -0,5047 | 1,27347E-27 |
| ACE2 | NUP85 | -0,5047 | 1,26841E-27 |
| ACE2 | TBK1 | -0,5048 | 1,23752E-27 |
| ACE2 | RP11-552F3.9 | -0,5048 | 1,22859E-27 |
| ACE2 | CTB-133G6.1 | -0,5048 | 1,23019E-27 |
| ACE2 | WASF2 | -0,5049 | 1,1795E-27 |
| ACE2 | RP11-418H16.1 | -0,5049 | 1,20298E-27 |
| ACE2 | NEK4 | -0,5049 | 1,19015E-27 |
| ACE2 | PARP9 | -0,5049 | 1,20612E-27 |
| ACE2 | RP3-406P24.5 | -0,5049 | 1,19743E-27 |
| ACE2 | BAP1 | -0,505 | 1,15217E-27 |
| ACE2 | WARS | -0,5051 | 1,14149E-27 |
| ACE2 | MEIS3P1 | -0,5051 | 1,11388E-27 |
| ACE2 | CATSPER2P1 | -0,5052 | 1,10075E-27 |
| ACE2 | HIST2H3DP1 | -0,5053 | 1,05665E-27 |
| ACE2 | COBLL1 | -0,5053 | 1,05711E-27 |
| ACE2 | DNAJA1 | -0,5053 | 1,07391E-27 |
| ACE2 | PPP6R2 | -0,5053 | 1,07791E-27 |
| ACE2 | RP11-1191J2.5 | -0,5054 | 1,03958E-27 |
| ACE2 | IWS1 | -0,5055 | 1,02149E-27 |
| ACE2 | CDKL2 | -0,5055 | 1,0199E-27 |
| ACE2 | FLJ20021 | -0,5055 | 1,01252E-27 |
| ACE2 | CXXC1 | -0,5055 | 1,0219E-27 |
| ACE2 | NAT10 | -0,5056 | 9,73171E-28 |
| ACE2 | DUXAP10 | -0,5056 | 9,73496E-28 |
| ACE2 | MARCH10 | -0,5056 | 9,8552E-28 |
| ACE2 | MYH9 | -0,5056 | 9,94359E-28 |
| ACE2 | FAM180B | -0,5057 | 9,52598E-28 |
| ACE2 | RP11-697E2.9 | -0,5057 | 9,44477E-28 |
| ACE2 | TMEM107 | -0,5057 | 9,61278E-28 |
| ACE2 | TGM2 | -0,5057 | 9,68315E-28 |
| ACE2 | ZFAND5 | -0,5058 | 9,41927E-28 |
| ACE2 | FCHSD1 | -0,5059 | 9,06805E-28 |
| ACE2 | RP1-37C10.3 | -0,506 | 8,91058E-28 |

| | | | |
|---|---|---|---|
| ACE2 | RPL12P16 | -0,506 | 8,77161E-28 |
| ACE2 | CTSH | -0,506 | 8,8984E-28 |
| ACE2 | DNAJA2 | -0,506 | 8,87668E-28 |
| ACE2 | C2orf73 | -0,5061 | 8,55346E-28 |
| ACE2 | BTN2A3P | -0,5061 | 8,6211E-28 |
| ACE2 | PPP1R3D | -0,5062 | 8,39126E-28 |
| ACE2 | CEP104 | -0,5063 | 8,1365E-28 |
| ACE2 | RARS2 | -0,5063 | 8,16868E-28 |
| ACE2 | PTCD1 | -0,5063 | 8,00627E-28 |
| ACE2 | RP11-754B17.1 | -0,5063 | 8,1096E-28 |
| ACE2 | GRB2 | -0,5063 | 8,00835E-28 |
| ACE2 | RP11-472N13.3 | -0,5064 | 7,8389E-28 |
| ACE2 | RP11-326C3.15 | -0,5064 | 7,95598E-28 |
| ACE2 | CTC-425O23.2 | -0,5064 | 7,84308E-28 |
| ACE2 | PAPPA2 | -0,5065 | 7,57837E-28 |
| ACE2 | POLL | -0,5066 | 7,40575E-28 |
| ACE2 | CTD-2517O10.6 | -0,5067 | 7,16519E-28 |
| ACE2 | RP11-656D10.3 | -0,5068 | 7,01516E-28 |
| ACE2 | TGFBRAP1 | -0,5069 | 6,93662E-28 |
| ACE2 | LTV1 | -0,5069 | 6,81634E-28 |
| ACE2 | ARFGAP2 | -0,5069 | 6,85449E-28 |
| ACE2 | LYSMD4 | -0,5069 | 6,89907E-28 |
| ACE2 | KANSL1 | -0,5069 | 6,94463E-28 |
| ACE2 | GPR4 | -0,507 | 6,63776E-28 |
| ACE2 | ARHGAP31-AS1 | -0,5072 | 6,30538E-28 |
| ACE2 | ZNF699 | -0,5072 | 6,36748E-28 |
| ACE2 | GLRX3 | -0,5073 | 6,19043E-28 |
| ACE2 | POU2F2 | -0,5073 | 6,14672E-28 |
| ACE2 | SLC39A1 | -0,5074 | 5,93901E-28 |
| ACE2 | TMEM267 | -0,5074 | 5,91894E-28 |
| ACE2 | EFNA5 | -0,5074 | 5,90399E-28 |
| ACE2 | HOXC9 | -0,5074 | 6,02655E-28 |
| ACE2 | ARL17B | -0,5074 | 6,0327E-28 |
| ACE2 | RP11-423P10.2 | -0,5075 | 5,84416E-28 |
| ACE2 | PKD2L2 | -0,5075 | 5,84394E-28 |
| ACE2 | SRL | -0,5075 | 5,7325E-28 |
| ACE2 | RP11-126K1.6 | -0,5076 | 5,59778E-28 |
| ACE2 | RP11-574O7.1 | -0,5076 | 5,60077E-28 |
| ACE2 | IRAK3 | -0,5078 | 5,39972E-28 |
| ACE2 | RP3-402G11.28 | -0,5078 | 5,28272E-28 |
| ACE2 | HRH4 | -0,5079 | 5,12514E-28 |
| ACE2 | RP11-15A1.4 | -0,5079 | 5,26367E-28 |
| ACE2 | TSC22D4 | -0,508 | 5,05082E-28 |
| ACE2 | RP11-509J21.4 | -0,508 | 5,07599E-28 |
| ACE2 | RP11-145P16.3 | -0,508 | 5,01225E-28 |
| ACE2 | RN7SL566P | -0,508 | 5,05391E-28 |
| ACE2 | ZNF226 | -0,508 | 5,12243E-28 |
| ACE2 | NAE1 | -0,5081 | 4,95086E-28 |
| ACE2 | USP16 | -0,5081 | 4,88383E-28 |
| ACE2 | NCOA7 | -0,5082 | 4,83774E-28 |
| ACE2 | MTRF1L | -0,5082 | 4,79633E-28 |
| ACE2 | FAM161B | -0,5082 | 4,80908E-28 |
| ACE2 | NDRG3 | -0,5082 | 4,74212E-28 |
| ACE2 | ADAM20 | -0,5083 | 4,64113E-28 |

| | | | |
|---|---|---|---|
| ACE2 | NFATC2 | -0,5083 | 4,68757E-28 |
| ACE2 | GCNT4 | -0,5084 | 4,51461E-28 |
| ACE2 | SPRY4 | -0,5084 | 4,52266E-28 |
| ACE2 | NPAP1 | -0,5084 | 4,5376E-28 |
| ACE2 | PLA2G6 | -0,5084 | 4,48657E-28 |
| ACE2 | HIPK1 | -0,5085 | 4,33559E-28 |
| ACE2 | FUT9 | -0,5085 | 4,35706E-28 |
| ACE2 | IFT80 | -0,5086 | 4,32523E-28 |
| ACE2 | RP11-234K19.1 | -0,5086 | 4,22946E-28 |
| ACE2 | SIK2 | -0,5086 | 4,26107E-28 |
| ACE2 | AATF | -0,5086 | 4,30835E-28 |
| ACE2 | EID2 | -0,5086 | 4,24019E-28 |
| ACE2 | RP4-694B14.8 | -0,5086 | 4,27803E-28 |
| ACE2 | CHMP3 | -0,5087 | 4,19828E-28 |
| ACE2 | PPP6R3 | -0,5087 | 4,13823E-28 |
| ACE2 | CDH24 | -0,5087 | 4,17598E-28 |
| ACE2 | LONP2 | -0,5087 | 4,18915E-28 |
| ACE2 | ZNF469 | -0,5087 | 4,14981E-28 |
| ACE2 | DIRAS1 | -0,5087 | 4,11174E-28 |
| ACE2 | SMG9 | -0,5087 | 4,17951E-28 |
| ACE2 | LRRN3 | -0,5088 | 3,99709E-28 |
| ACE2 | ASPH | -0,5089 | 3,91409E-28 |
| ACE2 | FAM168B | -0,509 | 3,78122E-28 |
| ACE2 | LRIT3 | -0,509 | 3,78817E-28 |
| ACE2 | RP11-278C7.4 | -0,509 | 3,8588E-28 |
| ACE2 | ATXN1L | -0,509 | 3,78581E-28 |
| ACE2 | NUP43 | -0,5091 | 3,71554E-28 |
| ACE2 | RP11-887P2.5 | -0,5091 | 3,69888E-28 |
| ACE2 | ADGRG6 | -0,5092 | 3,63509E-28 |
| ACE2 | PDE1C | -0,5092 | 3,57224E-28 |
| ACE2 | RP5-1029K10.2 | -0,5092 | 3,58611E-28 |
| ACE2 | MEF2B | -0,5092 | 3,59672E-28 |
| ACE2 | ACTN2 | -0,5093 | 3,51015E-28 |
| ACE2 | RP11-384K6.6 | -0,5094 | 3,37489E-28 |
| ACE2 | SNORD10 | -0,5094 | 3,45797E-28 |
| ACE2 | CTC-471J1.11 | -0,5094 | 3,43039E-28 |
| ACE2 | RAP1GDS1 | -0,5095 | 3,30824E-28 |
| ACE2 | RP11-482M8.3 | -0,5095 | 3,33145E-28 |
| ACE2 | RP11-230C9.2 | -0,5096 | 3,21571E-28 |
| ACE2 | AC010524.2 | -0,5096 | 3,26139E-28 |
| ACE2 | SERPINI2 | -0,5097 | 3,12778E-28 |
| ACE2 | LRPAP1 | -0,5098 | 3,05584E-28 |
| ACE2 | LINC00909 | -0,5098 | 3,02837E-28 |
| ACE2 | RP11-11N7.4 | -0,5099 | 2,99168E-28 |
| ACE2 | DYNC1I2 | -0,5099 | 2,95022E-28 |
| ACE2 | RP11-33B1.1 | -0,5099 | 2,9374E-28 |
| ACE2 | LINC00680 | -0,5099 | 2,94554E-28 |
| ACE2 | POLR2L | -0,5099 | 3,01343E-28 |
| ACE2 | PAPOLA | -0,5099 | 2,96835E-28 |
| ACE2 | POLH | -0,51 | 2,92787E-28 |
| ACE2 | VIT | -0,5101 | 2,81005E-28 |
| ACE2 | AC005330.2 | -0,5101 | 2,78661E-28 |
| ACE2 | SLC10A6 | -0,5102 | 2,75288E-28 |
| ACE2 | RP11-373N22.3 | -0,5102 | 2,73234E-28 |

| | | | |
|---|---|---|---|
| ACE2 | RN7SKP268 | -0,5102 | 2,70232E-28 |
| ACE2 | PPP2R5E | -0,5102 | 2,76776E-28 |
| ACE2 | ANGPT2 | -0,5103 | 2,65792E-28 |
| ACE2 | FSCN1P1 | -0,5103 | 2,67023E-28 |
| ACE2 | CTD-2666L21.1 | -0,5103 | 2,6842E-28 |
| ACE2 | LINC01858 | -0,5103 | 2,67838E-28 |
| ACE2 | PABPC1L2A | -0,5103 | 2,6587E-28 |
| ACE2 | LINC00921 | -0,5104 | 2,6114E-28 |
| ACE2 | JARID2 | -0,5106 | 2,42027E-28 |
| ACE2 | LAMA2 | -0,5106 | 2,42815E-28 |
| ACE2 | DLX3 | -0,5107 | 2,37538E-28 |
| ACE2 | AC009506.1 | -0,5108 | 2,32244E-28 |
| ACE2 | PLAG1 | -0,5108 | 2,28226E-28 |
| ACE2 | DMRT3 | -0,5108 | 2,29217E-28 |
| ACE2 | WDR76 | -0,5108 | 2,3206E-28 |
| ACE2 | RP11-705C15.2 | -0,5109 | 2,22509E-28 |
| ACE2 | BNIP3P28 | -0,5109 | 2,22445E-28 |
| ACE2 | AMMECR1L | -0,511 | 2,19516E-28 |
| ACE2 | ING3 | -0,511 | 2,21094E-28 |
| ACE2 | C8orf48 | -0,511 | 2,1711E-28 |
| ACE2 | PPWD1 | -0,5111 | 2,09444E-28 |
| ACE2 | MALRD1 | -0,5111 | 2,10234E-28 |
| ACE2 | TBCCD1 | -0,5112 | 2,07171E-28 |
| ACE2 | SART3 | -0,5112 | 2,06179E-28 |
| ACE2 | NDUFV3 | -0,5112 | 2,06131E-28 |
| ACE2 | SYT2 | -0,5113 | 1,97878E-28 |
| ACE2 | CBX3 | -0,5113 | 2,00715E-28 |
| ACE2 | MFAP1P1 | -0,5114 | 1,95835E-28 |
| ACE2 | ZC3H14 | -0,5114 | 1,95837E-28 |
| ACE2 | FAM89A | -0,5115 | 1,90404E-28 |
| ACE2 | RP5-826L7.1 | -0,5115 | 1,87353E-28 |
| ACE2 | ZRANB2 | -0,5116 | 1,86064E-28 |
| ACE2 | TSPAN10 | -0,5116 | 1,83751E-28 |
| ACE2 | BACH1 | -0,5116 | 1,82305E-28 |
| ACE2 | RP11-90L20.2 | -0,5117 | 1,78489E-28 |
| ACE2 | TTC27 | -0,5117 | 1,79445E-28 |
| ACE2 | NEMP2 | -0,5117 | 1,7916E-28 |
| ACE2 | KDM1A | -0,5118 | 1,76555E-28 |
| ACE2 | RP11-420L9.2 | -0,5118 | 1,76349E-28 |
| ACE2 | RP11-254F19.2 | -0,5118 | 1,75951E-28 |
| ACE2 | SEC63P1 | -0,5119 | 1,71362E-28 |
| ACE2 | RP11-477N3.1 | -0,5119 | 1,68952E-28 |
| ACE2 | CTD-3025N20.3 | -0,5119 | 1,68173E-28 |
| ACE2 | CTC-264K15.6 | -0,5119 | 1,69469E-28 |
| ACE2 | RP11-75A9.3 | -0,5119 | 1,67724E-28 |
| ACE2 | GAGE10 | -0,512 | 1,64355E-28 |
| ACE2 | RP6-24A23.3 | -0,512 | 1,65101E-28 |
| ACE2 | HIVEP1 | -0,5121 | 1,597E-28 |
| ACE2 | SMU1 | -0,5121 | 1,59221E-28 |
| ACE2 | ZNF627 | -0,5121 | 1,59669E-28 |
| ACE2 | ZBTB21 | -0,5121 | 1,61219E-28 |
| ACE2 | OTUD7B | -0,5122 | 1,55094E-28 |
| ACE2 | AC058791.1 | -0,5122 | 1,53629E-28 |
| ACE2 | HNRNPCP4 | -0,5122 | 1,55813E-28 |

| | | | |
|---|---|---|---|
| ACE2 | SH3D19 | -0,5123 | 1,49559E-28 |
| ACE2 | RP11-85B7.2 | -0,5123 | 1,50818E-28 |
| ACE2 | HAPLN2 | -0,5124 | 1,46646E-28 |
| ACE2 | LINC01021 | -0,5126 | 1,39105E-28 |
| ACE2 | TPT1P4 | -0,5126 | 1,39558E-28 |
| ACE2 | DBH | -0,5126 | 1,37884E-28 |
| ACE2 | RP5-1024N4.4 | -0,5127 | 1,35869E-28 |
| ACE2 | RP11-114M1.3 | -0,5128 | 1,29752E-28 |
| ACE2 | RP11-335L23.5 | -0,5128 | 1,31669E-28 |
| ACE2 | NOTCH1 | -0,5128 | 1,31387E-28 |
| ACE2 | NDUFS8 | -0,5128 | 1,3157E-28 |
| ACE2 | CITED1 | -0,5128 | 1,30352E-28 |
| ACE2 | RP4-669L17.10 | -0,5129 | 1,29324E-28 |
| ACE2 | TRIM33 | -0,5129 | 1,28876E-28 |
| ACE2 | JAK2 | -0,5129 | 1,27661E-28 |
| ACE2 | POC1B-AS1 | -0,5129 | 1,27045E-28 |
| ACE2 | RP11-483C6.1 | -0,5129 | 1,27247E-28 |
| ACE2 | ZNFX1 | -0,5129 | 1,27874E-28 |
| ACE2 | GAB3 | -0,5129 | 1,29344E-28 |
| ACE2 | LINC01788 | -0,513 | 1,23139E-28 |
| ACE2 | AC096574.5 | -0,513 | 1,23636E-28 |
| ACE2 | SRRT | -0,513 | 1,22785E-28 |
| ACE2 | KRT10 | -0,513 | 1,25623E-28 |
| ACE2 | RP1-198K11.5 | -0,513 | 1,22621E-28 |
| ACE2 | FPGS | -0,5131 | 1,20169E-28 |
| ACE2 | SMCO2 | -0,5133 | 1,14764E-28 |
| ACE2 | EEF1DP4 | -0,5134 | 1,10679E-28 |
| ACE2 | RPS3AP47 | -0,5134 | 1,09908E-28 |
| ACE2 | ACTG1P14 | -0,5135 | 1,07085E-28 |
| ACE2 | CBX2 | -0,5135 | 1,08226E-28 |
| ACE2 | OPRL1 | -0,5136 | 1,06157E-28 |
| ACE2 | NR0B1 | -0,5136 | 1,05485E-28 |
| ACE2 | RP11-342K6.2 | -0,5137 | 1,01396E-28 |
| ACE2 | RP11-266L9.8 | -0,5138 | 9,88594E-29 |
| ACE2 | LINC01529 | -0,5138 | 9,93629E-29 |
| ACE2 | UTRN | -0,5139 | 9,6344E-29 |
| ACE2 | DDX55 | -0,5139 | 9,74173E-29 |
| ACE2 | ZNF350-AS1 | -0,5139 | 9,68675E-29 |
| ACE2 | CHIC2 | -0,514 | 9,32355E-29 |
| ACE2 | RP11-379H18.1 | -0,514 | 9,22738E-29 |
| ACE2 | RNU6-8 | -0,514 | 9,23809E-29 |
| ACE2 | MAP3K7CL | -0,514 | 9,46158E-29 |
| ACE2 | RIMS3 | -0,5141 | 9,17295E-29 |
| ACE2 | PSMA1 | -0,5141 | 9,12743E-29 |
| ACE2 | RP11-843B15.4 | -0,5141 | 9,06334E-29 |
| ACE2 | TRAPPC13 | -0,5142 | 8,78854E-29 |
| ACE2 | HLCS | -0,5142 | 8,84116E-29 |
| ACE2 | ZSCAN31 | -0,5143 | 8,50783E-29 |
| ACE2 | FUS | -0,5143 | 8,62359E-29 |
| ACE2 | PCNT | -0,5143 | 8,56592E-29 |
| ACE2 | CACNA2D2 | -0,5144 | 8,29587E-29 |
| ACE2 | AP4E1 | -0,5144 | 8,42807E-29 |
| ACE2 | MED1 | -0,5144 | 8,25087E-29 |
| ACE2 | PARP4P3 | -0,5144 | 8,24204E-29 |

| | | | |
|---|---|---|---|
| ACE2 | RP1-224A6.9 | -0,5145 | 8,17143E-29 |
| ACE2 | NKTR | -0,5145 | 8,1157E-29 |
| ACE2 | SNCA-AS1 | -0,5145 | 8,05154E-29 |
| ACE2 | RP1-223B1.1 | -0,5145 | 8,10649E-29 |
| ACE2 | ABCB5 | -0,5145 | 8,21435E-29 |
| ACE2 | PLEKHA8 | -0,5145 | 8,05314E-29 |
| ACE2 | VGF | -0,5145 | 8,10268E-29 |
| ACE2 | MRPL50 | -0,5145 | 8,15219E-29 |
| ACE2 | PSMD6 | -0,5146 | 7,93489E-29 |
| ACE2 | IGSF10 | -0,5146 | 7,78881E-29 |
| ACE2 | AC106801.1 | -0,5146 | 7,80811E-29 |
| ACE2 | RMDN1 | -0,5146 | 7,81547E-29 |
| ACE2 | PRPF6 | -0,5146 | 7,83849E-29 |
| ACE2 | OLA1 | -0,5147 | 7,69984E-29 |
| ACE2 | TSSK5P | -0,5147 | 7,73876E-29 |
| ACE2 | AC112715.2 | -0,5148 | 7,3892E-29 |
| ACE2 | TRIM73 | -0,5148 | 7,50495E-29 |
| ACE2 | LIG3 | -0,5148 | 7,47495E-29 |
| ACE2 | SPRY2 | -0,5149 | 7,18333E-29 |
| ACE2 | RP11-547D13.1 | -0,5149 | 7,22784E-29 |
| ACE2 | RWDD4 | -0,515 | 6,98952E-29 |
| ACE2 | CTD-2026K11.4 | -0,515 | 7,01905E-29 |
| ACE2 | ERICH4 | -0,515 | 7,02319E-29 |
| ACE2 | OTUD3 | -0,5151 | 6,77744E-29 |
| ACE2 | NF2 | -0,5151 | 6,77562E-29 |
| ACE2 | MMS19 | -0,5152 | 6,71353E-29 |
| ACE2 | BEX5 | -0,5152 | 6,58581E-29 |
| ACE2 | ADIRF | -0,5153 | 6,45314E-29 |
| ACE2 | PAPLN | -0,5153 | 6,46009E-29 |
| ACE2 | ADSSL1 | -0,5153 | 6,38597E-29 |
| ACE2 | MIMT1 | -0,5153 | 6,54152E-29 |
| ACE2 | CAD | -0,5154 | 6,29679E-29 |
| ACE2 | RDH10-AS1 | -0,5154 | 6,3175E-29 |
| ACE2 | RP11-81K2.2 | -0,5154 | 6,19617E-29 |
| ACE2 | FAM156A | -0,5154 | 6,34934E-29 |
| ACE2 | IPO9-AS1 | -0,5155 | 6,08876E-29 |
| ACE2 | NUP35 | -0,5155 | 6,03854E-29 |
| ACE2 | RIC8A | -0,5155 | 6,11622E-29 |
| ACE2 | KLHL1 | -0,5155 | 6,10369E-29 |
| ACE2 | CREG2 | -0,5156 | 5,91775E-29 |
| ACE2 | RPL26P30 | -0,5156 | 5,87944E-29 |
| ACE2 | CTD-2047H16.5 | -0,5156 | 5,87293E-29 |
| ACE2 | ZNF284 | -0,5156 | 5,98279E-29 |
| ACE2 | PSMB5 | -0,5157 | 5,75405E-29 |
| ACE2 | TM6SF1 | -0,5157 | 5,78056E-29 |
| ACE2 | ATP1B1P1 | -0,5158 | 5,67219E-29 |
| ACE2 | ZFYVE26 | -0,5158 | 5,63839E-29 |
| ACE2 | LRRC37A16P | -0,5158 | 5,60351E-29 |
| ACE2 | RGS1 | -0,5159 | 5,45067E-29 |
| ACE2 | RP11-479O17.12 | -0,5159 | 5,36693E-29 |
| ACE2 | RP11-672A2.4 | -0,5159 | 5,4285E-29 |
| ACE2 | NAT6 | -0,516 | 5,29469E-29 |
| ACE2 | MB21D1 | -0,5161 | 5,20792E-29 |
| ACE2 | TRIP11 | -0,5161 | 5,12933E-29 |

| | | | |
|---|---|---|---|
| ACE2 | NPRL2 | -0,5162 | 5,03508E-29 |
| ACE2 | FAM120B | -0,5162 | 5,00426E-29 |
| ACE2 | MKX-AS1 | -0,5162 | 4,96844E-29 |
| ACE2 | CTDSP2 | -0,5162 | 4,97352E-29 |
| ACE2 | CTD-2196E14.6 | -0,5162 | 5,02803E-29 |
| ACE2 | ZNF850 | -0,5162 | 4,95641E-29 |
| ACE2 | NBPF19 | -0,5163 | 4,88985E-29 |
| ACE2 | SDHAP3 | -0,5163 | 4,85592E-29 |
| ACE2 | KCNK15 | -0,5163 | 4,91418E-29 |
| ACE2 | FMO1 | -0,5164 | 4,65399E-29 |
| ACE2 | KIAA1549 | -0,5164 | 4,66505E-29 |
| ACE2 | CA10 | -0,5165 | 4,58219E-29 |
| ACE2 | CACTIN-AS1 | -0,5165 | 4,59819E-29 |
| ACE2 | ZNF607 | -0,5165 | 4,61354E-29 |
| ACE2 | DNM1P46 | -0,5166 | 4,49777E-29 |
| ACE2 | PIH1D1 | -0,5166 | 4,38658E-29 |
| ACE2 | RP11-23J9.6 | -0,5167 | 4,32508E-29 |
| ACE2 | RP11-83B20.1 | -0,5167 | 4,35282E-29 |
| ACE2 | SUMO2P15 | -0,5167 | 4,34455E-29 |
| ACE2 | UBXN2B | -0,5168 | 4,24753E-29 |
| ACE2 | TMEM254-AS1 | -0,5168 | 4,24822E-29 |
| ACE2 | LAGE3 | -0,5168 | 4,22107E-29 |
| ACE2 | KIN | -0,5169 | 4,03795E-29 |
| ACE2 | AC007228.5 | -0,5169 | 4,135E-29 |
| ACE2 | ADHFE1 | -0,517 | 4,00783E-29 |
| ACE2 | ZIM2 | -0,517 | 3,96258E-29 |
| ACE2 | RP11-373D23.2 | -0,5171 | 3,81673E-29 |
| ACE2 | AMZ1 | -0,5171 | 3,87779E-29 |
| ACE2 | EEF1A1P6 | -0,5171 | 3,8502E-29 |
| ACE2 | LRRC63 | -0,5171 | 3,80745E-29 |
| ACE2 | RP1-79C4.4 | -0,5172 | 3,72891E-29 |
| ACE2 | NSL1 | -0,5172 | 3,73349E-29 |
| ACE2 | MTPN | -0,5172 | 3,70443E-29 |
| ACE2 | RP11-390G14.1 | -0,5174 | 3,53575E-29 |
| ACE2 | OTUD4 | -0,5174 | 3,52204E-29 |
| ACE2 | RP11-565F19.2 | -0,5174 | 3,52772E-29 |
| ACE2 | AK3P3 | -0,5175 | 3,48278E-29 |
| ACE2 | TPTE2P1 | -0,5175 | 3,46077E-29 |
| ACE2 | TRAPPC11 | -0,5176 | 3,32728E-29 |
| ACE2 | MGAT1 | -0,5176 | 3,32053E-29 |
| ACE2 | DICER1 | -0,5176 | 3,35479E-29 |
| ACE2 | MRPL9P1 | -0,5177 | 3,25576E-29 |
| ACE2 | CH25H | -0,5177 | 3,22667E-29 |
| ACE2 | BORCS8 | -0,5177 | 3,22134E-29 |
| ACE2 | SARS | -0,5178 | 3,16662E-29 |
| ACE2 | MCEE | -0,5178 | 3,17244E-29 |
| ACE2 | ATP6AP1L | -0,5178 | 3,15169E-29 |
| ACE2 | CTA-223H9.9 | -0,5178 | 3,10972E-29 |
| ACE2 | TATDN2 | -0,5179 | 3,0369E-29 |
| ACE2 | ZNF441 | -0,5179 | 3,09933E-29 |
| ACE2 | FAM65B | -0,5181 | 2,87364E-29 |
| ACE2 | AP3M1 | -0,5181 | 2,86244E-29 |
| ACE2 | RP11-672A2.5 | -0,5181 | 2,87E-29 |
| ACE2 | GRPR | -0,5181 | 2,92385E-29 |

| | | | |
|---|---|---|---|
| ACE2 | ADGRL3 | -0,5182 | 2,83602E-29 |
| ACE2 | MRPL38 | -0,5182 | 2,82657E-29 |
| ACE2 | ADGRL1 | -0,5183 | 2,76237E-29 |
| ACE2 | RP5-855D21.1 | -0,5184 | 2,62213E-29 |
| ACE2 | RAP1B | -0,5184 | 2,67786E-29 |
| ACE2 | PWAR1 | -0,5184 | 2,67023E-29 |
| ACE2 | SON | -0,5185 | 2,56592E-29 |
| ACE2 | LINC00316 | -0,5186 | 2,5245E-29 |
| ACE2 | RP11-307L14.1 | -0,5187 | 2,43667E-29 |
| ACE2 | RP11-154D17.1 | -0,5187 | 2,41228E-29 |
| ACE2 | RSAD1 | -0,5187 | 2,46278E-29 |
| ACE2 | LAMB4 | -0,5188 | 2,35354E-29 |
| ACE2 | SERBP1 | -0,5189 | 2,26321E-29 |
| ACE2 | RP11-100G15.7 | -0,5189 | 2,28686E-29 |
| ACE2 | RP5-906A24.1 | -0,5189 | 2,29229E-29 |
| ACE2 | ADM5 | -0,5189 | 2,28126E-29 |
| ACE2 | FNDC11 | -0,5189 | 2,26707E-29 |
| ACE2 | LEFTY2 | -0,519 | 2,22083E-29 |
| ACE2 | CCNH | -0,519 | 2,2419E-29 |
| ACE2 | SPINK13 | -0,519 | 2,25738E-29 |
| ACE2 | MPV17L | -0,5191 | 2,15403E-29 |
| ACE2 | MEIOC | -0,5191 | 2,15136E-29 |
| ACE2 | HPS4 | -0,5192 | 2,08274E-29 |
| ACE2 | SBF1 | -0,5192 | 2,12614E-29 |
| ACE2 | RP4-635E18.8 | -0,5193 | 2,06698E-29 |
| ACE2 | NELL1 | -0,5193 | 2,05085E-29 |
| ACE2 | ZGPAT | -0,5193 | 2,05844E-29 |
| ACE2 | RP11-46A10.6 | -0,5194 | 1,9861E-29 |
| ACE2 | LINC01625 | -0,5194 | 1,95961E-29 |
| ACE2 | LINC00683 | -0,5194 | 1,988E-29 |
| ACE2 | ZNF833P | -0,5194 | 1,99896E-29 |
| ACE2 | HMGXB4 | -0,5194 | 1,9992E-29 |
| ACE2 | IMPDH1P8 | -0,5195 | 1,93845E-29 |
| ACE2 | RP11-307O13.1 | -0,5195 | 1,91822E-29 |
| ACE2 | AF131216.6 | -0,5196 | 1,88636E-29 |
| ACE2 | SYNJ1 | -0,5197 | 1,80741E-29 |
| ACE2 | ANKRD20A17P | -0,5198 | 1,746E-29 |
| ACE2 | PEX14 | -0,52 | 1,68855E-29 |
| ACE2 | RP3-473L9.4 | -0,52 | 1,67661E-29 |
| ACE2 | MSL2 | -0,5201 | 1,64362E-29 |
| ACE2 | GRPEL2 | -0,5201 | 1,60318E-29 |
| ACE2 | RP3-331H24.6 | -0,5201 | 1,63149E-29 |
| ACE2 | RP11-88E10.5 | -0,5201 | 1,62283E-29 |
| ACE2 | ADORA2A-AS1 | -0,5201 | 1,61723E-29 |
| ACE2 | DDX56 | -0,5202 | 1,59701E-29 |
| ACE2 | SNRPA | -0,5202 | 1,56883E-29 |
| ACE2 | RP3-449M8.6 | -0,5203 | 1,51587E-29 |
| ACE2 | UBE2Q2P6 | -0,5203 | 1,53382E-29 |
| ACE2 | LGALS8 | -0,5204 | 1,46957E-29 |
| ACE2 | CACNA1C-IT2 | -0,5204 | 1,49452E-29 |
| ACE2 | RP11-425D17.1 | -0,5204 | 1,48088E-29 |
| ACE2 | GREB1L | -0,5204 | 1,48565E-29 |
| ACE2 | FOXO3 | -0,5205 | 1,45993E-29 |
| ACE2 | RP11-112L6.3 | -0,5205 | 1,42656E-29 |

| | | | |
|---|---|---|---|
| ACE2 | ZFX | -0,5205 | 1,43987E-29 |
| ACE2 | MRPS10 | -0,5206 | 1,40809E-29 |
| ACE2 | DRG1 | -0,5206 | 1,41976E-29 |
| ACE2 | RP11-192H23.6 | -0,5207 | 1,35579E-29 |
| ACE2 | AC004471.10 | -0,5207 | 1,36353E-29 |
| ACE2 | RP11-513M16.8 | -0,5208 | 1,33451E-29 |
| ACE2 | PLRG1 | -0,5209 | 1,28182E-29 |
| ACE2 | EPHA1-AS1 | -0,5209 | 1,27593E-29 |
| ACE2 | SIN3A | -0,5209 | 1,26778E-29 |
| ACE2 | AMOT | -0,5209 | 1,26895E-29 |
| ACE2 | SFPQ | -0,521 | 1,25667E-29 |
| ACE2 | RP11-1020A11.2 | -0,521 | 1,25501E-29 |
| ACE2 | RP11-295P9.3 | -0,521 | 1,23425E-29 |
| ACE2 | ALG10B | -0,521 | 1,25773E-29 |
| ACE2 | TBC1D16 | -0,521 | 1,25175E-29 |
| ACE2 | PLPPR3 | -0,521 | 1,25315E-29 |
| ACE2 | RP11-175B9.3 | -0,5211 | 1,20618E-29 |
| ACE2 | FNIP1 | -0,5211 | 1,23098E-29 |
| ACE2 | UBQLN1 | -0,5211 | 1,20979E-29 |
| ACE2 | HPS5 | -0,5211 | 1,19863E-29 |
| ACE2 | TAF7 | -0,5212 | 1,1629E-29 |
| ACE2 | RP11-291L22.9 | -0,5212 | 1,18095E-29 |
| ACE2 | CNPY3 | -0,5213 | 1,14089E-29 |
| ACE2 | EYA1 | -0,5213 | 1,13836E-29 |
| ACE2 | DPY19L4 | -0,5213 | 1,14753E-29 |
| ACE2 | RP11-423C15.3 | -0,5213 | 1,1419E-29 |
| ACE2 | ODF3L1 | -0,5213 | 1,1529E-29 |
| ACE2 | NPHP4 | -0,5214 | 1,11208E-29 |
| ACE2 | NT5DC2 | -0,5214 | 1,10183E-29 |
| ACE2 | HCG18 | -0,5214 | 1,09667E-29 |
| ACE2 | KIF27 | -0,5214 | 1,11445E-29 |
| ACE2 | RP11-676M6.1 | -0,5214 | 1,12049E-29 |
| ACE2 | LINC01436 | -0,5214 | 1,11105E-29 |
| ACE2 | CCND3 | -0,5216 | 1,05319E-29 |
| ACE2 | SNW1 | -0,5216 | 1,06232E-29 |
| ACE2 | GATA5 | -0,5216 | 1,05916E-29 |
| ACE2 | LIMS1-AS1 | -0,5217 | 1,02251E-29 |
| ACE2 | RP11-329B9.4 | -0,5218 | 9,94775E-30 |
| ACE2 | GABPA | -0,5218 | 9,8689E-30 |
| ACE2 | KCNH8 | -0,5219 | 9,73277E-30 |
| ACE2 | RP5-1159O4.1 | -0,5219 | 9,51593E-30 |
| ACE2 | HOXC10 | -0,5219 | 9,65025E-30 |
| ACE2 | RANBP2 | -0,522 | 9,31285E-30 |
| ACE2 | CDV3 | -0,522 | 9,21982E-30 |
| ACE2 | RP11-122C5.3 | -0,522 | 9,22303E-30 |
| ACE2 | LINC01679 | -0,522 | 9,38314E-30 |
| ACE2 | DNAJC19P5 | -0,5221 | 9,18222E-30 |
| ACE2 | CTC-471C19.1 | -0,5221 | 9,12749E-30 |
| ACE2 | HNRNPR | -0,5222 | 8,74916E-30 |
| ACE2 | PMS2 | -0,5222 | 8,76166E-30 |
| ACE2 | VN1R20P | -0,5223 | 8,59158E-30 |
| ACE2 | CDH23 | -0,5223 | 8,62878E-30 |
| ACE2 | ST3GAL1 | -0,5225 | 8,13219E-30 |
| ACE2 | COMP | -0,5226 | 7,95022E-30 |

| | | | |
|---|---|---|---|
| ACE2 | MMRN1 | -0,5227 | 7,50947E-30 |
| ACE2 | DDX46 | -0,5227 | 7,66075E-30 |
| ACE2 | GIMAP4 | -0,5227 | 7,65399E-30 |
| ACE2 | ARHGAP19 | -0,5227 | 7,57277E-30 |
| ACE2 | RAD23B | -0,5228 | 7,35909E-30 |
| ACE2 | ABCD2 | -0,5228 | 7,39771E-30 |
| ACE2 | YWHAH | -0,5228 | 7,32123E-30 |
| ACE2 | DMC1 | -0,5228 | 7,33756E-30 |
| ACE2 | PCSK1N | -0,5228 | 7,33895E-30 |
| ACE2 | PLSCR4 | -0,5229 | 7,17479E-30 |
| ACE2 | PRR18 | -0,5229 | 7,07881E-30 |
| ACE2 | ADARB2 | -0,5229 | 7,13236E-30 |
| ACE2 | FTSJ3 | -0,5229 | 7,10356E-30 |
| ACE2 | MAFG | -0,5229 | 7,15528E-30 |
| ACE2 | SLC2A1-AS1 | -0,523 | 6,94864E-30 |
| ACE2 | PCBP1-AS1 | -0,523 | 6,93133E-30 |
| ACE2 | POLR2D | -0,523 | 6,9603E-30 |
| ACE2 | RP11-1365D11.1 | -0,523 | 6,95008E-30 |
| ACE2 | ENOPH1 | -0,523 | 6,95627E-30 |
| ACE2 | RP11-567M16.6 | -0,523 | 6,87425E-30 |
| ACE2 | MAPK1IP1L | -0,5231 | 6,836E-30 |
| ACE2 | RP11-227G15.11 | -0,5231 | 6,86405E-30 |
| ACE2 | HHLA3 | -0,5232 | 6,63924E-30 |
| ACE2 | C10orf128 | -0,5232 | 6,56707E-30 |
| ACE2 | RP4-633O19__A.1 | -0,5232 | 6,63844E-30 |
| ACE2 | RTN4 | -0,5233 | 6,30963E-30 |
| ACE2 | FAM71F2 | -0,5233 | 6,31853E-30 |
| ACE2 | RP11-411A19.5 | -0,5233 | 6,44065E-30 |
| ACE2 | PSMA7 | -0,5233 | 6,43134E-30 |
| ACE2 | GLTSCR1L | -0,5234 | 6,26073E-30 |
| ACE2 | C8orf44 | -0,5234 | 6,25686E-30 |
| ACE2 | AC004156.3 | -0,5234 | 6,26464E-30 |
| ACE2 | RCN1P2 | -0,5235 | 6,01964E-30 |
| ACE2 | SUGT1 | -0,5235 | 5,96754E-30 |
| ACE2 | STARD7-AS1 | -0,5236 | 5,85003E-30 |
| ACE2 | FKBP9P1 | -0,5236 | 5,76226E-30 |
| ACE2 | PPRC1 | -0,5237 | 5,6827E-30 |
| ACE2 | NPIPB6 | -0,5237 | 5,63444E-30 |
| ACE2 | FBXL12 | -0,5237 | 5,68364E-30 |
| ACE2 | OSBPL9 | -0,5238 | 5,439E-30 |
| ACE2 | CDCA7L | -0,5238 | 5,47052E-30 |
| ACE2 | RP11-282I1.1 | -0,5238 | 5,47951E-30 |
| ACE2 | YPEL2 | -0,5239 | 5,42623E-30 |
| ACE2 | ZNF345 | -0,5239 | 5,41733E-30 |
| ACE2 | RP11-212D19.5 | -0,524 | 5,16897E-30 |
| ACE2 | RP11-1180F24.3 | -0,524 | 5,12585E-30 |
| ACE2 | SERINC3 | -0,524 | 5,17158E-30 |
| ACE2 | WBSCR22 | -0,5241 | 5,10781E-30 |
| ACE2 | UTP20 | -0,5241 | 5,05866E-30 |
| ACE2 | PPM1L | -0,5242 | 4,92987E-30 |
| ACE2 | RP11-615I2.6 | -0,5243 | 4,79476E-30 |
| ACE2 | POC5 | -0,5244 | 4,67848E-30 |
| ACE2 | CALCA | -0,5244 | 4,56349E-30 |
| ACE2 | WT1 | -0,5244 | 4,57854E-30 |

| | | | |
|---|---|---|---|
| ACE2 | HNRNPA1P16 | -0,5244 | 4,6421E-30 |
| ACE2 | SEMA3F | -0,5245 | 4,44802E-30 |
| ACE2 | KLHL24 | -0,5245 | 4,53958E-30 |
| ACE2 | BRIX1 | -0,5245 | 4,46757E-30 |
| ACE2 | UBN2 | -0,5245 | 4,53652E-30 |
| ACE2 | RP11-654A16.3 | -0,5245 | 4,46055E-30 |
| ACE2 | DNTTIP2 | -0,5246 | 4,32019E-30 |
| ACE2 | RNF214 | -0,5246 | 4,29812E-30 |
| ACE2 | STRAP | -0,5246 | 4,3902E-30 |
| ACE2 | CTD-2033D15.2 | -0,5246 | 4,40356E-30 |
| ACE2 | PRR34 | -0,5246 | 4,41949E-30 |
| ACE2 | RDH14 | -0,5247 | 4,2408E-30 |
| ACE2 | RP11-71E19.1 | -0,5247 | 4,23502E-30 |
| ACE2 | RP11-367J11.3 | -0,5247 | 4,19746E-30 |
| ACE2 | PCDHA7 | -0,5247 | 4,19136E-30 |
| ACE2 | ACR | -0,5247 | 4,18223E-30 |
| ACE2 | CDC26 | -0,5248 | 4,06801E-30 |
| ACE2 | RP11-339B21.8 | -0,5248 | 4,1525E-30 |
| ACE2 | SLC10A3 | -0,5248 | 4,10087E-30 |
| ACE2 | XXbac-BPG283O16.9 | -0,5249 | 4,04018E-30 |
| ACE2 | CDKN2A | -0,5249 | 4,03194E-30 |
| ACE2 | ZNF79 | -0,5249 | 3,97332E-30 |
| ACE2 | RP1-59D14.10 | -0,5249 | 3,9656E-30 |
| ACE2 | EXOSC2 | -0,525 | 3,92107E-30 |
| ACE2 | RNF2 | -0,5251 | 3,76916E-30 |
| ACE2 | TRIM52 | -0,5251 | 3,74954E-30 |
| ACE2 | ARHGAP5 | -0,5252 | 3,62238E-30 |
| ACE2 | LINC01024 | -0,5253 | 3,56789E-30 |
| ACE2 | RP11-2B6.3 | -0,5253 | 3,51487E-30 |
| ACE2 | ZCCHC4 | -0,5254 | 3,46165E-30 |
| ACE2 | SERPINB9 | -0,5254 | 3,43116E-30 |
| ACE2 | LINC00702 | -0,5254 | 3,44359E-30 |
| ACE2 | ZNF45 | -0,5254 | 3,41133E-30 |
| ACE2 | FAM90A1 | -0,5255 | 3,34548E-30 |
| ACE2 | RP11-1263C18.2 | -0,5256 | 3,24862E-30 |
| ACE2 | PPIL4 | -0,5256 | 3,26326E-30 |
| ACE2 | LINC00667 | -0,5256 | 3,24554E-30 |
| ACE2 | DICER1-AS1 | -0,5257 | 3,11371E-30 |
| ACE2 | MEG9 | -0,5257 | 3,14576E-30 |
| ACE2 | KRT25 | -0,5257 | 3,12688E-30 |
| ACE2 | AP2A1 | -0,5258 | 3,08725E-30 |
| ACE2 | CNOT3 | -0,5258 | 3,08102E-30 |
| ACE2 | SEC22B | -0,5259 | 2,94152E-30 |
| ACE2 | MORN2 | -0,526 | 2,92195E-30 |
| ACE2 | RP11-154H23.4 | -0,526 | 2,85152E-30 |
| ACE2 | MTIF3 | -0,526 | 2,84349E-30 |
| ACE2 | DUXAP8 | -0,526 | 2,8912E-30 |
| ACE2 | TRPV2 | -0,5261 | 2,8233E-30 |
| ACE2 | RP11-63G10.4 | -0,5262 | 2,68624E-30 |
| ACE2 | AGAP2 | -0,5263 | 2,65008E-30 |
| ACE2 | RPL7P9 | -0,5264 | 2,54219E-30 |
| ACE2 | SHC1 | -0,5264 | 2,55163E-30 |
| ACE2 | BRPF1 | -0,5264 | 2,54948E-30 |
| ACE2 | RP11-87H9.5 | -0,5264 | 2,57369E-30 |

| | | | |
|---|---|---|---|
| ACE2 | H2AFY2 | -0,5265 | 2,52073E-30 |
| ACE2 | CTA-445C9.14 | -0,5265 | 2,45854E-30 |
| ACE2 | CTD-2002J20.1 | -0,5266 | 2,38807E-30 |
| ACE2 | RAB18 | -0,5266 | 2,45042E-30 |
| ACE2 | ZNF674-AS1 | -0,5266 | 2,3953E-30 |
| ACE2 | MCCC1-AS1 | -0,5267 | 2,32134E-30 |
| ACE2 | CPSF6 | -0,5267 | 2,33927E-30 |
| ACE2 | SNX9 | -0,5268 | 2,24321E-30 |
| ACE2 | RP11-603J24.5 | -0,5268 | 2,29495E-30 |
| ACE2 | DYNLL1P4 | -0,5268 | 2,28646E-30 |
| ACE2 | KCTD11 | -0,5268 | 2,29705E-30 |
| ACE2 | RN7SKP208 | -0,5269 | 2,21054E-30 |
| ACE2 | ACKR4 | -0,5269 | 2,20967E-30 |
| ACE2 | HILPDA | -0,5269 | 2,2088E-30 |
| ACE2 | TUBGCP2 | -0,5269 | 2,23271E-30 |
| ACE2 | LYVE1 | -0,5269 | 2,18684E-30 |
| ACE2 | EHMT1 | -0,527 | 2,14334E-30 |
| ACE2 | DBIL5P | -0,527 | 2,13371E-30 |
| ACE2 | CCDC158 | -0,5271 | 2,05657E-30 |
| ACE2 | SAXO2 | -0,5271 | 2,09463E-30 |
| ACE2 | H3F3B | -0,5271 | 2,09126E-30 |
| ACE2 | FECH | -0,5271 | 2,10269E-30 |
| ACE2 | RP1-32I10.10 | -0,5271 | 2,08299E-30 |
| ACE2 | RHAG | -0,5272 | 2,01477E-30 |
| ACE2 | LINC01589 | -0,5272 | 2,01696E-30 |
| ACE2 | SZRD1 | -0,5273 | 1,96991E-30 |
| ACE2 | RP11-10N23.2 | -0,5273 | 1,95547E-30 |
| ACE2 | MAX | -0,5273 | 1,94815E-30 |
| ACE2 | YTHDF1 | -0,5273 | 1,94037E-30 |
| ACE2 | RP11-392O17.2 | -0,5274 | 1,89029E-30 |
| ACE2 | LINC01376 | -0,5274 | 1,89523E-30 |
| ACE2 | TMEM132E | -0,5274 | 1,91408E-30 |
| ACE2 | CCKBR | -0,5275 | 1,84075E-30 |
| ACE2 | CYYR1-AS1 | -0,5275 | 1,85765E-30 |
| ACE2 | RASGRF2-AS1 | -0,5276 | 1,77751E-30 |
| ACE2 | PFKFB3 | -0,5276 | 1,80575E-30 |
| ACE2 | NR4A1 | -0,5276 | 1,77797E-30 |
| ACE2 | ZNF394 | -0,5277 | 1,72413E-30 |
| ACE2 | PAFAH1B2 | -0,5277 | 1,71883E-30 |
| ACE2 | SNAP25-AS1 | -0,5277 | 1,76078E-30 |
| ACE2 | GLIS2-AS1 | -0,5278 | 1,66721E-30 |
| ACE2 | RP11-389C8.2 | -0,5279 | 1,62808E-30 |
| ACE2 | RHEBP2 | -0,5279 | 1,63549E-30 |
| ACE2 | SERTAD2 | -0,5281 | 1,55438E-30 |
| ACE2 | TANK | -0,5281 | 1,56839E-30 |
| ACE2 | TMEM140 | -0,5281 | 1,54729E-30 |
| ACE2 | CA9 | -0,5281 | 1,54042E-30 |
| ACE2 | SOCS2 | -0,5281 | 1,56852E-30 |
| ACE2 | C16orf46 | -0,5281 | 1,55655E-30 |
| ACE2 | RP1-234P15.4 | -0,5282 | 1,48819E-30 |
| ACE2 | TBX3 | -0,5282 | 1,48497E-30 |
| ACE2 | TAZ | -0,5282 | 1,48372E-30 |
| ACE2 | DYNC2LI1 | -0,5283 | 1,45549E-30 |
| ACE2 | PRRC2C | -0,5284 | 1,39778E-30 |

| | | | |
|---|---|---|---|
| ACE2 | MAPK8IP3 | -0,5284 | 1,43437E-30 |
| ACE2 | KAT14 | -0,5285 | 1,35924E-30 |
| ACE2 | THRB | -0,5287 | 1,28975E-30 |
| ACE2 | ATXN7 | -0,5287 | 1,29918E-30 |
| ACE2 | C16orf47 | -0,5287 | 1,31113E-30 |
| ACE2 | AC018890.6 | -0,5288 | 1,26371E-30 |
| ACE2 | CSPP1 | -0,5288 | 1,26508E-30 |
| ACE2 | ARMC3 | -0,5288 | 1,27016E-30 |
| ACE2 | RPL26P6 | -0,5288 | 1,27074E-30 |
| ACE2 | NABP2 | -0,5288 | 1,25788E-30 |
| ACE2 | RP11-463O9.9 | -0,5288 | 1,25882E-30 |
| ACE2 | C18orf25 | -0,5288 | 1,25005E-30 |
| ACE2 | ZNF280B | -0,5288 | 1,27406E-30 |
| ACE2 | RP11-436K8.1 | -0,5289 | 1,21502E-30 |
| ACE2 | RFTN1 | -0,5289 | 1,22958E-30 |
| ACE2 | C7orf25 | -0,5289 | 1,20843E-30 |
| ACE2 | TOX | -0,5289 | 1,20636E-30 |
| ACE2 | RP11-234G16.5 | -0,5289 | 1,21434E-30 |
| ACE2 | CDC23 | -0,5291 | 1,163E-30 |
| ACE2 | ZNF606 | -0,5291 | 1,15606E-30 |
| ACE2 | BSN-AS2 | -0,5292 | 1,1307E-30 |
| ACE2 | PCDHAC1 | -0,5292 | 1,12282E-30 |
| ACE2 | APBA1 | -0,5292 | 1,10925E-30 |
| ACE2 | TMEM132C | -0,5292 | 1,11848E-30 |
| ACE2 | RP11-243M5.5 | -0,5292 | 1,10671E-30 |
| ACE2 | CTC-524C5.2 | -0,5292 | 1,11447E-30 |
| ACE2 | FBN3 | -0,5292 | 1,12437E-30 |
| ACE2 | VAPB | -0,5292 | 1,10749E-30 |
| ACE2 | HUWE1 | -0,5292 | 1,10292E-30 |
| ACE2 | CLHC1 | -0,5293 | 1,07909E-30 |
| ACE2 | GAB1 | -0,5294 | 1,05141E-30 |
| ACE2 | RP11-247A12.7 | -0,5294 | 1,04448E-30 |
| ACE2 | SCLY | -0,5295 | 1,03022E-30 |
| ACE2 | KB-1042C11.5 | -0,5295 | 1,0102E-30 |
| ACE2 | PNPT1 | -0,5296 | 9,92994E-31 |
| ACE2 | RP11-526N18.1 | -0,5296 | 9,85019E-31 |
| ACE2 | OSBPL1A | -0,5296 | 1,00299E-30 |
| ACE2 | CH17-472G23.1 | -0,5297 | 9,64589E-31 |
| ACE2 | RP11-259K5.1 | -0,5298 | 9,22437E-31 |
| ACE2 | PRMT3 | -0,5298 | 9,19296E-31 |
| ACE2 | TMEM132A | -0,5298 | 9,37847E-31 |
| ACE2 | ZNF793-AS1 | -0,5298 | 9,28438E-31 |
| ACE2 | RP11-83N9.5 | -0,5299 | 8,9544E-31 |
| ACE2 | OMG | -0,5299 | 8,90502E-31 |
| ACE2 | RP11-213G2.5 | -0,53 | 8,69473E-31 |
| ACE2 | SLC12A1 | -0,53 | 8,79557E-31 |
| ACE2 | SF3B1 | -0,5301 | 8,51798E-31 |
| ACE2 | TBC1D2B | -0,5301 | 8,53207E-31 |
| ACE2 | RP11-46C24.7 | -0,5301 | 8,63742E-31 |
| ACE2 | CTC-204F22.1 | -0,5302 | 8,35756E-31 |
| ACE2 | SYP | -0,5302 | 8,38176E-31 |
| ACE2 | RASA1 | -0,5303 | 8,09553E-31 |
| ACE2 | RP11-712B9.5 | -0,5303 | 7,91613E-31 |
| ACE2 | AK4 | -0,5304 | 7,68699E-31 |

| | | | |
|---|---|---|---|
| ACE2 | GPR20 | -0,5304 | 7,82557E-31 |
| ACE2 | BCAS2 | -0,5305 | 7,55685E-31 |
| ACE2 | RP11-680F20.6 | -0,5305 | 7,49607E-31 |
| ACE2 | RP11-167B3.2 | -0,5305 | 7,44954E-31 |
| ACE2 | MAPT-IT1 | -0,5305 | 7,48629E-31 |
| ACE2 | HPD | -0,5306 | 7,33211E-31 |
| ACE2 | GABRB3 | -0,5306 | 7,38183E-31 |
| ACE2 | CTNS | -0,5306 | 7,28825E-31 |
| ACE2 | AC136289.1 | -0,5307 | 7,10714E-31 |
| ACE2 | CTB-158E9.2 | -0,5307 | 7,11706E-31 |
| ACE2 | CTD-3099C6.9 | -0,5307 | 7,12791E-31 |
| ACE2 | ACVR2B | -0,5309 | 6,65472E-31 |
| ACE2 | ORC5 | -0,5309 | 6,73144E-31 |
| ACE2 | RP11-286B14.1 | -0,531 | 6,41232E-31 |
| ACE2 | GNMT | -0,531 | 6,55369E-31 |
| ACE2 | PI15 | -0,531 | 6,52267E-31 |
| ACE2 | UTP23 | -0,531 | 6,5909E-31 |
| ACE2 | USP32 | -0,531 | 6,4899E-31 |
| ACE2 | RP11-383I23.2 | -0,5311 | 6,37064E-31 |
| ACE2 | ATR | -0,5311 | 6,27646E-31 |
| ACE2 | COL4A3BP | -0,5311 | 6,27176E-31 |
| ACE2 | TDH | -0,5312 | 6,02816E-31 |
| ACE2 | ASB14 | -0,5313 | 5,85943E-31 |
| ACE2 | CAPZA2 | -0,5313 | 6,01842E-31 |
| ACE2 | CCDC144NL | -0,5313 | 5,91499E-31 |
| ACE2 | C1orf140 | -0,5314 | 5,80201E-31 |
| ACE2 | NDST1 | -0,5314 | 5,69479E-31 |
| ACE2 | ZNF680 | -0,5314 | 5,70053E-31 |
| ACE2 | ZPR1 | -0,5314 | 5,77363E-31 |
| ACE2 | NLRP2B | -0,5314 | 5,69487E-31 |
| ACE2 | CDKN2AIPNL | -0,5315 | 5,54412E-31 |
| ACE2 | ZNF292 | -0,5316 | 5,48497E-31 |
| ACE2 | LINC00844 | -0,5316 | 5,43954E-31 |
| ACE2 | LINC01125 | -0,5317 | 5,30561E-31 |
| ACE2 | SCOC | -0,5317 | 5,33327E-31 |
| ACE2 | AC093627.10 | -0,5317 | 5,22636E-31 |
| ACE2 | RASL11B | -0,5318 | 5,11408E-31 |
| ACE2 | TXNDC16 | -0,5318 | 5,03137E-31 |
| ACE2 | L3MBTL4 | -0,5318 | 5,1198E-31 |
| ACE2 | APLN | -0,5318 | 5,1231E-31 |
| ACE2 | LRRC38 | -0,5319 | 4,91107E-31 |
| ACE2 | FAM212B-AS1 | -0,5319 | 4,88654E-31 |
| ACE2 | E2F5 | -0,5319 | 4,89489E-31 |
| ACE2 | KIAA0368 | -0,5319 | 4,97369E-31 |
| ACE2 | PSEN2 | -0,532 | 4,76176E-31 |
| ACE2 | RP11-195B17.1 | -0,532 | 4,84319E-31 |
| ACE2 | PELI2 | -0,532 | 4,78252E-31 |
| ACE2 | RP11-554D20.2 | -0,532 | 4,8584E-31 |
| ACE2 | PIK3C3 | -0,532 | 4,76107E-31 |
| ACE2 | ZNF157 | -0,532 | 4,86784E-31 |
| ACE2 | ELP3 | -0,5321 | 4,59934E-31 |
| ACE2 | LINC00487 | -0,5322 | 4,47836E-31 |
| ACE2 | LRRCC1 | -0,5322 | 4,58844E-31 |
| ACE2 | RP11-244E17.1 | -0,5322 | 4,54194E-31 |

| | | | |
|---|---|---|---|
| ACE2 | PRICKLE2-AS1 | -0,5323 | 4,35132E-31 |
| ACE2 | EXOC4 | -0,5323 | 4,34194E-31 |
| ACE2 | PQBP1 | -0,5323 | 4,3569E-31 |
| ACE2 | HAUS7 | -0,5323 | 4,40448E-31 |
| ACE2 | MRPS27 | -0,5324 | 4,20562E-31 |
| ACE2 | DISC1 | -0,5325 | 4,1148E-31 |
| ACE2 | AFF1 | -0,5325 | 4,18455E-31 |
| ACE2 | PSMB7 | -0,5325 | 4,08322E-31 |
| ACE2 | ASB16-AS1 | -0,5325 | 4,13962E-31 |
| ACE2 | FIG4 | -0,5326 | 3,97079E-31 |
| ACE2 | LINC00294 | -0,5326 | 4,04088E-31 |
| ACE2 | PPHLN1 | -0,5326 | 4,00382E-31 |
| ACE2 | CDC37 | -0,5326 | 4,0438E-31 |
| ACE2 | KCNE5 | -0,5326 | 4,06128E-31 |
| ACE2 | BCAT1 | -0,5327 | 3,91156E-31 |
| ACE2 | MPHOSPH9 | -0,5327 | 3,86698E-31 |
| ACE2 | C16orf71 | -0,5327 | 3,89814E-31 |
| ACE2 | HPN-AS1 | -0,5327 | 3,84755E-31 |
| ACE2 | PPIL1 | -0,5328 | 3,78513E-31 |
| ACE2 | CTD-2515H24.2 | -0,5328 | 3,80115E-31 |
| ACE2 | RP11-752L20.3 | -0,5329 | 3,68504E-31 |
| ACE2 | PACSIN1 | -0,5329 | 3,70515E-31 |
| ACE2 | ST6GALNAC6 | -0,5329 | 3,62869E-31 |
| ACE2 | NTMT1 | -0,5329 | 3,67248E-31 |
| ACE2 | HEXA | -0,5329 | 3,62436E-31 |
| ACE2 | RPL6P27 | -0,5329 | 3,67582E-31 |
| ACE2 | LINC00619 | -0,533 | 3,52027E-31 |
| ACE2 | RP11-60A24.3 | -0,533 | 3,56486E-31 |
| ACE2 | DBNDD2 | -0,533 | 3,52613E-31 |
| ACE2 | TADA3 | -0,5331 | 3,39761E-31 |
| ACE2 | WTAP | -0,5331 | 3,46995E-31 |
| ACE2 | OR13J1 | -0,5331 | 3,45037E-31 |
| ACE2 | THRAP3 | -0,5332 | 3,37717E-31 |
| ACE2 | CSNK2B | -0,5332 | 3,35819E-31 |
| ACE2 | LAG3 | -0,5332 | 3,3863E-31 |
| ACE2 | IARS | -0,5333 | 3,2426E-31 |
| ACE2 | RP5-1068E13.7 | -0,5333 | 3,26403E-31 |
| ACE2 | LINC00163 | -0,5333 | 3,2661E-31 |
| ACE2 | CCDC174 | -0,5334 | 3,11667E-31 |
| ACE2 | MBD4 | -0,5334 | 3,16546E-31 |
| ACE2 | GOT1L1 | -0,5334 | 3,18873E-31 |
| ACE2 | RP11-282O18.3 | -0,5334 | 3,12249E-31 |
| ACE2 | HAPLN3 | -0,5334 | 3,1781E-31 |
| ACE2 | C19orf43 | -0,5334 | 3,1666E-31 |
| ACE2 | NSUN4 | -0,5335 | 3,02165E-31 |
| ACE2 | RP11-449G16.1 | -0,5335 | 3,04056E-31 |
| ACE2 | ART5 | -0,5335 | 3,07239E-31 |
| ACE2 | MAP1LC3A | -0,5335 | 3,07479E-31 |
| ACE2 | CTC-510F12.3 | -0,5336 | 2,96235E-31 |
| ACE2 | TATDN3 | -0,5337 | 2,9029E-31 |
| ACE2 | LIPT1 | -0,5337 | 2,87917E-31 |
| ACE2 | CCDC121 | -0,5338 | 2,81732E-31 |
| ACE2 | RP11-1109F11.3 | -0,5338 | 2,81375E-31 |
| ACE2 | SPOPL | -0,534 | 2,63135E-31 |

| | | | |
|---|---|---|---|
| ACE2 | RP11-356B19.11 | -0,534 | 2,66364E-31 |
| ACE2 | SLITRK1 | -0,534 | 2,64755E-31 |
| ACE2 | AC012358.8 | -0,5341 | 2,53483E-31 |
| ACE2 | THAP2 | -0,5341 | 2,55894E-31 |
| ACE2 | MNS1 | -0,5341 | 2,53757E-31 |
| ACE2 | NDFIP1 | -0,5342 | 2,46909E-31 |
| ACE2 | MTRNR2L4 | -0,5342 | 2,48605E-31 |
| ACE2 | WDR13 | -0,5342 | 2,48243E-31 |
| ACE2 | RAP2A | -0,5343 | 2,4089E-31 |
| ACE2 | PRUNE1 | -0,5345 | 2,23487E-31 |
| ACE2 | RP11-574K11.5 | -0,5345 | 2,25389E-31 |
| ACE2 | NSD1 | -0,5346 | 2,18021E-31 |
| ACE2 | SFR1 | -0,5346 | 2,1599E-31 |
| ACE2 | C9orf172 | -0,5347 | 2,14557E-31 |
| ACE2 | DGKZ | -0,5347 | 2,09266E-31 |
| ACE2 | SUZ12 | -0,5347 | 2,14008E-31 |
| ACE2 | HTATSF1 | -0,5347 | 2,13928E-31 |
| ACE2 | RP11-307C18.1 | -0,5348 | 2,08387E-31 |
| ACE2 | FAM66A | -0,5348 | 2,06791E-31 |
| ACE2 | NXNL2 | -0,5348 | 2,06131E-31 |
| ACE2 | RP11-29H23.5 | -0,5349 | 1,98938E-31 |
| ACE2 | MTHFD1L | -0,5349 | 1,98583E-31 |
| ACE2 | NYNRIN | -0,5349 | 1,97149E-31 |
| ACE2 | WDR92 | -0,535 | 1,9122E-31 |
| ACE2 | RP1-95L4.4 | -0,535 | 1,91193E-31 |
| ACE2 | GNPTG | -0,535 | 1,92172E-31 |
| ACE2 | CXXC5 | -0,5351 | 1,89825E-31 |
| ACE2 | EXT2 | -0,5351 | 1,85318E-31 |
| ACE2 | CCND1 | -0,5351 | 1,8708E-31 |
| ACE2 | POLR2C | -0,5351 | 1,85188E-31 |
| ACE2 | ZSCAN22 | -0,5351 | 1,88793E-31 |
| ACE2 | SLC35G5 | -0,5352 | 1,84689E-31 |
| ACE2 | SMG6 | -0,5352 | 1,79403E-31 |
| ACE2 | STX11 | -0,5354 | 1,71856E-31 |
| ACE2 | WDR61 | -0,5354 | 1,71738E-31 |
| ACE2 | GSG1L | -0,5354 | 1,69173E-31 |
| ACE2 | YPEL3 | -0,5354 | 1,7094E-31 |
| ACE2 | ZNF275 | -0,5354 | 1,73665E-31 |
| ACE2 | AC114730.2 | -0,5355 | 1,64959E-31 |
| ACE2 | RAMP2-AS1 | -0,5356 | 1,60365E-31 |
| ACE2 | AC105760.2 | -0,5357 | 1,5408E-31 |
| ACE2 | ELFN2 | -0,5357 | 1,54348E-31 |
| ACE2 | ACBD4 | -0,5358 | 1,49525E-31 |
| ACE2 | LCA10 | -0,5358 | 1,49929E-31 |
| ACE2 | H6PD | -0,5359 | 1,48266E-31 |
| ACE2 | RP11-736K20.6 | -0,5359 | 1,48291E-31 |
| ACE2 | GOLGA8H | -0,5359 | 1,47979E-31 |
| ACE2 | KRT8P26 | -0,536 | 1,42813E-31 |
| ACE2 | KDM4D | -0,536 | 1,42051E-31 |
| ACE2 | CTD-2553L13.9 | -0,5362 | 1,36107E-31 |
| ACE2 | RPL23AP49 | -0,5363 | 1,31383E-31 |
| ACE2 | LINC01322 | -0,5363 | 1,32082E-31 |
| ACE2 | ZNF862 | -0,5363 | 1,31217E-31 |
| ACE2 | TNFAIP2 | -0,5363 | 1,31734E-31 |

| | | | |
|---|---|---|---|
| ACE2 | TMEM198 | -0,5364 | 1,27432E-31 |
| ACE2 | KDR | -0,5364 | 1,27486E-31 |
| ACE2 | SNAPC4 | -0,5364 | 1,27845E-31 |
| ACE2 | RP11-845M18.6 | -0,5364 | 1,27145E-31 |
| ACE2 | DNASE2 | -0,5364 | 1,24653E-31 |
| ACE2 | bP-21264C1.1 | -0,5364 | 1,26952E-31 |
| ACE2 | TESK1 | -0,5365 | 1,24146E-31 |
| ACE2 | RP11-371I20.2 | -0,5365 | 1,22492E-31 |
| ACE2 | ARL5A | -0,5366 | 1,186E-31 |
| ACE2 | NPIPB9 | -0,5366 | 1,1956E-31 |
| ACE2 | TGFB2-AS1 | -0,5367 | 1,16587E-31 |
| ACE2 | RP4-580N22.2 | -0,5367 | 1,15358E-31 |
| ACE2 | GOLGA2 | -0,5367 | 1,16844E-31 |
| ACE2 | PHF8 | -0,5367 | 1,16058E-31 |
| ACE2 | SYF2 | -0,5368 | 1,11652E-31 |
| ACE2 | FBXO25 | -0,5368 | 1,13216E-31 |
| ACE2 | NDP | -0,5368 | 1,13082E-31 |
| ACE2 | HTR1F | -0,5369 | 1,08631E-31 |
| ACE2 | RP11-1081M5.3 | -0,5369 | 1,08981E-31 |
| ACE2 | TIMM8AP1 | -0,537 | 1,04153E-31 |
| ACE2 | GOLPH3 | -0,537 | 1,05023E-31 |
| ACE2 | PRG2 | -0,5371 | 1,02624E-31 |
| ACE2 | RP11-524C21.1 | -0,5374 | 9,31359E-32 |
| ACE2 | AF186192.1 | -0,5374 | 9,19954E-32 |
| ACE2 | PDGFD | -0,5374 | 9,33414E-32 |
| ACE2 | UHRF1BP1L | -0,5374 | 9,43224E-32 |
| ACE2 | ACIN1 | -0,5375 | 9,12017E-32 |
| ACE2 | RNASEH1 | -0,5376 | 8,70672E-32 |
| ACE2 | SRSF7 | -0,5376 | 8,73676E-32 |
| ACE2 | PDCD1LG2 | -0,5376 | 8,63103E-32 |
| ACE2 | RP11-352M15.2 | -0,5376 | 8,81348E-32 |
| ACE2 | RP11-488C13.5 | -0,5376 | 8,82733E-32 |
| ACE2 | CACNA2D3-AS1 | -0,5377 | 8,42305E-32 |
| ACE2 | SMAD9-IT1 | -0,5377 | 8,3512E-32 |
| ACE2 | CAPS | -0,5377 | 8,41922E-32 |
| ACE2 | RP11-362K14.6 | -0,5378 | 8,26333E-32 |
| ACE2 | EEF1A1P5 | -0,5378 | 8,32517E-32 |
| ACE2 | CYP17A1-AS1 | -0,5379 | 8,08849E-32 |
| ACE2 | HNRNPLP2 | -0,5379 | 8,03949E-32 |
| ACE2 | TMA16 | -0,538 | 7,64521E-32 |
| ACE2 | ETF1 | -0,538 | 7,60877E-32 |
| ACE2 | PMS2P3 | -0,538 | 7,80546E-32 |
| ACE2 | CHST7 | -0,538 | 7,71613E-32 |
| ACE2 | RP11-416A14.1 | -0,5381 | 7,51168E-32 |
| ACE2 | SPIDR | -0,5381 | 7,48411E-32 |
| ACE2 | CCDC78 | -0,5381 | 7,5178E-32 |
| ACE2 | SDCBP | -0,5382 | 7,25509E-32 |
| ACE2 | C1QTNF9 | -0,5383 | 7,10966E-32 |
| ACE2 | C17orf107 | -0,5383 | 7,01869E-32 |
| ACE2 | SENP3 | -0,5383 | 6,97374E-32 |
| ACE2 | RP11-156E6.1 | -0,5383 | 7,06598E-32 |
| ACE2 | ARID2 | -0,5384 | 6,914E-32 |
| ACE2 | XXYLT1-AS2 | -0,5385 | 6,66781E-32 |
| ACE2 | LA16c-360A4.1 | -0,5385 | 6,59094E-32 |

| | | | |
|---|---|---|---|
| ACE2 | SERTAD3 | -0,5385 | 6,54383E-32 |
| ACE2 | BSCL2 | -0,5386 | 6,47227E-32 |
| ACE2 | TFPI2 | -0,5388 | 5,98881E-32 |
| ACE2 | DKFZP434A062 | -0,5388 | 5,98496E-32 |
| ACE2 | EPHX4 | -0,5389 | 5,89628E-32 |
| ACE2 | NUDT2 | -0,5389 | 5,88293E-32 |
| ACE2 | ZMIZ1 | -0,5389 | 5,8457E-32 |
| ACE2 | RP11-531A24.3 | -0,539 | 5,62988E-32 |
| ACE2 | RP11-399E6.1 | -0,5391 | 5,4529E-32 |
| ACE2 | PPP1CB | -0,5391 | 5,44593E-32 |
| ACE2 | PER1 | -0,5391 | 5,4996E-32 |
| ACE2 | CTC-525D6.1 | -0,5391 | 5,42294E-32 |
| ACE2 | RP5-1116H23.4 | -0,5391 | 5,53946E-32 |
| ACE2 | GPR1 | -0,5392 | 5,39152E-32 |
| ACE2 | RP11-464C19.3 | -0,5392 | 5,37837E-32 |
| ACE2 | ACTC1 | -0,5392 | 5,3464E-32 |
| ACE2 | GPATCH8 | -0,5392 | 5,3107E-32 |
| ACE2 | NBPF12 | -0,5393 | 5,22052E-32 |
| ACE2 | TNKS2 | -0,5393 | 5,18739E-32 |
| ACE2 | DHX33 | -0,5393 | 5,20184E-32 |
| ACE2 | TEKT3 | -0,5393 | 5,25037E-32 |
| ACE2 | LINC00894 | -0,5393 | 5,20766E-32 |
| ACE2 | FLG | -0,5394 | 5,06363E-32 |
| ACE2 | CH17-13I23.3 | -0,5395 | 4,82995E-32 |
| ACE2 | ZNF550 | -0,5395 | 4,88843E-32 |
| ACE2 | CNOT9 | -0,5396 | 4,75488E-32 |
| ACE2 | FKBP15 | -0,5396 | 4,74035E-32 |
| ACE2 | PCDHA11 | -0,5397 | 4,54804E-32 |
| ACE2 | BRAT1 | -0,5397 | 4,54651E-32 |
| ACE2 | ZNF623 | -0,5397 | 4,57489E-32 |
| ACE2 | GNAQ | -0,5397 | 4,62638E-32 |
| ACE2 | RNF208 | -0,5397 | 4,52954E-32 |
| ACE2 | RP11-139H15.5 | -0,5397 | 4,6411E-32 |
| ACE2 | RP11-411B10.8 | -0,5397 | 4,58962E-32 |
| ACE2 | ZNF530 | -0,5397 | 4,60247E-32 |
| ACE2 | LINC00315 | -0,5397 | 4,57509E-32 |
| ACE2 | MAPK1 | -0,5397 | 4,54204E-32 |
| ACE2 | RBBP4 | -0,5398 | 4,37697E-32 |
| ACE2 | RP11-472B18.1 | -0,5398 | 4,42864E-32 |
| ACE2 | RP5-874C20.7 | -0,5398 | 4,50184E-32 |
| ACE2 | ZNF655 | -0,5398 | 4,47678E-32 |
| ACE2 | MPHOSPH10 | -0,5399 | 4,29731E-32 |
| ACE2 | KB-431C1.5 | -0,5399 | 4,23614E-32 |
| ACE2 | RP11-345J18.2 | -0,5399 | 4,31466E-32 |
| ACE2 | CBFB | -0,5399 | 4,26528E-32 |
| ACE2 | GRM2 | -0,5401 | 4,05994E-32 |
| ACE2 | AC004540.5 | -0,5401 | 4,01072E-32 |
| ACE2 | UBQLNL | -0,5401 | 4,09503E-32 |
| ACE2 | RP11-123K19.1 | -0,5402 | 3,90786E-32 |
| ACE2 | SLC24A4 | -0,5403 | 3,8201E-32 |
| ACE2 | SMURF2P1 | -0,5403 | 3,80319E-32 |
| ACE2 | ZMYM1 | -0,5404 | 3,70248E-32 |
| ACE2 | FAM208A | -0,5404 | 3,64285E-32 |
| ACE2 | NDUFA4 | -0,5404 | 3,73663E-32 |

| | | | |
|---|---|---|---|
| ACE2 | PDDC1 | -0,5404 | 3,64778E-32 |
| ACE2 | PDCD2L | -0,5404 | 3,73676E-32 |
| ACE2 | RP11-227F19.1 | -0,5405 | 3,54328E-32 |
| ACE2 | ARID4A | -0,5405 | 3,56512E-32 |
| ACE2 | PTPN1 | -0,5405 | 3,5907E-32 |
| ACE2 | APCDD1L | -0,5405 | 3,57397E-32 |
| ACE2 | RCC1L | -0,5406 | 3,46107E-32 |
| ACE2 | LPL | -0,5406 | 3,43912E-32 |
| ACE2 | RP11-718O11.1 | -0,5406 | 3,47719E-32 |
| ACE2 | AC002398.13 | -0,5407 | 3,36372E-32 |
| ACE2 | TOGARAM2 | -0,5408 | 3,28756E-32 |
| ACE2 | TAF8 | -0,5408 | 3,20602E-32 |
| ACE2 | ZNF597 | -0,5409 | 3,11967E-32 |
| ACE2 | TMEM206 | -0,541 | 3,04946E-32 |
| ACE2 | IQCB1 | -0,5411 | 2,98581E-32 |
| ACE2 | ZNF740 | -0,5411 | 2,94705E-32 |
| ACE2 | ABCA1 | -0,5412 | 2,85136E-32 |
| ACE2 | CCPG1 | -0,5412 | 2,8658E-32 |
| ACE2 | MMP23B | -0,5413 | 2,75275E-32 |
| ACE2 | DIS3L2 | -0,5413 | 2,78504E-32 |
| ACE2 | CNOT6L | -0,5413 | 2,78108E-32 |
| ACE2 | RP11-1072C15.6 | -0,5413 | 2,77898E-32 |
| ACE2 | CCDC130 | -0,5413 | 2,78157E-32 |
| ACE2 | LINC01352 | -0,5414 | 2,68519E-32 |
| ACE2 | PPP3R1 | -0,5414 | 2,703E-32 |
| ACE2 | CADPS | -0,5414 | 2,69473E-32 |
| ACE2 | METTL23 | -0,5414 | 2,67225E-32 |
| ACE2 | TJP1 | -0,5415 | 2,65009E-32 |
| ACE2 | HMG20B | -0,5415 | 2,6014E-32 |
| ACE2 | UBXN10 | -0,5416 | 2,55015E-32 |
| ACE2 | CARD8 | -0,5416 | 2,53172E-32 |
| ACE2 | DHX34 | -0,5417 | 2,46394E-32 |
| ACE2 | RHBDD3 | -0,5417 | 2,42239E-32 |
| ACE2 | GPATCH11 | -0,5418 | 2,41494E-32 |
| ACE2 | PCDHA6 | -0,5418 | 2,34569E-32 |
| ACE2 | LRP11 | -0,5418 | 2,41561E-32 |
| ACE2 | RP11-1081M5.1 | -0,5418 | 2,3829E-32 |
| ACE2 | ADAMTSL1 | -0,5418 | 2,39129E-32 |
| ACE2 | RANBP10 | -0,5418 | 2,34927E-32 |
| ACE2 | STAG2 | -0,5418 | 2,36572E-32 |
| ACE2 | RBM44 | -0,5419 | 2,33588E-32 |
| ACE2 | CH17-360D5.2 | -0,5419 | 2,28935E-32 |
| ACE2 | RP1-197B17.3 | -0,5419 | 2,28956E-32 |
| ACE2 | RP11-426C22.4 | -0,5419 | 2,29106E-32 |
| ACE2 | FIZ1 | -0,5419 | 2,30604E-32 |
| ACE2 | AP000223.42 | -0,5419 | 2,33716E-32 |
| ACE2 | RAB7B | -0,542 | 2,24083E-32 |
| ACE2 | POLB | -0,542 | 2,23248E-32 |
| ACE2 | OAT | -0,542 | 2,23189E-32 |
| ACE2 | FAM86GP | -0,542 | 2,2412E-32 |
| ACE2 | RASD2 | -0,5421 | 2,15393E-32 |
| ACE2 | RSRC2 | -0,5422 | 2,13081E-32 |
| ACE2 | RP11-413H22.3 | -0,5422 | 2,09981E-32 |
| ACE2 | RP11-676J12.9 | -0,5422 | 2,07501E-32 |

| | | | |
|---|---|---|---|
| ACE2 | RP1-292L20.3 | -0,5423 | 2,04672E-32 |
| ACE2 | AC007969.5 | -0,5424 | 1,9808E-32 |
| ACE2 | TTN | -0,5424 | 1,95163E-32 |
| ACE2 | FLJ46284 | -0,5424 | 1,9549E-32 |
| ACE2 | CHD8 | -0,5424 | 2,00299E-32 |
| ACE2 | MAN2B2 | -0,5425 | 1,88793E-32 |
| ACE2 | LINC00242 | -0,5425 | 1,94335E-32 |
| ACE2 | AC073316.1 | -0,5425 | 1,90854E-32 |
| ACE2 | ZNF680P1 | -0,5425 | 1,90175E-32 |
| ACE2 | RP11-667K14.3 | -0,5425 | 1,92445E-32 |
| ACE2 | LMAN1 | -0,5425 | 1,94323E-32 |
| ACE2 | OR5K2 | -0,5426 | 1,83846E-32 |
| ACE2 | DLEU2L | -0,5427 | 1,8087E-32 |
| ACE2 | CSMD3 | -0,5427 | 1,77187E-32 |
| ACE2 | TAF2 | -0,5427 | 1,80812E-32 |
| ACE2 | MFGE8 | -0,5427 | 1,81476E-32 |
| ACE2 | ZNF551 | -0,5427 | 1,82654E-32 |
| ACE2 | FAM83D | -0,5427 | 1,82177E-32 |
| ACE2 | PRPF4 | -0,5428 | 1,77037E-32 |
| ACE2 | MINDY3 | -0,5428 | 1,74229E-32 |
| ACE2 | LINC01765 | -0,5429 | 1,66588E-32 |
| ACE2 | PNO1 | -0,5429 | 1,69775E-32 |
| ACE2 | TOX4 | -0,5429 | 1,66604E-32 |
| ACE2 | CRISPLD2 | -0,5429 | 1,70734E-32 |
| ACE2 | RP11-232D9.3 | -0,5429 | 1,66643E-32 |
| ACE2 | RAB3GAP2 | -0,543 | 1,65146E-32 |
| ACE2 | C4A | -0,543 | 1,62081E-32 |
| ACE2 | C18orf21 | -0,543 | 1,64722E-32 |
| ACE2 | SYS1 | -0,543 | 1,64134E-32 |
| ACE2 | RP11-47A8.5 | -0,5431 | 1,56523E-32 |
| ACE2 | FZD10-AS1 | -0,5431 | 1,59656E-32 |
| ACE2 | GNB1 | -0,5432 | 1,53772E-32 |
| ACE2 | RP11-153F10.2 | -0,5432 | 1,5315E-32 |
| ACE2 | CLCN3P1 | -0,5432 | 1,55346E-32 |
| ACE2 | GAS8 | -0,5432 | 1,54669E-32 |
| ACE2 | SPAG17 | -0,5433 | 1,49375E-32 |
| ACE2 | BOD1L1 | -0,5433 | 1,47909E-32 |
| ACE2 | KCTD21 | -0,5433 | 1,4976E-32 |
| ACE2 | RP11-161H23.9 | -0,5433 | 1,50659E-32 |
| ACE2 | LINC00398 | -0,5433 | 1,50377E-32 |
| ACE2 | EPOP | -0,5433 | 1,50267E-32 |
| ACE2 | RP11-21C4.1 | -0,5434 | 1,4413E-32 |
| ACE2 | HK1 | -0,5434 | 1,43819E-32 |
| ACE2 | KIF1B | -0,5435 | 1,38207E-32 |
| ACE2 | IKBKB | -0,5435 | 1,39663E-32 |
| ACE2 | VPS4A | -0,5435 | 1,41753E-32 |
| ACE2 | SLC47A1P1 | -0,5435 | 1,37834E-32 |
| ACE2 | AC115522.3 | -0,5436 | 1,3567E-32 |
| ACE2 | C2orf71 | -0,5438 | 1,27486E-32 |
| ACE2 | CNKSR3 | -0,5438 | 1,29367E-32 |
| ACE2 | CDK5RAP1 | -0,5438 | 1,27533E-32 |
| ACE2 | RFX8 | -0,5439 | 1,24853E-32 |
| ACE2 | FLI1 | -0,5439 | 1,24463E-32 |
| ACE2 | RP11-351M8.1 | -0,5439 | 1,21823E-32 |

| | | | |
|---|---|---|---|
| ACE2 | RP11-401P9.7 | -0,5439 | 1,23969E-32 |
| ACE2 | RP11-227G15.10 | -0,5439 | 1,23725E-32 |
| ACE2 | USP24 | -0,544 | 1,2053E-32 |
| ACE2 | AP006621.5 | -0,544 | 1,19332E-32 |
| ACE2 | ANKRD54 | -0,544 | 1,21334E-32 |
| ACE2 | RP11-169D4.2 | -0,5441 | 1,14181E-32 |
| ACE2 | RAB5B | -0,5441 | 1,16826E-32 |
| ACE2 | PAXBP1-AS1 | -0,5441 | 1,16559E-32 |
| ACE2 | CHTOP | -0,5442 | 1,11379E-32 |
| ACE2 | TAF1A | -0,5442 | 1,13987E-32 |
| ACE2 | NAA50 | -0,5442 | 1,11375E-32 |
| ACE2 | LARS | -0,5442 | 1,13342E-32 |
| ACE2 | GABBR2 | -0,5442 | 1,11588E-32 |
| ACE2 | KIAA0586 | -0,5442 | 1,12266E-32 |
| ACE2 | RP11-229O3.1 | -0,5442 | 1,14097E-32 |
| ACE2 | KLHL41 | -0,5443 | 1,07394E-32 |
| ACE2 | ALG1L15P | -0,5443 | 1,08374E-32 |
| ACE2 | RP5-940J5.8 | -0,5443 | 1,07717E-32 |
| ACE2 | ALDH1A1 | -0,5444 | 1,05606E-32 |
| ACE2 | ATG13 | -0,5444 | 1,05178E-32 |
| ACE2 | RP11-439H8.4 | -0,5445 | 1,01111E-32 |
| ACE2 | LBR | -0,5445 | 1,01377E-32 |
| ACE2 | OST4 | -0,5445 | 1,01252E-32 |
| ACE2 | TBX6 | -0,5445 | 1,01204E-32 |
| ACE2 | AC124789.1 | -0,5445 | 1,01094E-32 |
| ACE2 | RP11-541M12.6 | -0,5446 | 1,00179E-32 |
| ACE2 | RN7SL689P | -0,5447 | 9,73996E-33 |
| ACE2 | PCDHGB5 | -0,5447 | 9,46273E-33 |
| ACE2 | API5 | -0,5447 | 9,51122E-33 |
| ACE2 | SS18L1 | -0,5447 | 9,7475E-33 |
| ACE2 | LL0XNC01-237H1.2 | -0,5447 | 9,60904E-33 |
| ACE2 | KIAA1429 | -0,5448 | 9,37391E-33 |
| ACE2 | RP11-88G17.6 | -0,5449 | 9,00093E-33 |
| ACE2 | CA5BP1 | -0,545 | 8,64589E-33 |
| ACE2 | KCNN2 | -0,5451 | 8,51908E-33 |
| ACE2 | GPANK1 | -0,5451 | 8,41232E-33 |
| ACE2 | RP11-398E10.1 | -0,5451 | 8,34272E-33 |
| ACE2 | ZNF414 | -0,5451 | 8,34247E-33 |
| ACE2 | LOX | -0,5452 | 8,3027E-33 |
| ACE2 | HMBOX1 | -0,5452 | 8,09441E-33 |
| ACE2 | UBE2G1 | -0,5452 | 8,21456E-33 |
| ACE2 | RP11-6N17.3 | -0,5452 | 8,22925E-33 |
| ACE2 | RP11-714G18.1 | -0,5453 | 8,06927E-33 |
| ACE2 | C5orf56 | -0,5453 | 7,99411E-33 |
| ACE2 | ZNRF3 | -0,5453 | 8,03855E-33 |
| ACE2 | AC079584.3 | -0,5454 | 7,61252E-33 |
| ACE2 | RP11-379F4.8 | -0,5454 | 7,77827E-33 |
| ACE2 | EIF1AX | -0,5454 | 7,63545E-33 |
| ACE2 | PTGDS | -0,5455 | 7,48776E-33 |
| ACE2 | RFX7 | -0,5455 | 7,36134E-33 |
| ACE2 | CD300LG | -0,5455 | 7,57728E-33 |
| ACE2 | ZBTB11 | -0,5456 | 7,16066E-33 |
| ACE2 | RPL5P30 | -0,5456 | 7,1422E-33 |
| ACE2 | RP11-290L1.4 | -0,5456 | 7,26232E-33 |

| | | | |
|---|---|---|---|
| ACE2 | WDR7 | -0,5456 | 7,13336E-33 |
| ACE2 | FAM182A | -0,5458 | 6,88701E-33 |
| ACE2 | WDR44 | -0,5458 | 6,8359E-33 |
| ACE2 | RP11-188P20.3 | -0,5459 | 6,6022E-33 |
| ACE2 | RP5-855D21.2 | -0,5459 | 6,50704E-33 |
| ACE2 | ATG4C | -0,546 | 6,30378E-33 |
| ACE2 | UBE2L5P | -0,5461 | 6,22746E-33 |
| ACE2 | RP11-496N12.9 | -0,5462 | 5,8915E-33 |
| ACE2 | SRP72 | -0,5462 | 6,02026E-33 |
| ACE2 | AC068134.10 | -0,5463 | 5,74088E-33 |
| ACE2 | WWTR1-IT1 | -0,5463 | 5,78801E-33 |
| ACE2 | JADE2 | -0,5463 | 5,8621E-33 |
| ACE2 | ZFP36 | -0,5463 | 5,71049E-33 |
| ACE2 | PRRT3-AS1 | -0,5464 | 5,57717E-33 |
| ACE2 | AHCYP2 | -0,5464 | 5,65259E-33 |
| ACE2 | BTBD11 | -0,5464 | 5,69048E-33 |
| ACE2 | TNFSF15 | -0,5465 | 5,45597E-33 |
| ACE2 | ARL1 | -0,5465 | 5,51302E-33 |
| ACE2 | FRMD5 | -0,5465 | 5,50689E-33 |
| ACE2 | RP11-566E18.1 | -0,5466 | 5,27766E-33 |
| ACE2 | NIPBL | -0,5466 | 5,23133E-33 |
| ACE2 | SEM1 | -0,5466 | 5,27279E-33 |
| ACE2 | OAZ1 | -0,5466 | 5,22555E-33 |
| ACE2 | CHM | -0,5466 | 5,27827E-33 |
| ACE2 | RAB6C | -0,5467 | 5,07024E-33 |
| ACE2 | RP11-10L12.4 | -0,5467 | 5,05423E-33 |
| ACE2 | BISPR | -0,5467 | 5,13851E-33 |
| ACE2 | RAB6A | -0,5468 | 5,01982E-33 |
| ACE2 | SNHG19 | -0,5468 | 4,88343E-33 |
| ACE2 | RP11-211G3.2 | -0,5469 | 4,82949E-33 |
| ACE2 | CCL21 | -0,5469 | 4,83627E-33 |
| ACE2 | ADAMTSL2 | -0,5469 | 4,7682E-33 |
| ACE2 | RP11-2C24.5 | -0,5469 | 4,73165E-33 |
| ACE2 | NCBP2 | -0,547 | 4,62981E-33 |
| ACE2 | CTC-308K20.1 | -0,547 | 4,66087E-33 |
| ACE2 | NHP2 | -0,547 | 4,58819E-33 |
| ACE2 | SPOUT1 | -0,547 | 4,57507E-33 |
| ACE2 | PARN | -0,547 | 4,6409E-33 |
| ACE2 | TMEM14A | -0,5471 | 4,5485E-33 |
| ACE2 | IMPAD1 | -0,5471 | 4,5605E-33 |
| ACE2 | POLE3 | -0,5471 | 4,45259E-33 |
| ACE2 | CCAR1 | -0,5471 | 4,46032E-33 |
| ACE2 | EML3 | -0,5471 | 4,4645E-33 |
| ACE2 | RBMS3-AS2 | -0,5472 | 4,30769E-33 |
| ACE2 | RP11-735G4.1 | -0,5472 | 4,40968E-33 |
| ACE2 | ZNF225 | -0,5472 | 4,42633E-33 |
| ACE2 | GALNT2 | -0,5473 | 4,17808E-33 |
| ACE2 | KLHL2 | -0,5473 | 4,16113E-33 |
| ACE2 | UNC5A | -0,5474 | 4,04998E-33 |
| ACE2 | RP11-243M5.2 | -0,5474 | 4,11582E-33 |
| ACE2 | CA14 | -0,5475 | 3,90445E-33 |
| ACE2 | SPEF2 | -0,5475 | 3,98749E-33 |
| ACE2 | RP3-337H4.10 | -0,5475 | 3,92203E-33 |
| ACE2 | SCART1 | -0,5475 | 3,93227E-33 |

| | | | |
|---|---|---|---|
| ACE2 | RP11-281O15.2 | -0,5476 | 3,85921E-33 |
| ACE2 | GFRA2 | -0,5476 | 3,88639E-33 |
| ACE2 | AP000593.7 | -0,5476 | 3,84444E-33 |
| ACE2 | RPUSD4 | -0,5476 | 3,84717E-33 |
| ACE2 | CTD-2349P21.12 | -0,5476 | 3,81688E-33 |
| ACE2 | CACNB1 | -0,5476 | 3,85839E-33 |
| ACE2 | RP11-97N19.2 | -0,5476 | 3,8427E-33 |
| ACE2 | AASDH | -0,5477 | 3,69877E-33 |
| ACE2 | CREBZF | -0,5477 | 3,71191E-33 |
| ACE2 | CLEC2B | -0,5477 | 3,67513E-33 |
| ACE2 | RCN2 | -0,5477 | 3,74597E-33 |
| ACE2 | RAB11B | -0,5478 | 3,58742E-33 |
| ACE2 | NUPL2 | -0,5479 | 3,51492E-33 |
| ACE2 | TRIM4 | -0,548 | 3,3579E-33 |
| ACE2 | INO80 | -0,548 | 3,39509E-33 |
| ACE2 | NF1 | -0,548 | 3,39129E-33 |
| ACE2 | PPP4R1 | -0,548 | 3,37056E-33 |
| ACE2 | SLC9A3R2 | -0,5481 | 3,26191E-33 |
| ACE2 | WDR45B | -0,5481 | 3,23084E-33 |
| ACE2 | CTD-2583A14.1 | -0,5481 | 3,25644E-33 |
| ACE2 | TRDN | -0,5482 | 3,12749E-33 |
| ACE2 | OR52N4 | -0,5482 | 3,1912E-33 |
| ACE2 | BUD13 | -0,5482 | 3,18762E-33 |
| ACE2 | PPIEL | -0,5483 | 3,10745E-33 |
| ACE2 | LIMS4 | -0,5483 | 3,10216E-33 |
| ACE2 | SCN3A | -0,5483 | 3,03468E-33 |
| ACE2 | LINC00648 | -0,5483 | 3,11695E-33 |
| ACE2 | HAGHL | -0,5483 | 3,07401E-33 |
| ACE2 | TCAIM | -0,5484 | 2,93152E-33 |
| ACE2 | ZNF654 | -0,5484 | 2,96983E-33 |
| ACE2 | CRYBG3 | -0,5484 | 3,00246E-33 |
| ACE2 | PDP1 | -0,5484 | 3,01179E-33 |
| ACE2 | XXbac-BPG299F13.17 | -0,5485 | 2,91304E-33 |
| ACE2 | RP11-1081M5.2 | -0,5485 | 2,85542E-33 |
| ACE2 | NHLRC4 | -0,5485 | 2,91327E-33 |
| ACE2 | SOX13 | -0,5486 | 2,79547E-33 |
| ACE2 | SLN | -0,5486 | 2,78781E-33 |
| ACE2 | SNAI3-AS1 | -0,5486 | 2,80881E-33 |
| ACE2 | ZNF493 | -0,5486 | 2,83641E-33 |
| ACE2 | DMKN | -0,5486 | 2,82931E-33 |
| ACE2 | ACSL6 | -0,5487 | 2,69751E-33 |
| ACE2 | RAET1G | -0,5487 | 2,69588E-33 |
| ACE2 | KAT6A | -0,5487 | 2,67047E-33 |
| ACE2 | CEBPD | -0,5487 | 2,66301E-33 |
| ACE2 | SNHG9 | -0,5487 | 2,695E-33 |
| ACE2 | BCKDK | -0,5487 | 2,6733E-33 |
| ACE2 | ZNF233 | -0,5487 | 2,72513E-33 |
| ACE2 | TUSC2 | -0,5488 | 2,60723E-33 |
| ACE2 | HTR7P1 | -0,5488 | 2,59459E-33 |
| ACE2 | RP11-370I10.2 | -0,5488 | 2,64196E-33 |
| ACE2 | SLITRK3 | -0,5489 | 2,52552E-33 |
| ACE2 | LINC01394 | -0,5489 | 2,5414E-33 |
| ACE2 | FBXW2 | -0,5489 | 2,50851E-33 |
| ACE2 | RP11-258F1.1 | -0,5489 | 2,56478E-33 |

| | | | |
|---|---|---|---|
| ACE2 | RP13-131K19.6 | -0,549 | 2,47585E-33 |
| ACE2 | RP5-864K19.4 | -0,5491 | 2,3697E-33 |
| ACE2 | RNF181 | -0,5491 | 2,35618E-33 |
| ACE2 | LINC01300 | -0,5491 | 2,37516E-33 |
| ACE2 | ZNF804A | -0,5492 | 2,33586E-33 |
| ACE2 | TAS2R19 | -0,5492 | 2,32153E-33 |
| ACE2 | CCDC39 | -0,5493 | 2,21284E-33 |
| ACE2 | QDPR | -0,5493 | 2,23511E-33 |
| ACE2 | USO1 | -0,5493 | 2,24739E-33 |
| ACE2 | WNT2 | -0,5493 | 2,24614E-33 |
| ACE2 | QSER1 | -0,5493 | 2,24663E-33 |
| ACE2 | RB1 | -0,5493 | 2,26656E-33 |
| ACE2 | DHRS12 | -0,5493 | 2,19782E-33 |
| ACE2 | RP11-260M2.1 | -0,5494 | 2,14011E-33 |
| ACE2 | PRRX2 | -0,5494 | 2,17841E-33 |
| ACE2 | HOXC8 | -0,5494 | 2,17372E-33 |
| ACE2 | RP11-158H5.2 | -0,5494 | 2,17548E-33 |
| ACE2 | AC079354.3 | -0,5495 | 2,10993E-33 |
| ACE2 | SEC24B | -0,5495 | 2,12257E-33 |
| ACE2 | NAA15 | -0,5495 | 2,09742E-33 |
| ACE2 | CCDC113 | -0,5495 | 2,10123E-33 |
| ACE2 | ZNF491 | -0,5495 | 2,1259E-33 |
| ACE2 | RP11-295M18.6 | -0,5496 | 2,01378E-33 |
| ACE2 | EFHD1 | -0,5497 | 1,97401E-33 |
| ACE2 | SV2C | -0,5497 | 1,94742E-33 |
| ACE2 | RBAK | -0,5497 | 1,95744E-33 |
| ACE2 | DENND5B | -0,5497 | 1,94851E-33 |
| ACE2 | ARFGAP3 | -0,5497 | 1,94853E-33 |
| ACE2 | BTN2A2 | -0,5498 | 1,90844E-33 |
| ACE2 | EXOSC3 | -0,5499 | 1,84918E-33 |
| ACE2 | SOCS4 | -0,5499 | 1,85921E-33 |
| ACE2 | FAM110D | -0,55 | 1,77778E-33 |
| ACE2 | H1FX-AS1 | -0,5501 | 1,7445E-33 |
| ACE2 | ADGRA1 | -0,5501 | 1,71835E-33 |
| ACE2 | H1FNT | -0,5501 | 1,71065E-33 |
| ACE2 | IGFL2-AS1 | -0,5501 | 1,71363E-33 |
| ACE2 | RNF4 | -0,5502 | 1,65151E-33 |
| ACE2 | IK | -0,5502 | 1,6593E-33 |
| ACE2 | BARD1 | -0,5503 | 1,60773E-33 |
| ACE2 | LURAP1L | -0,5503 | 1,64558E-33 |
| ACE2 | RP3-414A15.11 | -0,5503 | 1,60652E-33 |
| ACE2 | HSD11B1 | -0,5504 | 1,55968E-33 |
| ACE2 | CPNE4 | -0,5504 | 1,55013E-33 |
| ACE2 | LSM3 | -0,5505 | 1,52043E-33 |
| ACE2 | HNRNPA0 | -0,5505 | 1,50031E-33 |
| ACE2 | KARS | -0,5505 | 1,49872E-33 |
| ACE2 | ZNHIT3 | -0,5505 | 1,49664E-33 |
| ACE2 | ZNF24 | -0,5505 | 1,50805E-33 |
| ACE2 | ZNF772 | -0,5505 | 1,53301E-33 |
| ACE2 | CLCF1 | -0,5506 | 1,49629E-33 |
| ACE2 | CNTN5 | -0,5506 | 1,47617E-33 |
| ACE2 | FAM185BP | -0,5507 | 1,42297E-33 |
| ACE2 | ANAPC5 | -0,5507 | 1,44772E-33 |
| ACE2 | FAM219B | -0,5507 | 1,44645E-33 |

| | | | |
|---|---|---|---|
| ACE2 | RP11-715J22.6 | -0,5507 | 1,42961E-33 |
| ACE2 | AC005197.2 | -0,5507 | 1,43197E-33 |
| ACE2 | AC020594.5 | -0,5508 | 1,38646E-33 |
| ACE2 | TSNARE1 | -0,5509 | 1,3273E-33 |
| ACE2 | FAM196A | -0,5509 | 1,33156E-33 |
| ACE2 | SIRT4 | -0,5509 | 1,3165E-33 |
| ACE2 | PIKFYVE | -0,551 | 1,28129E-33 |
| ACE2 | SSR3 | -0,551 | 1,27632E-33 |
| ACE2 | RPLP2 | -0,551 | 1,29513E-33 |
| ACE2 | MED13L | -0,551 | 1,31181E-33 |
| ACE2 | RP11-539G18.3 | -0,5511 | 1,25479E-33 |
| ACE2 | RP11-45A17.2 | -0,5511 | 1,27341E-33 |
| ACE2 | RP11-366L20.2 | -0,5511 | 1,26781E-33 |
| ACE2 | RP11-706P11.2 | -0,5511 | 1,25064E-33 |
| ACE2 | AC092641.2 | -0,5512 | 1,22198E-33 |
| ACE2 | TBC1D13 | -0,5512 | 1,22494E-33 |
| ACE2 | PSAP | -0,5512 | 1,22025E-33 |
| ACE2 | ASPA | -0,5512 | 1,22754E-33 |
| ACE2 | ZNF37CP | -0,5513 | 1,17917E-33 |
| ACE2 | UBP1 | -0,5514 | 1,12464E-33 |
| ACE2 | SMCO3 | -0,5514 | 1,1321E-33 |
| ACE2 | NR2C2AP | -0,5514 | 1,15086E-33 |
| ACE2 | DYRK1A | -0,5514 | 1,14539E-33 |
| ACE2 | GTF2A1L | -0,5515 | 1,09429E-33 |
| ACE2 | H1FX | -0,5515 | 1,11987E-33 |
| ACE2 | LAMA5-AS1 | -0,5515 | 1,09888E-33 |
| ACE2 | EMX2 | -0,5516 | 1,05876E-33 |
| ACE2 | NUCB2 | -0,5517 | 1,04558E-33 |
| ACE2 | WASHC4 | -0,5517 | 1,03637E-33 |
| ACE2 | RP4-730K3.3 | -0,5518 | 1,00914E-33 |
| ACE2 | RGPD8 | -0,5519 | 9,6693E-34 |
| ACE2 | RPL10AP6 | -0,5519 | 9,80957E-34 |
| ACE2 | IL6 | -0,5519 | 9,73293E-34 |
| ACE2 | GSN-AS1 | -0,5519 | 9,80936E-34 |
| ACE2 | MARCH9 | -0,5519 | 9,81505E-34 |
| ACE2 | RPLP1P6 | -0,552 | 9,33402E-34 |
| ACE2 | HMGB1P39 | -0,552 | 9,41826E-34 |
| ACE2 | RHOD | -0,552 | 9,52506E-34 |
| ACE2 | RP11-552F3.4 | -0,552 | 9,3676E-34 |
| ACE2 | REV1 | -0,5522 | 8,78516E-34 |
| ACE2 | UBC | -0,5522 | 8,68292E-34 |
| ACE2 | RNASEH1-AS1 | -0,5523 | 8,40698E-34 |
| ACE2 | LENG8 | -0,5523 | 8,53391E-34 |
| ACE2 | SCCPDH | -0,5524 | 8,17787E-34 |
| ACE2 | C16orf62 | -0,5524 | 8,13104E-34 |
| ACE2 | HIF3A | -0,5524 | 8,37169E-34 |
| ACE2 | AC009950.2 | -0,5525 | 7,90649E-34 |
| ACE2 | RP11-770J1.4 | -0,5525 | 8,00816E-34 |
| ACE2 | GPAT2P2 | -0,5526 | 7,79617E-34 |
| ACE2 | CCDC6 | -0,5526 | 7,83697E-34 |
| ACE2 | LINC00987 | -0,5527 | 7,51366E-34 |
| ACE2 | ESCO1 | -0,5527 | 7,48838E-34 |
| ACE2 | LINC01711 | -0,5527 | 7,56932E-34 |
| ACE2 | ANKRD7 | -0,5528 | 7,2608E-34 |

| | | | |
|---|---|---|---|
| ACE2 | STK36 | -0,5529 | 7,08577E-34 |
| ACE2 | PEBP1 | -0,5529 | 7,09593E-34 |
| ACE2 | LRRC43 | -0,5529 | 6,91966E-34 |
| ACE2 | GINS3 | -0,5529 | 7,09054E-34 |
| ACE2 | POLDIP3 | -0,5529 | 6,91824E-34 |
| ACE2 | PHTF2 | -0,553 | 6,8716E-34 |
| ACE2 | SMARCC2 | -0,553 | 6,87439E-34 |
| ACE2 | MYH7B | -0,553 | 6,73872E-34 |
| ACE2 | RP11-364B14.3 | -0,553 | 6,74968E-34 |
| ACE2 | AC004893.11 | -0,5531 | 6,57507E-34 |
| ACE2 | RP11-848P1.9 | -0,5531 | 6,63742E-34 |
| ACE2 | CEBPB | -0,5531 | 6,60982E-34 |
| ACE2 | MCMBP | -0,5532 | 6,47111E-34 |
| ACE2 | RP11-469N6.1 | -0,5532 | 6,42133E-34 |
| ACE2 | ACSBG1 | -0,5532 | 6,43309E-34 |
| ACE2 | RNF166 | -0,5532 | 6,31245E-34 |
| ACE2 | CTD-2528L19.6 | -0,5532 | 6,33173E-34 |
| ACE2 | CNTNAP5 | -0,5533 | 6,18907E-34 |
| ACE2 | TBPL1 | -0,5534 | 5,8754E-34 |
| ACE2 | MYOM2 | -0,5534 | 5,99147E-34 |
| ACE2 | HELZ | -0,5534 | 6,06283E-34 |
| ACE2 | AC006946.17 | -0,5534 | 5,93368E-34 |
| ACE2 | RP11-305K5.1 | -0,5535 | 5,69071E-34 |
| ACE2 | C21orf59 | -0,5535 | 5,70693E-34 |
| ACE2 | CTD-2201E18.5 | -0,5536 | 5,61194E-34 |
| ACE2 | TPTE2P3 | -0,5536 | 5,63474E-34 |
| ACE2 | RABGEF1 | -0,5537 | 5,34258E-34 |
| ACE2 | RP11-635N19.1 | -0,5537 | 5,32878E-34 |
| ACE2 | RP11-378A13.1 | -0,5538 | 5,22006E-34 |
| ACE2 | ZNF366 | -0,5538 | 5,23319E-34 |
| ACE2 | CPEB2 | -0,5539 | 5,14931E-34 |
| ACE2 | RP11-212I21.3 | -0,5539 | 5,03639E-34 |
| ACE2 | RP11-513I15.6 | -0,554 | 4,99142E-34 |
| ACE2 | RPL21P75 | -0,554 | 4,92454E-34 |
| ACE2 | SUGP1 | -0,554 | 4,8714E-34 |
| ACE2 | ZNF417 | -0,554 | 4,86667E-34 |
| ACE2 | CTD-3138B18.6 | -0,554 | 4,8447E-34 |
| ACE2 | ECE1 | -0,5541 | 4,82774E-34 |
| ACE2 | PLPP1 | -0,5541 | 4,77067E-34 |
| ACE2 | FOXO1 | -0,5541 | 4,72847E-34 |
| ACE2 | CLN5 | -0,5541 | 4,72901E-34 |
| ACE2 | RP11-6N17.4 | -0,5541 | 4,68643E-34 |
| ACE2 | LINC00652 | -0,5541 | 4,81372E-34 |
| ACE2 | ICK | -0,5542 | 4,55827E-34 |
| ACE2 | PTPMT1 | -0,5542 | 4,53489E-34 |
| ACE2 | SRSF5 | -0,5542 | 4,54817E-34 |
| ACE2 | RP11-344E13.4 | -0,5542 | 4,56919E-34 |
| ACE2 | MAFG-AS1 | -0,5542 | 4,54147E-34 |
| ACE2 | BCL11A | -0,5543 | 4,44161E-34 |
| ACE2 | ELFN1 | -0,5543 | 4,52569E-34 |
| ACE2 | RRAGA | -0,5543 | 4,4444E-34 |
| ACE2 | AIF1L | -0,5543 | 4,42578E-34 |
| ACE2 | MYOF | -0,5543 | 4,51471E-34 |
| ACE2 | RPF1 | -0,5544 | 4,25709E-34 |

| | | | |
|---|---|---|---|
| ACE2 | METTL14 | -0,5544 | 4,35446E-34 |
| ACE2 | DAG1 | -0,5545 | 4,16443E-34 |
| ACE2 | ZNF658 | -0,5545 | 4,12662E-34 |
| ACE2 | UFM1 | -0,5545 | 4,21273E-34 |
| ACE2 | AC007228.11 | -0,5545 | 4,20459E-34 |
| ACE2 | MYOC | -0,5546 | 4,04064E-34 |
| ACE2 | TRIM61 | -0,5546 | 4,06496E-34 |
| ACE2 | SNRPB2 | -0,5546 | 4,03317E-34 |
| ACE2 | RP11-490G2.2 | -0,5547 | 3,86557E-34 |
| ACE2 | LVRN | -0,5547 | 3,97126E-34 |
| ACE2 | ZNF335 | -0,5547 | 3,94893E-34 |
| ACE2 | RP11-338N10.1 | -0,5548 | 3,82477E-34 |
| ACE2 | KCTD18 | -0,5548 | 3,75945E-34 |
| ACE2 | GPR22 | -0,5548 | 3,79443E-34 |
| ACE2 | KLHL18 | -0,555 | 3,59406E-34 |
| ACE2 | RP11-480N24.4 | -0,555 | 3,58675E-34 |
| ACE2 | PROCR | -0,555 | 3,54409E-34 |
| ACE2 | DPY19L2P3 | -0,5551 | 3,46735E-34 |
| ACE2 | RAPGEF1 | -0,5551 | 3,41831E-34 |
| ACE2 | FAM35DP | -0,5551 | 3,43155E-34 |
| ACE2 | YY1 | -0,5551 | 3,41278E-34 |
| ACE2 | ADAMTS5 | -0,5551 | 3,46671E-34 |
| ACE2 | FUBP1 | -0,5552 | 3,31161E-34 |
| ACE2 | AC005154.6 | -0,5552 | 3,37222E-34 |
| ACE2 | PPME1 | -0,5552 | 3,30069E-34 |
| ACE2 | FNDC3A | -0,5552 | 3,34419E-34 |
| ACE2 | HAUS1 | -0,5552 | 3,33908E-34 |
| ACE2 | SYCE2 | -0,5552 | 3,2713E-34 |
| ACE2 | P2RY12 | -0,5553 | 3,21E-34 |
| ACE2 | CEP135 | -0,5553 | 3,23727E-34 |
| ACE2 | EIF2AK4 | -0,5553 | 3,2571E-34 |
| ACE2 | RP11-401P9.6 | -0,5553 | 3,18821E-34 |
| ACE2 | AC012613.2 | -0,5554 | 3,14944E-34 |
| ACE2 | KB-1208A12.3 | -0,5554 | 3,10673E-34 |
| ACE2 | RP11-411B10.2 | -0,5554 | 3,11268E-34 |
| ACE2 | RPL7L1 | -0,5555 | 3,02115E-34 |
| ACE2 | ATP6V1H | -0,5555 | 3,0064E-34 |
| ACE2 | CGRRF1 | -0,5555 | 3,00459E-34 |
| ACE2 | SLCO3A1 | -0,5555 | 3,0048E-34 |
| ACE2 | ZNF460 | -0,5555 | 3,04147E-34 |
| ACE2 | LINC01783 | -0,5556 | 2,88444E-34 |
| ACE2 | CLGN | -0,5556 | 2,94911E-34 |
| ACE2 | ZBTB2 | -0,5556 | 2,93436E-34 |
| ACE2 | RP11-809O17.1 | -0,5556 | 2,88171E-34 |
| ACE2 | RP11-403A3.3 | -0,5556 | 2,93602E-34 |
| ACE2 | TMEM114 | -0,5556 | 2,9508E-34 |
| ACE2 | PSMG2 | -0,5556 | 2,88559E-34 |
| ACE2 | LINC01289 | -0,5557 | 2,87042E-34 |
| ACE2 | KCNC2 | -0,5557 | 2,85679E-34 |
| ACE2 | RP11-1069G10.1 | -0,5557 | 2,78628E-34 |
| ACE2 | UBA3 | -0,5558 | 2,76927E-34 |
| ACE2 | TMEM30A | -0,5559 | 2,68257E-34 |
| ACE2 | SNHG5 | -0,5559 | 2,61462E-34 |
| ACE2 | MKLN1 | -0,5559 | 2,60708E-34 |

| | | | |
|---|---|---|---|
| ACE2 | GTF3C5 | -0,5559 | 2,67871E-34 |
| ACE2 | SLC35E3 | -0,5559 | 2,64168E-34 |
| ACE2 | RNF20 | -0,556 | 2,5454E-34 |
| ACE2 | TTC5 | -0,556 | 2,55835E-34 |
| ACE2 | RP11-66B24.2 | -0,556 | 2,59521E-34 |
| ACE2 | RP11-647K16.1 | -0,5561 | 2,45837E-34 |
| ACE2 | TMEM42 | -0,5561 | 2,47209E-34 |
| ACE2 | CTD-2135J3.3 | -0,5562 | 2,37716E-34 |
| ACE2 | FBXO33 | -0,5562 | 2,39644E-34 |
| ACE2 | ARHGEF9 | -0,5562 | 2,38013E-34 |
| ACE2 | RP11-177G23.2 | -0,5563 | 2,29628E-34 |
| ACE2 | CHRAC1 | -0,5563 | 2,35663E-34 |
| ACE2 | SENCR | -0,5563 | 2,33756E-34 |
| ACE2 | RP11-21G15.1 | -0,5563 | 2,29304E-34 |
| ACE2 | MED30 | -0,5564 | 2,218E-34 |
| ACE2 | CTC-296K1.4 | -0,5564 | 2,27435E-34 |
| ACE2 | PLIN5 | -0,5564 | 2,26693E-34 |
| ACE2 | LRRC4 | -0,5565 | 2,19971E-34 |
| ACE2 | UBE3A | -0,5565 | 2,14348E-34 |
| ACE2 | ZNF778 | -0,5565 | 2,18277E-34 |
| ACE2 | ITSN2 | -0,5566 | 2,07421E-34 |
| ACE2 | NSUN5 | -0,5566 | 2,08723E-34 |
| ACE2 | NPAT | -0,5566 | 2,08895E-34 |
| ACE2 | LA16c-312E8.4 | -0,5566 | 2,12987E-34 |
| ACE2 | RP11-1260E13.4 | -0,5566 | 2,11155E-34 |
| ACE2 | RP11-44N22.3 | -0,5567 | 2,0077E-34 |
| ACE2 | PICALM | -0,5567 | 2,05046E-34 |
| ACE2 | RP11-834C11.8 | -0,5567 | 2,03932E-34 |
| ACE2 | RP11-430G17.3 | -0,5568 | 1,94856E-34 |
| ACE2 | RDH16 | -0,5568 | 1,99127E-34 |
| ACE2 | HNRNPC | -0,5568 | 1,95922E-34 |
| ACE2 | GALR1 | -0,5568 | 1,93876E-34 |
| ACE2 | TPO | -0,5569 | 1,91523E-34 |
| ACE2 | RP11-104O19.4 | -0,5569 | 1,89853E-34 |
| ACE2 | CCDC192 | -0,557 | 1,8324E-34 |
| ACE2 | AC007319.1 | -0,5571 | 1,7899E-34 |
| ACE2 | HESX1 | -0,5571 | 1,79354E-34 |
| ACE2 | U91328.19 | -0,5571 | 1,78368E-34 |
| ACE2 | PGM5-AS1 | -0,5571 | 1,77393E-34 |
| ACE2 | PAPOLG | -0,5572 | 1,74665E-34 |
| ACE2 | PDE6B | -0,5572 | 1,74078E-34 |
| ACE2 | FABP4 | -0,5572 | 1,72957E-34 |
| ACE2 | WASH2P | -0,5573 | 1,68021E-34 |
| ACE2 | ERBB4 | -0,5573 | 1,66228E-34 |
| ACE2 | HEPACAM | -0,5573 | 1,67862E-34 |
| ACE2 | TCAP | -0,5573 | 1,66623E-34 |
| ACE2 | RPL18 | -0,5573 | 1,64717E-34 |
| ACE2 | PIGC | -0,5574 | 1,64371E-34 |
| ACE2 | PEX3 | -0,5574 | 1,6418E-34 |
| ACE2 | FAM200A | -0,5574 | 1,63003E-34 |
| ACE2 | POLG2 | -0,5574 | 1,60946E-34 |
| ACE2 | CBLN2 | -0,5575 | 1,56103E-34 |
| ACE2 | RP4-575N6.4 | -0,5576 | 1,53699E-34 |
| ACE2 | HECW2 | -0,5576 | 1,51278E-34 |

| | | | |
|---|---|---|---|
| ACE2 | RP11-43F13.3 | -0,5576 | 1,51464E-34 |
| ACE2 | DENR | -0,5576 | 1,51925E-34 |
| ACE2 | RP11-893F2.5 | -0,5576 | 1,53745E-34 |
| ACE2 | ATP5G2 | -0,5577 | 1,47503E-34 |
| ACE2 | RP3-508I15.20 | -0,5577 | 1,47752E-34 |
| ACE2 | KTN1-AS1 | -0,5578 | 1,40118E-34 |
| ACE2 | LINC01117 | -0,5579 | 1,38921E-34 |
| ACE2 | MTRR | -0,5579 | 1,36761E-34 |
| ACE2 | DBP | -0,5579 | 1,38804E-34 |
| ACE2 | CCDC191 | -0,558 | 1,32002E-34 |
| ACE2 | BHLHE40-AS1 | -0,5581 | 1,26735E-34 |
| ACE2 | RBL2 | -0,5581 | 1,30359E-34 |
| ACE2 | ANGPTL4 | -0,5581 | 1,29775E-34 |
| ACE2 | RP11-44D5.1 | -0,5582 | 1,2343E-34 |
| ACE2 | GHR | -0,5582 | 1,25172E-34 |
| ACE2 | PPP1R9B | -0,5582 | 1,2561E-34 |
| ACE2 | C5orf49 | -0,5583 | 1,19264E-34 |
| ACE2 | PNN | -0,5583 | 1,21457E-34 |
| ACE2 | TDRD9 | -0,5583 | 1,20331E-34 |
| ACE2 | CTB-66B24.1 | -0,5583 | 1,19552E-34 |
| ACE2 | XXbac-B562F10.11 | -0,5583 | 1,19036E-34 |
| ACE2 | AC093110.3 | -0,5584 | 1,14971E-34 |
| ACE2 | TTC33 | -0,5584 | 1,14696E-34 |
| ACE2 | C2CD3 | -0,5584 | 1,16582E-34 |
| ACE2 | TIMM44 | -0,5584 | 1,17997E-34 |
| ACE2 | GPR42 | -0,5584 | 1,18029E-34 |
| ACE2 | PYGO2 | -0,5585 | 1,14405E-34 |
| ACE2 | TMEM252 | -0,5585 | 1,14311E-34 |
| ACE2 | ALDH4A1 | -0,5588 | 1,01514E-34 |
| ACE2 | PSMD5 | -0,5588 | 1,0042E-34 |
| ACE2 | RP11-977G19.11 | -0,5589 | 9,95776E-35 |
| ACE2 | GADD45B | -0,5589 | 9,73027E-35 |
| ACE2 | AURKC | -0,5589 | 9,85558E-35 |
| ACE2 | RP3-402G11.27 | -0,5589 | 1,00068E-34 |
| ACE2 | RP4-598P13.1 | -0,559 | 9,40459E-35 |
| ACE2 | SRGAP2D | -0,559 | 9,63831E-35 |
| ACE2 | FEZ2 | -0,559 | 9,48508E-35 |
| ACE2 | ITGB1 | -0,559 | 9,61314E-35 |
| ACE2 | REEP3 | -0,559 | 9,49048E-35 |
| ACE2 | WNT3 | -0,559 | 9,45808E-35 |
| ACE2 | CNTD2 | -0,559 | 9,67983E-35 |
| ACE2 | ZNF514 | -0,5591 | 9,2359E-35 |
| ACE2 | SPHKAP | -0,5591 | 9,12282E-35 |
| ACE2 | RGS7BP | -0,5591 | 9,21015E-35 |
| ACE2 | LINC00847 | -0,5591 | 9,33066E-35 |
| ACE2 | DPP8 | -0,5591 | 9,32002E-35 |
| ACE2 | PRPF8 | -0,5591 | 9,25791E-35 |
| ACE2 | RP11-81A22.5 | -0,5591 | 9,34036E-35 |
| ACE2 | SNORD21 | -0,5592 | 8,97107E-35 |
| ACE2 | PPIG | -0,5592 | 8,90166E-35 |
| ACE2 | GIMAP8 | -0,5592 | 8,92799E-35 |
| ACE2 | AL590762.11 | -0,5592 | 8,80778E-35 |
| ACE2 | CUL9 | -0,5593 | 8,57816E-35 |
| ACE2 | MRPL35P2 | -0,5593 | 8,75558E-35 |

| | | | |
|---|---|---|---|
| ACE2 | ZNF581 | -0,5593 | 8,51329E-35 |
| ACE2 | ATP6AP2 | -0,5593 | 8,67743E-35 |
| ACE2 | RP11-490N5.3 | -0,5594 | 8,46739E-35 |
| ACE2 | PSMA3-AS1 | -0,5594 | 8,26627E-35 |
| ACE2 | GOLGA8K | -0,5595 | 8,12755E-35 |
| ACE2 | RP11-455I9.1 | -0,5597 | 7,58476E-35 |
| ACE2 | FLCN | -0,5597 | 7,50994E-35 |
| ACE2 | TXNL1 | -0,5597 | 7,69257E-35 |
| ACE2 | ASH1L | -0,5598 | 7,3978E-35 |
| ACE2 | CTC-512J14.7 | -0,56 | 6,82467E-35 |
| ACE2 | CTD-2195M15.1 | -0,56 | 6,78203E-35 |
| ACE2 | NUDT1 | -0,56 | 6,93271E-35 |
| ACE2 | PAM16 | -0,56 | 6,93223E-35 |
| ACE2 | WDR48 | -0,5601 | 6,59661E-35 |
| ACE2 | HDDC2 | -0,5601 | 6,54515E-35 |
| ACE2 | VPS37C | -0,5601 | 6,57346E-35 |
| ACE2 | RP11-728E14.3 | -0,5601 | 6,69708E-35 |
| ACE2 | AC008073.7 | -0,5602 | 6,36509E-35 |
| ACE2 | LRRFIP1 | -0,5602 | 6,3619E-35 |
| ACE2 | DLG5 | -0,5602 | 6,48233E-35 |
| ACE2 | ZCCHC8 | -0,5602 | 6,33613E-35 |
| ACE2 | GTF3C1 | -0,5602 | 6,49137E-35 |
| ACE2 | ZCCHC2 | -0,5602 | 6,45117E-35 |
| ACE2 | TMEM81 | -0,5603 | 6,29611E-35 |
| ACE2 | FAR2P2 | -0,5603 | 6,29139E-35 |
| ACE2 | SERPINE2 | -0,5603 | 6,18134E-35 |
| ACE2 | C10orf88 | -0,5604 | 5,98128E-35 |
| ACE2 | RBM4 | -0,5604 | 5,96982E-35 |
| ACE2 | ENOX1-AS1 | -0,5604 | 5,92997E-35 |
| ACE2 | SKINT1L | -0,5605 | 5,89046E-35 |
| ACE2 | AC093609.1 | -0,5605 | 5,86997E-35 |
| ACE2 | RBM27 | -0,5605 | 5,85603E-35 |
| ACE2 | RP11-272P10.2 | -0,5605 | 5,73724E-35 |
| ACE2 | ERH | -0,5605 | 5,90902E-35 |
| ACE2 | RBM8A | -0,5606 | 5,56318E-35 |
| ACE2 | CRTC1 | -0,5606 | 5,6132E-35 |
| ACE2 | SOS1 | -0,5607 | 5,4478E-35 |
| ACE2 | PARP14 | -0,5607 | 5,43859E-35 |
| ACE2 | SLC22A2 | -0,5607 | 5,3593E-35 |
| ACE2 | TMX3 | -0,5607 | 5,44428E-35 |
| ACE2 | ZRANB3 | -0,5609 | 5,17546E-35 |
| ACE2 | UBA5 | -0,5609 | 5,06997E-35 |
| ACE2 | RPL13A | -0,5609 | 5,05913E-35 |
| ACE2 | USP25 | -0,5609 | 5,05245E-35 |
| ACE2 | CUTA | -0,561 | 4,87528E-35 |
| ACE2 | ACTR10 | -0,561 | 4,99089E-35 |
| ACE2 | TLE3 | -0,561 | 4,95366E-35 |
| ACE2 | ZNF322 | -0,5611 | 4,82503E-35 |
| ACE2 | RP11-118K6.3 | -0,5611 | 4,78943E-35 |
| ACE2 | TMEM86B | -0,5611 | 4,78861E-35 |
| ACE2 | PI4KA | -0,5611 | 4,81277E-35 |
| ACE2 | RP11-392A14.9 | -0,5612 | 4,53495E-35 |
| ACE2 | AC007193.6 | -0,5612 | 4,6361E-35 |
| ACE2 | TIMM10B | -0,5613 | 4,47847E-35 |

| | | | |
|---|---|---|---|
| ACE2 | RP5-1063M23.3 | -0,5613 | 4,41957E-35 |
| ACE2 | CTC-444N24.6 | -0,5613 | 4,52418E-35 |
| ACE2 | TMEFF2 | -0,5614 | 4,29988E-35 |
| ACE2 | CINP | -0,5614 | 4,31507E-35 |
| ACE2 | CALB2 | -0,5614 | 4,32558E-35 |
| ACE2 | RP11-748H22.1 | -0,5615 | 4,19496E-35 |
| ACE2 | B3GLCT | -0,5615 | 4,23106E-35 |
| ACE2 | RP11-673E1.3 | -0,5616 | 4,00962E-35 |
| ACE2 | SIL1 | -0,5616 | 4,05265E-35 |
| ACE2 | RNF169 | -0,5617 | 3,89663E-35 |
| ACE2 | RP11-632L2.2 | -0,5617 | 3,88142E-35 |
| ACE2 | RP11-362F19.1 | -0,5618 | 3,75481E-35 |
| ACE2 | PSMB6 | -0,5618 | 3,82875E-35 |
| ACE2 | AC073410.1 | -0,562 | 3,4756E-35 |
| ACE2 | HUS1B | -0,562 | 3,5109E-35 |
| ACE2 | C12orf4 | -0,562 | 3,50869E-35 |
| ACE2 | CFAP20 | -0,562 | 3,58277E-35 |
| ACE2 | RP11-232D9.2 | -0,562 | 3,4913E-35 |
| ACE2 | BATF3 | -0,5621 | 3,42798E-35 |
| ACE2 | FBXO8 | -0,5621 | 3,38173E-35 |
| ACE2 | GFOD1 | -0,5621 | 3,36061E-35 |
| ACE2 | RP11-299L17.3 | -0,5621 | 3,42823E-35 |
| ACE2 | PITPNC1 | -0,5621 | 3,41002E-35 |
| ACE2 | KIAA0355 | -0,5622 | 3,31687E-35 |
| ACE2 | ERCC8 | -0,5623 | 3,23974E-35 |
| ACE2 | SYNCRIP | -0,5623 | 3,22763E-35 |
| ACE2 | MKL2 | -0,5623 | 3,17593E-35 |
| ACE2 | VN1R83P | -0,5623 | 3,14347E-35 |
| ACE2 | TAF1B | -0,5624 | 3,05232E-35 |
| ACE2 | CDC5L | -0,5624 | 3,10056E-35 |
| ACE2 | EEF2K | -0,5624 | 3,13075E-35 |
| ACE2 | ARL2BP | -0,5624 | 3,12996E-35 |
| ACE2 | STAU1 | -0,5624 | 3,12333E-35 |
| ACE2 | HP1BP3 | -0,5626 | 2,90003E-35 |
| ACE2 | ZNF32-AS2 | -0,5626 | 2,86172E-35 |
| ACE2 | UBTF | -0,5626 | 2,86409E-35 |
| ACE2 | RTCA-AS1 | -0,5627 | 2,79901E-35 |
| ACE2 | RPS18P9 | -0,5627 | 2,80958E-35 |
| ACE2 | CALB1 | -0,5627 | 2,80487E-35 |
| ACE2 | LINC01082 | -0,5627 | 2,79206E-35 |
| ACE2 | GPR183 | -0,5628 | 2,71264E-35 |
| ACE2 | ZNF780B | -0,5628 | 2,70088E-35 |
| ACE2 | UQCC1 | -0,5628 | 2,72059E-35 |
| ACE2 | ECHDC2 | -0,5629 | 2,63184E-35 |
| ACE2 | RP11-479O9.4 | -0,5629 | 2,58166E-35 |
| ACE2 | LINC01515 | -0,5629 | 2,59459E-35 |
| ACE2 | EIF4G2 | -0,5629 | 2,58983E-35 |
| ACE2 | LMF2 | -0,5629 | 2,63898E-35 |
| ACE2 | OSTN | -0,563 | 2,50885E-35 |
| ACE2 | CTD-3157E16.1 | -0,563 | 2,56108E-35 |
| ACE2 | RP11-395A13.2 | -0,5631 | 2,48401E-35 |
| ACE2 | PCDH8 | -0,5631 | 2,42645E-35 |
| ACE2 | U82695.9 | -0,5631 | 2,47716E-35 |
| ACE2 | SOHLH1 | -0,5633 | 2,32114E-35 |

| | | | |
|---|---|---|---|
| ACE2 | KCNIP2 | -0,5633 | 2,31923E-35 |
| ACE2 | P3H2-AS1 | -0,5634 | 2,22057E-35 |
| ACE2 | ZNF891 | -0,5634 | 2,23027E-35 |
| ACE2 | MAPK9 | -0,5635 | 2,11075E-35 |
| ACE2 | TCF25 | -0,5635 | 2,1661E-35 |
| ACE2 | CTB-131K11.1 | -0,5635 | 2,12595E-35 |
| ACE2 | AD001527.4 | -0,5635 | 2,15322E-35 |
| ACE2 | TBP | -0,5636 | 2,09798E-35 |
| ACE2 | FZD9 | -0,5636 | 2,07398E-35 |
| ACE2 | CTC-542B22.2 | -0,5636 | 2,05971E-35 |
| ACE2 | STARD4-AS1 | -0,5637 | 2,00459E-35 |
| ACE2 | ALDH1A2 | -0,5637 | 1,97819E-35 |
| ACE2 | PIK3R1 | -0,5638 | 1,95675E-35 |
| ACE2 | ITGA9-AS1 | -0,5639 | 1,88045E-35 |
| ACE2 | RP11-307C12.12 | -0,564 | 1,78952E-35 |
| ACE2 | CTDSPL2 | -0,564 | 1,81464E-35 |
| ACE2 | TP53TG5 | -0,564 | 1,83072E-35 |
| ACE2 | PIK3CD | -0,5641 | 1,72953E-35 |
| ACE2 | DHH | -0,5641 | 1,75507E-35 |
| ACE2 | KHDRBS1 | -0,5642 | 1,6885E-35 |
| ACE2 | RP11-530A18.1 | -0,5642 | 1,6854E-35 |
| ACE2 | ZNF720 | -0,5642 | 1,67835E-35 |
| ACE2 | STAT2 | -0,5643 | 1,61044E-35 |
| ACE2 | TRIL | -0,5644 | 1,57527E-35 |
| ACE2 | TAS2R20 | -0,5644 | 1,60094E-35 |
| ACE2 | KRT86 | -0,5644 | 1,57568E-35 |
| ACE2 | RP11-661O13.1 | -0,5644 | 1,57715E-35 |
| ACE2 | RP1-266L20.2 | -0,5645 | 1,53251E-35 |
| ACE2 | KRT17 | -0,5645 | 1,51672E-35 |
| ACE2 | GOSR2 | -0,5645 | 1,53669E-35 |
| ACE2 | SEC14L1 | -0,5645 | 1,51293E-35 |
| ACE2 | ZNF576 | -0,5645 | 1,54299E-35 |
| ACE2 | C1orf228 | -0,5646 | 1,49076E-35 |
| ACE2 | ELK4 | -0,5646 | 1,49668E-35 |
| ACE2 | ANKRD33B | -0,5646 | 1,45375E-35 |
| ACE2 | RP11-5C23.2 | -0,5646 | 1,49911E-35 |
| ACE2 | CARNS1 | -0,5646 | 1,47512E-35 |
| ACE2 | RPH3A | -0,5646 | 1,46589E-35 |
| ACE2 | ZNF230 | -0,5646 | 1,47461E-35 |
| ACE2 | HHEX | -0,5647 | 1,41496E-35 |
| ACE2 | PPP4R4 | -0,5647 | 1,43751E-35 |
| ACE2 | EIF2S1 | -0,5648 | 1,39235E-35 |
| ACE2 | YLPM1 | -0,5648 | 1,37872E-35 |
| ACE2 | RP11-666A8.7 | -0,5648 | 1,37001E-35 |
| ACE2 | RP11-579E24.2 | -0,5649 | 1,32555E-35 |
| ACE2 | RBFA | -0,5649 | 1,34108E-35 |
| ACE2 | RP11-285F7.2 | -0,565 | 1,27181E-35 |
| ACE2 | AC009967.3 | -0,565 | 1,30876E-35 |
| ACE2 | PTEN | -0,565 | 1,2969E-35 |
| ACE2 | CTD-2006H14.2 | -0,5651 | 1,24261E-35 |
| ACE2 | HRH3 | -0,5651 | 1,26015E-35 |
| ACE2 | BLZF1 | -0,5652 | 1,21285E-35 |
| ACE2 | LCNL1 | -0,5652 | 1,19905E-35 |
| ACE2 | SRPRA | -0,5652 | 1,20539E-35 |

| | | | |
|---|---|---|---|
| ACE2 | HS3ST4 | -0,5652 | 1,20921E-35 |
| ACE2 | CABIN1 | -0,5652 | 1,19384E-35 |
| ACE2 | MRNIP | -0,5653 | 1,15739E-35 |
| ACE2 | OVCH1 | -0,5653 | 1,16028E-35 |
| ACE2 | RP11-69I8.3 | -0,5654 | 1,13702E-35 |
| ACE2 | LINC01198 | -0,5654 | 1,14313E-35 |
| ACE2 | CASC4 | -0,5654 | 1,13632E-35 |
| ACE2 | SPESP1 | -0,5654 | 1,12565E-35 |
| ACE2 | RP11-585P4.6 | -0,5655 | 1,08931E-35 |
| ACE2 | RP11-840I19.5 | -0,5655 | 1,08347E-35 |
| ACE2 | MMP11 | -0,5655 | 1,10069E-35 |
| ACE2 | C16orf72 | -0,5656 | 1,0604E-35 |
| ACE2 | RP13-895J2.3 | -0,5657 | 1,02172E-35 |
| ACE2 | OGFR-AS1 | -0,5657 | 1,00809E-35 |
| ACE2 | RP3-339A18.6 | -0,5657 | 1,03281E-35 |
| ACE2 | RP11-372K14.2 | -0,5658 | 1,00084E-35 |
| ACE2 | MTO1 | -0,5658 | 9,81579E-36 |
| ACE2 | GJA1 | -0,5658 | 9,84211E-36 |
| ACE2 | JOSD2 | -0,5658 | 9,85927E-36 |
| ACE2 | PLD2 | -0,5659 | 9,5172E-36 |
| ACE2 | AADACL4 | -0,566 | 9,29506E-36 |
| ACE2 | AC012442.5 | -0,566 | 9,24768E-36 |
| ACE2 | AC097662.2 | -0,566 | 9,35336E-36 |
| ACE2 | NOS3 | -0,566 | 9,1455E-36 |
| ACE2 | ANP32B | -0,566 | 9,18423E-36 |
| ACE2 | MRPL23-AS1 | -0,566 | 9,32654E-36 |
| ACE2 | RP4-605O3.4 | -0,566 | 9,3346E-36 |
| ACE2 | MED12 | -0,566 | 9,11064E-36 |
| ACE2 | MXD4 | -0,5661 | 8,9302E-36 |
| ACE2 | LINC00674 | -0,5661 | 8,87397E-36 |
| ACE2 | EIF3I | -0,5662 | 8,53962E-36 |
| ACE2 | LINC02035 | -0,5662 | 8,56485E-36 |
| ACE2 | LINC00222 | -0,5662 | 8,53081E-36 |
| ACE2 | LOC100421166 | -0,5663 | 8,18295E-36 |
| ACE2 | RBM25 | -0,5664 | 8,13694E-36 |
| ACE2 | RASA4DP | -0,5665 | 7,78282E-36 |
| ACE2 | RP11-551L14.4 | -0,5665 | 7,71776E-36 |
| ACE2 | RP11-546D6.3 | -0,5665 | 7,75512E-36 |
| ACE2 | ZNF98 | -0,5665 | 7,70888E-36 |
| ACE2 | YES1P1 | -0,5666 | 7,44792E-36 |
| ACE2 | SNORD87 | -0,5667 | 7,36424E-36 |
| ACE2 | TTC17 | -0,5667 | 7,19023E-36 |
| ACE2 | CPZ | -0,5668 | 7,13901E-36 |
| ACE2 | CARD6 | -0,5668 | 6,9697E-36 |
| ACE2 | MYO7A | -0,5668 | 6,91646E-36 |
| ACE2 | RP11-834C11.15 | -0,5668 | 7,0656E-36 |
| ACE2 | AC084219.4 | -0,5668 | 6,96872E-36 |
| ACE2 | PKP4 | -0,5669 | 6,71187E-36 |
| ACE2 | BLOC1S5 | -0,5669 | 6,79353E-36 |
| ACE2 | RNY3P8 | -0,5669 | 6,68386E-36 |
| ACE2 | MSL3 | -0,5669 | 6,74048E-36 |
| ACE2 | PRG4 | -0,567 | 6,63225E-36 |
| ACE2 | EEF1G | -0,567 | 6,56681E-36 |
| ACE2 | ITGBL1 | -0,567 | 6,65984E-36 |

| | | | |
|---|---|---|---|
| ACE2 | LINC00607 | -0,5671 | 6,36441E-36 |
| ACE2 | SMARCC1 | -0,5671 | 6,32596E-36 |
| ACE2 | RP11-131H24.4 | -0,5671 | 6,44549E-36 |
| ACE2 | MEX3D | -0,5671 | 6,34126E-36 |
| ACE2 | TTI1 | -0,5671 | 6,30484E-36 |
| ACE2 | TFDP2 | -0,5673 | 5,8536E-36 |
| ACE2 | PPP1R9A | -0,5673 | 5,97185E-36 |
| ACE2 | NPIPA2 | -0,5673 | 5,99859E-36 |
| ACE2 | RP11-1094H24.4 | -0,5673 | 5,93955E-36 |
| ACE2 | MVB12B | -0,5674 | 5,74498E-36 |
| ACE2 | TEX2 | -0,5674 | 5,73956E-36 |
| ACE2 | AMH | -0,5674 | 5,71521E-36 |
| ACE2 | TARDBP | -0,5675 | 5,53808E-36 |
| ACE2 | RPAP1 | -0,5675 | 5,62639E-36 |
| ACE2 | ANKRD27 | -0,5675 | 5,50402E-36 |
| ACE2 | SATB1 | -0,5676 | 5,38443E-36 |
| ACE2 | EXOC7 | -0,5676 | 5,43079E-36 |
| ACE2 | GPAA1P1 | -0,5677 | 5,16678E-36 |
| ACE2 | SMARCD1 | -0,5677 | 5,12461E-36 |
| ACE2 | WHAMM | -0,5677 | 5,11483E-36 |
| ACE2 | C4B | -0,5678 | 4,99769E-36 |
| ACE2 | TRAM2 | -0,5678 | 5,01641E-36 |
| ACE2 | ALKBH3 | -0,5678 | 4,98404E-36 |
| ACE2 | MRPL57 | -0,5678 | 5,05918E-36 |
| ACE2 | RASGRF1 | -0,5678 | 4,93321E-36 |
| ACE2 | LSM14B | -0,5678 | 5,05211E-36 |
| ACE2 | ITSN1 | -0,5679 | 4,89983E-36 |
| ACE2 | CUX1 | -0,568 | 4,65294E-36 |
| ACE2 | FUBP3 | -0,568 | 4,62006E-36 |
| ACE2 | CDC42EP2 | -0,568 | 4,65717E-36 |
| ACE2 | MYO9A | -0,568 | 4,73221E-36 |
| ACE2 | SNAI1 | -0,568 | 4,5944E-36 |
| ACE2 | AC009299.5 | -0,5681 | 4,54699E-36 |
| ACE2 | CTB-33O18.3 | -0,5681 | 4,54094E-36 |
| ACE2 | ZNF317 | -0,5681 | 4,47265E-36 |
| ACE2 | AC074391.1 | -0,5682 | 4,42362E-36 |
| ACE2 | FAM86DP | -0,5682 | 4,40417E-36 |
| ACE2 | GAS2 | -0,5682 | 4,3922E-36 |
| ACE2 | FBXO15 | -0,5682 | 4,36812E-36 |
| ACE2 | SYMPK | -0,5682 | 4,40497E-36 |
| ACE2 | LINC00899 | -0,5682 | 4,3517E-36 |
| ACE2 | AC012146.7 | -0,5684 | 4,10887E-36 |
| ACE2 | MIR133A1HG | -0,5684 | 4,101E-36 |
| ACE2 | RP11-274H2.5 | -0,5685 | 3,87699E-36 |
| ACE2 | NMBR | -0,5685 | 3,97742E-36 |
| ACE2 | ZNF563 | -0,5685 | 3,91501E-36 |
| ACE2 | TRIM67 | -0,5686 | 3,767E-36 |
| ACE2 | YWHAG | -0,5686 | 3,79997E-36 |
| ACE2 | C12orf60 | -0,5686 | 3,77165E-36 |
| ACE2 | RP11-80I15.1 | -0,5687 | 3,6367E-36 |
| ACE2 | ADRB1 | -0,5687 | 3,69201E-36 |
| ACE2 | POLR3C | -0,5688 | 3,61557E-36 |
| ACE2 | ZNF75A | -0,5688 | 3,51044E-36 |
| ACE2 | NOV | -0,5689 | 3,49258E-36 |

| | | | |
|---|---|---|---|
| ACE2 | ARHGEF12 | -0,569 | 3,28424E-36 |
| ACE2 | RP11-541E12.1 | -0,5691 | 3,16187E-36 |
| ACE2 | RP11-517B11.7 | -0,5691 | 3,15885E-36 |
| ACE2 | DNHD1 | -0,5691 | 3,16727E-36 |
| ACE2 | NUMA1 | -0,5691 | 3,17785E-36 |
| ACE2 | C2orf27A | -0,5692 | 3,15559E-36 |
| ACE2 | MYNN | -0,5692 | 3,11753E-36 |
| ACE2 | BUD31 | -0,5692 | 3,14815E-36 |
| ACE2 | SLX4 | -0,5692 | 3,09362E-36 |
| ACE2 | RP11-174O3.3 | -0,5693 | 2,95517E-36 |
| ACE2 | POMT1 | -0,5693 | 2,95269E-36 |
| ACE2 | GABPB2 | -0,5694 | 2,8551E-36 |
| ACE2 | IL12A | -0,5694 | 2,88482E-36 |
| ACE2 | ZNF214 | -0,5694 | 2,87275E-36 |
| ACE2 | RP11-173M1.4 | -0,5694 | 2,85288E-36 |
| ACE2 | SOX18 | -0,5694 | 2,9187E-36 |
| ACE2 | RP5-1126H10.2 | -0,5695 | 2,81817E-36 |
| ACE2 | LINC01933 | -0,5695 | 2,81885E-36 |
| ACE2 | ARC | -0,5695 | 2,75373E-36 |
| ACE2 | LINC01503 | -0,5695 | 2,81748E-36 |
| ACE2 | TMEM72-AS1 | -0,5695 | 2,79433E-36 |
| ACE2 | WEE1 | -0,5695 | 2,82286E-36 |
| ACE2 | CTD-2292M16.8 | -0,5695 | 2,77153E-36 |
| ACE2 | RP3-453C12.15 | -0,5695 | 2,76568E-36 |
| ACE2 | BTBD2 | -0,5696 | 2,66464E-36 |
| ACE2 | NTRK2 | -0,5697 | 2,63527E-36 |
| ACE2 | BAHCC1 | -0,5697 | 2,57338E-36 |
| ACE2 | CEP350 | -0,5698 | 2,56076E-36 |
| ACE2 | ADRA2C | -0,5698 | 2,54206E-36 |
| ACE2 | CLOCK | -0,5698 | 2,51997E-36 |
| ACE2 | ZNF219 | -0,5699 | 2,43623E-36 |
| ACE2 | IRF9 | -0,57 | 2,38128E-36 |
| ACE2 | RP11-387H17.4 | -0,57 | 2,37743E-36 |
| ACE2 | PIAS4 | -0,57 | 2,34523E-36 |
| ACE2 | CTC-360J11.4 | -0,57 | 2,35106E-36 |
| ACE2 | FHL5 | -0,5701 | 2,30459E-36 |
| ACE2 | AF186192.5 | -0,5702 | 2,21544E-36 |
| ACE2 | HSPA13 | -0,5702 | 2,20416E-36 |
| ACE2 | SLC47A1 | -0,5703 | 2,12611E-36 |
| ACE2 | ASCC1 | -0,5704 | 2,08398E-36 |
| ACE2 | PPP2R2D | -0,5705 | 1,97301E-36 |
| ACE2 | MTMR6 | -0,5705 | 1,96439E-36 |
| ACE2 | CFD | -0,5705 | 2,01354E-36 |
| ACE2 | HRC | -0,5705 | 1,99504E-36 |
| ACE2 | ING1 | -0,5706 | 1,92146E-36 |
| ACE2 | SETD2 | -0,5707 | 1,8811E-36 |
| ACE2 | WI2-87327B8.1 | -0,5707 | 1,86009E-36 |
| ACE2 | SYN3 | -0,5707 | 1,85627E-36 |
| ACE2 | CAPN10 | -0,5708 | 1,78287E-36 |
| ACE2 | PSMC2 | -0,5708 | 1,76555E-36 |
| ACE2 | CTC-265F19.2 | -0,5708 | 1,8207E-36 |
| ACE2 | VGLL4 | -0,5709 | 1,72307E-36 |
| ACE2 | AK6 | -0,5709 | 1,75688E-36 |
| ACE2 | UBLCP1 | -0,5709 | 1,70639E-36 |

| | | | |
|---|---|---|---|
| ACE2 | TSPOAP1-AS1 | -0,5709 | 1,70882E-36 |
| ACE2 | TIMMDC1 | -0,571 | 1,68305E-36 |
| ACE2 | RP11-283I3.6 | -0,5711 | 1,64143E-36 |
| ACE2 | SLC25A30 | -0,5711 | 1,61145E-36 |
| ACE2 | CH17-408M7.1 | -0,5713 | 1,49722E-36 |
| ACE2 | PMVK | -0,5713 | 1,49737E-36 |
| ACE2 | PDE3B | -0,5713 | 1,48796E-36 |
| ACE2 | VWF | -0,5713 | 1,51648E-36 |
| ACE2 | PRRT2 | -0,5713 | 1,49358E-36 |
| ACE2 | COQ8B | -0,5713 | 1,49531E-36 |
| ACE2 | PIGP | -0,5713 | 1,50653E-36 |
| ACE2 | ANAPC4 | -0,5714 | 1,4765E-36 |
| ACE2 | SCG5 | -0,5714 | 1,47515E-36 |
| ACE2 | TMEM126A | -0,5715 | 1,39084E-36 |
| ACE2 | ICAM2 | -0,5715 | 1,39209E-36 |
| ACE2 | YIPF4 | -0,5716 | 1,36056E-36 |
| ACE2 | TOMM5 | -0,5716 | 1,34602E-36 |
| ACE2 | KPNB1 | -0,5716 | 1,36102E-36 |
| ACE2 | RP5-1116H23.3 | -0,5716 | 1,38338E-36 |
| ACE2 | LA16c-380F5.3 | -0,5717 | 1,33131E-36 |
| ACE2 | MTMR10 | -0,5718 | 1,25766E-36 |
| ACE2 | RP4-681N20.5 | -0,5718 | 1,26547E-36 |
| ACE2 | RP11-305F18.1 | -0,5718 | 1,28681E-36 |
| ACE2 | CREB1 | -0,5719 | 1,22405E-36 |
| ACE2 | RP11-127B20.2 | -0,5719 | 1,23727E-36 |
| ACE2 | DUSP1 | -0,5719 | 1,21877E-36 |
| ACE2 | EDRF1 | -0,5719 | 1,22401E-36 |
| ACE2 | DHPS | -0,5719 | 1,22255E-36 |
| ACE2 | IMPG2 | -0,572 | 1,18089E-36 |
| ACE2 | C9orf16 | -0,572 | 1,16989E-36 |
| ACE2 | PTRH1 | -0,5721 | 1,16231E-36 |
| ACE2 | MRPL43 | -0,5721 | 1,13258E-36 |
| ACE2 | CUL5 | -0,5721 | 1,15738E-36 |
| ACE2 | RP5-1142A6.9 | -0,5721 | 1,15876E-36 |
| ACE2 | AC003104.1 | -0,5721 | 1,15491E-36 |
| ACE2 | FBXW4P1 | -0,5721 | 1,14777E-36 |
| ACE2 | FAM3C | -0,5722 | 1,09864E-36 |
| ACE2 | AZIN1 | -0,5722 | 1,11653E-36 |
| ACE2 | SNX19 | -0,5722 | 1,10996E-36 |
| ACE2 | RP11-218E20.3 | -0,5722 | 1,12569E-36 |
| ACE2 | RP11-72I8.1 | -0,5723 | 1,07762E-36 |
| ACE2 | SPTBN1 | -0,5724 | 1,03426E-36 |
| ACE2 | SRP19 | -0,5724 | 1,04909E-36 |
| ACE2 | RP4-668J24.2 | -0,5724 | 1,03677E-36 |
| ACE2 | RPL24P8 | -0,5724 | 1,03523E-36 |
| ACE2 | NLRX1 | -0,5724 | 1,02369E-36 |
| ACE2 | DYNLL2 | -0,5724 | 1,03382E-36 |
| ACE2 | RP11-324I22.4 | -0,5725 | 9,86449E-37 |
| ACE2 | MRGPRE | -0,5725 | 9,83072E-37 |
| ACE2 | SPIN3 | -0,5725 | 9,85468E-37 |
| ACE2 | INSIG2 | -0,5726 | 9,48167E-37 |
| ACE2 | RP11-550I24.2 | -0,5726 | 9,72434E-37 |
| ACE2 | IFNAR2 | -0,5726 | 9,77612E-37 |
| ACE2 | LEPROT | -0,5727 | 9,31533E-37 |

| | | | |
|---|---|---|---|
| ACE2 | ADCY3 | -0,5727 | 9,40672E-37 |
| ACE2 | SLC9A7P1 | -0,5727 | 9,26834E-37 |
| ACE2 | CCIN | -0,5728 | 9,12927E-37 |
| ACE2 | RP11-266L9.3 | -0,5728 | 8,97889E-37 |
| ACE2 | FAM57B | -0,5728 | 8,91677E-37 |
| ACE2 | AC018766.6 | -0,5728 | 8,9405E-37 |
| ACE2 | AC002456.2 | -0,5729 | 8,74432E-37 |
| ACE2 | STAG3L4 | -0,573 | 8,40635E-37 |
| ACE2 | RP11-399K21.14 | -0,573 | 8,41816E-37 |
| ACE2 | MYO9B | -0,573 | 8,48749E-37 |
| ACE2 | XPC | -0,5731 | 7,99294E-37 |
| ACE2 | NPTX2 | -0,5731 | 8,02672E-37 |
| ACE2 | SMC2-AS1 | -0,5731 | 8,11521E-37 |
| ACE2 | JMJD1C | -0,5731 | 8,0738E-37 |
| ACE2 | CHORDC1 | -0,5731 | 7,9698E-37 |
| ACE2 | HSD17B1P1 | -0,5731 | 8,18506E-37 |
| ACE2 | NCAN | -0,5731 | 8,03431E-37 |
| ACE2 | RP11-489G11.3 | -0,5732 | 7,74614E-37 |
| ACE2 | RP3-368A4.6 | -0,5732 | 7,83741E-37 |
| ACE2 | LRRC3B | -0,5733 | 7,49386E-37 |
| ACE2 | SBDS | -0,5733 | 7,68243E-37 |
| ACE2 | SLC13A4 | -0,5733 | 7,5328E-37 |
| ACE2 | FSBP | -0,5733 | 7,66023E-37 |
| ACE2 | SETDB2 | -0,5733 | 7,53603E-37 |
| ACE2 | ZHX3 | -0,5733 | 7,53829E-37 |
| ACE2 | FXYD5 | -0,5734 | 7,19121E-37 |
| ACE2 | ZFAS1 | -0,5734 | 7,30242E-37 |
| ACE2 | PBRM1 | -0,5735 | 7,16962E-37 |
| ACE2 | TLE4 | -0,5735 | 7,17165E-37 |
| ACE2 | SP110 | -0,5736 | 6,72766E-37 |
| ACE2 | SERBP1P3 | -0,5736 | 6,80314E-37 |
| ACE2 | HSPA1L | -0,5736 | 6,78979E-37 |
| ACE2 | CNN2 | -0,5736 | 6,89756E-37 |
| ACE2 | CPLX1 | -0,5737 | 6,6543E-37 |
| ACE2 | TNPO1 | -0,5737 | 6,49657E-37 |
| ACE2 | KCNA5 | -0,5737 | 6,6145E-37 |
| ACE2 | CLCN7 | -0,5737 | 6,57337E-37 |
| ACE2 | AK9 | -0,5738 | 6,43357E-37 |
| ACE2 | BCL6B | -0,5738 | 6,2713E-37 |
| ACE2 | FBXO6 | -0,5739 | 6,10578E-37 |
| ACE2 | SRPX2 | -0,5739 | 6,17545E-37 |
| ACE2 | LY6G6D | -0,574 | 5,94227E-37 |
| ACE2 | THUMPD1 | -0,574 | 5,98324E-37 |
| ACE2 | CTD-3193O13.8 | -0,574 | 5,92742E-37 |
| ACE2 | MSANTD1 | -0,5741 | 5,73429E-37 |
| ACE2 | RP11-488C13.6 | -0,5741 | 5,75035E-37 |
| ACE2 | FOXN3-AS1 | -0,5741 | 5,70755E-37 |
| ACE2 | RP11-893F2.18 | -0,5741 | 5,68855E-37 |
| ACE2 | DACH2 | -0,5741 | 5,65832E-37 |
| ACE2 | RP11-730A19.9 | -0,5742 | 5,52779E-37 |
| ACE2 | KLF9 | -0,5743 | 5,37333E-37 |
| ACE2 | HMGN2P19 | -0,5744 | 5,24694E-37 |
| ACE2 | PROKR1 | -0,5744 | 5,13429E-37 |
| ACE2 | CCDC148 | -0,5744 | 5,15724E-37 |

| | | | |
|---|---|---|---|
| ACE2 | TATDN1 | -0,5745 | 5,00745E-37 |
| ACE2 | NUP160 | -0,5745 | 5,00634E-37 |
| ACE2 | SAFB2 | -0,5746 | 4,76923E-37 |
| ACE2 | AAR2 | -0,5746 | 4,76177E-37 |
| ACE2 | HNRNPH3 | -0,5747 | 4,64062E-37 |
| ACE2 | TRIM16L | -0,5747 | 4,69462E-37 |
| ACE2 | CTC-325H20.4 | -0,5747 | 4,57128E-37 |
| ACE2 | NFU1 | -0,5748 | 4,53986E-37 |
| ACE2 | GGCX | -0,5748 | 4,53378E-37 |
| ACE2 | RP11-256I23.2 | -0,5748 | 4,493E-37 |
| ACE2 | CFI | -0,5748 | 4,48832E-37 |
| ACE2 | CTD-3128G10.7 | -0,5748 | 4,47416E-37 |
| ACE2 | SCGN | -0,5749 | 4,28248E-37 |
| ACE2 | KLRA1P | -0,5749 | 4,32485E-37 |
| ACE2 | WI2-89031B12.1 | -0,5749 | 4,27754E-37 |
| ACE2 | OPRK1 | -0,575 | 4,25509E-37 |
| ACE2 | ZNF99 | -0,575 | 4,24049E-37 |
| ACE2 | JADE3 | -0,575 | 4,25246E-37 |
| ACE2 | WBP1L | -0,5751 | 4,08752E-37 |
| ACE2 | ATG2B | -0,5751 | 3,98297E-37 |
| ACE2 | RHOA | -0,5752 | 3,97179E-37 |
| ACE2 | ATXN7L1 | -0,5753 | 3,78461E-37 |
| ACE2 | RP11-319G6.1 | -0,5754 | 3,59459E-37 |
| ACE2 | RPL4P4 | -0,5755 | 3,50138E-37 |
| ACE2 | WISP3 | -0,5755 | 3,57703E-37 |
| ACE2 | CTB-152G17.6 | -0,5755 | 3,54184E-37 |
| ACE2 | EFR3A | -0,5755 | 3,48088E-37 |
| ACE2 | TRBV26OR9-2 | -0,5755 | 3,47607E-37 |
| ACE2 | TEX26-AS1 | -0,5755 | 3,54123E-37 |
| ACE2 | PIK3R3 | -0,5756 | 3,41203E-37 |
| ACE2 | RP4-597N16.4 | -0,5756 | 3,38047E-37 |
| ACE2 | GLIS3 | -0,5756 | 3,44882E-37 |
| ACE2 | WT1-AS | -0,5756 | 3,41338E-37 |
| ACE2 | TMEM257 | -0,5756 | 3,37304E-37 |
| ACE2 | TAF7L | -0,5757 | 3,2544E-37 |
| ACE2 | DBH-AS1 | -0,5758 | 3,17593E-37 |
| ACE2 | IGF1 | -0,5758 | 3,21195E-37 |
| ACE2 | CTD-2623N2.11 | -0,5758 | 3,21677E-37 |
| ACE2 | BEX2 | -0,5758 | 3,17253E-37 |
| ACE2 | BCYRN1 | -0,5759 | 3,11047E-37 |
| ACE2 | RP11-514P8.10 | -0,576 | 2,93065E-37 |
| ACE2 | PEX5 | -0,576 | 2,98864E-37 |
| ACE2 | RP11-64B16.4 | -0,576 | 2,94239E-37 |
| ACE2 | PTPRG | -0,5761 | 2,83224E-37 |
| ACE2 | AC003090.1 | -0,5761 | 2,88889E-37 |
| ACE2 | RGAG1 | -0,5761 | 2,83378E-37 |
| ACE2 | CTD-2287O16.1 | -0,5762 | 2,7376E-37 |
| ACE2 | RP11-981G7.6 | -0,5762 | 2,78136E-37 |
| ACE2 | RNASEK | -0,5762 | 2,7291E-37 |
| ACE2 | RCVRN | -0,5762 | 2,79675E-37 |
| ACE2 | RP11-861E21.3 | -0,5762 | 2,71627E-37 |
| ACE2 | MRGBP | -0,5762 | 2,76754E-37 |
| ACE2 | NAA30 | -0,5763 | 2,69472E-37 |
| ACE2 | LINC01310 | -0,5763 | 2,70307E-37 |

| | | | |
|---|---|---|---|
| ACE2 | RP11-375I20.6 | -0,5764 | 2,60126E-37 |
| ACE2 | PDGFRA | -0,5764 | 2,56608E-37 |
| ACE2 | RB1CC1 | -0,5764 | 2,55444E-37 |
| ACE2 | PITRM1 | -0,5764 | 2,56647E-37 |
| ACE2 | SCN4A | -0,5764 | 2,56847E-37 |
| ACE2 | MSTO1 | -0,5765 | 2,47771E-37 |
| ACE2 | PPEF1 | -0,5765 | 2,4924E-37 |
| ACE2 | RP11-367H1.1 | -0,5766 | 2,42463E-37 |
| ACE2 | PDE7B | -0,5766 | 2,38243E-37 |
| ACE2 | GLP2R | -0,5766 | 2,39467E-37 |
| ACE2 | ZNF234 | -0,5766 | 2,36981E-37 |
| ACE2 | CH17-469D17.1 | -0,5767 | 2,3089E-37 |
| ACE2 | GABRQ | -0,5767 | 2,29004E-37 |
| ACE2 | LRRC39 | -0,5768 | 2,2214E-37 |
| ACE2 | TSC22D2 | -0,5769 | 2,12345E-37 |
| ACE2 | FBLL1 | -0,5769 | 2,1538E-37 |
| ACE2 | CFL1P1 | -0,5769 | 2,15928E-37 |
| ACE2 | CTD-3234P18.6 | -0,5769 | 2,16089E-37 |
| ACE2 | IGFL1P1 | -0,5769 | 2,1524E-37 |
| ACE2 | RBM46 | -0,577 | 2,09871E-37 |
| ACE2 | GM2A | -0,5771 | 1,97414E-37 |
| ACE2 | RNGTT | -0,5771 | 2,00175E-37 |
| ACE2 | BNIP2 | -0,5771 | 1,97522E-37 |
| ACE2 | RP11-444D3.1 | -0,5772 | 1,90938E-37 |
| ACE2 | NT5DC3 | -0,5772 | 1,90929E-37 |
| ACE2 | MIPEPP3 | -0,5772 | 1,92243E-37 |
| ACE2 | FAM98A | -0,5773 | 1,84284E-37 |
| ACE2 | AZI2 | -0,5775 | 1,73171E-37 |
| ACE2 | C2orf80 | -0,5776 | 1,68695E-37 |
| ACE2 | CTD-2010I16.1 | -0,5776 | 1,66057E-37 |
| ACE2 | ARHGEF37 | -0,5777 | 1,60801E-37 |
| ACE2 | ZNF879 | -0,5777 | 1,63148E-37 |
| ACE2 | FRS2 | -0,5777 | 1,63689E-37 |
| ACE2 | ZNF596 | -0,5778 | 1,57371E-37 |
| ACE2 | AC009120.6 | -0,5778 | 1,59258E-37 |
| ACE2 | CYB5RL | -0,5779 | 1,53163E-37 |
| ACE2 | EIF2A | -0,5779 | 1,52634E-37 |
| ACE2 | ANKIB1 | -0,578 | 1,45296E-37 |
| ACE2 | PRKAG1 | -0,578 | 1,47926E-37 |
| ACE2 | RP11-759A24.1 | -0,578 | 1,47435E-37 |
| ACE2 | LINC00116 | -0,5781 | 1,38788E-37 |
| ACE2 | SPG20-AS1 | -0,5781 | 1,43302E-37 |
| ACE2 | ZNF8 | -0,5781 | 1,38927E-37 |
| ACE2 | CREG1 | -0,5782 | 1,3753E-37 |
| ACE2 | RP5-1098D14.1 | -0,5782 | 1,38547E-37 |
| ACE2 | RP11-98F14.12 | -0,5782 | 1,37425E-37 |
| ACE2 | RP11-227D13.1 | -0,5782 | 1,37003E-37 |
| ACE2 | ZNF264 | -0,5782 | 1,35392E-37 |
| ACE2 | RSU1 | -0,5783 | 1,29848E-37 |
| ACE2 | PLEKHA1 | -0,5783 | 1,2976E-37 |
| ACE2 | VPS26B | -0,5783 | 1,31533E-37 |
| ACE2 | NPTN | -0,5783 | 1,29851E-37 |
| ACE2 | ZNF181 | -0,5783 | 1,33104E-37 |
| ACE2 | AC006116.22 | -0,5783 | 1,30274E-37 |

| | | | |
|---|---|---|---|
| ACE2 | BEND7 | -0,5784 | 1,25888E-37 |
| ACE2 | TSPAN32 | -0,5784 | 1,25726E-37 |
| ACE2 | LIPE-AS1 | -0,5784 | 1,28784E-37 |
| ACE2 | MGMT | -0,5785 | 1,23856E-37 |
| ACE2 | ZNF518B | -0,5786 | 1,18099E-37 |
| ACE2 | UBOX5 | -0,5786 | 1,19507E-37 |
| ACE2 | ART3 | -0,5787 | 1,14966E-37 |
| ACE2 | FBXW7 | -0,5787 | 1,15893E-37 |
| ACE2 | DBT | -0,5788 | 1,11242E-37 |
| ACE2 | ANKRD36C | -0,5788 | 1,10684E-37 |
| ACE2 | PLK2 | -0,5788 | 1,0928E-37 |
| ACE2 | RUFY1 | -0,5788 | 1,0934E-37 |
| ACE2 | SARDH | -0,5788 | 1,09326E-37 |
| ACE2 | SLC9A7 | -0,5788 | 1,08918E-37 |
| ACE2 | RP11-1336O20.2 | -0,5789 | 1,06238E-37 |
| ACE2 | ANKRD31 | -0,5789 | 1,04933E-37 |
| ACE2 | INTS13 | -0,5789 | 1,08123E-37 |
| ACE2 | RP11-429B14.4 | -0,5789 | 1,04988E-37 |
| ACE2 | LINC00237 | -0,5789 | 1,05446E-37 |
| ACE2 | DGCR5 | -0,5789 | 1,08154E-37 |
| ACE2 | B3GAT3 | -0,5791 | 1,00421E-37 |
| ACE2 | ALKBH6 | -0,5791 | 9,94915E-38 |
| ACE2 | USP46 | -0,5792 | 9,68273E-38 |
| ACE2 | FNDC1-IT1 | -0,5792 | 9,52952E-38 |
| ACE2 | AP4M1 | -0,5792 | 9,74061E-38 |
| ACE2 | FAM58A | -0,5792 | 9,72221E-38 |
| ACE2 | PEX19 | -0,5793 | 9,20905E-38 |
| ACE2 | UBA6-AS1 | -0,5793 | 9,27309E-38 |
| ACE2 | ZRSR1 | -0,5793 | 9,20103E-38 |
| ACE2 | RP11-632K20.7 | -0,5793 | 9,11163E-38 |
| ACE2 | KCNK4 | -0,5794 | 8,99638E-38 |
| ACE2 | TSC2 | -0,5794 | 9,0553E-38 |
| ACE2 | PHF14 | -0,5795 | 8,51951E-38 |
| ACE2 | FBXO3 | -0,5795 | 8,63923E-38 |
| ACE2 | KCTD17 | -0,5795 | 8,5854E-38 |
| ACE2 | WDR64 | -0,5796 | 8,36351E-38 |
| ACE2 | RP11-38L15.3 | -0,5796 | 8,25076E-38 |
| ACE2 | NOG | -0,5796 | 8,45549E-38 |
| ACE2 | PANK2 | -0,5796 | 8,43682E-38 |
| ACE2 | RASSF2 | -0,5796 | 8,25273E-38 |
| ACE2 | PRH1 | -0,5798 | 7,62558E-38 |
| ACE2 | BDP1 | -0,58 | 7,23027E-38 |
| ACE2 | TUBBP1 | -0,58 | 7,26682E-38 |
| ACE2 | GRHPR | -0,58 | 7,18707E-38 |
| ACE2 | CAMTA2 | -0,58 | 7,12242E-38 |
| ACE2 | MICAL3 | -0,58 | 7,19595E-38 |
| ACE2 | RRP36 | -0,5801 | 6,91704E-38 |
| ACE2 | ERICH1 | -0,5801 | 6,84613E-38 |
| ACE2 | RP11-324I22.3 | -0,5801 | 6,90844E-38 |
| ACE2 | FOPNL | -0,5801 | 6,9454E-38 |
| ACE2 | SYNPR | -0,5803 | 6,58104E-38 |
| ACE2 | TLL2 | -0,5803 | 6,58159E-38 |
| ACE2 | SPIN2B | -0,5803 | 6,54687E-38 |
| ACE2 | RP1-315G1.3 | -0,5803 | 6,56051E-38 |

| | | | |
|---|---|---|---|
| ACE2 | RPL5P4 | -0,5804 | 6,31242E-38 |
| ACE2 | ANKRD42 | -0,5804 | 6,2812E-38 |
| ACE2 | MYCBP2-AS1 | -0,5804 | 6,23943E-38 |
| ACE2 | MIR137HG | -0,5805 | 6,02374E-38 |
| ACE2 | BRE | -0,5805 | 6,06805E-38 |
| ACE2 | BET1 | -0,5805 | 5,93598E-38 |
| ACE2 | GOLGA8R | -0,5805 | 6,05154E-38 |
| ACE2 | RP11-119F19.5 | -0,5806 | 5,91626E-38 |
| ACE2 | PSMF1 | -0,5806 | 5,8166E-38 |
| ACE2 | CELF3 | -0,5807 | 5,68187E-38 |
| ACE2 | BAZ2B | -0,5807 | 5,64781E-38 |
| ACE2 | HID1-AS1 | -0,5807 | 5,72438E-38 |
| ACE2 | ZNF148 | -0,5808 | 5,46003E-38 |
| ACE2 | FBXO32 | -0,5808 | 5,50391E-38 |
| ACE2 | ANAPC11 | -0,5808 | 5,35086E-38 |
| ACE2 | LMAN2L | -0,5809 | 5,27789E-38 |
| ACE2 | CUL2 | -0,5809 | 5,28626E-38 |
| ACE2 | FLT1 | -0,5809 | 5,2394E-38 |
| ACE2 | CHRNA5 | -0,5809 | 5,18492E-38 |
| ACE2 | CCDC90B | -0,581 | 5,14282E-38 |
| ACE2 | TRIM52-AS1 | -0,5811 | 4,81003E-38 |
| ACE2 | C12orf73 | -0,5811 | 4,90387E-38 |
| ACE2 | RAB3D | -0,5811 | 4,94249E-38 |
| ACE2 | DVL1 | -0,5812 | 4,70268E-38 |
| ACE2 | OSGEP | -0,5812 | 4,62443E-38 |
| ACE2 | ANKRD13C | -0,5813 | 4,54616E-38 |
| ACE2 | MCUB | -0,5813 | 4,53509E-38 |
| ACE2 | PLAT | -0,5813 | 4,57145E-38 |
| ACE2 | CLSTN3 | -0,5813 | 4,59681E-38 |
| ACE2 | DCTN4 | -0,5814 | 4,3124E-38 |
| ACE2 | LINC01578 | -0,5814 | 4,31356E-38 |
| ACE2 | LYAR | -0,5815 | 4,25295E-38 |
| ACE2 | BMS1P21 | -0,5815 | 4,30445E-38 |
| ACE2 | UBR3 | -0,5816 | 4,03013E-38 |
| ACE2 | LINC00887 | -0,5816 | 4,05241E-38 |
| ACE2 | AP001189.4 | -0,5816 | 4,02562E-38 |
| ACE2 | FRG1CP | -0,5816 | 4,02495E-38 |
| ACE2 | SLC25A17 | -0,5816 | 4,09309E-38 |
| ACE2 | FBXO11 | -0,5817 | 3,9228E-38 |
| ACE2 | RP11-290O12.2 | -0,5817 | 3,93008E-38 |
| ACE2 | REPS1 | -0,5817 | 3,9925E-38 |
| ACE2 | GIGYF1 | -0,5817 | 3,90585E-38 |
| ACE2 | CDC20P1 | -0,5817 | 3,92785E-38 |
| ACE2 | FNTB | -0,5817 | 3,88571E-38 |
| ACE2 | PHEX-AS1 | -0,5817 | 3,91552E-38 |
| ACE2 | CLVS1 | -0,5818 | 3,75239E-38 |
| ACE2 | MAD1L1 | -0,5819 | 3,65729E-38 |
| ACE2 | HOTAIR | -0,5819 | 3,68842E-38 |
| ACE2 | ZNF727 | -0,582 | 3,5168E-38 |
| ACE2 | PPP1R3B | -0,582 | 3,4815E-38 |
| ACE2 | KCNJ9 | -0,5821 | 3,41994E-38 |
| ACE2 | PCYOX1 | -0,5821 | 3,44744E-38 |
| ACE2 | MRPL36 | -0,5821 | 3,45612E-38 |
| ACE2 | TES | -0,5821 | 3,4644E-38 |

| | | | |
|---|---|---|---|
| ACE2 | AC074212.6 | -0,5821 | 3,47115E-38 |
| ACE2 | AC092667.2 | -0,5822 | 3,35339E-38 |
| ACE2 | ZNF487 | -0,5822 | 3,26634E-38 |
| ACE2 | AC015849.2 | -0,5822 | 3,25267E-38 |
| ACE2 | NPTN-IT1 | -0,5823 | 3,12913E-38 |
| ACE2 | CDC42EP4 | -0,5823 | 3,12674E-38 |
| ACE2 | GPSM2 | -0,5824 | 3,03974E-38 |
| ACE2 | FAM19A4 | -0,5824 | 3,06025E-38 |
| ACE2 | GLO1 | -0,5825 | 3,00582E-38 |
| ACE2 | ANKRD19P | -0,5825 | 2,95931E-38 |
| ACE2 | NOL9 | -0,5826 | 2,84284E-38 |
| ACE2 | STXBP5L | -0,5826 | 2,82803E-38 |
| ACE2 | CHD4 | -0,5826 | 2,80746E-38 |
| ACE2 | RP11-2N1.2 | -0,5826 | 2,87418E-38 |
| ACE2 | KB-1125A3.11 | -0,5826 | 2,89327E-38 |
| ACE2 | AP2A2 | -0,5827 | 2,72932E-38 |
| ACE2 | SCAMP1 | -0,5828 | 2,67754E-38 |
| ACE2 | EPHB6 | -0,5828 | 2,62414E-38 |
| ACE2 | RPL4P5 | -0,5828 | 2,63875E-38 |
| ACE2 | DCDC1 | -0,5828 | 2,68429E-38 |
| ACE2 | DRD2 | -0,5828 | 2,65041E-38 |
| ACE2 | CTD-3185P2.1 | -0,5828 | 2,62726E-38 |
| ACE2 | AQP4-AS1 | -0,5828 | 2,65504E-38 |
| ACE2 | AHR | -0,5829 | 2,58448E-38 |
| ACE2 | TWISTNB | -0,5829 | 2,58214E-38 |
| ACE2 | SCNM1 | -0,583 | 2,48833E-38 |
| ACE2 | ZNF697 | -0,5831 | 2,35178E-38 |
| ACE2 | RP11-111J6.2 | -0,5831 | 2,42356E-38 |
| ACE2 | LRRTM2 | -0,5831 | 2,35996E-38 |
| ACE2 | MAP2K5 | -0,5831 | 2,40745E-38 |
| ACE2 | ZBTB37 | -0,5832 | 2,26945E-38 |
| ACE2 | LINC01881 | -0,5832 | 2,33982E-38 |
| ACE2 | RP11-379F4.4 | -0,5832 | 2,26399E-38 |
| ACE2 | TSPAN19 | -0,5832 | 2,32919E-38 |
| ACE2 | ESD | -0,5832 | 2,29072E-38 |
| ACE2 | CFAP53 | -0,5832 | 2,29861E-38 |
| ACE2 | C1orf35 | -0,5833 | 2,25439E-38 |
| ACE2 | ARL8B | -0,5833 | 2,2419E-38 |
| ACE2 | JAKMIP1 | -0,5833 | 2,20073E-38 |
| ACE2 | RP11-696D21.2 | -0,5833 | 2,1959E-38 |
| ACE2 | TSR1 | -0,5833 | 2,23358E-38 |
| ACE2 | CSE1L | -0,5833 | 2,20352E-38 |
| ACE2 | RP3-449O17.1 | -0,5833 | 2,18398E-38 |
| ACE2 | NFE2L2 | -0,5834 | 2,15835E-38 |
| ACE2 | RNF13 | -0,5834 | 2,11901E-38 |
| ACE2 | BTAF1 | -0,5834 | 2,17986E-38 |
| ACE2 | RP5-1107A17.4 | -0,5834 | 2,16598E-38 |
| ACE2 | PAPSS1 | -0,5835 | 2,03744E-38 |
| ACE2 | AC006538.1 | -0,5835 | 2,0402E-38 |
| ACE2 | CCDC9 | -0,5835 | 2,0704E-38 |
| ACE2 | FAM124B | -0,5836 | 1,96963E-38 |
| ACE2 | MTG2 | -0,5837 | 1,95309E-38 |
| ACE2 | TBX1 | -0,5837 | 1,92365E-38 |
| ACE2 | RNF122 | -0,5838 | 1,84802E-38 |

| | | | |
|---|---|---|---|
| ACE2 | SYDE2 | -0,5839 | 1,80507E-38 |
| ACE2 | PAK5 | -0,5839 | 1,78271E-38 |
| ACE2 | MANBA | -0,584 | 1,73449E-38 |
| ACE2 | TSLP | -0,584 | 1,70746E-38 |
| ACE2 | RP4-650F12.2 | -0,5841 | 1,65221E-38 |
| ACE2 | PCBP2 | -0,5841 | 1,678E-38 |
| ACE2 | RP11-298D21.3 | -0,5841 | 1,66065E-38 |
| ACE2 | GATA2 | -0,5842 | 1,63194E-38 |
| ACE2 | UNC5D | -0,5842 | 1,6218E-38 |
| ACE2 | TOMM20 | -0,5843 | 1,55887E-38 |
| ACE2 | ZNF462 | -0,5843 | 1,55691E-38 |
| ACE2 | MSS51 | -0,5844 | 1,52227E-38 |
| ACE2 | SFTPD-AS1 | -0,5844 | 1,49199E-38 |
| ACE2 | CNIH2 | -0,5844 | 1,48544E-38 |
| ACE2 | RP1-159A19.4 | -0,5845 | 1,46498E-38 |
| ACE2 | RAB3GAP1 | -0,5845 | 1,44718E-38 |
| ACE2 | UCP3 | -0,5845 | 1,4509E-38 |
| ACE2 | CREBBP | -0,5845 | 1,45216E-38 |
| ACE2 | RANBP9 | -0,5846 | 1,39029E-38 |
| ACE2 | OPCML | -0,5846 | 1,40997E-38 |
| ACE2 | STX7 | -0,5847 | 1,32354E-38 |
| ACE2 | RP11-468H14.2 | -0,5847 | 1,34536E-38 |
| ACE2 | MEIS1-AS3 | -0,5848 | 1,29681E-38 |
| ACE2 | WDR24 | -0,5848 | 1,27478E-38 |
| ACE2 | NEURL4 | -0,5848 | 1,28655E-38 |
| ACE2 | RERE | -0,5849 | 1,23793E-38 |
| ACE2 | MRPS15 | -0,5849 | 1,24412E-38 |
| ACE2 | ITGB3BP | -0,5849 | 1,23305E-38 |
| ACE2 | MEF2D | -0,5849 | 1,27177E-38 |
| ACE2 | SCN1A | -0,5849 | 1,2393E-38 |
| ACE2 | SAP30L | -0,5849 | 1,24674E-38 |
| ACE2 | GTF2IRD2P1 | -0,5849 | 1,2679E-38 |
| ACE2 | CTD-2515A14.1 | -0,5849 | 1,25021E-38 |
| ACE2 | LINC01270 | -0,585 | 1,19552E-38 |
| ACE2 | KCNK15-AS1 | -0,5851 | 1,14571E-38 |
| ACE2 | EPHB1 | -0,5852 | 1,11254E-38 |
| ACE2 | NMNAT3 | -0,5852 | 1,10478E-38 |
| ACE2 | FAM200B | -0,5852 | 1,10403E-38 |
| ACE2 | HOXA10 | -0,5852 | 1,13774E-38 |
| ACE2 | GS1-259H13.2 | -0,5852 | 1,13326E-38 |
| ACE2 | GRIA4 | -0,5853 | 1,07116E-38 |
| ACE2 | APOLD1 | -0,5853 | 1,06779E-38 |
| ACE2 | ZSWIM4 | -0,5853 | 1,09717E-38 |
| ACE2 | LCA5L | -0,5853 | 1,06272E-38 |
| ACE2 | TAF1C | -0,5854 | 1,03456E-38 |
| ACE2 | RP11-159D12.11 | -0,5854 | 1,05706E-38 |
| ACE2 | MAP2K7 | -0,5854 | 1,02422E-38 |
| ACE2 | EIF2S2 | -0,5854 | 1,03242E-38 |
| ACE2 | BRD1 | -0,5854 | 1,02982E-38 |
| ACE2 | OSBPL10 | -0,5855 | 1,00058E-38 |
| ACE2 | ADAMTS16 | -0,5855 | 9,97075E-39 |
| ACE2 | RP5-1136G13.2 | -0,5855 | 1,02279E-38 |
| ACE2 | NAT9 | -0,5855 | 1,02003E-38 |
| ACE2 | C9orf72 | -0,5856 | 9,7694E-39 |

| | | | |
|---|---|---|---|
| ACE2 | RRM2P3 | -0,5856 | 9,71375E-39 |
| ACE2 | SLC8A1-AS1 | -0,5857 | 9,31224E-39 |
| ACE2 | GDF10 | -0,5858 | 8,89028E-39 |
| ACE2 | RP11-972P1.11 | -0,5858 | 9,06425E-39 |
| ACE2 | RIT2 | -0,5858 | 9,16958E-39 |
| ACE2 | CACNG8 | -0,5858 | 8,95887E-39 |
| ACE2 | GPR62 | -0,5859 | 8,78269E-39 |
| ACE2 | CCDC146 | -0,5859 | 8,72407E-39 |
| ACE2 | C7orf61 | -0,5859 | 8,56937E-39 |
| ACE2 | CNOT7 | -0,5859 | 8,67956E-39 |
| ACE2 | RBM17 | -0,5859 | 8,84469E-39 |
| ACE2 | RP11-326C3.12 | -0,5859 | 8,71902E-39 |
| ACE2 | GABRA3 | -0,5859 | 8,75335E-39 |
| ACE2 | TERF2 | -0,586 | 8,52882E-39 |
| ACE2 | MDM4 | -0,5861 | 8,07234E-39 |
| ACE2 | WDR31 | -0,5861 | 7,95863E-39 |
| ACE2 | LINC00493 | -0,5861 | 8,1162E-39 |
| ACE2 | FTX | -0,5861 | 7,95489E-39 |
| ACE2 | UTP14A | -0,5861 | 8,15314E-39 |
| ACE2 | AP001505.10 | -0,5862 | 7,9412E-39 |
| ACE2 | NCKAP1 | -0,5863 | 7,45845E-39 |
| ACE2 | URAHP | -0,5863 | 7,63504E-39 |
| ACE2 | RP4-565E6.1 | -0,5864 | 7,31272E-39 |
| ACE2 | CSMD2 | -0,5865 | 7,12683E-39 |
| ACE2 | AC073072.5 | -0,5865 | 7,10143E-39 |
| ACE2 | VPS13B | -0,5865 | 6,91591E-39 |
| ACE2 | ZNF436-AS1 | -0,5866 | 6,86152E-39 |
| ACE2 | RP11-1109F11.5 | -0,5866 | 6,79244E-39 |
| ACE2 | PXN | -0,5866 | 6,83316E-39 |
| ACE2 | MRPL30 | -0,5867 | 6,54402E-39 |
| ACE2 | RP11-1060G2.2 | -0,5867 | 6,58909E-39 |
| ACE2 | PRIMPOL | -0,5868 | 6,18017E-39 |
| ACE2 | TAF11 | -0,5868 | 6,31997E-39 |
| ACE2 | CEP78 | -0,5868 | 6,29392E-39 |
| ACE2 | ANO3 | -0,5868 | 6,28832E-39 |
| ACE2 | ZNF256 | -0,5868 | 6,33815E-39 |
| ACE2 | RANBP3L | -0,5869 | 6,02993E-39 |
| ACE2 | SEPHS1 | -0,5869 | 6,15278E-39 |
| ACE2 | CERS3-AS1 | -0,5869 | 6,13358E-39 |
| ACE2 | URB1 | -0,5869 | 6,1204E-39 |
| ACE2 | OR7E12P | -0,587 | 5,84302E-39 |
| ACE2 | MPO | -0,587 | 5,87096E-39 |
| ACE2 | RP11-338N10.3 | -0,5871 | 5,58429E-39 |
| ACE2 | CRIPT | -0,5871 | 5,60248E-39 |
| ACE2 | RP11-175K6.2 | -0,5871 | 5,70111E-39 |
| ACE2 | NEDD8 | -0,5871 | 5,60817E-39 |
| ACE2 | MBIP | -0,5871 | 5,66958E-39 |
| ACE2 | RP11-517P14.2 | -0,5872 | 5,3791E-39 |
| ACE2 | PUS3 | -0,5872 | 5,49412E-39 |
| ACE2 | TET2-AS1 | -0,5873 | 5,16845E-39 |
| ACE2 | LINC00840 | -0,5873 | 5,22528E-39 |
| ACE2 | IZUMO4 | -0,5873 | 5,22994E-39 |
| ACE2 | RP11-284E5.1 | -0,5874 | 5,10483E-39 |
| ACE2 | TM4SF4 | -0,5874 | 5,08898E-39 |

| | | | |
|---|---|---|---|
| ACE2 | LINC02104 | -0,5874 | 5,05922E-39 |
| ACE2 | TRIM32 | -0,5874 | 5,04579E-39 |
| ACE2 | PIGQ | -0,5874 | 4,96935E-39 |
| ACE2 | WDR83OS | -0,5874 | 5,05904E-39 |
| ACE2 | CEP250 | -0,5874 | 5,09506E-39 |
| ACE2 | ERAS | -0,5874 | 5,06323E-39 |
| ACE2 | LPGAT1 | -0,5875 | 4,91612E-39 |
| ACE2 | ENTPD3 | -0,5875 | 4,95187E-39 |
| ACE2 | TSC1 | -0,5875 | 4,86081E-39 |
| ACE2 | C1RL | -0,5875 | 4,89482E-39 |
| ACE2 | ANGPTL8 | -0,5875 | 4,86446E-39 |
| ACE2 | MTND6P21 | -0,5875 | 4,80525E-39 |
| ACE2 | PCDHA2 | -0,5876 | 4,68783E-39 |
| ACE2 | PALM2 | -0,5876 | 4,63973E-39 |
| ACE2 | MRPS26 | -0,5876 | 4,63696E-39 |
| ACE2 | SUSD2 | -0,5876 | 4,67074E-39 |
| ACE2 | DEPTOR | -0,5877 | 4,46122E-39 |
| ACE2 | MGA | -0,5877 | 4,58174E-39 |
| ACE2 | GRIPAP1 | -0,5877 | 4,60738E-39 |
| ACE2 | HS6ST1 | -0,5878 | 4,39462E-39 |
| ACE2 | MED15 | -0,5878 | 4,32012E-39 |
| ACE2 | CIR1 | -0,5879 | 4,21425E-39 |
| ACE2 | SECISBP2L | -0,5879 | 4,19214E-39 |
| ACE2 | AC026271.5 | -0,5879 | 4,18309E-39 |
| ACE2 | RP11-176H8.1 | -0,588 | 4,02925E-39 |
| ACE2 | RABAC1 | -0,588 | 4,1034E-39 |
| ACE2 | RNF185 | -0,588 | 3,99295E-39 |
| ACE2 | LRTOMT | -0,5881 | 3,8803E-39 |
| ACE2 | PIN4 | -0,5881 | 3,86745E-39 |
| ACE2 | MED6 | -0,5882 | 3,70436E-39 |
| ACE2 | ZNF416 | -0,5883 | 3,5978E-39 |
| ACE2 | BRD8 | -0,5884 | 3,46308E-39 |
| ACE2 | BACH2 | -0,5885 | 3,3298E-39 |
| ACE2 | MFAP5 | -0,5885 | 3,43805E-39 |
| ACE2 | LLNLR-268E12.1 | -0,5885 | 3,40057E-39 |
| ACE2 | CHMP7 | -0,5886 | 3,29845E-39 |
| ACE2 | RP11-479G22.8 | -0,5886 | 3,25567E-39 |
| ACE2 | SYCE1 | -0,5886 | 3,23528E-39 |
| ACE2 | ZNF283 | -0,5886 | 3,29706E-39 |
| ACE2 | UHRF1BP1 | -0,5887 | 3,18189E-39 |
| ACE2 | PHF3 | -0,5887 | 3,09096E-39 |
| ACE2 | RP11-330O11.3 | -0,5887 | 3,11107E-39 |
| ACE2 | CTC-296K1.3 | -0,5887 | 3,18952E-39 |
| ACE2 | RP11-79P5.9 | -0,5888 | 2,98453E-39 |
| ACE2 | AC073063.10 | -0,5888 | 3,04089E-39 |
| ACE2 | LRP6 | -0,5888 | 3,07928E-39 |
| ACE2 | GJC3 | -0,5889 | 2,94164E-39 |
| ACE2 | CLDN5 | -0,5889 | 2,94579E-39 |
| ACE2 | CCT6P3 | -0,589 | 2,86667E-39 |
| ACE2 | TAF4B | -0,589 | 2,76672E-39 |
| ACE2 | BDH2 | -0,5891 | 2,76123E-39 |
| ACE2 | TMEM14C | -0,5891 | 2,66644E-39 |
| ACE2 | LRRC47 | -0,5892 | 2,62463E-39 |
| ACE2 | DHX36 | -0,5892 | 2,6551E-39 |

| | | | |
|---|---|---|---|
| ACE2 | PSMB1 | -0,5892 | 2,62621E-39 |
| ACE2 | TSR3 | -0,5892 | 2,57648E-39 |
| ACE2 | AHSA2 | -0,5893 | 2,47898E-39 |
| ACE2 | ZNF602P | -0,5893 | 2,51371E-39 |
| ACE2 | ATP6V1G1 | -0,5893 | 2,53133E-39 |
| ACE2 | SF3B2 | -0,5893 | 2,52378E-39 |
| ACE2 | GLIPR1L2 | -0,5894 | 2,39982E-39 |
| ACE2 | AC007566.10 | -0,5895 | 2,38562E-39 |
| ACE2 | RIC1 | -0,5895 | 2,35604E-39 |
| ACE2 | PTPN5 | -0,5895 | 2,36728E-39 |
| ACE2 | RP11-686D22.4 | -0,5895 | 2,33936E-39 |
| ACE2 | RP5-1115A15.1 | -0,5896 | 2,24705E-39 |
| ACE2 | RP11-422P24.11 | -0,5896 | 2,25634E-39 |
| ACE2 | TTC1 | -0,5896 | 2,24614E-39 |
| ACE2 | IFT27 | -0,5896 | 2,2945E-39 |
| ACE2 | RP11-572M11.4 | -0,5897 | 2,19692E-39 |
| ACE2 | ADH1B | -0,5897 | 2,15819E-39 |
| ACE2 | GTF2E2 | -0,5897 | 2,20541E-39 |
| ACE2 | SLC1A2 | -0,5897 | 2,21794E-39 |
| ACE2 | RIOX2 | -0,5899 | 2,03662E-39 |
| ACE2 | RP11-139H14.5 | -0,5899 | 2,03217E-39 |
| ACE2 | RAD51B | -0,5899 | 2,054E-39 |
| ACE2 | SYPL1 | -0,59 | 1,96992E-39 |
| ACE2 | SNX16 | -0,59 | 1,96854E-39 |
| ACE2 | PRDX3P1 | -0,59 | 1,97917E-39 |
| ACE2 | TMEM167B | -0,5901 | 1,87285E-39 |
| ACE2 | PMF1 | -0,5901 | 1,90729E-39 |
| ACE2 | LINC00473 | -0,5901 | 1,90999E-39 |
| ACE2 | NHLRC2 | -0,5901 | 1,9047E-39 |
| ACE2 | ARMCX3-AS1 | -0,5901 | 1,9015E-39 |
| ACE2 | CCT4 | -0,5902 | 1,84558E-39 |
| ACE2 | HMGB1P6 | -0,5902 | 1,78916E-39 |
| ACE2 | MUM1 | -0,5902 | 1,82047E-39 |
| ACE2 | CLCN4 | -0,5902 | 1,82203E-39 |
| ACE2 | FAM217B | -0,5903 | 1,75051E-39 |
| ACE2 | LINC00323 | -0,5903 | 1,72684E-39 |
| ACE2 | DPY30 | -0,5904 | 1,69762E-39 |
| ACE2 | CENPC | -0,5904 | 1,6985E-39 |
| ACE2 | ADAL | -0,5904 | 1,68114E-39 |
| ACE2 | RP11-247C2.2 | -0,5904 | 1,66885E-39 |
| ACE2 | GEMIN2 | -0,5905 | 1,64064E-39 |
| ACE2 | ATXN2L | -0,5905 | 1,60829E-39 |
| ACE2 | CENPBD1P1 | -0,5905 | 1,62933E-39 |
| ACE2 | TUBGCP6 | -0,5905 | 1,60541E-39 |
| ACE2 | MUSTN1 | -0,5906 | 1,54295E-39 |
| ACE2 | HGF | -0,5906 | 1,58935E-39 |
| ACE2 | WWP1 | -0,5906 | 1,55194E-39 |
| ACE2 | F10 | -0,5906 | 1,57544E-39 |
| ACE2 | UNKL | -0,5906 | 1,58765E-39 |
| ACE2 | BCL7B | -0,5907 | 1,52876E-39 |
| ACE2 | RP11-365P13.5 | -0,5907 | 1,52066E-39 |
| ACE2 | HIST2H2BB | -0,5908 | 1,46589E-39 |
| ACE2 | S100A3 | -0,5908 | 1,46945E-39 |
| ACE2 | KIAA1614-AS1 | -0,5909 | 1,38662E-39 |

| | | | |
|---|---|---|---|
| ACE2 | AC092597.3 | -0,5909 | 1,39144E-39 |
| ACE2 | RP11-381K20.2 | -0,5909 | 1,37743E-39 |
| ACE2 | GPR158 | -0,5909 | 1,39114E-39 |
| ACE2 | HAUS2 | -0,5909 | 1,38379E-39 |
| ACE2 | RP11-861E21.1 | -0,5909 | 1,41567E-39 |
| ACE2 | OR51E1 | -0,591 | 1,32869E-39 |
| ACE2 | KCNC1 | -0,591 | 1,33705E-39 |
| ACE2 | MRE11 | -0,591 | 1,37124E-39 |
| ACE2 | AQP4 | -0,591 | 1,33261E-39 |
| ACE2 | ZNF558 | -0,591 | 1,34899E-39 |
| ACE2 | ATP7A | -0,5911 | 1,3057E-39 |
| ACE2 | RP11-722G7.1 | -0,5911 | 1,28605E-39 |
| ACE2 | ATOH8 | -0,5912 | 1,24422E-39 |
| ACE2 | FBXL13 | -0,5912 | 1,26737E-39 |
| ACE2 | NPTX1 | -0,5912 | 1,23899E-39 |
| ACE2 | RP11-108M9.3 | -0,5913 | 1,20325E-39 |
| ACE2 | MRGPRD | -0,5913 | 1,20415E-39 |
| ACE2 | ADAR | -0,5914 | 1,15925E-39 |
| ACE2 | PHC3 | -0,5914 | 1,15151E-39 |
| ACE2 | FAM169B | -0,5914 | 1,16945E-39 |
| ACE2 | FUT1 | -0,5914 | 1,14743E-39 |
| ACE2 | RP4-742C19.13 | -0,5914 | 1,14771E-39 |
| ACE2 | LINC01521 | -0,5915 | 1,10782E-39 |
| ACE2 | LARGE1 | -0,5915 | 1,12675E-39 |
| ACE2 | PEX5L | -0,5916 | 1,06503E-39 |
| ACE2 | RP11-342D11.2 | -0,5916 | 1,08218E-39 |
| ACE2 | DNAAF1 | -0,5916 | 1,08935E-39 |
| ACE2 | TPRG1 | -0,5917 | 1,0486E-39 |
| ACE2 | RP11-20I20.4 | -0,5917 | 1,02576E-39 |
| ACE2 | PGM2 | -0,5917 | 1,06137E-39 |
| ACE2 | ART4 | -0,5917 | 1,05553E-39 |
| ACE2 | C1orf122 | -0,5918 | 1,00261E-39 |
| ACE2 | PCDH12 | -0,5918 | 9,92563E-40 |
| ACE2 | ZNF318 | -0,5918 | 1,0197E-39 |
| ACE2 | ADO | -0,5918 | 1,00028E-39 |
| ACE2 | RP11-799M12.2 | -0,5919 | 9,57986E-40 |
| ACE2 | ANKRD37 | -0,5919 | 9,61899E-40 |
| ACE2 | TEX21P | -0,5919 | 9,64629E-40 |
| ACE2 | NDUFS5 | -0,592 | 9,48654E-40 |
| ACE2 | LHCGR | -0,592 | 9,32833E-40 |
| ACE2 | BRINP1 | -0,592 | 9,3065E-40 |
| ACE2 | DCTN1 | -0,5921 | 8,93189E-40 |
| ACE2 | AC005519.4 | -0,5921 | 9,02351E-40 |
| ACE2 | PLD5 | -0,5922 | 8,71827E-40 |
| ACE2 | CHD2 | -0,5922 | 8,58913E-40 |
| ACE2 | TCAM1P | -0,5922 | 8,54858E-40 |
| ACE2 | RBX1 | -0,5922 | 8,54476E-40 |
| ACE2 | GYS2 | -0,5923 | 8,47988E-40 |
| ACE2 | DKKL1 | -0,5923 | 8,21653E-40 |
| ACE2 | ZNF611 | -0,5923 | 8,31085E-40 |
| ACE2 | SEMA3G | -0,5924 | 8,01122E-40 |
| ACE2 | RP11-193M21.1 | -0,5924 | 7,9234E-40 |
| ACE2 | CYB5R3 | -0,5924 | 8,11348E-40 |
| ACE2 | TCEAL9 | -0,5924 | 8,1381E-40 |

| | | | |
|---|---|---|---|
| ACE2 | PRR16 | -0,5925 | 7,62006E-40 |
| ACE2 | RBM4B | -0,5925 | 7,80631E-40 |
| ACE2 | PPP1R15A | -0,5926 | 7,49451E-40 |
| ACE2 | ST6GAL2 | -0,5927 | 7,2702E-40 |
| ACE2 | RP11-452F19.3 | -0,5928 | 6,85111E-40 |
| ACE2 | RP11-438B23.2 | -0,5928 | 7,03017E-40 |
| ACE2 | RP13-39P12.3 | -0,5928 | 7,02287E-40 |
| ACE2 | ZYG11B | -0,5929 | 6,7993E-40 |
| ACE2 | NUP54 | -0,5929 | 6,61592E-40 |
| ACE2 | PAN3-AS1 | -0,5929 | 6,56798E-40 |
| ACE2 | SF3A2 | -0,5929 | 6,81123E-40 |
| ACE2 | RP11-313P22.1 | -0,5929 | 6,78156E-40 |
| ACE2 | TAF1 | -0,5929 | 6,61533E-40 |
| ACE2 | CRIM1 | -0,593 | 6,38661E-40 |
| ACE2 | RP11-664I21.6 | -0,593 | 6,36713E-40 |
| ACE2 | RGCC | -0,5931 | 6,18516E-40 |
| ACE2 | PAIP2B | -0,5932 | 6,07394E-40 |
| ACE2 | FBXO9 | -0,5932 | 5,91529E-40 |
| ACE2 | STARD3NL | -0,5932 | 5,96683E-40 |
| ACE2 | MTA3 | -0,5933 | 5,84571E-40 |
| ACE2 | AC098617.1 | -0,5933 | 5,81626E-40 |
| ACE2 | C9orf78 | -0,5933 | 5,8069E-40 |
| ACE2 | PRCP | -0,5933 | 5,75638E-40 |
| ACE2 | CREBL2 | -0,5933 | 5,7419E-40 |
| ACE2 | RP11-162A12.4 | -0,5933 | 5,71183E-40 |
| ACE2 | RBM42 | -0,5933 | 5,78344E-40 |
| ACE2 | AMER1 | -0,5933 | 5,70194E-40 |
| ACE2 | RPL10P7 | -0,5934 | 5,62335E-40 |
| ACE2 | ZFR | -0,5934 | 5,57082E-40 |
| ACE2 | AFF4 | -0,5934 | 5,50781E-40 |
| ACE2 | ADCY1 | -0,5934 | 5,6216E-40 |
| ACE2 | LINC00599 | -0,5934 | 5,58051E-40 |
| ACE2 | RP11-753H16.3 | -0,5934 | 5,61083E-40 |
| ACE2 | HSD17B6 | -0,5934 | 5,62573E-40 |
| ACE2 | WBP2 | -0,5934 | 5,55656E-40 |
| ACE2 | RPRD2 | -0,5935 | 5,39079E-40 |
| ACE2 | NKRF | -0,5935 | 5,38637E-40 |
| ACE2 | ZFAT | -0,5936 | 5,22199E-40 |
| ACE2 | EXOSC10 | -0,5937 | 4,90327E-40 |
| ACE2 | METAP1D | -0,5937 | 4,90226E-40 |
| ACE2 | CDK8 | -0,5937 | 4,92465E-40 |
| ACE2 | MEX3C | -0,5939 | 4,66371E-40 |
| ACE2 | STRN4 | -0,5939 | 4,69391E-40 |
| ACE2 | GRHL1 | -0,594 | 4,50732E-40 |
| ACE2 | THEGL | -0,594 | 4,43039E-40 |
| ACE2 | RP11-177G23.1 | -0,594 | 4,44962E-40 |
| ACE2 | KIF5B | -0,594 | 4,46066E-40 |
| ACE2 | FLRT1 | -0,594 | 4,4694E-40 |
| ACE2 | PIK3IP1-AS1 | -0,594 | 4,40235E-40 |
| ACE2 | SEPT3 | -0,594 | 4,51688E-40 |
| ACE2 | OSBPL6 | -0,5941 | 4,27709E-40 |
| ACE2 | INPP5K | -0,5941 | 4,29612E-40 |
| ACE2 | SNX17 | -0,5942 | 4,18589E-40 |
| ACE2 | STK25 | -0,5942 | 4,15301E-40 |

| | | | |
|---|---|---|---|
| ACE2 | RP11-299J3.8 | -0,5943 | 3,99235E-40 |
| ACE2 | PTPRQ | -0,5943 | 4,03579E-40 |
| ACE2 | ILF3 | -0,5943 | 3,90507E-40 |
| ACE2 | AC092835.2 | -0,5944 | 3,81745E-40 |
| ACE2 | DCAF16 | -0,5944 | 3,86403E-40 |
| ACE2 | INSC | -0,5944 | 3,7651E-40 |
| ACE2 | ZNF529-AS1 | -0,5944 | 3,84507E-40 |
| ACE2 | VBP1 | -0,5944 | 3,77932E-40 |
| ACE2 | MLF2 | -0,5945 | 3,70182E-40 |
| ACE2 | FAM65C | -0,5945 | 3,75598E-40 |
| ACE2 | ST6GALNAC3 | -0,5946 | 3,49216E-40 |
| ACE2 | HMGN4 | -0,5946 | 3,51754E-40 |
| ACE2 | GINM1 | -0,5946 | 3,49591E-40 |
| ACE2 | RINT1 | -0,5946 | 3,49146E-40 |
| ACE2 | RP11-70L8.4 | -0,5946 | 3,52503E-40 |
| ACE2 | RP11-817O13.9 | -0,5946 | 3,58702E-40 |
| ACE2 | SRR | -0,5946 | 3,49185E-40 |
| ACE2 | LRRC37A7P | -0,5946 | 3,55987E-40 |
| ACE2 | RP3-522J7.6 | -0,5946 | 3,57998E-40 |
| ACE2 | KCNH1 | -0,5947 | 3,36046E-40 |
| ACE2 | CD302 | -0,5947 | 3,42223E-40 |
| ACE2 | RYBP | -0,5947 | 3,4437E-40 |
| ACE2 | FAIM | -0,5947 | 3,48624E-40 |
| ACE2 | PIGH | -0,5947 | 3,42101E-40 |
| ACE2 | ZFYVE16 | -0,5948 | 3,27304E-40 |
| ACE2 | ERMARD | -0,5948 | 3,29452E-40 |
| ACE2 | MTDHP3 | -0,5948 | 3,23964E-40 |
| ACE2 | NORAD | -0,5948 | 3,27342E-40 |
| ACE2 | EDA2R | -0,5948 | 3,29807E-40 |
| ACE2 | LUC7L2 | -0,5949 | 3,20963E-40 |
| ACE2 | NTRK3 | -0,5949 | 3,19862E-40 |
| ACE2 | IL34 | -0,5949 | 3,18292E-40 |
| ACE2 | SLBP | -0,595 | 3,02476E-40 |
| ACE2 | POLR3D | -0,595 | 3,06191E-40 |
| ACE2 | SNURF | -0,595 | 3,03351E-40 |
| ACE2 | MTMR9LP | -0,5951 | 2,95932E-40 |
| ACE2 | HSPD1P4 | -0,5951 | 2,93016E-40 |
| ACE2 | EMC4 | -0,5951 | 2,96467E-40 |
| ACE2 | LRRC8C | -0,5952 | 2,83011E-40 |
| ACE2 | LINC00504 | -0,5952 | 2,84209E-40 |
| ACE2 | WDR25 | -0,5952 | 2,87129E-40 |
| ACE2 | SGF29 | -0,5952 | 2,85597E-40 |
| ACE2 | SUPT16H | -0,5953 | 2,75142E-40 |
| ACE2 | LINC01197 | -0,5953 | 2,71938E-40 |
| ACE2 | ZSCAN1 | -0,5953 | 2,72578E-40 |
| ACE2 | CCDC160 | -0,5953 | 2,71652E-40 |
| ACE2 | CDC42SE1 | -0,5955 | 2,51224E-40 |
| ACE2 | RP1-151F17.1 | -0,5955 | 2,53363E-40 |
| ACE2 | NARF | -0,5955 | 2,58394E-40 |
| ACE2 | TSHZ1 | -0,5955 | 2,57374E-40 |
| ACE2 | ESYT3 | -0,5956 | 2,40731E-40 |
| ACE2 | THAP6 | -0,5956 | 2,48612E-40 |
| ACE2 | LINC01554 | -0,5957 | 2,34775E-40 |
| ACE2 | IFITM4P | -0,5957 | 2,39866E-40 |

| | | | |
|---|---|---|---|
| ACE2 | NAP1L1P1 | -0,5957 | 2,33578E-40 |
| ACE2 | PPIL6 | -0,5958 | 2,29759E-40 |
| ACE2 | SCRT1 | -0,5958 | 2,29967E-40 |
| ACE2 | MTCYBP21 | -0,5958 | 2,29829E-40 |
| ACE2 | MALSU1 | -0,5959 | 2,20174E-40 |
| ACE2 | TRMT1 | -0,5959 | 2,15126E-40 |
| ACE2 | SRSF11 | -0,596 | 2,09768E-40 |
| ACE2 | RP3-428L16.1 | -0,596 | 2,11397E-40 |
| ACE2 | EIF3J | -0,596 | 2,09584E-40 |
| ACE2 | HNRNPU | -0,5961 | 2,00094E-40 |
| ACE2 | RGS9BP | -0,5961 | 2,03616E-40 |
| ACE2 | AMY2B | -0,5962 | 1,98466E-40 |
| ACE2 | LINC01355 | -0,5963 | 1,86781E-40 |
| ACE2 | SPAG16 | -0,5963 | 1,86425E-40 |
| ACE2 | LRRC77P | -0,5963 | 1,89489E-40 |
| ACE2 | NBAT1 | -0,5963 | 1,91528E-40 |
| ACE2 | MDFI | -0,5963 | 1,87722E-40 |
| ACE2 | DPY19L2P4 | -0,5963 | 1,88378E-40 |
| ACE2 | TRAFD1 | -0,5963 | 1,90949E-40 |
| ACE2 | CTD-3220F14.3 | -0,5963 | 1,88588E-40 |
| ACE2 | AC005740.6 | -0,5964 | 1,79565E-40 |
| ACE2 | OR2H2 | -0,5964 | 1,80265E-40 |
| ACE2 | ZNF674 | -0,5964 | 1,83705E-40 |
| ACE2 | MIR503HG | -0,5964 | 1,80531E-40 |
| ACE2 | HSP90AB1 | -0,5965 | 1,7305E-40 |
| ACE2 | ISCA1 | -0,5965 | 1,71654E-40 |
| ACE2 | RP5-1142A6.10 | -0,5965 | 1,74682E-40 |
| ACE2 | TIMM22 | -0,5965 | 1,72174E-40 |
| ACE2 | RP11-380B4.3 | -0,5966 | 1,66966E-40 |
| ACE2 | GBAS | -0,5967 | 1,60291E-40 |
| ACE2 | STX17 | -0,5968 | 1,53122E-40 |
| ACE2 | RP11-252A24.2 | -0,5968 | 1,58402E-40 |
| ACE2 | EFCAB13 | -0,5969 | 1,4767E-40 |
| ACE2 | TMEM92-AS1 | -0,5969 | 1,48643E-40 |
| ACE2 | RP11-578F21.9 | -0,597 | 1,42616E-40 |
| ACE2 | CTD-2527I21.14 | -0,597 | 1,4463E-40 |
| ACE2 | HDLBP | -0,5971 | 1,40917E-40 |
| ACE2 | DCP1A | -0,5971 | 1,3694E-40 |
| ACE2 | ATP6V1E1 | -0,5971 | 1,41262E-40 |
| ACE2 | SLC25A36 | -0,5972 | 1,31685E-40 |
| ACE2 | GALR2 | -0,5972 | 1,31846E-40 |
| ACE2 | WDTC1 | -0,5973 | 1,26762E-40 |
| ACE2 | ZSWIM8 | -0,5973 | 1,28192E-40 |
| ACE2 | AC141928.1 | -0,5974 | 1,22758E-40 |
| ACE2 | RPS14 | -0,5974 | 1,26211E-40 |
| ACE2 | CDAN1 | -0,5974 | 1,25393E-40 |
| ACE2 | CCL2 | -0,5974 | 1,2554E-40 |
| ACE2 | HNRNPUL1 | -0,5974 | 1,25978E-40 |
| ACE2 | RP4-686C3.7 | -0,5974 | 1,22876E-40 |
| ACE2 | CARF | -0,5975 | 1,19701E-40 |
| ACE2 | PCMT1 | -0,5975 | 1,1912E-40 |
| ACE2 | RGS6 | -0,5975 | 1,18845E-40 |
| ACE2 | RBM22P2 | -0,5976 | 1,13672E-40 |
| ACE2 | TNFAIP8L3 | -0,5976 | 1,15364E-40 |

| | | | |
|---|---|---|---|
| ACE2 | TRIM65 | -0,5976 | 1,1328E-40 |
| ACE2 | DESI2 | -0,5977 | 1,12497E-40 |
| ACE2 | YWHAQ | -0,5977 | 1,11503E-40 |
| ACE2 | FBXO36 | -0,5977 | 1,11986E-40 |
| ACE2 | RP11-54D18.2 | -0,5977 | 1,12516E-40 |
| ACE2 | RP11-573G6.6 | -0,5977 | 1,09324E-40 |
| ACE2 | NOL3 | -0,5977 | 1,09459E-40 |
| ACE2 | NR1D1 | -0,5977 | 1,09891E-40 |
| ACE2 | AC098614.2 | -0,5978 | 1,07342E-40 |
| ACE2 | UBR5 | -0,5978 | 1,08048E-40 |
| ACE2 | EVI5 | -0,5979 | 1,03523E-40 |
| ACE2 | ZC3H13 | -0,598 | 9,75781E-41 |
| ACE2 | SNX33 | -0,598 | 9,84043E-41 |
| ACE2 | RP11-322E11.5 | -0,598 | 9,95741E-41 |
| ACE2 | PCDHA1 | -0,5981 | 9,4998E-41 |
| ACE2 | ADNP | -0,5981 | 9,61144E-41 |
| ACE2 | CTA-268H5.14 | -0,5981 | 9,5314E-41 |
| ACE2 | C5orf64 | -0,5982 | 9,11384E-41 |
| ACE2 | WDR60 | -0,5982 | 9,21822E-41 |
| ACE2 | RP11-267M23.4 | -0,5982 | 9,24418E-41 |
| ACE2 | RP11-459F6.3 | -0,5982 | 9,22246E-41 |
| ACE2 | A2M | -0,5983 | 8,9363E-41 |
| ACE2 | GPR17 | -0,5984 | 8,39545E-41 |
| ACE2 | OSGEPL1 | -0,5984 | 8,62737E-41 |
| ACE2 | ACOX2 | -0,5984 | 8,59884E-41 |
| ACE2 | EIF3M | -0,5984 | 8,40212E-41 |
| ACE2 | PEX12 | -0,5985 | 8,35033E-41 |
| ACE2 | CTSL | -0,5986 | 7,81138E-41 |
| ACE2 | ZNF692 | -0,5987 | 7,6751E-41 |
| ACE2 | BCCIP | -0,5987 | 7,52484E-41 |
| ACE2 | OTUD6B-AS1 | -0,5988 | 7,27924E-41 |
| ACE2 | CYP4X1 | -0,5989 | 7,01999E-41 |
| ACE2 | AC105398.3 | -0,5989 | 7,11745E-41 |
| ACE2 | HOXD3 | -0,5989 | 7,09648E-41 |
| ACE2 | PCDHB19P | -0,5989 | 7,02165E-41 |
| ACE2 | RP11-475A13.1 | -0,5989 | 7,01668E-41 |
| ACE2 | GNRHR2 | -0,599 | 6,74232E-41 |
| ACE2 | ALMS1-IT1 | -0,599 | 6,72015E-41 |
| ACE2 | ACACB | -0,599 | 6,78159E-41 |
| ACE2 | GATA2-AS1 | -0,5991 | 6,57681E-41 |
| ACE2 | SERINC1 | -0,5991 | 6,53831E-41 |
| ACE2 | ARHGAP1 | -0,5991 | 6,48566E-41 |
| ACE2 | RP11-755F10.1 | -0,5991 | 6,63499E-41 |
| ACE2 | CBLN3 | -0,5991 | 6,47106E-41 |
| ACE2 | NCOA3 | -0,5991 | 6,44445E-41 |
| ACE2 | PRKAR2B | -0,5992 | 6,25762E-41 |
| ACE2 | CNTRL | -0,5992 | 6,28371E-41 |
| ACE2 | ANXA8L1 | -0,5992 | 6,37676E-41 |
| ACE2 | NSMCE4A | -0,5992 | 6,22E-41 |
| ACE2 | TRIM9 | -0,5992 | 6,4068E-41 |
| ACE2 | AL022476.2 | -0,5992 | 6,28289E-41 |
| ACE2 | NR1D2 | -0,5993 | 6,13766E-41 |
| ACE2 | LRRFIP1P1 | -0,5993 | 6,07268E-41 |
| ACE2 | MICA | -0,5993 | 6,04549E-41 |

| | | | |
|---|---|---|---|
| ACE2 | CCM2 | -0,5993 | 5,9747E-41 |
| ACE2 | VPS11 | -0,5993 | 5,94501E-41 |
| ACE2 | DUOX1 | -0,5993 | 5,95554E-41 |
| ACE2 | SLC17A7 | -0,5993 | 6,01553E-41 |
| ACE2 | BCDIN3D | -0,5994 | 5,8075E-41 |
| ACE2 | SOS1-IT1 | -0,5995 | 5,70452E-41 |
| ACE2 | C2orf49 | -0,5995 | 5,62182E-41 |
| ACE2 | CTTNBP2 | -0,5995 | 5,5339E-41 |
| ACE2 | NDUFB8 | -0,5995 | 5,58902E-41 |
| ACE2 | COPS3 | -0,5995 | 5,65214E-41 |
| ACE2 | PARL | -0,5996 | 5,32542E-41 |
| ACE2 | HNRNPD | -0,5996 | 5,50252E-41 |
| ACE2 | HSPB1P1 | -0,5996 | 5,34779E-41 |
| ACE2 | RP4-735C1.4 | -0,5998 | 5,08981E-41 |
| ACE2 | ZNF235 | -0,5998 | 5,01388E-41 |
| ACE2 | BCOR | -0,5998 | 5,06094E-41 |
| ACE2 | RPS4X | -0,5998 | 5,03203E-41 |
| ACE2 | RP11-1081L13.4 | -0,5999 | 4,85931E-41 |
| ACE2 | RP11-589M4.4 | -0,5999 | 4,90367E-41 |
| ACE2 | ZNF205 | -0,5999 | 4,82272E-41 |
| ACE2 | CH17-140K24.6 | -0,6 | 4,68631E-41 |
| ACE2 | EIF2S2P3 | -0,6001 | 4,51432E-41 |
| ACE2 | CCDC141 | -0,6002 | 4,29555E-41 |
| ACE2 | RIMS2 | -0,6002 | 4,33133E-41 |
| ACE2 | CTD-2530H12.4 | -0,6002 | 4,35249E-41 |
| ACE2 | JMJD7 | -0,6002 | 4,26849E-41 |
| ACE2 | AC006547.13 | -0,6002 | 4,24717E-41 |
| ACE2 | SDAD1 | -0,6004 | 4,03009E-41 |
| ACE2 | RP4-751H13.7 | -0,6004 | 4,02517E-41 |
| ACE2 | XPO7 | -0,6004 | 4,01495E-41 |
| ACE2 | PPM1E | -0,6004 | 3,93273E-41 |
| ACE2 | FEM1A | -0,6004 | 4,01727E-41 |
| ACE2 | ISG15 | -0,6005 | 3,7942E-41 |
| ACE2 | EIF2AK2 | -0,6005 | 3,85743E-41 |
| ACE2 | RP11-616M22.11 | -0,6005 | 3,89865E-41 |
| ACE2 | RPS9 | -0,6005 | 3,8087E-41 |
| ACE2 | WLS | -0,6006 | 3,73272E-41 |
| ACE2 | NKAIN3 | -0,6006 | 3,7629E-41 |
| ACE2 | DSCAML1 | -0,6006 | 3,66536E-41 |
| ACE2 | CTD-2547L24.3 | -0,6006 | 3,72159E-41 |
| ACE2 | SLC1A7 | -0,6007 | 3,55775E-41 |
| ACE2 | TFPI | -0,6007 | 3,53506E-41 |
| ACE2 | UBE4B | -0,6008 | 3,4756E-41 |
| ACE2 | RP11-259O2.1 | -0,6008 | 3,37362E-41 |
| ACE2 | CTD-2036P10.5 | -0,6008 | 3,46265E-41 |
| ACE2 | IGFBP5 | -0,6009 | 3,26605E-41 |
| ACE2 | RP11-339B21.10 | -0,6009 | 3,23824E-41 |
| ACE2 | RP11-644F5.11 | -0,6009 | 3,24575E-41 |
| ACE2 | GON4L | -0,601 | 3,20396E-41 |
| ACE2 | HIBADH | -0,601 | 3,17685E-41 |
| ACE2 | ACTR3B | -0,601 | 3,20272E-41 |
| ACE2 | RP11-658F2.8 | -0,601 | 3,22331E-41 |
| ACE2 | PCBP3-OT1 | -0,601 | 3,12502E-41 |
| ACE2 | AGO1 | -0,6011 | 3,02152E-41 |

| | | | |
|---|---|---|---|
| ACE2 | TFCP2 | -0,6011 | 3,09803E-41 |
| ACE2 | LINGO1 | -0,6011 | 3,05417E-41 |
| ACE2 | CRYZ | -0,6012 | 2,95908E-41 |
| ACE2 | CALCRL | -0,6012 | 2,91898E-41 |
| ACE2 | BTN2A1 | -0,6012 | 2,96677E-41 |
| ACE2 | NKAPP1 | -0,6012 | 2,8965E-41 |
| ACE2 | AGBL5-IT1 | -0,6013 | 2,8129E-41 |
| ACE2 | TANC1 | -0,6013 | 2,79025E-41 |
| ACE2 | VSX1 | -0,6013 | 2,87309E-41 |
| ACE2 | TRAM2-AS1 | -0,6014 | 2,71212E-41 |
| ACE2 | AC009228.1 | -0,6015 | 2,65185E-41 |
| ACE2 | SMARCA5 | -0,6015 | 2,59716E-41 |
| ACE2 | KATNBL1 | -0,6015 | 2,65291E-41 |
| ACE2 | UNC13A | -0,6015 | 2,60343E-41 |
| ACE2 | LGI3 | -0,6016 | 2,50129E-41 |
| ACE2 | MFSD14C | -0,6017 | 2,43015E-41 |
| ACE2 | CEP57 | -0,6017 | 2,39452E-41 |
| ACE2 | ALKBH1 | -0,6017 | 2,46941E-41 |
| ACE2 | RYR1 | -0,6017 | 2,39211E-41 |
| ACE2 | RP11-147L13.14 | -0,6018 | 2,35096E-41 |
| ACE2 | NDUFAF5 | -0,6018 | 2,34159E-41 |
| ACE2 | RP11-66N11.8 | -0,6018 | 2,37143E-41 |
| ACE2 | CREBRF | -0,6019 | 2,27921E-41 |
| ACE2 | ARMS2 | -0,6019 | 2,20615E-41 |
| ACE2 | GNS | -0,6019 | 2,28121E-41 |
| ACE2 | STAU2 | -0,602 | 2,14478E-41 |
| ACE2 | ITPR2 | -0,602 | 2,18435E-41 |
| ACE2 | KDM2A | -0,6021 | 2,09526E-41 |
| ACE2 | SAMHD1 | -0,6021 | 2,07471E-41 |
| ACE2 | CAPN6 | -0,6021 | 2,04968E-41 |
| ACE2 | MECR | -0,6022 | 1,99189E-41 |
| ACE2 | FAM193A | -0,6022 | 1,97698E-41 |
| ACE2 | CFAP58-AS1 | -0,6022 | 1,96989E-41 |
| ACE2 | RP11-834C11.6 | -0,6022 | 1,98549E-41 |
| ACE2 | DTD1 | -0,6022 | 2,00592E-41 |
| ACE2 | VASH2 | -0,6023 | 1,95983E-41 |
| ACE2 | PARP1 | -0,6023 | 1,93474E-41 |
| ACE2 | SMIM3 | -0,6023 | 1,92854E-41 |
| ACE2 | RP11-68I3.10 | -0,6023 | 1,95956E-41 |
| ACE2 | BBS7 | -0,6024 | 1,86073E-41 |
| ACE2 | SLC22A3 | -0,6024 | 1,88998E-41 |
| ACE2 | PSPC1-AS2 | -0,6024 | 1,83413E-41 |
| ACE2 | ZBTB1 | -0,6024 | 1,82141E-41 |
| ACE2 | ATP9B | -0,6024 | 1,83416E-41 |
| ACE2 | SERHL2 | -0,6024 | 1,85396E-41 |
| ACE2 | RP11-395L14.18 | -0,6025 | 1,79183E-41 |
| ACE2 | MED17 | -0,6025 | 1,78329E-41 |
| ACE2 | RP11-672L10.2 | -0,6025 | 1,819E-41 |
| ACE2 | PARD3 | -0,6026 | 1,73499E-41 |
| ACE2 | SPAG9 | -0,6026 | 1,73308E-41 |
| ACE2 | LINC00346 | -0,6027 | 1,65876E-41 |
| ACE2 | RP11-426C22.5 | -0,6028 | 1,57982E-41 |
| ACE2 | AC130469.2 | -0,6029 | 1,52718E-41 |
| ACE2 | APOL1 | -0,6029 | 1,53076E-41 |

| | | | |
|---|---|---|---|
| ACE2 | RP1-18D14.7 | -0,603 | 1,48315E-41 |
| ACE2 | SLC6A1 | -0,603 | 1,49896E-41 |
| ACE2 | BCLAF1 | -0,603 | 1,48641E-41 |
| ACE2 | RP11-157L3.9 | -0,603 | 1,48281E-41 |
| ACE2 | GLB1L3 | -0,6031 | 1,41694E-41 |
| ACE2 | ZNF268 | -0,6031 | 1,4124E-41 |
| ACE2 | FOXO3B | -0,6031 | 1,41698E-41 |
| ACE2 | CLTCL1 | -0,6031 | 1,42622E-41 |
| ACE2 | VPS41 | -0,6032 | 1,35249E-41 |
| ACE2 | RP11-462D18.4 | -0,6032 | 1,35708E-41 |
| ACE2 | SUMO1 | -0,6033 | 1,29069E-41 |
| ACE2 | RP11-44F14.6 | -0,6033 | 1,29152E-41 |
| ACE2 | WDR26 | -0,6034 | 1,24886E-41 |
| ACE2 | RP11-566K19.5 | -0,6034 | 1,24551E-41 |
| ACE2 | AP3S1 | -0,6035 | 1,23349E-41 |
| ACE2 | RP11-314C16.1 | -0,6035 | 1,22325E-41 |
| ACE2 | DPY19L2P2 | -0,6035 | 1,21531E-41 |
| ACE2 | WDR82 | -0,6036 | 1,15819E-41 |
| ACE2 | HERPUD2 | -0,6036 | 1,1612E-41 |
| ACE2 | TOPORS-AS1 | -0,6036 | 1,14952E-41 |
| ACE2 | RP4-647C14.2 | -0,6036 | 1,15127E-41 |
| ACE2 | NANOS3 | -0,6036 | 1,16706E-41 |
| ACE2 | DCAF6 | -0,6037 | 1,10768E-41 |
| ACE2 | CCNT2-AS1 | -0,6037 | 1,13331E-41 |
| ACE2 | CAMK2G | -0,6037 | 1,1047E-41 |
| ACE2 | FGF16 | -0,6037 | 1,11166E-41 |
| ACE2 | EPC2 | -0,6038 | 1,07714E-41 |
| ACE2 | RP11-956J14.1 | -0,6038 | 1,0759E-41 |
| ACE2 | TDRD3 | -0,6038 | 1,07608E-41 |
| ACE2 | EFCAB2 | -0,6039 | 1,04533E-41 |
| ACE2 | SAFB | -0,6039 | 1,0539E-41 |
| ACE2 | RP11-348F1.2 | -0,604 | 1,01813E-41 |
| ACE2 | SIRT1 | -0,6041 | 9,64115E-42 |
| ACE2 | CHRDL2 | -0,6041 | 9,76477E-42 |
| ACE2 | PRSS44 | -0,6042 | 9,42626E-42 |
| ACE2 | DDAH2 | -0,6042 | 9,17327E-42 |
| ACE2 | HECTD4 | -0,6042 | 9,14956E-42 |
| ACE2 | RP11-2N1.3 | -0,6042 | 9,26634E-42 |
| ACE2 | ZNF506 | -0,6042 | 9,43344E-42 |
| ACE2 | SCUBE3 | -0,6043 | 8,77432E-42 |
| ACE2 | RBFOX3 | -0,6043 | 8,93094E-42 |
| ACE2 | RPA2 | -0,6044 | 8,70509E-42 |
| ACE2 | GPR85 | -0,6044 | 8,56931E-42 |
| ACE2 | KAT6B | -0,6044 | 8,64626E-42 |
| ACE2 | MKRN3 | -0,6044 | 8,65472E-42 |
| ACE2 | C17orf75 | -0,6044 | 8,50281E-42 |
| ACE2 | TSPEAR-AS1 | -0,6044 | 8,49297E-42 |
| ACE2 | ZNF192P1 | -0,6045 | 8,1166E-42 |
| ACE2 | NAMA | -0,6045 | 8,34726E-42 |
| ACE2 | AKT1 | -0,6045 | 8,2432E-42 |
| ACE2 | ABHD17A | -0,6045 | 8,31153E-42 |
| ACE2 | RP11-129J12.1 | -0,6046 | 8,08729E-42 |
| ACE2 | RP11-867G23.1 | -0,6046 | 8,06775E-42 |
| ACE2 | ALOX15 | -0,6046 | 8,08124E-42 |

| | | | |
|---|---|---|---|
| ACE2 | KBTBD3 | -0,6047 | 7,56178E-42 |
| ACE2 | B3GAT1 | -0,6047 | 7,55542E-42 |
| ACE2 | CTD-2600O9.2 | -0,6047 | 7,64667E-42 |
| ACE2 | SRSF10 | -0,6048 | 7,32658E-42 |
| ACE2 | RP11-552M11.8 | -0,6048 | 7,49845E-42 |
| ACE2 | PLEKHG4B | -0,6048 | 7,44926E-42 |
| ACE2 | EIF3B | -0,6048 | 7,45145E-42 |
| ACE2 | ARHGAP33 | -0,6048 | 7,2715E-42 |
| ACE2 | IL1RAP | -0,6049 | 7,19461E-42 |
| ACE2 | ADAMTS1 | -0,6049 | 7,19584E-42 |
| ACE2 | MEGF8 | -0,605 | 6,80826E-42 |
| ACE2 | ATF6 | -0,6051 | 6,64634E-42 |
| ACE2 | RNF144B | -0,6051 | 6,66413E-42 |
| ACE2 | RC3H2 | -0,6051 | 6,55975E-42 |
| ACE2 | RP11-54G14.1 | -0,6051 | 6,65558E-42 |
| ACE2 | RIBC1 | -0,6051 | 6,47053E-42 |
| ACE2 | IL12A-AS1 | -0,6052 | 6,37748E-42 |
| ACE2 | EPHA7 | -0,6052 | 6,37987E-42 |
| ACE2 | CDK2 | -0,6052 | 6,41719E-42 |
| ACE2 | RP11-679B19.1 | -0,6052 | 6,23462E-42 |
| ACE2 | CTD-2349P21.9 | -0,6053 | 6,13066E-42 |
| ACE2 | RP11-377G16.2 | -0,6054 | 5,81735E-42 |
| ACE2 | HNRNPCP7 | -0,6054 | 5,76435E-42 |
| ACE2 | BPTF | -0,6054 | 5,74365E-42 |
| ACE2 | RP11-494K3.2 | -0,6055 | 5,64202E-42 |
| ACE2 | UBXN2A | -0,6055 | 5,60592E-42 |
| ACE2 | DAAM2 | -0,6055 | 5,5211E-42 |
| ACE2 | C10orf82 | -0,6055 | 5,66629E-42 |
| ACE2 | RP11-430H10.1 | -0,6055 | 5,58609E-42 |
| ACE2 | EXOSC9 | -0,6056 | 5,4252E-42 |
| ACE2 | GLTSCR1 | -0,6056 | 5,37732E-42 |
| ACE2 | RP11-87H9.4 | -0,6057 | 5,29347E-42 |
| ACE2 | PAN2 | -0,6057 | 5,21004E-42 |
| ACE2 | Z97634.3 | -0,6057 | 5,25547E-42 |
| ACE2 | AC005498.4 | -0,6057 | 5,28228E-42 |
| ACE2 | GPC3 | -0,6057 | 5,11075E-42 |
| ACE2 | PPP1R42 | -0,6058 | 4,96142E-42 |
| ACE2 | CD2BP2 | -0,6058 | 5,04949E-42 |
| ACE2 | ZNF232 | -0,6058 | 4,97615E-42 |
| ACE2 | KAZN | -0,6059 | 4,75478E-42 |
| ACE2 | POU2F1 | -0,6059 | 4,73089E-42 |
| ACE2 | APBB3 | -0,6059 | 4,75384E-42 |
| ACE2 | LRRC41 | -0,606 | 4,60981E-42 |
| ACE2 | PRKAB2 | -0,606 | 4,66371E-42 |
| ACE2 | HSPB11 | -0,6061 | 4,44584E-42 |
| ACE2 | RP11-154H23.3 | -0,6061 | 4,43333E-42 |
| ACE2 | HDAC9 | -0,6061 | 4,48803E-42 |
| ACE2 | ING4 | -0,6061 | 4,42709E-42 |
| ACE2 | RACK1 | -0,6062 | 4,24706E-42 |
| ACE2 | GRID1 | -0,6062 | 4,22918E-42 |
| ACE2 | UBQLN2 | -0,6062 | 4,22151E-42 |
| ACE2 | HPCAL4 | -0,6063 | 4,13942E-42 |
| ACE2 | LPIN1 | -0,6063 | 4,11271E-42 |
| ACE2 | SEMA3D | -0,6063 | 4,04494E-42 |

| | | | |
|---|---|---|---|
| ACE2 | FAM66B | -0,6063 | 4,06275E-42 |
| ACE2 | RP11-165A20.3 | -0,6063 | 4,14313E-42 |
| ACE2 | S100A2 | -0,6064 | 3,93662E-42 |
| ACE2 | HRAS | -0,6064 | 4,02424E-42 |
| ACE2 | NSMCE1 | -0,6064 | 3,96913E-42 |
| ACE2 | LYPLAL1 | -0,6065 | 3,83769E-42 |
| ACE2 | ATL2 | -0,6065 | 3,76158E-42 |
| ACE2 | CTD-2366F13.2 | -0,6065 | 3,85929E-42 |
| ACE2 | ACTN3 | -0,6065 | 3,82732E-42 |
| ACE2 | RPTOR | -0,6066 | 3,62739E-42 |
| ACE2 | CEP41 | -0,6067 | 3,5597E-42 |
| ACE2 | SETX | -0,6067 | 3,56609E-42 |
| ACE2 | BUD13P1 | -0,6067 | 3,54065E-42 |
| ACE2 | ITGAV | -0,6068 | 3,38075E-42 |
| ACE2 | CTD-2530H12.8 | -0,6068 | 3,42952E-42 |
| ACE2 | RPL7P38 | -0,6068 | 3,40789E-42 |
| ACE2 | KPNA3 | -0,6068 | 3,3828E-42 |
| ACE2 | LCMT1-AS1 | -0,6068 | 3,44148E-42 |
| ACE2 | RP1-137D17.1 | -0,6069 | 3,28484E-42 |
| ACE2 | TSNAX | -0,607 | 3,15885E-42 |
| ACE2 | FER1L5 | -0,607 | 3,09632E-42 |
| ACE2 | SSBP3 | -0,6071 | 2,97966E-42 |
| ACE2 | AC103563.8 | -0,6071 | 3,03113E-42 |
| ACE2 | PVRIG | -0,6071 | 2,98047E-42 |
| ACE2 | AC074289.1 | -0,6072 | 2,91995E-42 |
| ACE2 | ICE1 | -0,6072 | 2,85372E-42 |
| ACE2 | IMPDH1 | -0,6072 | 2,88711E-42 |
| ACE2 | GSTO1 | -0,6072 | 2,84396E-42 |
| ACE2 | ZSWIM7 | -0,6072 | 2,86448E-42 |
| ACE2 | KDM3A | -0,6073 | 2,73844E-42 |
| ACE2 | PI16 | -0,6074 | 2,68168E-42 |
| ACE2 | SACS-AS1 | -0,6074 | 2,65478E-42 |
| ACE2 | CTD-2536I1.2 | -0,6075 | 2,57381E-42 |
| ACE2 | CWC25 | -0,6075 | 2,54711E-42 |
| ACE2 | THRA | -0,6075 | 2,55515E-42 |
| ACE2 | ZNF846 | -0,6075 | 2,53435E-42 |
| ACE2 | MORF4L1P1 | -0,6076 | 2,46162E-42 |
| ACE2 | FAM218A | -0,6076 | 2,5111E-42 |
| ACE2 | GBGT1 | -0,6076 | 2,51756E-42 |
| ACE2 | FGF18 | -0,6077 | 2,37578E-42 |
| ACE2 | RP5-942I16.1 | -0,6077 | 2,36867E-42 |
| ACE2 | CNTNAP3B | -0,6077 | 2,39135E-42 |
| ACE2 | STX6 | -0,6078 | 2,2563E-42 |
| ACE2 | ARNTL2 | -0,6078 | 2,25746E-42 |
| ACE2 | RPL23A | -0,6078 | 2,33345E-42 |
| ACE2 | CRBN | -0,6079 | 2,20329E-42 |
| ACE2 | AC011239.2 | -0,608 | 2,11839E-42 |
| ACE2 | REEP5 | -0,608 | 2,12488E-42 |
| ACE2 | ZNF319 | -0,608 | 2,15985E-42 |
| ACE2 | GUCA1B | -0,6081 | 2,0553E-42 |
| ACE2 | CTD-2373N4.3 | -0,6081 | 2,00882E-42 |
| ACE2 | ZNF534 | -0,6081 | 2,04797E-42 |
| ACE2 | POGK | -0,6082 | 1,98125E-42 |
| ACE2 | AP001065.15 | -0,6082 | 1,96371E-42 |

| | | | |
|---|---|---|---|
| ACE2 | FAM193B | -0,6083 | 1,91858E-42 |
| ACE2 | FOXK2 | -0,6083 | 1,91713E-42 |
| ACE2 | EPG5 | -0,6083 | 1,92202E-42 |
| ACE2 | RP4-737E23.6 | -0,6083 | 1,91016E-42 |
| ACE2 | PAF1 | -0,6084 | 1,84285E-42 |
| ACE2 | RP11-1148L6.5 | -0,6084 | 1,81694E-42 |
| ACE2 | RP11-9G1.3 | -0,6085 | 1,71173E-42 |
| ACE2 | RPL12P30 | -0,6085 | 1,72504E-42 |
| ACE2 | LINC02001 | -0,6085 | 1,7543E-42 |
| ACE2 | CTBP1 | -0,6086 | 1,70118E-42 |
| ACE2 | ZSCAN9 | -0,6086 | 1,69318E-42 |
| ACE2 | RABGAP1 | -0,6086 | 1,70702E-42 |
| ACE2 | RICTOR | -0,6087 | 1,58027E-42 |
| ACE2 | CTC-471J1.2 | -0,6087 | 1,62824E-42 |
| ACE2 | RP11-362J17.1 | -0,6088 | 1,52313E-42 |
| ACE2 | RP11-258F22.1 | -0,6088 | 1,53047E-42 |
| ACE2 | NTF3 | -0,6088 | 1,57713E-42 |
| ACE2 | NUAK1 | -0,6088 | 1,52632E-42 |
| ACE2 | LTBP2 | -0,6088 | 1,5704E-42 |
| ACE2 | VSIG10L | -0,6088 | 1,54063E-42 |
| ACE2 | RP1-93H18.1 | -0,6089 | 1,46817E-42 |
| ACE2 | HAUS6 | -0,6089 | 1,47369E-42 |
| ACE2 | BCL9L | -0,6089 | 1,51021E-42 |
| ACE2 | CDKN2D | -0,6089 | 1,50596E-42 |
| ACE2 | LSM2 | -0,609 | 1,41023E-42 |
| ACE2 | MRVI1-AS1 | -0,609 | 1,44323E-42 |
| ACE2 | FAM92A1P1 | -0,609 | 1,4471E-42 |
| ACE2 | UTP4 | -0,609 | 1,46002E-42 |
| ACE2 | CFAP61 | -0,609 | 1,42306E-42 |
| ACE2 | MBTPS2 | -0,609 | 1,41209E-42 |
| ACE2 | RP11-385F5.5 | -0,6091 | 1,35946E-42 |
| ACE2 | CIB4 | -0,6091 | 1,39102E-42 |
| ACE2 | HOXD9 | -0,6091 | 1,36165E-42 |
| ACE2 | MT1L | -0,6091 | 1,36875E-42 |
| ACE2 | POLR2E | -0,6091 | 1,35451E-42 |
| ACE2 | CDH26 | -0,6091 | 1,37745E-42 |
| ACE2 | WDR6 | -0,6092 | 1,33655E-42 |
| ACE2 | TRA2B | -0,6092 | 1,33231E-42 |
| ACE2 | C9orf116 | -0,6092 | 1,33724E-42 |
| ACE2 | MDGA2 | -0,6092 | 1,34638E-42 |
| ACE2 | ZNF221 | -0,6092 | 1,34187E-42 |
| ACE2 | BHLHE40 | -0,6093 | 1,25155E-42 |
| ACE2 | CTBP1-AS2 | -0,6093 | 1,25434E-42 |
| ACE2 | C6orf106 | -0,6093 | 1,26596E-42 |
| ACE2 | SOCS7 | -0,6093 | 1,26059E-42 |
| ACE2 | ZBTB41 | -0,6094 | 1,213E-42 |
| ACE2 | RPL21P135 | -0,6094 | 1,23228E-42 |
| ACE2 | PYROXD2 | -0,6094 | 1,23506E-42 |
| ACE2 | NUDT21 | -0,6095 | 1,15615E-42 |
| ACE2 | CLEC3B | -0,6096 | 1,13553E-42 |
| ACE2 | TNRC18 | -0,6096 | 1,13704E-42 |
| ACE2 | ELMO1 | -0,6096 | 1,14542E-42 |
| ACE2 | PRRC2B | -0,6096 | 1,14865E-42 |
| ACE2 | ZNF605 | -0,6096 | 1,15393E-42 |

| | | | |
|---|---|---|---|
| ACE2 | LINC02145 | -0,6097 | 1,07578E-42 |
| ACE2 | LAMB1 | -0,6097 | 1,10281E-42 |
| ACE2 | HOXC6 | -0,6097 | 1,07516E-42 |
| ACE2 | TBXA2R | -0,6097 | 1,08043E-42 |
| ACE2 | C11orf57 | -0,6098 | 1,03391E-42 |
| ACE2 | NR1H2 | -0,6098 | 1,04108E-42 |
| ACE2 | APPL1 | -0,6099 | 1,00974E-42 |
| ACE2 | SLC16A12 | -0,6099 | 1,0079E-42 |
| ACE2 | TOB1-AS1 | -0,61 | 9,69394E-43 |
| ACE2 | DLGAP1-AS1 | -0,61 | 9,84968E-43 |
| ACE2 | LIPI | -0,61 | 9,84943E-43 |
| ACE2 | ESRRG | -0,6101 | 9,35433E-43 |
| ACE2 | UBR1 | -0,6101 | 9,18326E-43 |
| ACE2 | TOMM6 | -0,6102 | 9,04778E-43 |
| ACE2 | ZFAND3 | -0,6103 | 8,46712E-43 |
| ACE2 | GPIHBP1 | -0,6103 | 8,58348E-43 |
| ACE2 | MAGOHB | -0,6103 | 8,60217E-43 |
| ACE2 | CDH8 | -0,6103 | 8,67251E-43 |
| ACE2 | ZNF559-ZNF177 | -0,6103 | 8,50371E-43 |
| ACE2 | SH3TC2 | -0,6104 | 8,21445E-43 |
| ACE2 | TRHDE-AS1 | -0,6104 | 8,17566E-43 |
| ACE2 | TADA2A | -0,6104 | 8,18699E-43 |
| ACE2 | CTD-2587H19.2 | -0,6104 | 8,30024E-43 |
| ACE2 | ZKSCAN2 | -0,6105 | 7,85864E-43 |
| ACE2 | KIAA2012 | -0,6106 | 7,55655E-43 |
| ACE2 | LINC01489 | -0,6106 | 7,61407E-43 |
| ACE2 | DHX29 | -0,6107 | 7,34396E-43 |
| ACE2 | CBWD2 | -0,6108 | 7,01652E-43 |
| ACE2 | INTS12 | -0,6109 | 6,69581E-43 |
| ACE2 | RP11-393I2.4 | -0,6109 | 6,74775E-43 |
| ACE2 | RNF10 | -0,6109 | 6,71813E-43 |
| ACE2 | GLT1D1 | -0,6109 | 6,77916E-43 |
| ACE2 | KPTN | -0,6109 | 6,73427E-43 |
| ACE2 | LINC01697 | -0,6109 | 6,89461E-43 |
| ACE2 | LRRC75B | -0,6109 | 6,66039E-43 |
| ACE2 | NOL7 | -0,611 | 6,45312E-43 |
| ACE2 | OMD | -0,611 | 6,58537E-43 |
| ACE2 | LHX6 | -0,611 | 6,59105E-43 |
| ACE2 | PLXDC2 | -0,611 | 6,44286E-43 |
| ACE2 | NUCB1 | -0,611 | 6,5678E-43 |
| ACE2 | RP1-142L7.5 | -0,6111 | 6,16845E-43 |
| ACE2 | CCDC150P1 | -0,6111 | 6,14951E-43 |
| ACE2 | PDPN | -0,6112 | 6,08956E-43 |
| ACE2 | RRP15 | -0,6112 | 6,10864E-43 |
| ACE2 | CCNY | -0,6112 | 5,98378E-43 |
| ACE2 | USP12 | -0,6112 | 6,07394E-43 |
| ACE2 | CTB-47B11.3 | -0,6113 | 5,69814E-43 |
| ACE2 | BCL7A | -0,6113 | 5,67517E-43 |
| ACE2 | RAD1 | -0,6114 | 5,64593E-43 |
| ACE2 | ABCB4 | -0,6114 | 5,591E-43 |
| ACE2 | FZD6 | -0,6114 | 5,54119E-43 |
| ACE2 | KCNMA1-AS1 | -0,6114 | 5,58753E-43 |
| ACE2 | MARS | -0,6115 | 5,31995E-43 |
| ACE2 | RTEL1 | -0,6115 | 5,26482E-43 |

| | | | |
|---|---|---|---|
| ACE2 | LNPEP | -0,6116 | 5,08384E-43 |
| ACE2 | MEDAG | -0,6116 | 5,11455E-43 |
| ACE2 | BTBD19 | -0,6117 | 4,86581E-43 |
| ACE2 | HTR2B | -0,6117 | 4,9674E-43 |
| ACE2 | RPS2P32 | -0,6117 | 4,8589E-43 |
| ACE2 | CMB9-94B1.2 | -0,6117 | 4,89592E-43 |
| ACE2 | CABP7 | -0,6117 | 4,84896E-43 |
| ACE2 | BHMT | -0,6118 | 4,76667E-43 |
| ACE2 | RP11-11N9.4 | -0,6118 | 4,66471E-43 |
| ACE2 | SCYL1 | -0,6118 | 4,74927E-43 |
| ACE2 | SNUPN | -0,6118 | 4,72451E-43 |
| ACE2 | AC004951.5 | -0,6119 | 4,49307E-43 |
| ACE2 | RP11-130L8.2 | -0,612 | 4,33395E-43 |
| ACE2 | MMP2 | -0,612 | 4,32109E-43 |
| ACE2 | ZFP14 | -0,612 | 4,32956E-43 |
| ACE2 | CTD-2537I9.18 | -0,612 | 4,46201E-43 |
| ACE2 | RPL21P16 | -0,6121 | 4,15693E-43 |
| ACE2 | ATF7IP | -0,6121 | 4,14209E-43 |
| ACE2 | ZNF630 | -0,6121 | 4,19061E-43 |
| ACE2 | KBTBD2 | -0,6122 | 4,09613E-43 |
| ACE2 | MRPL24 | -0,6123 | 3,89828E-43 |
| ACE2 | LRP1B | -0,6123 | 3,8734E-43 |
| ACE2 | ZDHHC11 | -0,6124 | 3,76907E-43 |
| ACE2 | TDRD6 | -0,6124 | 3,80887E-43 |
| ACE2 | C11orf49 | -0,6124 | 3,71177E-43 |
| ACE2 | XAF1 | -0,6124 | 3,67677E-43 |
| ACE2 | CAPZB | -0,6125 | 3,54047E-43 |
| ACE2 | RP11-301L8.2 | -0,6125 | 3,5383E-43 |
| ACE2 | FRMPD4 | -0,6125 | 3,6043E-43 |
| ACE2 | LINC00629 | -0,6125 | 3,59241E-43 |
| ACE2 | STC2 | -0,6126 | 3,41943E-43 |
| ACE2 | ENG | -0,6126 | 3,41035E-43 |
| ACE2 | SPAG7 | -0,6126 | 3,42211E-43 |
| ACE2 | MRPL10 | -0,6126 | 3,45763E-43 |
| ACE2 | CCDC102B | -0,6126 | 3,50248E-43 |
| ACE2 | RP11-1166P10.1 | -0,6127 | 3,3288E-43 |
| ACE2 | PCMTD2 | -0,6127 | 3,28136E-43 |
| ACE2 | UBXN11 | -0,6128 | 3,20992E-43 |
| ACE2 | PIH1D2 | -0,6128 | 3,24118E-43 |
| ACE2 | FOXN4 | -0,6128 | 3,13428E-43 |
| ACE2 | ENSA | -0,6129 | 3,08533E-43 |
| ACE2 | FSTL5 | -0,6129 | 3,06327E-43 |
| ACE2 | LMO2 | -0,6129 | 3,02808E-43 |
| ACE2 | FAM53A | -0,613 | 2,92523E-43 |
| ACE2 | PTGES3P1 | -0,6131 | 2,87493E-43 |
| ACE2 | CNTNAP3P2 | -0,6131 | 2,84257E-43 |
| ACE2 | KCTD10 | -0,6131 | 2,808E-43 |
| ACE2 | LTBP4 | -0,6131 | 2,85445E-43 |
| ACE2 | EXOC6B | -0,6132 | 2,7672E-43 |
| ACE2 | NFATC3 | -0,6132 | 2,74743E-43 |
| ACE2 | AC005498.3 | -0,6132 | 2,68187E-43 |
| ACE2 | POLR3E | -0,6133 | 2,61064E-43 |
| ACE2 | ELAVL1 | -0,6133 | 2,6159E-43 |
| ACE2 | ZFYVE9 | -0,6134 | 2,53106E-43 |

| | | | |
|---|---|---|---|
| ACE2 | RPRM | -0,6134 | 2,48563E-43 |
| ACE2 | TRA2A | -0,6134 | 2,50439E-43 |
| ACE2 | ERV3-1 | -0,6134 | 2,53029E-43 |
| ACE2 | DNAJC24 | -0,6134 | 2,54958E-43 |
| ACE2 | DDX6 | -0,6135 | 2,41069E-43 |
| ACE2 | PCDHAC2 | -0,6136 | 2,28166E-43 |
| ACE2 | LINC00475 | -0,6136 | 2,29569E-43 |
| ACE2 | CFAP46 | -0,6136 | 2,33332E-43 |
| ACE2 | RAB21 | -0,6136 | 2,36287E-43 |
| ACE2 | SCAF1 | -0,6137 | 2,24224E-43 |
| ACE2 | RPP30 | -0,6138 | 2,12901E-43 |
| ACE2 | ZPLD1 | -0,6139 | 2,02908E-43 |
| ACE2 | ZNF410 | -0,6139 | 2,02761E-43 |
| ACE2 | GLS | -0,614 | 1,99972E-43 |
| ACE2 | PARP10 | -0,614 | 2,00211E-43 |
| ACE2 | CABYR | -0,614 | 1,95708E-43 |
| ACE2 | LINC00891 | -0,614 | 1,97559E-43 |
| ACE2 | ATP11A | -0,6141 | 1,87095E-43 |
| ACE2 | NKIRAS2 | -0,6141 | 1,905E-43 |
| ACE2 | C11orf88 | -0,6142 | 1,79931E-43 |
| ACE2 | TMTC3 | -0,6143 | 1,72026E-43 |
| ACE2 | TRIM47 | -0,6143 | 1,74169E-43 |
| ACE2 | C1orf229 | -0,6144 | 1,69294E-43 |
| ACE2 | ANK1 | -0,6144 | 1,67131E-43 |
| ACE2 | PTENP1 | -0,6144 | 1,70689E-43 |
| ACE2 | STMN2 | -0,6145 | 1,61741E-43 |
| ACE2 | RP11-967K21.1 | -0,6145 | 1,6442E-43 |
| ACE2 | CSF1 | -0,6146 | 1,56923E-43 |
| ACE2 | ACTR1B | -0,6146 | 1,58214E-43 |
| ACE2 | PJA2 | -0,6146 | 1,57162E-43 |
| ACE2 | LRRC29 | -0,6146 | 1,57513E-43 |
| ACE2 | GOLGA8B | -0,6147 | 1,49651E-43 |
| ACE2 | FAM20A | -0,6147 | 1,49749E-43 |
| ACE2 | RP11-64C12.8 | -0,6147 | 1,50472E-43 |
| ACE2 | FLJ37453 | -0,6148 | 1,41453E-43 |
| ACE2 | SPTAN1 | -0,6148 | 1,44519E-43 |
| ACE2 | RRP1B | -0,6148 | 1,45056E-43 |
| ACE2 | UPF3B | -0,6148 | 1,40757E-43 |
| ACE2 | ZNF589 | -0,6149 | 1,37864E-43 |
| ACE2 | TAF1D | -0,615 | 1,29965E-43 |
| ACE2 | HARS | -0,6151 | 1,28069E-43 |
| ACE2 | RP11-367F23.1 | -0,6151 | 1,2698E-43 |
| ACE2 | SYT6 | -0,6152 | 1,20548E-43 |
| ACE2 | CGGBP1 | -0,6152 | 1,24502E-43 |
| ACE2 | GOLT1B | -0,6152 | 1,21731E-43 |
| ACE2 | AC025171.1 | -0,6153 | 1,1923E-43 |
| ACE2 | RP11-1069G10.2 | -0,6153 | 1,15325E-43 |
| ACE2 | HINT3 | -0,6154 | 1,10705E-43 |
| ACE2 | NOM1 | -0,6154 | 1,13403E-43 |
| ACE2 | CDKN1B | -0,6154 | 1,10659E-43 |
| ACE2 | CTD-3018O17.5 | -0,6154 | 1,13384E-43 |
| ACE2 | IFRD1 | -0,6155 | 1,089E-43 |
| ACE2 | RCN1 | -0,6155 | 1,09215E-43 |
| ACE2 | RPL26 | -0,6156 | 1,03153E-43 |

| | | | |
|---|---|---|---|
| ACE2 | TBX18 | -0,6157 | 1,01664E-43 |
| ACE2 | RP11-242D8.1 | -0,6157 | 9,95554E-44 |
| ACE2 | RP11-693M3.1 | -0,6157 | 9,94282E-44 |
| ACE2 | BOLA1 | -0,6158 | 9,43051E-44 |
| ACE2 | NUP133 | -0,6158 | 9,59451E-44 |
| ACE2 | FMN2 | -0,6158 | 9,54075E-44 |
| ACE2 | RP11-395N3.2 | -0,6158 | 9,66695E-44 |
| ACE2 | WDR66 | -0,6158 | 9,46995E-44 |
| ACE2 | RASSF4 | -0,6159 | 9,3586E-44 |
| ACE2 | RP11-849H4.4 | -0,6159 | 9,24937E-44 |
| ACE2 | TPM4 | -0,6159 | 9,29573E-44 |
| ACE2 | RP3-467K16.2 | -0,616 | 9,00257E-44 |
| ACE2 | BTNL9 | -0,616 | 8,95266E-44 |
| ACE2 | CEP57L1 | -0,616 | 8,95449E-44 |
| ACE2 | LMLN | -0,6161 | 8,42425E-44 |
| ACE2 | RP11-148K1.12 | -0,6161 | 8,42481E-44 |
| ACE2 | CTD-3247H4.2 | -0,6161 | 8,48442E-44 |
| ACE2 | GOLGA8VP | -0,6161 | 8,43695E-44 |
| ACE2 | AP000318.2 | -0,6161 | 8,45153E-44 |
| ACE2 | MATN3 | -0,6162 | 8,04317E-44 |
| ACE2 | RP11-635L1.3 | -0,6162 | 8,21779E-44 |
| ACE2 | AJ011932.1 | -0,6162 | 8,1312E-44 |
| ACE2 | RP11-73M18.8 | -0,6163 | 7,91834E-44 |
| ACE2 | KIF26A | -0,6163 | 7,89488E-44 |
| ACE2 | UBL7 | -0,6163 | 7,76587E-44 |
| ACE2 | CILP2 | -0,6163 | 7,71207E-44 |
| ACE2 | C1orf101 | -0,6164 | 7,62462E-44 |
| ACE2 | ETAA1 | -0,6164 | 7,52855E-44 |
| ACE2 | FGF5 | -0,6164 | 7,51692E-44 |
| ACE2 | SLC2A8 | -0,6164 | 7,63809E-44 |
| ACE2 | HSPBAP1 | -0,6166 | 6,96544E-44 |
| ACE2 | PEBP4 | -0,6167 | 6,61881E-44 |
| ACE2 | CACNB2 | -0,6167 | 6,80721E-44 |
| ACE2 | ATP8A2 | -0,6167 | 6,78173E-44 |
| ACE2 | VIPAS39 | -0,6167 | 6,73703E-44 |
| ACE2 | NDUFV2-AS1 | -0,6167 | 6,57668E-44 |
| ACE2 | ZNF473 | -0,6167 | 6,57298E-44 |
| ACE2 | BNC1 | -0,6168 | 6,39528E-44 |
| ACE2 | UBE2Q2P1 | -0,6168 | 6,53554E-44 |
| ACE2 | SUPT4H1 | -0,6168 | 6,40798E-44 |
| ACE2 | BRD4 | -0,6168 | 6,52897E-44 |
| ACE2 | TMEM189 | -0,6168 | 6,30965E-44 |
| ACE2 | LINC01475 | -0,6169 | 6,08299E-44 |
| ACE2 | RP11-839D17.3 | -0,6169 | 6,22338E-44 |
| ACE2 | BBS2 | -0,6169 | 6,15238E-44 |
| ACE2 | ZNF132 | -0,6169 | 6,03752E-44 |
| ACE2 | SLAIN2 | -0,6171 | 5,57353E-44 |
| ACE2 | RP11-438D14.2 | -0,6171 | 5,59265E-44 |
| ACE2 | CEP128 | -0,6171 | 5,67768E-44 |
| ACE2 | ANKFY1 | -0,6171 | 5,5686E-44 |
| ACE2 | KIF28P | -0,6172 | 5,46673E-44 |
| ACE2 | PID1 | -0,6172 | 5,37369E-44 |
| ACE2 | SLC2A12 | -0,6173 | 5,13809E-44 |
| ACE2 | SREK1IP1 | -0,6174 | 5,03562E-44 |

| | | | |
|---|---|---|---|
| ACE2 | PRKAR1B | -0,6174 | 4,93744E-44 |
| ACE2 | HEYL | -0,6175 | 4,8802E-44 |
| ACE2 | FBXW11 | -0,6175 | 4,92296E-44 |
| ACE2 | FLOT2 | -0,6175 | 4,83181E-44 |
| ACE2 | CTC-241N9.1 | -0,6176 | 4,67991E-44 |
| ACE2 | TOGARAM1 | -0,6176 | 4,70864E-44 |
| ACE2 | RPS17 | -0,6176 | 4,7076E-44 |
| ACE2 | SAP18 | -0,6177 | 4,50277E-44 |
| ACE2 | ZNF175 | -0,6177 | 4,46201E-44 |
| ACE2 | GS1-124K5.10 | -0,6178 | 4,19466E-44 |
| ACE2 | PLXNA4 | -0,6178 | 4,24753E-44 |
| ACE2 | HCN1 | -0,6179 | 4,06004E-44 |
| ACE2 | CENPP | -0,6179 | 4,04762E-44 |
| ACE2 | CEP295 | -0,6179 | 4,14669E-44 |
| ACE2 | AMDHD1 | -0,6179 | 4,17767E-44 |
| ACE2 | AKAP11 | -0,6179 | 4,15186E-44 |
| ACE2 | SUPT6H | -0,6179 | 4,10994E-44 |
| ACE2 | TDRD10 | -0,618 | 3,89582E-44 |
| ACE2 | RP11-181K3.4 | -0,618 | 3,88536E-44 |
| ACE2 | PEX2 | -0,618 | 3,91967E-44 |
| ACE2 | CCDC144NL-AS1 | -0,618 | 3,87505E-44 |
| ACE2 | STK40 | -0,6181 | 3,83543E-44 |
| ACE2 | CHRM3 | -0,6181 | 3,81862E-44 |
| ACE2 | KLF11 | -0,6181 | 3,80455E-44 |
| ACE2 | MBNL1 | -0,6181 | 3,82983E-44 |
| ACE2 | PCDHGB3 | -0,6181 | 3,78166E-44 |
| ACE2 | ABCD1 | -0,6181 | 3,73823E-44 |
| ACE2 | CTB-113P19.5 | -0,6182 | 3,68471E-44 |
| ACE2 | TESC | -0,6182 | 3,57992E-44 |
| ACE2 | RP11-481J2.1 | -0,6182 | 3,56933E-44 |
| ACE2 | RP11-395G23.3 | -0,6183 | 3,50844E-44 |
| ACE2 | SHARPIN | -0,6183 | 3,50712E-44 |
| ACE2 | RERG-IT1 | -0,6183 | 3,47196E-44 |
| ACE2 | RP11-709D24.8 | -0,6183 | 3,5272E-44 |
| ACE2 | WDR70 | -0,6184 | 3,29549E-44 |
| ACE2 | NLRP1 | -0,6184 | 3,34067E-44 |
| ACE2 | RP11-462H3.2 | -0,6184 | 3,32954E-44 |
| ACE2 | NOTCH2 | -0,6185 | 3,23367E-44 |
| ACE2 | SCG2 | -0,6185 | 3,16963E-44 |
| ACE2 | MTERF4 | -0,6185 | 3,25826E-44 |
| ACE2 | ZCCHC9 | -0,6185 | 3,21753E-44 |
| ACE2 | GNAI1 | -0,6185 | 3,21715E-44 |
| ACE2 | FAM149B1 | -0,6185 | 3,22153E-44 |
| ACE2 | TUBGCP5 | -0,6185 | 3,28148E-44 |
| ACE2 | BCAS3 | -0,6185 | 3,18268E-44 |
| ACE2 | TAPT1 | -0,6186 | 3,06216E-44 |
| ACE2 | UBE2B | -0,6187 | 3,00085E-44 |
| ACE2 | NUDT9P1 | -0,6187 | 2,91485E-44 |
| ACE2 | CDC42BPA | -0,6188 | 2,83227E-44 |
| ACE2 | NELFE | -0,619 | 2,6717E-44 |
| ACE2 | CASP8AP2 | -0,619 | 2,60515E-44 |
| ACE2 | RCOR3 | -0,6191 | 2,56397E-44 |
| ACE2 | TMEM163 | -0,6191 | 2,52962E-44 |
| ACE2 | TFEB | -0,6191 | 2,56285E-44 |

| | | | |
|---|---|---|---|
| ACE2 | RP11-58A12.3 | -0,6191 | 2,5192E-44 |
| ACE2 | ZNF625 | -0,6191 | 2,49991E-44 |
| ACE2 | SERPINF1 | -0,6192 | 2,46127E-44 |
| ACE2 | SLITRK4 | -0,6192 | 2,46628E-44 |
| ACE2 | HNRNPLL | -0,6193 | 2,334E-44 |
| ACE2 | SENP2 | -0,6193 | 2,36493E-44 |
| ACE2 | FAM155A | -0,6193 | 2,33223E-44 |
| ACE2 | TNFRSF8 | -0,6194 | 2,19091E-44 |
| ACE2 | DNAJC10 | -0,6194 | 2,2632E-44 |
| ACE2 | PRSS45 | -0,6194 | 2,22547E-44 |
| ACE2 | GPBP1 | -0,6194 | 2,22623E-44 |
| ACE2 | RPS6 | -0,6194 | 2,23208E-44 |
| ACE2 | NUFIP1 | -0,6194 | 2,2264E-44 |
| ACE2 | FZD2 | -0,6194 | 2,19669E-44 |
| ACE2 | ST8SIA5 | -0,6194 | 2,20982E-44 |
| ACE2 | PACRG | -0,6195 | 2,14087E-44 |
| ACE2 | NELFCD | -0,6195 | 2,10927E-44 |
| ACE2 | UBE2F | -0,6196 | 2,0269E-44 |
| ACE2 | CTD-2013N24.2 | -0,6196 | 2,09144E-44 |
| ACE2 | KY | -0,6196 | 2,06451E-44 |
| ACE2 | FKBP9 | -0,6196 | 2,03022E-44 |
| ACE2 | AARD | -0,6196 | 2,02338E-44 |
| ACE2 | USP40 | -0,6197 | 1,94767E-44 |
| ACE2 | APBA3 | -0,6197 | 1,93918E-44 |
| ACE2 | ZNF337-AS1 | -0,6197 | 1,99864E-44 |
| ACE2 | RP11-316N24.2 | -0,6198 | 1,87433E-44 |
| ACE2 | KLHL42 | -0,6198 | 1,93055E-44 |
| ACE2 | RP11-622C24.2 | -0,6198 | 1,86023E-44 |
| ACE2 | RPA1 | -0,6198 | 1,92878E-44 |
| ACE2 | COMMD2 | -0,6199 | 1,8419E-44 |
| ACE2 | BAZ1B | -0,6199 | 1,79323E-44 |
| ACE2 | RP11-723D22.3 | -0,6199 | 1,83397E-44 |
| ACE2 | RP11-569G13.2 | -0,6199 | 1,82E-44 |
| ACE2 | RECQL | -0,6199 | 1,81335E-44 |
| ACE2 | MCHR1 | -0,6199 | 1,80669E-44 |
| ACE2 | CDK5R2 | -0,62 | 1,77828E-44 |
| ACE2 | RP11-930O11.3 | -0,62 | 1,77698E-44 |
| ACE2 | RP11-182J1.18 | -0,62 | 1,73069E-44 |
| ACE2 | KIAA2022 | -0,62 | 1,74458E-44 |
| ACE2 | BTC | -0,6201 | 1,70558E-44 |
| ACE2 | TVP23A | -0,6201 | 1,66658E-44 |
| ACE2 | ILDR2 | -0,6203 | 1,52169E-44 |
| ACE2 | RP11-274B21.9 | -0,6203 | 1,5496E-44 |
| ACE2 | ZFPM2-AS1 | -0,6203 | 1,5496E-44 |
| ACE2 | CFAP54 | -0,6203 | 1,56715E-44 |
| ACE2 | TAS2R15P | -0,6204 | 1,46248E-44 |
| ACE2 | VEZT | -0,6204 | 1,45722E-44 |
| ACE2 | VTN | -0,6204 | 1,50247E-44 |
| ACE2 | ADRA2B | -0,6205 | 1,45418E-44 |
| ACE2 | SELP | -0,6206 | 1,36664E-44 |
| ACE2 | LAPTM4A | -0,6206 | 1,36203E-44 |
| ACE2 | ENDOV | -0,6206 | 1,38641E-44 |
| ACE2 | BTF3 | -0,6207 | 1,30294E-44 |
| ACE2 | NAP1L4 | -0,6207 | 1,33012E-44 |

| | | | |
|---|---|---|---|
| ACE2 | RP11-438D14.3 | -0,6207 | 1,33143E-44 |
| ACE2 | SP4 | -0,6208 | 1,28112E-44 |
| ACE2 | STRN3 | -0,6208 | 1,26093E-44 |
| ACE2 | AKAP10 | -0,6208 | 1,24997E-44 |
| ACE2 | PFN1P2 | -0,6209 | 1,23251E-44 |
| ACE2 | UBE2W | -0,6209 | 1,22417E-44 |
| ACE2 | MBTPS1 | -0,6209 | 1,19754E-44 |
| ACE2 | TXLNG | -0,6209 | 1,22035E-44 |
| ACE2 | VAMP3 | -0,621 | 1,18464E-44 |
| ACE2 | RP4-564M11.2 | -0,621 | 1,18276E-44 |
| ACE2 | NIT2 | -0,621 | 1,14097E-44 |
| ACE2 | MT3 | -0,621 | 1,14615E-44 |
| ACE2 | CTB-25B13.5 | -0,621 | 1,17183E-44 |
| ACE2 | RP11-187C18.3 | -0,6211 | 1,12629E-44 |
| ACE2 | CACUL1 | -0,6211 | 1,09959E-44 |
| ACE2 | ARHGAP20 | -0,6212 | 1,09108E-44 |
| ACE2 | LINC01535 | -0,6212 | 1,05505E-44 |
| ACE2 | LRRC7 | -0,6213 | 1,02142E-44 |
| ACE2 | ZNF721 | -0,6213 | 1,03473E-44 |
| ACE2 | RWDD2A | -0,6213 | 1,04639E-44 |
| ACE2 | RP11-728F11.3 | -0,6213 | 1,03542E-44 |
| ACE2 | ZNF585B | -0,6213 | 1,03669E-44 |
| ACE2 | IP6K1 | -0,6214 | 9,8657E-45 |
| ACE2 | CRY1 | -0,6214 | 9,67669E-45 |
| ACE2 | TUBGCP3 | -0,6214 | 9,69236E-45 |
| ACE2 | MC1R | -0,6214 | 9,91191E-45 |
| ACE2 | PAFAH1B1 | -0,6214 | 9,88162E-45 |
| ACE2 | OBSCN | -0,6215 | 9,35796E-45 |
| ACE2 | CBX5 | -0,6215 | 9,34856E-45 |
| ACE2 | STARD3 | -0,6215 | 9,302E-45 |
| ACE2 | CYGB | -0,6215 | 9,4892E-45 |
| ACE2 | RP11-64C12.1 | -0,6215 | 9,57338E-45 |
| ACE2 | MIB1 | -0,6215 | 9,61942E-45 |
| ACE2 | TRIM28 | -0,6215 | 9,31702E-45 |
| ACE2 | MORC3 | -0,6215 | 9,60769E-45 |
| ACE2 | DNAJC21 | -0,6216 | 9,19571E-45 |
| ACE2 | ASGR1 | -0,6216 | 8,9492E-45 |
| ACE2 | PPP1R21 | -0,6217 | 8,78351E-45 |
| ACE2 | AC007403.3 | -0,6217 | 8,86756E-45 |
| ACE2 | RIOK1 | -0,6217 | 8,56317E-45 |
| ACE2 | ASNS | -0,6217 | 8,75075E-45 |
| ACE2 | ANKS6 | -0,6217 | 8,61868E-45 |
| ACE2 | RP11-497E19.1 | -0,6217 | 8,87554E-45 |
| ACE2 | AGAP2-AS1 | -0,6218 | 8,47182E-45 |
| ACE2 | ITM2A | -0,6218 | 8,4958E-45 |
| ACE2 | MTFR1L | -0,6219 | 7,87455E-45 |
| ACE2 | RP11-806K15.1 | -0,6219 | 8,05628E-45 |
| ACE2 | PMS2P1 | -0,6219 | 7,95292E-45 |
| ACE2 | NFIA-AS2 | -0,622 | 7,70558E-45 |
| ACE2 | ARAP3 | -0,622 | 7,66267E-45 |
| ACE2 | RP11-327J17.7 | -0,622 | 7,61228E-45 |
| ACE2 | RP11-563K23.1 | -0,6221 | 7,32069E-45 |
| ACE2 | MID1 | -0,6221 | 7,48959E-45 |
| ACE2 | MIR199A2 | -0,6222 | 6,99416E-45 |

| | | | |
|---|---|---|---|
| ACE2 | KPNA4 | -0,6222 | 7,06499E-45 |
| ACE2 | SYT15 | -0,6222 | 6,97847E-45 |
| ACE2 | RP11-15A1.8 | -0,6222 | 7,18752E-45 |
| ACE2 | RP11-17E13.2 | -0,6223 | 6,76654E-45 |
| ACE2 | ZNF32-AS1 | -0,6223 | 6,92197E-45 |
| ACE2 | TMEM159 | -0,6223 | 6,8165E-45 |
| ACE2 | SPNS1 | -0,6223 | 6,93513E-45 |
| ACE2 | ATP6V1B1 | -0,6224 | 6,42176E-45 |
| ACE2 | POMZP3 | -0,6224 | 6,41681E-45 |
| ACE2 | PRRT4 | -0,6224 | 6,64496E-45 |
| ACE2 | IPO13 | -0,6225 | 6,30691E-45 |
| ACE2 | FAM185A | -0,6225 | 6,36313E-45 |
| ACE2 | LMBR1 | -0,6225 | 6,18926E-45 |
| ACE2 | TBC1D15 | -0,6225 | 6,21186E-45 |
| ACE2 | SAMD11 | -0,6226 | 5,98583E-45 |
| ACE2 | FOXC1 | -0,6226 | 5,95558E-45 |
| ACE2 | HLA-F-AS1 | -0,6226 | 5,90121E-45 |
| ACE2 | ZC3HC1 | -0,6226 | 6,03289E-45 |
| ACE2 | WNT5B | -0,6226 | 5,92668E-45 |
| ACE2 | DEK | -0,6227 | 5,72336E-45 |
| ACE2 | NEXN-AS1 | -0,6228 | 5,54007E-45 |
| ACE2 | DNAJC19 | -0,6228 | 5,60653E-45 |
| ACE2 | RP11-586D19.1 | -0,6228 | 5,51974E-45 |
| ACE2 | RP11-127B20.3 | -0,6228 | 5,42771E-45 |
| ACE2 | SUB1 | -0,6228 | 5,47909E-45 |
| ACE2 | HBS1L | -0,6228 | 5,48349E-45 |
| ACE2 | RAMP3 | -0,6228 | 5,48904E-45 |
| ACE2 | ZNF746 | -0,6228 | 5,53117E-45 |
| ACE2 | BEAN1-AS1 | -0,6228 | 5,63621E-45 |
| ACE2 | SH2D5 | -0,6229 | 5,33901E-45 |
| ACE2 | TRAPPC12 | -0,6229 | 5,24197E-45 |
| ACE2 | C2orf42 | -0,623 | 5,0112E-45 |
| ACE2 | RP11-57H14.4 | -0,623 | 5,18068E-45 |
| ACE2 | PURA | -0,6231 | 4,79767E-45 |
| ACE2 | LINC02202 | -0,6231 | 4,90511E-45 |
| ACE2 | SPG21 | -0,6231 | 4,84939E-45 |
| ACE2 | LINC01581 | -0,6232 | 4,6122E-45 |
| ACE2 | RP11-795F19.5 | -0,6232 | 4,74066E-45 |
| ACE2 | DPY19L3 | -0,6232 | 4,7919E-45 |
| ACE2 | ANGEL2 | -0,6233 | 4,57709E-45 |
| ACE2 | RP11-586K2.1 | -0,6233 | 4,57423E-45 |
| ACE2 | KRTCAP2 | -0,6234 | 4,30838E-45 |
| ACE2 | CAPN7 | -0,6234 | 4,34424E-45 |
| ACE2 | C16orf89 | -0,6235 | 4,14796E-45 |
| ACE2 | GGA2 | -0,6235 | 4,12496E-45 |
| ACE2 | RNF187 | -0,6236 | 4,04946E-45 |
| ACE2 | RP11-366M4.11 | -0,6236 | 3,92686E-45 |
| ACE2 | KIF13A | -0,6236 | 3,94219E-45 |
| ACE2 | PIGK | -0,6237 | 3,77389E-45 |
| ACE2 | ACADL | -0,6237 | 3,85262E-45 |
| ACE2 | RPL7L1P9 | -0,6237 | 3,81351E-45 |
| ACE2 | FBXL5 | -0,6237 | 3,89228E-45 |
| ACE2 | MATN2 | -0,6237 | 3,77104E-45 |
| ACE2 | FEM1B | -0,6237 | 3,8111E-45 |

| | | | |
|---|---|---|---|
| ACE2 | MRPS14 | -0,6238 | 3,70252E-45 |
| ACE2 | RIF1 | -0,6238 | 3,68169E-45 |
| ACE2 | WDR49 | -0,6238 | 3,69785E-45 |
| ACE2 | BAG3 | -0,6238 | 3,70348E-45 |
| ACE2 | PSMC3IP | -0,6238 | 3,6722E-45 |
| ACE2 | NKAIN1 | -0,6239 | 3,52738E-45 |
| ACE2 | NRCAM | -0,6239 | 3,53895E-45 |
| ACE2 | HAUS3 | -0,624 | 3,34621E-45 |
| ACE2 | NEDD4 | -0,624 | 3,31846E-45 |
| ACE2 | ZNF652 | -0,624 | 3,33859E-45 |
| ACE2 | ELOC | -0,6241 | 3,2807E-45 |
| ACE2 | FRMD6-AS2 | -0,6241 | 3,20403E-45 |
| ACE2 | ATRAID | -0,6242 | 3,0526E-45 |
| ACE2 | HES7 | -0,6242 | 3,15609E-45 |
| ACE2 | SP100 | -0,6243 | 2,96723E-45 |
| ACE2 | ROBO2 | -0,6243 | 3,00343E-45 |
| ACE2 | RBMX | -0,6243 | 2,93176E-45 |
| ACE2 | RIT1 | -0,6244 | 2,80755E-45 |
| ACE2 | TEPP | -0,6244 | 2,85519E-45 |
| ACE2 | RP11-326C3.7 | -0,6246 | 2,61443E-45 |
| ACE2 | MNT | -0,6246 | 2,61874E-45 |
| ACE2 | LA16c-380F5.1 | -0,6247 | 2,51758E-45 |
| ACE2 | C3orf58 | -0,6248 | 2,41169E-45 |
| ACE2 | VPS37A | -0,6248 | 2,37647E-45 |
| ACE2 | MRPL20 | -0,6249 | 2,29636E-45 |
| ACE2 | PLPP3 | -0,6249 | 2,34099E-45 |
| ACE2 | NBPF20 | -0,6249 | 2,27926E-45 |
| ACE2 | KHDC1 | -0,6249 | 2,35301E-45 |
| ACE2 | RP11-864N7.4 | -0,6249 | 2,32514E-45 |
| ACE2 | RASSF3 | -0,6249 | 2,29458E-45 |
| ACE2 | RP11-489D6.2 | -0,6249 | 2,33871E-45 |
| ACE2 | GABPB1 | -0,6249 | 2,31102E-45 |
| ACE2 | SNTB2 | -0,6249 | 2,29481E-45 |
| ACE2 | GDF5 | -0,6249 | 2,27953E-45 |
| ACE2 | ARMC8 | -0,625 | 2,24011E-45 |
| ACE2 | ATP2B1-AS1 | -0,625 | 2,25246E-45 |
| ACE2 | TDRKH | -0,6251 | 2,14102E-45 |
| ACE2 | DDX59 | -0,6251 | 2,15712E-45 |
| ACE2 | RPS6KA2 | -0,6251 | 2,16113E-45 |
| ACE2 | AASS | -0,6251 | 2,10948E-45 |
| ACE2 | PHF20L1 | -0,6251 | 2,10063E-45 |
| ACE2 | DESI1 | -0,6251 | 2,17006E-45 |
| ACE2 | RP11-355F16.1 | -0,6252 | 2,06783E-45 |
| ACE2 | DOC2GP | -0,6252 | 2,03003E-45 |
| ACE2 | ZFYVE21 | -0,6252 | 2,08676E-45 |
| ACE2 | ERCC4 | -0,6252 | 2,02623E-45 |
| ACE2 | SF3A3 | -0,6253 | 1,9967E-45 |
| ACE2 | SIX2 | -0,6253 | 1,95773E-45 |
| ACE2 | PCDHGC4 | -0,6253 | 1,92917E-45 |
| ACE2 | AGK | -0,6254 | 1,84725E-45 |
| ACE2 | UBE2L6 | -0,6254 | 1,89082E-45 |
| ACE2 | HSPA2 | -0,6254 | 1,89615E-45 |
| ACE2 | IER5L | -0,6255 | 1,83372E-45 |
| ACE2 | IRF2BPL | -0,6255 | 1,79479E-45 |

| | | | |
|---|---|---|---|
| ACE2 | SH3RF3-AS1 | -0,6256 | 1,74367E-45 |
| ACE2 | FRMPD1 | -0,6256 | 1,73106E-45 |
| ACE2 | DCAF10 | -0,6256 | 1,74096E-45 |
| ACE2 | RP11-707G18.1 | -0,6256 | 1,73934E-45 |
| ACE2 | CTD-3092A11.2 | -0,6256 | 1,70654E-45 |
| ACE2 | DYNLRB1 | -0,6256 | 1,72134E-45 |
| ACE2 | UHRF2 | -0,6257 | 1,68095E-45 |
| ACE2 | HERC2P9 | -0,6257 | 1,64945E-45 |
| ACE2 | RP1-253P7.4 | -0,6257 | 1,63474E-45 |
| ACE2 | PRKCB | -0,6258 | 1,59683E-45 |
| ACE2 | RHBDL3 | -0,6258 | 1,57769E-45 |
| ACE2 | MICALL1 | -0,6258 | 1,57228E-45 |
| ACE2 | FTSJ1 | -0,6258 | 1,62875E-45 |
| ACE2 | C6orf89 | -0,6259 | 1,53408E-45 |
| ACE2 | BCDIN3D-AS1 | -0,6259 | 1,53224E-45 |
| ACE2 | MMP14 | -0,626 | 1,45921E-45 |
| ACE2 | PRSS46 | -0,6261 | 1,37906E-45 |
| ACE2 | AC108142.1 | -0,6261 | 1,43115E-45 |
| ACE2 | NUDCD1 | -0,6261 | 1,43088E-45 |
| ACE2 | ENY2 | -0,6261 | 1,40667E-45 |
| ACE2 | RAPSN | -0,6261 | 1,38405E-45 |
| ACE2 | ZNF222 | -0,6261 | 1,41742E-45 |
| ACE2 | ANOS1 | -0,6261 | 1,42639E-45 |
| ACE2 | ARSK | -0,6262 | 1,36483E-45 |
| ACE2 | PCDHGA1 | -0,6262 | 1,34385E-45 |
| ACE2 | KANSL2 | -0,6262 | 1,37618E-45 |
| ACE2 | BPI | -0,6262 | 1,34572E-45 |
| ACE2 | RP11-261C10.5 | -0,6263 | 1,28133E-45 |
| ACE2 | FAM180A | -0,6263 | 1,26904E-45 |
| ACE2 | HGSNAT | -0,6263 | 1,29664E-45 |
| ACE2 | RSF1 | -0,6263 | 1,30063E-45 |
| ACE2 | NKD2 | -0,6264 | 1,23602E-45 |
| ACE2 | NBN | -0,6264 | 1,26167E-45 |
| ACE2 | RP5-890O3.9 | -0,6265 | 1,16931E-45 |
| ACE2 | ECEL1 | -0,6265 | 1,20962E-45 |
| ACE2 | LINC00310 | -0,6265 | 1,21312E-45 |
| ACE2 | GOLGA7 | -0,6266 | 1,16373E-45 |
| ACE2 | CTD-2298J14.2 | -0,6266 | 1,14528E-45 |
| ACE2 | LINC01359 | -0,6267 | 1,10503E-45 |
| ACE2 | SP5 | -0,6267 | 1,1059E-45 |
| ACE2 | CNTROB | -0,6267 | 1,07652E-45 |
| ACE2 | NID1 | -0,6268 | 1,0453E-45 |
| ACE2 | CTB-108O6.2 | -0,6268 | 1,04062E-45 |
| ACE2 | C14orf166 | -0,6268 | 1,03692E-45 |
| ACE2 | AC016582.2 | -0,6268 | 1,0628E-45 |
| ACE2 | PLS3-AS1 | -0,6268 | 1,03966E-45 |
| ACE2 | ADAMTS6 | -0,6269 | 1,00469E-45 |
| ACE2 | KIZ-AS1 | -0,6269 | 9,94175E-46 |
| ACE2 | PITHD1 | -0,627 | 9,69753E-46 |
| ACE2 | RP11-266J6.2 | -0,627 | 9,79451E-46 |
| ACE2 | TAF6 | -0,627 | 9,63446E-46 |
| ACE2 | LINC00598 | -0,627 | 9,81915E-46 |
| ACE2 | CCM2L | -0,627 | 9,70268E-46 |
| ACE2 | NBAS | -0,6271 | 9,28091E-46 |

| | | | |
|---|---|---|---|
| ACE2 | HIGD1B | -0,6272 | 8,82138E-46 |
| ACE2 | FAM212B | -0,6273 | 8,50041E-46 |
| ACE2 | PRCC | -0,6273 | 8,6932E-46 |
| ACE2 | PFN4 | -0,6273 | 8,4946E-46 |
| ACE2 | RP11-320H14.1 | -0,6273 | 8,48267E-46 |
| ACE2 | TTC37 | -0,6273 | 8,45733E-46 |
| ACE2 | KRIT1 | -0,6273 | 8,58131E-46 |
| ACE2 | POU3F4 | -0,6273 | 8,35518E-46 |
| ACE2 | LIN37 | -0,6274 | 8,15735E-46 |
| ACE2 | PDRG1 | -0,6274 | 8,06286E-46 |
| ACE2 | TMEM126B | -0,6275 | 7,89982E-46 |
| ACE2 | MORF4L2 | -0,6275 | 7,86637E-46 |
| ACE2 | KCNQ1OT1 | -0,6276 | 7,40552E-46 |
| ACE2 | GPR180 | -0,6276 | 7,43158E-46 |
| ACE2 | RP4-616B8.5 | -0,6276 | 7,52196E-46 |
| ACE2 | DPH5 | -0,6277 | 7,33576E-46 |
| ACE2 | RP3-523C21.2 | -0,6277 | 7,08517E-46 |
| ACE2 | AGTR2 | -0,6277 | 7,09714E-46 |
| ACE2 | ESR1 | -0,6278 | 7,05105E-46 |
| ACE2 | CTB-55O6.12 | -0,6278 | 7,02554E-46 |
| ACE2 | STK38L | -0,6279 | 6,713E-46 |
| ACE2 | RP11-340F14.6 | -0,6279 | 6,55225E-46 |
| ACE2 | MED4 | -0,6279 | 6,56397E-46 |
| ACE2 | LINC01936 | -0,628 | 6,25305E-46 |
| ACE2 | PLEKHM3 | -0,628 | 6,35356E-46 |
| ACE2 | DYNC1LI1 | -0,628 | 6,2614E-46 |
| ACE2 | ATL3 | -0,628 | 6,38041E-46 |
| ACE2 | USE1 | -0,628 | 6,24472E-46 |
| ACE2 | ZNF507 | -0,628 | 6,47123E-46 |
| ACE2 | PPP4R1L | -0,628 | 6,23678E-46 |
| ACE2 | RP11-131L12.3 | -0,6281 | 5,99874E-46 |
| ACE2 | TUBB | -0,6282 | 5,93077E-46 |
| ACE2 | RAD54L2 | -0,6283 | 5,62591E-46 |
| ACE2 | GABRG2 | -0,6283 | 5,48817E-46 |
| ACE2 | AGTPBP1 | -0,6283 | 5,59314E-46 |
| ACE2 | ZFYVE1 | -0,6283 | 5,50651E-46 |
| ACE2 | RP11-520H14.7 | -0,6285 | 5,08513E-46 |
| ACE2 | ARID4B | -0,6285 | 5,20944E-46 |
| ACE2 | CDH10 | -0,6285 | 5,11125E-46 |
| ACE2 | SETD1B | -0,6285 | 5,14796E-46 |
| ACE2 | ZMYM2 | -0,6285 | 5,09092E-46 |
| ACE2 | AFG3L1P | -0,6285 | 5,22087E-46 |
| ACE2 | RASL10B | -0,6285 | 5,09533E-46 |
| ACE2 | METTL9 | -0,6286 | 4,84632E-46 |
| ACE2 | USP14 | -0,6286 | 4,83952E-46 |
| ACE2 | RP11-98G7.1 | -0,6287 | 4,76978E-46 |
| ACE2 | SEPT10 | -0,6287 | 4,63168E-46 |
| ACE2 | CWC22 | -0,6287 | 4,68345E-46 |
| ACE2 | RP11-613M10.6 | -0,6287 | 4,7718E-46 |
| ACE2 | GPAA1P2 | -0,6288 | 4,536E-46 |
| ACE2 | RP11-313J2.1 | -0,6288 | 4,57581E-46 |
| ACE2 | NONO | -0,6288 | 4,45617E-46 |
| ACE2 | CDKAL1 | -0,6289 | 4,40684E-46 |
| ACE2 | AC000403.4 | -0,629 | 4,10851E-46 |

| | | | |
|---|---|---|---|
| ACE2 | ABHD4 | -0,629 | 4,198E-46 |
| ACE2 | VPS39 | -0,629 | 4,16546E-46 |
| ACE2 | PRR29 | -0,629 | 4,09164E-46 |
| ACE2 | ABCA6 | -0,629 | 4,15689E-46 |
| ACE2 | MBD3 | -0,629 | 4,11598E-46 |
| ACE2 | CACTIN | -0,629 | 4,13968E-46 |
| ACE2 | KCTD3 | -0,6291 | 4,00153E-46 |
| ACE2 | MELTF-AS1 | -0,6291 | 4,01078E-46 |
| ACE2 | AP000648.5 | -0,6291 | 3,94603E-46 |
| ACE2 | C1QL1 | -0,6291 | 4,00684E-46 |
| ACE2 | TEX41 | -0,6292 | 3,85922E-46 |
| ACE2 | RSRC1 | -0,6292 | 3,77883E-46 |
| ACE2 | SLC7A2 | -0,6292 | 3,79563E-46 |
| ACE2 | SLF2 | -0,6292 | 3,82462E-46 |
| ACE2 | PAX9 | -0,6292 | 3,90239E-46 |
| ACE2 | RP11-857B24.5 | -0,6292 | 3,80619E-46 |
| ACE2 | RP11-680G24.4 | -0,6292 | 3,8882E-46 |
| ACE2 | HYDIN | -0,6292 | 3,8423E-46 |
| ACE2 | ZNF594 | -0,6292 | 3,83557E-46 |
| ACE2 | TMEM35B | -0,6293 | 3,69262E-46 |
| ACE2 | PAXIP1-AS2 | -0,6293 | 3,71944E-46 |
| ACE2 | C12orf65 | -0,6293 | 3,65601E-46 |
| ACE2 | MTMR1 | -0,6293 | 3,60061E-46 |
| ACE2 | LINC01140 | -0,6294 | 3,45926E-46 |
| ACE2 | RPL27A | -0,6294 | 3,50687E-46 |
| ACE2 | MYO15A | -0,6294 | 3,44769E-46 |
| ACE2 | MEOX2-AS1 | -0,6295 | 3,43951E-46 |
| ACE2 | KLF17 | -0,6296 | 3,30033E-46 |
| ACE2 | GLB1L | -0,6296 | 3,26477E-46 |
| ACE2 | GAPVD1 | -0,6296 | 3,18379E-46 |
| ACE2 | MPG | -0,6296 | 3,20319E-46 |
| ACE2 | RP11-404E16.1 | -0,6297 | 3,08572E-46 |
| ACE2 | RAF1 | -0,6297 | 3,05686E-46 |
| ACE2 | SVIL-AS1 | -0,6297 | 3,07701E-46 |
| ACE2 | LINC00900 | -0,6297 | 3,11746E-46 |
| ACE2 | LMCD1-AS1 | -0,6298 | 2,92505E-46 |
| ACE2 | SLC25A4 | -0,6298 | 2,96162E-46 |
| ACE2 | ZSCAN23 | -0,6298 | 2,95527E-46 |
| ACE2 | CPB2-AS1 | -0,6298 | 2,96827E-46 |
| ACE2 | PLPPR5 | -0,6299 | 2,84155E-46 |
| ACE2 | CTD-2287O16.5 | -0,6299 | 2,88314E-46 |
| ACE2 | CDC73 | -0,63 | 2,72966E-46 |
| ACE2 | KIDINS220 | -0,63 | 2,78766E-46 |
| ACE2 | AFF3 | -0,63 | 2,77344E-46 |
| ACE2 | CLDN18 | -0,63 | 2,75078E-46 |
| ACE2 | NADK2 | -0,63 | 2,75296E-46 |
| ACE2 | SH3PXD2B | -0,63 | 2,70665E-46 |
| ACE2 | SEMA3C | -0,63 | 2,76727E-46 |
| ACE2 | AC079630.4 | -0,63 | 2,73498E-46 |
| ACE2 | GABARAP | -0,63 | 2,686E-46 |
| ACE2 | ZNF30 | -0,63 | 2,74202E-46 |
| ACE2 | CTD-2619J13.17 | -0,63 | 2,70483E-46 |
| ACE2 | RN7SL861P | -0,6301 | 2,57035E-46 |
| ACE2 | MCTP1 | -0,6301 | 2,57434E-46 |

| | | | |
|---|---|---|---|
| ACE2 | CTD-3064M3.7 | -0,6301 | 2,56639E-46 |
| ACE2 | GIT1 | -0,6301 | 2,61258E-46 |
| ACE2 | LINC01097 | -0,6302 | 2,55443E-46 |
| ACE2 | TUBB2A | -0,6302 | 2,5148E-46 |
| ACE2 | PIK3C2A | -0,6302 | 2,55213E-46 |
| ACE2 | CEP126 | -0,6302 | 2,5434E-46 |
| ACE2 | RP11-650L12.2 | -0,6302 | 2,52552E-46 |
| ACE2 | RP11-161D15.3 | -0,6303 | 2,36651E-46 |
| ACE2 | EMC2 | -0,6303 | 2,40649E-46 |
| ACE2 | RP11-159N11.4 | -0,6303 | 2,40184E-46 |
| ACE2 | CACNG7 | -0,6303 | 2,41513E-46 |
| ACE2 | COPS4 | -0,6304 | 2,33299E-46 |
| ACE2 | ZNF398 | -0,6304 | 2,3457E-46 |
| ACE2 | ZNF189 | -0,6304 | 2,3E-46 |
| ACE2 | MID1IP1 | -0,6304 | 2,32611E-46 |
| ACE2 | DAP3 | -0,6305 | 2,20492E-46 |
| ACE2 | OXCT1 | -0,6305 | 2,1741E-46 |
| ACE2 | CDK4 | -0,6305 | 2,22252E-46 |
| ACE2 | KLHL33 | -0,6305 | 2,16857E-46 |
| ACE2 | STAM | -0,6306 | 2,11678E-46 |
| ACE2 | VENTX | -0,6306 | 2,14821E-46 |
| ACE2 | NPFF | -0,6306 | 2,11867E-46 |
| ACE2 | CTD-3193K9.3 | -0,6306 | 2,10035E-46 |
| ACE2 | RP11-694I15.7 | -0,6307 | 2,05684E-46 |
| ACE2 | SCAI | -0,6307 | 2,04574E-46 |
| ACE2 | CCDC91 | -0,6307 | 1,98984E-46 |
| ACE2 | ARNT | -0,6308 | 1,93565E-46 |
| ACE2 | THAP11 | -0,6308 | 1,94876E-46 |
| ACE2 | DPH1 | -0,6308 | 1,92417E-46 |
| ACE2 | SNU13 | -0,6308 | 1,90404E-46 |
| ACE2 | RP11-170N16.3 | -0,6309 | 1,85937E-46 |
| ACE2 | RP11-563J2.3 | -0,6309 | 1,85567E-46 |
| ACE2 | SYCP2 | -0,6309 | 1,84954E-46 |
| ACE2 | LMCD1 | -0,631 | 1,81965E-46 |
| ACE2 | RP11-341G23.4 | -0,631 | 1,80953E-46 |
| ACE2 | MED29 | -0,631 | 1,77594E-46 |
| ACE2 | QPCT | -0,6311 | 1,68978E-46 |
| ACE2 | RP5-1057I20.5 | -0,6311 | 1,70627E-46 |
| ACE2 | GAS6-AS2 | -0,6311 | 1,73453E-46 |
| ACE2 | TRIOBP | -0,6311 | 1,72925E-46 |
| ACE2 | RBPMS-AS1 | -0,6312 | 1,64736E-46 |
| ACE2 | SH2D3C | -0,6312 | 1,64707E-46 |
| ACE2 | ARHGAP5-AS1 | -0,6312 | 1,64676E-46 |
| ACE2 | SEC14L6 | -0,6312 | 1,67323E-46 |
| ACE2 | SUPT20H | -0,6313 | 1,57042E-46 |
| ACE2 | YPEL1 | -0,6313 | 1,56538E-46 |
| ACE2 | SMS | -0,6313 | 1,58087E-46 |
| ACE2 | USP21 | -0,6314 | 1,50811E-46 |
| ACE2 | RP11-809C18.3 | -0,6314 | 1,47673E-46 |
| ACE2 | OR51E2 | -0,6314 | 1,5082E-46 |
| ACE2 | ATM | -0,6314 | 1,50447E-46 |
| ACE2 | FARP1 | -0,6314 | 1,50775E-46 |
| ACE2 | DUSP12 | -0,6315 | 1,47188E-46 |
| ACE2 | RP11-434H6.7 | -0,6315 | 1,44519E-46 |

| | | | |
|---|---|---|---|
| ACE2 | ANKRD16 | -0,6315 | 1,44948E-46 |
| ACE2 | GTF2H1 | -0,6315 | 1,44196E-46 |
| ACE2 | CCDC82 | -0,6315 | 1,44225E-46 |
| ACE2 | MSANTD2 | -0,6315 | 1,46508E-46 |
| ACE2 | USPL1 | -0,6315 | 1,4234E-46 |
| ACE2 | HTRA2 | -0,6316 | 1,37223E-46 |
| ACE2 | CALCOCO1 | -0,6316 | 1,40283E-46 |
| ACE2 | RP11-171I2.3 | -0,6317 | 1,33884E-46 |
| ACE2 | TGFBI | -0,6317 | 1,32848E-46 |
| ACE2 | RPS24 | -0,6317 | 1,31244E-46 |
| ACE2 | HSF4 | -0,6317 | 1,3515E-46 |
| ACE2 | RPL29 | -0,6318 | 1,26319E-46 |
| ACE2 | OLFML1 | -0,6318 | 1,25474E-46 |
| ACE2 | METAP2 | -0,6318 | 1,25857E-46 |
| ACE2 | SCAPER | -0,6318 | 1,27817E-46 |
| ACE2 | RHBDF1 | -0,6318 | 1,26374E-46 |
| ACE2 | KIAA1191 | -0,6319 | 1,21192E-46 |
| ACE2 | ID4 | -0,6319 | 1,19874E-46 |
| ACE2 | VN1R108P | -0,6319 | 1,22784E-46 |
| ACE2 | COL20A1 | -0,6319 | 1,20132E-46 |
| ACE2 | SDCCAG3 | -0,6321 | 1,10024E-46 |
| ACE2 | RBM39 | -0,6321 | 1,11794E-46 |
| ACE2 | RPS19BP1 | -0,6321 | 1,10992E-46 |
| ACE2 | SERTAD4 | -0,6322 | 1,08917E-46 |
| ACE2 | AC073130.3 | -0,6322 | 1,07359E-46 |
| ACE2 | RP11-771K4.3 | -0,6322 | 1,05627E-46 |
| ACE2 | CTD-2547L24.4 | -0,6322 | 1,0661E-46 |
| ACE2 | RP11-6O2.4 | -0,6322 | 1,05498E-46 |
| ACE2 | FOXC2 | -0,6322 | 1,04992E-46 |
| ACE2 | CYP1B1 | -0,6323 | 1,01038E-46 |
| ACE2 | ANTXR2 | -0,6323 | 1,02703E-46 |
| ACE2 | RPS12 | -0,6323 | 1,02936E-46 |
| ACE2 | SGCZ | -0,6323 | 1,02504E-46 |
| ACE2 | PCDHGB1 | -0,6324 | 9,99367E-47 |
| ACE2 | NSD3 | -0,6324 | 9,64383E-47 |
| ACE2 | TIMM23B | -0,6324 | 9,99828E-47 |
| ACE2 | RP11-54O7.3 | -0,6325 | 9,45976E-47 |
| ACE2 | EDNRA | -0,6325 | 9,59886E-47 |
| ACE2 | CWF19L1 | -0,6325 | 9,24759E-47 |
| ACE2 | JMJD8 | -0,6325 | 9,40717E-47 |
| ACE2 | RP11-958N24.1 | -0,6325 | 9,28509E-47 |
| ACE2 | ERICH3 | -0,6326 | 9,091E-47 |
| ACE2 | DNAJB4 | -0,6326 | 9,10009E-47 |
| ACE2 | NBPF9 | -0,6326 | 9,09921E-47 |
| ACE2 | SLC4A7 | -0,6326 | 9,07462E-47 |
| ACE2 | RP11-713P17.4 | -0,6326 | 8,99582E-47 |
| ACE2 | RP11-455F5.3 | -0,6326 | 8,88761E-47 |
| ACE2 | MSL1 | -0,6326 | 9,20716E-47 |
| ACE2 | MAPK12 | -0,6326 | 9,0058E-47 |
| ACE2 | ARIH2 | -0,6327 | 8,81327E-47 |
| ACE2 | TAS2R14 | -0,6327 | 8,69693E-47 |
| ACE2 | MIA | -0,6327 | 8,81099E-47 |
| ACE2 | PSMB4 | -0,6328 | 8,26054E-47 |
| ACE2 | TBCE | -0,6328 | 8,15789E-47 |

| | | | |
|---|---|---|---|
| ACE2 | RP11-9N20.3 | -0,6328 | 8,2765E-47 |
| ACE2 | ABCA12 | -0,6329 | 7,82859E-47 |
| ACE2 | ETV5 | -0,6329 | 7,86218E-47 |
| ACE2 | RMND5B | -0,6329 | 7,84465E-47 |
| ACE2 | UPF2 | -0,6329 | 7,83813E-47 |
| ACE2 | GADD45GIP1 | -0,6329 | 7,80112E-47 |
| ACE2 | ZNF432 | -0,6329 | 7,78056E-47 |
| ACE2 | NNAT | -0,6329 | 8,01123E-47 |
| ACE2 | XXyac-YX65C7_A.2 | -0,633 | 7,46916E-47 |
| ACE2 | FOXK1 | -0,633 | 7,73071E-47 |
| ACE2 | ANKRD46 | -0,633 | 7,73508E-47 |
| ACE2 | RP11-195F19.9 | -0,633 | 7,48666E-47 |
| ACE2 | EIF2B1 | -0,633 | 7,51747E-47 |
| ACE2 | LINC01686 | -0,6331 | 7,33701E-47 |
| ACE2 | YTHDC1 | -0,6331 | 7,28682E-47 |
| ACE2 | SPIN1 | -0,6331 | 7,22382E-47 |
| ACE2 | CCDC59 | -0,6331 | 7,35701E-47 |
| ACE2 | TNFAIP1 | -0,6331 | 7,23543E-47 |
| ACE2 | DDX42 | -0,6331 | 7,34543E-47 |
| ACE2 | ATP6V1C1 | -0,6332 | 6,86404E-47 |
| ACE2 | ARGLU1 | -0,6332 | 7,03653E-47 |
| ACE2 | PRR36 | -0,6332 | 7,052E-47 |
| ACE2 | TSPEAR-AS2 | -0,6332 | 7,0582E-47 |
| ACE2 | C6orf226 | -0,6333 | 6,80958E-47 |
| ACE2 | ESYT2 | -0,6333 | 6,66143E-47 |
| ACE2 | SVIL | -0,6333 | 6,79017E-47 |
| ACE2 | RTBDN | -0,6333 | 6,74992E-47 |
| ACE2 | CTC-203F4.2 | -0,6334 | 6,41153E-47 |
| ACE2 | ZNF7 | -0,6334 | 6,46059E-47 |
| ACE2 | ZFP30 | -0,6334 | 6,43936E-47 |
| ACE2 | GNB1L | -0,6334 | 6,53293E-47 |
| ACE2 | ECM1 | -0,6335 | 6,12189E-47 |
| ACE2 | MAP4K3 | -0,6335 | 6,19904E-47 |
| ACE2 | RP11-753H16.5 | -0,6335 | 6,1679E-47 |
| ACE2 | FFAR3 | -0,6335 | 6,02655E-47 |
| ACE2 | WIPI2 | -0,6336 | 5,89381E-47 |
| ACE2 | IGF2-AS | -0,6336 | 5,83034E-47 |
| ACE2 | OTOF | -0,6337 | 5,6727E-47 |
| ACE2 | TXNRD2 | -0,6337 | 5,61845E-47 |
| ACE2 | EPB41L4A-AS1 | -0,6338 | 5,46899E-47 |
| ACE2 | ZNF511 | -0,6338 | 5,5129E-47 |
| ACE2 | KCNH5 | -0,6338 | 5,41285E-47 |
| ACE2 | SETD3 | -0,6338 | 5,42616E-47 |
| ACE2 | RP11-799B12.1 | -0,6338 | 5,29468E-47 |
| ACE2 | RASAL2 | -0,6339 | 5,18571E-47 |
| ACE2 | NICN1 | -0,6339 | 5,13166E-47 |
| ACE2 | ANKS1A | -0,6339 | 5,09917E-47 |
| ACE2 | RNH1 | -0,6339 | 5,09078E-47 |
| ACE2 | RP11-421F16.3 | -0,6339 | 5,19245E-47 |
| ACE2 | R3HDM2 | -0,6339 | 5,27907E-47 |
| ACE2 | STUB1 | -0,6339 | 5,17844E-47 |
| ACE2 | RP11-35G9.5 | -0,6339 | 5,19326E-47 |
| ACE2 | MYT1 | -0,6339 | 5,2304E-47 |
| ACE2 | SLFNL1 | -0,634 | 5,02959E-47 |

| | | | |
|---|---|---|---|
| ACE2 | EPHA4 | -0,634 | 4,91284E-47 |
| ACE2 | CSPG4P11 | -0,634 | 5,00647E-47 |
| ACE2 | DOC2B | -0,634 | 4,94599E-47 |
| ACE2 | MGAT5B | -0,634 | 4,86822E-47 |
| ACE2 | MAOB | -0,634 | 4,91409E-47 |
| ACE2 | NUP58 | -0,6341 | 4,7113E-47 |
| ACE2 | LINC01314 | -0,6341 | 4,80792E-47 |
| ACE2 | RP11-6O2.2 | -0,6341 | 4,79433E-47 |
| ACE2 | RTN4RL2 | -0,6342 | 4,45613E-47 |
| ACE2 | CLSTN2 | -0,6343 | 4,29728E-47 |
| ACE2 | CDHR3 | -0,6343 | 4,35065E-47 |
| ACE2 | ERCC6L2 | -0,6343 | 4,29622E-47 |
| ACE2 | ADCYAP1 | -0,6343 | 4,37422E-47 |
| ACE2 | MAPRE1 | -0,6343 | 4,32735E-47 |
| ACE2 | RP11-2C24.7 | -0,6344 | 4,21389E-47 |
| ACE2 | SMNDC1 | -0,6345 | 4,05303E-47 |
| ACE2 | PABPN1 | -0,6345 | 3,97736E-47 |
| ACE2 | ANGPTL1 | -0,6346 | 3,81628E-47 |
| ACE2 | RSPH9 | -0,6346 | 3,77614E-47 |
| ACE2 | SVOP | -0,6346 | 3,75232E-47 |
| ACE2 | CTB-36H16.2 | -0,6347 | 3,59583E-47 |
| ACE2 | TMC3 | -0,6347 | 3,71358E-47 |
| ACE2 | TMPRSS9 | -0,6347 | 3,63689E-47 |
| ACE2 | ZNF304 | -0,6347 | 3,69854E-47 |
| ACE2 | RP5-1139B12.3 | -0,6348 | 3,51446E-47 |
| ACE2 | RP11-173C1.1 | -0,6348 | 3,55772E-47 |
| ACE2 | WIPF3 | -0,6348 | 3,47871E-47 |
| ACE2 | IFIT3 | -0,635 | 3,24073E-47 |
| ACE2 | EIF2S3 | -0,635 | 3,26905E-47 |
| ACE2 | CIDECP | -0,6351 | 3,14265E-47 |
| ACE2 | BBIP1 | -0,6351 | 3,08172E-47 |
| ACE2 | CD248 | -0,6351 | 3,09551E-47 |
| ACE2 | TMPRSS5 | -0,6351 | 3,09253E-47 |
| ACE2 | TLR5 | -0,6352 | 2,94526E-47 |
| ACE2 | BCL6 | -0,6352 | 2,93759E-47 |
| ACE2 | RP11-124N14.3 | -0,6352 | 2,99113E-47 |
| ACE2 | SLC16A7 | -0,6352 | 3,00281E-47 |
| ACE2 | RP11-563J2.2 | -0,6353 | 2,87583E-47 |
| ACE2 | C17orf67 | -0,6353 | 2,88256E-47 |
| ACE2 | RNMT | -0,6354 | 2,71897E-47 |
| ACE2 | SUN1 | -0,6356 | 2,47735E-47 |
| ACE2 | RP11-309L24.2 | -0,6356 | 2,51833E-47 |
| ACE2 | PTGES3L | -0,6356 | 2,46566E-47 |
| ACE2 | C19orf47 | -0,6356 | 2,47847E-47 |
| ACE2 | TMEM201 | -0,6357 | 2,42048E-47 |
| ACE2 | RP11-474O21.5 | -0,6357 | 2,39741E-47 |
| ACE2 | NRDC | -0,6357 | 2,33497E-47 |
| ACE2 | NTNG2 | -0,6357 | 2,35723E-47 |
| ACE2 | KB-1460A1.5 | -0,6358 | 2,23261E-47 |
| ACE2 | ZNF16 | -0,6358 | 2,2791E-47 |
| ACE2 | NKD1 | -0,6358 | 2,27234E-47 |
| ACE2 | ADSL | -0,6358 | 2,2871E-47 |
| ACE2 | NAP1L2 | -0,6358 | 2,29794E-47 |
| ACE2 | GIN1 | -0,6359 | 2,21289E-47 |

| | | | |
|---|---|---|---|
| ACE2 | LINC00622 | -0,636 | 2,11606E-47 |
| ACE2 | POLR3G | -0,636 | 2,11242E-47 |
| ACE2 | SEC31A | -0,6361 | 2,00931E-47 |
| ACE2 | AKR1B1 | -0,6361 | 1,96271E-47 |
| ACE2 | GCSH | -0,6361 | 2,03186E-47 |
| ACE2 | ODAM | -0,6362 | 1,89845E-47 |
| ACE2 | IKZF5 | -0,6362 | 1,91564E-47 |
| ACE2 | ENDOU | -0,6362 | 1,87879E-47 |
| ACE2 | ATP2B2 | -0,6363 | 1,79799E-47 |
| ACE2 | MSRB2 | -0,6363 | 1,86631E-47 |
| ACE2 | AHNAK | -0,6363 | 1,82876E-47 |
| ACE2 | TRMT13 | -0,6364 | 1,72062E-47 |
| ACE2 | RP11-269F20.1 | -0,6364 | 1,76749E-47 |
| ACE2 | DNAJC27-AS1 | -0,6364 | 1,78866E-47 |
| ACE2 | UBXN10-AS1 | -0,6365 | 1,65538E-47 |
| ACE2 | HDAC3 | -0,6365 | 1,71564E-47 |
| ACE2 | CTD-3064M3.3 | -0,6365 | 1,66617E-47 |
| ACE2 | UBXN6 | -0,6365 | 1,70846E-47 |
| ACE2 | CTD-2527I21.15 | -0,6365 | 1,66572E-47 |
| ACE2 | TICAM2 | -0,6366 | 1,60423E-47 |
| ACE2 | USP27X | -0,6366 | 1,58674E-47 |
| ACE2 | IFT122 | -0,6367 | 1,53693E-47 |
| ACE2 | PTCD2 | -0,6367 | 1,53556E-47 |
| ACE2 | APOBEC3F | -0,6367 | 1,53741E-47 |
| ACE2 | RP11-351D16.3 | -0,6368 | 1,48039E-47 |
| ACE2 | SSB | -0,6369 | 1,42804E-47 |
| ACE2 | WDFY3 | -0,6369 | 1,39476E-47 |
| ACE2 | EXOC2 | -0,6369 | 1,43759E-47 |
| ACE2 | SURF2 | -0,6369 | 1,44553E-47 |
| ACE2 | PRMT7 | -0,6369 | 1,39792E-47 |
| ACE2 | XPNPEP3 | -0,6369 | 1,3856E-47 |
| ACE2 | PHEX | -0,6369 | 1,41937E-47 |
| ACE2 | RP11-266O8.1 | -0,637 | 1,35142E-47 |
| ACE2 | ZKSCAN3 | -0,6371 | 1,28356E-47 |
| ACE2 | LINC00957 | -0,6371 | 1,29448E-47 |
| ACE2 | SERPINH1 | -0,6371 | 1,27549E-47 |
| ACE2 | POMT2 | -0,6371 | 1,27844E-47 |
| ACE2 | RP11-848P1.5 | -0,6371 | 1,32287E-47 |
| ACE2 | UBQLN4 | -0,6372 | 1,24154E-47 |
| ACE2 | FAM114A1 | -0,6372 | 1,24788E-47 |
| ACE2 | FAM66D | -0,6372 | 1,22144E-47 |
| ACE2 | RP11-305E6.4 | -0,6372 | 1,22566E-47 |
| ACE2 | SEZ6 | -0,6372 | 1,24138E-47 |
| ACE2 | ZNF274 | -0,6372 | 1,24787E-47 |
| ACE2 | POLA1 | -0,6372 | 1,26074E-47 |
| ACE2 | LYSMD1 | -0,6373 | 1,19904E-47 |
| ACE2 | RP11-890B15.3 | -0,6373 | 1,20053E-47 |
| ACE2 | RBM19 | -0,6373 | 1,19897E-47 |
| ACE2 | TFAP2C | -0,6373 | 1,16436E-47 |
| ACE2 | TSPEAR | -0,6373 | 1,17865E-47 |
| ACE2 | GS1-124K5.4 | -0,6374 | 1,16266E-47 |
| ACE2 | GPS1 | -0,6375 | 1,09948E-47 |
| ACE2 | ZMAT5 | -0,6375 | 1,09373E-47 |
| ACE2 | ADGRD1 | -0,6376 | 1,04113E-47 |

| | | | |
|---|---|---|---|
| ACE2 | RPAIN | -0,6376 | 1,05537E-47 |
| ACE2 | RP11-156P1.3 | -0,6376 | 1,02613E-47 |
| ACE2 | PRKCA | -0,6376 | 1,03756E-47 |
| ACE2 | PLOD1 | -0,6377 | 1,01274E-47 |
| ACE2 | CPNE9 | -0,6377 | 9,95611E-48 |
| ACE2 | MON1A | -0,6377 | 9,89018E-48 |
| ACE2 | RBM24 | -0,6377 | 1,01839E-47 |
| ACE2 | NGDN | -0,6377 | 9,89919E-48 |
| ACE2 | TAOK1 | -0,6377 | 1,01802E-47 |
| ACE2 | SEPT9 | -0,6377 | 9,9829E-48 |
| ACE2 | GNPDA1 | -0,6378 | 9,62642E-48 |
| ACE2 | RSBN1L | -0,6378 | 9,35913E-48 |
| ACE2 | RP4-568B10.1 | -0,6378 | 9,57914E-48 |
| ACE2 | CLEC14A | -0,6378 | 9,68116E-48 |
| ACE2 | AGO2 | -0,6379 | 9,15599E-48 |
| ACE2 | RP11-23J9.7 | -0,6379 | 9,02973E-48 |
| ACE2 | STYX | -0,6379 | 8,96855E-48 |
| ACE2 | KLHL20 | -0,638 | 8,83834E-48 |
| ACE2 | B3GALNT2 | -0,638 | 8,60426E-48 |
| ACE2 | TAB2 | -0,638 | 8,83167E-48 |
| ACE2 | OTUD6B | -0,638 | 8,75459E-48 |
| ACE2 | KCNB1 | -0,638 | 8,93882E-48 |
| ACE2 | C4orf50 | -0,6381 | 8,42965E-48 |
| ACE2 | PTPRA | -0,6381 | 8,44817E-48 |
| ACE2 | KLHL15 | -0,6381 | 8,41909E-48 |
| ACE2 | SEPT7-AS1 | -0,6382 | 7,97983E-48 |
| ACE2 | PDAP1 | -0,6382 | 7,97574E-48 |
| ACE2 | RP11-441O15.3 | -0,6382 | 8,04542E-48 |
| ACE2 | ROCK1 | -0,6382 | 8,06169E-48 |
| ACE2 | ZNF865 | -0,6382 | 7,86957E-48 |
| ACE2 | CD93 | -0,6382 | 8,20004E-48 |
| ACE2 | CTSK | -0,6383 | 7,61072E-48 |
| ACE2 | RP11-168A11.4 | -0,6383 | 7,60166E-48 |
| ACE2 | KCTD16 | -0,6383 | 7,79034E-48 |
| ACE2 | ZNF618 | -0,6383 | 7,54321E-48 |
| ACE2 | LYL1 | -0,6383 | 7,6853E-48 |
| ACE2 | CTD-2175A23.1 | -0,6384 | 7,51755E-48 |
| ACE2 | DIRAS2 | -0,6384 | 7,35292E-48 |
| ACE2 | MUS81 | -0,6384 | 7,45117E-48 |
| ACE2 | AASDHPPT | -0,6384 | 7,24967E-48 |
| ACE2 | CACNA1C-AS1 | -0,6384 | 7,41009E-48 |
| ACE2 | EXOSC8 | -0,6384 | 7,31682E-48 |
| ACE2 | XXYLT1 | -0,6385 | 7,16358E-48 |
| ACE2 | PIAS2 | -0,6385 | 6,94195E-48 |
| ACE2 | KCNAB1 | -0,6386 | 6,855E-48 |
| ACE2 | LINC01638 | -0,6386 | 6,79171E-48 |
| ACE2 | C11orf96 | -0,6387 | 6,38146E-48 |
| ACE2 | LRRC32 | -0,6388 | 6,19026E-48 |
| ACE2 | AGTRAP | -0,6389 | 5,85327E-48 |
| ACE2 | RP11-522B15.3 | -0,639 | 5,6137E-48 |
| ACE2 | PM20D2 | -0,6391 | 5,47818E-48 |
| ACE2 | VCPKMT | -0,6391 | 5,38172E-48 |
| ACE2 | ULK4 | -0,6392 | 5,21751E-48 |
| ACE2 | FBXO24 | -0,6392 | 5,25979E-48 |

| | | | |
|---|---|---|---|
| ACE2 | GRIP1 | -0,6392 | 5,2011E-48 |
| ACE2 | MFSD14A | -0,6393 | 4,8709E-48 |
| ACE2 | ZNF330 | -0,6393 | 4,86916E-48 |
| ACE2 | CSAD | -0,6393 | 4,97835E-48 |
| ACE2 | APOBEC3G | -0,6393 | 5,06036E-48 |
| ACE2 | PRELP | -0,6394 | 4,8204E-48 |
| ACE2 | UBE2Q2L | -0,6394 | 4,68859E-48 |
| ACE2 | SFRP1 | -0,6395 | 4,60765E-48 |
| ACE2 | COL5A1-AS1 | -0,6395 | 4,49708E-48 |
| ACE2 | DNLZ | -0,6395 | 4,59118E-48 |
| ACE2 | CD59 | -0,6395 | 4,47143E-48 |
| ACE2 | BANF1 | -0,6395 | 4,5357E-48 |
| ACE2 | CTD-2008E3.1 | -0,6396 | 4,41402E-48 |
| ACE2 | TCAF1 | -0,6396 | 4,27796E-48 |
| ACE2 | SOX11 | -0,6398 | 4,04924E-48 |
| ACE2 | INTS4 | -0,6398 | 4,07366E-48 |
| ACE2 | GALK1 | -0,6398 | 3,99703E-48 |
| ACE2 | SGTA | -0,6398 | 3,90638E-48 |
| ACE2 | ARSB | -0,6399 | 3,81209E-48 |
| ACE2 | SNRPD2 | -0,6399 | 3,82901E-48 |
| ACE2 | RP11-171I2.1 | -0,64 | 3,66243E-48 |
| ACE2 | ZBTB49 | -0,64 | 3,62749E-48 |
| ACE2 | MMD | -0,64 | 3,68461E-48 |
| ACE2 | RP11-147L13.8 | -0,64 | 3,60395E-48 |
| ACE2 | PLK5 | -0,64 | 3,68839E-48 |
| ACE2 | AF001548.6 | -0,6401 | 3,43197E-48 |
| ACE2 | RP11-156L14.1 | -0,6401 | 3,42138E-48 |
| ACE2 | HNRNPA1 | -0,6402 | 3,3087E-48 |
| ACE2 | GOLGA6L5P | -0,6402 | 3,40846E-48 |
| ACE2 | ZNF431 | -0,6402 | 3,29132E-48 |
| ACE2 | MKNK1 | -0,6403 | 3,25594E-48 |
| ACE2 | CH17-472G23.4 | -0,6403 | 3,1318E-48 |
| ACE2 | SLC41A3 | -0,6403 | 3,22636E-48 |
| ACE2 | STOX2 | -0,6403 | 3,13165E-48 |
| ACE2 | USP39 | -0,6404 | 3,04469E-48 |
| ACE2 | RAB6C-AS1 | -0,6404 | 3,02063E-48 |
| ACE2 | FBLN2 | -0,6404 | 3,10521E-48 |
| ACE2 | GS1-358P8.4 | -0,6404 | 3,05608E-48 |
| ACE2 | RMST | -0,6405 | 2,95633E-48 |
| ACE2 | LINC01415 | -0,6405 | 2,98099E-48 |
| ACE2 | KRR1 | -0,6406 | 2,77926E-48 |
| ACE2 | AC079922.3 | -0,6407 | 2,7024E-48 |
| ACE2 | CLK4 | -0,6407 | 2,67015E-48 |
| ACE2 | BORCS7 | -0,6407 | 2,68617E-48 |
| ACE2 | KDM2B | -0,6407 | 2,69884E-48 |
| ACE2 | TEX30 | -0,6407 | 2,64437E-48 |
| ACE2 | RTFDC1 | -0,6407 | 2,70827E-48 |
| ACE2 | IPO11 | -0,6408 | 2,52484E-48 |
| ACE2 | PTPA | -0,6408 | 2,61186E-48 |
| ACE2 | RP11-441F2.5 | -0,6408 | 2,60168E-48 |
| ACE2 | BCAM | -0,6408 | 2,56921E-48 |
| ACE2 | RP11-307E17.8 | -0,6409 | 2,44321E-48 |
| ACE2 | SLC25A12 | -0,641 | 2,32565E-48 |
| ACE2 | RP11-630A13.4 | -0,641 | 2,36875E-48 |

| | | | |
|---|---|---|---|
| ACE2 | SPOCK3 | -0,641 | 2,32889E-48 |
| ACE2 | CTC-366B18.4 | -0,641 | 2,31412E-48 |
| ACE2 | USP28 | -0,641 | 2,36695E-48 |
| ACE2 | RAB11FIP5 | -0,6411 | 2,26534E-48 |
| ACE2 | WWC2-AS2 | -0,6411 | 2,26762E-48 |
| ACE2 | PAIP1 | -0,6411 | 2,20144E-48 |
| ACE2 | TEK | -0,6411 | 2,27347E-48 |
| ACE2 | PPFIA2 | -0,6411 | 2,29287E-48 |
| ACE2 | TNKS | -0,6412 | 2,18343E-48 |
| ACE2 | EMSY | -0,6412 | 2,17601E-48 |
| ACE2 | KSR1 | -0,6412 | 2,11819E-48 |
| ACE2 | DHX58 | -0,6412 | 2,10975E-48 |
| ACE2 | SEMA4F | -0,6413 | 2,02309E-48 |
| ACE2 | MYLK3 | -0,6413 | 2,02615E-48 |
| ACE2 | DRG2 | -0,6413 | 2,04197E-48 |
| ACE2 | MOSPD3 | -0,6414 | 1,94112E-48 |
| ACE2 | ADAM21 | -0,6414 | 2,00936E-48 |
| ACE2 | RBMXL1 | -0,6415 | 1,89118E-48 |
| ACE2 | NRBP1 | -0,6415 | 1,85873E-48 |
| ACE2 | KANK1 | -0,6415 | 1,85844E-48 |
| ACE2 | ZNF143 | -0,6415 | 1,91881E-48 |
| ACE2 | TMEM233 | -0,6415 | 1,90398E-48 |
| ACE2 | CTD-2325P2.4 | -0,6416 | 1,77004E-48 |
| ACE2 | CDC16 | -0,6417 | 1,69988E-48 |
| ACE2 | SKP1 | -0,6418 | 1,62921E-48 |
| ACE2 | ABCC1 | -0,6418 | 1,648E-48 |
| ACE2 | RP5-1039K5.16 | -0,6418 | 1,66431E-48 |
| ACE2 | NEK7 | -0,6419 | 1,55442E-48 |
| ACE2 | SYN2 | -0,6419 | 1,58034E-48 |
| ACE2 | DENND3 | -0,6419 | 1,5989E-48 |
| ACE2 | BCAN | -0,642 | 1,48585E-48 |
| ACE2 | RP11-1024P17.1 | -0,642 | 1,50298E-48 |
| ACE2 | PCDHB8 | -0,642 | 1,50467E-48 |
| ACE2 | UNC45A | -0,642 | 1,52351E-48 |
| ACE2 | ZNF547 | -0,6421 | 1,43321E-48 |
| ACE2 | KPNA1 | -0,6422 | 1,37988E-48 |
| ACE2 | PPP1R2 | -0,6422 | 1,405E-48 |
| ACE2 | TMEM151B | -0,6422 | 1,39011E-48 |
| ACE2 | SNX3 | -0,6422 | 1,38318E-48 |
| ACE2 | CELF1 | -0,6422 | 1,36103E-48 |
| ACE2 | IRS1 | -0,6423 | 1,32353E-48 |
| ACE2 | GEMIN5 | -0,6423 | 1,29381E-48 |
| ACE2 | RP11-51B23.3 | -0,6423 | 1,34296E-48 |
| ACE2 | AP001471.1 | -0,6423 | 1,33668E-48 |
| ACE2 | FGD5 | -0,6424 | 1,27598E-48 |
| ACE2 | CAMK2N2 | -0,6424 | 1,27636E-48 |
| ACE2 | LSM11 | -0,6424 | 1,27876E-48 |
| ACE2 | IGF2BP3 | -0,6424 | 1,27898E-48 |
| ACE2 | CTC-487M23.5 | -0,6425 | 1,18601E-48 |
| ACE2 | MEA1 | -0,6425 | 1,20274E-48 |
| ACE2 | SLC25A21-AS1 | -0,6425 | 1,23295E-48 |
| ACE2 | RP11-700H6.4 | -0,6425 | 1,19274E-48 |
| ACE2 | CCDC57 | -0,6425 | 1,2111E-48 |
| ACE2 | TFPT | -0,6425 | 1,19294E-48 |

| | | | |
|---|---|---|---|
| ACE2 | RP11-490M8.1 | -0,6426 | 1,1721E-48 |
| ACE2 | ANKAR | -0,6426 | 1,17918E-48 |
| ACE2 | SLMAP | -0,6426 | 1,17901E-48 |
| ACE2 | COPS7A | -0,6426 | 1,18388E-48 |
| ACE2 | ADCY4 | -0,6426 | 1,14873E-48 |
| ACE2 | ILF3-AS1 | -0,6426 | 1,14966E-48 |
| ACE2 | TULP2 | -0,6426 | 1,14691E-48 |
| ACE2 | UPB1 | -0,6426 | 1,14167E-48 |
| ACE2 | ATP6V0CP3 | -0,6427 | 1,1302E-48 |
| ACE2 | SNX31 | -0,6427 | 1,08357E-48 |
| ACE2 | NDST2 | -0,6427 | 1,09551E-48 |
| ACE2 | ZNF407 | -0,6427 | 1,09092E-48 |
| ACE2 | KCNIP4 | -0,6428 | 1,05233E-48 |
| ACE2 | RFESD | -0,6428 | 1,04798E-48 |
| ACE2 | SCAND2P | -0,6428 | 1,05853E-48 |
| ACE2 | PLCG1 | -0,6428 | 1,06525E-48 |
| ACE2 | HIST2H2BA | -0,6429 | 1,02877E-48 |
| ACE2 | RP4-561L24.3 | -0,643 | 9,5225E-49 |
| ACE2 | ITGB5 | -0,643 | 9,55523E-49 |
| ACE2 | ZSCAN12 | -0,643 | 9,5032E-49 |
| ACE2 | PTGFRN | -0,6431 | 9,23791E-49 |
| ACE2 | MRPL9 | -0,6431 | 9,43725E-49 |
| ACE2 | EFEMP1 | -0,6431 | 9,45795E-49 |
| ACE2 | TARID | -0,6431 | 9,09903E-49 |
| ACE2 | CEP95 | -0,6431 | 9,38517E-49 |
| ACE2 | MPP6 | -0,6432 | 8,83489E-49 |
| ACE2 | POLR1D | -0,6432 | 8,7309E-49 |
| ACE2 | DSCAM | -0,6432 | 8,95816E-49 |
| ACE2 | KCND1 | -0,6432 | 8,97957E-49 |
| ACE2 | RP1-278D1.2 | -0,6432 | 8,78534E-49 |
| ACE2 | RGS7 | -0,6433 | 8,33856E-49 |
| ACE2 | PWAR5 | -0,6433 | 8,53155E-49 |
| ACE2 | CCR10 | -0,6433 | 8,58894E-49 |
| ACE2 | RP11-225H22.4 | -0,6434 | 8,0179E-49 |
| ACE2 | SMARCE1 | -0,6434 | 7,94774E-49 |
| ACE2 | SSR4P1 | -0,6434 | 7,95266E-49 |
| ACE2 | S1PR1 | -0,6435 | 7,64322E-49 |
| ACE2 | FSCN1 | -0,6435 | 7,90749E-49 |
| ACE2 | C8orf58 | -0,6435 | 7,8226E-49 |
| ACE2 | ABC7-42404400C24.1 | -0,6435 | 7,7226E-49 |
| ACE2 | MPRIP | -0,6435 | 7,81667E-49 |
| ACE2 | TPGS2 | -0,6435 | 7,62536E-49 |
| ACE2 | TRAPPC2 | -0,6435 | 7,87638E-49 |
| ACE2 | ARPP21 | -0,6436 | 7,27702E-49 |
| ACE2 | SNRK | -0,6436 | 7,29955E-49 |
| ACE2 | RP11-671M22.4 | -0,6436 | 7,40018E-49 |
| ACE2 | ARFGAP1 | -0,6436 | 7,29556E-49 |
| ACE2 | ZNF365 | -0,6437 | 7,24169E-49 |
| ACE2 | CORO6 | -0,6437 | 7,24609E-49 |
| ACE2 | ZNF14 | -0,6437 | 7,02135E-49 |
| ACE2 | FTCD | -0,6437 | 7,13749E-49 |
| ACE2 | CCK | -0,6438 | 6,65186E-49 |
| ACE2 | TCF12 | -0,6438 | 6,76301E-49 |
| ACE2 | RP11-446H18.6 | -0,6439 | 6,4798E-49 |

| | | | |
|---|---|---|---|
| ACE2 | EFCC1 | -0,6439 | 6,59371E-49 |
| ACE2 | PAQR3 | -0,6439 | 6,40668E-49 |
| ACE2 | CPSF7 | -0,6439 | 6,48876E-49 |
| ACE2 | PTPN11 | -0,6439 | 6,52131E-49 |
| ACE2 | PRPF39 | -0,6439 | 6,43107E-49 |
| ACE2 | AVEN | -0,6439 | 6,39272E-49 |
| ACE2 | DUSP19 | -0,644 | 6,15003E-49 |
| ACE2 | ZNF142 | -0,644 | 6,25553E-49 |
| ACE2 | MAK16 | -0,644 | 6,10744E-49 |
| ACE2 | AC099850.1 | -0,644 | 6,08076E-49 |
| ACE2 | FMNL2 | -0,6441 | 5,87525E-49 |
| ACE2 | PCDHGA8 | -0,6441 | 5,97252E-49 |
| ACE2 | SMAD4 | -0,6441 | 6,05072E-49 |
| ACE2 | AMOTL2 | -0,6442 | 5,65699E-49 |
| ACE2 | MAP6D1 | -0,6442 | 5,68264E-49 |
| ACE2 | PDZRN4 | -0,6442 | 5,74262E-49 |
| ACE2 | CTD-2095E4.5 | -0,6442 | 5,75748E-49 |
| ACE2 | SP2-AS1 | -0,6442 | 5,79129E-49 |
| ACE2 | GRINA | -0,6443 | 5,4501E-49 |
| ACE2 | RABL2B | -0,6443 | 5,31933E-49 |
| ACE2 | DSE | -0,6444 | 5,12363E-49 |
| ACE2 | TIAL1 | -0,6444 | 5,09703E-49 |
| ACE2 | RP13-516M14.1 | -0,6444 | 5,09534E-49 |
| ACE2 | RPL18A | -0,6444 | 5,24449E-49 |
| ACE2 | ZNF2 | -0,6445 | 5,01328E-49 |
| ACE2 | DCUN1D5 | -0,6445 | 5,02177E-49 |
| ACE2 | CTD-2124B8.2 | -0,6446 | 4,68326E-49 |
| ACE2 | CTB-109A12.1 | -0,6446 | 4,82563E-49 |
| ACE2 | RP11-481A20.10 | -0,6446 | 4,84605E-49 |
| ACE2 | LGI1 | -0,6446 | 4,66685E-49 |
| ACE2 | KERA | -0,6446 | 4,80621E-49 |
| ACE2 | ADAMTS3 | -0,6447 | 4,54121E-49 |
| ACE2 | DNAJB2 | -0,6448 | 4,40788E-49 |
| ACE2 | RP11-346E8.1 | -0,6448 | 4,42288E-49 |
| ACE2 | PAPD7 | -0,6449 | 4,07315E-49 |
| ACE2 | CBL | -0,6449 | 4,06269E-49 |
| ACE2 | C1orf109 | -0,645 | 3,95448E-49 |
| ACE2 | ZDHHC4 | -0,645 | 4,0539E-49 |
| ACE2 | UQCRB | -0,645 | 4,00933E-49 |
| ACE2 | RPL8 | -0,645 | 4,02638E-49 |
| ACE2 | CTC1 | -0,645 | 4,00738E-49 |
| ACE2 | NATD1 | -0,645 | 3,93902E-49 |
| ACE2 | INO80B | -0,6451 | 3,86378E-49 |
| ACE2 | ZNF502 | -0,6451 | 3,77674E-49 |
| ACE2 | RP11-70P17.1 | -0,6452 | 3,66394E-49 |
| ACE2 | RP11-57H12.3 | -0,6452 | 3,64628E-49 |
| ACE2 | DMTF1 | -0,6452 | 3,57138E-49 |
| ACE2 | ZKSCAN5 | -0,6452 | 3,68543E-49 |
| ACE2 | SYNDIG1L | -0,6452 | 3,6081E-49 |
| ACE2 | KAT7 | -0,6452 | 3,58376E-49 |
| ACE2 | ZDHHC8 | -0,6452 | 3,61092E-49 |
| ACE2 | ERF | -0,6453 | 3,47951E-49 |
| ACE2 | CRYBB2P1 | -0,6453 | 3,49431E-49 |
| ACE2 | AGO4 | -0,6454 | 3,38099E-49 |

| | | | |
|---|---|---|---|
| ACE2 | POMGNT1 | -0,6454 | 3,25765E-49 |
| ACE2 | EMX2OS | -0,6454 | 3,36785E-49 |
| ACE2 | CALCB | -0,6454 | 3,31926E-49 |
| ACE2 | RP11-640N20.4 | -0,6454 | 3,35701E-49 |
| ACE2 | OR8T1P | -0,6455 | 3,12674E-49 |
| ACE2 | SETD1A | -0,6455 | 3,17675E-49 |
| ACE2 | LINC01563 | -0,6455 | 3,21426E-49 |
| ACE2 | RP11-159D12.8 | -0,6455 | 3,17888E-49 |
| ACE2 | KLHL26 | -0,6455 | 3,20256E-49 |
| ACE2 | APCDD1L-AS1 | -0,6455 | 3,11104E-49 |
| ACE2 | CD200 | -0,6456 | 3,02744E-49 |
| ACE2 | INTS6 | -0,6456 | 2,99915E-49 |
| ACE2 | TRAPPC3 | -0,6457 | 2,95554E-49 |
| ACE2 | RP5-1139B12.2 | -0,6457 | 2,93264E-49 |
| ACE2 | PCYOX1L | -0,6457 | 2,87615E-49 |
| ACE2 | ACTL6B | -0,6457 | 2,94171E-49 |
| ACE2 | CUL1 | -0,6457 | 2,92568E-49 |
| ACE2 | DENND4B | -0,6458 | 2,80468E-49 |
| ACE2 | EEFSEC | -0,6458 | 2,74018E-49 |
| ACE2 | PIK3CA | -0,6458 | 2,7251E-49 |
| ACE2 | VPS13C | -0,6458 | 2,78856E-49 |
| ACE2 | DOCK7 | -0,6459 | 2,63971E-49 |
| ACE2 | HSPA1A | -0,6459 | 2,67297E-49 |
| ACE2 | APOPT1 | -0,6459 | 2,61016E-49 |
| ACE2 | MTMR12 | -0,646 | 2,57643E-49 |
| ACE2 | ATRNL1 | -0,646 | 2,57303E-49 |
| ACE2 | C2CD4C | -0,646 | 2,51997E-49 |
| ACE2 | HAR1A | -0,646 | 2,5122E-49 |
| ACE2 | RP11-281O15.4 | -0,6461 | 2,38637E-49 |
| ACE2 | KB-1507C5.4 | -0,6461 | 2,40682E-49 |
| ACE2 | RP11-1391J7.1 | -0,6461 | 2,42369E-49 |
| ACE2 | GNPAT | -0,6462 | 2,27498E-49 |
| ACE2 | TRDMT1 | -0,6462 | 2,28574E-49 |
| ACE2 | SOCS2-AS1 | -0,6462 | 2,28247E-49 |
| ACE2 | RP11-318A15.8 | -0,6462 | 2,34463E-49 |
| ACE2 | MRPL40 | -0,6462 | 2,30868E-49 |
| ACE2 | DNM3OS | -0,6463 | 2,22456E-49 |
| ACE2 | RP11-334J6.6 | -0,6463 | 2,23673E-49 |
| ACE2 | CIRBP | -0,6463 | 2,16869E-49 |
| ACE2 | RP11-334A14.8 | -0,6464 | 2,12864E-49 |
| ACE2 | RELN | -0,6464 | 2,13664E-49 |
| ACE2 | RP11-267M23.1 | -0,6464 | 2,16387E-49 |
| ACE2 | TINF2 | -0,6464 | 2,14595E-49 |
| ACE2 | CTC-490E21.11 | -0,6464 | 2,12722E-49 |
| ACE2 | BCAP29 | -0,6465 | 2,00604E-49 |
| ACE2 | RNF220 | -0,6466 | 1,96781E-49 |
| ACE2 | BTRC | -0,6466 | 1,89911E-49 |
| ACE2 | MGEA5 | -0,6466 | 1,92343E-49 |
| ACE2 | DUSP3 | -0,6466 | 1,92574E-49 |
| ACE2 | LINC01534 | -0,6466 | 1,96186E-49 |
| ACE2 | SSX2IP | -0,6467 | 1,84918E-49 |
| ACE2 | STC1 | -0,6467 | 1,82589E-49 |
| ACE2 | BRD3 | -0,6467 | 1,85661E-49 |
| ACE2 | SRSF1 | -0,6467 | 1,86871E-49 |

| | | | |
|---|---|---|---|
| ACE2 | ATOX1 | -0,6468 | 1,75248E-49 |
| ACE2 | HEY1 | -0,6468 | 1,78231E-49 |
| ACE2 | NCOR2 | -0,6468 | 1,76454E-49 |
| ACE2 | BLMH | -0,6468 | 1,77564E-49 |
| ACE2 | AHCTF1 | -0,6469 | 1,68007E-49 |
| ACE2 | BCO2 | -0,6469 | 1,71666E-49 |
| ACE2 | CDR1 | -0,6469 | 1,67293E-49 |
| ACE2 | HSPG2 | -0,647 | 1,65172E-49 |
| ACE2 | ZRANB1 | -0,647 | 1,58154E-49 |
| ACE2 | SYT5 | -0,647 | 1,59259E-49 |
| ACE2 | LINC01715 | -0,6471 | 1,55754E-49 |
| ACE2 | RHOBTB3 | -0,6471 | 1,57229E-49 |
| ACE2 | DRD4 | -0,6471 | 1,57592E-49 |
| ACE2 | OGFOD1 | -0,6471 | 1,56669E-49 |
| ACE2 | WRB | -0,6471 | 1,52799E-49 |
| ACE2 | CTPS1 | -0,6472 | 1,47321E-49 |
| ACE2 | NAGK | -0,6472 | 1,48991E-49 |
| ACE2 | PHIP | -0,6472 | 1,45905E-49 |
| ACE2 | KL | -0,6472 | 1,45612E-49 |
| ACE2 | RTF1 | -0,6472 | 1,48825E-49 |
| ACE2 | DIS3L | -0,6472 | 1,44848E-49 |
| ACE2 | DLGAP4-AS1 | -0,6472 | 1,46732E-49 |
| ACE2 | SETMAR | -0,6473 | 1,4223E-49 |
| ACE2 | RBMS3-AS3 | -0,6473 | 1,44244E-49 |
| ACE2 | CYTL1 | -0,6473 | 1,39518E-49 |
| ACE2 | APOBEC2 | -0,6474 | 1,32042E-49 |
| ACE2 | HTR3A | -0,6474 | 1,35621E-49 |
| ACE2 | STOML1 | -0,6474 | 1,37105E-49 |
| ACE2 | MASP1 | -0,6475 | 1,29141E-49 |
| ACE2 | RP11-3B7.1 | -0,6476 | 1,26037E-49 |
| ACE2 | GLG1 | -0,6476 | 1,24501E-49 |
| ACE2 | UBE2O | -0,6476 | 1,25102E-49 |
| ACE2 | KIF17 | -0,6477 | 1,18666E-49 |
| ACE2 | RP11-553A21.3 | -0,6477 | 1,19701E-49 |
| ACE2 | ANGEL1 | -0,6477 | 1,15966E-49 |
| ACE2 | NSMCE3 | -0,6477 | 1,19404E-49 |
| ACE2 | NPR1 | -0,6478 | 1,11095E-49 |
| ACE2 | LYPLAL1-AS1 | -0,6478 | 1,14378E-49 |
| ACE2 | EHD3 | -0,6478 | 1,15152E-49 |
| ACE2 | LINC01990 | -0,6478 | 1,13047E-49 |
| ACE2 | CTC-325H20.8 | -0,6478 | 1,13564E-49 |
| ACE2 | CRTC2 | -0,6479 | 1,08441E-49 |
| ACE2 | AC005682.5 | -0,6479 | 1,09301E-49 |
| ACE2 | AMIGO2 | -0,6479 | 1,05671E-49 |
| ACE2 | ZNF140 | -0,6479 | 1,08385E-49 |
| ACE2 | GPX4 | -0,6479 | 1,07588E-49 |
| ACE2 | KLF2 | -0,6479 | 1,09065E-49 |
| ACE2 | NSMAF | -0,648 | 1,0125E-49 |
| ACE2 | BMS1P3 | -0,648 | 1,05085E-49 |
| ACE2 | LINC01138 | -0,6481 | 9,63026E-50 |
| ACE2 | RP11-203B9.4 | -0,6481 | 9,85184E-50 |
| ACE2 | WAC | -0,6481 | 1,00287E-49 |
| ACE2 | LINC00839 | -0,6481 | 9,81091E-50 |
| ACE2 | RP11-182L21.6 | -0,6481 | 9,67586E-50 |

| | | | |
|---|---|---|---|
| ACE2 | CCBE1 | -0,6481 | 9,9096E-50 |
| ACE2 | CYR61 | -0,6482 | 9,41322E-50 |
| ACE2 | CCDC107 | -0,6482 | 9,43206E-50 |
| ACE2 | NPRL3 | -0,6482 | 9,40368E-50 |
| ACE2 | OPRD1 | -0,6483 | 8,84522E-50 |
| ACE2 | ERICH6-AS1 | -0,6483 | 9,00017E-50 |
| ACE2 | ZDHHC11B | -0,6483 | 8,88938E-50 |
| ACE2 | GNG10 | -0,6483 | 8,83499E-50 |
| ACE2 | GPC5 | -0,6483 | 9,04011E-50 |
| ACE2 | ZNF106 | -0,6483 | 9,17478E-50 |
| ACE2 | PRUNE2 | -0,6484 | 8,65647E-50 |
| ACE2 | FXR2 | -0,6484 | 8,71802E-50 |
| ACE2 | LINC01694 | -0,6484 | 8,52557E-50 |
| ACE2 | CHRNB2 | -0,6485 | 8,0369E-50 |
| ACE2 | GPNMB | -0,6485 | 8,16211E-50 |
| ACE2 | TOM1L2 | -0,6485 | 8,07891E-50 |
| ACE2 | CYP2T1P | -0,6485 | 8,0359E-50 |
| ACE2 | MRPS6 | -0,6485 | 8,33615E-50 |
| ACE2 | C10orf10 | -0,6486 | 7,6753E-50 |
| ACE2 | NRIP2 | -0,6486 | 7,99134E-50 |
| ACE2 | EXOSC7 | -0,6487 | 7,56345E-50 |
| ACE2 | TLL1 | -0,6487 | 7,438E-50 |
| ACE2 | RP11-265E18.1 | -0,6487 | 7,594E-50 |
| ACE2 | NACC2 | -0,6487 | 7,65667E-50 |
| ACE2 | PI4K2A | -0,6487 | 7,36224E-50 |
| ACE2 | FAT3 | -0,6487 | 7,50764E-50 |
| ACE2 | FLYWCH1 | -0,6487 | 7,55935E-50 |
| ACE2 | FBXO7 | -0,6487 | 7,3397E-50 |
| ACE2 | RBBP7 | -0,6487 | 7,3687E-50 |
| ACE2 | POLR2G | -0,6488 | 7,27198E-50 |
| ACE2 | ALKBH8 | -0,6488 | 7,22024E-50 |
| ACE2 | LRP1 | -0,6488 | 7,32253E-50 |
| ACE2 | NOTCH3 | -0,6488 | 7,26362E-50 |
| ACE2 | NSFL1C | -0,6488 | 7,00762E-50 |
| ACE2 | RFPL1S | -0,6488 | 7,05795E-50 |
| ACE2 | RP4-758J18.7 | -0,6489 | 6,77225E-50 |
| ACE2 | MEMO1 | -0,6489 | 6,98776E-50 |
| ACE2 | RPL37A | -0,6489 | 6,90459E-50 |
| ACE2 | SLC35B4 | -0,6489 | 6,90746E-50 |
| ACE2 | KCNH3 | -0,6489 | 6,71865E-50 |
| ACE2 | CSRP2 | -0,649 | 6,44082E-50 |
| ACE2 | FRG1 | -0,6491 | 6,21813E-50 |
| ACE2 | TENM2 | -0,6491 | 6,13783E-50 |
| ACE2 | LATS1 | -0,6491 | 6,24515E-50 |
| ACE2 | DCUN1D2 | -0,6491 | 6,35851E-50 |
| ACE2 | DYNLRB2 | -0,6491 | 6,32796E-50 |
| ACE2 | RP3-422G23.4 | -0,6492 | 5,85392E-50 |
| ACE2 | SMARCA2 | -0,6492 | 6,04956E-50 |
| ACE2 | PDCD6IPP1 | -0,6492 | 5,8932E-50 |
| ACE2 | KLHL4 | -0,6492 | 5,95078E-50 |
| ACE2 | RP11-49C9.2 | -0,6492 | 6,0364E-50 |
| ACE2 | PYCR2 | -0,6493 | 5,63836E-50 |
| ACE2 | HCG11 | -0,6493 | 5,75584E-50 |
| ACE2 | FZD1 | -0,6493 | 5,78894E-50 |

| | | | |
|---|---|---|---|
| ACE2 | ATP6V1F | -0,6493 | 5,63028E-50 |
| ACE2 | VASN | -0,6493 | 5,81832E-50 |
| ACE2 | C3orf80 | -0,6494 | 5,48393E-50 |
| ACE2 | PRMT1 | -0,6494 | 5,56859E-50 |
| ACE2 | LBH | -0,6495 | 5,22849E-50 |
| ACE2 | LRRC8A | -0,6495 | 5,23625E-50 |
| ACE2 | PLBD2 | -0,6495 | 5,29075E-50 |
| ACE2 | LAMP5 | -0,6495 | 5,11522E-50 |
| ACE2 | OLFM3 | -0,6496 | 4,9025E-50 |
| ACE2 | MAEL | -0,6496 | 5,01655E-50 |
| ACE2 | MKRN2 | -0,6496 | 5,00057E-50 |
| ACE2 | MED12L | -0,6496 | 5,00107E-50 |
| ACE2 | BRD2 | -0,6496 | 4,96131E-50 |
| ACE2 | WDR73 | -0,6496 | 5,06093E-50 |
| ACE2 | PTCHD1 | -0,6496 | 5,03748E-50 |
| ACE2 | CHL1-AS1 | -0,6497 | 4,82541E-50 |
| ACE2 | INHBA | -0,6497 | 4,83363E-50 |
| ACE2 | CDC34 | -0,6497 | 4,69324E-50 |
| ACE2 | BNIP3P30 | -0,6497 | 4,69639E-50 |
| ACE2 | ENOX2 | -0,6497 | 4,78326E-50 |
| ACE2 | FAM46B | -0,6498 | 4,63108E-50 |
| ACE2 | CEP19 | -0,6498 | 4,61895E-50 |
| ACE2 | SORBS3 | -0,6498 | 4,57816E-50 |
| ACE2 | NOC3L | -0,6498 | 4,47739E-50 |
| ACE2 | TRIT1 | -0,6499 | 4,30662E-50 |
| ACE2 | SLC27A3 | -0,6499 | 4,38325E-50 |
| ACE2 | FAM131A | -0,6499 | 4,26346E-50 |
| ACE2 | SVEP1 | -0,6499 | 4,43043E-50 |
| ACE2 | RP11-359E3.4 | -0,6499 | 4,32658E-50 |
| ACE2 | BIVM | -0,6499 | 4,26671E-50 |
| ACE2 | RAB3A | -0,6499 | 4,27442E-50 |
| ACE2 | COMT | -0,6499 | 4,32053E-50 |
| ACE2 | RP11-654E17.2 | -0,65 | 4,22368E-50 |
| ACE2 | CLIC2 | -0,65 | 4,235E-50 |
| ACE2 | RSPO1 | -0,6501 | 3,93218E-50 |
| ACE2 | LINC01018 | -0,6501 | 3,97169E-50 |
| ACE2 | BCORL1 | -0,6501 | 3,9557E-50 |
| ACE2 | C4orf47 | -0,6502 | 3,85867E-50 |
| ACE2 | FBXO21 | -0,6502 | 3,7274E-50 |
| ACE2 | CENPB | -0,6502 | 3,77046E-50 |
| ACE2 | VEGFD | -0,6502 | 3,75176E-50 |
| ACE2 | GRIA3 | -0,6502 | 3,7623E-50 |
| ACE2 | AAMDC | -0,6503 | 3,6037E-50 |
| ACE2 | ESAM | -0,6503 | 3,54799E-50 |
| ACE2 | BTG3 | -0,6503 | 3,56827E-50 |
| ACE2 | HOOK3 | -0,6504 | 3,38279E-50 |
| ACE2 | RP11-498E2.9 | -0,6504 | 3,45985E-50 |
| ACE2 | APPBP2 | -0,6504 | 3,41727E-50 |
| ACE2 | LL0XNC01-240C2.1 | -0,6504 | 3,37665E-50 |
| ACE2 | RP11-326G21.1 | -0,6505 | 3,31238E-50 |
| ACE2 | EIF3J-AS1 | -0,6505 | 3,31007E-50 |
| ACE2 | TRMT2A | -0,6505 | 3,31352E-50 |
| ACE2 | TMEM50B | -0,6506 | 3,1193E-50 |
| ACE2 | RP3-467L1.4 | -0,6507 | 3,03139E-50 |

| | | | |
|---|---|---|---|
| ACE2 | RP11-403I13.8 | -0,6507 | 3,05039E-50 |
| ACE2 | AARS | -0,6507 | 2,96843E-50 |
| ACE2 | RGS4 | -0,6508 | 2,86658E-50 |
| ACE2 | KCNQ3 | -0,6508 | 2,90086E-50 |
| ACE2 | PTK2 | -0,6508 | 2,89991E-50 |
| ACE2 | TTLL5 | -0,6508 | 2,84801E-50 |
| ACE2 | ANKLE1 | -0,6508 | 2,832E-50 |
| ACE2 | RBBP9 | -0,6508 | 2,85386E-50 |
| ACE2 | SUPT7L | -0,6509 | 2,73217E-50 |
| ACE2 | HEIH | -0,6509 | 2,81028E-50 |
| ACE2 | RP11-428J1.5 | -0,6509 | 2,77927E-50 |
| ACE2 | AGBL3 | -0,6509 | 2,73834E-50 |
| ACE2 | CLIP1 | -0,6509 | 2,70255E-50 |
| ACE2 | TAL1 | -0,651 | 2,66311E-50 |
| ACE2 | ZBTB46 | -0,651 | 2,63208E-50 |
| ACE2 | RBP7 | -0,6511 | 2,55811E-50 |
| ACE2 | CARS2 | -0,6511 | 2,5162E-50 |
| ACE2 | DCAF8 | -0,6512 | 2,40129E-50 |
| ACE2 | PROS1 | -0,6512 | 2,38473E-50 |
| ACE2 | ZNF391 | -0,6513 | 2,33497E-50 |
| ACE2 | ELOVL5 | -0,6513 | 2,30232E-50 |
| ACE2 | ZACN | -0,6513 | 2,32815E-50 |
| ACE2 | YY2 | -0,6513 | 2,30188E-50 |
| ACE2 | CADM3-AS1 | -0,6514 | 2,17775E-50 |
| ACE2 | YPEL5 | -0,6514 | 2,14742E-50 |
| ACE2 | PODXL | -0,6514 | 2,18507E-50 |
| ACE2 | CHST8 | -0,6514 | 2,15049E-50 |
| ACE2 | CCDC117 | -0,6514 | 2,14696E-50 |
| ACE2 | EPHA6 | -0,6515 | 2,04593E-50 |
| ACE2 | UBXN7 | -0,6515 | 2,11836E-50 |
| ACE2 | LHPP | -0,6515 | 2,0948E-50 |
| ACE2 | MALAT1 | -0,6515 | 2,13507E-50 |
| ACE2 | NTN5 | -0,6515 | 2,11205E-50 |
| ACE2 | RP11-68I18.10 | -0,6516 | 1,96177E-50 |
| ACE2 | CSNK1A1 | -0,6516 | 2,02151E-50 |
| ACE2 | MAMDC2 | -0,6516 | 1,97661E-50 |
| ACE2 | COMMD3 | -0,6516 | 2,02705E-50 |
| ACE2 | AC108448.2 | -0,6516 | 2,00849E-50 |
| ACE2 | RPAP3 | -0,6516 | 1,94846E-50 |
| ACE2 | CSPG5 | -0,6517 | 1,86156E-50 |
| ACE2 | TULP1 | -0,6517 | 1,90723E-50 |
| ACE2 | EIF4B | -0,6517 | 1,94278E-50 |
| ACE2 | EEF2 | -0,6517 | 1,87402E-50 |
| ACE2 | UBE2M | -0,6517 | 1,91595E-50 |
| ACE2 | EIF1B-AS1 | -0,6518 | 1,85365E-50 |
| ACE2 | OSTM1 | -0,6518 | 1,80994E-50 |
| ACE2 | RP11-95O2.5 | -0,6518 | 1,81176E-50 |
| ACE2 | JMY | -0,6519 | 1,74496E-50 |
| ACE2 | RP11-350N15.3 | -0,6519 | 1,75795E-50 |
| ACE2 | NDUFAF8 | -0,6519 | 1,72822E-50 |
| ACE2 | AGPAT1 | -0,652 | 1,64175E-50 |
| ACE2 | NAV2 | -0,652 | 1,64017E-50 |
| ACE2 | STX5 | -0,652 | 1,68291E-50 |
| ACE2 | NMT1 | -0,652 | 1,65791E-50 |

| | | | |
|---|---|---|---|
| ACE2 | BCL9 | -0,6521 | 1,60034E-50 |
| ACE2 | FST | -0,6521 | 1,60781E-50 |
| ACE2 | AC025335.1 | -0,6521 | 1,60142E-50 |
| ACE2 | THSD7B | -0,6522 | 1,52205E-50 |
| ACE2 | FBXW8 | -0,6522 | 1,48586E-50 |
| ACE2 | ACKR3 | -0,6523 | 1,44622E-50 |
| ACE2 | DAB1 | -0,6524 | 1,40965E-50 |
| ACE2 | CTSO | -0,6524 | 1,37128E-50 |
| ACE2 | THAP5 | -0,6524 | 1,39823E-50 |
| ACE2 | TPBGL | -0,6524 | 1,40492E-50 |
| ACE2 | UBAP1 | -0,6525 | 1,32632E-50 |
| ACE2 | IPO7 | -0,6525 | 1,32328E-50 |
| ACE2 | ATP10A | -0,6526 | 1,26247E-50 |
| ACE2 | KCNMB3 | -0,6527 | 1,18925E-50 |
| ACE2 | LINC00377 | -0,6527 | 1,20485E-50 |
| ACE2 | TMEM39B | -0,6528 | 1,16265E-50 |
| ACE2 | PLXND1 | -0,6528 | 1,15402E-50 |
| ACE2 | NOL8 | -0,6528 | 1,16965E-50 |
| ACE2 | MED27 | -0,6528 | 1,1279E-50 |
| ACE2 | DNM1L | -0,6528 | 1,12902E-50 |
| ACE2 | MCPH1 | -0,6529 | 1,09613E-50 |
| ACE2 | VASH1 | -0,6529 | 1,11334E-50 |
| ACE2 | ISY1 | -0,653 | 1,04601E-50 |
| ACE2 | HAP1 | -0,653 | 1,06926E-50 |
| ACE2 | ZKSCAN8 | -0,6531 | 1,02147E-50 |
| ACE2 | ZNF484 | -0,6531 | 9,82905E-51 |
| ACE2 | RAMP2 | -0,6531 | 1,01094E-50 |
| ACE2 | CTD-3138B18.5 | -0,6531 | 1,01681E-50 |
| ACE2 | POLR3F | -0,6531 | 1,01509E-50 |
| ACE2 | MB21D2 | -0,6532 | 9,58915E-51 |
| ACE2 | EYA4 | -0,6532 | 9,76632E-51 |
| ACE2 | DHX40 | -0,6532 | 9,5375E-51 |
| ACE2 | METTL3 | -0,6533 | 9,14549E-51 |
| ACE2 | ZNF70 | -0,6533 | 8,94773E-51 |
| ACE2 | EXOSC1 | -0,6534 | 8,88417E-51 |
| ACE2 | FOLH1 | -0,6534 | 8,60686E-51 |
| ACE2 | NUTF2 | -0,6534 | 8,77039E-51 |
| ACE2 | ABHD17AP6 | -0,6534 | 8,75698E-51 |
| ACE2 | BABAM1 | -0,6534 | 8,69553E-51 |
| ACE2 | bP-2189O9.2 | -0,6534 | 8,51836E-51 |
| ACE2 | OPHN1 | -0,6534 | 8,65469E-51 |
| ACE2 | TRAPPC2L | -0,6535 | 8,44567E-51 |
| ACE2 | LL22NC03-80A10.6 | -0,6535 | 8,243E-51 |
| ACE2 | RPL11 | -0,6536 | 7,82223E-51 |
| ACE2 | RP11-124N19.4 | -0,6536 | 7,91677E-51 |
| ACE2 | MOV10L1 | -0,6536 | 8,03384E-51 |
| ACE2 | TMED8 | -0,6537 | 7,58694E-51 |
| ACE2 | CCP110 | -0,6537 | 7,72711E-51 |
| ACE2 | MIR637 | -0,6537 | 7,54883E-51 |
| ACE2 | ZNF728 | -0,6537 | 7,71994E-51 |
| ACE2 | RPAP2 | -0,6538 | 7,15872E-51 |
| ACE2 | THADA | -0,6538 | 7,23092E-51 |
| ACE2 | U2SURP | -0,6538 | 7,27387E-51 |
| ACE2 | CUTC | -0,6538 | 7,26518E-51 |

| | | | |
|---|---|---|---|
| ACE2 | SWAP70 | -0,6538 | 7,29049E-51 |
| ACE2 | KCNK7 | -0,6538 | 7,178E-51 |
| ACE2 | SRRM4 | -0,6538 | 7,19469E-51 |
| ACE2 | MIF4GD | -0,6538 | 7,34263E-51 |
| ACE2 | ZNF649 | -0,6538 | 7,21768E-51 |
| ACE2 | CECR7 | -0,6538 | 7,12408E-51 |
| ACE2 | TOP2B | -0,6539 | 7,04436E-51 |
| ACE2 | AZIN1-AS1 | -0,6539 | 6,9925E-51 |
| ACE2 | RP11-266K4.13 | -0,6539 | 7,02711E-51 |
| ACE2 | DDX24 | -0,6539 | 6,97302E-51 |
| ACE2 | C1GALT1C1L | -0,654 | 6,62729E-51 |
| ACE2 | ATRIP | -0,654 | 6,70716E-51 |
| ACE2 | AKR1E2 | -0,654 | 6,54119E-51 |
| ACE2 | TMEM255B | -0,654 | 6,58998E-51 |
| ACE2 | KIAA0430 | -0,654 | 6,60441E-51 |
| ACE2 | LZTR1 | -0,654 | 6,7398E-51 |
| ACE2 | TTC8 | -0,6541 | 6,42049E-51 |
| ACE2 | CDH4 | -0,6541 | 6,19832E-51 |
| ACE2 | IKBIP | -0,6542 | 6,14185E-51 |
| ACE2 | PCAT19 | -0,6542 | 6,01513E-51 |
| ACE2 | ZC3H12B | -0,6542 | 6,01941E-51 |
| ACE2 | RNF11 | -0,6543 | 5,64623E-51 |
| ACE2 | NMD3 | -0,6543 | 5,66263E-51 |
| ACE2 | ASB5 | -0,6543 | 5,84799E-51 |
| ACE2 | KIAA0895 | -0,6543 | 5,79438E-51 |
| ACE2 | DAAM1 | -0,6543 | 5,75283E-51 |
| ACE2 | GJA4 | -0,6544 | 5,58343E-51 |
| ACE2 | KLF3-AS1 | -0,6544 | 5,58841E-51 |
| ACE2 | SMIM8 | -0,6544 | 5,50565E-51 |
| ACE2 | KIAA2026 | -0,6544 | 5,57077E-51 |
| ACE2 | PRSS23 | -0,6544 | 5,49569E-51 |
| ACE2 | CLASRP | -0,6544 | 5,36511E-51 |
| ACE2 | GZF1 | -0,6544 | 5,4135E-51 |
| ACE2 | BGN | -0,6544 | 5,36153E-51 |
| ACE2 | SMYD1 | -0,6545 | 5,17315E-51 |
| ACE2 | NANOS1 | -0,6545 | 5,18519E-51 |
| ACE2 | ZNF449 | -0,6545 | 5,1609E-51 |
| ACE2 | LTBP1 | -0,6546 | 5,06884E-51 |
| ACE2 | FGL2 | -0,6546 | 5,04086E-51 |
| ACE2 | RP11-667F14.1 | -0,6546 | 4,87668E-51 |
| ACE2 | SPON2 | -0,6547 | 4,82275E-51 |
| ACE2 | KLHL3 | -0,6547 | 4,68437E-51 |
| ACE2 | RP4-751H13.5 | -0,6547 | 4,78139E-51 |
| ACE2 | RPL30 | -0,6547 | 4,79181E-51 |
| ACE2 | GACAT2 | -0,6547 | 4,83627E-51 |
| ACE2 | RP11-1191J2.2 | -0,6548 | 4,62321E-51 |
| ACE2 | ANGPT1 | -0,6548 | 4,64218E-51 |
| ACE2 | SHISA6 | -0,6548 | 4,64047E-51 |
| ACE2 | SLC25A42 | -0,6548 | 4,52442E-51 |
| ACE2 | LINC00205 | -0,6548 | 4,48256E-51 |
| ACE2 | CNKSR2 | -0,6548 | 4,64892E-51 |
| ACE2 | SDCCAG8 | -0,6549 | 4,26516E-51 |
| ACE2 | LINC02044 | -0,6549 | 4,43565E-51 |
| ACE2 | LINC00630 | -0,6549 | 4,24553E-51 |

| | | | |
|---|---|---|---|
| ACE2 | POM121C | -0,655 | 4,06482E-51 |
| ACE2 | ZNF773 | -0,655 | 4,19351E-51 |
| ACE2 | NTSR1 | -0,655 | 4,16351E-51 |
| ACE2 | ACSL4 | -0,655 | 4,17879E-51 |
| ACE2 | DPP6 | -0,6551 | 3,87932E-51 |
| ACE2 | GPR12 | -0,6551 | 4,0125E-51 |
| ACE2 | FAF1 | -0,6552 | 3,83544E-51 |
| ACE2 | AC074141.3 | -0,6552 | 3,69369E-51 |
| ACE2 | U52111.14 | -0,6552 | 3,81968E-51 |
| ACE2 | ANKRD26 | -0,6553 | 3,64503E-51 |
| ACE2 | GABARAPL2 | -0,6553 | 3,54631E-51 |
| ACE2 | COL3A1 | -0,6554 | 3,38923E-51 |
| ACE2 | TXLNA | -0,6555 | 3,30027E-51 |
| ACE2 | ZNF48 | -0,6555 | 3,34437E-51 |
| ACE2 | GBP1P1 | -0,6556 | 3,09426E-51 |
| ACE2 | ERICH2 | -0,6556 | 3,07328E-51 |
| ACE2 | MLH1 | -0,6556 | 3,09127E-51 |
| ACE2 | FAM160B1 | -0,6556 | 3,10341E-51 |
| ACE2 | RCBTB2 | -0,6556 | 3,18582E-51 |
| ACE2 | PFDN2 | -0,6557 | 2,97743E-51 |
| ACE2 | ZBTB20 | -0,6557 | 2,95175E-51 |
| ACE2 | FAM50B | -0,6557 | 3,05117E-51 |
| ACE2 | ZNF425 | -0,6557 | 3,00508E-51 |
| ACE2 | DAND5 | -0,6557 | 2,93357E-51 |
| ACE2 | PSMD4 | -0,6558 | 2,81997E-51 |
| ACE2 | KAT2B | -0,6558 | 2,89336E-51 |
| ACE2 | FGF10-AS1 | -0,6558 | 2,90454E-51 |
| ACE2 | FAM114A2 | -0,6558 | 2,83747E-51 |
| ACE2 | ZDHHC22 | -0,6558 | 2,82473E-51 |
| ACE2 | RP11-552F3.10 | -0,6558 | 2,80207E-51 |
| ACE2 | FABP3 | -0,6559 | 2,75734E-51 |
| ACE2 | PQLC3 | -0,6559 | 2,72102E-51 |
| ACE2 | HIVEP2 | -0,6559 | 2,70524E-51 |
| ACE2 | RP11-359E19.2 | -0,6559 | 2,7812E-51 |
| ACE2 | MARCH7 | -0,656 | 2,5745E-51 |
| ACE2 | C1QTNF2 | -0,656 | 2,64072E-51 |
| ACE2 | IFIT5 | -0,656 | 2,66206E-51 |
| ACE2 | TBC1D5 | -0,6561 | 2,46801E-51 |
| ACE2 | SMN1 | -0,6561 | 2,50393E-51 |
| ACE2 | CELF2 | -0,6561 | 2,49487E-51 |
| ACE2 | BMP7 | -0,6561 | 2,4747E-51 |
| ACE2 | GRIK1 | -0,6561 | 2,4582E-51 |
| ACE2 | ZFP69 | -0,6562 | 2,35541E-51 |
| ACE2 | SMYD3 | -0,6562 | 2,34638E-51 |
| ACE2 | RIPPLY2 | -0,6562 | 2,39824E-51 |
| ACE2 | SMTNL2 | -0,6562 | 2,32871E-51 |
| ACE2 | PAXBP1 | -0,6562 | 2,37307E-51 |
| ACE2 | SEZ6L | -0,6562 | 2,39802E-51 |
| ACE2 | MIIP | -0,6563 | 2,27777E-51 |
| ACE2 | ACTB | -0,6563 | 2,29464E-51 |
| ACE2 | TMEM245 | -0,6563 | 2,28192E-51 |
| ACE2 | ZNF22 | -0,6563 | 2,26327E-51 |
| ACE2 | BSN | -0,6564 | 2,10979E-51 |
| ACE2 | CTB-158E9.1 | -0,6564 | 2,20261E-51 |

| | | | |
|---|---|---|---|
| ACE2 | RP11-536O18.1 | -0,6564 | 2,12216E-51 |
| ACE2 | UBE2R2 | -0,6564 | 2,14526E-51 |
| ACE2 | LINC01537 | -0,6564 | 2,17675E-51 |
| ACE2 | CTA-212A2.1 | -0,6564 | 2,1893E-51 |
| ACE2 | CIART | -0,6565 | 2,0469E-51 |
| ACE2 | AC053503.12 | -0,6565 | 2,07253E-51 |
| ACE2 | CTC-487M23.6 | -0,6565 | 2,07557E-51 |
| ACE2 | ZNF287 | -0,6565 | 2,05142E-51 |
| ACE2 | RP11-753A21.1 | -0,6566 | 1,93552E-51 |
| ACE2 | PIK3CD-AS2 | -0,6568 | 1,78457E-51 |
| ACE2 | DNAJC27 | -0,6568 | 1,79829E-51 |
| ACE2 | WWTR1-AS1 | -0,6568 | 1,75244E-51 |
| ACE2 | IGF1R | -0,6568 | 1,75761E-51 |
| ACE2 | CDC25B | -0,6568 | 1,80545E-51 |
| ACE2 | COL25A1 | -0,6569 | 1,6965E-51 |
| ACE2 | SRP14 | -0,6569 | 1,71463E-51 |
| ACE2 | SNX1 | -0,6569 | 1,68339E-51 |
| ACE2 | ZNF500 | -0,6569 | 1,67131E-51 |
| ACE2 | RPL23AP7 | -0,657 | 1,62258E-51 |
| ACE2 | RP11-706J10.2 | -0,657 | 1,62394E-51 |
| ACE2 | LRRC10B | -0,657 | 1,59988E-51 |
| ACE2 | AJUBA | -0,657 | 1,60733E-51 |
| ACE2 | EXOC5 | -0,657 | 1,59432E-51 |
| ACE2 | ADAMTS9 | -0,6571 | 1,52608E-51 |
| ACE2 | LRRC70 | -0,6571 | 1,55344E-51 |
| ACE2 | CNGB1 | -0,6571 | 1,5276E-51 |
| ACE2 | RP11-894P9.1 | -0,6572 | 1,46883E-51 |
| ACE2 | ZNF444 | -0,6572 | 1,51112E-51 |
| ACE2 | PTGS2 | -0,6573 | 1,41473E-51 |
| ACE2 | GTF2H4 | -0,6573 | 1,42356E-51 |
| ACE2 | ZBTB26 | -0,6573 | 1,39702E-51 |
| ACE2 | TMEM132B | -0,6573 | 1,43714E-51 |
| ACE2 | PPP2R3C | -0,6573 | 1,43669E-51 |
| ACE2 | RP11-802O23.3 | -0,6574 | 1,33722E-51 |
| ACE2 | ANKRD50 | -0,6574 | 1,33357E-51 |
| ACE2 | RNF8 | -0,6574 | 1,32995E-51 |
| ACE2 | RP1-142L7.9 | -0,6574 | 1,37714E-51 |
| ACE2 | GDAP1 | -0,6574 | 1,35634E-51 |
| ACE2 | RP11-170K4.2 | -0,6575 | 1,26975E-51 |
| ACE2 | MSX1 | -0,6576 | 1,22212E-51 |
| ACE2 | RP11-216B9.6 | -0,6576 | 1,24785E-51 |
| ACE2 | RALGDS | -0,6576 | 1,25845E-51 |
| ACE2 | SBF2 | -0,6576 | 1,21645E-51 |
| ACE2 | CLDN11 | -0,6577 | 1,17986E-51 |
| ACE2 | VCAN | -0,6577 | 1,19558E-51 |
| ACE2 | ESYT1 | -0,6577 | 1,16518E-51 |
| ACE2 | PDS5B | -0,6577 | 1,19064E-51 |
| ACE2 | FGF13-AS1 | -0,6577 | 1,19685E-51 |
| ACE2 | RPP21 | -0,6578 | 1,14269E-51 |
| ACE2 | ATP5F1P5 | -0,6578 | 1,10619E-51 |
| ACE2 | CYLD | -0,6578 | 1,11114E-51 |
| ACE2 | LYPD5 | -0,6578 | 1,1182E-51 |
| ACE2 | CHSY3 | -0,6579 | 1,08248E-51 |
| ACE2 | RP11-946P6.4 | -0,6579 | 1,08579E-51 |

| | | | |
|---|---|---|---|
| ACE2 | CNTNAP4 | -0,6579 | 1,0663E-51 |
| ACE2 | LIN7A | -0,658 | 1,02435E-51 |
| ACE2 | HERC1 | -0,658 | 1,0358E-51 |
| ACE2 | AC004637.1 | -0,658 | 1,01612E-51 |
| ACE2 | DST | -0,6581 | 9,7275E-52 |
| ACE2 | C10orf90 | -0,6581 | 9,72493E-52 |
| ACE2 | PWP1 | -0,6581 | 9,56611E-52 |
| ACE2 | BEAN1 | -0,6581 | 9,64802E-52 |
| ACE2 | DACT3-AS1 | -0,6581 | 9,49864E-52 |
| ACE2 | RP11-536C12.1 | -0,6582 | 9,48496E-52 |
| ACE2 | TEAD1 | -0,6582 | 9,43096E-52 |
| ACE2 | LDHB | -0,6582 | 9,32533E-52 |
| ACE2 | TPT1-AS1 | -0,6582 | 9,1104E-52 |
| ACE2 | PARP11 | -0,6583 | 8,75631E-52 |
| ACE2 | DCLK1 | -0,6583 | 9,00095E-52 |
| ACE2 | PHF12 | -0,6583 | 8,74986E-52 |
| ACE2 | EP400NL | -0,6584 | 8,49866E-52 |
| ACE2 | RNF219 | -0,6584 | 8,39958E-52 |
| ACE2 | EXOG | -0,6585 | 8,01692E-52 |
| ACE2 | LINC00924 | -0,6585 | 8,0812E-52 |
| ACE2 | LRRC3 | -0,6585 | 8,20353E-52 |
| ACE2 | HS1BP3 | -0,6586 | 7,57271E-52 |
| ACE2 | NXF1 | -0,6586 | 7,83211E-52 |
| ACE2 | ZNF204P | -0,6587 | 7,43457E-52 |
| ACE2 | PHC1P1 | -0,6587 | 7,26312E-52 |
| ACE2 | PCNX4 | -0,6587 | 7,31567E-52 |
| ACE2 | SRPX | -0,6587 | 7,34097E-52 |
| ACE2 | MZT2B | -0,6588 | 6,87635E-52 |
| ACE2 | TMEM169 | -0,6588 | 6,84691E-52 |
| ACE2 | FOXRED2 | -0,6588 | 7,14231E-52 |
| ACE2 | USP48 | -0,6589 | 6,51947E-52 |
| ACE2 | MAN1A2 | -0,6589 | 6,51646E-52 |
| ACE2 | CH17-189H20.1 | -0,6589 | 6,67586E-52 |
| ACE2 | INSRR | -0,6589 | 6,62541E-52 |
| ACE2 | C1orf204 | -0,6589 | 6,78177E-52 |
| ACE2 | LINC00462 | -0,6589 | 6,57634E-52 |
| ACE2 | KCTD1 | -0,6589 | 6,65195E-52 |
| ACE2 | RBM5 | -0,659 | 6,30376E-52 |
| ACE2 | RP11-524C21.2 | -0,659 | 6,39022E-52 |
| ACE2 | PGM5P3-AS1 | -0,659 | 6,24217E-52 |
| ACE2 | GPR137C | -0,659 | 6,47443E-52 |
| ACE2 | RHOT1 | -0,659 | 6,37748E-52 |
| ACE2 | SPHK1 | -0,659 | 6,40869E-52 |
| ACE2 | SMAD2 | -0,659 | 6,39876E-52 |
| ACE2 | RP1-102E24.8 | -0,6591 | 5,95161E-52 |
| ACE2 | HSPA4L | -0,6592 | 5,86739E-52 |
| ACE2 | PTPDC1 | -0,6592 | 5,87207E-52 |
| ACE2 | UBA52 | -0,6592 | 5,76807E-52 |
| ACE2 | RPL21P28 | -0,6593 | 5,57713E-52 |
| ACE2 | HMX1 | -0,6593 | 5,57411E-52 |
| ACE2 | COL11A2 | -0,6593 | 5,59187E-52 |
| ACE2 | EFHC1 | -0,6593 | 5,63484E-52 |
| ACE2 | MMD2 | -0,6593 | 5,53876E-52 |
| ACE2 | RNF175 | -0,6594 | 5,32586E-52 |

| | | | |
|---|---|---|---|
| ACE2 | FLOT1 | -0,6594 | 5,34488E-52 |
| ACE2 | RP11-309L24.6 | -0,6594 | 5,21041E-52 |
| ACE2 | TACR2 | -0,6594 | 5,30881E-52 |
| ACE2 | PDE4A | -0,6594 | 5,3599E-52 |
| ACE2 | PCP4 | -0,6594 | 5,34127E-52 |
| ACE2 | RAD9B | -0,6595 | 5,0915E-52 |
| ACE2 | GPRC5B | -0,6595 | 4,93765E-52 |
| ACE2 | TNRC6A | -0,6595 | 5,00554E-52 |
| ACE2 | CX3CL1 | -0,6595 | 5,00476E-52 |
| ACE2 | LINC00908 | -0,6595 | 5,00582E-52 |
| ACE2 | AP001347.6 | -0,6595 | 4,98565E-52 |
| ACE2 | TUG1 | -0,6595 | 5,01822E-52 |
| ACE2 | PHC2 | -0,6596 | 4,82723E-52 |
| ACE2 | MROH2B | -0,6596 | 4,68316E-52 |
| ACE2 | DHX57 | -0,6598 | 4,26048E-52 |
| ACE2 | TFP1 | -0,6598 | 4,2653E-52 |
| ACE2 | OSBPL5 | -0,6598 | 4,34111E-52 |
| ACE2 | DUT | -0,6598 | 4,46175E-52 |
| ACE2 | CCDC159 | -0,6598 | 4,35268E-52 |
| ACE2 | RLIM | -0,6598 | 4,34019E-52 |
| ACE2 | BIN1 | -0,6599 | 4,17593E-52 |
| ACE2 | ZNF621 | -0,6599 | 4,10851E-52 |
| ACE2 | RP11-168P13.1 | -0,6599 | 4,07129E-52 |
| ACE2 | LINC00998 | -0,66 | 3,89653E-52 |
| ACE2 | FRMD6-AS1 | -0,66 | 4,00854E-52 |
| ACE2 | GREB1 | -0,6601 | 3,7641E-52 |
| ACE2 | ARID5A | -0,6601 | 3,86043E-52 |
| ACE2 | SH3BP5-AS1 | -0,6601 | 3,85609E-52 |
| ACE2 | TRMT11 | -0,6601 | 3,78366E-52 |
| ACE2 | RP1-111C20.4 | -0,6601 | 3,82662E-52 |
| ACE2 | GCK | -0,6601 | 3,70096E-52 |
| ACE2 | AC008992.1 | -0,6601 | 3,78113E-52 |
| ACE2 | TRMT44 | -0,6602 | 3,58986E-52 |
| ACE2 | SPOCK1 | -0,6602 | 3,66043E-52 |
| ACE2 | CWC15 | -0,6602 | 3,59995E-52 |
| ACE2 | RP11-64B16.2 | -0,6602 | 3,58657E-52 |
| ACE2 | NECAB2 | -0,6602 | 3,62796E-52 |
| ACE2 | GNG7 | -0,6602 | 3,6146E-52 |
| ACE2 | TMEM230 | -0,6602 | 3,54515E-52 |
| ACE2 | APOL3 | -0,6602 | 3,67339E-52 |
| ACE2 | PREPL | -0,6603 | 3,49763E-52 |
| ACE2 | AC006007.1 | -0,6603 | 3,38325E-52 |
| ACE2 | RP11-983P16.4 | -0,6603 | 3,50363E-52 |
| ACE2 | ALDH1L2 | -0,6603 | 3,42837E-52 |
| ACE2 | EPC1 | -0,6604 | 3,24654E-52 |
| ACE2 | LINC00959 | -0,6604 | 3,33118E-52 |
| ACE2 | LINC00671 | -0,6604 | 3,20233E-52 |
| ACE2 | AC139100.3 | -0,6604 | 3,27203E-52 |
| ACE2 | ADGRL4 | -0,6605 | 3,18153E-52 |
| ACE2 | RP11-417F21.1 | -0,6605 | 3,1943E-52 |
| ACE2 | SH3GL1 | -0,6605 | 3,17407E-52 |
| ACE2 | ECD | -0,6606 | 2,92799E-52 |
| ACE2 | RP11-102G14.1 | -0,6606 | 3,02994E-52 |
| ACE2 | ZNF578 | -0,6606 | 2,96682E-52 |

| | | | |
|---|---|---|---|
| ACE2 | CYYR1 | -0,6606 | 2,92068E-52 |
| ACE2 | LYST | -0,6607 | 2,91029E-52 |
| ACE2 | GALNT9 | -0,6607 | 2,79267E-52 |
| ACE2 | DOCK9 | -0,6607 | 2,79005E-52 |
| ACE2 | RTN1 | -0,6607 | 2,90777E-52 |
| ACE2 | AC113189.5 | -0,6607 | 2,84841E-52 |
| ACE2 | UTP6 | -0,6607 | 2,82749E-52 |
| ACE2 | SKA2 | -0,6607 | 2,78262E-52 |
| ACE2 | CEBPZOS | -0,6608 | 2,6854E-52 |
| ACE2 | ASPRV1 | -0,6608 | 2,67256E-52 |
| ACE2 | RP11-286H15.1 | -0,6608 | 2,67171E-52 |
| ACE2 | AL022344.7 | -0,6608 | 2,74825E-52 |
| ACE2 | CTNNBL1 | -0,6608 | 2,71608E-52 |
| ACE2 | VCAM1 | -0,6609 | 2,58697E-52 |
| ACE2 | SMPD4 | -0,6609 | 2,57535E-52 |
| ACE2 | RNF216-IT1 | -0,6609 | 2,60283E-52 |
| ACE2 | PECAM1 | -0,6609 | 2,54916E-52 |
| ACE2 | AC007750.5 | -0,661 | 2,45654E-52 |
| ACE2 | ZNF239 | -0,661 | 2,50313E-52 |
| ACE2 | RP5-858L17.1 | -0,661 | 2,42777E-52 |
| ACE2 | FLG-AS1 | -0,6611 | 2,38399E-52 |
| ACE2 | SNRNP27 | -0,6611 | 2,31397E-52 |
| ACE2 | LIFR | -0,6611 | 2,34057E-52 |
| ACE2 | AC018647.3 | -0,6611 | 2,35522E-52 |
| ACE2 | WDR59 | -0,6611 | 2,32853E-52 |
| ACE2 | LINC00304 | -0,6611 | 2,395E-52 |
| ACE2 | AC004448.5 | -0,6611 | 2,38301E-52 |
| ACE2 | PTGIR | -0,6611 | 2,37431E-52 |
| ACE2 | VMA21 | -0,6611 | 2,40024E-52 |
| ACE2 | LRIG1 | -0,6612 | 2,24078E-52 |
| ACE2 | GPM6A | -0,6612 | 2,19547E-52 |
| ACE2 | CDC37L1 | -0,6612 | 2,19113E-52 |
| ACE2 | RP11-114M1.1 | -0,6613 | 2,11784E-52 |
| ACE2 | CYHR1 | -0,6613 | 2,15193E-52 |
| ACE2 | RP11-304C12.5 | -0,6613 | 2,09671E-52 |
| ACE2 | KCNA1 | -0,6613 | 2,12775E-52 |
| ACE2 | ITGB3 | -0,6613 | 2,10662E-52 |
| ACE2 | SUGP2 | -0,6613 | 2,11549E-52 |
| ACE2 | LIX1L-AS1 | -0,6614 | 2,02525E-52 |
| ACE2 | GPR156 | -0,6614 | 2,01581E-52 |
| ACE2 | RP11-225B17.2 | -0,6614 | 2,00642E-52 |
| ACE2 | GATAD1 | -0,6614 | 2,08578E-52 |
| ACE2 | BMPR1A | -0,6614 | 2,04176E-52 |
| ACE2 | ZCCHC5 | -0,6615 | 1,93711E-52 |
| ACE2 | RP11-339B21.13 | -0,6616 | 1,83137E-52 |
| ACE2 | MOAP1 | -0,6616 | 1,84421E-52 |
| ACE2 | CXorf57 | -0,6616 | 1,84192E-52 |
| ACE2 | ZMAT4 | -0,6617 | 1,73077E-52 |
| ACE2 | RP11-573G6.4 | -0,6617 | 1,78794E-52 |
| ACE2 | KBTBD6 | -0,6617 | 1,72842E-52 |
| ACE2 | LINC01315 | -0,6617 | 1,76275E-52 |
| ACE2 | JPX | -0,6617 | 1,73368E-52 |
| ACE2 | PI4KB | -0,6618 | 1,72056E-52 |
| ACE2 | MEIS1-AS2 | -0,6618 | 1,66588E-52 |

| | | | |
|---|---|---|---|
| ACE2 | FAM175A | -0,6618 | 1,68901E-52 |
| ACE2 | BBS10 | -0,6618 | 1,67466E-52 |
| ACE2 | TPT1 | -0,6618 | 1,64597E-52 |
| ACE2 | GREM1 | -0,6618 | 1,67406E-52 |
| ACE2 | MEOX1 | -0,6618 | 1,71858E-52 |
| ACE2 | BCL2 | -0,6618 | 1,68817E-52 |
| ACE2 | ZNF404 | -0,6618 | 1,72345E-52 |
| ACE2 | NEFH | -0,6618 | 1,67252E-52 |
| ACE2 | LINC00982 | -0,6619 | 1,62592E-52 |
| ACE2 | KPNA6 | -0,6619 | 1,64299E-52 |
| ACE2 | ST3GAL5 | -0,6619 | 1,60632E-52 |
| ACE2 | CAMK2A | -0,6619 | 1,63278E-52 |
| ACE2 | HIRIP3 | -0,6619 | 1,59072E-52 |
| ACE2 | HOXB-AS3 | -0,6619 | 1,56812E-52 |
| ACE2 | ZNF182 | -0,6619 | 1,63644E-52 |
| ACE2 | ZNF662 | -0,662 | 1,55475E-52 |
| ACE2 | CCDC50 | -0,662 | 1,52703E-52 |
| ACE2 | PCDHA10 | -0,662 | 1,54304E-52 |
| ACE2 | SLC36A4 | -0,662 | 1,51208E-52 |
| ACE2 | CD276 | -0,662 | 1,5016E-52 |
| ACE2 | ZNF791 | -0,662 | 1,51158E-52 |
| ACE2 | RP11-569A11.1 | -0,6621 | 1,48822E-52 |
| ACE2 | RP11-34A14.3 | -0,6621 | 1,46104E-52 |
| ACE2 | LINC00989 | -0,6622 | 1,36432E-52 |
| ACE2 | RPS3A | -0,6622 | 1,39203E-52 |
| ACE2 | BNIP3L | -0,6622 | 1,38285E-52 |
| ACE2 | RPL13P5 | -0,6622 | 1,40243E-52 |
| ACE2 | MAGEE2 | -0,6623 | 1,3016E-52 |
| ACE2 | CTC-360G5.9 | -0,6624 | 1,29112E-52 |
| ACE2 | POFUT2 | -0,6624 | 1,24127E-52 |
| ACE2 | PRSS42 | -0,6625 | 1,2162E-52 |
| ACE2 | RBM18 | -0,6625 | 1,19536E-52 |
| ACE2 | RP11-333E1.1 | -0,6625 | 1,19037E-52 |
| ACE2 | SAE1 | -0,6625 | 1,1909E-52 |
| ACE2 | KIAA1143 | -0,6626 | 1,13533E-52 |
| ACE2 | PCDHGA11 | -0,6626 | 1,13764E-52 |
| ACE2 | KLHL7 | -0,6626 | 1,12787E-52 |
| ACE2 | DGKI | -0,6626 | 1,12572E-52 |
| ACE2 | RP11-88I18.2 | -0,6626 | 1,14656E-52 |
| ACE2 | AK8 | -0,6626 | 1,13921E-52 |
| ACE2 | TMEM221 | -0,6626 | 1,15355E-52 |
| ACE2 | NASP | -0,6627 | 1,08817E-52 |
| ACE2 | HTR2A | -0,6627 | 1,10337E-52 |
| ACE2 | C3 | -0,6627 | 1,09871E-52 |
| ACE2 | CEP44 | -0,6628 | 1,044E-52 |
| ACE2 | PRTFDC1 | -0,6628 | 1,06446E-52 |
| ACE2 | NMB | -0,6628 | 1,04016E-52 |
| ACE2 | RP11-231C14.6 | -0,6628 | 1,06648E-52 |
| ACE2 | HES4 | -0,6629 | 1,01626E-52 |
| ACE2 | SLFNL1-AS1 | -0,6629 | 1,01336E-52 |
| ACE2 | RP11-248N22.1 | -0,6629 | 1,00974E-52 |
| ACE2 | BRF1 | -0,6629 | 1,00157E-52 |
| ACE2 | ERG | -0,6629 | 9,99958E-53 |
| ACE2 | PRMT9 | -0,663 | 9,4446E-53 |

| | | | |
|---|---|---|---|
| ACE2 | MAST4 | -0,663 | 9,65979E-53 |
| ACE2 | RASGRP2 | -0,663 | 9,61189E-53 |
| ACE2 | DCC | -0,663 | 9,59751E-53 |
| ACE2 | WIZ | -0,663 | 9,44083E-53 |
| ACE2 | E2F6 | -0,6631 | 8,97033E-53 |
| ACE2 | ORC3 | -0,6631 | 9,21783E-53 |
| ACE2 | HDAC5 | -0,6631 | 8,91662E-53 |
| ACE2 | NXT1 | -0,6631 | 9,19081E-53 |
| ACE2 | ELN | -0,6632 | 8,63505E-53 |
| ACE2 | PXDNL | -0,6632 | 8,46998E-53 |
| ACE2 | NOS1 | -0,6632 | 8,4959E-53 |
| ACE2 | UBA2 | -0,6632 | 8,77352E-53 |
| ACE2 | CLCN6 | -0,6633 | 8,3005E-53 |
| ACE2 | PART1 | -0,6633 | 8,12874E-53 |
| ACE2 | PCOLCE | -0,6633 | 8,19825E-53 |
| ACE2 | BCL2L2 | -0,6633 | 8,33108E-53 |
| ACE2 | SLFN13 | -0,6633 | 8,18521E-53 |
| ACE2 | COX4I2 | -0,6633 | 8,18937E-53 |
| ACE2 | MKL1 | -0,6633 | 8,35916E-53 |
| ACE2 | TBC1D25 | -0,6633 | 8,37181E-53 |
| ACE2 | MOB3C | -0,6634 | 7,73861E-53 |
| ACE2 | TSSC4 | -0,6634 | 7,725E-53 |
| ACE2 | ZBTB44 | -0,6634 | 8,01215E-53 |
| ACE2 | USF2 | -0,6634 | 7,70032E-53 |
| ACE2 | ZNF474 | -0,6635 | 7,60729E-53 |
| ACE2 | MTCH1 | -0,6635 | 7,30065E-53 |
| ACE2 | FOXN3 | -0,6635 | 7,4094E-53 |
| ACE2 | ARMCX5-GPRASP2 | -0,6635 | 7,49084E-53 |
| ACE2 | ARMC1 | -0,6636 | 7,12187E-53 |
| ACE2 | LINC00202-1 | -0,6636 | 7,08585E-53 |
| ACE2 | RP11-452L6.7 | -0,6636 | 7,2763E-53 |
| ACE2 | CARTPT | -0,6637 | 6,81274E-53 |
| ACE2 | VIP | -0,6637 | 6,7966E-53 |
| ACE2 | RP11-392A14.8 | -0,6637 | 6,79073E-53 |
| ACE2 | USP36 | -0,6637 | 6,88417E-53 |
| ACE2 | RIIAD1 | -0,6638 | 6,59351E-53 |
| ACE2 | DEDD | -0,6638 | 6,48064E-53 |
| ACE2 | TP63 | -0,6638 | 6,37683E-53 |
| ACE2 | IGSF9B | -0,6638 | 6,46691E-53 |
| ACE2 | CTD-2026K11.6 | -0,6638 | 6,34925E-53 |
| ACE2 | CDR2 | -0,6638 | 6,31962E-53 |
| ACE2 | MYADM | -0,6638 | 6,58849E-53 |
| ACE2 | LINC01750 | -0,6639 | 6,11617E-53 |
| ACE2 | MSTO2P | -0,6639 | 6,26363E-53 |
| ACE2 | NAB1 | -0,6639 | 6,06464E-53 |
| ACE2 | RPL37 | -0,6639 | 6,02055E-53 |
| ACE2 | RPF2 | -0,6639 | 6,11495E-53 |
| ACE2 | PIP5K1A | -0,664 | 5,79978E-53 |
| ACE2 | MCFD2 | -0,664 | 5,9895E-53 |
| ACE2 | RP11-167N24.3 | -0,664 | 5,96291E-53 |
| ACE2 | CPS1 | -0,6641 | 5,57423E-53 |
| ACE2 | NFKBIL1 | -0,6641 | 5,50865E-53 |
| ACE2 | ACTR1A | -0,6641 | 5,62568E-53 |
| ACE2 | KCNJ5 | -0,6641 | 5,51689E-53 |

| | | | |
|---|---|---|---|
| ACE2 | VPS45 | -0,6642 | 5,39158E-53 |
| ACE2 | CREB3 | -0,6642 | 5,39709E-53 |
| ACE2 | RBBP6 | -0,6642 | 5,36465E-53 |
| ACE2 | SYT3 | -0,6642 | 5,36283E-53 |
| ACE2 | IBA57-AS1 | -0,6643 | 5,06199E-53 |
| ACE2 | DCLK3 | -0,6643 | 5,0101E-53 |
| ACE2 | SART1 | -0,6643 | 5,16377E-53 |
| ACE2 | TCP10L | -0,6643 | 4,96359E-53 |
| ACE2 | PHF21B | -0,6643 | 4,96302E-53 |
| ACE2 | CCDC30 | -0,6644 | 4,81781E-53 |
| ACE2 | CXXC4 | -0,6644 | 4,76079E-53 |
| ACE2 | ARID5B | -0,6644 | 4,93569E-53 |
| ACE2 | GHc-857G6.7 | -0,6644 | 4,92528E-53 |
| ACE2 | RP11-545G3.1 | -0,6645 | 4,60818E-53 |
| ACE2 | INPP5A | -0,6645 | 4,62155E-53 |
| ACE2 | DEAF1 | -0,6645 | 4,69611E-53 |
| ACE2 | PANX1 | -0,6645 | 4,54441E-53 |
| ACE2 | EGLN2 | -0,6645 | 4,71536E-53 |
| ACE2 | EVA1C | -0,6645 | 4,60228E-53 |
| ACE2 | HSPB7 | -0,6646 | 4,30262E-53 |
| ACE2 | EPM2AIP1 | -0,6646 | 4,45954E-53 |
| ACE2 | PCOLCE-AS1 | -0,6646 | 4,44932E-53 |
| ACE2 | SS18 | -0,6646 | 4,48966E-53 |
| ACE2 | MTND5P1 | -0,6646 | 4,40662E-53 |
| ACE2 | RP11-503P10.1 | -0,6647 | 4,28553E-53 |
| ACE2 | SLC26A4 | -0,6647 | 4,16261E-53 |
| ACE2 | SLC4A8 | -0,6647 | 4,18624E-53 |
| ACE2 | ABCA9 | -0,6647 | 4,19725E-53 |
| ACE2 | TROVE2 | -0,6648 | 3,92128E-53 |
| ACE2 | GPLD1 | -0,6648 | 3,90865E-53 |
| ACE2 | NEXN | -0,6649 | 3,76383E-53 |
| ACE2 | SHE | -0,6649 | 3,82721E-53 |
| ACE2 | ARRDC3 | -0,6649 | 3,79994E-53 |
| ACE2 | DNAJB6 | -0,6649 | 3,73774E-53 |
| ACE2 | SLC6A15 | -0,665 | 3,67858E-53 |
| ACE2 | RP11-701H24.7 | -0,665 | 3,55493E-53 |
| ACE2 | HYDIN2 | -0,6651 | 3,3865E-53 |
| ACE2 | PDLIM1P4 | -0,6651 | 3,44708E-53 |
| ACE2 | RFNG | -0,6651 | 3,39411E-53 |
| ACE2 | TGFBR3 | -0,6652 | 3,34748E-53 |
| ACE2 | SNCAIP | -0,6652 | 3,32978E-53 |
| ACE2 | C9orf170 | -0,6652 | 3,35605E-53 |
| ACE2 | RP11-96C23.11 | -0,6652 | 3,26161E-53 |
| ACE2 | DNM1P47 | -0,6652 | 3,30665E-53 |
| ACE2 | ABCA8 | -0,6652 | 3,21553E-53 |
| ACE2 | PRCD | -0,6652 | 3,34901E-53 |
| ACE2 | HGFAC | -0,6653 | 3,11523E-53 |
| ACE2 | RP11-16P20.3 | -0,6653 | 3,13088E-53 |
| ACE2 | FRG1HP | -0,6653 | 3,14005E-53 |
| ACE2 | PLEKHA8P1 | -0,6653 | 3,1171E-53 |
| ACE2 | RP3-414A15.12 | -0,6653 | 3,16767E-53 |
| ACE2 | ADAMTS7P3 | -0,6653 | 3,16303E-53 |
| ACE2 | RP4-800J21.3 | -0,6653 | 3,09015E-53 |
| ACE2 | VPS54 | -0,6654 | 2,96677E-53 |

| | | | |
|---|---|---|---|
| ACE2 | RFX3 | -0,6654 | 3,04324E-53 |
| ACE2 | GRIN2C | -0,6654 | 3,02784E-53 |
| ACE2 | RP11-760H22.2 | -0,6655 | 2,91138E-53 |
| ACE2 | AC006942.4 | -0,6655 | 2,82625E-53 |
| ACE2 | PMS1 | -0,6656 | 2,6821E-53 |
| ACE2 | PDE6D | -0,6656 | 2,75259E-53 |
| ACE2 | APBB2 | -0,6656 | 2,67801E-53 |
| ACE2 | PPP1R3G | -0,6656 | 2,68961E-53 |
| ACE2 | COLGALT1 | -0,6656 | 2,68152E-53 |
| ACE2 | ZNF543 | -0,6656 | 2,70646E-53 |
| ACE2 | AL590762.7 | -0,6656 | 2,76221E-53 |
| ACE2 | LINC01797 | -0,6657 | 2,62136E-53 |
| ACE2 | FAM138B | -0,6657 | 2,54532E-53 |
| ACE2 | GTF2IP1 | -0,6657 | 2,64274E-53 |
| ACE2 | DDX10 | -0,6657 | 2,56845E-53 |
| ACE2 | APOBEC3C | -0,6657 | 2,54871E-53 |
| ACE2 | SMPX | -0,6657 | 2,62001E-53 |
| ACE2 | RHOB | -0,6658 | 2,43686E-53 |
| ACE2 | CNOT4 | -0,6658 | 2,41959E-53 |
| ACE2 | ZNF793 | -0,6658 | 2,49857E-53 |
| ACE2 | ISM1 | -0,6659 | 2,30471E-53 |
| ACE2 | C21orf91 | -0,6659 | 2,33518E-53 |
| ACE2 | LPP | -0,666 | 2,21845E-53 |
| ACE2 | SNX25 | -0,666 | 2,28698E-53 |
| ACE2 | ZNF451 | -0,666 | 2,23705E-53 |
| ACE2 | SERPINI1 | -0,6661 | 2,08284E-53 |
| ACE2 | CUL4B | -0,6661 | 2,09811E-53 |
| ACE2 | SVBP | -0,6662 | 2,02149E-53 |
| ACE2 | PMS2CL | -0,6662 | 2,01369E-53 |
| ACE2 | GABARAPL1 | -0,6662 | 2,07239E-53 |
| ACE2 | CES1 | -0,6662 | 1,99739E-53 |
| ACE2 | FIS1 | -0,6663 | 1,94199E-53 |
| ACE2 | SOHLH2 | -0,6663 | 1,95449E-53 |
| ACE2 | RP11-1148L6.8 | -0,6663 | 1,96608E-53 |
| ACE2 | TBCA | -0,6664 | 1,83808E-53 |
| ACE2 | HOXA4 | -0,6664 | 1,79878E-53 |
| ACE2 | RHOBTB1 | -0,6664 | 1,8469E-53 |
| ACE2 | KITLG | -0,6664 | 1,84601E-53 |
| ACE2 | LUM | -0,6665 | 1,78917E-53 |
| ACE2 | LINC02094 | -0,6665 | 1,72816E-53 |
| ACE2 | MAU2 | -0,6665 | 1,72327E-53 |
| ACE2 | HAUS5 | -0,6665 | 1,76459E-53 |
| ACE2 | TSTD3 | -0,6666 | 1,67062E-53 |
| ACE2 | GNG3 | -0,6666 | 1,66315E-53 |
| ACE2 | RP11-674N23.4 | -0,6666 | 1,69349E-53 |
| ACE2 | C19orf68 | -0,6666 | 1,6532E-53 |
| ACE2 | C3orf70 | -0,6667 | 1,61874E-53 |
| ACE2 | HOXC5 | -0,6667 | 1,56476E-53 |
| ACE2 | RPS15A | -0,6667 | 1,58022E-53 |
| ACE2 | C1QTNF1 | -0,6667 | 1,59922E-53 |
| ACE2 | SLC6A17 | -0,6668 | 1,5147E-53 |
| ACE2 | CDC27 | -0,6668 | 1,48093E-53 |
| ACE2 | SGSM1 | -0,6668 | 1,50573E-53 |
| ACE2 | ZNF777 | -0,6669 | 1,44425E-53 |

| | | | |
|---|---|---|---|
| ACE2 | NDEL1 | -0,6669 | 1,4212E-53 |
| ACE2 | STARD8 | -0,6669 | 1,44344E-53 |
| ACE2 | ANKRD44 | -0,667 | 1,35327E-53 |
| ACE2 | CYP7B1 | -0,667 | 1,37987E-53 |
| ACE2 | GSN | -0,667 | 1,36145E-53 |
| ACE2 | GCGR | -0,667 | 1,36072E-53 |
| ACE2 | FRAS1 | -0,6671 | 1,32267E-53 |
| ACE2 | ITGA5 | -0,6671 | 1,28138E-53 |
| ACE2 | COMMD1 | -0,6672 | 1,27514E-53 |
| ACE2 | SMYD5 | -0,6672 | 1,2416E-53 |
| ACE2 | SLC24A2 | -0,6672 | 1,24608E-53 |
| ACE2 | MBD1 | -0,6672 | 1,26545E-53 |
| ACE2 | RRP7BP | -0,6672 | 1,25594E-53 |
| ACE2 | AGBL4 | -0,6673 | 1,16616E-53 |
| ACE2 | ACKR1 | -0,6673 | 1,21375E-53 |
| ACE2 | RP3-475N16.1 | -0,6673 | 1,18816E-53 |
| ACE2 | ITGB8 | -0,6673 | 1,20258E-53 |
| ACE2 | C10orf107 | -0,6673 | 1,18095E-53 |
| ACE2 | CASKIN2 | -0,6673 | 1,21513E-53 |
| ACE2 | DRC1 | -0,6674 | 1,10786E-53 |
| ACE2 | AC074286.1 | -0,6674 | 1,11859E-53 |
| ACE2 | B9D1 | -0,6675 | 1,09157E-53 |
| ACE2 | OGFRL1 | -0,6676 | 1,04309E-53 |
| ACE2 | MYOM1 | -0,6676 | 1,00957E-53 |
| ACE2 | RP11-470M17.2 | -0,6677 | 9,63616E-54 |
| ACE2 | RP11-690D19.3 | -0,6677 | 9,89211E-54 |
| ACE2 | RP11-1180F24.1 | -0,6677 | 9,92421E-54 |
| ACE2 | RP11-346C20.4 | -0,6677 | 9,98578E-54 |
| ACE2 | WDR75 | -0,6678 | 9,08756E-54 |
| ACE2 | KRBOX1 | -0,6678 | 9,38381E-54 |
| ACE2 | PFN2 | -0,6678 | 9,45923E-54 |
| ACE2 | MAB21L2 | -0,6678 | 9,20258E-54 |
| ACE2 | DOCK4 | -0,6678 | 9,21118E-54 |
| ACE2 | ZNF503-AS2 | -0,6678 | 9,34771E-54 |
| ACE2 | CCNA1 | -0,6678 | 9,37305E-54 |
| ACE2 | RP11-360F5.1 | -0,6679 | 8,92836E-54 |
| ACE2 | TMEM151A | -0,6679 | 9,05968E-54 |
| ACE2 | SAP30BP | -0,6679 | 8,93123E-54 |
| ACE2 | PANX2 | -0,6679 | 8,80884E-54 |
| ACE2 | DUSP10 | -0,668 | 8,63907E-54 |
| ACE2 | CYP21A1P | -0,668 | 8,34919E-54 |
| ACE2 | LINC00997 | -0,668 | 8,23598E-54 |
| ACE2 | AKNAD1 | -0,6681 | 7,84187E-54 |
| ACE2 | C5orf24 | -0,6681 | 7,91497E-54 |
| ACE2 | LINC00670 | -0,6681 | 7,8662E-54 |
| ACE2 | ZNF285 | -0,6681 | 7,86401E-54 |
| ACE2 | NTNG1 | -0,6682 | 7,77389E-54 |
| ACE2 | RAD23A | -0,6682 | 7,78964E-54 |
| ACE2 | POLR3H | -0,6682 | 7,61173E-54 |
| ACE2 | COL24A1 | -0,6683 | 7,38744E-54 |
| ACE2 | BMPR2 | -0,6683 | 7,37827E-54 |
| ACE2 | IL17RC | -0,6683 | 7,22181E-54 |
| ACE2 | RP11-155D18.13 | -0,6683 | 7,22418E-54 |
| ACE2 | CDIPT | -0,6683 | 7,34828E-54 |

| | | | |
|---|---|---|---|
| ACE2 | LZTFL1 | -0,6684 | 7,09781E-54 |
| ACE2 | DDB2 | -0,6684 | 7,07703E-54 |
| ACE2 | PSMC5 | -0,6684 | 6,82884E-54 |
| ACE2 | SERTM1 | -0,6685 | 6,44814E-54 |
| ACE2 | TANC2 | -0,6685 | 6,69937E-54 |
| ACE2 | SH3GL3 | -0,6686 | 6,21908E-54 |
| ACE2 | ZNF826P | -0,6686 | 6,22671E-54 |
| ACE2 | SRD5A3-AS1 | -0,6687 | 5,97403E-54 |
| ACE2 | PDE4D | -0,6687 | 5,94121E-54 |
| ACE2 | SRP68 | -0,6687 | 6,0076E-54 |
| ACE2 | NOTCH4 | -0,6688 | 5,71962E-54 |
| ACE2 | RNF38 | -0,6688 | 5,60469E-54 |
| ACE2 | STKLD1 | -0,6688 | 5,78078E-54 |
| ACE2 | PAGR1 | -0,6688 | 5,60166E-54 |
| ACE2 | NAP1L6 | -0,6688 | 5,72156E-54 |
| ACE2 | ALDH8A1 | -0,6689 | 5,33151E-54 |
| ACE2 | PLAC9 | -0,6689 | 5,38074E-54 |
| ACE2 | RIC8B | -0,6689 | 5,43541E-54 |
| ACE2 | NIN | -0,6689 | 5,52121E-54 |
| ACE2 | JDP2 | -0,6689 | 5,47908E-54 |
| ACE2 | IKZF4 | -0,669 | 5,09472E-54 |
| ACE2 | AC005618.6 | -0,6691 | 4,86262E-54 |
| ACE2 | ANAPC16 | -0,6691 | 4,85716E-54 |
| ACE2 | SCG3 | -0,6691 | 4,9027E-54 |
| ACE2 | TPM1 | -0,6691 | 4,97158E-54 |
| ACE2 | NPIPA1 | -0,6691 | 4,96385E-54 |
| ACE2 | HSD17B14 | -0,6691 | 4,92808E-54 |
| ACE2 | LSMEM2 | -0,6692 | 4,59138E-54 |
| ACE2 | FJX1 | -0,6692 | 4,65158E-54 |
| ACE2 | PPP2R5C | -0,6692 | 4,60061E-54 |
| ACE2 | ST7L | -0,6693 | 4,42946E-54 |
| ACE2 | ADRA1A | -0,6693 | 4,39283E-54 |
| ACE2 | PZP | -0,6693 | 4,55145E-54 |
| ACE2 | CSDC2 | -0,6693 | 4,43695E-54 |
| ACE2 | LPAR4 | -0,6693 | 4,51085E-54 |
| ACE2 | RP11-517P14.7 | -0,6695 | 4,073E-54 |
| ACE2 | COL13A1 | -0,6695 | 3,98497E-54 |
| ACE2 | PLA2G2C | -0,6696 | 3,91327E-54 |
| ACE2 | HAX1 | -0,6696 | 3,78723E-54 |
| ACE2 | LINC01799 | -0,6696 | 3,82426E-54 |
| ACE2 | ZEB2 | -0,6696 | 3,81919E-54 |
| ACE2 | RP11-326C3.11 | -0,6696 | 3,83863E-54 |
| ACE2 | RP11-161D15.1 | -0,6697 | 3,57755E-54 |
| ACE2 | PMS2P4 | -0,6697 | 3,62962E-54 |
| ACE2 | UCK1 | -0,6697 | 3,70468E-54 |
| ACE2 | HIPK3 | -0,6697 | 3,66098E-54 |
| ACE2 | RP3-412A9.16 | -0,6697 | 3,67449E-54 |
| ACE2 | PRKCE | -0,6698 | 3,50143E-54 |
| ACE2 | TCEA1 | -0,6698 | 3,50991E-54 |
| ACE2 | CHD6 | -0,6698 | 3,51891E-54 |
| ACE2 | PCIF1 | -0,6698 | 3,49104E-54 |
| ACE2 | JADE1 | -0,6699 | 3,27497E-54 |
| ACE2 | CTD-2368P22.1 | -0,6699 | 3,27205E-54 |
| ACE2 | USP27X-AS1 | -0,6699 | 3,34189E-54 |

| | | | |
|---|---|---|---|
| ACE2 | TMEM35A | -0,6699 | 3,3074E-54 |
| ACE2 | AC007743.1 | -0,67 | 3,11204E-54 |
| ACE2 | RP11-167N5.5 | -0,67 | 3,16944E-54 |
| ACE2 | FAM207A | -0,67 | 3,09153E-54 |
| ACE2 | CXorf67 | -0,67 | 3,16745E-54 |
| ACE2 | LINC00890 | -0,67 | 3,10647E-54 |
| ACE2 | AC103563.9 | -0,6701 | 3,04425E-54 |
| ACE2 | COPS9 | -0,6701 | 3,07788E-54 |
| ACE2 | RAB3C | -0,6701 | 2,98856E-54 |
| ACE2 | CKMT2 | -0,6701 | 3,0109E-54 |
| ACE2 | CAMK2B | -0,6701 | 3,00986E-54 |
| ACE2 | IGFBP6 | -0,6701 | 2,98035E-54 |
| ACE2 | PCNX1 | -0,6701 | 3,06625E-54 |
| ACE2 | ADRA1D | -0,6701 | 2,97176E-54 |
| ACE2 | ZCCHC12 | -0,6701 | 2,9466E-54 |
| ACE2 | PCOLCE2 | -0,6702 | 2,91112E-54 |
| ACE2 | MAP3K7 | -0,6702 | 2,82421E-54 |
| ACE2 | HMCN2 | -0,6702 | 2,8787E-54 |
| ACE2 | ZC3H7B | -0,6702 | 2,83461E-54 |
| ACE2 | DCUN1D4 | -0,6703 | 2,76134E-54 |
| ACE2 | OPN1SW | -0,6703 | 2,6787E-54 |
| ACE2 | PPP1R1A | -0,6703 | 2,69315E-54 |
| ACE2 | CTD-2006M22.2 | -0,6704 | 2,62187E-54 |
| ACE2 | POLR2K | -0,6704 | 2,57528E-54 |
| ACE2 | AP000436.4 | -0,6704 | 2,61465E-54 |
| ACE2 | AKTIP | -0,6704 | 2,5837E-54 |
| ACE2 | CDKN1C | -0,6705 | 2,41453E-54 |
| ACE2 | RP11-429A20.3 | -0,6705 | 2,46246E-54 |
| ACE2 | SPTB | -0,6705 | 2,45953E-54 |
| ACE2 | CHURC1 | -0,6705 | 2,4467E-54 |
| ACE2 | TFAP2E | -0,6706 | 2,33004E-54 |
| ACE2 | CITED4 | -0,6706 | 2,32585E-54 |
| ACE2 | POLN | -0,6706 | 2,33238E-54 |
| ACE2 | PCDHGB4 | -0,6706 | 2,40799E-54 |
| ACE2 | NSMCE2 | -0,6706 | 2,30332E-54 |
| ACE2 | UBXN1 | -0,6706 | 2,39307E-54 |
| ACE2 | RP11-269C23.5 | -0,6706 | 2,39055E-54 |
| ACE2 | LRRC37B | -0,6706 | 2,32336E-54 |
| ACE2 | PRKAR1A | -0,6706 | 2,33153E-54 |
| ACE2 | TTC3 | -0,6706 | 2,33435E-54 |
| ACE2 | C8orf59 | -0,6707 | 2,27636E-54 |
| ACE2 | MAPKAP1 | -0,6707 | 2,18723E-54 |
| ACE2 | HS6ST3 | -0,6707 | 2,24674E-54 |
| ACE2 | LINC01561 | -0,6708 | 2,13741E-54 |
| ACE2 | DCX | -0,6708 | 2,15759E-54 |
| ACE2 | CCL11 | -0,671 | 1,88256E-54 |
| ACE2 | TAF12 | -0,6711 | 1,84079E-54 |
| ACE2 | NPM1 | -0,6711 | 1,87804E-54 |
| ACE2 | HSPB1 | -0,6711 | 1,8166E-54 |
| ACE2 | ATP1A4 | -0,6713 | 1,6457E-54 |
| ACE2 | LAMC1 | -0,6713 | 1,63345E-54 |
| ACE2 | GPATCH2 | -0,6713 | 1,65784E-54 |
| ACE2 | ADAMTS19 | -0,6714 | 1,54469E-54 |
| ACE2 | RP11-616M22.1 | -0,6714 | 1,57885E-54 |

| | | | |
|---|---|---|---|
| ACE2 | STAT5A | -0,6714 | 1,59131E-54 |
| ACE2 | XYLT2 | -0,6714 | 1,56771E-54 |
| ACE2 | SLC27A5 | -0,6714 | 1,59082E-54 |
| ACE2 | DYNC1I1 | -0,6715 | 1,49819E-54 |
| ACE2 | LMBR1L | -0,6715 | 1,53877E-54 |
| ACE2 | ARPP19 | -0,6715 | 1,53064E-54 |
| ACE2 | CTD-2545G14.4 | -0,6715 | 1,53601E-54 |
| ACE2 | RPL34 | -0,6716 | 1,40884E-54 |
| ACE2 | SUFU | -0,6716 | 1,40605E-54 |
| ACE2 | PRDM11 | -0,6716 | 1,43234E-54 |
| ACE2 | EP400 | -0,6716 | 1,4222E-54 |
| ACE2 | NCOA1 | -0,6717 | 1,34555E-54 |
| ACE2 | RP11-666A8.9 | -0,6717 | 1,36248E-54 |
| ACE2 | TMEM242 | -0,6718 | 1,33003E-54 |
| ACE2 | RP11-302B13.1 | -0,6718 | 1,30493E-54 |
| ACE2 | AC010136.2 | -0,6719 | 1,25728E-54 |
| ACE2 | WDR53 | -0,6719 | 1,22959E-54 |
| ACE2 | LRRTM3 | -0,6719 | 1,21369E-54 |
| ACE2 | MPHOSPH8 | -0,6719 | 1,23779E-54 |
| ACE2 | ZNF331 | -0,6719 | 1,22205E-54 |
| ACE2 | PHGDH | -0,672 | 1,15668E-54 |
| ACE2 | RP11-532N4.2 | -0,672 | 1,16512E-54 |
| ACE2 | RP11-49G2.3 | -0,672 | 1,19437E-54 |
| ACE2 | DUSP7 | -0,6721 | 1,1138E-54 |
| ACE2 | ADAMTS9-AS1 | -0,6721 | 1,11394E-54 |
| ACE2 | RP4-584D14.7 | -0,6721 | 1,0913E-54 |
| ACE2 | TECPR2 | -0,6721 | 1,12745E-54 |
| ACE2 | DET1 | -0,6721 | 1,09733E-54 |
| ACE2 | TRAF3IP1 | -0,6722 | 1,06907E-54 |
| ACE2 | GOLGA7B | -0,6722 | 1,08727E-54 |
| ACE2 | UGGT2 | -0,6722 | 1,068E-54 |
| ACE2 | ZNF229 | -0,6722 | 1,07078E-54 |
| ACE2 | PFDN4 | -0,6722 | 1,05814E-54 |
| ACE2 | RP11-446H18.5 | -0,6723 | 1,01782E-54 |
| ACE2 | FNDC9 | -0,6723 | 1,02508E-54 |
| ACE2 | EHMT2 | -0,6723 | 1,002E-54 |
| ACE2 | SDAD1P1 | -0,6723 | 1,02164E-54 |
| ACE2 | AC093495.4 | -0,6724 | 9,55097E-55 |
| ACE2 | SEC22A | -0,6724 | 9,77839E-55 |
| ACE2 | AIG1 | -0,6724 | 9,86722E-55 |
| ACE2 | TMEM218 | -0,6724 | 9,43259E-55 |
| ACE2 | CTD-3157E16.2 | -0,6724 | 9,8253E-55 |
| ACE2 | MAPRE2 | -0,6724 | 9,61463E-55 |
| ACE2 | RPL23P2 | -0,6724 | 9,84796E-55 |
| ACE2 | TIMP1 | -0,6724 | 9,65165E-55 |
| ACE2 | MTERF3 | -0,6725 | 8,96683E-55 |
| ACE2 | TRIM44 | -0,6725 | 9,30288E-55 |
| ACE2 | ANO2 | -0,6725 | 8,96834E-55 |
| ACE2 | RTTN | -0,6726 | 8,56447E-55 |
| ACE2 | DENND6B | -0,6726 | 8,60267E-55 |
| ACE2 | RABL2A | -0,6727 | 8,42207E-55 |
| ACE2 | MTMR9 | -0,6727 | 8,30432E-55 |
| ACE2 | TCTN2 | -0,6727 | 8,41946E-55 |
| ACE2 | SLC4A5 | -0,6728 | 7,97039E-55 |

| | | | |
|---|---|---|---|
| ACE2 | RASSF1 | -0,6728 | 7,83761E-55 |
| ACE2 | ISYNA1 | -0,6728 | 8,04319E-55 |
| ACE2 | MYO16 | -0,6729 | 7,67316E-55 |
| ACE2 | ZNF595 | -0,673 | 7,21551E-55 |
| ACE2 | P4HA3 | -0,673 | 7,29608E-55 |
| ACE2 | YAP1 | -0,673 | 7,10485E-55 |
| ACE2 | GSTT2B | -0,673 | 7,30984E-55 |
| ACE2 | PABPC4 | -0,6731 | 6,95912E-55 |
| ACE2 | MYOZ2 | -0,6731 | 6,93666E-55 |
| ACE2 | HOMER1 | -0,6731 | 6,72585E-55 |
| ACE2 | LINC00472 | -0,6731 | 6,69784E-55 |
| ACE2 | CBARP | -0,6731 | 6,82196E-55 |
| ACE2 | GREM2 | -0,6732 | 6,36911E-55 |
| ACE2 | ABTB1 | -0,6732 | 6,49834E-55 |
| ACE2 | CCDC127 | -0,6732 | 6,61234E-55 |
| ACE2 | TPD52L1 | -0,6732 | 6,36421E-55 |
| ACE2 | C7orf26 | -0,6732 | 6,607E-55 |
| ACE2 | TRIM22 | -0,6732 | 6,44341E-55 |
| ACE2 | TBC1D17 | -0,6732 | 6,3616E-55 |
| ACE2 | GRIK1-AS1 | -0,6732 | 6,58665E-55 |
| ACE2 | FAM127B | -0,6732 | 6,39031E-55 |
| ACE2 | FAM228A | -0,6733 | 6,12803E-55 |
| ACE2 | UBE2E1 | -0,6733 | 6,0618E-55 |
| ACE2 | MYL3 | -0,6733 | 6,00169E-55 |
| ACE2 | DZIP3 | -0,6733 | 6,25835E-55 |
| ACE2 | SLC9B1 | -0,6733 | 6,23539E-55 |
| ACE2 | GDPD5 | -0,6733 | 6,16883E-55 |
| ACE2 | AVPR1A | -0,6733 | 6,12588E-55 |
| ACE2 | LINC00933 | -0,6733 | 6,03302E-55 |
| ACE2 | LINC01786 | -0,6734 | 5,87195E-55 |
| ACE2 | KCNQ5 | -0,6734 | 5,93638E-55 |
| ACE2 | CTB-12A17.2 | -0,6734 | 5,99074E-55 |
| ACE2 | MAGEF1 | -0,6735 | 5,55468E-55 |
| ACE2 | ALDH1A3 | -0,6735 | 5,48675E-55 |
| ACE2 | ANKRD36 | -0,6736 | 5,25456E-55 |
| ACE2 | SOGA3 | -0,6736 | 5,22285E-55 |
| ACE2 | GTF2IRD1 | -0,6736 | 5,2362E-55 |
| ACE2 | C17orf80 | -0,6736 | 5,41939E-55 |
| ACE2 | SLC8A2 | -0,6736 | 5,31232E-55 |
| ACE2 | UNC13C | -0,6737 | 4,97754E-55 |
| ACE2 | RP9 | -0,6738 | 4,80223E-55 |
| ACE2 | OR8S1 | -0,6738 | 4,727E-55 |
| ACE2 | C12orf29 | -0,6738 | 4,91196E-55 |
| ACE2 | ZNF10 | -0,6738 | 4,75634E-55 |
| ACE2 | TNPO2 | -0,6738 | 4,73424E-55 |
| ACE2 | RP3-428L16.2 | -0,6739 | 4,63653E-55 |
| ACE2 | WISP1 | -0,6739 | 4,6441E-55 |
| ACE2 | LINC01474 | -0,6739 | 4,60922E-55 |
| ACE2 | VCL | -0,6739 | 4,66869E-55 |
| ACE2 | SENP1 | -0,6739 | 4,63826E-55 |
| ACE2 | RP11-805I24.3 | -0,6739 | 4,49629E-55 |
| ACE2 | GMPS | -0,674 | 4,3608E-55 |
| ACE2 | CDO1 | -0,674 | 4,2466E-55 |
| ACE2 | MAN2A2 | -0,674 | 4,44122E-55 |

| | | | |
|---|---|---|---|
| ACE2 | CBR3 | -0,674 | 4,22986E-55 |
| ACE2 | TSR2 | -0,674 | 4,31301E-55 |
| ACE2 | RPL10 | -0,674 | 4,29335E-55 |
| ACE2 | RLF | -0,6741 | 4,21603E-55 |
| ACE2 | ABHD1 | -0,6741 | 4,17018E-55 |
| ACE2 | GLT8D1 | -0,6741 | 4,03523E-55 |
| ACE2 | C9orf50 | -0,6741 | 4,14009E-55 |
| ACE2 | FAM133A | -0,6741 | 4,13547E-55 |
| ACE2 | DDIT4L | -0,6742 | 3,91763E-55 |
| ACE2 | AC007796.1 | -0,6742 | 3,85614E-55 |
| ACE2 | METTL5 | -0,6743 | 3,64569E-55 |
| ACE2 | HCG23 | -0,6743 | 3,74491E-55 |
| ACE2 | CNTNAP3 | -0,6743 | 3,65198E-55 |
| ACE2 | HCN2 | -0,6743 | 3,67623E-55 |
| ACE2 | SUPT20HL2 | -0,6743 | 3,66692E-55 |
| ACE2 | ATP11C | -0,6743 | 3,77966E-55 |
| ACE2 | MCF2L2 | -0,6744 | 3,57291E-55 |
| ACE2 | VEGFC | -0,6744 | 3,52417E-55 |
| ACE2 | SNTG2 | -0,6745 | 3,3921E-55 |
| ACE2 | SAMD8 | -0,6745 | 3,36843E-55 |
| ACE2 | SYT14 | -0,6746 | 3,23432E-55 |
| ACE2 | RP11-532F6.3 | -0,6746 | 3,25643E-55 |
| ACE2 | TNC | -0,6746 | 3,13362E-55 |
| ACE2 | ACVR1 | -0,6747 | 2,97875E-55 |
| ACE2 | BCHE | -0,6747 | 2,98296E-55 |
| ACE2 | NEFM | -0,6747 | 3,07903E-55 |
| ACE2 | STAC2 | -0,6747 | 3,08025E-55 |
| ACE2 | CFLAR | -0,6748 | 2,88152E-55 |
| ACE2 | SEPT2 | -0,6748 | 2,92609E-55 |
| ACE2 | ARHGAP22 | -0,6748 | 2,92945E-55 |
| ACE2 | ZNF841 | -0,6748 | 2,93602E-55 |
| ACE2 | IFT52 | -0,6748 | 2,89363E-55 |
| ACE2 | C1orf56 | -0,675 | 2,57475E-55 |
| ACE2 | NEK10 | -0,675 | 2,67416E-55 |
| ACE2 | RP11-348P10.2 | -0,675 | 2,60996E-55 |
| ACE2 | PCDHB13 | -0,675 | 2,58614E-55 |
| ACE2 | SAR1A | -0,675 | 2,69104E-55 |
| ACE2 | ATP1A3 | -0,675 | 2,68516E-55 |
| ACE2 | IFI16 | -0,6751 | 2,4971E-55 |
| ACE2 | SLC7A14 | -0,6751 | 2,55966E-55 |
| ACE2 | DCTD | -0,6751 | 2,46958E-55 |
| ACE2 | PMP2 | -0,6751 | 2,47511E-55 |
| ACE2 | ZBED5 | -0,6751 | 2,46774E-55 |
| ACE2 | GTF2F2 | -0,6751 | 2,45039E-55 |
| ACE2 | DCAF12L2 | -0,6751 | 2,53707E-55 |
| ACE2 | PARK7 | -0,6752 | 2,3669E-55 |
| ACE2 | RP11-119F7.5 | -0,6752 | 2,42017E-55 |
| ACE2 | KLRG1 | -0,6752 | 2,40444E-55 |
| ACE2 | DCUN1D3 | -0,6752 | 2,37539E-55 |
| ACE2 | RFLNB | -0,6752 | 2,39276E-55 |
| ACE2 | FAT4 | -0,6753 | 2,27552E-55 |
| ACE2 | LINC00466 | -0,6754 | 2,15658E-55 |
| ACE2 | CTD-2591A1.1 | -0,6754 | 2,18076E-55 |
| ACE2 | COA1 | -0,6754 | 2,16316E-55 |

| | | | |
|---|---|---|---|
| ACE2 | RFX3-AS1 | -0,6754 | 2,12262E-55 |
| ACE2 | GRASP | -0,6754 | 2,13775E-55 |
| ACE2 | USHBP1 | -0,6754 | 2,16043E-55 |
| ACE2 | DNMT3A | -0,6755 | 2,086E-55 |
| ACE2 | PKD1P6 | -0,6755 | 2,07999E-55 |
| ACE2 | OSER1-AS1 | -0,6755 | 2,07974E-55 |
| ACE2 | PCYT1B | -0,6755 | 2,0385E-55 |
| ACE2 | C1QTNF7 | -0,6756 | 1,94185E-55 |
| ACE2 | TCF21 | -0,6756 | 1,90874E-55 |
| ACE2 | PARD6A | -0,6756 | 1,90884E-55 |
| ACE2 | TTC3P1 | -0,6756 | 1,9611E-55 |
| ACE2 | DPYD | -0,6757 | 1,87986E-55 |
| ACE2 | RGL1 | -0,6757 | 1,85518E-55 |
| ACE2 | FILIP1L | -0,6757 | 1,86972E-55 |
| ACE2 | DDHD1 | -0,6757 | 1,83334E-55 |
| ACE2 | AHNAK2 | -0,6757 | 1,84996E-55 |
| ACE2 | MOB3A | -0,6757 | 1,87549E-55 |
| ACE2 | PLIN4 | -0,6757 | 1,87537E-55 |
| ACE2 | HS3ST5 | -0,6758 | 1,73697E-55 |
| ACE2 | RP1-137D17.2 | -0,6758 | 1,73664E-55 |
| ACE2 | CAPS2 | -0,6758 | 1,77344E-55 |
| ACE2 | PCDHGC3 | -0,6759 | 1,70205E-55 |
| ACE2 | MYCBP2 | -0,6759 | 1,65713E-55 |
| ACE2 | DHX30 | -0,676 | 1,59627E-55 |
| ACE2 | PACRGL | -0,676 | 1,54911E-55 |
| ACE2 | UBE3D | -0,676 | 1,59009E-55 |
| ACE2 | PPP2R5B | -0,676 | 1,61788E-55 |
| ACE2 | TMEM88 | -0,676 | 1,55742E-55 |
| ACE2 | ZNF584 | -0,676 | 1,60514E-55 |
| ACE2 | SLC10A4 | -0,6761 | 1,51301E-55 |
| ACE2 | ZNF300P1 | -0,6761 | 1,47988E-55 |
| ACE2 | ARIH1 | -0,6761 | 1,50553E-55 |
| ACE2 | TMEM222 | -0,6762 | 1,4098E-55 |
| ACE2 | CH17-472G23.2 | -0,6762 | 1,41925E-55 |
| ACE2 | ACYP2 | -0,6762 | 1,45686E-55 |
| ACE2 | RP11-350N15.6 | -0,6762 | 1,43213E-55 |
| ACE2 | ST13 | -0,6762 | 1,44963E-55 |
| ACE2 | RP11-1379J22.2 | -0,6763 | 1,33836E-55 |
| ACE2 | NKAPL | -0,6763 | 1,33767E-55 |
| ACE2 | RPS10 | -0,6763 | 1,36992E-55 |
| ACE2 | GTF2IP4 | -0,6763 | 1,35912E-55 |
| ACE2 | RP11-174J11.1 | -0,6763 | 1,39695E-55 |
| ACE2 | CRY2 | -0,6763 | 1,35378E-55 |
| ACE2 | ZNF385A | -0,6763 | 1,34567E-55 |
| ACE2 | SLC25A21 | -0,6763 | 1,35197E-55 |
| ACE2 | EVL | -0,6764 | 1,27442E-55 |
| ACE2 | ZNF177 | -0,6764 | 1,27477E-55 |
| ACE2 | RP5-1198O20.4 | -0,6765 | 1,26023E-55 |
| ACE2 | VANGL2 | -0,6765 | 1,2225E-55 |
| ACE2 | RP11-95D17.1 | -0,6765 | 1,23339E-55 |
| ACE2 | PKNOX1 | -0,6765 | 1,208E-55 |
| ACE2 | WDR45 | -0,6765 | 1,25691E-55 |
| ACE2 | RP4-621F18.2 | -0,6766 | 1,15444E-55 |
| ACE2 | DCAF17 | -0,6766 | 1,14567E-55 |

| | | | |
|---|---|---|---|
| ACE2 | PAPPA | -0,6766 | 1,18417E-55 |
| ACE2 | ZNF174 | -0,6766 | 1,15715E-55 |
| ACE2 | CDS2 | -0,6766 | 1,14813E-55 |
| ACE2 | GBP1 | -0,6768 | 1,04476E-55 |
| ACE2 | ASNSD1 | -0,6768 | 1,05575E-55 |
| ACE2 | PCDHGA9 | -0,6768 | 1,03562E-55 |
| ACE2 | RELA | -0,6768 | 1,04551E-55 |
| ACE2 | AMN1 | -0,6768 | 1,0456E-55 |
| ACE2 | ZNF614 | -0,6769 | 9,90325E-56 |
| ACE2 | LANCL1 | -0,677 | 9,63665E-56 |
| ACE2 | WDR19 | -0,677 | 9,47039E-56 |
| ACE2 | DKK2 | -0,677 | 9,57658E-56 |
| ACE2 | ARSJ | -0,677 | 9,6341E-56 |
| ACE2 | RP1-151F17.2 | -0,677 | 9,81376E-56 |
| ACE2 | MAGEL2 | -0,677 | 9,5604E-56 |
| ACE2 | ZC3H7A | -0,677 | 9,77527E-56 |
| ACE2 | FMO2 | -0,6771 | 9,29101E-56 |
| ACE2 | AC053503.11 | -0,6771 | 9,02724E-56 |
| ACE2 | PSMC3 | -0,6771 | 9,04416E-56 |
| ACE2 | RENBP | -0,6771 | 8,99025E-56 |
| ACE2 | FBXO42 | -0,6772 | 8,78309E-56 |
| ACE2 | DISP1 | -0,6772 | 8,76888E-56 |
| ACE2 | TLN1 | -0,6772 | 8,68115E-56 |
| ACE2 | KCNMA1 | -0,6772 | 8,62115E-56 |
| ACE2 | OSBPL8 | -0,6772 | 8,5899E-56 |
| ACE2 | TBCEL | -0,6773 | 8,20949E-56 |
| ACE2 | SARM1 | -0,6773 | 8,04295E-56 |
| ACE2 | POLR2I | -0,6773 | 8,40234E-56 |
| ACE2 | RP11-389G6.3 | -0,6774 | 7,79501E-56 |
| ACE2 | PCDHGB2 | -0,6774 | 7,8508E-56 |
| ACE2 | FBXO30 | -0,6774 | 7,90154E-56 |
| ACE2 | NFIB | -0,6774 | 7,93742E-56 |
| ACE2 | FUT11 | -0,6774 | 7,69626E-56 |
| ACE2 | NUP188 | -0,6775 | 7,60912E-56 |
| ACE2 | COL4A2-AS1 | -0,6775 | 7,27308E-56 |
| ACE2 | SMYD4 | -0,6775 | 7,46215E-56 |
| ACE2 | AC016757.3 | -0,6776 | 6,88934E-56 |
| ACE2 | ZNF35 | -0,6776 | 7,00955E-56 |
| ACE2 | DPCD | -0,6776 | 7,18811E-56 |
| ACE2 | RP5-1039K5.12 | -0,6776 | 6,93893E-56 |
| ACE2 | TTC14 | -0,6777 | 6,61843E-56 |
| ACE2 | CNR1 | -0,6777 | 6,602E-56 |
| ACE2 | ARID1B | -0,6777 | 6,70109E-56 |
| ACE2 | MAFK | -0,6777 | 6,68336E-56 |
| ACE2 | TERF1 | -0,6777 | 6,59299E-56 |
| ACE2 | DDIT3 | -0,6777 | 6,88286E-56 |
| ACE2 | COMMD10 | -0,6778 | 6,35315E-56 |
| ACE2 | RP11-594N15.3 | -0,6778 | 6,40401E-56 |
| ACE2 | SMIM10L1 | -0,6778 | 6,29746E-56 |
| ACE2 | N4BP2L1 | -0,6778 | 6,40189E-56 |
| ACE2 | YEATS2-AS1 | -0,6779 | 6,19529E-56 |
| ACE2 | PNRC1 | -0,6779 | 5,99255E-56 |
| ACE2 | CHRM2 | -0,6779 | 6,08685E-56 |
| ACE2 | CTD-2245F17.6 | -0,6779 | 6,11541E-56 |

| | | | |
|---|---|---|---|
| ACE2 | AVPR2 | -0,6779 | 6,03375E-56 |
| ACE2 | PTPRG-AS1 | -0,678 | 5,64026E-56 |
| ACE2 | C3orf14 | -0,678 | 5,76414E-56 |
| ACE2 | PCDHGA12 | -0,678 | 5,78086E-56 |
| ACE2 | ROBO4 | -0,678 | 5,71801E-56 |
| ACE2 | VAMP2 | -0,678 | 5,88124E-56 |
| ACE2 | EIF3L | -0,678 | 5,82714E-56 |
| ACE2 | RP11-632P5.1 | -0,6781 | 5,49444E-56 |
| ACE2 | ADGRB1 | -0,6781 | 5,54615E-56 |
| ACE2 | SFRP5 | -0,6781 | 5,49653E-56 |
| ACE2 | IPO9 | -0,6782 | 5,24738E-56 |
| ACE2 | SOCS5 | -0,6782 | 5,30308E-56 |
| ACE2 | TTLL3 | -0,6782 | 5,1774E-56 |
| ACE2 | BRD9 | -0,6782 | 5,11629E-56 |
| ACE2 | RP11-154D6.1 | -0,6782 | 5,20066E-56 |
| ACE2 | GIT2 | -0,6782 | 5,27069E-56 |
| ACE2 | GPATCH2L | -0,6782 | 5,26331E-56 |
| ACE2 | SFRP2 | -0,6783 | 4,8808E-56 |
| ACE2 | NEFL | -0,6783 | 4,96511E-56 |
| ACE2 | CNNM1 | -0,6783 | 4,94739E-56 |
| ACE2 | INMT | -0,6784 | 4,63809E-56 |
| ACE2 | RP11-273G15.2 | -0,6784 | 4,69683E-56 |
| ACE2 | MOB2 | -0,6784 | 4,78756E-56 |
| ACE2 | RRAD | -0,6784 | 4,60398E-56 |
| ACE2 | MX2 | -0,6784 | 4,62426E-56 |
| ACE2 | CASQ2 | -0,6785 | 4,42906E-56 |
| ACE2 | PRDM6 | -0,6785 | 4,5836E-56 |
| ACE2 | SSPO | -0,6785 | 4,43964E-56 |
| ACE2 | XAB2 | -0,6785 | 4,49614E-56 |
| ACE2 | RPL23AP53 | -0,6786 | 4,29089E-56 |
| ACE2 | PIBF1 | -0,6786 | 4,14943E-56 |
| ACE2 | RP11-88E10.4 | -0,6786 | 4,27622E-56 |
| ACE2 | SAT2 | -0,6786 | 4,22784E-56 |
| ACE2 | DPF1 | -0,6786 | 4,19756E-56 |
| ACE2 | SIM2 | -0,6786 | 4,3398E-56 |
| ACE2 | AC006946.16 | -0,6786 | 4,18958E-56 |
| ACE2 | RBM6 | -0,6787 | 4,10274E-56 |
| ACE2 | CPSF4 | -0,6787 | 4,03214E-56 |
| ACE2 | AP000662.4 | -0,6787 | 4,10128E-56 |
| ACE2 | PIEZO1 | -0,6787 | 4,12216E-56 |
| ACE2 | ZIK1 | -0,6787 | 3,99309E-56 |
| ACE2 | EIF4A2 | -0,6788 | 3,83647E-56 |
| ACE2 | KATNA1 | -0,6788 | 3,75238E-56 |
| ACE2 | SPATS2 | -0,6788 | 3,83998E-56 |
| ACE2 | ZNF260 | -0,6788 | 3,78342E-56 |
| ACE2 | SEC63 | -0,6789 | 3,6324E-56 |
| ACE2 | RPS5 | -0,6789 | 3,64687E-56 |
| ACE2 | USP51 | -0,6789 | 3,58556E-56 |
| ACE2 | PHKA1 | -0,6789 | 3,61954E-56 |
| ACE2 | DAPL1 | -0,679 | 3,48979E-56 |
| ACE2 | RP11-496N12.6 | -0,6791 | 3,32522E-56 |
| ACE2 | AL133243.1 | -0,6791 | 3,34276E-56 |
| ACE2 | AC016995.3 | -0,6791 | 3,28819E-56 |
| ACE2 | PGM5 | -0,6791 | 3,27117E-56 |

| | | | |
|---|---|---|---|
| ACE2 | CORO1C | -0,6791 | 3,28582E-56 |
| ACE2 | SGCG | -0,6791 | 3,24065E-56 |
| ACE2 | EZH1 | -0,6791 | 3,24233E-56 |
| ACE2 | ZBTB8A | -0,6792 | 3,0587E-56 |
| ACE2 | PRRC2A | -0,6792 | 3,18464E-56 |
| ACE2 | MDGA1 | -0,6792 | 3,12442E-56 |
| ACE2 | DNM1P51 | -0,6792 | 3,178E-56 |
| ACE2 | ST8SIA3 | -0,6793 | 3,01724E-56 |
| ACE2 | GEMIN8 | -0,6793 | 2,99832E-56 |
| ACE2 | ZNF641 | -0,6794 | 2,77855E-56 |
| ACE2 | SDR39U1 | -0,6794 | 2,86592E-56 |
| ACE2 | BRD7 | -0,6794 | 2,80845E-56 |
| ACE2 | ASIC4 | -0,6795 | 2,71106E-56 |
| ACE2 | METTL21B | -0,6796 | 2,5556E-56 |
| ACE2 | RNPS1 | -0,6796 | 2,57311E-56 |
| ACE2 | CPO | -0,6797 | 2,41375E-56 |
| ACE2 | REC8 | -0,6797 | 2,47105E-56 |
| ACE2 | GDAP1L1 | -0,6797 | 2,36553E-56 |
| ACE2 | SLC2A11 | -0,6797 | 2,40499E-56 |
| ACE2 | HOXD-AS2 | -0,6798 | 2,31109E-56 |
| ACE2 | TMEM196 | -0,6798 | 2,31857E-56 |
| ACE2 | MTMR2 | -0,6798 | 2,27822E-56 |
| ACE2 | PHLDB1 | -0,6798 | 2,27354E-56 |
| ACE2 | RP11-686D22.3 | -0,6798 | 2,28929E-56 |
| ACE2 | KLHL10 | -0,6798 | 2,2823E-56 |
| ACE2 | PRKACA | -0,6798 | 2,34244E-56 |
| ACE2 | TRNAU1AP | -0,6799 | 2,16806E-56 |
| ACE2 | EPS15 | -0,6799 | 2,19419E-56 |
| ACE2 | ADD1 | -0,6799 | 2,17882E-56 |
| ACE2 | CLVS2 | -0,6799 | 2,19304E-56 |
| ACE2 | HDAC8 | -0,6799 | 2,23577E-56 |
| ACE2 | FAM13B | -0,68 | 2,11735E-56 |
| ACE2 | GPR146 | -0,68 | 2,10743E-56 |
| ACE2 | LATS2 | -0,68 | 2,07235E-56 |
| ACE2 | NID2 | -0,68 | 2,07258E-56 |
| ACE2 | SNHG16 | -0,68 | 2,10658E-56 |
| ACE2 | RYR2 | -0,6801 | 1,96564E-56 |
| ACE2 | RP11-597D13.9 | -0,6801 | 1,98761E-56 |
| ACE2 | FNBP4 | -0,6801 | 1,95101E-56 |
| ACE2 | AP1M1 | -0,6801 | 2,00864E-56 |
| ACE2 | DGCR14 | -0,6801 | 1,9808E-56 |
| ACE2 | CLASP1 | -0,6802 | 1,85994E-56 |
| ACE2 | N4BP3 | -0,6802 | 1,87081E-56 |
| ACE2 | CREM | -0,6802 | 1,87679E-56 |
| ACE2 | DDX5 | -0,6802 | 1,8959E-56 |
| ACE2 | HSD11B1L | -0,6802 | 1,87119E-56 |
| ACE2 | GRIP2 | -0,6803 | 1,81008E-56 |
| ACE2 | NEPRO | -0,6803 | 1,74199E-56 |
| ACE2 | THPO | -0,6803 | 1,80695E-56 |
| ACE2 | TMEM64 | -0,6803 | 1,82367E-56 |
| ACE2 | PTENP1-AS | -0,6803 | 1,74203E-56 |
| ACE2 | TRAPPC6B | -0,6803 | 1,74239E-56 |
| ACE2 | C19orf12 | -0,6803 | 1,76706E-56 |
| ACE2 | FOXF2 | -0,6804 | 1,7232E-56 |

| | | | |
|---|---|---|---|
| ACE2 | ZNF397 | -0,6804 | 1,72199E-56 |
| ACE2 | THAP8 | -0,6804 | 1,72711E-56 |
| ACE2 | FAXDC2 | -0,6805 | 1,57243E-56 |
| ACE2 | RALYL | -0,6805 | 1,61841E-56 |
| ACE2 | LSM14A | -0,6805 | 1,58721E-56 |
| ACE2 | UROD | -0,6806 | 1,53023E-56 |
| ACE2 | MYLK | -0,6806 | 1,55744E-56 |
| ACE2 | VAT1L | -0,6806 | 1,52251E-56 |
| ACE2 | NDUFA11 | -0,6806 | 1,5476E-56 |
| ACE2 | MDM1 | -0,6807 | 1,42746E-56 |
| ACE2 | EPS15L1 | -0,6807 | 1,4821E-56 |
| ACE2 | CACHD1 | -0,6808 | 1,37304E-56 |
| ACE2 | COA5 | -0,6808 | 1,36186E-56 |
| ACE2 | TMEM123 | -0,6808 | 1,36176E-56 |
| ACE2 | CEP290 | -0,6808 | 1,37512E-56 |
| ACE2 | PIRT | -0,6808 | 1,40062E-56 |
| ACE2 | TMEM100 | -0,6808 | 1,398E-56 |
| ACE2 | LRRC37A3 | -0,6808 | 1,34343E-56 |
| ACE2 | CRB1 | -0,6809 | 1,30287E-56 |
| ACE2 | TMEM45A | -0,6809 | 1,29208E-56 |
| ACE2 | PP12613 | -0,6809 | 1,30351E-56 |
| ACE2 | WBP4 | -0,6809 | 1,2817E-56 |
| ACE2 | SLC25A29 | -0,6809 | 1,29957E-56 |
| ACE2 | CLEC11A | -0,6809 | 1,2935E-56 |
| ACE2 | CCDC36 | -0,681 | 1,22336E-56 |
| ACE2 | PCDH18 | -0,681 | 1,24326E-56 |
| ACE2 | FLYWCH2 | -0,681 | 1,22516E-56 |
| ACE2 | IGLON5 | -0,681 | 1,23405E-56 |
| ACE2 | METTL8 | -0,6811 | 1,18347E-56 |
| ACE2 | SRSF3 | -0,6811 | 1,18137E-56 |
| ACE2 | RP11-531A24.5 | -0,6811 | 1,18936E-56 |
| ACE2 | RNF41 | -0,6811 | 1,15768E-56 |
| ACE2 | MTUS2 | -0,6811 | 1,20683E-56 |
| ACE2 | CTC-444N24.8 | -0,6811 | 1,20861E-56 |
| ACE2 | ZMYM3 | -0,6811 | 1,1613E-56 |
| ACE2 | RP11-31F15.2 | -0,6812 | 1,14663E-56 |
| ACE2 | VPS72 | -0,6812 | 1,12805E-56 |
| ACE2 | COLEC11 | -0,6812 | 1,14997E-56 |
| ACE2 | RBP1 | -0,6812 | 1,10241E-56 |
| ACE2 | SPCS3 | -0,6812 | 1,12489E-56 |
| ACE2 | RP11-549J18.1 | -0,6812 | 1,14604E-56 |
| ACE2 | SLC25A14 | -0,6812 | 1,14573E-56 |
| ACE2 | ANP32E | -0,6813 | 1,07078E-56 |
| ACE2 | PRPF3 | -0,6813 | 1,04318E-56 |
| ACE2 | IAH1 | -0,6813 | 1,04637E-56 |
| ACE2 | LINC00173 | -0,6813 | 1,08386E-56 |
| ACE2 | RNF24 | -0,6813 | 1,04356E-56 |
| ACE2 | PLA2G4A | -0,6814 | 1,03471E-56 |
| ACE2 | SESTD1 | -0,6814 | 1,00126E-56 |
| ACE2 | C8orf31 | -0,6814 | 9,90575E-57 |
| ACE2 | MAGI2 | -0,6815 | 9,51132E-57 |
| ACE2 | SEC14L2 | -0,6815 | 9,77932E-57 |
| ACE2 | BMI1 | -0,6816 | 9,25228E-57 |
| ACE2 | PTOV1-AS1 | -0,6816 | 8,9114E-57 |

| | | | |
|---|---|---|---|
| ACE2 | TTC21B | -0,6818 | 8,06838E-57 |
| ACE2 | FN1 | -0,6818 | 8,42991E-57 |
| ACE2 | NIPAL4 | -0,6818 | 8,32664E-57 |
| ACE2 | S1PR2 | -0,6818 | 8,16398E-57 |
| ACE2 | TRPC4 | -0,6819 | 7,9025E-57 |
| ACE2 | SPOP | -0,6819 | 7,91584E-57 |
| ACE2 | DONSON | -0,6819 | 7,69901E-57 |
| ACE2 | AC106786.1 | -0,682 | 7,43746E-57 |
| ACE2 | MYH3 | -0,682 | 7,32289E-57 |
| ACE2 | PLD3 | -0,682 | 7,47639E-57 |
| ACE2 | LRRC2 | -0,6821 | 7,08606E-57 |
| ACE2 | UFSP2 | -0,6821 | 6,95042E-57 |
| ACE2 | CSNK2A2 | -0,6821 | 6,95001E-57 |
| ACE2 | MRPS21 | -0,6822 | 6,77873E-57 |
| ACE2 | ANKRD53 | -0,6822 | 6,6075E-57 |
| ACE2 | MRFAP1L1 | -0,6822 | 6,71624E-57 |
| ACE2 | RP11-43F13.4 | -0,6822 | 6,66372E-57 |
| ACE2 | C8orf46 | -0,6822 | 6,77629E-57 |
| ACE2 | TRIM37 | -0,6822 | 6,75767E-57 |
| ACE2 | HEATR6 | -0,6822 | 6,52424E-57 |
| ACE2 | TTN-AS1 | -0,6823 | 6,46937E-57 |
| ACE2 | ZNF316 | -0,6823 | 6,309E-57 |
| ACE2 | CASC2 | -0,6823 | 6,21486E-57 |
| ACE2 | TIAM1 | -0,6823 | 6,35366E-57 |
| ACE2 | SASH1 | -0,6824 | 5,91006E-57 |
| ACE2 | DNM1 | -0,6824 | 5,89123E-57 |
| ACE2 | PDIA2 | -0,6824 | 6,19321E-57 |
| ACE2 | PRDM16 | -0,6825 | 5,69439E-57 |
| ACE2 | LINC01798 | -0,6825 | 5,8361E-57 |
| ACE2 | PTDSS2 | -0,6825 | 5,72171E-57 |
| ACE2 | AP000892.6 | -0,6825 | 5,72505E-57 |
| ACE2 | IFFO2 | -0,6826 | 5,51325E-57 |
| ACE2 | AGTR1 | -0,6826 | 5,50946E-57 |
| ACE2 | RAP2C-AS1 | -0,6826 | 5,40391E-57 |
| ACE2 | ATF2 | -0,6827 | 5,04197E-57 |
| ACE2 | XPOT | -0,6827 | 5,16865E-57 |
| ACE2 | TTLL1 | -0,6827 | 5,19191E-57 |
| ACE2 | LUC7L3 | -0,6828 | 4,8938E-57 |
| ACE2 | GPR27 | -0,6829 | 4,67569E-57 |
| ACE2 | AC073343.13 | -0,6829 | 4,63295E-57 |
| ACE2 | NR2C1 | -0,6829 | 4,78125E-57 |
| ACE2 | ANKRD40 | -0,6829 | 4,64946E-57 |
| ACE2 | ZC3H6 | -0,683 | 4,45589E-57 |
| ACE2 | U91319.1 | -0,683 | 4,34639E-57 |
| ACE2 | ZNF528 | -0,683 | 4,3127E-57 |
| ACE2 | HMGCLL1 | -0,6831 | 4,1327E-57 |
| ACE2 | GAS1 | -0,6831 | 4,29642E-57 |
| ACE2 | BBOF1 | -0,6831 | 4,15375E-57 |
| ACE2 | FAM133B | -0,6832 | 4,0649E-57 |
| ACE2 | TGFBR1 | -0,6833 | 3,74978E-57 |
| ACE2 | SFTPD | -0,6833 | 3,69852E-57 |
| ACE2 | KCTD2 | -0,6833 | 3,79397E-57 |
| ACE2 | SPRED3 | -0,6833 | 3,73248E-57 |
| ACE2 | ATRX | -0,6833 | 3,87762E-57 |

| | | | |
|---|---|---|---|
| ACE2 | VWCE | -0,6834 | 3,63745E-57 |
| ACE2 | CSDE1 | -0,6835 | 3,3874E-57 |
| ACE2 | COX20 | -0,6835 | 3,35703E-57 |
| ACE2 | IFITM2 | -0,6835 | 3,44263E-57 |
| ACE2 | ACSM5 | -0,6835 | 3,39335E-57 |
| ACE2 | ADPRM | -0,6835 | 3,40478E-57 |
| ACE2 | SNCB | -0,6836 | 3,28849E-57 |
| ACE2 | LIN52 | -0,6836 | 3,23135E-57 |
| ACE2 | SLC26A11 | -0,6836 | 3,23169E-57 |
| ACE2 | CTA-204B4.2 | -0,6837 | 3,0134E-57 |
| ACE2 | FAM122A | -0,6837 | 3,04226E-57 |
| ACE2 | POLR2M | -0,6837 | 3,02289E-57 |
| ACE2 | HDAC4 | -0,6838 | 2,93667E-57 |
| ACE2 | TNNT3 | -0,6838 | 2,85941E-57 |
| ACE2 | CHSY1 | -0,6838 | 2,90488E-57 |
| ACE2 | SAMD14 | -0,6838 | 2,84905E-57 |
| ACE2 | ARMCX6 | -0,6838 | 2,89745E-57 |
| ACE2 | CDC42EP3 | -0,6839 | 2,80967E-57 |
| ACE2 | RP11-162D16.2 | -0,6839 | 2,83812E-57 |
| ACE2 | RP11-230G5.2 | -0,6839 | 2,82385E-57 |
| ACE2 | CELF6 | -0,6839 | 2,70971E-57 |
| ACE2 | ZNF263 | -0,6839 | 2,74343E-57 |
| ACE2 | TEX14 | -0,6839 | 2,78535E-57 |
| ACE2 | ZNF426 | -0,6839 | 2,74816E-57 |
| ACE2 | ADIRF-AS1 | -0,684 | 2,59148E-57 |
| ACE2 | CRAMP1 | -0,684 | 2,6077E-57 |
| ACE2 | LRRC3-AS1 | -0,684 | 2,63697E-57 |
| ACE2 | CHRDL1 | -0,684 | 2,57912E-57 |
| ACE2 | YBX1 | -0,6841 | 2,48719E-57 |
| ACE2 | SCN11A | -0,6841 | 2,48922E-57 |
| ACE2 | RP11-731J8.2 | -0,6841 | 2,51525E-57 |
| ACE2 | RSPO3 | -0,6841 | 2,5445E-57 |
| ACE2 | SET | -0,6841 | 2,47909E-57 |
| ACE2 | PARK2 | -0,6842 | 2,32178E-57 |
| ACE2 | ZNF567 | -0,6842 | 2,32188E-57 |
| ACE2 | AR | -0,6842 | 2,3166E-57 |
| ACE2 | UTP11 | -0,6844 | 2,11796E-57 |
| ACE2 | CCDC3 | -0,6844 | 2,17619E-57 |
| ACE2 | RPLP1 | -0,6844 | 2,13693E-57 |
| ACE2 | CCDC154 | -0,6844 | 2,18445E-57 |
| ACE2 | C10orf25 | -0,6845 | 1,97552E-57 |
| ACE2 | PLPP4 | -0,6845 | 2,0442E-57 |
| ACE2 | MEGF9 | -0,6846 | 1,91483E-57 |
| ACE2 | TMEM185A | -0,6846 | 1,95983E-57 |
| ACE2 | MTHFD2P1 | -0,6847 | 1,82613E-57 |
| ACE2 | SOX7 | -0,6847 | 1,85366E-57 |
| ACE2 | RP11-400K9.4 | -0,6847 | 1,87433E-57 |
| ACE2 | RP11-147L13.15 | -0,6847 | 1,78163E-57 |
| ACE2 | SDK1 | -0,6848 | 1,72679E-57 |
| ACE2 | AAAS | -0,6848 | 1,76611E-57 |
| ACE2 | ADAMTSL5 | -0,6848 | 1,714E-57 |
| ACE2 | ELAVL4 | -0,6849 | 1,65012E-57 |
| ACE2 | TK2 | -0,6849 | 1,65099E-57 |
| ACE2 | RFTN2 | -0,685 | 1,5421E-57 |

| | | | |
|---|---|---|---|
| ACE2 | DLX6 | -0,685 | 1,56487E-57 |
| ACE2 | GPN3 | -0,685 | 1,57365E-57 |
| ACE2 | SYT4 | -0,685 | 1,60057E-57 |
| ACE2 | PARD3B | -0,6851 | 1,50577E-57 |
| ACE2 | ZNF300 | -0,6851 | 1,51924E-57 |
| ACE2 | NEDD1 | -0,6851 | 1,49993E-57 |
| ACE2 | ZNF709 | -0,6851 | 1,47343E-57 |
| ACE2 | ANAPC7 | -0,6852 | 1,3878E-57 |
| ACE2 | ZNF585A | -0,6852 | 1,44019E-57 |
| ACE2 | SH3BP5 | -0,6853 | 1,34933E-57 |
| ACE2 | RP11-230F18.5 | -0,6853 | 1,32628E-57 |
| ACE2 | CTU2 | -0,6853 | 1,33948E-57 |
| ACE2 | KRI1 | -0,6853 | 1,30247E-57 |
| ACE2 | ZNF383 | -0,6853 | 1,34603E-57 |
| ACE2 | FLJ27354 | -0,6854 | 1,29143E-57 |
| ACE2 | METTL6 | -0,6854 | 1,23812E-57 |
| ACE2 | HELQ | -0,6854 | 1,26584E-57 |
| ACE2 | SFXN3 | -0,6854 | 1,29736E-57 |
| ACE2 | AC022819.3 | -0,6854 | 1,28305E-57 |
| ACE2 | RP11-428J1.4 | -0,6855 | 1,22586E-57 |
| ACE2 | CACNA1B | -0,6855 | 1,18223E-57 |
| ACE2 | GLIPR1 | -0,6855 | 1,18405E-57 |
| ACE2 | MBNL1-AS1 | -0,6856 | 1,15471E-57 |
| ACE2 | AF001548.5 | -0,6856 | 1,12541E-57 |
| ACE2 | ERVK3-1 | -0,6856 | 1,1175E-57 |
| ACE2 | PLCXD3 | -0,6857 | 1,0902E-57 |
| ACE2 | FBXL22 | -0,6857 | 1,1029E-57 |
| ACE2 | ISLR2 | -0,6857 | 1,10471E-57 |
| ACE2 | LRRC58 | -0,6858 | 1,03824E-57 |
| ACE2 | PNISR | -0,6858 | 1,01752E-57 |
| ACE2 | TCHP | -0,6858 | 1,01515E-57 |
| ACE2 | CTB-50L17.10 | -0,6858 | 1,03218E-57 |
| ACE2 | OGFR | -0,6858 | 1,00077E-57 |
| ACE2 | SULT4A1 | -0,6858 | 1,00423E-57 |
| ACE2 | UBAP2L | -0,6859 | 9,88214E-58 |
| ACE2 | CLK1 | -0,6859 | 9,57335E-58 |
| ACE2 | PRRT3 | -0,6859 | 9,9031E-58 |
| ACE2 | LDLRAD3 | -0,6859 | 9,49419E-58 |
| ACE2 | KMT2A | -0,6859 | 9,58264E-58 |
| ACE2 | APC2 | -0,6859 | 9,85478E-58 |
| ACE2 | ZNF836 | -0,6859 | 9,87063E-58 |
| ACE2 | MACROD2 | -0,6859 | 9,71279E-58 |
| ACE2 | LA16c-385E7.1 | -0,686 | 9,30074E-58 |
| ACE2 | ZFHX3 | -0,686 | 9,14552E-58 |
| ACE2 | SLC9A9 | -0,6861 | 8,78969E-58 |
| ACE2 | RPS23 | -0,6861 | 8,96845E-58 |
| ACE2 | CAP2 | -0,6861 | 8,68734E-58 |
| ACE2 | RP11-452L6.6 | -0,6861 | 8,58707E-58 |
| ACE2 | RP11-401P9.5 | -0,6861 | 8,93112E-58 |
| ACE2 | GPC4 | -0,6861 | 8,82923E-58 |
| ACE2 | RP11-448G15.3 | -0,6862 | 8,21576E-58 |
| ACE2 | TBC1D1 | -0,6862 | 8,42609E-58 |
| ACE2 | NSA2 | -0,6862 | 8,52327E-58 |
| ACE2 | HAND1 | -0,6862 | 8,10907E-58 |

| | | | |
|---|---|---|---|
| ACE2 | KIAA0753 | -0,6862 | 8,41113E-58 |
| ACE2 | EBF1 | -0,6863 | 7,8951E-58 |
| ACE2 | N4BP2L2 | -0,6863 | 8,01921E-58 |
| ACE2 | OSR1 | -0,6864 | 7,42011E-58 |
| ACE2 | TMTC1 | -0,6864 | 7,54996E-58 |
| ACE2 | CNTN6 | -0,6865 | 7,17752E-58 |
| ACE2 | GPR179 | -0,6865 | 7,06107E-58 |
| ACE2 | GS1-590J6.3 | -0,6865 | 7,28013E-58 |
| ACE2 | DFFA | -0,6866 | 6,75412E-58 |
| ACE2 | ALCAM | -0,6866 | 6,57582E-58 |
| ACE2 | ANAPC13 | -0,6866 | 6,61067E-58 |
| ACE2 | PEG10 | -0,6866 | 6,61477E-58 |
| ACE2 | SRPK2 | -0,6866 | 6,7012E-58 |
| ACE2 | SH3PXD2A | -0,6866 | 6,77868E-58 |
| ACE2 | CASC1 | -0,6866 | 6,61558E-58 |
| ACE2 | ACTN1-AS1 | -0,6866 | 6,5994E-58 |
| ACE2 | STON1 | -0,6867 | 6,23475E-58 |
| ACE2 | SIK3 | -0,6867 | 6,3937E-58 |
| ACE2 | ATF7 | -0,6867 | 6,23701E-58 |
| ACE2 | RPL38 | -0,6867 | 6,39265E-58 |
| ACE2 | SRSF4 | -0,6868 | 6,04667E-58 |
| ACE2 | GUK1 | -0,6868 | 5,93265E-58 |
| ACE2 | DARS | -0,6868 | 6,01473E-58 |
| ACE2 | EOGT | -0,6868 | 6,03813E-58 |
| ACE2 | PCDHB3 | -0,6868 | 5,9541E-58 |
| ACE2 | SLC2A4 | -0,6868 | 6,15451E-58 |
| ACE2 | ZNF577 | -0,6868 | 6,17612E-58 |
| ACE2 | KCNQ2 | -0,6869 | 5,78471E-58 |
| ACE2 | MOSPD1 | -0,6869 | 5,74648E-58 |
| ACE2 | AMIGO1 | -0,687 | 5,49097E-58 |
| ACE2 | ZKSCAN7 | -0,687 | 5,39252E-58 |
| ACE2 | HOXA11 | -0,687 | 5,58718E-58 |
| ACE2 | ODF2 | -0,687 | 5,41253E-58 |
| ACE2 | RP11-147L13.11 | -0,687 | 5,56317E-58 |
| ACE2 | KCNK3 | -0,6871 | 5,24678E-58 |
| ACE2 | KLHL30-AS1 | -0,6871 | 5,23479E-58 |
| ACE2 | LINC00950 | -0,6871 | 5,21065E-58 |
| ACE2 | PITPNM2 | -0,6871 | 5,13087E-58 |
| ACE2 | COL5A3 | -0,6871 | 5,22347E-58 |
| ACE2 | S100A4 | -0,6872 | 4,94422E-58 |
| ACE2 | MYT1L | -0,6872 | 4,8672E-58 |
| ACE2 | MRPL33 | -0,6872 | 5,02096E-58 |
| ACE2 | SHISA5 | -0,6872 | 4,85927E-58 |
| ACE2 | SYCP2L | -0,6872 | 4,95218E-58 |
| ACE2 | PHYHIP | -0,6872 | 4,8673E-58 |
| ACE2 | SLC18A3 | -0,6872 | 5,00505E-58 |
| ACE2 | STEAP4 | -0,6873 | 4,55292E-58 |
| ACE2 | SCN4B | -0,6873 | 4,56028E-58 |
| ACE2 | TMEM158 | -0,6874 | 4,51835E-58 |
| ACE2 | SYT9 | -0,6874 | 4,38069E-58 |
| ACE2 | HMGB1 | -0,6874 | 4,39025E-58 |
| ACE2 | RP11-758N13.1 | -0,6874 | 4,48199E-58 |
| ACE2 | ORAI3 | -0,6874 | 4,36207E-58 |
| ACE2 | RP11-439E19.10 | -0,6875 | 4,24318E-58 |

| | | | |
|---|---|---|---|
| ACE2 | LRRC55 | -0,6875 | 4,26566E-58 |
| ACE2 | ADGRF5 | -0,6876 | 4,01976E-58 |
| ACE2 | POPDC3 | -0,6876 | 4,07434E-58 |
| ACE2 | COL4A2-AS2 | -0,6876 | 3,94667E-58 |
| ACE2 | ZFHX2 | -0,6876 | 4,01682E-58 |
| ACE2 | TNFAIP8L1 | -0,6876 | 3,97721E-58 |
| ACE2 | ZNF420 | -0,6877 | 3,81534E-58 |
| ACE2 | CWC27 | -0,6878 | 3,51741E-58 |
| ACE2 | NIPSNAP3B | -0,6879 | 3,4153E-58 |
| ACE2 | RPL7A | -0,6879 | 3,45111E-58 |
| ACE2 | ADA | -0,6879 | 3,4702E-58 |
| ACE2 | PTTG2 | -0,688 | 3,2184E-58 |
| ACE2 | ERGIC2 | -0,688 | 3,28034E-58 |
| ACE2 | SLC38A6 | -0,688 | 3,19292E-58 |
| ACE2 | SYNGR3 | -0,688 | 3,27483E-58 |
| ACE2 | SNRNP70 | -0,688 | 3,24727E-58 |
| ACE2 | DRP2 | -0,688 | 3,15077E-58 |
| ACE2 | ASB3 | -0,6881 | 3,11001E-58 |
| ACE2 | RHBDD1 | -0,6881 | 3,12293E-58 |
| ACE2 | HMGB3P22 | -0,6881 | 3,01293E-58 |
| ACE2 | RP11-286H14.8 | -0,6881 | 3,09896E-58 |
| ACE2 | KIAA0922 | -0,6882 | 2,8944E-58 |
| ACE2 | DIMT1 | -0,6882 | 2,83782E-58 |
| ACE2 | ZNF671 | -0,6882 | 2,89832E-58 |
| ACE2 | PALLD | -0,6883 | 2,68964E-58 |
| ACE2 | MKX | -0,6883 | 2,82199E-58 |
| ACE2 | ZNF236 | -0,6883 | 2,68228E-58 |
| ACE2 | C19orf18 | -0,6883 | 2,76538E-58 |
| ACE2 | EFCAB6 | -0,6883 | 2,75231E-58 |
| ACE2 | KIAA1644 | -0,6883 | 2,72534E-58 |
| ACE2 | WASHC3 | -0,6884 | 2,5667E-58 |
| ACE2 | CEP162 | -0,6885 | 2,48943E-58 |
| ACE2 | DAPK1 | -0,6885 | 2,43389E-58 |
| ACE2 | RNF182 | -0,6886 | 2,30448E-58 |
| ACE2 | ZNF783 | -0,6886 | 2,31747E-58 |
| ACE2 | ZNF250 | -0,6887 | 2,20468E-58 |
| ACE2 | RPS11 | -0,6887 | 2,24823E-58 |
| ACE2 | PRDM2 | -0,6888 | 2,09032E-58 |
| ACE2 | PENK | -0,6888 | 2,1559E-58 |
| ACE2 | PPP1R12B | -0,6889 | 1,96245E-58 |
| ACE2 | GALNT13 | -0,6889 | 1,99383E-58 |
| ACE2 | GUSBP4 | -0,6889 | 1,95326E-58 |
| ACE2 | STMN4 | -0,6889 | 1,96156E-58 |
| ACE2 | IFT140 | -0,6889 | 2,00497E-58 |
| ACE2 | CTD-3035K23.3 | -0,6889 | 1,95469E-58 |
| ACE2 | FKRP | -0,6889 | 2,02905E-58 |
| ACE2 | TCTEX1D1 | -0,689 | 1,88012E-58 |
| ACE2 | FMOD | -0,689 | 1,85362E-58 |
| ACE2 | KLHL30 | -0,689 | 1,87474E-58 |
| ACE2 | NPR3 | -0,689 | 1,86487E-58 |
| ACE2 | PCDHGA4 | -0,689 | 1,87989E-58 |
| ACE2 | SPX | -0,689 | 1,92018E-58 |
| ACE2 | GOLGA8N | -0,689 | 1,90408E-58 |
| ACE2 | GRK3 | -0,689 | 1,92238E-58 |

| | | | |
|---|---|---|---|
| ACE2 | SMIM1 | -0,6891 | 1,80546E-58 |
| ACE2 | ZNF146 | -0,6891 | 1,75405E-58 |
| ACE2 | NMNAT2 | -0,6892 | 1,72422E-58 |
| ACE2 | RP11-120J1.1 | -0,6892 | 1,72942E-58 |
| ACE2 | C15orf52 | -0,6892 | 1,67308E-58 |
| ACE2 | CIC | -0,6892 | 1,69638E-58 |
| ACE2 | RP11-92C4.6 | -0,6893 | 1,64946E-58 |
| ACE2 | MSANTD4 | -0,6893 | 1,57814E-58 |
| ACE2 | PPP1R12A | -0,6893 | 1,64247E-58 |
| ACE2 | RP11-822E23.6 | -0,6893 | 1,63078E-58 |
| ACE2 | ELOF1 | -0,6893 | 1,65613E-58 |
| ACE2 | TSPYL2 | -0,6893 | 1,62047E-58 |
| ACE2 | RPS27 | -0,6894 | 1,55979E-58 |
| ACE2 | RUFY2 | -0,6894 | 1,52079E-58 |
| ACE2 | PMEPA1 | -0,6894 | 1,56675E-58 |
| ACE2 | STUM | -0,6895 | 1,46158E-58 |
| ACE2 | NALCN | -0,6895 | 1,42225E-58 |
| ACE2 | NGB | -0,6895 | 1,42704E-58 |
| ACE2 | SIRT2 | -0,6895 | 1,4255E-58 |
| ACE2 | TMEM178A | -0,6896 | 1,35215E-58 |
| ACE2 | RP1-153G14.4 | -0,6896 | 1,38792E-58 |
| ACE2 | PDE5A | -0,6897 | 1,29789E-58 |
| ACE2 | ADAM20P1 | -0,6897 | 1,29072E-58 |
| ACE2 | XRCC1 | -0,6897 | 1,33932E-58 |
| ACE2 | USP4 | -0,6898 | 1,25972E-58 |
| ACE2 | HRASLS | -0,6898 | 1,21251E-58 |
| ACE2 | PLCG1-AS1 | -0,6898 | 1,22887E-58 |
| ACE2 | ASB1 | -0,6899 | 1,16268E-58 |
| ACE2 | AFAP1L1 | -0,6899 | 1,16302E-58 |
| ACE2 | RP5-1024C24.1 | -0,6899 | 1,15582E-58 |
| ACE2 | UBE2QL1 | -0,69 | 1,09197E-58 |
| ACE2 | CSGALNACT1 | -0,69 | 1,13016E-58 |
| ACE2 | RAB30 | -0,69 | 1,11216E-58 |
| ACE2 | PGPEP1L | -0,69 | 1,13362E-58 |
| ACE2 | TBL1X | -0,69 | 1,11142E-58 |
| ACE2 | AP2M1 | -0,6901 | 1,07667E-58 |
| ACE2 | RP11-481J2.4 | -0,6901 | 1,07419E-58 |
| ACE2 | NCAPH2 | -0,6901 | 1,05823E-58 |
| ACE2 | RNF25 | -0,6902 | 1,02405E-58 |
| ACE2 | FAM212A | -0,6902 | 9,82462E-59 |
| ACE2 | LINC02019 | -0,6902 | 9,88241E-59 |
| ACE2 | REV3L | -0,6902 | 1,02739E-58 |
| ACE2 | IQUB | -0,6902 | 9,9403E-59 |
| ACE2 | RP11-88I18.3 | -0,6902 | 9,86895E-59 |
| ACE2 | ATXN2 | -0,6902 | 9,99015E-59 |
| ACE2 | LRFN1 | -0,6902 | 9,7676E-59 |
| ACE2 | PEAR1 | -0,6903 | 9,29189E-59 |
| ACE2 | FLNC | -0,6903 | 9,71931E-59 |
| ACE2 | OSGIN2 | -0,6903 | 9,59863E-59 |
| ACE2 | RPL23AP1 | -0,6904 | 9,21387E-59 |
| ACE2 | RPS8 | -0,6905 | 8,74764E-59 |
| ACE2 | PLEKHA3 | -0,6905 | 8,41122E-59 |
| ACE2 | COL6A3 | -0,6905 | 8,69473E-59 |
| ACE2 | SEC62 | -0,6905 | 8,36985E-59 |

| | | | |
|---|---|---|---|
| ACE2 | PHKG1 | -0,6905 | 8,51491E-59 |
| ACE2 | KIAA1462 | -0,6905 | 8,43219E-59 |
| ACE2 | ACTA2 | -0,6905 | 8,70818E-59 |
| ACE2 | ZZZ3 | -0,6906 | 7,88753E-59 |
| ACE2 | SUMO2 | -0,6906 | 8,17277E-59 |
| ACE2 | ADARB1 | -0,6906 | 8,02858E-59 |
| ACE2 | MORN5 | -0,6908 | 7,18955E-59 |
| ACE2 | SLITRK5 | -0,6908 | 7,14132E-59 |
| ACE2 | MINDY2 | -0,6908 | 7,1033E-59 |
| ACE2 | NUBP2 | -0,6908 | 7,11489E-59 |
| ACE2 | PCDHB11 | -0,6909 | 6,96655E-59 |
| ACE2 | PINX1 | -0,6909 | 6,72441E-59 |
| ACE2 | METTL26 | -0,6909 | 7,04361E-59 |
| ACE2 | SEMA3E | -0,691 | 6,39285E-59 |
| ACE2 | RHOBTB2 | -0,691 | 6,51566E-59 |
| ACE2 | ZNF829 | -0,691 | 6,68426E-59 |
| ACE2 | GGT7 | -0,691 | 6,6173E-59 |
| ACE2 | DCTN3 | -0,6911 | 6,29582E-59 |
| ACE2 | SLC2A3 | -0,6911 | 6,17335E-59 |
| ACE2 | TCEAL4 | -0,6911 | 6,32138E-59 |
| ACE2 | CRELD1 | -0,6912 | 5,85603E-59 |
| ACE2 | DGKG | -0,6912 | 5,9383E-59 |
| ACE2 | SLC4A2 | -0,6912 | 5,79618E-59 |
| ACE2 | CH17-437K3.1 | -0,6913 | 5,6644E-59 |
| ACE2 | RP11-333I13.1 | -0,6913 | 5,42437E-59 |
| ACE2 | KLHL23 | -0,6913 | 5,60792E-59 |
| ACE2 | PFDN1 | -0,6913 | 5,43765E-59 |
| ACE2 | IFI27L2 | -0,6913 | 5,57261E-59 |
| ACE2 | CENPT | -0,6913 | 5,62329E-59 |
| ACE2 | AP000473.8 | -0,6913 | 5,5309E-59 |
| ACE2 | DIRC3 | -0,6914 | 5,13448E-59 |
| ACE2 | KCNE4 | -0,6914 | 5,32804E-59 |
| ACE2 | TNXA | -0,6914 | 5,28929E-59 |
| ACE2 | PHYHIPL | -0,6914 | 5,27248E-59 |
| ACE2 | TM2D3 | -0,6914 | 5,34216E-59 |
| ACE2 | ZNF579 | -0,6914 | 5,22551E-59 |
| ACE2 | SRM | -0,6915 | 5,1171E-59 |
| ACE2 | CC2D2B | -0,6915 | 5,03834E-59 |
| ACE2 | NME3 | -0,6916 | 4,63518E-59 |
| ACE2 | SNX29P2 | -0,6916 | 4,71477E-59 |
| ACE2 | STK32B | -0,6917 | 4,42453E-59 |
| ACE2 | TMEM43 | -0,6918 | 4,24356E-59 |
| ACE2 | AJAP1 | -0,6919 | 4,11351E-59 |
| ACE2 | RPS7 | -0,6919 | 4,06734E-59 |
| ACE2 | GRIA2 | -0,6919 | 4,09419E-59 |
| ACE2 | SNX18 | -0,6919 | 3,92738E-59 |
| ACE2 | IGFBP3 | -0,692 | 3,82588E-59 |
| ACE2 | PCGF6 | -0,692 | 3,79126E-59 |
| ACE2 | LRRIQ1 | -0,692 | 3,86532E-59 |
| ACE2 | CRABP1 | -0,692 | 3,90773E-59 |
| ACE2 | KRT222 | -0,692 | 3,82083E-59 |
| ACE2 | DFNA5 | -0,6921 | 3,60019E-59 |
| ACE2 | KANK3 | -0,6921 | 3,62783E-59 |
| ACE2 | FAM129A | -0,6922 | 3,46155E-59 |

| | | | |
|---|---|---|---|
| ACE2 | KLHDC3 | -0,6922 | 3,48261E-59 |
| ACE2 | MAPK4 | -0,6922 | 3,43903E-59 |
| ACE2 | NRTN | -0,6922 | 3,48184E-59 |
| ACE2 | PAIP2 | -0,6923 | 3,26712E-59 |
| ACE2 | MIR193BHG | -0,6923 | 3,32243E-59 |
| ACE2 | ZNF691 | -0,6924 | 3,06362E-59 |
| ACE2 | IKBKAP | -0,6924 | 3,1177E-59 |
| ACE2 | EIF2D | -0,6925 | 2,9099E-59 |
| ACE2 | LINC00578 | -0,6925 | 2,95908E-59 |
| ACE2 | TACC1 | -0,6925 | 2,94724E-59 |
| ACE2 | RP11-706C16.7 | -0,6925 | 2,9704E-59 |
| ACE2 | VSTM2B | -0,6925 | 2,89334E-59 |
| ACE2 | IL1RAPL1 | -0,6925 | 2,88282E-59 |
| ACE2 | CEP63 | -0,6926 | 2,79702E-59 |
| ACE2 | RP11-177H13.2 | -0,6926 | 2,76078E-59 |
| ACE2 | WNK1 | -0,6926 | 2,78098E-59 |
| ACE2 | VN1R85P | -0,6926 | 2,81757E-59 |
| ACE2 | ROM1 | -0,6927 | 2,61207E-59 |
| ACE2 | RPS16 | -0,6927 | 2,55743E-59 |
| ACE2 | KANSL3 | -0,6928 | 2,42729E-59 |
| ACE2 | SNHG7 | -0,6928 | 2,50699E-59 |
| ACE2 | MYL6 | -0,6928 | 2,41271E-59 |
| ACE2 | DRAM1 | -0,6928 | 2,51016E-59 |
| ACE2 | CDH5 | -0,6928 | 2,47661E-59 |
| ACE2 | WSB1 | -0,6928 | 2,49296E-59 |
| ACE2 | JMJD6 | -0,6928 | 2,45378E-59 |
| ACE2 | ACOT9 | -0,6928 | 2,43485E-59 |
| ACE2 | MSH6 | -0,6929 | 2,40917E-59 |
| ACE2 | C14orf28 | -0,6929 | 2,38735E-59 |
| ACE2 | KDSR | -0,6929 | 2,30619E-59 |
| ACE2 | SHROOM4 | -0,6929 | 2,34401E-59 |
| ACE2 | RPL9 | -0,693 | 2,1734E-59 |
| ACE2 | FNTA | -0,693 | 2,16718E-59 |
| ACE2 | LMO3 | -0,693 | 2,22636E-59 |
| ACE2 | RP11-567M16.1 | -0,6931 | 2,09287E-59 |
| ACE2 | DPYSL5 | -0,6932 | 1,95938E-59 |
| ACE2 | TF | -0,6932 | 1,96036E-59 |
| ACE2 | RP11-626E13.1 | -0,6932 | 2,02113E-59 |
| ACE2 | ATG12 | -0,6932 | 2,01519E-59 |
| ACE2 | SLC25A32 | -0,6932 | 2,02349E-59 |
| ACE2 | GATSL3 | -0,6932 | 2,03472E-59 |
| ACE2 | MAPK8IP2 | -0,6932 | 1,96709E-59 |
| ACE2 | EIF2B3 | -0,6933 | 1,88614E-59 |
| ACE2 | RAB13 | -0,6933 | 1,84684E-59 |
| ACE2 | PBX1 | -0,6933 | 1,92244E-59 |
| ACE2 | LINC01829 | -0,6933 | 1,9274E-59 |
| ACE2 | CCDC169 | -0,6933 | 1,90001E-59 |
| ACE2 | RP11-73M18.6 | -0,6933 | 1,88442E-59 |
| ACE2 | GSTM5 | -0,6934 | 1,76455E-59 |
| ACE2 | KLHL21 | -0,6935 | 1,71698E-59 |
| ACE2 | PHTF1 | -0,6935 | 1,68424E-59 |
| ACE2 | OLFML2B | -0,6935 | 1,71873E-59 |
| ACE2 | POPDC2 | -0,6935 | 1,684E-59 |
| ACE2 | FLT4 | -0,6935 | 1,71046E-59 |

| | | | |
|---|---|---|---|
| ACE2 | LUC7L | -0,6935 | 1,68985E-59 |
| ACE2 | TEKT2 | -0,6936 | 1,61889E-59 |
| ACE2 | RP11-228B15.4 | -0,6936 | 1,62283E-59 |
| ACE2 | GLMN | -0,6937 | 1,52099E-59 |
| ACE2 | RAP1A | -0,6937 | 1,49164E-59 |
| ACE2 | FAM189B | -0,6937 | 1,52841E-59 |
| ACE2 | IQCJ-SCHIP1 | -0,6937 | 1,52961E-59 |
| ACE2 | RPL7 | -0,6937 | 1,53914E-59 |
| ACE2 | CLCC1 | -0,6938 | 1,47756E-59 |
| ACE2 | SLC25A26 | -0,6938 | 1,44448E-59 |
| ACE2 | PURB | -0,6938 | 1,42328E-59 |
| ACE2 | CDK5RAP2 | -0,6938 | 1,42246E-59 |
| ACE2 | SOX1 | -0,6938 | 1,40933E-59 |
| ACE2 | MORC2 | -0,6938 | 1,42938E-59 |
| ACE2 | PTPRN | -0,6939 | 1,34318E-59 |
| ACE2 | SCN5A | -0,6939 | 1,38515E-59 |
| ACE2 | CTA-14H9.5 | -0,6939 | 1,39015E-59 |
| ACE2 | GS1-124K5.12 | -0,6939 | 1,33449E-59 |
| ACE2 | AP3M2 | -0,6939 | 1,36641E-59 |
| ACE2 | PARVA | -0,6939 | 1,39537E-59 |
| ACE2 | SYNPO2 | -0,694 | 1,32638E-59 |
| ACE2 | VWA3A | -0,694 | 1,31761E-59 |
| ACE2 | VEZF1 | -0,694 | 1,30764E-59 |
| ACE2 | CTD-2001J20.1 | -0,694 | 1,2815E-59 |
| ACE2 | SCGB1B2P | -0,694 | 1,27021E-59 |
| ACE2 | SMOX | -0,694 | 1,28653E-59 |
| ACE2 | SUSD4 | -0,6941 | 1,21329E-59 |
| ACE2 | WDR33 | -0,6941 | 1,24008E-59 |
| ACE2 | SAP30 | -0,6941 | 1,19578E-59 |
| ACE2 | C5 | -0,6941 | 1,2375E-59 |
| ACE2 | C12orf76 | -0,6941 | 1,20223E-59 |
| ACE2 | CDON | -0,6942 | 1,18115E-59 |
| ACE2 | WNT9A | -0,6943 | 1,09501E-59 |
| ACE2 | PPT2 | -0,6943 | 1,11147E-59 |
| ACE2 | EIF3E | -0,6943 | 1,07223E-59 |
| ACE2 | SWI5 | -0,6943 | 1,07506E-59 |
| ACE2 | DYNC2H1 | -0,6943 | 1,11199E-59 |
| ACE2 | NES | -0,6944 | 1,04178E-59 |
| ACE2 | OPTC | -0,6944 | 1,03016E-59 |
| ACE2 | KIT | -0,6944 | 1,05326E-59 |
| ACE2 | ANO1 | -0,6944 | 1,06714E-59 |
| ACE2 | SIVA1 | -0,6944 | 1,03641E-59 |
| ACE2 | TANGO6 | -0,6944 | 1,06116E-59 |
| ACE2 | PCP4L1 | -0,6945 | 9,84914E-60 |
| ACE2 | ZNF197 | -0,6945 | 9,90704E-60 |
| ACE2 | TRIM23 | -0,6945 | 9,96009E-60 |
| ACE2 | DCTN6 | -0,6945 | 1,00782E-59 |
| ACE2 | MEF2A | -0,6945 | 1,00063E-59 |
| ACE2 | MIR1-1HG | -0,6945 | 9,78758E-60 |
| ACE2 | KANTR | -0,6945 | 9,72804E-60 |
| ACE2 | INTS11 | -0,6946 | 9,16308E-60 |
| ACE2 | AC017104.6 | -0,6946 | 9,32263E-60 |
| ACE2 | SBDSP1 | -0,6946 | 9,49141E-60 |
| ACE2 | CALD1 | -0,6946 | 9,37976E-60 |

| | | | |
|---|---|---|---|
| ACE2 | RP11-62F24.2 | -0,6947 | 8,73001E-60 |
| ACE2 | UNC79 | -0,6947 | 8,73126E-60 |
| ACE2 | PTGS1 | -0,6948 | 8,35914E-60 |
| ACE2 | C20orf27 | -0,6948 | 8,58432E-60 |
| ACE2 | NAA10 | -0,6948 | 8,4777E-60 |
| ACE2 | SORBS2 | -0,6949 | 7,85315E-60 |
| ACE2 | NRF1 | -0,6949 | 8,13796E-60 |
| ACE2 | EIF1 | -0,6949 | 8,08307E-60 |
| ACE2 | TPRA1 | -0,695 | 7,40487E-60 |
| ACE2 | PPP3CC | -0,6951 | 7,16371E-60 |
| ACE2 | DAPK3 | -0,6951 | 7,03889E-60 |
| ACE2 | ATP2B3 | -0,6951 | 7,20057E-60 |
| ACE2 | PLA2G5 | -0,6952 | 6,79752E-60 |
| ACE2 | CPE | -0,6952 | 6,80946E-60 |
| ACE2 | IRAK1BP1 | -0,6952 | 6,77423E-60 |
| ACE2 | SNAPC1 | -0,6952 | 6,87946E-60 |
| ACE2 | NME5 | -0,6953 | 6,39086E-60 |
| ACE2 | PCMTD1 | -0,6953 | 6,4309E-60 |
| ACE2 | DYNLT3 | -0,6953 | 6,21498E-60 |
| ACE2 | RP11-384O8.1 | -0,6954 | 6,14918E-60 |
| ACE2 | PDZRN3 | -0,6954 | 6,10589E-60 |
| ACE2 | PIP4K2A | -0,6954 | 6,17541E-60 |
| ACE2 | IFIT1 | -0,6954 | 5,91295E-60 |
| ACE2 | TIE1 | -0,6955 | 5,75792E-60 |
| ACE2 | RFX5 | -0,6955 | 5,67241E-60 |
| ACE2 | CHMP4A | -0,6955 | 5,73216E-60 |
| ACE2 | TP53BP1 | -0,6955 | 5,73449E-60 |
| ACE2 | RP11-456K23.1 | -0,6955 | 5,82697E-60 |
| ACE2 | LLNLF-187D8.1 | -0,6955 | 5,84397E-60 |
| ACE2 | LRRC4B | -0,6955 | 5,63633E-60 |
| ACE2 | LRRC37BP1 | -0,6956 | 5,30054E-60 |
| ACE2 | DBF4B | -0,6956 | 5,47284E-60 |
| ACE2 | WFIKKN2 | -0,6956 | 5,30344E-60 |
| ACE2 | TUBB4A | -0,6956 | 5,56241E-60 |
| ACE2 | HNRNPA3 | -0,6957 | 4,99572E-60 |
| ACE2 | MFF | -0,6957 | 5,07722E-60 |
| ACE2 | JAZF1-AS1 | -0,6957 | 5,14156E-60 |
| ACE2 | LINGO2 | -0,6957 | 5,21955E-60 |
| ACE2 | MMRN2 | -0,6957 | 5,20266E-60 |
| ACE2 | COL4A1 | -0,6957 | 5,00414E-60 |
| ACE2 | C3orf62 | -0,6958 | 4,80664E-60 |
| ACE2 | PCDH7 | -0,6958 | 4,76543E-60 |
| ACE2 | GRP | -0,6958 | 4,83415E-60 |
| ACE2 | C22orf23 | -0,6958 | 4,8249E-60 |
| ACE2 | C6orf3 | -0,6959 | 4,63015E-60 |
| ACE2 | DENND4A | -0,6959 | 4,53212E-60 |
| ACE2 | LINC01081 | -0,6959 | 4,57551E-60 |
| ACE2 | CTC-543D15.8 | -0,6959 | 4,70842E-60 |
| ACE2 | TRIM39 | -0,696 | 4,35076E-60 |
| ACE2 | CYP21A2 | -0,696 | 4,33889E-60 |
| ACE2 | PLN | -0,696 | 4,38154E-60 |
| ACE2 | WDR91 | -0,696 | 4,34901E-60 |
| ACE2 | COL14A1 | -0,696 | 4,31909E-60 |
| ACE2 | SFSWAP | -0,696 | 4,33991E-60 |

| | | | |
|---|---|---|---|
| ACE2 | FBN1 | -0,6961 | 4,1765E-60 |
| ACE2 | DYX1C1 | -0,6961 | 4,16209E-60 |
| ACE2 | HSDL1 | -0,6961 | 4,21119E-60 |
| ACE2 | ASTN1 | -0,6962 | 3,94482E-60 |
| ACE2 | TMEM150C | -0,6962 | 3,89015E-60 |
| ACE2 | DLX2 | -0,6963 | 3,67461E-60 |
| ACE2 | KCTD20 | -0,6963 | 3,77605E-60 |
| ACE2 | ANXA1 | -0,6963 | 3,77365E-60 |
| ACE2 | VAMP1 | -0,6963 | 3,73767E-60 |
| ACE2 | KCNN1 | -0,6963 | 3,60126E-60 |
| ACE2 | RP11-358D17.2 | -0,6964 | 3,48281E-60 |
| ACE2 | TSPYL4 | -0,6964 | 3,51989E-60 |
| ACE2 | LOH12CR2 | -0,6964 | 3,53941E-60 |
| ACE2 | RSL1D1 | -0,6964 | 3,49453E-60 |
| ACE2 | WFDC1 | -0,6964 | 3,437E-60 |
| ACE2 | GID4 | -0,6964 | 3,45722E-60 |
| ACE2 | CTC-444N24.11 | -0,6964 | 3,45026E-60 |
| ACE2 | AC016747.3 | -0,6965 | 3,27197E-60 |
| ACE2 | INPP5E | -0,6965 | 3,36185E-60 |
| ACE2 | CALHM2 | -0,6965 | 3,39851E-60 |
| ACE2 | RP11-390P24.1 | -0,6965 | 3,32883E-60 |
| ACE2 | ZNF337 | -0,6965 | 3,37585E-60 |
| ACE2 | IL1RAPL2 | -0,6966 | 3,06818E-60 |
| ACE2 | DOCK11 | -0,6966 | 3,12981E-60 |
| ACE2 | LYPD1 | -0,6967 | 3,0512E-60 |
| ACE2 | KCNMB1 | -0,6967 | 2,89864E-60 |
| ACE2 | BHLHE22 | -0,6967 | 2,90814E-60 |
| ACE2 | RAPGEF3 | -0,6967 | 2,90446E-60 |
| ACE2 | NOVA2 | -0,6967 | 2,96414E-60 |
| ACE2 | CSRNP3 | -0,6968 | 2,8756E-60 |
| ACE2 | COL5A2 | -0,6968 | 2,86772E-60 |
| ACE2 | TTF1 | -0,6968 | 2,87128E-60 |
| ACE2 | RPGRIP1L | -0,6968 | 2,84275E-60 |
| ACE2 | SPEF1 | -0,6968 | 2,77488E-60 |
| ACE2 | MCOLN3 | -0,6969 | 2,71074E-60 |
| ACE2 | RGS16 | -0,6969 | 2,65482E-60 |
| ACE2 | TIAM2 | -0,6969 | 2,65096E-60 |
| ACE2 | CTD-2104P17.1 | -0,6969 | 2,67718E-60 |
| ACE2 | ZNF610 | -0,6969 | 2,62056E-60 |
| ACE2 | FBLN1 | -0,6969 | 2,7325E-60 |
| ACE2 | ARL9 | -0,697 | 2,55195E-60 |
| ACE2 | APH1B | -0,697 | 2,50683E-60 |
| ACE2 | NTAN1 | -0,697 | 2,56425E-60 |
| ACE2 | RP11-849N15.3 | -0,697 | 2,57031E-60 |
| ACE2 | ZNF555 | -0,697 | 2,4823E-60 |
| ACE2 | LINC00689 | -0,6971 | 2,42433E-60 |
| ACE2 | RAB8B | -0,6971 | 2,35936E-60 |
| ACE2 | PRMT2 | -0,6971 | 2,41987E-60 |
| ACE2 | C1R | -0,6972 | 2,30266E-60 |
| ACE2 | RCBTB1 | -0,6972 | 2,23122E-60 |
| ACE2 | ICE2 | -0,6972 | 2,28683E-60 |
| ACE2 | BAIAP2 | -0,6972 | 2,30035E-60 |
| ACE2 | ATXN1 | -0,6973 | 2,14707E-60 |
| ACE2 | STEAP1B | -0,6973 | 2,12997E-60 |

| | | | |
|---|---|---|---|
| ACE2 | EGFL7 | -0,6973 | 2,13786E-60 |
| ACE2 | DIXDC1 | -0,6973 | 2,15986E-60 |
| ACE2 | YAF2 | -0,6973 | 2,18328E-60 |
| ACE2 | SLFN11 | -0,6973 | 2,11302E-60 |
| ACE2 | PODN | -0,6974 | 2,04711E-60 |
| ACE2 | VAMP4 | -0,6974 | 2,02325E-60 |
| ACE2 | TRAM1L1 | -0,6974 | 2,022E-60 |
| ACE2 | DLGAP2 | -0,6974 | 1,98705E-60 |
| ACE2 | AQP6 | -0,6974 | 2,03112E-60 |
| ACE2 | ATXN3 | -0,6974 | 1,9995E-60 |
| ACE2 | TCEAL1 | -0,6974 | 2,01303E-60 |
| ACE2 | KIF9-AS1 | -0,6975 | 1,92196E-60 |
| ACE2 | PPP1R3C | -0,6975 | 1,90806E-60 |
| ACE2 | C12orf43 | -0,6975 | 1,96216E-60 |
| ACE2 | CAB39L | -0,6975 | 1,94326E-60 |
| ACE2 | ZNF134 | -0,6975 | 1,89018E-60 |
| ACE2 | FAM43B | -0,6976 | 1,8233E-60 |
| ACE2 | RP11-248J18.2 | -0,6976 | 1,81608E-60 |
| ACE2 | MOB1B | -0,6977 | 1,68841E-60 |
| ACE2 | MYCT1 | -0,6977 | 1,69133E-60 |
| ACE2 | FES | -0,6977 | 1,75649E-60 |
| ACE2 | RP11-686D22.7 | -0,6977 | 1,68677E-60 |
| ACE2 | VAX2 | -0,6978 | 1,60553E-60 |
| ACE2 | PCM1 | -0,6978 | 1,66275E-60 |
| ACE2 | IQSEC3 | -0,6978 | 1,61183E-60 |
| ACE2 | SMIM13 | -0,6979 | 1,49973E-60 |
| ACE2 | TRERF1 | -0,6979 | 1,54583E-60 |
| ACE2 | TCEAL8 | -0,6979 | 1,5151E-60 |
| ACE2 | RBM20 | -0,698 | 1,49281E-60 |
| ACE2 | EIF3F | -0,698 | 1,4334E-60 |
| ACE2 | IRGQ | -0,698 | 1,44805E-60 |
| ACE2 | CPNE1 | -0,698 | 1,47592E-60 |
| ACE2 | FZD7 | -0,6981 | 1,36374E-60 |
| ACE2 | TCTEX1D2 | -0,6981 | 1,37927E-60 |
| ACE2 | DSTNP2 | -0,6981 | 1,38296E-60 |
| ACE2 | IGFBP7-AS1 | -0,6982 | 1,2827E-60 |
| ACE2 | MAP3K14 | -0,6982 | 1,27239E-60 |
| ACE2 | TAS1R1 | -0,6983 | 1,21627E-60 |
| ACE2 | EVA1B | -0,6983 | 1,25629E-60 |
| ACE2 | R3HDM1 | -0,6983 | 1,25218E-60 |
| ACE2 | RXRB | -0,6983 | 1,20163E-60 |
| ACE2 | CYBRD1 | -0,6984 | 1,16838E-60 |
| ACE2 | PCDHGB7 | -0,6984 | 1,14773E-60 |
| ACE2 | ETFRF1 | -0,6984 | 1,1531E-60 |
| ACE2 | DCT | -0,6984 | 1,18392E-60 |
| ACE2 | CTA-38K21.2 | -0,6984 | 1,14377E-60 |
| ACE2 | DTNB | -0,6985 | 1,13339E-60 |
| ACE2 | CACNB4 | -0,6985 | 1,08731E-60 |
| ACE2 | DENND6A | -0,6985 | 1,11793E-60 |
| ACE2 | GPR135 | -0,6985 | 1,08909E-60 |
| ACE2 | RP11-666A8.13 | -0,6985 | 1,10002E-60 |
| ACE2 | ZNF790 | -0,6985 | 1,12326E-60 |
| ACE2 | SIRPA | -0,6985 | 1,09465E-60 |
| ACE2 | ALMS1 | -0,6986 | 1,07054E-60 |

| | | | |
|---|---|---|---|
| ACE2 | NRG2 | -0,6986 | 1,06024E-60 |
| ACE2 | PRR4 | -0,6987 | 1,01457E-60 |
| ACE2 | C14orf39 | -0,6987 | 9,60523E-61 |
| ACE2 | PARP16 | -0,6987 | 9,98388E-61 |
| ACE2 | TACR1 | -0,6988 | 9,16571E-61 |
| ACE2 | OSMR | -0,6988 | 9,3078E-61 |
| ACE2 | DCAKD | -0,6988 | 9,13834E-61 |
| ACE2 | WDR83 | -0,6988 | 9,16038E-61 |
| ACE2 | INPP5F | -0,6989 | 8,99374E-61 |
| ACE2 | SERPING1 | -0,6989 | 8,61775E-61 |
| ACE2 | C11orf68 | -0,6989 | 8,6817E-61 |
| ACE2 | CDK2AP1 | -0,6989 | 9,04185E-61 |
| ACE2 | ZNF154 | -0,6989 | 8,91013E-61 |
| ACE2 | MAP3K6 | -0,699 | 8,40028E-61 |
| ACE2 | PEF1 | -0,699 | 8,3447E-61 |
| ACE2 | LEPR | -0,699 | 8,50239E-61 |
| ACE2 | MED10 | -0,699 | 8,4864E-61 |
| ACE2 | UBE2V2 | -0,699 | 8,58767E-61 |
| ACE2 | ZNF615 | -0,699 | 8,41869E-61 |
| ACE2 | SLC43A1 | -0,6991 | 7,83374E-61 |
| ACE2 | PHOX2A | -0,6991 | 7,7476E-61 |
| ACE2 | PLEKHO2 | -0,6991 | 7,73367E-61 |
| ACE2 | ATP1A2 | -0,6992 | 7,62457E-61 |
| ACE2 | PALMD | -0,6993 | 7,21582E-61 |
| ACE2 | ARMC10 | -0,6993 | 7,10326E-61 |
| ACE2 | GGPS1 | -0,6994 | 6,73091E-61 |
| ACE2 | SEPT7 | -0,6994 | 6,64559E-61 |
| ACE2 | AMPH | -0,6994 | 6,53767E-61 |
| ACE2 | DIP2C | -0,6994 | 6,83666E-61 |
| ACE2 | ADAMTS8 | -0,6994 | 6,56387E-61 |
| ACE2 | RP11-46H11.3 | -0,6994 | 6,79309E-61 |
| ACE2 | FAM104B | -0,6994 | 6,72936E-61 |
| ACE2 | RP11-291I6.2 | -0,6995 | 6,30606E-61 |
| ACE2 | CDKN2C | -0,6996 | 5,90417E-61 |
| ACE2 | ERC1 | -0,6996 | 5,88825E-61 |
| ACE2 | LRFN5 | -0,6996 | 6,01848E-61 |
| ACE2 | SMURF2 | -0,6996 | 6,13408E-61 |
| ACE2 | GFPT2 | -0,6997 | 5,79108E-61 |
| ACE2 | PHF2 | -0,6997 | 5,78015E-61 |
| ACE2 | SYT1 | -0,6997 | 5,77889E-61 |
| ACE2 | RP11-6O2.3 | -0,6997 | 5,56559E-61 |
| ACE2 | DNER | -0,6998 | 5,47219E-61 |
| ACE2 | ZNF571 | -0,6998 | 5,2526E-61 |
| ACE2 | RBM11 | -0,6998 | 5,40935E-61 |
| ACE2 | LINC01354 | -0,6999 | 5,06953E-61 |
| ACE2 | TCOF1 | -0,6999 | 5,19432E-61 |
| ACE2 | FLRT2 | -0,6999 | 5,12293E-61 |
| ACE2 | RPS15 | -0,6999 | 5,09236E-61 |
| ACE2 | BVES | -0,7 | 4,93449E-61 |
| ACE2 | FAU | -0,7 | 4,93401E-61 |
| ACE2 | MORF4L1 | -0,7 | 4,71598E-61 |
| ACE2 | CDIP1 | -0,7 | 4,89739E-61 |
| ACE2 | SMIM10L2B | -0,7 | 4,67929E-61 |
| ACE2 | WDR54 | -0,7001 | 4,64862E-61 |

| | | | |
|---|---|---|---|
| ACE2 | ANKRD23 | -0,7001 | 4,49974E-61 |
| ACE2 | NR2F2 | -0,7002 | 4,41261E-61 |
| ACE2 | ZNF570 | -0,7002 | 4,32739E-61 |
| ACE2 | MACF1 | -0,7003 | 4,15619E-61 |
| ACE2 | GPR161 | -0,7003 | 4,14211E-61 |
| ACE2 | SPSB4 | -0,7003 | 4,11475E-61 |
| ACE2 | TTC9B | -0,7003 | 3,97819E-61 |
| ACE2 | RP11-33E12.2 | -0,7004 | 3,928E-61 |
| ACE2 | ZBED8 | -0,7004 | 3,8194E-61 |
| ACE2 | RAB23 | -0,7004 | 3,77688E-61 |
| ACE2 | TSSC2 | -0,7004 | 3,85859E-61 |
| ACE2 | DHX35 | -0,7004 | 3,90084E-61 |
| ACE2 | C2orf76 | -0,7005 | 3,54461E-61 |
| ACE2 | PLCD4 | -0,7005 | 3,54618E-61 |
| ACE2 | ZNF501 | -0,7005 | 3,68585E-61 |
| ACE2 | MGARP | -0,7005 | 3,70393E-61 |
| ACE2 | CTD-3162L10.1 | -0,7005 | 3,73529E-61 |
| ACE2 | IL6ST | -0,7006 | 3,42348E-61 |
| ACE2 | ARSI | -0,7006 | 3,42802E-61 |
| ACE2 | NAXD | -0,7006 | 3,42803E-61 |
| ACE2 | RFC1 | -0,7007 | 3,18263E-61 |
| ACE2 | WASF1 | -0,7007 | 3,25107E-61 |
| ACE2 | EBF2 | -0,7007 | 3,19714E-61 |
| ACE2 | RP11-244O19.1 | -0,7007 | 3,20989E-61 |
| ACE2 | LINC00632 | -0,7007 | 3,23432E-61 |
| ACE2 | ZFAND1 | -0,7008 | 3,0179E-61 |
| ACE2 | PRKG1 | -0,7008 | 3,05891E-61 |
| ACE2 | TPD52L2 | -0,7008 | 3,05367E-61 |
| ACE2 | FAM69A | -0,7009 | 2,86787E-61 |
| ACE2 | REEP1 | -0,7009 | 2,94996E-61 |
| ACE2 | RBM43 | -0,7009 | 2,93924E-61 |
| ACE2 | IQCA1 | -0,7009 | 2,89904E-61 |
| ACE2 | RP11-16P20.4 | -0,7009 | 2,83264E-61 |
| ACE2 | SORBS1 | -0,7009 | 2,96317E-61 |
| ACE2 | LOXL1-AS1 | -0,7009 | 2,92114E-61 |
| ACE2 | RN7SL417P | -0,7009 | 2,84266E-61 |
| ACE2 | URI1 | -0,7009 | 2,87156E-61 |
| ACE2 | DUSP15 | -0,7009 | 2,89754E-61 |
| ACE2 | SPATA4 | -0,701 | 2,68617E-61 |
| ACE2 | ALKBH2 | -0,701 | 2,76562E-61 |
| ACE2 | RP11-5C23.1 | -0,7011 | 2,53454E-61 |
| ACE2 | FADS2 | -0,7011 | 2,64721E-61 |
| ACE2 | GOLGA8A | -0,7011 | 2,547E-61 |
| ACE2 | TBC1D20 | -0,7011 | 2,6452E-61 |
| ACE2 | NTRK1 | -0,7012 | 2,43126E-61 |
| ACE2 | ZFHX4-AS1 | -0,7012 | 2,44173E-61 |
| ACE2 | UBE2V1 | -0,7012 | 2,41331E-61 |
| ACE2 | MAML1 | -0,7013 | 2,30456E-61 |
| ACE2 | RP11-37B2.1 | -0,7013 | 2,28337E-61 |
| ACE2 | HIPK4 | -0,7013 | 2,32418E-61 |
| ACE2 | PGM5P2 | -0,7014 | 2,16606E-61 |
| ACE2 | TRIM69 | -0,7014 | 2,23445E-61 |
| ACE2 | TBC1D26 | -0,7014 | 2,15404E-61 |
| ACE2 | CTC-359D24.3 | -0,7014 | 2,23179E-61 |

| | | | |
|---|---|---|---|
| ACE2 | RP11-208N14.5 | -0,7015 | 2,09781E-61 |
| ACE2 | MDC1 | -0,7016 | 1,98987E-61 |
| ACE2 | RP11-535M15.2 | -0,7016 | 1,91367E-61 |
| ACE2 | C10orf142 | -0,7016 | 1,92173E-61 |
| ACE2 | TUBB3 | -0,7016 | 1,99879E-61 |
| ACE2 | ARHGEF15 | -0,7016 | 1,94616E-61 |
| ACE2 | CEP131 | -0,7016 | 1,99692E-61 |
| ACE2 | LMOD1 | -0,7017 | 1,84696E-61 |
| ACE2 | AKAP12 | -0,7017 | 1,83583E-61 |
| ACE2 | NOX4 | -0,7017 | 1,89038E-61 |
| ACE2 | RABEP1 | -0,7017 | 1,87932E-61 |
| ACE2 | FAM227A | -0,7017 | 1,87997E-61 |
| ACE2 | SPP1 | -0,7018 | 1,74009E-61 |
| ACE2 | LPCAT1 | -0,7018 | 1,71607E-61 |
| ACE2 | DLG5-AS1 | -0,7018 | 1,76937E-61 |
| ACE2 | ZNF541 | -0,7018 | 1,72059E-61 |
| ACE2 | CAV1 | -0,7019 | 1,66215E-61 |
| ACE2 | SCN3B | -0,7019 | 1,64541E-61 |
| ACE2 | FRY | -0,7019 | 1,66628E-61 |
| ACE2 | WDFY2 | -0,7019 | 1,68172E-61 |
| ACE2 | TBC1D24 | -0,7019 | 1,70831E-61 |
| ACE2 | DNAJC6 | -0,702 | 1,58201E-61 |
| ACE2 | RPL32 | -0,702 | 1,55538E-61 |
| ACE2 | RNF150 | -0,702 | 1,54974E-61 |
| ACE2 | FAM20C | -0,702 | 1,57448E-61 |
| ACE2 | ZNF771 | -0,702 | 1,55754E-61 |
| ACE2 | PLD6 | -0,702 | 1,5693E-61 |
| ACE2 | SLC8A1 | -0,7021 | 1,5228E-61 |
| ACE2 | ADCYAP1R1 | -0,7021 | 1,47208E-61 |
| ACE2 | TUBB6 | -0,7021 | 1,51663E-61 |
| ACE2 | GAL | -0,7022 | 1,41156E-61 |
| ACE2 | RP11-66B24.1 | -0,7022 | 1,43669E-61 |
| ACE2 | ZNF766 | -0,7022 | 1,3871E-61 |
| ACE2 | ZNF835 | -0,7022 | 1,43411E-61 |
| ACE2 | ZIM2-AS1 | -0,7022 | 1,40261E-61 |
| ACE2 | GPR88 | -0,7023 | 1,294E-61 |
| ACE2 | MARK1 | -0,7023 | 1,33635E-61 |
| ACE2 | ACTG2 | -0,7023 | 1,34193E-61 |
| ACE2 | SCN2A | -0,7023 | 1,35313E-61 |
| ACE2 | LRRC34 | -0,7023 | 1,36485E-61 |
| ACE2 | GDF6 | -0,7023 | 1,31764E-61 |
| ACE2 | AOC4P | -0,7023 | 1,35038E-61 |
| ACE2 | BRAF | -0,7024 | 1,25645E-61 |
| ACE2 | FBLN5 | -0,7024 | 1,26993E-61 |
| ACE2 | LIMCH1 | -0,7025 | 1,18594E-61 |
| ACE2 | CFAP69 | -0,7025 | 1,16359E-61 |
| ACE2 | SOX17 | -0,7025 | 1,18363E-61 |
| ACE2 | CD44 | -0,7025 | 1,18405E-61 |
| ACE2 | SARNP | -0,7025 | 1,19526E-61 |
| ACE2 | RP11-611L7.2 | -0,7026 | 1,11338E-61 |
| ACE2 | SPIRE1 | -0,7026 | 1,10588E-61 |
| ACE2 | SENP7 | -0,7027 | 1,04901E-61 |
| ACE2 | MAF1 | -0,7027 | 1,06338E-61 |
| ACE2 | DPF2 | -0,7027 | 1,04987E-61 |

| | | | |
|---|---|---|---|
| ACE2 | ZBTB17 | -0,7028 | 1,01023E-61 |
| ACE2 | C11orf70 | -0,7028 | 1,01254E-61 |
| ACE2 | FERMT2 | -0,7028 | 9,84717E-62 |
| ACE2 | RPS6KL1 | -0,7028 | 1,00423E-61 |
| ACE2 | PTPRS | -0,7028 | 9,7861E-62 |
| ACE2 | TMCC1 | -0,7029 | 9,37859E-62 |
| ACE2 | NFIC | -0,7029 | 9,23575E-62 |
| ACE2 | ZNF302 | -0,7029 | 9,41464E-62 |
| ACE2 | CUBN | -0,703 | 8,95957E-62 |
| ACE2 | DACT1 | -0,703 | 8,72976E-62 |
| ACE2 | DGKE | -0,703 | 8,98778E-62 |
| ACE2 | ZNF527 | -0,703 | 8,72222E-62 |
| ACE2 | ZDBF2 | -0,7031 | 8,56825E-62 |
| ACE2 | RP11-20J15.3 | -0,7031 | 8,48628E-62 |
| ACE2 | UROS | -0,7031 | 8,59452E-62 |
| ACE2 | GYG1 | -0,7032 | 7,85727E-62 |
| ACE2 | SSH1 | -0,7032 | 8,22168E-62 |
| ACE2 | MDP1 | -0,7032 | 7,9584E-62 |
| ACE2 | PRRX1 | -0,7033 | 7,48219E-62 |
| ACE2 | SOX5 | -0,7033 | 7,55883E-62 |
| ACE2 | FDX2 | -0,7033 | 7,55979E-62 |
| ACE2 | TBCB | -0,7033 | 7,65283E-62 |
| ACE2 | ADPRH | -0,7034 | 7,27814E-62 |
| ACE2 | HCFC2 | -0,7034 | 7,17338E-62 |
| ACE2 | GATAD2B | -0,7035 | 6,58939E-62 |
| ACE2 | TOR1AIP1 | -0,7035 | 6,89347E-62 |
| ACE2 | TMEM130 | -0,7035 | 6,89176E-62 |
| ACE2 | TMTC2 | -0,7035 | 6,9034E-62 |
| ACE2 | SLFN12 | -0,7035 | 6,89014E-62 |
| ACE2 | ATCAY | -0,7035 | 6,71434E-62 |
| ACE2 | MANBAL | -0,7035 | 6,61381E-62 |
| ACE2 | EIF4ENIF1 | -0,7035 | 6,91854E-62 |
| ACE2 | COLGALT2 | -0,7036 | 6,24809E-62 |
| ACE2 | NDUFAF2 | -0,7036 | 6,55212E-62 |
| ACE2 | ZSCAN21 | -0,7036 | 6,26089E-62 |
| ACE2 | CELF4 | -0,7036 | 6,22144E-62 |
| ACE2 | PHLDB2 | -0,7037 | 6,07851E-62 |
| ACE2 | RP11-755F10.3 | -0,7037 | 6,17538E-62 |
| ACE2 | MED28 | -0,7038 | 5,84839E-62 |
| ACE2 | FADS3 | -0,7038 | 5,73109E-62 |
| ACE2 | ABCC9 | -0,7038 | 5,70095E-62 |
| ACE2 | CACNG4 | -0,7038 | 5,83337E-62 |
| ACE2 | WDCP | -0,7039 | 5,46124E-62 |
| ACE2 | ITGA9 | -0,7039 | 5,47663E-62 |
| ACE2 | THOC7 | -0,7039 | 5,31905E-62 |
| ACE2 | RP11-360F5.3 | -0,7039 | 5,32152E-62 |
| ACE2 | NKX3-1 | -0,7039 | 5,35463E-62 |
| ACE2 | ZNF74 | -0,7039 | 5,30232E-62 |
| ACE2 | TMEM150A | -0,704 | 5,03202E-62 |
| ACE2 | C3orf33 | -0,704 | 4,97547E-62 |
| ACE2 | OSMR-AS1 | -0,704 | 5,11028E-62 |
| ACE2 | ZNF273 | -0,704 | 4,97631E-62 |
| ACE2 | SNX29 | -0,704 | 5,05085E-62 |
| ACE2 | FAM172A | -0,7041 | 4,87057E-62 |

| | | | |
|---|---|---|---|
| ACE2 | CTD-2270F17.1 | -0,7041 | 4,82517E-62 |
| ACE2 | WNT16 | -0,7041 | 4,73998E-62 |
| ACE2 | ZNF467 | -0,7041 | 4,77366E-62 |
| ACE2 | MOK | -0,7041 | 4,72923E-62 |
| ACE2 | FAM50A | -0,7041 | 4,76248E-62 |
| ACE2 | XXbac-BPGBPG55C2( | -0,7042 | 4,62356E-62 |
| ACE2 | INA | -0,7042 | 4,65238E-62 |
| ACE2 | RP11-736K20.5 | -0,7042 | 4,63128E-62 |
| ACE2 | C4orf32 | -0,7043 | 4,29338E-62 |
| ACE2 | ADAM11 | -0,7043 | 4,19194E-62 |
| ACE2 | RAPGEF4 | -0,7044 | 4,07747E-62 |
| ACE2 | AC090044.2 | -0,7044 | 4,10135E-62 |
| ACE2 | ELAVL3 | -0,7044 | 4,04136E-62 |
| ACE2 | DSTN | -0,7044 | 4,15825E-62 |
| ACE2 | GTF2IRD2B | -0,7045 | 3,9059E-62 |
| ACE2 | DUSP26 | -0,7045 | 3,76909E-62 |
| ACE2 | CPLX2 | -0,7046 | 3,67226E-62 |
| ACE2 | PHF11 | -0,7046 | 3,5564E-62 |
| ACE2 | NLGN4X | -0,7046 | 3,65107E-62 |
| ACE2 | ADCY5 | -0,7047 | 3,33604E-62 |
| ACE2 | RP11-774O3.3 | -0,7047 | 3,37168E-62 |
| ACE2 | FAM162B | -0,7047 | 3,47068E-62 |
| ACE2 | LPAR1 | -0,7047 | 3,46024E-62 |
| ACE2 | RANP4 | -0,7047 | 3,40144E-62 |
| ACE2 | RGS2 | -0,7048 | 3,33482E-62 |
| ACE2 | RPL31 | -0,7048 | 3,23488E-62 |
| ACE2 | RP11-554A11.4 | -0,7048 | 3,15748E-62 |
| ACE2 | INAFM2 | -0,7048 | 3,16063E-62 |
| ACE2 | PIFO | -0,7049 | 3,03553E-62 |
| ACE2 | PTCHD4 | -0,7049 | 3,15031E-62 |
| ACE2 | EHBP1L1 | -0,7049 | 2,97982E-62 |
| ACE2 | RP11-582E3.6 | -0,7049 | 3,06064E-62 |
| ACE2 | ZNF136 | -0,7049 | 3,03735E-62 |
| ACE2 | SRRD | -0,7049 | 3,03351E-62 |
| ACE2 | CCDC85A | -0,705 | 2,82021E-62 |
| ACE2 | FAM161A | -0,705 | 2,89521E-62 |
| ACE2 | POLR3GL | -0,7051 | 2,78465E-62 |
| ACE2 | CTNNA2 | -0,7051 | 2,70961E-62 |
| ACE2 | WNT6 | -0,7051 | 2,67591E-62 |
| ACE2 | RHEB | -0,7051 | 2,74691E-62 |
| ACE2 | TMEM268 | -0,7051 | 2,67728E-62 |
| ACE2 | GPR176 | -0,7051 | 2,81064E-62 |
| ACE2 | VSTM2L | -0,7051 | 2,7657E-62 |
| ACE2 | LINC01778 | -0,7052 | 2,64251E-62 |
| ACE2 | RP11-819C21.1 | -0,7052 | 2,62056E-62 |
| ACE2 | STK19 | -0,7053 | 2,45194E-62 |
| ACE2 | NDUFA4L2 | -0,7053 | 2,40882E-62 |
| ACE2 | EHD2 | -0,7053 | 2,43904E-62 |
| ACE2 | WBP1 | -0,7054 | 2,36088E-62 |
| ACE2 | RP11-446H18.1 | -0,7054 | 2,36363E-62 |
| ACE2 | CPED1 | -0,7054 | 2,30746E-62 |
| ACE2 | LOXL2 | -0,7054 | 2,27135E-62 |
| ACE2 | WDYHV1 | -0,7054 | 2,35823E-62 |
| ACE2 | PCDHB1 | -0,7055 | 2,22381E-62 |

| | | | |
|---|---|---|---|
| ACE2 | ABL1 | -0,7055 | 2,12471E-62 |
| ACE2 | KAT8 | -0,7055 | 2,14753E-62 |
| ACE2 | RAB29 | -0,7056 | 2,07792E-62 |
| ACE2 | LINC00963 | -0,7056 | 2,04956E-62 |
| ACE2 | METTL24 | -0,7057 | 1,90306E-62 |
| ACE2 | SMOC2 | -0,7057 | 1,99981E-62 |
| ACE2 | FREM1 | -0,7057 | 1,9283E-62 |
| ACE2 | CCDC92 | -0,7057 | 1,94041E-62 |
| ACE2 | EMCN | -0,7058 | 1,78738E-62 |
| ACE2 | TTLL11 | -0,7058 | 1,81542E-62 |
| ACE2 | STAC | -0,7059 | 1,69894E-62 |
| ACE2 | NR2F1 | -0,7059 | 1,76414E-62 |
| ACE2 | DDX39B | -0,7059 | 1,71987E-62 |
| ACE2 | TRPM3 | -0,7059 | 1,76099E-62 |
| ACE2 | AP3B2 | -0,7059 | 1,7555E-62 |
| ACE2 | LONRF1 | -0,706 | 1,66494E-62 |
| ACE2 | LMF1 | -0,706 | 1,61744E-62 |
| ACE2 | ZNF423 | -0,706 | 1,59481E-62 |
| ACE2 | GBP2 | -0,7061 | 1,55665E-62 |
| ACE2 | EIF1B | -0,7061 | 1,53135E-62 |
| ACE2 | RNF217-AS1 | -0,7061 | 1,53346E-62 |
| ACE2 | KIAA0408 | -0,7061 | 1,57487E-62 |
| ACE2 | HSPB8 | -0,7061 | 1,56781E-62 |
| ACE2 | RBM38 | -0,7061 | 1,56013E-62 |
| ACE2 | L3MBTL2 | -0,7061 | 1,52345E-62 |
| ACE2 | CACNA1E | -0,7062 | 1,45231E-62 |
| ACE2 | ANXA5 | -0,7062 | 1,42607E-62 |
| ACE2 | SCRG1 | -0,7062 | 1,44111E-62 |
| ACE2 | COLEC12 | -0,7062 | 1,49073E-62 |
| ACE2 | SCML2 | -0,7062 | 1,47726E-62 |
| ACE2 | ZNF280C | -0,7062 | 1,49258E-62 |
| ACE2 | LDB2 | -0,7063 | 1,37485E-62 |
| ACE2 | C1S | -0,7063 | 1,41558E-62 |
| ACE2 | C12orf10 | -0,7063 | 1,41701E-62 |
| ACE2 | ZNF582 | -0,7063 | 1,34908E-62 |
| ACE2 | EMILIN3 | -0,7063 | 1,39209E-62 |
| ACE2 | CD81-AS1 | -0,7064 | 1,28728E-62 |
| ACE2 | RP11-21A7A.4 | -0,7064 | 1,31992E-62 |
| ACE2 | RPL6 | -0,7064 | 1,29112E-62 |
| ACE2 | ZNF271P | -0,7064 | 1,3117E-62 |
| ACE2 | TRAPPC2B | -0,7064 | 1,29188E-62 |
| ACE2 | RP11-557N21.1 | -0,7065 | 1,20895E-62 |
| ACE2 | BBS1 | -0,7065 | 1,23174E-62 |
| ACE2 | SDR9C7 | -0,7065 | 1,2208E-62 |
| ACE2 | EID3 | -0,7065 | 1,25325E-62 |
| ACE2 | ZSCAN30 | -0,7065 | 1,22843E-62 |
| ACE2 | CDH11 | -0,7066 | 1,15073E-62 |
| ACE2 | GJC1 | -0,7066 | 1,15828E-62 |
| ACE2 | GPR137B | -0,7067 | 1,0822E-62 |
| ACE2 | INPPL1 | -0,7067 | 1,10894E-62 |
| ACE2 | ZCWPW2 | -0,7068 | 1,06431E-62 |
| ACE2 | PER3 | -0,7069 | 9,78792E-63 |
| ACE2 | ATXN7L2 | -0,7069 | 9,84303E-63 |
| ACE2 | AGAP6 | -0,7069 | 9,57283E-63 |

| | | | |
|---|---|---|---|
| ACE2 | NPAS4 | -0,7069 | 9,99554E-63 |
| ACE2 | UBFD1 | -0,7069 | 9,71996E-63 |
| ACE2 | RP11-346C20.3 | -0,7069 | 9,70209E-63 |
| ACE2 | IFI6 | -0,707 | 9,03571E-63 |
| ACE2 | ATP6V1E2 | -0,707 | 9,23252E-63 |
| ACE2 | TNS1 | -0,707 | 9,17392E-63 |
| ACE2 | MORC4 | -0,707 | 9,11293E-63 |
| ACE2 | RYK | -0,7071 | 8,60411E-63 |
| ACE2 | KCNH2 | -0,7071 | 8,64969E-63 |
| ACE2 | STOM | -0,7071 | 8,6483E-63 |
| ACE2 | CIZ1 | -0,7071 | 8,5618E-63 |
| ACE2 | ZC3H10 | -0,7071 | 8,76181E-63 |
| ACE2 | SIX4 | -0,7071 | 8,63024E-63 |
| ACE2 | TWSG1 | -0,7071 | 8,97056E-63 |
| ACE2 | CEP89 | -0,7071 | 8,7911E-63 |
| ACE2 | DMWD | -0,7071 | 8,86171E-63 |
| ACE2 | RPL7P3 | -0,7071 | 8,89728E-63 |
| ACE2 | PTH1R | -0,7072 | 8,42366E-63 |
| ACE2 | CCDC116 | -0,7072 | 8,15438E-63 |
| ACE2 | TAB1 | -0,7072 | 8,19563E-63 |
| ACE2 | FHL1 | -0,7072 | 8,32472E-63 |
| ACE2 | SRGAP2B | -0,7073 | 7,72666E-63 |
| ACE2 | MAP3K20 | -0,7073 | 7,99242E-63 |
| ACE2 | TXNRD3 | -0,7073 | 7,80772E-63 |
| ACE2 | RP11-475B2.1 | -0,7073 | 7,66777E-63 |
| ACE2 | PNMA1 | -0,7073 | 7,81461E-63 |
| ACE2 | MARK3 | -0,7073 | 7,90515E-63 |
| ACE2 | ZFP1 | -0,7073 | 8,01436E-63 |
| ACE2 | MAST1 | -0,7073 | 7,5825E-63 |
| ACE2 | PLEKHF1 | -0,7073 | 7,66192E-63 |
| ACE2 | WDFY3-AS2 | -0,7074 | 7,32591E-63 |
| ACE2 | RASSF9 | -0,7074 | 7,4388E-63 |
| ACE2 | ULK1 | -0,7074 | 7,28952E-63 |
| ACE2 | SPACA6P-AS | -0,7074 | 7,41153E-63 |
| ACE2 | ZNF17 | -0,7074 | 7,16175E-63 |
| ACE2 | GPC1 | -0,7075 | 6,76448E-63 |
| ACE2 | HACD1 | -0,7075 | 7,11247E-63 |
| ACE2 | CSPG4 | -0,7075 | 7,09765E-63 |
| ACE2 | USP31 | -0,7075 | 6,8213E-63 |
| ACE2 | FAM20B | -0,7076 | 6,74018E-63 |
| ACE2 | MLF1 | -0,7076 | 6,61354E-63 |
| ACE2 | PCDHA3 | -0,7076 | 6,4837E-63 |
| ACE2 | CAMSAP1 | -0,7076 | 6,67428E-63 |
| ACE2 | ALG9 | -0,7076 | 6,67866E-63 |
| ACE2 | DTNBP1 | -0,7077 | 6,14239E-63 |
| ACE2 | ZNF117 | -0,7077 | 6,28239E-63 |
| ACE2 | IDS | -0,7077 | 6,09999E-63 |
| ACE2 | SPATA5 | -0,7078 | 5,77572E-63 |
| ACE2 | GTF2IRD2 | -0,7078 | 5,76618E-63 |
| ACE2 | RPL23AP82 | -0,7078 | 5,86585E-63 |
| ACE2 | ZCCHC7 | -0,7079 | 5,36832E-63 |
| ACE2 | LINC00856 | -0,7079 | 5,41855E-63 |
| ACE2 | TTC23 | -0,7079 | 5,40761E-63 |
| ACE2 | ZNF454 | -0,708 | 5,29761E-63 |

| | | | |
|---|---|---|---|
| ACE2 | HRASLS5 | -0,708 | 5,21673E-63 |
| ACE2 | ACTN1 | -0,708 | 5,31945E-63 |
| ACE2 | ZFP37 | -0,7081 | 4,84325E-63 |
| ACE2 | UBE2I | -0,7081 | 4,88618E-63 |
| ACE2 | CASC3 | -0,7081 | 4,8581E-63 |
| ACE2 | SIX5 | -0,7081 | 4,83265E-63 |
| ACE2 | CA5B | -0,7081 | 4,85203E-63 |
| ACE2 | AC010980.2 | -0,7082 | 4,68038E-63 |
| ACE2 | C19orf53 | -0,7082 | 4,56187E-63 |
| ACE2 | ZNF599 | -0,7082 | 4,69126E-63 |
| ACE2 | ZNF385B | -0,7083 | 4,51581E-63 |
| ACE2 | GTF2H5 | -0,7083 | 4,27556E-63 |
| ACE2 | TRMT61A | -0,7083 | 4,50528E-63 |
| ACE2 | LLGL1 | -0,7083 | 4,43372E-63 |
| ACE2 | SHANK3 | -0,7083 | 4,3761E-63 |
| ACE2 | PRKAA2 | -0,7084 | 4,05412E-63 |
| ACE2 | AC004985.12 | -0,7084 | 4,21165E-63 |
| ACE2 | CTD-2012K14.8 | -0,7084 | 4,13348E-63 |
| ACE2 | RP11-399E6.4 | -0,7085 | 3,82784E-63 |
| ACE2 | DVL3 | -0,7085 | 3,91936E-63 |
| ACE2 | TNS2 | -0,7085 | 4,01811E-63 |
| ACE2 | ACAP3 | -0,7086 | 3,75109E-63 |
| ACE2 | C8orf37 | -0,7086 | 3,76036E-63 |
| ACE2 | TMEM237 | -0,7087 | 3,47906E-63 |
| ACE2 | RPL35A | -0,7087 | 3,55852E-63 |
| ACE2 | TMEM170B | -0,7087 | 3,58706E-63 |
| ACE2 | GNA12 | -0,7087 | 3,55667E-63 |
| ACE2 | FOXP2 | -0,7087 | 3,39021E-63 |
| ACE2 | NINJ1 | -0,7087 | 3,44187E-63 |
| ACE2 | TMEM25 | -0,7087 | 3,50183E-63 |
| ACE2 | MYH11 | -0,7087 | 3,43761E-63 |
| ACE2 | RNF130 | -0,7088 | 3,20197E-63 |
| ACE2 | BLOC1S6 | -0,7088 | 3,31268E-63 |
| ACE2 | ZNF528-AS1 | -0,7088 | 3,36673E-63 |
| ACE2 | BTF3L4 | -0,7089 | 3,0786E-63 |
| ACE2 | MAP1B | -0,7089 | 3,13961E-63 |
| ACE2 | TXLNB | -0,7089 | 3,07168E-63 |
| ACE2 | FRRS1L | -0,7089 | 3,04654E-63 |
| ACE2 | PLA2G4B | -0,7089 | 3,19614E-63 |
| ACE2 | RPL22 | -0,709 | 2,9328E-63 |
| ACE2 | ATP8B2 | -0,709 | 2,98507E-63 |
| ACE2 | NCAM1-AS1 | -0,709 | 2,94741E-63 |
| ACE2 | CACNA1G | -0,709 | 2,85727E-63 |
| ACE2 | ETNK2 | -0,7091 | 2,6937E-63 |
| ACE2 | ANAPC10 | -0,7091 | 2,68864E-63 |
| ACE2 | ADRB3 | -0,7091 | 2,74013E-63 |
| ACE2 | ZHX2 | -0,7091 | 2,75536E-63 |
| ACE2 | RFXAP | -0,7091 | 2,78292E-63 |
| ACE2 | CHRNA3 | -0,7091 | 2,82848E-63 |
| ACE2 | VN1R1 | -0,7091 | 2,76675E-63 |
| ACE2 | DDX17 | -0,7091 | 2,83333E-63 |
| ACE2 | FAM198B | -0,7092 | 2,6589E-63 |
| ACE2 | STARD13 | -0,7092 | 2,58034E-63 |
| ACE2 | SNAPIN | -0,7093 | 2,46366E-63 |

| | | | |
|---|---|---|---|
| ACE2 | TRPC6 | -0,7093 | 2,40511E-63 |
| ACE2 | SRRM1 | -0,7094 | 2,36089E-63 |
| ACE2 | SNAP47 | -0,7094 | 2,37123E-63 |
| ACE2 | ZNF638 | -0,7094 | 2,34298E-63 |
| ACE2 | SH3RF3 | -0,7094 | 2,32153E-63 |
| ACE2 | SDPR | -0,7094 | 2,37631E-63 |
| ACE2 | LIFR-AS1 | -0,7094 | 2,39012E-63 |
| ACE2 | MOCS2 | -0,7094 | 2,32167E-63 |
| ACE2 | TEAD3 | -0,7094 | 2,36065E-63 |
| ACE2 | SF1 | -0,7094 | 2,26199E-63 |
| ACE2 | ANO6 | -0,7094 | 2,29212E-63 |
| ACE2 | SAMD4B | -0,7094 | 2,35893E-63 |
| ACE2 | SHISA7 | -0,7094 | 2,26598E-63 |
| ACE2 | NAA25 | -0,7095 | 2,16667E-63 |
| ACE2 | TNFRSF19 | -0,7095 | 2,20712E-63 |
| ACE2 | CCDC81 | -0,7096 | 2,03951E-63 |
| ACE2 | C16orf52 | -0,7096 | 2,03262E-63 |
| ACE2 | PIPOX | -0,7096 | 2,09958E-63 |
| ACE2 | TTL | -0,7097 | 1,97978E-63 |
| ACE2 | MSTN | -0,7097 | 1,93031E-63 |
| ACE2 | NSRP1 | -0,7097 | 1,93388E-63 |
| ACE2 | RPL17 | -0,7097 | 1,94777E-63 |
| ACE2 | RAMP1 | -0,7098 | 1,81967E-63 |
| ACE2 | HOPX | -0,7098 | 1,85968E-63 |
| ACE2 | HIP1 | -0,7098 | 1,79864E-63 |
| ACE2 | C8orf4 | -0,7098 | 1,89917E-63 |
| ACE2 | SPARCL1 | -0,7099 | 1,71288E-63 |
| ACE2 | GLRB | -0,7099 | 1,72996E-63 |
| ACE2 | NKAIN2 | -0,7099 | 1,73072E-63 |
| ACE2 | RIN3 | -0,7099 | 1,74682E-63 |
| ACE2 | TGFB1 | -0,7099 | 1,74527E-63 |
| ACE2 | LINC00240 | -0,71 | 1,64592E-63 |
| ACE2 | FAM13C | -0,71 | 1,68265E-63 |
| ACE2 | SLC39A6 | -0,71 | 1,63214E-63 |
| ACE2 | F8 | -0,71 | 1,6427E-63 |
| ACE2 | SNX27 | -0,7101 | 1,52226E-63 |
| ACE2 | POT1-AS1 | -0,7101 | 1,50767E-63 |
| ACE2 | RABEPK | -0,7101 | 1,59123E-63 |
| ACE2 | KCNIP3 | -0,7102 | 1,48001E-63 |
| ACE2 | NUDCD3 | -0,7102 | 1,47362E-63 |
| ACE2 | SNHG23 | -0,7102 | 1,4394E-63 |
| ACE2 | HOXB3 | -0,7102 | 1,48509E-63 |
| ACE2 | BST2 | -0,7102 | 1,44744E-63 |
| ACE2 | PPM1K | -0,7103 | 1,34912E-63 |
| ACE2 | NREP | -0,7103 | 1,34678E-63 |
| ACE2 | USP47 | -0,7103 | 1,39451E-63 |
| ACE2 | ZNF433 | -0,7103 | 1,34353E-63 |
| ACE2 | LINC01770 | -0,7104 | 1,2738E-63 |
| ACE2 | WDR47 | -0,7104 | 1,30961E-63 |
| ACE2 | ADRB2 | -0,7104 | 1,2838E-63 |
| ACE2 | FYCO1 | -0,7105 | 1,24131E-63 |
| ACE2 | ARHGAP31 | -0,7105 | 1,21637E-63 |
| ACE2 | RNF146 | -0,7105 | 1,24528E-63 |
| ACE2 | SETBP1 | -0,7105 | 1,24228E-63 |

| | | | |
|---|---|---|---|
| ACE2 | AES | -0,7105 | 1,20886E-63 |
| ACE2 | C8orf88 | -0,7106 | 1,15918E-63 |
| ACE2 | CD40 | -0,7106 | 1,18825E-63 |
| ACE2 | NRSN1 | -0,7107 | 1,10006E-63 |
| ACE2 | SH3BGR | -0,7107 | 1,09923E-63 |
| ACE2 | TIMP3 | -0,7107 | 1,11603E-63 |
| ACE2 | NAP1L3 | -0,7107 | 1,10497E-63 |
| ACE2 | GCFC2 | -0,7108 | 1,03167E-63 |
| ACE2 | TNFAIP6 | -0,7108 | 1,05329E-63 |
| ACE2 | DMGDH | -0,7108 | 1,00925E-63 |
| ACE2 | MTCL1 | -0,7108 | 1,05137E-63 |
| ACE2 | ATP2B4 | -0,7109 | 9,64845E-64 |
| ACE2 | CDHR4 | -0,7109 | 9,67088E-64 |
| ACE2 | ASPN | -0,7109 | 9,99385E-64 |
| ACE2 | CHST3 | -0,7109 | 9,8238E-64 |
| ACE2 | LRRC37A17P | -0,7109 | 9,76317E-64 |
| ACE2 | RP4-800G7.2 | -0,711 | 9,07666E-64 |
| ACE2 | CACNA1H | -0,711 | 9,38077E-64 |
| ACE2 | PRX | -0,711 | 8,95771E-64 |
| ACE2 | PHF13 | -0,7111 | 8,61693E-64 |
| ACE2 | CNGA3 | -0,7111 | 8,54714E-64 |
| ACE2 | LRRC17 | -0,7111 | 8,67527E-64 |
| ACE2 | CALY | -0,7111 | 8,51099E-64 |
| ACE2 | KIAA1549L | -0,7111 | 8,48653E-64 |
| ACE2 | NAA16 | -0,7111 | 8,46255E-64 |
| ACE2 | HEG1 | -0,7112 | 8,07263E-64 |
| ACE2 | RP11-535M15.1 | -0,7112 | 8,14768E-64 |
| ACE2 | C10orf11 | -0,7112 | 8,09345E-64 |
| ACE2 | FOXJ2 | -0,7112 | 8,27204E-64 |
| ACE2 | CLEC1A | -0,7112 | 8,42293E-64 |
| ACE2 | MAPKBP1 | -0,7112 | 8,11996E-64 |
| ACE2 | WWOX | -0,7112 | 8,02481E-64 |
| ACE2 | NAPB | -0,7112 | 8,03935E-64 |
| ACE2 | EXTL1 | -0,7113 | 7,74359E-64 |
| ACE2 | ZNF503 | -0,7113 | 7,54937E-64 |
| ACE2 | RP11-736K20.4 | -0,7113 | 7,53523E-64 |
| ACE2 | TAGLN | -0,7113 | 7,86665E-64 |
| ACE2 | JPH4 | -0,7113 | 7,83965E-64 |
| ACE2 | RBPMS2 | -0,7113 | 7,60709E-64 |
| ACE2 | ZNF565 | -0,7113 | 7,54538E-64 |
| ACE2 | PPP5D1 | -0,7113 | 7,53284E-64 |
| ACE2 | FAM149A | -0,7114 | 7,38122E-64 |
| ACE2 | RP11-91J19.3 | -0,7114 | 7,43851E-64 |
| ACE2 | RPS13 | -0,7114 | 7,41107E-64 |
| ACE2 | PPP1R32 | -0,7114 | 7,28144E-64 |
| ACE2 | TAGLN3 | -0,7115 | 6,95805E-64 |
| ACE2 | PDZD7 | -0,7115 | 6,75048E-64 |
| ACE2 | NRM | -0,7116 | 6,32911E-64 |
| ACE2 | SLIT1 | -0,7116 | 6,63306E-64 |
| ACE2 | HMGN1P3 | -0,7116 | 6,30081E-64 |
| ACE2 | FAM76A | -0,7117 | 6,05036E-64 |
| ACE2 | CRABP2 | -0,7117 | 6,16668E-64 |
| ACE2 | CLEC2L | -0,7117 | 6,00433E-64 |
| ACE2 | NRBP2 | -0,7117 | 6,22056E-64 |

| | | | |
|---|---|---|---|
| ACE2 | SNAPC5 | -0,7117 | 5,957E-64 |
| ACE2 | RP11-263K19.4 | -0,7118 | 5,73416E-64 |
| ACE2 | CEP120 | -0,7118 | 5,79813E-64 |
| ACE2 | ERICD | -0,7118 | 5,60628E-64 |
| ACE2 | FADS1 | -0,7118 | 5,63699E-64 |
| ACE2 | LINC01119 | -0,7119 | 5,36939E-64 |
| ACE2 | LGI2 | -0,7119 | 5,31276E-64 |
| ACE2 | RNF216P1 | -0,7119 | 5,30582E-64 |
| ACE2 | DPYSL4 | -0,7119 | 5,41677E-64 |
| ACE2 | NBEA | -0,7119 | 5,49322E-64 |
| ACE2 | CTIF | -0,7119 | 5,45388E-64 |
| ACE2 | BGLAP | -0,712 | 5,08931E-64 |
| ACE2 | PAQR6 | -0,712 | 5,07437E-64 |
| ACE2 | AFAP1 | -0,712 | 4,98833E-64 |
| ACE2 | PHOX2B | -0,712 | 5,1503E-64 |
| ACE2 | PDGFRB | -0,712 | 5,12972E-64 |
| ACE2 | GAS1RR | -0,712 | 5,25547E-64 |
| ACE2 | ZNF286B | -0,712 | 5,25511E-64 |
| ACE2 | PTOV1 | -0,712 | 5,20414E-64 |
| ACE2 | RBPJ | -0,7121 | 4,95646E-64 |
| ACE2 | NQO2 | -0,7121 | 4,70958E-64 |
| ACE2 | AKIP1 | -0,7121 | 4,71982E-64 |
| ACE2 | CACNA1C | -0,7121 | 4,82309E-64 |
| ACE2 | RBMS2 | -0,7121 | 4,85556E-64 |
| ACE2 | PATZ1 | -0,7121 | 4,82194E-64 |
| ACE2 | JAKMIP3 | -0,7122 | 4,5986E-64 |
| ACE2 | NDRG4 | -0,7122 | 4,48131E-64 |
| ACE2 | WBSCR17 | -0,7123 | 4,39216E-64 |
| ACE2 | LZTS1 | -0,7123 | 4,38596E-64 |
| ACE2 | ILK | -0,7123 | 4,36483E-64 |
| ACE2 | RP11-1055B8.3 | -0,7123 | 4,18295E-64 |
| ACE2 | EFCAB7 | -0,7124 | 3,9828E-64 |
| ACE2 | TGFBR2 | -0,7124 | 4,01507E-64 |
| ACE2 | ARHGAP10 | -0,7124 | 4,10046E-64 |
| ACE2 | NEK1 | -0,7124 | 4,12748E-64 |
| ACE2 | RPS20P22 | -0,7124 | 4,12532E-64 |
| ACE2 | OTUD5 | -0,7124 | 4,00227E-64 |
| ACE2 | TFE3 | -0,7124 | 3,9944E-64 |
| ACE2 | HMP19 | -0,7125 | 3,84435E-64 |
| ACE2 | RPL12 | -0,7125 | 3,76176E-64 |
| ACE2 | WSCD2 | -0,7125 | 3,73928E-64 |
| ACE2 | RP11-396B14.2 | -0,7125 | 3,73244E-64 |
| ACE2 | ZC4H2 | -0,7125 | 3,81633E-64 |
| ACE2 | C1QTNF3 | -0,7126 | 3,57844E-64 |
| ACE2 | MATR3 | -0,7126 | 3,59759E-64 |
| ACE2 | GSTA4 | -0,7126 | 3,57109E-64 |
| ACE2 | RBM28 | -0,7126 | 3,57158E-64 |
| ACE2 | CTD-2336O2.1 | -0,7126 | 3,54712E-64 |
| ACE2 | CDH6 | -0,7127 | 3,34437E-64 |
| ACE2 | TMEM8B | -0,7127 | 3,30923E-64 |
| ACE2 | WHRN | -0,7127 | 3,4561E-64 |
| ACE2 | BRSK2 | -0,7127 | 3,42191E-64 |
| ACE2 | CCDC84 | -0,7127 | 3,37118E-64 |
| ACE2 | KCNK2 | -0,7128 | 3,17548E-64 |

| | | | |
|---|---|---|---|
| ACE2 | RPSAP70 | -0,7128 | 3,2569E-64 |
| ACE2 | PCDHGA7 | -0,7128 | 3,14757E-64 |
| ACE2 | TMOD1 | -0,7128 | 3,27421E-64 |
| ACE2 | NFYB | -0,7128 | 3,1961E-64 |
| ACE2 | NTN1 | -0,7128 | 3,29465E-64 |
| ACE2 | RP11-481J13.1 | -0,7129 | 2,94645E-64 |
| ACE2 | ZNF160 | -0,7129 | 3,00618E-64 |
| ACE2 | KANSL1L | -0,713 | 2,90917E-64 |
| ACE2 | ITPR1-AS1 | -0,713 | 2,92164E-64 |
| ACE2 | ZNF76 | -0,713 | 2,93159E-64 |
| ACE2 | ZNF517 | -0,713 | 2,82427E-64 |
| ACE2 | DNAJB5 | -0,713 | 2,93898E-64 |
| ACE2 | UVRAG | -0,713 | 2,85035E-64 |
| ACE2 | MAMSTR | -0,713 | 2,7919E-64 |
| ACE2 | NKAP | -0,713 | 2,88047E-64 |
| ACE2 | MAP9 | -0,7131 | 2,69166E-64 |
| ACE2 | MLLT10 | -0,7131 | 2,73229E-64 |
| ACE2 | AIP | -0,7131 | 2,71497E-64 |
| ACE2 | ZNF843 | -0,7131 | 2,72438E-64 |
| ACE2 | CDKL5 | -0,7131 | 2,67871E-64 |
| ACE2 | ZCCHC18 | -0,7131 | 2,72057E-64 |
| ACE2 | NUDC | -0,7132 | 2,54027E-64 |
| ACE2 | TMEM145 | -0,7132 | 2,52833E-64 |
| ACE2 | FUZ | -0,7132 | 2,59498E-64 |
| ACE2 | ZNF583 | -0,7132 | 2,48491E-64 |
| ACE2 | WDR35 | -0,7133 | 2,38599E-64 |
| ACE2 | ZNF639 | -0,7134 | 2,32109E-64 |
| ACE2 | CASC10 | -0,7134 | 2,20875E-64 |
| ACE2 | MGAT4C | -0,7134 | 2,25624E-64 |
| ACE2 | ZNF227 | -0,7134 | 2,24388E-64 |
| ACE2 | MTCP1 | -0,7134 | 2,26309E-64 |
| ACE2 | ROR1 | -0,7135 | 2,11631E-64 |
| ACE2 | PPP2R2C | -0,7135 | 2,15576E-64 |
| ACE2 | EGFL8 | -0,7135 | 2,159E-64 |
| ACE2 | TRIM46 | -0,7136 | 1,99401E-64 |
| ACE2 | ZNF248 | -0,7136 | 2,03771E-64 |
| ACE2 | PYGM | -0,7136 | 2,01681E-64 |
| ACE2 | PSD2 | -0,7137 | 1,85672E-64 |
| ACE2 | PCDHB18P | -0,7137 | 1,86307E-64 |
| ACE2 | H2AFV | -0,7137 | 1,91591E-64 |
| ACE2 | TTC26 | -0,7137 | 1,84569E-64 |
| ACE2 | WDR86-AS1 | -0,7137 | 1,86374E-64 |
| ACE2 | HS3ST3A1 | -0,7138 | 1,75435E-64 |
| ACE2 | EFR3B | -0,7139 | 1,67978E-64 |
| ACE2 | INVS | -0,7139 | 1,65327E-64 |
| ACE2 | SGMS1 | -0,7139 | 1,64009E-64 |
| ACE2 | ZNF84 | -0,7139 | 1,71832E-64 |
| ACE2 | WASF3 | -0,7139 | 1,65901E-64 |
| ACE2 | TMEM179 | -0,7139 | 1,698E-64 |
| ACE2 | RP11-815I9.4 | -0,7139 | 1,70942E-64 |
| ACE2 | RPL19 | -0,7139 | 1,65826E-64 |
| ACE2 | RGS8 | -0,714 | 1,5735E-64 |
| ACE2 | TRAF5 | -0,714 | 1,60576E-64 |
| ACE2 | LINC00682 | -0,714 | 1,60283E-64 |

| | | | |
|---|---|---|---|
| ACE2 | MURC | -0,7141 | 1,50354E-64 |
| ACE2 | LINC00662 | -0,7141 | 1,49791E-64 |
| ACE2 | RP11-12M5.3 | -0,7142 | 1,37979E-64 |
| ACE2 | RP11-254F7.2 | -0,7142 | 1,4071E-64 |
| ACE2 | SETD5 | -0,7142 | 1,37947E-64 |
| ACE2 | RP11-635L1.2 | -0,7142 | 1,41341E-64 |
| ACE2 | COL1A2 | -0,7142 | 1,41718E-64 |
| ACE2 | BPGM | -0,7142 | 1,39843E-64 |
| ACE2 | RAD51D | -0,7142 | 1,39309E-64 |
| ACE2 | CFAP97 | -0,7143 | 1,34707E-64 |
| ACE2 | TSPYL5 | -0,7143 | 1,35748E-64 |
| ACE2 | UPF3A | -0,7143 | 1,35355E-64 |
| ACE2 | RP11-875O11.1 | -0,7144 | 1,23562E-64 |
| ACE2 | CHST15 | -0,7144 | 1,27759E-64 |
| ACE2 | THOC5 | -0,7144 | 1,22742E-64 |
| ACE2 | KLF8 | -0,7144 | 1,25795E-64 |
| ACE2 | FAM178B | -0,7145 | 1,20332E-64 |
| ACE2 | WDR17 | -0,7145 | 1,16854E-64 |
| ACE2 | AC005754.8 | -0,7145 | 1,18632E-64 |
| ACE2 | PABPC1P4 | -0,7145 | 1,19081E-64 |
| ACE2 | DCN | -0,7145 | 1,17236E-64 |
| ACE2 | MYOCD | -0,7145 | 1,18473E-64 |
| ACE2 | CTD-2541J13.1 | -0,7145 | 1,16581E-64 |
| ACE2 | ASXL1 | -0,7145 | 1,20133E-64 |
| ACE2 | RGN | -0,7145 | 1,15828E-64 |
| ACE2 | C1orf52 | -0,7146 | 1,13676E-64 |
| ACE2 | CCDC181 | -0,7146 | 1,1104E-64 |
| ACE2 | LIX1 | -0,7146 | 1,08429E-64 |
| ACE2 | EPM2A | -0,7147 | 1,03425E-64 |
| ACE2 | RASA4 | -0,7147 | 1,02798E-64 |
| ACE2 | IPO5 | -0,7147 | 1,02647E-64 |
| ACE2 | BRMS1L | -0,7147 | 1,0525E-64 |
| ACE2 | AOC2 | -0,7147 | 1,04376E-64 |
| ACE2 | SNAP25 | -0,7147 | 1,02518E-64 |
| ACE2 | RP11-1334A24.5 | -0,7148 | 9,70121E-65 |
| ACE2 | FBXO10 | -0,7148 | 9,84282E-65 |
| ACE2 | KB-68A7.1 | -0,7148 | 9,74149E-65 |
| ACE2 | PTPN14 | -0,7149 | 9,11821E-65 |
| ACE2 | COPS2 | -0,7149 | 9,12557E-65 |
| ACE2 | DDX19A | -0,7149 | 9,28841E-65 |
| ACE2 | SLC6A16 | -0,7149 | 9,34341E-65 |
| ACE2 | RP11-378J18.8 | -0,715 | 8,89198E-65 |
| ACE2 | NOP14-AS1 | -0,715 | 8,99186E-65 |
| ACE2 | GPX8 | -0,715 | 8,96737E-65 |
| ACE2 | MTX3 | -0,715 | 8,83127E-65 |
| ACE2 | C11orf58 | -0,715 | 9,03624E-65 |
| ACE2 | ATAD3C | -0,7151 | 8,46342E-65 |
| ACE2 | RXRG | -0,7151 | 8,37477E-65 |
| ACE2 | PCDHB17P | -0,7151 | 8,0608E-65 |
| ACE2 | TFAP2A-AS1 | -0,7151 | 8,38742E-65 |
| ACE2 | ZNF211 | -0,7151 | 8,2373E-65 |
| ACE2 | SUPT20HL1 | -0,7151 | 8,21825E-65 |
| ACE2 | TMEM47 | -0,7151 | 8,30546E-65 |
| ACE2 | GULP1 | -0,7152 | 7,59319E-65 |

| | | | |
|---|---|---|---|
| ACE2 | SRF | -0,7152 | 7,63234E-65 |
| ACE2 | RNF112 | -0,7152 | 7,62003E-65 |
| ACE2 | TLE2 | -0,7152 | 7,77446E-65 |
| ACE2 | PLEKHM2 | -0,7153 | 7,48906E-65 |
| ACE2 | CNBP | -0,7153 | 7,28822E-65 |
| ACE2 | KCTD7 | -0,7153 | 7,31365E-65 |
| ACE2 | PLA2G16 | -0,7153 | 7,40132E-65 |
| ACE2 | PFKM | -0,7153 | 7,22089E-65 |
| ACE2 | SLFN5 | -0,7153 | 7,37026E-65 |
| ACE2 | MPP3 | -0,7153 | 7,14034E-65 |
| ACE2 | ADAMTS12 | -0,7154 | 7,04286E-65 |
| ACE2 | ZCRB1 | -0,7154 | 7,03528E-65 |
| ACE2 | SNX21 | -0,7154 | 6,73811E-65 |
| ACE2 | BCAS4 | -0,7154 | 6,91804E-65 |
| ACE2 | FAM122C | -0,7154 | 7,01929E-65 |
| ACE2 | GNPDA2 | -0,7155 | 6,33305E-65 |
| ACE2 | SEMA3A | -0,7155 | 6,51107E-65 |
| ACE2 | LDB1 | -0,7155 | 6,55632E-65 |
| ACE2 | CPEB1-AS1 | -0,7155 | 6,60426E-65 |
| ACE2 | RPL23 | -0,7155 | 6,54827E-65 |
| ACE2 | KCNMB2 | -0,7156 | 6,15034E-65 |
| ACE2 | FYTTD1 | -0,7156 | 6,19484E-65 |
| ACE2 | ELOVL2 | -0,7156 | 6,23273E-65 |
| ACE2 | S1PR3 | -0,7156 | 6,08958E-65 |
| ACE2 | ST3GAL4-AS1 | -0,7156 | 6,25764E-65 |
| ACE2 | DNAH10OS | -0,7156 | 6,0973E-65 |
| ACE2 | FLJ16779 | -0,7156 | 6,20624E-65 |
| ACE2 | ITGB1BP2 | -0,7156 | 6,04755E-65 |
| ACE2 | IFI44L | -0,7157 | 5,83249E-65 |
| ACE2 | P2RX7 | -0,7157 | 5,79874E-65 |
| ACE2 | RP11-327J17.9 | -0,7157 | 5,76917E-65 |
| ACE2 | GNAZ | -0,7157 | 5,94798E-65 |
| ACE2 | ANKRD34A | -0,7158 | 5,44184E-65 |
| ACE2 | GRIA1 | -0,7158 | 5,40343E-65 |
| ACE2 | PLEKHA2 | -0,7158 | 5,53268E-65 |
| ACE2 | CPNE6 | -0,7158 | 5,32963E-65 |
| ACE2 | RSL24D1 | -0,7158 | 5,56925E-65 |
| ACE2 | NKILA | -0,7158 | 5,32387E-65 |
| ACE2 | RP11-420L9.5 | -0,7159 | 5,06054E-65 |
| ACE2 | COPS6 | -0,7159 | 5,00671E-65 |
| ACE2 | BNIP3 | -0,7159 | 5,15958E-65 |
| ACE2 | AOC3 | -0,7159 | 5,27486E-65 |
| ACE2 | ZNF677 | -0,7159 | 5,18027E-65 |
| ACE2 | ZNF549 | -0,7159 | 5,28057E-65 |
| ACE2 | HOXD8 | -0,716 | 4,95223E-65 |
| ACE2 | YBX3 | -0,716 | 4,74912E-65 |
| ACE2 | GPALPP1 | -0,7161 | 4,51257E-65 |
| ACE2 | TMEM204 | -0,7161 | 4,55347E-65 |
| ACE2 | CBX1 | -0,7161 | 4,58732E-65 |
| ACE2 | AHI1 | -0,7162 | 4,40377E-65 |
| ACE2 | ATP6V0E2 | -0,7162 | 4,36724E-65 |
| ACE2 | FZD4 | -0,7162 | 4,20055E-65 |
| ACE2 | PGR-AS1 | -0,7162 | 4,18897E-65 |
| ACE2 | PPP1R14A | -0,7162 | 4,25083E-65 |

| | | | |
|---|---|---|---|
| ACE2 | CTC-301O7.4 | -0,7162 | 4,21295E-65 |
| ACE2 | TRABD2B | -0,7163 | 4,16139E-65 |
| ACE2 | AC064875.2 | -0,7163 | 4,16416E-65 |
| ACE2 | FNDC4 | -0,7163 | 4,05304E-65 |
| ACE2 | AC053503.6 | -0,7163 | 4,12098E-65 |
| ACE2 | RAB33B | -0,7163 | 4,08849E-65 |
| ACE2 | MFAP3L | -0,7163 | 4,16917E-65 |
| ACE2 | AC010226.4 | -0,7163 | 4,1451E-65 |
| ACE2 | SNAP91 | -0,7163 | 4,1294E-65 |
| ACE2 | CTD-2054N24.2 | -0,7163 | 3,94412E-65 |
| ACE2 | RPS6KC1 | -0,7164 | 3,82162E-65 |
| ACE2 | ARHGDIG | -0,7164 | 3,75881E-65 |
| ACE2 | TTC39C | -0,7164 | 3,81036E-65 |
| ACE2 | PTGFR | -0,7165 | 3,69673E-65 |
| ACE2 | DOCK3 | -0,7165 | 3,49522E-65 |
| ACE2 | FAM220A | -0,7165 | 3,50776E-65 |
| ACE2 | RP11-867G23.10 | -0,7165 | 3,6511E-65 |
| ACE2 | CWF19L2 | -0,7165 | 3,56132E-65 |
| ACE2 | NEMP1 | -0,7165 | 3,50306E-65 |
| ACE2 | CCDC8 | -0,7165 | 3,49473E-65 |
| ACE2 | ABL2 | -0,7166 | 3,39313E-65 |
| ACE2 | FGF10 | -0,7166 | 3,37759E-65 |
| ACE2 | HOXB-AS1 | -0,7166 | 3,35825E-65 |
| ACE2 | ZNF559 | -0,7166 | 3,36022E-65 |
| ACE2 | CTC-510F12.2 | -0,7166 | 3,29301E-65 |
| ACE2 | SMIM17 | -0,7166 | 3,45477E-65 |
| ACE2 | DSTYK | -0,7167 | 3,28773E-65 |
| ACE2 | ATN1 | -0,7167 | 3,23802E-65 |
| ACE2 | FGF14-AS2 | -0,7167 | 3,14684E-65 |
| ACE2 | ZNF609 | -0,7167 | 3,15545E-65 |
| ACE2 | HLF | -0,7167 | 3,18667E-65 |
| ACE2 | MLLT11 | -0,7168 | 3,07854E-65 |
| ACE2 | ANGPTL7 | -0,7169 | 2,89912E-65 |
| ACE2 | BOK | -0,7169 | 2,90302E-65 |
| ACE2 | CHCHD6 | -0,7169 | 2,89949E-65 |
| ACE2 | DGKB | -0,7169 | 2,75365E-65 |
| ACE2 | ITPRIP | -0,7169 | 2,82888E-65 |
| ACE2 | MAP4K2 | -0,7169 | 2,87008E-65 |
| ACE2 | MED21 | -0,7169 | 2,91053E-65 |
| ACE2 | PTPRM | -0,7169 | 2,776E-65 |
| ACE2 | TMEM161A | -0,7169 | 2,84127E-65 |
| ACE2 | TMEM255A | -0,7169 | 2,91479E-65 |
| ACE2 | SGCB | -0,717 | 2,70841E-65 |
| ACE2 | SLC15A4 | -0,717 | 2,6832E-65 |
| ACE2 | MAP1A | -0,717 | 2,73618E-65 |
| ACE2 | ARHGAP6 | -0,717 | 2,63902E-65 |
| ACE2 | DNAJC8 | -0,7171 | 2,55362E-65 |
| ACE2 | ARMC9 | -0,7171 | 2,46681E-65 |
| ACE2 | CCL27 | -0,7171 | 2,46072E-65 |
| ACE2 | ZMAT1 | -0,7171 | 2,48971E-65 |
| ACE2 | C1orf54 | -0,7172 | 2,42936E-65 |
| ACE2 | KIF1A | -0,7172 | 2,41548E-65 |
| ACE2 | ABHD14A | -0,7172 | 2,43694E-65 |
| ACE2 | ZNF354C | -0,7172 | 2,41894E-65 |

| | | | |
|---|---|---|---|
| ACE2 | NRP1 | -0,7172 | 2,38267E-65 |
| ACE2 | FRA10AC1 | -0,7172 | 2,3085E-65 |
| ACE2 | C12orf57 | -0,7172 | 2,39168E-65 |
| ACE2 | PDE1B | -0,7172 | 2,30171E-65 |
| ACE2 | SAMD4A | -0,7172 | 2,37733E-65 |
| ACE2 | C14orf37 | -0,7172 | 2,35855E-65 |
| ACE2 | PTGIS | -0,7172 | 2,40609E-65 |
| ACE2 | MAPK11 | -0,7172 | 2,39135E-65 |
| ACE2 | LONRF2 | -0,7173 | 2,23079E-65 |
| ACE2 | LYRM1 | -0,7173 | 2,19234E-65 |
| ACE2 | NKAIN4 | -0,7173 | 2,23283E-65 |
| ACE2 | COPS8 | -0,7174 | 2,11911E-65 |
| ACE2 | BMT2 | -0,7174 | 2,0763E-65 |
| ACE2 | SLC43A3 | -0,7174 | 2,06946E-65 |
| ACE2 | CUX2 | -0,7174 | 2,14323E-65 |
| ACE2 | RRN3P2 | -0,7174 | 2,08586E-65 |
| ACE2 | CTXN1 | -0,7174 | 2,05773E-65 |
| ACE2 | THEM4 | -0,7175 | 1,97965E-65 |
| ACE2 | PDE8B | -0,7175 | 1,93182E-65 |
| ACE2 | ZCWPW1 | -0,7175 | 2,00504E-65 |
| ACE2 | TAF3 | -0,7175 | 1,93655E-65 |
| ACE2 | SLC15A3 | -0,7175 | 1,99803E-65 |
| ACE2 | STXBP4 | -0,7175 | 1,97417E-65 |
| ACE2 | PNCK | -0,7175 | 1,92345E-65 |
| ACE2 | ELF2 | -0,7176 | 1,81333E-65 |
| ACE2 | NR3C1 | -0,7176 | 1,84983E-65 |
| ACE2 | CNN1 | -0,7176 | 1,84812E-65 |
| ACE2 | RRAS | -0,7176 | 1,88442E-65 |
| ACE2 | TCEAL3 | -0,7176 | 1,89632E-65 |
| ACE2 | PTGER3 | -0,7177 | 1,76116E-65 |
| ACE2 | SERTAD4-AS1 | -0,7177 | 1,72593E-65 |
| ACE2 | CCNI | -0,7177 | 1,73695E-65 |
| ACE2 | RASA4B | -0,7177 | 1,73913E-65 |
| ACE2 | SKIDA1 | -0,7177 | 1,78807E-65 |
| ACE2 | HECTD2 | -0,7177 | 1,75879E-65 |
| ACE2 | SYNM | -0,7177 | 1,78143E-65 |
| ACE2 | UNK | -0,7177 | 1,80381E-65 |
| ACE2 | AKAP8L | -0,7177 | 1,80236E-65 |
| ACE2 | CRYZL1 | -0,7177 | 1,76203E-65 |
| ACE2 | LINC00334 | -0,7177 | 1,71408E-65 |
| ACE2 | PLPPR4 | -0,7178 | 1,70232E-65 |
| ACE2 | TMEM110 | -0,7179 | 1,57991E-65 |
| ACE2 | PNMA2 | -0,7179 | 1,55521E-65 |
| ACE2 | SPRYD3 | -0,7179 | 1,59462E-65 |
| ACE2 | AMER2 | -0,7179 | 1,52264E-65 |
| ACE2 | RORA | -0,7179 | 1,60099E-65 |
| ACE2 | PITPNB | -0,7179 | 1,57975E-65 |
| ACE2 | TENM1 | -0,7179 | 1,53902E-65 |
| ACE2 | CACNA2D1 | -0,718 | 1,50194E-65 |
| ACE2 | PRR14 | -0,718 | 1,49032E-65 |
| ACE2 | INHA | -0,7181 | 1,41957E-65 |
| ACE2 | FAM168A | -0,7181 | 1,38602E-65 |
| ACE2 | MPV17 | -0,7182 | 1,31062E-65 |
| ACE2 | CALU | -0,7182 | 1,32974E-65 |

| | | | |
|---|---|---|---|
| ACE2 | RAB2B | -0,7182 | 1,27537E-65 |
| ACE2 | MAPT | -0,7182 | 1,32999E-65 |
| ACE2 | DTNA | -0,7182 | 1,28905E-65 |
| ACE2 | LINC01273 | -0,7182 | 1,31665E-65 |
| ACE2 | RP3-333A15.2 | -0,7183 | 1,19215E-65 |
| ACE2 | EPN2 | -0,7183 | 1,19654E-65 |
| ACE2 | C22orf39 | -0,7183 | 1,24277E-65 |
| ACE2 | AC093673.5 | -0,7184 | 1,12797E-65 |
| ACE2 | CAMK1G | -0,7186 | 1,03117E-65 |
| ACE2 | LINC00506 | -0,7186 | 1,04511E-65 |
| ACE2 | RP11-689P11.2 | -0,7186 | 1,02671E-65 |
| ACE2 | CREB3L2 | -0,7186 | 1,05008E-65 |
| ACE2 | TSPAN18 | -0,7186 | 1,04365E-65 |
| ACE2 | SSSCA1 | -0,7186 | 1,02171E-65 |
| ACE2 | SACS | -0,7186 | 1,02272E-65 |
| ACE2 | LINC01579 | -0,7186 | 1,0366E-65 |
| ACE2 | SPEG | -0,7187 | 9,75539E-66 |
| ACE2 | RIPK2 | -0,7188 | 8,94177E-66 |
| ACE2 | RP11-508N22.12 | -0,7188 | 9,32489E-66 |
| ACE2 | ZNF516 | -0,7188 | 9,22467E-66 |
| ACE2 | DCBLD2 | -0,7189 | 8,39721E-66 |
| ACE2 | PDGFC | -0,7189 | 8,74611E-66 |
| ACE2 | FAM53C | -0,7189 | 8,30738E-66 |
| ACE2 | CREB5 | -0,7189 | 8,30416E-66 |
| ACE2 | CCNL2 | -0,719 | 8,11768E-66 |
| ACE2 | UCN | -0,719 | 8,06228E-66 |
| ACE2 | SFRP4 | -0,719 | 8,16976E-66 |
| ACE2 | B4GAT1 | -0,719 | 8,23289E-66 |
| ACE2 | KLHDC1 | -0,719 | 8,25273E-66 |
| ACE2 | LINC01096 | -0,7191 | 7,57289E-66 |
| ACE2 | FGFBP2 | -0,7191 | 7,47933E-66 |
| ACE2 | SENP6 | -0,7191 | 7,75017E-66 |
| ACE2 | PURG | -0,7191 | 7,37011E-66 |
| ACE2 | SBSPON | -0,7191 | 7,47534E-66 |
| ACE2 | RP11-629G13.1 | -0,7191 | 7,38172E-66 |
| ACE2 | CCNB1IP1 | -0,7191 | 7,71825E-66 |
| ACE2 | MAP4K5 | -0,7191 | 7,34924E-66 |
| ACE2 | RAD51C | -0,7191 | 7,47797E-66 |
| ACE2 | ZNF561-AS1 | -0,7191 | 7,47149E-66 |
| ACE2 | FAM210B | -0,7191 | 7,47785E-66 |
| ACE2 | CCDC80 | -0,7192 | 7,2854E-66 |
| ACE2 | PCDHGB6 | -0,7192 | 7,18823E-66 |
| ACE2 | IFITM3 | -0,7192 | 7,15263E-66 |
| ACE2 | KLF12 | -0,7192 | 7,01884E-66 |
| ACE2 | RPS6KB1 | -0,7192 | 6,92192E-66 |
| ACE2 | SPATA6 | -0,7193 | 6,61888E-66 |
| ACE2 | NKX3-2 | -0,7193 | 6,85987E-66 |
| ACE2 | PCDHGC5 | -0,7193 | 6,62452E-66 |
| ACE2 | PGR | -0,7193 | 6,68579E-66 |
| ACE2 | DVL2 | -0,7193 | 6,86232E-66 |
| ACE2 | KXD1 | -0,7193 | 6,66511E-66 |
| ACE2 | TSPY26P | -0,7193 | 6,55914E-66 |
| ACE2 | AGER | -0,7194 | 6,27488E-66 |
| ACE2 | TMEM74 | -0,7194 | 6,19211E-66 |

| | | | |
|---|---|---|---|
| ACE2 | LRRC27 | -0,7194 | 6,45366E-66 |
| ACE2 | HIC1 | -0,7194 | 6,15207E-66 |
| ACE2 | METTL16 | -0,7194 | 6,44978E-66 |
| ACE2 | FAM198A | -0,7195 | 5,93171E-66 |
| ACE2 | IQCG | -0,7195 | 5,88966E-66 |
| ACE2 | KCTD8 | -0,7195 | 5,78634E-66 |
| ACE2 | ECM2 | -0,7195 | 5,99565E-66 |
| ACE2 | MTERF2 | -0,7195 | 5,91424E-66 |
| ACE2 | LINC02210 | -0,7195 | 6,04101E-66 |
| ACE2 | COMMD7 | -0,7195 | 5,82862E-66 |
| ACE2 | ZNHIT6 | -0,7196 | 5,76525E-66 |
| ACE2 | RP11-410L14.2 | -0,7196 | 5,52595E-66 |
| ACE2 | PRKCDBP | -0,7196 | 5,50614E-66 |
| ACE2 | GLI2 | -0,7197 | 5,41948E-66 |
| ACE2 | SHISA3 | -0,7197 | 5,24522E-66 |
| ACE2 | HEY2 | -0,7197 | 5,33947E-66 |
| ACE2 | BNC2 | -0,7197 | 5,22623E-66 |
| ACE2 | RP11-359E10.1 | -0,7197 | 5,1626E-66 |
| ACE2 | TMX4 | -0,7197 | 5,31581E-66 |
| ACE2 | FILIP1 | -0,7198 | 4,93461E-66 |
| ACE2 | SMAD9 | -0,7198 | 5,02827E-66 |
| ACE2 | CNN3 | -0,7199 | 4,63128E-66 |
| ACE2 | FOXP1 | -0,7199 | 4,73295E-66 |
| ACE2 | TEX10 | -0,7199 | 4,77319E-66 |
| ACE2 | PCDHB5 | -0,72 | 4,44704E-66 |
| ACE2 | CNTNAP1 | -0,72 | 4,36296E-66 |
| ACE2 | GUSBP1 | -0,7201 | 4,15098E-66 |
| ACE2 | MAP3K4 | -0,7201 | 4,25567E-66 |
| ACE2 | ALKBH5 | -0,7201 | 4,07605E-66 |
| ACE2 | DEGS1 | -0,7202 | 3,87763E-66 |
| ACE2 | PRKRA | -0,7202 | 3,79612E-66 |
| ACE2 | ARHGEF26 | -0,7202 | 3,86534E-66 |
| ACE2 | PCDHGA3 | -0,7202 | 3,90053E-66 |
| ACE2 | NPC1L1 | -0,7202 | 3,82522E-66 |
| ACE2 | RANBP3 | -0,7202 | 3,82945E-66 |
| ACE2 | SLC24A3 | -0,7202 | 3,84943E-66 |
| ACE2 | JPH2 | -0,7202 | 3,89521E-66 |
| ACE2 | SMAD5 | -0,7203 | 3,55826E-66 |
| ACE2 | THSD7A | -0,7203 | 3,703E-66 |
| ACE2 | ZYX | -0,7203 | 3,73162E-66 |
| ACE2 | FBXO2 | -0,7204 | 3,3619E-66 |
| ACE2 | RBSN | -0,7204 | 3,51337E-66 |
| ACE2 | IL17B | -0,7204 | 3,5152E-66 |
| ACE2 | RP11-384P7.7 | -0,7204 | 3,40725E-66 |
| ACE2 | PDE2A | -0,7204 | 3,46188E-66 |
| ACE2 | ZNF660 | -0,7205 | 3,18084E-66 |
| ACE2 | CCDC136 | -0,7205 | 3,16218E-66 |
| ACE2 | SNAPC2 | -0,7205 | 3,15857E-66 |
| ACE2 | MRFAP1 | -0,7206 | 2,985E-66 |
| ACE2 | SAV1 | -0,7206 | 2,97772E-66 |
| ACE2 | PRSS27 | -0,7206 | 3,13955E-66 |
| ACE2 | TBX19 | -0,7207 | 2,83274E-66 |
| ACE2 | SOX2-OT | -0,7207 | 2,89442E-66 |
| ACE2 | PDZD2 | -0,7207 | 2,8131E-66 |

| | | | |
|---|---|---|---|
| ACE2 | MTAP | -0,7207 | 2,91307E-66 |
| ACE2 | AAED1 | -0,7207 | 2,86883E-66 |
| ACE2 | ZMYND11 | -0,7207 | 2,88545E-66 |
| ACE2 | ARMC4 | -0,7207 | 2,82345E-66 |
| ACE2 | ATXN10 | -0,7207 | 2,94194E-66 |
| ACE2 | PGM5P4-AS1 | -0,7208 | 2,70464E-66 |
| ACE2 | ROR2 | -0,7208 | 2,76591E-66 |
| ACE2 | TMEM120B | -0,7208 | 2,71711E-66 |
| ACE2 | THSD1 | -0,7208 | 2,74254E-66 |
| ACE2 | ERI3 | -0,7209 | 2,55877E-66 |
| ACE2 | PCDHB7 | -0,7209 | 2,56249E-66 |
| ACE2 | GMPR | -0,7209 | 2,61606E-66 |
| ACE2 | ZFHX4 | -0,7209 | 2,59045E-66 |
| ACE2 | DDX25 | -0,7209 | 2,55274E-66 |
| ACE2 | GATS | -0,721 | 2,42578E-66 |
| ACE2 | DDHD2 | -0,721 | 2,40391E-66 |
| ACE2 | ABCG4 | -0,721 | 2,35937E-66 |
| ACE2 | ZNF580 | -0,721 | 2,35583E-66 |
| ACE2 | LINC01278 | -0,721 | 2,38396E-66 |
| ACE2 | PCDHGA5 | -0,7211 | 2,20782E-66 |
| ACE2 | DPY19L1P1 | -0,7211 | 2,21356E-66 |
| ACE2 | POT1 | -0,7211 | 2,25696E-66 |
| ACE2 | TNNI3K | -0,7212 | 2,09452E-66 |
| ACE2 | EXTL3-AS1 | -0,7212 | 2,06743E-66 |
| ACE2 | EIF3H | -0,7212 | 2,16167E-66 |
| ACE2 | IGHMBP2 | -0,7212 | 2,10365E-66 |
| ACE2 | LINC01828 | -0,7213 | 1,95968E-66 |
| ACE2 | ECSCR | -0,7213 | 2,05723E-66 |
| ACE2 | LRRK2 | -0,7213 | 1,9395E-66 |
| ACE2 | RP11-977G19.5 | -0,7213 | 2,04822E-66 |
| ACE2 | CTD-2555O16.2 | -0,7213 | 1,97249E-66 |
| ACE2 | INTS3 | -0,7214 | 1,92214E-66 |
| ACE2 | DYRK3 | -0,7214 | 1,83803E-66 |
| ACE2 | KIRREL3 | -0,7214 | 1,8483E-66 |
| ACE2 | H19 | -0,7215 | 1,81516E-66 |
| ACE2 | RP11-182J1.14 | -0,7215 | 1,81139E-66 |
| ACE2 | RP11-893F2.13 | -0,7215 | 1,76028E-66 |
| ACE2 | NRK | -0,7215 | 1,81873E-66 |
| ACE2 | HERC5 | -0,7216 | 1,62757E-66 |
| ACE2 | MTA1 | -0,7216 | 1,61509E-66 |
| ACE2 | RAB31 | -0,7216 | 1,68878E-66 |
| ACE2 | ARL8A | -0,7217 | 1,60892E-66 |
| ACE2 | PWAR6 | -0,7217 | 1,57936E-66 |
| ACE2 | IFI44 | -0,7218 | 1,49092E-66 |
| ACE2 | FAM189A2 | -0,7218 | 1,46773E-66 |
| ACE2 | CCS | -0,7218 | 1,51652E-66 |
| ACE2 | DCAF5 | -0,7218 | 1,43292E-66 |
| ACE2 | MAGEE1 | -0,7218 | 1,48423E-66 |
| ACE2 | EPB41L2 | -0,7219 | 1,3712E-66 |
| ACE2 | AP000473.5 | -0,7219 | 1,34751E-66 |
| ACE2 | ARVCF | -0,7219 | 1,40255E-66 |
| ACE2 | S100PBP | -0,7221 | 1,2599E-66 |
| ACE2 | AEBP2 | -0,7221 | 1,24376E-66 |
| ACE2 | RBFADN | -0,7221 | 1,19628E-66 |

| | | | |
|---|---|---|---|
| ACE2 | OXCT1-AS1 | -0,7222 | 1,11783E-66 |
| ACE2 | ARL2 | -0,7222 | 1,18664E-66 |
| ACE2 | GPC6 | -0,7222 | 1,18469E-66 |
| ACE2 | SSBP4 | -0,7222 | 1,184E-66 |
| ACE2 | HSCB | -0,7222 | 1,15104E-66 |
| ACE2 | AC007405.6 | -0,7224 | 9,89568E-67 |
| ACE2 | NRP2 | -0,7224 | 1,00221E-66 |
| ACE2 | MAP2 | -0,7224 | 9,99372E-67 |
| ACE2 | RAI14 | -0,7224 | 9,89709E-67 |
| ACE2 | IFT20 | -0,7224 | 1,02245E-66 |
| ACE2 | PCBP3 | -0,7224 | 1,01518E-66 |
| ACE2 | FLNA | -0,7224 | 1,01348E-66 |
| ACE2 | GSTM2 | -0,7225 | 9,57172E-67 |
| ACE2 | CASQ1 | -0,7225 | 9,49208E-67 |
| ACE2 | DRC3 | -0,7225 | 9,52918E-67 |
| ACE2 | RFX2 | -0,7225 | 9,74637E-67 |
| ACE2 | HSPA12B | -0,7225 | 9,79615E-67 |
| ACE2 | CEBPB-AS1 | -0,7225 | 9,46894E-67 |
| ACE2 | ERC2 | -0,7226 | 9,05631E-67 |
| ACE2 | POLK | -0,7226 | 8,93574E-67 |
| ACE2 | CCSER2 | -0,7226 | 9,27053E-67 |
| ACE2 | GLTSCR2 | -0,7226 | 8,76294E-67 |
| ACE2 | NME7 | -0,7227 | 8,47867E-67 |
| ACE2 | MITF | -0,7227 | 8,68069E-67 |
| ACE2 | EEF1A1 | -0,7227 | 8,23901E-67 |
| ACE2 | GPR26 | -0,7227 | 8,36916E-67 |
| ACE2 | CFL2 | -0,7227 | 8,52591E-67 |
| ACE2 | ZNF519 | -0,7227 | 8,71704E-67 |
| ACE2 | LINC00869 | -0,7228 | 7,82245E-67 |
| ACE2 | RIMS1 | -0,7228 | 7,87687E-67 |
| ACE2 | AMOTL1 | -0,7228 | 7,93774E-67 |
| ACE2 | ITGA7 | -0,7228 | 7,988E-67 |
| ACE2 | LRCH2 | -0,7228 | 7,81388E-67 |
| ACE2 | LOXL3 | -0,7229 | 7,47513E-67 |
| ACE2 | RILPL2 | -0,7229 | 7,30643E-67 |
| ACE2 | SHANK1 | -0,7229 | 7,5505E-67 |
| ACE2 | METTL22 | -0,723 | 6,98015E-67 |
| ACE2 | PTPN12 | -0,7231 | 6,59197E-67 |
| ACE2 | AC025165.8 | -0,7231 | 6,68228E-67 |
| ACE2 | CHST11 | -0,7231 | 6,82536E-67 |
| ACE2 | MRPS31P4 | -0,7231 | 6,45516E-67 |
| ACE2 | RGS9 | -0,7231 | 6,50218E-67 |
| ACE2 | TNRC6C | -0,7231 | 6,55428E-67 |
| ACE2 | NYAP1 | -0,7232 | 6,36582E-67 |
| ACE2 | PDGFRL | -0,7232 | 6,38647E-67 |
| ACE2 | IFT43 | -0,7232 | 6,14301E-67 |
| ACE2 | ASB2 | -0,7232 | 6,15508E-67 |
| ACE2 | ZFP92 | -0,7232 | 6,08579E-67 |
| ACE2 | NBPF15 | -0,7233 | 5,91963E-67 |
| ACE2 | HOMER2 | -0,7233 | 5,99738E-67 |
| ACE2 | HSPB6 | -0,7233 | 5,71997E-67 |
| ACE2 | DENND2C | -0,7234 | 5,42954E-67 |
| ACE2 | MR1 | -0,7234 | 5,60039E-67 |
| ACE2 | IQSEC1 | -0,7234 | 5,48539E-67 |

| | | | |
|---|---|---|---|
| ACE2 | PHF20 | -0,7234 | 5,59751E-67 |
| ACE2 | AHDC1 | -0,7235 | 5,23279E-67 |
| ACE2 | PLCL1 | -0,7235 | 5,06995E-67 |
| ACE2 | FAM26E | -0,7235 | 5,33619E-67 |
| ACE2 | TMEM263 | -0,7235 | 5,23078E-67 |
| ACE2 | KCND3 | -0,7236 | 4,81887E-67 |
| ACE2 | ALS2 | -0,7236 | 4,9901E-67 |
| ACE2 | CTNND2 | -0,7236 | 4,97452E-67 |
| ACE2 | EXTL3 | -0,7236 | 4,89556E-67 |
| ACE2 | PHYHD1 | -0,7236 | 4,89258E-67 |
| ACE2 | COMMD6 | -0,7236 | 4,8131E-67 |
| ACE2 | RP11-478C6.6 | -0,7237 | 4,61937E-67 |
| ACE2 | CEP85L | -0,7237 | 4,69333E-67 |
| ACE2 | STX8 | -0,7237 | 4,48463E-67 |
| ACE2 | RP11-861E21.2 | -0,7237 | 4,60636E-67 |
| ACE2 | WDR41 | -0,7238 | 4,33948E-67 |
| ACE2 | RP11-359B12.2 | -0,7238 | 4,26006E-67 |
| ACE2 | HCFC1R1 | -0,7238 | 4,26891E-67 |
| ACE2 | NLGN1 | -0,7239 | 4,00636E-67 |
| ACE2 | GALNT11 | -0,7239 | 3,99264E-67 |
| ACE2 | ZNF280D | -0,7239 | 4,13057E-67 |
| ACE2 | YY1AP1 | -0,724 | 3,75193E-67 |
| ACE2 | PIP5KL1 | -0,724 | 3,91367E-67 |
| ACE2 | RP11-394B2.1 | -0,724 | 3,79624E-67 |
| ACE2 | ARHGAP29 | -0,7241 | 3,53995E-67 |
| ACE2 | AGBL5 | -0,7241 | 3,54232E-67 |
| ACE2 | TUBA1A | -0,7241 | 3,59798E-67 |
| ACE2 | RP3-508I15.9 | -0,7241 | 3,665E-67 |
| ACE2 | RASGRF2 | -0,7242 | 3,3263E-67 |
| ACE2 | LL21NC02-21A1.1 | -0,7242 | 3,46994E-67 |
| ACE2 | KIF26B | -0,7243 | 3,13583E-67 |
| ACE2 | EIF2B4 | -0,7243 | 3,22926E-67 |
| ACE2 | RPS27A | -0,7243 | 3,07979E-67 |
| ACE2 | SOX2 | -0,7243 | 3,21568E-67 |
| ACE2 | RP11-728F11.4 | -0,7243 | 3,2683E-67 |
| ACE2 | PRPH | -0,7243 | 3,09905E-67 |
| ACE2 | ZRSR2 | -0,7243 | 3,07829E-67 |
| ACE2 | CTB-1144G6.6 | -0,7244 | 2,95202E-67 |
| ACE2 | PRKD1 | -0,7244 | 3,07391E-67 |
| ACE2 | FBXO27 | -0,7244 | 3,03666E-67 |
| ACE2 | AKT2 | -0,7244 | 3,07154E-67 |
| ACE2 | PPM1F | -0,7244 | 3,05283E-67 |
| ACE2 | RPL21 | -0,7245 | 2,74136E-67 |
| ACE2 | RYR3 | -0,7245 | 2,81214E-67 |
| ACE2 | CHRNB4 | -0,7245 | 2,84435E-67 |
| ACE2 | PKD1 | -0,7245 | 2,88195E-67 |
| ACE2 | NCDN | -0,7246 | 2,64825E-67 |
| ACE2 | EGFLAM | -0,7246 | 2,63699E-67 |
| ACE2 | RORB | -0,7246 | 2,62845E-67 |
| ACE2 | MBNL2 | -0,7246 | 2,70982E-67 |
| ACE2 | DYM | -0,7246 | 2,70198E-67 |
| ACE2 | SYN1 | -0,7246 | 2,5677E-67 |
| ACE2 | CSRP1 | -0,7247 | 2,43863E-67 |
| ACE2 | ADAM23 | -0,7247 | 2,4337E-67 |

| | | | |
|---|---|---|---|
| ACE2 | HACD4 | -0,7247 | 2,51664E-67 |
| ACE2 | CCDC74B | -0,7248 | 2,31374E-67 |
| ACE2 | KDELC2 | -0,7248 | 2,26456E-67 |
| ACE2 | RP11-13P5.2 | -0,7249 | 2,25214E-67 |
| ACE2 | CSRNP2 | -0,7249 | 2,23843E-67 |
| ACE2 | IGDCC3 | -0,7249 | 2,19871E-67 |
| ACE2 | CBX6 | -0,7249 | 2,21857E-67 |
| ACE2 | TRNP1 | -0,725 | 2,03974E-67 |
| ACE2 | AGPAT4 | -0,725 | 2,02353E-67 |
| ACE2 | KCND2 | -0,725 | 2,02619E-67 |
| ACE2 | ZNF510 | -0,7251 | 1,90407E-67 |
| ACE2 | GRK5 | -0,7251 | 1,971E-67 |
| ACE2 | ST5 | -0,7251 | 1,93356E-67 |
| ACE2 | DCP1B | -0,7251 | 1,95724E-67 |
| ACE2 | ZNF548 | -0,7251 | 1,92838E-67 |
| ACE2 | RRAGB | -0,7251 | 1,91525E-67 |
| ACE2 | KNDC1 | -0,7252 | 1,85073E-67 |
| ACE2 | FTO | -0,7252 | 1,81265E-67 |
| ACE2 | DEFB124 | -0,7252 | 1,76809E-67 |
| ACE2 | MAP7D1 | -0,7253 | 1,72333E-67 |
| ACE2 | FAM163A | -0,7253 | 1,65947E-67 |
| ACE2 | PLXDC1 | -0,7253 | 1,66683E-67 |
| ACE2 | COPS7B | -0,7254 | 1,65704E-67 |
| ACE2 | ARMC2 | -0,7254 | 1,63639E-67 |
| ACE2 | FNBP1 | -0,7254 | 1,5801E-67 |
| ACE2 | PACS2 | -0,7254 | 1,63016E-67 |
| ACE2 | ZNF521 | -0,7254 | 1,60864E-67 |
| ACE2 | KIZ | -0,7254 | 1,57086E-67 |
| ACE2 | L3MBTL1 | -0,7254 | 1,64184E-67 |
| ACE2 | STX16 | -0,7254 | 1,56434E-67 |
| ACE2 | CPXM2 | -0,7255 | 1,54421E-67 |
| ACE2 | NRXN2 | -0,7255 | 1,526E-67 |
| ACE2 | ABI3BP | -0,7257 | 1,35432E-67 |
| ACE2 | FXR1 | -0,7257 | 1,35986E-67 |
| ACE2 | PTPN13 | -0,7257 | 1,32936E-67 |
| ACE2 | UBE2H | -0,7257 | 1,3349E-67 |
| ACE2 | USP13 | -0,7258 | 1,2239E-67 |
| ACE2 | CLUAP1 | -0,7258 | 1,25079E-67 |
| ACE2 | PARD6G | -0,7258 | 1,24355E-67 |
| ACE2 | PIP5K1C | -0,7258 | 1,22488E-67 |
| ACE2 | SOD3 | -0,7259 | 1,14612E-67 |
| ACE2 | MRGPRF | -0,7259 | 1,20917E-67 |
| ACE2 | POU3F3 | -0,726 | 1,11682E-67 |
| ACE2 | HPF1 | -0,726 | 1,10035E-67 |
| ACE2 | WWC2 | -0,726 | 1,10369E-67 |
| ACE2 | IL11RA | -0,726 | 1,12112E-67 |
| ACE2 | DUSP14 | -0,726 | 1,08741E-67 |
| ACE2 | NKIRAS1 | -0,7261 | 1,0279E-67 |
| ACE2 | HSF1 | -0,7261 | 1,05894E-67 |
| ACE2 | TBCD | -0,7261 | 1,02015E-67 |
| ACE2 | SCN1B | -0,7261 | 1,03424E-67 |
| ACE2 | EIF5B | -0,7262 | 9,92209E-68 |
| ACE2 | MBD5 | -0,7262 | 9,67066E-68 |
| ACE2 | IGIP | -0,7262 | 9,96092E-68 |

| | | | |
|---|---|---|---|
| ACE2 | TPPP3 | -0,7262 | 9,63415E-68 |
| ACE2 | FAM69C | -0,7262 | 9,71078E-68 |
| ACE2 | PPT1 | -0,7263 | 8,95779E-68 |
| ACE2 | FAP | -0,7263 | 8,92306E-68 |
| ACE2 | C10orf35 | -0,7263 | 8,93441E-68 |
| ACE2 | GPR83 | -0,7263 | 9,30114E-68 |
| ACE2 | ELMOD1 | -0,7263 | 9,26408E-68 |
| ACE2 | RPSAP18 | -0,7264 | 8,51518E-68 |
| ACE2 | RPL14 | -0,7264 | 8,59146E-68 |
| ACE2 | NCS1 | -0,7264 | 8,44282E-68 |
| ACE2 | DPH7 | -0,7264 | 8,61047E-68 |
| ACE2 | ZNF461 | -0,7265 | 8,21375E-68 |
| ACE2 | RP5-1085F17.3 | -0,7265 | 8,11195E-68 |
| ACE2 | MEAF6 | -0,7266 | 7,44622E-68 |
| ACE2 | OLFML3 | -0,7266 | 7,60558E-68 |
| ACE2 | RPL13 | -0,7266 | 7,78941E-68 |
| ACE2 | PDCD5 | -0,7266 | 7,45387E-68 |
| ACE2 | CYTH3 | -0,7267 | 6,98073E-68 |
| ACE2 | ASAP1 | -0,7267 | 6,99247E-68 |
| ACE2 | OLFM1 | -0,7267 | 6,94779E-68 |
| ACE2 | PNMAL1 | -0,7267 | 7,22722E-68 |
| ACE2 | C20orf203 | -0,7267 | 7,19881E-68 |
| ACE2 | GSAP | -0,7268 | 6,72734E-68 |
| ACE2 | ITPR1 | -0,7269 | 6,33319E-68 |
| ACE2 | CC2D2A | -0,7269 | 6,52316E-68 |
| ACE2 | SGTB | -0,7269 | 6,1674E-68 |
| ACE2 | R3HCC1 | -0,7269 | 6,27846E-68 |
| ACE2 | SALL2 | -0,7269 | 6,43385E-68 |
| ACE2 | ZBTB25 | -0,7269 | 6,21952E-68 |
| ACE2 | SLC16A4 | -0,727 | 5,87551E-68 |
| ACE2 | RP11-767C1.2 | -0,727 | 6,00876E-68 |
| ACE2 | ANXA6 | -0,727 | 6,08848E-68 |
| ACE2 | DCHS1 | -0,727 | 6,00151E-68 |
| ACE2 | RP11-817I4.1 | -0,727 | 5,81893E-68 |
| ACE2 | RP11-834C11.7 | -0,727 | 5,95312E-68 |
| ACE2 | PRPS1 | -0,727 | 6,0559E-68 |
| ACE2 | PBXIP1 | -0,7271 | 5,43413E-68 |
| ACE2 | CAMLG | -0,7271 | 5,66075E-68 |
| ACE2 | NALT1 | -0,7271 | 5,72161E-68 |
| ACE2 | NGFR | -0,7271 | 5,42098E-68 |
| ACE2 | AIDA | -0,7272 | 5,27001E-68 |
| ACE2 | AC012074.2 | -0,7272 | 5,13963E-68 |
| ACE2 | THBS4 | -0,7272 | 5,13717E-68 |
| ACE2 | PPP3CB | -0,7272 | 5,14453E-68 |
| ACE2 | SCN2B | -0,7272 | 5,19963E-68 |
| ACE2 | KLF7 | -0,7273 | 4,83932E-68 |
| ACE2 | RPL39L | -0,7273 | 5,06075E-68 |
| ACE2 | ARL4D | -0,7273 | 4,91788E-68 |
| ACE2 | PEG3 | -0,7273 | 4,9731E-68 |
| ACE2 | CEP97 | -0,7274 | 4,62551E-68 |
| ACE2 | ITGA1 | -0,7274 | 4,51224E-68 |
| ACE2 | RP3-425C14.4 | -0,7274 | 4,58264E-68 |
| ACE2 | FKBP14 | -0,7274 | 4,75299E-68 |
| ACE2 | LY6H | -0,7274 | 4,54971E-68 |

| | | | |
|---|---|---|---|
| ACE2 | LINC01505 | -0,7274 | 4,76392E-68 |
| ACE2 | LINC01341 | -0,7275 | 4,2439E-68 |
| ACE2 | AC007283.4 | -0,7275 | 4,48498E-68 |
| ACE2 | BRK1 | -0,7275 | 4,44753E-68 |
| ACE2 | SELENON | -0,7276 | 4,14954E-68 |
| ACE2 | B4GALT2 | -0,7276 | 4,11713E-68 |
| ACE2 | FIGN | -0,7276 | 4,03377E-68 |
| ACE2 | SIPA1 | -0,7276 | 4,07485E-68 |
| ACE2 | ISCU | -0,7276 | 3,99893E-68 |
| ACE2 | MLLT6 | -0,7276 | 4,06705E-68 |
| ACE2 | TCF15 | -0,7276 | 4,07507E-68 |
| ACE2 | CHODL | -0,7276 | 4,09932E-68 |
| ACE2 | FUNDC2 | -0,7276 | 4,11291E-68 |
| ACE2 | GSTM3 | -0,7277 | 3,87562E-68 |
| ACE2 | BACE1-AS | -0,7277 | 3,88665E-68 |
| ACE2 | MEG3 | -0,7277 | 3,91952E-68 |
| ACE2 | FNDC5 | -0,7278 | 3,60559E-68 |
| ACE2 | NDUFAF7 | -0,7278 | 3,63035E-68 |
| ACE2 | HLTF | -0,7278 | 3,59535E-68 |
| ACE2 | TPPP | -0,7278 | 3,56622E-68 |
| ACE2 | EPHA5 | -0,7279 | 3,36444E-68 |
| ACE2 | PIANP | -0,7279 | 3,46601E-68 |
| ACE2 | RASIP1 | -0,7279 | 3,4432E-68 |
| ACE2 | EPHA3 | -0,728 | 3,16213E-68 |
| ACE2 | NEIL2 | -0,728 | 3,2762E-68 |
| ACE2 | MED22 | -0,728 | 3,21913E-68 |
| ACE2 | CCDC124 | -0,728 | 3,2359E-68 |
| ACE2 | FOXO6 | -0,7281 | 3,01071E-68 |
| ACE2 | RUBCN | -0,7281 | 2,94995E-68 |
| ACE2 | FGFR1OP2 | -0,7281 | 3,07575E-68 |
| ACE2 | MSN | -0,7281 | 2,93548E-68 |
| ACE2 | BBX | -0,7282 | 2,86963E-68 |
| ACE2 | AC093323.3 | -0,7282 | 2,82458E-68 |
| ACE2 | TPST1 | -0,7282 | 2,74477E-68 |
| ACE2 | ENTPD1-AS1 | -0,7282 | 2,77723E-68 |
| ACE2 | LDB3 | -0,7283 | 2,66226E-68 |
| ACE2 | C14orf132 | -0,7283 | 2,58106E-68 |
| ACE2 | GNAO1 | -0,7283 | 2,5776E-68 |
| ACE2 | MECP2 | -0,7283 | 2,59011E-68 |
| ACE2 | PRR3 | -0,7284 | 2,48161E-68 |
| ACE2 | SRSF12 | -0,7284 | 2,41233E-68 |
| ACE2 | RAI1 | -0,7284 | 2,52738E-68 |
| ACE2 | RP11-893F2.14 | -0,7284 | 2,5381E-68 |
| ACE2 | NPEPL1 | -0,7284 | 2,41735E-68 |
| ACE2 | PXDN | -0,7285 | 2,25856E-68 |
| ACE2 | ZNF37A | -0,7285 | 2,31454E-68 |
| ACE2 | POLI | -0,7285 | 2,2762E-68 |
| ACE2 | WTIP | -0,7285 | 2,25596E-68 |
| ACE2 | PCSK2 | -0,7285 | 2,33578E-68 |
| ACE2 | INPP5B | -0,7286 | 2,17878E-68 |
| ACE2 | RAB28 | -0,7286 | 2,12363E-68 |
| ACE2 | CTC-463A16.1 | -0,7286 | 2,12157E-68 |
| ACE2 | THY1 | -0,7287 | 2,09636E-68 |
| ACE2 | GAP43 | -0,7288 | 1,93066E-68 |

| | | | |
|---|---|---|---|
| ACE2 | GKAP1 | -0,7288 | 1,97591E-68 |
| ACE2 | TMEM205 | -0,7288 | 1,95612E-68 |
| ACE2 | THAP7 | -0,7288 | 1,86676E-68 |
| ACE2 | SETD7 | -0,7289 | 1,78552E-68 |
| ACE2 | RP11-350N15.4 | -0,7289 | 1,79033E-68 |
| ACE2 | FGF13 | -0,7289 | 1,78273E-68 |
| ACE2 | CLIC4 | -0,729 | 1,69273E-68 |
| ACE2 | TTC4 | -0,729 | 1,67804E-68 |
| ACE2 | VGLL3 | -0,729 | 1,64302E-68 |
| ACE2 | RPL10A | -0,729 | 1,67168E-68 |
| ACE2 | LRRTM1 | -0,7291 | 1,55731E-68 |
| ACE2 | GDNF | -0,7291 | 1,59836E-68 |
| ACE2 | MORN4 | -0,7291 | 1,54935E-68 |
| ACE2 | RP11-356J5.12 | -0,7291 | 1,55448E-68 |
| ACE2 | NDN | -0,7291 | 1,58783E-68 |
| ACE2 | ZNF23 | -0,7291 | 1,58078E-68 |
| ACE2 | WWC3 | -0,7292 | 1,52798E-68 |
| ACE2 | ARHGEF33 | -0,7293 | 1,4277E-68 |
| ACE2 | ADAMTS9-AS2 | -0,7293 | 1,44573E-68 |
| ACE2 | RP3-331H24.7 | -0,7293 | 1,43788E-68 |
| ACE2 | PGM5P4 | -0,7294 | 1,27691E-68 |
| ACE2 | RPS25 | -0,7295 | 1,27116E-68 |
| ACE2 | RLBP1 | -0,7295 | 1,25969E-68 |
| ACE2 | CXorf36 | -0,7295 | 1,23317E-68 |
| ACE2 | NAV3 | -0,7296 | 1,16756E-68 |
| ACE2 | RAB11FIP3 | -0,7296 | 1,19259E-68 |
| ACE2 | FIBIN | -0,7297 | 1,0965E-68 |
| ACE2 | UNC80 | -0,7298 | 1,00343E-68 |
| ACE2 | SENP5 | -0,7298 | 1,00868E-68 |
| ACE2 | CNPY4 | -0,7298 | 1,03942E-68 |
| ACE2 | PCDH17 | -0,7298 | 1,00776E-68 |
| ACE2 | ZNF71 | -0,7298 | 1,05389E-68 |
| ACE2 | NUDT17 | -0,7299 | 9,68238E-69 |
| ACE2 | HACE1 | -0,7299 | 9,41705E-69 |
| ACE2 | MSRB3 | -0,7299 | 9,86404E-69 |
| ACE2 | DGCR6L | -0,7299 | 9,44566E-69 |
| ACE2 | RP11-148L24.1 | -0,73 | 8,79894E-69 |
| ACE2 | C7 | -0,73 | 8,8531E-69 |
| ACE2 | CD109 | -0,73 | 9,01327E-69 |
| ACE2 | FRMD4A | -0,73 | 8,9605E-69 |
| ACE2 | TRIP10 | -0,73 | 9,02175E-69 |
| ACE2 | SYNGAP1 | -0,7301 | 8,52156E-69 |
| ACE2 | FAXC | -0,7301 | 8,49808E-69 |
| ACE2 | RP11-662G23.1 | -0,7301 | 8,63902E-69 |
| ACE2 | STK33 | -0,7301 | 8,32635E-69 |
| ACE2 | LYRM9 | -0,7302 | 7,99688E-69 |
| ACE2 | ZNF208 | -0,7302 | 8,03502E-69 |
| ACE2 | CLASP2 | -0,7303 | 7,62059E-69 |
| ACE2 | REXO2 | -0,7304 | 7,08065E-69 |
| ACE2 | SLC48A1 | -0,7304 | 7,16827E-69 |
| ACE2 | NEK9 | -0,7304 | 7,12273E-69 |
| ACE2 | ITPKB | -0,7305 | 6,60838E-69 |
| ACE2 | TAC1 | -0,7305 | 6,7712E-69 |
| ACE2 | FAM131B | -0,7305 | 6,71385E-69 |

| | | | |
|---|---|---|---|
| ACE2 | ZNF880 | -0,7305 | 6,53293E-69 |
| ACE2 | CMTR1 | -0,7306 | 6,22719E-69 |
| ACE2 | BAG2 | -0,7306 | 6,0649E-69 |
| ACE2 | EBF3 | -0,7306 | 6,3075E-69 |
| ACE2 | ZNF546 | -0,7306 | 6,08528E-69 |
| ACE2 | TCEAL6 | -0,7306 | 6,03096E-69 |
| ACE2 | CRYGN | -0,7307 | 5,79019E-69 |
| ACE2 | USP35 | -0,7307 | 5,78409E-69 |
| ACE2 | EML1 | -0,7307 | 5,68453E-69 |
| ACE2 | C2CD2 | -0,7307 | 5,69308E-69 |
| ACE2 | ENAH | -0,7308 | 5,58452E-69 |
| ACE2 | GARNL3 | -0,7308 | 5,41484E-69 |
| ACE2 | RAB11FIP2 | -0,7308 | 5,29437E-69 |
| ACE2 | MEX3B | -0,7308 | 5,30321E-69 |
| ACE2 | LGI4 | -0,7308 | 5,37697E-69 |
| ACE2 | NFIX | -0,7309 | 5,14301E-69 |
| ACE2 | COL18A1 | -0,7309 | 5,20254E-69 |
| ACE2 | CHD5 | -0,731 | 4,65499E-69 |
| ACE2 | KIRREL | -0,731 | 4,7962E-69 |
| ACE2 | C5orf42 | -0,731 | 4,74975E-69 |
| ACE2 | STK3 | -0,731 | 4,638E-69 |
| ACE2 | AFAP1L2 | -0,731 | 4,76577E-69 |
| ACE2 | KIAA1586 | -0,7311 | 4,59609E-69 |
| ACE2 | RP11-1134I14.8 | -0,7311 | 4,4533E-69 |
| ACE2 | ISL2 | -0,7311 | 4,57464E-69 |
| ACE2 | PLEKHG2 | -0,7311 | 4,36763E-69 |
| ACE2 | SRPK3 | -0,7311 | 4,46045E-69 |
| ACE2 | PPOX | -0,7312 | 4,13812E-69 |
| ACE2 | C2orf40 | -0,7312 | 4,30649E-69 |
| ACE2 | CDK17 | -0,7312 | 4,21674E-69 |
| ACE2 | LRCH1 | -0,7312 | 4,13106E-69 |
| ACE2 | JPH3 | -0,7313 | 4,00348E-69 |
| ACE2 | SUGT1P1 | -0,7314 | 3,78923E-69 |
| ACE2 | SHC3 | -0,7314 | 3,59769E-69 |
| ACE2 | RWDD2B | -0,7314 | 3,60492E-69 |
| ACE2 | RBMX2 | -0,7314 | 3,73578E-69 |
| ACE2 | ZNF732 | -0,7315 | 3,53948E-69 |
| ACE2 | NKX2-3 | -0,7315 | 3,43669E-69 |
| ACE2 | SUZ12P1 | -0,7315 | 3,57895E-69 |
| ACE2 | ZNF788 | -0,7315 | 3,46234E-69 |
| ACE2 | LIMS1 | -0,7316 | 3,18343E-69 |
| ACE2 | RP11-517I3.2 | -0,7316 | 3,20359E-69 |
| ACE2 | TWIST1 | -0,7316 | 3,23894E-69 |
| ACE2 | NACA | -0,7316 | 3,31692E-69 |
| ACE2 | ITIH4 | -0,7317 | 3,15469E-69 |
| ACE2 | CCND2-AS1 | -0,7317 | 3,09532E-69 |
| ACE2 | HTRA3 | -0,7318 | 2,95841E-69 |
| ACE2 | NNT-AS1 | -0,7318 | 2,83723E-69 |
| ACE2 | PCDHB9 | -0,7318 | 2,9549E-69 |
| ACE2 | CSGALNACT2 | -0,7318 | 2,88899E-69 |
| ACE2 | DNM1P35 | -0,7318 | 2,95013E-69 |
| ACE2 | MDFIC | -0,7319 | 2,61947E-69 |
| ACE2 | OSCP1 | -0,732 | 2,54042E-69 |
| ACE2 | DCHS2 | -0,732 | 2,52864E-69 |

| | | | |
|---|---|---|---|
| ACE2 | RP11-545I5.3 | -0,732 | 2,54765E-69 |
| ACE2 | TRIM6 | -0,7321 | 2,36676E-69 |
| ACE2 | ZFR2 | -0,7321 | 2,30823E-69 |
| ACE2 | RPL3 | -0,7321 | 2,38913E-69 |
| ACE2 | AADAT | -0,7322 | 2,16737E-69 |
| ACE2 | MEF2C | -0,7322 | 2,27922E-69 |
| ACE2 | NUDT3 | -0,7322 | 2,25155E-69 |
| ACE2 | NFKB2 | -0,7322 | 2,28808E-69 |
| ACE2 | CTD-2047H16.4 | -0,7322 | 2,23575E-69 |
| ACE2 | DOK5 | -0,7322 | 2,23199E-69 |
| ACE2 | NISCH | -0,7323 | 2,12356E-69 |
| ACE2 | ELP2 | -0,7323 | 2,05781E-69 |
| ACE2 | ARL13B | -0,7324 | 2,01971E-69 |
| ACE2 | KLHL13 | -0,7324 | 1,96279E-69 |
| ACE2 | PCDHB16 | -0,7325 | 1,82824E-69 |
| ACE2 | CDH13 | -0,7325 | 1,89431E-69 |
| ACE2 | COL1A1 | -0,7325 | 1,8581E-69 |
| ACE2 | LINC01158 | -0,7326 | 1,68097E-69 |
| ACE2 | LMX1B | -0,7326 | 1,70795E-69 |
| ACE2 | KDELC1 | -0,7326 | 1,70705E-69 |
| ACE2 | EVI5L | -0,7326 | 1,69031E-69 |
| ACE2 | RAB9B | -0,7326 | 1,69822E-69 |
| ACE2 | ADAMTSL4 | -0,7327 | 1,59133E-69 |
| ACE2 | CYB5R1 | -0,7327 | 1,56837E-69 |
| ACE2 | TOMM7 | -0,7327 | 1,61898E-69 |
| ACE2 | TYW3 | -0,7328 | 1,50054E-69 |
| ACE2 | ZNF445 | -0,7328 | 1,52026E-69 |
| ACE2 | LINC01184 | -0,7329 | 1,39892E-69 |
| ACE2 | RP11-195F19.5 | -0,7329 | 1,44256E-69 |
| ACE2 | RNASEH2B | -0,7329 | 1,46061E-69 |
| ACE2 | SOX12 | -0,7329 | 1,39201E-69 |
| ACE2 | RAB36 | -0,7329 | 1,43726E-69 |
| ACE2 | RAI2 | -0,7329 | 1,38873E-69 |
| ACE2 | COL8A2 | -0,733 | 1,33224E-69 |
| ACE2 | DYSF | -0,733 | 1,32295E-69 |
| ACE2 | TET1 | -0,733 | 1,32065E-69 |
| ACE2 | TRIM8 | -0,733 | 1,32305E-69 |
| ACE2 | SORCS3 | -0,733 | 1,32555E-69 |
| ACE2 | RGMA | -0,733 | 1,35914E-69 |
| ACE2 | ZNF470 | -0,733 | 1,34761E-69 |
| ACE2 | ANKRD39 | -0,7331 | 1,26675E-69 |
| ACE2 | FSTL1 | -0,7331 | 1,27767E-69 |
| ACE2 | PSD | -0,7331 | 1,22059E-69 |
| ACE2 | STK32C | -0,7331 | 1,27145E-69 |
| ACE2 | ST3GAL6-AS1 | -0,7332 | 1,14748E-69 |
| ACE2 | AP4S1 | -0,7332 | 1,16201E-69 |
| ACE2 | SNHG14 | -0,7332 | 1,17128E-69 |
| ACE2 | PCDH10 | -0,7333 | 1,13564E-69 |
| ACE2 | PDLIM3 | -0,7333 | 1,12912E-69 |
| ACE2 | ZNF37BP | -0,7333 | 1,07195E-69 |
| ACE2 | DDX50 | -0,7333 | 1,07827E-69 |
| ACE2 | DRAP1 | -0,7333 | 1,07317E-69 |
| ACE2 | CCDC88A | -0,7334 | 1,02685E-69 |
| ACE2 | ARHGAP24 | -0,7334 | 1,02398E-69 |

| | | | |
|---|---|---|---|
| ACE2 | SLC29A1 | -0,7334 | 1,00712E-69 |
| ACE2 | MRVI1 | -0,7334 | 1,05164E-69 |
| ACE2 | WNK3 | -0,7334 | 1,03695E-69 |
| ACE2 | CNTN2 | -0,7335 | 9,68945E-70 |
| ACE2 | HLX | -0,7335 | 9,83393E-70 |
| ACE2 | BVES-AS1 | -0,7335 | 9,75318E-70 |
| ACE2 | TPCN2 | -0,7335 | 9,55613E-70 |
| ACE2 | TMEM59L | -0,7335 | 9,49337E-70 |
| ACE2 | MROH8 | -0,7335 | 9,45296E-70 |
| ACE2 | CHST12 | -0,7336 | 9,19048E-70 |
| ACE2 | C12orf54 | -0,7336 | 9,33047E-70 |
| ACE2 | DDX51 | -0,7336 | 9,32989E-70 |
| ACE2 | TGFB1I1 | -0,7336 | 9,28907E-70 |
| ACE2 | ZNF665 | -0,7336 | 8,93096E-70 |
| ACE2 | PRPF31 | -0,7336 | 9,33126E-70 |
| ACE2 | PORCN | -0,7336 | 9,16115E-70 |
| ACE2 | LINC01128 | -0,7337 | 8,49505E-70 |
| ACE2 | ST7 | -0,7337 | 8,69972E-70 |
| ACE2 | RAB3IL1 | -0,7337 | 8,67929E-70 |
| ACE2 | GRM6 | -0,7338 | 8,09537E-70 |
| ACE2 | AEBP1 | -0,7338 | 7,90546E-70 |
| ACE2 | SYT10 | -0,7338 | 7,99113E-70 |
| ACE2 | CTF1 | -0,7338 | 7,87921E-70 |
| ACE2 | BMPR1B | -0,7339 | 7,62614E-70 |
| ACE2 | TMEM200C | -0,7339 | 7,48881E-70 |
| ACE2 | ZNF83 | -0,7339 | 7,72082E-70 |
| ACE2 | NPTXR | -0,7339 | 7,46856E-70 |
| ACE2 | HDX | -0,7339 | 7,43541E-70 |
| ACE2 | CHADL | -0,734 | 7,01724E-70 |
| ACE2 | GRB14 | -0,7341 | 6,67896E-70 |
| ACE2 | ZCCHC17 | -0,7342 | 6,23518E-70 |
| ACE2 | VPS37D | -0,7344 | 5,35576E-70 |
| ACE2 | ZNF483 | -0,7344 | 5,44516E-70 |
| ACE2 | ARL14EP | -0,7344 | 5,338E-70 |
| ACE2 | FAM117A | -0,7344 | 5,41946E-70 |
| ACE2 | EEF1A2 | -0,7344 | 5,53738E-70 |
| ACE2 | LRSAM1 | -0,7345 | 5,07428E-70 |
| ACE2 | CBX7 | -0,7345 | 5,12054E-70 |
| ACE2 | CELSR2 | -0,7346 | 4,62666E-70 |
| ACE2 | SLC25A27 | -0,7346 | 4,90616E-70 |
| ACE2 | RPL27 | -0,7346 | 4,8139E-70 |
| ACE2 | NUDT10 | -0,7346 | 4,77792E-70 |
| ACE2 | FAM78B | -0,7347 | 4,43308E-70 |
| ACE2 | OBSL1 | -0,7347 | 4,35975E-70 |
| ACE2 | YEATS2 | -0,7347 | 4,54234E-70 |
| ACE2 | THBS2 | -0,7347 | 4,52351E-70 |
| ACE2 | CCDC180 | -0,7347 | 4,41143E-70 |
| ACE2 | EIF5A2 | -0,7348 | 4,06104E-70 |
| ACE2 | CDK9 | -0,7348 | 4,15201E-70 |
| ACE2 | SERP2 | -0,7348 | 4,1273E-70 |
| ACE2 | TSPAN2 | -0,7349 | 4,04004E-70 |
| ACE2 | NXPH3 | -0,7349 | 3,8629E-70 |
| ACE2 | TNXB | -0,735 | 3,6461E-70 |
| ACE2 | TRAF1 | -0,735 | 3,70759E-70 |

| | | | |
|---|---|---|---|
| ACE2 | GNAL | -0,735 | 3,61283E-70 |
| ACE2 | ZNF329 | -0,735 | 3,59314E-70 |
| ACE2 | DMD | -0,7351 | 3,35126E-70 |
| ACE2 | LINC01266 | -0,7352 | 3,30942E-70 |
| ACE2 | MCAM | -0,7352 | 3,21868E-70 |
| ACE2 | DES | -0,7353 | 3,09425E-70 |
| ACE2 | MPDZ | -0,7353 | 3,01831E-70 |
| ACE2 | AKAP6 | -0,7353 | 2,98307E-70 |
| ACE2 | TTLL7 | -0,7354 | 2,78976E-70 |
| ACE2 | MTR | -0,7354 | 2,90254E-70 |
| ACE2 | PKD2 | -0,7354 | 2,80752E-70 |
| ACE2 | CCND2 | -0,7354 | 2,7983E-70 |
| ACE2 | ZNF354A | -0,7355 | 2,73989E-70 |
| ACE2 | COL4A2 | -0,7355 | 2,63878E-70 |
| ACE2 | PIN1 | -0,7355 | 2,70465E-70 |
| ACE2 | PABPC5 | -0,7355 | 2,64746E-70 |
| ACE2 | ZMAT3 | -0,7356 | 2,51508E-70 |
| ACE2 | CXCL12 | -0,7356 | 2,54192E-70 |
| ACE2 | PCID2 | -0,7356 | 2,51846E-70 |
| ACE2 | AXL | -0,7356 | 2,44603E-70 |
| ACE2 | AC005301.9 | -0,7356 | 2,44947E-70 |
| ACE2 | SLC35F3 | -0,7357 | 2,26138E-70 |
| ACE2 | FER | -0,7357 | 2,30983E-70 |
| ACE2 | APLP1 | -0,7357 | 2,36864E-70 |
| ACE2 | NBPF8 | -0,7358 | 2,21561E-70 |
| ACE2 | MTURN | -0,7358 | 2,19625E-70 |
| ACE2 | FAM124A | -0,7358 | 2,17359E-70 |
| ACE2 | TMEM200B | -0,7359 | 2,09284E-70 |
| ACE2 | TRIM45 | -0,7359 | 2,10811E-70 |
| ACE2 | RP11-506B6.6 | -0,7359 | 2,07863E-70 |
| ACE2 | TSPAN4 | -0,7359 | 2,02912E-70 |
| ACE2 | OLFM2 | -0,736 | 1,9686E-70 |
| ACE2 | SCMH1 | -0,7361 | 1,75985E-70 |
| ACE2 | APLF | -0,7361 | 1,82123E-70 |
| ACE2 | SPG20 | -0,7361 | 1,77077E-70 |
| ACE2 | ANKRD10 | -0,7361 | 1,81776E-70 |
| ACE2 | CGNL1 | -0,7362 | 1,72903E-70 |
| ACE2 | TJAP1 | -0,7363 | 1,57498E-70 |
| ACE2 | RP11-461O7.1 | -0,7363 | 1,60864E-70 |
| ACE2 | ARMCX4 | -0,7363 | 1,59527E-70 |
| ACE2 | PRKD3 | -0,7364 | 1,51117E-70 |
| ACE2 | UIMC1 | -0,7364 | 1,4409E-70 |
| ACE2 | PREX2 | -0,7364 | 1,43942E-70 |
| ACE2 | CDADC1 | -0,7364 | 1,49074E-70 |
| ACE2 | TMEM231 | -0,7364 | 1,43626E-70 |
| ACE2 | SNTA1 | -0,7365 | 1,41663E-70 |
| ACE2 | SRRM3 | -0,7366 | 1,30184E-70 |
| ACE2 | C9orf47 | -0,7366 | 1,29221E-70 |
| ACE2 | AC006262.6 | -0,7366 | 1,26429E-70 |
| ACE2 | MAMLD1 | -0,7366 | 1,30337E-70 |
| ACE2 | MFAP2 | -0,7367 | 1,22257E-70 |
| ACE2 | MSANTD3 | -0,7367 | 1,23953E-70 |
| ACE2 | ZNF471 | -0,7367 | 1,22398E-70 |
| ACE2 | LSAMP | -0,7368 | 1,11002E-70 |

| | | | |
|---|---|---|---|
| ACE2 | ACTA2-AS1 | -0,7368 | 1,12008E-70 |
| ACE2 | LINC00327 | -0,7368 | 1,15272E-70 |
| ACE2 | DNALI1 | -0,7369 | 1,04273E-70 |
| ACE2 | CNIH3 | -0,7369 | 1,0718E-70 |
| ACE2 | SOX8 | -0,7369 | 1,08217E-70 |
| ACE2 | MEOX2 | -0,737 | 1,01665E-70 |
| ACE2 | DNAL1 | -0,737 | 9,97068E-71 |
| ACE2 | CD34 | -0,7371 | 9,31499E-71 |
| ACE2 | GUCY1A2 | -0,7371 | 9,64084E-71 |
| ACE2 | RDX | -0,7371 | 9,27647E-71 |
| ACE2 | KLHDC8B | -0,7372 | 8,60426E-71 |
| ACE2 | SPACA9 | -0,7372 | 8,6416E-71 |
| ACE2 | FXYD7 | -0,7372 | 8,74919E-71 |
| ACE2 | PARVB | -0,7372 | 8,5623E-71 |
| ACE2 | RNF115 | -0,7373 | 8,23765E-71 |
| ACE2 | FAM49A | -0,7373 | 8,28075E-71 |
| ACE2 | C7orf50 | -0,7373 | 8,09975E-71 |
| ACE2 | TIGD7 | -0,7373 | 8,22822E-71 |
| ACE2 | AC093838.4 | -0,7374 | 7,86391E-71 |
| ACE2 | MYO5A | -0,7374 | 7,60508E-71 |
| ACE2 | TUBD1 | -0,7374 | 7,83396E-71 |
| ACE2 | PPP1R18 | -0,7375 | 7,02958E-71 |
| ACE2 | KCNJ8 | -0,7375 | 7,13475E-71 |
| ACE2 | TIMM9 | -0,7376 | 6,86162E-71 |
| ACE2 | TFAP2A | -0,7377 | 6,45782E-71 |
| ACE2 | TMEM71 | -0,7377 | 6,14667E-71 |
| ACE2 | ZNF419 | -0,7377 | 6,31839E-71 |
| ACE2 | NXPE3 | -0,7378 | 6,02735E-71 |
| ACE2 | ZNF438 | -0,7378 | 5,97297E-71 |
| ACE2 | NGRN | -0,7378 | 5,94215E-71 |
| ACE2 | MTSS1L | -0,7378 | 5,75883E-71 |
| ACE2 | RGAG4 | -0,7378 | 5,79859E-71 |
| ACE2 | C19orf66 | -0,7379 | 5,41623E-71 |
| ACE2 | ZNF540 | -0,7379 | 5,40922E-71 |
| ACE2 | RP11-101E13.5 | -0,738 | 5,09412E-71 |
| ACE2 | CEP68 | -0,738 | 5,31666E-71 |
| ACE2 | RAB6B | -0,738 | 5,2033E-71 |
| ACE2 | FNDC1 | -0,738 | 5,12986E-71 |
| ACE2 | MYH10 | -0,738 | 5,35816E-71 |
| ACE2 | DSEL | -0,738 | 5,22259E-71 |
| ACE2 | TSEN15 | -0,7381 | 4,73799E-71 |
| ACE2 | IFT172 | -0,7381 | 4,74598E-71 |
| ACE2 | ZNF347 | -0,7381 | 5,02445E-71 |
| ACE2 | TAF9B | -0,7381 | 4,89429E-71 |
| ACE2 | PLS3 | -0,7381 | 4,83054E-71 |
| ACE2 | FCHSD2 | -0,7382 | 4,47525E-71 |
| ACE2 | COL12A1 | -0,7383 | 4,32798E-71 |
| ACE2 | CRTC3 | -0,7383 | 4,39037E-71 |
| ACE2 | ELMO2 | -0,7383 | 4,16628E-71 |
| ACE2 | ZNF354B | -0,7384 | 4,04263E-71 |
| ACE2 | PXDC1 | -0,7384 | 3,96935E-71 |
| ACE2 | AC005682.6 | -0,7384 | 3,89826E-71 |
| ACE2 | FBXW4 | -0,7384 | 3,93781E-71 |
| ACE2 | RP11-454H19.2 | -0,7384 | 3,95358E-71 |

| | | | |
|---|---|---|---|
| ACE2 | LMO4 | -0,7385 | 3,81858E-71 |
| ACE2 | PHF10 | -0,7385 | 3,81119E-71 |
| ACE2 | SEC61A2 | -0,7385 | 3,77545E-71 |
| ACE2 | ACSS3 | -0,7385 | 3,7994E-71 |
| ACE2 | PLEKHH3 | -0,7385 | 3,87071E-71 |
| ACE2 | SOGA1 | -0,7385 | 3,67861E-71 |
| ACE2 | NHS | -0,7385 | 3,72383E-71 |
| ACE2 | TWIST2 | -0,7386 | 3,48676E-71 |
| ACE2 | HILS1 | -0,7386 | 3,55319E-71 |
| ACE2 | ZFP28 | -0,7386 | 3,49446E-71 |
| ACE2 | LRCH3 | -0,7387 | 3,24413E-71 |
| ACE2 | RING1 | -0,7387 | 3,37301E-71 |
| ACE2 | CEND1 | -0,7387 | 3,35175E-71 |
| ACE2 | RP11-617F23.1 | -0,7387 | 3,25529E-71 |
| ACE2 | SNCG | -0,7388 | 3,01215E-71 |
| ACE2 | SGCA | -0,7388 | 3,13086E-71 |
| ACE2 | NECAB1 | -0,7389 | 2,81486E-71 |
| ACE2 | METRN | -0,7389 | 2,88652E-71 |
| ACE2 | IL33 | -0,739 | 2,6334E-71 |
| ACE2 | LINC00641 | -0,739 | 2,75981E-71 |
| ACE2 | ZNF566 | -0,739 | 2,73956E-71 |
| ACE2 | THAP3 | -0,7391 | 2,59505E-71 |
| ACE2 | ADGRB2 | -0,7391 | 2,47531E-71 |
| ACE2 | PCDHB14 | -0,7391 | 2,58492E-71 |
| ACE2 | CCT6P1 | -0,7391 | 2,5484E-71 |
| ACE2 | RTN2 | -0,7391 | 2,60029E-71 |
| ACE2 | DLGAP4 | -0,7391 | 2,49523E-71 |
| ACE2 | PALD1 | -0,7392 | 2,43549E-71 |
| ACE2 | FSTL3 | -0,7392 | 2,40502E-71 |
| ACE2 | RBFOX2 | -0,7392 | 2,41518E-71 |
| ACE2 | SLC35G2 | -0,7393 | 2,19761E-71 |
| ACE2 | GFRA3 | -0,7393 | 2,27802E-71 |
| ACE2 | HOXC4 | -0,7393 | 2,19129E-71 |
| ACE2 | RP11-789C17.1 | -0,7393 | 2,20725E-71 |
| ACE2 | ZNF569 | -0,7393 | 2,18657E-71 |
| ACE2 | GNAS | -0,7393 | 2,15546E-71 |
| ACE2 | SSBP2 | -0,7394 | 2,04482E-71 |
| ACE2 | RRP8 | -0,7394 | 2,11948E-71 |
| ACE2 | PAK3 | -0,7394 | 2,06856E-71 |
| ACE2 | PAMR1 | -0,7395 | 1,98202E-71 |
| ACE2 | ZNF529 | -0,7395 | 1,92478E-71 |
| ACE2 | ZDHHC1 | -0,7396 | 1,79084E-71 |
| ACE2 | ITGA2B | -0,7396 | 1,86093E-71 |
| ACE2 | MRC2 | -0,7396 | 1,78443E-71 |
| ACE2 | CTC-510F12.7 | -0,7396 | 1,82534E-71 |
| ACE2 | KIFC3 | -0,7397 | 1,69273E-71 |
| ACE2 | FAM204A | -0,7398 | 1,60221E-71 |
| ACE2 | ANKRD29 | -0,7398 | 1,62196E-71 |
| ACE2 | ZNF358 | -0,7398 | 1,64101E-71 |
| ACE2 | LINC00663 | -0,7398 | 1,59451E-71 |
| ACE2 | TTC25 | -0,7399 | 1,46241E-71 |
| ACE2 | PRELID3A | -0,7399 | 1,51355E-71 |
| ACE2 | KANK2 | -0,7399 | 1,5123E-71 |
| ACE2 | NBDY | -0,7399 | 1,53014E-71 |

| | | | |
|---|---|---|---|
| ACE2 | RPL5 | -0,74 | 1,40344E-71 |
| ACE2 | RET | -0,7401 | 1,33653E-71 |
| ACE2 | ZSWIM8-AS1 | -0,7402 | 1,19463E-71 |
| ACE2 | LRRC23 | -0,7402 | 1,22673E-71 |
| ACE2 | ALPK3 | -0,7402 | 1,21E-71 |
| ACE2 | TARSL2 | -0,7402 | 1,18669E-71 |
| ACE2 | CCDC40 | -0,7402 | 1,21729E-71 |
| ACE2 | RP5-1054A22.4 | -0,7402 | 1,25063E-71 |
| ACE2 | TMEM17 | -0,7403 | 1,13456E-71 |
| ACE2 | POGZ | -0,7404 | 1,07232E-71 |
| ACE2 | ARHGEF17 | -0,7404 | 1,07978E-71 |
| ACE2 | DLX5 | -0,7405 | 1,00651E-71 |
| ACE2 | UBALD1 | -0,7405 | 1,00079E-71 |
| ACE2 | PABPC4L | -0,7406 | 9,56736E-72 |
| ACE2 | GUCY1A3 | -0,7406 | 9,61104E-72 |
| ACE2 | LINC00094 | -0,7406 | 9,23607E-72 |
| ACE2 | CMTM3 | -0,7406 | 9,24991E-72 |
| ACE2 | PELP1 | -0,7406 | 9,38753E-72 |
| ACE2 | SIN3B | -0,7406 | 9,41466E-72 |
| ACE2 | SMTN | -0,7406 | 9,58027E-72 |
| ACE2 | SH3BGRL | -0,7406 | 9,30896E-72 |
| ACE2 | LINC00623 | -0,7407 | 8,50031E-72 |
| ACE2 | LDOC1 | -0,7407 | 8,94303E-72 |
| ACE2 | FAM182B | -0,7409 | 7,68927E-72 |
| ACE2 | PLXNB3 | -0,7409 | 7,88226E-72 |
| ACE2 | AKAP2 | -0,741 | 7,28103E-72 |
| ACE2 | NRIP3 | -0,741 | 7,19919E-72 |
| ACE2 | CAPRIN2 | -0,741 | 7,13328E-72 |
| ACE2 | TXNDC15 | -0,7411 | 6,83254E-72 |
| ACE2 | LYNX1 | -0,7411 | 6,86521E-72 |
| ACE2 | GALNT18 | -0,7411 | 6,72222E-72 |
| ACE2 | ARMCX3 | -0,7411 | 6,86342E-72 |
| ACE2 | GDNF-AS1 | -0,7412 | 6,2323E-72 |
| ACE2 | TPM2 | -0,7412 | 6,15556E-72 |
| ACE2 | NR2F1-AS1 | -0,7413 | 5,90353E-72 |
| ACE2 | SGCD | -0,7413 | 5,78364E-72 |
| ACE2 | RP3-333H23.9 | -0,7413 | 5,70637E-72 |
| ACE2 | RPL35 | -0,7414 | 5,39325E-72 |
| ACE2 | OTOGL | -0,7414 | 5,39512E-72 |
| ACE2 | PPP1R12C | -0,7414 | 5,3398E-72 |
| ACE2 | PACERR | -0,7415 | 5,12744E-72 |
| ACE2 | C2orf74 | -0,7415 | 5,26777E-72 |
| ACE2 | MARVELD1 | -0,7415 | 5,01154E-72 |
| ACE2 | RP11-817I4.2 | -0,7415 | 5,08996E-72 |
| ACE2 | ZNF676 | -0,7416 | 4,71696E-72 |
| ACE2 | TMEM184B | -0,7416 | 4,79246E-72 |
| ACE2 | ZMYM4 | -0,7417 | 4,35718E-72 |
| ACE2 | DLX6-AS1 | -0,7417 | 4,61473E-72 |
| ACE2 | NFATC1 | -0,7418 | 4,35134E-72 |
| ACE2 | ZBTB47 | -0,7419 | 3,8635E-72 |
| ACE2 | RPRD1A | -0,7419 | 4,01256E-72 |
| ACE2 | MARCH2 | -0,742 | 3,63405E-72 |
| ACE2 | PCDHB12 | -0,7421 | 3,38671E-72 |
| ACE2 | CAV2 | -0,7421 | 3,43466E-72 |

| | | | |
|---|---|---|---|
| ACE2 | TENM4 | -0,7421 | 3,37033E-72 |
| ACE2 | SPATA7 | -0,7421 | 3,4286E-72 |
| ACE2 | SPTBN4 | -0,7421 | 3,3621E-72 |
| ACE2 | AGO3 | -0,7422 | 3,12754E-72 |
| ACE2 | NAV1 | -0,7422 | 3,26424E-72 |
| ACE2 | THYN1 | -0,7422 | 3,19749E-72 |
| ACE2 | PRTG | -0,7422 | 3,13804E-72 |
| ACE2 | MAP4K4 | -0,7423 | 3,00843E-72 |
| ACE2 | NIFK-AS1 | -0,7423 | 2,92338E-72 |
| ACE2 | MAP3K20-AS1 | -0,7423 | 3,0526E-72 |
| ACE2 | RP11-798K23.5 | -0,7423 | 3,09992E-72 |
| ACE2 | KIF5A | -0,7423 | 3,05437E-72 |
| ACE2 | LYSMD2 | -0,7423 | 2,92944E-72 |
| ACE2 | PXK | -0,7424 | 2,82637E-72 |
| ACE2 | TUBB2B | -0,7424 | 2,85304E-72 |
| ACE2 | KHDRBS3 | -0,7424 | 2,83821E-72 |
| ACE2 | REM1 | -0,7424 | 2,8069E-72 |
| ACE2 | SKI | -0,7425 | 2,70629E-72 |
| ACE2 | COL11A1 | -0,7425 | 2,65914E-72 |
| ACE2 | DFNB59 | -0,7425 | 2,59106E-72 |
| ACE2 | PCDH15 | -0,7425 | 2,56408E-72 |
| ACE2 | AMZ2 | -0,7425 | 2,64523E-72 |
| ACE2 | MYL9 | -0,7425 | 2,71454E-72 |
| ACE2 | COL28A1 | -0,7426 | 2,49525E-72 |
| ACE2 | FGF7 | -0,7426 | 2,53755E-72 |
| ACE2 | VKORC1 | -0,7426 | 2,54225E-72 |
| ACE2 | LINC01160 | -0,7427 | 2,22964E-72 |
| ACE2 | GEM | -0,7427 | 2,3518E-72 |
| ACE2 | TERF2IP | -0,7427 | 2,35755E-72 |
| ACE2 | SLC23A2 | -0,7427 | 2,23361E-72 |
| ACE2 | ICA1L | -0,7429 | 1,9847E-72 |
| ACE2 | CAND2 | -0,7429 | 2,07086E-72 |
| ACE2 | CYS1 | -0,743 | 1,88652E-72 |
| ACE2 | EMILIN1 | -0,743 | 1,85461E-72 |
| ACE2 | UCHL1 | -0,743 | 1,84735E-72 |
| ACE2 | RUFY3 | -0,743 | 1,90446E-72 |
| ACE2 | TUSC3 | -0,743 | 1,84044E-72 |
| ACE2 | PDCD7 | -0,743 | 1,8726E-72 |
| ACE2 | RECK | -0,7431 | 1,77261E-72 |
| ACE2 | SYDE1 | -0,7431 | 1,80297E-72 |
| ACE2 | KIAA0930 | -0,7431 | 1,79723E-72 |
| ACE2 | BHLHB9 | -0,7431 | 1,77773E-72 |
| ACE2 | PTPRU | -0,7432 | 1,60113E-72 |
| ACE2 | COL21A1 | -0,7432 | 1,62892E-72 |
| ACE2 | RP11-158I9.5 | -0,7432 | 1,5973E-72 |
| ACE2 | GRK4 | -0,7433 | 1,56856E-72 |
| ACE2 | ZNF883 | -0,7433 | 1,56235E-72 |
| ACE2 | PELI3 | -0,7433 | 1,56107E-72 |
| ACE2 | MAP3K3 | -0,7433 | 1,54368E-72 |
| ACE2 | SNPH | -0,7433 | 1,53777E-72 |
| ACE2 | RPL24 | -0,7434 | 1,4078E-72 |
| ACE2 | PCDHGA6 | -0,7434 | 1,46973E-72 |
| ACE2 | CASKIN1 | -0,7434 | 1,42186E-72 |
| ACE2 | ATP10D | -0,7435 | 1,35785E-72 |

| | | | |
|---|---|---|---|
| ACE2 | BHMT2 | -0,7435 | 1,3161E-72 |
| ACE2 | FAM160B2 | -0,7435 | 1,3249E-72 |
| ACE2 | FAIM2 | -0,7435 | 1,34024E-72 |
| ACE2 | RNF215 | -0,7435 | 1,3913E-72 |
| ACE2 | FAM127C | -0,7435 | 1,37752E-72 |
| ACE2 | P3H1 | -0,7436 | 1,25353E-72 |
| ACE2 | MRAS | -0,7436 | 1,23517E-72 |
| ACE2 | JAKMIP2 | -0,7436 | 1,30086E-72 |
| ACE2 | TCEAL2 | -0,7436 | 1,26858E-72 |
| ACE2 | CAMSAP2 | -0,7437 | 1,21016E-72 |
| ACE2 | RP11-588K22.2 | -0,7437 | 1,19883E-72 |
| ACE2 | RNF217 | -0,7437 | 1,16776E-72 |
| ACE2 | ZDHHC17 | -0,7437 | 1,14236E-72 |
| ACE2 | UBAP1L | -0,7437 | 1,14261E-72 |
| ACE2 | PCNP | -0,7438 | 1,11688E-72 |
| ACE2 | RGS12 | -0,7438 | 1,13728E-72 |
| ACE2 | KCNT1 | -0,7438 | 1,07013E-72 |
| ACE2 | COPRS | -0,7438 | 1,09363E-72 |
| ACE2 | TMIE | -0,7439 | 1,01565E-72 |
| ACE2 | PPP2R2B | -0,7439 | 9,94422E-73 |
| ACE2 | SYNE1 | -0,7439 | 1,03657E-72 |
| ACE2 | GLI3 | -0,7439 | 9,97349E-73 |
| ACE2 | SCNN1D | -0,744 | 9,44361E-73 |
| ACE2 | GNB4 | -0,744 | 9,69959E-73 |
| ACE2 | C11orf74 | -0,744 | 9,8333E-73 |
| ACE2 | CRIP2 | -0,744 | 9,89704E-73 |
| ACE2 | ADAMTS7 | -0,744 | 9,55967E-73 |
| ACE2 | THCAT158 | -0,744 | 9,67557E-73 |
| ACE2 | ADGRL2 | -0,7441 | 8,93884E-73 |
| ACE2 | HFM1 | -0,7441 | 8,93959E-73 |
| ACE2 | SYPL2 | -0,7441 | 9,01562E-73 |
| ACE2 | WFS1 | -0,7441 | 9,27328E-73 |
| ACE2 | KCTD12 | -0,7441 | 9,08449E-73 |
| ACE2 | RP1-59D14.5 | -0,7441 | 8,71675E-73 |
| ACE2 | RP4-613B23.1 | -0,7442 | 8,50914E-73 |
| ACE2 | RRAGD | -0,7442 | 8,23486E-73 |
| ACE2 | ZNF333 | -0,7442 | 8,57981E-73 |
| ACE2 | MARK4 | -0,7442 | 8,18646E-73 |
| ACE2 | PELO | -0,7443 | 7,79241E-73 |
| ACE2 | CNP | -0,7443 | 7,69994E-73 |
| ACE2 | LAMA5 | -0,7443 | 7,88804E-73 |
| ACE2 | BOD1 | -0,7444 | 7,46953E-73 |
| ACE2 | CASP12 | -0,7444 | 7,10766E-73 |
| ACE2 | FAM171A2 | -0,7445 | 7,00295E-73 |
| ACE2 | TENM3 | -0,7447 | 6,11238E-73 |
| ACE2 | CTC-255N20.1 | -0,7447 | 6,06446E-73 |
| ACE2 | ZNF781 | -0,7447 | 6,06344E-73 |
| ACE2 | TRPS1 | -0,7448 | 5,78524E-73 |
| ACE2 | PARP6 | -0,7448 | 5,54268E-73 |
| ACE2 | FKBP8 | -0,7448 | 5,74667E-73 |
| ACE2 | PTBP2 | -0,7449 | 5,14237E-73 |
| ACE2 | PCDHB10 | -0,745 | 4,92468E-73 |
| ACE2 | IGF2 | -0,745 | 5,04058E-73 |
| ACE2 | NUPR1 | -0,745 | 4,73957E-73 |

| | | | |
|---|---|---|---|
| ACE2 | ERO1B | -0,7451 | 4,42695E-73 |
| ACE2 | C1QTNF4 | -0,7451 | 4,43368E-73 |
| ACE2 | GTF3A | -0,7451 | 4,51166E-73 |
| ACE2 | GNG8 | -0,7451 | 4,62844E-73 |
| ACE2 | PJA1 | -0,7451 | 4,44043E-73 |
| ACE2 | FRS3 | -0,7452 | 4,26953E-73 |
| ACE2 | ZC2HC1A | -0,7452 | 4,39119E-73 |
| ACE2 | PPFIBP1 | -0,7452 | 4,38622E-73 |
| ACE2 | CCDC69 | -0,7453 | 3,94086E-73 |
| ACE2 | ZSCAN26 | -0,7453 | 3,98415E-73 |
| ACE2 | OGN | -0,7453 | 3,99661E-73 |
| ACE2 | SELENOW | -0,7453 | 4,00993E-73 |
| ACE2 | THUMPD2 | -0,7455 | 3,54037E-73 |
| ACE2 | FAM234B | -0,7455 | 3,40237E-73 |
| ACE2 | ARHGAP23 | -0,7455 | 3,59355E-73 |
| ACE2 | PARP2 | -0,7456 | 3,17287E-73 |
| ACE2 | BBS4 | -0,7456 | 3,34104E-73 |
| ACE2 | DACT3 | -0,7456 | 3,28567E-73 |
| ACE2 | SLC2A4RG | -0,7456 | 3,25085E-73 |
| ACE2 | MAGED2 | -0,7457 | 3,04293E-73 |
| ACE2 | IL17RD | -0,7458 | 2,80295E-73 |
| ACE2 | KLHL22 | -0,7458 | 2,92802E-73 |
| ACE2 | LSM10 | -0,7459 | 2,7057E-73 |
| ACE2 | NUCKS1 | -0,7459 | 2,58218E-73 |
| ACE2 | CCDC149 | -0,7459 | 2,71602E-73 |
| ACE2 | XPA | -0,7459 | 2,64765E-73 |
| ACE2 | ULK2 | -0,7459 | 2,71712E-73 |
| ACE2 | ERCC1 | -0,7459 | 2,72439E-73 |
| ACE2 | LRRC20 | -0,746 | 2,44757E-73 |
| ACE2 | PAAF1 | -0,746 | 2,49014E-73 |
| ACE2 | ZMYM5 | -0,746 | 2,55115E-73 |
| ACE2 | FGF14 | -0,746 | 2,47684E-73 |
| ACE2 | PTRF | -0,746 | 2,53094E-73 |
| ACE2 | C20orf166-AS1 | -0,746 | 2,48777E-73 |
| ACE2 | UBE2G2 | -0,746 | 2,42496E-73 |
| ACE2 | DPY19L2 | -0,7461 | 2,28899E-73 |
| ACE2 | ENOX1 | -0,7461 | 2,28716E-73 |
| ACE2 | SPRY1 | -0,7462 | 2,17203E-73 |
| ACE2 | UBE2D4 | -0,7462 | 2,17875E-73 |
| ACE2 | RP11-266K4.14 | -0,7462 | 2,23578E-73 |
| ACE2 | ELK3 | -0,7462 | 2,21497E-73 |
| ACE2 | MPP2 | -0,7462 | 2,17203E-73 |
| ACE2 | FAM171B | -0,7463 | 2,02605E-73 |
| ACE2 | RARB | -0,7463 | 1,95749E-73 |
| ACE2 | ROBO3 | -0,7463 | 1,96277E-73 |
| ACE2 | CCDC184 | -0,7463 | 1,9974E-73 |
| ACE2 | MEIS2 | -0,7463 | 2,03611E-73 |
| ACE2 | SRGAP2 | -0,7465 | 1,72334E-73 |
| ACE2 | MAL | -0,7465 | 1,73881E-73 |
| ACE2 | PCDHGA2 | -0,7465 | 1,79936E-73 |
| ACE2 | IL17D | -0,7465 | 1,82056E-73 |
| ACE2 | TGFB3 | -0,7465 | 1,78897E-73 |
| ACE2 | CLIP3 | -0,7465 | 1,79171E-73 |
| ACE2 | SLC41A1 | -0,7466 | 1,63018E-73 |

| | | | |
|---|---|---|---|
| ACE2 | KCNIP1 | -0,7466 | 1,63308E-73 |
| ACE2 | ZCCHC24 | -0,7467 | 1,49947E-73 |
| ACE2 | CRYAB | -0,7467 | 1,58204E-73 |
| ACE2 | VWA1 | -0,7468 | 1,42591E-73 |
| ACE2 | CKMT2-AS1 | -0,7468 | 1,45185E-73 |
| ACE2 | RP11-210M15.2 | -0,7468 | 1,47612E-73 |
| ACE2 | FBF1 | -0,7468 | 1,42275E-73 |
| ACE2 | DDR2 | -0,7469 | 1,36251E-73 |
| ACE2 | LIMS2 | -0,7469 | 1,31494E-73 |
| ACE2 | KIF5C | -0,7469 | 1,38036E-73 |
| ACE2 | C19orf44 | -0,7469 | 1,30234E-73 |
| ACE2 | ZFP82 | -0,7469 | 1,38426E-73 |
| ACE2 | PCDHB6 | -0,747 | 1,27481E-73 |
| ACE2 | RNASEH2C | -0,747 | 1,28476E-73 |
| ACE2 | EID1 | -0,747 | 1,26356E-73 |
| ACE2 | FENDRR | -0,747 | 1,22907E-73 |
| ACE2 | FHOD3 | -0,747 | 1,26791E-73 |
| ACE2 | ZNF790-AS1 | -0,747 | 1,25947E-73 |
| ACE2 | S100A13 | -0,7471 | 1,20636E-73 |
| ACE2 | CACNA2D3 | -0,7471 | 1,19584E-73 |
| ACE2 | RNF212 | -0,7472 | 1,10411E-73 |
| ACE2 | MINDY4 | -0,7472 | 1,12564E-73 |
| ACE2 | CCDC106 | -0,7472 | 1,07179E-73 |
| ACE2 | HSPB2 | -0,7473 | 1,0165E-73 |
| ACE2 | KCNQ4 | -0,7474 | 9,27415E-74 |
| ACE2 | HAND2-AS1 | -0,7474 | 9,68046E-74 |
| ACE2 | CETN2 | -0,7474 | 9,3771E-74 |
| ACE2 | IFT22 | -0,7475 | 8,70239E-74 |
| ACE2 | HMGN3 | -0,7476 | 8,23462E-74 |
| ACE2 | FYN | -0,7476 | 8,45492E-74 |
| ACE2 | RP11-236L14.2 | -0,7476 | 8,58307E-74 |
| ACE2 | NUDT11 | -0,7476 | 8,11612E-74 |
| ACE2 | MRPL55 | -0,7477 | 7,72749E-74 |
| ACE2 | SCD5 | -0,7477 | 7,90252E-74 |
| ACE2 | ATF4 | -0,7477 | 7,98186E-74 |
| ACE2 | RP11-107N15.1 | -0,7478 | 7,26979E-74 |
| ACE2 | POMGNT2 | -0,7478 | 7,00108E-74 |
| ACE2 | AQP1 | -0,7479 | 6,97145E-74 |
| ACE2 | SNN | -0,7479 | 6,81365E-74 |
| ACE2 | SOX15 | -0,7479 | 6,62584E-74 |
| ACE2 | RP4-756G23.5 | -0,7479 | 6,98181E-74 |
| ACE2 | SLC9B2 | -0,748 | 6,22189E-74 |
| ACE2 | FMNL3 | -0,7481 | 5,97436E-74 |
| ACE2 | PCDHB15 | -0,7482 | 5,45067E-74 |
| ACE2 | MIER2 | -0,7482 | 5,68375E-74 |
| ACE2 | TMEM91 | -0,7482 | 5,35085E-74 |
| ACE2 | KCNT2 | -0,7483 | 4,97682E-74 |
| ACE2 | TRIP6 | -0,7483 | 5,09489E-74 |
| ACE2 | COL15A1 | -0,7483 | 5,27195E-74 |
| ACE2 | TRIB2 | -0,7484 | 4,69778E-74 |
| ACE2 | PTMS | -0,7484 | 4,80827E-74 |
| ACE2 | MGP | -0,7484 | 4,67582E-74 |
| ACE2 | LOXL1 | -0,7484 | 4,86215E-74 |
| ACE2 | KCNA2 | -0,7485 | 4,51192E-74 |

| | | | |
|---|---|---|---|
| ACE2 | PPFIA4 | -0,7485 | 4,37811E-74 |
| ACE2 | NAP1L5 | -0,7485 | 4,47477E-74 |
| ACE2 | PCDHB4 | -0,7486 | 4,16776E-74 |
| ACE2 | BEND6 | -0,7486 | 4,08062E-74 |
| ACE2 | RP11-128A17.2 | -0,7486 | 4,03779E-74 |
| ACE2 | DENND2A | -0,7487 | 3,91966E-74 |
| ACE2 | MPZ | -0,7488 | 3,64038E-74 |
| ACE2 | ARL6 | -0,7488 | 3,58751E-74 |
| ACE2 | SPARC | -0,7488 | 3,6428E-74 |
| ACE2 | AS3MT | -0,7488 | 3,67091E-74 |
| ACE2 | MEGF6 | -0,7489 | 3,4739E-74 |
| ACE2 | RP11-566K19.6 | -0,7489 | 3,50047E-74 |
| ACE2 | FAM120C | -0,7489 | 3,31834E-74 |
| ACE2 | STAT5B | -0,749 | 3,2687E-74 |
| ACE2 | CD99L2 | -0,749 | 3,12411E-74 |
| ACE2 | ASIC1 | -0,7491 | 2,91984E-74 |
| ACE2 | SMIM10L2A | -0,7491 | 2,9765E-74 |
| ACE2 | KLHL5 | -0,7492 | 2,7483E-74 |
| ACE2 | SLIT3 | -0,7492 | 2,757E-74 |
| ACE2 | BAALC | -0,7492 | 2,78616E-74 |
| ACE2 | ASTN2 | -0,7492 | 2,76667E-74 |
| ACE2 | TRMT112 | -0,7492 | 2,72084E-74 |
| ACE2 | SRCIN1 | -0,7492 | 2,82738E-74 |
| ACE2 | CAMTA1 | -0,7493 | 2,58288E-74 |
| ACE2 | ADD2 | -0,7493 | 2,63087E-74 |
| ACE2 | RP3-329A5.8 | -0,7493 | 2,58451E-74 |
| ACE2 | USP49 | -0,7493 | 2,51619E-74 |
| ACE2 | BACE1 | -0,7493 | 2,58505E-74 |
| ACE2 | RP11-342A23.2 | -0,7493 | 2,64172E-74 |
| ACE2 | TCFL5 | -0,7493 | 2,54168E-74 |
| ACE2 | LETM2 | -0,7494 | 2,36083E-74 |
| ACE2 | ZFPM2 | -0,7494 | 2,37123E-74 |
| ACE2 | RP4-639F20.1 | -0,7495 | 2,21653E-74 |
| ACE2 | CLMP | -0,7495 | 2,19595E-74 |
| ACE2 | RASL12 | -0,7495 | 2,20109E-74 |
| ACE2 | FGF12 | -0,7496 | 2,05052E-74 |
| ACE2 | CNTN1 | -0,7496 | 2,13357E-74 |
| ACE2 | MAP3K10 | -0,7496 | 2,14547E-74 |
| ACE2 | ARHGEF6 | -0,7496 | 2,04553E-74 |
| ACE2 | RP11-548H3.1 | -0,7497 | 1,89175E-74 |
| ACE2 | MAP7D3 | -0,7497 | 1,93067E-74 |
| ACE2 | AKT3 | -0,7498 | 1,7853E-74 |
| ACE2 | BICC1 | -0,7498 | 1,8242E-74 |
| ACE2 | THOC1 | -0,7498 | 1,84961E-74 |
| ACE2 | RPL15 | -0,7499 | 1,66189E-74 |
| ACE2 | RPS20 | -0,7499 | 1,68609E-74 |
| ACE2 | PCED1B | -0,7499 | 1,66528E-74 |
| ACE2 | GATM | -0,7499 | 1,74617E-74 |
| ACE2 | PLEKHH2 | -0,75 | 1,55493E-74 |
| ACE2 | ETV1 | -0,75 | 1,57319E-74 |
| ACE2 | ANKS1B | -0,75 | 1,6044E-74 |
| ACE2 | HNRNPA1L2 | -0,75 | 1,60678E-74 |
| ACE2 | CHD3 | -0,75 | 1,54638E-74 |
| ACE2 | GPX7 | -0,7501 | 1,44478E-74 |

| | | | |
|---|---|---|---|
| ACE2 | RWDD1 | -0,7501 | 1,45217E-74 |
| ACE2 | COQ10A | -0,7501 | 1,46553E-74 |
| ACE2 | EIF3G | -0,7501 | 1,43059E-74 |
| ACE2 | KCTD15 | -0,7501 | 1,48263E-74 |
| ACE2 | LINC01772 | -0,7502 | 1,40846E-74 |
| ACE2 | SCHIP1 | -0,7502 | 1,35999E-74 |
| ACE2 | TMEM109 | -0,7502 | 1,38587E-74 |
| ACE2 | ZNF532 | -0,7502 | 1,39987E-74 |
| ACE2 | TGFB2 | -0,7503 | 1,25053E-74 |
| ACE2 | CMSS1 | -0,7503 | 1,30026E-74 |
| ACE2 | HAND2 | -0,7503 | 1,30289E-74 |
| ACE2 | GRB10 | -0,7503 | 1,32364E-74 |
| ACE2 | GRIK2 | -0,7504 | 1,17772E-74 |
| ACE2 | VIPR2 | -0,7504 | 1,23323E-74 |
| ACE2 | FAM19A5 | -0,7504 | 1,19493E-74 |
| ACE2 | MID2 | -0,7504 | 1,19619E-74 |
| ACE2 | PNMA6A | -0,7504 | 1,16508E-74 |
| ACE2 | MAPK10 | -0,7505 | 1,1526E-74 |
| ACE2 | CHIC1 | -0,7505 | 1,15136E-74 |
| ACE2 | EXTL2 | -0,7507 | 9,55305E-75 |
| ACE2 | ADCY2 | -0,7507 | 9,86983E-75 |
| ACE2 | GRIK5 | -0,7507 | 9,84983E-75 |
| ACE2 | GPRASP1 | -0,7507 | 9,80804E-75 |
| ACE2 | SLC26A10 | -0,7508 | 8,90284E-75 |
| ACE2 | PSIP1 | -0,7509 | 8,64197E-75 |
| ACE2 | CYP2E1 | -0,7509 | 8,36833E-75 |
| ACE2 | PIEZO2 | -0,7509 | 8,50764E-75 |
| ACE2 | AC003973.3 | -0,7509 | 8,52747E-75 |
| ACE2 | SEC22C | -0,751 | 7,81437E-75 |
| ACE2 | SNHG6 | -0,751 | 8,12378E-75 |
| ACE2 | PLSCR3 | -0,751 | 8,16647E-75 |
| ACE2 | PRICKLE2 | -0,7511 | 7,16602E-75 |
| ACE2 | KATNAL1 | -0,7511 | 7,42597E-75 |
| ACE2 | TTBK2 | -0,7511 | 7,59764E-75 |
| ACE2 | ANKDD1A | -0,7511 | 7,14252E-75 |
| ACE2 | RP11-161M6.2 | -0,7511 | 7,15426E-75 |
| ACE2 | AARSD1 | -0,7511 | 7,18217E-75 |
| ACE2 | THBS3 | -0,7512 | 6,74233E-75 |
| ACE2 | BICD2 | -0,7512 | 6,65077E-75 |
| ACE2 | TMEM240 | -0,7513 | 6,46621E-75 |
| ACE2 | PRDM5 | -0,7513 | 6,29778E-75 |
| ACE2 | ADAM19 | -0,7513 | 6,47883E-75 |
| ACE2 | DLC1 | -0,7513 | 6,27884E-75 |
| ACE2 | BCL7C | -0,7513 | 6,38067E-75 |
| ACE2 | DMPK | -0,7513 | 6,37523E-75 |
| ACE2 | VIM | -0,7514 | 5,93556E-75 |
| ACE2 | RANGRF | -0,7514 | 6,05656E-75 |
| ACE2 | CLIP4 | -0,7515 | 5,50416E-75 |
| ACE2 | ST3GAL6 | -0,7515 | 5,71859E-75 |
| ACE2 | CMYA5 | -0,7516 | 5,05281E-75 |
| ACE2 | ABHD8 | -0,7517 | 4,88829E-75 |
| ACE2 | PDLIM7 | -0,7518 | 4,50533E-75 |
| ACE2 | MCC | -0,7519 | 4,12839E-75 |
| ACE2 | PRPH2 | -0,7519 | 4,25015E-75 |

| | | | |
|---|---|---|---|
| ACE2 | PRPF40B | -0,7519 | 4,14501E-75 |
| ACE2 | PDE10A | -0,752 | 4,03371E-75 |
| ACE2 | RARG | -0,752 | 3,94583E-75 |
| ACE2 | RNF216 | -0,7521 | 3,57694E-75 |
| ACE2 | NAALAD2 | -0,7521 | 3,76115E-75 |
| ACE2 | NXN | -0,7521 | 3,76348E-75 |
| ACE2 | ZNF542P | -0,7521 | 3,54982E-75 |
| ACE2 | ZNF385D | -0,7522 | 3,30066E-75 |
| ACE2 | POLR1E | -0,7522 | 3,48121E-75 |
| ACE2 | FXYD6 | -0,7522 | 3,378E-75 |
| ACE2 | MIR4697HG | -0,7522 | 3,35416E-75 |
| ACE2 | CTD-2525P14.5 | -0,7522 | 3,31934E-75 |
| ACE2 | C16orf86 | -0,7522 | 3,50834E-75 |
| ACE2 | CAMK1 | -0,7523 | 3,10604E-75 |
| ACE2 | RUNX1T1 | -0,7523 | 3,27504E-75 |
| ACE2 | TECTA | -0,7523 | 3,2133E-75 |
| ACE2 | EVC | -0,7524 | 2,91619E-75 |
| ACE2 | SCPEP1 | -0,7524 | 2,87851E-75 |
| ACE2 | KIF3C | -0,7525 | 2,7771E-75 |
| ACE2 | AGAP3 | -0,7525 | 2,73143E-75 |
| ACE2 | BFSP1 | -0,7525 | 2,83946E-75 |
| ACE2 | CECR2 | -0,7525 | 2,73742E-75 |
| ACE2 | KLHDC8A | -0,7526 | 2,5301E-75 |
| ACE2 | CLDND1 | -0,7526 | 2,55394E-75 |
| ACE2 | KIF3A | -0,7526 | 2,62933E-75 |
| ACE2 | TMEM121 | -0,7526 | 2,54118E-75 |
| ACE2 | ACBD6 | -0,7527 | 2,34933E-75 |
| ACE2 | PHF24 | -0,7527 | 2,38265E-75 |
| ACE2 | CUEDC1 | -0,7527 | 2,40005E-75 |
| ACE2 | NDE1 | -0,7528 | 2,26433E-75 |
| ACE2 | ST6GALNAC2 | -0,7528 | 2,27722E-75 |
| ACE2 | IFT46 | -0,7529 | 2,01934E-75 |
| ACE2 | ZNF512B | -0,7529 | 2,14223E-75 |
| ACE2 | BOC | -0,753 | 1,98992E-75 |
| ACE2 | TYRO3 | -0,753 | 1,99613E-75 |
| ACE2 | CTD-2036P10.6 | -0,753 | 1,88242E-75 |
| ACE2 | PLA2G4C | -0,7532 | 1,655E-75 |
| ACE2 | SLC45A1 | -0,7533 | 1,62564E-75 |
| ACE2 | SELENOM | -0,7533 | 1,56604E-75 |
| ACE2 | SEMA4C | -0,7534 | 1,46563E-75 |
| ACE2 | USP44 | -0,7534 | 1,43108E-75 |
| ACE2 | COX7A1 | -0,7534 | 1,47457E-75 |
| ACE2 | ZNF382 | -0,7534 | 1,50212E-75 |
| ACE2 | ROBO1 | -0,7535 | 1,3555E-75 |
| ACE2 | SOBP | -0,7535 | 1,41293E-75 |
| ACE2 | B4GALNT4 | -0,7535 | 1,32265E-75 |
| ACE2 | EEF1AKMT1 | -0,7535 | 1,32845E-75 |
| ACE2 | MDH1B | -0,7536 | 1,26937E-75 |
| ACE2 | RP11-620J15.3 | -0,7536 | 1,28254E-75 |
| ACE2 | C20orf194 | -0,7536 | 1,26512E-75 |
| ACE2 | ARHGEF3 | -0,7537 | 1,20972E-75 |
| ACE2 | ATP6V1G2 | -0,7537 | 1,1841E-75 |
| ACE2 | INAFM1 | -0,7537 | 1,15718E-75 |
| ACE2 | ZCCHC11 | -0,7538 | 1,11416E-75 |

| | | | |
|---|---|---|---|
| ACE2 | NGF | -0,7538 | 1,12155E-75 |
| ACE2 | ZNF767P | -0,7538 | 1,10419E-75 |
| ACE2 | AF001548.3 | -0,7538 | 1,12267E-75 |
| ACE2 | PRSS53 | -0,7538 | 1,11381E-75 |
| ACE2 | KLC2 | -0,7539 | 1,01533E-75 |
| ACE2 | CAMKK1 | -0,7539 | 1,0333E-75 |
| ACE2 | SORCS2 | -0,754 | 9,4233E-76 |
| ACE2 | CERCAM | -0,754 | 9,40859E-76 |
| ACE2 | CTD-3220F14.1 | -0,754 | 9,62174E-76 |
| ACE2 | TTC28-AS1 | -0,754 | 9,53232E-76 |
| ACE2 | TMEM108 | -0,7541 | 9,27248E-76 |
| ACE2 | TUBE1 | -0,7541 | 9,05072E-76 |
| ACE2 | PLEKHO1 | -0,7542 | 8,1206E-76 |
| ACE2 | CHL1-AS2 | -0,7542 | 8,1242E-76 |
| ACE2 | UBTD1 | -0,7542 | 8,64238E-76 |
| ACE2 | MEGF10 | -0,7543 | 7,55712E-76 |
| ACE2 | SEPT6 | -0,7543 | 7,94611E-76 |
| ACE2 | MEIS1 | -0,7544 | 7,4223E-76 |
| ACE2 | SCRN1 | -0,7544 | 7,11949E-76 |
| ACE2 | RP11-380L11.4 | -0,7544 | 7,1213E-76 |
| ACE2 | TEAD2 | -0,7544 | 7,31813E-76 |
| ACE2 | RAB28P5 | -0,7544 | 7,04327E-76 |
| ACE2 | CCDC66 | -0,7545 | 6,56262E-76 |
| ACE2 | ZEB1 | -0,7545 | 6,59091E-76 |
| ACE2 | RP11-429G19.3 | -0,7545 | 6,7584E-76 |
| ACE2 | PHF21A | -0,7545 | 6,74471E-76 |
| ACE2 | LRP2BP | -0,7547 | 6,03191E-76 |
| ACE2 | PACSIN3 | -0,7548 | 5,43747E-76 |
| ACE2 | UBE2Q2 | -0,7548 | 5,40253E-76 |
| ACE2 | SULT1C4 | -0,7549 | 5,01606E-76 |
| ACE2 | GUCY1B3 | -0,7549 | 5,11979E-76 |
| ACE2 | FAM227B | -0,7549 | 4,95958E-76 |
| ACE2 | LINC01116 | -0,755 | 4,66444E-76 |
| ACE2 | RNF165 | -0,755 | 4,59032E-76 |
| ACE2 | PCDHB2 | -0,7551 | 4,35766E-76 |
| ACE2 | ADAMTSL3 | -0,7551 | 4,46312E-76 |
| ACE2 | SV2B | -0,7552 | 4,14963E-76 |
| ACE2 | ERCC3 | -0,7553 | 3,79619E-76 |
| ACE2 | NRG3 | -0,7553 | 3,81193E-76 |
| ACE2 | TSNAXIP1 | -0,7553 | 3,71709E-76 |
| ACE2 | LAS1L | -0,7553 | 3,82574E-76 |
| ACE2 | B3GALNT1 | -0,7554 | 3,45867E-76 |
| ACE2 | AC156455.1 | -0,7554 | 3,60681E-76 |
| ACE2 | C1orf216 | -0,7555 | 3,36789E-76 |
| ACE2 | CARMN | -0,7555 | 3,29808E-76 |
| ACE2 | HSF2 | -0,7555 | 3,21344E-76 |
| ACE2 | SPAG8 | -0,7555 | 3,23056E-76 |
| ACE2 | CD27-AS1 | -0,7555 | 3,20406E-76 |
| ACE2 | MNAT1 | -0,7555 | 3,22563E-76 |
| ACE2 | FHL3 | -0,7556 | 3,19927E-76 |
| ACE2 | CFAP58 | -0,7556 | 3,05844E-76 |
| ACE2 | SV2A | -0,7557 | 2,94652E-76 |
| ACE2 | UST | -0,7557 | 2,90159E-76 |
| ACE2 | ARHGEF25 | -0,7557 | 2,95475E-76 |

| | | | |
|---|---|---|---|
| ACE2 | STRADA | -0,7557 | 2,80414E-76 |
| ACE2 | CHST9 | -0,7557 | 2,82709E-76 |
| ACE2 | RP11-412D9.4 | -0,7558 | 2,77423E-76 |
| ACE2 | B3GNT9 | -0,7558 | 2,68786E-76 |
| ACE2 | HMCN1 | -0,7559 | 2,53995E-76 |
| ACE2 | ATP6V0E2-AS1 | -0,7559 | 2,44048E-76 |
| ACE2 | HIKESHI | -0,7559 | 2,56614E-76 |
| ACE2 | BEX1 | -0,7559 | 2,57656E-76 |
| ACE2 | ZNF362 | -0,756 | 2,38731E-76 |
| ACE2 | PTHLH | -0,756 | 2,35174E-76 |
| ACE2 | ZFP36L1 | -0,756 | 2,31395E-76 |
| ACE2 | TBC1D19 | -0,7561 | 2,21551E-76 |
| ACE2 | RADIL | -0,7561 | 2,11614E-76 |
| ACE2 | FAM89B | -0,7562 | 2,07628E-76 |
| ACE2 | FAM127A | -0,7562 | 2,05496E-76 |
| ACE2 | WWTR1 | -0,7563 | 1,8249E-76 |
| ACE2 | JAZF1 | -0,7563 | 1,8666E-76 |
| ACE2 | LOXL4 | -0,7563 | 1,93741E-76 |
| ACE2 | ANKRD13D | -0,7564 | 1,71622E-76 |
| ACE2 | RASSF8 | -0,7564 | 1,77342E-76 |
| ACE2 | GAMT | -0,7564 | 1,73718E-76 |
| ACE2 | SDC3 | -0,7565 | 1,60427E-76 |
| ACE2 | RP11-398K22.12 | -0,7565 | 1,60886E-76 |
| ACE2 | RILPL1 | -0,7565 | 1,61985E-76 |
| ACE2 | CHST14 | -0,7566 | 1,5609E-76 |
| ACE2 | KAT2A | -0,7566 | 1,53413E-76 |
| ACE2 | ENTPD1 | -0,7567 | 1,42816E-76 |
| ACE2 | SSPN | -0,7567 | 1,35974E-76 |
| ACE2 | MN1 | -0,7567 | 1,43883E-76 |
| ACE2 | ABLIM3 | -0,7568 | 1,33425E-76 |
| ACE2 | KLC1 | -0,7568 | 1,30121E-76 |
| ACE2 | CTC-439O9.1 | -0,7568 | 1,29986E-76 |
| ACE2 | LINC00461 | -0,7569 | 1,23514E-76 |
| ACE2 | GNG2 | -0,7569 | 1,23986E-76 |
| ACE2 | C17orf51 | -0,7569 | 1,18436E-76 |
| ACE2 | PBX2 | -0,757 | 1,10557E-76 |
| ACE2 | FGFR1 | -0,757 | 1,12461E-76 |
| ACE2 | CTSF | -0,757 | 1,17595E-76 |
| ACE2 | LCA5 | -0,7571 | 1,08741E-76 |
| ACE2 | ITIH5 | -0,7571 | 1,07882E-76 |
| ACE2 | ZNF568 | -0,7571 | 1,07373E-76 |
| ACE2 | MIR99AHG | -0,7572 | 9,95082E-77 |
| ACE2 | STX2 | -0,7573 | 9,30242E-77 |
| ACE2 | EHBP1 | -0,7575 | 7,95346E-77 |
| ACE2 | RP11-324L17.1 | -0,7575 | 7,67001E-77 |
| ACE2 | COL8A1 | -0,7575 | 7,69994E-77 |
| ACE2 | ELOVL4 | -0,7575 | 7,80489E-77 |
| ACE2 | SERGEF | -0,7575 | 8,091E-77 |
| ACE2 | EMD | -0,7575 | 7,8216E-77 |
| ACE2 | RP11-73M7.6 | -0,7576 | 7,62044E-77 |
| ACE2 | DBN1 | -0,7576 | 7,45974E-77 |
| ACE2 | DXO | -0,7576 | 7,39254E-77 |
| ACE2 | PIAS3 | -0,7578 | 6,59714E-77 |
| ACE2 | SDC2 | -0,7578 | 6,51233E-77 |

| | | | |
|---|---|---|---|
| ACE2 | MAPK8IP1 | -0,7578 | 6,20562E-77 |
| ACE2 | PLCB1 | -0,7578 | 6,3194E-77 |
| ACE2 | PKIA | -0,7579 | 5,83016E-77 |
| ACE2 | RP11-454E5.4 | -0,7579 | 5,78938E-77 |
| ACE2 | MXRA7 | -0,7579 | 5,87204E-77 |
| ACE2 | DLX1 | -0,758 | 5,42603E-77 |
| ACE2 | PIP4K2B | -0,758 | 5,61E-77 |
| ACE2 | CROCC | -0,7581 | 5,0098E-77 |
| ACE2 | CYP27C1 | -0,7581 | 5,0041E-77 |
| ACE2 | GLIPR2 | -0,7581 | 5,00001E-77 |
| ACE2 | PYGO1 | -0,7581 | 5,02994E-77 |
| ACE2 | HKR1 | -0,7581 | 5,09401E-77 |
| ACE2 | LGALS1 | -0,7581 | 5,15211E-77 |
| ACE2 | SLC4A3 | -0,7582 | 4,69935E-77 |
| ACE2 | CYP2U1 | -0,7582 | 4,69096E-77 |
| ACE2 | FXYD2 | -0,7582 | 4,80807E-77 |
| ACE2 | TTYH1 | -0,7582 | 4,78363E-77 |
| ACE2 | CROCCP3 | -0,7583 | 4,4658E-77 |
| ACE2 | CHN1 | -0,7583 | 4,42531E-77 |
| ACE2 | IGSF11 | -0,7583 | 4,39989E-77 |
| ACE2 | C6orf48 | -0,7583 | 4,33788E-77 |
| ACE2 | IGFBP4 | -0,7583 | 4,48684E-77 |
| ACE2 | SCN7A | -0,7585 | 3,7328E-77 |
| ACE2 | ZNF876P | -0,7585 | 3,88449E-77 |
| ACE2 | GNL1 | -0,7585 | 3,95768E-77 |
| ACE2 | KPNA5 | -0,7586 | 3,55334E-77 |
| ACE2 | STARD9 | -0,7586 | 3,46242E-77 |
| ACE2 | C1orf123 | -0,7587 | 3,36253E-77 |
| ACE2 | VAT1 | -0,7587 | 3,23413E-77 |
| ACE2 | EID2B | -0,7587 | 3,36486E-77 |
| ACE2 | ILKAP | -0,7588 | 3,08441E-77 |
| ACE2 | FAM184A | -0,7588 | 3,05092E-77 |
| ACE2 | MICU3 | -0,7589 | 2,78554E-77 |
| ACE2 | MYEF2 | -0,759 | 2,69468E-77 |
| ACE2 | ARHGEF4 | -0,7591 | 2,54221E-77 |
| ACE2 | IGFBP7 | -0,7591 | 2,52973E-77 |
| ACE2 | GNG11 | -0,7591 | 2,46917E-77 |
| ACE2 | RBMS3 | -0,7592 | 2,3146E-77 |
| ACE2 | PRDM8 | -0,7592 | 2,32085E-77 |
| ACE2 | ASIC3 | -0,7592 | 2,34105E-77 |
| ACE2 | ZNF418 | -0,7592 | 2,3111E-77 |
| ACE2 | MAD2L2 | -0,7593 | 2,16413E-77 |
| ACE2 | NFASC | -0,7593 | 2,13056E-77 |
| ACE2 | KLHL29 | -0,7593 | 2,1909E-77 |
| ACE2 | LINC00882 | -0,7593 | 2,11073E-77 |
| ACE2 | BOLA3-AS1 | -0,7594 | 1,98217E-77 |
| ACE2 | F2RL2 | -0,7594 | 2,01922E-77 |
| ACE2 | BLVRA | -0,7594 | 2,01607E-77 |
| ACE2 | RP11-834C11.4 | -0,7594 | 2,01579E-77 |
| ACE2 | VAMP5 | -0,7597 | 1,60562E-77 |
| ACE2 | EIF3EP1 | -0,7597 | 1,63572E-77 |
| ACE2 | FAM69B | -0,7597 | 1,65073E-77 |
| ACE2 | DOK6 | -0,7597 | 1,65697E-77 |
| ACE2 | C3orf18 | -0,7598 | 1,45991E-77 |

| | | | |
|---|---|---|---|
| ACE2 | TSPOAP1 | -0,7598 | 1,48606E-77 |
| ACE2 | CTNNA3 | -0,7599 | 1,35571E-77 |
| ACE2 | RP11-713P17.3 | -0,7599 | 1,42961E-77 |
| ACE2 | DYRK4 | -0,7599 | 1,36706E-77 |
| ACE2 | ZNF536 | -0,7599 | 1,37862E-77 |
| ACE2 | SLC25A28 | -0,7602 | 1,10175E-77 |
| ACE2 | MYL6B | -0,7602 | 1,15441E-77 |
| ACE2 | SLAIN1 | -0,7602 | 1,155E-77 |
| ACE2 | SHF | -0,7602 | 1,14675E-77 |
| ACE2 | FOXF1 | -0,7602 | 1,1314E-77 |
| ACE2 | AC009948.5 | -0,7603 | 1,01565E-77 |
| ACE2 | MAP4 | -0,7603 | 1,02819E-77 |
| ACE2 | ABCA17P | -0,7603 | 1,03355E-77 |
| ACE2 | DEF8 | -0,7603 | 1,04981E-77 |
| ACE2 | RND2 | -0,7603 | 1,00895E-77 |
| ACE2 | ZNF343 | -0,7603 | 1,07605E-77 |
| ACE2 | NINL | -0,7603 | 1,01143E-77 |
| ACE2 | PRRG3 | -0,7603 | 1,03735E-77 |
| ACE2 | SEMA6C | -0,7604 | 9,80378E-78 |
| ACE2 | RUNDC3A | -0,7604 | 9,68929E-78 |
| ACE2 | CBY1 | -0,7604 | 9,57967E-78 |
| ACE2 | NRXN3 | -0,7605 | 9,2198E-78 |
| ACE2 | KRBOX4 | -0,7605 | 8,77513E-78 |
| ACE2 | C2orf81 | -0,7606 | 8,53463E-78 |
| ACE2 | PTP4A3 | -0,7606 | 8,67385E-78 |
| ACE2 | ST6GALNAC5 | -0,7607 | 7,63025E-78 |
| ACE2 | ITGB1BP1 | -0,7607 | 7,77564E-78 |
| ACE2 | BBS9 | -0,7607 | 7,99799E-78 |
| ACE2 | CPQ | -0,7607 | 7,71052E-78 |
| ACE2 | DENND5A | -0,7607 | 7,79648E-78 |
| ACE2 | MXRA8 | -0,7608 | 7,45998E-78 |
| ACE2 | TMEM67 | -0,7608 | 7,36267E-78 |
| ACE2 | GNB3 | -0,7608 | 7,15136E-78 |
| ACE2 | PPP1R3F | -0,7608 | 7,10278E-78 |
| ACE2 | ZNF512 | -0,7609 | 6,8954E-78 |
| ACE2 | GYPC | -0,7609 | 6,87154E-78 |
| ACE2 | KCTD6 | -0,7609 | 6,97772E-78 |
| ACE2 | PACS1 | -0,7609 | 6,5772E-78 |
| ACE2 | MAP6 | -0,7609 | 6,52955E-78 |
| ACE2 | NEGR1 | -0,761 | 6,25435E-78 |
| ACE2 | KIAA1614 | -0,761 | 6,25648E-78 |
| ACE2 | COL6A1 | -0,761 | 6,27474E-78 |
| ACE2 | PAQR7 | -0,7612 | 5,28846E-78 |
| ACE2 | COL27A1 | -0,7612 | 5,47706E-78 |
| ACE2 | CRTAC1 | -0,7613 | 4,85941E-78 |
| ACE2 | ISLR | -0,7613 | 5,12687E-78 |
| ACE2 | LRRC49 | -0,7614 | 4,82833E-78 |
| ACE2 | ZNF415 | -0,7614 | 4,6793E-78 |
| ACE2 | GJC2 | -0,7615 | 4,41029E-78 |
| ACE2 | RBPMS | -0,7615 | 4,20021E-78 |
| ACE2 | RP11-53O19.1 | -0,7616 | 3,92163E-78 |
| ACE2 | ARNT2 | -0,7616 | 4,17281E-78 |
| ACE2 | INPP4A | -0,7617 | 3,85447E-78 |
| ACE2 | EVC2 | -0,7617 | 3,64005E-78 |

| | | | |
|---|---|---|---|
| ACE2 | RP11-798M19.6 | -0,7618 | 3,43436E-78 |
| ACE2 | RP11-213G2.3 | -0,7618 | 3,58382E-78 |
| ACE2 | MAB21L1 | -0,7618 | 3,52963E-78 |
| ACE2 | WDPCP | -0,762 | 2,98121E-78 |
| ACE2 | FAM219A | -0,762 | 3,10341E-78 |
| ACE2 | ARL6IP4 | -0,762 | 3,10186E-78 |
| ACE2 | GDI1 | -0,762 | 3,08785E-78 |
| ACE2 | LRAT | -0,7621 | 2,73748E-78 |
| ACE2 | PRKRIP1 | -0,7621 | 2,7111E-78 |
| ACE2 | IGSF1 | -0,7621 | 2,70424E-78 |
| ACE2 | LINC00404 | -0,7622 | 2,6266E-78 |
| ACE2 | CHL1 | -0,7623 | 2,48827E-78 |
| ACE2 | TCEAL5 | -0,7623 | 2,48836E-78 |
| ACE2 | TMEM9 | -0,7624 | 2,24021E-78 |
| ACE2 | RHOQ | -0,7624 | 2,26226E-78 |
| ACE2 | DUSP22 | -0,7624 | 2,29785E-78 |
| ACE2 | GPRASP2 | -0,7624 | 2,2236E-78 |
| ACE2 | MRGPRF-AS1 | -0,7625 | 2,14409E-78 |
| ACE2 | FAM66C | -0,7625 | 2,08865E-78 |
| ACE2 | ACD | -0,7625 | 2,08789E-78 |
| ACE2 | STXBP1 | -0,7626 | 2,00093E-78 |
| ACE2 | C11orf63 | -0,7626 | 1,88763E-78 |
| ACE2 | STX1B | -0,7626 | 1,89422E-78 |
| ACE2 | DPYSL3 | -0,7627 | 1,78159E-78 |
| ACE2 | ZNF34 | -0,7627 | 1,82989E-78 |
| ACE2 | HHIPL1 | -0,7627 | 1,76155E-78 |
| ACE2 | ANKRD13B | -0,7627 | 1,75571E-78 |
| ACE2 | MEIS3 | -0,7627 | 1,76146E-78 |
| ACE2 | RNF180 | -0,7628 | 1,65933E-78 |
| ACE2 | UG0898H09 | -0,7629 | 1,57562E-78 |
| ACE2 | NCAM2 | -0,7629 | 1,49345E-78 |
| ACE2 | SEPT7P2 | -0,763 | 1,43735E-78 |
| ACE2 | RP11-177H2.1 | -0,763 | 1,40949E-78 |
| ACE2 | RAB12 | -0,763 | 1,48653E-78 |
| ACE2 | ANKRD65 | -0,7631 | 1,34838E-78 |
| ACE2 | PLEKHB1 | -0,7631 | 1,36455E-78 |
| ACE2 | IFT81 | -0,7631 | 1,35511E-78 |
| ACE2 | SLC27A1 | -0,7631 | 1,38248E-78 |
| ACE2 | RP4-665J23.1 | -0,7632 | 1,22405E-78 |
| ACE2 | FBXL7 | -0,7632 | 1,22714E-78 |
| ACE2 | PRNP | -0,7632 | 1,24774E-78 |
| ACE2 | AP1S2 | -0,7632 | 1,27571E-78 |
| ACE2 | YPEL4 | -0,7634 | 1,07098E-78 |
| ACE2 | NCAM1 | -0,7634 | 1,06469E-78 |
| ACE2 | CDH19 | -0,7634 | 1,04068E-78 |
| ACE2 | LDLRAD2 | -0,7635 | 1,01897E-78 |
| ACE2 | CEP170 | -0,7635 | 1,00652E-78 |
| ACE2 | LAMB2 | -0,7635 | 9,75456E-79 |
| ACE2 | CTD-2555O16.4 | -0,7635 | 9,58209E-79 |
| ACE2 | GNB5 | -0,7635 | 9,84387E-79 |
| ACE2 | MMP16 | -0,7636 | 9,1048E-79 |
| ACE2 | EEF1D | -0,7636 | 9,21802E-79 |
| ACE2 | LYRM4 | -0,7637 | 8,31784E-79 |
| ACE2 | QKI | -0,7637 | 8,63032E-79 |

| | | | |
|---|---|---|---|
| ACE2 | C11orf84 | -0,7637 | 8,76586E-79 |
| ACE2 | FAAP20 | -0,7638 | 7,66955E-79 |
| ACE2 | RBMS1 | -0,7638 | 8,2307E-79 |
| ACE2 | FZD3 | -0,7638 | 7,97312E-79 |
| ACE2 | LINC-PINT | -0,7639 | 7,48182E-79 |
| ACE2 | ZEB1-AS1 | -0,7639 | 7,33409E-79 |
| ACE2 | ABCA3 | -0,7639 | 7,14853E-79 |
| ACE2 | NCALD | -0,764 | 6,67044E-79 |
| ACE2 | CCDC85B | -0,764 | 6,96863E-79 |
| ACE2 | TPTEP1 | -0,764 | 7,08324E-79 |
| ACE2 | TRMT10B | -0,7641 | 6,33549E-79 |
| ACE2 | KYAT1 | -0,7641 | 6,15569E-79 |
| ACE2 | CTD-2554C21.2 | -0,7641 | 6,53347E-79 |
| ACE2 | AK5 | -0,7642 | 6,05344E-79 |
| ACE2 | PKMP3 | -0,7642 | 5,86498E-79 |
| ACE2 | DIRAS3 | -0,7643 | 5,569E-79 |
| ACE2 | TSPAN5 | -0,7643 | 5,61046E-79 |
| ACE2 | ZNF346 | -0,7643 | 5,3683E-79 |
| ACE2 | CLU | -0,7643 | 5,61508E-79 |
| ACE2 | PFDN5 | -0,7643 | 5,58794E-79 |
| ACE2 | SEPT4 | -0,7643 | 5,27926E-79 |
| ACE2 | RGS11 | -0,7644 | 5,27529E-79 |
| ACE2 | PDE1A | -0,7646 | 4,25161E-79 |
| ACE2 | WDR86 | -0,7646 | 4,29497E-79 |
| ACE2 | PHF1 | -0,7647 | 3,95478E-79 |
| ACE2 | RIC3 | -0,7647 | 3,97807E-79 |
| ACE2 | DCLK2 | -0,7648 | 3,70666E-79 |
| ACE2 | CYP46A1 | -0,7648 | 3,87238E-79 |
| ACE2 | LINC00888 | -0,765 | 3,36046E-79 |
| ACE2 | ZNF25 | -0,765 | 3,21938E-79 |
| ACE2 | RASA3 | -0,765 | 3,26482E-79 |
| ACE2 | CCDC74A | -0,7651 | 2,97824E-79 |
| ACE2 | INTU | -0,7651 | 3,00528E-79 |
| ACE2 | RGL2 | -0,7651 | 2,92831E-79 |
| ACE2 | RIMKLB | -0,7651 | 3,06428E-79 |
| ACE2 | ZNF133 | -0,7651 | 3,00407E-79 |
| ACE2 | TUB | -0,7652 | 2,86099E-79 |
| ACE2 | DTWD1 | -0,7652 | 2,86123E-79 |
| ACE2 | NENF | -0,7653 | 2,68335E-79 |
| ACE2 | LA16c-349E10.1 | -0,7653 | 2,69763E-79 |
| ACE2 | SCARF2 | -0,7653 | 2,54075E-79 |
| ACE2 | LRRTM4 | -0,7654 | 2,43339E-79 |
| ACE2 | EMP3 | -0,7654 | 2,4443E-79 |
| ACE2 | SHISA9 | -0,7655 | 2,3165E-79 |
| ACE2 | ZBTB4 | -0,7655 | 2,21548E-79 |
| ACE2 | CDR2L | -0,7655 | 2,17849E-79 |
| ACE2 | GPM6B | -0,7655 | 2,27153E-79 |
| ACE2 | PHLDA3 | -0,7656 | 2,10541E-79 |
| ACE2 | PDLIM4 | -0,7656 | 2,07815E-79 |
| ACE2 | C15orf41 | -0,7656 | 2,05324E-79 |
| ACE2 | ST3GAL2 | -0,7656 | 2,07109E-79 |
| ACE2 | AMZ2P1 | -0,7656 | 2,04377E-79 |
| ACE2 | CCDC102A | -0,7657 | 1,95039E-79 |
| ACE2 | MLLT1 | -0,7657 | 1,9109E-79 |

| | | | |
|---|---|---|---|
| ACE2 | UBE2E2 | -0,7658 | 1,80888E-79 |
| ACE2 | LHFP | -0,7659 | 1,68499E-79 |
| ACE2 | CORO2B | -0,7659 | 1,66273E-79 |
| ACE2 | IMMP2L | -0,7661 | 1,38865E-79 |
| ACE2 | ADGRA2 | -0,7661 | 1,41607E-79 |
| ACE2 | ST8SIA1 | -0,7661 | 1,37741E-79 |
| ACE2 | ANKRD35 | -0,7663 | 1,20833E-79 |
| ACE2 | AC009501.4 | -0,7663 | 1,24706E-79 |
| ACE2 | ANGPTL2 | -0,7663 | 1,2312E-79 |
| ACE2 | ARHGEF26-AS1 | -0,7664 | 1,16738E-79 |
| ACE2 | LRRN4CL | -0,7665 | 1,02524E-79 |
| ACE2 | LINC00665 | -0,7665 | 1,02441E-79 |
| ACE2 | FAM216A | -0,7666 | 1,01543E-79 |
| ACE2 | NPAS3 | -0,7666 | 1,01221E-79 |
| ACE2 | STX1A | -0,7667 | 9,39014E-80 |
| ACE2 | COPZ2 | -0,7667 | 9,01746E-80 |
| ACE2 | SMARCA1 | -0,7667 | 9,38005E-80 |
| ACE2 | LNP1 | -0,7668 | 8,73498E-80 |
| ACE2 | PDGFB | -0,7668 | 8,75216E-80 |
| ACE2 | NPHP1 | -0,7669 | 7,68054E-80 |
| ACE2 | LAMA4 | -0,7669 | 7,56282E-80 |
| ACE2 | S100B | -0,7669 | 7,84862E-80 |
| ACE2 | RP11-1114A5.4 | -0,767 | 7,39659E-80 |
| ACE2 | ZNF853 | -0,7671 | 6,97424E-80 |
| ACE2 | NPR2 | -0,7671 | 6,89771E-80 |
| ACE2 | SULF1 | -0,7672 | 6,30729E-80 |
| ACE2 | FGD1 | -0,7672 | 6,41548E-80 |
| ACE2 | VSTM4 | -0,7673 | 5,80688E-80 |
| ACE2 | MAP3K12 | -0,7673 | 5,61407E-80 |
| ACE2 | FAHD2B | -0,7674 | 5,23692E-80 |
| ACE2 | LY6E | -0,7674 | 5,43809E-80 |
| ACE2 | COL5A1 | -0,7674 | 5,26225E-80 |
| ACE2 | TTC7B | -0,7674 | 5,18367E-80 |
| ACE2 | B4GALNT1 | -0,7675 | 4,9436E-80 |
| ACE2 | PLEKHA4 | -0,7676 | 4,48258E-80 |
| ACE2 | MYOT | -0,7677 | 4,42268E-80 |
| ACE2 | HOXB4 | -0,7677 | 4,42518E-80 |
| ACE2 | ALK | -0,7678 | 3,99861E-80 |
| ACE2 | ZNF582-AS1 | -0,7678 | 4,08494E-80 |
| ACE2 | SLC13A3 | -0,7678 | 3,88325E-80 |
| ACE2 | CHRD | -0,7679 | 3,67269E-80 |
| ACE2 | C4orf3 | -0,7679 | 3,68095E-80 |
| ACE2 | CRISPLD1 | -0,7679 | 3,71215E-80 |
| ACE2 | MPPED2 | -0,7679 | 3,66465E-80 |
| ACE2 | DNAJC18 | -0,768 | 3,39915E-80 |
| ACE2 | PGBD1 | -0,768 | 3,41753E-80 |
| ACE2 | MAGI2-AS3 | -0,768 | 3,37081E-80 |
| ACE2 | TTC28 | -0,768 | 3,34598E-80 |
| ACE2 | PLPP7 | -0,7681 | 3,14868E-80 |
| ACE2 | ZNF428 | -0,7681 | 3,13877E-80 |
| ACE2 | PAM | -0,7682 | 2,97046E-80 |
| ACE2 | CPT1C | -0,7682 | 2,8606E-80 |
| ACE2 | RP11-157P1.4 | -0,7682 | 2,86097E-80 |
| ACE2 | FGF2 | -0,7683 | 2,75705E-80 |

| | | | |
|---|---|---|---|
| ACE2 | RP11-999E24.3 | -0,7685 | 2,30318E-80 |
| ACE2 | CRTAP | -0,7686 | 2,17289E-80 |
| ACE2 | SLC5A7 | -0,7688 | 1,82415E-80 |
| ACE2 | RUSC2 | -0,7688 | 1,89543E-80 |
| ACE2 | FAM92A | -0,7689 | 1,70499E-80 |
| ACE2 | TMEFF1 | -0,7689 | 1,70213E-80 |
| ACE2 | USP20 | -0,7689 | 1,77803E-80 |
| ACE2 | ZFP90 | -0,7689 | 1,77905E-80 |
| ACE2 | DPYSL2 | -0,769 | 1,57478E-80 |
| ACE2 | SMIM10 | -0,7691 | 1,53386E-80 |
| ACE2 | COL22A1 | -0,7692 | 1,3901E-80 |
| ACE2 | NPIPP1 | -0,7692 | 1,39465E-80 |
| ACE2 | TMEM178B | -0,7693 | 1,29482E-80 |
| ACE2 | ZNF32 | -0,7694 | 1,15514E-80 |
| ACE2 | ZNF667 | -0,7696 | 1,03703E-80 |
| ACE2 | PKIG | -0,7698 | 8,85974E-81 |
| ACE2 | JAM2 | -0,7698 | 8,50045E-81 |
| ACE2 | A2M-AS1 | -0,7699 | 8,1329E-81 |
| ACE2 | MEF2C-AS1 | -0,77 | 7,65897E-81 |
| ACE2 | PRICKLE1 | -0,7701 | 7,09153E-81 |
| ACE2 | FGF1 | -0,7702 | 6,46719E-81 |
| ACE2 | NFIA | -0,7703 | 6,08676E-81 |
| ACE2 | PNMAL2 | -0,7703 | 5,94752E-81 |
| ACE2 | COL9A3 | -0,7703 | 6,124E-81 |
| ACE2 | NHSL2 | -0,7704 | 5,33839E-81 |
| ACE2 | MAN1C1 | -0,7705 | 5,22575E-81 |
| ACE2 | ZSCAN16-AS1 | -0,7705 | 5,2473E-81 |
| ACE2 | RP3-525N10.2 | -0,7705 | 5,17569E-81 |
| ACE2 | SLITRK2 | -0,7705 | 5,06646E-81 |
| ACE2 | PEA15 | -0,7706 | 4,52346E-81 |
| ACE2 | DNM3 | -0,7706 | 4,86925E-81 |
| ACE2 | ARL3 | -0,7706 | 4,66053E-81 |
| ACE2 | WHAMMP3 | -0,7706 | 4,70591E-81 |
| ACE2 | NT5M | -0,7706 | 4,69154E-81 |
| ACE2 | ZDHHC15 | -0,7707 | 4,41994E-81 |
| ACE2 | SHISA4 | -0,771 | 3,43933E-81 |
| ACE2 | HNRNPDL | -0,771 | 3,56502E-81 |
| ACE2 | CDPF1 | -0,771 | 3,36825E-81 |
| ACE2 | LARP6 | -0,7711 | 3,14746E-81 |
| ACE2 | RERG | -0,7713 | 2,76537E-81 |
| ACE2 | ZNF626 | -0,7713 | 2,65238E-81 |
| ACE2 | ARHGEF10 | -0,7714 | 2,58884E-81 |
| ACE2 | CDH2 | -0,7714 | 2,4673E-81 |
| ACE2 | CNTLN | -0,7715 | 2,43652E-81 |
| ACE2 | NR2F2-AS1 | -0,7715 | 2,28397E-81 |
| ACE2 | SUPT3H | -0,7716 | 2,15857E-81 |
| ACE2 | TSPAN9 | -0,7717 | 2,07182E-81 |
| ACE2 | TIMP2 | -0,7717 | 2,02656E-81 |
| ACE2 | BBS5 | -0,7718 | 1,79526E-81 |
| ACE2 | CDK14 | -0,7718 | 1,8039E-81 |
| ACE2 | TULP3 | -0,7718 | 1,89483E-81 |
| ACE2 | ZNF286A | -0,7718 | 1,80453E-81 |
| ACE2 | PML | -0,7719 | 1,69762E-81 |
| ACE2 | PHC1 | -0,772 | 1,63391E-81 |

| | | | |
|---|---|---|---|
| ACE2 | AC108488.4 | -0,7721 | 1,4495E-81 |
| ACE2 | CEP164 | -0,7721 | 1,48624E-81 |
| ACE2 | DLG2 | -0,7722 | 1,38889E-81 |
| ACE2 | CAHM | -0,7724 | 1,17629E-81 |
| ACE2 | XKR4 | -0,7725 | 1,11489E-81 |
| ACE2 | LURAP1 | -0,7726 | 1,02023E-81 |
| ACE2 | DZIP1L | -0,7726 | 9,87151E-82 |
| ACE2 | NOVA1 | -0,7727 | 9,12632E-82 |
| ACE2 | PLP1 | -0,7727 | 8,92458E-82 |
| ACE2 | FRMD6 | -0,7729 | 8,12545E-82 |
| ACE2 | SRGAP2C | -0,7731 | 6,86866E-82 |
| ACE2 | FAM181B | -0,7731 | 6,80061E-82 |
| ACE2 | PRAF2 | -0,7731 | 6,77068E-82 |
| ACE2 | CFAP36 | -0,7732 | 6,20136E-82 |
| ACE2 | OLFML2A | -0,7733 | 6,0432E-82 |
| ACE2 | TCF4 | -0,7733 | 5,93137E-82 |
| ACE2 | ANTXR1 | -0,7734 | 5,5159E-82 |
| ACE2 | FKBP7 | -0,7734 | 5,31671E-82 |
| ACE2 | CDKL3 | -0,7736 | 4,5146E-82 |
| ACE2 | POLR2F | -0,7736 | 4,69415E-82 |
| ACE2 | FBXL2 | -0,7738 | 4,08183E-82 |
| ACE2 | NRSN2 | -0,7738 | 3,84646E-82 |
| ACE2 | CUEDC2 | -0,7739 | 3,52261E-82 |
| ACE2 | APBB1 | -0,774 | 3,46869E-82 |
| ACE2 | PMP22 | -0,774 | 3,47824E-82 |
| ACE2 | KCNN3 | -0,7741 | 3,09057E-82 |
| ACE2 | RGS10 | -0,7741 | 3,12273E-82 |
| ACE2 | NUMBL | -0,7741 | 3,08719E-82 |
| ACE2 | ARL10 | -0,7742 | 2,83814E-82 |
| ACE2 | GPR162 | -0,7742 | 2,86358E-82 |
| ACE2 | HDAC7 | -0,7742 | 2,90528E-82 |
| ACE2 | SSC5D | -0,7742 | 2,91604E-82 |
| ACE2 | HSPA12A | -0,7744 | 2,45329E-82 |
| ACE2 | CTD-2006C1.2 | -0,7744 | 2,54441E-82 |
| ACE2 | CA11 | -0,7744 | 2,55383E-82 |
| ACE2 | ENKD1 | -0,7745 | 2,27192E-82 |
| ACE2 | FKBP10 | -0,7746 | 2,10766E-82 |
| ACE2 | MFAP4 | -0,7748 | 1,83054E-82 |
| ACE2 | EBF4 | -0,7748 | 1,76033E-82 |
| ACE2 | CTNNAL1 | -0,7749 | 1,61758E-82 |
| ACE2 | TDRP | -0,775 | 1,59732E-82 |
| ACE2 | MIR497HG | -0,775 | 1,53078E-82 |
| ACE2 | NT5C3B | -0,7751 | 1,38772E-82 |
| ACE2 | GAS7 | -0,7752 | 1,35991E-82 |
| ACE2 | WIPF1 | -0,7753 | 1,2211E-82 |
| ACE2 | LAYN | -0,7754 | 1,11917E-82 |
| ACE2 | C11orf95 | -0,7755 | 1,04226E-82 |
| ACE2 | GRIK3 | -0,7756 | 9,7174E-83 |
| ACE2 | BEND5 | -0,7756 | 9,30572E-83 |
| ACE2 | TMEM136 | -0,7757 | 9,03534E-83 |
| ACE2 | DUBR | -0,7758 | 8,48553E-83 |
| ACE2 | PBX3 | -0,7759 | 7,64029E-83 |
| ACE2 | DPP7 | -0,7759 | 7,60615E-83 |
| ACE2 | VEGFB | -0,7759 | 7,34761E-83 |

| | | | |
|---|---|---|---|
| ACE2 | FAM228B | -0,776 | 7,2532E-83 |
| ACE2 | TNK2 | -0,776 | 6,77162E-83 |
| ACE2 | SLIT2 | -0,7761 | 6,22974E-83 |
| ACE2 | GLT8D2 | -0,7761 | 6,34311E-83 |
| ACE2 | CTB-193M12.5 | -0,7761 | 6,43237E-83 |
| ACE2 | NACAD | -0,7762 | 5,7627E-83 |
| ACE2 | SGIP1 | -0,7763 | 5,4689E-83 |
| ACE2 | DPT | -0,7763 | 5,40579E-83 |
| ACE2 | TGIF2 | -0,7763 | 5,54667E-83 |
| ACE2 | TMCC2 | -0,7765 | 4,55512E-83 |
| ACE2 | MAATS1 | -0,7765 | 4,72971E-83 |
| ACE2 | PHF19 | -0,7765 | 4,824E-83 |
| ACE2 | CHST10 | -0,7766 | 4,4297E-83 |
| ACE2 | HTRA1 | -0,7766 | 4,31518E-83 |
| ACE2 | KIAA1328 | -0,7768 | 3,58867E-83 |
| ACE2 | ST3GAL3 | -0,7769 | 3,41528E-83 |
| ACE2 | ZNF251 | -0,7769 | 3,41925E-83 |
| ACE2 | RP11-382A20.3 | -0,7769 | 3,45109E-83 |
| ACE2 | RP11-89K11.1 | -0,777 | 3,22166E-83 |
| ACE2 | PRIMA1 | -0,7771 | 2,83443E-83 |
| ACE2 | CSNK1E | -0,7771 | 2,9205E-83 |
| ACE2 | CMTM5 | -0,7772 | 2,76249E-83 |
| ACE2 | SLC12A4 | -0,7772 | 2,75089E-83 |
| ACE2 | TMEM55A | -0,7774 | 2,35871E-83 |
| ACE2 | CCDC28B | -0,7776 | 1,95658E-83 |
| ACE2 | MAGEH1 | -0,7776 | 1,93827E-83 |
| ACE2 | SLC16A1-AS1 | -0,7777 | 1,87205E-83 |
| ACE2 | ZFP2 | -0,778 | 1,42017E-83 |
| ACE2 | SLC39A13 | -0,778 | 1,4077E-83 |
| ACE2 | A4GALT | -0,778 | 1,45931E-83 |
| ACE2 | TSHZ2 | -0,7781 | 1,33986E-83 |
| ACE2 | CADM3 | -0,7782 | 1,18892E-83 |
| ACE2 | LAT | -0,7783 | 1,12078E-83 |
| ACE2 | ADAMTS10 | -0,7784 | 1,02896E-83 |
| ACE2 | TSHZ3 | -0,7784 | 1,06248E-83 |
| ACE2 | GDF11 | -0,7785 | 9,41774E-84 |
| ACE2 | RPL4 | -0,7785 | 9,64253E-84 |
| ACE2 | RP11-486G15.2 | -0,7786 | 9,14812E-84 |
| ACE2 | PALM | -0,7786 | 9,22726E-84 |
| ACE2 | GNAI2 | -0,7787 | 7,98914E-84 |
| ACE2 | TBKBP1 | -0,7787 | 8,46347E-84 |
| ACE2 | ZNF667-AS1 | -0,7788 | 7,31363E-84 |
| ACE2 | CRMP1 | -0,7789 | 6,81655E-84 |
| ACE2 | TTYH2 | -0,7791 | 6,01523E-84 |
| ACE2 | BEX4 | -0,7791 | 6,10947E-84 |
| ACE2 | FAM126A | -0,7792 | 5,30732E-84 |
| ACE2 | NAP1L1 | -0,7792 | 5,6388E-84 |
| ACE2 | NME4 | -0,7792 | 5,29032E-84 |
| ACE2 | CNRIP1 | -0,7793 | 4,95664E-84 |
| ACE2 | GFRA1 | -0,7793 | 4,91145E-84 |
| ACE2 | KIAA1755 | -0,7793 | 4,91284E-84 |
| ACE2 | LIX1L | -0,7794 | 4,85714E-84 |
| ACE2 | STK32A | -0,7796 | 3,95695E-84 |
| ACE2 | ARMCX1 | -0,7797 | 3,81806E-84 |

| | | | |
|---|---|---|---|
| ACE2 | CERS5 | -0,7799 | 3,17544E-84 |
| ACE2 | WHAMMP2 | -0,7799 | 3,12331E-84 |
| ACE2 | PKN1 | -0,7799 | 3,08543E-84 |
| ACE2 | SYT11 | -0,7801 | 2,60252E-84 |
| ACE2 | FXYD1 | -0,7801 | 2,69483E-84 |
| ACE2 | ZNF135 | -0,7801 | 2,55817E-84 |
| ACE2 | ADAM33 | -0,7802 | 2,54551E-84 |
| ACE2 | RP5-1057I20.4 | -0,7805 | 1,91394E-84 |
| ACE2 | C16orf45 | -0,7805 | 1,91971E-84 |
| ACE2 | ARMCX2 | -0,7805 | 1,89177E-84 |
| ACE2 | C15orf59 | -0,7808 | 1,49341E-84 |
| ACE2 | CPEB1 | -0,7808 | 1,50924E-84 |
| ACE2 | OAZ2 | -0,7809 | 1,40655E-84 |
| ACE2 | LINC00403 | -0,781 | 1,31274E-84 |
| ACE2 | CTD-3035K23.7 | -0,781 | 1,28989E-84 |
| ACE2 | FOXD3 | -0,7813 | 1,04346E-84 |
| ACE2 | TRPC1 | -0,7813 | 9,83353E-85 |
| ACE2 | PCDH9 | -0,7813 | 1,04179E-84 |
| ACE2 | FAM65A | -0,7813 | 9,9893E-85 |
| ACE2 | SGSM2 | -0,7815 | 8,70785E-85 |
| ACE2 | TMEM155 | -0,7816 | 7,93105E-85 |
| ACE2 | SEPT5 | -0,7816 | 7,60698E-85 |
| ACE2 | LZTS2 | -0,7817 | 7,39199E-85 |
| ACE2 | COL6A2 | -0,7817 | 7,57453E-85 |
| ACE2 | SPACA6 | -0,7818 | 6,6488E-85 |
| ACE2 | TCF7L1 | -0,7823 | 4,63729E-85 |
| ACE2 | LRP3 | -0,7827 | 3,27367E-85 |
| ACE2 | CADM2 | -0,7829 | 2,82461E-85 |
| ACE2 | COL16A1 | -0,783 | 2,54234E-85 |
| ACE2 | ADGRB3 | -0,7831 | 2,26637E-85 |
| ACE2 | SCARA3 | -0,7834 | 1,81387E-85 |
| ACE2 | C6orf1 | -0,7835 | 1,65393E-85 |
| ACE2 | DKK3 | -0,7835 | 1,61871E-85 |
| ACE2 | SMO | -0,7838 | 1,33936E-85 |
| ACE2 | DTX3 | -0,7841 | 1,02276E-85 |
| ACE2 | SGCE | -0,7842 | 9,19929E-86 |
| ACE2 | BEX3 | -0,7842 | 9,48005E-86 |
| ACE2 | KRBA1 | -0,7843 | 8,94618E-86 |
| ACE2 | USP11 | -0,7843 | 8,64914E-86 |
| ACE2 | FAM110B | -0,7844 | 8,14326E-86 |
| ACE2 | HOXB2 | -0,7844 | 7,73275E-86 |
| ACE2 | FAM229B | -0,7845 | 7,33778E-86 |
| ACE2 | KIF7 | -0,7845 | 7,28905E-86 |
| ACE2 | GRAMD1A | -0,7848 | 5,72174E-86 |
| ACE2 | NLGN3 | -0,7849 | 5,20549E-86 |
| ACE2 | SNCA | -0,785 | 4,93368E-86 |
| ACE2 | JAM3 | -0,7852 | 4,03384E-86 |
| ACE2 | TNFSF12 | -0,7852 | 4,05642E-86 |
| ACE2 | RP11-82L18.2 | -0,7855 | 3,34386E-86 |
| ACE2 | RAB34 | -0,7855 | 3,34497E-86 |
| ACE2 | SLC16A2 | -0,7855 | 3,26906E-86 |
| ACE2 | CEP112 | -0,7856 | 2,92639E-86 |
| ACE2 | RP5-1039K5.19 | -0,7856 | 2,92836E-86 |
| ACE2 | NRXN1 | -0,786 | 2,09993E-86 |

| | | | |
|---|---|---|---|
| ACE2 | COLQ | -0,786 | 2,13427E-86 |
| ACE2 | CADM1 | -0,786 | 2,0879E-86 |
| ACE2 | FAHD2CP | -0,7861 | 1,90173E-86 |
| ACE2 | GLIS2 | -0,7861 | 1,90943E-86 |
| ACE2 | NRN1 | -0,7862 | 1,74586E-86 |
| ACE2 | IFFO1 | -0,7862 | 1,79594E-86 |
| ACE2 | BICD1 | -0,7862 | 1,85907E-86 |
| ACE2 | PCBP4 | -0,7864 | 1,57056E-86 |
| ACE2 | TSPAN11 | -0,7865 | 1,40574E-86 |
| ACE2 | NAT14 | -0,7869 | 1,02311E-86 |
| ACE2 | NFATC4 | -0,7871 | 8,22967E-87 |
| ACE2 | ACAD11 | -0,7872 | 7,80123E-87 |
| ACE2 | C1orf198 | -0,7874 | 6,64219E-87 |
| ACE2 | ZNF496 | -0,7874 | 6,84454E-87 |
| ACE2 | PRRT1 | -0,7874 | 6,77399E-87 |
| ACE2 | ANK2 | -0,7877 | 5,21579E-87 |
| ACE2 | CD81 | -0,7878 | 4,58705E-87 |
| ACE2 | REEP2 | -0,7879 | 4,45193E-87 |
| ACE2 | NAB2 | -0,7879 | 4,46456E-87 |
| ACE2 | MOXD1 | -0,7881 | 3,592E-87 |
| ACE2 | FBXO44 | -0,7882 | 3,28035E-87 |
| ACE2 | TP73-AS1 | -0,7885 | 2,61464E-87 |
| ACE2 | GABBR1 | -0,7885 | 2,68674E-87 |
| ACE2 | PPM1M | -0,7887 | 2,19905E-87 |
| ACE2 | SMARCD3 | -0,7887 | 2,22802E-87 |
| ACE2 | HOMER3 | -0,7891 | 1,56862E-87 |
| ACE2 | ANKRD6 | -0,7894 | 1,19583E-87 |
| ACE2 | HABP4 | -0,7895 | 1,10571E-87 |
| ACE2 | ABI2 | -0,7896 | 9,9159E-88 |
| ACE2 | GPR173 | -0,7897 | 9,73523E-88 |
| ACE2 | RHOJ | -0,7903 | 5,52641E-88 |
| ACE2 | PTN | -0,7906 | 4,56455E-88 |
| ACE2 | NIM1K | -0,7907 | 3,87057E-88 |
| ACE2 | TMOD2 | -0,7908 | 3,74692E-88 |
| ACE2 | ATP1B2 | -0,7911 | 2,85573E-88 |
| ACE2 | C9orf3 | -0,7912 | 2,72472E-88 |
| ACE2 | MIR100HG | -0,7912 | 2,55674E-88 |
| ACE2 | ZSCAN18 | -0,7912 | 2,69946E-88 |
| ACE2 | NTM | -0,7918 | 1,59997E-88 |
| ACE2 | SHC4 | -0,7922 | 1,13284E-88 |
| ACE2 | GALNT16 | -0,7923 | 1,03139E-88 |
| ACE2 | L1CAM | -0,7928 | 6,62285E-89 |
| ACE2 | MCRIP1 | -0,793 | 5,75384E-89 |
| ACE2 | ANKS3 | -0,7933 | 4,47215E-89 |
| ACE2 | KCNMB4 | -0,7934 | 3,80206E-89 |
| ACE2 | CERS4 | -0,7937 | 3,03861E-89 |
| ACE2 | MMP17 | -0,7939 | 2,64512E-89 |
| ACE2 | PLPPR2 | -0,794 | 2,34648E-89 |
| ACE2 | PINLYP | -0,7942 | 1,95518E-89 |
| ACE2 | STMN3 | -0,7943 | 1,76599E-89 |
| ACE2 | PTCH2 | -0,7945 | 1,51698E-89 |
| ACE2 | TCTN1 | -0,7946 | 1,38175E-89 |
| ACE2 | RP9P | -0,7948 | 1,16362E-89 |
| ACE2 | CERS1 | -0,7949 | 1,05732E-89 |

| | | | |
|---|---|---|---|
| ACE2 | PTPRZ1 | -0,7955 | 6,10085E-90 |
| ACE2 | NLGN2 | -0,7956 | 5,63228E-90 |
| ACE2 | PDZD4 | -0,796 | 4,19458E-90 |
| ACE2 | FBXO17 | -0,7961 | 3,60768E-90 |
| ACE2 | TRAF3IP2-AS1 | -0,7967 | 2,29441E-90 |
| ACE2 | SHC2 | -0,797 | 1,71825E-90 |
| ACE2 | CASC15 | -0,7972 | 1,44142E-90 |
| ACE2 | LRRC4C | -0,7972 | 1,40662E-90 |
| ACE2 | DZIP1 | -0,7972 | 1,38955E-90 |
| ACE2 | LCAT | -0,7975 | 1,05963E-90 |
| ACE2 | DLG4 | -0,7975 | 1,10055E-90 |
| ACE2 | TRO | -0,7981 | 6,26211E-91 |
| ACE2 | RASGEF1C | -0,7982 | 5,85524E-91 |
| ACE2 | RASSF8-AS1 | -0,7987 | 3,7783E-91 |
| ACE2 | TCEAL7 | -0,7995 | 1,80445E-91 |
| ACE2 | NCKAP5L | -0,7998 | 1,40387E-91 |
| ACE2 | GPR155 | -0,8002 | 9,44702E-92 |
| ACE2 | CADM4 | -0,8005 | 7,63386E-92 |
| ACE2 | SYNGR1 | -0,8008 | 5,53728E-92 |
| ACE2 | BRSK1 | -0,8024 | 1,33554E-92 |
| ACE2 | LTBP3 | -0,8026 | 1,07192E-92 |
| ACE2 | SLC35F1 | -0,803 | 7,51585E-93 |
| ACE2 | FEZ1 | -0,803 | 7,91048E-93 |
| ACE2 | TUBG2 | -0,8031 | 7,23206E-93 |
| ACE2 | FOXD3-AS1 | -0,8035 | 5,06863E-93 |
| ACE2 | TCEA2 | -0,8035 | 4,90516E-93 |
| ACE2 | P3H3 | -0,8039 | 3,36699E-93 |
| ACE2 | EFS | -0,8047 | 1,55158E-93 |
| ACE2 | ENO2 | -0,8053 | 9,50895E-94 |
| ACE2 | PKNOX2 | -0,8054 | 8,68696E-94 |
| ACE2 | GPSM1 | -0,8057 | 6,17204E-94 |
| ACE2 | SNRPN | -0,8058 | 5,6017E-94 |
| ACE2 | EFEMP2 | -0,806 | 4,61723E-94 |
| ACE2 | SLC22A17 | -0,8088 | 3,50048E-95 |
| ACE2 | POU6F1 | -0,8107 | 5,9486E-96 |
| ACE2 | RARRES2 | -0,8111 | 4,05872E-96 |
| ACE2 | SORCS1 | -0,8137 | 3,17685E-97 |
| ACE2 | FLT3LG | -0,8161 | 3,05415E-98 |
