## Supplementary table 5 kidney for "Bioinformatic characterization of angiotensin-converting enzyme 2, the entry receptor for SARS-CoV-2"

| target_gene | correlated_gene | correlation | pval |
|---|---|---|---|
| ACE2 | ACE2 | 1 | 0 |
| ACE2 | TINAG | 0,9012 | 7,07074E-32 |
| ACE2 | METTL7B | 0,8705 | 2,78327E-27 |
| ACE2 | ANPEP | 0,8683 | 5,29795E-27 |
| ACE2 | LRP2 | 0,8671 | 7,58648E-27 |
| ACE2 | SLC13A1 | 0,8667 | 8,4994E-27 |
| ACE2 | UGT3A1 | 0,8608 | 4,51799E-26 |
| ACE2 | SLC27A2 | 0,8535 | 3,25692E-25 |
| ACE2 | TMEM27 | 0,8526 | 4,09779E-25 |
| ACE2 | CLRN3 | 0,8443 | 3,36078E-24 |
| ACE2 | ACP5 | 0,8367 | 2,05277E-23 |
| ACE2 | AMN | 0,8328 | 4,99318E-23 |
| ACE2 | ENTPD5 | 0,8258 | 2,35525E-22 |
| ACE2 | GGH | 0,8252 | 2,67986E-22 |
| ACE2 | TMEM82 | 0,8239 | 3,5992E-22 |
| ACE2 | SLC4A4 | 0,8235 | 3,88368E-22 |
| ACE2 | CDHR5 | 0,8207 | 7,06083E-22 |
| ACE2 | CUBN | 0,817 | 1,51319E-21 |
| ACE2 | SLC22A6 | 0,8111 | 4,9959E-21 |
| ACE2 | SLC7A9 | 0,8094 | 6,93524E-21 |
| ACE2 | USH1C | 0,8078 | 9,53485E-21 |
| ACE2 | ENPEP | 0,8062 | 1,30939E-20 |
| ACE2 | SLC47A1 | 0,8041 | 1,92972E-20 |
| ACE2 | GJB2 | 0,8018 | 2,9684E-20 |
| ACE2 | AZGP1 | 0,8009 | 3,5573E-20 |
| ACE2 | UGT2A3 | 0,8004 | 3,87934E-20 |
| ACE2 | CLDN2 | 0,7996 | 4,50646E-20 |
| ACE2 | IGSF11 | 0,798 | 6,08737E-20 |
| ACE2 | SLC17A1 | 0,7975 | 6,69226E-20 |
| ACE2 | SLC2A9 | 0,7973 | 6,84961E-20 |
| ACE2 | UGT1A9 | 0,7969 | 7,44895E-20 |
| ACE2 | MYO7B | 0,7939 | 1,2819E-19 |
| ACE2 | SLC23A1 | 0,792 | 1,79956E-19 |
| ACE2 | MTTP | 0,7916 | 1,91673E-19 |
| ACE2 | TCN2 | 0,7895 | 2,76585E-19 |
| ACE2 | SLC22A11 | 0,7893 | 2,87492E-19 |
| ACE2 | RHOBTB1 | 0,7888 | 3,11842E-19 |
| ACE2 | MGAM | 0,7882 | 3,494E-19 |
| ACE2 | AGMO | 0,7829 | 8,69422E-19 |
| ACE2 | ENPP3 | 0,7811 | 1,17011E-18 |
| ACE2 | PDZK1IP1 | 0,7808 | 1,23904E-18 |
| ACE2 | UGT2B7 | 0,7765 | 2,49618E-18 |
| ACE2 | GGT1 | 0,7746 | 3,44525E-18 |
| ACE2 | SLC17A3 | 0,7732 | 4,2989E-18 |
| ACE2 | AIFM1 | 0,7731 | 4,33017E-18 |
| ACE2 | SLC39A5 | 0,7728 | 4,54173E-18 |
| ACE2 | APOE | 0,7727 | 4,64981E-18 |
| ACE2 | ACE | 0,7718 | 5,32758E-18 |
| ACE2 | CDH9 | 0,7717 | 5,4198E-18 |
| ACE2 | ACSM2A | 0,7687 | 8,79806E-18 |
| ACE2 | SLC10A2 | 0,7671 | 1,1266E-17 |
| ACE2 | ANKS4B | 0,7668 | 1,18175E-17 |
| ACE2 | ENPP6 | 0,7659 | 1,35068E-17 |

| | | | |
|---|---|---|---|
| ACE2 | CHST13 | 0,763 | 2,12093E-17 |
| ACE2 | AQP11 | 0,7594 | 3,67423E-17 |
| ACE2 | SLC5A10 | 0,7591 | 3,80795E-17 |
| ACE2 | NAT8 | 0,757 | 5,21746E-17 |
| ACE2 | CTSL3P | 0,7568 | 5,4287E-17 |
| ACE2 | ACSM2B | 0,7556 | 6,43769E-17 |
| ACE2 | SLC13A3 | 0,7543 | 7,822E-17 |
| ACE2 | TM7SF3 | 0,7529 | 9,58218E-17 |
| ACE2 | NGEF | 0,7513 | 1,20566E-16 |
| ACE2 | SLC3A1 | 0,7508 | 1,29554E-16 |
| ACE2 | REEP6 | 0,7492 | 1,63982E-16 |
| ACE2 | PDZK1 | 0,7476 | 2,06217E-16 |
| ACE2 | HNF1A | 0,7462 | 2,50036E-16 |
| ACE2 | B3GNT4 | 0,7454 | 2,79505E-16 |
| ACE2 | CTSA | 0,7454 | 2,78214E-16 |
| ACE2 | CDHR2 | 0,7451 | 2,9125E-16 |
| ACE2 | SLC22A8 | 0,7448 | 3,03472E-16 |
| ACE2 | ABCC6 | 0,7432 | 3,80404E-16 |
| ACE2 | RP11-226B15.1 | 0,7432 | 3,79887E-16 |
| ACE2 | FUT6 | 0,7423 | 4,29427E-16 |
| ACE2 | KCNJ15 | 0,7416 | 4,72673E-16 |
| ACE2 | NOX4 | 0,7398 | 6,10652E-16 |
| ACE2 | LRRC19 | 0,7396 | 6,24694E-16 |
| ACE2 | ABCC6P1 | 0,7392 | 6,6283E-16 |
| ACE2 | SMIM24 | 0,7386 | 7,12884E-16 |
| ACE2 | PLA2G12B | 0,7379 | 7,91778E-16 |
| ACE2 | SLC9A3R1 | 0,7374 | 8,4265E-16 |
| ACE2 | SH3GL2 | 0,7368 | 9,15374E-16 |
| ACE2 | OIT3 | 0,7365 | 9,50737E-16 |
| ACE2 | ACOT7 | 0,7357 | 1,05387E-15 |
| ACE2 | UGT3A2 | 0,7354 | 1,10304E-15 |
| ACE2 | RNF157 | 0,7329 | 1,5483E-15 |
| ACE2 | GDA | 0,7328 | 1,55763E-15 |
| ACE2 | RP11-283I3.4 | 0,7325 | 1,63199E-15 |
| ACE2 | HAVCR2 | 0,7318 | 1,792E-15 |
| ACE2 | ENPP7 | 0,7315 | 1,86092E-15 |
| ACE2 | IL17RB | 0,7314 | 1,87263E-15 |
| ACE2 | SLC5A12 | 0,7302 | 2,20418E-15 |
| ACE2 | TTC38 | 0,7292 | 2,50163E-15 |
| ACE2 | DPEP1 | 0,7288 | 2,65946E-15 |
| ACE2 | ACSF2 | 0,7271 | 3,28705E-15 |
| ACE2 | SLC22A12 | 0,7264 | 3,62708E-15 |
| ACE2 | SLC16A4 | 0,7258 | 3,91286E-15 |
| ACE2 | SLC1A1 | 0,7258 | 3,88701E-15 |
| ACE2 | TST | 0,7258 | 3,92135E-15 |
| ACE2 | ABCC2 | 0,7252 | 4,22272E-15 |
| ACE2 | LGMN | 0,7245 | 4,59783E-15 |
| ACE2 | MYOM3 | 0,7244 | 4,63823E-15 |
| ACE2 | SLC7A7 | 0,7244 | 4,64773E-15 |
| ACE2 | GLIS1 | 0,7242 | 4,75553E-15 |
| ACE2 | TM6SF2 | 0,7241 | 4,83331E-15 |
| ACE2 | ALDH8A1 | 0,7238 | 5,02378E-15 |
| ACE2 | CLEC18B | 0,7235 | 5,20202E-15 |
| ACE2 | PEPD | 0,7232 | 5,45128E-15 |

| | | | |
|---|---|---|---|
| ACE2 | UGT1A1 | 0,7229 | 5,63967E-15 |
| ACE2 | HGD | 0,7226 | 5,83174E-15 |
| ACE2 | MIR194-2HG | 0,7225 | 5,96327E-15 |
| ACE2 | SLC28A1 | 0,722 | 6,31755E-15 |
| ACE2 | DPP4 | 0,7197 | 8,4765E-15 |
| ACE2 | CES2 | 0,7197 | 8,43882E-15 |
| ACE2 | MAP7D2 | 0,7193 | 8,92708E-15 |
| ACE2 | TM4SF5 | 0,7184 | 9,8977E-15 |
| ACE2 | CCL15 | 0,7175 | 1,10548E-14 |
| ACE2 | SLC16A9 | 0,7166 | 1,23622E-14 |
| ACE2 | CTAGE5 | 0,7166 | 1,24317E-14 |
| ACE2 | SLC37A4 | 0,7164 | 1,26796E-14 |
| ACE2 | SLC47A2 | 0,7163 | 1,29524E-14 |
| ACE2 | SLC30A2 | 0,7159 | 1,35756E-14 |
| ACE2 | AC009014.3 | 0,7155 | 1,4262E-14 |
| ACE2 | ERICH5 | 0,7155 | 1,43196E-14 |
| ACE2 | GLTPD2 | 0,7152 | 1,4845E-14 |
| ACE2 | FOLH1 | 0,7146 | 1,58189E-14 |
| ACE2 | SHMT1 | 0,7121 | 2,15764E-14 |
| ACE2 | AKR1A1 | 0,7116 | 2,29575E-14 |
| ACE2 | FCAMR | 0,7106 | 2,59235E-14 |
| ACE2 | GREB1 | 0,7064 | 4,2798E-14 |
| ACE2 | CTSB | 0,7061 | 4,402E-14 |
| ACE2 | CLEC18A | 0,7055 | 4,72306E-14 |
| ACE2 | GJB1 | 0,705 | 5,03761E-14 |
| ACE2 | GLB1L3 | 0,7041 | 5,5987E-14 |
| ACE2 | ADM2 | 0,7039 | 5,70648E-14 |
| ACE2 | LINC01843 | 0,7027 | 6,56979E-14 |
| ACE2 | RP5-856G1.2 | 0,7022 | 7,00597E-14 |
| ACE2 | SUSD2 | 0,7013 | 7,76367E-14 |
| ACE2 | KHDRBS2 | 0,7008 | 8,20297E-14 |
| ACE2 | FAH | 0,7008 | 8,2155E-14 |
| ACE2 | SLC22A4 | 0,6984 | 1,08784E-13 |
| ACE2 | SPAG5 | 0,698 | 1,14058E-13 |
| ACE2 | EHHADH | 0,6975 | 1,19486E-13 |
| ACE2 | MGST1 | 0,6951 | 1,57107E-13 |
| ACE2 | SLC17A4 | 0,6943 | 1,73244E-13 |
| ACE2 | FOLR1 | 0,6934 | 1,90828E-13 |
| ACE2 | SLC2A2 | 0,6932 | 1,95915E-13 |
| ACE2 | FAHD1 | 0,6931 | 1,96841E-13 |
| ACE2 | ENTPD2 | 0,6929 | 2,01717E-13 |
| ACE2 | KIAA1161 | 0,6926 | 2,09209E-13 |
| ACE2 | KHK | 0,6923 | 2,15002E-13 |
| ACE2 | CTSH | 0,6918 | 2,28956E-13 |
| ACE2 | PC | 0,691 | 2,50859E-13 |
| ACE2 | RASSF4 | 0,6899 | 2,80633E-13 |
| ACE2 | PRAP1 | 0,6897 | 2,87542E-13 |
| ACE2 | OSBPL6 | 0,6895 | 2,95521E-13 |
| ACE2 | CHDH | 0,6889 | 3,15832E-13 |
| ACE2 | XYLB | 0,6888 | 3,19713E-13 |
| ACE2 | TFEC | 0,6875 | 3,68386E-13 |
| ACE2 | MLXIPL | 0,6867 | 4,02568E-13 |
| ACE2 | GLB1L2 | 0,6866 | 4,05357E-13 |
| ACE2 | DEPDC7 | 0,6862 | 4,23709E-13 |

| | | | |
|---|---|---|---|
| ACE2 | NMRAL2P | 0,686 | 4,30532E-13 |
| ACE2 | PRLR | 0,686 | 4,31292E-13 |
| ACE2 | GPX3 | 0,6853 | 4,67584E-13 |
| ACE2 | RP5-856G1.1 | 0,6844 | 5,12471E-13 |
| ACE2 | DIO1 | 0,6839 | 5,44003E-13 |
| ACE2 | CES3 | 0,6825 | 6,33711E-13 |
| ACE2 | C11orf54 | 0,6816 | 6,92754E-13 |
| ACE2 | ACMSD | 0,6812 | 7,27139E-13 |
| ACE2 | PTPRD | 0,6812 | 7,2764E-13 |
| ACE2 | AC004485.3 | 0,6809 | 7,51519E-13 |
| ACE2 | EPB41L3 | 0,6808 | 7,5498E-13 |
| ACE2 | TUBAL3 | 0,6805 | 7,8602E-13 |
| ACE2 | VIL1 | 0,6801 | 8,1518E-13 |
| ACE2 | RAB29 | 0,6795 | 8,68202E-13 |
| ACE2 | CFI | 0,6795 | 8,67013E-13 |
| ACE2 | RNU1-70P | 0,6794 | 8,84371E-13 |
| ACE2 | XPNPEP2 | 0,6786 | 9,57654E-13 |
| ACE2 | WDR72 | 0,6784 | 9,80345E-13 |
| ACE2 | SLC6A19 | 0,6781 | 1,00633E-12 |
| ACE2 | UGT1A7 | 0,678 | 1,02376E-12 |
| ACE2 | ALPL | 0,6776 | 1,06796E-12 |
| ACE2 | SUSD3 | 0,6772 | 1,11473E-12 |
| ACE2 | NAPSA | 0,6762 | 1,23351E-12 |
| ACE2 | ECHS1 | 0,6759 | 1,27328E-12 |
| ACE2 | SLC22A18 | 0,6759 | 1,27893E-12 |
| ACE2 | DAB2 | 0,6751 | 1,38545E-12 |
| ACE2 | RENBP | 0,675 | 1,39301E-12 |
| ACE2 | AZGP1P1 | 0,6745 | 1,4658E-12 |
| ACE2 | HNF4A | 0,674 | 1,5564E-12 |
| ACE2 | LINC02027 | 0,6731 | 1,70506E-12 |
| ACE2 | ALDH1B1 | 0,6719 | 1,92789E-12 |
| ACE2 | DPYS | 0,6717 | 1,97372E-12 |
| ACE2 | ILDR2 | 0,6702 | 2,30739E-12 |
| ACE2 | NYX | 0,6693 | 2,52188E-12 |
| ACE2 | SYN2 | 0,6689 | 2,61921E-12 |
| ACE2 | PDZD3 | 0,6681 | 2,85582E-12 |
| ACE2 | TMEM106A | 0,6678 | 2,93044E-12 |
| ACE2 | DNAJC22 | 0,6673 | 3,10063E-12 |
| ACE2 | RP11-311F12.1 | 0,667 | 3,17301E-12 |
| ACE2 | SLC6A13 | 0,6654 | 3,73327E-12 |
| ACE2 | SERPINI1 | 0,6651 | 3,8537E-12 |
| ACE2 | PLG | 0,6649 | 3,94936E-12 |
| ACE2 | GPT2 | 0,6649 | 3,93905E-12 |
| ACE2 | ABCC6P2 | 0,6641 | 4,27803E-12 |
| ACE2 | ESPL1 | 0,6635 | 4,51404E-12 |
| ACE2 | CYP8B1 | 0,6626 | 4,9259E-12 |
| ACE2 | HECW1 | 0,6625 | 4,97401E-12 |
| ACE2 | ALDH4A1 | 0,6621 | 5,18609E-12 |
| ACE2 | KCNAB2 | 0,6609 | 5,83244E-12 |
| ACE2 | PDF | 0,6609 | 5,87205E-12 |
| ACE2 | CDH2 | 0,6609 | 5,8758E-12 |
| ACE2 | BBOX1 | 0,6608 | 5,88715E-12 |
| ACE2 | AHCY | 0,6607 | 5,99611E-12 |
| ACE2 | CDHR3 | 0,6591 | 6,95593E-12 |

| | | | |
|---|---|---|---|
| ACE2 | SHBG | 0,6587 | 7,30209E-12 |
| ACE2 | GIPC2 | 0,6584 | 7,48845E-12 |
| ACE2 | SLC5A2 | 0,6584 | 7,49561E-12 |
| ACE2 | CYP4A11 | 0,6583 | 7,58939E-12 |
| ACE2 | FUT3 | 0,6581 | 7,731E-12 |
| ACE2 | NLRP6 | 0,6573 | 8,35738E-12 |
| ACE2 | LINC01697 | 0,6572 | 8,45344E-12 |
| ACE2 | SLC36A2 | 0,6561 | 9,33167E-12 |
| ACE2 | STEAP1 | 0,656 | 9,46536E-12 |
| ACE2 | BAIAP2L2 | 0,656 | 9,50132E-12 |
| ACE2 | CYP4F3 | 0,6559 | 9,50737E-12 |
| ACE2 | STRIP2 | 0,6554 | 9,98092E-12 |
| ACE2 | PSAT1 | 0,6553 | 1,01396E-11 |
| ACE2 | TMEM176A | 0,6547 | 1,06837E-11 |
| ACE2 | CLEC18C | 0,6544 | 1,10109E-11 |
| ACE2 | ALDH1L1 | 0,6537 | 1,18163E-11 |
| ACE2 | QPRT | 0,6535 | 1,20178E-11 |
| ACE2 | TRIM71 | 0,6532 | 1,23379E-11 |
| ACE2 | F12 | 0,6526 | 1,31007E-11 |
| ACE2 | PDSS1 | 0,6522 | 1,36342E-11 |
| ACE2 | KCNH6 | 0,6521 | 1,37664E-11 |
| ACE2 | AMACR | 0,6518 | 1,4118E-11 |
| ACE2 | CKS2 | 0,6515 | 1,45187E-11 |
| ACE2 | MROH2A | 0,6512 | 1,50514E-11 |
| ACE2 | CRYM | 0,65 | 1,67644E-11 |
| ACE2 | CPN2 | 0,6498 | 1,71784E-11 |
| ACE2 | AGT | 0,6491 | 1,83617E-11 |
| ACE2 | NIT2 | 0,649 | 1,84126E-11 |
| ACE2 | SNX30 | 0,6487 | 1,89766E-11 |
| ACE2 | TSPAN18 | 0,6485 | 1,9425E-11 |
| ACE2 | SLC7A8 | 0,6483 | 1,96954E-11 |
| ACE2 | TMEM132E | 0,6476 | 2,11803E-11 |
| ACE2 | SLC5A9 | 0,6474 | 2,14939E-11 |
| ACE2 | NIPSNAP1 | 0,6469 | 2,2544E-11 |
| ACE2 | DNPH1 | 0,6457 | 2,51671E-11 |
| ACE2 | SLC3A2 | 0,6449 | 2,72433E-11 |
| ACE2 | RP11-556I14.1 | 0,6445 | 2,81093E-11 |
| ACE2 | HHLA2 | 0,6441 | 2,91548E-11 |
| ACE2 | AC026471.6 | 0,644 | 2,95455E-11 |
| ACE2 | LGALS2 | 0,644 | 2,94205E-11 |
| ACE2 | IL22RA1 | 0,6438 | 2,99588E-11 |
| ACE2 | LBX2-AS1 | 0,6438 | 3,00676E-11 |
| ACE2 | APOM | 0,6429 | 3,28547E-11 |
| ACE2 | SPAG5-AS1 | 0,6416 | 3,67382E-11 |
| ACE2 | MAOB | 0,6404 | 4,11339E-11 |
| ACE2 | ACAA1 | 0,6401 | 4,22406E-11 |
| ACE2 | MTNR1A | 0,6395 | 4,45706E-11 |
| ACE2 | TRIM10 | 0,6387 | 4,79565E-11 |
| ACE2 | NXNL2 | 0,6383 | 4,97363E-11 |
| ACE2 | ACOT4 | 0,6379 | 5,17482E-11 |
| ACE2 | UTP4 | 0,6378 | 5,20242E-11 |
| ACE2 | DDC | 0,6375 | 5,3678E-11 |
| ACE2 | SERPINF2 | 0,6367 | 5,77492E-11 |
| ACE2 | AC011738.4 | 0,6365 | 5,8681E-11 |

| | | | |
|---|---|---|---|
| ACE2 | DDO | 0,6364 | 5,91061E-11 |
| ACE2 | EXOC3L4 | 0,6364 | 5,93673E-11 |
| ACE2 | TMIGD1 | 0,636 | 6,14586E-11 |
| ACE2 | VSTM5 | 0,6357 | 6,31429E-11 |
| ACE2 | CGREF1 | 0,6355 | 6,43019E-11 |
| ACE2 | AOX1 | 0,6353 | 6,56147E-11 |
| ACE2 | CYP2C9 | 0,6352 | 6,58061E-11 |
| ACE2 | MUM1L1 | 0,6352 | 6,59218E-11 |
| ACE2 | HPN | 0,6351 | 6,67371E-11 |
| ACE2 | TYMS | 0,635 | 6,70895E-11 |
| ACE2 | NCKAP5 | 0,634 | 7,32959E-11 |
| ACE2 | DHTKD1 | 0,6337 | 7,55515E-11 |
| ACE2 | OCSTAMP | 0,6332 | 7,86349E-11 |
| ACE2 | CXCL14 | 0,6327 | 8,27444E-11 |
| ACE2 | GLDC | 0,6326 | 8,32499E-11 |
| ACE2 | SUCLG1 | 0,632 | 8,80913E-11 |
| ACE2 | TRIM14 | 0,6319 | 8,88551E-11 |
| ACE2 | ACAT1 | 0,6319 | 8,87011E-11 |
| ACE2 | KL | 0,6309 | 9,66434E-11 |
| ACE2 | MSRA | 0,6304 | 1,00795E-10 |
| ACE2 | SYT9 | 0,6304 | 1,01346E-10 |
| ACE2 | BIN1 | 0,6302 | 1,02904E-10 |
| ACE2 | GLB1L | 0,63 | 1,04994E-10 |
| ACE2 | CTC-224D3.1 | 0,629 | 1,14286E-10 |
| ACE2 | MIA2 | 0,6284 | 1,2113E-10 |
| ACE2 | SLC22A2 | 0,6273 | 1,32659E-10 |
| ACE2 | DGCR5 | 0,6273 | 1,32922E-10 |
| ACE2 | KB-1552D7.2 | 0,627 | 1,3625E-10 |
| ACE2 | HNF4G | 0,6268 | 1,38712E-10 |
| ACE2 | RAB11FIP3 | 0,6264 | 1,43114E-10 |
| ACE2 | GLYCTK | 0,6261 | 1,47351E-10 |
| ACE2 | UGT1A6 | 0,6259 | 1,49964E-10 |
| ACE2 | MPST | 0,6252 | 1,59181E-10 |
| ACE2 | TYMSOS | 0,6251 | 1,61217E-10 |
| ACE2 | OCIAD2 | 0,6244 | 1,71629E-10 |
| ACE2 | DPF3 | 0,6236 | 1,83764E-10 |
| ACE2 | TMEM150A | 0,6234 | 1,86159E-10 |
| ACE2 | BTD | 0,6232 | 1,89897E-10 |
| ACE2 | GLYAT | 0,6232 | 1,89847E-10 |
| ACE2 | CTD-3076M17.1 | 0,6231 | 1,915E-10 |
| ACE2 | SLC2A5 | 0,6229 | 1,944E-10 |
| ACE2 | ACY1 | 0,6225 | 2,00555E-10 |
| ACE2 | AGXT2 | 0,6219 | 2,11432E-10 |
| ACE2 | SLC13A2 | 0,6215 | 2,19913E-10 |
| ACE2 | GALM | 0,6214 | 2,20719E-10 |
| ACE2 | MSRB1 | 0,6209 | 2,30324E-10 |
| ACE2 | IYD | 0,6207 | 2,3563E-10 |
| ACE2 | SLC16A10 | 0,6201 | 2,47228E-10 |
| ACE2 | SLC22A24 | 0,62 | 2,4998E-10 |
| ACE2 | FGFR4 | 0,6199 | 2,51858E-10 |
| ACE2 | TRIP13 | 0,6191 | 2,68707E-10 |
| ACE2 | C4B | 0,6187 | 2,77833E-10 |
| ACE2 | RNF186 | 0,6182 | 2,91246E-10 |
| ACE2 | AK4 | 0,6182 | 2,89731E-10 |

| | | | |
|---|---|---|---|
| ACE2 | TTC29 | 0,618 | 2,94855E-10 |
| ACE2 | SUGCT | 0,618 | 2,96097E-10 |
| ACE2 | AP000445.1 | 0,6174 | 3,10274E-10 |
| ACE2 | QDPR | 0,6171 | 3,19349E-10 |
| ACE2 | RP11-94C24.13 | 0,6166 | 3,33302E-10 |
| ACE2 | GPR155 | 0,6163 | 3,41336E-10 |
| ACE2 | TRPM3 | 0,6158 | 3,54371E-10 |
| ACE2 | ANKRD33B | 0,6157 | 3,59229E-10 |
| ACE2 | TMEM206 | 0,6155 | 3,63371E-10 |
| ACE2 | PNMA6A | 0,6155 | 3,6516E-10 |
| ACE2 | RP13-942N8.1 | 0,6152 | 3,73928E-10 |
| ACE2 | GHRHR | 0,6149 | 3,83113E-10 |
| ACE2 | SOAT2 | 0,6147 | 3,90156E-10 |
| ACE2 | RP4-614C10.2 | 0,6145 | 3,96899E-10 |
| ACE2 | AMDHD1 | 0,6141 | 4,10285E-10 |
| ACE2 | PROZ | 0,6134 | 4,32676E-10 |
| ACE2 | SLC26A1 | 0,613 | 4,49145E-10 |
| ACE2 | SMIM10L2A | 0,613 | 4,48781E-10 |
| ACE2 | ESPN | 0,6128 | 4,57232E-10 |
| ACE2 | PCTP | 0,6126 | 4,62711E-10 |
| ACE2 | ACSM5 | 0,6124 | 4,73208E-10 |
| ACE2 | IGF2BP1 | 0,612 | 4,87831E-10 |
| ACE2 | SAMD5 | 0,6113 | 5,15374E-10 |
| ACE2 | CAPN12 | 0,6105 | 5,53234E-10 |
| ACE2 | RP1-154K9.2 | 0,6097 | 5,87918E-10 |
| ACE2 | SFXN2 | 0,6095 | 5,98529E-10 |
| ACE2 | ACSS2 | 0,6093 | 6,08199E-10 |
| ACE2 | TMED6 | 0,6088 | 6,36587E-10 |
| ACE2 | TACO1 | 0,6088 | 6,33386E-10 |
| ACE2 | LINC00939 | 0,6083 | 6,62001E-10 |
| ACE2 | SLC39A4 | 0,6077 | 6,97215E-10 |
| ACE2 | SEMA4G | 0,6077 | 6,95769E-10 |
| ACE2 | TPMT | 0,6076 | 7,03009E-10 |
| ACE2 | CALML4 | 0,6074 | 7,14635E-10 |
| ACE2 | RP11-146F11.4 | 0,6074 | 7,1381E-10 |
| ACE2 | ARSB | 0,6066 | 7,59724E-10 |
| ACE2 | HSD17B3 | 0,6066 | 7,60327E-10 |
| ACE2 | GALNT11 | 0,6065 | 7,63166E-10 |
| ACE2 | SERPINC1 | 0,6064 | 7,71874E-10 |
| ACE2 | TUBA4B | 0,6061 | 7,90447E-10 |
| ACE2 | RNF128 | 0,6059 | 8,01604E-10 |
| ACE2 | FMO1 | 0,6055 | 8,28886E-10 |
| ACE2 | MT1F | 0,6054 | 8,40099E-10 |
| ACE2 | BCHE | 0,6053 | 8,46762E-10 |
| ACE2 | P4HA2 | 0,6053 | 8,4409E-10 |
| ACE2 | ZYG11A | 0,6051 | 8,58205E-10 |
| ACE2 | ETNK2 | 0,6048 | 8,7786E-10 |
| ACE2 | PROC | 0,6037 | 9,5749E-10 |
| ACE2 | RBP5 | 0,6032 | 1,00052E-09 |
| ACE2 | GLRX | 0,6029 | 1,02754E-09 |
| ACE2 | ACO1 | 0,6026 | 1,05196E-09 |
| ACE2 | TMEM176B | 0,6013 | 1,15982E-09 |
| ACE2 | SLC22A18AS | 0,601 | 1,18909E-09 |
| ACE2 | AIG1 | 0,6004 | 1,25239E-09 |

| | | | |
|---|---|---|---|
| ACE2 | MTND4P20 | 0,6002 | 1,26912E-09 |
| ACE2 | XXyac-YM21GA2.7 | 0,6 | 1,28797E-09 |
| ACE2 | EAF2 | 0,5999 | 1,30527E-09 |
| ACE2 | ABHD6 | 0,5996 | 1,33183E-09 |
| ACE2 | ARSE | 0,5995 | 1,34115E-09 |
| ACE2 | GAMT | 0,5991 | 1,38331E-09 |
| ACE2 | CRAT | 0,598 | 1,5102E-09 |
| ACE2 | DNAJC12 | 0,5976 | 1,55778E-09 |
| ACE2 | TREH | 0,5975 | 1,57586E-09 |
| ACE2 | LGI3 | 0,5974 | 1,58193E-09 |
| ACE2 | ALKAL2 | 0,5969 | 1,65324E-09 |
| ACE2 | CYP2J2 | 0,5968 | 1,66763E-09 |
| ACE2 | MTCH2 | 0,5967 | 1,67957E-09 |
| ACE2 | NQO2 | 0,5964 | 1,70944E-09 |
| ACE2 | TRIM6 | 0,5964 | 1,71731E-09 |
| ACE2 | LINC00671 | 0,5961 | 1,75587E-09 |
| ACE2 | LINC01637 | 0,5961 | 1,75453E-09 |
| ACE2 | PIGR | 0,5959 | 1,78254E-09 |
| ACE2 | FTCDNL1 | 0,5956 | 1,82291E-09 |
| ACE2 | AC106876.2 | 0,5955 | 1,84438E-09 |
| ACE2 | BDH2 | 0,5955 | 1,8397E-09 |
| ACE2 | STRADB | 0,5949 | 1,92223E-09 |
| ACE2 | GBA3 | 0,5948 | 1,94654E-09 |
| ACE2 | SLC22A13 | 0,5946 | 1,9719E-09 |
| ACE2 | C4A | 0,5943 | 2,02237E-09 |
| ACE2 | RDH5 | 0,5938 | 2,10468E-09 |
| ACE2 | ACOX2 | 0,5937 | 2,11703E-09 |
| ACE2 | RP11-44F14.2 | 0,5935 | 2,14385E-09 |
| ACE2 | SLC25A10 | 0,5935 | 2,15363E-09 |
| ACE2 | DGCR9 | 0,5934 | 2,17283E-09 |
| ACE2 | LINC00526 | 0,5933 | 2,17827E-09 |
| ACE2 | PFN3 | 0,5923 | 2,36008E-09 |
| ACE2 | TMEM192 | 0,5922 | 2,3799E-09 |
| ACE2 | AC129492.6 | 0,5921 | 2,39432E-09 |
| ACE2 | SLC51A | 0,5918 | 2,45839E-09 |
| ACE2 | CTSV | 0,5912 | 2,56898E-09 |
| ACE2 | SORD | 0,5912 | 2,57536E-09 |
| ACE2 | MTFR1 | 0,5908 | 2,63868E-09 |
| ACE2 | SLC22A7 | 0,5906 | 2,68065E-09 |
| ACE2 | BHMT2 | 0,5901 | 2,79781E-09 |
| ACE2 | PROS1 | 0,5895 | 2,92358E-09 |
| ACE2 | MISP | 0,589 | 3,04926E-09 |
| ACE2 | DNAJC16 | 0,5887 | 3,10834E-09 |
| ACE2 | MCAT | 0,5876 | 3,38456E-09 |
| ACE2 | BNC2 | 0,5873 | 3,4648E-09 |
| ACE2 | LINC00955 | 0,5869 | 3,57857E-09 |
| ACE2 | SLC34A3 | 0,5865 | 3,667E-09 |
| ACE2 | RP11-238K6.1 | 0,5857 | 3,91774E-09 |
| ACE2 | LL22NC03-102D1.18 | 0,5857 | 3,92117E-09 |
| ACE2 | MAPT | 0,5854 | 3,98656E-09 |
| ACE2 | MAF | 0,5846 | 4,25276E-09 |
| ACE2 | LINC00494 | 0,5844 | 4,30087E-09 |
| ACE2 | TRIM15 | 0,5839 | 4,46322E-09 |
| ACE2 | SLC34A1 | 0,5838 | 4,49473E-09 |

| | | | |
|---|---|---|---|
| ACE2 | LACTB2 | 0,5838 | 4,51457E-09 |
| ACE2 | C16orf87 | 0,5838 | 4,52452E-09 |
| ACE2 | CYP4A22 | 0,5837 | 4,52785E-09 |
| ACE2 | TRPC7-AS1 | 0,5832 | 4,70381E-09 |
| ACE2 | TPP1 | 0,5832 | 4,72453E-09 |
| ACE2 | SLC51B | 0,5832 | 4,70037E-09 |
| ACE2 | ABAT | 0,583 | 4,80125E-09 |
| ACE2 | PCYOX1 | 0,5828 | 4,86742E-09 |
| ACE2 | UCN3 | 0,5827 | 4,89018E-09 |
| ACE2 | CTB-178M22.2 | 0,5824 | 4,99828E-09 |
| ACE2 | AARD | 0,582 | 5,15392E-09 |
| ACE2 | ALDH3B1 | 0,5818 | 5,23617E-09 |
| ACE2 | CYP4F2 | 0,5816 | 5,33244E-09 |
| ACE2 | DSEL | 0,5814 | 5,41181E-09 |
| ACE2 | ACO2 | 0,5814 | 5,38987E-09 |
| ACE2 | FUOM | 0,5808 | 5,64788E-09 |
| ACE2 | PTGR1 | 0,5805 | 5,78188E-09 |
| ACE2 | SMPD1 | 0,5799 | 6,03897E-09 |
| ACE2 | SMPDL3A | 0,5795 | 6,23499E-09 |
| ACE2 | DBI | 0,5791 | 6,39281E-09 |
| ACE2 | GOLGA2P5 | 0,5791 | 6,37897E-09 |
| ACE2 | ASL | 0,5788 | 6,52005E-09 |
| ACE2 | LIN52 | 0,5785 | 6,67574E-09 |
| ACE2 | TTLL6 | 0,5783 | 6,81041E-09 |
| ACE2 | HMGCL | 0,5782 | 6,84276E-09 |
| ACE2 | NPL | 0,5782 | 6,82165E-09 |
| ACE2 | ETFA | 0,5777 | 7,0818E-09 |
| ACE2 | TTYH3 | 0,5773 | 7,2831E-09 |
| ACE2 | LINC01833 | 0,5766 | 7,67857E-09 |
| ACE2 | SDHC | 0,5764 | 7,80052E-09 |
| ACE2 | UGT1A10 | 0,5763 | 7,84575E-09 |
| ACE2 | RP11-192H23.4 | 0,5758 | 8,14592E-09 |
| ACE2 | SFXN5 | 0,5755 | 8,35462E-09 |
| ACE2 | DMRTA1 | 0,5753 | 8,48422E-09 |
| ACE2 | HDHD3 | 0,5753 | 8,46435E-09 |
| ACE2 | FTCD | 0,5745 | 8,97532E-09 |
| ACE2 | ACAD11 | 0,5744 | 9,06143E-09 |
| ACE2 | MTHFS | 0,5742 | 9,19771E-09 |
| ACE2 | PCCB | 0,5728 | 1,01329E-08 |
| ACE2 | RP11-76I7.1 | 0,5727 | 1,01955E-08 |
| ACE2 | ABRACL | 0,5726 | 1,02627E-08 |
| ACE2 | LINC00907 | 0,5725 | 1,03355E-08 |
| ACE2 | PNPO | 0,5719 | 1,08394E-08 |
| ACE2 | AKR1C3 | 0,5716 | 1,10577E-08 |
| ACE2 | ASS1 | 0,5708 | 1,17121E-08 |
| ACE2 | FAM220CP | 0,5705 | 1,19747E-08 |
| ACE2 | ALDH2 | 0,5703 | 1,21269E-08 |
| ACE2 | RP11-71E19.1 | 0,57 | 1,2427E-08 |
| ACE2 | RP11-150O12.3 | 0,5695 | 1,29071E-08 |
| ACE2 | ENPP7P8 | 0,5695 | 1,2871E-08 |
| ACE2 | CMBL | 0,5694 | 1,29613E-08 |
| ACE2 | RNF152 | 0,5689 | 1,33888E-08 |
| ACE2 | C2 | 0,5686 | 1,37445E-08 |
| ACE2 | ASPG | 0,5685 | 1,37994E-08 |

| | | | |
|---|---|---|---|
| ACE2 | RP11-185E8.2 | 0,5684 | 1,39639E-08 |
| ACE2 | RP5-899B16.1 | 0,568 | 1,43257E-08 |
| ACE2 | GLUD1 | 0,568 | 1,42887E-08 |
| ACE2 | RP5-849H19.3 | 0,5677 | 1,46482E-08 |
| ACE2 | ATXN7L1 | 0,5677 | 1,45872E-08 |
| ACE2 | IQGAP2 | 0,5669 | 1,55036E-08 |
| ACE2 | LRRC43 | 0,5669 | 1,54376E-08 |
| ACE2 | GPR89B | 0,5662 | 1,62578E-08 |
| ACE2 | LINC00853 | 0,5658 | 1,67272E-08 |
| ACE2 | PNPLA3 | 0,5658 | 1,67402E-08 |
| ACE2 | FABP3 | 0,5652 | 1,74569E-08 |
| ACE2 | PLD1 | 0,5651 | 1,76054E-08 |
| ACE2 | GLOD5 | 0,5651 | 1,75414E-08 |
| ACE2 | AKR7A2 | 0,5645 | 1,83482E-08 |
| ACE2 | TMEM174 | 0,5644 | 1,84332E-08 |
| ACE2 | SYT6 | 0,5636 | 1,95246E-08 |
| ACE2 | KIAA1549 | 0,5635 | 1,97168E-08 |
| ACE2 | C10orf67 | 0,5635 | 1,96396E-08 |
| ACE2 | LDHD | 0,5633 | 1,99343E-08 |
| ACE2 | RP11-261N11.8 | 0,5633 | 1,99325E-08 |
| ACE2 | PAIP2B | 0,5632 | 2,01027E-08 |
| ACE2 | PLA2G16 | 0,5631 | 2,02509E-08 |
| ACE2 | ASLP1 | 0,5628 | 2,06813E-08 |
| ACE2 | SLC35D2 | 0,5622 | 2,16144E-08 |
| ACE2 | CYP4A22-AS1 | 0,5621 | 2,16587E-08 |
| ACE2 | PRSS55 | 0,5621 | 2,16763E-08 |
| ACE2 | AKR1C4 | 0,5618 | 2,21729E-08 |
| ACE2 | AHCYL2 | 0,5617 | 2,23254E-08 |
| ACE2 | AC027612.6 | 0,5615 | 2,26559E-08 |
| ACE2 | RP11-480I12.5 | 0,5614 | 2,27642E-08 |
| ACE2 | XXyac-YM21GA2.3 | 0,5613 | 2,30342E-08 |
| ACE2 | ALDH7A1 | 0,5612 | 2,32024E-08 |
| ACE2 | GCHFR | 0,561 | 2,34723E-08 |
| ACE2 | HAPLN4 | 0,561 | 2,34998E-08 |
| ACE2 | ABLIM3 | 0,5609 | 2,35522E-08 |
| ACE2 | GRHPR | 0,5609 | 2,35843E-08 |
| ACE2 | NKAIN4 | 0,5606 | 2,40819E-08 |
| ACE2 | RP1-276N6.2 | 0,5605 | 2,42492E-08 |
| ACE2 | MDH1 | 0,56 | 2,52069E-08 |
| ACE2 | SCN9A | 0,5588 | 2,74023E-08 |
| ACE2 | ETFB | 0,5588 | 2,7389E-08 |
| ACE2 | DECR2 | 0,5587 | 2,74127E-08 |
| ACE2 | C1QTNF12 | 0,5584 | 2,80867E-08 |
| ACE2 | KCNK5 | 0,5584 | 2,80814E-08 |
| ACE2 | FRMD5 | 0,5583 | 2,83477E-08 |
| ACE2 | CTSLP8 | 0,5579 | 2,91321E-08 |
| ACE2 | KCNK10 | 0,5577 | 2,95619E-08 |
| ACE2 | MT1E | 0,5577 | 2,94825E-08 |
| ACE2 | AC090286.4 | 0,5577 | 2,95105E-08 |
| ACE2 | GMNC | 0,5572 | 3,04494E-08 |
| ACE2 | GCH1 | 0,5571 | 3,07784E-08 |
| ACE2 | SCN8A | 0,557 | 3,09649E-08 |
| ACE2 | CHRNA4 | 0,5563 | 3,24251E-08 |
| ACE2 | CYP4F12 | 0,5562 | 3,27585E-08 |

| | | | |
|---|---|---|---|
| ACE2 | REEP1 | 0,5555 | 3,42367E-08 |
| ACE2 | RP11-598F7.4 | 0,5555 | 3,42367E-08 |
| ACE2 | LDHAL6EP | 0,5553 | 3,46766E-08 |
| ACE2 | GLMP | 0,5551 | 3,52507E-08 |
| ACE2 | CTC-457L16.1 | 0,5551 | 3,52978E-08 |
| ACE2 | GK | 0,5551 | 3,52102E-08 |
| ACE2 | ACY3 | 0,5549 | 3,57545E-08 |
| ACE2 | PBLD | 0,5548 | 3,5939E-08 |
| ACE2 | FTL | 0,5546 | 3,64696E-08 |
| ACE2 | CNTNAP3P2 | 0,5544 | 3,70638E-08 |
| ACE2 | PCBD1 | 0,5542 | 3,75353E-08 |
| ACE2 | APOC2 | 0,5542 | 3,74038E-08 |
| ACE2 | COL19A1 | 0,554 | 3,80055E-08 |
| ACE2 | PRELID1 | 0,5536 | 3,89325E-08 |
| ACE2 | MFSD12 | 0,5532 | 4,0006E-08 |
| ACE2 | AVP | 0,5529 | 4,08993E-08 |
| ACE2 | KYNU | 0,5528 | 4,12365E-08 |
| ACE2 | MYLK4 | 0,5528 | 4,11274E-08 |
| ACE2 | ARSF | 0,5527 | 4,1463E-08 |
| ACE2 | CDH6 | 0,5524 | 4,21747E-08 |
| ACE2 | DCSTAMP | 0,5522 | 4,28634E-08 |
| ACE2 | INHBC | 0,5518 | 4,41383E-08 |
| ACE2 | ZNRF2 | 0,5517 | 4,45006E-08 |
| ACE2 | CYP27A1 | 0,5512 | 4,578E-08 |
| ACE2 | GGT8P | 0,5507 | 4,74127E-08 |
| ACE2 | C5orf49 | 0,5506 | 4,78505E-08 |
| ACE2 | MAIP1 | 0,5505 | 4,79378E-08 |
| ACE2 | GRTP1 | 0,5495 | 5,13476E-08 |
| ACE2 | SH2D6 | 0,5487 | 5,44354E-08 |
| ACE2 | ASGR2 | 0,5485 | 5,48995E-08 |
| ACE2 | TXNDC17 | 0,5483 | 5,59019E-08 |
| ACE2 | SLC16A1 | 0,5481 | 5,64984E-08 |
| ACE2 | HIST1H2AA | 0,5481 | 5,64957E-08 |
| ACE2 | LINC01874 | 0,548 | 5,70163E-08 |
| ACE2 | RFFL | 0,5477 | 5,78394E-08 |
| ACE2 | RP11-100G15.7 | 0,5475 | 5,86356E-08 |
| ACE2 | ERBB3 | 0,5474 | 5,91616E-08 |
| ACE2 | GGT4P | 0,5464 | 6,32248E-08 |
| ACE2 | AC027612.4 | 0,5462 | 6,39276E-08 |
| ACE2 | PDSS2 | 0,5461 | 6,47056E-08 |
| ACE2 | CACNA1E | 0,5459 | 6,55181E-08 |
| ACE2 | HSPB8 | 0,5459 | 6,54334E-08 |
| ACE2 | NEFH | 0,5459 | 6,5391E-08 |
| ACE2 | CTD-2288F12.1 | 0,5458 | 6,59551E-08 |
| ACE2 | RP11-452I5.2 | 0,5449 | 6,98921E-08 |
| ACE2 | BCRP4 | 0,5449 | 7,0074E-08 |
| ACE2 | NT5DC1 | 0,5448 | 7,03443E-08 |
| ACE2 | PAH | 0,5446 | 7,11656E-08 |
| ACE2 | UPK1B | 0,5441 | 7,37792E-08 |
| ACE2 | ACADS | 0,5441 | 7,35422E-08 |
| ACE2 | SLC25A42 | 0,5441 | 7,35164E-08 |
| ACE2 | SLC5A8 | 0,5436 | 7,59453E-08 |
| ACE2 | ECHDC3 | 0,5432 | 7,82721E-08 |
| ACE2 | MARC2 | 0,543 | 7,93335E-08 |

| | | | |
|---|---|---|---|
| ACE2 | TMEM252 | 0,5428 | 8,04602E-08 |
| ACE2 | LRMP | 0,5427 | 8,077E-08 |
| ACE2 | NEK6 | 0,5426 | 8,14834E-08 |
| ACE2 | RNA5SP154 | 0,5423 | 8,29195E-08 |
| ACE2 | MYCNOS | 0,5422 | 8,35225E-08 |
| ACE2 | HAGH | 0,5421 | 8,38247E-08 |
| ACE2 | MT1X | 0,542 | 8,44017E-08 |
| ACE2 | CTD-2540F13.2 | 0,5418 | 8,56114E-08 |
| ACE2 | SSX2IP | 0,5417 | 8,64607E-08 |
| ACE2 | SCRN2 | 0,5417 | 8,62149E-08 |
| ACE2 | PRODH2 | 0,5416 | 8,69184E-08 |
| ACE2 | ATP6V1B2 | 0,5414 | 8,77695E-08 |
| ACE2 | SLC5A11 | 0,5411 | 8,94493E-08 |
| ACE2 | MYO7A | 0,5403 | 9,44384E-08 |
| ACE2 | METTL1 | 0,5392 | 1,01698E-07 |
| ACE2 | LINC01788 | 0,539 | 1,02846E-07 |
| ACE2 | POM121L1P | 0,5389 | 1,03453E-07 |
| ACE2 | RP11-60A24.3 | 0,5388 | 1,03863E-07 |
| ACE2 | PPP1R16B | 0,5386 | 1,05167E-07 |
| ACE2 | TMEM150B | 0,5383 | 1,07605E-07 |
| ACE2 | EVA1A | 0,5379 | 1,10129E-07 |
| ACE2 | RGS14 | 0,5375 | 1,12993E-07 |
| ACE2 | SLC12A6 | 0,5375 | 1,13174E-07 |
| ACE2 | MAJIN | 0,5372 | 1,15131E-07 |
| ACE2 | RP11-192H23.8 | 0,5363 | 1,22657E-07 |
| ACE2 | GLUD2 | 0,5359 | 1,25672E-07 |
| ACE2 | RP11-119J18.1 | 0,535 | 1,32891E-07 |
| ACE2 | SLC31A2 | 0,5348 | 1,34624E-07 |
| ACE2 | CFAP46 | 0,5347 | 1,35574E-07 |
| ACE2 | SLC25A44 | 0,5346 | 1,36525E-07 |
| ACE2 | AMOT | 0,5346 | 1,36678E-07 |
| ACE2 | PTER | 0,5345 | 1,3702E-07 |
| ACE2 | ANGPTL3 | 0,5344 | 1,38371E-07 |
| ACE2 | HCN3 | 0,5343 | 1,38567E-07 |
| ACE2 | RUNDC3B | 0,5343 | 1,38999E-07 |
| ACE2 | TRABD2A | 0,5341 | 1,40392E-07 |
| ACE2 | GSS | 0,5341 | 1,40568E-07 |
| ACE2 | FAM151A | 0,5339 | 1,42418E-07 |
| ACE2 | RP11-707M3.3 | 0,5339 | 1,4304E-07 |
| ACE2 | GCSH | 0,5339 | 1,42306E-07 |
| ACE2 | TMEM105 | 0,5338 | 1,43397E-07 |
| ACE2 | UGT2B10 | 0,5335 | 1,4656E-07 |
| ACE2 | RTKN | 0,5334 | 1,47443E-07 |
| ACE2 | TXN | 0,5334 | 1,47559E-07 |
| ACE2 | ST18 | 0,5333 | 1,47742E-07 |
| ACE2 | ISOC2 | 0,5333 | 1,4811E-07 |
| ACE2 | ZNF804B | 0,5332 | 1,48847E-07 |
| ACE2 | MTHFD1 | 0,5332 | 1,49404E-07 |
| ACE2 | LINC01060 | 0,5331 | 1,50459E-07 |
| ACE2 | LINCR-0001 | 0,5329 | 1,51666E-07 |
| ACE2 | CCDC150 | 0,5328 | 1,5316E-07 |
| ACE2 | HSD17B14 | 0,5327 | 1,53992E-07 |
| ACE2 | AC114812.5 | 0,5325 | 1,56204E-07 |
| ACE2 | LINC00871 | 0,5322 | 1,58806E-07 |

| | | | |
|---|---|---|---|
| ACE2 | MME | 0,532 | 1,60446E-07 |
| ACE2 | ADH6 | 0,532 | 1,60844E-07 |
| ACE2 | DOK4 | 0,532 | 1,60744E-07 |
| ACE2 | UNC79 | 0,5319 | 1,61442E-07 |
| ACE2 | GLS | 0,5317 | 1,64059E-07 |
| ACE2 | L2HGDH | 0,5317 | 1,64438E-07 |
| ACE2 | SLC17A2 | 0,5316 | 1,6456E-07 |
| ACE2 | MEI4 | 0,5316 | 1,64743E-07 |
| ACE2 | AC009403.2 | 0,5315 | 1,66483E-07 |
| ACE2 | FLJ22763 | 0,5313 | 1,67749E-07 |
| ACE2 | KIF21A | 0,5311 | 1,70091E-07 |
| ACE2 | PCCA | 0,5311 | 1,70329E-07 |
| ACE2 | CTD-2541J13.1 | 0,5311 | 1,69937E-07 |
| ACE2 | ACSM5P1 | 0,5309 | 1,72678E-07 |
| ACE2 | TRABD2B | 0,5307 | 1,74368E-07 |
| ACE2 | NPC1L1 | 0,5306 | 1,75907E-07 |
| ACE2 | RP1-95L4.4 | 0,5305 | 1,77074E-07 |
| ACE2 | NPY6R | 0,5303 | 1,78856E-07 |
| ACE2 | SLC22A5 | 0,5302 | 1,80263E-07 |
| ACE2 | SLC36A1 | 0,5301 | 1,81852E-07 |
| ACE2 | CYP27B1 | 0,5293 | 1,90277E-07 |
| ACE2 | CES4A | 0,5293 | 1,91127E-07 |
| ACE2 | SLC44A4 | 0,529 | 1,94437E-07 |
| ACE2 | AKR7A3 | 0,5289 | 1,95871E-07 |
| ACE2 | PDE7A | 0,5286 | 1,99227E-07 |
| ACE2 | FERMT1 | 0,5286 | 1,98893E-07 |
| ACE2 | RTCB | 0,5281 | 2,05195E-07 |
| ACE2 | GNPDA1 | 0,528 | 2,07338E-07 |
| ACE2 | GPD1 | 0,5274 | 2,1507E-07 |
| ACE2 | C3orf30 | 0,5273 | 2,15832E-07 |
| ACE2 | BPHL | 0,5272 | 2,17178E-07 |
| ACE2 | CRYL1 | 0,5271 | 2,1847E-07 |
| ACE2 | OGG1 | 0,5269 | 2,21381E-07 |
| ACE2 | BHMT | 0,5269 | 2,21992E-07 |
| ACE2 | THPO | 0,5268 | 2,23347E-07 |
| ACE2 | ACOX1 | 0,5266 | 2,26457E-07 |
| ACE2 | ACAA2 | 0,5266 | 2,25394E-07 |
| ACE2 | PMM1 | 0,5264 | 2,29264E-07 |
| ACE2 | EBNA1BP2 | 0,5262 | 2,31644E-07 |
| ACE2 | CNTNAP3B | 0,5261 | 2,33196E-07 |
| ACE2 | FAM189A1 | 0,5258 | 2,37299E-07 |
| ACE2 | CADM4 | 0,5258 | 2,37049E-07 |
| ACE2 | MT1H | 0,5255 | 2,41693E-07 |
| ACE2 | TNIK | 0,5252 | 2,46871E-07 |
| ACE2 | ATOH7 | 0,5252 | 2,46086E-07 |
| ACE2 | E2F3 | 0,5249 | 2,50786E-07 |
| ACE2 | NHEJ1 | 0,5248 | 2,53224E-07 |
| ACE2 | FAM20C | 0,5248 | 2,53224E-07 |
| ACE2 | RP5-984P4.6 | 0,5247 | 2,54621E-07 |
| ACE2 | SLC17A5 | 0,5246 | 2,55839E-07 |
| ACE2 | VRK3 | 0,5243 | 2,60359E-07 |
| ACE2 | RP11-63P12.7 | 0,5242 | 2,62658E-07 |
| ACE2 | NR1H4 | 0,5242 | 2,62844E-07 |
| ACE2 | KMO | 0,5239 | 2,66558E-07 |

| | | | |
|---|---|---|---|
| ACE2 | CTB-186H2.3 | 0,5239 | 2,67557E-07 |
| ACE2 | FADS6 | 0,5239 | 2,67497E-07 |
| ACE2 | SUCLG2 | 0,5238 | 2,68008E-07 |
| ACE2 | RP11-734K21.3 | 0,5236 | 2,72041E-07 |
| ACE2 | HADH | 0,5234 | 2,75704E-07 |
| ACE2 | FBP1 | 0,523 | 2,82418E-07 |
| ACE2 | GAS2L3 | 0,5229 | 2,83779E-07 |
| ACE2 | FN3K | 0,5227 | 2,86692E-07 |
| ACE2 | PFKFB2 | 0,5223 | 2,94365E-07 |
| ACE2 | MINPP1 | 0,5222 | 2,95557E-07 |
| ACE2 | NUTM2F | 0,5217 | 3,05639E-07 |
| ACE2 | TGFBR3L | 0,5217 | 3,05507E-07 |
| ACE2 | L3MBTL4 | 0,5216 | 3,08254E-07 |
| ACE2 | AP000347.2 | 0,5215 | 3,09001E-07 |
| ACE2 | RTCA | 0,5214 | 3,12013E-07 |
| ACE2 | HRASLS2 | 0,5213 | 3,13093E-07 |
| ACE2 | LAMTOR5 | 0,5212 | 3,15626E-07 |
| ACE2 | RBM47 | 0,5211 | 3,16291E-07 |
| ACE2 | CTD-3193O13.1 | 0,5207 | 3,25059E-07 |
| ACE2 | FAM135B | 0,5204 | 3,30969E-07 |
| ACE2 | RP11-644F5.15 | 0,5204 | 3,31578E-07 |
| ACE2 | PCDHAC1 | 0,5203 | 3,33702E-07 |
| ACE2 | C19orf12 | 0,5199 | 3,41275E-07 |
| ACE2 | SLC39A11 | 0,5198 | 3,43827E-07 |
| ACE2 | SDHB | 0,5196 | 3,46435E-07 |
| ACE2 | SLC16A13 | 0,5195 | 3,50225E-07 |
| ACE2 | LINC01781 | 0,5194 | 3,51476E-07 |
| ACE2 | COCH | 0,5191 | 3,58271E-07 |
| ACE2 | SFXN1 | 0,5189 | 3,61741E-07 |
| ACE2 | STON2 | 0,5189 | 3,62283E-07 |
| ACE2 | VNN1 | 0,5182 | 3,78748E-07 |
| ACE2 | PCK1 | 0,5179 | 3,85075E-07 |
| ACE2 | TIGD2 | 0,5178 | 3,86444E-07 |
| ACE2 | RHBDL1 | 0,5178 | 3,86672E-07 |
| ACE2 | ATP5G3 | 0,5176 | 3,91965E-07 |
| ACE2 | NUDT8 | 0,5173 | 3,9844E-07 |
| ACE2 | MROH3P | 0,5172 | 4,01509E-07 |
| ACE2 | KCNG2 | 0,5171 | 4,05379E-07 |
| ACE2 | TRPC7 | 0,517 | 4,05857E-07 |
| ACE2 | CYP2B6 | 0,5169 | 4,09642E-07 |
| ACE2 | GPHN | 0,5168 | 4,12855E-07 |
| ACE2 | TUBB4A | 0,5165 | 4,19099E-07 |
| ACE2 | AQP7 | 0,5162 | 4,26188E-07 |
| ACE2 | LBX2 | 0,5159 | 4,33521E-07 |
| ACE2 | PTH2R | 0,5156 | 4,41488E-07 |
| ACE2 | RP13-726E6.2 | 0,5156 | 4,41162E-07 |
| ACE2 | RP4-555D20.4 | 0,5152 | 4,54364E-07 |
| ACE2 | RP11-521M14.1 | 0,5149 | 4,61811E-07 |
| ACE2 | TDP2 | 0,5146 | 4,71096E-07 |
| ACE2 | HMOX1 | 0,5145 | 4,73927E-07 |
| ACE2 | RNU1-14P | 0,5144 | 4,7683E-07 |
| ACE2 | TSPAN1 | 0,5143 | 4,79506E-07 |
| ACE2 | ANK3 | 0,5143 | 4,78107E-07 |
| ACE2 | GCDH | 0,5143 | 4,77476E-07 |

| | | | |
|---|---|---|---|
| ACE2 | IDH1 | 0,5142 | 4,79995E-07 |
| ACE2 | CNDP2 | 0,5142 | 4,81118E-07 |
| ACE2 | RETSAT | 0,5138 | 4,93717E-07 |
| ACE2 | PAOX | 0,5133 | 5,07142E-07 |
| ACE2 | PRAME | 0,5131 | 5,14583E-07 |
| ACE2 | RGL1 | 0,512 | 5,48566E-07 |
| ACE2 | C10orf11 | 0,512 | 5,49358E-07 |
| ACE2 | AS3MT | 0,512 | 5,48247E-07 |
| ACE2 | RP11-21L23.4 | 0,5119 | 5,52957E-07 |
| ACE2 | GLYATL1 | 0,5116 | 5,62418E-07 |
| ACE2 | RP11-104J23.1 | 0,5116 | 5,62577E-07 |
| ACE2 | PPARA | 0,5116 | 5,61033E-07 |
| ACE2 | PLEKHB2 | 0,5115 | 5,66343E-07 |
| ACE2 | MMP24 | 0,5115 | 5,66587E-07 |
| ACE2 | MTMR11 | 0,5111 | 5,78273E-07 |
| ACE2 | LINC01014 | 0,5109 | 5,8424E-07 |
| ACE2 | NAT8B | 0,5108 | 5,88396E-07 |
| ACE2 | CYP21A2 | 0,5108 | 5,90012E-07 |
| ACE2 | OOSP1 | 0,5107 | 5,93837E-07 |
| ACE2 | ADORA2A-AS1 | 0,5107 | 5,92149E-07 |
| ACE2 | RP11-622A1.2 | 0,5104 | 6,04616E-07 |
| ACE2 | SOWAHB | 0,5102 | 6,0985E-07 |
| ACE2 | FUT5 | 0,5102 | 6,115E-07 |
| ACE2 | DOK6 | 0,51 | 6,1577E-07 |
| ACE2 | STX3 | 0,5095 | 6,35873E-07 |
| ACE2 | CPT2 | 0,5094 | 6,39176E-07 |
| ACE2 | RP1-239B22.5 | 0,509 | 6,56691E-07 |
| ACE2 | LAMP1 | 0,509 | 6,55752E-07 |
| ACE2 | GRIA3 | 0,5089 | 6,58774E-07 |
| ACE2 | CLPTM1L | 0,5087 | 6,66004E-07 |
| ACE2 | AGPAT3 | 0,5087 | 6,65907E-07 |
| ACE2 | BANF1P2 | 0,5086 | 6,69742E-07 |
| ACE2 | CNTN3 | 0,5084 | 6,76796E-07 |
| ACE2 | DMGDH | 0,5082 | 6,85291E-07 |
| ACE2 | ACSM3 | 0,5079 | 6,99679E-07 |
| ACE2 | CYB5A | 0,5078 | 7,02388E-07 |
| ACE2 | CARD10 | 0,5076 | 7,09361E-07 |
| ACE2 | PGPEP1 | 0,5072 | 7,26194E-07 |
| ACE2 | PKLR | 0,5069 | 7,40861E-07 |
| ACE2 | ERI3 | 0,5066 | 7,53977E-07 |
| ACE2 | ALCAM | 0,5065 | 7,60455E-07 |
| ACE2 | BX842568.2 | 0,5065 | 7,57426E-07 |
| ACE2 | B4GALT4 | 0,506 | 7,79431E-07 |
| ACE2 | AP000355.2 | 0,5058 | 7,90488E-07 |
| ACE2 | G6PC | 0,5057 | 7,92403E-07 |
| ACE2 | HIST1H2BA | 0,5056 | 7,97828E-07 |
| ACE2 | TMEM52 | 0,5055 | 8,06038E-07 |
| ACE2 | COX6A2 | 0,5052 | 8,18498E-07 |
| ACE2 | CYP17A1 | 0,505 | 8,28196E-07 |
| ACE2 | HYAL1 | 0,5047 | 8,411E-07 |
| ACE2 | HAAO | 0,5046 | 8,48765E-07 |
| ACE2 | HIST1H2APS1 | 0,5039 | 8,80004E-07 |
| ACE2 | LINC00854 | 0,5037 | 8,94132E-07 |
| ACE2 | IL32 | 0,5034 | 9,07746E-07 |

| | | | |
|---|---|---|---|
| ACE2 | SOBP | 0,5033 | 9,13016E-07 |
| ACE2 | PRSS51 | 0,5033 | 9,11856E-07 |
