## Supplementary table 5 testis for "Bioinformatic characterization of angiotensin-converting enzyme 2, the entry receptor for SARS-CoV-2"

| target_gene | correlated_gene | correlation | pval |
|---|---|---|---|
| ACE2 | ACE2 | 1 | 0 |
| ACE2 | PLP1 | 0,9092 | 1,0691E-138 |
| ACE2 | NDFIP1 | 0,8911 | 2,9095E-125 |
| ACE2 | SPARC | 0,8908 | 4,6743E-125 |
| ACE2 | SCP2 | 0,8886 | 1,3404E-123 |
| ACE2 | SMARCA1 | 0,8852 | 2,4395E-121 |
| ACE2 | ATP2B1 | 0,8829 | 6,4496E-120 |
| ACE2 | VSIG1 | 0,8815 | 4,6003E-119 |
| ACE2 | HTATSF1 | 0,8809 | 1,0949E-118 |
| ACE2 | ACO1 | 0,8808 | 1,2556E-118 |
| ACE2 | ENPP5 | 0,8802 | 3,0148E-118 |
| ACE2 | HPRT1 | 0,8802 | 3,0286E-118 |
| ACE2 | ALDH1A1 | 0,8799 | 4,3784E-118 |
| ACE2 | CPPED1 | 0,877 | 2,3897E-116 |
| ACE2 | SGCE | 0,8765 | 5,2518E-116 |
| ACE2 | GALNT1 | 0,876 | 9,5351E-116 |
| ACE2 | LAPTM4B | 0,8757 | 1,4604E-115 |
| ACE2 | FBXO21 | 0,8745 | 7,6319E-115 |
| ACE2 | TMEM87A | 0,874 | 1,4889E-114 |
| ACE2 | CLDN11 | 0,8727 | 8,1779E-114 |
| ACE2 | SEPT6 | 0,8722 | 1,6028E-113 |
| ACE2 | CARMIL1 | 0,8709 | 7,9579E-113 |
| ACE2 | MBNL3 | 0,8695 | 5,1442E-112 |
| ACE2 | MPZL1 | 0,8686 | 1,6326E-111 |
| ACE2 | HMGN4 | 0,8681 | 3,1968E-111 |
| ACE2 | MYO6 | 0,8675 | 6,2424E-111 |
| ACE2 | MANSC1 | 0,8675 | 6,4572E-111 |
| ACE2 | RBBP7 | 0,8671 | 1,0542E-110 |
| ACE2 | ATP1A1 | 0,8665 | 2,1683E-110 |
| ACE2 | KIAA1210 | 0,866 | 4,5401E-110 |
| ACE2 | PTN | 0,8656 | 6,6273E-110 |
| ACE2 | KLHL4 | 0,8653 | 1,0611E-109 |
| ACE2 | ACSL4 | 0,8632 | 1,3738E-108 |
| ACE2 | ITFG1 | 0,8623 | 3,9931E-108 |
| ACE2 | VCL | 0,8619 | 6,1761E-108 |
| ACE2 | EPHA4 | 0,8606 | 3,2233E-107 |
| ACE2 | SCRN1 | 0,8602 | 4,7881E-107 |
| ACE2 | AP3B1 | 0,8598 | 8,3966E-107 |
| ACE2 | MESDC2 | 0,8594 | 1,3456E-106 |
| ACE2 | LMBR1 | 0,859 | 1,9471E-106 |
| ACE2 | RNF13 | 0,8586 | 3,2621E-106 |
| ACE2 | NPTN | 0,8582 | 4,9654E-106 |
| ACE2 | ATP5C1 | 0,8577 | 9,552E-106 |
| ACE2 | TPM3 | 0,8574 | 1,3565E-105 |
| ACE2 | FEZ2 | 0,8574 | 1,394E-105 |
| ACE2 | ADH5 | 0,8572 | 1,6956E-105 |
| ACE2 | SERINC1 | 0,8566 | 3,3034E-105 |
| ACE2 | PRKAR2B | 0,8563 | 4,5015E-105 |
| ACE2 | HSPA9 | 0,8558 | 8,6927E-105 |
| ACE2 | C14orf37 | 0,8556 | 1,111E-104 |
| ACE2 | SERPINA5 | 0,8547 | 3,0176E-104 |
| ACE2 | VDAC1 | 0,8542 | 5,5483E-104 |
| ACE2 | LAMP2 | 0,8537 | 9,5532E-104 |

| | | | |
|---|---|---|---|
| ACE2 | ATP1B1 | 0,8536 | 1,0869E-103 |
| ACE2 | RAB5B | 0,8536 | 1,0777E-103 |
| ACE2 | AHCYL1 | 0,8535 | 1,1554E-103 |
| ACE2 | ME1 | 0,8534 | 1,3181E-103 |
| ACE2 | JAM2 | 0,8529 | 2,4129E-103 |
| ACE2 | GOT2 | 0,8524 | 3,8717E-103 |
| ACE2 | SLC30A9 | 0,8522 | 5,2833E-103 |
| ACE2 | PRDX3 | 0,8519 | 7,1489E-103 |
| ACE2 | PRNP | 0,8518 | 7,4264E-103 |
| ACE2 | APPL2 | 0,8516 | 9,5933E-103 |
| ACE2 | PREP | 0,8513 | 1,3272E-102 |
| ACE2 | RAB31 | 0,8494 | 1,1207E-101 |
| ACE2 | C1GALT1C1 | 0,8493 | 1,1933E-101 |
| ACE2 | NMT2 | 0,8491 | 1,5095E-101 |
| ACE2 | RDH11 | 0,849 | 1,7688E-101 |
| ACE2 | MBTPS2 | 0,849 | 1,6854E-101 |
| ACE2 | DKC1 | 0,8486 | 2,671E-101 |
| ACE2 | SH3BGRL | 0,8482 | 4,0136E-101 |
| ACE2 | SERINC5 | 0,848 | 5,4013E-101 |
| ACE2 | CAPZA2 | 0,8464 | 2,8279E-100 |
| ACE2 | KLHL15 | 0,8463 | 3,3114E-100 |
| ACE2 | APP | 0,846 | 4,1873E-100 |
| ACE2 | SLC30A5 | 0,8459 | 4,7143E-100 |
| ACE2 | MAPRE1 | 0,8459 | 4,8142E-100 |
| ACE2 | TMEM123 | 0,8457 | 6,0513E-100 |
| ACE2 | FAM199X | 0,8457 | 6,1949E-100 |
| ACE2 | NT5DC1 | 0,8453 | 9,2246E-100 |
| ACE2 | CALM1 | 0,8452 | 1,045E-99 |
| ACE2 | FAM49B | 0,8451 | 1,1645E-99 |
| ACE2 | ABCD3 | 0,845 | 1,221E-99 |
| ACE2 | HACD2 | 0,8448 | 1,5451E-99 |
| ACE2 | MDH1 | 0,8444 | 2,5343E-99 |
| ACE2 | MRPS35 | 0,844 | 3,8167E-99 |
| ACE2 | SNX6 | 0,8439 | 3,9551E-99 |
| ACE2 | AIFM1 | 0,8438 | 4,336E-99 |
| ACE2 | GPR37 | 0,8437 | 5,0687E-99 |
| ACE2 | GSR | 0,8436 | 5,715E-99 |
| ACE2 | CHODL | 0,8435 | 6,3922E-99 |
| ACE2 | TMED10 | 0,8431 | 9,114E-99 |
| ACE2 | CLINT1 | 0,843 | 1,09729E-98 |
| ACE2 | SC5D | 0,8427 | 1,44895E-98 |
| ACE2 | TRPC5OS | 0,8425 | 1,73535E-98 |
| ACE2 | TCEAL8 | 0,8424 | 2,04717E-98 |
| ACE2 | CDC42 | 0,8422 | 2,39616E-98 |
| ACE2 | ITGAV | 0,8416 | 4,41735E-98 |
| ACE2 | PRDX6 | 0,8415 | 5,10624E-98 |
| ACE2 | TMEM135 | 0,8415 | 4,91952E-98 |
| ACE2 | SPG21 | 0,8414 | 5,78329E-98 |
| ACE2 | HELZ | 0,8404 | 1,60045E-97 |
| ACE2 | GDI2 | 0,8395 | 3,81833E-97 |
| ACE2 | SGCB | 0,8393 | 4,82204E-97 |
| ACE2 | STAM | 0,8391 | 5,86936E-97 |
| ACE2 | UBR3 | 0,8387 | 8,98509E-97 |
| ACE2 | RDX | 0,8382 | 1,49499E-96 |

| | | | |
|---|---|---|---|
| ACE2 | C9orf64 | 0,838 | 1,75681E-96 |
| ACE2 | IPO7 | 0,8377 | 2,57544E-96 |
| ACE2 | DCBLD2 | 0,8372 | 3,98694E-96 |
| ACE2 | HLTF | 0,837 | 5,20978E-96 |
| ACE2 | EVI5 | 0,8369 | 5,34274E-96 |
| ACE2 | TJP1 | 0,8366 | 7,15716E-96 |
| ACE2 | SEMA3C | 0,8359 | 1,43846E-95 |
| ACE2 | PPA1 | 0,8358 | 1,71819E-95 |
| ACE2 | CMAS | 0,8358 | 1,61067E-95 |
| ACE2 | RAB7A | 0,835 | 3,64604E-95 |
| ACE2 | GATM | 0,8345 | 5,99895E-95 |
| ACE2 | TM9SF3 | 0,8341 | 8,67196E-95 |
| ACE2 | CTNND1 | 0,834 | 1,00498E-94 |
| ACE2 | NUDT10 | 0,8337 | 1,34434E-94 |
| ACE2 | ALDH9A1 | 0,8333 | 1,87208E-94 |
| ACE2 | CMTM6 | 0,8327 | 3,4773E-94 |
| ACE2 | HSD17B12 | 0,8327 | 3,47236E-94 |
| ACE2 | CERS6 | 0,8326 | 3,87384E-94 |
| ACE2 | GNAI1 | 0,8324 | 4,83865E-94 |
| ACE2 | CA5B | 0,8323 | 4,96175E-94 |
| ACE2 | MAPK1IP1L | 0,8319 | 7,32098E-94 |
| ACE2 | ARF4 | 0,8313 | 1,31128E-93 |
| ACE2 | SCARB2 | 0,8311 | 1,65538E-93 |
| ACE2 | SEC22B | 0,831 | 1,86E-93 |
| ACE2 | CPE | 0,831 | 1,74793E-93 |
| ACE2 | FAM213A | 0,8303 | 3,61385E-93 |
| ACE2 | RFT1 | 0,83 | 4,76452E-93 |
| ACE2 | MAGED1 | 0,8299 | 5,40508E-93 |
| ACE2 | SLC20A2 | 0,8296 | 7,21835E-93 |
| ACE2 | NCOA2 | 0,8296 | 7,13559E-93 |
| ACE2 | CCDC43 | 0,8292 | 9,9544E-93 |
| ACE2 | AP1S2 | 0,8292 | 1,0443E-92 |
| ACE2 | ATP6AP1 | 0,8292 | 1,00817E-92 |
| ACE2 | MCFD2 | 0,8289 | 1,41626E-92 |
| ACE2 | LMAN1 | 0,8282 | 2,66137E-92 |
| ACE2 | PLXDC2 | 0,8281 | 2,96584E-92 |
| ACE2 | TMEM192 | 0,8276 | 4,84472E-92 |
| ACE2 | TMED2 | 0,8271 | 7,37272E-92 |
| ACE2 | TTC8 | 0,827 | 8,05113E-92 |
| ACE2 | GNPDA1 | 0,8263 | 1,53214E-91 |
| ACE2 | DCAF12L2 | 0,8262 | 1,82752E-91 |
| ACE2 | BEX4 | 0,826 | 2,14534E-91 |
| ACE2 | ABI1 | 0,8259 | 2,37658E-91 |
| ACE2 | GJA1 | 0,8258 | 2,45114E-91 |
| ACE2 | EIF2AK4 | 0,8252 | 4,37646E-91 |
| ACE2 | SMAD4 | 0,8252 | 4,42903E-91 |
| ACE2 | ARL6IP5 | 0,825 | 5,59007E-91 |
| ACE2 | SLC15A2 | 0,825 | 5,19547E-91 |
| ACE2 | MOB3B | 0,8249 | 5,93036E-91 |
| ACE2 | LGR4 | 0,8249 | 5,85247E-91 |
| ACE2 | WSB2 | 0,8247 | 7,28991E-91 |
| ACE2 | HEG1 | 0,8245 | 8,55578E-91 |
| ACE2 | SLC25A5 | 0,8244 | 8,99004E-91 |
| ACE2 | PPT1 | 0,8243 | 1,03598E-90 |

| | | | |
|---|---|---|---|
| ACE2 | JAM3 | 0,8243 | 1,06408E-90 |
| ACE2 | OCRL | 0,8242 | 1,13942E-90 |
| ACE2 | PSAP | 0,8241 | 1,21819E-90 |
| ACE2 | CASC4 | 0,8238 | 1,60063E-90 |
| ACE2 | NUDT11 | 0,8237 | 1,73517E-90 |
| ACE2 | OSBPL11 | 0,8236 | 1,99339E-90 |
| ACE2 | ANO6 | 0,8235 | 2,15645E-90 |
| ACE2 | NDUFS1 | 0,8234 | 2,39411E-90 |
| ACE2 | FAM122B | 0,823 | 3,27051E-90 |
| ACE2 | ELOVL5 | 0,8229 | 3,59541E-90 |
| ACE2 | GLO1 | 0,8227 | 4,39934E-90 |
| ACE2 | CAMK2D | 0,8226 | 4,69359E-90 |
| ACE2 | ITGB8 | 0,8225 | 5,30719E-90 |
| ACE2 | LDHB | 0,8225 | 5,35096E-90 |
| ACE2 | TAF7 | 0,8222 | 7,02049E-90 |
| ACE2 | SLC25A4 | 0,8221 | 7,5465E-90 |
| ACE2 | CRK | 0,822 | 8,67129E-90 |
| ACE2 | AP3M1 | 0,8218 | 1,02604E-89 |
| ACE2 | SEC23IP | 0,8217 | 1,09213E-89 |
| ACE2 | LRRN3 | 0,821 | 2,03921E-89 |
| ACE2 | ERBB4 | 0,8208 | 2,47314E-89 |
| ACE2 | GBAS | 0,8208 | 2,40972E-89 |
| ACE2 | ATP6AP2 | 0,8207 | 2,66294E-89 |
| ACE2 | TBC1D8B | 0,8204 | 3,58989E-89 |
| ACE2 | GNAI3 | 0,8202 | 4,33299E-89 |
| ACE2 | CREG1 | 0,82 | 5,27865E-89 |
| ACE2 | EID1 | 0,8198 | 6,05162E-89 |
| ACE2 | TAOK3 | 0,8197 | 6,65152E-89 |
| ACE2 | B4GALT5 | 0,8197 | 6,82012E-89 |
| ACE2 | TUSC3 | 0,8193 | 9,6972E-89 |
| ACE2 | TAB3 | 0,8188 | 1,5302E-88 |
| ACE2 | SLC3A2 | 0,8187 | 1,64554E-88 |
| ACE2 | OTUD6B | 0,8186 | 1,73694E-88 |
| ACE2 | EXOC6 | 0,8184 | 2,14135E-88 |
| ACE2 | CFL2 | 0,8184 | 2,0879E-88 |
| ACE2 | MUT | 0,8183 | 2,3229E-88 |
| ACE2 | MAGT1 | 0,8182 | 2,52706E-88 |
| ACE2 | NBDY | 0,8179 | 3,28217E-88 |
| ACE2 | DAG1 | 0,8176 | 4,26117E-88 |
| ACE2 | SMIM10 | 0,8173 | 5,95926E-88 |
| ACE2 | B4GALT1 | 0,8172 | 6,07572E-88 |
| ACE2 | FUBP3 | 0,8172 | 6,03746E-88 |
| ACE2 | NXT2 | 0,8171 | 7,09114E-88 |
| ACE2 | MOSPD1 | 0,817 | 7,59181E-88 |
| ACE2 | RAB18 | 0,8169 | 8,30645E-88 |
| ACE2 | PGD | 0,8167 | 9,69938E-88 |
| ACE2 | RAB10 | 0,8165 | 1,21125E-87 |
| ACE2 | TCEAL9 | 0,8165 | 1,17652E-87 |
| ACE2 | AIG1 | 0,8164 | 1,27709E-87 |
| ACE2 | PTPRG | 0,8162 | 1,57543E-87 |
| ACE2 | TMEM106B | 0,8162 | 1,54084E-87 |
| ACE2 | ZYG11B | 0,8161 | 1,63565E-87 |
| ACE2 | NAP1L5 | 0,8157 | 2,26811E-87 |
| ACE2 | HPS3 | 0,8156 | 2,50664E-87 |

| | | | |
|---|---|---|---|
| ACE2 | TPMT | 0,8156 | 2,4744E-87 |
| ACE2 | GM2A | 0,8154 | 3,01854E-87 |
| ACE2 | FXN | 0,8152 | 3,65339E-87 |
| ACE2 | TFB2M | 0,8151 | 4,15044E-87 |
| ACE2 | CHM | 0,815 | 4,38453E-87 |
| ACE2 | MAP3K1 | 0,8147 | 5,7619E-87 |
| ACE2 | MYLK3 | 0,8145 | 6,57157E-87 |
| ACE2 | ANOS1 | 0,8142 | 8,57516E-87 |
| ACE2 | ARL8A | 0,814 | 1,01091E-86 |
| ACE2 | MTPN | 0,8138 | 1,22137E-86 |
| ACE2 | BTBD3 | 0,8136 | 1,46318E-86 |
| ACE2 | PEG10 | 0,8133 | 1,88374E-86 |
| ACE2 | SOX9 | 0,813 | 2,45436E-86 |
| ACE2 | LACC1 | 0,8129 | 2,58686E-86 |
| ACE2 | PPP3CA | 0,8128 | 3,00909E-86 |
| ACE2 | PPAT | 0,8126 | 3,47961E-86 |
| ACE2 | GRSF1 | 0,8121 | 5,3401E-86 |
| ACE2 | MCCC2 | 0,8114 | 9,44516E-86 |
| ACE2 | EZR | 0,8114 | 9,83318E-86 |
| ACE2 | MTMR4 | 0,8114 | 9,80468E-86 |
| ACE2 | ATP5A1 | 0,8113 | 1,05011E-85 |
| ACE2 | PHACTR2 | 0,8111 | 1,22233E-85 |
| ACE2 | TGOLN2 | 0,8109 | 1,53562E-85 |
| ACE2 | PPP2CA | 0,8109 | 1,4654E-85 |
| ACE2 | PRND | 0,8109 | 1,51366E-85 |
| ACE2 | ENDOD1 | 0,8108 | 1,60147E-85 |
| ACE2 | PLSCR4 | 0,8106 | 1,8982E-85 |
| ACE2 | NCKAP1 | 0,8105 | 2,00885E-85 |
| ACE2 | SETD7 | 0,8105 | 2,11048E-85 |
| ACE2 | USP13 | 0,8104 | 2,28219E-85 |
| ACE2 | BRWD3 | 0,8104 | 2,35697E-85 |
| ACE2 | REEP1 | 0,8103 | 2,45934E-85 |
| ACE2 | ARHGAP18 | 0,8103 | 2,46619E-85 |
| ACE2 | CXADR | 0,8103 | 2,41811E-85 |
| ACE2 | ZFAND5 | 0,8099 | 3,3541E-85 |
| ACE2 | CNIH1 | 0,8099 | 3,46074E-85 |
| ACE2 | GPRC5B | 0,8099 | 3,4343E-85 |
| ACE2 | WASL | 0,8093 | 5,56421E-85 |
| ACE2 | TMEM33 | 0,8087 | 9,29834E-85 |
| ACE2 | RP11-613D13.10 | 0,8087 | 9,51701E-85 |
| ACE2 | CD164 | 0,8084 | 1,26332E-84 |
| ACE2 | IDH1 | 0,8083 | 1,32055E-84 |
| ACE2 | ERP29 | 0,8082 | 1,39736E-84 |
| ACE2 | YIPF6 | 0,8078 | 2,09132E-84 |
| ACE2 | GUF1 | 0,8076 | 2,50332E-84 |
| ACE2 | ALDH6A1 | 0,8076 | 2,46037E-84 |
| ACE2 | ACOX1 | 0,8075 | 2,57924E-84 |
| ACE2 | PSMD10 | 0,8075 | 2,65805E-84 |
| ACE2 | SLC7A8 | 0,8072 | 3,4638E-84 |
| ACE2 | VEZT | 0,8071 | 3,80108E-84 |
| ACE2 | HNRNPH2 | 0,8071 | 3,5581E-84 |
| ACE2 | TMEM167A | 0,807 | 3,93702E-84 |
| ACE2 | MAP1B | 0,8068 | 4,55042E-84 |
| ACE2 | DNMBP | 0,8068 | 4,78364E-84 |

| | | | |
|---|---|---|---|
| ACE2 | ARPC5 | 0,8067 | 4,88569E-84 |
| ACE2 | CDS2 | 0,8067 | 5,24689E-84 |
| ACE2 | FUNDC1 | 0,8066 | 5,42037E-84 |
| ACE2 | MGST3 | 0,8064 | 6,5813E-84 |
| ACE2 | OSTC | 0,8064 | 6,27183E-84 |
| ACE2 | TOX | 0,8063 | 7,1899E-84 |
| ACE2 | AMIGO2 | 0,8056 | 1,3163E-83 |
| ACE2 | MCF2 | 0,8055 | 1,37406E-83 |
| ACE2 | DDX18 | 0,8054 | 1,45584E-83 |
| ACE2 | APOBEC3C | 0,8054 | 1,5457E-83 |
| ACE2 | DCAF8L2 | 0,8053 | 1,66985E-83 |
| ACE2 | DBT | 0,8051 | 1,84974E-83 |
| ACE2 | ATP9A | 0,8049 | 2,22244E-83 |
| ACE2 | GNE | 0,8047 | 2,697E-83 |
| ACE2 | SMIM15 | 0,8046 | 2,88964E-83 |
| ACE2 | ARMC10 | 0,8046 | 2,84833E-83 |
| ACE2 | KCTD3 | 0,8045 | 3,07126E-83 |
| ACE2 | SLC4A5 | 0,8043 | 3,74319E-83 |
| ACE2 | KIF1BP | 0,8039 | 5,2301E-83 |
| ACE2 | SLK | 0,8036 | 6,63733E-83 |
| ACE2 | HIBADH | 0,8034 | 7,51687E-83 |
| ACE2 | RBMXL1 | 0,8031 | 9,73083E-83 |
| ACE2 | TMED8 | 0,803 | 1,05065E-82 |
| ACE2 | SDHC | 0,8028 | 1,29041E-82 |
| ACE2 | SECISBP2L | 0,8028 | 1,26832E-82 |
| ACE2 | FAT1 | 0,8023 | 1,89526E-82 |
| ACE2 | MBNL2 | 0,8022 | 1,99542E-82 |
| ACE2 | BEX3 | 0,8019 | 2,53351E-82 |
| ACE2 | H2AFY | 0,8015 | 3,60953E-82 |
| ACE2 | WIPF3 | 0,8015 | 3,64038E-82 |
| ACE2 | DFNA5 | 0,8013 | 4,18198E-82 |
| ACE2 | TRIM35 | 0,801 | 5,28327E-82 |
| ACE2 | C3orf70 | 0,8009 | 5,77297E-82 |
| ACE2 | CHMP2B | 0,8007 | 6,92969E-82 |
| ACE2 | STK24 | 0,8007 | 7,03238E-82 |
| ACE2 | SSFA2 | 0,8005 | 7,89675E-82 |
| ACE2 | ELL2 | 0,8003 | 9,5354E-82 |
| ACE2 | RBM47 | 0,8 | 1,14748E-81 |
| ACE2 | POFUT1 | 0,7998 | 1,35418E-81 |
| ACE2 | VBP1 | 0,7998 | 1,4171E-81 |
| ACE2 | SUCLG2 | 0,7996 | 1,62074E-81 |
| ACE2 | C18orf25 | 0,7996 | 1,65627E-81 |
| ACE2 | RBM43 | 0,7995 | 1,74903E-81 |
| ACE2 | SKA2 | 0,7994 | 1,91531E-81 |
| ACE2 | TAF2 | 0,7993 | 2,13339E-81 |
| ACE2 | STARD7 | 0,799 | 2,68929E-81 |
| ACE2 | COX15 | 0,7989 | 2,95139E-81 |
| ACE2 | RRAGB | 0,7989 | 2,94644E-81 |
| ACE2 | FAM3C | 0,7986 | 3,51111E-81 |
| ACE2 | PREPL | 0,7985 | 3,85437E-81 |
| ACE2 | KLHL42 | 0,7983 | 4,78132E-81 |
| ACE2 | RAB6A | 0,7982 | 4,93969E-81 |
| ACE2 | PGM2 | 0,798 | 6,04351E-81 |
| ACE2 | GAB1 | 0,798 | 6,03078E-81 |

| | | | |
|---|---|---|---|
| ACE2 | GMPR | 0,798 | 5,94263E-81 |
| ACE2 | DPY19L4 | 0,7979 | 6,30108E-81 |
| ACE2 | BTG3 | 0,7979 | 6,49039E-81 |
| ACE2 | PANX1 | 0,7978 | 6,85939E-81 |
| ACE2 | IGF2R | 0,7976 | 8,29175E-81 |
| ACE2 | DHTKD1 | 0,7976 | 8,00839E-81 |
| ACE2 | APOOL | 0,7976 | 7,96766E-81 |
| ACE2 | SELENOT | 0,7975 | 8,95593E-81 |
| ACE2 | ARMCX3 | 0,7973 | 1,0471E-80 |
| ACE2 | CCDC6 | 0,7971 | 1,21505E-80 |
| ACE2 | PGAM1 | 0,797 | 1,28128E-80 |
| ACE2 | ARL15 | 0,7969 | 1,34573E-80 |
| ACE2 | HMGXB4 | 0,7966 | 1,81652E-80 |
| ACE2 | OCIAD1 | 0,7964 | 2,0618E-80 |
| ACE2 | LHFPL2 | 0,7963 | 2,20251E-80 |
| ACE2 | EIF2AK1 | 0,7961 | 2,57812E-80 |
| ACE2 | BMS1 | 0,7961 | 2,55783E-80 |
| ACE2 | CYB5B | 0,7954 | 4,54742E-80 |
| ACE2 | NUCKS1 | 0,7951 | 5,74614E-80 |
| ACE2 | CLTC | 0,795 | 6,19265E-80 |
| ACE2 | RHOU | 0,7948 | 7,02848E-80 |
| ACE2 | SNX18 | 0,7948 | 7,28556E-80 |
| ACE2 | TOR1A | 0,7944 | 9,51215E-80 |
| ACE2 | MAP3K5 | 0,7941 | 1,22896E-79 |
| ACE2 | VAPA | 0,794 | 1,31258E-79 |
| ACE2 | HDHD2 | 0,794 | 1,36094E-79 |
| ACE2 | PRKCI | 0,7938 | 1,5047E-79 |
| ACE2 | MGST2 | 0,7938 | 1,53284E-79 |
| ACE2 | SLF2 | 0,7934 | 2,06163E-79 |
| ACE2 | KITLG | 0,7934 | 2,06915E-79 |
| ACE2 | COA7 | 0,7932 | 2,4869E-79 |
| ACE2 | UMAD1 | 0,7931 | 2,6745E-79 |
| ACE2 | LEPROT | 0,7928 | 3,29803E-79 |
| ACE2 | SNX7 | 0,7927 | 3,65995E-79 |
| ACE2 | SERPINF1 | 0,7927 | 3,51431E-79 |
| ACE2 | CLU | 0,7925 | 4,24933E-79 |
| ACE2 | SEC62 | 0,7924 | 4,4401E-79 |
| ACE2 | DLD | 0,7923 | 4,90866E-79 |
| ACE2 | FAM98A | 0,7922 | 5,10763E-79 |
| ACE2 | UTP3 | 0,7921 | 5,72744E-79 |
| ACE2 | PFDN1 | 0,792 | 6,09208E-79 |
| ACE2 | SMARCA5 | 0,7918 | 7,40443E-79 |
| ACE2 | ATP11A | 0,7918 | 7,11674E-79 |
| ACE2 | TP53BP2 | 0,7917 | 8,06475E-79 |
| ACE2 | HIF1A | 0,7917 | 7,55929E-79 |
| ACE2 | YWHAB | 0,7916 | 8,48962E-79 |
| ACE2 | MAGEH1 | 0,7916 | 8,07947E-79 |
| ACE2 | PPIB | 0,7915 | 8,81951E-79 |
| ACE2 | HPS5 | 0,7912 | 1,18075E-78 |
| ACE2 | TMEM97 | 0,7911 | 1,20763E-78 |
| ACE2 | TXNDC15 | 0,7909 | 1,38476E-78 |
| ACE2 | ZXDB | 0,7909 | 1,40169E-78 |
| ACE2 | GNB1 | 0,79 | 2,86541E-78 |
| ACE2 | C1QBP | 0,7896 | 3,84455E-78 |

| | | | |
|---|---|---|---|
| ACE2 | DISP1 | 0,7895 | 4,08596E-78 |
| ACE2 | TACC1 | 0,7894 | 4,49665E-78 |
| ACE2 | OSTF1 | 0,7894 | 4,50331E-78 |
| ACE2 | SCAMP1 | 0,7893 | 4,61569E-78 |
| ACE2 | ATP6V1E1 | 0,7893 | 4,81977E-78 |
| ACE2 | ATF6 | 0,7891 | 5,63325E-78 |
| ACE2 | PCCA | 0,789 | 6,20348E-78 |
| ACE2 | NECTIN2 | 0,7889 | 6,46188E-78 |
| ACE2 | NAA50 | 0,7888 | 7,04572E-78 |
| ACE2 | CRIM1 | 0,7886 | 8,17456E-78 |
| ACE2 | GNAQ | 0,7886 | 8,34955E-78 |
| ACE2 | ENOX2 | 0,7886 | 8,20678E-78 |
| ACE2 | NAP1L3 | 0,7885 | 8,57872E-78 |
| ACE2 | KIAA1958 | 0,7884 | 9,13734E-78 |
| ACE2 | LRRC8D | 0,7879 | 1,35939E-77 |
| ACE2 | SHC4 | 0,7878 | 1,45621E-77 |
| ACE2 | TMEM14B | 0,7877 | 1,5971E-77 |
| ACE2 | SET | 0,7877 | 1,63694E-77 |
| ACE2 | KDSR | 0,7877 | 1,55888E-77 |
| ACE2 | TMED7 | 0,7876 | 1,66494E-77 |
| ACE2 | YES1 | 0,7876 | 1,74883E-77 |
| ACE2 | UCHL5 | 0,7874 | 1,96833E-77 |
| ACE2 | RGAG1 | 0,7873 | 2,10844E-77 |
| ACE2 | PNPLA4 | 0,7869 | 2,83119E-77 |
| ACE2 | ETFA | 0,7866 | 3,47491E-77 |
| ACE2 | PCSK6 | 0,786 | 5,53647E-77 |
| ACE2 | MRPL30 | 0,7858 | 6,65522E-77 |
| ACE2 | CTNNAL1 | 0,7856 | 7,63247E-77 |
| ACE2 | UBFD1 | 0,7856 | 7,44064E-77 |
| ACE2 | GCLC | 0,7853 | 9,53512E-77 |
| ACE2 | PON2 | 0,7851 | 1,07245E-76 |
| ACE2 | PEA15 | 0,785 | 1,14291E-76 |
| ACE2 | OPA1 | 0,785 | 1,20467E-76 |
| ACE2 | CUL5 | 0,7846 | 1,5472E-76 |
| ACE2 | PAFAH1B2 | 0,7846 | 1,54687E-76 |
| ACE2 | ALDH2 | 0,7845 | 1,70976E-76 |
| ACE2 | GTF2H3 | 0,7844 | 1,83101E-76 |
| ACE2 | GOLGA5 | 0,7843 | 2,03E-76 |
| ACE2 | NRAS | 0,7842 | 2,16318E-76 |
| ACE2 | KIF21A | 0,7842 | 2,07776E-76 |
| ACE2 | PDP2 | 0,7842 | 2,1452E-76 |
| ACE2 | TMEM30A | 0,7839 | 2,60496E-76 |
| ACE2 | EGLN1 | 0,7837 | 3,05359E-76 |
| ACE2 | DTD2 | 0,7836 | 3,40098E-76 |
| ACE2 | DAP3 | 0,7835 | 3,5573E-76 |
| ACE2 | ITM2B | 0,7834 | 3,91366E-76 |
| ACE2 | PRPS1 | 0,7834 | 3,8461E-76 |
| ACE2 | ANKRD17 | 0,7833 | 4,16629E-76 |
| ACE2 | CELF1 | 0,7828 | 5,87244E-76 |
| ACE2 | ANXA5 | 0,7827 | 6,16826E-76 |
| ACE2 | SLC31A1 | 0,7821 | 9,94191E-76 |
| ACE2 | ERO1A | 0,782 | 1,08379E-75 |
| ACE2 | LIMS1 | 0,7815 | 1,46391E-75 |
| ACE2 | CCDC47 | 0,7812 | 1,90956E-75 |

| | | | |
|---|---|---|---|
| ACE2 | PAICS | 0,7811 | 2,02218E-75 |
| ACE2 | WNK3 | 0,7811 | 1,9652E-75 |
| ACE2 | USP53 | 0,7809 | 2,30755E-75 |
| ACE2 | WT1 | 0,7808 | 2,40946E-75 |
| ACE2 | PCDHGB5 | 0,7806 | 2,92237E-75 |
| ACE2 | ZBED3 | 0,7803 | 3,54013E-75 |
| ACE2 | HEXA | 0,78 | 4,30148E-75 |
| ACE2 | ENOPH1 | 0,7797 | 5,45789E-75 |
| ACE2 | UBQLN2 | 0,7797 | 5,42693E-75 |
| ACE2 | POGK | 0,7796 | 5,89858E-75 |
| ACE2 | HEXB | 0,7796 | 5,8802E-75 |
| ACE2 | COX18 | 0,779 | 8,84783E-75 |
| ACE2 | VMP1 | 0,779 | 9,02848E-75 |
| ACE2 | HNRNPK | 0,7788 | 1,06198E-74 |
| ACE2 | KIAA1217 | 0,7788 | 1,05833E-74 |
| ACE2 | BMPR1A | 0,7788 | 1,06233E-74 |
| ACE2 | ZBTB33 | 0,7787 | 1,08676E-74 |
| ACE2 | SMIM20 | 0,7786 | 1,1936E-74 |
| ACE2 | RAP2C | 0,7785 | 1,29482E-74 |
| ACE2 | PHB | 0,7784 | 1,42918E-74 |
| ACE2 | ZMPSTE24 | 0,7782 | 1,54844E-74 |
| ACE2 | TADA1 | 0,7782 | 1,55775E-74 |
| ACE2 | NCK2 | 0,7782 | 1,56546E-74 |
| ACE2 | SORT1 | 0,7781 | 1,77222E-74 |
| ACE2 | SERPINE2 | 0,7781 | 1,75425E-74 |
| ACE2 | MAPK9 | 0,7781 | 1,71245E-74 |
| ACE2 | FCHSD2 | 0,778 | 1,81215E-74 |
| ACE2 | TTTY15 | 0,778 | 1,81666E-74 |
| ACE2 | STX12 | 0,7779 | 1,91043E-74 |
| ACE2 | FAM13A | 0,7779 | 2,0291E-74 |
| ACE2 | MXRA7 | 0,7779 | 1,98495E-74 |
| ACE2 | ATP2C1 | 0,7778 | 2,1935E-74 |
| ACE2 | PTPN14 | 0,7774 | 2,90773E-74 |
| ACE2 | CD2AP | 0,7773 | 2,95301E-74 |
| ACE2 | GNG10 | 0,777 | 3,61496E-74 |
| ACE2 | TAOK1 | 0,777 | 3,84241E-74 |
| ACE2 | ELMO2 | 0,777 | 3,61024E-74 |
| ACE2 | MORC4 | 0,777 | 3,67417E-74 |
| ACE2 | WASHC5 | 0,7768 | 4,3936E-74 |
| ACE2 | ERP44 | 0,7767 | 4,69298E-74 |
| ACE2 | DUSP16 | 0,7765 | 5,32448E-74 |
| ACE2 | EPPIN | 0,7764 | 5,5409E-74 |
| ACE2 | PHF6 | 0,7763 | 6,10928E-74 |
| ACE2 | DDAH1 | 0,7759 | 7,93478E-74 |
| ACE2 | CMPK1 | 0,7753 | 1,26108E-73 |
| ACE2 | ATP6V0E1 | 0,7753 | 1,26611E-73 |
| ACE2 | STK4 | 0,7753 | 1,23626E-73 |
| ACE2 | PGRMC1 | 0,7753 | 1,20721E-73 |
| ACE2 | AKAP7 | 0,7752 | 1,36765E-73 |
| ACE2 | TBL1X | 0,775 | 1,54691E-73 |
| ACE2 | ADAM17 | 0,7749 | 1,61284E-73 |
| ACE2 | PAPD4 | 0,7749 | 1,6116E-73 |
| ACE2 | CTBS | 0,7748 | 1,7117E-73 |
| ACE2 | CPNE3 | 0,7748 | 1,80943E-73 |

| | | | |
|---|---|---|---|
| ACE2 | XPR1 | 0,7747 | 1,88717E-73 |
| ACE2 | CDC42SE2 | 0,7746 | 2,03359E-73 |
| ACE2 | RPL7L1 | 0,7746 | 1,95736E-73 |
| ACE2 | GSTA4 | 0,7746 | 1,98014E-73 |
| ACE2 | AGL | 0,7743 | 2,45995E-73 |
| ACE2 | PPP1CB | 0,7741 | 2,78836E-73 |
| ACE2 | SUN2 | 0,774 | 3,04946E-73 |
| ACE2 | PPP2R3A | 0,7739 | 3,2252E-73 |
| ACE2 | SUSD1 | 0,7739 | 3,33376E-73 |
| ACE2 | LGMN | 0,7739 | 3,20453E-73 |
| ACE2 | ACTB | 0,7738 | 3,41873E-73 |
| ACE2 | JAZF1 | 0,7738 | 3,51933E-73 |
| ACE2 | STX3 | 0,7738 | 3,54693E-73 |
| ACE2 | DENND4A | 0,7738 | 3,62369E-73 |
| ACE2 | FECH | 0,7737 | 3,71939E-73 |
| ACE2 | QSOX1 | 0,7735 | 4,21215E-73 |
| ACE2 | WWP1 | 0,7735 | 4,45923E-73 |
| ACE2 | LPAR3 | 0,7734 | 4,54702E-73 |
| ACE2 | ME2 | 0,7734 | 4,60965E-73 |
| ACE2 | ATXN7L3B | 0,7733 | 5,01704E-73 |
| ACE2 | TWISTNB | 0,7732 | 5,26705E-73 |
| ACE2 | VAMP3 | 0,773 | 6,2573E-73 |
| ACE2 | ALDH3A2 | 0,7729 | 6,54518E-73 |
| ACE2 | ALDH7A1 | 0,7728 | 7,2145E-73 |
| ACE2 | ACVR1 | 0,7727 | 7,45608E-73 |
| ACE2 | RAB39B | 0,7726 | 8,12679E-73 |
| ACE2 | TCEAL5 | 0,7725 | 8,83229E-73 |
| ACE2 | GDF10 | 0,7724 | 9,25962E-73 |
| ACE2 | VTI1A | 0,7724 | 9,38001E-73 |
| ACE2 | IDH3A | 0,7723 | 1,01756E-72 |
| ACE2 | TCEAL3 | 0,7723 | 1,01378E-72 |
| ACE2 | PPCS | 0,7722 | 1,06952E-72 |
| ACE2 | SGPL1 | 0,7722 | 1,03723E-72 |
| ACE2 | DPY19L3 | 0,7722 | 1,05352E-72 |
| ACE2 | MAPK1 | 0,7722 | 1,05612E-72 |
| ACE2 | ITGA6 | 0,772 | 1,17509E-72 |
| ACE2 | NT5E | 0,772 | 1,19545E-72 |
| ACE2 | VGLL4 | 0,7718 | 1,35914E-72 |
| ACE2 | OCIAD2 | 0,7717 | 1,49505E-72 |
| ACE2 | UBE2H | 0,7717 | 1,46579E-72 |
| ACE2 | QTRT2 | 0,7716 | 1,6012E-72 |
| ACE2 | XIAP | 0,7716 | 1,59504E-72 |
| ACE2 | ADAM9 | 0,7714 | 1,84775E-72 |
| ACE2 | IBTK | 0,771 | 2,49286E-72 |
| ACE2 | FOXO1 | 0,771 | 2,4846E-72 |
| ACE2 | STRN | 0,7709 | 2,50155E-72 |
| ACE2 | DNAJC13 | 0,7709 | 2,58523E-72 |
| ACE2 | PFN2 | 0,7709 | 2,6581E-72 |
| ACE2 | PARVA | 0,7709 | 2,53775E-72 |
| ACE2 | QKI | 0,7708 | 2,8311E-72 |
| ACE2 | VHL | 0,7707 | 2,99361E-72 |
| ACE2 | EIF4G2 | 0,7706 | 3,09051E-72 |
| ACE2 | FUCA2 | 0,7705 | 3,3807E-72 |
| ACE2 | TMEM14C | 0,7704 | 3,58952E-72 |

| | | | |
|---|---|---|---|
| ACE2 | MRPL49 | 0,7704 | 3,54003E-72 |
| ACE2 | DHCR7 | 0,7704 | 3,68008E-72 |
| ACE2 | ZNF772 | 0,7704 | 3,70443E-72 |
| ACE2 | TMEM167B | 0,77 | 4,72773E-72 |
| ACE2 | REPS1 | 0,77 | 4,62241E-72 |
| ACE2 | UBE2E2 | 0,7699 | 5,17431E-72 |
| ACE2 | C5orf51 | 0,7699 | 5,04318E-72 |
| ACE2 | CLCN3 | 0,7698 | 5,27754E-72 |
| ACE2 | TPP1 | 0,7698 | 5,45738E-72 |
| ACE2 | BOD1L1 | 0,7697 | 5,83369E-72 |
| ACE2 | MAP3K21 | 0,7695 | 6,53946E-72 |
| ACE2 | HSD17B4 | 0,7695 | 6,69163E-72 |
| ACE2 | WDFY1 | 0,7694 | 7,28541E-72 |
| ACE2 | EMB | 0,7692 | 8,13289E-72 |
| ACE2 | MAP4K4 | 0,7691 | 8,83186E-72 |
| ACE2 | FBXO18 | 0,7687 | 1,13039E-71 |
| ACE2 | RAP1B | 0,7687 | 1,16508E-71 |
| ACE2 | PDIA5 | 0,7683 | 1,53427E-71 |
| ACE2 | NECAP1 | 0,7683 | 1,52239E-71 |
| ACE2 | SESN1 | 0,7682 | 1,64717E-71 |
| ACE2 | FOXO3 | 0,7681 | 1,69514E-71 |
| ACE2 | SSR1 | 0,7679 | 1,93455E-71 |
| ACE2 | ITGB1 | 0,7678 | 2,11252E-71 |
| ACE2 | USP9X | 0,7678 | 2,07242E-71 |
| ACE2 | ANP32A | 0,7677 | 2,28589E-71 |
| ACE2 | TAF1 | 0,7675 | 2,66789E-71 |
| ACE2 | PAK2 | 0,7672 | 3,13034E-71 |
| ACE2 | NHSL2 | 0,7672 | 3,18377E-71 |
| ACE2 | RAC1 | 0,767 | 3,64129E-71 |
| ACE2 | HEBP1 | 0,767 | 3,71294E-71 |
| ACE2 | PNPO | 0,7669 | 3,74575E-71 |
| ACE2 | NAB1 | 0,7667 | 4,53471E-71 |
| ACE2 | PCDHGA2 | 0,7667 | 4,55231E-71 |
| ACE2 | DENND4C | 0,7667 | 4,44088E-71 |
| ACE2 | PSMC6 | 0,7663 | 5,78872E-71 |
| ACE2 | ATP7A | 0,7656 | 8,9892E-71 |
| ACE2 | TMEM185A | 0,7656 | 9,1749E-71 |
| ACE2 | PSMA2 | 0,7655 | 1,01026E-70 |
| ACE2 | TIMP2 | 0,7653 | 1,12767E-70 |
| ACE2 | PRSS23 | 0,7652 | 1,22632E-70 |
| ACE2 | TMX1 | 0,7652 | 1,17042E-70 |
| ACE2 | PEBP1 | 0,7651 | 1,32339E-70 |
| ACE2 | MRPS23 | 0,765 | 1,38979E-70 |
| ACE2 | UBQLN1 | 0,7649 | 1,4591E-70 |
| ACE2 | SRSF8 | 0,7649 | 1,50605E-70 |
| ACE2 | NEU1 | 0,7647 | 1,67642E-70 |
| ACE2 | EMC1 | 0,7645 | 1,87758E-70 |
| ACE2 | MAP2K4 | 0,7645 | 1,92287E-70 |
| ACE2 | RBBP9 | 0,7645 | 1,86291E-70 |
| ACE2 | MUM1L1 | 0,7645 | 1,96025E-70 |
| ACE2 | NDRG4 | 0,7644 | 2,0317E-70 |
| ACE2 | WDR82 | 0,7642 | 2,27282E-70 |
| ACE2 | ARSD | 0,7642 | 2,28111E-70 |
| ACE2 | KIAA1462 | 0,764 | 2,67271E-70 |

| | | | |
|---|---|---|---|
| ACE2 | FAM120A | 0,7638 | 3,06009E-70 |
| ACE2 | NUP205 | 0,7637 | 3,26041E-70 |
| ACE2 | PRRC1 | 0,7635 | 3,73734E-70 |
| ACE2 | SLC41A1 | 0,7631 | 4,95061E-70 |
| ACE2 | CLEC12B | 0,7631 | 4,67171E-70 |
| ACE2 | RRN3 | 0,763 | 5,11785E-70 |
| ACE2 | ZNF711 | 0,763 | 5,26918E-70 |
| ACE2 | BCAR3 | 0,7629 | 5,49897E-70 |
| ACE2 | IARS2 | 0,7629 | 5,65591E-70 |
| ACE2 | MFSD2A | 0,7627 | 6,2887E-70 |
| ACE2 | PRKACB | 0,7626 | 6,46145E-70 |
| ACE2 | GBE1 | 0,7625 | 7,3077E-70 |
| ACE2 | CTSO | 0,7624 | 7,39441E-70 |
| ACE2 | EIF4E3 | 0,7623 | 8,36266E-70 |
| ACE2 | CD63 | 0,7623 | 7,88013E-70 |
| ACE2 | SYT11 | 0,7622 | 8,90971E-70 |
| ACE2 | RLIM | 0,7622 | 8,43977E-70 |
| ACE2 | KLHL9 | 0,7621 | 9,0714E-70 |
| ACE2 | POLR2K | 0,762 | 9,67648E-70 |
| ACE2 | POLE3 | 0,762 | 1,01929E-69 |
| ACE2 | EIF2S3 | 0,7619 | 1,05693E-69 |
| ACE2 | SPTSSA | 0,7618 | 1,09579E-69 |
| ACE2 | COMT | 0,7618 | 1,12025E-69 |
| ACE2 | CTNNB1 | 0,7616 | 1,24688E-69 |
| ACE2 | RHOBTB3 | 0,7616 | 1,26695E-69 |
| ACE2 | WDR44 | 0,7615 | 1,39575E-69 |
| ACE2 | HMGN3 | 0,7614 | 1,459E-69 |
| ACE2 | TLK1 | 0,7613 | 1,6051E-69 |
| ACE2 | TAX1BP1 | 0,7613 | 1,52655E-69 |
| ACE2 | MB21D2 | 0,761 | 1,92578E-69 |
| ACE2 | LAMTOR3 | 0,761 | 1,8966E-69 |
| ACE2 | MTMR1 | 0,761 | 1,89579E-69 |
| ACE2 | ZNF708 | 0,7608 | 2,18529E-69 |
| ACE2 | ZMYND8 | 0,7607 | 2,34484E-69 |
| ACE2 | SRPRA | 0,7606 | 2,52961E-69 |
| ACE2 | SLC25A36 | 0,7602 | 3,20996E-69 |
| ACE2 | YME1L1 | 0,7602 | 3,2402E-69 |
| ACE2 | GFM1 | 0,7601 | 3,33152E-69 |
| ACE2 | DUSP11 | 0,7598 | 4,16152E-69 |
| ACE2 | NUP133 | 0,7596 | 4,63542E-69 |
| ACE2 | ELF1 | 0,7596 | 4,69411E-69 |
| ACE2 | MYL12A | 0,7596 | 4,69748E-69 |
| ACE2 | ARCN1 | 0,7594 | 5,48399E-69 |
| ACE2 | SH3BP4 | 0,7592 | 6,06553E-69 |
| ACE2 | BIRC2 | 0,7591 | 6,35122E-69 |
| ACE2 | GABARAPL2 | 0,7591 | 6,60087E-69 |
| ACE2 | CLCN4 | 0,7591 | 6,57285E-69 |
| ACE2 | MOAP1 | 0,7589 | 7,58802E-69 |
| ACE2 | RIDA | 0,7587 | 8,66752E-69 |
| ACE2 | FAM127C | 0,7584 | 1,02167E-68 |
| ACE2 | ZEB2 | 0,7583 | 1,07216E-68 |
| ACE2 | PTPN4 | 0,7581 | 1,22924E-68 |
| ACE2 | CDS1 | 0,7581 | 1,25739E-68 |
| ACE2 | CDKN1B | 0,7581 | 1,19981E-68 |

| | | | |
|---|---|---|---|
| ACE2 | VTA1 | 0,758 | 1,27658E-68 |
| ACE2 | TMEM2 | 0,758 | 1,33089E-68 |
| ACE2 | TRMT2B | 0,7579 | 1,43778E-68 |
| ACE2 | HADHB | 0,7578 | 1,48156E-68 |
| ACE2 | GPAM | 0,7578 | 1,48742E-68 |
| ACE2 | HNRNPA1 | 0,7578 | 1,45446E-68 |
| ACE2 | GOLPH3 | 0,7577 | 1,55685E-68 |
| ACE2 | TOP1 | 0,7577 | 1,59758E-68 |
| ACE2 | MCCC1 | 0,7576 | 1,70839E-68 |
| ACE2 | MYL12B | 0,7575 | 1,81022E-68 |
| ACE2 | NAP1L2 | 0,7573 | 2,03565E-68 |
| ACE2 | ADAR | 0,7571 | 2,3426E-68 |
| ACE2 | VIPAS39 | 0,7571 | 2,37717E-68 |
| ACE2 | SPOCK1 | 0,757 | 2,42476E-68 |
| ACE2 | TMEM181 | 0,7569 | 2,69342E-68 |
| ACE2 | SSR3 | 0,7567 | 3,07044E-68 |
| ACE2 | ERGIC1 | 0,7566 | 3,16848E-68 |
| ACE2 | ZNF134 | 0,7566 | 3,123E-68 |
| ACE2 | NAXE | 0,7564 | 3,58986E-68 |
| ACE2 | SLC11A2 | 0,7563 | 3,8354E-68 |
| ACE2 | KLHL2 | 0,7561 | 4,45114E-68 |
| ACE2 | ADAM10 | 0,756 | 4,62151E-68 |
| ACE2 | PLAT | 0,7559 | 5,18234E-68 |
| ACE2 | AMD1 | 0,7557 | 5,58173E-68 |
| ACE2 | MSI2 | 0,7556 | 6,07888E-68 |
| ACE2 | MPDZ | 0,7555 | 6,6481E-68 |
| ACE2 | EPDR1 | 0,7553 | 7,55339E-68 |
| ACE2 | SLC25A30 | 0,7553 | 7,37213E-68 |
| ACE2 | AR | 0,7551 | 8,44043E-68 |
| ACE2 | ARL5A | 0,755 | 8,64271E-68 |
| ACE2 | PDE12 | 0,755 | 8,77904E-68 |
| ACE2 | NONO | 0,755 | 8,62775E-68 |
| ACE2 | PCSK5 | 0,7547 | 1,08279E-67 |
| ACE2 | FAM96A | 0,7547 | 1,07024E-67 |
| ACE2 | TXNL4B | 0,7546 | 1,17495E-67 |
| ACE2 | HMG20A | 0,7545 | 1,20183E-67 |
| ACE2 | CYB5R3 | 0,7544 | 1,26613E-67 |
| ACE2 | TUBB | 0,7543 | 1,34436E-67 |
| ACE2 | SLC52A3 | 0,7542 | 1,47352E-67 |
| ACE2 | SCML1 | 0,7542 | 1,51117E-67 |
| ACE2 | CALU | 0,7541 | 1,58292E-67 |
| ACE2 | HNRNPD | 0,754 | 1,67158E-67 |
| ACE2 | ZNF275 | 0,7539 | 1,72167E-67 |
| ACE2 | TCAIM | 0,7534 | 2,36404E-67 |
| ACE2 | PSEN2 | 0,7533 | 2,64695E-67 |
| ACE2 | GALNT7 | 0,7533 | 2,51283E-67 |
| ACE2 | SFXN1 | 0,7532 | 2,71918E-67 |
| ACE2 | APPL1 | 0,7531 | 2,89838E-67 |
| ACE2 | MYO10 | 0,7528 | 3,48729E-67 |
| ACE2 | SLC39A9 | 0,7528 | 3,52522E-67 |
| ACE2 | MAGEB1 | 0,7528 | 3,61809E-67 |
| ACE2 | SSB | 0,7527 | 3,86649E-67 |
| ACE2 | HOPX | 0,7527 | 3,76411E-67 |
| ACE2 | SEPT9 | 0,7526 | 4,04846E-67 |

| | | | |
|---|---|---|---|
| ACE2 | SCD | 0,7525 | 4,32067E-67 |
| ACE2 | TMED9 | 0,7524 | 4,47485E-67 |
| ACE2 | MEF2A | 0,7524 | 4,67097E-67 |
| ACE2 | GNG12 | 0,7522 | 5,13832E-67 |
| ACE2 | RNF2 | 0,7522 | 4,99851E-67 |
| ACE2 | RAB12 | 0,7522 | 5,11728E-67 |
| ACE2 | ZC3H13 | 0,752 | 5,72063E-67 |
| ACE2 | GSE1 | 0,7517 | 7,0977E-67 |
| ACE2 | CNST | 0,7514 | 8,66413E-67 |
| ACE2 | ADI1 | 0,7514 | 8,36839E-67 |
| ACE2 | DDX21 | 0,7514 | 8,53318E-67 |
| ACE2 | CTSD | 0,7513 | 8,9252E-67 |
| ACE2 | HCCS | 0,7511 | 1,02909E-66 |
| ACE2 | GATA4 | 0,751 | 1,07545E-66 |
| ACE2 | MARCKSL1 | 0,7508 | 1,24821E-66 |
| ACE2 | VPS35 | 0,7508 | 1,18996E-66 |
| ACE2 | PDHA1 | 0,7507 | 1,30791E-66 |
| ACE2 | VMA21 | 0,7505 | 1,44142E-66 |
| ACE2 | FOXO4 | 0,7504 | 1,54668E-66 |
| ACE2 | HSPD1 | 0,7503 | 1,70275E-66 |
| ACE2 | MBNL1 | 0,7503 | 1,71851E-66 |
| ACE2 | LMBRD1 | 0,7503 | 1,62694E-66 |
| ACE2 | SHTN1 | 0,7501 | 1,83986E-66 |
| ACE2 | STIM1 | 0,7501 | 1,86299E-66 |
| ACE2 | HOMER1 | 0,75 | 1,97998E-66 |
| ACE2 | ELK1 | 0,75 | 1,97321E-66 |
| ACE2 | KAT6B | 0,7499 | 2,07905E-66 |
| ACE2 | DHRS4-AS1 | 0,7499 | 2,11614E-66 |
| ACE2 | NFE2L1 | 0,7499 | 2,09575E-66 |
| ACE2 | DCTN4 | 0,7498 | 2,3184E-66 |
| ACE2 | USP27X | 0,7496 | 2,64078E-66 |
| ACE2 | CDYL2 | 0,7495 | 2,77778E-66 |
| ACE2 | GPHN | 0,7494 | 2,86055E-66 |
| ACE2 | ST3GAL5 | 0,7492 | 3,3688E-66 |
| ACE2 | SAMD8 | 0,7492 | 3,38216E-66 |
| ACE2 | SDHD | 0,749 | 3,81534E-66 |
| ACE2 | RNF149 | 0,7487 | 4,34362E-66 |
| ACE2 | MTM1 | 0,7487 | 4,45645E-66 |
| ACE2 | AFG3L2 | 0,7486 | 4,82606E-66 |
| ACE2 | SYK | 0,7485 | 5,10964E-66 |
| ACE2 | ANKRD46 | 0,7484 | 5,27016E-66 |
| ACE2 | MAML2 | 0,7484 | 5,49154E-66 |
| ACE2 | COQ2 | 0,748 | 6,76025E-66 |
| ACE2 | METTL8 | 0,7479 | 7,27118E-66 |
| ACE2 | ADPRH | 0,7479 | 7,23841E-66 |
| ACE2 | MRPL44 | 0,7478 | 7,89688E-66 |
| ACE2 | CISD2 | 0,7477 | 8,49291E-66 |
| ACE2 | ZNF430 | 0,7476 | 8,85928E-66 |
| ACE2 | PIK3C3 | 0,7473 | 1,05869E-65 |
| ACE2 | DIP2B | 0,7472 | 1,12336E-65 |
| ACE2 | KIDINS220 | 0,7471 | 1,18533E-65 |
| ACE2 | SLC7A2 | 0,7471 | 1,19158E-65 |
| ACE2 | UEVLD | 0,7471 | 1,22235E-65 |
| ACE2 | MRFAP1L1 | 0,747 | 1,2354E-65 |

| | | | |
|---|---|---|---|
| ACE2 | SLC35F6 | 0,7468 | 1,45062E-65 |
| ACE2 | TSPAN3 | 0,7468 | 1,46666E-65 |
| ACE2 | ZBTB14 | 0,7468 | 1,3877E-65 |
| ACE2 | FAM98B | 0,7467 | 1,51944E-65 |
| ACE2 | RBM12 | 0,7467 | 1,48196E-65 |
| ACE2 | GS1-358P8.4 | 0,7467 | 1,52985E-65 |
| ACE2 | TMED4 | 0,7466 | 1,65175E-65 |
| ACE2 | CDH2 | 0,7466 | 1,59971E-65 |
| ACE2 | SEL1L | 0,7465 | 1,74504E-65 |
| ACE2 | DCTN6 | 0,7464 | 1,84068E-65 |
| ACE2 | BLOC1S6 | 0,7464 | 1,83414E-65 |
| ACE2 | PPM1K | 0,7462 | 1,99676E-65 |
| ACE2 | PANK3 | 0,7462 | 2,08902E-65 |
| ACE2 | PSMD5 | 0,7462 | 2,0594E-65 |
| ACE2 | RP2 | 0,7461 | 2,24404E-65 |
| ACE2 | NAP1L1 | 0,746 | 2,31961E-65 |
| ACE2 | DSTN | 0,7459 | 2,38874E-65 |
| ACE2 | AK4 | 0,7458 | 2,57098E-65 |
| ACE2 | MOB1A | 0,7458 | 2,63496E-65 |
| ACE2 | NIPSNAP1 | 0,7458 | 2,55019E-65 |
| ACE2 | TTC37 | 0,7457 | 2,75105E-65 |
| ACE2 | CERS2 | 0,7456 | 2,93093E-65 |
| ACE2 | DMRT1 | 0,7456 | 3,01161E-65 |
| ACE2 | INIP | 0,7456 | 2,99544E-65 |
| ACE2 | RAB11FIP2 | 0,7456 | 2,88399E-65 |
| ACE2 | GALNT2 | 0,7455 | 3,08709E-65 |
| ACE2 | LPIN2 | 0,7455 | 3,1174E-65 |
| ACE2 | PUM2 | 0,7453 | 3,43638E-65 |
| ACE2 | ASNSD1 | 0,7453 | 3,61524E-65 |
| ACE2 | MYLK | 0,7451 | 3,99468E-65 |
| ACE2 | BCAT1 | 0,745 | 4,29467E-65 |
| ACE2 | PABPC5 | 0,7449 | 4,3939E-65 |
| ACE2 | KCTD20 | 0,7447 | 5,05055E-65 |
| ACE2 | TRERF1 | 0,7446 | 5,32669E-65 |
| ACE2 | ACBD3 | 0,7445 | 5,89707E-65 |
| ACE2 | DBR1 | 0,7445 | 5,58006E-65 |
| ACE2 | NOTCH2 | 0,7444 | 6,10931E-65 |
| ACE2 | BNIP2 | 0,7444 | 6,05551E-65 |
| ACE2 | DDX42 | 0,7442 | 6,79038E-65 |
| ACE2 | FAM69A | 0,7441 | 7,09893E-65 |
| ACE2 | MARK4 | 0,7441 | 7,23435E-65 |
| ACE2 | UBE2A | 0,7441 | 7,38035E-65 |
| ACE2 | PKP2 | 0,744 | 7,57159E-65 |
| ACE2 | SLC25A21 | 0,744 | 7,84748E-65 |
| ACE2 | MBTPS1 | 0,7439 | 8,39463E-65 |
| ACE2 | TAF9B | 0,7439 | 8,02874E-65 |
| ACE2 | HNRNPR | 0,7438 | 8,66618E-65 |
| ACE2 | NIPSNAP3A | 0,7437 | 9,11842E-65 |
| ACE2 | USP51 | 0,7437 | 9,32147E-65 |
| ACE2 | ARID5B | 0,7436 | 9,7142E-65 |
| ACE2 | MXI1 | 0,7435 | 1,06606E-64 |
| ACE2 | MAT2B | 0,7434 | 1,13945E-64 |
| ACE2 | UBE2E3 | 0,7431 | 1,3613E-64 |
| ACE2 | IREB2 | 0,7431 | 1,31981E-64 |

| | | | |
|---|---|---|---|
| ACE2 | MRPL32 | 0,743 | 1,44205E-64 |
| ACE2 | KIN | 0,7429 | 1,52111E-64 |
| ACE2 | MINDY3 | 0,7428 | 1,62661E-64 |
| ACE2 | TSEN15 | 0,7423 | 2,15988E-64 |
| ACE2 | ATXN1 | 0,7422 | 2,28055E-64 |
| ACE2 | FKBP5 | 0,7422 | 2,30527E-64 |
| ACE2 | M6PR | 0,7421 | 2,44545E-64 |
| ACE2 | EIF1AY | 0,7421 | 2,35878E-64 |
| ACE2 | LAMC1 | 0,742 | 2,52753E-64 |
| ACE2 | SUCO | 0,7418 | 2,80601E-64 |
| ACE2 | WBP4 | 0,7417 | 3,02411E-64 |
| ACE2 | PKN2 | 0,7416 | 3,20551E-64 |
| ACE2 | RYK | 0,7416 | 3,29781E-64 |
| ACE2 | FBXL3 | 0,7416 | 3,26345E-64 |
| ACE2 | SDCBP | 0,7414 | 3,59562E-64 |
| ACE2 | THRAP3 | 0,7413 | 3,89405E-64 |
| ACE2 | NKX3-1 | 0,7413 | 3,97532E-64 |
| ACE2 | ARNT | 0,7412 | 4,0071E-64 |
| ACE2 | SNX12 | 0,7411 | 4,35962E-64 |
| ACE2 | BANK1 | 0,741 | 4,48188E-64 |
| ACE2 | MORF4L2 | 0,7408 | 5,27345E-64 |
| ACE2 | GMFB | 0,7406 | 5,8103E-64 |
| ACE2 | PPP3R1 | 0,7404 | 6,76917E-64 |
| ACE2 | UMPS | 0,7404 | 6,52126E-64 |
| ACE2 | APMAP | 0,7404 | 6,57693E-64 |
| ACE2 | ID4 | 0,7403 | 7,0672E-64 |
| ACE2 | NUTF2 | 0,7402 | 7,33038E-64 |
| ACE2 | CXXC4 | 0,7397 | 9,91896E-64 |
| ACE2 | DARS | 0,7396 | 1,07358E-63 |
| ACE2 | SCRG1 | 0,7395 | 1,1113E-63 |
| ACE2 | LGALS3BP | 0,7393 | 1,23046E-63 |
| ACE2 | SOCS6 | 0,7393 | 1,23523E-63 |
| ACE2 | PELI1 | 0,7392 | 1,32039E-63 |
| ACE2 | NLN | 0,7392 | 1,34733E-63 |
| ACE2 | ATP6V1H | 0,7392 | 1,33157E-63 |
| ACE2 | ZFX | 0,7392 | 1,34724E-63 |
| ACE2 | PEX19 | 0,7391 | 1,39066E-63 |
| ACE2 | SACM1L | 0,7391 | 1,37108E-63 |
| ACE2 | RCBTB1 | 0,7391 | 1,37154E-63 |
| ACE2 | ARMT1 | 0,739 | 1,53576E-63 |
| ACE2 | CTNNA1 | 0,7389 | 1,57756E-63 |
| ACE2 | FBXL4 | 0,7389 | 1,61959E-63 |
| ACE2 | FAM45A | 0,7387 | 1,80142E-63 |
| ACE2 | SLAIN1 | 0,7386 | 1,89422E-63 |
| ACE2 | RTCA | 0,7385 | 2,04316E-63 |
| ACE2 | GLUD2 | 0,7385 | 1,98903E-63 |
| ACE2 | TMEM59 | 0,7383 | 2,26643E-63 |
| ACE2 | CPSF6 | 0,7383 | 2,19688E-63 |
| ACE2 | NDUFA4 | 0,7382 | 2,37389E-63 |
| ACE2 | NKAP | 0,7382 | 2,45532E-63 |
| ACE2 | ACTR2 | 0,738 | 2,68124E-63 |
| ACE2 | GNA13 | 0,7379 | 2,76781E-63 |
| ACE2 | ATP8B2 | 0,7378 | 3,09576E-63 |
| ACE2 | ANXA2 | 0,7378 | 3,07415E-63 |

| | | | |
|---|---|---|---|
| ACE2 | PAPSS1 | 0,7376 | 3,31572E-63 |
| ACE2 | BPTF | 0,7375 | 3,62807E-63 |
| ACE2 | TMEM50A | 0,7374 | 3,87393E-63 |
| ACE2 | TSPYL1 | 0,7373 | 4,13934E-63 |
| ACE2 | PIK3C2A | 0,7371 | 4,65559E-63 |
| ACE2 | MAMLD1 | 0,7371 | 4,62198E-63 |
| ACE2 | API5 | 0,737 | 4,93249E-63 |
| ACE2 | ADIPOR1 | 0,7367 | 5,68825E-63 |
| ACE2 | TMEM68 | 0,7367 | 5,72584E-63 |
| ACE2 | B3GNT2 | 0,7366 | 5,91099E-63 |
| ACE2 | SP3 | 0,7365 | 6,60901E-63 |
| ACE2 | FOXJ2 | 0,7365 | 6,28921E-63 |
| ACE2 | SNX1 | 0,7365 | 6,52474E-63 |
| ACE2 | ABCB7 | 0,7365 | 6,42331E-63 |
| ACE2 | AEBP2 | 0,7364 | 6,64968E-63 |
| ACE2 | MON2 | 0,7364 | 6,99937E-63 |
| ACE2 | PRDX1 | 0,7361 | 8,24171E-63 |
| ACE2 | HS2ST1 | 0,736 | 8,68403E-63 |
| ACE2 | LRPPRC | 0,736 | 8,4914E-63 |
| ACE2 | RWDD4 | 0,736 | 8,63373E-63 |
| ACE2 | GLTSCR1L | 0,7358 | 9,79212E-63 |
| ACE2 | OOSP2 | 0,7355 | 1,16887E-62 |
| ACE2 | GNB5 | 0,7355 | 1,15154E-62 |
| ACE2 | SLC5A3 | 0,7354 | 1,21494E-62 |
| ACE2 | PIK3CA | 0,7352 | 1,32703E-62 |
| ACE2 | VPS41 | 0,7352 | 1,34797E-62 |
| ACE2 | RETSAT | 0,7351 | 1,45433E-62 |
| ACE2 | LRRC58 | 0,7348 | 1,70863E-62 |
| ACE2 | RGL1 | 0,7346 | 1,90194E-62 |
| ACE2 | TEAD1 | 0,7345 | 2,03759E-62 |
| ACE2 | ADIPOR2 | 0,7345 | 2,01064E-62 |
| ACE2 | ZDHHC15 | 0,7343 | 2,32325E-62 |
| ACE2 | TMX2 | 0,7339 | 2,81716E-62 |
| ACE2 | EIF3M | 0,7338 | 3,03068E-62 |
| ACE2 | ATP2B4 | 0,7337 | 3,2678E-62 |
| ACE2 | SEC23A | 0,7336 | 3,46862E-62 |
| ACE2 | ACSL3 | 0,7335 | 3,59903E-62 |
| ACE2 | ARMCX6 | 0,7335 | 3,63885E-62 |
| ACE2 | F8 | 0,7335 | 3,59524E-62 |
| ACE2 | FLNB | 0,7334 | 3,91818E-62 |
| ACE2 | CACNA2D1 | 0,7332 | 4,33264E-62 |
| ACE2 | IARS | 0,7331 | 4,57722E-62 |
| ACE2 | TMEM43 | 0,7328 | 5,33991E-62 |
| ACE2 | ALAD | 0,7327 | 5,56591E-62 |
| ACE2 | TMLHE | 0,7327 | 5,63779E-62 |
| ACE2 | CDO1 | 0,7321 | 7,81002E-62 |
| ACE2 | AFDN | 0,7321 | 8,06576E-62 |
| ACE2 | PORCN | 0,7321 | 8,03295E-62 |
| ACE2 | MID1IP1 | 0,732 | 8,31658E-62 |
| ACE2 | MFSD14B | 0,7319 | 8,72407E-62 |
| ACE2 | KLHL23 | 0,7317 | 9,80033E-62 |
| ACE2 | B4GAT1 | 0,7315 | 1,12757E-61 |
| ACE2 | SNX25 | 0,7314 | 1,16729E-61 |
| ACE2 | TMEM242 | 0,7312 | 1,31569E-61 |

| | | | |
|---|---|---|---|
| ACE2 | SAR1A | 0,7312 | 1,37114E-61 |
| ACE2 | DENR | 0,7312 | 1,35229E-61 |
| ACE2 | EEF1A1 | 0,7311 | 1,44701E-61 |
| ACE2 | RRM2B | 0,7311 | 1,44268E-61 |
| ACE2 | WWC3 | 0,7311 | 1,44154E-61 |
| ACE2 | PJA2 | 0,7309 | 1,60843E-61 |
| ACE2 | UTY | 0,7307 | 1,77303E-61 |
| ACE2 | CTSA | 0,7306 | 1,92117E-61 |
| ACE2 | CDK2AP1 | 0,7305 | 2,00171E-61 |
| ACE2 | XPNPEP1 | 0,7303 | 2,25968E-61 |
| ACE2 | MTHFS | 0,7303 | 2,20302E-61 |
| ACE2 | ATRX | 0,7303 | 2,16354E-61 |
| ACE2 | SPTAN1 | 0,7301 | 2,51997E-61 |
| ACE2 | PRKD1 | 0,7301 | 2,53185E-61 |
| ACE2 | KCNK13 | 0,7301 | 2,46925E-61 |
| ACE2 | SUB1 | 0,7299 | 2,78561E-61 |
| ACE2 | CWC22 | 0,7298 | 2,92422E-61 |
| ACE2 | TMEM14A | 0,7298 | 2,93868E-61 |
| ACE2 | MPP5 | 0,7297 | 3,11912E-61 |
| ACE2 | MAP3K7 | 0,7295 | 3,42955E-61 |
| ACE2 | GANAB | 0,7295 | 3,56495E-61 |
| ACE2 | BICD1 | 0,7294 | 3,67288E-61 |
| ACE2 | MAIP1 | 0,7292 | 4,02533E-61 |
| ACE2 | CTSV | 0,7292 | 4,06318E-61 |
| ACE2 | COG3 | 0,7292 | 4,04699E-61 |
| ACE2 | GALNT18 | 0,7291 | 4,33123E-61 |
| ACE2 | TRIM44 | 0,729 | 4,58652E-61 |
| ACE2 | ZNF41 | 0,729 | 4,53768E-61 |
| ACE2 | EIF2A | 0,7289 | 4,77731E-61 |
| ACE2 | NSMAF | 0,7288 | 5,06825E-61 |
| ACE2 | RAB3GAP2 | 0,7286 | 5,87184E-61 |
| ACE2 | SEC24B | 0,7286 | 5,60033E-61 |
| ACE2 | ARL1 | 0,7286 | 5,77578E-61 |
| ACE2 | THOC2 | 0,7286 | 5,73173E-61 |
| ACE2 | SOAT1 | 0,7285 | 6,14864E-61 |
| ACE2 | HSD17B3 | 0,7285 | 6,02478E-61 |
| ACE2 | ANXA7 | 0,7284 | 6,27174E-61 |
| ACE2 | SLC39A7 | 0,7283 | 6,91229E-61 |
| ACE2 | ICMT | 0,7282 | 7,09729E-61 |
| ACE2 | DPH5 | 0,7282 | 7,08683E-61 |
| ACE2 | BEX1 | 0,7281 | 7,51036E-61 |
| ACE2 | FAM120C | 0,728 | 8,07113E-61 |
| ACE2 | SLC25A38 | 0,7278 | 9,02256E-61 |
| ACE2 | TXNDC5 | 0,7278 | 8,97191E-61 |
| ACE2 | SLC50A1 | 0,7277 | 9,7634E-61 |
| ACE2 | ING3 | 0,7277 | 9,4163E-61 |
| ACE2 | PURA | 0,7274 | 1,1256E-60 |
| ACE2 | DAZAP2 | 0,7274 | 1,09846E-60 |
| ACE2 | UPRT | 0,7273 | 1,20381E-60 |
| ACE2 | PDZD11 | 0,7272 | 1,28491E-60 |
| ACE2 | SLC35B4 | 0,7271 | 1,36224E-60 |
| ACE2 | CTAGE5 | 0,7271 | 1,29256E-60 |
| ACE2 | ANAPC13 | 0,7267 | 1,65053E-60 |
| ACE2 | TMEM55A | 0,7266 | 1,77344E-60 |

| | | | |
|---|---|---|---|
| ACE2 | HNRNPUL2 | 0,7265 | 1,82738E-60 |
| ACE2 | RASSF2 | 0,7265 | 1,89003E-60 |
| ACE2 | MTHFD1 | 0,7264 | 1,92706E-60 |
| ACE2 | YAP1 | 0,7262 | 2,15527E-60 |
| ACE2 | LRRC57 | 0,7261 | 2,30976E-60 |
| ACE2 | NEO1 | 0,7261 | 2,33862E-60 |
| ACE2 | MSMO1 | 0,726 | 2,39638E-60 |
| ACE2 | DCP1A | 0,7258 | 2,74649E-60 |
| ACE2 | SLC35D2 | 0,7258 | 2,66426E-60 |
| ACE2 | SLC25A3 | 0,7258 | 2,73586E-60 |
| ACE2 | TYRO3 | 0,7258 | 2,73204E-60 |
| ACE2 | KCTD12 | 0,7257 | 2,93215E-60 |
| ACE2 | RAB14 | 0,7256 | 2,99496E-60 |
| ACE2 | SH3RF1 | 0,7254 | 3,31017E-60 |
| ACE2 | YPEL2 | 0,7254 | 3,39913E-60 |
| ACE2 | IMPACT | 0,7254 | 3,33594E-60 |
| ACE2 | NCAM2 | 0,7253 | 3,59431E-60 |
| ACE2 | TMEM268 | 0,7252 | 3,83392E-60 |
| ACE2 | CYB5A | 0,7252 | 3,84641E-60 |
| ACE2 | PLIN2 | 0,7251 | 3,94593E-60 |
| ACE2 | ENAH | 0,725 | 4,31925E-60 |
| ACE2 | KBTBD6 | 0,725 | 4,30416E-60 |
| ACE2 | PRPF4B | 0,7249 | 4,36767E-60 |
| ACE2 | AXL | 0,7249 | 4,5426E-60 |
| ACE2 | XPO1 | 0,7246 | 5,16424E-60 |
| ACE2 | THEMIS2 | 0,7245 | 5,42355E-60 |
| ACE2 | LIPH | 0,7245 | 5,53857E-60 |
| ACE2 | TWF1 | 0,7244 | 5,85677E-60 |
| ACE2 | SND1 | 0,7243 | 6,11795E-60 |
| ACE2 | LCOR | 0,7243 | 6,17337E-60 |
| ACE2 | CBR4 | 0,7241 | 6,76783E-60 |
| ACE2 | TUBA1A | 0,7241 | 7,01487E-60 |
| ACE2 | ASB9 | 0,7241 | 6,81403E-60 |
| ACE2 | MBD2 | 0,724 | 7,35013E-60 |
| ACE2 | FNDC3B | 0,7239 | 7,74046E-60 |
| ACE2 | MAK16 | 0,7239 | 7,60644E-60 |
| ACE2 | FMR1 | 0,7239 | 7,66658E-60 |
| ACE2 | TNKS2 | 0,7238 | 8,37024E-60 |
| ACE2 | CLNS1A | 0,7237 | 8,46821E-60 |
| ACE2 | TGFBRAP1 | 0,7236 | 9,15612E-60 |
| ACE2 | TRAK1 | 0,7236 | 9,32199E-60 |
| ACE2 | FKBP14 | 0,7236 | 9,10741E-60 |
| ACE2 | CHD6 | 0,7236 | 9,23985E-60 |
| ACE2 | ECE1 | 0,7234 | 1,01024E-59 |
| ACE2 | MED28 | 0,7234 | 1,03465E-59 |
| ACE2 | TSPAN31 | 0,7232 | 1,12922E-59 |
| ACE2 | MPP1 | 0,7232 | 1,11888E-59 |
| ACE2 | TMEM267 | 0,7231 | 1,18769E-59 |
| ACE2 | RAB11FIP1 | 0,7229 | 1,30398E-59 |
| ACE2 | CEBPZOS | 0,7228 | 1,42082E-59 |
| ACE2 | PLEKHG1 | 0,7228 | 1,41697E-59 |
| ACE2 | NDRG1 | 0,7225 | 1,68927E-59 |
| ACE2 | CRTC3 | 0,7225 | 1,67272E-59 |
| ACE2 | GAS7 | 0,7225 | 1,67783E-59 |

| | | | |
|---|---|---|---|
| ACE2 | CTSZ | 0,7225 | 1,61831E-59 |
| ACE2 | GPM6B | 0,7224 | 1,7609E-59 |
| ACE2 | CLN5 | 0,7222 | 2,00719E-59 |
| ACE2 | YIPF5 | 0,722 | 2,17436E-59 |
| ACE2 | MYOZ2 | 0,7218 | 2,38318E-59 |
| ACE2 | SLC39A6 | 0,7218 | 2,48047E-59 |
| ACE2 | MAGEB3 | 0,7218 | 2,44587E-59 |
| ACE2 | DNAJB14 | 0,7217 | 2,621E-59 |
| ACE2 | USP33 | 0,7215 | 2,81017E-59 |
| ACE2 | TMEM56 | 0,7215 | 2,84159E-59 |
| ACE2 | USP9Y | 0,7215 | 2,85223E-59 |
| ACE2 | PHAX | 0,7212 | 3,28509E-59 |
| ACE2 | ASAH1 | 0,7212 | 3,3966E-59 |
| ACE2 | OSBP | 0,7212 | 3,33676E-59 |
| ACE2 | GXYLT1 | 0,7212 | 3,45502E-59 |
| ACE2 | SGK3 | 0,7211 | 3,54059E-59 |
| ACE2 | PTPN11 | 0,721 | 3,68529E-59 |
| ACE2 | FOXN3 | 0,7206 | 4,67952E-59 |
| ACE2 | NEK7 | 0,7205 | 4,7981E-59 |
| ACE2 | CYB5R1 | 0,7205 | 4,95917E-59 |
| ACE2 | ADK | 0,7205 | 4,79836E-59 |
| ACE2 | TSNAX | 0,7203 | 5,36495E-59 |
| ACE2 | RANBP6 | 0,7201 | 6,15578E-59 |
| ACE2 | PGK1 | 0,7201 | 6,22718E-59 |
| ACE2 | RABGAP1 | 0,72 | 6,49078E-59 |
| ACE2 | ARHGAP12 | 0,72 | 6,29308E-59 |
| ACE2 | COX7B | 0,72 | 6,56035E-59 |
| ACE2 | TBC1D9 | 0,7199 | 6,79908E-59 |
| ACE2 | SLC30A6 | 0,7198 | 7,24167E-59 |
| ACE2 | SNHG16 | 0,7198 | 7,30099E-59 |
| ACE2 | UBXN2A | 0,7197 | 7,48556E-59 |
| ACE2 | RIOX2 | 0,7197 | 7,58996E-59 |
| ACE2 | CASD1 | 0,7197 | 7,62772E-59 |
| ACE2 | MECP2 | 0,7196 | 7,94647E-59 |
| ACE2 | EBLN3P | 0,7195 | 8,2483E-59 |
| ACE2 | CSNK1A1 | 0,7194 | 8,69962E-59 |
| ACE2 | CAT | 0,7194 | 9,10973E-59 |
| ACE2 | MR1 | 0,7193 | 9,31227E-59 |
| ACE2 | GLUD1 | 0,7192 | 9,79525E-59 |
| ACE2 | AS3MT | 0,7191 | 1,04348E-58 |
| ACE2 | GSTA2 | 0,719 | 1,09464E-58 |
| ACE2 | CTBP2 | 0,7186 | 1,35974E-58 |
| ACE2 | FRG1BP | 0,7186 | 1,37561E-58 |
| ACE2 | LRRC42 | 0,7185 | 1,464E-58 |
| ACE2 | SNX4 | 0,7184 | 1,55494E-58 |
| ACE2 | CD81 | 0,7184 | 1,4796E-58 |
| ACE2 | DNAJC19 | 0,7183 | 1,55887E-58 |
| ACE2 | MRPL50 | 0,7183 | 1,63956E-58 |
| ACE2 | SLC33A1 | 0,7182 | 1,68766E-58 |
| ACE2 | ARAP2 | 0,7181 | 1,74937E-58 |
| ACE2 | RPS4Y1 | 0,718 | 1,89819E-58 |
| ACE2 | DNASE2 | 0,7179 | 2,01914E-58 |
| ACE2 | DKK3 | 0,7178 | 2,1224E-58 |
| ACE2 | FGD5-AS1 | 0,7176 | 2,28423E-58 |

| | | | |
|---|---|---|---|
| ACE2 | TGS1 | 0,7176 | 2,3208E-58 |
| ACE2 | CHAMP1 | 0,7176 | 2,30003E-58 |
| ACE2 | ACTR10 | 0,7174 | 2,59714E-58 |
| ACE2 | ATIC | 0,7173 | 2,78157E-58 |
| ACE2 | MRPL16 | 0,7173 | 2,75505E-58 |
| ACE2 | CHST9 | 0,7173 | 2,68342E-58 |
| ACE2 | FATE1 | 0,7173 | 2,73919E-58 |
| ACE2 | RMND5A | 0,7171 | 3,10092E-58 |
| ACE2 | DEFB119 | 0,7171 | 2,99655E-58 |
| ACE2 | PPIG | 0,7169 | 3,31762E-58 |
| ACE2 | COMMD2 | 0,7168 | 3,52538E-58 |
| ACE2 | TMEM41A | 0,7165 | 4,20108E-58 |
| ACE2 | NET1 | 0,7164 | 4,46307E-58 |
| ACE2 | RNASE4 | 0,7164 | 4,35289E-58 |
| ACE2 | TAF11 | 0,7162 | 4,83193E-58 |
| ACE2 | UST | 0,7162 | 4,8504E-58 |
| ACE2 | PPP2CB | 0,7162 | 4,85988E-58 |
| ACE2 | UBE4B | 0,7161 | 5,24456E-58 |
| ACE2 | MRS2 | 0,7161 | 5,0241E-58 |
| ACE2 | ALDH5A1 | 0,716 | 5,56283E-58 |
| ACE2 | LGALS3 | 0,716 | 5,54019E-58 |
| ACE2 | SMIM10L1 | 0,7159 | 5,76299E-58 |
| ACE2 | BCLAF1 | 0,7158 | 6,10164E-58 |
| ACE2 | PTK2 | 0,7158 | 6,17228E-58 |
| ACE2 | ACO2 | 0,7158 | 5,87714E-58 |
| ACE2 | TSR2 | 0,7158 | 6,03186E-58 |
| ACE2 | HIVEP1 | 0,7156 | 6,63054E-58 |
| ACE2 | GRB2 | 0,7155 | 7,21457E-58 |
| ACE2 | PHLDA1 | 0,7154 | 7,40093E-58 |
| ACE2 | SCYL2 | 0,7154 | 7,4753E-58 |
| ACE2 | REPS2 | 0,7153 | 7,76014E-58 |
| ACE2 | RECQL | 0,715 | 9,0088E-58 |
| ACE2 | KIAA1191 | 0,7149 | 9,79996E-58 |
| ACE2 | GPALPP1 | 0,7148 | 9,96921E-58 |
| ACE2 | CNBP | 0,7147 | 1,04964E-57 |
| ACE2 | UQCRC2 | 0,7145 | 1,19662E-57 |
| ACE2 | EHHADH | 0,7143 | 1,32164E-57 |
| ACE2 | SLC9A6 | 0,7142 | 1,37472E-57 |
| ACE2 | CLASP2 | 0,7141 | 1,48691E-57 |
| ACE2 | ASS1 | 0,7141 | 1,4849E-57 |
| ACE2 | CCDC115 | 0,714 | 1,54484E-57 |
| ACE2 | ALDH1B1 | 0,714 | 1,58489E-57 |
| ACE2 | SUSD3 | 0,7139 | 1,67566E-57 |
| ACE2 | ECHS1 | 0,7139 | 1,60781E-57 |
| ACE2 | RANBP3L | 0,7138 | 1,72702E-57 |
| ACE2 | SPCS2 | 0,7138 | 1,68544E-57 |
| ACE2 | DPH6 | 0,7138 | 1,70983E-57 |
| ACE2 | C11orf58 | 0,7135 | 2,04783E-57 |
| ACE2 | PIK3R1 | 0,7134 | 2,14149E-57 |
| ACE2 | ZC3HAV1 | 0,7134 | 2,10484E-57 |
| ACE2 | NPC2 | 0,7134 | 2,17757E-57 |
| ACE2 | ABHD4 | 0,7132 | 2,38766E-57 |
| ACE2 | ICK | 0,7131 | 2,49599E-57 |
| ACE2 | BCL2L13 | 0,7131 | 2,45219E-57 |

| | | | |
|---|---|---|---|
| ACE2 | APOBEC3G | 0,7131 | 2,52207E-57 |
| ACE2 | SEC31A | 0,7129 | 2,73909E-57 |
| ACE2 | SLC30A7 | 0,7128 | 2,89802E-57 |
| ACE2 | ZFAND1 | 0,7128 | 2,84283E-57 |
| ACE2 | COL1A2 | 0,7127 | 3,12396E-57 |
| ACE2 | IFNGR1 | 0,7126 | 3,23012E-57 |
| ACE2 | UQCRH | 0,7125 | 3,33904E-57 |
| ACE2 | ABCA8 | 0,7125 | 3,37119E-57 |
| ACE2 | DET1 | 0,7124 | 3,53062E-57 |
| ACE2 | SS18 | 0,7123 | 3,84267E-57 |
| ACE2 | ABCG2 | 0,7122 | 3,89955E-57 |
| ACE2 | NANP | 0,7121 | 4,21425E-57 |
| ACE2 | ILDR2 | 0,712 | 4,36918E-57 |
| ACE2 | NOP14 | 0,7119 | 4,71712E-57 |
| ACE2 | ZNF770 | 0,7119 | 4,74402E-57 |
| ACE2 | VASN | 0,7119 | 4,65966E-57 |
| ACE2 | MAP3K20 | 0,7118 | 4,86673E-57 |
| ACE2 | LRP6 | 0,7118 | 4,88159E-57 |
| ACE2 | BHLHB9 | 0,7118 | 4,98071E-57 |
| ACE2 | CUL4B | 0,7118 | 4,88069E-57 |
| ACE2 | PERP | 0,7117 | 5,1309E-57 |
| ACE2 | RAB30 | 0,7116 | 5,33773E-57 |
| ACE2 | GCH1 | 0,7115 | 5,82974E-57 |
| ACE2 | NDN | 0,7115 | 5,8454E-57 |
| ACE2 | PNRC2 | 0,7114 | 6,01811E-57 |
| ACE2 | METTL9 | 0,7113 | 6,49197E-57 |
| ACE2 | SMYD2 | 0,7111 | 6,87984E-57 |
| ACE2 | UBE2D1 | 0,7108 | 8,09037E-57 |
| ACE2 | CD58 | 0,7107 | 8,51335E-57 |
| ACE2 | B4GALT4 | 0,7106 | 9,33086E-57 |
| ACE2 | ZNF322 | 0,7106 | 9,22406E-57 |
| ACE2 | ARHGEF12 | 0,7106 | 8,93113E-57 |
| ACE2 | TMEM245 | 0,7104 | 1,01931E-56 |
| ACE2 | DHH | 0,7104 | 9,9236E-57 |
| ACE2 | NEDD9 | 0,7102 | 1,10247E-56 |
| ACE2 | DUBR | 0,7101 | 1,15622E-56 |
| ACE2 | SLC25A12 | 0,71 | 1,25438E-56 |
| ACE2 | LIPA | 0,7096 | 1,53996E-56 |
| ACE2 | TCF7L2 | 0,7096 | 1,54797E-56 |
| ACE2 | NARS | 0,7096 | 1,52946E-56 |
| ACE2 | NAA25 | 0,7095 | 1,62745E-56 |
| ACE2 | DDX6 | 0,7094 | 1,7292E-56 |
| ACE2 | STARD13 | 0,7092 | 1,84899E-56 |
| ACE2 | RIT1 | 0,709 | 2,12094E-56 |
| ACE2 | TATDN1 | 0,709 | 2,10302E-56 |
| ACE2 | SLC46A3 | 0,709 | 2,07436E-56 |
| ACE2 | MAP3K2 | 0,7089 | 2,19339E-56 |
| ACE2 | CAMK1 | 0,7089 | 2,20134E-56 |
| ACE2 | PCDHGA3 | 0,7089 | 2,18532E-56 |
| ACE2 | TPM4 | 0,7088 | 2,2727E-56 |
| ACE2 | HPF1 | 0,7086 | 2,61872E-56 |
| ACE2 | NCOA7 | 0,7086 | 2,5653E-56 |
| ACE2 | SLC25A13 | 0,7086 | 2,50655E-56 |
| ACE2 | PPP2R1A | 0,7086 | 2,5017E-56 |

| | | | |
|---|---|---|---|
| ACE2 | TRAM2-AS1 | 0,7085 | 2,74457E-56 |
| ACE2 | G6PD | 0,7085 | 2,70074E-56 |
| ACE2 | RAP1A | 0,7084 | 2,79012E-56 |
| ACE2 | DNAJC12 | 0,7084 | 2,83178E-56 |
| ACE2 | ZDHHC17 | 0,7083 | 2,9644E-56 |
| ACE2 | TRIB2 | 0,7082 | 3,06632E-56 |
| ACE2 | HNRNPA0 | 0,7082 | 3,15881E-56 |
| ACE2 | BCL2L2 | 0,7082 | 3,20755E-56 |
| ACE2 | YTHDF3 | 0,7081 | 3,341E-56 |
| ACE2 | GRN | 0,7081 | 3,2434E-56 |
| ACE2 | NOP10 | 0,708 | 3,43135E-56 |
| ACE2 | CLCC1 | 0,7079 | 3,63546E-56 |
| ACE2 | LCLAT1 | 0,7079 | 3,61798E-56 |
| ACE2 | ARHGAP20 | 0,7079 | 3,63384E-56 |
| ACE2 | HDAC8 | 0,7079 | 3,67615E-56 |
| ACE2 | RPS4X | 0,7078 | 3,88014E-56 |
| ACE2 | NCL | 0,7076 | 4,25566E-56 |
| ACE2 | COPG1 | 0,7076 | 4,24859E-56 |
| ACE2 | GLB1 | 0,7074 | 4,66593E-56 |
| ACE2 | ALG13 | 0,7074 | 4,76817E-56 |
| ACE2 | TRAM1L1 | 0,7073 | 5,08244E-56 |
| ACE2 | RAF1 | 0,7071 | 5,46152E-56 |
| ACE2 | PDS5A | 0,7068 | 6,51861E-56 |
| ACE2 | EIF2AK3 | 0,7067 | 6,82425E-56 |
| ACE2 | COPA | 0,7066 | 7,09162E-56 |
| ACE2 | PPP1R15B | 0,7065 | 7,57018E-56 |
| ACE2 | CCDC69 | 0,7065 | 7,39779E-56 |
| ACE2 | PPFIBP1 | 0,7065 | 7,62691E-56 |
| ACE2 | ITSN2 | 0,7064 | 7,83369E-56 |
| ACE2 | COPB2 | 0,7064 | 7,96891E-56 |
| ACE2 | NFIC | 0,7064 | 7,85289E-56 |
| ACE2 | ZNF776 | 0,7064 | 8,01294E-56 |
| ACE2 | ZMYND11 | 0,7063 | 8,11918E-56 |
| ACE2 | SPRYD4 | 0,7063 | 8,41649E-56 |
| ACE2 | PEX26 | 0,7063 | 8,41476E-56 |
| ACE2 | MERTK | 0,7062 | 8,87382E-56 |
| ACE2 | FAM110B | 0,7061 | 8,94746E-56 |
| ACE2 | CD300LG | 0,7061 | 9,24367E-56 |
| ACE2 | DEK | 0,706 | 9,81273E-56 |
| ACE2 | TMEM229A | 0,706 | 9,48807E-56 |
| ACE2 | OAT | 0,706 | 9,75687E-56 |
| ACE2 | CPQ | 0,7058 | 1,05173E-55 |
| ACE2 | ALG11 | 0,7057 | 1,09115E-55 |
| ACE2 | CADM1 | 0,7056 | 1,15764E-55 |
| ACE2 | GTDC1 | 0,7055 | 1,22554E-55 |
| ACE2 | CDC42EP4 | 0,7055 | 1,25339E-55 |
| ACE2 | TP53 | 0,7054 | 1,28202E-55 |
| ACE2 | DCAF12L1 | 0,7054 | 1,31863E-55 |
| ACE2 | TMSB10 | 0,7053 | 1,37595E-55 |
| ACE2 | DNMT3A | 0,7052 | 1,42152E-55 |
| ACE2 | PRKAB2 | 0,7051 | 1,53045E-55 |
| ACE2 | RAB9A | 0,7051 | 1,52193E-55 |
| ACE2 | NUP62CL | 0,7051 | 1,47914E-55 |
| ACE2 | MAPKAPK5 | 0,705 | 1,62787E-55 |

| | | | |
|---|---|---|---|
| ACE2 | ZNF45 | 0,705 | 1,58678E-55 |
| ACE2 | GFM2 | 0,7049 | 1,68784E-55 |
| ACE2 | SOGA1 | 0,7049 | 1,66839E-55 |
| ACE2 | VPS13C | 0,7047 | 1,84215E-55 |
| ACE2 | KCTD5 | 0,7047 | 1,84347E-55 |
| ACE2 | LYRM2 | 0,7046 | 1,91706E-55 |
| ACE2 | ZNF536 | 0,7045 | 2,03285E-55 |
| ACE2 | FAM102A | 0,7044 | 2,18826E-55 |
| ACE2 | TCEAL2 | 0,7044 | 2,19777E-55 |
| ACE2 | DCAF17 | 0,7043 | 2,27494E-55 |
| ACE2 | PHIP | 0,7041 | 2,47633E-55 |
| ACE2 | SMARCAD1 | 0,704 | 2,64319E-55 |
| ACE2 | PDCL | 0,704 | 2,7018E-55 |
| ACE2 | BMPR2 | 0,7039 | 2,72647E-55 |
| ACE2 | RAB5A | 0,7039 | 2,76469E-55 |
| ACE2 | PWWP2A | 0,7038 | 2,96019E-55 |
| ACE2 | FXYD6 | 0,7037 | 2,9896E-55 |
| ACE2 | TCEAL6 | 0,7037 | 3,07315E-55 |
| ACE2 | PDLIM5 | 0,7036 | 3,22774E-55 |
| ACE2 | CITED2 | 0,7036 | 3,24923E-55 |
| ACE2 | PPP1R3D | 0,7034 | 3,57729E-55 |
| ACE2 | TBX22 | 0,7033 | 3,78968E-55 |
| ACE2 | DMXL1 | 0,703 | 4,43347E-55 |
| ACE2 | IRS2 | 0,703 | 4,41708E-55 |
| ACE2 | USP22 | 0,703 | 4,32663E-55 |
| ACE2 | FAM210B | 0,703 | 4,25555E-55 |
| ACE2 | BLVRA | 0,7028 | 4,76997E-55 |
| ACE2 | BFAR | 0,7028 | 4,69301E-55 |
| ACE2 | PQLC3 | 0,7026 | 5,42474E-55 |
| ACE2 | NDUFA10 | 0,7026 | 5,27719E-55 |
| ACE2 | ACOT9 | 0,7026 | 5,20402E-55 |
| ACE2 | RHOA | 0,7025 | 5,61235E-55 |
| ACE2 | PHF24 | 0,7025 | 5,54264E-55 |
| ACE2 | FIG4 | 0,7024 | 5,95888E-55 |
| ACE2 | CNDP2 | 0,7024 | 5,99354E-55 |
| ACE2 | DOCK7 | 0,7023 | 6,29141E-55 |
| ACE2 | HEATR5B | 0,7023 | 6,30354E-55 |
| ACE2 | UTP20 | 0,7022 | 6,39477E-55 |
| ACE2 | TRIM26 | 0,7021 | 6,94338E-55 |
| ACE2 | MOB4 | 0,702 | 7,31104E-55 |
| ACE2 | DHRS7 | 0,702 | 7,33393E-55 |
| ACE2 | ANKRD13C | 0,7019 | 7,55812E-55 |
| ACE2 | ARF6 | 0,7017 | 8,45712E-55 |
| ACE2 | ABRACL | 0,7016 | 8,63919E-55 |
| ACE2 | KIF16B | 0,7016 | 8,58959E-55 |
| ACE2 | SERPINI1 | 0,7013 | 1,01145E-54 |
| ACE2 | AP001258.4 | 0,7013 | 1,00713E-54 |
| ACE2 | E2F6 | 0,7012 | 1,05563E-54 |
| ACE2 | FAM136A | 0,7012 | 1,07551E-54 |
| ACE2 | TEC | 0,7012 | 1,05206E-54 |
| ACE2 | CBR1 | 0,7012 | 1,04782E-54 |
| ACE2 | DCTN1 | 0,701 | 1,16327E-54 |
| ACE2 | FNBP1 | 0,701 | 1,1529E-54 |
| ACE2 | ZNF207 | 0,701 | 1,15197E-54 |

| | | | |
|---|---|---|---|
| ACE2 | GOLGA7 | 0,7009 | 1,2207E-54 |
| ACE2 | MED20 | 0,7008 | 1,3338E-54 |
| ACE2 | EPS8 | 0,7007 | 1,36594E-54 |
| ACE2 | DYRK2 | 0,7007 | 1,35082E-54 |
| ACE2 | PHACTR4 | 0,7006 | 1,40426E-54 |
| ACE2 | PPIP5K2 | 0,7006 | 1,41755E-54 |
| ACE2 | AGFG2 | 0,7005 | 1,50545E-54 |
| ACE2 | UBA3 | 0,7004 | 1,60383E-54 |
| ACE2 | CAND1 | 0,7004 | 1,60547E-54 |
| ACE2 | OSBPL9 | 0,7002 | 1,75914E-54 |
| ACE2 | COX8A | 0,7 | 1,88769E-54 |
| ACE2 | SELENOI | 0,6999 | 2,07636E-54 |
| ACE2 | DLGAP4 | 0,6999 | 2,03462E-54 |
| ACE2 | GNPDA2 | 0,6998 | 2,10653E-54 |
| ACE2 | USP47 | 0,6998 | 2,09377E-54 |
| ACE2 | PROS1 | 0,6997 | 2,26515E-54 |
| ACE2 | RPAP3 | 0,6997 | 2,23266E-54 |
| ACE2 | PPP2R2A | 0,6996 | 2,35728E-54 |
| ACE2 | PLXNC1 | 0,6995 | 2,525E-54 |
| ACE2 | PLEKHA2 | 0,6994 | 2,56548E-54 |
| ACE2 | PDIA6 | 0,6991 | 3,08609E-54 |
| ACE2 | OSGEPL1 | 0,6991 | 3,00683E-54 |
| ACE2 | TAF7L | 0,6991 | 2,96067E-54 |
| ACE2 | SLC35E1 | 0,699 | 3,18054E-54 |
| ACE2 | MFN1 | 0,6989 | 3,25889E-54 |
| ACE2 | KRR1 | 0,6989 | 3,31645E-54 |
| ACE2 | MVP | 0,6989 | 3,34535E-54 |
| ACE2 | MARC2 | 0,6988 | 3,45088E-54 |
| ACE2 | KRAS | 0,6988 | 3,51052E-54 |
| ACE2 | PHKA1 | 0,6988 | 3,56966E-54 |
| ACE2 | METTL2B | 0,6987 | 3,69729E-54 |
| ACE2 | DIRC2 | 0,6985 | 4,11627E-54 |
| ACE2 | C5orf24 | 0,6985 | 4,07724E-54 |
| ACE2 | RFC5 | 0,6984 | 4,18312E-54 |
| ACE2 | LMO7 | 0,6983 | 4,49901E-54 |
| ACE2 | HECTD1 | 0,6983 | 4,46896E-54 |
| ACE2 | APEX1 | 0,6982 | 4,67477E-54 |
| ACE2 | PUM3 | 0,6981 | 4,81733E-54 |
| ACE2 | LMO4 | 0,6979 | 5,38851E-54 |
| ACE2 | GPC5 | 0,6978 | 5,72667E-54 |
| ACE2 | STAU2 | 0,6977 | 6,04906E-54 |
| ACE2 | CLYBL | 0,6975 | 6,66689E-54 |
| ACE2 | UTP14C | 0,6974 | 6,79681E-54 |
| ACE2 | ZNF626 | 0,6974 | 6,97337E-54 |
| ACE2 | MTF2 | 0,6973 | 7,30757E-54 |
| ACE2 | C5orf22 | 0,697 | 8,39848E-54 |
| ACE2 | CDC23 | 0,697 | 8,40931E-54 |
| ACE2 | NDUFAF4 | 0,697 | 8,55921E-54 |
| ACE2 | FBXO9 | 0,6969 | 8,78608E-54 |
| ACE2 | FYN | 0,6969 | 8,80337E-54 |
| ACE2 | TUBA1C | 0,6968 | 9,41623E-54 |
| ACE2 | ARF3 | 0,6967 | 9,82251E-54 |
| ACE2 | PPIC | 0,6966 | 1,03063E-53 |
| ACE2 | TMEM168 | 0,6965 | 1,06122E-53 |

| | | | |
|---|---|---|---|
| ACE2 | NFE2L2 | 0,6964 | 1,1132E-53 |
| ACE2 | RICTOR | 0,6963 | 1,20331E-53 |
| ACE2 | TMEM64 | 0,6963 | 1,16181E-53 |
| ACE2 | ZZZ3 | 0,6962 | 1,27663E-53 |
| ACE2 | FAM120AOS | 0,6961 | 1,28697E-53 |
| ACE2 | FITM2 | 0,6961 | 1,3403E-53 |
| ACE2 | ANGEL2 | 0,696 | 1,34834E-53 |
| ACE2 | RABGGTB | 0,6958 | 1,5454E-53 |
| ACE2 | CDYL | 0,6958 | 1,52279E-53 |
| ACE2 | IKBKG | 0,6958 | 1,53949E-53 |
| ACE2 | LRRC47 | 0,6957 | 1,58298E-53 |
| ACE2 | SYNPO2 | 0,6957 | 1,59331E-53 |
| ACE2 | ANKRD40 | 0,6956 | 1,67938E-53 |
| ACE2 | ROCK2 | 0,6955 | 1,77634E-53 |
| ACE2 | DIAPH2 | 0,6955 | 1,75291E-53 |
| ACE2 | ATP6V1G1 | 0,6954 | 1,82119E-53 |
| ACE2 | USP12 | 0,6954 | 1,81171E-53 |
| ACE2 | SFPQ | 0,6953 | 1,89844E-53 |
| ACE2 | PLCD1 | 0,6953 | 1,92797E-53 |
| ACE2 | COMMD10 | 0,6953 | 1,92489E-53 |
| ACE2 | RAPGEF4 | 0,6952 | 2,03886E-53 |
| ACE2 | IGSF1 | 0,6952 | 2,06766E-53 |
| ACE2 | RBMX | 0,6952 | 2,03234E-53 |
| ACE2 | FTO | 0,6951 | 2,16159E-53 |
| ACE2 | MIER3 | 0,695 | 2,20317E-53 |
| ACE2 | GALNT11 | 0,6948 | 2,43869E-53 |
| ACE2 | CCDC122 | 0,6948 | 2,44085E-53 |
| ACE2 | RB1 | 0,6948 | 2,47639E-53 |
| ACE2 | HIGD1A | 0,6947 | 2,53173E-53 |
| ACE2 | SUSD6 | 0,6947 | 2,63801E-53 |
| ACE2 | PRMT1 | 0,6946 | 2,7467E-53 |
| ACE2 | IFIT5 | 0,6945 | 2,85526E-53 |
| ACE2 | RP11-705C15.3 | 0,6945 | 2,90153E-53 |
| ACE2 | LMCD1 | 0,6942 | 3,25805E-53 |
| ACE2 | JADE1 | 0,6942 | 3,22387E-53 |
| ACE2 | FAM214A | 0,6942 | 3,33509E-53 |
| ACE2 | REL | 0,6941 | 3,36604E-53 |
| ACE2 | SETMAR | 0,694 | 3,54976E-53 |
| ACE2 | BNIP3L | 0,694 | 3,69512E-53 |
| ACE2 | PSMA3 | 0,694 | 3,54538E-53 |
| ACE2 | HECTD2 | 0,6938 | 3,95108E-53 |
| ACE2 | PRKX | 0,6938 | 3,95459E-53 |
| ACE2 | EXOC8 | 0,6937 | 4,12085E-53 |
| ACE2 | GREB1 | 0,6937 | 4,26621E-53 |
| ACE2 | YAF2 | 0,6937 | 4,27615E-53 |
| ACE2 | MMD | 0,6936 | 4,44034E-53 |
| ACE2 | UBA1 | 0,6935 | 4,60609E-53 |
| ACE2 | PRPS2 | 0,6934 | 4,77728E-53 |
| ACE2 | SLC39A1 | 0,6933 | 4,94675E-53 |
| ACE2 | ANKS1A | 0,6933 | 4,97872E-53 |
| ACE2 | GLMP | 0,6932 | 5,36467E-53 |
| ACE2 | ZBTB41 | 0,6932 | 5,35401E-53 |
| ACE2 | POLA1 | 0,6932 | 5,40875E-53 |
| ACE2 | DHX15 | 0,6931 | 5,52068E-53 |

| | | | |
|---|---|---|---|
| ACE2 | H2AFY2 | 0,6929 | 6,10286E-53 |
| ACE2 | RP11-434D9.1 | 0,6928 | 6,4757E-53 |
| ACE2 | PPP1R21 | 0,6927 | 6,61609E-53 |
| ACE2 | ESYT1 | 0,6927 | 6,80619E-53 |
| ACE2 | RASA1 | 0,6926 | 7,17604E-53 |
| ACE2 | HNRNPC | 0,6926 | 7,07711E-53 |
| ACE2 | FBXW11 | 0,6925 | 7,6012E-53 |
| ACE2 | RP11-25K19.1 | 0,6923 | 8,10403E-53 |
| ACE2 | ADGRG1 | 0,6923 | 8,06901E-53 |
| ACE2 | LSR | 0,6922 | 8,49859E-53 |
| ACE2 | GPRASP2 | 0,6922 | 8,69068E-53 |
| ACE2 | ELMOD2 | 0,6921 | 9,00942E-53 |
| ACE2 | CLCN2 | 0,692 | 9,38201E-53 |
| ACE2 | RP11-573G6.4 | 0,692 | 9,65559E-53 |
| ACE2 | ACAD8 | 0,6919 | 9,97411E-53 |
| ACE2 | UGGT1 | 0,6918 | 1,04631E-52 |
| ACE2 | TRAF6 | 0,6918 | 1,02539E-52 |
| ACE2 | PRDM2 | 0,6917 | 1,0749E-52 |
| ACE2 | MTCH1 | 0,6917 | 1,09975E-52 |
| ACE2 | SCO1 | 0,6915 | 1,18263E-52 |
| ACE2 | IL1R1 | 0,6914 | 1,25834E-52 |
| ACE2 | TKT | 0,6914 | 1,23152E-52 |
| ACE2 | NUP54 | 0,6914 | 1,25545E-52 |
| ACE2 | MINPP1 | 0,6914 | 1,23009E-52 |
| ACE2 | SYTL4 | 0,6914 | 1,26689E-52 |
| ACE2 | RNF145 | 0,6913 | 1,30718E-52 |
| ACE2 | RNF170 | 0,6913 | 1,3474E-52 |
| ACE2 | VAPB | 0,6913 | 1,30256E-52 |
| ACE2 | PEX2 | 0,6912 | 1,39614E-52 |
| ACE2 | SLC9A7P1 | 0,6912 | 1,39759E-52 |
| ACE2 | SRXN1 | 0,6912 | 1,40947E-52 |
| ACE2 | CCNI | 0,6911 | 1,47172E-52 |
| ACE2 | ANP32E | 0,6909 | 1,58934E-52 |
| ACE2 | CRCP | 0,6909 | 1,61543E-52 |
| ACE2 | SVIL | 0,6909 | 1,61621E-52 |
| ACE2 | FADS1 | 0,6908 | 1,69886E-52 |
| ACE2 | FCF1 | 0,6908 | 1,64262E-52 |
| ACE2 | TRIM16 | 0,6908 | 1,65547E-52 |
| ACE2 | ANTXR2 | 0,6907 | 1,72167E-52 |
| ACE2 | ATXN10 | 0,6907 | 1,78725E-52 |
| ACE2 | ACP2 | 0,6905 | 1,89819E-52 |
| ACE2 | NOB1 | 0,6905 | 1,90848E-52 |
| ACE2 | STAG1 | 0,6904 | 2,05477E-52 |
| ACE2 | ADCY9 | 0,6904 | 2,06613E-52 |
| ACE2 | MASTL | 0,6902 | 2,2171E-52 |
| ACE2 | DDX59 | 0,69 | 2,47083E-52 |
| ACE2 | ICE2 | 0,69 | 2,49118E-52 |
| ACE2 | TCAF1 | 0,6898 | 2,7217E-52 |
| ACE2 | RIF1 | 0,6897 | 2,8254E-52 |
| ACE2 | CNP | 0,6897 | 2,88761E-52 |
| ACE2 | RNASE1 | 0,6896 | 2,91593E-52 |
| ACE2 | DAAM2 | 0,6895 | 3,13002E-52 |
| ACE2 | TAF4B | 0,6894 | 3,27264E-52 |
| ACE2 | PBDC1 | 0,6893 | 3,44998E-52 |

| | | | |
|---|---|---|---|
| ACE2 | STXBP3 | 0,6892 | 3,62281E-52 |
| ACE2 | CLN8 | 0,6891 | 3,77386E-52 |
| ACE2 | PTPRE | 0,6891 | 3,83884E-52 |
| ACE2 | ACADSB | 0,689 | 3,89174E-52 |
| ACE2 | ZNF644 | 0,6888 | 4,37096E-52 |
| ACE2 | PLOD2 | 0,6888 | 4,24488E-52 |
| ACE2 | HM13 | 0,6887 | 4,49541E-52 |
| ACE2 | STK10 | 0,6886 | 4,70618E-52 |
| ACE2 | SMNDC1 | 0,6886 | 4,68906E-52 |
| ACE2 | SH3D19 | 0,6883 | 5,41815E-52 |
| ACE2 | TERF2IP | 0,6883 | 5,48706E-52 |
| ACE2 | LRP8 | 0,688 | 6,44046E-52 |
| ACE2 | NFKB1 | 0,688 | 6,2729E-52 |
| ACE2 | R3HDM2 | 0,688 | 6,26988E-52 |
| ACE2 | HEATR6 | 0,6878 | 6,9903E-52 |
| ACE2 | SNX5 | 0,6878 | 6,99521E-52 |
| ACE2 | CALD1 | 0,6877 | 7,38063E-52 |
| ACE2 | ELP3 | 0,6877 | 7,30846E-52 |
| ACE2 | HNRNPDL | 0,6876 | 7,49671E-52 |
| ACE2 | IKZF5 | 0,6876 | 7,57785E-52 |
| ACE2 | STX6 | 0,6875 | 8,08336E-52 |
| ACE2 | CD99L2 | 0,6874 | 8,19623E-52 |
| ACE2 | SPOCK2 | 0,6873 | 8,75733E-52 |
| ACE2 | MARCH6 | 0,6872 | 9,34664E-52 |
| ACE2 | ZNF641 | 0,6872 | 9,10356E-52 |
| ACE2 | CEPT1 | 0,6871 | 9,44304E-52 |
| ACE2 | COPS2 | 0,6871 | 9,71544E-52 |
| ACE2 | DLST | 0,687 | 1,00246E-51 |
| ACE2 | SUPT6H | 0,687 | 1,00448E-51 |
| ACE2 | MAGED2 | 0,6869 | 1,07162E-51 |
| ACE2 | SMOC1 | 0,6867 | 1,14033E-51 |
| ACE2 | TMEM69 | 0,6866 | 1,22655E-51 |
| ACE2 | FAM114A1 | 0,6866 | 1,20843E-51 |
| ACE2 | ATE1 | 0,6865 | 1,26773E-51 |
| ACE2 | AVL9 | 0,6864 | 1,32209E-51 |
| ACE2 | ZYX | 0,6864 | 1,31793E-51 |
| ACE2 | PARP4 | 0,686 | 1,62737E-51 |
| ACE2 | EXT1 | 0,6859 | 1,65703E-51 |
| ACE2 | GPR107 | 0,6859 | 1,72111E-51 |
| ACE2 | TRPM7 | 0,6859 | 1,65465E-51 |
| ACE2 | MBOAT7 | 0,6859 | 1,65744E-51 |
| ACE2 | FNBP1L | 0,6856 | 1,90355E-51 |
| ACE2 | ABHD10 | 0,6855 | 2,01057E-51 |
| ACE2 | LSM1 | 0,6854 | 2,09483E-51 |
| ACE2 | MED12 | 0,6854 | 2,15948E-51 |
| ACE2 | KAT6A | 0,6852 | 2,36993E-51 |
| ACE2 | TXLNG | 0,6851 | 2,43392E-51 |
| ACE2 | ZFYVE16 | 0,685 | 2,6122E-51 |
| ACE2 | VPS39 | 0,685 | 2,59258E-51 |
| ACE2 | LNPEP | 0,6849 | 2,68788E-51 |
| ACE2 | PKD2 | 0,6846 | 3,05282E-51 |
| ACE2 | NOL11 | 0,6846 | 3,09957E-51 |
| ACE2 | SDF2 | 0,6845 | 3,26736E-51 |
| ACE2 | TMEM9B | 0,6844 | 3,45674E-51 |

| | | | |
|---|---|---|---|
| ACE2 | IGSF8 | 0,6843 | 3,55339E-51 |
| ACE2 | NFIA | 0,6842 | 3,78735E-51 |
| ACE2 | NMI | 0,6842 | 3,70627E-51 |
| ACE2 | GOLIM4 | 0,6842 | 3,69171E-51 |
| ACE2 | METTL5 | 0,6841 | 3,96586E-51 |
| ACE2 | RGAG4 | 0,6841 | 3,90646E-51 |
| ACE2 | ACAP2 | 0,684 | 4,12536E-51 |
| ACE2 | DDX39B | 0,684 | 4,16292E-51 |
| ACE2 | RALBP1 | 0,684 | 3,98175E-51 |
| ACE2 | C1orf109 | 0,6839 | 4,18528E-51 |
| ACE2 | SEC63 | 0,6839 | 4,30443E-51 |
| ACE2 | ENPP4 | 0,6837 | 4,74521E-51 |
| ACE2 | C8orf33 | 0,6836 | 4,8393E-51 |
| ACE2 | BROX | 0,6834 | 5,26697E-51 |
| ACE2 | PTCD2 | 0,6834 | 5,32284E-51 |
| ACE2 | PPIA | 0,6831 | 6,22167E-51 |
| ACE2 | EPAS1 | 0,683 | 6,59797E-51 |
| ACE2 | LINC00888 | 0,683 | 6,58215E-51 |
| ACE2 | TMCO3 | 0,683 | 6,48853E-51 |
| ACE2 | RORA | 0,683 | 6,33074E-51 |
| ACE2 | RPL39 | 0,683 | 6,52964E-51 |
| ACE2 | PBX2 | 0,6828 | 7,09432E-51 |
| ACE2 | SPTLC2 | 0,6828 | 7,20131E-51 |
| ACE2 | COPS7A | 0,6827 | 7,39928E-51 |
| ACE2 | TMEM39A | 0,6825 | 8,00735E-51 |
| ACE2 | PLEKHH1 | 0,6825 | 8,06168E-51 |
| ACE2 | ATF6B | 0,6824 | 8,53415E-51 |
| ACE2 | KIAA0141 | 0,6823 | 8,81433E-51 |
| ACE2 | ATXN2 | 0,6822 | 9,31094E-51 |
| ACE2 | SON | 0,6822 | 9,32631E-51 |
| ACE2 | LINC00998 | 0,6821 | 9,98331E-51 |
| ACE2 | ESD | 0,6821 | 9,98321E-51 |
| ACE2 | TMEM136 | 0,6819 | 1,05479E-50 |
| ACE2 | TMEM38A | 0,6819 | 1,0629E-50 |
| ACE2 | NDUFB11 | 0,6819 | 1,08046E-50 |
| ACE2 | VPS26A | 0,6818 | 1,13814E-50 |
| ACE2 | HEXIM1 | 0,6818 | 1,11452E-50 |
| ACE2 | PEG3 | 0,6818 | 1,12729E-50 |
| ACE2 | MPHOSPH10 | 0,6817 | 1,19782E-50 |
| ACE2 | ARMCX5 | 0,6817 | 1,1772E-50 |
| ACE2 | ATG5 | 0,6816 | 1,24107E-50 |
| ACE2 | TAGLN2 | 0,6815 | 1,26282E-50 |
| ACE2 | TMEM251 | 0,6815 | 1,29792E-50 |
| ACE2 | ITPRIPL2 | 0,6815 | 1,29856E-50 |
| ACE2 | SRSF9 | 0,6814 | 1,33059E-50 |
| ACE2 | BEX2 | 0,6814 | 1,36174E-50 |
| ACE2 | LARP1 | 0,6813 | 1,42325E-50 |
| ACE2 | GNL3L | 0,6813 | 1,39013E-50 |
| ACE2 | DIEXF | 0,6811 | 1,54079E-50 |
| ACE2 | BDH2 | 0,6811 | 1,56761E-50 |
| ACE2 | FAM110C | 0,681 | 1,58481E-50 |
| ACE2 | ASPH | 0,681 | 1,6265E-50 |
| ACE2 | DENND5A | 0,681 | 1,62501E-50 |
| ACE2 | SPTY2D1 | 0,6809 | 1,72922E-50 |

| | | | |
|---|---|---|---|
| ACE2 | WDR11 | 0,6807 | 1,8273E-50 |
| ACE2 | RAB5C | 0,6805 | 1,99612E-50 |
| ACE2 | CCT8 | 0,6805 | 2,0057E-50 |
| ACE2 | FAM133A | 0,6804 | 2,1061E-50 |
| ACE2 | NR0B1 | 0,6803 | 2,20035E-50 |
| ACE2 | CITED1 | 0,6803 | 2,19861E-50 |
| ACE2 | MFSD5 | 0,6802 | 2,31748E-50 |
| ACE2 | SPOP | 0,6802 | 2,28112E-50 |
| ACE2 | TUBA1B | 0,6801 | 2,40455E-50 |
| ACE2 | ZXDA | 0,6801 | 2,48794E-50 |
| ACE2 | TIMM17A | 0,68 | 2,5542E-50 |
| ACE2 | GLS | 0,6797 | 2,92566E-50 |
| ACE2 | ZNF532 | 0,6797 | 2,91139E-50 |
| ACE2 | EEF1B2 | 0,6796 | 3,12154E-50 |
| ACE2 | HDAC2 | 0,6796 | 3,0394E-50 |
| ACE2 | BIRC3 | 0,6796 | 3,09073E-50 |
| ACE2 | ZNF826P | 0,6796 | 3,04443E-50 |
| ACE2 | ACOT2 | 0,6795 | 3,17545E-50 |
| ACE2 | TTC19 | 0,6795 | 3,19794E-50 |
| ACE2 | PLD3 | 0,6795 | 3,24369E-50 |
| ACE2 | SLC35A2 | 0,6794 | 3,31712E-50 |
| ACE2 | CD151 | 0,6793 | 3,48662E-50 |
| ACE2 | METTL7A | 0,6792 | 3,71086E-50 |
| ACE2 | TMEM35B | 0,6791 | 3,89926E-50 |
| ACE2 | UBE2E1 | 0,6791 | 3,8017E-50 |
| ACE2 | ZFHX3 | 0,6791 | 3,89996E-50 |
| ACE2 | MCMBP | 0,679 | 4,01262E-50 |
| ACE2 | ST3GAL4 | 0,679 | 3,9577E-50 |
| ACE2 | OS9 | 0,679 | 3,94978E-50 |
| ACE2 | SKI | 0,6789 | 4,16396E-50 |
| ACE2 | LINC00847 | 0,6789 | 4,12547E-50 |
| ACE2 | MAN1A2 | 0,6787 | 4,58196E-50 |
| ACE2 | SMPDL3A | 0,6787 | 4,54668E-50 |
| ACE2 | MMP28 | 0,6787 | 4,64533E-50 |
| ACE2 | CLDN12 | 0,6786 | 4,84317E-50 |
| ACE2 | LSM10 | 0,6785 | 4,97953E-50 |
| ACE2 | GRHPR | 0,6785 | 5,08763E-50 |
| ACE2 | ITPR1 | 0,6783 | 5,50398E-50 |
| ACE2 | CNOT9 | 0,6782 | 5,83567E-50 |
| ACE2 | THAP12 | 0,6782 | 5,66325E-50 |
| ACE2 | PGGT1B | 0,6781 | 6,02872E-50 |
| ACE2 | SF3B1 | 0,678 | 6,33436E-50 |
| ACE2 | NFAT5 | 0,678 | 6,25643E-50 |
| ACE2 | SPTBN1 | 0,6779 | 6,54757E-50 |
| ACE2 | SLC35B2 | 0,6779 | 6,55705E-50 |
| ACE2 | WTAP | 0,6779 | 6,70809E-50 |
| ACE2 | MRPS10 | 0,6777 | 7,18175E-50 |
| ACE2 | TFCP2 | 0,6777 | 7,18942E-50 |
| ACE2 | ZC3H7B | 0,6777 | 7,41648E-50 |
| ACE2 | SSBP3 | 0,6776 | 7,61431E-50 |
| ACE2 | U2SURP | 0,6776 | 7,71296E-50 |
| ACE2 | TNRC6A | 0,6775 | 7,81572E-50 |
| ACE2 | TBL1XR1 | 0,6773 | 8,69856E-50 |
| ACE2 | CKAP4 | 0,6773 | 8,77357E-50 |

| | | | |
|---|---|---|---|
| ACE2 | FAM175B | 0,677 | 9,88981E-50 |
| ACE2 | TM2D3 | 0,677 | 9,87854E-50 |
| ACE2 | SCFD2 | 0,6768 | 1,07394E-49 |
| ACE2 | RP11-267M23.1 | 0,6768 | 1,07461E-49 |
| ACE2 | NEK9 | 0,6767 | 1,1641E-49 |
| ACE2 | MICAL2 | 0,6766 | 1,1653E-49 |
| ACE2 | EPM2AIP1 | 0,6764 | 1,30907E-49 |
| ACE2 | F2R | 0,6764 | 1,2915E-49 |
| ACE2 | ALKBH8 | 0,6763 | 1,34601E-49 |
| ACE2 | CACNB4 | 0,6762 | 1,45024E-49 |
| ACE2 | EXOSC6 | 0,6762 | 1,43368E-49 |
| ACE2 | SESTD1 | 0,6761 | 1,47303E-49 |
| ACE2 | STS | 0,6761 | 1,47962E-49 |
| ACE2 | N4BP1 | 0,676 | 1,52553E-49 |
| ACE2 | C17orf58 | 0,676 | 1,58892E-49 |
| ACE2 | ABI2 | 0,6758 | 1,69938E-49 |
| ACE2 | KLF3 | 0,6755 | 1,956E-49 |
| ACE2 | TMEM200B | 0,6754 | 2,0181E-49 |
| ACE2 | CD59 | 0,6754 | 2,06989E-49 |
| ACE2 | TRIO | 0,6752 | 2,27928E-49 |
| ACE2 | MOCS2 | 0,6752 | 2,26811E-49 |
| ACE2 | KYAT3 | 0,6751 | 2,28779E-49 |
| ACE2 | SETD5 | 0,6751 | 2,3626E-49 |
| ACE2 | RUNX1T1 | 0,675 | 2,46842E-49 |
| ACE2 | ARHGAP1 | 0,6749 | 2,51888E-49 |
| ACE2 | SLC38A2 | 0,6749 | 2,5435E-49 |
| ACE2 | SYNE3 | 0,6749 | 2,52671E-49 |
| ACE2 | LTBP1 | 0,6748 | 2,64296E-49 |
| ACE2 | MANBA | 0,6748 | 2,62047E-49 |
| ACE2 | ABCC4 | 0,6748 | 2,62806E-49 |
| ACE2 | RALB | 0,6746 | 2,92913E-49 |
| ACE2 | AKR1B1 | 0,6746 | 2,85879E-49 |
| ACE2 | PYGL | 0,6746 | 2,946E-49 |
| ACE2 | MTMR12 | 0,6745 | 3,0379E-49 |
| ACE2 | INTS13 | 0,6745 | 3,05084E-49 |
| ACE2 | LRCH3 | 0,6744 | 3,20713E-49 |
| ACE2 | WASHC4 | 0,6744 | 3,19371E-49 |
| ACE2 | COQ10B | 0,6743 | 3,2807E-49 |
| ACE2 | TTC21B | 0,6742 | 3,46362E-49 |
| ACE2 | STX7 | 0,6742 | 3,46023E-49 |
| ACE2 | RC3H2 | 0,6742 | 3,54643E-49 |
| ACE2 | PDIA3 | 0,6742 | 3,49405E-49 |
| ACE2 | HARS2 | 0,674 | 3,78629E-49 |
| ACE2 | CBX4 | 0,674 | 3,78366E-49 |
| ACE2 | SELENOF | 0,6739 | 3,99821E-49 |
| ACE2 | ARSK | 0,6739 | 4,05441E-49 |
| ACE2 | NECAB1 | 0,6738 | 4,2075E-49 |
| ACE2 | NAPG | 0,6738 | 4,24317E-49 |
| ACE2 | APOBEC3F | 0,6738 | 4,22666E-49 |
| ACE2 | RNASEK | 0,6737 | 4,26308E-49 |
| ACE2 | YTHDF2 | 0,6736 | 4,6279E-49 |
| ACE2 | CD276 | 0,6734 | 4,9918E-49 |
| ACE2 | RNMT | 0,6734 | 5,02842E-49 |
| ACE2 | PIM2 | 0,6734 | 5,02036E-49 |

| | | | |
|---|---|---|---|
| ACE2 | RP11-541N10.3 | 0,6731 | 5,73978E-49 |
| ACE2 | NIPAL3 | 0,673 | 6,04235E-49 |
| ACE2 | ANG | 0,673 | 5,9244E-49 |
| ACE2 | FKBP1A | 0,673 | 5,95792E-49 |
| ACE2 | ITPK1 | 0,6729 | 6,21096E-49 |
| ACE2 | SNX2 | 0,6728 | 6,58728E-49 |
| ACE2 | HSD17B11 | 0,6727 | 6,70373E-49 |
| ACE2 | MDN1 | 0,6726 | 7,08618E-49 |
| ACE2 | BLOC1S5 | 0,6725 | 7,48686E-49 |
| ACE2 | MID2 | 0,6725 | 7,31385E-49 |
| ACE2 | CCT6A | 0,6724 | 7,9133E-49 |
| ACE2 | EPCAM | 0,6723 | 7,9266E-49 |
| ACE2 | PSMB4 | 0,6722 | 8,58896E-49 |
| ACE2 | PLA2G4A | 0,6722 | 8,41738E-49 |
| ACE2 | ASB3 | 0,672 | 9,35669E-49 |
| ACE2 | PCYOX1 | 0,6719 | 9,65584E-49 |
| ACE2 | KDM5C | 0,6719 | 9,75815E-49 |
| ACE2 | MRPL35 | 0,6718 | 1,00048E-48 |
| ACE2 | IER3IP1 | 0,6718 | 1,01617E-48 |
| ACE2 | LYSMD3 | 0,6717 | 1,07013E-48 |
| ACE2 | GTF2I | 0,6716 | 1,12252E-48 |
| ACE2 | PCNP | 0,6715 | 1,14236E-48 |
| ACE2 | ZNF395 | 0,6715 | 1,15973E-48 |
| ACE2 | DEPDC5 | 0,6715 | 1,1372E-48 |
| ACE2 | APEX2 | 0,6715 | 1,15431E-48 |
| ACE2 | PTPN9 | 0,6714 | 1,22628E-48 |
| ACE2 | HIST1H2BK | 0,6713 | 1,27502E-48 |
| ACE2 | SEPT14 | 0,6713 | 1,2834E-48 |
| ACE2 | HEATR5A | 0,6712 | 1,31491E-48 |
| ACE2 | KHNYN | 0,6711 | 1,37743E-48 |
| ACE2 | HK2 | 0,671 | 1,42745E-48 |
| ACE2 | SYNPO | 0,6708 | 1,55488E-48 |
| ACE2 | DCTN5 | 0,6708 | 1,58522E-48 |
| ACE2 | HMGB1 | 0,6707 | 1,64437E-48 |
| ACE2 | PDCD6IP | 0,6706 | 1,74216E-48 |
| ACE2 | LAMA2 | 0,6706 | 1,70908E-48 |
| ACE2 | DOCK5 | 0,6706 | 1,7188E-48 |
| ACE2 | EIF4EBP2 | 0,6706 | 1,69017E-48 |
| ACE2 | PCBD1 | 0,6706 | 1,7139E-48 |
| ACE2 | IQGAP1 | 0,6705 | 1,77639E-48 |
| ACE2 | AFF1 | 0,6704 | 1,83523E-48 |
| ACE2 | AOX1 | 0,6703 | 1,98463E-48 |
| ACE2 | SLC44A1 | 0,6703 | 1,90953E-48 |
| ACE2 | KPNA4 | 0,6702 | 1,99748E-48 |
| ACE2 | IGFBPL1 | 0,6702 | 2,00779E-48 |
| ACE2 | KIAA0355 | 0,6701 | 2,09495E-48 |
| ACE2 | GNPTAB | 0,67 | 2,2051E-48 |
| ACE2 | TMEM126B | 0,6699 | 2,36606E-48 |
| ACE2 | HSD17B10 | 0,6699 | 2,32901E-48 |
| ACE2 | REV1 | 0,6698 | 2,3765E-48 |
| ACE2 | SLC39A14 | 0,6698 | 2,42114E-48 |
| ACE2 | PUS7L | 0,6698 | 2,38295E-48 |
| ACE2 | NCOA1 | 0,6697 | 2,52038E-48 |
| ACE2 | MBLAC2 | 0,6697 | 2,54225E-48 |

| | | | |
|---|---|---|---|
| ACE2 | ARPC4 | 0,6696 | 2,60174E-48 |
| ACE2 | CHPF2 | 0,6696 | 2,68332E-48 |
| ACE2 | BAG4 | 0,6696 | 2,64026E-48 |
| ACE2 | MACF1 | 0,6694 | 2,89332E-48 |
| ACE2 | SEMA7A | 0,6694 | 2,9511E-48 |
| ACE2 | CLCN5 | 0,6694 | 2,92744E-48 |
| ACE2 | DCAF8 | 0,6692 | 3,15174E-48 |
| ACE2 | FAM168B | 0,6692 | 3,09205E-48 |
| ACE2 | FYTTD1 | 0,6692 | 3,15624E-48 |
| ACE2 | NGRN | 0,6692 | 3,09217E-48 |
| ACE2 | SLC45A1 | 0,6688 | 3,76153E-48 |
| ACE2 | ECH1 | 0,6687 | 4,01025E-48 |
| ACE2 | MIS18BP1 | 0,6684 | 4,56626E-48 |
| ACE2 | ZNF326 | 0,6683 | 4,60251E-48 |
| ACE2 | UGT3A2 | 0,6682 | 4,8199E-48 |
| ACE2 | ELMSAN1 | 0,6682 | 4,86843E-48 |
| ACE2 | SEPT2 | 0,6678 | 5,71493E-48 |
| ACE2 | CASK | 0,6678 | 5,9175E-48 |
| ACE2 | NDST1 | 0,6677 | 5,94086E-48 |
| ACE2 | TNS3 | 0,6677 | 6,15629E-48 |
| ACE2 | FAM91A1 | 0,6677 | 6,12841E-48 |
| ACE2 | HDAC1 | 0,6674 | 6,98907E-48 |
| ACE2 | TRIB1 | 0,6674 | 7,01811E-48 |
| ACE2 | CDK17 | 0,6673 | 7,32688E-48 |
| ACE2 | HSPB8 | 0,6673 | 7,3565E-48 |
| ACE2 | DSCR3 | 0,6672 | 7,57119E-48 |
| ACE2 | GRAMD3 | 0,6671 | 7,70671E-48 |
| ACE2 | TGDS | 0,667 | 8,26526E-48 |
| ACE2 | DDHD2 | 0,6669 | 8,76487E-48 |
| ACE2 | VPS26B | 0,6669 | 8,59067E-48 |
| ACE2 | CYP39A1 | 0,6667 | 9,43682E-48 |
| ACE2 | MYO5C | 0,6667 | 9,49771E-48 |
| ACE2 | SEPHS2 | 0,6667 | 9,22757E-48 |
| ACE2 | PUDP | 0,6666 | 9,60725E-48 |
| ACE2 | AHSA1 | 0,6662 | 1,16239E-47 |
| ACE2 | TIMP1 | 0,6661 | 1,23319E-47 |
| ACE2 | USP27X-AS1 | 0,666 | 1,26351E-47 |
| ACE2 | DRAM2 | 0,6658 | 1,38252E-47 |
| ACE2 | LARS2 | 0,6658 | 1,38659E-47 |
| ACE2 | SPCS3 | 0,6658 | 1,38635E-47 |
| ACE2 | VAMP2 | 0,6658 | 1,3635E-47 |
| ACE2 | FRG1CP | 0,6658 | 1,40608E-47 |
| ACE2 | IFNGR2 | 0,6657 | 1,41744E-47 |
| ACE2 | ELMO1 | 0,6656 | 1,49244E-47 |
| ACE2 | ALG2 | 0,6655 | 1,56363E-47 |
| ACE2 | MID1 | 0,6655 | 1,53763E-47 |
| ACE2 | LARP4 | 0,6653 | 1,69856E-47 |
| ACE2 | C12orf66 | 0,6653 | 1,71581E-47 |
| ACE2 | ZNF791 | 0,6653 | 1,72409E-47 |
| ACE2 | PLEKHB2 | 0,6652 | 1,7711E-47 |
| ACE2 | AMOTL2 | 0,6652 | 1,8186E-47 |
| ACE2 | CCNG1 | 0,6652 | 1,78561E-47 |
| ACE2 | LRCH1 | 0,6651 | 1,84149E-47 |
| ACE2 | SENP7 | 0,665 | 1,94531E-47 |

| | | | |
|---|---|---|---|
| ACE2 | SASH1 | 0,665 | 1,92944E-47 |
| ACE2 | USP48 | 0,6648 | 2,16191E-47 |
| ACE2 | EPG5 | 0,6648 | 2,12575E-47 |
| ACE2 | TMPPE | 0,6647 | 2,24957E-47 |
| ACE2 | LRRC40 | 0,6646 | 2,29043E-47 |
| ACE2 | EIF4A2 | 0,6645 | 2,44448E-47 |
| ACE2 | PANK1 | 0,6645 | 2,42657E-47 |
| ACE2 | TCEAL4 | 0,6645 | 2,42024E-47 |
| ACE2 | SHISA4 | 0,6644 | 2,54093E-47 |
| ACE2 | BBS10 | 0,6644 | 2,56365E-47 |
| ACE2 | RP11-288C18.1 | 0,6643 | 2,64516E-47 |
| ACE2 | BCKDHB | 0,6643 | 2,62107E-47 |
| ACE2 | CHD9 | 0,6643 | 2,63928E-47 |
| ACE2 | ITM2C | 0,664 | 3,0093E-47 |
| ACE2 | ARAF | 0,664 | 3,00982E-47 |
| ACE2 | TIMM8A | 0,6638 | 3,19455E-47 |
| ACE2 | GNAI2 | 0,6637 | 3,37293E-47 |
| ACE2 | RPS6KB1 | 0,6637 | 3,422E-47 |
| ACE2 | PBX3 | 0,6636 | 3,5378E-47 |
| ACE2 | COPB1 | 0,6635 | 3,76045E-47 |
| ACE2 | TMEM35A | 0,6635 | 3,65313E-47 |
| ACE2 | KDM5D | 0,6635 | 3,64621E-47 |
| ACE2 | CAPN7 | 0,6634 | 3,91566E-47 |
| ACE2 | NAPB | 0,6634 | 3,89482E-47 |
| ACE2 | FBLN2 | 0,6631 | 4,47434E-47 |
| ACE2 | SKP2 | 0,6631 | 4,35508E-47 |
| ACE2 | UFM1 | 0,6631 | 4,30393E-47 |
| ACE2 | IL13RA1 | 0,6631 | 4,29975E-47 |
| ACE2 | DOCK9 | 0,6629 | 4,85414E-47 |
| ACE2 | FAM84A | 0,6628 | 4,9615E-47 |
| ACE2 | ARSG | 0,6628 | 5,06212E-47 |
| ACE2 | SEC61A1 | 0,6627 | 5,12537E-47 |
| ACE2 | POP7 | 0,6626 | 5,35127E-47 |
| ACE2 | RP11-57H14.4 | 0,6626 | 5,54135E-47 |
| ACE2 | HHAT | 0,6625 | 5,56631E-47 |
| ACE2 | RP11-240L7.4 | 0,6625 | 5,77951E-47 |
| ACE2 | RCOR1 | 0,6625 | 5,58745E-47 |
| ACE2 | DDX58 | 0,6624 | 5,85703E-47 |
| ACE2 | ARMCX1 | 0,6623 | 6,23888E-47 |
| ACE2 | SLC12A8 | 0,6622 | 6,41319E-47 |
| ACE2 | TAF5L | 0,662 | 7,13069E-47 |
| ACE2 | FNTA | 0,662 | 6,9222E-47 |
| ACE2 | IL1A | 0,6619 | 7,39807E-47 |
| ACE2 | RAP2B | 0,6619 | 7,20273E-47 |
| ACE2 | GRIPAP1 | 0,6619 | 7,1755E-47 |
| ACE2 | MED6 | 0,6617 | 7,97863E-47 |
| ACE2 | EIF2AK2 | 0,6616 | 8,33743E-47 |
| ACE2 | GNS | 0,6616 | 8,18381E-47 |
| ACE2 | FBXL20 | 0,6613 | 9,52029E-47 |
| ACE2 | LSM14A | 0,6613 | 9,27531E-47 |
| ACE2 | ANKRA2 | 0,6612 | 9,64613E-47 |
| ACE2 | ATG12 | 0,6612 | 9,67319E-47 |
| ACE2 | CCNJL | 0,6612 | 9,7517E-47 |
| ACE2 | TSPAN12 | 0,6611 | 1,03234E-46 |

| | | | |
|---|---|---|---|
| ACE2 | ZNF280D | 0,661 | 1,06121E-46 |
| ACE2 | FLRT3 | 0,6609 | 1,09858E-46 |
| ACE2 | LGALSL | 0,6607 | 1,21497E-46 |
| ACE2 | LONRF2 | 0,6607 | 1,19653E-46 |
| ACE2 | AKAP9 | 0,6607 | 1,21912E-46 |
| ACE2 | DERL1 | 0,6607 | 1,2251E-46 |
| ACE2 | ITPR2 | 0,6607 | 1,21173E-46 |
| ACE2 | ATP11C | 0,6607 | 1,1988E-46 |
| ACE2 | PPP4R2 | 0,6606 | 1,24284E-46 |
| ACE2 | STAT3 | 0,6605 | 1,30966E-46 |
| ACE2 | RHOC | 0,6604 | 1,38444E-46 |
| ACE2 | KALRN | 0,6604 | 1,35776E-46 |
| ACE2 | RNASEL | 0,6603 | 1,45742E-46 |
| ACE2 | PARM1 | 0,6603 | 1,45803E-46 |
| ACE2 | CSNK1G3 | 0,6603 | 1,45852E-46 |
| ACE2 | UNC5C | 0,6602 | 1,49348E-46 |
| ACE2 | SEPHS1 | 0,6602 | 1,48737E-46 |
| ACE2 | TJP2 | 0,6601 | 1,57034E-46 |
| ACE2 | MIER1 | 0,6599 | 1,70066E-46 |
| ACE2 | PPP1CA | 0,6599 | 1,6842E-46 |
| ACE2 | GSTA1 | 0,6598 | 1,76366E-46 |
| ACE2 | TRIP6 | 0,6598 | 1,77826E-46 |
| ACE2 | ARHGAP6 | 0,6598 | 1,74709E-46 |
| ACE2 | MYSM1 | 0,6597 | 1,8344E-46 |
| ACE2 | ZKSCAN8 | 0,6597 | 1,86935E-46 |
| ACE2 | NHLRC2 | 0,6597 | 1,83253E-46 |
| ACE2 | GPKOW | 0,6597 | 1,8809E-46 |
| ACE2 | FMNL3 | 0,6595 | 2,0085E-46 |
| ACE2 | COTL1 | 0,6594 | 2,12556E-46 |
| ACE2 | DDX24 | 0,6592 | 2,28659E-46 |
| ACE2 | MSANTD3 | 0,659 | 2,4922E-46 |
| ACE2 | STRADB | 0,6589 | 2,58411E-46 |
| ACE2 | TMEM87B | 0,6586 | 2,97404E-46 |
| ACE2 | PLGRKT | 0,6586 | 2,95203E-46 |
| ACE2 | CDK4 | 0,6586 | 2,99451E-46 |
| ACE2 | FRRS1 | 0,6584 | 3,16278E-46 |
| ACE2 | TSPAN5 | 0,6584 | 3,20929E-46 |
| ACE2 | AGK | 0,6584 | 3,17072E-46 |
| ACE2 | TRAPPC6B | 0,6584 | 3,22946E-46 |
| ACE2 | PDK2 | 0,6584 | 3,20369E-46 |
| ACE2 | PEPD | 0,6583 | 3,32498E-46 |
| ACE2 | ITSN1 | 0,6583 | 3,28949E-46 |
| ACE2 | SAMD5 | 0,6582 | 3,48285E-46 |
| ACE2 | SRC | 0,6582 | 3,47712E-46 |
| ACE2 | ITGB5 | 0,6581 | 3,56951E-46 |
| ACE2 | MTX3 | 0,6581 | 3,59872E-46 |
| ACE2 | KIAA1468 | 0,6581 | 3,5882E-46 |
| ACE2 | KLC1 | 0,658 | 3,84541E-46 |
| ACE2 | PLP2 | 0,6579 | 3,91771E-46 |
| ACE2 | RPL22 | 0,6577 | 4,22264E-46 |
| ACE2 | RUFY3 | 0,6577 | 4,2837E-46 |
| ACE2 | SETBP1 | 0,6576 | 4,56017E-46 |
| ACE2 | SDC4 | 0,6576 | 4,4747E-46 |
| ACE2 | MOSPD2 | 0,6576 | 4,38717E-46 |

| | | | |
|---|---|---|---|
| ACE2 | NT5C3A | 0,6575 | 4,59742E-46 |
| ACE2 | ADGRG2 | 0,6575 | 4,59853E-46 |
| ACE2 | PICALM | 0,6573 | 4,97824E-46 |
| ACE2 | CHST3 | 0,6572 | 5,17901E-46 |
| ACE2 | ARID4A | 0,6572 | 5,28767E-46 |
| ACE2 | AKT1 | 0,657 | 5,78159E-46 |
| ACE2 | PNRC1 | 0,6569 | 5,90448E-46 |
| ACE2 | PNP | 0,6568 | 6,18676E-46 |
| ACE2 | VSIR | 0,6567 | 6,37261E-46 |
| ACE2 | PRKCB | 0,6567 | 6,40528E-46 |
| ACE2 | CSRNP3 | 0,6566 | 6,81854E-46 |
| ACE2 | IRF2 | 0,6566 | 6,79975E-46 |
| ACE2 | CLN3 | 0,6566 | 6,72476E-46 |
| ACE2 | BET1 | 0,6564 | 7,43381E-46 |
| ACE2 | DNAJB5 | 0,6564 | 7,41615E-46 |
| ACE2 | SPAAR | 0,6564 | 7,39763E-46 |
| ACE2 | FAH | 0,6564 | 7,24393E-46 |
| ACE2 | DCK | 0,6562 | 8,04123E-46 |
| ACE2 | SLC19A2 | 0,6561 | 8,35876E-46 |
| ACE2 | GSTA3 | 0,656 | 8,70516E-46 |
| ACE2 | PRPF19 | 0,656 | 8,5771E-46 |
| ACE2 | SMAD5 | 0,6559 | 9,23936E-46 |
| ACE2 | TM9SF4 | 0,6559 | 9,0262E-46 |
| ACE2 | TEX10 | 0,6558 | 9,3521E-46 |
| ACE2 | IRF2BPL | 0,6558 | 9,30438E-46 |
| ACE2 | LARP7 | 0,6557 | 9,69329E-46 |
| ACE2 | GEMIN5 | 0,6557 | 9,84835E-46 |
| ACE2 | MLX | 0,6557 | 9,75766E-46 |
| ACE2 | PAK3 | 0,6557 | 9,77703E-46 |
| ACE2 | CXorf56 | 0,6557 | 9,70191E-46 |
| ACE2 | CALR | 0,6556 | 1,0439E-45 |
| ACE2 | DCTD | 0,6554 | 1,10239E-45 |
| ACE2 | BCL2 | 0,6554 | 1,11781E-45 |
| ACE2 | UBP1 | 0,6553 | 1,16049E-45 |
| ACE2 | TPT1 | 0,6552 | 1,232E-45 |
| ACE2 | HEYL | 0,6551 | 1,26843E-45 |
| ACE2 | GBA | 0,6551 | 1,25334E-45 |
| ACE2 | PDCL3 | 0,6551 | 1,26307E-45 |
| ACE2 | NEMF | 0,6551 | 1,25798E-45 |
| ACE2 | TIGD2 | 0,655 | 1,3066E-45 |
| ACE2 | UBE2L3 | 0,6548 | 1,43773E-45 |
| ACE2 | USP24 | 0,6547 | 1,51134E-45 |
| ACE2 | CS | 0,6546 | 1,52004E-45 |
| ACE2 | NDUFA1 | 0,6545 | 1,61387E-45 |
| ACE2 | EIF3I | 0,6544 | 1,66291E-45 |
| ACE2 | COPS8 | 0,6544 | 1,69932E-45 |
| ACE2 | DDX54 | 0,6542 | 1,80178E-45 |
| ACE2 | BCCIP | 0,6541 | 1,91993E-45 |
| ACE2 | PHB2 | 0,6541 | 1,90573E-45 |
| ACE2 | COG5 | 0,654 | 1,99086E-45 |
| ACE2 | NUBPL | 0,654 | 1,97895E-45 |
| ACE2 | SRP68 | 0,654 | 1,99463E-45 |
| ACE2 | LINC00909 | 0,6538 | 2,19542E-45 |
| ACE2 | PIGK | 0,6537 | 2,21162E-45 |

| | | | |
|---|---|---|---|
| ACE2 | INTS10 | 0,6537 | 2,28021E-45 |
| ACE2 | FERMT2 | 0,6537 | 2,29329E-45 |
| ACE2 | SLC25A53 | 0,6537 | 2,25369E-45 |
| ACE2 | GTF2B | 0,6536 | 2,36996E-45 |
| ACE2 | PABPC4 | 0,6535 | 2,44237E-45 |
| ACE2 | NUDT16P1 | 0,6535 | 2,42709E-45 |
| ACE2 | DFFA | 0,6533 | 2,63911E-45 |
| ACE2 | CDC123 | 0,6532 | 2,72556E-45 |
| ACE2 | SGTB | 0,6531 | 2,84593E-45 |
| ACE2 | ANXA6 | 0,6531 | 2,92582E-45 |
| ACE2 | FDXR | 0,6531 | 2,90926E-45 |
| ACE2 | TRAF3IP2 | 0,653 | 3,00197E-45 |
| ACE2 | ACVR1B | 0,653 | 2,96815E-45 |
| ACE2 | CCT8P1 | 0,6529 | 3,10826E-45 |
| ACE2 | DDX5 | 0,6529 | 3,12296E-45 |
| ACE2 | GPR27 | 0,6528 | 3,19748E-45 |
| ACE2 | TBC1D9B | 0,6528 | 3,26753E-45 |
| ACE2 | ARL6IP1 | 0,6527 | 3,45633E-45 |
| ACE2 | ABHD6 | 0,6526 | 3,59375E-45 |
| ACE2 | PEAK1 | 0,6526 | 3,59758E-45 |
| ACE2 | CHPF | 0,6525 | 3,73412E-45 |
| ACE2 | EPHX1 | 0,6524 | 3,87895E-45 |
| ACE2 | RASL12 | 0,6524 | 3,77196E-45 |
| ACE2 | TRIQK | 0,6523 | 4,00084E-45 |
| ACE2 | NHS | 0,6523 | 4,03781E-45 |
| ACE2 | AIMP1 | 0,6521 | 4,34291E-45 |
| ACE2 | ZSWIM5 | 0,652 | 4,42891E-45 |
| ACE2 | KLHL26 | 0,652 | 4,48204E-45 |
| ACE2 | RBSN | 0,6518 | 4,81473E-45 |
| ACE2 | USO1 | 0,6518 | 4,95493E-45 |
| ACE2 | PPM1L | 0,6516 | 5,41976E-45 |
| ACE2 | FAM20B | 0,6515 | 5,46003E-45 |
| ACE2 | OXSR1 | 0,6514 | 5,8286E-45 |
| ACE2 | TXN | 0,6514 | 5,66085E-45 |
| ACE2 | INTS4 | 0,6514 | 5,66412E-45 |
| ACE2 | ALG10B | 0,6514 | 5,66391E-45 |
| ACE2 | TANC2 | 0,6514 | 5,84913E-45 |
| ACE2 | FOXN2 | 0,6513 | 5,9012E-45 |
| ACE2 | ARHGAP10 | 0,6513 | 6,11749E-45 |
| ACE2 | GPX8 | 0,6513 | 5,95994E-45 |
| ACE2 | SLC10A7 | 0,6512 | 6,17727E-45 |
| ACE2 | YY1 | 0,6512 | 6,27876E-45 |
| ACE2 | ZNF738 | 0,6512 | 6,29694E-45 |
| ACE2 | SEC24A | 0,6511 | 6,64699E-45 |
| ACE2 | PTPDC1 | 0,6507 | 7,76216E-45 |
| ACE2 | RRAS2 | 0,6507 | 7,62606E-45 |
| ACE2 | POLR2G | 0,6507 | 7,51341E-45 |
| ACE2 | HSD11B1 | 0,6506 | 7,86577E-45 |
| ACE2 | FBXO4 | 0,6505 | 8,29175E-45 |
| ACE2 | TERF1 | 0,6505 | 8,17804E-45 |
| ACE2 | ZNF562 | 0,6505 | 8,37888E-45 |
| ACE2 | FAM32A | 0,6505 | 8,31101E-45 |
| ACE2 | NCSTN | 0,6503 | 9,05223E-45 |
| ACE2 | NR5A1 | 0,6501 | 9,69598E-45 |

| | | | |
|---|---|---|---|
| ACE2 | NHLRC3 | 0,6501 | 9,93447E-45 |
| ACE2 | GID8 | 0,65 | 1,0329E-44 |
| ACE2 | GNPTG | 0,6499 | 1,05705E-44 |
| ACE2 | FUBP1 | 0,6498 | 1,1139E-44 |
| ACE2 | RBP1 | 0,6498 | 1,08859E-44 |
| ACE2 | COPZ1 | 0,6497 | 1,13392E-44 |
| ACE2 | PRMT2 | 0,6497 | 1,14314E-44 |
| ACE2 | FYCO1 | 0,6496 | 1,20068E-44 |
| ACE2 | EXOSC2 | 0,6496 | 1,1824E-44 |
| ACE2 | ARL8B | 0,6495 | 1,25167E-44 |
| ACE2 | MSX2 | 0,6495 | 1,25062E-44 |
| ACE2 | FKBP9 | 0,6495 | 1,24127E-44 |
| ACE2 | DPYSL2 | 0,6495 | 1,26948E-44 |
| ACE2 | TMEM241 | 0,6495 | 1,24446E-44 |
| ACE2 | MED29 | 0,6495 | 1,24E-44 |
| ACE2 | P2RY1 | 0,6494 | 1,28812E-44 |
| ACE2 | LRIG1 | 0,6492 | 1,39608E-44 |
| ACE2 | WDFY3 | 0,6492 | 1,41588E-44 |
| ACE2 | IL6ST | 0,6492 | 1,4238E-44 |
| ACE2 | TUBB4A | 0,6492 | 1,3914E-44 |
| ACE2 | FAM127A | 0,6492 | 1,40678E-44 |
| ACE2 | ACADM | 0,6491 | 1,46911E-44 |
| ACE2 | PRMT6 | 0,6491 | 1,48495E-44 |
| ACE2 | MAML1 | 0,6491 | 1,48026E-44 |
| ACE2 | KANK1 | 0,6491 | 1,48319E-44 |
| ACE2 | SLBP | 0,649 | 1,53528E-44 |
| ACE2 | ISOC1 | 0,649 | 1,53715E-44 |
| ACE2 | C9orf66 | 0,649 | 1,50828E-44 |
| ACE2 | RPL21 | 0,649 | 1,5139E-44 |
| ACE2 | HMGCL | 0,6489 | 1,57869E-44 |
| ACE2 | C9orf72 | 0,6489 | 1,593E-44 |
| ACE2 | SF3B3 | 0,6486 | 1,80363E-44 |
| ACE2 | SAT1 | 0,6485 | 1,83896E-44 |
| ACE2 | GPC1 | 0,6484 | 1,97077E-44 |
| ACE2 | ALAS1 | 0,6484 | 1,95449E-44 |
| ACE2 | GCNA | 0,6484 | 1,95271E-44 |
| ACE2 | MEF2D | 0,6482 | 2,10383E-44 |
| ACE2 | HNRNPA2B1 | 0,648 | 2,29377E-44 |
| ACE2 | MYC | 0,648 | 2,31126E-44 |
| ACE2 | DDX46 | 0,6479 | 2,38975E-44 |
| ACE2 | MAPK14 | 0,6479 | 2,33836E-44 |
| ACE2 | CLSTN1 | 0,6476 | 2,71447E-44 |
| ACE2 | CSTB | 0,6476 | 2,63218E-44 |
| ACE2 | VAMP4 | 0,6475 | 2,76901E-44 |
| ACE2 | GPR137B | 0,6475 | 2,82572E-44 |
| ACE2 | CYFIP1 | 0,6475 | 2,81655E-44 |
| ACE2 | DDB1 | 0,6472 | 3,10856E-44 |
| ACE2 | SMS | 0,6472 | 3,21198E-44 |
| ACE2 | BPHL | 0,6471 | 3,23055E-44 |
| ACE2 | AFMID | 0,6471 | 3,26643E-44 |
| ACE2 | HMGN5 | 0,6471 | 3,22244E-44 |
| ACE2 | DVL3 | 0,6469 | 3,57366E-44 |
| ACE2 | GGCX | 0,6467 | 3,83577E-44 |
| ACE2 | MED23 | 0,6467 | 3,92627E-44 |

| | | | |
|---|---|---|---|
| ACE2 | PTPN12 | 0,6467 | 3,915E-44 |
| ACE2 | DYRK1A | 0,6467 | 3,84369E-44 |
| ACE2 | SDC2 | 0,6465 | 4,16472E-44 |
| ACE2 | SRI | 0,6464 | 4,42272E-44 |
| ACE2 | ZC3H7A | 0,6464 | 4,32993E-44 |
| ACE2 | PRDX2 | 0,6464 | 4,42395E-44 |
| ACE2 | BIRC6 | 0,6463 | 4,4692E-44 |
| ACE2 | UGDH | 0,6463 | 4,46343E-44 |
| ACE2 | MSRB2 | 0,6462 | 4,65059E-44 |
| ACE2 | MAF | 0,6462 | 4,71767E-44 |
| ACE2 | TSPAN13 | 0,6461 | 4,86257E-44 |
| ACE2 | ATP5G3 | 0,646 | 5,18271E-44 |
| ACE2 | AHR | 0,6459 | 5,35671E-44 |
| ACE2 | A2ML1 | 0,6459 | 5,39486E-44 |
| ACE2 | PPP1R12A | 0,6459 | 5,33306E-44 |
| ACE2 | LYSMD1 | 0,6457 | 5,68952E-44 |
| ACE2 | SNAPIN | 0,6457 | 5,76406E-44 |
| ACE2 | DPAGT1 | 0,6456 | 6,08819E-44 |
| ACE2 | SLC23A2 | 0,6456 | 5,99347E-44 |
| ACE2 | TFE3 | 0,6456 | 5,98208E-44 |
| ACE2 | KRT8P13 | 0,6454 | 6,36946E-44 |
| ACE2 | QDPR | 0,6451 | 7,21368E-44 |
| ACE2 | WT1-AS | 0,6451 | 7,40398E-44 |
| ACE2 | NRDE2 | 0,6451 | 7,25664E-44 |
| ACE2 | CHIC1 | 0,6451 | 7,21877E-44 |
| ACE2 | GAS6-AS2 | 0,645 | 7,56708E-44 |
| ACE2 | TOP2B | 0,6449 | 7,94547E-44 |
| ACE2 | PPIF | 0,6449 | 7,96348E-44 |
| ACE2 | TFG | 0,6446 | 8,80681E-44 |
| ACE2 | AF131215.2 | 0,6444 | 9,63828E-44 |
| ACE2 | IKBKAP | 0,6444 | 9,46631E-44 |
| ACE2 | SLC41A3 | 0,6441 | 1,06668E-43 |
| ACE2 | KBTBD2 | 0,6441 | 1,06719E-43 |
| ACE2 | HPS6 | 0,6441 | 1,08703E-43 |
| ACE2 | USP3 | 0,6441 | 1,07746E-43 |
| ACE2 | RHOT1 | 0,6441 | 1,09416E-43 |
| ACE2 | TAF13 | 0,644 | 1,13139E-43 |
| ACE2 | ZNF860 | 0,644 | 1,12817E-43 |
| ACE2 | ATF7 | 0,6439 | 1,17206E-43 |
| ACE2 | TXNDC11 | 0,6438 | 1,23473E-43 |
| ACE2 | DCAF16 | 0,6437 | 1,28759E-43 |
| ACE2 | CASP8AP2 | 0,6437 | 1,25894E-43 |
| ACE2 | LINC01152 | 0,6437 | 1,28018E-43 |
| ACE2 | RWDD3 | 0,6436 | 1,30128E-43 |
| ACE2 | CTSB | 0,6436 | 1,31615E-43 |
| ACE2 | SUMF2 | 0,6434 | 1,41649E-43 |
| ACE2 | CAPN5 | 0,6434 | 1,41865E-43 |
| ACE2 | MEDAG | 0,6434 | 1,41589E-43 |
| ACE2 | LRFN3 | 0,6434 | 1,41942E-43 |
| ACE2 | LSM5 | 0,6433 | 1,5E-43 |
| ACE2 | MRPL15 | 0,6433 | 1,50993E-43 |
| ACE2 | LRRC41 | 0,6432 | 1,5797E-43 |
| ACE2 | FKBP15 | 0,6432 | 1,55294E-43 |
| ACE2 | NDUFS2 | 0,643 | 1,70448E-43 |

| | | | |
|---|---|---|---|
| ACE2 | HECW2 | 0,643 | 1,67769E-43 |
| ACE2 | RFC1 | 0,6429 | 1,77111E-43 |
| ACE2 | TRIM38 | 0,6429 | 1,74356E-43 |
| ACE2 | APLP2 | 0,6429 | 1,72933E-43 |
| ACE2 | MRPL1 | 0,6427 | 1,87727E-43 |
| ACE2 | ZNF827 | 0,6427 | 1,85794E-43 |
| ACE2 | RPP14 | 0,6426 | 1,96733E-43 |
| ACE2 | SLC2A1 | 0,6425 | 2,03144E-43 |
| ACE2 | DTNA | 0,6425 | 2,04931E-43 |
| ACE2 | LIMD1 | 0,6423 | 2,21655E-43 |
| ACE2 | CGNL1 | 0,6423 | 2,20422E-43 |
| ACE2 | DOCK11 | 0,6423 | 2,18802E-43 |
| ACE2 | MMGT1 | 0,6423 | 2,17918E-43 |
| ACE2 | ITGB1BP1 | 0,6421 | 2,41694E-43 |
| ACE2 | PWAR6 | 0,6421 | 2,43934E-43 |
| ACE2 | TMTC2 | 0,642 | 2,49322E-43 |
| ACE2 | RAB40A | 0,642 | 2,49262E-43 |
| ACE2 | PDIA3P1 | 0,6419 | 2,57076E-43 |
| ACE2 | AKT3 | 0,6416 | 2,94289E-43 |
| ACE2 | MYO1B | 0,6416 | 2,88037E-43 |
| ACE2 | CRYZ | 0,6415 | 3,03896E-43 |
| ACE2 | BCL9L | 0,6413 | 3,25561E-43 |
| ACE2 | WIPF2 | 0,6411 | 3,50784E-43 |
| ACE2 | KREMEN1 | 0,641 | 3,69146E-43 |
| ACE2 | ZKSCAN1 | 0,6407 | 4,16889E-43 |
| ACE2 | STAG2 | 0,6406 | 4,29888E-43 |
| ACE2 | CTTN | 0,6405 | 4,40413E-43 |
| ACE2 | STRAP | 0,6405 | 4,56304E-43 |
| ACE2 | ZFR | 0,6404 | 4,7585E-43 |
| ACE2 | SLC35F5 | 0,6403 | 4,82915E-43 |
| ACE2 | UTP15 | 0,6403 | 4,893E-43 |
| ACE2 | FBXW2 | 0,6403 | 4,85713E-43 |
| ACE2 | USP10 | 0,6403 | 4,92249E-43 |
| ACE2 | GNB2 | 0,6402 | 5,05097E-43 |
| ACE2 | AMMECR1 | 0,6402 | 4,97732E-43 |
| ACE2 | PIR | 0,6401 | 5,32184E-43 |
| ACE2 | TSPYL2 | 0,6401 | 5,31013E-43 |
| ACE2 | AES | 0,64 | 5,38325E-43 |
| ACE2 | WLS | 0,6399 | 5,72041E-43 |
| ACE2 | EDEM1 | 0,6397 | 6,0987E-43 |
| ACE2 | MEAF6 | 0,6396 | 6,3567E-43 |
| ACE2 | AKIRIN2 | 0,6396 | 6,42335E-43 |
| ACE2 | NARS2 | 0,6396 | 6,35881E-43 |
| ACE2 | XYLT1 | 0,6396 | 6,42328E-43 |
| ACE2 | ARHGAP29 | 0,6394 | 6,8992E-43 |
| ACE2 | PARP1 | 0,6391 | 7,77557E-43 |
| ACE2 | SLC25A23 | 0,6391 | 7,78501E-43 |
| ACE2 | NPFFR1 | 0,639 | 8,11419E-43 |
| ACE2 | TSEN34 | 0,639 | 7,91663E-43 |
| ACE2 | ELP2 | 0,6389 | 8,34524E-43 |
| ACE2 | TCEAL1 | 0,6389 | 8,24757E-43 |
| ACE2 | EPC2 | 0,6388 | 8,7392E-43 |
| ACE2 | WDR75 | 0,6388 | 8,72667E-43 |
| ACE2 | RNF219 | 0,6388 | 8,55518E-43 |

| | | | |
|---|---|---|---|
| ACE2 | HMGCLL1 | 0,6387 | 8,99788E-43 |
| ACE2 | TRIM27 | 0,6385 | 9,96077E-43 |
| ACE2 | CMTM3 | 0,6385 | 9,84421E-43 |
| ACE2 | NUS1 | 0,6384 | 1,03697E-42 |
| ACE2 | CHCHD2 | 0,6384 | 1,01938E-42 |
| ACE2 | SERPINB9 | 0,6383 | 1,066E-42 |
| ACE2 | ST3GAL1 | 0,6383 | 1,04263E-42 |
| ACE2 | FBXW8 | 0,6382 | 1,10862E-42 |
| ACE2 | ACVR2A | 0,638 | 1,20047E-42 |
| ACE2 | UBA6 | 0,638 | 1,20349E-42 |
| ACE2 | MED17 | 0,638 | 1,1767E-42 |
| ACE2 | LPP-AS2 | 0,6379 | 1,21819E-42 |
| ACE2 | SRBD1 | 0,6378 | 1,28145E-42 |
| ACE2 | SERPINB8 | 0,6378 | 1,30901E-42 |
| ACE2 | S100A16 | 0,6377 | 1,36107E-42 |
| ACE2 | CCNG2 | 0,6377 | 1,34438E-42 |
| ACE2 | PIP4K2B | 0,6376 | 1,39712E-42 |
| ACE2 | ANAPC5 | 0,6375 | 1,43442E-42 |
| ACE2 | ILDR1 | 0,6374 | 1,50323E-42 |
| ACE2 | G3BP1 | 0,6374 | 1,47772E-42 |
| ACE2 | ZCCHC3 | 0,6374 | 1,4772E-42 |
| ACE2 | MED10 | 0,6373 | 1,58004E-42 |
| ACE2 | WNK1 | 0,6373 | 1,58695E-42 |
| ACE2 | SLC4A7 | 0,6372 | 1,6112E-42 |
| ACE2 | C3orf58 | 0,6371 | 1,67201E-42 |
| ACE2 | RUFY2 | 0,637 | 1,74776E-42 |
| ACE2 | C1orf74 | 0,6369 | 1,84894E-42 |
| ACE2 | AP1S1 | 0,6369 | 1,80465E-42 |
| ACE2 | PI4K2B | 0,6368 | 1,8662E-42 |
| ACE2 | BRIX1 | 0,6368 | 1,88618E-42 |
| ACE2 | BLMH | 0,6368 | 1,88609E-42 |
| ACE2 | SGSH | 0,6367 | 1,9974E-42 |
| ACE2 | ZNF680 | 0,6366 | 2,01889E-42 |
| ACE2 | IGSF3 | 0,6365 | 2,16225E-42 |
| ACE2 | DLG1 | 0,6363 | 2,30321E-42 |
| ACE2 | RP11-116O18.1 | 0,6363 | 2,31372E-42 |
| ACE2 | HLCS | 0,6363 | 2,29329E-42 |
| ACE2 | BCORL1 | 0,6362 | 2,42603E-42 |
| ACE2 | UBE2B | 0,6361 | 2,48083E-42 |
| ACE2 | ERV3-1 | 0,6361 | 2,45944E-42 |
| ACE2 | LARP4B | 0,6361 | 2,52447E-42 |
| ACE2 | CIPC | 0,636 | 2,53543E-42 |
| ACE2 | CDKN2AIPNL | 0,6359 | 2,72327E-42 |
| ACE2 | TAP1 | 0,6359 | 2,63658E-42 |
| ACE2 | GCLM | 0,6358 | 2,82628E-42 |
| ACE2 | CNOT6L | 0,6357 | 2,92943E-42 |
| ACE2 | NCAPD3 | 0,6357 | 2,8906E-42 |
| ACE2 | ULBP3 | 0,6356 | 2,96269E-42 |
| ACE2 | SPIN1 | 0,6355 | 3,10847E-42 |
| ACE2 | FBXO28 | 0,6354 | 3,2349E-42 |
| ACE2 | GDAP1 | 0,6354 | 3,18765E-42 |
| ACE2 | INO80D | 0,6353 | 3,31658E-42 |
| ACE2 | ZNF518A | 0,6351 | 3,62694E-42 |
| ACE2 | DCUN1D5 | 0,6351 | 3,67297E-42 |

| | | | |
|---|---|---|---|
| ACE2 | PRRG1 | 0,6351 | 3,62245E-42 |
| ACE2 | TXNRD1 | 0,6349 | 3,87003E-42 |
| ACE2 | SSH1 | 0,6349 | 3,87471E-42 |
| ACE2 | SIKE1 | 0,6348 | 4,12321E-42 |
| ACE2 | IKZF2 | 0,6348 | 4,08351E-42 |
| ACE2 | RALGAPA1 | 0,6348 | 4,04872E-42 |
| ACE2 | SERPINB1 | 0,6347 | 4,26815E-42 |
| ACE2 | RBM39 | 0,6347 | 4,17712E-42 |
| ACE2 | SENP6 | 0,6346 | 4,33155E-42 |
| ACE2 | KDELC1 | 0,6346 | 4,36551E-42 |
| ACE2 | CPT2 | 0,6345 | 4,49932E-42 |
| ACE2 | RASL11B | 0,6345 | 4,54409E-42 |
| ACE2 | SLC9A7 | 0,6345 | 4,52647E-42 |
| ACE2 | FAM127B | 0,6345 | 4,57151E-42 |
| ACE2 | EID2 | 0,6344 | 4,84857E-42 |
| ACE2 | LL0XNC01-221F2.2 | 0,6344 | 4,76066E-42 |
| ACE2 | HFE | 0,6343 | 4,9112E-42 |
| ACE2 | COQ10A | 0,6343 | 4,90073E-42 |
| ACE2 | CST9L | 0,6343 | 4,85688E-42 |
| ACE2 | NAA15 | 0,6342 | 5,13505E-42 |
| ACE2 | NTN4 | 0,6342 | 5,14746E-42 |
| ACE2 | VASP | 0,6339 | 5,73229E-42 |
| ACE2 | KDM6A | 0,6339 | 5,83425E-42 |
| ACE2 | KDM3B | 0,6338 | 5,92798E-42 |
| ACE2 | CORO2A | 0,6335 | 6,65236E-42 |
| ACE2 | MMS19 | 0,6335 | 6,72592E-42 |
| ACE2 | LRP10 | 0,6335 | 6,61277E-42 |
| ACE2 | SLC2A10 | 0,6335 | 6,82718E-42 |
| ACE2 | NPTX2 | 0,6334 | 6,96846E-42 |
| ACE2 | RPE | 0,6333 | 7,31625E-42 |
| ACE2 | YIPF3 | 0,6333 | 7,17361E-42 |
| ACE2 | TNFRSF1A | 0,6332 | 7,49768E-42 |
| ACE2 | CNEP1R1 | 0,6331 | 7,70449E-42 |
| ACE2 | BCOR | 0,6331 | 7,85267E-42 |
| ACE2 | DDX50 | 0,633 | 8,11676E-42 |
| ACE2 | LOXL3 | 0,6329 | 8,42068E-42 |
| ACE2 | MNAT1 | 0,6329 | 8,40622E-42 |
| ACE2 | STT3A | 0,6327 | 9,07927E-42 |
| ACE2 | SHMT2 | 0,6327 | 9,13723E-42 |
| ACE2 | TRAK2 | 0,6326 | 9,5992E-42 |
| ACE2 | ZNF616 | 0,6325 | 9,95428E-42 |
| ACE2 | GATAD1 | 0,6324 | 1,01832E-41 |
| ACE2 | ANXA11 | 0,6322 | 1,11482E-41 |
| ACE2 | RAB2A | 0,632 | 1,20476E-41 |
| ACE2 | PIK3CB | 0,6319 | 1,21475E-41 |
| ACE2 | MPRIP | 0,6319 | 1,24643E-41 |
| ACE2 | NNT-AS1 | 0,6318 | 1,25712E-41 |
| ACE2 | IDS | 0,6318 | 1,27377E-41 |
| ACE2 | PHC3 | 0,6316 | 1,39599E-41 |
| ACE2 | INSIG2 | 0,6314 | 1,49173E-41 |
| ACE2 | TCN1 | 0,6313 | 1,5489E-41 |
| ACE2 | ELK3 | 0,6313 | 1,53668E-41 |
| ACE2 | UTP14A | 0,6313 | 1,56234E-41 |
| ACE2 | MFAP1 | 0,6311 | 1,68041E-41 |

| | | | |
|---|---|---|---|
| ACE2 | NOM1 | 0,6309 | 1,80771E-41 |
| ACE2 | ROCK1 | 0,6308 | 1,88443E-41 |
| ACE2 | GABRQ | 0,6305 | 2,06188E-41 |
| ACE2 | USP28 | 0,6304 | 2,15358E-41 |
| ACE2 | MAP2K1 | 0,6303 | 2,25193E-41 |
| ACE2 | AGA | 0,6302 | 2,31105E-41 |
| ACE2 | LTBR | 0,6301 | 2,40151E-41 |
| ACE2 | SELL | 0,63 | 2,49573E-41 |
| ACE2 | POLR2B | 0,63 | 2,55122E-41 |
| ACE2 | PLRG1 | 0,63 | 2,49912E-41 |
| ACE2 | HMGN2 | 0,6299 | 2,63893E-41 |
| ACE2 | ARMCX5-GPRASP2 | 0,6298 | 2,70279E-41 |
| ACE2 | CERS4 | 0,6297 | 2,85768E-41 |
| ACE2 | PIK3R2 | 0,6297 | 2,85891E-41 |
| ACE2 | TTLL12 | 0,6296 | 2,94707E-41 |
| ACE2 | HDX | 0,6296 | 2,94634E-41 |
| ACE2 | TYW5 | 0,6295 | 3,07839E-41 |
| ACE2 | TMEM263 | 0,6295 | 3,06469E-41 |
| ACE2 | CHTOP | 0,6294 | 3,22177E-41 |
| ACE2 | FAM126B | 0,6294 | 3,15559E-41 |
| ACE2 | NELL2 | 0,6294 | 3,16102E-41 |
| ACE2 | PCMTD1 | 0,6293 | 3,31535E-41 |
| ACE2 | MARK3 | 0,6292 | 3,41752E-41 |
| ACE2 | BCAT2 | 0,6288 | 3,97032E-41 |
| ACE2 | MKLN1 | 0,6287 | 4,13185E-41 |
| ACE2 | FGF13 | 0,6286 | 4,28344E-41 |
| ACE2 | SLMAP | 0,6285 | 4,42358E-41 |
| ACE2 | CHSY1 | 0,6285 | 4,46666E-41 |
| ACE2 | DIRAS1 | 0,6285 | 4,49424E-41 |
| ACE2 | EMG1 | 0,6284 | 4,60317E-41 |
| ACE2 | YARS2 | 0,6284 | 4,64062E-41 |
| ACE2 | SYNE2 | 0,6284 | 4,63094E-41 |
| ACE2 | GLRB | 0,6283 | 4,78253E-41 |
| ACE2 | RBM22 | 0,6283 | 4,8235E-41 |
| ACE2 | DLAT | 0,6283 | 4,86745E-41 |
| ACE2 | UBE4A | 0,6283 | 4,78155E-41 |
| ACE2 | SLC10A3 | 0,6283 | 4,73633E-41 |
| ACE2 | NUDT9 | 0,6282 | 4,95847E-41 |
| ACE2 | PARD3 | 0,6282 | 5,03246E-41 |
| ACE2 | PIN4 | 0,6282 | 4,89561E-41 |
| ACE2 | RNF113A | 0,6282 | 5,02409E-41 |
| ACE2 | UBE2F | 0,6281 | 5,17143E-41 |
| ACE2 | TMEM165 | 0,6281 | 5,06883E-41 |
| ACE2 | TCEA1 | 0,6281 | 5,22559E-41 |
| ACE2 | NUDT19 | 0,6279 | 5,47091E-41 |
| ACE2 | SARS | 0,6278 | 5,69627E-41 |
| ACE2 | ZDHHC6 | 0,6278 | 5,666E-41 |
| ACE2 | C6orf47 | 0,6276 | 6,19842E-41 |
| ACE2 | SLC36A4 | 0,6275 | 6,44908E-41 |
| ACE2 | AREL1 | 0,6275 | 6,37577E-41 |
| ACE2 | LMTK2 | 0,6274 | 6,67284E-41 |
| ACE2 | TRIM56 | 0,6274 | 6,6976E-41 |
| ACE2 | NUCB1 | 0,6273 | 7,07894E-41 |
| ACE2 | PJA1 | 0,6272 | 7,12378E-41 |

| | | | |
|---|---|---|---|
| ACE2 | GATA6 | 0,6269 | 8,09766E-41 |
| ACE2 | SAMHD1 | 0,6269 | 8,09916E-41 |
| ACE2 | ZNF280C | 0,6268 | 8,25456E-41 |
| ACE2 | FRS2 | 0,6267 | 8,84383E-41 |
| ACE2 | DOCK10 | 0,6266 | 9,05681E-41 |
| ACE2 | SIRT5 | 0,6266 | 9,01478E-41 |
| ACE2 | MNDA | 0,6265 | 9,45956E-41 |
| ACE2 | ZNF678 | 0,6263 | 1,00562E-40 |
| ACE2 | CAND2 | 0,6262 | 1,05489E-40 |
| ACE2 | FSBP | 0,6262 | 1,03554E-40 |
| ACE2 | TMBIM1 | 0,6261 | 1,07379E-40 |
| ACE2 | TMEM218 | 0,6261 | 1,10589E-40 |
| ACE2 | PPP1R9A | 0,626 | 1,11352E-40 |
| ACE2 | BRMS1L | 0,626 | 1,14765E-40 |
| ACE2 | ZMYM3 | 0,626 | 1,14668E-40 |
| ACE2 | SEPSECS | 0,6259 | 1,17904E-40 |
| ACE2 | ADD3 | 0,6259 | 1,16456E-40 |
| ACE2 | USP31 | 0,6259 | 1,15691E-40 |
| ACE2 | GPR180 | 0,6258 | 1,19246E-40 |
| ACE2 | SPPL2A | 0,6256 | 1,29924E-40 |
| ACE2 | ARHGEF3 | 0,6255 | 1,37277E-40 |
| ACE2 | C1QTNF2 | 0,6255 | 1,36421E-40 |
| ACE2 | UBTF | 0,6255 | 1,3642E-40 |
| ACE2 | JADE2 | 0,6254 | 1,40258E-40 |
| ACE2 | RABEP1 | 0,6254 | 1,40338E-40 |
| ACE2 | SMARCE1 | 0,6254 | 1,42698E-40 |
| ACE2 | CRTAP | 0,6252 | 1,54183E-40 |
| ACE2 | SNX10 | 0,6252 | 1,51976E-40 |
| ACE2 | SEC11A | 0,6252 | 1,52773E-40 |
| ACE2 | NYAP1 | 0,6251 | 1,55345E-40 |
| ACE2 | TVP23B | 0,6251 | 1,58447E-40 |
| ACE2 | PPP6R3 | 0,625 | 1,61838E-40 |
| ACE2 | KIAA1551 | 0,625 | 1,65317E-40 |
| ACE2 | KCTD9 | 0,6249 | 1,68418E-40 |
| ACE2 | DNAJC3 | 0,6249 | 1,72153E-40 |
| ACE2 | KNOP1 | 0,6249 | 1,70079E-40 |
| ACE2 | SEZ6L2 | 0,6249 | 1,69565E-40 |
| ACE2 | POGLUT1 | 0,6248 | 1,76227E-40 |
| ACE2 | GLCE | 0,6247 | 1,80218E-40 |
| ACE2 | SH3TC2 | 0,6246 | 1,92458E-40 |
| ACE2 | REV3L | 0,6245 | 1,97898E-40 |
| ACE2 | KMT2E | 0,6244 | 2,0273E-40 |
| ACE2 | GABPA | 0,6244 | 2,06758E-40 |
| ACE2 | KLHL13 | 0,6244 | 2,0103E-40 |
| ACE2 | ABHD11 | 0,6243 | 2,13956E-40 |
| ACE2 | TMEM65 | 0,6243 | 2,12149E-40 |
| ACE2 | STAT1 | 0,6242 | 2,16031E-40 |
| ACE2 | MRPS27 | 0,6242 | 2,16017E-40 |
| ACE2 | CREBRF | 0,6242 | 2,21572E-40 |
| ACE2 | RHOB | 0,6241 | 2,29897E-40 |
| ACE2 | LAMB2 | 0,6241 | 2,27699E-40 |
| ACE2 | FGD6 | 0,6241 | 2,25047E-40 |
| ACE2 | LRIG2 | 0,624 | 2,38148E-40 |
| ACE2 | STAR | 0,624 | 2,40624E-40 |

| | | | |
|---|---|---|---|
| ACE2 | ZBED8 | 0,6239 | 2,43575E-40 |
| ACE2 | HTR7P1 | 0,6239 | 2,43716E-40 |
| ACE2 | FBXO5 | 0,6237 | 2,65691E-40 |
| ACE2 | NCOR2 | 0,6237 | 2,62409E-40 |
| ACE2 | PTPN2 | 0,6237 | 2,62934E-40 |
| ACE2 | SREBF2 | 0,6237 | 2,68438E-40 |
| ACE2 | GDI1 | 0,6237 | 2,59734E-40 |
| ACE2 | BCAP31 | 0,6235 | 2,87979E-40 |
| ACE2 | DUSP12 | 0,6234 | 2,92726E-40 |
| ACE2 | RBM25 | 0,6233 | 3,0597E-40 |
| ACE2 | AHDC1 | 0,6232 | 3,1657E-40 |
| ACE2 | FZD1 | 0,6232 | 3,12681E-40 |
| ACE2 | C14orf159 | 0,6232 | 3,21401E-40 |
| ACE2 | VAT1 | 0,623 | 3,44652E-40 |
| ACE2 | N6AMT1 | 0,623 | 3,4469E-40 |
| ACE2 | TRAPPC1 | 0,6229 | 3,59233E-40 |
| ACE2 | YWHAH | 0,6229 | 3,5525E-40 |
| ACE2 | PLOD1 | 0,6228 | 3,68871E-40 |
| ACE2 | UBLCP1 | 0,6228 | 3,71137E-40 |
| ACE2 | PRR12 | 0,6227 | 3,80505E-40 |
| ACE2 | LINC00641 | 0,6226 | 4,02336E-40 |
| ACE2 | BCKDK | 0,6226 | 4,02203E-40 |
| ACE2 | RP3-428L16.2 | 0,6225 | 4,13439E-40 |
| ACE2 | C16orf52 | 0,6225 | 4,05596E-40 |
| ACE2 | AMOT | 0,6225 | 4,11383E-40 |
| ACE2 | PDCD6 | 0,6223 | 4,47584E-40 |
| ACE2 | TDG | 0,6223 | 4,44648E-40 |
| ACE2 | ATP6V0C | 0,6223 | 4,36496E-40 |
| ACE2 | SKP1 | 0,6222 | 4,60027E-40 |
| ACE2 | DDR1 | 0,622 | 4,84877E-40 |
| ACE2 | NUP43 | 0,622 | 4,85193E-40 |
| ACE2 | BAZ1B | 0,622 | 4,97165E-40 |
| ACE2 | DNASE1L1 | 0,622 | 4,9981E-40 |
| ACE2 | XXyac-YR38GF2.1 | 0,6219 | 5,05628E-40 |
| ACE2 | SEPT7 | 0,6218 | 5,31683E-40 |
| ACE2 | GDF11 | 0,6218 | 5,27122E-40 |
| ACE2 | GABPB1 | 0,6218 | 5,2111E-40 |
| ACE2 | UGCG | 0,6217 | 5,52752E-40 |
| ACE2 | AP4E1 | 0,6217 | 5,46216E-40 |
| ACE2 | RP11-342K6.1 | 0,6216 | 5,67686E-40 |
| ACE2 | MYL9 | 0,6216 | 5,78297E-40 |
| ACE2 | SIGMAR1 | 0,6215 | 5,89536E-40 |
| ACE2 | ASB8 | 0,6215 | 5,83955E-40 |
| ACE2 | RASSF3 | 0,6215 | 5,84675E-40 |
| ACE2 | KIF13B | 0,6214 | 6,06387E-40 |
| ACE2 | FAM107B | 0,6214 | 6,09196E-40 |
| ACE2 | FAF2 | 0,6213 | 6,33482E-40 |
| ACE2 | TRIM8 | 0,6213 | 6,38327E-40 |
| ACE2 | NFATC2IP | 0,6213 | 6,47429E-40 |
| ACE2 | PLS3 | 0,6212 | 6,56775E-40 |
| ACE2 | SPOPL | 0,6211 | 6,76579E-40 |
| ACE2 | LAMTOR1 | 0,6211 | 6,7233E-40 |
| ACE2 | IL10RB | 0,6208 | 7,68157E-40 |
| ACE2 | ATP6V0B | 0,6207 | 7,91329E-40 |

| | | | |
|---|---|---|---|
| ACE2 | PTGFRN | 0,6207 | 7,88821E-40 |
| ACE2 | SLC40A1 | 0,6207 | 7,96245E-40 |
| ACE2 | AC000123.2 | 0,6207 | 8,05514E-40 |
| ACE2 | ME3 | 0,6207 | 7,98981E-40 |
| ACE2 | MTAP | 0,6204 | 8,7833E-40 |
| ACE2 | TMEM184C | 0,6203 | 9,21256E-40 |
| ACE2 | TRIM25 | 0,6203 | 8,99018E-40 |
| ACE2 | SUMF1 | 0,6201 | 9,75132E-40 |
| ACE2 | HYAL2 | 0,6201 | 9,99133E-40 |
| ACE2 | PRAF2 | 0,6201 | 9,69359E-40 |
| ACE2 | SHOC2 | 0,62 | 1,02134E-39 |
| ACE2 | PATZ1 | 0,62 | 1,03936E-39 |
| ACE2 | ZNF655 | 0,6199 | 1,07388E-39 |
| ACE2 | HS6ST1 | 0,6198 | 1,11368E-39 |
| ACE2 | MAML3 | 0,6198 | 1,10565E-39 |
| ACE2 | RP11-283I3.6 | 0,6197 | 1,15713E-39 |
| ACE2 | ATP2B1-AS1 | 0,6196 | 1,18683E-39 |
| ACE2 | LINC01560 | 0,6196 | 1,16973E-39 |
| ACE2 | ULBP1 | 0,6195 | 1,20964E-39 |
| ACE2 | FBP1 | 0,6195 | 1,21835E-39 |
| ACE2 | CREBZF | 0,6195 | 1,21523E-39 |
| ACE2 | TSPY26P | 0,6194 | 1,27644E-39 |
| ACE2 | CD9 | 0,6192 | 1,37784E-39 |
| ACE2 | MRPS31P4 | 0,6191 | 1,4366E-39 |
| ACE2 | TMTC4 | 0,6191 | 1,43695E-39 |
| ACE2 | PIK3R4 | 0,619 | 1,48855E-39 |
| ACE2 | KLF15 | 0,6189 | 1,53093E-39 |
| ACE2 | ATM | 0,6189 | 1,494E-39 |
| ACE2 | GTF2IP4 | 0,6187 | 1,61027E-39 |
| ACE2 | CALHM2 | 0,6186 | 1,69053E-39 |
| ACE2 | USP14 | 0,6186 | 1,68993E-39 |
| ACE2 | UNC119 | 0,6184 | 1,81234E-39 |
| ACE2 | NME2 | 0,6184 | 1,82474E-39 |
| ACE2 | BEX5 | 0,6184 | 1,85408E-39 |
| ACE2 | MFGE8 | 0,6183 | 1,91219E-39 |
| ACE2 | C16orf70 | 0,6183 | 1,88553E-39 |
| ACE2 | XPC | 0,6182 | 1,97835E-39 |
| ACE2 | HSP90AB1 | 0,6182 | 1,97341E-39 |
| ACE2 | DDX19A | 0,6181 | 2,04904E-39 |
| ACE2 | SFT2D2 | 0,618 | 2,06994E-39 |
| ACE2 | PROSC | 0,6179 | 2,15646E-39 |
| ACE2 | ABCC1 | 0,6179 | 2,21447E-39 |
| ACE2 | S100PBP | 0,6178 | 2,28177E-39 |
| ACE2 | TRRAP | 0,6178 | 2,24509E-39 |
| ACE2 | FAM3C2 | 0,6178 | 2,28658E-39 |
| ACE2 | RPS6KA3 | 0,6177 | 2,31522E-39 |
| ACE2 | PLSCR1 | 0,6176 | 2,40115E-39 |
| ACE2 | ORAI1 | 0,6176 | 2,42679E-39 |
| ACE2 | FOXRED2 | 0,6176 | 2,4591E-39 |
| ACE2 | PDCD4 | 0,6175 | 2,55549E-39 |
| ACE2 | HHLA3 | 0,6174 | 2,57865E-39 |
| ACE2 | CMBL | 0,6174 | 2,58712E-39 |
| ACE2 | HUS1 | 0,6174 | 2,60278E-39 |
| ACE2 | ZNF623 | 0,6174 | 2,57493E-39 |

| | | | |
|---|---|---|---|
| ACE2 | WBP1L | 0,6173 | 2,69848E-39 |
| ACE2 | IFI16 | 0,6172 | 2,85537E-39 |
| ACE2 | PLEKHA8 | 0,6171 | 2,92112E-39 |
| ACE2 | RP11-566E18.1 | 0,617 | 2,97471E-39 |
| ACE2 | STK38L | 0,617 | 3,06059E-39 |
| ACE2 | SMOC2 | 0,6169 | 3,15716E-39 |
| ACE2 | SMU1 | 0,6168 | 3,25593E-39 |
| ACE2 | C4orf33 | 0,6167 | 3,39993E-39 |
| ACE2 | LRRC8A | 0,6166 | 3,45312E-39 |
| ACE2 | VIM | 0,6166 | 3,46225E-39 |
| ACE2 | RP11-521C22.2 | 0,6166 | 3,45561E-39 |
| ACE2 | SEMA4B | 0,6166 | 3,44147E-39 |
| ACE2 | SHC1 | 0,6165 | 3,65505E-39 |
| ACE2 | ZRANB2 | 0,6164 | 3,6812E-39 |
| ACE2 | PCDHGB4 | 0,6164 | 3,76181E-39 |
| ACE2 | HMGCS1 | 0,6163 | 3,81793E-39 |
| ACE2 | ABCF3 | 0,6162 | 3,97288E-39 |
| ACE2 | FAM210A | 0,6161 | 4,13919E-39 |
| ACE2 | MAOB | 0,6161 | 4,16599E-39 |
| ACE2 | B4GALT6 | 0,616 | 4,32681E-39 |
| ACE2 | RBM3 | 0,616 | 4,24686E-39 |
| ACE2 | RP11-4O1.2 | 0,6158 | 4,69029E-39 |
| ACE2 | TBX3 | 0,6156 | 4,90826E-39 |
| ACE2 | DAAM1 | 0,6156 | 4,94893E-39 |
| ACE2 | STAU1 | 0,6156 | 5,03816E-39 |
| ACE2 | AL133243.1 | 0,6155 | 5,11537E-39 |
| ACE2 | TRIAP1 | 0,6155 | 5,07809E-39 |
| ACE2 | F3 | 0,6153 | 5,61623E-39 |
| ACE2 | VPS36 | 0,6153 | 5,62509E-39 |
| ACE2 | CAPN2 | 0,6151 | 5,9077E-39 |
| ACE2 | ADD1 | 0,6151 | 5,95263E-39 |
| ACE2 | SGIP1 | 0,615 | 6,08391E-39 |
| ACE2 | GATB | 0,615 | 6,19554E-39 |
| ACE2 | ILVBL | 0,615 | 6,22731E-39 |
| ACE2 | YY2 | 0,615 | 6,24998E-39 |
| ACE2 | MTFMT | 0,6149 | 6,35474E-39 |
| ACE2 | TRMT6 | 0,6149 | 6,39661E-39 |
| ACE2 | LRRN1 | 0,6148 | 6,54996E-39 |
| ACE2 | AASS | 0,6147 | 6,89588E-39 |
| ACE2 | SUCLA2 | 0,6147 | 6,99194E-39 |
| ACE2 | GPC3 | 0,6146 | 7,04998E-39 |
| ACE2 | LRIF1 | 0,6144 | 7,656E-39 |
| ACE2 | C19orf12 | 0,6144 | 7,75567E-39 |
| ACE2 | RP11-284E5.1 | 0,6143 | 7,7985E-39 |
| ACE2 | C1orf131 | 0,614 | 8,88497E-39 |
| ACE2 | ARNTL | 0,614 | 8,83539E-39 |
| ACE2 | BRWD1 | 0,6139 | 9,22767E-39 |
| ACE2 | AGO1 | 0,6138 | 9,46736E-39 |
| ACE2 | SHISA5 | 0,6138 | 9,48057E-39 |
| ACE2 | IMP3 | 0,6138 | 9,41413E-39 |
| ACE2 | TSPAN14 | 0,6137 | 9,72165E-39 |
| ACE2 | TIA1 | 0,6135 | 1,06958E-38 |
| ACE2 | TNPO1 | 0,6135 | 1,04277E-38 |
| ACE2 | CNKSR3 | 0,6135 | 1,06901E-38 |

| | | | |
|---|---|---|---|
| ACE2 | GRID1 | 0,6133 | 1,13048E-38 |
| ACE2 | LHFP | 0,6132 | 1,1576E-38 |
| ACE2 | KDM1B | 0,6131 | 1,19668E-38 |
| ACE2 | ACSS1 | 0,6131 | 1,21086E-38 |
| ACE2 | RP1-253P7.4 | 0,6128 | 1,35073E-38 |
| ACE2 | TMX3 | 0,6128 | 1,36705E-38 |
| ACE2 | TMEM47 | 0,6128 | 1,33511E-38 |
| ACE2 | CNPY4 | 0,6127 | 1,38191E-38 |
| ACE2 | RCAN3 | 0,6126 | 1,45883E-38 |
| ACE2 | HEPH | 0,6126 | 1,42224E-38 |
| ACE2 | LYN | 0,6125 | 1,50339E-38 |
| ACE2 | EIF4B | 0,6123 | 1,61603E-38 |
| ACE2 | SCN9A | 0,612 | 1,79723E-38 |
| ACE2 | PAXIP1-AS2 | 0,612 | 1,77592E-38 |
| ACE2 | CASP7 | 0,612 | 1,8092E-38 |
| ACE2 | CHMP4C | 0,6119 | 1,85488E-38 |
| ACE2 | TM9SF2 | 0,6119 | 1,87241E-38 |
| ACE2 | H6PD | 0,6118 | 1,90735E-38 |
| ACE2 | REST | 0,6117 | 1,97109E-38 |
| ACE2 | KDELC2 | 0,6117 | 1,9787E-38 |
| ACE2 | ZNF431 | 0,6117 | 1,97801E-38 |
| ACE2 | MATN3 | 0,6116 | 2,04293E-38 |
| ACE2 | SOX4 | 0,6116 | 2,03172E-38 |
| ACE2 | CCDC71L | 0,6116 | 2,06536E-38 |
| ACE2 | NHLRC1 | 0,6115 | 2,14677E-38 |
| ACE2 | LDHD | 0,6115 | 2,13539E-38 |
| ACE2 | SPIN3 | 0,6114 | 2,23273E-38 |
| ACE2 | CHURC1 | 0,6113 | 2,28606E-38 |
| ACE2 | FN3KRP | 0,6112 | 2,3658E-38 |
| ACE2 | MTR | 0,6111 | 2,49803E-38 |
| ACE2 | APOA1 | 0,6111 | 2,45306E-38 |
| ACE2 | SLC35A3 | 0,611 | 2,56187E-38 |
| ACE2 | SLC25A43 | 0,6108 | 2,71318E-38 |
| ACE2 | USP39 | 0,6105 | 3,06758E-38 |
| ACE2 | GNL3 | 0,6105 | 3,02485E-38 |
| ACE2 | MOB1B | 0,6105 | 3,061E-38 |
| ACE2 | ZNF665 | 0,6105 | 3,0721E-38 |
| ACE2 | NCK1 | 0,6104 | 3,08975E-38 |
| ACE2 | CHST12 | 0,6104 | 3,19537E-38 |
| ACE2 | PSAT1 | 0,6104 | 3,09336E-38 |
| ACE2 | RAD51B | 0,6104 | 3,18608E-38 |
| ACE2 | H1F0 | 0,6104 | 3,19704E-38 |
| ACE2 | UBR2 | 0,6103 | 3,28055E-38 |
| ACE2 | TARS | 0,6102 | 3,3455E-38 |
| ACE2 | RP11-690D19.3 | 0,6102 | 3,36696E-38 |
| ACE2 | SLC44A2 | 0,6102 | 3,32785E-38 |
| ACE2 | YTHDF1 | 0,6101 | 3,52582E-38 |
| ACE2 | GAB3 | 0,6101 | 3,47761E-38 |
| ACE2 | DGUOK | 0,6099 | 3,70171E-38 |
| ACE2 | ZNF704 | 0,6099 | 3,79494E-38 |
| ACE2 | UBE2Z | 0,6096 | 4,09667E-38 |
| ACE2 | MTDH | 0,6095 | 4,38199E-38 |
| ACE2 | LONP2 | 0,6095 | 4,33661E-38 |
| ACE2 | ZNF561 | 0,6095 | 4,23438E-38 |

| | | | |
|---|---|---|---|
| ACE2 | FSTL1 | 0,6094 | 4,47539E-38 |
| ACE2 | FOXC1 | 0,6094 | 4,43517E-38 |
| ACE2 | RP11-644F5.11 | 0,6094 | 4,45798E-38 |
| ACE2 | ARMC1 | 0,6093 | 4,64868E-38 |
| ACE2 | KMT2A | 0,6093 | 4,59321E-38 |
| ACE2 | CD46 | 0,6092 | 4,71311E-38 |
| ACE2 | SIAH2 | 0,6092 | 4,76117E-38 |
| ACE2 | GBP2 | 0,6091 | 4,92061E-38 |
| ACE2 | UVRAG | 0,6091 | 5,00891E-38 |
| ACE2 | GLG1 | 0,6091 | 5,01385E-38 |
| ACE2 | SLC39A8 | 0,609 | 5,16976E-38 |
| ACE2 | LDOC1 | 0,609 | 5,14983E-38 |
| ACE2 | HOMER2 | 0,6088 | 5,47044E-38 |
| ACE2 | ACSF2 | 0,6088 | 5,47625E-38 |
| ACE2 | PDE4DIP | 0,6087 | 5,74806E-38 |
| ACE2 | HNMT | 0,6087 | 5,78889E-38 |
| ACE2 | KIT | 0,6087 | 5,74517E-38 |
| ACE2 | PARN | 0,6087 | 5,74789E-38 |
| ACE2 | BAMBI | 0,6086 | 5,87577E-38 |
| ACE2 | TRPM3 | 0,6085 | 6,19157E-38 |
| ACE2 | CYBRD1 | 0,6083 | 6,52619E-38 |
| ACE2 | MYL6 | 0,6083 | 6,53063E-38 |
| ACE2 | KTN1-AS1 | 0,6083 | 6,44954E-38 |
| ACE2 | LIMK2 | 0,6083 | 6,54393E-38 |
| ACE2 | LINC00630 | 0,6083 | 6,4563E-38 |
| ACE2 | FAM13B | 0,6082 | 6,72935E-38 |
| ACE2 | LINC02035 | 0,6081 | 7,01318E-38 |
| ACE2 | TMEM164 | 0,6081 | 6,99309E-38 |
| ACE2 | XXYLT1 | 0,6078 | 7,86525E-38 |
| ACE2 | BLCAP | 0,6078 | 7,90762E-38 |
| ACE2 | SLC4A2 | 0,6077 | 7,98953E-38 |
| ACE2 | LETMD1 | 0,6077 | 8,09231E-38 |
| ACE2 | TXN2 | 0,6077 | 8,18849E-38 |
| ACE2 | NKAPP1 | 0,6077 | 8,07559E-38 |
| ACE2 | MTMR11 | 0,6076 | 8,26259E-38 |
| ACE2 | CHML | 0,6076 | 8,34671E-38 |
| ACE2 | ERCC6L2 | 0,6076 | 8,47782E-38 |
| ACE2 | PELI2 | 0,6076 | 8,44178E-38 |
| ACE2 | ZNF229 | 0,6076 | 8,41274E-38 |
| ACE2 | SHROOM2 | 0,6076 | 8,32194E-38 |
| ACE2 | SLC37A3 | 0,6075 | 8,58469E-38 |
| ACE2 | RHOBTB2 | 0,6075 | 8,77697E-38 |
| ACE2 | ITGA9 | 0,6074 | 8,99115E-38 |
| ACE2 | CSTF2 | 0,6074 | 8,87095E-38 |
| ACE2 | KDM4A | 0,607 | 1,01104E-37 |
| ACE2 | ZCCHC24 | 0,607 | 1,02666E-37 |
| ACE2 | PIGS | 0,607 | 1,01322E-37 |
| ACE2 | POLR3GL | 0,6068 | 1,10287E-37 |
| ACE2 | HMOX1 | 0,6068 | 1,11813E-37 |
| ACE2 | TMCO1 | 0,6067 | 1,16023E-37 |
| ACE2 | LRRFIP1 | 0,6067 | 1,13699E-37 |
| ACE2 | AAED1 | 0,6067 | 1,12761E-37 |
| ACE2 | PRR13 | 0,6067 | 1,15471E-37 |
| ACE2 | ATL2 | 0,6066 | 1,19156E-37 |

| | | | |
|---|---|---|---|
| ACE2 | B3GALNT1 | 0,6066 | 1,19314E-37 |
| ACE2 | ATP10D | 0,6066 | 1,18543E-37 |
| ACE2 | C2orf49 | 0,6064 | 1,2573E-37 |
| ACE2 | CBX6 | 0,6064 | 1,27585E-37 |
| ACE2 | GCC2 | 0,6063 | 1,29948E-37 |
| ACE2 | SIN3A | 0,6063 | 1,31142E-37 |
| ACE2 | ZNF792 | 0,6063 | 1,33248E-37 |
| ACE2 | ADNP | 0,6063 | 1,32277E-37 |
| ACE2 | KLF13 | 0,6062 | 1,37286E-37 |
| ACE2 | CTD-2001J20.1 | 0,6062 | 1,35648E-37 |
| ACE2 | TIMP3 | 0,6062 | 1,36318E-37 |
| ACE2 | UFL1 | 0,6061 | 1,41433E-37 |
| ACE2 | STRN3 | 0,6061 | 1,42183E-37 |
| ACE2 | MYZAP | 0,6061 | 1,40749E-37 |
| ACE2 | NPC1 | 0,6061 | 1,41308E-37 |
| ACE2 | FOXJ3 | 0,606 | 1,43583E-37 |
| ACE2 | SERPINH1 | 0,606 | 1,44322E-37 |
| ACE2 | MREG | 0,6059 | 1,48876E-37 |
| ACE2 | NUDT15 | 0,6059 | 1,51956E-37 |
| ACE2 | ABHD17C | 0,6059 | 1,49945E-37 |
| ACE2 | RAB22A | 0,6059 | 1,50005E-37 |
| ACE2 | PARVB | 0,6058 | 1,56572E-37 |
| ACE2 | C5orf15 | 0,6057 | 1,60431E-37 |
| ACE2 | LARS | 0,6057 | 1,6148E-37 |
| ACE2 | PAM | 0,6056 | 1,64186E-37 |
| ACE2 | IMPAD1 | 0,6056 | 1,66164E-37 |
| ACE2 | PTS | 0,6056 | 1,66895E-37 |
| ACE2 | EFNB2 | 0,6056 | 1,64311E-37 |
| ACE2 | LRRFIP1P1 | 0,6055 | 1,71918E-37 |
| ACE2 | ZDHHC21 | 0,6055 | 1,7479E-37 |
| ACE2 | ZNF674 | 0,6055 | 1,75242E-37 |
| ACE2 | TNKS1BP1 | 0,6054 | 1,81144E-37 |
| ACE2 | ZNF81 | 0,6054 | 1,79252E-37 |
| ACE2 | ITLN1 | 0,6052 | 1,89722E-37 |
| ACE2 | MAGEE1 | 0,6052 | 1,90552E-37 |
| ACE2 | PSAPL1 | 0,605 | 2,05536E-37 |
| ACE2 | PRIMPOL | 0,605 | 2,02282E-37 |
| ACE2 | ZNF658 | 0,6049 | 2,11171E-37 |
| ACE2 | CTSC | 0,6049 | 2,15799E-37 |
| ACE2 | TTYH3 | 0,6047 | 2,251E-37 |
| ACE2 | TPM1 | 0,6047 | 2,23985E-37 |
| ACE2 | PUS10 | 0,6046 | 2,39274E-37 |
| ACE2 | RP11-359E3.4 | 0,6046 | 2,31724E-37 |
| ACE2 | SMIM10L2B | 0,6045 | 2,47398E-37 |
| ACE2 | ARID1B | 0,6044 | 2,52657E-37 |
| ACE2 | FCHO2 | 0,6042 | 2,7098E-37 |
| ACE2 | DIMT1 | 0,6041 | 2,80717E-37 |
| ACE2 | ARPC3 | 0,6041 | 2,80741E-37 |
| ACE2 | SSPN | 0,6037 | 3,25386E-37 |
| ACE2 | DOPEY1 | 0,6036 | 3,3286E-37 |
| ACE2 | ZNF189 | 0,6036 | 3,26738E-37 |
| ACE2 | C11orf95 | 0,6035 | 3,39371E-37 |
| ACE2 | SART3 | 0,6035 | 3,39697E-37 |
| ACE2 | MSH6 | 0,6034 | 3,5521E-37 |

| | | | |
|---|---|---|---|
| ACE2 | RFC3 | 0,6033 | 3,63038E-37 |
| ACE2 | ASH2L | 0,6032 | 3,74932E-37 |
| ACE2 | VTN | 0,6032 | 3,82584E-37 |
| ACE2 | ZBTB22 | 0,6029 | 4,22959E-37 |
| ACE2 | GPD1L | 0,6028 | 4,31267E-37 |
| ACE2 | SHQ1 | 0,6028 | 4,29849E-37 |
| ACE2 | SLC35F2 | 0,6028 | 4,39968E-37 |
| ACE2 | CPT1A | 0,6027 | 4,54333E-37 |
| ACE2 | ETFBKMT | 0,6027 | 4,4824E-37 |
| ACE2 | ZNF512 | 0,6026 | 4,74497E-37 |
| ACE2 | PHF14 | 0,6026 | 4,68502E-37 |
| ACE2 | TSHZ3 | 0,6026 | 4,67072E-37 |
| ACE2 | NBN | 0,6024 | 5,0713E-37 |
| ACE2 | MAPRE2 | 0,6024 | 5,03031E-37 |
| ACE2 | RP3-425C14.4 | 0,6022 | 5,35859E-37 |
| ACE2 | SLFN5 | 0,6022 | 5,43995E-37 |
| ACE2 | UHMK1 | 0,6021 | 5,45516E-37 |
| ACE2 | ZNF292 | 0,6021 | 5,47143E-37 |
| ACE2 | LGALS1 | 0,602 | 5,76925E-37 |
| ACE2 | PTBP1 | 0,6019 | 5,88308E-37 |
| ACE2 | SCD5 | 0,6018 | 6,06828E-37 |
| ACE2 | GATA5 | 0,6018 | 6,04897E-37 |
| ACE2 | TNFSF4 | 0,6017 | 6,32907E-37 |
| ACE2 | KCNH2 | 0,6017 | 6,41105E-37 |
| ACE2 | POLR3A | 0,6017 | 6,28282E-37 |
| ACE2 | PTPN13 | 0,6016 | 6,51677E-37 |
| ACE2 | SCAF8 | 0,6016 | 6,58697E-37 |
| ACE2 | MRO | 0,6016 | 6,55204E-37 |
| ACE2 | UTP23 | 0,6015 | 6,78067E-37 |
| ACE2 | ARPC1B | 0,6014 | 7,11926E-37 |
| ACE2 | PSME1 | 0,6014 | 6,95384E-37 |
| ACE2 | ZFYVE21 | 0,6014 | 7,02811E-37 |
| ACE2 | HAUS3 | 0,6013 | 7,18466E-37 |
| ACE2 | ZUFSP | 0,6012 | 7,46143E-37 |
| ACE2 | ANKIB1 | 0,6012 | 7,44958E-37 |
| ACE2 | ST20-AS1 | 0,6012 | 7,46302E-37 |
| ACE2 | ZCCHC4 | 0,601 | 8,01338E-37 |
| ACE2 | DPP6 | 0,601 | 7,92938E-37 |
| ACE2 | PDGFRA | 0,6009 | 8,41995E-37 |
| ACE2 | PCBP2 | 0,6009 | 8,42822E-37 |
| ACE2 | TRAPPC11 | 0,6006 | 9,2687E-37 |
| ACE2 | MRC1 | 0,6006 | 9,13155E-37 |
| ACE2 | ZBTB7A | 0,6005 | 9,45755E-37 |
| ACE2 | TMEM187 | 0,6005 | 9,66597E-37 |
| ACE2 | SLC22A4 | 0,6003 | 1,02115E-36 |
| ACE2 | PPP3CB | 0,6003 | 1,02252E-36 |
| ACE2 | NOP58 | 0,6 | 1,14648E-36 |
| ACE2 | PRSS16 | 0,5997 | 1,26054E-36 |
| ACE2 | MSRB3 | 0,5997 | 1,27017E-36 |
| ACE2 | FBXO38 | 0,5995 | 1,34427E-36 |
| ACE2 | IFNAR2 | 0,5995 | 1,32949E-36 |
| ACE2 | SNIP1 | 0,5994 | 1,3733E-36 |
| ACE2 | TMEM200A | 0,5994 | 1,39631E-36 |
| ACE2 | TMEM177 | 0,5993 | 1,42518E-36 |

| | | | |
|---|---|---|---|
| ACE2 | PPARA | 0,5993 | 1,41198E-36 |
| ACE2 | SCML2 | 0,5993 | 1,41003E-36 |
| ACE2 | RBM24 | 0,5992 | 1,48422E-36 |
| ACE2 | ALDOC | 0,5992 | 1,50426E-36 |
| ACE2 | ABCD1 | 0,5992 | 1,48242E-36 |
| ACE2 | CDC37L1 | 0,5991 | 1,55353E-36 |
| ACE2 | CTD-2574D22.4 | 0,5991 | 1,52524E-36 |
| ACE2 | RNF135 | 0,5991 | 1,54179E-36 |
| ACE2 | ZADH2 | 0,5991 | 1,52259E-36 |
| ACE2 | SH3BP5 | 0,599 | 1,59782E-36 |
| ACE2 | TAPBP | 0,599 | 1,60226E-36 |
| ACE2 | PLEKHM3 | 0,5988 | 1,71922E-36 |
| ACE2 | EIF2B1 | 0,5987 | 1,75147E-36 |
| ACE2 | TSPYL4 | 0,5986 | 1,83763E-36 |
| ACE2 | RP11-400F19.6 | 0,5986 | 1,79706E-36 |
| ACE2 | SBDSP1 | 0,5984 | 1,95472E-36 |
| ACE2 | CST3 | 0,5984 | 1,91912E-36 |
| ACE2 | PCDH18 | 0,5983 | 2,00176E-36 |
| ACE2 | AUTS2 | 0,5983 | 1,98935E-36 |
| ACE2 | FZD3 | 0,5983 | 1,99791E-36 |
| ACE2 | RRAS | 0,5983 | 1,99173E-36 |
| ACE2 | DHX35 | 0,5982 | 2,09662E-36 |
| ACE2 | PRPF39 | 0,5981 | 2,15451E-36 |
| ACE2 | ABHD17B | 0,5979 | 2,29791E-36 |
| ACE2 | IPO11 | 0,5977 | 2,47638E-36 |
| ACE2 | VPS11 | 0,5976 | 2,50147E-36 |
| ACE2 | ZNF521 | 0,5976 | 2,57058E-36 |
| ACE2 | ZNF304 | 0,5976 | 2,56439E-36 |
| ACE2 | SNX15 | 0,5975 | 2,63234E-36 |
| ACE2 | PGLYRP2 | 0,5975 | 2,60529E-36 |
| ACE2 | OGT | 0,5974 | 2,67996E-36 |
| ACE2 | STK26 | 0,5973 | 2,78173E-36 |
| ACE2 | HS6ST2 | 0,5973 | 2,8077E-36 |
| ACE2 | PECR | 0,5972 | 2,89958E-36 |
| ACE2 | AK3 | 0,5972 | 2,90354E-36 |
| ACE2 | SUPV3L1 | 0,5972 | 2,88482E-36 |
| ACE2 | FBN1 | 0,5971 | 3,00164E-36 |
| ACE2 | CEACAM1 | 0,5971 | 3,02076E-36 |
| ACE2 | MTUS1 | 0,597 | 3,08627E-36 |
| ACE2 | TNIP1 | 0,5969 | 3,18475E-36 |
| ACE2 | YWHAG | 0,5969 | 3,2327E-36 |
| ACE2 | MTCP1 | 0,5968 | 3,35311E-36 |
| ACE2 | APH1A | 0,5966 | 3,49235E-36 |
| ACE2 | ZNF605 | 0,5965 | 3,71188E-36 |
| ACE2 | FGFR1OP2 | 0,5964 | 3,75107E-36 |
| ACE2 | CXorf38 | 0,5964 | 3,81316E-36 |
| ACE2 | PRRC2C | 0,5963 | 3,86769E-36 |
| ACE2 | HCFC1 | 0,5963 | 3,93028E-36 |
| ACE2 | MAN2B2 | 0,5962 | 4,06779E-36 |
| ACE2 | SH3KBP1 | 0,5962 | 4,11268E-36 |
| ACE2 | SYN1 | 0,5962 | 4,0403E-36 |
| ACE2 | WASHC2A | 0,5961 | 4,25873E-36 |
| ACE2 | CBFB | 0,5961 | 4,24171E-36 |
| ACE2 | HEATR1 | 0,596 | 4,26317E-36 |

| | | | |
|---|---|---|---|
| ACE2 | PBRM1 | 0,596 | 4,40144E-36 |
| ACE2 | TNFRSF19 | 0,596 | 4,28154E-36 |
| ACE2 | NPTN-IT1 | 0,5959 | 4,47694E-36 |
| ACE2 | CCNDBP1 | 0,5958 | 4,65455E-36 |
| ACE2 | TCEAL7 | 0,5958 | 4,57343E-36 |
| ACE2 | CERS5 | 0,5956 | 4,9469E-36 |
| ACE2 | KCNK15 | 0,5956 | 4,91097E-36 |
| ACE2 | NIPA2 | 0,5954 | 5,23659E-36 |
| ACE2 | LINC01852 | 0,5954 | 5,32336E-36 |
| ACE2 | RAD54B | 0,5953 | 5,47717E-36 |
| ACE2 | ACBD5 | 0,5953 | 5,53186E-36 |
| ACE2 | IRAK1 | 0,5952 | 5,60153E-36 |
| ACE2 | TFRC | 0,595 | 6,02504E-36 |
| ACE2 | SLC13A4 | 0,5949 | 6,23814E-36 |
| ACE2 | JADE3 | 0,5948 | 6,37226E-36 |
| ACE2 | TMEM163 | 0,5946 | 6,91035E-36 |
| ACE2 | AC074286.1 | 0,5946 | 6,978E-36 |
| ACE2 | FST | 0,5946 | 6,8755E-36 |
| ACE2 | HGSNAT | 0,5946 | 6,84268E-36 |
| ACE2 | JTB | 0,5944 | 7,39533E-36 |
| ACE2 | PDIA4 | 0,5944 | 7,42519E-36 |
| ACE2 | CTCF | 0,5944 | 7,2819E-36 |
| ACE2 | SPRED2 | 0,5943 | 7,7032E-36 |
| ACE2 | NIPBL | 0,5943 | 7,65685E-36 |
| ACE2 | HIST4H4 | 0,5943 | 7,69935E-36 |
| ACE2 | GTF2F2 | 0,5943 | 7,52104E-36 |
| ACE2 | EIF4A1 | 0,5943 | 7,74622E-36 |
| ACE2 | FNBP4 | 0,5942 | 7,992E-36 |
| ACE2 | SYNGR2 | 0,5941 | 8,1559E-36 |
| ACE2 | FAM102B | 0,594 | 8,39298E-36 |
| ACE2 | SLC16A1 | 0,594 | 8,32015E-36 |
| ACE2 | CTC-338M12.4 | 0,594 | 8,52983E-36 |
| ACE2 | FASTKD3 | 0,5939 | 8,79039E-36 |
| ACE2 | NMUR2 | 0,5939 | 8,73909E-36 |
| ACE2 | SYF2 | 0,5938 | 8,93751E-36 |
| ACE2 | BCR | 0,5938 | 8,96496E-36 |
| ACE2 | MAT2A | 0,5937 | 9,31488E-36 |
| ACE2 | AKR1A1 | 0,5936 | 9,51507E-36 |
| ACE2 | SPARCL1 | 0,5936 | 9,55985E-36 |
| ACE2 | MFHAS1 | 0,5936 | 9,70514E-36 |
| ACE2 | SMG8 | 0,5936 | 9,75153E-36 |
| ACE2 | STARD4 | 0,5935 | 9,95329E-36 |
| ACE2 | BMT2 | 0,5935 | 9,78479E-36 |
| ACE2 | TMEM229B | 0,5935 | 1,00466E-35 |
| ACE2 | FABP3 | 0,5934 | 1,01631E-35 |
| ACE2 | RBM8A | 0,5934 | 1,02859E-35 |
| ACE2 | XRCC5 | 0,5934 | 1,0157E-35 |
| ACE2 | EPB41L4B | 0,5934 | 1,02142E-35 |
| ACE2 | UNG | 0,5934 | 1,04224E-35 |
| ACE2 | CDC73 | 0,5933 | 1,05411E-35 |
| ACE2 | SRSF1 | 0,5933 | 1,07067E-35 |
| ACE2 | PPP4R3B | 0,5932 | 1,10407E-35 |
| ACE2 | CCDC125 | 0,5932 | 1,0834E-35 |
| ACE2 | MRPS6 | 0,5931 | 1,14751E-35 |

| | | | |
|---|---|---|---|
| ACE2 | GATSL3 | 0,5931 | 1,13633E-35 |
| ACE2 | ZNF718 | 0,593 | 1,1835E-35 |
| ACE2 | ANP32B | 0,593 | 1,16358E-35 |
| ACE2 | PAN3 | 0,593 | 1,15411E-35 |
| ACE2 | WSB1 | 0,5928 | 1,24282E-35 |
| ACE2 | NAMPT | 0,5927 | 1,30996E-35 |
| ACE2 | TEP1 | 0,5927 | 1,29655E-35 |
| ACE2 | HUNK | 0,5927 | 1,28883E-35 |
| ACE2 | UBQLN4 | 0,5926 | 1,34355E-35 |
| ACE2 | PCBP1 | 0,5926 | 1,35158E-35 |
| ACE2 | NCOA6 | 0,5926 | 1,35519E-35 |
| ACE2 | HECTD3 | 0,5925 | 1,36929E-35 |
| ACE2 | UGP2 | 0,5925 | 1,39695E-35 |
| ACE2 | PDZRN3 | 0,5924 | 1,43363E-35 |
| ACE2 | ATF1 | 0,5924 | 1,41144E-35 |
| ACE2 | SCPEP1 | 0,5924 | 1,43243E-35 |
| ACE2 | ZNF283 | 0,5924 | 1,43248E-35 |
| ACE2 | ZC3HAV1L | 0,5923 | 1,47292E-35 |
| ACE2 | RP11-264B17.2 | 0,5923 | 1,49709E-35 |
| ACE2 | PTPN1 | 0,5923 | 1,45177E-35 |
| ACE2 | NSG1 | 0,5921 | 1,57701E-35 |
| ACE2 | AC009784.3 | 0,5921 | 1,55287E-35 |
| ACE2 | TRMT10B | 0,5921 | 1,58316E-35 |
| ACE2 | STOM | 0,592 | 1,62171E-35 |
| ACE2 | MED14 | 0,592 | 1,62382E-35 |
| ACE2 | FKTN | 0,5919 | 1,66702E-35 |
| ACE2 | SETDB2 | 0,5919 | 1,67854E-35 |
| ACE2 | NFYB | 0,5917 | 1,80182E-35 |
| ACE2 | MAGEB4 | 0,5917 | 1,81477E-35 |
| ACE2 | HUWE1 | 0,5917 | 1,79862E-35 |
| ACE2 | ZMAT2 | 0,5916 | 1,87208E-35 |
| ACE2 | PHF12 | 0,5916 | 1,854E-35 |
| ACE2 | SLC12A7 | 0,5915 | 1,94885E-35 |
| ACE2 | ZNF527 | 0,5915 | 1,9188E-35 |
| ACE2 | RIMS4 | 0,5915 | 1,95028E-35 |
| ACE2 | TRIM4 | 0,5914 | 2,01089E-35 |
| ACE2 | RAD21 | 0,5914 | 1,95696E-35 |
| ACE2 | CNIH4 | 0,5912 | 2,14754E-35 |
| ACE2 | MAVS | 0,5912 | 2,13554E-35 |
| ACE2 | ARMC8 | 0,5911 | 2,208E-35 |
| ACE2 | ARPP19 | 0,591 | 2,22666E-35 |
| ACE2 | ZNF236 | 0,591 | 2,26968E-35 |
| ACE2 | ARHGAP17 | 0,5909 | 2,33271E-35 |
| ACE2 | PAK1IP1 | 0,5908 | 2,38658E-35 |
| ACE2 | PTAR1 | 0,5908 | 2,43285E-35 |
| ACE2 | RP11-1016B18.1 | 0,5907 | 2,47425E-35 |
| ACE2 | GLOD4 | 0,5906 | 2,61963E-35 |
| ACE2 | CHRM3-AS2 | 0,5905 | 2,66874E-35 |
| ACE2 | TIMM10B | 0,5905 | 2,6622E-35 |
| ACE2 | GPC4 | 0,5905 | 2,65321E-35 |
| ACE2 | RPIA | 0,5904 | 2,77449E-35 |
| ACE2 | ATP13A3 | 0,5904 | 2,71094E-35 |
| ACE2 | SEC14L6 | 0,5904 | 2,73355E-35 |
| ACE2 | ARHGEF26 | 0,5903 | 2,88943E-35 |

| | | | |
|---|---|---|---|
| ACE2 | ZNF845 | 0,5903 | 2,83871E-35 |
| ACE2 | EMD | 0,5903 | 2,84978E-35 |
| ACE2 | LUC7L2 | 0,5902 | 2,94917E-35 |
| ACE2 | AARD | 0,5902 | 2,96251E-35 |
| ACE2 | ECD | 0,5902 | 2,98602E-35 |
| ACE2 | COA1 | 0,5901 | 3,00252E-35 |
| ACE2 | PEMT | 0,5899 | 3,22628E-35 |
| ACE2 | RPL23 | 0,5899 | 3,28111E-35 |
| ACE2 | NFE2L3 | 0,5898 | 3,34192E-35 |
| ACE2 | PHKA2 | 0,5897 | 3,42157E-35 |
| ACE2 | ZNF654 | 0,5896 | 3,54567E-35 |
| ACE2 | FKRP | 0,5895 | 3,74602E-35 |
| ACE2 | HNRNPA3 | 0,5894 | 3,75861E-35 |
| ACE2 | CETN2 | 0,5894 | 3,77983E-35 |
| ACE2 | TET2 | 0,5893 | 3,91943E-35 |
| ACE2 | TOMM20 | 0,5892 | 4,05446E-35 |
| ACE2 | PIGW | 0,5892 | 4,06522E-35 |
| ACE2 | MCOLN1 | 0,5892 | 4,07655E-35 |
| ACE2 | ARMCX2 | 0,5892 | 4,09483E-35 |
| ACE2 | SEC14L4 | 0,5891 | 4,20318E-35 |
| ACE2 | LRRK2 | 0,589 | 4,30697E-35 |
| ACE2 | EFNB1 | 0,589 | 4,35281E-35 |
| ACE2 | ANAPC1 | 0,5889 | 4,54395E-35 |
| ACE2 | C5orf30 | 0,5889 | 4,54756E-35 |
| ACE2 | TMEM41B | 0,5889 | 4,50704E-35 |
| ACE2 | AC024560.3 | 0,5888 | 4,70066E-35 |
| ACE2 | PTER | 0,5888 | 4,60159E-35 |
| ACE2 | RAB35 | 0,5888 | 4,68071E-35 |
| ACE2 | NLGN4Y | 0,5888 | 4,59555E-35 |
| ACE2 | FAM219B | 0,5885 | 5,09743E-35 |
| ACE2 | ACVR2B | 0,5884 | 5,25133E-35 |
| ACE2 | FAXDC2 | 0,5882 | 5,68509E-35 |
| ACE2 | RPL41 | 0,5882 | 5,6251E-35 |
| ACE2 | KIRREL | 0,5881 | 5,87499E-35 |
| ACE2 | UBE2I | 0,5881 | 5,87654E-35 |
| ACE2 | PIGG | 0,588 | 6,04999E-35 |
| ACE2 | SEC24C | 0,5879 | 6,17653E-35 |
| ACE2 | TIMM17B | 0,5879 | 6,22834E-35 |
| ACE2 | ZNF117 | 0,5878 | 6,36385E-35 |
| ACE2 | ERVMER34-1 | 0,5876 | 6,93715E-35 |
| ACE2 | RP11-53O19.3 | 0,5876 | 6,88999E-35 |
| ACE2 | NKRF | 0,5875 | 7,17903E-35 |
| ACE2 | IMPDH2 | 0,5872 | 7,67481E-35 |
| ACE2 | GOLT1B | 0,5872 | 7,75249E-35 |
| ACE2 | RORC | 0,5871 | 7,99619E-35 |
| ACE2 | ARFGEF1 | 0,5871 | 8,03019E-35 |
| ACE2 | TLE1 | 0,5871 | 8,14785E-35 |
| ACE2 | MTA2 | 0,5871 | 7,92459E-35 |
| ACE2 | MAMDC2 | 0,587 | 8,21293E-35 |
| ACE2 | OMD | 0,587 | 8,42383E-35 |
| ACE2 | LMNA | 0,5869 | 8,63172E-35 |
| ACE2 | TBCA | 0,5869 | 8,6771E-35 |
| ACE2 | NAGLU | 0,5869 | 8,56344E-35 |
| ACE2 | UBL4A | 0,5869 | 8,57211E-35 |

| | | | |
|---|---|---|---|
| ACE2 | CDK13 | 0,5867 | 9,18834E-35 |
| ACE2 | HSCB | 0,5867 | 9,15452E-35 |
| ACE2 | CENPI | 0,5866 | 9,39938E-35 |
| ACE2 | MAGOHB | 0,5865 | 9,65091E-35 |
| ACE2 | RAP2A | 0,5865 | 9,67655E-35 |
| ACE2 | BRCC3 | 0,5865 | 9,81959E-35 |
| ACE2 | ABCB10 | 0,5863 | 1,03743E-34 |
| ACE2 | PCED1B | 0,5863 | 1,05802E-34 |
| ACE2 | PXN | 0,5863 | 1,04109E-34 |
| ACE2 | NFKBIA | 0,5863 | 1,04569E-34 |
| ACE2 | WDR81 | 0,5863 | 1,03853E-34 |
| ACE2 | RAB29 | 0,5862 | 1,08641E-34 |
| ACE2 | KMT2C | 0,5862 | 1,06032E-34 |
| ACE2 | EIF3C | 0,5862 | 1,08303E-34 |
| ACE2 | CLUH | 0,5862 | 1,07834E-34 |
| ACE2 | MSL3 | 0,5862 | 1,0649E-34 |
| ACE2 | IVNS1ABP | 0,5861 | 1,09879E-34 |
| ACE2 | PARP2 | 0,586 | 1,157E-34 |
| ACE2 | PPTC7 | 0,5859 | 1,17144E-34 |
| ACE2 | ENPP2 | 0,5858 | 1,22055E-34 |
| ACE2 | TSC22D4 | 0,5857 | 1,25368E-34 |
| ACE2 | CYB5RL | 0,5856 | 1,29178E-34 |
| ACE2 | ABCE1 | 0,5856 | 1,31673E-34 |
| ACE2 | DDX17 | 0,5856 | 1,30411E-34 |
| ACE2 | CDCA7L | 0,5855 | 1,32713E-34 |
| ACE2 | TMEM60 | 0,5855 | 1,34345E-34 |
| ACE2 | SLC25A25 | 0,5855 | 1,3426E-34 |
| ACE2 | MDM2 | 0,5854 | 1,38098E-34 |
| ACE2 | SNN | 0,5854 | 1,41083E-34 |
| ACE2 | RP11-244M2.1 | 0,5854 | 1,39975E-34 |
| ACE2 | NSUN2 | 0,5853 | 1,46092E-34 |
| ACE2 | ZBTB34 | 0,5853 | 1,45102E-34 |
| ACE2 | NAE1 | 0,5852 | 1,47203E-34 |
| ACE2 | SQSTM1 | 0,5851 | 1,53769E-34 |
| ACE2 | BTN2A2 | 0,5851 | 1,54669E-34 |
| ACE2 | RP11-440D17.3 | 0,585 | 1,56255E-34 |
| ACE2 | UXS1 | 0,585 | 1,60664E-34 |
| ACE2 | ARRDC4 | 0,585 | 1,56308E-34 |
| ACE2 | COX7A2L | 0,5849 | 1,62841E-34 |
| ACE2 | TRIM23 | 0,5849 | 1,65828E-34 |
| ACE2 | GAS1 | 0,5849 | 1,6188E-34 |
| ACE2 | RP11-421F16.3 | 0,5849 | 1,62853E-34 |
| ACE2 | TSHZ1 | 0,5849 | 1,65946E-34 |
| ACE2 | TSHZ2 | 0,5848 | 1,70477E-34 |
| ACE2 | OSBPL6 | 0,5847 | 1,73269E-34 |
| ACE2 | RP11-447D11.3 | 0,5847 | 1,72088E-34 |
| ACE2 | NIPAL2 | 0,5847 | 1,76619E-34 |
| ACE2 | CENPBD1P1 | 0,5847 | 1,73531E-34 |
| ACE2 | MME | 0,5846 | 1,77883E-34 |
| ACE2 | GTF2A1 | 0,5845 | 1,87496E-34 |
| ACE2 | RGS9 | 0,5845 | 1,85063E-34 |
| ACE2 | MAP3K4 | 0,5844 | 1,94202E-34 |
| ACE2 | SWAP70 | 0,5844 | 1,93582E-34 |
| ACE2 | SNRNP48 | 0,5843 | 1,98961E-34 |

| | | | |
|---|---|---|---|
| ACE2 | GPR157 | 0,5842 | 2,03098E-34 |
| ACE2 | TMCO4 | 0,5842 | 2,04562E-34 |
| ACE2 | PHKB | 0,5842 | 2,02925E-34 |
| ACE2 | SEPT11 | 0,5841 | 2,13646E-34 |
| ACE2 | ZKSCAN3 | 0,5841 | 2,10715E-34 |
| ACE2 | C6orf203 | 0,5841 | 2,14389E-34 |
| ACE2 | GRB14 | 0,584 | 2,14756E-34 |
| ACE2 | GPATCH8 | 0,584 | 2,1602E-34 |
| ACE2 | NOC3L | 0,5838 | 2,29755E-34 |
| ACE2 | MLF2 | 0,5838 | 2,29991E-34 |
| ACE2 | C15orf52 | 0,5838 | 2,2926E-34 |
| ACE2 | PPP1R14A | 0,5838 | 2,29168E-34 |
| ACE2 | EGFR | 0,5837 | 2,36462E-34 |
| ACE2 | RP3-368A4.6 | 0,5837 | 2,41647E-34 |
| ACE2 | TXNIP | 0,5835 | 2,56693E-34 |
| ACE2 | PTCD3 | 0,5835 | 2,56204E-34 |
| ACE2 | NAPEPLD | 0,5834 | 2,63784E-34 |
| ACE2 | ARHGEF40 | 0,5832 | 2,82098E-34 |
| ACE2 | MGAT5 | 0,5831 | 2,9096E-34 |
| ACE2 | PTPRK | 0,5831 | 2,87574E-34 |
| ACE2 | IST1 | 0,5831 | 2,92402E-34 |
| ACE2 | LAMB1 | 0,583 | 2,98418E-34 |
| ACE2 | SLC2A12 | 0,5829 | 3,05921E-34 |
| ACE2 | CA8 | 0,5829 | 3,09716E-34 |
| ACE2 | PTBP3 | 0,5827 | 3,26954E-34 |
| ACE2 | ABHD3 | 0,5827 | 3,33684E-34 |
| ACE2 | TMEM179B | 0,5826 | 3,44616E-34 |
| ACE2 | LNX2 | 0,5826 | 3,38831E-34 |
| ACE2 | ZNF765 | 0,5826 | 3,40989E-34 |
| ACE2 | PCYOX1L | 0,5825 | 3,47717E-34 |
| ACE2 | RP11-488L18.4 | 0,5824 | 3,64193E-34 |
| ACE2 | PPIL4 | 0,5824 | 3,65306E-34 |
| ACE2 | ZBTB6 | 0,5824 | 3,58789E-34 |
| ACE2 | A2M | 0,5824 | 3,61232E-34 |
| ACE2 | NIF3L1 | 0,5823 | 3,8102E-34 |
| ACE2 | MAP3K13 | 0,5821 | 3,96822E-34 |
| ACE2 | FZD4 | 0,5821 | 3,99715E-34 |
| ACE2 | GPR137C | 0,5821 | 4,00301E-34 |
| ACE2 | SRSF6 | 0,5821 | 3,94223E-34 |
| ACE2 | PNPLA8 | 0,582 | 4,14476E-34 |
| ACE2 | CPED1 | 0,582 | 4,10641E-34 |
| ACE2 | STOML2 | 0,582 | 4,13722E-34 |
| ACE2 | RNF150 | 0,5819 | 4,29339E-34 |
| ACE2 | AHCYL2 | 0,5819 | 4,19642E-34 |
| ACE2 | TWF2 | 0,5817 | 4,5124E-34 |
| ACE2 | ZNF557 | 0,5817 | 4,59617E-34 |
| ACE2 | SLFN12 | 0,5814 | 5,00401E-34 |
| ACE2 | IL15RA | 0,5813 | 5,1763E-34 |
| ACE2 | PYGB | 0,5813 | 5,12461E-34 |
| ACE2 | RCC1L | 0,5812 | 5,28541E-34 |
| ACE2 | MELK | 0,5812 | 5,32418E-34 |
| ACE2 | ENTPD4 | 0,5811 | 5,49489E-34 |
| ACE2 | TMEM30B | 0,581 | 5,64586E-34 |
| ACE2 | SUPT7L | 0,5809 | 5,86144E-34 |

| | | | |
|---|---|---|---|
| ACE2 | VKORC1 | 0,5809 | 5,79274E-34 |
| ACE2 | SRSF10 | 0,5807 | 6,22137E-34 |
| ACE2 | PDPN | 0,5806 | 6,36848E-34 |
| ACE2 | ARGLU1 | 0,5806 | 6,38853E-34 |
| ACE2 | FGFR2 | 0,5805 | 6,57585E-34 |
| ACE2 | ZNF445 | 0,5804 | 6,75145E-34 |
| ACE2 | ERRFI1 | 0,5802 | 7,38132E-34 |
| ACE2 | PTPRC | 0,5802 | 7,24419E-34 |
| ACE2 | GUCY1B3 | 0,5802 | 7,25646E-34 |
| ACE2 | LINC01549 | 0,5801 | 7,54751E-34 |
| ACE2 | GSN | 0,58 | 7,82842E-34 |
| ACE2 | RBBP5 | 0,5799 | 8,06664E-34 |
| ACE2 | LSM6 | 0,5799 | 7,98902E-34 |
| ACE2 | RAD50 | 0,5799 | 8,09187E-34 |
| ACE2 | MRFAP1 | 0,5798 | 8,24142E-34 |
| ACE2 | PTGR1 | 0,5798 | 8,15993E-34 |
| ACE2 | HMBS | 0,5798 | 8,18581E-34 |
| ACE2 | LAMA1 | 0,5798 | 8,22911E-34 |
| ACE2 | OTUD5 | 0,5796 | 8,82916E-34 |
| ACE2 | SLIT2 | 0,5795 | 9,22744E-34 |
| ACE2 | VSIG10 | 0,5794 | 9,47184E-34 |
| ACE2 | ACSM3 | 0,5794 | 9,39094E-34 |
| ACE2 | RPS6KA6 | 0,5794 | 9,48299E-34 |
| ACE2 | DDOST | 0,5793 | 9,57435E-34 |
| ACE2 | AC093323.3 | 0,5793 | 9,75367E-34 |
| ACE2 | MTMR3 | 0,5793 | 9,80539E-34 |
| ACE2 | PMM2 | 0,5792 | 9,90577E-34 |
| ACE2 | CD3EAP | 0,5792 | 9,98131E-34 |
| ACE2 | RBFOX2 | 0,5792 | 9,99219E-34 |
| ACE2 | DHDDS | 0,5791 | 1,0165E-33 |
| ACE2 | MCM3 | 0,5791 | 1,02784E-33 |
| ACE2 | PPP1R14C | 0,579 | 1,08114E-33 |
| ACE2 | IMPA1 | 0,579 | 1,06347E-33 |
| ACE2 | C11orf24 | 0,5789 | 1,09207E-33 |
| ACE2 | GMIP | 0,5787 | 1,18278E-33 |
| ACE2 | TRAPPC13 | 0,5786 | 1,22228E-33 |
| ACE2 | ORC4 | 0,5785 | 1,24584E-33 |
| ACE2 | PUS7 | 0,5785 | 1,22912E-33 |
| ACE2 | CPOX | 0,5784 | 1,27588E-33 |
| ACE2 | ABCC5 | 0,5782 | 1,38324E-33 |
| ACE2 | B3GAT3 | 0,5781 | 1,42347E-33 |
| ACE2 | NEPRO | 0,5779 | 1,48845E-33 |
| ACE2 | ANAPC4 | 0,5779 | 1,50713E-33 |
| ACE2 | LOH12CR2 | 0,5779 | 1,51042E-33 |
| ACE2 | JMJD1C | 0,5778 | 1,54522E-33 |
| ACE2 | ASAH2B | 0,5777 | 1,60629E-33 |
| ACE2 | TMEM246 | 0,5776 | 1,65758E-33 |
| ACE2 | TAF3 | 0,5774 | 1,74443E-33 |
| ACE2 | POLR2M | 0,5774 | 1,73627E-33 |
| ACE2 | RPL36A | 0,5774 | 1,75291E-33 |
| ACE2 | ADORA1 | 0,5773 | 1,79183E-33 |
| ACE2 | CTDSP1 | 0,5773 | 1,83418E-33 |
| ACE2 | SZRD1 | 0,5772 | 1,89821E-33 |
| ACE2 | TRIM5 | 0,5772 | 1,85801E-33 |

| | | | |
|---|---|---|---|
| ACE2 | TCEANC | 0,5772 | 1,89403E-33 |
| ACE2 | SCAP | 0,5771 | 1,90303E-33 |
| ACE2 | ZNF317 | 0,577 | 1,97321E-33 |
| ACE2 | PPT2 | 0,5769 | 2,08308E-33 |
| ACE2 | YIF1B | 0,5769 | 2,07921E-33 |
| ACE2 | ARFIP1 | 0,5767 | 2,22092E-33 |
| ACE2 | ABCA1 | 0,5767 | 2,2095E-33 |
| ACE2 | SLC39A10 | 0,5765 | 2,35827E-33 |
| ACE2 | RARS | 0,5765 | 2,30661E-33 |
| ACE2 | UBE3A | 0,5765 | 2,35525E-33 |
| ACE2 | ZNF470 | 0,5765 | 2,36362E-33 |
| ACE2 | ZXDC | 0,5763 | 2,50076E-33 |
| ACE2 | SUPT20HL2 | 0,5763 | 2,44442E-33 |
| ACE2 | INTS14 | 0,5762 | 2,58446E-33 |
| ACE2 | THUMPD1 | 0,576 | 2,69909E-33 |
| ACE2 | FCF1P2 | 0,5759 | 2,83679E-33 |
| ACE2 | MFSD8 | 0,5759 | 2,82863E-33 |
| ACE2 | ZNF740 | 0,5759 | 2,77256E-33 |
| ACE2 | SGPP1 | 0,5759 | 2,81955E-33 |
| ACE2 | UBE2Q2P1 | 0,5759 | 2,83138E-33 |
| ACE2 | ZMYM6 | 0,5758 | 2,90791E-33 |
| ACE2 | C7 | 0,5758 | 2,93144E-33 |
| ACE2 | PLEKHH3 | 0,5757 | 2,94278E-33 |
| ACE2 | PUM1 | 0,5755 | 3,16897E-33 |
| ACE2 | CENPQ | 0,5755 | 3,18326E-33 |
| ACE2 | ALDH18A1 | 0,5754 | 3,31732E-33 |
| ACE2 | SEH1L | 0,5754 | 3,26008E-33 |
| ACE2 | GTF2IRD1 | 0,5752 | 3,50534E-33 |
| ACE2 | NXPE3 | 0,5751 | 3,63184E-33 |
| ACE2 | RP11-762L8.6 | 0,5751 | 3,6343E-33 |
| ACE2 | TMEM159 | 0,5751 | 3,61643E-33 |
| ACE2 | C17orf49 | 0,5751 | 3,5543E-33 |
| ACE2 | TRIM65 | 0,5751 | 3,59878E-33 |
| ACE2 | NBPF15 | 0,575 | 3,72457E-33 |
| ACE2 | NID1 | 0,575 | 3,74524E-33 |
| ACE2 | RELA | 0,575 | 3,72282E-33 |
| ACE2 | FAM117B | 0,5749 | 3,87492E-33 |
| ACE2 | SNX27 | 0,5748 | 3,95312E-33 |
| ACE2 | CPNE5 | 0,5748 | 3,93463E-33 |
| ACE2 | RRS1 | 0,5748 | 3,93369E-33 |
| ACE2 | GAS6 | 0,5748 | 4,00589E-33 |
| ACE2 | NR1D2 | 0,5747 | 4,07407E-33 |
| ACE2 | NMD3 | 0,5747 | 4,05425E-33 |
| ACE2 | OSBPL8 | 0,5747 | 4,03154E-33 |
| ACE2 | ZNF12 | 0,5746 | 4,17285E-33 |
| ACE2 | MORC3 | 0,5746 | 4,22041E-33 |
| ACE2 | JAGN1 | 0,5745 | 4,28628E-33 |
| ACE2 | VWA5B2 | 0,5745 | 4,37924E-33 |
| ACE2 | RBAK | 0,5745 | 4,3686E-33 |
| ACE2 | ZNF320 | 0,5745 | 4,40025E-33 |
| ACE2 | RP11-284F21.7 | 0,5744 | 4,49552E-33 |
| ACE2 | ECHDC3 | 0,5743 | 4,56655E-33 |
| ACE2 | TPST2 | 0,5743 | 4,63995E-33 |
| ACE2 | IGIP | 0,5742 | 4,78566E-33 |

| | | | |
|---|---|---|---|
| ACE2 | PPP1R15A | 0,5742 | 4,82582E-33 |
| ACE2 | DLG3 | 0,5742 | 4,79491E-33 |
| ACE2 | DHX38 | 0,5741 | 4,98214E-33 |
| ACE2 | FAM198B | 0,574 | 5,01516E-33 |
| ACE2 | BIVM | 0,574 | 5,02304E-33 |
| ACE2 | MEGF10 | 0,5739 | 5,26635E-33 |
| ACE2 | SPRYD3 | 0,5739 | 5,20943E-33 |
| ACE2 | DLGAP1-AS1 | 0,5739 | 5,26909E-33 |
| ACE2 | FTH1 | 0,5737 | 5,54283E-33 |
| ACE2 | DCTPP1 | 0,5735 | 5,97829E-33 |
| ACE2 | PPP1R8 | 0,5733 | 6,3139E-33 |
| ACE2 | PLCL2 | 0,5733 | 6,36137E-33 |
| ACE2 | POLR1E | 0,5733 | 6,21072E-33 |
| ACE2 | CENPVL3 | 0,5732 | 6,53489E-33 |
| ACE2 | NINJ2 | 0,5731 | 6,62658E-33 |
| ACE2 | NKTR | 0,573 | 6,96054E-33 |
| ACE2 | CCNJ | 0,573 | 6,89347E-33 |
| ACE2 | SATB1 | 0,5729 | 7,1088E-33 |
| ACE2 | CLIP3 | 0,5729 | 7,20774E-33 |
| ACE2 | ADGRL2 | 0,5727 | 7,48183E-33 |
| ACE2 | SLC25A16 | 0,5725 | 7,99563E-33 |
| ACE2 | B2M | 0,5725 | 7,91641E-33 |
| ACE2 | NOD1 | 0,5724 | 8,39925E-33 |
| ACE2 | RPLP0 | 0,5724 | 8,4049E-33 |
| ACE2 | JPX | 0,5724 | 8,27728E-33 |
| ACE2 | PARD6B | 0,5723 | 8,55596E-33 |
| ACE2 | PSMA5 | 0,5721 | 9,10852E-33 |
| ACE2 | RP11-342K2.1 | 0,5721 | 9,19233E-33 |
| ACE2 | MSL2 | 0,572 | 9,47978E-33 |
| ACE2 | RP11-326I11.3 | 0,572 | 9,48346E-33 |
| ACE2 | RHEB | 0,5719 | 9,67407E-33 |
| ACE2 | TUFT1 | 0,5717 | 1,04117E-32 |
| ACE2 | PITPNC1 | 0,5717 | 1,02987E-32 |
| ACE2 | EBLN2 | 0,5715 | 1,09465E-32 |
| ACE2 | PALLD | 0,5715 | 1,0874E-32 |
| ACE2 | OCLM | 0,5714 | 1,11831E-32 |
| ACE2 | ATP2A2 | 0,5713 | 1,15484E-32 |
| ACE2 | SCFD1 | 0,5713 | 1,15205E-32 |
| ACE2 | RBM10 | 0,5713 | 1,15269E-32 |
| ACE2 | WWTR1 | 0,5712 | 1,19538E-32 |
| ACE2 | SELENOP | 0,5712 | 1,20729E-32 |
| ACE2 | ACLY | 0,5712 | 1,19792E-32 |
| ACE2 | THBD | 0,5712 | 1,17921E-32 |
| ACE2 | HS6ST1P1 | 0,5711 | 1,23897E-32 |
| ACE2 | GATAD2B | 0,5711 | 1,23599E-32 |
| ACE2 | LPAR1 | 0,5711 | 1,23396E-32 |
| ACE2 | HIPK3 | 0,5708 | 1,33668E-32 |
| ACE2 | DOK4 | 0,5708 | 1,33482E-32 |
| ACE2 | RP11-517B11.7 | 0,5706 | 1,42041E-32 |
| ACE2 | TRPC1 | 0,5706 | 1,42052E-32 |
| ACE2 | MBNL1-AS1 | 0,5706 | 1,43197E-32 |
| ACE2 | BTN2A1 | 0,5706 | 1,42623E-32 |
| ACE2 | OGFRL1 | 0,5705 | 1,4864E-32 |
| ACE2 | BORCS7 | 0,5705 | 1,50397E-32 |

| | | | |
|---|---|---|---|
| ACE2 | TERF2 | 0,5705 | 1,47102E-32 |
| ACE2 | NINL | 0,5705 | 1,47487E-32 |
| ACE2 | LINC01963 | 0,5704 | 1,54148E-32 |
| ACE2 | RHOG | 0,5704 | 1,55146E-32 |
| ACE2 | TRIM16L | 0,5703 | 1,59581E-32 |
| ACE2 | BOLA3-AS1 | 0,5702 | 1,61452E-32 |
| ACE2 | HERC6 | 0,5702 | 1,62507E-32 |
| ACE2 | RP11-872J21.5 | 0,5701 | 1,6928E-32 |
| ACE2 | SLC7A4 | 0,57 | 1,70465E-32 |
| ACE2 | TMEM161B | 0,5699 | 1,80236E-32 |
| ACE2 | MTHFD1L | 0,5698 | 1,86168E-32 |
| ACE2 | LDLRAD3 | 0,5698 | 1,8086E-32 |
| ACE2 | RBM33 | 0,5697 | 1,90864E-32 |
| ACE2 | DNAJA2 | 0,5697 | 1,87834E-32 |
| ACE2 | ANGPTL1 | 0,5696 | 1,97138E-32 |
| ACE2 | LONP1 | 0,5696 | 1,95225E-32 |
| ACE2 | MAGEF1 | 0,5695 | 2,01617E-32 |
| ACE2 | FAM133B | 0,5695 | 2,03226E-32 |
| ACE2 | ANGPTL3 | 0,5693 | 2,16877E-32 |
| ACE2 | KCTD2 | 0,5692 | 2,22663E-32 |
| ACE2 | KHK | 0,569 | 2,33987E-32 |
| ACE2 | AP2M1 | 0,569 | 2,3327E-32 |
| ACE2 | ATG14 | 0,569 | 2,33587E-32 |
| ACE2 | KCNIP3 | 0,5689 | 2,3905E-32 |
| ACE2 | NDUFS4 | 0,5689 | 2,43473E-32 |
| ACE2 | MCUR1 | 0,5689 | 2,4234E-32 |
| ACE2 | NUS1P1 | 0,5689 | 2,39651E-32 |
| ACE2 | PTOV1-AS1 | 0,5688 | 2,50512E-32 |
| ACE2 | GADD45A | 0,5687 | 2,54708E-32 |
| ACE2 | ADGRA3 | 0,5687 | 2,52849E-32 |
| ACE2 | B3GLCT | 0,5687 | 2,5915E-32 |
| ACE2 | SCNN1A | 0,5685 | 2,70513E-32 |
| ACE2 | XYLB | 0,5684 | 2,82251E-32 |
| ACE2 | CNOT6 | 0,5684 | 2,8262E-32 |
| ACE2 | LPAR2 | 0,5684 | 2,83457E-32 |
| ACE2 | SNRNP40 | 0,5683 | 2,88143E-32 |
| ACE2 | SMARCD2 | 0,5683 | 2,86948E-32 |
| ACE2 | RASGEF1B | 0,5682 | 2,97449E-32 |
| ACE2 | IFI27L2 | 0,5682 | 2,99541E-32 |
| ACE2 | NAAA | 0,5681 | 3,06773E-32 |
| ACE2 | KCNMB1 | 0,5681 | 3,12236E-32 |
| ACE2 | GPX7 | 0,568 | 3,19189E-32 |
| ACE2 | AL450992.2 | 0,568 | 3,19618E-32 |
| ACE2 | CUL7 | 0,568 | 3,17731E-32 |
| ACE2 | MAP4K5 | 0,568 | 3,20599E-32 |
| ACE2 | OIP5-AS1 | 0,568 | 3,19871E-32 |
| ACE2 | DIABLO | 0,5679 | 3,25378E-32 |
| ACE2 | HSPA4 | 0,5678 | 3,38444E-32 |
| ACE2 | KLF8 | 0,5678 | 3,41454E-32 |
| ACE2 | ZFYVE9 | 0,5677 | 3,44656E-32 |
| ACE2 | AMER1 | 0,5677 | 3,4874E-32 |
| ACE2 | N4BP2L2 | 0,5676 | 3,54438E-32 |
| ACE2 | MTHFD2 | 0,5675 | 3,63397E-32 |
| ACE2 | KIAA1161 | 0,5675 | 3,64722E-32 |

| | | | |
|---|---|---|---|
| ACE2 | FSHR | 0,5674 | 3,79312E-32 |
| ACE2 | TPRA1 | 0,5674 | 3,77473E-32 |
| ACE2 | HECA | 0,5674 | 3,77298E-32 |
| ACE2 | KLHL32 | 0,5673 | 3,88343E-32 |
| ACE2 | PIK3IP1 | 0,5672 | 3,98746E-32 |
| ACE2 | GDAP2 | 0,5671 | 4,18344E-32 |
| ACE2 | HBEGF | 0,5671 | 4,18953E-32 |
| ACE2 | ANKRD6 | 0,5671 | 4,21211E-32 |
| ACE2 | USF1 | 0,567 | 4,24409E-32 |
| ACE2 | ZNF22 | 0,5669 | 4,36903E-32 |
| ACE2 | KPNA1 | 0,5668 | 4,50268E-32 |
| ACE2 | CHMP4B | 0,5668 | 4,62375E-32 |
| ACE2 | NLGN4X | 0,5668 | 4,53384E-32 |
| ACE2 | RAI2 | 0,5668 | 4,59335E-32 |
| ACE2 | FAM26E | 0,5667 | 4,63636E-32 |
| ACE2 | DYNC1H1 | 0,5667 | 4,64472E-32 |
| ACE2 | IL4R | 0,5667 | 4,69916E-32 |
| ACE2 | TSPO | 0,5667 | 4,65455E-32 |
| ACE2 | ZNHIT6 | 0,5666 | 4,89695E-32 |
| ACE2 | RIPK1 | 0,5666 | 4,79303E-32 |
| ACE2 | PLEKHH2 | 0,5665 | 4,91492E-32 |
| ACE2 | OSBPL1A | 0,5665 | 4,97476E-32 |
| ACE2 | GNG7 | 0,5665 | 4,99971E-32 |
| ACE2 | FBXO11 | 0,5664 | 5,09223E-32 |
| ACE2 | TMEM25 | 0,5663 | 5,2589E-32 |
| ACE2 | SRP14 | 0,5663 | 5,34593E-32 |
| ACE2 | ADRA1D | 0,5663 | 5,31025E-32 |
| ACE2 | KIF1B | 0,5662 | 5,40873E-32 |
| ACE2 | PTMA | 0,5662 | 5,51295E-32 |
| ACE2 | TMCC1-AS1 | 0,5662 | 5,47041E-32 |
| ACE2 | ARHGEF9 | 0,5662 | 5,52937E-32 |
| ACE2 | GPRASP1 | 0,5662 | 5,45173E-32 |
| ACE2 | FSCN1 | 0,566 | 5,75444E-32 |
| ACE2 | PARP16 | 0,566 | 5,88239E-32 |
| ACE2 | DEFB123 | 0,5659 | 5,89657E-32 |
| ACE2 | MSN | 0,5659 | 5,95718E-32 |
| ACE2 | BCL9 | 0,5658 | 6,19208E-32 |
| ACE2 | SUPT16H | 0,5657 | 6,26727E-32 |
| ACE2 | CHST10 | 0,5656 | 6,62064E-32 |
| ACE2 | DOCK1 | 0,5656 | 6,58851E-32 |
| ACE2 | TMSB15B | 0,5656 | 6,45685E-32 |
| ACE2 | RPLP1 | 0,5655 | 6,66388E-32 |
| ACE2 | NRBF2 | 0,5654 | 6,92761E-32 |
| ACE2 | GLA | 0,5654 | 6,94974E-32 |
| ACE2 | DUSP7 | 0,5653 | 7,06336E-32 |
| ACE2 | SHPRH | 0,5651 | 7,66766E-32 |
| ACE2 | SBF2 | 0,5651 | 7,61923E-32 |
| ACE2 | IL7R | 0,5649 | 8,10883E-32 |
| ACE2 | CTD-2368P22.1 | 0,5649 | 8,06734E-32 |
| ACE2 | CTPS2 | 0,5649 | 8,04166E-32 |
| ACE2 | SPR | 0,5648 | 8,29287E-32 |
| ACE2 | WDR83OS | 0,5648 | 8,32836E-32 |
| ACE2 | ZBTB11 | 0,5647 | 8,48623E-32 |
| ACE2 | P4HB | 0,5647 | 8,66808E-32 |

| | | | |
|---|---|---|---|
| ACE2 | CHD8 | 0,5646 | 8,81435E-32 |
| ACE2 | UACA | 0,5646 | 8,88863E-32 |
| ACE2 | ERAL1 | 0,5646 | 8,86856E-32 |
| ACE2 | RP11-517I3.2 | 0,5645 | 9,06928E-32 |
| ACE2 | EDN1 | 0,5645 | 9,20938E-32 |
| ACE2 | PCLO | 0,5643 | 9,69714E-32 |
| ACE2 | TRNT1 | 0,5642 | 1,00586E-31 |
| ACE2 | CARD8-AS1 | 0,5642 | 9,8724E-32 |
| ACE2 | KIAA0368 | 0,5641 | 1,02214E-31 |
| ACE2 | METTL13 | 0,564 | 1,06622E-31 |
| ACE2 | LDB1 | 0,5639 | 1,09209E-31 |
| ACE2 | RBM26 | 0,5639 | 1,08214E-31 |
| ACE2 | ZC3H12B | 0,5639 | 1,08394E-31 |
| ACE2 | ANKH | 0,5638 | 1,12475E-31 |
| ACE2 | LIN7C | 0,5638 | 1,12815E-31 |
| ACE2 | GSS | 0,5638 | 1,13218E-31 |
| ACE2 | IGF1R | 0,5637 | 1,16876E-31 |
| ACE2 | COBLL1 | 0,5635 | 1,2221E-31 |
| ACE2 | GNB4 | 0,5635 | 1,22208E-31 |
| ACE2 | SLC41A2 | 0,5635 | 1,22703E-31 |
| ACE2 | KLHL5 | 0,5634 | 1,2634E-31 |
| ACE2 | RNF39 | 0,5634 | 1,25219E-31 |
| ACE2 | POLR3K | 0,5634 | 1,26537E-31 |
| ACE2 | AZGP1 | 0,5633 | 1,30066E-31 |
| ACE2 | HSPA8 | 0,5633 | 1,31893E-31 |
| ACE2 | SPSB1 | 0,5632 | 1,32341E-31 |
| ACE2 | CAMK2N1 | 0,5632 | 1,34996E-31 |
| ACE2 | PSMB2 | 0,5631 | 1,387E-31 |
| ACE2 | MFSD14C | 0,5629 | 1,46887E-31 |
| ACE2 | FOXN3-AS1 | 0,5629 | 1,44848E-31 |
| ACE2 | SIX1 | 0,5627 | 1,56441E-31 |
| ACE2 | SUSD5 | 0,5626 | 1,60367E-31 |
| ACE2 | SPRY2 | 0,5626 | 1,60513E-31 |
| ACE2 | ARSB | 0,5625 | 1,63185E-31 |
| ACE2 | NOL9 | 0,5624 | 1,72051E-31 |
| ACE2 | RPL18A | 0,5623 | 1,76508E-31 |
| ACE2 | IKBKB | 0,5622 | 1,78148E-31 |
| ACE2 | KANSL2 | 0,5621 | 1,86069E-31 |
| ACE2 | KIAA0232 | 0,562 | 1,89793E-31 |
| ACE2 | CASP6 | 0,562 | 1,92689E-31 |
| ACE2 | TMEM70 | 0,562 | 1,89959E-31 |
| ACE2 | NANS | 0,562 | 1,90545E-31 |
| ACE2 | SLC46A1 | 0,562 | 1,9136E-31 |
| ACE2 | APOO | 0,562 | 1,94196E-31 |
| ACE2 | CSGALNACT2 | 0,5618 | 2,02623E-31 |
| ACE2 | SORL1 | 0,5615 | 2,25105E-31 |
| ACE2 | CSDC2 | 0,5615 | 2,22191E-31 |
| ACE2 | CAV1 | 0,5614 | 2,27342E-31 |
| ACE2 | EBP | 0,5614 | 2,30208E-31 |
| ACE2 | CNOT4 | 0,5613 | 2,34477E-31 |
| ACE2 | PLLP | 0,5613 | 2,3345E-31 |
| ACE2 | PFKFB4 | 0,5611 | 2,47104E-31 |
| ACE2 | RIN3 | 0,5611 | 2,49463E-31 |
| ACE2 | FLNA | 0,5611 | 2,51769E-31 |

| | | | |
|---|---|---|---|
| ACE2 | NUDT12 | 0,561 | 2,56273E-31 |
| ACE2 | C8orf76 | 0,561 | 2,59221E-31 |
| ACE2 | LUZP4 | 0,561 | 2,59248E-31 |
| ACE2 | TOR3A | 0,5609 | 2,66198E-31 |
| ACE2 | C1orf21 | 0,5609 | 2,64033E-31 |
| ACE2 | AAMP | 0,5609 | 2,61742E-31 |
| ACE2 | F2RL3 | 0,5609 | 2,64572E-31 |
| ACE2 | RHBDL2 | 0,5608 | 2,75415E-31 |
| ACE2 | RP4-547N15.3 | 0,5608 | 2,72347E-31 |
| ACE2 | PCDHAC2 | 0,5608 | 2,69765E-31 |
| ACE2 | TMSB4Y | 0,5608 | 2,73087E-31 |
| ACE2 | MEF2C | 0,5607 | 2,82595E-31 |
| ACE2 | PDZRN4 | 0,5607 | 2,83575E-31 |
| ACE2 | CMTM4 | 0,5607 | 2,77668E-31 |
| ACE2 | LEPR | 0,5606 | 2,89929E-31 |
| ACE2 | SUZ12 | 0,5606 | 2,8928E-31 |
| ACE2 | TP73-AS1 | 0,5605 | 3,00319E-31 |
| ACE2 | GK5 | 0,5605 | 2,95633E-31 |
| ACE2 | HNRNPH3 | 0,5605 | 2,98504E-31 |
| ACE2 | RAB2B | 0,5605 | 2,97709E-31 |
| ACE2 | ZNF737 | 0,5605 | 3,02301E-31 |
| ACE2 | HSP90B1 | 0,5603 | 3,15727E-31 |
| ACE2 | OGFOD1 | 0,5603 | 3,21111E-31 |
| ACE2 | ABHD14B | 0,5602 | 3,25768E-31 |
| ACE2 | SAMD12 | 0,5601 | 3,3357E-31 |
| ACE2 | ZNF185 | 0,5601 | 3,36454E-31 |
| ACE2 | NAV1 | 0,56 | 3,42452E-31 |
| ACE2 | PRDM11 | 0,56 | 3,45225E-31 |
| ACE2 | RP11-174G6.5 | 0,56 | 3,42332E-31 |
| ACE2 | FAM122C | 0,56 | 3,43104E-31 |
| ACE2 | RP11-1020A11.2 | 0,5599 | 3,58635E-31 |
| ACE2 | ERICH5 | 0,5599 | 3,59082E-31 |
| ACE2 | MAFB | 0,5599 | 3,57687E-31 |
| ACE2 | SVEP1 | 0,5598 | 3,62325E-31 |
| ACE2 | ENSA | 0,5597 | 3,7554E-31 |
| ACE2 | CNGA2 | 0,5597 | 3,75632E-31 |
| ACE2 | CSTF2T | 0,5596 | 3,86751E-31 |
| ACE2 | CTSS | 0,5594 | 4,12207E-31 |
| ACE2 | LSG1 | 0,5594 | 4,10074E-31 |
| ACE2 | DICER1 | 0,5594 | 4,07449E-31 |
| ACE2 | OLMALINC | 0,5593 | 4,29577E-31 |
| ACE2 | MGRN1 | 0,5593 | 4,21052E-31 |
| ACE2 | PDLIM3 | 0,5589 | 4,71974E-31 |
| ACE2 | LSM8 | 0,5589 | 4,75283E-31 |
| ACE2 | AGO2 | 0,5589 | 4,84E-31 |
| ACE2 | GRTP1-AS1 | 0,5589 | 4,84353E-31 |
| ACE2 | JOSD1 | 0,5589 | 4,82454E-31 |
| ACE2 | FRY | 0,5588 | 4,92025E-31 |
| ACE2 | SAT2 | 0,5588 | 4,88035E-31 |
| ACE2 | NUAK2 | 0,5587 | 5,12017E-31 |
| ACE2 | ADAM23 | 0,5587 | 5,02663E-31 |
| ACE2 | KIAA2026 | 0,5587 | 5,07633E-31 |
| ACE2 | ISY1 | 0,5586 | 5,24291E-31 |
| ACE2 | LIX1L | 0,5585 | 5,36756E-31 |

| | | | |
|---|---|---|---|
| ACE2 | LRP1 | 0,5585 | 5,4032E-31 |
| ACE2 | GTF3C3 | 0,5584 | 5,60258E-31 |
| ACE2 | OXA1L | 0,5584 | 5,49541E-31 |
| ACE2 | CDK16 | 0,5584 | 5,59684E-31 |
| ACE2 | TNS1 | 0,558 | 6,26734E-31 |
| ACE2 | NUBP1 | 0,558 | 6,23208E-31 |
| ACE2 | PMAIP1 | 0,558 | 6,21978E-31 |
| ACE2 | DARS2 | 0,5579 | 6,42716E-31 |
| ACE2 | ZNF561-AS1 | 0,5579 | 6,40058E-31 |
| ACE2 | HILPDA | 0,5577 | 6,79524E-31 |
| ACE2 | CDK5 | 0,5576 | 6,9549E-31 |
| ACE2 | GATAD2A | 0,5576 | 7,06084E-31 |
| ACE2 | RXRA | 0,5574 | 7,43807E-31 |
| ACE2 | PIAS3 | 0,5572 | 7,85236E-31 |
| ACE2 | NME6 | 0,5572 | 7,91775E-31 |
| ACE2 | HERC3 | 0,5572 | 7,90037E-31 |
| ACE2 | FAM46A | 0,5572 | 7,7554E-31 |
| ACE2 | PLXNB3 | 0,5572 | 7,75198E-31 |
| ACE2 | PDIK1L | 0,5571 | 8,0387E-31 |
| ACE2 | ZBTB8A | 0,5571 | 7,99965E-31 |
| ACE2 | EARS2 | 0,5571 | 8,08727E-31 |
| ACE2 | FAM58A | 0,5571 | 8,09943E-31 |
| ACE2 | CTTNBP2NL | 0,557 | 8,28211E-31 |
| ACE2 | MARCH9 | 0,557 | 8,36219E-31 |
| ACE2 | HBS1L | 0,5569 | 8,68317E-31 |
| ACE2 | SLC35C2 | 0,5569 | 8,5214E-31 |
| ACE2 | IRF2BP2 | 0,5568 | 8,83042E-31 |
| ACE2 | FAT4 | 0,5568 | 8,79151E-31 |
| ACE2 | RAI1 | 0,5568 | 8,94906E-31 |
| ACE2 | POGZ | 0,5567 | 9,22577E-31 |
| ACE2 | CHD4 | 0,5567 | 9,02975E-31 |
| ACE2 | AC023481.1 | 0,5566 | 9,39552E-31 |
| ACE2 | FAM105A | 0,5566 | 9,46761E-31 |
| ACE2 | RP11-736K20.6 | 0,5566 | 9,42094E-31 |
| ACE2 | KRBOX4 | 0,5566 | 9,37896E-31 |
| ACE2 | FPGT | 0,5564 | 9,89269E-31 |
| ACE2 | MTMR14 | 0,5564 | 9,97824E-31 |
| ACE2 | CTSK | 0,5563 | 1,01736E-30 |
| ACE2 | ACPP | 0,5563 | 1,03422E-30 |
| ACE2 | NPHP3 | 0,5563 | 1,02661E-30 |
| ACE2 | STON1 | 0,5562 | 1,04378E-30 |
| ACE2 | RPS21 | 0,5562 | 1,06541E-30 |
| ACE2 | CELSR2 | 0,556 | 1,12259E-30 |
| ACE2 | RP5-1042I8.7 | 0,556 | 1,10309E-30 |
| ACE2 | DPYSL3 | 0,556 | 1,12569E-30 |
| ACE2 | RP11-507K2.6 | 0,5559 | 1,15969E-30 |
| ACE2 | DIS3L | 0,5559 | 1,15736E-30 |
| ACE2 | PFKL | 0,5559 | 1,15143E-30 |
| ACE2 | TMEM248 | 0,5558 | 1,19056E-30 |
| ACE2 | DVL2 | 0,5558 | 1,18577E-30 |
| ACE2 | CEP68 | 0,5556 | 1,27041E-30 |
| ACE2 | RPL10A | 0,5555 | 1,29677E-30 |
| ACE2 | FRMPD4 | 0,5555 | 1,27182E-30 |
| ACE2 | B4GALT3 | 0,5554 | 1,33978E-30 |

| | | | |
|---|---|---|---|
| ACE2 | ZNF658B | 0,5553 | 1,35826E-30 |
| ACE2 | TAP2 | 0,5552 | 1,38708E-30 |
| ACE2 | SNRNP27 | 0,555 | 1,50881E-30 |
| ACE2 | NCKIPSD | 0,555 | 1,48359E-30 |
| ACE2 | AUH | 0,555 | 1,49157E-30 |
| ACE2 | TYW3 | 0,5549 | 1,53123E-30 |
| ACE2 | STMND1 | 0,5548 | 1,5699E-30 |
| ACE2 | SGK1 | 0,5548 | 1,59522E-30 |
| ACE2 | PRUNE2 | 0,5548 | 1,59306E-30 |
| ACE2 | WDFY2 | 0,5548 | 1,56916E-30 |
| ACE2 | THAP11 | 0,5547 | 1,63882E-30 |
| ACE2 | TTC3P1 | 0,5547 | 1,60873E-30 |
| ACE2 | NAB2 | 0,5546 | 1,66379E-30 |
| ACE2 | RP11-230B22.1 | 0,5545 | 1,74694E-30 |
| ACE2 | ZNF501 | 0,5545 | 1,73793E-30 |
| ACE2 | SLC24A1 | 0,5545 | 1,73523E-30 |
| ACE2 | ZNF699 | 0,5545 | 1,74504E-30 |
| ACE2 | C4orf3 | 0,5544 | 1,78039E-30 |
| ACE2 | ASTN2 | 0,5544 | 1,75038E-30 |
| ACE2 | RP11-30K9.5 | 0,5543 | 1,84957E-30 |
| ACE2 | ANOS2P | 0,5543 | 1,85143E-30 |
| ACE2 | SLC47A1P1 | 0,554 | 1,99919E-30 |
| ACE2 | RHOQ | 0,5539 | 2,03734E-30 |
| ACE2 | RPS3 | 0,5539 | 2,02448E-30 |
| ACE2 | DCN | 0,5539 | 2,06645E-30 |
| ACE2 | RP11-278C7.5 | 0,5538 | 2,0927E-30 |
| ACE2 | IGFBP7 | 0,5537 | 2,15797E-30 |
| ACE2 | COPG2 | 0,5537 | 2,17294E-30 |
| ACE2 | HIPK2 | 0,5536 | 2,233E-30 |
| ACE2 | KCNN2 | 0,5535 | 2,33041E-30 |
| ACE2 | PHF13 | 0,5534 | 2,38994E-30 |
| ACE2 | KCNS3 | 0,5534 | 2,37277E-30 |
| ACE2 | RIN2 | 0,5534 | 2,34218E-30 |
| ACE2 | ITGA3 | 0,5533 | 2,45177E-30 |
| ACE2 | COMMD7 | 0,5532 | 2,50481E-30 |
| ACE2 | B3GALT2 | 0,5531 | 2,58893E-30 |
| ACE2 | NUMB | 0,5531 | 2,57594E-30 |
| ACE2 | GPT2 | 0,5531 | 2,59801E-30 |
| ACE2 | NIT1 | 0,553 | 2,62665E-30 |
| ACE2 | QPCT | 0,5529 | 2,76297E-30 |
| ACE2 | PIKFYVE | 0,5529 | 2,74359E-30 |
| ACE2 | TSTD3 | 0,5529 | 2,69578E-30 |
| ACE2 | RIMKLB | 0,5529 | 2,75962E-30 |
| ACE2 | SLC7A5 | 0,5529 | 2,69608E-30 |
| ACE2 | LINC01990 | 0,5528 | 2,7973E-30 |
| ACE2 | SLC16A3 | 0,5528 | 2,80625E-30 |
| ACE2 | HSPA13 | 0,5528 | 2,83099E-30 |
| ACE2 | RPRD2 | 0,5527 | 2,91629E-30 |
| ACE2 | CDC5L | 0,5527 | 2,92556E-30 |
| ACE2 | RASEF | 0,5526 | 2,99078E-30 |
| ACE2 | DCXR | 0,5525 | 3,07883E-30 |
| ACE2 | TIMM21 | 0,5525 | 3,08187E-30 |
| ACE2 | DOK1 | 0,5524 | 3,14596E-30 |
| ACE2 | HIST1H4I | 0,5524 | 3,16332E-30 |

| | | | |
|---|---|---|---|
| ACE2 | NEDD4 | 0,5524 | 3,16485E-30 |
| ACE2 | USP11 | 0,5524 | 3,17641E-30 |
| ACE2 | RP11-5C23.1 | 0,5523 | 3,26828E-30 |
| ACE2 | UQCC3 | 0,5523 | 3,28088E-30 |
| ACE2 | DCPS | 0,5522 | 3,35743E-30 |
| ACE2 | ATXN3 | 0,5522 | 3,35501E-30 |
| ACE2 | CMYA5 | 0,5521 | 3,4668E-30 |
| ACE2 | ZNF3 | 0,5521 | 3,41988E-30 |
| ACE2 | LZTS2 | 0,5518 | 3,78869E-30 |
| ACE2 | TGIF2 | 0,5517 | 3,81542E-30 |
| ACE2 | RPL10 | 0,5517 | 3,89949E-30 |
| ACE2 | SLC22A17 | 0,5516 | 3,96297E-30 |
| ACE2 | RP11-598P20.3 | 0,5515 | 4,04091E-30 |
| ACE2 | PRMT5 | 0,5515 | 4,04996E-30 |
| ACE2 | NYNRIN | 0,5514 | 4,16263E-30 |
| ACE2 | MIR99AHG | 0,5514 | 4,1438E-30 |
| ACE2 | KLF7 | 0,5512 | 4,43299E-30 |
| ACE2 | WFS1 | 0,5511 | 4,52764E-30 |
| ACE2 | SFT2D1 | 0,5511 | 4,54301E-30 |
| ACE2 | PHYHIP | 0,5511 | 4,54752E-30 |
| ACE2 | ABL1 | 0,5511 | 4,55931E-30 |
| ACE2 | SLC7A14 | 0,551 | 4,76351E-30 |
| ACE2 | SLC39A11 | 0,551 | 4,75847E-30 |
| ACE2 | MFSD1 | 0,5508 | 4,95112E-30 |
| ACE2 | PPARGC1B | 0,5508 | 4,92242E-30 |
| ACE2 | RP11-337C18.8 | 0,5507 | 5,11364E-30 |
| ACE2 | EVC | 0,5507 | 5,14048E-30 |
| ACE2 | SLC6A8 | 0,5507 | 5,12163E-30 |
| ACE2 | AGTR1 | 0,5506 | 5,24943E-30 |
| ACE2 | GSTP1 | 0,5506 | 5,26826E-30 |
| ACE2 | PPP1R26 | 0,5505 | 5,38752E-30 |
| ACE2 | TMEM62 | 0,5504 | 5,57624E-30 |
| ACE2 | CNKSR2 | 0,5504 | 5,61865E-30 |
| ACE2 | PGM1 | 0,5503 | 5,71295E-30 |
| ACE2 | RMND1 | 0,5503 | 5,75017E-30 |
| ACE2 | DSTNP1 | 0,5503 | 5,74949E-30 |
| ACE2 | UBR4 | 0,5502 | 5,97529E-30 |
| ACE2 | ABHD18 | 0,5502 | 5,95339E-30 |
| ACE2 | LINC00398 | 0,5502 | 5,93015E-30 |
| ACE2 | GJB1 | 0,5502 | 5,88593E-30 |
| ACE2 | SMN1 | 0,55 | 6,3242E-30 |
| ACE2 | ACTN4 | 0,55 | 6,19451E-30 |
| ACE2 | TST | 0,55 | 6,29251E-30 |
| ACE2 | NEGR1 | 0,5499 | 6,53581E-30 |
| ACE2 | UBE2D3 | 0,5499 | 6,4786E-30 |
| ACE2 | PTK2B | 0,5499 | 6,37227E-30 |
| ACE2 | THUMPD2 | 0,5497 | 6,91429E-30 |
| ACE2 | APAF1 | 0,5497 | 6,74294E-30 |
| ACE2 | TEFM | 0,5497 | 6,81211E-30 |
| ACE2 | NIFK | 0,5493 | 7,56298E-30 |
| ACE2 | CCDC68 | 0,5493 | 7,72045E-30 |
| ACE2 | ZNF182 | 0,5493 | 7,72956E-30 |
| ACE2 | RNASEH1 | 0,5492 | 7,82171E-30 |
| ACE2 | LDAH | 0,5492 | 7,94271E-30 |

| | | | |
|---|---|---|---|
| ACE2 | MGAT4B | 0,5491 | 8,10739E-30 |
| ACE2 | ADGRF1 | 0,5491 | 8,05348E-30 |
| ACE2 | PRIM2 | 0,5491 | 8,11129E-30 |
| ACE2 | DLEU7 | 0,5491 | 8,14854E-30 |
| ACE2 | SELENON | 0,5489 | 8,47326E-30 |
| ACE2 | NUDT5 | 0,5489 | 8,45794E-30 |
| ACE2 | MKL2 | 0,5489 | 8,6901E-30 |
| ACE2 | LPCAT2 | 0,5489 | 8,58845E-30 |
| ACE2 | TRAPPC2 | 0,5489 | 8,49907E-30 |
| ACE2 | CYP27A1 | 0,5488 | 8,70507E-30 |
| ACE2 | MANEA | 0,5486 | 9,28228E-30 |
| ACE2 | GOLGB1 | 0,5485 | 9,71242E-30 |
| ACE2 | HMGN3-AS1 | 0,5485 | 9,54434E-30 |
| ACE2 | SRP9 | 0,5484 | 9,78859E-30 |
| ACE2 | NME1 | 0,5483 | 1,01791E-29 |
| ACE2 | AFF4 | 0,5482 | 1,04982E-29 |
| ACE2 | COQ8A | 0,5481 | 1,08119E-29 |
| ACE2 | RP1-12G14.7 | 0,5479 | 1,13965E-29 |
| ACE2 | COL6A3 | 0,5478 | 1,16648E-29 |
| ACE2 | SKIV2L2 | 0,5478 | 1,18006E-29 |
| ACE2 | SLC18B1 | 0,5478 | 1,17344E-29 |
| ACE2 | GAMT | 0,5478 | 1,15755E-29 |
| ACE2 | ZNF662 | 0,5477 | 1,22099E-29 |
| ACE2 | TTC17 | 0,5477 | 1,20786E-29 |
| ACE2 | ABAT | 0,5476 | 1,24421E-29 |
| ACE2 | WBSCR22 | 0,5475 | 1,2618E-29 |
| ACE2 | CFL1 | 0,5475 | 1,27462E-29 |
| ACE2 | RAB11FIP1P1 | 0,5475 | 1,26384E-29 |
| ACE2 | TAX1BP3 | 0,5474 | 1,31259E-29 |
| ACE2 | ADGRG6 | 0,5472 | 1,37278E-29 |
| ACE2 | AASDH | 0,5471 | 1,43264E-29 |
| ACE2 | LINC01521 | 0,5471 | 1,42311E-29 |
| ACE2 | CD53 | 0,5469 | 1,49299E-29 |
| ACE2 | RMDN1 | 0,5469 | 1,52159E-29 |
| ACE2 | AC025335.1 | 0,5469 | 1,49772E-29 |
| ACE2 | GNA12 | 0,5468 | 1,53127E-29 |
| ACE2 | HACD4 | 0,5468 | 1,53544E-29 |
| ACE2 | CHST11 | 0,5468 | 1,55738E-29 |
| ACE2 | RNF185 | 0,5468 | 1,54393E-29 |
| ACE2 | FBXO30 | 0,5467 | 1,60856E-29 |
| ACE2 | SRPX | 0,5467 | 1,61838E-29 |
| ACE2 | ST6GALNAC6 | 0,5466 | 1,62009E-29 |
| ACE2 | PLA2G12A | 0,5465 | 1,67599E-29 |
| ACE2 | MBOAT1 | 0,5464 | 1,75904E-29 |
| ACE2 | SMG1 | 0,5464 | 1,741E-29 |
| ACE2 | SAFB2 | 0,5464 | 1,71373E-29 |
| ACE2 | SPIN4 | 0,5464 | 1,74831E-29 |
| ACE2 | ZDHHC14 | 0,5463 | 1,7946E-29 |
| ACE2 | SRP72 | 0,5462 | 1,85337E-29 |
| ACE2 | EFNA5 | 0,5462 | 1,81489E-29 |
| ACE2 | OMA1 | 0,5461 | 1,89887E-29 |
| ACE2 | RP11-545I5.3 | 0,5461 | 1,89638E-29 |
| ACE2 | KIAA0040 | 0,5459 | 1,97226E-29 |
| ACE2 | RP11-358D17.2 | 0,5459 | 2,02043E-29 |

| | | | |
|---|---|---|---|
| ACE2 | BTBD19 | 0,5458 | 2,03251E-29 |
| ACE2 | ANKRD10-IT1 | 0,5458 | 2,02882E-29 |
| ACE2 | BECN1 | 0,5458 | 2,055E-29 |
| ACE2 | APCDD1 | 0,5457 | 2,11859E-29 |
| ACE2 | OSTM1 | 0,5456 | 2,17118E-29 |
| ACE2 | PLOD3 | 0,5455 | 2,21289E-29 |
| ACE2 | ERF | 0,5453 | 2,34179E-29 |
| ACE2 | SNRPD3 | 0,5452 | 2,42798E-29 |
| ACE2 | NOCT | 0,5451 | 2,48805E-29 |
| ACE2 | SENP5 | 0,545 | 2,56513E-29 |
| ACE2 | RSL24D1 | 0,545 | 2,54397E-29 |
| ACE2 | FAM111A | 0,5449 | 2,62912E-29 |
| ACE2 | TRAP1 | 0,5449 | 2,62351E-29 |
| ACE2 | RP11-777F6.3 | 0,5448 | 2,75532E-29 |
| ACE2 | MED8 | 0,5447 | 2,77573E-29 |
| ACE2 | FRG1HP | 0,5447 | 2,83528E-29 |
| ACE2 | TMEM223 | 0,5447 | 2,7622E-29 |
| ACE2 | POR | 0,5446 | 2,87907E-29 |
| ACE2 | LUC7L3 | 0,5443 | 3,14583E-29 |
| ACE2 | RBX1 | 0,5442 | 3,22582E-29 |
| ACE2 | HACD1 | 0,5441 | 3,26676E-29 |
| ACE2 | USP45 | 0,544 | 3,40257E-29 |
| ACE2 | EPHX2 | 0,544 | 3,40937E-29 |
| ACE2 | DYNLRB1 | 0,5439 | 3,48798E-29 |
| ACE2 | RP11-364B14.3 | 0,5439 | 3,48849E-29 |
| ACE2 | PPP1R9B | 0,5438 | 3,61514E-29 |
| ACE2 | FHOD3 | 0,5438 | 3,56739E-29 |
| ACE2 | PLPP7 | 0,5437 | 3,73829E-29 |
| ACE2 | RP11-1148L6.5 | 0,5437 | 3,65679E-29 |
| ACE2 | GLIDR | 0,5436 | 3,84752E-29 |
| ACE2 | CYP11A1 | 0,5436 | 3,80148E-29 |
| ACE2 | MITF | 0,5434 | 3,99494E-29 |
| ACE2 | KCTD18 | 0,5433 | 4,10965E-29 |
| ACE2 | RNF121 | 0,5432 | 4,25847E-29 |
| ACE2 | RAPGEF1 | 0,5431 | 4,3292E-29 |
| ACE2 | ZDBF2 | 0,543 | 4,44719E-29 |
| ACE2 | UGDH-AS1 | 0,5429 | 4,65751E-29 |
| ACE2 | RSL1D1 | 0,5429 | 4,62286E-29 |
| ACE2 | HDHD3 | 0,5427 | 4,86347E-29 |
| ACE2 | EAPP | 0,5427 | 4,88247E-29 |
| ACE2 | ACAA1 | 0,5426 | 5,00692E-29 |
| ACE2 | CDK2 | 0,5426 | 5,08024E-29 |
| ACE2 | RP11-471B22.2 | 0,5426 | 5,01649E-29 |
| ACE2 | DNMT3B | 0,5426 | 5,03722E-29 |
| ACE2 | GPR155 | 0,5425 | 5,13704E-29 |
| ACE2 | SREK1 | 0,5425 | 5,2215E-29 |
| ACE2 | SEMA6A | 0,5425 | 5,17714E-29 |
| ACE2 | SAMD9 | 0,5425 | 5,1818E-29 |
| ACE2 | PFDN2 | 0,5424 | 5,2773E-29 |
| ACE2 | NPAT | 0,5424 | 5,32892E-29 |
| ACE2 | RCSD1 | 0,5423 | 5,50519E-29 |
| ACE2 | SMC5 | 0,5422 | 5,62699E-29 |
| ACE2 | DAPK1 | 0,5422 | 5,60583E-29 |
| ACE2 | RP11-63M22.2 | 0,5421 | 5,70833E-29 |

| | | | |
|---|---|---|---|
| ACE2 | ANXA9 | 0,5419 | 6,17763E-29 |
| ACE2 | CHCHD4 | 0,5419 | 6,05951E-29 |
| ACE2 | CASP3 | 0,5418 | 6,34044E-29 |
| ACE2 | KLF12 | 0,5416 | 6,64957E-29 |
| ACE2 | PPIAP22 | 0,5416 | 6,67001E-29 |
| ACE2 | FAM49A | 0,5415 | 6,83725E-29 |
| ACE2 | GYS1 | 0,5415 | 6,76745E-29 |
| ACE2 | SULF2 | 0,5415 | 6,74127E-29 |
| ACE2 | ZBTB5 | 0,5414 | 7,01407E-29 |
| ACE2 | MINK1 | 0,5414 | 6,97431E-29 |
| ACE2 | YEATS2 | 0,5412 | 7,50059E-29 |
| ACE2 | ARL10 | 0,5412 | 7,47045E-29 |
| ACE2 | FERMT1 | 0,5411 | 7,70714E-29 |
| ACE2 | SLC25A17 | 0,5411 | 7,61226E-29 |
| ACE2 | RPL6 | 0,541 | 7,83525E-29 |
| ACE2 | UROD | 0,5408 | 8,33631E-29 |
| ACE2 | MATR3 | 0,5407 | 8,60944E-29 |
| ACE2 | PARG | 0,5407 | 8,46042E-29 |
| ACE2 | DSC2 | 0,5407 | 8,53618E-29 |
| ACE2 | CCNT2 | 0,5406 | 8,85946E-29 |
| ACE2 | JAKMIP1 | 0,5405 | 8,97319E-29 |
| ACE2 | ETFDH | 0,5405 | 9,02665E-29 |
| ACE2 | S1PR3 | 0,5404 | 9,32545E-29 |
| ACE2 | PMP22 | 0,5404 | 9,15623E-29 |
| ACE2 | RP11-23N2.4 | 0,5403 | 9,4213E-29 |
| ACE2 | XPA | 0,5402 | 9,80584E-29 |
| ACE2 | SMIM17 | 0,5402 | 9,79882E-29 |
| ACE2 | SRGN | 0,5401 | 9,94476E-29 |
| ACE2 | IRS1 | 0,54 | 1,04464E-28 |
| ACE2 | IP6K2 | 0,5399 | 1,06625E-28 |
| ACE2 | CAPN1 | 0,5399 | 1,07514E-28 |
| ACE2 | ZNF836 | 0,5399 | 1,07024E-28 |
| ACE2 | RP11-710C12.1 | 0,5398 | 1,08197E-28 |
| ACE2 | HNRNPA3P9 | 0,5398 | 1,08671E-28 |
| ACE2 | SERPINF2 | 0,5398 | 1,09386E-28 |
| ACE2 | CTC-429P9.5 | 0,5398 | 1,08473E-28 |
| ACE2 | CTD-2537I9.12 | 0,5398 | 1,09056E-28 |
| ACE2 | SDHAF2 | 0,5397 | 1,10742E-28 |
| ACE2 | C1S | 0,5397 | 1,10748E-28 |
| ACE2 | ZNF564 | 0,5397 | 1,11426E-28 |
| ACE2 | ZNF154 | 0,5397 | 1,12998E-28 |
| ACE2 | RNF128 | 0,5396 | 1,16841E-28 |
| ACE2 | NXF3 | 0,5394 | 1,22363E-28 |
| ACE2 | ENC1 | 0,5392 | 1,2765E-28 |
| ACE2 | TGFBR2 | 0,5391 | 1,30916E-28 |
| ACE2 | RP11-679B19.1 | 0,539 | 1,37317E-28 |
| ACE2 | AKAP10 | 0,539 | 1,36007E-28 |
| ACE2 | ARHGEF10 | 0,5389 | 1,415E-28 |
| ACE2 | TNFAIP1 | 0,5389 | 1,40549E-28 |
| ACE2 | GRAMD1A | 0,5388 | 1,42259E-28 |
| ACE2 | GRWD1 | 0,5388 | 1,42772E-28 |
| ACE2 | WDR36 | 0,5387 | 1,48935E-28 |
| ACE2 | TANK | 0,5384 | 1,61489E-28 |
| ACE2 | HMGN1 | 0,5384 | 1,59832E-28 |

| | | | |
|---|---|---|---|
| ACE2 | AC092641.2 | 0,5383 | 1,6377E-28 |
| ACE2 | SMIM4 | 0,5383 | 1,66837E-28 |
| ACE2 | CTC-429P9.2 | 0,5383 | 1,66228E-28 |
| ACE2 | WDSUB1 | 0,5382 | 1,67174E-28 |
| ACE2 | INSIG1 | 0,5382 | 1,69597E-28 |
| ACE2 | ESYT2 | 0,5382 | 1,70586E-28 |
| ACE2 | PRRC2B | 0,5382 | 1,6982E-28 |
| ACE2 | RNF103 | 0,5381 | 1,7624E-28 |
| ACE2 | RP11-817I4.1 | 0,538 | 1,8055E-28 |
| ACE2 | SFRP1 | 0,5379 | 1,82437E-28 |
| ACE2 | EIF3L | 0,5379 | 1,85759E-28 |
| ACE2 | NAP1L6 | 0,5379 | 1,84008E-28 |
| ACE2 | NPY2R | 0,5378 | 1,90849E-28 |
| ACE2 | GRIK2 | 0,5378 | 1,8718E-28 |
| ACE2 | EP300 | 0,5378 | 1,88801E-28 |
| ACE2 | CBLB | 0,5377 | 1,93588E-28 |
| ACE2 | DMD | 0,5377 | 1,93496E-28 |
| ACE2 | PRLR | 0,5376 | 1,97788E-28 |
| ACE2 | CDKN1A | 0,5376 | 1,99112E-28 |
| ACE2 | KRIT1 | 0,5376 | 1,9971E-28 |
| ACE2 | NAALAD2 | 0,5376 | 2,00725E-28 |
| ACE2 | DDX1 | 0,5375 | 2,03163E-28 |
| ACE2 | TLN1 | 0,5375 | 2,07413E-28 |
| ACE2 | MOV10 | 0,5374 | 2,12555E-28 |
| ACE2 | PDCD2 | 0,5371 | 2,29772E-28 |
| ACE2 | RP13-554M15.2 | 0,5371 | 2,2603E-28 |
| ACE2 | PRAME | 0,537 | 2,34663E-28 |
| ACE2 | ZNF594 | 0,5369 | 2,42897E-28 |
| ACE2 | ZNF714 | 0,5369 | 2,4166E-28 |
| ACE2 | SALL4 | 0,5369 | 2,44464E-28 |
| ACE2 | FPGS | 0,5368 | 2,48559E-28 |
| ACE2 | RP11-493L12.4 | 0,5367 | 2,54427E-28 |
| ACE2 | AARS | 0,5367 | 2,52708E-28 |
| ACE2 | ZNF418 | 0,5367 | 2,58211E-28 |
| ACE2 | NDUFA5 | 0,5366 | 2,63932E-28 |
| ACE2 | LINC01003 | 0,5366 | 2,61181E-28 |
| ACE2 | CCDC85C | 0,5366 | 2,64602E-28 |
| ACE2 | METTL7B | 0,5365 | 2,7014E-28 |
| ACE2 | RPS2 | 0,5365 | 2,71031E-28 |
| ACE2 | LINC01278 | 0,5365 | 2,66721E-28 |
| ACE2 | ADCY5 | 0,5361 | 3,04229E-28 |
| ACE2 | OSMR | 0,5361 | 2,97166E-28 |
| ACE2 | RP11-96H19.1 | 0,5361 | 3,03615E-28 |
| ACE2 | TICAM1 | 0,5361 | 3,03694E-28 |
| ACE2 | UTP4 | 0,536 | 3,11187E-28 |
| ACE2 | LINC01496 | 0,536 | 3,11842E-28 |
| ACE2 | TMEM131 | 0,5359 | 3,20422E-28 |
| ACE2 | SLC25A20 | 0,5359 | 3,20708E-28 |
| ACE2 | LMBRD2 | 0,5359 | 3,15524E-28 |
| ACE2 | DNAAF5 | 0,5358 | 3,21911E-28 |
| ACE2 | NECTIN1 | 0,5358 | 3,26789E-28 |
| ACE2 | ZNF25 | 0,5357 | 3,35766E-28 |
| ACE2 | TTPAL | 0,5356 | 3,39742E-28 |
| ACE2 | PLCB3 | 0,5355 | 3,51837E-28 |

| | | | |
|---|---|---|---|
| ACE2 | WDR13 | 0,5355 | 3,52847E-28 |
| ACE2 | RPL14 | 0,5354 | 3,60467E-28 |
| ACE2 | TWIST1 | 0,5354 | 3,60834E-28 |
| ACE2 | RP11-439E19.10 | 0,5353 | 3,77813E-28 |
| ACE2 | LINC01378 | 0,5353 | 3,73662E-28 |
| ACE2 | FAM208B | 0,5353 | 3,76125E-28 |
| ACE2 | OAZ1 | 0,5353 | 3,73646E-28 |
| ACE2 | SLC7A6OS | 0,5352 | 3,83545E-28 |
| ACE2 | TMEM199 | 0,5352 | 3,8466E-28 |
| ACE2 | MRPS14 | 0,5351 | 3,90983E-28 |
| ACE2 | RAD1 | 0,5351 | 3,91569E-28 |
| ACE2 | MYH11 | 0,5351 | 3,9786E-28 |
| ACE2 | HID1 | 0,5351 | 3,92423E-28 |
| ACE2 | PITPNB | 0,5351 | 3,94525E-28 |
| ACE2 | RP11-819C21.1 | 0,535 | 4,01523E-28 |
| ACE2 | NSL1 | 0,5349 | 4,14413E-28 |
| ACE2 | KCTD6 | 0,5349 | 4,12851E-28 |
| ACE2 | R3HDM4 | 0,5349 | 4,1726E-28 |
| ACE2 | CTC-425O23.5 | 0,5348 | 4,24962E-28 |
| ACE2 | PTGER2 | 0,5347 | 4,36123E-28 |
| ACE2 | PSMD1 | 0,5346 | 4,53008E-28 |
| ACE2 | CTC-463A16.1 | 0,5346 | 4,54595E-28 |
| ACE2 | OLFM1 | 0,5346 | 4,48036E-28 |
| ACE2 | TMEM256 | 0,5346 | 4,47944E-28 |
| ACE2 | TRA2B | 0,5345 | 4,68576E-28 |
| ACE2 | NDUFB8 | 0,5344 | 4,75706E-28 |
| ACE2 | VWA1 | 0,5343 | 4,9495E-28 |
| ACE2 | BTBD2 | 0,5343 | 4,86721E-28 |
| ACE2 | BAX | 0,5343 | 4,8421E-28 |
| ACE2 | HIST3H2A | 0,5342 | 5,0412E-28 |
| ACE2 | CHP2 | 0,5342 | 4,99275E-28 |
| ACE2 | RP13-188A5.1 | 0,5342 | 5,02945E-28 |
| ACE2 | LINC00607 | 0,534 | 5,25129E-28 |
| ACE2 | ZNF629 | 0,534 | 5,25937E-28 |
| ACE2 | CIRBP | 0,5339 | 5,38247E-28 |
| ACE2 | FGD1 | 0,5339 | 5,39961E-28 |
| ACE2 | GSTM4 | 0,5338 | 5,63353E-28 |
| ACE2 | ST3GAL6 | 0,5338 | 5,64235E-28 |
| ACE2 | MIR22HG | 0,5337 | 5,71915E-28 |
| ACE2 | TTYH2 | 0,5337 | 5,70764E-28 |
| ACE2 | BLVRB | 0,5337 | 5,81642E-28 |
| ACE2 | HCG15 | 0,5336 | 5,83077E-28 |
| ACE2 | OPHN1 | 0,5336 | 5,88239E-28 |
| ACE2 | NOLC1 | 0,5335 | 6,14043E-28 |
| ACE2 | FBXO48 | 0,5332 | 6,6242E-28 |
| ACE2 | XPO4 | 0,5332 | 6,58887E-28 |
| ACE2 | WDR89 | 0,5331 | 6,8026E-28 |
| ACE2 | INTS6L | 0,5331 | 6,77633E-28 |
| ACE2 | NREP | 0,533 | 7,01619E-28 |
| ACE2 | AD000684.2 | 0,533 | 6,91084E-28 |
| ACE2 | IL20RA | 0,5329 | 7,06897E-28 |
| ACE2 | RP11-262H14.4 | 0,5328 | 7,31747E-28 |
| ACE2 | TIAM1 | 0,5328 | 7,27454E-28 |
| ACE2 | RAB9B | 0,5328 | 7,25299E-28 |

| | | | |
|---|---|---|---|
| ACE2 | ITFG2 | 0,5327 | 7,48679E-28 |
| ACE2 | RP11-900F13.3 | 0,5327 | 7,54145E-28 |
| ACE2 | RP11-96K19.2 | 0,5326 | 7,65584E-28 |
| ACE2 | RIOK2 | 0,5326 | 7,69329E-28 |
| ACE2 | FREM2 | 0,5326 | 7,82643E-28 |
| ACE2 | MCM6 | 0,5325 | 7,97795E-28 |
| ACE2 | ZNF843 | 0,5325 | 7,90667E-28 |
| ACE2 | SRSF4 | 0,5324 | 8,10834E-28 |
| ACE2 | PER2 | 0,5324 | 8,15253E-28 |
| ACE2 | GSAP | 0,5324 | 8,19431E-28 |
| ACE2 | SMAD9 | 0,5324 | 8,20446E-28 |
| ACE2 | RAB20 | 0,5324 | 8,08718E-28 |
| ACE2 | BRD7 | 0,5323 | 8,41815E-28 |
| ACE2 | MTCL1 | 0,5323 | 8,42822E-28 |
| ACE2 | YIPF4 | 0,5322 | 8,62891E-28 |
| ACE2 | RP11-736K20.5 | 0,5322 | 8,70399E-28 |
| ACE2 | NR2F2 | 0,5322 | 8,67712E-28 |
| ACE2 | PDGFD | 0,532 | 9,1319E-28 |
| ACE2 | GTF3A | 0,532 | 9,05866E-28 |
| ACE2 | ZSWIM3 | 0,532 | 8,97959E-28 |
| ACE2 | RBMX2 | 0,532 | 9,16016E-28 |
| ACE2 | STARD3 | 0,5319 | 9,41767E-28 |
| ACE2 | EDEM2 | 0,5319 | 9,31716E-28 |
| ACE2 | PTGES3P1 | 0,5318 | 9,48557E-28 |
| ACE2 | CCDC22 | 0,5318 | 9,60016E-28 |
| ACE2 | ATP1A2 | 0,5316 | 1,01232E-27 |
| ACE2 | TANC1 | 0,5316 | 1,01431E-27 |
| ACE2 | SLC46A2 | 0,5316 | 1,01879E-27 |
| ACE2 | HMGB1P6 | 0,5316 | 1,0233E-27 |
| ACE2 | MRPL4 | 0,5316 | 1,00998E-27 |
| ACE2 | NVL | 0,5315 | 1,04448E-27 |
| ACE2 | MAP9 | 0,5315 | 1,02641E-27 |
| ACE2 | STX11 | 0,5315 | 1,05249E-27 |
| ACE2 | RP11-295D4.3 | 0,5315 | 1,05088E-27 |
| ACE2 | ZSWIM4 | 0,5315 | 1,03644E-27 |
| ACE2 | CD93 | 0,5315 | 1,04104E-27 |
| ACE2 | GLRX | 0,5314 | 1,05914E-27 |
| ACE2 | PTCH1 | 0,5314 | 1,05825E-27 |
| ACE2 | TAPBPL | 0,5314 | 1,06521E-27 |
| ACE2 | RAPGEF2 | 0,5313 | 1,09567E-27 |
| ACE2 | KBTBD11 | 0,5313 | 1,10691E-27 |
| ACE2 | ADAMTSL5 | 0,5313 | 1,09376E-27 |
| ACE2 | FTCDNL1 | 0,5312 | 1,11305E-27 |
| ACE2 | COLGALT1 | 0,5312 | 1,11514E-27 |
| ACE2 | ZNF649 | 0,5312 | 1,11482E-27 |
| ACE2 | KCTD13 | 0,5311 | 1,15533E-27 |
| ACE2 | SMIM13 | 0,531 | 1,19486E-27 |
| ACE2 | NINJ1 | 0,531 | 1,17376E-27 |
| ACE2 | AC079630.4 | 0,531 | 1,19244E-27 |
| ACE2 | FTX | 0,531 | 1,18327E-27 |
| ACE2 | RP11-983P16.4 | 0,5309 | 1,23441E-27 |
| ACE2 | KIF3C | 0,5308 | 1,26551E-27 |
| ACE2 | TMEM51 | 0,5307 | 1,29581E-27 |
| ACE2 | FAM69C | 0,5307 | 1,28419E-27 |

| | | | |
|---|---|---|---|
| ACE2 | MPHOSPH8 | 0,5306 | 1,33622E-27 |
| ACE2 | SUPT4H1 | 0,5306 | 1,32816E-27 |
| ACE2 | DDX60 | 0,5305 | 1,34871E-27 |
| ACE2 | TRUB2 | 0,5305 | 1,35688E-27 |
| ACE2 | RP11-999E24.3 | 0,5305 | 1,36522E-27 |
| ACE2 | CDC42BPB | 0,5305 | 1,36239E-27 |
| ACE2 | ABT1 | 0,5304 | 1,37855E-27 |
| ACE2 | MSR1 | 0,5304 | 1,37796E-27 |
| ACE2 | MDFIC | 0,5303 | 1,41602E-27 |
| ACE2 | INPP5B | 0,5302 | 1,48655E-27 |
| ACE2 | PNKD | 0,5302 | 1,45575E-27 |
| ACE2 | BEND3P3 | 0,5302 | 1,48247E-27 |
| ACE2 | C10orf76 | 0,5301 | 1,49254E-27 |
| ACE2 | GLTSCR1 | 0,5301 | 1,51085E-27 |
| ACE2 | VSIG10L | 0,53 | 1,54251E-27 |
| ACE2 | PID1 | 0,5299 | 1,59166E-27 |
| ACE2 | NUDT16 | 0,5299 | 1,59198E-27 |
| ACE2 | GPBP1L1 | 0,5298 | 1,64102E-27 |
| ACE2 | RND3 | 0,5297 | 1,66802E-27 |
| ACE2 | UBASH3B | 0,5297 | 1,68498E-27 |
| ACE2 | ZNF550 | 0,5297 | 1,6746E-27 |
| ACE2 | MAGEB2 | 0,5297 | 1,69135E-27 |
| ACE2 | DCLRE1B | 0,5296 | 1,72726E-27 |
| ACE2 | C6orf141 | 0,5296 | 1,73463E-27 |
| ACE2 | RAP2C-AS1 | 0,5296 | 1,73682E-27 |
| ACE2 | PLPP3 | 0,5295 | 1,77629E-27 |
| ACE2 | ASB6 | 0,5295 | 1,75571E-27 |
| ACE2 | GSTO1 | 0,5295 | 1,75828E-27 |
| ACE2 | MAGEA4 | 0,5295 | 1,77859E-27 |
| ACE2 | RBBP4 | 0,5294 | 1,82914E-27 |
| ACE2 | ZNF619 | 0,5294 | 1,83899E-27 |
| ACE2 | STOX2 | 0,5294 | 1,81206E-27 |
| ACE2 | FAM65A | 0,5294 | 1,80407E-27 |
| ACE2 | FOXC2 | 0,5294 | 1,79758E-27 |
| ACE2 | ICE1 | 0,5293 | 1,88254E-27 |
| ACE2 | SH3PXD2B | 0,5293 | 1,87665E-27 |
| ACE2 | KCTD10 | 0,5293 | 1,85173E-27 |
| ACE2 | HSPE1 | 0,5292 | 1,93407E-27 |
| ACE2 | PCGF3 | 0,5292 | 1,89806E-27 |
| ACE2 | FAM35A | 0,5292 | 1,90331E-27 |
| ACE2 | RASSF10 | 0,5292 | 1,90125E-27 |
| ACE2 | TMEM132C | 0,5292 | 1,91776E-27 |
| ACE2 | RP11-10A14.4 | 0,5291 | 1,9681E-27 |
| ACE2 | NETO2 | 0,5291 | 1,99242E-27 |
| ACE2 | PDS5B | 0,529 | 2,01981E-27 |
| ACE2 | NUAK1 | 0,5288 | 2,11173E-27 |
| ACE2 | PTGES | 0,5286 | 2,24842E-27 |
| ACE2 | SOD1 | 0,5286 | 2,22521E-27 |
| ACE2 | ZCCHC17 | 0,5285 | 2,28922E-27 |
| ACE2 | SOS1 | 0,5285 | 2,30497E-27 |
| ACE2 | TGFB1 | 0,5285 | 2,29145E-27 |
| ACE2 | UCKL1 | 0,5285 | 2,28989E-27 |
| ACE2 | ARHGAP31 | 0,5284 | 2,39255E-27 |
| ACE2 | CCND1 | 0,5282 | 2,52397E-27 |

| | | | |
|---|---|---|---|
| ACE2 | CXorf23 | 0,5282 | 2,50422E-27 |
| ACE2 | LAPTM5 | 0,528 | 2,66374E-27 |
| ACE2 | POLR2C | 0,528 | 2,60599E-27 |
| ACE2 | TNIK | 0,5279 | 2,69345E-27 |
| ACE2 | FBLN5 | 0,5279 | 2,69895E-27 |
| ACE2 | RP11-57H12.5 | 0,5278 | 2,74846E-27 |
| ACE2 | GORAB | 0,5278 | 2,76464E-27 |
| ACE2 | FAM131A | 0,5278 | 2,75885E-27 |
| ACE2 | EBF3 | 0,5278 | 2,76604E-27 |
| ACE2 | SPON1 | 0,5278 | 2,81298E-27 |
| ACE2 | CREBL2 | 0,5278 | 2,80812E-27 |
| ACE2 | PALB2 | 0,5278 | 2,77187E-27 |
| ACE2 | CHRDL1 | 0,5278 | 2,77914E-27 |
| ACE2 | SFMBT2 | 0,5277 | 2,88922E-27 |
| ACE2 | WDR45 | 0,5277 | 2,87909E-27 |
| ACE2 | ADSL | 0,5276 | 2,91727E-27 |
| ACE2 | GPNMB | 0,5274 | 3,07107E-27 |
| ACE2 | SLC25A21-AS1 | 0,5274 | 3,08122E-27 |
| ACE2 | HSDL1 | 0,5274 | 3,10841E-27 |
| ACE2 | SPINK13 | 0,5273 | 3,15137E-27 |
| ACE2 | DNAJC25 | 0,5273 | 3,13687E-27 |
| ACE2 | RP11-348F1.2 | 0,5273 | 3,16736E-27 |
| ACE2 | ACTR1B | 0,5272 | 3,24861E-27 |
| ACE2 | RP11-47A8.5 | 0,5272 | 3,25542E-27 |
| ACE2 | SERPING1 | 0,5272 | 3,2317E-27 |
| ACE2 | CASP4 | 0,5271 | 3,32776E-27 |
| ACE2 | LINC01771 | 0,527 | 3,47394E-27 |
| ACE2 | RP11-135F9.3 | 0,5269 | 3,48275E-27 |
| ACE2 | RP11-426D19.1 | 0,5269 | 3,50083E-27 |
| ACE2 | RP11-284F21.10 | 0,5266 | 3,76965E-27 |
| ACE2 | COA6 | 0,5266 | 3,86062E-27 |
| ACE2 | KCTD21 | 0,5266 | 3,82594E-27 |
| ACE2 | SNRPE | 0,5264 | 4,07545E-27 |
| ACE2 | PPRC1 | 0,5264 | 3,99819E-27 |
| ACE2 | CSNK2A1 | 0,5264 | 4,03186E-27 |
| ACE2 | SSTR3 | 0,5264 | 4,075E-27 |
| ACE2 | ARSE | 0,5264 | 3,98947E-27 |
| ACE2 | ZFP36L2 | 0,5263 | 4,16598E-27 |
| ACE2 | TTC14 | 0,5263 | 4,102E-27 |
| ACE2 | ZBTB12 | 0,5263 | 4,17047E-27 |
| ACE2 | SIPA1L1 | 0,5263 | 4,11479E-27 |
| ACE2 | NSMCE4A | 0,5262 | 4,24921E-27 |
| ACE2 | CORO1B | 0,5262 | 4,19372E-27 |
| ACE2 | CCDC8 | 0,526 | 4,45062E-27 |
| ACE2 | RNF187 | 0,5259 | 4,54657E-27 |
| ACE2 | C4orf19 | 0,5259 | 4,56358E-27 |
| ACE2 | PLEK2 | 0,5259 | 4,61771E-27 |
| ACE2 | PIP5K1A | 0,5258 | 4,66936E-27 |
| ACE2 | TRADD | 0,5258 | 4,75767E-27 |
| ACE2 | NRG2 | 0,5257 | 4,78212E-27 |
| ACE2 | PCBD2 | 0,5256 | 4,96343E-27 |
| ACE2 | PFN1 | 0,5256 | 5,02496E-27 |
| ACE2 | PET117 | 0,5256 | 4,90492E-27 |
| ACE2 | MYBBP1A | 0,5255 | 5,13559E-27 |

| | | | |
|---|---|---|---|
| ACE2 | DCBLD1 | 0,5253 | 5,43188E-27 |
| ACE2 | RINL | 0,5253 | 5,43308E-27 |
| ACE2 | TRMT61B | 0,5252 | 5,46339E-27 |
| ACE2 | RASSF4 | 0,5251 | 5,64204E-27 |
| ACE2 | RP11-799D4.4 | 0,5251 | 5,60279E-27 |
| ACE2 | ARL6IP6 | 0,525 | 5,77053E-27 |
| ACE2 | PAIP1 | 0,525 | 5,88387E-27 |
| ACE2 | CPSF4 | 0,5249 | 6,01821E-27 |
| ACE2 | STAMBP | 0,5248 | 6,04802E-27 |
| ACE2 | PM20D2 | 0,5248 | 6,05761E-27 |
| ACE2 | ILF3 | 0,5248 | 6,09607E-27 |
| ACE2 | KAT14 | 0,5247 | 6,35692E-27 |
| ACE2 | C3orf33 | 0,5246 | 6,38847E-27 |
| ACE2 | SNAI2 | 0,5246 | 6,49058E-27 |
| ACE2 | ZNF71 | 0,5246 | 6,41696E-27 |
| ACE2 | TMEM230 | 0,5245 | 6,58281E-27 |
| ACE2 | SEPT8 | 0,5244 | 6,84766E-27 |
| ACE2 | DCUN1D2 | 0,5244 | 6,88697E-27 |
| ACE2 | CASKIN2 | 0,5244 | 6,84376E-27 |
| ACE2 | KIAA0087 | 0,5243 | 7,04934E-27 |
| ACE2 | SCARA5 | 0,5243 | 6,98529E-27 |
| ACE2 | CNPY2 | 0,5243 | 6,98657E-27 |
| ACE2 | ALDH4A1 | 0,5241 | 7,29601E-27 |
| ACE2 | BEND4 | 0,5241 | 7,44728E-27 |
| ACE2 | PAX9 | 0,5241 | 7,31588E-27 |
| ACE2 | CPEB4 | 0,524 | 7,62236E-27 |
| ACE2 | CLTB | 0,524 | 7,57465E-27 |
| ACE2 | PHYH | 0,524 | 7,63988E-27 |
| ACE2 | MTMR8 | 0,524 | 7,51778E-27 |
| ACE2 | ARHGAP21 | 0,5239 | 7,65174E-27 |
| ACE2 | GRTP1 | 0,5239 | 7,69342E-27 |
| ACE2 | RP11-44F14.6 | 0,5239 | 7,67681E-27 |
| ACE2 | MCTS1 | 0,5239 | 7,65322E-27 |
| ACE2 | C1RL | 0,5238 | 7,94131E-27 |
| ACE2 | SPATA5L1 | 0,5238 | 7,86255E-27 |
| ACE2 | LMOD1 | 0,5237 | 8,15904E-27 |
| ACE2 | TWNK | 0,5237 | 8,0856E-27 |
| ACE2 | FRYL | 0,5236 | 8,3954E-27 |
| ACE2 | ASCC1 | 0,5236 | 8,4709E-27 |
| ACE2 | MIR34AHG | 0,5235 | 8,58405E-27 |
| ACE2 | LAPTM4A | 0,5235 | 8,69295E-27 |
| ACE2 | RP11-930P14.2 | 0,5234 | 8,74814E-27 |
| ACE2 | PRPF4 | 0,5234 | 8,81182E-27 |
| ACE2 | LALBA | 0,5234 | 8,75716E-27 |
| ACE2 | FAM156A | 0,5234 | 8,74297E-27 |
| ACE2 | CHMP1B2P | 0,5233 | 9,01408E-27 |
| ACE2 | SRSF3 | 0,5232 | 9,30033E-27 |
| ACE2 | PRMT3 | 0,5232 | 9,25347E-27 |
| ACE2 | KIF13A | 0,5231 | 9,48858E-27 |
| ACE2 | FAM221A | 0,5231 | 9,46111E-27 |
| ACE2 | MIR100HG | 0,5231 | 9,66981E-27 |
| ACE2 | ZNF334 | 0,5231 | 9,4299E-27 |
| ACE2 | RP11-185E8.2 | 0,523 | 9,73077E-27 |
| ACE2 | SRL | 0,523 | 9,76023E-27 |

| | | | |
|---|---|---|---|
| ACE2 | TMEM127 | 0,5229 | 9,96004E-27 |
| ACE2 | ASF1A | 0,5229 | 1,00312E-26 |
| ACE2 | NSD3 | 0,5229 | 9,9397E-27 |
| ACE2 | AC069282.6 | 0,5228 | 1,03262E-26 |
| ACE2 | CH17-472G23.4 | 0,5227 | 1,06312E-26 |
| ACE2 | INPP4A | 0,5227 | 1,04876E-26 |
| ACE2 | PLAA | 0,5227 | 1,06069E-26 |
| ACE2 | SMAD3 | 0,5227 | 1,05408E-26 |
| ACE2 | RNPS1 | 0,5227 | 1,05918E-26 |
| ACE2 | ATRN | 0,5227 | 1,06123E-26 |
| ACE2 | EIF2B5 | 0,5225 | 1,123E-26 |
| ACE2 | GRPEL1 | 0,5225 | 1,10631E-26 |
| ACE2 | RP11-93B14.9 | 0,5225 | 1,12636E-26 |
| ACE2 | SYNCRIP | 0,5224 | 1,14304E-26 |
| ACE2 | TARDBP | 0,5223 | 1,17864E-26 |
| ACE2 | CHST7 | 0,5223 | 1,17271E-26 |
| ACE2 | TSPAN17 | 0,5222 | 1,19194E-26 |
| ACE2 | MLYCD | 0,5221 | 1,23308E-26 |
| ACE2 | CTA-363E19.2 | 0,522 | 1,28379E-26 |
| ACE2 | INO80 | 0,5219 | 1,28861E-26 |
| ACE2 | MGEA5 | 0,5218 | 1,32657E-26 |
| ACE2 | DCLRE1A | 0,5218 | 1,3306E-26 |
| ACE2 | LRRC75A-AS1 | 0,5218 | 1,34039E-26 |
| ACE2 | SMTN | 0,5218 | 1,34091E-26 |
| ACE2 | KAZN | 0,5217 | 1,38376E-26 |
| ACE2 | PPP1R3F | 0,5217 | 1,3581E-26 |
| ACE2 | RPN2 | 0,5214 | 1,48651E-26 |
| ACE2 | NUP37 | 0,5213 | 1,53114E-26 |
| ACE2 | PLEKHO2 | 0,5213 | 1,52542E-26 |
| ACE2 | RP11-81A1.6 | 0,5213 | 1,51892E-26 |
| ACE2 | CADPS2 | 0,5212 | 1,54527E-26 |
| ACE2 | MANSC4 | 0,5212 | 1,58181E-26 |
| ACE2 | CTDSPL | 0,5211 | 1,62264E-26 |
| ACE2 | ANKRD49 | 0,5211 | 1,62134E-26 |
| ACE2 | RPL23P2 | 0,5211 | 1,60872E-26 |
| ACE2 | ERLIN2 | 0,521 | 1,63514E-26 |
| ACE2 | SFXN3 | 0,5209 | 1,68226E-26 |
| ACE2 | AP1S3 | 0,5208 | 1,74843E-26 |
| ACE2 | MED4 | 0,5207 | 1,79651E-26 |
| ACE2 | ZC4H2 | 0,5206 | 1,83217E-26 |
| ACE2 | BOK | 0,5205 | 1,86419E-26 |
| ACE2 | KIAA1549 | 0,5205 | 1,87219E-26 |
| ACE2 | ANKFY1 | 0,5205 | 1,87631E-26 |
| ACE2 | CAMSAP3 | 0,5205 | 1,89604E-26 |
| ACE2 | SLC9A9 | 0,5204 | 1,92465E-26 |
| ACE2 | RP11-16E12.1 | 0,5204 | 1,90291E-26 |
| ACE2 | HPR | 0,5204 | 1,91395E-26 |
| ACE2 | PRKCA | 0,5204 | 1,92384E-26 |
| ACE2 | L1TD1 | 0,5203 | 1,95892E-26 |
| ACE2 | HSPD1P11 | 0,5202 | 2,03414E-26 |
| ACE2 | IFITM3 | 0,5202 | 2,00939E-26 |
| ACE2 | TATDN2P1 | 0,5202 | 2,01072E-26 |
| ACE2 | RIPK2 | 0,5201 | 2,07456E-26 |
| ACE2 | U2AF2 | 0,5201 | 2,09384E-26 |

| | | | |
|---|---|---|---|
| ACE2 | KLF10 | 0,5199 | 2,18268E-26 |
| ACE2 | CD84 | 0,5198 | 2,25288E-26 |
| ACE2 | MAP4K3 | 0,5198 | 2,21821E-26 |
| ACE2 | RALA | 0,5198 | 2,21612E-26 |
| ACE2 | GPSM1 | 0,5198 | 2,23647E-26 |
| ACE2 | NCOA3 | 0,5198 | 2,22333E-26 |
| ACE2 | MSH3 | 0,5197 | 2,32688E-26 |
| ACE2 | PNISR | 0,5197 | 2,31334E-26 |
| ACE2 | PMEPA1 | 0,5197 | 2,27305E-26 |
| ACE2 | ITGB2 | 0,5197 | 2,31564E-26 |
| ACE2 | LRIG3 | 0,5196 | 2,36338E-26 |
| ACE2 | USP18 | 0,5196 | 2,39062E-26 |
| ACE2 | GBP5 | 0,5195 | 2,41506E-26 |
| ACE2 | TSKU | 0,5195 | 2,45544E-26 |
| ACE2 | BARD1 | 0,5194 | 2,45751E-26 |
| ACE2 | RP3-438O4.4 | 0,5194 | 2,47168E-26 |
| ACE2 | RP11-38G5.4 | 0,5193 | 2,52406E-26 |
| ACE2 | ACOT1 | 0,5192 | 2,65049E-26 |
| ACE2 | NES | 0,5191 | 2,69226E-26 |
| ACE2 | MCUB | 0,5191 | 2,70517E-26 |
| ACE2 | LINC00844 | 0,5191 | 2,66057E-26 |
| ACE2 | NLRX1 | 0,5191 | 2,65491E-26 |
| ACE2 | ZMAT1 | 0,5191 | 2,70225E-26 |
| ACE2 | RNF130 | 0,519 | 2,73274E-26 |
| ACE2 | MICU1 | 0,519 | 2,76078E-26 |
| ACE2 | USP34 | 0,5187 | 2,96227E-26 |
| ACE2 | GGCT | 0,5187 | 2,95677E-26 |
| ACE2 | NOMO2 | 0,5187 | 3,00566E-26 |
| ACE2 | COA3 | 0,5187 | 2,95363E-26 |
| ACE2 | FIBIN | 0,5186 | 3,04795E-26 |
| ACE2 | LINC01270 | 0,5186 | 3,09006E-26 |
| ACE2 | SMC1A | 0,5186 | 3,0584E-26 |
| ACE2 | RPS23 | 0,5185 | 3,13154E-26 |
| ACE2 | ENPP1 | 0,5185 | 3,09543E-26 |
| ACE2 | HSD17B1 | 0,5185 | 3,14488E-26 |
| ACE2 | SNAPC2 | 0,5185 | 3,15676E-26 |
| ACE2 | CAPRIN2 | 0,5184 | 3,24915E-26 |
| ACE2 | SCN7A | 0,5182 | 3,38027E-26 |
| ACE2 | CHMP1B | 0,5182 | 3,40583E-26 |
| ACE2 | ZNF639 | 0,5181 | 3,44112E-26 |
| ACE2 | DHX40 | 0,5181 | 3,48446E-26 |
| ACE2 | ENTPD6 | 0,5181 | 3,44216E-26 |
| ACE2 | NOL8 | 0,518 | 3,52026E-26 |
| ACE2 | COL4A2 | 0,518 | 3,60387E-26 |
| ACE2 | BICD2 | 0,5179 | 3,68936E-26 |
| ACE2 | ADCY3 | 0,5178 | 3,73104E-26 |
| ACE2 | GLIPR2 | 0,5178 | 3,7824E-26 |
| ACE2 | PTPRH | 0,5178 | 3,76029E-26 |
| ACE2 | ITGA2 | 0,5177 | 3,85982E-26 |
| ACE2 | ZNF121 | 0,5177 | 3,81236E-26 |
| ACE2 | PQBP1 | 0,5177 | 3,86888E-26 |
| ACE2 | CNOT2 | 0,5176 | 3,92953E-26 |
| ACE2 | TSC22D1 | 0,5175 | 4,06851E-26 |
| ACE2 | RPS8 | 0,5174 | 4,13215E-26 |

| | | | |
|---|---|---|---|
| ACE2 | GPR55 | 0,5174 | 4,17525E-26 |
| ACE2 | C16orf91 | 0,5174 | 4,17188E-26 |
| ACE2 | BCAM | 0,5174 | 4,10358E-26 |
| ACE2 | SPICE1 | 0,5173 | 4,3148E-26 |
| ACE2 | ACTA2 | 0,5173 | 4,26149E-26 |
| ACE2 | SLC27A1 | 0,5173 | 4,22846E-26 |
| ACE2 | CXorf57 | 0,5173 | 4,21675E-26 |
| ACE2 | HP1BP3 | 0,5172 | 4,35252E-26 |
| ACE2 | GPN2 | 0,5172 | 4,31619E-26 |
| ACE2 | TRIM21 | 0,5172 | 4,37042E-26 |
| ACE2 | RP11-429P3.8 | 0,5172 | 4,35515E-26 |
| ACE2 | EMILIN3 | 0,5172 | 4,35637E-26 |
| ACE2 | ZNF419 | 0,517 | 4,58953E-26 |
| ACE2 | RACK1 | 0,5169 | 4,73858E-26 |
| ACE2 | GOPC | 0,5169 | 4,66944E-26 |
| ACE2 | RP11-455B3.1 | 0,5168 | 4,83824E-26 |
| ACE2 | MESDC1 | 0,5168 | 4,79785E-26 |
| ACE2 | LAMA3 | 0,5168 | 4,90269E-26 |
| ACE2 | NBPF8 | 0,5167 | 5,0058E-26 |
| ACE2 | SELENOH | 0,5166 | 5,07193E-26 |
| ACE2 | MAP4 | 0,5165 | 5,28601E-26 |
| ACE2 | GATC | 0,5165 | 5,22922E-26 |
| ACE2 | ZNF674-AS1 | 0,5165 | 5,2334E-26 |
| ACE2 | TMEM18 | 0,5164 | 5,3743E-26 |
| ACE2 | ITGA4 | 0,5164 | 5,3842E-26 |
| ACE2 | RP3-467N11.1 | 0,5164 | 5,32806E-26 |
| ACE2 | ZNF256 | 0,5162 | 5,64783E-26 |
| ACE2 | SCYL3 | 0,5161 | 5,79774E-26 |
| ACE2 | CLDN1 | 0,5161 | 5,74381E-26 |
| ACE2 | L3MBTL3 | 0,5161 | 5,79515E-26 |
| ACE2 | CDC25B | 0,5161 | 5,76279E-26 |
| ACE2 | INHA | 0,5159 | 6,12906E-26 |
| ACE2 | TCF12 | 0,5159 | 6,11537E-26 |
| ACE2 | CD2BP2 | 0,5158 | 6,28477E-26 |
| ACE2 | NR2C2 | 0,5156 | 6,56577E-26 |
| ACE2 | ISYNA1 | 0,5156 | 6,49879E-26 |
| ACE2 | RP11-15A1.3 | 0,5156 | 6,61982E-26 |
| ACE2 | AMMECR1L | 0,5153 | 7,16728E-26 |
| ACE2 | TCF21 | 0,5153 | 7,11698E-26 |
| ACE2 | DSCC1 | 0,5152 | 7,25746E-26 |
| ACE2 | ARHGAP5 | 0,515 | 7,55772E-26 |
| ACE2 | SDHAF4 | 0,5149 | 7,86641E-26 |
| ACE2 | CTD-2311M21.3 | 0,5149 | 7,94115E-26 |
| ACE2 | SNX30 | 0,5148 | 8,11062E-26 |
| ACE2 | AC004381.8 | 0,5148 | 8,0313E-26 |
| ACE2 | ZNF337 | 0,5148 | 7,95737E-26 |
| ACE2 | GADD45G | 0,5147 | 8,25213E-26 |
| ACE2 | ABCA2 | 0,5146 | 8,5438E-26 |
| ACE2 | ZBTB7C | 0,5146 | 8,55974E-26 |
| ACE2 | EHD2 | 0,5144 | 8,95804E-26 |
| ACE2 | HTT | 0,5143 | 9,16163E-26 |
| ACE2 | RP11-381K20.2 | 0,5143 | 9,16878E-26 |
| ACE2 | SLC25A18 | 0,5142 | 9,27073E-26 |
| ACE2 | CDCA7 | 0,514 | 9,76564E-26 |

| | | | |
|---|---|---|---|
| ACE2 | DTX3 | 0,5139 | 1,00781E-25 |
| ACE2 | RP11-474P2.6 | 0,5138 | 1,03441E-25 |
| ACE2 | BCDIN3D | 0,5138 | 1,03514E-25 |
| ACE2 | STUB1 | 0,5138 | 1,02997E-25 |
| ACE2 | ADCY6 | 0,5137 | 1,05194E-25 |
| ACE2 | IFIT3 | 0,5136 | 1,10018E-25 |
| ACE2 | WWC2 | 0,5134 | 1,13749E-25 |
| ACE2 | PFKFB3 | 0,5134 | 1,15549E-25 |
| ACE2 | HTRA1 | 0,5134 | 1,15737E-25 |
| ACE2 | CARD10 | 0,5134 | 1,14024E-25 |
| ACE2 | DPYD | 0,5133 | 1,1632E-25 |
| ACE2 | EXOSC7 | 0,5133 | 1,16221E-25 |
| ACE2 | RIC8B | 0,5133 | 1,18486E-25 |
| ACE2 | TP53INP1 | 0,5132 | 1,19685E-25 |
| ACE2 | FBLIM1 | 0,5131 | 1,24062E-25 |
| ACE2 | GTF2E1 | 0,5131 | 1,2251E-25 |
| ACE2 | ORAI3 | 0,513 | 1,26466E-25 |
| ACE2 | THSD7A | 0,5129 | 1,30072E-25 |
| ACE2 | RP4-742J24.2 | 0,5129 | 1,29651E-25 |
| ACE2 | FARSA | 0,5128 | 1,32362E-25 |
| ACE2 | ZNF773 | 0,5128 | 1,33117E-25 |
| ACE2 | RP4-758J24.6 | 0,5126 | 1,39515E-25 |
| ACE2 | TMEM115 | 0,5126 | 1,39827E-25 |
| ACE2 | DCLK2 | 0,5125 | 1,42306E-25 |
| ACE2 | SAMD9L | 0,5125 | 1,42325E-25 |
| ACE2 | CHTF8 | 0,5125 | 1,43191E-25 |
| ACE2 | PTPN23 | 0,5122 | 1,53959E-25 |
| ACE2 | RBPJ | 0,5122 | 1,5531E-25 |
| ACE2 | ZNF449 | 0,5122 | 1,5682E-25 |
| ACE2 | RP11-337N6.3 | 0,512 | 1,61996E-25 |
| ACE2 | MRE11 | 0,512 | 1,61909E-25 |
| ACE2 | ADCYAP1R1 | 0,5119 | 1,6687E-25 |
| ACE2 | SIRPA | 0,5118 | 1,69499E-25 |
| ACE2 | BHMT2 | 0,5116 | 1,82546E-25 |
| ACE2 | KIAA1429 | 0,5116 | 1,80913E-25 |
| ACE2 | MORF4L2-AS1 | 0,5116 | 1,79932E-25 |
| ACE2 | LAS1L | 0,5115 | 1,82993E-25 |
| ACE2 | SMIM10L2A | 0,5115 | 1,8689E-25 |
| ACE2 | MAP3K12 | 0,5114 | 1,8806E-25 |
| ACE2 | SNTA1 | 0,5114 | 1,89872E-25 |
| ACE2 | INPPL1 | 0,5113 | 1,92422E-25 |
| ACE2 | GCNT4 | 0,5112 | 1,97289E-25 |
| ACE2 | ROR2 | 0,5112 | 2,01325E-25 |
| ACE2 | C6orf1 | 0,5111 | 2,06421E-25 |
| ACE2 | VOPP1 | 0,5111 | 2,05516E-25 |
| ACE2 | CEP170B | 0,5111 | 2,04461E-25 |
| ACE2 | ATP6V1G2 | 0,511 | 2,09307E-25 |
| ACE2 | AC083843.1 | 0,511 | 2,08299E-25 |
| ACE2 | FAM201A | 0,5109 | 2,14357E-25 |
| ACE2 | USF2 | 0,5109 | 2,16672E-25 |
| ACE2 | MANEAL | 0,5108 | 2,219E-25 |
| ACE2 | SLC26A2 | 0,5108 | 2,20518E-25 |
| ACE2 | C6orf226 | 0,5108 | 2,19056E-25 |
| ACE2 | RALGDS | 0,5108 | 2,21167E-25 |

| | | | |
|---|---|---|---|
| ACE2 | LL0XNC01-36H8.1 | 0,5107 | 2,28388E-25 |
| ACE2 | CTC-565M22.1 | 0,5106 | 2,32743E-25 |
| ACE2 | EFEMP1 | 0,5105 | 2,3959E-25 |
| ACE2 | DYNC1I2 | 0,5105 | 2,38429E-25 |
| ACE2 | KLHL31 | 0,5105 | 2,40385E-25 |
| ACE2 | KIAA1456 | 0,5105 | 2,35013E-25 |
| ACE2 | EXOC2 | 0,5104 | 2,45207E-25 |
| ACE2 | CD82 | 0,5104 | 2,40713E-25 |
| ACE2 | KIAA0430 | 0,5103 | 2,51628E-25 |
| ACE2 | CHL1 | 0,5102 | 2,55591E-25 |
| ACE2 | ENPP6 | 0,5102 | 2,54599E-25 |
| ACE2 | MICU3 | 0,5102 | 2,54619E-25 |
| ACE2 | CDC27 | 0,5102 | 2,55533E-25 |
| ACE2 | CYFIP2 | 0,5101 | 2,64394E-25 |
| ACE2 | HNRNPAB | 0,5101 | 2,60859E-25 |
| ACE2 | NPM1 | 0,51 | 2,70643E-25 |
| ACE2 | ABO | 0,51 | 2,65923E-25 |
| ACE2 | PGAP3 | 0,51 | 2,69906E-25 |
| ACE2 | ZMAT3 | 0,5099 | 2,73991E-25 |
| ACE2 | EYA2 | 0,5099 | 2,75102E-25 |
| ACE2 | METTL16 | 0,5098 | 2,86264E-25 |
| ACE2 | MFAP3 | 0,5095 | 3,03715E-25 |
| ACE2 | LINC-PINT | 0,5095 | 3,03792E-25 |
| ACE2 | KLF9 | 0,5095 | 3,07304E-25 |
| ACE2 | DTWD1 | 0,5094 | 3,16074E-25 |
| ACE2 | GRAMD4P8 | 0,5093 | 3,2204E-25 |
| ACE2 | PARGP1 | 0,5093 | 3,1641E-25 |
| ACE2 | APOL2 | 0,5093 | 3,18643E-25 |
| ACE2 | STARD3NL | 0,5092 | 3,29843E-25 |
| ACE2 | RCBTB2 | 0,5091 | 3,38674E-25 |
| ACE2 | SLC12A4 | 0,5091 | 3,33964E-25 |
| ACE2 | SEMA6D | 0,509 | 3,48604E-25 |
| ACE2 | ERCC6L | 0,509 | 3,46591E-25 |
| ACE2 | POLQ | 0,5089 | 3,5202E-25 |
| ACE2 | DAB2 | 0,5089 | 3,56939E-25 |
| ACE2 | FAM171A1 | 0,5089 | 3,56707E-25 |
| ACE2 | PRR34-AS1 | 0,5089 | 3,53537E-25 |
| ACE2 | SLC16A4 | 0,5088 | 3,62171E-25 |
| ACE2 | BHLHE41 | 0,5088 | 3,61154E-25 |
| ACE2 | NEAT1 | 0,5087 | 3,76078E-25 |
| ACE2 | UBA6-AS1 | 0,5086 | 3,82937E-25 |
| ACE2 | TAMM41 | 0,5085 | 3,92125E-25 |
| ACE2 | TYK2 | 0,5084 | 4,04204E-25 |
| ACE2 | MRAS | 0,5083 | 4,05574E-25 |
| ACE2 | OSR2 | 0,5083 | 4,07349E-25 |
| ACE2 | ZNF549 | 0,5083 | 4,05735E-25 |
| ACE2 | ZNF362 | 0,5082 | 4,25856E-25 |
| ACE2 | SYT14 | 0,5082 | 4,17251E-25 |
| ACE2 | SLC16A13 | 0,5082 | 4,24436E-25 |
| ACE2 | EMP2 | 0,508 | 4,41682E-25 |
| ACE2 | SOCS4 | 0,5079 | 4,53993E-25 |
| ACE2 | TAF1A | 0,5078 | 4,66418E-25 |
| ACE2 | APBB2 | 0,5078 | 4,63504E-25 |
| ACE2 | EMC10 | 0,5078 | 4,61242E-25 |

| | | | |
|---|---|---|---|
| ACE2 | GPR75 | 0,5077 | 4,79104E-25 |
| ACE2 | IL17RB | 0,5076 | 4,91218E-25 |
| ACE2 | KLHL24 | 0,5076 | 4,85533E-25 |
| ACE2 | RP11-177C12.1 | 0,5076 | 4,91951E-25 |
| ACE2 | TBC1D17 | 0,5076 | 4,90514E-25 |
| ACE2 | BACE2 | 0,5076 | 4,84312E-25 |
| ACE2 | GNL2 | 0,5075 | 5,03593E-25 |
| ACE2 | MGAT4A | 0,5075 | 5,00921E-25 |
| ACE2 | EEF1A1P19 | 0,5075 | 5,02685E-25 |
| ACE2 | RP11-305E6.4 | 0,5075 | 5,04381E-25 |
| ACE2 | CPM | 0,5075 | 4,94609E-25 |
| ACE2 | MED9 | 0,5074 | 5,09882E-25 |
| ACE2 | LHX9 | 0,5073 | 5,21127E-25 |
| ACE2 | SLC29A1 | 0,5072 | 5,42101E-25 |
| ACE2 | SNX21 | 0,5072 | 5,33171E-25 |
| ACE2 | CMTR1 | 0,5071 | 5,57937E-25 |
| ACE2 | FMOD | 0,507 | 5,69064E-25 |
| ACE2 | RP11-177G23.1 | 0,5068 | 5,93026E-25 |
| ACE2 | AP000473.8 | 0,5068 | 5,92103E-25 |
| ACE2 | SLC25A32 | 0,5067 | 6,14347E-25 |
| ACE2 | MBIP | 0,5067 | 6,04877E-25 |
| ACE2 | CSNK1E | 0,5067 | 6,03374E-25 |
| ACE2 | C2orf27A | 0,5066 | 6,31984E-25 |
| ACE2 | C1R | 0,5066 | 6,2022E-25 |
| ACE2 | RP11-12D5.2 | 0,5066 | 6,30424E-25 |
| ACE2 | PPP2R5B | 0,5065 | 6,35116E-25 |
| ACE2 | ANO8 | 0,5065 | 6,43749E-25 |
| ACE2 | NEFH | 0,5065 | 6,37822E-25 |
| ACE2 | HIST2H2BB | 0,5064 | 6,55054E-25 |
| ACE2 | AK3P3 | 0,5064 | 6,49228E-25 |
| ACE2 | ZNF850 | 0,5063 | 6,80065E-25 |
| ACE2 | CA5BP1 | 0,5063 | 6,73431E-25 |
| ACE2 | AFF2 | 0,5063 | 6,76442E-25 |
| ACE2 | TMEM132A | 0,5062 | 6,82103E-25 |
| ACE2 | EMBP1 | 0,506 | 7,15878E-25 |
| ACE2 | HIVEP2 | 0,5059 | 7,35984E-25 |
| ACE2 | CTD-2292M16.8 | 0,5059 | 7,49736E-25 |
| ACE2 | C1D | 0,5057 | 7,81494E-25 |
| ACE2 | ACVR2B-AS1 | 0,5055 | 8,2004E-25 |
| ACE2 | TMX2P1 | 0,5055 | 8,28882E-25 |
| ACE2 | HIST1H2AC | 0,5054 | 8,48228E-25 |
| ACE2 | ARHGEF7 | 0,5054 | 8,29737E-25 |
| ACE2 | RPL13A | 0,5054 | 8,46463E-25 |
| ACE2 | MAGEC2 | 0,5053 | 8,62648E-25 |
| ACE2 | INTS5 | 0,5052 | 8,92338E-25 |
| ACE2 | TRIM2 | 0,5051 | 9,02975E-25 |
| ACE2 | SCAMP5 | 0,505 | 9,30291E-25 |
| ACE2 | STAT5A | 0,505 | 9,17667E-25 |
| ACE2 | IFI6 | 0,5049 | 9,49509E-25 |
| ACE2 | PLSCR3 | 0,5049 | 9,45058E-25 |
| ACE2 | CRYAB | 0,5047 | 9,99703E-25 |
| ACE2 | PRKAR1A | 0,5046 | 1,02618E-24 |
| ACE2 | PER3 | 0,5045 | 1,04932E-24 |
| ACE2 | AC104534.2 | 0,5045 | 1,05408E-24 |

| | | | |
|---|---|---|---|
| ACE2 | CTCFL | 0,5044 | 1,07224E-24 |
| ACE2 | LBR | 0,5043 | 1,10307E-24 |
| ACE2 | AC009237.8 | 0,5043 | 1,10826E-24 |
| ACE2 | EHD3 | 0,5042 | 1,11659E-24 |
| ACE2 | EXOC1 | 0,5042 | 1,11451E-24 |
| ACE2 | ZNF618 | 0,5042 | 1,14067E-24 |
| ACE2 | CD14 | 0,5041 | 1,15028E-24 |
| ACE2 | LEMD3 | 0,5041 | 1,1657E-24 |
| ACE2 | GCDH | 0,5041 | 1,14493E-24 |
| ACE2 | GTPBP3 | 0,5041 | 1,16009E-24 |
| ACE2 | RP5-907C10.3 | 0,504 | 1,18217E-24 |
| ACE2 | LRP5 | 0,504 | 1,1724E-24 |
| ACE2 | EML5 | 0,504 | 1,17598E-24 |
| ACE2 | RPL27 | 0,504 | 1,19317E-24 |
| ACE2 | RP11-890B15.3 | 0,5038 | 1,25656E-24 |
| ACE2 | FZD5 | 0,5037 | 1,2874E-24 |
| ACE2 | GFOD1 | 0,5037 | 1,28714E-24 |
| ACE2 | INMT | 0,5037 | 1,26362E-24 |
| ACE2 | DPPA4 | 0,5036 | 1,31734E-24 |
| ACE2 | PCDHB15 | 0,5035 | 1,34282E-24 |
| ACE2 | TNFRSF10B | 0,5035 | 1,32602E-24 |
| ACE2 | MCL1 | 0,5034 | 1,3775E-24 |
| ACE2 | BLOC1S3 | 0,5034 | 1,3679E-24 |
| ACE2 | SNHG8 | 0,5033 | 1,41286E-24 |
| ACE2 | MOCS1 | 0,5033 | 1,4015E-24 |
| ACE2 | FNDC3A | 0,5032 | 1,42298E-24 |
| ACE2 | CRISPLD2 | 0,5032 | 1,4298E-24 |
| ACE2 | NELFCD | 0,5032 | 1,44121E-24 |
| ACE2 | AP000432.1 | 0,5031 | 1,4922E-24 |
| ACE2 | CRNKL1 | 0,503 | 1,50402E-24 |
| ACE2 | PDK4 | 0,5029 | 1,53342E-24 |
| ACE2 | VSTM4 | 0,5029 | 1,54466E-24 |
| ACE2 | LPL | 0,5028 | 1,60027E-24 |
| ACE2 | P4HA1 | 0,5028 | 1,58362E-24 |
| ACE2 | PCNX4 | 0,5027 | 1,61964E-24 |
| ACE2 | C1orf115 | 0,5026 | 1,66888E-24 |
| ACE2 | UNC5B | 0,5026 | 1,66283E-24 |
| ACE2 | PLPPR5 | 0,5025 | 1,71237E-24 |
| ACE2 | CENPC | 0,5024 | 1,75324E-24 |
| ACE2 | ATP5G2 | 0,5024 | 1,75313E-24 |
| ACE2 | AC015849.19 | 0,5024 | 1,72924E-24 |
| ACE2 | DLSTP1 | 0,5023 | 1,77625E-24 |
| ACE2 | RP11-545E17.3 | 0,5023 | 1,81299E-24 |
| ACE2 | PHF8 | 0,5023 | 1,81307E-24 |
| ACE2 | ZCCHC18 | 0,5023 | 1,80181E-24 |
| ACE2 | TMEM133 | 0,5022 | 1,82319E-24 |
| ACE2 | XRN2 | 0,5022 | 1,83881E-24 |
| ACE2 | NSDHL | 0,5021 | 1,86277E-24 |
| ACE2 | OTUD3 | 0,502 | 1,9054E-24 |
| ACE2 | FHL2 | 0,5019 | 1,98981E-24 |
| ACE2 | SERP1 | 0,5019 | 1,95848E-24 |
| ACE2 | DNAJC7 | 0,5018 | 2,03452E-24 |
| ACE2 | ZNF219 | 0,5017 | 2,08534E-24 |
| ACE2 | SPTLC1 | 0,5016 | 2,10904E-24 |

| | | | |
|---|---|---|---|
| ACE2 | SLC43A3 | 0,5016 | 2,13823E-24 |
| ACE2 | TSPAN7 | 0,5016 | 2,14252E-24 |
| ACE2 | RP11-113K21.6 | 0,5014 | 2,23031E-24 |
| ACE2 | CERNA1 | 0,5014 | 2,212E-24 |
| ACE2 | CTD-2561J22.5 | 0,5014 | 2,20479E-24 |
| ACE2 | ZNF337-AS1 | 0,5014 | 2,233E-24 |
| ACE2 | FNIP1 | 0,5013 | 2,26537E-24 |
| ACE2 | COL14A1 | 0,5013 | 2,28338E-24 |
| ACE2 | FLJ20021 | 0,5012 | 2,33941E-24 |
| ACE2 | TXLNGY | 0,5012 | 2,31736E-24 |
| ACE2 | COL4A1 | 0,5011 | 2,40813E-24 |
| ACE2 | RNF115 | 0,501 | 2,46974E-24 |
| ACE2 | LCA5 | 0,501 | 2,45945E-24 |
| ACE2 | SLC7A1 | 0,5009 | 2,50622E-24 |
| ACE2 | NR3C1 | 0,5008 | 2,58734E-24 |
| ACE2 | ATP6V1C1 | 0,5008 | 2,58449E-24 |
| ACE2 | RP11-502I4.3 | 0,5008 | 2,58073E-24 |
| ACE2 | NCBP3 | 0,5008 | 2,6081E-24 |
| ACE2 | MINDY4 | 0,5006 | 2,68077E-24 |
| ACE2 | MEGF9 | 0,5006 | 2,72492E-24 |
| ACE2 | RP11-400K9.4 | 0,5005 | 2,78059E-24 |
| ACE2 | GAS5-AS1 | 0,5004 | 2,8089E-24 |
| ACE2 | ZNF195 | 0,5004 | 2,86933E-24 |
| ACE2 | ENPEP | 0,5003 | 2,88286E-24 |
| ACE2 | TRA2A | 0,5002 | 2,94597E-24 |
| ACE2 | ST5 | 0,5002 | 2,97888E-24 |
| ACE2 | GAPDH | 0,5002 | 2,99523E-24 |
| ACE2 | FAM50A | 0,5001 | 3,01833E-24 |
| ACE2 | SLC20A1 | 0,5 | 3,13258E-24 |
| ACE2 | WDR55 | 0,5 | 3,10847E-24 |
| ACE2 | DDX10 | 0,5 | 3,13429E-24 |
| ACE2 | MTMR10 | 0,5 | 3,11837E-24 |
| ACE2 | MPV17L | 0,5 | 3,12026E-24 |
| ACE2 | RGS17 | 0,4999 | 3,24152E-24 |
| ACE2 | METAP1D | 0,4997 | 3,33749E-24 |
| ACE2 | CD68 | 0,4997 | 3,39167E-24 |
| ACE2 | SDPR | 0,4996 | 3,48087E-24 |
| ACE2 | MLLT3 | 0,4996 | 3,45365E-24 |
| ACE2 | SSBP2 | 0,4995 | 3,55333E-24 |
| ACE2 | PLA2G4C | 0,4995 | 3,5279E-24 |
| ACE2 | IFIT1 | 0,4994 | 3,5778E-24 |
| ACE2 | SYTL2 | 0,4994 | 3,6247E-24 |
| ACE2 | WDHD1 | 0,4994 | 3,63429E-24 |
| ACE2 | CCND2 | 0,4993 | 3,72475E-24 |
| ACE2 | NAXD | 0,4993 | 3,67133E-24 |
| ACE2 | ARHGAP31-AS1 | 0,4992 | 3,80022E-24 |
| ACE2 | DAB2IP | 0,4992 | 3,74861E-24 |
| ACE2 | ZNF493 | 0,4992 | 3,82104E-24 |
| ACE2 | MRPL37 | 0,4991 | 3,90982E-24 |
| ACE2 | RP4-781B1.5 | 0,4991 | 3,92583E-24 |
| ACE2 | ETV3 | 0,499 | 3,9493E-24 |
| ACE2 | GIMAP4 | 0,4989 | 4,11386E-24 |
| ACE2 | AKAP13 | 0,4989 | 4,04194E-24 |
| ACE2 | OGG1 | 0,4987 | 4,26012E-24 |

| | | | |
|---|---|---|---|
| ACE2 | ABLIM1 | 0,4987 | 4,23542E-24 |
| ACE2 | RP11-454F8.2 | 0,4987 | 4,2505E-24 |
| ACE2 | BLACAT1 | 0,4986 | 4,37566E-24 |
| ACE2 | WASHC2C | 0,4986 | 4,4206E-24 |
| ACE2 | MAP2K3 | 0,4986 | 4,34521E-24 |
| ACE2 | MSANTD4 | 0,4985 | 4,44772E-24 |
| ACE2 | TATDN2P2 | 0,4984 | 4,6116E-24 |
| ACE2 | SMC3 | 0,4984 | 4,63257E-24 |
| ACE2 | ANKRD52 | 0,4984 | 4,64087E-24 |
| ACE2 | LXN | 0,4983 | 4,75295E-24 |
| ACE2 | ST6GALNAC4 | 0,4983 | 4,67155E-24 |
| ACE2 | SVIL-AS1 | 0,4983 | 4,7048E-24 |
| ACE2 | RP11-96D1.8 | 0,4983 | 4,72178E-24 |
| ACE2 | COL4A5 | 0,4983 | 4,65428E-24 |
| ACE2 | PODXL | 0,4982 | 4,85204E-24 |
| ACE2 | FSTL3 | 0,4982 | 4,7646E-24 |
| ACE2 | PPM1B | 0,4981 | 4,93345E-24 |
| ACE2 | YTHDC1 | 0,4981 | 4,95739E-24 |
| ACE2 | MGAT1 | 0,4981 | 4,88877E-24 |
| ACE2 | C9orf147 | 0,4981 | 4,91459E-24 |
| ACE2 | CFH | 0,498 | 5,04064E-24 |
| ACE2 | GYG2 | 0,498 | 5,11043E-24 |
| ACE2 | FAM84B | 0,4979 | 5,2396E-24 |
| ACE2 | SLC9A8 | 0,4979 | 5,1763E-24 |
| ACE2 | ZHX2 | 0,4977 | 5,44346E-24 |
| ACE2 | RGS10 | 0,4977 | 5,43789E-24 |
| ACE2 | FASN | 0,4977 | 5,44937E-24 |
| ACE2 | PDHX | 0,4976 | 5,6098E-24 |
| ACE2 | PHTF2 | 0,4975 | 5,66776E-24 |
| ACE2 | NACC1 | 0,4975 | 5,74522E-24 |
| ACE2 | ABTB2 | 0,4974 | 5,84107E-24 |
| ACE2 | ZBTB24 | 0,4973 | 6,00027E-24 |
| ACE2 | RP11-424G14.1 | 0,4973 | 5,92927E-24 |
| ACE2 | CGGBP1 | 0,4972 | 6,1041E-24 |
| ACE2 | ZNF559 | 0,4972 | 6,19593E-24 |
| ACE2 | FANCL | 0,4971 | 6,23358E-24 |
| ACE2 | RP11-210M15.2 | 0,4971 | 6,31216E-24 |
| ACE2 | PRKAA2 | 0,497 | 6,43874E-24 |
| ACE2 | CSRP1 | 0,497 | 6,42223E-24 |
| ACE2 | ADGRF5 | 0,497 | 6,38641E-24 |
| ACE2 | OSBPL3 | 0,497 | 6,39231E-24 |
| ACE2 | NKX2-8 | 0,497 | 6,40542E-24 |
| ACE2 | ST6GAL1 | 0,4969 | 6,65192E-24 |
| ACE2 | BMPR1B | 0,4969 | 6,55073E-24 |
| ACE2 | BMP6 | 0,4969 | 6,54494E-24 |
| ACE2 | ZHX1 | 0,4968 | 6,67246E-24 |
| ACE2 | CTDSP2 | 0,4968 | 6,81949E-24 |
| ACE2 | CYP4V2 | 0,4967 | 6,85206E-24 |
| ACE2 | INSR | 0,4967 | 6,98319E-24 |
| ACE2 | TPRG1-AS1 | 0,4966 | 7,10053E-24 |
| ACE2 | RINT1 | 0,4966 | 7,00235E-24 |
| ACE2 | ARL5B | 0,4965 | 7,28637E-24 |
| ACE2 | ZBTB25 | 0,4965 | 7,16628E-24 |
| ACE2 | VN1R85P | 0,4965 | 7,31974E-24 |

| | | | |
|---|---|---|---|
| ACE2 | NORAD | 0,4965 | 7,24798E-24 |
| ACE2 | POTEKP | 0,4964 | 7,41463E-24 |
| ACE2 | GPRIN3 | 0,4964 | 7,39352E-24 |
| ACE2 | IRGQ | 0,4964 | 7,49545E-24 |
| ACE2 | PLXND1 | 0,4963 | 7,5688E-24 |
| ACE2 | LINC00987 | 0,4963 | 7,62821E-24 |
| ACE2 | AC020951.1 | 0,4963 | 7,61547E-24 |
| ACE2 | HSPA5 | 0,4961 | 7,97799E-24 |
| ACE2 | DUSP8 | 0,4961 | 8,01375E-24 |
| ACE2 | CDH11 | 0,4961 | 7,94388E-24 |
| ACE2 | NEDD4L | 0,4961 | 7,87371E-24 |
| ACE2 | KHDRBS1 | 0,496 | 8,24459E-24 |
| ACE2 | POT1 | 0,496 | 8,09373E-24 |
| ACE2 | GVQW1 | 0,496 | 8,23058E-24 |
| ACE2 | IRF2BP1 | 0,496 | 8,1311E-24 |
| ACE2 | ERVK3-1 | 0,4959 | 8,4256E-24 |
| ACE2 | RP11-568K15.1 | 0,4956 | 8,95963E-24 |
| ACE2 | CTD-2308L22.1 | 0,4956 | 8,98372E-24 |
| ACE2 | ELOVL7 | 0,4956 | 8,92039E-24 |
| ACE2 | RPS12 | 0,4956 | 9,00542E-24 |
| ACE2 | BSPRY | 0,4955 | 9,13655E-24 |
| ACE2 | LINC00629 | 0,4955 | 9,27268E-24 |
| ACE2 | UBE2QL1 | 0,4954 | 9,49785E-24 |
| ACE2 | RP11-288H12.3 | 0,4954 | 9,4711E-24 |
| ACE2 | SH3PXD2A | 0,4954 | 9,31966E-24 |
| ACE2 | OSBPL5 | 0,4954 | 9,3549E-24 |
| ACE2 | TLDC2 | 0,4954 | 9,36966E-24 |
| ACE2 | NPTXR | 0,4954 | 9,32574E-24 |
| ACE2 | CD34 | 0,4953 | 9,5482E-24 |
| ACE2 | ZFYVE27 | 0,4953 | 9,60319E-24 |
| ACE2 | CHAD | 0,4953 | 9,67709E-24 |
| ACE2 | RBM38 | 0,4953 | 9,59359E-24 |
| ACE2 | RP3-414A15.10 | 0,4951 | 1,01136E-23 |
| ACE2 | ARHGAP25 | 0,495 | 1,02998E-23 |
| ACE2 | ZNF92 | 0,495 | 1,04142E-23 |
| ACE2 | POMK | 0,4949 | 1,05372E-23 |
| ACE2 | RPL26 | 0,4949 | 1,05397E-23 |
| ACE2 | DEGS1 | 0,4948 | 1,07574E-23 |
| ACE2 | RP1-152L7.5 | 0,4947 | 1,10662E-23 |
| ACE2 | RGMA | 0,4947 | 1,12237E-23 |
| ACE2 | ECHDC1 | 0,4946 | 1,14091E-23 |
| ACE2 | LTV1 | 0,4946 | 1,14158E-23 |
| ACE2 | CTC-490E21.10 | 0,4946 | 1,14806E-23 |
| ACE2 | PAPD7 | 0,4945 | 1,16169E-23 |
| ACE2 | TRIM9 | 0,4945 | 1,15821E-23 |
| ACE2 | WDR6 | 0,4944 | 1,19185E-23 |
| ACE2 | ETS1 | 0,4943 | 1,21326E-23 |
| ACE2 | TMSB15A | 0,4942 | 1,26349E-23 |
| ACE2 | ANKHD1 | 0,4941 | 1,29502E-23 |
| ACE2 | STAT6 | 0,4941 | 1,29505E-23 |
| ACE2 | IGFBP5 | 0,494 | 1,30461E-23 |
| ACE2 | CA11 | 0,494 | 1,32202E-23 |
| ACE2 | PLEKHA8P1 | 0,4939 | 1,3338E-23 |
| ACE2 | SDSL | 0,4939 | 1,33441E-23 |

| | | | |
|---|---|---|---|
| ACE2 | RASGRP1 | 0,4939 | 1,35319E-23 |
| ACE2 | ZFYVE26 | 0,4938 | 1,35887E-23 |
| ACE2 | LL0XNC01-7P3.1 | 0,4938 | 1,36569E-23 |
| ACE2 | RPS3A | 0,4937 | 1,41027E-23 |
| ACE2 | ARHGEF17 | 0,4937 | 1,39791E-23 |
| ACE2 | TINF2 | 0,4937 | 1,40076E-23 |
| ACE2 | APRT | 0,4937 | 1,40716E-23 |
| ACE2 | DSEL | 0,4936 | 1,45459E-23 |
| ACE2 | PGBD5 | 0,4935 | 1,4899E-23 |
| ACE2 | ST6GALNAC1 | 0,4935 | 1,47035E-23 |
| ACE2 | NPR2 | 0,4934 | 1,52226E-23 |
| ACE2 | RP11-30L15.4 | 0,4933 | 1,55992E-23 |
| ACE2 | RPL23AP64 | 0,4932 | 1,57353E-23 |
| ACE2 | ZNF726 | 0,4932 | 1,57983E-23 |
| ACE2 | CLIC2 | 0,4932 | 1,59437E-23 |
| ACE2 | PSMA1 | 0,4931 | 1,62683E-23 |
| ACE2 | SLC35E3 | 0,4931 | 1,6124E-23 |
| ACE2 | VPS13B | 0,493 | 1,65773E-23 |
| ACE2 | UBA2 | 0,493 | 1,67809E-23 |
| ACE2 | DDX60L | 0,4929 | 1,69644E-23 |
| ACE2 | RP11-705C15.2 | 0,4928 | 1,7605E-23 |
| ACE2 | SMARCD3 | 0,4927 | 1,79116E-23 |
| ACE2 | KRCC1 | 0,4926 | 1,81316E-23 |
| ACE2 | IFITM2 | 0,4926 | 1,82563E-23 |
| ACE2 | ANKDD1A | 0,4926 | 1,83828E-23 |
| ACE2 | ZNF831 | 0,4926 | 1,83533E-23 |
| ACE2 | ZNRF3 | 0,4926 | 1,80581E-23 |
| ACE2 | ZIC3 | 0,4925 | 1,88056E-23 |
| ACE2 | IMPG1 | 0,4924 | 1,91061E-23 |
| ACE2 | SUMO3 | 0,4924 | 1,9027E-23 |
| ACE2 | PAG1 | 0,4923 | 1,93934E-23 |
| ACE2 | TLR4 | 0,4923 | 1,96758E-23 |
| ACE2 | CHP1 | 0,4923 | 1,96486E-23 |
| ACE2 | STEAP4 | 0,4922 | 1,99391E-23 |
| ACE2 | INE1 | 0,4922 | 2,00816E-23 |
| ACE2 | RP11-119F19.5 | 0,492 | 2,08023E-23 |
| ACE2 | BAHD1 | 0,492 | 2,12294E-23 |
| ACE2 | SYS1 | 0,492 | 2,10741E-23 |
| ACE2 | TF | 0,4919 | 2,13903E-23 |
| ACE2 | FAM20C | 0,4919 | 2,13498E-23 |
| ACE2 | ALG10 | 0,4918 | 2,22237E-23 |
| ACE2 | RP11-1017G21.5 | 0,4918 | 2,19385E-23 |
| ACE2 | CTR9 | 0,4916 | 2,30414E-23 |
| ACE2 | CTC-325H20.7 | 0,4916 | 2,31219E-23 |
| ACE2 | ZNF589 | 0,4915 | 2,37884E-23 |
| ACE2 | LINC00092 | 0,4915 | 2,38085E-23 |
| ACE2 | SLC2A13 | 0,4915 | 2,34159E-23 |
| ACE2 | RHBDF2 | 0,4915 | 2,34979E-23 |
| ACE2 | STEAP3 | 0,4914 | 2,44396E-23 |
| ACE2 | HSPB3 | 0,4911 | 2,61249E-23 |
| ACE2 | IRAK3 | 0,4911 | 2,59603E-23 |
| ACE2 | POLR3B | 0,4911 | 2,61797E-23 |
| ACE2 | RP5-890O3.9 | 0,491 | 2,65381E-23 |
| ACE2 | SGCD | 0,4909 | 2,71502E-23 |

| | | | |
|---|---|---|---|
| ACE2 | URGCP | 0,4909 | 2,69298E-23 |
| ACE2 | RP11-68I18.10 | 0,4907 | 2,85545E-23 |
| ACE2 | TMEM170B | 0,4907 | 2,86024E-23 |
| ACE2 | SLC35C1 | 0,4907 | 2,82946E-23 |
| ACE2 | DYNC1LI2 | 0,4906 | 2,92351E-23 |
| ACE2 | MAN1C1 | 0,4905 | 3,02121E-23 |
| ACE2 | CAV2 | 0,4905 | 2,96099E-23 |
| ACE2 | DCP2 | 0,4903 | 3,12662E-23 |
| ACE2 | USP54 | 0,4903 | 3,09846E-23 |
| ACE2 | NMNAT2 | 0,4902 | 3,21595E-23 |
| ACE2 | AZI2 | 0,4902 | 3,20112E-23 |
| ACE2 | PTRF | 0,4902 | 3,21257E-23 |
| ACE2 | CICP14 | 0,4901 | 3,29592E-23 |
| ACE2 | MMP15 | 0,4901 | 3,2858E-23 |
| ACE2 | APC | 0,49 | 3,32424E-23 |
| ACE2 | ZNF785 | 0,49 | 3,34557E-23 |
| ACE2 | KANK2 | 0,49 | 3,3933E-23 |
| ACE2 | GPX3 | 0,4899 | 3,46709E-23 |
| ACE2 | EXT2 | 0,4899 | 3,47436E-23 |
| ACE2 | GLI2 | 0,4897 | 3,63939E-23 |
| ACE2 | CANX | 0,4897 | 3,57954E-23 |
| ACE2 | ELOVL4 | 0,4897 | 3,59587E-23 |
| ACE2 | BBS2 | 0,4897 | 3,63824E-23 |
| ACE2 | TRG-AS1 | 0,4896 | 3,6566E-23 |
| ACE2 | WBSCR17 | 0,4896 | 3,72355E-23 |
| ACE2 | ST6GAL2 | 0,4895 | 3,77837E-23 |
| ACE2 | CFI | 0,4895 | 3,79367E-23 |
| ACE2 | FBXO8 | 0,4895 | 3,7941E-23 |
| ACE2 | VPS52 | 0,4895 | 3,792E-23 |
| ACE2 | ZNF37BP | 0,4895 | 3,75128E-23 |
| ACE2 | FGR | 0,4894 | 3,83939E-23 |
| ACE2 | TAF1B | 0,4894 | 3,90367E-23 |
| ACE2 | SP110 | 0,4894 | 3,89216E-23 |
| ACE2 | PCDHGA7 | 0,4894 | 3,89594E-23 |
| ACE2 | CRTC1 | 0,4894 | 3,86276E-23 |
| ACE2 | EPHB1 | 0,4891 | 4,1376E-23 |
| ACE2 | TSPAN9 | 0,4891 | 4,14786E-23 |
| ACE2 | LEO1 | 0,4891 | 4,1608E-23 |
| ACE2 | BRD4 | 0,489 | 4,27979E-23 |
| ACE2 | RILPL2 | 0,4889 | 4,38664E-23 |
| ACE2 | LRP11 | 0,4888 | 4,40252E-23 |
| ACE2 | POM121C | 0,4886 | 4,61811E-23 |
| ACE2 | RP4-742C19.13 | 0,4886 | 4,60629E-23 |
| ACE2 | RBM28 | 0,4884 | 4,90472E-23 |
| ACE2 | PIANP | 0,4884 | 4,87713E-23 |
| ACE2 | CCDC149 | 0,4883 | 4,93738E-23 |
| ACE2 | COG6 | 0,4883 | 4,98648E-23 |
| ACE2 | CCDC51 | 0,4882 | 5,08113E-23 |
| ACE2 | EDNRA | 0,4882 | 5,15894E-23 |
| ACE2 | C1QTNF7 | 0,4881 | 5,22058E-23 |
| ACE2 | RP11-352M15.2 | 0,4881 | 5,2035E-23 |
| ACE2 | RPL7AP10 | 0,4881 | 5,17026E-23 |
| ACE2 | TRAF7 | 0,488 | 5,30553E-23 |
| ACE2 | ORMDL1 | 0,4879 | 5,45814E-23 |

| | | | |
|---|---|---|---|
| ACE2 | RCN1 | 0,4879 | 5,4756E-23 |
| ACE2 | SLC48A1 | 0,4879 | 5,42728E-23 |
| ACE2 | TMEM27 | 0,4879 | 5,47276E-23 |
| ACE2 | RP11-9E17.1 | 0,4878 | 5,65629E-23 |
| ACE2 | ZSCAN25 | 0,4877 | 5,69471E-23 |
| ACE2 | TRIM47 | 0,4877 | 5,76328E-23 |
| ACE2 | ZNF625 | 0,4877 | 5,71444E-23 |
| ACE2 | RP11-15A1.4 | 0,4877 | 5,7337E-23 |
| ACE2 | SIRPAP1 | 0,4877 | 5,77328E-23 |
| ACE2 | SGMS1 | 0,4876 | 5,91416E-23 |
| ACE2 | FRMPD3 | 0,4876 | 5,84919E-23 |
| ACE2 | ECM2 | 0,4874 | 6,11085E-23 |
| ACE2 | DDIT4 | 0,4874 | 6,13268E-23 |
| ACE2 | RP11-45M22.2 | 0,4874 | 6,099E-23 |
| ACE2 | KCND1 | 0,4874 | 6,08601E-23 |
| ACE2 | SPATS2L | 0,4873 | 6,2226E-23 |
| ACE2 | TLCD2 | 0,4873 | 6,22661E-23 |
| ACE2 | TOMM40 | 0,4873 | 6,31071E-23 |
| ACE2 | PTPN18 | 0,4871 | 6,51308E-23 |
| ACE2 | TP63 | 0,4871 | 6,64616E-23 |
| ACE2 | TAS2R5 | 0,4871 | 6,53692E-23 |
| ACE2 | ZNF703 | 0,4871 | 6,57608E-23 |
| ACE2 | ABHD14A | 0,487 | 6,76621E-23 |
| ACE2 | PLA2G5 | 0,4869 | 6,90075E-23 |
| ACE2 | PRR14L | 0,4869 | 6,90531E-23 |
| ACE2 | RGN | 0,4869 | 6,8721E-23 |
| ACE2 | BMP1 | 0,4868 | 7,02691E-23 |
| ACE2 | HLF | 0,4868 | 7,05065E-23 |
| ACE2 | TBC1D16 | 0,4868 | 6,98211E-23 |
| ACE2 | MGLL | 0,4867 | 7,25355E-23 |
| ACE2 | RP11-400F19.18 | 0,4867 | 7,24764E-23 |
| ACE2 | FBXL17 | 0,4865 | 7,62548E-23 |
| ACE2 | TXNDC16 | 0,4865 | 7,60522E-23 |
| ACE2 | ATP6V1D | 0,4865 | 7,58143E-23 |
| ACE2 | ZNF542P | 0,4865 | 7,63053E-23 |
| ACE2 | PCMTD2 | 0,4865 | 7,53957E-23 |
| ACE2 | ZBTB18 | 0,4864 | 7,75319E-23 |
| ACE2 | C8orf49 | 0,4864 | 7,76301E-23 |
| ACE2 | RP11-147L13.8 | 0,4864 | 7,73092E-23 |
| ACE2 | SUV39H1 | 0,4864 | 7,80589E-23 |
| ACE2 | ARMCX4 | 0,4863 | 7,94899E-23 |
| ACE2 | RN7SL574P | 0,4862 | 8,06593E-23 |
| ACE2 | TP53BP1 | 0,4861 | 8,29658E-23 |
| ACE2 | CHGA | 0,486 | 8,58377E-23 |
| ACE2 | VPS18 | 0,486 | 8,53172E-23 |
| ACE2 | THRA | 0,486 | 8,44038E-23 |
| ACE2 | APBA3 | 0,486 | 8,57742E-23 |
| ACE2 | AKAP8 | 0,486 | 8,58681E-23 |
| ACE2 | PINK1-AS | 0,4859 | 8,6022E-23 |
| ACE2 | HLA-E | 0,4859 | 8,7439E-23 |
| ACE2 | VEZF1 | 0,4859 | 8,60584E-23 |
| ACE2 | C6orf62 | 0,4856 | 9,21167E-23 |
| ACE2 | XKR8 | 0,4855 | 9,63067E-23 |
| ACE2 | BBS7 | 0,4855 | 9,61028E-23 |

| | | | |
|---|---|---|---|
| ACE2 | RPS6KA1 | 0,4853 | 9,94853E-23 |
| ACE2 | ZNF717 | 0,4853 | 9,90407E-23 |
| ACE2 | ADHFE1 | 0,4853 | 1,00189E-22 |
| ACE2 | RP11-359B12.2 | 0,4853 | 9,88906E-23 |
| ACE2 | RP11-359K18.4 | 0,4851 | 1,03586E-22 |
| ACE2 | SPIN2B | 0,4851 | 1,04282E-22 |
| ACE2 | SMYD1 | 0,485 | 1,06513E-22 |
| ACE2 | SNRNP200 | 0,485 | 1,07817E-22 |
| ACE2 | OGN | 0,485 | 1,05648E-22 |
| ACE2 | SFXN2 | 0,485 | 1,074E-22 |
| ACE2 | PATL1 | 0,485 | 1,05798E-22 |
| ACE2 | ITGA11 | 0,4849 | 1,09221E-22 |
| ACE2 | PHF5A | 0,4849 | 1,10367E-22 |
| ACE2 | BNIP3 | 0,4848 | 1,10626E-22 |
| ACE2 | KMT5B | 0,4848 | 1,10677E-22 |
| ACE2 | JMJD7 | 0,4848 | 1,11789E-22 |
| ACE2 | AC005785.5 | 0,4848 | 1,11653E-22 |
| ACE2 | EPB41 | 0,4847 | 1,14405E-22 |
| ACE2 | NAT8L | 0,4846 | 1,16166E-22 |
| ACE2 | ZBED5 | 0,4846 | 1,15856E-22 |
| ACE2 | NAA60 | 0,4846 | 1,17287E-22 |
| ACE2 | EXOC7 | 0,4846 | 1,17275E-22 |
| ACE2 | COL8A2 | 0,4845 | 1,18633E-22 |
| ACE2 | P4HA2 | 0,4845 | 1,1984E-22 |
| ACE2 | NAA16 | 0,4844 | 1,21761E-22 |
| ACE2 | RP11-680F8.4 | 0,4844 | 1,23452E-22 |
| ACE2 | DHX29 | 0,4842 | 1,27812E-22 |
| ACE2 | RP4-614C10.2 | 0,4839 | 1,37817E-22 |
| ACE2 | TACC2 | 0,4839 | 1,36101E-22 |
| ACE2 | SUMO1 | 0,4835 | 1,52281E-22 |
| ACE2 | CHST2 | 0,4835 | 1,50532E-22 |
| ACE2 | IFNAR1 | 0,4835 | 1,49747E-22 |
| ACE2 | MAGI2-AS3 | 0,4834 | 1,52916E-22 |
| ACE2 | IFI30 | 0,4834 | 1,541E-22 |
| ACE2 | MT-CO1 | 0,4834 | 1,54668E-22 |
| ACE2 | CD83 | 0,4833 | 1,58766E-22 |
| ACE2 | ARHGDIB | 0,4833 | 1,57507E-22 |
| ACE2 | SALL2 | 0,4833 | 1,58765E-22 |
| ACE2 | E2F3 | 0,4832 | 1,60481E-22 |
| ACE2 | RPS18 | 0,4832 | 1,62837E-22 |
| ACE2 | HEBP2 | 0,4832 | 1,62219E-22 |
| ACE2 | RAD18 | 0,4831 | 1,65147E-22 |
| ACE2 | MRPS26 | 0,4831 | 1,65121E-22 |
| ACE2 | TMEM184B | 0,4831 | 1,66981E-22 |
| ACE2 | ITM2A | 0,4831 | 1,6399E-22 |
| ACE2 | AP3S1 | 0,4828 | 1,78327E-22 |
| ACE2 | FAM83D | 0,4828 | 1,76564E-22 |
| ACE2 | PTGIS | 0,4828 | 1,75842E-22 |
| ACE2 | FBXL7 | 0,4827 | 1,82237E-22 |
| ACE2 | ATP1B2 | 0,4827 | 1,8044E-22 |
| ACE2 | ZNF888 | 0,4827 | 1,8209E-22 |
| ACE2 | CNRIP1 | 0,4826 | 1,86436E-22 |
| ACE2 | NPY1R | 0,4826 | 1,84708E-22 |
| ACE2 | SMAD2 | 0,4826 | 1,8426E-22 |

| | | | |
|---|---|---|---|
| ACE2 | XK | 0,4826 | 1,83536E-22 |
| ACE2 | AGAP1 | 0,4825 | 1,89492E-22 |
| ACE2 | C10orf10 | 0,4825 | 1,87509E-22 |
| ACE2 | BMX | 0,4825 | 1,87498E-22 |
| ACE2 | SOX13 | 0,4824 | 1,93924E-22 |
| ACE2 | EPB41L1 | 0,4824 | 1,93473E-22 |
| ACE2 | TBC1D22B | 0,4823 | 1,99926E-22 |
| ACE2 | LINC00963 | 0,4823 | 1,99915E-22 |
| ACE2 | CYP4F11 | 0,4823 | 1,96877E-22 |
| ACE2 | PTPRA | 0,4823 | 1,97491E-22 |
| ACE2 | AHNAK | 0,4822 | 2,00846E-22 |
| ACE2 | NUPR1 | 0,4822 | 2,03201E-22 |
| ACE2 | ARL13B | 0,4821 | 2,07559E-22 |
| ACE2 | RASD1 | 0,4821 | 2,06848E-22 |
| ACE2 | MRI1 | 0,4821 | 2,07282E-22 |
| ACE2 | MAGOH | 0,482 | 2,14189E-22 |
| ACE2 | TRAV8-3 | 0,4819 | 2,14703E-22 |
| ACE2 | AC004893.11 | 0,4818 | 2,21229E-22 |
| ACE2 | RP4-657E11.10 | 0,4817 | 2,27582E-22 |
| ACE2 | CTC-498J12.1 | 0,4816 | 2,34812E-22 |
| ACE2 | TEAD3 | 0,4816 | 2,34189E-22 |
| ACE2 | TIRAP | 0,4816 | 2,31186E-22 |
| ACE2 | HIST2H2BE | 0,4815 | 2,38654E-22 |
| ACE2 | KMT2D | 0,4815 | 2,37289E-22 |
| ACE2 | MDM4 | 0,4814 | 2,41559E-22 |
| ACE2 | PEX7 | 0,4814 | 2,45027E-22 |
| ACE2 | POLE2 | 0,4814 | 2,44898E-22 |
| ACE2 | CP | 0,4813 | 2,50363E-22 |
| ACE2 | ATP5SL | 0,4813 | 2,47191E-22 |
| ACE2 | LGI3 | 0,4812 | 2,53208E-22 |
| ACE2 | KLHDC1 | 0,4812 | 2,52066E-22 |
| ACE2 | PGPEP1 | 0,4811 | 2,59593E-22 |
| ACE2 | GARNL3 | 0,481 | 2,63419E-22 |
| ACE2 | HEY1 | 0,4809 | 2,6941E-22 |
| ACE2 | USP8 | 0,4809 | 2,72782E-22 |
| ACE2 | MYOCD | 0,4809 | 2,71076E-22 |
| ACE2 | DMXL2 | 0,4808 | 2,78121E-22 |
| ACE2 | ZNF823 | 0,4808 | 2,76224E-22 |
| ACE2 | NGEF | 0,4807 | 2,837E-22 |
| ACE2 | LDHA | 0,4807 | 2,86815E-22 |
| ACE2 | RP11-867O8.5 | 0,4807 | 2,8709E-22 |
| ACE2 | AC000403.4 | 0,4807 | 2,84415E-22 |
| ACE2 | AKT2 | 0,4807 | 2,8246E-22 |
| ACE2 | COL4A4 | 0,4806 | 2,90145E-22 |
| ACE2 | PCDHGA4 | 0,4806 | 2,88322E-22 |
| ACE2 | ZNF33A | 0,4806 | 2,91568E-22 |
| ACE2 | ZNF333 | 0,4806 | 2,88713E-22 |
| ACE2 | BMP2 | 0,4806 | 2,8896E-22 |
| ACE2 | C2CD2 | 0,4805 | 2,97842E-22 |
| ACE2 | THBS2 | 0,4804 | 3,04521E-22 |
| ACE2 | NAPA | 0,4803 | 3,14342E-22 |
| ACE2 | ST6GALNAC3 | 0,4802 | 3,17715E-22 |
| ACE2 | PRPF38B | 0,4802 | 3,2104E-22 |
| ACE2 | PCDHGB6 | 0,4802 | 3,214E-22 |

| | | | |
|---|---|---|---|
| ACE2 | ELOVL2 | 0,4802 | 3,1666E-22 |
| ACE2 | NKIRAS2 | 0,4802 | 3,16441E-22 |
| ACE2 | GOSR1 | 0,4801 | 3,22698E-22 |
| ACE2 | SERTAD1 | 0,4801 | 3,27439E-22 |
| ACE2 | AC007285.6 | 0,48 | 3,35527E-22 |
| ACE2 | IGSF6 | 0,48 | 3,30335E-22 |
| ACE2 | CACNB2 | 0,4799 | 3,41062E-22 |
| ACE2 | SYCN | 0,4799 | 3,41469E-22 |
| ACE2 | APOL1 | 0,4799 | 3,44999E-22 |
| ACE2 | CTHRC1 | 0,4798 | 3,4764E-22 |
| ACE2 | RP3-394A18.1 | 0,4798 | 3,494E-22 |
| ACE2 | FADS2 | 0,4797 | 3,57952E-22 |
| ACE2 | PIWIL4 | 0,4797 | 3,58849E-22 |
| ACE2 | FGF7 | 0,4797 | 3,5482E-22 |
| ACE2 | C16orf58 | 0,4797 | 3,57113E-22 |
| ACE2 | SLC26A11 | 0,4797 | 3,58599E-22 |
| ACE2 | RP11-448A19.1 | 0,4796 | 3,68422E-22 |
| ACE2 | RP11-848P1.2 | 0,4796 | 3,62672E-22 |
| ACE2 | KCNB1 | 0,4796 | 3,6879E-22 |
| ACE2 | FAM122A | 0,4795 | 3,7581E-22 |
| ACE2 | EED | 0,4795 | 3,74696E-22 |
| ACE2 | MMP14 | 0,4795 | 3,7093E-22 |
| ACE2 | NIPSNAP3B | 0,4794 | 3,82655E-22 |
| ACE2 | SETD6 | 0,4794 | 3,86263E-22 |
| ACE2 | RP4-681N20.5 | 0,4794 | 3,78431E-22 |
| ACE2 | GPR161 | 0,4793 | 3,87769E-22 |
| ACE2 | RAB11FIP3 | 0,4793 | 3,91231E-22 |
| ACE2 | FMN2 | 0,4792 | 3,95678E-22 |
| ACE2 | EFR3A | 0,4792 | 3,98149E-22 |
| ACE2 | TRMT112 | 0,4791 | 4,0679E-22 |
| ACE2 | DUT | 0,4788 | 4,37433E-22 |
| ACE2 | OR7A5 | 0,4787 | 4,46444E-22 |
| ACE2 | PSMA7 | 0,4787 | 4,46328E-22 |
| ACE2 | VPS13D | 0,4786 | 4,61343E-22 |
| ACE2 | C1QC | 0,4786 | 4,59522E-22 |
| ACE2 | NTRK2 | 0,4786 | 4,57197E-22 |
| ACE2 | FN1 | 0,4784 | 4,82317E-22 |
| ACE2 | RBM12B | 0,4784 | 4,7874E-22 |
| ACE2 | ATF7IP2 | 0,4784 | 4,78394E-22 |
| ACE2 | ITPKC | 0,4784 | 4,83724E-22 |
| ACE2 | COX16 | 0,4783 | 4,86131E-22 |
| ACE2 | TMEM109 | 0,4782 | 4,97819E-22 |
| ACE2 | LRRN4CL | 0,478 | 5,1899E-22 |
| ACE2 | ZNF271P | 0,478 | 5,23011E-22 |
| ACE2 | PODN | 0,4779 | 5,33869E-22 |
| ACE2 | TMCC1 | 0,4779 | 5,4091E-22 |
| ACE2 | EDNRB | 0,4779 | 5,4009E-22 |
| ACE2 | ZNF486 | 0,4779 | 5,31681E-22 |
| ACE2 | PASD1 | 0,4779 | 5,30353E-22 |
| ACE2 | RNASE6 | 0,4778 | 5,47343E-22 |
| ACE2 | SYNE1 | 0,4777 | 5,64944E-22 |
| ACE2 | EMP1 | 0,4777 | 5,637E-22 |
| ACE2 | WDR59 | 0,4777 | 5,55404E-22 |
| ACE2 | CTD-3220F14.2 | 0,4776 | 5,68894E-22 |

| | | | |
|---|---|---|---|
| ACE2 | FAM129A | 0,4775 | 5,82222E-22 |
| ACE2 | TMEM206 | 0,4774 | 5,9345E-22 |
| ACE2 | ATN1 | 0,4774 | 6,05686E-22 |
| ACE2 | ACTG1 | 0,4773 | 6,18917E-22 |
| ACE2 | KIAA0907 | 0,4772 | 6,20803E-22 |
| ACE2 | RP11-70L8.4 | 0,4771 | 6,42803E-22 |
| ACE2 | LINC01128 | 0,4769 | 6,74605E-22 |
| ACE2 | TMEM203 | 0,4769 | 6,76329E-22 |
| ACE2 | RAB6C | 0,4768 | 6,80329E-22 |
| ACE2 | CALCRL | 0,4768 | 6,90934E-22 |
| ACE2 | CKMT2 | 0,4768 | 6,80791E-22 |
| ACE2 | SKIV2L | 0,4767 | 7,05091E-22 |
| ACE2 | RP11-203M5.7 | 0,4767 | 7,06622E-22 |
| ACE2 | ZNF441 | 0,4767 | 7,04264E-22 |
| ACE2 | SLC25A40 | 0,4766 | 7,09357E-22 |
| ACE2 | ARHGAP26 | 0,4765 | 7,40224E-22 |
| ACE2 | UBE2G2 | 0,4765 | 7,32059E-22 |
| ACE2 | MT-ND5 | 0,4765 | 7,37135E-22 |
| ACE2 | SYPL2 | 0,4763 | 7,66983E-22 |
| ACE2 | TTC13 | 0,4763 | 7,64486E-22 |
| ACE2 | VCAM1 | 0,4762 | 7,89562E-22 |
| ACE2 | HHIP | 0,476 | 8,13047E-22 |
| ACE2 | SMIM3 | 0,476 | 8,15402E-22 |
| ACE2 | MANEA-AS1 | 0,476 | 8,15282E-22 |
| ACE2 | IDNK | 0,476 | 8,25373E-22 |
| ACE2 | RP5-965F6.2 | 0,4759 | 8,44065E-22 |
| ACE2 | DDR2 | 0,4759 | 8,34635E-22 |
| ACE2 | DOCK4 | 0,4759 | 8,41363E-22 |
| ACE2 | RHOJ | 0,4758 | 8,51034E-22 |
| ACE2 | LBH | 0,4757 | 8,82911E-22 |
| ACE2 | RPN1 | 0,4757 | 8,80393E-22 |
| ACE2 | DDB2 | 0,4757 | 8,79376E-22 |
| ACE2 | ASCC3 | 0,4756 | 8,91274E-22 |
| ACE2 | ZNF503 | 0,4756 | 8,93746E-22 |
| ACE2 | SLC14A2 | 0,4756 | 8,93083E-22 |
| ACE2 | RENBP | 0,4756 | 8,977E-22 |
| ACE2 | PLXNA1 | 0,4755 | 9,10116E-22 |
| ACE2 | PLAGL1 | 0,4755 | 9,24454E-22 |
| ACE2 | CYB561D1 | 0,4754 | 9,41442E-22 |
| ACE2 | RPP25 | 0,4754 | 9,36159E-22 |
| ACE2 | RP11-513M16.7 | 0,4753 | 9,59467E-22 |
| ACE2 | ALOX5AP | 0,4753 | 9,54937E-22 |
| ACE2 | MT-ND4 | 0,4753 | 9,60866E-22 |
| ACE2 | RNF4 | 0,4751 | 1,01222E-21 |
| ACE2 | AC004540.5 | 0,4751 | 1,01026E-21 |
| ACE2 | MYO5A | 0,475 | 1,02476E-21 |
| ACE2 | ZSCAN21 | 0,4749 | 1,05388E-21 |
| ACE2 | CWF19L2 | 0,4749 | 1,05295E-21 |
| ACE2 | LINC00920 | 0,4749 | 1,05259E-21 |
| ACE2 | UBE2G1 | 0,4749 | 1,03672E-21 |
| ACE2 | EPB41L2 | 0,4748 | 1,06761E-21 |
| ACE2 | STXBP1 | 0,4748 | 1,07132E-21 |
| ACE2 | RPL23A | 0,4748 | 1,07593E-21 |
| ACE2 | RP11-156E6.1 | 0,4748 | 1,0739E-21 |

| | | | |
|---|---|---|---|
| ACE2 | ENG | 0,4746 | 1,11948E-21 |
| ACE2 | CILP | 0,4746 | 1,11788E-21 |
| ACE2 | SIN3B | 0,4746 | 1,12347E-21 |
| ACE2 | KTI12 | 0,4745 | 1,15186E-21 |
| ACE2 | ARRB2 | 0,4745 | 1,14988E-21 |
| ACE2 | ZNF579 | 0,4744 | 1,16744E-21 |
| ACE2 | FCER1G | 0,4743 | 1,19725E-21 |
| ACE2 | HSPB1 | 0,4743 | 1,19601E-21 |
| ACE2 | LTBP2 | 0,4743 | 1,19263E-21 |
| ACE2 | TMEM182 | 0,4742 | 1,22254E-21 |
| ACE2 | DIP2C | 0,4742 | 1,21706E-21 |
| ACE2 | RP11-894P9.1 | 0,4741 | 1,26139E-21 |
| ACE2 | UBE2Q2 | 0,474 | 1,28138E-21 |
| ACE2 | DRG2 | 0,474 | 1,28075E-21 |
| ACE2 | CTB-88F18.5 | 0,4739 | 1,30377E-21 |
| ACE2 | ZNF786 | 0,4739 | 1,30152E-21 |
| ACE2 | ASH1L | 0,4738 | 1,32708E-21 |
| ACE2 | GPRC5A | 0,4738 | 1,33219E-21 |
| ACE2 | CDK12 | 0,4738 | 1,33098E-21 |
| ACE2 | SLX4IP | 0,4738 | 1,33637E-21 |
| ACE2 | MXRA5 | 0,4738 | 1,3445E-21 |
| ACE2 | GPR173 | 0,4738 | 1,33435E-21 |
| ACE2 | RP11-552M11.8 | 0,4737 | 1,35576E-21 |
| ACE2 | SLC25A11 | 0,4737 | 1,36936E-21 |
| ACE2 | NXPH3 | 0,4737 | 1,35941E-21 |
| ACE2 | GON4L | 0,4736 | 1,38412E-21 |
| ACE2 | TNIP2 | 0,4736 | 1,3863E-21 |
| ACE2 | HIST1H1C | 0,4736 | 1,38707E-21 |
| ACE2 | SSBP3-AS1 | 0,4735 | 1,43194E-21 |
| ACE2 | SNX17 | 0,4735 | 1,42809E-21 |
| ACE2 | ZNF620 | 0,4735 | 1,42655E-21 |
| ACE2 | ADAMTS5 | 0,4735 | 1,43367E-21 |
| ACE2 | MS4A4A | 0,4733 | 1,47648E-21 |
| ACE2 | NOP56 | 0,4733 | 1,50056E-21 |
| ACE2 | RUSC1 | 0,4732 | 1,53673E-21 |
| ACE2 | RBL1 | 0,4732 | 1,53186E-21 |
| ACE2 | RPAIN | 0,4731 | 1,55286E-21 |
| ACE2 | SP100 | 0,473 | 1,57951E-21 |
| ACE2 | TMOD1 | 0,473 | 1,58213E-21 |
| ACE2 | RPL27A | 0,473 | 1,59153E-21 |
| ACE2 | ZNF606 | 0,4729 | 1,61401E-21 |
| ACE2 | EBI3 | 0,4728 | 1,68176E-21 |
| ACE2 | NFIB | 0,4727 | 1,69865E-21 |
| ACE2 | RP3-461P17.6 | 0,4727 | 1,70077E-21 |
| ACE2 | CLIC4 | 0,4726 | 1,74166E-21 |
| ACE2 | TMEM98 | 0,4726 | 1,74856E-21 |
| ACE2 | YRDC | 0,4725 | 1,76963E-21 |
| ACE2 | SOS1-IT1 | 0,4724 | 1,81339E-21 |
| ACE2 | DBP | 0,4724 | 1,82652E-21 |
| ACE2 | NFATC2 | 0,4724 | 1,82595E-21 |
| ACE2 | LUM | 0,4722 | 1,91158E-21 |
| ACE2 | ASB13 | 0,4721 | 1,96331E-21 |
| ACE2 | SLC22A15 | 0,472 | 1,97115E-21 |
| ACE2 | IQSEC1 | 0,472 | 1,9695E-21 |

| | | | |
|---|---|---|---|
| ACE2 | RP11-582E3.6 | 0,4719 | 2,01901E-21 |
| ACE2 | ELOCP19 | 0,4718 | 2,06645E-21 |
| ACE2 | RP1-151F17.1 | 0,4718 | 2,06522E-21 |
| ACE2 | CHUK | 0,4718 | 2,07879E-21 |
| ACE2 | CTD-2537I9.13 | 0,4718 | 2,09208E-21 |
| ACE2 | SNRPGP10 | 0,4717 | 2,11317E-21 |
| ACE2 | ATP5B | 0,4717 | 2,14052E-21 |
| ACE2 | NRIP1 | 0,4717 | 2,13605E-21 |
| ACE2 | ACOX2 | 0,4716 | 2,17991E-21 |
| ACE2 | ALDH1L1 | 0,4716 | 2,16085E-21 |
| ACE2 | CAP2 | 0,4716 | 2,15032E-21 |
| ACE2 | ARRDC3 | 0,4715 | 2,21805E-21 |
| ACE2 | SYVN1 | 0,4715 | 2,21964E-21 |
| ACE2 | VPS29 | 0,4715 | 2,22709E-21 |
| ACE2 | CREB3L4 | 0,4713 | 2,31577E-21 |
| ACE2 | RP1-80N2.3 | 0,4713 | 2,31646E-21 |
| ACE2 | IGFBP3 | 0,4713 | 2,33638E-21 |
| ACE2 | MYOF | 0,4713 | 2,34061E-21 |
| ACE2 | MIR17HG | 0,4713 | 2,30277E-21 |
| ACE2 | FZD6 | 0,4712 | 2,38273E-21 |
| ACE2 | RAB7B | 0,4711 | 2,41212E-21 |
| ACE2 | SERPINA3 | 0,4711 | 2,41392E-21 |
| ACE2 | PCDHGC3 | 0,471 | 2,47749E-21 |
| ACE2 | TMCC3 | 0,471 | 2,49514E-21 |
| ACE2 | TOB2 | 0,471 | 2,47149E-21 |
| ACE2 | PLXNB2 | 0,471 | 2,46971E-21 |
| ACE2 | ACKR3 | 0,4709 | 2,5089E-21 |
| ACE2 | MYO7A | 0,4708 | 2,56271E-21 |
| ACE2 | ZFP69B | 0,4706 | 2,68239E-21 |
| ACE2 | AC003090.1 | 0,4706 | 2,69527E-21 |
| ACE2 | HERC5 | 0,4705 | 2,73818E-21 |
| ACE2 | FURIN | 0,4705 | 2,75616E-21 |
| ACE2 | ATP6V0A2 | 0,4704 | 2,84857E-21 |
| ACE2 | RTFDC1 | 0,4704 | 2,81098E-21 |
| ACE2 | COLEC12 | 0,4703 | 2,91419E-21 |
| ACE2 | NETO1 | 0,4703 | 2,88021E-21 |
| ACE2 | IFIT2 | 0,4701 | 3,00853E-21 |
| ACE2 | SLC38A1 | 0,4701 | 2,99887E-21 |
| ACE2 | FZD2 | 0,4701 | 3,00246E-21 |
| ACE2 | FN3K | 0,4701 | 3,02209E-21 |
| ACE2 | HCLS1 | 0,4699 | 3,1326E-21 |
| ACE2 | BTRC | 0,4698 | 3,2164E-21 |
| ACE2 | MMP2 | 0,4698 | 3,25108E-21 |
| ACE2 | RNF26 | 0,4697 | 3,303E-21 |
| ACE2 | TBC1D24 | 0,4697 | 3,25641E-21 |
| ACE2 | CTD-2574D22.3 | 0,4697 | 3,29556E-21 |
| ACE2 | TMEM37 | 0,4696 | 3,37461E-21 |
| ACE2 | TLR1 | 0,4696 | 3,35713E-21 |
| ACE2 | CTC-425O23.2 | 0,4696 | 3,38323E-21 |
| ACE2 | PIGA | 0,4696 | 3,36531E-21 |
| ACE2 | VCP | 0,4695 | 3,43054E-21 |
| ACE2 | GIT2 | 0,4695 | 3,45846E-21 |
| ACE2 | LINC01902 | 0,4695 | 3,40856E-21 |
| ACE2 | FKBP9P1 | 0,4694 | 3,55195E-21 |

| | | | |
|---|---|---|---|
| ACE2 | TGFB1I1 | 0,4694 | 3,49503E-21 |
| ACE2 | IPO9 | 0,4693 | 3,55368E-21 |
| ACE2 | LAYN | 0,4693 | 3,60436E-21 |
| ACE2 | EIF3J-AS1 | 0,4691 | 3,79209E-21 |
| ACE2 | SLC16A9 | 0,4688 | 4,02826E-21 |
| ACE2 | PLEKHF1 | 0,4688 | 3,96209E-21 |
| ACE2 | PLXNA3 | 0,4688 | 4,00625E-21 |
| ACE2 | PLCXD3 | 0,4687 | 4,05414E-21 |
| ACE2 | STX4 | 0,4687 | 4,06758E-21 |
| ACE2 | MYO9B | 0,4687 | 4,07326E-21 |
| ACE2 | STIM2 | 0,4686 | 4,1768E-21 |
| ACE2 | LAGE3 | 0,4686 | 4,17097E-21 |
| ACE2 | RP11-513I15.6 | 0,4685 | 4,29321E-21 |
| ACE2 | RP11-111K18.2 | 0,4685 | 4,31843E-21 |
| ACE2 | RP11-500G22.5 | 0,4685 | 4,31648E-21 |
| ACE2 | ZNF518B | 0,4684 | 4,40762E-21 |
| ACE2 | RP11-73M18.6 | 0,4684 | 4,3814E-21 |
| ACE2 | SCAMP4 | 0,4683 | 4,49147E-21 |
| ACE2 | ACAP2-IT1 | 0,4681 | 4,61854E-21 |
| ACE2 | KEL | 0,4681 | 4,6815E-21 |
| ACE2 | JAG1 | 0,468 | 4,74092E-21 |
| ACE2 | L1CAM | 0,468 | 4,77367E-21 |
| ACE2 | BTC | 0,4679 | 4,82529E-21 |
| ACE2 | FRMPD1 | 0,4678 | 4,98781E-21 |
| ACE2 | SLC29A3 | 0,4678 | 4,92692E-21 |
| ACE2 | HAUS2 | 0,4678 | 5,00985E-21 |
| ACE2 | KAT2B | 0,4677 | 5,05993E-21 |
| ACE2 | ADAT2 | 0,4677 | 5,0799E-21 |
| ACE2 | CLDN23 | 0,4675 | 5,31044E-21 |
| ACE2 | NQO1 | 0,4675 | 5,34656E-21 |
| ACE2 | TRMT44 | 0,4674 | 5,39357E-21 |
| ACE2 | ANXA2P2 | 0,4674 | 5,426E-21 |
| ACE2 | PDDC1 | 0,4674 | 5,44158E-21 |
| ACE2 | PLEKHG3 | 0,4674 | 5,4328E-21 |
| ACE2 | SLF1 | 0,4673 | 5,5711E-21 |
| ACE2 | SPIDR | 0,4673 | 5,56654E-21 |
| ACE2 | BEND2 | 0,4673 | 5,52391E-21 |
| ACE2 | NDUFS5 | 0,4672 | 5,70118E-21 |
| ACE2 | SOD2 | 0,4671 | 5,74904E-21 |
| ACE2 | KIAA1147 | 0,4671 | 5,84401E-21 |
| ACE2 | RP11-158H5.7 | 0,4671 | 5,78339E-21 |
| ACE2 | TYW1 | 0,4669 | 5,99088E-21 |
| ACE2 | LYZ | 0,4669 | 6,07785E-21 |
| ACE2 | EZH1 | 0,4668 | 6,17511E-21 |
| ACE2 | PTPRM | 0,4668 | 6,22963E-21 |
| ACE2 | KIF5C | 0,4667 | 6,30496E-21 |
| ACE2 | RP11-313J2.1 | 0,4667 | 6,29259E-21 |
| ACE2 | CGREF1 | 0,4666 | 6,39862E-21 |
| ACE2 | KRT8P33 | 0,4666 | 6,46984E-21 |
| ACE2 | SLC35D1 | 0,4664 | 6,80523E-21 |
| ACE2 | ZNF112 | 0,4664 | 6,78896E-21 |
| ACE2 | ZNF354B | 0,4663 | 6,90648E-21 |
| ACE2 | NUFIP2 | 0,4663 | 6,82683E-21 |
| ACE2 | MAGEA12 | 0,4663 | 6,91766E-21 |

| | | | |
|---|---|---|---|
| ACE2 | MARVELD1 | 0,4662 | 6,99912E-21 |
| ACE2 | ZYG11A | 0,4661 | 7,20055E-21 |
| ACE2 | LINC00284 | 0,466 | 7,29012E-21 |
| ACE2 | FLCN | 0,466 | 7,27479E-21 |
| ACE2 | PRKCSH | 0,466 | 7,29176E-21 |
| ACE2 | AC000120.7 | 0,4658 | 7,7303E-21 |
| ACE2 | IGF2 | 0,4658 | 7,74821E-21 |
| ACE2 | RBMS2 | 0,4658 | 7,66006E-21 |
| ACE2 | SARM1 | 0,4658 | 7,66461E-21 |
| ACE2 | RP11-348F1.3 | 0,4658 | 7,68782E-21 |
| ACE2 | BLOC1S4 | 0,4657 | 7,8137E-21 |
| ACE2 | TCERG1 | 0,4657 | 7,92197E-21 |
| ACE2 | TKTL1 | 0,4657 | 7,88256E-21 |
| ACE2 | NACC2 | 0,4656 | 8,00796E-21 |
| ACE2 | LTBP4 | 0,4656 | 8,05366E-21 |
| ACE2 | CDC42EP1 | 0,4656 | 7,97598E-21 |
| ACE2 | GSPT2 | 0,4655 | 8,27106E-21 |
| ACE2 | SEC13 | 0,4654 | 8,41822E-21 |
| ACE2 | NRBP2 | 0,4654 | 8,35591E-21 |
| ACE2 | PHF23 | 0,4654 | 8,37809E-21 |
| ACE2 | SLC16A2 | 0,4654 | 8,27792E-21 |
| ACE2 | DROSHA | 0,4653 | 8,47564E-21 |
| ACE2 | RP11-359J14.2 | 0,4653 | 8,61696E-21 |
| ACE2 | PARP3 | 0,4652 | 8,63733E-21 |
| ACE2 | RP11-290L1.4 | 0,4652 | 8,71571E-21 |
| ACE2 | ZNF697 | 0,4651 | 8,858E-21 |
| ACE2 | ZNF14 | 0,4651 | 8,96215E-21 |
| ACE2 | LITAF | 0,4649 | 9,35798E-21 |
| ACE2 | KCNK15-AS1 | 0,4647 | 9,71438E-21 |
| ACE2 | TLN2 | 0,4646 | 9,9219E-21 |
| ACE2 | RP11-390G14.1 | 0,4645 | 1,02407E-20 |
| ACE2 | CD27-AS1 | 0,4645 | 1,02455E-20 |
| ACE2 | C1QA | 0,4644 | 1,03498E-20 |
| ACE2 | C6orf89 | 0,4644 | 1,02825E-20 |
| ACE2 | PHEX | 0,4644 | 1,03785E-20 |
| ACE2 | KCNE3 | 0,4643 | 1,06511E-20 |
| ACE2 | CDC16 | 0,4643 | 1,05031E-20 |
| ACE2 | HIST2H2BA | 0,4642 | 1,0825E-20 |
| ACE2 | DLK1 | 0,4642 | 1,0746E-20 |
| ACE2 | LINC01197 | 0,4642 | 1,07398E-20 |
| ACE2 | KCNA2 | 0,4641 | 1,09537E-20 |
| ACE2 | POLE4 | 0,4641 | 1,10246E-20 |
| ACE2 | NOTCH1 | 0,4641 | 1,11182E-20 |
| ACE2 | ZNF347 | 0,4641 | 1,09605E-20 |
| ACE2 | DSTYK | 0,464 | 1,13907E-20 |
| ACE2 | PRELID1 | 0,464 | 1,12119E-20 |
| ACE2 | EPOR | 0,4639 | 1,14406E-20 |
| ACE2 | EMX2OS | 0,4638 | 1,17919E-20 |
| ACE2 | PYM1 | 0,4638 | 1,17809E-20 |
| ACE2 | FAM180A | 0,4637 | 1,20712E-20 |
| ACE2 | MOCOS | 0,4637 | 1,1973E-20 |
| ACE2 | GTF2IP1 | 0,4636 | 1,22506E-20 |
| ACE2 | TNS2 | 0,4635 | 1,26887E-20 |
| ACE2 | TRIP11 | 0,4635 | 1,27003E-20 |

| | | | |
|---|---|---|---|
| ACE2 | TFAP4 | 0,4635 | 1,26804E-20 |
| ACE2 | MICA | 0,4634 | 1,29188E-20 |
| ACE2 | ADGRE5 | 0,4634 | 1,29023E-20 |
| ACE2 | C22orf39 | 0,4633 | 1,30354E-20 |
| ACE2 | WDR45B | 0,4632 | 1,33829E-20 |
| ACE2 | FAM96B | 0,4631 | 1,3612E-20 |
| ACE2 | RP4-545K15.5 | 0,4631 | 1,38373E-20 |
| ACE2 | SORBS2 | 0,463 | 1,40099E-20 |
| ACE2 | LRRC17 | 0,4629 | 1,44146E-20 |
| ACE2 | NRF1 | 0,4629 | 1,42165E-20 |
| ACE2 | ST3GAL2 | 0,4629 | 1,43696E-20 |
| ACE2 | RP11-346E8.1 | 0,4629 | 1,43355E-20 |
| ACE2 | CD163 | 0,4628 | 1,45017E-20 |
| ACE2 | RNF144A | 0,4627 | 1,48399E-20 |
| ACE2 | PMS2P3 | 0,4627 | 1,48483E-20 |
| ACE2 | TLR2 | 0,4626 | 1,52443E-20 |
| ACE2 | AZIN1 | 0,4626 | 1,50945E-20 |
| ACE2 | LINC00346 | 0,4626 | 1,53112E-20 |
| ACE2 | ALDH1A3 | 0,4626 | 1,5276E-20 |
| ACE2 | ZNF566 | 0,4626 | 1,5274E-20 |
| ACE2 | ARX | 0,4626 | 1,53673E-20 |
| ACE2 | RPS6KA5 | 0,4625 | 1,54622E-20 |
| ACE2 | MLLT6 | 0,4625 | 1,54282E-20 |
| ACE2 | PTP4A2 | 0,4624 | 1,60506E-20 |
| ACE2 | RP11-230F18.5 | 0,4624 | 1,58745E-20 |
| ACE2 | RP11-887P2.5 | 0,4623 | 1,63795E-20 |
| ACE2 | TMEM100 | 0,4623 | 1,63432E-20 |
| ACE2 | MEX3A | 0,4622 | 1,6558E-20 |
| ACE2 | CAMK2G | 0,4621 | 1,69323E-20 |
| ACE2 | CTD-2515A14.1 | 0,462 | 1,74331E-20 |
| ACE2 | RP11-873E20.1 | 0,462 | 1,72904E-20 |
| ACE2 | SGMS1-AS1 | 0,4619 | 1,76227E-20 |
| ACE2 | CTD-2005H7.1 | 0,4619 | 1,76575E-20 |
| ACE2 | NEMP1 | 0,4619 | 1,7724E-20 |
| ACE2 | RP11-793J2.1 | 0,4619 | 1,76906E-20 |
| ACE2 | CTC-523E23.3 | 0,4619 | 1,75196E-20 |
| ACE2 | RP11-700J17.1 | 0,4618 | 1,81314E-20 |
| ACE2 | WDYHV1 | 0,4618 | 1,82538E-20 |
| ACE2 | SEMA4C | 0,4617 | 1,8587E-20 |
| ACE2 | KBTBD8 | 0,4617 | 1,83116E-20 |
| ACE2 | NDRG2 | 0,4617 | 1,86121E-20 |
| ACE2 | AC004540.4 | 0,4614 | 1,96809E-20 |
| ACE2 | RP11-817I4.2 | 0,4614 | 1,98101E-20 |
| ACE2 | GLYR1 | 0,4614 | 1,94817E-20 |
| ACE2 | STXBP4 | 0,4614 | 1,96149E-20 |
| ACE2 | PARP9 | 0,4613 | 2,00701E-20 |
| ACE2 | ESCO2 | 0,4613 | 2,0197E-20 |
| ACE2 | CTD-2002H8.2 | 0,4613 | 2,02194E-20 |
| ACE2 | MTERF3 | 0,4612 | 2,05271E-20 |
| ACE2 | STEAP1 | 0,4611 | 2,10469E-20 |
| ACE2 | CALM2P2 | 0,4611 | 2,10888E-20 |
| ACE2 | C11orf54 | 0,4611 | 2,08493E-20 |
| ACE2 | MTRNR2L4 | 0,4611 | 2,1163E-20 |
| ACE2 | AOC3 | 0,4611 | 2,07658E-20 |

| | | | |
|---|---|---|---|
| ACE2 | HELB | 0,461 | 2,15754E-20 |
| ACE2 | RPS16 | 0,461 | 2,1505E-20 |
| ACE2 | PRPF38A | 0,4608 | 2,23817E-20 |
| ACE2 | EML4 | 0,4608 | 2,24966E-20 |
| ACE2 | UNC93B1 | 0,4608 | 2,25211E-20 |
| ACE2 | ARHGEF1 | 0,4608 | 2,2552E-20 |
| ACE2 | DHX9 | 0,4607 | 2,28661E-20 |
| ACE2 | EMX2 | 0,4607 | 2,30758E-20 |
| ACE2 | CRIPT | 0,4606 | 2,31209E-20 |
| ACE2 | KRT18 | 0,4606 | 2,32222E-20 |
| ACE2 | CTD-2623N2.3 | 0,4606 | 2,31321E-20 |
| ACE2 | ANKRD27 | 0,4606 | 2,31322E-20 |
| ACE2 | PROK1 | 0,4605 | 2,40381E-20 |
| ACE2 | RP11-686D22.7 | 0,4604 | 2,42391E-20 |
| ACE2 | FBXO42 | 0,4603 | 2,4851E-20 |
| ACE2 | LINC01087 | 0,4602 | 2,54539E-20 |
| ACE2 | RP1-30M3.6 | 0,46 | 2,64802E-20 |
| ACE2 | OPRK1 | 0,46 | 2,66954E-20 |
| ACE2 | IRF9 | 0,46 | 2,62248E-20 |
| ACE2 | PRELP | 0,4599 | 2,69004E-20 |
| ACE2 | DANCR | 0,4599 | 2,70569E-20 |
| ACE2 | CTC-241N9.1 | 0,4599 | 2,70468E-20 |
| ACE2 | FAM3A | 0,4598 | 2,75698E-20 |
| ACE2 | KCTD7 | 0,4597 | 2,81043E-20 |
| ACE2 | ZDHHC7 | 0,4597 | 2,82334E-20 |
| ACE2 | RPS27L | 0,4596 | 2,88064E-20 |
| ACE2 | IL6R | 0,4594 | 3,02789E-20 |
| ACE2 | PITRM1 | 0,4594 | 3,03144E-20 |
| ACE2 | GTPBP10 | 0,4593 | 3,04609E-20 |
| ACE2 | RP11-144I2.1 | 0,4593 | 3,06328E-20 |
| ACE2 | CTC-548K16.6 | 0,4593 | 3,05544E-20 |
| ACE2 | RBM45 | 0,4592 | 3,10744E-20 |
| ACE2 | LSM7 | 0,4592 | 3,14995E-20 |
| ACE2 | AC093673.5 | 0,4591 | 3,21344E-20 |
| ACE2 | RP11-180M15.7 | 0,4591 | 3,22045E-20 |
| ACE2 | PLCD3 | 0,4591 | 3,22934E-20 |
| ACE2 | MAGEA6 | 0,459 | 3,29473E-20 |
| ACE2 | MT-ND6 | 0,459 | 3,28105E-20 |
| ACE2 | PSMG4 | 0,4589 | 3,3289E-20 |
| ACE2 | RP11-155G14.5 | 0,4589 | 3,32464E-20 |
| ACE2 | MIR497HG | 0,4589 | 3,35588E-20 |
| ACE2 | OLFML1 | 0,4588 | 3,3792E-20 |
| ACE2 | SEMA6C | 0,4587 | 3,46534E-20 |
| ACE2 | RP11-379F4.7 | 0,4586 | 3,53859E-20 |
| ACE2 | SRD5A1 | 0,4586 | 3,52839E-20 |
| ACE2 | CLIP2 | 0,4586 | 3,53569E-20 |
| ACE2 | EHBP1L1 | 0,4586 | 3,5499E-20 |
| ACE2 | CLDN20 | 0,4583 | 3,78729E-20 |
| ACE2 | RP11-467H10.1 | 0,4583 | 3,78769E-20 |
| ACE2 | FAM129B | 0,4582 | 3,84145E-20 |
| ACE2 | RP11-573M3.3 | 0,4582 | 3,87084E-20 |
| ACE2 | NCBP2-AS2 | 0,4581 | 3,92665E-20 |
| ACE2 | CDH23 | 0,4581 | 3,97812E-20 |
| ACE2 | TMEM204 | 0,4581 | 3,96006E-20 |

| | | | |
|---|---|---|---|
| ACE2 | PAQR8 | 0,458 | 4,0163E-20 |
| ACE2 | RP11-73M18.8 | 0,458 | 4,02483E-20 |
| ACE2 | ASB7 | 0,458 | 4,04053E-20 |
| ACE2 | AXIN2 | 0,458 | 4,06649E-20 |
| ACE2 | SLC8B1 | 0,4579 | 4,15466E-20 |
| ACE2 | LL0XNC01-116E7.4 | 0,4579 | 4,08068E-20 |
| ACE2 | SLC16A11 | 0,4578 | 4,17118E-20 |
| ACE2 | INHBB | 0,4577 | 4,33664E-20 |
| ACE2 | ID2 | 0,4576 | 4,3874E-20 |
| ACE2 | NPR3 | 0,4576 | 4,39712E-20 |
| ACE2 | CH17-340M24.3 | 0,4576 | 4,4012E-20 |
| ACE2 | CLIC1 | 0,4575 | 4,44074E-20 |
| ACE2 | PKN1 | 0,4575 | 4,45644E-20 |
| ACE2 | IGHV3-74 | 0,4574 | 4,55526E-20 |
| ACE2 | LINC00891 | 0,4573 | 4,70678E-20 |
| ACE2 | GBP3 | 0,4572 | 4,75794E-20 |
| ACE2 | UBE3C | 0,4572 | 4,7642E-20 |
| ACE2 | ITIH5 | 0,4572 | 4,78193E-20 |
| ACE2 | GREB1L | 0,4572 | 4,81353E-20 |
| ACE2 | PRORSD1P | 0,4571 | 4,84043E-20 |
| ACE2 | FNTB | 0,4571 | 4,90739E-20 |
| ACE2 | TMEM143 | 0,457 | 4,98105E-20 |
| ACE2 | APOBEC3D | 0,457 | 5,02204E-20 |
| ACE2 | PTPRD-AS1 | 0,4569 | 5,09931E-20 |
| ACE2 | ZNF37A | 0,4569 | 5,04799E-20 |
| ACE2 | CMTM7 | 0,4568 | 5,18132E-20 |
| ACE2 | RP11-61I13.3 | 0,4568 | 5,165E-20 |
| ACE2 | CLEC4C | 0,4568 | 5,15927E-20 |
| ACE2 | RPS17 | 0,4568 | 5,18472E-20 |
| ACE2 | CTC-439O9.1 | 0,4568 | 5,15593E-20 |
| ACE2 | NBPF26 | 0,4567 | 5,3028E-20 |
| ACE2 | PAPOLG | 0,4566 | 5,39169E-20 |
| ACE2 | NCS1 | 0,4566 | 5,45741E-20 |
| ACE2 | TMEM19 | 0,4566 | 5,46058E-20 |
| ACE2 | ADNP2 | 0,4566 | 5,46897E-20 |
| ACE2 | SUPT20HL1 | 0,4566 | 5,404E-20 |
| ACE2 | ITPR3 | 0,4565 | 5,55142E-20 |
| ACE2 | SLC1A3 | 0,4564 | 5,61988E-20 |
| ACE2 | VPS4A | 0,4564 | 5,70577E-20 |
| ACE2 | CNN2 | 0,4564 | 5,70569E-20 |
| ACE2 | SLAMF8 | 0,4563 | 5,76829E-20 |
| ACE2 | RP11-124N2.1 | 0,4563 | 5,78123E-20 |
| ACE2 | AGAP3 | 0,4563 | 5,77774E-20 |
| ACE2 | AKR1C1 | 0,4563 | 5,77068E-20 |
| ACE2 | RP11-680F8.1 | 0,4563 | 5,76329E-20 |
| ACE2 | CLN6 | 0,4563 | 5,81668E-20 |
| ACE2 | NANOS1 | 0,4562 | 5,89335E-20 |
| ACE2 | WFIKKN2 | 0,4562 | 5,94119E-20 |
| ACE2 | MITD1 | 0,4561 | 6,0547E-20 |
| ACE2 | ZNF841 | 0,4561 | 6,04753E-20 |
| ACE2 | GLIPR1 | 0,456 | 6,19368E-20 |
| ACE2 | CYP20A1 | 0,4559 | 6,28051E-20 |
| ACE2 | MVB12B | 0,4559 | 6,31161E-20 |
| ACE2 | RP11-73M18.7 | 0,4559 | 6,2531E-20 |

| | | | |
|---|---|---|---|
| ACE2 | CLK1 | 0,4558 | 6,42497E-20 |
| ACE2 | PSMB8 | 0,4557 | 6,59885E-20 |
| ACE2 | BRPF3 | 0,4557 | 6,52227E-20 |
| ACE2 | PABPC1L | 0,4557 | 6,58316E-20 |
| ACE2 | LINC00886 | 0,4556 | 6,70108E-20 |
| ACE2 | PTGDS | 0,4556 | 6,6624E-20 |
| ACE2 | STARD5 | 0,4556 | 6,72752E-20 |
| ACE2 | RUNDC3B | 0,4555 | 6,76115E-20 |
| ACE2 | RP11-757G1.6 | 0,4555 | 6,80058E-20 |
| ACE2 | LL22NC03-2H8.4 | 0,4555 | 6,76549E-20 |
| ACE2 | RP11-395I14.2 | 0,4553 | 7,14483E-20 |
| ACE2 | SLITRK4 | 0,4553 | 7,13707E-20 |
| ACE2 | MYOZ1 | 0,4552 | 7,18069E-20 |
| ACE2 | RP11-226L15.5 | 0,4551 | 7,3548E-20 |
| ACE2 | MTERF1 | 0,4551 | 7,39623E-20 |
| ACE2 | IL18BP | 0,4551 | 7,46853E-20 |
| ACE2 | TUBD1 | 0,455 | 7,59575E-20 |
| ACE2 | BRF2 | 0,4549 | 7,68398E-20 |
| ACE2 | SLCO2B1 | 0,4549 | 7,76642E-20 |
| ACE2 | VDAC1P8 | 0,4548 | 7,84571E-20 |
| ACE2 | SP1 | 0,4548 | 7,86013E-20 |
| ACE2 | CISD3 | 0,4548 | 7,89226E-20 |
| ACE2 | GGT1 | 0,4548 | 7,84895E-20 |
| ACE2 | NNT | 0,4547 | 7,97797E-20 |
| ACE2 | RPS2P46 | 0,4547 | 8,11902E-20 |
| ACE2 | TSC22D3 | 0,4546 | 8,13529E-20 |
| ACE2 | RP11-177C12.5 | 0,4545 | 8,3521E-20 |
| ACE2 | AIDA | 0,4544 | 8,51039E-20 |
| ACE2 | NFKB2 | 0,4544 | 8,49819E-20 |
| ACE2 | ARHGDIA | 0,4544 | 8,49235E-20 |
| ACE2 | DOPEY2 | 0,4544 | 8,59015E-20 |
| ACE2 | CTB-152G17.6 | 0,4543 | 8,81656E-20 |
| ACE2 | RIMKLBP2 | 0,4542 | 9,01943E-20 |
| ACE2 | FGFRL1 | 0,4541 | 9,09562E-20 |
| ACE2 | OBP2B | 0,4541 | 9,07727E-20 |
| ACE2 | VAC14 | 0,454 | 9,39955E-20 |
| ACE2 | CTD-2267D19.3 | 0,454 | 9,3009E-20 |
| ACE2 | KLK10 | 0,454 | 9,29583E-20 |
| ACE2 | ZNF263 | 0,4539 | 9,59151E-20 |
| ACE2 | RAB33B | 0,4538 | 9,68039E-20 |
| ACE2 | RP1-80N2.4 | 0,4538 | 9,74661E-20 |
| ACE2 | TACR2 | 0,4538 | 9,67765E-20 |
| ACE2 | BSDC1 | 0,4537 | 9,84913E-20 |
| ACE2 | TOM1L2 | 0,4536 | 1,0088E-19 |
| ACE2 | SLC35E2 | 0,4535 | 1,03774E-19 |
| ACE2 | ZNF852 | 0,4535 | 1,0366E-19 |
| ACE2 | KDR | 0,4535 | 1,02433E-19 |
| ACE2 | HRH1 | 0,4533 | 1,08313E-19 |
| ACE2 | CCDC25 | 0,4533 | 1,07081E-19 |
| ACE2 | CTC-444N24.7 | 0,4533 | 1,07751E-19 |
| ACE2 | RP3-522J7.7 | 0,4531 | 1,11467E-19 |
| ACE2 | RP11-204M4.3 | 0,453 | 1,15292E-19 |
| ACE2 | IL17RC | 0,4529 | 1,18058E-19 |
| ACE2 | SPTB | 0,4529 | 1,17564E-19 |

| | | | |
|---|---|---|---|
| ACE2 | ACOT4 | 0,4529 | 1,16217E-19 |
| ACE2 | TRPV2 | 0,4529 | 1,17542E-19 |
| ACE2 | TNFRSF10C | 0,4528 | 1,20474E-19 |
| ACE2 | HTR3B | 0,4528 | 1,19707E-19 |
| ACE2 | OSGEP | 0,4528 | 1,1955E-19 |
| ACE2 | LINC01012 | 0,4526 | 1,24476E-19 |
| ACE2 | HOXB7 | 0,4525 | 1,26454E-19 |
| ACE2 | PDE7B | 0,4524 | 1,28443E-19 |
| ACE2 | RP11-10N23.2 | 0,4524 | 1,28825E-19 |
| ACE2 | CEMIP | 0,4524 | 1,30435E-19 |
| ACE2 | HDAC5 | 0,4524 | 1,30865E-19 |
| ACE2 | GALNT4 | 0,4523 | 1,3193E-19 |
| ACE2 | PTGR2 | 0,4523 | 1,31307E-19 |
| ACE2 | RPRD1B | 0,4521 | 1,3757E-19 |
| ACE2 | ZNF638 | 0,4519 | 1,4271E-19 |
| ACE2 | PLEC | 0,4519 | 1,43736E-19 |
| ACE2 | DSG2 | 0,4519 | 1,45062E-19 |
| ACE2 | POU2F1 | 0,4518 | 1,47604E-19 |
| ACE2 | CLDN3 | 0,4518 | 1,46767E-19 |
| ACE2 | RP11-977G19.14 | 0,4518 | 1,47611E-19 |
| ACE2 | AKAP11 | 0,4518 | 1,48052E-19 |
| ACE2 | NME4 | 0,4518 | 1,46528E-19 |
| ACE2 | C6 | 0,4517 | 1,48403E-19 |
| ACE2 | RP11-1079K10.3 | 0,4517 | 1,49774E-19 |
| ACE2 | SGSM1 | 0,4517 | 1,48468E-19 |
| ACE2 | PHF3 | 0,4516 | 1,5369E-19 |
| ACE2 | RP11-251G23.5 | 0,4516 | 1,51587E-19 |
| ACE2 | YWHAZ | 0,4515 | 1,55255E-19 |
| ACE2 | SRGAP2B | 0,4514 | 1,57942E-19 |
| ACE2 | NRP1 | 0,4514 | 1,5943E-19 |
| ACE2 | C14orf132 | 0,4513 | 1,62112E-19 |
| ACE2 | RP11-826N14.2 | 0,4512 | 1,67195E-19 |
| ACE2 | EMILIN1 | 0,4511 | 1,70051E-19 |
| ACE2 | PIGX | 0,4511 | 1,69332E-19 |
| ACE2 | ZNF761 | 0,4511 | 1,68332E-19 |
| ACE2 | RCHY1 | 0,451 | 1,73582E-19 |
| ACE2 | BID | 0,451 | 1,71839E-19 |
| ACE2 | RP11-110I1.5 | 0,4509 | 1,7758E-19 |
| ACE2 | ZFP14 | 0,4509 | 1,75968E-19 |
| ACE2 | C8orf37 | 0,4508 | 1,7989E-19 |
| ACE2 | ZNF835 | 0,4508 | 1,81522E-19 |
| ACE2 | XXbac-BPGBPG55C20. | 0,4506 | 1,87842E-19 |
| ACE2 | COQ3 | 0,4506 | 1,87473E-19 |
| ACE2 | DPPA2 | 0,4505 | 1,9353E-19 |
| ACE2 | CTF1 | 0,4505 | 1,90493E-19 |
| ACE2 | HKR1 | 0,4505 | 1,89845E-19 |
| ACE2 | NAA10 | 0,4505 | 1,89871E-19 |
| ACE2 | CTD-2227E11.1 | 0,4504 | 1,93962E-19 |
| ACE2 | MAP1A | 0,4504 | 1,95638E-19 |
| ACE2 | RP1-228H13.5 | 0,4503 | 2,0002E-19 |
| ACE2 | RP11-967K21.1 | 0,4503 | 1,98368E-19 |
| ACE2 | DIO3 | 0,4503 | 2,00149E-19 |
| ACE2 | ZNF407 | 0,4503 | 1,99655E-19 |
| ACE2 | MAGEA3 | 0,4502 | 2,03041E-19 |

| | | | |
|---|---|---|---|
| ACE2 | KANK4 | 0,45 | 2,13447E-19 |
| ACE2 | RP4-675C20.4 | 0,45 | 2,10182E-19 |
| ACE2 | PRCC | 0,45 | 2,12523E-19 |
| ACE2 | ACSS2 | 0,4499 | 2,17403E-19 |
| ACE2 | RP11-449J21.3 | 0,4498 | 2,23229E-19 |
| ACE2 | KCNJ3 | 0,4497 | 2,25937E-19 |
| ACE2 | CHRAC1 | 0,4497 | 2,24093E-19 |
| ACE2 | RP11-503E24.2 | 0,4496 | 2,28647E-19 |
| ACE2 | LY6E | 0,4496 | 2,32029E-19 |
| ACE2 | SEMA3A | 0,4494 | 2,38132E-19 |
| ACE2 | GGT5 | 0,4494 | 2,42186E-19 |
| ACE2 | SPAST | 0,4493 | 2,44411E-19 |
| ACE2 | ADAMTS2 | 0,4493 | 2,43823E-19 |
| ACE2 | FBXO27 | 0,4493 | 2,4684E-19 |
| ACE2 | C1QB | 0,4492 | 2,51702E-19 |
| ACE2 | SLA | 0,4492 | 2,48785E-19 |
| ACE2 | CLUHP3 | 0,4492 | 2,49818E-19 |
| ACE2 | HIST1H4H | 0,4491 | 2,55106E-19 |
| ACE2 | TMSB4X | 0,4489 | 2,6442E-19 |
| ACE2 | RP11-536K7.3 | 0,4488 | 2,703E-19 |
| ACE2 | CTPS1 | 0,4487 | 2,76111E-19 |
| ACE2 | ZNF484 | 0,4487 | 2,75716E-19 |
| ACE2 | RAB34 | 0,4487 | 2,78441E-19 |
| ACE2 | RPS11 | 0,4486 | 2,83386E-19 |
| ACE2 | RP4-598P13.1 | 0,4485 | 2,85812E-19 |
| ACE2 | NIPA1 | 0,4485 | 2,89144E-19 |
| ACE2 | LIPG | 0,4485 | 2,8632E-19 |
| ACE2 | THUMPD3-AS1 | 0,4484 | 2,96844E-19 |
| ACE2 | LINC02001 | 0,4484 | 2,92886E-19 |
| ACE2 | BCAS4 | 0,4484 | 2,9191E-19 |
| ACE2 | RP11-404O13.4 | 0,4483 | 3,00818E-19 |
| ACE2 | APOBEC3H | 0,4483 | 3,01306E-19 |
| ACE2 | VEGFD | 0,4483 | 2,99389E-19 |
| ACE2 | MT-CYB | 0,4481 | 3,15026E-19 |
| ACE2 | TMEM128 | 0,4479 | 3,28291E-19 |
| ACE2 | HNRNPA3P16 | 0,4479 | 3,22588E-19 |
| ACE2 | TMX4 | 0,4479 | 3,22929E-19 |
| ACE2 | MAGEC1 | 0,4479 | 3,27194E-19 |
| ACE2 | NT5C3AP1 | 0,4478 | 3,3458E-19 |
| ACE2 | GALNT10 | 0,4478 | 3,31917E-19 |
| ACE2 | AF131215.3 | 0,4478 | 3,30863E-19 |
| ACE2 | ATR | 0,4477 | 3,37211E-19 |
| ACE2 | CH17-431G21.1 | 0,4476 | 3,47615E-19 |
| ACE2 | XIST | 0,4476 | 3,46831E-19 |
| ACE2 | CPNE4 | 0,4475 | 3,53072E-19 |
| ACE2 | CHMP5 | 0,4475 | 3,5357E-19 |
| ACE2 | SORD | 0,4475 | 3,52704E-19 |
| ACE2 | DIP2A | 0,4475 | 3,53534E-19 |
| ACE2 | HAT1 | 0,4474 | 3,59718E-19 |
| ACE2 | RP11-250B2.6 | 0,4474 | 3,62339E-19 |
| ACE2 | RP11-16E12.2 | 0,4474 | 3,61377E-19 |
| ACE2 | UBE2D4 | 0,4473 | 3,67413E-19 |
| ACE2 | FIGNL1 | 0,4473 | 3,67634E-19 |
| ACE2 | MAGEA1 | 0,4473 | 3,68462E-19 |

| | | | |
|---|---|---|---|
| ACE2 | KCNC4 | 0,4472 | 3,7919E-19 |
| ACE2 | CREB5 | 0,4472 | 3,73543E-19 |
| ACE2 | WIF1 | 0,4472 | 3,77321E-19 |
| ACE2 | ITPA | 0,4472 | 3,77236E-19 |
| ACE2 | KLHL33 | 0,4471 | 3,85113E-19 |
| ACE2 | LRP3 | 0,447 | 3,91543E-19 |
| ACE2 | GGT7 | 0,447 | 3,92184E-19 |
| ACE2 | MT-ND4L | 0,447 | 3,88097E-19 |
| ACE2 | LIMS4 | 0,4469 | 3,96444E-19 |
| ACE2 | HOOK3 | 0,4469 | 3,98947E-19 |
| ACE2 | CEBPB | 0,4469 | 4,00995E-19 |
| ACE2 | ARID1A | 0,4468 | 4,08854E-19 |
| ACE2 | DCUN1D4 | 0,4468 | 4,0447E-19 |
| ACE2 | SCHIP1 | 0,4467 | 4,1906E-19 |
| ACE2 | DSE | 0,4467 | 4,17823E-19 |
| ACE2 | CHMP7 | 0,4467 | 4,16802E-19 |
| ACE2 | FRMD8 | 0,4467 | 4,14246E-19 |
| ACE2 | RP11-554K11.2 | 0,4466 | 4,25823E-19 |
| ACE2 | NAA35 | 0,4466 | 4,23946E-19 |
| ACE2 | RP11-362A9.3 | 0,4465 | 4,34779E-19 |
| ACE2 | RP11-159D12.8 | 0,4465 | 4,29231E-19 |
| ACE2 | ZNF554 | 0,4465 | 4,36345E-19 |
| ACE2 | RBMS3 | 0,4464 | 4,40669E-19 |
| ACE2 | ZNF830 | 0,4463 | 4,46502E-19 |
| ACE2 | RP11-1148L6.9 | 0,4463 | 4,46289E-19 |
| ACE2 | SRSF11 | 0,4462 | 4,58238E-19 |
| ACE2 | RP11-519C12.1 | 0,4461 | 4,71939E-19 |
| ACE2 | VSTM2L | 0,446 | 4,79421E-19 |
| ACE2 | ARHGAP32 | 0,4459 | 4,93621E-19 |
| ACE2 | FOSL2 | 0,4458 | 4,95685E-19 |
| ACE2 | GABRB2 | 0,4457 | 5,06103E-19 |
| ACE2 | ABCC10 | 0,4457 | 5,08821E-19 |
| ACE2 | RPL11P3 | 0,4456 | 5,19587E-19 |
| ACE2 | LINC00324 | 0,4456 | 5,23855E-19 |
| ACE2 | APOBEC3B | 0,4456 | 5,18309E-19 |
| ACE2 | GALM | 0,4455 | 5,28471E-19 |
| ACE2 | BCAR1 | 0,4455 | 5,3446E-19 |
| ACE2 | MAGEB16 | 0,4455 | 5,26275E-19 |
| ACE2 | ZFP1 | 0,4453 | 5,54793E-19 |
| ACE2 | AC093110.3 | 0,4452 | 5,58623E-19 |
| ACE2 | ACAD9 | 0,4452 | 5,6161E-19 |
| ACE2 | ALOX5 | 0,4452 | 5,65389E-19 |
| ACE2 | RP11-884K10.7 | 0,4451 | 5,74419E-19 |
| ACE2 | RPL17P46 | 0,4451 | 5,7433E-19 |
| ACE2 | RP11-115H13.1 | 0,445 | 5,83735E-19 |
| ACE2 | RGS7BP | 0,4449 | 5,93998E-19 |
| ACE2 | PTGES2 | 0,4447 | 6,23189E-19 |
| ACE2 | RP11-326C3.13 | 0,4447 | 6,28321E-19 |
| ACE2 | RP11-110I1.11 | 0,4447 | 6,17055E-19 |
| ACE2 | SMAD6 | 0,4447 | 6,21536E-19 |
| ACE2 | FAM172A | 0,4446 | 6,36453E-19 |
| ACE2 | KCNIP1 | 0,4446 | 6,29525E-19 |
| ACE2 | TMEM9B-AS1 | 0,4446 | 6,30903E-19 |
| ACE2 | TAF1D | 0,4446 | 6,35236E-19 |

| | | | |
|---|---|---|---|
| ACE2 | RNF213 | 0,4446 | 6,37775E-19 |
| ACE2 | ZDHHC9 | 0,4446 | 6,33199E-19 |
| ACE2 | CRYL1 | 0,4445 | 6,50841E-19 |
| ACE2 | F8A1 | 0,4445 | 6,47967E-19 |
| ACE2 | IL27RA | 0,4444 | 6,59874E-19 |
| ACE2 | TTF2 | 0,4443 | 6,76718E-19 |
| ACE2 | PRDM8 | 0,4443 | 6,71781E-19 |
| ACE2 | RPS14 | 0,4443 | 6,78307E-19 |
| ACE2 | LFNG | 0,4442 | 6,9006E-19 |
| ACE2 | AKAP2 | 0,4442 | 6,85395E-19 |
| ACE2 | CORO1C | 0,4442 | 6,89621E-19 |
| ACE2 | LYVE1 | 0,4441 | 7,00637E-19 |
| ACE2 | PRRG4 | 0,4441 | 7,07288E-19 |
| ACE2 | JPH4 | 0,4441 | 7,07058E-19 |
| ACE2 | ATG16L1 | 0,444 | 7,17124E-19 |
| ACE2 | ADGRA2 | 0,444 | 7,23345E-19 |
| ACE2 | TAS2R31 | 0,4439 | 7,24456E-19 |
| ACE2 | CCDC12 | 0,4438 | 7,42646E-19 |
| ACE2 | FAM86C1 | 0,4438 | 7,39367E-19 |
| ACE2 | SRSF5 | 0,4438 | 7,43633E-19 |
| ACE2 | ZNF595 | 0,4437 | 7,61917E-19 |
| ACE2 | MAP3K15 | 0,4437 | 7,54238E-19 |
| ACE2 | CCR1 | 0,4436 | 7,80211E-19 |
| ACE2 | STARD4-AS1 | 0,4436 | 7,70832E-19 |
| ACE2 | LINC01625 | 0,4436 | 7,7804E-19 |
| ACE2 | CCAR1 | 0,4436 | 7,7385E-19 |
| ACE2 | CLMP | 0,4436 | 7,79176E-19 |
| ACE2 | HIST2H3PS2 | 0,4435 | 7,90834E-19 |
| ACE2 | RPL10P16 | 0,4435 | 7,92183E-19 |
| ACE2 | SNCA | 0,4434 | 8,098E-19 |
| ACE2 | ITGA1 | 0,4434 | 8,10272E-19 |
| ACE2 | FGF10 | 0,4433 | 8,28436E-19 |
| ACE2 | RP11-348N5.7 | 0,4433 | 8,32167E-19 |
| ACE2 | LDLRAP1 | 0,4432 | 8,45387E-19 |
| ACE2 | RP11-134G8.7 | 0,4431 | 8,65857E-19 |
| ACE2 | LINC00667 | 0,4431 | 8,55225E-19 |
| ACE2 | RP3-342P20.2 | 0,443 | 8,69306E-19 |
| ACE2 | ZNF226 | 0,443 | 8,73498E-19 |
| ACE2 | PLA2R1 | 0,4429 | 8,88567E-19 |
| ACE2 | ZNF597 | 0,4429 | 8,84745E-19 |
| ACE2 | BTG2 | 0,4428 | 9,18755E-19 |
| ACE2 | CTTNBP2 | 0,4427 | 9,37649E-19 |
| ACE2 | MSC-AS1 | 0,4427 | 9,32227E-19 |
| ACE2 | NLGN2 | 0,4426 | 9,4762E-19 |
| ACE2 | PSMB9 | 0,4425 | 9,63372E-19 |
| ACE2 | UQCRHL | 0,4424 | 9,78718E-19 |
| ACE2 | PARL | 0,4423 | 9,9867E-19 |
| ACE2 | ANAPC16 | 0,4423 | 9,99843E-19 |
| ACE2 | CTBP1-AS2 | 0,4422 | 1,03297E-18 |
| ACE2 | TPM2 | 0,4422 | 1,02336E-18 |
| ACE2 | ZNF844 | 0,4422 | 1,01935E-18 |
| ACE2 | FTL | 0,4422 | 1,01764E-18 |
| ACE2 | RP1-286D6.5 | 0,4421 | 1,04783E-18 |
| ACE2 | C2 | 0,4421 | 1,03905E-18 |

| | | | |
|---|---|---|---|
| ACE2 | BBX | 0,442 | 1,066E-18 |
| ACE2 | EMP3 | 0,442 | 1,068E-18 |
| ACE2 | WDR17 | 0,4419 | 1,09229E-18 |
| ACE2 | FER | 0,4419 | 1,09443E-18 |
| ACE2 | PMPCB | 0,4419 | 1,08502E-18 |
| ACE2 | TAS2R4 | 0,4419 | 1,09261E-18 |
| ACE2 | EEF1A1P5 | 0,4419 | 1,08793E-18 |
| ACE2 | CCBE1 | 0,4419 | 1,09651E-18 |
| ACE2 | NGF | 0,4418 | 1,10357E-18 |
| ACE2 | TPD52L2 | 0,4418 | 1,11712E-18 |
| ACE2 | CTD-2530H12.8 | 0,4417 | 1,14495E-18 |
| ACE2 | POSTN | 0,4417 | 1,14215E-18 |
| ACE2 | BNC2 | 0,4416 | 1,16913E-18 |
| ACE2 | RPS2P5 | 0,4416 | 1,14967E-18 |
| ACE2 | RP11-192M23.1 | 0,4416 | 1,16447E-18 |
| ACE2 | LPCAT1 | 0,4415 | 1,18709E-18 |
| ACE2 | NCALD | 0,4415 | 1,19188E-18 |
| ACE2 | SEMA4G | 0,4415 | 1,17039E-18 |
| ACE2 | C12orf75 | 0,4415 | 1,17542E-18 |
| ACE2 | AC009237.11 | 0,4414 | 1,19486E-18 |
| ACE2 | HAND2 | 0,4414 | 1,21045E-18 |
| ACE2 | RASL11A | 0,4414 | 1,19456E-18 |
| ACE2 | CTB-75G16.1 | 0,4413 | 1,21743E-18 |
| ACE2 | CEP120 | 0,4412 | 1,2487E-18 |
| ACE2 | EPHB2 | 0,441 | 1,30725E-18 |
| ACE2 | RP11-102N12.3 | 0,441 | 1,31391E-18 |
| ACE2 | LINC00519 | 0,441 | 1,29188E-18 |
| ACE2 | EFTUD2 | 0,441 | 1,29581E-18 |
| ACE2 | RP11-723D22.3 | 0,4409 | 1,32737E-18 |
| ACE2 | ZNF638-IT1 | 0,4408 | 1,35336E-18 |
| ACE2 | CYP26B1 | 0,4408 | 1,35989E-18 |
| ACE2 | RP1-151F17.2 | 0,4407 | 1,38315E-18 |
| ACE2 | MMRN2 | 0,4407 | 1,38853E-18 |
| ACE2 | NISCH | 0,4406 | 1,40685E-18 |
| ACE2 | GIMAP8 | 0,4406 | 1,41535E-18 |
| ACE2 | VWF | 0,4406 | 1,41552E-18 |
| ACE2 | FXYD5 | 0,4405 | 1,44969E-18 |
| ACE2 | RP11-254A24.2 | 0,4404 | 1,47182E-18 |
| ACE2 | PPP5C | 0,4404 | 1,46371E-18 |
| ACE2 | LSS | 0,4404 | 1,47184E-18 |
| ACE2 | TMEM129 | 0,4403 | 1,49276E-18 |
| ACE2 | NBEAL2 | 0,4402 | 1,53515E-18 |
| ACE2 | TMEM130 | 0,4402 | 1,53417E-18 |
| ACE2 | RP11-244H3.1 | 0,4401 | 1,54746E-18 |
| ACE2 | RPS15AP10 | 0,4401 | 1,56424E-18 |
| ACE2 | AKR1C3 | 0,44 | 1,58575E-18 |
| ACE2 | PFAS | 0,44 | 1,59007E-18 |
| ACE2 | ARHGAP23 | 0,44 | 1,59871E-18 |
| ACE2 | RRAGC | 0,4399 | 1,62982E-18 |
| ACE2 | LY75 | 0,4399 | 1,61931E-18 |
| ACE2 | CACNB3 | 0,4399 | 1,62276E-18 |
| ACE2 | RP11-120K18.2 | 0,4398 | 1,65245E-18 |
| ACE2 | CCDC28B | 0,4397 | 1,68021E-18 |
| ACE2 | RP11-162A12.2 | 0,4397 | 1,70228E-18 |

| | | | |
|---|---|---|---|
| ACE2 | AC141928.1 | 0,4396 | 1,7364E-18 |
| ACE2 | ACSM5 | 0,4396 | 1,72828E-18 |
| ACE2 | CFDP1 | 0,4396 | 1,70629E-18 |
| ACE2 | DTNB | 0,4395 | 1,76837E-18 |
| ACE2 | GLIS3 | 0,4395 | 1,77323E-18 |
| ACE2 | GLS2 | 0,4395 | 1,75564E-18 |
| ACE2 | SSC5D | 0,4395 | 1,76542E-18 |
| ACE2 | TBL1Y | 0,4395 | 1,746E-18 |
| ACE2 | WNT2B | 0,4394 | 1,79029E-18 |
| ACE2 | DDX41 | 0,4394 | 1,79742E-18 |
| ACE2 | CPVL | 0,4394 | 1,80033E-18 |
| ACE2 | LINC01184 | 0,4392 | 1,87994E-18 |
| ACE2 | VSIG4 | 0,4392 | 1,87802E-18 |
| ACE2 | THNSL2 | 0,4391 | 1,90666E-18 |
| ACE2 | RP11-231L11.3 | 0,4391 | 1,91307E-18 |
| ACE2 | GPRC5C | 0,4391 | 1,89748E-18 |
| ACE2 | PDE4B | 0,439 | 1,92269E-18 |
| ACE2 | HNRNPKP4 | 0,439 | 1,9357E-18 |
| ACE2 | STK32A | 0,439 | 1,95324E-18 |
| ACE2 | PCP4L1 | 0,4388 | 2,01502E-18 |
| ACE2 | RP5-1007F24.1 | 0,4388 | 2,011E-18 |
| ACE2 | DLX5 | 0,4388 | 2,00005E-18 |
| ACE2 | FADD | 0,4388 | 1,99887E-18 |
| ACE2 | PLD2 | 0,4388 | 2,03341E-18 |
| ACE2 | HSPG2 | 0,4387 | 2,06115E-18 |
| ACE2 | CDKAL1 | 0,4387 | 2,06739E-18 |
| ACE2 | KCNE4 | 0,4386 | 2,08691E-18 |
| ACE2 | FADS3 | 0,4384 | 2,18558E-18 |
| ACE2 | YWHAQ | 0,4383 | 2,21247E-18 |
| ACE2 | REG3G | 0,4383 | 2,22287E-18 |
| ACE2 | RNF43 | 0,4383 | 2,21522E-18 |
| ACE2 | SHKBP1 | 0,4382 | 2,261E-18 |
| ACE2 | SRGAP3 | 0,4381 | 2,33341E-18 |
| ACE2 | LDB2 | 0,4381 | 2,33673E-18 |
| ACE2 | MRPL48 | 0,4381 | 2,32123E-18 |
| ACE2 | ZNF70 | 0,4381 | 2,31075E-18 |
| ACE2 | PBX1 | 0,438 | 2,34476E-18 |
| ACE2 | C7orf25 | 0,438 | 2,33976E-18 |
| ACE2 | MET | 0,438 | 2,3698E-18 |
| ACE2 | RARRES3 | 0,438 | 2,35204E-18 |
| ACE2 | PAPLN | 0,438 | 2,35899E-18 |
| ACE2 | PBLD | 0,4379 | 2,4152E-18 |
| ACE2 | PDCD7 | 0,4379 | 2,41798E-18 |
| ACE2 | ZNF211 | 0,4379 | 2,38589E-18 |
| ACE2 | PXDC1 | 0,4378 | 2,4528E-18 |
| ACE2 | PRTFDC1 | 0,4378 | 2,47965E-18 |
| ACE2 | RNLS | 0,4378 | 2,46411E-18 |
| ACE2 | ETNK1 | 0,4378 | 2,45456E-18 |
| ACE2 | ARFGEF2 | 0,4377 | 2,52329E-18 |
| ACE2 | DGCR2 | 0,4376 | 2,53084E-18 |
| ACE2 | HPCAL1 | 0,4375 | 2,62285E-18 |
| ACE2 | RP11-730G20.2 | 0,4374 | 2,63938E-18 |
| ACE2 | RP11-4L24.4 | 0,4374 | 2,63147E-18 |
| ACE2 | RP11-186N15.3 | 0,4374 | 2,67164E-18 |

| | | | |
|---|---|---|---|
| ACE2 | RRBP1 | 0,4374 | 2,66806E-18 |
| ACE2 | FBRSL1 | 0,4373 | 2,68698E-18 |
| ACE2 | ZNF581 | 0,4373 | 2,7098E-18 |
| ACE2 | AC009299.5 | 0,4372 | 2,7794E-18 |
| ACE2 | FCGR3A | 0,4371 | 2,79975E-18 |
| ACE2 | LINC00954 | 0,4371 | 2,82707E-18 |
| ACE2 | RP11-1277A3.2 | 0,4371 | 2,79116E-18 |
| ACE2 | FAM111B | 0,4371 | 2,83574E-18 |
| ACE2 | ALPK3 | 0,4371 | 2,82273E-18 |
| ACE2 | SRRM1 | 0,437 | 2,86677E-18 |
| ACE2 | RP11-531A24.5 | 0,437 | 2,88886E-18 |
| ACE2 | CDCA4P1 | 0,437 | 2,90004E-18 |
| ACE2 | BTF3L4 | 0,4369 | 2,93847E-18 |
| ACE2 | C5orf34 | 0,4369 | 2,95562E-18 |
| ACE2 | MTCO1P28 | 0,4369 | 2,93146E-18 |
| ACE2 | MPND | 0,4369 | 2,93375E-18 |
| ACE2 | SMIM7 | 0,4369 | 2,90505E-18 |
| ACE2 | HSF2 | 0,4368 | 2,96275E-18 |
| ACE2 | PML | 0,4368 | 2,99415E-18 |
| ACE2 | MANBAL | 0,4368 | 2,9858E-18 |
| ACE2 | LYNX1 | 0,4367 | 3,03919E-18 |
| ACE2 | CD27 | 0,4367 | 3,0384E-18 |
| ACE2 | TMBIM6 | 0,4367 | 3,05538E-18 |
| ACE2 | LAMC3 | 0,4365 | 3,17072E-18 |
| ACE2 | DUSP5 | 0,4365 | 3,185E-18 |
| ACE2 | MARK2 | 0,4365 | 3,17853E-18 |
| ACE2 | CTD-2104P17.1 | 0,4364 | 3,26035E-18 |
| ACE2 | CYYR1 | 0,4364 | 3,23606E-18 |
| ACE2 | RNF146 | 0,4363 | 3,32657E-18 |
| ACE2 | NAT16 | 0,4363 | 3,29269E-18 |
| ACE2 | MED22 | 0,4363 | 3,28299E-18 |
| ACE2 | ST3GAL4-AS1 | 0,4363 | 3,29327E-18 |
| ACE2 | IGSF9B | 0,4363 | 3,30176E-18 |
| ACE2 | TFPI | 0,4362 | 3,35567E-18 |
| ACE2 | POP5 | 0,4361 | 3,4313E-18 |
| ACE2 | MRPL45 | 0,4361 | 3,43048E-18 |
| ACE2 | SMIM1 | 0,436 | 3,50983E-18 |
| ACE2 | PDCD10 | 0,436 | 3,49298E-18 |
| ACE2 | AP000640.10 | 0,4359 | 3,58383E-18 |
| ACE2 | PAK4 | 0,4359 | 3,5657E-18 |
| ACE2 | HIF3A | 0,4359 | 3,53221E-18 |
| ACE2 | RP11-715F3.2 | 0,4358 | 3,65926E-18 |
| ACE2 | HIST2H3DP1 | 0,4355 | 3,86203E-18 |
| ACE2 | RHPN2 | 0,4355 | 3,81612E-18 |
| ACE2 | PGBD1 | 0,4354 | 3,91169E-18 |
| ACE2 | APBB1IP | 0,4354 | 3,90909E-18 |
| ACE2 | HERC2 | 0,4354 | 3,95274E-18 |
| ACE2 | FTH1P8 | 0,4354 | 3,90164E-18 |
| ACE2 | RP5-1139B12.2 | 0,4353 | 3,98147E-18 |
| ACE2 | BMPR1APS1 | 0,4353 | 3,98609E-18 |
| ACE2 | FAM185BP | 0,4352 | 4,11066E-18 |
| ACE2 | EPHB6 | 0,4352 | 4,12447E-18 |
| ACE2 | FAM196A | 0,4352 | 4,05966E-18 |
| ACE2 | CABLES1 | 0,4352 | 4,06959E-18 |

| | | | |
|---|---|---|---|
| ACE2 | NOP9 | 0,4351 | 4,20452E-18 |
| ACE2 | EPRS | 0,435 | 4,23562E-18 |
| ACE2 | AC093724.2 | 0,435 | 4,28649E-18 |
| ACE2 | RNF152 | 0,4349 | 4,34379E-18 |
| ACE2 | RN7SL244P | 0,4348 | 4,40981E-18 |
| ACE2 | RP11-384P7.7 | 0,4348 | 4,45339E-18 |
| ACE2 | FBXL8 | 0,4348 | 4,40374E-18 |
| ACE2 | RP3-413H6.2 | 0,4347 | 4,51353E-18 |
| ACE2 | CNOT7 | 0,4346 | 4,58944E-18 |
| ACE2 | WNT11 | 0,4345 | 4,71433E-18 |
| ACE2 | DAPP1 | 0,4344 | 4,80884E-18 |
| ACE2 | PDE8B | 0,4343 | 4,88808E-18 |
| ACE2 | SLC25A27 | 0,4343 | 4,86136E-18 |
| ACE2 | PURB | 0,4343 | 4,87622E-18 |
| ACE2 | ACVRL1 | 0,4342 | 4,96915E-18 |
| ACE2 | EBPL | 0,4342 | 5,00764E-18 |
| ACE2 | FOXI3 | 0,4341 | 5,04494E-18 |
| ACE2 | IGFBP2 | 0,4341 | 5,06337E-18 |
| ACE2 | RXRB | 0,4341 | 5,08436E-18 |
| ACE2 | TSG101 | 0,4341 | 5,10803E-18 |
| ACE2 | CBR3 | 0,4341 | 5,1044E-18 |
| ACE2 | KLHL21 | 0,434 | 5,16464E-18 |
| ACE2 | TIGD5 | 0,434 | 5,20877E-18 |
| ACE2 | TAGLN | 0,4339 | 5,23085E-18 |
| ACE2 | C2CD4D | 0,4338 | 5,33781E-18 |
| ACE2 | APIP | 0,4338 | 5,39024E-18 |
| ACE2 | COX19 | 0,4337 | 5,47231E-18 |
| ACE2 | RP11-268G12.1 | 0,4337 | 5,45155E-18 |
| ACE2 | RASGEF1A | 0,4336 | 5,62606E-18 |
| ACE2 | FZD10 | 0,4336 | 5,5994E-18 |
| ACE2 | GMPPA | 0,4335 | 5,69085E-18 |
| ACE2 | TRO | 0,4335 | 5,71133E-18 |
| ACE2 | RP11-378J18.8 | 0,4334 | 5,81946E-18 |
| ACE2 | LMO2 | 0,4334 | 5,83949E-18 |
| ACE2 | A2M-AS1 | 0,4334 | 5,84294E-18 |
| ACE2 | PPCDC | 0,4334 | 5,84506E-18 |
| ACE2 | LINC01948 | 0,4332 | 5,98251E-18 |
| ACE2 | DYNC1I1 | 0,4332 | 6,04184E-18 |
| ACE2 | RTF1 | 0,4332 | 5,96948E-18 |
| ACE2 | TFDP3 | 0,4331 | 6,1253E-18 |
| ACE2 | CROT | 0,433 | 6,26343E-18 |
| ACE2 | RP5-1180D12.1 | 0,433 | 6,28426E-18 |
| ACE2 | ZNF630 | 0,433 | 6,26183E-18 |
| ACE2 | MRPS28 | 0,4329 | 6,33047E-18 |
| ACE2 | LPCAT3 | 0,4329 | 6,4331E-18 |
| ACE2 | LINC00863 | 0,4328 | 6,47623E-18 |
| ACE2 | PDCD1LG2 | 0,4327 | 6,70272E-18 |
| ACE2 | LOXL4 | 0,4327 | 6,68361E-18 |
| ACE2 | NCKAP5L | 0,4327 | 6,64419E-18 |
| ACE2 | IGFBP6 | 0,4326 | 6,71969E-18 |
| ACE2 | HELZ2 | 0,4326 | 6,7552E-18 |
| ACE2 | ACTG1P20 | 0,4325 | 6,89584E-18 |
| ACE2 | TIFA | 0,4325 | 6,87719E-18 |
| ACE2 | GIMAP1 | 0,4323 | 7,19843E-18 |

| | | | |
|---|---|---|---|
| ACE2 | TAS2R19 | 0,4322 | 7,38245E-18 |
| ACE2 | PRELID3A | 0,4322 | 7,2836E-18 |
| ACE2 | RP11-49K24.8 | 0,4321 | 7,43436E-18 |
| ACE2 | RP11-490K7.4 | 0,4319 | 7,78469E-18 |
| ACE2 | SLC25A24 | 0,4318 | 7,92899E-18 |
| ACE2 | RP11-262H14.5 | 0,4318 | 7,94258E-18 |
| ACE2 | TBC1D13 | 0,4318 | 7,8309E-18 |
| ACE2 | ZNF710 | 0,4318 | 7,96996E-18 |
| ACE2 | LANCL3 | 0,4318 | 7,83094E-18 |
| ACE2 | ZSWIM1 | 0,4317 | 8,05059E-18 |
| ACE2 | GIMAP7 | 0,4316 | 8,17323E-18 |
| ACE2 | CTD-2017C7.2 | 0,4315 | 8,36161E-18 |
| ACE2 | RBM5-AS1 | 0,4314 | 8,52474E-18 |
| ACE2 | FAIM | 0,4314 | 8,47248E-18 |
| ACE2 | OR7E7P | 0,4314 | 8,56462E-18 |
| ACE2 | GDE1 | 0,4314 | 8,5873E-18 |
| ACE2 | TIMM44 | 0,4314 | 8,50038E-18 |
| ACE2 | LINC01126 | 0,4313 | 8,66813E-18 |
| ACE2 | SHPK | 0,4313 | 8,70432E-18 |
| ACE2 | CTB-55O6.12 | 0,4312 | 8,88331E-18 |
| ACE2 | RSRP1 | 0,4311 | 9,08267E-18 |
| ACE2 | GTF2H4 | 0,4311 | 9,02469E-18 |
| ACE2 | BZW2 | 0,431 | 9,16097E-18 |
| ACE2 | PDZK1IP1 | 0,4309 | 9,34852E-18 |
| ACE2 | TNFSF10 | 0,4309 | 9,44264E-18 |
| ACE2 | RBM23 | 0,4309 | 9,33828E-18 |
| ACE2 | SRSF2 | 0,4309 | 9,31647E-18 |
| ACE2 | TNFAIP8 | 0,4307 | 9,7807E-18 |
| ACE2 | GSPT1 | 0,4307 | 9,70643E-18 |
| ACE2 | SOWAHD | 0,4307 | 9,7476E-18 |
| ACE2 | GGA2 | 0,4306 | 9,9597E-18 |
| ACE2 | SLC1A6 | 0,4306 | 9,85535E-18 |
| ACE2 | LRCH2 | 0,4306 | 9,87632E-18 |
| ACE2 | RP11-277P12.20 | 0,4305 | 1,00791E-17 |
| ACE2 | RP11-258C19.4 | 0,4305 | 1,01126E-17 |
| ACE2 | NEIL3 | 0,4304 | 1,03767E-17 |
| ACE2 | MYLIP | 0,4304 | 1,04068E-17 |
| ACE2 | ZNF720 | 0,4304 | 1,04213E-17 |
| ACE2 | TPRG1L | 0,4303 | 1,05745E-17 |
| ACE2 | CH17-472G23.2 | 0,4302 | 1,06809E-17 |
| ACE2 | FBXO32 | 0,4302 | 1,07982E-17 |
| ACE2 | CNR1 | 0,4301 | 1,09398E-17 |
| ACE2 | RP1-266L20.9 | 0,4301 | 1,08643E-17 |
| ACE2 | KCNJ8 | 0,4301 | 1,10406E-17 |
| ACE2 | BRSK2 | 0,43 | 1,11865E-17 |
| ACE2 | ABCC9 | 0,43 | 1,11626E-17 |
| ACE2 | RNU6-1160P | 0,4299 | 1,12795E-17 |
| ACE2 | AP3S2 | 0,4299 | 1,14459E-17 |
| ACE2 | SERTAD3 | 0,4299 | 1,13455E-17 |
| ACE2 | CSF1 | 0,4298 | 1,15933E-17 |
| ACE2 | RPL12 | 0,4298 | 1,15518E-17 |
| ACE2 | MRPL19 | 0,4297 | 1,18301E-17 |
| ACE2 | MFAP4 | 0,4297 | 1,18061E-17 |
| ACE2 | RNF169 | 0,4296 | 1,19537E-17 |

| | | | |
|---|---|---|---|
| ACE2 | LINC00940 | 0,4294 | 1,24892E-17 |
| ACE2 | HLA-C | 0,4293 | 1,28161E-17 |
| ACE2 | RP3-425C14.6 | 0,4293 | 1,27627E-17 |
| ACE2 | RSF1 | 0,4293 | 1,28228E-17 |
| ACE2 | ZSCAN32 | 0,4293 | 1,2759E-17 |
| ACE2 | RP11-257O5.2 | 0,4292 | 1,29897E-17 |
| ACE2 | CC2D2A | 0,4291 | 1,3275E-17 |
| ACE2 | MTND1P8 | 0,4291 | 1,32118E-17 |
| ACE2 | TCP11L2 | 0,429 | 1,3635E-17 |
| ACE2 | DBX2 | 0,4289 | 1,36761E-17 |
| ACE2 | PMF1 | 0,4288 | 1,41545E-17 |
| ACE2 | GNAS | 0,4288 | 1,41388E-17 |
| ACE2 | SETD2 | 0,4286 | 1,45414E-17 |
| ACE2 | RIC3 | 0,4286 | 1,47288E-17 |
| ACE2 | C1RL-AS1 | 0,4286 | 1,46104E-17 |
| ACE2 | CFLAR | 0,4285 | 1,48727E-17 |
| ACE2 | PDGFRB | 0,4285 | 1,48E-17 |
| ACE2 | CTDNEP1 | 0,4285 | 1,47664E-17 |
| ACE2 | CMB9-55A18.1 | 0,4284 | 1,50888E-17 |
| ACE2 | FLI1 | 0,4284 | 1,5052E-17 |
| ACE2 | TLR8 | 0,4284 | 1,52951E-17 |
| ACE2 | TAS2R6P | 0,4283 | 1,54245E-17 |
| ACE2 | HEATR3 | 0,4283 | 1,55197E-17 |
| ACE2 | PPP4R3CP | 0,4283 | 1,5326E-17 |
| ACE2 | NGFR | 0,4282 | 1,56606E-17 |
| ACE2 | PYGO2 | 0,4281 | 1,59302E-17 |
| ACE2 | TCAF1P1 | 0,4281 | 1,60595E-17 |
| ACE2 | UTP6 | 0,4281 | 1,61922E-17 |
| ACE2 | MSH2 | 0,428 | 1,64937E-17 |
| ACE2 | TRPM6 | 0,428 | 1,64418E-17 |
| ACE2 | NUMA1 | 0,428 | 1,62898E-17 |
| ACE2 | FUT4 | 0,428 | 1,622E-17 |
| ACE2 | RP11-326C3.11 | 0,4279 | 1,65415E-17 |
| ACE2 | RPL18 | 0,4278 | 1,69483E-17 |
| ACE2 | TRABD2B | 0,4277 | 1,73936E-17 |
| ACE2 | CUTA | 0,4277 | 1,73732E-17 |
| ACE2 | KLRG1 | 0,4277 | 1,72504E-17 |
| ACE2 | CBX3 | 0,4276 | 1,77766E-17 |
| ACE2 | WAPL | 0,4276 | 1,76137E-17 |
| ACE2 | SS18L1 | 0,4276 | 1,76604E-17 |
| ACE2 | HERC1 | 0,4275 | 1,79814E-17 |
| ACE2 | VIT | 0,4274 | 1,81859E-17 |
| ACE2 | LGSN | 0,4274 | 1,83491E-17 |
| ACE2 | PLA2G15 | 0,4274 | 1,83287E-17 |
| ACE2 | TMEM74 | 0,4273 | 1,86355E-17 |
| ACE2 | GALK1 | 0,4273 | 1,86813E-17 |
| ACE2 | LINC01340 | 0,4272 | 1,90091E-17 |
| ACE2 | RP11-355O1.11 | 0,4272 | 1,90112E-17 |
| ACE2 | RP11-730A19.9 | 0,4272 | 1,90384E-17 |
| ACE2 | PSRC1 | 0,4271 | 1,94763E-17 |
| ACE2 | AKIP1 | 0,4271 | 1,95425E-17 |
| ACE2 | SPRY1 | 0,427 | 1,9755E-17 |
| ACE2 | KCNMB4 | 0,427 | 1,98156E-17 |
| ACE2 | C14orf93 | 0,427 | 1,97434E-17 |

| | | | |
|---|---|---|---|
| ACE2 | PHLDB2 | 0,4269 | 2,02439E-17 |
| ACE2 | UPK3BP1 | 0,4269 | 2,03482E-17 |
| ACE2 | PANK4 | 0,4268 | 2,04564E-17 |
| ACE2 | RC3H1 | 0,4268 | 2,05557E-17 |
| ACE2 | CTA-253N17.1 | 0,4268 | 2,03863E-17 |
| ACE2 | RP11-736K20.4 | 0,4268 | 2,04309E-17 |
| ACE2 | MFN2 | 0,4267 | 2,09488E-17 |
| ACE2 | TUBGCP2 | 0,4267 | 2,08564E-17 |
| ACE2 | CTC-490E21.11 | 0,4267 | 2,10611E-17 |
| ACE2 | TOMM34 | 0,4267 | 2,0953E-17 |
| ACE2 | RBM20 | 0,4264 | 2,20449E-17 |
| ACE2 | CDKN2C | 0,4263 | 2,24265E-17 |
| ACE2 | BOLA3 | 0,4263 | 2,26319E-17 |
| ACE2 | KCNAB1 | 0,4263 | 2,24985E-17 |
| ACE2 | IMPA1P1 | 0,4263 | 2,26793E-17 |
| ACE2 | IYD | 0,4262 | 2,29809E-17 |
| ACE2 | ADGRG4 | 0,4262 | 2,30505E-17 |
| ACE2 | RP11-307E17.8 | 0,4261 | 2,33284E-17 |
| ACE2 | SNX33 | 0,4261 | 2,36237E-17 |
| ACE2 | PTPRZ1 | 0,426 | 2,37989E-17 |
| ACE2 | FAM224B | 0,426 | 2,38405E-17 |
| ACE2 | PLTP | 0,4259 | 2,43852E-17 |
| ACE2 | ATG4A | 0,4259 | 2,44138E-17 |
| ACE2 | KLHL29 | 0,4258 | 2,45989E-17 |
| ACE2 | LINC01336 | 0,4258 | 2,48825E-17 |
| ACE2 | ITLN2 | 0,4257 | 2,51502E-17 |
| ACE2 | CLASP1 | 0,4257 | 2,5505E-17 |
| ACE2 | SLC4A3 | 0,4257 | 2,53574E-17 |
| ACE2 | VN1R1 | 0,4257 | 2,5312E-17 |
| ACE2 | RBBP8P1 | 0,4257 | 2,55186E-17 |
| ACE2 | NT5DC2 | 0,4256 | 2,55742E-17 |
| ACE2 | LMF1 | 0,4256 | 2,59192E-17 |
| ACE2 | CENPB | 0,4256 | 2,5787E-17 |
| ACE2 | SPIN2A | 0,4256 | 2,56724E-17 |
| ACE2 | RP11-373E16.3 | 0,4255 | 2,64415E-17 |
| ACE2 | ATP8A1 | 0,4255 | 2,60621E-17 |
| ACE2 | STAT2 | 0,4255 | 2,62953E-17 |
| ACE2 | EDN3 | 0,4255 | 2,6345E-17 |
| ACE2 | CPB2 | 0,4254 | 2,67607E-17 |
| ACE2 | TAF15 | 0,4254 | 2,67428E-17 |
| ACE2 | CTB-50E14.5 | 0,4254 | 2,67702E-17 |
| ACE2 | RP11-250B2.4 | 0,4253 | 2,72642E-17 |
| ACE2 | LINC01285 | 0,4253 | 2,75357E-17 |
| ACE2 | FGD5 | 0,4252 | 2,79689E-17 |
| ACE2 | SCML2P2 | 0,4252 | 2,77931E-17 |
| ACE2 | DHX8 | 0,4252 | 2,79528E-17 |
| ACE2 | GHc-857G6.7 | 0,4252 | 2,77526E-17 |
| ACE2 | SLC25A5-AS1 | 0,4252 | 2,75529E-17 |
| ACE2 | DNAJB11 | 0,4251 | 2,85337E-17 |
| ACE2 | HNRNPH1 | 0,4251 | 2,81358E-17 |
| ACE2 | UBAP2L | 0,425 | 2,91302E-17 |
| ACE2 | WFDC2 | 0,425 | 2,86124E-17 |
| ACE2 | RP11-278C7.3 | 0,4249 | 2,92842E-17 |
| ACE2 | ADAL | 0,4249 | 2,94053E-17 |

| | | | |
|---|---|---|---|
| ACE2 | CITF22-92A6.1 | 0,4249 | 2,91699E-17 |
| ACE2 | RP11-158K1.3 | 0,4247 | 3,03292E-17 |
| ACE2 | MAGEB10 | 0,4247 | 3,03631E-17 |
| ACE2 | TOR4A | 0,4246 | 3,09992E-17 |
| ACE2 | FHL3 | 0,4245 | 3,1923E-17 |
| ACE2 | CAB39 | 0,4245 | 3,18821E-17 |
| ACE2 | GUCY2C | 0,4245 | 3,1492E-17 |
| ACE2 | LMX1B | 0,4244 | 3,23046E-17 |
| ACE2 | PTGES3 | 0,4244 | 3,206E-17 |
| ACE2 | PRR11 | 0,4244 | 3,22442E-17 |
| ACE2 | CPAMD8 | 0,4244 | 3,2443E-17 |
| ACE2 | FBLN1 | 0,4244 | 3,25025E-17 |
| ACE2 | RP11-677M14.8 | 0,4243 | 3,30861E-17 |
| ACE2 | PARP11 | 0,4243 | 3,27311E-17 |
| ACE2 | AADAT | 0,4242 | 3,36101E-17 |
| ACE2 | GJA4 | 0,4241 | 3,44511E-17 |
| ACE2 | TMEM63A | 0,4241 | 3,40772E-17 |
| ACE2 | C18orf8 | 0,4241 | 3,45207E-17 |
| ACE2 | TTI1 | 0,4241 | 3,43723E-17 |
| ACE2 | MCHR1 | 0,4241 | 3,43314E-17 |
| ACE2 | CXCR3 | 0,4241 | 3,44972E-17 |
| ACE2 | TBC1D5 | 0,424 | 3,4768E-17 |
| ACE2 | RP11-589M4.4 | 0,424 | 3,50131E-17 |
| ACE2 | DDX23 | 0,4239 | 3,55761E-17 |
| ACE2 | CBLN3 | 0,4238 | 3,59957E-17 |
| ACE2 | RP11-70D24.3 | 0,4238 | 3,62581E-17 |
| ACE2 | GIMAP6 | 0,4237 | 3,67895E-17 |
| ACE2 | RP11-398C13.8 | 0,4237 | 3,67044E-17 |
| ACE2 | RPL18AP3 | 0,4237 | 3,65764E-17 |
| ACE2 | CYP4B1 | 0,4236 | 3,78466E-17 |
| ACE2 | CAPN13 | 0,4236 | 3,72524E-17 |
| ACE2 | SLC8A1 | 0,4236 | 3,75868E-17 |
| ACE2 | CLSTN3 | 0,4236 | 3,72753E-17 |
| ACE2 | CAMKK2 | 0,4236 | 3,77273E-17 |
| ACE2 | DCLK1 | 0,4236 | 3,78583E-17 |
| ACE2 | FAM107A | 0,4235 | 3,85893E-17 |
| ACE2 | BMF | 0,4235 | 3,83265E-17 |
| ACE2 | PRKD2 | 0,4234 | 3,92298E-17 |
| ACE2 | KLF3-AS1 | 0,4232 | 4,06891E-17 |
| ACE2 | RP11-343J18.1 | 0,4231 | 4,11594E-17 |
| ACE2 | ZBTB43 | 0,4231 | 4,16264E-17 |
| ACE2 | RPS6KA4 | 0,4231 | 4,0961E-17 |
| ACE2 | HDAC7 | 0,4231 | 4,12376E-17 |
| ACE2 | AK4P1 | 0,4231 | 4,09729E-17 |
| ACE2 | COG1 | 0,423 | 4,17682E-17 |
| ACE2 | AC005519.4 | 0,4228 | 4,36505E-17 |
| ACE2 | RP11-92A5.2 | 0,4227 | 4,45669E-17 |
| ACE2 | HEIH | 0,4227 | 4,43307E-17 |
| ACE2 | RAB3GAP1 | 0,4226 | 4,51489E-17 |
| ACE2 | HELLS | 0,4226 | 4,52631E-17 |
| ACE2 | RAVER2 | 0,4225 | 4,57859E-17 |
| ACE2 | DNM3OS | 0,4225 | 4,6232E-17 |
| ACE2 | TMEM106C | 0,4225 | 4,62008E-17 |
| ACE2 | PREX1 | 0,4225 | 4,639E-17 |

| | | | |
|---|---|---|---|
| ACE2 | TTPA | 0,4224 | 4,66807E-17 |
| ACE2 | PTCHD1 | 0,4224 | 4,7303E-17 |
| ACE2 | MUL1 | 0,4223 | 4,81694E-17 |
| ACE2 | UHRF2 | 0,4223 | 4,81024E-17 |
| ACE2 | PAPSS2 | 0,4223 | 4,74958E-17 |
| ACE2 | ZFP36L1 | 0,4223 | 4,80125E-17 |
| ACE2 | MAGOH2P | 0,4223 | 4,76301E-17 |
| ACE2 | RASGRP4 | 0,4223 | 4,77883E-17 |
| ACE2 | CICP16 | 0,4222 | 4,88336E-17 |
| ACE2 | CELF2 | 0,4222 | 4,92458E-17 |
| ACE2 | HHEX | 0,4222 | 4,84018E-17 |
| ACE2 | LTBP3 | 0,4222 | 4,88914E-17 |
| ACE2 | CTC-591M7.1 | 0,4222 | 4,91325E-17 |
| ACE2 | SLC2A4RG | 0,4222 | 4,84681E-17 |
| ACE2 | RAB38 | 0,4221 | 4,95502E-17 |
| ACE2 | NBL1 | 0,4218 | 5,24721E-17 |
| ACE2 | MMP19 | 0,4218 | 5,24825E-17 |
| ACE2 | ZDHHC18 | 0,4217 | 5,3924E-17 |
| ACE2 | RWDD4P2 | 0,4217 | 5,38715E-17 |
| ACE2 | RP11-12G12.7 | 0,4217 | 5,38033E-17 |
| ACE2 | DCAF8L1 | 0,4217 | 5,38826E-17 |
| ACE2 | RP11-767N6.2 | 0,4216 | 5,432E-17 |
| ACE2 | SLC25A15 | 0,4216 | 5,44722E-17 |
| ACE2 | EIF4A3 | 0,4216 | 5,48394E-17 |
| ACE2 | ELF4 | 0,4216 | 5,42887E-17 |
| ACE2 | TBC1D23 | 0,4215 | 5,60384E-17 |
| ACE2 | GRAMD4 | 0,4215 | 5,51869E-17 |
| ACE2 | GFPT2 | 0,4213 | 5,77711E-17 |
| ACE2 | RFWD3 | 0,4213 | 5,73721E-17 |
| ACE2 | MT-TS1 | 0,4213 | 5,81013E-17 |
| ACE2 | SART1 | 0,4212 | 5,87644E-17 |
| ACE2 | ELOVL1 | 0,4209 | 6,17345E-17 |
| ACE2 | LETM1 | 0,4209 | 6,23793E-17 |
| ACE2 | ANXA1 | 0,4209 | 6,20522E-17 |
| ACE2 | ALPL | 0,4207 | 6,43079E-17 |
| ACE2 | AKR7A2P1 | 0,4207 | 6,4268E-17 |
| ACE2 | RPL30 | 0,4207 | 6,45463E-17 |
| ACE2 | RP11-248J18.2 | 0,4207 | 6,48851E-17 |
| ACE2 | LLNLR-245B6.1 | 0,4207 | 6,41574E-17 |
| ACE2 | LINC01659 | 0,4207 | 6,43192E-17 |
| ACE2 | CD4 | 0,4206 | 6,61177E-17 |
| ACE2 | SOCS2 | 0,4206 | 6,59175E-17 |
| ACE2 | EMSY | 0,4205 | 6,64019E-17 |
| ACE2 | AVPR1A | 0,4205 | 6,75653E-17 |
| ACE2 | BRS3 | 0,4204 | 6,81415E-17 |
| ACE2 | GABRB3 | 0,4203 | 6,90526E-17 |
| ACE2 | CTD-2536I1.2 | 0,4203 | 6,97604E-17 |
| ACE2 | ID1 | 0,4203 | 6,98747E-17 |
| ACE2 | STARD8 | 0,4203 | 6,91147E-17 |
| ACE2 | MKNK1 | 0,4202 | 7,03929E-17 |
| ACE2 | CAD | 0,4202 | 7,0365E-17 |
| ACE2 | GANC | 0,4202 | 7,11336E-17 |
| ACE2 | RAB3D | 0,4202 | 7,02216E-17 |
| ACE2 | HARS | 0,4201 | 7,15536E-17 |

| | | | |
|---|---|---|---|
| ACE2 | GIPC2 | 0,42 | 7,41805E-17 |
| ACE2 | RP11-124N14.3 | 0,4199 | 7,46535E-17 |
| ACE2 | RP11-524F11.1 | 0,4199 | 7,51568E-17 |
| ACE2 | CBX7 | 0,4198 | 7,67576E-17 |
| ACE2 | RAB21 | 0,4197 | 7,74836E-17 |
| ACE2 | MED16 | 0,4197 | 7,72836E-17 |
| ACE2 | CCDC117 | 0,4197 | 7,84025E-17 |
| ACE2 | AC139103.1 | 0,4196 | 7,89391E-17 |
| ACE2 | FAM189A2 | 0,4196 | 7,8567E-17 |
| ACE2 | RP11-752L20.3 | 0,4195 | 8,01504E-17 |
| ACE2 | CTD-2010I16.1 | 0,4195 | 7,99987E-17 |
| ACE2 | PDPK1 | 0,4194 | 8,14272E-17 |
| ACE2 | ZNF519 | 0,4194 | 8,25487E-17 |
| ACE2 | RAB23 | 0,4193 | 8,37735E-17 |
| ACE2 | PDCL3P5 | 0,4193 | 8,36385E-17 |
| ACE2 | CHEK2 | 0,4193 | 8,43954E-17 |
| ACE2 | TEAD4 | 0,4192 | 8,58629E-17 |
| ACE2 | ZNF480 | 0,4192 | 8,60406E-17 |
| ACE2 | ZNF528 | 0,4192 | 8,5093E-17 |
| ACE2 | CROCCP3 | 0,4191 | 8,61455E-17 |
| ACE2 | KB-431C1.5 | 0,4191 | 8,63658E-17 |
| ACE2 | RP13-216E22.4 | 0,4191 | 8,71345E-17 |
| ACE2 | BTN2A3P | 0,419 | 8,85098E-17 |
| ACE2 | PCDHGA5 | 0,4188 | 9,11889E-17 |
| ACE2 | HDAC3 | 0,4188 | 9,21977E-17 |
| ACE2 | MOB3A | 0,4187 | 9,39532E-17 |
| ACE2 | RILPL1 | 0,4186 | 9,49899E-17 |
| ACE2 | RP11-395A13.2 | 0,4185 | 9,69568E-17 |
| ACE2 | PRR3 | 0,4185 | 9,71754E-17 |
| ACE2 | PNN | 0,4185 | 9,66243E-17 |
| ACE2 | RP11-90B9.2 | 0,4185 | 9,66384E-17 |
| ACE2 | IL18R1 | 0,4184 | 9,94766E-17 |
| ACE2 | HCG16 | 0,4184 | 9,84202E-17 |
| ACE2 | MGP | 0,4184 | 9,94349E-17 |
| ACE2 | LINC00506 | 0,4183 | 1,01035E-16 |
| ACE2 | CPN2 | 0,4183 | 1,01225E-16 |
| ACE2 | IRF6 | 0,4182 | 1,02456E-16 |
| ACE2 | MAP3K8 | 0,4182 | 1,01771E-16 |
| ACE2 | TENM1 | 0,4182 | 1,01653E-16 |
| ACE2 | KATNBL1 | 0,418 | 1,06239E-16 |
| ACE2 | NOMO1 | 0,418 | 1,05436E-16 |
| ACE2 | DLGAP1-AS2 | 0,4179 | 1,08619E-16 |
| ACE2 | SARDH | 0,4178 | 1,10489E-16 |
| ACE2 | HES1 | 0,4177 | 1,11581E-16 |
| ACE2 | GABRA6 | 0,4177 | 1,11766E-16 |
| ACE2 | ZNF462 | 0,4177 | 1,11614E-16 |
| ACE2 | MT1X | 0,4177 | 1,12418E-16 |
| ACE2 | PLK3 | 0,4176 | 1,15364E-16 |
| ACE2 | CREB3L1 | 0,4176 | 1,1547E-16 |
| ACE2 | U52111.14 | 0,4176 | 1,1396E-16 |
| ACE2 | RARB | 0,4175 | 1,16367E-16 |
| ACE2 | RXFP1 | 0,4175 | 1,17711E-16 |
| ACE2 | MATN2 | 0,4175 | 1,16328E-16 |
| ACE2 | CLEC2B | 0,4175 | 1,17159E-16 |

| | | | |
|---|---|---|---|
| ACE2 | ING4 | 0,4174 | 1,18536E-16 |
| ACE2 | PTPRO | 0,4173 | 1,21718E-16 |
| ACE2 | ADAM1B | 0,4173 | 1,21674E-16 |
| ACE2 | RILP | 0,4173 | 1,20312E-16 |
| ACE2 | HHIP-AS1 | 0,4172 | 1,22708E-16 |
| ACE2 | CAAP1 | 0,4172 | 1,22355E-16 |
| ACE2 | PDE8A | 0,4172 | 1,24232E-16 |
| ACE2 | DLGAP1 | 0,4172 | 1,23745E-16 |
| ACE2 | MIB1 | 0,4172 | 1,23891E-16 |
| ACE2 | TLR6 | 0,4171 | 1,24862E-16 |
| ACE2 | PMS2 | 0,4171 | 1,25931E-16 |
| ACE2 | SAP30BP | 0,4171 | 1,25617E-16 |
| ACE2 | C1orf27 | 0,417 | 1,27177E-16 |
| ACE2 | XX-FW83563B9.5 | 0,417 | 1,27585E-16 |
| ACE2 | MCM2 | 0,4169 | 1,31395E-16 |
| ACE2 | ABTB1 | 0,4169 | 1,29512E-16 |
| ACE2 | FRG1DP | 0,4169 | 1,30253E-16 |
| ACE2 | FBXL5 | 0,4167 | 1,35527E-16 |
| ACE2 | IRF4 | 0,4166 | 1,37545E-16 |
| ACE2 | RP11-351J23.1 | 0,4166 | 1,36881E-16 |
| ACE2 | GATS | 0,4166 | 1,36826E-16 |
| ACE2 | POU3F1 | 0,4165 | 1,39863E-16 |
| ACE2 | RP5-849H19.3 | 0,4165 | 1,4107E-16 |
| ACE2 | RP11-576D8.4 | 0,4165 | 1,40815E-16 |
| ACE2 | ACVR1C | 0,4165 | 1,39849E-16 |
| ACE2 | TOMM6 | 0,4165 | 1,41259E-16 |
| ACE2 | OR7C1 | 0,4165 | 1,39333E-16 |
| ACE2 | AP000473.5 | 0,4165 | 1,41097E-16 |
| ACE2 | ZFP92 | 0,4165 | 1,39805E-16 |
| ACE2 | MASP2 | 0,4164 | 1,41806E-16 |
| ACE2 | RP1-296G17.3 | 0,4164 | 1,42222E-16 |
| ACE2 | RP11-390E23.6 | 0,4163 | 1,46237E-16 |
| ACE2 | RP11-181K3.4 | 0,4162 | 1,4832E-16 |
| ACE2 | CTD-2283N19.1 | 0,4161 | 1,50041E-16 |
| ACE2 | WRNIP1 | 0,4161 | 1,51644E-16 |
| ACE2 | ARF5 | 0,4161 | 1,51423E-16 |
| ACE2 | ZNF417 | 0,4161 | 1,51929E-16 |
| ACE2 | CGN | 0,416 | 1,53657E-16 |
| ACE2 | TLE2 | 0,416 | 1,54653E-16 |
| ACE2 | INHBA | 0,4159 | 1,57402E-16 |
| ACE2 | NEDD8 | 0,4159 | 1,57344E-16 |
| ACE2 | PTPN7 | 0,4158 | 1,58567E-16 |
| ACE2 | CEBPD | 0,4158 | 1,60461E-16 |
| ACE2 | PECAM1 | 0,4158 | 1,59127E-16 |
| ACE2 | HNRNPA1P49 | 0,4156 | 1,66401E-16 |
| ACE2 | TNFRSF10D | 0,4156 | 1,64692E-16 |
| ACE2 | RAB43 | 0,4155 | 1,69561E-16 |
| ACE2 | TEK | 0,4155 | 1,68322E-16 |
| ACE2 | KRTAP5-AS1 | 0,4155 | 1,67414E-16 |
| ACE2 | SPRY4 | 0,4154 | 1,70149E-16 |
| ACE2 | SCRN2 | 0,4154 | 1,70996E-16 |
| ACE2 | ANO10 | 0,4153 | 1,76055E-16 |
| ACE2 | ZNF516 | 0,4153 | 1,73932E-16 |
| ACE2 | DRP2 | 0,4153 | 1,75724E-16 |

| | | | |
|---|---|---|---|
| ACE2 | SNAP25 | 0,4152 | 1,785E-16 |
| ACE2 | SNORA2 | 0,4151 | 1,81561E-16 |
| ACE2 | ZNF287 | 0,4151 | 1,82567E-16 |
| ACE2 | TPTE2P5 | 0,415 | 1,85874E-16 |
| ACE2 | ZNF468 | 0,4148 | 1,90507E-16 |
| ACE2 | TBC1D8 | 0,4147 | 1,93563E-16 |
| ACE2 | THYN1 | 0,4147 | 1,94049E-16 |
| ACE2 | RP11-527N22.1 | 0,4145 | 2,0129E-16 |
| ACE2 | CALCOCO1 | 0,4145 | 2,02671E-16 |
| ACE2 | LINC00924 | 0,4144 | 2,04529E-16 |
| ACE2 | WNK2 | 0,4143 | 2,08279E-16 |
| ACE2 | MYCBP2 | 0,4143 | 2,09935E-16 |
| ACE2 | SOX1 | 0,4143 | 2,09435E-16 |
| ACE2 | ZNF260 | 0,4143 | 2,09935E-16 |
| ACE2 | VN1R81P | 0,4142 | 2,14596E-16 |
| ACE2 | NR2C1 | 0,4141 | 2,15625E-16 |
| ACE2 | FANCB | 0,4141 | 2,17469E-16 |
| ACE2 | SYT2 | 0,414 | 2,23135E-16 |
| ACE2 | RP13-487P22.1 | 0,414 | 2,21787E-16 |
| ACE2 | AC004019.13 | 0,414 | 2,19279E-16 |
| ACE2 | AC005336.4 | 0,4139 | 2,24814E-16 |
| ACE2 | COG2 | 0,4138 | 2,27801E-16 |
| ACE2 | DTX2P1 | 0,4138 | 2,283E-16 |
| ACE2 | LZTS1 | 0,4138 | 2,27328E-16 |
| ACE2 | XXbac-B562F10.11 | 0,4138 | 2,27854E-16 |
| ACE2 | UCHL1 | 0,4137 | 2,32724E-16 |
| ACE2 | PGAM1P2 | 0,4136 | 2,35993E-16 |
| ACE2 | GGACT | 0,4136 | 2,36031E-16 |
| ACE2 | RGS2 | 0,4135 | 2,40977E-16 |
| ACE2 | CYP1B1 | 0,4135 | 2,41857E-16 |
| ACE2 | MRPS5 | 0,4135 | 2,40116E-16 |
| ACE2 | CLPTM1L | 0,4135 | 2,43619E-16 |
| ACE2 | RASAL1 | 0,4135 | 2,42889E-16 |
| ACE2 | AGRP | 0,4135 | 2,41052E-16 |
| ACE2 | CYTIP | 0,4134 | 2,45813E-16 |
| ACE2 | AC010975.1 | 0,4133 | 2,51098E-16 |
| ACE2 | GBGT1 | 0,4133 | 2,50709E-16 |
| ACE2 | PRPF8 | 0,4132 | 2,54523E-16 |
| ACE2 | NONOP2 | 0,4131 | 2,58863E-16 |
| ACE2 | DKFZP434I0714 | 0,4131 | 2,62209E-16 |
| ACE2 | RP11-273G15.2 | 0,4131 | 2,62094E-16 |
| ACE2 | AP1AR | 0,413 | 2,65261E-16 |
| ACE2 | RNASET2 | 0,413 | 2,64569E-16 |
| ACE2 | RPUSD2 | 0,413 | 2,6447E-16 |
| ACE2 | INTS2 | 0,413 | 2,66235E-16 |
| ACE2 | FOXF2 | 0,4129 | 2,68354E-16 |
| ACE2 | PPDPF | 0,4129 | 2,67672E-16 |
| ACE2 | RP11-1348G14.4 | 0,4128 | 2,74081E-16 |
| ACE2 | CCDC106 | 0,4128 | 2,74028E-16 |
| ACE2 | CCM2 | 0,4127 | 2,82506E-16 |
| ACE2 | AC024084.1 | 0,4126 | 2,86165E-16 |
| ACE2 | ASPHD1 | 0,4126 | 2,8491E-16 |
| ACE2 | ZNF208 | 0,4126 | 2,87642E-16 |
| ACE2 | TIMP4 | 0,4125 | 2,91859E-16 |

| | | | |
|---|---|---|---|
| ACE2 | ERN1 | 0,4125 | 2,88316E-16 |
| ACE2 | AL133243.2 | 0,4124 | 2,96776E-16 |
| ACE2 | FBL | 0,4124 | 2,98171E-16 |
| ACE2 | PARP14 | 0,4123 | 2,99551E-16 |
| ACE2 | F10 | 0,4123 | 3,01476E-16 |
| ACE2 | B3GAT2 | 0,4122 | 3,06801E-16 |
| ACE2 | RPUSD4 | 0,4122 | 3,06704E-16 |
| ACE2 | C14orf119 | 0,4122 | 3,05193E-16 |
| ACE2 | RPL7AP2 | 0,4122 | 3,04879E-16 |
| ACE2 | SYTL5 | 0,4122 | 3,05199E-16 |
| ACE2 | PHYHD1 | 0,4121 | 3,12421E-16 |
| ACE2 | NPDC1 | 0,4121 | 3,13041E-16 |
| ACE2 | NPNT | 0,412 | 3,1577E-16 |
| ACE2 | P2RX7 | 0,412 | 3,19814E-16 |
| ACE2 | AC093495.4 | 0,4119 | 3,24025E-16 |
| ACE2 | PRKDC | 0,4119 | 3,25349E-16 |
| ACE2 | NUDT22 | 0,4118 | 3,31672E-16 |
| ACE2 | RAP1GAP2 | 0,4118 | 3,32332E-16 |
| ACE2 | CTGF | 0,4117 | 3,36259E-16 |
| ACE2 | RP5-855D21.2 | 0,4117 | 3,33548E-16 |
| ACE2 | LINC01232 | 0,4117 | 3,33608E-16 |
| ACE2 | SPNS1 | 0,4117 | 3,32526E-16 |
| ACE2 | RP11-45M22.3 | 0,4116 | 3,39654E-16 |
| ACE2 | TCF23 | 0,4114 | 3,55231E-16 |
| ACE2 | ACBD4 | 0,4114 | 3,55378E-16 |
| ACE2 | CCNL1 | 0,4113 | 3,59681E-16 |
| ACE2 | FBXW7 | 0,4113 | 3,59943E-16 |
| ACE2 | TBX18 | 0,4113 | 3,5973E-16 |
| ACE2 | CUL4A | 0,4113 | 3,57483E-16 |
| ACE2 | VAT1L | 0,4113 | 3,58011E-16 |
| ACE2 | RIC1 | 0,4112 | 3,65671E-16 |
| ACE2 | AQR | 0,4112 | 3,68681E-16 |
| ACE2 | FLNC | 0,4111 | 3,75542E-16 |
| ACE2 | C5 | 0,4111 | 3,747E-16 |
| ACE2 | EPHA2 | 0,411 | 3,83124E-16 |
| ACE2 | GBP1 | 0,411 | 3,82168E-16 |
| ACE2 | CTD-2538G9.5 | 0,411 | 3,83398E-16 |
| ACE2 | S1PR1 | 0,4109 | 3,84958E-16 |
| ACE2 | FAM43A | 0,4109 | 3,84975E-16 |
| ACE2 | VEGFB | 0,4109 | 3,84182E-16 |
| ACE2 | ERC1 | 0,4109 | 3,86353E-16 |
| ACE2 | ST13 | 0,4109 | 3,83976E-16 |
| ACE2 | GDA | 0,4107 | 4,01797E-16 |
| ACE2 | KANTR | 0,4107 | 4,0189E-16 |
| ACE2 | NIT2 | 0,4106 | 4,0959E-16 |
| ACE2 | SDC1 | 0,4105 | 4,1565E-16 |
| ACE2 | VAMP5 | 0,4105 | 4,14869E-16 |
| ACE2 | ITPR1-AS1 | 0,4105 | 4,14443E-16 |
| ACE2 | NEIL2 | 0,4105 | 4,14038E-16 |
| ACE2 | ULK1 | 0,4105 | 4,16128E-16 |
| ACE2 | RRP15 | 0,4104 | 4,21342E-16 |
| ACE2 | HLA-DMB | 0,4104 | 4,21588E-16 |
| ACE2 | RP11-676M6.1 | 0,4104 | 4,22022E-16 |
| ACE2 | RP11-120M18.2 | 0,4104 | 4,21446E-16 |

| | | | |
|---|---|---|---|
| ACE2 | FLG | 0,4103 | 4,28454E-16 |
| ACE2 | ZNF621 | 0,4103 | 4,28327E-16 |
| ACE2 | AC116366.6 | 0,4103 | 4,27954E-16 |
| ACE2 | RP11-609N14.4 | 0,4103 | 4,35327E-16 |
| ACE2 | SURF6 | 0,4102 | 4,38289E-16 |
| ACE2 | DPP9 | 0,4102 | 4,40047E-16 |
| ACE2 | RP11-93B14.10 | 0,4102 | 4,40765E-16 |
| ACE2 | UXT-AS1 | 0,4102 | 4,38855E-16 |
| ACE2 | TLE1P1 | 0,4101 | 4,48357E-16 |
| ACE2 | SMYD5 | 0,41 | 4,5781E-16 |
| ACE2 | C4orf50 | 0,41 | 4,58217E-16 |
| ACE2 | RP11-166O4.6 | 0,41 | 4,52794E-16 |
| ACE2 | XPOT | 0,41 | 4,56768E-16 |
| ACE2 | HNRNPA1P59 | 0,4099 | 4,62113E-16 |
| ACE2 | CAMK1G | 0,4099 | 4,67303E-16 |
| ACE2 | BOC | 0,4099 | 4,64086E-16 |
| ACE2 | ELN | 0,4099 | 4,66744E-16 |
| ACE2 | SMO | 0,4099 | 4,60826E-16 |
| ACE2 | NOMO3 | 0,4099 | 4,67308E-16 |
| ACE2 | PKDCC | 0,4098 | 4,71733E-16 |
| ACE2 | ETAA1 | 0,4098 | 4,70172E-16 |
| ACE2 | RP11-632K20.7 | 0,4098 | 4,75798E-16 |
| ACE2 | PRMT7 | 0,4098 | 4,74877E-16 |
| ACE2 | KIAA0319L | 0,4097 | 4,80606E-16 |
| ACE2 | RP11-326C3.16 | 0,4097 | 4,78761E-16 |
| ACE2 | RP11-46H11.11 | 0,4097 | 4,83629E-16 |
| ACE2 | NOL3 | 0,4097 | 4,82772E-16 |
| ACE2 | PCNX1 | 0,4096 | 4,8936E-16 |
| ACE2 | CEP192 | 0,4096 | 4,88387E-16 |
| ACE2 | RP11-284F21.9 | 0,4095 | 4,97027E-16 |
| ACE2 | RBM17 | 0,4095 | 4,95775E-16 |
| ACE2 | SOX3 | 0,4095 | 4,98961E-16 |
| ACE2 | SNORA55 | 0,4094 | 5,09816E-16 |
| ACE2 | RPAP2 | 0,4094 | 5,08096E-16 |
| ACE2 | SLC23A1 | 0,4093 | 5,18109E-16 |
| ACE2 | LCMT2 | 0,4093 | 5,17226E-16 |
| ACE2 | ITGB4 | 0,4093 | 5,15364E-16 |
| ACE2 | RTCB | 0,4093 | 5,15962E-16 |
| ACE2 | MCM10 | 0,4092 | 5,27551E-16 |
| ACE2 | C11orf57 | 0,4091 | 5,37632E-16 |
| ACE2 | TMEM186 | 0,4091 | 5,34668E-16 |
| ACE2 | ESX1 | 0,4091 | 5,3601E-16 |
| ACE2 | CDON | 0,409 | 5,4712E-16 |
| ACE2 | RPA1 | 0,409 | 5,40105E-16 |
| ACE2 | FGFR1 | 0,4089 | 5,58345E-16 |
| ACE2 | HNRNPA1L2 | 0,4089 | 5,57357E-16 |
| ACE2 | HSBP1L1 | 0,4089 | 5,56896E-16 |
| ACE2 | CYBB | 0,4089 | 5,58951E-16 |
| ACE2 | PEX14 | 0,4088 | 5,64554E-16 |
| ACE2 | IFITM1 | 0,4088 | 5,67293E-16 |
| ACE2 | RP4-657D16.3 | 0,4087 | 5,75549E-16 |
| ACE2 | NRIP2 | 0,4087 | 5,79195E-16 |
| ACE2 | SASH3 | 0,4086 | 5,79876E-16 |
| ACE2 | CNTNAP2 | 0,4085 | 5,99455E-16 |

| | | | |
|---|---|---|---|
| ACE2 | IRF8 | 0,4085 | 5,90189E-16 |
| ACE2 | ACSF3 | 0,4084 | 6,03135E-16 |
| ACE2 | TBC1D20 | 0,4084 | 6,10091E-16 |
| ACE2 | RP11-82K18.2 | 0,4083 | 6,12985E-16 |
| ACE2 | AP000560.3 | 0,4083 | 6,14829E-16 |
| ACE2 | HMGB1P14 | 0,4082 | 6,2687E-16 |
| ACE2 | PYCARD | 0,4082 | 6,27517E-16 |
| ACE2 | ZNF813 | 0,4082 | 6,27284E-16 |
| ACE2 | ZNF587 | 0,4082 | 6,27876E-16 |
| ACE2 | HS3ST1 | 0,408 | 6,55961E-16 |
| ACE2 | DCAF4L1 | 0,4079 | 6,60327E-16 |
| ACE2 | EPC1 | 0,4079 | 6,59275E-16 |
| ACE2 | RP13-192B19.2 | 0,4079 | 6,61824E-16 |
| ACE2 | TNRC18P1 | 0,4078 | 6,69417E-16 |
| ACE2 | RP11-473O4.5 | 0,4078 | 6,7684E-16 |
| ACE2 | HNRNPA1P14 | 0,4077 | 6,81519E-16 |
| ACE2 | CDKL5 | 0,4077 | 6,81314E-16 |
| ACE2 | FRMD6-AS1 | 0,4075 | 7,06549E-16 |
| ACE2 | EIF1AX | 0,4075 | 7,17637E-16 |
| ACE2 | CTC-303L1.2 | 0,4074 | 7,18091E-16 |
| ACE2 | RPL13AP5 | 0,4074 | 7,21485E-16 |
| ACE2 | PDLIM1 | 0,4073 | 7,43297E-16 |
| ACE2 | PLCB4 | 0,4073 | 7,36143E-16 |
| ACE2 | CHRNB1 | 0,4072 | 7,55225E-16 |
| ACE2 | SMG1P3 | 0,4069 | 7,94019E-16 |
| ACE2 | KANSL1 | 0,4069 | 7,86482E-16 |
| ACE2 | CHERP | 0,4069 | 7,9091E-16 |
| ACE2 | SAMD4B | 0,4069 | 7,96835E-16 |
| ACE2 | AF127936.7 | 0,4069 | 7,90305E-16 |
| ACE2 | RP11-345P4.6 | 0,4068 | 8,1262E-16 |
| ACE2 | FLRT2 | 0,4068 | 8,03328E-16 |
| ACE2 | DDX50P1 | 0,4067 | 8,15899E-16 |
| ACE2 | PPIL1 | 0,4067 | 8,13431E-16 |
| ACE2 | TTC9 | 0,4067 | 8,25245E-16 |
| ACE2 | RP11-767N6.7 | 0,4065 | 8,5527E-16 |
| ACE2 | FAXC | 0,4064 | 8,70055E-16 |
| ACE2 | PTGS1 | 0,4064 | 8,65855E-16 |
| ACE2 | LINC01415 | 0,4064 | 8,6518E-16 |
| ACE2 | DGUOK-AS1 | 0,4063 | 8,76423E-16 |
| ACE2 | SOS2 | 0,4063 | 8,73439E-16 |
| ACE2 | GLIS2 | 0,4063 | 8,86123E-16 |
| ACE2 | INSM1 | 0,4063 | 8,76061E-16 |
| ACE2 | WFDC6 | 0,4063 | 8,78349E-16 |
| ACE2 | LRRC3 | 0,4063 | 8,8258E-16 |
| ACE2 | RP11-226B15.1 | 0,4063 | 8,78823E-16 |
| ACE2 | SALL1 | 0,4062 | 9,0209E-16 |
| ACE2 | COX10-AS1 | 0,4062 | 9,00497E-16 |
| ACE2 | PTPRJ | 0,4061 | 9,19721E-16 |
| ACE2 | SLC22A3 | 0,406 | 9,34517E-16 |
| ACE2 | MT-ATP6 | 0,406 | 9,29263E-16 |
| ACE2 | MBD4 | 0,4059 | 9,36933E-16 |
| ACE2 | LINC01091 | 0,4059 | 9,50389E-16 |
| ACE2 | RPA4 | 0,4059 | 9,50113E-16 |
| ACE2 | GNG11 | 0,4057 | 9,74484E-16 |

| | | | |
|---|---|---|---|
| ACE2 | AC025918.2 | 0,4057 | 9,76828E-16 |
| ACE2 | RPL7P57 | 0,4057 | 9,73378E-16 |
| ACE2 | SLC45A3 | 0,4056 | 9,94794E-16 |
| ACE2 | RP11-134O21.1 | 0,4056 | 1,00111E-15 |
| ACE2 | MS4A6A | 0,4056 | 1,0015E-15 |
| ACE2 | RP11-431N15.2 | 0,4056 | 9,9985E-16 |
| ACE2 | ACTN1 | 0,4055 | 1,00777E-15 |
| ACE2 | FAM103A1 | 0,4055 | 1,01412E-15 |
| ACE2 | ZNF17 | 0,4055 | 1,01297E-15 |
| ACE2 | SFTPD-AS1 | 0,4054 | 1,02664E-15 |
| ACE2 | STXBP5L | 0,4053 | 1,04861E-15 |
| ACE2 | PRDM4 | 0,4053 | 1,05031E-15 |
| ACE2 | MLH3 | 0,4053 | 1,05888E-15 |
| ACE2 | RP11-49I11.1 | 0,4053 | 1,0499E-15 |
| ACE2 | GAN | 0,4052 | 1,07308E-15 |
| ACE2 | SHISA6 | 0,405 | 1,09808E-15 |
| ACE2 | NFYC-AS1 | 0,4049 | 1,12003E-15 |
| ACE2 | MTSS1 | 0,4048 | 1,13787E-15 |
| ACE2 | RP11-466H18.1 | 0,4048 | 1,15112E-15 |
| ACE2 | RP11-762H8.2 | 0,4048 | 1,14143E-15 |
| ACE2 | FBXL19 | 0,4048 | 1,14136E-15 |
| ACE2 | RET | 0,4047 | 1,16489E-15 |
| ACE2 | CD248 | 0,4047 | 1,16508E-15 |
| ACE2 | FTH1P16 | 0,4047 | 1,16394E-15 |
| ACE2 | PTPRS | 0,4046 | 1,17958E-15 |
| ACE2 | ZNF808 | 0,4046 | 1,19569E-15 |
| ACE2 | SLC12A5 | 0,4045 | 1,2165E-15 |
| ACE2 | RP1-102K2.9 | 0,4045 | 1,21622E-15 |
| ACE2 | CACNA1E | 0,4044 | 1,22409E-15 |
| ACE2 | LATS1 | 0,4044 | 1,24162E-15 |
| ACE2 | INF2 | 0,4044 | 1,22934E-15 |
| ACE2 | PPIL2 | 0,4044 | 1,22708E-15 |
| ACE2 | TMEM150C | 0,4042 | 1,26436E-15 |
| ACE2 | ZNF771 | 0,4042 | 1,26798E-15 |
| ACE2 | HPS4 | 0,4042 | 1,27418E-15 |
| ACE2 | TM2D1 | 0,4041 | 1,29409E-15 |
| ACE2 | TC2N | 0,4041 | 1,28967E-15 |
| ACE2 | SIRPB2 | 0,4041 | 1,29092E-15 |
| ACE2 | MAGEA10 | 0,4041 | 1,30562E-15 |
| ACE2 | ZNF804A | 0,404 | 1,32316E-15 |
| ACE2 | RSU1 | 0,4039 | 1,34636E-15 |
| ACE2 | NACA | 0,4039 | 1,34227E-15 |
| ACE2 | EML3 | 0,4038 | 1,37879E-15 |
| ACE2 | CXorf40A | 0,4038 | 1,37126E-15 |
| ACE2 | MTCO1P12 | 0,4034 | 1,47681E-15 |
| ACE2 | EIF3J | 0,4034 | 1,47362E-15 |
| ACE2 | USP36 | 0,4034 | 1,47446E-15 |
| ACE2 | AGMO | 0,4033 | 1,48792E-15 |
| ACE2 | PARD6G | 0,4033 | 1,49614E-15 |
| ACE2 | SIGLEC7 | 0,4033 | 1,49691E-15 |
| ACE2 | GIMAP2 | 0,4032 | 1,5215E-15 |
| ACE2 | KCNJ2 | 0,4032 | 1,51902E-15 |
| ACE2 | MPEG1 | 0,4031 | 1,55968E-15 |
| ACE2 | TBCEL | 0,4031 | 1,55654E-15 |

| | | | |
|---|---|---|---|
| ACE2 | TMTC3 | 0,4031 | 1,54189E-15 |
| ACE2 | LILRB4 | 0,4031 | 1,53727E-15 |
| ACE2 | RP11-1134I14.8 | 0,403 | 1,57029E-15 |
| ACE2 | MCM5 | 0,4029 | 1,59724E-15 |
| ACE2 | CDK2AP2 | 0,4028 | 1,64144E-15 |
| ACE2 | SALL3 | 0,4028 | 1,62892E-15 |
| ACE2 | TBXAS1 | 0,4026 | 1,68321E-15 |
| ACE2 | RP5-1107A17.4 | 0,4026 | 1,67461E-15 |
| ACE2 | RP11-511P7.6 | 0,4025 | 1,73158E-15 |
| ACE2 | IFI27 | 0,4025 | 1,72457E-15 |
| ACE2 | DNAJC16 | 0,4024 | 1,7427E-15 |
| ACE2 | PEX11B | 0,4024 | 1,76331E-15 |
| ACE2 | RP11-417L19.5 | 0,4024 | 1,74478E-15 |
| ACE2 | JAK1 | 0,4023 | 1,77635E-15 |
| ACE2 | RP11-362F19.1 | 0,4023 | 1,78417E-15 |
| ACE2 | LCP2 | 0,4023 | 1,7883E-15 |
| ACE2 | MGST1 | 0,4023 | 1,77649E-15 |
| ACE2 | RP11-417O11.5 | 0,4022 | 1,80351E-15 |
| ACE2 | SCN8A | 0,4022 | 1,80857E-15 |
| ACE2 | RAMP2-AS1 | 0,4022 | 1,79553E-15 |
| ACE2 | RAC2 | 0,4022 | 1,80832E-15 |
| ACE2 | SFSWAP | 0,4021 | 1,84449E-15 |
| ACE2 | WDR3 | 0,402 | 1,86429E-15 |
| ACE2 | SHROOM4 | 0,402 | 1,85925E-15 |
| ACE2 | DLC1 | 0,4019 | 1,89139E-15 |
| ACE2 | NDP | 0,4019 | 1,90568E-15 |
| ACE2 | DNPH1 | 0,4017 | 1,98486E-15 |
| ACE2 | FMR1-IT1 | 0,4017 | 1,9765E-15 |
| ACE2 | SLC35E2B | 0,4016 | 2,02025E-15 |
| ACE2 | SNRPGP2 | 0,4016 | 1,99497E-15 |
| ACE2 | PRDM16 | 0,4015 | 2,06024E-15 |
| ACE2 | ACOX3 | 0,4015 | 2,0494E-15 |
| ACE2 | DMRTA1 | 0,4015 | 2,04746E-15 |
| ACE2 | CTD-2591A1.1 | 0,4014 | 2,07406E-15 |
| ACE2 | SLC24A4 | 0,4014 | 2,0969E-15 |
| ACE2 | YWHAE | 0,4012 | 2,13895E-15 |
| ACE2 | ZNF365 | 0,4011 | 2,21094E-15 |
| ACE2 | MFSD4A | 0,401 | 2,2426E-15 |
| ACE2 | RP11-290F5.2 | 0,401 | 2,21649E-15 |
| ACE2 | HIST2H2BF | 0,4009 | 2,27176E-15 |
| ACE2 | AC093642.1 | 0,4009 | 2,25466E-15 |
| ACE2 | DHX33 | 0,4009 | 2,28952E-15 |
| ACE2 | COL4A3 | 0,4008 | 2,3081E-15 |
| ACE2 | TRIM32 | 0,4008 | 2,31712E-15 |
| ACE2 | HSD17B1P1 | 0,4008 | 2,29282E-15 |
| ACE2 | RP11-283G6.3 | 0,4006 | 2,39633E-15 |
| ACE2 | GCN1 | 0,4005 | 2,44263E-15 |
| ACE2 | RN7SL381P | 0,4005 | 2,42388E-15 |
| ACE2 | TRIP10 | 0,4005 | 2,41836E-15 |
| ACE2 | PHLDB1 | 0,4004 | 2,49752E-15 |
| ACE2 | IL32 | 0,4003 | 2,50335E-15 |
| ACE2 | SNHG20 | 0,4002 | 2,55844E-15 |
| ACE2 | LENG9 | 0,4002 | 2,55458E-15 |
| ACE2 | LINC01748 | 0,4001 | 2,6187E-15 |

| | | | |
|---|---|---|---|
| ACE2 | HNRNPU | 0,4001 | 2,61503E-15 |
| ACE2 | CTD-2353F22.2 | 0,4001 | 2,61953E-15 |
| ACE2 | STOX1 | 0,4001 | 2,62045E-15 |
| ACE2 | NRK | 0,4001 | 2,61536E-15 |
| ACE2 | COMMD6 | 0,4 | 2,63606E-15 |
| ACE2 | RP11-70D24.2 | 0,4 | 2,67742E-15 |
| ACE2 | RP11-259K5.2 | 0,3999 | 2,70813E-15 |
| ACE2 | AC058791.1 | 0,3999 | 2,71494E-15 |
| ACE2 | FLYWCH1 | 0,3999 | 2,70163E-15 |
| ACE2 | ZNF512B | 0,3999 | 2,70546E-15 |
| ACE2 | RHOA-IT1 | 0,3998 | 2,74006E-15 |
| ACE2 | TPD52 | 0,3998 | 2,76632E-15 |
| ACE2 | GABBR1 | 0,3997 | 2,7969E-15 |
| ACE2 | CYP2U1 | 0,3996 | 2,84959E-15 |
| ACE2 | TFEC | 0,3996 | 2,86192E-15 |
| ACE2 | FBXL12 | 0,3996 | 2,86294E-15 |
| ACE2 | KCNQ1 | 0,3995 | 2,90778E-15 |
| ACE2 | MGA | 0,3995 | 2,89865E-15 |
| ACE2 | HERPUD1 | 0,3995 | 2,87275E-15 |
| ACE2 | COX7A1 | 0,3995 | 2,87564E-15 |
| ACE2 | TMEM63B | 0,3994 | 2,96393E-15 |
| ACE2 | HNRNPA1P48 | 0,3994 | 2,92249E-15 |
| ACE2 | NOTCH3 | 0,3994 | 2,92074E-15 |
| ACE2 | KIAA1671 | 0,3994 | 2,96448E-15 |
| ACE2 | RP3-355L5.4 | 0,3993 | 2,98329E-15 |
| ACE2 | MYRF | 0,3993 | 3,00582E-15 |
| ACE2 | TRIP4 | 0,3993 | 2,99823E-15 |
| ACE2 | COL6A2 | 0,3993 | 2,97476E-15 |
| ACE2 | RP5-1000K24.2 | 0,3993 | 3,0097E-15 |
| ACE2 | FIGN | 0,3992 | 3,06983E-15 |
| ACE2 | ARHGEF28 | 0,3992 | 3,053E-15 |
| ACE2 | PEX1 | 0,3992 | 3,05494E-15 |
| ACE2 | PPP2R2D | 0,3992 | 3,03825E-15 |
| ACE2 | DANT2 | 0,3992 | 3,05161E-15 |
| ACE2 | RANP4 | 0,3992 | 3,05309E-15 |
| ACE2 | C9orf170 | 0,3991 | 3,09913E-15 |
| ACE2 | LRP4 | 0,3991 | 3,09603E-15 |
| ACE2 | WISP2 | 0,3991 | 3,07981E-15 |
| ACE2 | PMS2P4 | 0,399 | 3,17235E-15 |
| ACE2 | ZNF106 | 0,399 | 3,15177E-15 |
| ACE2 | C1QTNF1 | 0,399 | 3,14428E-15 |
| ACE2 | VN1R108P | 0,399 | 3,13822E-15 |
| ACE2 | MRPS2 | 0,3989 | 3,21983E-15 |
| ACE2 | PNO1 | 0,3988 | 3,25446E-15 |
| ACE2 | COPS4 | 0,3987 | 3,34032E-15 |
| ACE2 | MYOT | 0,3987 | 3,30423E-15 |
| ACE2 | ZNF90 | 0,3987 | 3,30114E-15 |
| ACE2 | ITGAM | 0,3985 | 3,41815E-15 |
| ACE2 | TNFRSF11A | 0,3985 | 3,47091E-15 |
| ACE2 | TNRC6B | 0,3983 | 3,54169E-15 |
| ACE2 | SMC4 | 0,3982 | 3,63711E-15 |
| ACE2 | PDLIM2 | 0,3982 | 3,6526E-15 |
| ACE2 | WAC | 0,3981 | 3,70795E-15 |
| ACE2 | PEX12 | 0,3981 | 3,69446E-15 |

| | | | |
|---|---|---|---|
| ACE2 | NELFB | 0,398 | 3,72149E-15 |
| ACE2 | AJ006998.2 | 0,398 | 3,75588E-15 |
| ACE2 | LAMTOR5 | 0,3979 | 3,80934E-15 |
| ACE2 | USP49 | 0,3979 | 3,82667E-15 |
| ACE2 | CENPBD1 | 0,3979 | 3,84346E-15 |
| ACE2 | MAMSTR | 0,3979 | 3,83862E-15 |
| ACE2 | NADK | 0,3978 | 3,84951E-15 |
| ACE2 | RBM12B-AS1 | 0,3978 | 3,87696E-15 |
| ACE2 | PHF21A | 0,3978 | 3,85398E-15 |
| ACE2 | RP11-338K17.5 | 0,3978 | 3,88224E-15 |
| ACE2 | PCOLCE2 | 0,3976 | 3,99942E-15 |
| ACE2 | H1FX | 0,3975 | 4,10881E-15 |
| ACE2 | PRAG1 | 0,3975 | 4,1009E-15 |
| ACE2 | CTD-2006H14.2 | 0,3975 | 4,06327E-15 |
| ACE2 | RSPO2 | 0,3975 | 4,05683E-15 |
| ACE2 | SPTLC3 | 0,3975 | 4,05361E-15 |
| ACE2 | PNMA3 | 0,3975 | 4,07194E-15 |
| ACE2 | HNRNPM | 0,3974 | 4,13639E-15 |
| ACE2 | HRC | 0,3974 | 4,16251E-15 |
| ACE2 | FIBCD1 | 0,3973 | 4,19941E-15 |
| ACE2 | PRR31 | 0,3973 | 4,22606E-15 |
| ACE2 | THY1 | 0,3973 | 4,19891E-15 |
| ACE2 | IL17RA | 0,3973 | 4,19925E-15 |
| ACE2 | GBP4 | 0,3972 | 4,33887E-15 |
| ACE2 | FSTL5 | 0,3972 | 4,32196E-15 |
| ACE2 | DECR1 | 0,3972 | 4,32626E-15 |
| ACE2 | IGFBP4 | 0,3972 | 4,30819E-15 |
| ACE2 | PLEK | 0,3971 | 4,38689E-15 |
| ACE2 | RP11-175K6.2 | 0,3971 | 4,34945E-15 |
| ACE2 | NUDT7 | 0,3971 | 4,37525E-15 |
| ACE2 | LINC01979 | 0,3971 | 4,41607E-15 |
| ACE2 | DNAJC5 | 0,3971 | 4,39027E-15 |
| ACE2 | HPCAL4 | 0,397 | 4,45916E-15 |
| ACE2 | CYP4A22-AS1 | 0,397 | 4,45244E-15 |
| ACE2 | INTU | 0,397 | 4,43931E-15 |
| ACE2 | RP11-253E3.3 | 0,3969 | 4,55359E-15 |
| ACE2 | RP11-774O3.3 | 0,3968 | 4,5884E-15 |
| ACE2 | RP11-330M2.4 | 0,3967 | 4,6722E-15 |
| ACE2 | RNU6-26P | 0,3966 | 4,77544E-15 |
| ACE2 | CXCL9 | 0,3966 | 4,7529E-15 |
| ACE2 | IL17B | 0,3966 | 4,78498E-15 |
| ACE2 | ZNF572 | 0,3966 | 4,76258E-15 |
| ACE2 | RP11-762I7.4 | 0,3966 | 4,79087E-15 |
| ACE2 | RP11-230F18.6 | 0,3966 | 4,80048E-15 |
| ACE2 | PTTG1IP | 0,3966 | 4,74214E-15 |
| ACE2 | LYPLAL1 | 0,3965 | 4,89463E-15 |
| ACE2 | KCNG1 | 0,3965 | 4,8759E-15 |
| ACE2 | NDC1 | 0,3964 | 4,89629E-15 |
| ACE2 | SEMA4F | 0,3964 | 4,94929E-15 |
| ACE2 | PDPK2P | 0,3964 | 4,93913E-15 |
| ACE2 | MAG | 0,3963 | 5,03339E-15 |
| ACE2 | FAM171B | 0,3962 | 5,08418E-15 |
| ACE2 | SEMA4D | 0,3962 | 5,09928E-15 |
| ACE2 | PXMP4 | 0,3962 | 5,14403E-15 |

| | | | |
|---|---|---|---|
| ACE2 | RP11-646I6.3 | 0,3959 | 5,37553E-15 |
| ACE2 | CRMP1 | 0,3958 | 5,4824E-15 |
| ACE2 | RP11-486O12.2 | 0,3958 | 5,44333E-15 |
| ACE2 | CFD | 0,3958 | 5,44613E-15 |
| ACE2 | PTBP2 | 0,3956 | 5,69126E-15 |
| ACE2 | PLAC9P1 | 0,3956 | 5,69485E-15 |
| ACE2 | PDHB | 0,3956 | 5,7044E-15 |
| ACE2 | PRIM1 | 0,3956 | 5,64006E-15 |
| ACE2 | PCDHGB2 | 0,3955 | 5,76364E-15 |
| ACE2 | RP11-122K13.12 | 0,3955 | 5,75334E-15 |
| ACE2 | RASA4 | 0,3954 | 5,84266E-15 |
| ACE2 | MTMR9 | 0,3954 | 5,89933E-15 |
| ACE2 | CDKL1 | 0,3954 | 5,88019E-15 |
| ACE2 | ENOSF1 | 0,3954 | 5,83855E-15 |
| ACE2 | LRRC75B | 0,3954 | 5,81387E-15 |
| ACE2 | HOXA1 | 0,3953 | 5,92761E-15 |
| ACE2 | ALG6 | 0,3952 | 6,09904E-15 |
| ACE2 | DPT | 0,3952 | 6,07275E-15 |
| ACE2 | MICB | 0,3952 | 6,06518E-15 |
| ACE2 | EPB41L4A | 0,395 | 6,26531E-15 |
| ACE2 | DUXAP8 | 0,395 | 6,30005E-15 |
| ACE2 | SLC16A12 | 0,3949 | 6,38189E-15 |
| ACE2 | DENND6A | 0,3948 | 6,46906E-15 |
| ACE2 | HLA-A | 0,3948 | 6,52385E-15 |
| ACE2 | ZCCHC10 | 0,3947 | 6,64727E-15 |
| ACE2 | CHRD | 0,3946 | 6,72865E-15 |
| ACE2 | ANKRD29 | 0,3946 | 6,68947E-15 |
| ACE2 | DHPS | 0,3944 | 6,93673E-15 |
| ACE2 | SERPINE1 | 0,3942 | 7,19377E-15 |
| ACE2 | MMP1 | 0,3942 | 7,15413E-15 |
| ACE2 | DPEP1 | 0,3942 | 7,19511E-15 |
| ACE2 | GADD45B | 0,3942 | 7,18737E-15 |
| ACE2 | ZBTB12BP | 0,3941 | 7,3181E-15 |
| ACE2 | SRR | 0,3941 | 7,25504E-15 |
| ACE2 | APOC1 | 0,3941 | 7,31033E-15 |
| ACE2 | PPIEL | 0,394 | 7,48924E-15 |
| ACE2 | ARHGEF26-AS1 | 0,394 | 7,49647E-15 |
| ACE2 | ARHGEF6 | 0,394 | 7,42795E-15 |
| ACE2 | C2CD5 | 0,3939 | 7,5948E-15 |
| ACE2 | DNAJC8 | 0,3938 | 7,62885E-15 |
| ACE2 | ADRA2C | 0,3938 | 7,70461E-15 |
| ACE2 | ITGB1P1 | 0,3938 | 7,62844E-15 |
| ACE2 | LPAR4 | 0,3938 | 7,69269E-15 |
| ACE2 | ITPKB | 0,3937 | 7,7867E-15 |
| ACE2 | ERBIN | 0,3937 | 7,77448E-15 |
| ACE2 | RASSF9 | 0,3937 | 7,77076E-15 |
| ACE2 | SMCR8 | 0,3937 | 7,81796E-15 |
| ACE2 | PPARG | 0,3936 | 7,97071E-15 |
| ACE2 | ATP6V0E2 | 0,3936 | 7,97774E-15 |
| ACE2 | RP11-90B9.3 | 0,3936 | 8,01356E-15 |
| ACE2 | AP001469.5 | 0,3936 | 7,98699E-15 |
| ACE2 | C22orf29 | 0,3933 | 8,36071E-15 |
| ACE2 | NENF | 0,3932 | 8,5449E-15 |
| ACE2 | FAM160B2 | 0,3932 | 8,46089E-15 |

| | | | |
|---|---|---|---|
| ACE2 | GAS5 | 0,3931 | 8,62821E-15 |
| ACE2 | KDM7A | 0,3931 | 8,72961E-15 |
| ACE2 | RP11-887P2.3 | 0,393 | 8,80404E-15 |
| ACE2 | RP11-21K12.2 | 0,393 | 8,77E-15 |
| ACE2 | RP11-384F7.2 | 0,3929 | 8,95625E-15 |
| ACE2 | RBM18 | 0,3929 | 8,93044E-15 |
| ACE2 | SYT17 | 0,3929 | 8,90844E-15 |
| ACE2 | SLC1A7 | 0,3928 | 9,16784E-15 |
| ACE2 | WNT9A | 0,3927 | 9,2133E-15 |
| ACE2 | AGTRAP | 0,3926 | 9,45993E-15 |
| ACE2 | CDC42SE1 | 0,3926 | 9,3737E-15 |
| ACE2 | THBS1 | 0,3926 | 9,46587E-15 |
| ACE2 | GYPC | 0,3925 | 9,55491E-15 |
| ACE2 | CARD6 | 0,3925 | 9,5286E-15 |
| ACE2 | RP11-875O11.1 | 0,3925 | 9,53414E-15 |
| ACE2 | RSPRY1 | 0,3925 | 9,65054E-15 |
| ACE2 | SETDB1 | 0,3923 | 9,87211E-15 |
| ACE2 | DYSF | 0,3923 | 9,98572E-15 |
| ACE2 | SMIM14 | 0,3923 | 9,87638E-15 |
| ACE2 | C5orf42 | 0,3923 | 9,87058E-15 |
| ACE2 | TSPAN4 | 0,3922 | 1,00945E-14 |
| ACE2 | CTC-550B14.7 | 0,3922 | 1,01533E-14 |
| ACE2 | IGF1 | 0,3921 | 1,03403E-14 |
| ACE2 | RP11-739P1.3 | 0,392 | 1,03581E-14 |
| ACE2 | UPF3AP3 | 0,392 | 1,03804E-14 |
| ACE2 | DNAJC17 | 0,392 | 1,03932E-14 |
| ACE2 | FBXO40 | 0,3919 | 1,06047E-14 |
| ACE2 | ZNF488 | 0,3919 | 1,06161E-14 |
| ACE2 | LA16c-360H6.3 | 0,3919 | 1,05565E-14 |
| ACE2 | PCNX2 | 0,3918 | 1,07959E-14 |
| ACE2 | RP11-769O8.1 | 0,3918 | 1,08719E-14 |
| ACE2 | DOK2 | 0,3917 | 1,09877E-14 |
| ACE2 | CRTAC1 | 0,3916 | 1,11765E-14 |
| ACE2 | RP11-390K5.6 | 0,3916 | 1,12386E-14 |
| ACE2 | B3GALT5 | 0,3916 | 1,1129E-14 |
| ACE2 | AGRN | 0,3913 | 1,16772E-14 |
| ACE2 | MAGEA11 | 0,3913 | 1,16484E-14 |
| ACE2 | RP11-347E10.1 | 0,3911 | 1,21003E-14 |
| ACE2 | RP11-218M22.1 | 0,3909 | 1,2635E-14 |
| ACE2 | EXOC3L2 | 0,3909 | 1,25496E-14 |
| ACE2 | RNF8 | 0,3908 | 1,27307E-14 |
| ACE2 | MRGPRF | 0,3907 | 1,30812E-14 |
| ACE2 | CYP27C1 | 0,3906 | 1,32829E-14 |
| ACE2 | RP11-573G6.6 | 0,3906 | 1,32075E-14 |
| ACE2 | COMMD9 | 0,3906 | 1,31689E-14 |
| ACE2 | APOA1-AS | 0,3906 | 1,32268E-14 |
| ACE2 | TTC7B | 0,3906 | 1,31916E-14 |
| ACE2 | RP11-797A18.5 | 0,3906 | 1,32873E-14 |
| ACE2 | TPM3P9 | 0,3905 | 1,35274E-14 |
| ACE2 | ACTBP2 | 0,3904 | 1,36305E-14 |
| ACE2 | BICC1 | 0,3904 | 1,36539E-14 |
| ACE2 | SSSCA1-AS1 | 0,3904 | 1,3594E-14 |
| ACE2 | ACAN | 0,3904 | 1,3564E-14 |
| ACE2 | COL18A1 | 0,3903 | 1,39243E-14 |

| | | | |
|---|---|---|---|
| ACE2 | PKD1P6 | 0,3902 | 1,4091E-14 |
| ACE2 | AP5Z1 | 0,3901 | 1,42699E-14 |
| ACE2 | ATP5HP4 | 0,3901 | 1,44403E-14 |
| ACE2 | PHLPP2 | 0,3901 | 1,43296E-14 |
| ACE2 | RPL5 | 0,39 | 1,46297E-14 |
| ACE2 | SOCS5 | 0,39 | 1,45286E-14 |
| ACE2 | TRUB1 | 0,39 | 1,47146E-14 |
| ACE2 | AC005517.3 | 0,39 | 1,45259E-14 |
| ACE2 | ALDH16A1 | 0,39 | 1,45807E-14 |
| ACE2 | WDR12 | 0,3899 | 1,47493E-14 |
| ACE2 | GPIHBP1 | 0,3899 | 1,49245E-14 |
| ACE2 | SNRPGP15 | 0,3898 | 1,5006E-14 |
| ACE2 | MIR133A1HG | 0,3896 | 1,56428E-14 |
| ACE2 | SLC25A14 | 0,3896 | 1,57354E-14 |
| ACE2 | MTATP6P1 | 0,3895 | 1,59916E-14 |
| ACE2 | PLPPR4 | 0,3895 | 1,58675E-14 |
| ACE2 | RP1-178F15.4 | 0,3895 | 1,58525E-14 |
| ACE2 | RP11-182L21.6 | 0,3895 | 1,58629E-14 |
| ACE2 | HSP90AB3P | 0,3894 | 1,6171E-14 |
| ACE2 | C20orf196 | 0,3894 | 1,60151E-14 |
| ACE2 | THEM5 | 0,3893 | 1,64665E-14 |
| ACE2 | ELAVL2 | 0,3893 | 1,63038E-14 |
| ACE2 | AC027612.6 | 0,3892 | 1,65592E-14 |
| ACE2 | FZD7 | 0,3892 | 1,67995E-14 |
| ACE2 | RP11-179B2.2 | 0,3892 | 1,65995E-14 |
| ACE2 | NGDN | 0,389 | 1,73389E-14 |
| ACE2 | RP11-60A24.3 | 0,389 | 1,71807E-14 |
| ACE2 | TMEM169 | 0,3889 | 1,75809E-14 |
| ACE2 | FGL2 | 0,3889 | 1,75212E-14 |
| ACE2 | SPG11 | 0,3889 | 1,74799E-14 |
| ACE2 | SESN2 | 0,3888 | 1,79606E-14 |
| ACE2 | ASL | 0,3888 | 1,79877E-14 |
| ACE2 | RP11-382A20.3 | 0,3888 | 1,77758E-14 |
| ACE2 | ZNF530 | 0,3888 | 1,7794E-14 |
| ACE2 | LSM2 | 0,3887 | 1,80731E-14 |
| ACE2 | TRHDE-AS1 | 0,3887 | 1,80498E-14 |
| ACE2 | TGIF1 | 0,3887 | 1,82222E-14 |
| ACE2 | TRPV3 | 0,3886 | 1,85044E-14 |
| ACE2 | ST6GALNAC2 | 0,3886 | 1,83811E-14 |
| ACE2 | BTBD8 | 0,3885 | 1,86198E-14 |
| ACE2 | DLEU2L | 0,3884 | 1,89405E-14 |
| ACE2 | TATDN2 | 0,3884 | 1,90672E-14 |
| ACE2 | TUBGCP5 | 0,3884 | 1,90819E-14 |
| ACE2 | RP11-769O8.3 | 0,3884 | 1,91758E-14 |
| ACE2 | ARL4AP4 | 0,3883 | 1,93784E-14 |
| ACE2 | CFB | 0,3883 | 1,94515E-14 |
| ACE2 | QPRT | 0,3883 | 1,92797E-14 |
| ACE2 | KLHL8 | 0,3882 | 1,97061E-14 |
| ACE2 | FAM78B | 0,3881 | 2,00629E-14 |
| ACE2 | HSPB6 | 0,3881 | 1,99091E-14 |
| ACE2 | NIPAL1 | 0,388 | 2,02425E-14 |
| ACE2 | RP11-213H15.1 | 0,388 | 2,04958E-14 |
| ACE2 | UBE3B | 0,388 | 2,05082E-14 |
| ACE2 | RP11-1000B6.3 | 0,388 | 2,03128E-14 |

| | | | |
|---|---|---|---|
| ACE2 | CTD-2329K10.1 | 0,388 | 2,03855E-14 |
| ACE2 | APOL3 | 0,388 | 2,02751E-14 |
| ACE2 | RP11-344P13.6 | 0,3879 | 2,089E-14 |
| ACE2 | NSF | 0,3879 | 2,07354E-14 |
| ACE2 | ANTXR1 | 0,3878 | 2,12082E-14 |
| ACE2 | RP11-370I10.12 | 0,3878 | 2,10352E-14 |
| ACE2 | RP11-432J24.5 | 0,3877 | 2,15525E-14 |
| ACE2 | TCF3 | 0,3877 | 2,15139E-14 |
| ACE2 | SLC38A11 | 0,3876 | 2,18631E-14 |
| ACE2 | DEFB122 | 0,3876 | 2,19317E-14 |
| ACE2 | CTB-36H16.2 | 0,3874 | 2,26696E-14 |
| ACE2 | GRPEL2 | 0,3874 | 2,26151E-14 |
| ACE2 | COL27A1 | 0,3874 | 2,24901E-14 |
| ACE2 | GPR34 | 0,3874 | 2,25476E-14 |
| ACE2 | PDLIM1P4 | 0,3873 | 2,28617E-14 |
| ACE2 | BEND6 | 0,3873 | 2,27853E-14 |
| ACE2 | MS4A4E | 0,3873 | 2,30118E-14 |
| ACE2 | RTP4 | 0,3872 | 2,33451E-14 |
| ACE2 | RBM48 | 0,3872 | 2,32506E-14 |
| ACE2 | SAFB | 0,3872 | 2,32621E-14 |
| ACE2 | CDPF1 | 0,3872 | 2,3408E-14 |
| ACE2 | PCDH19 | 0,3872 | 2,3208E-14 |
| ACE2 | LINC00493 | 0,3871 | 2,36141E-14 |
| ACE2 | ZRSR2 | 0,3871 | 2,36524E-14 |
| ACE2 | LINGO4 | 0,387 | 2,42037E-14 |
| ACE2 | IFT80 | 0,387 | 2,40786E-14 |
| ACE2 | ATG16L2 | 0,387 | 2,38851E-14 |
| ACE2 | COX6B1P5 | 0,3869 | 2,45987E-14 |
| ACE2 | PLIN5 | 0,3869 | 2,45842E-14 |
| ACE2 | TXNRD2 | 0,3869 | 2,44871E-14 |
| ACE2 | TRIM22 | 0,3868 | 2,48556E-14 |
| ACE2 | SRGAP2C | 0,3867 | 2,53618E-14 |
| ACE2 | AC097724.3 | 0,3867 | 2,51156E-14 |
| ACE2 | KLF6 | 0,3867 | 2,54427E-14 |
| ACE2 | PLEKHA5 | 0,3867 | 2,5296E-14 |
| ACE2 | PIGC | 0,3866 | 2,55485E-14 |
| ACE2 | AC074117.10 | 0,3866 | 2,56914E-14 |
| ACE2 | RPRM | 0,3866 | 2,59274E-14 |
| ACE2 | ICAM1 | 0,3866 | 2,57889E-14 |
| ACE2 | CTD-3131K8.2 | 0,3866 | 2,58539E-14 |
| ACE2 | CENPA | 0,3865 | 2,61605E-14 |
| ACE2 | PSEN1 | 0,3865 | 2,61448E-14 |
| ACE2 | CTD-3138B18.5 | 0,3865 | 2,63283E-14 |
| ACE2 | APOL6 | 0,3864 | 2,67599E-14 |
| ACE2 | WTH3DI | 0,3863 | 2,69647E-14 |
| ACE2 | ADH1B | 0,3863 | 2,70153E-14 |
| ACE2 | KNDC1 | 0,3863 | 2,70947E-14 |
| ACE2 | FDX1 | 0,3859 | 2,86758E-14 |
| ACE2 | TNFRSF1B | 0,3858 | 2,94303E-14 |
| ACE2 | GALNT13 | 0,3858 | 2,93242E-14 |
| ACE2 | GFRA1 | 0,3858 | 2,94813E-14 |
| ACE2 | RP11-361L15.5 | 0,3858 | 2,92869E-14 |
| ACE2 | NBEAL1 | 0,3857 | 2,98745E-14 |
| ACE2 | TMEM176A | 0,3857 | 2,96449E-14 |

| | | | |
|---|---|---|---|
| ACE2 | CNTNAP3B | 0,3857 | 3,00743E-14 |
| ACE2 | PIGCP1 | 0,3857 | 2,99233E-14 |
| ACE2 | RP11-135N5.3 | 0,3857 | 2,9962E-14 |
| ACE2 | SPP1 | 0,3856 | 3,01985E-14 |
| ACE2 | RP11-389C8.2 | 0,3856 | 3,01296E-14 |
| ACE2 | SSH3 | 0,3856 | 3,02953E-14 |
| ACE2 | OTUD6A | 0,3854 | 3,15279E-14 |
| ACE2 | KCNE1B | 0,3853 | 3,17666E-14 |
| ACE2 | SERTAD2 | 0,3851 | 3,30459E-14 |
| ACE2 | USP19 | 0,3851 | 3,30683E-14 |
| ACE2 | PCDHGA1 | 0,3851 | 3,27455E-14 |
| ACE2 | ACCS | 0,3851 | 3,27541E-14 |
| ACE2 | MS4A7 | 0,385 | 3,37621E-14 |
| ACE2 | KCNK3 | 0,3848 | 3,44389E-14 |
| ACE2 | AC093162.5 | 0,3847 | 3,54667E-14 |
| ACE2 | RP11-114M5.1 | 0,3847 | 3,54719E-14 |
| ACE2 | MIR4697HG | 0,3847 | 3,52044E-14 |
| ACE2 | RP3-323P24.3 | 0,3847 | 3,54966E-14 |
| ACE2 | SRPX2 | 0,3847 | 3,53584E-14 |
| ACE2 | PAICSP1 | 0,3846 | 3,58398E-14 |
| ACE2 | MRPS34 | 0,3846 | 3,58721E-14 |
| ACE2 | IL1R2 | 0,3845 | 3,61356E-14 |
| ACE2 | TCEA1P4 | 0,3845 | 3,61062E-14 |
| ACE2 | FGGY | 0,3844 | 3,69622E-14 |
| ACE2 | FNBP1P1 | 0,3844 | 3,67897E-14 |
| ACE2 | XRN1 | 0,3844 | 3,72421E-14 |
| ACE2 | RP11-462H3.2 | 0,3844 | 3,72534E-14 |
| ACE2 | APOD | 0,3843 | 3,76845E-14 |
| ACE2 | RP11-588K22.2 | 0,3843 | 3,76807E-14 |
| ACE2 | ST3GAL6-AS1 | 0,3842 | 3,79791E-14 |
| ACE2 | AC093627.10 | 0,3842 | 3,81808E-14 |
| ACE2 | EEF1A1P6 | 0,3842 | 3,79984E-14 |
| ACE2 | SCART1 | 0,3842 | 3,81826E-14 |
| ACE2 | PTTG4P | 0,3842 | 3,83894E-14 |
| ACE2 | AC010761.13 | 0,3842 | 3,83743E-14 |
| ACE2 | GCAT | 0,3842 | 3,79741E-14 |
| ACE2 | RTKN | 0,3841 | 3,86349E-14 |
| ACE2 | PITPNA | 0,3841 | 3,90413E-14 |
| ACE2 | GABRA3 | 0,3841 | 3,90423E-14 |
| ACE2 | ZNF816 | 0,384 | 3,92553E-14 |
| ACE2 | C20orf166-AS1 | 0,384 | 3,93019E-14 |
| ACE2 | PAK6 | 0,3839 | 4,02876E-14 |
| ACE2 | PON3 | 0,3838 | 4,06092E-14 |
| ACE2 | PIGT | 0,3838 | 4,0732E-14 |
| ACE2 | PRPF6 | 0,3838 | 4,10799E-14 |
| ACE2 | MIXL1 | 0,3836 | 4,20054E-14 |
| ACE2 | IFRD1 | 0,3836 | 4,22882E-14 |
| ACE2 | CHCHD7 | 0,3836 | 4,20886E-14 |
| ACE2 | CTB-193M12.5 | 0,3836 | 4,2308E-14 |
| ACE2 | C19orf35 | 0,3836 | 4,21425E-14 |
| ACE2 | RBM41 | 0,3836 | 4,19904E-14 |
| ACE2 | MT-ATP8 | 0,3836 | 4,22964E-14 |
| ACE2 | TNFSF12 | 0,3835 | 4,30145E-14 |
| ACE2 | TRMT13 | 0,3834 | 4,36333E-14 |

| | | | |
|---|---|---|---|
| ACE2 | NOSTRIN | 0,3834 | 4,36095E-14 |
| ACE2 | MYO1D | 0,3834 | 4,38796E-14 |
| ACE2 | AC003682.16 | 0,3834 | 4,38558E-14 |
| ACE2 | RP11-354M20.3 | 0,3833 | 4,40848E-14 |
| ACE2 | EXTL1 | 0,3832 | 4,53501E-14 |
| ACE2 | RP11-266L9.6 | 0,3832 | 4,51738E-14 |
| ACE2 | RBFOX3 | 0,3832 | 4,49417E-14 |
| ACE2 | ZC3H15 | 0,3831 | 4,57773E-14 |
| ACE2 | LAMA5 | 0,3831 | 4,5594E-14 |
| ACE2 | MGARP | 0,383 | 4,68804E-14 |
| ACE2 | RTL1 | 0,3829 | 4,72405E-14 |
| ACE2 | SLC47A1 | 0,3829 | 4,73946E-14 |
| ACE2 | JPH1 | 0,3828 | 4,82886E-14 |
| ACE2 | FUT1 | 0,3827 | 4,86798E-14 |
| ACE2 | AC004383.3 | 0,3827 | 4,90513E-14 |
| ACE2 | RFX6 | 0,3826 | 4,93618E-14 |
| ACE2 | LINC01474 | 0,3826 | 4,97906E-14 |
| ACE2 | OGDHL | 0,3826 | 4,95546E-14 |
| ACE2 | RP11-253M7.4 | 0,3826 | 4,97359E-14 |
| ACE2 | IDH2 | 0,3826 | 4,95123E-14 |
| ACE2 | ALMS1-IT1 | 0,3825 | 5,0382E-14 |
| ACE2 | RP11-7F17.8 | 0,3825 | 5,03081E-14 |
| ACE2 | FOXL1 | 0,3825 | 5,04007E-14 |
| ACE2 | FLT4 | 0,3824 | 5,15777E-14 |
| ACE2 | SLFN13 | 0,3824 | 5,10063E-14 |
| ACE2 | RP11-17P16.2 | 0,3823 | 5,24807E-14 |
| ACE2 | CTD-2621I17.3 | 0,3823 | 5,23278E-14 |
| ACE2 | PUF60 | 0,3822 | 5,26628E-14 |
| ACE2 | MAP3K11 | 0,3822 | 5,34502E-14 |
| ACE2 | BGN | 0,3822 | 5,27111E-14 |
| ACE2 | RP11-130F10.1 | 0,3821 | 5,35312E-14 |
| ACE2 | ZNF75A | 0,3821 | 5,3749E-14 |
| ACE2 | CENPV | 0,3821 | 5,42329E-14 |
| ACE2 | ID3 | 0,382 | 5,45116E-14 |
| ACE2 | CMKLR1 | 0,382 | 5,52114E-14 |
| ACE2 | CORO1A | 0,382 | 5,45204E-14 |
| ACE2 | POLR2A | 0,3819 | 5,61475E-14 |
| ACE2 | KCTD1 | 0,3819 | 5,61598E-14 |
| ACE2 | RP11-332H14.2 | 0,3818 | 5,64515E-14 |
| ACE2 | RP11-173A16.2 | 0,3818 | 5,68497E-14 |
| ACE2 | ZNF66 | 0,3818 | 5,67217E-14 |
| ACE2 | SNX9 | 0,3817 | 5,72766E-14 |
| ACE2 | RP11-454L9.2 | 0,3817 | 5,72499E-14 |
| ACE2 | HSD17B14 | 0,3817 | 5,79987E-14 |
| ACE2 | PGM3 | 0,3816 | 5,80981E-14 |
| ACE2 | GLI1 | 0,3816 | 5,86301E-14 |
| ACE2 | IGF2BP1 | 0,3816 | 5,883E-14 |
| ACE2 | URB1 | 0,3816 | 5,87531E-14 |
| ACE2 | AL590762.11 | 0,3816 | 5,81902E-14 |
| ACE2 | SERTAD4 | 0,3815 | 5,91506E-14 |
| ACE2 | AC012485.2 | 0,3815 | 5,97637E-14 |
| ACE2 | SMAD7 | 0,3815 | 5,9682E-14 |
| ACE2 | ELOVL3 | 0,3814 | 6,04711E-14 |
| ACE2 | TPH1 | 0,3814 | 6,00661E-14 |

| | | | |
|---|---|---|---|
| ACE2 | RHBDF1 | 0,3814 | 6,05044E-14 |
| ACE2 | COL6A1 | 0,3814 | 6,03391E-14 |
| ACE2 | KB-1732A1.1 | 0,3813 | 6,15509E-14 |
| ACE2 | BAZ2A | 0,3813 | 6,10329E-14 |
| ACE2 | GATA6-AS1 | 0,3813 | 6,16915E-14 |
| ACE2 | HTRA3 | 0,3812 | 6,28977E-14 |
| ACE2 | DEAF1 | 0,3812 | 6,22562E-14 |
| ACE2 | SRM | 0,3811 | 6,29737E-14 |
| ACE2 | S100A13 | 0,3811 | 6,33235E-14 |
| ACE2 | C15orf62 | 0,3811 | 6,31262E-14 |
| ACE2 | CD47 | 0,381 | 6,41554E-14 |
| ACE2 | CDKN1C | 0,381 | 6,4252E-14 |
| ACE2 | CNTNAP3 | 0,3809 | 6,50355E-14 |
| ACE2 | TTC9C | 0,3809 | 6,52086E-14 |
| ACE2 | KIAA1644 | 0,3809 | 6,53787E-14 |
| ACE2 | GTF3C2 | 0,3808 | 6,66238E-14 |
| ACE2 | RP11-81A22.4 | 0,3808 | 6,68916E-14 |
| ACE2 | RALGAPB | 0,3808 | 6,69804E-14 |
| ACE2 | SPEN | 0,3807 | 6,78375E-14 |
| ACE2 | FCF1P1 | 0,3807 | 6,73748E-14 |
| ACE2 | CNTNAP1 | 0,3807 | 6,78308E-14 |
| ACE2 | SIPA1L2 | 0,3806 | 6,84581E-14 |
| ACE2 | HMGB1P5 | 0,3806 | 6,87537E-14 |
| ACE2 | EP400 | 0,3806 | 6,89909E-14 |
| ACE2 | HOXD4 | 0,3805 | 6,95432E-14 |
| ACE2 | NXPH4 | 0,3804 | 7,064E-14 |
| ACE2 | RPS11P5 | 0,3804 | 7,05311E-14 |
| ACE2 | TMEM170A | 0,3804 | 7,16351E-14 |
| ACE2 | BHLHE22 | 0,3803 | 7,24219E-14 |
| ACE2 | NOV | 0,3803 | 7,23996E-14 |
| ACE2 | SARNP | 0,3803 | 7,28435E-14 |
| ACE2 | BEND5 | 0,3802 | 7,40252E-14 |
| ACE2 | MALT1 | 0,3802 | 7,4011E-14 |
| ACE2 | RAMP1 | 0,3801 | 7,47848E-14 |
| ACE2 | P2RY4 | 0,3801 | 7,51901E-14 |
| ACE2 | AMPD2 | 0,38 | 7,61638E-14 |
| ACE2 | RP11-440D17.4 | 0,38 | 7,57138E-14 |
| ACE2 | MYO1E | 0,38 | 7,64099E-14 |
| ACE2 | BX842568.2 | 0,38 | 7,56342E-14 |
| ACE2 | RP11-462G12.1 | 0,3799 | 7,69833E-14 |
| ACE2 | ZNF676 | 0,3799 | 7,66553E-14 |
| ACE2 | RP13-650J16.1 | 0,3798 | 7,83587E-14 |
| ACE2 | RP11-576C2.1 | 0,3798 | 7,8691E-14 |
| ACE2 | PEAR1 | 0,3797 | 7,99896E-14 |
| ACE2 | RP11-16N11.2 | 0,3797 | 7,97954E-14 |
| ACE2 | PLA2G7 | 0,3797 | 7,95471E-14 |
| ACE2 | CRYZP1 | 0,3797 | 8,02798E-14 |
| ACE2 | GAS1RR | 0,3797 | 7,90502E-14 |
| ACE2 | MYCLP1 | 0,3796 | 8,06959E-14 |
| ACE2 | CPZ | 0,3795 | 8,24123E-14 |
| ACE2 | CASP2 | 0,3795 | 8,23563E-14 |
| ACE2 | ZNF24 | 0,3795 | 8,19615E-14 |
| ACE2 | TRIM52 | 0,3794 | 8,38847E-14 |
| ACE2 | ADH5P4 | 0,3794 | 8,38092E-14 |

| | | | |
|---|---|---|---|
| ACE2 | HS3ST2 | 0,3794 | 8,36084E-14 |
| ACE2 | E2F6P4 | 0,3793 | 8,52189E-14 |
| ACE2 | NCMAP | 0,3792 | 8,57664E-14 |
| ACE2 | CCDC93 | 0,3792 | 8,6827E-14 |
| ACE2 | TRIM71 | 0,3792 | 8,63002E-14 |
| ACE2 | TREM2 | 0,3792 | 8,68255E-14 |
| ACE2 | SMIM8 | 0,3792 | 8,6368E-14 |
| ACE2 | SNED1 | 0,3791 | 8,83962E-14 |
| ACE2 | RP11-410L14.1 | 0,3791 | 8,72137E-14 |
| ACE2 | NLRP5 | 0,3791 | 8,72138E-14 |
| ACE2 | BCL6 | 0,379 | 8,94946E-14 |
| ACE2 | ZNF793 | 0,379 | 8,93495E-14 |
| ACE2 | TAF8 | 0,3789 | 9,09153E-14 |
| ACE2 | SH3BGRL2 | 0,3789 | 9,11887E-14 |
| ACE2 | AKAP5 | 0,3789 | 9,04149E-14 |
| ACE2 | ARNT2 | 0,3789 | 9,05293E-14 |
| ACE2 | RAG1 | 0,3788 | 9,25069E-14 |
| ACE2 | TRIM62 | 0,3786 | 9,50808E-14 |
| ACE2 | OST4 | 0,3786 | 9,55045E-14 |
| ACE2 | RSAD1 | 0,3786 | 9,57959E-14 |
| ACE2 | GIPC1 | 0,3786 | 9,46615E-14 |
| ACE2 | TMEM158 | 0,3785 | 9,65144E-14 |
| ACE2 | TTC33 | 0,3785 | 9,63559E-14 |
| ACE2 | RP11-486G15.2 | 0,3784 | 9,87646E-14 |
| ACE2 | RP11-849H4.4 | 0,3784 | 9,90231E-14 |
| ACE2 | RPL37 | 0,3783 | 9,95541E-14 |
| ACE2 | RP11-520H14.7 | 0,3782 | 1,0181E-13 |
| ACE2 | PVT1 | 0,3782 | 1,01324E-13 |
| ACE2 | CCDC3 | 0,3782 | 1,01785E-13 |
| ACE2 | ARRDC2 | 0,3781 | 1,02662E-13 |
| ACE2 | EPB41L5 | 0,378 | 1,05199E-13 |
| ACE2 | LINC02198 | 0,378 | 1,04477E-13 |
| ACE2 | ADAM22 | 0,3779 | 1,06881E-13 |
| ACE2 | FXYD1 | 0,3778 | 1,08431E-13 |
| ACE2 | NCBP2 | 0,3776 | 1,12621E-13 |
| ACE2 | SRD5A3 | 0,3776 | 1,12471E-13 |
| ACE2 | SCGN | 0,3776 | 1,1126E-13 |
| ACE2 | RP11-338K17.6 | 0,3776 | 1,12487E-13 |
| ACE2 | AC002310.12 | 0,3776 | 1,11409E-13 |
| ACE2 | RP13-210D15.8 | 0,3776 | 1,10941E-13 |
| ACE2 | SYT1 | 0,3775 | 1,14159E-13 |
| ACE2 | AC007193.10 | 0,3775 | 1,14066E-13 |
| ACE2 | EFNA1 | 0,3774 | 1,15638E-13 |
| ACE2 | AC004980.7 | 0,3773 | 1,17157E-13 |
| ACE2 | FAM234B | 0,3773 | 1,1646E-13 |
| ACE2 | SRRM2 | 0,3773 | 1,17703E-13 |
| ACE2 | CORO6 | 0,3773 | 1,16609E-13 |
| ACE2 | TWIST2 | 0,3772 | 1,19941E-13 |
| ACE2 | ASIC1 | 0,3772 | 1,18409E-13 |
| ACE2 | SLC27A2 | 0,3772 | 1,19195E-13 |
| ACE2 | DTX4 | 0,3771 | 1,21549E-13 |
| ACE2 | LRFN1 | 0,3771 | 1,21998E-13 |
| ACE2 | IQCB1 | 0,377 | 1,22787E-13 |
| ACE2 | SH2B1 | 0,377 | 1,23884E-13 |

| | | | |
|---|---|---|---|
| ACE2 | TAZ | 0,377 | 1,2362E-13 |
| ACE2 | RMDN3 | 0,3769 | 1,25422E-13 |
| ACE2 | RP11-667M19.9 | 0,3768 | 1,27282E-13 |
| ACE2 | ANKRD26 | 0,3767 | 1,29792E-13 |
| ACE2 | RHNO1 | 0,3767 | 1,29943E-13 |
| ACE2 | CHMP4A | 0,3767 | 1,28937E-13 |
| ACE2 | FAM208A | 0,3765 | 1,32622E-13 |
| ACE2 | SLC7A10 | 0,3765 | 1,3382E-13 |
| ACE2 | LINC01237 | 0,3764 | 1,3437E-13 |
| ACE2 | LINC01085 | 0,3764 | 1,35594E-13 |
| ACE2 | RABGGTA | 0,3763 | 1,36633E-13 |
| ACE2 | RP11-188D8.1 | 0,3762 | 1,39356E-13 |
| ACE2 | KCTD14 | 0,3762 | 1,39691E-13 |
| ACE2 | DIRAS3 | 0,3761 | 1,41409E-13 |
| ACE2 | AC005618.6 | 0,3761 | 1,42199E-13 |
| ACE2 | RP11-20D14.6 | 0,3761 | 1,42715E-13 |
| ACE2 | ARHGAP5-AS1 | 0,376 | 1,43602E-13 |
| ACE2 | PTMS | 0,3759 | 1,46851E-13 |
| ACE2 | KLHL22 | 0,3759 | 1,45855E-13 |
| ACE2 | RP11-181K12.1 | 0,3758 | 1,50248E-13 |
| ACE2 | RP11-338K17.10 | 0,3758 | 1,4826E-13 |
| ACE2 | LSAMP | 0,3757 | 1,50811E-13 |
| ACE2 | ERCC6 | 0,3757 | 1,50452E-13 |
| ACE2 | RP11-554D14.2 | 0,3757 | 1,5181E-13 |
| ACE2 | ZNF559-ZNF177 | 0,3756 | 1,55047E-13 |
| ACE2 | CYP2A6 | 0,3756 | 1,54163E-13 |
| ACE2 | AC124861.1 | 0,3755 | 1,55431E-13 |
| ACE2 | HIST1H3H | 0,3755 | 1,56445E-13 |
| ACE2 | RP5-1139B12.3 | 0,3754 | 1,60039E-13 |
| ACE2 | VANGL2 | 0,3753 | 1,60463E-13 |
| ACE2 | P3H3 | 0,3753 | 1,61896E-13 |
| ACE2 | TRPC4AP | 0,3753 | 1,61212E-13 |
| ACE2 | PFN1P2 | 0,3752 | 1,64249E-13 |
| ACE2 | RNU6-103P | 0,3752 | 1,64704E-13 |
| ACE2 | SCARF2 | 0,3752 | 1,6517E-13 |
| ACE2 | PVRIG | 0,3751 | 1,6679E-13 |
| ACE2 | RPL24P8 | 0,3751 | 1,6545E-13 |
| ACE2 | AP001372.2 | 0,3751 | 1,67029E-13 |
| ACE2 | TTC5 | 0,3751 | 1,67937E-13 |
| ACE2 | PRPS1P2 | 0,3749 | 1,72762E-13 |
| ACE2 | SOCS5P4 | 0,3749 | 1,72563E-13 |
| ACE2 | RP5-997D16.2 | 0,3748 | 1,74725E-13 |
| ACE2 | RP11-767C1.2 | 0,3748 | 1,7506E-13 |
| ACE2 | RP11-252I14.2 | 0,3748 | 1,74799E-13 |
| ACE2 | TXNDC12 | 0,3747 | 1,78592E-13 |
| ACE2 | PIK3C2B | 0,3747 | 1,76644E-13 |
| ACE2 | IRF5 | 0,3747 | 1,78351E-13 |
| ACE2 | SLCO5A1 | 0,3747 | 1,79029E-13 |
| ACE2 | CAMK1D | 0,3747 | 1,77518E-13 |
| ACE2 | TOX2 | 0,3747 | 1,77008E-13 |
| ACE2 | POU3F2 | 0,3746 | 1,79307E-13 |
| ACE2 | BLOC1S2 | 0,3746 | 1,81121E-13 |
| ACE2 | CNTN1 | 0,3746 | 1,7961E-13 |
| ACE2 | LINC01545 | 0,3746 | 1,80851E-13 |

| | | | |
|---|---|---|---|
| ACE2 | AC013480.2 | 0,3745 | 1,84087E-13 |
| ACE2 | AC007365.3 | 0,3745 | 1,84355E-13 |
| ACE2 | LINC01354 | 0,3743 | 1,90735E-13 |
| ACE2 | ZBTB46 | 0,3743 | 1,8925E-13 |
| ACE2 | OR3A1 | 0,3742 | 1,93315E-13 |
| ACE2 | CTD-2349P21.3 | 0,3742 | 1,91422E-13 |
| ACE2 | GATSL2 | 0,3741 | 1,95195E-13 |
| ACE2 | ITGA8 | 0,3741 | 1,94454E-13 |
| ACE2 | DNAJC1 | 0,3741 | 1,9654E-13 |
| ACE2 | CUEDC1 | 0,374 | 1,98682E-13 |
| ACE2 | FAM87A | 0,3739 | 2,00165E-13 |
| ACE2 | TRIM34 | 0,3739 | 2,03123E-13 |
| ACE2 | KLHL1 | 0,3739 | 2,01372E-13 |
| ACE2 | ZNF385B | 0,3738 | 2,0567E-13 |
| ACE2 | LIMS2 | 0,3737 | 2,08907E-13 |
| ACE2 | ZFP37 | 0,3737 | 2,08738E-13 |
| ACE2 | CTD-2049O4.1 | 0,3736 | 2,11931E-13 |
| ACE2 | AP1G1 | 0,3736 | 2,11976E-13 |
| ACE2 | MRPL10 | 0,3736 | 2,12771E-13 |
| ACE2 | ZNF592 | 0,3735 | 2,15331E-13 |
| ACE2 | JOSD2 | 0,3735 | 2,16293E-13 |
| ACE2 | CHD3 | 0,3734 | 2,19543E-13 |
| ACE2 | GNRHR | 0,3733 | 2,21986E-13 |
| ACE2 | ZNF205 | 0,3733 | 2,21823E-13 |
| ACE2 | ERMAP | 0,3732 | 2,25487E-13 |
| ACE2 | HMGA1 | 0,3732 | 2,23988E-13 |
| ACE2 | LINC00957 | 0,3732 | 2,25166E-13 |
| ACE2 | RP11-481C4.2 | 0,3732 | 2,24455E-13 |
| ACE2 | ADAM15 | 0,3731 | 2,27849E-13 |
| ACE2 | RP11-499E18.1 | 0,3731 | 2,29534E-13 |
| ACE2 | TFPI2 | 0,3731 | 2,27516E-13 |
| ACE2 | RP11-212I21.4 | 0,3731 | 2,29657E-13 |
| ACE2 | NPEPPS | 0,3731 | 2,28746E-13 |
| ACE2 | RPS19P3 | 0,373 | 2,34435E-13 |
| ACE2 | RP5-966M1.7 | 0,3729 | 2,3451E-13 |
| ACE2 | RP11-613F22.8 | 0,3729 | 2,38109E-13 |
| ACE2 | LL0XNC01-116E7.2 | 0,3729 | 2,37649E-13 |
| ACE2 | RP11-346C20.3 | 0,3728 | 2,3912E-13 |
| ACE2 | ZNF460 | 0,3728 | 2,38645E-13 |
| ACE2 | POLR2J4 | 0,3727 | 2,42212E-13 |
| ACE2 | KCTD15 | 0,3727 | 2,44362E-13 |
| ACE2 | RP11-325E14.5 | 0,3727 | 2,44282E-13 |
| ACE2 | DCAF13P3 | 0,3725 | 2,52516E-13 |
| ACE2 | RP11-968O1.5 | 0,3724 | 2,57523E-13 |
| ACE2 | TRAF3IP2-AS1 | 0,3723 | 2,59529E-13 |
| ACE2 | VSTM5 | 0,3723 | 2,60916E-13 |
| ACE2 | C15orf59 | 0,3723 | 2,5821E-13 |
| ACE2 | RP11-147L13.15 | 0,3723 | 2,58734E-13 |
| ACE2 | RHOXF1 | 0,3722 | 2,63365E-13 |
| ACE2 | ETV5 | 0,3721 | 2,67333E-13 |
| ACE2 | GIN1 | 0,3721 | 2,68664E-13 |
| ACE2 | AC016722.4 | 0,372 | 2,71408E-13 |
| ACE2 | TRANK1 | 0,372 | 2,72369E-13 |
| ACE2 | TYROBP | 0,3719 | 2,77303E-13 |

| | | | |
|---|---|---|---|
| ACE2 | ZNF586 | 0,3719 | 2,78369E-13 |
| ACE2 | RGS14 | 0,3718 | 2,80985E-13 |
| ACE2 | NFYC | 0,3717 | 2,8779E-13 |
| ACE2 | AC011239.1 | 0,3717 | 2,84752E-13 |
| ACE2 | FASTKD2 | 0,3717 | 2,86836E-13 |
| ACE2 | LINC01554 | 0,3717 | 2,85112E-13 |
| ACE2 | RP11-274B21.1 | 0,3717 | 2,86208E-13 |
| ACE2 | NUTM2A-AS1 | 0,3717 | 2,8721E-13 |
| ACE2 | AP3D1 | 0,3717 | 2,83569E-13 |
| ACE2 | RP11-767I20.1 | 0,3716 | 2,92121E-13 |
| ACE2 | CTD-2184D3.5 | 0,3715 | 2,95399E-13 |
| ACE2 | SLCO4A1-AS1 | 0,3715 | 2,96349E-13 |
| ACE2 | CCDC85A | 0,3714 | 2,98146E-13 |
| ACE2 | RPLP2 | 0,3714 | 3,00507E-13 |
| ACE2 | N4BP2L2-IT2 | 0,3713 | 3,04996E-13 |
| ACE2 | PLK5 | 0,3713 | 3,04603E-13 |
| ACE2 | C7orf50 | 0,3712 | 3,09479E-13 |
| ACE2 | SURF4 | 0,3711 | 3,16274E-13 |
| ACE2 | PSMD13 | 0,3711 | 3,12088E-13 |
| ACE2 | PHRF1 | 0,3711 | 3,16135E-13 |
| ACE2 | RP11-269G24.3 | 0,3711 | 3,13941E-13 |
| ACE2 | PLEKHA4 | 0,3711 | 3,14621E-13 |
| ACE2 | RP11-977G19.12 | 0,371 | 3,1751E-13 |
| ACE2 | ARHGEF25 | 0,371 | 3,1869E-13 |
| ACE2 | RP11-549B18.1 | 0,371 | 3,18404E-13 |
| ACE2 | VGLL3 | 0,3709 | 3,24827E-13 |
| ACE2 | CDR2L | 0,3709 | 3,24938E-13 |
| ACE2 | LINC01116 | 0,3708 | 3,29635E-13 |
| ACE2 | RIMKLBP1 | 0,3708 | 3,30425E-13 |
| ACE2 | COMMD5 | 0,3707 | 3,34919E-13 |
| ACE2 | ZNF587B | 0,3707 | 3,32407E-13 |
| ACE2 | DUSP1 | 0,3706 | 3,39543E-13 |
| ACE2 | RP11-284H18.1 | 0,3706 | 3,3897E-13 |
| ACE2 | TTF1 | 0,3706 | 3,41226E-13 |
| ACE2 | RP11-285G1.14 | 0,3706 | 3,37275E-13 |
| ACE2 | GDNF | 0,3704 | 3,52668E-13 |
| ACE2 | SCAF4 | 0,3704 | 3,50764E-13 |
| ACE2 | HTR2B | 0,3703 | 3,55897E-13 |
| ACE2 | RP11-285A1.1 | 0,3702 | 3,59082E-13 |
| ACE2 | RP11-351C21.2 | 0,3701 | 3,66414E-13 |
| ACE2 | TBC1D2B | 0,3701 | 3,65287E-13 |
| ACE2 | RP11-686D22.3 | 0,3701 | 3,65241E-13 |
| ACE2 | AC019097.7 | 0,37 | 3,7276E-13 |
| ACE2 | STX16 | 0,37 | 3,75516E-13 |
| ACE2 | GLYATL1P3 | 0,3699 | 3,76533E-13 |
| ACE2 | RP11-244F12.3 | 0,3698 | 3,81984E-13 |
| ACE2 | MIR186 | 0,3697 | 3,89661E-13 |
| ACE2 | FAM53B-AS1 | 0,3697 | 3,92111E-13 |
| ACE2 | KIAA1549L | 0,3697 | 3,89155E-13 |
| ACE2 | RP11-536L3.4 | 0,3696 | 3,99167E-13 |
| ACE2 | RP4-671O14.6 | 0,3696 | 3,98891E-13 |
| ACE2 | TLL1 | 0,3695 | 4,0165E-13 |
| ACE2 | PPP1R1B | 0,3695 | 4,0173E-13 |
| ACE2 | JUN | 0,3694 | 4,12499E-13 |

| | | | |
|---|---|---|---|
| ACE2 | SLC16A14 | 0,3694 | 4,12353E-13 |
| ACE2 | KCTD21-AS1 | 0,3694 | 4,1015E-13 |
| ACE2 | RP11-603J24.7 | 0,3694 | 4,09169E-13 |
| ACE2 | ZNF529 | 0,3694 | 4,06816E-13 |
| ACE2 | REEP2 | 0,3693 | 4,18685E-13 |
| ACE2 | UGGT2 | 0,3693 | 4,16018E-13 |
| ACE2 | POM121 | 0,3692 | 4,2168E-13 |
| ACE2 | ADAMDEC1 | 0,3692 | 4,2214E-13 |
| ACE2 | RBMY2FP | 0,3692 | 4,2262E-13 |
| ACE2 | RPP40 | 0,3691 | 4,29001E-13 |
| ACE2 | CTB-25B13.5 | 0,3691 | 4,28586E-13 |
| ACE2 | RP11-678G14.3 | 0,3691 | 4,32394E-13 |
| ACE2 | PLCH2 | 0,369 | 4,3748E-13 |
| ACE2 | S100A11 | 0,369 | 4,39137E-13 |
| ACE2 | ADGRL1 | 0,3689 | 4,40578E-13 |
| ACE2 | RBMS1 | 0,3687 | 4,57469E-13 |
| ACE2 | IMPA2 | 0,3687 | 4,59203E-13 |
| ACE2 | VNN1 | 0,3686 | 4,60434E-13 |
| ACE2 | ZNF469 | 0,3685 | 4,70693E-13 |
| ACE2 | CTD-3222D19.8 | 0,3685 | 4,69838E-13 |
| ACE2 | MPV17L2 | 0,3685 | 4,73004E-13 |
| ACE2 | ZBTB7B | 0,3684 | 4,76053E-13 |
| ACE2 | RFWD2 | 0,3683 | 4,87E-13 |
| ACE2 | MNT | 0,3683 | 4,8813E-13 |
| ACE2 | KCNK1 | 0,3682 | 4,91858E-13 |
| ACE2 | ATP6V0CP3 | 0,3682 | 4,96211E-13 |
| ACE2 | PRR14 | 0,3682 | 4,96111E-13 |
| ACE2 | CTB-92J24.2 | 0,3682 | 4,93614E-13 |
| ACE2 | BLOC1S1 | 0,3681 | 5,04799E-13 |
| ACE2 | CXorf36 | 0,3681 | 4,98161E-13 |
| ACE2 | RP1-283E3.4 | 0,368 | 5,10825E-13 |
| ACE2 | NDUFB1P2 | 0,368 | 5,07009E-13 |
| ACE2 | LL0XNC01-237H1.2 | 0,3679 | 5,1575E-13 |
| ACE2 | ZC3H6 | 0,3678 | 5,22138E-13 |
| ACE2 | AIF1 | 0,3678 | 5,22879E-13 |
| ACE2 | SCUBE1 | 0,3678 | 5,24719E-13 |
| ACE2 | SOX12 | 0,3677 | 5,36946E-13 |
| ACE2 | MAGEB6P1 | 0,3676 | 5,40687E-13 |
| ACE2 | GPLD1 | 0,3675 | 5,52896E-13 |
| ACE2 | RP11-274B21.13 | 0,3675 | 5,49144E-13 |
| ACE2 | SLC18A2 | 0,3675 | 5,50316E-13 |
| ACE2 | RP11-504I13.3 | 0,3675 | 5,50457E-13 |
| ACE2 | LPIN3 | 0,3675 | 5,54751E-13 |
| ACE2 | CAPN6 | 0,3675 | 5,49891E-13 |
| ACE2 | RPSA | 0,3674 | 5,59739E-13 |
| ACE2 | RP1-95L4.4 | 0,3674 | 5,54978E-13 |
| ACE2 | C1orf140 | 0,3673 | 5,64681E-13 |
| ACE2 | RPS15A | 0,3673 | 5,6366E-13 |
| ACE2 | FUT8-AS1 | 0,3672 | 5,7621E-13 |
| ACE2 | ARSJ | 0,3671 | 5,85746E-13 |
| ACE2 | NRBF2P5 | 0,3671 | 5,82623E-13 |
| ACE2 | FOLR2 | 0,3671 | 5,90173E-13 |
| ACE2 | NFIX | 0,367 | 5,98797E-13 |
| ACE2 | CTC-429P9.3 | 0,367 | 5,99295E-13 |

| | | | |
|---|---|---|---|
| ACE2 | STRIP1 | 0,3669 | 6,07353E-13 |
| ACE2 | H19 | 0,3669 | 6,07107E-13 |
| ACE2 | RPL39P3 | 0,3668 | 6,16423E-13 |
| ACE2 | NNMT | 0,3668 | 6,13672E-13 |
| ACE2 | ERCC3 | 0,3667 | 6,25935E-13 |
| ACE2 | CTD-3157E16.2 | 0,3667 | 6,1897E-13 |
| ACE2 | TMEM40 | 0,3666 | 6,34915E-13 |
| ACE2 | TAS2R20 | 0,3666 | 6,31824E-13 |
| ACE2 | JRK | 0,3665 | 6,46802E-13 |
| ACE2 | RPL4 | 0,3665 | 6,41035E-13 |
| ACE2 | CYBA | 0,3665 | 6,4594E-13 |
| ACE2 | CTD-2245F17.3 | 0,3665 | 6,45533E-13 |
| ACE2 | RP11-245J9.5 | 0,3664 | 6,49302E-13 |
| ACE2 | TMPO | 0,3664 | 6,53792E-13 |
| ACE2 | RP11-234A1.1 | 0,3663 | 6,58336E-13 |
| ACE2 | IPCEF1 | 0,3663 | 6,62843E-13 |
| ACE2 | GCHFR | 0,3663 | 6,65064E-13 |
| ACE2 | MEGF6 | 0,3662 | 6,75559E-13 |
| ACE2 | UBE2W | 0,3662 | 6,72187E-13 |
| ACE2 | RASD2 | 0,3662 | 6,73296E-13 |
| ACE2 | TNFSF13 | 0,3661 | 6,86515E-13 |
| ACE2 | KRT222 | 0,3661 | 6,81707E-13 |
| ACE2 | RBM5 | 0,366 | 6,95746E-13 |
| ACE2 | RARRES2 | 0,366 | 6,89345E-13 |
| ACE2 | COMTD1 | 0,366 | 6,93291E-13 |
| ACE2 | SOX15 | 0,366 | 6,99034E-13 |
| ACE2 | TLR5 | 0,3659 | 7,09231E-13 |
| ACE2 | CXCL12 | 0,3659 | 7,09331E-13 |
| ACE2 | SMUG1 | 0,3659 | 7,0901E-13 |
| ACE2 | PRDM13 | 0,3658 | 7,21412E-13 |
| ACE2 | RAB6C-AS1 | 0,3657 | 7,28638E-13 |
| ACE2 | PRPF40A | 0,3657 | 7,22708E-13 |
| ACE2 | ANGPTL4 | 0,3657 | 7,23806E-13 |
| ACE2 | RP11-311P8.2 | 0,3657 | 7,23938E-13 |
| ACE2 | ACAA2 | 0,3656 | 7,43105E-13 |
| ACE2 | MXRA8 | 0,3655 | 7,48885E-13 |
| ACE2 | TSPAN2 | 0,3655 | 7,48698E-13 |
| ACE2 | RP11-177H2.1 | 0,3655 | 7,54795E-13 |
| ACE2 | TRAF1 | 0,3655 | 7,55776E-13 |
| ACE2 | RP11-513G19.1 | 0,3655 | 7,55778E-13 |
| ACE2 | SNHG21 | 0,3655 | 7,53178E-13 |
| ACE2 | TRIM72 | 0,3655 | 7,55497E-13 |
| ACE2 | DHCR24 | 0,3654 | 7,59725E-13 |
| ACE2 | AC010733.5 | 0,3654 | 7,65126E-13 |
| ACE2 | TNFRSF10A | 0,3653 | 7,71738E-13 |
| ACE2 | SCYL1 | 0,3653 | 7,71904E-13 |
| ACE2 | RPL7P8 | 0,3652 | 7,81856E-13 |
| ACE2 | AQP10 | 0,3652 | 7,84933E-13 |
| ACE2 | RP11-166B2.3 | 0,3652 | 7,88919E-13 |
| ACE2 | CD300C | 0,3652 | 7,80842E-13 |
| ACE2 | RP3-467D16.3 | 0,365 | 8,13031E-13 |
| ACE2 | RP11-1114A5.4 | 0,365 | 8,16188E-13 |
| ACE2 | DUSP2 | 0,3649 | 8,27554E-13 |
| ACE2 | GABPB1-AS1 | 0,3649 | 8,25063E-13 |

| | | | |
|---|---|---|---|
| ACE2 | RP11-1082L8.3 | 0,3648 | 8,30828E-13 |
| ACE2 | RP5-1042K10.13 | 0,3648 | 8,3165E-13 |
| ACE2 | LINC01186 | 0,3648 | 8,32327E-13 |
| ACE2 | CKM | 0,3647 | 8,52277E-13 |
| ACE2 | MEOX2 | 0,3646 | 8,55381E-13 |
| ACE2 | GUCY1A2 | 0,3646 | 8,57248E-13 |
| ACE2 | RP5-1014D13.2 | 0,3646 | 8,65133E-13 |
| ACE2 | PAIP1P1 | 0,3645 | 8,75636E-13 |
| ACE2 | SNRPGP14 | 0,3645 | 8,77606E-13 |
| ACE2 | MAP3K6 | 0,3644 | 8,92473E-13 |
| ACE2 | TAS2R3 | 0,3644 | 8,8691E-13 |
| ACE2 | CTD-2284J15.1 | 0,3644 | 8,84845E-13 |
| ACE2 | MACROD2 | 0,3644 | 8,84674E-13 |
| ACE2 | LINC00899 | 0,3644 | 8,89859E-13 |
| ACE2 | SYNC | 0,3643 | 9,0261E-13 |
| ACE2 | TGFBI | 0,3643 | 9,00432E-13 |
| ACE2 | HSD17B7 | 0,3641 | 9,29791E-13 |
| ACE2 | BCL7A | 0,3641 | 9,29536E-13 |
| ACE2 | RP11-317B17.4 | 0,364 | 9,51674E-13 |
| ACE2 | RNF24 | 0,364 | 9,51633E-13 |
| ACE2 | RP5-1099D15.1 | 0,3639 | 9,64433E-13 |
| ACE2 | SEC22C | 0,3637 | 9,88988E-13 |
| ACE2 | MRVI1 | 0,3637 | 9,92485E-13 |
| ACE2 | HAAO | 0,3636 | 1,00631E-12 |
| ACE2 | C12orf76 | 0,3636 | 1,00124E-12 |
| ACE2 | THSD4 | 0,3636 | 1,00214E-12 |
| ACE2 | MPHOSPH6 | 0,3635 | 1,02519E-12 |
| ACE2 | CYGB | 0,3635 | 1,02411E-12 |
| ACE2 | DFFBP1 | 0,3634 | 1,03369E-12 |
| ACE2 | KLHL12 | 0,3633 | 1,05562E-12 |
| ACE2 | AC064850.4 | 0,3633 | 1,05264E-12 |
| ACE2 | GART | 0,3633 | 1,04354E-12 |
| ACE2 | RP11-270C12.3 | 0,3632 | 1,0755E-12 |
| ACE2 | CXCR4 | 0,3631 | 1,08046E-12 |
| ACE2 | ZNF227 | 0,3631 | 1,0852E-12 |
| ACE2 | LIMK1 | 0,363 | 1,09901E-12 |
| ACE2 | DNM1P35 | 0,363 | 1,1084E-12 |
| ACE2 | DECR2 | 0,3629 | 1,12264E-12 |
| ACE2 | RP3-332B22.1 | 0,3628 | 1,13645E-12 |
| ACE2 | SPN | 0,3628 | 1,14012E-12 |
| ACE2 | TMEM161B-AS1 | 0,3627 | 1,15366E-12 |
| ACE2 | RBPMS2 | 0,3627 | 1,14906E-12 |
| ACE2 | SLCO4A1 | 0,3627 | 1,15578E-12 |
| ACE2 | SERPINB9P1 | 0,3626 | 1,17412E-12 |
| ACE2 | RP11-225N10.3 | 0,3624 | 1,20403E-12 |
| ACE2 | BTF3 | 0,3624 | 1,20042E-12 |
| ACE2 | TET1 | 0,3624 | 1,19709E-12 |
| ACE2 | PCGF6 | 0,3624 | 1,20981E-12 |
| ACE2 | LAG3 | 0,3624 | 1,20698E-12 |
| ACE2 | LINC00665 | 0,3624 | 1,21408E-12 |
| ACE2 | SCN2B | 0,3623 | 1,22315E-12 |
| ACE2 | HIST1H2BH | 0,3622 | 1,24163E-12 |
| ACE2 | RP11-14N7.2 | 0,3621 | 1,25454E-12 |
| ACE2 | MZT1 | 0,3621 | 1,2711E-12 |

| | | | |
|---|---|---|---|
| ACE2 | RP11-936I5.1 | 0,3621 | 1,2551E-12 |
| ACE2 | RP11-199H2.2 | 0,362 | 1,2869E-12 |
| ACE2 | SRSF7 | 0,362 | 1,28987E-12 |
| ACE2 | GUCY1A3 | 0,362 | 1,27265E-12 |
| ACE2 | BTNL8 | 0,362 | 1,29101E-12 |
| ACE2 | RP11-23J9.7 | 0,362 | 1,28184E-12 |
| ACE2 | ZFAS1 | 0,362 | 1,28172E-12 |
| ACE2 | TRGV6 | 0,3618 | 1,31333E-12 |
| ACE2 | VPS50 | 0,3618 | 1,32528E-12 |
| ACE2 | ZNF136 | 0,3617 | 1,33482E-12 |
| ACE2 | FMO4 | 0,3616 | 1,35363E-12 |
| ACE2 | CTD-2287O16.5 | 0,3616 | 1,3686E-12 |
| ACE2 | LINC02204 | 0,3616 | 1,37025E-12 |
| ACE2 | RP11-433A19.2 | 0,3615 | 1,38389E-12 |
| ACE2 | RP11-380B4.3 | 0,3615 | 1,38851E-12 |
| ACE2 | RPAP1 | 0,3614 | 1,40779E-12 |
| ACE2 | LEMD1-AS1 | 0,3613 | 1,43004E-12 |
| ACE2 | C2orf72 | 0,3613 | 1,43571E-12 |
| ACE2 | SLC39A4 | 0,3613 | 1,43507E-12 |
| ACE2 | MCAM | 0,3613 | 1,42154E-12 |
| ACE2 | MYOSLID | 0,3612 | 1,44275E-12 |
| ACE2 | HIST1H3E | 0,3612 | 1,4536E-12 |
| ACE2 | RP11-299G20.5 | 0,3612 | 1,43921E-12 |
| ACE2 | EHD4 | 0,3611 | 1,47354E-12 |
| ACE2 | RP11-575L7.8 | 0,361 | 1,4868E-12 |
| ACE2 | VN2R19P | 0,361 | 1,48816E-12 |
| ACE2 | AF129075.5 | 0,361 | 1,49434E-12 |
| ACE2 | PDE5A | 0,3609 | 1,51708E-12 |
| ACE2 | TSC22D2 | 0,3608 | 1,52939E-12 |
| ACE2 | TMEM173 | 0,3608 | 1,53073E-12 |
| ACE2 | HIST2H3D | 0,3607 | 1,56843E-12 |
| ACE2 | AC015933.2 | 0,3607 | 1,56946E-12 |
| ACE2 | HOMER3 | 0,3607 | 1,55379E-12 |
| ACE2 | HMGA1P2 | 0,3606 | 1,58869E-12 |
| ACE2 | TTYH1 | 0,3606 | 1,59089E-12 |
| ACE2 | SULT2A1 | 0,3605 | 1,61803E-12 |
| ACE2 | RGS1 | 0,3604 | 1,62795E-12 |
| ACE2 | MAP3K20-AS1 | 0,3604 | 1,62952E-12 |
| ACE2 | TMEM26 | 0,3604 | 1,62701E-12 |
| ACE2 | PIM1 | 0,3603 | 1,66122E-12 |
| ACE2 | RP11-326C3.7 | 0,3603 | 1,66286E-12 |
| ACE2 | EBF4 | 0,3603 | 1,65084E-12 |
| ACE2 | PACSIN2 | 0,3602 | 1,68661E-12 |
| ACE2 | SELENBP1 | 0,3601 | 1,71905E-12 |
| ACE2 | ABI3BP | 0,3601 | 1,721E-12 |
| ACE2 | EPB41L4A-AS2 | 0,3601 | 1,71614E-12 |
| ACE2 | TM2D2 | 0,3601 | 1,72051E-12 |
| ACE2 | RARG | 0,3601 | 1,70656E-12 |
| ACE2 | RGS19 | 0,3601 | 1,7194E-12 |
| ACE2 | JDP2 | 0,36 | 1,72755E-12 |
| ACE2 | RFK | 0,3598 | 1,79648E-12 |
| ACE2 | HSPA8P1 | 0,3597 | 1,80746E-12 |
| ACE2 | BCS1L | 0,3596 | 1,83853E-12 |
| ACE2 | SP4 | 0,3596 | 1,84276E-12 |

| | | | |
|---|---|---|---|
| ACE2 | CES4A | 0,3596 | 1,83696E-12 |
| ACE2 | AC002128.5 | 0,3596 | 1,85187E-12 |
| ACE2 | HSPB2 | 0,3595 | 1,86485E-12 |
| ACE2 | MYEF2 | 0,3595 | 1,88197E-12 |
| ACE2 | PCAT14 | 0,3595 | 1,8644E-12 |
| ACE2 | HLA-DRA | 0,3594 | 1,91454E-12 |
| ACE2 | ALKBH1 | 0,3594 | 1,8941E-12 |
| ACE2 | RP5-855D21.1 | 0,3593 | 1,94224E-12 |
| ACE2 | RP11-81K2.2 | 0,3593 | 1,91931E-12 |
| ACE2 | MIRLET7BHG | 0,3593 | 1,92205E-12 |
| ACE2 | ZFPM1 | 0,3592 | 1,96059E-12 |
| ACE2 | FAM53B | 0,3591 | 1,97902E-12 |
| ACE2 | CX3CL1 | 0,3591 | 1,97741E-12 |
| ACE2 | XBP1 | 0,3591 | 1,9741E-12 |
| ACE2 | BHLHE40 | 0,3589 | 2,05806E-12 |
| ACE2 | BAK1 | 0,3589 | 2,06248E-12 |
| ACE2 | IGDCC4 | 0,3589 | 2,03611E-12 |
| ACE2 | SLC26A6 | 0,3588 | 2,07524E-12 |
| ACE2 | CTA-204B4.2 | 0,3588 | 2,07133E-12 |
| ACE2 | AC010084.1 | 0,3588 | 2,08836E-12 |
| ACE2 | PTPN5 | 0,3587 | 2,12077E-12 |
| ACE2 | DLX3 | 0,3587 | 2,11829E-12 |
| ACE2 | IFT52 | 0,3587 | 2,11242E-12 |
| ACE2 | ATP2A3 | 0,3586 | 2,15244E-12 |
| ACE2 | MAP3K7CL | 0,3585 | 2,16434E-12 |
| ACE2 | RP11-493E12.2 | 0,3584 | 2,21941E-12 |
| ACE2 | CTD-2184D3.7 | 0,3584 | 2,20841E-12 |
| ACE2 | CYP2B7P | 0,3584 | 2,2075E-12 |
| ACE2 | ZNF281 | 0,3583 | 2,23357E-12 |
| ACE2 | ANKRD39 | 0,3583 | 2,25868E-12 |
| ACE2 | ANK3 | 0,3583 | 2,25378E-12 |
| ACE2 | RAD54L2 | 0,3582 | 2,28109E-12 |
| ACE2 | FABP4 | 0,3582 | 2,29378E-12 |
| ACE2 | ERCC1 | 0,3581 | 2,30689E-12 |
| ACE2 | PARK7 | 0,358 | 2,34064E-12 |
| ACE2 | RYBP | 0,358 | 2,35185E-12 |
| ACE2 | RERG | 0,358 | 2,34934E-12 |
| ACE2 | MED13L | 0,358 | 2,33657E-12 |
| ACE2 | WDR76 | 0,358 | 2,34994E-12 |
| ACE2 | DNAJA3 | 0,3579 | 2,39465E-12 |
| ACE2 | RP11-75A9.3 | 0,3579 | 2,36984E-12 |
| ACE2 | RP11-30K9.1 | 0,3577 | 2,45769E-12 |
| ACE2 | EVC2 | 0,3576 | 2,47672E-12 |
| ACE2 | GSTM2 | 0,3575 | 2,53404E-12 |
| ACE2 | KB-1208A12.3 | 0,3574 | 2,55176E-12 |
| ACE2 | SIX5 | 0,3574 | 2,57947E-12 |
| ACE2 | LRFN5 | 0,3573 | 2,61312E-12 |
| ACE2 | KCNJ16 | 0,3573 | 2,59915E-12 |
| ACE2 | RP11-31F15.2 | 0,3572 | 2,63498E-12 |
| ACE2 | RP11-174N3.4 | 0,3572 | 2,66087E-12 |
| ACE2 | MCM9 | 0,3572 | 2,65314E-12 |
| ACE2 | POLG | 0,3572 | 2,65748E-12 |
| ACE2 | ADAMTS1 | 0,3572 | 2,64686E-12 |
| ACE2 | UXT | 0,3572 | 2,64759E-12 |

| | | | |
|---|---|---|---|
| ACE2 | AJAP1 | 0,3571 | 2,69104E-12 |
| ACE2 | TIMMDC1 | 0,3571 | 2,67036E-12 |
| ACE2 | HMGB3P14 | 0,3571 | 2,68477E-12 |
| ACE2 | RP1-102E24.10 | 0,3571 | 2,68992E-12 |
| ACE2 | MCF2L | 0,3571 | 2,6872E-12 |
| ACE2 | NOG | 0,3571 | 2,68627E-12 |
| ACE2 | GAL3ST4 | 0,357 | 2,71023E-12 |
| ACE2 | NAA30 | 0,357 | 2,71731E-12 |
| ACE2 | ZNF267 | 0,357 | 2,73323E-12 |
| ACE2 | ZNF397 | 0,357 | 2,706E-12 |
| ACE2 | B3GALT6 | 0,3569 | 2,75134E-12 |
| ACE2 | SLC44A3-AS1 | 0,3569 | 2,75643E-12 |
| ACE2 | NMT1 | 0,3569 | 2,75277E-12 |
| ACE2 | PRR26 | 0,3568 | 2,8196E-12 |
| ACE2 | RP11-142E9.1 | 0,3568 | 2,8091E-12 |
| ACE2 | IL10RA | 0,3567 | 2,85901E-12 |
| ACE2 | RP11-521B24.4 | 0,3567 | 2,83377E-12 |
| ACE2 | PCDHGA6 | 0,3566 | 2,88617E-12 |
| ACE2 | DUSP22 | 0,3566 | 2,87792E-12 |
| ACE2 | PM20D1 | 0,3565 | 2,92947E-12 |
| ACE2 | TULP4 | 0,3565 | 2,93956E-12 |
| ACE2 | PCDHGA12 | 0,3564 | 2,98871E-12 |
| ACE2 | MLEC | 0,3564 | 2,9926E-12 |
| ACE2 | RP11-78F17.1 | 0,3564 | 2,98594E-12 |
| ACE2 | TRDMT1 | 0,3563 | 3,02174E-12 |
| ACE2 | CPEB1 | 0,3563 | 3,03771E-12 |
| ACE2 | N4BP2 | 0,3562 | 3,08095E-12 |
| ACE2 | RPS3AP38 | 0,3562 | 3,05219E-12 |
| ACE2 | PTRH2 | 0,3562 | 3,05275E-12 |
| ACE2 | RP11-661O13.1 | 0,3561 | 3,1379E-12 |
| ACE2 | CD36 | 0,356 | 3,18739E-12 |
| ACE2 | TCTN3 | 0,356 | 3,18158E-12 |
| ACE2 | DDX47 | 0,356 | 3,15849E-12 |
| ACE2 | HMGN2P3 | 0,356 | 3,17315E-12 |
| ACE2 | RP11-635N19.1 | 0,356 | 3,15553E-12 |
| ACE2 | PQLC2 | 0,3559 | 3,2329E-12 |
| ACE2 | IQCJ-SCHIP1 | 0,3559 | 3,19852E-12 |
| ACE2 | ECT2 | 0,3559 | 3,23203E-12 |
| ACE2 | RP11-388P9.2 | 0,3559 | 3,21446E-12 |
| ACE2 | ZFY-AS1 | 0,3559 | 3,2221E-12 |
| ACE2 | PRDM6 | 0,3558 | 3,27343E-12 |
| ACE2 | CTD-2555A7.3 | 0,3558 | 3,24333E-12 |
| ACE2 | RBM4 | 0,3557 | 3,3175E-12 |
| ACE2 | TCF19 | 0,3556 | 3,36148E-12 |
| ACE2 | RP11-278C7.1 | 0,3556 | 3,38048E-12 |
| ACE2 | RP11-20I23.5 | 0,3556 | 3,37291E-12 |
| ACE2 | LINC00857 | 0,3554 | 3,43528E-12 |
| ACE2 | AC141586.5 | 0,3554 | 3,44801E-12 |
| ACE2 | NOBOX | 0,3552 | 3,56082E-12 |
| ACE2 | RP11-292E2.2 | 0,3551 | 3,62784E-12 |
| ACE2 | MAP7D2 | 0,3551 | 3,63443E-12 |
| ACE2 | ZNF502 | 0,355 | 3,66301E-12 |
| ACE2 | RP11-350J20.5 | 0,355 | 3,69527E-12 |
| ACE2 | DAGLA | 0,355 | 3,67444E-12 |

| | | | |
|---|---|---|---|
| ACE2 | VPS33B | 0,355 | 3,67416E-12 |
| ACE2 | ATAD5 | 0,355 | 3,67439E-12 |
| ACE2 | RPL5P30 | 0,3549 | 3,74124E-12 |
| ACE2 | LINC00216 | 0,3549 | 3,72473E-12 |
| ACE2 | CBLN4 | 0,3549 | 3,73062E-12 |
| ACE2 | MTMR9LP | 0,3548 | 3,79571E-12 |
| ACE2 | MYOZ3 | 0,3548 | 3,79362E-12 |
| ACE2 | LCN6 | 0,3548 | 3,77747E-12 |
| ACE2 | RP11-843P14.1 | 0,3547 | 3,85183E-12 |
| ACE2 | PNPLA3 | 0,3547 | 3,85382E-12 |
| ACE2 | MBOAT2 | 0,3546 | 3,91573E-12 |
| ACE2 | TOPAZ1 | 0,3546 | 3,876E-12 |
| ACE2 | AEBP1 | 0,3546 | 3,87927E-12 |
| ACE2 | IKZF1 | 0,3545 | 3,94966E-12 |
| ACE2 | LRP12 | 0,3545 | 3,92722E-12 |
| ACE2 | TPTE2P1 | 0,3545 | 3,97695E-12 |
| ACE2 | DTX2 | 0,3544 | 3,99276E-12 |
| ACE2 | RP11-121A8.1 | 0,3543 | 4,07725E-12 |
| ACE2 | LINC01816 | 0,3542 | 4,10936E-12 |
| ACE2 | CNTNAP3P2 | 0,3542 | 4,11929E-12 |
| ACE2 | VAV2 | 0,3542 | 4,1183E-12 |
| ACE2 | TIMM23B | 0,3542 | 4,10507E-12 |
| ACE2 | FAM155B | 0,3542 | 4,14132E-12 |
| ACE2 | SNHG1 | 0,3541 | 4,17679E-12 |
| ACE2 | COCH | 0,3541 | 4,21556E-12 |
| ACE2 | CCNB3 | 0,3541 | 4,21495E-12 |
| ACE2 | FZD10-AS1 | 0,354 | 4,23609E-12 |
| ACE2 | MAP2K7 | 0,354 | 4,24896E-12 |
| ACE2 | CRKL | 0,354 | 4,28973E-12 |
| ACE2 | TMEFF2 | 0,3539 | 4,32522E-12 |
| ACE2 | PHF20L1 | 0,3539 | 4,32435E-12 |
| ACE2 | FCN2 | 0,3539 | 4,30662E-12 |
| ACE2 | KIF4A | 0,3539 | 4,33616E-12 |
| ACE2 | AC007228.11 | 0,3538 | 4,38558E-12 |
| ACE2 | MT-CO2 | 0,3538 | 4,35628E-12 |
| ACE2 | COPS6 | 0,3536 | 4,53439E-12 |
| ACE2 | RP11-616K22.1 | 0,3536 | 4,49914E-12 |
| ACE2 | KCNE5 | 0,3536 | 4,53661E-12 |
| ACE2 | RP11-484D2.4 | 0,3535 | 4,61508E-12 |
| ACE2 | RP11-1026M7.2 | 0,3534 | 4,64256E-12 |
| ACE2 | SORBS1 | 0,3534 | 4,68417E-12 |
| ACE2 | MAPK8IP2 | 0,3534 | 4,64182E-12 |
| ACE2 | RP11-629O1.2 | 0,3533 | 4,72279E-12 |
| ACE2 | MS4A2 | 0,3533 | 4,69445E-12 |
| ACE2 | PALM | 0,3533 | 4,74444E-12 |
| ACE2 | SV2A | 0,3532 | 4,79695E-12 |
| ACE2 | RP11-1148L6.8 | 0,3532 | 4,76799E-12 |
| ACE2 | STAB1 | 0,3531 | 4,84846E-12 |
| ACE2 | ROBO2 | 0,3531 | 4,8919E-12 |
| ACE2 | BOD1 | 0,353 | 4,95539E-12 |
| ACE2 | EEF1B2P3 | 0,353 | 4,96684E-12 |
| ACE2 | RP11-67L14.1 | 0,3529 | 4,97498E-12 |
| ACE2 | PDGFA | 0,3529 | 4,99366E-12 |
| ACE2 | FEM1A | 0,3529 | 4,98932E-12 |

| | | | |
|---|---|---|---|
| ACE2 | TOR1AIP2 | 0,3528 | 5,05775E-12 |
| ACE2 | TAF6L | 0,3528 | 5,08606E-12 |
| ACE2 | TIMELESS | 0,3528 | 5,12211E-12 |
| ACE2 | SPRED1 | 0,3528 | 5,07188E-12 |
| ACE2 | PRDX3P1 | 0,3528 | 5,11843E-12 |
| ACE2 | RPLP0P6 | 0,3527 | 5,17134E-12 |
| ACE2 | SATB2 | 0,3527 | 5,17579E-12 |
| ACE2 | PDLIM7 | 0,3527 | 5,17795E-12 |
| ACE2 | RP11-284B18.3 | 0,3527 | 5,19571E-12 |
| ACE2 | SRPK3 | 0,3527 | 5,17866E-12 |
| ACE2 | DEPTOR | 0,3526 | 5,2405E-12 |
| ACE2 | ZNF160 | 0,3526 | 5,23805E-12 |
| ACE2 | PLA2G2A | 0,3525 | 5,29866E-12 |
| ACE2 | CCR2 | 0,3525 | 5,34022E-12 |
| ACE2 | ZNF343 | 0,3525 | 5,33696E-12 |
| ACE2 | MYH9 | 0,3525 | 5,29687E-12 |
| ACE2 | UBXN4 | 0,3524 | 5,38505E-12 |
| ACE2 | RP11-398J10.2 | 0,3524 | 5,41901E-12 |
| ACE2 | RP11-397P13.7 | 0,3523 | 5,44984E-12 |
| ACE2 | GIGYF1 | 0,3523 | 5,43675E-12 |
| ACE2 | RP11-87H9.5 | 0,3523 | 5,50778E-12 |
| ACE2 | RP11-46D6.1 | 0,3523 | 5,45388E-12 |
| ACE2 | CTC-448F2.4 | 0,3523 | 5,47042E-12 |
| ACE2 | LINC02202 | 0,3522 | 5,56622E-12 |
| ACE2 | CTD-2555O16.4 | 0,3522 | 5,55862E-12 |
| ACE2 | HMGB3P9 | 0,3521 | 5,67398E-12 |
| ACE2 | RP11-701H24.3 | 0,352 | 5,75998E-12 |
| ACE2 | MFNG | 0,352 | 5,73727E-12 |
| ACE2 | ARHGAP36 | 0,352 | 5,75651E-12 |
| ACE2 | RP11-101E13.5 | 0,3519 | 5,8412E-12 |
| ACE2 | FAM96AP2 | 0,3519 | 5,8427E-12 |
| ACE2 | CTB-127C13.1 | 0,3519 | 5,8388E-12 |
| ACE2 | C14orf1 | 0,3519 | 5,76923E-12 |
| ACE2 | LOC100421166 | 0,3519 | 5,82058E-12 |
| ACE2 | NASP | 0,3518 | 5,85901E-12 |
| ACE2 | TTC3 | 0,3518 | 5,86403E-12 |
| ACE2 | GHDC | 0,3517 | 5,98198E-12 |
| ACE2 | EIF4A2P1 | 0,3517 | 6,00756E-12 |
| ACE2 | PNKP | 0,3517 | 5,93682E-12 |
| ACE2 | FERP1 | 0,3517 | 6,01925E-12 |
| ACE2 | MARK1 | 0,3516 | 6,06627E-12 |
| ACE2 | ARHGAP19 | 0,3516 | 6,06542E-12 |
| ACE2 | RP11-260M19.2 | 0,3516 | 6,10756E-12 |
| ACE2 | RP11-488L18.10 | 0,3515 | 6,1623E-12 |
| ACE2 | GPR146 | 0,3515 | 6,17935E-12 |
| ACE2 | RP11-98F14.12 | 0,3515 | 6,17245E-12 |
| ACE2 | MGAT2 | 0,3515 | 6,17238E-12 |
| ACE2 | CTD-2012K14.5 | 0,3515 | 6,11593E-12 |
| ACE2 | FBXO47 | 0,3515 | 6,16347E-12 |
| ACE2 | FASTK | 0,3514 | 6,24844E-12 |
| ACE2 | ZNF280B | 0,3514 | 6,26976E-12 |
| ACE2 | IGF2-AS | 0,3513 | 6,31382E-12 |
| ACE2 | PDE9A | 0,3513 | 6,32772E-12 |
| ACE2 | FOXR2 | 0,3513 | 6,29463E-12 |

| | | | |
|---|---|---|---|
| ACE2 | TFP1 | 0,3512 | 6,40954E-12 |
| ACE2 | EFNA4 | 0,3511 | 6,56156E-12 |
| ACE2 | BANF1 | 0,3511 | 6,57531E-12 |
| ACE2 | NOD2 | 0,3511 | 6,55672E-12 |
| ACE2 | EEF1A1P30 | 0,3511 | 6,56703E-12 |
| ACE2 | GRIA3 | 0,3511 | 6,50577E-12 |
| ACE2 | MYH10 | 0,351 | 6,62794E-12 |
| ACE2 | FLOT2 | 0,351 | 6,66204E-12 |
| ACE2 | ZNF224 | 0,351 | 6,65055E-12 |
| ACE2 | LINC00173 | 0,3508 | 6,85289E-12 |
| ACE2 | PNMAL2 | 0,3508 | 6,8291E-12 |
| ACE2 | RP11-386G11.11 | 0,3507 | 6,90251E-12 |
| ACE2 | AC009120.3 | 0,3507 | 6,93189E-12 |
| ACE2 | SDAD1 | 0,3506 | 7,06197E-12 |
| ACE2 | RTN4IP1 | 0,3506 | 7,04503E-12 |
| ACE2 | METTL25 | 0,3506 | 6,98101E-12 |
| ACE2 | MIPEP | 0,3506 | 7,03194E-12 |
| ACE2 | CNPPD1 | 0,3505 | 7,10155E-12 |
| ACE2 | PGF | 0,3505 | 7,17273E-12 |
| ACE2 | S100A6 | 0,3504 | 7,25206E-12 |
| ACE2 | S100A1 | 0,3504 | 7,2261E-12 |
| ACE2 | VPS54 | 0,3504 | 7,24957E-12 |
| ACE2 | RP11-480I12.5 | 0,3503 | 7,38353E-12 |
| ACE2 | RP11-5C23.2 | 0,3503 | 7,37864E-12 |
| ACE2 | RPL12P9 | 0,3503 | 7,33946E-12 |
| ACE2 | RP11-513M16.8 | 0,3502 | 7,48971E-12 |
| ACE2 | SLC25A25-AS1 | 0,3502 | 7,42511E-12 |
| ACE2 | VPS37D | 0,3501 | 7,52858E-12 |
| ACE2 | ZNF580 | 0,3501 | 7,58127E-12 |
| ACE2 | CTB-131K11.1 | 0,35 | 7,64558E-12 |
| ACE2 | ZNF615 | 0,35 | 7,70759E-12 |
| ACE2 | GHR | 0,3499 | 7,80741E-12 |
| ACE2 | MARCH5 | 0,3499 | 7,80439E-12 |
| ACE2 | LINC00371 | 0,3499 | 7,78579E-12 |
| ACE2 | ASB16-AS1 | 0,3499 | 7,79602E-12 |
| ACE2 | SS18L2 | 0,3498 | 7,88589E-12 |
| ACE2 | CD74 | 0,3497 | 8,03575E-12 |
| ACE2 | TMEM9 | 0,3496 | 8,11675E-12 |
| ACE2 | CHRNA5 | 0,3496 | 8,08901E-12 |
| ACE2 | KIFAP3 | 0,3495 | 8,21545E-12 |
| ACE2 | RP11-195B17.1 | 0,3495 | 8,23133E-12 |
| ACE2 | MARS2 | 0,3495 | 8,27751E-12 |
| ACE2 | RP11-227H15.4 | 0,3495 | 8,28674E-12 |
| ACE2 | HLA-DPB1 | 0,3494 | 8,31227E-12 |
| ACE2 | MCOLN3 | 0,3493 | 8,51532E-12 |
| ACE2 | RP11-350G8.5 | 0,3493 | 8,54572E-12 |
| ACE2 | HOXA4 | 0,3493 | 8,46418E-12 |
| ACE2 | AC008746.5 | 0,3493 | 8,49361E-12 |
| ACE2 | MPV17 | 0,3492 | 8,62969E-12 |
| ACE2 | FABP6 | 0,3492 | 8,58498E-12 |
| ACE2 | CTD-2270F17.1 | 0,3492 | 8,652E-12 |
| ACE2 | RP11-231C14.6 | 0,3492 | 8,666E-12 |
| ACE2 | A4GALT | 0,3492 | 8,60197E-12 |
| ACE2 | CTD-2517O10.5 | 0,3491 | 8,7034E-12 |

| | | | |
|---|---|---|---|
| ACE2 | RP4-800G7.2 | 0,3491 | 8,79825E-12 |
| ACE2 | CSTA | 0,3489 | 9,05691E-12 |
| ACE2 | CXorf40B | 0,3488 | 9,11572E-12 |
| ACE2 | C1orf122 | 0,3487 | 9,20639E-12 |
| ACE2 | CES3 | 0,3487 | 9,25792E-12 |
| ACE2 | ZBED6CL | 0,3486 | 9,37895E-12 |
| ACE2 | RP11-180I4.4 | 0,3486 | 9,43216E-12 |
| ACE2 | RPS19 | 0,3486 | 9,34242E-12 |
| ACE2 | RP11-29H23.7 | 0,3485 | 9,48992E-12 |
| ACE2 | HIGD2A | 0,3485 | 9,59122E-12 |
| ACE2 | PSMB8-AS1 | 0,3485 | 9,5862E-12 |
| ACE2 | WNT5B | 0,3484 | 9,68269E-12 |
| ACE2 | HOXC9 | 0,3484 | 9,73203E-12 |
| ACE2 | LINC01719 | 0,3483 | 9,87501E-12 |
| ACE2 | PI4KB | 0,3483 | 9,88729E-12 |
| ACE2 | MED27 | 0,3483 | 9,85918E-12 |
| ACE2 | RNGTTP1 | 0,3483 | 9,76903E-12 |
| ACE2 | MARCH1 | 0,3482 | 9,96733E-12 |
| ACE2 | TTTY14 | 0,3482 | 9,91126E-12 |
| ACE2 | RASSF5 | 0,3481 | 1,01298E-11 |
| ACE2 | TRBC2 | 0,3481 | 1,00775E-11 |
| ACE2 | ABHD15 | 0,3481 | 1,01005E-11 |
| ACE2 | RP11-178C3.6 | 0,3481 | 1,00629E-11 |
| ACE2 | CTC-325H20.8 | 0,348 | 1,02249E-11 |
| ACE2 | ARFGAP3 | 0,348 | 1,02113E-11 |
| ACE2 | COBL | 0,3479 | 1,04499E-11 |
| ACE2 | ITGA5 | 0,3479 | 1,04715E-11 |
| ACE2 | LINC00485 | 0,3479 | 1,0469E-11 |
| ACE2 | PDK3 | 0,3479 | 1,04699E-11 |
| ACE2 | HOXD10 | 0,3477 | 1,06636E-11 |
| ACE2 | ZNF280A | 0,3477 | 1,07062E-11 |
| ACE2 | OLFML3 | 0,3476 | 1,08525E-11 |
| ACE2 | BRCA2 | 0,3476 | 1,08965E-11 |
| ACE2 | SIAH1P1 | 0,3476 | 1,0894E-11 |
| ACE2 | RP11-490M8.1 | 0,3475 | 1,09846E-11 |
| ACE2 | DTD1 | 0,3475 | 1,10862E-11 |
| ACE2 | TMEM240 | 0,3474 | 1,12327E-11 |
| ACE2 | LGMNP1 | 0,3473 | 1,13965E-11 |
| ACE2 | ORAOV1 | 0,3472 | 1,14826E-11 |
| ACE2 | RP11-728F11.4 | 0,3471 | 1,16606E-11 |
| ACE2 | SAMSN1 | 0,3471 | 1,17634E-11 |
| ACE2 | GTF2H2C | 0,347 | 1,18228E-11 |
| ACE2 | LIN52 | 0,347 | 1,18231E-11 |
| ACE2 | RHOXF2 | 0,347 | 1,19025E-11 |
| ACE2 | TOMM70 | 0,3469 | 1,20237E-11 |
| ACE2 | GSC2 | 0,3468 | 1,22182E-11 |
| ACE2 | RP11-537E18.1 | 0,3467 | 1,2384E-11 |
| ACE2 | GFI1 | 0,3466 | 1,26096E-11 |
| ACE2 | RP11-448G15.3 | 0,3465 | 1,28294E-11 |
| ACE2 | TBX21 | 0,3465 | 1,26587E-11 |
| ACE2 | VPS72 | 0,3464 | 1,29662E-11 |
| ACE2 | RP11-549J18.1 | 0,3464 | 1,29567E-11 |
| ACE2 | GGTA1P | 0,3464 | 1,28458E-11 |
| ACE2 | TRIT1 | 0,3463 | 1,31912E-11 |

| | | | |
|---|---|---|---|
| ACE2 | INTS3 | 0,3463 | 1,31369E-11 |
| ACE2 | RUFY1 | 0,3463 | 1,31105E-11 |
| ACE2 | CH17-13I23.3 | 0,3462 | 1,32737E-11 |
| ACE2 | PHF19 | 0,3462 | 1,33075E-11 |
| ACE2 | ABHD13 | 0,3461 | 1,34535E-11 |
| ACE2 | HDHD5 | 0,3461 | 1,35351E-11 |
| ACE2 | RGS18 | 0,346 | 1,36563E-11 |
| ACE2 | ENPP7P10 | 0,346 | 1,37724E-11 |
| ACE2 | ZRSR1 | 0,346 | 1,36364E-11 |
| ACE2 | TRAF3 | 0,346 | 1,36085E-11 |
| ACE2 | FAM19A5 | 0,346 | 1,36912E-11 |
| ACE2 | POLM | 0,3459 | 1,39433E-11 |
| ACE2 | GPR183 | 0,3459 | 1,38926E-11 |
| ACE2 | JAG2 | 0,3458 | 1,41831E-11 |
| ACE2 | LINC01160 | 0,3457 | 1,43079E-11 |
| ACE2 | AIF1L | 0,3457 | 1,42925E-11 |
| ACE2 | CHMP6 | 0,3457 | 1,42359E-11 |
| ACE2 | ZNF444P1 | 0,3456 | 1,45298E-11 |
| ACE2 | ZNF524 | 0,3456 | 1,458E-11 |
| ACE2 | LINC01647 | 0,3455 | 1,47373E-11 |
| ACE2 | ELFN1 | 0,3455 | 1,46594E-11 |
| ACE2 | LDLR | 0,3454 | 1,48978E-11 |
| ACE2 | BPNT1 | 0,3453 | 1,50773E-11 |
| ACE2 | MTERF4 | 0,3452 | 1,54132E-11 |
| ACE2 | RP11-215P8.3 | 0,3452 | 1,54493E-11 |
| ACE2 | PACSIN3 | 0,3452 | 1,54538E-11 |
| ACE2 | MED18 | 0,3451 | 1,56895E-11 |
| ACE2 | FAM185A | 0,3451 | 1,5532E-11 |
| ACE2 | SQLE | 0,3451 | 1,55752E-11 |
| ACE2 | ZNF385A | 0,3451 | 1,55157E-11 |
| ACE2 | ENTPD3 | 0,345 | 1,57779E-11 |
| ACE2 | POLR2H | 0,345 | 1,5831E-11 |
| ACE2 | STON2 | 0,3449 | 1,61378E-11 |
| ACE2 | RP6-24A23.7 | 0,3449 | 1,61235E-11 |
| ACE2 | RP11-342K6.2 | 0,3448 | 1,62641E-11 |
| ACE2 | TOGARAM1 | 0,3448 | 1,63465E-11 |
| ACE2 | RP11-1070N10.3 | 0,3448 | 1,62399E-11 |
| ACE2 | ADRA2B | 0,3447 | 1,64051E-11 |
| ACE2 | SRCIN1 | 0,3447 | 1,65076E-11 |
| ACE2 | C18orf63 | 0,3447 | 1,65762E-11 |
| ACE2 | AP000692.9 | 0,3447 | 1,64226E-11 |
| ACE2 | BTK | 0,3447 | 1,65052E-11 |
| ACE2 | LRRC32 | 0,3446 | 1,66987E-11 |
| ACE2 | CSPG4 | 0,3445 | 1,68826E-11 |
| ACE2 | ZNF682 | 0,3445 | 1,70115E-11 |
| ACE2 | ZFAND2B | 0,3443 | 1,75537E-11 |
| ACE2 | U91328.19 | 0,3443 | 1,75164E-11 |
| ACE2 | AKR1C8P | 0,3443 | 1,74517E-11 |
| ACE2 | PALMD | 0,3442 | 1,76467E-11 |
| ACE2 | RP11-660L16.2 | 0,3442 | 1,76668E-11 |
| ACE2 | RP11-77K12.3 | 0,3442 | 1,75947E-11 |
| ACE2 | LINC01793 | 0,3441 | 1,80156E-11 |
| ACE2 | EIF3E | 0,3441 | 1,80126E-11 |
| ACE2 | RP11-262A16.1 | 0,3441 | 1,79469E-11 |

| | | | |
|---|---|---|---|
| ACE2 | NECAP2 | 0,344 | 1,83533E-11 |
| ACE2 | SNHG5 | 0,344 | 1,83359E-11 |
| ACE2 | PALD1 | 0,344 | 1,82952E-11 |
| ACE2 | PNPLA2 | 0,344 | 1,81937E-11 |
| ACE2 | SECTM1 | 0,344 | 1,81867E-11 |
| ACE2 | RP11-218C14.8 | 0,3439 | 1,8596E-11 |
| ACE2 | C17orf62 | 0,3438 | 1,88121E-11 |
| ACE2 | CTA-228A9.3 | 0,3438 | 1,86822E-11 |
| ACE2 | EDEM3 | 0,3437 | 1,9154E-11 |
| ACE2 | FZD8 | 0,3437 | 1,90689E-11 |
| ACE2 | PPP4R3A | 0,3437 | 1,90227E-11 |
| ACE2 | RP11-35G9.5 | 0,3437 | 1,91197E-11 |
| ACE2 | RP1-290I10.5 | 0,3436 | 1,91712E-11 |
| ACE2 | SAGE1 | 0,3436 | 1,93994E-11 |
| ACE2 | SNRPF | 0,3435 | 1,96486E-11 |
| ACE2 | FMNL2 | 0,3434 | 1,98151E-11 |
| ACE2 | RP11-81K2.1 | 0,3434 | 1,97922E-11 |
| ACE2 | TGFBR3 | 0,3433 | 2,01531E-11 |
| ACE2 | RP5-827C21.2 | 0,3433 | 2,02048E-11 |
| ACE2 | RP11-266K4.13 | 0,3433 | 2,00743E-11 |
| ACE2 | TNFSF13B | 0,3432 | 2,0544E-11 |
| ACE2 | NME3 | 0,3432 | 2,03619E-11 |
| ACE2 | RTCA-AS1 | 0,3431 | 2,06832E-11 |
| ACE2 | FABP7P2 | 0,3431 | 2,06214E-11 |
| ACE2 | SPOCD1 | 0,343 | 2,10032E-11 |
| ACE2 | PRPF3 | 0,343 | 2,09433E-11 |
| ACE2 | RP11-121C2.2 | 0,343 | 2,1053E-11 |
| ACE2 | BTBD7P1 | 0,343 | 2,10746E-11 |
| ACE2 | CREB3L2 | 0,3429 | 2,14047E-11 |
| ACE2 | C8orf46 | 0,3429 | 2,11829E-11 |
| ACE2 | RP11-17G12.2 | 0,3429 | 2,12865E-11 |
| ACE2 | RP11-57H12.3 | 0,3428 | 2,15763E-11 |
| ACE2 | EEF1A1P11 | 0,3428 | 2,16712E-11 |
| ACE2 | ZNF425 | 0,3428 | 2,17633E-11 |
| ACE2 | HOXC8 | 0,3428 | 2,16404E-11 |
| ACE2 | AC009095.4 | 0,3428 | 2,15795E-11 |
| ACE2 | AC083899.3 | 0,3426 | 2,21991E-11 |
| ACE2 | RPS2P7 | 0,3426 | 2,21685E-11 |
| ACE2 | STYK1 | 0,3425 | 2,24199E-11 |
| ACE2 | RP11-565F19.2 | 0,3425 | 2,25753E-11 |
| ACE2 | RP11-497G19.3 | 0,3424 | 2,28888E-11 |
| ACE2 | CTB-25B13.12 | 0,3424 | 2,28895E-11 |
| ACE2 | CTD-2231E14.2 | 0,3424 | 2,27751E-11 |
| ACE2 | CTC-448F2.6 | 0,3424 | 2,297E-11 |
| ACE2 | ZNF492 | 0,3423 | 2,3354E-11 |
| ACE2 | XKRX | 0,3423 | 2,30821E-11 |
| ACE2 | TAGAP | 0,3422 | 2,3562E-11 |
| ACE2 | CTD-3236F5.1 | 0,3422 | 2,35302E-11 |
| ACE2 | RP11-172H24.4 | 0,3422 | 2,35282E-11 |
| ACE2 | RP11-498C9.15 | 0,3422 | 2,35683E-11 |
| ACE2 | ATF4 | 0,3421 | 2,38307E-11 |
| ACE2 | HMGCR | 0,342 | 2,42572E-11 |
| ACE2 | TFAP2C | 0,3419 | 2,45271E-11 |
| ACE2 | MIA | 0,3418 | 2,49983E-11 |

| | | | |
|---|---|---|---|
| ACE2 | PNMA6A | 0,3417 | 2,52603E-11 |
| ACE2 | BARHL2 | 0,3415 | 2,60272E-11 |
| ACE2 | RP11-259N19.1 | 0,3415 | 2,60455E-11 |
| ACE2 | RN7SL689P | 0,3415 | 2,59575E-11 |
| ACE2 | RN7SL481P | 0,3415 | 2,59031E-11 |
| ACE2 | MIR202HG | 0,3415 | 2,59624E-11 |
| ACE2 | AC025165.8 | 0,3415 | 2,60657E-11 |
| ACE2 | EEF1AKMT1 | 0,3415 | 2,61422E-11 |
| ACE2 | CTB-134H23.3 | 0,3415 | 2,58862E-11 |
| ACE2 | RPS28 | 0,3415 | 2,6184E-11 |
| ACE2 | RSBN1 | 0,3414 | 2,62854E-11 |
| ACE2 | VLDLR | 0,3414 | 2,65739E-11 |
| ACE2 | HMGB1P1 | 0,3414 | 2,62886E-11 |
| ACE2 | ADCY2 | 0,3413 | 2,67864E-11 |
| ACE2 | AP000648.5 | 0,3413 | 2,68248E-11 |
| ACE2 | SPAG7 | 0,3413 | 2,66879E-11 |
| ACE2 | MT-RNR2 | 0,3413 | 2,69084E-11 |
| ACE2 | ARHGEF19 | 0,3412 | 2,72973E-11 |
| ACE2 | WARS2-IT1 | 0,3412 | 2,70896E-11 |
| ACE2 | RP11-818O24.3 | 0,3411 | 2,73973E-11 |
| ACE2 | CXorf67 | 0,3411 | 2,75707E-11 |
| ACE2 | LINC01587 | 0,341 | 2,79454E-11 |
| ACE2 | KDELR3 | 0,341 | 2,80582E-11 |
| ACE2 | RP11-177B4.1 | 0,3409 | 2,8485E-11 |
| ACE2 | COLCA1 | 0,3409 | 2,81504E-11 |
| ACE2 | LPCAT4 | 0,3409 | 2,85167E-11 |
| ACE2 | CTD-2547G23.4 | 0,3409 | 2,81273E-11 |
| ACE2 | NAP1L4P1 | 0,3408 | 2,87264E-11 |
| ACE2 | ARRB1 | 0,3408 | 2,87444E-11 |
| ACE2 | GAGE1 | 0,3408 | 2,85405E-11 |
| ACE2 | RP3-337H4.8 | 0,3407 | 2,92579E-11 |
| ACE2 | SGCG | 0,3407 | 2,9141E-11 |
| ACE2 | RP4-613B23.1 | 0,3406 | 2,93681E-11 |
| ACE2 | CSK | 0,3406 | 2,96552E-11 |
| ACE2 | PLAUR | 0,3406 | 2,94183E-11 |
| ACE2 | RHOXF1P3 | 0,3406 | 2,93568E-11 |
| ACE2 | ADAMTSL4 | 0,3402 | 3,1274E-11 |
| ACE2 | EFNA3 | 0,3402 | 3,14231E-11 |
| ACE2 | SCARA3 | 0,3402 | 3,14017E-11 |
| ACE2 | CPSF2 | 0,3402 | 3,13241E-11 |
| ACE2 | ADA | 0,3402 | 3,13991E-11 |
| ACE2 | E2F5 | 0,3401 | 3,16332E-11 |
| ACE2 | GACAT2 | 0,3401 | 3,17269E-11 |
| ACE2 | DNTTIP2 | 0,34 | 3,22023E-11 |
| ACE2 | MIR199A2 | 0,34 | 3,19396E-11 |
| ACE2 | RP11-297J22.1 | 0,34 | 3,20604E-11 |
| ACE2 | PRSS35 | 0,34 | 3,19615E-11 |
| ACE2 | ACBD3-AS1 | 0,3399 | 3,26528E-11 |
| ACE2 | TNFSF8 | 0,3399 | 3,2522E-11 |
| ACE2 | CTD-2213F21.2 | 0,3399 | 3,27133E-11 |
| ACE2 | RNPS1P1 | 0,3398 | 3,3092E-11 |
| ACE2 | RP11-861E21.2 | 0,3398 | 3,29089E-11 |
| ACE2 | KCNS1 | 0,3398 | 3,32002E-11 |
| ACE2 | STEAP2 | 0,3397 | 3,33891E-11 |

| | | | |
|---|---|---|---|
| ACE2 | ANKRD10 | 0,3397 | 3,36166E-11 |
| ACE2 | FKBP10 | 0,3397 | 3,34918E-11 |
| ACE2 | LINC01909 | 0,3397 | 3,34013E-11 |
| ACE2 | CDC42BPA | 0,3396 | 3,41983E-11 |
| ACE2 | ERI2 | 0,3396 | 3,39796E-11 |
| ACE2 | RPL3 | 0,3396 | 3,39041E-11 |
| ACE2 | SLC11A1 | 0,3395 | 3,43342E-11 |
| ACE2 | FAM13A-AS1 | 0,3395 | 3,47325E-11 |
| ACE2 | CCDC152 | 0,3395 | 3,44603E-11 |
| ACE2 | URM1 | 0,3394 | 3,50826E-11 |
| ACE2 | LA16c-316G12.2 | 0,3394 | 3,48995E-11 |
| ACE2 | LIN7B | 0,3394 | 3,49682E-11 |
| ACE2 | RPS5P2 | 0,3394 | 3,47772E-11 |
| ACE2 | RFXANK | 0,3393 | 3,52611E-11 |
| ACE2 | AC007193.9 | 0,3393 | 3,55086E-11 |
| ACE2 | FAM236A | 0,3393 | 3,5423E-11 |
| ACE2 | PAN3-AS1 | 0,3392 | 3,61563E-11 |
| ACE2 | PSMD12 | 0,3392 | 3,58217E-11 |
| ACE2 | SIGLEC1 | 0,3392 | 3,59557E-11 |
| ACE2 | SV2C | 0,3391 | 3,66945E-11 |
| ACE2 | ZNF84 | 0,3391 | 3,64182E-11 |
| ACE2 | AP003068.23 | 0,339 | 3,7024E-11 |
| ACE2 | TBXA2R | 0,339 | 3,69859E-11 |
| ACE2 | AC093609.1 | 0,3389 | 3,72894E-11 |
| ACE2 | RP11-1102P22.1 | 0,3389 | 3,73915E-11 |
| ACE2 | SERTAD4-AS1 | 0,3388 | 3,81377E-11 |
| ACE2 | MEIS2 | 0,3388 | 3,81808E-11 |
| ACE2 | OAF | 0,3387 | 3,87682E-11 |
| ACE2 | RP1-197B17.5 | 0,3387 | 3,87779E-11 |
| ACE2 | PTOV1-AS2 | 0,3387 | 3,85184E-11 |
| ACE2 | NFKBIL1 | 0,3386 | 3,91081E-11 |
| ACE2 | MT1M | 0,3386 | 3,89876E-11 |
| ACE2 | UBXN7 | 0,3385 | 3,97382E-11 |
| ACE2 | RP11-344N10.5 | 0,3385 | 3,97878E-11 |
| ACE2 | IRX3 | 0,3385 | 3,95185E-11 |
| ACE2 | DUTP6 | 0,3384 | 4,00116E-11 |
| ACE2 | PSD4 | 0,3384 | 4,00897E-11 |
| ACE2 | HIST1H2BN | 0,3384 | 4,05188E-11 |
| ACE2 | CTD-2516F10.2 | 0,3384 | 4,01677E-11 |
| ACE2 | LINC01697 | 0,3384 | 4,00008E-11 |
| ACE2 | PRKAG2-AS1 | 0,3383 | 4,1054E-11 |
| ACE2 | PSIP1 | 0,3383 | 4,06012E-11 |
| ACE2 | HERC2P9 | 0,3383 | 4,09891E-11 |
| ACE2 | TCF4 | 0,3383 | 4,10225E-11 |
| ACE2 | RP11-148L24.1 | 0,3382 | 4,16873E-11 |
| ACE2 | ZNF503-AS2 | 0,3382 | 4,13776E-11 |
| ACE2 | PTK6 | 0,3382 | 4,1233E-11 |
| ACE2 | CTD-2377O17.1 | 0,3381 | 4,17363E-11 |
| ACE2 | HLA-B | 0,3381 | 4,21046E-11 |
| ACE2 | CDC6 | 0,3381 | 4,21985E-11 |
| ACE2 | RP3-468K18.6 | 0,338 | 4,28647E-11 |
| ACE2 | P2RY14 | 0,3379 | 4,28851E-11 |
| ACE2 | SLC7A11 | 0,3379 | 4,33388E-11 |
| ACE2 | GALT | 0,3379 | 4,33404E-11 |

| | | | |
|---|---|---|---|
| ACE2 | RP11-1084A12.1 | 0,3379 | 4,31308E-11 |
| ACE2 | BNIP3P25 | 0,3379 | 4,32722E-11 |
| ACE2 | AC093818.1 | 0,3378 | 4,39459E-11 |
| ACE2 | TIMM22 | 0,3378 | 4,37545E-11 |
| ACE2 | KAT2A | 0,3378 | 4,3766E-11 |
| ACE2 | BCL3 | 0,3378 | 4,36887E-11 |
| ACE2 | TINAGL1 | 0,3377 | 4,4472E-11 |
| ACE2 | CTB-33G10.1 | 0,3377 | 4,43323E-11 |
| ACE2 | LINC01694 | 0,3376 | 4,52571E-11 |
| ACE2 | RP11-84A19.4 | 0,3375 | 4,56234E-11 |
| ACE2 | KYAT1 | 0,3375 | 4,56918E-11 |
| ACE2 | CTD-2179L22.1 | 0,3374 | 4,59966E-11 |
| ACE2 | UGT8 | 0,3373 | 4,66313E-11 |
| ACE2 | RP11-1134I14.4 | 0,3373 | 4,68071E-11 |
| ACE2 | RCC2 | 0,3372 | 4,74064E-11 |
| ACE2 | EXO5 | 0,3371 | 4,85145E-11 |
| ACE2 | SP8 | 0,3371 | 4,79513E-11 |
| ACE2 | KDM8 | 0,3371 | 4,81043E-11 |
| ACE2 | PROB1 | 0,3369 | 4,98192E-11 |
| ACE2 | ERICH1 | 0,3368 | 5,0668E-11 |
| ACE2 | RP11-727A23.4 | 0,3368 | 5,06707E-11 |
| ACE2 | SNORA71B | 0,3368 | 5,01991E-11 |
| ACE2 | HOXC13 | 0,3367 | 5,09164E-11 |
| ACE2 | SLC30A4 | 0,3367 | 5,12649E-11 |
| ACE2 | POLR3G | 0,3366 | 5,14134E-11 |
| ACE2 | DCAF4L2 | 0,3365 | 5,21872E-11 |
| ACE2 | LINC00461 | 0,3364 | 5,29226E-11 |
| ACE2 | AMHR2 | 0,3364 | 5,33135E-11 |
| ACE2 | RBM11 | 0,3364 | 5,3179E-11 |
| ACE2 | FCN3 | 0,3363 | 5,40607E-11 |
| ACE2 | HMGN2P5 | 0,3363 | 5,36321E-11 |
| ACE2 | MAFF | 0,3362 | 5,4761E-11 |
| ACE2 | AC004691.5 | 0,3361 | 5,5808E-11 |
| ACE2 | RP11-552F3.9 | 0,3361 | 5,52133E-11 |
| ACE2 | RP11-1365D11.1 | 0,336 | 5,60795E-11 |
| ACE2 | VEGFC | 0,336 | 5,60449E-11 |
| ACE2 | RNF34 | 0,336 | 5,66127E-11 |
| ACE2 | IL2RA | 0,3359 | 5,72499E-11 |
| ACE2 | CARD16 | 0,3358 | 5,75281E-11 |
| ACE2 | RP11-274B21.10 | 0,3357 | 5,90131E-11 |
| ACE2 | SLC13A5 | 0,3357 | 5,84233E-11 |
| ACE2 | CTSL | 0,3356 | 5,91715E-11 |
| ACE2 | FOXF1 | 0,3356 | 5,90586E-11 |
| ACE2 | CTD-2095E4.4 | 0,3356 | 5,92551E-11 |
| ACE2 | CD302 | 0,3355 | 6,00887E-11 |
| ACE2 | DBNL | 0,3355 | 6,00408E-11 |
| ACE2 | NOX4 | 0,3354 | 6,08308E-11 |
| ACE2 | LINC01148 | 0,3353 | 6,18044E-11 |
| ACE2 | CCDC102A | 0,3353 | 6,1626E-11 |
| ACE2 | ZNF497 | 0,3352 | 6,31317E-11 |
| ACE2 | RNPC3 | 0,3351 | 6,36804E-11 |
| ACE2 | SPRR2F | 0,3351 | 6,4012E-11 |
| ACE2 | FBXO17 | 0,3351 | 6,40914E-11 |
| ACE2 | RP11-250B2.5 | 0,3349 | 6,50981E-11 |

| | | | |
|---|---|---|---|
| ACE2 | IMPDH1 | 0,3349 | 6,59622E-11 |
| ACE2 | LINC01028 | 0,3349 | 6,56847E-11 |
| ACE2 | IL1RAPL2 | 0,3349 | 6,55121E-11 |
| ACE2 | RP11-264B17.4 | 0,3348 | 6,64564E-11 |
| ACE2 | LINC00894 | 0,3348 | 6,62447E-11 |
| ACE2 | FKBP2 | 0,3347 | 6,74038E-11 |
| ACE2 | VPS4B | 0,3347 | 6,75796E-11 |
| ACE2 | SCN1B | 0,3347 | 6,74923E-11 |
| ACE2 | VPS16 | 0,3347 | 6,73053E-11 |
| ACE2 | ADARB1 | 0,3347 | 6,74147E-11 |
| ACE2 | RP11-855A2.2 | 0,3346 | 6,78317E-11 |
| ACE2 | TBC1D14 | 0,3345 | 6,90886E-11 |
| ACE2 | RP11-502M1.2 | 0,3345 | 6,9612E-11 |
| ACE2 | TCIRG1 | 0,3344 | 7,03366E-11 |
| ACE2 | OR7E47P | 0,3344 | 7,0271E-11 |
| ACE2 | RP11-484D2.3 | 0,3342 | 7,22164E-11 |
| ACE2 | L3MBTL4-AS1 | 0,3342 | 7,21124E-11 |
| ACE2 | SRF | 0,3341 | 7,27387E-11 |
| ACE2 | STAM-AS1 | 0,3341 | 7,28616E-11 |
| ACE2 | RNU6-1016P | 0,334 | 7,37459E-11 |
| ACE2 | RP11-475I24.3 | 0,334 | 7,37431E-11 |
| ACE2 | RP11-294J22.7 | 0,334 | 7,37148E-11 |
| ACE2 | FAM65C | 0,334 | 7,38763E-11 |
| ACE2 | SH3RF3 | 0,3339 | 7,48711E-11 |
| ACE2 | ANKS6 | 0,3339 | 7,52957E-11 |
| ACE2 | ETV6 | 0,3339 | 7,51558E-11 |
| ACE2 | CTD-2325P2.4 | 0,3339 | 7,51397E-11 |
| ACE2 | PISD | 0,3339 | 7,56969E-11 |
| ACE2 | ILKAP | 0,3337 | 7,78256E-11 |
| ACE2 | ROBO1 | 0,3337 | 7,68628E-11 |
| ACE2 | RP11-571I18.4 | 0,3337 | 7,72889E-11 |
| ACE2 | SDR39U1 | 0,3337 | 7,69652E-11 |
| ACE2 | RTN2 | 0,3337 | 7,68403E-11 |
| ACE2 | CLDN18 | 0,3336 | 7,81352E-11 |
| ACE2 | MARCO | 0,3335 | 7,92228E-11 |
| ACE2 | HNRNPA1P35 | 0,3335 | 7,9473E-11 |
| ACE2 | GMDS | 0,3335 | 7,99748E-11 |
| ACE2 | DCTN3 | 0,3335 | 7,89697E-11 |
| ACE2 | SLC27A6 | 0,3334 | 8,04859E-11 |
| ACE2 | NTAN1P2 | 0,3334 | 8,10573E-11 |
| ACE2 | WHAMM | 0,3334 | 8,08805E-11 |
| ACE2 | ZFP36 | 0,3334 | 8,01584E-11 |
| ACE2 | ESYT3 | 0,3332 | 8,31028E-11 |
| ACE2 | MFSD3 | 0,3332 | 8,2271E-11 |
| ACE2 | PWAR5 | 0,333 | 8,51763E-11 |
| ACE2 | ATP6V0A1 | 0,333 | 8,56579E-11 |
| ACE2 | RPL6P27 | 0,333 | 8,52832E-11 |
| ACE2 | RP11-283G6.4 | 0,3329 | 8,66188E-11 |
| ACE2 | LL22NC03-2H8.5 | 0,3329 | 8,60033E-11 |
| ACE2 | FMO2 | 0,3328 | 8,74282E-11 |
| ACE2 | ABL2 | 0,3328 | 8,76567E-11 |
| ACE2 | TMEM8A | 0,3328 | 8,78469E-11 |
| ACE2 | SREBF1 | 0,3328 | 8,76018E-11 |
| ACE2 | C21orf91 | 0,3327 | 8,93158E-11 |

| | | | |
|---|---|---|---|
| ACE2 | TMEM155 | 0,3326 | 8,97885E-11 |
| ACE2 | CRYM-AS1 | 0,3326 | 8,97176E-11 |
| ACE2 | NEXN | 0,3325 | 9,16368E-11 |
| ACE2 | MRGPRE | 0,3325 | 9,14122E-11 |
| ACE2 | ICAM2 | 0,3325 | 9,10866E-11 |
| ACE2 | ZFYVE28 | 0,3324 | 9,19486E-11 |
| ACE2 | NPM1P27 | 0,3324 | 9,21172E-11 |
| ACE2 | TSTD2 | 0,3324 | 9,26452E-11 |
| ACE2 | CLIC6 | 0,3324 | 9,27796E-11 |
| ACE2 | B3GAT1 | 0,3323 | 9,31793E-11 |
| ACE2 | SUMO2 | 0,3323 | 9,35389E-11 |
| ACE2 | RAB8A | 0,3323 | 9,42214E-11 |
| ACE2 | HSPA12A | 0,3321 | 9,68651E-11 |
| ACE2 | CTD-2371O3.2 | 0,3321 | 9,59891E-11 |
| ACE2 | PARPBP | 0,3321 | 9,57396E-11 |
| ACE2 | LRRC4B | 0,3321 | 9,59446E-11 |
| ACE2 | TBC1D8-AS1 | 0,332 | 9,70467E-11 |
| ACE2 | ZFP62 | 0,332 | 9,75577E-11 |
| ACE2 | DUSP9 | 0,332 | 9,82092E-11 |
| ACE2 | AGPS | 0,3319 | 9,86431E-11 |
| ACE2 | RP11-342L8.2 | 0,3319 | 9,90084E-11 |
| ACE2 | CARNS1 | 0,3319 | 9,96343E-11 |
| ACE2 | PPIP5K1 | 0,3318 | 1,00269E-10 |
| ACE2 | SIX2 | 0,3317 | 1,0135E-10 |
| ACE2 | RP11-65D17.1 | 0,3316 | 1,02903E-10 |
| ACE2 | SPSB3 | 0,3316 | 1,02563E-10 |
| ACE2 | KCNN1 | 0,3316 | 1,02748E-10 |
| ACE2 | BHLHE40-AS1 | 0,3315 | 1,04586E-10 |
| ACE2 | LA16c-358B7.3 | 0,3315 | 1,04031E-10 |
| ACE2 | ETS2 | 0,3315 | 1,04627E-10 |
| ACE2 | RBM34 | 0,3314 | 1,0599E-10 |
| ACE2 | AMER3 | 0,3314 | 1,06122E-10 |
| ACE2 | UBXN2B | 0,3314 | 1,0568E-10 |
| ACE2 | HNRNPA1P10 | 0,3314 | 1,05438E-10 |
| ACE2 | UPF1 | 0,3314 | 1,06456E-10 |
| ACE2 | RUNX1 | 0,3314 | 1,05606E-10 |
| ACE2 | EDA2R | 0,3314 | 1,06395E-10 |
| ACE2 | RP11-30P6.6 | 0,3313 | 1,07595E-10 |
| ACE2 | SLITRK6 | 0,3313 | 1,08114E-10 |
| ACE2 | CTD-2104P17.2 | 0,3313 | 1,07715E-10 |
| ACE2 | TTC38 | 0,3313 | 1,07398E-10 |
| ACE2 | SLC6A11 | 0,3312 | 1,08681E-10 |
| ACE2 | B4GALT7 | 0,3312 | 1,09615E-10 |
| ACE2 | ADM | 0,3312 | 1,08364E-10 |
| ACE2 | CTD-2555O16.2 | 0,3312 | 1,08254E-10 |
| ACE2 | IKBIP | 0,3311 | 1,11093E-10 |
| ACE2 | RN7SKP70 | 0,331 | 1,11834E-10 |
| ACE2 | OAS2 | 0,331 | 1,12525E-10 |
| ACE2 | IFI27L1 | 0,331 | 1,12149E-10 |
| ACE2 | PROCR | 0,331 | 1,12205E-10 |
| ACE2 | TRGV9 | 0,3309 | 1,13396E-10 |
| ACE2 | HNRNPA1P36 | 0,3309 | 1,12769E-10 |
| ACE2 | RCE1 | 0,3309 | 1,13626E-10 |
| ACE2 | HOXB9 | 0,3309 | 1,12786E-10 |

| | | | |
|---|---|---|---|
| ACE2 | TAS2R14 | 0,3308 | 1,14948E-10 |
| ACE2 | AP000254.8 | 0,3308 | 1,15023E-10 |
| ACE2 | LINC00680 | 0,3307 | 1,1735E-10 |
| ACE2 | RP11-631N16.2 | 0,3307 | 1,16995E-10 |
| ACE2 | GSC | 0,3307 | 1,16082E-10 |
| ACE2 | ZNF366 | 0,3306 | 1,18923E-10 |
| ACE2 | PCDH1 | 0,3306 | 1,17842E-10 |
| ACE2 | CLEC4M | 0,3305 | 1,2E-10 |
| ACE2 | CEBPA | 0,3305 | 1,19734E-10 |
| ACE2 | ZNF548 | 0,3305 | 1,20347E-10 |
| ACE2 | TTTY10 | 0,3305 | 1,20252E-10 |
| ACE2 | FAM175A | 0,3304 | 1,20833E-10 |
| ACE2 | PAXIP1 | 0,3304 | 1,21924E-10 |
| ACE2 | BCL11B | 0,3304 | 1,20899E-10 |
| ACE2 | STC2 | 0,3303 | 1,2354E-10 |
| ACE2 | SPANXA2-OT1 | 0,3303 | 1,23785E-10 |
| ACE2 | LINC01119 | 0,3302 | 1,24921E-10 |
| ACE2 | VCPIP1 | 0,3302 | 1,24623E-10 |
| ACE2 | EFNA2 | 0,3302 | 1,25206E-10 |
| ACE2 | CTD-3222D19.9 | 0,3302 | 1,25415E-10 |
| ACE2 | NPPB | 0,3301 | 1,27176E-10 |
| ACE2 | KIF7 | 0,3301 | 1,26304E-10 |
| ACE2 | PFKFB1 | 0,3301 | 1,25871E-10 |
| ACE2 | C1orf52 | 0,33 | 1,27767E-10 |
| ACE2 | ITGA9-AS1 | 0,33 | 1,27614E-10 |
| ACE2 | LIFR | 0,33 | 1,27461E-10 |
| ACE2 | ANGPTL2 | 0,33 | 1,27715E-10 |
| ACE2 | ALKAL1 | 0,3299 | 1,2966E-10 |
| ACE2 | RP11-217B7.2 | 0,3299 | 1,29815E-10 |
| ACE2 | SENCR | 0,3298 | 1,32387E-10 |
| ACE2 | RP11-5G9.5 | 0,3298 | 1,31972E-10 |
| ACE2 | RAB17 | 0,3297 | 1,32929E-10 |
| ACE2 | FGFBP3 | 0,3297 | 1,33924E-10 |
| ACE2 | RP1-199J3.5 | 0,3296 | 1,35123E-10 |
| ACE2 | CDH24 | 0,3296 | 1,34753E-10 |
| ACE2 | SNRPB2 | 0,3296 | 1,34847E-10 |
| ACE2 | GHET1 | 0,3295 | 1,36384E-10 |
| ACE2 | AQP3 | 0,3295 | 1,38077E-10 |
| ACE2 | TSEN2 | 0,3294 | 1,38456E-10 |
| ACE2 | RP11-630A13.4 | 0,3294 | 1,3841E-10 |
| ACE2 | AC007796.1 | 0,3294 | 1,39369E-10 |
| ACE2 | CHIT1 | 0,3293 | 1,40975E-10 |
| ACE2 | ZNF284 | 0,3292 | 1,4261E-10 |
| ACE2 | RP11-490D19.8 | 0,3291 | 1,45663E-10 |
| ACE2 | ZNF865 | 0,3291 | 1,44228E-10 |
| ACE2 | MYOM3 | 0,329 | 1,46272E-10 |
| ACE2 | HOXD9 | 0,329 | 1,46171E-10 |
| ACE2 | RP11-351J23.2 | 0,329 | 1,46918E-10 |
| ACE2 | ZNF252P-AS1 | 0,329 | 1,46184E-10 |
| ACE2 | CTD-2528L19.6 | 0,329 | 1,46231E-10 |
| ACE2 | AC026150.8 | 0,3289 | 1,49097E-10 |
| ACE2 | TNFRSF12A | 0,3289 | 1,48206E-10 |
| ACE2 | APPBP2 | 0,3289 | 1,4829E-10 |
| ACE2 | RP11-1252D15.1 | 0,3288 | 1,50198E-10 |

| | | | |
|---|---|---|---|
| ACE2 | OTULIN | 0,3288 | 1,50735E-10 |
| ACE2 | PLPP1 | 0,3288 | 1,49997E-10 |
| ACE2 | SNCG | 0,3288 | 1,50167E-10 |
| ACE2 | RP3-467L1.6 | 0,3287 | 1,53484E-10 |
| ACE2 | TFCP2L1 | 0,3286 | 1,55833E-10 |
| ACE2 | NRN1 | 0,3286 | 1,55639E-10 |
| ACE2 | CNTLN | 0,3285 | 1,56795E-10 |
| ACE2 | GRK2 | 0,3285 | 1,57516E-10 |
| ACE2 | RP11-340F14.6 | 0,3285 | 1,56884E-10 |
| ACE2 | RP11-290D2.6 | 0,3285 | 1,5759E-10 |
| ACE2 | CTD-2319I12.10 | 0,3285 | 1,57449E-10 |
| ACE2 | GPR143 | 0,3285 | 1,57387E-10 |
| ACE2 | VPS37A | 0,3284 | 1,6014E-10 |
| ACE2 | RP1-267D11.6 | 0,3284 | 1,59592E-10 |
| ACE2 | LINC01770 | 0,3283 | 1,61099E-10 |
| ACE2 | AC074289.1 | 0,3283 | 1,61828E-10 |
| ACE2 | XXbac-BPG283O16.9 | 0,3283 | 1,60846E-10 |
| ACE2 | DDX12P | 0,3283 | 1,61631E-10 |
| ACE2 | FUT8 | 0,3283 | 1,61929E-10 |
| ACE2 | MX1 | 0,3283 | 1,61891E-10 |
| ACE2 | APOL4 | 0,3283 | 1,60784E-10 |
| ACE2 | AC073063.10 | 0,328 | 1,68562E-10 |
| ACE2 | MT-CO3 | 0,3279 | 1,69741E-10 |
| ACE2 | MRPS18B | 0,3278 | 1,71851E-10 |
| ACE2 | XXyac-YX65C7_A.2 | 0,3277 | 1,75475E-10 |
| ACE2 | MYCL | 0,3276 | 1,78229E-10 |
| ACE2 | LINC01266 | 0,3276 | 1,77796E-10 |
| ACE2 | EOMES | 0,3276 | 1,78322E-10 |
| ACE2 | RP11-142A22.4 | 0,3276 | 1,7839E-10 |
| ACE2 | AC007000.10 | 0,3276 | 1,76535E-10 |
| ACE2 | CYCSP34 | 0,3276 | 1,76285E-10 |
| ACE2 | ADRA1B | 0,3275 | 1,80456E-10 |
| ACE2 | GNG4 | 0,3273 | 1,8571E-10 |
| ACE2 | CDH6 | 0,3273 | 1,85623E-10 |
| ACE2 | RP11-568G11.4 | 0,3272 | 1,87428E-10 |
| ACE2 | MXD4 | 0,3272 | 1,86122E-10 |
| ACE2 | CTC-529P8.1 | 0,3271 | 1,90083E-10 |
| ACE2 | LINC01234 | 0,327 | 1,92765E-10 |
| ACE2 | SAMD1 | 0,3269 | 1,93855E-10 |
| ACE2 | USP26 | 0,3269 | 1,94903E-10 |
| ACE2 | TRGV3 | 0,3268 | 1,97193E-10 |
| ACE2 | HMGB1P31 | 0,3267 | 1,99331E-10 |
| ACE2 | LINC02147 | 0,3267 | 1,99978E-10 |
| ACE2 | RP11-617F23.1 | 0,3267 | 2,00174E-10 |
| ACE2 | RP11-466F5.3 | 0,3266 | 2,01771E-10 |
| ACE2 | RP11-79N23.1 | 0,3266 | 2,03574E-10 |
| ACE2 | TUBG2 | 0,3266 | 2,03754E-10 |
| ACE2 | TRNP1 | 0,3265 | 2,0539E-10 |
| ACE2 | RN7SL145P | 0,3265 | 2,0597E-10 |
| ACE2 | MT-TE | 0,3265 | 2,06408E-10 |
| ACE2 | PABPC4L | 0,3264 | 2,09551E-10 |
| ACE2 | TSFM | 0,3264 | 2,07775E-10 |
| ACE2 | HCFC1R1 | 0,3264 | 2,08175E-10 |
| ACE2 | TRPM4 | 0,3264 | 2,07316E-10 |

| | | | |
|---|---|---|---|
| ACE2 | ANKZF1 | 0,3263 | 2,12026E-10 |
| ACE2 | HOXC6 | 0,3263 | 2,11009E-10 |
| ACE2 | SNX20 | 0,3263 | 2,11487E-10 |
| ACE2 | CTD-3222D19.12 | 0,3263 | 2,11111E-10 |
| ACE2 | MAPK12 | 0,3263 | 2,11661E-10 |
| ACE2 | ADH1C | 0,3262 | 2,12855E-10 |
| ACE2 | ATG2A | 0,3262 | 2,15175E-10 |
| ACE2 | RABEP2 | 0,3262 | 2,15551E-10 |
| ACE2 | ZBTB45 | 0,3262 | 2,15014E-10 |
| ACE2 | ELOA-AS1 | 0,3261 | 2,17406E-10 |
| ACE2 | DDI2 | 0,326 | 2,19663E-10 |
| ACE2 | AC068580.6 | 0,326 | 2,18697E-10 |
| ACE2 | KCNAB3 | 0,326 | 2,2113E-10 |
| ACE2 | ZNF558 | 0,326 | 2,19375E-10 |
| ACE2 | IDH3B | 0,3259 | 2,23634E-10 |
| ACE2 | LINC01448 | 0,3258 | 2,24769E-10 |
| ACE2 | RP11-863H1.1 | 0,3257 | 2,29696E-10 |
| ACE2 | TRPC6 | 0,3256 | 2,33357E-10 |
| ACE2 | PAGE2 | 0,3256 | 2,32129E-10 |
| ACE2 | GPATCH3 | 0,3254 | 2,38675E-10 |
| ACE2 | GUK1 | 0,3254 | 2,37927E-10 |
| ACE2 | PITX2 | 0,3254 | 2,37686E-10 |
| ACE2 | ELAVL3 | 0,3254 | 2,39125E-10 |
| ACE2 | NCK1-AS1 | 0,3253 | 2,40094E-10 |
| ACE2 | RP5-855D21.3 | 0,3253 | 2,4217E-10 |
| ACE2 | MAGIX | 0,3253 | 2,42159E-10 |
| ACE2 | PDSS2 | 0,3252 | 2,44688E-10 |
| ACE2 | TOX4 | 0,3252 | 2,46339E-10 |
| ACE2 | STAP2 | 0,3252 | 2,4407E-10 |
| ACE2 | PIWIL2 | 0,3251 | 2,48735E-10 |
| ACE2 | DPM2 | 0,3251 | 2,49087E-10 |
| ACE2 | OAS1 | 0,3251 | 2,47493E-10 |
| ACE2 | TNFRSF14 | 0,325 | 2,52101E-10 |
| ACE2 | UBE2Q1 | 0,3248 | 2,59821E-10 |
| ACE2 | COL15A1 | 0,3248 | 2,57165E-10 |
| ACE2 | LINC02176 | 0,3248 | 2,58705E-10 |
| ACE2 | KCNJ12 | 0,3248 | 2,5684E-10 |
| ACE2 | RP13-346H10.2 | 0,3248 | 2,59776E-10 |
| ACE2 | MTCO1P53 | 0,3248 | 2,57407E-10 |
| ACE2 | PIRT | 0,3247 | 2,62318E-10 |
| ACE2 | FAM110D | 0,3246 | 2,64072E-10 |
| ACE2 | CT55 | 0,3246 | 2,63852E-10 |
| ACE2 | PCDHGA10 | 0,3245 | 2,69975E-10 |
| ACE2 | CTD-3234P18.6 | 0,3245 | 2,69073E-10 |
| ACE2 | RRM2 | 0,3244 | 2,71073E-10 |
| ACE2 | POLDIP2 | 0,3244 | 2,70558E-10 |
| ACE2 | FTSJ3 | 0,3243 | 2,76271E-10 |
| ACE2 | ZNF346-IT1 | 0,3242 | 2,81123E-10 |
| ACE2 | SCARF1 | 0,3242 | 2,8099E-10 |
| ACE2 | MEIS3P2 | 0,3242 | 2,80189E-10 |
| ACE2 | ATP5G1 | 0,3242 | 2,79256E-10 |
| ACE2 | AP000355.2 | 0,3242 | 2,79512E-10 |
| ACE2 | TDP2 | 0,3241 | 2,82594E-10 |
| ACE2 | HEXDC-IT1 | 0,3241 | 2,82361E-10 |

| | | | |
|---|---|---|---|
| ACE2 | CHI3L1 | 0,324 | 2,8595E-10 |
| ACE2 | PYCR2 | 0,3239 | 2,89393E-10 |
| ACE2 | ASCL1 | 0,3239 | 2,89932E-10 |
| ACE2 | RP11-162D9.3 | 0,3238 | 2,96478E-10 |
| ACE2 | IL1RAPL1 | 0,3238 | 2,9459E-10 |
| ACE2 | RP11-483I13.6 | 0,3237 | 3,00789E-10 |
| ACE2 | ORC5 | 0,3237 | 2,98409E-10 |
| ACE2 | TNFRSF11B | 0,3237 | 2,97256E-10 |
| ACE2 | ZNF213-AS1 | 0,3236 | 3,04409E-10 |
| ACE2 | BCL6B | 0,3236 | 3,04759E-10 |
| ACE2 | SFRP4 | 0,3235 | 3,07024E-10 |
| ACE2 | RTN1 | 0,3235 | 3,0588E-10 |
| ACE2 | SNHG14 | 0,3234 | 3,12888E-10 |
| ACE2 | PPP1R1A | 0,3233 | 3,16281E-10 |
| ACE2 | PPL | 0,3232 | 3,19834E-10 |
| ACE2 | RP5-882C2.2 | 0,3232 | 3,19717E-10 |
| ACE2 | PAGE2B | 0,3232 | 3,18187E-10 |
| ACE2 | RP11-503N18.5 | 0,3231 | 3,22905E-10 |
| ACE2 | CTD-2517O10.6 | 0,3231 | 3,23833E-10 |
| ACE2 | RP11-23J9.6 | 0,3231 | 3,25522E-10 |
| ACE2 | TMOD3 | 0,3231 | 3,23104E-10 |
| ACE2 | RAB43P1 | 0,3231 | 3,21696E-10 |
| ACE2 | RNU6-850P | 0,323 | 3,25848E-10 |
| ACE2 | SLC16A6 | 0,323 | 3,25874E-10 |
| ACE2 | GPR1 | 0,3229 | 3,33195E-10 |
| ACE2 | POP1 | 0,3229 | 3,33635E-10 |
| ACE2 | MTBP | 0,3229 | 3,32652E-10 |
| ACE2 | RP11-662B19.2 | 0,3228 | 3,3664E-10 |
| ACE2 | TYRO3P | 0,3227 | 3,40986E-10 |
| ACE2 | TVP23A | 0,3227 | 3,39921E-10 |
| ACE2 | FTH1P24 | 0,3226 | 3,45502E-10 |
| ACE2 | RP5-1021I20.1 | 0,3226 | 3,47921E-10 |
| ACE2 | RP11-441F2.5 | 0,3226 | 3,43984E-10 |
| ACE2 | PDXP | 0,3226 | 3,43601E-10 |
| ACE2 | BTN3A1 | 0,3225 | 3,51167E-10 |
| ACE2 | CYP2G1P | 0,3225 | 3,49224E-10 |
| ACE2 | RP11-21C4.1 | 0,3224 | 3,54465E-10 |
| ACE2 | ZNF157 | 0,3224 | 3,5271E-10 |
| ACE2 | NUP107 | 0,3223 | 3,61528E-10 |
| ACE2 | SPINT1 | 0,3223 | 3,61061E-10 |
| ACE2 | DOCK6 | 0,3223 | 3,6112E-10 |
| ACE2 | ERG | 0,3223 | 3,58498E-10 |
| ACE2 | RP11-184M15.1 | 0,3222 | 3,64379E-10 |
| ACE2 | RAD52 | 0,3222 | 3,62529E-10 |
| ACE2 | AFP | 0,3221 | 3,68544E-10 |
| ACE2 | FDX2 | 0,3221 | 3,69984E-10 |
| ACE2 | TRNAU1AP | 0,322 | 3,74956E-10 |
| ACE2 | CXCL6 | 0,322 | 3,74194E-10 |
| ACE2 | PDAP1 | 0,322 | 3,72522E-10 |
| ACE2 | SULT1B1 | 0,3219 | 3,78177E-10 |
| ACE2 | HNRNPCP7 | 0,3219 | 3,78582E-10 |
| ACE2 | CTC-308K20.3 | 0,3218 | 3,84046E-10 |
| ACE2 | CYP3A5 | 0,3218 | 3,84662E-10 |
| ACE2 | CYP4F3 | 0,3218 | 3,82829E-10 |

| | | | |
|---|---|---|---|
| ACE2 | RP11-690I21.2 | 0,3217 | 3,89588E-10 |
| ACE2 | ECSCR | 0,3217 | 3,88946E-10 |
| ACE2 | SLC24A2 | 0,3217 | 3,87058E-10 |
| ACE2 | CTC-459F4.9 | 0,3217 | 3,90488E-10 |
| ACE2 | VN1R107P | 0,3217 | 3,91325E-10 |
| ACE2 | GYS2 | 0,3216 | 3,93115E-10 |
| ACE2 | FAM213B | 0,3215 | 4,00019E-10 |
| ACE2 | RASGRP3 | 0,3215 | 3,98429E-10 |
| ACE2 | SNHG18 | 0,3215 | 3,98671E-10 |
| ACE2 | C9orf106 | 0,3215 | 3,99742E-10 |
| ACE2 | EEF1A1P14 | 0,3214 | 4,05464E-10 |
| ACE2 | ADPGK | 0,3214 | 4,04448E-10 |
| ACE2 | FGF19 | 0,3213 | 4,12839E-10 |
| ACE2 | LINC00310 | 0,3213 | 4,09704E-10 |
| ACE2 | AC005301.9 | 0,3213 | 4,11756E-10 |
| ACE2 | MAGEE2 | 0,3212 | 4,17888E-10 |
| ACE2 | MLLT10P1 | 0,3211 | 4,20541E-10 |
| ACE2 | GPX1 | 0,321 | 4,26591E-10 |
| ACE2 | AC005150.1 | 0,321 | 4,29595E-10 |
| ACE2 | CTD-2651B20.1 | 0,321 | 4,25084E-10 |
| ACE2 | CLSPN | 0,3209 | 4,35152E-10 |
| ACE2 | TUBGCP3 | 0,3209 | 4,31862E-10 |
| ACE2 | LINC00870 | 0,3208 | 4,35604E-10 |
| ACE2 | HEXDC | 0,3208 | 4,39508E-10 |
| ACE2 | GCFC2 | 0,3207 | 4,42353E-10 |
| ACE2 | RP11-700H6.1 | 0,3207 | 4,44636E-10 |
| ACE2 | AC114730.2 | 0,3206 | 4,51687E-10 |
| ACE2 | DAP | 0,3206 | 4,50276E-10 |
| ACE2 | RP11-771K4.3 | 0,3206 | 4,47966E-10 |
| ACE2 | RP11-117D22.2 | 0,3205 | 4,56765E-10 |
| ACE2 | LINC01124 | 0,3204 | 4,63276E-10 |
| ACE2 | ESAM | 0,3203 | 4,64976E-10 |
| ACE2 | MICU2 | 0,3203 | 4,69573E-10 |
| ACE2 | DES | 0,3202 | 4,71507E-10 |
| ACE2 | FAM124B | 0,3202 | 4,71719E-10 |
| ACE2 | SBNO1 | 0,3202 | 4,74868E-10 |
| ACE2 | RHOXF1-AS1 | 0,3201 | 4,80189E-10 |
| ACE2 | MUTYH | 0,32 | 4,87264E-10 |
| ACE2 | RP11-173M1.5 | 0,32 | 4,84678E-10 |
| ACE2 | TLR9 | 0,3199 | 4,94265E-10 |
| ACE2 | TRBV25-1 | 0,3199 | 4,94061E-10 |
| ACE2 | RP11-380M21.1 | 0,3198 | 4,98019E-10 |
| ACE2 | TRIB3 | 0,3198 | 5,00777E-10 |
| ACE2 | SYNDIG1 | 0,3198 | 5,01845E-10 |
| ACE2 | R3HDM1 | 0,3196 | 5,1516E-10 |
| ACE2 | FHOD1 | 0,3196 | 5,14183E-10 |
| ACE2 | AC005154.6 | 0,3195 | 5,22712E-10 |
| ACE2 | ZIC1 | 0,3194 | 5,28231E-10 |
| ACE2 | TMED1 | 0,3194 | 5,24645E-10 |
| ACE2 | KLHL18 | 0,3193 | 5,3092E-10 |
| ACE2 | RPL9 | 0,3193 | 5,33016E-10 |
| ACE2 | KIAA1109 | 0,3193 | 5,34041E-10 |
| ACE2 | KLHL34 | 0,3193 | 5,32524E-10 |
| ACE2 | ACTN2 | 0,3192 | 5,42577E-10 |

| | | | |
|---|---|---|---|
| ACE2 | SLC12A9 | 0,3192 | 5,36994E-10 |
| ACE2 | RN7SL726P | 0,3192 | 5,41207E-10 |
| ACE2 | TPCN1 | 0,3192 | 5,42221E-10 |
| ACE2 | ZNF235 | 0,3192 | 5,43405E-10 |
| ACE2 | AAR2 | 0,3191 | 5,50778E-10 |
| ACE2 | CLK2 | 0,319 | 5,52535E-10 |
| ACE2 | NR2F1 | 0,319 | 5,55394E-10 |
| ACE2 | NUDT3 | 0,3189 | 5,60343E-10 |
| ACE2 | RP11-508N22.12 | 0,3189 | 5,59092E-10 |
| ACE2 | TRIM3 | 0,3189 | 5,60598E-10 |
| ACE2 | COPE | 0,3189 | 5,61007E-10 |
| ACE2 | NR5A2 | 0,3188 | 5,71223E-10 |
| ACE2 | RNASE2 | 0,3188 | 5,73113E-10 |
| ACE2 | SUGT1P2 | 0,3187 | 5,76327E-10 |
| ACE2 | CAMKK1 | 0,3187 | 5,76083E-10 |
| ACE2 | RP11-355F16.1 | 0,3185 | 5,93594E-10 |
| ACE2 | AF131215.9 | 0,3185 | 5,94744E-10 |
| ACE2 | TPGS1 | 0,3185 | 5,89921E-10 |
| ACE2 | RP11-75C10.9 | 0,3184 | 5,97652E-10 |
| ACE2 | COL8A1 | 0,3183 | 6,11082E-10 |
| ACE2 | RP11-95I19.1 | 0,3183 | 6,05458E-10 |
| ACE2 | NPR1 | 0,3182 | 6,18428E-10 |
| ACE2 | HIST1H2AD | 0,3182 | 6,19456E-10 |
| ACE2 | C1GALT1 | 0,3182 | 6,13786E-10 |
| ACE2 | LINC00900 | 0,3182 | 6,12327E-10 |
| ACE2 | RP11-680C21.1 | 0,3182 | 6,18632E-10 |
| ACE2 | NCF2 | 0,318 | 6,29727E-10 |
| ACE2 | RP11-342K6.3 | 0,318 | 6,30141E-10 |
| ACE2 | LINC01410 | 0,318 | 6,27915E-10 |
| ACE2 | DCLRE1C | 0,318 | 6,29706E-10 |
| ACE2 | LDHAP4 | 0,3179 | 6,36331E-10 |
| ACE2 | TOP3B | 0,3179 | 6,38152E-10 |
| ACE2 | UNKL | 0,3178 | 6,4906E-10 |
| ACE2 | CH507-254M2.1 | 0,3178 | 6,45116E-10 |
| ACE2 | POMT2 | 0,3177 | 6,55942E-10 |
| ACE2 | RP11-638I2.2 | 0,3177 | 6,53318E-10 |
| ACE2 | FLT3LG | 0,3177 | 6,58479E-10 |
| ACE2 | PLAC1 | 0,3177 | 6,56858E-10 |
| ACE2 | VIPR2 | 0,3176 | 6,62776E-10 |
| ACE2 | MAN1B1 | 0,3176 | 6,69066E-10 |
| ACE2 | AC008592.3 | 0,3175 | 6,7787E-10 |
| ACE2 | RP11-888D10.4 | 0,3175 | 6,73407E-10 |
| ACE2 | CTC-510F12.3 | 0,3175 | 6,78353E-10 |
| ACE2 | TEX13B | 0,3175 | 6,70092E-10 |
| ACE2 | PSMB1 | 0,3174 | 6,80111E-10 |
| ACE2 | WRN | 0,3174 | 6,79753E-10 |
| ACE2 | AKAP17BP | 0,3174 | 6,82804E-10 |
| ACE2 | ARHGEF16 | 0,3173 | 6,88292E-10 |
| ACE2 | CTD-2154I11.2 | 0,3173 | 6,88371E-10 |
| ACE2 | AQP1 | 0,3173 | 6,94134E-10 |
| ACE2 | ZNF321P | 0,3173 | 6,94099E-10 |
| ACE2 | UPP1 | 0,3172 | 7,04055E-10 |
| ACE2 | FRG1JP | 0,3172 | 7,02272E-10 |
| ACE2 | ZNF404 | 0,3172 | 7,02241E-10 |

| | | | |
|---|---|---|---|
| ACE2 | FHIT | 0,3171 | 7,0874E-10 |
| ACE2 | RP11-119F7.5 | 0,317 | 7,18163E-10 |
| ACE2 | CHEK1 | 0,317 | 7,18126E-10 |
| ACE2 | DCAKD | 0,317 | 7,21656E-10 |
| ACE2 | RP11-50B3.4 | 0,3169 | 7,28003E-10 |
| ACE2 | HOXB8 | 0,3169 | 7,24811E-10 |
| ACE2 | NRTN | 0,3169 | 7,27382E-10 |
| ACE2 | RP4-735C1.4 | 0,3168 | 7,36759E-10 |
| ACE2 | ACAD11 | 0,3168 | 7,38168E-10 |
| ACE2 | TJAP1 | 0,3168 | 7,35747E-10 |
| ACE2 | SPX | 0,3168 | 7,37838E-10 |
| ACE2 | MILR1 | 0,3168 | 7,42445E-10 |
| ACE2 | ARHGAP4 | 0,3168 | 7,33721E-10 |
| ACE2 | MAEA | 0,3167 | 7,49382E-10 |
| ACE2 | DNAJB6 | 0,3167 | 7,5197E-10 |
| ACE2 | IVD | 0,3167 | 7,52379E-10 |
| ACE2 | PSTPIP2 | 0,3167 | 7,43166E-10 |
| ACE2 | PDE3A | 0,3165 | 7,71101E-10 |
| ACE2 | SNRPGP4 | 0,3165 | 7,6707E-10 |
| ACE2 | SRY | 0,3164 | 7,75359E-10 |
| ACE2 | LINC00941 | 0,3163 | 7,90302E-10 |
| ACE2 | RP11-454F8.4 | 0,3163 | 7,88152E-10 |
| ACE2 | SPNS2 | 0,3163 | 7,91043E-10 |
| ACE2 | HCG11 | 0,3162 | 7,96596E-10 |
| ACE2 | GUSBP12 | 0,3162 | 7,99022E-10 |
| ACE2 | C1orf204 | 0,3161 | 8,08165E-10 |
| ACE2 | SCRN3 | 0,3161 | 8,12506E-10 |
| ACE2 | SMARCC2 | 0,3161 | 8,05172E-10 |
| ACE2 | FAM168A | 0,316 | 8,22618E-10 |
| ACE2 | GPATCH11 | 0,3158 | 8,44269E-10 |
| ACE2 | MTA3 | 0,3158 | 8,37756E-10 |
| ACE2 | GIMAP5 | 0,3158 | 8,41911E-10 |
| ACE2 | SEC61B | 0,3158 | 8,41707E-10 |
| ACE2 | LINC01252 | 0,3158 | 8,41832E-10 |
| ACE2 | PDZRN3-AS1 | 0,3157 | 8,5517E-10 |
| ACE2 | HNRNPKP1 | 0,3157 | 8,45839E-10 |
| ACE2 | AC000078.5 | 0,3157 | 8,50769E-10 |
| ACE2 | HIKESHI | 0,3156 | 8,57153E-10 |
| ACE2 | RP3-405J10.3 | 0,3156 | 8,67596E-10 |
| ACE2 | CCDC88C | 0,3156 | 8,61631E-10 |
| ACE2 | C15orf56 | 0,3156 | 8,62044E-10 |
| ACE2 | F13A1 | 0,3155 | 8,733E-10 |
| ACE2 | HLA-DPA1 | 0,3155 | 8,75236E-10 |
| ACE2 | RP11-522B15.3 | 0,3155 | 8,69801E-10 |
| ACE2 | LRRC75A | 0,3155 | 8,76083E-10 |
| ACE2 | TEF | 0,3155 | 8,68729E-10 |
| ACE2 | RP11-731C17.2 | 0,3154 | 8,84648E-10 |
| ACE2 | ENDOU | 0,3154 | 8,87463E-10 |
| ACE2 | GCSH | 0,3154 | 8,81337E-10 |
| ACE2 | RP11-60A14.1 | 0,3153 | 8,94022E-10 |
| ACE2 | SLC35E1P1 | 0,3153 | 8,92821E-10 |
| ACE2 | IGLON5 | 0,3153 | 9,00823E-10 |
| ACE2 | SARAF | 0,3152 | 9,04011E-10 |
| ACE2 | TBX2 | 0,3152 | 9,1304E-10 |

| | | | |
|---|---|---|---|
| ACE2 | LINC01132 | 0,3151 | 9,16623E-10 |
| ACE2 | SNX29P2 | 0,3151 | 9,15618E-10 |
| ACE2 | GRPR | 0,3151 | 9,21633E-10 |
| ACE2 | FGFR3 | 0,315 | 9,25901E-10 |
| ACE2 | CCDC18-AS1 | 0,3149 | 9,40695E-10 |
| ACE2 | NUS1P2 | 0,3149 | 9,45968E-10 |
| ACE2 | ZFP3 | 0,3149 | 9,40057E-10 |
| ACE2 | RP11-53I6.4 | 0,3149 | 9,38944E-10 |
| ACE2 | TCEAL3-AS1 | 0,3149 | 9,39806E-10 |
| ACE2 | TMEM196 | 0,3148 | 9,50018E-10 |
| ACE2 | RP11-34A14.3 | 0,3148 | 9,58491E-10 |
| ACE2 | DNM1L | 0,3148 | 9,58097E-10 |
| ACE2 | NDUFAB1 | 0,3148 | 9,50733E-10 |
| ACE2 | WNK4 | 0,3148 | 9,59774E-10 |
| ACE2 | TEX13C | 0,3148 | 9,61079E-10 |
| ACE2 | AP1G2 | 0,3147 | 9,71177E-10 |
| ACE2 | MDGA1 | 0,3146 | 9,82198E-10 |
| ACE2 | LOX | 0,3145 | 9,97604E-10 |
| ACE2 | RP11-463O12.5 | 0,3145 | 9,92565E-10 |
| ACE2 | MYT1 | 0,3145 | 9,99363E-10 |
| ACE2 | SAMD11 | 0,3143 | 1,02248E-09 |
| ACE2 | CYS1 | 0,3143 | 1,01421E-09 |
| ACE2 | ZNF442 | 0,3143 | 1,01899E-09 |
| ACE2 | GAPDHP42 | 0,3142 | 1,03491E-09 |
| ACE2 | C10orf128 | 0,3142 | 1,03071E-09 |
| ACE2 | MRNIP | 0,3141 | 1,04556E-09 |
| ACE2 | KIF26A | 0,3141 | 1,04237E-09 |
| ACE2 | AC006128.2 | 0,3141 | 1,04445E-09 |
| ACE2 | RP11-659P15.1 | 0,314 | 1,06238E-09 |
| ACE2 | MS4A3 | 0,3139 | 1,07067E-09 |
| ACE2 | GJA3 | 0,3139 | 1,0709E-09 |
| ACE2 | GREM1 | 0,3139 | 1,07565E-09 |
| ACE2 | RP11-227D13.1 | 0,3139 | 1,07275E-09 |
| ACE2 | HACD3 | 0,3139 | 1,07128E-09 |
| ACE2 | CH17-262O2.1 | 0,3138 | 1,09321E-09 |
| ACE2 | RP11-517P14.2 | 0,3138 | 1,08318E-09 |
| ACE2 | CHORDC1 | 0,3138 | 1,0913E-09 |
| ACE2 | HCN2 | 0,3138 | 1,08477E-09 |
| ACE2 | MPPED2 | 0,3137 | 1,0971E-09 |
| ACE2 | SNRNP70 | 0,3137 | 1,10427E-09 |
| ACE2 | SPO11 | 0,3137 | 1,09969E-09 |
| ACE2 | USP40 | 0,3136 | 1,11785E-09 |
| ACE2 | GTF3C1 | 0,3136 | 1,10831E-09 |
| ACE2 | BCLAF1P2 | 0,3136 | 1,11489E-09 |
| ACE2 | CTD-2619J13.9 | 0,3136 | 1,11555E-09 |
| ACE2 | DOK7 | 0,3134 | 1,14112E-09 |
| ACE2 | CTB-78O21.1 | 0,3133 | 1,15511E-09 |
| ACE2 | SYT15 | 0,3133 | 1,16117E-09 |
| ACE2 | RP11-325K4.2 | 0,3133 | 1,15823E-09 |
| ACE2 | TAB1 | 0,3133 | 1,16431E-09 |
| ACE2 | FTOP1 | 0,3132 | 1,16601E-09 |
| ACE2 | C6orf3 | 0,3132 | 1,1662E-09 |
| ACE2 | MIR181A1HG | 0,3131 | 1,18477E-09 |
| ACE2 | RP11-708J19.3 | 0,313 | 1,20204E-09 |

| | | | |
|---|---|---|---|
| ACE2 | TMOD2 | 0,313 | 1,20533E-09 |
| ACE2 | PANX2 | 0,313 | 1,20919E-09 |
| ACE2 | EMCN | 0,3129 | 1,21902E-09 |
| ACE2 | RBBP8 | 0,3129 | 1,2252E-09 |
| ACE2 | MIS18A-AS1 | 0,3129 | 1,2218E-09 |
| ACE2 | MT1A | 0,3128 | 1,22939E-09 |
| ACE2 | SOD3 | 0,3127 | 1,24504E-09 |
| ACE2 | RP11-243M5.5 | 0,3127 | 1,25242E-09 |
| ACE2 | ULK3 | 0,3127 | 1,24794E-09 |
| ACE2 | ANKRD65 | 0,3126 | 1,27291E-09 |
| ACE2 | ALDH3A1 | 0,3126 | 1,26048E-09 |
| ACE2 | HMGB1P21 | 0,3125 | 1,2906E-09 |
| ACE2 | CTD-2562J17.6 | 0,3125 | 1,28919E-09 |
| ACE2 | SETP14 | 0,3123 | 1,31967E-09 |
| ACE2 | N4BP2L1 | 0,3123 | 1,30855E-09 |
| ACE2 | LINC01271 | 0,3123 | 1,32358E-09 |
| ACE2 | CCDC120 | 0,3123 | 1,30982E-09 |
| ACE2 | GPR63 | 0,3122 | 1,32866E-09 |
| ACE2 | CLDN10 | 0,3122 | 1,33459E-09 |
| ACE2 | MCC | 0,312 | 1,36481E-09 |
| ACE2 | CMAHP | 0,3119 | 1,37926E-09 |
| ACE2 | SNORA2A | 0,3119 | 1,37932E-09 |
| ACE2 | C17orf75 | 0,3119 | 1,37663E-09 |
| ACE2 | NBPF10 | 0,3118 | 1,39931E-09 |
| ACE2 | RABL3 | 0,3118 | 1,40973E-09 |
| ACE2 | RPS13 | 0,3118 | 1,39879E-09 |
| ACE2 | MT-TC | 0,3118 | 1,40986E-09 |
| ACE2 | CCL26 | 0,3117 | 1,42211E-09 |
| ACE2 | CBWD1 | 0,3117 | 1,4226E-09 |
| ACE2 | IGHV3-73 | 0,3117 | 1,42406E-09 |
| ACE2 | FAM89A | 0,3116 | 1,44292E-09 |
| ACE2 | AC079250.1 | 0,3116 | 1,4431E-09 |
| ACE2 | DAZL | 0,3116 | 1,44059E-09 |
| ACE2 | NUP93 | 0,3116 | 1,43765E-09 |
| ACE2 | UBE2L4 | 0,3116 | 1,42909E-09 |
| ACE2 | JAK2 | 0,3115 | 1,46466E-09 |
| ACE2 | SALRNA1 | 0,3115 | 1,45771E-09 |
| ACE2 | RP11-902B17.1 | 0,3114 | 1,47723E-09 |
| ACE2 | RP11-781P6.1 | 0,3114 | 1,48111E-09 |
| ACE2 | ZCCHC12 | 0,3114 | 1,47979E-09 |
| ACE2 | PDE2A | 0,3113 | 1,48445E-09 |
| ACE2 | GREM2 | 0,3111 | 1,52407E-09 |
| ACE2 | IL1B | 0,3111 | 1,52813E-09 |
| ACE2 | RARRES2P1 | 0,3111 | 1,5287E-09 |
| ACE2 | TSPAN33 | 0,3111 | 1,53273E-09 |
| ACE2 | VAMP1 | 0,311 | 1,54816E-09 |
| ACE2 | ZNF358 | 0,311 | 1,54181E-09 |
| ACE2 | CERS1 | 0,311 | 1,54449E-09 |
| ACE2 | RP11-578F21.10 | 0,3109 | 1,56777E-09 |
| ACE2 | COG7 | 0,3109 | 1,56314E-09 |
| ACE2 | RP11-17E13.2 | 0,3108 | 1,58342E-09 |
| ACE2 | RP11-326C3.14 | 0,3108 | 1,58354E-09 |
| ACE2 | CES5A | 0,3108 | 1,60118E-09 |
| ACE2 | RP4-536B24.2 | 0,3108 | 1,59926E-09 |

| | | | |
|---|---|---|---|
| ACE2 | SLC13A2 | 0,3108 | 1,59713E-09 |
| ACE2 | CYSLTR1 | 0,3108 | 1,58602E-09 |
| ACE2 | PABPC5-AS1 | 0,3106 | 1,62621E-09 |
| ACE2 | PRDM1 | 0,3105 | 1,64354E-09 |
| ACE2 | RP11-1217F2.22 | 0,3105 | 1,64749E-09 |
| ACE2 | RP3-395M20.8 | 0,3104 | 1,66448E-09 |
| ACE2 | RNPEPL1 | 0,3104 | 1,67787E-09 |
| ACE2 | MTIF2 | 0,3102 | 1,71084E-09 |
| ACE2 | RP3-395M20.3 | 0,3101 | 1,74707E-09 |
| ACE2 | LINC01503 | 0,3101 | 1,73243E-09 |
| ACE2 | PYROXD2 | 0,31 | 1,76514E-09 |
| ACE2 | GAPDHP71 | 0,3098 | 1,80183E-09 |
| ACE2 | CTD-2609K8.3 | 0,3098 | 1,81284E-09 |
| ACE2 | ZMIZ1 | 0,3096 | 1,85532E-09 |
| ACE2 | ADAMTS20 | 0,3096 | 1,85931E-09 |
| ACE2 | SEC62-AS1 | 0,3095 | 1,88517E-09 |
| ACE2 | HSF4 | 0,3095 | 1,87668E-09 |
| ACE2 | DTWD2 | 0,3094 | 1,90237E-09 |
| ACE2 | ITGB1BP2 | 0,3094 | 1,89029E-09 |
| ACE2 | RP5-1126H10.2 | 0,3093 | 1,91762E-09 |
| ACE2 | RP11-855A2.1 | 0,3093 | 1,91983E-09 |
| ACE2 | CLCA1 | 0,3092 | 1,95077E-09 |
| ACE2 | AKR1C2 | 0,3092 | 1,93812E-09 |
| ACE2 | SBF2-AS1 | 0,3092 | 1,9403E-09 |
| ACE2 | KCNA5 | 0,3091 | 1,97537E-09 |
| ACE2 | KDM2A | 0,309 | 1,987E-09 |
| ACE2 | RPS3AP26 | 0,3089 | 2,01073E-09 |
| ACE2 | RPL14P1 | 0,3089 | 2,03159E-09 |
| ACE2 | CTD-3126B10.2 | 0,3089 | 2,02968E-09 |
| ACE2 | DBNDD1 | 0,3089 | 2,02321E-09 |
| ACE2 | RP4-758J18.13 | 0,3088 | 2,05594E-09 |
| ACE2 | RP3-507I15.2 | 0,3088 | 2,03967E-09 |
| ACE2 | CCL2 | 0,3087 | 2,06808E-09 |
| ACE2 | CYP2A7 | 0,3087 | 2,07733E-09 |
| ACE2 | RP6-109B7.4 | 0,3087 | 2,06589E-09 |
| ACE2 | RP11-89C3.3 | 0,3086 | 2,11093E-09 |
| ACE2 | TMEM237 | 0,3085 | 2,11918E-09 |
| ACE2 | KRTAP5-8 | 0,3085 | 2,13504E-09 |
| ACE2 | IKBKGP1 | 0,3085 | 2,13363E-09 |
| ACE2 | LINC02158 | 0,3082 | 2,21359E-09 |
| ACE2 | RBMS3-AS3 | 0,3081 | 2,2485E-09 |
| ACE2 | BTF3P12 | 0,3081 | 2,24862E-09 |
| ACE2 | IPO9-AS1 | 0,308 | 2,27232E-09 |
| ACE2 | AC006033.22 | 0,308 | 2,25781E-09 |
| ACE2 | CLEC1A | 0,308 | 2,26485E-09 |
| ACE2 | ESF1 | 0,3079 | 2,29253E-09 |
| ACE2 | RP11-320A16.1 | 0,3078 | 2,3097E-09 |
| ACE2 | LINC01630 | 0,3078 | 2,31493E-09 |
| ACE2 | RP11-308D16.2 | 0,3078 | 2,32704E-09 |
| ACE2 | RP11-164P12.3 | 0,3077 | 2,35168E-09 |
| ACE2 | LAMA4 | 0,3077 | 2,34132E-09 |
| ACE2 | RP11-672A2.4 | 0,3077 | 2,35314E-09 |
| ACE2 | TSSK5P | 0,3076 | 2,37727E-09 |
| ACE2 | CDH3 | 0,3076 | 2,39063E-09 |

| | | | |
|---|---|---|---|
| ACE2 | PDZD2 | 0,3075 | 2,42305E-09 |
| ACE2 | RP11-215P8.2 | 0,3075 | 2,41198E-09 |
| ACE2 | HSP90B2P | 0,3075 | 2,41262E-09 |
| ACE2 | SGK2 | 0,3075 | 2,41731E-09 |
| ACE2 | RP1-197B17.7 | 0,3074 | 2,44145E-09 |
| ACE2 | EMC4 | 0,3074 | 2,42605E-09 |
| ACE2 | UBR1 | 0,3074 | 2,433E-09 |
| ACE2 | COL1A1 | 0,3074 | 2,43547E-09 |
| ACE2 | NCAN | 0,3074 | 2,43064E-09 |
| ACE2 | FPR3 | 0,3074 | 2,45428E-09 |
| ACE2 | RP11-118B18.2 | 0,3073 | 2,45604E-09 |
| ACE2 | RBFA | 0,3073 | 2,4657E-09 |
| ACE2 | SLC30A3 | 0,3071 | 2,52923E-09 |
| ACE2 | MT1F | 0,3071 | 2,54071E-09 |
| ACE2 | SLC7A6 | 0,307 | 2,5739E-09 |
| ACE2 | CTD-2647L4.4 | 0,3069 | 2,59634E-09 |
| ACE2 | ACOT6 | 0,3069 | 2,60381E-09 |
| ACE2 | SOX8 | 0,3068 | 2,61614E-09 |
| ACE2 | GIPC3 | 0,3068 | 2,63513E-09 |
| ACE2 | SETD1A | 0,3067 | 2,66405E-09 |
| ACE2 | RSL24D1P6 | 0,3067 | 2,67836E-09 |
| ACE2 | TBC1D4 | 0,3066 | 2,70915E-09 |
| ACE2 | EIF3CL | 0,3066 | 2,71179E-09 |
| ACE2 | MAGEB6 | 0,3066 | 2,68007E-09 |
| ACE2 | DNAH5 | 0,3065 | 2,71785E-09 |
| ACE2 | LRRC25 | 0,3065 | 2,71467E-09 |
| ACE2 | EPS15 | 0,3064 | 2,75063E-09 |
| ACE2 | RP11-139I14.2 | 0,3064 | 2,74861E-09 |
| ACE2 | ZC3H12C | 0,3064 | 2,76685E-09 |
| ACE2 | LINC01772 | 0,3063 | 2,80244E-09 |
| ACE2 | CH17-472G23.1 | 0,3063 | 2,79376E-09 |
| ACE2 | C4orf32 | 0,3063 | 2,80072E-09 |
| ACE2 | ACADL | 0,3062 | 2,8459E-09 |
| ACE2 | MYADM | 0,3062 | 2,83608E-09 |
| ACE2 | RPS20P4 | 0,3061 | 2,87767E-09 |
| ACE2 | TMEM59L | 0,3061 | 2,86675E-09 |
| ACE2 | CTD-2620I22.3 | 0,3061 | 2,87201E-09 |
| ACE2 | CTC-512J12.7 | 0,306 | 2,90125E-09 |
| ACE2 | MIR1254-1 | 0,3059 | 2,94531E-09 |
| ACE2 | CA7 | 0,3059 | 2,92791E-09 |
| ACE2 | GPR50 | 0,3059 | 2,95269E-09 |
| ACE2 | IWS1 | 0,3057 | 3,02286E-09 |
| ACE2 | RNF5 | 0,3057 | 3,01778E-09 |
| ACE2 | ISLR | 0,3057 | 3,01764E-09 |
| ACE2 | RPL36 | 0,3057 | 3,02228E-09 |
| ACE2 | NECAP1P2 | 0,3056 | 3,04851E-09 |
| ACE2 | NT5C2 | 0,3056 | 3,06255E-09 |
| ACE2 | DPF2 | 0,3056 | 3,053E-09 |
| ACE2 | PTMAP5 | 0,3056 | 3,07196E-09 |
| ACE2 | MT1E | 0,3056 | 3,05701E-09 |
| ACE2 | TNFAIP8L2 | 0,3055 | 3,10103E-09 |
| ACE2 | ECEL1 | 0,3055 | 3,09203E-09 |
| ACE2 | DYNLT1 | 0,3055 | 3,08738E-09 |
| ACE2 | RP4-605O3.4 | 0,3055 | 3,09828E-09 |

| | | | |
|---|---|---|---|
| ACE2 | RP11-305K5.1 | 0,3054 | 3,14184E-09 |
| ACE2 | RP11-489G11.3 | 0,3054 | 3,13553E-09 |
| ACE2 | SLC6A19 | 0,3054 | 3,12603E-09 |
| ACE2 | CENPH | 0,3054 | 3,12827E-09 |
| ACE2 | UFSP1 | 0,3054 | 3,11101E-09 |
| ACE2 | PDE7A | 0,3054 | 3,11536E-09 |
| ACE2 | FKBP1C | 0,3053 | 3,18497E-09 |
| ACE2 | AC004067.5 | 0,3052 | 3,2134E-09 |
| ACE2 | MARCKS | 0,3052 | 3,19395E-09 |
| ACE2 | TRBV24-1 | 0,3052 | 3,21498E-09 |
| ACE2 | MMP24 | 0,3052 | 3,18957E-09 |
| ACE2 | AAAS | 0,3051 | 3,26085E-09 |
| ACE2 | RP11-793H13.3 | 0,3051 | 3,24849E-09 |
| ACE2 | RP11-398E10.1 | 0,3051 | 3,23816E-09 |
| ACE2 | NR2C2AP | 0,3051 | 3,26505E-09 |
| ACE2 | ADAM12 | 0,305 | 3,30802E-09 |
| ACE2 | UBE2CP2 | 0,3049 | 3,34549E-09 |
| ACE2 | RP11-496D24.2 | 0,3047 | 3,39372E-09 |
| ACE2 | MEIS3P1 | 0,3047 | 3,40234E-09 |
| ACE2 | CTD-2515H24.2 | 0,3046 | 3,46871E-09 |
| ACE2 | CTD-3162L10.5 | 0,3046 | 3,46219E-09 |
| ACE2 | LINC00652 | 0,3046 | 3,45891E-09 |
| ACE2 | GAS2L1 | 0,3046 | 3,45104E-09 |
| ACE2 | CCNL2 | 0,3045 | 3,50348E-09 |
| ACE2 | PDE4DIPP1 | 0,3045 | 3,50502E-09 |
| ACE2 | DLGAP1-AS4 | 0,3045 | 3,4863E-09 |
| ACE2 | SLC19A1 | 0,3045 | 3,50002E-09 |
| ACE2 | EEF1A1P12 | 0,3044 | 3,56294E-09 |
| ACE2 | AP002954.3 | 0,3044 | 3,5577E-09 |
| ACE2 | CTC-444N24.11 | 0,3044 | 3,55562E-09 |
| ACE2 | MDFI | 0,3043 | 3,57194E-09 |
| ACE2 | RP3-415N12.1 | 0,3043 | 3,59294E-09 |
| ACE2 | OR10V2P | 0,3043 | 3,57763E-09 |
| ACE2 | CH17-408M7.1 | 0,3042 | 3,65081E-09 |
| ACE2 | LGALS12 | 0,3042 | 3,63158E-09 |
| ACE2 | ART4 | 0,3042 | 3,65281E-09 |
| ACE2 | FXR2 | 0,3042 | 3,64867E-09 |
| ACE2 | ARHGEF10L | 0,3041 | 3,67956E-09 |
| ACE2 | CASP8 | 0,3041 | 3,69383E-09 |
| ACE2 | MYRIP | 0,3041 | 3,6747E-09 |
| ACE2 | ACKR2 | 0,3041 | 3,65627E-09 |
| ACE2 | RASGRP2 | 0,3041 | 3,69661E-09 |
| ACE2 | DTX3L | 0,304 | 3,70231E-09 |
| ACE2 | FAM26F | 0,304 | 3,71208E-09 |
| ACE2 | RP11-760H22.2 | 0,304 | 3,70401E-09 |
| ACE2 | CTC-429P9.1 | 0,304 | 3,7201E-09 |
| ACE2 | AC097382.5 | 0,3039 | 3,75419E-09 |
| ACE2 | ADPRHL1 | 0,3039 | 3,74643E-09 |
| ACE2 | CEACAM4 | 0,3039 | 3,76483E-09 |
| ACE2 | CAMK2B | 0,3038 | 3,81924E-09 |
| ACE2 | RP11-43F13.3 | 0,3037 | 3,87136E-09 |
| ACE2 | HNRNPA1P28 | 0,3036 | 3,93116E-09 |
| ACE2 | S100A4 | 0,3035 | 3,94331E-09 |
| ACE2 | FTH1P2 | 0,3035 | 3,94629E-09 |

| | | | |
|---|---|---|---|
| ACE2 | NEFL | 0,3035 | 3,94947E-09 |
| ACE2 | NCBP1 | 0,3035 | 3,95763E-09 |
| ACE2 | THOC6 | 0,3035 | 3,96068E-09 |
| ACE2 | GALC | 0,3034 | 4,00821E-09 |
| ACE2 | BFSP1 | 0,3034 | 3,99504E-09 |
| ACE2 | RP11-1101H11.1 | 0,3033 | 4,07521E-09 |
| ACE2 | TAC1 | 0,3033 | 4,04731E-09 |
| ACE2 | CCR10 | 0,3033 | 4,06923E-09 |
| ACE2 | LINC01623 | 0,3032 | 4,08601E-09 |
| ACE2 | CTD-2033A16.2 | 0,3032 | 4,08522E-09 |
| ACE2 | MYH8 | 0,3032 | 4,12937E-09 |
| ACE2 | WDR77 | 0,3031 | 4,13531E-09 |
| ACE2 | PTPN21 | 0,3031 | 4,17588E-09 |
| ACE2 | AC004985.12 | 0,303 | 4,20417E-09 |
| ACE2 | STK3 | 0,303 | 4,21943E-09 |
| ACE2 | CTD-2026K11.4 | 0,3028 | 4,32323E-09 |
| ACE2 | RP11-286N22.10 | 0,3027 | 4,35291E-09 |
| ACE2 | ZBTB40 | 0,3026 | 4,41452E-09 |
| ACE2 | SNAI1P1 | 0,3026 | 4,39787E-09 |
| ACE2 | RP11-383F6.1 | 0,3026 | 4,43984E-09 |
| ACE2 | TMEM260 | 0,3026 | 4,40495E-09 |
| ACE2 | SLC25A29 | 0,3026 | 4,42658E-09 |
| ACE2 | RP11-284F21.11 | 0,3025 | 4,47029E-09 |
| ACE2 | CTA-315H11.2 | 0,3025 | 4,47204E-09 |
| ACE2 | ROM1 | 0,3025 | 4,46899E-09 |
| ACE2 | RP11-21L23.2 | 0,3025 | 4,48437E-09 |
| ACE2 | HLA-H | 0,3024 | 4,53091E-09 |
| ACE2 | RNU1-73P | 0,3023 | 4,57346E-09 |
| ACE2 | ASPM | 0,3022 | 4,65218E-09 |
| ACE2 | RP11-496H1.1 | 0,3022 | 4,64721E-09 |
| ACE2 | CTB-113I20.2 | 0,3022 | 4,61739E-09 |
| ACE2 | DACT2 | 0,3021 | 4,7131E-09 |
| ACE2 | RP11-64B16.2 | 0,3021 | 4,71105E-09 |
| ACE2 | AC005197.2 | 0,3021 | 4,70139E-09 |
| ACE2 | TCEA3 | 0,302 | 4,73737E-09 |
| ACE2 | MT2A | 0,302 | 4,74624E-09 |
| ACE2 | PRKACA | 0,3019 | 4,82997E-09 |
| ACE2 | CHGB | 0,3019 | 4,8074E-09 |
| ACE2 | LINC01750 | 0,3018 | 4,88054E-09 |
| ACE2 | HESX1 | 0,3018 | 4,85147E-09 |
| ACE2 | CTD-2011F17.2 | 0,3018 | 4,87996E-09 |
| ACE2 | LIMA1 | 0,3018 | 4,90362E-09 |
| ACE2 | RPL21P10 | 0,3018 | 4,90306E-09 |
| ACE2 | RP11-49I11.4 | 0,3018 | 4,87819E-09 |
| ACE2 | LCMT1-AS2 | 0,3017 | 4,93027E-09 |
| ACE2 | NR2F2-AS1 | 0,3016 | 5,00047E-09 |
| ACE2 | BNIP3P30 | 0,3016 | 5,02844E-09 |
| ACE2 | CTD-2515H24.4 | 0,3015 | 5,05395E-09 |
| ACE2 | MIR503HG | 0,3015 | 5,07418E-09 |
| ACE2 | AGGF1 | 0,3014 | 5,12738E-09 |
| ACE2 | RP11-307C19.1 | 0,3014 | 5,13567E-09 |
| ACE2 | RP11-1166P10.1 | 0,3014 | 5,14053E-09 |
| ACE2 | SDCCAG8 | 0,3012 | 5,23522E-09 |
| ACE2 | LINC00839 | 0,3012 | 5,22095E-09 |

| | | | |
|---|---|---|---|
| ACE2 | RN7SL809P | 0,3012 | 5,24025E-09 |
| ACE2 | SIX4 | 0,3012 | 5,23932E-09 |
| ACE2 | SNRPD1 | 0,3012 | 5,27789E-09 |
| ACE2 | JPH2 | 0,3012 | 5,27216E-09 |
| ACE2 | ARVCF | 0,3012 | 5,27321E-09 |
| ACE2 | H3F3A | 0,301 | 5,36328E-09 |
| ACE2 | MYOC | 0,3009 | 5,46392E-09 |
| ACE2 | AC013275.2 | 0,3009 | 5,44504E-09 |
| ACE2 | CTC-287O8.1 | 0,3009 | 5,44469E-09 |
| ACE2 | CTB-58E17.3 | 0,3009 | 5,43751E-09 |
| ACE2 | LL22NC03-80A10.6 | 0,3009 | 5,45287E-09 |
| ACE2 | PDZD4 | 0,3009 | 5,41375E-09 |
| ACE2 | GCNT2 | 0,3008 | 5,53276E-09 |
| ACE2 | RP11-381E24.1 | 0,3007 | 5,56754E-09 |
| ACE2 | BST2 | 0,3007 | 5,5524E-09 |
| ACE2 | MID1IP1-AS1 | 0,3007 | 5,56102E-09 |
| ACE2 | H3F3AP6 | 0,3006 | 5,66002E-09 |
| ACE2 | RP4-800G7.3 | 0,3006 | 5,65827E-09 |
| ACE2 | RP4-562J12.1 | 0,3006 | 5,61234E-09 |
| ACE2 | GRIK3 | 0,3005 | 5,69528E-09 |
| ACE2 | EGR4 | 0,3005 | 5,74177E-09 |
| ACE2 | TMEM255B | 0,3005 | 5,72448E-09 |
| ACE2 | AC007256.5 | 0,3004 | 5,77354E-09 |
| ACE2 | C19orf60 | 0,3004 | 5,79731E-09 |
| ACE2 | HIST1H1D | 0,3003 | 5,85633E-09 |
| ACE2 | AC007743.1 | 0,3002 | 5,93647E-09 |
| ACE2 | CBWD2 | 0,3002 | 5,94329E-09 |
| ACE2 | CTD-2006M22.2 | 0,3002 | 5,90986E-09 |
| ACE2 | EIF4EP1 | 0,3002 | 5,90265E-09 |
| ACE2 | ACTR3 | 0,3001 | 6,01542E-09 |
| ACE2 | PTGES2-AS1 | 0,3001 | 6,03603E-09 |
| ACE2 | ERBB3 | 0,3001 | 5,97667E-09 |
| ACE2 | RFXAP | 0,3001 | 6,01326E-09 |
| ACE2 | FAM159B | 0,3 | 6,11065E-09 |
| ACE2 | LY86 | 0,3 | 6,06665E-09 |
| ACE2 | TCF24 | 0,3 | 6,05945E-09 |
| ACE2 | RRM1 | 0,3 | 6,09715E-09 |
| ACE2 | LL0XNC01-116E7.1 | 0,3 | 6,1037E-09 |
| ACE2 | SLC27A3 | 0,2999 | 6,15528E-09 |
| ACE2 | RAMP2 | 0,2999 | 6,1296E-09 |
| ACE2 | CTD-2373N4.3 | 0,2998 | 6,24321E-09 |
| ACE2 | CRTC2 | 0,2996 | 6,34524E-09 |
| ACE2 | ZNF384 | 0,2996 | 6,33853E-09 |
| ACE2 | SLC19A3 | 0,2995 | 6,41528E-09 |
| ACE2 | SELENOK | 0,2995 | 6,42587E-09 |
| ACE2 | HMGCS2 | 0,2994 | 6,56345E-09 |
| ACE2 | LINC01960 | 0,2994 | 6,50697E-09 |
| ACE2 | RP11-806O11.1 | 0,2994 | 6,54581E-09 |
| ACE2 | TMEM50B | 0,2994 | 6,54305E-09 |
| ACE2 | COX5B | 0,2993 | 6,64064E-09 |
| ACE2 | AC011899.9 | 0,2993 | 6,63251E-09 |
| ACE2 | PRELID3B | 0,2993 | 6,59488E-09 |
| ACE2 | ADAMTS9-AS1 | 0,2992 | 6,72436E-09 |
| ACE2 | CTC-325J23.2 | 0,2992 | 6,70294E-09 |

| | | | |
|---|---|---|---|
| ACE2 | MLXIP | 0,2992 | 6,66944E-09 |
| ACE2 | PLPP2 | 0,2992 | 6,71021E-09 |
| ACE2 | MT-ND1 | 0,2992 | 6,70277E-09 |
| ACE2 | FAM198A | 0,2991 | 6,77458E-09 |
| ACE2 | TMEM261 | 0,2991 | 6,73885E-09 |
| ACE2 | KMT5A | 0,2991 | 6,81212E-09 |
| ACE2 | ARGFXP2 | 0,2991 | 6,81246E-09 |
| ACE2 | LINC00339 | 0,299 | 6,89173E-09 |
| ACE2 | HAND2-AS1 | 0,299 | 6,82141E-09 |
| ACE2 | RN7SKP287 | 0,299 | 6,87528E-09 |
| ACE2 | RP11-13N13.5 | 0,299 | 6,83127E-09 |
| ACE2 | AC004951.5 | 0,2989 | 6,95984E-09 |
| ACE2 | WDR72 | 0,2989 | 6,90395E-09 |
| ACE2 | CNN3 | 0,2988 | 7,02174E-09 |
| ACE2 | NCLP1 | 0,2987 | 7,12025E-09 |
| ACE2 | TXNL1 | 0,2987 | 7,13257E-09 |
| ACE2 | ZNF74 | 0,2987 | 7,10882E-09 |
| ACE2 | PNMA2 | 0,2986 | 7,15926E-09 |
| ACE2 | DFFB | 0,2985 | 7,29182E-09 |
| ACE2 | RP11-307L14.1 | 0,2985 | 7,2678E-09 |
| ACE2 | BYSL | 0,2985 | 7,31585E-09 |
| ACE2 | TCERG1L | 0,2985 | 7,26643E-09 |
| ACE2 | SIGIRR | 0,2985 | 7,3248E-09 |
| ACE2 | RP11-490H24.5 | 0,2985 | 7,27594E-09 |
| ACE2 | RP11-387H17.6 | 0,2985 | 7,26657E-09 |
| ACE2 | TADA2B | 0,2984 | 7,36783E-09 |
| ACE2 | FGF1 | 0,2984 | 7,33465E-09 |
| ACE2 | CREBBP | 0,2984 | 7,35511E-09 |
| ACE2 | RN7SL573P | 0,2984 | 7,35673E-09 |
| ACE2 | CUX1 | 0,2983 | 7,49459E-09 |
| ACE2 | SNORD22 | 0,2983 | 7,44984E-09 |
| ACE2 | ACP6 | 0,2982 | 7,59682E-09 |
| ACE2 | MEIS1 | 0,2982 | 7,51814E-09 |
| ACE2 | DRD5 | 0,2981 | 7,66109E-09 |
| ACE2 | RP11-764K9.1 | 0,2981 | 7,60601E-09 |
| ACE2 | KIAA2022 | 0,298 | 7,71415E-09 |
| ACE2 | NDUFB4P6 | 0,2979 | 7,7991E-09 |
| ACE2 | PIP5K1B | 0,2979 | 7,8279E-09 |
| ACE2 | CLRN1 | 0,2978 | 7,93809E-09 |
| ACE2 | KB-1517D11.4 | 0,2978 | 7,96895E-09 |
| ACE2 | FTLP3 | 0,2978 | 7,9735E-09 |
| ACE2 | HLA-F-AS1 | 0,2977 | 7,98416E-09 |
| ACE2 | C11orf96 | 0,2977 | 8,00069E-09 |
| ACE2 | RP11-573M3.2 | 0,2977 | 8,01271E-09 |
| ACE2 | RP11-1113L8.2 | 0,2977 | 8,05561E-09 |
| ACE2 | GPATCH1 | 0,2976 | 8,11399E-09 |
| ACE2 | LILRB5 | 0,2976 | 8,09248E-09 |
| ACE2 | RP11-65L3.4 | 0,2975 | 8,18021E-09 |
| ACE2 | VCAN | 0,2975 | 8,24574E-09 |
| ACE2 | UBR5-AS1 | 0,2975 | 8,26329E-09 |
| ACE2 | RP3-414A15.12 | 0,2975 | 8,16993E-09 |
| ACE2 | RP11-96D1.3 | 0,2975 | 8,23165E-09 |
| ACE2 | APOE | 0,2975 | 8,23638E-09 |
| ACE2 | WAS | 0,2975 | 8,25669E-09 |

| | | | |
|---|---|---|---|
| ACE2 | OR8D4 | 0,2974 | 8,34903E-09 |
| ACE2 | RP1-78O14.1 | 0,2974 | 8,2971E-09 |
| ACE2 | RP11-34F13.2 | 0,2974 | 8,31321E-09 |
| ACE2 | ZSCAN16 | 0,2973 | 8,39995E-09 |
| ACE2 | NPM1P37 | 0,2973 | 8,40544E-09 |
| ACE2 | CRISPLD1 | 0,2973 | 8,44398E-09 |
| ACE2 | SLC2A3P2 | 0,2971 | 8,64837E-09 |
| ACE2 | CHL1-AS1 | 0,2971 | 8,64147E-09 |
| ACE2 | RP11-532F6.3 | 0,2971 | 8,6692E-09 |
| ACE2 | ZSCAN5B | 0,2971 | 8,65211E-09 |
| ACE2 | RP11-16E18.3 | 0,297 | 8,77146E-09 |
| ACE2 | RP11-66N24.3 | 0,2969 | 8,8397E-09 |
| ACE2 | RNF217 | 0,2968 | 8,98895E-09 |
| ACE2 | RP11-147L13.12 | 0,2968 | 8,97085E-09 |
| ACE2 | RUSC1-AS1 | 0,2967 | 9,07761E-09 |
| ACE2 | RP11-271C24.3 | 0,2967 | 9,00475E-09 |
| ACE2 | GATA3 | 0,2966 | 9,18589E-09 |
| ACE2 | RCAN2 | 0,2965 | 9,22744E-09 |
| ACE2 | GABRG2 | 0,2964 | 9,42808E-09 |
| ACE2 | CCND2-AS2 | 0,2963 | 9,47743E-09 |
| ACE2 | RP11-977G19.11 | 0,2963 | 9,50447E-09 |
| ACE2 | CDK5RAP1 | 0,2963 | 9,49798E-09 |
| ACE2 | MKRN7P | 0,2963 | 9,52295E-09 |
| ACE2 | LINC01419 | 0,2962 | 9,64406E-09 |
| ACE2 | CTD-3064M3.3 | 0,2962 | 9,65478E-09 |
| ACE2 | TVP23C | 0,2962 | 9,56059E-09 |
| ACE2 | SNRPEP4 | 0,2962 | 9,59686E-09 |
| ACE2 | QSER1 | 0,2961 | 9,75918E-09 |
| ACE2 | AATK | 0,2961 | 9,66882E-09 |
| ACE2 | VN1R83P | 0,2961 | 9,72346E-09 |
| ACE2 | GPER1 | 0,296 | 9,81534E-09 |
| ACE2 | RP11-326I11.5 | 0,2959 | 1,00164E-08 |
| ACE2 | RP11-6O2.3 | 0,2959 | 9,93561E-09 |
| ACE2 | GGT6 | 0,2959 | 9,97728E-09 |
| ACE2 | GAPDHP1 | 0,2959 | 9,98589E-09 |
| ACE2 | RPSAP72 | 0,2958 | 1,00248E-08 |
| ACE2 | CTC-542B22.2 | 0,2958 | 1,01245E-08 |
| ACE2 | LILRB2 | 0,2958 | 1,00543E-08 |
| ACE2 | CELA2A | 0,2957 | 1,01844E-08 |
| ACE2 | RP11-497H17.1 | 0,2957 | 1,02525E-08 |
| ACE2 | HMGB3P24 | 0,2956 | 1,03442E-08 |
| ACE2 | RP11-477D19.2 | 0,2955 | 1,0494E-08 |
| ACE2 | SLC45A4 | 0,2955 | 1,04244E-08 |
| ACE2 | PTPRD | 0,2954 | 1,05523E-08 |
| ACE2 | BMPR1APS2 | 0,2953 | 1,06949E-08 |
| ACE2 | RP11-51F16.9 | 0,2953 | 1,07239E-08 |
| ACE2 | RP1-122K4.2 | 0,2953 | 1,07458E-08 |
| ACE2 | RN7SL262P | 0,2953 | 1,07489E-08 |
| ACE2 | RP11-325K4.3 | 0,2952 | 1,08248E-08 |
| ACE2 | S100B | 0,2952 | 1,07896E-08 |
| ACE2 | RP5-1050E16.2 | 0,2951 | 1,09678E-08 |
| ACE2 | ENTPD2 | 0,295 | 1,10324E-08 |
| ACE2 | FSD2 | 0,295 | 1,11412E-08 |
| ACE2 | CTA-38K21.2 | 0,295 | 1,1087E-08 |

| | | | |
|---|---|---|---|
| ACE2 | TAF5 | 0,2949 | 1,11615E-08 |
| ACE2 | PRKAG1 | 0,2949 | 1,1214E-08 |
| ACE2 | RP11-567L7.3 | 0,2949 | 1,11905E-08 |
| ACE2 | RP11-1149O23.3 | 0,2948 | 1,13603E-08 |
| ACE2 | LRSAM1 | 0,2947 | 1,14806E-08 |
| ACE2 | RP11-346C4.3 | 0,2947 | 1,14863E-08 |
| ACE2 | ZDHHC8 | 0,2947 | 1,14688E-08 |
| ACE2 | RHOXF2B | 0,2947 | 1,14673E-08 |
| ACE2 | RP11-474P2.4 | 0,2946 | 1,15717E-08 |
| ACE2 | EVL | 0,2946 | 1,16864E-08 |
| ACE2 | SYDE2 | 0,2945 | 1,18053E-08 |
| ACE2 | ZNF575 | 0,2945 | 1,17991E-08 |
| ACE2 | LLNLF-187D8.1 | 0,2945 | 1,17037E-08 |
| ACE2 | TMOD4 | 0,2944 | 1,19072E-08 |
| ACE2 | AC010226.4 | 0,2944 | 1,19765E-08 |
| ACE2 | U62631.5 | 0,2944 | 1,19527E-08 |
| ACE2 | AC113404.1 | 0,2943 | 1,20495E-08 |
| ACE2 | RP3-380E11.2 | 0,2943 | 1,20522E-08 |
| ACE2 | RP11-475C16.1 | 0,2943 | 1,2119E-08 |
| ACE2 | RPS28P7 | 0,2943 | 1,20316E-08 |
| ACE2 | HIST1H1PS1 | 0,2942 | 1,22109E-08 |
| ACE2 | RP11-1072C15.4 | 0,2942 | 1,21682E-08 |
| ACE2 | SCG2 | 0,2941 | 1,23208E-08 |
| ACE2 | RP11-44F21.3 | 0,2941 | 1,24119E-08 |
| ACE2 | CKS1B | 0,294 | 1,24677E-08 |
| ACE2 | C1orf186 | 0,294 | 1,24449E-08 |
| ACE2 | RBBP4P1 | 0,294 | 1,24237E-08 |
| ACE2 | RP6-74O6.6 | 0,2939 | 1,26257E-08 |
| ACE2 | RP11-399B17.1 | 0,2939 | 1,26792E-08 |
| ACE2 | RP11-697N18.3 | 0,2939 | 1,25805E-08 |
| ACE2 | TOMM5 | 0,2939 | 1,26943E-08 |
| ACE2 | RP11-152N13.16 | 0,2939 | 1,25998E-08 |
| ACE2 | GAPDHP70 | 0,2939 | 1,26948E-08 |
| ACE2 | CTD-2012K14.8 | 0,2939 | 1,25785E-08 |
| ACE2 | CELA2B | 0,2938 | 1,28295E-08 |
| ACE2 | TRIM45 | 0,2938 | 1,27654E-08 |
| ACE2 | GPCPD1 | 0,2938 | 1,28237E-08 |
| ACE2 | ARNTL2 | 0,2936 | 1,31E-08 |
| ACE2 | SPHK1 | 0,2936 | 1,31039E-08 |
| ACE2 | USHBP1 | 0,2936 | 1,31076E-08 |
| ACE2 | LSAMP-AS1 | 0,2935 | 1,33288E-08 |
| ACE2 | CTD-2666L21.2 | 0,2935 | 1,32056E-08 |
| ACE2 | KB-1995A5.4 | 0,2935 | 1,32362E-08 |
| ACE2 | RP11-692D12.1 | 0,2934 | 1,34385E-08 |
| ACE2 | RP11-36C20.1 | 0,2934 | 1,34448E-08 |
| ACE2 | LINC01376 | 0,2933 | 1,35002E-08 |
| ACE2 | GJC1 | 0,2933 | 1,35339E-08 |
| ACE2 | HIGD1B | 0,2933 | 1,36362E-08 |
| ACE2 | RP6-159A1.4 | 0,2933 | 1,35296E-08 |
| ACE2 | CTC-497E21.3 | 0,2932 | 1,37411E-08 |
| ACE2 | RP11-492D6.3 | 0,2932 | 1,37578E-08 |
| ACE2 | SNORD89 | 0,2931 | 1,39405E-08 |
| ACE2 | IMP4 | 0,2931 | 1,38287E-08 |
| ACE2 | WWC1 | 0,2931 | 1,38945E-08 |

| | | | |
|---|---|---|---|
| ACE2 | APBB1 | 0,293 | 1,40471E-08 |
| ACE2 | TMSB4XP8 | 0,2929 | 1,41758E-08 |
| ACE2 | RP11-568J23.5 | 0,2929 | 1,41749E-08 |
| ACE2 | ZDHHC8P1 | 0,2929 | 1,41686E-08 |
| ACE2 | RP4-761J14.8 | 0,2928 | 1,44405E-08 |
| ACE2 | RP11-568A7.2 | 0,2927 | 1,45414E-08 |
| ACE2 | TRBV26OR9-2 | 0,2927 | 1,45457E-08 |
| ACE2 | RTN3 | 0,2927 | 1,46349E-08 |
| ACE2 | RP11-74J13.9 | 0,2927 | 1,4528E-08 |
| ACE2 | RPL34 | 0,2925 | 1,48757E-08 |
| ACE2 | CTD-2081C10.7 | 0,2925 | 1,49452E-08 |
| ACE2 | HR | 0,2925 | 1,49053E-08 |
| ACE2 | RP13-923O23.7 | 0,2925 | 1,49894E-08 |
| ACE2 | UPF2 | 0,2925 | 1,49416E-08 |
| ACE2 | KRTAP5-1 | 0,2925 | 1,48756E-08 |
| ACE2 | RP11-697E2.9 | 0,2925 | 1,48317E-08 |
| ACE2 | RP11-364L4.1 | 0,2924 | 1,50094E-08 |
| ACE2 | RP11-248B24.1 | 0,2924 | 1,50173E-08 |
| ACE2 | RP11-104C4.6 | 0,2924 | 1,50909E-08 |
| ACE2 | BANCR | 0,2923 | 1,52152E-08 |
| ACE2 | KMT5C | 0,2923 | 1,53075E-08 |
| ACE2 | RAC1P2 | 0,2921 | 1,55533E-08 |
| ACE2 | CTD-2332E11.2 | 0,2921 | 1,57188E-08 |
| ACE2 | WNT5A-AS1 | 0,292 | 1,58824E-08 |
| ACE2 | NEK6 | 0,292 | 1,57567E-08 |
| ACE2 | PRDM15 | 0,292 | 1,58579E-08 |
| ACE2 | RN7SL333P | 0,2919 | 1,59718E-08 |
| ACE2 | TUBGCP6 | 0,2919 | 1,60336E-08 |
| ACE2 | INTS6L-AS1 | 0,2919 | 1,59235E-08 |
| ACE2 | TAPT1-AS1 | 0,2918 | 1,62397E-08 |
| ACE2 | CTA-276F8.1 | 0,2918 | 1,62476E-08 |
| ACE2 | HRCT1 | 0,2917 | 1,64348E-08 |
| ACE2 | RP11-521B24.5 | 0,2917 | 1,63942E-08 |
| ACE2 | LINS1 | 0,2917 | 1,63442E-08 |
| ACE2 | LGI4 | 0,2917 | 1,63627E-08 |
| ACE2 | SNAP25-AS1 | 0,2917 | 1,63117E-08 |
| ACE2 | CCR5 | 0,2916 | 1,66271E-08 |
| ACE2 | RP11-219B17.3 | 0,2916 | 1,66513E-08 |
| ACE2 | MAGEA8 | 0,2916 | 1,66453E-08 |
| ACE2 | LINC00499 | 0,2915 | 1,67333E-08 |
| ACE2 | RP11-315D13.1 | 0,2915 | 1,67225E-08 |
| ACE2 | UPB1 | 0,2915 | 1,68261E-08 |
| ACE2 | RP11-312J18.5 | 0,2914 | 1,69054E-08 |
| ACE2 | TOP2A | 0,2914 | 1,69286E-08 |
| ACE2 | RP13-128O4.3 | 0,2914 | 1,69899E-08 |
| ACE2 | CDC42-IT1 | 0,2913 | 1,71015E-08 |
| ACE2 | RP11-275F13.3 | 0,2913 | 1,71992E-08 |
| ACE2 | HNRNPKP2 | 0,2913 | 1,72259E-08 |
| ACE2 | WDCP | 0,2912 | 1,73188E-08 |
| ACE2 | CCDC58 | 0,2912 | 1,74E-08 |
| ACE2 | OSBPL7 | 0,2912 | 1,73433E-08 |
| ACE2 | MAB21L1 | 0,2911 | 1,75471E-08 |
| ACE2 | RP11-737O24.3 | 0,291 | 1,78141E-08 |
| ACE2 | KLHL20 | 0,2909 | 1,80872E-08 |

| | | | |
|---|---|---|---|
| ACE2 | RP11-452L6.6 | 0,2909 | 1,80073E-08 |
| ACE2 | HIBCH | 0,2908 | 1,82282E-08 |
| ACE2 | NEURL1B | 0,2908 | 1,81269E-08 |
| ACE2 | HSPD1P21 | 0,2908 | 1,83211E-08 |
| ACE2 | MAP2 | 0,2907 | 1,84403E-08 |
| ACE2 | RPL7AP26 | 0,2907 | 1,84859E-08 |
| ACE2 | APOLD1 | 0,2907 | 1,83608E-08 |
| ACE2 | SLC9B2 | 0,2906 | 1,87531E-08 |
| ACE2 | PPIAP31 | 0,2906 | 1,86004E-08 |
| ACE2 | ELMOD1 | 0,2906 | 1,8687E-08 |
| ACE2 | RIPPLY2 | 0,2905 | 1,89155E-08 |
| ACE2 | RP11-498P14.5 | 0,2905 | 1,89417E-08 |
| ACE2 | RP11-23E10.4 | 0,2904 | 1,90426E-08 |
| ACE2 | HSPH1 | 0,2903 | 1,9259E-08 |
| ACE2 | RP11-38M8.1 | 0,2902 | 1,96061E-08 |
| ACE2 | RP11-724N1.1 | 0,2902 | 1,96264E-08 |
| ACE2 | RP11-343B18.2 | 0,2902 | 1,95505E-08 |
| ACE2 | EIF4EBP1 | 0,29 | 1,994E-08 |
| ACE2 | LCN10 | 0,29 | 1,99167E-08 |
| ACE2 | RP11-30K9.6 | 0,29 | 1,9918E-08 |
| ACE2 | FTH1P1 | 0,2899 | 2,01926E-08 |
| ACE2 | MAN1A2P1 | 0,2899 | 2,0202E-08 |
| ACE2 | VDAC1P1 | 0,2899 | 2,01359E-08 |
| ACE2 | RP11-434P11.2 | 0,2898 | 2,04134E-08 |
| ACE2 | RP11-325L7.2 | 0,2898 | 2,04105E-08 |
| ACE2 | RP11-510M2.4 | 0,2898 | 2,05815E-08 |
| ACE2 | TPD52L1 | 0,2897 | 2,06403E-08 |
| ACE2 | DLL1 | 0,2897 | 2,06415E-08 |
| ACE2 | LDLRAD4-AS1 | 0,2897 | 2,08282E-08 |
| ACE2 | ARHGEF37 | 0,2896 | 2,10762E-08 |
| ACE2 | FCGR2B | 0,2895 | 2,11574E-08 |
| ACE2 | DUOXA1 | 0,2895 | 2,13096E-08 |
| ACE2 | NOS2 | 0,2895 | 2,11602E-08 |
| ACE2 | COLEC11 | 0,2894 | 2,1522E-08 |
| ACE2 | IL1RL1 | 0,2894 | 2,15737E-08 |
| ACE2 | SFRP5 | 0,2894 | 2,14248E-08 |
| ACE2 | RNF207 | 0,2893 | 2,16094E-08 |
| ACE2 | MSX1 | 0,2893 | 2,18281E-08 |
| ACE2 | ATAD2 | 0,2893 | 2,17199E-08 |
| ACE2 | EIF4EBP2P2 | 0,2892 | 2,20455E-08 |
| ACE2 | TTC31 | 0,2891 | 2,22998E-08 |
| ACE2 | DPP3P2 | 0,2891 | 2,23197E-08 |
| ACE2 | ZNF840P | 0,2891 | 2,21229E-08 |
| ACE2 | RBM44 | 0,289 | 2,25861E-08 |
| ACE2 | TGM2 | 0,289 | 2,24138E-08 |
| ACE2 | FMO9P | 0,2889 | 2,28676E-08 |
| ACE2 | RP11-260M2.1 | 0,2889 | 2,28373E-08 |
| ACE2 | PMS2P9 | 0,2889 | 2,26449E-08 |
| ACE2 | RP13-216E22.5 | 0,2889 | 2,28667E-08 |
| ACE2 | RP5-998N21.4 | 0,2888 | 2,29904E-08 |
| ACE2 | ADPRM | 0,2888 | 2,31388E-08 |
| ACE2 | NIFK-AS1 | 0,2887 | 2,32932E-08 |
| ACE2 | AOAH | 0,2887 | 2,32597E-08 |
| ACE2 | RP11-428G5.7 | 0,2887 | 2,32674E-08 |

| | | | |
|---|---|---|---|
| ACE2 | PCYT1A | 0,2886 | 2,35108E-08 |
| ACE2 | INHBE | 0,2886 | 2,35872E-08 |
| ACE2 | XAB2 | 0,2885 | 2,38952E-08 |
| ACE2 | EGR1 | 0,2883 | 2,44828E-08 |
| ACE2 | PTGER4P2 | 0,2883 | 2,45268E-08 |
| ACE2 | RPS6P25 | 0,2883 | 2,44894E-08 |
| ACE2 | DAZ2 | 0,2883 | 2,4284E-08 |
| ACE2 | FAM167B | 0,2882 | 2,4694E-08 |
| ACE2 | MAN2A1 | 0,2882 | 2,47898E-08 |
| ACE2 | PLAC9 | 0,2882 | 2,46206E-08 |
| ACE2 | NLRP8 | 0,2882 | 2,46129E-08 |
| ACE2 | BMS1P22 | 0,2882 | 2,47783E-08 |
| ACE2 | CA14 | 0,2881 | 2,48811E-08 |
| ACE2 | RP11-164P12.5 | 0,288 | 2,53063E-08 |
| ACE2 | RPS24 | 0,288 | 2,51668E-08 |
| ACE2 | RP11-384C4.2 | 0,2879 | 2,54607E-08 |
| ACE2 | CPA3 | 0,2879 | 2,56288E-08 |
| ACE2 | RASA4B | 0,2879 | 2,55481E-08 |
| ACE2 | LINC00702 | 0,2879 | 2,5443E-08 |
| ACE2 | MYCBP2-AS1 | 0,2879 | 2,54279E-08 |
| ACE2 | CLEC10A | 0,2879 | 2,54329E-08 |
| ACE2 | TFAP2A | 0,2878 | 2,582E-08 |
| ACE2 | LINC01537 | 0,2878 | 2,58961E-08 |
| ACE2 | LINC00278 | 0,2878 | 2,59567E-08 |
| ACE2 | RP3-425C14.5 | 0,2877 | 2,61642E-08 |
| ACE2 | MATN1-AS1 | 0,2876 | 2,63106E-08 |
| ACE2 | RP11-404O13.5 | 0,2874 | 2,7181E-08 |
| ACE2 | TBC1D25 | 0,2874 | 2,70839E-08 |
| ACE2 | HIST1H2AL | 0,2873 | 2,7522E-08 |
| ACE2 | KLRA1P | 0,2873 | 2,73486E-08 |
| ACE2 | TIPRL | 0,2872 | 2,77017E-08 |
| ACE2 | WT1-AS_2 | 0,2872 | 2,78372E-08 |
| ACE2 | C5orf56 | 0,2871 | 2,80282E-08 |
| ACE2 | CES2 | 0,2871 | 2,80767E-08 |
| ACE2 | SSR4 | 0,2871 | 2,79357E-08 |
| ACE2 | RP11-378A13.1 | 0,287 | 2,83402E-08 |
| ACE2 | VPS25 | 0,287 | 2,82456E-08 |
| ACE2 | ABCA9-AS1 | 0,287 | 2,83034E-08 |
| ACE2 | UAP1L1 | 0,2869 | 2,88169E-08 |
| ACE2 | RP11-474P2.7 | 0,2869 | 2,86574E-08 |
| ACE2 | FRMD4B | 0,2868 | 2,88657E-08 |
| ACE2 | GPR162 | 0,2868 | 2,91264E-08 |
| ACE2 | FCMR | 0,2867 | 2,92889E-08 |
| ACE2 | C8orf31 | 0,2867 | 2,93957E-08 |
| ACE2 | CTC-487M23.5 | 0,2866 | 2,96335E-08 |
| ACE2 | HCRTR2 | 0,2866 | 2,97792E-08 |
| ACE2 | METTL15 | 0,2866 | 2,97429E-08 |
| ACE2 | RP11-483C6.1 | 0,2866 | 2,96773E-08 |
| ACE2 | PIGHP1 | 0,2865 | 2,9994E-08 |
| ACE2 | CTD-2527I21.5 | 0,2865 | 2,99157E-08 |
| ACE2 | CTD-2336O2.1 | 0,2864 | 3,04498E-08 |
| ACE2 | RP11-85I17.2 | 0,2864 | 3,05467E-08 |
| ACE2 | PHOX2A | 0,2864 | 3,02243E-08 |
| ACE2 | WIPI1 | 0,2864 | 3,03016E-08 |

| | | | |
|---|---|---|---|
| ACE2 | AC000068.9 | 0,2864 | 3,0505E-08 |
| ACE2 | RAB41 | 0,2864 | 3,02451E-08 |
| ACE2 | DPH2 | 0,2863 | 3,06508E-08 |
| ACE2 | HSPA8P15 | 0,2863 | 3,07302E-08 |
| ACE2 | TINCR | 0,2863 | 3,09092E-08 |
| ACE2 | RP11-243M5.2 | 0,2862 | 3,11443E-08 |
| ACE2 | SMG1P5 | 0,2862 | 3,12351E-08 |
| ACE2 | SIRT6 | 0,2862 | 3,11614E-08 |
| ACE2 | DPPA3 | 0,2861 | 3,12853E-08 |
| ACE2 | ANO9 | 0,286 | 3,18668E-08 |
| ACE2 | HYDIN2 | 0,2859 | 3,22734E-08 |
| ACE2 | TSPY11P | 0,2859 | 3,20839E-08 |
| ACE2 | TRAF5 | 0,2858 | 3,25996E-08 |
| ACE2 | ANKRD31 | 0,2858 | 3,25049E-08 |
| ACE2 | IGHV3-72 | 0,2858 | 3,27365E-08 |
| ACE2 | SERINC2 | 0,2857 | 3,28769E-08 |
| ACE2 | MISP3 | 0,2856 | 3,34484E-08 |
| ACE2 | CASZ1 | 0,2855 | 3,3636E-08 |
| ACE2 | LRRN2 | 0,2855 | 3,36175E-08 |
| ACE2 | RP11-77P6.2 | 0,2855 | 3,35535E-08 |
| ACE2 | RP11-847H18.2 | 0,2855 | 3,37533E-08 |
| ACE2 | LINC02074 | 0,2855 | 3,35167E-08 |
| ACE2 | ALG1L13P | 0,2854 | 3,42719E-08 |
| ACE2 | POF1B | 0,2853 | 3,43879E-08 |
| ACE2 | ZNF467 | 0,2852 | 3,48912E-08 |
| ACE2 | RBM42 | 0,2852 | 3,48985E-08 |
| ACE2 | PTCD1 | 0,2851 | 3,54383E-08 |
| ACE2 | LYST | 0,285 | 3,58955E-08 |
| ACE2 | AC073343.13 | 0,285 | 3,58834E-08 |
| ACE2 | CADM2 | 0,2849 | 3,59784E-08 |
| ACE2 | SNX11 | 0,2849 | 3,6313E-08 |
| ACE2 | XXbac-BPG299F13.17 | 0,2848 | 3,63297E-08 |
| ACE2 | KCNA4 | 0,2848 | 3,65326E-08 |
| ACE2 | CTD-2147F2.1 | 0,2848 | 3,66102E-08 |
| ACE2 | RP11-147L13.13 | 0,2848 | 3,63295E-08 |
| ACE2 | RNA5SP187 | 0,2846 | 3,73825E-08 |
| ACE2 | ESRP1 | 0,2846 | 3,74044E-08 |
| ACE2 | ZNF780B | 0,2845 | 3,76397E-08 |
| ACE2 | KCNJ13 | 0,2844 | 3,81048E-08 |
| ACE2 | AC092597.3 | 0,2844 | 3,83186E-08 |
| ACE2 | RP11-358B23.6 | 0,2844 | 3,80635E-08 |
| ACE2 | SEPT7P4 | 0,2843 | 3,87792E-08 |
| ACE2 | RP11-349F21.5 | 0,2843 | 3,88731E-08 |
| ACE2 | NTHL1 | 0,2843 | 3,86522E-08 |
| ACE2 | ZNF663P | 0,2843 | 3,86771E-08 |
| ACE2 | PLPP4 | 0,2842 | 3,89995E-08 |
| ACE2 | CNOT3 | 0,2842 | 3,89731E-08 |
| ACE2 | CH507-42P11.8 | 0,2842 | 3,89004E-08 |
| ACE2 | GABRE | 0,2842 | 3,9241E-08 |
| ACE2 | SPEG | 0,2841 | 3,94269E-08 |
| ACE2 | RP11-61K12.2 | 0,2841 | 3,97271E-08 |
| ACE2 | CNOT8 | 0,2841 | 3,93842E-08 |
| ACE2 | SMG1P1 | 0,2841 | 3,93929E-08 |
| ACE2 | RP11-93O14.2 | 0,2841 | 3,95696E-08 |

| | | | |
|---|---|---|---|
| ACE2 | RP11-131M11.2 | 0,284 | 3,99399E-08 |
| ACE2 | LINC00621 | 0,284 | 4,01072E-08 |
| ACE2 | RP11-506M13.3 | 0,2839 | 4,04298E-08 |
| ACE2 | ISG15 | 0,2838 | 4,07237E-08 |
| ACE2 | AC009404.2 | 0,2838 | 4,11419E-08 |
| ACE2 | ABCA5 | 0,2838 | 4,09695E-08 |
| ACE2 | RELB | 0,2838 | 4,09696E-08 |
| ACE2 | BIK | 0,2838 | 4,10558E-08 |
| ACE2 | RP11-284A20.1 | 0,2837 | 4,13582E-08 |
| ACE2 | KIAA0391 | 0,2837 | 4,14773E-08 |
| ACE2 | RP11-849N15.3 | 0,2837 | 4,11996E-08 |
| ACE2 | KLF2 | 0,2837 | 4,15397E-08 |
| ACE2 | ERVWE2 | 0,2837 | 4,12519E-08 |
| ACE2 | SMARCAL1 | 0,2836 | 4,19785E-08 |
| ACE2 | AC011380.8 | 0,2836 | 4,18861E-08 |
| ACE2 | RP11-797H7.1 | 0,2836 | 4,20398E-08 |
| ACE2 | DUOX1 | 0,2836 | 4,17471E-08 |
| ACE2 | ZSCAN30 | 0,2836 | 4,19024E-08 |
| ACE2 | HPGD | 0,2835 | 4,24555E-08 |
| ACE2 | CLEC7A | 0,2834 | 4,30671E-08 |
| ACE2 | LY86-AS1 | 0,2833 | 4,32618E-08 |
| ACE2 | AC005154.8 | 0,2833 | 4,33177E-08 |
| ACE2 | ACTG1P14 | 0,2833 | 4,31192E-08 |
| ACE2 | AMIGO1 | 0,2832 | 4,38601E-08 |
| ACE2 | SCAF11 | 0,2832 | 4,38721E-08 |
| ACE2 | DLX4 | 0,2832 | 4,38009E-08 |
| ACE2 | MSL1 | 0,2831 | 4,44326E-08 |
| ACE2 | RNF14 | 0,283 | 4,50902E-08 |
| ACE2 | SCARNA12 | 0,283 | 4,46018E-08 |
| ACE2 | GACAT3 | 0,2828 | 4,56734E-08 |
| ACE2 | COL5A2 | 0,2828 | 4,57561E-08 |
| ACE2 | RP11-15L13.4 | 0,2828 | 4,57555E-08 |
| ACE2 | CDK6 | 0,2828 | 4,59212E-08 |
| ACE2 | DDIAS | 0,2828 | 4,5655E-08 |
| ACE2 | RP13-20L14.10 | 0,2828 | 4,56959E-08 |
| ACE2 | HHIPL1 | 0,2827 | 4,65696E-08 |
| ACE2 | GAREM2 | 0,2826 | 4,70956E-08 |
| ACE2 | NFKBID | 0,2826 | 4,68178E-08 |
| ACE2 | RP11-181C21.4 | 0,2825 | 4,73075E-08 |
| ACE2 | SEC61G | 0,2825 | 4,73871E-08 |
| ACE2 | CBWD5 | 0,2825 | 4,75506E-08 |
| ACE2 | RBMS1P1 | 0,2825 | 4,73325E-08 |
| ACE2 | JAK3 | 0,2825 | 4,76832E-08 |
| ACE2 | ARID5A | 0,2824 | 4,792E-08 |
| ACE2 | FAM181B | 0,2824 | 4,82837E-08 |
| ACE2 | IDH1-AS1 | 0,2823 | 4,8495E-08 |
| ACE2 | CTD-3126B10.5 | 0,2823 | 4,87635E-08 |
| ACE2 | TULP3 | 0,2822 | 4,93912E-08 |
| ACE2 | PSMA3-AS1 | 0,2822 | 4,89557E-08 |
| ACE2 | CALB2 | 0,2822 | 4,92891E-08 |
| ACE2 | RP5-1142A6.10 | 0,2822 | 4,91203E-08 |
| ACE2 | ACP5 | 0,2822 | 4,89405E-08 |
| ACE2 | AC011290.5 | 0,2821 | 4,95632E-08 |
| ACE2 | LONRF3 | 0,2821 | 4,98574E-08 |

| | | | |
|---|---|---|---|
| ACE2 | RP11-575H3.1 | 0,282 | 5,01643E-08 |
| ACE2 | LINC00540 | 0,282 | 5,05319E-08 |
| ACE2 | RP4-751H13.5 | 0,2819 | 5,07967E-08 |
| ACE2 | AC010136.2 | 0,2818 | 5,11321E-08 |
| ACE2 | ZNF777 | 0,2818 | 5,12748E-08 |
| ACE2 | NUFIP1 | 0,2818 | 5,13133E-08 |
| ACE2 | RP11-83B20.3 | 0,2817 | 5,17336E-08 |
| ACE2 | AF127577.13 | 0,2817 | 5,19735E-08 |
| ACE2 | MEN1 | 0,2816 | 5,23595E-08 |
| ACE2 | NALCN | 0,2816 | 5,26658E-08 |
| ACE2 | RP1-56J10.8 | 0,2816 | 5,23648E-08 |
| ACE2 | CAP1 | 0,2815 | 5,33441E-08 |
| ACE2 | ORC6 | 0,2815 | 5,2913E-08 |
| ACE2 | AC069257.6 | 0,2814 | 5,37264E-08 |
| ACE2 | RNA5SP317 | 0,2814 | 5,40461E-08 |
| ACE2 | RAPGEF3 | 0,2813 | 5,44198E-08 |
| ACE2 | RP11-439A17.9 | 0,2812 | 5,52953E-08 |
| ACE2 | TNRC18 | 0,2812 | 5,49845E-08 |
| ACE2 | TWSG1 | 0,2811 | 5,59335E-08 |
| ACE2 | ZNF92P3 | 0,2809 | 5,70299E-08 |
| ACE2 | THBS4 | 0,2808 | 5,73252E-08 |
| ACE2 | TIGD7 | 0,2808 | 5,7309E-08 |
| ACE2 | C16orf54 | 0,2808 | 5,7754E-08 |
| ACE2 | INSL3 | 0,2808 | 5,77735E-08 |
| ACE2 | TSPY13P | 0,2808 | 5,77985E-08 |
| ACE2 | RELL1 | 0,2807 | 5,82073E-08 |
| ACE2 | LYSMD4 | 0,2807 | 5,83497E-08 |
| ACE2 | SNORD10 | 0,2807 | 5,78868E-08 |
| ACE2 | IP6K3 | 0,2806 | 5,87629E-08 |
| ACE2 | ETV7 | 0,2806 | 5,90865E-08 |
| ACE2 | UBE3D | 0,2806 | 5,90776E-08 |
| ACE2 | RN7SL81P | 0,2804 | 6,03573E-08 |
| ACE2 | TNNT3 | 0,2804 | 6,04518E-08 |
| ACE2 | RP11-225H22.4 | 0,2803 | 6,10699E-08 |
| ACE2 | CTD-2349P21.10 | 0,2803 | 6,11438E-08 |
| ACE2 | RP11-465B22.3 | 0,2802 | 6,16073E-08 |
| ACE2 | RP11-367F23.1 | 0,2802 | 6,12741E-08 |
| ACE2 | RP11-197M22.2 | 0,2802 | 6,15244E-08 |
| ACE2 | CTD-3032J10.3 | 0,2802 | 6,12504E-08 |
| ACE2 | TMEM161A | 0,2801 | 6,19262E-08 |
| ACE2 | ADCY1 | 0,28 | 6,28922E-08 |
| ACE2 | RP11-481J2.4 | 0,28 | 6,26617E-08 |
| ACE2 | NTRK1 | 0,2799 | 6,38932E-08 |
| ACE2 | COX4I2 | 0,2799 | 6,38141E-08 |
| ACE2 | NFASC | 0,2798 | 6,45229E-08 |
| ACE2 | RP11-778D9.4 | 0,2798 | 6,44681E-08 |
| ACE2 | LINC01511 | 0,2798 | 6,44765E-08 |
| ACE2 | LINC01011 | 0,2797 | 6,48249E-08 |
| ACE2 | AC003005.2 | 0,2797 | 6,54956E-08 |
| ACE2 | SLC31A2 | 0,2796 | 6,59128E-08 |
| ACE2 | Z69666.2 | 0,2796 | 6,57509E-08 |
| ACE2 | CARD8 | 0,2796 | 6,56294E-08 |
| ACE2 | RP11-798M19.6 | 0,2795 | 6,64749E-08 |
| ACE2 | RHEBL1 | 0,2795 | 6,64316E-08 |

| | | | |
|---|---|---|---|
| ACE2 | RP11-897M7.3 | 0,2795 | 6,63722E-08 |
| ACE2 | RP11-745A24.1 | 0,2794 | 6,76041E-08 |
| ACE2 | ELL2P1 | 0,2792 | 6,8574E-08 |
| ACE2 | TMEM17 | 0,2792 | 6,88081E-08 |
| ACE2 | NSMF | 0,2792 | 6,91383E-08 |
| ACE2 | CKAP2 | 0,2792 | 6,86218E-08 |
| ACE2 | ABCD4 | 0,2792 | 6,89033E-08 |
| ACE2 | HERC2P4 | 0,2792 | 6,88193E-08 |
| ACE2 | AC137932.4 | 0,2792 | 6,86582E-08 |
| ACE2 | RP1-40E16.11 | 0,2791 | 6,94858E-08 |
| ACE2 | TMEM214 | 0,279 | 7,08417E-08 |
| ACE2 | CTC-338M12.9 | 0,279 | 7,04834E-08 |
| ACE2 | NR1D1 | 0,279 | 7,0094E-08 |
| ACE2 | TSPAN11 | 0,2789 | 7,14878E-08 |
| ACE2 | TMEM51-AS1 | 0,2788 | 7,19026E-08 |
| ACE2 | SNORA66 | 0,2788 | 7,21588E-08 |
| ACE2 | TTL | 0,2787 | 7,28778E-08 |
| ACE2 | DHFR2 | 0,2787 | 7,25973E-08 |
| ACE2 | RP11-478C1.7 | 0,2787 | 7,26938E-08 |
| ACE2 | PPP1R3B | 0,2787 | 7,24751E-08 |
| ACE2 | ZNF784 | 0,2787 | 7,27555E-08 |
| ACE2 | CHCHD10 | 0,2787 | 7,31808E-08 |
| ACE2 | AC114973.1 | 0,2786 | 7,40266E-08 |
| ACE2 | MICE | 0,2786 | 7,37835E-08 |
| ACE2 | HGF | 0,2786 | 7,37677E-08 |
| ACE2 | TAGLN2P1 | 0,2786 | 7,3297E-08 |
| ACE2 | APBA1 | 0,2786 | 7,39299E-08 |
| ACE2 | RP11-950C14.10 | 0,2786 | 7,33794E-08 |
| ACE2 | CYP4F22 | 0,2786 | 7,35567E-08 |
| ACE2 | CD52 | 0,2784 | 7,57421E-08 |
| ACE2 | NTSR2 | 0,2784 | 7,56294E-08 |
| ACE2 | PTGER4P2-CDK2AP2F | 0,2784 | 7,50166E-08 |
| ACE2 | RP1-128M12.3 | 0,2783 | 7,62818E-08 |
| ACE2 | RP11-976B16.1 | 0,2783 | 7,61775E-08 |
| ACE2 | DMPK | 0,2783 | 7,58402E-08 |
| ACE2 | SORCS1 | 0,2781 | 7,75113E-08 |
| ACE2 | RP11-524F11.3 | 0,2781 | 7,75577E-08 |
| ACE2 | FRZB | 0,278 | 7,90211E-08 |
| ACE2 | RP11-11C20.3 | 0,278 | 7,88174E-08 |
| ACE2 | C1QL3 | 0,278 | 7,89641E-08 |
| ACE2 | GBP1P1 | 0,2779 | 7,9568E-08 |
| ACE2 | RP11-589N15.2 | 0,2779 | 7,9744E-08 |
| ACE2 | AC009120.5 | 0,2779 | 7,98496E-08 |
| ACE2 | NBPF9 | 0,2778 | 8,05507E-08 |
| ACE2 | MLPH | 0,2778 | 8,03037E-08 |
| ACE2 | LOXL2 | 0,2778 | 8,06017E-08 |
| ACE2 | RP11-350N15.4 | 0,2778 | 8,08583E-08 |
| ACE2 | RP11-138H8.6 | 0,2778 | 8,08766E-08 |
| ACE2 | CPSF7 | 0,2777 | 8,18957E-08 |
| ACE2 | AAK1 | 0,2776 | 8,24802E-08 |
| ACE2 | ENY2 | 0,2776 | 8,20231E-08 |
| ACE2 | OR5BK1P | 0,2776 | 8,26758E-08 |
| ACE2 | MB21D1 | 0,2775 | 8,33841E-08 |
| ACE2 | ZNF252P | 0,2775 | 8,33839E-08 |

| | | | |
|---|---|---|---|
| ACE2 | TRPC3 | 0,2774 | 8,37946E-08 |
| ACE2 | ITGB3 | 0,2774 | 8,42357E-08 |
| ACE2 | FOXRED1 | 0,2773 | 8,53657E-08 |
| ACE2 | RN7SL225P | 0,2772 | 8,61722E-08 |
| ACE2 | CYP2F2P | 0,2772 | 8,60509E-08 |
| ACE2 | BEST4 | 0,2771 | 8,66167E-08 |
| ACE2 | JAZF1-AS1 | 0,2771 | 8,6844E-08 |
| ACE2 | IGHV5-78 | 0,2771 | 8,74517E-08 |
| ACE2 | RP11-122G18.12 | 0,277 | 8,79128E-08 |
| ACE2 | RPS12P26 | 0,277 | 8,81563E-08 |
| ACE2 | RNF166 | 0,277 | 8,76923E-08 |
| ACE2 | MIR635 | 0,277 | 8,82806E-08 |
| ACE2 | SH3RF3-AS1 | 0,2769 | 8,94864E-08 |
| ACE2 | CH17-296N19.1 | 0,2769 | 8,85636E-08 |
| ACE2 | MKNK2 | 0,2769 | 8,89204E-08 |
| ACE2 | DMC1 | 0,2769 | 8,85765E-08 |
| ACE2 | MCCC1-AS1 | 0,2768 | 9,01087E-08 |
| ACE2 | GSDMD | 0,2768 | 8,98867E-08 |
| ACE2 | RPS3AP34 | 0,2767 | 9,15109E-08 |
| ACE2 | GOLM1 | 0,2767 | 9,15322E-08 |
| ACE2 | RPL29P3 | 0,2767 | 9,10959E-08 |
| ACE2 | RP11-1260E13.4 | 0,2767 | 9,11333E-08 |
| ACE2 | CD22 | 0,2767 | 9,07158E-08 |
| ACE2 | EIF4E1B | 0,2766 | 9,23703E-08 |
| ACE2 | TMPOP1 | 0,2765 | 9,32524E-08 |
| ACE2 | CSTF3 | 0,2765 | 9,27096E-08 |
| ACE2 | RP11-430E17.1 | 0,2765 | 9,2976E-08 |
| ACE2 | NKILA | 0,2765 | 9,29784E-08 |
| ACE2 | LINC01262 | 0,2764 | 9,38416E-08 |
| ACE2 | CHEK2P2 | 0,2764 | 9,44781E-08 |
| ACE2 | MIR3142HG | 0,2763 | 9,48379E-08 |
| ACE2 | GSTM5 | 0,2762 | 9,66936E-08 |
| ACE2 | RP11-480I12.9 | 0,2762 | 9,66076E-08 |
| ACE2 | ST8SIA4 | 0,2762 | 9,63877E-08 |
| ACE2 | ZBTB10 | 0,2762 | 9,61521E-08 |
| ACE2 | RP11-13K12.1 | 0,2762 | 9,6022E-08 |
| ACE2 | CTD-2527I21.14 | 0,2762 | 9,65513E-08 |
| ACE2 | RP4-549F15.1 | 0,2761 | 9,77695E-08 |
| ACE2 | SH3TC1 | 0,2761 | 9,77145E-08 |
| ACE2 | ANKRD11 | 0,2761 | 9,70302E-08 |
| ACE2 | A1BG-AS1 | 0,2761 | 9,7409E-08 |
| ACE2 | RP11-516A11.1 | 0,2761 | 9,67762E-08 |
| ACE2 | HAPLN3 | 0,276 | 9,84602E-08 |
| ACE2 | CTC-293G12.1 | 0,2759 | 9,89571E-08 |
| ACE2 | OCA2 | 0,2759 | 9,9048E-08 |
| ACE2 | HMGN2P15 | 0,2759 | 9,91978E-08 |
| ACE2 | ANXA3 | 0,2758 | 1,00159E-07 |
| ACE2 | MTFP1 | 0,2758 | 1,00695E-07 |
| ACE2 | SHROOM2P1 | 0,2758 | 1,00523E-07 |
| ACE2 | LINC01914 | 0,2757 | 1,01202E-07 |
| ACE2 | SNHG7 | 0,2757 | 1,01173E-07 |
| ACE2 | CTD-2245F17.6 | 0,2757 | 1,01588E-07 |
| ACE2 | ZNF250 | 0,2756 | 1,03128E-07 |
| ACE2 | RP11-403A3.3 | 0,2756 | 1,03136E-07 |

| | | | |
|---|---|---|---|
| ACE2 | RASA3 | 0,2756 | 1,03318E-07 |
| ACE2 | CTC-529I10.1 | 0,2756 | 1,02748E-07 |
| ACE2 | RP1-290F12.3 | 0,2756 | 1,02347E-07 |
| ACE2 | ARL4C | 0,2755 | 1,03437E-07 |
| ACE2 | RP11-844P9.4 | 0,2755 | 1,03642E-07 |
| ACE2 | RP11-373D7.1 | 0,2755 | 1,04335E-07 |
| ACE2 | CTD-2547L16.3 | 0,2755 | 1,04243E-07 |
| ACE2 | RP11-80H5.7 | 0,2755 | 1,03945E-07 |
| ACE2 | RP11-123O10.4 | 0,2755 | 1,0398E-07 |
| ACE2 | RP11-261C10.8 | 0,2754 | 1,05243E-07 |
| ACE2 | MTCO1P40 | 0,2754 | 1,05698E-07 |
| ACE2 | RP11-855A2.3 | 0,2754 | 1,05319E-07 |
| ACE2 | BNIP3P22 | 0,2754 | 1,05008E-07 |
| ACE2 | CTD-2233K9.1 | 0,2754 | 1,05187E-07 |
| ACE2 | PNMAL1 | 0,2752 | 1,0767E-07 |
| ACE2 | LL22NC03-86G7.1 | 0,2752 | 1,07978E-07 |
| ACE2 | RP3-331H24.7 | 0,2751 | 1,08407E-07 |
| ACE2 | MINCR | 0,2751 | 1,08759E-07 |
| ACE2 | RP3-414A15.11 | 0,2751 | 1,08875E-07 |
| ACE2 | RPL11 | 0,275 | 1,10309E-07 |
| ACE2 | RP1-179N16.3 | 0,275 | 1,09633E-07 |
| ACE2 | RP5-965G21.6 | 0,275 | 1,10159E-07 |
| ACE2 | CSF1R | 0,2749 | 1,11296E-07 |
| ACE2 | RP11-21L23.3 | 0,2749 | 1,11132E-07 |
| ACE2 | LTA4H | 0,2749 | 1,11576E-07 |
| ACE2 | LINC01101 | 0,2748 | 1,12168E-07 |
| ACE2 | RP11-713P17.3 | 0,2748 | 1,11775E-07 |
| ACE2 | PXN-AS1 | 0,2748 | 1,11731E-07 |
| ACE2 | DPY30 | 0,2747 | 1,14116E-07 |
| ACE2 | RP1-249I4.2 | 0,2747 | 1,13046E-07 |
| ACE2 | TRPC5 | 0,2747 | 1,12995E-07 |
| ACE2 | RP11-65L3.2 | 0,2746 | 1,1461E-07 |
| ACE2 | RPL5P1 | 0,2746 | 1,15332E-07 |
| ACE2 | CTD-2224J9.8 | 0,2746 | 1,15106E-07 |
| ACE2 | MGME1 | 0,2746 | 1,14856E-07 |
| ACE2 | TEX11 | 0,2745 | 1,16107E-07 |
| ACE2 | LLPH | 0,2744 | 1,17027E-07 |
| ACE2 | RP11-621L6.2 | 0,2744 | 1,17926E-07 |
| ACE2 | RNA5SP479 | 0,2744 | 1,1693E-07 |
| ACE2 | HNRNPA1P22 | 0,2743 | 1,18578E-07 |
| ACE2 | RP11-104G3.2 | 0,2743 | 1,19286E-07 |
| ACE2 | ASAH2 | 0,2743 | 1,1901E-07 |
| ACE2 | PER1 | 0,2743 | 1,18105E-07 |
| ACE2 | FGF13-AS1 | 0,2743 | 1,18913E-07 |
| ACE2 | PAPPA2 | 0,2742 | 1,20115E-07 |
| ACE2 | B3GNT8 | 0,2742 | 1,19575E-07 |
| ACE2 | RP11-747H7.3 | 0,2741 | 1,21927E-07 |
| ACE2 | CTC-325H20.4 | 0,2741 | 1,20692E-07 |
| ACE2 | GAP43 | 0,274 | 1,22239E-07 |
| ACE2 | RP11-39H3.2 | 0,274 | 1,2306E-07 |
| ACE2 | EIF5 | 0,274 | 1,22836E-07 |
| ACE2 | CTD-3157E16.1 | 0,274 | 1,22182E-07 |
| ACE2 | ELANE | 0,274 | 1,22182E-07 |
| ACE2 | RP1-138B7.4 | 0,274 | 1,21996E-07 |

| | | | |
|---|---|---|---|
| ACE2 | MAP2K4P1 | 0,274 | 1,23248E-07 |
| ACE2 | LINC01447 | 0,2739 | 1,23753E-07 |
| ACE2 | RP11-626G11.6 | 0,2739 | 1,24522E-07 |
| ACE2 | AC016739.2 | 0,2738 | 1,24738E-07 |
| ACE2 | OR3A2 | 0,2738 | 1,25374E-07 |
| ACE2 | RP11-17P16.1 | 0,2736 | 1,27677E-07 |
| ACE2 | PKN3 | 0,2736 | 1,27995E-07 |
| ACE2 | GAREM1 | 0,2736 | 1,28193E-07 |
| ACE2 | LINC01901 | 0,2736 | 1,28676E-07 |
| ACE2 | NXF1 | 0,2735 | 1,29279E-07 |
| ACE2 | RP11-334C17.5 | 0,2735 | 1,29319E-07 |
| ACE2 | bP-2171C21.5 | 0,2735 | 1,30192E-07 |
| ACE2 | EBF1 | 0,2734 | 1,30774E-07 |
| ACE2 | GIF | 0,2734 | 1,30324E-07 |
| ACE2 | RP11-269G24.7 | 0,2734 | 1,30951E-07 |
| ACE2 | AC002454.1 | 0,2733 | 1,32933E-07 |
| ACE2 | RP11-187A9.3 | 0,2733 | 1,3301E-07 |
| ACE2 | NLRC5 | 0,2733 | 1,31787E-07 |
| ACE2 | RNU6-1048P | 0,2731 | 1,34647E-07 |
| ACE2 | AC005534.6 | 0,2731 | 1,36093E-07 |
| ACE2 | RN7SL566P | 0,2731 | 1,34896E-07 |
| ACE2 | RP11-476C8.2 | 0,273 | 1,36508E-07 |
| ACE2 | ZBED3-AS1 | 0,273 | 1,36492E-07 |
| ACE2 | RPL23AP66 | 0,273 | 1,36218E-07 |
| ACE2 | HMGXB3 | 0,2729 | 1,38408E-07 |
| ACE2 | RP11-38L15.3 | 0,2729 | 1,38224E-07 |
| ACE2 | TMEM221 | 0,2729 | 1,3872E-07 |
| ACE2 | MKRN4P | 0,2729 | 1,38235E-07 |
| ACE2 | ICOS | 0,2728 | 1,39171E-07 |
| ACE2 | QRSL1P3 | 0,2728 | 1,3976E-07 |
| ACE2 | RP11-384M15.3 | 0,2728 | 1,40417E-07 |
| ACE2 | FAM83G | 0,2728 | 1,39425E-07 |
| ACE2 | TNRC6C-AS1 | 0,2728 | 1,40577E-07 |
| ACE2 | CTD-2541J13.1 | 0,2728 | 1,40314E-07 |
| ACE2 | RP3-412A9.16 | 0,2728 | 1,39228E-07 |
| ACE2 | CITF22-92A6.2 | 0,2728 | 1,40554E-07 |
| ACE2 | DENND2A | 0,2727 | 1,40934E-07 |
| ACE2 | CCS | 0,2727 | 1,41189E-07 |
| ACE2 | RP11-351M8.1 | 0,2727 | 1,41967E-07 |
| ACE2 | RP11-230C9.4 | 0,2726 | 1,43312E-07 |
| ACE2 | SSRP1 | 0,2725 | 1,44715E-07 |
| ACE2 | PWP1 | 0,2725 | 1,44936E-07 |
| ACE2 | ZNF146 | 0,2725 | 1,44157E-07 |
| ACE2 | NMU | 0,2724 | 1,45508E-07 |
| ACE2 | ERP27 | 0,2723 | 1,47947E-07 |
| ACE2 | CTD-2574D22.2 | 0,2723 | 1,46996E-07 |
| ACE2 | ZC2HC1A | 0,2722 | 1,4867E-07 |
| ACE2 | PTENP1-AS | 0,2722 | 1,49108E-07 |
| ACE2 | EEF1A1P4 | 0,2722 | 1,49719E-07 |
| ACE2 | LAMP1 | 0,2722 | 1,49943E-07 |
| ACE2 | AC091177.1 | 0,2722 | 1,48886E-07 |
| ACE2 | TAF4 | 0,2721 | 1,51436E-07 |
| ACE2 | WWTR1-AS1 | 0,272 | 1,53092E-07 |
| ACE2 | GAPDHP62 | 0,272 | 1,52845E-07 |

| | | | |
|---|---|---|---|
| ACE2 | MAPKAPK5-AS1 | 0,272 | 1,53335E-07 |
| ACE2 | RN7SL49P | 0,272 | 1,52193E-07 |
| ACE2 | PAGE5 | 0,272 | 1,52091E-07 |
| ACE2 | FRMD7 | 0,272 | 1,5232E-07 |
| ACE2 | ZSWIM8 | 0,2719 | 1,54013E-07 |
| ACE2 | RP11-133K1.9 | 0,2719 | 1,54745E-07 |
| ACE2 | CFHR1 | 0,2718 | 1,56566E-07 |
| ACE2 | RP11-88I21.1 | 0,2718 | 1,56631E-07 |
| ACE2 | ABC7-42404400C24.1 | 0,2718 | 1,56592E-07 |
| ACE2 | RP11-59N23.3 | 0,2718 | 1,56167E-07 |
| ACE2 | AC002368.4 | 0,2718 | 1,55729E-07 |
| ACE2 | TM4SF1 | 0,2717 | 1,57654E-07 |
| ACE2 | TSPOAP1 | 0,2717 | 1,57491E-07 |
| ACE2 | PATJ | 0,2716 | 1,59774E-07 |
| ACE2 | PRRT1 | 0,2716 | 1,59954E-07 |
| ACE2 | AC145124.2 | 0,2716 | 1,58572E-07 |
| ACE2 | CKAP5 | 0,2716 | 1,59481E-07 |
| ACE2 | RP11-124N19.4 | 0,2716 | 1,5877E-07 |
| ACE2 | THSD1 | 0,2716 | 1,59514E-07 |
| ACE2 | RP11-161M6.6 | 0,2716 | 1,59074E-07 |
| ACE2 | TMEM171 | 0,2715 | 1,61648E-07 |
| ACE2 | WNT9B | 0,2715 | 1,6076E-07 |
| ACE2 | ZNF30-AS1 | 0,2715 | 1,61069E-07 |
| ACE2 | AC012668.2 | 0,2714 | 1,62232E-07 |
| ACE2 | BEND3P1 | 0,2714 | 1,62936E-07 |
| ACE2 | DMAP1 | 0,2713 | 1,63941E-07 |
| ACE2 | RPL32P3 | 0,2713 | 1,64004E-07 |
| ACE2 | EMC2 | 0,2713 | 1,63846E-07 |
| ACE2 | RP11-770J1.3 | 0,2713 | 1,64282E-07 |
| ACE2 | SNAP23 | 0,2713 | 1,65157E-07 |
| ACE2 | AC092066.1 | 0,2713 | 1,65515E-07 |
| ACE2 | RP11-850A17.1 | 0,2712 | 1,65958E-07 |
| ACE2 | SNAI1 | 0,2711 | 1,67448E-07 |
| ACE2 | EEF1A1P9 | 0,271 | 1,70309E-07 |
| ACE2 | TMEM154 | 0,271 | 1,70714E-07 |
| ACE2 | PMS2P11 | 0,271 | 1,70741E-07 |
| ACE2 | SOCS3 | 0,271 | 1,70483E-07 |
| ACE2 | RP3-449O17.1 | 0,271 | 1,70018E-07 |
| ACE2 | GAD2 | 0,2709 | 1,71692E-07 |
| ACE2 | CBFA2T3 | 0,2708 | 1,731E-07 |
| ACE2 | XIRP1 | 0,2707 | 1,76402E-07 |
| ACE2 | CTD-2530N21.5 | 0,2707 | 1,75226E-07 |
| ACE2 | CASP12 | 0,2707 | 1,75994E-07 |
| ACE2 | CD226 | 0,2707 | 1,76452E-07 |
| ACE2 | CTD-3193K9.3 | 0,2706 | 1,77402E-07 |
| ACE2 | ABCA6 | 0,2706 | 1,77521E-07 |
| ACE2 | UQCC1 | 0,2706 | 1,77848E-07 |
| ACE2 | ADAM8 | 0,2705 | 1,79201E-07 |
| ACE2 | C14orf39 | 0,2705 | 1,80548E-07 |
| ACE2 | KRT27 | 0,2705 | 1,80556E-07 |
| ACE2 | NELFA | 0,2704 | 1,82366E-07 |
| ACE2 | F2RL1 | 0,2704 | 1,80636E-07 |
| ACE2 | PCDHGA8 | 0,2704 | 1,81544E-07 |
| ACE2 | MRPL58 | 0,2704 | 1,81541E-07 |

| | | | |
|---|---|---|---|
| ACE2 | TSPY1 | 0,2703 | 1,8441E-07 |
| ACE2 | AC133785.1 | 0,2702 | 1,85491E-07 |
| ACE2 | SYNGAP1 | 0,2702 | 1,85185E-07 |
| ACE2 | RP11-148K1.10 | 0,2702 | 1,84531E-07 |
| ACE2 | AUP1 | 0,2701 | 1,87754E-07 |
| ACE2 | COL5A1 | 0,2701 | 1,87975E-07 |
| ACE2 | RP11-326C3.12 | 0,2701 | 1,87621E-07 |
| ACE2 | EEA1 | 0,2701 | 1,88399E-07 |
| ACE2 | ABCC3 | 0,2701 | 1,87978E-07 |
| ACE2 | UNC119B | 0,27 | 1,89105E-07 |
| ACE2 | MT-TY | 0,27 | 1,90277E-07 |
| ACE2 | KCNV1 | 0,2699 | 1,90913E-07 |
| ACE2 | COX6A2 | 0,2699 | 1,91582E-07 |
| ACE2 | RP11-54O7.18 | 0,2698 | 1,94699E-07 |
| ACE2 | HOXA10 | 0,2698 | 1,9432E-07 |
| ACE2 | AC002543.2 | 0,2698 | 1,93495E-07 |
| ACE2 | LINC00543 | 0,2698 | 1,9426E-07 |
| ACE2 | BOLA2P2 | 0,2697 | 1,95323E-07 |
| ACE2 | TAS2R10 | 0,2697 | 1,94773E-07 |
| ACE2 | ZMAT5 | 0,2697 | 1,9572E-07 |
| ACE2 | SH2D1B | 0,2696 | 1,97364E-07 |
| ACE2 | PPP2R2DP1 | 0,2696 | 1,97174E-07 |
| ACE2 | OR2S1P | 0,2696 | 1,97149E-07 |
| ACE2 | FAM219A | 0,2695 | 2,00906E-07 |
| ACE2 | HKDC1 | 0,2695 | 2,00483E-07 |
| ACE2 | RP11-23F23.3 | 0,2695 | 2,00145E-07 |
| ACE2 | RP11-61A14.1 | 0,2694 | 2,02981E-07 |
| ACE2 | RP11-331F4.5 | 0,2694 | 2,02067E-07 |
| ACE2 | METTL4 | 0,2694 | 2,02838E-07 |
| ACE2 | ZNF69 | 0,2694 | 2,02059E-07 |
| ACE2 | CAPZA1 | 0,2693 | 2,04891E-07 |
| ACE2 | RGL3 | 0,2692 | 2,07616E-07 |
| ACE2 | CTD-2325A15.5 | 0,2691 | 2,09093E-07 |
| ACE2 | RP11-59C5.3 | 0,2691 | 2,09029E-07 |
| ACE2 | ERVFRD-3 | 0,269 | 2,1039E-07 |
| ACE2 | PGM5 | 0,269 | 2,10884E-07 |
| ACE2 | PHLDA2 | 0,269 | 2,10089E-07 |
| ACE2 | OR9A3P | 0,2689 | 2,12844E-07 |
| ACE2 | ZGRF1 | 0,2688 | 2,15154E-07 |
| ACE2 | PDE4D | 0,2688 | 2,14957E-07 |
| ACE2 | GS1-124K5.10 | 0,2688 | 2,15816E-07 |
| ACE2 | DLEU7-AS1 | 0,2688 | 2,14635E-07 |
| ACE2 | ALG12 | 0,2688 | 2,16473E-07 |
| ACE2 | RERE | 0,2687 | 2,18627E-07 |
| ACE2 | SLC51A | 0,2687 | 2,1795E-07 |
| ACE2 | STMN4 | 0,2686 | 2,21348E-07 |
| ACE2 | A1CF | 0,2686 | 2,19475E-07 |
| ACE2 | NPM1P26 | 0,2686 | 2,19486E-07 |
| ACE2 | UNC13A | 0,2686 | 2,20313E-07 |
| ACE2 | ANKRD36C | 0,2685 | 2,22482E-07 |
| ACE2 | PRKAG2 | 0,2685 | 2,21899E-07 |
| ACE2 | RNVU1-1 | 0,2684 | 2,25843E-07 |
| ACE2 | KCNC3 | 0,2684 | 2,25101E-07 |
| ACE2 | FAM9A | 0,2684 | 2,24812E-07 |

| | | | |
|---|---|---|---|
| ACE2 | PIGP | 0,2683 | 2,28598E-07 |
| ACE2 | WNT7B | 0,2683 | 2,2668E-07 |
| ACE2 | HNRNPA1P68 | 0,2682 | 2,3051E-07 |
| ACE2 | RP11-371M22.1 | 0,2682 | 2,29978E-07 |
| ACE2 | SACS | 0,2682 | 2,31218E-07 |
| ACE2 | RP11-78O7.2 | 0,2682 | 2,30656E-07 |
| ACE2 | FBXO41 | 0,2681 | 2,32492E-07 |
| ACE2 | ZBTB20-AS4 | 0,2681 | 2,31807E-07 |
| ACE2 | MTCO2P3 | 0,2681 | 2,32385E-07 |
| ACE2 | SETP8 | 0,2681 | 2,32708E-07 |
| ACE2 | RP11-71H17.1 | 0,268 | 2,35305E-07 |
| ACE2 | RP11-547D13.1 | 0,268 | 2,35464E-07 |
| ACE2 | PPM1F | 0,268 | 2,34374E-07 |
| ACE2 | BDH2P1 | 0,2679 | 2,37664E-07 |
| ACE2 | ERVK13-1 | 0,2679 | 2,38199E-07 |
| ACE2 | MRPS12 | 0,2679 | 2,36752E-07 |
| ACE2 | HPDL | 0,2678 | 2,39857E-07 |
| ACE2 | UBE2FP1 | 0,2678 | 2,39826E-07 |
| ACE2 | FAM86HP | 0,2678 | 2,39855E-07 |
| ACE2 | AC003986.5 | 0,2678 | 2,41458E-07 |
| ACE2 | LTB4R2 | 0,2678 | 2,40326E-07 |
| ACE2 | RP1-37M3.8 | 0,2678 | 2,38948E-07 |
| ACE2 | CACNA2D2 | 0,2677 | 2,42301E-07 |
| ACE2 | EEF1A1P13 | 0,2677 | 2,42507E-07 |
| ACE2 | ILK | 0,2677 | 2,41907E-07 |
| ACE2 | DPP3 | 0,2677 | 2,43873E-07 |
| ACE2 | SHC2 | 0,2677 | 2,4291E-07 |
| ACE2 | RN7SL674P | 0,2676 | 2,45262E-07 |
| ACE2 | TMSB10P1 | 0,2676 | 2,46146E-07 |
| ACE2 | CYTH3 | 0,2676 | 2,44546E-07 |
| ACE2 | ADAP2 | 0,2676 | 2,451E-07 |
| ACE2 | ENGASE | 0,2676 | 2,4527E-07 |
| ACE2 | OCEL1 | 0,2676 | 2,44808E-07 |
| ACE2 | RP11-214J9.1 | 0,2675 | 2,48729E-07 |
| ACE2 | RPL10AP6 | 0,2674 | 2,49894E-07 |
| ACE2 | LINC01947 | 0,2674 | 2,50204E-07 |
| ACE2 | ANKRD63 | 0,2674 | 2,49356E-07 |
| ACE2 | RUBCN | 0,2673 | 2,53966E-07 |
| ACE2 | DDX18P1 | 0,2673 | 2,54514E-07 |
| ACE2 | SERPINA1 | 0,2673 | 2,53464E-07 |
| ACE2 | RP11-622C24.2 | 0,2673 | 2,52998E-07 |
| ACE2 | PSMD10P1 | 0,2673 | 2,52482E-07 |
| ACE2 | SLC2A1-AS1 | 0,2672 | 2,56257E-07 |
| ACE2 | G0S2 | 0,2672 | 2,55697E-07 |
| ACE2 | RPL21P32 | 0,2672 | 2,55257E-07 |
| ACE2 | RP11-775G23.1 | 0,2672 | 2,55305E-07 |
| ACE2 | TPSB2 | 0,2671 | 2,5823E-07 |
| ACE2 | RP5-858L17.1 | 0,2671 | 2,58702E-07 |
| ACE2 | FOXQ1 | 0,267 | 2,60374E-07 |
| ACE2 | CASP1 | 0,267 | 2,60774E-07 |
| ACE2 | ATP5H | 0,267 | 2,60317E-07 |
| ACE2 | ARHGEF18 | 0,267 | 2,6288E-07 |
| ACE2 | ETHE1 | 0,267 | 2,61014E-07 |
| ACE2 | CCND3 | 0,2669 | 2,63678E-07 |

| | | | |
|---|---|---|---|
| ACE2 | MTND4P4 | 0,2669 | 2,65642E-07 |
| ACE2 | GS1-5L10.1 | 0,2669 | 2,63411E-07 |
| ACE2 | CLEC12A | 0,2668 | 2,66654E-07 |
| ACE2 | RP11-81A1.4 | 0,2668 | 2,68033E-07 |
| ACE2 | RP11-153F10.2 | 0,2667 | 2,69489E-07 |
| ACE2 | TCOF1 | 0,2667 | 2,69835E-07 |
| ACE2 | BTNL9 | 0,2667 | 2,71436E-07 |
| ACE2 | TFAP2A-AS1 | 0,2666 | 2,73104E-07 |
| ACE2 | MFAP5 | 0,2666 | 2,72699E-07 |
| ACE2 | RP11-263K4.1 | 0,2666 | 2,74336E-07 |
| ACE2 | SENP3 | 0,2666 | 2,72907E-07 |
| ACE2 | ZNF80 | 0,2665 | 2,75548E-07 |
| ACE2 | RP5-1049N15.2 | 0,2665 | 2,761E-07 |
| ACE2 | PRSS53 | 0,2665 | 2,76472E-07 |
| ACE2 | CYP4X1 | 0,2664 | 2,78929E-07 |
| ACE2 | FGD5P1 | 0,2663 | 2,83242E-07 |
| ACE2 | OCLN | 0,2663 | 2,81892E-07 |
| ACE2 | BTAF1 | 0,2663 | 2,82039E-07 |
| ACE2 | RN7SL23P | 0,2663 | 2,8121E-07 |
| ACE2 | RP11-426C22.6 | 0,2663 | 2,82852E-07 |
| ACE2 | CA4 | 0,2663 | 2,81767E-07 |
| ACE2 | SLC17A9 | 0,2663 | 2,81243E-07 |
| ACE2 | MRAP | 0,2663 | 2,80956E-07 |
| ACE2 | HIST1H2AI | 0,2662 | 2,85518E-07 |
| ACE2 | RP11-676J12.7 | 0,2662 | 2,85601E-07 |
| ACE2 | CCDC144NL | 0,2662 | 2,84041E-07 |
| ACE2 | GUCD1 | 0,2662 | 2,8358E-07 |
| ACE2 | RAPH1 | 0,2661 | 2,86578E-07 |
| ACE2 | MACROD1 | 0,2661 | 2,88197E-07 |
| ACE2 | RP11-51L5.3 | 0,2661 | 2,87089E-07 |
| ACE2 | WTIP | 0,2661 | 2,89207E-07 |
| ACE2 | ARFGAP2 | 0,266 | 2,90367E-07 |
| ACE2 | RP11-114F3.5 | 0,266 | 2,91083E-07 |
| ACE2 | RP11-930O11.3 | 0,266 | 2,8994E-07 |
| ACE2 | DHX36 | 0,2659 | 2,92886E-07 |
| ACE2 | CTD-2235C13.1 | 0,2659 | 2,9524E-07 |
| ACE2 | RP11-511H9.3 | 0,2659 | 2,95527E-07 |
| ACE2 | RP11-74E22.3 | 0,2659 | 2,95571E-07 |
| ACE2 | RP11-4B16.4 | 0,2659 | 2,92792E-07 |
| ACE2 | LINC01194 | 0,2658 | 2,97517E-07 |
| ACE2 | RP1-223B1.1 | 0,2658 | 2,97614E-07 |
| ACE2 | TMEM220-AS1 | 0,2658 | 2,96036E-07 |
| ACE2 | RP11-299J3.6 | 0,2657 | 2,99159E-07 |
| ACE2 | CEND1 | 0,2657 | 3,01783E-07 |
| ACE2 | RP11-408A13.4 | 0,2656 | 3,03223E-07 |
| ACE2 | FAR1 | 0,2656 | 3,0299E-07 |
| ACE2 | RP11-417N10.3 | 0,2656 | 3,05089E-07 |
| ACE2 | PPP1R3G | 0,2655 | 3,05457E-07 |
| ACE2 | RP11-342K6.4 | 0,2654 | 3,11023E-07 |
| ACE2 | TAF1C | 0,2654 | 3,10977E-07 |
| ACE2 | RP5-890E16.5 | 0,2654 | 3,08786E-07 |
| ACE2 | MT-TN | 0,2654 | 3,09609E-07 |
| ACE2 | CCRL2 | 0,2653 | 3,14988E-07 |
| ACE2 | LEP | 0,2653 | 3,12073E-07 |

| | | | |
|---|---|---|---|
| ACE2 | NSFL1C | 0,2653 | 3,12401E-07 |
| ACE2 | SASS6 | 0,2652 | 3,18372E-07 |
| ACE2 | MYD88 | 0,2652 | 3,16369E-07 |
| ACE2 | RP11-228M15.1 | 0,2652 | 3,15925E-07 |
| ACE2 | CTB-17P3.4 | 0,2651 | 3,18576E-07 |
| ACE2 | RP11-228B15.4 | 0,2651 | 3,18678E-07 |
| ACE2 | ZSCAN23 | 0,265 | 3,24262E-07 |
| ACE2 | ALG1L9P | 0,265 | 3,25191E-07 |
| ACE2 | MEG3 | 0,265 | 3,2232E-07 |
| ACE2 | PDGFC | 0,2649 | 3,26763E-07 |
| ACE2 | RP11-477J21.6 | 0,2649 | 3,25751E-07 |
| ACE2 | ARHGEF5 | 0,2648 | 3,32185E-07 |
| ACE2 | TBCD | 0,2647 | 3,33337E-07 |
| ACE2 | LINC01736 | 0,2646 | 3,38159E-07 |
| ACE2 | PCDHGB7 | 0,2646 | 3,35837E-07 |
| ACE2 | ABHD17AP5 | 0,2646 | 3,37116E-07 |
| ACE2 | BAHCC1 | 0,2644 | 3,4369E-07 |
| ACE2 | CBS | 0,2644 | 3,44179E-07 |
| ACE2 | CICP3 | 0,2643 | 3,48375E-07 |
| ACE2 | bP-2171C21.6 | 0,2643 | 3,50021E-07 |
| ACE2 | MT-ND3 | 0,2643 | 3,47599E-07 |
| ACE2 | RP11-382D12.2 | 0,2642 | 3,53725E-07 |
| ACE2 | RP11-408P14.1 | 0,2642 | 3,53712E-07 |
| ACE2 | MIR4458HG | 0,2641 | 3,55627E-07 |
| ACE2 | GPBP1 | 0,2641 | 3,56185E-07 |
| ACE2 | RP11-567J20.2 | 0,2641 | 3,54607E-07 |
| ACE2 | RP11-1079K10.2 | 0,2641 | 3,5446E-07 |
| ACE2 | SLC35B1 | 0,2641 | 3,54032E-07 |
| ACE2 | B4GALNT4 | 0,264 | 3,59635E-07 |
| ACE2 | RP11-981P6.1 | 0,264 | 3,60669E-07 |
| ACE2 | AC093616.4 | 0,2639 | 3,62319E-07 |
| ACE2 | CTBP1 | 0,2639 | 3,6168E-07 |
| ACE2 | SHB | 0,2639 | 3,638E-07 |
| ACE2 | SLITRK1 | 0,2638 | 3,65485E-07 |
| ACE2 | RP11-346C20.4 | 0,2638 | 3,68059E-07 |
| ACE2 | AC005329.7 | 0,2638 | 3,68041E-07 |
| ACE2 | COL6A4P1 | 0,2637 | 3,69869E-07 |
| ACE2 | ALG1L2 | 0,2637 | 3,73085E-07 |
| ACE2 | KB-1836B5.1 | 0,2636 | 3,74634E-07 |
| ACE2 | RN7SL181P | 0,2636 | 3,74574E-07 |
| ACE2 | RP3-395M20.2 | 0,2635 | 3,78203E-07 |
| ACE2 | RP11-791G15.2 | 0,2635 | 3,77574E-07 |
| ACE2 | CASS4 | 0,2635 | 3,77594E-07 |
| ACE2 | SLC23A3 | 0,2634 | 3,83366E-07 |
| ACE2 | FAM86B3P | 0,2634 | 3,81751E-07 |
| ACE2 | TONSL | 0,2634 | 3,83148E-07 |
| ACE2 | ITGB7 | 0,2634 | 3,81843E-07 |
| ACE2 | MTUS2 | 0,2634 | 3,8449E-07 |
| ACE2 | KLRF1 | 0,2633 | 3,8798E-07 |
| ACE2 | CCL28 | 0,2632 | 3,90719E-07 |
| ACE2 | SNORA29 | 0,2632 | 3,90423E-07 |
| ACE2 | DSC3 | 0,2631 | 3,95213E-07 |
| ACE2 | PNMA5 | 0,2631 | 3,97295E-07 |
| ACE2 | RP11-305L7.7 | 0,263 | 3,99237E-07 |

| | | | |
|---|---|---|---|
| ACE2 | ADCY4 | 0,263 | 4,00695E-07 |
| ACE2 | RP11-71L14.3 | 0,263 | 4,00346E-07 |
| ACE2 | RP11-252K23.2 | 0,263 | 3,99223E-07 |
| ACE2 | PPP1R13L | 0,263 | 3,98621E-07 |
| ACE2 | RP11-127B16.1 | 0,2629 | 4,03777E-07 |
| ACE2 | ZNF282 | 0,2629 | 4,04344E-07 |
| ACE2 | ARMC10P1 | 0,2628 | 4,06846E-07 |
| ACE2 | RP11-351D16.3 | 0,2628 | 4,09379E-07 |
| ACE2 | LAX1 | 0,2627 | 4,10429E-07 |
| ACE2 | RP11-818O24.2 | 0,2627 | 4,14281E-07 |
| ACE2 | ELAC2 | 0,2627 | 4,11919E-07 |
| ACE2 | YWHAEP7 | 0,2627 | 4,10632E-07 |
| ACE2 | SSX3 | 0,2627 | 4,14409E-07 |
| ACE2 | AP006621.5 | 0,2626 | 4,17334E-07 |
| ACE2 | DGCR12 | 0,2626 | 4,15431E-07 |
| ACE2 | RP3-333H23.9 | 0,2626 | 4,14568E-07 |
| ACE2 | CTSH | 0,2625 | 4,19508E-07 |
| ACE2 | SNX18P1Y | 0,2625 | 4,22302E-07 |
| ACE2 | CHN1 | 0,2624 | 4,23453E-07 |
| ACE2 | HMGB2 | 0,2624 | 4,25054E-07 |
| ACE2 | SMN2 | 0,2624 | 4,26961E-07 |
| ACE2 | TSR3 | 0,2624 | 4,24528E-07 |
| ACE2 | RBMXL3 | 0,2624 | 4,24914E-07 |
| ACE2 | DENND3 | 0,2623 | 4,31939E-07 |
| ACE2 | RP11-8P13.5 | 0,2623 | 4,29374E-07 |
| ACE2 | SMAD9-IT1 | 0,2622 | 4,35065E-07 |
| ACE2 | CTDSPL2 | 0,2622 | 4,35786E-07 |
| ACE2 | RP11-535A5.1 | 0,2622 | 4,33091E-07 |
| ACE2 | SLC10A5 | 0,2621 | 4,40626E-07 |
| ACE2 | RP11-87H9.4 | 0,2621 | 4,4083E-07 |
| ACE2 | AC004538.3 | 0,262 | 4,42492E-07 |
| ACE2 | AQP2 | 0,262 | 4,41619E-07 |
| ACE2 | FANCI | 0,262 | 4,45139E-07 |
| ACE2 | NMRAL1 | 0,262 | 4,43714E-07 |
| ACE2 | RP1-56K13.2 | 0,262 | 4,42383E-07 |
| ACE2 | HPS1 | 0,2619 | 4,49201E-07 |
| ACE2 | PPHLN1 | 0,2619 | 4,47755E-07 |
| ACE2 | RPL31P53 | 0,2619 | 4,46585E-07 |
| ACE2 | GPR132 | 0,2619 | 4,49822E-07 |
| ACE2 | AC002400.1 | 0,2619 | 4,48778E-07 |
| ACE2 | AC090587.4 | 0,2618 | 4,55367E-07 |
| ACE2 | AC003104.1 | 0,2618 | 4,5327E-07 |
| ACE2 | PAGE1 | 0,2618 | 4,51408E-07 |
| ACE2 | HNRNPH1P1 | 0,2617 | 4,57228E-07 |
| ACE2 | RP11-629G13.1 | 0,2617 | 4,58955E-07 |
| ACE2 | AC105053.3 | 0,2616 | 4,63251E-07 |
| ACE2 | CTD-2060C23.1 | 0,2616 | 4,6404E-07 |
| ACE2 | EEPD1 | 0,2616 | 4,64516E-07 |
| ACE2 | CABP4 | 0,2616 | 4,6392E-07 |
| ACE2 | ROBO4 | 0,2616 | 4,63198E-07 |
| ACE2 | ABCD2 | 0,2616 | 4,62863E-07 |
| ACE2 | RP11-749H20.2 | 0,2616 | 4,62255E-07 |
| ACE2 | DAND5 | 0,2616 | 4,63581E-07 |
| ACE2 | RP11-314N13.3 | 0,2615 | 4,65814E-07 |

| | | | |
|---|---|---|---|
| ACE2 | RN7SL431P | 0,2614 | 4,72647E-07 |
| ACE2 | POU6F1 | 0,2614 | 4,713E-07 |
| ACE2 | LA16c-381G6.1 | 0,2614 | 4,73125E-07 |
| ACE2 | RP11-407G23.7 | 0,2614 | 4,72417E-07 |
| ACE2 | AC104058.1 | 0,2613 | 4,7548E-07 |
| ACE2 | CYCS | 0,2612 | 4,83898E-07 |
| ACE2 | C9orf47 | 0,2611 | 4,88719E-07 |
| ACE2 | NDUFB1 | 0,2611 | 4,87124E-07 |
| ACE2 | RP11-563J2.3 | 0,261 | 4,94538E-07 |
| ACE2 | H3F3AP4 | 0,2609 | 4,95341E-07 |
| ACE2 | PACSIN1 | 0,2609 | 4,97205E-07 |
| ACE2 | VGF | 0,2609 | 4,96638E-07 |
| ACE2 | KRBA1 | 0,2609 | 4,95583E-07 |
| ACE2 | PTP4A3 | 0,2609 | 4,96479E-07 |
| ACE2 | ZNF33B | 0,2609 | 4,9794E-07 |
| ACE2 | RP11-49K24.5 | 0,2609 | 4,96126E-07 |
| ACE2 | RP11-49C9.2 | 0,2609 | 4,97819E-07 |
| ACE2 | FOXI2 | 0,2608 | 5,02986E-07 |
| ACE2 | CCDC85B | 0,2608 | 5,03321E-07 |
| ACE2 | OGFOD3 | 0,2608 | 5,00416E-07 |
| ACE2 | ZNF551 | 0,2608 | 5,04276E-07 |
| ACE2 | INPP5J | 0,2608 | 5,0447E-07 |
| ACE2 | AVPR2 | 0,2608 | 5,04616E-07 |
| ACE2 | RP11-342I1.2 | 0,2607 | 5,08939E-07 |
| ACE2 | RP11-243M5.3 | 0,2607 | 5,07179E-07 |
| ACE2 | FRG1EP | 0,2607 | 5,08356E-07 |
| ACE2 | RAB13 | 0,2606 | 5,13725E-07 |
| ACE2 | TM4SF18 | 0,2606 | 5,15369E-07 |
| ACE2 | FAM214B | 0,2606 | 5,12023E-07 |
| ACE2 | AC022819.3 | 0,2606 | 5,1081E-07 |
| ACE2 | TRMT10A | 0,2605 | 5,18467E-07 |
| ACE2 | RMI2 | 0,2605 | 5,19421E-07 |
| ACE2 | JUNB | 0,2605 | 5,17574E-07 |
| ACE2 | IGLV7-46 | 0,2605 | 5,17088E-07 |
| ACE2 | GTF2H1 | 0,2604 | 5,21635E-07 |
| ACE2 | RP11-449G16.1 | 0,2603 | 5,28359E-07 |
| ACE2 | RP11-250B2.3 | 0,2603 | 5,26903E-07 |
| ACE2 | UPK3B | 0,2603 | 5,29661E-07 |
| ACE2 | ANGPT2 | 0,2603 | 5,29384E-07 |
| ACE2 | MTCO1P2 | 0,2603 | 5,27135E-07 |
| ACE2 | RP11-467L19.14 | 0,2603 | 5,29905E-07 |
| ACE2 | RP11-686D22.4 | 0,2603 | 5,29611E-07 |
| ACE2 | RP3-467N11.2 | 0,2602 | 5,33716E-07 |
| ACE2 | CHI3L2 | 0,2601 | 5,42561E-07 |
| ACE2 | SLC35A5 | 0,2601 | 5,42001E-07 |
| ACE2 | RPS29 | 0,2601 | 5,43108E-07 |
| ACE2 | RP11-162I7.1 | 0,2601 | 5,39678E-07 |
| ACE2 | LINC00205 | 0,2601 | 5,41652E-07 |
| ACE2 | PVALB | 0,2601 | 5,40101E-07 |
| ACE2 | STX1A | 0,26 | 5,49155E-07 |
| ACE2 | RP11-571M6.17 | 0,26 | 5,46313E-07 |
| ACE2 | MT1G | 0,26 | 5,47831E-07 |
| ACE2 | RP6-24A23.3 | 0,26 | 5,45143E-07 |
| ACE2 | CLSTN2 | 0,2599 | 5,51134E-07 |

| | | | |
|---|---|---|---|
| ACE2 | RP11-530C5.4 | 0,2599 | 5,51093E-07 |
| ACE2 | RP13-150K15.1 | 0,2599 | 5,53855E-07 |
| ACE2 | LINC01936 | 0,2597 | 5,65665E-07 |
| ACE2 | RP11-173A8.2 | 0,2597 | 5,65157E-07 |
| ACE2 | NAGPA-AS1 | 0,2597 | 5,61158E-07 |
| ACE2 | RP3-514P16.1 | 0,2597 | 5,60961E-07 |
| ACE2 | MAST4 | 0,2596 | 5,71213E-07 |
| ACE2 | RP11-276H19.2 | 0,2596 | 5,68661E-07 |
| ACE2 | PFN1P6 | 0,2595 | 5,77421E-07 |
| ACE2 | THADA | 0,2595 | 5,7548E-07 |
| ACE2 | CCDC141 | 0,2595 | 5,74835E-07 |
| ACE2 | HCG23 | 0,2595 | 5,72655E-07 |
| ACE2 | HNRNPA1P4 | 0,2595 | 5,73394E-07 |
| ACE2 | LRP1-AS | 0,2595 | 5,78211E-07 |
| ACE2 | CDC42P6 | 0,2594 | 5,83731E-07 |
| ACE2 | MTCO3P10 | 0,2594 | 5,78468E-07 |
| ACE2 | CD81-AS1 | 0,2594 | 5,83738E-07 |
| ACE2 | RPS4XP14 | 0,2593 | 5,87274E-07 |
| ACE2 | CSAD | 0,2593 | 5,8679E-07 |
| ACE2 | ZFHX2-AS1 | 0,2593 | 5,88443E-07 |
| ACE2 | GPRIN1 | 0,2592 | 5,94464E-07 |
| ACE2 | AL022393.9 | 0,2592 | 5,91138E-07 |
| ACE2 | MTND4P26 | 0,2591 | 5,985E-07 |
| ACE2 | IDI1 | 0,2591 | 5,98831E-07 |
| ACE2 | PCBP2-OT1 | 0,2591 | 6,00878E-07 |
| ACE2 | GSDMA | 0,2591 | 5,98235E-07 |
| ACE2 | AC005363.9 | 0,259 | 6,03045E-07 |
| ACE2 | CTD-3193K9.1 | 0,259 | 6,03552E-07 |
| ACE2 | MTND5P11 | 0,2589 | 6,0983E-07 |
| ACE2 | RP11-405O10.2 | 0,2589 | 6,11187E-07 |
| ACE2 | AC004076.5 | 0,2589 | 6,12866E-07 |
| ACE2 | FAAH2 | 0,2589 | 6,13696E-07 |
| ACE2 | AL133243.4 | 0,2588 | 6,15677E-07 |
| ACE2 | RP11-301L8.2 | 0,2588 | 6,20972E-07 |
| ACE2 | RFC2 | 0,2588 | 6,15545E-07 |
| ACE2 | LINC01692 | 0,2587 | 6,23645E-07 |
| ACE2 | CTC-338M12.13 | 0,2586 | 6,30238E-07 |
| ACE2 | SOCS5P5 | 0,2586 | 6,2924E-07 |
| ACE2 | MALAT1 | 0,2586 | 6,3465E-07 |
| ACE2 | USP21 | 0,2585 | 6,40826E-07 |
| ACE2 | TMUB1 | 0,2585 | 6,39639E-07 |
| ACE2 | SUMO2P19 | 0,2585 | 6,37767E-07 |
| ACE2 | AC068580.5 | 0,2585 | 6,38185E-07 |
| ACE2 | RP11-474C8.8 | 0,2585 | 6,35792E-07 |
| ACE2 | RB1CC1 | 0,2584 | 6,45565E-07 |
| ACE2 | RP11-693M3.1 | 0,2584 | 6,45948E-07 |
| ACE2 | NCOA4 | 0,2583 | 6,52141E-07 |
| ACE2 | RP11-182L21.5 | 0,2583 | 6,54045E-07 |
| ACE2 | LHFPL1 | 0,2583 | 6,50032E-07 |
| ACE2 | TSPY15P | 0,2583 | 6,51778E-07 |
| ACE2 | RP11-480A16.1 | 0,2581 | 6,62809E-07 |
| ACE2 | IL33 | 0,2581 | 6,63671E-07 |
| ACE2 | TEKT4P2 | 0,2581 | 6,65056E-07 |
| ACE2 | RP11-864N7.2 | 0,258 | 6,72489E-07 |

| | | | |
|---|---|---|---|
| ACE2 | DLG5 | 0,2579 | 6,78394E-07 |
| ACE2 | NAV3 | 0,2579 | 6,8052E-07 |
| ACE2 | LMO7DN | 0,2579 | 6,75665E-07 |
| ACE2 | KIAA0100 | 0,2579 | 6,75197E-07 |
| ACE2 | RP11-95O2.1 | 0,2579 | 6,80332E-07 |
| ACE2 | RN7SL559P | 0,2578 | 6,82449E-07 |
| ACE2 | LINC00989 | 0,2578 | 6,86297E-07 |
| ACE2 | RP11-142C4.6 | 0,2578 | 6,86246E-07 |
| ACE2 | PYGM | 0,2577 | 6,89382E-07 |
| ACE2 | RP11-8L8.2 | 0,2577 | 6,95601E-07 |
| ACE2 | RP5-1029K10.2 | 0,2577 | 6,93112E-07 |
| ACE2 | NHP2 | 0,2576 | 7,03068E-07 |
| ACE2 | PHKG1P1 | 0,2576 | 6,98306E-07 |
| ACE2 | AKR1B10 | 0,2576 | 6,97486E-07 |
| ACE2 | ASPN | 0,2576 | 6,99804E-07 |
| ACE2 | DUXAP9 | 0,2576 | 6,97221E-07 |
| ACE2 | RP11-613M10.6 | 0,2575 | 7,10151E-07 |
| ACE2 | GLIS2-AS1 | 0,2575 | 7,0796E-07 |
| ACE2 | NCOR1 | 0,2575 | 7,10429E-07 |
| ACE2 | TBC1D10A | 0,2575 | 7,08914E-07 |
| ACE2 | PGGHG | 0,2574 | 7,13668E-07 |
| ACE2 | RP5-1031D4.3 | 0,2573 | 7,22209E-07 |
| ACE2 | RP11-733O18.1 | 0,2573 | 7,24293E-07 |
| ACE2 | TRMT10C | 0,2572 | 7,30984E-07 |
| ACE2 | FCF1P3 | 0,2572 | 7,27262E-07 |
| ACE2 | RP11-989F5.1 | 0,2572 | 7,26097E-07 |
| ACE2 | LINC00926 | 0,2572 | 7,31552E-07 |
| ACE2 | CTIF | 0,2572 | 7,28725E-07 |
| ACE2 | RP1-79C4.1 | 0,2571 | 7,33996E-07 |
| ACE2 | HOXD1 | 0,2571 | 7,40003E-07 |
| ACE2 | WDR86 | 0,2571 | 7,3792E-07 |
| ACE2 | TMEM253 | 0,2571 | 7,34904E-07 |
| ACE2 | DSG3 | 0,2571 | 7,33349E-07 |
| ACE2 | AC107081.5 | 0,257 | 7,41719E-07 |
| ACE2 | ALKBH2 | 0,257 | 7,45178E-07 |
| ACE2 | RP11-435J9.2 | 0,257 | 7,44651E-07 |
| ACE2 | LDHAP2 | 0,2569 | 7,48659E-07 |
| ACE2 | RP11-44F14.1 | 0,2569 | 7,50447E-07 |
| ACE2 | FAM156B | 0,2569 | 7,53765E-07 |
| ACE2 | HADHA | 0,2568 | 7,61381E-07 |
| ACE2 | IRF7 | 0,2568 | 7,63333E-07 |
| ACE2 | RP11-104N10.2 | 0,2567 | 7,67263E-07 |
| ACE2 | RPL4P6 | 0,2567 | 7,67083E-07 |
| ACE2 | ESPN | 0,2566 | 7,74343E-07 |
| ACE2 | ZNF862 | 0,2566 | 7,77665E-07 |
| ACE2 | ABCB10P1 | 0,2566 | 7,77563E-07 |
| ACE2 | GLP2R | 0,2566 | 7,73256E-07 |
| ACE2 | FBXW4 | 0,2565 | 7,83415E-07 |
| ACE2 | GABRA5 | 0,2565 | 7,87014E-07 |
| ACE2 | TNMD | 0,2565 | 7,80266E-07 |
| ACE2 | GSTM1 | 0,2564 | 7,90192E-07 |
| ACE2 | RP5-1142A6.9 | 0,2564 | 7,93574E-07 |
| ACE2 | CBX1 | 0,2564 | 7,93902E-07 |
| ACE2 | RP11-120D5.1 | 0,2564 | 7,91459E-07 |

| | | | |
|---|---|---|---|
| ACE2 | PRDM14 | 0,2563 | 7,96869E-07 |
| ACE2 | SPTBN4 | 0,2563 | 7,97478E-07 |
| ACE2 | CTA-212A2.1 | 0,2563 | 7,97621E-07 |
| ACE2 | RPL31 | 0,2562 | 8,10149E-07 |
| ACE2 | C3AR1 | 0,2562 | 8,07694E-07 |
| ACE2 | RP11-394B2.5 | 0,2562 | 8,08424E-07 |
| ACE2 | RP11-1109F11.3 | 0,2561 | 8,16484E-07 |
| ACE2 | ELOVL6 | 0,256 | 8,2504E-07 |
| ACE2 | AC012078.2 | 0,256 | 8,21688E-07 |
| ACE2 | HIST1H2AG | 0,2559 | 8,30271E-07 |
| ACE2 | RP11-579D7.8 | 0,2559 | 8,28798E-07 |
| ACE2 | RAB11B | 0,2559 | 8,35923E-07 |
| ACE2 | GALK2 | 0,2558 | 8,37951E-07 |
| ACE2 | RAB11A | 0,2557 | 8,46007E-07 |
| ACE2 | S1PR4 | 0,2557 | 8,51946E-07 |
| ACE2 | ADGRE3 | 0,2557 | 8,46478E-07 |
| ACE2 | RP11-110G21.1 | 0,2556 | 8,61915E-07 |
| ACE2 | POU4F1 | 0,2556 | 8,54616E-07 |
| ACE2 | RGS16 | 0,2555 | 8,65998E-07 |
| ACE2 | ECI1 | 0,2555 | 8,71156E-07 |
| ACE2 | AC002550.6 | 0,2555 | 8,65839E-07 |
| ACE2 | HAUS1 | 0,2555 | 8,64368E-07 |
| ACE2 | CTD-2017D11.2 | 0,2555 | 8,68121E-07 |
| ACE2 | CXCL5 | 0,2554 | 8,75069E-07 |
| ACE2 | MSC | 0,2554 | 8,74597E-07 |
| ACE2 | RP11-45A16.4 | 0,2554 | 8,80079E-07 |
| ACE2 | RANBP2 | 0,2553 | 8,87247E-07 |
| ACE2 | RP11-466G12.2 | 0,2553 | 8,88404E-07 |
| ACE2 | TMEM147 | 0,2553 | 8,85411E-07 |
| ACE2 | RP11-656D10.3 | 0,2552 | 8,95106E-07 |
| ACE2 | NAT10 | 0,2551 | 9,05168E-07 |
| ACE2 | RP11-151N17.1 | 0,2551 | 9,06725E-07 |
| ACE2 | MAP1LC3C | 0,255 | 9,08738E-07 |
| ACE2 | NARF-IT1 | 0,255 | 9,08799E-07 |
| ACE2 | BSG | 0,255 | 9,09304E-07 |
| ACE2 | IVL | -0,255 | 9,10748E-07 |
| ACE2 | CTD-2258A20.3 | -0,255 | 9,12705E-07 |
| ACE2 | LINC01780 | -0,2551 | 9,05602E-07 |
| ACE2 | AC139712.4 | -0,2551 | 8,99212E-07 |
| ACE2 | CDC20B | -0,2551 | 9,07758E-07 |
| ACE2 | RP11-1I2.1 | -0,2551 | 9,0202E-07 |
| ACE2 | ZNF806 | -0,2552 | 8,94754E-07 |
| ACE2 | RP11-356I2.4 | -0,2552 | 8,90471E-07 |
| ACE2 | RP11-168P13.1 | -0,2552 | 8,90317E-07 |
| ACE2 | AC073257.1 | -0,2553 | 8,8953E-07 |
| ACE2 | NIPAL4 | -0,2553 | 8,82613E-07 |
| ACE2 | RP11-278A23.1 | -0,2553 | 8,81023E-07 |
| ACE2 | RP11-286N3.1 | -0,2553 | 8,87896E-07 |
| ACE2 | RP11-569G9.7 | -0,2554 | 8,79196E-07 |
| ACE2 | MYL3 | -0,2554 | 8,75754E-07 |
| ACE2 | ETF1P1 | -0,2554 | 8,77641E-07 |
| ACE2 | CTC-537E7.1 | -0,2555 | 8,71082E-07 |
| ACE2 | RP11-815N9.2 | -0,2556 | 8,58066E-07 |
| ACE2 | SBDS | -0,2556 | 8,55069E-07 |

| | | | |
|---|---|---|---|
| ACE2 | RP11-324F11.1 | -0,2556 | 8,57499E-07 |
| ACE2 | LINC01133 | -0,2557 | 8,50754E-07 |
| ACE2 | RP11-192H6.2 | -0,2557 | 8,4819E-07 |
| ACE2 | UCP2 | -0,2557 | 8,52744E-07 |
| ACE2 | RP11-214K3.20 | -0,2557 | 8,48405E-07 |
| ACE2 | RNU5F-1 | -0,2558 | 8,44624E-07 |
| ACE2 | TEDDM2P | -0,2558 | 8,41076E-07 |
| ACE2 | E2F8 | -0,2559 | 8,30256E-07 |
| ACE2 | MANF | -0,256 | 8,24799E-07 |
| ACE2 | RP11-501E14.1 | -0,256 | 8,25332E-07 |
| ACE2 | FAM187B2P | -0,256 | 8,27069E-07 |
| ACE2 | RP11-608O21.1 | -0,2561 | 8,19499E-07 |
| ACE2 | RP11-455F5.4 | -0,2561 | 8,18366E-07 |
| ACE2 | RP11-96L14.8 | -0,2562 | 8,09529E-07 |
| ACE2 | KAT8 | -0,2562 | 8,10774E-07 |
| ACE2 | DUXAP4 | -0,2563 | 7,9863E-07 |
| ACE2 | RP11-183E24.1 | -0,2564 | 7,89833E-07 |
| ACE2 | RP11-1072C15.6 | -0,2564 | 7,92986E-07 |
| ACE2 | TUBB8P8 | -0,2565 | 7,83405E-07 |
| ACE2 | RP11-461L13.4 | -0,2565 | 7,8242E-07 |
| ACE2 | RP11-16D22.2 | -0,2565 | 7,8E-07 |
| ACE2 | POTED | -0,2565 | 7,81533E-07 |
| ACE2 | ARG2 | -0,2566 | 7,76208E-07 |
| ACE2 | RP11-108K3.3 | -0,2566 | 7,73519E-07 |
| ACE2 | CLK4 | -0,2567 | 7,6544E-07 |
| ACE2 | PPP1R26-AS1 | -0,2567 | 7,67432E-07 |
| ACE2 | HIF1A-AS2 | -0,2567 | 7,67964E-07 |
| ACE2 | RP11-354P11.4 | -0,2567 | 7,69529E-07 |
| ACE2 | CITED4 | -0,2568 | 7,60631E-07 |
| ACE2 | AC010872.1 | -0,2568 | 7,58834E-07 |
| ACE2 | RP11-495P10.9 | -0,2569 | 7,52137E-07 |
| ACE2 | XXyac-YRM2039.3 | -0,2569 | 7,52845E-07 |
| ACE2 | RP11-463M16.5 | -0,2569 | 7,54599E-07 |
| ACE2 | RP11-184C23.4 | -0,257 | 7,45636E-07 |
| ACE2 | RP11-753D20.1 | -0,257 | 7,47318E-07 |
| ACE2 | RP11-275I4.1 | -0,257 | 7,47117E-07 |
| ACE2 | PIGZ | -0,2571 | 7,35878E-07 |
| ACE2 | GCM2 | -0,2571 | 7,34056E-07 |
| ACE2 | LINC01996 | -0,2571 | 7,3675E-07 |
| ACE2 | AIFM3 | -0,2571 | 7,35835E-07 |
| ACE2 | RPL12P25 | -0,2572 | 7,26347E-07 |
| ACE2 | EIF3FP1 | -0,2572 | 7,2971E-07 |
| ACE2 | CACNA2D4 | -0,2573 | 7,20151E-07 |
| ACE2 | RP11-81H3.2 | -0,2573 | 7,20156E-07 |
| ACE2 | ISL2 | -0,2573 | 7,2002E-07 |
| ACE2 | RPL18P13 | -0,2573 | 7,24763E-07 |
| ACE2 | CELSR3 | -0,2574 | 7,14701E-07 |
| ACE2 | TMEM212-IT1 | -0,2574 | 7,17213E-07 |
| ACE2 | KB-1683C8.1 | -0,2574 | 7,17614E-07 |
| ACE2 | RP11-230L22.4 | -0,2574 | 7,14938E-07 |
| ACE2 | FAM149B1 | -0,2574 | 7,13837E-07 |
| ACE2 | RP11-315O6.2 | -0,2574 | 7,17723E-07 |
| ACE2 | LINC01226 | -0,2575 | 7,04183E-07 |
| ACE2 | RP1-140J1.1 | -0,2575 | 7,06335E-07 |

| | | | |
|---|---|---|---|
| ACE2 | RP11-605P22.1 | -0,2575 | 7,06395E-07 |
| ACE2 | RP11-432M24.4 | -0,2575 | 7,09456E-07 |
| ACE2 | ANKRD30BP2 | -0,2575 | 7,09901E-07 |
| ACE2 | NPAP1P6 | -0,2576 | 6,96432E-07 |
| ACE2 | RP11-69M1.4 | -0,2576 | 7,02928E-07 |
| ACE2 | RP11-285M22.3 | -0,2576 | 7,00049E-07 |
| ACE2 | RP5-1004I9.2 | -0,2576 | 7,03188E-07 |
| ACE2 | TRBV7-4 | -0,2577 | 6,9443E-07 |
| ACE2 | PARP4P3 | -0,2577 | 6,91693E-07 |
| ACE2 | CDH10 | -0,2578 | 6,82192E-07 |
| ACE2 | RP11-168P6.2 | -0,2578 | 6,89076E-07 |
| ACE2 | BNIP3P13 | -0,2578 | 6,86379E-07 |
| ACE2 | SPHKAP | -0,2579 | 6,75845E-07 |
| ACE2 | C8orf87 | -0,2579 | 6,76011E-07 |
| ACE2 | ARF4P3 | -0,2579 | 6,81369E-07 |
| ACE2 | PDCD5 | -0,2579 | 6,77351E-07 |
| ACE2 | CTC-360J11.4 | -0,2579 | 6,79262E-07 |
| ACE2 | FAM118A | -0,2579 | 6,79695E-07 |
| ACE2 | LINC01558 | -0,258 | 6,73065E-07 |
| ACE2 | RHOBTB1 | -0,258 | 6,6988E-07 |
| ACE2 | IGHG4 | -0,258 | 6,71957E-07 |
| ACE2 | RP11-434D2.8 | -0,258 | 6,7138E-07 |
| ACE2 | RN7SKP169 | -0,258 | 6,74825E-07 |
| ACE2 | RP11-619J20.1 | -0,2581 | 6,62525E-07 |
| ACE2 | ST18 | -0,2581 | 6,63444E-07 |
| ACE2 | SPIC | -0,2581 | 6,65435E-07 |
| ACE2 | RP11-357N13.2 | -0,2582 | 6,57976E-07 |
| ACE2 | GS1-388B5.1 | -0,2583 | 6,51614E-07 |
| ACE2 | AC073071.1 | -0,2583 | 6,50129E-07 |
| ACE2 | SH2B3 | -0,2583 | 6,51337E-07 |
| ACE2 | RP11-342M21.2 | -0,2583 | 6,47914E-07 |
| ACE2 | SPINT2 | -0,2583 | 6,49197E-07 |
| ACE2 | FMO11P | -0,2584 | 6,44507E-07 |
| ACE2 | AOX2P | -0,2584 | 6,45066E-07 |
| ACE2 | AC020743.3 | -0,2584 | 6,42098E-07 |
| ACE2 | RP11-252P19.1 | -0,2585 | 6,34846E-07 |
| ACE2 | DEFA4 | -0,2585 | 6,35393E-07 |
| ACE2 | CCDC26 | -0,2585 | 6,39997E-07 |
| ACE2 | CKBP1 | -0,2585 | 6,40015E-07 |
| ACE2 | RABGAP1L | -0,2586 | 6,31622E-07 |
| ACE2 | RP11-630D6.5 | -0,2586 | 6,30511E-07 |
| ACE2 | BNIP1 | -0,2586 | 6,332E-07 |
| ACE2 | TRIM53CP | -0,2586 | 6,31088E-07 |
| ACE2 | A2ML1-AS1 | -0,2586 | 6,31518E-07 |
| ACE2 | RP11-122G18.11 | -0,2587 | 6,2434E-07 |
| ACE2 | RP11-378I6.1 | -0,2587 | 6,21907E-07 |
| ACE2 | OR52E5 | -0,2587 | 6,24935E-07 |
| ACE2 | RP11-11K13.1 | -0,2587 | 6,23166E-07 |
| ACE2 | MTSS1L | -0,2587 | 6,25548E-07 |
| ACE2 | GS1-304P7.2 | -0,2588 | 6,18788E-07 |
| ACE2 | AC006003.3 | -0,2588 | 6,18126E-07 |
| ACE2 | PSME4 | -0,2589 | 6,14973E-07 |
| ACE2 | CTSW | -0,2589 | 6,1125E-07 |
| ACE2 | RP11-800A3.3 | -0,2589 | 6,09681E-07 |

| | | | |
|---|---|---|---|
| ACE2 | OPCML-IT1 | -0,2589 | 6,15046E-07 |
| ACE2 | RP11-215P8.4 | -0,2591 | 5,99349E-07 |
| ACE2 | MPP6 | -0,2591 | 6,01293E-07 |
| ACE2 | CTC-398G3.1 | -0,2591 | 5,98572E-07 |
| ACE2 | DCLRE1CP1 | -0,2592 | 5,93776E-07 |
| ACE2 | TBC1D32 | -0,2593 | 5,8845E-07 |
| ACE2 | AC005304.1 | -0,2593 | 5,87971E-07 |
| ACE2 | BTNL10 | -0,2594 | 5,82436E-07 |
| ACE2 | AC009478.1 | -0,2594 | 5,803E-07 |
| ACE2 | PPP1R3E | -0,2594 | 5,79999E-07 |
| ACE2 | CDC25A | -0,2595 | 5,76432E-07 |
| ACE2 | C18orf54 | -0,2596 | 5,69381E-07 |
| ACE2 | RECQL4 | -0,2597 | 5,6575E-07 |
| ACE2 | EXOSC8 | -0,2597 | 5,62681E-07 |
| ACE2 | TEX30 | -0,2597 | 5,63303E-07 |
| ACE2 | ABCC13 | -0,2597 | 5,64857E-07 |
| ACE2 | RP11-366M4.11 | -0,2598 | 5,55665E-07 |
| ACE2 | ARV1 | -0,2599 | 5,50808E-07 |
| ACE2 | RP11-507D14.4 | -0,2599 | 5,4958E-07 |
| ACE2 | LINC01220 | -0,2599 | 5,53666E-07 |
| ACE2 | PPIHP1 | -0,26 | 5,45992E-07 |
| ACE2 | MARK3P3 | -0,2601 | 5,42872E-07 |
| ACE2 | RP11-17E2.2 | -0,2601 | 5,41906E-07 |
| ACE2 | ZNF454 | -0,2601 | 5,41814E-07 |
| ACE2 | AC007327.5 | -0,2601 | 5,39175E-07 |
| ACE2 | KCNK18 | -0,2601 | 5,39453E-07 |
| ACE2 | POTEB3 | -0,2601 | 5,43374E-07 |
| ACE2 | MAPK8IP3 | -0,2601 | 5,41367E-07 |
| ACE2 | TADA3 | -0,2603 | 5,31316E-07 |
| ACE2 | SLC4A4 | -0,2603 | 5,30684E-07 |
| ACE2 | NUDT13 | -0,2603 | 5,27085E-07 |
| ACE2 | RP11-37N22.1 | -0,2604 | 5,21623E-07 |
| ACE2 | TRBV25OR9-2 | -0,2604 | 5,21762E-07 |
| ACE2 | RP11-432J22.2 | -0,2604 | 5,22454E-07 |
| ACE2 | RP11-434D2.9 | -0,2604 | 5,21606E-07 |
| ACE2 | SRP72P1 | -0,2604 | 5,26839E-07 |
| ACE2 | RP4-784A16.3 | -0,2605 | 5,21367E-07 |
| ACE2 | ACTN4P1 | -0,2605 | 5,17047E-07 |
| ACE2 | AC129778.2 | -0,2605 | 5,16437E-07 |
| ACE2 | MC3R | -0,2605 | 5,20059E-07 |
| ACE2 | RP11-307C12.12 | -0,2606 | 5,14487E-07 |
| ACE2 | CLVS2 | -0,2606 | 5,11373E-07 |
| ACE2 | RP11-478K15.6 | -0,2606 | 5,12254E-07 |
| ACE2 | CTC-543D15.1 | -0,2606 | 5,15827E-07 |
| ACE2 | RP4-669L17.10 | -0,2607 | 5,05419E-07 |
| ACE2 | PHACTR1 | -0,2607 | 5,07205E-07 |
| ACE2 | SAP18 | -0,2607 | 5,08874E-07 |
| ACE2 | AC005780.1 | -0,2607 | 5,10447E-07 |
| ACE2 | CTB-60E11.9 | -0,2607 | 5,10448E-07 |
| ACE2 | OR5H7P | -0,2608 | 5,03634E-07 |
| ACE2 | POC5 | -0,2608 | 5,03743E-07 |
| ACE2 | TRAV41 | -0,2608 | 5,00857E-07 |
| ACE2 | AC092295.4 | -0,2608 | 5,01672E-07 |
| ACE2 | CDC42EP3P1 | -0,2609 | 4,96856E-07 |

| | | | |
|---|---|---|---|
| ACE2 | HMGB3P15 | -0,2611 | 4,89454E-07 |
| ACE2 | GUCA1B | -0,2611 | 4,88566E-07 |
| ACE2 | LINC02075 | -0,2611 | 4,8796E-07 |
| ACE2 | NUTF2P6 | -0,2612 | 4,80428E-07 |
| ACE2 | AC016735.1 | -0,2613 | 4,7666E-07 |
| ACE2 | ZNF705D | -0,2613 | 4,78707E-07 |
| ACE2 | GCNT1 | -0,2613 | 4,78814E-07 |
| ACE2 | EIF4BP3 | -0,2613 | 4,7909E-07 |
| ACE2 | TRAF6P1 | -0,2613 | 4,76365E-07 |
| ACE2 | TFDP2 | -0,2614 | 4,7255E-07 |
| ACE2 | CTD-2269E23.2 | -0,2614 | 4,72032E-07 |
| ACE2 | CTD-2014N11.1 | -0,2614 | 4,72407E-07 |
| ACE2 | SHF | -0,2614 | 4,7294E-07 |
| ACE2 | PPA2 | -0,2615 | 4,66476E-07 |
| ACE2 | RNMTL1P2 | -0,2615 | 4,66932E-07 |
| ACE2 | RP11-680F20.10 | -0,2615 | 4,69777E-07 |
| ACE2 | LINC00407 | -0,2615 | 4,66535E-07 |
| ACE2 | RP1-313L4.3 | -0,2616 | 4,63308E-07 |
| ACE2 | AC141930.3 | -0,2616 | 4,63479E-07 |
| ACE2 | MTCO3P30 | -0,2616 | 4,61803E-07 |
| ACE2 | AC090952.5 | -0,2617 | 4,58586E-07 |
| ACE2 | SRSF9P1 | -0,2617 | 4,57742E-07 |
| ACE2 | COX7A2P2 | -0,2618 | 4,53685E-07 |
| ACE2 | LINC00599 | -0,2618 | 4,54856E-07 |
| ACE2 | KB-1930G5.4 | -0,2618 | 4,52647E-07 |
| ACE2 | PTPN20CP | -0,2618 | 4,52373E-07 |
| ACE2 | RP11-96C23.11 | -0,2618 | 4,51745E-07 |
| ACE2 | PELI3 | -0,2618 | 4,52475E-07 |
| ACE2 | RP11-234B24.2 | -0,2618 | 4,53245E-07 |
| ACE2 | PPP1CC | -0,2618 | 4,50751E-07 |
| ACE2 | RP11-429O2.1 | -0,2619 | 4,46591E-07 |
| ACE2 | COL7A1 | -0,262 | 4,42245E-07 |
| ACE2 | RP11-517C16.2 | -0,262 | 4,44191E-07 |
| ACE2 | HCG25 | -0,2621 | 4,37426E-07 |
| ACE2 | AC092610.12 | -0,2621 | 4,37644E-07 |
| ACE2 | RP11-20L24.1 | -0,2621 | 4,384E-07 |
| ACE2 | MTCO1P23 | -0,2622 | 4,34969E-07 |
| ACE2 | TBX5 | -0,2622 | 4,34737E-07 |
| ACE2 | RP11-589M4.3 | -0,2622 | 4,35725E-07 |
| ACE2 | ZNF232 | -0,2622 | 4,35121E-07 |
| ACE2 | LINC01674 | -0,2622 | 4,32243E-07 |
| ACE2 | RP1-101G11.3 | -0,2622 | 4,36468E-07 |
| ACE2 | RNU4-57P | -0,2623 | 4,28334E-07 |
| ACE2 | KLF3P1 | -0,2623 | 4,30254E-07 |
| ACE2 | RP11-113E21.3 | -0,2623 | 4,32105E-07 |
| ACE2 | FSCN2 | -0,2623 | 4,30453E-07 |
| ACE2 | SPAG16 | -0,2624 | 4,23261E-07 |
| ACE2 | AC064853.3 | -0,2624 | 4,23803E-07 |
| ACE2 | AC004471.9 | -0,2624 | 4,24408E-07 |
| ACE2 | FBXO44 | -0,2625 | 4,22531E-07 |
| ACE2 | TUBA3E | -0,2625 | 4,22202E-07 |
| ACE2 | RP11-80P20.3 | -0,2625 | 4,20244E-07 |
| ACE2 | GUSBP8 | -0,2625 | 4,19459E-07 |
| ACE2 | CTC-507E12.1 | -0,2625 | 4,22687E-07 |

| | | | |
|---|---|---|---|
| ACE2 | CTB-32P11.1 | -0,2625 | 4,21162E-07 |
| ACE2 | RP1-266L20.4 | -0,2626 | 4,18613E-07 |
| ACE2 | HNF4G | -0,2626 | 4,18087E-07 |
| ACE2 | RP11-799N11.1 | -0,2626 | 4,18556E-07 |
| ACE2 | Z69890.1 | -0,2627 | 4,11689E-07 |
| ACE2 | RP11-740P5.2 | -0,2628 | 4,06738E-07 |
| ACE2 | RP11-614F17.1 | -0,2628 | 4,07797E-07 |
| ACE2 | AP001255.2 | -0,2628 | 4,0851E-07 |
| ACE2 | FOXD4L3 | -0,2629 | 4,04644E-07 |
| ACE2 | TACC1P1 | -0,2629 | 4,02512E-07 |
| ACE2 | RP11-180P8.1 | -0,2629 | 4,04916E-07 |
| ACE2 | AC092835.2 | -0,263 | 3,98431E-07 |
| ACE2 | LRRC49 | -0,263 | 4,01297E-07 |
| ACE2 | RP11-40E6.1 | -0,2631 | 3,94473E-07 |
| ACE2 | RP11-1069G10.2 | -0,2631 | 3,93962E-07 |
| ACE2 | CTD-2342I9.1 | -0,2632 | 3,90017E-07 |
| ACE2 | ENOX1-AS1 | -0,2632 | 3,89907E-07 |
| ACE2 | RP11-66B24.2 | -0,2632 | 3,92813E-07 |
| ACE2 | LINC01473 | -0,2633 | 3,8719E-07 |
| ACE2 | LTF | -0,2633 | 3,8698E-07 |
| ACE2 | SBSPON | -0,2633 | 3,87771E-07 |
| ACE2 | HHLA1 | -0,2633 | 3,87733E-07 |
| ACE2 | RP11-490B18.5 | -0,2633 | 3,87796E-07 |
| ACE2 | AC020594.5 | -0,2634 | 3,847E-07 |
| ACE2 | AC008937.3 | -0,2634 | 3,83714E-07 |
| ACE2 | SLC4A8 | -0,2634 | 3,82113E-07 |
| ACE2 | LINC01254 | -0,2634 | 3,82056E-07 |
| ACE2 | TRBV5-7 | -0,2635 | 3,78883E-07 |
| ACE2 | RP11-678G14.5 | -0,2635 | 3,79893E-07 |
| ACE2 | AC010891.2 | -0,2635 | 3,80533E-07 |
| ACE2 | LINC02149 | -0,2636 | 3,76988E-07 |
| ACE2 | RP11-387M24.5 | -0,2636 | 3,76414E-07 |
| ACE2 | LINGO2 | -0,2636 | 3,73499E-07 |
| ACE2 | LINC01813 | -0,2637 | 3,714E-07 |
| ACE2 | AC006372.4 | -0,2637 | 3,72938E-07 |
| ACE2 | UBE2V1P5 | -0,2637 | 3,71778E-07 |
| ACE2 | MFAP1P1 | -0,2637 | 3,71683E-07 |
| ACE2 | ZNF774 | -0,2637 | 3,72027E-07 |
| ACE2 | RP11-700H6.4 | -0,2637 | 3,70586E-07 |
| ACE2 | RPP21 | -0,2638 | 3,66528E-07 |
| ACE2 | RP11-941H19.1 | -0,2638 | 3,68941E-07 |
| ACE2 | RP11-295I5.4 | -0,2638 | 3,68019E-07 |
| ACE2 | RP11-632B21.1 | -0,2638 | 3,66976E-07 |
| ACE2 | RP11-407A16.6 | -0,2638 | 3,65424E-07 |
| ACE2 | RN7SKP146 | -0,2638 | 3,66198E-07 |
| ACE2 | CALY | -0,2639 | 3,62391E-07 |
| ACE2 | GMPSP1 | -0,264 | 3,58816E-07 |
| ACE2 | TAS2R2P | -0,264 | 3,5804E-07 |
| ACE2 | SNHG11 | -0,264 | 3,60636E-07 |
| ACE2 | RP11-241G9.3 | -0,2641 | 3,5741E-07 |
| ACE2 | RPL35 | -0,2641 | 3,56997E-07 |
| ACE2 | HNRNPRP2 | -0,2641 | 3,5625E-07 |
| ACE2 | RP13-122B23.8 | -0,2642 | 3,50487E-07 |
| ACE2 | LL22NC03-27C5.1 | -0,2644 | 3,43235E-07 |

| | | | |
|---|---|---|---|
| ACE2 | S100A7L2 | -0,2645 | 3,4267E-07 |
| ACE2 | RP11-574K11.29 | -0,2645 | 3,39852E-07 |
| ACE2 | RP11-59K5.1 | -0,2645 | 3,4198E-07 |
| ACE2 | PCBP3-OT1 | -0,2645 | 3,40038E-07 |
| ACE2 | TTTY16 | -0,2645 | 3,4167E-07 |
| ACE2 | MGAT4EP | -0,2646 | 3,35992E-07 |
| ACE2 | RBM7 | -0,2646 | 3,38583E-07 |
| ACE2 | AE000658.30 | -0,2646 | 3,38692E-07 |
| ACE2 | KRTAP17-1 | -0,2646 | 3,36668E-07 |
| ACE2 | MGAM | -0,2647 | 3,32357E-07 |
| ACE2 | AC011995.3 | -0,2648 | 3,31168E-07 |
| ACE2 | RPS6KA2-IT1 | -0,2648 | 3,29053E-07 |
| ACE2 | RPL27AP5 | -0,2649 | 3,27072E-07 |
| ACE2 | IL6RP1 | -0,2649 | 3,2694E-07 |
| ACE2 | RP4-594L9.2 | -0,265 | 3,24803E-07 |
| ACE2 | GRIK4 | -0,265 | 3,24835E-07 |
| ACE2 | TECTA | -0,265 | 3,23436E-07 |
| ACE2 | NDUFAF5 | -0,265 | 3,25E-07 |
| ACE2 | RNU6-766P | -0,2651 | 3,19959E-07 |
| ACE2 | SEMA5A | -0,2651 | 3,1894E-07 |
| ACE2 | NDE1P2 | -0,2651 | 3,21775E-07 |
| ACE2 | MIAT | -0,2651 | 3,19098E-07 |
| ACE2 | MXD1 | -0,2652 | 3,15919E-07 |
| ACE2 | RP11-648M2.1 | -0,2652 | 3,17799E-07 |
| ACE2 | CTD-2544H17.1 | -0,2652 | 3,15677E-07 |
| ACE2 | LINC01526 | -0,2652 | 3,16675E-07 |
| ACE2 | MTRF1L | -0,2652 | 3,1743E-07 |
| ACE2 | CTAGE15 | -0,2652 | 3,15522E-07 |
| ACE2 | POM121L6P | -0,2653 | 3,12608E-07 |
| ACE2 | AC007003.1 | -0,2653 | 3,14717E-07 |
| ACE2 | SPANXA1 | -0,2653 | 3,12044E-07 |
| ACE2 | HOXA6 | -0,2654 | 3,11574E-07 |
| ACE2 | TMPRSS11GP | -0,2655 | 3,06723E-07 |
| ACE2 | VPS37B | -0,2655 | 3,08478E-07 |
| ACE2 | RP11-190D6.1 | -0,2655 | 3,06906E-07 |
| ACE2 | SOX14 | -0,2656 | 3,04032E-07 |
| ACE2 | RP11-1191J2.2 | -0,2656 | 3,04787E-07 |
| ACE2 | TRBV6-4 | -0,2658 | 2,98636E-07 |
| ACE2 | PGBD4P5 | -0,2659 | 2,94016E-07 |
| ACE2 | RP11-219J21.2 | -0,2659 | 2,93763E-07 |
| ACE2 | FAM27D1 | -0,2659 | 2,94376E-07 |
| ACE2 | RPL18P11 | -0,2659 | 2,93855E-07 |
| ACE2 | RP11-468N14.1 | -0,266 | 2,90791E-07 |
| ACE2 | LYPD5 | -0,266 | 2,91913E-07 |
| ACE2 | AP000347.4 | -0,266 | 2,91902E-07 |
| ACE2 | ATP5G2P3 | -0,2661 | 2,88815E-07 |
| ACE2 | CTB-76P12.1 | -0,2662 | 2,83523E-07 |
| ACE2 | MTND4LP24 | -0,2662 | 2,86206E-07 |
| ACE2 | HNRNPCP2 | -0,2663 | 2,83008E-07 |
| ACE2 | RP11-453O5.1 | -0,2663 | 2,81775E-07 |
| ACE2 | UQCRB | -0,2663 | 2,82726E-07 |
| ACE2 | BUD31 | -0,2664 | 2,77657E-07 |
| ACE2 | RALGPS1 | -0,2664 | 2,77878E-07 |
| ACE2 | AP000593.7 | -0,2664 | 2,7847E-07 |

| | | | |
|---|---|---|---|
| ACE2 | CCDC190 | -0,2666 | 2,72963E-07 |
| ACE2 | RPS6P4 | -0,2666 | 2,73441E-07 |
| ACE2 | LINC02199 | -0,2666 | 2,7191E-07 |
| ACE2 | EME1 | -0,2666 | 2,72422E-07 |
| ACE2 | RP11-863P13.6 | -0,2667 | 2,7005E-07 |
| ACE2 | THAP8 | -0,2668 | 2,6856E-07 |
| ACE2 | UGT1A1 | -0,2669 | 2,64597E-07 |
| ACE2 | RP11-93O17.1 | -0,2669 | 2,64462E-07 |
| ACE2 | ABALON | -0,2669 | 2,65494E-07 |
| ACE2 | RP11-400D2.3 | -0,267 | 2,60969E-07 |
| ACE2 | RANBP9 | -0,267 | 2,62566E-07 |
| ACE2 | CTD-2210P24.1 | -0,267 | 2,62712E-07 |
| ACE2 | CEP57 | -0,267 | 2,60501E-07 |
| ACE2 | AMN | -0,267 | 2,6141E-07 |
| ACE2 | REEP6 | -0,267 | 2,61954E-07 |
| ACE2 | TTTY11 | -0,2671 | 2,5993E-07 |
| ACE2 | AC090018.1 | -0,2672 | 2,57393E-07 |
| ACE2 | RP11-225B17.2 | -0,2672 | 2,55487E-07 |
| ACE2 | KLHDC10 | -0,2672 | 2,55127E-07 |
| ACE2 | FGF8 | -0,2672 | 2,56328E-07 |
| ACE2 | KIAA0895L | -0,2672 | 2,56124E-07 |
| ACE2 | OR1AB1P | -0,2672 | 2,54846E-07 |
| ACE2 | KRTAP10-2 | -0,2672 | 2,57295E-07 |
| ACE2 | ZNF648 | -0,2673 | 2,53806E-07 |
| ACE2 | AC093732.1 | -0,2673 | 2,53538E-07 |
| ACE2 | RP11-65F13.3 | -0,2673 | 2,53912E-07 |
| ACE2 | RP11-116N8.1 | -0,2673 | 2,53864E-07 |
| ACE2 | DENND5B-AS1 | -0,2674 | 2,49642E-07 |
| ACE2 | WDR45BP1 | -0,2674 | 2,50908E-07 |
| ACE2 | RP11-385M4.2 | -0,2675 | 2,49219E-07 |
| ACE2 | CTC-573M9.1 | -0,2675 | 2,48607E-07 |
| ACE2 | RP11-545M17.1 | -0,2675 | 2,49144E-07 |
| ACE2 | KPNA2 | -0,2675 | 2,48192E-07 |
| ACE2 | BRAT1 | -0,2676 | 2,45669E-07 |
| ACE2 | CFTR | -0,2676 | 2,45624E-07 |
| ACE2 | RP11-597G23.1 | -0,2676 | 2,44174E-07 |
| ACE2 | NOC2LP1 | -0,2677 | 2,42978E-07 |
| ACE2 | RP11-560G2.2 | -0,2677 | 2,4353E-07 |
| ACE2 | IDO1 | -0,2678 | 2,40884E-07 |
| ACE2 | CSPG4P13 | -0,2678 | 2,40752E-07 |
| ACE2 | UBIAD1 | -0,2679 | 2,36774E-07 |
| ACE2 | HNRNPCL1 | -0,2679 | 2,38408E-07 |
| ACE2 | LINC01804 | -0,2679 | 2,37354E-07 |
| ACE2 | ZNF398 | -0,2679 | 2,38098E-07 |
| ACE2 | ANKRD36 | -0,2681 | 2,31375E-07 |
| ACE2 | SOWAHC | -0,2681 | 2,31618E-07 |
| ACE2 | PTPRN2 | -0,2681 | 2,32835E-07 |
| ACE2 | LINC01163 | -0,2681 | 2,32483E-07 |
| ACE2 | RP11-315D16.4 | -0,2681 | 2,31513E-07 |
| ACE2 | MORC1-AS1 | -0,2682 | 2,29302E-07 |
| ACE2 | FGG | -0,2682 | 2,3127E-07 |
| ACE2 | KRT18P45 | -0,2682 | 2,29215E-07 |
| ACE2 | RP11-279L11.1 | -0,2682 | 2,30019E-07 |
| ACE2 | ZNF750 | -0,2682 | 2,30686E-07 |

| | | | |
|---|---|---|---|
| ACE2 | MTF1 | -0,2684 | 2,25338E-07 |
| ACE2 | BCL11A | -0,2684 | 2,24925E-07 |
| ACE2 | RP11-405F3.5 | -0,2684 | 2,24047E-07 |
| ACE2 | ZNF402P | -0,2684 | 2,24102E-07 |
| ACE2 | CXXC4-AS1 | -0,2685 | 2,22583E-07 |
| ACE2 | CTB-138E5.1 | -0,2685 | 2,21632E-07 |
| ACE2 | PRRC2A | -0,2685 | 2,2312E-07 |
| ACE2 | HMGA1P8 | -0,2686 | 2,21294E-07 |
| ACE2 | RP11-319G6.3 | -0,2686 | 2,20848E-07 |
| ACE2 | RP11-1006G14.5 | -0,2686 | 2,20868E-07 |
| ACE2 | RP11-626G11.4 | -0,2686 | 2,19889E-07 |
| ACE2 | LINC01098 | -0,2687 | 2,17775E-07 |
| ACE2 | CDK5R2 | -0,2688 | 2,16867E-07 |
| ACE2 | KB-1184D12.1 | -0,2688 | 2,15642E-07 |
| ACE2 | LINC01006 | -0,2689 | 2,12735E-07 |
| ACE2 | RP11-363E7.4 | -0,2689 | 2,12401E-07 |
| ACE2 | OR8B7P | -0,2689 | 2,13939E-07 |
| ACE2 | BCAR4 | -0,2689 | 2,14278E-07 |
| ACE2 | C19orf48 | -0,2689 | 2,1359E-07 |
| ACE2 | HOTTIP | -0,269 | 2,11454E-07 |
| ACE2 | AC108039.1 | -0,2691 | 2,09506E-07 |
| ACE2 | GPR6 | -0,2691 | 2,08214E-07 |
| ACE2 | SCARNA13 | -0,2691 | 2,09263E-07 |
| ACE2 | CTD-2022H16.1 | -0,2691 | 2,09327E-07 |
| ACE2 | RP11-14I2.2 | -0,2692 | 2,07108E-07 |
| ACE2 | DMRT3 | -0,2692 | 2,07477E-07 |
| ACE2 | RPL17P1 | -0,2692 | 2,06114E-07 |
| ACE2 | IGLVVII-41-1 | -0,2692 | 2,07226E-07 |
| ACE2 | ITIH1 | -0,2693 | 2,0435E-07 |
| ACE2 | RP11-359P18.1 | -0,2693 | 2,04344E-07 |
| ACE2 | KRTAP2-5P | -0,2693 | 2,04527E-07 |
| ACE2 | RP11-182I21.2 | -0,2695 | 1,99272E-07 |
| ACE2 | AC005077.7 | -0,2695 | 2,00683E-07 |
| ACE2 | LHX4 | -0,2696 | 1,98498E-07 |
| ACE2 | SNX18P4 | -0,2696 | 1,98382E-07 |
| ACE2 | CARD17 | -0,2696 | 1,98783E-07 |
| ACE2 | PPIAP25 | -0,2696 | 1,9869E-07 |
| ACE2 | KRTAP29-1 | -0,2696 | 1,97484E-07 |
| ACE2 | AP000350.7 | -0,2696 | 1,98577E-07 |
| ACE2 | TTTY19 | -0,2696 | 1,9867E-07 |
| ACE2 | RN7SL837P | -0,2697 | 1,96808E-07 |
| ACE2 | CUZD1 | -0,2697 | 1,96161E-07 |
| ACE2 | ANKRD20A9P | -0,2697 | 1,96614E-07 |
| ACE2 | LINC01875 | -0,2698 | 1,94604E-07 |
| ACE2 | LINC01853 | -0,2698 | 1,93735E-07 |
| ACE2 | POLD4 | -0,2698 | 1,94634E-07 |
| ACE2 | RP11-2I17.4 | -0,2698 | 1,93126E-07 |
| ACE2 | RP11-469H8.6 | -0,2699 | 1,91875E-07 |
| ACE2 | CMTR2 | -0,2699 | 1,92467E-07 |
| ACE2 | RP11-1060G2.1 | -0,27 | 1,90305E-07 |
| ACE2 | NHEJ1 | -0,2701 | 1,8742E-07 |
| ACE2 | RUVBL1-AS1 | -0,2701 | 1,87064E-07 |
| ACE2 | RP11-659O3.1 | -0,2701 | 1,87894E-07 |
| ACE2 | RP11-443C10.1 | -0,2701 | 1,87244E-07 |

| | | | |
|---|---|---|---|
| ACE2 | RP11-221N13.2 | -0,2702 | 1,86196E-07 |
| ACE2 | MARK2P6 | -0,2703 | 1,84381E-07 |
| ACE2 | RP5-1006K12.1 | -0,2703 | 1,84526E-07 |
| ACE2 | RP11-432F4.2 | -0,2703 | 1,84477E-07 |
| ACE2 | AC104532.3 | -0,2703 | 1,82992E-07 |
| ACE2 | CTC-453G23.7 | -0,2703 | 1,83319E-07 |
| ACE2 | SPANXN4 | -0,2703 | 1,8369E-07 |
| ACE2 | RP11-227F8.2 | -0,2704 | 1,8127E-07 |
| ACE2 | DDX49 | -0,2705 | 1,79414E-07 |
| ACE2 | CYP2T3P | -0,2705 | 1,79887E-07 |
| ACE2 | RP11-508M8.1 | -0,2706 | 1,78293E-07 |
| ACE2 | RP11-203L2.4 | -0,2706 | 1,77306E-07 |
| ACE2 | E2F3P1 | -0,2706 | 1,77803E-07 |
| ACE2 | AL589986.2 | -0,2707 | 1,75733E-07 |
| ACE2 | CNTN4-AS2 | -0,2707 | 1,75913E-07 |
| ACE2 | NFYA | -0,2707 | 1,75607E-07 |
| ACE2 | CTD-2105E13.16 | -0,2707 | 1,75381E-07 |
| ACE2 | AC010731.3 | -0,2708 | 1,73743E-07 |
| ACE2 | RP11-324I22.3 | -0,2708 | 1,74344E-07 |
| ACE2 | PPIAP32 | -0,2708 | 1,72996E-07 |
| ACE2 | RP11-731F5.2 | -0,2708 | 1,74384E-07 |
| ACE2 | CACNG8 | -0,2708 | 1,73391E-07 |
| ACE2 | bP-21201H5.2 | -0,2708 | 1,74595E-07 |
| ACE2 | RP11-336A10.7 | -0,2709 | 1,72718E-07 |
| ACE2 | AC092628.3 | -0,271 | 1,6953E-07 |
| ACE2 | NAMPTP3 | -0,271 | 1,6949E-07 |
| ACE2 | FAN1 | -0,2711 | 1,68173E-07 |
| ACE2 | RP11-157P23.2 | -0,2711 | 1,6812E-07 |
| ACE2 | RP11-375H19.2 | -0,2712 | 1,66096E-07 |
| ACE2 | RYR3 | -0,2712 | 1,67129E-07 |
| ACE2 | DBF4P1 | -0,2713 | 1,65319E-07 |
| ACE2 | RP13-714J12.1 | -0,2713 | 1,64539E-07 |
| ACE2 | RASGRF1 | -0,2713 | 1,65416E-07 |
| ACE2 | MIR6807 | -0,2713 | 1,6467E-07 |
| ACE2 | RP11-366M4.12 | -0,2715 | 1,61788E-07 |
| ACE2 | LRRC14B | -0,2715 | 1,61732E-07 |
| ACE2 | RAD51AP1 | -0,2715 | 1,61286E-07 |
| ACE2 | CTD-2373N4.5 | -0,2716 | 1,58674E-07 |
| ACE2 | ZNF23 | -0,2716 | 1,59895E-07 |
| ACE2 | ENDOV | -0,2716 | 1,58618E-07 |
| ACE2 | RP11-10N16.2 | -0,2717 | 1,57531E-07 |
| ACE2 | RP11-5O24.1 | -0,2717 | 1,5844E-07 |
| ACE2 | RP5-1103G7.10 | -0,2717 | 1,56995E-07 |
| ACE2 | RP5-837I24.2 | -0,2718 | 1,55345E-07 |
| ACE2 | CTA-246H3.11 | -0,2718 | 1,55578E-07 |
| ACE2 | RP11-855C21.1 | -0,2719 | 1,54176E-07 |
| ACE2 | PARP4P2 | -0,2719 | 1,54831E-07 |
| ACE2 | TP53TG3D | -0,2719 | 1,54639E-07 |
| ACE2 | AC005786.3 | -0,2719 | 1,54562E-07 |
| ACE2 | KCND3 | -0,272 | 1,51811E-07 |
| ACE2 | TM4SF19 | -0,272 | 1,52177E-07 |
| ACE2 | RP11-476M19.3 | -0,272 | 1,52891E-07 |
| ACE2 | ADH1A | -0,2721 | 1,50923E-07 |
| ACE2 | LHFPL3-AS1 | -0,2721 | 1,50153E-07 |

| | | | |
|---|---|---|---|
| ACE2 | RP11-143J24.1 | -0,2721 | 1,5126E-07 |
| ACE2 | AC009228.1 | -0,2722 | 1,48616E-07 |
| ACE2 | CEP57L1 | -0,2723 | 1,48019E-07 |
| ACE2 | RP11-395B7.2 | -0,2723 | 1,47351E-07 |
| ACE2 | OTUD6B-AS1 | -0,2723 | 1,48336E-07 |
| ACE2 | ALDH1A2 | -0,2723 | 1,48507E-07 |
| ACE2 | RP11-1094M14.7 | -0,2723 | 1,48133E-07 |
| ACE2 | CYP2D7 | -0,2723 | 1,48259E-07 |
| ACE2 | RP11-1198D22.1 | -0,2724 | 1,45488E-07 |
| ACE2 | SOBP | -0,2724 | 1,45818E-07 |
| ACE2 | FRRS1L | -0,2724 | 1,46488E-07 |
| ACE2 | CDK5RAP2 | -0,2724 | 1,45319E-07 |
| ACE2 | ACTBP14 | -0,2724 | 1,45989E-07 |
| ACE2 | CTC-448F2.7 | -0,2724 | 1,46017E-07 |
| ACE2 | DGCR8 | -0,2724 | 1,45438E-07 |
| ACE2 | SELP | -0,2725 | 1,43902E-07 |
| ACE2 | FAM134C | -0,2725 | 1,45199E-07 |
| ACE2 | CD180 | -0,2726 | 1,43391E-07 |
| ACE2 | PRELID3BP11 | -0,2726 | 1,4219E-07 |
| ACE2 | RPL30P5 | -0,2727 | 1,41336E-07 |
| ACE2 | NUDT6 | -0,2727 | 1,41226E-07 |
| ACE2 | RP11-1084J3.1 | -0,2727 | 1,42144E-07 |
| ACE2 | ADORA2B | -0,2727 | 1,42099E-07 |
| ACE2 | FGF7P1 | -0,2727 | 1,41104E-07 |
| ACE2 | MIR6801 | -0,2727 | 1,41352E-07 |
| ACE2 | AC017048.2 | -0,2728 | 1,40419E-07 |
| ACE2 | RP11-170K4.2 | -0,2728 | 1,40459E-07 |
| ACE2 | ACTR6 | -0,2728 | 1,40353E-07 |
| ACE2 | PPIAP28 | -0,2728 | 1,39547E-07 |
| ACE2 | SNHG10 | -0,2728 | 1,39307E-07 |
| ACE2 | CTC-559E9.8 | -0,2728 | 1,3918E-07 |
| ACE2 | AP000290.7 | -0,2728 | 1,40129E-07 |
| ACE2 | RP11-69E11.4 | -0,2729 | 1,37808E-07 |
| ACE2 | C1GALT1P3 | -0,2729 | 1,38528E-07 |
| ACE2 | PAX5 | -0,2729 | 1,38283E-07 |
| ACE2 | RCN1P2 | -0,2729 | 1,38749E-07 |
| ACE2 | AC020956.3 | -0,2729 | 1,37742E-07 |
| ACE2 | SENP2 | -0,273 | 1,36309E-07 |
| ACE2 | SLC25A51 | -0,273 | 1,37557E-07 |
| ACE2 | RP11-9M16.2 | -0,273 | 1,36816E-07 |
| ACE2 | RP11-927P21.6 | -0,273 | 1,36487E-07 |
| ACE2 | MYL5 | -0,2732 | 1,33786E-07 |
| ACE2 | AC073635.5 | -0,2732 | 1,33206E-07 |
| ACE2 | FABP5P11 | -0,2732 | 1,33422E-07 |
| ACE2 | RP11-494O16.4 | -0,2732 | 1,34324E-07 |
| ACE2 | ROS1 | -0,2733 | 1,32335E-07 |
| ACE2 | RP11-379L18.3 | -0,2733 | 1,32386E-07 |
| ACE2 | RP11-697H9.2 | -0,2734 | 1,31559E-07 |
| ACE2 | RP11-408E5.6 | -0,2734 | 1,30603E-07 |
| ACE2 | RP11-569D9.6 | -0,2734 | 1,31512E-07 |
| ACE2 | PDCD2L | -0,2734 | 1,30427E-07 |
| ACE2 | KLF2P2 | -0,2735 | 1,29796E-07 |
| ACE2 | CDC27P1 | -0,2735 | 1,29722E-07 |
| ACE2 | HTR2A | -0,2735 | 1,29875E-07 |

| | | | |
|---|---|---|---|
| ACE2 | ANKRD20A18P | -0,2735 | 1,29617E-07 |
| ACE2 | ZDHHC23 | -0,2736 | 1,28721E-07 |
| ACE2 | RP11-278J6.4 | -0,2736 | 1,28439E-07 |
| ACE2 | STIP1 | -0,2736 | 1,28102E-07 |
| ACE2 | LINC00504 | -0,2737 | 1,26358E-07 |
| ACE2 | HSPD1P12 | -0,2737 | 1,26521E-07 |
| ACE2 | KRTAP20-4 | -0,2737 | 1,26805E-07 |
| ACE2 | RP11-98J23.2 | -0,2738 | 1,25422E-07 |
| ACE2 | SLC5A4P1 | -0,2738 | 1,25839E-07 |
| ACE2 | NANOG | -0,2738 | 1,24789E-07 |
| ACE2 | AC007271.3 | -0,2739 | 1,24632E-07 |
| ACE2 | PCDHA13 | -0,2739 | 1,24378E-07 |
| ACE2 | DIAPH1 | -0,2739 | 1,2461E-07 |
| ACE2 | GPR89P | -0,274 | 1,22852E-07 |
| ACE2 | OR56A3 | -0,274 | 1,22584E-07 |
| ACE2 | UPK1A | -0,274 | 1,22153E-07 |
| ACE2 | CPSF3 | -0,2741 | 1,2141E-07 |
| ACE2 | ZSWIM5P3 | -0,2741 | 1,21158E-07 |
| ACE2 | SMIM5 | -0,2741 | 1,21602E-07 |
| ACE2 | AVPI1 | -0,2743 | 1,18546E-07 |
| ACE2 | FILIP1 | -0,2744 | 1,17933E-07 |
| ACE2 | RP11-293D9.2 | -0,2744 | 1,16946E-07 |
| ACE2 | RP11-268I9.3 | -0,2744 | 1,17032E-07 |
| ACE2 | AC010729.1 | -0,2745 | 1,15885E-07 |
| ACE2 | SLC9A9-AS1 | -0,2745 | 1,15604E-07 |
| ACE2 | TRBV12-3 | -0,2745 | 1,16347E-07 |
| ACE2 | ANKRD20A8P | -0,2746 | 1,14818E-07 |
| ACE2 | RP11-336N8.2 | -0,2746 | 1,14816E-07 |
| ACE2 | RP11-151A10.3 | -0,2746 | 1,15238E-07 |
| ACE2 | RP1-41C23.1 | -0,2746 | 1,14693E-07 |
| ACE2 | HIST1H2BA | -0,2747 | 1,13924E-07 |
| ACE2 | RP11-90K17.2 | -0,2748 | 1,1247E-07 |
| ACE2 | CNTNAP5 | -0,2749 | 1,11496E-07 |
| ACE2 | TTLL4 | -0,2749 | 1,1132E-07 |
| ACE2 | RP11-155G14.1 | -0,2749 | 1,11392E-07 |
| ACE2 | SLC6A17 | -0,275 | 1,09893E-07 |
| ACE2 | AL591704.9 | -0,275 | 1,09872E-07 |
| ACE2 | ZIC4 | -0,275 | 1,10424E-07 |
| ACE2 | METAP1 | -0,2751 | 1,09243E-07 |
| ACE2 | CNGB3 | -0,2751 | 1,0807E-07 |
| ACE2 | DEPDC4 | -0,2751 | 1,09036E-07 |
| ACE2 | RP11-62L18.3 | -0,2752 | 1,07886E-07 |
| ACE2 | RP11-715H19.2 | -0,2752 | 1,07685E-07 |
| ACE2 | CYP4F29P | -0,2752 | 1,07307E-07 |
| ACE2 | LL22NC01-81G9.3 | -0,2752 | 1,07754E-07 |
| ACE2 | EIF5B | -0,2753 | 1,06592E-07 |
| ACE2 | PLK4 | -0,2753 | 1,06549E-07 |
| ACE2 | RP11-447H19.1 | -0,2753 | 1,06631E-07 |
| ACE2 | GPSM3 | -0,2753 | 1,0635E-07 |
| ACE2 | CCL13 | -0,2753 | 1,0589E-07 |
| ACE2 | RP11-10J5.1 | -0,2754 | 1,05024E-07 |
| ACE2 | GNPAT | -0,2755 | 1,04218E-07 |
| ACE2 | AC107079.1 | -0,2755 | 1,04197E-07 |
| ACE2 | AC013402.4 | -0,2756 | 1,02682E-07 |

| | | | |
|---|---|---|---|
| ACE2 | GLE1 | -0,2756 | 1,02361E-07 |
| ACE2 | DHRS4L2 | -0,2756 | 1,03188E-07 |
| ACE2 | MIIP | -0,2757 | 1,01804E-07 |
| ACE2 | RP11-114G11.5 | -0,2757 | 1,01241E-07 |
| ACE2 | RP11-548K12.3 | -0,2757 | 1,01475E-07 |
| ACE2 | GRIA4 | -0,2757 | 1,02242E-07 |
| ACE2 | RP11-625L16.1 | -0,2757 | 1,01536E-07 |
| ACE2 | RP11-316I3.1 | -0,2758 | 1,00717E-07 |
| ACE2 | RP11-463H12.2 | -0,2758 | 1,00119E-07 |
| ACE2 | HNRNPH1P3 | -0,2758 | 1,00981E-07 |
| ACE2 | RPL21P18 | -0,2758 | 1,00299E-07 |
| ACE2 | RP3-461P17.9 | -0,2758 | 1,00689E-07 |
| ACE2 | PRDX2P4 | -0,2759 | 9,93781E-08 |
| ACE2 | PFN1P3 | -0,276 | 9,87696E-08 |
| ACE2 | AC106053.1 | -0,276 | 9,88435E-08 |
| ACE2 | LHFPL4 | -0,276 | 9,83008E-08 |
| ACE2 | KATNBL1P4 | -0,276 | 9,82491E-08 |
| ACE2 | RP5-1106E3.1 | -0,276 | 9,88982E-08 |
| ACE2 | TSC22D1-AS1 | -0,276 | 9,84724E-08 |
| ACE2 | MYH1 | -0,276 | 9,83035E-08 |
| ACE2 | MAST3 | -0,276 | 9,80609E-08 |
| ACE2 | OR2T11 | -0,2761 | 9,77135E-08 |
| ACE2 | ATP1B3 | -0,2761 | 9,73675E-08 |
| ACE2 | RP11-235G24.3 | -0,2761 | 9,71315E-08 |
| ACE2 | C9orf40 | -0,2761 | 9,67816E-08 |
| ACE2 | RP11-6O2.2 | -0,2761 | 9,74029E-08 |
| ACE2 | RP5-1050D4.4 | -0,2761 | 9,76256E-08 |
| ACE2 | LGALS16 | -0,2762 | 9,60732E-08 |
| ACE2 | GPR89B | -0,2763 | 9,53141E-08 |
| ACE2 | PLD5 | -0,2763 | 9,48628E-08 |
| ACE2 | AC006028.11 | -0,2763 | 9,56831E-08 |
| ACE2 | RP11-1C8.5 | -0,2763 | 9,46933E-08 |
| ACE2 | FAM86DP | -0,2764 | 9,44006E-08 |
| ACE2 | NPY | -0,2764 | 9,3835E-08 |
| ACE2 | CTC-430J12.2 | -0,2765 | 9,33368E-08 |
| ACE2 | RP11-474C8.3 | -0,2765 | 9,32584E-08 |
| ACE2 | ERICH6-AS1 | -0,2766 | 9,20316E-08 |
| ACE2 | IDE | -0,2766 | 9,24165E-08 |
| ACE2 | RP11-282I1.2 | -0,2766 | 9,22681E-08 |
| ACE2 | PCDHA8 | -0,2767 | 9,15018E-08 |
| ACE2 | LINC00933 | -0,2767 | 9,1507E-08 |
| ACE2 | RP11-613C6.2 | -0,2767 | 9,1219E-08 |
| ACE2 | LINC01840 | -0,2768 | 9,05121E-08 |
| ACE2 | RP11-236F9.2 | -0,2768 | 9,03193E-08 |
| ACE2 | OR51H2P | -0,2768 | 9,03219E-08 |
| ACE2 | CTD-3184A7.4 | -0,2768 | 9,05215E-08 |
| ACE2 | GPN1 | -0,2769 | 8,86433E-08 |
| ACE2 | AL137067.2 | -0,2769 | 8,89003E-08 |
| ACE2 | RPL21P11 | -0,2769 | 8,87676E-08 |
| ACE2 | TMUB2 | -0,2769 | 8,91393E-08 |
| ACE2 | LINC00307 | -0,2769 | 8,89481E-08 |
| ACE2 | RP11-46E17.6 | -0,2769 | 8,9274E-08 |
| ACE2 | RP11-333O1.1 | -0,277 | 8,76259E-08 |
| ACE2 | PTF1A | -0,277 | 8,85427E-08 |

| | | | |
|---|---|---|---|
| ACE2 | CCDC88A | -0,2771 | 8,69358E-08 |
| ACE2 | RP11-415C15.2 | -0,2771 | 8,66696E-08 |
| ACE2 | RP11-119D18.1 | -0,2773 | 8,53585E-08 |
| ACE2 | RP11-769N22.2 | -0,2774 | 8,44076E-08 |
| ACE2 | GRM4 | -0,2774 | 8,3992E-08 |
| ACE2 | IGHV1OR16-2 | -0,2774 | 8,41029E-08 |
| ACE2 | LILRB1 | -0,2774 | 8,45158E-08 |
| ACE2 | DNAJA1P6 | -0,2774 | 8,4477E-08 |
| ACE2 | RN7SL73P | -0,2776 | 8,20284E-08 |
| ACE2 | MYO1H | -0,2776 | 8,22256E-08 |
| ACE2 | GP2 | -0,2776 | 8,26113E-08 |
| ACE2 | CTC-457E21.1 | -0,2776 | 8,19849E-08 |
| ACE2 | RP4-669P10.16 | -0,2776 | 8,22934E-08 |
| ACE2 | AC068137.13 | -0,2778 | 8,08437E-08 |
| ACE2 | RP11-415C15.1 | -0,2778 | 8,09776E-08 |
| ACE2 | METTL14 | -0,2778 | 8,0185E-08 |
| ACE2 | RP5-1136A10.1 | -0,2778 | 8,01598E-08 |
| ACE2 | FENDRR | -0,2778 | 8,06339E-08 |
| ACE2 | ERO1B | -0,2779 | 7,93467E-08 |
| ACE2 | RP11-424M21.1 | -0,2779 | 7,92345E-08 |
| ACE2 | RP13-46H24.1 | -0,2779 | 8,00806E-08 |
| ACE2 | RP11-586K12.13 | -0,2779 | 7,96635E-08 |
| ACE2 | LAD1 | -0,278 | 7,85554E-08 |
| ACE2 | RP11-403I13.9 | -0,2781 | 7,7774E-08 |
| ACE2 | PROKR1 | -0,2781 | 7,82366E-08 |
| ACE2 | TAGLN3 | -0,2781 | 7,78871E-08 |
| ACE2 | CTD-3035K23.6 | -0,2781 | 7,79915E-08 |
| ACE2 | RPL21P41 | -0,2782 | 7,68065E-08 |
| ACE2 | RP3-425P12.1 | -0,2782 | 7,71353E-08 |
| ACE2 | RP11-503C24.6 | -0,2782 | 7,69612E-08 |
| ACE2 | RP4-745K6.1 | -0,2782 | 7,69169E-08 |
| ACE2 | RP11-204E4.4 | -0,2782 | 7,71821E-08 |
| ACE2 | CEP19 | -0,2783 | 7,61412E-08 |
| ACE2 | RP13-577H12.2 | -0,2783 | 7,62374E-08 |
| ACE2 | RP11-645N11.3 | -0,2783 | 7,57927E-08 |
| ACE2 | RP11-381I15.1 | -0,2783 | 7,58568E-08 |
| ACE2 | MC5R | -0,2784 | 7,49546E-08 |
| ACE2 | WBP1LP1 | -0,2785 | 7,40937E-08 |
| ACE2 | PGLYRP1 | -0,2785 | 7,44895E-08 |
| ACE2 | RP11-647K16.1 | -0,2786 | 7,3342E-08 |
| ACE2 | KB-1568E2.1 | -0,2786 | 7,38483E-08 |
| ACE2 | CTD-2288F12.1 | -0,2786 | 7,39653E-08 |
| ACE2 | RP11-443B20.1 | -0,2787 | 7,29047E-08 |
| ACE2 | CTD-3064H18.2 | -0,2787 | 7,2735E-08 |
| ACE2 | DCAF6 | -0,2788 | 7,18171E-08 |
| ACE2 | UGT1A2P | -0,2788 | 7,23063E-08 |
| ACE2 | CTD-2288O8.1 | -0,2788 | 7,21431E-08 |
| ACE2 | RN7SL352P | -0,2788 | 7,20837E-08 |
| ACE2 | BEND3 | -0,2788 | 7,16495E-08 |
| ACE2 | SLC25A24P1 | -0,2789 | 7,14189E-08 |
| ACE2 | RP11-556G22.3 | -0,2789 | 7,15564E-08 |
| ACE2 | OR7E39P | -0,2789 | 7,16159E-08 |
| ACE2 | CTB-161M19.1 | -0,279 | 7,0683E-08 |
| ACE2 | RP11-344B5.2 | -0,279 | 7,06931E-08 |

| | | | |
|---|---|---|---|
| ACE2 | LINC02181 | -0,279 | 7,01918E-08 |
| ACE2 | LINC00683 | -0,279 | 7,02975E-08 |
| ACE2 | RPL21P67 | -0,2791 | 6,97496E-08 |
| ACE2 | RP11-144G7.2 | -0,2792 | 6,8679E-08 |
| ACE2 | RP11-466F5.6 | -0,2793 | 6,79582E-08 |
| ACE2 | VTRNA1-1 | -0,2793 | 6,83334E-08 |
| ACE2 | TBC1D30 | -0,2793 | 6,79563E-08 |
| ACE2 | AP4B1-AS1 | -0,2794 | 6,73522E-08 |
| ACE2 | RP1-117P20.3 | -0,2794 | 6,73137E-08 |
| ACE2 | RP11-576N17.3 | -0,2794 | 6,771E-08 |
| ACE2 | FAM27C | -0,2794 | 6,75434E-08 |
| ACE2 | NAP1L1P1 | -0,2794 | 6,76395E-08 |
| ACE2 | RP11-299H22.3 | -0,2794 | 6,76157E-08 |
| ACE2 | AC013439.4 | -0,2795 | 6,67815E-08 |
| ACE2 | RP11-654D12.2 | -0,2795 | 6,6891E-08 |
| ACE2 | RP11-543H23.1 | -0,2795 | 6,65683E-08 |
| ACE2 | PDZK1 | -0,2796 | 6,55347E-08 |
| ACE2 | NPY5R | -0,2796 | 6,56293E-08 |
| ACE2 | DEFB109P3 | -0,2796 | 6,58797E-08 |
| ACE2 | RP11-248E9.6 | -0,2796 | 6,58427E-08 |
| ACE2 | PRDX2P1 | -0,2796 | 6,58597E-08 |
| ACE2 | RP11-57P19.1 | -0,2796 | 6,61693E-08 |
| ACE2 | RP11-297C4.1 | -0,2796 | 6,6204E-08 |
| ACE2 | SGSM3 | -0,2796 | 6,59368E-08 |
| ACE2 | RP11-402P6.13 | -0,2796 | 6,61881E-08 |
| ACE2 | FOXD4 | -0,2797 | 6,4996E-08 |
| ACE2 | RP1-288L1.4 | -0,2797 | 6,51729E-08 |
| ACE2 | IGSF21 | -0,2798 | 6,40828E-08 |
| ACE2 | AC004988.1 | -0,2798 | 6,42126E-08 |
| ACE2 | RP11-956E11.1 | -0,2798 | 6,43346E-08 |
| ACE2 | VAV3-AS1 | -0,2799 | 6,39423E-08 |
| ACE2 | DAPK3 | -0,2799 | 6,35314E-08 |
| ACE2 | EFHC2 | -0,2799 | 6,39876E-08 |
| ACE2 | AC016903.2 | -0,28 | 6,28614E-08 |
| ACE2 | FGF9 | -0,28 | 6,26918E-08 |
| ACE2 | OR10G3 | -0,28 | 6,27788E-08 |
| ACE2 | GS1-531I17.3 | -0,28 | 6,29616E-08 |
| ACE2 | LINC01657 | -0,2801 | 6,25326E-08 |
| ACE2 | LRRC10 | -0,2801 | 6,21361E-08 |
| ACE2 | LINC00396 | -0,2801 | 6,23196E-08 |
| ACE2 | LRRCC1 | -0,2802 | 6,13118E-08 |
| ACE2 | RP13-895J2.8 | -0,2802 | 6,18308E-08 |
| ACE2 | PTP4A1P1 | -0,2803 | 6,12253E-08 |
| ACE2 | AC018890.6 | -0,2803 | 6,06306E-08 |
| ACE2 | MYLKP1 | -0,2803 | 6,08697E-08 |
| ACE2 | CSH2 | -0,2803 | 6,08717E-08 |
| ACE2 | RP11-174O3.4 | -0,2804 | 6,02111E-08 |
| ACE2 | GADL1 | -0,2805 | 5,93987E-08 |
| ACE2 | EEF1DP5 | -0,2805 | 5,94341E-08 |
| ACE2 | RP11-463C14.1 | -0,2805 | 5,92114E-08 |
| ACE2 | CTD-2210P24.2 | -0,2805 | 5,97513E-08 |
| ACE2 | RP4-646N3.1 | -0,2805 | 5,97487E-08 |
| ACE2 | AKAP6 | -0,2807 | 5,84205E-08 |
| ACE2 | AC005532.5 | -0,2808 | 5,73208E-08 |

| | | | |
|---|---|---|---|
| ACE2 | SPDYE2B | -0,2808 | 5,76857E-08 |
| ACE2 | FEM1AP4 | -0,2808 | 5,76024E-08 |
| ACE2 | COX6A1 | -0,2809 | 5,71582E-08 |
| ACE2 | HMGN3P1 | -0,281 | 5,62007E-08 |
| ACE2 | OR7E91P | -0,281 | 5,6241E-08 |
| ACE2 | RP11-79H23.3 | -0,281 | 5,65619E-08 |
| ACE2 | RNU2-60P | -0,281 | 5,64588E-08 |
| ACE2 | RP11-582E3.4 | -0,281 | 5,63023E-08 |
| ACE2 | RP11-638L3.1 | -0,281 | 5,64688E-08 |
| ACE2 | GOLGA6L11P | -0,2811 | 5,57397E-08 |
| ACE2 | CTC-351M12.1 | -0,2812 | 5,48919E-08 |
| ACE2 | BACH1 | -0,2812 | 5,52429E-08 |
| ACE2 | RP11-209D14.2 | -0,2813 | 5,43386E-08 |
| ACE2 | PGBD4P8 | -0,2814 | 5,36083E-08 |
| ACE2 | MT1HL1 | -0,2814 | 5,40356E-08 |
| ACE2 | RNA5SP260 | -0,2814 | 5,407E-08 |
| ACE2 | RP11-70J12.1 | -0,2814 | 5,36841E-08 |
| ACE2 | RP11-407G23.1 | -0,2815 | 5,29229E-08 |
| ACE2 | RP11-763E3.1 | -0,2815 | 5,33954E-08 |
| ACE2 | RP11-214O1.3 | -0,2816 | 5,23984E-08 |
| ACE2 | ATP1A1-AS1 | -0,2817 | 5,21847E-08 |
| ACE2 | HMHB1 | -0,2817 | 5,20228E-08 |
| ACE2 | AP000233.4 | -0,2817 | 5,19169E-08 |
| ACE2 | SCNN1D | -0,2818 | 5,13529E-08 |
| ACE2 | CTD-2529P6.4 | -0,2818 | 5,1585E-08 |
| ACE2 | CTD-2291D10.3 | -0,2818 | 5,11391E-08 |
| ACE2 | PCDHGA11 | -0,2819 | 5,11026E-08 |
| ACE2 | RP3-330M21.5 | -0,2819 | 5,05503E-08 |
| ACE2 | GRK5 | -0,2819 | 5,07836E-08 |
| ACE2 | LINC00353 | -0,2819 | 5,10688E-08 |
| ACE2 | CTD-3187F8.2 | -0,2819 | 5,07319E-08 |
| ACE2 | NDUFB6 | -0,282 | 5,01565E-08 |
| ACE2 | POM121L10P | -0,282 | 5,0211E-08 |
| ACE2 | HAGLROS | -0,2821 | 4,96004E-08 |
| ACE2 | WRAP53 | -0,2821 | 4,96454E-08 |
| ACE2 | RP4-781K5.9 | -0,2822 | 4,91286E-08 |
| ACE2 | LA16c-329F2.1 | -0,2823 | 4,84037E-08 |
| ACE2 | ZNF274 | -0,2823 | 4,85532E-08 |
| ACE2 | SNRPCP1 | -0,2824 | 4,82233E-08 |
| ACE2 | RP11-196G11.4 | -0,2824 | 4,77854E-08 |
| ACE2 | FAM159A | -0,2825 | 4,72607E-08 |
| ACE2 | WFDC10A | -0,2825 | 4,7489E-08 |
| ACE2 | AC092660.1 | -0,2826 | 4,68496E-08 |
| ACE2 | RP11-481E4.2 | -0,2826 | 4,69756E-08 |
| ACE2 | BRI3BP | -0,2827 | 4,64186E-08 |
| ACE2 | KIF18B | -0,2827 | 4,6362E-08 |
| ACE2 | CTC-543D15.3 | -0,2827 | 4,62212E-08 |
| ACE2 | CDC42EP5 | -0,2827 | 4,63456E-08 |
| ACE2 | KCNK4 | -0,2828 | 4,57017E-08 |
| ACE2 | RP11-846F4.11 | -0,2828 | 4,60608E-08 |
| ACE2 | CTD-2363C16.2 | -0,2829 | 4,56202E-08 |
| ACE2 | RP5-933K21.3 | -0,2829 | 4,56144E-08 |
| ACE2 | ZDHHC24 | -0,283 | 4,47554E-08 |
| ACE2 | RP1-172H20.4 | -0,283 | 4,47591E-08 |

| | | | |
|---|---|---|---|
| ACE2 | RP1-47M23.3 | -0,2831 | 4,4384E-08 |
| ACE2 | CDC40 | -0,2831 | 4,43874E-08 |
| ACE2 | CXADRP1 | -0,2831 | 4,43932E-08 |
| ACE2 | ASLP1 | -0,2831 | 4,42581E-08 |
| ACE2 | DEC1 | -0,2832 | 4,36675E-08 |
| ACE2 | UBE2MP1 | -0,2832 | 4,40332E-08 |
| ACE2 | GNGT2 | -0,2832 | 4,38196E-08 |
| ACE2 | EID3 | -0,2833 | 4,31519E-08 |
| ACE2 | RP5-1024G6.7 | -0,2834 | 4,29424E-08 |
| ACE2 | RPL34P6 | -0,2834 | 4,27911E-08 |
| ACE2 | GLT8D1 | -0,2834 | 4,26792E-08 |
| ACE2 | NUPL2 | -0,2834 | 4,27673E-08 |
| ACE2 | RP11-176H8.1 | -0,2834 | 4,2676E-08 |
| ACE2 | KCNN4 | -0,2834 | 4,2988E-08 |
| ACE2 | AP001628.6 | -0,2834 | 4,28369E-08 |
| ACE2 | WASHC5-AS1 | -0,2835 | 4,21605E-08 |
| ACE2 | RP1-59D14.10 | -0,2835 | 4,22816E-08 |
| ACE2 | RP11-430L17.1 | -0,2836 | 4,18806E-08 |
| ACE2 | AC060834.2 | -0,2836 | 4,20887E-08 |
| ACE2 | NPM1P31 | -0,2836 | 4,17621E-08 |
| ACE2 | RP11-172C16.4 | -0,2836 | 4,17634E-08 |
| ACE2 | KRT17P8 | -0,2836 | 4,18822E-08 |
| ACE2 | NACAP3 | -0,2836 | 4,21287E-08 |
| ACE2 | AC009120.9 | -0,2836 | 4,20615E-08 |
| ACE2 | CTD-2521M24.4 | -0,2836 | 4,16634E-08 |
| ACE2 | RP11-946L16.1 | -0,2837 | 4,13197E-08 |
| ACE2 | RP11-973N13.5 | -0,2837 | 4,14047E-08 |
| ACE2 | RP11-757F18.5 | -0,2838 | 4,09607E-08 |
| ACE2 | PTCHD3 | -0,2838 | 4,11594E-08 |
| ACE2 | RNF41 | -0,2838 | 4,09099E-08 |
| ACE2 | RP11-756H6.1 | -0,2838 | 4,0954E-08 |
| ACE2 | CHDH | -0,2839 | 4,04294E-08 |
| ACE2 | OR7E109P | -0,2839 | 4,05095E-08 |
| ACE2 | CEACAM7 | -0,2839 | 4,06862E-08 |
| ACE2 | AP000253.1 | -0,284 | 4,01011E-08 |
| ACE2 | AC005915.1 | -0,2841 | 3,96945E-08 |
| ACE2 | LINC01752 | -0,2841 | 3,9639E-08 |
| ACE2 | AC002056.5 | -0,2841 | 3,96397E-08 |
| ACE2 | CENPF | -0,2842 | 3,93232E-08 |
| ACE2 | MCTP2 | -0,2842 | 3,91558E-08 |
| ACE2 | VN2R17P | -0,2842 | 3,91737E-08 |
| ACE2 | DPP4 | -0,2843 | 3,86681E-08 |
| ACE2 | RP11-168O22.1 | -0,2843 | 3,85633E-08 |
| ACE2 | RP4-710L4.2 | -0,2843 | 3,87744E-08 |
| ACE2 | TRIM50 | -0,2844 | 3,81476E-08 |
| ACE2 | MTND1P20 | -0,2844 | 3,80497E-08 |
| ACE2 | AUNIP | -0,2845 | 3,79725E-08 |
| ACE2 | LINC01326 | -0,2845 | 3,78221E-08 |
| ACE2 | CTAGE8 | -0,2845 | 3,76566E-08 |
| ACE2 | ZFP91 | -0,2845 | 3,80017E-08 |
| ACE2 | DACT1 | -0,2845 | 3,77541E-08 |
| ACE2 | LINC00589 | -0,2846 | 3,71848E-08 |
| ACE2 | RP11-118F2.2 | -0,2846 | 3,75501E-08 |
| ACE2 | TMEM132B | -0,2847 | 3,70489E-08 |

| | | | |
|---|---|---|---|
| ACE2 | KCNH5 | -0,2847 | 3,69284E-08 |
| ACE2 | BRD8 | -0,285 | 3,58568E-08 |
| ACE2 | LRRC4 | -0,285 | 3,56834E-08 |
| ACE2 | GAPDHP54 | -0,2852 | 3,50338E-08 |
| ACE2 | LINC01176 | -0,2853 | 3,44684E-08 |
| ACE2 | RP4-739H11.3 | -0,2854 | 3,41973E-08 |
| ACE2 | PCM1 | -0,2854 | 3,41932E-08 |
| ACE2 | RP5-943J3.1 | -0,2855 | 3,36787E-08 |
| ACE2 | DSG1 | -0,2855 | 3,36403E-08 |
| ACE2 | CTA-992D9.6 | -0,2855 | 3,37522E-08 |
| ACE2 | RP3-428L16.1 | -0,2856 | 3,3142E-08 |
| ACE2 | ZAR1L | -0,2856 | 3,31793E-08 |
| ACE2 | RP11-520P18.1 | -0,2857 | 3,29094E-08 |
| ACE2 | LPAR5 | -0,2857 | 3,30794E-08 |
| ACE2 | LINC01075 | -0,2857 | 3,29815E-08 |
| ACE2 | DNM1P47 | -0,2858 | 3,25261E-08 |
| ACE2 | CTC-471C19.2 | -0,2859 | 3,22988E-08 |
| ACE2 | CTD-2201E9.2 | -0,2859 | 3,21561E-08 |
| ACE2 | DCUN1D2-AS | -0,2859 | 3,22611E-08 |
| ACE2 | TTC39C | -0,2859 | 3,20734E-08 |
| ACE2 | RP11-448G15.1 | -0,286 | 3,17524E-08 |
| ACE2 | SEC24B-AS1 | -0,286 | 3,18306E-08 |
| ACE2 | RP11-325L7.1 | -0,286 | 3,18104E-08 |
| ACE2 | CHID1 | -0,286 | 3,1843E-08 |
| ACE2 | CTD-2021H9.3 | -0,286 | 3,18247E-08 |
| ACE2 | AC114273.3 | -0,286 | 3,183E-08 |
| ACE2 | RP4-811M8.1 | -0,286 | 3,18305E-08 |
| ACE2 | RP11-163G10.4 | -0,2861 | 3,16124E-08 |
| ACE2 | AC002480.5 | -0,2861 | 3,16141E-08 |
| ACE2 | NDUFAF8 | -0,2861 | 3,14008E-08 |
| ACE2 | RP11-1280N14.3 | -0,2862 | 3,113E-08 |
| ACE2 | LINC00240 | -0,2862 | 3,10647E-08 |
| ACE2 | TRGJP2 | -0,2862 | 3,09744E-08 |
| ACE2 | PNPLA5 | -0,2862 | 3,11276E-08 |
| ACE2 | RP11-536O18.1 | -0,2863 | 3,08419E-08 |
| ACE2 | LRRK1 | -0,2863 | 3,05767E-08 |
| ACE2 | GNA15 | -0,2863 | 3,08667E-08 |
| ACE2 | RP4-564O4.1 | -0,2863 | 3,06143E-08 |
| ACE2 | AL109763.1 | -0,2863 | 3,08464E-08 |
| ACE2 | REEP3 | -0,2864 | 3,03594E-08 |
| ACE2 | DMRTB1 | -0,2865 | 2,99984E-08 |
| ACE2 | FSD1L | -0,2865 | 3,01419E-08 |
| ACE2 | MEI1 | -0,2865 | 3,01547E-08 |
| ACE2 | PTMAP2 | -0,2867 | 2,92051E-08 |
| ACE2 | GOLGA8DP | -0,2867 | 2,92294E-08 |
| ACE2 | RIMS3 | -0,2869 | 2,85786E-08 |
| ACE2 | ANKRD18CP | -0,2869 | 2,85384E-08 |
| ACE2 | G2E3 | -0,2869 | 2,85878E-08 |
| ACE2 | RP11-501O2.3 | -0,287 | 2,85089E-08 |
| ACE2 | TKTL2 | -0,287 | 2,82374E-08 |
| ACE2 | SCUBE3 | -0,287 | 2,83733E-08 |
| ACE2 | SVBP | -0,2871 | 2,78926E-08 |
| ACE2 | RP11-363D14.1 | -0,2871 | 2,81482E-08 |
| ACE2 | SPATA31A6 | -0,2871 | 2,80596E-08 |

| | | | |
|---|---|---|---|
| ACE2 | NPSR1-AS1 | -0,2872 | 2,76711E-08 |
| ACE2 | RAB11B-AS1 | -0,2872 | 2,78436E-08 |
| ACE2 | UBA52P7 | -0,2873 | 2,74614E-08 |
| ACE2 | PCDHA10 | -0,2874 | 2,70412E-08 |
| ACE2 | PKIA | -0,2874 | 2,70902E-08 |
| ACE2 | LINC01018 | -0,2875 | 2,67965E-08 |
| ACE2 | ASB10 | -0,2875 | 2,69152E-08 |
| ACE2 | RP11-321L2.2 | -0,2875 | 2,67452E-08 |
| ACE2 | WNT8B | -0,2875 | 2,67924E-08 |
| ACE2 | CTD-2135D7.3 | -0,2875 | 2,666E-08 |
| ACE2 | AC093063.2 | -0,2875 | 2,6789E-08 |
| ACE2 | OR2T27 | -0,2876 | 2,64835E-08 |
| ACE2 | CTB-70G10.1 | -0,2876 | 2,64425E-08 |
| ACE2 | LENG8-AS1 | -0,2876 | 2,65014E-08 |
| ACE2 | RP11-1J7.1 | -0,2877 | 2,62861E-08 |
| ACE2 | RP11-1379J22.2 | -0,2877 | 2,61379E-08 |
| ACE2 | RP11-401E14.2 | -0,2878 | 2,59772E-08 |
| ACE2 | LL22NC03-23C6.12 | -0,2878 | 2,57875E-08 |
| ACE2 | KMO | -0,2879 | 2,54699E-08 |
| ACE2 | PSCA | -0,2879 | 2,55873E-08 |
| ACE2 | RP11-61F12.1 | -0,2879 | 2,54709E-08 |
| ACE2 | LINC01529 | -0,2879 | 2,56477E-08 |
| ACE2 | DDRGK1 | -0,2879 | 2,54448E-08 |
| ACE2 | MYCNUT | -0,288 | 2,53295E-08 |
| ACE2 | RFTN2 | -0,288 | 2,51971E-08 |
| ACE2 | RP11-697K23.3 | -0,288 | 2,52315E-08 |
| ACE2 | CTC-205M6.1 | -0,288 | 2,51535E-08 |
| ACE2 | RPSAP43 | -0,288 | 2,53781E-08 |
| ACE2 | BSNDP4 | -0,288 | 2,53984E-08 |
| ACE2 | MEST | -0,288 | 2,51738E-08 |
| ACE2 | KB-1396H2.2 | -0,288 | 2,53773E-08 |
| ACE2 | ADGRA1 | -0,288 | 2,53347E-08 |
| ACE2 | CTD-2034I21.1 | -0,288 | 2,5207E-08 |
| ACE2 | FBXO46 | -0,288 | 2,52618E-08 |
| ACE2 | USP41 | -0,288 | 2,52235E-08 |
| ACE2 | TRAF3IP3 | -0,2881 | 2,50112E-08 |
| ACE2 | ZNF184 | -0,2881 | 2,49462E-08 |
| ACE2 | ZNF192P2 | -0,2881 | 2,48931E-08 |
| ACE2 | RP4-705F19.1 | -0,2882 | 2,46144E-08 |
| ACE2 | HCG4P8 | -0,2882 | 2,46027E-08 |
| ACE2 | ATP6V1E1P1 | -0,2883 | 2,43554E-08 |
| ACE2 | AC114752.1 | -0,2883 | 2,44351E-08 |
| ACE2 | AC006160.5 | -0,2883 | 2,43985E-08 |
| ACE2 | RP11-80H5.6 | -0,2883 | 2,42919E-08 |
| ACE2 | RP11-486L19.2 | -0,2883 | 2,44146E-08 |
| ACE2 | AP000357.4 | -0,2883 | 2,42894E-08 |
| ACE2 | AC068138.1 | -0,2884 | 2,42377E-08 |
| ACE2 | SLC8A2 | -0,2884 | 2,41212E-08 |
| ACE2 | RP11-457P14.5 | -0,2885 | 2,38902E-08 |
| ACE2 | TRHDE | -0,2885 | 2,37078E-08 |
| ACE2 | MPI | -0,2885 | 2,38502E-08 |
| ACE2 | RP11-142G1.1 | -0,2885 | 2,3928E-08 |
| ACE2 | RP1-274L7.4 | -0,2885 | 2,38492E-08 |
| ACE2 | P3H2 | -0,2886 | 2,3556E-08 |

| | | | |
|---|---|---|---|
| ACE2 | FRMD6-AS2 | -0,2886 | 2,35975E-08 |
| ACE2 | AC098795.1 | -0,2887 | 2,33948E-08 |
| ACE2 | RP11-20I20.2 | -0,2888 | 2,29299E-08 |
| ACE2 | AC005772.2 | -0,2888 | 2,31193E-08 |
| ACE2 | RP11-159M11.2 | -0,2889 | 2,2728E-08 |
| ACE2 | PCGF5 | -0,2889 | 2,26876E-08 |
| ACE2 | TOM1L1 | -0,2889 | 2,28398E-08 |
| ACE2 | ZNF133 | -0,2889 | 2,28622E-08 |
| ACE2 | RP11-848G14.2 | -0,289 | 2,24864E-08 |
| ACE2 | RP11-403N16.2 | -0,289 | 2,23652E-08 |
| ACE2 | LINP1 | -0,289 | 2,24309E-08 |
| ACE2 | RP1-288H2.2 | -0,289 | 2,24209E-08 |
| ACE2 | LRRC37BP1 | -0,289 | 2,25058E-08 |
| ACE2 | CCDC191 | -0,2891 | 2,20982E-08 |
| ACE2 | RPSAP47 | -0,2892 | 2,20709E-08 |
| ACE2 | RP11-160E2.6 | -0,2892 | 2,19092E-08 |
| ACE2 | OR6M2P | -0,2893 | 2,16113E-08 |
| ACE2 | KATNAL1 | -0,2893 | 2,1693E-08 |
| ACE2 | RP11-329J18.3 | -0,2893 | 2,17379E-08 |
| ACE2 | SFMBT1 | -0,2894 | 2,14876E-08 |
| ACE2 | RP11-387A1.5 | -0,2895 | 2,11512E-08 |
| ACE2 | DUSP19 | -0,2895 | 2,11599E-08 |
| ACE2 | TAF1L | -0,2895 | 2,1227E-08 |
| ACE2 | LINC00911 | -0,2895 | 2,11246E-08 |
| ACE2 | CTA-363E6.2 | -0,2895 | 2,11423E-08 |
| ACE2 | RP11-102J14.1 | -0,2895 | 2,12979E-08 |
| ACE2 | ETV3L | -0,2896 | 2,09848E-08 |
| ACE2 | LINC01985 | -0,2896 | 2,10054E-08 |
| ACE2 | AC091729.7 | -0,2896 | 2,08487E-08 |
| ACE2 | RP11-274A11.4 | -0,2896 | 2,09352E-08 |
| ACE2 | ZNF730 | -0,2896 | 2,09476E-08 |
| ACE2 | ZFP30 | -0,2896 | 2,09077E-08 |
| ACE2 | RP11-231G15.3 | -0,2898 | 2,03973E-08 |
| ACE2 | AC096579.13 | -0,2899 | 2,02066E-08 |
| ACE2 | RP11-309M7.1 | -0,2899 | 2,01942E-08 |
| ACE2 | RPL35P4 | -0,2899 | 2,01569E-08 |
| ACE2 | AC092625.1 | -0,29 | 2,00964E-08 |
| ACE2 | SLC9C1 | -0,29 | 2,00762E-08 |
| ACE2 | RP5-1068H6.6 | -0,2901 | 1,97289E-08 |
| ACE2 | AF067845.3 | -0,2902 | 1,94841E-08 |
| ACE2 | CTD-3023L14.3 | -0,2902 | 1,94392E-08 |
| ACE2 | RP11-23E10.5 | -0,2902 | 1,96129E-08 |
| ACE2 | TTC39DP | -0,2903 | 1,92121E-08 |
| ACE2 | GPR141 | -0,2903 | 1,92953E-08 |
| ACE2 | RNF20 | -0,2903 | 1,92921E-08 |
| ACE2 | MTND5P42 | -0,2903 | 1,93132E-08 |
| ACE2 | MARCH4 | -0,2904 | 1,91076E-08 |
| ACE2 | RP11-18H21.3 | -0,2904 | 1,89857E-08 |
| ACE2 | HDAC9 | -0,2904 | 1,89914E-08 |
| ACE2 | RP11-432I5.2 | -0,2904 | 1,91307E-08 |
| ACE2 | CNN2P10 | -0,2905 | 1,89345E-08 |
| ACE2 | MED12L | -0,2905 | 1,87645E-08 |
| ACE2 | LINC01327 | -0,2906 | 1,86606E-08 |
| ACE2 | FAM149A | -0,2906 | 1,87517E-08 |

| | | | |
|---|---|---|---|
| ACE2 | RP11-138I18.2 | -0,2906 | 1,86691E-08 |
| ACE2 | RP11-629N8.3 | -0,2906 | 1,86275E-08 |
| ACE2 | AC010091.1 | -0,2907 | 1,84957E-08 |
| ACE2 | RP11-662J14.1 | -0,2907 | 1,84773E-08 |
| ACE2 | RP13-577H12.3 | -0,2908 | 1,82633E-08 |
| ACE2 | KRT17P1 | -0,2908 | 1,81744E-08 |
| ACE2 | RP11-350G24.1 | -0,2909 | 1,79172E-08 |
| ACE2 | RP11-661D19.3 | -0,2909 | 1,7903E-08 |
| ACE2 | RP11-807E13.2 | -0,2909 | 1,79207E-08 |
| ACE2 | RP11-358H12.1 | -0,291 | 1,77093E-08 |
| ACE2 | ZSCAN16-AS1 | -0,291 | 1,77589E-08 |
| ACE2 | TNRC18P3 | -0,291 | 1,78078E-08 |
| ACE2 | OR4N2 | -0,291 | 1,77731E-08 |
| ACE2 | LINC00907 | -0,291 | 1,78627E-08 |
| ACE2 | POP4 | -0,291 | 1,77311E-08 |
| ACE2 | PCDHB19P | -0,2911 | 1,76734E-08 |
| ACE2 | EIF4H | -0,2911 | 1,75642E-08 |
| ACE2 | OR13D1 | -0,2911 | 1,75672E-08 |
| ACE2 | RP11-323P17.2 | -0,2911 | 1,7645E-08 |
| ACE2 | KIAA0586 | -0,2912 | 1,73343E-08 |
| ACE2 | USP43 | -0,2912 | 1,72894E-08 |
| ACE2 | ZNF225 | -0,2912 | 1,74473E-08 |
| ACE2 | AF212831.2 | -0,2912 | 1,7408E-08 |
| ACE2 | SAR1P1 | -0,2913 | 1,72432E-08 |
| ACE2 | GTF2IRD2P1 | -0,2913 | 1,72159E-08 |
| ACE2 | TBC1D3J | -0,2913 | 1,72224E-08 |
| ACE2 | OR7A1P | -0,2913 | 1,7109E-08 |
| ACE2 | RNA5SP174 | -0,2915 | 1,67573E-08 |
| ACE2 | HMMR | -0,2916 | 1,66201E-08 |
| ACE2 | RN7SL22P | -0,2916 | 1,66211E-08 |
| ACE2 | SPCS2P2 | -0,2916 | 1,66784E-08 |
| ACE2 | RP11-529A4.10 | -0,2916 | 1,65367E-08 |
| ACE2 | RP11-297M9.2 | -0,2916 | 1,6665E-08 |
| ACE2 | CTD-3194G12.2 | -0,2916 | 1,64903E-08 |
| ACE2 | ZNF540 | -0,2916 | 1,6636E-08 |
| ACE2 | KRT18P23 | -0,2916 | 1,6582E-08 |
| ACE2 | PCDHA11 | -0,2917 | 1,6483E-08 |
| ACE2 | RP11-941H19.3 | -0,2917 | 1,64046E-08 |
| ACE2 | RP11-108E14.1 | -0,2918 | 1,6255E-08 |
| ACE2 | RP11-298D21.1 | -0,2918 | 1,61469E-08 |
| ACE2 | CTD-3145H4.1 | -0,2919 | 1,60807E-08 |
| ACE2 | CYP4F60P | -0,2919 | 1,59216E-08 |
| ACE2 | RP11-755E23.3 | -0,2919 | 1,59208E-08 |
| ACE2 | RP11-45P15.4 | -0,2919 | 1,60329E-08 |
| ACE2 | RP11-666O2.4 | -0,2919 | 1,60961E-08 |
| ACE2 | RNU6ATAC38P | -0,2919 | 1,60179E-08 |
| ACE2 | KANSL3 | -0,292 | 1,57319E-08 |
| ACE2 | FBXO33 | -0,292 | 1,575E-08 |
| ACE2 | RP11-109D20.1 | -0,2921 | 1,56799E-08 |
| ACE2 | BPIFB2 | -0,2921 | 1,55734E-08 |
| ACE2 | C1orf220 | -0,2922 | 1,54418E-08 |
| ACE2 | RP11-78B10.2 | -0,2922 | 1,54672E-08 |
| ACE2 | FANCM | -0,2922 | 1,5506E-08 |
| ACE2 | GRB10 | -0,2923 | 1,52909E-08 |

| | | | |
|---|---|---|---|
| ACE2 | RP11-44N11.1 | -0,2923 | 1,53438E-08 |
| ACE2 | CACNA1B | -0,2923 | 1,52382E-08 |
| ACE2 | LGALS9 | -0,2923 | 1,52343E-08 |
| ACE2 | LINC02111 | -0,2924 | 1,51391E-08 |
| ACE2 | RP3-510L9.1 | -0,2924 | 1,50128E-08 |
| ACE2 | AC002064.4 | -0,2924 | 1,50488E-08 |
| ACE2 | FAM66E | -0,2924 | 1,51085E-08 |
| ACE2 | RLN1 | -0,2924 | 1,50042E-08 |
| ACE2 | ADRB1 | -0,2924 | 1,5098E-08 |
| ACE2 | ZNF414 | -0,2924 | 1,50338E-08 |
| ACE2 | UCK1 | -0,2925 | 1,49249E-08 |
| ACE2 | RP11-77K12.10 | -0,2925 | 1,49131E-08 |
| ACE2 | ZNF90P2 | -0,2926 | 1,47052E-08 |
| ACE2 | KLHL7-AS1 | -0,2926 | 1,47741E-08 |
| ACE2 | RP11-547C5.1 | -0,2926 | 1,47617E-08 |
| ACE2 | RP11-473C19.1 | -0,2926 | 1,47982E-08 |
| ACE2 | UPK1B | -0,2927 | 1,45517E-08 |
| ACE2 | PELO | -0,2927 | 1,45094E-08 |
| ACE2 | RP11-819M15.2 | -0,2927 | 1,45665E-08 |
| ACE2 | RP11-244E17.1 | -0,2927 | 1,46156E-08 |
| ACE2 | AC103563.7 | -0,2928 | 1,4429E-08 |
| ACE2 | RP11-59O6.3 | -0,2928 | 1,43569E-08 |
| ACE2 | COG8 | -0,2928 | 1,44326E-08 |
| ACE2 | SNX29 | -0,2929 | 1,41536E-08 |
| ACE2 | XPO6 | -0,2929 | 1,42073E-08 |
| ACE2 | RP4-713A8.1 | -0,293 | 1,40377E-08 |
| ACE2 | LIN54 | -0,293 | 1,40694E-08 |
| ACE2 | RP11-388K12.1 | -0,293 | 1,40484E-08 |
| ACE2 | NLRP10 | -0,293 | 1,40286E-08 |
| ACE2 | RP11-25L3.3 | -0,293 | 1,40383E-08 |
| ACE2 | RP11-720L8.1 | -0,2931 | 1,38474E-08 |
| ACE2 | AC090044.2 | -0,2932 | 1,36625E-08 |
| ACE2 | TRIM40 | -0,2932 | 1,366E-08 |
| ACE2 | TNRC18P2 | -0,2932 | 1,36832E-08 |
| ACE2 | RP11-736E3.3 | -0,2932 | 1,37939E-08 |
| ACE2 | REXO4 | -0,2932 | 1,37929E-08 |
| ACE2 | RNA5SP301 | -0,2932 | 1,38065E-08 |
| ACE2 | MS4A1 | -0,2932 | 1,37056E-08 |
| ACE2 | RP11-234B24.4 | -0,2932 | 1,37254E-08 |
| ACE2 | SDF2L1 | -0,2932 | 1,37306E-08 |
| ACE2 | RP5-1125M8.2 | -0,2934 | 1,34734E-08 |
| ACE2 | LRRC2-AS1 | -0,2934 | 1,34291E-08 |
| ACE2 | RP11-334E6.2 | -0,2934 | 1,34642E-08 |
| ACE2 | AC007394.3 | -0,2936 | 1,31155E-08 |
| ACE2 | CSN2 | -0,2936 | 1,30905E-08 |
| ACE2 | AC034228.4 | -0,2936 | 1,31503E-08 |
| ACE2 | KIZ | -0,2936 | 1,31707E-08 |
| ACE2 | AC008073.9 | -0,2937 | 1,29126E-08 |
| ACE2 | RP11-673E1.4 | -0,2937 | 1,30132E-08 |
| ACE2 | ANAPC2 | -0,2937 | 1,29639E-08 |
| ACE2 | PHBP17 | -0,2937 | 1,2896E-08 |
| ACE2 | AC011525.4 | -0,2937 | 1,28681E-08 |
| ACE2 | UBE2D2 | -0,2938 | 1,27533E-08 |
| ACE2 | DPY19L2P1 | -0,2938 | 1,27308E-08 |

| | | | |
|---|---|---|---|
| ACE2 | RP11-881M11.2 | -0,2938 | 1,27178E-08 |
| ACE2 | VPS9D1-AS1 | -0,2938 | 1,27829E-08 |
| ACE2 | LINC02162 | -0,2939 | 1,26379E-08 |
| ACE2 | RPL36AP39 | -0,2939 | 1,25748E-08 |
| ACE2 | BAZ1A | -0,2939 | 1,26958E-08 |
| ACE2 | TSPAN10 | -0,2939 | 1,26044E-08 |
| ACE2 | NPL | -0,294 | 1,24403E-08 |
| ACE2 | RP11-385E5.5 | -0,294 | 1,2445E-08 |
| ACE2 | AC010969.1 | -0,2941 | 1,2332E-08 |
| ACE2 | CHL1-AS2 | -0,2941 | 1,24072E-08 |
| ACE2 | WISP1 | -0,2941 | 1,23916E-08 |
| ACE2 | PGBD4 | -0,2941 | 1,23226E-08 |
| ACE2 | XCR1 | -0,2942 | 1,21261E-08 |
| ACE2 | RP11-118N24.2 | -0,2942 | 1,21985E-08 |
| ACE2 | ADAM3A | -0,2942 | 1,21897E-08 |
| ACE2 | NUDCD3 | -0,2943 | 1,20806E-08 |
| ACE2 | AC012322.1 | -0,2943 | 1,21185E-08 |
| ACE2 | KRT19P3 | -0,2944 | 1,19351E-08 |
| ACE2 | RP11-692C24.1 | -0,2944 | 1,19039E-08 |
| ACE2 | RP11-44I10.3 | -0,2945 | 1,17078E-08 |
| ACE2 | TNK1 | -0,2945 | 1,1723E-08 |
| ACE2 | EYA4 | -0,2946 | 1,16944E-08 |
| ACE2 | DGKA | -0,2946 | 1,16125E-08 |
| ACE2 | FDPSP7 | -0,2947 | 1,1471E-08 |
| ACE2 | RP11-790J24.1 | -0,2947 | 1,14892E-08 |
| ACE2 | AC013271.3 | -0,2948 | 1,1422E-08 |
| ACE2 | GALNT6 | -0,2948 | 1,13107E-08 |
| ACE2 | RP11-496B10.3 | -0,2949 | 1,12771E-08 |
| ACE2 | RP11-661C8.2 | -0,2949 | 1,12512E-08 |
| ACE2 | RP11-72K17.1 | -0,2949 | 1,12685E-08 |
| ACE2 | COL26A1 | -0,2949 | 1,12718E-08 |
| ACE2 | OR9N1P | -0,2949 | 1,12003E-08 |
| ACE2 | RP4-735N21.1 | -0,295 | 1,11208E-08 |
| ACE2 | LINC01019 | -0,295 | 1,11384E-08 |
| ACE2 | C14orf28 | -0,295 | 1,11202E-08 |
| ACE2 | RP11-222N13.1 | -0,2951 | 1,09197E-08 |
| ACE2 | RP11-99C10.1 | -0,2951 | 1,10068E-08 |
| ACE2 | LINC02183 | -0,2951 | 1,09957E-08 |
| ACE2 | MST1L | -0,2952 | 1,0841E-08 |
| ACE2 | KRT18P5 | -0,2952 | 1,08202E-08 |
| ACE2 | TRGC1 | -0,2953 | 1,07124E-08 |
| ACE2 | RP5-945I17.2 | -0,2953 | 1,07231E-08 |
| ACE2 | APOC3 | -0,2953 | 1,07465E-08 |
| ACE2 | RP11-463O9.9 | -0,2953 | 1,0668E-08 |
| ACE2 | AC004004.2 | -0,2953 | 1,07525E-08 |
| ACE2 | TRIML2 | -0,2954 | 1,05106E-08 |
| ACE2 | RP5-1120P11.3 | -0,2954 | 1,05522E-08 |
| ACE2 | PPP1R17 | -0,2954 | 1,05845E-08 |
| ACE2 | TRBV28 | -0,2954 | 1,0545E-08 |
| ACE2 | RNF215 | -0,2954 | 1,05503E-08 |
| ACE2 | CLK3 | -0,2955 | 1,05034E-08 |
| ACE2 | MKL1 | -0,2955 | 1,03987E-08 |
| ACE2 | RP11-480D4.1 | -0,2956 | 1,03377E-08 |
| ACE2 | RP1 | -0,2956 | 1,03334E-08 |

| | | | |
|---|---|---|---|
| ACE2 | P4HA3 | -0,2956 | 1,0272E-08 |
| ACE2 | KIRREL3 | -0,2956 | 1,03153E-08 |
| ACE2 | RP11-814P5.1 | -0,2956 | 1,02757E-08 |
| ACE2 | MYO5B | -0,2956 | 1,03234E-08 |
| ACE2 | RP11-674N23.4 | -0,2957 | 1,01916E-08 |
| ACE2 | RP11-122L4.2 | -0,2958 | 1,01041E-08 |
| ACE2 | RP11-114H24.6 | -0,2958 | 1,00473E-08 |
| ACE2 | AP000146.2 | -0,2958 | 1,00484E-08 |
| ACE2 | KCNAB1-AS2 | -0,2959 | 9,97469E-09 |
| ACE2 | RP11-737F9.2 | -0,2959 | 9,95602E-09 |
| ACE2 | LINC00411 | -0,2959 | 9,93923E-09 |
| ACE2 | CD200 | -0,296 | 9,78312E-09 |
| ACE2 | NAIP | -0,296 | 9,85769E-09 |
| ACE2 | RP13-580B18.1 | -0,296 | 9,86632E-09 |
| ACE2 | CD3G | -0,296 | 9,79239E-09 |
| ACE2 | OPCML-IT2 | -0,296 | 9,80299E-09 |
| ACE2 | PDE4C | -0,296 | 9,85992E-09 |
| ACE2 | ZNF568 | -0,296 | 9,86842E-09 |
| ACE2 | AGBL4 | -0,2961 | 9,67985E-09 |
| ACE2 | RP11-385M4.3 | -0,2961 | 9,70387E-09 |
| ACE2 | AC007179.1 | -0,2961 | 9,6747E-09 |
| ACE2 | GRIFIN | -0,2961 | 9,75917E-09 |
| ACE2 | LINC00959 | -0,2961 | 9,67067E-09 |
| ACE2 | RP11-108P20.2 | -0,2961 | 9,66398E-09 |
| ACE2 | AC008937.2 | -0,2962 | 9,59036E-09 |
| ACE2 | MRPS18CP6 | -0,2962 | 9,63609E-09 |
| ACE2 | RP11-451G4.3 | -0,2963 | 9,49098E-09 |
| ACE2 | RP11-79C6.2 | -0,2963 | 9,43862E-09 |
| ACE2 | GLULP4 | -0,2963 | 9,48834E-09 |
| ACE2 | OR8G2P | -0,2963 | 9,45263E-09 |
| ACE2 | OR2T8 | -0,2964 | 9,42412E-09 |
| ACE2 | FLT1 | -0,2964 | 9,35301E-09 |
| ACE2 | CXADRP2 | -0,2964 | 9,32774E-09 |
| ACE2 | RP11-479A21.1 | -0,2965 | 9,28686E-09 |
| ACE2 | RP11-15I11.3 | -0,2966 | 9,15824E-09 |
| ACE2 | RP11-296L22.8 | -0,2966 | 9,18592E-09 |
| ACE2 | RP11-25H11.1 | -0,2966 | 9,12564E-09 |
| ACE2 | RP5-881L22.6 | -0,2966 | 9,20415E-09 |
| ACE2 | GAPDHP26 | -0,2967 | 9,09724E-09 |
| ACE2 | SLC22A10 | -0,2967 | 9,01114E-09 |
| ACE2 | DDT | -0,2967 | 8,99884E-09 |
| ACE2 | NOTCH2NL | -0,2968 | 8,9391E-09 |
| ACE2 | PPIAL4G | -0,2968 | 8,98696E-09 |
| ACE2 | RP4-609E1.2 | -0,2969 | 8,78811E-09 |
| ACE2 | AC079610.2 | -0,2969 | 8,82435E-09 |
| ACE2 | RP11-20F24.2 | -0,2969 | 8,84057E-09 |
| ACE2 | RP11-326L17.1 | -0,2969 | 8,84698E-09 |
| ACE2 | CBX3P8 | -0,297 | 8,74745E-09 |
| ACE2 | DHRS4 | -0,297 | 8,76497E-09 |
| ACE2 | RP11-94B19.7 | -0,297 | 8,71749E-09 |
| ACE2 | TUBB8P9 | -0,2971 | 8,57954E-09 |
| ACE2 | OR2A2 | -0,2971 | 8,58543E-09 |
| ACE2 | ARID3C | -0,2971 | 8,61756E-09 |
| ACE2 | LDB3 | -0,2971 | 8,58219E-09 |

| | | | |
|---|---|---|---|
| ACE2 | AP000797.4 | -0,2972 | 8,53572E-09 |
| ACE2 | RP11-507B12.2 | -0,2972 | 8,48436E-09 |
| ACE2 | LINC01063 | -0,2973 | 8,43225E-09 |
| ACE2 | VN1R10P | -0,2974 | 8,32245E-09 |
| ACE2 | CHORDC2P | -0,2974 | 8,35863E-09 |
| ACE2 | LINC01494 | -0,2975 | 8,20734E-09 |
| ACE2 | RP11-46H11.3 | -0,2975 | 8,26469E-09 |
| ACE2 | NOL10 | -0,2976 | 8,12444E-09 |
| ACE2 | IGKV1-12 | -0,2976 | 8,09631E-09 |
| ACE2 | LINC02045 | -0,2976 | 8,0745E-09 |
| ACE2 | SEL1L3 | -0,2976 | 8,11663E-09 |
| ACE2 | MTO1 | -0,2976 | 8,10294E-09 |
| ACE2 | COPZ2 | -0,2976 | 8,16194E-09 |
| ACE2 | TSHB | -0,2977 | 8,04208E-09 |
| ACE2 | STON1-GTF2A1L | -0,2977 | 7,97658E-09 |
| ACE2 | RP11-26H16.4 | -0,2978 | 7,9267E-09 |
| ACE2 | AC079305.11 | -0,2978 | 7,90501E-09 |
| ACE2 | HELT | -0,2978 | 7,93408E-09 |
| ACE2 | CACNA1H | -0,2978 | 7,9516E-09 |
| ACE2 | MARVELD3 | -0,2978 | 7,92976E-09 |
| ACE2 | RPL21P122 | -0,2978 | 7,88033E-09 |
| ACE2 | RP11-430C7.5 | -0,2979 | 7,87768E-09 |
| ACE2 | RP11-656A15.1 | -0,2979 | 7,863E-09 |
| ACE2 | LINC01839 | -0,2979 | 7,81289E-09 |
| ACE2 | BTF3P13 | -0,2979 | 7,83258E-09 |
| ACE2 | KB-1980E6.3 | -0,2979 | 7,8351E-09 |
| ACE2 | RP11-555J4.3 | -0,2979 | 7,79889E-09 |
| ACE2 | RP11-353M9.1 | -0,2979 | 7,82289E-09 |
| ACE2 | ZFPL1 | -0,2979 | 7,87071E-09 |
| ACE2 | MTDHP4 | -0,2979 | 7,78597E-09 |
| ACE2 | RP11-15F12.1 | -0,298 | 7,71138E-09 |
| ACE2 | CTC-510F12.4 | -0,2981 | 7,64944E-09 |
| ACE2 | RP5-1003J2.4 | -0,2982 | 7,55868E-09 |
| ACE2 | RASSF8-AS1 | -0,2982 | 7,54919E-09 |
| ACE2 | RP4-684O24.5 | -0,2982 | 7,51034E-09 |
| ACE2 | LINC00498 | -0,2983 | 7,46896E-09 |
| ACE2 | PRDX2P2 | -0,2983 | 7,47789E-09 |
| ACE2 | SLC15A5 | -0,2983 | 7,44774E-09 |
| ACE2 | RP11-167H9.5 | -0,2984 | 7,39406E-09 |
| ACE2 | RP11-89C3.2 | -0,2984 | 7,33519E-09 |
| ACE2 | RP11-569A11.1 | -0,2985 | 7,25158E-09 |
| ACE2 | ZNF876P | -0,2985 | 7,30376E-09 |
| ACE2 | LINC02028 | -0,2986 | 7,20695E-09 |
| ACE2 | RP11-703C10.1 | -0,2986 | 7,17108E-09 |
| ACE2 | RP11-1060J15.7 | -0,2986 | 7,23855E-09 |
| ACE2 | LINC01481 | -0,2986 | 7,15893E-09 |
| ACE2 | RP11-809H16.3 | -0,2986 | 7,19608E-09 |
| ACE2 | RP11-561C5.1 | -0,2987 | 7,06962E-09 |
| ACE2 | RP11-1099M24.8 | -0,2987 | 7,09192E-09 |
| ACE2 | RP11-548K12.9 | -0,2988 | 7,04726E-09 |
| ACE2 | LINC01249 | -0,2989 | 6,92725E-09 |
| ACE2 | DUSP8P5 | -0,2989 | 6,97854E-09 |
| ACE2 | LINC02129 | -0,2989 | 6,97658E-09 |
| ACE2 | XRCC2 | -0,299 | 6,83625E-09 |

| | | | |
|---|---|---|---|
| ACE2 | RP11-1112C15.2 | -0,299 | 6,88237E-09 |
| ACE2 | IGHV3-29 | -0,299 | 6,89468E-09 |
| ACE2 | CYMP-AS1 | -0,2991 | 6,73514E-09 |
| ACE2 | RN7SKP281 | -0,2991 | 6,74221E-09 |
| ACE2 | PCDHA7 | -0,2991 | 6,76902E-09 |
| ACE2 | CTD-2054N24.2 | -0,2991 | 6,79155E-09 |
| ACE2 | RP11-123C5.5 | -0,2991 | 6,75537E-09 |
| ACE2 | CAMP | -0,2992 | 6,70831E-09 |
| ACE2 | LINC00397 | -0,2992 | 6,68952E-09 |
| ACE2 | OR4N3P | -0,2994 | 6,49463E-09 |
| ACE2 | NANOGP1 | -0,2995 | 6,45862E-09 |
| ACE2 | LINC01836 | -0,2995 | 6,41851E-09 |
| ACE2 | RP5-1033E15.3 | -0,2995 | 6,47013E-09 |
| ACE2 | RP11-34P13.14 | -0,2996 | 6,35765E-09 |
| ACE2 | TNFRSF13C | -0,2996 | 6,39022E-09 |
| ACE2 | LINC00662 | -0,2997 | 6,31575E-09 |
| ACE2 | CTD-2320J21.1 | -0,2998 | 6,18932E-09 |
| ACE2 | NANOGP6 | -0,2998 | 6,21463E-09 |
| ACE2 | PTHLH | -0,2998 | 6,25189E-09 |
| ACE2 | RP11-624L4.1 | -0,2998 | 6,24174E-09 |
| ACE2 | ARHGAP45 | -0,2998 | 6,20427E-09 |
| ACE2 | FOXD3 | -0,2999 | 6,14478E-09 |
| ACE2 | HSP90AA4P | -0,2999 | 6,18461E-09 |
| ACE2 | RP11-321A17.3 | -0,2999 | 6,11292E-09 |
| ACE2 | ANKRD30BL | -0,3 | 6,05121E-09 |
| ACE2 | RP11-700E23.1 | -0,3 | 6,04685E-09 |
| ACE2 | RP11-460B17.3 | -0,3 | 6,10847E-09 |
| ACE2 | RP11-106D4.2 | -0,3 | 6,09753E-09 |
| ACE2 | EPHX3 | -0,3 | 6,04082E-09 |
| ACE2 | FCF1P6 | -0,3001 | 5,97388E-09 |
| ACE2 | AC073464.11 | -0,3001 | 5,97784E-09 |
| ACE2 | RPL7AP49 | -0,3001 | 5,98057E-09 |
| ACE2 | NPAP1P4 | -0,3001 | 6,03473E-09 |
| ACE2 | LLNLR-249E10.1 | -0,3001 | 6,00559E-09 |
| ACE2 | RP11-707M1.1 | -0,3002 | 5,92824E-09 |
| ACE2 | UBBP2 | -0,3003 | 5,85422E-09 |
| ACE2 | RP11-157E16.1 | -0,3003 | 5,82237E-09 |
| ACE2 | RP11-738G5.2 | -0,3003 | 5,84861E-09 |
| ACE2 | RP11-333A23.2 | -0,3003 | 5,83384E-09 |
| ACE2 | SP7 | -0,3003 | 5,8509E-09 |
| ACE2 | CRYM | -0,3003 | 5,85843E-09 |
| ACE2 | HAO1 | -0,3003 | 5,87678E-09 |
| ACE2 | WDR20 | -0,3004 | 5,8135E-09 |
| ACE2 | AC139712.2 | -0,3005 | 5,72265E-09 |
| ACE2 | CLGN | -0,3005 | 5,71009E-09 |
| ACE2 | OR6V1 | -0,3005 | 5,71724E-09 |
| ACE2 | RP11-115D19.4 | -0,3006 | 5,63588E-09 |
| ACE2 | AC005740.4 | -0,3006 | 5,63026E-09 |
| ACE2 | AC004920.3 | -0,3006 | 5,67477E-09 |
| ACE2 | CEACAM20 | -0,3006 | 5,62084E-09 |
| ACE2 | NRXN1 | -0,3007 | 5,60083E-09 |
| ACE2 | CTB-113I20.1 | -0,3007 | 5,54505E-09 |
| ACE2 | LEF1 | -0,3008 | 5,49274E-09 |
| ACE2 | RP11-63E5.6 | -0,3008 | 5,52848E-09 |

| | | | |
|---|---|---|---|
| ACE2 | ZNF101P1 | -0,3008 | 5,53553E-09 |
| ACE2 | RP11-12J10.4 | -0,3008 | 5,50444E-09 |
| ACE2 | EAF2 | -0,3009 | 5,45184E-09 |
| ACE2 | RP11-212P7.2 | -0,3009 | 5,43873E-09 |
| ACE2 | ATP5A1P2 | -0,301 | 5,40206E-09 |
| ACE2 | CTAGE6 | -0,301 | 5,40865E-09 |
| ACE2 | RP11-23E19.1 | -0,301 | 5,39134E-09 |
| ACE2 | RP11-462L8.1 | -0,3011 | 5,28956E-09 |
| ACE2 | RP11-233G1.4 | -0,3011 | 5,29163E-09 |
| ACE2 | ARHGAP27 | -0,3011 | 5,3136E-09 |
| ACE2 | UQCRC2P1 | -0,3012 | 5,2565E-09 |
| ACE2 | RP11-393N21.1 | -0,3014 | 5,10983E-09 |
| ACE2 | HMGB3P16 | -0,3014 | 5,13803E-09 |
| ACE2 | LINC01928 | -0,3014 | 5,14594E-09 |
| ACE2 | LINC01618 | -0,3016 | 4,99231E-09 |
| ACE2 | DDX3P3 | -0,3016 | 5,02315E-09 |
| ACE2 | RPS27P16 | -0,3016 | 5,00056E-09 |
| ACE2 | RP11-215D10.1 | -0,3016 | 5,01946E-09 |
| ACE2 | AC011406.2 | -0,3017 | 4,94124E-09 |
| ACE2 | NOL7 | -0,3017 | 4,95009E-09 |
| ACE2 | FAM74A6 | -0,3017 | 4,95896E-09 |
| ACE2 | NADSYN1 | -0,3017 | 4,91014E-09 |
| ACE2 | LINC01925 | -0,3017 | 4,92994E-09 |
| ACE2 | RP4-657M3.2 | -0,3018 | 4,87936E-09 |
| ACE2 | TTC39B | -0,3018 | 4,85724E-09 |
| ACE2 | RP1-92C8.3 | -0,3019 | 4,8426E-09 |
| ACE2 | CTD-2235C13.2 | -0,302 | 4,74055E-09 |
| ACE2 | PLEKHF2 | -0,302 | 4,77947E-09 |
| ACE2 | KLRC2 | -0,302 | 4,77832E-09 |
| ACE2 | COL21A1 | -0,3021 | 4,71343E-09 |
| ACE2 | ZNF44 | -0,3021 | 4,72805E-09 |
| ACE2 | CSMD3 | -0,3022 | 4,61974E-09 |
| ACE2 | RASL10A | -0,3022 | 4,66839E-09 |
| ACE2 | CDH26 | -0,3023 | 4,61564E-09 |
| ACE2 | GRAP | -0,3024 | 4,5094E-09 |
| ACE2 | CTC-296K1.3 | -0,3024 | 4,54568E-09 |
| ACE2 | LRRC2 | -0,3025 | 4,451E-09 |
| ACE2 | RP11-497D6.5 | -0,3025 | 4,49158E-09 |
| ACE2 | CTC-457E21.9 | -0,3025 | 4,47287E-09 |
| ACE2 | OR2T32P | -0,3026 | 4,40572E-09 |
| ACE2 | LINC00862 | -0,3027 | 4,37724E-09 |
| ACE2 | SPAG1 | -0,3027 | 4,35317E-09 |
| ACE2 | OR7E104P | -0,3027 | 4,38694E-09 |
| ACE2 | GOSR2 | -0,3027 | 4,39001E-09 |
| ACE2 | TUBB8P12 | -0,3027 | 4,34562E-09 |
| ACE2 | CHADL | -0,3027 | 4,35467E-09 |
| ACE2 | HS1BP3 | -0,3028 | 4,33541E-09 |
| ACE2 | ALG3 | -0,3029 | 4,25912E-09 |
| ACE2 | RP11-125O18.1 | -0,3029 | 4,27976E-09 |
| ACE2 | AC105342.1 | -0,303 | 4,22439E-09 |
| ACE2 | LINC02017 | -0,303 | 4,21538E-09 |
| ACE2 | RANBP17 | -0,303 | 4,20119E-09 |
| ACE2 | RP11-350E12.4 | -0,303 | 4,20605E-09 |
| ACE2 | SETP16 | -0,303 | 4,21697E-09 |

| | | | |
|---|---|---|---|
| ACE2 | RP11-160C18.4 | -0,303 | 4,20032E-09 |
| ACE2 | AC016735.3 | -0,3031 | 4,13753E-09 |
| ACE2 | RP11-529P9.1 | -0,3031 | 4,17317E-09 |
| ACE2 | ANKRD34B | -0,3031 | 4,17448E-09 |
| ACE2 | CNOT10 | -0,3032 | 4,12356E-09 |
| ACE2 | FLJ31104 | -0,3032 | 4,10697E-09 |
| ACE2 | UQCRHP1 | -0,3032 | 4,12889E-09 |
| ACE2 | PEX11A | -0,3033 | 4,05879E-09 |
| ACE2 | NBPF6 | -0,3034 | 4,01442E-09 |
| ACE2 | C1orf162 | -0,3034 | 3,98991E-09 |
| ACE2 | RP11-42L13.3 | -0,3034 | 3,98842E-09 |
| ACE2 | SPDEF | -0,3034 | 3,98515E-09 |
| ACE2 | RP11-473J6.1 | -0,3034 | 3,98557E-09 |
| ACE2 | RP11-187C18.4 | -0,3034 | 4,01458E-09 |
| ACE2 | RP11-362A1.1 | -0,3034 | 4,00291E-09 |
| ACE2 | LINC01801 | -0,3034 | 4,0175E-09 |
| ACE2 | CTD-3096P4.1 | -0,3035 | 3,96813E-09 |
| ACE2 | OR4N4 | -0,3035 | 3,97029E-09 |
| ACE2 | C2orf54 | -0,3036 | 3,91076E-09 |
| ACE2 | CTD-2534J5.1 | -0,3037 | 3,85535E-09 |
| ACE2 | RP11-62L10.1 | -0,3038 | 3,80439E-09 |
| ACE2 | RP11-11H9.2 | -0,3038 | 3,83324E-09 |
| ACE2 | MESP2 | -0,3038 | 3,81654E-09 |
| ACE2 | RP11-552F3.4 | -0,3038 | 3,81446E-09 |
| ACE2 | RPL12P37 | -0,3038 | 3,81476E-09 |
| ACE2 | AC005387.3 | -0,3038 | 3,81989E-09 |
| ACE2 | ENPP7P2 | -0,3039 | 3,75857E-09 |
| ACE2 | RP11-702F3.3 | -0,3039 | 3,75645E-09 |
| ACE2 | AC108868.6 | -0,304 | 3,71529E-09 |
| ACE2 | ZNF137P | -0,304 | 3,72452E-09 |
| ACE2 | RP11-1319K7.1 | -0,3041 | 3,66661E-09 |
| ACE2 | OPTN | -0,3041 | 3,65592E-09 |
| ACE2 | MAPT | -0,3041 | 3,66518E-09 |
| ACE2 | RP11-543H23.2 | -0,3041 | 3,68624E-09 |
| ACE2 | PLEKHG2 | -0,3041 | 3,66149E-09 |
| ACE2 | AC024592.9 | -0,3042 | 3,64511E-09 |
| ACE2 | XCL1 | -0,3043 | 3,58763E-09 |
| ACE2 | RP11-248G5.9 | -0,3043 | 3,57345E-09 |
| ACE2 | AC090505.5 | -0,3044 | 3,53005E-09 |
| ACE2 | DKFZp779M0652 | -0,3044 | 3,52322E-09 |
| ACE2 | L3MBTL2 | -0,3044 | 3,52673E-09 |
| ACE2 | RP11-574M7.2 | -0,3045 | 3,48716E-09 |
| ACE2 | RP11-185H22.1 | -0,3045 | 3,52047E-09 |
| ACE2 | RP11-455F5.6 | -0,3045 | 3,51263E-09 |
| ACE2 | METRN | -0,3046 | 3,47559E-09 |
| ACE2 | AC010740.1 | -0,3047 | 3,41755E-09 |
| ACE2 | TRIM41 | -0,3047 | 3,40853E-09 |
| ACE2 | SPDYE2 | -0,3048 | 3,37944E-09 |
| ACE2 | AC009487.6 | -0,3049 | 3,34438E-09 |
| ACE2 | CTC-548H10.2 | -0,3049 | 3,33504E-09 |
| ACE2 | GS1-124K5.12 | -0,3049 | 3,31465E-09 |
| ACE2 | CTD-2014E2.7 | -0,3049 | 3,33806E-09 |
| ACE2 | AC092162.1 | -0,305 | 3,27988E-09 |
| ACE2 | BNIP3P33 | -0,305 | 3,30896E-09 |

| | | | |
|---|---|---|---|
| ACE2 | ZNF578 | -0,3051 | 3,249E-09 |
| ACE2 | RP11-379F4.8 | -0,3052 | 3,19691E-09 |
| ACE2 | RP11-122M14.3 | -0,3053 | 3,15114E-09 |
| ACE2 | R3HCC1L | -0,3053 | 3,18014E-09 |
| ACE2 | TPT1P12 | -0,3053 | 3,16746E-09 |
| ACE2 | MYRFL | -0,3053 | 3,17837E-09 |
| ACE2 | CTD-2228K2.7 | -0,3054 | 3,1495E-09 |
| ACE2 | ASS1P10 | -0,3054 | 3,11517E-09 |
| ACE2 | CTB-51J22.1 | -0,3054 | 3,12527E-09 |
| ACE2 | OR11H6 | -0,3054 | 3,12712E-09 |
| ACE2 | TRIM39 | -0,3055 | 3,08835E-09 |
| ACE2 | GJD2 | -0,3055 | 3,08936E-09 |
| ACE2 | LA16c-358B7.4 | -0,3056 | 3,07032E-09 |
| ACE2 | IL17RD | -0,3057 | 2,99958E-09 |
| ACE2 | FCRL2 | -0,3058 | 2,99267E-09 |
| ACE2 | PCDHB18P | -0,3058 | 2,99665E-09 |
| ACE2 | SOGA3 | -0,3058 | 2,98534E-09 |
| ACE2 | CNR2 | -0,3059 | 2,94364E-09 |
| ACE2 | RP4-533D7.5 | -0,3059 | 2,96012E-09 |
| ACE2 | LINC01221 | -0,3059 | 2,94922E-09 |
| ACE2 | LINC01814 | -0,3059 | 2,95425E-09 |
| ACE2 | AC010149.4 | -0,3059 | 2,93994E-09 |
| ACE2 | RP5-958B11.2 | -0,3059 | 2,9383E-09 |
| ACE2 | HAR1A | -0,3059 | 2,94637E-09 |
| ACE2 | LINC00837 | -0,306 | 2,89292E-09 |
| ACE2 | RP11-867G2.4 | -0,306 | 2,90027E-09 |
| ACE2 | CTC-428H11.2 | -0,3061 | 2,85209E-09 |
| ACE2 | RAET1E-AS1 | -0,3061 | 2,87661E-09 |
| ACE2 | LPGAT1 | -0,3063 | 2,81134E-09 |
| ACE2 | TRBV21OR9-2 | -0,3063 | 2,80618E-09 |
| ACE2 | RP5-1185I7.1 | -0,3064 | 2,76836E-09 |
| ACE2 | FTH1P11 | -0,3064 | 2,78133E-09 |
| ACE2 | STK39 | -0,3065 | 2,72704E-09 |
| ACE2 | TRBV20OR9-2 | -0,3065 | 2,73894E-09 |
| ACE2 | RP11-116D17.2 | -0,3065 | 2,72643E-09 |
| ACE2 | RP11-326L9.1 | -0,3065 | 2,74352E-09 |
| ACE2 | CTD-2651B20.2 | -0,3065 | 2,74574E-09 |
| ACE2 | ACSM1 | -0,3065 | 2,74412E-09 |
| ACE2 | CHCHD3P3 | -0,3066 | 2,69716E-09 |
| ACE2 | KRT79 | -0,3066 | 2,70254E-09 |
| ACE2 | EVPLL | -0,3066 | 2,69135E-09 |
| ACE2 | RP5-875O13.6 | -0,3067 | 2,66014E-09 |
| ACE2 | INSL5 | -0,3067 | 2,65595E-09 |
| ACE2 | RP11-113D6.9 | -0,3067 | 2,65907E-09 |
| ACE2 | BCRP9 | -0,3067 | 2,67412E-09 |
| ACE2 | RP11-358N2.2 | -0,3067 | 2,65597E-09 |
| ACE2 | CLUAP1 | -0,3068 | 2,63232E-09 |
| ACE2 | HOXA13 | -0,3069 | 2,58652E-09 |
| ACE2 | CCDC90B | -0,3069 | 2,60664E-09 |
| ACE2 | CBY1 | -0,3069 | 2,5975E-09 |
| ACE2 | RP11-297K8.2 | -0,307 | 2,56283E-09 |
| ACE2 | RN7SL776P | -0,307 | 2,5643E-09 |
| ACE2 | RP11-84C10.3 | -0,307 | 2,5725E-09 |
| ACE2 | LINC01666 | -0,307 | 2,56513E-09 |

| | | | |
|---|---|---|---|
| ACE2 | GK | -0,307 | 2,56904E-09 |
| ACE2 | AC097662.2 | -0,3072 | 2,50529E-09 |
| ACE2 | LIMCH1 | -0,3072 | 2,51503E-09 |
| ACE2 | GRIN3A | -0,3072 | 2,50614E-09 |
| ACE2 | RP11-1157N2__B.2 | -0,3072 | 2,50256E-09 |
| ACE2 | RP11-190J23.1 | -0,3073 | 2,4659E-09 |
| ACE2 | CTB-35F21.3 | -0,3073 | 2,48476E-09 |
| ACE2 | RN7SL68P | -0,3073 | 2,46184E-09 |
| ACE2 | SPTBN5 | -0,3073 | 2,4802E-09 |
| ACE2 | RP4-714D9.4 | -0,3074 | 2,44388E-09 |
| ACE2 | SOCS5P3 | -0,3074 | 2,45122E-09 |
| ACE2 | AC139887.4 | -0,3074 | 2,4479E-09 |
| ACE2 | RP11-526F3.1 | -0,3074 | 2,43037E-09 |
| ACE2 | TP53TG1 | -0,3074 | 2,43166E-09 |
| ACE2 | KLRC3 | -0,3074 | 2,44407E-09 |
| ACE2 | LINC01053 | -0,3074 | 2,44799E-09 |
| ACE2 | C2orf69P2 | -0,3074 | 2,43076E-09 |
| ACE2 | RP1-127L4.7 | -0,3074 | 2,45022E-09 |
| ACE2 | KCNMB3P1 | -0,3075 | 2,40506E-09 |
| ACE2 | ATP13A5-AS1 | -0,3076 | 2,3916E-09 |
| ACE2 | TMEM151A | -0,3076 | 2,36711E-09 |
| ACE2 | TMBIM4 | -0,3076 | 2,36983E-09 |
| ACE2 | IL22 | -0,3076 | 2,39362E-09 |
| ACE2 | RABL2B | -0,3076 | 2,38697E-09 |
| ACE2 | RP11-145G20.1 | -0,3077 | 2,36157E-09 |
| ACE2 | IFT74 | -0,3077 | 2,3518E-09 |
| ACE2 | AKR1C4 | -0,3077 | 2,34773E-09 |
| ACE2 | TPTE2P2 | -0,3077 | 2,35549E-09 |
| ACE2 | SULT1A1 | -0,3077 | 2,33627E-09 |
| ACE2 | RP11-388K12.2 | -0,3078 | 2,31793E-09 |
| ACE2 | MOK | -0,3078 | 2,31762E-09 |
| ACE2 | LINC01805 | -0,3079 | 2,28117E-09 |
| ACE2 | AC106874.1 | -0,3079 | 2,28765E-09 |
| ACE2 | RP11-83C7.1 | -0,3079 | 2,30195E-09 |
| ACE2 | RP1-168P16.1 | -0,3079 | 2,29683E-09 |
| ACE2 | FBXO7 | -0,3079 | 2,3053E-09 |
| ACE2 | RSPO1 | -0,308 | 2,26211E-09 |
| ACE2 | BCL2L15 | -0,308 | 2,27264E-09 |
| ACE2 | POLR2E | -0,308 | 2,25387E-09 |
| ACE2 | RP11-338C15.3 | -0,3081 | 2,22394E-09 |
| ACE2 | AC097499.2 | -0,3081 | 2,24864E-09 |
| ACE2 | NUP98 | -0,3081 | 2,2382E-09 |
| ACE2 | ACER1 | -0,3081 | 2,23229E-09 |
| ACE2 | RP11-114H24.4 | -0,3082 | 2,19763E-09 |
| ACE2 | RP3-514A23.2 | -0,3083 | 2,17379E-09 |
| ACE2 | RP11-294C11.11 | -0,3083 | 2,17942E-09 |
| ACE2 | RP1-125I3.4 | -0,3084 | 2,15398E-09 |
| ACE2 | HHATL-AS1 | -0,3084 | 2,16585E-09 |
| ACE2 | RP13-516M14.2 | -0,3084 | 2,15444E-09 |
| ACE2 | CTD-2613O8.1 | -0,3085 | 2,13142E-09 |
| ACE2 | RP11-379I19.2 | -0,3085 | 2,11627E-09 |
| ACE2 | VSTM1 | -0,3085 | 2,1393E-09 |
| ACE2 | CCR9 | -0,3086 | 2,08862E-09 |
| ACE2 | HMGN2P27 | -0,3086 | 2,10211E-09 |

| | | | |
|---|---|---|---|
| ACE2 | DAPK2 | -0,3086 | 2,10909E-09 |
| ACE2 | RP11-352B15.2 | -0,3086 | 2,09202E-09 |
| ACE2 | POM121L8P | -0,3086 | 2,09513E-09 |
| ACE2 | RP11-213G6.2 | -0,3087 | 2,06694E-09 |
| ACE2 | PLAU | -0,3087 | 2,08566E-09 |
| ACE2 | RP5-1092A3.5 | -0,3088 | 2,05244E-09 |
| ACE2 | AC092431.3 | -0,3088 | 2,04759E-09 |
| ACE2 | RP11-367E12.4 | -0,3088 | 2,04115E-09 |
| ACE2 | RP11-40F6.1 | -0,3088 | 2,03815E-09 |
| ACE2 | KLF17P1 | -0,3088 | 2,04228E-09 |
| ACE2 | LINC01269 | -0,3088 | 2,03794E-09 |
| ACE2 | CIAPIN1P | -0,3088 | 2,04932E-09 |
| ACE2 | CRYZL1 | -0,3088 | 2,05211E-09 |
| ACE2 | BCRP7 | -0,3088 | 2,03618E-09 |
| ACE2 | ZIK1P1 | -0,3088 | 2,0394E-09 |
| ACE2 | CCR3 | -0,3089 | 2,01873E-09 |
| ACE2 | PCDHA1 | -0,3089 | 2,02549E-09 |
| ACE2 | TMEM198B | -0,3089 | 2,02839E-09 |
| ACE2 | WIZ | -0,3089 | 2,01934E-09 |
| ACE2 | RP11-227F19.1 | -0,309 | 1,98719E-09 |
| ACE2 | SLC10A4 | -0,309 | 1,9966E-09 |
| ACE2 | RP11-403A21.3 | -0,309 | 2,00208E-09 |
| ACE2 | TFAP2E | -0,3091 | 1,9761E-09 |
| ACE2 | ZRANB2-AS2 | -0,3091 | 1,9655E-09 |
| ACE2 | LGALS8-AS1 | -0,3091 | 1,97736E-09 |
| ACE2 | TMEM86A | -0,3091 | 1,98088E-09 |
| ACE2 | RP11-505K9.1 | -0,3091 | 1,96779E-09 |
| ACE2 | CYP4Z1 | -0,3093 | 1,91266E-09 |
| ACE2 | EEF1A1P37 | -0,3093 | 1,92091E-09 |
| ACE2 | ADAMTSL1 | -0,3093 | 1,917E-09 |
| ACE2 | CTD-2102P23.1 | -0,3093 | 1,92794E-09 |
| ACE2 | IL20 | -0,3094 | 1,90014E-09 |
| ACE2 | IGKV2-4 | -0,3094 | 1,89493E-09 |
| ACE2 | RP11-680B3.2 | -0,3094 | 1,88931E-09 |
| ACE2 | KCNQ1-AS1 | -0,3095 | 1,88379E-09 |
| ACE2 | RP1-205F14P.1 | -0,3095 | 1,87791E-09 |
| ACE2 | OTOL1 | -0,3097 | 1,81878E-09 |
| ACE2 | RWDD2A | -0,3097 | 1,82322E-09 |
| ACE2 | SEM1 | -0,3097 | 1,81916E-09 |
| ACE2 | GGTLC4P | -0,3097 | 1,83652E-09 |
| ACE2 | RP11-314C16.1 | -0,3098 | 1,79728E-09 |
| ACE2 | RP11-510I5.4 | -0,3098 | 1,80533E-09 |
| ACE2 | MTCYBP21 | -0,3098 | 1,80936E-09 |
| ACE2 | RP11-86H7.6 | -0,3099 | 1,78957E-09 |
| ACE2 | RP11-268P4.2 | -0,3099 | 1,78491E-09 |
| ACE2 | RP11-1299A16.1 | -0,31 | 1,76721E-09 |
| ACE2 | SMURF2P1 | -0,3101 | 1,74538E-09 |
| ACE2 | RNU4-74P | -0,3102 | 1,71105E-09 |
| ACE2 | RN7SKP264 | -0,3102 | 1,7111E-09 |
| ACE2 | RP11-164N3.2 | -0,3102 | 1,71773E-09 |
| ACE2 | UBL7-AS1 | -0,3102 | 1,71732E-09 |
| ACE2 | EMX1 | -0,3103 | 1,68968E-09 |
| ACE2 | ADAMTS6 | -0,3103 | 1,70047E-09 |
| ACE2 | PCDHGC5 | -0,3103 | 1,68794E-09 |

| | | | |
|---|---|---|---|
| ACE2 | CKLF | -0,3103 | 1,70389E-09 |
| ACE2 | RPS23P7 | -0,3103 | 1,69852E-09 |
| ACE2 | AC012363.13 | -0,3104 | 1,66892E-09 |
| ACE2 | CTD-2227I18.1 | -0,3104 | 1,66711E-09 |
| ACE2 | RP5-991C6.2 | -0,3104 | 1,68175E-09 |
| ACE2 | RPL7P29 | -0,3104 | 1,66438E-09 |
| ACE2 | RP11-57A19.7 | -0,3104 | 1,68114E-09 |
| ACE2 | RP11-290H9.5 | -0,3105 | 1,6448E-09 |
| ACE2 | SPCS1 | -0,3106 | 1,63637E-09 |
| ACE2 | RP11-366M4.13 | -0,3106 | 1,62959E-09 |
| ACE2 | LCTL | -0,3106 | 1,63419E-09 |
| ACE2 | PCAT29 | -0,3106 | 1,63069E-09 |
| ACE2 | RHCG | -0,3106 | 1,62296E-09 |
| ACE2 | PYCR1 | -0,3106 | 1,62771E-09 |
| ACE2 | RP11-1376P16.1 | -0,3106 | 1,64108E-09 |
| ACE2 | FTH1P22 | -0,3107 | 1,62155E-09 |
| ACE2 | RP11-618M23.3 | -0,3107 | 1,62132E-09 |
| ACE2 | RP11-165I9.9 | -0,3107 | 1,60466E-09 |
| ACE2 | RP11-61A14.4 | -0,3108 | 1,58637E-09 |
| ACE2 | LLNLR-276E7.1 | -0,3108 | 1,59974E-09 |
| ACE2 | CTD-2195M15.3 | -0,3109 | 1,56643E-09 |
| ACE2 | RP11-641A6.3 | -0,311 | 1,55077E-09 |
| ACE2 | EFCAB13 | -0,311 | 1,56063E-09 |
| ACE2 | OSBPL9P5 | -0,3111 | 1,52776E-09 |
| ACE2 | CTD-2033D15.3 | -0,3111 | 1,53279E-09 |
| ACE2 | RPL35AP3 | -0,3112 | 1,50476E-09 |
| ACE2 | RP11-822E23.6 | -0,3112 | 1,51524E-09 |
| ACE2 | AC092168.4 | -0,3113 | 1,49994E-09 |
| ACE2 | DMP1 | -0,3113 | 1,49774E-09 |
| ACE2 | RP11-1360M22.10 | -0,3113 | 1,50062E-09 |
| ACE2 | RP11-588H7.1 | -0,3114 | 1,46705E-09 |
| ACE2 | RP11-374M1.3 | -0,3114 | 1,47112E-09 |
| ACE2 | FAM216A | -0,3114 | 1,48E-09 |
| ACE2 | RPS6KL1 | -0,3114 | 1,46581E-09 |
| ACE2 | PIK3CDP1 | -0,3117 | 1,42822E-09 |
| ACE2 | GGT8P | -0,3117 | 1,41562E-09 |
| ACE2 | AC079305.5 | -0,3117 | 1,42611E-09 |
| ACE2 | ARHGEF38 | -0,3117 | 1,428E-09 |
| ACE2 | MAP1LC3BP1 | -0,3117 | 1,42751E-09 |
| ACE2 | PHPT1 | -0,3117 | 1,42486E-09 |
| ACE2 | SLC2A14 | -0,3117 | 1,42262E-09 |
| ACE2 | RP11-265N7.1 | -0,3117 | 1,41262E-09 |
| ACE2 | AC010967.3 | -0,3118 | 1,40044E-09 |
| ACE2 | CTAGE7P | -0,3118 | 1,40624E-09 |
| ACE2 | USP12-AS1 | -0,3118 | 1,39885E-09 |
| ACE2 | ISCA1P4 | -0,3118 | 1,40267E-09 |
| ACE2 | RP1-102D24.5 | -0,3118 | 1,40114E-09 |
| ACE2 | LINC01151 | -0,3119 | 1,38303E-09 |
| ACE2 | RPS26P21 | -0,312 | 1,36942E-09 |
| ACE2 | OFCC1 | -0,312 | 1,3733E-09 |
| ACE2 | TOB2P1 | -0,312 | 1,36149E-09 |
| ACE2 | HTR1B | -0,312 | 1,3718E-09 |
| ACE2 | ESPL1 | -0,312 | 1,36424E-09 |
| ACE2 | RP11-368J21.3 | -0,312 | 1,35869E-09 |

| | | | |
|---|---|---|---|
| ACE2 | DNAJC15 | -0,3121 | 1,34293E-09 |
| ACE2 | UBALD2 | -0,3121 | 1,34394E-09 |
| ACE2 | RP1-265C24.8 | -0,3122 | 1,33424E-09 |
| ACE2 | MGC27382 | -0,3123 | 1,30728E-09 |
| ACE2 | RP11-745L13.2 | -0,3123 | 1,32083E-09 |
| ACE2 | RP1-28O17.1 | -0,3123 | 1,31778E-09 |
| ACE2 | FRG2HP | -0,3123 | 1,3092E-09 |
| ACE2 | DUSP10 | -0,3124 | 1,30588E-09 |
| ACE2 | F12 | -0,3125 | 1,27842E-09 |
| ACE2 | MRPL42 | -0,3125 | 1,27597E-09 |
| ACE2 | RP11-296O14.3 | -0,3126 | 1,25977E-09 |
| ACE2 | RP11-644F5.16 | -0,3126 | 1,26641E-09 |
| ACE2 | HCFC2 | -0,3126 | 1,26317E-09 |
| ACE2 | RP11-989E6.8 | -0,3126 | 1,26301E-09 |
| ACE2 | LYRM4 | -0,3127 | 1,25155E-09 |
| ACE2 | AC005488.11 | -0,3127 | 1,24337E-09 |
| ACE2 | RP4-568B10.1 | -0,3127 | 1,25522E-09 |
| ACE2 | AC123023.1 | -0,3128 | 1,22903E-09 |
| ACE2 | IGLC7 | -0,3128 | 1,23479E-09 |
| ACE2 | TBC1D1 | -0,3129 | 1,21202E-09 |
| ACE2 | RP11-566K11.4 | -0,3129 | 1,22275E-09 |
| ACE2 | CTB-78F1.1 | -0,313 | 1,19937E-09 |
| ACE2 | FAM177B | -0,3131 | 1,18506E-09 |
| ACE2 | YWHAEP1 | -0,3131 | 1,18418E-09 |
| ACE2 | SLC26A3 | -0,3131 | 1,18425E-09 |
| ACE2 | CSNK1A1L | -0,3131 | 1,18427E-09 |
| ACE2 | RP11-297C4.2 | -0,3131 | 1,18171E-09 |
| ACE2 | ARMC6 | -0,3132 | 1,1739E-09 |
| ACE2 | MOGS | -0,3133 | 1,1549E-09 |
| ACE2 | RPS27AP10 | -0,3133 | 1,15957E-09 |
| ACE2 | CLIP1 | -0,3133 | 1,15132E-09 |
| ACE2 | RP1-163M9.7 | -0,3134 | 1,13762E-09 |
| ACE2 | RP11-955H22.4 | -0,3134 | 1,13893E-09 |
| ACE2 | CXADRP3 | -0,3134 | 1,13868E-09 |
| ACE2 | RP11-451H23.2 | -0,3135 | 1,12284E-09 |
| ACE2 | LINC00350 | -0,3135 | 1,1304E-09 |
| ACE2 | ZNF525 | -0,3135 | 1,13465E-09 |
| ACE2 | RP11-809N15.3 | -0,3136 | 1,10826E-09 |
| ACE2 | RP11-108A14.1 | -0,3136 | 1,10893E-09 |
| ACE2 | ZWINT | -0,3136 | 1,11052E-09 |
| ACE2 | RP11-488C13.4 | -0,3136 | 1,117E-09 |
| ACE2 | C1orf50 | -0,3137 | 1,10162E-09 |
| ACE2 | RP11-612J15.1 | -0,3137 | 1,1051E-09 |
| ACE2 | DOHH | -0,3137 | 1,09446E-09 |
| ACE2 | ECEL1P1 | -0,3138 | 1,09028E-09 |
| ACE2 | RP11-167N5.5 | -0,3138 | 1,08941E-09 |
| ACE2 | TMEM231 | -0,3138 | 1,09112E-09 |
| ACE2 | RN7SL364P | -0,3138 | 1,0926E-09 |
| ACE2 | RP11-561P12.5 | -0,3139 | 1,0788E-09 |
| ACE2 | NKD1 | -0,3139 | 1,07549E-09 |
| ACE2 | RP11-153F1.1 | -0,314 | 1,0583E-09 |
| ACE2 | CEP128 | -0,314 | 1,0531E-09 |
| ACE2 | RP11-1029J19.4 | -0,3141 | 1,04363E-09 |
| ACE2 | USP32 | -0,3141 | 1,04839E-09 |

| | | | |
|---|---|---|---|
| ACE2 | WDR7 | -0,3141 | 1,04652E-09 |
| ACE2 | AC010504.2 | -0,3141 | 1,04101E-09 |
| ACE2 | RP3-492P14.2 | -0,3142 | 1,03659E-09 |
| ACE2 | RP5-905N1.2 | -0,3143 | 1,01386E-09 |
| ACE2 | DNAJB2 | -0,3144 | 1,01205E-09 |
| ACE2 | ELOC | -0,3144 | 1,0107E-09 |
| ACE2 | AQP11 | -0,3144 | 1,00553E-09 |
| ACE2 | MRPL12 | -0,3144 | 1,00159E-09 |
| ACE2 | LDHAL6A | -0,3145 | 9,88108E-10 |
| ACE2 | C4orf26 | -0,3146 | 9,80993E-10 |
| ACE2 | RP11-409I10.2 | -0,3146 | 9,81623E-10 |
| ACE2 | RP11-182J1.14 | -0,3147 | 9,64929E-10 |
| ACE2 | AC019178.3 | -0,3148 | 9,52284E-10 |
| ACE2 | RP11-770E5.1 | -0,3148 | 9,5844E-10 |
| ACE2 | RP11-232A1.2 | -0,3149 | 9,39447E-10 |
| ACE2 | RP11-867G2.8 | -0,3149 | 9,47154E-10 |
| ACE2 | AC018641.7 | -0,3151 | 9,19466E-10 |
| ACE2 | AF186192.1 | -0,3151 | 9,16535E-10 |
| ACE2 | HMGB3 | -0,3151 | 9,24514E-10 |
| ACE2 | AC009518.3 | -0,3152 | 9,07112E-10 |
| ACE2 | AOC1 | -0,3152 | 9,0323E-10 |
| ACE2 | RP11-298E9.6 | -0,3152 | 9,08266E-10 |
| ACE2 | LINC01734 | -0,3152 | 9,05928E-10 |
| ACE2 | SAA3P | -0,3153 | 8,98092E-10 |
| ACE2 | MTA1 | -0,3153 | 8,99063E-10 |
| ACE2 | GOLGA8EP | -0,3153 | 8,94792E-10 |
| ACE2 | THAP5 | -0,3154 | 8,84141E-10 |
| ACE2 | RP11-420N3.2 | -0,3154 | 8,86836E-10 |
| ACE2 | SPATA5 | -0,3155 | 8,74798E-10 |
| ACE2 | MCF2L-AS1 | -0,3155 | 8,68651E-10 |
| ACE2 | C7orf43 | -0,3157 | 8,53495E-10 |
| ACE2 | LINC01507 | -0,3157 | 8,47449E-10 |
| ACE2 | CIZ1 | -0,3157 | 8,54447E-10 |
| ACE2 | GOLGA8CP | -0,3159 | 8,32181E-10 |
| ACE2 | BSN | -0,316 | 8,23642E-10 |
| ACE2 | RP11-10N16.3 | -0,3161 | 8,13248E-10 |
| ACE2 | TBC1D19 | -0,3161 | 8,07466E-10 |
| ACE2 | RP1-28C20.1 | -0,3161 | 8,10527E-10 |
| ACE2 | ZNF679 | -0,3162 | 8,02405E-10 |
| ACE2 | PCDH20 | -0,3162 | 7,986E-10 |
| ACE2 | SLX1A | -0,3162 | 8,01514E-10 |
| ACE2 | RP5-1028K7.2 | -0,3162 | 8,02675E-10 |
| ACE2 | SNX24 | -0,3163 | 7,91697E-10 |
| ACE2 | PCOLCE-AS1 | -0,3163 | 7,8474E-10 |
| ACE2 | TRIM28 | -0,3163 | 7,87422E-10 |
| ACE2 | RASA4CP | -0,3164 | 7,77974E-10 |
| ACE2 | PRSS3P4 | -0,3164 | 7,77353E-10 |
| ACE2 | RP11-946P6.6 | -0,3165 | 7,70169E-10 |
| ACE2 | AC009134.1 | -0,3165 | 7,66173E-10 |
| ACE2 | EIF4G3 | -0,3167 | 7,4994E-10 |
| ACE2 | RAB4A | -0,3167 | 7,50644E-10 |
| ACE2 | AC078883.3 | -0,3167 | 7,43893E-10 |
| ACE2 | OR7E29P | -0,3167 | 7,49468E-10 |
| ACE2 | RP13-635I23.3 | -0,3167 | 7,48057E-10 |

| | | | |
|---|---|---|---|
| ACE2 | SLC25A1P4 | -0,3167 | 7,48353E-10 |
| ACE2 | LRRC37A2 | -0,3167 | 7,45241E-10 |
| ACE2 | AC019084.7 | -0,3168 | 7,34861E-10 |
| ACE2 | RP11-128P10.1 | -0,3168 | 7,37914E-10 |
| ACE2 | DISP2 | -0,3168 | 7,39994E-10 |
| ACE2 | GS1-115G20.1 | -0,3169 | 7,29618E-10 |
| ACE2 | RP11-567E21.3 | -0,3169 | 7,29117E-10 |
| ACE2 | ZCRB1 | -0,3169 | 7,30404E-10 |
| ACE2 | ZNF544 | -0,3169 | 7,25992E-10 |
| ACE2 | NOTCH2P1 | -0,3171 | 7,14167E-10 |
| ACE2 | KCNC1 | -0,3171 | 7,12373E-10 |
| ACE2 | RP11-313I2.7 | -0,3171 | 7,0807E-10 |
| ACE2 | CTB-114C7.4 | -0,3172 | 7,03885E-10 |
| ACE2 | RP1-253B10.1 | -0,3172 | 6,97311E-10 |
| ACE2 | RP11-566H8.1 | -0,3172 | 7,04082E-10 |
| ACE2 | FGD4 | -0,3173 | 6,91111E-10 |
| ACE2 | RP11-258F1.2 | -0,3173 | 6,92915E-10 |
| ACE2 | CTD-2026D20.2 | -0,3173 | 6,94049E-10 |
| ACE2 | RP11-343D2.11 | -0,3174 | 6,81894E-10 |
| ACE2 | PPM1N | -0,3174 | 6,79754E-10 |
| ACE2 | OR7E21P | -0,3175 | 6,7722E-10 |
| ACE2 | PHB2P1 | -0,3175 | 6,78669E-10 |
| ACE2 | AC005625.1 | -0,3175 | 6,76022E-10 |
| ACE2 | RP4-806M20.4 | -0,3175 | 6,73852E-10 |
| ACE2 | AP001439.2 | -0,3175 | 6,73073E-10 |
| ACE2 | LACTB2 | -0,3176 | 6,62752E-10 |
| ACE2 | CTD-3074O7.7 | -0,3176 | 6,65602E-10 |
| ACE2 | SLC35G4 | -0,3176 | 6,63787E-10 |
| ACE2 | KRT8P30 | -0,3177 | 6,59214E-10 |
| ACE2 | AC008691.1 | -0,3177 | 6,58101E-10 |
| ACE2 | RP11-945C19.4 | -0,3177 | 6,55783E-10 |
| ACE2 | CTD-2616J11.2 | -0,3178 | 6,47703E-10 |
| ACE2 | AC093843.1 | -0,3179 | 6,43177E-10 |
| ACE2 | NDUFAF1 | -0,3179 | 6,43447E-10 |
| ACE2 | MGAT3 | -0,3179 | 6,37078E-10 |
| ACE2 | ANK2 | -0,318 | 6,29365E-10 |
| ACE2 | RP11-1049H7.2 | -0,318 | 6,34048E-10 |
| ACE2 | RP11-612B6.1 | -0,318 | 6,3374E-10 |
| ACE2 | SPDYA | -0,3181 | 6,2589E-10 |
| ACE2 | PDK1 | -0,3181 | 6,25009E-10 |
| ACE2 | AC006000.5 | -0,3181 | 6,27799E-10 |
| ACE2 | SKAP2 | -0,3182 | 6,17605E-10 |
| ACE2 | DNAJC30 | -0,3182 | 6,19248E-10 |
| ACE2 | SHMT1 | -0,3182 | 6,12522E-10 |
| ACE2 | LRRC37A14P | -0,3182 | 6,12059E-10 |
| ACE2 | RP11-410C4.5 | -0,3183 | 6,11518E-10 |
| ACE2 | FAM161A | -0,3183 | 6,06179E-10 |
| ACE2 | RP11-149F8.3 | -0,3183 | 6,09786E-10 |
| ACE2 | SAMD13 | -0,3184 | 5,96613E-10 |
| ACE2 | ARHGAP33 | -0,3184 | 5,96429E-10 |
| ACE2 | CTD-2263F21.1 | -0,3185 | 5,95103E-10 |
| ACE2 | PICSAR | -0,3185 | 5,93728E-10 |
| ACE2 | SATL1 | -0,3185 | 5,94928E-10 |
| ACE2 | RP11-551G24.3 | -0,3186 | 5,83645E-10 |

| | | | |
|---|---|---|---|
| ACE2 | ASB12 | -0,3186 | 5,87461E-10 |
| ACE2 | IL36RN | -0,3187 | 5,78668E-10 |
| ACE2 | NDUFC1 | -0,3187 | 5,78107E-10 |
| ACE2 | RPL7P23 | -0,3187 | 5,77579E-10 |
| ACE2 | RP11-464C19.2 | -0,3187 | 5,80445E-10 |
| ACE2 | PSKH1 | -0,3188 | 5,72246E-10 |
| ACE2 | GEMIN7 | -0,3188 | 5,69151E-10 |
| ACE2 | DAP3P1 | -0,3189 | 5,61193E-10 |
| ACE2 | RAB32 | -0,3189 | 5,61591E-10 |
| ACE2 | HS3ST6 | -0,3189 | 5,61863E-10 |
| ACE2 | ARL6IP1P1 | -0,319 | 5,56617E-10 |
| ACE2 | UNC13D | -0,319 | 5,57568E-10 |
| ACE2 | OXNAD1 | -0,3191 | 5,49549E-10 |
| ACE2 | NKAIN3 | -0,3191 | 5,44593E-10 |
| ACE2 | AC114755.3 | -0,3192 | 5,40702E-10 |
| ACE2 | SERPINI2 | -0,3192 | 5,40545E-10 |
| ACE2 | NUGGC | -0,3192 | 5,3946E-10 |
| ACE2 | FAM230A | -0,3192 | 5,43196E-10 |
| ACE2 | LINC01819 | -0,3193 | 5,31284E-10 |
| ACE2 | RP11-192C21.2 | -0,3193 | 5,35677E-10 |
| ACE2 | RP11-6L6.7 | -0,3193 | 5,36118E-10 |
| ACE2 | CTB-11I22.1 | -0,3193 | 5,34587E-10 |
| ACE2 | RP11-631F7.1 | -0,3193 | 5,34653E-10 |
| ACE2 | RP11-463I20.3 | -0,3193 | 5,34863E-10 |
| ACE2 | HES2 | -0,3194 | 5,24646E-10 |
| ACE2 | GCSAML | -0,3194 | 5,24423E-10 |
| ACE2 | OR2T7 | -0,3194 | 5,27291E-10 |
| ACE2 | KRTAP20-2 | -0,3195 | 5,22445E-10 |
| ACE2 | NME8 | -0,3196 | 5,14637E-10 |
| ACE2 | RP11-63B13.1 | -0,3197 | 5,05883E-10 |
| ACE2 | GUSBP4 | -0,3197 | 5,08505E-10 |
| ACE2 | IL11 | -0,3197 | 5,04219E-10 |
| ACE2 | RP1-140K8.1 | -0,3198 | 4,99131E-10 |
| ACE2 | XXbac-BPG249D20.9 | -0,3198 | 5,02215E-10 |
| ACE2 | CACNA1C-IT2 | -0,3198 | 5,01285E-10 |
| ACE2 | RP11-80B9.1 | -0,3199 | 4,9234E-10 |
| ACE2 | PDE10A | -0,3199 | 4,95848E-10 |
| ACE2 | RP11-17G12.3 | -0,3199 | 4,95263E-10 |
| ACE2 | RHOT1P1 | -0,3199 | 4,91156E-10 |
| ACE2 | TMEM38B | -0,32 | 4,84212E-10 |
| ACE2 | AC005786.6 | -0,32 | 4,84057E-10 |
| ACE2 | HORMAD2-AS1 | -0,32 | 4,89016E-10 |
| ACE2 | OR7E90P | -0,3201 | 4,77485E-10 |
| ACE2 | HERPUD2 | -0,3201 | 4,78714E-10 |
| ACE2 | RP11-34H11.6 | -0,3201 | 4,77805E-10 |
| ACE2 | CTD-2562G15.3 | -0,3201 | 4,8111E-10 |
| ACE2 | STXBP2 | -0,3201 | 4,8173E-10 |
| ACE2 | RP11-339F13.2 | -0,3202 | 4,7304E-10 |
| ACE2 | RP11-685M7.3 | -0,3202 | 4,77224E-10 |
| ACE2 | KB-1615E4.2 | -0,3203 | 4,66165E-10 |
| ACE2 | MKX-AS1 | -0,3203 | 4,69823E-10 |
| ACE2 | CLEC16A | -0,3203 | 4,65849E-10 |
| ACE2 | AC092159.3 | -0,3204 | 4,62958E-10 |
| ACE2 | IMMP1LP1 | -0,3204 | 4,60434E-10 |

| | | | |
|---|---|---|---|
| ACE2 | SLC1A5 | -0,3204 | 4,614E-10 |
| ACE2 | MRPL55 | -0,3205 | 4,55086E-10 |
| ACE2 | RFX4 | -0,3205 | 4,54179E-10 |
| ACE2 | RPL10P14 | -0,3205 | 4,55112E-10 |
| ACE2 | MB | -0,3205 | 4,55653E-10 |
| ACE2 | CER1 | -0,3206 | 4,47497E-10 |
| ACE2 | KLF2P3 | -0,3207 | 4,44515E-10 |
| ACE2 | ZNF713 | -0,3207 | 4,46616E-10 |
| ACE2 | TRBV13 | -0,3207 | 4,41714E-10 |
| ACE2 | TGM5 | -0,3207 | 4,4143E-10 |
| ACE2 | SFTPD | -0,3208 | 4,40031E-10 |
| ACE2 | CSDAP1 | -0,3208 | 4,41092E-10 |
| ACE2 | AC010761.14 | -0,3208 | 4,36338E-10 |
| ACE2 | RP11-191L17.1 | -0,3209 | 4,32751E-10 |
| ACE2 | AC024028.1 | -0,3209 | 4,3295E-10 |
| ACE2 | RNY1P9 | -0,3209 | 4,33333E-10 |
| ACE2 | FOXL2NB | -0,321 | 4,24607E-10 |
| ACE2 | WDR1 | -0,321 | 4,25363E-10 |
| ACE2 | RP11-440G9.1 | -0,321 | 4,28579E-10 |
| ACE2 | OR7E36P | -0,321 | 4,24933E-10 |
| ACE2 | PSG8 | -0,321 | 4,2641E-10 |
| ACE2 | CTD-2066L21.3 | -0,3211 | 4,20234E-10 |
| ACE2 | RP11-216P16.4 | -0,3211 | 4,18895E-10 |
| ACE2 | NKG7 | -0,3211 | 4,22431E-10 |
| ACE2 | RP11-36D19.9 | -0,3212 | 4,16216E-10 |
| ACE2 | PDXDC1 | -0,3212 | 4,14127E-10 |
| ACE2 | RP4-680D5.2 | -0,3213 | 4,11627E-10 |
| ACE2 | MYO5BP3 | -0,3213 | 4,09555E-10 |
| ACE2 | PRB4 | -0,3213 | 4,1095E-10 |
| ACE2 | SEPHS1P1 | -0,3214 | 4,05199E-10 |
| ACE2 | ZNF725P | -0,3214 | 4,05184E-10 |
| ACE2 | NABP1 | -0,3215 | 4,02328E-10 |
| ACE2 | LINC01612 | -0,3215 | 4,00029E-10 |
| ACE2 | POLG2 | -0,3215 | 3,99791E-10 |
| ACE2 | RP11-128P17.4 | -0,3215 | 4,01785E-10 |
| ACE2 | TCFL5 | -0,3215 | 3,99326E-10 |
| ACE2 | RP11-342M1.3 | -0,3216 | 3,93282E-10 |
| ACE2 | PHTF1 | -0,3216 | 3,94806E-10 |
| ACE2 | NXPE2 | -0,3216 | 3,93015E-10 |
| ACE2 | PTBP1P | -0,3216 | 3,95301E-10 |
| ACE2 | PROK2 | -0,3217 | 3,88165E-10 |
| ACE2 | DNAJC2 | -0,3217 | 3,888E-10 |
| ACE2 | AGGF1P4 | -0,3217 | 3,88125E-10 |
| ACE2 | RP11-351A20.2 | -0,3217 | 3,9182E-10 |
| ACE2 | KRT25 | -0,3217 | 3,87043E-10 |
| ACE2 | AP000704.5 | -0,3217 | 3,88731E-10 |
| ACE2 | PGAM1P3 | -0,3218 | 3,83007E-10 |
| ACE2 | ITGB3BP | -0,3219 | 3,79056E-10 |
| ACE2 | RP11-275A14.1 | -0,3219 | 3,78656E-10 |
| ACE2 | MUC15 | -0,3219 | 3,80135E-10 |
| ACE2 | CD3E | -0,3219 | 3,79373E-10 |
| ACE2 | RPS20P2 | -0,3219 | 3,79092E-10 |
| ACE2 | MCHR2 | -0,322 | 3,75038E-10 |
| ACE2 | RP3-340H11.2 | -0,3221 | 3,70095E-10 |

| | | | |
|---|---|---|---|
| ACE2 | C3orf14 | -0,3222 | 3,64262E-10 |
| ACE2 | CTC-311G1.1 | -0,3222 | 3,66854E-10 |
| ACE2 | CTD-2647L4.1 | -0,3222 | 3,64791E-10 |
| ACE2 | RN7SL241P | -0,3222 | 3,6537E-10 |
| ACE2 | AL022341.3 | -0,3222 | 3,64866E-10 |
| ACE2 | RP11-321E2.12 | -0,3223 | 3,57532E-10 |
| ACE2 | DSTNP2 | -0,3223 | 3,61437E-10 |
| ACE2 | CLDN9 | -0,3223 | 3,61788E-10 |
| ACE2 | CTB-186G2.4 | -0,3223 | 3,59394E-10 |
| ACE2 | RP11-274B21.2 | -0,3224 | 3,57205E-10 |
| ACE2 | ERGIC3 | -0,3224 | 3,54191E-10 |
| ACE2 | LINC01104 | -0,3225 | 3,48201E-10 |
| ACE2 | RP11-240E2.2 | -0,3225 | 3,51495E-10 |
| ACE2 | NEUROG2 | -0,3226 | 3,43778E-10 |
| ACE2 | AGPAT1 | -0,3226 | 3,45694E-10 |
| ACE2 | TRBV12-4 | -0,3226 | 3,44055E-10 |
| ACE2 | LINC01257 | -0,3226 | 3,46401E-10 |
| ACE2 | RP11-294C11.7 | -0,3226 | 3,45725E-10 |
| ACE2 | RP5-1170K4.7 | -0,3226 | 3,47081E-10 |
| ACE2 | ZDHHC20P1 | -0,3227 | 3,43208E-10 |
| ACE2 | KRT89P | -0,3227 | 3,38997E-10 |
| ACE2 | RP11-527D15.1 | -0,3228 | 3,38819E-10 |
| ACE2 | RP11-315I20.3 | -0,3229 | 3,32451E-10 |
| ACE2 | MTND5P31 | -0,3229 | 3,32152E-10 |
| ACE2 | LARP1B | -0,3229 | 3,31634E-10 |
| ACE2 | RP3-443C4.2 | -0,3229 | 3,32806E-10 |
| ACE2 | ALKBH3 | -0,3229 | 3,33954E-10 |
| ACE2 | SERPINC1 | -0,323 | 3,28178E-10 |
| ACE2 | CAPN14 | -0,323 | 3,29746E-10 |
| ACE2 | MPRIPP1 | -0,323 | 3,26268E-10 |
| ACE2 | RP11-662M24.2 | -0,323 | 3,2596E-10 |
| ACE2 | CTD-2396E7.11 | -0,323 | 3,27954E-10 |
| ACE2 | CCER2 | -0,323 | 3,29943E-10 |
| ACE2 | TRIM55 | -0,3231 | 3,25593E-10 |
| ACE2 | GINS3 | -0,3231 | 3,25268E-10 |
| ACE2 | AL133247.2 | -0,3232 | 3,18888E-10 |
| ACE2 | UGT2B10 | -0,3232 | 3,20332E-10 |
| ACE2 | RP11-672A2.3 | -0,3232 | 3,20336E-10 |
| ACE2 | CTC-297N7.7 | -0,3232 | 3,18604E-10 |
| ACE2 | CTD-3064M3.4 | -0,3233 | 3,15675E-10 |
| ACE2 | CTD-2325B11.1 | -0,3234 | 3,09534E-10 |
| ACE2 | CTD-2122P11.1 | -0,3234 | 3,11812E-10 |
| ACE2 | RP11-589F5.4 | -0,3234 | 3,12771E-10 |
| ACE2 | OR10A4 | -0,3234 | 3,12787E-10 |
| ACE2 | RP11-480C22.1 | -0,3234 | 3,11954E-10 |
| ACE2 | NPAS3 | -0,3234 | 3,10304E-10 |
| ACE2 | HNRNPLP2 | -0,3234 | 3,11858E-10 |
| ACE2 | RP11-510C10.2 | -0,3235 | 3,08056E-10 |
| ACE2 | RP11-370I10.10 | -0,3235 | 3,07295E-10 |
| ACE2 | LA16c-313D11.12 | -0,3235 | 3,05028E-10 |
| ACE2 | CTB-99A3.1 | -0,3236 | 3,04523E-10 |
| ACE2 | LINC01713 | -0,3236 | 3,0302E-10 |
| ACE2 | TOR1B | -0,3237 | 2,98067E-10 |
| ACE2 | FAM216B | -0,3237 | 3,00633E-10 |

| | | | |
|---|---|---|---|
| ACE2 | CDCA2 | -0,3238 | 2,93019E-10 |
| ACE2 | RP11-539L10.5 | -0,3239 | 2,91501E-10 |
| ACE2 | GRXCR1 | -0,3239 | 2,8979E-10 |
| ACE2 | EEF1E1 | -0,3239 | 2,91419E-10 |
| ACE2 | RP11-599B13.7 | -0,324 | 2,8895E-10 |
| ACE2 | METTL21A | -0,3241 | 2,82616E-10 |
| ACE2 | PANO1 | -0,3241 | 2,84483E-10 |
| ACE2 | CDH5 | -0,3241 | 2,83405E-10 |
| ACE2 | DNM1P28 | -0,3242 | 2,79328E-10 |
| ACE2 | LINC01023 | -0,3243 | 2,7589E-10 |
| ACE2 | FAM35DP | -0,3243 | 2,74332E-10 |
| ACE2 | HOXA-AS3 | -0,3244 | 2,71065E-10 |
| ACE2 | OR2M1P | -0,3245 | 2,6876E-10 |
| ACE2 | RP11-399F4.2 | -0,3245 | 2,68949E-10 |
| ACE2 | RP11-249C24.10 | -0,3245 | 2,67608E-10 |
| ACE2 | RP11-627G18.4 | -0,3246 | 2,6364E-10 |
| ACE2 | ACOT8 | -0,3246 | 2,66417E-10 |
| ACE2 | RP11-579D7.4 | -0,3247 | 2,62619E-10 |
| ACE2 | RP11-603J24.5 | -0,3247 | 2,61645E-10 |
| ACE2 | RAX | -0,3247 | 2,60905E-10 |
| ACE2 | LINC01005 | -0,3248 | 2,59054E-10 |
| ACE2 | CTB-107G13.1 | -0,3248 | 2,57047E-10 |
| ACE2 | LINC00404 | -0,3248 | 2,56606E-10 |
| ACE2 | RP11-15A1.7 | -0,3248 | 2,56802E-10 |
| ACE2 | RP11-526I2.1 | -0,3249 | 2,53385E-10 |
| ACE2 | AC011523.2 | -0,3249 | 2,56434E-10 |
| ACE2 | AC093642.4 | -0,325 | 2,52509E-10 |
| ACE2 | RBMS3-AS1 | -0,3251 | 2,48527E-10 |
| ACE2 | OR2H4P | -0,3251 | 2,47402E-10 |
| ACE2 | OR8C1P | -0,3251 | 2,48525E-10 |
| ACE2 | RP11-109N23.6 | -0,3252 | 2,46092E-10 |
| ACE2 | RPL5P6 | -0,3253 | 2,39913E-10 |
| ACE2 | RP4-705D16.3 | -0,3253 | 2,42472E-10 |
| ACE2 | FLJ37453 | -0,3254 | 2,37569E-10 |
| ACE2 | RP11-7F17.5 | -0,3254 | 2,39525E-10 |
| ACE2 | POLR1B | -0,3255 | 2,36086E-10 |
| ACE2 | CCSER1 | -0,3255 | 2,34086E-10 |
| ACE2 | LINC01332 | -0,3255 | 2,35856E-10 |
| ACE2 | RP11-367G6.3 | -0,3255 | 2,35442E-10 |
| ACE2 | RP11-632L2.2 | -0,3255 | 2,34748E-10 |
| ACE2 | DNM1P34 | -0,3255 | 2,33826E-10 |
| ACE2 | AC067968.3 | -0,3255 | 2,33793E-10 |
| ACE2 | CLDN14 | -0,3255 | 2,33752E-10 |
| ACE2 | LINC02082 | -0,3256 | 2,30482E-10 |
| ACE2 | PLPPR1 | -0,3256 | 2,33355E-10 |
| ACE2 | SIGLEC14 | -0,3256 | 2,31506E-10 |
| ACE2 | CTD-2537I9.16 | -0,3256 | 2,30901E-10 |
| ACE2 | RP11-386M24.9 | -0,3257 | 2,29522E-10 |
| ACE2 | TBC1D3L | -0,3257 | 2,28529E-10 |
| ACE2 | RP1-298M8.1 | -0,3257 | 2,29492E-10 |
| ACE2 | RP11-717H13.1 | -0,3258 | 2,25131E-10 |
| ACE2 | MIR4526 | -0,3258 | 2,25294E-10 |
| ACE2 | IL23R | -0,3259 | 2,24144E-10 |
| ACE2 | CYP3A43 | -0,3259 | 2,22388E-10 |

| | | | |
|---|---|---|---|
| ACE2 | RPS19P1 | -0,3259 | 2,22E-10 |
| ACE2 | LINC02200 | -0,326 | 2,19799E-10 |
| ACE2 | PIGV | -0,3261 | 2,15615E-10 |
| ACE2 | RP11-715C4.1 | -0,3261 | 2,16006E-10 |
| ACE2 | LINC02069 | -0,3262 | 2,15242E-10 |
| ACE2 | GLIPR1L1 | -0,3262 | 2,14642E-10 |
| ACE2 | HYKK | -0,3262 | 2,13298E-10 |
| ACE2 | MRPS9 | -0,3263 | 2,1026E-10 |
| ACE2 | P2RY6 | -0,3263 | 2,10207E-10 |
| ACE2 | RP11-438H24.2 | -0,3263 | 2,11295E-10 |
| ACE2 | RP11-417L19.6 | -0,3266 | 2,02796E-10 |
| ACE2 | RP11-568J23.4 | -0,3266 | 2,04055E-10 |
| ACE2 | C7orf57 | -0,3267 | 2,01322E-10 |
| ACE2 | RP11-468H14.2 | -0,3267 | 2,00874E-10 |
| ACE2 | SLC6A12 | -0,3267 | 2,00519E-10 |
| ACE2 | RP11-121J20.1 | -0,3267 | 1,98933E-10 |
| ACE2 | NSUN6 | -0,3268 | 1,96757E-10 |
| ACE2 | SCOCP1 | -0,3268 | 1,96635E-10 |
| ACE2 | THAP7-AS1 | -0,3268 | 1,98155E-10 |
| ACE2 | CTNNA2 | -0,3269 | 1,94881E-10 |
| ACE2 | TRBV29OR9-2 | -0,3269 | 1,94766E-10 |
| ACE2 | AP005135.2 | -0,3269 | 1,95702E-10 |
| ACE2 | RP1-74B13.2 | -0,3269 | 1,94847E-10 |
| ACE2 | CTD-2139B15.1 | -0,327 | 1,91822E-10 |
| ACE2 | RP11-15G8.1 | -0,327 | 1,92282E-10 |
| ACE2 | XKR6 | -0,327 | 1,9267E-10 |
| ACE2 | PRR21 | -0,3271 | 1,89852E-10 |
| ACE2 | LINC01932 | -0,3271 | 1,88543E-10 |
| ACE2 | KLC4 | -0,3271 | 1,88843E-10 |
| ACE2 | RP1-228P16.5 | -0,3271 | 1,90302E-10 |
| ACE2 | MAP3K9 | -0,3271 | 1,89472E-10 |
| ACE2 | MTTP | -0,3272 | 1,86126E-10 |
| ACE2 | SRPK1 | -0,3272 | 1,86666E-10 |
| ACE2 | RP11-309L24.2 | -0,3272 | 1,87496E-10 |
| ACE2 | GALR3 | -0,3272 | 1,88084E-10 |
| ACE2 | ESRRG | -0,3273 | 1,84373E-10 |
| ACE2 | RP11-512G4.1 | -0,3273 | 1,84232E-10 |
| ACE2 | RP11-208N14.4 | -0,3273 | 1,85267E-10 |
| ACE2 | RP11-183I6.2 | -0,3273 | 1,85628E-10 |
| ACE2 | SPATA31A5 | -0,3273 | 1,84652E-10 |
| ACE2 | STK24-AS1 | -0,3273 | 1,84416E-10 |
| ACE2 | RPL36AP29 | -0,3274 | 1,82696E-10 |
| ACE2 | LINC01559 | -0,3274 | 1,8229E-10 |
| ACE2 | RP11-155I9.1 | -0,3274 | 1,83019E-10 |
| ACE2 | CTC-552D5.1 | -0,3275 | 1,7937E-10 |
| ACE2 | RP11-1167A19.2 | -0,3275 | 1,79767E-10 |
| ACE2 | PSG10P | -0,3275 | 1,80524E-10 |
| ACE2 | CTD-2639E6.9 | -0,3275 | 1,80049E-10 |
| ACE2 | AC114730.11 | -0,3276 | 1,77813E-10 |
| ACE2 | VN1R40P | -0,3276 | 1,76945E-10 |
| ACE2 | RP11-165H23.1 | -0,3276 | 1,76796E-10 |
| ACE2 | PPP2R1B | -0,3276 | 1,76346E-10 |
| ACE2 | AC108004.3 | -0,3276 | 1,78545E-10 |
| ACE2 | RP11-91H12.3 | -0,3277 | 1,75931E-10 |

| | | | |
|---|---|---|---|
| ACE2 | RP11-677O4.2 | -0,3277 | 1,74524E-10 |
| ACE2 | LINC01258 | -0,3278 | 1,72447E-10 |
| ACE2 | FKBP6 | -0,3278 | 1,72175E-10 |
| ACE2 | MMP10 | -0,3279 | 1,69792E-10 |
| ACE2 | ZNF331 | -0,3279 | 1,6932E-10 |
| ACE2 | RP11-126K1.9 | -0,328 | 1,67483E-10 |
| ACE2 | UQCRBP2 | -0,328 | 1,68957E-10 |
| ACE2 | CFL1P1 | -0,328 | 1,69101E-10 |
| ACE2 | PSMC1P8 | -0,328 | 1,6842E-10 |
| ACE2 | RP11-50D16.4 | -0,328 | 1,6726E-10 |
| ACE2 | LINC00449 | -0,328 | 1,66995E-10 |
| ACE2 | RP11-478H13.4 | -0,3281 | 1,66131E-10 |
| ACE2 | ETFRF1 | -0,3281 | 1,66571E-10 |
| ACE2 | TESMIN | -0,3282 | 1,63155E-10 |
| ACE2 | SPANXN3 | -0,3282 | 1,64471E-10 |
| ACE2 | AC027119.1 | -0,3283 | 1,61258E-10 |
| ACE2 | BCHE | -0,3283 | 1,60474E-10 |
| ACE2 | EPB41L4A-AS1 | -0,3283 | 1,61139E-10 |
| ACE2 | KATNBL1P6 | -0,3283 | 1,61331E-10 |
| ACE2 | RP11-44N12.4 | -0,3283 | 1,62132E-10 |
| ACE2 | RP11-1223D19.2 | -0,3284 | 1,5927E-10 |
| ACE2 | RP1-97J1.2 | -0,3284 | 1,58717E-10 |
| ACE2 | PROSER2 | -0,3284 | 1,59857E-10 |
| ACE2 | LINC00453 | -0,3284 | 1,59357E-10 |
| ACE2 | AC005357.1 | -0,3284 | 1,58627E-10 |
| ACE2 | LINC00489 | -0,3284 | 1,58679E-10 |
| ACE2 | ERVH48-1 | -0,3284 | 1,60181E-10 |
| ACE2 | CTA-351J1.1 | -0,3285 | 1,57508E-10 |
| ACE2 | TCF4-AS1 | -0,3285 | 1,56476E-10 |
| ACE2 | RP11-560A15.3 | -0,3285 | 1,56488E-10 |
| ACE2 | RP11-203F10.5 | -0,3286 | 1,56051E-10 |
| ACE2 | C4BPAP1 | -0,3286 | 1,54186E-10 |
| ACE2 | SYT9 | -0,3286 | 1,55962E-10 |
| ACE2 | BLZF2P | -0,3286 | 1,55316E-10 |
| ACE2 | RPL10P15 | -0,3286 | 1,54843E-10 |
| ACE2 | THAP12P6 | -0,3287 | 1,52147E-10 |
| ACE2 | AC007682.1 | -0,3288 | 1,50768E-10 |
| ACE2 | RP1-153P14.5 | -0,3288 | 1,50507E-10 |
| ACE2 | RP11-349P19.1 | -0,3288 | 1,50386E-10 |
| ACE2 | ATP10A | -0,3288 | 1,50805E-10 |
| ACE2 | RP11-495P10.3 | -0,3289 | 1,47896E-10 |
| ACE2 | RP4-580N22.2 | -0,3289 | 1,48986E-10 |
| ACE2 | RP11-278J20.2 | -0,3289 | 1,49216E-10 |
| ACE2 | RP11-760D2.4 | -0,3289 | 1,48616E-10 |
| ACE2 | EIF4A1P6 | -0,3289 | 1,48392E-10 |
| ACE2 | CTD-2382E5.2 | -0,3289 | 1,4844E-10 |
| ACE2 | LINC01754 | -0,3289 | 1,49507E-10 |
| ACE2 | GNAQP1 | -0,3291 | 1,44936E-10 |
| ACE2 | RP11-520A21.1 | -0,3291 | 1,44091E-10 |
| ACE2 | CACNA1C-AS3 | -0,3291 | 1,44725E-10 |
| ACE2 | RP11-158M2.2 | -0,3291 | 1,45569E-10 |
| ACE2 | RP11-672L10.3 | -0,3291 | 1,44584E-10 |
| ACE2 | GDPD4 | -0,3292 | 1,42533E-10 |
| ACE2 | ADAMTS7P4 | -0,3292 | 1,42329E-10 |

| | | | |
|---|---|---|---|
| ACE2 | NEFHP1 | -0,3294 | 1,3995E-10 |
| ACE2 | AP000648.6 | -0,3294 | 1,39615E-10 |
| ACE2 | HSF5 | -0,3294 | 1,3937E-10 |
| ACE2 | HMGB2P1 | -0,3294 | 1,40052E-10 |
| ACE2 | CMTM8 | -0,3295 | 1,36436E-10 |
| ACE2 | PCDHA2 | -0,3295 | 1,36428E-10 |
| ACE2 | RP11-620J15.4 | -0,3295 | 1,37839E-10 |
| ACE2 | IGLV3-22 | -0,3295 | 1,3685E-10 |
| ACE2 | AC008277.1 | -0,3296 | 1,36051E-10 |
| ACE2 | KLF4 | -0,3296 | 1,35223E-10 |
| ACE2 | RP11-279N8.1 | -0,3296 | 1,35729E-10 |
| ACE2 | ZFP64 | -0,3296 | 1,35204E-10 |
| ACE2 | GOLGA6B | -0,3297 | 1,33236E-10 |
| ACE2 | IL2RB | -0,3297 | 1,34452E-10 |
| ACE2 | RP11-574H6.1 | -0,3298 | 1,32432E-10 |
| ACE2 | SPATA31A7 | -0,3298 | 1,31103E-10 |
| ACE2 | RP11-196H14.4 | -0,3298 | 1,31766E-10 |
| ACE2 | RNU6-430P | -0,3298 | 1,31814E-10 |
| ACE2 | RP11-530N7.3 | -0,3298 | 1,32117E-10 |
| ACE2 | RP11-261N11.8 | -0,3298 | 1,32397E-10 |
| ACE2 | APOA2 | -0,3299 | 1,29185E-10 |
| ACE2 | CSN3 | -0,3299 | 1,30105E-10 |
| ACE2 | RP11-222G7.2 | -0,3299 | 1,29124E-10 |
| ACE2 | TMEM116 | -0,3299 | 1,30818E-10 |
| ACE2 | CTC-312O10.2 | -0,3299 | 1,30461E-10 |
| ACE2 | H2AFB3 | -0,3299 | 1,30256E-10 |
| ACE2 | LSP1P3 | -0,33 | 1,28286E-10 |
| ACE2 | SUGCT | -0,33 | 1,2865E-10 |
| ACE2 | RPSAP66 | -0,33 | 1,28612E-10 |
| ACE2 | FOXN1 | -0,3301 | 1,26703E-10 |
| ACE2 | C1orf210 | -0,3302 | 1,25422E-10 |
| ACE2 | RP11-742B18.1 | -0,3302 | 1,24514E-10 |
| ACE2 | RP11-548K12.10 | -0,3302 | 1,25472E-10 |
| ACE2 | PPFIBP2 | -0,3302 | 1,24299E-10 |
| ACE2 | MYO1A | -0,3302 | 1,24201E-10 |
| ACE2 | LINC00463 | -0,3302 | 1,24176E-10 |
| ACE2 | LINC01946 | -0,3303 | 1,23237E-10 |
| ACE2 | CTD-2247C11.2 | -0,3303 | 1,22405E-10 |
| ACE2 | RP11-425D10.1 | -0,3303 | 1,23419E-10 |
| ACE2 | RP11-267M23.7 | -0,3303 | 1,23868E-10 |
| ACE2 | ASCL4 | -0,3303 | 1,23191E-10 |
| ACE2 | EDDM3A | -0,3303 | 1,23381E-10 |
| ACE2 | NBPF25P | -0,3306 | 1,18469E-10 |
| ACE2 | AC159540.1 | -0,3306 | 1,18704E-10 |
| ACE2 | RP11-661C8.3 | -0,3306 | 1,1788E-10 |
| ACE2 | RP11-45H22.3 | -0,3306 | 1,18478E-10 |
| ACE2 | RP11-150O12.5 | -0,3306 | 1,18258E-10 |
| ACE2 | SAMD12-AS1 | -0,3306 | 1,18421E-10 |
| ACE2 | CRIP1P4 | -0,3306 | 1,17669E-10 |
| ACE2 | CCDC18 | -0,3307 | 1,16142E-10 |
| ACE2 | RP4-712E4.2 | -0,3307 | 1,15828E-10 |
| ACE2 | TMPRSS11E | -0,3307 | 1,17324E-10 |
| ACE2 | NDUFB11P1 | -0,3307 | 1,17286E-10 |
| ACE2 | RP11-722P11.4 | -0,3307 | 1,16114E-10 |

| | | | |
|---|---|---|---|
| ACE2 | CTD-2021H9.1 | -0,3307 | 1,17285E-10 |
| ACE2 | ATP6AP1L | -0,3308 | 1,14827E-10 |
| ACE2 | RP11-403N16.5 | -0,3308 | 1,15001E-10 |
| ACE2 | NRGN | -0,3308 | 1,14302E-10 |
| ACE2 | RP11-72M17.1 | -0,3308 | 1,15656E-10 |
| ACE2 | ZNF200 | -0,3309 | 1,14124E-10 |
| ACE2 | LLNLR-285B5.1 | -0,3309 | 1,14218E-10 |
| ACE2 | EEF1A1P34 | -0,3309 | 1,12762E-10 |
| ACE2 | RP11-404I7.1 | -0,331 | 1,11929E-10 |
| ACE2 | RP11-367G18.1 | -0,331 | 1,11751E-10 |
| ACE2 | RP11-1082L8.2 | -0,331 | 1,12156E-10 |
| ACE2 | TMEM254-AS1 | -0,331 | 1,11679E-10 |
| ACE2 | IPO5 | -0,331 | 1,11536E-10 |
| ACE2 | DIRC3 | -0,3311 | 1,11116E-10 |
| ACE2 | CTB-1144G6.6 | -0,3311 | 1,10795E-10 |
| ACE2 | RP11-588P7.2 | -0,3311 | 1,10808E-10 |
| ACE2 | SEC14L1 | -0,3311 | 1,11057E-10 |
| ACE2 | AC018832.1 | -0,3312 | 1,09295E-10 |
| ACE2 | STAG3L2 | -0,3313 | 1,07902E-10 |
| ACE2 | DBF4 | -0,3313 | 1,07337E-10 |
| ACE2 | RP11-57C13.4 | -0,3313 | 1,07042E-10 |
| ACE2 | ZNF646P1 | -0,3313 | 1,07854E-10 |
| ACE2 | KRTAP24-1 | -0,3313 | 1,07583E-10 |
| ACE2 | CREB1 | -0,3314 | 1,05386E-10 |
| ACE2 | XXbac-BPG154L12.5 | -0,3314 | 1,05896E-10 |
| ACE2 | BRI3BPP1 | -0,3315 | 1,04979E-10 |
| ACE2 | TEX9 | -0,3316 | 1,03526E-10 |
| ACE2 | RP11-89N17.1 | -0,3317 | 1,01121E-10 |
| ACE2 | ZNF79 | -0,3317 | 1,01056E-10 |
| ACE2 | RP11-186F10.2 | -0,3317 | 1,01553E-10 |
| ACE2 | RP11-88E10.5 | -0,3317 | 1,01423E-10 |
| ACE2 | RP11-191G24.1 | -0,3318 | 1,00993E-10 |
| ACE2 | PCDHA3 | -0,3319 | 9,94037E-11 |
| ACE2 | RP11-125M16.1 | -0,3319 | 9,8634E-11 |
| ACE2 | RP3-446N13.2 | -0,3319 | 9,8593E-11 |
| ACE2 | ZNF100 | -0,3319 | 9,94043E-11 |
| ACE2 | NAPSB | -0,3319 | 9,9023E-11 |
| ACE2 | LINC00640 | -0,332 | 9,74442E-11 |
| ACE2 | CCL8 | -0,332 | 9,80816E-11 |
| ACE2 | WDFY3-AS1 | -0,3321 | 9,6813E-11 |
| ACE2 | LINC02118 | -0,3321 | 9,59656E-11 |
| ACE2 | ALOX15 | -0,3321 | 9,62236E-11 |
| ACE2 | MTFR1L | -0,3323 | 9,42812E-11 |
| ACE2 | LRRC14 | -0,3323 | 9,35362E-11 |
| ACE2 | KRT16P6 | -0,3323 | 9,3564E-11 |
| ACE2 | RP11-403I13.7 | -0,3324 | 9,27146E-11 |
| ACE2 | SMG7 | -0,3324 | 9,27023E-11 |
| ACE2 | CTD-2196P11.2 | -0,3324 | 9,29319E-11 |
| ACE2 | EHF | -0,3324 | 9,20177E-11 |
| ACE2 | CTAGE3P | -0,3324 | 9,19713E-11 |
| ACE2 | AC009120.8 | -0,3324 | 9,21647E-11 |
| ACE2 | MEP1B | -0,3324 | 9,30052E-11 |
| ACE2 | LMLN-AS1 | -0,3326 | 8,98247E-11 |
| ACE2 | C1orf101 | -0,3327 | 8,81873E-11 |

| | | | |
|---|---|---|---|
| ACE2 | TMEM244 | -0,3327 | 8,87568E-11 |
| ACE2 | LINC02119 | -0,3328 | 8,7646E-11 |
| ACE2 | POM121L2 | -0,3328 | 8,69521E-11 |
| ACE2 | NCR3LG1 | -0,3328 | 8,79189E-11 |
| ACE2 | RP11-295G12.1 | -0,3328 | 8,69376E-11 |
| ACE2 | TUBGCP4 | -0,3328 | 8,73018E-11 |
| ACE2 | ARL11 | -0,3329 | 8,58967E-11 |
| ACE2 | MRPL38 | -0,3329 | 8,65547E-11 |
| ACE2 | C1orf195 | -0,333 | 8,52437E-11 |
| ACE2 | AC002480.3 | -0,333 | 8,4846E-11 |
| ACE2 | ONECUT1 | -0,333 | 8,5027E-11 |
| ACE2 | AC098614.2 | -0,3331 | 8,40643E-11 |
| ACE2 | RP11-284M14.1 | -0,3331 | 8,36375E-11 |
| ACE2 | RAC1P6 | -0,3331 | 8,35901E-11 |
| ACE2 | RNASE7 | -0,3331 | 8,3805E-11 |
| ACE2 | RP11-260G13.1 | -0,3331 | 8,43193E-11 |
| ACE2 | LINC00951 | -0,3332 | 8,24168E-11 |
| ACE2 | RP11-72I2.1 | -0,3332 | 8,26289E-11 |
| ACE2 | CTD-2060L22.1 | -0,3332 | 8,31861E-11 |
| ACE2 | PKD1L1 | -0,3332 | 8,25635E-11 |
| ACE2 | RP11-98D18.17 | -0,3333 | 8,13628E-11 |
| ACE2 | AARSP1 | -0,3333 | 8,19125E-11 |
| ACE2 | NOL8P1 | -0,3334 | 8,02687E-11 |
| ACE2 | LINC01608 | -0,3334 | 8,08729E-11 |
| ACE2 | CTD-3035K23.3 | -0,3335 | 7,90254E-11 |
| ACE2 | DNAH8 | -0,3336 | 7,84894E-11 |
| ACE2 | ZNRD1-AS1_2 | -0,3336 | 7,88921E-11 |
| ACE2 | KIAA0753 | -0,3336 | 7,88901E-11 |
| ACE2 | RP11-379B18.5 | -0,3337 | 7,72541E-11 |
| ACE2 | RP11-959I15.1 | -0,3337 | 7,78458E-11 |
| ACE2 | RP11-424M22.3 | -0,3337 | 7,73266E-11 |
| ACE2 | FLJ21408 | -0,3337 | 7,6827E-11 |
| ACE2 | GDPD1 | -0,3337 | 7,75313E-11 |
| ACE2 | OR6R1P | -0,3338 | 7,62345E-11 |
| ACE2 | TUBG1P | -0,3338 | 7,65597E-11 |
| ACE2 | KB-1299A7.2 | -0,3338 | 7,59247E-11 |
| ACE2 | RP4-760C5.3 | -0,3338 | 7,63209E-11 |
| ACE2 | MTX1P1 | -0,3339 | 7,48332E-11 |
| ACE2 | CTC-209H22.3 | -0,3339 | 7,50274E-11 |
| ACE2 | RP11-3K24.3 | -0,3339 | 7,53844E-11 |
| ACE2 | RABAC1 | -0,3339 | 7,55816E-11 |
| ACE2 | POTEI | -0,334 | 7,44815E-11 |
| ACE2 | SLC2A2 | -0,334 | 7,38026E-11 |
| ACE2 | RP11-529E10.7 | -0,334 | 7,44497E-11 |
| ACE2 | CTC-332L22.1 | -0,334 | 7,41439E-11 |
| ACE2 | HIST1H2AB | -0,334 | 7,42689E-11 |
| ACE2 | KRTAP3-2 | -0,334 | 7,39328E-11 |
| ACE2 | RP11-496N12.6 | -0,3341 | 7,28949E-11 |
| ACE2 | AC017002.4 | -0,3341 | 7,36257E-11 |
| ACE2 | CASC21 | -0,3341 | 7,28957E-11 |
| ACE2 | SMARCA4 | -0,3341 | 7,28494E-11 |
| ACE2 | CTC-523E23.5 | -0,3341 | 7,32643E-11 |
| ACE2 | GGT2 | -0,3341 | 7,34139E-11 |
| ACE2 | RP11-407H12.8 | -0,3342 | 7,20487E-11 |

| | | | |
|---|---|---|---|
| ACE2 | LLNLR-263F3.1 | -0,3342 | 7,1958E-11 |
| ACE2 | PRUNE1 | -0,3343 | 7,15754E-11 |
| ACE2 | LINC01923 | -0,3343 | 7,13773E-11 |
| ACE2 | RP11-34H11.3 | -0,3343 | 7,14815E-11 |
| ACE2 | ATXN2L | -0,3343 | 7,11312E-11 |
| ACE2 | LINC01639 | -0,3343 | 7,07772E-11 |
| ACE2 | OR10J9P | -0,3344 | 7,01626E-11 |
| ACE2 | AC106870.1 | -0,3344 | 6,99199E-11 |
| ACE2 | PLPP6 | -0,3344 | 6,97294E-11 |
| ACE2 | SESN3 | -0,3344 | 7,04285E-11 |
| ACE2 | OR7E159P | -0,3344 | 7,01829E-11 |
| ACE2 | SENP8 | -0,3344 | 7,03244E-11 |
| ACE2 | RP4-654H19.2 | -0,3345 | 6,92828E-11 |
| ACE2 | RP11-543E8.2 | -0,3345 | 6,94141E-11 |
| ACE2 | RP11-3B7.1 | -0,3345 | 6,95188E-11 |
| ACE2 | RP11-292D4.4 | -0,3345 | 6,94458E-11 |
| ACE2 | RP5-851M4.1 | -0,3345 | 6,8853E-11 |
| ACE2 | SNAP91 | -0,3346 | 6,86989E-11 |
| ACE2 | DPY19L2P4 | -0,3346 | 6,79725E-11 |
| ACE2 | FAM27E2 | -0,3346 | 6,84976E-11 |
| ACE2 | Z83851.4 | -0,3346 | 6,86644E-11 |
| ACE2 | ATMIN | -0,3347 | 6,73365E-11 |
| ACE2 | CTC-539A10.7 | -0,3347 | 6,69604E-11 |
| ACE2 | RP5-1097P24.1 | -0,3347 | 6,73678E-11 |
| ACE2 | LINC01651 | -0,3348 | 6,64467E-11 |
| ACE2 | CTD-2247C11.4 | -0,3349 | 6,5592E-11 |
| ACE2 | RP13-968A2.1 | -0,3349 | 6,56793E-11 |
| ACE2 | RNA5SP252 | -0,3349 | 6,51943E-11 |
| ACE2 | RP11-88I18.2 | -0,3349 | 6,58662E-11 |
| ACE2 | RP11-428C19.4 | -0,3349 | 6,52112E-11 |
| ACE2 | IL12RB1 | -0,3349 | 6,53709E-11 |
| ACE2 | GPR32 | -0,3349 | 6,50783E-11 |
| ACE2 | KRT18P13 | -0,335 | 6,45167E-11 |
| ACE2 | RP11-134E15.1 | -0,335 | 6,47465E-11 |
| ACE2 | RP11-812E19.3 | -0,335 | 6,42015E-11 |
| ACE2 | RPL21P126 | -0,335 | 6,41796E-11 |
| ACE2 | TPRKB | -0,3351 | 6,34809E-11 |
| ACE2 | AC073465.1 | -0,3351 | 6,34866E-11 |
| ACE2 | GUSBP9 | -0,3351 | 6,36013E-11 |
| ACE2 | CTA-331P3.1 | -0,3351 | 6,33714E-11 |
| ACE2 | RP11-396B14.2 | -0,3351 | 6,39428E-11 |
| ACE2 | ELOCP29 | -0,3351 | 6,35605E-11 |
| ACE2 | SHISA7 | -0,3351 | 6,36542E-11 |
| ACE2 | AC017006.2 | -0,3352 | 6,3089E-11 |
| ACE2 | MCIDAS | -0,3352 | 6,29755E-11 |
| ACE2 | TTTY22 | -0,3352 | 6,27232E-11 |
| ACE2 | RANBP3 | -0,3353 | 6,17491E-11 |
| ACE2 | HLA-K | -0,3354 | 6,13665E-11 |
| ACE2 | RP11-732M18.2 | -0,3354 | 6,07607E-11 |
| ACE2 | AC005281.1 | -0,3354 | 6,12494E-11 |
| ACE2 | RP11-366L20.4 | -0,3354 | 6,12523E-11 |
| ACE2 | CGB3 | -0,3354 | 6,11836E-11 |
| ACE2 | SLC22A18AS | -0,3355 | 5,99599E-11 |
| ACE2 | NAP1L4 | -0,3355 | 6,07149E-11 |

| | | | |
|---|---|---|---|
| ACE2 | IL27 | -0,3355 | 6,07064E-11 |
| ACE2 | OR2AK2 | -0,3356 | 5,97316E-11 |
| ACE2 | SNORD54 | -0,3356 | 5,94789E-11 |
| ACE2 | CTC-508F8.1 | -0,3356 | 5,91034E-11 |
| ACE2 | RP11-392B6.1 | -0,3357 | 5,88217E-11 |
| ACE2 | CXorf58 | -0,3357 | 5,84155E-11 |
| ACE2 | AC007040.8 | -0,3358 | 5,76074E-11 |
| ACE2 | PTP4A1 | -0,3358 | 5,79283E-11 |
| ACE2 | RP5-937E21.8 | -0,3358 | 5,76293E-11 |
| ACE2 | RP11-395L14.13 | -0,3359 | 5,68524E-11 |
| ACE2 | AC009961.3 | -0,3359 | 5,7159E-11 |
| ACE2 | RP4-625H18.2 | -0,3359 | 5,7026E-11 |
| ACE2 | SOHLH2 | -0,3359 | 5,68242E-11 |
| ACE2 | ZG16B | -0,3359 | 5,67813E-11 |
| ACE2 | CTD-3193O13.8 | -0,3359 | 5,66916E-11 |
| ACE2 | RPL36AP45 | -0,336 | 5,59259E-11 |
| ACE2 | RP11-69I13.1 | -0,3361 | 5,53402E-11 |
| ACE2 | OR4E1 | -0,3361 | 5,55832E-11 |
| ACE2 | RP11-586K12.11 | -0,3361 | 5,57701E-11 |
| ACE2 | AC019185.4 | -0,3362 | 5,50385E-11 |
| ACE2 | RP11-300A12.2 | -0,3362 | 5,48799E-11 |
| ACE2 | ZPBP2 | -0,3362 | 5,43744E-11 |
| ACE2 | THOC5 | -0,3363 | 5,3689E-11 |
| ACE2 | RP11-335L23.4 | -0,3364 | 5,32504E-11 |
| ACE2 | LINC02105 | -0,3364 | 5,32752E-11 |
| ACE2 | RP11-204E9.3 | -0,3364 | 5,32773E-11 |
| ACE2 | RP11-444D13.1 | -0,3365 | 5,22897E-11 |
| ACE2 | C1orf54 | -0,3366 | 5,18076E-11 |
| ACE2 | AC108039.3 | -0,3366 | 5,17087E-11 |
| ACE2 | DNAH7 | -0,3366 | 5,20412E-11 |
| ACE2 | CNDP1 | -0,3366 | 5,18526E-11 |
| ACE2 | RP4-671G15.2 | -0,3367 | 5,12655E-11 |
| ACE2 | RP11-462B18.1 | -0,3367 | 5,0818E-11 |
| ACE2 | CTAGE16P | -0,3367 | 5,09631E-11 |
| ACE2 | CEP131 | -0,3367 | 5,09356E-11 |
| ACE2 | RNU6-1165P | -0,3368 | 5,04567E-11 |
| ACE2 | SLN | -0,3368 | 5,02139E-11 |
| ACE2 | CYB5D2 | -0,3368 | 5,0388E-11 |
| ACE2 | OR9Q2 | -0,3369 | 4,96661E-11 |
| ACE2 | RNU1-11P | -0,337 | 4,90793E-11 |
| ACE2 | RP11-488I20.9 | -0,337 | 4,86597E-11 |
| ACE2 | FKBP1AP1 | -0,337 | 4,90726E-11 |
| ACE2 | LINC01907 | -0,3371 | 4,85248E-11 |
| ACE2 | RP11-111F5.2 | -0,3371 | 4,80849E-11 |
| ACE2 | HOGA1 | -0,3371 | 4,82097E-11 |
| ACE2 | RP11-64C12.6 | -0,3371 | 4,84546E-11 |
| ACE2 | LARP6 | -0,3372 | 4,78047E-11 |
| ACE2 | RP11-112L7.1 | -0,3373 | 4,69032E-11 |
| ACE2 | FAM166AP6 | -0,3374 | 4,65551E-11 |
| ACE2 | RIOK3 | -0,3374 | 4,64526E-11 |
| ACE2 | RP11-195B3.1 | -0,3375 | 4,58456E-11 |
| ACE2 | SLC25A3P1 | -0,3376 | 4,48638E-11 |
| ACE2 | PSMD6-AS1 | -0,3376 | 4,51273E-11 |
| ACE2 | CTD-2532K18.1 | -0,3376 | 4,48922E-11 |

| | | | |
|---|---|---|---|
| ACE2 | RP11-191A15.2 | -0,3376 | 4,4965E-11 |
| ACE2 | RP1-166H1.2 | -0,3376 | 4,52569E-11 |
| ACE2 | RP11-676B18.2 | -0,3376 | 4,50337E-11 |
| ACE2 | RP11-160N1.8 | -0,3377 | 4,41673E-11 |
| ACE2 | HSPE1P4 | -0,3377 | 4,41722E-11 |
| ACE2 | BVES | -0,3378 | 4,3856E-11 |
| ACE2 | RP11-36D19.8 | -0,3378 | 4,36259E-11 |
| ACE2 | TPTE2P6 | -0,3378 | 4,35108E-11 |
| ACE2 | ACTR5 | -0,3378 | 4,35637E-11 |
| ACE2 | SVOPL | -0,3379 | 4,29806E-11 |
| ACE2 | RP11-419K12.1 | -0,3379 | 4,31251E-11 |
| ACE2 | SNAPC4 | -0,3379 | 4,31298E-11 |
| ACE2 | CAPS2 | -0,3379 | 4,29463E-11 |
| ACE2 | BAIAP3 | -0,3379 | 4,28777E-11 |
| ACE2 | ZBTB3 | -0,338 | 4,23272E-11 |
| ACE2 | ATP4A | -0,338 | 4,27344E-11 |
| ACE2 | RP4-639J15.3 | -0,3381 | 4,19966E-11 |
| ACE2 | AC130464.1 | -0,3381 | 4,20306E-11 |
| ACE2 | RP1-72A23.1 | -0,3382 | 4,15894E-11 |
| ACE2 | ZC3H18 | -0,3382 | 4,15562E-11 |
| ACE2 | TTC30A | -0,3383 | 4,06622E-11 |
| ACE2 | RAD21L1 | -0,3383 | 4,06197E-11 |
| ACE2 | CDC45 | -0,3383 | 4,07107E-11 |
| ACE2 | KMT2E-AS1 | -0,3384 | 4,01587E-11 |
| ACE2 | KB-1458E12.1 | -0,3384 | 4,0532E-11 |
| ACE2 | GOLGA8O | -0,3384 | 4,00661E-11 |
| ACE2 | GTSCR1 | -0,3385 | 3,95108E-11 |
| ACE2 | LIX1-AS1 | -0,3386 | 3,93551E-11 |
| ACE2 | STRBP | -0,3386 | 3,89073E-11 |
| ACE2 | MS4A10 | -0,3386 | 3,91398E-11 |
| ACE2 | RP11-180M15.4 | -0,3386 | 3,90514E-11 |
| ACE2 | MIR8078 | -0,3386 | 3,91546E-11 |
| ACE2 | GP1BA | -0,3387 | 3,87613E-11 |
| ACE2 | STX18-IT1 | -0,3388 | 3,80231E-11 |
| ACE2 | P4HA2-AS1 | -0,3389 | 3,73692E-11 |
| ACE2 | RP11-534L6.5 | -0,3389 | 3,73823E-11 |
| ACE2 | RP11-650K20.2 | -0,3389 | 3,7478E-11 |
| ACE2 | KIF1C | -0,3389 | 3,74858E-11 |
| ACE2 | LINC00469 | -0,3389 | 3,74184E-11 |
| ACE2 | ALPK1 | -0,339 | 3,71706E-11 |
| ACE2 | ZEB1 | -0,339 | 3,70945E-11 |
| ACE2 | RPL10P13 | -0,339 | 3,72684E-11 |
| ACE2 | EEF1DP3 | -0,339 | 3,69985E-11 |
| ACE2 | AC006116.12 | -0,339 | 3,70476E-11 |
| ACE2 | SLC4A11 | -0,3391 | 3,62724E-11 |
| ACE2 | TPP2 | -0,3392 | 3,58843E-11 |
| ACE2 | ZNF19 | -0,3392 | 3,57821E-11 |
| ACE2 | RCAN1 | -0,3392 | 3,60349E-11 |
| ACE2 | ACTRT1 | -0,3392 | 3,60483E-11 |
| ACE2 | VIP | -0,3393 | 3,56774E-11 |
| ACE2 | RP11-881M11.8 | -0,3393 | 3,54054E-11 |
| ACE2 | LINC01912 | -0,3393 | 3,56785E-11 |
| ACE2 | CRYGD | -0,3394 | 3,49358E-11 |
| ACE2 | POLH | -0,3394 | 3,50358E-11 |

| | | | |
|---|---|---|---|
| ACE2 | MTCO3P24 | -0,3394 | 3,4926E-11 |
| ACE2 | GALNT3 | -0,3395 | 3,42915E-11 |
| ACE2 | TRBV7-9 | -0,3395 | 3,45192E-11 |
| ACE2 | ANO5 | -0,3395 | 3,43646E-11 |
| ACE2 | AP000462.3 | -0,3395 | 3,43113E-11 |
| ACE2 | FAM110A | -0,3395 | 3,46269E-11 |
| ACE2 | LINC01954 | -0,3396 | 3,39959E-11 |
| ACE2 | RP11-142L1.1 | -0,3397 | 3,36328E-11 |
| ACE2 | RP11-714G18.1 | -0,3398 | 3,28978E-11 |
| ACE2 | RP11-5N11.2 | -0,3398 | 3,30673E-11 |
| ACE2 | RP11-164H13.1 | -0,3398 | 3,32354E-11 |
| ACE2 | MECR | -0,3399 | 3,26175E-11 |
| ACE2 | RELN | -0,3399 | 3,24263E-11 |
| ACE2 | GLULP5 | -0,3399 | 3,26644E-11 |
| ACE2 | CTD-2046I8.1 | -0,34 | 3,19957E-11 |
| ACE2 | TRIM36-IT1 | -0,34 | 3,20768E-11 |
| ACE2 | BCRP5 | -0,34 | 3,23475E-11 |
| ACE2 | IGDCC3 | -0,3401 | 3,18823E-11 |
| ACE2 | RP11-276H19.4 | -0,3402 | 3,12505E-11 |
| ACE2 | CTD-2302E22.4 | -0,3402 | 3,1195E-11 |
| ACE2 | CTD-2014B16.1 | -0,3402 | 3,12211E-11 |
| ACE2 | PITX1 | -0,3403 | 3,08843E-11 |
| ACE2 | CTC-359M8.3 | -0,3403 | 3,08516E-11 |
| ACE2 | CH507-42P11.7 | -0,3403 | 3,06663E-11 |
| ACE2 | AC007161.5 | -0,3404 | 3,05139E-11 |
| ACE2 | COX20 | -0,3405 | 3,01664E-11 |
| ACE2 | NPM1P38 | -0,3405 | 2,98249E-11 |
| ACE2 | RP11-320N21.2 | -0,3405 | 2,98939E-11 |
| ACE2 | OR7E111P | -0,3405 | 2,98637E-11 |
| ACE2 | RP11-255P5.3 | -0,3405 | 2,98201E-11 |
| ACE2 | MTG1 | -0,3406 | 2,96862E-11 |
| ACE2 | RP11-98J9.2 | -0,3406 | 2,97216E-11 |
| ACE2 | FAM76B | -0,3407 | 2,91996E-11 |
| ACE2 | CTD-2354A18.1 | -0,3407 | 2,9146E-11 |
| ACE2 | KRT18P32 | -0,3408 | 2,88217E-11 |
| ACE2 | NIM1K | -0,3408 | 2,8682E-11 |
| ACE2 | RP11-570H19.2 | -0,3408 | 2,86123E-11 |
| ACE2 | RP11-541G9.1 | -0,3408 | 2,87074E-11 |
| ACE2 | TMEM156 | -0,3409 | 2,84867E-11 |
| ACE2 | LINC00707 | -0,3409 | 2,84566E-11 |
| ACE2 | RP11-1099M24.6 | -0,3409 | 2,82402E-11 |
| ACE2 | DBF4B | -0,3409 | 2,8353E-11 |
| ACE2 | AC073218.1 | -0,341 | 2,784E-11 |
| ACE2 | RP11-696F12.1 | -0,341 | 2,79777E-11 |
| ACE2 | RP11-1084A12.2 | -0,341 | 2,80649E-11 |
| ACE2 | CTD-2583P5.1 | -0,341 | 2,80341E-11 |
| ACE2 | THUMPD3 | -0,3411 | 2,74427E-11 |
| ACE2 | RP11-196O2.1 | -0,3411 | 2,75495E-11 |
| ACE2 | MRPL11 | -0,3411 | 2,75571E-11 |
| ACE2 | CHRM5 | -0,3411 | 2,7505E-11 |
| ACE2 | DNAH3 | -0,3411 | 2,7495E-11 |
| ACE2 | MRPL39 | -0,3411 | 2,73999E-11 |
| ACE2 | MIR219A2 | -0,3412 | 2,71689E-11 |
| ACE2 | BEND7 | -0,3412 | 2,71398E-11 |

| | | | |
|---|---|---|---|
| ACE2 | ART5 | -0,3412 | 2,70262E-11 |
| ACE2 | RP11-718O11.1 | -0,3412 | 2,72772E-11 |
| ACE2 | IGFLR1 | -0,3412 | 2,7064E-11 |
| ACE2 | MKKS | -0,3412 | 2,71616E-11 |
| ACE2 | CTB-133G6.1 | -0,3413 | 2,68316E-11 |
| ACE2 | LL22NC03-13G6.2 | -0,3413 | 2,67646E-11 |
| ACE2 | AC010127.3 | -0,3414 | 2,64881E-11 |
| ACE2 | RP11-43F13.1 | -0,3414 | 2,63827E-11 |
| ACE2 | RP11-72L22.1 | -0,3414 | 2,64372E-11 |
| ACE2 | RP11-14K2.1 | -0,3414 | 2,64303E-11 |
| ACE2 | RP5-827L5.2 | -0,3414 | 2,65348E-11 |
| ACE2 | PKNOX1 | -0,3414 | 2,63572E-11 |
| ACE2 | HYI-AS1 | -0,3416 | 2,55977E-11 |
| ACE2 | RP11-356C4.3 | -0,3416 | 2,55875E-11 |
| ACE2 | ALDOAP1 | -0,3417 | 2,52089E-11 |
| ACE2 | XXbac-BPG252P9.10 | -0,3417 | 2,51693E-11 |
| ACE2 | GOLGA8S | -0,3417 | 2,54626E-11 |
| ACE2 | AP3B2 | -0,3417 | 2,53736E-11 |
| ACE2 | ADGRV1 | -0,3418 | 2,47696E-11 |
| ACE2 | KRT75 | -0,3418 | 2,4938E-11 |
| ACE2 | CTC-264K15.6 | -0,3418 | 2,50991E-11 |
| ACE2 | RPL21P129 | -0,3418 | 2,49178E-11 |
| ACE2 | RP11-492A10.1 | -0,3419 | 2,44725E-11 |
| ACE2 | LCN2 | -0,3419 | 2,45544E-11 |
| ACE2 | CNTROB | -0,3419 | 2,45902E-11 |
| ACE2 | SAGE4P | -0,3419 | 2,47454E-11 |
| ACE2 | ZNF746 | -0,342 | 2,43376E-11 |
| ACE2 | CERS3 | -0,342 | 2,42716E-11 |
| ACE2 | RP11-484D4.3 | -0,3421 | 2,38966E-11 |
| ACE2 | RP11-164C12.1 | -0,3421 | 2,3804E-11 |
| ACE2 | RP11-91K8.5 | -0,3422 | 2,36212E-11 |
| ACE2 | ADAM5 | -0,3422 | 2,35281E-11 |
| ACE2 | AQP12B | -0,3423 | 2,328E-11 |
| ACE2 | RP11-351O2.2 | -0,3423 | 2,32591E-11 |
| ACE2 | RP11-51F16.1 | -0,3423 | 2,32132E-11 |
| ACE2 | AC104623.2 | -0,3424 | 2,27609E-11 |
| ACE2 | AC092570.3 | -0,3424 | 2,27684E-11 |
| ACE2 | OR2N1P | -0,3424 | 2,29337E-11 |
| ACE2 | WDR82P2 | -0,3425 | 2,25027E-11 |
| ACE2 | RP11-364B6.2 | -0,3426 | 2,24055E-11 |
| ACE2 | RP11-347I19.7 | -0,3426 | 2,22829E-11 |
| ACE2 | RP11-889D3.1 | -0,3428 | 2,15792E-11 |
| ACE2 | RP3-402G11.26 | -0,3428 | 2,17807E-11 |
| ACE2 | SPANXA2 | -0,3428 | 2,17881E-11 |
| ACE2 | RP13-436F16.1 | -0,3429 | 2,1422E-11 |
| ACE2 | AC079163.1 | -0,3429 | 2,13287E-11 |
| ACE2 | DUSP28 | -0,3429 | 2,14534E-11 |
| ACE2 | LINC00491 | -0,3429 | 2,1288E-11 |
| ACE2 | MRRFP1 | -0,3429 | 2,13965E-11 |
| ACE2 | GIGYF2 | -0,343 | 2,09266E-11 |
| ACE2 | LINC00293 | -0,343 | 2,09014E-11 |
| ACE2 | RP11-599B13.9 | -0,343 | 2,11653E-11 |
| ACE2 | SLC9A4 | -0,3431 | 2,06969E-11 |
| ACE2 | RP11-731K22.1 | -0,3431 | 2,08478E-11 |

| | | | |
|---|---|---|---|
| ACE2 | PENK | -0,3431 | 2,07286E-11 |
| ACE2 | RP11-885L14.1 | -0,3431 | 2,05914E-11 |
| ACE2 | CCDC144CP | -0,3431 | 2,07154E-11 |
| ACE2 | SMIM19 | -0,3432 | 2,03166E-11 |
| ACE2 | RP4-616B8.5 | -0,3433 | 2,00567E-11 |
| ACE2 | CSF2RBP1 | -0,3433 | 2,0134E-11 |
| ACE2 | RP1-171K16.5 | -0,3433 | 2,0201E-11 |
| ACE2 | CTB-113D17.1 | -0,3434 | 1,99562E-11 |
| ACE2 | RP3-412A9.15 | -0,3434 | 1,9774E-11 |
| ACE2 | LINC01675 | -0,3435 | 1,96141E-11 |
| ACE2 | RP11-269C4.2 | -0,3435 | 1,96855E-11 |
| ACE2 | RP11-403B2.7 | -0,3435 | 1,96449E-11 |
| ACE2 | DDX53 | -0,3435 | 1,95992E-11 |
| ACE2 | RP11-756G20.1 | -0,3436 | 1,91811E-11 |
| ACE2 | BCRP2 | -0,3436 | 1,93177E-11 |
| ACE2 | MKRN1 | -0,3437 | 1,89303E-11 |
| ACE2 | FXYD2 | -0,3437 | 1,91427E-11 |
| ACE2 | AC005522.6 | -0,3438 | 1,86897E-11 |
| ACE2 | LINC01518 | -0,3438 | 1,86526E-11 |
| ACE2 | IQCK | -0,3438 | 1,86362E-11 |
| ACE2 | RP11-98G7.1 | -0,3439 | 1,85162E-11 |
| ACE2 | KY | -0,3439 | 1,85179E-11 |
| ACE2 | LINC01483 | -0,3439 | 1,86224E-11 |
| ACE2 | PTCD2P2 | -0,344 | 1,82423E-11 |
| ACE2 | VPS35P1 | -0,3441 | 1,79436E-11 |
| ACE2 | VN1R35P | -0,3442 | 1,76966E-11 |
| ACE2 | PRB2 | -0,3442 | 1,76374E-11 |
| ACE2 | RP11-714M23.2 | -0,3442 | 1,76523E-11 |
| ACE2 | DIO1 | -0,3443 | 1,74059E-11 |
| ACE2 | RP4-814D15.1 | -0,3443 | 1,73457E-11 |
| ACE2 | RP11-687D19.1 | -0,3443 | 1,73772E-11 |
| ACE2 | MECOM | -0,3444 | 1,72E-11 |
| ACE2 | HOXB-AS2 | -0,3444 | 1,72282E-11 |
| ACE2 | ZNF285B | -0,3444 | 1,73317E-11 |
| ACE2 | LETM1P2 | -0,3444 | 1,72353E-11 |
| ACE2 | TRIM31 | -0,3445 | 1,7038E-11 |
| ACE2 | RP11-624D11.2 | -0,3445 | 1,68949E-11 |
| ACE2 | CTD-2308G16.1 | -0,3445 | 1,68947E-11 |
| ACE2 | RXRG | -0,3446 | 1,68345E-11 |
| ACE2 | GNA14-AS1 | -0,3446 | 1,66997E-11 |
| ACE2 | CTD-2021H9.2 | -0,3446 | 1,67422E-11 |
| ACE2 | SETP9 | -0,3447 | 1,65171E-11 |
| ACE2 | RP11-360P21.2 | -0,3447 | 1,64672E-11 |
| ACE2 | RP1-200K18.1 | -0,3447 | 1,66109E-11 |
| ACE2 | AP005273.1 | -0,3447 | 1,6547E-11 |
| ACE2 | SEC14L3 | -0,3447 | 1,6611E-11 |
| ACE2 | RP11-355B11.2 | -0,3448 | 1,61778E-11 |
| ACE2 | TSLP | -0,3448 | 1,62554E-11 |
| ACE2 | RP11-734K23.9 | -0,3449 | 1,5983E-11 |
| ACE2 | OR2F1 | -0,3449 | 1,60331E-11 |
| ACE2 | BARX1-AS1 | -0,3449 | 1,60057E-11 |
| ACE2 | CTC-261N6.1 | -0,345 | 1,57385E-11 |
| ACE2 | RP11-953B20.2 | -0,345 | 1,56939E-11 |
| ACE2 | RP11-108K3.2 | -0,345 | 1,58479E-11 |

| | | | |
|---|---|---|---|
| ACE2 | AC084149.2 | -0,3451 | 1,56655E-11 |
| ACE2 | ABHD16A | -0,3451 | 1,55194E-11 |
| ACE2 | RP11-350J20.13 | -0,3451 | 1,56062E-11 |
| ACE2 | TIMM23 | -0,3451 | 1,55114E-11 |
| ACE2 | CTSLP4 | -0,3451 | 1,54792E-11 |
| ACE2 | RP11-1028N23.2 | -0,3451 | 1,55993E-11 |
| ACE2 | CYP2A13 | -0,3451 | 1,56042E-11 |
| ACE2 | PLEKHA3P1 | -0,3451 | 1,56382E-11 |
| ACE2 | RPL10P5 | -0,3452 | 1,53795E-11 |
| ACE2 | RP11-372H2.1 | -0,3452 | 1,53403E-11 |
| ACE2 | LINC01097 | -0,3452 | 1,54663E-11 |
| ACE2 | RP11-556O9.2 | -0,3452 | 1,5278E-11 |
| ACE2 | TYMS | -0,3452 | 1,5315E-11 |
| ACE2 | MCOLN2 | -0,3453 | 1,50504E-11 |
| ACE2 | HRG | -0,3453 | 1,51986E-11 |
| ACE2 | RP11-110H1.9 | -0,3453 | 1,50554E-11 |
| ACE2 | LINC01663 | -0,3453 | 1,51832E-11 |
| ACE2 | RP11-740P5.3 | -0,3454 | 1,49729E-11 |
| ACE2 | LINC01935 | -0,3454 | 1,48437E-11 |
| ACE2 | CEP55 | -0,3454 | 1,50144E-11 |
| ACE2 | RP11-804F13.1 | -0,3454 | 1,49042E-11 |
| ACE2 | RP11-282O18.3 | -0,3454 | 1,49673E-11 |
| ACE2 | RPL5P2 | -0,3454 | 1,49864E-11 |
| ACE2 | RP1-128O3.6 | -0,3455 | 1,47872E-11 |
| ACE2 | RN7SL51P | -0,3456 | 1,4485E-11 |
| ACE2 | RP11-1149M10.2 | -0,3456 | 1,45342E-11 |
| ACE2 | TMEM126A | -0,3456 | 1,44056E-11 |
| ACE2 | COX6B1 | -0,3456 | 1,45831E-11 |
| ACE2 | TGM3 | -0,3456 | 1,44102E-11 |
| ACE2 | XIRP2 | -0,3457 | 1,42212E-11 |
| ACE2 | SLAIN2 | -0,3457 | 1,42624E-11 |
| ACE2 | RP11-793B23.1 | -0,3457 | 1,42131E-11 |
| ACE2 | KIAA0408 | -0,3457 | 1,43221E-11 |
| ACE2 | AHCY | -0,3457 | 1,43741E-11 |
| ACE2 | DNAJC22 | -0,3458 | 1,4123E-11 |
| ACE2 | CHIAP3 | -0,3459 | 1,38906E-11 |
| ACE2 | RP11-533E19.7 | -0,3459 | 1,38446E-11 |
| ACE2 | ID2-AS1 | -0,3459 | 1,38359E-11 |
| ACE2 | GABRA4 | -0,3459 | 1,39798E-11 |
| ACE2 | RP11-138A9.1 | -0,3459 | 1,38599E-11 |
| ACE2 | ST13P13 | -0,3459 | 1,3899E-11 |
| ACE2 | SLC10A1 | -0,3459 | 1,38125E-11 |
| ACE2 | RP1-32I10.10 | -0,3459 | 1,39184E-11 |
| ACE2 | RP11-293A21.1 | -0,346 | 1,37015E-11 |
| ACE2 | G6PC | -0,346 | 1,36805E-11 |
| ACE2 | AC096574.4 | -0,3461 | 1,3421E-11 |
| ACE2 | PANX3 | -0,3461 | 1,34848E-11 |
| ACE2 | RP11-61O1.1 | -0,3461 | 1,34957E-11 |
| ACE2 | RP11-62C3.6 | -0,3462 | 1,32186E-11 |
| ACE2 | RPL27P12 | -0,3462 | 1,32888E-11 |
| ACE2 | UBL3 | -0,3462 | 1,32845E-11 |
| ACE2 | POM121L3P | -0,3462 | 1,32661E-11 |
| ACE2 | IQCC | -0,3463 | 1,30301E-11 |
| ACE2 | RP11-34F13.3 | -0,3463 | 1,31428E-11 |

| | | | |
|---|---|---|---|
| ACE2 | RP11-465I4.3 | -0,3463 | 1,31897E-11 |
| ACE2 | ACTBL2 | -0,3464 | 1,29671E-11 |
| ACE2 | AP006216.10 | -0,3464 | 1,29248E-11 |
| ACE2 | CTD-3099C6.9 | -0,3464 | 1,28387E-11 |
| ACE2 | CUL3 | -0,3465 | 1,28205E-11 |
| ACE2 | ZDHHC11 | -0,3465 | 1,2737E-11 |
| ACE2 | OR8G3P | -0,3465 | 1,28184E-11 |
| ACE2 | CTD-2055G21.1 | -0,3465 | 1,26676E-11 |
| ACE2 | TCF15 | -0,3465 | 1,28024E-11 |
| ACE2 | RP11-29A19.2 | -0,3466 | 1,25463E-11 |
| ACE2 | AIFM2 | -0,3466 | 1,25935E-11 |
| ACE2 | RP11-20D14.4 | -0,3466 | 1,25725E-11 |
| ACE2 | SLC26A4-AS1 | -0,3467 | 1,22895E-11 |
| ACE2 | KCNG2 | -0,3467 | 1,24637E-11 |
| ACE2 | KRT80 | -0,3468 | 1,21165E-11 |
| ACE2 | AC002117.1 | -0,3468 | 1,22645E-11 |
| ACE2 | RP11-64C12.4 | -0,3468 | 1,2214E-11 |
| ACE2 | AC087430.1 | -0,3469 | 1,20548E-11 |
| ACE2 | RP11-102M11.2 | -0,3469 | 1,20126E-11 |
| ACE2 | AC009014.3 | -0,3469 | 1,2043E-11 |
| ACE2 | LINC00282 | -0,3471 | 1,17351E-11 |
| ACE2 | RP5-881L22.5 | -0,3471 | 1,17491E-11 |
| ACE2 | CRACR2B | -0,3472 | 1,15423E-11 |
| ACE2 | CACNA1C-IT1 | -0,3472 | 1,14576E-11 |
| ACE2 | MRPS7 | -0,3472 | 1,14633E-11 |
| ACE2 | RP11-677O4.4 | -0,3472 | 1,15472E-11 |
| ACE2 | MMP9 | -0,3472 | 1,14975E-11 |
| ACE2 | RP11-109P6.2 | -0,3473 | 1,1423E-11 |
| ACE2 | CTD-2058B24.3 | -0,3473 | 1,129E-11 |
| ACE2 | GOLGA8R | -0,3473 | 1,12861E-11 |
| ACE2 | RP11-127I20.8 | -0,3473 | 1,13765E-11 |
| ACE2 | SIM2 | -0,3473 | 1,13563E-11 |
| ACE2 | SNU13 | -0,3473 | 1,13293E-11 |
| ACE2 | NEUROG1 | -0,3474 | 1,12636E-11 |
| ACE2 | ASB2 | -0,3474 | 1,12046E-11 |
| ACE2 | AC006116.14 | -0,3474 | 1,11618E-11 |
| ACE2 | RP11-351N4.4 | -0,3475 | 1,09727E-11 |
| ACE2 | RP11-775C24.5 | -0,3475 | 1,09741E-11 |
| ACE2 | CTD-2525I3.8 | -0,3475 | 1,10567E-11 |
| ACE2 | FAM189B | -0,3476 | 1,08732E-11 |
| ACE2 | RP11-295K2.3 | -0,3476 | 1,08157E-11 |
| ACE2 | CD83P1 | -0,3476 | 1,08852E-11 |
| ACE2 | MRPS17 | -0,3476 | 1,08695E-11 |
| ACE2 | RP11-589C21.1 | -0,3476 | 1,08302E-11 |
| ACE2 | ZNF385C | -0,3476 | 1,08593E-11 |
| ACE2 | RPS27AP13 | -0,3477 | 1,06784E-11 |
| ACE2 | CTD-2544M6.1 | -0,3477 | 1,07604E-11 |
| ACE2 | HNRNPA1P11 | -0,3477 | 1,0783E-11 |
| ACE2 | FRG2JP | -0,3478 | 1,06023E-11 |
| ACE2 | EIF4ENIF1 | -0,3478 | 1,05192E-11 |
| ACE2 | LINC01566 | -0,3479 | 1,03986E-11 |
| ACE2 | RP11-53L24.1 | -0,3479 | 1,03388E-11 |
| ACE2 | RP4-568F9.3 | -0,3479 | 1,04453E-11 |
| ACE2 | PABPC1L2A | -0,3479 | 1,04091E-11 |

| | | | |
|---|---|---|---|
| ACE2 | CCDC148-AS1 | -0,348 | 1,02961E-11 |
| ACE2 | ANKAR | -0,348 | 1,01829E-11 |
| ACE2 | RETNLB | -0,348 | 1,0323E-11 |
| ACE2 | CRAT37 | -0,348 | 1,0215E-11 |
| ACE2 | TBC1D3G | -0,348 | 1,02114E-11 |
| ACE2 | AC005008.3 | -0,3481 | 1,01693E-11 |
| ACE2 | RP11-310E22.6 | -0,3481 | 1,00652E-11 |
| ACE2 | RP11-432I5.8 | -0,3481 | 1,01041E-11 |
| ACE2 | LZIC | -0,3482 | 9,93049E-12 |
| ACE2 | FNDC5 | -0,3482 | 9,99553E-12 |
| ACE2 | AC007312.3 | -0,3482 | 1,00145E-11 |
| ACE2 | KRT8P10 | -0,3483 | 9,76251E-12 |
| ACE2 | FAM184A | -0,3483 | 9,74877E-12 |
| ACE2 | TBC1D3B | -0,3483 | 9,78201E-12 |
| ACE2 | DCTN1-AS1 | -0,3484 | 9,64184E-12 |
| ACE2 | ZIM3 | -0,3484 | 9,67564E-12 |
| ACE2 | AC023469.1 | -0,3485 | 9,58891E-12 |
| ACE2 | IMMP1L | -0,3485 | 9,51261E-12 |
| ACE2 | RP11-314D7.1 | -0,3485 | 9,5339E-12 |
| ACE2 | FLJ30679 | -0,3485 | 9,56395E-12 |
| ACE2 | CCKAR | -0,3486 | 9,38568E-12 |
| ACE2 | RP11-323I15.2 | -0,3486 | 9,42372E-12 |
| ACE2 | AC073987.1 | -0,3487 | 9,20944E-12 |
| ACE2 | UPP2 | -0,3487 | 9,31228E-12 |
| ACE2 | OR1D2 | -0,3487 | 9,2047E-12 |
| ACE2 | RP3-483K16.4 | -0,3488 | 9,17983E-12 |
| ACE2 | GUSB | -0,3488 | 9,16973E-12 |
| ACE2 | RP11-18H21.2 | -0,3489 | 8,94784E-12 |
| ACE2 | RPL31P44 | -0,3489 | 9,03313E-12 |
| ACE2 | AC091814.2 | -0,3489 | 9,03509E-12 |
| ACE2 | AP002856.7 | -0,349 | 8,87538E-12 |
| ACE2 | LINC01203 | -0,349 | 8,89474E-12 |
| ACE2 | RP11-459F6.3 | -0,3491 | 8,75569E-12 |
| ACE2 | GALE | -0,3492 | 8,60746E-12 |
| ACE2 | LINC01137 | -0,3492 | 8,58764E-12 |
| ACE2 | LINC00570 | -0,3492 | 8,56596E-12 |
| ACE2 | RP11-989E6.13 | -0,3492 | 8,66766E-12 |
| ACE2 | RP11-61J19.2 | -0,3493 | 8,43104E-12 |
| ACE2 | DMWD | -0,3493 | 8,45243E-12 |
| ACE2 | RP11-3L21.2 | -0,3495 | 8,23765E-12 |
| ACE2 | CHRNA10 | -0,3495 | 8,29712E-12 |
| ACE2 | RBFOX1 | -0,3495 | 8,2658E-12 |
| ACE2 | RP11-740C1.2 | -0,3496 | 8,11669E-12 |
| ACE2 | IMPDH1P5 | -0,3496 | 8,16822E-12 |
| ACE2 | RP11-438P9.2 | -0,3497 | 8,04212E-12 |
| ACE2 | PHKBP2 | -0,3497 | 8,00756E-12 |
| ACE2 | TEX2 | -0,3497 | 8,00668E-12 |
| ACE2 | SLA2 | -0,3497 | 8,04043E-12 |
| ACE2 | AP000688.14 | -0,3497 | 8,02502E-12 |
| ACE2 | PAF1 | -0,3498 | 7,93137E-12 |
| ACE2 | AC006116.15 | -0,3498 | 7,94245E-12 |
| ACE2 | RP11-178L8.1 | -0,3499 | 7,77168E-12 |
| ACE2 | PCDHB5 | -0,35 | 7,69774E-12 |
| ACE2 | ALG1 | -0,35 | 7,63799E-12 |

| | | | |
|---|---|---|---|
| ACE2 | TPTE2P4 | -0,35 | 7,70904E-12 |
| ACE2 | RP11-101K23.1 | -0,3501 | 7,52916E-12 |
| ACE2 | ZSCAN5A | -0,3501 | 7,54179E-12 |
| ACE2 | GARSP1 | -0,3502 | 7,45458E-12 |
| ACE2 | FBXO15 | -0,3502 | 7,39843E-12 |
| ACE2 | RP11-278H7.3 | -0,3503 | 7,3569E-12 |
| ACE2 | OR1F12 | -0,3503 | 7,35294E-12 |
| ACE2 | RP11-368P15.1 | -0,3503 | 7,30751E-12 |
| ACE2 | RP11-78A19.2 | -0,3503 | 7,29065E-12 |
| ACE2 | HNRNPFP1 | -0,3504 | 7,1919E-12 |
| ACE2 | RP11-342F21.1 | -0,3504 | 7,24192E-12 |
| ACE2 | PRSS45 | -0,3505 | 7,15244E-12 |
| ACE2 | RER1 | -0,3506 | 7,07E-12 |
| ACE2 | RP11-552E20.4 | -0,3506 | 7,01933E-12 |
| ACE2 | RP5-986I17.2 | -0,3506 | 7,00592E-12 |
| ACE2 | COA4 | -0,3507 | 6,89531E-12 |
| ACE2 | RP11-592N21.2 | -0,3507 | 6,96251E-12 |
| ACE2 | RP11-31I22.3 | -0,3507 | 6,95748E-12 |
| ACE2 | NAALADL2 | -0,3508 | 6,82622E-12 |
| ACE2 | KARSP3 | -0,3509 | 6,71824E-12 |
| ACE2 | MTND4P29 | -0,351 | 6,64467E-12 |
| ACE2 | THOC1 | -0,351 | 6,58914E-12 |
| ACE2 | CDC42EP3 | -0,3511 | 6,4875E-12 |
| ACE2 | HSD17B2 | -0,3511 | 6,49251E-12 |
| ACE2 | CLPP | -0,3511 | 6,54451E-12 |
| ACE2 | CNNM4 | -0,3512 | 6,41901E-12 |
| ACE2 | DNAJC10 | -0,3512 | 6,43881E-12 |
| ACE2 | PACRG-AS3 | -0,3512 | 6,41582E-12 |
| ACE2 | KRTAP13-2 | -0,3512 | 6,42418E-12 |
| ACE2 | RP11-168E17.1 | -0,3513 | 6,2941E-12 |
| ACE2 | RP11-83J21.3 | -0,3513 | 6,36387E-12 |
| ACE2 | FAM215A | -0,3513 | 6,32828E-12 |
| ACE2 | OR4D1 | -0,3513 | 6,30425E-12 |
| ACE2 | RP11-651P23.4 | -0,3514 | 6,20543E-12 |
| ACE2 | RP11-526P5.1 | -0,3514 | 6,22217E-12 |
| ACE2 | NAA20 | -0,3514 | 6,26207E-12 |
| ACE2 | CA15P1 | -0,3514 | 6,28835E-12 |
| ACE2 | CPSF1P1 | -0,3514 | 6,26885E-12 |
| ACE2 | CD109 | -0,3516 | 6,10379E-12 |
| ACE2 | AC006373.1 | -0,3516 | 6,09034E-12 |
| ACE2 | MRPS33 | -0,3516 | 6,02826E-12 |
| ACE2 | RP11-39M21.2 | -0,3516 | 6,0974E-12 |
| ACE2 | KRT18P18 | -0,3516 | 6,06688E-12 |
| ACE2 | PPP1R12BP1 | -0,3516 | 6,07554E-12 |
| ACE2 | PCBP4 | -0,3517 | 5,9506E-12 |
| ACE2 | TPRX1P1 | -0,3517 | 5,96233E-12 |
| ACE2 | LINC01001 | -0,3517 | 6,00035E-12 |
| ACE2 | OR10H1 | -0,3517 | 6,0055E-12 |
| ACE2 | NXPH2 | -0,3518 | 5,85715E-12 |
| ACE2 | RPLP1P6 | -0,3518 | 5,89197E-12 |
| ACE2 | C16orf72 | -0,3518 | 5,87963E-12 |
| ACE2 | SPRR2G | -0,3519 | 5,76657E-12 |
| ACE2 | PGLYRP3 | -0,3519 | 5,78906E-12 |
| ACE2 | SPDYE12P | -0,3519 | 5,8469E-12 |

| | | | |
|---|---|---|---|
| ACE2 | RP4-584D14.5 | -0,3519 | 5,79504E-12 |
| ACE2 | WEE2-AS1 | -0,352 | 5,70976E-12 |
| ACE2 | RPL13 | -0,352 | 5,68951E-12 |
| ACE2 | RP11-431M7.2 | -0,3521 | 5,63909E-12 |
| ACE2 | TMEM71 | -0,3521 | 5,59858E-12 |
| ACE2 | MAPK8IP1 | -0,3521 | 5,61573E-12 |
| ACE2 | LINC00664 | -0,3521 | 5,61433E-12 |
| ACE2 | GOLGA2P3Y | -0,3522 | 5,54396E-12 |
| ACE2 | RP11-321E2.13 | -0,3523 | 5,50153E-12 |
| ACE2 | RP11-503C24.3 | -0,3523 | 5,45181E-12 |
| ACE2 | RP11-402L1.11 | -0,3523 | 5,51261E-12 |
| ACE2 | TRPM8 | -0,3524 | 5,37657E-12 |
| ACE2 | RP11-175D17.3 | -0,3524 | 5,40735E-12 |
| ACE2 | CH17-140K24.3 | -0,3526 | 5,20934E-12 |
| ACE2 | RP11-108K3.1 | -0,3526 | 5,2083E-12 |
| ACE2 | NNAT | -0,3526 | 5,23722E-12 |
| ACE2 | CD48 | -0,353 | 4,9673E-12 |
| ACE2 | RP11-479J7.1 | -0,353 | 4,94672E-12 |
| ACE2 | CTD-2186M15.1 | -0,353 | 4,91984E-12 |
| ACE2 | IGFALS | -0,353 | 4,93492E-12 |
| ACE2 | RP5-1029F21.3 | -0,353 | 4,93347E-12 |
| ACE2 | KCMF1 | -0,3531 | 4,84068E-12 |
| ACE2 | RP11-214K3.19 | -0,3532 | 4,79235E-12 |
| ACE2 | RP11-355I22.2 | -0,3532 | 4,76349E-12 |
| ACE2 | RP11-700E23.2 | -0,3533 | 4,7317E-12 |
| ACE2 | CTAGE11P | -0,3533 | 4,73423E-12 |
| ACE2 | ACTN1-AS1 | -0,3533 | 4,69898E-12 |
| ACE2 | ASS1P1 | -0,3534 | 4,63784E-12 |
| ACE2 | RP4-668E10.4 | -0,3534 | 4,63708E-12 |
| ACE2 | RP11-69C17.4 | -0,3534 | 4,64764E-12 |
| ACE2 | ICAM3 | -0,3534 | 4,6533E-12 |
| ACE2 | MPL | -0,3535 | 4,57139E-12 |
| ACE2 | AC000111.6 | -0,3535 | 4,5937E-12 |
| ACE2 | KIF24 | -0,3535 | 4,56847E-12 |
| ACE2 | TNPO1P1 | -0,3535 | 4,6091E-12 |
| ACE2 | RP1-89D4.1 | -0,3535 | 4,56371E-12 |
| ACE2 | RP11-345J4.3 | -0,3535 | 4,57073E-12 |
| ACE2 | FHP1 | -0,3536 | 4,54032E-12 |
| ACE2 | TRERNA1 | -0,3536 | 4,48883E-12 |
| ACE2 | TRAPPC3 | -0,3537 | 4,46333E-12 |
| ACE2 | AC097468.4 | -0,3537 | 4,47965E-12 |
| ACE2 | CTD-2083E4.6 | -0,3537 | 4,47403E-12 |
| ACE2 | FAM153A | -0,3537 | 4,43113E-12 |
| ACE2 | RP11-314D7.2 | -0,3537 | 4,42382E-12 |
| ACE2 | ATG10 | -0,3538 | 4,39835E-12 |
| ACE2 | SEZ6L | -0,3538 | 4,40341E-12 |
| ACE2 | RP11-528N21.1 | -0,3539 | 4,3451E-12 |
| ACE2 | RP11-91J19.2 | -0,3539 | 4,33463E-12 |
| ACE2 | TOMM22P5 | -0,3539 | 4,29466E-12 |
| ACE2 | RP11-161H23.8 | -0,3539 | 4,29246E-12 |
| ACE2 | RP11-37L2.1 | -0,3539 | 4,33624E-12 |
| ACE2 | LINC01049 | -0,3539 | 4,32914E-12 |
| ACE2 | RP11-109N23.4 | -0,3539 | 4,33657E-12 |
| ACE2 | GDF5OS | -0,3539 | 4,35114E-12 |

| | | | |
|---|---|---|---|
| ACE2 | RP11-545A16.4 | -0,354 | 4,27396E-12 |
| ACE2 | RP11-105N14.1 | -0,354 | 4,26451E-12 |
| ACE2 | TSPO2 | -0,354 | 4,23183E-12 |
| ACE2 | RP11-375N15.1 | -0,354 | 4,28479E-12 |
| ACE2 | KRTAP13-1 | -0,354 | 4,25082E-12 |
| ACE2 | CLNK | -0,3541 | 4,1715E-12 |
| ACE2 | WNT8A | -0,3541 | 4,19449E-12 |
| ACE2 | AC099342.1 | -0,3541 | 4,20377E-12 |
| ACE2 | AC142119.1 | -0,3542 | 4,15432E-12 |
| ACE2 | ZFPM2-AS1 | -0,3542 | 4,16357E-12 |
| ACE2 | RP11-680F8.3 | -0,3542 | 4,12619E-12 |
| ACE2 | AADACL4 | -0,3543 | 4,0572E-12 |
| ACE2 | PMS1 | -0,3543 | 4,05697E-12 |
| ACE2 | EEF1GP8 | -0,3543 | 4,09249E-12 |
| ACE2 | RP11-173M1.4 | -0,3543 | 4,08784E-12 |
| ACE2 | DHX30 | -0,3544 | 4,00262E-12 |
| ACE2 | FAM204A | -0,3544 | 4,02083E-12 |
| ACE2 | RP11-98D18.3 | -0,3545 | 3,93224E-12 |
| ACE2 | RP11-48O20.5 | -0,3545 | 3,97996E-12 |
| ACE2 | LINC00342 | -0,3545 | 3,94798E-12 |
| ACE2 | HBZP1 | -0,3545 | 3,9558E-12 |
| ACE2 | RNF11P2 | -0,3546 | 3,91418E-12 |
| ACE2 | MIF | -0,3546 | 3,87234E-12 |
| ACE2 | GCSAM | -0,3547 | 3,85989E-12 |
| ACE2 | XXbac-BPGBPG24O18. | -0,3548 | 3,79196E-12 |
| ACE2 | VSTM2A-OT1 | -0,3548 | 3,77869E-12 |
| ACE2 | RP11-986G18.3 | -0,3548 | 3,80035E-12 |
| ACE2 | GLIS1 | -0,3549 | 3,72308E-12 |
| ACE2 | RP11-83A16.1 | -0,3549 | 3,71437E-12 |
| ACE2 | CTD-3212A4.2 | -0,3549 | 3,7202E-12 |
| ACE2 | RP11-661A12.12 | -0,3549 | 3,72758E-12 |
| ACE2 | RP11-69M1.8 | -0,3549 | 3,74905E-12 |
| ACE2 | FAM90A27P | -0,3549 | 3,71992E-12 |
| ACE2 | OPN3 | -0,355 | 3,67786E-12 |
| ACE2 | RP1-242N11.1 | -0,355 | 3,67372E-12 |
| ACE2 | RP11-513O17.1 | -0,355 | 3,65629E-12 |
| ACE2 | CD6 | -0,355 | 3,67212E-12 |
| ACE2 | CD40LG | -0,355 | 3,64564E-12 |
| ACE2 | CUL9 | -0,3551 | 3,62152E-12 |
| ACE2 | OR52L1 | -0,3551 | 3,64057E-12 |
| ACE2 | RP11-362K2.2 | -0,3551 | 3,60627E-12 |
| ACE2 | GOLGA8G | -0,3551 | 3,62662E-12 |
| ACE2 | RP11-56B16.5 | -0,3551 | 3,6151E-12 |
| ACE2 | PPP4R1 | -0,3551 | 3,60824E-12 |
| ACE2 | XRCC1 | -0,3551 | 3,64432E-12 |
| ACE2 | AC013448.1 | -0,3552 | 3,54802E-12 |
| ACE2 | OR7E101P | -0,3553 | 3,50573E-12 |
| ACE2 | PPM1A | -0,3553 | 3,49768E-12 |
| ACE2 | BRK1 | -0,3554 | 3,43857E-12 |
| ACE2 | RP11-242P2.1 | -0,3554 | 3,46937E-12 |
| ACE2 | LINC01959 | -0,3555 | 3,41691E-12 |
| ACE2 | RP1-287H17.1 | -0,3555 | 3,42787E-12 |
| ACE2 | RP4-593H12.1 | -0,3555 | 3,42898E-12 |
| ACE2 | POU2F2 | -0,3555 | 3,41211E-12 |

| | | | |
|---|---|---|---|
| ACE2 | AC097721.2 | -0,3556 | 3,37888E-12 |
| ACE2 | EEF1GP2 | -0,3556 | 3,34517E-12 |
| ACE2 | RP11-602N24.3 | -0,3557 | 3,33073E-12 |
| ACE2 | SLC6A4 | -0,3557 | 3,30009E-12 |
| ACE2 | CTC-559E9.2 | -0,3557 | 3,29346E-12 |
| ACE2 | NDUFA3P4 | -0,3558 | 3,26756E-12 |
| ACE2 | HOXA11 | -0,3559 | 3,21725E-12 |
| ACE2 | RP11-998D10.4 | -0,3559 | 3,2138E-12 |
| ACE2 | RP1-35C21.2 | -0,356 | 3,16455E-12 |
| ACE2 | CTD-2140G10.2 | -0,356 | 3,14186E-12 |
| ACE2 | RP4-680D5.9 | -0,3561 | 3,13727E-12 |
| ACE2 | RP11-203F10.1 | -0,3561 | 3,10138E-12 |
| ACE2 | RP11-71B7.1 | -0,3561 | 3,13489E-12 |
| ACE2 | OR2K2 | -0,3561 | 3,10955E-12 |
| ACE2 | NDUFA9 | -0,3561 | 3,13643E-12 |
| ACE2 | RP11-115J16.1 | -0,3563 | 3,01346E-12 |
| ACE2 | PRH2 | -0,3563 | 3,01314E-12 |
| ACE2 | GOLGA8F | -0,3563 | 3,02223E-12 |
| ACE2 | DYTN | -0,3564 | 2,98425E-12 |
| ACE2 | TTLL11-IT1 | -0,3564 | 2,98922E-12 |
| ACE2 | RP11-49L2.1 | -0,3564 | 2,9696E-12 |
| ACE2 | NUBP2 | -0,3565 | 2,95784E-12 |
| ACE2 | NBPF13P | -0,3566 | 2,88466E-12 |
| ACE2 | RNU1-70P | -0,3566 | 2,88089E-12 |
| ACE2 | RP11-420M1.2 | -0,3566 | 2,8934E-12 |
| ACE2 | GDPGP1 | -0,3566 | 2,89474E-12 |
| ACE2 | FAR2P2 | -0,3567 | 2,83021E-12 |
| ACE2 | FLJ13224 | -0,3567 | 2,86502E-12 |
| ACE2 | FGF14 | -0,3567 | 2,8567E-12 |
| ACE2 | XPNPEP3 | -0,3567 | 2,84249E-12 |
| ACE2 | CNGA3 | -0,3568 | 2,80038E-12 |
| ACE2 | CDR2 | -0,3568 | 2,81896E-12 |
| ACE2 | TRAM2 | -0,3569 | 2,78685E-12 |
| ACE2 | RP3-486I3.7 | -0,3569 | 2,75278E-12 |
| ACE2 | AC004009.3 | -0,3569 | 2,74798E-12 |
| ACE2 | CHRNB3 | -0,357 | 2,72288E-12 |
| ACE2 | PHF2 | -0,3571 | 2,6718E-12 |
| ACE2 | CADM3-AS1 | -0,3572 | 2,65991E-12 |
| ACE2 | LINC00636 | -0,3572 | 2,63123E-12 |
| ACE2 | GNA11 | -0,3572 | 2,63251E-12 |
| ACE2 | DPRX | -0,3572 | 2,64262E-12 |
| ACE2 | MROH2A | -0,3573 | 2,60299E-12 |
| ACE2 | RP11-542K23.10 | -0,3573 | 2,6236E-12 |
| ACE2 | RP11-486A14.2 | -0,3573 | 2,62336E-12 |
| ACE2 | RP11-626G11.1 | -0,3573 | 2,59786E-12 |
| ACE2 | ANKRD20A5P | -0,3573 | 2,6206E-12 |
| ACE2 | IGLV1-41 | -0,3573 | 2,60763E-12 |
| ACE2 | CDK15 | -0,3574 | 2,56894E-12 |
| ACE2 | RN7SL28P | -0,3574 | 2,57547E-12 |
| ACE2 | AC011516.1 | -0,3574 | 2,55247E-12 |
| ACE2 | NR1I3 | -0,3575 | 2,53704E-12 |
| ACE2 | NAF1 | -0,3575 | 2,53314E-12 |
| ACE2 | TMA16 | -0,3576 | 2,50513E-12 |
| ACE2 | WAC-AS1 | -0,3576 | 2,49898E-12 |

| | | | |
|---|---|---|---|
| ACE2 | EIF4HP2 | -0,3576 | 2,47425E-12 |
| ACE2 | FTH1P6 | -0,3577 | 2,4703E-12 |
| ACE2 | LINC02220 | -0,3577 | 2,4599E-12 |
| ACE2 | RP11-752G15.8 | -0,3577 | 2,47245E-12 |
| ACE2 | RP11-558O12.1 | -0,3577 | 2,45722E-12 |
| ACE2 | GNG5 | -0,3578 | 2,41941E-12 |
| ACE2 | RP11-122G18.7 | -0,3578 | 2,40321E-12 |
| ACE2 | CCDC174 | -0,3578 | 2,41206E-12 |
| ACE2 | RP11-469J4.3 | -0,3578 | 2,41246E-12 |
| ACE2 | LINC01182 | -0,3578 | 2,41081E-12 |
| ACE2 | KIAA2013 | -0,3579 | 2,37242E-12 |
| ACE2 | AC109815.2 | -0,3579 | 2,39821E-12 |
| ACE2 | TNNT2 | -0,358 | 2,34447E-12 |
| ACE2 | DLX2 | -0,358 | 2,35022E-12 |
| ACE2 | KAAG1 | -0,358 | 2,36215E-12 |
| ACE2 | RP11-1415C14.4 | -0,3581 | 2,31155E-12 |
| ACE2 | RP1-111D6.5 | -0,3581 | 2,29815E-12 |
| ACE2 | SIK3 | -0,3581 | 2,30813E-12 |
| ACE2 | RP11-356K23.2 | -0,3581 | 2,32474E-12 |
| ACE2 | RN7SKP275 | -0,3582 | 2,27713E-12 |
| ACE2 | HIST1H2BPS1 | -0,3582 | 2,26769E-12 |
| ACE2 | TRIM73 | -0,3582 | 2,28393E-12 |
| ACE2 | ATP5S | -0,3582 | 2,29086E-12 |
| ACE2 | AC112518.3 | -0,3583 | 2,22661E-12 |
| ACE2 | ABHD12B | -0,3583 | 2,24827E-12 |
| ACE2 | RP5-1103B4.3 | -0,3584 | 2,19746E-12 |
| ACE2 | RP11-67M9.1 | -0,3584 | 2,19359E-12 |
| ACE2 | LINC01606 | -0,3584 | 2,20332E-12 |
| ACE2 | RP11-115J23.1 | -0,3584 | 2,19774E-12 |
| ACE2 | PGAM1P5 | -0,3584 | 2,20052E-12 |
| ACE2 | BOLL | -0,3585 | 2,18876E-12 |
| ACE2 | LRP2BP | -0,3585 | 2,16367E-12 |
| ACE2 | KIAA0319 | -0,3585 | 2,18192E-12 |
| ACE2 | UBE2R2 | -0,3585 | 2,16996E-12 |
| ACE2 | ANGPT4 | -0,3585 | 2,18559E-12 |
| ACE2 | RP11-505P4.7 | -0,3586 | 2,12862E-12 |
| ACE2 | ZNF541 | -0,3586 | 2,13102E-12 |
| ACE2 | RP5-1119A7.17 | -0,3586 | 2,15718E-12 |
| ACE2 | SLC7A3 | -0,3586 | 2,1482E-12 |
| ACE2 | RP5-1092L12.2 | -0,3587 | 2,12246E-12 |
| ACE2 | ANKRD1 | -0,3587 | 2,12319E-12 |
| ACE2 | ATG7 | -0,3588 | 2,08021E-12 |
| ACE2 | FXR1 | -0,3588 | 2,07047E-12 |
| ACE2 | RP11-505C13.1 | -0,3588 | 2,07613E-12 |
| ACE2 | LINC01520 | -0,3588 | 2,0793E-12 |
| ACE2 | TKFC | -0,3588 | 2,0941E-12 |
| ACE2 | ACYP1 | -0,3588 | 2,0783E-12 |
| ACE2 | BMP10 | -0,3589 | 2,04391E-12 |
| ACE2 | AC005102.1 | -0,3589 | 2,06075E-12 |
| ACE2 | TLK1P1 | -0,3589 | 2,03931E-12 |
| ACE2 | RP11-29P20.1 | -0,3589 | 2,04128E-12 |
| ACE2 | KCNJ15 | -0,3591 | 1,97488E-12 |
| ACE2 | IGKV1-8 | -0,3592 | 1,95171E-12 |
| ACE2 | CCNH | -0,3592 | 1,96262E-12 |

| | | | |
|---|---|---|---|
| ACE2 | CTD-2382H12.1 | -0,3592 | 1,95473E-12 |
| ACE2 | IK | -0,3594 | 1,88894E-12 |
| ACE2 | TUBBP9 | -0,3594 | 1,89648E-12 |
| ACE2 | RP11-874J12.3 | -0,3595 | 1,88104E-12 |
| ACE2 | LINC01858 | -0,3595 | 1,86811E-12 |
| ACE2 | RP4-561L24.3 | -0,3596 | 1,85662E-12 |
| ACE2 | RP11-156E8.1 | -0,3596 | 1,85133E-12 |
| ACE2 | RP11-647P12.1 | -0,3596 | 1,83489E-12 |
| ACE2 | KB-1460A1.3 | -0,3596 | 1,83085E-12 |
| ACE2 | RP11-245A18.1 | -0,3596 | 1,83382E-12 |
| ACE2 | RP11-323A7.1 | -0,3596 | 1,84174E-12 |
| ACE2 | GOLGA2P11 | -0,3596 | 1,84319E-12 |
| ACE2 | CXorf66 | -0,3596 | 1,85481E-12 |
| ACE2 | CEP70 | -0,3597 | 1,81088E-12 |
| ACE2 | ARHGAP24 | -0,3597 | 1,80706E-12 |
| ACE2 | ING2 | -0,3597 | 1,82401E-12 |
| ACE2 | RTN4RL2 | -0,3597 | 1,81435E-12 |
| ACE2 | CDIP1 | -0,3597 | 1,81393E-12 |
| ACE2 | ZNF861P | -0,3597 | 1,8128E-12 |
| ACE2 | PTOV1 | -0,3597 | 1,82491E-12 |
| ACE2 | RP11-848G14.5 | -0,3598 | 1,79786E-12 |
| ACE2 | RP11-342M3.2 | -0,3598 | 1,79048E-12 |
| ACE2 | RP11-122G18.10 | -0,3599 | 1,75463E-12 |
| ACE2 | CCDC167 | -0,3599 | 1,76343E-12 |
| ACE2 | RP11-728K20.1 | -0,3599 | 1,76521E-12 |
| ACE2 | LINC01955 | -0,36 | 1,73211E-12 |
| ACE2 | RP11-30P16.1 | -0,36 | 1,72847E-12 |
| ACE2 | CTAGE1 | -0,36 | 1,7428E-12 |
| ACE2 | PCDH10 | -0,3601 | 1,70383E-12 |
| ACE2 | CTD-2296D1.1 | -0,3601 | 1,70554E-12 |
| ACE2 | RP11-285G1.2 | -0,3601 | 1,71507E-12 |
| ACE2 | HTATIP2 | -0,3601 | 1,70152E-12 |
| ACE2 | RP11-66B24.4 | -0,3601 | 1,69739E-12 |
| ACE2 | GJA5 | -0,3602 | 1,69166E-12 |
| ACE2 | PRICKLE2-AS2 | -0,3602 | 1,68594E-12 |
| ACE2 | AC107399.2 | -0,3602 | 1,67609E-12 |
| ACE2 | RP11-799D4.3 | -0,3602 | 1,69229E-12 |
| ACE2 | SAMD10 | -0,3602 | 1,69416E-12 |
| ACE2 | DDX11L2 | -0,3603 | 1,64798E-12 |
| ACE2 | RP11-760D2.5 | -0,3603 | 1,65627E-12 |
| ACE2 | ATP5J2 | -0,3603 | 1,65927E-12 |
| ACE2 | FGF17 | -0,3603 | 1,65277E-12 |
| ACE2 | RP11-17A4.4 | -0,3603 | 1,66255E-12 |
| ACE2 | SLC35F4 | -0,3603 | 1,65441E-12 |
| ACE2 | PLA2G10 | -0,3603 | 1,64942E-12 |
| ACE2 | CARD14 | -0,3603 | 1,6669E-12 |
| ACE2 | FAM46B | -0,3604 | 1,63325E-12 |
| ACE2 | CIR1 | -0,3604 | 1,64533E-12 |
| ACE2 | SCARNA22 | -0,3604 | 1,64538E-12 |
| ACE2 | RP5-1096D14.6 | -0,3604 | 1,62307E-12 |
| ACE2 | RP11-304F15.6 | -0,3604 | 1,63963E-12 |
| ACE2 | GTF2IP5 | -0,3605 | 1,61208E-12 |
| ACE2 | RP11-294C11.1 | -0,3605 | 1,60333E-12 |
| ACE2 | RP11-97N19.3 | -0,3605 | 1,60054E-12 |

| | | | |
|---|---|---|---|
| ACE2 | RP11-727F15.13 | -0,3606 | 1,59375E-12 |
| ACE2 | NPSR1 | -0,3607 | 1,55478E-12 |
| ACE2 | ATE1-AS1 | -0,3607 | 1,55227E-12 |
| ACE2 | ZC3H3 | -0,3608 | 1,54064E-12 |
| ACE2 | RNASEH1P2 | -0,3608 | 1,54573E-12 |
| ACE2 | PLAGL2 | -0,3608 | 1,5361E-12 |
| ACE2 | PCDHA5 | -0,3609 | 1,51841E-12 |
| ACE2 | RP11-251G23.2 | -0,3609 | 1,51814E-12 |
| ACE2 | RP11-285G1.9 | -0,3609 | 1,52056E-12 |
| ACE2 | RP11-770G2.2 | -0,3609 | 1,52522E-12 |
| ACE2 | GGH | -0,361 | 1,48836E-12 |
| ACE2 | INHBC | -0,361 | 1,48195E-12 |
| ACE2 | SIRT3 | -0,3611 | 1,46936E-12 |
| ACE2 | RP11-413N10.3 | -0,3611 | 1,4781E-12 |
| ACE2 | RP11-133K1.6 | -0,3611 | 1,46251E-12 |
| ACE2 | PPP2R2C | -0,3612 | 1,44013E-12 |
| ACE2 | TRBV4-2 | -0,3612 | 1,44198E-12 |
| ACE2 | MSRB1 | -0,3612 | 1,44997E-12 |
| ACE2 | SLCO4C1 | -0,3613 | 1,41913E-12 |
| ACE2 | RP11-386M24.3 | -0,3613 | 1,43238E-12 |
| ACE2 | AC025171.1 | -0,3614 | 1,40245E-12 |
| ACE2 | TCEA1P3 | -0,3614 | 1,40077E-12 |
| ACE2 | FAM166AP7 | -0,3614 | 1,41331E-12 |
| ACE2 | IGLVI-42 | -0,3614 | 1,40294E-12 |
| ACE2 | EML2 | -0,3615 | 1,3873E-12 |
| ACE2 | RP11-543C4.3 | -0,3616 | 1,3556E-12 |
| ACE2 | RP11-802D6.1 | -0,3616 | 1,36622E-12 |
| ACE2 | CTAGE13P | -0,3617 | 1,34675E-12 |
| ACE2 | SMPD1 | -0,3617 | 1,34718E-12 |
| ACE2 | MRPS11 | -0,3617 | 1,34509E-12 |
| ACE2 | SERPINB4 | -0,3617 | 1,33785E-12 |
| ACE2 | KIF9-AS1 | -0,3618 | 1,31701E-12 |
| ACE2 | RP11-755B10.3 | -0,3618 | 1,32456E-12 |
| ACE2 | CEP63 | -0,3618 | 1,32266E-12 |
| ACE2 | SLC6A7 | -0,3618 | 1,32974E-12 |
| ACE2 | MRPL17 | -0,3618 | 1,31326E-12 |
| ACE2 | RP11-103J8.2 | -0,3618 | 1,3305E-12 |
| ACE2 | POPDC3 | -0,3619 | 1,29889E-12 |
| ACE2 | ATG13 | -0,3619 | 1,30744E-12 |
| ACE2 | RP11-297M9.1 | -0,3619 | 1,29837E-12 |
| ACE2 | IFT74-AS1 | -0,362 | 1,28122E-12 |
| ACE2 | RPL13P5 | -0,362 | 1,27426E-12 |
| ACE2 | RP11-583F2.1 | -0,362 | 1,29022E-12 |
| ACE2 | SORCS3 | -0,3621 | 1,26637E-12 |
| ACE2 | RP11-432J3.4 | -0,3621 | 1,27213E-12 |
| ACE2 | LINC00552 | -0,3621 | 1,26481E-12 |
| ACE2 | FAAP24 | -0,3621 | 1,2707E-12 |
| ACE2 | RP11-325I22.3 | -0,3622 | 1,25169E-12 |
| ACE2 | RP11-503C24.4 | -0,3622 | 1,24574E-12 |
| ACE2 | MIR9-3HG | -0,3622 | 1,23847E-12 |
| ACE2 | CD55 | -0,3623 | 1,23232E-12 |
| ACE2 | CDH23-AS1 | -0,3623 | 1,21993E-12 |
| ACE2 | ADAM2 | -0,3624 | 1,20266E-12 |
| ACE2 | ODF2-AS1 | -0,3624 | 1,20761E-12 |

| | | | |
|---|---|---|---|
| ACE2 | CTB-52I2.4 | -0,3624 | 1,1972E-12 |
| ACE2 | RP11-158D2.2 | -0,3625 | 1,18821E-12 |
| ACE2 | RP11-203P2.2 | -0,3625 | 1,17913E-12 |
| ACE2 | CYP4A22 | -0,3626 | 1,16815E-12 |
| ACE2 | AC091633.3 | -0,3626 | 1,1664E-12 |
| ACE2 | RAB8B | -0,3626 | 1,1752E-12 |
| ACE2 | MIR4432HG | -0,3627 | 1,14998E-12 |
| ACE2 | SH2D3C | -0,3627 | 1,15058E-12 |
| ACE2 | RP11-456O19.5 | -0,3627 | 1,15972E-12 |
| ACE2 | MIR3180-4 | -0,3628 | 1,13698E-12 |
| ACE2 | RP11-356C4.5 | -0,3628 | 1,14123E-12 |
| ACE2 | AC114765.2 | -0,3629 | 1,11996E-12 |
| ACE2 | ELOCP2 | -0,3629 | 1,1105E-12 |
| ACE2 | ZNF543 | -0,3629 | 1,11503E-12 |
| ACE2 | IGLVIV-65 | -0,3629 | 1,11223E-12 |
| ACE2 | RP11-745C15.2 | -0,363 | 1,10629E-12 |
| ACE2 | RP11-465L8.1 | -0,363 | 1,10028E-12 |
| ACE2 | PDE6G | -0,363 | 1,10234E-12 |
| ACE2 | MIR3945HG | -0,3631 | 1,08935E-12 |
| ACE2 | RP11-548K12.7 | -0,3631 | 1,08841E-12 |
| ACE2 | HOXB3 | -0,3631 | 1,09122E-12 |
| ACE2 | DCAF15 | -0,3631 | 1,08665E-12 |
| ACE2 | CELA3B | -0,3632 | 1,07388E-12 |
| ACE2 | LINC01994 | -0,3632 | 1,06928E-12 |
| ACE2 | ATP11A-AS1 | -0,3633 | 1,0486E-12 |
| ACE2 | RP11-114H24.7 | -0,3633 | 1,05382E-12 |
| ACE2 | AP000692.10 | -0,3633 | 1,051E-12 |
| ACE2 | TRIM58 | -0,3634 | 1,0328E-12 |
| ACE2 | FAM227B | -0,3634 | 1,04228E-12 |
| ACE2 | RP11-214N1.1 | -0,3635 | 1,02024E-12 |
| ACE2 | LINC01859 | -0,3635 | 1,02319E-12 |
| ACE2 | RP11-448G4.2 | -0,3636 | 1,00482E-12 |
| ACE2 | MGAT4D | -0,3636 | 9,99681E-13 |
| ACE2 | RP11-513O17.3 | -0,3636 | 1,00237E-12 |
| ACE2 | C1orf106 | -0,3638 | 9,71043E-13 |
| ACE2 | LDHAP1 | -0,3638 | 9,72531E-13 |
| ACE2 | ZNF438 | -0,3638 | 9,78028E-13 |
| ACE2 | SLC38A8 | -0,3638 | 9,68569E-13 |
| ACE2 | AC091705.1 | -0,3639 | 9,62975E-13 |
| ACE2 | FBXO10 | -0,3639 | 9,53369E-13 |
| ACE2 | RANBP20P | -0,3639 | 9,59206E-13 |
| ACE2 | BMP15 | -0,364 | 9,51139E-13 |
| ACE2 | RP11-407A16.5 | -0,3641 | 9,30387E-13 |
| ACE2 | CADM2-AS1 | -0,3642 | 9,15572E-13 |
| ACE2 | RP11-676F20.1 | -0,3642 | 9,18935E-13 |
| ACE2 | RP11-61O1.2 | -0,3642 | 9,20606E-13 |
| ACE2 | GGT3P | -0,3642 | 9,15527E-13 |
| ACE2 | RP11-124B13.1 | -0,3643 | 9,01271E-13 |
| ACE2 | SCN3B | -0,3643 | 9,0135E-13 |
| ACE2 | CUX2 | -0,3643 | 9,02638E-13 |
| ACE2 | RNF19B | -0,3644 | 8,82462E-13 |
| ACE2 | CNTRL | -0,3644 | 8,87733E-13 |
| ACE2 | CRACR2A | -0,3644 | 8,90641E-13 |
| ACE2 | REXO1 | -0,3644 | 8,85749E-13 |

| | | | |
|---|---|---|---|
| ACE2 | CTA-929C8.7 | -0,3644 | 8,84922E-13 |
| ACE2 | WDR60 | -0,3645 | 8,80654E-13 |
| ACE2 | RP11-460M2.1 | -0,3647 | 8,52386E-13 |
| ACE2 | TP53TG3HP | -0,3647 | 8,50852E-13 |
| ACE2 | RP11-321A17.5 | -0,3647 | 8,54503E-13 |
| ACE2 | RP5-901A4.1 | -0,3648 | 8,31272E-13 |
| ACE2 | ASTE1 | -0,3649 | 8,16314E-13 |
| ACE2 | RP11-506K19.2 | -0,3649 | 8,20858E-13 |
| ACE2 | CTNNA3 | -0,365 | 8,10056E-13 |
| ACE2 | STT3A-AS1 | -0,365 | 8,14847E-13 |
| ACE2 | RP11-352M15.1 | -0,365 | 8,15322E-13 |
| ACE2 | AC006116.21 | -0,365 | 8,11941E-13 |
| ACE2 | ESRG | -0,3651 | 7,9407E-13 |
| ACE2 | KCTD9P6 | -0,3651 | 8,03297E-13 |
| ACE2 | SPAG6 | -0,3651 | 7,92525E-13 |
| ACE2 | ASXL1 | -0,3653 | 7,70646E-13 |
| ACE2 | CTA-342B11.1 | -0,3653 | 7,77368E-13 |
| ACE2 | AC093698.4 | -0,3654 | 7,61489E-13 |
| ACE2 | KB-1562D12.1 | -0,3654 | 7,65183E-13 |
| ACE2 | DAOA-AS1 | -0,3654 | 7,59292E-13 |
| ACE2 | RP11-297L17.3 | -0,3655 | 7,51896E-13 |
| ACE2 | RN7SL664P | -0,3656 | 7,39217E-13 |
| ACE2 | OR5H6 | -0,3656 | 7,36762E-13 |
| ACE2 | RP11-1024P17.1 | -0,3657 | 7,2291E-13 |
| ACE2 | RPS26P38 | -0,3658 | 7,13306E-13 |
| ACE2 | CTA-276F8.2 | -0,3658 | 7,18157E-13 |
| ACE2 | RP11-1O2.1 | -0,3658 | 7,16292E-13 |
| ACE2 | KCTD10P1 | -0,3659 | 7,05087E-13 |
| ACE2 | RP11-897M7.4 | -0,3659 | 7,0838E-13 |
| ACE2 | POTEC | -0,3659 | 7,0614E-13 |
| ACE2 | RP5-1056L3.3 | -0,366 | 6,9016E-13 |
| ACE2 | ZFAND3 | -0,366 | 6,97798E-13 |
| ACE2 | RP11-463C8.5 | -0,366 | 6,98674E-13 |
| ACE2 | TSHR | -0,366 | 6,99044E-13 |
| ACE2 | RP11-169F17.1 | -0,366 | 6,97916E-13 |
| ACE2 | AC012451.1 | -0,3661 | 6,81321E-13 |
| ACE2 | STK32B | -0,3661 | 6,874E-13 |
| ACE2 | RP11-429B11.2 | -0,3661 | 6,84908E-13 |
| ACE2 | LINC02213 | -0,3661 | 6,86154E-13 |
| ACE2 | AC004869.3 | -0,3661 | 6,79252E-13 |
| ACE2 | COQ5 | -0,3661 | 6,8471E-13 |
| ACE2 | AL132796.1 | -0,3661 | 6,84115E-13 |
| ACE2 | RP11-666A20.4 | -0,3662 | 6,70324E-13 |
| ACE2 | RP11-803P9.1 | -0,3662 | 6,7402E-13 |
| ACE2 | LINC01076 | -0,3662 | 6,7086E-13 |
| ACE2 | RXFP2 | -0,3662 | 6,7626E-13 |
| ACE2 | LRRC37A12P | -0,3663 | 6,65541E-13 |
| ACE2 | RP11-194G10.3 | -0,3663 | 6,64075E-13 |
| ACE2 | OSTN-AS1 | -0,3663 | 6,66831E-13 |
| ACE2 | RP11-304M2.3 | -0,3664 | 6,54756E-13 |
| ACE2 | RP11-274M17.3 | -0,3664 | 6,56533E-13 |
| ACE2 | RP11-692C24.2 | -0,3664 | 6,50017E-13 |
| ACE2 | NPIPA2 | -0,3664 | 6,5589E-13 |
| ACE2 | CRHR2 | -0,3665 | 6,41714E-13 |

| | | | |
|---|---|---|---|
| ACE2 | MGAT3-AS1 | -0,3665 | 6,38511E-13 |
| ACE2 | RP11-76C10.3 | -0,3666 | 6,30784E-13 |
| ACE2 | TOE1 | -0,3667 | 6,18425E-13 |
| ACE2 | EML6 | -0,3667 | 6,24173E-13 |
| ACE2 | TRBV10-1 | -0,3667 | 6,24467E-13 |
| ACE2 | RORB | -0,3667 | 6,25463E-13 |
| ACE2 | CTC-304I17.2 | -0,3667 | 6,24612E-13 |
| ACE2 | KRTAP16-1 | -0,3667 | 6,25706E-13 |
| ACE2 | C6orf58 | -0,3668 | 6,17474E-13 |
| ACE2 | AC004837.5 | -0,3668 | 6,11301E-13 |
| ACE2 | SMIM24 | -0,3668 | 6,10035E-13 |
| ACE2 | CTA-384D8.35 | -0,3668 | 6,14796E-13 |
| ACE2 | RP3-467K16.2 | -0,3669 | 6,04674E-13 |
| ACE2 | TRIM59 | -0,3669 | 6,08573E-13 |
| ACE2 | POM121L14P | -0,3669 | 6,05079E-13 |
| ACE2 | RP11-1360M22.2 | -0,3669 | 6,05112E-13 |
| ACE2 | AC136896.2 | -0,3669 | 6,03132E-13 |
| ACE2 | RP1-56K13.5 | -0,3669 | 6,04039E-13 |
| ACE2 | LINC01597 | -0,3669 | 6,08622E-13 |
| ACE2 | RP11-78A18.2 | -0,367 | 5,95112E-13 |
| ACE2 | CTD-2630F21.1 | -0,367 | 5,9347E-13 |
| ACE2 | EFR3B | -0,3671 | 5,86162E-13 |
| ACE2 | LINC01826 | -0,3671 | 5,84113E-13 |
| ACE2 | RP11-557H15.5 | -0,3671 | 5,83949E-13 |
| ACE2 | RP11-455J15.1 | -0,3671 | 5,85491E-13 |
| ACE2 | RP11-386I23.1 | -0,3672 | 5,74564E-13 |
| ACE2 | C11orf52 | -0,3673 | 5,67381E-13 |
| ACE2 | SRPK2 | -0,3674 | 5,57341E-13 |
| ACE2 | PEBP4 | -0,3674 | 5,5913E-13 |
| ACE2 | RP11-402F9.3 | -0,3674 | 5,54963E-13 |
| ACE2 | PLSCR5-AS1 | -0,3675 | 5,47965E-13 |
| ACE2 | TRBV6-7 | -0,3675 | 5,52092E-13 |
| ACE2 | AC004381.6 | -0,3676 | 5,42487E-13 |
| ACE2 | IGFL3 | -0,3676 | 5,45193E-13 |
| ACE2 | SPANXB1 | -0,3676 | 5,40597E-13 |
| ACE2 | XX-FW81066F1.2 | -0,3676 | 5,38478E-13 |
| ACE2 | CTC-305H11.1 | -0,3677 | 5,31207E-13 |
| ACE2 | AC090587.5 | -0,3677 | 5,30295E-13 |
| ACE2 | RP11-680E19.2 | -0,3677 | 5,34984E-13 |
| ACE2 | RP11-451N19.3 | -0,3677 | 5,34224E-13 |
| ACE2 | CA15P2 | -0,3677 | 5,3325E-13 |
| ACE2 | GS1-542M4.4 | -0,3678 | 5,28444E-13 |
| ACE2 | ZBTB42 | -0,3679 | 5,20287E-13 |
| ACE2 | TP73 | -0,368 | 5,13086E-13 |
| ACE2 | FECHP1 | -0,368 | 5,07241E-13 |
| ACE2 | ATF5 | -0,368 | 5,05925E-13 |
| ACE2 | RP11-15B17.1 | -0,3681 | 4,99444E-13 |
| ACE2 | MNX1 | -0,3681 | 5,01567E-13 |
| ACE2 | BAG2 | -0,3682 | 4,93995E-13 |
| ACE2 | RP11-430H10.2 | -0,3682 | 4,95255E-13 |
| ACE2 | DEDD2 | -0,3682 | 4,96057E-13 |
| ACE2 | PRRT3 | -0,3683 | 4,86261E-13 |
| ACE2 | RP3-401D24.1 | -0,3684 | 4,78851E-13 |
| ACE2 | RP11-327L9.1 | -0,3684 | 4,80976E-13 |

| | | | |
|---|---|---|---|
| ACE2 | OR52L2P | -0,3684 | 4,81264E-13 |
| ACE2 | RPL31P61 | -0,3684 | 4,80192E-13 |
| ACE2 | TMEM86B | -0,3684 | 4,802E-13 |
| ACE2 | LGALS8 | -0,3685 | 4,70423E-13 |
| ACE2 | FARS2 | -0,3685 | 4,74083E-13 |
| ACE2 | IQUB | -0,3685 | 4,68801E-13 |
| ACE2 | LINC01838 | -0,3685 | 4,6944E-13 |
| ACE2 | RP11-98D18.7 | -0,3686 | 4,62131E-13 |
| ACE2 | EAF1-AS1 | -0,3686 | 4,65568E-13 |
| ACE2 | RP11-386B13.4 | -0,3686 | 4,64329E-13 |
| ACE2 | HLA-P | -0,3686 | 4,60882E-13 |
| ACE2 | MAPK13 | -0,3686 | 4,6412E-13 |
| ACE2 | CTB-3M24.3 | -0,3687 | 4,5351E-13 |
| ACE2 | EXOSC3 | -0,3687 | 4,54794E-13 |
| ACE2 | RP11-403N16.3 | -0,3687 | 4,58888E-13 |
| ACE2 | CUBNP1 | -0,3688 | 4,52574E-13 |
| ACE2 | RP11-719K4.7 | -0,3688 | 4,49961E-13 |
| ACE2 | TMA16P2 | -0,3689 | 4,40392E-13 |
| ACE2 | F5 | -0,3689 | 4,42483E-13 |
| ACE2 | OR6B1 | -0,3689 | 4,39588E-13 |
| ACE2 | RP11-402G3.4 | -0,3689 | 4,42165E-13 |
| ACE2 | ZNF461 | -0,3689 | 4,40034E-13 |
| ACE2 | LINC01876 | -0,369 | 4,37429E-13 |
| ACE2 | ARF1P3 | -0,369 | 4,38526E-13 |
| ACE2 | RP11-843A23.1 | -0,369 | 4,33468E-13 |
| ACE2 | SPRYD7 | -0,369 | 4,35447E-13 |
| ACE2 | TBX4 | -0,369 | 4,38278E-13 |
| ACE2 | KCNB2 | -0,3691 | 4,26461E-13 |
| ACE2 | C18orf12 | -0,3691 | 4,2789E-13 |
| ACE2 | KB-1042C11.5 | -0,3692 | 4,21502E-13 |
| ACE2 | RP11-429J17.5 | -0,3693 | 4,1605E-13 |
| ACE2 | UBE2C | -0,3693 | 4,13944E-13 |
| ACE2 | SMC4P1 | -0,3694 | 4,10722E-13 |
| ACE2 | PCED1CP | -0,3694 | 4,10129E-13 |
| ACE2 | SP2 | -0,3694 | 4,06378E-13 |
| ACE2 | RP11-321E2.6 | -0,3696 | 3,95478E-13 |
| ACE2 | DDC | -0,3696 | 3,95782E-13 |
| ACE2 | RP11-462B18.2 | -0,3696 | 3,95574E-13 |
| ACE2 | RP11-436A7.1 | -0,3697 | 3,91978E-13 |
| ACE2 | RP11-554I8.1 | -0,3697 | 3,91336E-13 |
| ACE2 | RP11-158L12.6 | -0,3697 | 3,88313E-13 |
| ACE2 | SYCE2 | -0,3697 | 3,90019E-13 |
| ACE2 | CTD-2516F10.4 | -0,3698 | 3,83571E-13 |
| ACE2 | RP11-165D7.5 | -0,3698 | 3,85541E-13 |
| ACE2 | DNAJB1P1 | -0,3699 | 3,79126E-13 |
| ACE2 | CNTN6 | -0,3699 | 3,76278E-13 |
| ACE2 | RP11-173P15.10 | -0,3699 | 3,79859E-13 |
| ACE2 | RP11-89K22.1 | -0,3699 | 3,76717E-13 |
| ACE2 | RP11-624J12.1 | -0,3699 | 3,79777E-13 |
| ACE2 | SPACA6P-AS | -0,3699 | 3,79978E-13 |
| ACE2 | AF067845.1 | -0,37 | 3,73494E-13 |
| ACE2 | RP11-299D14.2 | -0,37 | 3,72067E-13 |
| ACE2 | CTC-281F24.1 | -0,37 | 3,71102E-13 |
| ACE2 | RP11-879F14.1 | -0,37 | 3,75278E-13 |

| | | | |
|---|---|---|---|
| ACE2 | RP11-388C12.8 | -0,3701 | 3,65709E-13 |
| ACE2 | RPL13AP7 | -0,3701 | 3,66796E-13 |
| ACE2 | OTUD7B | -0,3702 | 3,60782E-13 |
| ACE2 | TMF1 | -0,3702 | 3,64138E-13 |
| ACE2 | RP11-597D13.9 | -0,3702 | 3,62938E-13 |
| ACE2 | SPINK5 | -0,3702 | 3,6223E-13 |
| ACE2 | STXBP5 | -0,3702 | 3,58916E-13 |
| ACE2 | RP11-537H15.4 | -0,3703 | 3,55234E-13 |
| ACE2 | PITRM1-AS1 | -0,3703 | 3,5473E-13 |
| ACE2 | ZNHIT1 | -0,3704 | 3,52013E-13 |
| ACE2 | RP11-618M23.4 | -0,3704 | 3,4752E-13 |
| ACE2 | RP11-219O3.2 | -0,3704 | 3,49165E-13 |
| ACE2 | SCGB2A2 | -0,3704 | 3,50903E-13 |
| ACE2 | FRMD5 | -0,3704 | 3,48618E-13 |
| ACE2 | RPL21P116 | -0,3704 | 3,50729E-13 |
| ACE2 | CH17-53B9.4 | -0,3704 | 3,4943E-13 |
| ACE2 | AC012354.6 | -0,3705 | 3,46529E-13 |
| ACE2 | AC073257.2 | -0,3705 | 3,42118E-13 |
| ACE2 | KIF4B | -0,3705 | 3,43168E-13 |
| ACE2 | RP11-102N11.1 | -0,3705 | 3,46596E-13 |
| ACE2 | TRBV17 | -0,3705 | 3,42861E-13 |
| ACE2 | AC025016.1 | -0,3705 | 3,42787E-13 |
| ACE2 | TUBB8P4 | -0,3705 | 3,46609E-13 |
| ACE2 | RP11-318A15.2 | -0,3705 | 3,45786E-13 |
| ACE2 | TRBV7-7 | -0,3706 | 3,36847E-13 |
| ACE2 | LINC00964 | -0,3706 | 3,38099E-13 |
| ACE2 | SNF8 | -0,3707 | 3,31976E-13 |
| ACE2 | BARHL1 | -0,3708 | 3,27201E-13 |
| ACE2 | RP13-735L24.1 | -0,3708 | 3,26821E-13 |
| ACE2 | RP11-123M21.2 | -0,3708 | 3,29751E-13 |
| ACE2 | CHPT1 | -0,3708 | 3,271E-13 |
| ACE2 | MTIF3 | -0,3708 | 3,29448E-13 |
| ACE2 | RP11-509A17.3 | -0,3708 | 3,27404E-13 |
| ACE2 | CTA-243E7.2 | -0,3708 | 3,27734E-13 |
| ACE2 | CARM1 | -0,3709 | 3,22358E-13 |
| ACE2 | ANKEF1 | -0,3709 | 3,21519E-13 |
| ACE2 | AC067945.4 | -0,371 | 3,20521E-13 |
| ACE2 | RP11-555E9.1 | -0,371 | 3,18977E-13 |
| ACE2 | CTB-179K24.3 | -0,371 | 3,21202E-13 |
| ACE2 | AC004112.4 | -0,3711 | 3,15965E-13 |
| ACE2 | UBN2 | -0,3711 | 3,14136E-13 |
| ACE2 | CSRP2 | -0,3711 | 3,14446E-13 |
| ACE2 | KLHDC7A | -0,3712 | 3,08422E-13 |
| ACE2 | AC097532.2 | -0,3712 | 3,07843E-13 |
| ACE2 | CTC-276P9.2 | -0,3712 | 3,09016E-13 |
| ACE2 | RP11-386M24.10 | -0,3712 | 3,10823E-13 |
| ACE2 | RP11-271K11.5 | -0,3712 | 3,08307E-13 |
| ACE2 | GNB1L | -0,3712 | 3,10656E-13 |
| ACE2 | ADORA2BP1 | -0,3713 | 3,05253E-13 |
| ACE2 | AC011525.2 | -0,3713 | 3,03258E-13 |
| ACE2 | SGMS2 | -0,3714 | 3,01539E-13 |
| ACE2 | CTB-58E17.1 | -0,3714 | 2,97808E-13 |
| ACE2 | LINC01832 | -0,3715 | 2,95748E-13 |
| ACE2 | RP3-468B3.2 | -0,3716 | 2,92301E-13 |

| | | | |
|---|---|---|---|
| ACE2 | RP11-806L2.6 | -0,3716 | 2,89235E-13 |
| ACE2 | POU6F2-AS1 | -0,3717 | 2,84242E-13 |
| ACE2 | RCC2P3 | -0,3717 | 2,83497E-13 |
| ACE2 | KRT18P46 | -0,3718 | 2,81335E-13 |
| ACE2 | RP11-93I21.3 | -0,3718 | 2,78938E-13 |
| ACE2 | RAE1 | -0,3718 | 2,8148E-13 |
| ACE2 | EBNA1BP2 | -0,3719 | 2,77798E-13 |
| ACE2 | AC013727.2 | -0,3719 | 2,78062E-13 |
| ACE2 | NYAP2 | -0,3719 | 2,77974E-13 |
| ACE2 | AK4P5 | -0,3719 | 2,78064E-13 |
| ACE2 | AC006978.6 | -0,3719 | 2,77096E-13 |
| ACE2 | CASC8 | -0,3719 | 2,76159E-13 |
| ACE2 | RP11-60E8.2 | -0,3719 | 2,77877E-13 |
| ACE2 | RP11-49G10.3 | -0,3719 | 2,77079E-13 |
| ACE2 | LINC01845 | -0,372 | 2,73744E-13 |
| ACE2 | CTD-2254N19.1 | -0,3721 | 2,67461E-13 |
| ACE2 | OR1L8 | -0,3721 | 2,66306E-13 |
| ACE2 | RAB40B | -0,3721 | 2,68554E-13 |
| ACE2 | MROH8 | -0,3721 | 2,68857E-13 |
| ACE2 | RP11-269F21.2 | -0,3722 | 2,64558E-13 |
| ACE2 | RP11-374F3.5 | -0,3723 | 2,60631E-13 |
| ACE2 | NHLRC4 | -0,3723 | 2,58547E-13 |
| ACE2 | VN1R51P | -0,3724 | 2,57331E-13 |
| ACE2 | RABEPK | -0,3724 | 2,55351E-13 |
| ACE2 | LINC01416 | -0,3724 | 2,55285E-13 |
| ACE2 | EEF1GP5 | -0,3724 | 2,54911E-13 |
| ACE2 | TMEFF1 | -0,3725 | 2,49964E-13 |
| ACE2 | IFT140 | -0,3725 | 2,50013E-13 |
| ACE2 | CTC-218B8.3 | -0,3725 | 2,5365E-13 |
| ACE2 | LIM2 | -0,3725 | 2,5307E-13 |
| ACE2 | MPPED1 | -0,3725 | 2,5217E-13 |
| ACE2 | RP11-716O23.2 | -0,3726 | 2,47051E-13 |
| ACE2 | AC011290.4 | -0,3726 | 2,47501E-13 |
| ACE2 | MZF1-AS1 | -0,3726 | 2,46012E-13 |
| ACE2 | RP11-390F4.3 | -0,3727 | 2,42824E-13 |
| ACE2 | LINC00856 | -0,3727 | 2,458E-13 |
| ACE2 | LINC02088 | -0,3727 | 2,44842E-13 |
| ACE2 | RP11-672L10.6 | -0,3727 | 2,43871E-13 |
| ACE2 | PRODH2 | -0,3727 | 2,45721E-13 |
| ACE2 | CLDN17 | -0,3727 | 2,44917E-13 |
| ACE2 | MYCBP | -0,3728 | 2,39113E-13 |
| ACE2 | RP11-29H23.5 | -0,3728 | 2,41827E-13 |
| ACE2 | MAPKAPK2 | -0,3728 | 2,41944E-13 |
| ACE2 | RP11-362I1.1 | -0,3728 | 2,38706E-13 |
| ACE2 | PCDHB3 | -0,3728 | 2,39887E-13 |
| ACE2 | RP11-848D3.4 | -0,3728 | 2,40926E-13 |
| ACE2 | POLR2F | -0,3728 | 2,39027E-13 |
| ACE2 | RP11-1055B8.2 | -0,3729 | 2,34886E-13 |
| ACE2 | SLC16A1-AS1 | -0,373 | 2,31178E-13 |
| ACE2 | HNRNPA1P2 | -0,373 | 2,30839E-13 |
| ACE2 | PPP3CC | -0,373 | 2,34423E-13 |
| ACE2 | ATP8B5P | -0,373 | 2,32554E-13 |
| ACE2 | HJURP | -0,3731 | 2,29069E-13 |
| ACE2 | CTB-49A3.2 | -0,3731 | 2,29944E-13 |

| | | | |
|---|---|---|---|
| ACE2 | BACH2 | -0,3731 | 2,30756E-13 |
| ACE2 | WNT10B | -0,3731 | 2,27552E-13 |
| ACE2 | CEP350 | -0,3732 | 2,2661E-13 |
| ACE2 | AC092684.1 | -0,3732 | 2,25601E-13 |
| ACE2 | TMEM5-AS1 | -0,3732 | 2,24412E-13 |
| ACE2 | RLBP1 | -0,3732 | 2,24949E-13 |
| ACE2 | VSX2 | -0,3733 | 2,21716E-13 |
| ACE2 | RPSAP31 | -0,3734 | 2,18294E-13 |
| ACE2 | PPATP1 | -0,3735 | 2,14041E-13 |
| ACE2 | CTD-2194D22.5 | -0,3735 | 2,15843E-13 |
| ACE2 | RP11-220D10.1 | -0,3735 | 2,14891E-13 |
| ACE2 | WASF1 | -0,3736 | 2,10611E-13 |
| ACE2 | RP11-366H4.3 | -0,3737 | 2,08373E-13 |
| ACE2 | RP5-906C1.1 | -0,3737 | 2,06683E-13 |
| ACE2 | SLAMF6 | -0,3738 | 2,05741E-13 |
| ACE2 | RP11-109E24.1 | -0,3738 | 2,04883E-13 |
| ACE2 | CTD-3065J16.9 | -0,3738 | 2,06183E-13 |
| ACE2 | NAA11 | -0,3739 | 2,03184E-13 |
| ACE2 | RP11-264E18.1 | -0,3739 | 2,03212E-13 |
| ACE2 | RP1-177I10.1 | -0,374 | 2,00076E-13 |
| ACE2 | LINC02175 | -0,374 | 1,99107E-13 |
| ACE2 | RP11-498M5.2 | -0,3741 | 1,94438E-13 |
| ACE2 | ZNF815P | -0,3741 | 1,94341E-13 |
| ACE2 | GEMIN2 | -0,3741 | 1,96271E-13 |
| ACE2 | MZT1P2 | -0,3742 | 1,92047E-13 |
| ACE2 | RP11-1D12.1 | -0,3742 | 1,92062E-13 |
| ACE2 | RP11-144G16.1 | -0,3742 | 1,93284E-13 |
| ACE2 | RP11-299H22.6 | -0,3742 | 1,93508E-13 |
| ACE2 | RPL12P12 | -0,3742 | 1,91124E-13 |
| ACE2 | AC108051.2 | -0,3743 | 1,88478E-13 |
| ACE2 | SUPT3H | -0,3743 | 1,90436E-13 |
| ACE2 | RP11-834C11.6 | -0,3743 | 1,8881E-13 |
| ACE2 | LINC00363 | -0,3743 | 1,88863E-13 |
| ACE2 | RP11-346J10.3 | -0,3744 | 1,86095E-13 |
| ACE2 | C17orf51 | -0,3744 | 1,87171E-13 |
| ACE2 | CTC-360P9.1 | -0,3744 | 1,85708E-13 |
| ACE2 | GRHL3 | -0,3745 | 1,83544E-13 |
| ACE2 | KLRC4 | -0,3745 | 1,83145E-13 |
| ACE2 | RP11-403P17.4 | -0,3745 | 1,83157E-13 |
| ACE2 | GFRA2 | -0,3746 | 1,80228E-13 |
| ACE2 | RP11-732A21.2 | -0,3746 | 1,80494E-13 |
| ACE2 | AP001601.2 | -0,3746 | 1,80453E-13 |
| ACE2 | OR5AC2 | -0,3748 | 1,75956E-13 |
| ACE2 | SEMA5A-AS1 | -0,3748 | 1,76158E-13 |
| ACE2 | FAM74A3 | -0,3748 | 1,73859E-13 |
| ACE2 | KCNIP2 | -0,3748 | 1,74441E-13 |
| ACE2 | CTD-2014E2.2 | -0,3748 | 1,7439E-13 |
| ACE2 | BMS1P4 | -0,375 | 1,68017E-13 |
| ACE2 | RP11-744N12.3 | -0,375 | 1,68257E-13 |
| ACE2 | RPL21P103 | -0,3751 | 1,67832E-13 |
| ACE2 | RP11-359N5.1 | -0,3751 | 1,67657E-13 |
| ACE2 | RP11-442O18.1 | -0,3752 | 1,64998E-13 |
| ACE2 | AP1M1 | -0,3752 | 1,6486E-13 |
| ACE2 | RP11-340F16.2 | -0,3753 | 1,60955E-13 |

| | | | |
|---|---|---|---|
| ACE2 | HCG14 | -0,3753 | 1,61735E-13 |
| ACE2 | GS1-259H13.7 | -0,3753 | 1,6147E-13 |
| ACE2 | RP11-84O12.3 | -0,3754 | 1,57986E-13 |
| ACE2 | AC004899.3 | -0,3754 | 1,5901E-13 |
| ACE2 | OR11K2P | -0,3754 | 1,58326E-13 |
| ACE2 | AC005863.1 | -0,3754 | 1,59326E-13 |
| ACE2 | RN7SKP208 | -0,3755 | 1,56763E-13 |
| ACE2 | CTAGE9 | -0,3755 | 1,5602E-13 |
| ACE2 | SPDYE20P | -0,3755 | 1,56547E-13 |
| ACE2 | RP11-178L8.6 | -0,3755 | 1,57287E-13 |
| ACE2 | RP1-193H18.3 | -0,3755 | 1,55516E-13 |
| ACE2 | RP11-99H20.1 | -0,3756 | 1,52716E-13 |
| ACE2 | GALNT16 | -0,3756 | 1,528E-13 |
| ACE2 | RP13-270P17.2 | -0,3756 | 1,54906E-13 |
| ACE2 | PPP4R1L | -0,3756 | 1,54746E-13 |
| ACE2 | BRMS1 | -0,3757 | 1,52169E-13 |
| ACE2 | P2RY2 | -0,3757 | 1,50357E-13 |
| ACE2 | PSMC1P11 | -0,3758 | 1,48795E-13 |
| ACE2 | RP11-575B7.3 | -0,376 | 1,43805E-13 |
| ACE2 | ARF4-AS1 | -0,376 | 1,44069E-13 |
| ACE2 | KPNA5 | -0,3761 | 1,4266E-13 |
| ACE2 | RP11-428L9.1 | -0,3762 | 1,4057E-13 |
| ACE2 | RP1-65P5.5 | -0,3762 | 1,40833E-13 |
| ACE2 | AGPAT3 | -0,3762 | 1,38939E-13 |
| ACE2 | NCOA4P3 | -0,3763 | 1,38107E-13 |
| ACE2 | CDH12 | -0,3763 | 1,37314E-13 |
| ACE2 | DNAJB1 | -0,3763 | 1,36799E-13 |
| ACE2 | RPL21P131 | -0,3763 | 1,36831E-13 |
| ACE2 | RP11-505O17.1 | -0,3764 | 1,35432E-13 |
| ACE2 | ASAP2 | -0,3764 | 1,35195E-13 |
| ACE2 | INO80B | -0,3764 | 1,3534E-13 |
| ACE2 | RP4-669L17.8 | -0,3765 | 1,34356E-13 |
| ACE2 | RP11-324O2.3 | -0,3765 | 1,3299E-13 |
| ACE2 | TRIM7 | -0,3766 | 1,32077E-13 |
| ACE2 | AC006042.7 | -0,3766 | 1,30158E-13 |
| ACE2 | HINT2 | -0,3766 | 1,30424E-13 |
| ACE2 | ANKRD26P3 | -0,3766 | 1,3058E-13 |
| ACE2 | CTB-129P6.11 | -0,3766 | 1,31827E-13 |
| ACE2 | PRAMEF1 | -0,3767 | 1,29825E-13 |
| ACE2 | GOLPH3L | -0,3767 | 1,29396E-13 |
| ACE2 | RP11-278H7.1 | -0,3767 | 1,29638E-13 |
| ACE2 | ELL3 | -0,3767 | 1,2961E-13 |
| ACE2 | RP11-978I15.10 | -0,3768 | 1,26036E-13 |
| ACE2 | RP11-51J9.5 | -0,3768 | 1,26497E-13 |
| ACE2 | RP13-16H11.7 | -0,3768 | 1,26204E-13 |
| ACE2 | GSG2 | -0,3768 | 1,26278E-13 |
| ACE2 | RP11-749I16.3 | -0,3768 | 1,26189E-13 |
| ACE2 | AC007948.1 | -0,3768 | 1,27284E-13 |
| ACE2 | TRIM29 | -0,3769 | 1,2575E-13 |
| ACE2 | PLS1 | -0,377 | 1,22506E-13 |
| ACE2 | RPS17P11 | -0,377 | 1,22252E-13 |
| ACE2 | CTB-113P19.1 | -0,377 | 1,23151E-13 |
| ACE2 | RP11-457I16.4 | -0,377 | 1,2346E-13 |
| ACE2 | RP11-605F20.1 | -0,377 | 1,23425E-13 |

| | | | |
|---|---|---|---|
| ACE2 | LINC01660 | -0,377 | 1,23515E-13 |
| ACE2 | PLA2G3 | -0,377 | 1,23007E-13 |
| ACE2 | RP4-799P18.4 | -0,3771 | 1,20226E-13 |
| ACE2 | PLCH1 | -0,3771 | 1,20843E-13 |
| ACE2 | FEM1B | -0,3771 | 1,20953E-13 |
| ACE2 | RP11-319G9.3 | -0,3771 | 1,22023E-13 |
| ACE2 | CCDC77 | -0,3772 | 1,20086E-13 |
| ACE2 | KLK6 | -0,3772 | 1,18945E-13 |
| ACE2 | LINC01740 | -0,3773 | 1,16825E-13 |
| ACE2 | GRIA1 | -0,3773 | 1,16468E-13 |
| ACE2 | WDR90 | -0,3773 | 1,16888E-13 |
| ACE2 | LIFR-AS1 | -0,3774 | 1,15799E-13 |
| ACE2 | GPC6-AS2 | -0,3774 | 1,15172E-13 |
| ACE2 | AP001042.1 | -0,3774 | 1,15907E-13 |
| ACE2 | SPANXN2 | -0,3774 | 1,15368E-13 |
| ACE2 | RP11-327I22.5 | -0,3775 | 1,14346E-13 |
| ACE2 | NTS | -0,3775 | 1,12858E-13 |
| ACE2 | GLYAT | -0,3776 | 1,11325E-13 |
| ACE2 | RP11-164C12.2 | -0,3776 | 1,11429E-13 |
| ACE2 | LINC00869 | -0,3777 | 1,10215E-13 |
| ACE2 | SMAD1-AS1 | -0,3777 | 1,09723E-13 |
| ACE2 | KB-1000E4.2 | -0,3777 | 1,10136E-13 |
| ACE2 | SWT1 | -0,3778 | 1,07713E-13 |
| ACE2 | RP3-406P24.4 | -0,3778 | 1,08141E-13 |
| ACE2 | C7orf69 | -0,3778 | 1,08195E-13 |
| ACE2 | RPL7P51 | -0,3778 | 1,07539E-13 |
| ACE2 | RGMB | -0,3779 | 1,06646E-13 |
| ACE2 | FTH1P10 | -0,378 | 1,04311E-13 |
| ACE2 | EIF3KP1 | -0,378 | 1,04891E-13 |
| ACE2 | ZBED5-AS1 | -0,378 | 1,04333E-13 |
| ACE2 | RP11-126K1.2 | -0,3781 | 1,03157E-13 |
| ACE2 | RP11-319G9.5 | -0,3781 | 1,03932E-13 |
| ACE2 | ZNF667-AS1 | -0,3781 | 1,02824E-13 |
| ACE2 | ANKRD23 | -0,3782 | 1,01792E-13 |
| ACE2 | TRGC2 | -0,3782 | 1,01833E-13 |
| ACE2 | RP11-455I9.1 | -0,3782 | 1,0133E-13 |
| ACE2 | COQ4 | -0,3782 | 1,01911E-13 |
| ACE2 | AHCYP4 | -0,3783 | 1,00597E-13 |
| ACE2 | ITGA10 | -0,3784 | 9,8957E-14 |
| ACE2 | RP11-677O4.6 | -0,3784 | 9,82746E-14 |
| ACE2 | ACOT7 | -0,3785 | 9,69061E-14 |
| ACE2 | OR51S1 | -0,3785 | 9,64943E-14 |
| ACE2 | CIAPIN1 | -0,3785 | 9,65614E-14 |
| ACE2 | TULP3P1 | -0,3786 | 9,4911E-14 |
| ACE2 | RP11-562L8.1 | -0,3786 | 9,47626E-14 |
| ACE2 | RP11-96A1.4 | -0,3787 | 9,41047E-14 |
| ACE2 | AC007556.3 | -0,3789 | 9,05082E-14 |
| ACE2 | TDGF1P2 | -0,3789 | 9,04114E-14 |
| ACE2 | IGSF22 | -0,3789 | 9,06435E-14 |
| ACE2 | KIF3B | -0,3789 | 9,10889E-14 |
| ACE2 | CPXCR1 | -0,3789 | 9,1074E-14 |
| ACE2 | RP11-245P10.4 | -0,379 | 8,94832E-14 |
| ACE2 | RP11-23P13.6 | -0,379 | 8,97281E-14 |
| ACE2 | RP11-671M22.4 | -0,379 | 8,89064E-14 |

| | | | |
|---|---|---|---|
| ACE2 | RP11-525K10.2 | -0,379 | 8,93543E-14 |
| ACE2 | BCRP6 | -0,379 | 8,96461E-14 |
| ACE2 | RP11-359K18.3 | -0,3791 | 8,82934E-14 |
| ACE2 | RP11-170N16.3 | -0,3792 | 8,58415E-14 |
| ACE2 | RP3-406A7.1 | -0,3792 | 8,61337E-14 |
| ACE2 | RP4-601P9.1 | -0,3792 | 8,66333E-14 |
| ACE2 | U51244.2 | -0,3793 | 8,49181E-14 |
| ACE2 | ARHGAP11B | -0,3793 | 8,44111E-14 |
| ACE2 | RP11-152L20.3 | -0,3793 | 8,45588E-14 |
| ACE2 | RNA5SP258 | -0,3794 | 8,29549E-14 |
| ACE2 | AC007773.2 | -0,3794 | 8,41292E-14 |
| ACE2 | RP11-254A17.1 | -0,3795 | 8,24672E-14 |
| ACE2 | ZC3H14 | -0,3795 | 8,1824E-14 |
| ACE2 | CCL23 | -0,3795 | 8,18045E-14 |
| ACE2 | NLGN1 | -0,3796 | 8,13276E-14 |
| ACE2 | POU2AF1 | -0,3797 | 7,97745E-14 |
| ACE2 | RBKS | -0,3798 | 7,81878E-14 |
| ACE2 | CTC-311G1.2 | -0,3798 | 7,83516E-14 |
| ACE2 | KRT7 | -0,3798 | 7,86846E-14 |
| ACE2 | RPL24P7 | -0,3799 | 7,72117E-14 |
| ACE2 | LINC01195 | -0,3799 | 7,74929E-14 |
| ACE2 | LINC01360 | -0,38 | 7,64254E-14 |
| ACE2 | POLK | -0,38 | 7,52567E-14 |
| ACE2 | RIMBP2 | -0,38 | 7,61222E-14 |
| ACE2 | DLGAP1-AS3 | -0,3801 | 7,50566E-14 |
| ACE2 | LINC00518 | -0,3802 | 7,36858E-14 |
| ACE2 | RP11-2F20.1 | -0,3802 | 7,31271E-14 |
| ACE2 | DUX4L27 | -0,3802 | 7,34478E-14 |
| ACE2 | AC006050.2 | -0,3802 | 7,36097E-14 |
| ACE2 | AC012123.1 | -0,3802 | 7,28996E-14 |
| ACE2 | ZNRD1ASP | -0,3803 | 7,17088E-14 |
| ACE2 | OR6B2 | -0,3804 | 7,11796E-14 |
| ACE2 | AC007091.1 | -0,3804 | 7,10031E-14 |
| ACE2 | AC083884.8 | -0,3804 | 7,15532E-14 |
| ACE2 | RP11-266E6.3 | -0,3804 | 7,07928E-14 |
| ACE2 | RP5-1061H20.4 | -0,3805 | 6,95541E-14 |
| ACE2 | RP11-405K6.1 | -0,3805 | 7,02404E-14 |
| ACE2 | RP11-885B4.2 | -0,3805 | 6,96434E-14 |
| ACE2 | RP11-104J23.2 | -0,3805 | 7,03499E-14 |
| ACE2 | RP11-595B24.2 | -0,3805 | 6,97011E-14 |
| ACE2 | RP11-153N17.1 | -0,3806 | 6,8583E-14 |
| ACE2 | DENND2D | -0,3809 | 6,58509E-14 |
| ACE2 | ACBD6 | -0,3809 | 6,59758E-14 |
| ACE2 | CTD-2234N14.2 | -0,3809 | 6,58533E-14 |
| ACE2 | RP11-264M12.2 | -0,3809 | 6,60242E-14 |
| ACE2 | UBBP4 | -0,3809 | 6,54729E-14 |
| ACE2 | OR7D1P | -0,3809 | 6,52618E-14 |
| ACE2 | AURKA | -0,3809 | 6,56931E-14 |
| ACE2 | CTD-3074O7.2 | -0,381 | 6,50131E-14 |
| ACE2 | RP3-348I23.2 | -0,3812 | 6,29441E-14 |
| ACE2 | RP11-53I6.3 | -0,3812 | 6,19955E-14 |
| ACE2 | TRAPPC2B | -0,3813 | 6,12438E-14 |
| ACE2 | CEP104 | -0,3814 | 6,01764E-14 |
| ACE2 | EEF1DP4 | -0,3814 | 6,02766E-14 |

| | | | |
|---|---|---|---|
| ACE2 | RP3-445O10.1 | -0,3815 | 5,93492E-14 |
| ACE2 | LRRC8C | -0,3815 | 5,9362E-14 |
| ACE2 | RP11-556N4.1 | -0,3815 | 5,97582E-14 |
| ACE2 | CIT | -0,3815 | 5,95037E-14 |
| ACE2 | RP11-368I7.6 | -0,3815 | 5,92577E-14 |
| ACE2 | RPS12P28 | -0,3815 | 5,92606E-14 |
| ACE2 | RNF225 | -0,3815 | 5,91019E-14 |
| ACE2 | TEX28 | -0,3815 | 5,96563E-14 |
| ACE2 | 7SK | -0,3816 | 5,81622E-14 |
| ACE2 | HGH1 | -0,3816 | 5,8675E-14 |
| ACE2 | AC016582.2 | -0,3816 | 5,81469E-14 |
| ACE2 | RP1-92O14.6 | -0,3817 | 5,80088E-14 |
| ACE2 | AC093166.2 | -0,3817 | 5,79999E-14 |
| ACE2 | AC122136.2 | -0,3817 | 5,7915E-14 |
| ACE2 | RP11-7C6.1 | -0,3817 | 5,70985E-14 |
| ACE2 | RP3-461F17.1 | -0,3817 | 5,78913E-14 |
| ACE2 | ASNSP1 | -0,3818 | 5,65074E-14 |
| ACE2 | RP11-401H2.1 | -0,3818 | 5,61953E-14 |
| ACE2 | RP11-500G22.4 | -0,3818 | 5,67056E-14 |
| ACE2 | RP11-669E14.6 | -0,3818 | 5,61826E-14 |
| ACE2 | LRRC37A17P | -0,3818 | 5,65285E-14 |
| ACE2 | AC079613.1 | -0,3819 | 5,57536E-14 |
| ACE2 | CTD-2516K3.3 | -0,3819 | 5,53655E-14 |
| ACE2 | RP3-403L10.3 | -0,3819 | 5,61081E-14 |
| ACE2 | C20orf202 | -0,3819 | 5,60273E-14 |
| ACE2 | POU4F2 | -0,382 | 5,44108E-14 |
| ACE2 | RP11-540B6.6 | -0,382 | 5,5028E-14 |
| ACE2 | TAAR2 | -0,3821 | 5,38278E-14 |
| ACE2 | CTD-2555I5.1 | -0,3821 | 5,37669E-14 |
| ACE2 | CCDC175 | -0,3821 | 5,4357E-14 |
| ACE2 | RP11-212E8.1 | -0,3821 | 5,40945E-14 |
| ACE2 | TRAIP | -0,3822 | 5,27423E-14 |
| ACE2 | CCDC183-AS1 | -0,3823 | 5,19311E-14 |
| ACE2 | VCPKMT | -0,3823 | 5,22605E-14 |
| ACE2 | OR5H14 | -0,3824 | 5,14538E-14 |
| ACE2 | VPS51 | -0,3824 | 5,16254E-14 |
| ACE2 | HSPBAP1 | -0,3825 | 5,02525E-14 |
| ACE2 | DMBT1P1 | -0,3825 | 5,04687E-14 |
| ACE2 | RP11-359E8.5 | -0,3826 | 4,93769E-14 |
| ACE2 | AC104781.1 | -0,3826 | 4,97268E-14 |
| ACE2 | LNPK | -0,3826 | 4,99235E-14 |
| ACE2 | GDF9 | -0,3826 | 4,94262E-14 |
| ACE2 | CFAP54 | -0,3826 | 4,9298E-14 |
| ACE2 | RP11-299H22.1 | -0,3826 | 4,96174E-14 |
| ACE2 | FAM47C | -0,3826 | 4,9709E-14 |
| ACE2 | TNFRSF8 | -0,3827 | 4,91621E-14 |
| ACE2 | SLC25A5P6 | -0,3827 | 4,86161E-14 |
| ACE2 | RNU4-72P | -0,3827 | 4,85251E-14 |
| ACE2 | RP11-319G9.1 | -0,3827 | 4,91776E-14 |
| ACE2 | RP11-22C11.2 | -0,3828 | 4,77841E-14 |
| ACE2 | SGO1P2 | -0,3829 | 4,76561E-14 |
| ACE2 | RP11-81H14.2 | -0,3829 | 4,74582E-14 |
| ACE2 | BRIP1 | -0,3829 | 4,76487E-14 |
| ACE2 | RP13-143G15.4 | -0,383 | 4,64782E-14 |

| | | | |
|---|---|---|---|
| ACE2 | WNT2 | -0,383 | 4,61907E-14 |
| ACE2 | LDLRAD4 | -0,383 | 4,65319E-14 |
| ACE2 | bP-21201H5.1 | -0,3831 | 4,54998E-14 |
| ACE2 | RP11-76E17.4 | -0,3832 | 4,50901E-14 |
| ACE2 | BAD | -0,3833 | 4,42314E-14 |
| ACE2 | XX-CR54.1 | -0,3834 | 4,32758E-14 |
| ACE2 | LINC02011 | -0,3834 | 4,35381E-14 |
| ACE2 | GNL1 | -0,3834 | 4,35537E-14 |
| ACE2 | LRRC1 | -0,3834 | 4,39439E-14 |
| ACE2 | CTB-176F20.3 | -0,3834 | 4,34414E-14 |
| ACE2 | RPL36AL | -0,3835 | 4,31946E-14 |
| ACE2 | DNAL1 | -0,3835 | 4,30454E-14 |
| ACE2 | RP11-138E16.1 | -0,3835 | 4,28447E-14 |
| ACE2 | NPHS1 | -0,3835 | 4,2702E-14 |
| ACE2 | CTD-2033D15.1 | -0,3836 | 4,18969E-14 |
| ACE2 | RP11-133N21.10 | -0,3837 | 4,16618E-14 |
| ACE2 | FAP | -0,3838 | 4,10816E-14 |
| ACE2 | NPIPB15 | -0,3838 | 4,107E-14 |
| ACE2 | RPL29P32 | -0,3838 | 4,11495E-14 |
| ACE2 | AC012442.6 | -0,3839 | 4,03817E-14 |
| ACE2 | LINC00620 | -0,3839 | 4,02935E-14 |
| ACE2 | TLE4 | -0,3839 | 4,02092E-14 |
| ACE2 | CTC-344H19.4 | -0,3839 | 4,03466E-14 |
| ACE2 | RP11-452J21.2 | -0,384 | 3,96658E-14 |
| ACE2 | RP11-216P16.6 | -0,384 | 3,92472E-14 |
| ACE2 | CCDC178 | -0,384 | 3,97396E-14 |
| ACE2 | AP000695.4 | -0,384 | 3,94163E-14 |
| ACE2 | RP4-660H19.1 | -0,3841 | 3,91243E-14 |
| ACE2 | AC090505.1 | -0,3841 | 3,90701E-14 |
| ACE2 | RP11-286E11.2 | -0,3841 | 3,89343E-14 |
| ACE2 | TXNP1 | -0,3841 | 3,89237E-14 |
| ACE2 | RP11-285A18.2 | -0,3841 | 3,9096E-14 |
| ACE2 | RP11-488I20.8 | -0,3841 | 3,85785E-14 |
| ACE2 | RP11-679B19.2 | -0,3841 | 3,85869E-14 |
| ACE2 | AC005176.2 | -0,3842 | 3,80413E-14 |
| ACE2 | GVQW2 | -0,3843 | 3,78475E-14 |
| ACE2 | TMPRSS11A | -0,3844 | 3,67133E-14 |
| ACE2 | RP11-624A4.1 | -0,3844 | 3,67509E-14 |
| ACE2 | RP11-395P13.7 | -0,3844 | 3,69023E-14 |
| ACE2 | GS1-251I9.2 | -0,3844 | 3,69788E-14 |
| ACE2 | MS4A8 | -0,3845 | 3,63917E-14 |
| ACE2 | OACYLP | -0,3845 | 3,65367E-14 |
| ACE2 | RP11-522M21.2 | -0,3847 | 3,5401E-14 |
| ACE2 | FAM53C | -0,3847 | 3,54581E-14 |
| ACE2 | FAM157A | -0,3848 | 3,48624E-14 |
| ACE2 | RP11-544L8__B.4 | -0,3848 | 3,45842E-14 |
| ACE2 | RP11-60C6.5 | -0,3849 | 3,40873E-14 |
| ACE2 | RP11-146F11.4 | -0,3849 | 3,41191E-14 |
| ACE2 | RP5-888M10.2 | -0,385 | 3,37496E-14 |
| ACE2 | LINC00211 | -0,385 | 3,34414E-14 |
| ACE2 | RP11-386B13.1 | -0,385 | 3,37363E-14 |
| ACE2 | FAM166AP2 | -0,385 | 3,3578E-14 |
| ACE2 | U2AF1L4 | -0,3851 | 3,30197E-14 |
| ACE2 | C1orf216 | -0,3852 | 3,21687E-14 |

| | | | |
|---|---|---|---|
| ACE2 | KCNA10 | -0,3853 | 3,17995E-14 |
| ACE2 | VN1R2 | -0,3853 | 3,20869E-14 |
| ACE2 | AC006116.13 | -0,3853 | 3,17487E-14 |
| ACE2 | OOEP | -0,3854 | 3,14953E-14 |
| ACE2 | ZNF239 | -0,3854 | 3,1203E-14 |
| ACE2 | LINC00668 | -0,3855 | 3,10211E-14 |
| ACE2 | LBP | -0,3855 | 3,08557E-14 |
| ACE2 | NUP210L | -0,3856 | 3,02351E-14 |
| ACE2 | ZBTB20-AS3 | -0,3856 | 3,01418E-14 |
| ACE2 | RP11-150O12.4 | -0,3856 | 3,01674E-14 |
| ACE2 | TNFSF11 | -0,3856 | 3,01575E-14 |
| ACE2 | SERAC1 | -0,3857 | 2,96963E-14 |
| ACE2 | KCNH3 | -0,3857 | 2,99439E-14 |
| ACE2 | FRG2KP | -0,3857 | 2,97801E-14 |
| ACE2 | RP11-663P9.1 | -0,3858 | 2,91941E-14 |
| ACE2 | LINC01607 | -0,3858 | 2,94658E-14 |
| ACE2 | RP11-676J12.8 | -0,3858 | 2,93158E-14 |
| ACE2 | AC013448.2 | -0,3859 | 2,90726E-14 |
| ACE2 | GMNN | -0,3859 | 2,89847E-14 |
| ACE2 | MAPKBP1 | -0,3859 | 2,88691E-14 |
| ACE2 | PLEKHG6 | -0,386 | 2,84104E-14 |
| ACE2 | CTD-2196E14.5 | -0,386 | 2,8569E-14 |
| ACE2 | C4BPAP2 | -0,3861 | 2,77818E-14 |
| ACE2 | TEX261 | -0,3861 | 2,78253E-14 |
| ACE2 | LIMD1-AS1 | -0,3861 | 2,81728E-14 |
| ACE2 | RP11-378I6.2 | -0,3861 | 2,81224E-14 |
| ACE2 | RP11-834C11.15 | -0,3861 | 2,77971E-14 |
| ACE2 | CCNG2P1 | -0,3861 | 2,80588E-14 |
| ACE2 | RP11-386G11.10 | -0,3862 | 2,74123E-14 |
| ACE2 | AC140725.4 | -0,3862 | 2,74245E-14 |
| ACE2 | C5orf58 | -0,3863 | 2,71835E-14 |
| ACE2 | RP11-6F6.1 | -0,3863 | 2,72547E-14 |
| ACE2 | KDM4D | -0,3863 | 2,69513E-14 |
| ACE2 | CCDC144NL-AS1 | -0,3863 | 2,72099E-14 |
| ACE2 | OR7D4 | -0,3863 | 2,69044E-14 |
| ACE2 | RP5-1121H13.1 | -0,3863 | 2,69869E-14 |
| ACE2 | RP11-266N13.2 | -0,3864 | 2,67885E-14 |
| ACE2 | LINC00566 | -0,3864 | 2,67539E-14 |
| ACE2 | RPL12P45 | -0,3865 | 2,59911E-14 |
| ACE2 | RP11-567C20.3 | -0,3865 | 2,61986E-14 |
| ACE2 | LRRIQ3 | -0,3865 | 2,61549E-14 |
| ACE2 | RP11-544D21.2 | -0,3865 | 2,63103E-14 |
| ACE2 | RP11-495O11.1 | -0,3865 | 2,62066E-14 |
| ACE2 | LINC00615 | -0,3865 | 2,61164E-14 |
| ACE2 | FAM47A | -0,3865 | 2,6081E-14 |
| ACE2 | RP5-1182A14.5 | -0,3866 | 2,56702E-14 |
| ACE2 | LINC01710 | -0,3866 | 2,58537E-14 |
| ACE2 | AC012358.4 | -0,3866 | 2,56314E-14 |
| ACE2 | DIRC3-AS1 | -0,3866 | 2,5884E-14 |
| ACE2 | AC068580.7 | -0,3866 | 2,55945E-14 |
| ACE2 | SERPINB3 | -0,3866 | 2,56468E-14 |
| ACE2 | TFB1M | -0,3867 | 2,54158E-14 |
| ACE2 | RP11-325L12.7 | -0,3867 | 2,51935E-14 |
| ACE2 | RP11-989E6.3 | -0,3867 | 2,51853E-14 |

| | | | |
|---|---|---|---|
| ACE2 | HOXB1 | -0,3867 | 2,54533E-14 |
| ACE2 | SPDYE18 | -0,3868 | 2,47046E-14 |
| ACE2 | RP11-125H8.1 | -0,3868 | 2,47152E-14 |
| ACE2 | ABCC11 | -0,3868 | 2,47946E-14 |
| ACE2 | DOC2B | -0,3868 | 2,49303E-14 |
| ACE2 | ZNF859P | -0,3869 | 2,43987E-14 |
| ACE2 | CTD-2530H12.2 | -0,3869 | 2,46383E-14 |
| ACE2 | SF3B5 | -0,387 | 2,40961E-14 |
| ACE2 | FLJ42969 | -0,387 | 2,41888E-14 |
| ACE2 | MRPL46 | -0,387 | 2,41757E-14 |
| ACE2 | ALDH1L1-AS2 | -0,3871 | 2,36857E-14 |
| ACE2 | RP11-77B22.2 | -0,3871 | 2,35597E-14 |
| ACE2 | NPIPB9 | -0,3871 | 2,36145E-14 |
| ACE2 | CHKB | -0,3871 | 2,35855E-14 |
| ACE2 | RP5-1193P9.4 | -0,3872 | 2,34009E-14 |
| ACE2 | LINC00708 | -0,3872 | 2,33316E-14 |
| ACE2 | CFAP73 | -0,3872 | 2,34566E-14 |
| ACE2 | OR7E33P | -0,3872 | 2,34354E-14 |
| ACE2 | USP42 | -0,3873 | 2,29469E-14 |
| ACE2 | CYSRT1 | -0,3873 | 2,29157E-14 |
| ACE2 | LINC01465 | -0,3873 | 2,28406E-14 |
| ACE2 | RP11-417L19.2 | -0,3874 | 2,25184E-14 |
| ACE2 | OR4M1 | -0,3874 | 2,24358E-14 |
| ACE2 | RP11-104O19.4 | -0,3875 | 2,21379E-14 |
| ACE2 | ZNF222 | -0,3875 | 2,20314E-14 |
| ACE2 | CADM4 | -0,3876 | 2,17079E-14 |
| ACE2 | ELFN2 | -0,3876 | 2,18106E-14 |
| ACE2 | KLHDC7B | -0,3876 | 2,16797E-14 |
| ACE2 | RP4-740C4.5 | -0,3877 | 2,15828E-14 |
| ACE2 | RP11-521C10.1 | -0,3877 | 2,13982E-14 |
| ACE2 | AQP9 | -0,3877 | 2,14463E-14 |
| ACE2 | LINC02002 | -0,3877 | 2,15529E-14 |
| ACE2 | AC139099.7 | -0,3877 | 2,14778E-14 |
| ACE2 | RP11-814M22.2 | -0,3878 | 2,092E-14 |
| ACE2 | EPPK1 | -0,3878 | 2,09754E-14 |
| ACE2 | CTAGE10P | -0,3878 | 2,12307E-14 |
| ACE2 | METTL21C | -0,3878 | 2,10746E-14 |
| ACE2 | LLNLR-275H8.1 | -0,3878 | 2,10196E-14 |
| ACE2 | AC097461.4 | -0,3879 | 2,06725E-14 |
| ACE2 | TLR3 | -0,3879 | 2,06024E-14 |
| ACE2 | UNC5B-AS1 | -0,3879 | 2,08391E-14 |
| ACE2 | OR4D6 | -0,3879 | 2,05936E-14 |
| ACE2 | ERCC4 | -0,3879 | 2,08856E-14 |
| ACE2 | BPIFB5P | -0,3879 | 2,071E-14 |
| ACE2 | ZSCAN9 | -0,388 | 2,05218E-14 |
| ACE2 | AC112715.2 | -0,3882 | 1,97095E-14 |
| ACE2 | PCDHB8 | -0,3882 | 1,9823E-14 |
| ACE2 | KRT8P51 | -0,3882 | 1,95895E-14 |
| ACE2 | KIF27 | -0,3882 | 1,96937E-14 |
| ACE2 | ZNF782 | -0,3883 | 1,93159E-14 |
| ACE2 | SNAPC5 | -0,3883 | 1,93765E-14 |
| ACE2 | SLC25A33 | -0,3885 | 1,87515E-14 |
| ACE2 | TBC1D3H | -0,3885 | 1,87431E-14 |
| ACE2 | RP13-266I11.1 | -0,3885 | 1,862E-14 |

| | | | |
|---|---|---|---|
| ACE2 | RP11-142J21.1 | -0,3886 | 1,84466E-14 |
| ACE2 | PTGER3 | -0,3887 | 1,82607E-14 |
| ACE2 | RP4-621B10.8 | -0,3887 | 1,80872E-14 |
| ACE2 | ADCY8 | -0,3887 | 1,82775E-14 |
| ACE2 | RP11-280K24.1 | -0,3887 | 1,80571E-14 |
| ACE2 | RP11-665C16.6 | -0,3887 | 1,82048E-14 |
| ACE2 | KRBOX1-AS1 | -0,3888 | 1,77739E-14 |
| ACE2 | CTC-558O2.1 | -0,3888 | 1,78308E-14 |
| ACE2 | RP11-587H10.2 | -0,3888 | 1,79859E-14 |
| ACE2 | RP11-115D19.1 | -0,3889 | 1,75072E-14 |
| ACE2 | TRBV8-2 | -0,3889 | 1,76241E-14 |
| ACE2 | GOLGA6L3 | -0,3889 | 1,75354E-14 |
| ACE2 | RP11-451L19.1 | -0,3889 | 1,76663E-14 |
| ACE2 | HSP90AB2P | -0,389 | 1,72489E-14 |
| ACE2 | ODC1 | -0,3891 | 1,69769E-14 |
| ACE2 | GS1-122H1.1 | -0,3892 | 1,67866E-14 |
| ACE2 | LINC00879 | -0,3892 | 1,66972E-14 |
| ACE2 | BTBD9-AS1 | -0,3892 | 1,65943E-14 |
| ACE2 | RP11-321M21.1 | -0,3892 | 1,66589E-14 |
| ACE2 | KCNJ14 | -0,3892 | 1,66967E-14 |
| ACE2 | MYLPF | -0,3893 | 1,63742E-14 |
| ACE2 | RP11-1148O4.2 | -0,3893 | 1,65407E-14 |
| ACE2 | LACTBL1 | -0,3894 | 1,62443E-14 |
| ACE2 | OR8A1 | -0,3894 | 1,61399E-14 |
| ACE2 | RP11-335I12.2 | -0,3895 | 1,58933E-14 |
| ACE2 | AB019440.50 | -0,3895 | 1,5859E-14 |
| ACE2 | RP11-314N14.1 | -0,3896 | 1,5575E-14 |
| ACE2 | OR2C3 | -0,3897 | 1,53452E-14 |
| ACE2 | KRT8P43 | -0,3897 | 1,54362E-14 |
| ACE2 | CFAP69 | -0,3897 | 1,53272E-14 |
| ACE2 | RP11-558A11.2 | -0,3897 | 1,52968E-14 |
| ACE2 | CCDC155 | -0,3897 | 1,54211E-14 |
| ACE2 | C2orf78 | -0,3898 | 1,51626E-14 |
| ACE2 | CTB-73N10.1 | -0,3898 | 1,50466E-14 |
| ACE2 | RPL21P90 | -0,3898 | 1,51653E-14 |
| ACE2 | OR51A9P | -0,3898 | 1,50249E-14 |
| ACE2 | RP11-1197K16.2 | -0,3898 | 1,5041E-14 |
| ACE2 | CTD-2313F11.1 | -0,3899 | 1,48202E-14 |
| ACE2 | CTB-113P19.3 | -0,3899 | 1,47362E-14 |
| ACE2 | RP11-552O4.1 | -0,3899 | 1,49208E-14 |
| ACE2 | AC006486.10 | -0,3899 | 1,4738E-14 |
| ACE2 | RP11-78C3.1 | -0,3901 | 1,43479E-14 |
| ACE2 | RAB36 | -0,3901 | 1,44383E-14 |
| ACE2 | RP11-563K23.1 | -0,3902 | 1,41428E-14 |
| ACE2 | AC079779.4 | -0,3903 | 1,37951E-14 |
| ACE2 | RP11-18H7.1 | -0,3903 | 1,38319E-14 |
| ACE2 | OR7E161P | -0,3903 | 1,38382E-14 |
| ACE2 | INTS11 | -0,3905 | 1,33212E-14 |
| ACE2 | LINC01850 | -0,3905 | 1,35091E-14 |
| ACE2 | TPI1P3 | -0,3905 | 1,34372E-14 |
| ACE2 | SDHAF3 | -0,3905 | 1,33783E-14 |
| ACE2 | CNTF | -0,3905 | 1,34012E-14 |
| ACE2 | RP11-21M24.2 | -0,3905 | 1,34416E-14 |
| ACE2 | RP11-207C16.4 | -0,3906 | 1,3172E-14 |

| | | | |
|---|---|---|---|
| ACE2 | RP11-298I3.4 | -0,3906 | 1,31382E-14 |
| ACE2 | AC006116.20 | -0,3906 | 1,31275E-14 |
| ACE2 | LINC01358 | -0,3907 | 1,30205E-14 |
| ACE2 | AC018359.3 | -0,3907 | 1,29504E-14 |
| ACE2 | EGFLAM | -0,3908 | 1,2782E-14 |
| ACE2 | RFTN1 | -0,3909 | 1,24743E-14 |
| ACE2 | RP11-603J24.4 | -0,3909 | 1,24867E-14 |
| ACE2 | MLST8 | -0,3909 | 1,26581E-14 |
| ACE2 | OR2G6 | -0,391 | 1,22544E-14 |
| ACE2 | AC011753.6 | -0,391 | 1,22993E-14 |
| ACE2 | BBS12 | -0,391 | 1,22889E-14 |
| ACE2 | RP11-945A11.1 | -0,391 | 1,2338E-14 |
| ACE2 | BOLA2B | -0,391 | 1,24133E-14 |
| ACE2 | KB-68A7.2 | -0,391 | 1,23155E-14 |
| ACE2 | AGR2 | -0,3911 | 1,21689E-14 |
| ACE2 | CTD-2007A10.1 | -0,3911 | 1,21199E-14 |
| ACE2 | TDGF1P7 | -0,3911 | 1,21046E-14 |
| ACE2 | RP11-651P23.2 | -0,3912 | 1,18373E-14 |
| ACE2 | RP11-386I8.4 | -0,3912 | 1,20113E-14 |
| ACE2 | MIA2 | -0,3912 | 1,19732E-14 |
| ACE2 | USP12-AS2 | -0,3913 | 1,1656E-14 |
| ACE2 | SYT14P1 | -0,3914 | 1,15272E-14 |
| ACE2 | KLK3 | -0,3914 | 1,15963E-14 |
| ACE2 | CHIA | -0,3915 | 1,13869E-14 |
| ACE2 | RP11-93H12.4 | -0,3915 | 1,14102E-14 |
| ACE2 | COL20A1 | -0,3915 | 1,13031E-14 |
| ACE2 | LINC01707 | -0,3916 | 1,11734E-14 |
| ACE2 | CCDC39-AS1 | -0,3916 | 1,11965E-14 |
| ACE2 | CTD-3023L14.1 | -0,3916 | 1,1142E-14 |
| ACE2 | VTI1B | -0,3916 | 1,11868E-14 |
| ACE2 | RP11-147L13.7 | -0,3916 | 1,10961E-14 |
| ACE2 | RP1-118J21.25 | -0,3917 | 1,08809E-14 |
| ACE2 | RP11-315A16.1 | -0,3917 | 1,09932E-14 |
| ACE2 | AC060834.3 | -0,3917 | 1,09291E-14 |
| ACE2 | NXPE2P1 | -0,3917 | 1,09213E-14 |
| ACE2 | UBE2Q2L | -0,3917 | 1,10526E-14 |
| ACE2 | COX11P1 | -0,3918 | 1,08507E-14 |
| ACE2 | RP11-71J4.2 | -0,3918 | 1,08727E-14 |
| ACE2 | IGLV1-36 | -0,3918 | 1,07432E-14 |
| ACE2 | AC114814.4 | -0,3919 | 1,05492E-14 |
| ACE2 | RP11-371A19.2 | -0,3919 | 1,0607E-14 |
| ACE2 | LINC01224 | -0,3919 | 1,06236E-14 |
| ACE2 | AC007182.6 | -0,392 | 1,04635E-14 |
| ACE2 | RP11-1096D5.2 | -0,392 | 1,04065E-14 |
| ACE2 | UBE2Q2P9 | -0,3921 | 1,02992E-14 |
| ACE2 | ACAT1 | -0,3922 | 1,01225E-14 |
| ACE2 | RP11-442J21.2 | -0,3922 | 1,00136E-14 |
| ACE2 | RP4-777L9.3 | -0,3922 | 1,00469E-14 |
| ACE2 | RHBDD3 | -0,3922 | 1,01266E-14 |
| ACE2 | WDR63 | -0,3923 | 9,92212E-15 |
| ACE2 | RP11-177F15.1 | -0,3923 | 9,87635E-15 |
| ACE2 | EFCAB2 | -0,3923 | 9,88925E-15 |
| ACE2 | NICN1 | -0,3923 | 9,87009E-15 |
| ACE2 | AFAP1L1 | -0,3923 | 9,91213E-15 |

| | | | |
|---|---|---|---|
| ACE2 | AC012146.7 | -0,3923 | 9,88704E-15 |
| ACE2 | PRR9 | -0,3924 | 9,7154E-15 |
| ACE2 | WBP1 | -0,3924 | 9,80088E-15 |
| ACE2 | RNF7 | -0,3924 | 9,79885E-15 |
| ACE2 | RP11-501O2.1 | -0,3924 | 9,7675E-15 |
| ACE2 | LINC02197 | -0,3924 | 9,6968E-15 |
| ACE2 | CDKN2B | -0,3924 | 9,79834E-15 |
| ACE2 | RP11-673F18.1 | -0,3924 | 9,80934E-15 |
| ACE2 | AC013402.5 | -0,3927 | 9,24615E-15 |
| ACE2 | RP5-1132H15.3 | -0,3927 | 9,33632E-15 |
| ACE2 | RP11-75L1.1 | -0,3927 | 9,27783E-15 |
| ACE2 | RP5-1057B20.3 | -0,3927 | 9,20405E-15 |
| ACE2 | AC002472.11 | -0,3927 | 9,32224E-15 |
| ACE2 | MARCH3 | -0,3928 | 9,11786E-15 |
| ACE2 | RGPD1 | -0,3929 | 9,01364E-15 |
| ACE2 | LINC01818 | -0,393 | 8,85499E-15 |
| ACE2 | POMZP3 | -0,393 | 8,88142E-15 |
| ACE2 | ZFHX4 | -0,393 | 8,79457E-15 |
| ACE2 | ZCCHC8 | -0,3931 | 8,6648E-15 |
| ACE2 | PAPD5 | -0,3931 | 8,68394E-15 |
| ACE2 | AC010468.2 | -0,3932 | 8,44685E-15 |
| ACE2 | OR8G7P | -0,3932 | 8,55268E-15 |
| ACE2 | RP13-401N8.3 | -0,3932 | 8,56599E-15 |
| ACE2 | IL31RA | -0,3933 | 8,3924E-15 |
| ACE2 | OR7E31P | -0,3933 | 8,44118E-15 |
| ACE2 | CTC-559E9.12 | -0,3933 | 8,31958E-15 |
| ACE2 | RP4-656G21.1 | -0,3934 | 8,26165E-15 |
| ACE2 | YBX1 | -0,3934 | 8,21991E-15 |
| ACE2 | AC016751.3 | -0,3934 | 8,21719E-15 |
| ACE2 | RP11-293F5.8 | -0,3935 | 8,14141E-15 |
| ACE2 | KIF6 | -0,3936 | 7,97934E-15 |
| ACE2 | RP11-613D13.8 | -0,3936 | 7,96061E-15 |
| ACE2 | LINC00904 | -0,3936 | 7,93079E-15 |
| ACE2 | TSPAN6 | -0,3936 | 8,01838E-15 |
| ACE2 | RP4-614N24.1 | -0,3937 | 7,86558E-15 |
| ACE2 | RP5-1086K13.1 | -0,3937 | 7,78708E-15 |
| ACE2 | PAX3 | -0,3937 | 7,88399E-15 |
| ACE2 | YAP1P1 | -0,3937 | 7,82016E-15 |
| ACE2 | RP11-114F3.4 | -0,3937 | 7,83569E-15 |
| ACE2 | RP11-512N21.3 | -0,3937 | 7,86106E-15 |
| ACE2 | RP11-269C23.5 | -0,3937 | 7,80374E-15 |
| ACE2 | LINC01926 | -0,3937 | 7,88618E-15 |
| ACE2 | IFNLR1 | -0,3938 | 7,65017E-15 |
| ACE2 | GAK | -0,3939 | 7,61423E-15 |
| ACE2 | CTD-2376I20.1 | -0,3939 | 7,52948E-15 |
| ACE2 | GOLGA8K | -0,3939 | 7,5583E-15 |
| ACE2 | AC034228.2 | -0,394 | 7,49136E-15 |
| ACE2 | CTD-2517M22.14 | -0,394 | 7,38706E-15 |
| ACE2 | CHRM4 | -0,394 | 7,45398E-15 |
| ACE2 | RP11-474C8.5 | -0,394 | 7,41871E-15 |
| ACE2 | CTA-929C8.8 | -0,3941 | 7,31796E-15 |
| ACE2 | TRBV7-6 | -0,3942 | 7,13097E-15 |
| ACE2 | SZT2 | -0,3943 | 7,05067E-15 |
| ACE2 | GRAMD1C | -0,3943 | 7,10056E-15 |

| | | | |
|---|---|---|---|
| ACE2 | CTD-2003C8.2 | -0,3943 | 7,05728E-15 |
| ACE2 | PLB1 | -0,3944 | 6,99123E-15 |
| ACE2 | CTC-270D5.1 | -0,3944 | 6,93149E-15 |
| ACE2 | ZNF354A | -0,3944 | 6,89734E-15 |
| ACE2 | RP11-697E22.2 | -0,3945 | 6,83436E-15 |
| ACE2 | LINC01767 | -0,3946 | 6,74931E-15 |
| ACE2 | ARNTL2-AS1 | -0,3946 | 6,7202E-15 |
| ACE2 | CTD-2012K14.7 | -0,3946 | 6,67147E-15 |
| ACE2 | AC011193.1 | -0,3946 | 6,70801E-15 |
| ACE2 | JUP | -0,3946 | 6,75821E-15 |
| ACE2 | RP11-794M8.1 | -0,3946 | 6,76355E-15 |
| ACE2 | RP11-364B14.1 | -0,3946 | 6,68878E-15 |
| ACE2 | TRIM54 | -0,3947 | 6,57358E-15 |
| ACE2 | CCDC13-AS1 | -0,3947 | 6,62247E-15 |
| ACE2 | LIAS | -0,3947 | 6,6216E-15 |
| ACE2 | RP11-43F13.4 | -0,3947 | 6,55208E-15 |
| ACE2 | EYA1 | -0,3947 | 6,60453E-15 |
| ACE2 | AP001425.14 | -0,3947 | 6,57909E-15 |
| ACE2 | DDX6P1 | -0,3948 | 6,4501E-15 |
| ACE2 | RASSF7 | -0,3948 | 6,48086E-15 |
| ACE2 | DYNLL1P1 | -0,3948 | 6,49294E-15 |
| ACE2 | RP11-481F24.3 | -0,3948 | 6,54312E-15 |
| ACE2 | AC106900.6 | -0,3949 | 6,332E-15 |
| ACE2 | MPP4 | -0,3949 | 6,35523E-15 |
| ACE2 | RP11-857B24.1 | -0,3949 | 6,42854E-15 |
| ACE2 | PRNT | -0,3949 | 6,37234E-15 |
| ACE2 | RPLP0P1 | -0,3949 | 6,37426E-15 |
| ACE2 | AC105393.2 | -0,395 | 6,2631E-15 |
| ACE2 | DARS-AS1 | -0,395 | 6,22409E-15 |
| ACE2 | DPPA5P4 | -0,395 | 6,2369E-15 |
| ACE2 | CBX2 | -0,395 | 6,23617E-15 |
| ACE2 | GSTTP1 | -0,395 | 6,26569E-15 |
| ACE2 | TIMM10 | -0,3951 | 6,19136E-15 |
| ACE2 | OR5AZ1P | -0,3951 | 6,19087E-15 |
| ACE2 | CATSPERB | -0,3951 | 6,11783E-15 |
| ACE2 | RP13-152O15.5 | -0,3952 | 6,06418E-15 |
| ACE2 | DEFB135 | -0,3953 | 6,00537E-15 |
| ACE2 | LINC02039 | -0,3954 | 5,81892E-15 |
| ACE2 | RP11-738B7.1 | -0,3954 | 5,83713E-15 |
| ACE2 | RP11-150O12.3 | -0,3954 | 5,90333E-15 |
| ACE2 | RP11-304F15.3 | -0,3954 | 5,88803E-15 |
| ACE2 | DHX34 | -0,3954 | 5,81106E-15 |
| ACE2 | DNASE2B | -0,3955 | 5,7945E-15 |
| ACE2 | RP11-438F14.3 | -0,3955 | 5,71868E-15 |
| ACE2 | LINC01596 | -0,3955 | 5,77671E-15 |
| ACE2 | RP3-393K13.1 | -0,3955 | 5,7492E-15 |
| ACE2 | RP11-603J24.6 | -0,3956 | 5,68474E-15 |
| ACE2 | LINC00905 | -0,3956 | 5,65664E-15 |
| ACE2 | CHRNB2 | -0,3957 | 5,60703E-15 |
| ACE2 | RP11-578F21.4 | -0,3957 | 5,52731E-15 |
| ACE2 | MYADML2 | -0,3957 | 5,56627E-15 |
| ACE2 | RPL12P39 | -0,3957 | 5,56906E-15 |
| ACE2 | RN7SL313P | -0,3958 | 5,45982E-15 |
| ACE2 | LINC00159 | -0,3958 | 5,42553E-15 |

| | | | |
|---|---|---|---|
| ACE2 | MST1P2 | -0,3959 | 5,393E-15 |
| ACE2 | SNRPFP2 | -0,3959 | 5,34641E-15 |
| ACE2 | RP11-466F5.8 | -0,3959 | 5,39855E-15 |
| ACE2 | CFL1P8 | -0,3959 | 5,3514E-15 |
| ACE2 | LA16c-312E8.2 | -0,3959 | 5,38431E-15 |
| ACE2 | CCL14 | -0,3959 | 5,33677E-15 |
| ACE2 | RP11-123M21.1 | -0,396 | 5,31225E-15 |
| ACE2 | RP4-651E10.4 | -0,3961 | 5,18498E-15 |
| ACE2 | LGI1 | -0,3961 | 5,17636E-15 |
| ACE2 | PTH2 | -0,3961 | 5,17219E-15 |
| ACE2 | RP11-292D4.3 | -0,3962 | 5,1331E-15 |
| ACE2 | RN7SKP77 | -0,3962 | 5,09234E-15 |
| ACE2 | PIK3CD-AS2 | -0,3963 | 5,02712E-15 |
| ACE2 | STAP1 | -0,3963 | 5,04582E-15 |
| ACE2 | RP11-713N11.4 | -0,3963 | 4,98343E-15 |
| ACE2 | RP11-263E1.1 | -0,3963 | 4,99141E-15 |
| ACE2 | HOMER2P1 | -0,3963 | 4,99946E-15 |
| ACE2 | MRPS25 | -0,3964 | 4,96665E-15 |
| ACE2 | CA5AP1 | -0,3964 | 4,92595E-15 |
| ACE2 | KLK2 | -0,3964 | 4,96234E-15 |
| ACE2 | RP11-354H21.2 | -0,3965 | 4,82628E-15 |
| ACE2 | RP11-355E10.1 | -0,3965 | 4,89317E-15 |
| ACE2 | LRRC3C | -0,3965 | 4,86546E-15 |
| ACE2 | ZNF233 | -0,3966 | 4,77971E-15 |
| ACE2 | PTH1R | -0,3967 | 4,71069E-15 |
| ACE2 | RP11-586D19.1 | -0,3967 | 4,68033E-15 |
| ACE2 | UOX | -0,3968 | 4,60033E-15 |
| ACE2 | ANKRD34C | -0,3968 | 4,59927E-15 |
| ACE2 | AC008088.4 | -0,3968 | 4,57574E-15 |
| ACE2 | CTD-2337I7.1 | -0,3969 | 4,5459E-15 |
| ACE2 | RP11-734K21.2 | -0,397 | 4,48867E-15 |
| ACE2 | ZNF766 | -0,397 | 4,46091E-15 |
| ACE2 | OSGIN2 | -0,3971 | 4,39557E-15 |
| ACE2 | ZNF839 | -0,3971 | 4,34355E-15 |
| ACE2 | GOLGA6L17P | -0,3971 | 4,3604E-15 |
| ACE2 | APOBR | -0,3971 | 4,4015E-15 |
| ACE2 | LINC00466 | -0,3972 | 4,33186E-15 |
| ACE2 | OCM | -0,3972 | 4,29552E-15 |
| ACE2 | PATE4 | -0,3972 | 4,31081E-15 |
| ACE2 | RP11-401F2.4 | -0,3972 | 4,31419E-15 |
| ACE2 | KRTAP10-6 | -0,3972 | 4,33439E-15 |
| ACE2 | RP11-316M21.6 | -0,3973 | 4,23827E-15 |
| ACE2 | IGKV6D-21 | -0,3974 | 4,16839E-15 |
| ACE2 | ANKRD13A | -0,3974 | 4,15046E-15 |
| ACE2 | OR7E23P | -0,3974 | 4,12639E-15 |
| ACE2 | RP1-90G24.8 | -0,3974 | 4,14156E-15 |
| ACE2 | CTC-350I8.1 | -0,3975 | 4,05714E-15 |
| ACE2 | OGFRP1 | -0,3975 | 4,0932E-15 |
| ACE2 | HNRNPA1P54 | -0,3976 | 4,04846E-15 |
| ACE2 | LINC00486 | -0,3976 | 3,99334E-15 |
| ACE2 | EHBP1 | -0,3976 | 4,0017E-15 |
| ACE2 | RP11-174O3.1 | -0,3976 | 4,05257E-15 |
| ACE2 | AC087501.1 | -0,3976 | 4,00994E-15 |
| ACE2 | CTD-2619J13.3 | -0,3976 | 4,04701E-15 |

| | | | |
|---|---|---|---|
| ACE2 | LINC01824 | -0,3977 | 3,98056E-15 |
| ACE2 | RSPH3 | -0,3977 | 3,98143E-15 |
| ACE2 | ANXA2R | -0,3978 | 3,89515E-15 |
| ACE2 | RP11-479O9.4 | -0,3978 | 3,86505E-15 |
| ACE2 | CYP4F25P | -0,3979 | 3,83889E-15 |
| ACE2 | GPRC5D | -0,3979 | 3,81992E-15 |
| ACE2 | ZIC2 | -0,3979 | 3,81082E-15 |
| ACE2 | RP3-322G13.7 | -0,3979 | 3,84883E-15 |
| ACE2 | RP11-302I18.3 | -0,398 | 3,77779E-15 |
| ACE2 | INPP5D | -0,398 | 3,72543E-15 |
| ACE2 | RP11-478P10.1 | -0,398 | 3,76272E-15 |
| ACE2 | ANKK1 | -0,398 | 3,78118E-15 |
| ACE2 | RP11-521B24.3 | -0,398 | 3,76832E-15 |
| ACE2 | MC1R | -0,398 | 3,7398E-15 |
| ACE2 | TTN | -0,3981 | 3,69188E-15 |
| ACE2 | TRAF3IP1 | -0,3981 | 3,69128E-15 |
| ACE2 | LINC00572 | -0,3981 | 3,70011E-15 |
| ACE2 | RP5-1119A7.14 | -0,3981 | 3,6776E-15 |
| ACE2 | RP1-47A17.1 | -0,3981 | 3,66942E-15 |
| ACE2 | AC009263.2 | -0,3982 | 3,59671E-15 |
| ACE2 | CTD-2552K11.2 | -0,3982 | 3,59759E-15 |
| ACE2 | TUBB8 | -0,3982 | 3,65222E-15 |
| ACE2 | RP11-403I13.4 | -0,3983 | 3,58944E-15 |
| ACE2 | HIST1H2AA | -0,3983 | 3,55106E-15 |
| ACE2 | HLA-J | -0,3983 | 3,55826E-15 |
| ACE2 | RPL12P29 | -0,3983 | 3,54639E-15 |
| ACE2 | RP11-715J22.3 | -0,3983 | 3,53443E-15 |
| ACE2 | AC104984.4 | -0,3983 | 3,55032E-15 |
| ACE2 | CTB-32O4.3 | -0,3983 | 3,5555E-15 |
| ACE2 | SLC6A6P1 | -0,3983 | 3,55574E-15 |
| ACE2 | IGLV3-21 | -0,3983 | 3,58498E-15 |
| ACE2 | SAP18P2 | -0,3984 | 3,5028E-15 |
| ACE2 | CTC-308K20.1 | -0,3984 | 3,47645E-15 |
| ACE2 | RP5-1043L3.1 | -0,3985 | 3,44497E-15 |
| ACE2 | CRYGS | -0,3985 | 3,44121E-15 |
| ACE2 | KLF7P1 | -0,3986 | 3,36143E-15 |
| ACE2 | AC108056.1 | -0,3986 | 3,37704E-15 |
| ACE2 | RPSAP52 | -0,3986 | 3,37785E-15 |
| ACE2 | RP11-96D1.6 | -0,3986 | 3,37419E-15 |
| ACE2 | RP11-766H1.1 | -0,3986 | 3,39301E-15 |
| ACE2 | TPT1P11 | -0,3987 | 3,33974E-15 |
| ACE2 | EIF1B-AS1 | -0,3987 | 3,34601E-15 |
| ACE2 | KRT18P34 | -0,3987 | 3,32754E-15 |
| ACE2 | KIAA1324L | -0,3987 | 3,31176E-15 |
| ACE2 | GOLGA7B | -0,3987 | 3,3018E-15 |
| ACE2 | CTD-2292M16.7 | -0,3988 | 3,27531E-15 |
| ACE2 | RP11-804M7.1 | -0,3988 | 3,25674E-15 |
| ACE2 | STX1B | -0,3988 | 3,25737E-15 |
| ACE2 | ALMS1 | -0,3989 | 3,20679E-15 |
| ACE2 | AC119403.1 | -0,3989 | 3,21263E-15 |
| ACE2 | FAM182A | -0,3989 | 3,19269E-15 |
| ACE2 | RP4-669P10.20 | -0,3989 | 3,19616E-15 |
| ACE2 | DAZAP2P1 | -0,399 | 3,18088E-15 |
| ACE2 | CCDC62 | -0,399 | 3,14123E-15 |

| | | | |
|---|---|---|---|
| ACE2 | PAFAH1B1 | -0,399 | 3,14039E-15 |
| ACE2 | CEP250 | -0,399 | 3,17316E-15 |
| ACE2 | C3orf38 | -0,3991 | 3,12763E-15 |
| ACE2 | CCDC146 | -0,3991 | 3,07869E-15 |
| ACE2 | RP11-244N9.6 | -0,3991 | 3,11157E-15 |
| ACE2 | CFAP58-AS1 | -0,3991 | 3,12556E-15 |
| ACE2 | RP11-475A13.2 | -0,3991 | 3,11605E-15 |
| ACE2 | RP4-633O19__A.1 | -0,3991 | 3,08296E-15 |
| ACE2 | HAX1 | -0,3992 | 3,06817E-15 |
| ACE2 | RP11-524K14.1 | -0,3992 | 3,06095E-15 |
| ACE2 | ANGEL1 | -0,3992 | 3,04931E-15 |
| ACE2 | LINC01706 | -0,3992 | 3,07281E-15 |
| ACE2 | UTS2B | -0,3993 | 2,99376E-15 |
| ACE2 | IL21-AS1 | -0,3993 | 3,00297E-15 |
| ACE2 | RP11-141O11.1 | -0,3993 | 3,00991E-15 |
| ACE2 | SMPD4P1 | -0,3993 | 3,00726E-15 |
| ACE2 | AC007405.4 | -0,3994 | 2,93687E-15 |
| ACE2 | RP11-775H9.2 | -0,3994 | 2,95029E-15 |
| ACE2 | RP4-639J15.2 | -0,3994 | 2,93659E-15 |
| ACE2 | TLX1NB | -0,3994 | 2,9353E-15 |
| ACE2 | CIB3 | -0,3994 | 2,94492E-15 |
| ACE2 | RP11-324C10.1 | -0,3995 | 2,89328E-15 |
| ACE2 | AOC2 | -0,3995 | 2,89431E-15 |
| ACE2 | LINC01687 | -0,3995 | 2,90193E-15 |
| ACE2 | SERBP1P5 | -0,3996 | 2,84595E-15 |
| ACE2 | FAM83B | -0,3996 | 2,83264E-15 |
| ACE2 | RP11-114H23.1 | -0,3996 | 2,85761E-15 |
| ACE2 | RP11-517A5.6 | -0,3996 | 2,84573E-15 |
| ACE2 | DHRS3 | -0,3998 | 2,73701E-15 |
| ACE2 | LINC01641 | -0,3998 | 2,72763E-15 |
| ACE2 | RP11-433A10.2 | -0,3998 | 2,76261E-15 |
| ACE2 | CTD-2035E11.5 | -0,3998 | 2,76999E-15 |
| ACE2 | AC006130.3 | -0,3998 | 2,74629E-15 |
| ACE2 | OR7E89P | -0,4 | 2,66999E-15 |
| ACE2 | DCHS1 | -0,4 | 2,67276E-15 |
| ACE2 | ZNF420 | -0,4 | 2,64555E-15 |
| ACE2 | DDC-AS1 | -0,4001 | 2,5989E-15 |
| ACE2 | RP11-402L5.1 | -0,4001 | 2,62393E-15 |
| ACE2 | ZBTB20-AS5 | -0,4002 | 2,55628E-15 |
| ACE2 | PLEKHA1 | -0,4002 | 2,56537E-15 |
| ACE2 | PIGBOS1 | -0,4002 | 2,55408E-15 |
| ACE2 | CTC-265F19.3 | -0,4002 | 2,57568E-15 |
| ACE2 | AP001468.58 | -0,4002 | 2,54541E-15 |
| ACE2 | CRELD2 | -0,4002 | 2,54328E-15 |
| ACE2 | RP11-190P13.2 | -0,4003 | 2,5018E-15 |
| ACE2 | MIR583HG | -0,4003 | 2,53866E-15 |
| ACE2 | OR5V1 | -0,4003 | 2,53053E-15 |
| ACE2 | RP3-495K2.1 | -0,4004 | 2,49101E-15 |
| ACE2 | FOXD3-AS1 | -0,4005 | 2,45423E-15 |
| ACE2 | RP11-815M8.1 | -0,4005 | 2,42099E-15 |
| ACE2 | RP11-98C1.2 | -0,4005 | 2,42325E-15 |
| ACE2 | CTD-2260A17.2 | -0,4005 | 2,44801E-15 |
| ACE2 | RPL19P16 | -0,4005 | 2,45462E-15 |
| ACE2 | RDH8 | -0,4005 | 2,42379E-15 |

| | | | |
|---|---|---|---|
| ACE2 | PRAMEF14 | -0,4006 | 2,37438E-15 |
| ACE2 | FBXW12 | -0,4006 | 2,38535E-15 |
| ACE2 | RP3-400B16.1 | -0,4006 | 2,3756E-15 |
| ACE2 | RP3-365O12.2 | -0,4006 | 2,40884E-15 |
| ACE2 | RNU1-65P | -0,4006 | 2,39216E-15 |
| ACE2 | RP11-300G22.2 | -0,4006 | 2,39279E-15 |
| ACE2 | DHRS11 | -0,4006 | 2,40304E-15 |
| ACE2 | GJB3 | -0,4007 | 2,3642E-15 |
| ACE2 | RP11-291B21.2 | -0,4007 | 2,35334E-15 |
| ACE2 | TRIM69 | -0,4007 | 2,36911E-15 |
| ACE2 | ZNF971P | -0,4007 | 2,34137E-15 |
| ACE2 | ZFAT | -0,4008 | 2,30172E-15 |
| ACE2 | LINC01309 | -0,4008 | 2,29328E-15 |
| ACE2 | LINC02130 | -0,4009 | 2,26517E-15 |
| ACE2 | AC006076.1 | -0,4009 | 2,27862E-15 |
| ACE2 | CTD-2154B17.4 | -0,401 | 2,23056E-15 |
| ACE2 | LINC01002 | -0,401 | 2,22194E-15 |
| ACE2 | TBRG4 | -0,4011 | 2,18685E-15 |
| ACE2 | GPR12 | -0,4011 | 2,17873E-15 |
| ACE2 | PBK | -0,4012 | 2,14442E-15 |
| ACE2 | CALCOCO2 | -0,4012 | 2,15701E-15 |
| ACE2 | XKR7 | -0,4012 | 2,13998E-15 |
| ACE2 | RP11-565A3.1 | -0,4013 | 2,12434E-15 |
| ACE2 | NPIPB7 | -0,4013 | 2,10497E-15 |
| ACE2 | SNAP23P1 | -0,4015 | 2,03102E-15 |
| ACE2 | TCEANC2 | -0,4016 | 2,02176E-15 |
| ACE2 | STX18 | -0,4016 | 2,01763E-15 |
| ACE2 | C14orf178 | -0,4016 | 1,99402E-15 |
| ACE2 | NPRL3 | -0,4016 | 2,00271E-15 |
| ACE2 | CD177P1 | -0,4016 | 2,00311E-15 |
| ACE2 | PHF20 | -0,4016 | 1,99397E-15 |
| ACE2 | RP11-61L23.2 | -0,4017 | 1,9922E-15 |
| ACE2 | SPOCK3 | -0,4018 | 1,94124E-15 |
| ACE2 | RP11-646J21.2 | -0,4018 | 1,94807E-15 |
| ACE2 | SLC4A1 | -0,4018 | 1,95109E-15 |
| ACE2 | CTNNBL1 | -0,4018 | 1,92635E-15 |
| ACE2 | PUSL1 | -0,4019 | 1,89766E-15 |
| ACE2 | CLCN3P1 | -0,4019 | 1,91848E-15 |
| ACE2 | CDKN2D | -0,4019 | 1,90811E-15 |
| ACE2 | RN7SL583P | -0,402 | 1,87074E-15 |
| ACE2 | KCNT2 | -0,402 | 1,88985E-15 |
| ACE2 | CPNE9 | -0,402 | 1,88884E-15 |
| ACE2 | HIST1H2BB | -0,402 | 1,86372E-15 |
| ACE2 | RP4-673D20.4 | -0,402 | 1,88239E-15 |
| ACE2 | DNAJC24 | -0,4021 | 1,83948E-15 |
| ACE2 | LINC01193 | -0,4021 | 1,84061E-15 |
| ACE2 | ZNF668 | -0,4021 | 1,85079E-15 |
| ACE2 | PPP2R5A | -0,4022 | 1,79674E-15 |
| ACE2 | PNOC | -0,4022 | 1,79976E-15 |
| ACE2 | CTA-797E19.2 | -0,4022 | 1,81828E-15 |
| ACE2 | RP11-1074O12.1 | -0,4022 | 1,80525E-15 |
| ACE2 | NARSP2 | -0,4023 | 1,77593E-15 |
| ACE2 | ZBTB47 | -0,4024 | 1,73528E-15 |
| ACE2 | RP11-368M16.6 | -0,4024 | 1,7567E-15 |

| | | | |
|---|---|---|---|
| ACE2 | ARMC9 | -0,4025 | 1,7096E-15 |
| ACE2 | UTRN | -0,4025 | 1,72548E-15 |
| ACE2 | CARD11 | -0,4025 | 1,73191E-15 |
| ACE2 | C3orf62 | -0,4026 | 1,68877E-15 |
| ACE2 | TPT1P3 | -0,4026 | 1,69145E-15 |
| ACE2 | PCDHA12 | -0,4027 | 1,65433E-15 |
| ACE2 | TRIL | -0,4028 | 1,63008E-15 |
| ACE2 | RP1-116K23.1 | -0,4028 | 1,62107E-15 |
| ACE2 | UMOD | -0,4028 | 1,63896E-15 |
| ACE2 | DYNAP | -0,4029 | 1,59783E-15 |
| ACE2 | RP11-420K14.8 | -0,4029 | 1,60028E-15 |
| ACE2 | AP5S1 | -0,4029 | 1,59876E-15 |
| ACE2 | CATSPER2P1 | -0,403 | 1,57931E-15 |
| ACE2 | RPL31P57 | -0,403 | 1,57961E-15 |
| ACE2 | RP11-398B16.2 | -0,4032 | 1,53095E-15 |
| ACE2 | LINC00400 | -0,4032 | 1,51377E-15 |
| ACE2 | C18orf21 | -0,4032 | 1,50873E-15 |
| ACE2 | KLK11 | -0,4032 | 1,50906E-15 |
| ACE2 | ZSCAN4 | -0,4032 | 1,51027E-15 |
| ACE2 | IGKV6-21 | -0,4033 | 1,49378E-15 |
| ACE2 | RP11-189E14.3 | -0,4033 | 1,49436E-15 |
| ACE2 | MC2R | -0,4033 | 1,50501E-15 |
| ACE2 | AC073094.4 | -0,4034 | 1,46681E-15 |
| ACE2 | LPXN | -0,4034 | 1,46707E-15 |
| ACE2 | TRAV39 | -0,4034 | 1,47705E-15 |
| ACE2 | LRRC37A4P | -0,4034 | 1,47284E-15 |
| ACE2 | RP11-10N23.5 | -0,4035 | 1,4314E-15 |
| ACE2 | DCAF10 | -0,4035 | 1,45057E-15 |
| ACE2 | RP11-540N6.1 | -0,4035 | 1,44355E-15 |
| ACE2 | LINC01320 | -0,4036 | 1,41951E-15 |
| ACE2 | RP11-445N20.3 | -0,4036 | 1,42851E-15 |
| ACE2 | RP11-321F8.4 | -0,4036 | 1,4134E-15 |
| ACE2 | IRX6 | -0,4036 | 1,42625E-15 |
| ACE2 | RP11-495P10.4 | -0,4037 | 1,39299E-15 |
| ACE2 | ACOT12 | -0,4038 | 1,36084E-15 |
| ACE2 | RP5-1099C19.2 | -0,4038 | 1,35783E-15 |
| ACE2 | FANCC | -0,4038 | 1,36322E-15 |
| ACE2 | RP11-36B15.1 | -0,4039 | 1,344E-15 |
| ACE2 | NUP155 | -0,4039 | 1,33379E-15 |
| ACE2 | HAVCR1 | -0,4039 | 1,35124E-15 |
| ACE2 | RP11-959I15.4 | -0,404 | 1,32667E-15 |
| ACE2 | CTD-2319I12.4 | -0,404 | 1,32505E-15 |
| ACE2 | ATAD1 | -0,4041 | 1,30119E-15 |
| ACE2 | RP11-883G14.3 | -0,4041 | 1,2926E-15 |
| ACE2 | RP1-10C16.1 | -0,4042 | 1,26975E-15 |
| ACE2 | FABP5P10 | -0,4042 | 1,27009E-15 |
| ACE2 | EIF4E2 | -0,4042 | 1,26463E-15 |
| ACE2 | AC114755.2 | -0,4043 | 1,25944E-15 |
| ACE2 | ARFGEF3 | -0,4043 | 1,24675E-15 |
| ACE2 | HOXA5 | -0,4043 | 1,24553E-15 |
| ACE2 | RPL30P11 | -0,4043 | 1,25032E-15 |
| ACE2 | CTD-2125J1.1 | -0,4043 | 1,2539E-15 |
| ACE2 | RP11-306I1.2 | -0,4044 | 1,23309E-15 |
| ACE2 | OR9H1P | -0,4044 | 1,23926E-15 |

| | | | |
|---|---|---|---|
| ACE2 | SELPLG | -0,4044 | 1,23044E-15 |
| ACE2 | RP11-823P9.3 | -0,4045 | 1,21E-15 |
| ACE2 | LINC01556 | -0,4045 | 1,19981E-15 |
| ACE2 | AC005703.2 | -0,4045 | 1,20634E-15 |
| ACE2 | MRPS10P1 | -0,4046 | 1,18825E-15 |
| ACE2 | RP11-368M16.9 | -0,4046 | 1,18421E-15 |
| ACE2 | CHRM1 | -0,4046 | 1,19371E-15 |
| ACE2 | OR8D1 | -0,4046 | 1,18716E-15 |
| ACE2 | CCDC36 | -0,4047 | 1,16803E-15 |
| ACE2 | GPX5 | -0,4047 | 1,1739E-15 |
| ACE2 | RP5-916L7.2 | -0,4047 | 1,17778E-15 |
| ACE2 | FAM169B | -0,4048 | 1,13882E-15 |
| ACE2 | RP11-742D12.2 | -0,4048 | 1,14131E-15 |
| ACE2 | CDKL2 | -0,4049 | 1,1283E-15 |
| ACE2 | SLC44A4 | -0,4049 | 1,13185E-15 |
| ACE2 | TRBV23OR9-2 | -0,4049 | 1,1319E-15 |
| ACE2 | PPP1R13B | -0,4049 | 1,13131E-15 |
| ACE2 | CTD-2012B7.1 | -0,405 | 1,10952E-15 |
| ACE2 | RP11-304C12.5 | -0,405 | 1,10434E-15 |
| ACE2 | LINC00426 | -0,405 | 1,1101E-15 |
| ACE2 | RP11-304L19.1 | -0,405 | 1,116E-15 |
| ACE2 | RP3-388N13.5 | -0,4051 | 1,08386E-15 |
| ACE2 | RP11-845M18.6 | -0,4051 | 1,08208E-15 |
| ACE2 | LINC02046 | -0,4052 | 1,07441E-15 |
| ACE2 | UFL1-AS1 | -0,4052 | 1,0754E-15 |
| ACE2 | CENPU | -0,4053 | 1,05932E-15 |
| ACE2 | RP11-646J21.4 | -0,4053 | 1,04855E-15 |
| ACE2 | LRRC37B | -0,4053 | 1,05497E-15 |
| ACE2 | ANKRD30B | -0,4053 | 1,0486E-15 |
| ACE2 | RP11-339A11.2 | -0,4054 | 1,02874E-15 |
| ACE2 | RP3-454G6.2 | -0,4054 | 1,03329E-15 |
| ACE2 | PACRGL | -0,4054 | 1,04032E-15 |
| ACE2 | ENO4 | -0,4054 | 1,02755E-15 |
| ACE2 | RP1-68D18.4 | -0,4054 | 1,0299E-15 |
| ACE2 | ESPNP | -0,4055 | 1,0172E-15 |
| ACE2 | LVCAT1 | -0,4055 | 1,00949E-15 |
| ACE2 | RP11-118B18.3 | -0,4055 | 1,0214E-15 |
| ACE2 | CSTP1 | -0,4055 | 1,01242E-15 |
| ACE2 | LINC02026 | -0,4056 | 1,00467E-15 |
| ACE2 | RP11-356O9.1 | -0,4056 | 9,92861E-16 |
| ACE2 | GNAS-AS1 | -0,4056 | 9,87712E-16 |
| ACE2 | NOL4L | -0,4057 | 9,86118E-16 |
| ACE2 | IQGAP2 | -0,4059 | 9,50776E-16 |
| ACE2 | C8orf88 | -0,4059 | 9,43395E-16 |
| ACE2 | RP11-494O16.3 | -0,4059 | 9,49314E-16 |
| ACE2 | TGFA | -0,406 | 9,35418E-16 |
| ACE2 | SPESP1 | -0,406 | 9,34035E-16 |
| ACE2 | RNF139 | -0,4061 | 9,11407E-16 |
| ACE2 | RP11-508N22.6 | -0,4061 | 9,08444E-16 |
| ACE2 | RP11-655M14.12 | -0,4061 | 9,12782E-16 |
| ACE2 | ATP6V0D1 | -0,4061 | 9,17612E-16 |
| ACE2 | RP11-208N20.1 | -0,4062 | 9,02195E-16 |
| ACE2 | KRT18P1 | -0,4062 | 8,98338E-16 |
| ACE2 | RP11-274B18.4 | -0,4062 | 8,91904E-16 |

| | | | |
|---|---|---|---|
| ACE2 | LINC01640 | -0,4062 | 8,9493E-16 |
| ACE2 | FAM19A4 | -0,4063 | 8,76577E-16 |
| ACE2 | LINC02211 | -0,4063 | 8,73817E-16 |
| ACE2 | CTB-37A13.1 | -0,4063 | 8,84548E-16 |
| ACE2 | METTL24 | -0,4063 | 8,8414E-16 |
| ACE2 | AP000936.4 | -0,4063 | 8,73632E-16 |
| ACE2 | GNG8 | -0,4063 | 8,75148E-16 |
| ACE2 | BSNDP2 | -0,4063 | 8,73687E-16 |
| ACE2 | AC092155.1 | -0,4064 | 8,70614E-16 |
| ACE2 | KYNU | -0,4064 | 8,67503E-16 |
| ACE2 | RP11-818F20.4 | -0,4064 | 8,63769E-16 |
| ACE2 | RP11-461F11.2 | -0,4064 | 8,71151E-16 |
| ACE2 | BCRP1 | -0,4064 | 8,6246E-16 |
| ACE2 | CTC-441N14.1 | -0,4065 | 8,47779E-16 |
| ACE2 | OR1K1 | -0,4065 | 8,5409E-16 |
| ACE2 | RP11-60C6.6 | -0,4065 | 8,52666E-16 |
| ACE2 | RP11-522B15.1 | -0,4065 | 8,46633E-16 |
| ACE2 | LL09NC01-251B2.3 | -0,4066 | 8,36757E-16 |
| ACE2 | RNF6P1 | -0,4066 | 8,318E-16 |
| ACE2 | RP11-867G23.8 | -0,4067 | 8,26803E-16 |
| ACE2 | CTD-3162L10.3 | -0,4067 | 8,13094E-16 |
| ACE2 | BCRP4 | -0,4067 | 8,16394E-16 |
| ACE2 | SFXN5 | -0,4068 | 8,06842E-16 |
| ACE2 | AC005041.11 | -0,4068 | 8,11314E-16 |
| ACE2 | LINC00882 | -0,4068 | 8,07707E-16 |
| ACE2 | RNF168 | -0,4068 | 8,10476E-16 |
| ACE2 | RBM17P4 | -0,4068 | 8,08088E-16 |
| ACE2 | MSLN | -0,4068 | 8,01483E-16 |
| ACE2 | RP11-20G6.2 | -0,4068 | 8,10713E-16 |
| ACE2 | SNRPD2 | -0,4068 | 8,03476E-16 |
| ACE2 | AC009518.8 | -0,4069 | 7,97742E-16 |
| ACE2 | AQP7P1 | -0,4069 | 7,8492E-16 |
| ACE2 | BOD1P2 | -0,4069 | 7,85046E-16 |
| ACE2 | LIPT1 | -0,407 | 7,72815E-16 |
| ACE2 | RP11-729M20.1 | -0,407 | 7,79363E-16 |
| ACE2 | RP11-273B19.1 | -0,4071 | 7,63864E-16 |
| ACE2 | CTD-2024P10.2 | -0,4071 | 7,70262E-16 |
| ACE2 | ZNF396 | -0,4071 | 7,64029E-16 |
| ACE2 | C2CD3 | -0,4072 | 7,55852E-16 |
| ACE2 | IL17D | -0,4072 | 7,44095E-16 |
| ACE2 | BBOF1 | -0,4072 | 7,46139E-16 |
| ACE2 | GOLGA6L22 | -0,4073 | 7,35839E-16 |
| ACE2 | NBPF4 | -0,4074 | 7,18334E-16 |
| ACE2 | LCT | -0,4074 | 7,28877E-16 |
| ACE2 | RPS6P20 | -0,4074 | 7,19788E-16 |
| ACE2 | ASIC2 | -0,4074 | 7,24835E-16 |
| ACE2 | LINC01015 | -0,4075 | 7,11091E-16 |
| ACE2 | SPSB2 | -0,4075 | 7,17238E-16 |
| ACE2 | CRYGB | -0,4076 | 6,98555E-16 |
| ACE2 | RP5-991O23.1 | -0,4077 | 6,91902E-16 |
| ACE2 | RP11-699F16.2 | -0,4077 | 6,82841E-16 |
| ACE2 | RP11-39M21.1 | -0,4077 | 6,86733E-16 |
| ACE2 | LA16c-380F5.3 | -0,4077 | 6,81846E-16 |
| ACE2 | RP11-374M1.10 | -0,4078 | 6,69609E-16 |

| | | | |
|---|---|---|---|
| ACE2 | CTC-457E21.4 | -0,4078 | 6,74217E-16 |
| ACE2 | RP11-705O24.3 | -0,4079 | 6,57815E-16 |
| ACE2 | VTCN1 | -0,408 | 6,46527E-16 |
| ACE2 | ZSWIM2 | -0,408 | 6,50376E-16 |
| ACE2 | RP11-429B14.4 | -0,408 | 6,48385E-16 |
| ACE2 | HFE2 | -0,4081 | 6,35478E-16 |
| ACE2 | RP11-157L3.11 | -0,4081 | 6,34221E-16 |
| ACE2 | CTD-2514C3.1 | -0,4081 | 6,34003E-16 |
| ACE2 | KCTD16 | -0,4082 | 6,28678E-16 |
| ACE2 | NCAPH2 | -0,4082 | 6,2528E-16 |
| ACE2 | FYB | -0,4083 | 6,21239E-16 |
| ACE2 | R3HCC1 | -0,4083 | 6,18493E-16 |
| ACE2 | RP11-292D4.1 | -0,4084 | 6,08575E-16 |
| ACE2 | RP11-528M18.2 | -0,4084 | 6,08463E-16 |
| ACE2 | ZNF585A | -0,4084 | 6,0637E-16 |
| ACE2 | NAALADL2-AS1 | -0,4085 | 5,93906E-16 |
| ACE2 | RP11-89H19.2 | -0,4085 | 5,94928E-16 |
| ACE2 | ELK2BP | -0,4085 | 6,00304E-16 |
| ACE2 | RP11-304L19.2 | -0,4085 | 5,99858E-16 |
| ACE2 | RP11-114L10.2 | -0,4086 | 5,87909E-16 |
| ACE2 | RP11-328C8.4 | -0,4086 | 5,83077E-16 |
| ACE2 | SLC24A5 | -0,4086 | 5,89579E-16 |
| ACE2 | AE000662.92 | -0,4087 | 5,74952E-16 |
| ACE2 | ANKS4B | -0,4087 | 5,71793E-16 |
| ACE2 | RP3-518E13.2 | -0,4088 | 5,66456E-16 |
| ACE2 | EMC3 | -0,4088 | 5,68011E-16 |
| ACE2 | CTC-228N24.2 | -0,4088 | 5,60623E-16 |
| ACE2 | RP1-232N11.2 | -0,4088 | 5,59775E-16 |
| ACE2 | FBXO36 | -0,4089 | 5,52691E-16 |
| ACE2 | RP11-100L22.4 | -0,4089 | 5,52533E-16 |
| ACE2 | PSME2P6 | -0,4089 | 5,58957E-16 |
| ACE2 | OIT3 | -0,4089 | 5,56632E-16 |
| ACE2 | CTD-2530H12.7 | -0,4089 | 5,58349E-16 |
| ACE2 | SPAG5 | -0,4089 | 5,52344E-16 |
| ACE2 | AP001347.6 | -0,4089 | 5,52789E-16 |
| ACE2 | RP11-789C7.1 | -0,409 | 5,44448E-16 |
| ACE2 | RP11-268F1.3 | -0,4091 | 5,31227E-16 |
| ACE2 | BRDT | -0,4092 | 5,24281E-16 |
| ACE2 | ACTBP7 | -0,4092 | 5,22603E-16 |
| ACE2 | AC007787.2 | -0,4092 | 5,27624E-16 |
| ACE2 | BCKDHA | -0,4092 | 5,22566E-16 |
| ACE2 | RP11-787D18.1 | -0,4093 | 5,17748E-16 |
| ACE2 | RP5-857K21.4 | -0,4094 | 5,03584E-16 |
| ACE2 | EIF4E | -0,4094 | 5,02933E-16 |
| ACE2 | SLCO3A1 | -0,4094 | 5,05052E-16 |
| ACE2 | ATP5J2P5 | -0,4095 | 4,97393E-16 |
| ACE2 | ACTG1P12 | -0,4096 | 4,85516E-16 |
| ACE2 | LINC02123 | -0,4097 | 4,76758E-16 |
| ACE2 | RP11-321L2.1 | -0,4097 | 4,82211E-16 |
| ACE2 | THTPA | -0,4097 | 4,8003E-16 |
| ACE2 | LINC02184 | -0,4097 | 4,83235E-16 |
| ACE2 | HMGB1P45 | -0,4098 | 4,75483E-16 |
| ACE2 | SLC1A2 | -0,4098 | 4,70522E-16 |
| ACE2 | C11orf70 | -0,4098 | 4,73918E-16 |

| | | | |
|---|---|---|---|
| ACE2 | RP11-495K9.6 | -0,4098 | 4,74303E-16 |
| ACE2 | RGS9BP | -0,4098 | 4,69444E-16 |
| ACE2 | AC017053.1 | -0,4099 | 4,64088E-16 |
| ACE2 | AC013733.4 | -0,4099 | 4,64045E-16 |
| ACE2 | AC009473.1 | -0,4099 | 4,66086E-16 |
| ACE2 | TMEM99 | -0,4099 | 4,60007E-16 |
| ACE2 | CACNA1I | -0,4099 | 4,63261E-16 |
| ACE2 | SHCBP1L | -0,41 | 4,55583E-16 |
| ACE2 | NMBR | -0,41 | 4,58272E-16 |
| ACE2 | TMBIM7P | -0,41 | 4,55683E-16 |
| ACE2 | TP53AIP1 | -0,41 | 4,55944E-16 |
| ACE2 | RP11-471B22.3 | -0,41 | 4,57263E-16 |
| ACE2 | RP11-80F22.4 | -0,41 | 4,54316E-16 |
| ACE2 | CTC-459F4.5 | -0,41 | 4,55355E-16 |
| ACE2 | GGPS1 | -0,4101 | 4,50965E-16 |
| ACE2 | AC008067.2 | -0,4101 | 4,4964E-16 |
| ACE2 | AC005593.2 | -0,4101 | 4,46797E-16 |
| ACE2 | RP11-24P4.1 | -0,4101 | 4,50866E-16 |
| ACE2 | RP11-187C18.3 | -0,4101 | 4,44874E-16 |
| ACE2 | ADAMTS7P1 | -0,4101 | 4,45977E-16 |
| ACE2 | RP11-640N20.1 | -0,4101 | 4,4977E-16 |
| ACE2 | ZNF57 | -0,4101 | 4,45235E-16 |
| ACE2 | ATG3 | -0,4102 | 4,36634E-16 |
| ACE2 | RP11-257I8.2 | -0,4102 | 4,36427E-16 |
| ACE2 | AC005009.1 | -0,4102 | 4,38181E-16 |
| ACE2 | RANBP1 | -0,4103 | 4,28868E-16 |
| ACE2 | PAIP2B | -0,4104 | 4,20219E-16 |
| ACE2 | GLIPR1L2 | -0,4104 | 4,25059E-16 |
| ACE2 | RAI1-AS1 | -0,4104 | 4,23108E-16 |
| ACE2 | C17orf53 | -0,4104 | 4,26748E-16 |
| ACE2 | CWC27 | -0,4105 | 4,19606E-16 |
| ACE2 | CPLX3 | -0,4105 | 4,17871E-16 |
| ACE2 | NRG4 | -0,4105 | 4,15781E-16 |
| ACE2 | KSR1 | -0,4105 | 4,12801E-16 |
| ACE2 | ATXN3L | -0,4105 | 4,14415E-16 |
| ACE2 | RP5-1073O3.2 | -0,4106 | 4,06065E-16 |
| ACE2 | TBC1D27 | -0,4106 | 4,05684E-16 |
| ACE2 | RASL10B | -0,4106 | 4,10935E-16 |
| ACE2 | HSD52 | -0,4107 | 3,98688E-16 |
| ACE2 | NBR1 | -0,4107 | 4,03962E-16 |
| ACE2 | EPS8L3 | -0,4108 | 3,91746E-16 |
| ACE2 | RP3-382I10.3 | -0,4108 | 3,92063E-16 |
| ACE2 | WDR78 | -0,4109 | 3,90607E-16 |
| ACE2 | DEDD | -0,4109 | 3,90271E-16 |
| ACE2 | IL17F | -0,4109 | 3,84834E-16 |
| ACE2 | NRG3 | -0,4109 | 3,89773E-16 |
| ACE2 | CDK5R1 | -0,4109 | 3,8876E-16 |
| ACE2 | ZNF610 | -0,4109 | 3,88081E-16 |
| ACE2 | AC026202.3 | -0,411 | 3,78716E-16 |
| ACE2 | RP11-497H16.7 | -0,411 | 3,79564E-16 |
| ACE2 | SPTLC1P2 | -0,411 | 3,8355E-16 |
| ACE2 | RP11-306G20.1 | -0,411 | 3,78751E-16 |
| ACE2 | COX10 | -0,411 | 3,80539E-16 |
| ACE2 | AC138517.6 | -0,4111 | 3,74396E-16 |

| | | | |
|---|---|---|---|
| ACE2 | LRRC3-AS1 | -0,4111 | 3,70866E-16 |
| ACE2 | RP11-157J24.1 | -0,4112 | 3,67905E-16 |
| ACE2 | INTS6-AS1 | -0,4112 | 3,6632E-16 |
| ACE2 | LINC01103 | -0,4113 | 3,6302E-16 |
| ACE2 | RP11-237D3.1 | -0,4113 | 3,63512E-16 |
| ACE2 | TUBA4B | -0,4114 | 3,52762E-16 |
| ACE2 | GS1-18A18.2 | -0,4114 | 3,55992E-16 |
| ACE2 | COQ6 | -0,4114 | 3,54318E-16 |
| ACE2 | RP11-430L3.1 | -0,4115 | 3,46121E-16 |
| ACE2 | RNU6-725P | -0,4116 | 3,40568E-16 |
| ACE2 | RP11-153M7.5 | -0,4116 | 3,4073E-16 |
| ACE2 | RP11-930P14.1 | -0,4116 | 3,41649E-16 |
| ACE2 | ANKRD42 | -0,4116 | 3,4279E-16 |
| ACE2 | TBR1 | -0,4117 | 3,33542E-16 |
| ACE2 | RP11-1415C14.3 | -0,4117 | 3,35154E-16 |
| ACE2 | RP11-18B16.2 | -0,4117 | 3,33064E-16 |
| ACE2 | GRIN1 | -0,4117 | 3,35179E-16 |
| ACE2 | CTXN1 | -0,4117 | 3,37227E-16 |
| ACE2 | BCRP3 | -0,4117 | 3,3579E-16 |
| ACE2 | RP3-332O11.2 | -0,4118 | 3,3213E-16 |
| ACE2 | GHRLOS | -0,4118 | 3,27822E-16 |
| ACE2 | RPL31P28 | -0,4118 | 3,30552E-16 |
| ACE2 | LINC02091 | -0,4118 | 3,26716E-16 |
| ACE2 | LINC00908 | -0,4118 | 3,26992E-16 |
| ACE2 | RN7SKP214 | -0,4118 | 3,29729E-16 |
| ACE2 | RP11-63G10.3 | -0,4119 | 3,21889E-16 |
| ACE2 | FOLR3 | -0,4119 | 3,24516E-16 |
| ACE2 | LINC02205 | -0,4119 | 3,21676E-16 |
| ACE2 | LINC01477 | -0,4119 | 3,21202E-16 |
| ACE2 | SLC5A1 | -0,4119 | 3,22333E-16 |
| ACE2 | KLF14 | -0,412 | 3,17871E-16 |
| ACE2 | MRGPRG | -0,412 | 3,19594E-16 |
| ACE2 | RP11-313F23.3 | -0,412 | 3,16954E-16 |
| ACE2 | DLX2-AS1 | -0,4122 | 3,06117E-16 |
| ACE2 | RHPN1 | -0,4122 | 3,05825E-16 |
| ACE2 | RP11-461K13.1 | -0,4122 | 3,0774E-16 |
| ACE2 | SKOR1 | -0,4122 | 3,053E-16 |
| ACE2 | RP11-31I22.4 | -0,4123 | 2,9903E-16 |
| ACE2 | RP11-253D19.1 | -0,4124 | 2,94774E-16 |
| ACE2 | CWF19L1 | -0,4124 | 2,957E-16 |
| ACE2 | GOLGA8T | -0,4124 | 2,96652E-16 |
| ACE2 | ZNF768 | -0,4124 | 2,9369E-16 |
| ACE2 | KRTDAP | -0,4124 | 2,94341E-16 |
| ACE2 | TMEM150A | -0,4125 | 2,90128E-16 |
| ACE2 | IL20RB-AS1 | -0,4125 | 2,88186E-16 |
| ACE2 | RPL7AP57 | -0,4125 | 2,91539E-16 |
| ACE2 | AC017104.6 | -0,4126 | 2,84128E-16 |
| ACE2 | RP11-157L3.10 | -0,4126 | 2,86906E-16 |
| ACE2 | RP11-575F12.1 | -0,4126 | 2,84006E-16 |
| ACE2 | RP11-428O18.6 | -0,4127 | 2,78928E-16 |
| ACE2 | CTD-2008P7.1 | -0,4127 | 2,77988E-16 |
| ACE2 | RP11-115D7.3 | -0,4128 | 2,73826E-16 |
| ACE2 | FOXO6 | -0,4128 | 2,74277E-16 |
| ACE2 | ANKRD55 | -0,4128 | 2,74137E-16 |

| | | | |
|---|---|---|---|
| ACE2 | RP11-314A20.2 | -0,4128 | 2,77394E-16 |
| ACE2 | CYLC1 | -0,4128 | 2,7542E-16 |
| ACE2 | OR10AH1P | -0,4129 | 2,70649E-16 |
| ACE2 | AC004895.4 | -0,4129 | 2,69628E-16 |
| ACE2 | LINC00654 | -0,4129 | 2,70149E-16 |
| ACE2 | PINK1 | -0,413 | 2,67365E-16 |
| ACE2 | SLC44A3 | -0,413 | 2,66033E-16 |
| ACE2 | LINC01847 | -0,413 | 2,65275E-16 |
| ACE2 | AC003075.4 | -0,413 | 2,66661E-16 |
| ACE2 | RP11-863K10.4 | -0,413 | 2,65942E-16 |
| ACE2 | ZNF350-AS1 | -0,413 | 2,64468E-16 |
| ACE2 | AC008132.13 | -0,413 | 2,67311E-16 |
| ACE2 | AC007050.17 | -0,413 | 2,63138E-16 |
| ACE2 | COPS5 | -0,4131 | 2,58518E-16 |
| ACE2 | RP11-1082L8.1 | -0,4131 | 2,59864E-16 |
| ACE2 | GLDN | -0,4131 | 2,61623E-16 |
| ACE2 | RP11-643M14.1 | -0,4132 | 2,55635E-16 |
| ACE2 | CD300LD | -0,4132 | 2,54538E-16 |
| ACE2 | RP11-685A21.1 | -0,4132 | 2,54596E-16 |
| ACE2 | RP11-438D8.2 | -0,4133 | 2,51207E-16 |
| ACE2 | GPANK1 | -0,4133 | 2,52909E-16 |
| ACE2 | RP1-40E16.2 | -0,4134 | 2,45284E-16 |
| ACE2 | TBPL1 | -0,4134 | 2,46503E-16 |
| ACE2 | ADAM21P1 | -0,4134 | 2,46104E-16 |
| ACE2 | ARHGAP28 | -0,4137 | 2,34401E-16 |
| ACE2 | POLR3C | -0,4138 | 2,2739E-16 |
| ACE2 | RP11-381O7.3 | -0,4138 | 2,31431E-16 |
| ACE2 | CTB-31O20.8 | -0,4138 | 2,27361E-16 |
| ACE2 | TUSC2 | -0,4139 | 2,23691E-16 |
| ACE2 | RP11-518I13.1 | -0,4139 | 2,26192E-16 |
| ACE2 | ZNF415 | -0,4139 | 2,24009E-16 |
| ACE2 | TNC | -0,414 | 2,22538E-16 |
| ACE2 | RP11-167N4.2 | -0,414 | 2,21735E-16 |
| ACE2 | GAA | -0,414 | 2,21993E-16 |
| ACE2 | DGCR10 | -0,414 | 2,23162E-16 |
| ACE2 | DAW1 | -0,4141 | 2,18302E-16 |
| ACE2 | LRIT3 | -0,4141 | 2,1647E-16 |
| ACE2 | CACNA1S | -0,4142 | 2,14863E-16 |
| ACE2 | LINC00945 | -0,4142 | 2,13184E-16 |
| ACE2 | TRAPPC3L | -0,4143 | 2,08009E-16 |
| ACE2 | PDE11A | -0,4144 | 2,04113E-16 |
| ACE2 | RP11-202G18.1 | -0,4144 | 2,06553E-16 |
| ACE2 | CTD-2035E11.3 | -0,4145 | 2,02411E-16 |
| ACE2 | UBDP1 | -0,4145 | 2,01395E-16 |
| ACE2 | RP11-379I19.3 | -0,4145 | 2,00356E-16 |
| ACE2 | FOXD1 | -0,4146 | 1,98896E-16 |
| ACE2 | RPL12P1 | -0,4146 | 1,98532E-16 |
| ACE2 | RP11-744I24.2 | -0,4146 | 1,98043E-16 |
| ACE2 | THAP7 | -0,4146 | 1,98231E-16 |
| ACE2 | IQCJ | -0,4147 | 1,93373E-16 |
| ACE2 | CTD-2035E11.4 | -0,4147 | 1,94055E-16 |
| ACE2 | FAM47B | -0,4147 | 1,95841E-16 |
| ACE2 | RP11-40G16.1 | -0,4148 | 1,90761E-16 |
| ACE2 | RP11-96A15.1 | -0,4148 | 1,91974E-16 |

| | | | |
|---|---|---|---|
| ACE2 | RP1-63G5.7 | -0,4148 | 1,91382E-16 |
| ACE2 | CTD-2256P15.2 | -0,4149 | 1,87402E-16 |
| ACE2 | KATNA1 | -0,4149 | 1,87327E-16 |
| ACE2 | AC073850.6 | -0,4149 | 1,87144E-16 |
| ACE2 | SSX2IP | -0,415 | 1,84789E-16 |
| ACE2 | RP11-464F9.22 | -0,415 | 1,85979E-16 |
| ACE2 | CTC-268N12.2 | -0,415 | 1,83608E-16 |
| ACE2 | LINC02081 | -0,415 | 1,84854E-16 |
| ACE2 | KRT8P48 | -0,4151 | 1,80878E-16 |
| ACE2 | TMEM72-AS1 | -0,4151 | 1,82531E-16 |
| ACE2 | SMG6-IT1 | -0,4151 | 1,811E-16 |
| ACE2 | LRRC3B | -0,4152 | 1,78731E-16 |
| ACE2 | CTSLP6 | -0,4152 | 1,76644E-16 |
| ACE2 | RP11-310E22.5 | -0,4152 | 1,7813E-16 |
| ACE2 | AC139143.2 | -0,4152 | 1,77054E-16 |
| ACE2 | RP11-17M15.2 | -0,4152 | 1,76176E-16 |
| ACE2 | RP11-381P6.1 | -0,4152 | 1,7854E-16 |
| ACE2 | PSMD6 | -0,4153 | 1,73859E-16 |
| ACE2 | FTH1P18 | -0,4153 | 1,76138E-16 |
| ACE2 | RP11-136C24.2 | -0,4155 | 1,6899E-16 |
| ACE2 | LINC01396 | -0,4155 | 1,67163E-16 |
| ACE2 | RP11-1023L17.2 | -0,4155 | 1,69108E-16 |
| ACE2 | RP11-140A10.3 | -0,4155 | 1,69085E-16 |
| ACE2 | STMN2 | -0,4156 | 1,6626E-16 |
| ACE2 | RP11-715J22.2 | -0,4156 | 1,64093E-16 |
| ACE2 | RP11-180D21.3 | -0,4157 | 1,63241E-16 |
| ACE2 | CYP2AB1P | -0,4157 | 1,61508E-16 |
| ACE2 | BAG6 | -0,4157 | 1,62877E-16 |
| ACE2 | CTA-392C11.1 | -0,4157 | 1,63722E-16 |
| ACE2 | CNTD1 | -0,4157 | 1,61808E-16 |
| ACE2 | MRPL40 | -0,4157 | 1,629E-16 |
| ACE2 | SEMA3B | -0,4158 | 1,58554E-16 |
| ACE2 | SFTPC | -0,4158 | 1,6055E-16 |
| ACE2 | AGAP9 | -0,4158 | 1,58463E-16 |
| ACE2 | FICD | -0,4158 | 1,58026E-16 |
| ACE2 | PTGER4 | -0,4159 | 1,57727E-16 |
| ACE2 | FTH1P12 | -0,4159 | 1,55435E-16 |
| ACE2 | RPL21P111 | -0,4159 | 1,55018E-16 |
| ACE2 | C19orf81 | -0,4159 | 1,56998E-16 |
| ACE2 | CYCSP10 | -0,416 | 1,53348E-16 |
| ACE2 | COLQ | -0,416 | 1,5317E-16 |
| ACE2 | RP11-230G5.2 | -0,416 | 1,52249E-16 |
| ACE2 | DPYSL5 | -0,4161 | 1,49475E-16 |
| ACE2 | PRR16 | -0,4161 | 1,49502E-16 |
| ACE2 | HLA-G | -0,4161 | 1,51667E-16 |
| ACE2 | RADIL | -0,4161 | 1,5036E-16 |
| ACE2 | IGF2BP3 | -0,4161 | 1,50398E-16 |
| ACE2 | SLC27A4 | -0,4161 | 1,4978E-16 |
| ACE2 | RPS17P1 | -0,4163 | 1,45176E-16 |
| ACE2 | HTR7 | -0,4163 | 1,46064E-16 |
| ACE2 | PWWP2B | -0,4163 | 1,46458E-16 |
| ACE2 | BDNF | -0,4163 | 1,45691E-16 |
| ACE2 | LGR6 | -0,4164 | 1,44038E-16 |
| ACE2 | CUTC | -0,4164 | 1,43463E-16 |

| | | | |
|---|---|---|---|
| ACE2 | LRRC37A9P | -0,4164 | 1,41676E-16 |
| ACE2 | C17orf78 | -0,4164 | 1,42131E-16 |
| ACE2 | KRT42P | -0,4164 | 1,44051E-16 |
| ACE2 | RP11-290K4.2 | -0,4165 | 1,39214E-16 |
| ACE2 | AP000351.13 | -0,4165 | 1,41339E-16 |
| ACE2 | FAM183BP | -0,4166 | 1,38729E-16 |
| ACE2 | MOS | -0,4166 | 1,38492E-16 |
| ACE2 | DNAJC5G | -0,4167 | 1,3389E-16 |
| ACE2 | THAP12P7 | -0,4167 | 1,35234E-16 |
| ACE2 | PROSER3 | -0,4167 | 1,3561E-16 |
| ACE2 | RP11-543C4.1 | -0,4168 | 1,32002E-16 |
| ACE2 | CNGB1 | -0,4168 | 1,32908E-16 |
| ACE2 | RP11-550I24.2 | -0,4169 | 1,29087E-16 |
| ACE2 | RP11-23F23.2 | -0,4169 | 1,3136E-16 |
| ACE2 | RP11-304F15.4 | -0,4169 | 1,29118E-16 |
| ACE2 | RP1-191L6.2 | -0,4169 | 1,30439E-16 |
| ACE2 | CAMTA1-IT1 | -0,417 | 1,27566E-16 |
| ACE2 | CTD-2139B15.2 | -0,417 | 1,29038E-16 |
| ACE2 | RP11-437J19.1 | -0,417 | 1,28328E-16 |
| ACE2 | RP11-794P6.6 | -0,417 | 1,27803E-16 |
| ACE2 | DNM1P46 | -0,417 | 1,26839E-16 |
| ACE2 | XXbac-B476C20.13 | -0,417 | 1,27566E-16 |
| ACE2 | CCDC138 | -0,4171 | 1,25767E-16 |
| ACE2 | RP11-358M3.1 | -0,4171 | 1,25715E-16 |
| ACE2 | LINC01585 | -0,4171 | 1,25492E-16 |
| ACE2 | IL21 | -0,4172 | 1,22352E-16 |
| ACE2 | AC026954.6 | -0,4172 | 1,23473E-16 |
| ACE2 | SOX9-AS1 | -0,4172 | 1,22971E-16 |
| ACE2 | RMDN2-AS1 | -0,4173 | 1,21177E-16 |
| ACE2 | MAPK6PS2 | -0,4173 | 1,20577E-16 |
| ACE2 | RP11-817O13.9 | -0,4174 | 1,19028E-16 |
| ACE2 | TFR2 | -0,4175 | 1,16403E-16 |
| ACE2 | RP11-552E10.1 | -0,4175 | 1,17734E-16 |
| ACE2 | NME9 | -0,4176 | 1,1402E-16 |
| ACE2 | RP11-69C17.2 | -0,4177 | 1,12799E-16 |
| ACE2 | SNPH | -0,4177 | 1,1328E-16 |
| ACE2 | RP11-384O8.1 | -0,4178 | 1,11111E-16 |
| ACE2 | RP5-894D12.4 | -0,4178 | 1,11128E-16 |
| ACE2 | CTC-304I17.3 | -0,4178 | 1,09606E-16 |
| ACE2 | RP4-595K12.2 | -0,4179 | 1,0746E-16 |
| ACE2 | VAV3 | -0,4179 | 1,08944E-16 |
| ACE2 | RP11-384F7.1 | -0,4179 | 1,08163E-16 |
| ACE2 | RP11-501M7.1 | -0,4179 | 1,08732E-16 |
| ACE2 | MMP7 | -0,4179 | 1,07784E-16 |
| ACE2 | RP11-112N19.1 | -0,4179 | 1,08025E-16 |
| ACE2 | RP11-56N19.5 | -0,418 | 1,06751E-16 |
| ACE2 | FRMD1 | -0,418 | 1,05798E-16 |
| ACE2 | RPL21P120 | -0,418 | 1,05564E-16 |
| ACE2 | AP000472.3 | -0,418 | 1,05999E-16 |
| ACE2 | TMEM31 | -0,418 | 1,06725E-16 |
| ACE2 | RN7SL371P | -0,4181 | 1,05195E-16 |
| ACE2 | TESK2 | -0,4181 | 1,0541E-16 |
| ACE2 | RP11-115J16.2 | -0,4181 | 1,04699E-16 |
| ACE2 | ASCC2 | -0,4181 | 1,04253E-16 |

| | | | |
|---|---|---|---|
| ACE2 | FER1L6-AS2 | -0,4182 | 1,03318E-16 |
| ACE2 | CTC-339O9.1 | -0,4182 | 1,02202E-16 |
| ACE2 | GJB4 | -0,4183 | 1,00253E-16 |
| ACE2 | OPRPN | -0,4184 | 9,8898E-17 |
| ACE2 | CTD-2374C24.1 | -0,4184 | 9,9623E-17 |
| ACE2 | RP11-433M22.2 | -0,4184 | 9,83738E-17 |
| ACE2 | RP11-344J7.2 | -0,4185 | 9,64732E-17 |
| ACE2 | NFKBIE | -0,4185 | 9,64946E-17 |
| ACE2 | RPS26P52 | -0,4185 | 9,75457E-17 |
| ACE2 | EPHA8 | -0,4186 | 9,60382E-17 |
| ACE2 | RMDN2 | -0,4186 | 9,57616E-17 |
| ACE2 | RP11-325O24.6 | -0,4186 | 9,5083E-17 |
| ACE2 | RP11-58A18.1 | -0,4186 | 9,58801E-17 |
| ACE2 | CTD-3099C6.5 | -0,4186 | 9,52148E-17 |
| ACE2 | SPANXN5 | -0,4186 | 9,51612E-17 |
| ACE2 | UBE2J1 | -0,4188 | 9,21006E-17 |
| ACE2 | RP11-363J20.1 | -0,4189 | 9,0149E-17 |
| ACE2 | CTD-2243E23.1 | -0,4189 | 9,00452E-17 |
| ACE2 | RP11-739N10.1 | -0,4189 | 9,01028E-17 |
| ACE2 | CDH20 | -0,419 | 8,85911E-17 |
| ACE2 | RP11-393I23.4 | -0,4191 | 8,75559E-17 |
| ACE2 | RP11-281O15.4 | -0,4191 | 8,68763E-17 |
| ACE2 | RP11-429J17.4 | -0,4191 | 8,73993E-17 |
| ACE2 | EVA1C | -0,4191 | 8,70924E-17 |
| ACE2 | CTA-398F10.2 | -0,4192 | 8,48788E-17 |
| ACE2 | RP11-53B5.1 | -0,4192 | 8,54502E-17 |
| ACE2 | PROSER2-AS1 | -0,4192 | 8,57551E-17 |
| ACE2 | MKRN2 | -0,4193 | 8,32651E-17 |
| ACE2 | RP13-539F13.3 | -0,4193 | 8,41218E-17 |
| ACE2 | IL4 | -0,4193 | 8,31164E-17 |
| ACE2 | AGAP6 | -0,4193 | 8,34422E-17 |
| ACE2 | RP11-520B13.8 | -0,4194 | 8,19242E-17 |
| ACE2 | RP11-232C2.2 | -0,4194 | 8,26017E-17 |
| ACE2 | RP11-111E14.1 | -0,4194 | 8,15037E-17 |
| ACE2 | NPHP3-AS1 | -0,4195 | 8,07834E-17 |
| ACE2 | LINC01933 | -0,4195 | 8,03841E-17 |
| ACE2 | SHBG | -0,4195 | 8,09523E-17 |
| ACE2 | RP11-1137G4.5 | -0,4195 | 8,09227E-17 |
| ACE2 | DPY19L1P2 | -0,4196 | 7,97422E-17 |
| ACE2 | RP11-458F8.1 | -0,4196 | 7,92098E-17 |
| ACE2 | SNTG1 | -0,4196 | 7,96757E-17 |
| ACE2 | RP11-188C12.2 | -0,4196 | 7,95543E-17 |
| ACE2 | RP11-598C10.1 | -0,4196 | 7,92408E-17 |
| ACE2 | RP11-361I14.2 | -0,4196 | 7,90414E-17 |
| ACE2 | CTD-3253I12.1 | -0,4197 | 7,83468E-17 |
| ACE2 | HGS | -0,4197 | 7,75376E-17 |
| ACE2 | LARGE-IT1 | -0,4197 | 7,74992E-17 |
| ACE2 | RP11-644C3.1 | -0,4198 | 7,64225E-17 |
| ACE2 | THG1L | -0,4198 | 7,69112E-17 |
| ACE2 | GABRR2 | -0,4198 | 7,59917E-17 |
| ACE2 | RP11-375N15.2 | -0,4198 | 7,69208E-17 |
| ACE2 | SMARCE1P5 | -0,4198 | 7,6809E-17 |
| ACE2 | CTD-2540L5.9 | -0,4198 | 7,59925E-17 |
| ACE2 | AC005324.7 | -0,4198 | 7,57084E-17 |

| | | | |
|---|---|---|---|
| ACE2 | WDR87 | -0,4198 | 7,67521E-17 |
| ACE2 | AC009487.5 | -0,4199 | 7,49359E-17 |
| ACE2 | KIF15 | -0,4199 | 7,49554E-17 |
| ACE2 | AC004906.3 | -0,4199 | 7,55211E-17 |
| ACE2 | RP11-447B18.1 | -0,42 | 7,33635E-17 |
| ACE2 | RP1-167A14.2 | -0,42 | 7,31175E-17 |
| ACE2 | RP11-862L9.3 | -0,42 | 7,32698E-17 |
| ACE2 | ACSL1 | -0,4201 | 7,23649E-17 |
| ACE2 | PGBD4P1 | -0,4201 | 7,2133E-17 |
| ACE2 | MSMB | -0,4201 | 7,23945E-17 |
| ACE2 | CCT4P1 | -0,4202 | 7,08046E-17 |
| ACE2 | RP11-213G2.1 | -0,4202 | 7,0567E-17 |
| ACE2 | CALML4 | -0,4202 | 7,13304E-17 |
| ACE2 | ANXA10 | -0,4203 | 6,94596E-17 |
| ACE2 | CTB-33O18.2 | -0,4203 | 6,91183E-17 |
| ACE2 | SFTA2 | -0,4203 | 6,93088E-17 |
| ACE2 | RP11-499O7.7 | -0,4203 | 6,9204E-17 |
| ACE2 | NDUFB2-AS1 | -0,4204 | 6,79119E-17 |
| ACE2 | AC007463.2 | -0,4205 | 6,73237E-17 |
| ACE2 | RP11-775B15.2 | -0,4205 | 6,66023E-17 |
| ACE2 | STK33 | -0,4205 | 6,74581E-17 |
| ACE2 | TMEM52B | -0,4205 | 6,65826E-17 |
| ACE2 | UNC13C | -0,4205 | 6,64846E-17 |
| ACE2 | PARP6 | -0,4205 | 6,74965E-17 |
| ACE2 | ZNF807 | -0,4205 | 6,71913E-17 |
| ACE2 | B3GAT3P1 | -0,4206 | 6,54261E-17 |
| ACE2 | CDHR4 | -0,4207 | 6,44507E-17 |
| ACE2 | LINC00032 | -0,4207 | 6,4615E-17 |
| ACE2 | ARHGAP42 | -0,4207 | 6,51124E-17 |
| ACE2 | RP11-333E1.2 | -0,4207 | 6,47987E-17 |
| ACE2 | DIAPH3-AS2 | -0,4208 | 6,28293E-17 |
| ACE2 | RP1-101G11.2 | -0,4208 | 6,33166E-17 |
| ACE2 | FCGR1A | -0,4209 | 6,21133E-17 |
| ACE2 | CCDC142 | -0,4209 | 6,25212E-17 |
| ACE2 | XXbac-BPGBPG34I8.1 | -0,4209 | 6,19375E-17 |
| ACE2 | LINC01686 | -0,421 | 6,13336E-17 |
| ACE2 | FAM167A | -0,4211 | 5,9612E-17 |
| ACE2 | CYP2R1 | -0,4211 | 5,96187E-17 |
| ACE2 | LA16c-313D11.9 | -0,4211 | 6,03114E-17 |
| ACE2 | CTD-3037G24.4 | -0,4211 | 5,9868E-17 |
| ACE2 | YBX1P2 | -0,4212 | 5,91498E-17 |
| ACE2 | ORMDL2 | -0,4212 | 5,91413E-17 |
| ACE2 | RP11-164N3.3 | -0,4213 | 5,82071E-17 |
| ACE2 | RP11-707P17.1 | -0,4213 | 5,76615E-17 |
| ACE2 | RP11-46A10.2 | -0,4214 | 5,69975E-17 |
| ACE2 | RP11-15K3.1 | -0,4214 | 5,67641E-17 |
| ACE2 | SCEL-AS1 | -0,4214 | 5,62679E-17 |
| ACE2 | WARS | -0,4214 | 5,70019E-17 |
| ACE2 | MYL6P5 | -0,4214 | 5,64803E-17 |
| ACE2 | CH17-118O6.3 | -0,4215 | 5,60568E-17 |
| ACE2 | ENTPD1-AS1 | -0,4215 | 5,56972E-17 |
| ACE2 | GGT4P | -0,4215 | 5,51959E-17 |
| ACE2 | LINC01680 | -0,4216 | 5,45646E-17 |
| ACE2 | SRSF12 | -0,4216 | 5,43604E-17 |

| | | | |
|---|---|---|---|
| ACE2 | LCA5L | -0,4216 | 5,45369E-17 |
| ACE2 | ZNF890P | -0,4217 | 5,3287E-17 |
| ACE2 | LINC02155 | -0,4217 | 5,31593E-17 |
| ACE2 | LHPP | -0,4217 | 5,40035E-17 |
| ACE2 | FAM90A1 | -0,4217 | 5,37825E-17 |
| ACE2 | GSKIP | -0,4217 | 5,35968E-17 |
| ACE2 | RP5-837I24.6 | -0,4218 | 5,26858E-17 |
| ACE2 | RP11-400N9.1 | -0,4218 | 5,22838E-17 |
| ACE2 | RP11-294C11.3 | -0,4218 | 5,21978E-17 |
| ACE2 | TUBBP8 | -0,4218 | 5,30317E-17 |
| ACE2 | NEK8 | -0,4219 | 5,16347E-17 |
| ACE2 | AC037445.1 | -0,4219 | 5,14405E-17 |
| ACE2 | RP11-430G17.1 | -0,422 | 5,09348E-17 |
| ACE2 | RP11-240A16.1 | -0,422 | 5,04176E-17 |
| ACE2 | RP4-753D10.3 | -0,422 | 5,04705E-17 |
| ACE2 | PCDHB12 | -0,4221 | 4,98023E-17 |
| ACE2 | FOXG1 | -0,4222 | 4,91317E-17 |
| ACE2 | RP11-107D24.2 | -0,4222 | 4,84082E-17 |
| ACE2 | SPANXD | -0,4222 | 4,90343E-17 |
| ACE2 | EXD1 | -0,4223 | 4,82643E-17 |
| ACE2 | RP11-91H12.1 | -0,4224 | 4,68226E-17 |
| ACE2 | RP11-274B21.4 | -0,4224 | 4,67121E-17 |
| ACE2 | RP11-272D20.2 | -0,4225 | 4,62608E-17 |
| ACE2 | XXbac-BPG181B23.6 | -0,4225 | 4,62385E-17 |
| ACE2 | GAS8 | -0,4226 | 4,5455E-17 |
| ACE2 | GGNBP2 | -0,4226 | 4,55595E-17 |
| ACE2 | RNF38 | -0,4227 | 4,41406E-17 |
| ACE2 | RP11-314D7.3 | -0,4227 | 4,4839E-17 |
| ACE2 | RP11-591L14.2 | -0,4227 | 4,4233E-17 |
| ACE2 | SCNM1 | -0,4228 | 4,36559E-17 |
| ACE2 | PSMC1P10 | -0,4228 | 4,3486E-17 |
| ACE2 | CHRND | -0,4228 | 4,34152E-17 |
| ACE2 | PHYHIPL | -0,4228 | 4,35496E-17 |
| ACE2 | TTR | -0,4228 | 4,40514E-17 |
| ACE2 | RP11-98J23.1 | -0,4229 | 4,24756E-17 |
| ACE2 | EDARADD | -0,423 | 4,2439E-17 |
| ACE2 | RP11-674E16.4 | -0,423 | 4,23397E-17 |
| ACE2 | RP11-567J20.1 | -0,423 | 4,22406E-17 |
| ACE2 | RP11-221N13.3 | -0,423 | 4,182E-17 |
| ACE2 | LINC00934 | -0,423 | 4,21824E-17 |
| ACE2 | DNM1P51 | -0,423 | 4,1944E-17 |
| ACE2 | AP000320.6 | -0,423 | 4,21696E-17 |
| ACE2 | RP11-456P18.2 | -0,4231 | 4,15716E-17 |
| ACE2 | LRRC34 | -0,4231 | 4,11261E-17 |
| ACE2 | GS1-251I9.3 | -0,4231 | 4,11329E-17 |
| ACE2 | HNRNPUP1 | -0,4231 | 4,14413E-17 |
| ACE2 | LINC00235 | -0,4231 | 4,09614E-17 |
| ACE2 | CTC-297N7.9 | -0,4231 | 4,15228E-17 |
| ACE2 | LINC02057 | -0,4232 | 4,05257E-17 |
| ACE2 | SMAP1 | -0,4232 | 4,06365E-17 |
| ACE2 | LINC01855 | -0,4233 | 3,97524E-17 |
| ACE2 | RP11-804A23.1 | -0,4234 | 3,89396E-17 |
| ACE2 | SH2D3A | -0,4234 | 3,9358E-17 |
| ACE2 | SUV39H2 | -0,4235 | 3,82037E-17 |

| | | | |
|---|---|---|---|
| ACE2 | RP11-537A6.9 | -0,4235 | 3,82051E-17 |
| ACE2 | NPTX1 | -0,4235 | 3,79521E-17 |
| ACE2 | MFSD4B | -0,4236 | 3,7346E-17 |
| ACE2 | NUP214 | -0,4236 | 3,78649E-17 |
| ACE2 | STK17A | -0,4238 | 3,63112E-17 |
| ACE2 | RP11-317N8.4 | -0,4238 | 3,62339E-17 |
| ACE2 | CDRT15 | -0,4238 | 3,60106E-17 |
| ACE2 | LINC01800 | -0,4239 | 3,51872E-17 |
| ACE2 | GTF2IP7 | -0,4239 | 3,52366E-17 |
| ACE2 | RP11-488L1.1 | -0,424 | 3,51323E-17 |
| ACE2 | RP5-991B18.1 | -0,424 | 3,46743E-17 |
| ACE2 | CYMP | -0,4241 | 3,3919E-17 |
| ACE2 | ATP2B2 | -0,4241 | 3,41038E-17 |
| ACE2 | RP11-3G20.1 | -0,4241 | 3,44983E-17 |
| ACE2 | RP11-139B1.1 | -0,4241 | 3,41473E-17 |
| ACE2 | CPNE6 | -0,4241 | 3,44829E-17 |
| ACE2 | PSMB6 | -0,4241 | 3,39678E-17 |
| ACE2 | LINC01441 | -0,4241 | 3,42359E-17 |
| ACE2 | FAM230B | -0,4241 | 3,39452E-17 |
| ACE2 | LINC02018 | -0,4242 | 3,37806E-17 |
| ACE2 | RP11-458K10.2 | -0,4242 | 3,38348E-17 |
| ACE2 | DENND6A-AS1 | -0,4243 | 3,32009E-17 |
| ACE2 | PEX10 | -0,4244 | 3,26345E-17 |
| ACE2 | UBXN10-AS1 | -0,4244 | 3,23333E-17 |
| ACE2 | RP11-839G9.1 | -0,4244 | 3,24816E-17 |
| ACE2 | AC005518.2 | -0,4245 | 3,18527E-17 |
| ACE2 | LINC01164 | -0,4245 | 3,18313E-17 |
| ACE2 | RP11-71C5.2 | -0,4245 | 3,16215E-17 |
| ACE2 | SEMG2 | -0,4245 | 3,19046E-17 |
| ACE2 | NSUN7 | -0,4246 | 3,11406E-17 |
| ACE2 | RP11-142M10.2 | -0,4246 | 3,09472E-17 |
| ACE2 | ZNF569 | -0,4246 | 3,12585E-17 |
| ACE2 | NXNL2 | -0,4247 | 3,02703E-17 |
| ACE2 | INA | -0,4247 | 3,08157E-17 |
| ACE2 | SLC25A35 | -0,4247 | 3,07626E-17 |
| ACE2 | ANKRD61 | -0,4248 | 2,9903E-17 |
| ACE2 | RP11-403D15.2 | -0,4248 | 2,98332E-17 |
| ACE2 | TMPRSS4-AS1 | -0,4248 | 3,00033E-17 |
| ACE2 | DCP1B | -0,4248 | 2,98642E-17 |
| ACE2 | CD163L1 | -0,4248 | 3,01534E-17 |
| ACE2 | GPATCH2L | -0,4248 | 2,99912E-17 |
| ACE2 | SNAP29 | -0,4249 | 2,92019E-17 |
| ACE2 | VN1R20P | -0,425 | 2,89633E-17 |
| ACE2 | RP11-55L3.1 | -0,425 | 2,87654E-17 |
| ACE2 | C6orf48 | -0,425 | 2,90569E-17 |
| ACE2 | LINC02055 | -0,425 | 2,89126E-17 |
| ACE2 | RP11-120E11.2 | -0,425 | 2,91488E-17 |
| ACE2 | RP11-77K12.5 | -0,425 | 2,87977E-17 |
| ACE2 | XIRP2-AS1 | -0,4251 | 2,84131E-17 |
| ACE2 | CTB-91J4.1 | -0,4251 | 2,82937E-17 |
| ACE2 | CDC14C | -0,4252 | 2,80143E-17 |
| ACE2 | PPP2R3C | -0,4252 | 2,8028E-17 |
| ACE2 | OR11A1 | -0,4253 | 2,74548E-17 |
| ACE2 | RP11-991C1.2 | -0,4253 | 2,71695E-17 |

| | | | |
|---|---|---|---|
| ACE2 | NRADDP | -0,4254 | 2,67501E-17 |
| ACE2 | ZNF705G | -0,4254 | 2,66212E-17 |
| ACE2 | DCAF5 | -0,4254 | 2,67081E-17 |
| ACE2 | FAM149B1P1 | -0,4254 | 2,66894E-17 |
| ACE2 | SPIRE1 | -0,4254 | 2,69771E-17 |
| ACE2 | RIMBP3 | -0,4254 | 2,66796E-17 |
| ACE2 | RP11-367B6.2 | -0,4255 | 2,60847E-17 |
| ACE2 | RP11-89K11.1 | -0,4255 | 2,61486E-17 |
| ACE2 | RP1-288L1.5 | -0,4255 | 2,63764E-17 |
| ACE2 | RP5-837M10.1 | -0,4256 | 2,56416E-17 |
| ACE2 | C11orf68 | -0,4256 | 2,58672E-17 |
| ACE2 | RFFL | -0,4256 | 2,58317E-17 |
| ACE2 | CTB-60E11.12 | -0,4257 | 2,53264E-17 |
| ACE2 | AC012361.1 | -0,4258 | 2,47427E-17 |
| ACE2 | PRKAR2A | -0,4258 | 2,47167E-17 |
| ACE2 | LINC01388 | -0,4258 | 2,46302E-17 |
| ACE2 | AC092620.3 | -0,4259 | 2,45322E-17 |
| ACE2 | LINC01848 | -0,4259 | 2,43557E-17 |
| ACE2 | HINT1 | -0,4259 | 2,41658E-17 |
| ACE2 | CASC6 | -0,4259 | 2,41688E-17 |
| ACE2 | SAMD15 | -0,4259 | 2,43866E-17 |
| ACE2 | FAR2P3 | -0,426 | 2,38549E-17 |
| ACE2 | EGFEM1P | -0,4261 | 2,33932E-17 |
| ACE2 | RPL5P24 | -0,4261 | 2,35974E-17 |
| ACE2 | C9orf57 | -0,4261 | 2,3669E-17 |
| ACE2 | CASC2 | -0,4261 | 2,35426E-17 |
| ACE2 | CTD-2213F21.3 | -0,4261 | 2,34273E-17 |
| ACE2 | RP11-552C15.1 | -0,4261 | 2,3601E-17 |
| ACE2 | PARP10 | -0,4262 | 2,28998E-17 |
| ACE2 | OR7E105P | -0,4262 | 2,30053E-17 |
| ACE2 | RP11-1072A3.3 | -0,4262 | 2,29901E-17 |
| ACE2 | ZHX3 | -0,4262 | 2,32304E-17 |
| ACE2 | RP11-294L11.1 | -0,4263 | 2,2747E-17 |
| ACE2 | TSGA10 | -0,4263 | 2,25745E-17 |
| ACE2 | RP11-756P10.4 | -0,4263 | 2,24388E-17 |
| ACE2 | PHYKPL | -0,4263 | 2,26998E-17 |
| ACE2 | RP11-307P5.2 | -0,4263 | 2,2727E-17 |
| ACE2 | VN1R25P | -0,4263 | 2,24285E-17 |
| ACE2 | RP11-574K11.32 | -0,4263 | 2,25368E-17 |
| ACE2 | DRAM1 | -0,4263 | 2,26321E-17 |
| ACE2 | CTC-523E23.15 | -0,4263 | 2,27528E-17 |
| ACE2 | RP1-309H15.2 | -0,4264 | 2,232E-17 |
| ACE2 | RPL7P33 | -0,4264 | 2,19938E-17 |
| ACE2 | RP11-475D12.2 | -0,4264 | 2,20875E-17 |
| ACE2 | BIN2 | -0,4264 | 2,19886E-17 |
| ACE2 | LINC01684 | -0,4264 | 2,22018E-17 |
| ACE2 | CTC-1337H24.3 | -0,4265 | 2,16141E-17 |
| ACE2 | KLRK1 | -0,4265 | 2,15459E-17 |
| ACE2 | SCCPDH | -0,4266 | 2,13082E-17 |
| ACE2 | RP11-576E20.1 | -0,4266 | 2,11658E-17 |
| ACE2 | POC1B | -0,4266 | 2,12925E-17 |
| ACE2 | PRAMEF2 | -0,4267 | 2,10768E-17 |
| ACE2 | RP11-734K21.5 | -0,4267 | 2,10177E-17 |
| ACE2 | UCN2 | -0,4267 | 2,11096E-17 |

| | | | |
|---|---|---|---|
| ACE2 | ASB15 | -0,4267 | 2,08809E-17 |
| ACE2 | TTC41P | -0,4268 | 2,03699E-17 |
| ACE2 | LINC01667 | -0,4268 | 2,04916E-17 |
| ACE2 | GSTO3P | -0,4269 | 1,99912E-17 |
| ACE2 | AADACP1 | -0,4269 | 2,00284E-17 |
| ACE2 | SEMA6A-AS1 | -0,4269 | 2,027E-17 |
| ACE2 | COQ9 | -0,4269 | 2,01998E-17 |
| ACE2 | RP5-983L19.2 | -0,4269 | 2,01367E-17 |
| ACE2 | LINC01844 | -0,427 | 1,96841E-17 |
| ACE2 | RP11-508N12.2 | -0,427 | 1,98992E-17 |
| ACE2 | RP11-482E14.2 | -0,427 | 1,97737E-17 |
| ACE2 | CACNA1C-AS2 | -0,427 | 1,97214E-17 |
| ACE2 | ZNF391 | -0,4271 | 1,9369E-17 |
| ACE2 | RP11-436P7.1 | -0,4271 | 1,95922E-17 |
| ACE2 | FAR2 | -0,4271 | 1,92592E-17 |
| ACE2 | TANGO2 | -0,4271 | 1,93577E-17 |
| ACE2 | RP11-51G5.1 | -0,4272 | 1,88969E-17 |
| ACE2 | RP13-638C3.5 | -0,4272 | 1,90865E-17 |
| ACE2 | MTNR1A | -0,4273 | 1,88312E-17 |
| ACE2 | AC072061.2 | -0,4273 | 1,87011E-17 |
| ACE2 | FOXM1 | -0,4273 | 1,88335E-17 |
| ACE2 | MAPK8IP1P2 | -0,4273 | 1,86347E-17 |
| ACE2 | C17orf47 | -0,4273 | 1,86244E-17 |
| ACE2 | RP11-629B11.5 | -0,4274 | 1,82222E-17 |
| ACE2 | AF131216.5 | -0,4274 | 1,83328E-17 |
| ACE2 | AC062020.2 | -0,4275 | 1,7931E-17 |
| ACE2 | C7orf62 | -0,4276 | 1,7567E-17 |
| ACE2 | GRIK5 | -0,4276 | 1,76672E-17 |
| ACE2 | RP11-539L10.3 | -0,4277 | 1,74152E-17 |
| ACE2 | PSMD8 | -0,4277 | 1,73079E-17 |
| ACE2 | AC008073.7 | -0,4278 | 1,69761E-17 |
| ACE2 | RP11-96B2.1 | -0,4278 | 1,70495E-17 |
| ACE2 | RP11-510J16.5 | -0,4278 | 1,6875E-17 |
| ACE2 | ZNF763 | -0,4278 | 1,68806E-17 |
| ACE2 | IGSF10 | -0,4279 | 1,65731E-17 |
| ACE2 | NPM1P30 | -0,4279 | 1,68038E-17 |
| ACE2 | RP11-697H9.3 | -0,4279 | 1,67848E-17 |
| ACE2 | RPL13AP3 | -0,4279 | 1,6526E-17 |
| ACE2 | OLFM2 | -0,4279 | 1,66032E-17 |
| ACE2 | RN7SKP66 | -0,428 | 1,62487E-17 |
| ACE2 | BNC1 | -0,428 | 1,62183E-17 |
| ACE2 | AC003958.2 | -0,428 | 1,64204E-17 |
| ACE2 | RP11-120A1.1 | -0,4281 | 1,59143E-17 |
| ACE2 | LINC01861 | -0,4281 | 1,59316E-17 |
| ACE2 | SCG5 | -0,4281 | 1,59402E-17 |
| ACE2 | NPM1P29 | -0,4282 | 1,57573E-17 |
| ACE2 | MKRN9P | -0,4282 | 1,58331E-17 |
| ACE2 | CSNK2A2 | -0,4282 | 1,56754E-17 |
| ACE2 | RP11-91I8.3 | -0,4282 | 1,58922E-17 |
| ACE2 | BACH1-IT2 | -0,4282 | 1,5641E-17 |
| ACE2 | AP000345.2 | -0,4282 | 1,56824E-17 |
| ACE2 | UCK2 | -0,4283 | 1,53538E-17 |
| ACE2 | RP11-429P3.5 | -0,4283 | 1,55078E-17 |
| ACE2 | RP13-415G19.2 | -0,4283 | 1,55275E-17 |

| | | | |
|---|---|---|---|
| ACE2 | FAM212B-AS1 | -0,4284 | 1,51469E-17 |
| ACE2 | GRK7 | -0,4284 | 1,50862E-17 |
| ACE2 | B3GNT5 | -0,4284 | 1,52902E-17 |
| ACE2 | RN7SKP293 | -0,4284 | 1,52301E-17 |
| ACE2 | RP11-368M16.7 | -0,4284 | 1,50373E-17 |
| ACE2 | TGIF2LY | -0,4284 | 1,5158E-17 |
| ACE2 | AC004063.1 | -0,4285 | 1,47706E-17 |
| ACE2 | RP11-290O12.2 | -0,4285 | 1,48187E-17 |
| ACE2 | C17orf80 | -0,4285 | 1,47731E-17 |
| ACE2 | RP11-709D24.5 | -0,4286 | 1,46607E-17 |
| ACE2 | RP11-297C4.3 | -0,4286 | 1,45912E-17 |
| ACE2 | RP11-8H2.1 | -0,4286 | 1,46743E-17 |
| ACE2 | LINC00895 | -0,4286 | 1,45958E-17 |
| ACE2 | RP11-196H14.2 | -0,4287 | 1,43928E-17 |
| ACE2 | CPNE2 | -0,4287 | 1,42819E-17 |
| ACE2 | TBC1D3I | -0,4287 | 1,43059E-17 |
| ACE2 | SF3A2 | -0,4287 | 1,41943E-17 |
| ACE2 | LINC02216 | -0,4288 | 1,39543E-17 |
| ACE2 | RP11-728K20.2 | -0,4288 | 1,40291E-17 |
| ACE2 | RP11-157P23.1 | -0,4288 | 1,41003E-17 |
| ACE2 | PGCP1 | -0,4289 | 1,38572E-17 |
| ACE2 | RGPD8 | -0,4289 | 1,38603E-17 |
| ACE2 | UST-AS1 | -0,4289 | 1,39022E-17 |
| ACE2 | RP11-77E14.2 | -0,4289 | 1,3739E-17 |
| ACE2 | ZNF85 | -0,4289 | 1,3866E-17 |
| ACE2 | RP11-465B22.8 | -0,429 | 1,34794E-17 |
| ACE2 | SH2D6 | -0,429 | 1,34473E-17 |
| ACE2 | RP11-806J6.1 | -0,429 | 1,36146E-17 |
| ACE2 | MYBPC1 | -0,429 | 1,35271E-17 |
| ACE2 | SAG | -0,4291 | 1,31727E-17 |
| ACE2 | RP11-263J14.1 | -0,4291 | 1,32039E-17 |
| ACE2 | GAPDHP22 | -0,4291 | 1,3349E-17 |
| ACE2 | RP11-24M17.6 | -0,4291 | 1,33671E-17 |
| ACE2 | ANKRD45 | -0,4292 | 1,29958E-17 |
| ACE2 | SFXN4 | -0,4292 | 1,30965E-17 |
| ACE2 | RP11-159D23.2 | -0,4292 | 1,29279E-17 |
| ACE2 | S100A8 | -0,4293 | 1,27391E-17 |
| ACE2 | RP1-90J20.8 | -0,4293 | 1,28274E-17 |
| ACE2 | CFAP206 | -0,4293 | 1,26895E-17 |
| ACE2 | TMC1 | -0,4293 | 1,27949E-17 |
| ACE2 | MYH3 | -0,4293 | 1,27658E-17 |
| ACE2 | RNF138 | -0,4293 | 1,2782E-17 |
| ACE2 | PLVAP | -0,4293 | 1,27724E-17 |
| ACE2 | RP11-536P6.3 | -0,4293 | 1,27928E-17 |
| ACE2 | ADGRL3-AS1 | -0,4294 | 1,25703E-17 |
| ACE2 | RP11-522B15.7 | -0,4294 | 1,25284E-17 |
| ACE2 | RP11-266L9.1 | -0,4294 | 1,2563E-17 |
| ACE2 | LINC00303 | -0,4295 | 1,2367E-17 |
| ACE2 | LINC01613 | -0,4295 | 1,22029E-17 |
| ACE2 | RP11-672F9.1 | -0,4295 | 1,22629E-17 |
| ACE2 | ZNRF1 | -0,4295 | 1,23753E-17 |
| ACE2 | RASSF1 | -0,4296 | 1,21618E-17 |
| ACE2 | RP11-446F3.2 | -0,4296 | 1,19803E-17 |
| ACE2 | ARHGAP15 | -0,4297 | 1,19297E-17 |

| | | | |
|---|---|---|---|
| ACE2 | SULF1 | -0,4297 | 1,18945E-17 |
| ACE2 | CFAP58 | -0,4297 | 1,17837E-17 |
| ACE2 | LINC00644 | -0,4297 | 1,1869E-17 |
| ACE2 | RP11-77H9.8 | -0,4297 | 1,17903E-17 |
| ACE2 | DOT1L | -0,4297 | 1,18979E-17 |
| ACE2 | TAF1A-AS1 | -0,4299 | 1,13599E-17 |
| ACE2 | TPRG1 | -0,4299 | 1,14404E-17 |
| ACE2 | TM4SF19-AS1 | -0,4299 | 1,13728E-17 |
| ACE2 | RP11-317O24.1 | -0,4299 | 1,14204E-17 |
| ACE2 | RP11-428L9.2 | -0,43 | 1,12014E-17 |
| ACE2 | PPIAP3 | -0,43 | 1,12536E-17 |
| ACE2 | RP11-495P10.1 | -0,4302 | 1,07621E-17 |
| ACE2 | NPM1P52 | -0,4302 | 1,07347E-17 |
| ACE2 | BTBD17 | -0,4303 | 1,04501E-17 |
| ACE2 | RP11-29B9.1 | -0,4304 | 1,03268E-17 |
| ACE2 | TMEM105 | -0,4304 | 1,03617E-17 |
| ACE2 | CABLES2 | -0,4304 | 1,04328E-17 |
| ACE2 | PKDREJ | -0,4304 | 1,04272E-17 |
| ACE2 | CITF22-49D8.1 | -0,4304 | 1,04059E-17 |
| ACE2 | LINC01867 | -0,4305 | 1,0054E-17 |
| ACE2 | AC016738.3 | -0,4305 | 1,02291E-17 |
| ACE2 | CDHR5 | -0,4305 | 1,01688E-17 |
| ACE2 | CTD-2350J17.1 | -0,4307 | 9,70384E-18 |
| ACE2 | LINC01708 | -0,4308 | 9,51759E-18 |
| ACE2 | LINC00210 | -0,4308 | 9,60931E-18 |
| ACE2 | GNA14 | -0,4308 | 9,50361E-18 |
| ACE2 | AKR1C6P | -0,4308 | 9,53391E-18 |
| ACE2 | PCGF1 | -0,4309 | 9,4656E-18 |
| ACE2 | E2F7 | -0,4309 | 9,42532E-18 |
| ACE2 | NABP2 | -0,431 | 9,19431E-18 |
| ACE2 | RP11-785F11.1 | -0,4311 | 9,03629E-18 |
| ACE2 | ATP5F1P7 | -0,4311 | 8,96344E-18 |
| ACE2 | RP11-47G4.2 | -0,4311 | 8,95791E-18 |
| ACE2 | CT83 | -0,4311 | 9,05472E-18 |
| ACE2 | NDUFB4 | -0,4312 | 8,89429E-18 |
| ACE2 | RP11-380J14.4 | -0,4313 | 8,65881E-18 |
| ACE2 | DPYD-AS2 | -0,4313 | 8,76031E-18 |
| ACE2 | RP11-85O21.2 | -0,4313 | 8,77968E-18 |
| ACE2 | RP11-33E24.3 | -0,4314 | 8,53439E-18 |
| ACE2 | CUBN | -0,4314 | 8,50072E-18 |
| ACE2 | IGKV1-13 | -0,4315 | 8,32881E-18 |
| ACE2 | NGLY1 | -0,4315 | 8,41566E-18 |
| ACE2 | RP11-447M12.2 | -0,4315 | 8,44541E-18 |
| ACE2 | SPG20-AS1 | -0,4315 | 8,30291E-18 |
| ACE2 | RP11-540B6.3 | -0,4315 | 8,43847E-18 |
| ACE2 | RP5-1085F17.4 | -0,4315 | 8,34974E-18 |
| ACE2 | TCTEX1D2 | -0,4316 | 8,27668E-18 |
| ACE2 | RP11-385N23.1 | -0,4316 | 8,18007E-18 |
| ACE2 | RPL29P30 | -0,4316 | 8,27776E-18 |
| ACE2 | LINC02097 | -0,4316 | 8,24967E-18 |
| ACE2 | RP11-141A19.1 | -0,4317 | 7,98616E-18 |
| ACE2 | ST7L | -0,4317 | 8,00946E-18 |
| ACE2 | AC105402.4 | -0,4317 | 8,10475E-18 |
| ACE2 | CTD-2576D5.4 | -0,4317 | 8,01315E-18 |

| | | | |
|---|---|---|---|
| ACE2 | CTD-2582D11.1 | -0,4317 | 8,10457E-18 |
| ACE2 | SDR16C6P | -0,4318 | 7,82821E-18 |
| ACE2 | MPP7 | -0,4318 | 7,82638E-18 |
| ACE2 | MRPL21 | -0,4318 | 7,89106E-18 |
| ACE2 | LINC01427 | -0,4318 | 7,92705E-18 |
| ACE2 | RPS15P5 | -0,4319 | 7,81996E-18 |
| ACE2 | TMEM144 | -0,4319 | 7,79616E-18 |
| ACE2 | NPIPB14P | -0,432 | 7,60978E-18 |
| ACE2 | CH17-195P21.1 | -0,4321 | 7,48609E-18 |
| ACE2 | RP11-46H11.12 | -0,4321 | 7,46298E-18 |
| ACE2 | A3GALT2 | -0,4322 | 7,37156E-18 |
| ACE2 | FAM138B | -0,4322 | 7,27518E-18 |
| ACE2 | AC004066.3 | -0,4322 | 7,26234E-18 |
| ACE2 | PCDHGC4 | -0,4322 | 7,3564E-18 |
| ACE2 | PPP1R16A | -0,4322 | 7,24342E-18 |
| ACE2 | RP11-85L21.4 | -0,4322 | 7,27384E-18 |
| ACE2 | GPHA2 | -0,4323 | 7,22839E-18 |
| ACE2 | AC092664.1 | -0,4324 | 7,02773E-18 |
| ACE2 | RPS26P2 | -0,4324 | 7,08686E-18 |
| ACE2 | RP11-89C3.4 | -0,4324 | 7,0744E-18 |
| ACE2 | DACH1 | -0,4324 | 7,08999E-18 |
| ACE2 | CTD-2545H1.2 | -0,4324 | 6,97712E-18 |
| ACE2 | GTSE1-AS1 | -0,4324 | 7,01968E-18 |
| ACE2 | AC010894.5 | -0,4325 | 6,92557E-18 |
| ACE2 | KHDRBS3 | -0,4325 | 6,95053E-18 |
| ACE2 | ZNF570 | -0,4325 | 6,93075E-18 |
| ACE2 | PIN1P1 | -0,4326 | 6,71028E-18 |
| ACE2 | C1orf229 | -0,4326 | 6,77225E-18 |
| ACE2 | IL37 | -0,4326 | 6,71052E-18 |
| ACE2 | SLC39A12-AS1 | -0,4326 | 6,74523E-18 |
| ACE2 | RP11-42H13.1 | -0,4326 | 6,82296E-18 |
| ACE2 | AC007795.1 | -0,4326 | 6,75197E-18 |
| ACE2 | RN7SL555P | -0,4326 | 6,81695E-18 |
| ACE2 | ZAN | -0,4327 | 6,61416E-18 |
| ACE2 | CEP76 | -0,4327 | 6,63562E-18 |
| ACE2 | RP11-419L4.1 | -0,4328 | 6,52009E-18 |
| ACE2 | GPR65 | -0,4328 | 6,4682E-18 |
| ACE2 | RP11-332H18.3 | -0,4328 | 6,49716E-18 |
| ACE2 | LINC01900 | -0,4328 | 6,45979E-18 |
| ACE2 | RPL12P42 | -0,4328 | 6,53352E-18 |
| ACE2 | PINLYP | -0,4328 | 6,49392E-18 |
| ACE2 | SPATA2L | -0,4329 | 6,41542E-18 |
| ACE2 | LINC01635 | -0,433 | 6,2252E-18 |
| ACE2 | CTA-360L10.1 | -0,433 | 6,32508E-18 |
| ACE2 | RP11-103B5.4 | -0,433 | 6,29977E-18 |
| ACE2 | RP11-64B16.3 | -0,433 | 6,28311E-18 |
| ACE2 | CTD-2525I3.3 | -0,433 | 6,22498E-18 |
| ACE2 | CCDC39 | -0,4331 | 6,15413E-18 |
| ACE2 | NDUFB2 | -0,4331 | 6,08777E-18 |
| ACE2 | RUBCNL | -0,4331 | 6,16861E-18 |
| ACE2 | RP11-156P1.3 | -0,4331 | 6,14145E-18 |
| ACE2 | CYB561D2 | -0,4332 | 6,01028E-18 |
| ACE2 | LRP1B | -0,4333 | 5,94108E-18 |
| ACE2 | RPL32P9 | -0,4334 | 5,82861E-18 |

| | | | |
|---|---|---|---|
| ACE2 | RP11-356K23.1 | -0,4334 | 5,79401E-18 |
| ACE2 | CINP | -0,4334 | 5,82701E-18 |
| ACE2 | GOLGA6L6 | -0,4334 | 5,74651E-18 |
| ACE2 | CTD-3037G24.3 | -0,4334 | 5,7544E-18 |
| ACE2 | SLC32A1 | -0,4334 | 5,79011E-18 |
| ACE2 | RP4-669H2.1 | -0,4334 | 5,749E-18 |
| ACE2 | AP000302.58 | -0,4334 | 5,82326E-18 |
| ACE2 | RP11-730N24.1 | -0,4335 | 5,67708E-18 |
| ACE2 | KB-1205A7.1 | -0,4335 | 5,68181E-18 |
| ACE2 | SMTNL1 | -0,4335 | 5,64104E-18 |
| ACE2 | PRKCA-AS1 | -0,4335 | 5,64722E-18 |
| ACE2 | RP11-389O22.5 | -0,4336 | 5,55934E-18 |
| ACE2 | ZNF596 | -0,4336 | 5,5266E-18 |
| ACE2 | AC004870.3 | -0,4337 | 5,43499E-18 |
| ACE2 | RP11-700P18.1 | -0,4337 | 5,41832E-18 |
| ACE2 | RP11-810K23.6 | -0,4337 | 5,43819E-18 |
| ACE2 | PKMYT1 | -0,4337 | 5,45172E-18 |
| ACE2 | RP11-574O16.1 | -0,4338 | 5,33789E-18 |
| ACE2 | GLT6D1 | -0,4338 | 5,32151E-18 |
| ACE2 | UMODL1 | -0,4338 | 5,33072E-18 |
| ACE2 | IGSF11 | -0,4339 | 5,30055E-18 |
| ACE2 | PRMT1P1 | -0,4339 | 5,26073E-18 |
| ACE2 | MROH1 | -0,4339 | 5,25698E-18 |
| ACE2 | TUBB4B | -0,4339 | 5,22218E-18 |
| ACE2 | RP13-820C6.4 | -0,4339 | 5,2209E-18 |
| ACE2 | CYP4F35P | -0,4339 | 5,27985E-18 |
| ACE2 | MST1 | -0,434 | 5,17424E-18 |
| ACE2 | GS1-39E22.1 | -0,434 | 5,2095E-18 |
| ACE2 | DPY19L1P1 | -0,434 | 5,17721E-18 |
| ACE2 | LINC01673 | -0,434 | 5,13953E-18 |
| ACE2 | RP1-225E12.2 | -0,4341 | 5,04202E-18 |
| ACE2 | RP11-525A16.4 | -0,4341 | 5,0282E-18 |
| ACE2 | RP1-122P22.4 | -0,4341 | 5,07894E-18 |
| ACE2 | EEF1A1P1 | -0,4341 | 5,02138E-18 |
| ACE2 | PRKCE | -0,4342 | 4,95737E-18 |
| ACE2 | CYP4F30P | -0,4342 | 4,96754E-18 |
| ACE2 | OR5B21 | -0,4342 | 4,98804E-18 |
| ACE2 | RASSF8 | -0,4342 | 4,94837E-18 |
| ACE2 | RP11-279O17.2 | -0,4342 | 4,93503E-18 |
| ACE2 | RPL21P50 | -0,4343 | 4,8537E-18 |
| ACE2 | RP11-890B15.2 | -0,4343 | 4,85439E-18 |
| ACE2 | LINC01863 | -0,4344 | 4,75106E-18 |
| ACE2 | RP11-211C9.1 | -0,4344 | 4,79599E-18 |
| ACE2 | RP11-769N19.2 | -0,4344 | 4,75719E-18 |
| ACE2 | MIMT1 | -0,4344 | 4,76736E-18 |
| ACE2 | RP11-488L18.6 | -0,4345 | 4,65653E-18 |
| ACE2 | LRR1 | -0,4345 | 4,69973E-18 |
| ACE2 | GPD2 | -0,4346 | 4,59194E-18 |
| ACE2 | DCAF1 | -0,4346 | 4,59187E-18 |
| ACE2 | RNF125 | -0,4346 | 4,59726E-18 |
| ACE2 | OR2L13 | -0,4347 | 4,46174E-18 |
| ACE2 | CMPK2 | -0,4347 | 4,51203E-18 |
| ACE2 | CTD-2006C1.2 | -0,4347 | 4,53104E-18 |
| ACE2 | LINC01605 | -0,4348 | 4,41822E-18 |

| | | | |
|---|---|---|---|
| ACE2 | MAPT-IT1 | -0,4348 | 4,39648E-18 |
| ACE2 | IGLVIV-66-1 | -0,4348 | 4,43065E-18 |
| ACE2 | RP11-319E12.2 | -0,4349 | 4,34452E-18 |
| ACE2 | RP11-98J9.1 | -0,4349 | 4,34762E-18 |
| ACE2 | ARL6IP4 | -0,4349 | 4,30116E-18 |
| ACE2 | RP11-69J7.1 | -0,435 | 4,24182E-18 |
| ACE2 | RP11-219B4.6 | -0,435 | 4,22505E-18 |
| ACE2 | RP11-85G21.3 | -0,4351 | 4,19095E-18 |
| ACE2 | C10orf126 | -0,4351 | 4,14471E-18 |
| ACE2 | RP11-325D8.1 | -0,4351 | 4,19423E-18 |
| ACE2 | TDP1 | -0,4351 | 4,18038E-18 |
| ACE2 | CTD-2196E14.3 | -0,4351 | 4,19326E-18 |
| ACE2 | CTD-2023N9.2 | -0,4352 | 4,05025E-18 |
| ACE2 | CISD1 | -0,4353 | 4,01062E-18 |
| ACE2 | YBX3 | -0,4353 | 4,03977E-18 |
| ACE2 | NFAM1 | -0,4353 | 4,01918E-18 |
| ACE2 | VIL1 | -0,4354 | 3,89163E-18 |
| ACE2 | RP11-249L12.1 | -0,4354 | 3,95575E-18 |
| ACE2 | RP11-397H6.1 | -0,4355 | 3,84607E-18 |
| ACE2 | ADGRF3 | -0,4356 | 3,7761E-18 |
| ACE2 | AL022344.7 | -0,4356 | 3,74288E-18 |
| ACE2 | RBBP6 | -0,4356 | 3,76046E-18 |
| ACE2 | AC008079.10 | -0,4356 | 3,78814E-18 |
| ACE2 | LINC00207 | -0,4356 | 3,78139E-18 |
| ACE2 | RP5-1065P14.2 | -0,4357 | 3,69021E-18 |
| ACE2 | RP11-1277A3.1 | -0,4357 | 3,70265E-18 |
| ACE2 | SYNJ2BP | -0,4357 | 3,69722E-18 |
| ACE2 | ZNF295-AS1 | -0,4357 | 3,67687E-18 |
| ACE2 | VPREB1 | -0,4357 | 3,69875E-18 |
| ACE2 | TFAM | -0,4358 | 3,59928E-18 |
| ACE2 | RP11-131L12.2 | -0,4358 | 3,61956E-18 |
| ACE2 | B4GALNT2 | -0,4358 | 3,65234E-18 |
| ACE2 | LINC01756 | -0,4359 | 3,54593E-18 |
| ACE2 | AC013472.4 | -0,4359 | 3,56121E-18 |
| ACE2 | FAM19A1 | -0,4359 | 3,57502E-18 |
| ACE2 | RN7SKP230 | -0,4359 | 3,59085E-18 |
| ACE2 | FAM25E | -0,4359 | 3,55734E-18 |
| ACE2 | RP11-436F21.1 | -0,436 | 3,47734E-18 |
| ACE2 | RP4-796I8.1 | -0,436 | 3,47559E-18 |
| ACE2 | CECR1 | -0,436 | 3,51633E-18 |
| ACE2 | MORC2-AS1 | -0,436 | 3,51521E-18 |
| ACE2 | RP1-68D18.3 | -0,4361 | 3,39473E-18 |
| ACE2 | RP11-1086I4.2 | -0,4361 | 3,43128E-18 |
| ACE2 | CTD-2168K21.1 | -0,4362 | 3,3612E-18 |
| ACE2 | AMER2 | -0,4362 | 3,34416E-18 |
| ACE2 | KIAA0556 | -0,4362 | 3,34282E-18 |
| ACE2 | CFAP52 | -0,4362 | 3,36833E-18 |
| ACE2 | RP11-342D14.1 | -0,4362 | 3,38822E-18 |
| ACE2 | RP11-499F19.3 | -0,4363 | 3,28046E-18 |
| ACE2 | ATP5O | -0,4363 | 3,29749E-18 |
| ACE2 | RP1-17K7.1 | -0,4364 | 3,22007E-18 |
| ACE2 | NIFKP3 | -0,4364 | 3,23706E-18 |
| ACE2 | LINC01570 | -0,4364 | 3,24194E-18 |
| ACE2 | AC009892.10 | -0,4364 | 3,20462E-18 |

| | | | |
|---|---|---|---|
| ACE2 | RP11-74E24.2 | -0,4365 | 3,19271E-18 |
| ACE2 | RNF141 | -0,4365 | 3,17966E-18 |
| ACE2 | RP11-153K16.1 | -0,4365 | 3,19266E-18 |
| ACE2 | CD207 | -0,4366 | 3,0945E-18 |
| ACE2 | XXbac-BPG157A10.21 | -0,4366 | 3,13527E-18 |
| ACE2 | RP1-38C16.2 | -0,4366 | 3,08594E-18 |
| ACE2 | MAF1 | -0,4366 | 3,11568E-18 |
| ACE2 | FLJ42102 | -0,4366 | 3,11364E-18 |
| ACE2 | SERPINB11 | -0,4366 | 3,11488E-18 |
| ACE2 | RP11-77G23.2 | -0,4367 | 3,06699E-18 |
| ACE2 | RP11-361H10.7 | -0,4367 | 3,0279E-18 |
| ACE2 | FAR2P4 | -0,4368 | 2,9982E-18 |
| ACE2 | CASC15 | -0,4368 | 2,97649E-18 |
| ACE2 | RP11-439H9.1 | -0,4368 | 2,97683E-18 |
| ACE2 | RP11-563N12.2 | -0,4368 | 2,98458E-18 |
| ACE2 | ADAM18 | -0,4368 | 3,00801E-18 |
| ACE2 | TLE3 | -0,4368 | 2,98418E-18 |
| ACE2 | RP11-3M1.1 | -0,4368 | 2,96846E-18 |
| ACE2 | RP1-66N13.4 | -0,4368 | 2,98324E-18 |
| ACE2 | BRD9 | -0,4369 | 2,90625E-18 |
| ACE2 | RP11-34N19.1 | -0,4369 | 2,92995E-18 |
| ACE2 | RP11-255G12.2 | -0,4369 | 2,90808E-18 |
| ACE2 | TES | -0,437 | 2,8782E-18 |
| ACE2 | RP11-589C21.6 | -0,437 | 2,84967E-18 |
| ACE2 | LINC02148 | -0,4371 | 2,81683E-18 |
| ACE2 | RP11-588P7.1 | -0,4371 | 2,82852E-18 |
| ACE2 | AC010492.2 | -0,4371 | 2,81E-18 |
| ACE2 | CPO | -0,4372 | 2,77789E-18 |
| ACE2 | FAM170B | -0,4372 | 2,73714E-18 |
| ACE2 | SDR9C7 | -0,4372 | 2,74913E-18 |
| ACE2 | HSPA4L | -0,4373 | 2,72313E-18 |
| ACE2 | JAKMIP2 | -0,4373 | 2,69158E-18 |
| ACE2 | RP3-446N13.4 | -0,4373 | 2,71225E-18 |
| ACE2 | RP11-383C5.5 | -0,4374 | 2,6551E-18 |
| ACE2 | TENM4 | -0,4374 | 2,67546E-18 |
| ACE2 | AC007326.10 | -0,4374 | 2,6359E-18 |
| ACE2 | RP4-565E6.1 | -0,4375 | 2,61908E-18 |
| ACE2 | SLC5A10 | -0,4376 | 2,53378E-18 |
| ACE2 | MAEL | -0,4377 | 2,50996E-18 |
| ACE2 | RP11-588P8.1 | -0,4377 | 2,48489E-18 |
| ACE2 | RP11-58B2.1 | -0,4377 | 2,49746E-18 |
| ACE2 | ZNF775 | -0,4377 | 2,5271E-18 |
| ACE2 | CTD-2270L9.4 | -0,4377 | 2,52638E-18 |
| ACE2 | LRRC15 | -0,4378 | 2,47417E-18 |
| ACE2 | RP11-335O4.1 | -0,4378 | 2,46493E-18 |
| ACE2 | RP4-794I6.4 | -0,4378 | 2,45533E-18 |
| ACE2 | RP11-440J4.2 | -0,4379 | 2,42977E-18 |
| ACE2 | AC002519.6 | -0,4379 | 2,4014E-18 |
| ACE2 | LINC01568 | -0,4379 | 2,4077E-18 |
| ACE2 | TMEM160 | -0,4379 | 2,3979E-18 |
| ACE2 | AC068489.1 | -0,438 | 2,36964E-18 |
| ACE2 | RNU1-96P | -0,438 | 2,33915E-18 |
| ACE2 | CCNO | -0,438 | 2,38E-18 |
| ACE2 | TCP10 | -0,438 | 2,33856E-18 |

| | | | |
|---|---|---|---|
| ACE2 | AC005014.5 | -0,438 | 2,37154E-18 |
| ACE2 | C10orf67 | -0,438 | 2,35787E-18 |
| ACE2 | LINC01039 | -0,438 | 2,37833E-18 |
| ACE2 | RP11-420K14.1 | -0,438 | 2,34522E-18 |
| ACE2 | C20orf96 | -0,438 | 2,37616E-18 |
| ACE2 | RP4-726N1.2 | -0,438 | 2,37078E-18 |
| ACE2 | CTC-340I23.2 | -0,4381 | 2,33071E-18 |
| ACE2 | STRCP1 | -0,4381 | 2,3349E-18 |
| ACE2 | FAF1 | -0,4382 | 2,28825E-18 |
| ACE2 | AC097381.1 | -0,4382 | 2,28106E-18 |
| ACE2 | RP11-620J15.3 | -0,4382 | 2,26619E-18 |
| ACE2 | AC053503.4 | -0,4383 | 2,21457E-18 |
| ACE2 | RP11-339B21.8 | -0,4383 | 2,22174E-18 |
| ACE2 | CTSLP1 | -0,4383 | 2,22615E-18 |
| ACE2 | LINC01885 | -0,4384 | 2,1676E-18 |
| ACE2 | OR7E155P | -0,4384 | 2,17085E-18 |
| ACE2 | GPR18 | -0,4384 | 2,17157E-18 |
| ACE2 | LRRC53 | -0,4385 | 2,12114E-18 |
| ACE2 | MAGI2-AS2 | -0,4385 | 2,14322E-18 |
| ACE2 | HELLPAR | -0,4385 | 2,12698E-18 |
| ACE2 | MIR3180-1 | -0,4385 | 2,1292E-18 |
| ACE2 | KRT73 | -0,4386 | 2,09371E-18 |
| ACE2 | RP11-711C17.1 | -0,4386 | 2,10196E-18 |
| ACE2 | POMP | -0,4387 | 2,04641E-18 |
| ACE2 | MTHFSD | -0,4387 | 2,05699E-18 |
| ACE2 | MS4A13 | -0,4388 | 2,01921E-18 |
| ACE2 | SEPT3 | -0,4388 | 2,00696E-18 |
| ACE2 | LINC00958 | -0,4389 | 1,95779E-18 |
| ACE2 | RP11-507B12.1 | -0,4389 | 1,95806E-18 |
| ACE2 | IQCH | -0,4389 | 1,97434E-18 |
| ACE2 | CRTC3-AS1 | -0,4389 | 1,96814E-18 |
| ACE2 | HCG4P3 | -0,439 | 1,95598E-18 |
| ACE2 | RP11-863N1.4 | -0,439 | 1,94724E-18 |
| ACE2 | GLTSCR2-AS1 | -0,439 | 1,93594E-18 |
| ACE2 | D21S2088E | -0,4391 | 1,9052E-18 |
| ACE2 | PDZPH1P | -0,4392 | 1,85404E-18 |
| ACE2 | LINC02079 | -0,4392 | 1,87667E-18 |
| ACE2 | AC145343.2 | -0,4392 | 1,87921E-18 |
| ACE2 | RABL2A | -0,4393 | 1,81579E-18 |
| ACE2 | PSMA2P2 | -0,4393 | 1,83471E-18 |
| ACE2 | RP11-380D11.2 | -0,4393 | 1,81424E-18 |
| ACE2 | AIMP2 | -0,4394 | 1,79721E-18 |
| ACE2 | HNRNPRP1 | -0,4394 | 1,79591E-18 |
| ACE2 | RP11-567C2.1 | -0,4394 | 1,7922E-18 |
| ACE2 | AC144833.1 | -0,4394 | 1,77979E-18 |
| ACE2 | RP11-483E23.2 | -0,4394 | 1,79149E-18 |
| ACE2 | ZNF574 | -0,4394 | 1,79878E-18 |
| ACE2 | GAPDHP74 | -0,4395 | 1,77003E-18 |
| ACE2 | SMR3B | -0,4395 | 1,76249E-18 |
| ACE2 | TCP10L2 | -0,4395 | 1,73975E-18 |
| ACE2 | RP11-324P9.1 | -0,4395 | 1,77092E-18 |
| ACE2 | PROZ | -0,4396 | 1,73108E-18 |
| ACE2 | RORA-AS1 | -0,4396 | 1,70968E-18 |
| ACE2 | ZNF567 | -0,4396 | 1,70994E-18 |

| | | | |
|---|---|---|---|
| ACE2 | NEUROD1 | -0,4397 | 1,69326E-18 |
| ACE2 | SLC38A9 | -0,4397 | 1,67436E-18 |
| ACE2 | AP000470.2 | -0,4397 | 1,70371E-18 |
| ACE2 | PMCHL2 | -0,4398 | 1,63965E-18 |
| ACE2 | RP11-136K7.1 | -0,4399 | 1,62515E-18 |
| ACE2 | TRIM60P16 | -0,4399 | 1,62629E-18 |
| ACE2 | RP11-359J6.1 | -0,4399 | 1,60799E-18 |
| ACE2 | WDR66 | -0,4399 | 1,61016E-18 |
| ACE2 | RP11-159D12.10 | -0,4399 | 1,61515E-18 |
| ACE2 | UQCRC1 | -0,44 | 1,57574E-18 |
| ACE2 | ARMC2 | -0,44 | 1,58805E-18 |
| ACE2 | ZNF273 | -0,44 | 1,58268E-18 |
| ACE2 | RP11-626H12.3 | -0,44 | 1,60428E-18 |
| ACE2 | PIAS2 | -0,44 | 1,60479E-18 |
| ACE2 | LINC01741 | -0,4401 | 1,55074E-18 |
| ACE2 | LINC02077 | -0,4401 | 1,55522E-18 |
| ACE2 | RP11-539I5.1 | -0,4401 | 1,57017E-18 |
| ACE2 | GRM7-AS2 | -0,4402 | 1,52522E-18 |
| ACE2 | CTC-218H9.1 | -0,4402 | 1,54232E-18 |
| ACE2 | RP1-34L19.1 | -0,4402 | 1,5206E-18 |
| ACE2 | AC003985.1 | -0,4402 | 1,54022E-18 |
| ACE2 | DIAPH3 | -0,4402 | 1,53354E-18 |
| ACE2 | CAGE1 | -0,4403 | 1,50111E-18 |
| ACE2 | LINC02210 | -0,4403 | 1,48906E-18 |
| ACE2 | HSF2BP | -0,4403 | 1,4904E-18 |
| ACE2 | AC009264.1 | -0,4404 | 1,46171E-18 |
| ACE2 | RP5-951N9.1 | -0,4404 | 1,46781E-18 |
| ACE2 | MIR600HG | -0,4405 | 1,44054E-18 |
| ACE2 | LINC01788 | -0,4406 | 1,42146E-18 |
| ACE2 | UBTD2 | -0,4406 | 1,40129E-18 |
| ACE2 | SRGAP3-AS1 | -0,4407 | 1,37658E-18 |
| ACE2 | LINC01359 | -0,4408 | 1,35299E-18 |
| ACE2 | AF178030.2 | -0,4408 | 1,35144E-18 |
| ACE2 | RP11-49O14.3 | -0,4408 | 1,34797E-18 |
| ACE2 | RP11-720D4.2 | -0,4408 | 1,34527E-18 |
| ACE2 | COIL | -0,4408 | 1,35005E-18 |
| ACE2 | CTD-2542C24.8 | -0,4408 | 1,37033E-18 |
| ACE2 | PPP1R2P2 | -0,4408 | 1,35287E-18 |
| ACE2 | RP11-256L11.3 | -0,4409 | 1,33749E-18 |
| ACE2 | E2F2 | -0,441 | 1,30807E-18 |
| ACE2 | MTCH2 | -0,441 | 1,30668E-18 |
| ACE2 | DNAH10 | -0,441 | 1,30856E-18 |
| ACE2 | LINC02125 | -0,441 | 1,312E-18 |
| ACE2 | CCDC80 | -0,4412 | 1,25494E-18 |
| ACE2 | RP11-136K7.3 | -0,4412 | 1,24877E-18 |
| ACE2 | RP3-406P24.3 | -0,4412 | 1,2499E-18 |
| ACE2 | RP5-1093G23.1 | -0,4412 | 1,25962E-18 |
| ACE2 | RP11-219B4.7 | -0,4412 | 1,26111E-18 |
| ACE2 | C9orf129 | -0,4412 | 1,26579E-18 |
| ACE2 | SLC12A1 | -0,4412 | 1,24343E-18 |
| ACE2 | RP11-152O14.3 | -0,4412 | 1,2419E-18 |
| ACE2 | CFAP20 | -0,4412 | 1,24256E-18 |
| ACE2 | MAP3K10 | -0,4412 | 1,24418E-18 |
| ACE2 | KB-1592A4.15 | -0,4412 | 1,25333E-18 |

| | | | |
|---|---|---|---|
| ACE2 | TAT | -0,4413 | 1,21759E-18 |
| ACE2 | MUM1 | -0,4413 | 1,22555E-18 |
| ACE2 | RP5-837J1.6 | -0,4414 | 1,20167E-18 |
| ACE2 | UBE2DNL | -0,4414 | 1,19955E-18 |
| ACE2 | RP11-327F22.1 | -0,4415 | 1,1753E-18 |
| ACE2 | DYM | -0,4415 | 1,19029E-18 |
| ACE2 | CTD-2314G24.2 | -0,4416 | 1,16001E-18 |
| ACE2 | TRBV5-3 | -0,4416 | 1,16096E-18 |
| ACE2 | NARF | -0,4416 | 1,15102E-18 |
| ACE2 | LINC00633 | -0,4416 | 1,1496E-18 |
| ACE2 | CC2D2B | -0,4417 | 1,13218E-18 |
| ACE2 | SLX4 | -0,4417 | 1,12448E-18 |
| ACE2 | DND1P1 | -0,4417 | 1,13456E-18 |
| ACE2 | CTD-2532D12.5 | -0,4417 | 1,14506E-18 |
| ACE2 | AC006273.4 | -0,4417 | 1,13726E-18 |
| ACE2 | AF038458.3 | -0,4418 | 1,11114E-18 |
| ACE2 | DDX20 | -0,4419 | 1,09739E-18 |
| ACE2 | VN1R48P | -0,4419 | 1,08391E-18 |
| ACE2 | SLC25A30-AS1 | -0,4419 | 1,08576E-18 |
| ACE2 | RP11-307C19.2 | -0,4419 | 1,09432E-18 |
| ACE2 | APLF | -0,442 | 1,07631E-18 |
| ACE2 | CTD-2342N23.3 | -0,442 | 1,07187E-18 |
| ACE2 | ST8SIA6-AS1 | -0,442 | 1,07321E-18 |
| ACE2 | RP11-522B15.5 | -0,442 | 1,06748E-18 |
| ACE2 | LL22NC03-24A12.9 | -0,442 | 1,06822E-18 |
| ACE2 | RP11-410L14.2 | -0,4421 | 1,05767E-18 |
| ACE2 | KRT72 | -0,4421 | 1,04104E-18 |
| ACE2 | IGHEP2 | -0,4422 | 1,01919E-18 |
| ACE2 | RP11-442O18.2 | -0,4422 | 1,01839E-18 |
| ACE2 | BPIFA1 | -0,4422 | 1,02753E-18 |
| ACE2 | MICALL1 | -0,4422 | 1,01789E-18 |
| ACE2 | RP1-85F18.6 | -0,4422 | 1,03147E-18 |
| ACE2 | RP11-404K5.1 | -0,4423 | 1,00045E-18 |
| ACE2 | SHH | -0,4423 | 1,01633E-18 |
| ACE2 | TRPA1 | -0,4423 | 1,01369E-18 |
| ACE2 | MAP2K6 | -0,4423 | 1,01157E-18 |
| ACE2 | LINC01924 | -0,4423 | 1,00884E-18 |
| ACE2 | FAM3B | -0,4423 | 9,98461E-19 |
| ACE2 | AC007326.9 | -0,4423 | 1,0046E-18 |
| ACE2 | LINC01980 | -0,4424 | 9,91303E-19 |
| ACE2 | RP11-497K21.1 | -0,4424 | 9,94626E-19 |
| ACE2 | CTD-2561F5.1 | -0,4424 | 9,78637E-19 |
| ACE2 | RP4-550H1.4 | -0,4424 | 9,92712E-19 |
| ACE2 | AP000275.64 | -0,4424 | 9,82143E-19 |
| ACE2 | AC007078.4 | -0,4425 | 9,64934E-19 |
| ACE2 | CCDC158 | -0,4426 | 9,57732E-19 |
| ACE2 | ZER1 | -0,4426 | 9,48675E-19 |
| ACE2 | OR4N1P | -0,4426 | 9,55279E-19 |
| ACE2 | EEF1A1P2 | -0,4426 | 9,517E-19 |
| ACE2 | RP11-133L19.1 | -0,4426 | 9,51279E-19 |
| ACE2 | ZNF485 | -0,4427 | 9,23075E-19 |
| ACE2 | RP11-98J9.3 | -0,4427 | 9,28035E-19 |
| ACE2 | KLKP1 | -0,4427 | 9,2369E-19 |
| ACE2 | RP11-114H23.2 | -0,4428 | 9,02556E-19 |

| | | | |
|---|---|---|---|
| ACE2 | LATS2 | -0,4428 | 9,04643E-19 |
| ACE2 | CTD-2260A17.3 | -0,4429 | 8,93159E-19 |
| ACE2 | SMIM9 | -0,4429 | 8,90892E-19 |
| ACE2 | LINC01312 | -0,443 | 8,69717E-19 |
| ACE2 | PWRN3 | -0,443 | 8,72591E-19 |
| ACE2 | LINC00661 | -0,443 | 8,77164E-19 |
| ACE2 | OSBPL10 | -0,4431 | 8,51279E-19 |
| ACE2 | IFT122 | -0,4431 | 8,62014E-19 |
| ACE2 | RP11-369E15.2 | -0,4431 | 8,53808E-19 |
| ACE2 | RP11-652G5.1 | -0,4431 | 8,5564E-19 |
| ACE2 | NEPNP | -0,4432 | 8,33281E-19 |
| ACE2 | ARMC4P1 | -0,4432 | 8,4868E-19 |
| ACE2 | RP11-444A22.1 | -0,4433 | 8,31341E-19 |
| ACE2 | LINC00513 | -0,4433 | 8,30319E-19 |
| ACE2 | PKIA-AS1 | -0,4433 | 8,23759E-19 |
| ACE2 | IL25 | -0,4433 | 8,23949E-19 |
| ACE2 | SNUPN | -0,4433 | 8,22589E-19 |
| ACE2 | RP11-98D18.1 | -0,4434 | 8,06753E-19 |
| ACE2 | RP11-418H16.1 | -0,4434 | 8,15397E-19 |
| ACE2 | CCDC110 | -0,4434 | 8,01292E-19 |
| ACE2 | TP53I11 | -0,4434 | 8,10219E-19 |
| ACE2 | GOLGA6L9 | -0,4434 | 8,11052E-19 |
| ACE2 | KRT40 | -0,4434 | 8,0039E-19 |
| ACE2 | AC064871.3 | -0,4435 | 7,96572E-19 |
| ACE2 | WDR49 | -0,4435 | 7,86925E-19 |
| ACE2 | RP11-108B14.5 | -0,4435 | 7,89332E-19 |
| ACE2 | HSD3B7 | -0,4435 | 7,92158E-19 |
| ACE2 | GNRH2 | -0,4435 | 7,84297E-19 |
| ACE2 | TMEM54 | -0,4436 | 7,68706E-19 |
| ACE2 | CSRNP1 | -0,4436 | 7,79229E-19 |
| ACE2 | ZEB1-AS1 | -0,4436 | 7,8371E-19 |
| ACE2 | ITIH4-AS1 | -0,4437 | 7,63988E-19 |
| ACE2 | MCF2L2 | -0,4437 | 7,53919E-19 |
| ACE2 | PSORS1C2 | -0,4437 | 7,64493E-19 |
| ACE2 | RP11-587H10.1 | -0,4437 | 7,5642E-19 |
| ACE2 | RPL5P14 | -0,4438 | 7,41653E-19 |
| ACE2 | RP11-17A1.3 | -0,4438 | 7,41105E-19 |
| ACE2 | RP11-331H13.1 | -0,4438 | 7,48674E-19 |
| ACE2 | SNAP47 | -0,4439 | 7,33904E-19 |
| ACE2 | IAH1 | -0,4439 | 7,33728E-19 |
| ACE2 | PCDHA9 | -0,4439 | 7,27533E-19 |
| ACE2 | RP11-216P16.7 | -0,4439 | 7,30303E-19 |
| ACE2 | RP11-120E13.1 | -0,4439 | 7,31957E-19 |
| ACE2 | LRRC8B | -0,444 | 7,21119E-19 |
| ACE2 | RP11-419C19.1 | -0,444 | 7,09343E-19 |
| ACE2 | PITPNM1 | -0,444 | 7,11828E-19 |
| ACE2 | CYP19A1 | -0,444 | 7,22086E-19 |
| ACE2 | FAM8A1 | -0,4441 | 7,02287E-19 |
| ACE2 | CCT8L1P | -0,4441 | 7,03574E-19 |
| ACE2 | CTD-2292P10.4 | -0,4441 | 7,07438E-19 |
| ACE2 | RP11-519G16.3 | -0,4441 | 7,01216E-19 |
| ACE2 | AC009505.2 | -0,4442 | 6,89607E-19 |
| ACE2 | LINC02042 | -0,4442 | 6,87295E-19 |
| ACE2 | EFHC1 | -0,4442 | 6,86725E-19 |

| | | | |
|---|---|---|---|
| ACE2 | MIP | -0,4442 | 6,95145E-19 |
| ACE2 | NOVA1-AS1 | -0,4442 | 6,86335E-19 |
| ACE2 | FAM192A | -0,4442 | 6,92399E-19 |
| ACE2 | TUBB3 | -0,4442 | 6,91295E-19 |
| ACE2 | ZNF101 | -0,4442 | 6,89746E-19 |
| ACE2 | CTA-929C8.6 | -0,4442 | 6,83469E-19 |
| ACE2 | SLC9B1 | -0,4443 | 6,8125E-19 |
| ACE2 | GS1-124K5.9 | -0,4443 | 6,76114E-19 |
| ACE2 | NFE4 | -0,4443 | 6,7399E-19 |
| ACE2 | ENDOG | -0,4443 | 6,68057E-19 |
| ACE2 | LINC01165 | -0,4443 | 6,70239E-19 |
| ACE2 | OR52N2 | -0,4443 | 6,77517E-19 |
| ACE2 | PCSK6-AS1 | -0,4443 | 6,73503E-19 |
| ACE2 | KB-1125A3.10 | -0,4443 | 6,73458E-19 |
| ACE2 | ADAM30 | -0,4444 | 6,63737E-19 |
| ACE2 | AC010745.1 | -0,4444 | 6,65733E-19 |
| ACE2 | RP11-460H9.1 | -0,4444 | 6,5696E-19 |
| ACE2 | RP11-549K20.1 | -0,4444 | 6,58133E-19 |
| ACE2 | RAB3IP | -0,4444 | 6,58891E-19 |
| ACE2 | PTCHD3P2 | -0,4445 | 6,42797E-19 |
| ACE2 | RP5-1157M23.2 | -0,4445 | 6,52498E-19 |
| ACE2 | ZCCHC6 | -0,4445 | 6,53842E-19 |
| ACE2 | EIF5AL1 | -0,4445 | 6,46605E-19 |
| ACE2 | CFAP43 | -0,4445 | 6,52277E-19 |
| ACE2 | LYPD6 | -0,4446 | 6,37577E-19 |
| ACE2 | RP11-223C24.2 | -0,4446 | 6,37914E-19 |
| ACE2 | GTF2IP23 | -0,4446 | 6,39589E-19 |
| ACE2 | RP11-523O18.7 | -0,4446 | 6,33121E-19 |
| ACE2 | OR13G1 | -0,4447 | 6,24311E-19 |
| ACE2 | LIMS1-AS1 | -0,4447 | 6,19097E-19 |
| ACE2 | AC017048.1 | -0,4447 | 6,21489E-19 |
| ACE2 | RP11-654C22.2 | -0,4447 | 6,21179E-19 |
| ACE2 | RN7SKP30 | -0,4447 | 6,21279E-19 |
| ACE2 | CTA-392C11.2 | -0,4447 | 6,19751E-19 |
| ACE2 | CTD-2515H24.1 | -0,4447 | 6,21448E-19 |
| ACE2 | CLHC1 | -0,4448 | 6,07083E-19 |
| ACE2 | C1orf168 | -0,4449 | 5,99544E-19 |
| ACE2 | RP11-271K21.12 | -0,4449 | 5,94078E-19 |
| ACE2 | CTD-2050E21.2 | -0,4449 | 5,97057E-19 |
| ACE2 | WNT16 | -0,4449 | 6,02424E-19 |
| ACE2 | RP11-1084E5.1 | -0,4449 | 6,03376E-19 |
| ACE2 | ARL2BPP3 | -0,4449 | 5,9975E-19 |
| ACE2 | RP5-928E24.2 | -0,4449 | 5,93009E-19 |
| ACE2 | REN | -0,4451 | 5,72016E-19 |
| ACE2 | DPY19L2P3 | -0,4451 | 5,79226E-19 |
| ACE2 | AC139099.4 | -0,4451 | 5,71287E-19 |
| ACE2 | RP11-459E5.1 | -0,4452 | 5,63576E-19 |
| ACE2 | MTNR1B | -0,4452 | 5,67795E-19 |
| ACE2 | RP11-443B7.1 | -0,4453 | 5,54825E-19 |
| ACE2 | HHATL | -0,4453 | 5,4639E-19 |
| ACE2 | CTD-2194L12.3 | -0,4453 | 5,5117E-19 |
| ACE2 | TCAM1P | -0,4453 | 5,55927E-19 |
| ACE2 | AC012314.20 | -0,4453 | 5,46161E-19 |
| ACE2 | RP11-59J16.2 | -0,4454 | 5,43443E-19 |

| | | | |
|---|---|---|---|
| ACE2 | CTD-3075F15.1 | -0,4454 | 5,4352E-19 |
| ACE2 | RP11-805I24.3 | -0,4454 | 5,45876E-19 |
| ACE2 | ZNF473 | -0,4454 | 5,46026E-19 |
| ACE2 | SPTLC1P5 | -0,4455 | 5,33737E-19 |
| ACE2 | PLA2G2C | -0,4456 | 5,21476E-19 |
| ACE2 | AC073333.8 | -0,4456 | 5,20516E-19 |
| ACE2 | CTD-3065B20.2 | -0,4456 | 5,21025E-19 |
| ACE2 | C2orf42 | -0,4457 | 5,05406E-19 |
| ACE2 | RP11-44N11.3 | -0,4457 | 5,05993E-19 |
| ACE2 | FAM74A4 | -0,4457 | 5,09618E-19 |
| ACE2 | LINC00474 | -0,4457 | 5,13739E-19 |
| ACE2 | LRRC37A6P | -0,4457 | 5,06287E-19 |
| ACE2 | OR56A1 | -0,4457 | 5,1341E-19 |
| ACE2 | DLEC1 | -0,4458 | 5,03218E-19 |
| ACE2 | RP11-1D12.2 | -0,4458 | 4,96185E-19 |
| ACE2 | RP11-382N13.6 | -0,4458 | 4,98532E-19 |
| ACE2 | KLK15 | -0,4458 | 5,00062E-19 |
| ACE2 | TTC4 | -0,4459 | 4,90335E-19 |
| ACE2 | AC073043.1 | -0,4459 | 4,84358E-19 |
| ACE2 | RP3-449H6.1 | -0,4459 | 4,93415E-19 |
| ACE2 | RP1-29G21.1 | -0,4459 | 4,91922E-19 |
| ACE2 | C11orf63 | -0,446 | 4,83388E-19 |
| ACE2 | RP11-26L16.1 | -0,446 | 4,78645E-19 |
| ACE2 | KRT18P55 | -0,446 | 4,83599E-19 |
| ACE2 | DSG4 | -0,446 | 4,74699E-19 |
| ACE2 | RP11-431M7.3 | -0,4461 | 4,68531E-19 |
| ACE2 | CEP85L | -0,4461 | 4,69125E-19 |
| ACE2 | RP11-849F2.4 | -0,4461 | 4,69235E-19 |
| ACE2 | LINC01920 | -0,4463 | 4,46833E-19 |
| ACE2 | SPEF2 | -0,4463 | 4,50712E-19 |
| ACE2 | LINC01335 | -0,4463 | 4,54204E-19 |
| ACE2 | GOT1 | -0,4463 | 4,55066E-19 |
| ACE2 | HPN | -0,4463 | 4,51717E-19 |
| ACE2 | RP11-2A4.4 | -0,4464 | 4,44687E-19 |
| ACE2 | RP1-124C6.1 | -0,4464 | 4,38585E-19 |
| ACE2 | RP11-308N19.3 | -0,4464 | 4,40896E-19 |
| ACE2 | RP11-111M22.2 | -0,4464 | 4,3877E-19 |
| ACE2 | RP13-444H2.1 | -0,4464 | 4,45288E-19 |
| ACE2 | OR14J1 | -0,4465 | 4,33575E-19 |
| ACE2 | RP11-62F24.2 | -0,4465 | 4,34293E-19 |
| ACE2 | CAB39L | -0,4465 | 4,33651E-19 |
| ACE2 | PPP1R2P9 | -0,4465 | 4,35059E-19 |
| ACE2 | RP11-547I7.1 | -0,4466 | 4,23917E-19 |
| ACE2 | GPBAR1 | -0,4466 | 4,25458E-19 |
| ACE2 | LNX1 | -0,4466 | 4,27995E-19 |
| ACE2 | CYB5R4 | -0,4466 | 4,20755E-19 |
| ACE2 | PAPOLB | -0,4466 | 4,24211E-19 |
| ACE2 | AP000462.1 | -0,4466 | 4,27589E-19 |
| ACE2 | RP11-219E7.3 | -0,4466 | 4,20396E-19 |
| ACE2 | SPDYE6 | -0,4467 | 4,14267E-19 |
| ACE2 | AC008278.2 | -0,4468 | 4,0701E-19 |
| ACE2 | RP11-863K10.7 | -0,4468 | 4,09448E-19 |
| ACE2 | MIRLET7DHG | -0,4468 | 4,03909E-19 |
| ACE2 | EME2 | -0,4468 | 4,10835E-19 |

| | | | |
|---|---|---|---|
| ACE2 | FAM74A1 | -0,4469 | 3,99308E-19 |
| ACE2 | RP11-1030E3.1 | -0,4469 | 4,02809E-19 |
| ACE2 | AC073316.2 | -0,447 | 3,89151E-19 |
| ACE2 | AP000474.1 | -0,447 | 3,90039E-19 |
| ACE2 | RP11-389G6.3 | -0,4471 | 3,85315E-19 |
| ACE2 | KB-1410C5.3 | -0,4472 | 3,76287E-19 |
| ACE2 | DNM1P17 | -0,4472 | 3,78638E-19 |
| ACE2 | RP11-57G10.8 | -0,4472 | 3,7415E-19 |
| ACE2 | SIX3 | -0,4473 | 3,64635E-19 |
| ACE2 | RP11-94M14.2 | -0,4473 | 3,64433E-19 |
| ACE2 | LINC01695 | -0,4474 | 3,63757E-19 |
| ACE2 | PPEF1 | -0,4474 | 3,6125E-19 |
| ACE2 | AC007040.6 | -0,4475 | 3,52746E-19 |
| ACE2 | CTD-2631K10.1 | -0,4475 | 3,51392E-19 |
| ACE2 | RP11-675M1.2 | -0,4475 | 3,5122E-19 |
| ACE2 | RBM26-AS1 | -0,4475 | 3,50697E-19 |
| ACE2 | KCTD19 | -0,4475 | 3,56509E-19 |
| ACE2 | AC064852.4 | -0,4476 | 3,49076E-19 |
| ACE2 | LRCH4 | -0,4476 | 3,49379E-19 |
| ACE2 | DEFB134 | -0,4476 | 3,45693E-19 |
| ACE2 | AC009505.4 | -0,4477 | 3,38213E-19 |
| ACE2 | EPHA5 | -0,4477 | 3,3675E-19 |
| ACE2 | RP13-631K18.3 | -0,4477 | 3,37646E-19 |
| ACE2 | GPR68 | -0,4477 | 3,36344E-19 |
| ACE2 | CAPZB | -0,4478 | 3,32645E-19 |
| ACE2 | RP11-816J6.3 | -0,4478 | 3,34815E-19 |
| ACE2 | C5orf64 | -0,4478 | 3,30589E-19 |
| ACE2 | AC093627.9 | -0,4479 | 3,23707E-19 |
| ACE2 | SDK1 | -0,4479 | 3,28641E-19 |
| ACE2 | TRMT2A | -0,4479 | 3,23823E-19 |
| ACE2 | RP11-211G3.2 | -0,448 | 3,18113E-19 |
| ACE2 | ITK | -0,448 | 3,19541E-19 |
| ACE2 | RP11-386I8.6 | -0,448 | 3,21501E-19 |
| ACE2 | KLK7 | -0,448 | 3,17979E-19 |
| ACE2 | PKD2L2 | -0,4481 | 3,11506E-19 |
| ACE2 | RP4-742N3.1 | -0,4482 | 3,09097E-19 |
| ACE2 | RP4-712E4.1 | -0,4483 | 3,00528E-19 |
| ACE2 | ZNF192P1 | -0,4483 | 2,97961E-19 |
| ACE2 | ATP8A2P1 | -0,4483 | 2,98479E-19 |
| ACE2 | RP11-266L9.3 | -0,4483 | 3,01296E-19 |
| ACE2 | GFY | -0,4483 | 3,02125E-19 |
| ACE2 | RP11-358D14.3 | -0,4483 | 2,99857E-19 |
| ACE2 | KLB | -0,4484 | 2,95983E-19 |
| ACE2 | LINC01038 | -0,4484 | 2,9718E-19 |
| ACE2 | KRI1 | -0,4484 | 2,94543E-19 |
| ACE2 | SPATC1L | -0,4484 | 2,93428E-19 |
| ACE2 | NBPF7 | -0,4485 | 2,88912E-19 |
| ACE2 | MAPK10 | -0,4485 | 2,89879E-19 |
| ACE2 | GPR150 | -0,4485 | 2,86022E-19 |
| ACE2 | RP11-545A16.1 | -0,4486 | 2,81963E-19 |
| ACE2 | RP11-141B14.1 | -0,4486 | 2,84915E-19 |
| ACE2 | MROH2B | -0,4486 | 2,85037E-19 |
| ACE2 | SLC6A5 | -0,4486 | 2,80528E-19 |
| ACE2 | CTD-2022H16.3 | -0,4486 | 2,8535E-19 |

| | | | |
|---|---|---|---|
| ACE2 | RP11-74K19.1 | -0,4487 | 2,75762E-19 |
| ACE2 | EEF1B2P7 | -0,4487 | 2,75342E-19 |
| ACE2 | AC003991.3 | -0,4487 | 2,79634E-19 |
| ACE2 | RP11-313C15.1 | -0,4487 | 2,76852E-19 |
| ACE2 | RP11-262K1.1 | -0,4488 | 2,71567E-19 |
| ACE2 | MDH1B | -0,4488 | 2,73839E-19 |
| ACE2 | LINC00415 | -0,4488 | 2,69028E-19 |
| ACE2 | RP11-557L19.1 | -0,4489 | 2,65595E-19 |
| ACE2 | AC008278.3 | -0,4489 | 2,63858E-19 |
| ACE2 | OR51A7 | -0,4489 | 2,64857E-19 |
| ACE2 | LA16c-380A1.2 | -0,4489 | 2,67152E-19 |
| ACE2 | RP11-394O4.3 | -0,4491 | 2,52842E-19 |
| ACE2 | ACAT2 | -0,4491 | 2,56142E-19 |
| ACE2 | RP11-677I18.4 | -0,4491 | 2,54671E-19 |
| ACE2 | LINC02191 | -0,4491 | 2,52834E-19 |
| ACE2 | KIAA1328 | -0,4491 | 2,5283E-19 |
| ACE2 | RP11-768G7.3 | -0,4492 | 2,49274E-19 |
| ACE2 | IGFL2 | -0,4492 | 2,4838E-19 |
| ACE2 | AC092159.2 | -0,4493 | 2,43332E-19 |
| ACE2 | RP11-328K15.1 | -0,4493 | 2,46193E-19 |
| ACE2 | FBXW9 | -0,4493 | 2,4705E-19 |
| ACE2 | LINC01654 | -0,4494 | 2,38342E-19 |
| ACE2 | RP11-488P3.1 | -0,4494 | 2,40178E-19 |
| ACE2 | RP11-91H12.4 | -0,4494 | 2,38321E-19 |
| ACE2 | EAF1 | -0,4495 | 2,36524E-19 |
| ACE2 | LINC01229 | -0,4495 | 2,33007E-19 |
| ACE2 | CTB-35F21.1 | -0,4496 | 2,30447E-19 |
| ACE2 | RP11-114G22.1 | -0,4496 | 2,32132E-19 |
| ACE2 | RP1-118J21.5 | -0,4497 | 2,23757E-19 |
| ACE2 | PSMC1P7 | -0,4497 | 2,25293E-19 |
| ACE2 | DDX25 | -0,4497 | 2,28037E-19 |
| ACE2 | RP11-543H12.1 | -0,4497 | 2,27404E-19 |
| ACE2 | RP13-638C3.3 | -0,4497 | 2,26013E-19 |
| ACE2 | FAR2P1 | -0,4498 | 2,21159E-19 |
| ACE2 | UQCR10 | -0,4498 | 2,22639E-19 |
| ACE2 | FAM53A | -0,4499 | 2,16268E-19 |
| ACE2 | RP11-756P10.5 | -0,4499 | 2,14486E-19 |
| ACE2 | RP11-796I2.1 | -0,4499 | 2,17346E-19 |
| ACE2 | RP11-320N21.1 | -0,4499 | 2,15383E-19 |
| ACE2 | RP11-370I10.2 | -0,4499 | 2,17704E-19 |
| ACE2 | MTATP6P24 | -0,4499 | 2,172E-19 |
| ACE2 | RP1-249H1.2 | -0,45 | 2,129E-19 |
| ACE2 | RP11-799O21.2 | -0,45 | 2,13202E-19 |
| ACE2 | RP11-121G22.3 | -0,45 | 2,10409E-19 |
| ACE2 | CEP170 | -0,4501 | 2,07119E-19 |
| ACE2 | AC007390.4 | -0,4501 | 2,09257E-19 |
| ACE2 | AC013733.3 | -0,4501 | 2,06365E-19 |
| ACE2 | RP3-425P12.2 | -0,4501 | 2,07008E-19 |
| ACE2 | SNRPC | -0,4501 | 2,09486E-19 |
| ACE2 | RN7SL272P | -0,4501 | 2,06408E-19 |
| ACE2 | KLHL11 | -0,4501 | 2,08126E-19 |
| ACE2 | CTA-796E4.3 | -0,4501 | 2,08235E-19 |
| ACE2 | AC009480.3 | -0,4502 | 2,0436E-19 |
| ACE2 | RP11-380O24.1 | -0,4502 | 2,04776E-19 |

| | | | |
|---|---|---|---|
| ACE2 | CTB-131B5.2 | -0,4502 | 2,04459E-19 |
| ACE2 | ALS2CR11 | -0,4503 | 1,98576E-19 |
| ACE2 | CTD-2057J6.1 | -0,4503 | 1,97686E-19 |
| ACE2 | VN1R64P | -0,4503 | 1,98512E-19 |
| ACE2 | AC093074.1 | -0,4503 | 1,98895E-19 |
| ACE2 | ROPN1L-AS1 | -0,4504 | 1,95731E-19 |
| ACE2 | GS1-166A23.2 | -0,4504 | 1,9428E-19 |
| ACE2 | ACSL5 | -0,4504 | 1,9738E-19 |
| ACE2 | ZNF970P | -0,4504 | 1,94486E-19 |
| ACE2 | ANKLE2 | -0,4504 | 1,94414E-19 |
| ACE2 | RP11-24D15.1 | -0,4504 | 1,96861E-19 |
| ACE2 | ZNF622 | -0,4505 | 1,91737E-19 |
| ACE2 | LINC01298 | -0,4505 | 1,92789E-19 |
| ACE2 | RP11-347H15.4 | -0,4505 | 1,93092E-19 |
| ACE2 | LINC01138 | -0,4506 | 1,8838E-19 |
| ACE2 | MELTF-AS1 | -0,4506 | 1,87825E-19 |
| ACE2 | PIEZO2 | -0,4506 | 1,8781E-19 |
| ACE2 | CTC-454I21.4 | -0,4506 | 1,8898E-19 |
| ACE2 | ISX-AS1 | -0,4506 | 1,86291E-19 |
| ACE2 | AC073218.3 | -0,4507 | 1,83718E-19 |
| ACE2 | RP11-809O17.1 | -0,4507 | 1,83422E-19 |
| ACE2 | RP11-16B13.1 | -0,4507 | 1,85102E-19 |
| ACE2 | LINC00523 | -0,4507 | 1,84351E-19 |
| ACE2 | INO80E | -0,4507 | 1,85304E-19 |
| ACE2 | RP11-879F14.3 | -0,4507 | 1,82982E-19 |
| ACE2 | LHB | -0,4507 | 1,83712E-19 |
| ACE2 | TMEM212 | -0,4508 | 1,8051E-19 |
| ACE2 | AC009133.12 | -0,4508 | 1,7997E-19 |
| ACE2 | GFRA4 | -0,4508 | 1,81234E-19 |
| ACE2 | RP11-126K1.6 | -0,4509 | 1,77186E-19 |
| ACE2 | RP11-342A1.1 | -0,4509 | 1,76933E-19 |
| ACE2 | RP11-776A13.3 | -0,4509 | 1,76163E-19 |
| ACE2 | B4GALT4-AS1 | -0,451 | 1,72257E-19 |
| ACE2 | RP11-5P22.3 | -0,451 | 1,7258E-19 |
| ACE2 | RP11-481C4.1 | -0,451 | 1,7434E-19 |
| ACE2 | TMEM234 | -0,4511 | 1,691E-19 |
| ACE2 | KLHL30-AS1 | -0,4511 | 1,67944E-19 |
| ACE2 | LINC02110 | -0,4511 | 1,68922E-19 |
| ACE2 | URAD | -0,4511 | 1,70383E-19 |
| ACE2 | TMEM42 | -0,4512 | 1,6439E-19 |
| ACE2 | AC008592.7 | -0,4512 | 1,65982E-19 |
| ACE2 | RBPMS2P1 | -0,4512 | 1,66633E-19 |
| ACE2 | ZNF503-AS1 | -0,4512 | 1,66031E-19 |
| ACE2 | RP11-43N22.1 | -0,4512 | 1,66031E-19 |
| ACE2 | RP11-214K3.21 | -0,4512 | 1,67048E-19 |
| ACE2 | RP11-2C15.1 | -0,4512 | 1,64316E-19 |
| ACE2 | C22orf15 | -0,4512 | 1,64951E-19 |
| ACE2 | RP11-102N12.2 | -0,4513 | 1,6157E-19 |
| ACE2 | RP11-91J3.2 | -0,4513 | 1,6196E-19 |
| ACE2 | TUBA8 | -0,4513 | 1,62935E-19 |
| ACE2 | LINC01997 | -0,4514 | 1,60048E-19 |
| ACE2 | RP11-63G10.4 | -0,4515 | 1,55985E-19 |
| ACE2 | PGLYRP4 | -0,4515 | 1,54543E-19 |
| ACE2 | EFHB | -0,4515 | 1,55505E-19 |

| | | | |
|---|---|---|---|
| ACE2 | MEX3C | -0,4515 | 1,56067E-19 |
| ACE2 | DAZAP1 | -0,4515 | 1,5596E-19 |
| ACE2 | LINC00114 | -0,4515 | 1,54772E-19 |
| ACE2 | CTD-2252P21.1 | -0,4516 | 1,53551E-19 |
| ACE2 | HSPD1P10 | -0,4516 | 1,53831E-19 |
| ACE2 | RP1-121G13.2 | -0,4516 | 1,52098E-19 |
| ACE2 | AC074389.7 | -0,4516 | 1,53816E-19 |
| ACE2 | RP11-6D1.3 | -0,4516 | 1,51823E-19 |
| ACE2 | RP5-944M2.2 | -0,4516 | 1,51589E-19 |
| ACE2 | LINC00359 | -0,4516 | 1,52691E-19 |
| ACE2 | LINC01581 | -0,4516 | 1,52595E-19 |
| ACE2 | RP11-521D12.2 | -0,4517 | 1,48701E-19 |
| ACE2 | AC017006.3 | -0,4517 | 1,50877E-19 |
| ACE2 | RP11-895K13.2 | -0,4517 | 1,49506E-19 |
| ACE2 | TTTY18 | -0,4517 | 1,50747E-19 |
| ACE2 | RP5-875O13.7 | -0,4518 | 1,46632E-19 |
| ACE2 | RP11-570K4.1 | -0,4518 | 1,4613E-19 |
| ACE2 | RP11-503C24.1 | -0,4518 | 1,47752E-19 |
| ACE2 | RP11-685B13.2 | -0,4518 | 1,47326E-19 |
| ACE2 | RP11-783L4.1 | -0,4518 | 1,46984E-19 |
| ACE2 | RP11-285E9.5 | -0,4518 | 1,46051E-19 |
| ACE2 | DZANK1 | -0,4518 | 1,48013E-19 |
| ACE2 | AC144449.1 | -0,4519 | 1,43587E-19 |
| ACE2 | NBPF22P | -0,4519 | 1,42523E-19 |
| ACE2 | RP11-44M6.1 | -0,4519 | 1,43653E-19 |
| ACE2 | ABCC2 | -0,4519 | 1,43891E-19 |
| ACE2 | LOXL1-AS1 | -0,4519 | 1,43591E-19 |
| ACE2 | RP5-1029F21.4 | -0,4519 | 1,42784E-19 |
| ACE2 | TEDDM1 | -0,452 | 1,41895E-19 |
| ACE2 | RP11-121E16.1 | -0,452 | 1,39935E-19 |
| ACE2 | GOLGA8H | -0,452 | 1,40056E-19 |
| ACE2 | VCAN-AS1 | -0,4521 | 1,37495E-19 |
| ACE2 | AC005481.5 | -0,4521 | 1,37885E-19 |
| ACE2 | RP11-473E2.4 | -0,4521 | 1,37253E-19 |
| ACE2 | CTD-2363C16.1 | -0,4522 | 1,33809E-19 |
| ACE2 | ANKRD20A10P | -0,4522 | 1,36231E-19 |
| ACE2 | RP11-394G3.2 | -0,4523 | 1,33353E-19 |
| ACE2 | RP11-10C8.2 | -0,4523 | 1,31262E-19 |
| ACE2 | RP11-599B13.3 | -0,4524 | 1,29828E-19 |
| ACE2 | CLIP4 | -0,4525 | 1,27709E-19 |
| ACE2 | NLGN1-AS1 | -0,4525 | 1,25899E-19 |
| ACE2 | USPL1 | -0,4525 | 1,25991E-19 |
| ACE2 | USP44 | -0,4526 | 1,25117E-19 |
| ACE2 | PLA2G6 | -0,4526 | 1,23176E-19 |
| ACE2 | C3orf67-AS1 | -0,4527 | 1,22971E-19 |
| ACE2 | CNN2P9 | -0,4527 | 1,23004E-19 |
| ACE2 | SMCO2 | -0,4527 | 1,21727E-19 |
| ACE2 | RP11-752G15.10 | -0,4527 | 1,2126E-19 |
| ACE2 | RP5-998H6.2 | -0,4527 | 1,22345E-19 |
| ACE2 | RP11-404F10.2 | -0,4528 | 1,19762E-19 |
| ACE2 | FOXD4L1 | -0,4528 | 1,18596E-19 |
| ACE2 | RP11-710F7.2 | -0,4528 | 1,20556E-19 |
| ACE2 | RP11-492L8.2 | -0,4528 | 1,18267E-19 |
| ACE2 | AC005162.4 | -0,4528 | 1,20462E-19 |

| | | | |
|---|---|---|---|
| ACE2 | CTD-2529O21.1 | -0,4528 | 1,20102E-19 |
| ACE2 | GPR45 | -0,4529 | 1,16009E-19 |
| ACE2 | CCDC126 | -0,4529 | 1,17394E-19 |
| ACE2 | AP001462.6 | -0,4529 | 1,1596E-19 |
| ACE2 | CTD-2650P22.1 | -0,4529 | 1,16169E-19 |
| ACE2 | RP3-448I9.1 | -0,453 | 1,14793E-19 |
| ACE2 | RP11-3B12.2 | -0,453 | 1,14504E-19 |
| ACE2 | SLC16A7 | -0,453 | 1,14476E-19 |
| ACE2 | RP11-927P21.11 | -0,453 | 1,15271E-19 |
| ACE2 | SIRPG-AS1 | -0,453 | 1,13715E-19 |
| ACE2 | PROKR2 | -0,453 | 1,14129E-19 |
| ACE2 | TAF9 | -0,4531 | 1,1157E-19 |
| ACE2 | BIN3 | -0,4531 | 1,12334E-19 |
| ACE2 | LINC00967 | -0,4531 | 1,1291E-19 |
| ACE2 | FAM205C | -0,4531 | 1,11846E-19 |
| ACE2 | ARHGEF7-IT1 | -0,4531 | 1,11958E-19 |
| ACE2 | AC099684.1 | -0,4531 | 1,12159E-19 |
| ACE2 | TMEM232 | -0,4532 | 1,10516E-19 |
| ACE2 | FOXK1 | -0,4532 | 1,10121E-19 |
| ACE2 | LACTB2-AS1 | -0,4533 | 1,06861E-19 |
| ACE2 | MYO3A | -0,4533 | 1,06884E-19 |
| ACE2 | RP11-19J5.2 | -0,4533 | 1,07562E-19 |
| ACE2 | RP11-96K19.5 | -0,4534 | 1,05439E-19 |
| ACE2 | RP11-814E24.1 | -0,4534 | 1,04748E-19 |
| ACE2 | SYT13 | -0,4534 | 1,0641E-19 |
| ACE2 | RP5-890E16.2 | -0,4534 | 1,05977E-19 |
| ACE2 | AC096559.2 | -0,4535 | 1,02385E-19 |
| ACE2 | DHRS7B | -0,4535 | 1,0404E-19 |
| ACE2 | RP11-64C12.3 | -0,4535 | 1,02964E-19 |
| ACE2 | DNTTIP1 | -0,4535 | 1,02615E-19 |
| ACE2 | RP11-533E19.5 | -0,4536 | 1,01296E-19 |
| ACE2 | SLC25A47 | -0,4536 | 1,01894E-19 |
| ACE2 | LINC00993 | -0,4537 | 9,94737E-20 |
| ACE2 | RP11-107C16.2 | -0,4537 | 9,95994E-20 |
| ACE2 | RP11-794P6.1 | -0,4537 | 9,99855E-20 |
| ACE2 | RP11-330M19.1 | -0,4538 | 9,70069E-20 |
| ACE2 | AC103563.9 | -0,4538 | 9,74803E-20 |
| ACE2 | RP11-5P15.1 | -0,4538 | 9,79469E-20 |
| ACE2 | RP11-437J2.4 | -0,4538 | 9,60736E-20 |
| ACE2 | MMP20 | -0,4538 | 9,76565E-20 |
| ACE2 | LINC01648 | -0,4539 | 9,60184E-20 |
| ACE2 | RP11-167P20.1 | -0,4539 | 9,54089E-20 |
| ACE2 | AL031601.5 | -0,4539 | 9,50738E-20 |
| ACE2 | FANCF | -0,4539 | 9,5239E-20 |
| ACE2 | RP13-895J2.7 | -0,4539 | 9,42667E-20 |
| ACE2 | RP5-994D16.9 | -0,454 | 9,40553E-20 |
| ACE2 | RP11-350F16.2 | -0,454 | 9,21868E-20 |
| ACE2 | RP11-281N10.1 | -0,454 | 9,32873E-20 |
| ACE2 | KCNMA1-AS2 | -0,454 | 9,36826E-20 |
| ACE2 | SNTN | -0,4541 | 9,19119E-20 |
| ACE2 | RP11-433C9.3 | -0,4541 | 9,05637E-20 |
| ACE2 | AC084082.3 | -0,4541 | 9,12954E-20 |
| ACE2 | RP11-100M12.1 | -0,4541 | 9,14812E-20 |
| ACE2 | C1orf226 | -0,4542 | 8,91802E-20 |

| | | | |
|---|---|---|---|
| ACE2 | SNRPCP9 | -0,4542 | 9,00096E-20 |
| ACE2 | FAAHP1 | -0,4543 | 8,77958E-20 |
| ACE2 | SEMA4A | -0,4543 | 8,76407E-20 |
| ACE2 | RPS18P1 | -0,4543 | 8,75507E-20 |
| ACE2 | WDTC1 | -0,4544 | 8,57909E-20 |
| ACE2 | PHGDH | -0,4544 | 8,48187E-20 |
| ACE2 | CCDC66 | -0,4544 | 8,65366E-20 |
| ACE2 | RP11-87E22.2 | -0,4544 | 8,50555E-20 |
| ACE2 | SNX18P26 | -0,4544 | 8,63249E-20 |
| ACE2 | LINC01021 | -0,4545 | 8,38704E-20 |
| ACE2 | RPL26L1 | -0,4545 | 8,31662E-20 |
| ACE2 | RP11-756K15.2 | -0,4545 | 8,38559E-20 |
| ACE2 | HPN-AS1 | -0,4545 | 8,34036E-20 |
| ACE2 | OR10J1 | -0,4546 | 8,22075E-20 |
| ACE2 | RP1-111B22.3 | -0,4546 | 8,14521E-20 |
| ACE2 | TUSC1 | -0,4546 | 8,1579E-20 |
| ACE2 | RP11-154P18.2 | -0,4546 | 8,14219E-20 |
| ACE2 | PPP1R2P10 | -0,4546 | 8,17902E-20 |
| ACE2 | RP11-720D4.3 | -0,4547 | 8,11769E-20 |
| ACE2 | AC005775.2 | -0,4547 | 8,06928E-20 |
| ACE2 | LAIR2 | -0,4547 | 8,06179E-20 |
| ACE2 | BIRC6-AS2 | -0,4548 | 7,85833E-20 |
| ACE2 | EGFLAM-AS4 | -0,4548 | 7,87827E-20 |
| ACE2 | PCDHGB8P | -0,4548 | 7,82711E-20 |
| ACE2 | DOCK4-AS1 | -0,4548 | 7,8239E-20 |
| ACE2 | RP11-76C10.2 | -0,4548 | 7,9236E-20 |
| ACE2 | RP11-159F24.2 | -0,4549 | 7,68989E-20 |
| ACE2 | LINC01435 | -0,4549 | 7,75054E-20 |
| ACE2 | TRPM5 | -0,4549 | 7,78798E-20 |
| ACE2 | ST8SIA1 | -0,4549 | 7,68862E-20 |
| ACE2 | LINC01177 | -0,4549 | 7,75621E-20 |
| ACE2 | BIRC5 | -0,4549 | 7,65057E-20 |
| ACE2 | COX6C | -0,455 | 7,57316E-20 |
| ACE2 | RP11-157J13.1 | -0,455 | 7,52105E-20 |
| ACE2 | KARSP2 | -0,455 | 7,61639E-20 |
| ACE2 | RP11-56L13.1 | -0,455 | 7,52983E-20 |
| ACE2 | MRGBP | -0,455 | 7,57583E-20 |
| ACE2 | ARRDC3-AS1 | -0,4551 | 7,47331E-20 |
| ACE2 | GOLGA6C | -0,4551 | 7,45221E-20 |
| ACE2 | LYPD6B | -0,4552 | 7,1926E-20 |
| ACE2 | RP11-100F15.1 | -0,4552 | 7,20551E-20 |
| ACE2 | SBF1 | -0,4552 | 7,23928E-20 |
| ACE2 | FAM86JP | -0,4553 | 7,09112E-20 |
| ACE2 | RP11-1191J2.4 | -0,4553 | 7,17777E-20 |
| ACE2 | LINC02113 | -0,4553 | 7,07174E-20 |
| ACE2 | CTD-2236F14.1 | -0,4554 | 6,92293E-20 |
| ACE2 | CTB-111H14.1 | -0,4554 | 6,97273E-20 |
| ACE2 | CPA6 | -0,4554 | 7,00378E-20 |
| ACE2 | RP11-3G20.2 | -0,4554 | 6,90301E-20 |
| ACE2 | LRRTM3 | -0,4554 | 6,91966E-20 |
| ACE2 | RP11-368P15.2 | -0,4554 | 6,99089E-20 |
| ACE2 | RP11-443G13.2 | -0,4554 | 6,94722E-20 |
| ACE2 | RPL23AP82 | -0,4554 | 6,91558E-20 |
| ACE2 | NDUFB3 | -0,4555 | 6,84807E-20 |

| | | | |
|---|---|---|---|
| ACE2 | NFE2 | -0,4555 | 6,77509E-20 |
| ACE2 | RP11-466A19.6 | -0,4555 | 6,86484E-20 |
| ACE2 | TTTY12 | -0,4555 | 6,84568E-20 |
| ACE2 | ARTN | -0,4557 | 6,56622E-20 |
| ACE2 | LINC00309 | -0,4557 | 6,59931E-20 |
| ACE2 | LINC01574 | -0,4557 | 6,56942E-20 |
| ACE2 | DMBT1 | -0,4557 | 6,57531E-20 |
| ACE2 | RP11-613F22.7 | -0,4557 | 6,49287E-20 |
| ACE2 | RP11-259O18.5 | -0,4557 | 6,53191E-20 |
| ACE2 | LINC00382 | -0,4557 | 6,48036E-20 |
| ACE2 | RP11-385M4.1 | -0,4558 | 6,39503E-20 |
| ACE2 | RP11-332M2.1 | -0,4558 | 6,38737E-20 |
| ACE2 | RP11-227F19.5 | -0,4558 | 6,37474E-20 |
| ACE2 | RP11-152P17.3 | -0,4558 | 6,37646E-20 |
| ACE2 | MFF | -0,4559 | 6,28202E-20 |
| ACE2 | CARTPT | -0,4559 | 6,23179E-20 |
| ACE2 | DKK4 | -0,4559 | 6,2244E-20 |
| ACE2 | LINC00867 | -0,4559 | 6,28636E-20 |
| ACE2 | AC010907.5 | -0,456 | 6,11294E-20 |
| ACE2 | LINC01966 | -0,456 | 6,08305E-20 |
| ACE2 | POTEA | -0,456 | 6,0911E-20 |
| ACE2 | RP11-135D11.2 | -0,456 | 6,20059E-20 |
| ACE2 | LINC00445 | -0,456 | 6,201E-20 |
| ACE2 | USP29 | -0,456 | 6,15715E-20 |
| ACE2 | LINC00601 | -0,4561 | 5,96589E-20 |
| ACE2 | GRIP1 | -0,4561 | 6,07219E-20 |
| ACE2 | CTD-3022G6.1 | -0,4561 | 6,07398E-20 |
| ACE2 | LGALS13 | -0,4562 | 5,90427E-20 |
| ACE2 | TSPEAR-AS1 | -0,4562 | 5,93626E-20 |
| ACE2 | GPATCH2 | -0,4563 | 5,7683E-20 |
| ACE2 | AC113617.1 | -0,4563 | 5,70801E-20 |
| ACE2 | RP1-168P16.3 | -0,4563 | 5,81634E-20 |
| ACE2 | RP11-775D22.3 | -0,4564 | 5,70479E-20 |
| ACE2 | FAM78A | -0,4564 | 5,61984E-20 |
| ACE2 | TBX15 | -0,4565 | 5,56402E-20 |
| ACE2 | RP4-669B10.3 | -0,4565 | 5,57177E-20 |
| ACE2 | RP11-180I4.2 | -0,4565 | 5,53885E-20 |
| ACE2 | RP11-327P2.5 | -0,4565 | 5,52812E-20 |
| ACE2 | RP11-219E7.4 | -0,4565 | 5,53605E-20 |
| ACE2 | ZNHIT3 | -0,4565 | 5,53558E-20 |
| ACE2 | CTD-2553L13.7 | -0,4565 | 5,52618E-20 |
| ACE2 | RP11-51L5.2 | -0,4566 | 5,38752E-20 |
| ACE2 | RP11-768G7.1 | -0,4567 | 5,35384E-20 |
| ACE2 | RP11-79E3.3 | -0,4567 | 5,29108E-20 |
| ACE2 | AC074389.6 | -0,4567 | 5,35852E-20 |
| ACE2 | ADAM21 | -0,4567 | 5,26543E-20 |
| ACE2 | PROM2 | -0,4568 | 5,15346E-20 |
| ACE2 | HCP5 | -0,4568 | 5,1617E-20 |
| ACE2 | KRTAP5-10 | -0,4568 | 5,21721E-20 |
| ACE2 | LINC00609 | -0,4569 | 5,03806E-20 |
| ACE2 | RP11-876N24.1 | -0,4569 | 5,04407E-20 |
| ACE2 | CTB-66B24.1 | -0,4569 | 5,04391E-20 |
| ACE2 | LINC02146 | -0,457 | 4,96097E-20 |
| ACE2 | CTD-2233C11.2 | -0,457 | 5,00486E-20 |

| | | | |
|---|---|---|---|
| ACE2 | PPP4R1-AS1 | -0,457 | 5,01281E-20 |
| ACE2 | RP4-671O14.7 | -0,457 | 4,96417E-20 |
| ACE2 | DRD2 | -0,4571 | 4,86175E-20 |
| ACE2 | RP11-521E5.1 | -0,4572 | 4,81875E-20 |
| ACE2 | AC004593.3 | -0,4572 | 4,74934E-20 |
| ACE2 | FOLH1B | -0,4572 | 4,74149E-20 |
| ACE2 | RNASEH2A | -0,4572 | 4,81372E-20 |
| ACE2 | RP11-78O9.1 | -0,4573 | 4,6589E-20 |
| ACE2 | RPSAP11 | -0,4573 | 4,70886E-20 |
| ACE2 | WDR27 | -0,4573 | 4,6715E-20 |
| ACE2 | RP11-280O24.3 | -0,4573 | 4,70776E-20 |
| ACE2 | AP000439.5 | -0,4573 | 4,63814E-20 |
| ACE2 | EFCAB5 | -0,4573 | 4,71392E-20 |
| ACE2 | TISP43 | -0,4574 | 4,59238E-20 |
| ACE2 | GNAT3 | -0,4574 | 4,61689E-20 |
| ACE2 | OR7E15P | -0,4574 | 4,56274E-20 |
| ACE2 | RP11-567M16.2 | -0,4574 | 4,59228E-20 |
| ACE2 | ABCA4 | -0,4575 | 4,47077E-20 |
| ACE2 | MCCD1P2 | -0,4575 | 4,4924E-20 |
| ACE2 | ACTL6B | -0,4575 | 4,52968E-20 |
| ACE2 | PSD3 | -0,4576 | 4,3672E-20 |
| ACE2 | AMBP | -0,4576 | 4,43115E-20 |
| ACE2 | DEUP1 | -0,4576 | 4,39777E-20 |
| ACE2 | ARHGAP35 | -0,4576 | 4,36422E-20 |
| ACE2 | LY6G6F | -0,4577 | 4,28307E-20 |
| ACE2 | MYB-AS1 | -0,4577 | 4,30033E-20 |
| ACE2 | RP11-548K12.13 | -0,4577 | 4,34555E-20 |
| ACE2 | AB015752.3 | -0,4577 | 4,30626E-20 |
| ACE2 | SCUBE2 | -0,4577 | 4,28337E-20 |
| ACE2 | HS3ST3B1 | -0,4577 | 4,3456E-20 |
| ACE2 | RP3-333B15.6 | -0,4577 | 4,33352E-20 |
| ACE2 | NDUFA6-AS1 | -0,4577 | 4,28806E-20 |
| ACE2 | RP5-1101C3.1 | -0,4578 | 4,20542E-20 |
| ACE2 | PLXNA4 | -0,4578 | 4,24916E-20 |
| ACE2 | CDK20 | -0,4578 | 4,19572E-20 |
| ACE2 | TMPRSS12 | -0,4578 | 4,20495E-20 |
| ACE2 | WFDC11 | -0,4578 | 4,25389E-20 |
| ACE2 | GLRA3 | -0,4579 | 4,10109E-20 |
| ACE2 | STK11 | -0,4579 | 4,11719E-20 |
| ACE2 | AJ011932.1 | -0,4579 | 4,11997E-20 |
| ACE2 | CTB-174D11.2 | -0,458 | 4,03654E-20 |
| ACE2 | RP11-21B23.3 | -0,458 | 4,02834E-20 |
| ACE2 | CCT6B | -0,458 | 4,01694E-20 |
| ACE2 | CTC-523E23.14 | -0,458 | 4,04083E-20 |
| ACE2 | CTB-60E11.11 | -0,458 | 4,0703E-20 |
| ACE2 | RP11-34P13.16 | -0,4581 | 3,95056E-20 |
| ACE2 | CTB-164N12.1 | -0,4582 | 3,87984E-20 |
| ACE2 | RP11-295D22.1 | -0,4582 | 3,88197E-20 |
| ACE2 | RP11-216N14.9 | -0,4583 | 3,76663E-20 |
| ACE2 | RP11-526P5.2 | -0,4583 | 3,7811E-20 |
| ACE2 | LINC00529 | -0,4584 | 3,7464E-20 |
| ACE2 | RP11-1070N10.4 | -0,4584 | 3,70811E-20 |
| ACE2 | RP11-62C7.2 | -0,4584 | 3,74626E-20 |
| ACE2 | OR1F2P | -0,4584 | 3,72624E-20 |

| | | | |
|---|---|---|---|
| ACE2 | AC009158.1 | -0,4584 | 3,72536E-20 |
| ACE2 | RP11-325K19.2 | -0,4584 | 3,74021E-20 |
| ACE2 | AC004070.1 | -0,4584 | 3,69624E-20 |
| ACE2 | CTD-2066L21.1 | -0,4585 | 3,60185E-20 |
| ACE2 | PNLIPRP3 | -0,4585 | 3,63117E-20 |
| ACE2 | AP001092.4 | -0,4585 | 3,67138E-20 |
| ACE2 | TAS2R12 | -0,4585 | 3,6363E-20 |
| ACE2 | IGFBP7-AS1 | -0,4586 | 3,5401E-20 |
| ACE2 | CTB-23I7.1 | -0,4586 | 3,5312E-20 |
| ACE2 | CSMD1 | -0,4587 | 3,5065E-20 |
| ACE2 | CACFD1 | -0,4587 | 3,48373E-20 |
| ACE2 | TRBV5-1 | -0,4588 | 3,44256E-20 |
| ACE2 | GKAP1 | -0,4588 | 3,41998E-20 |
| ACE2 | RP11-341G23.3 | -0,4588 | 3,43208E-20 |
| ACE2 | PRAMEF11 | -0,4589 | 3,37333E-20 |
| ACE2 | LINC01865 | -0,4589 | 3,34774E-20 |
| ACE2 | RP3-359N14.2 | -0,4589 | 3,33604E-20 |
| ACE2 | NAV2-AS4 | -0,4589 | 3,33832E-20 |
| ACE2 | RPS2P6 | -0,4589 | 3,33765E-20 |
| ACE2 | LINC00994 | -0,459 | 3,30052E-20 |
| ACE2 | MFSD13B | -0,459 | 3,23978E-20 |
| ACE2 | LLNLR-304G9.1 | -0,459 | 3,29485E-20 |
| ACE2 | NR4A2 | -0,4591 | 3,23542E-20 |
| ACE2 | DNHD1 | -0,4591 | 3,18806E-20 |
| ACE2 | CCDC57 | -0,4591 | 3,21086E-20 |
| ACE2 | NPRL2 | -0,4592 | 3,16157E-20 |
| ACE2 | GPC2 | -0,4592 | 3,11611E-20 |
| ACE2 | OR8G5 | -0,4592 | 3,15297E-20 |
| ACE2 | CTD-2128A3.3 | -0,4592 | 3,16808E-20 |
| ACE2 | RP11-189E14.5 | -0,4592 | 3,13559E-20 |
| ACE2 | RP11-118E18.4 | -0,4592 | 3,13652E-20 |
| ACE2 | TMEM108 | -0,4593 | 3,04743E-20 |
| ACE2 | AC006150.1 | -0,4594 | 2,9809E-20 |
| ACE2 | EMC6 | -0,4594 | 2,9998E-20 |
| ACE2 | CTB-32H22.1 | -0,4595 | 2,96291E-20 |
| ACE2 | GS1-278J22.2 | -0,4595 | 2,91586E-20 |
| ACE2 | KRT18P59 | -0,4595 | 2,9693E-20 |
| ACE2 | ADAT1 | -0,4595 | 2,93647E-20 |
| ACE2 | YIPF2 | -0,4595 | 2,9243E-20 |
| ACE2 | LINC00487 | -0,4596 | 2,89819E-20 |
| ACE2 | TNIP3 | -0,4596 | 2,89277E-20 |
| ACE2 | RP11-756P10.2 | -0,4596 | 2,90901E-20 |
| ACE2 | CCDC34 | -0,4596 | 2,86028E-20 |
| ACE2 | MDGA2 | -0,4596 | 2,87601E-20 |
| ACE2 | RP11-770E5.2 | -0,4597 | 2,79357E-20 |
| ACE2 | RP11-164D18.2 | -0,4597 | 2,81751E-20 |
| ACE2 | CTA-989H11.1 | -0,4597 | 2,83362E-20 |
| ACE2 | AFAP1-AS1 | -0,4598 | 2,7683E-20 |
| ACE2 | BRCC3P1 | -0,4598 | 2,78511E-20 |
| ACE2 | ANKRD7 | -0,46 | 2,62532E-20 |
| ACE2 | RP11-94H18.1 | -0,46 | 2,667E-20 |
| ACE2 | C11orf49 | -0,46 | 2,62293E-20 |
| ACE2 | CTC-448F2.3 | -0,4601 | 2,5956E-20 |
| ACE2 | RP11-478J18.2 | -0,4602 | 2,51427E-20 |

| | | | |
|---|---|---|---|
| ACE2 | RP11-634B7.4 | -0,4602 | 2,55493E-20 |
| ACE2 | AC016644.1 | -0,4602 | 2,51976E-20 |
| ACE2 | AJ003147.9 | -0,4602 | 2,55408E-20 |
| ACE2 | DNAH2 | -0,4602 | 2,51562E-20 |
| ACE2 | RP11-126O1.5 | -0,4602 | 2,51653E-20 |
| ACE2 | NBPF5P | -0,4603 | 2,46338E-20 |
| ACE2 | RP11-545A16.3 | -0,4604 | 2,44028E-20 |
| ACE2 | ATAT1 | -0,4605 | 2,4066E-20 |
| ACE2 | KRT8P41 | -0,4605 | 2,36243E-20 |
| ACE2 | DUXB | -0,4605 | 2,39338E-20 |
| ACE2 | RP1-97G4.1 | -0,4606 | 2,34301E-20 |
| ACE2 | HMGB1P33 | -0,4606 | 2,34655E-20 |
| ACE2 | RP11-357N13.1 | -0,4606 | 2,31054E-20 |
| ACE2 | LINC01869 | -0,4606 | 2,31363E-20 |
| ACE2 | TREML3P | -0,4607 | 2,26531E-20 |
| ACE2 | CTD-3239E11.2 | -0,4607 | 2,28398E-20 |
| ACE2 | RP11-268P4.5 | -0,4607 | 2,29048E-20 |
| ACE2 | FBXW10 | -0,4607 | 2,28771E-20 |
| ACE2 | RP11-17A4.2 | -0,4608 | 2,22599E-20 |
| ACE2 | LINC02101 | -0,4609 | 2,19892E-20 |
| ACE2 | RP11-340I6.3 | -0,4609 | 2,18515E-20 |
| ACE2 | RP11-465K16.1 | -0,4609 | 2,16672E-20 |
| ACE2 | ZFHX4-AS1 | -0,4609 | 2,17641E-20 |
| ACE2 | TMC2 | -0,4609 | 2,19508E-20 |
| ACE2 | ZC3H11B | -0,461 | 2,15629E-20 |
| ACE2 | RP11-161D15.2 | -0,461 | 2,12936E-20 |
| ACE2 | CYSTM1 | -0,461 | 2,1573E-20 |
| ACE2 | SAYSD1 | -0,461 | 2,14735E-20 |
| ACE2 | AC005160.3 | -0,461 | 2,14004E-20 |
| ACE2 | RP11-218M22.2 | -0,461 | 2,15829E-20 |
| ACE2 | PPP1R8P1 | -0,4611 | 2,08768E-20 |
| ACE2 | RP11-176D17.3 | -0,4611 | 2,11162E-20 |
| ACE2 | RP11-577G20.1 | -0,4611 | 2,0839E-20 |
| ACE2 | BVES-AS1 | -0,4611 | 2,10901E-20 |
| ACE2 | IL19 | -0,4612 | 2,06652E-20 |
| ACE2 | LINC02020 | -0,4612 | 2,07203E-20 |
| ACE2 | LINC01007 | -0,4612 | 2,05091E-20 |
| ACE2 | CARM1P1 | -0,4612 | 2,04261E-20 |
| ACE2 | RP1-29C18.8 | -0,4612 | 2,06683E-20 |
| ACE2 | LINC01777 | -0,4613 | 1,98913E-20 |
| ACE2 | CCDC180 | -0,4613 | 2,00142E-20 |
| ACE2 | BAG5 | -0,4613 | 2,01115E-20 |
| ACE2 | RP11-416N4.1 | -0,4613 | 2,02544E-20 |
| ACE2 | DHX57 | -0,4614 | 1,96875E-20 |
| ACE2 | RP11-23D5.1 | -0,4614 | 1,98246E-20 |
| ACE2 | CTC-338M12.5 | -0,4615 | 1,90966E-20 |
| ACE2 | RP11-685M7.5 | -0,4615 | 1,91311E-20 |
| ACE2 | CFAP161 | -0,4615 | 1,92472E-20 |
| ACE2 | AC111155.1 | -0,4615 | 1,91494E-20 |
| ACE2 | RP11-134L10.1 | -0,4616 | 1,89865E-20 |
| ACE2 | GRHL2 | -0,4616 | 1,89512E-20 |
| ACE2 | ANKRD13B | -0,4616 | 1,8862E-20 |
| ACE2 | RP4-640H8.2 | -0,4616 | 1,88517E-20 |
| ACE2 | CABS1 | -0,4617 | 1,82764E-20 |

| | | | |
|---|---|---|---|
| ACE2 | MYPOP | -0,4617 | 1,83469E-20 |
| ACE2 | AC006547.13 | -0,4617 | 1,84496E-20 |
| ACE2 | RP11-79E3.2 | -0,4618 | 1,80265E-20 |
| ACE2 | RP11-255I10.2 | -0,4618 | 1,81872E-20 |
| ACE2 | RP11-94H18.2 | -0,4618 | 1,8116E-20 |
| ACE2 | RP11-358H18.3 | -0,4618 | 1,807E-20 |
| ACE2 | RP11-611O2.2 | -0,4618 | 1,82301E-20 |
| ACE2 | LINC01893 | -0,4618 | 1,80803E-20 |
| ACE2 | RP3-508I15.14 | -0,4618 | 1,79594E-20 |
| ACE2 | SLC22A16 | -0,4619 | 1,754E-20 |
| ACE2 | KCNQ1DN | -0,462 | 1,72727E-20 |
| ACE2 | AQP6 | -0,462 | 1,73777E-20 |
| ACE2 | RP11-620J15.2 | -0,462 | 1,72265E-20 |
| ACE2 | LINC00944 | -0,462 | 1,73709E-20 |
| ACE2 | LRRC39 | -0,4621 | 1,68986E-20 |
| ACE2 | RSPH10B2 | -0,4621 | 1,68342E-20 |
| ACE2 | RP11-554F20.1 | -0,4621 | 1,70879E-20 |
| ACE2 | NOXRED1 | -0,4621 | 1,6851E-20 |
| ACE2 | RP5-1042K10.10 | -0,4621 | 1,70368E-20 |
| ACE2 | RP11-340A13.2 | -0,4622 | 1,6485E-20 |
| ACE2 | BBIP1 | -0,4622 | 1,67177E-20 |
| ACE2 | LARGE2 | -0,4622 | 1,6491E-20 |
| ACE2 | CTD-2066L21.2 | -0,4623 | 1,63907E-20 |
| ACE2 | SLC8A1-AS1 | -0,4624 | 1,60034E-20 |
| ACE2 | LINC01986 | -0,4624 | 1,58952E-20 |
| ACE2 | PP2D1 | -0,4624 | 1,59226E-20 |
| ACE2 | PLS1-AS1 | -0,4624 | 1,59937E-20 |
| ACE2 | SOX6 | -0,4624 | 1,57741E-20 |
| ACE2 | SNRNP35 | -0,4624 | 1,60022E-20 |
| ACE2 | SMIM21 | -0,4624 | 1,59739E-20 |
| ACE2 | AC011551.3 | -0,4624 | 1,58998E-20 |
| ACE2 | PCDHB2 | -0,4625 | 1,56744E-20 |
| ACE2 | CNBD1 | -0,4625 | 1,56297E-20 |
| ACE2 | RP11-476B13.2 | -0,4625 | 1,55031E-20 |
| ACE2 | RP1-76B20.11 | -0,4626 | 1,51941E-20 |
| ACE2 | CACNA2D3 | -0,4627 | 1,50484E-20 |
| ACE2 | TRIM36 | -0,4627 | 1,49694E-20 |
| ACE2 | RP11-64B16.4 | -0,4627 | 1,50058E-20 |
| ACE2 | ERICH6B | -0,4627 | 1,50762E-20 |
| ACE2 | FGFR1OP | -0,4628 | 1,47515E-20 |
| ACE2 | CCL27 | -0,4628 | 1,4748E-20 |
| ACE2 | RP11-363J20.2 | -0,4628 | 1,46837E-20 |
| ACE2 | IGHV3-49 | -0,4628 | 1,46194E-20 |
| ACE2 | RFX2 | -0,4628 | 1,45913E-20 |
| ACE2 | SNX18P14 | -0,4629 | 1,42734E-20 |
| ACE2 | PXDNL | -0,4629 | 1,41578E-20 |
| ACE2 | SPATA31A3 | -0,4629 | 1,41605E-20 |
| ACE2 | NRDC | -0,463 | 1,39143E-20 |
| ACE2 | POMT1 | -0,463 | 1,40646E-20 |
| ACE2 | LDHC | -0,463 | 1,40743E-20 |
| ACE2 | PAQR5 | -0,463 | 1,39507E-20 |
| ACE2 | SYNGR4 | -0,463 | 1,39838E-20 |
| ACE2 | AC008074.3 | -0,4631 | 1,3707E-20 |
| ACE2 | CAMLG | -0,4631 | 1,35582E-20 |

| | | | |
|---|---|---|---|
| ACE2 | RP11-1028N23.3 | -0,4631 | 1,37875E-20 |
| ACE2 | PSMD10P2 | -0,4633 | 1,29877E-20 |
| ACE2 | RP11-655C2.3 | -0,4633 | 1,31249E-20 |
| ACE2 | GUCY2D | -0,4633 | 1,31021E-20 |
| ACE2 | RP4-682C21.5 | -0,4634 | 1,2945E-20 |
| ACE2 | RP11-44K6.3 | -0,4634 | 1,28362E-20 |
| ACE2 | LINC00898 | -0,4634 | 1,27726E-20 |
| ACE2 | LINC01282 | -0,4634 | 1,28095E-20 |
| ACE2 | OR5AC4P | -0,4635 | 1,24984E-20 |
| ACE2 | RP11-23J9.3 | -0,4635 | 1,25484E-20 |
| ACE2 | RP11-57C13.6 | -0,4635 | 1,26374E-20 |
| ACE2 | RP11-676J12.9 | -0,4635 | 1,27051E-20 |
| ACE2 | RP5-1033H22.2 | -0,4636 | 1,22038E-20 |
| ACE2 | GJB7 | -0,4636 | 1,24246E-20 |
| ACE2 | LINC00408 | -0,4636 | 1,2323E-20 |
| ACE2 | RP5-963E22.5 | -0,4636 | 1,22794E-20 |
| ACE2 | PSMG1 | -0,4636 | 1,22057E-20 |
| ACE2 | GLB1L | -0,4637 | 1,19967E-20 |
| ACE2 | RP11-473M20.11 | -0,4637 | 1,19914E-20 |
| ACE2 | LINC00992 | -0,4638 | 1,16748E-20 |
| ACE2 | LRRC43 | -0,4638 | 1,17503E-20 |
| ACE2 | GALNT5 | -0,4639 | 1,16376E-20 |
| ACE2 | RESP18 | -0,4639 | 1,14226E-20 |
| ACE2 | AC074019.1 | -0,4639 | 1,14787E-20 |
| ACE2 | RP11-35J1.2 | -0,4639 | 1,15452E-20 |
| ACE2 | RORB-AS1 | -0,4639 | 1,14775E-20 |
| ACE2 | OSBPL9P4 | -0,4639 | 1,15294E-20 |
| ACE2 | OCSTAMP | -0,4639 | 1,16139E-20 |
| ACE2 | ERICH3-AS1 | -0,464 | 1,12829E-20 |
| ACE2 | RPL39P18 | -0,464 | 1,13681E-20 |
| ACE2 | CTB-191D16.1 | -0,464 | 1,11938E-20 |
| ACE2 | RP11-476F14.1 | -0,464 | 1,12876E-20 |
| ACE2 | RPL21P17 | -0,4641 | 1,11021E-20 |
| ACE2 | LINC00884 | -0,4641 | 1,11181E-20 |
| ACE2 | LINC01470 | -0,4641 | 1,09899E-20 |
| ACE2 | ROR1-AS1 | -0,4642 | 1,07842E-20 |
| ACE2 | EEF1B2P8 | -0,4642 | 1,08194E-20 |
| ACE2 | AC005013.5 | -0,4642 | 1,08326E-20 |
| ACE2 | SNX32 | -0,4642 | 1,08958E-20 |
| ACE2 | LRRC9 | -0,4642 | 1,07568E-20 |
| ACE2 | VN1R6P | -0,4642 | 1,07429E-20 |
| ACE2 | DNAJA1P5 | -0,4643 | 1,05566E-20 |
| ACE2 | RP11-109G23.1 | -0,4643 | 1,04978E-20 |
| ACE2 | ASNS | -0,4643 | 1,06261E-20 |
| ACE2 | RP11-813F20.4 | -0,4643 | 1,05359E-20 |
| ACE2 | AC012354.8 | -0,4644 | 1,04292E-20 |
| ACE2 | BBS5 | -0,4644 | 1,03938E-20 |
| ACE2 | ESR1 | -0,4644 | 1,03772E-20 |
| ACE2 | CETN4P | -0,4645 | 1,01789E-20 |
| ACE2 | OR2B4P | -0,4645 | 1,02086E-20 |
| ACE2 | RP11-125D12.2 | -0,4645 | 1,01728E-20 |
| ACE2 | RP11-760D2.10 | -0,4645 | 1,02556E-20 |
| ACE2 | SPDYE7P | -0,4645 | 1,01762E-20 |
| ACE2 | RP11-640N20.6 | -0,4645 | 1,02434E-20 |

| | | | |
|---|---|---|---|
| ACE2 | C3orf67 | -0,4646 | 9,87824E-21 |
| ACE2 | RASGRF2-AS1 | -0,4646 | 9,94143E-21 |
| ACE2 | SUGT1P1 | -0,4646 | 9,95578E-21 |
| ACE2 | RP1-186E20.1 | -0,4646 | 9,95636E-21 |
| ACE2 | TNR-IT1 | -0,4647 | 9,69098E-21 |
| ACE2 | ABCB6 | -0,4647 | 9,63852E-21 |
| ACE2 | CAP1P2 | -0,4647 | 9,62983E-21 |
| ACE2 | CTD-2384A14.1 | -0,4647 | 9,72719E-21 |
| ACE2 | LINC01017 | -0,4648 | 9,45499E-21 |
| ACE2 | CTD-2033C11.1 | -0,4648 | 9,47433E-21 |
| ACE2 | CLEC9A | -0,4648 | 9,43986E-21 |
| ACE2 | RP4-669K10.8 | -0,4649 | 9,30875E-21 |
| ACE2 | RP11-500C11.3 | -0,4649 | 9,28385E-21 |
| ACE2 | RP11-4C20.3 | -0,4649 | 9,37472E-21 |
| ACE2 | FLNB-AS1 | -0,465 | 9,14575E-21 |
| ACE2 | RNU1-30P | -0,465 | 9,12773E-21 |
| ACE2 | PLA2G1B | -0,465 | 9,05791E-21 |
| ACE2 | KRT8P5 | -0,465 | 9,17737E-21 |
| ACE2 | LINC01810 | -0,4651 | 8,9655E-21 |
| ACE2 | FBXL2 | -0,4651 | 8,82605E-21 |
| ACE2 | RPL7AP28 | -0,4651 | 8,83861E-21 |
| ACE2 | RP11-419C19.3 | -0,4651 | 8,93396E-21 |
| ACE2 | ASIC3 | -0,4651 | 8,8621E-21 |
| ACE2 | RP11-313F23.4 | -0,4651 | 8,97069E-21 |
| ACE2 | RP11-483P21.2 | -0,4651 | 9,00106E-21 |
| ACE2 | CTD-2006O16.2 | -0,4652 | 8,64969E-21 |
| ACE2 | GRK3 | -0,4652 | 8,70665E-21 |
| ACE2 | RP11-525K10.3 | -0,4653 | 8,46085E-21 |
| ACE2 | LINC02071 | -0,4653 | 8,45735E-21 |
| ACE2 | PADI2 | -0,4654 | 8,34486E-21 |
| ACE2 | RP11-389G6.4 | -0,4654 | 8,43249E-21 |
| ACE2 | PCNAP1 | -0,4654 | 8,36033E-21 |
| ACE2 | RP11-436D23.1 | -0,4654 | 8,43559E-21 |
| ACE2 | RP11-162N7.1 | -0,4654 | 8,34846E-21 |
| ACE2 | FTCD | -0,4654 | 8,28175E-21 |
| ACE2 | RP11-701I24.3 | -0,4655 | 8,11764E-21 |
| ACE2 | UPK2 | -0,4655 | 8,13667E-21 |
| ACE2 | CTD-2554C21.2 | -0,4655 | 8,25668E-21 |
| ACE2 | AL121578.7 | -0,4655 | 8,26196E-21 |
| ACE2 | PLCH1-AS2 | -0,4656 | 8,07658E-21 |
| ACE2 | AC006014.8 | -0,4656 | 8,04942E-21 |
| ACE2 | RP11-627K11.1 | -0,4656 | 7,97159E-21 |
| ACE2 | RN7SKP113 | -0,4657 | 7,88702E-21 |
| ACE2 | CTD-2074D8.1 | -0,4657 | 7,84662E-21 |
| ACE2 | RNU1-83P | -0,4658 | 7,64367E-21 |
| ACE2 | BEST3 | -0,4658 | 7,71824E-21 |
| ACE2 | HAGH | -0,4658 | 7,59642E-21 |
| ACE2 | LINC00251 | -0,4659 | 7,53174E-21 |
| ACE2 | C4orf22 | -0,466 | 7,32151E-21 |
| ACE2 | RP11-351M16.4 | -0,466 | 7,28172E-21 |
| ACE2 | RP11-278A23.4 | -0,466 | 7,42106E-21 |
| ACE2 | RP11-277J6.3 | -0,466 | 7,34808E-21 |
| ACE2 | RP11-420K14.6 | -0,466 | 7,3557E-21 |
| ACE2 | RP5-828H9.1 | -0,466 | 7,37521E-21 |

| | | | |
|---|---|---|---|
| ACE2 | RP11-16L9.2 | -0,4661 | 7,14949E-21 |
| ACE2 | RP11-474I11.7 | -0,4661 | 7,23049E-21 |
| ACE2 | LINC01317 | -0,4662 | 7,02963E-21 |
| ACE2 | TACC3 | -0,4662 | 7,02376E-21 |
| ACE2 | ZFP42 | -0,4662 | 7,09384E-21 |
| ACE2 | RP11-461L18.1 | -0,4662 | 7,10446E-21 |
| ACE2 | RPL7AP7 | -0,4662 | 7,06402E-21 |
| ACE2 | ZPBP | -0,4662 | 7,01848E-21 |
| ACE2 | PSMC4 | -0,4662 | 6,97284E-21 |
| ACE2 | PLCG1-AS1 | -0,4662 | 7,09959E-21 |
| ACE2 | TIAM2 | -0,4663 | 6,93183E-21 |
| ACE2 | LINC00930 | -0,4663 | 6,85263E-21 |
| ACE2 | LINC02140 | -0,4663 | 6,83496E-21 |
| ACE2 | CFAP44 | -0,4664 | 6,72053E-21 |
| ACE2 | TCEA1P1 | -0,4664 | 6,76642E-21 |
| ACE2 | LINC00477 | -0,4664 | 6,80818E-21 |
| ACE2 | RP11-117L5.1 | -0,4664 | 6,76636E-21 |
| ACE2 | TOX4P1 | -0,4665 | 6,56332E-21 |
| ACE2 | OR51A6P | -0,4665 | 6,63026E-21 |
| ACE2 | MGAT5B | -0,4665 | 6,54992E-21 |
| ACE2 | RP11-529E15.1 | -0,4666 | 6,51549E-21 |
| ACE2 | NUP35 | -0,4666 | 6,39447E-21 |
| ACE2 | RP11-54C4.3 | -0,4666 | 6,44314E-21 |
| ACE2 | DGKI | -0,4666 | 6,50813E-21 |
| ACE2 | NPAP1P2 | -0,4666 | 6,41427E-21 |
| ACE2 | AP000640.2 | -0,4666 | 6,45297E-21 |
| ACE2 | RP11-632K5.2 | -0,4666 | 6,38651E-21 |
| ACE2 | RP11-181C3.1 | -0,4666 | 6,50456E-21 |
| ACE2 | LINC01216 | -0,4667 | 6,33483E-21 |
| ACE2 | WDR24 | -0,4667 | 6,37084E-21 |
| ACE2 | CHMP1A | -0,4667 | 6,31213E-21 |
| ACE2 | AC007563.5 | -0,4668 | 6,16583E-21 |
| ACE2 | RP11-379K17.4 | -0,4668 | 6,21207E-21 |
| ACE2 | ELDR | -0,4668 | 6,24038E-21 |
| ACE2 | RP11-335F8.2 | -0,4668 | 6,23612E-21 |
| ACE2 | CDKL3 | -0,4669 | 6,06796E-21 |
| ACE2 | DSG2-AS1 | -0,4669 | 6,05948E-21 |
| ACE2 | LRRFIP2 | -0,467 | 5,95542E-21 |
| ACE2 | FILIP1L | -0,467 | 5,89787E-21 |
| ACE2 | FOSL1P1 | -0,467 | 5,97965E-21 |
| ACE2 | RP11-257P3.3 | -0,467 | 5,9271E-21 |
| ACE2 | CTC-465D4.1 | -0,467 | 5,9817E-21 |
| ACE2 | SLC25A52 | -0,467 | 5,91612E-21 |
| ACE2 | PRKCQ-AS1 | -0,4671 | 5,80695E-21 |
| ACE2 | TRBVB | -0,4672 | 5,70624E-21 |
| ACE2 | CHST5 | -0,4672 | 5,72551E-21 |
| ACE2 | RP11-219A15.2 | -0,4672 | 5,6306E-21 |
| ACE2 | FAM65B | -0,4673 | 5,54871E-21 |
| ACE2 | RFX3 | -0,4673 | 5,57784E-21 |
| ACE2 | RP11-690D19.1 | -0,4673 | 5,56449E-21 |
| ACE2 | ALX1 | -0,4673 | 5,59962E-21 |
| ACE2 | FAM228B | -0,4674 | 5,3787E-21 |
| ACE2 | LINC00575 | -0,4674 | 5,48472E-21 |
| ACE2 | SMIM15-AS1 | -0,4674 | 5,42563E-21 |

| | | | |
|---|---|---|---|
| ACE2 | RP11-93O17.2 | -0,4674 | 5,4599E-21 |
| ACE2 | SPATA7 | -0,4674 | 5,38531E-21 |
| ACE2 | RARA-AS1 | -0,4675 | 5,32448E-21 |
| ACE2 | AP1M2 | -0,4675 | 5,31586E-21 |
| ACE2 | CST11 | -0,4675 | 5,28976E-21 |
| ACE2 | DCDC2B | -0,4676 | 5,14918E-21 |
| ACE2 | DNTT | -0,4676 | 5,15585E-21 |
| ACE2 | ASRGL1 | -0,4676 | 5,20792E-21 |
| ACE2 | OR8B12 | -0,4676 | 5,19363E-21 |
| ACE2 | FAM92A1P1 | -0,4676 | 5,23882E-21 |
| ACE2 | RP11-849N15.4 | -0,4676 | 5,17591E-21 |
| ACE2 | AC104451.2 | -0,4677 | 5,08668E-21 |
| ACE2 | SNX18P27 | -0,4677 | 5,11415E-21 |
| ACE2 | PA2G4 | -0,4678 | 4,96363E-21 |
| ACE2 | SLC2A4 | -0,4678 | 4,9962E-21 |
| ACE2 | AC005009.2 | -0,4679 | 4,8183E-21 |
| ACE2 | SCAANT1 | -0,468 | 4,76086E-21 |
| ACE2 | FAM192BP | -0,468 | 4,75295E-21 |
| ACE2 | RP11-370A5.1 | -0,468 | 4,76895E-21 |
| ACE2 | UFD1L | -0,468 | 4,72007E-21 |
| ACE2 | ASH2LP1 | -0,468 | 4,72107E-21 |
| ACE2 | SYT6 | -0,4681 | 4,70819E-21 |
| ACE2 | AC104653.1 | -0,4682 | 4,60555E-21 |
| ACE2 | CCDC171 | -0,4682 | 4,61172E-21 |
| ACE2 | LINC00588 | -0,4683 | 4,43762E-21 |
| ACE2 | ARAP1-AS2 | -0,4683 | 4,50142E-21 |
| ACE2 | LINC01058 | -0,4683 | 4,50577E-21 |
| ACE2 | MIR3681HG | -0,4684 | 4,36618E-21 |
| ACE2 | RP11-152D8.1 | -0,4684 | 4,34108E-21 |
| ACE2 | RP11-345F18.1 | -0,4684 | 4,32991E-21 |
| ACE2 | WBSCR27 | -0,4684 | 4,3466E-21 |
| ACE2 | RP11-1018N14.4 | -0,4684 | 4,39873E-21 |
| ACE2 | RP11-243E13.2 | -0,4684 | 4,32372E-21 |
| ACE2 | LINC00334 | -0,4685 | 4,31945E-21 |
| ACE2 | FAM200A | -0,4686 | 4,16023E-21 |
| ACE2 | RP11-2C24.7 | -0,4686 | 4,22458E-21 |
| ACE2 | ABCB11 | -0,4687 | 4,07806E-21 |
| ACE2 | RP11-40A13.2 | -0,4687 | 4,04869E-21 |
| ACE2 | MRPL34 | -0,4687 | 4,09E-21 |
| ACE2 | AC079779.5 | -0,4688 | 3,98383E-21 |
| ACE2 | UGT1A6 | -0,4688 | 3,98711E-21 |
| ACE2 | RP11-255E6.6 | -0,4688 | 4,01344E-21 |
| ACE2 | C3orf30 | -0,4688 | 4,02471E-21 |
| ACE2 | FEZF1 | -0,4688 | 4,02364E-21 |
| ACE2 | PSMA4 | -0,4688 | 4,01673E-21 |
| ACE2 | AA06 | -0,4688 | 4,02805E-21 |
| ACE2 | PDCL3P4 | -0,4689 | 3,92992E-21 |
| ACE2 | RP11-360K13.1 | -0,4689 | 3,90546E-21 |
| ACE2 | ATP8A2P3 | -0,4689 | 3,91561E-21 |
| ACE2 | RNF175 | -0,469 | 3,8235E-21 |
| ACE2 | RP1-90J20.2 | -0,469 | 3,83618E-21 |
| ACE2 | CA3 | -0,469 | 3,81239E-21 |
| ACE2 | RP11-315O6.1 | -0,469 | 3,829E-21 |
| ACE2 | RP11-390P24.1 | -0,469 | 3,8707E-21 |

| | | | |
|---|---|---|---|
| ACE2 | CTC-525D6.5 | -0,469 | 3,79313E-21 |
| ACE2 | LCE5A | -0,4691 | 3,72908E-21 |
| ACE2 | RP11-713D19.1 | -0,4691 | 3,72063E-21 |
| ACE2 | AC100830.3 | -0,4691 | 3,72074E-21 |
| ACE2 | CTD-2589H19.6 | -0,4692 | 3,70789E-21 |
| ACE2 | SAA2 | -0,4692 | 3,70926E-21 |
| ACE2 | ITGAE | -0,4692 | 3,66004E-21 |
| ACE2 | ANKRD24 | -0,4692 | 3,69876E-21 |
| ACE2 | SULT6B1 | -0,4693 | 3,57005E-21 |
| ACE2 | RP11-169E6.1 | -0,4693 | 3,61331E-21 |
| ACE2 | MAP3K19 | -0,4694 | 3,53539E-21 |
| ACE2 | LINC01524 | -0,4694 | 3,52793E-21 |
| ACE2 | TTN-AS1 | -0,4695 | 3,46951E-21 |
| ACE2 | RP11-499P1.2 | -0,4695 | 3,43981E-21 |
| ACE2 | TRIM42 | -0,4695 | 3,43891E-21 |
| ACE2 | LEKR1 | -0,4695 | 3,46881E-21 |
| ACE2 | RP11-472K22.2 | -0,4695 | 3,43802E-21 |
| ACE2 | HTR5A | -0,4695 | 3,44278E-21 |
| ACE2 | AHNAK2 | -0,4695 | 3,44968E-21 |
| ACE2 | VN1R3 | -0,4695 | 3,42775E-21 |
| ACE2 | NEK10 | -0,4696 | 3,35835E-21 |
| ACE2 | APTR | -0,4696 | 3,36374E-21 |
| ACE2 | RP11-754N21.1 | -0,4696 | 3,39352E-21 |
| ACE2 | RP4-799D16.1 | -0,4697 | 3,28274E-21 |
| ACE2 | RP11-4N23.1 | -0,4698 | 3,2088E-21 |
| ACE2 | RP1-90K10.4 | -0,4699 | 3,12869E-21 |
| ACE2 | AC013436.6 | -0,4699 | 3,13286E-21 |
| ACE2 | CPNE8 | -0,4699 | 3,12935E-21 |
| ACE2 | RP11-244B22.10 | -0,4699 | 3,13791E-21 |
| ACE2 | AC114808.3 | -0,47 | 3,11153E-21 |
| ACE2 | ATP5G1P1 | -0,47 | 3,09652E-21 |
| ACE2 | KRT18P28 | -0,4701 | 3,04568E-21 |
| ACE2 | FSIP2 | -0,4701 | 3,01159E-21 |
| ACE2 | OXSM | -0,4701 | 3,00264E-21 |
| ACE2 | C6orf52 | -0,4701 | 3,00435E-21 |
| ACE2 | RP11-753D20.4 | -0,4701 | 3,00697E-21 |
| ACE2 | RP11-605F22.2 | -0,4701 | 2,99285E-21 |
| ACE2 | RNF126 | -0,4701 | 3,04158E-21 |
| ACE2 | RP11-286E11.1 | -0,4702 | 2,92212E-21 |
| ACE2 | DDX43P3 | -0,4702 | 2,91812E-21 |
| ACE2 | TNFAIP2 | -0,4702 | 2,96725E-21 |
| ACE2 | USP3-AS1 | -0,4702 | 2,94994E-21 |
| ACE2 | LGALS7B | -0,4702 | 2,94961E-21 |
| ACE2 | TEX264 | -0,4703 | 2,9123E-21 |
| ACE2 | CTD-2201E18.3 | -0,4704 | 2,80541E-21 |
| ACE2 | LINC00906 | -0,4704 | 2,83128E-21 |
| ACE2 | AC007563.3 | -0,4705 | 2,7901E-21 |
| ACE2 | CTD-2341M24.1 | -0,4705 | 2,78369E-21 |
| ACE2 | CDRT1 | -0,4705 | 2,73881E-21 |
| ACE2 | SPACA6 | -0,4705 | 2,73676E-21 |
| ACE2 | RP11-107E5.4 | -0,4706 | 2,69558E-21 |
| ACE2 | PCDH9-AS4 | -0,4706 | 2,73149E-21 |
| ACE2 | RP11-2C24.5 | -0,4706 | 2,71683E-21 |
| ACE2 | CTD-2008L17.2 | -0,4706 | 2,69374E-21 |

| | | | |
|---|---|---|---|
| ACE2 | RP11-107M16.2 | -0,4707 | 2,63688E-21 |
| ACE2 | PPP1R42 | -0,4707 | 2,65389E-21 |
| ACE2 | LINC00960 | -0,4708 | 2,56412E-21 |
| ACE2 | RP11-24H2.3 | -0,4708 | 2,58933E-21 |
| ACE2 | C15orf39 | -0,4708 | 2,60409E-21 |
| ACE2 | LINC01365 | -0,4709 | 2,53312E-21 |
| ACE2 | CIB2 | -0,4709 | 2,52876E-21 |
| ACE2 | KRT8P45 | -0,471 | 2,46706E-21 |
| ACE2 | TRAPPC12 | -0,471 | 2,45558E-21 |
| ACE2 | CCDC169 | -0,471 | 2,48735E-21 |
| ACE2 | TMEM106A | -0,471 | 2,46217E-21 |
| ACE2 | SIRPB3P | -0,471 | 2,49475E-21 |
| ACE2 | RP11-292D4.2 | -0,4711 | 2,41722E-21 |
| ACE2 | NHEG1 | -0,4711 | 2,42506E-21 |
| ACE2 | RP11-760D2.11 | -0,4711 | 2,43719E-21 |
| ACE2 | RAG2 | -0,4711 | 2,40968E-21 |
| ACE2 | RP11-676F20.5 | -0,4711 | 2,42427E-21 |
| ACE2 | RP11-567M21.3 | -0,4711 | 2,40595E-21 |
| ACE2 | RP5-968J1.1 | -0,4711 | 2,44335E-21 |
| ACE2 | FHDC1 | -0,4712 | 2,39346E-21 |
| ACE2 | RP11-210G22.1 | -0,4712 | 2,38658E-21 |
| ACE2 | CLIP1-AS1 | -0,4712 | 2,38276E-21 |
| ACE2 | RPS2P1 | -0,4712 | 2,37566E-21 |
| ACE2 | BNIPL | -0,4713 | 2,30518E-21 |
| ACE2 | COL6A5 | -0,4713 | 2,30239E-21 |
| ACE2 | AC004744.3 | -0,4713 | 2,29691E-21 |
| ACE2 | BMS1P19 | -0,4714 | 2,25514E-21 |
| ACE2 | SOSTDC1 | -0,4714 | 2,28595E-21 |
| ACE2 | RP11-242P2.2 | -0,4714 | 2,28569E-21 |
| ACE2 | RORA-AS2 | -0,4714 | 2,24294E-21 |
| ACE2 | RP11-410C4.4 | -0,4715 | 2,24066E-21 |
| ACE2 | AC018643.4 | -0,4715 | 2,20986E-21 |
| ACE2 | RP11-359E19.2 | -0,4715 | 2,21178E-21 |
| ACE2 | SPTA1 | -0,4716 | 2,15043E-21 |
| ACE2 | RP11-420L9.4 | -0,4716 | 2,16258E-21 |
| ACE2 | CTD-2373H9.5 | -0,4717 | 2,12288E-21 |
| ACE2 | RTKN2 | -0,4718 | 2,06113E-21 |
| ACE2 | ADAM20 | -0,4718 | 2,08991E-21 |
| ACE2 | CTD-2126E3.4 | -0,4718 | 2,05947E-21 |
| ACE2 | HDAC4 | -0,4719 | 2,04687E-21 |
| ACE2 | PILRB | -0,4719 | 2,02578E-21 |
| ACE2 | POM121L9P | -0,4719 | 2,05029E-21 |
| ACE2 | RP11-473M10.3 | -0,472 | 1,9728E-21 |
| ACE2 | RTP5 | -0,4722 | 1,90665E-21 |
| ACE2 | AKAP3 | -0,4722 | 1,88644E-21 |
| ACE2 | HMGN2P46 | -0,4722 | 1,8937E-21 |
| ACE2 | LYG1 | -0,4723 | 1,86345E-21 |
| ACE2 | DNAH12 | -0,4723 | 1,87064E-21 |
| ACE2 | RP11-444P10.1 | -0,4723 | 1,87256E-21 |
| ACE2 | ADO | -0,4724 | 1,82325E-21 |
| ACE2 | AC004637.1 | -0,4724 | 1,81514E-21 |
| ACE2 | EIF6 | -0,4724 | 1,80636E-21 |
| ACE2 | AC098872.3 | -0,4725 | 1,78075E-21 |
| ACE2 | WNT10A | -0,4725 | 1,77605E-21 |

| | | | |
|---|---|---|---|
| ACE2 | RP11-478H13.1 | -0,4725 | 1,78769E-21 |
| ACE2 | RP11-432J24.3 | -0,4725 | 1,77094E-21 |
| ACE2 | MS4A14 | -0,4725 | 1,79388E-21 |
| ACE2 | CDCA3 | -0,4725 | 1,78151E-21 |
| ACE2 | FAM222B | -0,4725 | 1,77638E-21 |
| ACE2 | PPM1E | -0,4725 | 1,77461E-21 |
| ACE2 | GINS1 | -0,4725 | 1,78872E-21 |
| ACE2 | RFPL1 | -0,4725 | 1,79917E-21 |
| ACE2 | KRBOX1 | -0,4726 | 1,76009E-21 |
| ACE2 | NDUFA2 | -0,4726 | 1,73605E-21 |
| ACE2 | RP11-53B2.2 | -0,4726 | 1,75715E-21 |
| ACE2 | RP11-495P10.5 | -0,4727 | 1,70082E-21 |
| ACE2 | RNF32 | -0,4727 | 1,70579E-21 |
| ACE2 | RBM22P2 | -0,4727 | 1,70816E-21 |
| ACE2 | NREP-AS1 | -0,4728 | 1,65363E-21 |
| ACE2 | CCDC102B | -0,4728 | 1,65619E-21 |
| ACE2 | RP4-555D20.3 | -0,4729 | 1,62426E-21 |
| ACE2 | CTD-2176I21.2 | -0,4729 | 1,62303E-21 |
| ACE2 | XXbac-BCX196D17.5 | -0,4729 | 1,6389E-21 |
| ACE2 | PCK2 | -0,4729 | 1,62277E-21 |
| ACE2 | RP11-74E22.6 | -0,4729 | 1,63382E-21 |
| ACE2 | RP4-639F20.1 | -0,473 | 1,58786E-21 |
| ACE2 | RP11-227F19.2 | -0,473 | 1,58187E-21 |
| ACE2 | RP4-630C24.3 | -0,473 | 1,59801E-21 |
| ACE2 | TRAPPC2P2 | -0,473 | 1,58041E-21 |
| ACE2 | ATXN2-AS | -0,473 | 1,58563E-21 |
| ACE2 | AC002115.9 | -0,473 | 1,60087E-21 |
| ACE2 | RP1-92O14.3 | -0,4731 | 1,5613E-21 |
| ACE2 | SCIN | -0,4731 | 1,54741E-21 |
| ACE2 | CLK2P1 | -0,4731 | 1,57251E-21 |
| ACE2 | RP11-20L19.1 | -0,4731 | 1,57177E-21 |
| ACE2 | AC003973.4 | -0,4731 | 1,55237E-21 |
| ACE2 | RP3-510H16.3 | -0,4731 | 1,57447E-21 |
| ACE2 | RP11-148B18.1 | -0,4732 | 1,53587E-21 |
| ACE2 | EGF | -0,4732 | 1,53734E-21 |
| ACE2 | SRP54-AS1 | -0,4732 | 1,51116E-21 |
| ACE2 | CA3-AS1 | -0,4733 | 1,49463E-21 |
| ACE2 | CTD-3118D11.3 | -0,4733 | 1,49389E-21 |
| ACE2 | RP4-553F4.2 | -0,4733 | 1,47932E-21 |
| ACE2 | AATBC | -0,4733 | 1,49955E-21 |
| ACE2 | FLAD1 | -0,4734 | 1,46902E-21 |
| ACE2 | TNR | -0,4734 | 1,45431E-21 |
| ACE2 | ABT1P1 | -0,4734 | 1,46014E-21 |
| ACE2 | LINC02114 | -0,4734 | 1,45268E-21 |
| ACE2 | LRRC63 | -0,4734 | 1,46471E-21 |
| ACE2 | LSM4 | -0,4735 | 1,4136E-21 |
| ACE2 | MAGEB5 | -0,4735 | 1,42101E-21 |
| ACE2 | SCN10A | -0,4736 | 1,40526E-21 |
| ACE2 | RDH10 | -0,4736 | 1,39507E-21 |
| ACE2 | GPR83 | -0,4736 | 1,38894E-21 |
| ACE2 | LINC02132 | -0,4736 | 1,40448E-21 |
| ACE2 | RP6-102O10.1 | -0,4737 | 1,36032E-21 |
| ACE2 | TUBA3D | -0,4737 | 1,3753E-21 |
| ACE2 | LYAR | -0,4737 | 1,35276E-21 |

| | | | |
|---|---|---|---|
| ACE2 | EGR3 | -0,4737 | 1,35712E-21 |
| ACE2 | RP11-267M23.4 | -0,4737 | 1,36305E-21 |
| ACE2 | RP11-150D5.2 | -0,4737 | 1,37182E-21 |
| ACE2 | RP11-98I9.4 | -0,4738 | 1,34269E-21 |
| ACE2 | C10orf35 | -0,4738 | 1,32734E-21 |
| ACE2 | TMEM233 | -0,4738 | 1,34762E-21 |
| ACE2 | RP11-151A6.6 | -0,4738 | 1,32477E-21 |
| ACE2 | LINC00596 | -0,4738 | 1,33654E-21 |
| ACE2 | RP11-404K5.4 | -0,4739 | 1,31626E-21 |
| ACE2 | CTC-265N9.1 | -0,4739 | 1,29454E-21 |
| ACE2 | DCDC2 | -0,4739 | 1,30008E-21 |
| ACE2 | FAM95C | -0,4739 | 1,31865E-21 |
| ACE2 | RP11-403N16.4 | -0,4739 | 1,29361E-21 |
| ACE2 | RP11-213G2.3 | -0,4739 | 1,31501E-21 |
| ACE2 | RP11-178F10.3 | -0,4739 | 1,31123E-21 |
| ACE2 | LINC01938 | -0,474 | 1,2844E-21 |
| ACE2 | CCSER2 | -0,474 | 1,26607E-21 |
| ACE2 | RP11-899L11.1 | -0,474 | 1,28484E-21 |
| ACE2 | DDA1 | -0,474 | 1,28552E-21 |
| ACE2 | MRPL47 | -0,4742 | 1,22109E-21 |
| ACE2 | GLP1R | -0,4742 | 1,21181E-21 |
| ACE2 | AC009518.2 | -0,4742 | 1,23307E-21 |
| ACE2 | C15orf41 | -0,4742 | 1,21794E-21 |
| ACE2 | FRG2DP | -0,4742 | 1,232E-21 |
| ACE2 | CTC-304I17.5 | -0,4742 | 1,22135E-21 |
| ACE2 | RP11-666A8.9 | -0,4742 | 1,22646E-21 |
| ACE2 | LINC00399 | -0,4743 | 1,18893E-21 |
| ACE2 | LINC01180 | -0,4743 | 1,18958E-21 |
| ACE2 | LINC01397 | -0,4744 | 1,17065E-21 |
| ACE2 | LVRN | -0,4744 | 1,18061E-21 |
| ACE2 | DNAJB5-AS1 | -0,4744 | 1,17235E-21 |
| ACE2 | ASNSP3 | -0,4744 | 1,17587E-21 |
| ACE2 | ADNP-AS1 | -0,4744 | 1,17305E-21 |
| ACE2 | C15orf48 | -0,4745 | 1,13637E-21 |
| ACE2 | WBP11P1 | -0,4745 | 1,1419E-21 |
| ACE2 | GDF5 | -0,4745 | 1,14538E-21 |
| ACE2 | RP11-24I21.1 | -0,4746 | 1,12399E-21 |
| ACE2 | AL162151.4 | -0,4746 | 1,1187E-21 |
| ACE2 | OR2L2 | -0,4747 | 1,0934E-21 |
| ACE2 | RP11-552M14.1 | -0,4747 | 1,10194E-21 |
| ACE2 | RP11-445N20.2 | -0,4747 | 1,08658E-21 |
| ACE2 | CAMTA2 | -0,4747 | 1,08589E-21 |
| ACE2 | RP11-120M18.5 | -0,4747 | 1,09869E-21 |
| ACE2 | PAGE3 | -0,4747 | 1,10141E-21 |
| ACE2 | OR14L1P | -0,4748 | 1,07631E-21 |
| ACE2 | RP4-673D20.6 | -0,4748 | 1,06485E-21 |
| ACE2 | RP11-437J2.3 | -0,4749 | 1,04821E-21 |
| ACE2 | RP11-244B22.6 | -0,4749 | 1,03528E-21 |
| ACE2 | C19orf68 | -0,4749 | 1,04372E-21 |
| ACE2 | CFL1P2 | -0,475 | 1,01242E-21 |
| ACE2 | HMCN1 | -0,475 | 1,02447E-21 |
| ACE2 | MYH15 | -0,475 | 1,01281E-21 |
| ACE2 | RP11-454P21.1 | -0,475 | 1,01541E-21 |
| ACE2 | RP11-530C5.1 | -0,475 | 1,02297E-21 |

| | | | |
|---|---|---|---|
| ACE2 | SLC6A2 | -0,475 | 1,0176E-21 |
| ACE2 | PAXBP1-AS1 | -0,475 | 1,02099E-21 |
| ACE2 | RP11-335L23.5 | -0,4751 | 1,004E-21 |
| ACE2 | RSPH4A | -0,4751 | 1,00532E-21 |
| ACE2 | LCP1 | -0,4751 | 1,00067E-21 |
| ACE2 | LYRM1 | -0,4751 | 9,97863E-22 |
| ACE2 | HOOK1 | -0,4752 | 9,6906E-22 |
| ACE2 | AC007362.3 | -0,4752 | 9,88655E-22 |
| ACE2 | CTD-2410N18.3 | -0,4752 | 9,78176E-22 |
| ACE2 | RP3-400B16.4 | -0,4752 | 9,69369E-22 |
| ACE2 | ECT2L | -0,4752 | 9,79933E-22 |
| ACE2 | CTA-398F10.1 | -0,4752 | 9,88346E-22 |
| ACE2 | FAM83H-AS1 | -0,4752 | 9,82638E-22 |
| ACE2 | SLCO1C1 | -0,4752 | 9,72215E-22 |
| ACE2 | AMN1 | -0,4752 | 9,70886E-22 |
| ACE2 | RP11-434D2.12 | -0,4752 | 9,7561E-22 |
| ACE2 | GUSBP1 | -0,4753 | 9,47467E-22 |
| ACE2 | RDH10-AS1 | -0,4753 | 9,47302E-22 |
| ACE2 | LINC00502 | -0,4753 | 9,47566E-22 |
| ACE2 | HIGD1C | -0,4753 | 9,62735E-22 |
| ACE2 | CENPJ | -0,4753 | 9,6653E-22 |
| ACE2 | NEK5 | -0,4753 | 9,49483E-22 |
| ACE2 | RP11-22D3.2 | -0,4753 | 9,49265E-22 |
| ACE2 | RP11-472M19.2 | -0,4754 | 9,30168E-22 |
| ACE2 | ZNF317P1 | -0,4754 | 9,39598E-22 |
| ACE2 | LINC01044 | -0,4754 | 9,40036E-22 |
| ACE2 | LINC02171 | -0,4755 | 9,1546E-22 |
| ACE2 | RP11-159D8.1 | -0,4755 | 9,08596E-22 |
| ACE2 | RP11-618M23.1 | -0,4756 | 8,95971E-22 |
| ACE2 | ADAM29 | -0,4757 | 8,8502E-22 |
| ACE2 | RP4-591C20.9 | -0,4757 | 8,77928E-22 |
| ACE2 | RP11-29H23.4 | -0,4758 | 8,60869E-22 |
| ACE2 | RP11-179H18.8 | -0,4758 | 8,57738E-22 |
| ACE2 | TPTE2 | -0,4758 | 8,61075E-22 |
| ACE2 | FAM92B | -0,4758 | 8,62595E-22 |
| ACE2 | TIMM13 | -0,4758 | 8,64848E-22 |
| ACE2 | LINC01070 | -0,4759 | 8,30015E-22 |
| ACE2 | GUCY2F | -0,4759 | 8,46817E-22 |
| ACE2 | RP11-397C12.1 | -0,476 | 8,2588E-22 |
| ACE2 | AC007875.2 | -0,476 | 8,23621E-22 |
| ACE2 | CTC-301O7.4 | -0,476 | 8,24344E-22 |
| ACE2 | RP5-858B6.3 | -0,4761 | 7,964E-22 |
| ACE2 | MYEOV | -0,4761 | 7,98792E-22 |
| ACE2 | RP11-361H10.3 | -0,4761 | 8,00481E-22 |
| ACE2 | RP11-127O4.2 | -0,4762 | 7,85342E-22 |
| ACE2 | CNGA4 | -0,4762 | 7,82657E-22 |
| ACE2 | ZFAND4 | -0,4763 | 7,64081E-22 |
| ACE2 | AGAP12P | -0,4763 | 7,64162E-22 |
| ACE2 | ZAR1 | -0,4764 | 7,51198E-22 |
| ACE2 | RP11-506N2.1 | -0,4764 | 7,49473E-22 |
| ACE2 | NRG1-IT3 | -0,4764 | 7,44363E-22 |
| ACE2 | RP1-302D9.1 | -0,4764 | 7,54589E-22 |
| ACE2 | SH3D21 | -0,4765 | 7,39343E-22 |
| ACE2 | CCDC121 | -0,4765 | 7,3186E-22 |

| | | | |
|---|---|---|---|
| ACE2 | AC013727.1 | -0,4765 | 7,25869E-22 |
| ACE2 | PDE1A | -0,4765 | 7,2748E-22 |
| ACE2 | AC006960.7 | -0,4765 | 7,2654E-22 |
| ACE2 | RP5-884M6.1 | -0,4765 | 7,29683E-22 |
| ACE2 | LINC01598 | -0,4765 | 7,30802E-22 |
| ACE2 | ZKSCAN4 | -0,4766 | 7,09274E-22 |
| ACE2 | RP11-472N13.2 | -0,4766 | 7,2449E-22 |
| ACE2 | SLC25A19 | -0,4766 | 7,21196E-22 |
| ACE2 | RP11-122M14.1 | -0,4767 | 6,96579E-22 |
| ACE2 | RP11-463D19.1 | -0,4767 | 7,03975E-22 |
| ACE2 | ODF3 | -0,4767 | 6,95428E-22 |
| ACE2 | C11orf16 | -0,4767 | 6,93871E-22 |
| ACE2 | CTB-41I6.1 | -0,4767 | 6,94471E-22 |
| ACE2 | TECR | -0,4767 | 7,03793E-22 |
| ACE2 | AP001048.4 | -0,4767 | 7,01897E-22 |
| ACE2 | MYADML | -0,4768 | 6,90553E-22 |
| ACE2 | LINC00838 | -0,4768 | 6,77803E-22 |
| ACE2 | CCDC65 | -0,4768 | 6,88349E-22 |
| ACE2 | RP11-817O13.8 | -0,4768 | 6,84508E-22 |
| ACE2 | METTL6 | -0,477 | 6,59741E-22 |
| ACE2 | RP1-111D6.3 | -0,477 | 6,51116E-22 |
| ACE2 | GBX1 | -0,477 | 6,50171E-22 |
| ACE2 | RP11-521O16.2 | -0,4771 | 6,39383E-22 |
| ACE2 | RP11-172C16.5 | -0,4771 | 6,46712E-22 |
| ACE2 | RP4-734C18.1 | -0,4771 | 6,3654E-22 |
| ACE2 | HTR1D | -0,4772 | 6,29026E-22 |
| ACE2 | C8B | -0,4772 | 6,21963E-22 |
| ACE2 | TMPRSS7 | -0,4772 | 6,23034E-22 |
| ACE2 | RP11-423H2.5 | -0,4772 | 6,28549E-22 |
| ACE2 | C19orf18 | -0,4772 | 6,2728E-22 |
| ACE2 | DISC1 | -0,4773 | 6,16082E-22 |
| ACE2 | RP11-36N20.1 | -0,4773 | 6,14455E-22 |
| ACE2 | RBMXL2 | -0,4773 | 6,12077E-22 |
| ACE2 | CDADC1 | -0,4773 | 6,17812E-22 |
| ACE2 | ACOXL | -0,4774 | 5,96269E-22 |
| ACE2 | PPP2R2B-IT1 | -0,4774 | 6,04407E-22 |
| ACE2 | CTD-3247F14.2 | -0,4774 | 5,97133E-22 |
| ACE2 | SCGB1D4 | -0,4774 | 6,00148E-22 |
| ACE2 | RP11-76E17.2 | -0,4774 | 6,04202E-22 |
| ACE2 | KCNJ18 | -0,4774 | 6,04334E-22 |
| ACE2 | RP11-61L19.2 | -0,4774 | 6,04881E-22 |
| ACE2 | RNF144A-AS1 | -0,4775 | 5,85109E-22 |
| ACE2 | RP11-73B2.6 | -0,4775 | 5,82371E-22 |
| ACE2 | RP11-489E7.4 | -0,4775 | 5,81942E-22 |
| ACE2 | IARS2P1 | -0,4775 | 5,8729E-22 |
| ACE2 | RAB3IL1 | -0,4775 | 5,81196E-22 |
| ACE2 | RP11-418I22.3 | -0,4775 | 5,79642E-22 |
| ACE2 | RP11-204E4.3 | -0,4775 | 5,79658E-22 |
| ACE2 | ANO7L1 | -0,4776 | 5,70923E-22 |
| ACE2 | RP11-467P9.1 | -0,4776 | 5,79154E-22 |
| ACE2 | CTD-2210P24.3 | -0,4776 | 5,68687E-22 |
| ACE2 | CASC1 | -0,4776 | 5,78455E-22 |
| ACE2 | RP11-348H3.2 | -0,4777 | 5,60414E-22 |
| ACE2 | LINC01499 | -0,4777 | 5,65942E-22 |

| | | | |
|---|---|---|---|
| ACE2 | RP11-697E22.3 | -0,4777 | 5,54442E-22 |
| ACE2 | RP11-378J18.9 | -0,4778 | 5,42869E-22 |
| ACE2 | C3orf20 | -0,4778 | 5,47605E-22 |
| ACE2 | RP11-748C4.1 | -0,4778 | 5,52927E-22 |
| ACE2 | RP11-1007G5.2 | -0,4778 | 5,44504E-22 |
| ACE2 | DHX40P1 | -0,4778 | 5,52139E-22 |
| ACE2 | AC007386.2 | -0,4779 | 5,39243E-22 |
| ACE2 | LINC01964 | -0,4779 | 5,35437E-22 |
| ACE2 | RP11-650J17.2 | -0,4779 | 5,36544E-22 |
| ACE2 | RP11-548H18.2 | -0,4779 | 5,37868E-22 |
| ACE2 | CTB-167B5.1 | -0,478 | 5,28231E-22 |
| ACE2 | RP11-539E19.2 | -0,478 | 5,21169E-22 |
| ACE2 | RP11-484D2.2 | -0,478 | 5,21605E-22 |
| ACE2 | RP11-187E13.1 | -0,478 | 5,21472E-22 |
| ACE2 | CCNB2 | -0,478 | 5,26088E-22 |
| ACE2 | LINC00305 | -0,478 | 5,27181E-22 |
| ACE2 | OR7E25P | -0,478 | 5,19831E-22 |
| ACE2 | DNAJC21 | -0,4781 | 5,16481E-22 |
| ACE2 | RP11-91K8.4 | -0,4782 | 4,98443E-22 |
| ACE2 | GRID2 | -0,4782 | 5,05031E-22 |
| ACE2 | PMCHL1 | -0,4782 | 4,96238E-22 |
| ACE2 | RP11-344L13.2 | -0,4782 | 4,98583E-22 |
| ACE2 | SYCE1 | -0,4782 | 5,032E-22 |
| ACE2 | ABC14-1188822O7.1 | -0,4783 | 4,93909E-22 |
| ACE2 | SPATA16 | -0,4783 | 4,92653E-22 |
| ACE2 | LINC01513 | -0,4783 | 4,93022E-22 |
| ACE2 | RP11-15B24.5 | -0,4783 | 4,87148E-22 |
| ACE2 | STAMBPL1 | -0,4783 | 4,89441E-22 |
| ACE2 | RP11-702B10.2 | -0,4783 | 4,93811E-22 |
| ACE2 | WDR7-OT1 | -0,4783 | 4,88772E-22 |
| ACE2 | NPHP1 | -0,4784 | 4,80788E-22 |
| ACE2 | AC009506.1 | -0,4784 | 4,83314E-22 |
| ACE2 | LHFPL3-AS2 | -0,4784 | 4,76757E-22 |
| ACE2 | ERICH6 | -0,4785 | 4,63917E-22 |
| ACE2 | PABPC3 | -0,4785 | 4,70573E-22 |
| ACE2 | RP11-466A19.3 | -0,4785 | 4,63276E-22 |
| ACE2 | IGF2BP2 | -0,4786 | 4,55746E-22 |
| ACE2 | RP3-388N13.3 | -0,4786 | 4,5853E-22 |
| ACE2 | RP11-778O17.4 | -0,4786 | 4,59223E-22 |
| ACE2 | CYTH2 | -0,4786 | 4,59292E-22 |
| ACE2 | AC019048.1 | -0,4787 | 4,49748E-22 |
| ACE2 | PPP1R14BP2 | -0,4787 | 4,46915E-22 |
| ACE2 | RP11-617I14.1 | -0,4787 | 4,52179E-22 |
| ACE2 | RP11-503L19.1 | -0,4787 | 4,44032E-22 |
| ACE2 | AC063976.3 | -0,4787 | 4,50413E-22 |
| ACE2 | ZSCAN31 | -0,4787 | 4,52383E-22 |
| ACE2 | CCDC83 | -0,4787 | 4,44548E-22 |
| ACE2 | LLNLR-271C9.1 | -0,4787 | 4,52245E-22 |
| ACE2 | RP11-65D24.2 | -0,4788 | 4,39444E-22 |
| ACE2 | RN7SKP108 | -0,4788 | 4,34306E-22 |
| ACE2 | CDC25C | -0,4789 | 4,28426E-22 |
| ACE2 | PPP1R11 | -0,4789 | 4,24949E-22 |
| ACE2 | RP11-219E7.2 | -0,4789 | 4,32323E-22 |
| ACE2 | RP11-313A24.1 | -0,479 | 4,2098E-22 |

| | | | |
|---|---|---|---|
| ACE2 | DLGAP2-AS1 | -0,479 | 4,22536E-22 |
| ACE2 | CBX3P4 | -0,479 | 4,21563E-22 |
| ACE2 | DEPDC7 | -0,4791 | 4,1149E-22 |
| ACE2 | RP11-707A18.1 | -0,4792 | 4,02996E-22 |
| ACE2 | RP11-133K1.8 | -0,4792 | 4,02589E-22 |
| ACE2 | PBX4 | -0,4792 | 4,02543E-22 |
| ACE2 | LINC01934 | -0,4793 | 3,89669E-22 |
| ACE2 | RP11-126H7.4 | -0,4793 | 3,91542E-22 |
| ACE2 | RP4-799G3.2 | -0,4794 | 3,86189E-22 |
| ACE2 | TTLL5 | -0,4794 | 3,83676E-22 |
| ACE2 | DCLK3 | -0,4795 | 3,7626E-22 |
| ACE2 | KCCAT333 | -0,4795 | 3,73193E-22 |
| ACE2 | RP11-112J1.3 | -0,4795 | 3,71094E-22 |
| ACE2 | CPHXL | -0,4795 | 3,70586E-22 |
| ACE2 | C12orf56 | -0,4796 | 3,64362E-22 |
| ACE2 | UBE2N | -0,4796 | 3,67786E-22 |
| ACE2 | SERTM1 | -0,4796 | 3,67827E-22 |
| ACE2 | CCDC61 | -0,4796 | 3,69349E-22 |
| ACE2 | CTD-2186M15.3 | -0,4797 | 3,58091E-22 |
| ACE2 | RP11-286H14.4 | -0,4797 | 3,59503E-22 |
| ACE2 | OR7E26P | -0,4797 | 3,57339E-22 |
| ACE2 | RP11-517P14.7 | -0,4797 | 3,58174E-22 |
| ACE2 | SPDYE22P | -0,4797 | 3,5423E-22 |
| ACE2 | SVIP | -0,4798 | 3,47049E-22 |
| ACE2 | RPS6P21 | -0,4798 | 3,506E-22 |
| ACE2 | GOLGA8UP | -0,4798 | 3,52369E-22 |
| ACE2 | AMZ2P1 | -0,4798 | 3,45763E-22 |
| ACE2 | AGBL5 | -0,4799 | 3,43403E-22 |
| ACE2 | HK1 | -0,4799 | 3,43164E-22 |
| ACE2 | RP11-500M8.4 | -0,4799 | 3,438E-22 |
| ACE2 | DZIP1 | -0,4799 | 3,39167E-22 |
| ACE2 | CCDC134 | -0,4799 | 3,40648E-22 |
| ACE2 | CLEC20A | -0,48 | 3,35679E-22 |
| ACE2 | LINC01391 | -0,48 | 3,30662E-22 |
| ACE2 | SPTLC1P1 | -0,48 | 3,30261E-22 |
| ACE2 | RP11-702F3.4 | -0,48 | 3,36357E-22 |
| ACE2 | S100P | -0,4801 | 3,23536E-22 |
| ACE2 | C6orf118 | -0,4801 | 3,25143E-22 |
| ACE2 | RP1-266L20.2 | -0,4801 | 3,25857E-22 |
| ACE2 | RP11-383M4.6 | -0,4801 | 3,23557E-22 |
| ACE2 | RP11-20D14.3 | -0,4801 | 3,29607E-22 |
| ACE2 | NUCB1-AS1 | -0,4801 | 3,24794E-22 |
| ACE2 | AC006116.17 | -0,4801 | 3,26043E-22 |
| ACE2 | RP11-430C7.2 | -0,4802 | 3,2158E-22 |
| ACE2 | CPS1 | -0,4802 | 3,20128E-22 |
| ACE2 | HIST1H2APS5 | -0,4802 | 3,19971E-22 |
| ACE2 | MEX3B | -0,4802 | 3,19558E-22 |
| ACE2 | RP11-44L9.2 | -0,4802 | 3,1595E-22 |
| ACE2 | LAIR1 | -0,4802 | 3,20166E-22 |
| ACE2 | RP5-821D11.7 | -0,4802 | 3,18828E-22 |
| ACE2 | LMNB1 | -0,4803 | 3,15255E-22 |
| ACE2 | ZNF124 | -0,4804 | 3,03657E-22 |
| ACE2 | RP11-115J16.3 | -0,4804 | 3,02123E-22 |
| ACE2 | RP11-33N16.2 | -0,4804 | 3,06973E-22 |

| | | | |
|---|---|---|---|
| ACE2 | FSIP1 | -0,4804 | 3,03344E-22 |
| ACE2 | GOLGA6L10 | -0,4804 | 3,01694E-22 |
| ACE2 | SIGLECL1 | -0,4805 | 3,01307E-22 |
| ACE2 | RP5-872K7.7 | -0,4805 | 2,96301E-22 |
| ACE2 | CHCHD2P6 | -0,4806 | 2,88425E-22 |
| ACE2 | CLVS1 | -0,4806 | 2,93419E-22 |
| ACE2 | RP11-420O16.1 | -0,4807 | 2,82176E-22 |
| ACE2 | PRSS51 | -0,4807 | 2,85619E-22 |
| ACE2 | PTPRT | -0,4807 | 2,87486E-22 |
| ACE2 | RP5-968D22.3 | -0,4808 | 2,78242E-22 |
| ACE2 | RP11-425A23.1 | -0,4808 | 2,77692E-22 |
| ACE2 | RP11-124O11.1 | -0,4808 | 2,77863E-22 |
| ACE2 | RMST | -0,4808 | 2,75662E-22 |
| ACE2 | RP11-572C21.1 | -0,4808 | 2,77708E-22 |
| ACE2 | RP11-517O13.3 | -0,4808 | 2,76197E-22 |
| ACE2 | RP11-62G11.2 | -0,4809 | 2,71787E-22 |
| ACE2 | C14orf79 | -0,4809 | 2,72147E-22 |
| ACE2 | ST8SIA2 | -0,4809 | 2,7234E-22 |
| ACE2 | DLG4 | -0,4809 | 2,74696E-22 |
| ACE2 | LINC01953 | -0,481 | 2,65154E-22 |
| ACE2 | SPATA9 | -0,481 | 2,66457E-22 |
| ACE2 | UBR7 | -0,481 | 2,64367E-22 |
| ACE2 | OR7D2 | -0,481 | 2,63483E-22 |
| ACE2 | RRNAD1 | -0,4811 | 2,58801E-22 |
| ACE2 | CNTN4-AS1 | -0,4811 | 2,63185E-22 |
| ACE2 | RP11-103H7.3 | -0,4811 | 2,57758E-22 |
| ACE2 | CWC25 | -0,4811 | 2,59317E-22 |
| ACE2 | AC004535.2 | -0,4812 | 2,57181E-22 |
| ACE2 | HMBOX1 | -0,4812 | 2,5336E-22 |
| ACE2 | VRTN | -0,4812 | 2,53874E-22 |
| ACE2 | RP11-272D12.1 | -0,4812 | 2,52605E-22 |
| ACE2 | RP5-1121H13.3 | -0,4812 | 2,54392E-22 |
| ACE2 | CCDC30 | -0,4813 | 2,48795E-22 |
| ACE2 | LINC01958 | -0,4813 | 2,48301E-22 |
| ACE2 | OR5E1P | -0,4813 | 2,50285E-22 |
| ACE2 | SLC25A1P1 | -0,4813 | 2,47921E-22 |
| ACE2 | RP11-37C7.2 | -0,4813 | 2,48662E-22 |
| ACE2 | RP11-293E1.1 | -0,4813 | 2,50952E-22 |
| ACE2 | RP11-202G11.2 | -0,4814 | 2,44194E-22 |
| ACE2 | C21orf59 | -0,4814 | 2,43068E-22 |
| ACE2 | CYP2J2 | -0,4815 | 2,365E-22 |
| ACE2 | SPATA1 | -0,4815 | 2,35577E-22 |
| ACE2 | RP11-540E16.2 | -0,4815 | 2,3768E-22 |
| ACE2 | AC083862.6 | -0,4815 | 2,3526E-22 |
| ACE2 | IFITM8P | -0,4815 | 2,37857E-22 |
| ACE2 | LINC02098 | -0,4815 | 2,35504E-22 |
| ACE2 | OTOA | -0,4815 | 2,39628E-22 |
| ACE2 | RP11-805I24.1 | -0,4815 | 2,39862E-22 |
| ACE2 | ANKRD54P1 | -0,4816 | 2,33479E-22 |
| ACE2 | RP11-126O1.4 | -0,4816 | 2,33938E-22 |
| ACE2 | ATP1A3 | -0,4816 | 2,31654E-22 |
| ACE2 | DRG1 | -0,4816 | 2,31475E-22 |
| ACE2 | RP11-165F24.3 | -0,4817 | 2,25191E-22 |
| ACE2 | FAM95B1 | -0,4817 | 2,26464E-22 |

| | | | |
|---|---|---|---|
| ACE2 | LINC01493 | -0,4817 | 2,28159E-22 |
| ACE2 | CCNA1 | -0,4817 | 2,28112E-22 |
| ACE2 | MIPOL1 | -0,4817 | 2,27816E-22 |
| ACE2 | RP11-566K19.9 | -0,4817 | 2,27827E-22 |
| ACE2 | RP11-193H5.1 | -0,4818 | 2,23113E-22 |
| ACE2 | RP1-155D22.1 | -0,4818 | 2,20571E-22 |
| ACE2 | RP11-124G5.3 | -0,4818 | 2,22474E-22 |
| ACE2 | RP11-16B9.1 | -0,4818 | 2,21854E-22 |
| ACE2 | RP1-159A19.4 | -0,4819 | 2,16813E-22 |
| ACE2 | THPO | -0,4819 | 2,17019E-22 |
| ACE2 | RP11-350F16.1 | -0,4819 | 2,15278E-22 |
| ACE2 | FRMPD2B | -0,4819 | 2,18935E-22 |
| ACE2 | RP11-407A16.3 | -0,4819 | 2,18578E-22 |
| ACE2 | RP11-226P1.2 | -0,4819 | 2,19351E-22 |
| ACE2 | LINC02207 | -0,4819 | 2,17016E-22 |
| ACE2 | TFAP2B | -0,482 | 2,13658E-22 |
| ACE2 | CLMN | -0,482 | 2,12806E-22 |
| ACE2 | RP11-157B13.7 | -0,482 | 2,1451E-22 |
| ACE2 | MEP1A | -0,4821 | 2,07339E-22 |
| ACE2 | SPRR4 | -0,4822 | 2,01207E-22 |
| ACE2 | AC012462.3 | -0,4822 | 2,02162E-22 |
| ACE2 | AC091633.2 | -0,4822 | 2,04609E-22 |
| ACE2 | RP11-292B1.2 | -0,4822 | 2,00666E-22 |
| ACE2 | CDKN2A-AS1 | -0,4822 | 2,02013E-22 |
| ACE2 | NTRK3 | -0,4822 | 2,01509E-22 |
| ACE2 | KRT16P2 | -0,4822 | 2,04424E-22 |
| ACE2 | RP1-41P2.7 | -0,4822 | 2,023E-22 |
| ACE2 | AC091770.3 | -0,4823 | 1,99321E-22 |
| ACE2 | PTPRQ | -0,4823 | 1,99304E-22 |
| ACE2 | DICER1-AS1 | -0,4823 | 1,96243E-22 |
| ACE2 | MIR181C | -0,4823 | 2,00371E-22 |
| ACE2 | RP11-598C10.2 | -0,4824 | 1,92866E-22 |
| ACE2 | OR10D3 | -0,4824 | 1,95386E-22 |
| ACE2 | CTD-2135D7.2 | -0,4824 | 1,93485E-22 |
| ACE2 | TBC1D3F | -0,4824 | 1,93777E-22 |
| ACE2 | RP11-29H23.6 | -0,4825 | 1,90346E-22 |
| ACE2 | CCER1 | -0,4825 | 1,89316E-22 |
| ACE2 | AC008753.6 | -0,4825 | 1,87404E-22 |
| ACE2 | RP11-483F11.7 | -0,4826 | 1,84774E-22 |
| ACE2 | RP5-823G15.5 | -0,4826 | 1,83349E-22 |
| ACE2 | RP11-154F14.3 | -0,4827 | 1,82571E-22 |
| ACE2 | TRBV9 | -0,4827 | 1,80708E-22 |
| ACE2 | RPL7P59 | -0,4827 | 1,8064E-22 |
| ACE2 | MYO5BP1 | -0,4827 | 1,81885E-22 |
| ACE2 | RP11-69H14.6 | -0,4827 | 1,78992E-22 |
| ACE2 | RP11-691H4.4 | -0,4827 | 1,79303E-22 |
| ACE2 | RP11-680N20.1 | -0,4827 | 1,80096E-22 |
| ACE2 | DLX6 | -0,4828 | 1,78552E-22 |
| ACE2 | CTD-3023L14.2 | -0,4828 | 1,7846E-22 |
| ACE2 | LINC01874 | -0,4829 | 1,74294E-22 |
| ACE2 | LIN28B-AS1 | -0,4829 | 1,7437E-22 |
| ACE2 | HDAC1P1 | -0,4829 | 1,72866E-22 |
| ACE2 | AC008268.1 | -0,483 | 1,67587E-22 |
| ACE2 | RP11-613D13.4 | -0,483 | 1,68325E-22 |

| | | | |
|---|---|---|---|
| ACE2 | RGCC | -0,483 | 1,68158E-22 |
| ACE2 | MESP1 | -0,483 | 1,68483E-22 |
| ACE2 | AC010746.3 | -0,4832 | 1,59869E-22 |
| ACE2 | RP11-342M3.1 | -0,4832 | 1,6065E-22 |
| ACE2 | AP006216.5 | -0,4832 | 1,60238E-22 |
| ACE2 | RP11-452D21.1 | -0,4832 | 1,62115E-22 |
| ACE2 | PPP6R2 | -0,4832 | 1,63094E-22 |
| ACE2 | SLC9A1 | -0,4833 | 1,57132E-22 |
| ACE2 | RP11-60L3.2 | -0,4833 | 1,56014E-22 |
| ACE2 | GNG12-AS1 | -0,4834 | 1,54005E-22 |
| ACE2 | RP11-367N14.2 | -0,4834 | 1,53864E-22 |
| ACE2 | RP11-542F9.1 | -0,4834 | 1,55655E-22 |
| ACE2 | UBXN8 | -0,4834 | 1,54603E-22 |
| ACE2 | CTD-2501E16.2 | -0,4834 | 1,54335E-22 |
| ACE2 | LINC02092 | -0,4834 | 1,53271E-22 |
| ACE2 | REG4 | -0,4835 | 1,51E-22 |
| ACE2 | RP11-158I3.3 | -0,4835 | 1,50076E-22 |
| ACE2 | AC002401.1 | -0,4835 | 1,49718E-22 |
| ACE2 | ZNF565 | -0,4835 | 1,50864E-22 |
| ACE2 | RP11-469L4.1 | -0,4836 | 1,45792E-22 |
| ACE2 | RP11-566H8.3 | -0,4836 | 1,4727E-22 |
| ACE2 | TRBV7-3 | -0,4837 | 1,42425E-22 |
| ACE2 | RP11-151A6.4 | -0,4837 | 1,45295E-22 |
| ACE2 | OR7E106P | -0,4837 | 1,44875E-22 |
| ACE2 | CDK5RAP3 | -0,4837 | 1,42375E-22 |
| ACE2 | RP1-317E23.3 | -0,4838 | 1,39156E-22 |
| ACE2 | SNAP47-AS1 | -0,4838 | 1,4027E-22 |
| ACE2 | RP11-467K18.2 | -0,4838 | 1,40613E-22 |
| ACE2 | RP11-547D23.1 | -0,4838 | 1,40497E-22 |
| ACE2 | RP11-993B23.3 | -0,4839 | 1,38632E-22 |
| ACE2 | RP11-119B16.2 | -0,4839 | 1,38943E-22 |
| ACE2 | LINC01643 | -0,4839 | 1,37359E-22 |
| ACE2 | TBC1D22A-AS1 | -0,4839 | 1,37544E-22 |
| ACE2 | RP5-1024G6.5 | -0,484 | 1,3447E-22 |
| ACE2 | CTC-264O10.2 | -0,484 | 1,33912E-22 |
| ACE2 | RP11-360A18.1 | -0,484 | 1,34069E-22 |
| ACE2 | RPLP0P2 | -0,484 | 1,34577E-22 |
| ACE2 | RP13-379L11.2 | -0,484 | 1,35648E-22 |
| ACE2 | PRELID3BP7 | -0,4841 | 1,32785E-22 |
| ACE2 | RP11-417L14.1 | -0,4841 | 1,31042E-22 |
| ACE2 | RP11-484D2.5 | -0,4841 | 1,31147E-22 |
| ACE2 | CWC15 | -0,4841 | 1,3194E-22 |
| ACE2 | EEF2KMT | -0,4841 | 1,30063E-22 |
| ACE2 | C21orf58 | -0,4841 | 1,32127E-22 |
| ACE2 | RP1-67A8.3 | -0,4842 | 1,27259E-22 |
| ACE2 | LINC01450 | -0,4842 | 1,29163E-22 |
| ACE2 | RP11-192H23.7 | -0,4842 | 1,28497E-22 |
| ACE2 | RP11-705O1.8 | -0,4842 | 1,28121E-22 |
| ACE2 | LINC01347 | -0,4843 | 1,24324E-22 |
| ACE2 | RP11-65D24.1 | -0,4843 | 1,26175E-22 |
| ACE2 | RP3-449M8.6 | -0,4843 | 1,24959E-22 |
| ACE2 | CTD-2071N1.1 | -0,4843 | 1,25646E-22 |
| ACE2 | LRRC28 | -0,4843 | 1,24689E-22 |
| ACE2 | RP11-302L19.1 | -0,4844 | 1,22864E-22 |

| | | | |
|---|---|---|---|
| ACE2 | TTC6 | -0,4844 | 1,2349E-22 |
| ACE2 | ENKD1 | -0,4844 | 1,22718E-22 |
| ACE2 | BCL2L11 | -0,4845 | 1,20237E-22 |
| ACE2 | LINC01937 | -0,4845 | 1,20158E-22 |
| ACE2 | CCDC54 | -0,4845 | 1,20967E-22 |
| ACE2 | RP11-73B2.4 | -0,4845 | 1,20376E-22 |
| ACE2 | BRAP | -0,4845 | 1,20225E-22 |
| ACE2 | RP11-210M15.1 | -0,4845 | 1,20095E-22 |
| ACE2 | LRRC7 | -0,4846 | 1,15806E-22 |
| ACE2 | PPM1M | -0,4846 | 1,17296E-22 |
| ACE2 | THAP9 | -0,4846 | 1,17225E-22 |
| ACE2 | RP11-351M16.3 | -0,4846 | 1,16184E-22 |
| ACE2 | CTB-147C22.8 | -0,4846 | 1,1829E-22 |
| ACE2 | GDAP1L1 | -0,4846 | 1,16224E-22 |
| ACE2 | NKAPL | -0,4847 | 1,15594E-22 |
| ACE2 | RP11-153K11.3 | -0,4847 | 1,14119E-22 |
| ACE2 | RP11-408J6.1 | -0,4847 | 1,15306E-22 |
| ACE2 | SLC26A4 | -0,4848 | 1,11064E-22 |
| ACE2 | CCDC81 | -0,4848 | 1,1162E-22 |
| ACE2 | DERL2 | -0,4848 | 1,11736E-22 |
| ACE2 | LINC01823 | -0,4849 | 1,08476E-22 |
| ACE2 | AC009495.2 | -0,4849 | 1,09234E-22 |
| ACE2 | RP4-753M9.1 | -0,4849 | 1,09191E-22 |
| ACE2 | LINC01364 | -0,485 | 1,07092E-22 |
| ACE2 | CTB-178M22.2 | -0,485 | 1,07929E-22 |
| ACE2 | LINC00929 | -0,485 | 1,07747E-22 |
| ACE2 | MTFR1 | -0,4851 | 1,03384E-22 |
| ACE2 | LLNLF-173C4.1 | -0,4851 | 1,03383E-22 |
| ACE2 | DUS3L | -0,4851 | 1,05016E-22 |
| ACE2 | SELENOV | -0,4851 | 1,04255E-22 |
| ACE2 | RP11-180O5.2 | -0,4852 | 1,01472E-22 |
| ACE2 | VDAC2P3 | -0,4852 | 1,0323E-22 |
| ACE2 | RP11-299J3.8 | -0,4852 | 1,02963E-22 |
| ACE2 | RP11-696N14.1 | -0,4852 | 1,0258E-22 |
| ACE2 | CTD-3088G3.6 | -0,4852 | 1,02575E-22 |
| ACE2 | RP1-158P9.1 | -0,4853 | 9,93525E-23 |
| ACE2 | ATP2B2-IT1 | -0,4853 | 9,88212E-23 |
| ACE2 | ATXN7L1 | -0,4853 | 1,00644E-22 |
| ACE2 | CTD-2023N9.1 | -0,4854 | 9,64435E-23 |
| ACE2 | FANK1-AS1 | -0,4854 | 9,8313E-23 |
| ACE2 | MIR762HG | -0,4854 | 9,68353E-23 |
| ACE2 | AF096876.1 | -0,4854 | 9,69251E-23 |
| ACE2 | CCL20 | -0,4855 | 9,60888E-23 |
| ACE2 | C12orf54 | -0,4855 | 9,5071E-23 |
| ACE2 | CATSPER2 | -0,4855 | 9,43258E-23 |
| ACE2 | RP4-635E18.6 | -0,4856 | 9,34399E-23 |
| ACE2 | CALR4P | -0,4856 | 9,26166E-23 |
| ACE2 | KRTCAP2 | -0,4856 | 9,39292E-23 |
| ACE2 | RP5-832C2.5 | -0,4857 | 9,07902E-23 |
| ACE2 | PKLR | -0,4857 | 9,04974E-23 |
| ACE2 | CTD-2247C11.1 | -0,4857 | 9,04655E-23 |
| ACE2 | SPAM1 | -0,4857 | 9,08403E-23 |
| ACE2 | POLB | -0,4857 | 9,11812E-23 |
| ACE2 | FAM225B | -0,4857 | 9,02546E-23 |

| | | | |
|---|---|---|---|
| ACE2 | RP11-897M7.1 | -0,4857 | 9,17276E-23 |
| ACE2 | PFDN4 | -0,4857 | 9,06569E-23 |
| ACE2 | RP3-448I9.2 | -0,4858 | 8,83921E-23 |
| ACE2 | POT1-AS1 | -0,4858 | 8,90187E-23 |
| ACE2 | C8orf48 | -0,4858 | 8,82575E-23 |
| ACE2 | RGS22 | -0,4858 | 8,94257E-23 |
| ACE2 | PDZD8 | -0,4858 | 8,86838E-23 |
| ACE2 | RAPSN | -0,4858 | 8,90362E-23 |
| ACE2 | NDRG3 | -0,4858 | 8,92269E-23 |
| ACE2 | PADI1 | -0,4859 | 8,74139E-23 |
| ACE2 | MDS2 | -0,4859 | 8,78523E-23 |
| ACE2 | HOXA3 | -0,4859 | 8,65319E-23 |
| ACE2 | RP11-359E19.1 | -0,4859 | 8,64224E-23 |
| ACE2 | TRIM14 | -0,4859 | 8,63387E-23 |
| ACE2 | SLC3A1 | -0,486 | 8,53474E-23 |
| ACE2 | AC013444.1 | -0,486 | 8,57662E-23 |
| ACE2 | AADACL2-AS1 | -0,486 | 8,57775E-23 |
| ACE2 | ZBBX | -0,486 | 8,58089E-23 |
| ACE2 | ATP23 | -0,486 | 8,57685E-23 |
| ACE2 | RP11-856M7.1 | -0,486 | 8,54765E-23 |
| ACE2 | TPGS2 | -0,486 | 8,50595E-23 |
| ACE2 | GJA9 | -0,4861 | 8,23835E-23 |
| ACE2 | LINC02013 | -0,4861 | 8,37344E-23 |
| ACE2 | SCOC-AS1 | -0,4861 | 8,28498E-23 |
| ACE2 | RP11-203B7.2 | -0,4861 | 8,36612E-23 |
| ACE2 | RP11-613E4.4 | -0,4861 | 8,34339E-23 |
| ACE2 | AC022153.1 | -0,4861 | 8,22038E-23 |
| ACE2 | RPSAP10 | -0,4862 | 8,03027E-23 |
| ACE2 | INSC | -0,4862 | 8,09725E-23 |
| ACE2 | SAXO2 | -0,4862 | 8,03503E-23 |
| ACE2 | HTR5BP | -0,4863 | 7,87943E-23 |
| ACE2 | TTC29 | -0,4863 | 7,96034E-23 |
| ACE2 | RP11-327O17.2 | -0,4863 | 7,91833E-23 |
| ACE2 | RP11-712L6.5 | -0,4863 | 7,89254E-23 |
| ACE2 | NTRK3-AS1 | -0,4863 | 7,83604E-23 |
| ACE2 | NDNF | -0,4864 | 7,81475E-23 |
| ACE2 | LINC01742 | -0,4864 | 7,73692E-23 |
| ACE2 | AC105398.3 | -0,4865 | 7,58282E-23 |
| ACE2 | PFDN6 | -0,4865 | 7,48339E-23 |
| ACE2 | RP11-27M9.1 | -0,4865 | 7,54032E-23 |
| ACE2 | RP11-367J7.3 | -0,4866 | 7,32766E-23 |
| ACE2 | RP11-5O17.1 | -0,4866 | 7,44258E-23 |
| ACE2 | CTD-2116N20.1 | -0,4866 | 7,33831E-23 |
| ACE2 | RP11-945A11.2 | -0,4866 | 7,4632E-23 |
| ACE2 | LINC00443 | -0,4866 | 7,32601E-23 |
| ACE2 | LINC00368 | -0,4866 | 7,34729E-23 |
| ACE2 | RPL23AP8 | -0,4866 | 7,36916E-23 |
| ACE2 | GABRA1 | -0,4867 | 7,30285E-23 |
| ACE2 | RP11-407N17.4 | -0,4867 | 7,28497E-23 |
| ACE2 | NOS2P4 | -0,4867 | 7,2666E-23 |
| ACE2 | RP11-231L11.1 | -0,4868 | 7,05039E-23 |
| ACE2 | RP11-284A20.2 | -0,4868 | 7,10136E-23 |
| ACE2 | CTC-471J1.11 | -0,4868 | 7,09137E-23 |
| ACE2 | RP11-171A24.2 | -0,4869 | 6,83551E-23 |

| | | | |
|---|---|---|---|
| ACE2 | ACSL6 | -0,487 | 6,68813E-23 |
| ACE2 | HDGFL1 | -0,487 | 6,81332E-23 |
| ACE2 | PINX1 | -0,487 | 6,7729E-23 |
| ACE2 | RP11-379L18.1 | -0,487 | 6,78185E-23 |
| ACE2 | PIAS4 | -0,487 | 6,73443E-23 |
| ACE2 | ZNF653 | -0,487 | 6,69455E-23 |
| ACE2 | CASP14 | -0,487 | 6,81427E-23 |
| ACE2 | LAMTOR5-AS1 | -0,4871 | 6,56805E-23 |
| ACE2 | LINC01142 | -0,4871 | 6,60262E-23 |
| ACE2 | C4BPB | -0,4871 | 6,52773E-23 |
| ACE2 | AC092839.3 | -0,4871 | 6,53278E-23 |
| ACE2 | TMEM225 | -0,4871 | 6,54633E-23 |
| ACE2 | SLC9A3R1 | -0,4871 | 6,62906E-23 |
| ACE2 | SIRT7 | -0,4871 | 6,60219E-23 |
| ACE2 | ZNF385D-AS2 | -0,4872 | 6,40567E-23 |
| ACE2 | RP11-215P8.1 | -0,4872 | 6,46058E-23 |
| ACE2 | MED7 | -0,4872 | 6,43291E-23 |
| ACE2 | AP000593.6 | -0,4872 | 6,36718E-23 |
| ACE2 | CCDC150P1 | -0,4872 | 6,45593E-23 |
| ACE2 | DNAH17 | -0,4872 | 6,38032E-23 |
| ACE2 | TPT1P5 | -0,4873 | 6,34307E-23 |
| ACE2 | SNX18P2 | -0,4873 | 6,28137E-23 |
| ACE2 | RP11-332M4.1 | -0,4873 | 6,27843E-23 |
| ACE2 | ARMC4 | -0,4873 | 6,2219E-23 |
| ACE2 | ABHD2 | -0,4873 | 6,26651E-23 |
| ACE2 | AC008079.9 | -0,4873 | 6,26258E-23 |
| ACE2 | RP11-261C10.4 | -0,4874 | 6,09342E-23 |
| ACE2 | RP11-482E14.1 | -0,4874 | 6,1118E-23 |
| ACE2 | CTC-378H22.2 | -0,4874 | 6,19596E-23 |
| ACE2 | LINC01665 | -0,4875 | 6,03775E-23 |
| ACE2 | RP11-206L10.9 | -0,4876 | 5,91246E-23 |
| ACE2 | ERICH3 | -0,4876 | 5,85191E-23 |
| ACE2 | XXyac-YX155B6.5 | -0,4876 | 5,8161E-23 |
| ACE2 | RP11-295M18.6 | -0,4876 | 5,90945E-23 |
| ACE2 | WDR64 | -0,4876 | 5,89711E-23 |
| ACE2 | RP3-416J7.5 | -0,4876 | 5,8812E-23 |
| ACE2 | YTHDF3-AS1 | -0,4876 | 5,84974E-23 |
| ACE2 | LINC00544 | -0,4876 | 5,90898E-23 |
| ACE2 | RP11-439E19.7 | -0,4877 | 5,75383E-23 |
| ACE2 | CLDN16 | -0,4877 | 5,78164E-23 |
| ACE2 | LINC02215 | -0,4877 | 5,74649E-23 |
| ACE2 | MIR670HG | -0,4877 | 5,71338E-23 |
| ACE2 | MIR5572 | -0,4877 | 5,7031E-23 |
| ACE2 | RP11-410N8.4 | -0,4877 | 5,78352E-23 |
| ACE2 | EEF1GP4 | -0,4878 | 5,60894E-23 |
| ACE2 | RP11-204P2.3 | -0,4878 | 5,55338E-23 |
| ACE2 | JRKL | -0,4878 | 5,56642E-23 |
| ACE2 | KRT33B | -0,4878 | 5,59554E-23 |
| ACE2 | RP11-159D12.6 | -0,4878 | 5,61243E-23 |
| ACE2 | LINC02080 | -0,4878 | 5,56046E-23 |
| ACE2 | CNN1 | -0,4878 | 5,61875E-23 |
| ACE2 | RP11-174O3.3 | -0,4879 | 5,43354E-23 |
| ACE2 | LINC01208 | -0,4879 | 5,41357E-23 |
| ACE2 | RP11-277K23.1 | -0,4879 | 5,41101E-23 |

| | | | |
|---|---|---|---|
| ACE2 | RP11-240G22.5 | -0,4879 | 5,47929E-23 |
| ACE2 | LOXL1 | -0,4879 | 5,5142E-23 |
| ACE2 | FAM134A | -0,488 | 5,32875E-23 |
| ACE2 | KIF2B | -0,488 | 5,34061E-23 |
| ACE2 | STK35 | -0,488 | 5,3213E-23 |
| ACE2 | CALML3-AS1 | -0,4881 | 5,22904E-23 |
| ACE2 | LINC00454 | -0,4881 | 5,17854E-23 |
| ACE2 | RP11-128P17.2 | -0,4881 | 5,27675E-23 |
| ACE2 | C6orf120 | -0,4882 | 5,05852E-23 |
| ACE2 | RP11-458K10.3 | -0,4882 | 5,06476E-23 |
| ACE2 | RP11-797M17.1 | -0,4882 | 5,1436E-23 |
| ACE2 | CTD-2532K18.2 | -0,4883 | 5,03671E-23 |
| ACE2 | RP11-351C8.1 | -0,4883 | 4,99882E-23 |
| ACE2 | RP11-73M7.9 | -0,4885 | 4,74683E-23 |
| ACE2 | AGT | -0,4885 | 4,75888E-23 |
| ACE2 | RP11-434H14.1 | -0,4885 | 4,81049E-23 |
| ACE2 | CNTN4 | -0,4885 | 4,74869E-23 |
| ACE2 | SLCO6A1 | -0,4885 | 4,72238E-23 |
| ACE2 | IFNWP2 | -0,4885 | 4,75441E-23 |
| ACE2 | LINC02167 | -0,4885 | 4,77627E-23 |
| ACE2 | FRG2C | -0,4886 | 4,67487E-23 |
| ACE2 | RP11-16N2.1 | -0,4886 | 4,67228E-23 |
| ACE2 | CTC-436K13.3 | -0,4886 | 4,65549E-23 |
| ACE2 | CTC-229L21.1 | -0,4886 | 4,66754E-23 |
| ACE2 | NAPSA | -0,4886 | 4,68366E-23 |
| ACE2 | ZCWPW2 | -0,4887 | 4,57276E-23 |
| ACE2 | PKIB | -0,4887 | 4,57112E-23 |
| ACE2 | TRBV16 | -0,4887 | 4,59108E-23 |
| ACE2 | RP11-759A9.1 | -0,4887 | 4,59714E-23 |
| ACE2 | RP11-649G15.2 | -0,4887 | 4,54983E-23 |
| ACE2 | PSPC1P1 | -0,4887 | 4,54699E-23 |
| ACE2 | RP11-463I20.2 | -0,4887 | 4,59017E-23 |
| ACE2 | LINC02036 | -0,4888 | 4,44447E-23 |
| ACE2 | RP11-346J10.1 | -0,4888 | 4,42387E-23 |
| ACE2 | MBOAT4 | -0,4888 | 4,48167E-23 |
| ACE2 | CARS-AS1 | -0,4888 | 4,45324E-23 |
| ACE2 | RP11-1148O4.1 | -0,4888 | 4,42761E-23 |
| ACE2 | SLC2A5 | -0,4889 | 4,34368E-23 |
| ACE2 | LINC01784 | -0,4889 | 4,33825E-23 |
| ACE2 | FAM166AP4 | -0,4889 | 4,31478E-23 |
| ACE2 | RP11-481J13.1 | -0,4889 | 4,30723E-23 |
| ACE2 | RP11-480O10.1 | -0,4889 | 4,3859E-23 |
| ACE2 | CHCHD2P8 | -0,4889 | 4,30898E-23 |
| ACE2 | LINC02025 | -0,489 | 4,22172E-23 |
| ACE2 | SLC45A2 | -0,489 | 4,21175E-23 |
| ACE2 | MAK | -0,4891 | 4,18703E-23 |
| ACE2 | ASB4 | -0,4891 | 4,18732E-23 |
| ACE2 | FUT11 | -0,4891 | 4,18883E-23 |
| ACE2 | LINC01389 | -0,4892 | 4,09156E-23 |
| ACE2 | RP11-354I13.2 | -0,4892 | 4,07066E-23 |
| ACE2 | MRPL54 | -0,4892 | 4,06277E-23 |
| ACE2 | RP11-690I21.1 | -0,4893 | 3,98451E-23 |
| ACE2 | RP3-466I7.1 | -0,4893 | 3,92257E-23 |
| ACE2 | RP11-360L9.4 | -0,4893 | 3,94022E-23 |

| | | | |
|---|---|---|---|
| ACE2 | RP11-90L1.8 | -0,4893 | 3,98714E-23 |
| ACE2 | AC016757.3 | -0,4894 | 3,83754E-23 |
| ACE2 | RP11-384J4.2 | -0,4894 | 3,90141E-23 |
| ACE2 | RP11-666O2.2 | -0,4894 | 3,89902E-23 |
| ACE2 | ZNRD1 | -0,4895 | 3,76726E-23 |
| ACE2 | DPYS | -0,4895 | 3,81446E-23 |
| ACE2 | GOLGA8VP | -0,4895 | 3,78404E-23 |
| ACE2 | RP11-206L10.2 | -0,4896 | 3,67523E-23 |
| ACE2 | KBTBD12 | -0,4896 | 3,71943E-23 |
| ACE2 | SLITRK3 | -0,4896 | 3,66579E-23 |
| ACE2 | FBXL13 | -0,4896 | 3,68199E-23 |
| ACE2 | FBXO25 | -0,4896 | 3,72077E-23 |
| ACE2 | SLC7A13 | -0,4896 | 3,7126E-23 |
| ACE2 | PPFIA2 | -0,4896 | 3,72413E-23 |
| ACE2 | STRADA | -0,4896 | 3,70856E-23 |
| ACE2 | RP11-718B12.2 | -0,4897 | 3,61729E-23 |
| ACE2 | RP11-415J8.3 | -0,4898 | 3,52403E-23 |
| ACE2 | ARRDC1 | -0,4898 | 3,4978E-23 |
| ACE2 | LINC00841 | -0,4898 | 3,51203E-23 |
| ACE2 | RP11-219F10.1 | -0,4898 | 3,52823E-23 |
| ACE2 | GPC6-AS1 | -0,4898 | 3,54067E-23 |
| ACE2 | ZNF563 | -0,4899 | 3,42104E-23 |
| ACE2 | CTB-55O6.13 | -0,4899 | 3,46512E-23 |
| ACE2 | RP11-211G3.3 | -0,49 | 3,32403E-23 |
| ACE2 | RP11-280G9.1 | -0,49 | 3,37969E-23 |
| ACE2 | RNF139-AS1 | -0,49 | 3,34295E-23 |
| ACE2 | MIR6786 | -0,49 | 3,35974E-23 |
| ACE2 | SPHK2 | -0,49 | 3,32296E-23 |
| ACE2 | CLTCL1 | -0,49 | 3,35359E-23 |
| ACE2 | UTAT33 | -0,4901 | 3,26482E-23 |
| ACE2 | LINC01968 | -0,4902 | 3,18071E-23 |
| ACE2 | CCL24 | -0,4902 | 3,19607E-23 |
| ACE2 | IAPP | -0,4902 | 3,22929E-23 |
| ACE2 | LINC02076 | -0,4902 | 3,2131E-23 |
| ACE2 | LINC02052 | -0,4903 | 3,15453E-23 |
| ACE2 | RP11-654D12.3 | -0,4903 | 3,10554E-23 |
| ACE2 | RPGRIP1 | -0,4903 | 3,16091E-23 |
| ACE2 | OR1R1P | -0,4903 | 3,14661E-23 |
| ACE2 | TG | -0,4904 | 3,03028E-23 |
| ACE2 | TGIF2LX | -0,4904 | 3,08491E-23 |
| ACE2 | LINC01846 | -0,4905 | 3,00868E-23 |
| ACE2 | HEATR4 | -0,4905 | 2,96756E-23 |
| ACE2 | CTB-147C22.6 | -0,4905 | 3,00397E-23 |
| ACE2 | C9orf135-AS1 | -0,4906 | 2,88951E-23 |
| ACE2 | RP11-300J18.2 | -0,4906 | 2,92843E-23 |
| ACE2 | SOCS7 | -0,4906 | 2,89763E-23 |
| ACE2 | RP11-376O6.2 | -0,4907 | 2,81803E-23 |
| ACE2 | HMGB3P18 | -0,4907 | 2,84708E-23 |
| ACE2 | HAS2 | -0,4907 | 2,87723E-23 |
| ACE2 | ZPR1 | -0,4907 | 2,85346E-23 |
| ACE2 | RP1-228P16.3 | -0,4907 | 2,83889E-23 |
| ACE2 | ALS2CR12 | -0,4908 | 2,80174E-23 |
| ACE2 | SOX5 | -0,4908 | 2,78214E-23 |
| ACE2 | LINC02169 | -0,4908 | 2,78325E-23 |

| | | | |
|---|---|---|---|
| ACE2 | RP11-822E23.2 | -0,4908 | 2,77053E-23 |
| ACE2 | MED26 | -0,4908 | 2,8031E-23 |
| ACE2 | ISPD-AS1 | -0,4909 | 2,70327E-23 |
| ACE2 | AL022344.5 | -0,4909 | 2,68889E-23 |
| ACE2 | TMPRSS5 | -0,4909 | 2,7465E-23 |
| ACE2 | RP11-61E11.2 | -0,4909 | 2,73463E-23 |
| ACE2 | AC083864.3 | -0,491 | 2,64958E-23 |
| ACE2 | RP11-280K24.4 | -0,491 | 2,6868E-23 |
| ACE2 | RP11-114G11.4 | -0,4911 | 2,6117E-23 |
| ACE2 | RP11-370B11.4 | -0,4911 | 2,59349E-23 |
| ACE2 | KRT18P40 | -0,4911 | 2,56583E-23 |
| ACE2 | RP11-174O3.6 | -0,4912 | 2,56009E-23 |
| ACE2 | TRIM66 | -0,4912 | 2,51084E-23 |
| ACE2 | RP11-380N8.7 | -0,4912 | 2,54051E-23 |
| ACE2 | LINC00331 | -0,4912 | 2,53328E-23 |
| ACE2 | LINC01661 | -0,4913 | 2,49419E-23 |
| ACE2 | AC073130.1 | -0,4913 | 2,45939E-23 |
| ACE2 | RP11-89K10.1 | -0,4913 | 2,45422E-23 |
| ACE2 | RP11-386G21.2 | -0,4914 | 2,40703E-23 |
| ACE2 | RP11-555H7.2 | -0,4914 | 2,42968E-23 |
| ACE2 | EIF4A1P9 | -0,4914 | 2,40618E-23 |
| ACE2 | LINC00901 | -0,4915 | 2,37404E-23 |
| ACE2 | RP11-109G23.3 | -0,4915 | 2,35321E-23 |
| ACE2 | RP11-52J3.3 | -0,4915 | 2,35845E-23 |
| ACE2 | GAS6-AS1 | -0,4915 | 2,3622E-23 |
| ACE2 | RDM1 | -0,4915 | 2,38447E-23 |
| ACE2 | RP11-134J21.1 | -0,4916 | 2,28571E-23 |
| ACE2 | FGA | -0,4916 | 2,31247E-23 |
| ACE2 | MGAT4C | -0,4916 | 2,30101E-23 |
| ACE2 | RP11-563E2.2 | -0,4917 | 2,24999E-23 |
| ACE2 | RP11-796G6.1 | -0,4917 | 2,24643E-23 |
| ACE2 | C21orf91-OT1 | -0,4917 | 2,26757E-23 |
| ACE2 | ZNF35 | -0,4918 | 2,20205E-23 |
| ACE2 | RP11-503L23.1 | -0,4918 | 2,18781E-23 |
| ACE2 | MGAM2 | -0,4919 | 2,16513E-23 |
| ACE2 | RP11-358H18.2 | -0,4919 | 2,13953E-23 |
| ACE2 | RP11-4B16.3 | -0,4919 | 2,12615E-23 |
| ACE2 | AP000266.7 | -0,4919 | 2,17106E-23 |
| ACE2 | OXCT2P1 | -0,492 | 2,09769E-23 |
| ACE2 | RP11-758I14.3 | -0,492 | 2,07892E-23 |
| ACE2 | LSMEM1 | -0,492 | 2,11059E-23 |
| ACE2 | GMNC | -0,4921 | 2,06082E-23 |
| ACE2 | TRIM64EP | -0,4921 | 2,05302E-23 |
| ACE2 | GPC5-IT1 | -0,4921 | 2,03673E-23 |
| ACE2 | CMC2 | -0,4921 | 2,03465E-23 |
| ACE2 | RP11-803A13.1 | -0,4922 | 1,98553E-23 |
| ACE2 | LINC01345 | -0,4923 | 1,9354E-23 |
| ACE2 | RP3-340N1.2 | -0,4923 | 1,94091E-23 |
| ACE2 | RSPH10B | -0,4923 | 1,96988E-23 |
| ACE2 | EVX1-AS | -0,4923 | 1,97608E-23 |
| ACE2 | C11orf74 | -0,4923 | 1,95145E-23 |
| ACE2 | IKZF3 | -0,4923 | 1,94534E-23 |
| ACE2 | RP11-541P9.3 | -0,4924 | 1,89981E-23 |
| ACE2 | MVK | -0,4924 | 1,89984E-23 |

| | | | |
|---|---|---|---|
| ACE2 | OR11H7 | -0,4924 | 1,90332E-23 |
| ACE2 | TUBG1 | -0,4924 | 1,89797E-23 |
| ACE2 | C19orf67 | -0,4924 | 1,91465E-23 |
| ACE2 | SPAG17 | -0,4925 | 1,88036E-23 |
| ACE2 | LINC01774 | -0,4926 | 1,81771E-23 |
| ACE2 | RP11-161D15.3 | -0,4926 | 1,82195E-23 |
| ACE2 | RP11-128M1.1 | -0,4926 | 1,82961E-23 |
| ACE2 | RP11-675F6.3 | -0,4927 | 1,79197E-23 |
| ACE2 | PIH1D2 | -0,4927 | 1,79077E-23 |
| ACE2 | RP11-104J23.1 | -0,4927 | 1,77483E-23 |
| ACE2 | CTD-2126E3.3 | -0,4927 | 1,79148E-23 |
| ACE2 | AC096554.1 | -0,4929 | 1,68036E-23 |
| ACE2 | PLCXD2 | -0,4929 | 1,71137E-23 |
| ACE2 | RP11-168L22.2 | -0,4929 | 1,71991E-23 |
| ACE2 | DCDC1 | -0,4929 | 1,6969E-23 |
| ACE2 | RP11-426C22.7 | -0,4929 | 1,68258E-23 |
| ACE2 | RPSAP73 | -0,4931 | 1,64037E-23 |
| ACE2 | C9orf43 | -0,4931 | 1,61298E-23 |
| ACE2 | DYDC2 | -0,4931 | 1,62552E-23 |
| ACE2 | RP11-272B17.2 | -0,4931 | 1,60298E-23 |
| ACE2 | TUBA3C | -0,4931 | 1,61199E-23 |
| ACE2 | RP11-473E2.2 | -0,4932 | 1,58311E-23 |
| ACE2 | POLD2 | -0,4933 | 1,54344E-23 |
| ACE2 | FABP12 | -0,4933 | 1,56115E-23 |
| ACE2 | CHKA | -0,4933 | 1,55136E-23 |
| ACE2 | RP11-739B23.1 | -0,4933 | 1,53126E-23 |
| ACE2 | CRAT8 | -0,4933 | 1,5465E-23 |
| ACE2 | FLJ22447 | -0,4933 | 1,54558E-23 |
| ACE2 | GTF2A2 | -0,4933 | 1,53172E-23 |
| ACE2 | HHIPL2 | -0,4934 | 1,52581E-23 |
| ACE2 | RP11-534P6.1 | -0,4934 | 1,49333E-23 |
| ACE2 | CTD-2006K23.1 | -0,4934 | 1,50351E-23 |
| ACE2 | SEL1L2 | -0,4934 | 1,52586E-23 |
| ACE2 | VIPR1-AS1 | -0,4935 | 1,48219E-23 |
| ACE2 | RP11-713P17.5 | -0,4935 | 1,47996E-23 |
| ACE2 | RP11-899L11.3 | -0,4935 | 1,45958E-23 |
| ACE2 | SLC16A6P1 | -0,4935 | 1,48347E-23 |
| ACE2 | DNAJC28 | -0,4935 | 1,4834E-23 |
| ACE2 | RP11-73B2.2 | -0,4936 | 1,44824E-23 |
| ACE2 | KB-1552D7.2 | -0,4936 | 1,43723E-23 |
| ACE2 | LINC01543 | -0,4936 | 1,44438E-23 |
| ACE2 | CACNA1A | -0,4936 | 1,44239E-23 |
| ACE2 | ARHGEF33 | -0,4937 | 1,39886E-23 |
| ACE2 | LINC01941 | -0,4937 | 1,39731E-23 |
| ACE2 | OR8B2 | -0,4937 | 1,39329E-23 |
| ACE2 | KXD1 | -0,4937 | 1,40476E-23 |
| ACE2 | LINC00308 | -0,4937 | 1,4161E-23 |
| ACE2 | LINC00315 | -0,4937 | 1,41285E-23 |
| ACE2 | LNP1 | -0,4938 | 1,3682E-23 |
| ACE2 | TM4SF1-AS1 | -0,4938 | 1,38074E-23 |
| ACE2 | RP11-543D10.2 | -0,4938 | 1,38895E-23 |
| ACE2 | AF186192.6 | -0,4938 | 1,37651E-23 |
| ACE2 | AC068483.1 | -0,494 | 1,31233E-23 |
| ACE2 | RP11-785H20.1 | -0,494 | 1,29934E-23 |

| | | | |
|---|---|---|---|
| ACE2 | RP11-272P10.2 | -0,494 | 1,30813E-23 |
| ACE2 | RP13-16H11.8 | -0,494 | 1,32471E-23 |
| ACE2 | LINC02124 | -0,494 | 1,30919E-23 |
| ACE2 | ATP2C2 | -0,494 | 1,3142E-23 |
| ACE2 | PTGES3L | -0,494 | 1,30842E-23 |
| ACE2 | ITGB6 | -0,4941 | 1,26806E-23 |
| ACE2 | CTB-78H18.1 | -0,4941 | 1,28171E-23 |
| ACE2 | HSPBP1 | -0,4942 | 1,25963E-23 |
| ACE2 | VN1R34P | -0,4943 | 1,21427E-23 |
| ACE2 | ARRDC1-AS1 | -0,4943 | 1,22845E-23 |
| ACE2 | RP11-676F20.2 | -0,4943 | 1,22317E-23 |
| ACE2 | C1orf94 | -0,4944 | 1,18246E-23 |
| ACE2 | LINC01717 | -0,4944 | 1,19427E-23 |
| ACE2 | AC113167.1 | -0,4944 | 1,20395E-23 |
| ACE2 | GCM1 | -0,4944 | 1,18793E-23 |
| ACE2 | KCNU1 | -0,4944 | 1,204E-23 |
| ACE2 | C10orf25 | -0,4944 | 1,20524E-23 |
| ACE2 | LAMP5-AS1 | -0,4944 | 1,19128E-23 |
| ACE2 | RP11-363E6.4 | -0,4945 | 1,17548E-23 |
| ACE2 | RP1-18D14.7 | -0,4946 | 1,13709E-23 |
| ACE2 | RP11-5A1.1 | -0,4946 | 1,13339E-23 |
| ACE2 | NDUFAF2 | -0,4947 | 1,10454E-23 |
| ACE2 | TRDN | -0,4947 | 1,12063E-23 |
| ACE2 | RP11-380F14.2 | -0,4947 | 1,12177E-23 |
| ACE2 | GPRIN2 | -0,4947 | 1,11273E-23 |
| ACE2 | FETUB | -0,4948 | 1,08017E-23 |
| ACE2 | HIST1H2APS1 | -0,4948 | 1,07742E-23 |
| ACE2 | HOMEZ | -0,4948 | 1,08154E-23 |
| ACE2 | CLIC5 | -0,4949 | 1,05115E-23 |
| ACE2 | AP000289.6 | -0,4949 | 1,06919E-23 |
| ACE2 | LINC01704 | -0,495 | 1,04293E-23 |
| ACE2 | RP11-145H9.3 | -0,495 | 1,03423E-23 |
| ACE2 | AK1 | -0,495 | 1,03E-23 |
| ACE2 | KRT23 | -0,495 | 1,02696E-23 |
| ACE2 | RP5-1059H15.1 | -0,4951 | 1,00677E-23 |
| ACE2 | SYNJ2 | -0,4951 | 1,01675E-23 |
| ACE2 | RP11-379P1.4 | -0,4951 | 1,01935E-23 |
| ACE2 | C11orf84 | -0,4951 | 1,00664E-23 |
| ACE2 | RP11-822E23.7 | -0,4951 | 1,01984E-23 |
| ACE2 | RP11-293E1.2 | -0,4951 | 1,00301E-23 |
| ACE2 | PSMD4 | -0,4952 | 9,92149E-24 |
| ACE2 | RP11-576I22.2 | -0,4952 | 9,94487E-24 |
| ACE2 | LINC01830 | -0,4952 | 9,78532E-24 |
| ACE2 | FKBP7 | -0,4952 | 9,85576E-24 |
| ACE2 | MS4A5 | -0,4952 | 9,88008E-24 |
| ACE2 | DNAI1 | -0,4953 | 9,5954E-24 |
| ACE2 | TP53I13 | -0,4954 | 9,45368E-24 |
| ACE2 | SYNJ2-IT1 | -0,4955 | 9,23347E-24 |
| ACE2 | RP11-588G21.2 | -0,4955 | 9,15596E-24 |
| ACE2 | RP11-411G9.2 | -0,4955 | 9,19753E-24 |
| ACE2 | RNASE12 | -0,4955 | 9,1168E-24 |
| ACE2 | RP3-436N22.3 | -0,4956 | 9,01285E-24 |
| ACE2 | RP11-171N4.3 | -0,4956 | 8,96222E-24 |
| ACE2 | MAS1LP1 | -0,4956 | 8,96992E-24 |

| | | | |
|---|---|---|---|
| ACE2 | CCDC92 | -0,4956 | 8,86723E-24 |
| ACE2 | RP11-452B18.2 | -0,4957 | 8,73822E-24 |
| ACE2 | RP11-53L24.2 | -0,4957 | 8,75641E-24 |
| ACE2 | KATNAL2 | -0,4957 | 8,72725E-24 |
| ACE2 | AP000351.3 | -0,4957 | 8,82784E-24 |
| ACE2 | RP11-267A15.3 | -0,4958 | 8,49779E-24 |
| ACE2 | TRBV6-8 | -0,4958 | 8,581E-24 |
| ACE2 | HEMGN | -0,4958 | 8,46772E-24 |
| ACE2 | RP11-876F14.1 | -0,4958 | 8,48479E-24 |
| ACE2 | MYO16-AS2 | -0,4958 | 8,51305E-24 |
| ACE2 | PRSS54 | -0,4958 | 8,54906E-24 |
| ACE2 | RP11-700H13.1 | -0,4958 | 8,51112E-24 |
| ACE2 | AC017060.1 | -0,4959 | 8,32118E-24 |
| ACE2 | RP11-809H16.5 | -0,4959 | 8,31669E-24 |
| ACE2 | RP11-2L4.1 | -0,4959 | 8,28644E-24 |
| ACE2 | CST12P | -0,4959 | 8,34183E-24 |
| ACE2 | ADRM1 | -0,4959 | 8,40938E-24 |
| ACE2 | RP11-413E1.4 | -0,496 | 8,1617E-24 |
| ACE2 | HLCS-IT1 | -0,496 | 8,07007E-24 |
| ACE2 | AP001625.5 | -0,496 | 8,08668E-24 |
| ACE2 | LINC02117 | -0,4961 | 7,94159E-24 |
| ACE2 | RP11-179A9.3 | -0,4961 | 7,94207E-24 |
| ACE2 | KB-1183D5.13 | -0,4961 | 7,91618E-24 |
| ACE2 | RP11-10F11.2 | -0,4962 | 7,70254E-24 |
| ACE2 | GOLGA6L1 | -0,4963 | 7,59506E-24 |
| ACE2 | OVOL3 | -0,4963 | 7,52091E-24 |
| ACE2 | PROX1 | -0,4964 | 7,43873E-24 |
| ACE2 | CENPT | -0,4964 | 7,50215E-24 |
| ACE2 | AACS | -0,4965 | 7,20388E-24 |
| ACE2 | RP11-480G7.2 | -0,4965 | 7,17277E-24 |
| ACE2 | AMZ2 | -0,4965 | 7,32209E-24 |
| ACE2 | RP11-269F19.2 | -0,4966 | 7,07377E-24 |
| ACE2 | NDUFAF7 | -0,4966 | 7,06092E-24 |
| ACE2 | AC104113.3 | -0,4966 | 7,13458E-24 |
| ACE2 | SYN3 | -0,4966 | 7,09419E-24 |
| ACE2 | UBAC1 | -0,4967 | 6,88667E-24 |
| ACE2 | CTC-412M14.5 | -0,4967 | 6,90219E-24 |
| ACE2 | BRINP3 | -0,4968 | 6,6949E-24 |
| ACE2 | RP11-428J1.5 | -0,4968 | 6,79033E-24 |
| ACE2 | XXbac-BPG258E24.10 | -0,4968 | 6,69174E-24 |
| ACE2 | RP11-555J4.4 | -0,4968 | 6,72715E-24 |
| ACE2 | SLC7A11-AS1 | -0,4969 | 6,64807E-24 |
| ACE2 | LINC01449 | -0,4969 | 6,58434E-24 |
| ACE2 | CYP4F62P | -0,497 | 6,39801E-24 |
| ACE2 | ADH4 | -0,497 | 6,44703E-24 |
| ACE2 | RP11-475C16.2 | -0,497 | 6,44474E-24 |
| ACE2 | PCDH8P1 | -0,497 | 6,40518E-24 |
| ACE2 | GTF2A1L | -0,4971 | 6,19937E-24 |
| ACE2 | RP11-1338A24.1 | -0,4971 | 6,30952E-24 |
| ACE2 | RP11-480G3.1 | -0,4971 | 6,33558E-24 |
| ACE2 | RP11-586L23.1 | -0,4971 | 6,24405E-24 |
| ACE2 | RP11-459F6.1 | -0,4971 | 6,21135E-24 |
| ACE2 | LINC01733 | -0,4972 | 6,17259E-24 |
| ACE2 | LAMP3 | -0,4973 | 5,99072E-24 |

| | | | |
|---|---|---|---|
| ACE2 | RP11-428K3.1 | -0,4974 | 5,83155E-24 |
| ACE2 | RP11-180C1.1 | -0,4974 | 5,80797E-24 |
| ACE2 | RP11-532M24.1 | -0,4974 | 5,87127E-24 |
| ACE2 | RP11-394A14.4 | -0,4974 | 5,86156E-24 |
| ACE2 | RP11-507K2.2 | -0,4974 | 5,84615E-24 |
| ACE2 | TXNDC12-AS1 | -0,4976 | 5,61649E-24 |
| ACE2 | RFX8 | -0,4976 | 5,60676E-24 |
| ACE2 | RP11-298C2.1 | -0,4976 | 5,60104E-24 |
| ACE2 | EIF4BP8 | -0,4976 | 5,57169E-24 |
| ACE2 | USP32P3 | -0,4976 | 5,53612E-24 |
| ACE2 | LINC00326 | -0,4977 | 5,41056E-24 |
| ACE2 | RP11-15B24.3 | -0,4977 | 5,49847E-24 |
| ACE2 | RP11-295P9.6 | -0,4977 | 5,46558E-24 |
| ACE2 | C20orf27 | -0,4977 | 5,46741E-24 |
| ACE2 | RP11-492O8.1 | -0,4977 | 5,48321E-24 |
| ACE2 | MYLK-AS2 | -0,4978 | 5,31677E-24 |
| ACE2 | ROCK1P1 | -0,4978 | 5,26435E-24 |
| ACE2 | TSPEAR | -0,4978 | 5,3545E-24 |
| ACE2 | GPR156 | -0,498 | 5,00643E-24 |
| ACE2 | COMMD3 | -0,498 | 5,01252E-24 |
| ACE2 | RP11-56I23.1 | -0,498 | 5,04234E-24 |
| ACE2 | NKPD1 | -0,498 | 5,0172E-24 |
| ACE2 | AC130709.1 | -0,4981 | 4,89071E-24 |
| ACE2 | CTC-359M8.1 | -0,4981 | 4,95917E-24 |
| ACE2 | EFCAB9 | -0,4981 | 4,98402E-24 |
| ACE2 | RP3-377H17.2 | -0,4981 | 4,9934E-24 |
| ACE2 | RP11-69M1.6 | -0,4981 | 4,89131E-24 |
| ACE2 | LPAR6 | -0,4981 | 4,99646E-24 |
| ACE2 | AC100802.3 | -0,4982 | 4,77223E-24 |
| ACE2 | RP6-91H8.2 | -0,4982 | 4,80643E-24 |
| ACE2 | AF001548.5 | -0,4982 | 4,79362E-24 |
| ACE2 | RP11-441F2.2 | -0,4982 | 4,76448E-24 |
| ACE2 | RP5-971N18.3 | -0,4982 | 4,86048E-24 |
| ACE2 | KCNQ5-IT1 | -0,4983 | 4,7268E-24 |
| ACE2 | AE000662.93 | -0,4983 | 4,67807E-24 |
| ACE2 | HES7 | -0,4983 | 4,65314E-24 |
| ACE2 | RP11-905K4.1 | -0,4983 | 4,66815E-24 |
| ACE2 | USP25 | -0,4983 | 4,73437E-24 |
| ACE2 | RP11-662M24.1 | -0,4984 | 4,62357E-24 |
| ACE2 | RP11-283I3.1 | -0,4985 | 4,49768E-24 |
| ACE2 | AC002066.1 | -0,4986 | 4,34571E-24 |
| ACE2 | LINC01153 | -0,4986 | 4,38167E-24 |
| ACE2 | HYLS1 | -0,4986 | 4,37945E-24 |
| ACE2 | RNASE13 | -0,4986 | 4,34355E-24 |
| ACE2 | MDP1 | -0,4986 | 4,35244E-24 |
| ACE2 | CTD-2538C1.2 | -0,4986 | 4,42788E-24 |
| ACE2 | B9D2 | -0,4986 | 4,37999E-24 |
| ACE2 | RP11-103C3.2 | -0,4987 | 4,29146E-24 |
| ACE2 | LINC01792 | -0,4987 | 4,32356E-24 |
| ACE2 | RP11-6N13.4 | -0,4987 | 4,32411E-24 |
| ACE2 | SNX31 | -0,4987 | 4,23516E-24 |
| ACE2 | TTLL7-IT1 | -0,4988 | 4,17814E-24 |
| ACE2 | CTD-2655K5.1 | -0,4988 | 4,18077E-24 |
| ACE2 | HEXIM2 | -0,4988 | 4,12738E-24 |

| | | | |
|---|---|---|---|
| ACE2 | COX6CP13 | -0,4989 | 4,05544E-24 |
| ACE2 | TIGD4 | -0,4989 | 4,12207E-24 |
| ACE2 | AC006372.5 | -0,4989 | 4,11998E-24 |
| ACE2 | MBD1 | -0,4989 | 4,08862E-24 |
| ACE2 | EFCAB6-AS1 | -0,4989 | 4,09696E-24 |
| ACE2 | CTC-336P14.1 | -0,499 | 4,01778E-24 |
| ACE2 | KLHL41 | -0,499 | 4,02089E-24 |
| ACE2 | RP11-497D6.3 | -0,499 | 3,94004E-24 |
| ACE2 | FAM161B | -0,499 | 3,99119E-24 |
| ACE2 | SMDT1 | -0,499 | 3,96924E-24 |
| ACE2 | LINC01981 | -0,4991 | 3,91822E-24 |
| ACE2 | AC004692.5 | -0,4991 | 3,90824E-24 |
| ACE2 | CALCR | -0,4991 | 3,85922E-24 |
| ACE2 | IRX1P1 | -0,4991 | 3,90549E-24 |
| ACE2 | ZMYND12 | -0,4992 | 3,75633E-24 |
| ACE2 | RP11-161I10.1 | -0,4992 | 3,77964E-24 |
| ACE2 | IFT22 | -0,4992 | 3,7512E-24 |
| ACE2 | RP11-15B24.1 | -0,4992 | 3,77144E-24 |
| ACE2 | RP11-61K9.3 | -0,4992 | 3,75535E-24 |
| ACE2 | RP11-47I22.2 | -0,4992 | 3,82817E-24 |
| ACE2 | RP11-76E12.1 | -0,4992 | 3,80028E-24 |
| ACE2 | AC093230.2 | -0,4992 | 3,77145E-24 |
| ACE2 | KIAA1257 | -0,4993 | 3,70258E-24 |
| ACE2 | RP3-335N17.2 | -0,4993 | 3,72192E-24 |
| ACE2 | RP11-354A14.1 | -0,4993 | 3,73982E-24 |
| ACE2 | CYP4F34P | -0,4993 | 3,69332E-24 |
| ACE2 | EXOC3L1 | -0,4993 | 3,70073E-24 |
| ACE2 | AC013460.1 | -0,4994 | 3,61267E-24 |
| ACE2 | TRIM74 | -0,4994 | 3,64124E-24 |
| ACE2 | BORCS6 | -0,4994 | 3,57457E-24 |
| ACE2 | OSER1-AS1 | -0,4994 | 3,63245E-24 |
| ACE2 | TTC21A | -0,4995 | 3,56612E-24 |
| ACE2 | RP11-81K13.1 | -0,4995 | 3,55944E-24 |
| ACE2 | CTD-2555O16.1 | -0,4995 | 3,54145E-24 |
| ACE2 | TTBK2 | -0,4996 | 3,46745E-24 |
| ACE2 | RP11-1223D19.1 | -0,4997 | 3,39793E-24 |
| ACE2 | LINC02107 | -0,4997 | 3,32604E-24 |
| ACE2 | AC008427.2 | -0,4997 | 3,37957E-24 |
| ACE2 | RP11-437F6.1 | -0,4997 | 3,33342E-24 |
| ACE2 | RP11-799D4.1 | -0,4997 | 3,32922E-24 |
| ACE2 | LINC02003 | -0,4997 | 3,39148E-24 |
| ACE2 | RP11-35O7.1 | -0,4998 | 3,30344E-24 |
| ACE2 | ZMAT4 | -0,4998 | 3,29786E-24 |
| ACE2 | ARHGEF7-AS1 | -0,4998 | 3,27676E-24 |
| ACE2 | RP11-99L13.1 | -0,4998 | 3,29289E-24 |
| ACE2 | C14orf177 | -0,4998 | 3,25589E-24 |
| ACE2 | TTC12 | -0,4999 | 3,21892E-24 |
| ACE2 | RP11-318G21.4 | -0,4999 | 3,23464E-24 |
| ACE2 | FPR2 | -0,4999 | 3,23722E-24 |
| ACE2 | RP1-101D8.1 | -0,4999 | 3,23734E-24 |
| ACE2 | RP11-8L2.1 | -0,5 | 3,16126E-24 |
| ACE2 | HVCN1 | -0,5 | 3,09937E-24 |
| ACE2 | RP11-666A8.13 | -0,5 | 3,15335E-24 |
| ACE2 | RP11-72I2.2 | -0,5001 | 3,08906E-24 |

| | | | |
|---|---|---|---|
| ACE2 | LINC00343 | -0,5002 | 2,97295E-24 |
| ACE2 | LINC01583 | -0,5002 | 2,94758E-24 |
| ACE2 | RP11-357H3.1 | -0,5002 | 3,00267E-24 |
| ACE2 | LRRIQ1 | -0,5003 | 2,90178E-24 |
| ACE2 | LINC00421 | -0,5003 | 2,93979E-24 |
| ACE2 | AC025811.3 | -0,5003 | 2,91506E-24 |
| ACE2 | LINC02173 | -0,5004 | 2,83946E-24 |
| ACE2 | RP11-347L18.1 | -0,5004 | 2,85562E-24 |
| ACE2 | RP5-1048B16.1 | -0,5004 | 2,86321E-24 |
| ACE2 | HCAR2 | -0,5004 | 2,86271E-24 |
| ACE2 | UBE2S | -0,5004 | 2,86807E-24 |
| ACE2 | RPS7P5 | -0,5005 | 2,79679E-24 |
| ACE2 | LINC01248 | -0,5005 | 2,73791E-24 |
| ACE2 | RP11-257I14.1 | -0,5005 | 2,77207E-24 |
| ACE2 | RP11-973N13.4 | -0,5005 | 2,74592E-24 |
| ACE2 | KCNH1 | -0,5006 | 2,72641E-24 |
| ACE2 | ADAD1 | -0,5006 | 2,71708E-24 |
| ACE2 | TMEM26-AS1 | -0,5006 | 2,73654E-24 |
| ACE2 | LINC02180 | -0,5006 | 2,73215E-24 |
| ACE2 | RP11-369J9.1 | -0,5007 | 2,65248E-24 |
| ACE2 | ADGB | -0,5007 | 2,61707E-24 |
| ACE2 | GSG1L | -0,5007 | 2,65277E-24 |
| ACE2 | RPS2P45 | -0,5007 | 2,66283E-24 |
| ACE2 | LL22NC03-104C7.1 | -0,5007 | 2,63365E-24 |
| ACE2 | RP11-210L7.3 | -0,5008 | 2,57288E-24 |
| ACE2 | LINC00381 | -0,5008 | 2,60438E-24 |
| ACE2 | RP11-737O24.1 | -0,5008 | 2,59258E-24 |
| ACE2 | RP11-336K24.5 | -0,5009 | 2,503E-24 |
| ACE2 | FCRL3 | -0,5009 | 2,50536E-24 |
| ACE2 | AC072062.3 | -0,5009 | 2,51477E-24 |
| ACE2 | RP11-80H8.4 | -0,5009 | 2,53102E-24 |
| ACE2 | RP3-442L6.4 | -0,5009 | 2,51632E-24 |
| ACE2 | KRT19P1 | -0,5009 | 2,52165E-24 |
| ACE2 | RP11-87E22.1 | -0,5009 | 2,52687E-24 |
| ACE2 | RP11-100G15.12 | -0,5009 | 2,54518E-24 |
| ACE2 | AC012494.1 | -0,501 | 2,47245E-24 |
| ACE2 | RP11-511H23.2 | -0,501 | 2,43721E-24 |
| ACE2 | SPATA45 | -0,5011 | 2,41935E-24 |
| ACE2 | SDCCAG3 | -0,5011 | 2,40729E-24 |
| ACE2 | SIX3-AS1 | -0,5012 | 2,31716E-24 |
| ACE2 | PCAT2 | -0,5012 | 2,35198E-24 |
| ACE2 | FLT3 | -0,5012 | 2,33174E-24 |
| ACE2 | RP11-10K17.6 | -0,5012 | 2,33977E-24 |
| ACE2 | RP11-452L6.7 | -0,5012 | 2,33163E-24 |
| ACE2 | RP11-124I4.2 | -0,5013 | 2,26349E-24 |
| ACE2 | RP11-653J6.1 | -0,5013 | 2,29241E-24 |
| ACE2 | RP11-593F5.2 | -0,5014 | 2,24867E-24 |
| ACE2 | RP11-760D2.7 | -0,5014 | 2,2213E-24 |
| ACE2 | RSU1P1 | -0,5014 | 2,20256E-24 |
| ACE2 | LINC01069 | -0,5014 | 2,20518E-24 |
| ACE2 | RP11-548M13.1 | -0,5014 | 2,20911E-24 |
| ACE2 | ZNF18 | -0,5014 | 2,21309E-24 |
| ACE2 | ERICH4 | -0,5014 | 2,22993E-24 |
| ACE2 | DUSP21 | -0,5014 | 2,24498E-24 |

| | | | |
|---|---|---|---|
| ACE2 | TUBA4A | -0,5015 | 2,20098E-24 |
| ACE2 | ITIH3 | -0,5015 | 2,19239E-24 |
| ACE2 | RP11-27P7.1 | -0,5015 | 2,19133E-24 |
| ACE2 | RP11-481G8.2 | -0,5015 | 2,1892E-24 |
| ACE2 | CFAP61 | -0,5015 | 2,17382E-24 |
| ACE2 | EYS | -0,5016 | 2,10367E-24 |
| ACE2 | CCDC7 | -0,5016 | 2,12901E-24 |
| ACE2 | RP11-788H18.1 | -0,5016 | 2,13982E-24 |
| ACE2 | CAPN15 | -0,5016 | 2,11872E-24 |
| ACE2 | ZNF689 | -0,5016 | 2,1246E-24 |
| ACE2 | LINC02131 | -0,5016 | 2,13413E-24 |
| ACE2 | LINC01428 | -0,5016 | 2,13705E-24 |
| ACE2 | SPATA4 | -0,5017 | 2,06561E-24 |
| ACE2 | TTLL2 | -0,5017 | 2,06521E-24 |
| ACE2 | RP11-384G23.1 | -0,5017 | 2,07007E-24 |
| ACE2 | FAM207A | -0,5017 | 2,08933E-24 |
| ACE2 | KB-1995A5.5 | -0,5018 | 2,01826E-24 |
| ACE2 | RP5-1139B12.4 | -0,5019 | 1,96111E-24 |
| ACE2 | CTD-2185K10.1 | -0,5019 | 1,9719E-24 |
| ACE2 | RP11-340A13.1 | -0,5019 | 1,97869E-24 |
| ACE2 | RP11-432J9.6 | -0,5019 | 1,95396E-24 |
| ACE2 | TPH2 | -0,5019 | 1,95848E-24 |
| ACE2 | RP1-37N7.1 | -0,5019 | 1,98128E-24 |
| ACE2 | LINC01525 | -0,502 | 1,94639E-24 |
| ACE2 | NPM1P32 | -0,502 | 1,92537E-24 |
| ACE2 | RP11-384K6.6 | -0,502 | 1,90455E-24 |
| ACE2 | KCNV2 | -0,502 | 1,936E-24 |
| ACE2 | LINC01516 | -0,502 | 1,92291E-24 |
| ACE2 | CMB9-22P13.2 | -0,502 | 1,90503E-24 |
| ACE2 | CACNA1C-AS4 | -0,502 | 1,93279E-24 |
| ACE2 | RP11-175P13.2 | -0,502 | 1,92157E-24 |
| ACE2 | RP11-433J20.1 | -0,5021 | 1,89966E-24 |
| ACE2 | RP11-168C9.1 | -0,5021 | 1,88927E-24 |
| ACE2 | LINC00332 | -0,5021 | 1,87949E-24 |
| ACE2 | TMC7 | -0,5021 | 1,86518E-24 |
| ACE2 | RP11-527L4.9 | -0,5021 | 1,89686E-24 |
| ACE2 | LL22NC03-102D1.18 | -0,5021 | 1,86811E-24 |
| ACE2 | GRM6 | -0,5022 | 1,82108E-24 |
| ACE2 | RP13-884E18.4 | -0,5023 | 1,79457E-24 |
| ACE2 | CTD-2154B17.1 | -0,5023 | 1,79574E-24 |
| ACE2 | C6orf201 | -0,5023 | 1,80969E-24 |
| ACE2 | UNC5D | -0,5023 | 1,77145E-24 |
| ACE2 | CTD-2554C21.1 | -0,5023 | 1,80659E-24 |
| ACE2 | LINC01548 | -0,5023 | 1,78727E-24 |
| ACE2 | HYPM | -0,5023 | 1,80111E-24 |
| ACE2 | BFSP2-AS1 | -0,5024 | 1,74599E-24 |
| ACE2 | RP1-199J3.7 | -0,5024 | 1,72893E-24 |
| ACE2 | RP11-2E11.5 | -0,5024 | 1,75201E-24 |
| ACE2 | ACADS | -0,5024 | 1,7568E-24 |
| ACE2 | ELL | -0,5024 | 1,75878E-24 |
| ACE2 | RP1-45N11.1 | -0,5025 | 1,69695E-24 |
| ACE2 | DPY19L2P2 | -0,5025 | 1,72244E-24 |
| ACE2 | MFSD11 | -0,5025 | 1,72086E-24 |
| ACE2 | CTD-3096M3.2 | -0,5025 | 1,70752E-24 |

| | | | |
|---|---|---|---|
| ACE2 | FBLL1 | -0,5026 | 1,65608E-24 |
| ACE2 | TRY2P | -0,5026 | 1,65506E-24 |
| ACE2 | ADAM32 | -0,5026 | 1,64697E-24 |
| ACE2 | KB-1930G5.3 | -0,5026 | 1,64578E-24 |
| ACE2 | UBE2SP1 | -0,5026 | 1,64813E-24 |
| ACE2 | LINC00446 | -0,5027 | 1,63524E-24 |
| ACE2 | PRR15L | -0,5027 | 1,61862E-24 |
| ACE2 | RP11-647F2.2 | -0,5027 | 1,63913E-24 |
| ACE2 | CCDC181 | -0,5028 | 1,58957E-24 |
| ACE2 | ICE2P2 | -0,5028 | 1,57772E-24 |
| ACE2 | IL23A | -0,5028 | 1,59087E-24 |
| ACE2 | LINC01672 | -0,5029 | 1,54382E-24 |
| ACE2 | CDC20 | -0,5029 | 1,55268E-24 |
| ACE2 | DYRK3 | -0,5029 | 1,5428E-24 |
| ACE2 | CTD-2376I4.1 | -0,5029 | 1,54628E-24 |
| ACE2 | RP11-727F15.12 | -0,5029 | 1,53498E-24 |
| ACE2 | INCA1 | -0,5029 | 1,54398E-24 |
| ACE2 | CTD-2666L21.3 | -0,5029 | 1,56135E-24 |
| ACE2 | RFTN1P1 | -0,5029 | 1,56681E-24 |
| ACE2 | ZNF607 | -0,503 | 1,51568E-24 |
| ACE2 | RP11-753N8.1 | -0,5031 | 1,47778E-24 |
| ACE2 | HYAL3 | -0,5032 | 1,43142E-24 |
| ACE2 | LINC02032 | -0,5032 | 1,42221E-24 |
| ACE2 | CTD-2318H23.1 | -0,5032 | 1,43669E-24 |
| ACE2 | HMMR-AS1 | -0,5032 | 1,42604E-24 |
| ACE2 | LEMD2 | -0,5032 | 1,43645E-24 |
| ACE2 | C2orf73 | -0,5033 | 1,39583E-24 |
| ACE2 | IL1RN | -0,5033 | 1,42049E-24 |
| ACE2 | RP11-359D24.1 | -0,5033 | 1,42096E-24 |
| ACE2 | RP11-826N14.4 | -0,5033 | 1,41034E-24 |
| ACE2 | MOXD2P | -0,5033 | 1,39575E-24 |
| ACE2 | FRMD4A | -0,5033 | 1,39051E-24 |
| ACE2 | LINC00563 | -0,5033 | 1,40332E-24 |
| ACE2 | RP1-140A9.1 | -0,5034 | 1,37369E-24 |
| ACE2 | RP5-1024N4.6 | -0,5034 | 1,36243E-24 |
| ACE2 | RP4-613A2.1 | -0,5034 | 1,37988E-24 |
| ACE2 | AC010096.1 | -0,5034 | 1,36922E-24 |
| ACE2 | KCNH7 | -0,5034 | 1,36547E-24 |
| ACE2 | RP11-123J14.4 | -0,5034 | 1,3716E-24 |
| ACE2 | TRPV6 | -0,5034 | 1,35932E-24 |
| ACE2 | KB-1639H6.2 | -0,5034 | 1,35372E-24 |
| ACE2 | AC009542.2 | -0,5035 | 1,34112E-24 |
| ACE2 | NOL6 | -0,5035 | 1,32498E-24 |
| ACE2 | RP11-711C17.2 | -0,5035 | 1,34349E-24 |
| ACE2 | KCNF1 | -0,5036 | 1,29347E-24 |
| ACE2 | RP11-456N14.6 | -0,5036 | 1,30322E-24 |
| ACE2 | RP11-110H1.8 | -0,5036 | 1,31676E-24 |
| ACE2 | INPP4B | -0,5037 | 1,26934E-24 |
| ACE2 | RP11-35J1.1 | -0,5037 | 1,26021E-24 |
| ACE2 | RP11-390F4.10 | -0,5037 | 1,25991E-24 |
| ACE2 | RP11-439K3.3 | -0,5037 | 1,27794E-24 |
| ACE2 | RP11-511B23.2 | -0,5037 | 1,26546E-24 |
| ACE2 | ZNF440 | -0,5037 | 1,26422E-24 |
| ACE2 | RP5-1028L10.1 | -0,5038 | 1,22869E-24 |

| | | | |
|---|---|---|---|
| ACE2 | AC010880.1 | -0,5038 | 1,2557E-24 |
| ACE2 | CTD-3007L5.1 | -0,5038 | 1,23446E-24 |
| ACE2 | RP11-38F22.1 | -0,5038 | 1,23014E-24 |
| ACE2 | CH17-342O10.1 | -0,5038 | 1,25468E-24 |
| ACE2 | RP11-367J11.3 | -0,5039 | 1,22352E-24 |
| ACE2 | C11orf1 | -0,5039 | 1,20636E-24 |
| ACE2 | AHSP | -0,5039 | 1,21405E-24 |
| ACE2 | RP13-210D15.4 | -0,5039 | 1,21777E-24 |
| ACE2 | RP11-131H24.4 | -0,504 | 1,19017E-24 |
| ACE2 | CTD-2576F9.1 | -0,504 | 1,18896E-24 |
| ACE2 | ALOX12P2 | -0,504 | 1,19274E-24 |
| ACE2 | LUZP1 | -0,5041 | 1,14969E-24 |
| ACE2 | GAPDHP72 | -0,5041 | 1,15922E-24 |
| ACE2 | ST14 | -0,5041 | 1,16604E-24 |
| ACE2 | RP11-482D24.2 | -0,5041 | 1,16545E-24 |
| ACE2 | NEK11 | -0,5042 | 1,12099E-24 |
| ACE2 | SMC2-AS1 | -0,5042 | 1,13609E-24 |
| ACE2 | RP11-45F15.1 | -0,5042 | 1,13423E-24 |
| ACE2 | C3orf52 | -0,5043 | 1,10271E-24 |
| ACE2 | SMCO1 | -0,5043 | 1,11218E-24 |
| ACE2 | CTD-2275D24.1 | -0,5043 | 1,107E-24 |
| ACE2 | FAM99B | -0,5043 | 1,0952E-24 |
| ACE2 | RP11-705C15.4 | -0,5044 | 1,06618E-24 |
| ACE2 | MUC16 | -0,5044 | 1,07618E-24 |
| ACE2 | RP11-473A10.2 | -0,5045 | 1,04901E-24 |
| ACE2 | RP11-411G2.2 | -0,5045 | 1,05398E-24 |
| ACE2 | AC011196.3 | -0,5045 | 1,05913E-24 |
| ACE2 | CTD-2050I18.1 | -0,5045 | 1,04484E-24 |
| ACE2 | EEFSEC | -0,5046 | 1,0328E-24 |
| ACE2 | AFAP1 | -0,5046 | 1,03058E-24 |
| ACE2 | RP3-522P13.3 | -0,5046 | 1,01246E-24 |
| ACE2 | AC093716.1 | -0,5046 | 1,02636E-24 |
| ACE2 | AC079781.7 | -0,5046 | 1,01033E-24 |
| ACE2 | WTAPP1 | -0,5046 | 1,01461E-24 |
| ACE2 | RP11-80F22.15 | -0,5046 | 1,02599E-24 |
| ACE2 | ITIH4 | -0,5047 | 9,98493E-25 |
| ACE2 | C4orf17 | -0,5047 | 9,87083E-25 |
| ACE2 | RP11-369E15.1 | -0,5047 | 9,99512E-25 |
| ACE2 | RP11-49K4.2 | -0,5047 | 1,00773E-24 |
| ACE2 | C12orf71 | -0,5047 | 1,00126E-24 |
| ACE2 | RP11-456O19.3 | -0,5047 | 1,00697E-24 |
| ACE2 | LINC01594 | -0,5048 | 9,7274E-25 |
| ACE2 | RP1-60O19.1 | -0,5048 | 9,75984E-25 |
| ACE2 | RP11-276H1.2 | -0,5048 | 9,69605E-25 |
| ACE2 | RFPL4AP6 | -0,5048 | 9,67159E-25 |
| ACE2 | KCND3-AS1 | -0,5049 | 9,38036E-25 |
| ACE2 | GS1-72M22.1 | -0,5049 | 9,47648E-25 |
| ACE2 | LINC00442 | -0,5049 | 9,59952E-25 |
| ACE2 | RP5-864K19.4 | -0,505 | 9,27974E-25 |
| ACE2 | CDC37L1-AS1 | -0,505 | 9,16659E-25 |
| ACE2 | SYP-AS1 | -0,505 | 9,27669E-25 |
| ACE2 | RP11-263K19.6 | -0,5051 | 8,97685E-25 |
| ACE2 | LINC02174 | -0,5051 | 9,08626E-25 |
| ACE2 | MIR3943 | -0,5051 | 9,05107E-25 |

| | | | |
|---|---|---|---|
| ACE2 | LRGUK | -0,5051 | 8,98119E-25 |
| ACE2 | HSPA8P11 | -0,5051 | 9,04212E-25 |
| ACE2 | RP1-239B22.5 | -0,5051 | 9,11777E-25 |
| ACE2 | CEACAM21 | -0,5051 | 8,98648E-25 |
| ACE2 | MUC4 | -0,5052 | 8,87384E-25 |
| ACE2 | RP11-114J13.1 | -0,5052 | 8,8874E-25 |
| ACE2 | RAD21-AS1 | -0,5052 | 8,88774E-25 |
| ACE2 | RP11-379C10.4 | -0,5052 | 8,86794E-25 |
| ACE2 | RP11-25G10.2 | -0,5052 | 8,83281E-25 |
| ACE2 | CUBNP3 | -0,5052 | 8,83572E-25 |
| ACE2 | AC064874.1 | -0,5053 | 8,6672E-25 |
| ACE2 | CTD-2192J16.26 | -0,5053 | 8,59022E-25 |
| ACE2 | AC034243.1 | -0,5054 | 8,47253E-25 |
| ACE2 | DENND1A | -0,5054 | 8,47189E-25 |
| ACE2 | RP11-544P1.3 | -0,5054 | 8,48275E-25 |
| ACE2 | RP6-99M1.1 | -0,5054 | 8,39051E-25 |
| ACE2 | RP11-150O12.6 | -0,5055 | 8,29201E-25 |
| ACE2 | RPS17P15 | -0,5055 | 8,22672E-25 |
| ACE2 | RP11-356J5.12 | -0,5055 | 8,1725E-25 |
| ACE2 | CACNA1C | -0,5055 | 8,15531E-25 |
| ACE2 | TYMSOS | -0,5055 | 8,28681E-25 |
| ACE2 | RP11-276E15.4 | -0,5056 | 7,98133E-25 |
| ACE2 | BSX | -0,5056 | 8,08739E-25 |
| ACE2 | KIF21B | -0,5057 | 7,82978E-25 |
| ACE2 | CCNYL2 | -0,5057 | 7,81494E-25 |
| ACE2 | APOF | -0,5057 | 7,87403E-25 |
| ACE2 | SKAP1 | -0,5057 | 7,79187E-25 |
| ACE2 | CCDC170 | -0,5058 | 7,64107E-25 |
| ACE2 | PFKP | -0,5058 | 7,6417E-25 |
| ACE2 | NACAP1 | -0,5059 | 7,49707E-25 |
| ACE2 | LUZP2 | -0,5059 | 7,40431E-25 |
| ACE2 | RP11-22P4.2 | -0,5059 | 7,40533E-25 |
| ACE2 | RP5-1077H22.1 | -0,506 | 7,1616E-25 |
| ACE2 | RP11-255A11.2 | -0,506 | 7,24904E-25 |
| ACE2 | CISTR | -0,506 | 7,21394E-25 |
| ACE2 | EP400NL | -0,506 | 7,33164E-25 |
| ACE2 | AKAP4 | -0,506 | 7,20042E-25 |
| ACE2 | EHHADH-AS1 | -0,5061 | 7,1223E-25 |
| ACE2 | ZNF793-AS1 | -0,5061 | 7,1166E-25 |
| ACE2 | LINC02059 | -0,5062 | 6,92004E-25 |
| ACE2 | ARHGAP26-IT1 | -0,5062 | 6,89248E-25 |
| ACE2 | SLC26A8 | -0,5062 | 6,92052E-25 |
| ACE2 | SDAD1P1 | -0,5062 | 6,92268E-25 |
| ACE2 | CARS | -0,5062 | 6,91876E-25 |
| ACE2 | RP11-566K19.6 | -0,5062 | 6,97023E-25 |
| ACE2 | ZNF790 | -0,5062 | 6,83422E-25 |
| ACE2 | MACROD2-IT1 | -0,5062 | 6,9222E-25 |
| ACE2 | RP11-530C5.2 | -0,5063 | 6,74168E-25 |
| ACE2 | LINC00376 | -0,5063 | 6,69362E-25 |
| ACE2 | RP1-300I2.2 | -0,5063 | 6,7126E-25 |
| ACE2 | RP11-346J10.2 | -0,5064 | 6,62342E-25 |
| ACE2 | TMEM217 | -0,5064 | 6,5848E-25 |
| ACE2 | RP11-203B9.4 | -0,5064 | 6,49085E-25 |
| ACE2 | RP5-1148A21.3 | -0,5064 | 6,60725E-25 |

| | | | |
|---|---|---|---|
| ACE2 | KCNMA1-AS3 | -0,5064 | 6,55368E-25 |
| ACE2 | ZNF573 | -0,5064 | 6,54468E-25 |
| ACE2 | RP11-495K9.2 | -0,5065 | 6,33417E-25 |
| ACE2 | ADPGK-AS1 | -0,5065 | 6,36511E-25 |
| ACE2 | EFTUD1P1 | -0,5065 | 6,44636E-25 |
| ACE2 | LINC00254 | -0,5065 | 6,42028E-25 |
| ACE2 | RP11-12K11.2 | -0,5066 | 6,20331E-25 |
| ACE2 | LINC00327 | -0,5066 | 6,3177E-25 |
| ACE2 | AC006277.2 | -0,5066 | 6,27562E-25 |
| ACE2 | AC005262.3 | -0,5066 | 6,32606E-25 |
| ACE2 | PMM1 | -0,5066 | 6,29524E-25 |
| ACE2 | PAX4 | -0,5067 | 6,09715E-25 |
| ACE2 | WASHC3 | -0,5067 | 6,03976E-25 |
| ACE2 | CHRNE | -0,5067 | 6,057E-25 |
| ACE2 | RP11-98D18.15 | -0,5068 | 5,95509E-25 |
| ACE2 | ENAM | -0,5068 | 5,96682E-25 |
| ACE2 | CTD-2161F6.2 | -0,5068 | 5,90885E-25 |
| ACE2 | SNCAIP | -0,5068 | 6,02271E-25 |
| ACE2 | HRAT17 | -0,5069 | 5,73958E-25 |
| ACE2 | RP11-84C10.1 | -0,5069 | 5,77957E-25 |
| ACE2 | CHST1 | -0,507 | 5,61174E-25 |
| ACE2 | GPA33 | -0,5071 | 5,48152E-25 |
| ACE2 | HSPA1L | -0,5071 | 5,4908E-25 |
| ACE2 | C7orf31 | -0,5071 | 5,4737E-25 |
| ACE2 | STPG3-AS1 | -0,5071 | 5,5099E-25 |
| ACE2 | TMEM63C | -0,5071 | 5,5356E-25 |
| ACE2 | SRD5A2 | -0,5072 | 5,44159E-25 |
| ACE2 | RP11-379F4.4 | -0,5072 | 5,41662E-25 |
| ACE2 | ADCY10P1 | -0,5072 | 5,45238E-25 |
| ACE2 | RP1-45I4.2 | -0,5072 | 5,43961E-25 |
| ACE2 | RP11-618M23.2 | -0,5072 | 5,40442E-25 |
| ACE2 | SLIT1 | -0,5072 | 5,38852E-25 |
| ACE2 | ANO1 | -0,5072 | 5,38919E-25 |
| ACE2 | AC015923.1 | -0,5072 | 5,38144E-25 |
| ACE2 | RP11-760L24.1 | -0,5072 | 5,4415E-25 |
| ACE2 | RP11-412P11.1 | -0,5073 | 5,2022E-25 |
| ACE2 | SOX30 | -0,5073 | 5,19919E-25 |
| ACE2 | C6orf222 | -0,5073 | 5,25365E-25 |
| ACE2 | ZNF479 | -0,5073 | 5,24853E-25 |
| ACE2 | LMNTD1 | -0,5073 | 5,23443E-25 |
| ACE2 | RP11-76C10.4 | -0,5073 | 5,30256E-25 |
| ACE2 | PCLAF | -0,5073 | 5,30675E-25 |
| ACE2 | OR1C1 | -0,5074 | 5,15453E-25 |
| ACE2 | AC009305.1 | -0,5074 | 5,12967E-25 |
| ACE2 | AC007557.3 | -0,5074 | 5,11812E-25 |
| ACE2 | CPHL1P | -0,5074 | 5,09875E-25 |
| ACE2 | UQCC2 | -0,5074 | 5,1846E-25 |
| ACE2 | RP13-60M5.2 | -0,5074 | 5,13227E-25 |
| ACE2 | RP11-100F15.2 | -0,5074 | 5,1447E-25 |
| ACE2 | RP11-1086F11.1 | -0,5075 | 4,95707E-25 |
| ACE2 | ADD2 | -0,5075 | 5,06301E-25 |
| ACE2 | CDH16 | -0,5075 | 5,01554E-25 |
| ACE2 | RP11-122D10.1 | -0,5076 | 4,91659E-25 |
| ACE2 | RP11-411B10.2 | -0,5076 | 4,83034E-25 |

| | | | |
|---|---|---|---|
| ACE2 | XKR3 | -0,5076 | 4,93661E-25 |
| ACE2 | AC079354.5 | -0,5077 | 4,79775E-25 |
| ACE2 | AC116609.2 | -0,5078 | 4,70214E-25 |
| ACE2 | RP11-328K2.1 | -0,5078 | 4,64422E-25 |
| ACE2 | FLJ27354 | -0,5079 | 4,58528E-25 |
| ACE2 | GATA3-AS1 | -0,5079 | 4,47875E-25 |
| ACE2 | CCDC86 | -0,5079 | 4,55428E-25 |
| ACE2 | SPATA8-AS1 | -0,508 | 4,44745E-25 |
| ACE2 | TMEM200C | -0,508 | 4,40791E-25 |
| ACE2 | CNTN2 | -0,5081 | 4,26326E-25 |
| ACE2 | DNAH6 | -0,5081 | 4,32501E-25 |
| ACE2 | RP11-452D21.2 | -0,5081 | 4,29908E-25 |
| ACE2 | RP11-353P15.1 | -0,5081 | 4,34217E-25 |
| ACE2 | MLIP | -0,5082 | 4,17543E-25 |
| ACE2 | DHRS7C | -0,5082 | 4,25763E-25 |
| ACE2 | CTD-2213F21.4 | -0,5083 | 4,07793E-25 |
| ACE2 | TMEM132E | -0,5083 | 4,05521E-25 |
| ACE2 | SCGB2B2 | -0,5083 | 4,06287E-25 |
| ACE2 | KRTAP21-2 | -0,5083 | 4,06532E-25 |
| ACE2 | SCN3A | -0,5084 | 4,04362E-25 |
| ACE2 | ARMC3 | -0,5084 | 4,00894E-25 |
| ACE2 | CTD-2071N1.2 | -0,5084 | 3,99406E-25 |
| ACE2 | CTA-363E6.6 | -0,5084 | 3,9647E-25 |
| ACE2 | CTA-963H5.5 | -0,5084 | 4,01334E-25 |
| ACE2 | AC092155.4 | -0,5085 | 3,87624E-25 |
| ACE2 | C2orf69P1 | -0,5085 | 3,89367E-25 |
| ACE2 | TNNI3 | -0,5085 | 3,94791E-25 |
| ACE2 | LINC01626 | -0,5086 | 3,80042E-25 |
| ACE2 | OR52D1 | -0,5086 | 3,8352E-25 |
| ACE2 | RP11-379J13.2 | -0,5086 | 3,81722E-25 |
| ACE2 | RP11-131H24.5 | -0,5086 | 3,79934E-25 |
| ACE2 | CILP2 | -0,5086 | 3,78621E-25 |
| ACE2 | CTD-3224I3.3 | -0,5087 | 3,73042E-25 |
| ACE2 | OR10AE3P | -0,5087 | 3,69412E-25 |
| ACE2 | METTL23 | -0,5087 | 3,72608E-25 |
| ACE2 | LINC01046 | -0,5088 | 3,65769E-25 |
| ACE2 | RP11-1000B6.2 | -0,5088 | 3,62639E-25 |
| ACE2 | RP11-624L4.2 | -0,5088 | 3,63953E-25 |
| ACE2 | RP11-330L19.2 | -0,5088 | 3,59261E-25 |
| ACE2 | PPIL6 | -0,5089 | 3,5436E-25 |
| ACE2 | RP11-64K12.9 | -0,5089 | 3,50838E-25 |
| ACE2 | RP11-103J17.1 | -0,509 | 3,47836E-25 |
| ACE2 | RP11-713M15.1 | -0,509 | 3,4928E-25 |
| ACE2 | FCAMR | -0,5091 | 3,3958E-25 |
| ACE2 | AC093326.3 | -0,5091 | 3,32813E-25 |
| ACE2 | TEX37 | -0,5091 | 3,32483E-25 |
| ACE2 | SNTG2 | -0,5092 | 3,31369E-25 |
| ACE2 | CCDC173 | -0,5092 | 3,26311E-25 |
| ACE2 | RP11-178F10.2 | -0,5092 | 3,27283E-25 |
| ACE2 | DMTN | -0,5093 | 3,20628E-25 |
| ACE2 | C8orf89 | -0,5093 | 3,217E-25 |
| ACE2 | PRKCG | -0,5093 | 3,18568E-25 |
| ACE2 | CLSTN2-AS1 | -0,5094 | 3,11921E-25 |
| ACE2 | SLC17A4 | -0,5094 | 3,1612E-25 |

| | | | |
|---|---|---|---|
| ACE2 | IGLVIV-64 | -0,5094 | 3,10369E-25 |
| ACE2 | LETM2 | -0,5095 | 3,0499E-25 |
| ACE2 | LINC01609 | -0,5095 | 3,02258E-25 |
| ACE2 | PHBP18 | -0,5095 | 3,02685E-25 |
| ACE2 | DCC | -0,5096 | 2,97236E-25 |
| ACE2 | TRPM1 | -0,5097 | 2,874E-25 |
| ACE2 | RP11-68I3.7 | -0,5097 | 2,89756E-25 |
| ACE2 | LINC01883 | -0,5098 | 2,81811E-25 |
| ACE2 | BSN-AS1 | -0,5098 | 2,83232E-25 |
| ACE2 | RP11-242F11.2 | -0,5098 | 2,8417E-25 |
| ACE2 | EXPH5 | -0,5098 | 2,83259E-25 |
| ACE2 | RP11-627G18.2 | -0,5098 | 2,80765E-25 |
| ACE2 | C1QL4 | -0,5099 | 2,72839E-25 |
| ACE2 | RP11-718I15.1 | -0,5099 | 2,76199E-25 |
| ACE2 | GS1-24F4.3 | -0,51 | 2,68715E-25 |
| ACE2 | SDCBP2 | -0,51 | 2,68164E-25 |
| ACE2 | ZIM2 | -0,5101 | 2,62539E-25 |
| ACE2 | RP1-127D3.4 | -0,5102 | 2,56351E-25 |
| ACE2 | CTD-2244C20.2 | -0,5102 | 2,53005E-25 |
| ACE2 | RUSC2 | -0,5102 | 2,57326E-25 |
| ACE2 | TMC5 | -0,5102 | 2,5385E-25 |
| ACE2 | RP11-45I20.1 | -0,5103 | 2,50801E-25 |
| ACE2 | RP11-15B24.4 | -0,5103 | 2,52229E-25 |
| ACE2 | OLFM5P | -0,5103 | 2,52077E-25 |
| ACE2 | UBAC2-AS1 | -0,5103 | 2,47838E-25 |
| ACE2 | RP11-1078H9.1 | -0,5103 | 2,50408E-25 |
| ACE2 | CTC-524C5.5 | -0,5103 | 2,48659E-25 |
| ACE2 | RP11-795H16.3 | -0,5103 | 2,5029E-25 |
| ACE2 | MYL12BP2 | -0,5104 | 2,42299E-25 |
| ACE2 | CTD-2001E22.1 | -0,5104 | 2,45258E-25 |
| ACE2 | RP11-168K9.1 | -0,5104 | 2,43304E-25 |
| ACE2 | LINC00452 | -0,5104 | 2,42679E-25 |
| ACE2 | RP11-255M2.3 | -0,5104 | 2,45663E-25 |
| ACE2 | RP11-527H14.4 | -0,5104 | 2,45579E-25 |
| ACE2 | AC073636.1 | -0,5105 | 2,37126E-25 |
| ACE2 | RP11-265B7.1 | -0,5105 | 2,35122E-25 |
| ACE2 | SPATA31B1P | -0,5105 | 2,37517E-25 |
| ACE2 | RP11-152H18.4 | -0,5105 | 2,38813E-25 |
| ACE2 | LINC00939 | -0,5105 | 2,36757E-25 |
| ACE2 | TRIM13 | -0,5105 | 2,37714E-25 |
| ACE2 | CSNK1G1 | -0,5105 | 2,35237E-25 |
| ACE2 | LINC01628 | -0,5106 | 2,30288E-25 |
| ACE2 | CTD-2587H24.5 | -0,5106 | 2,30888E-25 |
| ACE2 | ARL13A | -0,5106 | 2,32777E-25 |
| ACE2 | RP11-135A1.3 | -0,5107 | 2,27205E-25 |
| ACE2 | GATA1 | -0,5107 | 2,25805E-25 |
| ACE2 | RP11-168O16.2 | -0,5108 | 2,19067E-25 |
| ACE2 | RPL7AP25 | -0,5108 | 2,17574E-25 |
| ACE2 | RP11-574O7.1 | -0,5108 | 2,1829E-25 |
| ACE2 | CTD-2534I21.8 | -0,5109 | 2,16711E-25 |
| ACE2 | CTC-537E7.2 | -0,511 | 2,11003E-25 |
| ACE2 | SAMD3 | -0,511 | 2,11901E-25 |
| ACE2 | SLC25A1P3 | -0,511 | 2,10657E-25 |
| ACE2 | NRARP | -0,511 | 2,11953E-25 |

| | | | |
|---|---|---|---|
| ACE2 | CTD-2583P5.3 | -0,511 | 2,10008E-25 |
| ACE2 | LINC02102 | -0,5111 | 2,04691E-25 |
| ACE2 | KRT81 | -0,5112 | 1,98787E-25 |
| ACE2 | AC007731.1 | -0,5112 | 2,00965E-25 |
| ACE2 | IQSEC2 | -0,5112 | 1,98022E-25 |
| ACE2 | RP4-803A2.2 | -0,5113 | 1,93669E-25 |
| ACE2 | AXDND1 | -0,5113 | 1,96256E-25 |
| ACE2 | KCNQ5 | -0,5113 | 1,93655E-25 |
| ACE2 | SLC6A16 | -0,5113 | 1,96679E-25 |
| ACE2 | RP3-430A16.1 | -0,5114 | 1,88985E-25 |
| ACE2 | RP11-678G14.4 | -0,5114 | 1,89498E-25 |
| ACE2 | VPS9D1 | -0,5115 | 1,87082E-25 |
| ACE2 | RP11-105N14.2 | -0,5116 | 1,79101E-25 |
| ACE2 | SEPT4-AS1 | -0,5116 | 1,79348E-25 |
| ACE2 | C2orf16 | -0,5117 | 1,7489E-25 |
| ACE2 | AC007163.6 | -0,5117 | 1,76867E-25 |
| ACE2 | NUP210P1 | -0,5117 | 1,75673E-25 |
| ACE2 | RP11-577G20.2 | -0,5117 | 1,75953E-25 |
| ACE2 | RP11-730G20.1 | -0,5117 | 1,74469E-25 |
| ACE2 | RLN2 | -0,5117 | 1,76814E-25 |
| ACE2 | AP000442.4 | -0,5117 | 1,73659E-25 |
| ACE2 | AC079354.3 | -0,5118 | 1,69625E-25 |
| ACE2 | IRF1 | -0,5118 | 1,71253E-25 |
| ACE2 | RP11-138E9.2 | -0,5118 | 1,71808E-25 |
| ACE2 | RP11-1E4.1 | -0,5119 | 1,69251E-25 |
| ACE2 | NOL4 | -0,5119 | 1,6914E-25 |
| ACE2 | RP11-344A5.1 | -0,512 | 1,647E-25 |
| ACE2 | CEP152 | -0,512 | 1,64412E-25 |
| ACE2 | C16orf45 | -0,512 | 1,64214E-25 |
| ACE2 | RP11-65N13.8 | -0,5121 | 1,60629E-25 |
| ACE2 | LIN37 | -0,5121 | 1,59929E-25 |
| ACE2 | CALM3 | -0,5121 | 1,58033E-25 |
| ACE2 | RP11-506B6.6 | -0,5122 | 1,55109E-25 |
| ACE2 | RP11-157I4.4 | -0,5122 | 1,53434E-25 |
| ACE2 | GPR158 | -0,5122 | 1,55181E-25 |
| ACE2 | TRMT5 | -0,5122 | 1,54428E-25 |
| ACE2 | ZC2HC1C | -0,5122 | 1,55476E-25 |
| ACE2 | RPL3L | -0,5122 | 1,56233E-25 |
| ACE2 | RP3-400B16.3 | -0,5123 | 1,52956E-25 |
| ACE2 | INTS1 | -0,5123 | 1,50242E-25 |
| ACE2 | NOS2P2 | -0,5123 | 1,50935E-25 |
| ACE2 | NUP62 | -0,5123 | 1,51137E-25 |
| ACE2 | FCRL5 | -0,5124 | 1,48008E-25 |
| ACE2 | ZNF474 | -0,5124 | 1,46137E-25 |
| ACE2 | SLC5A8 | -0,5124 | 1,46067E-25 |
| ACE2 | RIMBP3C | -0,5124 | 1,46283E-25 |
| ACE2 | SI | -0,5125 | 1,43042E-25 |
| ACE2 | IGF2BP2-AS1 | -0,5125 | 1,44932E-25 |
| ACE2 | RP11-196D18.1 | -0,5125 | 1,43229E-25 |
| ACE2 | RP11-498P14.4 | -0,5125 | 1,43392E-25 |
| ACE2 | TOPORSLP1 | -0,5125 | 1,4498E-25 |
| ACE2 | LINC01056 | -0,5125 | 1,44482E-25 |
| ACE2 | RGPD4 | -0,5126 | 1,39242E-25 |
| ACE2 | RP11-713M6.2 | -0,5126 | 1,40729E-25 |

| | | | |
|---|---|---|---|
| ACE2 | EMC8 | -0,5126 | 1,41787E-25 |
| ACE2 | AC015849.12 | -0,5126 | 1,3907E-25 |
| ACE2 | SCAND1 | -0,5126 | 1,41669E-25 |
| ACE2 | FIS1 | -0,5127 | 1,37528E-25 |
| ACE2 | GRM5 | -0,5127 | 1,37123E-25 |
| ACE2 | NAGS | -0,5127 | 1,37449E-25 |
| ACE2 | LINC01337 | -0,5128 | 1,34557E-25 |
| ACE2 | PSMB7 | -0,5128 | 1,32611E-25 |
| ACE2 | ANO3 | -0,5128 | 1,34591E-25 |
| ACE2 | RP11-348B17.1 | -0,5128 | 1,31801E-25 |
| ACE2 | QPCTL | -0,5128 | 1,32436E-25 |
| ACE2 | SPATA17-AS1 | -0,5129 | 1,3066E-25 |
| ACE2 | RP11-563N6.6 | -0,5129 | 1,31381E-25 |
| ACE2 | RP11-840I19.3 | -0,5129 | 1,31375E-25 |
| ACE2 | CCDC154 | -0,5129 | 1,29702E-25 |
| ACE2 | RP11-515E23.1 | -0,5129 | 1,28725E-25 |
| ACE2 | RP11-697B24.1 | -0,513 | 1,26246E-25 |
| ACE2 | MGMT | -0,513 | 1,26161E-25 |
| ACE2 | ABCG4 | -0,513 | 1,25418E-25 |
| ACE2 | LINC00448 | -0,513 | 1,27133E-25 |
| ACE2 | RP11-79P5.9 | -0,5131 | 1,23222E-25 |
| ACE2 | RP11-121A14.2 | -0,5131 | 1,22448E-25 |
| ACE2 | RP1-251M9.2 | -0,5131 | 1,23683E-25 |
| ACE2 | CTD-3028N15.1 | -0,5131 | 1,24157E-25 |
| ACE2 | PCBP3 | -0,5131 | 1,24057E-25 |
| ACE2 | RP11-390B4.3 | -0,5132 | 1,21007E-25 |
| ACE2 | RP11-338O1.2 | -0,5132 | 1,22018E-25 |
| ACE2 | RP11-99L13.2 | -0,5132 | 1,19147E-25 |
| ACE2 | ABCC12 | -0,5132 | 1,20141E-25 |
| ACE2 | RPL34-AS1 | -0,5133 | 1,19067E-25 |
| ACE2 | LRRC37A5P | -0,5133 | 1,18447E-25 |
| ACE2 | STX5 | -0,5133 | 1,16964E-25 |
| ACE2 | AC118653.2 | -0,5133 | 1,18345E-25 |
| ACE2 | LL22NC03-23C6.13 | -0,5133 | 1,16303E-25 |
| ACE2 | LINC01398 | -0,5134 | 1,13631E-25 |
| ACE2 | DNM3-IT1 | -0,5134 | 1,14488E-25 |
| ACE2 | CPEB2-AS1 | -0,5134 | 1,13454E-25 |
| ACE2 | LINC01950 | -0,5134 | 1,15054E-25 |
| ACE2 | RP1-140C12.2 | -0,5134 | 1,14227E-25 |
| ACE2 | LINC02127 | -0,5134 | 1,13691E-25 |
| ACE2 | FHAD1 | -0,5135 | 1,11393E-25 |
| ACE2 | MYT1L-AS1 | -0,5135 | 1,109E-25 |
| ACE2 | LRRC6 | -0,5135 | 1,10918E-25 |
| ACE2 | SVOP | -0,5135 | 1,1195E-25 |
| ACE2 | TBX5-AS1 | -0,5135 | 1,11229E-25 |
| ACE2 | RP11-357K6.2 | -0,5135 | 1,10751E-25 |
| ACE2 | APOBEC4 | -0,5136 | 1,07695E-25 |
| ACE2 | LINC01276 | -0,5136 | 1,08525E-25 |
| ACE2 | FAM230C | -0,5136 | 1,08866E-25 |
| ACE2 | RP11-507K2.3 | -0,5136 | 1,08475E-25 |
| ACE2 | CTD-2319I12.2 | -0,5137 | 1,05232E-25 |
| ACE2 | USH2A | -0,5138 | 1,04427E-25 |
| ACE2 | RP11-244K5.6 | -0,5138 | 1,02444E-25 |
| ACE2 | BCL2L1 | -0,5138 | 1,0275E-25 |

| | | | |
|---|---|---|---|
| ACE2 | CFAP36 | -0,5139 | 1,02177E-25 |
| ACE2 | RP11-14D22.1 | -0,5139 | 1,00665E-25 |
| ACE2 | RP11-753D20.3 | -0,5139 | 1,01772E-25 |
| ACE2 | ZNF821 | -0,5139 | 1,00004E-25 |
| ACE2 | RP11-396N11.1 | -0,5139 | 1,01658E-25 |
| ACE2 | C3P1 | -0,5139 | 1,0136E-25 |
| ACE2 | ZNF571-AS1 | -0,5139 | 1,02041E-25 |
| ACE2 | RP11-798K23.1 | -0,514 | 9,92554E-26 |
| ACE2 | LINC01505 | -0,514 | 9,82926E-26 |
| ACE2 | TSPAN19 | -0,514 | 9,73303E-26 |
| ACE2 | BIRC8 | -0,514 | 9,75435E-26 |
| ACE2 | ARF4P2 | -0,514 | 9,93599E-26 |
| ACE2 | NCBP2L | -0,514 | 9,82124E-26 |
| ACE2 | CCDC148 | -0,5141 | 9,53597E-26 |
| ACE2 | RP11-350E12.6 | -0,5141 | 9,70616E-26 |
| ACE2 | PLCZ1 | -0,5141 | 9,69053E-26 |
| ACE2 | RP11-609L23.1 | -0,5141 | 9,65021E-26 |
| ACE2 | RP11-960B9.2 | -0,5141 | 9,52868E-26 |
| ACE2 | RP13-131K19.1 | -0,5142 | 9,32404E-26 |
| ACE2 | RP11-561O23.5 | -0,5142 | 9,33072E-26 |
| ACE2 | LINC00619 | -0,5142 | 9,397E-26 |
| ACE2 | LINC2194 | -0,5142 | 9,40675E-26 |
| ACE2 | DRD3 | -0,5143 | 9,13354E-26 |
| ACE2 | ZP1 | -0,5143 | 9,19613E-26 |
| ACE2 | PIP5K1C | -0,5143 | 9,06505E-26 |
| ACE2 | CFAP74 | -0,5144 | 8,84615E-26 |
| ACE2 | RP13-39P12.3 | -0,5144 | 8,85481E-26 |
| ACE2 | RP11-469N6.2 | -0,5144 | 8,83299E-26 |
| ACE2 | CDKN3 | -0,5144 | 8,91909E-26 |
| ACE2 | RP5-991G20.2 | -0,5144 | 8,89668E-26 |
| ACE2 | AC013470.6 | -0,5145 | 8,64318E-26 |
| ACE2 | AC073342.12 | -0,5145 | 8,60315E-26 |
| ACE2 | RP11-713M15.2 | -0,5145 | 8,75027E-26 |
| ACE2 | RP11-114O8.1 | -0,5146 | 8,54358E-26 |
| ACE2 | GRIN2B | -0,5146 | 8,37945E-26 |
| ACE2 | RP1-4G17.2 | -0,5146 | 8,55461E-26 |
| ACE2 | SAP18P3 | -0,5147 | 8,2161E-26 |
| ACE2 | C12orf50 | -0,5147 | 8,31085E-26 |
| ACE2 | RP11-1217F2.15 | -0,5148 | 8,06381E-26 |
| ACE2 | SLC25A51P1 | -0,5149 | 7,8844E-26 |
| ACE2 | RP11-738G5.1 | -0,5149 | 7,79852E-26 |
| ACE2 | DDX19B | -0,5149 | 7,86678E-26 |
| ACE2 | LYPD4 | -0,5149 | 7,75357E-26 |
| ACE2 | ANKRD60 | -0,5149 | 7,93856E-26 |
| ACE2 | DNAH14 | -0,515 | 7,67122E-26 |
| ACE2 | NT5C1B | -0,515 | 7,61583E-26 |
| ACE2 | RP11-526A4.1 | -0,515 | 7,70646E-26 |
| ACE2 | LECT2 | -0,515 | 7,73266E-26 |
| ACE2 | AC000124.1 | -0,515 | 7,63749E-26 |
| ACE2 | COX5A | -0,515 | 7,6214E-26 |
| ACE2 | ATP8B3 | -0,515 | 7,69066E-26 |
| ACE2 | RP3-522D1.1 | -0,5151 | 7,39534E-26 |
| ACE2 | GPR87 | -0,5151 | 7,51519E-26 |
| ACE2 | MAP6D1 | -0,5151 | 7,48151E-26 |

| | | | |
|---|---|---|---|
| ACE2 | RP11-164O23.7 | -0,5152 | 7,34629E-26 |
| ACE2 | LINC02060 | -0,5152 | 7,31947E-26 |
| ACE2 | TYSND1 | -0,5152 | 7,21256E-26 |
| ACE2 | NR0B2 | -0,5153 | 7,11316E-26 |
| ACE2 | LINC01688 | -0,5153 | 7,04846E-26 |
| ACE2 | ACTBP13 | -0,5153 | 7,06038E-26 |
| ACE2 | LINC00567 | -0,5153 | 7,07114E-26 |
| ACE2 | RP11-486O13.4 | -0,5153 | 7,15722E-26 |
| ACE2 | GPI | -0,5153 | 7,07353E-26 |
| ACE2 | TM4SF20 | -0,5154 | 6,8538E-26 |
| ACE2 | RP11-651P23.5 | -0,5154 | 6,97338E-26 |
| ACE2 | RAI14 | -0,5154 | 6,83241E-26 |
| ACE2 | RP3-399L15.3 | -0,5154 | 6,94258E-26 |
| ACE2 | CDH13 | -0,5154 | 6,85922E-26 |
| ACE2 | AADACL3 | -0,5155 | 6,69137E-26 |
| ACE2 | RP11-350N15.5 | -0,5155 | 6,82146E-26 |
| ACE2 | CDCA5 | -0,5155 | 6,76187E-26 |
| ACE2 | LINC01068 | -0,5155 | 6,74066E-26 |
| ACE2 | RP11-68L1.2 | -0,5156 | 6,64855E-26 |
| ACE2 | CTC-340A15.2 | -0,5156 | 6,62823E-26 |
| ACE2 | CAPN11 | -0,5156 | 6,56986E-26 |
| ACE2 | RP11-535C21.3 | -0,5156 | 6,64411E-26 |
| ACE2 | ACRV1 | -0,5156 | 6,60983E-26 |
| ACE2 | ZC3H12A | -0,5157 | 6,37202E-26 |
| ACE2 | ZBTB17 | -0,5158 | 6,24802E-26 |
| ACE2 | LINC02067 | -0,5159 | 6,05405E-26 |
| ACE2 | UBE2R2-AS1 | -0,5159 | 6,10881E-26 |
| ACE2 | LINC00378 | -0,5159 | 6,13594E-26 |
| ACE2 | RP5-1044H5.1 | -0,5161 | 5,73715E-26 |
| ACE2 | LINC01209 | -0,5161 | 5,83401E-26 |
| ACE2 | RP11-631F7.2 | -0,5161 | 5,81454E-26 |
| ACE2 | GS1-259H13.2 | -0,5161 | 5,84401E-26 |
| ACE2 | CTD-3088G3.4 | -0,5161 | 5,71596E-26 |
| ACE2 | AVP | -0,5161 | 5,82159E-26 |
| ACE2 | MSANTD1 | -0,5162 | 5,68451E-26 |
| ACE2 | LINC00533 | -0,5163 | 5,45708E-26 |
| ACE2 | RP11-440G5.2 | -0,5163 | 5,54597E-26 |
| ACE2 | RP5-944M2.3 | -0,5163 | 5,46574E-26 |
| ACE2 | RP11-463J10.2 | -0,5163 | 5,4477E-26 |
| ACE2 | RP13-270P17.1 | -0,5163 | 5,55777E-26 |
| ACE2 | GALR1 | -0,5163 | 5,50412E-26 |
| ACE2 | PIN1 | -0,5163 | 5,53949E-26 |
| ACE2 | DNAH1 | -0,5164 | 5,36874E-26 |
| ACE2 | RP11-428L21.1 | -0,5164 | 5,39457E-26 |
| ACE2 | CRADD | -0,5164 | 5,33013E-26 |
| ACE2 | LINC00524 | -0,5164 | 5,29689E-26 |
| ACE2 | RP11-63A1.1 | -0,5164 | 5,31893E-26 |
| ACE2 | TUBAP | -0,5164 | 5,34762E-26 |
| ACE2 | RP11-375O18.2 | -0,5165 | 5,17887E-26 |
| ACE2 | RP11-284G10.1 | -0,5165 | 5,187E-26 |
| ACE2 | SRRM4 | -0,5165 | 5,23131E-26 |
| ACE2 | RP11-527N12.2 | -0,5165 | 5,18937E-26 |
| ACE2 | CCL1 | -0,5165 | 5,20169E-26 |
| ACE2 | RP11-605B16.1 | -0,5166 | 5,06485E-26 |

| | | | |
|---|---|---|---|
| ACE2 | KLHL40 | -0,5166 | 5,10974E-26 |
| ACE2 | ZNF846 | -0,5166 | 5,04457E-26 |
| ACE2 | ELK2AP | -0,5167 | 4,93078E-26 |
| ACE2 | RN7SL645P | -0,5167 | 4,96778E-26 |
| ACE2 | AC016292.1 | -0,5167 | 4,92881E-26 |
| ACE2 | FAM182B | -0,5167 | 5,02305E-26 |
| ACE2 | LINC01744 | -0,5168 | 4,90048E-26 |
| ACE2 | IL5RA | -0,5168 | 4,90281E-26 |
| ACE2 | LINC02103 | -0,5168 | 4,88763E-26 |
| ACE2 | IMP3P2 | -0,5168 | 4,89775E-26 |
| ACE2 | STX17-AS1 | -0,5168 | 4,79962E-26 |
| ACE2 | RP11-384M20.1 | -0,5168 | 4,81781E-26 |
| ACE2 | CTB-50L17.16 | -0,5168 | 4,84569E-26 |
| ACE2 | AC006126.3 | -0,5168 | 4,89712E-26 |
| ACE2 | RP11-264J4.10 | -0,5169 | 4,74384E-26 |
| ACE2 | LINC01539 | -0,5169 | 4,70599E-26 |
| ACE2 | ISM1-AS1 | -0,5169 | 4,69651E-26 |
| ACE2 | RP1-65J11.5 | -0,517 | 4,5512E-26 |
| ACE2 | FNDC4 | -0,517 | 4,64964E-26 |
| ACE2 | RP11-287D1.2 | -0,517 | 4,57221E-26 |
| ACE2 | RP11-326K13.4 | -0,517 | 4,60567E-26 |
| ACE2 | RP11-44N11.2 | -0,5171 | 4,50641E-26 |
| ACE2 | RP11-111M22.3 | -0,5171 | 4,52769E-26 |
| ACE2 | DACT3-AS1 | -0,5171 | 4,48627E-26 |
| ACE2 | AC104777.4 | -0,5172 | 4,34773E-26 |
| ACE2 | RP11-418I22.2 | -0,5172 | 4,32363E-26 |
| ACE2 | CELF4 | -0,5172 | 4,34102E-26 |
| ACE2 | RP11-33B1.1 | -0,5173 | 4,24364E-26 |
| ACE2 | SEPT7P2 | -0,5174 | 4,16932E-26 |
| ACE2 | RP5-1099E6.3 | -0,5175 | 4,09468E-26 |
| ACE2 | AL121656.5 | -0,5175 | 4,06943E-26 |
| ACE2 | AC147651.5 | -0,5175 | 4,07505E-26 |
| ACE2 | LHX2 | -0,5175 | 3,99915E-26 |
| ACE2 | SLC16A12-AS1 | -0,5175 | 4,00414E-26 |
| ACE2 | RP11-765K14.1 | -0,5176 | 3,99469E-26 |
| ACE2 | RP11-554L12.1 | -0,5176 | 3,9023E-26 |
| ACE2 | TESC-AS1 | -0,5176 | 3,92219E-26 |
| ACE2 | RP11-278A23.2 | -0,5176 | 3,93086E-26 |
| ACE2 | RP11-243J16.7 | -0,5176 | 3,96961E-26 |
| ACE2 | MRPL53 | -0,5177 | 3,82608E-26 |
| ACE2 | RP11-54O15.3 | -0,5177 | 3,82843E-26 |
| ACE2 | STAB2 | -0,5177 | 3,8301E-26 |
| ACE2 | AC091199.1 | -0,5177 | 3,8474E-26 |
| ACE2 | EEF1GP7 | -0,5178 | 3,79405E-26 |
| ACE2 | CTD-2530H12.1 | -0,5178 | 3,78249E-26 |
| ACE2 | RP11-1029J19.2 | -0,5178 | 3,72152E-26 |
| ACE2 | CTD-2555C10.3 | -0,5178 | 3,73307E-26 |
| ACE2 | RP11-313P18.1 | -0,5178 | 3,76675E-26 |
| ACE2 | ZNF677 | -0,5178 | 3,72766E-26 |
| ACE2 | CFAP221 | -0,5179 | 3,68598E-26 |
| ACE2 | MPIG6B | -0,5179 | 3,63517E-26 |
| ACE2 | OVOL1-AS1 | -0,5179 | 3,67825E-26 |
| ACE2 | ISG20 | -0,5179 | 3,68304E-26 |
| ACE2 | PRAC2 | -0,5179 | 3,67025E-26 |

| | | | |
|---|---|---|---|
| ACE2 | SIGLEC15 | -0,5179 | 3,62227E-26 |
| ACE2 | TYRL | -0,518 | 3,59941E-26 |
| ACE2 | LRRC71 | -0,5181 | 3,48518E-26 |
| ACE2 | AC016738.4 | -0,5181 | 3,50213E-26 |
| ACE2 | AGAP7P | -0,5181 | 3,43656E-26 |
| ACE2 | RPL12P38 | -0,5181 | 3,49255E-26 |
| ACE2 | RP11-19O2.2 | -0,5182 | 3,38052E-26 |
| ACE2 | LINC01022 | -0,5182 | 3,40715E-26 |
| ACE2 | C10orf113 | -0,5182 | 3,3903E-26 |
| ACE2 | ZDHHC1 | -0,5182 | 3,35155E-26 |
| ACE2 | ASB17 | -0,5183 | 3,29273E-26 |
| ACE2 | RP11-504P24.3 | -0,5183 | 3,33472E-26 |
| ACE2 | ZFR2 | -0,5183 | 3,29147E-26 |
| ACE2 | MYCBPAP | -0,5184 | 3,21162E-26 |
| ACE2 | LINC01815 | -0,5185 | 3,10832E-26 |
| ACE2 | CDHR3 | -0,5185 | 3,11462E-26 |
| ACE2 | RP11-514D23.3 | -0,5185 | 3,09534E-26 |
| ACE2 | WFDC3 | -0,5185 | 3,1453E-26 |
| ACE2 | LINC01315 | -0,5185 | 3,14419E-26 |
| ACE2 | POMC | -0,5186 | 3,04511E-26 |
| ACE2 | RP11-875H7.2 | -0,5186 | 3,08978E-26 |
| ACE2 | AC099344.3 | -0,5187 | 2,97378E-26 |
| ACE2 | LINC00362 | -0,5187 | 3,00398E-26 |
| ACE2 | LRRC37A3 | -0,5187 | 2,9697E-26 |
| ACE2 | AC092566.1 | -0,5187 | 2,96672E-26 |
| ACE2 | C4orf47 | -0,5188 | 2,86669E-26 |
| ACE2 | RP11-66N24.7 | -0,5189 | 2,83181E-26 |
| ACE2 | CCDC40 | -0,5189 | 2,81195E-26 |
| ACE2 | LINC02049 | -0,519 | 2,75428E-26 |
| ACE2 | AC002463.3 | -0,519 | 2,78641E-26 |
| ACE2 | RP11-527L4.6 | -0,519 | 2,72861E-26 |
| ACE2 | TTC3-AS1 | -0,519 | 2,76854E-26 |
| ACE2 | CCDC91 | -0,5191 | 2,678E-26 |
| ACE2 | RP11-642M2.1 | -0,5191 | 2,7001E-26 |
| ACE2 | RP11-580I16.2 | -0,5191 | 2,67801E-26 |
| ACE2 | RP11-58O15.1 | -0,5192 | 2,6047E-26 |
| ACE2 | SEPSECS-AS1 | -0,5192 | 2,61788E-26 |
| ACE2 | LINC02109 | -0,5192 | 2,59425E-26 |
| ACE2 | AC074389.5 | -0,5192 | 2,6374E-26 |
| ACE2 | EMC9 | -0,5192 | 2,59776E-26 |
| ACE2 | RP11-485M7.3 | -0,5193 | 2,52799E-26 |
| ACE2 | C10orf142 | -0,5193 | 2,56847E-26 |
| ACE2 | RGPD3 | -0,5194 | 2,46336E-26 |
| ACE2 | RP11-1055B8.9 | -0,5194 | 2,49042E-26 |
| ACE2 | AP001596.6 | -0,5194 | 2,48613E-26 |
| ACE2 | RP11-706F1.1 | -0,5195 | 2,44071E-26 |
| ACE2 | RP11-178H8.1 | -0,5195 | 2,43746E-26 |
| ACE2 | PAH | -0,5195 | 2,44763E-26 |
| ACE2 | RP11-470M1.2 | -0,5195 | 2,43524E-26 |
| ACE2 | LINC02134 | -0,5195 | 2,43118E-26 |
| ACE2 | RP11-453A12.1 | -0,5195 | 2,44552E-26 |
| ACE2 | SLC5A9 | -0,5196 | 2,3354E-26 |
| ACE2 | ATP2B2-IT2 | -0,5196 | 2,38356E-26 |
| ACE2 | RP11-304C12.3 | -0,5196 | 2,35903E-26 |

| | | | |
|---|---|---|---|
| ACE2 | FAM155A-IT1 | -0,5196 | 2,3659E-26 |
| ACE2 | IPO4 | -0,5196 | 2,38661E-26 |
| ACE2 | NPIPB8 | -0,5196 | 2,33807E-26 |
| ACE2 | AC012501.2 | -0,5197 | 2,31351E-26 |
| ACE2 | PPP1R1C | -0,5197 | 2,32744E-26 |
| ACE2 | CARD18 | -0,5197 | 2,28659E-26 |
| ACE2 | RP11-523O18.5 | -0,5198 | 2,2713E-26 |
| ACE2 | AC009411.1 | -0,5199 | 2,21212E-26 |
| ACE2 | LINC02050 | -0,5199 | 2,19834E-26 |
| ACE2 | AHRR | -0,5199 | 2,2059E-26 |
| ACE2 | RP11-958J22.3 | -0,5199 | 2,17883E-26 |
| ACE2 | CTC-499B15.7 | -0,5199 | 2,20641E-26 |
| ACE2 | AP001442.2 | -0,5199 | 2,18926E-26 |
| ACE2 | RP11-391M7.3 | -0,52 | 2,14799E-26 |
| ACE2 | HLA-V | -0,5201 | 2,07175E-26 |
| ACE2 | RP11-7F17.4 | -0,5202 | 2,04594E-26 |
| ACE2 | LINC01913 | -0,5203 | 1,99549E-26 |
| ACE2 | SLC22A7 | -0,5203 | 1,98248E-26 |
| ACE2 | NDFIP2-AS1 | -0,5203 | 1,95964E-26 |
| ACE2 | RP4-539M6.22 | -0,5203 | 1,96956E-26 |
| ACE2 | AC004911.2 | -0,5204 | 1,92662E-26 |
| ACE2 | GLTP | -0,5204 | 1,9249E-26 |
| ACE2 | C17orf107 | -0,5204 | 1,90857E-26 |
| ACE2 | LINC01895 | -0,5204 | 1,9447E-26 |
| ACE2 | DEFB124 | -0,5204 | 1,92083E-26 |
| ACE2 | AC007238.1 | -0,5205 | 1,88746E-26 |
| ACE2 | RP11-422J15.1 | -0,5205 | 1,88335E-26 |
| ACE2 | RP11-258F1.1 | -0,5205 | 1,89511E-26 |
| ACE2 | RP11-757O6.6 | -0,5205 | 1,8655E-26 |
| ACE2 | RP11-383G10.5 | -0,5206 | 1,82808E-26 |
| ACE2 | RP11-686G23.2 | -0,5206 | 1,81191E-26 |
| ACE2 | FUT3 | -0,5206 | 1,83083E-26 |
| ACE2 | RP11-220I1.2 | -0,5207 | 1,7829E-26 |
| ACE2 | POLR1D | -0,5208 | 1,72407E-26 |
| ACE2 | RP11-829H16.3 | -0,5208 | 1,72661E-26 |
| ACE2 | STARD6 | -0,5208 | 1,73483E-26 |
| ACE2 | RP11-80K6.2 | -0,5208 | 1,75358E-26 |
| ACE2 | DSCR8 | -0,5208 | 1,75265E-26 |
| ACE2 | NME5 | -0,5209 | 1,70501E-26 |
| ACE2 | RP11-40A13.1 | -0,5209 | 1,67836E-26 |
| ACE2 | AP005530.1 | -0,5209 | 1,70064E-26 |
| ACE2 | CHRM2 | -0,521 | 1,62705E-26 |
| ACE2 | KLF5 | -0,521 | 1,64791E-26 |
| ACE2 | COMP | -0,5211 | 1,60439E-26 |
| ACE2 | LINC00536 | -0,5212 | 1,56641E-26 |
| ACE2 | UNC79 | -0,5212 | 1,55206E-26 |
| ACE2 | KB-173C10.1 | -0,5213 | 1,52548E-26 |
| ACE2 | RP1-149A16.12 | -0,5213 | 1,50795E-26 |
| ACE2 | STAT4 | -0,5214 | 1,47446E-26 |
| ACE2 | GSTA8P | -0,5214 | 1,49519E-26 |
| ACE2 | RP3-399L15.2 | -0,5214 | 1,48005E-26 |
| ACE2 | C7orf66 | -0,5214 | 1,491E-26 |
| ACE2 | RP11-162D16.2 | -0,5214 | 1,47264E-26 |
| ACE2 | AP000997.2 | -0,5214 | 1,47194E-26 |

| | | | |
|---|---|---|---|
| ACE2 | PCOTH | -0,5214 | 1,4834E-26 |
| ACE2 | AC017104.2 | -0,5215 | 1,44785E-26 |
| ACE2 | BMPR1B-AS1 | -0,5215 | 1,45355E-26 |
| ACE2 | PDHA2 | -0,5215 | 1,46483E-26 |
| ACE2 | SPATA31D4 | -0,5215 | 1,42971E-26 |
| ACE2 | RP11-58G13.1 | -0,5215 | 1,46414E-26 |
| ACE2 | AC010967.2 | -0,5216 | 1,41157E-26 |
| ACE2 | RP11-92B11.4 | -0,5216 | 1,39676E-26 |
| ACE2 | RP11-644K8.1 | -0,5217 | 1,38967E-26 |
| ACE2 | OR7E122P | -0,5217 | 1,38957E-26 |
| ACE2 | RP11-606D9.1 | -0,5217 | 1,37256E-26 |
| ACE2 | LINC00403 | -0,5217 | 1,3738E-26 |
| ACE2 | FSCB | -0,5217 | 1,37144E-26 |
| ACE2 | PPM1G | -0,5218 | 1,34204E-26 |
| ACE2 | UBA52P6 | -0,5218 | 1,34E-26 |
| ACE2 | PRMT5-AS1 | -0,5218 | 1,32119E-26 |
| ACE2 | RP11-313P18.2 | -0,5218 | 1,32369E-26 |
| ACE2 | RP11-573D15.8 | -0,5219 | 1,31485E-26 |
| ACE2 | RP11-319E12.1 | -0,5219 | 1,31117E-26 |
| ACE2 | AC006380.3 | -0,5219 | 1,29989E-26 |
| ACE2 | RP11-20P5.2 | -0,5219 | 1,30061E-26 |
| ACE2 | RP11-216P16.5 | -0,5219 | 1,28797E-26 |
| ACE2 | RP11-271M24.2 | -0,5219 | 1,31478E-26 |
| ACE2 | RP11-6N17.3 | -0,5219 | 1,30442E-26 |
| ACE2 | BMP8A | -0,522 | 1,27795E-26 |
| ACE2 | RP11-1008C21.1 | -0,522 | 1,27231E-26 |
| ACE2 | NWD1 | -0,522 | 1,26825E-26 |
| ACE2 | RP1-102G20.5 | -0,5221 | 1,24207E-26 |
| ACE2 | RP11-85I21.1 | -0,5221 | 1,24947E-26 |
| ACE2 | SPATA31E1 | -0,5221 | 1,2228E-26 |
| ACE2 | OR2AT1P | -0,5221 | 1,23065E-26 |
| ACE2 | CYP2D8P | -0,5221 | 1,23778E-26 |
| ACE2 | AL672294.1 | -0,5222 | 1,22088E-26 |
| ACE2 | ACSM6 | -0,5222 | 1,20755E-26 |
| ACE2 | WDR62 | -0,5223 | 1,16899E-26 |
| ACE2 | RP1-100J12.1 | -0,5223 | 1,16096E-26 |
| ACE2 | DGCR9 | -0,5223 | 1,17077E-26 |
| ACE2 | RP11-518L10.5 | -0,5224 | 1,15598E-26 |
| ACE2 | LINC01797 | -0,5224 | 1,15947E-26 |
| ACE2 | FAM76A | -0,5225 | 1,10204E-26 |
| ACE2 | TRIM46 | -0,5225 | 1,12592E-26 |
| ACE2 | AHSG | -0,5225 | 1,12344E-26 |
| ACE2 | OR51H1 | -0,5225 | 1,10937E-26 |
| ACE2 | ST8SIA5 | -0,5225 | 1,12532E-26 |
| ACE2 | C2orf74 | -0,5226 | 1,10006E-26 |
| ACE2 | RP3-495K2.3 | -0,5226 | 1,09047E-26 |
| ACE2 | RP11-670N15.1 | -0,5226 | 1,09272E-26 |
| ACE2 | RAB37 | -0,5226 | 1,0995E-26 |
| ACE2 | CXorf65 | -0,5226 | 1,09008E-26 |
| ACE2 | CLCA2 | -0,5227 | 1,06403E-26 |
| ACE2 | SLC36A3 | -0,5227 | 1,04657E-26 |
| ACE2 | RP1-207H1.3 | -0,5228 | 1,02467E-26 |
| ACE2 | RP11-361D14.2 | -0,5229 | 9,92714E-27 |
| ACE2 | CTD-2015H6.3 | -0,5229 | 1,01492E-26 |

| | | | |
|---|---|---|---|
| ACE2 | FBXO39 | -0,5229 | 1,01324E-26 |
| ACE2 | PGAM5P1 | -0,523 | 9,68107E-27 |
| ACE2 | RP11-118M9.3 | -0,523 | 9,87992E-27 |
| ACE2 | MRGPRG-AS1 | -0,523 | 9,78258E-27 |
| ACE2 | FBF1 | -0,523 | 9,75939E-27 |
| ACE2 | RP11-415J8.5 | -0,5231 | 9,61825E-27 |
| ACE2 | LINC01081 | -0,5231 | 9,64083E-27 |
| ACE2 | ATP7B | -0,5232 | 9,22825E-27 |
| ACE2 | ADAM11 | -0,5232 | 9,32203E-27 |
| ACE2 | SLC34A2 | -0,5233 | 9,15101E-27 |
| ACE2 | RP5-897D18.1 | -0,5233 | 9,05452E-27 |
| ACE2 | RP5-1115A15.1 | -0,5234 | 8,74429E-27 |
| ACE2 | RP5-855F14.2 | -0,5234 | 8,77027E-27 |
| ACE2 | MON1A | -0,5234 | 8,89024E-27 |
| ACE2 | CCDC162P | -0,5234 | 8,84782E-27 |
| ACE2 | RP11-680H20.2 | -0,5234 | 8,91655E-27 |
| ACE2 | FOXB1 | -0,5234 | 8,79143E-27 |
| ACE2 | LINC01709 | -0,5235 | 8,50669E-27 |
| ACE2 | HMP19 | -0,5236 | 8,35811E-27 |
| ACE2 | RP11-20A20.2 | -0,5236 | 8,3888E-27 |
| ACE2 | RP11-317J10.4 | -0,5237 | 8,08203E-27 |
| ACE2 | BAALC-AS1 | -0,5237 | 8,18978E-27 |
| ACE2 | RP11-626G11.5 | -0,5237 | 8,11482E-27 |
| ACE2 | CDC14A | -0,5238 | 7,87384E-27 |
| ACE2 | LINC01961 | -0,5238 | 7,96735E-27 |
| ACE2 | PHOSPHO2 | -0,5238 | 7,91534E-27 |
| ACE2 | GRK6 | -0,5238 | 8,01952E-27 |
| ACE2 | RP11-470M17.2 | -0,5238 | 8,00653E-27 |
| ACE2 | ORM2 | -0,5238 | 7,88525E-27 |
| ACE2 | ARPC5L | -0,5238 | 8,05382E-27 |
| ACE2 | CTD-2020K17.4 | -0,5238 | 7,99035E-27 |
| ACE2 | RP11-171N4.2 | -0,5239 | 7,84856E-27 |
| ACE2 | RP11-60A8.2 | -0,5239 | 7,78689E-27 |
| ACE2 | RP11-770E5.3 | -0,5239 | 7,80701E-27 |
| ACE2 | RP11-100L22.3 | -0,5239 | 7,8188E-27 |
| ACE2 | RP11-845M18.7 | -0,5239 | 7,84191E-27 |
| ACE2 | CTB-60B18.10 | -0,5239 | 7,83879E-27 |
| ACE2 | SPATA17 | -0,524 | 7,60981E-27 |
| ACE2 | CTB-33O18.1 | -0,524 | 7,55172E-27 |
| ACE2 | LRRD1 | -0,524 | 7,61139E-27 |
| ACE2 | RP11-646J21.3 | -0,524 | 7,46675E-27 |
| ACE2 | RP5-1021I20.6 | -0,524 | 7,53353E-27 |
| ACE2 | MRPS21 | -0,5241 | 7,31651E-27 |
| ACE2 | RP11-263K19.4 | -0,5241 | 7,37719E-27 |
| ACE2 | LINC01020 | -0,5241 | 7,39227E-27 |
| ACE2 | GOLGA2P5 | -0,5241 | 7,35303E-27 |
| ACE2 | PCDH9-AS3 | -0,5241 | 7,29997E-27 |
| ACE2 | RP11-666E17.1 | -0,5241 | 7,34925E-27 |
| ACE2 | RP11-139H15.6 | -0,5241 | 7,26829E-27 |
| ACE2 | LRRC37A11P | -0,5241 | 7,30239E-27 |
| ACE2 | KIAA1324 | -0,5242 | 7,17148E-27 |
| ACE2 | OR14A2 | -0,5242 | 7,24558E-27 |
| ACE2 | TCTE3 | -0,5242 | 7,15053E-27 |
| ACE2 | ARSA | -0,5242 | 7,13478E-27 |

| | | | |
|---|---|---|---|
| ACE2 | RP11-133L14.5 | -0,5243 | 7,02789E-27 |
| ACE2 | ACD | -0,5243 | 6,91207E-27 |
| ACE2 | CTD-3162L10.1 | -0,5243 | 7,07144E-27 |
| ACE2 | MPP2 | -0,5244 | 6,78145E-27 |
| ACE2 | RPSAP28 | -0,5245 | 6,60163E-27 |
| ACE2 | GDPD5 | -0,5245 | 6,67876E-27 |
| ACE2 | LINC00029 | -0,5245 | 6,58679E-27 |
| ACE2 | RGPD4-AS1 | -0,5246 | 6,45517E-27 |
| ACE2 | C3orf49 | -0,5246 | 6,43738E-27 |
| ACE2 | CCDC113 | -0,5246 | 6,43514E-27 |
| ACE2 | RP11-23I7.1 | -0,5247 | 6,20682E-27 |
| ACE2 | RP1-81P11.1 | -0,5247 | 6,26052E-27 |
| ACE2 | KB-1572G7.4 | -0,5247 | 6,24017E-27 |
| ACE2 | CFAP57 | -0,5248 | 6,13251E-27 |
| ACE2 | CTD-2020K17.3 | -0,5248 | 6,1372E-27 |
| ACE2 | DDX39B-AS1 | -0,525 | 5,74685E-27 |
| ACE2 | RP11-262H14.3 | -0,525 | 5,86508E-27 |
| ACE2 | FLJ36000 | -0,525 | 5,81882E-27 |
| ACE2 | RP11-167H9.4 | -0,5251 | 5,69313E-27 |
| ACE2 | RP5-856G1.2 | -0,5251 | 5,59102E-27 |
| ACE2 | RP11-981G7.6 | -0,5252 | 5,55656E-27 |
| ACE2 | AC005614.3 | -0,5252 | 5,46814E-27 |
| ACE2 | GK2 | -0,5253 | 5,31702E-27 |
| ACE2 | RP11-461O7.2 | -0,5253 | 5,33246E-27 |
| ACE2 | LINC01478 | -0,5253 | 5,3082E-27 |
| ACE2 | RP11-99H8.1 | -0,5254 | 5,19564E-27 |
| ACE2 | OPN5 | -0,5254 | 5,18463E-27 |
| ACE2 | RPS3AP29 | -0,5254 | 5,26373E-27 |
| ACE2 | RP11-113D6.10 | -0,5254 | 5,20964E-27 |
| ACE2 | KCNE2 | -0,5254 | 5,24932E-27 |
| ACE2 | DSCR10 | -0,5254 | 5,1885E-27 |
| ACE2 | PZP | -0,5255 | 5,16391E-27 |
| ACE2 | RP11-512C24.3 | -0,5255 | 5,10674E-27 |
| ACE2 | DCHS2 | -0,5256 | 4,91843E-27 |
| ACE2 | PACRG | -0,5256 | 4,92978E-27 |
| ACE2 | FSCN1P1 | -0,5256 | 4,92039E-27 |
| ACE2 | ERFE | -0,5257 | 4,88825E-27 |
| ACE2 | RP11-789F5.1 | -0,5257 | 4,81367E-27 |
| ACE2 | LINC00336 | -0,5257 | 4,78728E-27 |
| ACE2 | FAM177A1 | -0,5257 | 4,81495E-27 |
| ACE2 | EFCAB6 | -0,5257 | 4,8253E-27 |
| ACE2 | AC012506.1 | -0,5258 | 4,74509E-27 |
| ACE2 | BAK1P2 | -0,5259 | 4,61611E-27 |
| ACE2 | SH2D7 | -0,5259 | 4,57724E-27 |
| ACE2 | ZNF345 | -0,5259 | 4,64808E-27 |
| ACE2 | LINC01759 | -0,526 | 4,43939E-27 |
| ACE2 | DRC7 | -0,526 | 4,5194E-27 |
| ACE2 | LINC01992 | -0,526 | 4,43305E-27 |
| ACE2 | RP11-258B16.1 | -0,526 | 4,45232E-27 |
| ACE2 | TP53RK | -0,526 | 4,48791E-27 |
| ACE2 | AC019064.1 | -0,5261 | 4,33807E-27 |
| ACE2 | CACNG3 | -0,5261 | 4,38508E-27 |
| ACE2 | RP11-26N15.2 | -0,5261 | 4,39878E-27 |
| ACE2 | FAM151A | -0,5262 | 4,21925E-27 |

| | | | |
|---|---|---|---|
| ACE2 | RP11-79O8.1 | -0,5262 | 4,27491E-27 |
| ACE2 | AC102953.4 | -0,5262 | 4,19866E-27 |
| ACE2 | SRRT | -0,5262 | 4,22388E-27 |
| ACE2 | IQSEC3P1 | -0,5262 | 4,22411E-27 |
| ACE2 | OVCH1-AS1 | -0,5262 | 4,23605E-27 |
| ACE2 | CTC-276P9.3 | -0,5263 | 4,15238E-27 |
| ACE2 | PRPS1L1 | -0,5263 | 4,101E-27 |
| ACE2 | ETNK2 | -0,5264 | 4,05006E-27 |
| ACE2 | RAET1E | -0,5264 | 4,00627E-27 |
| ACE2 | DNAJC9-AS1 | -0,5264 | 4,06836E-27 |
| ACE2 | RP11-223P11.3 | -0,5264 | 4,04513E-27 |
| ACE2 | RP11-457D20.2 | -0,5264 | 4,04326E-27 |
| ACE2 | RP5-994D16.3 | -0,5265 | 3,9405E-27 |
| ACE2 | LINC01753 | -0,5265 | 3,90989E-27 |
| ACE2 | RPL23AP25 | -0,5265 | 3,89406E-27 |
| ACE2 | NKX3-2 | -0,5265 | 3,89878E-27 |
| ACE2 | RP11-379B8.1 | -0,5265 | 3,93019E-27 |
| ACE2 | DUSP13 | -0,5265 | 3,9633E-27 |
| ACE2 | RP11-467D18.2 | -0,5266 | 3,78105E-27 |
| ACE2 | RP1-111D6.4 | -0,5266 | 3,81539E-27 |
| ACE2 | IL12RB2 | -0,5267 | 3,73589E-27 |
| ACE2 | IL10 | -0,5267 | 3,6941E-27 |
| ACE2 | RP5-978I12.1 | -0,5267 | 3,72434E-27 |
| ACE2 | AC078842.4 | -0,5267 | 3,70568E-27 |
| ACE2 | RP11-787D18.2 | -0,5268 | 3,63415E-27 |
| ACE2 | RP11-202I11.2 | -0,5268 | 3,61544E-27 |
| ACE2 | BTBD16 | -0,5268 | 3,65264E-27 |
| ACE2 | ANKS1B | -0,5268 | 3,58861E-27 |
| ACE2 | CTD-2200P10.1 | -0,5268 | 3,65286E-27 |
| ACE2 | F11-AS1 | -0,5269 | 3,53615E-27 |
| ACE2 | FER1L6 | -0,5269 | 3,57104E-27 |
| ACE2 | ENKUR | -0,5269 | 3,51252E-27 |
| ACE2 | RP11-580I1.1 | -0,527 | 3,42861E-27 |
| ACE2 | RP11-805I24.4 | -0,527 | 3,43706E-27 |
| ACE2 | NEBL | -0,5271 | 3,3328E-27 |
| ACE2 | LA16c-361A3.3 | -0,5271 | 3,37993E-27 |
| ACE2 | CENPM | -0,5271 | 3,35486E-27 |
| ACE2 | RP11-395P13.4 | -0,5272 | 3,25084E-27 |
| ACE2 | UBE2L5P | -0,5272 | 3,28338E-27 |
| ACE2 | RPS20P35 | -0,5272 | 3,24588E-27 |
| ACE2 | RP11-319E16.1 | -0,5273 | 3,19394E-27 |
| ACE2 | KLK13 | -0,5273 | 3,21527E-27 |
| ACE2 | RP11-543B16.3 | -0,5274 | 3,0736E-27 |
| ACE2 | IQCG | -0,5274 | 3,12199E-27 |
| ACE2 | C4orf36 | -0,5274 | 3,05005E-27 |
| ACE2 | RP11-123O10.3 | -0,5274 | 3,05672E-27 |
| ACE2 | RP11-482M8.1 | -0,5274 | 3,0994E-27 |
| ACE2 | RP11-94C24.6 | -0,5274 | 3,12177E-27 |
| ACE2 | LINC01362 | -0,5275 | 2,99142E-27 |
| ACE2 | TRIM52-AS1 | -0,5275 | 2,99302E-27 |
| ACE2 | C7orf72 | -0,5275 | 3,00068E-27 |
| ACE2 | RP11-363N22.2 | -0,5275 | 3,008E-27 |
| ACE2 | RP11-893F2.13 | -0,5275 | 3,00309E-27 |
| ACE2 | ZNF165 | -0,5276 | 2,95298E-27 |

| | | | |
|---|---|---|---|
| ACE2 | STYXL1 | -0,5276 | 2,95825E-27 |
| ACE2 | RNF183 | -0,5276 | 2,9142E-27 |
| ACE2 | AP000442.1 | -0,5276 | 2,92065E-27 |
| ACE2 | RP1-53O8.2 | -0,5277 | 2,85304E-27 |
| ACE2 | RP11-1060J15.4 | -0,5277 | 2,82171E-27 |
| ACE2 | GOLGA6L2 | -0,5278 | 2,74494E-27 |
| ACE2 | NXPE1 | -0,5279 | 2,71176E-27 |
| ACE2 | GJB6 | -0,5279 | 2,70799E-27 |
| ACE2 | RP11-486O13.2 | -0,5279 | 2,73181E-27 |
| ACE2 | LINC01971 | -0,5279 | 2,71371E-27 |
| ACE2 | CTD-2373J6.2 | -0,528 | 2,63124E-27 |
| ACE2 | RP11-83N9.5 | -0,528 | 2,64593E-27 |
| ACE2 | RP11-94P11.4 | -0,528 | 2,65158E-27 |
| ACE2 | CTD-3046C4.1 | -0,5281 | 2,54775E-27 |
| ACE2 | MRLN | -0,5281 | 2,58281E-27 |
| ACE2 | RP1-17K7.2 | -0,5281 | 2,59168E-27 |
| ACE2 | CLEC4A | -0,5281 | 2,54994E-27 |
| ACE2 | LINC01952 | -0,5282 | 2,50731E-27 |
| ACE2 | BRWD1-AS1 | -0,5282 | 2,47045E-27 |
| ACE2 | MYO5BP2 | -0,5283 | 2,43502E-27 |
| ACE2 | ZNF213 | -0,5283 | 2,4358E-27 |
| ACE2 | AC027612.4 | -0,5284 | 2,34537E-27 |
| ACE2 | HSPA9P1 | -0,5284 | 2,36041E-27 |
| ACE2 | CTC-558O19.1 | -0,5284 | 2,36979E-27 |
| ACE2 | EFCAB1 | -0,5284 | 2,34466E-27 |
| ACE2 | PRB3 | -0,5284 | 2,3807E-27 |
| ACE2 | RP11-357K6.1 | -0,5284 | 2,34571E-27 |
| ACE2 | RP11-562A8.5 | -0,5284 | 2,36601E-27 |
| ACE2 | KIF17 | -0,5285 | 2,3056E-27 |
| ACE2 | RP11-686D16.1 | -0,5285 | 2,30211E-27 |
| ACE2 | TXLNB | -0,5285 | 2,28431E-27 |
| ACE2 | RP11-421M1.8 | -0,5286 | 2,26112E-27 |
| ACE2 | SPATA6L | -0,5286 | 2,26752E-27 |
| ACE2 | RP1-17K7.3 | -0,5286 | 2,25471E-27 |
| ACE2 | C11orf98 | -0,5286 | 2,2615E-27 |
| ACE2 | C16orf96 | -0,5286 | 2,22499E-27 |
| ACE2 | SLC16A5 | -0,5286 | 2,23652E-27 |
| ACE2 | AC116609.1 | -0,5287 | 2,20799E-27 |
| ACE2 | RP11-334G22.1 | -0,5287 | 2,19308E-27 |
| ACE2 | LINC01375 | -0,5287 | 2,17956E-27 |
| ACE2 | RP11-133L19.2 | -0,5287 | 2,16435E-27 |
| ACE2 | RP11-386G11.8 | -0,5289 | 2,08839E-27 |
| ACE2 | TEX26 | -0,5289 | 2,08708E-27 |
| ACE2 | GTF2IP13 | -0,529 | 2,00859E-27 |
| ACE2 | NRG1 | -0,529 | 2,04254E-27 |
| ACE2 | RP11-689E3.2 | -0,529 | 2,02461E-27 |
| ACE2 | LINC01862 | -0,529 | 2,00997E-27 |
| ACE2 | MUC1 | -0,5291 | 1,96274E-27 |
| ACE2 | RP11-779O18.3 | -0,5291 | 1,9453E-27 |
| ACE2 | RP11-73O6.3 | -0,5291 | 1,99099E-27 |
| ACE2 | CYP26C1 | -0,5291 | 1,95238E-27 |
| ACE2 | PIPSL | -0,5291 | 1,95587E-27 |
| ACE2 | RPL12P35 | -0,5291 | 1,97478E-27 |
| ACE2 | ZNF546 | -0,5291 | 1,98815E-27 |

| | | | |
|---|---|---|---|
| ACE2 | RP11-849F2.5 | -0,5292 | 1,9274E-27 |
| ACE2 | DNAH9 | -0,5292 | 1,91469E-27 |
| ACE2 | RP11-34P13.15 | -0,5293 | 1,84832E-27 |
| ACE2 | PPP3R2 | -0,5293 | 1,87259E-27 |
| ACE2 | RP4-550H1.7 | -0,5293 | 1,86818E-27 |
| ACE2 | LINC02201 | -0,5294 | 1,81414E-27 |
| ACE2 | CTC-338M12.6 | -0,5294 | 1,80744E-27 |
| ACE2 | NEUROD4 | -0,5294 | 1,83977E-27 |
| ACE2 | CTD-2647E9.3 | -0,5294 | 1,83926E-27 |
| ACE2 | CTD-2562J15.6 | -0,5294 | 1,79882E-27 |
| ACE2 | DESI1 | -0,5295 | 1,7918E-27 |
| ACE2 | SMIM12 | -0,5296 | 1,72983E-27 |
| ACE2 | AIM2 | -0,5296 | 1,74242E-27 |
| ACE2 | AC012456.3 | -0,5296 | 1,72991E-27 |
| ACE2 | LINC01854 | -0,5296 | 1,74433E-27 |
| ACE2 | RP11-572O17.1 | -0,5296 | 1,72599E-27 |
| ACE2 | MAL2 | -0,5296 | 1,70747E-27 |
| ACE2 | RP11-535C7.1 | -0,5297 | 1,66737E-27 |
| ACE2 | PPEF2 | -0,5297 | 1,69703E-27 |
| ACE2 | TTC23L | -0,5297 | 1,66087E-27 |
| ACE2 | CTD-2023N9.3 | -0,5297 | 1,68383E-27 |
| ACE2 | FGL1 | -0,5297 | 1,69237E-27 |
| ACE2 | GRK6P1 | -0,5297 | 1,6703E-27 |
| ACE2 | NOX5 | -0,5297 | 1,67601E-27 |
| ACE2 | CDRT8 | -0,5297 | 1,67248E-27 |
| ACE2 | AARSD1 | -0,5297 | 1,69679E-27 |
| ACE2 | MAP3K14 | -0,5297 | 1,66414E-27 |
| ACE2 | AKAP14 | -0,5297 | 1,693E-27 |
| ACE2 | RP11-313P13.5 | -0,5298 | 1,65053E-27 |
| ACE2 | RP11-855O10.2 | -0,5298 | 1,63178E-27 |
| ACE2 | STK16 | -0,5299 | 1,58445E-27 |
| ACE2 | RP11-264J4.8 | -0,5299 | 1,59927E-27 |
| ACE2 | LINC01533 | -0,5299 | 1,60064E-27 |
| ACE2 | RP1-27K12.4 | -0,53 | 1,53896E-27 |
| ACE2 | LINC02164 | -0,53 | 1,54865E-27 |
| ACE2 | C16orf46 | -0,53 | 1,56743E-27 |
| ACE2 | RP11-1151B14.3 | -0,53 | 1,54862E-27 |
| ACE2 | CLDN34 | -0,53 | 1,5666E-27 |
| ACE2 | HNRNPA1P8 | -0,5301 | 1,52822E-27 |
| ACE2 | RP11-408E5.5 | -0,5301 | 1,49597E-27 |
| ACE2 | AP000281.2 | -0,5301 | 1,51212E-27 |
| ACE2 | CCDC179 | -0,5302 | 1,46724E-27 |
| ACE2 | PIK3R5 | -0,5302 | 1,46463E-27 |
| ACE2 | ZNF645 | -0,5302 | 1,46224E-27 |
| ACE2 | RP11-274B21.12 | -0,5303 | 1,43025E-27 |
| ACE2 | PDZD9 | -0,5303 | 1,41992E-27 |
| ACE2 | PITPNM3 | -0,5305 | 1,35107E-27 |
| ACE2 | AKR1D1 | -0,5306 | 1,32904E-27 |
| ACE2 | KRT7-AS | -0,5306 | 1,32267E-27 |
| ACE2 | RP11-26M5.2 | -0,5307 | 1,28836E-27 |
| ACE2 | CTD-2210P24.6 | -0,5307 | 1,28804E-27 |
| ACE2 | RP11-25I15.2 | -0,5307 | 1,2966E-27 |
| ACE2 | AC108025.2 | -0,5308 | 1,26674E-27 |
| ACE2 | AC005522.7 | -0,5308 | 1,26619E-27 |

| | | | |
|---|---|---|---|
| ACE2 | FAM205A | -0,5308 | 1,24975E-27 |
| ACE2 | AP000770.1 | -0,5308 | 1,24509E-27 |
| ACE2 | RP11-809H16.4 | -0,5308 | 1,25326E-27 |
| ACE2 | RP11-315L6.1 | -0,5308 | 1,24874E-27 |
| ACE2 | AC100848.1 | -0,5308 | 1,25211E-27 |
| ACE2 | MMP24-AS1 | -0,5308 | 1,25936E-27 |
| ACE2 | TGM4 | -0,5309 | 1,22237E-27 |
| ACE2 | OR1F1 | -0,5309 | 1,21797E-27 |
| ACE2 | CTA-125H2.2 | -0,5309 | 1,2128E-27 |
| ACE2 | KISS1 | -0,5311 | 1,16883E-27 |
| ACE2 | TMEM72 | -0,5311 | 1,16096E-27 |
| ACE2 | KRT85 | -0,5311 | 1,1419E-27 |
| ACE2 | RP11-127I20.5 | -0,5311 | 1,16281E-27 |
| ACE2 | LINC01256 | -0,5312 | 1,12373E-27 |
| ACE2 | RP11-284J1.1 | -0,5312 | 1,12491E-27 |
| ACE2 | TROAP | -0,5312 | 1,12569E-27 |
| ACE2 | RP11-983C2.2 | -0,5312 | 1,14051E-27 |
| ACE2 | TAS1R1 | -0,5313 | 1,08511E-27 |
| ACE2 | AC064853.2 | -0,5313 | 1,09804E-27 |
| ACE2 | STK19 | -0,5313 | 1,08767E-27 |
| ACE2 | RHBDD2 | -0,5313 | 1,10477E-27 |
| ACE2 | ENOX1 | -0,5313 | 1,08404E-27 |
| ACE2 | TP53INP2 | -0,5313 | 1,10738E-27 |
| ACE2 | MAD2L2 | -0,5314 | 1,07394E-27 |
| ACE2 | AC012501.3 | -0,5314 | 1,07032E-27 |
| ACE2 | LINC02143 | -0,5314 | 1,07981E-27 |
| ACE2 | RP5-1198O20.4 | -0,5315 | 1,0503E-27 |
| ACE2 | AC068042.1 | -0,5315 | 1,03097E-27 |
| ACE2 | TRIM61 | -0,5315 | 1,04846E-27 |
| ACE2 | RP11-546M21.6 | -0,5315 | 1,03134E-27 |
| ACE2 | KNCN | -0,5316 | 1,02054E-27 |
| ACE2 | SLIRP | -0,5316 | 1,00829E-27 |
| ACE2 | RP11-1017G21.4 | -0,5316 | 1,01595E-27 |
| ACE2 | RP11-109P11.1 | -0,5316 | 1,00625E-27 |
| ACE2 | AC114814.3 | -0,5317 | 9,87109E-28 |
| ACE2 | POU5F2 | -0,5317 | 9,78227E-28 |
| ACE2 | RP11-379P15.1 | -0,5318 | 9,5743E-28 |
| ACE2 | ANO1-AS2 | -0,5318 | 9,52068E-28 |
| ACE2 | GOLGA6D | -0,5318 | 9,6708E-28 |
| ACE2 | LINC01440 | -0,5318 | 9,6425E-28 |
| ACE2 | THBS3 | -0,5319 | 9,35753E-28 |
| ACE2 | AC097467.2 | -0,5319 | 9,40408E-28 |
| ACE2 | RP11-146E23.2 | -0,5319 | 9,22701E-28 |
| ACE2 | CSNK1G2P1 | -0,5319 | 9,353E-28 |
| ACE2 | RP11-677I18.3 | -0,5319 | 9,34022E-28 |
| ACE2 | CTA-246H3.8 | -0,5319 | 9,21794E-28 |
| ACE2 | SNORA73 | -0,5321 | 8,91555E-28 |
| ACE2 | SPDYE4 | -0,5321 | 8,86121E-28 |
| ACE2 | DMRTA2 | -0,5322 | 8,50868E-28 |
| ACE2 | ARHGEF3-AS1 | -0,5322 | 8,55679E-28 |
| ACE2 | VWA2 | -0,5322 | 8,53188E-28 |
| ACE2 | PDCD6IPP2 | -0,5323 | 8,32531E-28 |
| ACE2 | RP11-520D19.1 | -0,5324 | 8,20292E-28 |
| ACE2 | LINC02150 | -0,5324 | 8,1784E-28 |

| | | | |
|---|---|---|---|
| ACE2 | SLC36A2 | -0,5324 | 8,11875E-28 |
| ACE2 | RP11-117L5.4 | -0,5324 | 8,131E-28 |
| ACE2 | RP11-468I15.1 | -0,5324 | 8,09262E-28 |
| ACE2 | MIR663AHG | -0,5324 | 8,20459E-28 |
| ACE2 | AC021188.4 | -0,5325 | 7,88186E-28 |
| ACE2 | RP11-319G6.1 | -0,5325 | 7,83869E-28 |
| ACE2 | RP11-69H7.2 | -0,5325 | 7,9984E-28 |
| ACE2 | DACT3 | -0,5325 | 7,88995E-28 |
| ACE2 | CTA-221G9.7 | -0,5325 | 7,98956E-28 |
| ACE2 | LINC01778 | -0,5326 | 7,65228E-28 |
| ACE2 | ACCSL | -0,5326 | 7,67909E-28 |
| ACE2 | RP11-57A19.2 | -0,5326 | 7,82244E-28 |
| ACE2 | INPP5K | -0,5326 | 7,6416E-28 |
| ACE2 | CTD-3035K23.7 | -0,5326 | 7,64122E-28 |
| ACE2 | XXbac-BPGBPG55C20. | -0,5327 | 7,48757E-28 |
| ACE2 | PCDH15 | -0,5327 | 7,59906E-28 |
| ACE2 | RP11-955H22.3 | -0,5327 | 7,6217E-28 |
| ACE2 | GAPDHP52 | -0,5327 | 7,48319E-28 |
| ACE2 | AC017104.4 | -0,5328 | 7,29103E-28 |
| ACE2 | CPEB3 | -0,5328 | 7,42588E-28 |
| ACE2 | RP11-384K6.8 | -0,5329 | 7,07582E-28 |
| ACE2 | RP11-417J1.1 | -0,5329 | 7,0969E-28 |
| ACE2 | CCDC136 | -0,5329 | 7,07018E-28 |
| ACE2 | FLJ40288 | -0,5329 | 7,12176E-28 |
| ACE2 | LINC02209 | -0,5329 | 7,17714E-28 |
| ACE2 | RP11-326E22.1 | -0,5329 | 7,22014E-28 |
| ACE2 | FAM71C | -0,5329 | 7,21045E-28 |
| ACE2 | LINC02188 | -0,5329 | 7,18883E-28 |
| ACE2 | FAM134B | -0,533 | 6,93873E-28 |
| ACE2 | FAM225A | -0,533 | 7,03502E-28 |
| ACE2 | LINC00560 | -0,533 | 7,01043E-28 |
| ACE2 | C16orf78 | -0,533 | 6,98373E-28 |
| ACE2 | CTD-2369P2.5 | -0,533 | 6,95563E-28 |
| ACE2 | LINC01304 | -0,5331 | 6,77147E-28 |
| ACE2 | PAQR9 | -0,5331 | 6,70699E-28 |
| ACE2 | TRAC | -0,5331 | 6,72703E-28 |
| ACE2 | AC006538.1 | -0,5331 | 6,71247E-28 |
| ACE2 | SLC30A10 | -0,5332 | 6,58524E-28 |
| ACE2 | RP11-254I22.3 | -0,5332 | 6,62555E-28 |
| ACE2 | RP11-96D1.5 | -0,5332 | 6,58683E-28 |
| ACE2 | RP11-396C23.4 | -0,5333 | 6,4006E-28 |
| ACE2 | HERC2P10 | -0,5333 | 6,38699E-28 |
| ACE2 | ZNF385D | -0,5334 | 6,25845E-28 |
| ACE2 | CRB2 | -0,5334 | 6,22698E-28 |
| ACE2 | RP11-723O4.6 | -0,5335 | 6,04418E-28 |
| ACE2 | RP11-586D19.2 | -0,5335 | 6,04682E-28 |
| ACE2 | TMPOP2 | -0,5335 | 6,00819E-28 |
| ACE2 | LINC01970 | -0,5335 | 6,0458E-28 |
| ACE2 | FAM27B | -0,5336 | 5,84153E-28 |
| ACE2 | RP11-128B16.5 | -0,5336 | 5,84781E-28 |
| ACE2 | GJB5 | -0,5337 | 5,75118E-28 |
| ACE2 | CASQ1 | -0,5337 | 5,82485E-28 |
| ACE2 | LRRC77P | -0,5337 | 5,70137E-28 |
| ACE2 | WWC2-AS2 | -0,5337 | 5,70549E-28 |

| | | | |
|---|---|---|---|
| ACE2 | C8orf58 | -0,5337 | 5,75426E-28 |
| ACE2 | RP11-861E23.2 | -0,5337 | 5,82083E-28 |
| ACE2 | VARS | -0,5338 | 5,52716E-28 |
| ACE2 | AC007349.5 | -0,5338 | 5,57262E-28 |
| ACE2 | LINC01289 | -0,5338 | 5,65114E-28 |
| ACE2 | LINC01501 | -0,5338 | 5,62813E-28 |
| ACE2 | LINC00358 | -0,5338 | 5,52736E-28 |
| ACE2 | LINC00221 | -0,5338 | 5,55298E-28 |
| ACE2 | RP11-568N6.1 | -0,5339 | 5,38314E-28 |
| ACE2 | LINC02040 | -0,5339 | 5,42206E-28 |
| ACE2 | LAMB4 | -0,5339 | 5,51345E-28 |
| ACE2 | LINC00208 | -0,5339 | 5,48696E-28 |
| ACE2 | CALML3 | -0,5339 | 5,4309E-28 |
| ACE2 | TMPRSS4 | -0,5339 | 5,41632E-28 |
| ACE2 | GABRG3 | -0,5339 | 5,46377E-28 |
| ACE2 | KNSTRN | -0,5339 | 5,44256E-28 |
| ACE2 | ASXL3 | -0,5339 | 5,4291E-28 |
| ACE2 | RP11-446J8.1 | -0,534 | 5,25867E-28 |
| ACE2 | GTF2IP14 | -0,534 | 5,28288E-28 |
| ACE2 | RP11-157L3.12 | -0,534 | 5,28893E-28 |
| ACE2 | LINC01495 | -0,534 | 5,28776E-28 |
| ACE2 | LINC00539 | -0,534 | 5,36721E-28 |
| ACE2 | MOV10L1 | -0,534 | 5,31914E-28 |
| ACE2 | ACOT13 | -0,5341 | 5,13333E-28 |
| ACE2 | RP11-121A14.3 | -0,5341 | 5,19314E-28 |
| ACE2 | RP11-70F11.11 | -0,5341 | 5,21405E-28 |
| ACE2 | MVD | -0,5341 | 5,136E-28 |
| ACE2 | LRRIQ4 | -0,5342 | 5,04325E-28 |
| ACE2 | GPR19 | -0,5342 | 5,07877E-28 |
| ACE2 | RP11-78O7.3 | -0,5342 | 5,02881E-28 |
| ACE2 | LRP4-AS1 | -0,5343 | 4,83803E-28 |
| ACE2 | CTD-2520I13.1 | -0,5343 | 4,90923E-28 |
| ACE2 | DENND6B | -0,5343 | 4,8425E-28 |
| ACE2 | RP4-799P18.3 | -0,5344 | 4,78176E-28 |
| ACE2 | AC012594.1 | -0,5345 | 4,59546E-28 |
| ACE2 | AC004549.6 | -0,5345 | 4,62273E-28 |
| ACE2 | FREM1 | -0,5345 | 4,68356E-28 |
| ACE2 | RP11-4C20.4 | -0,5345 | 4,60976E-28 |
| ACE2 | OVCH1 | -0,5345 | 4,64067E-28 |
| ACE2 | AP000282.3 | -0,5345 | 4,61572E-28 |
| ACE2 | LINC00635 | -0,5346 | 4,52099E-28 |
| ACE2 | SAXO1 | -0,5346 | 4,53029E-28 |
| ACE2 | PHACTR2-AS1 | -0,5347 | 4,37031E-28 |
| ACE2 | SPDYE3 | -0,5347 | 4,4262E-28 |
| ACE2 | TEX36 | -0,5347 | 4,37696E-28 |
| ACE2 | CTD-2506J14.1 | -0,5347 | 4,36702E-28 |
| ACE2 | RP1-138B7.7 | -0,5347 | 4,37559E-28 |
| ACE2 | UBXN10 | -0,5348 | 4,26328E-28 |
| ACE2 | KHDC1L | -0,5348 | 4,31293E-28 |
| ACE2 | CELF2-AS2 | -0,5348 | 4,24808E-28 |
| ACE2 | LINC01776 | -0,5349 | 4,14933E-28 |
| ACE2 | RP11-43N5.1 | -0,5349 | 4,19847E-28 |
| ACE2 | CELF3 | -0,535 | 4,00414E-28 |
| ACE2 | RP11-118K6.3 | -0,535 | 4,06911E-28 |

| | | | |
|---|---|---|---|
| ACE2 | RP11-399E6.1 | -0,5351 | 3,96945E-28 |
| ACE2 | MYLK-AS1 | -0,5351 | 3,90363E-28 |
| ACE2 | SLC35G5 | -0,5351 | 3,90954E-28 |
| ACE2 | FAM170B-AS1 | -0,5351 | 3,92723E-28 |
| ACE2 | RP11-109A6.2 | -0,5351 | 3,93434E-28 |
| ACE2 | RP11-683O4.1 | -0,5351 | 3,89695E-28 |
| ACE2 | AC055764.1 | -0,5352 | 3,85648E-28 |
| ACE2 | TRBV5-6 | -0,5352 | 3,78124E-28 |
| ACE2 | RP11-45F15.2 | -0,5352 | 3,81459E-28 |
| ACE2 | RP11-1O10.1 | -0,5352 | 3,80936E-28 |
| ACE2 | LIN28A | -0,5353 | 3,69395E-28 |
| ACE2 | CTD-2130F23.1 | -0,5353 | 3,70898E-28 |
| ACE2 | LIPT2 | -0,5353 | 3,75161E-28 |
| ACE2 | KSR2 | -0,5353 | 3,78109E-28 |
| ACE2 | FNDC8 | -0,5353 | 3,74778E-28 |
| ACE2 | RP11-426J5.2 | -0,5353 | 3,71995E-28 |
| ACE2 | RP4-697P8.3 | -0,5353 | 3,70714E-28 |
| ACE2 | PXYLP1 | -0,5354 | 3,65865E-28 |
| ACE2 | CEP170P1 | -0,5354 | 3,67433E-28 |
| ACE2 | PTP4A1P2 | -0,5354 | 3,6552E-28 |
| ACE2 | CSNK1A1P1 | -0,5354 | 3,62882E-28 |
| ACE2 | OSER1 | -0,5354 | 3,62538E-28 |
| ACE2 | RP11-259P1.1 | -0,5354 | 3,58589E-28 |
| ACE2 | ADAMTSL4-AS1 | -0,5355 | 3,50203E-28 |
| ACE2 | RP11-305P14.1 | -0,5355 | 3,54079E-28 |
| ACE2 | RP5-856G1.1 | -0,5355 | 3,5283E-28 |
| ACE2 | C15orf65 | -0,5355 | 3,52009E-28 |
| ACE2 | RP11-293N14.1 | -0,5355 | 3,51014E-28 |
| ACE2 | STX8 | -0,5355 | 3,56005E-28 |
| ACE2 | RP11-282C5.1 | -0,5356 | 3,40302E-28 |
| ACE2 | C10orf107 | -0,5356 | 3,39737E-28 |
| ACE2 | LBX1-AS1 | -0,5356 | 3,45824E-28 |
| ACE2 | RP11-115F18.1 | -0,5356 | 3,39965E-28 |
| ACE2 | CYP2G2P | -0,5356 | 3,41383E-28 |
| ACE2 | RP4-621N11.2 | -0,5356 | 3,47231E-28 |
| ACE2 | TRIM11 | -0,5357 | 3,39105E-28 |
| ACE2 | PHF7 | -0,5357 | 3,37105E-28 |
| ACE2 | CTD-2001C12.1 | -0,5357 | 3,37407E-28 |
| ACE2 | RP11-326N17.1 | -0,5357 | 3,35076E-28 |
| ACE2 | CTD-2650P22.2 | -0,5357 | 3,33345E-28 |
| ACE2 | DNAJA4 | -0,5357 | 3,34324E-28 |
| ACE2 | OVOL2 | -0,5357 | 3,30347E-28 |
| ACE2 | LINC01646 | -0,5358 | 3,24441E-28 |
| ACE2 | AC116609.3 | -0,5358 | 3,26978E-28 |
| ACE2 | LINC01799 | -0,5358 | 3,29483E-28 |
| ACE2 | LY6G6D | -0,5358 | 3,27069E-28 |
| ACE2 | RP11-798K3.2 | -0,5358 | 3,26441E-28 |
| ACE2 | RP11-49P4.7 | -0,5359 | 3,1621E-28 |
| ACE2 | CAPZA3 | -0,5359 | 3,1919E-28 |
| ACE2 | LINC01906 | -0,5359 | 3,13553E-28 |
| ACE2 | AC000095.12 | -0,5359 | 3,16964E-28 |
| ACE2 | TMEM30CP | -0,536 | 3,05214E-28 |
| ACE2 | WISP3 | -0,536 | 3,08544E-28 |
| ACE2 | AMOTL1 | -0,536 | 3,05383E-28 |

| | | | |
|---|---|---|---|
| ACE2 | RP11-857B24.5 | -0,536 | 3,08761E-28 |
| ACE2 | LINC00691 | -0,5361 | 2,96832E-28 |
| ACE2 | RP11-731J8.3 | -0,5361 | 3,00618E-28 |
| ACE2 | RP11-728F11.3 | -0,5361 | 3,0386E-28 |
| ACE2 | AP002954.4 | -0,5361 | 2,98952E-28 |
| ACE2 | LA16c-380A1.1 | -0,5361 | 3,02924E-28 |
| ACE2 | RP11-67K19.3 | -0,5362 | 2,91168E-28 |
| ACE2 | RP11-168P8.5 | -0,5362 | 2,94667E-28 |
| ACE2 | ZSCAN29 | -0,5362 | 2,92929E-28 |
| ACE2 | NPIPB2 | -0,5362 | 2,91345E-28 |
| ACE2 | RP11-1137G4.4 | -0,5362 | 2,91573E-28 |
| ACE2 | ZNF709 | -0,5362 | 2,90416E-28 |
| ACE2 | CHAF1B | -0,5362 | 2,89861E-28 |
| ACE2 | CCIN | -0,5363 | 2,84614E-28 |
| ACE2 | RP5-1125A11.4 | -0,5363 | 2,83104E-28 |
| ACE2 | RP3-388M5.9 | -0,5363 | 2,85415E-28 |
| ACE2 | RP11-434D11.4 | -0,5364 | 2,75219E-28 |
| ACE2 | ADGRD2 | -0,5364 | 2,75077E-28 |
| ACE2 | GCNT3 | -0,5364 | 2,79723E-28 |
| ACE2 | INO80C | -0,5364 | 2,77338E-28 |
| ACE2 | HAR1B | -0,5364 | 2,76533E-28 |
| ACE2 | GRIK1 | -0,5364 | 2,72998E-28 |
| ACE2 | CTD-2281E23.1 | -0,5365 | 2,70786E-28 |
| ACE2 | HPSE2 | -0,5365 | 2,65929E-28 |
| ACE2 | PLA2G4E-AS1 | -0,5365 | 2,67884E-28 |
| ACE2 | ORC1 | -0,5366 | 2,59083E-28 |
| ACE2 | PPP6R1 | -0,5366 | 2,62129E-28 |
| ACE2 | MYO3B | -0,5367 | 2,5261E-28 |
| ACE2 | RP13-580B18.4 | -0,5367 | 2,52042E-28 |
| ACE2 | ASTN2-AS1 | -0,5367 | 2,5265E-28 |
| ACE2 | RP11-314P15.1 | -0,5367 | 2,52786E-28 |
| ACE2 | AC005324.6 | -0,5368 | 2,50954E-28 |
| ACE2 | GNAT1 | -0,5369 | 2,42962E-28 |
| ACE2 | RP11-439C8.2 | -0,5369 | 2,43063E-28 |
| ACE2 | CTD-2126E3.1 | -0,5369 | 2,42244E-28 |
| ACE2 | LINC01995 | -0,537 | 2,37103E-28 |
| ACE2 | RP11-386B13.3 | -0,537 | 2,32486E-28 |
| ACE2 | C6orf163 | -0,537 | 2,32971E-28 |
| ACE2 | OLAH | -0,537 | 2,3435E-28 |
| ACE2 | YPEL4 | -0,537 | 2,3204E-28 |
| ACE2 | NENFP2 | -0,537 | 2,35233E-28 |
| ACE2 | LINC01746 | -0,537 | 2,37203E-28 |
| ACE2 | C22orf23 | -0,537 | 2,37697E-28 |
| ACE2 | RP11-161H23.9 | -0,5371 | 2,26546E-28 |
| ACE2 | RP11-52L5.6 | -0,5371 | 2,25958E-28 |
| ACE2 | LINC01043 | -0,5371 | 2,25577E-28 |
| ACE2 | RP4-697K14.3 | -0,5371 | 2,29602E-28 |
| ACE2 | VEPH1 | -0,5372 | 2,23285E-28 |
| ACE2 | SETD9 | -0,5372 | 2,22783E-28 |
| ACE2 | MEPCE | -0,5372 | 2,20209E-28 |
| ACE2 | LINC01241 | -0,5372 | 2,24971E-28 |
| ACE2 | RP1-292L20.3 | -0,5372 | 2,20254E-28 |
| ACE2 | RP11-324J13.2 | -0,5374 | 2,1144E-28 |
| ACE2 | SPACA1 | -0,5374 | 2,13324E-28 |

| | | | |
|---|---|---|---|
| ACE2 | MYH16 | -0,5374 | 2,09957E-28 |
| ACE2 | PITX3 | -0,5374 | 2,08872E-28 |
| ACE2 | CTD-2384A14.2 | -0,5374 | 2,11807E-28 |
| ACE2 | RP11-856M7.2 | -0,5374 | 2,09219E-28 |
| ACE2 | LINC00656 | -0,5374 | 2,10758E-28 |
| ACE2 | IGHMBP2 | -0,5375 | 2,03716E-28 |
| ACE2 | MIR1539 | -0,5375 | 2,03894E-28 |
| ACE2 | RP11-268G13.1 | -0,5375 | 2,02875E-28 |
| ACE2 | RP11-536C5.7 | -0,5376 | 1,97334E-28 |
| ACE2 | CELF6 | -0,5376 | 1,97568E-28 |
| ACE2 | TTC39A | -0,5377 | 1,93675E-28 |
| ACE2 | AC005008.2 | -0,5377 | 1,93155E-28 |
| ACE2 | AC068858.1 | -0,5377 | 1,9383E-28 |
| ACE2 | BEAN1 | -0,5377 | 1,95568E-28 |
| ACE2 | DGCR6 | -0,5377 | 1,94503E-28 |
| ACE2 | LINC01653 | -0,5378 | 1,86905E-28 |
| ACE2 | CTD-2296D1.4 | -0,5378 | 1,90231E-28 |
| ACE2 | DEFB121 | -0,5378 | 1,87591E-28 |
| ACE2 | AC004237.1 | -0,5379 | 1,8207E-28 |
| ACE2 | GUCA1A | -0,5379 | 1,84014E-28 |
| ACE2 | LINC00301 | -0,5379 | 1,83259E-28 |
| ACE2 | LRRC23 | -0,5379 | 1,85203E-28 |
| ACE2 | PLEKHM2 | -0,538 | 1,77951E-28 |
| ACE2 | AC000099.1 | -0,538 | 1,76792E-28 |
| ACE2 | GPX2 | -0,538 | 1,8116E-28 |
| ACE2 | UCA1 | -0,538 | 1,8099E-28 |
| ACE2 | PDE6D | -0,5381 | 1,76137E-28 |
| ACE2 | PRSS58 | -0,5381 | 1,74308E-28 |
| ACE2 | FAM205BP | -0,5381 | 1,76065E-28 |
| ACE2 | ABC13-47488600E17.1 | -0,5381 | 1,75443E-28 |
| ACE2 | SP140 | -0,5382 | 1,67308E-28 |
| ACE2 | LINC02030 | -0,5382 | 1,69511E-28 |
| ACE2 | SMAD1-AS2 | -0,5382 | 1,69052E-28 |
| ACE2 | LINC01333 | -0,5382 | 1,67456E-28 |
| ACE2 | PTK7 | -0,5382 | 1,67617E-28 |
| ACE2 | LINC01927 | -0,5382 | 1,70209E-28 |
| ACE2 | MLLT1 | -0,5382 | 1,67615E-28 |
| ACE2 | GS1-279B7.1 | -0,5383 | 1,66704E-28 |
| ACE2 | RP11-406A20.1 | -0,5383 | 1,62592E-28 |
| ACE2 | ANKRD30BP3 | -0,5383 | 1,6323E-28 |
| ACE2 | RP11-727A23.10 | -0,5383 | 1,6624E-28 |
| ACE2 | THAP2 | -0,5383 | 1,62699E-28 |
| ACE2 | EFCAB14-AS1 | -0,5384 | 1,60949E-28 |
| ACE2 | C1orf87 | -0,5384 | 1,60331E-28 |
| ACE2 | KIAA1211 | -0,5384 | 1,61225E-28 |
| ACE2 | FAM92A | -0,5384 | 1,60567E-28 |
| ACE2 | RP11-479G22.8 | -0,5384 | 1,60527E-28 |
| ACE2 | HRASLS5 | -0,5384 | 1,60583E-28 |
| ACE2 | THCAT158 | -0,5384 | 1,58686E-28 |
| ACE2 | LINC01555 | -0,5385 | 1,54162E-28 |
| ACE2 | GSTCD | -0,5385 | 1,56705E-28 |
| ACE2 | RP11-613E4.5 | -0,5385 | 1,56608E-28 |
| ACE2 | EML2-AS1 | -0,5385 | 1,56185E-28 |
| ACE2 | FAM81B | -0,5386 | 1,50022E-28 |

| | | | |
|---|---|---|---|
| ACE2 | CTB-131B5.5 | -0,5386 | 1,50265E-28 |
| ACE2 | LINC01519 | -0,5386 | 1,52343E-28 |
| ACE2 | RP11-379K22.3 | -0,5386 | 1,53652E-28 |
| ACE2 | SYT5 | -0,5386 | 1,51969E-28 |
| ACE2 | ADCY10 | -0,5387 | 1,49522E-28 |
| ACE2 | RP11-344J7.3 | -0,5387 | 1,47742E-28 |
| ACE2 | RP11-12A20.7 | -0,5387 | 1,47329E-28 |
| ACE2 | MRPL52 | -0,5387 | 1,45596E-28 |
| ACE2 | RP11-270M14.4 | -0,5387 | 1,45643E-28 |
| ACE2 | RP11-519M16.1 | -0,5388 | 1,44684E-28 |
| ACE2 | RP11-955H22.1 | -0,5388 | 1,42054E-28 |
| ACE2 | CTC-457E21.10 | -0,5388 | 1,45501E-28 |
| ACE2 | AKR1E2 | -0,5389 | 1,41055E-28 |
| ACE2 | C16orf97 | -0,5389 | 1,41511E-28 |
| ACE2 | RP11-625N16.1 | -0,539 | 1,3596E-28 |
| ACE2 | RP11-527F13.1 | -0,539 | 1,35003E-28 |
| ACE2 | LINC00366 | -0,539 | 1,35127E-28 |
| ACE2 | SERP2 | -0,539 | 1,35405E-28 |
| ACE2 | SYMPK | -0,539 | 1,36411E-28 |
| ACE2 | RP3-366N23.4 | -0,5391 | 1,31869E-28 |
| ACE2 | MIR3663HG | -0,5391 | 1,32772E-28 |
| ACE2 | RP11-299L17.3 | -0,5391 | 1,31512E-28 |
| ACE2 | PRPH | -0,5392 | 1,29348E-28 |
| ACE2 | LINC01696 | -0,5393 | 1,25208E-28 |
| ACE2 | AC016716.2 | -0,5393 | 1,25224E-28 |
| ACE2 | RP11-395C17.1 | -0,5393 | 1,23652E-28 |
| ACE2 | RP11-673E11.2 | -0,5393 | 1,25362E-28 |
| ACE2 | RP11-328C8.2 | -0,5393 | 1,26758E-28 |
| ACE2 | LINC01589 | -0,5393 | 1,24492E-28 |
| ACE2 | RP5-892K4.1 | -0,5394 | 1,207E-28 |
| ACE2 | DIS3L2 | -0,5394 | 1,21427E-28 |
| ACE2 | DLG1-AS1 | -0,5394 | 1,21826E-28 |
| ACE2 | RP11-300M6.1 | -0,5394 | 1,22031E-28 |
| ACE2 | C6orf10 | -0,5394 | 1,22191E-28 |
| ACE2 | PEX16 | -0,5394 | 1,20241E-28 |
| ACE2 | PLUT | -0,5395 | 1,18107E-28 |
| ACE2 | WASF3-AS1 | -0,5396 | 1,14623E-28 |
| ACE2 | LINC01531 | -0,5396 | 1,14042E-28 |
| ACE2 | AL591704.7 | -0,5397 | 1,1084E-28 |
| ACE2 | RP11-475D12.1 | -0,5397 | 1,12605E-28 |
| ACE2 | RP11-285E23.2 | -0,5397 | 1,11342E-28 |
| ACE2 | RP11-966I7.3 | -0,5397 | 1,11511E-28 |
| ACE2 | CTC-490G23.4 | -0,5397 | 1,12297E-28 |
| ACE2 | RP4-639J15.1 | -0,5398 | 1,08453E-28 |
| ACE2 | HTR2A-AS1 | -0,5398 | 1,07872E-28 |
| ACE2 | PKMP1 | -0,5399 | 1,06197E-28 |
| ACE2 | TMCO5A | -0,5399 | 1,05895E-28 |
| ACE2 | NOS2P1 | -0,5399 | 1,05471E-28 |
| ACE2 | RPL12P3 | -0,5399 | 1,06536E-28 |
| ACE2 | RP11-416K24.1 | -0,54 | 1,04644E-28 |
| ACE2 | RP11-201E8.1 | -0,54 | 1,04591E-28 |
| ACE2 | KRT8P36 | -0,54 | 1,03682E-28 |
| ACE2 | CPLX2 | -0,54 | 1,02713E-28 |
| ACE2 | RP11-179A10.1 | -0,54 | 1,02996E-28 |

| | | | |
|---|---|---|---|
| ACE2 | SPA17 | -0,54 | 1,02081E-28 |
| ACE2 | ARMC7 | -0,54 | 1,03447E-28 |
| ACE2 | RP3-522J7.6 | -0,54 | 1,04487E-28 |
| ACE2 | RP11-22M7.2 | -0,5401 | 1,0167E-28 |
| ACE2 | ARL9 | -0,5401 | 1,00729E-28 |
| ACE2 | OLR1 | -0,5401 | 1,01697E-28 |
| ACE2 | RP11-364P2.2 | -0,5402 | 9,64515E-29 |
| ACE2 | RP11-123O1.1 | -0,5402 | 9,77842E-29 |
| ACE2 | LINC00710 | -0,5402 | 9,81138E-29 |
| ACE2 | POLL | -0,5402 | 9,88822E-29 |
| ACE2 | CTB-41I6.2 | -0,5402 | 9,79317E-29 |
| ACE2 | CTD-2526M8.3 | -0,5402 | 9,88439E-29 |
| ACE2 | CCDC9 | -0,5403 | 9,3846E-29 |
| ACE2 | TBL2 | -0,5404 | 9,29764E-29 |
| ACE2 | NRXN3 | -0,5404 | 9,3648E-29 |
| ACE2 | RPL36AP48 | -0,5404 | 9,36404E-29 |
| ACE2 | LINC00229 | -0,5404 | 9,23719E-29 |
| ACE2 | AF146191.4 | -0,5405 | 9,03099E-29 |
| ACE2 | PSMC3 | -0,5405 | 8,89472E-29 |
| ACE2 | COLCA2 | -0,5405 | 9,07128E-29 |
| ACE2 | KRT18P17 | -0,5406 | 8,77868E-29 |
| ACE2 | FAM217A | -0,5406 | 8,74982E-29 |
| ACE2 | VPS28 | -0,5406 | 8,81397E-29 |
| ACE2 | AC124997.1 | -0,5406 | 8,67091E-29 |
| ACE2 | CABYR | -0,5406 | 8,7214E-29 |
| ACE2 | LINC01722 | -0,5406 | 8,75154E-29 |
| ACE2 | FGF12-AS1 | -0,5407 | 8,50183E-29 |
| ACE2 | FRMD3 | -0,5407 | 8,48336E-29 |
| ACE2 | AC009518.4 | -0,5408 | 8,37689E-29 |
| ACE2 | RP11-254F7.2 | -0,5409 | 8,00505E-29 |
| ACE2 | LINC00550 | -0,5409 | 7,95746E-29 |
| ACE2 | LINC01685 | -0,541 | 7,91955E-29 |
| ACE2 | LINC01066 | -0,541 | 7,8546E-29 |
| ACE2 | LINC01065 | -0,541 | 7,82443E-29 |
| ACE2 | AC005062.2 | -0,5411 | 7,6054E-29 |
| ACE2 | TMEM120A | -0,5411 | 7,59118E-29 |
| ACE2 | RP11-783K16.13 | -0,5411 | 7,51986E-29 |
| ACE2 | LINC02189 | -0,5411 | 7,65736E-29 |
| ACE2 | CCDC42 | -0,5411 | 7,597E-29 |
| ACE2 | AC104695.4 | -0,5412 | 7,47682E-29 |
| ACE2 | AC009313.1 | -0,5412 | 7,3537E-29 |
| ACE2 | CTD-2523D13.1 | -0,5412 | 7,50024E-29 |
| ACE2 | RP11-401F2.3 | -0,5412 | 7,43372E-29 |
| ACE2 | RP11-433J20.2 | -0,5413 | 7,18336E-29 |
| ACE2 | RN7SL176P | -0,5413 | 7,29611E-29 |
| ACE2 | CCL4 | -0,5413 | 7,17255E-29 |
| ACE2 | IGSF23 | -0,5413 | 7,25165E-29 |
| ACE2 | PDE6A | -0,5414 | 7,08244E-29 |
| ACE2 | RP5-866L20.1 | -0,5415 | 6,82427E-29 |
| ACE2 | LINC01851 | -0,5415 | 6,84724E-29 |
| ACE2 | GNG3 | -0,5415 | 6,87902E-29 |
| ACE2 | SLC25A5P1 | -0,5415 | 6,90039E-29 |
| ACE2 | RP3-406A7.3 | -0,5416 | 6,56805E-29 |
| ACE2 | CALR3 | -0,5416 | 6,59123E-29 |

| | | | |
|---|---|---|---|
| ACE2 | RP11-1102P16.1 | -0,5417 | 6,51792E-29 |
| ACE2 | PPP1R36 | -0,5417 | 6,52322E-29 |
| ACE2 | CTD-2184D3.3 | -0,5417 | 6,44063E-29 |
| ACE2 | NPPA | -0,5418 | 6,20439E-29 |
| ACE2 | RP11-17A4.3 | -0,5418 | 6,27998E-29 |
| ACE2 | CBLN1 | -0,5418 | 6,30274E-29 |
| ACE2 | LOXHD1 | -0,5418 | 6,20889E-29 |
| ACE2 | CTD-2260A17.1 | -0,5419 | 6,14194E-29 |
| ACE2 | RP3-399L15.1 | -0,5419 | 6,15978E-29 |
| ACE2 | ANKRD19P | -0,5419 | 6,15746E-29 |
| ACE2 | RP11-375D13.4 | -0,5419 | 6,09295E-29 |
| ACE2 | RP11-612A1.1 | -0,5419 | 6,09359E-29 |
| ACE2 | RP4-683M8.2 | -0,542 | 5,98145E-29 |
| ACE2 | FAM170A | -0,542 | 5,86751E-29 |
| ACE2 | LINC01467 | -0,542 | 6,01913E-29 |
| ACE2 | RP11-820I16.1 | -0,542 | 5,97401E-29 |
| ACE2 | LINC02053 | -0,5421 | 5,72723E-29 |
| ACE2 | LPP-AS1 | -0,5421 | 5,72261E-29 |
| ACE2 | MIR4307HG | -0,5421 | 5,72802E-29 |
| ACE2 | CD200R1L | -0,5422 | 5,66192E-29 |
| ACE2 | KLK12 | -0,5422 | 5,58161E-29 |
| ACE2 | RP11-152O14.4 | -0,5423 | 5,51439E-29 |
| ACE2 | FAM71A | -0,5424 | 5,37489E-29 |
| ACE2 | OPRM1 | -0,5424 | 5,3758E-29 |
| ACE2 | EDNRB-AS1 | -0,5424 | 5,24585E-29 |
| ACE2 | FAM71D | -0,5424 | 5,30469E-29 |
| ACE2 | RP11-94B19.5 | -0,5424 | 5,26176E-29 |
| ACE2 | LINC01522 | -0,5424 | 5,29962E-29 |
| ACE2 | TPI1P2 | -0,5425 | 5,10764E-29 |
| ACE2 | LINC00441 | -0,5425 | 5,12096E-29 |
| ACE2 | DIO2-AS1 | -0,5425 | 5,15078E-29 |
| ACE2 | THOP1 | -0,5425 | 5,20271E-29 |
| ACE2 | TMIGD3 | -0,5426 | 5,04681E-29 |
| ACE2 | IQCJ-SCHIP1-AS1 | -0,5426 | 5,06827E-29 |
| ACE2 | PGK2 | -0,5426 | 5,04261E-29 |
| ACE2 | TMEM125 | -0,5427 | 4,83691E-29 |
| ACE2 | IFT172 | -0,5427 | 4,90498E-29 |
| ACE2 | FAHD2A | -0,5427 | 4,8377E-29 |
| ACE2 | RP13-254B10.1 | -0,5427 | 4,94609E-29 |
| ACE2 | HAS2-AS1 | -0,5427 | 4,85088E-29 |
| ACE2 | RP11-871F6.3 | -0,5428 | 4,72587E-29 |
| ACE2 | LIN7A | -0,5428 | 4,70997E-29 |
| ACE2 | LINC01048 | -0,5428 | 4,75256E-29 |
| ACE2 | KCNRG | -0,5428 | 4,73445E-29 |
| ACE2 | CLEC4F | -0,5429 | 4,62804E-29 |
| ACE2 | CCDC74A | -0,5429 | 4,6224E-29 |
| ACE2 | RP11-760D2.1 | -0,543 | 4,51843E-29 |
| ACE2 | RFX3-AS1 | -0,543 | 4,45784E-29 |
| ACE2 | RP11-148I19.1 | -0,543 | 4,45851E-29 |
| ACE2 | RP11-141M1.1 | -0,543 | 4,43695E-29 |
| ACE2 | RP11-428L21.2 | -0,5431 | 4,43354E-29 |
| ACE2 | MEF2C-AS1 | -0,5431 | 4,31868E-29 |
| ACE2 | RPL12P27 | -0,5431 | 4,42609E-29 |
| ACE2 | RP11-184I16.4 | -0,5432 | 4,26258E-29 |

| | | | |
|---|---|---|---|
| ACE2 | RNF133 | -0,5432 | 4,29631E-29 |
| ACE2 | RP11-680E19.1 | -0,5432 | 4,30071E-29 |
| ACE2 | RP11-74M13.5 | -0,5432 | 4,31074E-29 |
| ACE2 | SCLY | -0,5433 | 4,15671E-29 |
| ACE2 | YBX2P1 | -0,5434 | 3,99285E-29 |
| ACE2 | MS4A18 | -0,5434 | 4,06091E-29 |
| ACE2 | RP11-28O3.2 | -0,5434 | 4,01789E-29 |
| ACE2 | LINC01649 | -0,5435 | 3,88359E-29 |
| ACE2 | LINC01032 | -0,5435 | 3,86245E-29 |
| ACE2 | RP11-572N21.1 | -0,5435 | 3,89574E-29 |
| ACE2 | C9orf135 | -0,5435 | 3,86814E-29 |
| ACE2 | BSCL2 | -0,5435 | 3,9498E-29 |
| ACE2 | RP11-478K15.7 | -0,5435 | 3,86345E-29 |
| ACE2 | AC010649.1 | -0,5435 | 3,9239E-29 |
| ACE2 | KLHL6-AS1 | -0,5436 | 3,85052E-29 |
| ACE2 | ASB5 | -0,5436 | 3,85232E-29 |
| ACE2 | RP5-1077I2.3 | -0,5436 | 3,80459E-29 |
| ACE2 | DCUN1D1 | -0,5437 | 3,65723E-29 |
| ACE2 | RP5-908M14.10 | -0,5437 | 3,68184E-29 |
| ACE2 | C10orf90 | -0,5438 | 3,57869E-29 |
| ACE2 | IGLVIVOR22-2 | -0,5438 | 3,5513E-29 |
| ACE2 | RP11-285E9.6 | -0,5439 | 3,52748E-29 |
| ACE2 | RP11-746B8.1 | -0,5439 | 3,51613E-29 |
| ACE2 | RP4-784A16.2 | -0,544 | 3,38082E-29 |
| ACE2 | AC009411.2 | -0,544 | 3,39201E-29 |
| ACE2 | AC018766.6 | -0,544 | 3,42344E-29 |
| ACE2 | RP11-357P18.3 | -0,5441 | 3,27532E-29 |
| ACE2 | AC009303.1 | -0,5441 | 3,30754E-29 |
| ACE2 | CAMKV | -0,5441 | 3,28485E-29 |
| ACE2 | RP11-778H2.1 | -0,5441 | 3,27893E-29 |
| ACE2 | RP3-446N13.3 | -0,5441 | 3,34204E-29 |
| ACE2 | RP11-626P14.2 | -0,5441 | 3,299E-29 |
| ACE2 | GOLGA6A | -0,5441 | 3,34117E-29 |
| ACE2 | ANTXRLP1 | -0,5442 | 3,2583E-29 |
| ACE2 | RP5-1066H13.4 | -0,5443 | 3,10507E-29 |
| ACE2 | C3orf80 | -0,5443 | 3,09889E-29 |
| ACE2 | CTB-113P19.4 | -0,5443 | 3,16633E-29 |
| ACE2 | RP11-1080G15.2 | -0,5443 | 3,13782E-29 |
| ACE2 | ARHGEF39 | -0,5443 | 3,09E-29 |
| ACE2 | SPZ1 | -0,5444 | 3,06039E-29 |
| ACE2 | NDUFA8 | -0,5444 | 3,0308E-29 |
| ACE2 | LINGO1-AS2 | -0,5444 | 3,074E-29 |
| ACE2 | LRRC29 | -0,5444 | 3,01334E-29 |
| ACE2 | KB-1440D3.13 | -0,5444 | 3,02873E-29 |
| ACE2 | SKINT1L | -0,5445 | 2,93549E-29 |
| ACE2 | INPP1 | -0,5445 | 2,96904E-29 |
| ACE2 | DNAJC5B | -0,5445 | 2,92511E-29 |
| ACE2 | LINC01251 | -0,5445 | 2,97881E-29 |
| ACE2 | AGAP14 | -0,5445 | 2,99341E-29 |
| ACE2 | RP11-179A16.2 | -0,5445 | 2,96782E-29 |
| ACE2 | PRR19 | -0,5445 | 2,98631E-29 |
| ACE2 | CTC-503K11.2 | -0,5446 | 2,84262E-29 |
| ACE2 | RP11-141M1.3 | -0,5446 | 2,86529E-29 |
| ACE2 | RP11-872J21.3 | -0,5446 | 2,88082E-29 |

| | | | |
|---|---|---|---|
| ACE2 | ADSSL1 | -0,5446 | 2,86566E-29 |
| ACE2 | SDCCAG3P1 | -0,5446 | 2,90894E-29 |
| ACE2 | TBX1 | -0,5446 | 2,89572E-29 |
| ACE2 | RP11-67L3.4 | -0,5447 | 2,81181E-29 |
| ACE2 | GRM8 | -0,5447 | 2,80088E-29 |
| ACE2 | FAM135B | -0,5447 | 2,7856E-29 |
| ACE2 | HARBI1 | -0,5447 | 2,81492E-29 |
| ACE2 | GOLGA8IP | -0,5447 | 2,76144E-29 |
| ACE2 | KRT34 | -0,5447 | 2,76696E-29 |
| ACE2 | CTB-180A7.8 | -0,5447 | 2,76259E-29 |
| ACE2 | HYDIN | -0,5448 | 2,72455E-29 |
| ACE2 | RP4-777L9.2 | -0,5448 | 2,69828E-29 |
| ACE2 | LINC01650 | -0,5449 | 2,6767E-29 |
| ACE2 | CRISP1 | -0,5449 | 2,6129E-29 |
| ACE2 | RP11-85A1.3 | -0,5449 | 2,67496E-29 |
| ACE2 | RP11-535A19.1 | -0,5449 | 2,63672E-29 |
| ACE2 | RP11-65M17.3 | -0,5449 | 2,67757E-29 |
| ACE2 | SLFNL1-AS1 | -0,545 | 2,54573E-29 |
| ACE2 | RP11-12M5.4 | -0,545 | 2,55695E-29 |
| ACE2 | RP11-281P23.2 | -0,545 | 2,57204E-29 |
| ACE2 | CTC-347C20.1 | -0,545 | 2,57032E-29 |
| ACE2 | ZNF204P | -0,545 | 2,5552E-29 |
| ACE2 | OGFOD2 | -0,545 | 2,55556E-29 |
| ACE2 | LINC02135 | -0,545 | 2,56294E-29 |
| ACE2 | VAMP8 | -0,5451 | 2,53302E-29 |
| ACE2 | FGF14-IT1 | -0,5451 | 2,51497E-29 |
| ACE2 | LINC00919 | -0,5451 | 2,53081E-29 |
| ACE2 | RP11-323H21.3 | -0,5452 | 2,43217E-29 |
| ACE2 | LINC00649 | -0,5452 | 2,40097E-29 |
| ACE2 | RP11-73E17.2 | -0,5453 | 2,38148E-29 |
| ACE2 | CCDC105 | -0,5453 | 2,36598E-29 |
| ACE2 | GLRX2 | -0,5454 | 2,30532E-29 |
| ACE2 | COMMD1 | -0,5454 | 2,30564E-29 |
| ACE2 | RP11-548K12.12 | -0,5454 | 2,2719E-29 |
| ACE2 | AF067845.2 | -0,5454 | 2,27313E-29 |
| ACE2 | PSD | -0,5454 | 2,33015E-29 |
| ACE2 | STRC | -0,5454 | 2,29922E-29 |
| ACE2 | RP5-1024N4.2 | -0,5455 | 2,24574E-29 |
| ACE2 | FAM183DP | -0,5455 | 2,2227E-29 |
| ACE2 | AC002064.7 | -0,5455 | 2,23526E-29 |
| ACE2 | RP11-712B9.2 | -0,5455 | 2,23327E-29 |
| ACE2 | AC015849.16 | -0,5455 | 2,2553E-29 |
| ACE2 | C5AR2 | -0,5455 | 2,2208E-29 |
| ACE2 | RIIAD1 | -0,5456 | 2,1992E-29 |
| ACE2 | MLF1 | -0,5456 | 2,17912E-29 |
| ACE2 | AC007405.6 | -0,5458 | 2,05097E-29 |
| ACE2 | RP11-149I2.4 | -0,5458 | 2,07734E-29 |
| ACE2 | RP11-20J15.3 | -0,5458 | 2,07456E-29 |
| ACE2 | RP13-81N3.2 | -0,5458 | 2,05451E-29 |
| ACE2 | KLHL10 | -0,5458 | 2,05002E-29 |
| ACE2 | TUBA3GP | -0,5458 | 2,06127E-29 |
| ACE2 | AC012462.2 | -0,546 | 1,94621E-29 |
| ACE2 | RP11-415D17.4 | -0,546 | 1,92914E-29 |
| ACE2 | TPD52L3 | -0,546 | 1,939E-29 |

| | | | |
|---|---|---|---|
| ACE2 | RP11-168K11.3 | -0,546 | 1,96556E-29 |
| ACE2 | CEP112 | -0,546 | 1,95008E-29 |
| ACE2 | BOD1L2 | -0,546 | 1,96E-29 |
| ACE2 | GRXCR2 | -0,5461 | 1,89375E-29 |
| ACE2 | CALHM3 | -0,5461 | 1,88148E-29 |
| ACE2 | IQCE | -0,5462 | 1,85875E-29 |
| ACE2 | SUN3 | -0,5462 | 1,83088E-29 |
| ACE2 | PWRN2 | -0,5463 | 1,76798E-29 |
| ACE2 | AC072062.1 | -0,5464 | 1,7305E-29 |
| ACE2 | PC | -0,5464 | 1,73328E-29 |
| ACE2 | RP11-525K10.1 | -0,5464 | 1,73466E-29 |
| ACE2 | LRRC74B | -0,5464 | 1,74483E-29 |
| ACE2 | AC011738.4 | -0,5465 | 1,70719E-29 |
| ACE2 | RPS19P7 | -0,5465 | 1,68592E-29 |
| ACE2 | UNC45B | -0,5465 | 1,67679E-29 |
| ACE2 | RP11-2N1.1 | -0,5466 | 1,63479E-29 |
| ACE2 | CCDC150 | -0,5467 | 1,5828E-29 |
| ACE2 | RP11-571L19.7 | -0,5467 | 1,58775E-29 |
| ACE2 | RP11-1085N6.2 | -0,5467 | 1,6114E-29 |
| ACE2 | NAALADL1 | -0,5468 | 1,55414E-29 |
| ACE2 | RP11-676F20.4 | -0,5468 | 1,56514E-29 |
| ACE2 | RP11-279O9.4 | -0,5469 | 1,50447E-29 |
| ACE2 | TMEM215 | -0,5469 | 1,52254E-29 |
| ACE2 | RP1-7G5.6 | -0,5469 | 1,4949E-29 |
| ACE2 | EIF3G | -0,5469 | 1,50193E-29 |
| ACE2 | RP5-905H7.11 | -0,547 | 1,46988E-29 |
| ACE2 | HNF4A | -0,547 | 1,46015E-29 |
| ACE2 | RP11-236J17.6 | -0,5471 | 1,41887E-29 |
| ACE2 | ZNF649-AS1 | -0,5471 | 1,43626E-29 |
| ACE2 | LINC00700 | -0,5472 | 1,38957E-29 |
| ACE2 | DGAT2 | -0,5472 | 1,4007E-29 |
| ACE2 | LINC00458 | -0,5472 | 1,39731E-29 |
| ACE2 | RP11-288L9.4 | -0,5473 | 1,36449E-29 |
| ACE2 | RP11-364C11.3 | -0,5473 | 1,36387E-29 |
| ACE2 | TSSK4 | -0,5473 | 1,33837E-29 |
| ACE2 | RP1-168P16.2 | -0,5473 | 1,34784E-29 |
| ACE2 | TIFAB | -0,5474 | 1,31089E-29 |
| ACE2 | RP3-449H6.2 | -0,5474 | 1,31317E-29 |
| ACE2 | DDI1 | -0,5474 | 1,32835E-29 |
| ACE2 | RP3-503A6.2 | -0,5475 | 1,26298E-29 |
| ACE2 | PLEKHJ1 | -0,5475 | 1,2777E-29 |
| ACE2 | RP11-473M14.3 | -0,5476 | 1,24553E-29 |
| ACE2 | ZNF20 | -0,5476 | 1,23556E-29 |
| ACE2 | GK3P | -0,5477 | 1,19393E-29 |
| ACE2 | RP11-779O18.2 | -0,5477 | 1,22054E-29 |
| ACE2 | AC006372.6 | -0,5477 | 1,21694E-29 |
| ACE2 | CTD-3088G3.8 | -0,5477 | 1,22076E-29 |
| ACE2 | CTD-2588J1.1 | -0,5478 | 1,16414E-29 |
| ACE2 | LINC01579 | -0,5478 | 1,16769E-29 |
| ACE2 | RASIP1 | -0,5478 | 1,17364E-29 |
| ACE2 | LINC00427 | -0,5479 | 1,14421E-29 |
| ACE2 | CRNDE | -0,5479 | 1,13475E-29 |
| ACE2 | DRC3 | -0,5479 | 1,15386E-29 |
| ACE2 | TMEM191A | -0,5479 | 1,14448E-29 |

| | | | |
|---|---|---|---|
| ACE2 | RTP1 | -0,548 | 1,09941E-29 |
| ACE2 | RP11-297L17.4 | -0,548 | 1,0991E-29 |
| ACE2 | CTD-2086O20.1 | -0,548 | 1,09567E-29 |
| ACE2 | MIR646HG | -0,548 | 1,09528E-29 |
| ACE2 | LINC01231 | -0,5481 | 1,06559E-29 |
| ACE2 | ZDHHC12 | -0,5481 | 1,0749E-29 |
| ACE2 | C10orf111 | -0,5481 | 1,08917E-29 |
| ACE2 | CCDC38 | -0,5481 | 1,06137E-29 |
| ACE2 | LINC01739 | -0,5482 | 1,03118E-29 |
| ACE2 | RP11-728G15.1 | -0,5482 | 1,05957E-29 |
| ACE2 | C22orf46 | -0,5482 | 1,04314E-29 |
| ACE2 | RP11-118M12.1 | -0,5483 | 1,03047E-29 |
| ACE2 | REM2 | -0,5483 | 1,00637E-29 |
| ACE2 | RP11-504G3.4 | -0,5484 | 9,93931E-30 |
| ACE2 | EWSAT1 | -0,5484 | 9,8704E-30 |
| ACE2 | AC092652.1 | -0,5485 | 9,5737E-30 |
| ACE2 | RP11-650J17.1 | -0,5485 | 9,6468E-30 |
| ACE2 | RP11-379I19.1 | -0,5485 | 9,61823E-30 |
| ACE2 | FAM181A-AS1 | -0,5485 | 9,46617E-30 |
| ACE2 | LRRC37A16P | -0,5485 | 9,47652E-30 |
| ACE2 | TSSK6 | -0,5485 | 9,51457E-30 |
| ACE2 | ZNF599 | -0,5485 | 9,69579E-30 |
| ACE2 | AC006042.6 | -0,5486 | 9,4117E-30 |
| ACE2 | RP11-6B19.1 | -0,5486 | 9,46327E-30 |
| ACE2 | TNP2 | -0,5486 | 9,28166E-30 |
| ACE2 | SLC25A39 | -0,5486 | 9,38354E-30 |
| ACE2 | ASPSCR1 | -0,5486 | 9,34219E-30 |
| ACE2 | AC002511.2 | -0,5486 | 9,30858E-30 |
| ACE2 | NECAB3 | -0,5486 | 9,35015E-30 |
| ACE2 | RABIF | -0,5487 | 8,96762E-30 |
| ACE2 | MIR7515HG | -0,5487 | 9,03827E-30 |
| ACE2 | AC073464.7 | -0,5487 | 9,01089E-30 |
| ACE2 | PPP2R2B | -0,5487 | 9,151E-30 |
| ACE2 | CTD-2647E9.1 | -0,5487 | 9,15168E-30 |
| ACE2 | SLC37A1 | -0,5487 | 9,09995E-30 |
| ACE2 | CTB-49A3.5 | -0,5488 | 8,85167E-30 |
| ACE2 | RP11-283I3.4 | -0,5488 | 8,82804E-30 |
| ACE2 | TMEM102 | -0,5488 | 8,87377E-30 |
| ACE2 | RP11-314N13.9 | -0,5488 | 8,77228E-30 |
| ACE2 | RP11-378I13.1 | -0,5489 | 8,46923E-30 |
| ACE2 | AC096732.2 | -0,5489 | 8,56775E-30 |
| ACE2 | HIST1H2BJ | -0,5489 | 8,56898E-30 |
| ACE2 | ENO1-AS1 | -0,549 | 8,31998E-30 |
| ACE2 | AC007422.1 | -0,549 | 8,24041E-30 |
| ACE2 | AC093901.1 | -0,549 | 8,31475E-30 |
| ACE2 | LINC00692 | -0,549 | 8,28459E-30 |
| ACE2 | C4orf45 | -0,549 | 8,39387E-30 |
| ACE2 | EDRF1-AS1 | -0,549 | 8,23193E-30 |
| ACE2 | RPL23AP75 | -0,549 | 8,37481E-30 |
| ACE2 | RP11-166P13.4 | -0,549 | 8,37783E-30 |
| ACE2 | RP5-945F2.3 | -0,5492 | 7,8108E-30 |
| ACE2 | GFOD2 | -0,5492 | 7,88427E-30 |
| ACE2 | RP4-673D20.3 | -0,5492 | 7,88418E-30 |
| ACE2 | RP11-779P15.2 | -0,5493 | 7,59328E-30 |

| | | | |
|---|---|---|---|
| ACE2 | COMMD4 | -0,5493 | 7,69508E-30 |
| ACE2 | ATP5G1P4 | -0,5494 | 7,42833E-30 |
| ACE2 | RP11-185E12.2 | -0,5494 | 7,46619E-30 |
| ACE2 | LINC01300 | -0,5494 | 7,36914E-30 |
| ACE2 | RP11-293M10.2 | -0,5494 | 7,36354E-30 |
| ACE2 | RP11-403P17.6 | -0,5494 | 7,40643E-30 |
| ACE2 | RP11-108M9.3 | -0,5495 | 7,26976E-30 |
| ACE2 | LINC01999 | -0,5496 | 7,0021E-30 |
| ACE2 | ZNF341-AS1 | -0,5496 | 6,97106E-30 |
| ACE2 | RP3-333H23.8 | -0,5496 | 6,96302E-30 |
| ACE2 | RP11-727F15.9 | -0,5497 | 6,90897E-30 |
| ACE2 | MAST1 | -0,5497 | 6,78148E-30 |
| ACE2 | WDR88 | -0,5497 | 6,89156E-30 |
| ACE2 | RP11-546B15.1 | -0,5498 | 6,70131E-30 |
| ACE2 | PLSCR2 | -0,5499 | 6,51676E-30 |
| ACE2 | NPAP1P3 | -0,5499 | 6,46814E-30 |
| ACE2 | AC011752.1 | -0,55 | 6,22685E-30 |
| ACE2 | LINC01210 | -0,5501 | 6,07648E-30 |
| ACE2 | RP11-100N20.1 | -0,5501 | 6,13385E-30 |
| ACE2 | CRH | -0,5501 | 6,10158E-30 |
| ACE2 | FRMPD2 | -0,5501 | 6,0869E-30 |
| ACE2 | DRAP1 | -0,5502 | 5,87781E-30 |
| ACE2 | UBE2U | -0,5503 | 5,77192E-30 |
| ACE2 | RNASE3 | -0,5503 | 5,83584E-30 |
| ACE2 | RP11-481J2.1 | -0,5503 | 5,81076E-30 |
| ACE2 | DUSP18 | -0,5503 | 5,81196E-30 |
| ACE2 | CENPS | -0,5504 | 5,64528E-30 |
| ACE2 | TBL3 | -0,5504 | 5,62368E-30 |
| ACE2 | RP11-275I14.2 | -0,5505 | 5,48751E-30 |
| ACE2 | RP11-818C3.1 | -0,5505 | 5,42718E-30 |
| ACE2 | RP11-393K12.2 | -0,5505 | 5,46732E-30 |
| ACE2 | AC007557.4 | -0,5506 | 5,35174E-30 |
| ACE2 | OR7E14P | -0,5506 | 5,29025E-30 |
| ACE2 | RP11-15F12.3 | -0,5506 | 5,23038E-30 |
| ACE2 | RP11-866E20.3 | -0,5507 | 5,17943E-30 |
| ACE2 | PRAL | -0,5508 | 4,97083E-30 |
| ACE2 | AC002398.13 | -0,5508 | 5,05024E-30 |
| ACE2 | LARGE-AS1 | -0,5508 | 5,02457E-30 |
| ACE2 | GRM7 | -0,5509 | 4,87838E-30 |
| ACE2 | NKX1-1 | -0,5509 | 4,80412E-30 |
| ACE2 | RP11-63K6.5 | -0,5509 | 4,80373E-30 |
| ACE2 | NPHS2 | -0,551 | 4,7083E-30 |
| ACE2 | LINC01377 | -0,551 | 4,66792E-30 |
| ACE2 | AC006153.3 | -0,551 | 4,70242E-30 |
| ACE2 | RP11-1103G16.1 | -0,551 | 4,76415E-30 |
| ACE2 | NAGK | -0,5511 | 4,59618E-30 |
| ACE2 | RP11-778J15.1 | -0,5511 | 4,62467E-30 |
| ACE2 | RP11-286O18.1 | -0,5511 | 4,55225E-30 |
| ACE2 | RP1-117O3.2 | -0,5512 | 4,3901E-30 |
| ACE2 | GABBR2 | -0,5512 | 4,40214E-30 |
| ACE2 | C12orf65 | -0,5512 | 4,46761E-30 |
| ACE2 | LINC01584 | -0,5512 | 4,50964E-30 |
| ACE2 | DBI | -0,5513 | 4,35193E-30 |
| ACE2 | RP11-103H7.2 | -0,5513 | 4,33698E-30 |

| | | | |
|---|---|---|---|
| ACE2 | RP1-191J18.66 | -0,5514 | 4,26025E-30 |
| ACE2 | FAM46C | -0,5515 | 4,06428E-30 |
| ACE2 | RP11-506H20.1 | -0,5515 | 4,12283E-30 |
| ACE2 | PTCHD3P3 | -0,5515 | 4,10761E-30 |
| ACE2 | KB-1043D8.6 | -0,5515 | 4,08588E-30 |
| ACE2 | RP11-64D24.4 | -0,5515 | 4,13368E-30 |
| ACE2 | RP11-83B20.1 | -0,5515 | 4,10164E-30 |
| ACE2 | RP11-265N7.2 | -0,5515 | 4,08625E-30 |
| ACE2 | LINC02093 | -0,5515 | 4,10483E-30 |
| ACE2 | KIAA1683 | -0,5515 | 4,08445E-30 |
| ACE2 | RAB11FIP5 | -0,5516 | 4,0013E-30 |
| ACE2 | ST7-OT4 | -0,5516 | 3,92115E-30 |
| ACE2 | AC004941.3 | -0,5516 | 3,97051E-30 |
| ACE2 | RP3-508I15.21 | -0,5516 | 3,91318E-30 |
| ACE2 | RP11-680A11.5 | -0,5517 | 3,84078E-30 |
| ACE2 | LINC01491 | -0,5517 | 3,8079E-30 |
| ACE2 | RP5-850O15.3 | -0,5518 | 3,73547E-30 |
| ACE2 | RP11-45K10.2 | -0,5518 | 3,72391E-30 |
| ACE2 | RP11-867G2.5 | -0,5518 | 3,7483E-30 |
| ACE2 | RP11-328C8.5 | -0,5518 | 3,79602E-30 |
| ACE2 | FAM163B | -0,5519 | 3,63605E-30 |
| ACE2 | TREH | -0,5519 | 3,64562E-30 |
| ACE2 | RP11-768F21.1 | -0,5519 | 3,62183E-30 |
| ACE2 | PDILT | -0,5519 | 3,68804E-30 |
| ACE2 | CTNS | -0,5519 | 3,68258E-30 |
| ACE2 | CTD-2376I4.2 | -0,552 | 3,53074E-30 |
| ACE2 | GS1-24F4.2 | -0,552 | 3,57506E-30 |
| ACE2 | RP11-643G16.4 | -0,552 | 3,55217E-30 |
| ACE2 | RP11-303E16.7 | -0,552 | 3,51579E-30 |
| ACE2 | GEMIN4 | -0,552 | 3,52427E-30 |
| ACE2 | RGSL1 | -0,5521 | 3,43626E-30 |
| ACE2 | ULK4 | -0,5521 | 3,42717E-30 |
| ACE2 | SRRM3 | -0,5521 | 3,42038E-30 |
| ACE2 | RP4-778K6.1 | -0,5521 | 3,47291E-30 |
| ACE2 | FIBP | -0,5521 | 3,39981E-30 |
| ACE2 | ZNF433 | -0,5521 | 3,44649E-30 |
| ACE2 | DRC1 | -0,5522 | 3,35134E-30 |
| ACE2 | CDKL4 | -0,5522 | 3,29557E-30 |
| ACE2 | TEX35 | -0,5523 | 3,24057E-30 |
| ACE2 | C2orf83 | -0,5523 | 3,23398E-30 |
| ACE2 | OSBPL10-AS1 | -0,5523 | 3,27314E-30 |
| ACE2 | RP11-63L7.5 | -0,5523 | 3,25581E-30 |
| ACE2 | REEP4 | -0,5523 | 3,25111E-30 |
| ACE2 | RP11-697E14.2 | -0,5523 | 3,26669E-30 |
| ACE2 | ESRRB | -0,5523 | 3,2333E-30 |
| ACE2 | LRRC69 | -0,5524 | 3,12736E-30 |
| ACE2 | RP11-407A16.7 | -0,5524 | 3,18406E-30 |
| ACE2 | SPATA6 | -0,5525 | 3,0841E-30 |
| ACE2 | EBLN1 | -0,5525 | 3,03325E-30 |
| ACE2 | RSPH14 | -0,5525 | 3,06031E-30 |
| ACE2 | APOL5 | -0,5525 | 3,03306E-30 |
| ACE2 | HNRNPA1P15 | -0,5526 | 2,97094E-30 |
| ACE2 | RP11-322I2.1 | -0,5526 | 2,94137E-30 |
| ACE2 | LIPJ | -0,5527 | 2,86593E-30 |

| | | | |
|---|---|---|---|
| ACE2 | RP11-300J18.1 | -0,5527 | 2,91677E-30 |
| ACE2 | AC104088.1 | -0,5528 | 2,81113E-30 |
| ACE2 | RP11-600L4.1 | -0,5528 | 2,85299E-30 |
| ACE2 | RP11-402J6.1 | -0,5528 | 2,80092E-30 |
| ACE2 | EEF1D | -0,5528 | 2,77576E-30 |
| ACE2 | MIR5188 | -0,5528 | 2,77575E-30 |
| ACE2 | TTLL13P | -0,5528 | 2,81511E-30 |
| ACE2 | CCHCR1 | -0,5529 | 2,75061E-30 |
| ACE2 | RP11-363L24.3 | -0,5529 | 2,75829E-30 |
| ACE2 | BTBD6 | -0,5529 | 2,72303E-30 |
| ACE2 | HIGD2B | -0,5529 | 2,72547E-30 |
| ACE2 | C16orf71 | -0,5529 | 2,74985E-30 |
| ACE2 | BACH1-IT1 | -0,5529 | 2,69714E-30 |
| ACE2 | TMEM269 | -0,553 | 2,68187E-30 |
| ACE2 | LINC01884 | -0,553 | 2,66994E-30 |
| ACE2 | LINC02063 | -0,553 | 2,66002E-30 |
| ACE2 | HNRNPA1P38 | -0,553 | 2,67998E-30 |
| ACE2 | RP11-449J1.1 | -0,5531 | 2,57831E-30 |
| ACE2 | RP11-297A16.2 | -0,5531 | 2,59823E-30 |
| ACE2 | HAGHL | -0,5531 | 2,59459E-30 |
| ACE2 | RP11-344J7.4 | -0,5532 | 2,47251E-30 |
| ACE2 | RP4-536B24.4 | -0,5532 | 2,53002E-30 |
| ACE2 | CFAP53 | -0,5532 | 2,50399E-30 |
| ACE2 | AC007365.1 | -0,5533 | 2,41172E-30 |
| ACE2 | KBTBD3 | -0,5533 | 2,41687E-30 |
| ACE2 | CTD-2600O9.2 | -0,5533 | 2,42721E-30 |
| ACE2 | RP11-353K11.1 | -0,5534 | 2,39106E-30 |
| ACE2 | HHLA2 | -0,5534 | 2,33874E-30 |
| ACE2 | SSMEM1 | -0,5534 | 2,34776E-30 |
| ACE2 | RP11-819M15.1 | -0,5534 | 2,39439E-30 |
| ACE2 | SUGP1 | -0,5534 | 2,37863E-30 |
| ACE2 | GUCA1C | -0,5535 | 2,3129E-30 |
| ACE2 | RP11-944C7.1 | -0,5535 | 2,27944E-30 |
| ACE2 | CCDC189 | -0,5535 | 2,33037E-30 |
| ACE2 | ZBTB20 | -0,5536 | 2,25225E-30 |
| ACE2 | RP11-344G13.1 | -0,5536 | 2,26501E-30 |
| ACE2 | MSRA | -0,5536 | 2,23629E-30 |
| ACE2 | RP11-837J7.3 | -0,5536 | 2,21854E-30 |
| ACE2 | SEC1P | -0,5536 | 2,23519E-30 |
| ACE2 | AGBL2 | -0,5537 | 2,16658E-30 |
| ACE2 | IL21R | -0,5537 | 2,18715E-30 |
| ACE2 | PLCB1-IT1 | -0,5537 | 2,14461E-30 |
| ACE2 | RP11-174G17 | -0,5538 | 2,12635E-30 |
| ACE2 | C2orf40 | -0,5538 | 2,10132E-30 |
| ACE2 | RP11-527L4.5 | -0,5538 | 2,1237E-30 |
| ACE2 | LPO | -0,5538 | 2,11882E-30 |
| ACE2 | CTD-2588C8.8 | -0,5538 | 2,09621E-30 |
| ACE2 | RP13-467H17.1 | -0,5539 | 2,04983E-30 |
| ACE2 | POC1A | -0,554 | 1,98364E-30 |
| ACE2 | AC005076.5 | -0,554 | 2,00659E-30 |
| ACE2 | C21orf2 | -0,554 | 1,9975E-30 |
| ACE2 | LINC01911 | -0,5541 | 1,92885E-30 |
| ACE2 | MOBP | -0,5541 | 1,95042E-30 |
| ACE2 | ACOXL-AS1 | -0,5542 | 1,88413E-30 |

| | | | |
|---|---|---|---|
| ACE2 | FXYD4 | -0,5542 | 1,86805E-30 |
| ACE2 | RP11-1006G14.1 | -0,5542 | 1,87838E-30 |
| ACE2 | RP11-343K8.3 | -0,5542 | 1,86115E-30 |
| ACE2 | SLC25A2 | -0,5543 | 1,8327E-30 |
| ACE2 | RP11-867G2.6 | -0,5543 | 1,80999E-30 |
| ACE2 | PSMA6 | -0,5543 | 1,81073E-30 |
| ACE2 | ANKRD44-IT1 | -0,5544 | 1,77216E-30 |
| ACE2 | PIK3R6 | -0,5544 | 1,75829E-30 |
| ACE2 | OSCP1 | -0,5545 | 1,73195E-30 |
| ACE2 | MUC12 | -0,5545 | 1,73494E-30 |
| ACE2 | RP11-112L6.3 | -0,5545 | 1,71888E-30 |
| ACE2 | RP11-358D14.2 | -0,5545 | 1,70229E-30 |
| ACE2 | RP11-138I18.1 | -0,5546 | 1,66093E-30 |
| ACE2 | SPATA13 | -0,5546 | 1,66734E-30 |
| ACE2 | RP11-96D24.1 | -0,5546 | 1,67451E-30 |
| ACE2 | MAGI1 | -0,5547 | 1,64926E-30 |
| ACE2 | DNAJC18 | -0,5547 | 1,62333E-30 |
| ACE2 | RP11-40H20.4 | -0,5548 | 1,57054E-30 |
| ACE2 | ALPK2 | -0,5548 | 1,57742E-30 |
| ACE2 | LINC01972 | -0,5549 | 1,51396E-30 |
| ACE2 | OVOS2 | -0,5549 | 1,54661E-30 |
| ACE2 | NLRC3 | -0,5549 | 1,55368E-30 |
| ACE2 | PVRL3-AS1 | -0,555 | 1,51039E-30 |
| ACE2 | C11orf65 | -0,555 | 1,47312E-30 |
| ACE2 | RP11-684B21.1 | -0,555 | 1,47909E-30 |
| ACE2 | AC006942.4 | -0,555 | 1,48236E-30 |
| ACE2 | PDCL2 | -0,5551 | 1,44148E-30 |
| ACE2 | GRP | -0,5551 | 1,44522E-30 |
| ACE2 | CECR6 | -0,5552 | 1,3941E-30 |
| ACE2 | LINC01763 | -0,5553 | 1,37786E-30 |
| ACE2 | MRRF | -0,5553 | 1,36388E-30 |
| ACE2 | LINC00444 | -0,5553 | 1,35516E-30 |
| ACE2 | LINC00479 | -0,5553 | 1,34823E-30 |
| ACE2 | MNX1-AS1 | -0,5554 | 1,33111E-30 |
| ACE2 | RP4-816N1.6 | -0,5554 | 1,32581E-30 |
| ACE2 | TPK1 | -0,5555 | 1,28861E-30 |
| ACE2 | CTD-2626G11.2 | -0,5555 | 1,29503E-30 |
| ACE2 | RP11-162K11.4 | -0,5556 | 1,26303E-30 |
| ACE2 | C2orf88 | -0,5557 | 1,22778E-30 |
| ACE2 | FABP5P3 | -0,5557 | 1,2324E-30 |
| ACE2 | KB-1083B1.1 | -0,5557 | 1,22644E-30 |
| ACE2 | RP11-447M12.3 | -0,5557 | 1,20145E-30 |
| ACE2 | CELF2-AS1 | -0,5557 | 1,20779E-30 |
| ACE2 | CTD-2270P14.5 | -0,5557 | 1,2083E-30 |
| ACE2 | RP11-466A19.7 | -0,5557 | 1,20702E-30 |
| ACE2 | RP11-634B7.5 | -0,5558 | 1,19744E-30 |
| ACE2 | LINC01100 | -0,5558 | 1,17613E-30 |
| ACE2 | ZC2HC1B | -0,5558 | 1,18468E-30 |
| ACE2 | AP4M1 | -0,5558 | 1,17494E-30 |
| ACE2 | CA2 | -0,5559 | 1,14327E-30 |
| ACE2 | IZUMO3 | -0,5559 | 1,15467E-30 |
| ACE2 | HAGLR | -0,556 | 1,12646E-30 |
| ACE2 | PSMF1 | -0,556 | 1,10512E-30 |
| ACE2 | C20orf187 | -0,556 | 1,13147E-30 |

| | | | |
|---|---|---|---|
| ACE2 | AGPAT4 | -0,5561 | 1,07735E-30 |
| ACE2 | RP11-715G15.1 | -0,5562 | 1,03952E-30 |
| ACE2 | ACR | -0,5562 | 1,04091E-30 |
| ACE2 | SPSB4 | -0,5563 | 1,02998E-30 |
| ACE2 | LINC01277 | -0,5563 | 1,02539E-30 |
| ACE2 | RP11-347C18.3 | -0,5563 | 1,0192E-30 |
| ACE2 | CEP295NL | -0,5563 | 1,01989E-30 |
| ACE2 | CACTIN-AS1 | -0,5563 | 1,03233E-30 |
| ACE2 | EPGN | -0,5564 | 1,0031E-30 |
| ACE2 | MRM3 | -0,5564 | 9,80382E-31 |
| ACE2 | AC069363.1 | -0,5564 | 9,87031E-31 |
| ACE2 | RP11-523J2.1 | -0,5565 | 9,77791E-31 |
| ACE2 | MRPL43 | -0,5565 | 9,5277E-31 |
| ACE2 | RP11-114H24.2 | -0,5565 | 9,58007E-31 |
| ACE2 | RP11-390E23.3 | -0,5566 | 9,29619E-31 |
| ACE2 | RP11-763K15.1 | -0,5566 | 9,30588E-31 |
| ACE2 | EFCAB10 | -0,5567 | 9,18628E-31 |
| ACE2 | PURG | -0,5567 | 9,07661E-31 |
| ACE2 | BTG4 | -0,5567 | 9,15487E-31 |
| ACE2 | RP11-125D12.1 | -0,5568 | 8,77639E-31 |
| ACE2 | OPRL1 | -0,5568 | 8,76366E-31 |
| ACE2 | C1orf189 | -0,5569 | 8,59192E-31 |
| ACE2 | ZNF646 | -0,5569 | 8,66277E-31 |
| ACE2 | NTN1 | -0,5569 | 8,59588E-31 |
| ACE2 | CYLC2 | -0,557 | 8,22406E-31 |
| ACE2 | RP11-762H8.3 | -0,557 | 8,23075E-31 |
| ACE2 | AC010731.4 | -0,5571 | 8,13554E-31 |
| ACE2 | SLC17A1 | -0,5571 | 8,173E-31 |
| ACE2 | NT5C3B | -0,5571 | 8,0127E-31 |
| ACE2 | CTD-2623N2.11 | -0,5571 | 8,11311E-31 |
| ACE2 | LINC01192 | -0,5572 | 7,82094E-31 |
| ACE2 | AGPAT2 | -0,5572 | 7,94051E-31 |
| ACE2 | BCL2L14 | -0,5572 | 7,88336E-31 |
| ACE2 | FZD9 | -0,5573 | 7,74014E-31 |
| ACE2 | RP11-147I3.1 | -0,5573 | 7,54961E-31 |
| ACE2 | LINC00354 | -0,5573 | 7,58648E-31 |
| ACE2 | AP001630.5 | -0,5573 | 7,74495E-31 |
| ACE2 | CCDC192 | -0,5574 | 7,41728E-31 |
| ACE2 | PPP1R2P3 | -0,5574 | 7,3303E-31 |
| ACE2 | AC004009.1 | -0,5574 | 7,42067E-31 |
| ACE2 | RAB27B | -0,5574 | 7,49689E-31 |
| ACE2 | LINC01899 | -0,5574 | 7,50655E-31 |
| ACE2 | CTD-2666L21.1 | -0,5574 | 7,43241E-31 |
| ACE2 | RP11-404P21.6 | -0,5575 | 7,19157E-31 |
| ACE2 | RP11-38C18.3 | -0,5576 | 7,06113E-31 |
| ACE2 | LRRC52 | -0,5576 | 6,93219E-31 |
| ACE2 | LPIN1 | -0,5576 | 7,04282E-31 |
| ACE2 | RP11-161D15.1 | -0,5576 | 6,91277E-31 |
| ACE2 | LINC02112 | -0,5576 | 7,06094E-31 |
| ACE2 | RP11-510M2.6 | -0,5576 | 7,08846E-31 |
| ACE2 | LINC01984 | -0,5576 | 6,98746E-31 |
| ACE2 | TTC25 | -0,5576 | 6,92782E-31 |
| ACE2 | SPATA32 | -0,5576 | 6,90658E-31 |
| ACE2 | RP11-927P21.4 | -0,5576 | 7,04886E-31 |

| | | | |
|---|---|---|---|
| ACE2 | MIEF1 | -0,5576 | 6,9503E-31 |
| ACE2 | LINC00854 | -0,5577 | 6,82118E-31 |
| ACE2 | RP11-655H13.2 | -0,5578 | 6,61576E-31 |
| ACE2 | LINC01146 | -0,5578 | 6,6301E-31 |
| ACE2 | LINC01872 | -0,5578 | 6,65896E-31 |
| ACE2 | AC015971.2 | -0,5579 | 6,32062E-31 |
| ACE2 | CCDC74B | -0,5579 | 6,36918E-31 |
| ACE2 | PROM1 | -0,5579 | 6,33073E-31 |
| ACE2 | RP11-87C7.3 | -0,5579 | 6,44095E-31 |
| ACE2 | PITPNA-AS1 | -0,5579 | 6,43263E-31 |
| ACE2 | DCDC2C | -0,558 | 6,23511E-31 |
| ACE2 | AC008697.1 | -0,558 | 6,29434E-31 |
| ACE2 | PLEKHS1 | -0,558 | 6,30153E-31 |
| ACE2 | CTC-774J1.1 | -0,558 | 6,15797E-31 |
| ACE2 | CCDC63 | -0,558 | 6,19049E-31 |
| ACE2 | RNF113B | -0,558 | 6,24969E-31 |
| ACE2 | STPG1 | -0,5581 | 6,00801E-31 |
| ACE2 | RP4-591L5.1 | -0,5581 | 6,09666E-31 |
| ACE2 | NUTM2F | -0,5581 | 6,00478E-31 |
| ACE2 | AKNAD1 | -0,5583 | 5,76062E-31 |
| ACE2 | RP11-96A1.5 | -0,5583 | 5,61989E-31 |
| ACE2 | CTC-329D1.2 | -0,5583 | 5,73789E-31 |
| ACE2 | OR8T1P | -0,5583 | 5,63764E-31 |
| ACE2 | NPIPB10P | -0,5583 | 5,63177E-31 |
| ACE2 | KRT17P3 | -0,5583 | 5,65325E-31 |
| ACE2 | RIMBP3B | -0,5583 | 5,72886E-31 |
| ACE2 | CAV3 | -0,5584 | 5,61348E-31 |
| ACE2 | UBALD1 | -0,5584 | 5,49845E-31 |
| ACE2 | AL022476.2 | -0,5584 | 5,46302E-31 |
| ACE2 | RP1-37J18.1 | -0,5585 | 5,36658E-31 |
| ACE2 | TNNI3K | -0,5585 | 5,3731E-31 |
| ACE2 | TP53I3 | -0,5585 | 5,4014E-31 |
| ACE2 | RP11-561E1.1 | -0,5585 | 5,40477E-31 |
| ACE2 | RP5-1021I20.5 | -0,5585 | 5,33547E-31 |
| ACE2 | SATB2-AS1 | -0,5586 | 5,26282E-31 |
| ACE2 | RP11-407B7.3 | -0,5586 | 5,16573E-31 |
| ACE2 | RP11-580I1.2 | -0,5586 | 5,21314E-31 |
| ACE2 | PLAC8L1 | -0,5587 | 5,13948E-31 |
| ACE2 | MPC1L | -0,5587 | 5,11804E-31 |
| ACE2 | RP11-231E19.1 | -0,5588 | 4,95466E-31 |
| ACE2 | PRSS3 | -0,5588 | 4,97419E-31 |
| ACE2 | LINC00433 | -0,5588 | 4,85163E-31 |
| ACE2 | RP11-1079H9.1 | -0,5588 | 4,90677E-31 |
| ACE2 | FAM83C | -0,5588 | 4,86408E-31 |
| ACE2 | CLCNKB | -0,5589 | 4,80878E-31 |
| ACE2 | MYCNOS | -0,5589 | 4,81959E-31 |
| ACE2 | AC016768.1 | -0,5589 | 4,77626E-31 |
| ACE2 | CAMKMT | -0,5589 | 4,84051E-31 |
| ACE2 | RP11-112J3.16 | -0,5589 | 4,82201E-31 |
| ACE2 | SHANK2-AS1 | -0,5589 | 4,82729E-31 |
| ACE2 | SUMO1P1 | -0,5589 | 4,79416E-31 |
| ACE2 | CT62 | -0,559 | 4,57593E-31 |
| ACE2 | CTA-299D3.8 | -0,559 | 4,70442E-31 |
| ACE2 | RP5-1056L3.1 | -0,5591 | 4,48544E-31 |

| | | | |
|---|---|---|---|
| ACE2 | RP11-280H21.1 | -0,5591 | 4,46314E-31 |
| ACE2 | RP11-424I19.1 | -0,5591 | 4,46443E-31 |
| ACE2 | GIT1 | -0,5591 | 4,55E-31 |
| ACE2 | PTRHD1 | -0,5592 | 4,4273E-31 |
| ACE2 | CASC22 | -0,5592 | 4,43175E-31 |
| ACE2 | RP11-568J23.6 | -0,5592 | 4,4357E-31 |
| ACE2 | CTC-281F24.3 | -0,5592 | 4,3697E-31 |
| ACE2 | CTC-304I17.4 | -0,5592 | 4,32765E-31 |
| ACE2 | CFAP44-AS1 | -0,5593 | 4,19061E-31 |
| ACE2 | ARHGAP26-AS1 | -0,5593 | 4,2953E-31 |
| ACE2 | SHANK2-AS2 | -0,5594 | 4,13629E-31 |
| ACE2 | LINC01431 | -0,5594 | 4,14308E-31 |
| ACE2 | KIF2C | -0,5595 | 3,98185E-31 |
| ACE2 | MAP6 | -0,5595 | 4,0055E-31 |
| ACE2 | TTC39C-AS1 | -0,5595 | 4,04286E-31 |
| ACE2 | PARD6G-AS1 | -0,5595 | 4,03508E-31 |
| ACE2 | RP11-589C21.5 | -0,5596 | 3,87855E-31 |
| ACE2 | RP11-83B20.2 | -0,5596 | 3,90992E-31 |
| ACE2 | PSMD6-AS2 | -0,5597 | 3,77449E-31 |
| ACE2 | CTD-2161F6.1 | -0,5597 | 3,7805E-31 |
| ACE2 | SIL1 | -0,5597 | 3,76666E-31 |
| ACE2 | RP1-166H4.2 | -0,5597 | 3,817E-31 |
| ACE2 | MCHR2-AS1 | -0,5597 | 3,83024E-31 |
| ACE2 | ZCCHC13 | -0,5597 | 3,78725E-31 |
| ACE2 | FAM92A1P2 | -0,5598 | 3,70761E-31 |
| ACE2 | RP11-383C5.3 | -0,5598 | 3,65256E-31 |
| ACE2 | LINC00559 | -0,5598 | 3,635E-31 |
| ACE2 | PSMB11 | -0,5598 | 3,62908E-31 |
| ACE2 | RP5-908M14.5 | -0,5598 | 3,63643E-31 |
| ACE2 | DNAJC27-AS1 | -0,5599 | 3,54175E-31 |
| ACE2 | XXbac-BPG254F23.7 | -0,5599 | 3,61069E-31 |
| ACE2 | RP11-392P7.6 | -0,5599 | 3,55118E-31 |
| ACE2 | WDR37 | -0,56 | 3,4938E-31 |
| ACE2 | DYDC1 | -0,56 | 3,43867E-31 |
| ACE2 | LINC01892 | -0,56 | 3,46051E-31 |
| ACE2 | SPPL2B | -0,56 | 3,4136E-31 |
| ACE2 | NMNAT3 | -0,5601 | 3,37765E-31 |
| ACE2 | RP11-393I2.4 | -0,5601 | 3,38234E-31 |
| ACE2 | FAM220A | -0,5601 | 3,32469E-31 |
| ACE2 | SEPT10P1 | -0,5601 | 3,32926E-31 |
| ACE2 | AP001625.4 | -0,5601 | 3,36976E-31 |
| ACE2 | AC007740.1 | -0,5602 | 3,28307E-31 |
| ACE2 | ANKRD26P1 | -0,5602 | 3,22728E-31 |
| ACE2 | RP11-923I11.4 | -0,5603 | 3,13656E-31 |
| ACE2 | LINC00158 | -0,5603 | 3,18025E-31 |
| ACE2 | TUSC7 | -0,5604 | 3,11084E-31 |
| ACE2 | ANKUB1 | -0,5604 | 3,07931E-31 |
| ACE2 | LINC01082 | -0,5604 | 3,03588E-31 |
| ACE2 | RP11-169N13.4 | -0,5605 | 3,02194E-31 |
| ACE2 | RP11-79C6.3 | -0,5605 | 2,9859E-31 |
| ACE2 | RP11-368M16.3 | -0,5605 | 2,97231E-31 |
| ACE2 | PIK3IP1-AS1 | -0,5605 | 2,94738E-31 |
| ACE2 | RP11-74C13.3 | -0,5606 | 2,90677E-31 |
| ACE2 | IQCF5-AS1 | -0,5607 | 2,85462E-31 |

| | | | |
|---|---|---|---|
| ACE2 | PFDN5 | -0,5607 | 2,8038E-31 |
| ACE2 | UBE2E2-AS1 | -0,5608 | 2,72921E-31 |
| ACE2 | RP11-324I22.2 | -0,5608 | 2,71571E-31 |
| ACE2 | AP006285.6 | -0,5608 | 2,75308E-31 |
| ACE2 | RP11-671P2.1 | -0,5608 | 2,7218E-31 |
| ACE2 | EIF4EBP3 | -0,5609 | 2,64276E-31 |
| ACE2 | RP4-568F9.6 | -0,5609 | 2,65001E-31 |
| ACE2 | TUBB1 | -0,5609 | 2,65223E-31 |
| ACE2 | ALDH7A1P1 | -0,561 | 2,53768E-31 |
| ACE2 | METTL22 | -0,561 | 2,57343E-31 |
| ACE2 | AC010894.3 | -0,5611 | 2,46782E-31 |
| ACE2 | RPS27AP12 | -0,5611 | 2,5169E-31 |
| ACE2 | NELL1 | -0,5611 | 2,51135E-31 |
| ACE2 | RP11-650L12.4 | -0,5611 | 2,50799E-31 |
| ACE2 | PSMC5 | -0,5611 | 2,51215E-31 |
| ACE2 | ZNF487 | -0,5612 | 2,41753E-31 |
| ACE2 | PDYN | -0,5612 | 2,43835E-31 |
| ACE2 | PKIG | -0,5612 | 2,46116E-31 |
| ACE2 | GGA1 | -0,5613 | 2,35556E-31 |
| ACE2 | RP11-573D15.9 | -0,5614 | 2,31988E-31 |
| ACE2 | SORCS3-AS1 | -0,5614 | 2,30023E-31 |
| ACE2 | RP11-749H17.2 | -0,5614 | 2,28123E-31 |
| ACE2 | RP11-10A14.3 | -0,5615 | 2,21791E-31 |
| ACE2 | RP11-219J21.1 | -0,5615 | 2,24579E-31 |
| ACE2 | TMEM262 | -0,5615 | 2,20598E-31 |
| ACE2 | RP11-227H15.5 | -0,5616 | 2,13864E-31 |
| ACE2 | RP5-1027O15.1 | -0,5616 | 2,17357E-31 |
| ACE2 | C12orf60 | -0,5617 | 2,08791E-31 |
| ACE2 | GABRG3-AS1 | -0,5617 | 2,09672E-31 |
| ACE2 | KB-1269D1.8 | -0,5617 | 2,11006E-31 |
| ACE2 | SHOX2 | -0,5618 | 2,01729E-31 |
| ACE2 | HMGB1P20 | -0,5618 | 2,01823E-31 |
| ACE2 | HMOX2 | -0,5619 | 2,00003E-31 |
| ACE2 | PLEKHM1P1 | -0,5619 | 1,94432E-31 |
| ACE2 | PIH1D3 | -0,5619 | 1,96139E-31 |
| ACE2 | TTC7A | -0,562 | 1,91514E-31 |
| ACE2 | FOXP1-IT1 | -0,562 | 1,88711E-31 |
| ACE2 | RP11-456H18.1 | -0,562 | 1,91015E-31 |
| ACE2 | MKNK2P1 | -0,562 | 1,9412E-31 |
| ACE2 | LINC00395 | -0,562 | 1,89976E-31 |
| ACE2 | SRGAP3-AS3 | -0,5621 | 1,83308E-31 |
| ACE2 | RP11-436H11.1 | -0,5621 | 1,83295E-31 |
| ACE2 | LINC00271 | -0,5621 | 1,88283E-31 |
| ACE2 | MROH9 | -0,5622 | 1,81947E-31 |
| ACE2 | ZSCAN2 | -0,5622 | 1,81086E-31 |
| ACE2 | XXbac-B444P24.8 | -0,5622 | 1,8247E-31 |
| ACE2 | RP11-619L12.4 | -0,5623 | 1,74595E-31 |
| ACE2 | RP11-217B7.3 | -0,5623 | 1,76875E-31 |
| ACE2 | RP13-941N14.1 | -0,5623 | 1,73576E-31 |
| ACE2 | RP11-452I5.2 | -0,5623 | 1,7452E-31 |
| ACE2 | OVGP1 | -0,5624 | 1,69156E-31 |
| ACE2 | GSTK1 | -0,5624 | 1,72133E-31 |
| ACE2 | CCDC88B | -0,5624 | 1,68656E-31 |
| ACE2 | TMEM132D | -0,5624 | 1,71667E-31 |

| | | | |
|---|---|---|---|
| ACE2 | AC018693.6 | -0,5625 | 1,64165E-31 |
| ACE2 | UBQLN3 | -0,5625 | 1,65259E-31 |
| ACE2 | RP11-867G23.1 | -0,5625 | 1,65291E-31 |
| ACE2 | RP11-111A22.1 | -0,5625 | 1,67184E-31 |
| ACE2 | KATNB1 | -0,5625 | 1,63942E-31 |
| ACE2 | SYCE1L | -0,5625 | 1,65041E-31 |
| ACE2 | OR2U2P | -0,5626 | 1,62091E-31 |
| ACE2 | S1PR5 | -0,5626 | 1,61571E-31 |
| ACE2 | LINC01725 | -0,5627 | 1,56507E-31 |
| ACE2 | LYG2 | -0,5627 | 1,55654E-31 |
| ACE2 | RASGEF1C | -0,5627 | 1,54355E-31 |
| ACE2 | RP11-368M16.8 | -0,5627 | 1,56392E-31 |
| ACE2 | FBXL16 | -0,5627 | 1,56564E-31 |
| ACE2 | MIEN1 | -0,5627 | 1,57155E-31 |
| ACE2 | CFAP65 | -0,5628 | 1,51114E-31 |
| ACE2 | RP11-389G6.5 | -0,5628 | 1,52413E-31 |
| ACE2 | CTD-2024I7.13 | -0,5628 | 1,49132E-31 |
| ACE2 | RP11-502N13.2 | -0,5628 | 1,51611E-31 |
| ACE2 | NACA2 | -0,5628 | 1,5122E-31 |
| ACE2 | CNTFR-AS1 | -0,5629 | 1,46419E-31 |
| ACE2 | CSNK1G2-AS1 | -0,5629 | 1,48488E-31 |
| ACE2 | RP11-774D14.1 | -0,563 | 1,42553E-31 |
| ACE2 | LINC01442 | -0,563 | 1,42941E-31 |
| ACE2 | WDR93 | -0,563 | 1,42336E-31 |
| ACE2 | RP11-343H19.1 | -0,5631 | 1,36173E-31 |
| ACE2 | AC062032.1 | -0,5632 | 1,34219E-31 |
| ACE2 | HIP1 | -0,5632 | 1,33054E-31 |
| ACE2 | RP4-737E23.7 | -0,5632 | 1,32467E-31 |
| ACE2 | LINC01712 | -0,5633 | 1,2978E-31 |
| ACE2 | WNT7A | -0,5633 | 1,28687E-31 |
| ACE2 | C15orf32 | -0,5633 | 1,31686E-31 |
| ACE2 | RP11-28O3.1 | -0,5633 | 1,29444E-31 |
| ACE2 | CTC-312O10.3 | -0,5633 | 1,29725E-31 |
| ACE2 | PRAMENP | -0,5633 | 1,28349E-31 |
| ACE2 | ZBTB48 | -0,5634 | 1,27657E-31 |
| ACE2 | ADAM6 | -0,5634 | 1,2588E-31 |
| ACE2 | RP11-461O7.1 | -0,5634 | 1,25993E-31 |
| ACE2 | LINC01682 | -0,5635 | 1,23886E-31 |
| ACE2 | DPP10-AS3 | -0,5635 | 1,22393E-31 |
| ACE2 | ORMDL3 | -0,5635 | 1,21757E-31 |
| ACE2 | LINC02072 | -0,5635 | 1,24151E-31 |
| ACE2 | MFAP3L | -0,5636 | 1,17561E-31 |
| ACE2 | WBP1LP2 | -0,5636 | 1,2027E-31 |
| ACE2 | ODF4 | -0,5636 | 1,18316E-31 |
| ACE2 | RP11-68L18.1 | -0,5637 | 1,14572E-31 |
| ACE2 | OR8B3 | -0,5637 | 1,14638E-31 |
| ACE2 | LRRC37A7P | -0,5637 | 1,15527E-31 |
| ACE2 | LINC02142 | -0,5638 | 1,11017E-31 |
| ACE2 | RP11-498B4.5 | -0,5638 | 1,12515E-31 |
| ACE2 | GOLGA8J | -0,5638 | 1,12595E-31 |
| ACE2 | RP11-131L23.1 | -0,5639 | 1,09464E-31 |
| ACE2 | RP11-38C18.2 | -0,5639 | 1,07513E-31 |
| ACE2 | GRID1-AS1 | -0,5639 | 1,07425E-31 |
| ACE2 | RP11-310E22.4 | -0,5639 | 1,07191E-31 |

| | | | |
|---|---|---|---|
| ACE2 | DSG1-AS1 | -0,5639 | 1,08922E-31 |
| ACE2 | RPP38 | -0,5641 | 1,03539E-31 |
| ACE2 | SGTA | -0,5641 | 1,0279E-31 |
| ACE2 | AC016909.1 | -0,5642 | 9,89502E-32 |
| ACE2 | CTB-33O18.3 | -0,5642 | 1,00404E-31 |
| ACE2 | LINC01392 | -0,5642 | 9,82505E-32 |
| ACE2 | NRG3-AS1 | -0,5642 | 9,81158E-32 |
| ACE2 | AP006285.7 | -0,5642 | 9,86887E-32 |
| ACE2 | FZR1 | -0,5642 | 9,97291E-32 |
| ACE2 | GAPDHS | -0,5642 | 1,00395E-31 |
| ACE2 | RP5-991C6.4 | -0,5643 | 9,70368E-32 |
| ACE2 | RP4-566L20.1 | -0,5643 | 9,62854E-32 |
| ACE2 | ATG9A | -0,5644 | 9,4529E-32 |
| ACE2 | RP11-716O23.1 | -0,5644 | 9,49976E-32 |
| ACE2 | POM121L12 | -0,5644 | 9,29265E-32 |
| ACE2 | MESTIT1 | -0,5644 | 9,40496E-32 |
| ACE2 | GOT1L1 | -0,5644 | 9,46739E-32 |
| ACE2 | RP11-434I12.4 | -0,5644 | 9,23973E-32 |
| ACE2 | C10orf53 | -0,5645 | 9,1361E-32 |
| ACE2 | FLJ40194 | -0,5645 | 9,15403E-32 |
| ACE2 | RP11-95O2.5 | -0,5645 | 8,99389E-32 |
| ACE2 | RP11-575L7.2 | -0,5646 | 8,75308E-32 |
| ACE2 | GLT1D1 | -0,5646 | 8,87709E-32 |
| ACE2 | CTD-2553L13.10 | -0,5646 | 8,79911E-32 |
| ACE2 | CTB-161M19.4 | -0,5647 | 8,60995E-32 |
| ACE2 | RP11-305A4.3 | -0,5647 | 8,48436E-32 |
| ACE2 | RP5-1002M8.4 | -0,5647 | 8,55283E-32 |
| ACE2 | RP11-265D19.6 | -0,5647 | 8,68108E-32 |
| ACE2 | LINC02190 | -0,5648 | 8,25692E-32 |
| ACE2 | RP11-347D21.1 | -0,5648 | 8,21073E-32 |
| ACE2 | LZTS1-AS1 | -0,5649 | 8,11795E-32 |
| ACE2 | SAA4 | -0,5649 | 7,97281E-32 |
| ACE2 | C17orf112 | -0,5649 | 8,00379E-32 |
| ACE2 | ADIPOQ-AS1 | -0,5651 | 7,60586E-32 |
| ACE2 | AF038458.5 | -0,5651 | 7,64958E-32 |
| ACE2 | LAMC2 | -0,5652 | 7,42951E-32 |
| ACE2 | RP11-80B9.4 | -0,5652 | 7,4797E-32 |
| ACE2 | AC073254.1 | -0,5652 | 7,30743E-32 |
| ACE2 | OR2H2 | -0,5652 | 7,37835E-32 |
| ACE2 | CYP2C8 | -0,5652 | 7,30751E-32 |
| ACE2 | ALK | -0,5653 | 7,10928E-32 |
| ACE2 | LINC01374 | -0,5653 | 7,05089E-32 |
| ACE2 | LINC01890 | -0,5654 | 6,92784E-32 |
| ACE2 | RP11-272J7.4 | -0,5654 | 6,96544E-32 |
| ACE2 | RP11-348J12.2 | -0,5654 | 6,8863E-32 |
| ACE2 | PLK1 | -0,5654 | 6,86417E-32 |
| ACE2 | ODF3B | -0,5654 | 6,98149E-32 |
| ACE2 | GSTO2 | -0,5655 | 6,639E-32 |
| ACE2 | CSF2RB | -0,5655 | 6,82511E-32 |
| ACE2 | ACE3P | -0,5656 | 6,49102E-32 |
| ACE2 | SPRR2C | -0,5657 | 6,42599E-32 |
| ACE2 | APOA5 | -0,5657 | 6,274E-32 |
| ACE2 | PLEKHD1 | -0,5657 | 6,37597E-32 |
| ACE2 | CTC-304I17.6 | -0,5657 | 6,38392E-32 |

| | | | |
|---|---|---|---|
| ACE2 | FAM228A | -0,5658 | 6,09253E-32 |
| ACE2 | FHL5 | -0,5658 | 6,19156E-32 |
| ACE2 | TEKT1 | -0,5658 | 6,08586E-32 |
| ACE2 | CEP41 | -0,5659 | 6,04182E-32 |
| ACE2 | NDUFA13 | -0,5659 | 5,96953E-32 |
| ACE2 | RP11-550H2.2 | -0,566 | 5,86601E-32 |
| ACE2 | C6orf229 | -0,566 | 5,87191E-32 |
| ACE2 | CCDC75P1 | -0,5661 | 5,5699E-32 |
| ACE2 | LINC01553 | -0,5661 | 5,69834E-32 |
| ACE2 | UBXN1 | -0,5661 | 5,60106E-32 |
| ACE2 | DIAPH3-AS1 | -0,5661 | 5,64211E-32 |
| ACE2 | GPAT3 | -0,5662 | 5,46515E-32 |
| ACE2 | WI2-1959D15.3 | -0,5662 | 5,39138E-32 |
| ACE2 | CTD-3076O17.2 | -0,5662 | 5,50872E-32 |
| ACE2 | RP4-726F1.2 | -0,5663 | 5,22961E-32 |
| ACE2 | RP11-10A14.7 | -0,5663 | 5,26358E-32 |
| ACE2 | AC011513.4 | -0,5663 | 5,25905E-32 |
| ACE2 | TDRKH-AS1 | -0,5664 | 5,11365E-32 |
| ACE2 | MAT1A | -0,5664 | 5,09147E-32 |
| ACE2 | EVI5L | -0,5664 | 5,20998E-32 |
| ACE2 | LIPE-AS1 | -0,5664 | 5,14221E-32 |
| ACE2 | GGTLC1 | -0,5664 | 5,08459E-32 |
| ACE2 | C1orf185 | -0,5665 | 4,96648E-32 |
| ACE2 | LINC01677 | -0,5665 | 4,9487E-32 |
| ACE2 | RP11-318I4.1 | -0,5665 | 5,02669E-32 |
| ACE2 | RP3-470L22.1 | -0,5665 | 4,93785E-32 |
| ACE2 | CSNK2B | -0,5665 | 5,05538E-32 |
| ACE2 | CTD-2568P8.1 | -0,5665 | 4,96215E-32 |
| ACE2 | SLAMF9 | -0,5666 | 4,8372E-32 |
| ACE2 | RP11-347H15.5 | -0,5666 | 4,8538E-32 |
| ACE2 | RP11-262A12.1 | -0,5666 | 4,91056E-32 |
| ACE2 | NRIR | -0,5667 | 4,75677E-32 |
| ACE2 | CTB-16J4.1 | -0,5667 | 4,75781E-32 |
| ACE2 | CYHR1 | -0,5667 | 4,72537E-32 |
| ACE2 | PDF | -0,5667 | 4,66034E-32 |
| ACE2 | AURKC | -0,5667 | 4,72629E-32 |
| ACE2 | RP11-167N24.3 | -0,5668 | 4,50484E-32 |
| ACE2 | RP11-725G5.2 | -0,5668 | 4,62442E-32 |
| ACE2 | LINC01227 | -0,5668 | 4,5434E-32 |
| ACE2 | SH3BGR | -0,5668 | 4,55813E-32 |
| ACE2 | RP11-416L21.2 | -0,5669 | 4,36768E-32 |
| ACE2 | RP11-610J23.1 | -0,5669 | 4,44708E-32 |
| ACE2 | RP11-495L19.1 | -0,5669 | 4,48352E-32 |
| ACE2 | DHRS4L1 | -0,5669 | 4,39723E-32 |
| ACE2 | REC8 | -0,5669 | 4,45274E-32 |
| ACE2 | CTD-3220F14.1 | -0,5669 | 4,37312E-32 |
| ACE2 | RP4-715N11.2 | -0,5669 | 4,42969E-32 |
| ACE2 | PEX5L-AS2 | -0,567 | 4,30371E-32 |
| ACE2 | GALNTL5 | -0,567 | 4,31708E-32 |
| ACE2 | WDR38 | -0,567 | 4,25994E-32 |
| ACE2 | ADIRF-AS1 | -0,567 | 4,27177E-32 |
| ACE2 | CATSPER1 | -0,567 | 4,32471E-32 |
| ACE2 | AC000036.4 | -0,567 | 4,31822E-32 |
| ACE2 | DAB1-AS1 | -0,5671 | 4,19691E-32 |

| | | | |
|---|---|---|---|
| ACE2 | R3HDM2P2 | -0,5671 | 4,22E-32 |
| ACE2 | RP3-333B15.4 | -0,5671 | 4,12671E-32 |
| ACE2 | LEXM | -0,5672 | 3,99197E-32 |
| ACE2 | OR6F1 | -0,5672 | 4,01638E-32 |
| ACE2 | RP11-203B7.1 | -0,5672 | 4,07005E-32 |
| ACE2 | RP11-58O3.2 | -0,5672 | 4,05719E-32 |
| ACE2 | LINC01517 | -0,5672 | 4,00554E-32 |
| ACE2 | CTB-174D11.3 | -0,5673 | 3,94838E-32 |
| ACE2 | NRG1-IT1 | -0,5673 | 3,9247E-32 |
| ACE2 | CTB-186H2.3 | -0,5673 | 3,89478E-32 |
| ACE2 | AC011747.3 | -0,5674 | 3,75816E-32 |
| ACE2 | TTLL3 | -0,5674 | 3,79137E-32 |
| ACE2 | EEF1GP3 | -0,5674 | 3,84196E-32 |
| ACE2 | CTC-471F3.5 | -0,5674 | 3,78024E-32 |
| ACE2 | LINC02064 | -0,5675 | 3,65836E-32 |
| ACE2 | RP4-591N18.2 | -0,5675 | 3,63966E-32 |
| ACE2 | ERI3-IT1 | -0,5676 | 3,55337E-32 |
| ACE2 | RP11-2O17.2 | -0,5676 | 3,59298E-32 |
| ACE2 | NEFM | -0,5676 | 3,6034E-32 |
| ACE2 | RP11-552I14.1 | -0,5676 | 3,60093E-32 |
| ACE2 | RP3-496C20.1 | -0,5676 | 3,55128E-32 |
| ACE2 | BAG1 | -0,5677 | 3,45117E-32 |
| ACE2 | RP11-75N6.3 | -0,5677 | 3,5043E-32 |
| ACE2 | HSF1 | -0,5678 | 3,383E-32 |
| ACE2 | RP11-1078H9.6 | -0,5678 | 3,34728E-32 |
| ACE2 | RP11-439E19.9 | -0,5679 | 3,23687E-32 |
| ACE2 | TAS2R1 | -0,5679 | 3,28928E-32 |
| ACE2 | CRYBA2 | -0,568 | 3,20373E-32 |
| ACE2 | NAV2-AS5 | -0,568 | 3,12794E-32 |
| ACE2 | OTOP2 | -0,568 | 3,21709E-32 |
| ACE2 | RP11-574M7.1 | -0,5681 | 3,10567E-32 |
| ACE2 | LINC00238 | -0,5681 | 3,11593E-32 |
| ACE2 | RP11-244B22.11 | -0,5681 | 3,09194E-32 |
| ACE2 | USH1G | -0,5681 | 3,08575E-32 |
| ACE2 | HIPK4 | -0,5681 | 3,06696E-32 |
| ACE2 | RP11-547D24.3 | -0,5682 | 2,98708E-32 |
| ACE2 | RP5-968P14.2 | -0,5682 | 2,96918E-32 |
| ACE2 | AC004869.2 | -0,5682 | 3,01928E-32 |
| ACE2 | PRSS55 | -0,5682 | 3,00447E-32 |
| ACE2 | LINC02186 | -0,5682 | 2,99872E-32 |
| ACE2 | EFCAB3 | -0,5682 | 2,98549E-32 |
| ACE2 | JSRP1 | -0,5682 | 2,98227E-32 |
| ACE2 | HPCA | -0,5683 | 2,87721E-32 |
| ACE2 | S100A7A | -0,5683 | 2,87584E-32 |
| ACE2 | C1orf105 | -0,5683 | 2,92502E-32 |
| ACE2 | RP11-343P9.1 | -0,5683 | 2,91769E-32 |
| ACE2 | SEPT7P9 | -0,5685 | 2,71273E-32 |
| ACE2 | ROPN1B | -0,5686 | 2,60756E-32 |
| ACE2 | CTC-459M5.2 | -0,5686 | 2,6149E-32 |
| ACE2 | CLYBL-AS1 | -0,5686 | 2,66475E-32 |
| ACE2 | LINC01745 | -0,5688 | 2,50616E-32 |
| ACE2 | CTC-461F20.1 | -0,5688 | 2,5036E-32 |
| ACE2 | NRIP3 | -0,5688 | 2,51714E-32 |
| ACE2 | CTB-174D11.1 | -0,5689 | 2,41748E-32 |

| | | | |
|---|---|---|---|
| ACE2 | CAPSL | -0,569 | 2,31175E-32 |
| ACE2 | PKD2L1 | -0,569 | 2,3594E-32 |
| ACE2 | RP11-7M8.2 | -0,569 | 2,31898E-32 |
| ACE2 | RP11-10L12.4 | -0,5691 | 2,23805E-32 |
| ACE2 | FAM222A-AS1 | -0,5691 | 2,29748E-32 |
| ACE2 | HCAR1 | -0,5691 | 2,29589E-32 |
| ACE2 | SIVA1 | -0,5691 | 2,27472E-32 |
| ACE2 | HEATR9 | -0,5691 | 2,27445E-32 |
| ACE2 | CEP164 | -0,5692 | 2,1911E-32 |
| ACE2 | RP11-383C6.2 | -0,5693 | 2,1267E-32 |
| ACE2 | TMEM74B | -0,5693 | 2,15329E-32 |
| ACE2 | LINC01096 | -0,5694 | 2,05583E-32 |
| ACE2 | CTD-2506P8.6 | -0,5694 | 2,06256E-32 |
| ACE2 | OR7E19P | -0,5694 | 2,06501E-32 |
| ACE2 | H1FX-AS1 | -0,5696 | 1,93692E-32 |
| ACE2 | RP4-718D20.3 | -0,5696 | 1,92523E-32 |
| ACE2 | LINC01931 | -0,5697 | 1,88143E-32 |
| ACE2 | SPINK2 | -0,5697 | 1,868E-32 |
| ACE2 | KCNMA1-AS1 | -0,5699 | 1,79396E-32 |
| ACE2 | ARMS2 | -0,5699 | 1,80442E-32 |
| ACE2 | CFAP46 | -0,5699 | 1,77757E-32 |
| ACE2 | CDHR2 | -0,57 | 1,70744E-32 |
| ACE2 | CEP162 | -0,57 | 1,72605E-32 |
| ACE2 | C10orf120 | -0,57 | 1,74458E-32 |
| ACE2 | RP11-1028N23.4 | -0,5701 | 1,65837E-32 |
| ACE2 | LINC01755 | -0,5702 | 1,62414E-32 |
| ACE2 | KCP | -0,5702 | 1,63533E-32 |
| ACE2 | INSL6 | -0,5703 | 1,58049E-32 |
| ACE2 | KIRREL3-AS2 | -0,5703 | 1,59246E-32 |
| ACE2 | RP11-121M22.1 | -0,5704 | 1,51823E-32 |
| ACE2 | KRT73-AS1 | -0,5704 | 1,54775E-32 |
| ACE2 | CTA-243E7.1 | -0,5704 | 1,51802E-32 |
| ACE2 | EVX2 | -0,5705 | 1,4958E-32 |
| ACE2 | LINC02087 | -0,5705 | 1,48364E-32 |
| ACE2 | RP11-499E14.1 | -0,5706 | 1,44277E-32 |
| ACE2 | RP11-755O11.2 | -0,5706 | 1,42589E-32 |
| ACE2 | FRAT2 | -0,5707 | 1,40171E-32 |
| ACE2 | RP11-783K16.14 | -0,5707 | 1,37586E-32 |
| ACE2 | RP4-601P9.2 | -0,5707 | 1,37671E-32 |
| ACE2 | RP11-163M18.1 | -0,5707 | 1,41224E-32 |
| ACE2 | RP11-168L7.3 | -0,5707 | 1,40278E-32 |
| ACE2 | VWA3B | -0,5708 | 1,36551E-32 |
| ACE2 | RP11-308K2.1 | -0,5708 | 1,35973E-32 |
| ACE2 | PDE1C | -0,5708 | 1,3576E-32 |
| ACE2 | C9orf153 | -0,5708 | 1,33692E-32 |
| ACE2 | RP11-781A6.1 | -0,5708 | 1,36326E-32 |
| ACE2 | IGLL1 | -0,5708 | 1,34982E-32 |
| ACE2 | RP11-147C23.1 | -0,5709 | 1,32334E-32 |
| ACE2 | WDR53 | -0,5709 | 1,30805E-32 |
| ACE2 | RP11-642D21.1 | -0,5709 | 1,31627E-32 |
| ACE2 | ARHGAP22 | -0,5709 | 1,32785E-32 |
| ACE2 | NODAL | -0,5709 | 1,31823E-32 |
| ACE2 | TAC3 | -0,5709 | 1,32446E-32 |
| ACE2 | HMGA2 | -0,571 | 1,29058E-32 |

| | | | |
|---|---|---|---|
| ACE2 | GRK4 | -0,5711 | 1,24398E-32 |
| ACE2 | CEACAM19 | -0,5711 | 1,2491E-32 |
| ACE2 | LINC00880 | -0,5712 | 1,1796E-32 |
| ACE2 | THEGL | -0,5712 | 1,18993E-32 |
| ACE2 | TTLL9 | -0,5712 | 1,20945E-32 |
| ACE2 | RP4-753D4.2 | -0,5713 | 1,15578E-32 |
| ACE2 | ELAVL4 | -0,5714 | 1,13641E-32 |
| ACE2 | ACSBG2 | -0,5715 | 1,08595E-32 |
| ACE2 | RP11-353N4.5 | -0,5716 | 1,05339E-32 |
| ACE2 | LINC02029 | -0,5716 | 1,04946E-32 |
| ACE2 | TOPORS-AS1 | -0,5716 | 1,0688E-32 |
| ACE2 | CES1P2 | -0,5716 | 1,07321E-32 |
| ACE2 | CTD-2538C1.3 | -0,5716 | 1,05815E-32 |
| ACE2 | RP11-654C20.1 | -0,5717 | 1,01855E-32 |
| ACE2 | DNAAF1 | -0,5717 | 1,02289E-32 |
| ACE2 | LOC401913 | -0,5717 | 1,02731E-32 |
| ACE2 | COX7B2 | -0,5718 | 9,89664E-33 |
| ACE2 | TRIML1 | -0,5718 | 1,00084E-32 |
| ACE2 | RP11-981G7.3 | -0,5718 | 9,89186E-33 |
| ACE2 | RP11-547C13.1 | -0,5718 | 1,00492E-32 |
| ACE2 | MIR4300HG | -0,5718 | 9,98343E-33 |
| ACE2 | EIF5A2 | -0,5719 | 9,5609E-33 |
| ACE2 | RP11-421L21.3 | -0,572 | 9,23059E-33 |
| ACE2 | AC010729.2 | -0,572 | 9,2896E-33 |
| ACE2 | ACTN3 | -0,572 | 9,43322E-33 |
| ACE2 | SCGB1B2P | -0,572 | 9,49158E-33 |
| ACE2 | SYT16 | -0,5721 | 9,07656E-33 |
| ACE2 | RP11-1379J22.7 | -0,5722 | 8,71141E-33 |
| ACE2 | RP11-646J21.6 | -0,5722 | 8,72229E-33 |
| ACE2 | AKAP1 | -0,5722 | 8,72288E-33 |
| ACE2 | SLC44A5 | -0,5723 | 8,45776E-33 |
| ACE2 | RP11-296E7.1 | -0,5723 | 8,46677E-33 |
| ACE2 | RP11-321G12.1 | -0,5723 | 8,42145E-33 |
| ACE2 | RP11-709D24.6 | -0,5723 | 8,45414E-33 |
| ACE2 | RP11-701P16.2 | -0,5724 | 8,36888E-33 |
| ACE2 | AC002465.2 | -0,5724 | 8,25372E-33 |
| ACE2 | LINC01782 | -0,5724 | 8,39011E-33 |
| ACE2 | PIGQ | -0,5725 | 8,1256E-33 |
| ACE2 | CA10 | -0,5725 | 8,09149E-33 |
| ACE2 | FITM1 | -0,5726 | 7,84413E-33 |
| ACE2 | RP11-247C2.2 | -0,5726 | 7,82835E-33 |
| ACE2 | CTD-2525I3.6 | -0,5726 | 7,77375E-33 |
| ACE2 | GS1-279B7.2 | -0,5727 | 7,56188E-33 |
| ACE2 | C5orf49 | -0,5727 | 7,44315E-33 |
| ACE2 | CTA-292E10.6 | -0,5727 | 7,60741E-33 |
| ACE2 | SERBP1P3 | -0,5728 | 7,30719E-33 |
| ACE2 | AC004692.4 | -0,5728 | 7,21907E-33 |
| ACE2 | LINC00840 | -0,5728 | 7,24818E-33 |
| ACE2 | MAATS1 | -0,5729 | 7,01239E-33 |
| ACE2 | CAHM | -0,5729 | 7,14992E-33 |
| ACE2 | RP11-626I20.3 | -0,5729 | 7,09939E-33 |
| ACE2 | RP11-303E16.3 | -0,573 | 6,79593E-33 |
| ACE2 | RP11-566H8.2 | -0,5731 | 6,7359E-33 |
| ACE2 | RP11-165F24.5 | -0,5731 | 6,58503E-33 |

| | | | |
|---|---|---|---|
| ACE2 | FOXN4 | -0,5731 | 6,69288E-33 |
| ACE2 | LINC00299 | -0,5732 | 6,39324E-33 |
| ACE2 | REC114 | -0,5732 | 6,3789E-33 |
| ACE2 | HIRIP3 | -0,5732 | 6,39148E-33 |
| ACE2 | LINC00112 | -0,5732 | 6,40649E-33 |
| ACE2 | RP11-467L13.7 | -0,5733 | 6,35364E-33 |
| ACE2 | RP11-335K5.2 | -0,5733 | 6,25478E-33 |
| ACE2 | LINC01681 | -0,5734 | 6,18306E-33 |
| ACE2 | RP11-521I2.3 | -0,5734 | 6,04369E-33 |
| ACE2 | RP11-711K1.7 | -0,5734 | 6,14055E-33 |
| ACE2 | FGF14-AS1 | -0,5734 | 6,0008E-33 |
| ACE2 | RP11-496H1.2 | -0,5735 | 5,962E-33 |
| ACE2 | ICA1 | -0,5735 | 5,92743E-33 |
| ACE2 | AC005077.14 | -0,5735 | 5,89255E-33 |
| ACE2 | CTC-559E9.6 | -0,5735 | 5,91006E-33 |
| ACE2 | AC079807.2 | -0,5736 | 5,71529E-33 |
| ACE2 | HECW1-IT1 | -0,5736 | 5,686E-33 |
| ACE2 | PRR29-AS1 | -0,5736 | 5,75328E-33 |
| ACE2 | LINC01905 | -0,5736 | 5,63837E-33 |
| ACE2 | RP11-112L6.2 | -0,5736 | 5,66127E-33 |
| ACE2 | LINC01676 | -0,5737 | 5,48667E-33 |
| ACE2 | SCML4 | -0,5737 | 5,48975E-33 |
| ACE2 | SPACA7 | -0,5737 | 5,60969E-33 |
| ACE2 | PRKCZ | -0,5738 | 5,36509E-33 |
| ACE2 | NOS1AP | -0,5738 | 5,4057E-33 |
| ACE2 | PPP1R7 | -0,5738 | 5,45449E-33 |
| ACE2 | CRISP2 | -0,5738 | 5,46041E-33 |
| ACE2 | MICALCL | -0,5738 | 5,35841E-33 |
| ACE2 | RP11-64B16.5 | -0,5738 | 5,43873E-33 |
| ACE2 | AC013472.3 | -0,5739 | 5,20265E-33 |
| ACE2 | RP11-338L18.1 | -0,5739 | 5,24467E-33 |
| ACE2 | RP11-661G16.2 | -0,5739 | 5,18907E-33 |
| ACE2 | USP50 | -0,5739 | 5,2561E-33 |
| ACE2 | AC009299.2 | -0,574 | 4,98683E-33 |
| ACE2 | RPL36P20 | -0,574 | 5,10896E-33 |
| ACE2 | RP11-758H9.2 | -0,574 | 4,99741E-33 |
| ACE2 | TTC22 | -0,5741 | 4,97137E-33 |
| ACE2 | DGAT1 | -0,5741 | 4,88126E-33 |
| ACE2 | RP11-514F8.2 | -0,5741 | 4,89866E-33 |
| ACE2 | RIN1 | -0,5741 | 4,90728E-33 |
| ACE2 | RP11-271K11.1 | -0,5741 | 4,9116E-33 |
| ACE2 | CTD-2561B21.5 | -0,5741 | 4,84479E-33 |
| ACE2 | MEGF11 | -0,5742 | 4,74859E-33 |
| ACE2 | SLC22A31 | -0,5742 | 4,71276E-33 |
| ACE2 | VRK3 | -0,5742 | 4,78372E-33 |
| ACE2 | MPC2 | -0,5744 | 4,41526E-33 |
| ACE2 | RP11-393B14.2 | -0,5744 | 4,43707E-33 |
| ACE2 | GAS2L1P2 | -0,5744 | 4,42706E-33 |
| ACE2 | CFAP77 | -0,5744 | 4,49144E-33 |
| ACE2 | GTSF1 | -0,5744 | 4,42743E-33 |
| ACE2 | SLPI | -0,5744 | 4,42058E-33 |
| ACE2 | RP11-667F14.1 | -0,5745 | 4,39312E-33 |
| ACE2 | FAM186B | -0,5745 | 4,34863E-33 |
| ACE2 | RP11-252M21.6 | -0,5745 | 4,27945E-33 |

| | | | |
|---|---|---|---|
| ACE2 | PMFBP1 | -0,5745 | 4,36566E-33 |
| ACE2 | RP1-28H20.3 | -0,5745 | 4,32625E-33 |
| ACE2 | RP11-61G19.2 | -0,5746 | 4,24557E-33 |
| ACE2 | FLJ12825 | -0,5746 | 4,17187E-33 |
| ACE2 | LINC01134 | -0,5747 | 4,11373E-33 |
| ACE2 | RP11-709A23.2 | -0,5747 | 4,02799E-33 |
| ACE2 | CTC-441N14.4 | -0,5748 | 3,94473E-33 |
| ACE2 | C10orf105 | -0,5749 | 3,88736E-33 |
| ACE2 | MRPS15 | -0,575 | 3,73328E-33 |
| ACE2 | LINC01445 | -0,575 | 3,71333E-33 |
| ACE2 | AC002467.7 | -0,575 | 3,69484E-33 |
| ACE2 | NRM | -0,5751 | 3,61411E-33 |
| ACE2 | ITPKA | -0,5751 | 3,60762E-33 |
| ACE2 | RSPH6A | -0,5751 | 3,619E-33 |
| ACE2 | LINC01509 | -0,5752 | 3,51441E-33 |
| ACE2 | APOOP5 | -0,5752 | 3,44846E-33 |
| ACE2 | ROPN1 | -0,5753 | 3,39097E-33 |
| ACE2 | SERGEF | -0,5753 | 3,42614E-33 |
| ACE2 | RP11-637A17.2 | -0,5753 | 3,34434E-33 |
| ACE2 | MTPAP | -0,5754 | 3,30969E-33 |
| ACE2 | RP11-559M23.1 | -0,5755 | 3,181E-33 |
| ACE2 | CXXC5 | -0,5755 | 3,15828E-33 |
| ACE2 | CDKN2B-AS1 | -0,5755 | 3,15091E-33 |
| ACE2 | FAM227A | -0,5755 | 3,13075E-33 |
| ACE2 | AF015720.3 | -0,5756 | 3,10887E-33 |
| ACE2 | TNFAIP6 | -0,5757 | 2,94338E-33 |
| ACE2 | RP11-483A20.3 | -0,5757 | 2,96373E-33 |
| ACE2 | AC005487.2 | -0,5757 | 3,00566E-33 |
| ACE2 | LINC00364 | -0,5757 | 3,00724E-33 |
| ACE2 | CENPW | -0,5758 | 2,9347E-33 |
| ACE2 | RP11-423C15.3 | -0,5758 | 2,86953E-33 |
| ACE2 | CCT8L2 | -0,5758 | 2,90406E-33 |
| ACE2 | EPHA6 | -0,5759 | 2,77374E-33 |
| ACE2 | ACTRT3 | -0,5759 | 2,76708E-33 |
| ACE2 | TCP11 | -0,5759 | 2,78004E-33 |
| ACE2 | AC007557.2 | -0,576 | 2,75039E-33 |
| ACE2 | RP11-361M10.3 | -0,576 | 2,68811E-33 |
| ACE2 | CD2 | -0,5762 | 2,55156E-33 |
| ACE2 | RP11-480I12.10 | -0,5762 | 2,58868E-33 |
| ACE2 | RP11-445O3.2 | -0,5762 | 2,56468E-33 |
| ACE2 | HECW1 | -0,5762 | 2,52501E-33 |
| ACE2 | CIRBP-AS1 | -0,5762 | 2,57203E-33 |
| ACE2 | RP11-417F21.1 | -0,5763 | 2,46562E-33 |
| ACE2 | FER1L6-AS1 | -0,5763 | 2,47743E-33 |
| ACE2 | RP11-29B9.2 | -0,5763 | 2,47979E-33 |
| ACE2 | NEURL1-AS1 | -0,5763 | 2,49605E-33 |
| ACE2 | RP11-99A1.2 | -0,5763 | 2,51637E-33 |
| ACE2 | LINC01125 | -0,5764 | 2,39884E-33 |
| ACE2 | SLC10A6 | -0,5764 | 2,43599E-33 |
| ACE2 | RP11-369E15.4 | -0,5764 | 2,41538E-33 |
| ACE2 | RP11-236J17.7 | -0,5764 | 2,42524E-33 |
| ACE2 | RP11-983C2.3 | -0,5764 | 2,4148E-33 |
| ACE2 | RP11-1109M24.5 | -0,5764 | 2,36575E-33 |
| ACE2 | RP11-6N17.4 | -0,5764 | 2,38305E-33 |

| | | | |
|---|---|---|---|
| ACE2 | LACE1 | -0,5765 | 2,3561E-33 |
| ACE2 | PLA2G12B | -0,5766 | 2,25526E-33 |
| ACE2 | COX8C | -0,5766 | 2,22569E-33 |
| ACE2 | CYB5R2 | -0,5767 | 2,17341E-33 |
| ACE2 | CTD-2545H1.1 | -0,5767 | 2,21331E-33 |
| ACE2 | EQTN | -0,5768 | 2,13658E-33 |
| ACE2 | RP4-706G24.1 | -0,5769 | 2,04745E-33 |
| ACE2 | RP11-298H24.1 | -0,5769 | 2,05514E-33 |
| ACE2 | LINC01917 | -0,5769 | 2,03888E-33 |
| ACE2 | IPO13 | -0,577 | 1,98325E-33 |
| ACE2 | AGBL3 | -0,577 | 1,98221E-33 |
| ACE2 | C11orf97 | -0,577 | 2,00487E-33 |
| ACE2 | LINC01486 | -0,577 | 1,9662E-33 |
| ACE2 | RP3-446N13.1 | -0,577 | 1,99668E-33 |
| ACE2 | LRRC72 | -0,5771 | 1,92942E-33 |
| ACE2 | B4GALT1-AS1 | -0,5771 | 1,95903E-33 |
| ACE2 | RP11-65L19.4 | -0,5771 | 1,91408E-33 |
| ACE2 | LINC01791 | -0,5771 | 1,92584E-33 |
| ACE2 | RP11-371I20.2 | -0,5772 | 1,88293E-33 |
| ACE2 | HN1 | -0,5772 | 1,8563E-33 |
| ACE2 | RP11-584P21.2 | -0,5773 | 1,83764E-33 |
| ACE2 | CLPB | -0,5774 | 1,75162E-33 |
| ACE2 | TSGA13 | -0,5775 | 1,6943E-33 |
| ACE2 | RP11-686F15.2 | -0,5775 | 1,71752E-33 |
| ACE2 | CTB-50L17.10 | -0,5775 | 1,70018E-33 |
| ACE2 | WFDC5 | -0,5775 | 1,68774E-33 |
| ACE2 | CTD-2228K2.2 | -0,5776 | 1,66573E-33 |
| ACE2 | RP5-942I16.1 | -0,5776 | 1,64272E-33 |
| ACE2 | LINC02165 | -0,5776 | 1,66091E-33 |
| ACE2 | CLEC17A | -0,5776 | 1,67435E-33 |
| ACE2 | LINC00971 | -0,5777 | 1,57715E-33 |
| ACE2 | SPATA8 | -0,5777 | 1,57671E-33 |
| ACE2 | RP11-347C12.3 | -0,5777 | 1,58607E-33 |
| ACE2 | RP11-310I24.1 | -0,5778 | 1,55598E-33 |
| ACE2 | RP11-227G15.12 | -0,5778 | 1,54572E-33 |
| ACE2 | RP11-343H5.6 | -0,5779 | 1,5136E-33 |
| ACE2 | MAD2L1BP | -0,5779 | 1,49252E-33 |
| ACE2 | RP11-203J24.8 | -0,5779 | 1,48214E-33 |
| ACE2 | PFN3 | -0,578 | 1,43958E-33 |
| ACE2 | MEIG1 | -0,578 | 1,47965E-33 |
| ACE2 | CNPY3 | -0,5781 | 1,43331E-33 |
| ACE2 | GJA10 | -0,5781 | 1,40993E-33 |
| ACE2 | PRKACG | -0,5782 | 1,34984E-33 |
| ACE2 | MRPL41 | -0,5782 | 1,37E-33 |
| ACE2 | RP11-69E11.8 | -0,5783 | 1,34328E-33 |
| ACE2 | RP11-359I18.5 | -0,5783 | 1,30665E-33 |
| ACE2 | SLIT1-AS1 | -0,5783 | 1,33365E-33 |
| ACE2 | MAP1S | -0,5783 | 1,324E-33 |
| ACE2 | CTD-2525I3.2 | -0,5783 | 1,3284E-33 |
| ACE2 | RP11-12M5.1 | -0,5784 | 1,27257E-33 |
| ACE2 | TEX29 | -0,5784 | 1,28227E-33 |
| ACE2 | RP11-846C15.2 | -0,5784 | 1,29394E-33 |
| ACE2 | CDC34 | -0,5784 | 1,29117E-33 |
| ACE2 | AGBL1 | -0,5785 | 1,23173E-33 |

| | | | |
|---|---|---|---|
| ACE2 | ZNF829 | -0,5785 | 1,24414E-33 |
| ACE2 | CLCN1 | -0,5786 | 1,19199E-33 |
| ACE2 | LINC00592 | -0,5786 | 1,21425E-33 |
| ACE2 | GABARAP | -0,5786 | 1,20089E-33 |
| ACE2 | RP11-448P19.1 | -0,5787 | 1,15595E-33 |
| ACE2 | EFCAB11 | -0,5788 | 1,13157E-33 |
| ACE2 | RP11-417O18.2 | -0,5789 | 1,09536E-33 |
| ACE2 | LINC01919 | -0,5789 | 1,09531E-33 |
| ACE2 | IGSF11-AS1 | -0,579 | 1,07074E-33 |
| ACE2 | RP11-79M19.2 | -0,579 | 1,06967E-33 |
| ACE2 | RP11-109E10.1 | -0,579 | 1,07931E-33 |
| ACE2 | LINC01989 | -0,579 | 1,06302E-33 |
| ACE2 | LINC01349 | -0,5791 | 1,03757E-33 |
| ACE2 | OSGIN1 | -0,5792 | 9,99403E-34 |
| ACE2 | C2orf66 | -0,5793 | 9,55759E-34 |
| ACE2 | TEX36-AS1 | -0,5793 | 9,60939E-34 |
| ACE2 | RP4-650F12.2 | -0,5794 | 9,43265E-34 |
| ACE2 | LINC01796 | -0,5794 | 9,25683E-34 |
| ACE2 | RP11-752L20.5 | -0,5794 | 9,44715E-34 |
| ACE2 | RP11-297A16.4 | -0,5794 | 9,38518E-34 |
| ACE2 | RP11-687M24.8 | -0,5795 | 9,16491E-34 |
| ACE2 | PRH1 | -0,5796 | 8,70505E-34 |
| ACE2 | RP11-268P4.4 | -0,5796 | 8,81658E-34 |
| ACE2 | RP11-218M11.3 | -0,5797 | 8,49115E-34 |
| ACE2 | ADAD2 | -0,5798 | 8,23065E-34 |
| ACE2 | MAFG-AS1 | -0,5798 | 8,19382E-34 |
| ACE2 | SBK2 | -0,5798 | 8,27967E-34 |
| ACE2 | TEX13A | -0,5798 | 8,22747E-34 |
| ACE2 | BRINP2 | -0,5799 | 8,05264E-34 |
| ACE2 | SPATA18 | -0,5799 | 8,00244E-34 |
| ACE2 | MS4A6E | -0,5799 | 8,04481E-34 |
| ACE2 | RP1-261D10.2 | -0,5799 | 8,00261E-34 |
| ACE2 | CTC-273B12.5 | -0,5799 | 7,93846E-34 |
| ACE2 | NAALADL2-AS3 | -0,58 | 7,69804E-34 |
| ACE2 | ADAMTS7P3 | -0,58 | 7,7302E-34 |
| ACE2 | SPPL2C | -0,58 | 7,82508E-34 |
| ACE2 | RP4-738P15.1 | -0,58 | 7,77767E-34 |
| ACE2 | CTC-441N14.2 | -0,5801 | 7,59461E-34 |
| ACE2 | RP11-30L15.6 | -0,5801 | 7,43714E-34 |
| ACE2 | MRPL28 | -0,5801 | 7,46149E-34 |
| ACE2 | C17orf105 | -0,5801 | 7,57203E-34 |
| ACE2 | CCDC87 | -0,5802 | 7,29476E-34 |
| ACE2 | C22orf31 | -0,5802 | 7,21066E-34 |
| ACE2 | MORN1 | -0,5803 | 6,97633E-34 |
| ACE2 | RP1-155D22.2 | -0,5803 | 6,96918E-34 |
| ACE2 | RP11-493E12.1 | -0,5804 | 6,85511E-34 |
| ACE2 | AC004862.6 | -0,5804 | 6,79627E-34 |
| ACE2 | TTLL6 | -0,5804 | 6,75636E-34 |
| ACE2 | KIAA2012 | -0,5805 | 6,65153E-34 |
| ACE2 | RP11-556O15.1 | -0,5805 | 6,7122E-34 |
| ACE2 | AC099552.2 | -0,5805 | 6,61337E-34 |
| ACE2 | LINC02166 | -0,5805 | 6,69732E-34 |
| ACE2 | RNF114 | -0,5805 | 6,58213E-34 |
| ACE2 | RP11-804N13.1 | -0,5806 | 6,41975E-34 |

| | | | |
|---|---|---|---|
| ACE2 | RP11-669N7.3 | -0,5806 | 6,35958E-34 |
| ACE2 | ODF3L1 | -0,5806 | 6,48018E-34 |
| ACE2 | RP11-464F9.20 | -0,5807 | 6,1503E-34 |
| ACE2 | HBZ | -0,5807 | 6,21134E-34 |
| ACE2 | CTD-2050B12.1 | -0,5807 | 6,23217E-34 |
| ACE2 | GPR135 | -0,5808 | 6,08674E-34 |
| ACE2 | RP11-848P1.5 | -0,5808 | 6,08503E-34 |
| ACE2 | CCDC151 | -0,5808 | 6,08479E-34 |
| ACE2 | RP11-415D17.3 | -0,581 | 5,64003E-34 |
| ACE2 | ZSCAN1 | -0,581 | 5,70545E-34 |
| ACE2 | LINC00616 | -0,5811 | 5,44477E-34 |
| ACE2 | C11orf88 | -0,5811 | 5,40386E-34 |
| ACE2 | RP11-557N21.1 | -0,5812 | 5,29648E-34 |
| ACE2 | AC007563.1 | -0,5812 | 5,38116E-34 |
| ACE2 | TGM7 | -0,5812 | 5,24008E-34 |
| ACE2 | C5orf46 | -0,5813 | 5,1853E-34 |
| ACE2 | MAMDC2-AS1 | -0,5813 | 5,15816E-34 |
| ACE2 | LINC00942 | -0,5813 | 5,14808E-34 |
| ACE2 | GFER | -0,5814 | 5,04452E-34 |
| ACE2 | YJEFN3 | -0,5815 | 4,81686E-34 |
| ACE2 | SLC4A10 | -0,5816 | 4,73099E-34 |
| ACE2 | RP11-734K21.3 | -0,5817 | 4,49665E-34 |
| ACE2 | RP5-1022J11.2 | -0,5817 | 4,50842E-34 |
| ACE2 | BAP1 | -0,5818 | 4,4043E-34 |
| ACE2 | RP11-25O10.2 | -0,5819 | 4,31299E-34 |
| ACE2 | MBD3L1 | -0,5819 | 4,23732E-34 |
| ACE2 | CTD-2623N2.5 | -0,5819 | 4,25271E-34 |
| ACE2 | PPP1R14B | -0,582 | 4,08072E-34 |
| ACE2 | PRMT8 | -0,582 | 4,07438E-34 |
| ACE2 | RSPO4 | -0,582 | 4,10211E-34 |
| ACE2 | ARMC12 | -0,5821 | 4,03523E-34 |
| ACE2 | RPS26P8 | -0,5821 | 4,04172E-34 |
| ACE2 | ZDHHC3 | -0,5822 | 3,91552E-34 |
| ACE2 | HABP4 | -0,5822 | 3,84855E-34 |
| ACE2 | BPIFB1 | -0,5822 | 3,83739E-34 |
| ACE2 | ZMIZ2 | -0,5823 | 3,70773E-34 |
| ACE2 | ARL4A | -0,5824 | 3,61943E-34 |
| ACE2 | RP11-235C23.5 | -0,5824 | 3,63704E-34 |
| ACE2 | LINC01886 | -0,5825 | 3,49102E-34 |
| ACE2 | KIRREL3-AS3 | -0,5825 | 3,47245E-34 |
| ACE2 | NPM2 | -0,5826 | 3,40249E-34 |
| ACE2 | CREM | -0,5826 | 3,38248E-34 |
| ACE2 | FBXO3-AS1 | -0,5826 | 3,38779E-34 |
| ACE2 | CCL18 | -0,5826 | 3,40215E-34 |
| ACE2 | RP11-32D16.1 | -0,5827 | 3,28716E-34 |
| ACE2 | RP11-613H2.1 | -0,5827 | 3,30043E-34 |
| ACE2 | RP11-1129I3.1 | -0,5827 | 3,32E-34 |
| ACE2 | RP11-379J5.5 | -0,5827 | 3,30674E-34 |
| ACE2 | CST13P | -0,5827 | 3,33861E-34 |
| ACE2 | ADRA2A | -0,5828 | 3,19716E-34 |
| ACE2 | CLIC1P1 | -0,5828 | 3,1797E-34 |
| ACE2 | PBXIP1 | -0,5829 | 3,13431E-34 |
| ACE2 | HMGB3P22 | -0,5829 | 3,08573E-34 |
| ACE2 | SPATS1 | -0,583 | 2,98682E-34 |

| | | | |
|---|---|---|---|
| ACE2 | MMEL1 | -0,5831 | 2,93967E-34 |
| ACE2 | RP11-304L19.4 | -0,5831 | 2,90918E-34 |
| ACE2 | XXbac-BPG308J9.3 | -0,5832 | 2,79146E-34 |
| ACE2 | RP1-69D17.4 | -0,5833 | 2,72359E-34 |
| ACE2 | AC007391.2 | -0,5834 | 2,64011E-34 |
| ACE2 | EXOC3-AS1 | -0,5834 | 2,62187E-34 |
| ACE2 | ANO2 | -0,5834 | 2,64468E-34 |
| ACE2 | RP11-607P23.1 | -0,5834 | 2,65816E-34 |
| ACE2 | LINC02152 | -0,5834 | 2,61234E-34 |
| ACE2 | LINC01718 | -0,5834 | 2,61456E-34 |
| ACE2 | RP11-474B12.1 | -0,5835 | 2,55805E-34 |
| ACE2 | RP11-73G16.3 | -0,5836 | 2,45337E-34 |
| ACE2 | GPS2 | -0,5836 | 2,4528E-34 |
| ACE2 | HDAC11-AS1 | -0,5837 | 2,42963E-34 |
| ACE2 | DTHD1 | -0,5837 | 2,39967E-34 |
| ACE2 | RP11-806K15.1 | -0,5837 | 2,41299E-34 |
| ACE2 | EGFLAM-AS2 | -0,5837 | 2,40105E-34 |
| ACE2 | PRSS37 | -0,5837 | 2,36715E-34 |
| ACE2 | OTOG | -0,5837 | 2,37073E-34 |
| ACE2 | NDUFS8 | -0,5837 | 2,41319E-34 |
| ACE2 | RP11-73M7.6 | -0,5838 | 2,32624E-34 |
| ACE2 | ZNF32-AS2 | -0,5838 | 2,3203E-34 |
| ACE2 | RP11-177H2.2 | -0,5839 | 2,23371E-34 |
| ACE2 | RP11-364C11.2 | -0,5839 | 2,28332E-34 |
| ACE2 | RP11-245D16.4 | -0,584 | 2,15269E-34 |
| ACE2 | LINC01620 | -0,584 | 2,2023E-34 |
| ACE2 | ZNRF3-AS1 | -0,584 | 2,17419E-34 |
| ACE2 | CCR6 | -0,5841 | 2,12793E-34 |
| ACE2 | TMEM81 | -0,5842 | 2,06862E-34 |
| ACE2 | RUNX2 | -0,5842 | 2,04324E-34 |
| ACE2 | LYZL6 | -0,5842 | 2,04851E-34 |
| ACE2 | AKNA | -0,5843 | 1,99312E-34 |
| ACE2 | RP11-274B21.3 | -0,5844 | 1,93329E-34 |
| ACE2 | LINC00705 | -0,5844 | 1,9488E-34 |
| ACE2 | IGHGP | -0,5844 | 1,93764E-34 |
| ACE2 | RP4-781K5.4 | -0,5846 | 1,81099E-34 |
| ACE2 | RP11-522M21.3 | -0,5847 | 1,7211E-34 |
| ACE2 | CTD-2201I18.1 | -0,5847 | 1,76101E-34 |
| ACE2 | CCDC89 | -0,5847 | 1,73394E-34 |
| ACE2 | NKX6-1 | -0,5848 | 1,70222E-34 |
| ACE2 | BX255923.3 | -0,5848 | 1,70794E-34 |
| ACE2 | PRR29 | -0,5848 | 1,71272E-34 |
| ACE2 | TRABD2A | -0,5849 | 1,65484E-34 |
| ACE2 | LKAAEAR1 | -0,5849 | 1,65729E-34 |
| ACE2 | C1orf167 | -0,585 | 1,56246E-34 |
| ACE2 | LINC00690 | -0,585 | 1,5585E-34 |
| ACE2 | RP11-262I2.2 | -0,585 | 1,58675E-34 |
| ACE2 | AC010974.3 | -0,5851 | 1,51564E-34 |
| ACE2 | EIF2B4 | -0,5852 | 1,47016E-34 |
| ACE2 | LINC02034 | -0,5852 | 1,48762E-34 |
| ACE2 | FUT10 | -0,5854 | 1,39901E-34 |
| ACE2 | MCM8-AS1 | -0,5854 | 1,41343E-34 |
| ACE2 | TP53TG5 | -0,5854 | 1,37077E-34 |
| ACE2 | TPT1-AS1 | -0,5855 | 1,33351E-34 |

| | | | |
|---|---|---|---|
| ACE2 | RP11-542P2.2 | -0,5856 | 1,31799E-34 |
| ACE2 | PPP1R35 | -0,5856 | 1,2983E-34 |
| ACE2 | RP11-688I9.4 | -0,5856 | 1,32161E-34 |
| ACE2 | SIDT1 | -0,5857 | 1,27178E-34 |
| ACE2 | MLIP-AS1 | -0,5857 | 1,2662E-34 |
| ACE2 | RP11-87C12.6 | -0,5857 | 1,27575E-34 |
| ACE2 | RP11-215E13.2 | -0,5857 | 1,25237E-34 |
| ACE2 | RP5-930J4.2 | -0,5858 | 1,23583E-34 |
| ACE2 | RP11-320M2.1 | -0,5858 | 1,20972E-34 |
| ACE2 | AC005077.5 | -0,5858 | 1,22384E-34 |
| ACE2 | RP11-533E19.2 | -0,5859 | 1,18007E-34 |
| ACE2 | RP11-260E18.1 | -0,5859 | 1,17373E-34 |
| ACE2 | GRM3 | -0,5859 | 1,17743E-34 |
| ACE2 | RPL12P33 | -0,5859 | 1,18722E-34 |
| ACE2 | RGS6 | -0,5859 | 1,17659E-34 |
| ACE2 | PCDHB7 | -0,586 | 1,15687E-34 |
| ACE2 | NLRP1 | -0,586 | 1,1538E-34 |
| ACE2 | FAM187B | -0,586 | 1,15439E-34 |
| ACE2 | MAST2 | -0,5861 | 1,11876E-34 |
| ACE2 | TSPAN18 | -0,5861 | 1,09399E-34 |
| ACE2 | DNAI2 | -0,5863 | 1,04104E-34 |
| ACE2 | PACRG-AS2 | -0,5864 | 1,00617E-34 |
| ACE2 | RP11-961A15.3 | -0,5864 | 1,00414E-34 |
| ACE2 | RP11-146I2.1 | -0,5865 | 9,8362E-35 |
| ACE2 | CTC-232P5.3 | -0,5865 | 9,64795E-35 |
| ACE2 | PLEKHA6 | -0,5866 | 9,57135E-35 |
| ACE2 | APOBEC2 | -0,5866 | 9,38681E-35 |
| ACE2 | RPL17P43 | -0,5866 | 9,47336E-35 |
| ACE2 | GP9 | -0,5867 | 9,25826E-35 |
| ACE2 | RP11-489P6.1 | -0,5867 | 9,01686E-35 |
| ACE2 | NRL | -0,5868 | 8,8524E-35 |
| ACE2 | LINC01808 | -0,5869 | 8,6744E-35 |
| ACE2 | CCDC37-AS1 | -0,5869 | 8,60885E-35 |
| ACE2 | KLK5 | -0,5869 | 8,46751E-35 |
| ACE2 | FUNDC2P2 | -0,587 | 8,2895E-35 |
| ACE2 | ABRA | -0,587 | 8,21879E-35 |
| ACE2 | LINC01868 | -0,5871 | 7,95439E-35 |
| ACE2 | RP11-393J16.4 | -0,5871 | 7,97149E-35 |
| ACE2 | AL450226.2 | -0,5871 | 7,92609E-35 |
| ACE2 | LINC01141 | -0,5873 | 7,45479E-35 |
| ACE2 | RPSAP9 | -0,5873 | 7,61966E-35 |
| ACE2 | TMEM219 | -0,5873 | 7,62397E-35 |
| ACE2 | RP11-543N12.1 | -0,5873 | 7,46972E-35 |
| ACE2 | LINC00608 | -0,5874 | 7,31979E-35 |
| ACE2 | RP11-374P20.4 | -0,5874 | 7,3078E-35 |
| ACE2 | ARIH2OS | -0,5875 | 6,96023E-35 |
| ACE2 | RP11-45L9.1 | -0,5875 | 7,06087E-35 |
| ACE2 | FAM57A | -0,5875 | 6,98977E-35 |
| ACE2 | ANKFN1 | -0,5875 | 7,05013E-35 |
| ACE2 | LINC01010 | -0,5876 | 6,88481E-35 |
| ACE2 | TXNDC8 | -0,5876 | 6,935E-35 |
| ACE2 | ZNF555 | -0,5876 | 6,88631E-35 |
| ACE2 | BPIFC | -0,5876 | 6,91182E-35 |
| ACE2 | CAMK4 | -0,5877 | 6,68186E-35 |

| | | | |
|---|---|---|---|
| ACE2 | RP1-292B18.4 | -0,5877 | 6,58106E-35 |
| ACE2 | RP3-441A12.1 | -0,5879 | 6,27656E-35 |
| ACE2 | RP11-165M1.3 | -0,5879 | 6,17096E-35 |
| ACE2 | KCNH6 | -0,5879 | 6,19152E-35 |
| ACE2 | TMEM53 | -0,588 | 6,04273E-35 |
| ACE2 | PRR27 | -0,5881 | 5,87055E-35 |
| ACE2 | C11orf80 | -0,5881 | 5,72916E-35 |
| ACE2 | FBXL14 | -0,5881 | 5,74688E-35 |
| ACE2 | PRC1-AS1 | -0,5881 | 5,83977E-35 |
| ACE2 | GPAT2P2 | -0,5882 | 5,72693E-35 |
| ACE2 | LINC01500 | -0,5882 | 5,68345E-35 |
| ACE2 | GRM7-AS3 | -0,5883 | 5,45915E-35 |
| ACE2 | RP11-544A12.4 | -0,5883 | 5,51745E-35 |
| ACE2 | RP11-810P12.5 | -0,5883 | 5,42219E-35 |
| ACE2 | SLC38A7 | -0,5883 | 5,42667E-35 |
| ACE2 | ARHGAP40 | -0,5883 | 5,49184E-35 |
| ACE2 | CTD-2560E9.5 | -0,5884 | 5,25245E-35 |
| ACE2 | RP11-560G2.1 | -0,5884 | 5,27534E-35 |
| ACE2 | RP11-321C24.1 | -0,5884 | 5,23804E-35 |
| ACE2 | RP11-96D1.7 | -0,5884 | 5,33327E-35 |
| ACE2 | ITGA2B | -0,5884 | 5,28881E-35 |
| ACE2 | RP11-567M16.3 | -0,5884 | 5,3237E-35 |
| ACE2 | RP11-214D15.2 | -0,5885 | 5,18212E-35 |
| ACE2 | RP11-397C18.2 | -0,5885 | 5,16781E-35 |
| ACE2 | PRSS30P | -0,5885 | 5,035E-35 |
| ACE2 | LINC01255 | -0,5885 | 5,03181E-35 |
| ACE2 | NT5DC4 | -0,5886 | 4,98542E-35 |
| ACE2 | RP11-435M3.2 | -0,5886 | 4,87291E-35 |
| ACE2 | SAMD4A | -0,5886 | 4,89333E-35 |
| ACE2 | C20orf144 | -0,5886 | 5,01886E-35 |
| ACE2 | LINC01072 | -0,5887 | 4,72957E-35 |
| ACE2 | RPL39L | -0,5888 | 4,68012E-35 |
| ACE2 | CTD-2296D1.3 | -0,5888 | 4,66272E-35 |
| ACE2 | C11orf91 | -0,5888 | 4,58006E-35 |
| ACE2 | IFT57 | -0,5889 | 4,42277E-35 |
| ACE2 | RSU1P2 | -0,5889 | 4,54747E-35 |
| ACE2 | LINC01351 | -0,589 | 4,32358E-35 |
| ACE2 | TSSK1B | -0,589 | 4,34047E-35 |
| ACE2 | TRBV7-1 | -0,589 | 4,34212E-35 |
| ACE2 | FAIM2 | -0,589 | 4,27831E-35 |
| ACE2 | RP11-669B18.1 | -0,589 | 4,41462E-35 |
| ACE2 | RP11-388E23.2 | -0,589 | 4,2922E-35 |
| ACE2 | AC005703.3 | -0,589 | 4,30125E-35 |
| ACE2 | CASC17 | -0,589 | 4,32023E-35 |
| ACE2 | ARRDC5 | -0,5891 | 4,17156E-35 |
| ACE2 | LINC00687 | -0,5891 | 4,1888E-35 |
| ACE2 | DCST1 | -0,5893 | 3,9701E-35 |
| ACE2 | PRSS56 | -0,5893 | 3,93377E-35 |
| ACE2 | ADTRP | -0,5893 | 3,90578E-35 |
| ACE2 | RP3-325F22.3 | -0,5893 | 3,90401E-35 |
| ACE2 | AC002383.2 | -0,5893 | 3,99387E-35 |
| ACE2 | RP11-744I24.3 | -0,5893 | 3,94459E-35 |
| ACE2 | IQCA1L | -0,5893 | 3,95852E-35 |
| ACE2 | RP11-540O11.1 | -0,5893 | 3,89171E-35 |

| | | | |
|---|---|---|---|
| ACE2 | UBL7 | -0,5893 | 3,88773E-35 |
| ACE2 | RP11-165M1.2 | -0,5893 | 3,93539E-35 |
| ACE2 | LINC00868 | -0,5893 | 3,89711E-35 |
| ACE2 | LINC00558 | -0,5894 | 3,80029E-35 |
| ACE2 | LINC01664 | -0,5894 | 3,8095E-35 |
| ACE2 | RP11-508N22.9 | -0,5895 | 3,67712E-35 |
| ACE2 | RP11-388C12.1 | -0,5895 | 3,68165E-35 |
| ACE2 | UBE2M | -0,5895 | 3,69614E-35 |
| ACE2 | LINC00659 | -0,5895 | 3,68156E-35 |
| ACE2 | NEXN-AS1 | -0,5896 | 3,54309E-35 |
| ACE2 | MORN2 | -0,5896 | 3,58758E-35 |
| ACE2 | SMG9 | -0,5896 | 3,59048E-35 |
| ACE2 | RP11-517B11.4 | -0,5898 | 3,32185E-35 |
| ACE2 | RP11-528I4.2 | -0,5898 | 3,30479E-35 |
| ACE2 | MRPS24 | -0,5898 | 3,3791E-35 |
| ACE2 | NTMT1 | -0,5898 | 3,29863E-35 |
| ACE2 | LINC02156 | -0,5898 | 3,33885E-35 |
| ACE2 | RP11-526P6.1 | -0,5898 | 3,39174E-35 |
| ACE2 | RP11-168O16.1 | -0,5899 | 3,21191E-35 |
| ACE2 | CYP51A1-AS1 | -0,5899 | 3,20395E-35 |
| ACE2 | RP11-94B19.3 | -0,59 | 3,14964E-35 |
| ACE2 | CDNF | -0,5901 | 2,99257E-35 |
| ACE2 | RP11-620J15.1 | -0,5901 | 3,03099E-35 |
| ACE2 | ZFHX2 | -0,5901 | 3,05871E-35 |
| ACE2 | RP11-893F2.14 | -0,5901 | 3,06291E-35 |
| ACE2 | ZNF330 | -0,5902 | 2,98335E-35 |
| ACE2 | ADAMTS13 | -0,5902 | 2,93912E-35 |
| ACE2 | USP2 | -0,5903 | 2,81185E-35 |
| ACE2 | LNX1-AS1 | -0,5904 | 2,78868E-35 |
| ACE2 | ALKBH3-AS1 | -0,5904 | 2,70828E-35 |
| ACE2 | LINC01490 | -0,5904 | 2,76375E-35 |
| ACE2 | UBA52 | -0,5904 | 2,77014E-35 |
| ACE2 | RP5-1017F8.2 | -0,5904 | 2,76436E-35 |
| ACE2 | LINC00943 | -0,5905 | 2,69648E-35 |
| ACE2 | OTUB2 | -0,5905 | 2,65833E-35 |
| ACE2 | LA16c-306E5.1 | -0,5905 | 2,64812E-35 |
| ACE2 | RP11-749H17.1 | -0,5905 | 2,68026E-35 |
| ACE2 | MOCS3 | -0,5905 | 2,66529E-35 |
| ACE2 | SPTLC1P4 | -0,5906 | 2,58166E-35 |
| ACE2 | RP11-102C16.3 | -0,5906 | 2,60518E-35 |
| ACE2 | ACRBP | -0,5906 | 2,61414E-35 |
| ACE2 | H3F3C | -0,5906 | 2,53656E-35 |
| ACE2 | RP4-753D10.5 | -0,5906 | 2,58726E-35 |
| ACE2 | FAM71F2 | -0,5907 | 2,52144E-35 |
| ACE2 | AANAT | -0,5907 | 2,5349E-35 |
| ACE2 | RP11-67L3.2 | -0,5908 | 2,38137E-35 |
| ACE2 | RP11-675F6.4 | -0,5908 | 2,391E-35 |
| ACE2 | KIFC3 | -0,5908 | 2,41284E-35 |
| ACE2 | LINC00467 | -0,5909 | 2,33378E-35 |
| ACE2 | ATP2A1-AS1 | -0,5909 | 2,35091E-35 |
| ACE2 | DPP7 | -0,591 | 2,28801E-35 |
| ACE2 | LINC00928 | -0,591 | 2,25714E-35 |
| ACE2 | ESRP2 | -0,591 | 2,27557E-35 |
| ACE2 | SGO1-AS1 | -0,5911 | 2,18683E-35 |

| | | | |
|---|---|---|---|
| ACE2 | CDH18 | -0,5911 | 2,15491E-35 |
| ACE2 | MNS1 | -0,5911 | 2,19047E-35 |
| ACE2 | RP11-347D21.2 | -0,5911 | 2,17508E-35 |
| ACE2 | FAM24A | -0,5912 | 2,09886E-35 |
| ACE2 | C8G | -0,5913 | 2,06955E-35 |
| ACE2 | ELOBP2 | -0,5913 | 2,01695E-35 |
| ACE2 | RP11-143E21.6 | -0,5913 | 2,06726E-35 |
| ACE2 | MACROD2-AS1 | -0,5913 | 2,03762E-35 |
| ACE2 | RP11-359G22.4 | -0,5913 | 2,05681E-35 |
| ACE2 | AC012506.3 | -0,5914 | 1,98025E-35 |
| ACE2 | NUTM2G | -0,5914 | 2,0111E-35 |
| ACE2 | WDR54 | -0,5915 | 1,94404E-35 |
| ACE2 | NPC1L1 | -0,5915 | 1,9438E-35 |
| ACE2 | ADRA1A | -0,5915 | 1,8893E-35 |
| ACE2 | SPAG8 | -0,5915 | 1,93385E-35 |
| ACE2 | CCNY | -0,5915 | 1,93596E-35 |
| ACE2 | RP11-557J10.5 | -0,5916 | 1,88226E-35 |
| ACE2 | FAM186A | -0,5916 | 1,85365E-35 |
| ACE2 | RGL4 | -0,5916 | 1,87214E-35 |
| ACE2 | ANTXRL | -0,5917 | 1,77457E-35 |
| ACE2 | RP11-474L23.3 | -0,5917 | 1,81146E-35 |
| ACE2 | SH3GL3 | -0,5917 | 1,8155E-35 |
| ACE2 | C19orf47 | -0,5917 | 1,82103E-35 |
| ACE2 | LINC02054 | -0,5918 | 1,74274E-35 |
| ACE2 | TBX18-AS1 | -0,5918 | 1,71811E-35 |
| ACE2 | EFNB3 | -0,5918 | 1,71704E-35 |
| ACE2 | LINC00658 | -0,5918 | 1,76209E-35 |
| ACE2 | RP11-273B19.2 | -0,5919 | 1,70155E-35 |
| ACE2 | RP11-565A3.2 | -0,5919 | 1,70304E-35 |
| ACE2 | RP11-463J10.4 | -0,5919 | 1,65663E-35 |
| ACE2 | TEKT5 | -0,5919 | 1,71036E-35 |
| ACE2 | C17orf100 | -0,5919 | 1,66436E-35 |
| ACE2 | RP11-1112J20.2 | -0,592 | 1,65209E-35 |
| ACE2 | RP11-101E7.2 | -0,592 | 1,62884E-35 |
| ACE2 | RP1-228P16.4 | -0,5921 | 1,56366E-35 |
| ACE2 | RNASE11 | -0,5921 | 1,57965E-35 |
| ACE2 | SLC22A14 | -0,5922 | 1,54646E-35 |
| ACE2 | RP11-634H22.1 | -0,5922 | 1,54997E-35 |
| ACE2 | CCDC183 | -0,5922 | 1,53709E-35 |
| ACE2 | FAM83C-AS1 | -0,5922 | 1,50358E-35 |
| ACE2 | RIBC2 | -0,5922 | 1,53024E-35 |
| ACE2 | AC092687.5 | -0,5923 | 1,46127E-35 |
| ACE2 | RP11-457M11.7 | -0,5923 | 1,45915E-35 |
| ACE2 | RP11-95M15.1 | -0,5923 | 1,45735E-35 |
| ACE2 | VN1R32P | -0,5924 | 1,40622E-35 |
| ACE2 | TMEM139 | -0,5924 | 1,43776E-35 |
| ACE2 | C9orf131 | -0,5924 | 1,42383E-35 |
| ACE2 | CCDC168 | -0,5925 | 1,36044E-35 |
| ACE2 | RP11-387H17.4 | -0,5926 | 1,35439E-35 |
| ACE2 | LINC01783 | -0,5927 | 1,30338E-35 |
| ACE2 | AK8 | -0,5927 | 1,27763E-35 |
| ACE2 | C9orf163 | -0,5927 | 1,27968E-35 |
| ACE2 | CTD-3105H18.4 | -0,5927 | 1,30491E-35 |
| ACE2 | G6PC2 | -0,5928 | 1,26396E-35 |

| | | | |
|---|---|---|---|
| ACE2 | ST7-AS2 | -0,5928 | 1,25858E-35 |
| ACE2 | RP11-539L10.2 | -0,5929 | 1,1986E-35 |
| ACE2 | VASH2 | -0,593 | 1,16351E-35 |
| ACE2 | RNF181 | -0,593 | 1,1557E-35 |
| ACE2 | LINC02024 | -0,593 | 1,15588E-35 |
| ACE2 | RP11-597D13.7 | -0,593 | 1,18466E-35 |
| ACE2 | RP11-632K5.3 | -0,593 | 1,16954E-35 |
| ACE2 | NSMCE1 | -0,593 | 1,15688E-35 |
| ACE2 | PPP1R12C | -0,5931 | 1,12151E-35 |
| ACE2 | RP11-554D14.1 | -0,5932 | 1,09975E-35 |
| ACE2 | RP11-297L17.2 | -0,5932 | 1,1108E-35 |
| ACE2 | RP11-181B18.1 | -0,5933 | 1,06657E-35 |
| ACE2 | AC012506.4 | -0,5933 | 1,07025E-35 |
| ACE2 | TOM1 | -0,5933 | 1,07697E-35 |
| ACE2 | RP11-111A21.1 | -0,5934 | 1,0142E-35 |
| ACE2 | ZNF513 | -0,5935 | 1,00072E-35 |
| ACE2 | TESC | -0,5935 | 9,95182E-36 |
| ACE2 | ADAM20P1 | -0,5935 | 1,005E-35 |
| ACE2 | RP11-349E4.1 | -0,5936 | 9,49747E-36 |
| ACE2 | AGBL1-AS1 | -0,5936 | 9,53903E-36 |
| ACE2 | GEMIN6 | -0,5937 | 9,22556E-36 |
| ACE2 | RP11-486O13.5 | -0,5937 | 9,22081E-36 |
| ACE2 | RP11-1007O24.3 | -0,5937 | 9,22542E-36 |
| ACE2 | WSCD1 | -0,5937 | 9,31661E-36 |
| ACE2 | PLCD4 | -0,5938 | 8,93077E-36 |
| ACE2 | CRIP3 | -0,5938 | 8,97215E-36 |
| ACE2 | AC004471.10 | -0,5938 | 8,91357E-36 |
| ACE2 | RP11-550H2.1 | -0,5939 | 8,66628E-36 |
| ACE2 | RP3-523C21.3 | -0,5939 | 8,83582E-36 |
| ACE2 | MROH7 | -0,5941 | 8,03155E-36 |
| ACE2 | SERPINB6 | -0,5941 | 8,08884E-36 |
| ACE2 | MIR193BHG | -0,5941 | 8,15582E-36 |
| ACE2 | ZMYND15 | -0,5941 | 8,27556E-36 |
| ACE2 | IBA57 | -0,5942 | 7,8873E-36 |
| ACE2 | RP11-539G18.3 | -0,5942 | 7,90279E-36 |
| ACE2 | RP11-227D13.5 | -0,5942 | 7,84807E-36 |
| ACE2 | ATP1A4 | -0,5943 | 7,54163E-36 |
| ACE2 | RP11-688I9.2 | -0,5943 | 7,74224E-36 |
| ACE2 | RP11-25I15.3 | -0,5943 | 7,73448E-36 |
| ACE2 | TIMM50 | -0,5943 | 7,71189E-36 |
| ACE2 | FAM19A3 | -0,5946 | 6,91974E-36 |
| ACE2 | RP11-536G4.2 | -0,5946 | 6,91437E-36 |
| ACE2 | CTD-2184C24.2 | -0,5946 | 6,87287E-36 |
| ACE2 | TEX21P | -0,5946 | 6,92991E-36 |
| ACE2 | ARID3A | -0,5947 | 6,72241E-36 |
| ACE2 | NF2 | -0,5947 | 6,73273E-36 |
| ACE2 | RIBC1 | -0,5947 | 6,76016E-36 |
| ACE2 | CLCNKA | -0,5948 | 6,42328E-36 |
| ACE2 | RP11-67C2.2 | -0,5948 | 6,36221E-36 |
| ACE2 | SIRT2 | -0,5948 | 6,36973E-36 |
| ACE2 | EP300-AS1 | -0,5948 | 6,48895E-36 |
| ACE2 | LINC01798 | -0,5949 | 6,15539E-36 |
| ACE2 | RP11-725M22.1 | -0,5949 | 6,15477E-36 |
| ACE2 | RP4-584D14.6 | -0,5949 | 6,2153E-36 |

| | | | |
|---|---|---|---|
| ACE2 | TMPO-AS1 | -0,5949 | 6,27028E-36 |
| ACE2 | RP11-103J8.1 | -0,5949 | 6,19046E-36 |
| ACE2 | RP3-333B15.5 | -0,5949 | 6,21754E-36 |
| ACE2 | CTD-2330K9.2 | -0,595 | 6,06005E-36 |
| ACE2 | LINC00494 | -0,595 | 6,04735E-36 |
| ACE2 | RP11-259K15.2 | -0,5951 | 5,82886E-36 |
| ACE2 | LSMEM2 | -0,5952 | 5,61711E-36 |
| ACE2 | LINC01040 | -0,5952 | 5,62145E-36 |
| ACE2 | TSPAN1 | -0,5953 | 5,53926E-36 |
| ACE2 | ATP6V1E2 | -0,5953 | 5,4927E-36 |
| ACE2 | RP11-486A14.1 | -0,5953 | 5,46472E-36 |
| ACE2 | RP11-522N14.2 | -0,5953 | 5,4291E-36 |
| ACE2 | KLHDC4 | -0,5953 | 5,51737E-36 |
| ACE2 | RP11-316O14.1 | -0,5954 | 5,216E-36 |
| ACE2 | PRR7-AS1 | -0,5954 | 5,28829E-36 |
| ACE2 | SH2D5 | -0,5957 | 4,75707E-36 |
| ACE2 | STX18-AS1 | -0,5957 | 4,72579E-36 |
| ACE2 | CEP164P1 | -0,5957 | 4,74293E-36 |
| ACE2 | TMCO2 | -0,5958 | 4,66488E-36 |
| ACE2 | CTD-2530N21.4 | -0,5958 | 4,67763E-36 |
| ACE2 | GPR25 | -0,596 | 4,26764E-36 |
| ACE2 | LRWD1 | -0,596 | 4,36151E-36 |
| ACE2 | RP11-575L7.4 | -0,596 | 4,27558E-36 |
| ACE2 | NUTM1 | -0,596 | 4,30438E-36 |
| ACE2 | TNFRSF21 | -0,5961 | 4,13642E-36 |
| ACE2 | SMIM2-AS1 | -0,5961 | 4,12876E-36 |
| ACE2 | RP11-1078H9.5 | -0,5961 | 4,18403E-36 |
| ACE2 | LYSMD2 | -0,5961 | 4,1478E-36 |
| ACE2 | LINC01888 | -0,5962 | 4,04474E-36 |
| ACE2 | PTPMT1 | -0,5963 | 3,928E-36 |
| ACE2 | CHD5 | -0,5964 | 3,80914E-36 |
| ACE2 | LINC01811 | -0,5964 | 3,79217E-36 |
| ACE2 | PSORS1C1 | -0,5964 | 3,82271E-36 |
| ACE2 | RP11-34D15.2 | -0,5964 | 3,76413E-36 |
| ACE2 | UBQLNL | -0,5964 | 3,8029E-36 |
| ACE2 | RALY-AS1 | -0,5964 | 3,73489E-36 |
| ACE2 | AY269186.2 | -0,5965 | 3,7015E-36 |
| ACE2 | RP11-446F17.3 | -0,5965 | 3,67269E-36 |
| ACE2 | TRBV27 | -0,5965 | 3,67462E-36 |
| ACE2 | RP11-1137G4.3 | -0,5965 | 3,67209E-36 |
| ACE2 | LINC01803 | -0,5966 | 3,51243E-36 |
| ACE2 | PLCXD2-AS1 | -0,5966 | 3,53971E-36 |
| ACE2 | LINC01339 | -0,5966 | 3,48973E-36 |
| ACE2 | RP11-15E18.1 | -0,5966 | 3,5905E-36 |
| ACE2 | UGT3A1 | -0,5967 | 3,38921E-36 |
| ACE2 | IL7 | -0,5967 | 3,40569E-36 |
| ACE2 | CTC-527H23.2 | -0,5967 | 3,40377E-36 |
| ACE2 | LINC01016 | -0,5968 | 3,27026E-36 |
| ACE2 | UNC5CL | -0,5968 | 3,31334E-36 |
| ACE2 | JHDM1D-AS1 | -0,5968 | 3,33654E-36 |
| ACE2 | LINC01592 | -0,5968 | 3,36274E-36 |
| ACE2 | MFSD6L | -0,5968 | 3,2971E-36 |
| ACE2 | FAM27E3 | -0,5969 | 3,22932E-36 |
| ACE2 | SGF29 | -0,5969 | 3,2547E-36 |

| | | | |
|---|---|---|---|
| ACE2 | NFIA-AS1 | -0,597 | 3,05374E-36 |
| ACE2 | TEX43 | -0,597 | 3,07412E-36 |
| ACE2 | LINC00643 | -0,597 | 3,14586E-36 |
| ACE2 | IFRD2 | -0,5971 | 2,96476E-36 |
| ACE2 | NAPRT | -0,5971 | 3,0335E-36 |
| ACE2 | GGN | -0,5971 | 3,01709E-36 |
| ACE2 | TAT-AS1 | -0,5972 | 2,90668E-36 |
| ACE2 | OR7E24 | -0,5972 | 2,9468E-36 |
| ACE2 | PRICKLE2-AS1 | -0,5973 | 2,77095E-36 |
| ACE2 | RP11-264K23.1 | -0,5973 | 2,85156E-36 |
| ACE2 | RP11-202D18.2 | -0,5974 | 2,73378E-36 |
| ACE2 | LMNTD2 | -0,5974 | 2,70478E-36 |
| ACE2 | RANGAP1 | -0,5974 | 2,68892E-36 |
| ACE2 | CATSPERD | -0,5975 | 2,59743E-36 |
| ACE2 | C19orf44 | -0,5975 | 2,61238E-36 |
| ACE2 | AC005754.7 | -0,5976 | 2,54154E-36 |
| ACE2 | FAM83E | -0,5976 | 2,53716E-36 |
| ACE2 | RP4-665N4.4 | -0,5978 | 2,37252E-36 |
| ACE2 | CFAP99 | -0,5978 | 2,38771E-36 |
| ACE2 | LINC00167 | -0,5978 | 2,3622E-36 |
| ACE2 | TEX26-AS1 | -0,5978 | 2,37054E-36 |
| ACE2 | RP11-486O13.3 | -0,5978 | 2,33781E-36 |
| ACE2 | CCDC114 | -0,5979 | 2,28411E-36 |
| ACE2 | LINC01350 | -0,598 | 2,19974E-36 |
| ACE2 | STARD13-AS | -0,5981 | 2,16204E-36 |
| ACE2 | RP11-80H8.3 | -0,5982 | 2,04327E-36 |
| ACE2 | LINC00347 | -0,5982 | 2,04173E-36 |
| ACE2 | NPPA-AS1 | -0,5983 | 2,02858E-36 |
| ACE2 | DUS2 | -0,5984 | 1,93337E-36 |
| ACE2 | ARHGEF7-AS2 | -0,5987 | 1,73388E-36 |
| ACE2 | LINC00226 | -0,5989 | 1,65721E-36 |
| ACE2 | IZUMO1 | -0,5989 | 1,61463E-36 |
| ACE2 | ODF2 | -0,599 | 1,58196E-36 |
| ACE2 | RP13-653N12.1 | -0,599 | 1,60016E-36 |
| ACE2 | TBC1D29 | -0,599 | 1,59289E-36 |
| ACE2 | DALRD3 | -0,5991 | 1,55685E-36 |
| ACE2 | MEA1 | -0,5991 | 1,55213E-36 |
| ACE2 | TCP10L | -0,5991 | 1,53077E-36 |
| ACE2 | LIX1L-AS1 | -0,5992 | 1,49735E-36 |
| ACE2 | GPR39 | -0,5992 | 1,46285E-36 |
| ACE2 | C2orf82 | -0,5992 | 1,45683E-36 |
| ACE2 | RPS6KA2-AS1 | -0,5993 | 1,45574E-36 |
| ACE2 | ANKRD9 | -0,5993 | 1,41009E-36 |
| ACE2 | TMEM202 | -0,5993 | 1,42773E-36 |
| ACE2 | SPATA31D5P | -0,5994 | 1,39476E-36 |
| ACE2 | SLFNL1 | -0,5995 | 1,32988E-36 |
| ACE2 | RP11-183E24.2 | -0,5995 | 1,32776E-36 |
| ACE2 | DKFZP434H168 | -0,5995 | 1,32097E-36 |
| ACE2 | LINC00272 | -0,5996 | 1,29877E-36 |
| ACE2 | CACNA1C-IT3 | -0,5996 | 1,2989E-36 |
| ACE2 | MRPL36 | -0,5997 | 1,23412E-36 |
| ACE2 | LINC00265 | -0,5997 | 1,25803E-36 |
| ACE2 | SSNA1 | -0,5997 | 1,25241E-36 |
| ACE2 | AC005391.2 | -0,5997 | 1,2552E-36 |

| | | | |
|---|---|---|---|
| ACE2 | GGNBP1 | -0,5998 | 1,19561E-36 |
| ACE2 | RP11-542C10.1 | -0,6 | 1,1453E-36 |
| ACE2 | C15orf61 | -0,6 | 1,14606E-36 |
| ACE2 | ABHD8 | -0,6001 | 1,10235E-36 |
| ACE2 | LINC01728 | -0,6001 | 1,0847E-36 |
| ACE2 | C1orf158 | -0,6002 | 1,06717E-36 |
| ACE2 | RP11-11N7.4 | -0,6002 | 1,06898E-36 |
| ACE2 | AC110769.3 | -0,6002 | 1,06572E-36 |
| ACE2 | RPL23AP26 | -0,6002 | 1,07156E-36 |
| ACE2 | RP11-474P2.2 | -0,6002 | 1,06508E-36 |
| ACE2 | LINC00424 | -0,6002 | 1,06905E-36 |
| ACE2 | RP11-566K19.5 | -0,6002 | 1,05425E-36 |
| ACE2 | LINC01580 | -0,6002 | 1,04004E-36 |
| ACE2 | SRD5A3-AS1 | -0,6003 | 1,03117E-36 |
| ACE2 | RP11-313E19.2 | -0,6003 | 1,00643E-36 |
| ACE2 | SPTBN2 | -0,6003 | 1,01831E-36 |
| ACE2 | MKNK1-AS1 | -0,6004 | 9,85529E-37 |
| ACE2 | RP11-158I23.1 | -0,6004 | 9,91233E-37 |
| ACE2 | FAM99A | -0,6005 | 9,53704E-37 |
| ACE2 | ELOBP1 | -0,6005 | 9,53556E-37 |
| ACE2 | RP11-572O6.1 | -0,6006 | 9,1179E-37 |
| ACE2 | ATP6V1F | -0,6006 | 9,22996E-37 |
| ACE2 | RP11-483P21.6 | -0,6006 | 9,14419E-37 |
| ACE2 | CITF22-62D4.1 | -0,6006 | 9,19448E-37 |
| ACE2 | LINC01962 | -0,6007 | 9,03311E-37 |
| ACE2 | GTF2IP20 | -0,6008 | 8,66471E-37 |
| ACE2 | PRKAR2A-AS1 | -0,6009 | 8,37308E-37 |
| ACE2 | RNF148 | -0,6009 | 8,34718E-37 |
| ACE2 | CD37 | -0,6009 | 8,43121E-37 |
| ACE2 | LINC01149 | -0,6011 | 7,73123E-37 |
| ACE2 | CTD-2021A8.3 | -0,6011 | 7,66373E-37 |
| ACE2 | GID4 | -0,6011 | 7,8057E-37 |
| ACE2 | RP11-160H12.2 | -0,6012 | 7,40364E-37 |
| ACE2 | AZIN2 | -0,6013 | 7,26532E-37 |
| ACE2 | LINC01655 | -0,6013 | 7,31703E-37 |
| ACE2 | LINC00677 | -0,6013 | 7,37138E-37 |
| ACE2 | LINC01729 | -0,6013 | 7,31782E-37 |
| ACE2 | LINC02056 | -0,6014 | 7,03436E-37 |
| ACE2 | LINC01514 | -0,6014 | 7,11196E-37 |
| ACE2 | RPS11P6 | -0,6014 | 6,92165E-37 |
| ACE2 | RP11-213G2.2 | -0,6017 | 6,34907E-37 |
| ACE2 | C11orf42 | -0,6017 | 6,24743E-37 |
| ACE2 | METTL26 | -0,6017 | 6,40071E-37 |
| ACE2 | CHST8 | -0,6017 | 6,4056E-37 |
| ACE2 | KIF9 | -0,6019 | 5,83852E-37 |
| ACE2 | TEKT3 | -0,6019 | 5,97515E-37 |
| ACE2 | ANKRD53 | -0,602 | 5,66581E-37 |
| ACE2 | RP11-369E15.3 | -0,602 | 5,69495E-37 |
| ACE2 | OPLAH | -0,602 | 5,68847E-37 |
| ACE2 | STK32C | -0,602 | 5,81968E-37 |
| ACE2 | FOXP1-AS1 | -0,6021 | 5,50911E-37 |
| ACE2 | PCAT6 | -0,6022 | 5,31663E-37 |
| ACE2 | RP11-331G2.7 | -0,6022 | 5,33031E-37 |
| ACE2 | RP3-417L20.4 | -0,6022 | 5,35493E-37 |

| | | | |
|---|---|---|---|
| ACE2 | ZC3H10 | -0,6022 | 5,29641E-37 |
| ACE2 | LINC01052 | -0,6022 | 5,30514E-37 |
| ACE2 | RND2 | -0,6022 | 5,44326E-37 |
| ACE2 | RP1-198K11.5 | -0,6022 | 5,28435E-37 |
| ACE2 | GTF2IRD1P1 | -0,6023 | 5,21073E-37 |
| ACE2 | FAM71B | -0,6024 | 4,96839E-37 |
| ACE2 | RP11-16C18.3 | -0,6025 | 4,91565E-37 |
| ACE2 | TDRD10 | -0,6026 | 4,60327E-37 |
| ACE2 | RP11-338C15.5 | -0,6026 | 4,67889E-37 |
| ACE2 | RP11-1080G15.1 | -0,6026 | 4,63361E-37 |
| ACE2 | RP11-190D6.2 | -0,6026 | 4,64988E-37 |
| ACE2 | TH2LCRR | -0,6027 | 4,55695E-37 |
| ACE2 | CA12 | -0,6027 | 4,55542E-37 |
| ACE2 | FBRS | -0,6027 | 4,46025E-37 |
| ACE2 | RP11-538P18.2 | -0,6028 | 4,43068E-37 |
| ACE2 | RP11-185P18.2 | -0,6028 | 4,39742E-37 |
| ACE2 | AGAP5 | -0,6029 | 4,18696E-37 |
| ACE2 | RP11-340I6.7 | -0,603 | 4,09446E-37 |
| ACE2 | SDR42E2 | -0,6031 | 4,00618E-37 |
| ACE2 | WBP2NL | -0,6031 | 3,91583E-37 |
| ACE2 | RP11-557H15.3 | -0,6032 | 3,81228E-37 |
| ACE2 | ZNF705A | -0,6032 | 3,76287E-37 |
| ACE2 | C21orf62-AS1 | -0,6032 | 3,77735E-37 |
| ACE2 | SIRPD | -0,6033 | 3,67681E-37 |
| ACE2 | AC022007.5 | -0,6035 | 3,47959E-37 |
| ACE2 | DNAJB13 | -0,6035 | 3,43545E-37 |
| ACE2 | RP11-307O10.1 | -0,6035 | 3,44787E-37 |
| ACE2 | HSFY1P1 | -0,6035 | 3,45505E-37 |
| ACE2 | LINC01394 | -0,6037 | 3,24022E-37 |
| ACE2 | ADAM7 | -0,6037 | 3,22554E-37 |
| ACE2 | C5orf52 | -0,6038 | 3,12889E-37 |
| ACE2 | RP11-291L22.9 | -0,6038 | 3,06048E-37 |
| ACE2 | COX6B2 | -0,6038 | 3,09283E-37 |
| ACE2 | FTMT | -0,6039 | 3,02252E-37 |
| ACE2 | SCN4A | -0,6039 | 2,95455E-37 |
| ACE2 | FAT2 | -0,6041 | 2,7974E-37 |
| ACE2 | WDR97 | -0,6041 | 2,78741E-37 |
| ACE2 | RP11-128P17.3 | -0,6041 | 2,75363E-37 |
| ACE2 | MIR3976HG | -0,6042 | 2,68373E-37 |
| ACE2 | CPA5 | -0,6044 | 2,54318E-37 |
| ACE2 | RP11-403F21.4 | -0,6044 | 2,50844E-37 |
| ACE2 | TTLL8 | -0,6044 | 2,53856E-37 |
| ACE2 | NSUN4 | -0,6045 | 2,39644E-37 |
| ACE2 | SNRPD2P2 | -0,6045 | 2,41474E-37 |
| ACE2 | RAB11FIP4 | -0,6045 | 2,42658E-37 |
| ACE2 | LINC01837 | -0,6045 | 2,44224E-37 |
| ACE2 | RP11-168P8.3 | -0,6046 | 2,36724E-37 |
| ACE2 | CNIH2 | -0,6046 | 2,39082E-37 |
| ACE2 | TOMM20L | -0,6046 | 2,39264E-37 |
| ACE2 | CNBD2 | -0,6046 | 2,36524E-37 |
| ACE2 | ATRIP | -0,6047 | 2,24743E-37 |
| ACE2 | SPERT | -0,6048 | 2,19656E-37 |
| ACE2 | FSCN3 | -0,6049 | 2,09958E-37 |
| ACE2 | RP11-496I9.1 | -0,6049 | 2,13262E-37 |

| | | | |
|---|---|---|---|
| ACE2 | RP4-737E23.2 | -0,605 | 2,06182E-37 |
| ACE2 | AC010980.2 | -0,6051 | 1,98284E-37 |
| ACE2 | LINC01233 | -0,6051 | 1,95209E-37 |
| ACE2 | RP11-109M19.1 | -0,6052 | 1,88434E-37 |
| ACE2 | RP11-46O21.2 | -0,6053 | 1,8651E-37 |
| ACE2 | RP11-646J21.7 | -0,6054 | 1,7621E-37 |
| ACE2 | PCCA-AS1 | -0,6054 | 1,80578E-37 |
| ACE2 | RP11-426L16.9 | -0,6055 | 1,71605E-37 |
| ACE2 | LEMD1 | -0,6055 | 1,7239E-37 |
| ACE2 | LINC02168 | -0,6056 | 1,67101E-37 |
| ACE2 | CTB-32O4.2 | -0,6056 | 1,64273E-37 |
| ACE2 | FAM209A | -0,6056 | 1,65403E-37 |
| ACE2 | CHCHD6 | -0,6057 | 1,60028E-37 |
| ACE2 | RP11-404K5.2 | -0,6057 | 1,5967E-37 |
| ACE2 | TLCD1 | -0,6057 | 1,60478E-37 |
| ACE2 | ODF1 | -0,6058 | 1,54305E-37 |
| ACE2 | SEPT12 | -0,6058 | 1,54697E-37 |
| ACE2 | SGSM2 | -0,6058 | 1,57342E-37 |
| ACE2 | MAP3K14-AS1 | -0,6059 | 1,49049E-37 |
| ACE2 | LRRC18 | -0,6061 | 1,41478E-37 |
| ACE2 | SLC35G6 | -0,6061 | 1,4163E-37 |
| ACE2 | MYO15A | -0,6061 | 1,40203E-37 |
| ACE2 | LINC01107 | -0,6062 | 1,33959E-37 |
| ACE2 | CCDC184 | -0,6062 | 1,37627E-37 |
| ACE2 | RP11-474D1.2 | -0,6062 | 1,37181E-37 |
| ACE2 | SPATA46 | -0,6063 | 1,3301E-37 |
| ACE2 | AC007131.2 | -0,6063 | 1,31039E-37 |
| ACE2 | RP11-61N20.3 | -0,6063 | 1,30966E-37 |
| ACE2 | LINC01974 | -0,6063 | 1,29301E-37 |
| ACE2 | LRRC10B | -0,6064 | 1,26253E-37 |
| ACE2 | CTD-3194G12.1 | -0,6064 | 1,26382E-37 |
| ACE2 | CTA-407F11.8 | -0,6064 | 1,26984E-37 |
| ACE2 | RP3-465N24.6 | -0,6065 | 1,22478E-37 |
| ACE2 | RP11-769N22.1 | -0,6065 | 1,24288E-37 |
| ACE2 | RP11-2C7.1 | -0,6065 | 1,21608E-37 |
| ACE2 | EPHA10 | -0,6066 | 1,20158E-37 |
| ACE2 | RP11-173A6.4 | -0,6066 | 1,1696E-37 |
| ACE2 | ZBTB20-AS1 | -0,6067 | 1,1452E-37 |
| ACE2 | SCRT1 | -0,6067 | 1,15255E-37 |
| ACE2 | RP11-295P9.3 | -0,6067 | 1,13631E-37 |
| ACE2 | KRT33A | -0,6067 | 1,15222E-37 |
| ACE2 | RP11-797E24.3 | -0,6067 | 1,13081E-37 |
| ACE2 | TSPAN16 | -0,6067 | 1,12363E-37 |
| ACE2 | RP11-19P22.7 | -0,6068 | 1,11976E-37 |
| ACE2 | PDE4A | -0,6068 | 1,09058E-37 |
| ACE2 | CST8 | -0,6071 | 9,84379E-38 |
| ACE2 | KB-1410C5.2 | -0,6072 | 9,69198E-38 |
| ACE2 | PLCE1-AS2 | -0,6072 | 9,48203E-38 |
| ACE2 | H1FOO | -0,6073 | 9,25679E-38 |
| ACE2 | RPH3A | -0,6073 | 9,35208E-38 |
| ACE2 | RP11-755F10.3 | -0,6075 | 8,73206E-38 |
| ACE2 | RP11-485O10.3 | -0,6075 | 8,51724E-38 |
| ACE2 | CSE1L-AS1 | -0,6075 | 8,69903E-38 |
| ACE2 | AC009487.4 | -0,6077 | 8,18339E-38 |

| | | | |
|---|---|---|---|
| ACE2 | SAC3D1 | -0,6077 | 8,03607E-38 |
| ACE2 | GORASP1 | -0,6078 | 7,91084E-38 |
| ACE2 | RMND5B | -0,6078 | 7,74055E-38 |
| ACE2 | AC004448.5 | -0,6078 | 7,89648E-38 |
| ACE2 | WDR83 | -0,6078 | 7,77851E-38 |
| ACE2 | MYH7B | -0,6078 | 7,81379E-38 |
| ACE2 | CRAT | -0,6079 | 7,50949E-38 |
| ACE2 | TXNDC2 | -0,6079 | 7,62402E-38 |
| ACE2 | ZNF692 | -0,608 | 7,20083E-38 |
| ACE2 | RP11-732A19.9 | -0,608 | 7,32176E-38 |
| ACE2 | CCDC33 | -0,608 | 7,35214E-38 |
| ACE2 | SLIT2-IT1 | -0,6081 | 7,07917E-38 |
| ACE2 | SLC35G3 | -0,6081 | 6,91096E-38 |
| ACE2 | RGMB-AS1 | -0,6082 | 6,7993E-38 |
| ACE2 | MRVI1-AS1 | -0,6082 | 6,85772E-38 |
| ACE2 | AC096649.2 | -0,6083 | 6,48049E-38 |
| ACE2 | UBE2J2 | -0,6084 | 6,27719E-38 |
| ACE2 | JMJD4 | -0,6084 | 6,32414E-38 |
| ACE2 | LY6G6C | -0,6084 | 6,41314E-38 |
| ACE2 | WDFY4 | -0,6084 | 6,23725E-38 |
| ACE2 | C3orf35 | -0,6085 | 6,1357E-38 |
| ACE2 | ROPN1L | -0,6085 | 6,03474E-38 |
| ACE2 | RP1-170O19.17 | -0,6085 | 6,08901E-38 |
| ACE2 | SPDYE1 | -0,6085 | 6,03576E-38 |
| ACE2 | GPR85 | -0,6085 | 6,17123E-38 |
| ACE2 | DFNB59 | -0,6086 | 5,96748E-38 |
| ACE2 | CASC16 | -0,6086 | 5,85184E-38 |
| ACE2 | CD320 | -0,6086 | 5,98682E-38 |
| ACE2 | AQP7 | -0,6088 | 5,59577E-38 |
| ACE2 | RP11-3L23.2 | -0,6088 | 5,56215E-38 |
| ACE2 | GJA8 | -0,6089 | 5,27587E-38 |
| ACE2 | AC007009.1 | -0,6089 | 5,29331E-38 |
| ACE2 | SIRT4 | -0,6089 | 5,2663E-38 |
| ACE2 | DYNLRB2 | -0,6089 | 5,39892E-38 |
| ACE2 | ING1 | -0,609 | 5,16572E-38 |
| ACE2 | RSPH1 | -0,609 | 5,07752E-38 |
| ACE2 | CYB5D1 | -0,6092 | 4,78756E-38 |
| ACE2 | HCG2040054 | -0,6093 | 4,58039E-38 |
| ACE2 | ERVW-1 | -0,6093 | 4,68616E-38 |
| ACE2 | RP11-429A20.3 | -0,6093 | 4,56337E-38 |
| ACE2 | CTD-2329C7.2 | -0,6093 | 4,5638E-38 |
| ACE2 | FLJ46284 | -0,6095 | 4,23757E-38 |
| ACE2 | AC114271.2 | -0,6096 | 4,2063E-38 |
| ACE2 | KCNMA1 | -0,6097 | 4,00974E-38 |
| ACE2 | MCRIP1 | -0,6097 | 4,034E-38 |
| ACE2 | CFAP45 | -0,6098 | 3,87334E-38 |
| ACE2 | C4orf51 | -0,6098 | 3,88592E-38 |
| ACE2 | RP11-716D16.1 | -0,6098 | 3,90607E-38 |
| ACE2 | CD19 | -0,6098 | 3,85157E-38 |
| ACE2 | HEMK1 | -0,6099 | 3,80082E-38 |
| ACE2 | TREML2 | -0,6099 | 3,69605E-38 |
| ACE2 | RP11-154D6.1 | -0,6099 | 3,72389E-38 |
| ACE2 | AC005808.3 | -0,6101 | 3,50084E-38 |
| ACE2 | RFPL3S | -0,6101 | 3,45661E-38 |

| | | | |
|---|---|---|---|
| ACE2 | RP11-318C2.1 | -0,6102 | 3,31327E-38 |
| ACE2 | KRTAP5-7 | -0,6104 | 3,15952E-38 |
| ACE2 | RP11-178L8.5 | -0,6104 | 3,14142E-38 |
| ACE2 | CDH22 | -0,6104 | 3,16785E-38 |
| ACE2 | RP11-428F8.2 | -0,6106 | 2,96731E-38 |
| ACE2 | RP11-628O18.1 | -0,6107 | 2,87166E-38 |
| ACE2 | KIZ-AS1 | -0,6107 | 2,81276E-38 |
| ACE2 | IFT27 | -0,6107 | 2,83292E-38 |
| ACE2 | PTTG1 | -0,6108 | 2,77659E-38 |
| ACE2 | DBIL5P | -0,6108 | 2,75809E-38 |
| ACE2 | TOB1-AS1 | -0,6108 | 2,71362E-38 |
| ACE2 | RP11-143M1.3 | -0,6109 | 2,66536E-38 |
| ACE2 | PSD2-AS1 | -0,6112 | 2,33479E-38 |
| ACE2 | LINC00598 | -0,6113 | 2,27517E-38 |
| ACE2 | HOMER2P2 | -0,6113 | 2,31579E-38 |
| ACE2 | LIPE | -0,6113 | 2,30534E-38 |
| ACE2 | LINC01273 | -0,6113 | 2,29915E-38 |
| ACE2 | SLC6A13 | -0,6114 | 2,22212E-38 |
| ACE2 | C12orf10 | -0,6114 | 2,18947E-38 |
| ACE2 | VWA5B1 | -0,6115 | 2,11137E-38 |
| ACE2 | RP5-991B18.2 | -0,6115 | 2,13344E-38 |
| ACE2 | ITPRIPL1 | -0,6117 | 1,98957E-38 |
| ACE2 | IQCF2 | -0,6117 | 2,00594E-38 |
| ACE2 | LINC00704 | -0,6117 | 1,96783E-38 |
| ACE2 | LINC00606 | -0,6118 | 1,91555E-38 |
| ACE2 | GPR182 | -0,6118 | 1,92081E-38 |
| ACE2 | APH1B | -0,6118 | 1,91504E-38 |
| ACE2 | PPP1R27 | -0,6119 | 1,86466E-38 |
| ACE2 | RP11-630C16.1 | -0,6121 | 1,7177E-38 |
| ACE2 | LINC01013 | -0,6122 | 1,66629E-38 |
| ACE2 | LINC00703 | -0,6123 | 1,63277E-38 |
| ACE2 | LINC00202-1 | -0,6124 | 1,53592E-38 |
| ACE2 | RP11-327F22.4 | -0,6124 | 1,55436E-38 |
| ACE2 | TNP1 | -0,6125 | 1,50181E-38 |
| ACE2 | C10orf82 | -0,6125 | 1,5111E-38 |
| ACE2 | PLIN1 | -0,6125 | 1,50825E-38 |
| ACE2 | ETV2 | -0,6125 | 1,48482E-38 |
| ACE2 | C3orf22 | -0,6127 | 1,40831E-38 |
| ACE2 | CBY3 | -0,6127 | 1,40621E-38 |
| ACE2 | RP11-705O24.1 | -0,6127 | 1,38488E-38 |
| ACE2 | RP11-201M22.1 | -0,6127 | 1,39722E-38 |
| ACE2 | FKBP8 | -0,6127 | 1,41612E-38 |
| ACE2 | TMEM75 | -0,6128 | 1,36393E-38 |
| ACE2 | NR1H3 | -0,6128 | 1,33708E-38 |
| ACE2 | RP1-58B11.1 | -0,6129 | 1,29833E-38 |
| ACE2 | HIPK1-AS1 | -0,6129 | 1,30529E-38 |
| ACE2 | GPT | -0,6129 | 1,31388E-38 |
| ACE2 | RP11-168K9.2 | -0,6129 | 1,311E-38 |
| ACE2 | RP11-476H16.1 | -0,6129 | 1,31032E-38 |
| ACE2 | RFX1 | -0,6129 | 1,29776E-38 |
| ACE2 | DNAJB3 | -0,613 | 1,25866E-38 |
| ACE2 | C9orf50 | -0,6131 | 1,23013E-38 |
| ACE2 | AC009120.4 | -0,6131 | 1,1986E-38 |
| ACE2 | FAM104A | -0,6131 | 1,22706E-38 |

| | | | |
|---|---|---|---|
| ACE2 | ARL2 | -0,6132 | 1,17248E-38 |
| ACE2 | RCC1 | -0,6133 | 1,13856E-38 |
| ACE2 | IGSF9 | -0,6133 | 1,12299E-38 |
| ACE2 | AC012370.3 | -0,6135 | 1,06404E-38 |
| ACE2 | LINC01471 | -0,6135 | 1,03815E-38 |
| ACE2 | CYP4F26P | -0,6135 | 1,05673E-38 |
| ACE2 | RP11-85O21.5 | -0,6135 | 1,04865E-38 |
| ACE2 | LINC01764 | -0,6135 | 1,04556E-38 |
| ACE2 | C5orf64-AS1 | -0,6137 | 9,63627E-39 |
| ACE2 | CYP11B1 | -0,6137 | 9,77989E-39 |
| ACE2 | PHC2 | -0,6138 | 9,51192E-39 |
| ACE2 | RP11-400L8.2 | -0,6138 | 9,58164E-39 |
| ACE2 | CCDC74BP1 | -0,6138 | 9,50215E-39 |
| ACE2 | ADD3-AS1 | -0,6139 | 9,0767E-39 |
| ACE2 | C20orf173 | -0,6139 | 9,20904E-39 |
| ACE2 | RP11-334J6.6 | -0,614 | 8,89457E-39 |
| ACE2 | AP003774.5 | -0,614 | 8,91692E-39 |
| ACE2 | RP11-273B20.3 | -0,6141 | 8,44078E-39 |
| ACE2 | RP11-62H20.1 | -0,6141 | 8,42718E-39 |
| ACE2 | RP11-217B1.2 | -0,6141 | 8,421E-39 |
| ACE2 | LINCR-0001 | -0,6142 | 8,25771E-39 |
| ACE2 | RP4-583K8.1 | -0,6142 | 8,10055E-39 |
| ACE2 | RP11-156L14.1 | -0,6143 | 7,95124E-39 |
| ACE2 | NPIPB1P | -0,6143 | 8,04156E-39 |
| ACE2 | LINC01424 | -0,6143 | 7,84413E-39 |
| ACE2 | LINC01487 | -0,6144 | 7,75E-39 |
| ACE2 | FAM71E2 | -0,6144 | 7,6229E-39 |
| ACE2 | C2orf71 | -0,6145 | 7,39921E-39 |
| ACE2 | ARHGAP22-IT1 | -0,6145 | 7,28364E-39 |
| ACE2 | GCKR | -0,6146 | 7,00563E-39 |
| ACE2 | RPUSD3 | -0,6146 | 7,08931E-39 |
| ACE2 | LINC00593 | -0,6146 | 7,19809E-39 |
| ACE2 | LINC01760 | -0,6147 | 6,84665E-39 |
| ACE2 | AC002064.5 | -0,6149 | 6,36181E-39 |
| ACE2 | HMCN2 | -0,6149 | 6,44604E-39 |
| ACE2 | IQCD | -0,6149 | 6,40707E-39 |
| ACE2 | RP11-526N18.1 | -0,6149 | 6,35781E-39 |
| ACE2 | CDKN2A | -0,615 | 6,09294E-39 |
| ACE2 | DHRS9 | -0,6151 | 5,85548E-39 |
| ACE2 | FAM163A | -0,6152 | 5,78603E-39 |
| ACE2 | SLC22A23 | -0,6152 | 5,83613E-39 |
| ACE2 | TSNAXIP1 | -0,6152 | 5,76794E-39 |
| ACE2 | CELSR3-AS1 | -0,6154 | 5,38574E-39 |
| ACE2 | AC114783.1 | -0,6155 | 5,08682E-39 |
| ACE2 | MYBPC3 | -0,6155 | 5,19822E-39 |
| ACE2 | RP11-94B19.2 | -0,6155 | 5,09915E-39 |
| ACE2 | RPL29P7 | -0,6156 | 5,05273E-39 |
| ACE2 | MOXD1 | -0,6156 | 5,00599E-39 |
| ACE2 | NOS3 | -0,6156 | 4,98239E-39 |
| ACE2 | MYO18B | -0,6156 | 4,92465E-39 |
| ACE2 | LMX1A | -0,6157 | 4,79768E-39 |
| ACE2 | TEKT4 | -0,6157 | 4,79987E-39 |
| ACE2 | MEX3D | -0,6157 | 4,81034E-39 |
| ACE2 | ATG9B | -0,6158 | 4,68379E-39 |

| | | | |
|---|---|---|---|
| ACE2 | LINC00535 | -0,6158 | 4,5909E-39 |
| ACE2 | STAC3 | -0,6158 | 4,5822E-39 |
| ACE2 | AIM1L | -0,616 | 4,35221E-39 |
| ACE2 | SCP2D1 | -0,616 | 4,24179E-39 |
| ACE2 | RP11-261C10.7 | -0,6161 | 4,24027E-39 |
| ACE2 | XRRA1 | -0,6161 | 4,15043E-39 |
| ACE2 | CTSF | -0,6162 | 4,05598E-39 |
| ACE2 | LINC00634 | -0,6162 | 3,99166E-39 |
| ACE2 | RP11-864J10.4 | -0,6163 | 3,87237E-39 |
| ACE2 | AIPL1 | -0,6163 | 3,91433E-39 |
| ACE2 | COL2A1 | -0,6165 | 3,59005E-39 |
| ACE2 | PHF1 | -0,6168 | 3,21118E-39 |
| ACE2 | RP11-505K9.4 | -0,6169 | 3,08268E-39 |
| ACE2 | RP11-6N13.1 | -0,617 | 3,02209E-39 |
| ACE2 | DYRK4 | -0,617 | 3,0173E-39 |
| ACE2 | C6orf99 | -0,6171 | 2,85941E-39 |
| ACE2 | RP3-446N13.5 | -0,6171 | 2,8725E-39 |
| ACE2 | LINC00113 | -0,6171 | 2,91316E-39 |
| ACE2 | MAPT-AS1 | -0,6172 | 2,77048E-39 |
| ACE2 | ATPIF1 | -0,6175 | 2,56293E-39 |
| ACE2 | RP11-17M16.2 | -0,6175 | 2,53718E-39 |
| ACE2 | LINC01523 | -0,6175 | 2,53653E-39 |
| ACE2 | BMP7-AS1 | -0,6175 | 2,53843E-39 |
| ACE2 | MZT2B | -0,6176 | 2,46451E-39 |
| ACE2 | NAMA | -0,6177 | 2,32441E-39 |
| ACE2 | DIO2 | -0,6177 | 2,33446E-39 |
| ACE2 | RP5-1004I9.1 | -0,6177 | 2,37474E-39 |
| ACE2 | ADORA2A-AS1 | -0,6177 | 2,35881E-39 |
| ACE2 | HRASLS | -0,6178 | 2,27758E-39 |
| ACE2 | NPIPB6 | -0,6178 | 2,23185E-39 |
| ACE2 | MEIS3 | -0,6178 | 2,24645E-39 |
| ACE2 | ERI3 | -0,618 | 2,07927E-39 |
| ACE2 | ZNF706 | -0,618 | 2,13944E-39 |
| ACE2 | RP13-653N12.2 | -0,618 | 2,13435E-39 |
| ACE2 | NFKBIB | -0,618 | 2,13181E-39 |
| ACE2 | RP1-69D17.3 | -0,6181 | 2,01139E-39 |
| ACE2 | IL4I1 | -0,6181 | 2,01657E-39 |
| ACE2 | TULP2 | -0,6183 | 1,86025E-39 |
| ACE2 | LINC01849 | -0,6184 | 1,79612E-39 |
| ACE2 | RP5-956O18.2 | -0,6185 | 1,77755E-39 |
| ACE2 | AC079305.10 | -0,6186 | 1,67789E-39 |
| ACE2 | RP11-863K10.2 | -0,6186 | 1,69902E-39 |
| ACE2 | PGAP2 | -0,6186 | 1,70216E-39 |
| ACE2 | OXCT2 | -0,6187 | 1,63038E-39 |
| ACE2 | GSG1 | -0,6187 | 1,60957E-39 |
| ACE2 | LINC00637 | -0,6187 | 1,62278E-39 |
| ACE2 | CDRT4 | -0,6187 | 1,6213E-39 |
| ACE2 | NDUFS7 | -0,6188 | 1,55763E-39 |
| ACE2 | RP11-73G16.1 | -0,6189 | 1,53504E-39 |
| ACE2 | RP11-491H19.1 | -0,6189 | 1,53684E-39 |
| ACE2 | IFT43 | -0,6189 | 1,52086E-39 |
| ACE2 | LRRC20 | -0,6193 | 1,3041E-39 |
| ACE2 | RBMX2P5 | -0,6194 | 1,28887E-39 |
| ACE2 | TMEM249 | -0,6194 | 1,26077E-39 |

| | | | |
|---|---|---|---|
| ACE2 | ACPT | -0,6195 | 1,20233E-39 |
| ACE2 | FAM183A | -0,6196 | 1,18754E-39 |
| ACE2 | AC009365.4 | -0,6197 | 1,11971E-39 |
| ACE2 | RP11-665J16.1 | -0,6197 | 1,12955E-39 |
| ACE2 | CTB-147C22.9 | -0,6197 | 1,1315E-39 |
| ACE2 | SLC13A3 | -0,6197 | 1,14351E-39 |
| ACE2 | FBXO6 | -0,6199 | 1,05102E-39 |
| ACE2 | COL11A1 | -0,62 | 1,03704E-39 |
| ACE2 | LINC01878 | -0,62 | 1,02127E-39 |
| ACE2 | JMJD1C-AS1 | -0,62 | 1,01671E-39 |
| ACE2 | COL22A1 | -0,6201 | 9,84245E-40 |
| ACE2 | HSPB7 | -0,6202 | 9,5793E-40 |
| ACE2 | DZIP1L | -0,6203 | 9,0941E-40 |
| ACE2 | ZNF483 | -0,6204 | 8,93316E-40 |
| ACE2 | RGS8 | -0,6205 | 8,45281E-40 |
| ACE2 | CMTM2 | -0,6205 | 8,42415E-40 |
| ACE2 | EPHA1 | -0,6206 | 8,07426E-40 |
| ACE2 | C1orf234 | -0,6207 | 8,00811E-40 |
| ACE2 | FANCD2OS | -0,6208 | 7,53538E-40 |
| ACE2 | RP11-109E24.2 | -0,6208 | 7,64629E-40 |
| ACE2 | PKHD1 | -0,6208 | 7,51207E-40 |
| ACE2 | CCDC185 | -0,621 | 7,19842E-40 |
| ACE2 | RP11-536N17.1 | -0,621 | 7,18291E-40 |
| ACE2 | CCDC60 | -0,621 | 7,07631E-40 |
| ACE2 | ASGR1 | -0,621 | 7,22424E-40 |
| ACE2 | COL9A1 | -0,6211 | 6,94908E-40 |
| ACE2 | LRRC36 | -0,6211 | 6,86134E-40 |
| ACE2 | DNAJB7 | -0,6211 | 6,87336E-40 |
| ACE2 | EFCAB12 | -0,6212 | 6,56563E-40 |
| ACE2 | RP11-359E10.1 | -0,6212 | 6,67628E-40 |
| ACE2 | PIFO | -0,6213 | 6,45226E-40 |
| ACE2 | CCDC13 | -0,6213 | 6,3747E-40 |
| ACE2 | AC145123.2 | -0,6213 | 6,38077E-40 |
| ACE2 | FRAT1 | -0,6213 | 6,35489E-40 |
| ACE2 | KCNIP2-AS1 | -0,6213 | 6,29407E-40 |
| ACE2 | RP11-664D1.1 | -0,6214 | 6,08861E-40 |
| ACE2 | MRPL51 | -0,6215 | 6,01061E-40 |
| ACE2 | RPL10L | -0,6215 | 5,96932E-40 |
| ACE2 | ST3GAL3 | -0,6216 | 5,70055E-40 |
| ACE2 | C1orf100 | -0,6216 | 5,65145E-40 |
| ACE2 | SORBS3 | -0,6216 | 5,67106E-40 |
| ACE2 | ACE | -0,6216 | 5,60076E-40 |
| ACE2 | RP4-665J23.1 | -0,6217 | 5,42543E-40 |
| ACE2 | RP1-93H18.1 | -0,6217 | 5,46153E-40 |
| ACE2 | LINC01766 | -0,6218 | 5,29763E-40 |
| ACE2 | CTD-2330K9.3 | -0,6219 | 5,07636E-40 |
| ACE2 | FGD3 | -0,6219 | 5,0372E-40 |
| ACE2 | FAM19A2 | -0,6219 | 5,06636E-40 |
| ACE2 | NDUFB10 | -0,6219 | 5,07571E-40 |
| ACE2 | GP6 | -0,622 | 4,83913E-40 |
| ACE2 | NOL12 | -0,622 | 4,86514E-40 |
| ACE2 | RP11-794G24.1 | -0,6221 | 4,73027E-40 |
| ACE2 | SPAG5-AS1 | -0,6221 | 4,7639E-40 |
| ACE2 | MURC | -0,6223 | 4,38026E-40 |

| | | | |
|---|---|---|---|
| ACE2 | SPEF1 | -0,6223 | 4,33576E-40 |
| ACE2 | LINC01214 | -0,6224 | 4,19933E-40 |
| ACE2 | MROH5 | -0,6224 | 4,29587E-40 |
| ACE2 | SPATA31C1 | -0,6224 | 4,25834E-40 |
| ACE2 | LINC00642 | -0,6225 | 4,04536E-40 |
| ACE2 | XYLT2 | -0,6227 | 3,82958E-40 |
| ACE2 | CHRNA1 | -0,6228 | 3,67355E-40 |
| ACE2 | LINC01939 | -0,6231 | 3,32928E-40 |
| ACE2 | HCN4 | -0,6231 | 3,29718E-40 |
| ACE2 | RP5-1027G4.3 | -0,6232 | 3,14005E-40 |
| ACE2 | MARCH11 | -0,6235 | 2,88782E-40 |
| ACE2 | AL591893.1 | -0,6236 | 2,73094E-40 |
| ACE2 | RP1-297M16.2 | -0,6237 | 2,60907E-40 |
| ACE2 | LRRC74A | -0,6237 | 2,66717E-40 |
| ACE2 | APOPT1 | -0,6237 | 2,67875E-40 |
| ACE2 | RP13-401N8.1 | -0,6237 | 2,61337E-40 |
| ACE2 | AC062017.1 | -0,6239 | 2,44418E-40 |
| ACE2 | RP11-231D20.2 | -0,624 | 2,3258E-40 |
| ACE2 | DOCK9-AS2 | -0,624 | 2,38223E-40 |
| ACE2 | C1orf116 | -0,6241 | 2,25128E-40 |
| ACE2 | ELOA2 | -0,6244 | 2,07525E-40 |
| ACE2 | OASL | -0,6245 | 1,9701E-40 |
| ACE2 | LINC01603 | -0,6247 | 1,79671E-40 |
| ACE2 | RP11-20B24.5 | -0,6247 | 1,8489E-40 |
| ACE2 | RP11-219B4.5 | -0,6248 | 1,76843E-40 |
| ACE2 | SRRM2-AS1 | -0,6248 | 1,77778E-40 |
| ACE2 | SLC38A6 | -0,6249 | 1,68996E-40 |
| ACE2 | C20orf85 | -0,6249 | 1,67302E-40 |
| ACE2 | RP11-503N18.4 | -0,6252 | 1,49558E-40 |
| ACE2 | CXCL16 | -0,6252 | 1,49891E-40 |
| ACE2 | CYP17A1-AS1 | -0,6254 | 1,43198E-40 |
| ACE2 | RP11-12M5.3 | -0,6256 | 1,28449E-40 |
| ACE2 | RP11-15J22.8 | -0,6256 | 1,32588E-40 |
| ACE2 | RP11-160H22.3 | -0,6259 | 1,16287E-40 |
| ACE2 | COX7A2 | -0,626 | 1,13993E-40 |
| ACE2 | CFAP157 | -0,626 | 1,13595E-40 |
| ACE2 | PFN4 | -0,6261 | 1,09716E-40 |
| ACE2 | PHLDB3 | -0,6261 | 1,08896E-40 |
| ACE2 | RP5-1069C8.3 | -0,6262 | 1,05653E-40 |
| ACE2 | DLGAP4-AS1 | -0,6262 | 1,04002E-40 |
| ACE2 | SPATA12 | -0,6263 | 1,01386E-40 |
| ACE2 | UBXN6 | -0,6263 | 1,02785E-40 |
| ACE2 | SLC7A7 | -0,6265 | 9,44746E-41 |
| ACE2 | RP11-365F18.1 | -0,6267 | 8,64673E-41 |
| ACE2 | UCP3 | -0,6267 | 8,76052E-41 |
| ACE2 | RNF157-AS1 | -0,6267 | 8,75404E-41 |
| ACE2 | BPI | -0,6267 | 8,63822E-41 |
| ACE2 | ZNRF4 | -0,6268 | 8,26007E-41 |
| ACE2 | NDUFV3 | -0,6269 | 8,11148E-41 |
| ACE2 | MSTO2P | -0,627 | 7,77046E-41 |
| ACE2 | AP003025.2 | -0,6271 | 7,60131E-41 |
| ACE2 | CATSPERG | -0,6272 | 7,27359E-41 |
| ACE2 | PLA2G4E | -0,6273 | 6,96576E-41 |
| ACE2 | LINGO1-AS1 | -0,6273 | 6,84783E-41 |

| | | | |
|---|---|---|---|
| ACE2 | BCL7C | -0,6273 | 6,95227E-41 |
| ACE2 | IL12A-AS1 | -0,6275 | 6,49474E-41 |
| ACE2 | DGCR6L | -0,6277 | 5,89445E-41 |
| ACE2 | CCNT2-AS1 | -0,6279 | 5,64702E-41 |
| ACE2 | RP11-834C11.4 | -0,6279 | 5,63068E-41 |
| ACE2 | TCTE1 | -0,628 | 5,3402E-41 |
| ACE2 | TBC1D3P5 | -0,628 | 5,31659E-41 |
| ACE2 | C2orf70 | -0,6281 | 5,22423E-41 |
| ACE2 | RNF126P1 | -0,6281 | 5,24926E-41 |
| ACE2 | CYP4F32P | -0,6282 | 4,9857E-41 |
| ACE2 | KLHDC3 | -0,6282 | 5,01723E-41 |
| ACE2 | TTLL10-AS1 | -0,6283 | 4,71699E-41 |
| ACE2 | FAM229B | -0,6283 | 4,70716E-41 |
| ACE2 | AC016907.3 | -0,6284 | 4,68347E-41 |
| ACE2 | FAM166A | -0,6284 | 4,54017E-41 |
| ACE2 | QRICH2 | -0,6284 | 4,69286E-41 |
| ACE2 | DGCR14 | -0,6284 | 4,54352E-41 |
| ACE2 | LURAP1 | -0,6285 | 4,44786E-41 |
| ACE2 | EVX1 | -0,6285 | 4,52453E-41 |
| ACE2 | RP11-156K13.3 | -0,6286 | 4,29949E-41 |
| ACE2 | CTD-2572N17.1 | -0,6286 | 4,27822E-41 |
| ACE2 | NAT6 | -0,6288 | 3,9826E-41 |
| ACE2 | DDX39A | -0,6288 | 4,03965E-41 |
| ACE2 | YPEL1 | -0,6289 | 3,83037E-41 |
| ACE2 | DCST2 | -0,629 | 3,68209E-41 |
| ACE2 | RP11-700N1.1 | -0,629 | 3,68188E-41 |
| ACE2 | ZNF32-AS3 | -0,6292 | 3,47567E-41 |
| ACE2 | RP11-70F11.8 | -0,6292 | 3,38024E-41 |
| ACE2 | TBC1D10B | -0,6292 | 3,39197E-41 |
| ACE2 | COPRS | -0,6292 | 3,35785E-41 |
| ACE2 | LINC01976 | -0,6294 | 3,15955E-41 |
| ACE2 | CEP83-AS1 | -0,6295 | 3,05766E-41 |
| ACE2 | CATSPER3 | -0,6299 | 2,59163E-41 |
| ACE2 | RAD51D | -0,63 | 2,57391E-41 |
| ACE2 | PRM1 | -0,6301 | 2,40632E-41 |
| ACE2 | RP11-227D13.4 | -0,6302 | 2,32165E-41 |
| ACE2 | RP11-554D20.1 | -0,6302 | 2,3294E-41 |
| ACE2 | U73166.2 | -0,6303 | 2,27741E-41 |
| ACE2 | APOM | -0,6303 | 2,21734E-41 |
| ACE2 | LINC01723 | -0,6305 | 2,08703E-41 |
| ACE2 | CTC-457E21.5 | -0,6306 | 1,98266E-41 |
| ACE2 | LYZL4 | -0,6308 | 1,89649E-41 |
| ACE2 | GTF2F1 | -0,6308 | 1,89977E-41 |
| ACE2 | DNAAF3 | -0,6308 | 1,88971E-41 |
| ACE2 | CIB4 | -0,6309 | 1,77971E-41 |
| ACE2 | RP11-188C12.3 | -0,6309 | 1,78023E-41 |
| ACE2 | HMGB4 | -0,6311 | 1,65906E-41 |
| ACE2 | UBE2Q1-AS1 | -0,6312 | 1,58685E-41 |
| ACE2 | ZNF688 | -0,6312 | 1,59701E-41 |
| ACE2 | LINC00935 | -0,6313 | 1,57202E-41 |
| ACE2 | RP11-396F22.1 | -0,6314 | 1,50473E-41 |
| ACE2 | MUC19 | -0,6314 | 1,47718E-41 |
| ACE2 | RP11-394A14.2 | -0,6314 | 1,50232E-41 |
| ACE2 | CHRNB4 | -0,6314 | 1,47394E-41 |

| | | | |
|---|---|---|---|
| ACE2 | ZNF683 | -0,6315 | 1,43555E-41 |
| ACE2 | P2RX3 | -0,6315 | 1,44526E-41 |
| ACE2 | TIMD4 | -0,6316 | 1,39836E-41 |
| ACE2 | RP11-628E19.2 | -0,6316 | 1,40633E-41 |
| ACE2 | RP11-160C18.2 | -0,6316 | 1,36708E-41 |
| ACE2 | LINC01492 | -0,6317 | 1,32652E-41 |
| ACE2 | CTD-2649C14.3 | -0,6317 | 1,3133E-41 |
| ACE2 | C20orf78 | -0,6317 | 1,35545E-41 |
| ACE2 | OR2H1 | -0,6319 | 1,22774E-41 |
| ACE2 | RTP2 | -0,632 | 1,16881E-41 |
| ACE2 | ELP5 | -0,6321 | 1,12167E-41 |
| ACE2 | PSENEN | -0,6321 | 1,16231E-41 |
| ACE2 | MORN3 | -0,6322 | 1,10743E-41 |
| ACE2 | SLC14A2-AS1 | -0,6322 | 1,09795E-41 |
| ACE2 | CTD-3074O7.5 | -0,6323 | 1,05225E-41 |
| ACE2 | ODF3L2 | -0,6323 | 1,05542E-41 |
| ACE2 | RP11-665I14.1 | -0,6324 | 1,02553E-41 |
| ACE2 | SLC25A37 | -0,6324 | 1,03743E-41 |
| ACE2 | C16orf95 | -0,6325 | 9,86174E-42 |
| ACE2 | CTC-203F4.2 | -0,6326 | 9,55031E-42 |
| ACE2 | SLC51B | -0,6326 | 9,53546E-42 |
| ACE2 | RP11-309L24.4 | -0,6327 | 8,94278E-42 |
| ACE2 | RP11-274J16.5 | -0,6327 | 9,01165E-42 |
| ACE2 | LINC00521 | -0,6328 | 8,80891E-42 |
| ACE2 | TBC1D21 | -0,6329 | 8,3959E-42 |
| ACE2 | RP13-131K19.2 | -0,6331 | 7,69382E-42 |
| ACE2 | TSSK1A | -0,6331 | 7,94783E-42 |
| ACE2 | MST1R | -0,6335 | 6,65978E-42 |
| ACE2 | SMIM2 | -0,6335 | 6,62696E-42 |
| ACE2 | KRT15 | -0,6335 | 6,74514E-42 |
| ACE2 | FMC1 | -0,6336 | 6,56739E-42 |
| ACE2 | RP11-554E23.2 | -0,6337 | 6,2883E-42 |
| ACE2 | LINC01344 | -0,6338 | 5,94216E-42 |
| ACE2 | RP11-462P6.1 | -0,6339 | 5,84936E-42 |
| ACE2 | S1PR2 | -0,6339 | 5,71145E-42 |
| ACE2 | EGLN2 | -0,6339 | 5,80472E-42 |
| ACE2 | KRTAP5-9 | -0,634 | 5,61691E-42 |
| ACE2 | B4GALNT3 | -0,634 | 5,53086E-42 |
| ACE2 | RP11-862G15.1 | -0,634 | 5,60625E-42 |
| ACE2 | DYRK1B | -0,6341 | 5,38132E-42 |
| ACE2 | LMNB2 | -0,6343 | 4,98862E-42 |
| ACE2 | CCDC96 | -0,6344 | 4,83302E-42 |
| ACE2 | NOS2P3 | -0,6344 | 4,73938E-42 |
| ACE2 | SIRPG | -0,6344 | 4,76405E-42 |
| ACE2 | GTF3C2-AS1 | -0,6346 | 4,40929E-42 |
| ACE2 | ELOB | -0,6346 | 4,40297E-42 |
| ACE2 | SPATA31A1 | -0,6347 | 4,17221E-42 |
| ACE2 | YBEY | -0,6347 | 4,17851E-42 |
| ACE2 | LYZL1 | -0,6348 | 4,13312E-42 |
| ACE2 | RP11-24M17.5 | -0,6348 | 4,1415E-42 |
| ACE2 | DPP10-AS1 | -0,6349 | 3,86052E-42 |
| ACE2 | LINC01988 | -0,6349 | 3,88705E-42 |
| ACE2 | C1QTNF3 | -0,6349 | 3,8831E-42 |
| ACE2 | FAAP20 | -0,635 | 3,81967E-42 |

| | | | |
|---|---|---|---|
| ACE2 | RP11-535A19.2 | -0,6351 | 3,61833E-42 |
| ACE2 | PNPLA6 | -0,6351 | 3,69506E-42 |
| ACE2 | RP11-360D2.1 | -0,6352 | 3,4554E-42 |
| ACE2 | PDSS1P2 | -0,6352 | 3,51592E-42 |
| ACE2 | CSMD2-AS1 | -0,6353 | 3,40981E-42 |
| ACE2 | RP3-467K16.4 | -0,6354 | 3,2947E-42 |
| ACE2 | FAM57B | -0,6354 | 3,27789E-42 |
| ACE2 | RP11-253M7.1 | -0,6355 | 3,08426E-42 |
| ACE2 | C8orf74 | -0,6361 | 2,45906E-42 |
| ACE2 | IQCF6 | -0,6362 | 2,35046E-42 |
| ACE2 | RP11-119J18.1 | -0,6362 | 2,35099E-42 |
| ACE2 | FOXJ1 | -0,6363 | 2,27342E-42 |
| ACE2 | HDAC11 | -0,6364 | 2,18347E-42 |
| ACE2 | AL157902.3 | -0,6365 | 2,0871E-42 |
| ACE2 | CTD-3185P2.1 | -0,6365 | 2,08754E-42 |
| ACE2 | RP11-486I11.2 | -0,6365 | 2,15176E-42 |
| ACE2 | TBATA | -0,6366 | 2,05227E-42 |
| ACE2 | RBBP8NL | -0,6366 | 2,01544E-42 |
| ACE2 | KRT8P12 | -0,6368 | 1,85948E-42 |
| ACE2 | RP11-398C13.6 | -0,6368 | 1,86515E-42 |
| ACE2 | ECE2 | -0,6369 | 1,80552E-42 |
| ACE2 | FAM221B | -0,6369 | 1,81409E-42 |
| ACE2 | PWRN1 | -0,637 | 1,75248E-42 |
| ACE2 | RP11-470L19.6 | -0,6371 | 1,66775E-42 |
| ACE2 | FFAR4 | -0,6371 | 1,6533E-42 |
| ACE2 | SPATA25 | -0,6371 | 1,67819E-42 |
| ACE2 | HNF1B | -0,6373 | 1,56822E-42 |
| ACE2 | SPATA20 | -0,6373 | 1,54475E-42 |
| ACE2 | RP3-476K8.3 | -0,6374 | 1,5228E-42 |
| ACE2 | RP4-736L20.3 | -0,6375 | 1,43768E-42 |
| ACE2 | KCNQ5-AS1 | -0,6375 | 1,42704E-42 |
| ACE2 | AC005523.3 | -0,6375 | 1,46575E-42 |
| ACE2 | ALLC | -0,6376 | 1,37545E-42 |
| ACE2 | SMIM23 | -0,6376 | 1,37214E-42 |
| ACE2 | RP11-402G3.3 | -0,6376 | 1,36642E-42 |
| ACE2 | IL20RB | -0,6377 | 1,32084E-42 |
| ACE2 | NEURL1 | -0,6377 | 1,31375E-42 |
| ACE2 | RP11-796E2.4 | -0,6377 | 1,35025E-42 |
| ACE2 | C12orf42 | -0,6377 | 1,32113E-42 |
| ACE2 | IL13 | -0,6378 | 1,29319E-42 |
| ACE2 | AMZ1 | -0,6378 | 1,30359E-42 |
| ACE2 | SLC22A18 | -0,6378 | 1,27013E-42 |
| ACE2 | AF131216.7 | -0,6379 | 1,24684E-42 |
| ACE2 | TPPP2 | -0,6379 | 1,24492E-42 |
| ACE2 | CARHSP1 | -0,6379 | 1,22231E-42 |
| ACE2 | SRRM5 | -0,6379 | 1,22246E-42 |
| ACE2 | RP11-218M11.7 | -0,638 | 1,21099E-42 |
| ACE2 | CTDP1 | -0,638 | 1,17555E-42 |
| ACE2 | BANF2 | -0,638 | 1,1847E-42 |
| ACE2 | RP11-140K17.3 | -0,6382 | 1,09897E-42 |
| ACE2 | ISG20L2 | -0,6384 | 1,02637E-42 |
| ACE2 | ADIG | -0,6384 | 1,01522E-42 |
| ACE2 | RNFT2 | -0,6386 | 9,51715E-43 |
| ACE2 | LINC01747 | -0,6386 | 9,30002E-43 |

| | | | |
|---|---|---|---|
| ACE2 | LYZL2 | -0,6388 | 8,75175E-43 |
| ACE2 | SLC39A3 | -0,6388 | 8,80131E-43 |
| ACE2 | EHD1 | -0,6389 | 8,46885E-43 |
| ACE2 | LINC00174 | -0,639 | 8,026E-43 |
| ACE2 | CRIP2 | -0,639 | 8,05685E-43 |
| ACE2 | C15orf59-AS1 | -0,6391 | 7,74696E-43 |
| ACE2 | CATIP | -0,6392 | 7,51584E-43 |
| ACE2 | IQCF4 | -0,6392 | 7,44085E-43 |
| ACE2 | LINC01820 | -0,6393 | 7,18192E-43 |
| ACE2 | LINC01095 | -0,6393 | 7,16485E-43 |
| ACE2 | SLC39A13 | -0,6393 | 7,0525E-43 |
| ACE2 | CTC-1337H24.4 | -0,6393 | 7,1168E-43 |
| ACE2 | RP11-53B2.1 | -0,6394 | 6,96997E-43 |
| ACE2 | TEKT2 | -0,6397 | 6,02076E-43 |
| ACE2 | RP3-486I3.4 | -0,6397 | 6,19488E-43 |
| ACE2 | MPST | -0,6397 | 6,22424E-43 |
| ACE2 | ANAPC11 | -0,6398 | 5,83164E-43 |
| ACE2 | TTLL11 | -0,6399 | 5,59191E-43 |
| ACE2 | CTD-2162K18.4 | -0,6399 | 5,58758E-43 |
| ACE2 | SPATA19 | -0,64 | 5,46885E-43 |
| ACE2 | C1orf194 | -0,6401 | 5,18942E-43 |
| ACE2 | RP11-524O1.4 | -0,6403 | 4,84584E-43 |
| ACE2 | SPATA42 | -0,6404 | 4,65962E-43 |
| ACE2 | ISLR2 | -0,6404 | 4,60357E-43 |
| ACE2 | SPATC1 | -0,6405 | 4,40886E-43 |
| ACE2 | RP11-195L15.2 | -0,6405 | 4,56667E-43 |
| ACE2 | SGCA | -0,6405 | 4,47528E-43 |
| ACE2 | LY6G5C | -0,6406 | 4,27594E-43 |
| ACE2 | C11orf53 | -0,6407 | 4,16387E-43 |
| ACE2 | RP5-1057J7.1 | -0,6408 | 4,05906E-43 |
| ACE2 | DKFZP434K028 | -0,6408 | 4,01975E-43 |
| ACE2 | SLC27A5 | -0,6408 | 3,96693E-43 |
| ACE2 | RP11-687M24.7 | -0,6409 | 3,8798E-43 |
| ACE2 | LINC01498 | -0,6409 | 3,86853E-43 |
| ACE2 | RANGRF | -0,6409 | 3,85007E-43 |
| ACE2 | IQCF5 | -0,641 | 3,72653E-43 |
| ACE2 | LEF1-AS1 | -0,6411 | 3,60516E-43 |
| ACE2 | CPT1B | -0,6413 | 3,26765E-43 |
| ACE2 | ACTL7B | -0,6415 | 3,04489E-43 |
| ACE2 | ZBTB32 | -0,6417 | 2,83278E-43 |
| ACE2 | TTC16 | -0,642 | 2,48322E-43 |
| ACE2 | CMTM1 | -0,6421 | 2,4231E-43 |
| ACE2 | FER1L5 | -0,6422 | 2,28036E-43 |
| ACE2 | ACAP1 | -0,6422 | 2,32403E-43 |
| ACE2 | OSBP2 | -0,6422 | 2,30479E-43 |
| ACE2 | RP11-126O1.2 | -0,6424 | 2,14216E-43 |
| ACE2 | CSTL1 | -0,6424 | 2,15725E-43 |
| ACE2 | LRRC27 | -0,6425 | 2,08152E-43 |
| ACE2 | CETN1 | -0,6425 | 2,06228E-43 |
| ACE2 | SPATA24 | -0,6426 | 1,93309E-43 |
| ACE2 | RP4-569M23.5 | -0,6426 | 1,97842E-43 |
| ACE2 | HK3 | -0,6427 | 1,88848E-43 |
| ACE2 | COL11A2 | -0,6427 | 1,86076E-43 |
| ACE2 | RP11-340I6.8 | -0,6427 | 1,89158E-43 |

| | | | |
|---|---|---|---|
| ACE2 | AGAP10P | -0,6428 | 1,85066E-43 |
| ACE2 | DNALI1 | -0,6431 | 1,58373E-43 |
| ACE2 | RP5-1051H14.2 | -0,6433 | 1,4964E-43 |
| ACE2 | NRAV | -0,6433 | 1,46648E-43 |
| ACE2 | RP11-783K16.5 | -0,6434 | 1,44183E-43 |
| ACE2 | CCDC116 | -0,6435 | 1,36264E-43 |
| ACE2 | RP11-779O18.1 | -0,6438 | 1,24246E-43 |
| ACE2 | SMIM22 | -0,6439 | 1,19403E-43 |
| ACE2 | SEPT4 | -0,6439 | 1,16558E-43 |
| ACE2 | PARD6A | -0,6441 | 1,09125E-43 |
| ACE2 | MFSD7 | -0,6442 | 1,03147E-43 |
| ACE2 | VPS13A-AS1 | -0,6442 | 1,0362E-43 |
| ACE2 | TSGA10IP | -0,6442 | 1,03913E-43 |
| ACE2 | FAHD2B | -0,6444 | 9,75604E-44 |
| ACE2 | ELMO3 | -0,6445 | 9,35059E-44 |
| ACE2 | HOXA-AS2 | -0,6449 | 7,86711E-44 |
| ACE2 | C10orf55 | -0,6449 | 7,75819E-44 |
| ACE2 | RP4-676J13.2 | -0,645 | 7,70399E-44 |
| ACE2 | PRM3 | -0,6451 | 7,30492E-44 |
| ACE2 | PHOSPHO1 | -0,6451 | 7,32944E-44 |
| ACE2 | RHBG | -0,6454 | 6,52677E-44 |
| ACE2 | RGR | -0,6454 | 6,49409E-44 |
| ACE2 | ARHGAP9 | -0,6456 | 5,95994E-44 |
| ACE2 | LIMD2 | -0,6458 | 5,49199E-44 |
| ACE2 | RP11-94B19.4 | -0,6458 | 5,49865E-44 |
| ACE2 | AC144838.3 | -0,6459 | 5,40253E-44 |
| ACE2 | GOLGA6L7P | -0,646 | 5,16355E-44 |
| ACE2 | C16orf82 | -0,646 | 5,09225E-44 |
| ACE2 | ACTL7A | -0,6461 | 4,90478E-44 |
| ACE2 | LINC01827 | -0,6462 | 4,69266E-44 |
| ACE2 | LINC01743 | -0,6464 | 4,42772E-44 |
| ACE2 | XX-FYM637E10_5.1 | -0,6464 | 4,35952E-44 |
| ACE2 | AC019055.1 | -0,6465 | 4,25735E-44 |
| ACE2 | TMEM95 | -0,6465 | 4,17407E-44 |
| ACE2 | SPATA3 | -0,6466 | 4,08877E-44 |
| ACE2 | FANK1 | -0,6467 | 3,81929E-44 |
| ACE2 | SLC35E4 | -0,6467 | 3,90866E-44 |
| ACE2 | TRAPPC2L | -0,6468 | 3,7703E-44 |
| ACE2 | MRPL14 | -0,6469 | 3,57531E-44 |
| ACE2 | FBXO24 | -0,647 | 3,42088E-44 |
| ACE2 | C17orf64 | -0,647 | 3,46809E-44 |
| ACE2 | DRICH1 | -0,647 | 3,46226E-44 |
| ACE2 | ZNF628 | -0,6472 | 3,09966E-44 |
| ACE2 | PAOX | -0,6473 | 3,08891E-44 |
| ACE2 | CTC-453G23.4 | -0,6475 | 2,824E-44 |
| ACE2 | ERICH2 | -0,6477 | 2,6098E-44 |
| ACE2 | RP11-798K23.3 | -0,6477 | 2,53662E-44 |
| ACE2 | LINC00982 | -0,6479 | 2,42817E-44 |
| ACE2 | C9orf116 | -0,6479 | 2,42715E-44 |
| ACE2 | DNPEP | -0,648 | 2,24258E-44 |
| ACE2 | AC074391.1 | -0,6481 | 2,1844E-44 |
| ACE2 | TMEM205 | -0,6481 | 2,20657E-44 |
| ACE2 | PRADC1 | -0,6485 | 1,85221E-44 |
| ACE2 | FAM71F1 | -0,6485 | 1,86171E-44 |

| | | | |
|---|---|---|---|
| ACE2 | SPATA3-AS1 | -0,6486 | 1,80021E-44 |
| ACE2 | RP11-92C4.3 | -0,6486 | 1,82296E-44 |
| ACE2 | CCDC70 | -0,6486 | 1,79418E-44 |
| ACE2 | FAHD2CP | -0,6487 | 1,69927E-44 |
| ACE2 | DISP3 | -0,6489 | 1,59036E-44 |
| ACE2 | AC079922.3 | -0,6489 | 1,56682E-44 |
| ACE2 | LINC01551 | -0,6489 | 1,56895E-44 |
| ACE2 | LRIT1 | -0,6494 | 1,3178E-44 |
| ACE2 | RP13-942N8.1 | -0,6494 | 1,30702E-44 |
| ACE2 | RDH16 | -0,6495 | 1,24137E-44 |
| ACE2 | PCP2 | -0,6495 | 1,23002E-44 |
| ACE2 | LENG1 | -0,6495 | 1,23037E-44 |
| ACE2 | TSACC | -0,6496 | 1,1877E-44 |
| ACE2 | LINC00384 | -0,6496 | 1,19295E-44 |
| ACE2 | CDIPT-AS1 | -0,6496 | 1,2091E-44 |
| ACE2 | C1orf228 | -0,6497 | 1,15625E-44 |
| ACE2 | WI2-2373I1.1 | -0,6497 | 1,16386E-44 |
| ACE2 | BPIFA3 | -0,6498 | 1,09432E-44 |
| ACE2 | TRAPPC5 | -0,6499 | 1,0465E-44 |
| ACE2 | SWI5 | -0,65 | 1,00531E-44 |
| ACE2 | LINC00694 | -0,6502 | 9,50837E-45 |
| ACE2 | CDRT15P1 | -0,6503 | 9,00781E-45 |
| ACE2 | GS1-124K5.4 | -0,6504 | 8,82526E-45 |
| ACE2 | SPACA4 | -0,6508 | 7,27956E-45 |
| ACE2 | ACTL9 | -0,651 | 6,8624E-45 |
| ACE2 | RP3-453P22.2 | -0,6511 | 6,64755E-45 |
| ACE2 | C10orf62 | -0,6512 | 6,36943E-45 |
| ACE2 | SMKR1 | -0,6513 | 6,05594E-45 |
| ACE2 | RP11-282I1.1 | -0,6514 | 5,75005E-45 |
| ACE2 | RHBDF1P1 | -0,6515 | 5,55544E-45 |
| ACE2 | RP4-816N1.8 | -0,6515 | 5,5403E-45 |
| ACE2 | AC002310.14 | -0,6515 | 5,47959E-45 |
| ACE2 | RP11-44N22.3 | -0,6516 | 5,34382E-45 |
| ACE2 | LINC01198 | -0,6517 | 5,07818E-45 |
| ACE2 | TMEM191B | -0,6519 | 4,61595E-45 |
| ACE2 | TEX44 | -0,652 | 4,52647E-45 |
| ACE2 | C19orf45 | -0,652 | 4,54317E-45 |
| ACE2 | CCDC24 | -0,6523 | 4,03462E-45 |
| ACE2 | RP11-552D8.1 | -0,6523 | 3,94237E-45 |
| ACE2 | ABCA17P | -0,6523 | 3,92794E-45 |
| ACE2 | BRWD1-AS2 | -0,6525 | 3,64058E-45 |
| ACE2 | CATSPER4 | -0,6526 | 3,50712E-45 |
| ACE2 | RP11-462G2.2 | -0,6526 | 3,5727E-45 |
| ACE2 | DPF3 | -0,6526 | 3,60114E-45 |
| ACE2 | LRFN4 | -0,6527 | 3,44123E-45 |
| ACE2 | SPTY2D1-AS1 | -0,6528 | 3,21863E-45 |
| ACE2 | KLF17 | -0,6529 | 3,14831E-45 |
| ACE2 | GFI1B | -0,653 | 2,95003E-45 |
| ACE2 | SPATA31D3 | -0,6531 | 2,9259E-45 |
| ACE2 | GTSF1L | -0,6531 | 2,85311E-45 |
| ACE2 | SMG7-AS1 | -0,6532 | 2,7496E-45 |
| ACE2 | CTD-2033D24.2 | -0,6534 | 2,54174E-45 |
| ACE2 | C11orf71 | -0,6536 | 2,33599E-45 |
| ACE2 | FAM209B | -0,6539 | 2,07909E-45 |

| | | | |
|---|---|---|---|
| ACE2 | TIPARP-AS1 | -0,6542 | 1,79326E-45 |
| ACE2 | RP11-881M11.1 | -0,6542 | 1,86187E-45 |
| ACE2 | CCDC166 | -0,6544 | 1,69674E-45 |
| ACE2 | ADAMTS17 | -0,6545 | 1,61743E-45 |
| ACE2 | IQCF1 | -0,6546 | 1,55766E-45 |
| ACE2 | SPATA31D1 | -0,6546 | 1,57296E-45 |
| ACE2 | LINC01121 | -0,6548 | 1,43831E-45 |
| ACE2 | RP11-661A12.8 | -0,655 | 1,31773E-45 |
| ACE2 | DPCD | -0,6552 | 1,21422E-45 |
| ACE2 | RP5-908M14.9 | -0,6552 | 1,23273E-45 |
| ACE2 | RP11-104H15.12 | -0,6553 | 1,1715E-45 |
| ACE2 | HSPB9 | -0,6554 | 1,09437E-45 |
| ACE2 | C9orf24 | -0,6555 | 1,05751E-45 |
| ACE2 | PPP1R32 | -0,6555 | 1,05033E-45 |
| ACE2 | RP11-810P12.7 | -0,6555 | 1,05325E-45 |
| ACE2 | THEG | -0,6555 | 1,08837E-45 |
| ACE2 | RP11-614F17.2 | -0,6557 | 1,00343E-45 |
| ACE2 | DKKL1 | -0,6558 | 9,26053E-46 |
| ACE2 | AC114812.5 | -0,6559 | 8,87748E-46 |
| ACE2 | GALNT8 | -0,6559 | 9,20052E-46 |
| ACE2 | TPPP3 | -0,656 | 8,82036E-46 |
| ACE2 | C17orf97 | -0,656 | 8,59323E-46 |
| ACE2 | TTC34 | -0,6561 | 8,4156E-46 |
| ACE2 | DLGAP2 | -0,6563 | 7,80237E-46 |
| ACE2 | RP11-1006G14.2 | -0,6565 | 6,93334E-46 |
| ACE2 | RP5-1022P6.7 | -0,6565 | 6,9217E-46 |
| ACE2 | C1orf61 | -0,6566 | 6,8674E-46 |
| ACE2 | DXO | -0,6566 | 6,91615E-46 |
| ACE2 | C12orf80 | -0,6569 | 5,90898E-46 |
| ACE2 | MTX1 | -0,657 | 5,74271E-46 |
| ACE2 | RUNDC3A | -0,6571 | 5,52045E-46 |
| ACE2 | ITFG1-AS1 | -0,6572 | 5,31702E-46 |
| ACE2 | WDR34 | -0,6573 | 5,11697E-46 |
| ACE2 | AC007249.3 | -0,6574 | 4,87149E-46 |
| ACE2 | CTD-2568A17.1 | -0,6575 | 4,62704E-46 |
| ACE2 | COL25A1 | -0,6576 | 4,49483E-46 |
| ACE2 | DNAJB8 | -0,6577 | 4,2026E-46 |
| ACE2 | RP11-14J7.6 | -0,6577 | 4,22873E-46 |
| ACE2 | RP11-114H24.3 | -0,6577 | 4,2888E-46 |
| ACE2 | KLHDC9 | -0,6579 | 3,87575E-46 |
| ACE2 | RP11-173A6.3 | -0,6579 | 3,96629E-46 |
| ACE2 | C7orf34 | -0,6579 | 3,91984E-46 |
| ACE2 | RP11-77H9.1 | -0,6579 | 3,89553E-46 |
| ACE2 | RP11-886P16.10 | -0,6581 | 3,69318E-46 |
| ACE2 | CDRT15P2 | -0,6582 | 3,45577E-46 |
| ACE2 | CTB-31O20.4 | -0,6582 | 3,50429E-46 |
| ACE2 | VWA3A | -0,6584 | 3,18246E-46 |
| ACE2 | TRPT1 | -0,6586 | 2,94472E-46 |
| ACE2 | NUDT1 | -0,659 | 2,52158E-46 |
| ACE2 | ZSWIM8-AS1 | -0,659 | 2,44058E-46 |
| ACE2 | PRR30 | -0,6595 | 1,99015E-46 |
| ACE2 | RP11-203H19.2 | -0,6595 | 2,05535E-46 |
| ACE2 | TEX40 | -0,6595 | 2,00942E-46 |
| ACE2 | TEPP | -0,6596 | 1,90033E-46 |

| | | | |
|---|---|---|---|
| ACE2 | TESK1 | -0,6597 | 1,85881E-46 |
| ACE2 | RP11-322N21.2 | -0,6598 | 1,75025E-46 |
| ACE2 | RSPH9 | -0,6598 | 1,74279E-46 |
| ACE2 | OAZ3 | -0,6599 | 1,73177E-46 |
| ACE2 | COL9A3 | -0,6601 | 1,53914E-46 |
| ACE2 | DSCR9 | -0,6602 | 1,48484E-46 |
| ACE2 | DNAL4 | -0,6602 | 1,52826E-46 |
| ACE2 | LINC00877 | -0,6608 | 1,15266E-46 |
| ACE2 | CCDC187 | -0,6608 | 1,18581E-46 |
| ACE2 | ISOC2 | -0,6609 | 1,12946E-46 |
| ACE2 | DERL3 | -0,6611 | 1,02531E-46 |
| ACE2 | RP1-313I6.12 | -0,6613 | 9,59812E-47 |
| ACE2 | PRSS46 | -0,6614 | 8,92134E-47 |
| ACE2 | AC015849.2 | -0,6614 | 8,99208E-47 |
| ACE2 | TXNL4A | -0,6614 | 8,85956E-47 |
| ACE2 | C5orf60 | -0,6617 | 7,80316E-47 |
| ACE2 | TSSC4 | -0,6617 | 8,09528E-47 |
| ACE2 | IZUMO2 | -0,6617 | 7,86729E-47 |
| ACE2 | FNDC11 | -0,6617 | 8,00394E-47 |
| ACE2 | SLC25A10 | -0,6619 | 7,34876E-47 |
| ACE2 | SUN5 | -0,6621 | 6,85866E-47 |
| ACE2 | C22orf42 | -0,6621 | 6,69473E-47 |
| ACE2 | TEX33 | -0,6621 | 6,60689E-47 |
| ACE2 | SLC2A8 | -0,6622 | 6,41065E-47 |
| ACE2 | DOC2A | -0,6622 | 6,40161E-47 |
| ACE2 | KCTD17 | -0,6626 | 5,42031E-47 |
| ACE2 | CHST13 | -0,6627 | 5,15336E-47 |
| ACE2 | C3orf18 | -0,6628 | 4,95346E-47 |
| ACE2 | OVOL1 | -0,6628 | 4,93408E-47 |
| ACE2 | LINC02219 | -0,6629 | 4,78809E-47 |
| ACE2 | SH3RF2 | -0,6633 | 4,10502E-47 |
| ACE2 | RP11-19P22.5 | -0,6636 | 3,56756E-47 |
| ACE2 | CFAP100 | -0,6637 | 3,39969E-47 |
| ACE2 | RP11-526D8.7 | -0,6639 | 3,17377E-47 |
| ACE2 | AC024619.2 | -0,664 | 2,92657E-47 |
| ACE2 | UBOX5-AS1 | -0,6642 | 2,75558E-47 |
| ACE2 | MAPRE3 | -0,6645 | 2,37276E-47 |
| ACE2 | MAD1L1 | -0,6645 | 2,37127E-47 |
| ACE2 | MSI1 | -0,6646 | 2,34772E-47 |
| ACE2 | ACTRT2 | -0,665 | 1,9801E-47 |
| ACE2 | TMEM91 | -0,6651 | 1,8795E-47 |
| ACE2 | CTD-2308N23.2 | -0,6652 | 1,79323E-47 |
| ACE2 | CTD-2201G3.1 | -0,6653 | 1,71334E-47 |
| ACE2 | RP11-758P17.3 | -0,6653 | 1,67673E-47 |
| ACE2 | SMIM2-IT1 | -0,6653 | 1,72469E-47 |
| ACE2 | CCDC159 | -0,6653 | 1,70505E-47 |
| ACE2 | LELP1 | -0,6654 | 1,64184E-47 |
| ACE2 | B3GALT4 | -0,6654 | 1,62872E-47 |
| ACE2 | DNAJB8-AS1 | -0,6657 | 1,41531E-47 |
| ACE2 | STKLD1 | -0,6662 | 1,16903E-47 |
| ACE2 | NDUFA3 | -0,6662 | 1,17543E-47 |
| ACE2 | TMPRSS6 | -0,6662 | 1,17899E-47 |
| ACE2 | CTD-2267D19.1 | -0,6663 | 1,09366E-47 |
| ACE2 | LINC00927 | -0,6665 | 1,02289E-47 |

| | | | |
|---|---|---|---|
| ACE2 | RP11-379C10.1 | -0,6667 | 9,43439E-48 |
| ACE2 | TMCO5B | -0,6667 | 9,28491E-48 |
| ACE2 | CSNK1G2 | -0,6667 | 9,55487E-48 |
| ACE2 | GOLGA2P6 | -0,6668 | 9,09169E-48 |
| ACE2 | FAM83F | -0,6668 | 9,14071E-48 |
| ACE2 | CCDC153 | -0,667 | 8,24096E-48 |
| ACE2 | RP11-342M1.7 | -0,6671 | 7,99118E-48 |
| ACE2 | SPACA9 | -0,6671 | 7,88946E-48 |
| ACE2 | CASC23 | -0,6672 | 7,48648E-48 |
| ACE2 | LINC00390 | -0,6673 | 7,30168E-48 |
| ACE2 | NELFE | -0,6676 | 6,37735E-48 |
| ACE2 | LRRC46 | -0,6677 | 5,94321E-48 |
| ACE2 | RP11-545G3.1 | -0,6682 | 4,92401E-48 |
| ACE2 | TBC1D3P2 | -0,6685 | 4,3352E-48 |
| ACE2 | CCDC157 | -0,6687 | 3,91395E-48 |
| ACE2 | IGHV1-24 | -0,6689 | 3,52646E-48 |
| ACE2 | CCDC129 | -0,6691 | 3,26987E-48 |
| ACE2 | MAPK15 | -0,6692 | 3,16105E-48 |
| ACE2 | LINC01356 | -0,6693 | 2,95949E-48 |
| ACE2 | LINC01940 | -0,6695 | 2,73727E-48 |
| ACE2 | PDXK | -0,6695 | 2,79319E-48 |
| ACE2 | RP11-402G3.5 | -0,6697 | 2,56287E-48 |
| ACE2 | CCDC27 | -0,67 | 2,23807E-48 |
| ACE2 | PRM2 | -0,6701 | 2,11105E-48 |
| ACE2 | RP11-311H10.4 | -0,6706 | 1,69412E-48 |
| ACE2 | RP11-452K12.4 | -0,6708 | 1,56756E-48 |
| ACE2 | SPATA41 | -0,6708 | 1,57874E-48 |
| ACE2 | TJP3 | -0,671 | 1,43967E-48 |
| ACE2 | SMCP | -0,6711 | 1,35318E-48 |
| ACE2 | C19orf70 | -0,6713 | 1,26995E-48 |
| ACE2 | LRRC56 | -0,6721 | 8,66603E-49 |
| ACE2 | TOR2A | -0,6723 | 8,16475E-49 |
| ACE2 | GPR137 | -0,6723 | 8,24787E-49 |
| ACE2 | STXBP5-AS1 | -0,6724 | 7,68253E-49 |
| ACE2 | PRKCDBP | -0,6724 | 7,77627E-49 |
| ACE2 | PHKG2 | -0,6725 | 7,35076E-49 |
| ACE2 | RP11-295G20.2 | -0,6726 | 7,08887E-49 |
| ACE2 | RP11-218M11.1 | -0,6726 | 6,96693E-49 |
| ACE2 | SPAG4 | -0,6726 | 7,21042E-49 |
| ACE2 | ZMYND10 | -0,6731 | 5,70913E-49 |
| ACE2 | CPTP | -0,6733 | 5,27285E-49 |
| ACE2 | TTLL10 | -0,6734 | 4,99073E-49 |
| ACE2 | ZMIZ1-AS1 | -0,6735 | 4,74838E-49 |
| ACE2 | ARHGEF2 | -0,6737 | 4,38579E-49 |
| ACE2 | ANKMY1 | -0,6737 | 4,44592E-49 |
| ACE2 | RP11-363E6.3 | -0,674 | 3,7376E-49 |
| ACE2 | SPATA31C2 | -0,6741 | 3,64075E-49 |
| ACE2 | PROCA1 | -0,6741 | 3,64122E-49 |
| ACE2 | TBC1D28 | -0,6743 | 3,35128E-49 |
| ACE2 | CREB3 | -0,6744 | 3,12383E-49 |
| ACE2 | DEF6 | -0,6746 | 2,97535E-49 |
| ACE2 | RP11-197N18.7 | -0,6746 | 2,87652E-49 |
| ACE2 | EFCAB8 | -0,6747 | 2,84444E-49 |
| ACE2 | FAM181A | -0,6751 | 2,37999E-49 |

| | | | |
|---|---|---|---|
| ACE2 | RP11-307F22.2 | -0,6752 | 2,18909E-49 |
| ACE2 | C17orf98 | -0,6752 | 2,26688E-49 |
| ACE2 | TNFAIP8L1 | -0,6755 | 1,97166E-49 |
| ACE2 | LINC02153 | -0,6763 | 1,33996E-49 |
| ACE2 | CIB1 | -0,6763 | 1,38094E-49 |
| ACE2 | SLC5A2 | -0,6764 | 1,31117E-49 |
| ACE2 | LRTOMT | -0,6765 | 1,23197E-49 |
| ACE2 | B9D1 | -0,6768 | 1,10366E-49 |
| ACE2 | LINC00917 | -0,6773 | 8,56149E-50 |
| ACE2 | PRRX2 | -0,6775 | 7,82434E-50 |
| ACE2 | CTD-2021A8.2 | -0,6776 | 7,47437E-50 |
| ACE2 | LINC00851 | -0,6777 | 7,11026E-50 |
| ACE2 | SSUH2 | -0,6779 | 6,48832E-50 |
| ACE2 | DUS1L | -0,678 | 6,45921E-50 |
| ACE2 | TMEM247 | -0,6782 | 5,83996E-50 |
| ACE2 | TMEM190 | -0,6788 | 4,39717E-50 |
| ACE2 | AC138647.1 | -0,6792 | 3,69672E-50 |
| ACE2 | C7orf71 | -0,6793 | 3,53273E-50 |
| ACE2 | NDUFS6 | -0,6799 | 2,72114E-50 |
| ACE2 | FBXW5 | -0,6802 | 2,29217E-50 |
| ACE2 | H2AFJ | -0,6805 | 2,07742E-50 |
| ACE2 | USP2-AS1 | -0,6808 | 1,77702E-50 |
| ACE2 | TSKS | -0,6809 | 1,66256E-50 |
| ACE2 | TEAD2 | -0,6811 | 1,55519E-50 |
| ACE2 | SPEM1 | -0,6816 | 1,20711E-50 |
| ACE2 | RP3-477O4.5 | -0,6819 | 1,09506E-50 |
| ACE2 | AQP5 | -0,682 | 1,03147E-50 |
| ACE2 | BSN-AS2 | -0,6823 | 8,99653E-51 |
| ACE2 | GSTZ1 | -0,6824 | 8,50443E-51 |
| ACE2 | ABHD1 | -0,6825 | 8,13396E-51 |
| ACE2 | C17orf74 | -0,6827 | 7,47901E-51 |
| ACE2 | SYCE3 | -0,6829 | 6,79924E-51 |
| ACE2 | PGP | -0,6832 | 5,86817E-51 |
| ACE2 | STPG3 | -0,6834 | 5,41174E-51 |
| ACE2 | RP11-24M17.4 | -0,6837 | 4,70111E-51 |
| ACE2 | RP11-146F11.1 | -0,6837 | 4,63415E-51 |
| ACE2 | C2orf81 | -0,6839 | 4,23752E-51 |
| ACE2 | LMO1 | -0,6842 | 3,63274E-51 |
| ACE2 | NDUFV2-AS1 | -0,6842 | 3,75006E-51 |
| ACE2 | C3orf84 | -0,6844 | 3,3199E-51 |
| ACE2 | GOLGA2P9 | -0,6846 | 3,12415E-51 |
| ACE2 | LINC01343 | -0,6847 | 2,92722E-51 |
| ACE2 | PTDSS2 | -0,6852 | 2,37009E-51 |
| ACE2 | CORT | -0,6854 | 2,08044E-51 |
| ACE2 | SPATA21 | -0,6857 | 1,84915E-51 |
| ACE2 | C1orf56 | -0,6857 | 1,83106E-51 |
| ACE2 | MCRIP2 | -0,6857 | 1,86948E-51 |
| ACE2 | MOSPD3 | -0,6858 | 1,77248E-51 |
| ACE2 | C20orf141 | -0,6871 | 9,70028E-52 |
| ACE2 | RUVBL2 | -0,6872 | 9,24396E-52 |
| ACE2 | JARID2-AS1 | -0,6877 | 7,38564E-52 |
| ACE2 | AZIN1-AS1 | -0,6881 | 5,94499E-52 |
| ACE2 | ZDHHC19 | -0,6882 | 5,61835E-52 |
| ACE2 | AC068831.6 | -0,6883 | 5,55224E-52 |

| | | | |
|---|---|---|---|
| ACE2 | LINC01658 | -0,6883 | 5,55634E-52 |
| ACE2 | C17orf50 | -0,6886 | 4,84202E-52 |
| ACE2 | TBC1D3P1 | -0,6886 | 4,84965E-52 |
| ACE2 | RNF151 | -0,6889 | 4,16374E-52 |
| ACE2 | LINC01191 | -0,6897 | 2,84119E-52 |
| ACE2 | AC116614.1 | -0,6899 | 2,51569E-52 |
| ACE2 | PPM1J | -0,6905 | 1,94827E-52 |
| ACE2 | POC1B-AS1 | -0,6906 | 1,82402E-52 |
| ACE2 | HPD | -0,6909 | 1,60303E-52 |
| ACE2 | AC006019.3 | -0,6911 | 1,47094E-52 |
| ACE2 | TCEA2 | -0,6911 | 1,46951E-52 |
| ACE2 | C7orf61 | -0,6913 | 1,34408E-52 |
| ACE2 | PCSK4 | -0,6913 | 1,2994E-52 |
| ACE2 | SLC25A28 | -0,6916 | 1,16913E-52 |
| ACE2 | OXLD1 | -0,692 | 9,64038E-53 |
| ACE2 | ARL4D | -0,6921 | 9,14229E-53 |
| ACE2 | LINC00264 | -0,6922 | 8,48452E-53 |
| ACE2 | SMPD2 | -0,6924 | 7,92953E-53 |
| ACE2 | C1orf159 | -0,6927 | 6,59958E-53 |
| ACE2 | SMIM6 | -0,6927 | 6,7845E-53 |
| ACE2 | TOGARAM2 | -0,693 | 5,8087E-53 |
| ACE2 | ACTL10 | -0,6933 | 5,11231E-53 |
| ACE2 | PLEKHO1 | -0,6934 | 4,80247E-53 |
| ACE2 | FAM71E1 | -0,6934 | 4,82511E-53 |
| ACE2 | MSTO1 | -0,6935 | 4,5366E-53 |
| ACE2 | NUPR2 | -0,6935 | 4,49862E-53 |
| ACE2 | RP11-727F15.14 | -0,6935 | 4,68965E-53 |
| ACE2 | RP11-831A10.2 | -0,6935 | 4,50131E-53 |
| ACE2 | IDO2 | -0,6937 | 4,15498E-53 |
| ACE2 | CA9 | -0,6938 | 3,99207E-53 |
| ACE2 | TSSK3 | -0,694 | 3,55485E-53 |
| ACE2 | TEX38 | -0,6942 | 3,27553E-53 |
| ACE2 | RP5-1023B21.1 | -0,6946 | 2,71768E-53 |
| ACE2 | IZUMO4 | -0,6948 | 2,45482E-53 |
| ACE2 | AC105339.1 | -0,695 | 2,2323E-53 |
| ACE2 | PICK1 | -0,6951 | 2,16983E-53 |
| ACE2 | SSSCA1 | -0,6957 | 1,61948E-53 |
| ACE2 | UBXN11 | -0,696 | 1,34409E-53 |
| ACE2 | C16orf92 | -0,6965 | 1,05138E-53 |
| ACE2 | IQCF3 | -0,6966 | 1,01292E-53 |
| ACE2 | ATPAF2 | -0,6968 | 9,41264E-54 |
| ACE2 | CCDC188 | -0,6969 | 8,8173E-54 |
| ACE2 | C16orf86 | -0,697 | 8,64692E-54 |
| ACE2 | CLDND2 | -0,6979 | 5,56002E-54 |
| ACE2 | MARCH10 | -0,698 | 5,30173E-54 |
| ACE2 | FUZ | -0,6983 | 4,42633E-54 |
| ACE2 | NDUFAF3 | -0,6989 | 3,39157E-54 |
| ACE2 | LINC00202-2 | -0,6997 | 2,26315E-54 |
| ACE2 | MORN5 | -0,7002 | 1,78783E-54 |
| ACE2 | TMEM89 | -0,7005 | 1,51376E-54 |
| ACE2 | OR7E156P | -0,7006 | 1,45951E-54 |
| ACE2 | YBX2 | -0,7006 | 1,4634E-54 |
| ACE2 | PTRH1 | -0,7008 | 1,28883E-54 |
| ACE2 | ATG4D | -0,7028 | 4,75156E-55 |

| | | | |
|---|---|---|---|
| ACE2 | POLR2I | -0,7029 | 4,47128E-55 |
| ACE2 | IBA57-AS1 | -0,7032 | 3,92582E-55 |
| ACE2 | GRINA | -0,7033 | 3,7531E-55 |
| ACE2 | FAM173A | -0,7034 | 3,50748E-55 |
| ACE2 | TRIM17 | -0,7043 | 2,26667E-55 |
| ACE2 | CHCHD5 | -0,705 | 1,59479E-55 |
| ACE2 | CCDC182 | -0,7055 | 1,23911E-55 |
| ACE2 | TBC1D26 | -0,7062 | 8,53174E-56 |
| ACE2 | PRR22 | -0,7068 | 6,39629E-56 |
| ACE2 | C19orf71 | -0,7069 | 6,18083E-56 |
| ACE2 | IRGC | -0,707 | 5,73946E-56 |
| ACE2 | AMDHD2 | -0,7074 | 4,74934E-56 |
| ACE2 | SURF2 | -0,7102 | 1,11786E-56 |
| ACE2 | RP11-326A19.3 | -0,7105 | 9,44347E-57 |
| ACE2 | RP11-280O1.2 | -0,7109 | 7,90579E-57 |
| ACE2 | LRRC73 | -0,7117 | 5,03592E-57 |
| ACE2 | TMEM239 | -0,7124 | 3,52212E-57 |
| ACE2 | CDRT15L2 | -0,7128 | 2,95986E-57 |
| ACE2 | TEX22 | -0,7134 | 2,08001E-57 |
| ACE2 | SPATA33 | -0,7134 | 2,16243E-57 |
| ACE2 | LYL1 | -0,7135 | 2,01226E-57 |
| ACE2 | AC026471.6 | -0,714 | 1,59352E-57 |
| ACE2 | MAP2K2 | -0,714 | 1,56095E-57 |
| ACE2 | RP1-149M18.3 | -0,7147 | 1,10084E-57 |
| ACE2 | CAPN3 | -0,7154 | 7,3199E-58 |
| ACE2 | DUSP15 | -0,7159 | 5,78244E-58 |
| ACE2 | CTC-479C5.6 | -0,7167 | 3,8225E-58 |
| ACE2 | RP11-458F8.4 | -0,7173 | 2,74397E-58 |
| ACE2 | ELOF1 | -0,7182 | 1,70901E-58 |
| ACE2 | TAF10 | -0,72 | 6,28328E-59 |
| ACE2 | FAM229A | -0,7209 | 3,93841E-59 |
| ACE2 | TMEM210 | -0,7209 | 3,96963E-59 |
| ACE2 | RP11-266L9.8 | -0,7214 | 3,00108E-59 |
| ACE2 | UBL4B | -0,7215 | 2,89983E-59 |
| ACE2 | SHARPIN | -0,7235 | 9,37909E-60 |
| ACE2 | PEX11G | -0,7238 | 8,22216E-60 |
| ACE2 | H1FNT | -0,7242 | 6,47367E-60 |
| ACE2 | HILS1 | -0,7255 | 3,18459E-60 |
| ACE2 | NOSIP | -0,7265 | 1,88157E-60 |
| ACE2 | ALKBH7 | -0,7281 | 7,62989E-61 |
| ACE2 | DNAJC4 | -0,7283 | 6,98832E-61 |
| ACE2 | SPACA3 | -0,7285 | 6,16131E-61 |
| ACE2 | KRT86 | -0,7286 | 5,70875E-61 |
| ACE2 | PRRX2-AS1 | -0,7298 | 2,97073E-61 |
| ACE2 | PCYT2 | -0,7303 | 2,17398E-61 |
| ACE2 | FAAH | -0,7307 | 1,7222E-61 |
| ACE2 | NAA38 | -0,7307 | 1,73638E-61 |
| ACE2 | C11orf94 | -0,7328 | 5,47079E-62 |
| ACE2 | ZNHIT2 | -0,7338 | 3,12013E-62 |
| ACE2 | RBAKDN | -0,7346 | 1,88781E-62 |
| ACE2 | STARD10 | -0,7359 | 8,96596E-63 |
| ACE2 | AC008694.3 | -0,7373 | 4,11375E-63 |
| ACE2 | SLC25A41 | -0,7418 | 2,90032E-64 |
| ACE2 | GPX4 | -0,7432 | 1,26852E-64 |

| | | | |
|---|---|---|---|
| ACE2 | TMEM191C | -0,7442 | 7,06166E-65 |
| ACE2 | WBSCR28 | -0,7446 | 5,40527E-65 |
| ACE2 | RP11-195F19.5 | -0,7463 | 1,87605E-65 |
| ACE2 | EXOSC4 | -0,7464 | 1,85209E-65 |
| ACE2 | PQLC1 | -0,7466 | 1,56596E-65 |
| ACE2 | C16orf90 | -0,7503 | 1,64278E-66 |
| ACE2 | AURKAIP1 | -0,7645 | 1,85839E-70 |
