## Supplementary table 6 lung for "Bioinformatic characterization of angiotensin-converting enzyme 2, the entry receptor for SARS-CoV-2"

| | source | native | name | p_value | query | significar | term_size | query_size | intersection_si | effective_domain_si | precisior | recall | parents |
|---|---|---|---|---|---|---|---|---|---|---|---|---|---|
| 0 | KEGG | KEGG:04614 | Renin-angiotensin system | 0,045858097 | positive_corr | True | 23 | 1 | 1 | 7788 | 1 | 0,043478261 | ['KEGG:00000'] |
