## Supplementary table 6 intestine for "Bioinformatic characterization of angiotensin-converting enzyme 2, the entry receptor for SARS-CoV-2"

| | source | native | name | p_value | significar | query | term_size | query_size | intersection_si | effective_domain_si | precision | recall | parents |
|---|---|---|---|---|---|---|---|---|---|---|---|---|---|
| 1184 | GO:BP | GO:0006629 | lipid metabolic process | 1,09029E-32 | True | positive_corr | 1414 | 1037 | 204 | 17906 | 0,196721311 | 0,14427157 | ['GO:0044238', 'GO:0071704'] |
| 1454 | GO:BP | GO:0044281 | small molecule metabolic process | 1,24794E-31 | True | positive_corr | 1987 | 1037 | 251 | 17906 | 0,242044359 | 0,126321087 | ['GO:0008152'] |
| 1024 | GO:BP | GO:0006082 | organic acid metabolic process | 4,91712E-24 | True | positive_corr | 1177 | 1037 | 165 | 17906 | 0,159112825 | 0,140186916 | ['GO:0044237', 'GO:0044281', 'GO:0071704'] |
| 1019 | GO:BP | GO:0044255 | cellular lipid metabolic process | 2,1196E-23 | True | positive_corr | 1070 | 1037 | 154 | 17906 | 0,148505304 | 0,143925234 | ['GO:0006629', 'GO:0044237'] |
| 1016 | GO:BP | GO:0043436 | oxoacid metabolic process | 1,59456E-22 | True | positive_corr | 1157 | 1037 | 160 | 17906 | 0,154291225 | 0,138288678 | ['GO:0006082'] |
| 1455 | GO:BP | GO:0006820 | anion transport | 1,07636E-20 | True | positive_corr | 654 | 1037 | 109 | 17906 | 0,105110897 | 0,166666667 | ['GO:0006811'] |
| 1456 | GO:BP | GO:0006805 | xenobiotic metabolic process | 1,27263E-20 | True | positive_corr | 127 | 1037 | 45 | 17906 | 0,043394407 | 0,354330709 | ['GO:0044237', 'GO:0071466'] |
| 1458 | GO:BP | GO:0044242 | cellular lipid catabolic process | 2,8506E-20 | True | positive_corr | 215 | 1037 | 58 | 17906 | 0,055930569 | 0,269767442 | ['GO:0016042', 'GO:0044248', 'GO:0044255'] |
| 1465 | GO:BP | GO:0019752 | carboxylic acid metabolic process | 2,92346E-20 | True | positive_corr | 1066 | 1037 | 147 | 17906 | 0,141755063 | 0,137898687 | ['GO:0043436'] |
| 1009 | GO:BP | GO:0032787 | monocarboxylic acid metabolic process | 9,07954E-20 | True | positive_corr | 661 | 1037 | 108 | 17906 | 0,104146577 | 0,163388805 | ['GO:0019752'] |
| 1028 | GO:BP | GO:0016042 | lipid catabolic process | 4,24582E-19 | True | positive_corr | 337 | 1037 | 72 | 17906 | 0,069431051 | 0,213649852 | ['GO:0006629', 'GO:1901575'] |
| 1472 | GO:BP | GO:0006811 | ion transport | 1,44278E-18 | True | positive_corr | 1723 | 1037 | 198 | 17906 | 0,190935391 | 0,114915844 | ['GO:0006810'] |
| 999 | GO:BP | GO:0015711 | organic anion transport | 2,57933E-17 | True | positive_corr | 508 | 1037 | 88 | 17906 | 0,084860174 | 0,173228346 | ['GO:0006820', 'GO:0071702'] |
| 997 | GO:BP | GO:0007586 | digestion | 5,66746E-17 | True | positive_corr | 139 | 1037 | 43 | 17906 | 0,041465767 | 0,309352518 | ['GO:0032501'] |
| 1488 | GO:BP | GO:0071466 | cellular response to xenobiotic stimulus | 1,61105E-16 | True | positive_corr | 184 | 1037 | 49 | 17906 | 0,047251688 | 0,266304348 | ['GO:0009410', 'GO:0070887'] |
| 1493 | GO:BP | GO:0044282 | small molecule catabolic process | 7,98178E-16 | True | positive_corr | 447 | 1037 | 79 | 17906 | 0,076181292 | 0,176733781 | ['GO:0009056', 'GO:0044281'] |
| 992 | GO:BP | GO:0071840 | cellular component organization or biogenesis | 4,53775E-15 | True | negative_corr | 6667 | 2460 | 1112 | 17906 | 0,45203252 | 0,16679166 | ['GO:0008150'] |
| 1494 | GO:BP | GO:0008202 | steroid metabolic process | 6,25904E-15 | True | positive_corr | 332 | 1037 | 65 | 17906 | 0,06268081 | 0,195783133 | ['GO:0006629', 'GO:1901360'] |
| 1496 | GO:BP | GO:0030030 | cell projection organization | 6,31859E-15 | True | negative_corr | 1578 | 2460 | 337 | 17906 | 0,13699187 | 0,21356147 | ['GO:0016043'] |
| 982 | GO:BP | GO:0120036 | plasma membrane bounded cell projection organization | 8,05706E-15 | True | negative_corr | 1539 | 2460 | 330 | 17906 | 0,134146341 | 0,214424951 | ['GO:0030030'] |
| 981 | GO:BP | GO:0005975 | carbohydrate metabolic process | 1,13186E-13 | True | positive_corr | 640 | 1037 | 94 | 17906 | 0,090646095 | 0,146875 | ['GO:0044238', 'GO:0071704'] |
| 1000 | GO:BP | GO:0006631 | fatty acid metabolic process | 1,40011E-13 | True | positive_corr | 380 | 1037 | 68 | 17906 | 0,06557377 | 0,178947368 | ['GO:0032787', 'GO:0044255'] |
| 1497 | GO:BP | GO:0009653 | anatomical structure morphogenesis | 1,60507E-13 | True | negative_corr | 2820 | 2460 | 532 | 17906 | 0,216260163 | 0,188652482 | ['GO:0032502', 'GO:0048856'] |
| 1451 | GO:BP | GO:0006066 | alcohol metabolic process | 9,69653E-13 | True | positive_corr | 375 | 1037 | 66 | 17906 | 0,06364513 | 0,176 | ['GO:0044281', 'GO:1901615'] |
| 1031 | GO:BP | GO:0010817 | regulation of hormone levels | 2,32891E-12 | True | positive_corr | 542 | 1037 | 82 | 17906 | 0,079074253 | 0,151291513 | ['GO:0065008'] |
| 1417 | GO:BP | GO:1901615 | organic hydroxy compound metabolic process | 2,46717E-12 | True | positive_corr | 553 | 1037 | 83 | 17906 | 0,080038573 | 0,150090416 | ['GO:0071704'] |
| 1419 | GO:BP | GO:0016043 | cellular component organization | 4,42708E-12 | True | negative_corr | 6481 | 2460 | 1067 | 17906 | 0,433739837 | 0,164635087 | ['GO:0009987', 'GO:0071840'] |
| 1425 | GO:BP | GO:0050892 | intestinal absorption | 5,18772E-12 | True | positive_corr | 38 | 1037 | 20 | 17906 | 0,019286403 | 0,526315789 | ['GO:0006810', 'GO:0022600'] |
| 1426 | GO:BP | GO:0008610 | lipid biosynthetic process | 1,28675E-11 | True | positive_corr | 711 | 1037 | 96 | 17906 | 0,092574735 | 0,135021097 | ['GO:0006629', 'GO:1901576'] |
| 1061 | GO:BP | GO:0006413 | translational initiation | 1,56041E-11 | True | negative_corr | 196 | 2460 | 70 | 17906 | 0,028455285 | 0,357142857 | ['GO:0006412', 'GO:0044237'] |
| 1060 | GO:BP | GO:0044085 | cellular component biogenesis | 3,28801E-11 | True | negative_corr | 3241 | 2460 | 585 | 17906 | 0,237804878 | 0,180499846 | ['GO:0071840'] |
| 1427 | GO:BP | GO:0055085 | transmembrane transport | 6,78637E-11 | True | positive_corr | 1634 | 1037 | 169 | 17906 | 0,162970106 | 0,103427173 | ['GO:0006810'] |
| 1428 | GO:BP | GO:0072329 | monocarboxylic acid catabolic process | 1,44872E-10 | True | positive_corr | 130 | 1037 | 34 | 17906 | 0,032786885 | 0,261538462 | ['GO:0032787', 'GO:0046395'] |
| 1437 | GO:BP | GO:0042445 | hormone metabolic process | 1,55791E-10 | True | positive_corr | 244 | 1037 | 48 | 17906 | 0,046287367 | 0,196721311 | ['GO:0008152', 'GO:0010817'] |
| 1030 | GO:BP | GO:0046942 | carboxylic acid transport | 2,56745E-10 | True | positive_corr | 340 | 1037 | 58 | 17906 | 0,055930569 | 0,170588235 | ['GO:0015711', 'GO:0015849'] |
| 1053 | GO:BP | GO:0120031 | plasma membrane bounded cell projection assembly | 2,65923E-10 | True | negative_corr | 562 | 2460 | 142 | 17906 | 0,057723577 | 0,252669039 | ['GO:0030031', 'GO:0120036'] |
| 1440 | GO:BP | GO:0015849 | organic acid transport | 3,33098E-10 | True | positive_corr | 342 | 1037 | 58 | 17906 | 0,055930569 | 0,169590643 | ['GO:0071702'] |
| 1442 | GO:BP | GO:0060271 | cilium assembly | 4,42185E-10 | True | negative_corr | 369 | 2460 | 104 | 17906 | 0,042276423 | 0,281842818 | ['GO:0044782', 'GO:0070925', 'GO:0120031'] |
| 1044 | GO:BP | GO:0022600 | digestive system process | 5,87686E-10 | True | positive_corr | 100 | 1037 | 29 | 17906 | 0,027965284 | 0,29 | ['GO:0003008', 'GO:0007586'] |
| 1043 | GO:BP | GO:0044283 | small molecule biosynthetic process | 7,26284E-10 | True | positive_corr | 724 | 1037 | 93 | 17906 | 0,089681774 | 0,128453039 | ['GO:0009058', 'GO:0044281'] |
| 1042 | GO:BP | GO:0044782 | cilium organization | 7,67532E-10 | True | negative_corr | 387 | 2460 | 107 | 17906 | 0,043495935 | 0,276485788 | ['GO:0006996', 'GO:0120036'] |
| 1041 | GO:BP | GO:0030031 | cell projection assembly | 8,18395E-10 | True | negative_corr | 575 | 2460 | 143 | 17906 | 0,058130081 | 0,248695652 | ['GO:0022607', 'GO:0030030'] |
| 1448 | GO:BP | GO:0007399 | nervous system development | 9,36317E-10 | True | negative_corr | 2432 | 2460 | 452 | 17906 | 0,183739837 | 0,185855263 | ['GO:0048731'] |
| 1449 | GO:BP | GO:0045047 | protein targeting to ER | 1,05414E-09 | True | negative_corr | 110 | 2460 | 46 | 17906 | 0,018699187 | 0,418181818 | ['GO:0006605', 'GO:0072599'] |
| 1450 | GO:BP | GO:0006614 | SRP-dependent cotranslational protein targeting to membrane | 1,75833E-09 | True | negative_corr | 96 | 2460 | 42 | 17906 | 0,017073171 | 0,4375 | ['GO:0006613', 'GO:0045047'] |
| 1050 | GO:BP | GO:0006721 | terpenoid metabolic process | 2,11622E-09 | True | positive_corr | 119 | 1037 | 31 | 17906 | 0,029893925 | 0,260504202 | ['GO:0006720'] |
| 975 | GO:BP | GO:0009062 | fatty acid catabolic process | 2,32188E-09 | True | positive_corr | 105 | 1037 | 29 | 17906 | 0,027965284 | 0,276190476 | ['GO:0006631', 'GO:0044242', 'GO:0072329'] |
| 974 | GO:BP | GO:0007224 | smoothened signaling pathway | 2,8343E-09 | True | negative_corr | 145 | 2460 | 54 | 17906 | 0,02195122 | 0,372413793 | ['GO:0007166'] |
| 973 | GO:BP | GO:0006613 | cotranslational protein targeting to membrane | 2,92656E-09 | True | negative_corr | 101 | 2460 | 43 | 17906 | 0,017479675 | 0,425742574 | ['GO:0006612'] |
| 1528 | GO:BP | GO:0072599 | establishment of protein localization to endoplasmic reticulum | 4,90272E-09 | True | negative_corr | 114 | 2460 | 46 | 17906 | 0,018699187 | 0,403508772 | ['GO:0070972', 'GO:0072594'] |
| 909 | GO:BP | GO:0044260 | cellular macromolecule metabolic process | 5,53226E-09 | True | negative_corr | 8429 | 2460 | 1318 | 17906 | 0,535772358 | 0,156364931 | ['GO:0043170', 'GO:0044237'] |
| 908 | GO:BP | GO:0055114 | oxidation-reduction process | 5,54073E-09 | True | positive_corr | 1028 | 1037 | 116 | 17906 | 0,111861138 | 0,112840467 | ['GO:0008152'] |
| 907 | GO:BP | GO:0006720 | isoprenoid metabolic process | 7,97188E-09 | True | positive_corr | 140 | 1037 | 33 | 17906 | 0,031822565 | 0,235714286 | ['GO:0044255'] |
| 902 | GO:BP | GO:0009410 | response to xenobiotic stimulus | 8,25655E-09 | True | positive_corr | 299 | 1037 | 51 | 17906 | 0,049180328 | 0,170568562 | ['GO:0042221'] |
| 1529 | GO:BP | GO:0005996 | monosaccharide metabolic process | 1,07068E-08 | True | positive_corr | 301 | 1037 | 51 | 17906 | 0,049180328 | 0,169435216 | ['GO:0005975', 'GO:0044281'] |
| 897 | GO:BP | GO:0016070 | RNA metabolic process | 1,15218E-08 | True | negative_corr | 4714 | 2460 | 789 | 17906 | 0,320731707 | 0,16737378 | ['GO:0090304'] |
| 1545 | GO:BP | GO:0000902 | cell morphogenesis | 2,16722E-08 | True | negative_corr | 1068 | 2460 | 225 | 17906 | 0,091463415 | 0,210674157 | ['GO:0032989'] |
| 888 | GO:BP | GO:0034754 | cellular hormone metabolic process | 2,63537E-08 | True | positive_corr | 130 | 1037 | 31 | 17906 | 0,029893925 | 0,238461538 | ['GO:0042445', 'GO:0044237'] |
| 919 | GO:BP | GO:0022607 | cellular component assembly | 2,9534E-08 | True | negative_corr | 3000 | 2460 | 531 | 17906 | 0,215853659 | 0,177 | ['GO:0016043', 'GO:0044085'] |
| 885 | GO:BP | GO:0051641 | cellular localization | 3,04717E-08 | True | negative_corr | 2889 | 2460 | 514 | 17906 | 0,208943089 | 0,177916234 | ['GO:0051179'] |
| 1546 | GO:BP | GO:0017144 | drug metabolic process | 3,75465E-08 | True | positive_corr | 600 | 1037 | 78 | 17906 | 0,075216972 | 0,13 | ['GO:0044237'] |
| 882 | GO:BP | GO:0016101 | diterpenoid metabolic process | 3,92516E-08 | True | positive_corr | 109 | 1037 | 28 | 17906 | 0,027000964 | 0,256880734 | ['GO:0006721'] |
| 877 | GO:BP | GO:0032989 | cellular component morphogenesis | 4,45808E-08 | True | negative_corr | 1177 | 2460 | 242 | 17906 | 0,098373984 | 0,205607477 | ['GO:0009653', 'GO:0016043', 'GO:0048869'] |
| 875 | GO:BP | GO:0007155 | cell adhesion | 5,38391E-08 | True | negative_corr | 1440 | 2460 | 285 | 17906 | 0,115853659 | 0,197916667 | ['GO:0022610'] |

| | | | | | | | | | | | | | |
|---|---|---|---|---|---|---|---|---|---|---|---|---|---|
| 1279 | GO:BP | GO:0001523 | retinoid metabolic process | 5,43931E-08 | True | positive_corr | 103 | 1037 | 27 | 17906 | 0,026036644 | 0,262135922 | ['GO:0016101'] |
| 1549 | GO:BP | GO:0022610 | biological adhesion | 5,77115E-08 | True | negative_corr | 1447 | 2460 | 286 | 17906 | 0,116260163 | 0,197650311 | ['GO:0008150'] |
| 868 | GO:BP | GO:0046395 | carboxylic acid catabolic process | 6,30133E-08 | True | positive_corr | 276 | 1037 | 47 | 17906 | 0,045323047 | 0,170289855 | ['GO:0016054', 'GO:0019752'] |
| 1552 | GO:BP | GO:0016054 | organic acid catabolic process | 6,30133E-08 | True | positive_corr | 276 | 1037 | 47 | 17906 | 0,045323047 | 0,170289855 | ['GO:0006082', 'GO:0044248', 'GO:0044282', 'GO:1901575'] |
| 1554 | GO:BP | GO:0090304 | nucleic acid metabolic process | 7,16419E-08 | True | negative_corr | 5207 | 2460 | 855 | 17906 | 0,347560976 | 0,164202036 | ['GO:0006139', 'GO:0043170'] |
| 884 | GO:BP | GO:0000184 | nuclear-transcribed mRNA catabolic process, nonsense-mediated dec | 8,0697E-08 | True | negative_corr | 122 | 2460 | 46 | 17906 | 0,018699187 | 0,37704918 | ['GO:0000956'] |
| 1526 | GO:BP | GO:0070972 | protein localization to endoplasmic reticulum | 8,49851E-08 | True | negative_corr | 139 | 2460 | 50 | 17906 | 0,020325203 | 0,35971223 | ['GO:0033365'] |
| 1522 | GO:BP | GO:0006810 | transport | 9,5773E-08 | True | positive_corr | 5337 | 1037 | 405 | 17906 | 0,390549662 | 0,075885329 | ['GO:0051234'] |
| 922 | GO:BP | GO:0000904 | cell morphogenesis involved in differentiation | 1,03806E-07 | True | negative_corr | 769 | 2460 | 171 | 17906 | 0,069512195 | 0,22236671 | ['GO:0000902', 'GO:0048468'] |
| 972 | GO:BP | GO:0070727 | cellular macromolecule localization | 1,16851E-07 | True | negative_corr | 1994 | 2460 | 372 | 17906 | 0,151219512 | 0,186559679 | ['GO:0033036', 'GO:0051641'] |
| 1500 | GO:BP | GO:0034613 | cellular protein localization | 1,30923E-07 | True | negative_corr | 1983 | 2460 | 370 | 17906 | 0,150406504 | 0,186585981 | ['GO:0008104', 'GO:0070727'] |
| 1503 | GO:BP | GO:0001676 | long-chain fatty acid metabolic process | 1,41939E-07 | True | positive_corr | 107 | 1037 | 27 | 17906 | 0,026036644 | 0,252336449 | ['GO:0006631'] |
| 1504 | GO:BP | GO:0051128 | regulation of cellular component organization | 1,82008E-07 | True | negative_corr | 2541 | 2460 | 456 | 17906 | 0,185365854 | 0,179456907 | ['GO:0016043', 'GO:0050794'] |
| 1505 | GO:BP | GO:0055088 | lipid homeostasis | 1,8231E-07 | True | positive_corr | 156 | 1037 | 33 | 17906 | 0,031822565 | 0,211538462 | ['GO:0048878'] |
| 1507 | GO:BP | GO:0022008 | neurogenesis | 1,82356E-07 | True | negative_corr | 1679 | 2460 | 321 | 17906 | 0,130487805 | 0,191185229 | ['GO:0007399', 'GO:0030154'] |
| 1511 | GO:BP | GO:0048731 | system development | 2,41656E-07 | True | negative_corr | 5020 | 2460 | 824 | 17906 | 0,33495935 | 0,164143426 | ['GO:0007275', 'GO:0048856'] |
| 963 | GO:BP | GO:0032530 | regulation of microvillus organization | 2,75596E-07 | True | positive_corr | 13 | 1037 | 10 | 17906 | 0,009643202 | 0,769230769 | ['GO:0032528', 'GO:0120035'] |
| 962 | GO:BP | GO:0019585 | glucuronate metabolic process | 2,92189E-07 | True | positive_corr | 24 | 1037 | 13 | 17906 | 0,012536162 | 0,541666667 | ['GO:0006063'] |
| 1512 | GO:BP | GO:0006063 | uronic acid metabolic process | 2,92189E-07 | True | positive_corr | 24 | 1037 | 13 | 17906 | 0,012536162 | 0,541666667 | ['GO:0005996', 'GO:0032787'] |
| 953 | GO:BP | GO:0032528 | microvillus organization | 2,92189E-07 | True | positive_corr | 24 | 1037 | 13 | 17906 | 0,012536162 | 0,541666667 | ['GO:0120036'] |
| 948 | GO:BP | GO:0048468 | cell development | 3,38762E-07 | True | negative_corr | 2215 | 2460 | 404 | 17906 | 0,164227642 | 0,182392777 | ['GO:0030154', 'GO:0048856', 'GO:0048869'] |
| 947 | GO:BP | GO:0007275 | multicellular organism development | 3,44549E-07 | True | negative_corr | 5604 | 2460 | 907 | 17906 | 0,368699187 | 0,16184868 | ['GO:0032501', 'GO:0048856'] |
| 945 | GO:BP | GO:0030182 | neuron differentiation | 3,65697E-07 | True | negative_corr | 1409 | 2460 | 276 | 17906 | 0,112195122 | 0,195883605 | ['GO:0030154', 'GO:0048699'] |
| 1519 | GO:BP | GO:0051234 | establishment of localization | 4,71418E-07 | True | positive_corr | 5460 | 1037 | 409 | 17906 | 0,394406943 | 0,074908425 | ['GO:0051179'] |
| 939 | GO:BP | GO:0034308 | primary alcohol metabolic process | 4,82805E-07 | True | positive_corr | 90 | 1037 | 24 | 17906 | 0,023143684 | 0,266666667 | ['GO:0006066'] |
| 938 | GO:BP | GO:0007017 | microtubule-based process | 5,05612E-07 | True | negative_corr | 778 | 2460 | 170 | 17906 | 0,069105691 | 0,218508997 | ['GO:0009987'] |
| 937 | GO:BP | GO:0071702 | organic substance transport | 6,89869E-07 | True | positive_corr | 2937 | 1037 | 246 | 17906 | 0,237222758 | 0,083758938 | ['GO:0006810'] |
| 935 | GO:BP | GO:0006641 | triglyceride metabolic process | 7,9388E-07 | True | positive_corr | 107 | 1037 | 26 | 17906 | 0,025072324 | 0,242990654 | ['GO:0006639'] |
| 931 | GO:BP | GO:0006690 | icosanoid metabolic process | 8,35493E-07 | True | positive_corr | 115 | 1037 | 27 | 17906 | 0,026036644 | 0,234782609 | ['GO:0019752', 'GO:1901568'] |
| 925 | GO:BP | GO:0070925 | organelle assembly | 8,46841E-07 | True | negative_corr | 847 | 2460 | 181 | 17906 | 0,073577236 | 0,213695396 | ['GO:0006996', 'GO:0022607'] |
| 1074 | GO:BP | GO:0048699 | generation of neurons | 1,19966E-06 | True | negative_corr | 1574 | 2460 | 300 | 17906 | 0,12195122 | 0,190597205 | ['GO:0022008'] |
| 1561 | GO:BP | GO:0006139 | nucleobase-containing compound metabolic process | 1,42971E-06 | True | negative_corr | 5736 | 2460 | 921 | 17906 | 0,374390244 | 0,160564854 | ['GO:0006725', 'GO:0034641', 'GO:0044238', 'GO:0046483', 'GO:1901360'] |
| 1414 | GO:BP | GO:0019080 | viral gene expression | 1,47219E-06 | True | negative_corr | 195 | 2460 | 60 | 17906 | 0,024390244 | 0,307692308 | ['GO:0016032'] |
| 1408 | GO:BP | GO:0032502 | developmental process | 1,8638E-06 | True | negative_corr | 6630 | 2460 | 1047 | 17906 | 0,425609756 | 0,157918552 | ['GO:0008150'] |
| 1298 | GO:BP | GO:0035295 | tube development | 2,0225E-06 | True | negative_corr | 1143 | 2460 | 229 | 17906 | 0,093089431 | 0,200349956 | ['GO:0007275', 'GO:0048856'] |
| 1299 | GO:BP | GO:0046394 | carboxylic acid biosynthetic process | 2,14882E-06 | True | positive_corr | 347 | 1037 | 51 | 17906 | 0,049180328 | 0,146974063 | ['GO:0016053', 'GO:0019752'] |
| 1213 | GO:BP | GO:0007010 | cytoskeleton organization | 2,22619E-06 | True | negative_corr | 1371 | 2460 | 266 | 17906 | 0,108130081 | 0,194018964 | ['GO:0006996'] |
| 1212 | GO:BP | GO:0033365 | protein localization to organelle | 2,3106E-06 | True | negative_corr | 964 | 2460 | 199 | 17906 | 0,080894309 | 0,206431535 | ['GO:0034613'] |
| 1211 | GO:BP | GO:0009790 | embryo development | 2,32629E-06 | True | negative_corr | 1060 | 2460 | 215 | 17906 | 0,087398374 | 0,202830189 | ['GO:0007275'] |
| 1210 | GO:BP | GO:0016053 | organic acid biosynthetic process | 2,38049E-06 | True | positive_corr | 348 | 1037 | 51 | 17906 | 0,049180328 | 0,146551724 | ['GO:0006082', 'GO:0044249', 'GO:0044283', 'GO:1901576'] |
| 1208 | GO:BP | GO:0098656 | anion transmembrane transport | 2,95325E-06 | True | positive_corr | 298 | 1037 | 46 | 17906 | 0,044358727 | 0,154362416 | ['GO:0006820', 'GO:0034220'] |
| 1206 | GO:BP | GO:0048729 | tissue morphogenesis | 3,19385E-06 | True | negative_corr | 686 | 2460 | 151 | 17906 | 0,061382114 | 0,220116618 | ['GO:0009653', 'GO:0009888'] |
| 1203 | GO:BP | GO:0035239 | tube morphogenesis | 3,26427E-06 | True | negative_corr | 944 | 2460 | 195 | 17906 | 0,079268293 | 0,206567797 | ['GO:0009653', 'GO:0035295'] |
| 1293 | GO:BP | GO:0006639 | acylglycerol metabolic process | 3,27834E-06 | True | positive_corr | 130 | 1037 | 28 | 17906 | 0,027000964 | 0,215384615 | ['GO:0006638', 'GO:0046486'] |
| 1199 | GO:BP | GO:0015748 | organophosphate ester transport | 3,42628E-06 | True | positive_corr | 122 | 1037 | 27 | 17906 | 0,026036644 | 0,221311475 | ['GO:0071702'] |
| 1303 | GO:BP | GO:0006638 | neutral lipid metabolic process | 3,94305E-06 | True | positive_corr | 131 | 1037 | 28 | 17906 | 0,027000964 | 0,213740458 | ['GO:0044255'] |
| 1304 | GO:BP | GO:0034220 | ion transmembrane transport | 4,55275E-06 | True | positive_corr | 1184 | 1037 | 119 | 17906 | 0,114754098 | 0,100506757 | ['GO:0006811', 'GO:0055085'] |
| 1192 | GO:BP | GO:0042632 | cholesterol homeostasis | 5,01649E-06 | True | positive_corr | 100 | 1037 | 24 | 17906 | 0,023143684 | 0,24 | ['GO:0055092'] |
| 1191 | GO:BP | GO:0061512 | protein localization to cilium | 5,53787E-06 | True | negative_corr | 55 | 2460 | 26 | 17906 | 0,010569106 | 0,472727273 | ['GO:0033365'] |
| 1190 | GO:BP | GO:0006869 | lipid transport | 6,14473E-06 | True | positive_corr | 390 | 1037 | 54 | 17906 | 0,052073288 | 0,138461538 | ['GO:0010876', 'GO:0071702'] |
| 1189 | GO:BP | GO:0055092 | sterol homeostasis | 6,22375E-06 | True | positive_corr | 101 | 1037 | 24 | 17906 | 0,023143684 | 0,237623762 | ['GO:0055088'] |
| 1898 | GO:BP | GO:0031344 | regulation of cell projection organization | 6,29508E-06 | True | negative_corr | 704 | 2460 | 153 | 17906 | 0,062195122 | 0,217329545 | ['GO:0030030', 'GO:0051128'] |
| 1305 | GO:BP | GO:0061448 | connective tissue development | 6,30849E-06 | True | negative_corr | 281 | 2460 | 76 | 17906 | 0,030894309 | 0,270462633 | ['GO:0009888'] |
| 1182 | GO:BP | GO:0048856 | anatomical structure development | 6,69851E-06 | True | negative_corr | 6105 | 2460 | 968 | 17906 | 0,393495935 | 0,158558559 | ['GO:0032502'] |
| 1198 | GO:BP | GO:1901568 | fatty acid derivative metabolic process | 6,77744E-06 | True | positive_corr | 169 | 1037 | 32 | 17906 | 0,030858245 | 0,189349112 | ['GO:0071704'] |
| 1308 | GO:BP | GO:0048666 | neuron development | 6,90546E-06 | True | negative_corr | 1146 | 2460 | 227 | 17906 | 0,092276423 | 0,198080279 | ['GO:0030182', 'GO:0048468'] |
| 1218 | GO:BP | GO:0120035 | regulation of plasma membrane bounded cell projection organization | 7,39925E-06 | True | negative_corr | 694 | 2460 | 151 | 17906 | 0,061382114 | 0,217579251 | ['GO:0031344', 'GO:0120036'] |
| 1225 | GO:BP | GO:0006694 | steroid biosynthetic process | 9,45578E-06 | True | positive_corr | 199 | 1037 | 35 | 17906 | 0,033751205 | 0,175879397 | ['GO:0008202', 'GO:0008610', 'GO:1901362'] |
| 1275 | GO:BP | GO:0033559 | unsaturated fatty acid metabolic process | 9,53054E-06 | True | positive_corr | 111 | 1037 | 25 | 17906 | 0,024108004 | 0,225225225 | ['GO:0006631'] |
| 1273 | GO:BP | GO:0048519 | negative regulation of biological process | 1,04106E-05 | True | negative_corr | 5681 | 2460 | 906 | 17906 | 0,368292683 | 0,159478965 | ['GO:0008150', 'GO:0050789'] |
| 1272 | GO:BP | GO:0009059 | macromolecule biosynthetic process | 1,14065E-05 | True | negative_corr | 5167 | 2460 | 832 | 17906 | 0,338211382 | 0,16102187 | ['GO:0043170', 'GO:1901576'] |
| 1280 | GO:BP | GO:0044271 | cellular nitrogen compound biosynthetic process | 1,17173E-05 | True | negative_corr | 5008 | 2460 | 809 | 17906 | 0,328861789 | 0,161541534 | ['GO:0034641', 'GO:0044249'] |
| 1270 | GO:BP | GO:0043170 | macromolecule metabolic process | 1,17542E-05 | True | negative_corr | 9592 | 2460 | 1450 | 17906 | 0,589430894 | 0,15116764 | ['GO:0071704'] |
| 1269 | GO:BP | GO:0007156 | homophilic cell adhesion via plasma membrane adhesion molecules | 1,2911E-05 | True | negative_corr | 167 | 2460 | 52 | 17906 | 0,021138211 | 0,311377246 | ['GO:0098742'] |
| 1265 | GO:BP | GO:0098742 | cell-cell adhesion via plasma-membrane adhesion molecules | 1,29715E-05 | True | negative_corr | 275 | 2460 | 74 | 17906 | 0,030081301 | 0,269090909 | ['GO:0098609'] |

| | | | | | | | | | | | | | |
|---|---|---|---|---|---|---|---|---|---|---|---|---|---|
| 1262 | GO:BP | GO:0015718 | monocarboxylic acid transport | 1,43422E-05 | True | positive_corr | 174 | 1037 | 32 | 17906 | 0,030858245 | 0,183908046 | ['GO:0046942'] |
| 1261 | GO:BP | GO:0009887 | animal organ morphogenesis | 1,53071E-05 | True | negative_corr | 1095 | 2460 | 217 | 17906 | 0,088211382 | 0,198173516 | ['GO:0009653', 'GO:0048513'] |
| 1220 | GO:BP | GO:0030258 | lipid modification | 1,60858E-05 | True | positive_corr | 242 | 1037 | 39 | 17906 | 0,037608486 | 0,161157025 | ['GO:0044255'] |
| 1259 | GO:BP | GO:0051179 | localization | 1,6737E-05 | True | positive_corr | 6889 | 1037 | 487 | 17906 | 0,469623915 | 0,070692408 | ['GO:0008150'] |
| 1251 | GO:BP | GO:0034645 | cellular macromolecule biosynthetic process | 2,03735E-05 | True | negative_corr | 5008 | 2460 | 807 | 17906 | 0,32804878 | 0,161142173 | ['GO:0009059', 'GO:0044249', 'GO:0044260'] |
| 1289 | GO:BP | GO:0030855 | epithelial cell differentiation | 2,1752E-05 | True | positive_corr | 786 | 1037 | 86 | 17906 | 0,082931533 | 0,109414758 | ['GO:0030154', 'GO:0060429'] |
| 1291 | GO:BP | GO:0065008 | regulation of biological quality | 2,20015E-05 | True | positive_corr | 4156 | 1037 | 318 | 17906 | 0,306653809 | 0,076515881 | ['GO:0065007'] |
| 1243 | GO:BP | GO:0002009 | morphogenesis of an epithelium | 2,39253E-05 | True | negative_corr | 556 | 2460 | 125 | 17906 | 0,050813008 | 0,224820144 | ['GO:0048729', 'GO:0060429'] |
| 1240 | GO:BP | GO:0048646 | anatomical structure formation involved in morphogenesis | 2,55727E-05 | True | negative_corr | 1206 | 2460 | 234 | 17906 | 0,095121951 | 0,194029851 | ['GO:0009653', 'GO:0032502'] |
| 1239 | GO:BP | GO:0008210 | estrogen metabolic process | 2,63135E-05 | True | positive_corr | 32 | 1037 | 13 | 17906 | 0,012536162 | 0,40625 | ['GO:0008202', 'GO:0034754'] |
| 1292 | GO:BP | GO:0031175 | neuron projection development | 2,72684E-05 | True | negative_corr | 1017 | 2460 | 203 | 17906 | 0,082520325 | 0,199606686 | ['GO:0048666', 'GO:0120036'] |
| 1236 | GO:BP | GO:0010876 | lipid localization | 3,17236E-05 | True | positive_corr | 431 | 1037 | 56 | 17906 | 0,054001929 | 0,129930394 | ['GO:0033036'] |
| 1234 | GO:BP | GO:0048878 | chemical homeostasis | 3,17458E-05 | True | positive_corr | 1224 | 1037 | 119 | 17906 | 0,114754098 | 0,097222222 | ['GO:0042592'] |
| 1286 | GO:BP | GO:0048858 | cell projection morphogenesis | 3,18112E-05 | True | negative_corr | 691 | 2460 | 148 | 17906 | 0,060162602 | 0,214182344 | ['GO:0000902', 'GO:0030030', 'GO:0032990'] |
| 1309 | GO:BP | GO:0019637 | organophosphate metabolic process | 3,22831E-05 | True | positive_corr | 1051 | 1037 | 106 | 17906 | 0,102217936 | 0,100856327 | ['GO:0006793', 'GO:0071704'] |
| 1310 | GO:BP | GO:0031589 | cell-substrate adhesion | 3,68046E-05 | True | negative_corr | 355 | 2460 | 88 | 17906 | 0,035772358 | 0,247887324 | ['GO:0007155'] |
| 1312 | GO:BP | GO:0120039 | plasma membrane bounded cell projection morphogenesis | 3,81193E-05 | True | negative_corr | 687 | 2460 | 147 | 17906 | 0,059756098 | 0,213973799 | ['GO:0048858'] |
| 1117 | GO:BP | GO:0000956 | nuclear-transcribed mRNA catabolic process | 4,09621E-05 | True | negative_corr | 211 | 2460 | 60 | 17906 | 0,024390244 | 0,28436019 | ['GO:0006402'] |
| 1114 | GO:BP | GO:0008643 | carbohydrate transport | 4,29272E-05 | True | positive_corr | 154 | 1037 | 29 | 17906 | 0,027965284 | 0,188311688 | ['GO:0071702'] |
| 1113 | GO:BP | GO:0048812 | neuron projection morphogenesis | 5,38083E-05 | True | negative_corr | 673 | 2460 | 144 | 17906 | 0,058536585 | 0,213967311 | ['GO:0031175', 'GO:0120039'] |
| 1111 | GO:BP | GO:0006814 | sodium ion transport | 5,89374E-05 | True | positive_corr | 223 | 1037 | 36 | 17906 | 0,034715526 | 0,161434978 | ['GO:0015672', 'GO:0030001'] |
| 1389 | GO:BP | GO:0043009 | chordate embryonic development | 6,2278E-05 | True | negative_corr | 657 | 2460 | 141 | 17906 | 0,057317073 | 0,214611872 | ['GO:0009792'] |
| 1106 | GO:BP | GO:0060562 | epithelial tube morphogenesis | 6,27693E-05 | True | negative_corr | 332 | 2460 | 83 | 17906 | 0,033739837 | 0,25 | ['GO:0002009', 'GO:0035239'] |
| 1105 | GO:BP | GO:0052695 | cellular glucuronidation | 6,45933E-05 | True | positive_corr | 19 | 1037 | 10 | 17906 | 0,009643202 | 0,526315789 | ['GO:0019585'] |
| 1104 | GO:BP | GO:0009792 | embryo development ending in birth or egg hatching | 6,5494E-05 | True | negative_corr | 675 | 2460 | 144 | 17906 | 0,058536585 | 0,213333333 | ['GO:0009790'] |
| 1102 | GO:BP | GO:0045216 | cell-cell junction organization | 7,04008E-05 | True | positive_corr | 195 | 1037 | 33 | 17906 | 0,031822565 | 0,169230769 | ['GO:0034330'] |
| 1118 | GO:BP | GO:0032990 | cell part morphogenesis | 7,10751E-05 | True | negative_corr | 711 | 2460 | 150 | 17906 | 0,06097561 | 0,210970464 | ['GO:0032989'] |
| 1098 | GO:BP | GO:0030154 | cell differentiation | 7,15814E-05 | True | negative_corr | 4335 | 2460 | 705 | 17906 | 0,286585366 | 0,162629758 | ['GO:0048869'] |
| 1096 | GO:BP | GO:0043062 | extracellular structure organization | 7,17538E-05 | True | negative_corr | 376 | 2460 | 91 | 17906 | 0,03699187 | 0,242021277 | ['GO:0016043'] |
| 1094 | GO:BP | GO:0072657 | protein localization to membrane | 7,1936E-05 | True | negative_corr | 641 | 2460 | 138 | 17906 | 0,056097561 | 0,215288612 | ['GO:0034613'] |
| 1093 | GO:BP | GO:0010467 | gene expression | 7,95152E-05 | True | negative_corr | 5672 | 2460 | 897 | 17906 | 0,364634146 | 0,158145275 | ['GO:0043170'] |
| 1088 | GO:BP | GO:0048869 | cellular developmental process | 8,10478E-05 | True | negative_corr | 4531 | 2460 | 733 | 17906 | 0,29796748 | 0,161774443 | ['GO:0009987', 'GO:0032502'] |
| 1087 | GO:BP | GO:0034330 | cell junction organization | 8,52323E-05 | True | negative_corr | 297 | 2460 | 76 | 17906 | 0,030894309 | 0,255892256 | ['GO:0016043'] |
| 1393 | GO:BP | GO:0045664 | regulation of neuron differentiation | 8,8901E-05 | True | negative_corr | 684 | 2460 | 145 | 17906 | 0,058943089 | 0,211988304 | ['GO:0030182', 'GO:0050767'] |
| 1407 | GO:BP | GO:0008104 | protein localization | 9,24835E-05 | True | negative_corr | 2923 | 2460 | 496 | 17906 | 0,201626016 | 0,169688676 | ['GO:0033036'] |
| 1084 | GO:BP | GO:0051252 | regulation of RNA metabolic process | 9,62819E-05 | True | negative_corr | 3856 | 2460 | 634 | 17906 | 0,257723577 | 0,164419087 | ['GO:0016070', 'GO:0019219', 'GO:0060255'] |
| 1083 | GO:BP | GO:0046483 | heterocycle metabolic process | 9,88201E-05 | True | negative_corr | 5910 | 2460 | 930 | 17906 | 0,37804878 | 0,157360406 | ['GO:0044237'] |
| 1097 | GO:BP | GO:0106001 | intestinal hexose absorption | 0,000103383 | True | positive_corr | 6 | 1037 | 6 | 17906 | 0,005785921 | 1 | ['GO:0008643', 'GO:0050892'] |
| 1385 | GO:BP | GO:0019369 | arachidonic acid metabolic process | 0,000103843 | True | positive_corr | 54 | 1037 | 16 | 17906 | 0,015429122 | 0,296296296 | ['GO:0001676', 'GO:0006690', 'GO:0033559'] |
| 1123 | GO:BP | GO:0002181 | cytoplasmic translation | 0,000111138 | True | negative_corr | 103 | 2460 | 36 | 17906 | 0,014634146 | 0,349514563 | ['GO:0006412'] |
| 1378 | GO:BP | GO:0019373 | epoxygenase P450 pathway | 0,000122473 | True | positive_corr | 20 | 1037 | 10 | 17906 | 0,009643202 | 0,5 | ['GO:0019369'] |
| 1315 | GO:BP | GO:0030198 | extracellular matrix organization | 0,000129768 | True | negative_corr | 375 | 2460 | 90 | 17906 | 0,036585366 | 0,24 | ['GO:0043062'] |
| 1169 | GO:BP | GO:0021915 | neural tube development | 0,000129977 | True | negative_corr | 168 | 2460 | 50 | 17906 | 0,020325203 | 0,297619048 | ['GO:0007399', 'GO:0035295', 'GO:0043009', 'GO:0060429'] |
| 1167 | GO:BP | GO:0019083 | viral transcription | 0,000139273 | True | negative_corr | 178 | 2460 | 52 | 17906 | 0,021138211 | 0,292134831 | ['GO:0016032', 'GO:0019080'] |
| 1166 | GO:BP | GO:0048523 | negative regulation of cellular process | 0,000141906 | True | negative_corr | 4996 | 2460 | 798 | 17906 | 0,324390244 | 0,159727782 | ['GO:0009987', 'GO:0048519', 'GO:0050794'] |
| 1321 | GO:BP | GO:0048598 | embryonic morphogenesis | 0,000154973 | True | negative_corr | 608 | 2460 | 131 | 17906 | 0,053252033 | 0,215460526 | ['GO:0009653', 'GO:0009790'] |
| 1326 | GO:BP | GO:0006725 | cellular aromatic compound metabolic process | 0,000185554 | True | negative_corr | 5956 | 2460 | 934 | 17906 | 0,379674797 | 0,156816655 | ['GO:0044237'] |
| 1162 | GO:BP | GO:0060255 | regulation of macromolecule metabolic process | 0,000192915 | True | negative_corr | 6438 | 2460 | 1002 | 17906 | 0,407317073 | 0,155638397 | ['GO:0019222', 'GO:0043170'] |
| 1336 | GO:BP | GO:0000226 | microtubule cytoskeleton organization | 0,000204844 | True | negative_corr | 576 | 2460 | 125 | 17906 | 0,050813008 | 0,217013889 | ['GO:0007010', 'GO:0007017'] |
| 1159 | GO:BP | GO:0048667 | cell morphogenesis involved in neuron differentiation | 0,00020906 | True | negative_corr | 611 | 2460 | 131 | 17906 | 0,053252033 | 0,214402619 | ['GO:0000904', 'GO:0048666'] |
| 1340 | GO:BP | GO:0010975 | regulation of neuron projection development | 0,000214939 | True | negative_corr | 519 | 2460 | 115 | 17906 | 0,046747967 | 0,221579961 | ['GO:0031175', 'GO:0045664', 'GO:0120035'] |
| 1350 | GO:BP | GO:0010468 | regulation of gene expression | 0,000255223 | True | negative_corr | 4727 | 2460 | 757 | 17906 | 0,307723577 | 0,160143854 | ['GO:0010467', 'GO:0060255'] |
| 1354 | GO:BP | GO:0006635 | fatty acid beta-oxidation | 0,000271392 | True | positive_corr | 72 | 1037 | 18 | 17906 | 0,017357763 | 0,25 | ['GO:0009062', 'GO:0019395'] |
| 1150 | GO:BP | GO:0044267 | cellular protein metabolic process | 0,000284515 | True | negative_corr | 5357 | 2460 | 847 | 17906 | 0,344308943 | 0,158110883 | ['GO:0019538', 'GO:0044260'] |
| 1143 | GO:BP | GO:0042573 | retinoic acid metabolic process | 0,000284531 | True | positive_corr | 32 | 1037 | 12 | 17906 | 0,011571842 | 0,375 | ['GO:0001523', 'GO:0032787', 'GO:0034754'] |
| 1138 | GO:BP | GO:0006812 | cation transport | 0,000288227 | True | positive_corr | 1205 | 1037 | 114 | 17906 | 0,109932498 | 0,094605809 | ['GO:0006811'] |
| 1359 | GO:BP | GO:0019219 | regulation of nucleobase-containing compound metabolic process | 0,000288858 | True | negative_corr | 4116 | 2460 | 668 | 17906 | 0,271544715 | 0,162293489 | ['GO:0006139', 'GO:0031323', 'GO:0051171', 'GO:0080090'] |
| 1131 | GO:BP | GO:0072358 | cardiovascular system development | 0,000314259 | True | negative_corr | 829 | 2460 | 167 | 17906 | 0,067886179 | 0,201447527 | ['GO:0048731', 'GO:0072359'] |
| 1130 | GO:BP | GO:0016266 | O-glycan processing | 0,000316097 | True | positive_corr | 58 | 1037 | 16 | 17906 | 0,015429122 | 0,275862069 | ['GO:0006493'] |
| 1368 | GO:BP | GO:0007389 | pattern specification process | 0,000325665 | True | negative_corr | 449 | 2460 | 102 | 17906 | 0,041463415 | 0,227171492 | ['GO:0007275', 'GO:0032501'] |
| 1376 | GO:BP | GO:2001141 | regulation of RNA biosynthetic process | 0,000331194 | True | negative_corr | 3593 | 2460 | 591 | 17906 | 0,240243902 | 0,164486502 | ['GO:0010556', 'GO:0031326', 'GO:0032774', 'GO:0051252'] |
| 1125 | GO:BP | GO:0019627 | urea metabolic process | 0,000339376 | True | positive_corr | 13 | 1037 | 8 | 17906 | 0,007714561 | 0,615384615 | ['GO:0043603', 'GO:0044281', 'GO:0071941', 'GO:1901564'] |
| 1077 | GO:BP | GO:0001501 | skeletal system development | 0,000348211 | True | negative_corr | 535 | 2460 | 117 | 17906 | 0,047560976 | 0,218691589 | ['GO:0048731'] |
| 859 | GO:BP | GO:0072001 | renal system development | 0,000356008 | True | negative_corr | 296 | 2460 | 74 | 17906 | 0,030081301 | 0,25 | ['GO:0001655', 'GO:0048731'] |
| 1548 | GO:BP | GO:1903506 | regulation of nucleic acid-templated transcription | 0,000356598 | True | negative_corr | 3588 | 2460 | 590 | 17906 | 0,239837398 | 0,164437012 | ['GO:0097659', 'GO:2001141'] |

| | | | | | | | | | | | | | |
|---|---|---|---|---|---|---|---|---|---|---|---|---|---|
| 1562 | GO:BP | GO:0022613 | ribonucleoprotein complex biogenesis | 0,000359724 | True | negative_corr | 433 | 2460 | 99 | 17906 | 0,040243902 | 0,228637413 | ['GO:0044085'] |
| 1744 | GO:BP | GO:0046486 | glycerolipid metabolic process | 0,000375812 | True | positive_corr | 416 | 1037 | 52 | 17906 | 0,050144648 | 0,125 | ['GO:0044255'] |
| 523 | GO:BP | GO:0001944 | vasculature development | 0,000404933 | True | negative_corr | 820 | 2460 | 165 | 17906 | 0,067073171 | 0,201219512 | ['GO:0048731', 'GO:0072358'] |
| 1748 | GO:BP | GO:0006355 | regulation of transcription, DNA-templated | 0,000416884 | True | negative_corr | 3517 | 2460 | 579 | 17906 | 0,235365854 | 0,164628945 | ['GO:0006351', 'GO:0010468', 'GO:1903506', 'GO:2000112'] |
| 500 | GO:BP | GO:0015914 | phospholipid transport | 0,00044089 | True | positive_corr | 90 | 1037 | 20 | 17906 | 0,019286403 | 0,222222222 | ['GO:0006869', 'GO:0015711', 'GO:0015748'] |
| 488 | GO:BP | GO:0042493 | response to drug | 0,00045582 | True | positive_corr | 1093 | 1037 | 105 | 17906 | 0,101253616 | 0,096065874 | ['GO:0042221'] |
| 484 | GO:BP | GO:0044087 | regulation of cellular component biogenesis | 0,000471918 | True | negative_corr | 968 | 2460 | 189 | 17906 | 0,076829268 | 0,195247934 | ['GO:0044085', 'GO:0050789'] |
| 1763 | GO:BP | GO:0042737 | drug catabolic process | 0,000489854 | True | positive_corr | 143 | 1037 | 26 | 17906 | 0,025072324 | 0,181818182 | ['GO:0017144', 'GO:0044248'] |
| 1768 | GO:BP | GO:0006790 | sulfur compound metabolic process | 0,000516018 | True | positive_corr | 374 | 1037 | 48 | 17906 | 0,046287367 | 0,128342246 | ['GO:0044237'] |
| 474 | GO:BP | GO:1902652 | secondary alcohol metabolic process | 0,000575846 | True | positive_corr | 163 | 1037 | 28 | 17906 | 0,027000964 | 0,171779141 | ['GO:0006066'] |
| 1740 | GO:BP | GO:0032774 | RNA biosynthetic process | 0,000604426 | True | negative_corr | 3758 | 2460 | 613 | 17906 | 0,249186992 | 0,16311868 | ['GO:0009059', 'GO:0016070', 'GO:0034654'] |
| 473 | GO:BP | GO:0097435 | supramolecular fiber organization | 0,000609467 | True | negative_corr | 693 | 2460 | 143 | 17906 | 0,058130081 | 0,206349206 | ['GO:0016043'] |
| 461 | GO:BP | GO:0034378 | chylomicron assembly | 0,000612039 | True | positive_corr | 10 | 1037 | 7 | 17906 | 0,006750241 | 0,7 | ['GO:0034377'] |
| 1769 | GO:BP | GO:0051216 | cartilage development | 0,000623798 | True | negative_corr | 216 | 2460 | 58 | 17906 | 0,023577236 | 0,268518519 | ['GO:0001501', 'GO:0048513', 'GO:0061448'] |
| 1775 | GO:BP | GO:0001568 | blood vessel development | 0,000676614 | True | negative_corr | 784 | 2460 | 158 | 17906 | 0,064227642 | 0,201530612 | ['GO:0001944', 'GO:0048856'] |
| 1779 | GO:BP | GO:0023051 | regulation of signaling | 0,000679613 | True | negative_corr | 3679 | 2460 | 601 | 17906 | 0,244308943 | 0,163359609 | ['GO:0023052', 'GO:0050789'] |
| 452 | GO:BP | GO:0032532 | regulation of microvillus length | 0,000687953 | True | positive_corr | 7 | 1037 | 6 | 17906 | 0,005785921 | 0,857142857 | ['GO:0032530', 'GO:0032536'] |
| 1784 | GO:BP | GO:0016052 | carbohydrate catabolic process | 0,000698483 | True | positive_corr | 204 | 1037 | 32 | 17906 | 0,030858245 | 0,156862745 | ['GO:0005975', 'GO:1901575'] |
| 1787 | GO:BP | GO:0019395 | fatty acid oxidation | 0,000723414 | True | positive_corr | 101 | 1037 | 21 | 17906 | 0,020250723 | 0,207920792 | ['GO:0006631', 'GO:0034440'] |
| 1792 | GO:BP | GO:0071941 | nitrogen cycle metabolic process | 0,000751538 | True | positive_corr | 14 | 1037 | 8 | 17906 | 0,007714561 | 0,571428571 | ['GO:0006807'] |
| 426 | GO:BP | GO:0009812 | flavonoid metabolic process | 0,000751538 | True | positive_corr | 14 | 1037 | 8 | 17906 | 0,007714561 | 0,571428571 | ['GO:0071704'] |
| 462 | GO:BP | GO:0061564 | axon development | 0,000757564 | True | negative_corr | 531 | 2460 | 115 | 17906 | 0,046747967 | 0,216572505 | ['GO:0031175'] |
| 425 | GO:BP | GO:0097659 | nucleic acid-templated transcription | 0,000780328 | True | negative_corr | 3744 | 2460 | 610 | 17906 | 0,24796748 | 0,16292735 | ['GO:0032774'] |
| 1732 | GO:BP | GO:0006612 | protein targeting to membrane | 0,000783961 | True | negative_corr | 197 | 2460 | 54 | 17906 | 0,02195122 | 0,274111675 | ['GO:0006605', 'GO:0090150'] |
| 551 | GO:BP | GO:0051960 | regulation of nervous system development | 0,000813338 | True | negative_corr | 963 | 2460 | 187 | 17906 | 0,07601626 | 0,194184839 | ['GO:0007399', 'GO:2000026'] |
| 1687 | GO:BP | GO:0046461 | neutral lipid catabolic process | 0,00081896 | True | positive_corr | 41 | 1037 | 13 | 17906 | 0,012536162 | 0,317073171 | ['GO:0006638', 'GO:0044242'] |
| 611 | GO:BP | GO:0046464 | acylglycerol catabolic process | 0,00081896 | True | positive_corr | 41 | 1037 | 13 | 17906 | 0,012536162 | 0,317073171 | ['GO:0006639', 'GO:0046461', 'GO:0046503'] |
| 1697 | GO:BP | GO:0001822 | kidney development | 0,000846738 | True | negative_corr | 286 | 2460 | 71 | 17906 | 0,028861789 | 0,248251748 | ['GO:0048513', 'GO:0072001'] |
| 602 | GO:BP | GO:0034377 | plasma lipoprotein particle assembly | 0,000852711 | True | positive_corr | 29 | 1037 | 11 | 17906 | 0,010607522 | 0,379310345 | ['GO:0065005', 'GO:0071827', 'GO:0097006'] |
| 597 | GO:BP | GO:0097164 | ammonium ion metabolic process | 0,000876304 | True | positive_corr | 206 | 1037 | 32 | 17906 | 0,030858245 | 0,155339806 | ['GO:0006807'] |
| 858 | GO:BP | GO:0001655 | urogenital system development | 0,000884146 | True | negative_corr | 335 | 2460 | 80 | 17906 | 0,032520325 | 0,23880597 | ['GO:0048731'] |
| 592 | GO:BP | GO:0046466 | membrane lipid catabolic process | 0,000894351 | True | positive_corr | 35 | 1037 | 12 | 17906 | 0,011571842 | 0,342857143 | ['GO:0006643', 'GO:0044242'] |
| 586 | GO:BP | GO:0050793 | regulation of developmental process | 0,00089689 | True | negative_corr | 2779 | 2460 | 467 | 17906 | 0,189837398 | 0,16804606 | ['GO:0032502', 'GO:0050789'] |
| 1704 | GO:BP | GO:2000112 | regulation of cellular macromolecule biosynthetic process | 0,000911656 | True | negative_corr | 4036 | 2460 | 652 | 17906 | 0,26504065 | 0,161546085 | ['GO:0010556', 'GO:0031326', 'GO:0034645'] |
| 550 | GO:BP | GO:0033036 | macromolecule localization | 0,000923013 | True | negative_corr | 3292 | 2460 | 543 | 17906 | 0,220731707 | 0,164945322 | ['GO:0051179'] |
| 577 | GO:BP | GO:0007409 | axonogenesis | 0,000939219 | True | negative_corr | 487 | 2460 | 107 | 17906 | 0,043495935 | 0,219712526 | ['GO:0048667', 'GO:0048812', 'GO:0061564'] |
| 571 | GO:BP | GO:0042273 | ribosomal large subunit biogenesis | 0,000996327 | True | negative_corr | 76 | 2460 | 28 | 17906 | 0,011382114 | 0,368421053 | ['GO:0022613', 'GO:0042254'] |
| 1711 | GO:BP | GO:0034440 | lipid oxidation | 0,001024405 | True | positive_corr | 103 | 1037 | 21 | 17906 | 0,020250723 | 0,203883495 | ['GO:0030258', 'GO:0055114'] |
| 566 | GO:BP | GO:0044241 | lipid digestion | 0,001071681 | True | positive_corr | 19 | 1037 | 9 | 17906 | 0,008678881 | 0,473684211 | ['GO:0007586'] |
| 563 | GO:BP | GO:0006575 | cellular modified amino acid metabolic process | 0,001128226 | True | positive_corr | 188 | 1037 | 30 | 17906 | 0,028929605 | 0,159574468 | ['GO:0044237', 'GO:1901564'] |
| 1718 | GO:BP | GO:0010646 | regulation of cell communication | 0,001160491 | True | negative_corr | 3638 | 2460 | 593 | 17906 | 0,241056911 | 0,163001649 | ['GO:0007154', 'GO:0050794'] |
| 1722 | GO:BP | GO:0045595 | regulation of cell differentiation | 0,001267606 | True | negative_corr | 1932 | 2460 | 338 | 17906 | 0,137398374 | 0,17494824 | ['GO:0030154', 'GO:0050793', 'GO:0050794'] |
| 1725 | GO:BP | GO:0008203 | cholesterol metabolic process | 0,001296458 | True | positive_corr | 150 | 1037 | 26 | 17906 | 0,025072324 | 0,173333333 | ['GO:0016125', 'GO:1902652'] |
| 1726 | GO:BP | GO:0072359 | circulatory system development | 0,001315021 | True | negative_corr | 1194 | 2460 | 223 | 17906 | 0,090650407 | 0,186767169 | ['GO:0048731'] |
| 1728 | GO:BP | GO:0006996 | organelle organization | 0,001456624 | True | negative_corr | 3966 | 2460 | 640 | 17906 | 0,260162602 | 0,161371659 | ['GO:0016043'] |
| 1705 | GO:BP | GO:0003002 | regionalization | 0,001560804 | True | negative_corr | 356 | 2460 | 83 | 17906 | 0,033739837 | 0,233146067 | ['GO:0007389'] |
| 1676 | GO:BP | GO:0048589 | developmental growth | 0,001584396 | True | negative_corr | 698 | 2460 | 142 | 17906 | 0,057723577 | 0,203438395 | ['GO:0032502', 'GO:0040007'] |
| 419 | GO:BP | GO:0051171 | regulation of nitrogen compound metabolic process | 0,001765194 | True | negative_corr | 6009 | 2460 | 932 | 17906 | 0,378861789 | 0,155100682 | ['GO:0006807', 'GO:0019222'] |
| 1797 | GO:BP | GO:1904970 | brush border assembly | 0,001793275 | True | positive_corr | 5 | 1037 | 5 | 17906 | 0,004821601 | 1 | ['GO:0022607'] |
| 1838 | GO:BP | GO:1901616 | organic hydroxy compound catabolic process | 0,001834729 | True | positive_corr | 73 | 1037 | 17 | 17906 | 0,016393443 | 0,232876712 | ['GO:1901575', 'GO:1901615'] |
| 251 | GO:BP | GO:0070085 | glycosylation | 0,002031626 | True | positive_corr | 267 | 1037 | 37 | 17906 | 0,035679846 | 0,138576779 | ['GO:0008152'] |
| 250 | GO:BP | GO:0051130 | positive regulation of cellular component organization | 0,002034677 | True | negative_corr | 1245 | 2460 | 230 | 17906 | 0,093495935 | 0,184738956 | ['GO:0016043', 'GO:0048522', 'GO:0051128'] |
| 243 | GO:BP | GO:0040007 | growth | 0,002163628 | True | negative_corr | 1020 | 2460 | 194 | 17906 | 0,078861789 | 0,190196078 | ['GO:0008150'] |
| 239 | GO:BP | GO:0007043 | cell-cell junction assembly | 0,002210093 | True | positive_corr | 135 | 1037 | 24 | 17906 | 0,023143684 | 0,177777778 | ['GO:0034329', 'GO:0045216'] |
| 1854 | GO:BP | GO:1901575 | organic substance catabolic process | 0,002269587 | True | positive_corr | 2239 | 1037 | 182 | 17906 | 0,175506268 | 0,081286289 | ['GO:0009056', 'GO:0071704'] |
| 195 | GO:BP | GO:0034329 | cell junction assembly | 0,002308671 | True | negative_corr | 245 | 2460 | 62 | 17906 | 0,025203252 | 0,253061224 | ['GO:0022607', 'GO:0034330'] |
| 1855 | GO:BP | GO:0006821 | chloride transport | 0,002343874 | True | positive_corr | 108 | 1037 | 21 | 17906 | 0,020250723 | 0,194444444 | ['GO:0015698'] |
| 1856 | GO:BP | GO:0034641 | cellular nitrogen compound metabolic process | 0,002392306 | True | negative_corr | 6508 | 2460 | 1001 | 17906 | 0,406910569 | 0,153810695 | ['GO:0006807', 'GO:0044237'] |
| 265 | GO:BP | GO:0006351 | transcription, DNA-templated | 0,00239858 | True | negative_corr | 3698 | 2460 | 599 | 17906 | 0,243495935 | 0,161979448 | ['GO:0010467', 'GO:0034645', 'GO:0097659'] |
| 1857 | GO:BP | GO:0042908 | xenobiotic transport | 0,002557579 | True | positive_corr | 26 | 1037 | 10 | 17906 | 0,009643202 | 0,384615385 | ['GO:0006810'] |
| 179 | GO:BP | GO:0042572 | retinol metabolic process | 0,002732243 | True | positive_corr | 45 | 1037 | 13 | 17906 | 0,012536162 | 0,288888889 | ['GO:0001523', 'GO:0034308', 'GO:0034754'] |
| 167 | GO:BP | GO:0022406 | membrane docking | 0,002812803 | True | negative_corr | 184 | 2460 | 50 | 17906 | 0,020325203 | 0,27173913 | ['GO:0009987'] |
| 1866 | GO:BP | GO:0015698 | inorganic anion transport | 0,002888875 | True | positive_corr | 166 | 1037 | 27 | 17906 | 0,026036644 | 0,162650602 | ['GO:0006820'] |
| 1871 | GO:BP | GO:0070293 | renal absorption | 0,002901385 | True | positive_corr | 16 | 1037 | 8 | 17906 | 0,007714561 | 0,5 | ['GO:0003014'] |
| 1881 | GO:BP | GO:0030033 | microvillus assembly | 0,002901385 | True | positive_corr | 16 | 1037 | 8 | 17906 | 0,007714561 | 0,5 | ['GO:0032528', 'GO:0120031'] |

| | | | | | | | | | | | | | |
|---|---|---|---|---|---|---|---|---|---|---|---|---|---|
| 1886 | GO:BP | GO:0072330 | monocarboxylic acid biosynthetic process | 0,003000355 | True | positive_corr | 228 | 1037 | 33 | 17906 | 0,031822565 | 0,144736842 | ['GO:0032787', 'GO:0046394'] |
| 85 | GO:BP | GO:0051129 | negative regulation of cellular component organization | 0,003024485 | True | negative_corr | 748 | 2460 | 149 | 17906 | 0,060569106 | 0,199197861 | ['GO:0016043', 'GO:0048523', 'GO:0051128'] |
| 80 | GO:BP | GO:0010556 | regulation of macromolecule biosynthetic process | 0,003034528 | True | negative_corr | 4186 | 2460 | 669 | 17906 | 0,27195122 | 0,159818442 | ['GO:0009059', 'GO:0009889', 'GO:0060255'] |
| 1892 | GO:BP | GO:0006886 | intracellular protein transport | 0,003065059 | True | negative_corr | 1182 | 2460 | 219 | 17906 | 0,08902439 | 0,185279188 | ['GO:0015031', 'GO:0034613', 'GO:0046907'] |
| 1865 | GO:BP | GO:0006807 | nitrogen compound metabolic process | 0,003102995 | True | negative_corr | 10111 | 2460 | 1497 | 17906 | 0,608536585 | 0,148056572 | ['GO:0008152'] |
| 1794 | GO:BP | GO:0006633 | fatty acid biosynthetic process | 0,003206154 | True | positive_corr | 157 | 1037 | 26 | 17906 | 0,025072324 | 0,165605096 | ['GO:0006631', 'GO:0008610', 'GO:0072330'] |
| 1837 | GO:BP | GO:0046365 | monosaccharide catabolic process | 0,003360135 | True | positive_corr | 68 | 1037 | 16 | 17906 | 0,015429122 | 0,235294118 | ['GO:0005996', 'GO:0016052', 'GO:0044282'] |
| 1836 | GO:BP | GO:0034654 | nucleobase-containing compound biosynthetic process | 0,00367713 | True | negative_corr | 4226 | 2460 | 674 | 17906 | 0,27398374 | 0,159488878 | ['GO:0006139', 'GO:0018130', 'GO:0019438', 'GO:0044271', 'GO:1901362'] |
| 1798 | GO:BP | GO:0019433 | triglyceride catabolic process | 0,003849656 | True | positive_corr | 33 | 1037 | 11 | 17906 | 0,010607522 | 0,333333333 | ['GO:0006641', 'GO:0046464'] |
| 383 | GO:BP | GO:0065005 | protein-lipid complex assembly | 0,003849656 | True | positive_corr | 33 | 1037 | 11 | 17906 | 0,010607522 | 0,333333333 | ['GO:0065003', 'GO:0071825'] |
| 382 | GO:BP | GO:0008589 | regulation of smoothened signaling pathway | 0,003955245 | True | negative_corr | 85 | 2460 | 29 | 17906 | 0,011788618 | 0,341176471 | ['GO:0007224', 'GO:0009966'] |
| 381 | GO:BP | GO:0048880 | sensory system development | 0,003957898 | True | negative_corr | 375 | 2460 | 85 | 17906 | 0,034552846 | 0,226666667 | ['GO:0048731'] |
| 1799 | GO:BP | GO:1901264 | carbohydrate derivative transport | 0,004088713 | True | positive_corr | 77 | 1037 | 17 | 17906 | 0,016393443 | 0,220779221 | ['GO:0071702'] |
| 1800 | GO:BP | GO:0150063 | visual system development | 0,00425452 | True | negative_corr | 370 | 2460 | 84 | 17906 | 0,034146341 | 0,227027027 | ['GO:0048880'] |
| 378 | GO:BP | GO:0099118 | microtubule-based protein transport | 0,004333185 | True | negative_corr | 68 | 2460 | 25 | 17906 | 0,010162602 | 0,367647059 | ['GO:0015031', 'GO:0099111'] |
| 1810 | GO:BP | GO:0098840 | protein transport along microtubule | 0,004333185 | True | negative_corr | 68 | 2460 | 25 | 17906 | 0,010162602 | 0,367647059 | ['GO:0006886', 'GO:0010970', 'GO:0099118'] |
| 1811 | GO:BP | GO:0006749 | glutathione metabolic process | 0,0044602 | True | positive_corr | 54 | 1037 | 14 | 17906 | 0,013500482 | 0,259259259 | ['GO:0006518', 'GO:0006575', 'GO:0006790', 'GO:0051186'] |
| 276 | GO:BP | GO:0006486 | protein glycosylation | 0,004596604 | True | positive_corr | 254 | 1037 | 35 | 17906 | 0,033751205 | 0,137795276 | ['GO:0006464', 'GO:0009101', 'GO:0043413'] |
| 1816 | GO:BP | GO:0043413 | macromolecule glycosylation | 0,004596604 | True | positive_corr | 254 | 1037 | 35 | 17906 | 0,033751205 | 0,137795276 | ['GO:0043412', 'GO:0070085'] |
| 1818 | GO:BP | GO:0050794 | regulation of cellular process | 0,004820866 | True | negative_corr | 11187 | 2460 | 1640 | 17906 | 0,666666667 | 0,146598731 | ['GO:0009987', 'GO:0050789'] |
| 1819 | GO:BP | GO:0046503 | glycerolipid catabolic process | 0,004944312 | True | positive_corr | 62 | 1037 | 15 | 17906 | 0,014464802 | 0,241935484 | ['GO:0044242', 'GO:0046486'] |
| 332 | GO:BP | GO:0001654 | eye development | 0,005130955 | True | negative_corr | 366 | 2460 | 83 | 17906 | 0,033739837 | 0,226775956 | ['GO:0007423', 'GO:0150063'] |
| 324 | GO:BP | GO:0097711 | ciliary basal body-plasma membrane docking | 0,005181175 | True | negative_corr | 95 | 2460 | 31 | 17906 | 0,012601626 | 0,326315789 | ['GO:0060271', 'GO:0140056'] |
| 1820 | GO:BP | GO:0050789 | regulation of biological process | 0,005401568 | True | negative_corr | 11940 | 2460 | 1740 | 17906 | 0,707317073 | 0,145728643 | ['GO:0008150', 'GO:0065007'] |
| 311 | GO:BP | GO:0043010 | camera-type eye development | 0,006302085 | True | negative_corr | 323 | 2460 | 75 | 17906 | 0,030487805 | 0,232198142 | ['GO:0001654'] |
| 1821 | GO:BP | GO:0042254 | ribosome biogenesis | 0,006393955 | True | negative_corr | 301 | 2460 | 71 | 17906 | 0,028861789 | 0,235880399 | ['GO:0022613'] |
| 1833 | GO:BP | GO:0050767 | regulation of neurogenesis | 0,006534552 | True | negative_corr | 856 | 2460 | 165 | 17906 | 0,067073171 | 0,192757009 | ['GO:0022008', 'GO:0048699', 'GO:0051960', 'GO:0060284'] |
| 288 | GO:BP | GO:0019318 | hexose metabolic process | 0,006610002 | True | positive_corr | 258 | 1037 | 35 | 17906 | 0,033751205 | 0,135658915 | ['GO:0005996'] |
| 353 | GO:BP | GO:1901135 | carbohydrate derivative metabolic process | 0,006970768 | True | positive_corr | 1130 | 1037 | 103 | 17906 | 0,099324976 | 0,091150442 | ['GO:0071704'] |
| 1675 | GO:BP | GO:0006766 | vitamin metabolic process | 0,006989565 | True | positive_corr | 134 | 1037 | 23 | 17906 | 0,022179364 | 0,171641791 | ['GO:0044281'] |
| 595 | GO:BP | GO:0030199 | collagen fibril organization | 0,007222828 | True | negative_corr | 53 | 2460 | 21 | 17906 | 0,008536585 | 0,396226415 | ['GO:0030198', 'GO:0097435'] |
| 624 | GO:BP | GO:0042073 | intraciliary transport | 0,007222828 | True | negative_corr | 53 | 2460 | 21 | 17906 | 0,008536585 | 0,396226415 | ['GO:0031503', 'GO:0044782', 'GO:0098840'] |
| 798 | GO:BP | GO:0032536 | regulation of cell projection size | 0,007499382 | True | positive_corr | 13 | 1037 | 7 | 17906 | 0,006750241 | 0,538461538 | ['GO:0032535'] |
| 797 | GO:BP | GO:0022604 | regulation of cell morphogenesis | 0,007660025 | True | negative_corr | 508 | 2460 | 107 | 17906 | 0,043495935 | 0,210629921 | ['GO:0000902', 'GO:0022603', 'GO:0051128'] |
| 1602 | GO:BP | GO:0048041 | focal adhesion assembly | 0,007682439 | True | negative_corr | 83 | 2460 | 28 | 17906 | 0,011382114 | 0,337349398 | ['GO:0007044', 'GO:0007160'] |
| 783 | GO:BP | GO:0009056 | catabolic process | 0,00805604 | True | positive_corr | 2693 | 1037 | 209 | 17906 | 0,201542912 | 0,077608615 | ['GO:0008152'] |
| 781 | GO:BP | GO:0015908 | fatty acid transport | 0,008087039 | True | positive_corr | 98 | 1037 | 19 | 17906 | 0,018322083 | 0,193877551 | ['GO:0006869', 'GO:0015718'] |
| 773 | GO:BP | GO:0140056 | organelle localization by membrane tethering | 0,008316214 | True | negative_corr | 175 | 2460 | 47 | 17906 | 0,019105691 | 0,268571429 | ['GO:0022406', 'GO:0051640'] |
| 1604 | GO:BP | GO:0098609 | cell-cell adhesion | 0,008583228 | True | negative_corr | 866 | 2460 | 166 | 17906 | 0,067479675 | 0,191685912 | ['GO:0007155'] |
| 1606 | GO:BP | GO:1901360 | organic cyclic compound metabolic process | 0,00984551 | True | negative_corr | 6185 | 2460 | 949 | 17906 | 0,385772358 | 0,153435732 | ['GO:0071704'] |
| 767 | GO:BP | GO:1905515 | non-motile cilium assembly | 0,009912903 | True | negative_corr | 58 | 2460 | 22 | 17906 | 0,008943089 | 0,379310345 | ['GO:0060271'] |
| 799 | GO:BP | GO:0031346 | positive regulation of cell projection organization | 0,010300971 | True | negative_corr | 395 | 2460 | 87 | 17906 | 0,035365854 | 0,220253165 | ['GO:0030030', 'GO:0031344', 'GO:0051130'] |
| 1609 | GO:BP | GO:1905039 | carboxylic acid transmembrane transport | 0,010320531 | True | positive_corr | 137 | 1037 | 23 | 17906 | 0,022179364 | 0,167883212 | ['GO:0046942', 'GO:0098656', 'GO:1903825'] |
| 762 | GO:BP | GO:1903825 | organic acid transmembrane transport | 0,010320531 | True | positive_corr | 137 | 1037 | 23 | 17906 | 0,022179364 | 0,167883212 | ['GO:0015849', 'GO:0055085'] |
| 1610 | GO:BP | GO:0007267 | cell-cell signaling | 0,010357507 | True | negative_corr | 1699 | 2460 | 296 | 17906 | 0,120325203 | 0,174220129 | ['GO:0007154', 'GO:0023052'] |
| 1611 | GO:BP | GO:0016125 | sterol metabolic process | 0,010523056 | True | positive_corr | 167 | 1037 | 26 | 17906 | 0,025072324 | 0,155688623 | ['GO:0008202', 'GO:1901615'] |
| 758 | GO:BP | GO:0006402 | mRNA catabolic process | 0,011254523 | True | negative_corr | 373 | 2460 | 83 | 17906 | 0,033739837 | 0,222520107 | ['GO:0006401', 'GO:0010629', 'GO:0016071'] |
| 620 | GO:BP | GO:0090287 | regulation of cellular response to growth factor stimulus | 0,011262267 | True | negative_corr | 300 | 2460 | 70 | 17906 | 0,028455285 | 0,233333333 | ['GO:0048583', 'GO:0050794', 'GO:0071363'] |
| 751 | GO:BP | GO:0033043 | regulation of organelle organization | 0,011378518 | True | negative_corr | 1325 | 2460 | 238 | 17906 | 0,096747967 | 0,179622642 | ['GO:0006996', 'GO:0051128'] |
| 1622 | GO:BP | GO:0042738 | exogenous drug catabolic process | 0,011652686 | True | positive_corr | 24 | 1037 | 9 | 17906 | 0,008678881 | 0,375 | ['GO:0042493', 'GO:0042737'] |
| 745 | GO:BP | GO:0006691 | leukotriene metabolic process | 0,011695565 | True | positive_corr | 30 | 1037 | 10 | 17906 | 0,009643202 | 0,333333333 | ['GO:0006690'] |
| 744 | GO:BP | GO:2000026 | regulation of multicellular organismal development | 0,011879694 | True | negative_corr | 2210 | 2460 | 373 | 17906 | 0,151626016 | 0,168778281 | ['GO:0007275', 'GO:0050793', 'GO:0051239'] |
| 763 | GO:BP | GO:0031323 | regulation of cellular metabolic process | 0,012829612 | True | negative_corr | 6393 | 2460 | 977 | 17906 | 0,397154472 | 0,152823401 | ['GO:0019222', 'GO:0044237', 'GO:0050794'] |
| 1623 | GO:BP | GO:0090109 | regulation of cell-substrate junction assembly | 0,012847668 | True | negative_corr | 63 | 2460 | 23 | 17906 | 0,009349593 | 0,365079365 | ['GO:0007044', 'GO:0150116', 'GO:1901888'] |
| 803 | GO:BP | GO:0150116 | regulation of cell-substrate junction organization | 0,012847668 | True | negative_corr | 63 | 2460 | 23 | 17906 | 0,009349593 | 0,365079365 | ['GO:0051128', 'GO:0150115'] |
| 807 | GO:BP | GO:0051893 | regulation of focal adhesion assembly | 0,012847668 | True | negative_corr | 63 | 2460 | 23 | 17906 | 0,009349593 | 0,365079365 | ['GO:0001952', 'GO:0048041', 'GO:0090109'] |
| 851 | GO:BP | GO:1905114 | cell surface receptor signaling pathway involved in cell-cell signaling | 0,012920489 | True | negative_corr | 633 | 2460 | 127 | 17906 | 0,051626016 | 0,200631912 | ['GO:0007166', 'GO:0007267'] |
| 1567 | GO:BP | GO:0060284 | regulation of cell development | 0,013362173 | True | negative_corr | 991 | 2460 | 185 | 17906 | 0,075203252 | 0,186680121 | ['GO:0045595', 'GO:0048468'] |
| 1570 | GO:BP | GO:0006464 | cellular protein modification process | 0,01367605 | True | negative_corr | 4216 | 2460 | 667 | 17906 | 0,271138211 | 0,158206831 | ['GO:0036211', 'GO:0044267'] |
| 837 | GO:BP | GO:0036211 | protein modification process | 0,01367605 | True | negative_corr | 4216 | 2460 | 667 | 17906 | 0,271138211 | 0,158206831 | ['GO:0019538', 'GO:0043412'] |
| 836 | GO:BP | GO:0019222 | regulation of metabolic process | 0,014283525 | True | negative_corr | 6982 | 2460 | 1059 | 17906 | 0,430487805 | 0,151675738 | ['GO:0008152', 'GO:0050789'] |
| 833 | GO:BP | GO:0060485 | mesenchyme development | 0,014622493 | True | negative_corr | 280 | 2460 | 66 | 17906 | 0,026829268 | 0,235714286 | ['GO:0009888', 'GO:0048513'] |
| 1572 | GO:BP | GO:0030029 | actin filament-based process | 0,0147654 | True | negative_corr | 781 | 2460 | 151 | 17906 | 0,061382114 | 0,193341869 | ['GO:0009987'] |
| 831 | GO:BP | GO:0019438 | aromatic compound biosynthetic process | 0,014965162 | True | negative_corr | 4302 | 2460 | 679 | 17906 | 0,27601626 | 0,157833566 | ['GO:0006725', 'GO:0044249'] |
| 1578 | GO:BP | GO:0032270 | positive regulation of cellular protein metabolic process | 0,015694557 | True | negative_corr | 1642 | 2460 | 286 | 17906 | 0,116260163 | 0,174177832 | ['GO:0031325', 'GO:0032268', 'GO:0044267', 'GO:0051247'] |

| | | | | | | | | | | | | | |
|---|---|---|---|---|---|---|---|---|---|---|---|---|---|
| 805 | GO:BP | GO:0030149 | sphingolipid catabolic process | 0,016372167 | True | positive_corr | 31 | 1037 | 10 | 17906 | 0,009643202 | 0,322580645 | ['GO:0006665', 'GO:0046466', 'GO:1901565'] |
| 824 | GO:BP | GO:0018130 | heterocycle biosynthetic process | 0,016995235 | True | negative_corr | 4292 | 2460 | 677 | 17906 | 0,275203252 | 0,157735322 | ['GO:0044249', 'GO:0046483'] |
| 1583 | GO:BP | GO:0031326 | regulation of cellular biosynthetic process | 0,017171823 | True | negative_corr | 4334 | 2460 | 683 | 17906 | 0,277642276 | 0,15759114 | ['GO:0009889', 'GO:0031323', 'GO:0044249'] |
| 821 | GO:BP | GO:0051640 | organelle localization | 0,017785163 | True | negative_corr | 643 | 2460 | 128 | 17906 | 0,05203252 | 0,199066874 | ['GO:0051641'] |
| 820 | GO:BP | GO:0048522 | positive regulation of cellular process | 0,018762807 | True | negative_corr | 5605 | 2460 | 864 | 17906 | 0,351219512 | 0,154148082 | ['GO:0009987', 'GO:0048518', 'GO:0050794'] |
| 1588 | GO:BP | GO:0050808 | synapse organization | 0,019361095 | True | negative_corr | 430 | 2460 | 92 | 17906 | 0,037398374 | 0,213953488 | ['GO:0016043'] |
| 812 | GO:BP | GO:0022603 | regulation of anatomical structure morphogenesis | 0,019971656 | True | negative_corr | 1194 | 2460 | 216 | 17906 | 0,087804878 | 0,180904523 | ['GO:0009653', 'GO:0050793'] |
| 811 | GO:BP | GO:0030001 | metal ion transport | 0,020488893 | True | positive_corr | 922 | 1037 | 86 | 17906 | 0,082931533 | 0,093275488 | ['GO:0006812'] |
| 810 | GO:BP | GO:0080090 | regulation of primary metabolic process | 0,020626578 | True | negative_corr | 6189 | 2460 | 946 | 17906 | 0,384552846 | 0,152851834 | ['GO:0019222', 'GO:0044238'] |
| 809 | GO:BP | GO:0035735 | intraciliary transport involved in cilium assembly | 0,020919074 | True | negative_corr | 40 | 2460 | 17 | 17906 | 0,006910569 | 0,425 | ['GO:0042073', 'GO:0060271'] |
| 808 | GO:BP | GO:0002062 | chondrocyte differentiation | 0,021299394 | True | negative_corr | 125 | 2460 | 36 | 17906 | 0,014634146 | 0,288 | ['GO:0030154', 'GO:0051216'] |
| 1579 | GO:BP | GO:0072006 | nephron development | 0,021818216 | True | negative_corr | 145 | 2460 | 40 | 17906 | 0,016260163 | 0,275862069 | ['GO:0001822', 'GO:0048856'] |
| 736 | GO:BP | GO:0015721 | bile acid and bile salt transport | 0,022583314 | True | positive_corr | 32 | 1037 | 10 | 17906 | 0,009643202 | 0,3125 | ['GO:0006869', 'GO:0015718', 'GO:0015850'] |
| 1614 | GO:BP | GO:0001570 | vasculogenesis | 0,024847868 | True | negative_corr | 83 | 2460 | 27 | 17906 | 0,01097561 | 0,325301205 | ['GO:0030154', 'GO:0048514'] |
| 1653 | GO:BP | GO:0034310 | primary alcohol catabolic process | 0,025376155 | True | positive_corr | 15 | 1037 | 7 | 17906 | 0,006750241 | 0,466666667 | ['GO:0034308', 'GO:0046164'] |
| 1652 | GO:BP | GO:0006000 | fructose metabolic process | 0,025376155 | True | positive_corr | 15 | 1037 | 7 | 17906 | 0,006750241 | 0,466666667 | ['GO:0019318'] |
| 674 | GO:BP | GO:0007160 | cell-matrix adhesion | 0,026685798 | True | negative_corr | 230 | 2460 | 56 | 17906 | 0,022764228 | 0,243478261 | ['GO:0031589'] |
| 1625 | GO:BP | GO:0009966 | regulation of signal transduction | 0,027012792 | True | negative_corr | 3244 | 2460 | 523 | 17906 | 0,212601626 | 0,161220715 | ['GO:0007165', 'GO:0010646', 'GO:0023051', 'GO:0048583'] |
| 672 | GO:BP | GO:0035924 | cellular response to vascular endothelial growth factor stimulus | 0,028147888 | True | negative_corr | 70 | 2460 | 24 | 17906 | 0,009756098 | 0,342857143 | ['GO:0071363'] |
| 1654 | GO:BP | GO:0031503 | protein-containing complex localization | 0,028642693 | True | negative_corr | 291 | 2460 | 67 | 17906 | 0,027235772 | 0,23024055 | ['GO:0008104'] |
| 666 | GO:BP | GO:0048705 | skeletal system morphogenesis | 0,029299662 | True | negative_corr | 247 | 2460 | 59 | 17906 | 0,02398374 | 0,238866397 | ['GO:0001501', 'GO:0009887'] |
| 665 | GO:BP | GO:0043412 | macromolecule modification | 0,029993797 | True | negative_corr | 4428 | 2460 | 694 | 17906 | 0,282113821 | 0,156729901 | ['GO:0043170'] |
| 1655 | GO:BP | GO:0090150 | establishment of protein localization to membrane | 0,030754952 | True | negative_corr | 331 | 2460 | 74 | 17906 | 0,030081301 | 0,223564955 | ['GO:0045184', 'GO:0072657'] |
| 1657 | GO:BP | GO:0036101 | leukotriene B4 catabolic process | 0,031077646 | True | positive_corr | 4 | 1037 | 4 | 17906 | 0,003857281 | 1 | ['GO:0036100', 'GO:0036102', 'GO:0042758', 'GO:1901616'] |
| 1634 | GO:BP | GO:0036100 | leukotriene catabolic process | 0,031077646 | True | positive_corr | 4 | 1037 | 4 | 17906 | 0,003857281 | 1 | ['GO:0006691', 'GO:1901523'] |
| 654 | GO:BP | GO:0036102 | leukotriene B4 metabolic process | 0,031077646 | True | positive_corr | 4 | 1037 | 4 | 17906 | 0,003857281 | 1 | ['GO:0001676', 'GO:0006691', 'GO:0033559', 'GO:1901615'] |
| 649 | GO:BP | GO:0071407 | cellular response to organic cyclic compound | 0,032745385 | True | positive_corr | 624 | 1037 | 63 | 17906 | 0,06075217 | 0,100961538 | ['GO:0014070', 'GO:0071310'] |
| 1662 | GO:BP | GO:0009889 | regulation of biosynthetic process | 0,032880948 | True | negative_corr | 4417 | 2460 | 692 | 17906 | 0,281300813 | 0,156667421 | ['GO:0009058', 'GO:0019222'] |
| 1664 | GO:BP | GO:0015672 | monovalent inorganic cation transport | 0,034072557 | True | positive_corr | 547 | 1037 | 57 | 17906 | 0,054966249 | 0,104204753 | ['GO:0006812'] |
| 1665 | GO:BP | GO:0030300 | regulation of intestinal cholesterol absorption | 0,034129687 | True | positive_corr | 7 | 1037 | 5 | 17906 | 0,004821601 | 0,714285714 | ['GO:0030299', 'GO:0032374', 'GO:1904729'] |
| 1671 | GO:BP | GO:0052696 | flavonoid glucuronidation | 0,034129687 | True | positive_corr | 7 | 1037 | 5 | 17906 | 0,004821601 | 0,714285714 | ['GO:0009812', 'GO:0052695'] |
| 629 | GO:BP | GO:0086042 | cardiac muscle cell-cardiac muscle cell adhesion | 0,034129687 | True | positive_corr | 7 | 1037 | 5 | 17906 | 0,004821601 | 0,714285714 | ['GO:0034109'] |
| 628 | GO:BP | GO:1904729 | regulation of intestinal lipid absorption | 0,034129687 | True | positive_corr | 7 | 1037 | 5 | 17906 | 0,004821601 | 0,714285714 | ['GO:0032368', 'GO:0098856', 'GO:1904478'] |
| 1674 | GO:BP | GO:0000470 | maturation of LSU-rRNA | 0,036019723 | True | negative_corr | 30 | 2460 | 14 | 17906 | 0,005691057 | 0,466666667 | ['GO:0006364', 'GO:0042273'] |
| 625 | GO:BP | GO:0061061 | muscle structure development | 0,037098571 | True | negative_corr | 683 | 2460 | 133 | 17906 | 0,054065041 | 0,194729136 | ['GO:0048856'] |
| 653 | GO:BP | GO:0000050 | urea cycle | 0,037100984 | True | positive_corr | 11 | 1037 | 6 | 17906 | 0,005785921 | 0,545454545 | ['GO:0019627', 'GO:0043604', 'GO:1901566'] |
| 1631 | GO:BP | GO:0034379 | very-low-density lipoprotein particle assembly | 0,037100984 | True | positive_corr | 11 | 1037 | 6 | 17906 | 0,005785921 | 0,545454545 | ['GO:0034377'] |
| 1278 | GO:BP | GO:0050770 | regulation of axonogenesis | 0,039838466 | True | negative_corr | 190 | 2460 | 48 | 17906 | 0,019512195 | 0,252631579 | ['GO:0007409', 'GO:0010769', 'GO:0010975'] |
| 695 | GO:BP | GO:0150115 | cell-substrate junction organization | 0,0398484 | True | negative_corr | 99 | 2460 | 30 | 17906 | 0,012195122 | 0,303030303 | ['GO:0034330'] |
| 1630 | GO:BP | GO:0007044 | cell-substrate junction assembly | 0,0398484 | True | negative_corr | 99 | 2460 | 30 | 17906 | 0,012195122 | 0,303030303 | ['GO:0034329', 'GO:0150115'] |
| 718 | GO:BP | GO:0051169 | nuclear transport | 0,0400075 | True | negative_corr | 362 | 2460 | 79 | 17906 | 0,032113821 | 0,218232044 | ['GO:0046907'] |
| 719 | GO:BP | GO:0010769 | regulation of cell morphogenesis involved in differentiation | 0,041347939 | True | negative_corr | 311 | 2460 | 70 | 17906 | 0,028455285 | 0,225080386 | ['GO:0000904', 'GO:0022604', 'GO:0060284'] |
| 706 | GO:BP | GO:0006364 | rRNA processing | 0,041458629 | True | negative_corr | 217 | 2460 | 53 | 17906 | 0,021544715 | 0,244239631 | ['GO:0016072', 'GO:0034470', 'GO:0042254'] |
| 732 | GO:BP | GO:0046907 | intracellular transport | 0,042311484 | True | negative_corr | 1781 | 2460 | 304 | 17906 | 0,123577236 | 0,170690623 | ['GO:0006810', 'GO:0051641'] |
| 726 | GO:BP | GO:0030036 | actin cytoskeleton organization | 0,043443393 | True | negative_corr | 679 | 2460 | 132 | 17906 | 0,053658537 | 0,194403535 | ['GO:0007010', 'GO:0030029'] |
| 733 | GO:BP | GO:0051493 | regulation of cytoskeleton organization | 0,045370584 | True | negative_corr | 564 | 2460 | 113 | 17906 | 0,045934959 | 0,20035461 | ['GO:0007010', 'GO:0033043'] |
| 702 | GO:BP | GO:0051181 | cofactor transport | 0,046560564 | True | positive_corr | 49 | 1037 | 12 | 17906 | 0,011571842 | 0,244897959 | ['GO:0006810'] |
| 707 | GO:BP | GO:0016072 | rRNA metabolic process | 0,048508334 | True | negative_corr | 229 | 2460 | 55 | 17906 | 0,022357724 | 0,240174672 | ['GO:0034660'] |
| 1876 | GO:CC | GO:0045177 | apical part of cell | 1,40868E-35 | True | positive_corr | 400 | 1107 | 102 | 18856 | 0,092140921 | 0,255 | ['GO:0110165'] |
| 1872 | GO:CC | GO:0016324 | apical plasma membrane | 3,62706E-35 | True | positive_corr | 331 | 1107 | 92 | 18856 | 0,083107498 | 0,277945619 | ['GO:0045177', 'GO:0098590'] |
| 1698 | GO:CC | GO:0031224 | intrinsic component of membrane | 2,26768E-31 | True | positive_corr | 5842 | 1107 | 530 | 18856 | 0,478771454 | 0,090722355 | ['GO:0016020', 'GO:0110165'] |
| 1553 | GO:CC | GO:0005622 | intracellular | 5,20674E-31 | True | negative_corr | 14626 | 2576 | 2224 | 18856 | 0,863354037 | 0,152057979 | ['GO:0005575'] |
| 1789 | GO:CC | GO:0016021 | integral component of membrane | 2,66808E-30 | True | positive_corr | 5692 | 1107 | 517 | 18856 | 0,467028004 | 0,090829234 | ['GO:0031224'] |
| 1346 | GO:CC | GO:0005903 | brush border | 5,60193E-27 | True | positive_corr | 108 | 1107 | 47 | 18856 | 0,042457091 | 0,435185185 | ['GO:0098862'] |
| 1699 | GO:CC | GO:0016020 | membrane | 6,16756E-25 | True | positive_corr | 9731 | 1107 | 744 | 18856 | 0,672086721 | 0,076456685 | ['GO:0110165'] |
| 1661 | GO:CC | GO:0070062 | extracellular exosome | 2,91956E-23 | True | positive_corr | 2144 | 1107 | 244 | 18856 | 0,220415537 | 0,11380597 | ['GO:0005615', 'GO:1903561'] |
| 1320 | GO:CC | GO:1903561 | extracellular vesicle | 5,87364E-23 | True | positive_corr | 2167 | 1107 | 245 | 18856 | 0,22131888 | 0,113059529 | ['GO:0031982', 'GO:0043230'] |
| 1757 | GO:CC | GO:0043230 | extracellular organelle | 8,21875E-23 | True | positive_corr | 2172 | 1107 | 245 | 18856 | 0,22131888 | 0,112799263 | ['GO:0005576', 'GO:0043226'] |
| 1780 | GO:CC | GO:0043226 | organelle | 9,58606E-23 | True | negative_corr | 13720 | 2576 | 2085 | 18856 | 0,80939441 | 0,15196793 | ['GO:0110165'] |
| 1541 | GO:CC | GO:0043229 | intracellular organelle | 7,73468E-22 | True | negative_corr | 12840 | 2576 | 1972 | 18856 | 0,76552795 | 0,153582555 | ['GO:0005622', 'GO:0043226'] |
| 1603 | GO:CC | GO:0098862 | cluster of actin-based cell projections | 8,22918E-21 | True | positive_corr | 161 | 1107 | 50 | 18856 | 0,045167118 | 0,310559006 | ['GO:0110165'] |
| 1520 | GO:CC | GO:0098590 | plasma membrane region | 3,70911E-20 | True | positive_corr | 1153 | 1107 | 154 | 18856 | 0,139114724 | 0,133564614 | ['GO:0005886', 'GO:0016020'] |
| 1466 | GO:CC | GO:0005634 | nucleus | 2,97467E-18 | True | negative_corr | 7486 | 2576 | 1239 | 18856 | 0,480978261 | 0,16550895 | ['GO:0043231'] |
| 1521 | GO:CC | GO:0031982 | vesicle | 4,14044E-17 | True | positive_corr | 3914 | 1107 | 354 | 18856 | 0,319783198 | 0,090444558 | ['GO:0043227'] |
| 1639 | GO:CC | GO:0005829 | cytosol | 1,83152E-16 | True | negative_corr | 5154 | 2576 | 894 | 18856 | 0,347049689 | 0,173457509 | ['GO:0005737', 'GO:0110165'] |

| | | | | | | | | | | | | | |
|---|---|---|---|---|---|---|---|---|---|---|---|---|---|
| 1486 | GO:CC | GO:0030054 | cell junction | 1,75784E-15 | True | negative_corr | 1327 | 2576 | 292 | 18856 | 0,113354037 | 0,220045215 | ['GO:0110165'] |
| 1755 | GO:CC | GO:0043232 | intracellular non-membrane-bounded organelle | 3,92908E-15 | True | negative_corr | 4954 | 2576 | 857 | 18856 | 0,332686335 | 0,172991522 | ['GO:0043228', 'GO:0043229'] |
| 1517 | GO:CC | GO:0043228 | non-membrane-bounded organelle | 4,7389E-15 | True | negative_corr | 4964 | 2576 | 858 | 18856 | 0,333074534 | 0,17284448 | ['GO:0043226'] |
| 1840 | GO:CC | GO:0031526 | brush border membrane | 4,10285E-14 | True | positive_corr | 56 | 1107 | 25 | 18856 | 0,022583559 | 0,446428571 | ['GO:0005903', 'GO:0031253'] |
| 1830 | GO:CC | GO:0070161 | anchoring junction | 5,73464E-13 | True | negative_corr | 785 | 2576 | 188 | 18856 | 0,072981366 | 0,239490446 | ['GO:0030054'] |
| 1589 | GO:CC | GO:0015630 | microtubule cytoskeleton | 6,38318E-13 | True | negative_corr | 1242 | 2576 | 268 | 18856 | 0,104037267 | 0,215780998 | ['GO:0005856'] |
| 1527 | GO:CC | GO:0005856 | cytoskeleton | 7,66492E-13 | True | negative_corr | 2263 | 2576 | 435 | 18856 | 0,16886646 | 0,192222713 | ['GO:0043232'] |
| 1516 | GO:CC | GO:0005737 | cytoplasm | 9,82696E-13 | True | negative_corr | 11634 | 2576 | 1767 | 18856 | 0,685947205 | 0,151882414 | ['GO:0005622', 'GO:0110165'] |
| 1388 | GO:CC | GO:0043227 | membrane-bounded organelle | 7,31002E-12 | True | negative_corr | 12584 | 2576 | 1885 | 18856 | 0,731754658 | 0,149793388 | ['GO:0043226'] |
| 1390 | GO:CC | GO:0042995 | cell projection | 1,25352E-11 | True | negative_corr | 2288 | 2576 | 433 | 18856 | 0,168090062 | 0,189248252 | ['GO:0110165'] |
| 1313 | GO:CC | GO:0043231 | intracellular membrane-bounded organelle | 4,44039E-11 | True | negative_corr | 11067 | 2576 | 1681 | 18856 | 0,652562112 | 0,151893015 | ['GO:0043227', 'GO:0043229'] |
| 1717 | GO:CC | GO:0005925 | focal adhesion | 4,86717E-11 | True | negative_corr | 410 | 2576 | 112 | 18856 | 0,043478261 | 0,273170732 | ['GO:0030055'] |
| 1756 | GO:CC | GO:0005815 | microtubule organizing center | 5,16217E-11 | True | negative_corr | 770 | 2576 | 179 | 18856 | 0,069487578 | 0,232467532 | ['GO:0015630', 'GO:0110165'] |
| 1498 | GO:CC | GO:0120025 | plasma membrane bounded cell projection | 8,17643E-11 | True | negative_corr | 2200 | 2576 | 415 | 18856 | 0,161102484 | 0,188636364 | ['GO:0042995'] |
| 37 | GO:CC | GO:0031090 | organelle membrane | 8,57375E-11 | True | positive_corr | 3531 | 1107 | 303 | 18856 | 0,273712737 | 0,085811385 | ['GO:0016020', 'GO:0043227'] |
| 1071 | GO:CC | GO:0045202 | synapse | 1,23816E-10 | True | negative_corr | 1362 | 2576 | 279 | 18856 | 0,108307453 | 0,204845815 | ['GO:0110165'] |
| 1013 | GO:CC | GO:0030055 | cell-substrate junction | 1,64924E-10 | True | negative_corr | 417 | 2576 | 112 | 18856 | 0,043478261 | 0,268585132 | ['GO:0070161'] |
| 637 | GO:CC | GO:0022626 | cytosolic ribosome | 2,44646E-10 | True | negative_corr | 108 | 2576 | 45 | 18856 | 0,017468944 | 0,416666667 | ['GO:0005829', 'GO:0005840'] |
| 617 | GO:CC | GO:0012505 | endomembrane system | 2,44923E-10 | True | positive_corr | 4539 | 1107 | 368 | 18856 | 0,332429991 | 0,081075127 | ['GO:0110165'] |
| 1025 | GO:CC | GO:0071944 | cell periphery | 3,17566E-10 | True | positive_corr | 5732 | 1107 | 444 | 18856 | 0,401084011 | 0,077459874 | ['GO:0110165'] |
| 613 | GO:CC | GO:0005902 | microvillus | 4,33994E-10 | True | positive_corr | 85 | 1107 | 26 | 18856 | 0,023486902 | 0,305882353 | ['GO:0098858'] |
| 590 | GO:CC | GO:0005615 | extracellular space | 6,36967E-10 | True | positive_corr | 3543 | 1107 | 300 | 18856 | 0,27100271 | 0,084674005 | ['GO:0005576', 'GO:0110165'] |
| 576 | GO:CC | GO:0005886 | plasma membrane | 1,20904E-09 | True | positive_corr | 5604 | 1107 | 433 | 18856 | 0,391147245 | 0,077266238 | ['GO:0016020', 'GO:0071944'] |
| 569 | GO:CC | GO:0005813 | centrosome | 1,55998E-09 | True | negative_corr | 586 | 2576 | 141 | 18856 | 0,054736025 | 0,240614334 | ['GO:0005815'] |
| 561 | GO:CC | GO:0022625 | cytosolic large ribosomal subunit | 3,60036E-09 | True | negative_corr | 59 | 2576 | 30 | 18856 | 0,011645963 | 0,508474576 | ['GO:0015934', 'GO:0022626'] |
| 560 | GO:CC | GO:0016323 | basolateral plasma membrane | 3,70878E-09 | True | positive_corr | 221 | 1107 | 42 | 18856 | 0,037940379 | 0,190045249 | ['GO:0098590'] |
| 555 | GO:CC | GO:0005911 | cell-cell junction | 4,62192E-09 | True | positive_corr | 439 | 1107 | 64 | 18856 | 0,057813911 | 0,145785877 | ['GO:0070161'] |
| 546 | GO:CC | GO:0043296 | apical junction complex | 7,14947E-09 | True | positive_corr | 148 | 1107 | 33 | 18856 | 0,029810298 | 0,222972973 | ['GO:0005911'] |
| 1072 | GO:CC | GO:0005654 | nucleoplasm | 9,26798E-09 | True | negative_corr | 4012 | 2576 | 678 | 18856 | 0,263198758 | 0,168993021 | ['GO:0031981', 'GO:0110165'] |
| 534 | GO:CC | GO:0005887 | integral component of plasma membrane | 5,68727E-08 | True | positive_corr | 1624 | 1107 | 157 | 18856 | 0,141824752 | 0,096674877 | ['GO:0016021', 'GO:0031226'] |
| 531 | GO:CC | GO:0005789 | endoplasmic reticulum membrane | 6,39563E-08 | True | positive_corr | 1075 | 1107 | 115 | 18856 | 0,103884372 | 0,106976744 | ['GO:0031090', 'GO:0042175'] |
| 512 | GO:CC | GO:0036064 | ciliary basal body | 7,3484E-08 | True | negative_corr | 133 | 2576 | 47 | 18856 | 0,018245342 | 0,353383459 | ['GO:0005815', 'GO:0005929'] |
| 509 | GO:CC | GO:0031226 | intrinsic component of plasma membrane | 8,22743E-08 | True | positive_corr | 1700 | 1107 | 162 | 18856 | 0,146341463 | 0,095294118 | ['GO:0005886', 'GO:0031224'] |
| 508 | GO:CC | GO:0042175 | nuclear outer membrane-endoplasmic reticulum membrane network | 1,13533E-07 | True | positive_corr | 1098 | 1107 | 116 | 18856 | 0,104787715 | 0,10564663 | ['GO:0012505', 'GO:0016020'] |
| 483 | GO:CC | GO:0031253 | cell projection membrane | 1,20852E-07 | True | positive_corr | 335 | 1107 | 51 | 18856 | 0,046070461 | 0,152238806 | ['GO:0098590', 'GO:0120025'] |
| 477 | GO:CC | GO:0031981 | nuclear lumen | 1,69481E-07 | True | negative_corr | 4780 | 2576 | 781 | 18856 | 0,30318323 | 0,163389121 | ['GO:0005634', 'GO:0070013'] |
| 1119 | GO:CC | GO:1990904 | ribonucleoprotein complex | 2,83447E-07 | True | negative_corr | 697 | 2576 | 153 | 18856 | 0,05939441 | 0,219512195 | ['GO:0032991'] |
| 639 | GO:CC | GO:0031528 | microvillus membrane | 7,42839E-07 | True | positive_corr | 24 | 1107 | 12 | 18856 | 0,010840108 | 0,5 | ['GO:0005902', 'GO:0031253'] |
| 1012 | GO:CC | GO:0005576 | extracellular region | 1,58111E-06 | True | positive_corr | 4548 | 1107 | 349 | 18856 | 0,315266486 | 0,076737027 | ['GO:0110165'] |
| 645 | GO:CC | GO:0098858 | actin-based cell projection | 2,30189E-06 | True | positive_corr | 211 | 1107 | 36 | 18856 | 0,032520325 | 0,170616114 | ['GO:0120025'] |
| 664 | GO:CC | GO:0070160 | tight junction | 3,44976E-06 | True | positive_corr | 132 | 1107 | 27 | 18856 | 0,024390244 | 0,204545455 | ['GO:0005911'] |
| 861 | GO:CC | GO:0005783 | endoplasmic reticulum | 4,34009E-06 | True | positive_corr | 1810 | 1107 | 163 | 18856 | 0,147244806 | 0,090055249 | ['GO:0005737', 'GO:0012505', 'GO:0043231'] |
| 832 | GO:CC | GO:0035869 | ciliary transition zone | 7,3624E-06 | True | negative_corr | 72 | 2576 | 29 | 18856 | 0,011257764 | 0,402777778 | ['GO:0005929', 'GO:0110165'] |
| 910 | GO:CC | GO:0031974 | membrane-enclosed lumen | 1,03604E-05 | True | negative_corr | 5898 | 2576 | 926 | 18856 | 0,35947205 | 0,157002374 | ['GO:0110165'] |
| 920 | GO:CC | GO:0043233 | organelle lumen | 1,03604E-05 | True | negative_corr | 5898 | 2576 | 926 | 18856 | 0,35947205 | 0,157002374 | ['GO:0031974', 'GO:0043226'] |
| 806 | GO:CC | GO:0070013 | intracellular organelle lumen | 1,03604E-05 | True | negative_corr | 5898 | 2576 | 926 | 18856 | 0,35947205 | 0,157002374 | ['GO:0043229', 'GO:0043233'] |
| 1277 | GO:CC | GO:0062023 | collagen-containing extracellular matrix | 1,19684E-05 | True | negative_corr | 407 | 2576 | 96 | 18856 | 0,037267081 | 0,235872236 | ['GO:0031012'] |
| 952 | GO:CC | GO:0044391 | ribosomal subunit | 1,35839E-05 | True | negative_corr | 192 | 2576 | 55 | 18856 | 0,021350932 | 0,286458333 | ['GO:0005840', 'GO:1990904'] |
| 795 | GO:CC | GO:0036477 | somatodendritic compartment | 1,4938E-05 | True | negative_corr | 863 | 2576 | 174 | 18856 | 0,067546584 | 0,201622248 | ['GO:0110165'] |
| 777 | GO:CC | GO:0005923 | bicellular tight junction | 1,92173E-05 | True | positive_corr | 125 | 1107 | 25 | 18856 | 0,022583559 | 0,2 | ['GO:0043296', 'GO:0070160'] |
| 770 | GO:CC | GO:0016328 | lateral plasma membrane | 2,07758E-05 | True | positive_corr | 62 | 1107 | 17 | 18856 | 0,01535682 | 0,274193548 | ['GO:0005886', 'GO:0110165'] |
| 453 | GO:CC | GO:0005840 | ribosome | 2,14429E-05 | True | negative_corr | 245 | 2576 | 65 | 18856 | 0,025232919 | 0,265306122 | ['GO:0043232'] |
| 969 | GO:CC | GO:0005929 | cilium | 2,45709E-05 | True | negative_corr | 642 | 2576 | 136 | 18856 | 0,052795031 | 0,211838006 | ['GO:0043226', 'GO:0120025'] |
| 760 | GO:CC | GO:0097542 | ciliary tip | 5,04676E-05 | True | negative_corr | 46 | 2576 | 21 | 18856 | 0,008152174 | 0,456521739 | ['GO:0005929', 'GO:0110165'] |
| 971 | GO:CC | GO:0005730 | nucleolus | 5,13021E-05 | True | negative_corr | 927 | 2576 | 182 | 18856 | 0,070652174 | 0,196332255 | ['GO:0031981', 'GO:0043232'] |
| 755 | GO:CC | GO:0030027 | lamellipodium | 0,000105299 | True | negative_corr | 198 | 2576 | 54 | 18856 | 0,020962733 | 0,272727273 | ['GO:0031252', 'GO:0120025'] |
| 977 | GO:CC | GO:0070993 | translation preinitiation complex | 0,000120983 | True | negative_corr | 18 | 2576 | 12 | 18856 | 0,004658385 | 0,666666667 | ['GO:0005737', 'GO:1990904'] |
| 986 | GO:CC | GO:0070161 | anchoring junction | 0,000163611 | True | positive_corr | 785 | 1107 | 81 | 18856 | 0,073170732 | 0,103184713 | ['GO:0030054'] |
| 746 | GO:CC | GO:0015934 | large ribosomal subunit | 0,000248367 | True | negative_corr | 119 | 2576 | 37 | 18856 | 0,014363354 | 0,31092437 | ['GO:0044391'] |
| 735 | GO:CC | GO:0016327 | apicolateral plasma membrane | 0,000250811 | True | positive_corr | 20 | 1107 | 9 | 18856 | 0,008130081 | 0,45 | ['GO:0098590'] |
| 995 | GO:CC | GO:0030425 | dendrite | 0,000310025 | True | negative_corr | 622 | 2576 | 128 | 18856 | 0,049689441 | 0,205787781 | ['GO:0043005', 'GO:0097447'] |
| 1002 | GO:CC | GO:0005737 | cytoplasm | 0,000362834 | True | positive_corr | 11634 | 1107 | 757 | 18856 | 0,683830172 | 0,065067904 | ['GO:0005622', 'GO:0110165'] |
| 667 | GO:CC | GO:0097447 | dendritic tree | 0,000370047 | True | negative_corr | 624 | 2576 | 128 | 18856 | 0,049689441 | 0,205128205 | ['GO:0036477', 'GO:0043005'] |
| 970 | GO:CC | GO:0005874 | microtubule | 0,000464347 | True | negative_corr | 427 | 2576 | 94 | 18856 | 0,036490683 | 0,220140515 | ['GO:0015630', 'GO:0099513'] |
| 450 | GO:CC | GO:0016282 | eukaryotic 43S preinitiation complex | 0,000596137 | True | negative_corr | 17 | 2576 | 11 | 18856 | 0,004270186 | 0,647058824 | ['GO:0070993'] |

| | | | | | | | | | | | | | |
|---|---|---|---|---|---|---|---|---|---|---|---|---|---|
| 883 | GO:CC | GO:0005788 | endoplasmic reticulum lumen | 0,000748922 | True | negative_corr | 307 | 2576 | 72 | 18856 | 0,027950311 | 0,234527687 | ['GO:0005783', 'GO:0070013'] |
| 320 | GO:CC | GO:0097546 | ciliary base | 0,00094517 | True | negative_corr | 34 | 2576 | 16 | 18856 | 0,00621118 | 0,470588235 | ['GO:0005929', 'GO:0110165'] |
| 185 | GO:CC | GO:0043005 | neuron projection | 0,001020264 | True | negative_corr | 1376 | 2576 | 246 | 18856 | 0,095496894 | 0,17877907 | ['GO:0120025'] |
| 86 | GO:CC | GO:0033290 | eukaryotic 48S preinitiation complex | 0,001220161 | True | negative_corr | 15 | 2576 | 10 | 18856 | 0,003881988 | 0,666666667 | ['GO:0070993'] |
| 1157 | GO:CC | GO:0005814 | centriole | 0,001769203 | True | negative_corr | 143 | 2576 | 40 | 18856 | 0,01552795 | 0,27972028 | ['GO:0005815', 'GO:0043232'] |
| 1215 | GO:CC | GO:0032391 | photoreceptor connecting cilium | 0,001820444 | True | negative_corr | 43 | 2576 | 18 | 18856 | 0,006987578 | 0,418604651 | ['GO:0035869', 'GO:0097733'] |
| 1214 | GO:CC | GO:0098794 | postsynapse | 0,003240652 | True | negative_corr | 638 | 2576 | 126 | 18856 | 0,048913043 | 0,197492163 | ['GO:0045202', 'GO:0110165'] |
| 90 | GO:CC | GO:0099081 | supramolecular polymer | 0,003653623 | True | negative_corr | 1001 | 2576 | 184 | 18856 | 0,071428571 | 0,183816184 | ['GO:0099080'] |
| 292 | GO:CC | GO:0044297 | cell body | 0,003992246 | True | negative_corr | 592 | 2576 | 118 | 18856 | 0,045807453 | 0,199324324 | ['GO:0110165'] |
| 1196 | GO:CC | GO:0034385 | triglyceride-rich plasma lipoprotein particle | 0,004723757 | True | positive_corr | 21 | 1107 | 8 | 18856 | 0,007226739 | 0,380952381 | ['GO:0034358'] |
| 333 | GO:CC | GO:0034361 | very-low-density lipoprotein particle | 0,004723757 | True | positive_corr | 21 | 1107 | 8 | 18856 | 0,007226739 | 0,380952381 | ['GO:0034385'] |
| 101 | GO:CC | GO:0031012 | extracellular matrix | 0,004854845 | True | negative_corr | 534 | 2576 | 108 | 18856 | 0,041925466 | 0,202247191 | ['GO:0005576', 'GO:0110165'] |
| 130 | GO:CC | GO:0099512 | supramolecular fiber | 0,004874021 | True | negative_corr | 993 | 2576 | 182 | 18856 | 0,070652174 | 0,183282981 | ['GO:0099081'] |
| 252 | GO:CC | GO:0032991 | protein-containing complex | 0,005141135 | True | negative_corr | 5484 | 2576 | 840 | 18856 | 0,326086957 | 0,153172867 | ['GO:0005575'] |
| 236 | GO:CC | GO:0005852 | eukaryotic translation initiation factor 3 complex | 0,006078027 | True | negative_corr | 17 | 2576 | 10 | 18856 | 0,003881988 | 0,588235294 | ['GO:0005737', 'GO:0032991'] |
| 1179 | GO:CC | GO:0099080 | supramolecular complex | 0,006985953 | True | negative_corr | 1308 | 2576 | 230 | 18856 | 0,089285714 | 0,175840979 | ['GO:0110165'] |
| 151 | GO:CC | GO:0005911 | cell-cell junction | 0,008588881 | True | negative_corr | 439 | 2576 | 91 | 18856 | 0,035326087 | 0,207289294 | ['GO:0070161'] |
| 204 | GO:CC | GO:0030990 | intraciliary transport particle | 0,00989627 | True | negative_corr | 28 | 2576 | 13 | 18856 | 0,005046584 | 0,464285714 | ['GO:0032991'] |
| 189 | GO:CC | GO:0098984 | neuron to neuron synapse | 0,01213713 | True | negative_corr | 372 | 2576 | 79 | 18856 | 0,030667702 | 0,212365591 | ['GO:0045202'] |
| 180 | GO:CC | GO:0042383 | sarcolemma | 0,013346836 | True | negative_corr | 134 | 2576 | 36 | 18856 | 0,013975155 | 0,268656716 | ['GO:0005886'] |
| 1181 | GO:CC | GO:0005777 | peroxisome | 0,016443418 | True | positive_corr | 137 | 1107 | 21 | 18856 | 0,01897019 | 0,153284672 | ['GO:0042579'] |
| 364 | GO:CC | GO:0042579 | microbody | 0,016443418 | True | positive_corr | 137 | 1107 | 21 | 18856 | 0,01897019 | 0,153284672 | ['GO:0005737', 'GO:0043231'] |
| 187 | GO:CC | GO:0030054 | cell junction | 0,018421156 | True | positive_corr | 1327 | 1107 | 112 | 18856 | 0,101174345 | 0,084400904 | ['GO:0110165'] |
| 1238 | GO:CC | GO:0005915 | zonula adherens | 0,023954046 | True | positive_corr | 9 | 1107 | 5 | 18856 | 0,004516712 | 0,555555556 | ['GO:0005912', 'GO:0043296'] |
| 446 | GO:CC | GO:0005844 | polysome | 0,023995723 | True | negative_corr | 73 | 2576 | 23 | 18856 | 0,008928571 | 0,315068493 | ['GO:1990904'] |
| 374 | GO:CC | GO:0005581 | collagen trimer | 0,024099606 | True | negative_corr | 97 | 2576 | 28 | 18856 | 0,010869565 | 0,288659794 | ['GO:0032991'] |
| 46 | GO:CC | GO:0042627 | chylomicron | 0,027053272 | True | positive_corr | 14 | 1107 | 6 | 18856 | 0,005420054 | 0,428571429 | ['GO:0034358'] |
| 1124 | GO:CC | GO:0031252 | cell leading edge | 0,02908999 | True | negative_corr | 417 | 2576 | 85 | 18856 | 0,032996894 | 0,20383693 | ['GO:0110165'] |
| 443 | GO:CC | GO:0032279 | asymmetric synapse | 0,035575784 | True | negative_corr | 348 | 2576 | 73 | 18856 | 0,028338509 | 0,209770115 | ['GO:0098984'] |
| 379 | GO:CC | GO:0043025 | neuronal cell body | 0,037147128 | True | negative_corr | 523 | 2576 | 102 | 18856 | 0,039596273 | 0,195028681 | ['GO:0036477', 'GO:0044297'] |
| 1255 | GO:CC | GO:0014069 | postsynaptic density | 0,042506103 | True | negative_corr | 344 | 2576 | 72 | 18856 | 0,027950311 | 0,209302326 | ['GO:0032279', 'GO:0099572'] |
| 405 | GO:CC | GO:0034451 | centriolar satellite | 0,043067756 | True | negative_corr | 95 | 2576 | 27 | 18856 | 0,010481366 | 0,284210526 | ['GO:0005813', 'GO:0110165'] |
| 1597 | GO:MF | GO:0005515 | protein binding | 3,71655E-19 | True | negative_corr | 12743 | 2548 | 1991 | 18126 | 0,781397174 | 0,156242643 | ['GO:0005488'] |
| 1615 | GO:MF | GO:0005215 | transporter activity | 6,09623E-19 | True | positive_corr | 1223 | 1059 | 157 | 18126 | 0,148253069 | 0,128372854 | ['GO:0003674'] |
| 871 | GO:MF | GO:0022857 | transmembrane transporter activity | 2,00293E-17 | True | positive_corr | 1068 | 1059 | 140 | 18126 | 0,132200189 | 0,131086142 | ['GO:0005215'] |
| 1564 | GO:MF | GO:0005488 | binding | 5,48923E-15 | True | negative_corr | 15944 | 2548 | 2364 | 18126 | 0,927786499 | 0,148268941 | ['GO:0003674'] |
| 1565 | GO:MF | GO:0015075 | ion transmembrane transporter activity | 2,15305E-14 | True | positive_corr | 887 | 1059 | 117 | 18126 | 0,110481586 | 0,131905299 | ['GO:0022857'] |
| 844 | GO:MF | GO:0008509 | anion transmembrane transporter activity | 8,30308E-13 | True | positive_corr | 336 | 1059 | 61 | 18126 | 0,057601511 | 0,181547619 | ['GO:0015075'] |
| 1891 | GO:MF | GO:0003824 | catalytic activity | 1,77097E-12 | True | positive_corr | 5853 | 1059 | 460 | 18126 | 0,434372049 | 0,078592175 | ['GO:0003674'] |
| 1601 | GO:MF | GO:0015318 | inorganic molecular entity transmembrane transporter activity | 6,7066E-12 | True | positive_corr | 830 | 1059 | 106 | 18126 | 0,100094429 | 0,127710843 | ['GO:0022857'] |
| 761 | GO:MF | GO:0008514 | organic anion transmembrane transporter activity | 2,26875E-10 | True | positive_corr | 211 | 1059 | 43 | 18126 | 0,040604344 | 0,203791469 | ['GO:0008509'] |
| 764 | GO:MF | GO:0022804 | active transmembrane transporter activity | 1,05016E-09 | True | positive_corr | 352 | 1059 | 57 | 18126 | 0,053824363 | 0,161931818 | ['GO:0022857'] |
| 769 | GO:MF | GO:0015291 | secondary active transmembrane transporter activity | 4,0221E-09 | True | positive_corr | 238 | 1059 | 44 | 18126 | 0,041548631 | 0,18487395 | ['GO:0022804'] |
| 823 | GO:MF | GO:0048037 | cofactor binding | 7,04507E-09 | True | positive_corr | 517 | 1059 | 71 | 18126 | 0,067044381 | 0,137330754 | ['GO:0005488'] |
| 1890 | GO:MF | GO:0015370 | solute:sodium symporter activity | 1,10793E-08 | True | positive_corr | 75 | 1059 | 23 | 18126 | 0,021718602 | 0,306666667 | ['GO:0015081', 'GO:0015294'] |
| 822 | GO:MF | GO:0046943 | carboxylic acid transmembrane transporter activity | 1,38931E-08 | True | positive_corr | 150 | 1059 | 33 | 18126 | 0,031161473 | 0,22 | ['GO:0005342', 'GO:0008514', 'GO:0015318'] |
| 1889 | GO:MF | GO:0005342 | organic acid transmembrane transporter activity | 1,38931E-08 | True | positive_corr | 150 | 1059 | 33 | 18126 | 0,031161473 | 0,22 | ['GO:0022857'] |
| 1874 | GO:MF | GO:0016491 | oxidoreductase activity | 8,41472E-08 | True | positive_corr | 768 | 1059 | 90 | 18126 | 0,084985836 | 0,1171875 | ['GO:0003824'] |
| 451 | GO:MF | GO:0015293 | symporter activity | 9,89386E-08 | True | positive_corr | 144 | 1059 | 31 | 18126 | 0,029272899 | 0,215277778 | ['GO:0015291'] |
| 663 | GO:MF | GO:0015294 | solute:cation symporter activity | 1,12924E-07 | True | positive_corr | 105 | 1059 | 26 | 18126 | 0,024551464 | 0,247619048 | ['GO:0015293', 'GO:0022853', 'GO:0022890'] |
| 676 | GO:MF | GO:0008194 | UDP-glycosyltransferase activity | 4,86032E-07 | True | positive_corr | 153 | 1059 | 31 | 18126 | 0,029272899 | 0,202614379 | ['GO:0016757'] |
| 1773 | GO:MF | GO:0022853 | active ion transmembrane transporter activity | 4,93539E-07 | True | positive_corr | 245 | 1059 | 41 | 18126 | 0,03871577 | 0,167346939 | ['GO:0015075', 'GO:0022804'] |
| 457 | GO:MF | GO:0008324 | cation transmembrane transporter activity | 1,01139E-06 | True | positive_corr | 660 | 1059 | 78 | 18126 | 0,073654391 | 0,118181818 | ['GO:0015075'] |
| 475 | GO:MF | GO:0005496 | steroid binding | 1,97563E-06 | True | positive_corr | 95 | 1059 | 23 | 18126 | 0,021718602 | 0,242105263 | ['GO:0008289', 'GO:0097159'] |
| 1762 | GO:MF | GO:0015081 | sodium ion transmembrane transporter activity | 2,80902E-06 | True | positive_corr | 155 | 1059 | 30 | 18126 | 0,028328612 | 0,193548387 | ['GO:0015077', 'GO:0046873'] |
| 1796 | GO:MF | GO:0005319 | lipid transporter activity | 3,75777E-06 | True | positive_corr | 148 | 1059 | 29 | 18126 | 0,027384325 | 0,195945946 | ['GO:0005215'] |
| 392 | GO:MF | GO:0016712 | oxidoreductase activity, acting on paired donors, with incorporation or | 4,35528E-06 | True | positive_corr | 37 | 1059 | 14 | 18126 | 0,013220019 | 0,378378378 | ['GO:0004497', 'GO:0016705'] |
| 380 | GO:MF | GO:0016616 | oxidoreductase activity, acting on the CH-OH group of donors, NAD | 6,11214E-06 | True | positive_corr | 125 | 1059 | 26 | 18126 | 0,024551464 | 0,208 | ['GO:0016614'] |
| 1801 | GO:MF | GO:0016614 | oxidoreductase activity, acting on CH-OH group of donors | 6,60144E-06 | True | positive_corr | 134 | 1059 | 27 | 18126 | 0,025495751 | 0,201492537 | ['GO:0016491'] |
| 1807 | GO:MF | GO:0005198 | structural molecule activity | 8,00574E-06 | True | negative_corr | 683 | 2548 | 149 | 18126 | 0,058477237 | 0,218155198 | ['GO:0003674'] |
| 1547 | GO:MF | GO:0016709 | oxidoreductase activity, acting on paired donors, with incorporation or | 9,58803E-06 | True | positive_corr | 45 | 1059 | 15 | 18126 | 0,014164306 | 0,333333333 | ['GO:0004497', 'GO:0016705'] |
| 1731 | GO:MF | GO:0008092 | cytoskeletal protein binding | 1,19027E-05 | True | negative_corr | 991 | 2548 | 201 | 18126 | 0,0788854 | 0,202825429 | ['GO:0005515'] |
| 1812 | GO:MF | GO:0050662 | coenzyme binding | 1,40492E-05 | True | positive_corr | 295 | 1059 | 43 | 18126 | 0,040604344 | 0,145762712 | ['GO:0048037'] |
| 558 | GO:MF | GO:0004497 | monooxygenase activity | 1,52024E-05 | True | positive_corr | 105 | 1059 | 23 | 18126 | 0,021718602 | 0,219047619 | ['GO:0016491'] |
| 1723 | GO:MF | GO:0022890 | inorganic cation transmembrane transporter activity | 2,15393E-05 | True | positive_corr | 609 | 1059 | 70 | 18126 | 0,066100094 | 0,114942529 | ['GO:0008324', 'GO:0015318'] |

| | | | | | | | | | | | | | |
|---|---|---|---|---|---|---|---|---|---|---|---|---|---|
| 1628 | GO:MF | GO:0016758 | transferase activity, transferring hexosyl groups | 2,16452E-05 | True | positive_corr | 217 | 1059 | 35 | 18126 | 0,033050047 | 0,161290323 | ['GO:0016757'] |
| 1707 | GO:MF | GO:0036094 | small molecule binding | 2,89063E-05 | True | positive_corr | 2574 | 1059 | 212 | 18126 | 0,200188857 | 0,082362082 | ['GO:0005488'] |
| 607 | GO:MF | GO:0008392 | arachidonic acid epoxygenase activity | 3,08301E-05 | True | positive_corr | 16 | 1059 | 9 | 18126 | 0,008498584 | 0,5625 | ['GO:0008391'] |
| 1693 | GO:MF | GO:0008289 | lipid binding | 3,57916E-05 | True | positive_corr | 741 | 1059 | 80 | 18126 | 0,075542965 | 0,107962213 | ['GO:0005488'] |
| 1829 | GO:MF | GO:0015631 | tubulin binding | 4,04352E-05 | True | negative_corr | 343 | 2548 | 85 | 18126 | 0,033359498 | 0,247813411 | ['GO:0008092'] |
| 1678 | GO:MF | GO:0008391 | arachidonic acid monooxygenase activity | 6,21092E-05 | True | positive_corr | 17 | 1059 | 9 | 18126 | 0,008498584 | 0,529411765 | ['GO:0004497', 'GO:0016705'] |
| 1677 | GO:MF | GO:0046873 | metal ion transmembrane transporter activity | 6,36213E-05 | True | positive_corr | 446 | 1059 | 55 | 18126 | 0,051935788 | 0,123318386 | ['GO:0022890'] |
| 1834 | GO:MF | GO:0019899 | enzyme binding | 6,92592E-05 | True | negative_corr | 2246 | 2548 | 398 | 18126 | 0,156200942 | 0,177203918 | ['GO:0005515'] |
| 1835 | GO:MF | GO:0003735 | structural constituent of ribosome | 0,000107945 | True | negative_corr | 171 | 2548 | 50 | 18126 | 0,019623234 | 0,292397661 | ['GO:0005198'] |
| 281 | GO:MF | GO:0043168 | anion binding | 0,000234963 | True | positive_corr | 2841 | 1059 | 225 | 18126 | 0,212464589 | 0,079197466 | ['GO:0043167'] |
| 273 | GO:MF | GO:0003707 | steroid hormone receptor activity | 0,000246322 | True | positive_corr | 56 | 1059 | 15 | 18126 | 0,014164306 | 0,267857143 | ['GO:0038023'] |
| 641 | GO:MF | GO:0033764 | steroid dehydrogenase activity, acting on the CH-OH group of donors | 0,000256 | True | positive_corr | 30 | 1059 | 11 | 18126 | 0,010387158 | 0,366666667 | ['GO:0016229', 'GO:0016616'] |
| 1785 | GO:MF | GO:0120013 | lipid transfer activity | 0,000309822 | True | positive_corr | 43 | 1059 | 13 | 18126 | 0,012275732 | 0,302325581 | ['GO:0005319'] |
| 1844 | GO:MF | GO:0030020 | extracellular matrix structural constituent conferring tensile strength | 0,000374558 | True | negative_corr | 41 | 2548 | 19 | 18126 | 0,007456829 | 0,463414634 | ['GO:0005201'] |
| 673 | GO:MF | GO:0015077 | monovalent inorganic cation transmembrane transporter activity | 0,000532038 | True | positive_corr | 393 | 1059 | 48 | 18126 | 0,045325779 | 0,122137405 | ['GO:0022890'] |
| 675 | GO:MF | GO:0015144 | carbohydrate transmembrane transporter activity | 0,000681143 | True | positive_corr | 39 | 1059 | 12 | 18126 | 0,011331445 | 0,307692308 | ['GO:0022857'] |
| 309 | GO:MF | GO:0070330 | aromatase activity | 0,000766605 | True | positive_corr | 27 | 1059 | 10 | 18126 | 0,009442871 | 0,37037037 | ['GO:0016712'] |
| 1271 | GO:MF | GO:0015020 | glucuronosyltransferase activity | 0,001080007 | True | positive_corr | 34 | 1059 | 11 | 18126 | 0,010387158 | 0,323529412 | ['GO:0008194', 'GO:0016758'] |
| 1899 | GO:MF | GO:0016229 | steroid dehydrogenase activity | 0,001080007 | True | positive_corr | 34 | 1059 | 11 | 18126 | 0,010387158 | 0,323529412 | ['GO:0016491'] |
| 1420 | GO:MF | GO:0046872 | metal ion binding | 0,001184671 | True | negative_corr | 4227 | 2548 | 687 | 18126 | 0,269623234 | 0,162526615 | ['GO:0043169'] |
| 1040 | GO:MF | GO:0005201 | extracellular matrix structural constituent | 0,001415961 | True | negative_corr | 165 | 2548 | 46 | 18126 | 0,018053375 | 0,278787879 | ['GO:0005198'] |
| 1037 | GO:MF | GO:0017016 | Ras GTPase binding | 0,001559851 | True | negative_corr | 428 | 2548 | 95 | 18126 | 0,037284144 | 0,221962617 | ['GO:0031267'] |
| 1515 | GO:MF | GO:0004177 | aminopeptidase activity | 0,001602839 | True | positive_corr | 49 | 1059 | 13 | 18126 | 0,012275732 | 0,265306122 | ['GO:0008238'] |
| 956 | GO:MF | GO:0016705 | oxidoreductase activity, acting on paired donors, with incorporation or | 0,001610052 | True | positive_corr | 164 | 1059 | 26 | 18126 | 0,024551464 | 0,158536585 | ['GO:0016491'] |
| 964 | GO:MF | GO:0031267 | small GTPase binding | 0,002280004 | True | negative_corr | 443 | 2548 | 97 | 18126 | 0,038069074 | 0,218961625 | ['GO:0051020'] |
| 1329 | GO:MF | GO:0016757 | transferase activity, transferring glycosyl groups | 0,00283677 | True | positive_corr | 288 | 1059 | 37 | 18126 | 0,034938621 | 0,128472222 | ['GO:0016740'] |
| 1330 | GO:MF | GO:0008017 | microtubule binding | 0,003359157 | True | negative_corr | 253 | 2548 | 62 | 18126 | 0,02433281 | 0,245059289 | ['GO:0015631'] |
| 1092 | GO:MF | GO:0015103 | inorganic anion transmembrane transporter activity | 0,003611066 | True | positive_corr | 141 | 1059 | 23 | 18126 | 0,021718602 | 0,163120567 | ['GO:0008509', 'GO:0015318'] |
| 976 | GO:MF | GO:0008395 | steroid hydroxylase activity | 0,00366047 | True | positive_corr | 38 | 1059 | 11 | 18126 | 0,010387158 | 0,289473684 | ['GO:0004497'] |
| 988 | GO:MF | GO:0008238 | exopeptidase activity | 0,004630359 | True | positive_corr | 114 | 1059 | 20 | 18126 | 0,018885741 | 0,175438596 | ['GO:0070011'] |
| 1344 | GO:MF | GO:0004769 | steroid delta-isomerase activity | 0,005912897 | True | positive_corr | 4 | 1059 | 4 | 18126 | 0,003777148 | 1 | ['GO:0016863'] |
| 1487 | GO:MF | GO:0050051 | leukotriene-B4 20-monooxygenase activity | 0,005912897 | True | positive_corr | 4 | 1059 | 4 | 18126 | 0,003777148 | 1 | ['GO:0016709'] |
| 1122 | GO:MF | GO:0050294 | steroid sulfotransferase activity | 0,005912897 | True | positive_corr | 4 | 1059 | 4 | 18126 | 0,003777148 | 1 | ['GO:0008146'] |
| 1015 | GO:MF | GO:0005506 | iron ion binding | 0,006425789 | True | positive_corr | 156 | 1059 | 24 | 18126 | 0,02266289 | 0,153846154 | ['GO:0046914'] |
| 1382 | GO:MF | GO:0033293 | monocarboxylic acid binding | 0,00726623 | True | positive_corr | 72 | 1059 | 15 | 18126 | 0,014164306 | 0,208333333 | ['GO:0031406'] |
| 996 | GO:MF | GO:0008376 | acetylgalactosaminyltransferase activity | 0,007445742 | True | positive_corr | 48 | 1059 | 12 | 18126 | 0,011331445 | 0,25 | ['GO:0008194', 'GO:0016758'] |
| 1485 | GO:MF | GO:0019904 | protein domain specific binding | 0,009369746 | True | negative_corr | 712 | 2548 | 140 | 18126 | 0,054945055 | 0,196629213 | ['GO:0005515'] |
| 1366 | GO:MF | GO:0050431 | transforming growth factor beta binding | 0,009831636 | True | negative_corr | 23 | 2548 | 12 | 18126 | 0,004709576 | 0,52173913 | ['GO:0019838', 'GO:0019955'] |
| 1471 | GO:MF | GO:0043169 | cation binding | 0,010796244 | True | negative_corr | 4321 | 2548 | 691 | 18126 | 0,271193093 | 0,159916686 | ['GO:0043167'] |
| 1374 | GO:MF | GO:1901618 | organic hydroxy compound transmembrane transporter activity | 0,011642757 | True | positive_corr | 50 | 1059 | 12 | 18126 | 0,011331445 | 0,24 | ['GO:0015318'] |
| 1381 | GO:MF | GO:0140103 | catalytic activity, acting on a glycoprotein | 0,015027463 | True | positive_corr | 23 | 1059 | 8 | 18126 | 0,007554297 | 0,347826087 | ['GO:0140096'] |
| 1014 | GO:MF | GO:0051020 | GTPase binding | 0,015244602 | True | negative_corr | 551 | 2548 | 112 | 18126 | 0,043956044 | 0,203266788 | ['GO:0019899'] |
| 1445 | GO:MF | GO:0031406 | carboxylic acid binding | 0,016187611 | True | positive_corr | 219 | 1059 | 29 | 18126 | 0,027384325 | 0,132420091 | ['GO:0043168', 'GO:0043177'] |
| 944 | GO:MF | GO:0043177 | organic acid binding | 0,017307219 | True | positive_corr | 231 | 1059 | 30 | 18126 | 0,028328612 | 0,12987013 | ['GO:0036094'] |
| 1532 | GO:MF | GO:0030165 | PDZ domain binding | 0,019625206 | True | positive_corr | 87 | 1059 | 16 | 18126 | 0,015108593 | 0,183908046 | ['GO:0019904'] |
| 1217 | GO:MF | GO:0042910 | xenobiotic transmembrane transporter activity | 0,020917252 | True | positive_corr | 18 | 1059 | 7 | 18126 | 0,006610009 | 0,388888889 | ['GO:0022857'] |
| 1049 | GO:MF | GO:0015108 | chloride transmembrane transporter activity | 0,02478481 | True | positive_corr | 98 | 1059 | 17 | 18126 | 0,01605288 | 0,173469388 | ['GO:0015103'] |
| 1195 | GO:MF | GO:0008374 | O-acyltransferase activity | 0,026383081 | True | positive_corr | 46 | 1059 | 11 | 18126 | 0,010387158 | 0,239130435 | ['GO:0016747'] |
| 1183 | GO:MF | GO:0086083 | cell adhesive protein binding involved in bundle of His cell-Purkinje my | 0,028187572 | True | positive_corr | 5 | 1059 | 4 | 18126 | 0,003777148 | 0,8 | ['GO:0086080'] |
| 1086 | GO:MF | GO:0046906 | tetrapyrrole binding | 0,030739675 | True | positive_corr | 150 | 1059 | 22 | 18126 | 0,020774315 | 0,146666667 | ['GO:0097159', 'GO:1901363'] |
| 1534 | GO:MF | GO:0120020 | cholesterol transfer activity | 0,031452017 | True | positive_corr | 19 | 1059 | 7 | 18126 | 0,006610009 | 0,368421053 | ['GO:0120015'] |
| 1282 | GO:MF | GO:0020037 | heme binding | 0,032154289 | True | positive_corr | 140 | 1059 | 21 | 18126 | 0,019830028 | 0,15 | ['GO:0046906', 'GO:0048037'] |
| 1533 | GO:MF | GO:0008233 | peptidase activity | 0,033024351 | True | positive_corr | 641 | 1059 | 62 | 18126 | 0,058545798 | 0,096723869 | ['GO:0016787', 'GO:0140096'] |
| 1246 | GO:MF | GO:0008401 | retinoic acid 4-hydroxylase activity | 0,035570877 | True | positive_corr | 9 | 1059 | 5 | 18126 | 0,004721435 | 0,555555556 | ['GO:0004497', 'GO:0016705'] |
| 893 | GO:MF | GO:0003676 | nucleic acid binding | 0,036073287 | True | negative_corr | 4361 | 2548 | 691 | 18126 | 0,271193093 | 0,158449897 | ['GO:0097159', 'GO:1901363'] |
| 1178 | GO:MF | GO:0005254 | chloride channel activity | 0,03667209 | True | positive_corr | 73 | 1059 | 14 | 18126 | 0,013220019 | 0,191780822 | ['GO:0005253', 'GO:0015108'] |
| 1082 | GO:MF | GO:0030170 | pyridoxal phosphate binding | 0,038717091 | True | positive_corr | 56 | 1059 | 12 | 18126 | 0,011331445 | 0,214285714 | ['GO:0043168', 'GO:0050662', 'GO:0070279'] |
| 913 | GO:MF | GO:0016782 | transferase activity, transferring sulfur-containing groups | 0,042830488 | True | positive_corr | 74 | 1059 | 14 | 18126 | 0,013220019 | 0,189189189 | ['GO:0016740'] |
| 921 | GO:MF | GO:0019900 | kinase binding | 0,045644983 | True | negative_corr | 741 | 2548 | 141 | 18126 | 0,05533752 | 0,190283401 | ['GO:0019899'] |
| 1172 | GO:MF | GO:0120015 | sterol transfer activity | 0,045954432 | True | positive_corr | 20 | 1059 | 7 | 18126 | 0,006610009 | 0,35 | ['GO:0015248', 'GO:0120013'] |
| 932 | GO:MF | GO:0070279 | vitamin B6 binding | 0,046464529 | True | positive_corr | 57 | 1059 | 12 | 18126 | 0,011331445 | 0,210526316 | ['GO:0008144', 'GO:0019842', 'GO:0097159', 'GO:1901363'] |
| 1059 | GO:MF | GO:0005548 | phospholipid transporter activity | 0,049048376 | True | positive_corr | 49 | 1059 | 11 | 18126 | 0,010387158 | 0,224489796 | ['GO:0005319'] |
| 1782 | HP | HP:0011297 | Abnormal digit morphology | 3,7025E-07 | True | negative_corr | 1133 | 666 | 246 | 4225 | 0,369369369 | 0,217122683 | ['HP:0002813'] |
| 1057 | HP | HP:0001155 | Abnormality of the hand | 5,85216E-07 | True | negative_corr | 1178 | 666 | 253 | 4225 | 0,37987988 | 0,214770798 | ['HP:0002817'] |
| 1761 | HP | HP:0001162 | Postaxial hand polydactyly | 9,04199E-07 | True | negative_corr | 92 | 666 | 39 | 4225 | 0,058558559 | 0,423913043 | ['HP:0001161', 'HP:0004207', 'HP:0100259'] |

| | | | | | | | | | | | | | |
|---|---|---|---|---|---|---|---|---|---|---|---|---|---|
| 1774 | HP | HP:0001159 | Syndactyly | 1,57011E-06 | True | negative_corr | 328 | 666 | 93 | 4225 | 0,13963964 | 0,283536585 | ['HP:0011297'] |
| 1068 | HP | HP:0001167 | Abnormality of finger | 1,60448E-06 | True | negative_corr | 926 | 666 | 207 | 4225 | 0,310810811 | 0,223542117 | ['HP:0001155', 'HP:0011297'] |
| 1767 | HP | HP:0002817 | Abnormality of the upper limb | 1,61253E-06 | True | negative_corr | 1282 | 666 | 269 | 4225 | 0,403903904 | 0,209828393 | ['HP:0040064'] |
| 1070 | HP | HP:0100259 | Postaxial polydactyly | 1,99871E-06 | True | negative_corr | 125 | 666 | 47 | 4225 | 0,070570571 | 0,376 | ['HP:0010442'] |
| 892 | HP | HP:0040064 | Abnormality of limbs | 2,72542E-06 | True | negative_corr | 1790 | 666 | 352 | 4225 | 0,528528529 | 0,196648045 | ['HP:0000118'] |
| 1751 | HP | HP:0040068 | Abnormality of limb bone | 9,64685E-06 | True | negative_corr | 1296 | 666 | 268 | 4225 | 0,402402402 | 0,206790123 | ['HP:0000924', 'HP:0040064'] |
| 1781 | HP | HP:0002813 | Abnormality of limb bone morphology | 9,64685E-06 | True | negative_corr | 1296 | 666 | 268 | 4225 | 0,402402402 | 0,206790123 | ['HP:0011844', 'HP:0040068'] |
| 1786 | HP | HP:0010442 | Polydactyly | 1,45934E-05 | True | negative_corr | 214 | 666 | 66 | 4225 | 0,099099099 | 0,308411215 | ['HP:0011297'] |
| 1349 | HP | HP:0005918 | Abnormal finger phalanx morphology | 2,02435E-05 | True | negative_corr | 557 | 666 | 135 | 4225 | 0,202702703 | 0,242369838 | ['HP:0001167'] |
| 1377 | HP | HP:0001760 | Abnormality of the foot | 2,58107E-05 | True | negative_corr | 1091 | 666 | 231 | 4225 | 0,346846847 | 0,211732356 | ['HP:0002814'] |
| 1897 | HP | HP:0001161 | Hand polydactyly | 4,01946E-05 | True | negative_corr | 161 | 666 | 53 | 4225 | 0,07957958 | 0,329192547 | ['HP:0009997', 'HP:0010442'] |
| 1893 | HP | HP:0010902 | Abnormal circulating glutamine family amino acid concentration | 4,64683E-05 | True | positive_corr | 17 | 294 | 10 | 4225 | 0,034013605 | 0,588235294 | ['HP:0003112'] |
| 1249 | HP | HP:0011844 | Abnormal appendicular skeleton morphology | 4,6878E-05 | True | negative_corr | 1398 | 666 | 282 | 4225 | 0,423423423 | 0,201716738 | ['HP:0011842'] |
| 1884 | HP | HP:0002814 | Abnormality of the lower limb | 0,000126906 | True | negative_corr | 1303 | 666 | 264 | 4225 | 0,396396396 | 0,202609363 | ['HP:0040064'] |
| 1870 | HP | HP:0000787 | Nephrolithiasis | 0,000135414 | True | positive_corr | 96 | 294 | 23 | 4225 | 0,078231293 | 0,239583333 | ['HP:0012210'] |
| 1311 | HP | HP:0009997 | Duplication of phalanx of hand | 0,000165116 | True | negative_corr | 185 | 666 | 57 | 4225 | 0,085585586 | 0,308108108 | ['HP:0004275', 'HP:0005918'] |
| 1864 | HP | HP:0032180 | Abnormal circulating metabolite concentration | 0,000201844 | True | positive_corr | 963 | 294 | 105 | 4225 | 0,357142857 | 0,109034268 | ['HP:0001939'] |
| 1314 | HP | HP:0009142 | Duplication of bones involving the upper extremities | 0,000203198 | True | negative_corr | 186 | 666 | 57 | 4225 | 0,085585586 | 0,306451613 | ['HP:0002817'] |
| 1165 | HP | HP:0004275 | Duplication of hand bones | 0,000203198 | True | negative_corr | 186 | 666 | 57 | 4225 | 0,085585586 | 0,306451613 | ['HP:0001155', 'HP:0009142'] |
| 1850 | HP | HP:0002157 | Azotemia | 0,000538773 | True | positive_corr | 103 | 294 | 23 | 4225 | 0,078231293 | 0,223300971 | ['HP:0004364'] |
| 1788 | HP | HP:0004364 | Abnormal circulating nitrogen compound concentration | 0,000932961 | True | positive_corr | 106 | 294 | 23 | 4225 | 0,078231293 | 0,216981132 | ['HP:0032180'] |
| 1849 | HP | HP:0012072 | Aciduria | 0,000957567 | True | positive_corr | 182 | 294 | 32 | 4225 | 0,108843537 | 0,175824176 | ['HP:0003110'] |
| 1163 | HP | HP:0001780 | Abnormality of toe | 0,001182928 | True | negative_corr | 539 | 666 | 125 | 4225 | 0,187687688 | 0,231910946 | ['HP:0001760', 'HP:0011297'] |
| 1158 | HP | HP:0003110 | Abnormality of urine homeostasis | 0,001267 | True | positive_corr | 555 | 294 | 68 | 4225 | 0,231292517 | 0,122522523 | ['HP:0001939', 'HP:0011277'] |
| 1343 | HP | HP:0000974 | Hyperextensible skin | 0,001299508 | True | negative_corr | 52 | 666 | 23 | 4225 | 0,034534535 | 0,442307692 | ['HP:0008067'] |
| 1720 | HP | HP:0001156 | Brachydactyly | 0,001397615 | True | negative_corr | 302 | 666 | 79 | 4225 | 0,118618619 | 0,261589404 | ['HP:0011927'] |
| 1355 | HP | HP:0006101 | Finger syndactyly | 0,001567811 | True | negative_corr | 178 | 666 | 53 | 4225 | 0,07957958 | 0,297752809 | ['HP:0001159'] |
| 1135 | HP | HP:0001829 | Foot polydactyly | 0,002601748 | True | negative_corr | 101 | 666 | 35 | 4225 | 0,052552553 | 0,346534653 | ['HP:0001780', 'HP:0009136', 'HP:0010442'] |
| 1365 | HP | HP:0001259 | Coma | 0,002670896 | True | positive_corr | 81 | 294 | 19 | 4225 | 0,06462585 | 0,234567901 | ['HP:0004372'] |
| 1806 | HP | HP:0009136 | Duplication involving bones of the feet | 0,003195324 | True | negative_corr | 106 | 666 | 36 | 4225 | 0,054054054 | 0,339622642 | ['HP:0001760', 'HP:0040069'] |
| 1126 | HP | HP:0009121 | Abnormal axial skeleton morphology | 0,003285601 | True | negative_corr | 2121 | 666 | 389 | 4225 | 0,584084084 | 0,183404055 | ['HP:0011842'] |
| 1793 | HP | HP:0005916 | Abnormal metacarpal morphology | 0,003348068 | True | negative_corr | 141 | 666 | 44 | 4225 | 0,066066066 | 0,312056738 | ['HP:0001163'] |
| 1164 | HP | HP:0000090 | Nephronophthisis | 0,003777428 | True | negative_corr | 37 | 666 | 18 | 4225 | 0,027027027 | 0,486486486 | ['HP:0100957'] |
| 1438 | HP | HP:0001987 | Hyperammonemia | 0,004117777 | True | positive_corr | 48 | 294 | 14 | 4225 | 0,047619048 | 0,291666667 | ['HP:0002157'] |
| 872 | HP | HP:0011927 | Short digit | 0,006744422 | True | negative_corr | 459 | 666 | 107 | 4225 | 0,160660661 | 0,233115468 | ['HP:0011297'] |
| 1590 | HP | HP:0002014 | Diarrhea | 0,007209878 | True | positive_corr | 277 | 294 | 40 | 4225 | 0,136054422 | 0,144404332 | ['HP:0011458'] |
| 813 | HP | HP:0009115 | Aplasia/hypoplasia involving the skeleton | 0,007262605 | True | negative_corr | 933 | 666 | 192 | 4225 | 0,288288288 | 0,205787781 | ['HP:0011842'] |
| 1591 | HP | HP:0010981 | Hypolipoproteinemia | 0,007625709 | True | positive_corr | 21 | 294 | 9 | 4225 | 0,030612245 | 0,428571429 | ['HP:0010979'] |
| 1592 | HP | HP:0000692 | Misalignment of teeth | 0,00860343 | True | negative_corr | 260 | 666 | 68 | 4225 | 0,102102102 | 0,261538462 | ['HP:0000164'] |
| 1029 | HP | HP:0002084 | Encephalocele | 0,008608405 | True | negative_corr | 93 | 666 | 32 | 4225 | 0,048048048 | 0,344086022 | ['HP:0002011', 'HP:0011815'] |
| 965 | HP | HP:0011815 | Cephalocele | 0,008608405 | True | negative_corr | 93 | 666 | 32 | 4225 | 0,048048048 | 0,344086022 | ['HP:0000929'] |
| 966 | HP | HP:0003119 | Abnormal circulating lipid concentration | 0,012556394 | True | positive_corr | 204 | 294 | 32 | 4225 | 0,108843537 | 0,156862745 | ['HP:0032180'] |
| 1673 | HP | HP:0001939 | Abnormality of metabolism/homeostasis | 0,013994031 | True | positive_corr | 2027 | 294 | 177 | 4225 | 0,602040816 | 0,087321164 | ['HP:0000118'] |
| 1501 | HP | HP:0005920 | Abnormal epiphysis morphology of the phalanges of the hand | 0,014536024 | True | negative_corr | 33 | 666 | 16 | 4225 | 0,024024024 | 0,484848485 | ['HP:0005918', 'HP:0005924'] |
| 1670 | HP | HP:0002418 | Abnormality of midbrain morphology | 0,015496999 | True | negative_corr | 51 | 666 | 21 | 4225 | 0,031531532 | 0,411764706 | ['HP:0012443'] |
| 994 | HP | HP:0003111 | Abnormal blood ion concentration | 0,016790962 | True | positive_corr | 246 | 294 | 36 | 4225 | 0,12244898 | 0,146341463 | ['HP:0032180'] |
| 1489 | HP | HP:0000977 | Soft skin | 0,017207756 | True | negative_corr | 30 | 666 | 15 | 4225 | 0,022522523 | 0,5 | ['HP:0010647'] |
| 1464 | HP | HP:0001395 | Hepatic fibrosis | 0,01841767 | True | negative_corr | 109 | 666 | 35 | 4225 | 0,052552553 | 0,321100917 | ['HP:0410042'] |
| 1008 | HP | HP:0040070 | Abnormal upper limb bone morphology | 0,019200608 | True | negative_corr | 363 | 666 | 87 | 4225 | 0,130630631 | 0,239669421 | ['HP:0002813', 'HP:0002817'] |
| 1468 | HP | HP:0002419 | Molar tooth sign on MRI | 0,023497572 | True | negative_corr | 34 | 666 | 16 | 4225 | 0,024024024 | 0,470588235 | ['HP:0002418'] |
| 1470 | HP | HP:0004422 | Biparietal narrowing | 0,024382123 | True | negative_corr | 60 | 666 | 23 | 4225 | 0,034534535 | 0,383333333 | ['HP:0002648'] |
| 1646 | HP | HP:0006494 | Aplasia/Hypoplasia involving bones of the feet | 0,025327052 | True | negative_corr | 223 | 666 | 59 | 4225 | 0,088588589 | 0,264573991 | ['HP:0001760', 'HP:0006493'] |
| 1683 | HP | HP:0002612 | Congenital hepatic fibrosis | 0,026599297 | True | negative_corr | 45 | 666 | 19 | 4225 | 0,028528529 | 0,422222222 | ['HP:0001395'] |
| 827 | HP | HP:0000174 | Abnormal palate morphology | 0,027494872 | True | negative_corr | 864 | 666 | 177 | 4225 | 0,265765766 | 0,204861111 | ['HP:0000163'] |
| 1560 | HP | HP:0001163 | Abnormality of the metacarpal bones | 0,027815135 | True | negative_corr | 180 | 666 | 50 | 4225 | 0,075075075 | 0,277777778 | ['HP:0001155', 'HP:0040070'] |
| 1716 | HP | HP:0006706 | Cystic liver disease | 0,028545952 | True | negative_corr | 31 | 666 | 15 | 4225 | 0,022522523 | 0,483870968 | ['HP:0410042'] |
| 1550 | HP | HP:0003563 | Decreased LDL cholesterol concentration | 0,028816152 | True | positive_corr | 10 | 294 | 6 | 4225 | 0,020408163 | 0,6 | ['HP:0010981', 'HP:0031886'] |
| 863 | HP | HP:0010049 | Short metacarpal | 0,029603931 | True | negative_corr | 98 | 666 | 32 | 4225 | 0,048048048 | 0,326530612 | ['HP:0005914'] |
| 862 | HP | HP:0100797 | Toenail dysplasia | 0,032082673 | True | negative_corr | 13 | 666 | 9 | 4225 | 0,013513514 | 0,692307692 | ['HP:0002164', 'HP:0008388'] |
| 1706 | HP | HP:0005914 | Aplasia/Hypoplasia involving the metacarpal bones | 0,039168912 | True | negative_corr | 108 | 666 | 34 | 4225 | 0,051051051 | 0,314814815 | ['HP:0005916', 'HP:0005927'] |
| 838 | HP | HP:0011277 | Abnormality of the urinary system physiology | 0,039390411 | True | positive_corr | 878 | 294 | 90 | 4225 | 0,306122449 | 0,102505695 | ['HP:0000079'] |
| 860 | HP | HP:0003355 | Aminoaciduria | 0,041189064 | True | positive_corr | 131 | 294 | 23 | 4225 | 0,078231293 | 0,175572519 | ['HP:0012072'] |
| 1563 | HP | HP:0000309 | Abnormality of the midface | 0,041778599 | True | negative_corr | 466 | 666 | 105 | 4225 | 0,157657658 | 0,225321888 | ['HP:0000271'] |
| 1439 | HP | HP:0004207 | Abnormal 5th finger morphology | 0,044718725 | True | negative_corr | 360 | 666 | 85 | 4225 | 0,127627628 | 0,236111111 | ['HP:0001167'] |
| 1569 | HP | HP:0010903 | Abnormal circulating glutamine concentration | 0,047600404 | True | positive_corr | 7 | 294 | 5 | 4225 | 0,017006803 | 0,714285714 | ['HP:0010902'] |

| | | | | | | | | | | | | | |
|---|---|---|---|---|---|---|---|---|---|---|---|---|---|
| 1701 | HP | HP:0001397 | Hepatic steatosis | 0,048066937 | True | positive_corr | 106 | 294 | 20 | 4225 | 0,068027211 | 0,188679245 | ['HP:0006561'] |
| 1877 | KEGG | KEGG:01100 | Metabolic pathways | 1,5222E-26 | True | positive_corr | 1482 | 544 | 210 | 7788 | 0,386029412 | 0,141700405 | ['KEGG:00000'] |
| 1577 | KEGG | KEGG:05204 | Chemical carcinogenesis | 4,11398E-14 | True | positive_corr | 80 | 544 | 31 | 7788 | 0,056985294 | 0,3875 | ['KEGG:00000'] |
| 690 | KEGG | KEGG:00982 | Drug metabolism - cytochrome P450 | 2,79137E-13 | True | positive_corr | 69 | 544 | 28 | 7788 | 0,051470588 | 0,405797101 | ['KEGG:00000'] |
| 1542 | KEGG | KEGG:00980 | Metabolism of xenobiotics by cytochrome P450 | 1,56316E-12 | True | positive_corr | 73 | 544 | 28 | 7788 | 0,051470588 | 0,383561644 | ['KEGG:00000'] |
| 703 | KEGG | KEGG:03010 | Ribosome | 2,13097E-10 | True | negative_corr | 134 | 1003 | 49 | 7788 | 0,04885344 | 0,365671642 | ['KEGG:00000'] |
| 1861 | KEGG | KEGG:00830 | Retinol metabolism | 4,87197E-10 | True | positive_corr | 66 | 544 | 24 | 7788 | 0,044117647 | 0,363636364 | ['KEGG:00000'] |
| 1153 | KEGG | KEGG:00140 | Steroid hormone biosynthesis | 3,19826E-09 | True | positive_corr | 60 | 544 | 22 | 7788 | 0,040441176 | 0,366666667 | ['KEGG:00000'] |
| 1281 | KEGG | KEGG:00040 | Pentose and glucuronate interconversions | 1,97947E-08 | True | positive_corr | 34 | 544 | 16 | 7788 | 0,029411765 | 0,470588235 | ['KEGG:00000'] |
| 741 | KEGG | KEGG:00983 | Drug metabolism - other enzymes | 2,77904E-08 | True | positive_corr | 78 | 544 | 24 | 7788 | 0,044117647 | 0,307692308 | ['KEGG:00000'] |
| 1824 | KEGG | KEGG:04975 | Fat digestion and absorption | 6,20835E-08 | True | positive_corr | 41 | 544 | 17 | 7788 | 0,03125 | 0,414634146 | ['KEGG:00000'] |
| 1858 | KEGG | KEGG:03320 | PPAR signaling pathway | 3,16422E-06 | True | positive_corr | 76 | 544 | 21 | 7788 | 0,038602941 | 0,276315789 | ['KEGG:00000'] |
| 1557 | KEGG | KEGG:04976 | Bile secretion | 6,1633E-06 | True | positive_corr | 72 | 544 | 20 | 7788 | 0,036764706 | 0,277777778 | ['KEGG:00000'] |
| 135 | KEGG | KEGG:04950 | Maturity onset diabetes of the young | 5,80437E-05 | True | positive_corr | 26 | 544 | 11 | 7788 | 0,020220588 | 0,423076923 | ['KEGG:00000'] |
| 1161 | KEGG | KEGG:00053 | Ascorbate and aldarate metabolism | 9,18306E-05 | True | positive_corr | 27 | 544 | 11 | 7788 | 0,020220588 | 0,407407407 | ['KEGG:00000'] |
| 936 | KEGG | KEGG:04530 | Tight junction | 0,000160611 | True | positive_corr | 168 | 544 | 30 | 7788 | 0,055147059 | 0,178571429 | ['KEGG:00000'] |
| 1300 | KEGG | KEGG:04974 | Protein digestion and absorption | 0,000189129 | True | positive_corr | 95 | 544 | 21 | 7788 | 0,038602941 | 0,221052632 | ['KEGG:00000'] |
| 1518 | KEGG | KEGG:04973 | Carbohydrate digestion and absorption | 0,000249697 | True | positive_corr | 47 | 544 | 14 | 7788 | 0,025735294 | 0,29787234 | ['KEGG:00000'] |
| 1180 | KEGG | KEGG:04978 | Mineral absorption | 0,000764732 | True | positive_corr | 58 | 544 | 15 | 7788 | 0,027573529 | 0,25862069 | ['KEGG:00000'] |
| 901 | KEGG | KEGG:03013 | RNA transport | 0,000853706 | True | negative_corr | 159 | 1003 | 41 | 7788 | 0,040877368 | 0,257861635 | ['KEGG:00000'] |
| 1719 | KEGG | KEGG:00600 | Sphingolipid metabolism | 0,001424017 | True | positive_corr | 47 | 544 | 13 | 7788 | 0,023897059 | 0,276595745 | ['KEGG:00000'] |
| 1367 | KEGG | KEGG:04010 | MAPK signaling pathway | 0,001560441 | True | negative_corr | 295 | 1003 | 64 | 7788 | 0,063808574 | 0,216949153 | ['KEGG:00000'] |
| 1110 | KEGG | KEGG:01200 | Carbon metabolism | 0,001767039 | True | positive_corr | 117 | 544 | 22 | 7788 | 0,040441176 | 0,188034188 | ['KEGG:00000'] |
| 1386 | KEGG | KEGG:04977 | Vitamin digestion and absorption | 0,002114399 | True | positive_corr | 24 | 544 | 9 | 7788 | 0,016544118 | 0,375 | ['KEGG:00000'] |
| 456 | KEGG | KEGG:00860 | Porphyrin and chlorophyll metabolism | 0,00214314 | True | positive_corr | 42 | 544 | 12 | 7788 | 0,022058824 | 0,285714286 | ['KEGG:00000'] |
| 1411 | KEGG | KEGG:04015 | Rap1 signaling pathway | 0,004802978 | True | negative_corr | 210 | 1003 | 48 | 7788 | 0,047856431 | 0,228571429 | ['KEGG:00000'] |
| 1776 | KEGG | KEGG:00051 | Fructose and mannose metabolism | 0,006121397 | True | positive_corr | 33 | 544 | 10 | 7788 | 0,018382353 | 0,303030303 | ['KEGG:00000'] |
| 1413 | KEGG | KEGG:00480 | Glutathione metabolism | 0,007127081 | True | positive_corr | 54 | 544 | 13 | 7788 | 0,023897059 | 0,240740741 | ['KEGG:00000'] |
| 1743 | KEGG | KEGG:00590 | Arachidonic acid metabolism | 0,008004624 | True | positive_corr | 62 | 544 | 14 | 7788 | 0,025735294 | 0,225806452 | ['KEGG:00000'] |
| 636 | KEGG | KEGG:00071 | Fatty acid degradation | 0,018127568 | True | positive_corr | 44 | 544 | 11 | 7788 | 0,020220588 | 0,25 | ['KEGG:00000'] |
| 1734 | KEGG | KEGG:04360 | Axon guidance | 0,018834807 | True | negative_corr | 180 | 1003 | 41 | 7788 | 0,040877368 | 0,227777778 | ['KEGG:00000'] |
| 1733 | KEGG | KEGG:01230 | Biosynthesis of amino acids | 0,01934179 | True | positive_corr | 75 | 544 | 15 | 7788 | 0,027573529 | 0,2 | ['KEGG:00000'] |
| 1076 | KEGG | KEGG:00052 | Galactose metabolism | 0,020950915 | True | positive_corr | 31 | 544 | 9 | 7788 | 0,016544118 | 0,290322581 | ['KEGG:00000'] |
| 556 | KEGG | KEGG:00512 | Mucin type O-glycan biosynthesis | 0,020950915 | True | positive_corr | 31 | 544 | 9 | 7788 | 0,016544118 | 0,290322581 | ['KEGG:00000'] |
| 430 | KEGG | KEGG:05168 | Herpes simplex virus 1 infection | 0,0217429 | True | negative_corr | 487 | 1003 | 90 | 7788 | 0,089730808 | 0,184804928 | ['KEGG:00000'] |
| 406 | KEGG | KEGG:00561 | Glycerolipid metabolism | 0,026863585 | True | positive_corr | 61 | 544 | 13 | 7788 | 0,023897059 | 0,213114754 | ['KEGG:00000'] |
| 1715 | KEGG | KEGG:00920 | Sulfur metabolism | 0,034573733 | True | positive_corr | 10 | 544 | 5 | 7788 | 0,009191176 | 0,5 | ['KEGG:00000'] |
| 1375 | KEGG | KEGG:04146 | Peroxisome | 0,040676329 | True | positive_corr | 80 | 544 | 15 | 7788 | 0,027573529 | 0,1875 | ['KEGG:00000'] |
| 1721 | KEGG | KEGG:00601 | Glycosphingolipid biosynthesis - lacto and neolacto series | 0,041078243 | True | positive_corr | 27 | 544 | 8 | 7788 | 0,014705882 | 0,296296296 | ['KEGG:00000'] |
| 579 | KEGG | KEGG:00220 | Arginine biosynthesis | 0,043184737 | True | positive_corr | 21 | 544 | 7 | 7788 | 0,012867647 | 0,333333333 | ['KEGG:00000'] |
| 1441 | WP | WP:WP2882 | Nuclear Receptors Meta-Pathway | 5,89646E-18 | True | positive_corr | 318 | 491 | 74 | 6646 | 0,150712831 | 0,232704403 | ['WP:000000'] |
| 1048 | WP | WP:WP702 | Metapathway biotransformation Phase I and II | 2,85022E-16 | True | positive_corr | 182 | 491 | 52 | 6646 | 0,105906314 | 0,285714286 | ['WP:000000'] |
| 857 | WP | WP:WP4536 | Genes related to primary cilium development (based on CRISPR) | 2,94238E-15 | True | negative_corr | 103 | 962 | 51 | 6646 | 0,053014553 | 0,495145631 | ['WP:000000'] |
| 570 | WP | WP:WP477 | Cytoplasmic Ribosomal Proteins | 3,18702E-10 | True | negative_corr | 88 | 962 | 40 | 6646 | 0,041580042 | 0,454545455 | ['WP:000000'] |
| 1058 | WP | WP:WP2884 | NRF2 pathway | 1,0496E-09 | True | positive_corr | 144 | 491 | 37 | 6646 | 0,075356415 | 0,256944444 | ['WP:000000'] |
| 1703 | WP | WP:WP691 | Tamoxifen metabolism | 1,26124E-09 | True | positive_corr | 21 | 491 | 14 | 6646 | 0,028513238 | 0,666666667 | ['WP:000000'] |
| 1085 | WP | WP:WP2876 | Pregnane X Receptor pathway | 2,48877E-09 | True | positive_corr | 33 | 491 | 17 | 6646 | 0,034623218 | 0,515151515 | ['WP:000000'] |
| 1400 | WP | WP:WP2875 | Constitutive Androstane Receptor Pathway | 1,79311E-08 | True | positive_corr | 32 | 491 | 16 | 6646 | 0,032586558 | 0,5 | ['WP:000000'] |
| 606 | WP | WP:WP1604 | Codeine and Morphine Metabolism | 1,38095E-06 | True | positive_corr | 15 | 491 | 10 | 6646 | 0,020366599 | 0,666666667 | ['WP:000000'] |
| 1129 | WP | WP:WP2289 | Drug Induction of Bile Acid Pathway | 7,80634E-06 | True | positive_corr | 17 | 491 | 10 | 6646 | 0,020366599 | 0,588235294 | ['WP:000000'] |
| 1173 | WP | WP:WP3942 | PPAR signaling pathway | 2,79833E-05 | True | positive_corr | 67 | 491 | 19 | 6646 | 0,038696538 | 0,28358209 | ['WP:000000'] |
| 687 | WP | WP:WP299 | Nuclear Receptors in Lipid Metabolism and Toxicity | 3,5226E-05 | True | positive_corr | 33 | 491 | 13 | 6646 | 0,026476578 | 0,393939394 | ['WP:000000'] |
| 369 | WP | WP:WP4656 | Joubert Syndrome | 4,033E-05 | True | negative_corr | 76 | 962 | 29 | 6646 | 0,03014553 | 0,381578947 | ['WP:000000'] |
| 341 | WP | WP:WP692 | Sulfation Biotransformation Reaction | 6,34941E-05 | True | positive_corr | 16 | 491 | 9 | 6646 | 0,018329939 | 0,5625 | ['WP:000000'] |
| 1156 | WP | WP:WP698 | Glucuronidation | 0,000129701 | True | positive_corr | 26 | 491 | 11 | 6646 | 0,022403259 | 0,423076923 | ['WP:000000'] |
| 1663 | WP | WP:WP43 | Oxidation by Cytochrome P450 | 0,000162723 | True | positive_corr | 61 | 491 | 17 | 6646 | 0,034623218 | 0,278688525 | ['WP:000000'] |
| 1379 | WP | WP:WP697 | Estrogen metabolism | 0,000235875 | True | positive_corr | 18 | 491 | 9 | 6646 | 0,018329939 | 0,5 | ['WP:000000'] |
| 1463 | WP | WP:WP2878 | PPAR Alpha Pathway | 0,00115836 | True | positive_corr | 26 | 491 | 10 | 6646 | 0,020366599 | 0,384615385 | ['WP:000000'] |
| 1380 | WP | WP:WP4595 | Urea cycle and associated pathways | 0,001165724 | True | positive_corr | 21 | 491 | 9 | 6646 | 0,018329939 | 0,428571429 | ['WP:000000'] |
| 297 | WP | WP:WP4720 | Eicosanoid metabolism via Cytochrome P450 Mono-Oxygenases (CY | 0,001589847 | True | positive_corr | 9 | 491 | 6 | 6646 | 0,012219959 | 0,666666667 | ['WP:000000'] |
| 1341 | WP | WP:WP4571 | Urea cycle and related diseases | 0,001589847 | True | positive_corr | 9 | 491 | 6 | 6646 | 0,012219959 | 0,666666667 | ['WP:000000'] |
| 1339 | WP | WP:WP694 | Arylamine metabolism | 0,001768571 | True | positive_corr | 6 | 491 | 5 | 6646 | 0,010183299 | 0,833333333 | ['WP:000000'] |
| 626 | WP | WP:WP229 | Irinotecan Pathway | 0,001940716 | True | positive_corr | 13 | 491 | 7 | 6646 | 0,014256619 | 0,538461538 | ['WP:000000'] |
| 614 | WP | WP:WP143 | Fatty Acid Beta Oxidation | 0,002785776 | True | positive_corr | 34 | 491 | 11 | 6646 | 0,022403259 | 0,323529412 | ['WP:000000'] |
| 954 | WP | WP:WP4331 | Neovascularisation processes | 0,003168014 | True | negative_corr | 37 | 962 | 16 | 6646 | 0,016632017 | 0,432432432 | ['WP:000000'] |

| | | | | | | | | | | | | | |
|---|---|---|---|---|---|---|---|---|---|---|---|---|---|
| 1387 | WP | WP:WP430 | Statin Pathway | 0,003566665 | True | positive_corr | 29 | 491 | 10 | 6646 | 0,020366599 | 0,344827586 | ['WP:000000'] |
| 1481 | WP | WP:WP3925 | Amino Acid metabolism | 0,013553752 | True | positive_corr | 91 | 491 | 18 | 6646 | 0,036659878 | 0,197802198 | ['WP:000000'] |
| 1766 | WP | WP:WP368 | Mitochondrial LC-Fatty Acid Beta-Oxidation | 0,016931503 | True | positive_corr | 17 | 491 | 7 | 6646 | 0,014256619 | 0,411764706 | ['WP:000000'] |
| 1764 | WP | WP:WP3871 | Valproic acid pathway | 0,02511628 | True | positive_corr | 13 | 491 | 6 | 6646 | 0,012219959 | 0,461538462 | ['WP:000000'] |
| 1032 | WP | WP:WP4532 | Intraflagellar transport proteins binding to dynein | 0,025161833 | True | negative_corr | 27 | 962 | 12 | 6646 | 0,012474012 | 0,444444444 | ['WP:000000'] |
