## Supplementary table 6 colon for "Bioinformatic characterization of angiotensin-converting enzyme 2, the entry receptor for SARS-CoV-2"

| | source | native | name | p_value | query | significar | term_size | query_size | intersection_si | effective_domain_si | precision | recall | parents |
|---|---|---|---|---|---|---|---|---|---|---|---|---|---|
| 926 | GO:BF | GO:0007399 | nervous system development | 6,06871E-29 | negative_corr | True | 2432 | 2072 | 466 | 17906 | 0,224903475 | 0,191611842 | ['GO:0048731'] |
| 932 | GO:BF | GO:0120036 | plasma membrane bounded cell projection organization | 1,30767E-27 | negative_corr | True | 1539 | 2072 | 329 | 17906 | 0,158783784 | 0,213775179 | ['GO:0030030'] |
| 931 | GO:BF | GO:0030030 | cell projection organization | 1,34252E-27 | negative_corr | True | 1578 | 2072 | 335 | 17906 | 0,161679537 | 0,212294043 | ['GO:0016043'] |
| 1590 | GO:BF | GO:0032989 | cellular component morphogenesis | 3,6E-23 | negative_corr | True | 1177 | 2072 | 260 | 17906 | 0,125482625 | 0,220900595 | ['GO:0009653', 'GO:0016043', 'GO:0048869'] |
| 929 | GO:BF | GO:0022008 | neurogenesis | 9,79719E-23 | negative_corr | True | 1679 | 2072 | 336 | 17906 | 0,162162162 | 0,200119119 | ['GO:0007399', 'GO:0030154'] |
| 928 | GO:BF | GO:0031175 | neuron projection development | 1,06855E-22 | negative_corr | True | 1017 | 2072 | 233 | 17906 | 0,112451737 | 0,229105211 | ['GO:0048666', 'GO:0120036'] |
| 927 | GO:BF | GO:0000902 | cell morphogenesis | 1,35625E-22 | negative_corr | True | 1068 | 2072 | 241 | 17906 | 0,116312741 | 0,225655431 | ['GO:0032989'] |
| 1220 | GO:BF | GO:0000904 | cell morphogenesis involved in differentiation | 1,88437E-22 | negative_corr | True | 769 | 2072 | 191 | 17906 | 0,092181467 | 0,248374512 | ['GO:0000902', 'GO:0048468'] |
| 1591 | GO:BF | GO:0048666 | neuron development | 4,87675E-22 | negative_corr | True | 1146 | 2072 | 252 | 17906 | 0,121621622 | 0,219895288 | ['GO:0030182', 'GO:0048468'] |
| 922 | GO:BF | GO:0030182 | neuron differentiation | 4,96345E-22 | negative_corr | True | 1409 | 2072 | 293 | 17906 | 0,141409266 | 0,2079489 | ['GO:0030154', 'GO:0048699'] |
| 1597 | GO:BF | GO:0006629 | lipid metabolic process | 8,30197E-22 | positive_corr | True | 1414 | 998 | 176 | 17906 | 0,176352705 | 0,12446959 | ['GO:0044238', 'GO:0071704'] |
| 914 | GO:BF | GO:0048699 | generation of neurons | 1,10261E-21 | negative_corr | True | 1574 | 2072 | 317 | 17906 | 0,152992278 | 0,201397713 | ['GO:0022008'] |
| 1221 | GO:BF | GO:0009653 | anatomical structure morphogenesis | 1,30701E-21 | negative_corr | True | 2820 | 2072 | 495 | 17906 | 0,238899614 | 0,175531915 | ['GO:0032502', 'GO:0048856'] |
| 912 | GO:BF | GO:0048812 | neuron projection morphogenesis | 4,34652E-20 | negative_corr | True | 673 | 2072 | 169 | 17906 | 0,081563707 | 0,251114413 | ['GO:0031175', 'GO:0120039'] |
| 1599 | GO:BF | GO:0048667 | cell morphogenesis involved in neuron differentiation | 5,18658E-20 | negative_corr | True | 611 | 2072 | 158 | 17906 | 0,076254826 | 0,258592471 | ['GO:0000904', 'GO:0048666'] |
| 905 | GO:BF | GO:0120039 | plasma membrane bounded cell projection morphogenesis | 6,57361E-20 | negative_corr | True | 687 | 2072 | 171 | 17906 | 0,082528958 | 0,248908297 | ['GO:0048858'] |
| 903 | GO:BF | GO:0044281 | small molecule metabolic process | 7,8813E-20 | positive_corr | True | 1987 | 998 | 216 | 17906 | 0,216432866 | 0,108706593 | ['GO:0008152'] |
| 900 | GO:BF | GO:0048858 | cell projection morphogenesis | 1,29344E-19 | negative_corr | True | 691 | 2072 | 171 | 17906 | 0,082528958 | 0,247467438 | ['GO:0000902', 'GO:0030030', 'GO:0032990'] |
| 1589 | GO:BF | GO:0048468 | cell development | 2,87751E-19 | negative_corr | True | 2215 | 2072 | 402 | 17906 | 0,194015444 | 0,181489842 | ['GO:0030154', 'GO:0048856', 'GO:0048869'] |
| 1224 | GO:BF | GO:0048731 | system development | 3,23906E-18 | negative_corr | True | 5020 | 2072 | 768 | 17906 | 0,370656371 | 0,152988048 | ['GO:0007275', 'GO:0048856'] |
| 936 | GO:BF | GO:0032990 | cell part morphogenesis | 3,4057E-18 | negative_corr | True | 711 | 2072 | 171 | 17906 | 0,082528958 | 0,240506329 | ['GO:0032989'] |
| 1588 | GO:BF | GO:0007275 | multicellular organism development | 1,15623E-17 | negative_corr | True | 5604 | 2072 | 838 | 17906 | 0,404440154 | 0,149536046 | ['GO:0032501', 'GO:0048856'] |
| 1200 | GO:BF | GO:0048856 | anatomical structure development | 5,55573E-17 | negative_corr | True | 6105 | 2072 | 896 | 17906 | 0,432432432 | 0,146764947 | ['GO:0032502'] |
| 1201 | GO:BF | GO:0032502 | developmental process | 1,4818E-15 | negative_corr | True | 6630 | 2072 | 952 | 17906 | 0,459459459 | 0,143589744 | ['GO:0008150'] |
| 979 | GO:BF | GO:0061564 | axon development | 3,59791E-15 | negative_corr | True | 531 | 2072 | 133 | 17906 | 0,064189189 | 0,25047081 | ['GO:0031175'] |
| 978 | GO:BF | GO:0044255 | cellular lipid metabolic process | 4,1184E-15 | positive_corr | True | 1070 | 998 | 132 | 17906 | 0,132264529 | 0,123364486 | ['GO:0006629', 'GO:0044237'] |
| 974 | GO:BF | GO:0007409 | axonogenesis | 1,35141E-14 | negative_corr | True | 487 | 2072 | 124 | 17906 | 0,05984556 | 0,254620123 | ['GO:0048667', 'GO:0048812', 'GO:0061564'] |
| 970 | GO:BF | GO:0006811 | ion transport | 2,6408E-13 | positive_corr | True | 1723 | 998 | 178 | 17906 | 0,178356713 | 0,103308183 | ['GO:0006810'] |
| 966 | GO:BF | GO:0010975 | regulation of neuron projection development | 4,21886E-13 | negative_corr | True | 519 | 2072 | 126 | 17906 | 0,060810811 | 0,242774566 | ['GO:0031175', 'GO:0045664', 'GO:0120035'] |
| 965 | GO:BF | GO:0031344 | regulation of cell projection organization | 7,25274E-13 | negative_corr | True | 704 | 2072 | 156 | 17906 | 0,075289575 | 0,221590909 | ['GO:0030030', 'GO:0051128'] |
| 1582 | GO:BF | GO:0120035 | regulation of plasma membrane bounded cell projection organization | 9,99833E-13 | negative_corr | True | 694 | 2072 | 154 | 17906 | 0,074324324 | 0,221902017 | ['GO:0031344', 'GO:0120036'] |
| 1499 | GO:BF | GO:0030154 | cell differentiation | 1,30141E-12 | negative_corr | True | 4335 | 2072 | 653 | 17906 | 0,31515444 | 0,150634371 | ['GO:0048869'] |
| 953 | GO:BF | GO:0048869 | cellular developmental process | 2,64737E-12 | negative_corr | True | 4531 | 2072 | 676 | 17906 | 0,326254826 | 0,149194438 | ['GO:0009987', 'GO:0032502'] |
| 950 | GO:BF | GO:0007267 | cell-cell signaling | 6,19358E-12 | negative_corr | True | 1699 | 2072 | 301 | 17906 | 0,14527027 | 0,177163037 | ['GO:0007154', 'GO:0023052'] |
| 949 | GO:BF | GO:0006820 | anion transport | 8,05926E-12 | positive_corr | True | 654 | 998 | 89 | 17906 | 0,089178357 | 0,136085627 | ['GO:0006811'] |
| 1496 | GO:BF | GO:0007268 | chemical synaptic transmission | 8,87189E-12 | negative_corr | True | 717 | 2072 | 155 | 17906 | 0,07480695 | 0,216178522 | ['GO:0098916'] |
| 1215 | GO:BF | GO:0098916 | anterograde trans-synaptic signaling | 8,87189E-12 | negative_corr | True | 717 | 2072 | 155 | 17906 | 0,07480695 | 0,216178522 | ['GO:0099537'] |
| 1217 | GO:BF | GO:0045664 | regulation of neuron differentiation | 1,60434E-11 | negative_corr | True | 684 | 2072 | 149 | 17906 | 0,071911197 | 0,217836257 | ['GO:0030182', 'GO:0050767'] |
| 942 | GO:BF | GO:0099536 | synaptic signaling | 2,7175E-11 | negative_corr | True | 745 | 2072 | 158 | 17906 | 0,076254826 | 0,212080537 | ['GO:0007267'] |
| 937 | GO:BF | GO:0099537 | trans-synaptic signaling | 2,73002E-11 | negative_corr | True | 726 | 2072 | 155 | 17906 | 0,07480695 | 0,213498623 | ['GO:0099536'] |
| 1581 | GO:BF | GO:0061061 | muscle structure development | 3,1464E-11 | negative_corr | True | 683 | 2072 | 148 | 17906 | 0,071428571 | 0,216691069 | ['GO:0048856'] |
| 898 | GO:BF | GO:0051960 | regulation of nervous system development | 4,11499E-11 | negative_corr | True | 963 | 2072 | 191 | 17906 | 0,092181467 | 0,198338525 | ['GO:0007399', 'GO:2000026'] |
| 1601 | GO:BF | GO:0015711 | organic anion transport | 5,88633E-11 | positive_corr | True | 508 | 998 | 74 | 17906 | 0,074148297 | 0,145669291 | ['GO:0006820', 'GO:0071702'] |
| 1615 | GO:BF | GO:0023051 | regulation of signaling | 1,64767E-10 | negative_corr | True | 3679 | 2072 | 558 | 17906 | 0,269305019 | 0,15167165 | ['GO:0023052', 'GO:0050789'] |
| 830 | GO:BF | GO:0006082 | organic acid metabolic process | 6,12283E-10 | positive_corr | True | 1177 | 998 | 127 | 17906 | 0,127254509 | 0,107901444 | ['GO:0044237', 'GO:0044281', 'GO:0071704'] |
| 1238 | GO:BF | GO:0007155 | cell adhesion | 7,27196E-10 | negative_corr | True | 1440 | 2072 | 256 | 17906 | 0,123552124 | 0,177777778 | ['GO:0022610'] |
| 822 | GO:BF | GO:0016043 | cellular component organization | 7,79904E-10 | negative_corr | True | 6481 | 2072 | 900 | 17906 | 0,434362934 | 0,138867459 | ['GO:0009987', 'GO:0071840'] |
| 1622 | GO:BF | GO:0006810 | transport | 1,04907E-09 | positive_corr | True | 5337 | 998 | 401 | 17906 | 0,401803607 | 0,075135844 | ['GO:0051234'] |
| 1488 | GO:BF | GO:0022610 | biological adhesion | 1,2682E-09 | negative_corr | True | 1447 | 2072 | 256 | 17906 | 0,123552124 | 0,176917761 | ['GO:0008150'] |
| 1637 | GO:BF | GO:0007010 | cytoskeleton organization | 1,43894E-09 | negative_corr | True | 1371 | 2072 | 245 | 17906 | 0,118243243 | 0,178701678 | ['GO:0006996'] |
| 815 | GO:BF | GO:0050808 | synapse organization | 1,6128E-09 | negative_corr | True | 430 | 2072 | 102 | 17906 | 0,049227799 | 0,237209302 | ['GO:0016043'] |
| 810 | GO:BF | GO:0010646 | regulation of cell communication | 1,69916E-09 | negative_corr | True | 3638 | 2072 | 547 | 17906 | 0,263996139 | 0,150357339 | ['GO:0007154', 'GO:0050794'] |
| 1243 | GO:BF | GO:0043436 | oxoacid metabolic process | 1,96528E-09 | positive_corr | True | 1157 | 998 | 124 | 17906 | 0,124248497 | 0,107173725 | ['GO:0006082'] |
| 1244 | GO:BF | GO:0050767 | regulation of neurogenesis | 2,27048E-09 | negative_corr | True | 856 | 2072 | 169 | 17906 | 0,081563707 | 0,197429907 | ['GO:0022008', 'GO:0048699', 'GO:0051960', 'GO:0060284'] |
| 803 | GO:BF | GO:0022604 | regulation of cell morphogenesis | 3,6016E-09 | negative_corr | True | 508 | 2072 | 114 | 17906 | 0,055019305 | 0,224409449 | ['GO:0000902', 'GO:0022603', 'GO:0051128'] |
| 1245 | GO:BF | GO:0050793 | regulation of developmental process | 3,94707E-09 | negative_corr | True | 2779 | 2072 | 434 | 17906 | 0,209459459 | 0,156171285 | ['GO:0032502', 'GO:0050789'] |
| 800 | GO:BF | GO:0006805 | xenobiotic metabolic process | 4,55827E-09 | positive_corr | True | 127 | 998 | 31 | 17906 | 0,031062124 | 0,244094488 | ['GO:0044237', 'GO:0071466'] |
| 1483 | GO:BF | GO:0005975 | carbohydrate metabolic process | 7,55922E-09 | positive_corr | True | 640 | 998 | 81 | 17906 | 0,081162325 | 0,1265625 | ['GO:0044238', 'GO:0071704'] |
| 798 | GO:BF | GO:0071840 | cellular component organization or biogenesis | 9,24789E-09 | negative_corr | True | 6667 | 2072 | 915 | 17906 | 0,441602317 | 0,137243138 | ['GO:0008150'] |
| 1147 | GO:BF | GO:0050804 | modulation of chemical synaptic transmission | 1,25851E-08 | negative_corr | True | 456 | 2072 | 104 | 17906 | 0,05019305 | 0,228070175 | ['GO:0007268', 'GO:0099177'] |
| 1614 | GO:BF | GO:0099177 | regulation of trans-synaptic signaling | 1,44857E-08 | negative_corr | True | 457 | 2072 | 104 | 17906 | 0,05019305 | 0,227571116 | ['GO:0010646', 'GO:0023051', 'GO:0099537'] |
| 1225 | GO:BF | GO:0055085 | transmembrane transport | 2,03179E-08 | positive_corr | True | 1634 | 998 | 156 | 17906 | 0,156312625 | 0,095471236 | ['GO:0006810'] |
| 833 | GO:BF | GO:0019752 | carboxylic acid metabolic process | 2,12614E-08 | positive_corr | True | 1066 | 998 | 114 | 17906 | 0,114228457 | 0,106941839 | ['GO:0043436'] |
| 835 | GO:BF | GO:0051234 | establishment of localization | 2,26049E-08 | positive_corr | True | 5460 | 998 | 402 | 17906 | 0,402805611 | 0,073626374 | ['GO:0051179'] |

| | | | | | | | | | | | | |
|---|---|---|---|---|---|---|---|---|---|---|---|---|
| 885 | GO:BF | GO:0045595 regulation of cell differentiation | 2,75372E-08 | negative_corr | True | 1932 | 2072 | 317 | 17906 | 0,152992278 | 0,164078675 | ['GO:0030154', 'GO:0050793', 'GO:0050794'] |
| 884 | GO:BF | GO:0048589 developmental growth | 3,54385E-08 | negative_corr | True | 698 | 2072 | 141 | 17906 | 0,068050193 | 0,202005731 | ['GO:0032502', 'GO:0040007'] |
| 1493 | GO:BF | GO:0007586 digestion | 5,71318E-08 | positive_corr | True | 139 | 998 | 31 | 17906 | 0,031062124 | 0,223021583 | ['GO:0032501'] |
| 1231 | GO:BF | GO:2000026 regulation of multicellular organismal development | 7,86207E-08 | negative_corr | True | 2210 | 2072 | 352 | 17906 | 0,16988417 | 0,159276018 | ['GO:0007275', 'GO:0050793', 'GO:0051239'] |
| 870 | GO:BF | GO:0008202 steroid metabolic process | 9,957E-08 | positive_corr | True | 332 | 998 | 51 | 17906 | 0,051102204 | 0,153614458 | ['GO:0006629', 'GO:1901360'] |
| 867 | GO:BF | GO:0060284 regulation of cell development | 1,05286E-07 | negative_corr | True | 991 | 2072 | 183 | 17906 | 0,088320463 | 0,184661958 | ['GO:0045595', 'GO:0048468'] |
| 862 | GO:BF | GO:0032530 regulation of microvillus organization | 1,69829E-07 | positive_corr | True | 13 | 998 | 10 | 17906 | 0,01002004 | 0,769230769 | ['GO:0032528', 'GO:0120035'] |
| 861 | GO:BF | GO:1901615 organic hydroxy compound metabolic process | 2,40878E-07 | positive_corr | True | 553 | 998 | 70 | 17906 | 0,070140281 | 0,126582278 | ['GO:0071704'] |
| 851 | GO:BF | GO:0051128 regulation of cellular component organization | 2,42163E-07 | negative_corr | True | 2541 | 2072 | 393 | 17906 | 0,189671815 | 0,154663518 | ['GO:0016043', 'GO:0050794'] |
| 847 | GO:BF | GO:0032787 monocarboxylic acid metabolic process | 2,47439E-07 | positive_corr | True | 661 | 998 | 79 | 17906 | 0,079158317 | 0,119515885 | ['GO:0019752'] |
| 846 | GO:BF | GO:0010769 regulation of cell morphogenesis involved in differentiation | 3,28653E-07 | negative_corr | True | 311 | 2072 | 76 | 17906 | 0,036679537 | 0,24437299 | ['GO:0000904', 'GO:0022604', 'GO:0060284'] |
| 845 | GO:BF | GO:0071466 cellular response to xenobiotic stimulus | 3,42569E-07 | positive_corr | True | 184 | 998 | 35 | 17906 | 0,03507014 | 0,190217391 | ['GO:0009410', 'GO:0070887'] |
| 1232 | GO:BF | GO:0051179 localization | 4,78889E-07 | positive_corr | True | 6889 | 998 | 479 | 17906 | 0,47995992 | 0,069531137 | ['GO:0008150'] |
| 1235 | GO:BF | GO:0044282 small molecule catabolic process | 5,41935E-07 | positive_corr | True | 447 | 998 | 60 | 17906 | 0,06012024 | 0,134228188 | ['GO:0009056', 'GO:0044281'] |
| 1612 | GO:BF | GO:1905114 cell surface receptor signaling pathway involved in cell-cell signaling | 6,02817E-07 | negative_corr | True | 633 | 2072 | 127 | 17906 | 0,061293436 | 0,200631912 | ['GO:0007166', 'GO:0007267'] |
| 838 | GO:BF | GO:0120031 plasma membrane bounded cell projection assembly | 6,2595E-07 | negative_corr | True | 562 | 2072 | 116 | 17906 | 0,055984556 | 0,206405694 | ['GO:0030031', 'GO:0120036'] |
| 1613 | GO:BF | GO:0008610 lipid biosynthetic process | 6,29983E-07 | positive_corr | True | 711 | 998 | 82 | 17906 | 0,082164329 | 0,11533052 | ['GO:0006629', 'GO:1901576'] |
| 834 | GO:BF | GO:0007224 smoothened signaling pathway | 6,56404E-07 | negative_corr | True | 145 | 2072 | 45 | 17906 | 0,021718147 | 0,310344828 | ['GO:0007166'] |
| 985 | GO:BF | GO:0007610 behavior | 7,14223E-07 | negative_corr | True | 615 | 2072 | 124 | 17906 | 0,05984556 | 0,201626016 | ['GO:0008150'] |
| 986 | GO:BF | GO:0034220 ion transmembrane transport | 7,90415E-07 | positive_corr | True | 1184 | 998 | 118 | 17906 | 0,118236473 | 0,099662162 | ['GO:0006811', 'GO:0055085'] |
| 987 | GO:BF | GO:0007156 homophilic cell adhesion via plasma membrane adhesion molecules | 9,51343E-07 | negative_corr | True | 167 | 2072 | 49 | 17906 | 0,023648649 | 0,293413174 | ['GO:0098742'] |
| 1115 | GO:BF | GO:0006066 alcohol metabolic process | 9,63811E-07 | positive_corr | True | 375 | 998 | 53 | 17906 | 0,053106212 | 0,141333333 | ['GO:0044281', 'GO:1901615'] |
| 1114 | GO:BF | GO:0030031 cell projection assembly | 1,29864E-06 | negative_corr | True | 575 | 2072 | 117 | 17906 | 0,056467181 | 0,203478261 | ['GO:0022607', 'GO:0030030'] |
| 1113 | GO:BF | GO:0050770 regulation of axonogenesis | 1,42615E-06 | negative_corr | True | 190 | 2072 | 53 | 17906 | 0,025579151 | 0,278947368 | ['GO:0007409', 'GO:0010769', 'GO:0010975'] |
| 1112 | GO:BF | GO:0023052 signaling | 1,64261E-06 | negative_corr | True | 6772 | 2072 | 911 | 17906 | 0,439671815 | 0,134524513 | ['GO:0008150'] |
| 1529 | GO:BF | GO:0007423 sensory organ development | 2,63131E-06 | negative_corr | True | 562 | 2072 | 114 | 17906 | 0,055019305 | 0,202846975 | ['GO:0048513'] |
| 1107 | GO:BF | GO:0072657 protein localization to membrane | 2,73208E-06 | negative_corr | True | 641 | 2072 | 126 | 17906 | 0,060810811 | 0,196567863 | ['GO:0034613'] |
| 1106 | GO:BF | GO:0032528 microvillus organization | 3,05274E-06 | positive_corr | True | 24 | 998 | 12 | 17906 | 0,012024048 | 0,5 | ['GO:0120036'] |
| 1105 | GO:BF | GO:0016266 O-glycan processing | 3,12181E-06 | positive_corr | True | 58 | 998 | 18 | 17906 | 0,018036072 | 0,310344828 | ['GO:0006493'] |
| 1104 | GO:BF | GO:0034754 cellular hormone metabolic process | 5,93009E-06 | positive_corr | True | 130 | 998 | 27 | 17906 | 0,027054108 | 0,207692308 | ['GO:0042445', 'GO:0044237'] |
| 1530 | GO:BF | GO:0032501 multicellular organismal process | 6,49766E-06 | negative_corr | True | 7873 | 2072 | 1036 | 17906 | 0,5 | 0,131588975 | ['GO:0008150'] |
| 1532 | GO:BF | GO:0030855 epithelial cell differentiation | 6,92787E-06 | positive_corr | True | 786 | 998 | 85 | 17906 | 0,085170341 | 0,108142494 | ['GO:0030154', 'GO:0060429'] |
| 1512 | GO:BF | GO:0030029 actin filament-based process | 8,62617E-06 | negative_corr | True | 781 | 2072 | 145 | 17906 | 0,069980695 | 0,185659411 | ['GO:0009987'] |
| 1100 | GO:BF | GO:0009887 animal organ morphogenesis | 8,65722E-06 | negative_corr | True | 1095 | 2072 | 190 | 17906 | 0,091698842 | 0,173515982 | ['GO:0009653', 'GO:0048513'] |
| 1533 | GO:BF | GO:0031346 positive regulation of cell projection organization | 9,87226E-06 | negative_corr | True | 395 | 2072 | 86 | 17906 | 0,041505792 | 0,217721519 | ['GO:0030030', 'GO:0031344', 'GO:0051130'] |
| 1097 | GO:BF | GO:0097485 neuron projection guidance | 1,039E-05 | negative_corr | True | 286 | 2072 | 68 | 17906 | 0,032818533 | 0,237762238 | ['GO:0006928', 'GO:0006935', 'GO:0048812'] |
| 1538 | GO:BF | GO:0040007 growth | 1,0748E-05 | negative_corr | True | 1020 | 2072 | 179 | 17906 | 0,086389961 | 0,175490196 | ['GO:0008150'] |
| 1093 | GO:BF | GO:0009966 regulation of signal transduction | 1,16642E-05 | negative_corr | True | 3244 | 2072 | 473 | 17906 | 0,228281853 | 0,145807645 | ['GO:0007165', 'GO:0010646', 'GO:0023051', 'GO:0048583'] |
| 1116 | GO:BF | GO:0051239 regulation of multicellular organismal process | 1,23232E-05 | negative_corr | True | 3356 | 2072 | 487 | 17906 | 0,23503861 | 0,14511323 | ['GO:0032501', 'GO:0050789'] |
| 1539 | GO:BF | GO:1901135 carbohydrate derivative metabolic process | 1,26916E-05 | positive_corr | True | 1130 | 998 | 110 | 17906 | 0,110220441 | 0,097345133 | ['GO:0071704'] |
| 1119 | GO:BF | GO:0050789 regulation of biological process | 1,35429E-05 | negative_corr | True | 11940 | 2072 | 1496 | 17906 | 0,722007722 | 0,125293132 | ['GO:0008150', 'GO:0065007'] |
| 1121 | GO:BF | GO:0016042 lipid catabolic process | 1,47929E-05 | positive_corr | True | 337 | 998 | 47 | 17906 | 0,047094188 | 0,139465875 | ['GO:0006629', 'GO:1901575'] |
| 1519 | GO:BF | GO:0007411 axon guidance | 1,88666E-05 | negative_corr | True | 284 | 2072 | 67 | 17906 | 0,032335907 | 0,235915493 | ['GO:0007409', 'GO:0097485'] |
| 1144 | GO:BF | GO:0046942 carboxylic acid transport | 1,9694E-05 | positive_corr | True | 340 | 998 | 47 | 17906 | 0,047094188 | 0,138235294 | ['GO:0015711', 'GO:0015849'] |
| 1152 | GO:BF | GO:0043413 macromolecule glycosylation | 2,02143E-05 | positive_corr | True | 254 | 998 | 39 | 17906 | 0,039078156 | 0,153543307 | ['GO:0043412', 'GO:0070085'] |
| 1142 | GO:BF | GO:0006486 protein glycosylation | 2,02143E-05 | positive_corr | True | 254 | 998 | 39 | 17906 | 0,039078156 | 0,153543307 | ['GO:0006464', 'GO:0009101', 'GO:0043413'] |
| 1141 | GO:BF | GO:0051093 negative regulation of developmental process | 2,16195E-05 | negative_corr | True | 1072 | 2072 | 185 | 17906 | 0,089285714 | 0,172574627 | ['GO:0032502', 'GO:0048519', 'GO:0050793'] |
| 1514 | GO:BF | GO:0015849 organic acid transport | 2,37773E-05 | positive_corr | True | 342 | 998 | 47 | 17906 | 0,047094188 | 0,137426901 | ['GO:0071702'] |
| 1139 | GO:BF | GO:0045216 cell-cell junction organization | 2,51187E-05 | positive_corr | True | 195 | 998 | 33 | 17906 | 0,033066132 | 0,169230769 | ['GO:0034330'] |
| 1158 | GO:BF | GO:0065008 regulation of biological quality | 2,57665E-05 | positive_corr | True | 4156 | 998 | 307 | 17906 | 0,30761523 | 0,073869105 | ['GO:0065007'] |
| 1135 | GO:BF | GO:0070085 glycosylation | 2,66237E-05 | positive_corr | True | 267 | 998 | 40 | 17906 | 0,04008016 | 0,149812734 | ['GO:0008152'] |
| 1160 | GO:BF | GO:0061512 protein localization to cilium | 2,9395E-05 | negative_corr | True | 55 | 2072 | 23 | 17906 | 0,011100386 | 0,418181818 | ['GO:0033365'] |
| 1524 | GO:BF | GO:0010976 positive regulation of neuron projection development | 2,95965E-05 | negative_corr | True | 293 | 2072 | 68 | 17906 | 0,032818533 | 0,232081911 | ['GO:0010975', 'GO:0031175', 'GO:0031346', 'GO:0045666'] |
| 1513 | GO:BF | GO:0042391 regulation of membrane potential | 2,99345E-05 | negative_corr | True | 442 | 2072 | 92 | 17906 | 0,044401544 | 0,208144796 | ['GO:0065008'] |
| 1528 | GO:BF | GO:0031589 cell-substrate adhesion | 3,42192E-05 | negative_corr | True | 355 | 2072 | 78 | 17906 | 0,037644788 | 0,21971831 | ['GO:0007155'] |
| 1129 | GO:BF | GO:0044057 regulation of system process | 4,4734E-05 | negative_corr | True | 623 | 2072 | 119 | 17906 | 0,057432432 | 0,191011236 | ['GO:0003008', 'GO:0051239'] |
| 1127 | GO:BF | GO:0051962 positive regulation of nervous system development | 4,48663E-05 | negative_corr | True | 563 | 2072 | 110 | 17906 | 0,053088803 | 0,195381883 | ['GO:0007399', 'GO:0051094', 'GO:0051240', 'GO:0051960'] |
| 1125 | GO:BF | GO:0022600 digestive system process | 5,9005E-05 | positive_corr | True | 100 | 998 | 22 | 17906 | 0,022044088 | 0,22 | ['GO:0003008', 'GO:0007586'] |
| 1124 | GO:BF | GO:0010817 regulation of hormone levels | 6,05896E-05 | positive_corr | True | 542 | 998 | 63 | 17906 | 0,063126253 | 0,116236162 | ['GO:0065008'] |
| 1120 | GO:BF | GO:0050794 regulation of cellular process | 6,17705E-05 | negative_corr | True | 11187 | 2072 | 1407 | 17906 | 0,679054054 | 0,125770984 | ['GO:0009987', 'GO:0050789'] |
| 1079 | GO:BF | GO:0006928 movement of cell or subcellular component | 6,36611E-05 | negative_corr | True | 2252 | 2072 | 341 | 17906 | 0,16457529 | 0,151420959 | ['GO:0009987'] |
| 1078 | GO:BF | GO:0007017 microtubule-based process | 6,92399E-05 | negative_corr | True | 778 | 2072 | 141 | 17906 | 0,068050193 | 0,181233933 | ['GO:0009987'] |
| 1541 | GO:BF | GO:0098656 anion transmembrane transport | 7,02556E-05 | positive_corr | True | 298 | 998 | 42 | 17906 | 0,042084168 | 0,140939597 | ['GO:0006820', 'GO:0034220'] |
| 1508 | GO:BF | GO:0050769 positive regulation of neurogenesis | 7,81045E-05 | negative_corr | True | 496 | 2072 | 99 | 17906 | 0,047779923 | 0,199596774 | ['GO:0010720', 'GO:0022008', 'GO:0050767', 'GO:0051962'] |
| 1014 | GO:BF | GO:0007154 cell communication | 8,48802E-05 | negative_corr | True | 6798 | 2072 | 900 | 17906 | 0,434362934 | 0,13239188 | ['GO:0009987'] |

| | | | | | | | | | | | |
|---|---|---|---|---|---|---|---|---|---|---|---|
| 1013 | GO:BF GO:0030198 extracellular matrix organization | 8,81143E-05 | negative_corr | True | 375 | 2072 | 80 | 17906 | 0,038610039 | 0,213333333 | ['GO:0043062'] |
| 1183 | GO:BF GO:0006631 fatty acid metabolic process | 9,21604E-05 | positive_corr | True | 380 | 998 | 49 | 17906 | 0,049098196 | 0,128947368 | ['GO:0032787', 'GO:0044255'] |
| 1010 | GO:BF GO:0045666 positive regulation of neuron differentiation | 9,22031E-05 | negative_corr | True | 388 | 2072 | 82 | 17906 | 0,03957529 | 0,211340206 | ['GO:0030182', 'GO:0045664', 'GO:0050769'] |
| 1009 | GO:BF GO:0005996 monosaccharide metabolic process | 9,39814E-05 | positive_corr | True | 301 | 998 | 42 | 17906 | 0,042084168 | 0,139534884 | ['GO:0005975', 'GO:0044281'] |
| 1573 | GO:BF GO:0007264 small GTPase mediated signal transduction | 9,466E-05 | negative_corr | True | 584 | 2072 | 112 | 17906 | 0,054054054 | 0,191780822 | ['GO:0035556'] |
| 1007 | GO:BF GO:0043062 extracellular structure organization | 9,94983E-05 | negative_corr | True | 376 | 2072 | 80 | 17906 | 0,038610039 | 0,212765957 | ['GO:0016043'] |
| 1005 | GO:BF GO:0042692 muscle cell differentiation | 9,97141E-05 | negative_corr | True | 395 | 2072 | 83 | 17906 | 0,040057915 | 0,210126582 | ['GO:0030154', 'GO:0061061'] |
| 1507 | GO:BF GO:0019637 organophosphate metabolic process | 0,000109382 | positive_corr | True | 1051 | 998 | 101 | 17906 | 0,101202405 | 0,096098953 | ['GO:0006793', 'GO:0071704'] |
| 1576 | GO:BF GO:0071363 cellular response to growth factor stimulus | 0,000113765 | negative_corr | True | 715 | 2072 | 131 | 17906 | 0,063223938 | 0,183216783 | ['GO:0070848', 'GO:0071310'] |
| 1579 | GO:BF GO:0009790 embryo development | 0,000115241 | negative_corr | True | 1060 | 2072 | 180 | 17906 | 0,086872587 | 0,169811321 | ['GO:0007275'] |
| 1580 | GO:BF GO:0070848 response to growth factor | 0,000118079 | negative_corr | True | 743 | 2072 | 135 | 17906 | 0,06515444 | 0,181695828 | ['GO:0010033'] |
| 1187 | GO:BF GO:1901137 carbohydrate derivative biosynthetic process | 0,000145348 | positive_corr | True | 669 | 998 | 72 | 17906 | 0,072144289 | 0,107623318 | ['GO:1901135', 'GO:1901576'] |
| 993 | GO:BF GO:0043010 camera-type eye development | 0,000148932 | negative_corr | True | 323 | 2072 | 71 | 17906 | 0,034266409 | 0,219814241 | ['GO:0001654'] |
| 992 | GO:BF GO:0098742 cell-cell adhesion via plasma-membrane adhesion molecules | 0,000168381 | negative_corr | True | 275 | 2072 | 63 | 17906 | 0,030405405 | 0,229090909 | ['GO:0098609'] |
| 988 | GO:BF GO:0050892 intestinal absorption | 0,000170233 | positive_corr | True | 38 | 998 | 13 | 17906 | 0,013026052 | 0,342105263 | ['GO:0006810', 'GO:0022600'] |
| 1023 | GO:BF GO:0098815 modulation of excitatory postsynaptic potential | 0,000188215 | negative_corr | True | 43 | 2072 | 19 | 17906 | 0,009169884 | 0,441860465 | ['GO:0009966', 'GO:0031644', 'GO:0042391', 'GO:0050804', 'GO:0060079'] |
| 1028 | GO:BF GO:0048880 sensory system development | 0,000188252 | negative_corr | True | 375 | 2072 | 79 | 17906 | 0,038127413 | 0,210666667 | ['GO:0048731'] |
| 1029 | GO:BF GO:0042445 hormone metabolic process | 0,000213885 | positive_corr | True | 244 | 998 | 36 | 17906 | 0,036072144 | 0,147540984 | ['GO:0008152', 'GO:0010817'] |
| 1175 | GO:BF GO:0016358 dendrite development | 0,000225646 | negative_corr | True | 253 | 2072 | 59 | 17906 | 0,028474903 | 0,233201581 | ['GO:0031175', 'GO:0048856'] |
| 1072 | GO:BF GO:0031345 negative regulation of cell projection organization | 0,000231896 | negative_corr | True | 189 | 2072 | 48 | 17906 | 0,023166023 | 0,253968254 | ['GO:0030030', 'GO:0031344', 'GO:0051129'] |
| 1170 | GO:BF GO:0022603 regulation of anatomical structure morphogenesis | 0,000236396 | negative_corr | True | 1194 | 2072 | 197 | 17906 | 0,09507722 | 0,164991625 | ['GO:0009653', 'GO:0050793'] |
| 1553 | GO:BF GO:0060348 bone development | 0,000246706 | negative_corr | True | 224 | 2072 | 54 | 17906 | 0,026061776 | 0,241071429 | ['GO:0001501', 'GO:0048513'] |
| 1173 | GO:BF GO:0006821 chloride transport | 0,0002556 | positive_corr | True | 108 | 998 | 22 | 17906 | 0,022044088 | 0,203703704 | ['GO:0015698'] |
| 1060 | GO:BF GO:0044242 cellular lipid catabolic process | 0,000285267 | positive_corr | True | 215 | 998 | 33 | 17906 | 0,033066132 | 0,153488372 | ['GO:0016042', 'GO:0044248', 'GO:0044255'] |
| 1564 | GO:BF GO:0001654 eye development | 0,000291041 | negative_corr | True | 366 | 2072 | 77 | 17906 | 0,037162162 | 0,210382514 | ['GO:0007423', 'GO:0150063'] |
| 1057 | GO:BF GO:0065007 biological regulation | 0,000309135 | negative_corr | True | 12628 | 2072 | 1561 | 17906 | 0,753378378 | 0,123614191 | ['GO:0008150'] |
| 1056 | GO:BF GO:0048523 negative regulation of cellular process | 0,000311074 | negative_corr | True | 4996 | 2072 | 679 | 17906 | 0,327702703 | 0,135908727 | ['GO:0009987', 'GO:0048519', 'GO:0050794'] |
| 794 | GO:BF GO:0097435 supramolecular fiber organization | 0,000321941 | negative_corr | True | 693 | 2072 | 126 | 17906 | 0,060810811 | 0,181818182 | ['GO:0016043'] |
| 1565 | GO:BF GO:0048167 regulation of synaptic plasticity | 0,0003278 | negative_corr | True | 191 | 2072 | 48 | 17906 | 0,023166023 | 0,251308901 | ['GO:0050804', 'GO:0065008'] |
| 1051 | GO:BF GO:0015698 inorganic anion transport | 0,000349895 | positive_corr | True | 166 | 998 | 28 | 17906 | 0,028056112 | 0,168674699 | ['GO:0006820'] |
| 1050 | GO:BF GO:0050890 cognition | 0,00035785 | negative_corr | True | 305 | 2072 | 67 | 17906 | 0,032335907 | 0,219672131 | ['GO:0050877'] |
| 1048 | GO:BF GO:0044782 cilium organization | 0,000361549 | negative_corr | True | 387 | 2072 | 80 | 17906 | 0,038610039 | 0,206718346 | ['GO:0006996', 'GO:0120036'] |
| 1570 | GO:BF GO:0032879 regulation of localization | 0,000405655 | negative_corr | True | 2942 | 2072 | 424 | 17906 | 0,204633205 | 0,144119646 | ['GO:0050789', 'GO:0051179'] |
| 1040 | GO:BF GO:0060271 cilium assembly | 0,0004135 | negative_corr | True | 369 | 2072 | 77 | 17906 | 0,037162162 | 0,208672087 | ['GO:0044782', 'GO:0070925', 'GO:0120031'] |
| 1039 | GO:BF GO:0048513 animal organ development | 0,000418224 | negative_corr | True | 3677 | 2072 | 516 | 17906 | 0,249034749 | 0,140331792 | ['GO:0048731', 'GO:0048856'] |
| 1038 | GO:BF GO:0150063 visual system development | 0,000464191 | negative_corr | True | 370 | 2072 | 77 | 17906 | 0,037162162 | 0,208108108 | ['GO:0048880'] |
| 1174 | GO:BF GO:0007417 central nervous system development | 0,000487912 | negative_corr | True | 1031 | 2072 | 173 | 17906 | 0,083494208 | 0,167798254 | ['GO:0007399', 'GO:0048731'] |
| 1052 | GO:BF GO:0032532 regulation of microvillus length | 0,000492627 | positive_corr | True | 7 | 998 | 6 | 17906 | 0,006012024 | 0,857142857 | ['GO:0032530', 'GO:0032536'] |
| 1257 | GO:BF GO:0086042 cardiac muscle cell-cardiac muscle cell adhesion | 0,000492627 | positive_corr | True | 7 | 998 | 6 | 17906 | 0,006012024 | 0,857142857 | ['GO:0034109'] |
| 796 | GO:BF GO:0071702 organic substance transport | 0,00052118 | positive_corr | True | 2937 | 998 | 224 | 17906 | 0,224448898 | 0,076268301 | ['GO:0006810'] |
| 786 | GO:BF GO:0007611 learning or memory | 0,000530131 | negative_corr | True | 265 | 2072 | 60 | 17906 | 0,028957529 | 0,226415094 | ['GO:0007610', 'GO:0050890'] |
| 478 | GO:BF GO:0060560 developmental growth involved in morphogenesis | 0,000539916 | negative_corr | True | 241 | 2072 | 56 | 17906 | 0,027027027 | 0,232365145 | ['GO:0009653', 'GO:0048589'] |
| 1349 | GO:BF GO:0006355 regulation of transcription, DNA-templated | 0,00057665 | negative_corr | True | 3517 | 2072 | 495 | 17906 | 0,238899614 | 0,140744953 | ['GO:0006351', 'GO:0010468', 'GO:1903506', 'GO:2000112'] |
| 468 | GO:BF GO:0045665 negative regulation of neuron differentiation | 0,000620028 | negative_corr | True | 230 | 2072 | 54 | 17906 | 0,026061776 | 0,234782609 | ['GO:0030182', 'GO:0045664', 'GO:0050768'] |
| 467 | GO:BF GO:0009410 response to xenobiotic stimulus | 0,000621856 | positive_corr | True | 299 | 998 | 40 | 17906 | 0,04008016 | 0,133779264 | ['GO:0042221'] |
| 466 | GO:BF GO:0006493 protein O-linked glycosylation | 0,000694535 | positive_corr | True | 114 | 998 | 22 | 17906 | 0,022044088 | 0,192982456 | ['GO:0006486'] |
| 1351 | GO:BF GO:0010977 negative regulation of neuron projection development | 0,000731888 | negative_corr | True | 156 | 2072 | 41 | 17906 | 0,019787645 | 0,262820513 | ['GO:0010975', 'GO:0031175', 'GO:0031345', 'GO:0045665'] |
| 463 | GO:BF GO:0061387 regulation of extent of cell growth | 0,00076311 | negative_corr | True | 113 | 2072 | 33 | 17906 | 0,015926641 | 0,292035398 | ['GO:0001558', 'GO:0008361'] |
| 451 | GO:BF GO:0035556 intracellular signal transduction | 0,000807213 | negative_corr | True | 2896 | 2072 | 416 | 17906 | 0,200772201 | 0,143646409 | ['GO:0007165'] |
| 443 | GO:BF GO:0051961 negative regulation of nervous system development | 0,000811445 | negative_corr | True | 324 | 2072 | 69 | 17906 | 0,033301158 | 0,212962963 | ['GO:0007399', 'GO:0051093', 'GO:0051241', 'GO:0051960'] |
| 1441 | GO:BF GO:0060079 excitatory postsynaptic potential | 0,000826916 | negative_corr | True | 103 | 2072 | 31 | 17906 | 0,01496139 | 0,300970874 | ['GO:0060078', 'GO:0099565'] |
| 438 | GO:BF GO:0021915 neural tube development | 0,000843915 | negative_corr | True | 168 | 2072 | 43 | 17906 | 0,020752896 | 0,255952381 | ['GO:0007399', 'GO:0035295', 'GO:0043009', 'GO:0060429'] |
| 426 | GO:BF GO:0061448 connective tissue development | 0,000879595 | negative_corr | True | 281 | 2072 | 62 | 17906 | 0,02992278 | 0,220640569 | ['GO:0009888'] |
| 1355 | GO:BF GO:2001141 regulation of RNA biosynthetic process | 0,000881747 | negative_corr | True | 3593 | 2072 | 503 | 17906 | 0,242760618 | 0,139994434 | ['GO:0010556', 'GO:0031326', 'GO:0032774', 'GO:0051252'] |
| 422 | GO:BF GO:0090287 regulation of cellular response to growth factor stimulus | 0,000925485 | negative_corr | True | 300 | 2072 | 65 | 17906 | 0,031370656 | 0,216666667 | ['GO:0048583', 'GO:0050794', 'GO:0071363'] |
| 1360 | GO:BF GO:0046903 secretion | 0,000935842 | positive_corr | True | 1709 | 998 | 143 | 17906 | 0,143286573 | 0,083674664 | ['GO:0006810'] |
| 1363 | GO:BF GO:0048638 regulation of developmental growth | 0,000941967 | negative_corr | True | 357 | 2072 | 74 | 17906 | 0,035714286 | 0,207282913 | ['GO:0040008', 'GO:0048589', 'GO:0050793'] |
| 418 | GO:BF GO:0001501 skeletal system development | 0,000964783 | negative_corr | True | 535 | 2072 | 101 | 17906 | 0,048745174 | 0,188785047 | ['GO:0048731'] |
| 483 | GO:BF GO:0097479 synaptic vesicle localization | 0,000981257 | negative_corr | True | 60 | 2072 | 22 | 17906 | 0,010617761 | 0,366666667 | ['GO:0051648'] |
| 484 | GO:BF GO:1903506 regulation of nucleic acid-templated transcription | 0,000983393 | negative_corr | True | 3588 | 2072 | 502 | 17906 | 0,242277992 | 0,139910814 | ['GO:0097659', 'GO:2001141'] |
| 490 | GO:BF GO:0009101 glycoprotein biosynthetic process | 0,000995656 | positive_corr | True | 339 | 998 | 43 | 17906 | 0,043086172 | 0,126843658 | ['GO:0009100', 'GO:0034645', 'GO:1901137', 'GO:1901566'] |
| 1670 | GO:BF GO:0007043 cell-cell junction assembly | 0,001000048 | positive_corr | True | 135 | 998 | 24 | 17906 | 0,024048096 | 0,177777778 | ['GO:0034329', 'GO:0045216'] |
| 569 | GO:BF GO:0007517 muscle organ development | 0,001085658 | negative_corr | True | 410 | 2072 | 82 | 17906 | 0,03957529 | 0,2 | ['GO:0048513', 'GO:0061061'] |
| 787 | GO:BF GO:0017144 drug metabolic process | 0,001108364 | positive_corr | True | 600 | 998 | 64 | 17906 | 0,064128257 | 0,106666667 | ['GO:0044237'] |
| 566 | GO:BF GO:0019373 epoxygenase P450 pathway | 0,001203384 | positive_corr | True | 20 | 998 | 9 | 17906 | 0,009018036 | 0,45 | ['GO:0019369'] |

| | | | | | | | | | | | | |
|---|---|---|---|---|---|---|---|---|---|---|---|---|
| 1333 | GO:BF | GO:1990778 protein localization to cell periphery | 0,001257866 | negative_corr | True | 334 | 2072 | 70 | 17906 | 0,033783784 | 0,209580838 | ['GO:0034613'] |
| 556 | GO:BF | GO:0048675 axon extension | 0,001283318 | negative_corr | True | 126 | 2072 | 35 | 17906 | 0,016891892 | 0,277777778 | ['GO:0007409', 'GO:1990138'] |
| 555 | GO:BF | GO:0010720 positive regulation of cell development | 0,001289778 | negative_corr | True | 579 | 2072 | 107 | 17906 | 0,051640927 | 0,184801382 | ['GO:0045597', 'GO:0048468', 'GO:0060284'] |
| 1334 | GO:BF | GO:1904970 brush border assembly | 0,001331939 | positive_corr | True | 5 | 998 | 5 | 17906 | 0,00501002 | 1 | ['GO:0022607'] |
| 1658 | GO:BF | GO:0048519 negative regulation of biological process | 0,001354326 | negative_corr | True | 5681 | 2072 | 756 | 17906 | 0,364864865 | 0,133075163 | ['GO:0008150', 'GO:0050789'] |
| 1366 | GO:BF | GO:0034330 cell junction organization | 0,001420831 | positive_corr | True | 297 | 998 | 39 | 17906 | 0,039078156 | 0,131313131 | ['GO:0016043'] |
| 551 | GO:BF | GO:0035637 multicellular organismal signaling | 0,001456234 | negative_corr | True | 206 | 2072 | 49 | 17906 | 0,023648649 | 0,237864078 | ['GO:0023052', 'GO:0032501'] |
| 519 | GO:BF | GO:0048598 embryonic morphogenesis | 0,001476641 | negative_corr | True | 608 | 2072 | 111 | 17906 | 0,053571429 | 0,182565789 | ['GO:0009653', 'GO:0009790'] |
| 1336 | GO:BF | GO:0030036 actin cytoskeleton organization | 0,00170324 | negative_corr | True | 679 | 2072 | 121 | 17906 | 0,058397683 | 0,17820324 | ['GO:0007010', 'GO:0030029'] |
| 1668 | GO:BF | GO:0034032 purine nucleoside bisphosphate metabolic process | 0,001740325 | positive_corr | True | 139 | 998 | 24 | 17906 | 0,024048096 | 0,172661871 | ['GO:0033865', 'GO:0072521'] |
| 1442 | GO:BF | GO:0033875 ribonucleoside bisphosphate metabolic process | 0,001740325 | positive_corr | True | 139 | 998 | 24 | 17906 | 0,024048096 | 0,172661871 | ['GO:0033865'] |
| 498 | GO:BF | GO:0033865 nucleoside bisphosphate metabolic process | 0,001740325 | positive_corr | True | 139 | 998 | 24 | 17906 | 0,024048096 | 0,172661871 | ['GO:0006753'] |
| 1340 | GO:BF | GO:0000226 microtubule cytoskeleton organization | 0,001793135 | negative_corr | True | 576 | 2072 | 106 | 17906 | 0,051158301 | 0,184027778 | ['GO:0007010', 'GO:0007017'] |
| 495 | GO:BF | GO:0048813 dendrite morphogenesis | 0,001902943 | negative_corr | True | 150 | 2072 | 39 | 17906 | 0,018822394 | 0,26 | ['GO:0016358', 'GO:0048667', 'GO:0048812'] |
| 1669 | GO:BF | GO:0006814 sodium ion transport | 0,002078583 | positive_corr | True | 223 | 998 | 32 | 17906 | 0,032064128 | 0,143497758 | ['GO:0015672', 'GO:0030001'] |
| 1335 | GO:BF | GO:0060429 epithelium development | 0,002081025 | positive_corr | True | 1319 | 998 | 115 | 17906 | 0,115230461 | 0,087187263 | ['GO:0009888'] |
| 570 | GO:BF | GO:0019369 arachidonic acid metabolic process | 0,002546392 | positive_corr | True | 54 | 998 | 14 | 17906 | 0,014028056 | 0,259259259 | ['GO:0001676', 'GO:0006690', 'GO:0033559'] |
| 1367 | GO:BF | GO:0001764 neuron migration | 0,002566291 | negative_corr | True | 163 | 2072 | 41 | 17906 | 0,019787645 | 0,251533742 | ['GO:0016477', 'GO:0048699'] |
| 395 | GO:BF | GO:0048588 developmental cell growth | 0,002595075 | negative_corr | True | 240 | 2072 | 54 | 17906 | 0,026061776 | 0,225 | ['GO:0016049', 'GO:0048468', 'GO:0048589'] |
| 254 | GO:BF | GO:0051129 negative regulation of cellular component organization | 0,002645727 | negative_corr | True | 748 | 2072 | 130 | 17906 | 0,062741313 | 0,173796791 | ['GO:0016043', 'GO:0048523', 'GO:0051128'] |
| 1396 | GO:BF | GO:0099643 signal release from synapse | 0,002788883 | negative_corr | True | 175 | 2072 | 43 | 17906 | 0,020752896 | 0,245714286 | ['GO:0023061', 'GO:0099536'] |
| 248 | GO:BF | GO:0007269 neurotransmitter secretion | 0,002788883 | negative_corr | True | 175 | 2072 | 43 | 17906 | 0,020752896 | 0,245714286 | ['GO:0001505', 'GO:0006836', 'GO:0007268', 'GO:0099643'] |
| 247 | GO:BF | GO:0048646 anatomical structure formation involved in morphogenesis | 0,002826924 | negative_corr | True | 1206 | 2072 | 193 | 17906 | 0,093146718 | 0,160033167 | ['GO:0009653', 'GO:0032502'] |
| 1397 | GO:BF | GO:0051641 cellular localization | 0,002835568 | negative_corr | True | 2889 | 2072 | 411 | 17906 | 0,198359073 | 0,142263759 | ['GO:0051179'] |
| 1401 | GO:BF | GO:0034330 cell junction organization | 0,003001999 | negative_corr | True | 297 | 2072 | 63 | 17906 | 0,030405405 | 0,212121212 | ['GO:0016043'] |
| 238 | GO:BF | GO:0051252 regulation of RNA metabolic process | 0,003123975 | negative_corr | True | 3856 | 2072 | 531 | 17906 | 0,256274131 | 0,137707469 | ['GO:0016070', 'GO:0019219', 'GO:0060255'] |
| 233 | GO:BF | GO:0007167 enzyme linked receptor protein signaling pathway | 0,003316641 | negative_corr | True | 1076 | 2072 | 175 | 17906 | 0,084459459 | 0,162639405 | ['GO:0007166'] |
| 227 | GO:BF | GO:0008366 axon ensheathment | 0,003420921 | negative_corr | True | 142 | 2072 | 37 | 17906 | 0,017857143 | 0,26056338 | ['GO:0007272'] |
| 226 | GO:BF | GO:0007272 ensheathment of neurons | 0,003420921 | negative_corr | True | 142 | 2072 | 37 | 17906 | 0,017857143 | 0,26056338 | ['GO:0007399', 'GO:0009987'] |
| 221 | GO:BF | GO:0051668 localization within membrane | 0,003805717 | negative_corr | True | 94 | 2072 | 28 | 17906 | 0,013513514 | 0,29787234 | ['GO:0051641'] |
| 1420 | GO:BF | GO:0006812 cation transport | 0,003874504 | positive_corr | True | 1205 | 998 | 106 | 17906 | 0,106212425 | 0,087966805 | ['GO:0006811'] |
| 1408 | GO:BF | GO:0008361 regulation of cell size | 0,003952396 | negative_corr | True | 183 | 2072 | 44 | 17906 | 0,021235521 | 0,240437158 | ['GO:0032535'] |
| 1409 | GO:BF | GO:0001676 long-chain fatty acid metabolic process | 0,004056321 | positive_corr | True | 107 | 998 | 20 | 17906 | 0,02004008 | 0,186915888 | ['GO:0006631'] |
| 164 | GO:BF | GO:0007265 Ras protein signal transduction | 0,004296899 | negative_corr | True | 457 | 2072 | 87 | 17906 | 0,041988417 | 0,190371991 | ['GO:0007264'] |
| 163 | GO:BF | GO:0007612 learning | 0,004379298 | negative_corr | True | 149 | 2072 | 38 | 17906 | 0,018339768 | 0,255033557 | ['GO:0007611'] |
| 144 | GO:BF | GO:0048878 chemical homeostasis | 0,004605994 | positive_corr | True | 1224 | 998 | 107 | 17906 | 0,107214429 | 0,087418301 | ['GO:0042592'] |
| 255 | GO:BF | GO:0034613 cellular protein localization | 0,004704773 | negative_corr | True | 1983 | 2072 | 294 | 17906 | 0,141891892 | 0,148260212 | ['GO:0008104', 'GO:0070727'] |
| 257 | GO:BF | GO:0006694 steroid biosynthetic process | 0,004899701 | positive_corr | True | 199 | 998 | 29 | 17906 | 0,029058116 | 0,145728643 | ['GO:0008202', 'GO:0008610', 'GO:1901362'] |
| 1391 | GO:BF | GO:0015701 bicarbonate transport | 0,004906314 | positive_corr | True | 42 | 998 | 12 | 17906 | 0,012024048 | 0,285714286 | ['GO:0015711'] |
| 1387 | GO:BF | GO:0098660 inorganic ion transmembrane transport | 0,00498508 | positive_corr | True | 871 | 998 | 82 | 17906 | 0,082164329 | 0,094144661 | ['GO:0034220'] |
| 379 | GO:BF | GO:0099173 postsynapse organization | 0,005176331 | negative_corr | True | 173 | 2072 | 42 | 17906 | 0,02027027 | 0,242774566 | ['GO:0016043', 'GO:0050808'] |
| 378 | GO:BF | GO:0070727 cellular macromolecule localization | 0,005418713 | negative_corr | True | 1994 | 2072 | 295 | 17906 | 0,142374517 | 0,147943831 | ['GO:0033036', 'GO:0051641'] |
| 372 | GO:BF | GO:0042552 myelination | 0,005509768 | negative_corr | True | 139 | 2072 | 36 | 17906 | 0,017374517 | 0,258992806 | ['GO:0008366'] |
| 1369 | GO:BF | GO:0098609 cell-cell adhesion | 0,005599286 | negative_corr | True | 866 | 2072 | 145 | 17906 | 0,069980695 | 0,16743649 | ['GO:0007155'] |
| 342 | GO:BF | GO:1901568 fatty acid derivative metabolic process | 0,00591052 | positive_corr | True | 169 | 998 | 26 | 17906 | 0,026052104 | 0,153846154 | ['GO:0071704'] |
| 335 | GO:BF | GO:0044283 small molecule biosynthetic process | 0,005974707 | positive_corr | True | 724 | 998 | 71 | 17906 | 0,071142285 | 0,098066298 | ['GO:0009058', 'GO:0044281'] |
| 327 | GO:BF | GO:1901617 organic hydroxy compound biosynthetic process | 0,006064698 | positive_corr | True | 268 | 998 | 35 | 17906 | 0,03507014 | 0,130597015 | ['GO:1901576', 'GO:1901615'] |
| 1370 | GO:BF | GO:0050768 negative regulation of neurogenesis | 0,00608713 | negative_corr | True | 303 | 2072 | 63 | 17906 | 0,030405405 | 0,207920792 | ['GO:0010721', 'GO:0022008', 'GO:0050767', 'GO:0051961'] |
| 1368 | GO:BF | GO:0099565 chemical synaptic transmission, postsynaptic | 0,006121872 | negative_corr | True | 112 | 2072 | 31 | 17906 | 0,01496139 | 0,276785714 | ['GO:0007268', 'GO:0050877', 'GO:1905114'] |
| 325 | GO:BF | GO:0045596 negative regulation of cell differentiation | 0,006174261 | negative_corr | True | 767 | 2072 | 131 | 17906 | 0,063223938 | 0,170795306 | ['GO:0030154', 'GO:0045595', 'GO:0048523', 'GO:0051093'] |
| 309 | GO:BF | GO:0051216 cartilage development | 0,00629919 | negative_corr | True | 216 | 2072 | 49 | 17906 | 0,023648649 | 0,226851852 | ['GO:0001501', 'GO:0048513', 'GO:0061448'] |
| 299 | GO:BF | GO:2000463 positive regulation of excitatory postsynaptic potential | 0,006692271 | negative_corr | True | 27 | 2072 | 13 | 17906 | 0,006274131 | 0,481481481 | ['GO:0009967', 'GO:0031646', 'GO:0050806', 'GO:0060079', 'GO:0098815'] |
| 292 | GO:BF | GO:0045597 positive regulation of cell differentiation | 0,006810331 | negative_corr | True | 1022 | 2072 | 166 | 17906 | 0,08011583 | 0,162426614 | ['GO:0030154', 'GO:0045595', 'GO:0048522', 'GO:0051094'] |
| 289 | GO:BF | GO:0072329 monocarboxylic acid catabolic process | 0,007080858 | positive_corr | True | 130 | 998 | 22 | 17906 | 0,022044088 | 0,169230769 | ['GO:0032787', 'GO:0046395'] |
| 285 | GO:BF | GO:0033559 unsaturated fatty acid metabolic process | 0,00735377 | positive_corr | True | 111 | 998 | 20 | 17906 | 0,02004008 | 0,18018018 | ['GO:0006631'] |
| 284 | GO:BF | GO:0072359 circulatory system development | 0,007595994 | negative_corr | True | 1194 | 2072 | 189 | 17906 | 0,091216216 | 0,158291457 | ['GO:0048731'] |
| 1431 | GO:BF | GO:0051094 positive regulation of developmental process | 0,007612832 | negative_corr | True | 1457 | 2072 | 224 | 17906 | 0,108108108 | 0,153740563 | ['GO:0032502', 'GO:0048518', 'GO:0050793'] |
| 1429 | GO:BF | GO:0051552 flavone metabolic process | 0,007622206 | positive_corr | True | 6 | 998 | 5 | 17906 | 0,00501002 | 0,833333333 | ['GO:0009812', 'GO:0042440'] |
| 1372 | GO:BF | GO:0086073 bundle of His cell-Purkinje myocyte adhesion involved in cell commu | 0,007622206 | positive_corr | True | 6 | 998 | 5 | 17906 | 0,00501002 | 0,833333333 | ['GO:0034113', 'GO:0086042', 'GO:0086069'] |
| 571 | GO:BF | GO:0019585 glucuronate metabolic process | 0,007648795 | positive_corr | True | 24 | 998 | 9 | 17906 | 0,009018036 | 0,375 | ['GO:0006063'] |
| 1657 | GO:BF | GO:0006063 uronic acid metabolic process | 0,007648795 | positive_corr | True | 24 | 998 | 9 | 17906 | 0,009018036 | 0,375 | ['GO:0005996', 'GO:0032787'] |
| 579 | GO:BF | GO:0007507 heart development | 0,007842958 | negative_corr | True | 594 | 2072 | 106 | 17906 | 0,051158301 | 0,178451178 | ['GO:0048513', 'GO:0072359'] |
| 725 | GO:BF | GO:0060562 epithelial tube morphogenesis | 0,008745808 | negative_corr | True | 332 | 2072 | 67 | 17906 | 0,032335907 | 0,201807229 | ['GO:0002009', 'GO:0035239'] |
| 1289 | GO:BF | GO:1904062 regulation of cation transmembrane transport | 0,009171227 | negative_corr | True | 352 | 2072 | 70 | 17906 | 0,033783784 | 0,198863636 | ['GO:0034765', 'GO:0098655'] |
| 722 | GO:BF | GO:0009812 flavonoid metabolic process | 0,009934725 | positive_corr | True | 14 | 998 | 7 | 17906 | 0,007014028 | 0,5 | ['GO:0071704'] |

| | | | | | | | | | | | | | |
|---|---|---|---|---|---|---|---|---|---|---|---|---|---|
| 721 | GO:BF | GO:0043113 | receptor clustering | 0,009939114 | negative_corr | True | 58 | 2072 | 20 | 17906 | 0,00965251 | 0,344827586 | ['GO:0051668', 'GO:0072657'] |
| 1292 | GO:BF | GO:0015718 | monocarboxylic acid transport | 0,01027188 | positive_corr | True | 174 | 998 | 26 | 17906 | 0,026052104 | 0,149425287 | ['GO:0046942'] |
| 713 | GO:BF | GO:0002009 | morphogenesis of an epithelium | 0,010482991 | negative_corr | True | 556 | 2072 | 100 | 17906 | 0,048262548 | 0,179856115 | ['GO:0048729', 'GO:0060429'] |
| 712 | GO:BF | GO:0048568 | embryonic organ development | 0,010949533 | negative_corr | True | 447 | 2072 | 84 | 17906 | 0,040540541 | 0,187919463 | ['GO:0009790', 'GO:0048513'] |
| 1288 | GO:BF | GO:0072001 | renal system development | 0,011786386 | negative_corr | True | 296 | 2072 | 61 | 17906 | 0,029440154 | 0,206081081 | ['GO:0001655', 'GO:0048731'] |
| 1462 | GO:BF | GO:0006643 | membrane lipid metabolic process | 0,012036832 | positive_corr | True | 208 | 998 | 29 | 17906 | 0,029058116 | 0,139423077 | ['GO:0044255'] |
| 709 | GO:BF | GO:0009062 | fatty acid catabolic process | 0,012113243 | positive_corr | True | 105 | 998 | 19 | 17906 | 0,019038076 | 0,180952381 | ['GO:0006631', 'GO:0044242', 'GO:0072329'] |
| 706 | GO:BF | GO:0002934 | desmosome organization | 0,012775011 | positive_corr | True | 10 | 998 | 6 | 17906 | 0,006012024 | 0,6 | ['GO:0045216'] |
| 705 | GO:BF | GO:0098661 | inorganic anion transmembrane transport | 0,01292143 | positive_corr | True | 115 | 998 | 20 | 17906 | 0,02004008 | 0,173913043 | ['GO:0015698', 'GO:0098656', 'GO:0098660'] |
| 704 | GO:BF | GO:0043009 | chordate embryonic development | 0,013205586 | negative_corr | True | 657 | 2072 | 114 | 17906 | 0,055019305 | 0,173515982 | ['GO:0009792'] |
| 573 | GO:BF | GO:0048705 | skeletal system morphogenesis | 0,014456191 | negative_corr | True | 247 | 2072 | 53 | 17906 | 0,025579151 | 0,214574899 | ['GO:0001501', 'GO:0009887'] |
| 702 | GO:BF | GO:0042573 | retinoic acid metabolic process | 0,014472833 | positive_corr | True | 32 | 998 | 10 | 17906 | 0,01002004 | 0,3125 | ['GO:0001523', 'GO:0032787', 'GO:0034754'] |
| 697 | GO:BF | GO:0009100 | glycoprotein metabolic process | 0,015100156 | positive_corr | True | 413 | 998 | 46 | 17906 | 0,046092184 | 0,111380145 | ['GO:0019538', 'GO:0044260', 'GO:1901135'] |
| 1298 | GO:BF | GO:0007416 | synapse assembly | 0,015226284 | negative_corr | True | 180 | 2072 | 42 | 17906 | 0,02027027 | 0,233333333 | ['GO:0007399', 'GO:0022607', 'GO:0050808'] |
| 1647 | GO:BF | GO:0030277 | maintenance of gastrointestinal epithelium | 0,015627387 | positive_corr | True | 20 | 998 | 8 | 17906 | 0,008016032 | 0,4 | ['GO:0010669', 'GO:0022600'] |
| 1469 | GO:BF | GO:0051130 | positive regulation of cellular component organization | 0,01567434 | negative_corr | True | 1245 | 2072 | 194 | 17906 | 0,093629344 | 0,155823293 | ['GO:0016043', 'GO:0048522', 'GO:0051128'] |
| 736 | GO:BF | GO:0016055 | Wnt signaling pathway | 0,016464069 | negative_corr | True | 527 | 2072 | 95 | 17906 | 0,045849421 | 0,180265655 | ['GO:0198738', 'GO:1905114'] |
| 1258 | GO:BF | GO:0001508 | action potential | 0,016969952 | negative_corr | True | 134 | 2072 | 34 | 17906 | 0,016409266 | 0,253731343 | ['GO:0042391'] |
| 1264 | GO:BF | GO:0007626 | locomotory behavior | 0,017159235 | negative_corr | True | 205 | 2072 | 46 | 17906 | 0,022200772 | 0,224390244 | ['GO:0007610'] |
| 1266 | GO:BF | GO:0055086 | nucleobase-containing small molecule metabolic process | 0,017879574 | positive_corr | True | 638 | 998 | 63 | 17906 | 0,063126253 | 0,098746082 | ['GO:0006139', 'GO:0044281'] |
| 778 | GO:BF | GO:0035295 | tube development | 0,018172351 | negative_corr | True | 1143 | 2072 | 180 | 17906 | 0,086872587 | 0,157480315 | ['GO:0007275', 'GO:0048856'] |
| 1479 | GO:BF | GO:0009888 | tissue development | 0,019163569 | negative_corr | True | 2118 | 2072 | 307 | 17906 | 0,148166023 | 0,144948064 | ['GO:0048856'] |
| 775 | GO:BF | GO:0198738 | cell-cell signaling by wnt | 0,019337081 | negative_corr | True | 529 | 2072 | 95 | 17906 | 0,045849421 | 0,179584121 | ['GO:0007267'] |
| 771 | GO:BF | GO:0098693 | regulation of synaptic vesicle cycle | 0,019916401 | negative_corr | True | 118 | 2072 | 31 | 17906 | 0,01496139 | 0,262711864 | ['GO:0060627', 'GO:0099504'] |
| 1477 | GO:BF | GO:2000807 | regulation of synaptic vesicle clustering | 0,020032615 | negative_corr | True | 9 | 2072 | 7 | 17906 | 0,003378378 | 0,777777778 | ['GO:0060341', 'GO:0097091', 'GO:0098693'] |
| 1471 | GO:BF | GO:0006793 | phosphorus metabolic process | 0,020222811 | positive_corr | True | 3393 | 998 | 243 | 17906 | 0,243486974 | 0,071618037 | ['GO:0044237'] |
| 1643 | GO:BF | GO:0034308 | primary alcohol metabolic process | 0,020858319 | positive_corr | True | 90 | 998 | 17 | 17906 | 0,017034068 | 0,188888889 | ['GO:0006066'] |
| 745 | GO:BF | GO:0090407 | organophosphate biosynthetic process | 0,021126841 | positive_corr | True | 601 | 998 | 60 | 17906 | 0,06012024 | 0,099833611 | ['GO:0019637', 'GO:1901576'] |
| 1275 | GO:BF | GO:0009888 | tissue development | 0,021640229 | positive_corr | True | 2118 | 998 | 163 | 17906 | 0,163326653 | 0,076959396 | ['GO:0048856'] |
| 743 | GO:BF | GO:0061337 | cardiac conduction | 0,021906433 | negative_corr | True | 147 | 2072 | 36 | 17906 | 0,017374517 | 0,244897959 | ['GO:0008016', 'GO:0035637'] |
| 1277 | GO:BF | GO:0060078 | regulation of postsynaptic membrane potential | 0,021906433 | negative_corr | True | 147 | 2072 | 36 | 17906 | 0,017374517 | 0,244897959 | ['GO:0042391'] |
| 739 | GO:BF | GO:0002028 | regulation of sodium ion transport | 0,021979336 | negative_corr | True | 86 | 2072 | 25 | 17906 | 0,012065637 | 0,290697674 | ['GO:0006814', 'GO:0010959'] |
| 1285 | GO:BF | GO:0048522 | positive regulation of cellular process | 0,02242391 | negative_corr | True | 5605 | 2072 | 735 | 17906 | 0,35472973 | 0,131132917 | ['GO:0009987', 'GO:0048518', 'GO:0050794'] |
| 1286 | GO:BF | GO:0051241 | negative regulation of multicellular organismal process | 0,022687352 | negative_corr | True | 1358 | 2072 | 208 | 17906 | 0,1003861 | 0,153166421 | ['GO:0032501', 'GO:0048519', 'GO:0051239'] |
| 696 | GO:BF | GO:0006720 | isoprenoid metabolic process | 0,024510546 | positive_corr | True | 140 | 998 | 22 | 17906 | 0,022044088 | 0,157142857 | ['GO:0044255'] |
| 681 | GO:BF | GO:0051056 | regulation of small GTPase mediated signal transduction | 0,024702636 | negative_corr | True | 342 | 2072 | 67 | 17906 | 0,032335907 | 0,195906433 | ['GO:0007264', 'GO:1902531'] |
| 703 | GO:BF | GO:0006936 | muscle contraction | 0,02503158 | negative_corr | True | 362 | 2072 | 70 | 17906 | 0,033783784 | 0,193370166 | ['GO:0003012'] |
| 679 | GO:BF | GO:0097659 | nucleic acid-templated transcription | 0,026238478 | negative_corr | True | 3744 | 2072 | 509 | 17906 | 0,245656371 | 0,135950855 | ['GO:0032774'] |
| 640 | GO:BF | GO:0010721 | negative regulation of cell development | 0,028347751 | negative_corr | True | 350 | 2072 | 68 | 17906 | 0,032818533 | 0,194285714 | ['GO:0045596', 'GO:0048468', 'GO:0060284'] |
| 632 | GO:BF | GO:0009792 | embryo development ending in birth or egg hatching | 0,028546386 | negative_corr | True | 675 | 2072 | 115 | 17906 | 0,055501931 | 0,17037037 | ['GO:0009790'] |
| 630 | GO:BF | GO:0014706 | striated muscle tissue development | 0,028961666 | negative_corr | True | 397 | 2072 | 75 | 17906 | 0,036196911 | 0,188916877 | ['GO:0060537'] |
| 629 | GO:BF | GO:0010771 | negative regulation of cell morphogenesis involved in differentiation | 0,029325033 | negative_corr | True | 98 | 2072 | 27 | 17906 | 0,013030888 | 0,275510204 | ['GO:0000904', 'GO:0010721', 'GO:0010769', 'GO:0051129'] |
| 626 | GO:BF | GO:0003015 | heart process | 0,029872104 | negative_corr | True | 298 | 2072 | 60 | 17906 | 0,028957529 | 0,201342282 | ['GO:0003013'] |
| 622 | GO:BF | GO:0030033 | microvillus assembly | 0,030002783 | positive_corr | True | 16 | 998 | 7 | 17906 | 0,007014028 | 0,4375 | ['GO:0032528', 'GO:0120031'] |
| 616 | GO:BF | GO:0055114 | oxidation-reduction process | 0,030225206 | positive_corr | True | 1028 | 998 | 90 | 17906 | 0,090180361 | 0,087548638 | ['GO:0008152'] |
| 612 | GO:BF | GO:0016054 | organic acid catabolic process | 0,03025335 | positive_corr | True | 276 | 998 | 34 | 17906 | 0,034068136 | 0,123188406 | ['GO:0006082', 'GO:0044248', 'GO:0044282', 'GO:1901575'] |
| 1461 | GO:BF | GO:0046395 | carboxylic acid catabolic process | 0,03025335 | positive_corr | True | 276 | 998 | 34 | 17906 | 0,034068136 | 0,123188406 | ['GO:0016054', 'GO:0019752'] |
| 606 | GO:BF | GO:0006790 | sulfur compound metabolic process | 0,031096784 | positive_corr | True | 374 | 998 | 42 | 17906 | 0,042084168 | 0,112299465 | ['GO:0044237'] |
| 603 | GO:BF | GO:0032774 | RNA biosynthetic process | 0,031331759 | negative_corr | True | 3758 | 2072 | 510 | 17906 | 0,246138996 | 0,135710484 | ['GO:0009059', 'GO:0016070', 'GO:0034654'] |
| 1445 | GO:BF | GO:2000112 | regulation of cellular macromolecule biosynthetic process | 0,031490819 | negative_corr | True | 4036 | 2072 | 544 | 17906 | 0,262548263 | 0,134786918 | ['GO:0010556', 'GO:0031326', 'GO:0034645'] |
| 595 | GO:BF | GO:0046165 | alcohol biosynthetic process | 0,031514459 | positive_corr | True | 174 | 998 | 25 | 17906 | 0,0250501 | 0,143678161 | ['GO:0006066', 'GO:0044283', 'GO:1901617'] |
| 594 | GO:BF | GO:0010669 | epithelial structure maintenance | 0,033005137 | positive_corr | True | 28 | 998 | 9 | 17906 | 0,009018036 | 0,321428571 | ['GO:0001894'] |
| 593 | GO:BF | GO:0001822 | kidney development | 0,033154618 | negative_corr | True | 286 | 2072 | 58 | 17906 | 0,027992278 | 0,202797203 | ['GO:0048513', 'GO:0072001'] |
| 1332 | GO:BF | GO:0051493 | regulation of cytoskeleton organization | 0,033544326 | negative_corr | True | 564 | 2072 | 99 | 17906 | 0,047779923 | 0,175531915 | ['GO:0007010', 'GO:0033043'] |
| 1656 | GO:BF | GO:0048041 | focal adhesion assembly | 0,036463072 | negative_corr | True | 83 | 2072 | 24 | 17906 | 0,011583012 | 0,289156627 | ['GO:0007044', 'GO:0007160'] |
| 582 | GO:BF | GO:1990138 | neuron projection extension | 0,036513332 | negative_corr | True | 174 | 2072 | 40 | 17906 | 0,019305019 | 0,229885057 | ['GO:0048588', 'GO:0048812', 'GO:0060560'] |
| 1328 | GO:BF | GO:0006351 | transcription, DNA-templated | 0,036651493 | negative_corr | True | 3698 | 2072 | 502 | 17906 | 0,242277992 | 0,135749054 | ['GO:0010467', 'GO:0034645', 'GO:0097659'] |
| 1314 | GO:BF | GO:0040008 | regulation of growth | 0,036686612 | negative_corr | True | 693 | 2072 | 117 | 17906 | 0,056467181 | 0,168831169 | ['GO:0040007', 'GO:0050789'] |
| 1518 | GO:BF | GO:1902476 | chloride transmembrane transport | 0,038092813 | positive_corr | True | 94 | 998 | 17 | 17906 | 0,017034068 | 0,180851064 | ['GO:0006821', 'GO:0098661'] |
| 672 | GO:BF | GO:0034329 | cell junction assembly | 0,042612507 | positive_corr | True | 245 | 998 | 31 | 17906 | 0,031062124 | 0,126530612 | ['GO:0022607', 'GO:0034330'] |
| 649 | GO:BF | GO:0044262 | cellular carbohydrate metabolic process | 0,044228735 | positive_corr | True | 293 | 998 | 35 | 17906 | 0,03507014 | 0,119453925 | ['GO:0005975', 'GO:0044237'] |
| 671 | GO:BF | GO:0035239 | tube morphogenesis | 0,044766965 | negative_corr | True | 944 | 2072 | 151 | 17906 | 0,072876448 | 0,159957627 | ['GO:0009653', 'GO:0035295'] |
| 1313 | GO:BF | GO:0010959 | regulation of metal ion transport | 0,046302784 | negative_corr | True | 409 | 2072 | 76 | 17906 | 0,036679537 | 0,185819071 | ['GO:0030001', 'GO:0043269'] |
| 673 | GO:BF | GO:0060537 | muscle tissue development | 0,047108634 | negative_corr | True | 416 | 2072 | 77 | 17906 | 0,037162162 | 0,185096154 | ['GO:0009888'] |
| 653 | GO:BF | GO:0019318 | hexose metabolic process | 0,047117446 | positive_corr | True | 258 | 998 | 32 | 17906 | 0,032064128 | 0,124031008 | ['GO:0005996'] |

| | | | | | | | | | | | | |
|---|---|---|---|---|---|---|---|---|---|---|---|---|
| 654 | GO:BF | GO:0006690 icosanoid metabolic process | 0,047215058 | positive_corr | True | 115 | 998 | 19 | 17906 | 0,019038076 | 0,165217391 | ['GO:0019752', 'GO:1901568'] |
| 1654 | GO:BF | GO:0021535 cell migration in hindbrain | 0,048803014 | negative_corr | True | 16 | 2072 | 9 | 17906 | 0,004343629 | 0,5625 | ['GO:0016477', 'GO:0030902'] |
| 1310 | GO:BF | GO:0007158 neuron cell-cell adhesion | 0,048803014 | negative_corr | True | 16 | 2072 | 9 | 17906 | 0,004343629 | 0,5625 | ['GO:0098609'] |
| 663 | GO:BF | GO:0010970 transport along microtubule | 0,048925441 | negative_corr | True | 164 | 2072 | 38 | 17906 | 0,018339768 | 0,231707317 | ['GO:0030705', 'GO:0099111'] |
| 1230 | GO:CC | GO:0042995 cell projection | 3,95156E-29 | negative_corr | True | 2288 | 2182 | 444 | 18856 | 0,203483043 | 0,194055944 | ['GO:0110165'] |
| 1491 | GO:CC | GO:0120025 plasma membrane bounded cell projection | 6,81227E-27 | negative_corr | True | 2200 | 2182 | 424 | 18856 | 0,19431714 | 0,192727273 | ['GO:0042995'] |
| 1353 | GO:CC | GO:0045202 synapse | 1,04994E-26 | negative_corr | True | 1362 | 2182 | 297 | 18856 | 0,136113657 | 0,218061674 | ['GO:0110165'] |
| 1172 | GO:CC | GO:0045177 apical part of cell | 7,68883E-26 | positive_corr | True | 400 | 1064 | 87 | 18856 | 0,081766917 | 0,2175 | ['GO:0110165'] |
| 1255 | GO:CC | GO:0016324 apical plasma membrane | 3,17606E-23 | positive_corr | True | 331 | 1064 | 75 | 18856 | 0,070488722 | 0,226586103 | ['GO:0045177', 'GO:0098590'] |
| 1302 | GO:CC | GO:0070062 extracellular exosome | 2,09536E-22 | positive_corr | True | 2144 | 1064 | 235 | 18856 | 0,220864662 | 0,109608209 | ['GO:0005615', 'GO:1903561'] |
| 1407 | GO:CC | GO:1903561 extracellular vesicle | 3,97346E-22 | positive_corr | True | 2167 | 1064 | 236 | 18856 | 0,221804511 | 0,108906322 | ['GO:0031982', 'GO:0043230'] |
| 1222 | GO:CC | GO:0043230 extracellular organelle | 5,49028E-22 | positive_corr | True | 2172 | 1064 | 236 | 18856 | 0,221804511 | 0,108655617 | ['GO:0005576', 'GO:0043226'] |
| 1274 | GO:CC | GO:0016020 membrane | 1,81213E-21 | positive_corr | True | 9731 | 1064 | 707 | 18856 | 0,664473684 | 0,072654403 | ['GO:0110165'] |
| 1216 | GO:CC | GO:0031224 intrinsic component of membrane | 3,00657E-21 | positive_corr | True | 5842 | 1064 | 481 | 18856 | 0,452067669 | 0,082334817 | ['GO:0016020', 'GO:0110165'] |
| 1350 | GO:CC | GO:0016021 integral component of membrane | 3,31721E-20 | positive_corr | True | 5692 | 1064 | 468 | 18856 | 0,439849624 | 0,082220661 | ['GO:0031224'] |
| 1239 | GO:CC | GO:0030054 cell junction | 1,31952E-19 | negative_corr | True | 1327 | 2182 | 271 | 18856 | 0,124197984 | 0,204220045 | ['GO:0110165'] |
| 1311 | GO:CC | GO:0036477 somatodendritic compartment | 6,88569E-19 | negative_corr | True | 863 | 2182 | 196 | 18856 | 0,089825848 | 0,227114716 | ['GO:0110165'] |
| 1492 | GO:CC | GO:0043005 neuron projection | 3,32061E-18 | negative_corr | True | 1376 | 2182 | 274 | 18856 | 0,125572869 | 0,199127907 | ['GO:0120025'] |
| 1382 | GO:CC | GO:0031982 vesicle | 8,0847E-17 | positive_corr | True | 3914 | 1064 | 342 | 18856 | 0,321428571 | 0,087378641 | ['GO:0043227'] |
| 1463 | GO:CC | GO:0030425 dendrite | 3,56981E-14 | negative_corr | True | 622 | 2182 | 144 | 18856 | 0,0659945 | 0,231511254 | ['GO:0043005', 'GO:0097447'] |
| 1352 | GO:CC | GO:0097447 dendritic tree | 4,77628E-14 | negative_corr | True | 624 | 2182 | 144 | 18856 | 0,0659945 | 0,230769231 | ['GO:0036477', 'GO:0043005'] |
| 1315 | GO:CC | GO:0005622 intracellular | 2,00801E-13 | negative_corr | True | 14626 | 2182 | 1835 | 18856 | 0,840971586 | 0,125461507 | ['GO:0005575'] |
| 140 | GO:CC | GO:0071944 cell periphery | 3,13245E-13 | positive_corr | True | 5732 | 1064 | 443 | 18856 | 0,416353383 | 0,077285415 | ['GO:0110165'] |
| 1143 | GO:CC | GO:0098590 plasma membrane region | 3,68058E-13 | positive_corr | True | 1153 | 1064 | 133 | 18856 | 0,125 | 0,115351258 | ['GO:0005886', 'GO:0016020'] |
| 499 | GO:CC | GO:0005856 cytoskeleton | 4,29071E-13 | negative_corr | True | 2263 | 2182 | 381 | 18856 | 0,174610449 | 0,168360583 | ['GO:0043232'] |
| 503 | GO:CC | GO:0005903 brush border | 1,09033E-12 | positive_corr | True | 108 | 1064 | 32 | 18856 | 0,030075188 | 0,296296296 | ['GO:0098862'] |
| 1662 | GO:CC | GO:0030424 axon | 1,18639E-12 | negative_corr | True | 647 | 2182 | 144 | 18856 | 0,0659945 | 0,222565688 | ['GO:0043005'] |
| 554 | GO:CC | GO:0098794 postsynapse | 1,86047E-12 | negative_corr | True | 638 | 2182 | 142 | 18856 | 0,06507791 | 0,222570533 | ['GO:0045202', 'GO:0110165'] |
| 557 | GO:CC | GO:0044297 cell body | 6,64902E-12 | negative_corr | True | 592 | 2182 | 133 | 18856 | 0,060953254 | 0,224662162 | ['GO:0110165'] |
| 585 | GO:CC | GO:0005886 plasma membrane | 7,60833E-12 | positive_corr | True | 5604 | 1064 | 429 | 18856 | 0,403195489 | 0,076552463 | ['GO:0016020', 'GO:0071944'] |
| 601 | GO:CC | GO:0070161 anchoring junction | 4,95558E-11 | negative_corr | True | 785 | 2182 | 161 | 18856 | 0,073785518 | 0,205095541 | ['GO:0030054'] |
| 604 | GO:CC | GO:0043025 neuronal cell body | 6,57828E-11 | negative_corr | True | 523 | 2182 | 119 | 18856 | 0,054537122 | 0,227533461 | ['GO:0036477', 'GO:0044297'] |
| 1655 | GO:CC | GO:0043296 apical junction complex | 9,85093E-11 | positive_corr | True | 148 | 1064 | 35 | 18856 | 0,032894737 | 0,236486486 | ['GO:0005911'] |
| 643 | GO:CC | GO:0062023 collagen-containing extracellular matrix | 6,9112E-10 | negative_corr | True | 407 | 2182 | 97 | 18856 | 0,044454629 | 0,238329238 | ['GO:0031012'] |
| 646 | GO:CC | GO:0005615 extracellular space | 8,1366E-10 | positive_corr | True | 3543 | 1064 | 290 | 18856 | 0,272556391 | 0,081851538 | ['GO:0005576', 'GO:0110165'] |
| 650 | GO:CC | GO:0012505 endomembrane system | 2,83494E-09 | positive_corr | True | 4539 | 1064 | 351 | 18856 | 0,329887218 | 0,077329808 | ['GO:0110165'] |
| 660 | GO:CC | GO:0098862 cluster of actin-based cell projections | 6,4548E-09 | positive_corr | True | 161 | 1064 | 34 | 18856 | 0,031954887 | 0,211180124 | ['GO:0110165'] |
| 674 | GO:CC | GO:0005911 cell-cell junction | 7,85618E-09 | positive_corr | True | 439 | 1064 | 62 | 18856 | 0,058270677 | 0,141230068 | ['GO:0070161'] |
| 680 | GO:CC | GO:0005902 microvillus | 6,63277E-08 | positive_corr | True | 85 | 1064 | 23 | 18856 | 0,021616541 | 0,270588235 | ['GO:0098858'] |
| 710 | GO:CC | GO:0070160 tight junction | 7,2547E-08 | positive_corr | True | 132 | 1064 | 29 | 18856 | 0,027255639 | 0,21969697 | ['GO:0005911'] |
| 711 | GO:CC | GO:0015630 microtubule cytoskeleton | 8,62107E-08 | negative_corr | True | 1242 | 2182 | 216 | 18856 | 0,098991751 | 0,173913043 | ['GO:0005856'] |
| 720 | GO:CC | GO:0031090 organelle membrane | 1,32724E-07 | positive_corr | True | 3531 | 1064 | 279 | 18856 | 0,262218045 | 0,079014444 | ['GO:0016020', 'GO:0043227'] |
| 1650 | GO:CC | GO:0016323 basolateral plasma membrane | 2,52188E-07 | positive_corr | True | 221 | 1064 | 38 | 18856 | 0,035714286 | 0,171945701 | ['GO:0098590'] |
| 724 | GO:CC | GO:0005576 extracellular region | 3,41045E-07 | positive_corr | True | 4548 | 1064 | 341 | 18856 | 0,320488722 | 0,074978012 | ['GO:0110165'] |
| 726 | GO:CC | GO:0098984 neuron to neuron synapse | 8,43018E-07 | negative_corr | True | 372 | 2182 | 83 | 18856 | 0,038038497 | 0,22311828 | ['GO:0045202'] |
| 731 | GO:CC | GO:0032279 asymmetric synapse | 8,92213E-07 | negative_corr | True | 348 | 2182 | 79 | 18856 | 0,036205316 | 0,227011494 | ['GO:0098984'] |
| 1149 | GO:CC | GO:0014069 postsynaptic density | 1,19045E-06 | negative_corr | True | 344 | 2182 | 78 | 18856 | 0,035747021 | 0,226744186 | ['GO:0032279', 'GO:0099572'] |
| 496 | GO:CC | GO:0099572 postsynaptic specialization | 1,46482E-06 | negative_corr | True | 370 | 2182 | 82 | 18856 | 0,037580202 | 0,221621622 | ['GO:0043226', 'GO:0098794'] |
| 738 | GO:CC | GO:0030427 site of polarized growth | 1,74094E-06 | negative_corr | True | 195 | 2182 | 52 | 18856 | 0,023831347 | 0,266666667 | ['GO:0110165'] |
| 465 | GO:CC | GO:0005923 bicellular tight junction | 2,13827E-06 | positive_corr | True | 125 | 1064 | 26 | 18856 | 0,02443609 | 0,208 | ['GO:0043296', 'GO:0070160'] |
| 448 | GO:CC | GO:0005887 integral component of plasma membrane | 2,67079E-06 | positive_corr | True | 1624 | 1064 | 146 | 18856 | 0,137218045 | 0,089901478 | ['GO:0016021', 'GO:0031226'] |
| 174 | GO:CC | GO:0031012 extracellular matrix | 2,67499E-06 | negative_corr | True | 534 | 2182 | 107 | 18856 | 0,04903758 | 0,200374532 | ['GO:0005576', 'GO:0110165'] |
| 175 | GO:CC | GO:0030426 growth cone | 3,11003E-06 | negative_corr | True | 187 | 2182 | 50 | 18856 | 0,022914757 | 0,267379679 | ['GO:0030427', 'GO:0150034'] |
| 184 | GO:CC | GO:0043229 intracellular organelle | 3,41331E-06 | negative_corr | True | 12840 | 2182 | 1600 | 18856 | 0,733272227 | 0,124610592 | ['GO:0005622', 'GO:0043226'] |
| 220 | GO:CC | GO:0031226 intrinsic component of plasma membrane | 5,56065E-06 | positive_corr | True | 1700 | 1064 | 150 | 18856 | 0,140977444 | 0,088235294 | ['GO:0005886', 'GO:0031224'] |
| 239 | GO:CC | GO:0098978 glutamatergic synapse | 5,62485E-06 | negative_corr | True | 368 | 2182 | 80 | 18856 | 0,036663611 | 0,217391304 | ['GO:0045202'] |
| 244 | GO:CC | GO:0098590 plasma membrane region | 6,62669E-06 | negative_corr | True | 1153 | 2182 | 195 | 18856 | 0,089367553 | 0,169124024 | ['GO:0005886', 'GO:0016020'] |
| 249 | GO:CC | GO:0005737 cytoplasm | 8,30451E-06 | positive_corr | True | 11634 | 1064 | 740 | 18856 | 0,695488722 | 0,06360667 | ['GO:0005622', 'GO:0110165'] |
| 261 | GO:CC | GO:0097060 synaptic membrane | 8,37187E-06 | negative_corr | True | 390 | 2182 | 83 | 18856 | 0,038038497 | 0,212820513 | ['GO:0045202', 'GO:0098590'] |
| 266 | GO:CC | GO:0043204 perikaryon | 8,55747E-06 | negative_corr | True | 154 | 2182 | 43 | 18856 | 0,019706691 | 0,279220779 | ['GO:0043025', 'GO:0110165'] |
| 1674 | GO:CC | GO:0031252 cell leading edge | 1,01728E-05 | negative_corr | True | 417 | 2182 | 87 | 18856 | 0,039871677 | 0,208633094 | ['GO:0110165'] |
| 280 | GO:CC | GO:0042383 sarcolemma | 1,10621E-05 | negative_corr | True | 134 | 2182 | 39 | 18856 | 0,017873511 | 0,291044776 | ['GO:0005886'] |
| 300 | GO:CC | GO:0043226 organelle | 1,24356E-05 | negative_corr | True | 13720 | 2182 | 1692 | 18856 | 0,77543538 | 0,123323615 | ['GO:0110165'] |
| 318 | GO:CC | GO:0016328 lateral plasma membrane | 1,25991E-05 | positive_corr | True | 62 | 1064 | 17 | 18856 | 0,015977444 | 0,274193548 | ['GO:0005886', 'GO:0110165'] |
| 326 | GO:CC | GO:0150034 distal axon | 1,37982E-05 | negative_corr | True | 307 | 2182 | 69 | 18856 | 0,031622365 | 0,2247557 | ['GO:0030424', 'GO:0110165'] |

| | | | | | | | | | | | | |
|---|---|---|---|---|---|---|---|---|---|---|---|---|
| 343 | GO:CC | GO:0031526 brush border membrane | 1,67837E-05 | positive_corr | True | 56 | 1064 | 16 | 18856 | 0,015037594 | 0,285714286 | ['GO:0005903', 'GO:0031253'] |
| 397 | GO:CC | GO:0022625 cytosolic large ribosomal subunit | 2,14447E-05 | negative_corr | True | 59 | 2182 | 23 | 18856 | 0,010540788 | 0,389830508 | ['GO:0015934', 'GO:0022626'] |
| 400 | GO:CC | GO:0005737 cytoplasm | 2,18153E-05 | negative_corr | True | 11634 | 2182 | 1459 | 18856 | 0,668652612 | 0,125408286 | ['GO:0005622', 'GO:0110165'] |
| 417 | GO:CC | GO:0030027 lamellipodium | 2,31507E-05 | negative_corr | True | 198 | 2182 | 50 | 18856 | 0,022914757 | 0,252525253 | ['GO:0031252', 'GO:0120025'] |
| 419 | GO:CC | GO:0098858 actin-based cell projection | 3,53515E-05 | positive_corr | True | 211 | 1064 | 33 | 18856 | 0,031015038 | 0,156398104 | ['GO:0120025'] |
| 421 | GO:CC | GO:0005925 focal adhesion | 4,06903E-05 | negative_corr | True | 410 | 2182 | 84 | 18856 | 0,038496792 | 0,204878049 | ['GO:0030055'] |
| 423 | GO:CC | GO:0030055 cell-substrate junction | 4,32481E-05 | negative_corr | True | 417 | 2182 | 85 | 18856 | 0,038955087 | 0,20383693 | ['GO:0070161'] |
| 439 | GO:CC | GO:0043232 intracellular non-membrane-bounded organelle | 5,99607E-05 | negative_corr | True | 4954 | 2182 | 674 | 18856 | 0,308890926 | 0,136051675 | ['GO:0043228', 'GO:0043229'] |
| 440 | GO:CC | GO:0043228 non-membrane-bounded organelle | 6,35346E-05 | negative_corr | True | 4964 | 2182 | 675 | 18856 | 0,309349221 | 0,135979049 | ['GO:0043226'] |
| 464 | GO:CC | GO:0070161 anchoring junction | 6,93616E-05 | positive_corr | True | 785 | 1064 | 80 | 18856 | 0,07518797 | 0,101910828 | ['GO:0030054'] |
| 1645 | GO:CC | GO:0005911 cell-cell junction | 0,000112264 | negative_corr | True | 439 | 2182 | 87 | 18856 | 0,039871677 | 0,198177677 | ['GO:0070161'] |
| 737 | GO:CC | GO:0098793 presynapse | 0,00018935 | negative_corr | True | 531 | 2182 | 100 | 18856 | 0,045829514 | 0,188323917 | ['GO:0045202', 'GO:0110165'] |
| 747 | GO:CC | GO:0016327 apicolateral plasma membrane | 0,000193869 | positive_corr | True | 20 | 1064 | 9 | 18856 | 0,008458647 | 0,45 | ['GO:0098590'] |
| 883 | GO:CC | GO:0005815 microtubule organizing center | 0,000257922 | negative_corr | True | 770 | 2182 | 134 | 18856 | 0,061411549 | 0,174025974 | ['GO:0015630', 'GO:0110165'] |
| 899 | GO:CC | GO:0035869 ciliary transition zone | 0,000336165 | negative_corr | True | 72 | 2182 | 24 | 18856 | 0,010999083 | 0,333333333 | ['GO:0005929', 'GO:0110165'] |
| 904 | GO:CC | GO:0031253 cell projection membrane | 0,000376285 | positive_corr | True | 335 | 1064 | 42 | 18856 | 0,039473684 | 0,125373134 | ['GO:0098590', 'GO:0120025'] |
| 744 | GO:CC | GO:0005929 cilium | 0,000379895 | negative_corr | True | 642 | 2182 | 115 | 18856 | 0,052703941 | 0,179127726 | ['GO:0043226', 'GO:0120025'] |
| 1587 | GO:CC | GO:0036064 ciliary basal body | 0,000766687 | negative_corr | True | 133 | 2182 | 35 | 18856 | 0,01604033 | 0,263157895 | ['GO:0005815', 'GO:0005929'] |
| 1586 | GO:CC | GO:0045211 postsynaptic membrane | 0,001172289 | negative_corr | True | 286 | 2182 | 60 | 18856 | 0,027497709 | 0,20979021 | ['GO:0097060', 'GO:0098794'] |
| 1585 | GO:CC | GO:0031528 microvillus membrane | 0,001228959 | positive_corr | True | 24 | 1064 | 9 | 18856 | 0,008458647 | 0,375 | ['GO:0005902', 'GO:0031253'] |
| 948 | GO:CC | GO:0030054 cell junction | 0,001315711 | positive_corr | True | 1327 | 1064 | 114 | 18856 | 0,107142857 | 0,085908063 | ['GO:0110165'] |
| 980 | GO:CC | GO:0005604 basement membrane | 0,001414622 | negative_corr | True | 98 | 2182 | 28 | 18856 | 0,012832264 | 0,285714286 | ['GO:0062023'] |
| 981 | GO:CC | GO:0099081 supramolecular polymer | 0,001729493 | negative_corr | True | 1001 | 2182 | 162 | 18856 | 0,074243813 | 0,161838162 | ['GO:0099080'] |
| 994 | GO:CC | GO:0099512 supramolecular fiber | 0,002555366 | negative_corr | True | 993 | 2182 | 160 | 18856 | 0,073327223 | 0,161127895 | ['GO:0099081'] |
| 1572 | GO:CC | GO:0097542 ciliary tip | 0,002689489 | negative_corr | True | 46 | 2182 | 17 | 18856 | 0,007791017 | 0,369565217 | ['GO:0005929', 'GO:0110165'] |
| 1021 | GO:CC | GO:0044304 main axon | 0,00282429 | negative_corr | True | 70 | 2182 | 22 | 18856 | 0,010082493 | 0,314285714 | ['GO:0030424', 'GO:0110165'] |
| 1033 | GO:CC | GO:0005874 microtubule | 0,003124628 | negative_corr | True | 427 | 2182 | 80 | 18856 | 0,036663611 | 0,18735363 | ['GO:0015630', 'GO:0099513'] |
| 1036 | GO:CC | GO:0061689 tricellular tight junction | 0,003653807 | positive_corr | True | 4 | 1064 | 4 | 18856 | 0,003759398 | 1 | ['GO:0070160'] |
| 1555 | GO:CC | GO:0005783 endoplasmic reticulum | 0,004255315 | positive_corr | True | 1810 | 1064 | 144 | 18856 | 0,135338346 | 0,079558011 | ['GO:0005737', 'GO:0012505', 'GO:0043231'] |
| 1101 | GO:CC | GO:0031410 cytoplasmic vesicle | 0,005247964 | positive_corr | True | 2372 | 1064 | 180 | 18856 | 0,169172932 | 0,075885329 | ['GO:0005737', 'GO:0097708'] |
| 1133 | GO:CC | GO:0005813 centrosome | 0,005455148 | negative_corr | True | 586 | 2182 | 102 | 18856 | 0,046746104 | 0,174061433 | ['GO:0005815'] |
| 1136 | GO:CC | GO:0097708 intracellular vesicle | 0,00565289 | positive_corr | True | 2375 | 1064 | 180 | 18856 | 0,169172932 | 0,075789474 | ['GO:0031982', 'GO:0043229'] |
| 878 | GO:CC | GO:0042175 nuclear outer membrane-endoplasmic reticulum membrane network | 0,006778808 | positive_corr | True | 1098 | 1064 | 95 | 18856 | 0,089285714 | 0,086520947 | ['GO:0012505', 'GO:0016020'] |
| 873 | GO:CC | GO:0034703 cation channel complex | 0,006898001 | negative_corr | True | 225 | 2182 | 48 | 18856 | 0,021998167 | 0,213333333 | ['GO:0034702'] |
| 913 | GO:CC | GO:0015629 actin cytoskeleton | 0,007022827 | positive_corr | True | 517 | 1064 | 53 | 18856 | 0,04981203 | 0,102514507 | ['GO:0005856'] |
| 1603 | GO:CC | GO:0005789 endoplasmic reticulum membrane | 0,008360712 | positive_corr | True | 1075 | 1064 | 93 | 18856 | 0,087406015 | 0,086511628 | ['GO:0031090', 'GO:0042175'] |
| 763 | GO:CC | GO:0022626 cytosolic ribosome | 0,010469595 | negative_corr | True | 108 | 2182 | 28 | 18856 | 0,012832264 | 0,259259259 | ['GO:0005829', 'GO:0005840'] |
| 764 | GO:CC | GO:0098588 bounding membrane of organelle | 0,011381759 | positive_corr | True | 2092 | 1064 | 160 | 18856 | 0,15037594 | 0,076481836 | ['GO:0031090'] |
| 776 | GO:CC | GO:0005634 nucleus | 0,014618706 | negative_corr | True | 7486 | 2182 | 952 | 18856 | 0,436296975 | 0,127170719 | ['GO:0043231'] |
| 783 | GO:CC | GO:0030315 T-tubule | 0,017089392 | negative_corr | True | 52 | 2182 | 17 | 18856 | 0,007791017 | 0,326923077 | ['GO:0042383', 'GO:0110165'] |
| 784 | GO:CC | GO:0014704 intercalated disc | 0,017089392 | negative_corr | True | 52 | 2182 | 17 | 18856 | 0,007791017 | 0,326923077 | ['GO:0044291'] |
| 785 | GO:CC | GO:0098805 whole membrane | 0,026817483 | positive_corr | True | 1694 | 1064 | 132 | 18856 | 0,12406015 | 0,077922078 | ['GO:0016020'] |
| 1642 | GO:CC | GO:0042581 specific granule | 0,02903237 | positive_corr | True | 158 | 1064 | 22 | 18856 | 0,020676692 | 0,139240506 | ['GO:0030141'] |
| 795 | GO:CC | GO:0044291 cell-cell contact zone | 0,030805391 | negative_corr | True | 75 | 2182 | 21 | 18856 | 0,009624198 | 0,28 | ['GO:0005911'] |
| 842 | GO:CC | GO:0005777 peroxisome | 0,031051016 | positive_corr | True | 137 | 1064 | 20 | 18856 | 0,018796992 | 0,145985401 | ['GO:0042579'] |
| 799 | GO:CC | GO:0042579 microbody | 0,031051016 | positive_corr | True | 137 | 1064 | 20 | 18856 | 0,018796992 | 0,145985401 | ['GO:0005737', 'GO:0043231'] |
| 1414 | GO:CC | GO:0031234 extrinsic component of cytoplasmic side of plasma membrane | 0,034246864 | negative_corr | True | 92 | 2182 | 24 | 18856 | 0,010999083 | 0,260869565 | ['GO:0005622', 'GO:0009898', 'GO:0019897'] |
| 840 | GO:CC | GO:0005912 adherens junction | 0,043636594 | negative_corr | True | 99 | 2182 | 25 | 18856 | 0,011457379 | 0,252525253 | ['GO:0005911'] |
| 807 | GO:CC | GO:0099568 cytoplasmic region | 0,044357353 | negative_corr | True | 248 | 2182 | 49 | 18856 | 0,022456462 | 0,197580645 | ['GO:0005737'] |
| 836 | GO:CC | GO:0055037 recycling endosome | 0,045255795 | positive_corr | True | 174 | 1064 | 23 | 18856 | 0,021616541 | 0,132183908 | ['GO:0005768'] |
| 818 | GO:CC | GO:0015629 actin cytoskeleton | 0,046495273 | negative_corr | True | 517 | 2182 | 88 | 18856 | 0,040329973 | 0,170212766 | ['GO:0005856'] |
| 828 | GO:CC | GO:0005614 interstitial matrix | 0,047914641 | negative_corr | True | 12 | 2182 | 7 | 18856 | 0,003208066 | 0,583333333 | ['GO:0062023'] |
| 806 | GO:CC | GO:0008328 ionotropic glutamate receptor complex | 0,049418654 | negative_corr | True | 51 | 2182 | 16 | 18856 | 0,007332722 | 0,31372549 | ['GO:0005887', 'GO:0034702', 'GO:0098878'] |
| 1583 | GO:MF | GO:0005515 protein binding | 1,03007E-11 | negative_corr | True | 12743 | 2154 | 1662 | 18126 | 0,771587744 | 0,130424547 | ['GO:0005488'] |
| 1522 | GO:MF | GO:0022857 transmembrane transporter activity | 3,14834E-11 | positive_corr | True | 1068 | 1033 | 122 | 18126 | 0,118102614 | 0,11423221 | ['GO:0005215'] |
| 1600 | GO:MF | GO:0008092 cytoskeletal protein binding | 3,47811E-11 | negative_corr | True | 991 | 2154 | 197 | 18126 | 0,091457753 | 0,198789102 | ['GO:0005515'] |
| 1526 | GO:MF | GO:0005488 binding | 7,88547E-11 | negative_corr | True | 15944 | 2154 | 1993 | 18126 | 0,925255339 | 0,125 | ['GO:0003674'] |
| 1546 | GO:MF | GO:0005215 transporter activity | 1,89285E-10 | positive_corr | True | 1223 | 1033 | 132 | 18126 | 0,127783156 | 0,107931316 | ['GO:0003674'] |
| 1671 | GO:MF | GO:0003824 catalytic activity | 6,90756E-10 | positive_corr | True | 5853 | 1033 | 438 | 18126 | 0,424007744 | 0,074833419 | ['GO:0003674'] |
| 1423 | GO:MF | GO:0015075 ion transmembrane transporter activity | 1,2594E-09 | positive_corr | True | 887 | 1033 | 103 | 18126 | 0,099709584 | 0,116121759 | ['GO:0022857'] |
| 1428 | GO:MF | GO:0008509 anion transmembrane transporter activity | 6,35351E-08 | positive_corr | True | 336 | 1033 | 51 | 18126 | 0,049370765 | 0,151785714 | ['GO:0015075'] |
| 1556 | GO:MF | GO:0015318 inorganic molecular entity transmembrane transporter activity | 2,39932E-07 | positive_corr | True | 830 | 1033 | 92 | 18126 | 0,089060987 | 0,110843373 | ['GO:0022857'] |
| 1618 | GO:MF | GO:0008514 organic anion transmembrane transporter activity | 1,42786E-06 | positive_corr | True | 211 | 1033 | 36 | 18126 | 0,034849952 | 0,170616114 | ['GO:0008509'] |
| 1415 | GO:MF | GO:0015631 tubulin binding | 1,94741E-06 | negative_corr | True | 343 | 2154 | 79 | 18126 | 0,036675952 | 0,2303207 | ['GO:0008092'] |
| 1575 | GO:MF | GO:0046872 metal ion binding | 2,80421E-06 | negative_corr | True | 4227 | 2154 | 610 | 18126 | 0,283194058 | 0,144310386 | ['GO:0043169'] |

| | | | | | | | | | | | | |
|---|---|---|---|---|---|---|---|---|---|---|---|---|
| 1449 | GO:MF | GO:0008194 | UDP-glycosyltransferase activity | 4,27184E-06 | positive_corr | True | 153 | 1033 | 29 | 18126 | 0,028073572 | 0,189542484 ['GO:0016757'] |
| 1482 | GO:MF | GO:0005201 | extracellular matrix structural constituent | 8,91621E-06 | negative_corr | True | 165 | 2154 | 46 | 18126 | 0,021355617 | 0,278787879 ['GO:0005198'] |
| 1639 | GO:MF | GO:0005198 | structural molecule activity | 1,21661E-05 | negative_corr | True | 683 | 2154 | 130 | 18126 | 0,060352832 | 0,19033675 ['GO:0003674'] |
| 1500 | GO:MF | GO:0022804 | active transmembrane transporter activity | 2,14981E-05 | positive_corr | True | 352 | 1033 | 47 | 18126 | 0,045498548 | 0,133522727 ['GO:0022857'] |
| 1427 | GO:MF | GO:0043169 | cation binding | 2,18948E-05 | negative_corr | True | 4321 | 2154 | 615 | 18126 | 0,28551532 | 0,142328165 ['GO:0043167'] |
| 1501 | GO:MF | GO:0050662 | coenzyme binding | 5,13833E-05 | positive_corr | True | 295 | 1033 | 41 | 18126 | 0,039690223 | 0,138983051 ['GO:0048037'] |
| 1067 | GO:MF | GO:0051015 | actin filament binding | 0,000103622 | positive_corr | True | 196 | 1033 | 31 | 18126 | 0,030009681 | 0,158163265 ['GO:0003779', 'GO:0044877'] |
| 1059 | GO:MF | GO:0016758 | transferase activity, transferring hexosyl groups | 0,000109412 | positive_corr | True | 217 | 1033 | 33 | 18126 | 0,031945789 | 0,152073733 ['GO:0016757'] |
| 940 | GO:MF | GO:0015291 | secondary active transmembrane transporter activity | 0,00010972 | positive_corr | True | 238 | 1033 | 35 | 18126 | 0,033881897 | 0,147058824 ['GO:0022804'] |
| 943 | GO:MF | GO:0043167 | ion binding | 0,00013232 | negative_corr | True | 6308 | 2154 | 855 | 18126 | 0,396935933 | 0,135542169 ['GO:0005488'] |
| 944 | GO:MF | GO:0008017 | microtubule binding | 0,000261128 | negative_corr | True | 253 | 2154 | 58 | 18126 | 0,026926648 | 0,229249012 ['GO:0015631'] |
| 964 | GO:MF | GO:0086083 | cell adhesive protein binding involved in bundle of His cell-Purkinje m | 0,000273253 | positive_corr | True | 5 | 1033 | 5 | 18126 | 0,004840271 | 1 ['GO:0086080'] |
| 999 | GO:MF | GO:0048037 | cofactor binding | 0,000282782 | positive_corr | True | 517 | 1033 | 58 | 18126 | 0,056147144 | 0,112185687 ['GO:0005488'] |
| 1002 | GO:MF | GO:0008392 | arachidonic acid epoxygenase activity | 0,000424149 | positive_corr | True | 16 | 1033 | 8 | 18126 | 0,007744434 | 0,5 ['GO:0008391'] |
| 892 | GO:MF | GO:0043168 | anion binding | 0,000536607 | positive_corr | True | 2841 | 1033 | 218 | 18126 | 0,211035818 | 0,076733545 ['GO:0043167'] |
| 1004 | GO:MF | GO:0046943 | carboxylic acid transmembrane transporter activity | 0,000561802 | positive_corr | True | 150 | 1033 | 25 | 18126 | 0,024201355 | 0,166666667 ['GO:0005342', 'GO:0008514', 'GO:0015318'] |
| 1066 | GO:MF | GO:0005342 | organic acid transmembrane transporter activity | 0,000561802 | positive_corr | True | 150 | 1033 | 25 | 18126 | 0,024201355 | 0,166666667 ['GO:0022857'] |
| 1675 | GO:MF | GO:0008391 | arachidonic acid monooxygenase activity | 0,000761085 | positive_corr | True | 17 | 1033 | 8 | 18126 | 0,007744434 | 0,470588235 ['GO:0004497', 'GO:0016705'] |
| 1099 | GO:MF | GO:0030020 | extracellular matrix structural constituent conferring tensile strength | 0,00079264 | negative_corr | True | 41 | 2154 | 17 | 18126 | 0,007892293 | 0,414634146 ['GO:0005201'] |
| 1103 | GO:MF | GO:0008376 | acetylgalactosaminyltransferase activity | 0,000848215 | positive_corr | True | 48 | 1033 | 13 | 18126 | 0,012584705 | 0,270833333 ['GO:0008194', 'GO:0016758'] |
| 1108 | GO:MF | GO:0016247 | channel regulator activity | 0,001282509 | negative_corr | True | 142 | 2154 | 37 | 18126 | 0,017177344 | 0,26056338 ['GO:0098772'] |
| 1140 | GO:MF | GO:0016757 | transferase activity, transferring glycosyl groups | 0,00144535 | positive_corr | True | 288 | 1033 | 37 | 18126 | 0,035818006 | 0,128472222 ['GO:0016740'] |
| 1041 | GO:MF | GO:0098632 | cell-cell adhesion mediator activity | 0,001785141 | positive_corr | True | 51 | 1033 | 13 | 18126 | 0,012584705 | 0,254901961 ['GO:0005515', 'GO:0098631'] |
| 1155 | GO:MF | GO:0015103 | inorganic anion transmembrane transporter activity | 0,002158163 | positive_corr | True | 141 | 1033 | 23 | 18126 | 0,022265247 | 0,163120567 ['GO:0008509', 'GO:0015318'] |
| 841 | GO:MF | GO:0016614 | oxidoreductase activity, acting on CH-OH group of donors | 0,003100519 | positive_corr | True | 134 | 1033 | 22 | 18126 | 0,021297193 | 0,164179104 ['GO:0016491'] |
| 801 | GO:MF | GO:0036094 | small molecule binding | 0,003592087 | positive_corr | True | 2574 | 1033 | 196 | 18126 | 0,189738625 | 0,076146076 ['GO:0005488'] |
| 469 | GO:MF | GO:0033293 | monocarboxylic acid binding | 0,004861617 | positive_corr | True | 72 | 1033 | 15 | 18126 | 0,014520813 | 0,208333333 ['GO:0031406'] |
| 491 | GO:MF | GO:0019904 | protein domain specific binding | 0,005138263 | negative_corr | True | 712 | 2154 | 123 | 18126 | 0,057103064 | 0,172752809 ['GO:0005515'] |
| 494 | GO:MF | GO:0016936 | galactoside binding | 0,005208932 | positive_corr | True | 7 | 1033 | 5 | 18126 | 0,004840271 | 0,714285714 ['GO:0097367'] |
| 502 | GO:MF | GO:0003924 | GTPase activity | 0,005469071 | negative_corr | True | 328 | 2154 | 66 | 18126 | 0,030640669 | 0,201219512 ['GO:0017111'] |
| 530 | GO:MF | GO:0005509 | calcium ion binding | 0,011453405 | negative_corr | True | 717 | 2154 | 122 | 18126 | 0,056638812 | 0,170153417 ['GO:0046872'] |
| 553 | GO:MF | GO:0016616 | oxidoreductase activity, acting on the CH-OH group of donors, NAI | 0,011827593 | positive_corr | True | 125 | 1033 | 20 | 18126 | 0,019361084 | 0,16 ['GO:0016614'] |
| 837 | GO:MF | GO:0098631 | cell adhesion mediator activity | 0,012012121 | positive_corr | True | 60 | 1033 | 13 | 18126 | 0,012584705 | 0,216666667 ['GO:0005488'] |
| 568 | GO:MF | GO:0022853 | active ion transmembrane transporter activity | 0,012027934 | positive_corr | True | 245 | 1033 | 31 | 18126 | 0,030009681 | 0,126530612 ['GO:0015075', 'GO:0022804'] |
| 670 | GO:MF | GO:0016712 | oxidoreductase activity, acting on paired donors, with incorporation | 0,01346522 | positive_corr | True | 37 | 1033 | 10 | 18126 | 0,009680542 | 0,27027027 ['GO:0004497', 'GO:0016705'] |
| 723 | GO:MF | GO:0015108 | chloride transmembrane transporter activity | 0,016315882 | positive_corr | True | 98 | 1033 | 17 | 18126 | 0,016456922 | 0,173469388 ['GO:0015103'] |
| 729 | GO:MF | GO:0008324 | cation transmembrane transporter activity | 0,018514288 | positive_corr | True | 660 | 1033 | 63 | 18126 | 0,060987415 | 0,095454545 ['GO:0015075'] |
| 740 | GO:MF | GO:0086080 | protein binding involved in heterotypic cell-cell adhesion | 0,018792169 | positive_corr | True | 13 | 1033 | 6 | 18126 | 0,005808325 | 0,461538462 ['GO:0098632'] |
| 755 | GO:MF | GO:0022890 | inorganic cation transmembrane transporter activity | 0,021324228 | positive_corr | True | 609 | 1033 | 59 | 18126 | 0,057115198 | 0,096880131 ['GO:0008324', 'GO:0015318'] |
| 790 | GO:MF | GO:0099106 | ion channel regulator activity | 0,021659874 | negative_corr | True | 119 | 2154 | 30 | 18126 | 0,013927577 | 0,25210084 ['GO:0016247'] |
| 642 | GO:MF | GO:0005254 | chloride channel activity | 0,02521687 | positive_corr | True | 73 | 1033 | 14 | 18126 | 0,013552759 | 0,191780822 ['GO:0005253', 'GO:0015108'] |
| 1182 | GO:MF | GO:0003707 | steroid hormone receptor activity | 0,02740048 | positive_corr | True | 56 | 1033 | 12 | 18126 | 0,011616651 | 0,214285714 ['GO:0038023'] |
| 1676 | GO:MF | GO:0030695 | GTPase regulator activity | 0,029510642 | negative_corr | True | 306 | 2154 | 60 | 18126 | 0,027855153 | 0,196078431 ['GO:0060589'] |
| 1287 | GO:MF | GO:0045296 | cadherin binding | 0,030489426 | positive_corr | True | 330 | 1033 | 37 | 18126 | 0,035818006 | 0,112121212 ['GO:0050839'] |
| 1251 | GO:MF | GO:0003779 | actin binding | 0,03351234 | positive_corr | True | 433 | 1033 | 45 | 18126 | 0,043562439 | 0,103926097 ['GO:0008092'] |
| 1295 | GO:MF | GO:0005253 | anion channel activity | 0,037803609 | positive_corr | True | 85 | 1033 | 15 | 18126 | 0,014520813 | 0,176470588 ['GO:0005216', 'GO:0015103'] |
| 1237 | GO:MF | GO:0017124 | SH3 domain binding | 0,04332756 | negative_corr | True | 129 | 2154 | 31 | 18126 | 0,014391829 | 0,240310078 ['GO:0019904'] |
| 1203 | GO:MF | GO:0050839 | cell adhesion molecule binding | 0,049047886 | positive_corr | True | 506 | 1033 | 50 | 18126 | 0,048402711 | 0,098814229 ['GO:0005515'] |
| 1301 | GO:MF | GO:0004089 | carbonate dehydratase activity | 0,049654269 | positive_corr | True | 15 | 1033 | 6 | 18126 | 0,005808325 | 0,4 ['GO:0016836'] |
| 1312 | GO:MF | GO:0030165 | PDZ domain binding | 0,049738485 | positive_corr | True | 87 | 1033 | 15 | 18126 | 0,014520813 | 0,172413793 ['GO:0019904'] |
| 1468 | HP | HP:0040064 | Abnormality of limbs | 1,42615E-05 | negative_corr | True | 1790 | 566 | 302 | 4225 | 0,533568905 | 0,168715084 ['HP:0000118'] |
| 894 | HP | HP:0001760 | Abnormality of the foot | 1,93887E-05 | negative_corr | True | 1091 | 566 | 202 | 4225 | 0,356890459 | 0,185151237 ['HP:0002814'] |
| 919 | HP | HP:0001155 | Abnormality of the hand | 5,2422E-05 | negative_corr | True | 1178 | 566 | 213 | 4225 | 0,376325088 | 0,180814941 ['HP:0002817'] |
| 935 | HP | HP:0011844 | Abnormal appendicular skeleton morphology | 7,75107E-05 | negative_corr | True | 1398 | 566 | 244 | 4225 | 0,431095406 | 0,17453505 ['HP:0011842'] |
| 1602 | HP | HP:0002817 | Abnormality of the upper limb | 9,37903E-05 | negative_corr | True | 1282 | 566 | 227 | 4225 | 0,401060071 | 0,177067083 ['HP:0040064'] |
| 945 | HP | HP:0002814 | Abnormality of the lower limb | 9,50573E-05 | negative_corr | True | 1303 | 566 | 230 | 4225 | 0,406360424 | 0,176515733 ['HP:0040064'] |
| 954 | HP | HP:0011297 | Abnormal digit morphology | 0,000110347 | negative_corr | True | 1133 | 566 | 205 | 4225 | 0,362190813 | 0,180935569 ['HP:0002813'] |
| 982 | HP | HP:0001167 | Abnormality of finger | 0,000121015 | negative_corr | True | 926 | 566 | 174 | 4225 | 0,307420495 | 0,187904968 ['HP:0001155', 'HP:0011297'] |
| 1011 | HP | HP:0000925 | Abnormality of the vertebral column | 0,000202422 | negative_corr | True | 1182 | 566 | 211 | 4225 | 0,372791519 | 0,178510998 ['HP:0009121'] |
| 930 | HP | HP:0000974 | Hyperextensible skin | 0,000311117 | negative_corr | True | 52 | 566 | 22 | 4225 | 0,038869258 | 0,423076923 ['HP:0008067'] |
| 1291 | HP | HP:0005918 | Abnormal finger phalanx morphology | 0,000542174 | negative_corr | True | 557 | 566 | 114 | 4225 | 0,201413428 | 0,204667864 ['HP:0001167'] |
| 819 | HP | HP:0100259 | Postaxial polydactyly | 0,000551519 | negative_corr | True | 125 | 566 | 38 | 4225 | 0,067137809 | 0,304 ['HP:0010442'] |
| 1571 | HP | HP:0040068 | Abnormality of limb bone | 0,001113828 | negative_corr | True | 1296 | 566 | 224 | 4225 | 0,395759717 | 0,172839506 ['HP:0000924', 'HP:0040064'] |
| 1628 | HP | HP:0002813 | Abnormality of limb bone morphology | 0,001113828 | negative_corr | True | 1296 | 566 | 224 | 4225 | 0,395759717 | 0,172839506 ['HP:0011844', 'HP:0040068'] |
| 1634 | HP | HP:0010442 | Polydactyly | 0,001797925 | negative_corr | True | 214 | 566 | 54 | 4225 | 0,09540636 | 0,252336449 ['HP:0011297'] |

| | | | | | | | | | | | | | |
|---|---|---|---|---|---|---|---|---|---|---|---|---|---|
| 1635 | HP | HP:0001162 | Postaxial hand polydactyly | 0,001865757 | negative_corr | True | 92 | 566 | 30 | 4225 | 0,053003534 | 0,326086957 | ['HP:0001161', 'HP:0004207', 'HP:0100259'] |
| 1636 | HP | HP:0001288 | Gait disturbance | 0,002363396 | negative_corr | True | 857 | 566 | 158 | 4225 | 0,279151943 | 0,184364061 | ['HP:0100022'] |
| 1339 | HP | HP:0001942 | Metabolic acidosis | 0,004964734 | positive_corr | True | 89 | 254 | 18 | 4225 | 0,070866142 | 0,202247191 | ['HP:0001941'] |
| 1653 | HP | HP:0010674 | Abnormality of the curvature of the vertebral column | 0,006266506 | negative_corr | True | 937 | 566 | 168 | 4225 | 0,296819788 | 0,179295624 | ['HP:0000925'] |
| 1354 | HP | HP:0005622 | Broad long bones | 0,007023269 | negative_corr | True | 112 | 566 | 33 | 4225 | 0,058303887 | 0,294642857 | ['HP:0011314'] |
| 1659 | HP | HP:0002650 | Scoliosis | 0,009482844 | negative_corr | True | 869 | 566 | 157 | 4225 | 0,277385159 | 0,180667434 | ['HP:0010674'] |
| 1392 | HP | HP:0003111 | Abnormal blood ion concentration | 0,0096378 | positive_corr | True | 246 | 254 | 33 | 4225 | 0,12992126 | 0,134146341 | ['HP:0032180'] |
| 1424 | HP | HP:0001161 | Hand polydactyly | 0,011832805 | negative_corr | True | 161 | 566 | 42 | 4225 | 0,074204947 | 0,260869565 | ['HP:0009997', 'HP:0010442'] |
| 1394 | HP | HP:0006817 | Aplasia/Hypoplasia of the cerebellar vermis | 0,014938963 | negative_corr | True | 206 | 566 | 50 | 4225 | 0,088339223 | 0,242718447 | ['HP:0002334'] |
| 1672 | HP | HP:0004299 | Hernia of the abdominal wall | 0,017000391 | negative_corr | True | 364 | 566 | 77 | 4225 | 0,136042403 | 0,211538462 | ['HP:0010866', 'HP:0100790'] |
| 1673 | HP | HP:0030877 | Reduced FEV1/FVC ratio | 0,017262625 | positive_corr | True | 4 | 254 | 4 | 4225 | 0,015748031 | 1 | ['HP:0006536', 'HP:0032340'] |
| 831 | HP | HP:0009997 | Duplication of phalanx of hand | 0,017447474 | negative_corr | True | 185 | 566 | 46 | 4225 | 0,081272085 | 0,248648649 | ['HP:0004275', 'HP:0005918'] |
| 1489 | HP | HP:0001539 | Omphalocele | 0,019041453 | negative_corr | True | 64 | 566 | 22 | 4225 | 0,038869258 | 0,34375 | ['HP:0004299'] |
| 1186 | HP | HP:0011949 | Acute infectious pneumonia | 0,019412642 | positive_corr | True | 7 | 254 | 5 | 4225 | 0,019685039 | 0,714285714 | ['HP:0011948'] |
| 1130 | HP | HP:0000888 | Horizontal ribs | 0,019444041 | negative_corr | True | 14 | 566 | 9 | 4225 | 0,01590106 | 0,642857143 | ['HP:0000772'] |
| 1131 | HP | HP:0004275 | Duplication of hand bones | 0,020306715 | negative_corr | True | 186 | 566 | 46 | 4225 | 0,081272085 | 0,247311828 | ['HP:0001155', 'HP:0009142'] |
| 1495 | HP | HP:0009142 | Duplication of bones involving the upper extremities | 0,020306715 | negative_corr | True | 186 | 566 | 46 | 4225 | 0,081272085 | 0,247311828 | ['HP:0002817'] |
| 1537 | HP | HP:0006009 | Broad phalanx | 0,024426517 | negative_corr | True | 103 | 566 | 30 | 4225 | 0,053003534 | 0,291262136 | ['HP:0005622', 'HP:0011297'] |
| 1516 | HP | HP:0010866 | Abdominal wall defect | 0,025813516 | negative_corr | True | 368 | 566 | 77 | 4225 | 0,136042403 | 0,20923913 | ['HP:0004298'] |
| 1515 | HP | HP:0000782 | Abnormality of the scapula | 0,029004456 | negative_corr | True | 119 | 566 | 33 | 4225 | 0,058303887 | 0,277310924 | ['HP:0000765'] |
| 1510 | HP | HP:0001780 | Abnormality of toe | 0,033622283 | negative_corr | True | 539 | 566 | 104 | 4225 | 0,183745583 | 0,192949907 | ['HP:0001760', 'HP:0011297'] |
| 1550 | HP | HP:0002438 | Cerebellar malformation | 0,039289831 | negative_corr | True | 294 | 566 | 64 | 4225 | 0,113074205 | 0,217687075 | ['HP:0001317'] |
| 1509 | HP | HP:0031865 | Abnormal liver physiology | 0,040906671 | positive_corr | True | 12 | 254 | 6 | 4225 | 0,023622047 | 0,5 | ['HP:0001392'] |
| 1494 | HP | HP:0001406 | Intrahepatic cholestasis | 0,040906671 | positive_corr | True | 12 | 254 | 6 | 4225 | 0,023622047 | 0,5 | ['HP:0001396', 'HP:0031865'] |
| 1065 | HP | HP:0001317 | Abnormal cerebellum morphology | 0,041915817 | negative_corr | True | 717 | 566 | 131 | 4225 | 0,231448763 | 0,182705718 | ['HP:0011283'] |
| 1184 | HP | HP:0002334 | Abnormality of the cerebellar vermis | 0,046284356 | negative_corr | True | 237 | 566 | 54 | 4225 | 0,09540636 | 0,227848101 | ['HP:0002438'] |
| 1061 | HP | HP:0003691 | Scapular winging | 0,047303246 | negative_corr | True | 72 | 566 | 23 | 4225 | 0,040636042 | 0,319444444 | ['HP:0000782', 'HP:0001435'] |
| 1504 | HP | HP:0000841 | Hyperactive renin-angiotensin system | 0,049228169 | positive_corr | True | 8 | 254 | 5 | 4225 | 0,019685039 | 0,625 | ['HP:0000847'] |
| 1211 | HP | HP:0008242 | Pseudohypoaldosteronism | 0,049228169 | positive_corr | True | 8 | 254 | 5 | 4225 | 0,019685039 | 0,625 | ['HP:0011733'] |
| 1527 | KEGG | KEGG:0110 | Metabolic pathways | 5,94225E-21 | positive_corr | True | 1482 | 508 | 188 | 7788 | 0,37007874 | 0,126855601 | ['KEGG:00000'] |
| 1627 | KEGG | KEGG:0083 | Retinol metabolism | 1,36333E-05 | positive_corr | True | 66 | 508 | 18 | 7788 | 0,035433071 | 0,272727273 | ['KEGG:00000'] |
| 269 | KEGG | KEGG:0520 | Chemical carcinogenesis | 6,52087E-05 | positive_corr | True | 80 | 508 | 19 | 7788 | 0,037401575 | 0,2375 | ['KEGG:00000'] |
| 1132 | KEGG | KEGG:0004 | Pentose and glucuronate interconversions | 8,43058E-05 | positive_corr | True | 34 | 508 | 12 | 7788 | 0,023622047 | 0,352941176 | ['KEGG:00000'] |
| 1145 | KEGG | KEGG:0497 | Bile secretion | 0,000279873 | positive_corr | True | 72 | 508 | 17 | 7788 | 0,033464567 | 0,236111111 | ['KEGG:00000'] |
| 816 | KEGG | KEGG:0436 | Axon guidance | 0,000293836 | negative_corr | True | 180 | 814 | 40 | 7788 | 0,049140049 | 0,222222222 | ['KEGG:00000'] |
| 1503 | KEGG | KEGG:0014 | Steroid hormone biosynthesis | 0,00051125 | positive_corr | True | 60 | 508 | 15 | 7788 | 0,029527559 | 0,25 | ['KEGG:00000'] |
| 1148 | KEGG | KEGG:0451 | Focal adhesion | 0,000586709 | negative_corr | True | 198 | 814 | 42 | 7788 | 0,051597052 | 0,212121212 | ['KEGG:00000'] |
| 1454 | KEGG | KEGG:0098 | Drug metabolism - cytochrome P450 | 0,000710378 | positive_corr | True | 69 | 508 | 16 | 7788 | 0,031496063 | 0,231884058 | ['KEGG:00000'] |
| 1421 | KEGG | KEGG:0496 | Proximal tubule bicarbonate reclamation | 0,000790372 | positive_corr | True | 23 | 508 | 9 | 7788 | 0,017716535 | 0,391304348 | ['KEGG:00000'] |
| 1214 | KEGG | KEGG:0453 | Tight junction | 0,001138987 | positive_corr | True | 168 | 508 | 27 | 7788 | 0,053149606 | 0,160714286 | ['KEGG:00000'] |
| 1569 | KEGG | KEGG:0472 | Glutamatergic synapse | 0,001333509 | negative_corr | True | 114 | 814 | 28 | 7788 | 0,034398034 | 0,245614035 | ['KEGG:00000'] |
| 1436 | KEGG | KEGG:0092 | Sulfur metabolism | 0,001389843 | positive_corr | True | 10 | 508 | 6 | 7788 | 0,011811024 | 0,6 | ['KEGG:00000'] |
| 1385 | KEGG | KEGG:0098 | Metabolism of xenobiotics by cytochrome P450 | 0,001534067 | positive_corr | True | 73 | 508 | 16 | 7788 | 0,031496063 | 0,219178082 | ['KEGG:00000'] |
| 1386 | KEGG | KEGG:0495 | Maturity onset diabetes of the young | 0,002532055 | positive_corr | True | 26 | 508 | 9 | 7788 | 0,017716535 | 0,346153846 | ['KEGG:00000'] |
| 1638 | KEGG | KEGG:0401 | MAPK signaling pathway | 0,002799037 | negative_corr | True | 295 | 814 | 54 | 7788 | 0,066339066 | 0,183050847 | ['KEGG:00000'] |
| 1162 | KEGG | KEGG:0005 | Ascorbate and aldarate metabolism | 0,003580872 | positive_corr | True | 27 | 508 | 9 | 7788 | 0,017716535 | 0,333333333 | ['KEGG:00000'] |
| 1053 | KEGG | KEGG:0451 | ECM-receptor interaction | 0,008592287 | negative_corr | True | 88 | 814 | 22 | 7788 | 0,027027027 | 0,25 | ['KEGG:00000'] |
| 1402 | KEGG | KEGG:0402 | cGMP-PKG signaling pathway | 0,008617059 | negative_corr | True | 165 | 814 | 34 | 7788 | 0,041769042 | 0,206060606 | ['KEGG:00000'] |
| 1412 | KEGG | KEGG:0332 | PPAR signaling pathway | 0,010337165 | positive_corr | True | 76 | 508 | 15 | 7788 | 0,029527559 | 0,197368421 | ['KEGG:00000'] |
| 1008 | KEGG | KEGG:0513 | Pathogenic Escherichia coli infection | 0,011304031 | positive_corr | True | 201 | 508 | 28 | 7788 | 0,05511811 | 0,139303483 | ['KEGG:00000'] |
| 1073 | KEGG | KEGG:0439 | Hippo signaling pathway | 0,011801373 | negative_corr | True | 154 | 814 | 32 | 7788 | 0,039312039 | 0,207792208 | ['KEGG:00000'] |
| 1003 | KEGG | KEGG:0492 | Relaxin signaling pathway | 0,012539336 | negative_corr | True | 128 | 814 | 28 | 7788 | 0,034398034 | 0,21875 | ['KEGG:00000'] |
| 1250 | KEGG | KEGG:0098 | Drug metabolism - other enzymes | 0,014081738 | positive_corr | True | 78 | 508 | 15 | 7788 | 0,029527559 | 0,192307692 | ['KEGG:00000'] |
| 1226 | KEGG | KEGG:0414 | Peroxisome | 0,018954478 | positive_corr | True | 80 | 508 | 15 | 7788 | 0,029527559 | 0,1875 | ['KEGG:00000'] |
| 1102 | KEGG | KEGG:0472 | Dopaminergic synapse | 0,021902455 | negative_corr | True | 132 | 814 | 28 | 7788 | 0,034398034 | 0,212121212 | ['KEGG:00000'] |
| 1547 | KEGG | KEGG:0060 | Glycosphingolipid biosynthesis - lacto and neolacto series | 0,025227796 | positive_corr | True | 27 | 508 | 8 | 7788 | 0,015748031 | 0,296296296 | ['KEGG:00000'] |
| 907 | KEGG | KEGG:0541 | Dilated cardiomyopathy (DCM) | 0,028651396 | negative_corr | True | 95 | 814 | 22 | 7788 | 0,027027027 | 0,231578947 | ['KEGG:00000'] |
| 1593 | KEGG | KEGG:0541 | Arrhythmogenic right ventricular cardiomyopathy (ARVC) | 0,031522982 | negative_corr | True | 77 | 814 | 19 | 7788 | 0,023341523 | 0,246753247 | ['KEGG:00000'] |
| 1592 | KEGG | KEGG:0497 | Mineral absorption | 0,032460975 | positive_corr | True | 58 | 508 | 12 | 7788 | 0,023622047 | 0,206896552 | ['KEGG:00000'] |
| 925 | KEGG | KEGG:0471 | Circadian entrainment | 0,039257716 | negative_corr | True | 97 | 814 | 22 | 7788 | 0,027027027 | 0,226804124 | ['KEGG:00000'] |
| 1094 | KEGG | KEGG:0451 | Cell adhesion molecules (CAMs) | 0,043115286 | negative_corr | True | 144 | 814 | 29 | 7788 | 0,035626536 | 0,201388889 | ['KEGG:00000'] |
| 1584 | WP | WP:WP453 | Genes related to primary cilium development (based on CRISPR) | 2,69076E-09 | negative_corr | True | 103 | 810 | 39 | 6646 | 0,048148148 | 0,378640777 | ['WP:000000'] |
| 1269 | WP | WP:WP465 | Joubert Syndrome | 1,76694E-05 | negative_corr | True | 76 | 810 | 27 | 6646 | 0,033333333 | 0,355263158 | ['WP:000000'] |
| 1270 | WP | WP:WP288 | Nuclear Receptors Meta-Pathway | 1,9547E-05 | positive_corr | True | 318 | 429 | 46 | 6646 | 0,107226107 | 0,144654088 | ['WP:000000'] |
| 1490 | WP | WP:WP702 | Metapathway biotransformation Phase I and II | 2,16478E-05 | positive_corr | True | 182 | 429 | 32 | 6646 | 0,074592075 | 0,175824176 | ['WP:000000'] |

| | | | | | | | | | | | | | |
|---|---|---|---|---|---|---|---|---|---|---|---|---|---|
| 1240 | WP | WP:WP2289 | Drug Induction of Bile Acid Pathway | 0,000629718 | positive_corr | True | 17 | 429 | 8 | 6646 | 0,018648019 | 0,470588235 | ['WP:000000'] |
| 1390 | WP | WP:WP477 | Cytoplasmic Ribosomal Proteins | 0,001606209 | negative_corr | True | 88 | 810 | 26 | 6646 | 0,032098765 | 0,295454545 | ['WP:000000'] |
| 584 | WP | WP:WP2875 | Constitutive Androstane Receptor Pathway | 0,003093164 | positive_corr | True | 32 | 429 | 10 | 6646 | 0,023310023 | 0,3125 | ['WP:000000'] |
| 839 | WP | WP:WP698 | Glucuronidation | 0,003202395 | positive_corr | True | 26 | 429 | 9 | 6646 | 0,020979021 | 0,346153846 | ['WP:000000'] |
| 1164 | WP | WP:WP299 | Nuclear Receptors in Lipid Metabolism and Toxicity | 0,00418564 | positive_corr | True | 33 | 429 | 10 | 6646 | 0,023310023 | 0,303030303 | ['WP:000000'] |
| 832 | WP | WP:WP2876 | Pregnane X Receptor pathway | 0,00418564 | positive_corr | True | 33 | 429 | 10 | 6646 | 0,023310023 | 0,303030303 | ['WP:000000'] |
| 1329 | WP | WP:WP691 | Tamoxifen metabolism | 0,004186087 | positive_corr | True | 21 | 429 | 8 | 6646 | 0,018648019 | 0,380952381 | ['WP:000000'] |
| 1646 | WP | WP:WP382 | MAPK Signaling Pathway | 0,005942023 | negative_corr | True | 246 | 810 | 52 | 6646 | 0,064197531 | 0,211382114 | ['WP:000000'] |
| 1596 | WP | WP:WP3967 | miR-509-3p alteration of YAP1/ECM axis | 0,009098353 | negative_corr | True | 17 | 810 | 9 | 6646 | 0,011111111 | 0,529411765 | ['WP:000000'] |
| 1236 | WP | WP:WP536 | Calcium Regulation in the Cardiac Cell | 0,014670412 | negative_corr | True | 150 | 810 | 35 | 6646 | 0,043209877 | 0,233333333 | ['WP:000000'] |
| 1464 | WP | WP:WP4720 | Eicosanoid metabolism via Cytochrome P450 Mono-Oxygenases (C | 0,017087516 | positive_corr | True | 9 | 429 | 5 | 6646 | 0,011655012 | 0,555555556 | ['WP:000000'] |
| 1610 | WP | WP:WP3925 | Amino Acid metabolism | 0,030682146 | positive_corr | True | 91 | 429 | 16 | 6646 | 0,037296037 | 0,175824176 | ['WP:000000'] |
| 1611 | WP | WP:WP1604 | Codeine and Morphine Metabolism | 0,032592925 | positive_corr | True | 15 | 429 | 6 | 6646 | 0,013986014 | 0,4 | ['WP:000000'] |
| 1621 | WP | WP:WP694 | Arylamine metabolism | 0,035602137 | positive_corr | True | 6 | 429 | 4 | 6646 | 0,009324009 | 0,666666667 | ['WP:000000'] |
| 829 | WP | WP:WP1603 | Nicotine Activity on Chromaffin Cells | 0,0359368 | negative_corr | True | 4 | 810 | 4 | 6646 | 0,004938272 | 1 | ['WP:000000'] |
| 1271 | WP | WP:WP2858 | Ectoderm Differentiation | 0,037892055 | negative_corr | True | 139 | 810 | 32 | 6646 | 0,039506173 | 0,230215827 | ['WP:000000'] |
| 1080 | WP | WP:WP3942 | PPAR signaling pathway | 0,044358773 | positive_corr | True | 67 | 429 | 13 | 6646 | 0,03030303 | 0,194029851 | ['WP:000000'] |
