## Supplementary table 6 kidney for "Bioinformatic characterization of angiotensin-converting enzyme 2, the entry receptor for SARS-CoV-2"

| | source | native | name | p_value | query | significar | term_size | query_size | intersection_si | effective_domain_si | precision | recall | parents |
|---|---|---|---|---|---|---|---|---|---|---|---|---|---|
| 130 | GO:BF | GO:0015711 | organic anion transport | 2,7837E-17 | positive_corr | True | 508 | 118 | 30 | 17906 | 0,254237288 | 0,059055118 | ['GO:0006820', 'GO:0071702'] |
| 34 | GO:BF | GO:0006820 | anion transport | 3,27017E-16 | positive_corr | True | 654 | 118 | 32 | 17906 | 0,271186441 | 0,048929664 | ['GO:0006811'] |
| 38 | GO:BF | GO:0046942 | carboxylic acid transport | 1,22018E-11 | positive_corr | True | 340 | 118 | 21 | 17906 | 0,177966102 | 0,061764706 | ['GO:0015711', 'GO:0015849'] |
| 92 | GO:BF | GO:0015849 | organic acid transport | 1,37081E-11 | positive_corr | True | 342 | 118 | 21 | 17906 | 0,177966102 | 0,061403509 | ['GO:0071702'] |
| 46 | GO:BF | GO:0044281 | small molecule metabolic process | 2,95325E-11 | positive_corr | True | 1987 | 118 | 45 | 17906 | 0,381355932 | 0,022647207 | ['GO:0008152'] |
| 48 | GO:BF | GO:0006082 | organic acid metabolic process | 1,59101E-10 | positive_corr | True | 1177 | 118 | 34 | 17906 | 0,288135593 | 0,028887001 | ['GO:0044237', 'GO:0044281', 'GO:0071704'] |
| 49 | GO:BF | GO:0055085 | transmembrane transport | 3,20883E-09 | positive_corr | True | 1634 | 118 | 38 | 17906 | 0,322033898 | 0,023255814 | ['GO:0006810'] |
| 129 | GO:BF | GO:0006811 | ion transport | 1,61087E-08 | positive_corr | True | 1723 | 118 | 38 | 17906 | 0,322033898 | 0,022054556 | ['GO:0006810'] |
| 54 | GO:BF | GO:1904970 | brush border assembly | 2,03793E-08 | positive_corr | True | 5 | 118 | 5 | 17906 | 0,042372881 | 1 | ['GO:0022607'] |
| 56 | GO:BF | GO:0046415 | urate metabolic process | 2,126E-07 | positive_corr | True | 13 | 118 | 6 | 17906 | 0,050847458 | 0,461538462 | ['GO:0006082', 'GO:0072521'] |
| 57 | GO:BF | GO:0043436 | oxoacid metabolic process | 2,32733E-06 | positive_corr | True | 1157 | 118 | 28 | 17906 | 0,237288136 | 0,024200519 | ['GO:0006082'] |
| 126 | GO:BF | GO:0071702 | organic substance transport | 5,87321E-06 | positive_corr | True | 2937 | 118 | 46 | 17906 | 0,389830508 | 0,01566224 | ['GO:0006810'] |
| 124 | GO:BF | GO:0019752 | carboxylic acid metabolic process | 8,73823E-06 | positive_corr | True | 1066 | 118 | 26 | 17906 | 0,220338983 | 0,024390244 | ['GO:0043436'] |
| 62 | GO:BF | GO:0015718 | monocarboxylic acid transport | 3,69396E-05 | positive_corr | True | 174 | 118 | 11 | 17906 | 0,093220339 | 0,063218391 | ['GO:0046942'] |
| 120 | GO:BF | GO:0015747 | urate transport | 4,75163E-05 | positive_corr | True | 6 | 118 | 4 | 17906 | 0,033898305 | 0,666666667 | ['GO:0015711', 'GO:0071705'] |
| 119 | GO:BF | GO:0051181 | cofactor transport | 5,5028E-05 | positive_corr | True | 49 | 118 | 7 | 17906 | 0,059322034 | 0,142857143 | ['GO:0006810'] |
| 65 | GO:BF | GO:0006766 | vitamin metabolic process | 0,000450345 | positive_corr | True | 134 | 118 | 9 | 17906 | 0,076271186 | 0,067164179 | ['GO:0044281'] |
| 66 | GO:BF | GO:0051186 | cofactor metabolic process | 0,000526587 | positive_corr | True | 453 | 118 | 15 | 17906 | 0,127118644 | 0,033112583 | ['GO:0044237'] |
| 67 | GO:BF | GO:1903825 | organic acid transmembrane transport | 0,000544003 | positive_corr | True | 137 | 118 | 9 | 17906 | 0,076271186 | 0,065693431 | ['GO:0015849', 'GO:0055085'] |
| 118 | GO:BF | GO:1905039 | carboxylic acid transmembrane transport | 0,000544003 | positive_corr | True | 137 | 118 | 9 | 17906 | 0,076271186 | 0,065693431 | ['GO:0046942', 'GO:0098656', 'GO:1903825'] |
| 116 | GO:BF | GO:0072337 | modified amino acid transport | 0,000974459 | positive_corr | True | 25 | 118 | 5 | 17906 | 0,042372881 | 0,2 | ['GO:0071702', 'GO:0071705'] |
| 76 | GO:BF | GO:0006855 | drug transmembrane transport | 0,001104093 | positive_corr | True | 75 | 118 | 7 | 17906 | 0,059322034 | 0,093333333 | ['GO:0015893', 'GO:0055085'] |
| 77 | GO:BF | GO:0098656 | anion transmembrane transport | 0,001145431 | positive_corr | True | 298 | 118 | 12 | 17906 | 0,101694915 | 0,040268456 | ['GO:0006820', 'GO:0034220'] |
| 78 | GO:BF | GO:0042908 | xenobiotic transport | 0,00120014 | positive_corr | True | 26 | 118 | 5 | 17906 | 0,042372881 | 0,192307692 | ['GO:0006810'] |
| 79 | GO:BF | GO:0015893 | drug transport | 0,002017239 | positive_corr | True | 207 | 118 | 10 | 17906 | 0,084745763 | 0,048309179 | ['GO:0006810', 'GO:0042493'] |
| 84 | GO:BF | GO:0016264 | gap junction assembly | 0,002185462 | positive_corr | True | 13 | 118 | 4 | 17906 | 0,033898305 | 0,307692308 | ['GO:0007043'] |
| 86 | GO:BF | GO:0006767 | water-soluble vitamin metabolic process | 0,003045606 | positive_corr | True | 87 | 118 | 7 | 17906 | 0,059322034 | 0,08045977 | ['GO:0006766'] |
| 98 | GO:BF | GO:0006814 | sodium ion transport | 0,00395232 | positive_corr | True | 223 | 118 | 10 | 17906 | 0,084745763 | 0,044843049 | ['GO:0015672', 'GO:0030001'] |
| 33 | GO:BF | GO:0006810 | transport | 0,005608485 | positive_corr | True | 5337 | 118 | 59 | 17906 | 0,5 | 0,0110549 | ['GO:0051234'] |
| 136 | GO:BF | GO:0006790 | sulfur compound metabolic process | 0,012051095 | positive_corr | True | 374 | 118 | 12 | 17906 | 0,101694915 | 0,032085561 | ['GO:0044237'] |
| 94 | GO:BF | GO:0051234 | establishment of localization | 0,012386551 | positive_corr | True | 5460 | 118 | 59 | 17906 | 0,5 | 0,010805861 | ['GO:0051179'] |
| 142 | GO:BF | GO:0032787 | monocarboxylic acid metabolic process | 0,012449654 | positive_corr | True | 661 | 118 | 16 | 17906 | 0,13559322 | 0,024205749 | ['GO:0019752'] |
| 137 | GO:BF | GO:0051180 | vitamin transport | 0,012635539 | positive_corr | True | 41 | 118 | 5 | 17906 | 0,042372881 | 0,12195122 | ['GO:0006810'] |
| 19 | GO:BF | GO:0051179 | localization | 0,01439641 | positive_corr | True | 6889 | 118 | 69 | 17906 | 0,584745763 | 0,010015967 | ['GO:0008150'] |
| 20 | GO:BF | GO:0042493 | response to drug | 0,015137469 | positive_corr | True | 1093 | 118 | 21 | 17906 | 0,177966102 | 0,019213175 | ['GO:0042221'] |
| 3 | GO:BF | GO:0032532 | regulation of microvillus length | 0,017103977 | positive_corr | True | 7 | 118 | 3 | 17906 | 0,025423729 | 0,428571429 | ['GO:0032530', 'GO:0032536'] |
| 28 | GO:BF | GO:0000101 | sulfur amino acid transport | 0,017103977 | positive_corr | True | 7 | 118 | 3 | 17906 | 0,025423729 | 0,428571429 | ['GO:0006865', 'GO:0072348'] |
| 143 | GO:BF | GO:0072348 | sulfur compound transport | 0,022513053 | positive_corr | True | 46 | 118 | 5 | 17906 | 0,042372881 | 0,108695652 | ['GO:0006810'] |
| 147 | GO:BF | GO:0072521 | purine-containing compound metabolic process | 0,024656044 | positive_corr | True | 471 | 118 | 13 | 17906 | 0,110169492 | 0,027600849 | ['GO:0006725', 'GO:0034641', 'GO:0046483', 'GO:1901360', 'GO:1901564'] |
| 5 | GO:BF | GO:0015698 | inorganic anion transport | 0,02535017 | positive_corr | True | 166 | 118 | 8 | 17906 | 0,06779661 | 0,048192771 | ['GO:0006820'] |
| 148 | GO:BF | GO:1990961 | xenobiotic detoxification by transmembrane export across the plasma membra | 0,02723467 | positive_corr | True | 8 | 118 | 3 | 17906 | 0,025423729 | 0,375 | ['GO:0042908', 'GO:0098754', 'GO:0140115'] |
| 22 | GO:BF | GO:0015889 | cobalamin transport | 0,02723467 | positive_corr | True | 8 | 118 | 3 | 17906 | 0,025423729 | 0,375 | ['GO:0015893', 'GO:0051180', 'GO:0051181', 'GO:0071702', 'GO:0071705'] |
| 18 | GO:BF | GO:0006063 | uronic acid metabolic process | 0,03070898 | positive_corr | True | 24 | 118 | 4 | 17906 | 0,033898305 | 0,166666667 | ['GO:0005996', 'GO:0032787'] |
| 149 | GO:BF | GO:0019585 | glucuronate metabolic process | 0,03070898 | positive_corr | True | 24 | 118 | 4 | 17906 | 0,033898305 | 0,166666667 | ['GO:0006063'] |
| 140 | GO:BF | GO:0032528 | microvillus organization | 0,03070898 | positive_corr | True | 24 | 118 | 4 | 17906 | 0,033898305 | 0,166666667 | ['GO:0120036'] |
| 10 | GO:CC | GO:0045177 | apical part of cell | 4,91579E-22 | positive_corr | True | 400 | 125 | 31 | 18856 | 0,248 | 0,0775 | ['GO:0110165'] |
| 15 | GO:CC | GO:0016324 | apical plasma membrane | 1,82888E-19 | positive_corr | True | 331 | 125 | 27 | 18856 | 0,216 | 0,081570997 | ['GO:0045177', 'GO:0098590'] |
| 107 | GO:CC | GO:0005903 | brush border | 4,99743E-18 | positive_corr | True | 108 | 125 | 18 | 18856 | 0,144 | 0,166666667 | ['GO:0098862'] |
| 7 | GO:CC | GO:0031526 | brush border membrane | 2,35568E-16 | positive_corr | True | 56 | 125 | 14 | 18856 | 0,112 | 0,25 | ['GO:0005903', 'GO:0031253'] |
| 6 | GO:CC | GO:0098862 | cluster of actin-based cell projections | 8,2835E-15 | positive_corr | True | 161 | 125 | 18 | 18856 | 0,144 | 0,111801242 | ['GO:0110165'] |
| 4 | GO:CC | GO:0070062 | extracellular exosome | 5,41568E-14 | positive_corr | True | 2144 | 125 | 50 | 18856 | 0,4 | 0,023320896 | ['GO:0005615', 'GO:1903561'] |
| 105 | GO:CC | GO:1903561 | extracellular vesicle | 8,39996E-14 | positive_corr | True | 2167 | 125 | 50 | 18856 | 0,4 | 0,023073373 | ['GO:0031982', 'GO:0043230'] |
| 2 | GO:CC | GO:0043230 | extracellular organelle | 9,23373E-14 | positive_corr | True | 2172 | 125 | 50 | 18856 | 0,4 | 0,023020258 | ['GO:0005576', 'GO:0043226'] |
| 99 | GO:CC | GO:0098590 | plasma membrane region | 4,06274E-12 | positive_corr | True | 1153 | 125 | 35 | 18856 | 0,28 | 0,030355594 | ['GO:0005886', 'GO:0016020'] |
| 1 | GO:CC | GO:0031224 | intrinsic component of membrane | 1,35123E-10 | positive_corr | True | 5842 | 125 | 78 | 18856 | 0,624 | 0,013351592 | ['GO:0016020', 'GO:0110165'] |
| 9 | GO:CC | GO:0016021 | integral component of membrane | 4,233E-10 | positive_corr | True | 5692 | 125 | 76 | 18856 | 0,608 | 0,013352073 | ['GO:0031224'] |
| 115 | GO:CC | GO:0031982 | vesicle | 8,52587E-10 | positive_corr | True | 3914 | 125 | 61 | 18856 | 0,488 | 0,015585079 | ['GO:0043227'] |
| 0 | GO:CC | GO:0005615 | extracellular space | 2,11602E-09 | positive_corr | True | 3543 | 125 | 57 | 18856 | 0,456 | 0,016088061 | ['GO:0005576', 'GO:0110165'] |
| 58 | GO:CC | GO:0005902 | microvillus | 3,3543E-09 | positive_corr | True | 85 | 125 | 11 | 18856 | 0,088 | 0,129411765 | ['GO:0098858'] |
| 29 | GO:CC | GO:0031253 | cell projection membrane | 2,55545E-07 | positive_corr | True | 335 | 125 | 16 | 18856 | 0,128 | 0,047761194 | ['GO:0098590', 'GO:0120025'] |
| 133 | GO:CC | GO:0005886 | plasma membrane | 2,66821E-07 | positive_corr | True | 5604 | 125 | 70 | 18856 | 0,56 | 0,012491078 | ['GO:0016020', 'GO:0071944'] |
| 132 | GO:CC | GO:0005576 | extracellular region | 5,79763E-07 | positive_corr | True | 4548 | 125 | 61 | 18856 | 0,488 | 0,013412489 | ['GO:0110165'] |
| 39 | GO:CC | GO:0071944 | cell periphery | 7,78695E-07 | positive_corr | True | 5732 | 125 | 70 | 18856 | 0,56 | 0,012212142 | ['GO:0110165'] |
| 40 | GO:CC | GO:0031528 | microvillus membrane | 3,14391E-06 | positive_corr | True | 24 | 125 | 6 | 18856 | 0,048 | 0,25 | ['GO:0005902', 'GO:0031253'] |

| | | | | | | | | | | | | | |
|---|---|---|---|---|---|---|---|---|---|---|---|---|---|
| 125 | GO:CC | GO:0016020 | membrane | 1,66472E-05 | positive_corr | True | 9731 | 125 | 94 | 18856 | 0,752 | 0,00965985 | ['GO:0110165'] |
| 43 | GO:CC | GO:0098858 | actin-based cell projection | 5,53012E-05 | positive_corr | True | 211 | 125 | 11 | 18856 | 0,088 | 0,052132701 | ['GO:0120025'] |
| 27 | GO:CC | GO:0016323 | basolateral plasma membrane | 0,005726826 | positive_corr | True | 221 | 125 | 9 | 18856 | 0,072 | 0,040723982 | ['GO:0098590'] |
| 42 | GO:CC | GO:0005887 | integral component of plasma membrane | 0,006621174 | positive_corr | True | 1624 | 125 | 26 | 18856 | 0,208 | 0,016009852 | ['GO:0016021', 'GO:0031226'] |
| 47 | GO:CC | GO:0005773 | vacuole | 0,007012194 | positive_corr | True | 797 | 125 | 17 | 18856 | 0,136 | 0,021329987 | ['GO:0005737', 'GO:0043231'] |
| 25 | GO:CC | GO:0009986 | cell surface | 0,011971149 | positive_corr | True | 918 | 125 | 18 | 18856 | 0,144 | 0,019607843 | ['GO:0110165'] |
| 52 | GO:CC | GO:0031226 | intrinsic component of plasma membrane | 0,01451719 | positive_corr | True | 1700 | 125 | 26 | 18856 | 0,208 | 0,015294118 | ['GO:0005886', 'GO:0031224'] |
| 26 | GO:CC | GO:0000323 | lytic vacuole | 0,020230574 | positive_corr | True | 697 | 125 | 15 | 18856 | 0,12 | 0,021520803 | ['GO:0005773'] |
| 21 | GO:CC | GO:0005764 | lysosome | 0,020230574 | positive_corr | True | 697 | 125 | 15 | 18856 | 0,12 | 0,021520803 | ['GO:0000323'] |
| 114 | GO:MF | GO:0008514 | organic anion transmembrane transporter activity | 8,1459E-14 | positive_corr | True | 211 | 120 | 19 | 18126 | 0,158333333 | 0,090047393 | ['GO:0008509'] |
| 95 | GO:MF | GO:0005215 | transporter activity | 1,11271E-13 | positive_corr | True | 1223 | 120 | 38 | 18126 | 0,316666667 | 0,031071137 | ['GO:0003674'] |
| 96 | GO:MF | GO:0015291 | secondary active transmembrane transporter activity | 7,65621E-13 | positive_corr | True | 238 | 120 | 19 | 18126 | 0,158333333 | 0,079831933 | ['GO:0022804'] |
| 127 | GO:MF | GO:0008509 | anion transmembrane transporter activity | 3,00446E-12 | positive_corr | True | 336 | 120 | 21 | 18126 | 0,175 | 0,0625 | ['GO:0015075'] |
| 135 | GO:MF | GO:0022857 | transmembrane transporter activity | 3,19275E-12 | positive_corr | True | 1068 | 120 | 34 | 18126 | 0,283333333 | 0,031835206 | ['GO:0005215'] |
| 146 | GO:MF | GO:0022804 | active transmembrane transporter activity | 7,55156E-12 | positive_corr | True | 352 | 120 | 21 | 18126 | 0,175 | 0,059659091 | ['GO:0022857'] |
| 102 | GO:MF | GO:0015293 | symporter activity | 3,38962E-10 | positive_corr | True | 144 | 120 | 14 | 18126 | 0,116666667 | 0,097222222 | ['GO:0015291'] |
| 104 | GO:MF | GO:0046943 | carboxylic acid transmembrane transporter activity | 1,02346E-08 | positive_corr | True | 150 | 120 | 13 | 18126 | 0,108333333 | 0,086666667 | ['GO:0005342', 'GO:0008514', 'GO:0015318'] |
| 128 | GO:MF | GO:0005342 | organic acid transmembrane transporter activity | 1,02346E-08 | positive_corr | True | 150 | 120 | 13 | 18126 | 0,108333333 | 0,086666667 | ['GO:0022857'] |
| 131 | GO:MF | GO:0015318 | inorganic molecular entity transmembrane transporter activity | 1,29058E-08 | positive_corr | True | 830 | 120 | 26 | 18126 | 0,216666667 | 0,031325301 | ['GO:0022857'] |
| 117 | GO:MF | GO:0015370 | solute:sodium symporter activity | 3,15812E-08 | positive_corr | True | 75 | 120 | 10 | 18126 | 0,083333333 | 0,133333333 | ['GO:0015081', 'GO:0015294'] |
| 100 | GO:MF | GO:0015294 | solute:cation symporter activity | 4,90972E-08 | positive_corr | True | 105 | 120 | 11 | 18126 | 0,091666667 | 0,104761905 | ['GO:0015293', 'GO:0022853', 'GO:0022890'] |
| 139 | GO:MF | GO:0015075 | ion transmembrane transporter activity | 5,53334E-08 | positive_corr | True | 887 | 120 | 26 | 18126 | 0,216666667 | 0,029312289 | ['GO:0022857'] |
| 69 | GO:MF | GO:0022853 | active ion transmembrane transporter activity | 4,35646E-07 | positive_corr | True | 245 | 120 | 14 | 18126 | 0,116666667 | 0,057142857 | ['GO:0015075', 'GO:0022804'] |
| 91 | GO:MF | GO:0008238 | exopeptidase activity | 2,0894E-06 | positive_corr | True | 114 | 120 | 10 | 18126 | 0,083333333 | 0,087719298 | ['GO:0070011'] |
| 53 | GO:MF | GO:0004177 | aminopeptidase activity | 1,5782E-05 | positive_corr | True | 49 | 120 | 7 | 18126 | 0,058333333 | 0,142857143 | ['GO:0008238'] |
| 51 | GO:MF | GO:0005343 | organic acid:sodium symporter activity | 1,91625E-05 | positive_corr | True | 30 | 120 | 6 | 18126 | 0,05 | 0,2 | ['GO:0005342', 'GO:0015370'] |
| 45 | GO:MF | GO:0015081 | sodium ion transmembrane transporter activity | 4,01207E-05 | positive_corr | True | 155 | 120 | 10 | 18126 | 0,083333333 | 0,064516129 | ['GO:0015077', 'GO:0046873'] |
| 41 | GO:MF | GO:0008235 | metalloexopeptidase activity | 6,68182E-05 | positive_corr | True | 60 | 120 | 7 | 18126 | 0,058333333 | 0,116666667 | ['GO:0008237', 'GO:0008238'] |
| 37 | GO:MF | GO:0030247 | polysaccharide binding | 0,000415969 | positive_corr | True | 27 | 120 | 5 | 18126 | 0,041666667 | 0,185185185 | ['GO:0030246'] |
| 36 | GO:MF | GO:0015143 | urate transmembrane transporter activity | 0,000557669 | positive_corr | True | 4 | 120 | 3 | 18126 | 0,025 | 0,75 | ['GO:0008514', 'GO:1901702'] |
| 35 | GO:MF | GO:1901702 | salt transmembrane transporter activity | 0,000557669 | positive_corr | True | 4 | 120 | 3 | 18126 | 0,025 | 0,75 | ['GO:0008324', 'GO:0008509'] |
| 32 | GO:MF | GO:0005452 | inorganic anion exchanger activity | 0,000617495 | positive_corr | True | 13 | 120 | 4 | 18126 | 0,033333333 | 0,307692308 | ['GO:0015103', 'GO:0015291', 'GO:0022853'] |
| 31 | GO:MF | GO:0016805 | dipeptidase activity | 0,000860072 | positive_corr | True | 14 | 120 | 4 | 18126 | 0,033333333 | 0,285714286 | ['GO:0008238'] |
| 30 | GO:MF | GO:0008239 | dipeptidyl-peptidase activity | 0,000860072 | positive_corr | True | 14 | 120 | 4 | 18126 | 0,033333333 | 0,285714286 | ['GO:0004177'] |
| 24 | GO:MF | GO:0015020 | glucuronosyltransferase activity | 0,001381635 | positive_corr | True | 34 | 120 | 5 | 18126 | 0,041666667 | 0,147058824 | ['GO:0008194', 'GO:0016758'] |
| 17 | GO:MF | GO:0003996 | acyl-CoA ligase activity | 0,001547822 | positive_corr | True | 16 | 120 | 4 | 18126 | 0,033333333 | 0,25 | ['GO:0016405', 'GO:0016878'] |
| 14 | GO:MF | GO:0015103 | inorganic anion transmembrane transporter activity | 0,002195201 | positive_corr | True | 141 | 120 | 8 | 18126 | 0,066666667 | 0,056737589 | ['GO:0008509', 'GO:0015318'] |
| 13 | GO:MF | GO:0042910 | xenobiotic transmembrane transporter activity | 0,002575853 | positive_corr | True | 18 | 120 | 4 | 18126 | 0,033333333 | 0,222222222 | ['GO:0022857'] |
| 12 | GO:MF | GO:0046873 | metal ion transmembrane transporter activity | 0,004044951 | positive_corr | True | 446 | 120 | 13 | 18126 | 0,108333333 | 0,029147982 | ['GO:0022890'] |
| 11 | GO:MF | GO:0000099 | sulfur amino acid transmembrane transporter activity | 0,00480911 | positive_corr | True | 7 | 120 | 3 | 18126 | 0,025 | 0,428571429 | ['GO:0015171', 'GO:1901682'] |
| 8 | GO:MF | GO:0015077 | monovalent inorganic cation transmembrane transporter activity | 0,005793158 | positive_corr | True | 393 | 120 | 12 | 18126 | 0,1 | 0,030534351 | ['GO:0022890'] |
| 55 | GO:MF | GO:0015645 | fatty acid ligase activity | 0,006032801 | positive_corr | True | 22 | 120 | 4 | 18126 | 0,033333333 | 0,181818182 | ['GO:0016878'] |
| 59 | GO:MF | GO:0004180 | carboxypeptidase activity | 0,006388181 | positive_corr | True | 46 | 120 | 5 | 18126 | 0,041666667 | 0,108695652 | ['GO:0008238'] |
| 154 | GO:MF | GO:0017153 | sodium:dicarboxylate symporter activity | 0,011430506 | positive_corr | True | 9 | 120 | 3 | 18126 | 0,025 | 0,333333333 | ['GO:0005310', 'GO:0005343'] |
| 74 | GO:MF | GO:0016405 | CoA-ligase activity | 0,012079725 | positive_corr | True | 26 | 120 | 4 | 18126 | 0,033333333 | 0,153846154 | ['GO:0016877'] |
| 87 | GO:MF | GO:0008324 | cation transmembrane transporter activity | 0,015173862 | positive_corr | True | 660 | 120 | 15 | 18126 | 0,125 | 0,022727273 | ['GO:0015075'] |
| 68 | GO:MF | GO:0047760 | butyrate-CoA ligase activity | 0,016250362 | positive_corr | True | 10 | 120 | 3 | 18126 | 0,025 | 0,3 | ['GO:0016405', 'GO:0016878'] |
| 70 | GO:MF | GO:0008028 | monocarboxylic acid transmembrane transporter activity | 0,020044258 | positive_corr | True | 58 | 120 | 5 | 18126 | 0,041666667 | 0,086206897 | ['GO:0046943'] |
| 82 | GO:MF | GO:0016878 | acid-thiol ligase activity | 0,02169517 | positive_corr | True | 30 | 120 | 4 | 18126 | 0,033333333 | 0,133333333 | ['GO:0016877'] |
| 61 | GO:MF | GO:0022890 | inorganic cation transmembrane transporter activity | 0,025141825 | positive_corr | True | 609 | 120 | 14 | 18126 | 0,116666667 | 0,022988506 | ['GO:0008324', 'GO:0015318'] |
| 81 | GO:MF | GO:0070011 | peptidase activity, acting on L-amino acid peptides | 0,027938192 | positive_corr | True | 615 | 120 | 14 | 18126 | 0,116666667 | 0,022764228 | ['GO:0008233'] |
| 75 | GO:MF | GO:0008233 | peptidase activity | 0,043448224 | positive_corr | True | 641 | 120 | 14 | 18126 | 0,116666667 | 0,021840874 | ['GO:0016787', 'GO:0140096'] |
| 90 | HP | HP:0003532 | Ornithinuria | 0,000521062 | positive_corr | True | 3 | 35 | 3 | 4225 | 0,085714286 | 1 | ['HP:0003355'] |
| 88 | HP | HP:0003268 | Argininuria | 0,000521062 | positive_corr | True | 3 | 35 | 3 | 4225 | 0,085714286 | 1 | ['HP:0003355'] |
| 63 | HP | HP:0000787 | Nephrolithiasis | 0,000752665 | positive_corr | True | 96 | 35 | 8 | 4225 | 0,228571429 | 0,083333333 | ['HP:0012210'] |
| 64 | HP | HP:0003297 | Hyperlysinuria | 0,002072399 | positive_corr | True | 4 | 35 | 3 | 4225 | 0,085714286 | 0,75 | ['HP:0003355'] |
| 85 | HP | HP:0003110 | Abnormality of urine homeostasis | 0,002316255 | positive_corr | True | 555 | 35 | 16 | 4225 | 0,457142857 | 0,028828829 | ['HP:0001939', 'HP:0011277'] |
| 72 | HP | HP:0011277 | Abnormality of the urinary system physiology | 0,002434347 | positive_corr | True | 878 | 35 | 20 | 4225 | 0,571428571 | 0,022779043 | ['HP:0000079'] |
| 93 | KEGG | KEGG:0461 | Renin-angiotensin system | 7,94809E-05 | positive_corr | True | 23 | 65 | 5 | 7788 | 0,076923077 | 0,217391304 | ['KEGG:00000'] |
| 121 | KEGG | KEGG:0497 | Vitamin digestion and absorption | 0,003172051 | positive_corr | True | 24 | 65 | 4 | 7788 | 0,061538462 | 0,166666667 | ['KEGG:00000'] |
| 111 | KEGG | KEGG:0497 | Protein digestion and absorption | 0,009955364 | positive_corr | True | 95 | 65 | 6 | 7788 | 0,092307692 | 0,063157895 | ['KEGG:00000'] |
| 60 | KEGG | KEGG:0004 | Pentose and glucuronate interconversions | 0,013003474 | positive_corr | True | 34 | 65 | 4 | 7788 | 0,061538462 | 0,117647059 | ['KEGG:00000'] |
| 134 | WP | WP:WP288 | NRF2 pathway | 0,001563774 | positive_corr | True | 144 | 53 | 8 | 6646 | 0,150943396 | 0,055555556 | ['WP:000000'] |
| 83 | WP | WP:WP4271 | Vitamin B12 Disorders | 0,01302529 | positive_corr | True | 13 | 53 | 3 | 6646 | 0,056603774 | 0,230769231 | ['WP:000000'] |
| 153 | WP | WP:WP229 | Irinotecan Pathway | 0,01302529 | positive_corr | True | 13 | 53 | 3 | 6646 | 0,056603774 | 0,230769231 | ['WP:000000'] |

| | | | | | | | | | | | | | |
|---|---|---|---|---|---|---|---|---|---|---|---|---|---|
| 122 | WP | WP:WP1604 | Codeine and Morphine Metabolism | 0,020489397 | positive_corr | True | 15 | 53 | 3 | 6646 | 0,056603774 | 0,2 | ['WP:000000'] |
| 123 | WP | WP:WP4506 | Tyrosine Metabolism | 0,037284954 | positive_corr | True | 4 | 53 | 2 | 6646 | 0,037735849 | 0,5 | ['WP:000000'] |
