## Supplementary table 6 testis for "Bioinformatic characterization of angiotensin-converting enzyme 2, the entry receptor for SARS-CoV-2"

| | source | native | name | p_value | query | significar | term_size | query_size | intersection_si | effective_domain_si | precision | recall | parents |
|---|---|---|---|---|---|---|---|---|---|---|---|---|---|
| 779 | GO:BF | GO:0016192 | vesicle-mediated transport | 2,38933E-13 | positive_corr | True | 2146 | 1185 | 238 | 17906 | 0,200843882 | 0,110904007 | ['GO:0006810'] |
| 476 | GO:BF | GO:0046907 | intracellular transport | 4,68179E-12 | positive_corr | True | 1781 | 1185 | 203 | 17906 | 0,171308017 | 0,11398091 | ['GO:0006810', 'GO:0051641'] |
| 481 | GO:BF | GO:0051641 | cellular localization | 9,88701E-12 | positive_corr | True | 2889 | 1185 | 292 | 17906 | 0,246413502 | 0,101073036 | ['GO:0051179'] |
| 484 | GO:BF | GO:0044237 | cellular metabolic process | 1,05706E-11 | positive_corr | True | 10860 | 1185 | 843 | 17906 | 0,711392405 | 0,077624309 | ['GO:0008152', 'GO:0009987'] |
| 500 | GO:BF | GO:0006810 | transport | 2,50106E-11 | positive_corr | True | 5337 | 1185 | 473 | 17906 | 0,399156118 | 0,088626569 | ['GO:0051234'] |
| 502 | GO:BF | GO:1901564 | organonitrogen compound metabolic process | 8,78541E-11 | positive_corr | True | 6889 | 1185 | 579 | 17906 | 0,488607595 | 0,084047031 | ['GO:0006807', 'GO:0071704'] |
| 979 | GO:BF | GO:0051234 | establishment of localization | 9,5424E-11 | positive_corr | True | 5460 | 1185 | 479 | 17906 | 0,404219409 | 0,087728938 | ['GO:0051179'] |
| 519 | GO:BF | GO:0065008 | regulation of biological quality | 1,86559E-10 | positive_corr | True | 4156 | 1185 | 383 | 17906 | 0,323206751 | 0,092155919 | ['GO:0065007'] |
| 982 | GO:BF | GO:0051179 | localization | 2,19419E-10 | positive_corr | True | 6889 | 1185 | 577 | 17906 | 0,486919831 | 0,083756714 | ['GO:0008150'] |
| 521 | GO:BF | GO:0034613 | cellular protein localization | 4,30158E-10 | positive_corr | True | 1983 | 1185 | 213 | 17906 | 0,179746835 | 0,107413011 | ['GO:0008104', 'GO:0070727'] |
| 965 | GO:BF | GO:0033036 | macromolecule localization | 5,34663E-10 | positive_corr | True | 3292 | 1185 | 316 | 17906 | 0,266666667 | 0,095990279 | ['GO:0051179'] |
| 956 | GO:BF | GO:0008104 | protein localization | 7,35413E-10 | positive_corr | True | 2923 | 1185 | 287 | 17906 | 0,242194093 | 0,098186794 | ['GO:0033036'] |
| 534 | GO:BF | GO:0070727 | cellular macromolecule localization | 7,53643E-10 | positive_corr | True | 1994 | 1185 | 213 | 17906 | 0,179746835 | 0,106820461 | ['GO:0033036', 'GO:0051641'] |
| 933 | GO:BF | GO:0044248 | cellular catabolic process | 2,32774E-09 | positive_corr | True | 2314 | 1185 | 237 | 17906 | 0,2 | 0,102420052 | ['GO:0009056', 'GO:0044237'] |
| 541 | GO:BF | GO:0044281 | small molecule metabolic process | 3,31669E-09 | positive_corr | True | 1987 | 1185 | 210 | 17906 | 0,17721519 | 0,105686965 | ['GO:0008152'] |
| 561 | GO:BF | GO:0043312 | neutrophil degranulation | 9,17407E-09 | positive_corr | True | 484 | 1185 | 75 | 17906 | 0,063291139 | 0,154958678 | ['GO:0002283', 'GO:0002446', 'GO:0043299'] |
| 908 | GO:BF | GO:0002283 | neutrophil activation involved in immune response | 1,25029E-08 | positive_corr | True | 487 | 1185 | 75 | 17906 | 0,063291139 | 0,154004107 | ['GO:0002275', 'GO:0042119'] |
| 970 | GO:BF | GO:0002446 | neutrophil mediated immunity | 1,41951E-08 | positive_corr | True | 498 | 1185 | 76 | 17906 | 0,064135021 | 0,152610442 | ['GO:0002444'] |
| 574 | GO:BF | GO:0009056 | catabolic process | 3,29354E-08 | positive_corr | True | 2693 | 1185 | 262 | 17906 | 0,221097046 | 0,097289268 | ['GO:0008152'] |
| 983 | GO:BF | GO:0042119 | neutrophil activation | 3,78089E-08 | positive_corr | True | 498 | 1185 | 75 | 17906 | 0,063291139 | 0,15060241 | ['GO:0036230'] |
| 458 | GO:BF | GO:0036230 | granulocyte activation | 7,47468E-08 | positive_corr | True | 505 | 1185 | 75 | 17906 | 0,063291139 | 0,148514851 | ['GO:0002274'] |
| 1015 | GO:BF | GO:0008152 | metabolic process | 1,19247E-07 | positive_corr | True | 11681 | 1185 | 874 | 17906 | 0,737552743 | 0,074822361 | ['GO:0008150'] |
| 371 | GO:BF | GO:0016043 | cellular component organization | 1,32481E-07 | positive_corr | True | 6481 | 1185 | 534 | 17906 | 0,450632911 | 0,082394692 | ['GO:0009987', 'GO:0071840'] |
| 775 | GO:BF | GO:0045055 | regulated exocytosis | 1,33032E-07 | positive_corr | True | 793 | 1185 | 102 | 17906 | 0,086075949 | 0,128625473 | ['GO:0006887'] |
| 384 | GO:BF | GO:0043603 | cellular amide metabolic process | 2,0334E-07 | positive_corr | True | 1181 | 1185 | 136 | 17906 | 0,114767932 | 0,115156647 | ['GO:0034641'] |
| 1014 | GO:BF | GO:0002444 | myeloid leukocyte mediated immunity | 2,67276E-07 | positive_corr | True | 549 | 1185 | 78 | 17906 | 0,065822785 | 0,142076503 | ['GO:0002443'] |
| 1010 | GO:BF | GO:0071840 | cellular component organization or biogenesis | 2,71929E-07 | positive_corr | True | 6667 | 1185 | 545 | 17906 | 0,459915612 | 0,081745913 | ['GO:0008150'] |
| 1009 | GO:BF | GO:0006082 | organic acid metabolic process | 3,16227E-07 | positive_corr | True | 1177 | 1185 | 135 | 17906 | 0,113924051 | 0,114698386 | ['GO:0044237', 'GO:0044281', 'GO:0071704'] |
| 467 | GO:BF | GO:0043299 | leukocyte degranulation | 3,60297E-07 | positive_corr | True | 532 | 1185 | 76 | 17906 | 0,064135021 | 0,142857143 | ['GO:0002252', 'GO:0045055'] |
| 392 | GO:BF | GO:0002274 | myeloid leukocyte activation | 3,79553E-07 | positive_corr | True | 657 | 1185 | 88 | 17906 | 0,074261603 | 0,133942161 | ['GO:0045321'] |
| 397 | GO:BF | GO:1901566 | organonitrogen compound biosynthetic process | 4,2463E-07 | positive_corr | True | 1905 | 1185 | 195 | 17906 | 0,164556962 | 0,102362205 | ['GO:1901564', 'GO:1901576'] |
| 426 | GO:BF | GO:0043436 | oxoacid metabolic process | 7,42141E-07 | positive_corr | True | 1157 | 1185 | 132 | 17906 | 0,111392405 | 0,114088159 | ['GO:0006082'] |
| 427 | GO:BF | GO:0006887 | exocytosis | 7,95016E-07 | positive_corr | True | 907 | 1185 | 110 | 17906 | 0,092827004 | 0,121278942 | ['GO:0016192', 'GO:0032940'] |
| 444 | GO:BF | GO:0002275 | myeloid cell activation involved in immune response | 8,68328E-07 | positive_corr | True | 542 | 1185 | 76 | 17906 | 0,064135021 | 0,140221402 | ['GO:0002274', 'GO:0002366'] |
| 445 | GO:BF | GO:0045184 | establishment of protein localization | 1,39544E-06 | positive_corr | True | 2235 | 1185 | 219 | 17906 | 0,184810127 | 0,097986577 | ['GO:0008104', 'GO:0051234'] |
| 987 | GO:BF | GO:0019752 | carboxylic acid metabolic process | 1,58875E-06 | positive_corr | True | 1066 | 1185 | 123 | 17906 | 0,103797468 | 0,115384615 | ['GO:0043436'] |
| 986 | GO:BF | GO:0071702 | organic substance transport | 2,36719E-06 | positive_corr | True | 2937 | 1185 | 272 | 17906 | 0,229535865 | 0,092611508 | ['GO:0006810'] |
| 395 | GO:BF | GO:0015031 | protein transport | 2,37474E-06 | positive_corr | True | 2121 | 1185 | 209 | 17906 | 0,176371308 | 0,098538425 | ['GO:0015833', 'GO:0045184'] |
| 902 | GO:BF | GO:0006886 | intracellular protein transport | 2,99594E-06 | positive_corr | True | 1182 | 1185 | 132 | 17906 | 0,111392405 | 0,111675127 | ['GO:0015031', 'GO:0034613', 'GO:0046907'] |
| 891 | GO:BF | GO:0048193 | Golgi vesicle transport | 3,5442E-06 | positive_corr | True | 377 | 1185 | 58 | 17906 | 0,048945148 | 0,153846154 | ['GO:0016192'] |
| 886 | GO:BF | GO:1901575 | organic substance catabolic process | 4,50118E-06 | positive_corr | True | 2239 | 1185 | 217 | 17906 | 0,183122363 | 0,096918267 | ['GO:0009056', 'GO:0071704'] |
| 825 | GO:BF | GO:0071704 | organic substance metabolic process | 4,82767E-06 | positive_corr | True | 11103 | 1185 | 829 | 17906 | 0,699578059 | 0,074664505 | ['GO:0008152'] |
| 824 | GO:BF | GO:0031175 | neuron projection development | 5,15263E-06 | positive_corr | True | 1017 | 1185 | 117 | 17906 | 0,098734177 | 0,115044248 | ['GO:0048666', 'GO:0120036'] |
| 700 | GO:BF | GO:0015833 | peptide transport | 7,05761E-06 | positive_corr | True | 2161 | 1185 | 210 | 17906 | 0,17721519 | 0,097177233 | ['GO:0042886', 'GO:0071702'] |
| 705 | GO:BF | GO:0009894 | regulation of catabolic process | 7,52048E-06 | positive_corr | True | 1012 | 1185 | 116 | 17906 | 0,097890295 | 0,114624506 | ['GO:0009056', 'GO:0019222'] |
| 716 | GO:BF | GO:0071705 | nitrogen compound transport | 1,01382E-05 | positive_corr | True | 2506 | 1185 | 236 | 17906 | 0,199156118 | 0,094173982 | ['GO:0006810'] |
| 815 | GO:BF | GO:0042886 | amide transport | 1,01674E-05 | positive_corr | True | 2196 | 1185 | 212 | 17906 | 0,178902954 | 0,096539162 | ['GO:0071705'] |
| 805 | GO:BF | GO:0000902 | cell morphogenesis | 1,28433E-05 | positive_corr | True | 1068 | 1185 | 120 | 17906 | 0,101265823 | 0,112359551 | ['GO:0032989'] |
| 685 | GO:BF | GO:0051128 | regulation of cellular component organization | 1,39574E-05 | positive_corr | True | 2541 | 1185 | 238 | 17906 | 0,200843882 | 0,093663912 | ['GO:0016043', 'GO:0050794'] |
| 725 | GO:BF | GO:0046903 | secretion | 2,19602E-05 | positive_corr | True | 1709 | 1185 | 172 | 17906 | 0,145147679 | 0,100643651 | ['GO:0006810'] |
| 797 | GO:BF | GO:0032989 | cellular component morphogenesis | 2,78559E-05 | positive_corr | True | 1177 | 1185 | 128 | 17906 | 0,108016878 | 0,108751062 | ['GO:0009653', 'GO:0016043', 'GO:0048869'] |
| 796 | GO:BF | GO:1901565 | organonitrogen compound catabolic process | 4,92843E-05 | positive_corr | True | 1333 | 1185 | 140 | 17906 | 0,11814346 | 0,105026257 | ['GO:1901564', 'GO:1901575'] |
| 742 | GO:BF | GO:0140352 | export from cell | 4,95313E-05 | positive_corr | True | 1616 | 1185 | 163 | 17906 | 0,137552743 | 0,100866337 | ['GO:0006810', 'GO:0009987'] |
| 783 | GO:BF | GO:0007167 | enzyme linked receptor protein signaling pathway | 6,91772E-05 | positive_corr | True | 1076 | 1185 | 118 | 17906 | 0,099578059 | 0,109665428 | ['GO:0007166'] |
| 753 | GO:BF | GO:0044238 | primary metabolic process | 8,29238E-05 | positive_corr | True | 10594 | 1185 | 789 | 17906 | 0,665822785 | 0,074476119 | ['GO:0008152'] |
| 761 | GO:BF | GO:0061024 | membrane organization | 8,65392E-05 | positive_corr | True | 893 | 1185 | 102 | 17906 | 0,086075949 | 0,114221725 | ['GO:0016043'] |
| 1327 | GO:BF | GO:0032940 | secretion by cell | 8,67736E-05 | positive_corr | True | 1567 | 1185 | 158 | 17906 | 0,133333333 | 0,100829611 | ['GO:0046903', 'GO:0140352'] |
| 727 | GO:BF | GO:0051186 | cofactor metabolic process | 9,44585E-05 | positive_corr | True | 453 | 1185 | 62 | 17906 | 0,052320675 | 0,136865342 | ['GO:0044237'] |
| 679 | GO:BF | GO:0007034 | vacuolar transport | 0,000148958 | positive_corr | True | 144 | 1185 | 29 | 17906 | 0,024472574 | 0,201388889 | ['GO:0006810'] |
| 827 | GO:BF | GO:0007169 | transmembrane receptor protein tyrosine kinase signaling pathway | 0,000166062 | positive_corr | True | 744 | 1185 | 88 | 17906 | 0,074261603 | 0,11827957 | ['GO:0007167'] |
| 833 | GO:BF | GO:0031329 | regulation of cellular catabolic process | 0,00019968 | positive_corr | True | 850 | 1185 | 97 | 17906 | 0,08185654 | 0,114117647 | ['GO:0009894', 'GO:0031323', 'GO:0044248'] |
| 884 | GO:BF | GO:0044267 | cellular protein metabolic process | 0,00021677 | positive_corr | True | 5357 | 1185 | 436 | 17906 | 0,367932489 | 0,081388833 | ['GO:0019538', 'GO:0044260'] |
| 879 | GO:BF | GO:0006796 | phosphate-containing compound metabolic process | 0,00025878 | positive_corr | True | 3367 | 1185 | 293 | 17906 | 0,247257384 | 0,087021087 | ['GO:0006793'] |
| 875 | GO:BF | GO:0002263 | cell activation involved in immune response | 0,000291955 | positive_corr | True | 708 | 1185 | 84 | 17906 | 0,070886076 | 0,118644068 | ['GO:0001775', 'GO:0002252', 'GO:0006955'] |

| | | | | | | | | | | | | | |
|---|---|---|---|---|---|---|---|---|---|---|---|---|---|
| 873 | GO:BF | GO:0006790 | sulfur compound metabolic process | 0,000298305 | positive_corr | True | 374 | 1185 | 53 | 17906 | 0,044725738 | 0,14171123 | ['GO:0044237'] |
| 605 | GO:BF | GO:0006793 | phosphorus metabolic process | 0,000367345 | positive_corr | True | 3393 | 1185 | 294 | 17906 | 0,248101266 | 0,086648983 | ['GO:0044237'] |
| 606 | GO:BF | GO:0002366 | leukocyte activation involved in immune response | 0,000456439 | positive_corr | True | 704 | 1185 | 83 | 17906 | 0,070042194 | 0,117897727 | ['GO:0002263', 'GO:0045321'] |
| 870 | GO:BF | GO:0007041 | lysosomal transport | 0,000489026 | positive_corr | True | 103 | 1185 | 23 | 17906 | 0,019409283 | 0,223300971 | ['GO:0007034'] |
| 868 | GO:BF | GO:1901698 | response to nitrogen compound | 0,0005945 | positive_corr | True | 1168 | 1185 | 122 | 17906 | 0,102953586 | 0,104452055 | ['GO:0042221'] |
| 625 | GO:BF | GO:0006099 | tricarboxylic acid cycle | 0,000601995 | positive_corr | True | 36 | 1185 | 13 | 17906 | 0,010970464 | 0,361111111 | ['GO:0009060', 'GO:0044238'] |
| 866 | GO:BF | GO:0007399 | nervous system development | 0,000656635 | positive_corr | True | 2432 | 1185 | 221 | 17906 | 0,18649789 | 0,090871711 | ['GO:0048731'] |
| 863 | GO:BF | GO:0055114 | oxidation-reduction process | 0,000743251 | positive_corr | True | 1028 | 1185 | 110 | 17906 | 0,092827004 | 0,107003891 | ['GO:0008152'] |
| 645 | GO:BF | GO:0009060 | aerobic respiration | 0,000751631 | positive_corr | True | 90 | 1185 | 21 | 17906 | 0,017721519 | 0,233333333 | ['GO:0045333'] |
| 647 | GO:BF | GO:0072659 | protein localization to plasma membrane | 0,000802572 | positive_corr | True | 275 | 1185 | 42 | 17906 | 0,035443038 | 0,152727273 | ['GO:0072657', 'GO:1990778'] |
| 648 | GO:BF | GO:0016482 | cytosolic transport | 0,000939081 | positive_corr | True | 165 | 1185 | 30 | 17906 | 0,025316456 | 0,181818182 | ['GO:0046907'] |
| 649 | GO:BF | GO:0010243 | response to organonitrogen compound | 0,001133794 | positive_corr | True | 1085 | 1185 | 114 | 17906 | 0,096202532 | 0,105069124 | ['GO:0010033', 'GO:1901698'] |
| 858 | GO:BF | GO:0048666 | neuron development | 0,001150712 | positive_corr | True | 1146 | 1185 | 119 | 17906 | 0,100421941 | 0,103839442 | ['GO:0030182', 'GO:0048468'] |
| 840 | GO:BF | GO:0016050 | vesicle organization | 0,001200253 | positive_corr | True | 339 | 1185 | 48 | 17906 | 0,040506329 | 0,14159292 | ['GO:0006996'] |
| 367 | GO:BF | GO:0016032 | viral process | 0,001324502 | positive_corr | True | 838 | 1185 | 93 | 17906 | 0,078481013 | 0,11097852 | ['GO:0044403'] |
| 362 | GO:BF | GO:0034332 | adherens junction organization | 0,001406515 | positive_corr | True | 71 | 1185 | 18 | 17906 | 0,015189873 | 0,253521127 | ['GO:0045216'] |
| 373 | GO:BF | GO:0120036 | plasma membrane bounded cell projection organization | 0,001608948 | positive_corr | True | 1539 | 1185 | 150 | 17906 | 0,126582278 | 0,097465887 | ['GO:0030030'] |
| 1037 | GO:BF | GO:0006629 | lipid metabolic process | 0,001657604 | positive_corr | True | 1414 | 1185 | 140 | 17906 | 0,11814346 | 0,099009901 | ['GO:0044238', 'GO:0071704'] |
| 1199 | GO:BF | GO:0030030 | cell projection organization | 0,001674015 | positive_corr | True | 1578 | 1185 | 153 | 17906 | 0,129113924 | 0,096958175 | ['GO:0016043'] |
| 1192 | GO:BF | GO:0006665 | sphingolipid metabolic process | 0,00178986 | positive_corr | True | 161 | 1185 | 29 | 17906 | 0,024472574 | 0,180124224 | ['GO:0006643', 'GO:1901564'] |
| 136 | GO:BF | GO:0044403 | symbiotic process | 0,001797593 | positive_corr | True | 891 | 1185 | 97 | 17906 | 0,08185654 | 0,108866442 | ['GO:0044419'] |
| 145 | GO:BF | GO:0010256 | endomembrane system organization | 0,00185383 | positive_corr | True | 449 | 1185 | 58 | 17906 | 0,048945148 | 0,129175947 | ['GO:0016043'] |
| 1179 | GO:BF | GO:1990778 | protein localization to cell periphery | 0,001887683 | positive_corr | True | 334 | 1185 | 47 | 17906 | 0,039662447 | 0,140718563 | ['GO:0034613'] |
| 149 | GO:BF | GO:0048468 | cell development | 0,001904542 | positive_corr | True | 2215 | 1185 | 202 | 17906 | 0,170464135 | 0,091196388 | ['GO:0030154', 'GO:0048856', 'GO:0048869'] |
| 158 | GO:BF | GO:0006996 | organelle organization | 0,001930105 | positive_corr | True | 3966 | 1185 | 331 | 17906 | 0,279324895 | 0,083459405 | ['GO:0016043'] |
| 1178 | GO:BF | GO:0031647 | regulation of protein stability | 0,001939149 | positive_corr | True | 284 | 1185 | 42 | 17906 | 0,035443038 | 0,147887324 | ['GO:0065008'] |
| 176 | GO:BF | GO:0050821 | protein stabilization | 0,002071613 | positive_corr | True | 180 | 1185 | 31 | 17906 | 0,026160338 | 0,172222222 | ['GO:0031647'] |
| 180 | GO:BF | GO:0005975 | carbohydrate metabolic process | 0,002278615 | positive_corr | True | 640 | 1185 | 75 | 17906 | 0,063291139 | 0,1171875 | ['GO:0044238', 'GO:0071704'] |
| 189 | GO:BF | GO:0019538 | protein metabolic process | 0,002496615 | positive_corr | True | 5940 | 1185 | 469 | 17906 | 0,395780591 | 0,078956229 | ['GO:0043170', 'GO:0044238', 'GO:1901564'] |
| 190 | GO:BF | GO:0010288 | response to lead ion | 0,003230586 | positive_corr | True | 24 | 1185 | 10 | 17906 | 0,008438819 | 0,416666667 | ['GO:0010038'] |
| 1169 | GO:BF | GO:0044272 | sulfur compound biosynthetic process | 0,003289402 | positive_corr | True | 193 | 1185 | 32 | 17906 | 0,027004219 | 0,165803109 | ['GO:0006790', 'GO:0044249'] |
| 202 | GO:BF | GO:0051130 | positive regulation of cellular component organization | 0,003432483 | positive_corr | True | 1245 | 1185 | 125 | 17906 | 0,105485232 | 0,100401606 | ['GO:0016043', 'GO:0048522', 'GO:0051128'] |
| 1163 | GO:BF | GO:0030182 | neuron differentiation | 0,003658586 | positive_corr | True | 1409 | 1185 | 138 | 17906 | 0,116455696 | 0,097941803 | ['GO:0030154', 'GO:0048699'] |
| 208 | GO:BF | GO:0043412 | macromolecule modification | 0,003733106 | positive_corr | True | 4428 | 1185 | 362 | 17906 | 0,305485232 | 0,081752484 | ['GO:0043170'] |
| 361 | GO:BF | GO:0061919 | process utilizing autophagic mechanism | 0,003864223 | positive_corr | True | 514 | 1185 | 63 | 17906 | 0,053164557 | 0,122568093 | ['GO:0009987'] |
| 111 | GO:BF | GO:0006914 | autophagy | 0,003864223 | positive_corr | True | 514 | 1185 | 63 | 17906 | 0,053164557 | 0,122568093 | ['GO:0044248', 'GO:0061919'] |
| 212 | GO:BF | GO:0006807 | nitrogen compound metabolic process | 0,00395753 | positive_corr | True | 10111 | 1185 | 746 | 17906 | 0,629535865 | 0,073781031 | ['GO:0008152'] |
| 1213 | GO:BF | GO:0055076 | transition metal ion homeostasis | 0,003974657 | positive_corr | True | 132 | 1185 | 25 | 17906 | 0,021097046 | 0,189393939 | ['GO:0055065'] |
| 1223 | GO:BF | GO:1901135 | carbohydrate derivative metabolic process | 0,004907847 | positive_corr | True | 1130 | 1185 | 115 | 17906 | 0,097046414 | 0,101769912 | ['GO:0071704'] |
| 1313 | GO:BF | GO:0010506 | regulation of autophagy | 0,004914163 | positive_corr | True | 335 | 1185 | 46 | 17906 | 0,038818565 | 0,137313433 | ['GO:0006914', 'GO:0031329'] |
| 1296 | GO:BF | GO:0006464 | cellular protein modification process | 0,005060736 | positive_corr | True | 4216 | 1185 | 346 | 17906 | 0,291983122 | 0,082068311 | ['GO:0036211', 'GO:0044267'] |
| 1291 | GO:BF | GO:0036211 | protein modification process | 0,005060736 | positive_corr | True | 4216 | 1185 | 346 | 17906 | 0,291983122 | 0,082068311 | ['GO:0019538', 'GO:0043412'] |
| 1280 | GO:BF | GO:0046395 | carboxylic acid catabolic process | 0,005919132 | positive_corr | True | 276 | 1185 | 40 | 17906 | 0,033755274 | 0,144927536 | ['GO:0016054', 'GO:0019752'] |
| 1278 | GO:BF | GO:0016054 | organic acid catabolic process | 0,005919132 | positive_corr | True | 276 | 1185 | 40 | 17906 | 0,033755274 | 0,144927536 | ['GO:0006082', 'GO:0044248', 'GO:0044282', 'GO:1901575'] |
| 1269 | GO:BF | GO:0046916 | cellular transition metal ion homeostasis | 0,006487576 | positive_corr | True | 110 | 1185 | 22 | 17906 | 0,018565401 | 0,2 | ['GO:0006875', 'GO:0055076'] |
| 1265 | GO:BF | GO:0044282 | small molecule catabolic process | 0,007510064 | positive_corr | True | 447 | 1185 | 56 | 17906 | 0,047257384 | 0,125279642 | ['GO:0009056', 'GO:0044281'] |
| 1263 | GO:BF | GO:0044419 | interspecies interaction between organisms | 0,007683952 | positive_corr | True | 945 | 1185 | 99 | 17906 | 0,083544304 | 0,104761905 | ['GO:0051704'] |
| 1259 | GO:BF | GO:0002252 | immune effector process | 0,008846881 | positive_corr | True | 1281 | 1185 | 126 | 17906 | 0,106329114 | 0,098360656 | ['GO:0002376'] |
| 1258 | GO:BF | GO:0072657 | protein localization to membrane | 0,0090953 | positive_corr | True | 641 | 1185 | 73 | 17906 | 0,061603376 | 0,113884555 | ['GO:0034613'] |
| 1257 | GO:BF | GO:0007264 | small GTPase mediated signal transduction | 0,009215046 | positive_corr | True | 584 | 1185 | 68 | 17906 | 0,057383966 | 0,116438356 | ['GO:0035556'] |
| 1255 | GO:BF | GO:0006900 | vesicle budding from membrane | 0,0092831 | positive_corr | True | 104 | 1185 | 21 | 17906 | 0,017721519 | 0,201923077 | ['GO:0016050', 'GO:0016192', 'GO:0061024'] |
| 1252 | GO:BF | GO:0007033 | vacuole organization | 0,009351728 | positive_corr | True | 165 | 1185 | 28 | 17906 | 0,023628692 | 0,16969697 | ['GO:0006996'] |
| 1247 | GO:BF | GO:1901699 | cellular response to nitrogen compound | 0,011638414 | positive_corr | True | 727 | 1185 | 80 | 17906 | 0,067510549 | 0,110041265 | ['GO:0070887', 'GO:1901698'] |
| 1246 | GO:BF | GO:0045216 | cell-cell junction organization | 0,01182698 | positive_corr | True | 195 | 1185 | 31 | 17906 | 0,026160338 | 0,158974359 | ['GO:0034330'] |
| 1243 | GO:BF | GO:0010033 | response to organic substance | 0,01343952 | positive_corr | True | 3421 | 1185 | 286 | 17906 | 0,241350211 | 0,083601286 | ['GO:0042221'] |
| 1224 | GO:BF | GO:0046148 | pigment biosynthetic process | 0,014741808 | positive_corr | True | 60 | 1185 | 15 | 17906 | 0,012658228 | 0,25 | ['GO:0042440'] |
| 1222 | GO:BF | GO:0071417 | cellular response to organonitrogen compound | 0,01474685 | positive_corr | True | 673 | 1185 | 75 | 17906 | 0,063291139 | 0,111441308 | ['GO:0010243', 'GO:0071310', 'GO:0071495', 'GO:1901699'] |
| 215 | GO:BF | GO:0000186 | activation of MAPKK activity | 0,014986161 | positive_corr | True | 53 | 1185 | 14 | 17906 | 0,011814346 | 0,264150943 | ['GO:0032147', 'GO:0043406'] |
| 1162 | GO:BF | GO:0044093 | positive regulation of molecular function | 0,015439232 | positive_corr | True | 1822 | 1185 | 167 | 17906 | 0,14092827 | 0,091657519 | ['GO:0065009'] |
| 1151 | GO:BF | GO:0000904 | cell morphogenesis involved in differentiation | 0,016240431 | positive_corr | True | 769 | 1185 | 83 | 17906 | 0,070042194 | 0,10793238 | ['GO:0000902', 'GO:0048468'] |
| 262 | GO:BF | GO:0009057 | macromolecule catabolic process | 0,016796759 | positive_corr | True | 1450 | 1185 | 138 | 17906 | 0,116455696 | 0,095172414 | ['GO:0043170', 'GO:1901575'] |
| 354 | GO:BF | GO:0030163 | protein catabolic process | 0,018824124 | positive_corr | True | 965 | 1185 | 99 | 17906 | 0,083544304 | 0,102590674 | ['GO:0009057', 'GO:0019538', 'GO:1901565'] |
| 288 | GO:BF | GO:0043085 | positive regulation of catalytic activity | 0,018828008 | positive_corr | True | 1466 | 1185 | 139 | 17906 | 0,117299578 | 0,094815825 | ['GO:0044093', 'GO:0050790'] |
| 291 | GO:BF | GO:0019725 | cellular homeostasis | 0,019464185 | positive_corr | True | 978 | 1185 | 100 | 17906 | 0,084388186 | 0,102249489 | ['GO:0009987', 'GO:0042592'] |
| 1045 | GO:BF | GO:0010035 | response to inorganic substance | 0,020575187 | positive_corr | True | 575 | 1185 | 66 | 17906 | 0,055696203 | 0,114782609 | ['GO:0042221'] |

| | | | | | | | | | | | | | |
|---|---|---|---|---|---|---|---|---|---|---|---|---|---|
| 1056 | GO:BF | GO:0006888 | endoplasmic reticulum to Golgi vesicle-mediated transport | 0,020769964 | positive_corr | True | 210 | 1185 | 32 | 17906 | 0,027004219 | 0,152380952 | ['GO:0046907', 'GO:0048193'] |
| 1095 | GO:BF | GO:0048699 | generation of neurons | 0,022552654 | positive_corr | True | 1574 | 1185 | 147 | 17906 | 0,124050633 | 0,09339263 | ['GO:0022008'] |
| 1063 | GO:BF | GO:0022008 | neurogenesis | 0,023664353 | positive_corr | True | 1679 | 1185 | 155 | 17906 | 0,130801688 | 0,092316855 | ['GO:0007399', 'GO:0030154'] |
| 1082 | GO:BF | GO:0009893 | positive regulation of metabolic process | 0,026479227 | positive_corr | True | 3742 | 1185 | 307 | 17906 | 0,25907173 | 0,082041689 | ['GO:0008152', 'GO:0019222', 'GO:0048518'] |
| 312 | GO:BF | GO:0001775 | cell activation | 0,028416942 | positive_corr | True | 1465 | 1185 | 138 | 17906 | 0,116455696 | 0,094197952 | ['GO:0009987'] |
| 1081 | GO:BF | GO:1904375 | regulation of protein localization to cell periphery | 0,028479801 | positive_corr | True | 120 | 1185 | 22 | 17906 | 0,018565401 | 0,183333333 | ['GO:1903827', 'GO:1990778'] |
| 329 | GO:BF | GO:0016197 | endosomal transport | 0,028545615 | positive_corr | True | 233 | 1185 | 34 | 17906 | 0,028691983 | 0,145922747 | ['GO:0016192', 'GO:0046907'] |
| 1079 | GO:BF | GO:0033365 | protein localization to organelle | 0,030549886 | positive_corr | True | 964 | 1185 | 98 | 17906 | 0,082700422 | 0,101659751 | ['GO:0034613'] |
| 337 | GO:BF | GO:0072350 | tricarboxylic acid metabolic process | 0,031012456 | positive_corr | True | 14 | 1185 | 7 | 17906 | 0,005907173 | 0,5 | ['GO:0019752'] |
| 1083 | GO:BF | GO:0008360 | regulation of cell shape | 0,031075781 | positive_corr | True | 166 | 1185 | 27 | 17906 | 0,02278481 | 0,162650602 | ['GO:0022604', 'GO:0065008'] |
| 1078 | GO:BF | GO:0002443 | leukocyte mediated immunity | 0,032617989 | positive_corr | True | 880 | 1185 | 91 | 17906 | 0,076793249 | 0,103409091 | ['GO:0002252'] |
| 259 | GO:BF | GO:0070887 | cellular response to chemical stimulus | 0,033442645 | positive_corr | True | 3418 | 1185 | 283 | 17906 | 0,238818565 | 0,082796957 | ['GO:0042221', 'GO:0051716'] |
| 1096 | GO:BF | GO:0007163 | establishment or maintenance of cell polarity | 0,034313349 | positive_corr | True | 225 | 1185 | 33 | 17906 | 0,027848101 | 0,146666667 | ['GO:0009987'] |
| 1038 | GO:BF | GO:0051188 | cofactor biosynthetic process | 0,034342574 | positive_corr | True | 235 | 1185 | 34 | 17906 | 0,028691983 | 0,144680851 | ['GO:0044249', 'GO:0051186'] |
| 1145 | GO:BF | GO:0010998 | regulation of translational initiation by eIF2 alpha phosphorylation | 0,034559846 | positive_corr | True | 10 | 1185 | 6 | 17906 | 0,005063291 | 0,6 | ['GO:0006468', 'GO:0043558'] |
| 1141 | GO:BF | GO:0045321 | leukocyte activation | 0,035067287 | positive_corr | True | 1305 | 1185 | 125 | 17906 | 0,105485232 | 0,095785441 | ['GO:0001775', 'GO:0002376'] |
| 233 | GO:BF | GO:1903827 | regulation of cellular protein localization | 0,035135492 | positive_corr | True | 550 | 1185 | 63 | 17906 | 0,053164557 | 0,114545455 | ['GO:0032880', 'GO:0034613', 'GO:0060341'] |
| 1129 | GO:BF | GO:0034330 | cell junction organization | 0,035847274 | positive_corr | True | 297 | 1185 | 40 | 17906 | 0,033755274 | 0,134680135 | ['GO:0016043'] |
| 1134 | GO:BF | GO:1901605 | alpha-amino acid metabolic process | 0,036905954 | positive_corr | True | 206 | 1185 | 31 | 17906 | 0,026160338 | 0,150485437 | ['GO:0006520'] |
| 1128 | GO:BF | GO:0048194 | Golgi vesicle budding | 0,037886672 | positive_corr | True | 80 | 1185 | 17 | 17906 | 0,014345992 | 0,2125 | ['GO:0006900', 'GO:0048193'] |
| 1116 | GO:BF | GO:0006732 | coenzyme metabolic process | 0,040121735 | positive_corr | True | 257 | 1185 | 36 | 17906 | 0,030379747 | 0,140077821 | ['GO:0051186'] |
| 253 | GO:BF | GO:0040007 | growth | 0,040329833 | positive_corr | True | 1020 | 1185 | 102 | 17906 | 0,086075949 | 0,1 | ['GO:0008150'] |
| 238 | GO:BF | GO:0006643 | membrane lipid metabolic process | 0,04487448 | positive_corr | True | 208 | 1185 | 31 | 17906 | 0,026160338 | 0,149038462 | ['GO:0044255'] |
| 336 | GO:BF | GO:0048812 | neuron projection morphogenesis | 0,049665357 | positive_corr | True | 673 | 1185 | 73 | 17906 | 0,061603376 | 0,108469539 | ['GO:0031175', 'GO:0120039'] |
| 1058 | GO:CC | GO:0005737 | cytoplasm | 4,54796E-46 | positive_corr | True | 11634 | 1225 | 986 | 18856 | 0,804897959 | 0,08475159 | ['GO:0005622', 'GO:0110165'] |
| 795 | GO:CC | GO:0005622 | intracellular | 4,58826E-39 | positive_corr | True | 14626 | 1225 | 1123 | 18856 | 0,916734694 | 0,076781075 | ['GO:0005575'] |
| 1042 | GO:CC | GO:0043227 | membrane-bounded organelle | 3,71461E-38 | positive_corr | True | 12584 | 1225 | 1019 | 18856 | 0,831836735 | 0,080975842 | ['GO:0043226'] |
| 1018 | GO:CC | GO:0043226 | organelle | 4,05786E-32 | positive_corr | True | 13720 | 1225 | 1064 | 18856 | 0,868571429 | 0,07755102 | ['GO:0110165'] |
| 1029 | GO:CC | GO:0031090 | organelle membrane | 4,48789E-30 | positive_corr | True | 3531 | 1225 | 397 | 18856 | 0,324081633 | 0,112432739 | ['GO:0016020', 'GO:0043227'] |
| 790 | GO:CC | GO:0043231 | intracellular membrane-bounded organelle | 3,0211E-29 | positive_corr | True | 11067 | 1225 | 909 | 18856 | 0,742040816 | 0,08213608 | ['GO:0043227', 'GO:0043229'] |
| 1030 | GO:CC | GO:0043229 | intracellular organelle | 6,67736E-29 | positive_corr | True | 12840 | 1225 | 1009 | 18856 | 0,823673469 | 0,078582555 | ['GO:0005622', 'GO:0043226'] |
| 1200 | GO:CC | GO:0031982 | vesicle | 5,33335E-27 | positive_corr | True | 3914 | 1225 | 418 | 18856 | 0,34122449 | 0,106796117 | ['GO:0043227'] |
| 1075 | GO:CC | GO:0070062 | extracellular exosome | 3,05491E-25 | positive_corr | True | 2144 | 1225 | 268 | 18856 | 0,21877551 | 0,125 | ['GO:0005615', 'GO:1903561'] |
| 991 | GO:CC | GO:1903561 | extracellular vesicle | 3,06074E-25 | positive_corr | True | 2167 | 1225 | 270 | 18856 | 0,220408163 | 0,124596216 | ['GO:0031982', 'GO:0043230'] |
| 845 | GO:CC | GO:0043230 | extracellular organelle | 4,43938E-25 | positive_corr | True | 2172 | 1225 | 270 | 18856 | 0,220408163 | 0,124309392 | ['GO:0005576', 'GO:0043226'] |
| 931 | GO:CC | GO:0012505 | endomembrane system | 2,54322E-24 | positive_corr | True | 4539 | 1225 | 457 | 18856 | 0,373061224 | 0,10068297 | ['GO:0110165'] |
| 984 | GO:CC | GO:0098805 | whole membrane | 2,59313E-23 | positive_corr | True | 1694 | 1225 | 223 | 18856 | 0,182040816 | 0,131641086 | ['GO:0016020'] |
| 1194 | GO:CC | GO:0005829 | cytosol | 1,4611E-20 | positive_corr | True | 5154 | 1225 | 489 | 18856 | 0,399183673 | 0,094877765 | ['GO:0005737', 'GO:0110165'] |
| 1093 | GO:CC | GO:0031410 | cytoplasmic vesicle | 1,6677E-20 | positive_corr | True | 2372 | 1225 | 274 | 18856 | 0,223673469 | 0,115514334 | ['GO:0005737', 'GO:0097708'] |
| 869 | GO:CC | GO:0097708 | intracellular vesicle | 2,01876E-20 | positive_corr | True | 2375 | 1225 | 274 | 18856 | 0,223673469 | 0,115368421 | ['GO:0031982', 'GO:0043229'] |
| 872 | GO:CC | GO:0098588 | bounding membrane of organelle | 4,79899E-20 | positive_corr | True | 2092 | 1225 | 249 | 18856 | 0,203265306 | 0,119024857 | ['GO:0031090'] |
| 892 | GO:CC | GO:0005794 | Golgi apparatus | 7,10798E-17 | positive_corr | True | 1586 | 1225 | 196 | 18856 | 0,16 | 0,123581337 | ['GO:0005737', 'GO:0012505', 'GO:0043231'] |
| 1156 | GO:CC | GO:0042175 | nuclear outer membrane-endoplasmic reticulum membrane network | 6,10905E-14 | positive_corr | True | 1098 | 1225 | 144 | 18856 | 0,11755102 | 0,131147541 | ['GO:0012505', 'GO:0016020'] |
| 1146 | GO:CC | GO:0005789 | endoplasmic reticulum membrane | 7,03594E-13 | positive_corr | True | 1075 | 1225 | 139 | 18856 | 0,113469388 | 0,129302326 | ['GO:0031090', 'GO:0042175'] |
| 929 | GO:CC | GO:0005764 | lysosome | 2,16466E-12 | positive_corr | True | 697 | 1225 | 102 | 18856 | 0,083265306 | 0,146341463 | ['GO:0000323'] |
| 930 | GO:CC | GO:0000323 | lytic vacuole | 2,16466E-12 | positive_corr | True | 697 | 1225 | 102 | 18856 | 0,083265306 | 0,146341463 | ['GO:0005773'] |
| 822 | GO:CC | GO:0005773 | vacuole | 4,20809E-12 | positive_corr | True | 797 | 1225 | 111 | 18856 | 0,090612245 | 0,139272271 | ['GO:0005737', 'GO:0043231'] |
| 774 | GO:CC | GO:0005739 | mitochondrion | 1,03548E-11 | positive_corr | True | 1631 | 1225 | 184 | 18856 | 0,150204082 | 0,112814224 | ['GO:0005737', 'GO:0043231'] |
| 0 | GO:CC | GO:0030141 | secretory granule | 5,85097E-11 | positive_corr | True | 839 | 1225 | 112 | 18856 | 0,091428571 | 0,133492253 | ['GO:0012505', 'GO:0099503'] |
| 346 | GO:CC | GO:0030135 | coated vesicle | 3,24086E-10 | positive_corr | True | 294 | 1225 | 55 | 18856 | 0,044897959 | 0,18707483 | ['GO:0031410'] |
| 209 | GO:CC | GO:0005798 | Golgi-associated vesicle | 4,61095E-10 | positive_corr | True | 181 | 1225 | 41 | 18856 | 0,033469388 | 0,226519337 | ['GO:0031410'] |
| 528 | GO:CC | GO:0005768 | endosome | 6,05703E-10 | positive_corr | True | 935 | 1225 | 118 | 18856 | 0,096326531 | 0,126203209 | ['GO:0012505', 'GO:0031410'] |
| 536 | GO:CC | GO:0000139 | Golgi membrane | 6,85016E-10 | positive_corr | True | 752 | 1225 | 101 | 18856 | 0,08244898 | 0,134308511 | ['GO:0005794', 'GO:0098588'] |
| 204 | GO:CC | GO:0099503 | secretory vesicle | 9,9885E-10 | positive_corr | True | 1009 | 1225 | 124 | 18856 | 0,10122449 | 0,122893954 | ['GO:0031410'] |
| 197 | GO:CC | GO:0005783 | endoplasmic reticulum | 3,01611E-09 | positive_corr | True | 1810 | 1225 | 190 | 18856 | 0,155102041 | 0,104972376 | ['GO:0005737', 'GO:0012505', 'GO:0043231'] |
| 569 | GO:CC | GO:0070161 | anchoring junction | 8,93578E-09 | positive_corr | True | 785 | 1225 | 101 | 18856 | 0,08244898 | 0,12866242 | ['GO:0030054'] |
| 169 | GO:CC | GO:0010008 | endosome membrane | 2,27487E-08 | positive_corr | True | 481 | 1225 | 71 | 18856 | 0,057959184 | 0,147609148 | ['GO:0005768', 'GO:0098588', 'GO:0098805'] |
| 146 | GO:CC | GO:0042470 | melanosome | 1,24177E-07 | positive_corr | True | 104 | 1225 | 27 | 18856 | 0,022040816 | 0,259615385 | ['GO:0048770'] |
| 135 | GO:CC | GO:0048770 | pigment granule | 1,24177E-07 | positive_corr | True | 104 | 1225 | 27 | 18856 | 0,022040816 | 0,259615385 | ['GO:0031410'] |
| 570 | GO:CC | GO:0005925 | focal adhesion | 1,41782E-07 | positive_corr | True | 410 | 1225 | 62 | 18856 | 0,050612245 | 0,151219512 | ['GO:0030055'] |
| 127 | GO:CC | GO:0031975 | envelope | 1,56032E-07 | positive_corr | True | 1219 | 1225 | 135 | 18856 | 0,110204082 | 0,110746514 | ['GO:0110165'] |
| 577 | GO:CC | GO:0031967 | organelle envelope | 1,56032E-07 | positive_corr | True | 1219 | 1225 | 135 | 18856 | 0,110204082 | 0,110746514 | ['GO:0031975', 'GO:0043227', 'GO:0043229'] |
| 119 | GO:CC | GO:0098852 | lytic vacuole membrane | 2,12736E-07 | positive_corr | True | 365 | 1225 | 57 | 18856 | 0,046530612 | 0,156164384 | ['GO:0005774'] |
| 99 | GO:CC | GO:0005765 | lysosomal membrane | 2,12736E-07 | positive_corr | True | 365 | 1225 | 57 | 18856 | 0,046530612 | 0,156164384 | ['GO:0005764', 'GO:0098852'] |
| 593 | GO:CC | GO:0030055 | cell-substrate junction | 2,83111E-07 | positive_corr | True | 417 | 1225 | 62 | 18856 | 0,050612245 | 0,148681055 | ['GO:0070161'] |

| | | | | | | | | | | | | |
|---|---|---|---|---|---|---|---|---|---|---|---|---|
| 527 | GO:CC | GO:0042582 azurophil granule | 3,3368E-07 | positive_corr | True | 154 | 1225 | 33 | 18856 | 0,026938776 | 0,214285714 | ['GO:0005766', 'GO:0030141'] |
| 522 | GO:CC | GO:0005766 primary lysosome | 3,3368E-07 | positive_corr | True | 154 | 1225 | 33 | 18856 | 0,026938776 | 0,214285714 | ['GO:0005764'] |
| 211 | GO:CC | GO:0005774 vacuolar membrane | 4,58151E-07 | positive_corr | True | 422 | 1225 | 62 | 18856 | 0,050612245 | 0,146919431 | ['GO:0005773', 'GO:0098588', 'GO:0098805'] |
| 220 | GO:CC | GO:0016020 membrane | 4,87715E-07 | positive_corr | True | 9731 | 1225 | 733 | 18856 | 0,598367347 | 0,075326277 | ['GO:0110165'] |
| 333 | GO:CC | GO:0005759 mitochondrial matrix | 6,08507E-07 | positive_corr | True | 476 | 1225 | 67 | 18856 | 0,054693878 | 0,140756303 | ['GO:0005739', 'GO:0070013'] |
| 332 | GO:CC | GO:0012506 vesicle membrane | 1,28434E-06 | positive_corr | True | 801 | 1225 | 96 | 18856 | 0,078367347 | 0,119850187 | ['GO:0031090', 'GO:0031982'] |
| 347 | GO:CC | GO:0030659 cytoplasmic vesicle membrane | 2,95645E-06 | positive_corr | True | 780 | 1225 | 93 | 18856 | 0,075918367 | 0,119230769 | ['GO:0012506', 'GO:0031410'] |
| 355 | GO:CC | GO:0005740 mitochondrial envelope | 1,04662E-05 | positive_corr | True | 766 | 1225 | 90 | 18856 | 0,073469388 | 0,117493473 | ['GO:0005739', 'GO:0031967'] |
| 356 | GO:CC | GO:0031252 cell leading edge | 1,17744E-05 | positive_corr | True | 417 | 1225 | 58 | 18856 | 0,047346939 | 0,139088729 | ['GO:0110165'] |
| 273 | GO:CC | GO:0005938 cell cortex | 1,20587E-05 | positive_corr | True | 326 | 1225 | 49 | 18856 | 0,04 | 0,150306748 | ['GO:0005737', 'GO:0071944'] |
| 363 | GO:CC | GO:0048471 perinuclear region of cytoplasm | 1,31177E-05 | positive_corr | True | 724 | 1225 | 86 | 18856 | 0,070204082 | 0,11878453 | ['GO:0005737', 'GO:0110165'] |
| 597 | GO:CC | GO:0031966 mitochondrial membrane | 2,50709E-05 | positive_corr | True | 723 | 1225 | 85 | 18856 | 0,069387755 | 0,117565698 | ['GO:0005740', 'GO:0031090'] |
| 252 | GO:CC | GO:0005775 vacuolar lumen | 2,51045E-05 | positive_corr | True | 173 | 1225 | 32 | 18856 | 0,026122449 | 0,184971098 | ['GO:0005773', 'GO:0070013'] |
| 235 | GO:CC | GO:0045335 phagocytic vesicle | 3,10145E-05 | positive_corr | True | 132 | 1225 | 27 | 18856 | 0,022040816 | 0,204545455 | ['GO:0030139'] |
| 370 | GO:CC | GO:0031984 organelle subcompartment | 3,85974E-05 | positive_corr | True | 379 | 1225 | 53 | 18856 | 0,043265306 | 0,139841689 | ['GO:0043227', 'GO:0043229', 'GO:0110165'] |
| 387 | GO:CC | GO:0030134 COPII-coated ER to Golgi transport vesicle | 4,02221E-05 | positive_corr | True | 94 | 1225 | 22 | 18856 | 0,017959184 | 0,234042553 | ['GO:0005798', 'GO:0030135'] |
| 388 | GO:CC | GO:0030667 secretory granule membrane | 4,19953E-05 | positive_corr | True | 299 | 1225 | 45 | 18856 | 0,036734694 | 0,150501672 | ['GO:0030141', 'GO:0030659', 'GO:0098588', 'GO:0098805'] |
| 390 | GO:CC | GO:0034774 secretory granule lumen | 4,24127E-05 | positive_corr | True | 319 | 1225 | 47 | 18856 | 0,038367347 | 0,147335423 | ['GO:0030141', 'GO:0060205'] |
| 420 | GO:CC | GO:0060205 cytoplasmic vesicle lumen | 6,1927E-05 | positive_corr | True | 323 | 1225 | 47 | 18856 | 0,038367347 | 0,145510836 | ['GO:0031410', 'GO:0031983', 'GO:0070013'] |
| 457 | GO:CC | GO:0030027 lamellipodium | 6,44969E-05 | positive_corr | True | 198 | 1225 | 34 | 18856 | 0,027755102 | 0,171717172 | ['GO:0031252', 'GO:0120025'] |
| 369 | GO:CC | GO:0098791 Golgi subcompartment | 6,46753E-05 | positive_corr | True | 354 | 1225 | 50 | 18856 | 0,040816327 | 0,141242938 | ['GO:0005794', 'GO:0031984'] |
| 601 | GO:CC | GO:0031983 vesicle lumen | 7,45934E-05 | positive_corr | True | 325 | 1225 | 47 | 18856 | 0,038367347 | 0,144615385 | ['GO:0031982', 'GO:0043233'] |
| 126 | GO:CC | GO:0005615 extracellular space | 9,87361E-05 | positive_corr | True | 3543 | 1225 | 299 | 18856 | 0,244081633 | 0,084391758 | ['GO:0005576', 'GO:0110165'] |
| 340 | GO:CC | GO:0005770 late endosome | 0,00010797 | positive_corr | True | 269 | 1225 | 41 | 18856 | 0,033469388 | 0,152416357 | ['GO:0005768'] |
| 7 | GO:CC | GO:0045121 membrane raft | 0,000119678 | positive_corr | True | 320 | 1225 | 46 | 18856 | 0,03755102 | 0,14375 | ['GO:0098857'] |
| 665 | GO:CC | GO:0098857 membrane microdomain | 0,000131116 | positive_corr | True | 321 | 1225 | 46 | 18856 | 0,03755102 | 0,143302181 | ['GO:0098589'] |
| 26 | GO:CC | GO:0098589 membrane region | 0,000168051 | positive_corr | True | 334 | 1225 | 47 | 18856 | 0,038367347 | 0,140718563 | ['GO:0016020', 'GO:0098805'] |
| 670 | GO:CC | GO:0005623 cell | 0,000286164 | positive_corr | True | 1116 | 1225 | 114 | 18856 | 0,093061224 | 0,102150538 | ['GO:0005575'] |
| 741 | GO:CC | GO:0030175 filopodium | 0,000312059 | positive_corr | True | 105 | 1225 | 22 | 18856 | 0,017959184 | 0,20952381 | ['GO:0098858'] |
| 686 | GO:CC | GO:0005911 cell-cell junction | 0,000363377 | positive_corr | True | 439 | 1225 | 56 | 18856 | 0,045714286 | 0,127562642 | ['GO:0070161'] |
| 29 | GO:CC | GO:0030660 Golgi-associated vesicle membrane | 0,000516137 | positive_corr | True | 108 | 1225 | 22 | 18856 | 0,017959184 | 0,203703704 | ['GO:0005798', 'GO:0098588', 'GO:0098805'] |
| 25 | GO:CC | GO:0005802 trans-Golgi network | 0,000552081 | positive_corr | True | 236 | 1225 | 36 | 18856 | 0,029387755 | 0,152542373 | ['GO:0098791'] |
| 602 | GO:CC | GO:0030133 transport vesicle | 0,000681712 | positive_corr | True | 415 | 1225 | 53 | 18856 | 0,043265306 | 0,127710843 | ['GO:0012505', 'GO:0031410'] |
| 12 | GO:CC | GO:0030054 cell junction | 0,000735733 | positive_corr | True | 1327 | 1225 | 129 | 18856 | 0,105306122 | 0,097211756 | ['GO:0110165'] |
| 11 | GO:CC | GO:0030662 coated vesicle membrane | 0,001068506 | positive_corr | True | 175 | 1225 | 29 | 18856 | 0,023673469 | 0,165714286 | ['GO:0030135', 'GO:0030659', 'GO:0098588', 'GO:0098805'] |
| 10 | GO:CC | GO:0005769 early endosome | 0,001529431 | positive_corr | True | 361 | 1225 | 47 | 18856 | 0,038367347 | 0,130193906 | ['GO:0005768'] |
| 730 | GO:CC | GO:0030136 clathrin-coated vesicle | 0,001790005 | positive_corr | True | 189 | 1225 | 30 | 18856 | 0,024489796 | 0,158730159 | ['GO:0030135'] |
| 2 | GO:CC | GO:0035578 azurophil granule lumen | 0,001816768 | positive_corr | True | 91 | 1225 | 19 | 18856 | 0,015510204 | 0,208791209 | ['GO:0005775', 'GO:0034774', 'GO:0042582'] |
| 18 | GO:CC | GO:1902494 catalytic complex | 0,002075725 | positive_corr | True | 1393 | 1225 | 132 | 18856 | 0,107755102 | 0,094759512 | ['GO:0032991'] |
| 659 | GO:CC | GO:0043233 organelle lumen | 0,00317303 | positive_corr | True | 5898 | 1225 | 452 | 18856 | 0,368979592 | 0,076636148 | ['GO:0031974', 'GO:0043226'] |
| 27 | GO:CC | GO:0070013 intracellular organelle lumen | 0,00317303 | positive_corr | True | 5898 | 1225 | 452 | 18856 | 0,368979592 | 0,076636148 | ['GO:0043229', 'GO:0043233'] |
| 1 | GO:CC | GO:0031974 membrane-enclosed lumen | 0,00317303 | positive_corr | True | 5898 | 1225 | 452 | 18856 | 0,368979592 | 0,076636148 | ['GO:0110165'] |
| 69 | GO:CC | GO:0019866 organelle inner membrane | 0,003879559 | positive_corr | True | 543 | 1225 | 62 | 18856 | 0,050612245 | 0,114180479 | ['GO:0031090', 'GO:0031967'] |
| 619 | GO:CC | GO:0043202 lysosomal lumen | 0,00412206 | positive_corr | True | 96 | 1225 | 19 | 18856 | 0,015510204 | 0,197916667 | ['GO:0005764', 'GO:0005775'] |
| 622 | GO:CC | GO:0032991 protein-containing complex | 0,00531786 | positive_corr | True | 5484 | 1225 | 422 | 18856 | 0,344489796 | 0,076951131 | ['GO:0005575'] |
| 766 | GO:CC | GO:0012507 ER to Golgi transport vesicle membrane | 0,005585016 | positive_corr | True | 58 | 1225 | 14 | 18856 | 0,011428571 | 0,24137931 | ['GO:0030134', 'GO:0030658', 'GO:0030660', 'GO:0030662'] |
| 30 | GO:CC | GO:0031901 early endosome membrane | 0,005779982 | positive_corr | True | 152 | 1225 | 25 | 18856 | 0,020408163 | 0,164473684 | ['GO:0005769', 'GO:0010008'] |
| 623 | GO:CC | GO:0098858 actin-based cell projection | 0,006235759 | positive_corr | True | 211 | 1225 | 31 | 18856 | 0,025306122 | 0,146919431 | ['GO:0120025'] |
| 722 | GO:CC | GO:0044291 cell-cell contact zone | 0,007535799 | positive_corr | True | 75 | 1225 | 16 | 18856 | 0,013061224 | 0,213333333 | ['GO:0005911'] |
| 48 | GO:CC | GO:0030139 endocytic vesicle | 0,008735468 | positive_corr | True | 298 | 1225 | 39 | 18856 | 0,031836735 | 0,130872483 | ['GO:0031410'] |
| 653 | GO:CC | GO:0030140 trans-Golgi network transport vesicle | 0,011146298 | positive_corr | True | 33 | 1225 | 10 | 18856 | 0,008163265 | 0,303030303 | ['GO:0005798', 'GO:0030133', 'GO:0030136'] |
| 45 | GO:CC | GO:0030137 COPI-coated vesicle | 0,01656906 | positive_corr | True | 28 | 1225 | 9 | 18856 | 0,007346939 | 0,321428571 | ['GO:0005798', 'GO:0030135'] |
| 41 | GO:CC | GO:0032009 early phagosome | 0,018084572 | positive_corr | True | 12 | 1225 | 6 | 18856 | 0,004897959 | 0,5 | ['GO:0045335'] |
| 40 | GO:CC | GO:0005912 adherens junction | 0,022825622 | positive_corr | True | 99 | 1225 | 18 | 18856 | 0,014693878 | 0,181818182 | ['GO:0005911'] |
| 633 | GO:CC | GO:0005743 mitochondrial inner membrane | 0,023731775 | positive_corr | True | 482 | 1225 | 54 | 18856 | 0,044081633 | 0,112033195 | ['GO:0019866', 'GO:0031966'] |
| 330 | GO:MF | GO:0003824 catalytic activity | 1,69813E-14 | positive_corr | True | 5853 | 1206 | 524 | 18126 | 0,434494196 | 0,089526738 | ['GO:0003674'] |
| 1085 | GO:MF | GO:0005515 protein binding | 1,58796E-13 | positive_corr | True | 12743 | 1206 | 968 | 18126 | 0,8026534 | 0,075963274 | ['GO:0005488'] |
| 1319 | GO:MF | GO:0000166 nucleotide binding | 1,03073E-09 | positive_corr | True | 2167 | 1206 | 225 | 18126 | 0,186567164 | 0,10383018 | ['GO:0036094', 'GO:1901265'] |
| 1310 | GO:MF | GO:1901265 nucleoside phosphate binding | 1,08018E-09 | positive_corr | True | 2168 | 1206 | 225 | 18126 | 0,186567164 | 0,103782288 | ['GO:0097159', 'GO:1901363'] |
| 261 | GO:MF | GO:0019899 enzyme binding | 2,2539E-09 | positive_corr | True | 2246 | 1206 | 230 | 18126 | 0,190713101 | 0,102404274 | ['GO:0005515'] |
| 319 | GO:MF | GO:0045296 cadherin binding | 1,35467E-08 | positive_corr | True | 330 | 1206 | 57 | 18126 | 0,047263682 | 0,172727273 | ['GO:0050839'] |
| 1100 | GO:MF | GO:0050839 cell adhesion molecule binding | 5,87607E-08 | positive_corr | True | 506 | 1206 | 74 | 18126 | 0,061359867 | 0,146245059 | ['GO:0005515'] |
| 307 | GO:MF | GO:0036094 small molecule binding | 8,95985E-08 | positive_corr | True | 2574 | 1206 | 249 | 18126 | 0,206467662 | 0,096736597 | ['GO:0005488'] |
| 96 | GO:MF | GO:0032555 purine ribonucleotide binding | 1,23375E-07 | positive_corr | True | 1913 | 1206 | 196 | 18126 | 0,16252073 | 0,102456874 | ['GO:0017076', 'GO:0032553'] |
| 1290 | GO:MF | GO:0032553 ribonucleotide binding | 1,4486E-07 | positive_corr | True | 1929 | 1206 | 197 | 18126 | 0,163349917 | 0,102125454 | ['GO:0000166', 'GO:0097367'] |

| | | | | | | | | | | | | | |
|---|---|---|---|---|---|---|---|---|---|---|---|---|---|
| 1284 | GO:MF | GO:0017076 | purine nucleotide binding | 2,19153E-07 | positive_corr | True | 1926 | 1206 | 196 | 18126 | 0,16252073 | 0,101765317 | ['GO:0000166'] |
| 1283 | GO:MF | GO:0035639 | purine ribonucleoside triphosphate binding | 3,22209E-07 | positive_corr | True | 1848 | 1206 | 189 | 18126 | 0,156716418 | 0,102272727 | ['GO:0043168', 'GO:1901265'] |
| 1130 | GO:MF | GO:0019904 | protein domain specific binding | 2,82304E-06 | positive_corr | True | 712 | 1206 | 89 | 18126 | 0,073797678 | 0,125 | ['GO:0005515'] |
| 232 | GO:MF | GO:0043168 | anion binding | 7,5687E-06 | positive_corr | True | 2841 | 1206 | 260 | 18126 | 0,215588723 | 0,091517071 | ['GO:0043167'] |
| 224 | GO:MF | GO:0097367 | carbohydrate derivative binding | 2,01885E-05 | positive_corr | True | 2269 | 1206 | 214 | 18126 | 0,177446103 | 0,094314676 | ['GO:0005488'] |
| 1159 | GO:MF | GO:0019901 | protein kinase binding | 2,26835E-05 | positive_corr | True | 655 | 1206 | 81 | 18126 | 0,067164179 | 0,123664122 | ['GO:0019900'] |
| 1176 | GO:MF | GO:0019900 | kinase binding | 3,81591E-05 | positive_corr | True | 741 | 1206 | 88 | 18126 | 0,072968491 | 0,118758435 | ['GO:0019899'] |
| 1253 | GO:MF | GO:0044877 | protein-containing complex binding | 5,7817E-05 | positive_corr | True | 1241 | 1206 | 130 | 18126 | 0,107794362 | 0,10475423 | ['GO:0005488'] |
| 1070 | GO:MF | GO:0003924 | GTPase activity | 0,000112119 | positive_corr | True | 328 | 1206 | 48 | 18126 | 0,039800995 | 0,146341463 | ['GO:0017111'] |
| 1212 | GO:MF | GO:0005525 | GTP binding | 0,000302796 | positive_corr | True | 380 | 1206 | 52 | 18126 | 0,043117745 | 0,136842105 | ['GO:0032550', 'GO:0032561', 'GO:0035639'] |
| 109 | GO:MF | GO:0032550 | purine ribonucleoside binding | 0,00041886 | positive_corr | True | 384 | 1206 | 52 | 18126 | 0,043117745 | 0,135416667 | ['GO:0001883', 'GO:0032549'] |
| 1123 | GO:MF | GO:0019003 | GDP binding | 0,00051021 | positive_corr | True | 73 | 1206 | 18 | 18126 | 0,014925373 | 0,246575342 | ['GO:0032550', 'GO:0032561', 'GO:0043168'] |
| 221 | GO:MF | GO:0032549 | ribonucleoside binding | 0,00053199 | positive_corr | True | 387 | 1206 | 52 | 18126 | 0,043117745 | 0,134366925 | ['GO:0001882'] |
| 769 | GO:MF | GO:0001883 | purine nucleoside binding | 0,00053199 | positive_corr | True | 387 | 1206 | 52 | 18126 | 0,043117745 | 0,134366925 | ['GO:0001882'] |
| 341 | GO:MF | GO:0001882 | nucleoside binding | 0,000916565 | positive_corr | True | 394 | 1206 | 52 | 18126 | 0,043117745 | 0,131979695 | ['GO:0036094', 'GO:0097159', 'GO:0097367', 'GO:1901363'] |
| 566 | GO:MF | GO:0032559 | adenyl ribonucleotide binding | 0,00105336 | positive_corr | True | 1560 | 1206 | 150 | 18126 | 0,124378109 | 0,096153846 | ['GO:0030554', 'GO:0032555'] |
| 556 | GO:MF | GO:0008092 | cytoskeletal protein binding | 0,001093609 | positive_corr | True | 991 | 1206 | 104 | 18126 | 0,086235489 | 0,104944501 | ['GO:0005515'] |
| 901 | GO:MF | GO:0050662 | coenzyme binding | 0,001317498 | positive_corr | True | 295 | 1206 | 42 | 18126 | 0,034825871 | 0,142372881 | ['GO:0048037'] |
| 932 | GO:MF | GO:0032561 | guanyl ribonucleotide binding | 0,001549724 | positive_corr | True | 401 | 1206 | 52 | 18126 | 0,043117745 | 0,12967581 | ['GO:0019001', 'GO:0032555'] |
| 505 | GO:MF | GO:0019001 | guanyl nucleotide binding | 0,001549724 | positive_corr | True | 401 | 1206 | 52 | 18126 | 0,043117745 | 0,12967581 | ['GO:0017076'] |
| 592 | GO:MF | GO:0030554 | adenyl nucleotide binding | 0,001566381 | positive_corr | True | 1571 | 1206 | 150 | 18126 | 0,124378109 | 0,095480586 | ['GO:0017076'] |
| 612 | GO:MF | GO:0016491 | oxidoreductase activity | 0,0022697 | positive_corr | True | 768 | 1206 | 84 | 18126 | 0,069651741 | 0,109375 | ['GO:0003824'] |
| 985 | GO:MF | GO:0005488 | binding | 0,003209033 | positive_corr | True | 15944 | 1206 | 1107 | 18126 | 0,917910448 | 0,069430507 | ['GO:0003674'] |
| 455 | GO:MF | GO:0043878 | glyceraldehyde-3-phosphate dehydrogenase (NAD+) (non-phosphorylating) activity | 0,003942982 | positive_corr | True | 6 | 1206 | 5 | 18126 | 0,004145937 | 0,833333333 | ['GO:0016620'] |
| 1013 | GO:MF | GO:0005524 | ATP binding | 0,004987036 | positive_corr | True | 1502 | 1206 | 142 | 18126 | 0,11774461 | 0,094540613 | ['GO:0008144', 'GO:0032559', 'GO:0035639'] |
| 848 | GO:MF | GO:0016787 | hydrolase activity | 0,008102716 | positive_corr | True | 2544 | 1206 | 220 | 18126 | 0,182421227 | 0,086477987 | ['GO:0003824'] |
| 847 | GO:MF | GO:0016620 | oxidoreductase activity, acting on the aldehyde or oxo group of donors, NAD or NADP as acc | 0,009931231 | positive_corr | True | 37 | 1206 | 11 | 18126 | 0,009121061 | 0,297297297 | ['GO:0016903'] |
| 662 | GO:MF | GO:0043167 | ion binding | 0,012191093 | positive_corr | True | 6308 | 1206 | 486 | 18126 | 0,402985075 | 0,077045022 | ['GO:0005488'] |
| 368 | GO:MF | GO:0016903 | oxidoreductase activity, acting on the aldehyde or oxo group of donors | 0,01410502 | positive_corr | True | 45 | 1206 | 12 | 18126 | 0,009950249 | 0,266666667 | ['GO:0016491'] |
| 849 | GO:MF | GO:0004709 | MAP kinase kinase kinase activity | 0,01479972 | positive_corr | True | 26 | 1206 | 9 | 18126 | 0,007462687 | 0,346153846 | ['GO:0004674'] |
| 814 | GO:MF | GO:0048037 | cofactor binding | 0,018360932 | positive_corr | True | 517 | 1206 | 59 | 18126 | 0,048922056 | 0,114119923 | ['GO:0005488'] |
| 765 | GO:MF | GO:0004029 | aldehyde dehydrogenase (NAD+) activity | 0,020461478 | positive_corr | True | 16 | 1206 | 7 | 18126 | 0,005804312 | 0,4375 | ['GO:0004030'] |
| 823 | GO:MF | GO:0051287 | NAD binding | 0,026433088 | positive_corr | True | 55 | 1206 | 13 | 18126 | 0,010779436 | 0,236363636 | ['GO:0000166', 'GO:0050662'] |
| 357 | GO:MF | GO:0004030 | aldehyde dehydrogenase [NAD(P)+] activity | 0,032788001 | positive_corr | True | 17 | 1206 | 7 | 18126 | 0,005804312 | 0,411764706 | ['GO:0016620'] |
| 751 | GO:MF | GO:0008144 | drug binding | 0,03866353 | positive_corr | True | 1762 | 1206 | 157 | 18126 | 0,130182421 | 0,089103292 | ['GO:0005488'] |
| 770 | GO:MF | GO:0016614 | oxidoreductase activity, acting on CH-OH group of donors | 0,038837036 | positive_corr | True | 134 | 1206 | 22 | 18126 | 0,018242123 | 0,164179104 | ['GO:0016491'] |
| 821 | GO:MF | GO:0016616 | oxidoreductase activity, acting on the CH-OH group of donors, NAD or NADP as acceptor | 0,039935235 | positive_corr | True | 125 | 1206 | 21 | 18126 | 0,017412935 | 0,168 | ['GO:0016614'] |
| 728 | GO:MF | GO:0016740 | transferase activity | 0,043609446 | positive_corr | True | 2381 | 1206 | 203 | 18126 | 0,168325041 | 0,085258295 | ['GO:0003824'] |
| 832 | GO:MF | GO:0140096 | catalytic activity, acting on a protein | 0,047458055 | positive_corr | True | 2236 | 1206 | 192 | 18126 | 0,15920398 | 0,085867621 | ['GO:0003824'] |
| 907 | HP | HP:0001417 | X-linked inheritance | 6,48469E-09 | positive_corr | True | 237 | 397 | 57 | 4225 | 0,143576826 | 0,240506329 | ['HP:0010985'] |
| 731 | HP | HP:0010985 | Gonosomal inheritance | 1,20133E-08 | positive_corr | True | 247 | 397 | 58 | 4225 | 0,146095718 | 0,234817814 | ['HP:0000005'] |
| 1209 | HP | HP:0001419 | X-linked recessive inheritance | 3,68157E-05 | positive_corr | True | 172 | 397 | 40 | 4225 | 0,100755668 | 0,23255814 | ['HP:0001417'] |
| 586 | HP | HP:0001290 | Generalized hypotonia | 7,8015E-05 | positive_corr | True | 909 | 397 | 129 | 4225 | 0,324937028 | 0,141914191 | ['HP:0001252'] |
| 1220 | HP | HP:0011804 | Abnormal muscle physiology | 0,000191163 | positive_corr | True | 2011 | 397 | 238 | 4225 | 0,599496222 | 0,11834908 | ['HP:0003011'] |
| 694 | HP | HP:0002167 | Neurological speech impairment | 0,000217125 | positive_corr | True | 961 | 397 | 133 | 4225 | 0,335012594 | 0,138397503 | ['HP:0011446'] |
| 1285 | HP | HP:0003808 | Abnormal muscle tone | 0,000329111 | positive_corr | True | 1675 | 397 | 205 | 4225 | 0,516372796 | 0,12238806 | ['HP:0011804'] |
| 1242 | HP | HP:0001276 | Hypertonia | 0,000868373 | positive_corr | True | 834 | 397 | 117 | 4225 | 0,294710327 | 0,14028777 | ['HP:0002493', 'HP:0003808'] |
| 640 | HP | HP:0001250 | Seizures | 0,001409857 | positive_corr | True | 1529 | 397 | 188 | 4225 | 0,473551637 | 0,122956181 | ['HP:0012638'] |
| 855 | HP | HP:0002493 | Upper motor neuron dysfunction | 0,001602894 | positive_corr | True | 1059 | 397 | 140 | 4225 | 0,352644836 | 0,132200189 | ['HP:0011442'] |
| 1251 | HP | HP:0010729 | Cherry red spot of the macula | 0,002466402 | positive_corr | True | 9 | 397 | 7 | 4225 | 0,017632242 | 0,777777778 | ['HP:0000630'] |
| 706 | HP | HP:0001252 | Muscular hypotonia | 0,003049432 | positive_corr | True | 1413 | 397 | 175 | 4225 | 0,440806045 | 0,123849965 | ['HP:0003808'] |
| 1193 | HP | HP:0001268 | Mental deterioration | 0,005204161 | positive_corr | True | 289 | 397 | 51 | 4225 | 0,128463476 | 0,176470588 | ['HP:0100543'] |
| 841 | HP | HP:0003011 | Abnormality of the musculature | 0,005947951 | positive_corr | True | 2298 | 397 | 258 | 4225 | 0,649874055 | 0,11227154 | ['HP:0000118'] |
| 811 | HP | HP:0011446 | Abnormality of higher mental function | 0,006124259 | positive_corr | True | 2146 | 397 | 244 | 4225 | 0,614609572 | 0,113699907 | ['HP:0012638'] |
| 1188 | HP | HP:0000726 | Dementia | 0,008364545 | positive_corr | True | 158 | 397 | 33 | 4225 | 0,083123426 | 0,208860759 | ['HP:0001268'] |
| 1328 | HP | HP:0011442 | Abnormality of central motor function | 0,012081245 | positive_corr | True | 1552 | 397 | 186 | 4225 | 0,468513854 | 0,119845361 | ['HP:0012638'] |
| 248 | HP | HP:0008715 | Testicular dysgenesis | 0,019108583 | positive_corr | True | 11 | 397 | 7 | 4225 | 0,017632242 | 0,636363636 | ['HP:0000035'] |
| 1072 | HP | HP:0002060 | Abnormality of the cerebrum | 0,021842116 | positive_corr | True | 1524 | 397 | 182 | 4225 | 0,458438287 | 0,119422572 | ['HP:0100547'] |
| 925 | HP | HP:0001423 | X-linked dominant inheritance | 0,022809862 | positive_corr | True | 65 | 397 | 18 | 4225 | 0,04534005 | 0,276923077 | ['HP:0001417'] |
| 1165 | HP | HP:0009023 | Abdominal wall muscle weakness | 0,025969884 | negative_corr | True | 11 | 4 | 2 | 4225 | 0,5 | 0,181818182 | ['HP:0001324'] |
| 1017 | HP | HP:0002071 | Abnormality of extrapyramidal motor function | 0,027318163 | positive_corr | True | 219 | 397 | 40 | 4225 | 0,100755668 | 0,182648402 | ['HP:0011442'] |
| 1041 | HP | HP:0010936 | Abnormality of the lower urinary tract | 0,030415669 | positive_corr | True | 616 | 397 | 87 | 4225 | 0,219143577 | 0,141233766 | ['HP:0000079'] |
| 1099 | HP | HP:0012759 | Neurodevelopmental abnormality | 0,031879577 | positive_corr | True | 2216 | 397 | 247 | 4225 | 0,622166247 | 0,111462094 | ['HP:0012638'] |
| 258 | HP | HP:0100547 | Abnormality of forebrain morphology | 0,038997239 | positive_corr | True | 1547 | 397 | 183 | 4225 | 0,460957179 | 0,118293471 | ['HP:0012443'] |
| 1039 | HP | HP:0001300 | Parkinsonism | 0,049997187 | positive_corr | True | 94 | 397 | 22 | 4225 | 0,055415617 | 0,234042553 | ['HP:0002071'] |

| | | | | | | | | | | | | | |
|---|---|---|---|---|---|---|---|---|---|---|---|---|---|
| 1127 | KEGG | KEGG:0414 | Lysosome | 1,75808E-09 | positive_corr | True | 123 | 594 | 34 | 7788 | 0,057239057 | 0,276422764 | ['KEGG:00000'] |
| 1023 | KEGG | KEGG:0110 | Metabolic pathways | 2,68766E-06 | positive_corr | True | 1482 | 594 | 166 | 7788 | 0,279461279 | 0,112010796 | ['KEGG:00000'] |
| 1031 | KEGG | KEGG:0120 | Carbon metabolism | 3,73262E-06 | positive_corr | True | 117 | 594 | 28 | 7788 | 0,047138047 | 0,239316239 | ['KEGG:00000'] |
| 819 | KEGG | KEGG:0028 | Valine, leucine and isoleucine degradation | 4,09131E-06 | positive_corr | True | 48 | 594 | 17 | 7788 | 0,028619529 | 0,354166667 | ['KEGG:00000'] |
| 1183 | KEGG | KEGG:0510 | Bacterial invasion of epithelial cells | 3,37105E-05 | positive_corr | True | 73 | 594 | 20 | 7788 | 0,033670034 | 0,273972603 | ['KEGG:00000'] |
| 721 | KEGG | KEGG:0472 | Neurotrophin signaling pathway | 0,000278162 | positive_corr | True | 119 | 594 | 25 | 7788 | 0,042087542 | 0,210084034 | ['KEGG:00000'] |
| 1052 | KEGG | KEGG:0521 | Renal cell carcinoma | 0,001021758 | positive_corr | True | 68 | 594 | 17 | 7788 | 0,028619529 | 0,25 | ['KEGG:00000'] |
| 1055 | KEGG | KEGG:0467 | Leukocyte transendothelial migration | 0,001047171 | positive_corr | True | 112 | 594 | 23 | 7788 | 0,038720539 | 0,205357143 | ['KEGG:00000'] |
| 1073 | KEGG | KEGG:0451 | Focal adhesion | 0,001577077 | positive_corr | True | 198 | 594 | 33 | 7788 | 0,055555556 | 0,166666667 | ['KEGG:00000'] |
| 820 | KEGG | KEGG:0062 | Pyruvate metabolism | 0,00225874 | positive_corr | True | 39 | 594 | 12 | 7788 | 0,02020202 | 0,307692308 | ['KEGG:00000'] |
| 1016 | KEGG | KEGG:0453 | Tight junction | 0,002659865 | positive_corr | True | 168 | 594 | 29 | 7788 | 0,048821549 | 0,172619048 | ['KEGG:00000'] |
| 924 | KEGG | KEGG:0407 | Sphingolipid signaling pathway | 0,002958932 | positive_corr | True | 119 | 594 | 23 | 7788 | 0,038720539 | 0,193277311 | ['KEGG:00000'] |
| 851 | KEGG | KEGG:0002 | Citrate cycle (TCA cycle) | 0,005170173 | positive_corr | True | 30 | 594 | 10 | 7788 | 0,016835017 | 0,333333333 | ['KEGG:00000'] |
| 1067 | KEGG | KEGG:0401 | ErbB signaling pathway | 0,005388599 | positive_corr | True | 84 | 594 | 18 | 7788 | 0,03030303 | 0,214285714 | ['KEGG:00000'] |
| 899 | KEGG | KEGG:0461 | Platelet activation | 0,005849643 | positive_corr | True | 124 | 594 | 23 | 7788 | 0,038720539 | 0,185483871 | ['KEGG:00000'] |
| 1207 | KEGG | KEGG:0041 | beta-Alanine metabolism | 0,007117044 | positive_corr | True | 31 | 594 | 10 | 7788 | 0,016835017 | 0,322580645 | ['KEGG:00000'] |
| 449 | KEGG | KEGG:0452 | Adherens junction | 0,0074205 | positive_corr | True | 71 | 594 | 16 | 7788 | 0,026936027 | 0,225352113 | ['KEGG:00000'] |
| 1160 | KEGG | KEGG:0121 | Fatty acid metabolism | 0,007772032 | positive_corr | True | 57 | 594 | 14 | 7788 | 0,023569024 | 0,245614035 | ['KEGG:00000'] |
| 471 | KEGG | KEGG:0007 | Fatty acid degradation | 0,00855174 | positive_corr | True | 44 | 594 | 12 | 7788 | 0,02020202 | 0,272727273 | ['KEGG:00000'] |
| 583 | KEGG | KEGG:0414 | Protein processing in endoplasmic reticulum | 0,014523542 | positive_corr | True | 166 | 594 | 27 | 7788 | 0,045454545 | 0,162650602 | ['KEGG:00000'] |
| 916 | KEGG | KEGG:0481 | Regulation of actin cytoskeleton | 0,015450171 | positive_corr | True | 212 | 594 | 32 | 7788 | 0,053872054 | 0,150943396 | ['KEGG:00000'] |
| 470 | KEGG | KEGG:0064 | Propanoate metabolism | 0,01706713 | positive_corr | True | 34 | 594 | 10 | 7788 | 0,016835017 | 0,294117647 | ['KEGG:00000'] |
| 226 | KEGG | KEGG:0513 | Salmonella infection | 0,018474665 | positive_corr | True | 214 | 594 | 32 | 7788 | 0,053872054 | 0,14953271 | ['KEGG:00000'] |
| 1142 | KEGG | KEGG:0027 | Cysteine and methionine metabolism | 0,021299463 | positive_corr | True | 48 | 594 | 12 | 7788 | 0,02020202 | 0,25 | ['KEGG:00000'] |
| 1152 | KEGG | KEGG:0414 | Autophagy - animal | 0,025141963 | positive_corr | True | 136 | 594 | 23 | 7788 | 0,038720539 | 0,169117647 | ['KEGG:00000'] |
| 1324 | WP | WP:WP388 | VEGFA-VEGFR2 Signaling | 5,46469E-08 | positive_corr | True | 428 | 563 | 75 | 6646 | 0,13321492 | 0,175233645 | ['WP:000000'] |
| 1139 | WP | WP:WP313 | Signaling of Hepatocyte Growth Factor Receptor | 0,001813037 | positive_corr | True | 34 | 563 | 12 | 6646 | 0,021314387 | 0,352941176 | ['WP:000000'] |
| 1314 | WP | WP:WP465 | Gastrin Signaling Pathway | 0,003363592 | positive_corr | True | 114 | 563 | 24 | 6646 | 0,042628774 | 0,210526316 | ['WP:000000'] |
| 1168 | WP | WP:WP673 | ErbB Signaling Pathway | 0,007207058 | positive_corr | True | 90 | 563 | 20 | 6646 | 0,035523979 | 0,222222222 | ['WP:000000'] |
| 1089 | WP | WP:WP420 | MET in type 1 papillary renal cell carcinoma | 0,010220419 | positive_corr | True | 58 | 563 | 15 | 6646 | 0,026642984 | 0,25862069 | ['WP:000000'] |
| 962 | WP | WP:WP306 | Focal Adhesion | 0,015334927 | positive_corr | True | 197 | 563 | 33 | 6646 | 0,058614565 | 0,16751269 | ['WP:000000'] |
| 304 | WP | WP:WP429 | Metabolic reprogramming in colon cancer | 0,019455493 | positive_corr | True | 42 | 563 | 12 | 6646 | 0,021314387 | 0,285714286 | ['WP:000000'] |
| 871 | WP | WP:WP392 | Amino Acid metabolism | 0,026889184 | positive_corr | True | 91 | 563 | 19 | 6646 | 0,03374778 | 0,208791209 | ['WP:000000'] |
| 1215 | WP | WP:WP245 | TCA Cycle and Deficiency of Pyruvate Dehydrogenase complex (PDHc) | 0,027096987 | positive_corr | True | 16 | 563 | 7 | 6646 | 0,012433393 | 0,4375 | ['WP:000000'] |
| 1219 | WP | WP:WP244 | Alpha 6 Beta 4 signaling pathway | 0,041903846 | positive_corr | True | 33 | 563 | 10 | 6646 | 0,017761989 | 0,303030303 | ['WP:000000'] |
| 1300 | WP | WP:WP328 | Copper homeostasis | 0,044943287 | positive_corr | True | 52 | 563 | 13 | 6646 | 0,023090586 | 0,25 | ['WP:000000'] |
| 1053 | WP | WP:WP368 | Association Between Physico-Chemical Features and Toxicity Associated Pathways | 0,048774641 | positive_corr | True | 66 | 563 | 15 | 6646 | 0,026642984 | 0,227272727 | ['WP:000000'] |
